# Unroasted and Roasted Coffee-Derived Extracellular Vesicles Inhibit Proliferation and Migration of SK-MEL-28 Melanoma Cells via Distinct Molecular Pathways

**DOI:** 10.1101/2025.10.27.684740

**Authors:** Ela Doruk Korkmaz, Seren Kucuk Vardari, Ilgın Işıltan, Benan Temizci

## Abstract

Melanoma is among the most common cancers in both men and women, and it can spread earlier and more aggressively than other skin cancers, driving demand for innovative and biocompatible therapeutic strategies. Coffee, one of the most widely consumed beverages worldwide, has been epidemiologically linked to a reduced risk of several cancers, including melanoma. It contains bioactive compounds with reported anti-cancer properties; however, the molecular mechanisms underlying its potential protective effects against melanoma remain insufficiently explored. Plant-derived extracellular vesicles (EVs) have emerged as promising natural nanocarriers due to their high stability, cellular uptake efficiency, and low toxicity. Here, we report the first comparative investigation of EVs isolated from unroasted and roasted *Coffea arabica* beans and their anti-cancer effects on SK-MEL-28 melanoma cells. Both EV types selectively reduced SK-MEL-28 cell viability, induced apoptosis, suppressed tumor spheroid growth, and impaired cellular migration. Mechanistically, unroasted coffee EVs downregulated *SERPINA1* and inhibited the PI3K/AKT pathway, whereas roasted coffee EVs attenuated MAPK signaling by reducing BRAF phosphorylation. These findings demonstrate that coffee-derived EVs exhibit potent anti-melanoma activities through distinct oncogenic pathways and identify them as edible, naturally occurring nanocarriers with therapeutic potential against skin cancer.

## 1. Introduction

Cancer is driven by genetic and epigenetic changes influenced by diet, environment, and lifestyle. Diet is the most significant modifiable factor affecting cancer risk, capable of inducing epigenetic changes through DNA methylation, histone modification, and microRNA regulation [1]. Evidence shows that sustainable, plant-based diets are associated with lower mortality and reduced risks of cancer, likely due to their high content of fiber, vitamins, antioxidants, and phytochemicals, which reduce inflammation and protect against chronic diseases [2,3]. Natural chemopreventive compounds, especially phytochemicals, have shown potential in reversing cancer-related epigenetic changes and contributing antioxidant, anti-inflammatory, and anti-cancer effects [4–6]. Studies on post-diagnostic diets in cancer patients show that eating patterns rich in fruits, vegetables, whole grains, legumes, nuts, and low-fat dairy are linked to better outcomes, including lower all-cause and cancer-specific mortality [7–9].

Bitter-tasting foods like beans, nuts, and coffee have proven biological activities, including antihyperlipidemic, antihypertensive, anti-inflammatory, antioxidant, and anti-tumor effects [10]. Coffee, globally the most widely consumed bitter and stimulant beverage, is also a functional food rich in over 1,000 bioactive compounds, including chlorogenic acids, caffeine, and polyphenols [11]. These compounds contribute to its antioxidant, anti-inflammatory, and anti-cancer properties. Epidemiological evidence links coffee consumption, including decaffeinated varieties, to reduced risk of several cancers such as liver, endometrial, prostate, colorectal, melanoma, and breast [12].

Despite coffee’s potential health benefits in preventing cancer and other chronic diseases, the low bioavailability of active phytochemicals of coffee, the fraction reaching circulation in active form, limits its therapeutic efficacy. Plant-derived bioactive compounds, such as polyphenols, flavonoids, and alkaloids found in coffee, suffer from poor oral bioavailability, rapid degradation in the gastrointestinal tract, and low intestinal permeability, resulting in insufficient systemic exposure to exert therapeutic effects [6,13,14]. Some of the current studies have focused on innovative delivery systems such as liposome-based encapsulations of coffee extracts to enhance bioavailability of phytochemicals [15,16]. While artificial encapsulations are limited by concerns related to safety, cost, and scalability, plant-derived extracellular vesicles (EVs) have recently emerged as safe, biocompatible, and cost-effective alternatives to the delivery of bioactive compounds and pharmaceuticals, addressing a major challenge that limits the clinical impact of many phytochemicals [17].

Plant-derived EVs, lipid bilayer vesicles naturally secreted by plants, biochemically resemble their parent cells, and contain RNAs, proteins, lipids, and many other biological components that the parental cells have. They also contain higher concentrations of bioactive compounds compared to the extract forms, and confer enhanced bioavailability of natural bioactive phytochemicals. Plant-derived EVs offer an innovative solution by functioning as biocompatible, stable and having the potential to infiltrate mammalian cells to regulate biological activities. They convey immense potential as cancer therapeutics due to their low immunogenicity, high intracellular intake rate, and low toxicity [18–23].

While habitual coffee consumption has been shown to have protective effects and an inverse correlation with different types of skin cancers such as basal cell carcinoma, squamous cell carcinoma, and melanoma in many cohort studies [24–27], topical use of coffee compounds has also been investigated in skin cancers. Topical application of caffeine in UV-pretreated high-risk mice was shown to inhibit UV-induced squamous cell carcinomas in hairless mice, indicating caffeine’s capacity to eliminate UV-damaged cells through apoptosis in prevention of UV-induced skin cancers [28,29]. Another study found that topical application of spent coffee ground extracts in solution prevented UVB-induced photoaging by down-regulating matrix metalloproteinases; therefore, upholding caffeine’s potential as an anti-photoaging and anti-skin cancer agent [30], while another study indicated the potential of non-caffeine metabolites, including anti-inflammatory phytochemicals as the key mediators in decreasing non-melanoma skin cancer risk [31]. Likewise, coffee extracts and its isolated active biocompounds have been investigated related to other dermo-cosmetic applications, such as incorporation into anti-pollution film-forming facial sprays; inclusion in cosmetic products aimed at preventing chronic inflammatory skin conditions like eczema and telangiectasia, pivotal identifying feature of basal cell carcinoma; and use in skin-whitening products through the inhibition of melanin production [32–35]. All these findings highlight the potential of coffee as a multifunctional, sustainable cosmeceutical ingredient, offering skin protection, making it a promising candidate for skin health applications.

Although cohort studies have suggested that coffee consumption reduces the incidence of melanoma, molecular investigations into its anti-cancer mechanisms remain limited. Building on the extensive epidemiologic data linking coffee consumption with reduced skin cancer risk, as well as findings suggesting that topical application of coffee may help protect skin health and prevent skin cancers, this study aimed to investigate the anti-skin cancer effects of coffee by examining coffee-derived extracellular vesicles as nano-carriers of bioactive compounds in SK-MEL-28 melanoma cells, a widely used model for studying cutaneous melanoma, particularly regarding the effects of coffee, which remains relatively underexplored in the literature. While coffee is typically consumed roasted, research shows that roasting levels influence anti-cancer activity. Green coffee demonstrated the strongest effect against oral squamous cell carcinoma [36], while lightly roasted coffee was more effective in colon cancer cells [37]. Therefore, we used EVs from both unroasted and roasted beans of *Coffea arabica*, which is one of the most consumed coffee types [38], to investigate their anti-cancer effects comparatively on SK-MEL-28 melanoma cells.

This study aimed to investigate the anti–skin cancer effects of coffee by examining coffee-derived extracellular vesicles as nano-carriers of bioactive compounds in SK-MEL-28 melanoma cells, a widely used model for studying cutaneous melanoma that remains relatively underexplored in the literature.

In this study, we demonstrated for the first time that extracellular vesicles from both unroasted and roasted coffee exhibit anti-cancer effects in SK-MEL-28 melanoma cells, including reduced cell viability through apoptosis induction, suppression of tumor growth, and decreased cell migration, although via distinct molecular mechanisms.

## 2. Materials and Methods

### 2.1. Isolation and Characterization of Coffee Extracellular Vesicles

Unroasted and roasted coffee Arabica beans (Arifoglu, Istanbul Turkey) were ground and incubated in a two-fold volume of 1X Dulbecco’s Phosphate-Buffered Saline (DPBS) (Gibco Life Technologies) for 1 hour at 4 °C. After filtration through gauze and glass filters to remove debris, the extracts were sequentially centrifuged at 300 x g for 10 minutes, 3000 x g for 15 minutes, and finally at 10,000 x g for 30 minutes, all at 4 °C. The supernatants were passed through a 0.45 µm filter (Millipore). Coffee extracellular vesicles (EVs) were then purified and concentrated using a tangential flow filtration system (TFF-EVs-S, Hansa Biomed Life Sciences). TFF was performed by the peristaltic pump Masterflex L/S, model 7535-04, operating with a flow velocity of 80 ml/min. The retentate, containing EVs, was washed 3 times with 50 ml 1X PBS, and then was recovered in 1X PBS buffer.

Particle size and zeta potential of the coffee EVs were measured by nanoparticle tracking analysis (NTA) using a ZetaView PMX-120 instrument (Particle Metrix) as described in Fortunato et al. (2021) [39]. The system was calibrated with 100 nm polystyrene beads, and diluted samples (in 1X PBS) were analyzed across 11 positions under standardized conditions (Camera sensitivity 85, shutter speed 100, high video quality (capturing 60 frames) at 30 frames/s (1 cycle), minimal area 10, maximal area 1000, and brightness 25). The ZetaView PMX-120 determines particle size and concentration by monitoring the Brownian motion of individual particles in video sequences, deriving their diffusion coefficients, and converting these into diameters via the Stokes–Einstein equation. Concentration is calculated by counting the particles detected within the instrument’s calibrated viewing volume and correcting for dilution, while size distribution is obtained by compiling the measured diameters into a histogram.

To further characterize the coffee EVs, a typical plant EV marker TET8 (Tetraspanin 8) protein levels were quantified by ELISA. Briefly, coffee EVs were bound to high-binding plates (Hansa Biomed Life Sciences) overnight at 37 °C in a humid chamber. Unbound material was washed with PBS-T (0.5% Tween in PBS), and wells were blocked with 3% bovine serum albumin (Capricorn Scientific) in PBS for 2 hours at room temperature. After washing, wells were incubated with rabbit anti-TET8 antibody (1:1000, PhytoAB, PAB-PHY1490S) for 2 hours, rinsed with PBS, and treated with HRP-conjugated anti-rabbit secondary antibody (1:5000, Bio-Rad, #1706515) for 1 hour. Following three PBS-T washes, 100 µL TMB (3,3’,5,5’-tetramethylbenzidine) substrate (Abcam) was added for 20 minutes, stopped with 100 µL H₂SO₄ (Sigma-Aldrich), and read at 450 nm on a Varioskan LUX reader. PBS served as blank, and results were expressed as fold change over background (OD₄₅₀ sample / OD₄₅₀ blank).

EVs were prepared for electron microscopy (EM) as in Puhka et al. (2017) [40] by loading onto carbon coated and glow discharged 200 mesh copper grids with pioloform support membrane. EVs were fixed with 2.0% paraformaldehyde (PFA) in sodium phosphate (NaPO_4_) buffer, stained with 2% neutral uranyl acetate, further stained, and embedded in uranyl acetate and methyl cellulose mixture (1.8/0.4%). EVs were viewed with transmission EM using TEM Hitachi HT7800 operating at 100 kV. Images were taken with Gatan Rio9 bottom mounted CMOS-camera, model 1809 (Gatan Inc., USA) with 3072 x 3072 pixels image size and no binning.

### 2.2. Cell Culture

MEL-ST melanocytes [41,42] and SK-MEL-28 melanoma cells (RRID:CVCL_0526, HTB-72) (both kindly provided by Prof. Dr. N.C. Tolga Emre) were cultured at 37°C with 5% CO_2_ in Dulbecco’s Modified Eagle Medium (DMEM; Gibco Life Technologies) supplied with 10% Fetal Bovine Serum (FBS; Capricorn Scientific) and 1% penicillin-streptomycin (Capricorn Scientific). When the cells achieved around 70% confluence, they were passaged using 2.5% trypsin (Gibco Life Technologies).

### 2.3. Cellular Uptake

The cellular uptake of coffee EVs was examined by confocal microscopy using DIOC18 lipophilic carbocyanine dye. For this aim, SK-MEL-28 cells were seeded at 5×10^4^ cells/coverslip onto coverslips coated with 0.01% poly-L-Lysine in 12-well plates. After overnight incubation, unroasted and roasted coffee EVs were stained with DIOC18 dye (Biotium), a green fluorescent dye, at 37°C for 45 minutes. Excessive dye was removed using 100-kDa Amicon ultracentrifugal filters. Then, cells were treated with fluorescently-labelled unroasted or roasted coffee EVs for 24 hours. Untreated cells were used as a negative control. After 24 hours incubation, the cells were fixed with 4% paraformaldehyde for 15 minutes, and washed three times with 1X PBS. Then, the cells were blocked with 3% (w/v) bovine serum albumin in 1X PBS containing 0.1% Triton X-100 for 1 hour at room temperature. Following, the cells were stained with phalloidin (1:40, Sigma Aldrich) to label actin cytoskeleton for 1 hour at room temperature, and then washed with 1X PBS. Finally, the coverslips were mounted with PBS on slides, and cells were visualized by TCS SP2 SE Confocal Microscope (Leica) with a 63X oil immersion objective. All images were captured in a single focal plane using Leica Application Suite X (LAS X) software (version 1.4.5, RRID:SCR_013673).

### 2.4. MTT Cell Viability Assay

To investigate the cytotoxic effects of coffee EVs on melanocytes and melanoma cancer cells, both MEL-ST and SK-MEL-28 cells were seeded at a density of 5×10^3^ cells/well in DMEM containing 5% FBS into 96-well plates. After overnight incubation to allow adherence, cells were treated with varying concentrations of unroasted or roasted coffee EVs (0 - 8 x 10^8^ particles/µl) for 24 hours. Following, 10 µl of MTT reagent (3-(4, 5-dimethylthiazolyl-2)-2,5-diphenyltetrazolium bromide; Millipore) at 5 mg/ml prepared in 1X PBS was added to each well and incubated at 37°C for 4 hours. Formazan salts were then dissolved in 200 µL of DMSO (dimethyl sulfoxide), and absorbance was measured at 570 nm with a reference wavelength of 655 nm using a Bio-Rad Benchmark Plus microplate reader (Bio-Rad Laboratories).

### 2.5. DAPI Staining of Nuclei

MEL-ST and SK-MEL-28 cells were seeded at a density of 5×10^4^ cells/coverslip onto 0.01% poly-L-Lysine (Sigma-Aldrich) coated coverslips in 24-well plates. After overnight incubation, the cells were treated with 4×10^8^ particles/µl unroasted or roasted coffee EVs for 24 hours. Next, the cells were fixed with 4% paraformaldehyde for 15 minutes and washed with 1X PBS. They were then incubated with DAPI (at a 1:1000 dilution in PBS, Invitrogen) for 10 minutes at room temperature. After washing with 1X PBS, ProLong™ Diamond Antifade Mounting Medium (Invitrogen) was added onto slides, and coverslips were placed onto the mounting medium. Prepared samples were observed using a TCS SP2 SE Confocal Microscope (Leica) with a 63X oil immersion objective. All images were captured in a single focal plane using Leica Application Suite X (LAS X) software (version 1.4.5, RRID:SCR_013673).

### 2.6. 7-AAD and Annexin-V Apoptosis Detection Assay

Flow cytometry analysis was performed after dual-staining with 7-AAD (7-Aminoactinomycin D) and Annexin-V to determine whether unroasted or roasted coffee EVs induced apoptosis in melanoma cells. SK-MEL-28 cells were seeded into 24-well plates at 5×10^4^ cells/well density and incubated overnight. The cells were then treated with 4×10^8^ particles/µl unroasted or roasted coffee EVs. After 24 hours, the cells were trypsinized and centrifuged at 370 x g for 5 minutes. Cell pellets were washed twice with 200 µl ice-cold FACS (Fluorescence-Activated Cell Sorting) buffer (2% FBS in 1X PBS) and stained sequentially with 7-AAD (IM3422, Beckman Coulter) and/or Alexa Fluor 647-conjugated Annexin-V (422201, BioLegend). For 7-AAD staining, the cell pellets were dissolved in 50 µl 7-AAD staining solution (prepared by mixing 5 µL 7-AAD with 45 µL FACS buffer) and incubated in the dark at 4°C for 15 minutes. After incubation, 150 µl of FACS buffer was added to each tube, and the cells were centrifuged. Subsequently, the cells were stained with 100 µl Annexin-V solution (1 µl Alexa Fluor 647-conjugated Annexin-V diluted in 99 µl Annexin-V binding buffer) and incubated in the dark at 4°C for 20 minutes. After centrifugation, cell pellets were resuspended in 200 µl Annexin-V binding buffer (BioLegend). Unstained cells were used as negative controls, while the cells stained only with 7-AAD or Annexin-V served as positive controls. Finally, the cells were analyzed with flow cytometry (BD Accuri C6), and the percentage of apoptotic cells was calculated using FlowJo software (version 10, RRID:SCR_008520).

### 2.7. 3D Spheroid Formation

To determine whether coffee EVs inhibit tumor growth, 3D spheroids were generated using SK-MEL-28 melanoma cells. For this purpose, SK-MEL-28 melanoma cells were harvested from monolayer cultures, and 7.5×10^3^ cells were seeded into each well of 96-well ultra-low attachment plates (Thermo Fisher Scientific). The plates were centrifuged at 500 x g for 5 minutes to facilitate spheroid formation and incubated at 37°C, 5% CO_2_. After approximately 3 days, when the spheroids had formed, SK-MEL-28 spheroids were treated with 4×10^8^ particles/µl unroasted or roasted coffee EVs. Seeding of cells was designated as day -3, and the treatment as day 0. Spheroids were imaged daily under a Zeiss Axiovert A1 fluorescence microscope using a 5x objective until the fourth day after treatment. To assess how EV treatment affected spheroid size, the area of spheroids was measured using Image J software (version 1.54, RRID:SCR_003070).

### 2.8. Wound Healing Assay

To examine whether coffee EVs affect the migratory ability of SK-MEL-28 melanoma cells, cells were seeded at 8×10^5^ cells/well density in a 6-well plate. After the cells reached 90-95% confluency, the cell layers were scraped using a 10 µl pipette tip to create a straight line, and the wells were washed with 1X PBS. After 24 hours, cells were treated with unroasted or roasted coffee EVs at 4×10^8^ particles/µl, and all cells were incubated for up to 48 hours. Scratch closure was monitored daily under a Zeiss Axiovert A1 fluorescence microscope using a 5x objective. The scratch closure area was calculated by using the wound healing size tool plugin [43] of the Image J software (version 1.54, RRID:SCR_003070). Closure percentages in treated cells were compared to those in control cells to determine the effect of coffee EVs application on migration.

### 2.9. RNA Isolation and Whole Transcriptome RNA Sequencing

RNA-sequencing-based whole transcriptome analysis was performed to identify differentially expressed genes after treatment with unroasted coffee EV in SK-MEL-28 melanoma cells. For this aim, first, total RNA extraction was performed. Briefly, SK-MEL-28 cells were seeded at 8×10^5^ cells/dish in a 100 mm cell culture dish and treated with 4×10^8^ particles/μL unroasted coffee EV for 24 hours. Untreated cells were used as the control group. Then, cells were harvested, and total RNA from control and unroasted coffee EV-treated SK-MEL-28 cells was isolated using MN Nucleospin RNA kit (Macherey-Nage) according to the manufacturer’s instructions. RNA concentrations were measured using a Nanodrop 2000 (Thermo Fisher Scientific). For transcriptome analysis, 1 μg of RNA from each sample was prepared in a total volume of 30 μl and dried in GenTegra RNA tubes for 24 hours using a biosafety hood. Transcriptome analysis, including library preparation, sequencing, and differential gene expression analysis, was performed by Macrogen. Briefly, libraries were prepared using the TruSeq stranded mRNA kit following the manufacturer’s instructions and sequenced on the NovaSeq 6000 platform to generate paired-end reads (2×100 bp). Standard bioinformatic and differential gene expression (DGE) analyses were performed.

### 2.10. Antibodies

Primary antibodies used for immunoblotting are as follows: SERPINA1 rabbit polyclonal antibody (1:800, Affinity Biosciences Cat#DF6182, RRID:AB_2838148), PTEN rabbit polyclonal antibody (1:500, Santa Cruz Biotechnology Cat#6817-R, RRID:AB_654892), phospho-Akt (Thr308) rabbit polyclonal antibody (1:500, Santa Cruz Biotechnology Cat#16646-R, RRID:AB_667742), Bcl-2 rabbit monoclonal antibody (1:1000, Cell Signaling Technology Cat#4223, RRID:AB_1903909), phospho-BRAF (Ser445) rabbit polyclonal antibody (1:1000, Cell Signaling Technology Cat#2696, RRID:AB_390721), phospho-MEK1/2 (Ser217/221) rabbit polyclonal antibody (1:1000, Cell Signaling Technology Cat#9121, RRID:AB_331648), Cyclin D1 rabbit polyclonal antibody (1:500, Santa Cruz Biotechnology Cat#sc-717, RRID:AB_631336), and β-actin mouse mAb (1:1000, Cell Signaling Technology Cat#3700, RRID: AB_2242334). Secondary antibodies used for immunoblotting are as follows: Anti-mouse IgG (H+L) (DyLight™ 800 4X PEG Conjugate) (1:15000, Cell Signaling Technology Cat#5257, RRID: AB_10693543), anti-rabbit IgG (H+L) (DyLight 800 Conjugate) (1:15000, Cell Signaling Technology Cat#5151, RRID: AB_10697505), anti-rabbit IgG (H+L) (DyLight 680 Conjugate) (1:15000, Cell Signaling Technology Cat#5366, RRID:AB_10693812), anti-mouse IgG (H+L) (DyLight 680 Conjugate) (1:15000, Cell Signaling Technology Cat#5470, RRID:AB_10696895).

### 2.11. Protein Isolation and Immunoblotting

Total proteins were isolated from SK-MEL-28 melanoma cells treated with unroasted or roasted coffee EVs. After 24 hours, cells were lysed using 1% NP40 buffer (150 mM NaCl, 1% NP-40, 50 mM pH 8 Tris-Cl), supplemented with protease and phosphatase inhibitor cocktails (Roche). Lysates were centrifuged at 12000 x g for 25 minutes at 4 °C, and the resulting supernatants were collected as total protein extracts. Protein concentrations were quantified using the Pierce™ BCA Protein Assay Kit (Thermo Fisher Scientific). 30 µg of total protein per sample was separated on a 12% SDS gel and transferred onto nitrocellulose membranes (Thermo Fisher Scientific). Membranes were blocked with 5% non-fat dry milk in 1X TBS-T for 1 hour at room temperature. The membranes were then incubated with the indicated primary antibodies overnight at 4°C and then with appropriate secondary antibodies at room temperature for 1 hour. After each primary and secondary antibody incubation, membranes were washed three times with 1X TBS-T. Protein bands were visualized using the LI-COR Odyssey CLx Near-Infrared Fluorescence Imaging System. Densitometric analyses were performed using Image Studio Lite (version 5.2, RRID: SCR_013715) and Adobe Photoshop CS5 (version 12.0, RRID: SCR_014199).

### 2.12. Statistical Analysis

GraphPad Prism software (version 8.0.1, RRID:SCR_002798) was used for the statistical analyses. The one-way or two-way ANOVA was utilized for multiple group comparisons; Student’s t-test was utilized for two-group comparisons. Error bars in the graphs were generated using ± standard error of the mean (SEM). Statistical significance was set at p<0.05 and defined as follows: n.s. (non-significant):p>0.05, *p<0.05, **p<0.01, ***p<0.001, ****p<0.0001.

## 3. Results

### 3.1. Characterization of Coffee EVs

To investigate the effects of coffee EVs on melanoma cells, EVs from unroasted and roasted beans of *Coffea arabica* were isolated. After isolation, the size distribution and surface zeta potentials of the EVs were analyzed using the NTA ZetaView PMX-120 NTA instrument (Particle Metrix) to verify their biophysical characteristics. The size of the coffee EVs ranged from 50 – 200 nm for unroasted beans (mean size 128.1 nm) and 50–150 nm for roasted beans (mean size 102.3 nm) (Fig. 1A). The zeta potential, an indicator of colloidal stability, was measured at approximately - 60 for unroasted and -65 mV for roasted EVs (in 0.1% PBS diluent), indicating that stable EVs were isolated (Fig. 1B). In addition, TET8 expression, a common marker of plant extracellular vesicles, was positively detected in both samples (Fig. 1C), confirming the successful isolation of EVs from coffee beans, and TEM images confirmed the presence of vesicular structures (Fig. 1D). All together these findings demonstrated that stable EV particles were successfully isolated from both types of coffee beans.

**Fig. 1.**
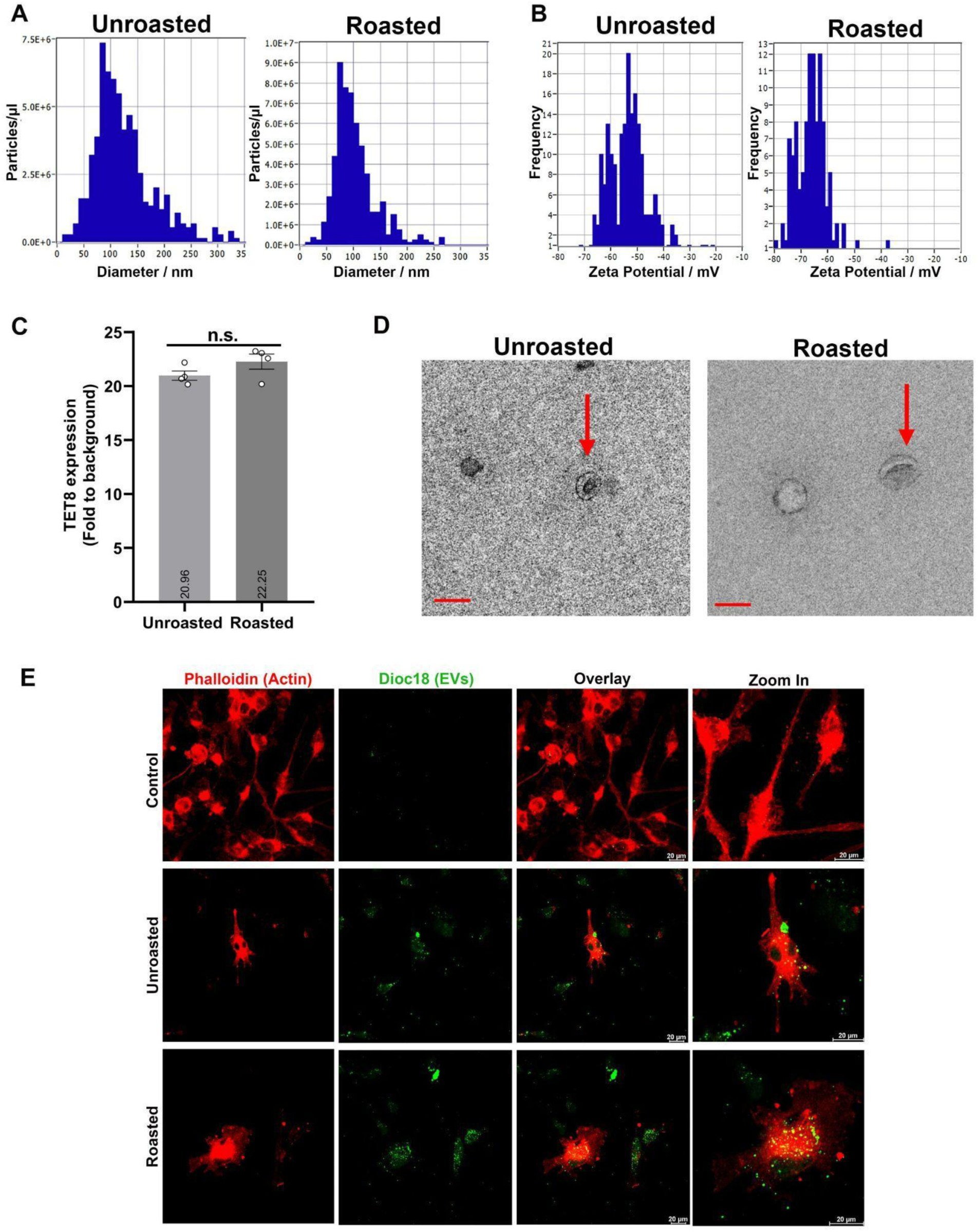
Characterization of unroasted and roasted coffee EVs. **A, B)** Particle size distribution (**A**) and surface zeta potential (**B**) were analyzed with the NTA ZetaView PMX-120 NTA instrument. **C)** TET8 expression, an EV marker of plant cells, was analyzed by ELISA, and represented as a bar graph (n=4). Statistical analysis was performed using a Student’s t-test. Data are presented as mean ± SEM. n.s. p>0.05. **D)** Representative TEM images of unroasted and roasted coffee EVs showing preserved vesicle morphology. Scale bar: 100 nm. **E)** Cellular uptake of unroasted and roasted coffee EVs was confirmed in SK-MEL-28 melanoma cells. Control cells were not treated with coffee EVs. DIOC18 dye (green) was used for EVs staining, and phalloidin (red) was used for labelling actin cytoskeleton. Cells were visualized using a TCS SP2 SE Confocal Microscope (Leica) with a 63X oil immersion objective. In each panel, the right images show higher magnification views. Scale bar: 20 μm.

Next, we examined whether coffee EVs were taken up by SK-MEL-28 cells to assess their potential effects. For this purpose, coffee EVs were stained with DiOC18, and then the cells were treated with DiOC18-labeled EVs for 24 hours. Immunocytochemistry results confirmed the cellular uptake of coffee EVs by SK-MEL-28 cells, as indicated by the intracellular presence of DiOC18-labeled EVs (Fig. 1E).

### 3.2. Coffee EVs Decrease Viability and Trigger Apoptosis in SK-MEL-28 Melanoma Cells

To investigate the effects of unroasted and roasted coffee EVs on SK-MEL-28 melanoma cells, the potential cytotoxic effects of coffee EVs were first examined on normal melanocytes. For this purpose, MEL-ST melanocytes were treated with unroasted and roasted coffee EVs across a range of doses (0 to 8×10^8^ particles/µl). 24 hours after treatment, the viability of melanocytes was assessed by an MTT assay. The results showed that unroasted or roasted coffee EVs did not affect the viability of healthy melanocyte MEL-ST cells at 2×10^8^ and 4×10^8^ particles/μl doses. In contrast, doses of 6×10^8^ and 8×10^8^ particles/μl doses of both unroasted and roasted coffee EVs significantly reduced MEL-ST melanocyte cell viability (Fig. 2A).

**Fig. 2.**
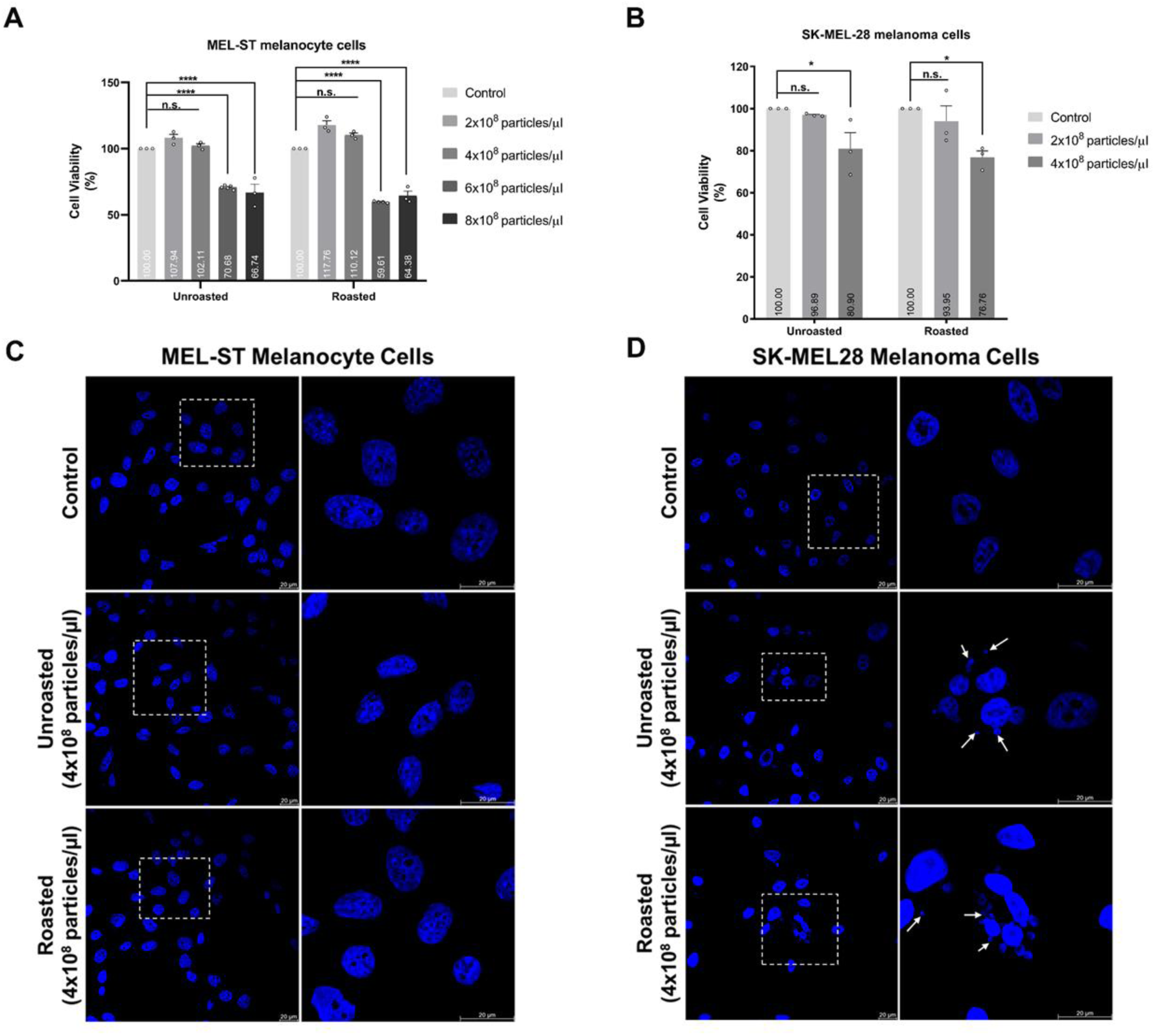
Coffee EVs decrease cell viability and disrupt nuclear morphology in SK-MEL-28 melanoma cells without affecting healthy MEL-ST melanocytes. **A, B)** MTT cell viability analysis of MEL-ST (**A**) and SK-MEL-28 (**B**) cells was performed following treatment with unroasted or roasted coffee EVs. Experiments were performed with three biological replicates, each containing five independent technical replicates. Statistical analysis was performed using a two-way ANOVA test. Data are presented as mean ± SEM. n.s. p>0.05, *p<0.05, ****p<0.0001. **C, D)** DAPI staining of the nuclei of MEL-ST (**C**) and SK-MEL-28 (**D**) cells was conducted to analyze nuclear morphology after treatment with unroasted or roasted coffee EVs. Cells were visualized using a TCS SP2 SE Confocal Microscope (Leica) with a 63X oil immersion objective. In each panel, the right images show higher magnification views of the boxed areas indicated in the images. Scale bar: 20 μm.

Given that healthy cells should remain unaffected during treatment with anti-cancer agents [44], selected concentrations up to 4×10⁸ particles/µL were chosen to assess their effects on SK-MEL-28 melanoma cells. MTT analysis revealed that treatment with unroasted and roasted coffee EVs at 2×10^8^ particles/μl did not alter the viability of SK-MEL-28 melanoma cells (Fig. 2B), similar to the result observed in MEL-ST melanocyte cells (Fig. 2A). However, treatment with either type of EVs at 4×10⁸ particles/µL resulted in a statistically significant decrease in SK-MEL-28 melanoma cell viability (Fig. 2B). Furthermore, there was no significant difference between unroasted and roasted coffee EVs in their effects on SK-MEL-28 melanoma cell viability (Fig. 2B).

After determining that 4×10^8^ particles/µl of both types of EVs did not affect MEL-ST melanocytes but did reduce SK-MEL-28 melanoma cell viability, possible alterations in the nuclei of MEL-ST melanocytes and SK-MEL-28 melanoma cells were analyzed using DAPI staining. As shown in Fig. 2C, no changes were observed in the nuclear structure of MEL-ST melanocytes with either type of coffee EVs. In contrast, unroasted and roasted coffee EVs altered the nuclear structure of SK-MEL-28 melanoma cells, forming apoptotic bodies, including membrane blebbing and cell shrinkage (Fig. 2D). These results indicated that unroasted and roasted coffee EVs triggered apoptosis in SK-MEL-28 melanoma cells.

Flow cytometry analysis utilizing 7-AAD and Annexin-V dual staining was performed to investigate the stages of apoptosis observed in SK-MEL-28 melanoma cells upon coffee EVs treatment. The results revealed that both types of coffee EVs significantly enhanced the percentage of late apoptotic SK-MEL-28 melanoma cells, with no significant difference in the effects of either coffee EVs (Fig. 3). In contrast, treatment with roasted coffee EVs application raised the early apoptotic cell population to a higher level than unroasted coffee EVs (Fig. 3), indicating that the roasted coffee EVs can increase the apoptosis earlier. Altogether, these results indicated that both types of coffee EVs reduced cell viability by triggering apoptosis in SK-MEL-28 melanoma cancer cells without affecting MEL-ST melanocytes.

**Fig. 3.**
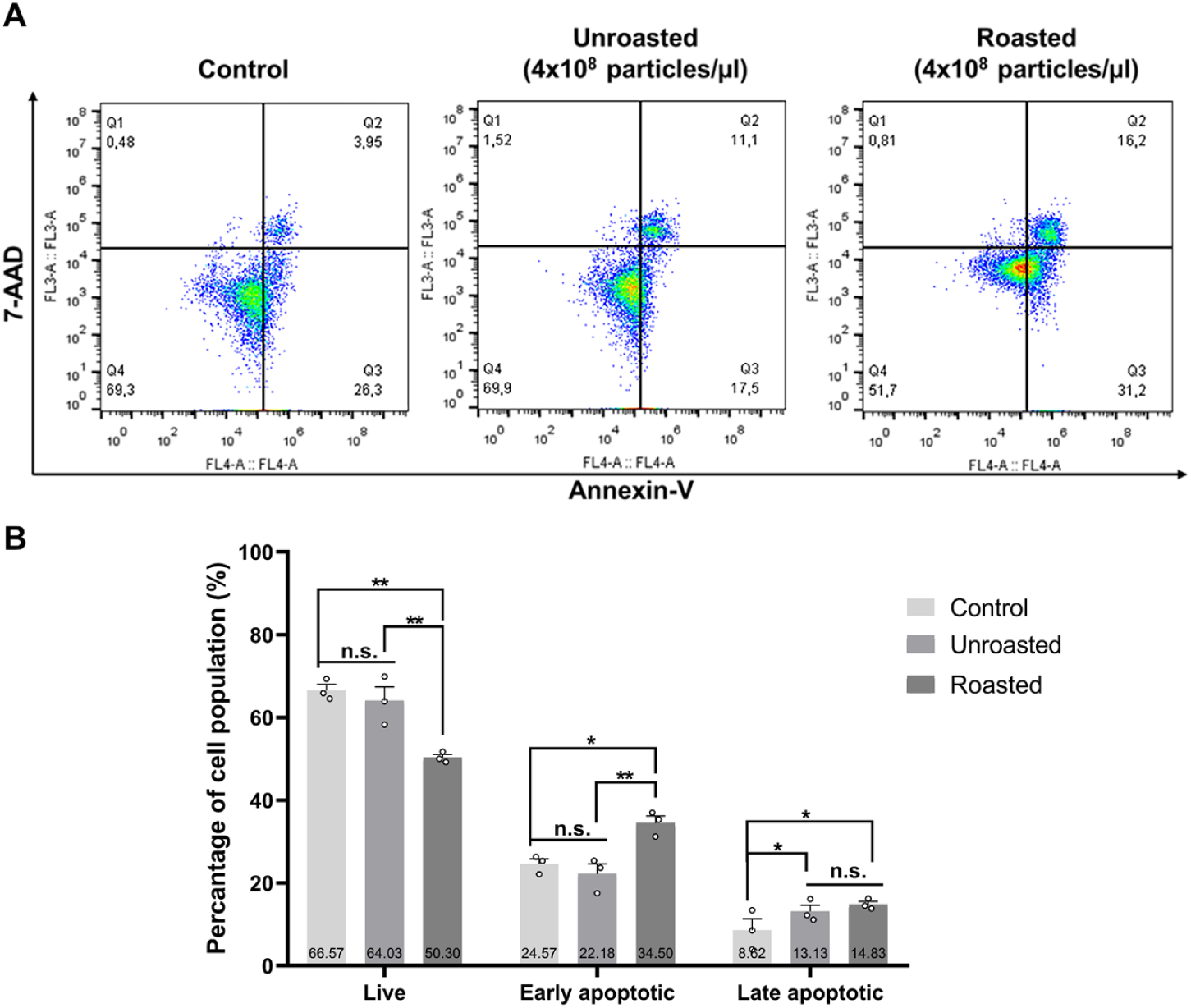
Coffee EV treatment induces apoptotic cell death in SK-MEL-28 melanoma cells. **A)** Flow cytometry analysis was performed to determine the stages of apoptosis following treatment with unroasted or roasted coffee EVs. Representative histograms from flow cytometry analysis are shown in the upper panel, with the Q1, Q2, Q3, and Q4 quadrants representing necrotic, late apoptotic, early apoptotic, and live cells, respectively. The experiment was performed with three biological replicates. **B)** The percentages of live, early apoptotic, and late apoptotic cells were quantified and are presented as a bar graph, as shown in the lower panel. Statistical analysis was performed by a one-way ANOVA test. Data are presented as mean ± SEM. n.s. p>0.05, *p<0.05, **p<0.01

### 3.3. Coffee EVs Reduce Tumor Size in 3D Spheroid Models of SK-MEL-28 Melanoma Cells

Following the finding that coffee EVs induce apoptosis in SK-MEL-28 melanoma cells, we investigated their effects on the growth of SK-MEL-28 melanoma tumors using a 3D spheroid formation assay. For this purpose, SK-MEL-28 melanoma spheroids were treated with either unroasted or roasted coffee EVs, and the images of control and treated spheroids were captured daily to compare their growth (Fig. 4A). The spheroid areas were quantitatively analyzed following the treatment. The results showed that the growth of SK-MEL-28 melanoma spheroids treated with unroasted coffee EVs was significantly slower than that of the control spheroids starting from day 2 after treatment (Fig. 4B). In contrast, the growth of spheroids treated with roasted coffee EVs was suppressed from day 1, with their area remaining constant for two days post-treatment. Although there was a slight increase in size after day 2, it remained significantly smaller than the control spheroids (Fig. 4B). These findings indicate that while both types of coffee EVs had similar inhibitory effects on SK-MEL-28 spheroid growth at later time points, roasted coffee EVs suppressed spheroid growth more rapidly, whereas unroasted coffee EVs required longer time to elicit a comparable response.

**Fig. 4.**
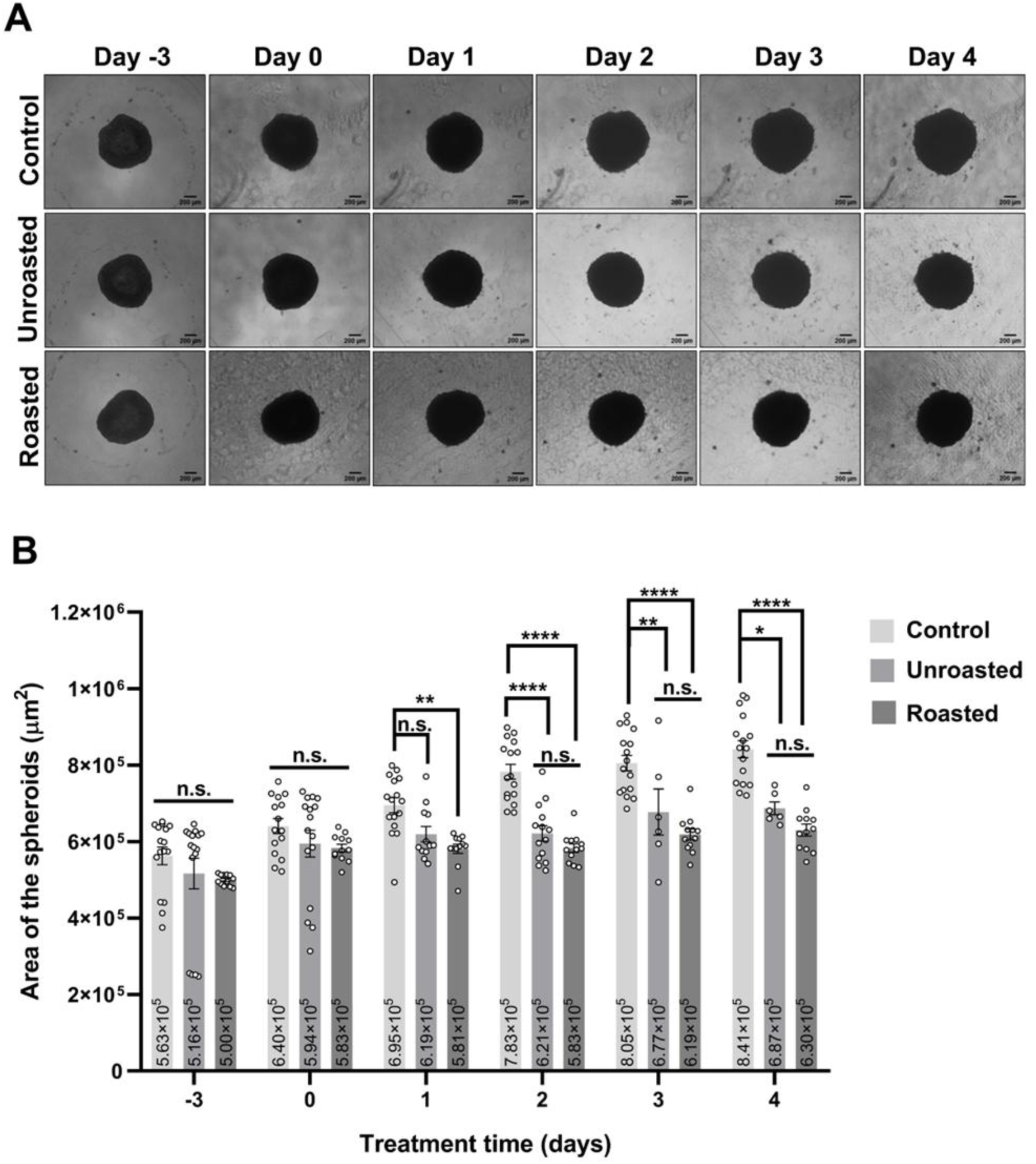
Treatment with coffee EVs mitigates the growth of SK-MEL-28 spheroids. **A)** Representative images of the control, unroasted coffee EV-treated and roasted coffee EV-treated SK-MEL-28 melanoma spheroids from day -3 to day 4. Day -3: Spheroids after formation; Day 0: Spheroids just before treatment; Day 1, 2, 3, and 4: Spheroids after treatment. Scale bar: 200 μm. The experiment was performed with two biological replicates, each containing at least three independent technical replicates. **B)** The areas of spheroids were quantified and represented in the bar graph (n≥6). Statistical analysis was performed by a two-way ANOVA test. Data are presented as mean ± SEM. n.s. p>0.05, *p<0.05, **p<0.01, ****p<0.0001.

### 3.4. Coffee EVs Decrease the Migration Capacity of SK-MEL-28 Melanoma Cells

After demonstrating that both coffee EVs hinder SK-MEL-28 melanoma spheroid growth, we investigated their impact on the migratory capacity of SK-MEL-28 melanoma cells. To assess the effect of coffee EVs on cell migration, a wound healing assay was performed on SK-MEL-28 melanoma cells treated for 24 hours with either type of coffee EVs, alongside untreated controls. Images of the scratch area were taken over a 48-hour period, and the percentage of closure in cells treated with coffee EVs was compared to that in control cells using ImageJ software. The results showed that while both coffee EVs reduced cell migration, the roasted coffee EV had a greater inhibitory effect at 24 hours (Fig. 5). However, by 48 hours, the level of migration inhibition was comparable for the two coffee EVs (Fig. 5B). Although both coffee EVs exhibited similar inhibitory effects on cell migration at 48 hours, the difference observed at the 24-hour time point suggests that unroasted and roasted coffee EVs may exert their effects through distinct molecular pathways.

**Fig. 5.**
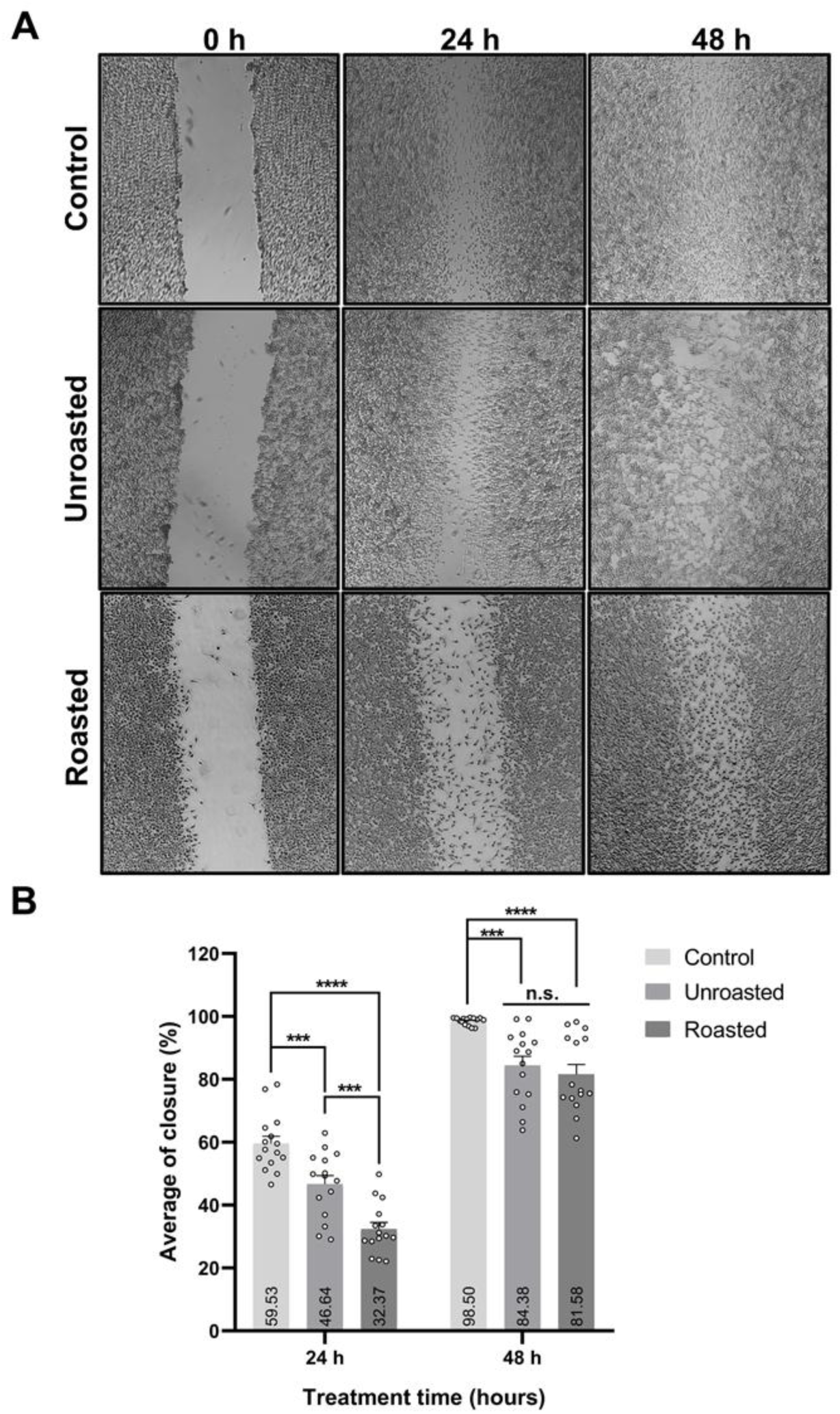
SK-MEL-28 melanoma cell migration capability is decreased upon coffee EV treatment. **A)** Representative images from in vitro scratch wound healing assays of control, unroasted coffee EV-treated, or roasted coffee EV-treated SK-MEL-28 melanoma cells from 0 to 48 hours. The experiment was performed with two biological replicates, each containing at least seven independent technical replicates. **B)** The percentage of closure was quantified and represented in the bar graph (n=15). Statistical analysis was performed by a two-way ANOVA test. Data are presented as mean ± SEM. n.s. p>0.05, ***p<0.001, ****p<0.0001.

### 3.5. Unroasted and Roasted Coffee EVs Exhibit Their Effects on SK-MEL-28 Cells by Activating Different Signaling Pathways

While roasted coffee EVs induced early apoptosis to a greater extent than unroasted EVs, both types exhibited comparable suppressive effects on spheroid tumor growth in long-term analyses. Similarly, while roasted coffee EVs more effectively impaired SK-MEL-28 melanoma cell migration during the initial 24-hour period, this difference diminished over time, and the effects of both EV types became comparable at later time points. These observations led us to hypothesize that exosomes isolated from roasted or unroasted coffee may exert their anti-cancer effects on SK-MEL-28 melanoma cells through different mechanisms.

To investigate transcriptional changes induced by exosome treatment, transcriptome profiling was performed on SK-MEL-28 melanoma cells treated with unroasted coffee EVs to identify gene expression alterations that could explain the observed effects. Notably, unroasted coffee EV treatment caused the most pronounced decrease (30-fold) in *SERPINA1* mRNA levels in SK-MEL-28 cells (Fig.S1).

SerpinA1 is a serine protease inhibitor known to promote tumorigenesis by suppressing PTEN, a negative regulator of the PI3K/AKT pathway [45]. Therefore, we examined the impact of decreased *SERPINA1* mRNA expression level on PI3K/AKT signaling. The results showed that SerpinA1 levels decreased significantly only in SK-MEL-28 cells treated with unroasted coffee EVs (Fig. 6A, B). This reduction in SerpinA1 expression led to a significant increase in PTEN expression upon unroasted coffee EV treatment, while there was no change in roasted coffee EVs-treated cells (Fig. 6A, C). Supporting these findings, only cells treated with unroasted coffee EVs showed a reduction in phosphorylated AKT at the Thr308 residue (Fig. 6A, D), which is triggered by active PI3K [46]. In contrast, no change in p-AKT levels was observed in roasted coffee EV-treated cells (Fig. 6A, D). Given these results, we further analyzed the expression of Bcl-2, an anti-apoptotic protein downstream of the PI3K/AKT [47]. Consistent with the p-AKT findings, Bcl-2 expression was markedly reduced only in the unroasted EV-treated group, while roasted EV treatment had no effect (Fig. 6A, E). Together, these results suggest that unroasted coffee EVs may exert anti-proliferative effects on SK-MEL-28 melanoma cells by suppressing the PI3K/AKT signaling pathway through downregulation of *SERPINA1*.

**Fig. 6.**
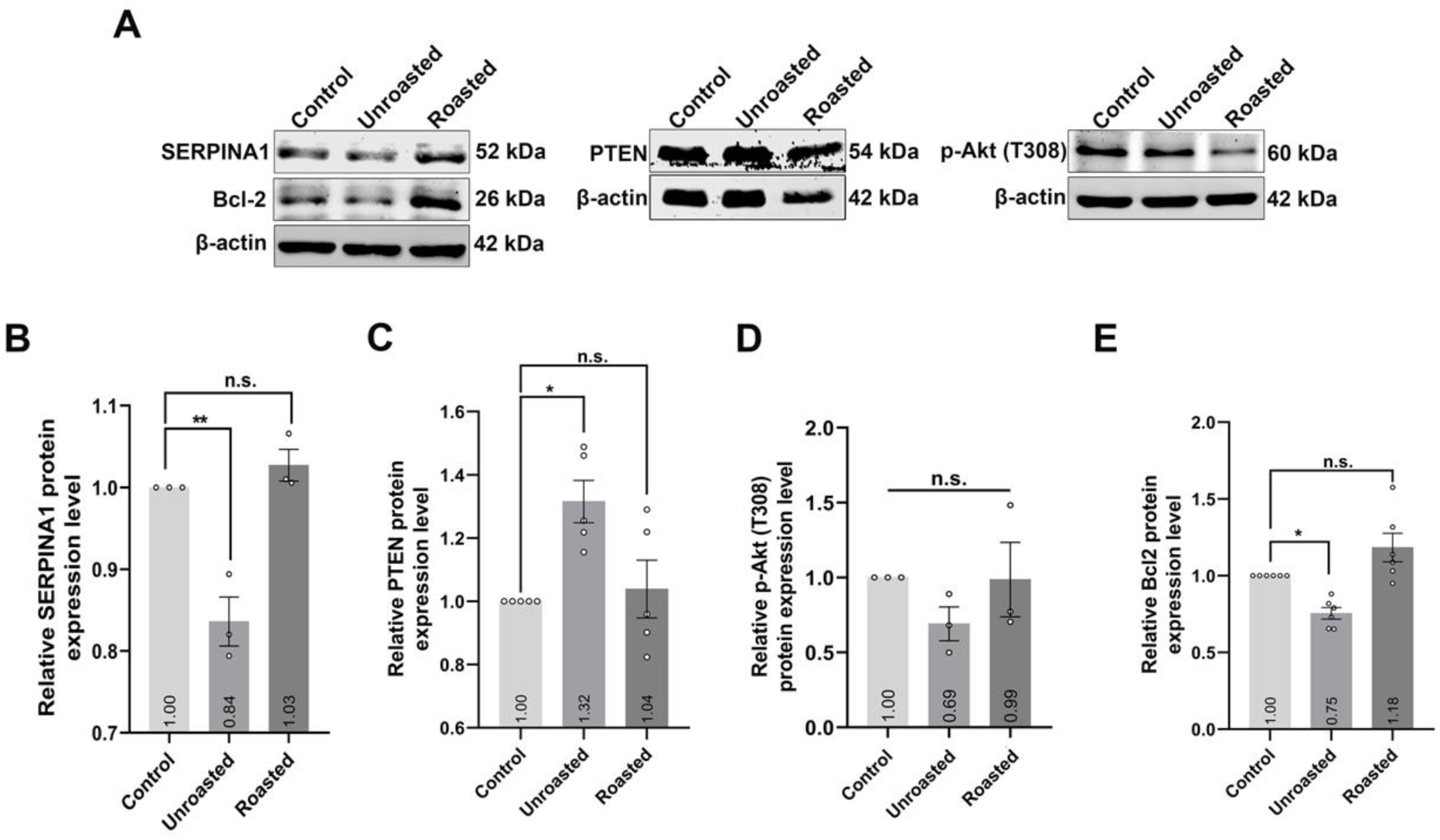
PI3K/Akt signaling pathway is suppressed only in unroasted coffee EV-treated SK-MEL-28 melanoma cells. **A)** The expression levels of SERPINA1, PTEN, p-Akt (T308), and Bcl-2 proteins in SK-MEL-28 melanoma cells upon unroasted or roasted coffee EV treatment. β-actin was used as an internal control. The experiments were performed at least three biological replicates. **B, C, D, E)** The quantification of the band densities is represented in the bar graph. Data were normalized to β-actin and the control experiment group. Statistical analysis was performed by a one-way ANOVA test. Data are presented as mean ± SEM. n.s. p>0.05, *p<0.05, **p<0.01.

Compounds found in coffee, such as caffeic acid and chlorogenic acid, have been shown to exert anti-cancer effects by inhibiting the mitogen-activated protein kinase (MAPK) signaling pathway [48,49]. To investigate whether coffee EVs exert MAPK signaling pathway-dependent anti-cancer effects on SK-MEL-28 melanoma cells, we first examined how each type of coffee EV treatment affected the phosphorylation level of BRAF, a key component of the MAPK pathway [50], known to be overactive in SK-MEL-28 cells [51]. The results showed that phosphorylated BRAF (p-BRAF) levels were significantly reduced in cells treated with roasted coffee EVs, while no change was observed with unroasted coffee EV treatment (Fig. 7A, B). Following the observed decrease in the p-BRAF levels with roasted coffee EVs, we also examined the level of phosphorylated MEK1/2 (p-MEK1/2), a downstream effector of the MAPK pathway, and the Cyclin D1, whose expression is regulated by MAPK signaling [52] (Fig. 7). Interestingly, p-MEK1/2 levels were significantly decreased only in SK-MEL-28 melanoma cells treated with roasted coffee EVs (Fig. 7A, C). However, Cyclin D1 expression was significantly reduced in cells treated with either type of coffee EV (Fig. 7A, D). These findings suggest that roasted coffee EVs may exert anti-proliferative effects on SK-MEL-28 melanoma cells by attenuating BRAF activity and inhibiting the MAPK pathway signaling.

**Fig. 7.**
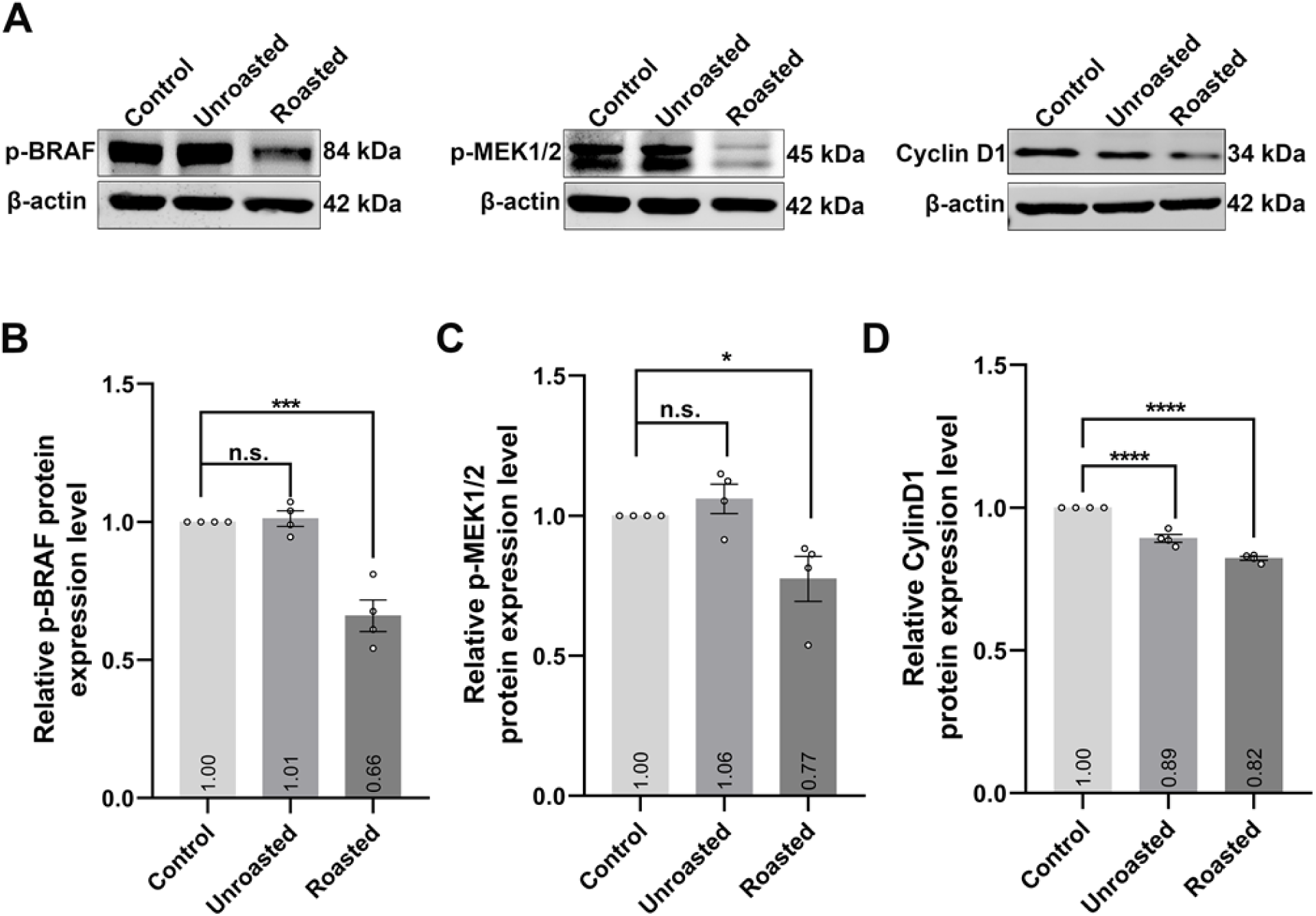
MAPK signaling pathway is suppressed in roasted coffee EV-treated SK-MEL-28 melanoma cells. **A)** The expression levels of p-BRAF, p-MEK1/2, and Cyclin D1 proteins in SK-MEL-28 melanoma cells upon unroasted or roasted coffee EV treatment. β-actin was used as an internal control. The experiments were performed at least three biological replicates. **B, C, D)** The quantification of the band densities is represented in the bar graph. Data were normalized to β-actin and the control experiment group. Statistical analysis was performed by a one-way ANOVA test. Data are presented as mean ± SEM. n.s. p>0.05, *p<0.05, ***p<0.001, ****p<0.0001.

## 4. Discussion

Nutrition plays a significant role in both carcinogenesis, and prevention of cancer. Eating habits have been directly linked to certain cancers, such as stomach, colon, and esophagus [53–55]. Coffee is one of the most widely consumed beverages worldwide, and it was once categorized by the IARC (International Agency for Research on Cancer) as potentially carcinogenic (Group 2B) to humans due to its high temperature consumption [56]. However, coffee was reclassified after accumulating evidence from numerous studies revealed that it may actually have protective effects against certain types of cancer rather than contributing to their development [57].

Several epidemiological studies have demonstrated an inverse relationship between coffee consumption and various cancer incidences [58–60]. In particular, melanoma has shown a significant inverse correlation with coffee intake and associated risk [24,26]. While cohort studies suggest a potential protective association between coffee consumption and melanoma incidence, the molecular mechanisms underlying this relationship remain poorly understood; instead, studies have focused on the benefits of isolated compounds of coffee such as caffeic acid and caffeine [61–63]. While these findings highlight the potential of coffee and its constituents’ anti-cancer effects, emerging interest has shifted toward exploring more complex, naturally encapsulated forms of coffee bioactive compounds, specifically, coffee-derived extracellular vesicles (EVs) [64,65]. Kantarcioglu et al. (2023) demonstrated potential therapeutic effects of roasted coffee EVs on liver fibrosis, suggesting anti-cancer potential in hepatocellular carcinoma [64]. Despite the fact that coffee is typically consumed in roasted form, research using coffee extracts has demonstrated that the roasting affects the cellular anti-cancer responses that coffee elicits [36,37]. Therefore, it was essential to investigate the potential differences in possible anti-cancer effects of unroasted and roasted coffee-derived EVs on melanoma cells, in order to further support the proposed preventive benefits of coffee consumption against melanoma.

In this study, to elucidate the molecular mechanisms underlying the coffee’s potential anti-cancer effects on melanoma, we chose to use EVs isolated from unroasted and roasted *Coffea arabica* beans to comparatively examine the potential differential effects of coffee EVs on SK-MEL-28 melanoma cells, a suitable model for studying cutaneous melanoma [66]. We first investigated how coffee EVs, which were isolated from unroasted and roasted coffee beans and confirmed to have plant EV properties (Fig. 1), affect the viability of normal MEL-ST melanocyte cells and SK-MEL-28 melanoma cells. At a dose of 4×10⁸ particles/μl, neither type of EVs impacted MEL-ST melanocyte viability (Fig. 2A), whereas both significantly reduced the viability of SK-MEL-28 cells (Fig. 2B). Furthermore, we analyzed nuclear morphology in both cell types and observed that unlike MEL-ST melanocytes, both treatments induced apoptotic body formation in SK-MEL-28 melanoma cells (Fig. 2C, D). These results suggest that both types of coffee EVs have the potential to induce apoptosis in SK-MEL-28 melanoma cells. This was further supported by flow cytometry analysis, which showed a significant increase in the percentage of late-apoptotic SK-MEL-28 melanoma cells following treatment with either type of coffee EVs (Fig. 3). Although both coffee EVs were found to induce late apoptosis in SK-MEL-28 melanoma cells to a similar extent in flow cytometry analyses, roasted coffee EVs triggered early apoptosis to a greater extent than unroasted coffee EVs (Fig. 3).

These findings suggest that extending the treatment duration may further enhance late apoptosis, particularly with roasted coffee EVs, since coffee components such as phenolic acids and caffeic acid, which increase during the roasting of coffee beans, are known to exert apoptotic effects [67–70]. However, 3D tumor spheroid analysis contradicted the assumption that roasted coffee EVs would show a stronger apoptotic effect over time than unroasted coffee EVs, since in long-term analyses, both types of coffee EVs exhibited similar growth-inhibitory effects on SK-MEL-28 melanoma spheroids (Fig. 4). Likewise, wound healing analyses demonstrated that although roasted coffee EVs elicited an earlier response, both EV types comparably reduced the migration capacity of SK-MEL-28 melanoma cells (Fig. 5). Melanoidins, compounds formed during roasting, are known to suppress the activity of several matrix metalloproteinases (MMP-1, MMP-2, and MMP-9), which play an active role in tumor growth and metastasis [71–73]. Thus, it is reasonable to expect roasted coffee EVs to induce earlier cell death and migration inhibition.

There are two possible explanations for why both EV types eventually produced similar long-term outcomes in spheroid growth inhibition and wound closure, despite differing short-term responses: Firstly, cells may have adapted to the effects of roasted coffee EVs over time, and/or certain compounds in the EVs may have only transient activity. Nevertheless, since the growth-inhibitory effect of roasted coffee EVs remained consistent over time in the long-term tumor spheroid analysis (Fig. 4), it is unlikely that the compounds lost their efficacy or that the cells developed resistance. Instead, as an alternative approach, it is more plausible that unroasted coffee EVs exert their effects more gradually. We propose that unlike the rapid cellular response triggered by roasted coffee EVs, unroasted coffee EVs may act through slower mechanisms such as transcriptional regulation, ultimately leading to anti-cancer effects similar to that of roasted coffee EVs over the long term.

Since cellular responses via RNA-based regulation or silencing requires more time [74], we reasoned that the delayed effects of unroasted coffee EVs could be attributed to RNA based regulation. Therefore, the transcriptome profile of SK-MEL-28 melanoma cells treated with unroasted coffee EVs was investigated, and the transcription of 80 genes were found to be significantly altered upon unroasted coffee EV treatment (Fig S1 and supplementary document). We noticed that the most significant transcriptional change detected in SK-MEL-28 melanoma cells with unroasted coffee EV treatment was on the *SERPINA1* gene, whose mRNA levels lowered by about 30-fold (Fig. S1). SerpinA1, also known as alpha-1 antitrypsin, is a serine protease inhibitor, and elevated SerpinA1 expression was observed in breast cancer, prostate cancer, papillary thyroid carcinoma and colorectal adenocarcinoma [75–78]. Recently, *SERPINA1* was found to be upregulated in cutaneous melanoma (CM) tissues and was significantly associated with tumor aggressiveness during CM progression and poor prognosis, indicating as a possible biomarker for CM diagnosis, as well as its therapeutic target [79]. Therefore, the decrease in *SERPINA1* gene expression following treatment with unroasted coffee EVs is noteworthy, further supporting the importance of SerpinA1 in cutaneous melanoma. Since SerpinA1 has been shown to stimulate cell proliferation and migration by activating the PI3K signaling pathway and suppressing PTEN [45], we hypothesized that the tumor growth inhibitory and migration impairing effect of unroasted coffee EVs could be due to decreased SerpinA1 protein expression and therefore, we investigated the effects of both coffee EVs on PI3K/AKT pathway. The results indicated that unroasted coffee EVs suppress the PI3K/AKT signaling pathway by downregulating SerpinA1 expression (Fig. 6). To understand more about the mechanism by which unroasted coffee EVs reduce *SERPINA1* mRNA levels, we consulted a study that identified 15 novel miRNAs in coffee EVs [64]. Using the miRDB database custom prediction tool, we found that among these newly discovered miRNAs, novel_9 is predicted to target the 3′UTR of *SERPINA1* mRNA, with a very high target score of 95%. This finding suggests that the suppression of SK-MEL-28 cell proliferation and migration caused by unroasted coffee EV treatment may be controlled by miRNA-based suppression of *SERPINA1* transcription.

Spent coffee ground extracts were shown to attenuate melanogenesis in B16F10 melanoma cells by downregulating the protein kinase A (PKA), phosphatidylinositol 3-kinase (PI3K)/Akt, and MAPK signaling pathways, highlighting the potential of coffee as an inhibitor of melanoma formation [80]. Likewise, isolated coffee compounds such as chlorogenic acid and caffeic acid, which are intensified during the roasting process, have been demonstrated to exhibit anti-cancer effects on melanoma cells by inducing apoptosis [61–63]. Additionally, it is known that SK-MEL-28 melanoma cells harbor mutations of BRAF, a serine/threonine kinase central to the MAPK signaling pathway and have an overactive MAPK pathway [51]. Thereupon, we investigated the effects of both types of coffee EVs on the MAPK kinase signaling pathway in SK-MEL-28 melanoma cells. Unlike unroasted coffee EVs, only roasted coffee EV treatment was found to reduce p-BRAF levels and p-MEK1/2 kinases located downstream of p-BRAF (Fig.7A, B, C). In contrast, Cyclin D1 expression levels were found to be decreased in both coffee EV treatments (Fig.7A, D). Cyclin D1 transcription is driven by ERK signaling and enhanced by AKT-mediated β-catenin stabilization, while AKT also boosts Cyclin D1 protein stability by inhibiting GSK-3β [81–83]. These results indicate that roasted coffee EVs suppress tumor growth mainly by blocking MAPK signaling in SK-MEL-28 melanoma cells, whereas unroasted coffee EVs reduce Cyclin D1 levels indirectly through PI3K/AKT signaling.

As the first study investigating the molecular mechanisms underlying the apoptotic effects of EVs derived from unroasted and roasted coffee beans comparatively, our findings revealed that unroasted and roasted coffee EVs may exhibit anti-cancer effects on SK-MEL-28 melanoma cells, via reducing cell viability by triggering apoptosis, suppressing tumor growth, and decreasing migration capability, through different mechanisms: unroasted coffee EV suppresses the PI3K/AKT signaling pathway by downregulating *SERPINA1* expression, while roasted coffee EV inhibits the MAPK pathway by attenuating BRAF activity.

Plant-derived EVs demonstrate anti-cancer properties not only due to their intrinsic bioactive components but also because their edibility and biocompatibility allow them to function as carriers in combination therapies without introducing additional toxicity or side effects. Collectively, our study revealing the anti-cancer effects of coffee EVs suggest that coffee EVs have the potential to be used as both natural therapeutic agents and vehicles to carry the conventional chemotherapeutics for delivery in combinatorial cancer treatments. Unlike conventional strategies that rely on liposome-based encapsulation, the use of intrinsic coffee-derived EVs could eliminate the need for synthetic delivery systems.

## Conclusion

Given its global consumption and accessibility, while coffee represents a readily available dietary source and offers cancer prevention in a cup, coffee-derived exosomes enriched in bioactive compounds and phytochemicals may serve as chemotherapeutic agents and natural nanocarriers. Advancements in multidisciplinary research that explore molecular pathways and therapeutic actions, alongside the enhancement of delivery methods, are anticipated to accelerate the transformation of coffee into next-generation treatments, paving the way for new possibilities in the prevention and management of skin cancers.

## Author Contributions

Ela Doruk Korkmaz and Seren Kucukvardar contributed to the publication equally (co-first authorship). Ela Doruk Korkmaz performed exosome isolation and characterization. Ela Doruk Korkmaz and Ilgin Isiltan performed MTT cell viability and DAPI staining analyses. Ela Doruk Korkmaz and Seren Kucukvardar performed all other experiments including flow cytometry, 3D-spheroid formation, wound healing, and western blot analysis. Seren Kucukvardar and Benan Temizci analyzed all data and wrote the original draft. Benan Temizci designed the study, supervised the research process. All authors approved and contributed to the final manuscript.

## Author Approvals

All authors have seen and approved the manuscript, and that it hasn’t been accepted or published elsewhere

## Funding

Transcriptome profiling analysis was supported by TUBITAK 2204A-1689B012405831 to Ela Doruk Korkmaz

## Data Availability

All data supporting the results of this investigation are accessible either within the publication, supplementary material, or from the corresponding author on reasonable request.

## Conflict of Interest

The authors declare that they have no conflict of interest.

---

**Supplementary Fig. 1.**
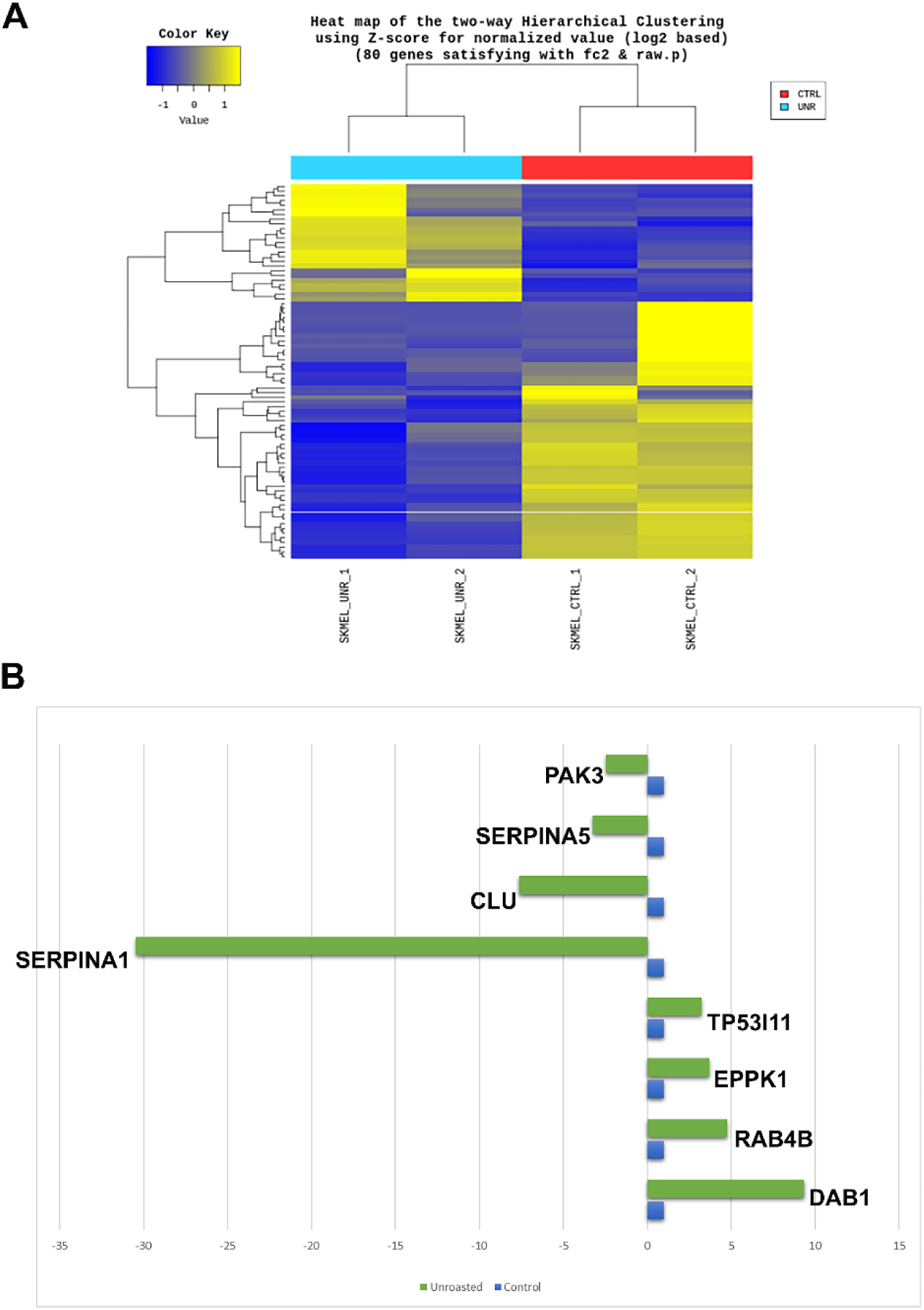
Transcriptome profiling analysis of the RNA-sequencing data distinguishing the control and unroasted coffee EV-treated SK-MEL-28 cells. **A)** The hierarchical clustering analysis representing the similarity of expression patterns between samples and genes. Experiment was performed with two biological replicates (SKMEL_UR_1 and SKMEL_UR_2 are RNA samples from unroasted coffee EV-treated SK-MEL-28 cells; SKMEL_CTRL_1 and SKMEL_CTRL_2 are RNA samples from control SK-MEL-28 cells). **B)** Graphical representation of selected genes showing decreased or increased expression following treatment.

## Notes

### Competing Interest Statement

The authors have declared no competing interest.

## References

1. Gerhauser, C. Cancer chemoprevention and nutriepigenetics: state of the art and future challenges. Top. Curr. Chem. 2013, 329, 73–132.

2. Thompson, A.S.; Tresserra-Rimbau, A.; Karavasiloglou, N.; Jennings, A.; Cantwell, M.; Hill, C.; Perez-Cornago, A., Bondonno, N.P.; Murphy, N.; Rohrmann, S., et al. Association of Healthful Plant-based Diet Adherence With Risk of Mortality and Major Chronic Diseases Among Adults in the UK. JAMA Netw. Open. 2023, 6, e234714.

3. Kasper, M.; Al Masri, M.; Kühn, T.; Rohrmann, S.; Wirnitzer, K.; Leitzmann, M.; Jochem, C. Sustainable diets and cancer: a systematic review and meta-analysis. EClinicalMedicine 2025, 83, 103215.

4. Flabouraris, G.; Karikas, G.A. Nutri-epigenetics and synthetic analogs in cancer chemoprevention. J. BUON. 2016, 21, 4–16.

5. Del Saz-Lara, A.; López de Las Hazas, M.C.; Visioli, F.; Dávalos, A. Nutri-Epigenetic Effects of Phenolic Compounds from Extra Virgin Olive Oil: A Systematic Review. Adv. Nutr. 2022, 13, 2039–2060.

6. Hossain, M.S.; Wazed, M.A.; Asha S, Amin, M.R.; Shimul, I.M. Dietary Phytochemicals in Health and Disease: Mechanisms, Clinical Evidence, and Applications-A Comprehensive Review. Food Sci. Nutr. 2025, 13, e70101.

7. Aune, D. Plant Foods, Antioxidant Biomarkers, and the Risk of Cardiovascular Disease, Cancer, and Mortality: A Review of the Evidence. Adv. Nutr. 2019, 10, 404–421.

8. Rinninella, E.; Mele, M.C.; Cintoni, M.; Raoul, P.; Ianiro, G.; Salerno, L.; Pozzo, C.; Bria E; Muscaritoli, M.; Molfino, A., et al. The Facts about Food after Cancer Diagnosis: A Systematic Review of Prospective Cohort Studies. Nutrients 2020, 12, 2345.

9. Jabbari, M.; Pourmoradian, S.; Eini-Zinab, H.; Mosharkesh, E.; Hosseini Balam, F.; Yaghmaei, Y.; Yadegari, A.; Amini, B.; Arman Moghadam, D.; Barati, M.;, et al. Levels of evidence for the association between different food groups/items consumption and the risk of various cancer sites: an umbrella review. Int. J. Food Sci. Nutr. 2022, 73, 861–874.

10. Qiao, K.; Zhao, M.; Huang, Y.; Liang, L.; Zhang, Y. Bitter Perception and Effects of Foods Rich in Bitter Compounds on Human Health: A Comprehensive Review. Foods. 2024, 13, 3747.

11. de Melo Pereira, G.V.; de Carvalho Neto, D.P.; Magalhães Júnior, AI.; do Prado, F.G.; Pagnoncelli, M.G.B.; Karp, S.G.; Soccol, C.R. Chemical composition and health properties of coffee and coffee by-products. Adv. Food Nutr. Res. 2020, 91, 65–96.

12. Nigra, A.D.; Teodoro, A.J.; Gil, G.A. A Decade of Research on Coffee as an Anticarcinogenic Beverage. Oxid. Med. Cell. Longev. 2021, 2021, 4420479.

13. Lachance, J.C.; Radhakrishnan, S.; Madiwale, G.; Guerrier, S.; Vanamala, J.K.P. Targeting hallmarks of cancer with a food-system-based approach. Nutrition 2020, 69, 110563.

14. Ciupei, D.; Colişar, A.; Leopold, L.; Stănilă, A.; Diaconeasa, Z.M. Polyphenols: From Classification to Therapeutic Potential and Bioavailability. Foods 2024, 13, 4131.

15. Pavelkova, R.; Matouskova, P.; Hoova, J.; Porizka, J.; Marova, I. Preparation and characterisation of organic UV filters based on combined PHB/liposomes with natural phenolic compounds. J. Biotechnol. 2020, *324S*, 100021.

16. Saewan, N.; Jimtaisong, A.; Panyachariwat, N.; Chaiwut, P. In Vitro and In Vivo Anti-Aging Effect of Coffee Berry Nanoliposomes. Molecules 2023, 28, 6830.

17. Madhan, S.; Dhar, R.; Devi, A. Plant-derived exosomes: a green approach for cancer drug delivery. J. Mater. Chem. B. 2024, 12, 2236–2252.

18. Zhang, G.; Huang, X.; Xiu, H.; Sun, Y.; Chen, J.; Cheng, G.; Song, Z.; Peng, Y.; Shen, Y.; Wang, J. et al. Extracellular vesicles: Natural liver-accumulating drug delivery vehicles for the treatment of liver diseases. J. Extracell. Vesicles 2020, 10, e12030.

19. Zhang, Z.; Yu, Y.; Zhu, G.; Zeng, L.; Xu, S.; Cheng, H.; Ouyang, Z.; Chen, J.; Pathak, J.L.; Wu, L. et al. The Emerging Role of Plant-Derived Exosomes-Like Nanoparticles in Immune Regulation and Periodontitis Treatment. Front. Immunol. 2022, 13, 896745.

20. Langellotto, M.D.; Rassu, G.; Serri, C.; Demartis, S.; Giunchedi, P.; Gavini, E. Plant-derived extracellular vesicles: a synergetic combination of a drug delivery system and a source of natural bioactive compounds. Drug Deliv. Transl. Res. 2025, 15, 831–845.

21. Nemati, M.; Singh, B.; Mir, R.A.; Nemati, M.; Babaei, A.; Ahmadi, M.; Rasmi, Y.; Golezani, A.G.; Rezaie, J. Plant-derived extracellular vesicles: a novel nanomedicine approach with advantages and challenges. Cell Commun. Signal. 2022, 20, 69.

22. Sha, A.; Luo, Y.; Xiao, W.; He, J.; Chen, X.; Xiong, Z.; Peng, L.; Zou, L.; Liu, B.; Li, Q. Plant-Derived Exosome-like Nanoparticles: A Comprehensive Overview of Their Composition, Biogenesis, Isolation, and Biological Applications. Int. J. Mol. Sci. 2024, 25, 12092.

23. Sah, N.K.; Arora, S.; Sahu, R.C.; Kumar, D.; Agrawal, A.K. Plant-based exosome-like extracellular vesicles as encapsulation vehicles for enhanced bioavailability and breast cancer therapy: recent advances and challenges. Med. Oncol. 2025, 42, 184.

24. Di Maso, M.; Boffetta, P.; Negri, E.; La Vecchia, C.; Bravi, F. Caffeinated Coffee Consumption and Health Outcomes in the US Population: A Dose-Response Meta-Analysis and Estimation of Disease Cases and Deaths Avoided. Adv. Nutr. 2021, 12, 1160–1176.

25. Navarro-Bielsa, A.; Gracia-Cazaña, T.; Almagro, M.; De la Fuente-Meira, S.; Flórez, Á.; Yélamos, O.; Montero-Vilchez, T.; González-Cruz, C.; Diago, A.; Abadías-Granado, I. et al. The Influence of the Exposome in the Cutaneous Squamous Cell Carcinoma, a Multicenter Case-Control Study. Cancers (Basel) 2023, 15, 5376.

26. Paiva, M.; Yumeen, S.; Kahn, B.J.; Nan, H.; Cho, E.; Saliba, E.; Qureshi, A. Coffee, Citrus, and Alcohol: A Review of What We Drink and How it May Affect our Risk for Skin Cancer. Yale J. Biol. Med. 2023, 96, 205–210.

27. Navarro-Bielsa, A.; Gracia-Cazaña, T.; Almagro, M.; De-la-Fuente-Meira, S.; Florez, Á.; Yélamos, O.; Montero-Vilchez, T.; González-Cruz, C.; Diago, A.; Abadías-Granado, I. et al. Exposome and basal cell carcinoma: a multicenter case-control study. Int. J. Dermatol. 2024, 63, 907–915.

28. Lu, Y.P.; Lou, Y.R.; Xie, J.G.; Peng, Q.Y.; Liao, J.; Yang, C.S.; Huang, M.T.; Conney, A.H. Topical applications of caffeine or (-)-epigallocatechin gallate (EGCG) inhibit carcinogenesis and selectively increase apoptosis in UVB-induced skin tumors in mice. Proc. Natl. Acad. Sci. USA 2002, 99, 12455–60.

29. Koo, S.W.; Hirakawa, S.; Fujii, S.; Kawasumi, M.; Nghiem, P. Protection from photodamage by topical application of caffeine after ultraviolet irradiation. Br. J. Dermatol. 2007, 156, 957–64.

30. Choi, H.S.; Park, E.D.; Park, Y.; Han, S.H.; Hong, K.B.; Suh, H.J. Topical application of spent coffee ground extracts protects skin from ultraviolet B-induced photoaging in hairless mice. Photochem. Photobiol. Sci. 2016, 15, 779–90.

31. Bray, E.R.; Kirsner, R.S.; Issa, N.T. Coffee and skin-Considerations beyond the caffeine perspective. J. Am. Acad. Dermatol. 2020, 82, e63.

32. de Mello, V.; de Mesquita Júnior, G.A.; Alvim, J.G.E.; Costa, J.C.D.; Vilela, F.M.P. Recent patent applications for coffee and coffee by-products as active ingredients in cosmetics. Int. J. Cosmet. Sci. 2023, 45, 267–287.

33. Elias, M.L.; Israeli, A.F.; Madan, R. Caffeine in Skincare: Its Role in Skin Cancer, Sun Protection, and Cosmetics. Indian J. Dermatol. 2023, 68, 546–550.

34. Preedalikit, W.; Chittasupho, C.; Leelapornpisid, P.; Qi, S.; Kiattisin, K. Development and Evaluation of Anti-Pollution Film-Forming Facial Spray Containing Coffee Cherry Pulp Extract. Pharmaceutics 2025, 17, 360.

35. Ruse, G.; Jîjie, A.R.; Moacă, E.A.; Pătrașcu, D.; Ardelean, F.; Jojic, A.A.; Ardelean, S.; Tchiakpe-Antal, D.S. *Coffea arabica*: An Emerging Active Ingredient in Dermato-Cosmetic Applications. Pharmaceuticals (Basel) 2025, 18, 171.

36. Rashad, A.E.E.M.; Abdelaziz, M.A.M.; Abdulaziz, M.A.M.; The impact of altering the concentration of coffee constituents on their anticancer effect on oral squamous cell carcinoma cell line - *in vitro* study. Contemp. Oncol. (Pozn) 2024, 28, 63–70.

37. Mojica, B.E.; Fong, L.E.; Biju, D.; Muharram, A.; Davis, I.M.; Vela, K.O.; Rios, D.; Osorio-Camacena, E.; Kaur, B.; Rojas, S.M. et al. The Impact of the Roast Levels of Coffee Extracts on their Potential Anticancer Activities. J. Food Sci. 2018, 83, 1125–1130.

38. Freitas, V.V.; Borges, L.L.; Vidigal, M.C.; dos Santos, M.H.; Stringheta, P.C. Coffee: A comprehensive overview of origin, market, and the quality process. Trends Food Sci. Technol. 2024, 146, 104411.

39. Fortunato, D.; Mladenović, D.; Criscuoli, M.; Loria, F.; Veiman, K.L.; Zocco, D.; Koort, K.; Zarovni, N. Opportunities and Pitfalls of Fluorescent Labeling Methodologies for Extracellular Vesicle Profiling on High-Resolution Single-Particle Platforms. Int. J. Mol. Sci. 2021, 22, 10510.

40. Puhka, M.; Takatalo, M.; Nordberg, M.E.; Valkonen, S.; Nandania, J.; Aatonen, M.; Yliperttula, M.; Laitinen, S.; Velagapudi, V.; Mirtti, T.; Kallioniemi, O.; Rannikko, A.; Siljander, P.R.; Af Hällström, T.M. Metabolomic Profiling of Extracellular Vesicles and Alternative Normalization Methods Reveal Enriched Metabolites and Strategies to Study Prostate Cancer-Related Changes. Theranostics. 2017, 7, 3824–3841.

41. Gupta, P.B.; Kuperwasser, C.; Brunet, J.P.; Ramaswamy, S.; Kuo, W.L.; Gray, J.W.; Naber, S.P.; Weinberg, R.A. The melanocyte differentiation program predisposes to metastasis after neoplastic transformation. Nat. Genet. 2005, 37, 1047–54.

42. Sobhiafshar, U.; Çakici, B.; Yilmaz, E.; Yildiz Ayhan, N.; Hedaya, L.; Ayhan, M.C.; Yerinde, C.; Alankuş, Y.B.; Gürkaşlar, H.K.; Firat-Karalar, E.N. et al. Interferon regulatory factor 4 modulates epigenetic silencing and cancer-critical pathways in melanoma cells. Mol. Oncol. 2024, 18, 2423–2448.

43. Suarez-Arnedo, A.; Torres Figueroa, F.; Clavijo, C.; Arbeláez, P.; Cruz, J.C.; Muñoz-Camargo, C. An image J plugin for the high throughput image analysis of in vitro scratch wound healing assays. PLoS One 2020, 15, e0232565.

44. Blagosklonny, M.V. Selective protection of normal cells from chemotherapy, while killing drug-resistant cancer cells. Oncotarget 2023, 14, 193–206.

45. Xiubing, C.; Huazhen, L.; Xueyan, W.; Jing, N.; Qing, L.; Haixing, J.; Shanyu, Q.; Jiefu, L. SERPINA1 promotes the invasion, metastasis, and proliferation of pancreatic ductal adenocarcinoma via the PI3K/Akt/NF-κB pathway. Biochem. Pharmacol. 2024, 230, 116580.

46. Shi, X.; Wang, J.; Lei,Y.; Cong, C.; Tan, D.; Zhou, X. Research progress on the PI3K/AKT signaling pathway in gynecological cancer (Review). Mol. Med. Rep. 2019, 19, 4529–4535.

47. Jiang, Y.; Fang, B.; Xu, B.; Chen, L. The RAS-PI3K-AKT-NF-κB pathway transcriptionally regulates the expression of BCL2 family and IAP family genes and inhibits apoptosis in fibrous epulis. J. Clin. Lab. Anal. 2020, 34, e23102.

48. Alam, M.; Ashraf, G.M.; Sheikh, K.; Khan, A.; Ali, S.; Ansari, M.M.; Adnan, M.; Pasupuleti, V.R.; Hassan, M.I. Potential Therapeutic Implications of Caffeic Acid in Cancer Signaling: Past, Present, and Future. Front. Pharmacol. 2022, 13, 845871.

49. Ontawong, A.; Duangjai, A.; Vaddhanaphuti, C.S.; Amornlerdpison, D.; Pengnet, S.; Kamkaew, N. Chlorogenic acid rich in coffee pulp extract suppresses inflammatory status by inhibiting the p38, MAPK, and NF-κB pathways. Heliyon 2023, 9, e13917.

50. Sumimoto, H.; Imabayashi, F.; Iwata, T.; Kawakami, Y. The BRAF-MAPK signaling pathway is essential for cancer-immune evasion in human melanoma cells. J. Exp. Med. 2006, 203, 1651–6.

51. Castellani, G.; Buccarelli, M.; Arasi, M.B.; Rossi, S.; Pisanu, M.E.; Bellenghi, M.; Lintas, C.; Tabolacci, C. BRAF Mutations in Melanoma: Biological Aspects, Therapeutic Implications, and Circulating Biomarkers. Cancers (Basel) 2023, 15, 4026.

52. Bhatt, K.V.; Spofford, L.S.; Aram, G.; McMullen, M.; Pumiglia, K.; Aplin, A.E. Adhesion control of cyclin D1 and p27Kip1 levels is deregulated in melanoma cells through BRAF-MEK-ERK signaling. Oncogene 2005, 24, 3459–71.

53. Abnet, C.C.; Corley, D.A.; Freedman, N.D.; Kamangar, F. Diet and upper gastrointestinal malignancies. Gastroenterology 2015, 148, 1234–1243.e4.

54. Zhang, F.X.; Miao, Y.; Ruan, J.G.; Meng, S.P.; Dong, J.D.; Yin, H.; Huang, Y.; Chen, F.R.; Wang, Z.C.; Lai, Y.F. Association Between Nitrite and Nitrate Intake and Risk of Gastric Cancer: A Systematic Review and Meta-Analysis. Med. Sci. Monit. 2019, 25, 1788–1799.

55. Nucci, D.; Marino, A.; Realdon, S.; Nardi, M.; Fatigoni, C.; Gianfredi, V. Lifestyle, WCRF/AICR Recommendations, and Esophageal Adenocarcinoma Risk: A Systematic Review of the Literature. Nutrients 2021, 13, 3525.

56. IARC Working Group on the Evaluation of Carcinogenic Risks to Humans. Coffee, tea, mate, methylxanthines and methylglyoxal. Lyon (FR): International Agency for Research on Cancer; 1991.

57. IARC Working Group on the Evaluation of Carcinogenic Risks to Humans. Drinking Coffee, Mate, and Very Hot Beverages. Lyon (FR): International Agency for Research on Cancer; 2018.

58. Torres-Collado, L.; Compañ-Gabucio, L.M.; González-Palacios, S.; Notario-Barandiaran, L.; Oncina-Cánovas, A.; Vioque, J.; García-de la Hera, M. Coffee Consumption and All-Cause, Cardiovascular, and Cancer Mortality in an Adult Mediterranean Population. Nutrients 2021, 13, 1241.

59. Carter, P.; Yuan, S.; Kar, S.; Vithayathil, M.; Mason, A.M.; Burgess, S.; Larsson, S.C. Coffee consumption and cancer risk: a Mendelian randomisation study. Clin. Nutr. 2022, 41, 2113–2123.

60. Shin, S.; Lee, J.E.; Loftfield, E.; Shu, X.O.; Abe, S.K.; Rahman, M.S.; Saito, E.; Islam, M.R.; Tsugane, S.; Sawada, N. et al. Coffee and tea consumption and mortality from all causes, cardiovascular disease and cancer: a pooled analysis of prospective studies from the Asia Cohort Consortium. Int. J. Epidemiol. 2022, 51, 626–640.

61. Kudugunti, S.K.; Vad, N.M.; Whiteside, A.J.; Naik, B.U.; Yusuf, M.A.; Srivenugopal, K.S.; Moridani, M.Y. Biochemical mechanism of caffeic acid phenylethyl ester (CAPE) selective toxicity towards melanoma cell lines. Chem. Biol. Interact. 2010, 188, 1–14.

62. Pelinson, L.P.; Assmann, C.E.; Palma, T.V.; da Cruz, I.B.M.; Pillat, M.M.; Mânica, A.; Stefanello, N.; Weis, G.C.C.; de Oliveira Alves, A.; de Andrade, C.M. et al. Antiproliferative and apoptotic effects of caffeic acid on SK-Mel-28 human melanoma cancer cells. Mol. Biol. Rep. 2019, 46, 2085–2092.

63. Manica, D.; da Silva, G.B.; de Lima, J.; Cassol, J.; Dallagnol, P.; Narzetti, R.A.; Moreno, M.; Bagatini, M.D. Caffeine reduces viability, induces apoptosis, inhibits migration and modulates the CD39/CD73 axis in metastatic cutaneous melanoma cells. Purinergic Signal. 2024, 20, 385–397.

64. Kantarcıoğlu, M.; Yıldırım, G.; Akpınar Oktar, P.; Yanbakan, S.; Özer, Z.B.; Yurtsever Sarıca, D.; Taşdelen, S.; Bayrak, E.; Akın Balı, D.F.; Öztürk, S. et al. Coffee-Derived Exosome-Like Nanoparticles: Are They the Secret Heroes? Turk. J. Gastroenterol. 2023, 34, 161–169.

65. Esmekaya, M.A.; Ertekin, B. Neuroprotective effects of coffee-derived exosome-like nanoparticles against Aβ-induced neurotoxicity. Gen. Physiol. Biophys. 2024, 43, 535–543.

66. Haridas, P.; McGovern, J.A.; McElwain, S.D.L.; Simpson, M.J. Quantitative comparison of the spreading and invasion of radial growth phase and metastatic melanoma cells in a three-dimensional human skin equivalent model. PeerJ. 2017, 5, e3754.

67. Chang, W.C.; Hsieh, C.H.; Hsiao, M.W.; Lin, W.C.; Hung, Y.C.; Ye, J.C. Caffeic acid induces apoptosis in human cervical cancer cells through the mitochondrial pathway. Taiwan J. Obstet. Gynecol. 2010, 49, 419–24.

68. Somporn, C.; Kamtuo, A.; Theerakulpisut, P.; Siriamornpun, S. Effects of roasting degree on radical scavenging activity, phenolics and volatile compounds of Arabica coffee beans (Coffea arabica L. cv. Catimor). Int. J. Food. Sci. Technol. 2011, 46, 2287–96.

69. Eroğlu, C.; Seçme, M.; Bağcı, G.; Dodurga, Y. Assessment of the anticancer mechanism of ferulic acid via cell cycle and apoptotic pathways in human prostate cancer cell lines. Tumour Biol. 2015, 36, 9437–46.

70. Surya, S.; Sampathkumar, P.; Sivasankaran, S.M.; Pethanasamy, M.; Elanchezhiyan, C.; Deepa, B.; Manoharan, S. Vanillic acid exhibits potent antiproliferative and free radical scavenging effects under in vitro conditions. In. J. Nutr. Pharmacol. Neurol. Dis. 2023, 13, 188–98.

71. De Marco, L.M.; Fischer, S.; Henle, T. High molecular weight coffee melanoidins are inhibitors for matrix metalloproteases. J. Agric. Food Chem. 2011, 59, 11417–23.

72. Gialeli, C.; Theocharis, A.D.; Karamanos, N.K. Roles of matrix metalloproteinases in cancer progression and their pharmacological targeting. FEBS J. 2011, 278, 16–27.

73. Moreira, A.S.; Nunes, F.M.; Domingues, M.R.; Coimbra, M.A. Coffee melanoidins: structures, mechanisms of formation and potential health impacts. Food Funct. 2012, 3, 903–15.

74. Kurreck, J. RNA interference: from basic research to therapeutic applications. Angew. Chem. Int. Ed. Engl. 2009, 48, 1378–98.

75. Tahara, E.; Ito, H.; Taniyama, K.; Yokozaki, H.; Hata, J. Alpha 1-antitrypsin, alpha 1-antichymotrypsin, and alpha 2-macroglobulin in human gastric carcinomas: a retrospective immunohistochemical study. Hum. Pathol. 1984, 15, 957–64.

76. Karashima, S.; Kataoka, H.; Itoh, H.; Maruyama, R.; Koono, M. Prognostic significance of alpha-1-antitrypsin in early stage of colorectal carcinomas. Int. J. Cancer. 1990, 45, 244–50.

77. Jarzab, B.; Wiench, M.; Fujarewicz, K.; Simek, K.; Jarzab, M.; Oczko-Wojciechowska, M.; Wloch, J.; Czarniecka, A.; Chmielik, E.; Lange, D. et al. Gene expression profile of papillary thyroid cancer: sources of variability and diagnostic implications. Cancer Res. 2005, 65, 1587–97.

78. El-Akawi, Z.J.; Al-Hindawi, F.K.; Bashir, N.A. Alpha-1 antitrypsin (alpha1-AT) plasma levels in lung, prostate and breast cancer patients. Neuro. Endocrinol. Lett. 2008, 29, 482–4.

79. Hu, F.; Mei, R.; Zhang, H.; Hao, D.; Li, W. Bioinformatics analysis of prognostic value and immune cell infiltration of *SERPINA1* gene in cutaneous melanoma. Ann. Transl. Med. 2022, 10, 966.

80. Huang, H.C.; Wei, C.M.; Siao, J.H.; Tsai, T.C.; Ko, W.P.; Chang, K.J.; Hii, C.H.; Chang, T.M. Supercritical Fluid Extract of Spent Coffee Grounds Attenuates Melanogenesis through Downregulation of the PKA, PI3K/Akt, and MAPK Signaling Pathways. Evid. Based Complement Alternat. Med. 2016, 2016, 5860296.

81. Zhang, W.; Liu, H.T. MAPK signal pathways in the regulation of cell proliferation in mammalian cells. Cell Res. 2002, 12, 9–18.

82. Fang, D.; Hawke, D.; Zheng, Y.; Xia, Y.; Meisenhelder, J.; Nika, H.; Mills, G.B.; Kobayashi, R.; Hunter, T.; Lu, Z. Phosphorylation of beta-catenin by AKT promotes beta-catenin transcriptional activity. J. Biol. Chem. 2007, 282, 11221–9.

83. Alao, J.P. The regulation of cyclin D1 degradation: roles in cancer development and the potential for therapeutic invention. Mol. Cancer. 2007, 6, 24.

