## Supplemental Document for "Unroasted and Roasted Coffee-Derived Extracellular Vesicles Inhibit Proliferation and Migration of SK-MEL-28 Melanoma Cells via Distinct Molecular Pathways"

A

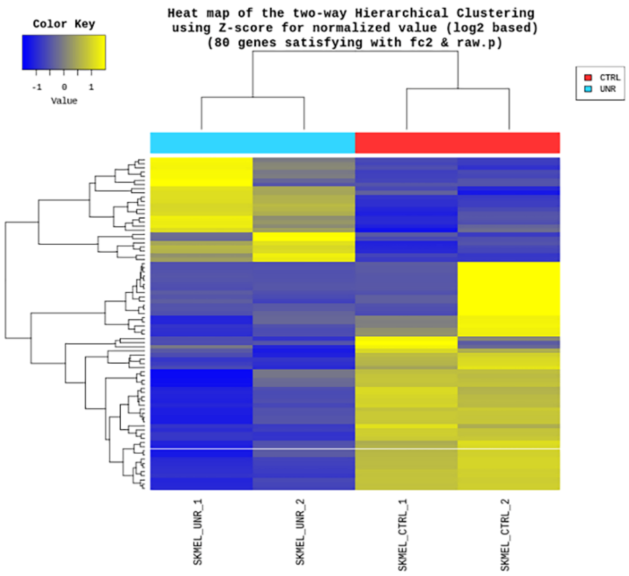


B

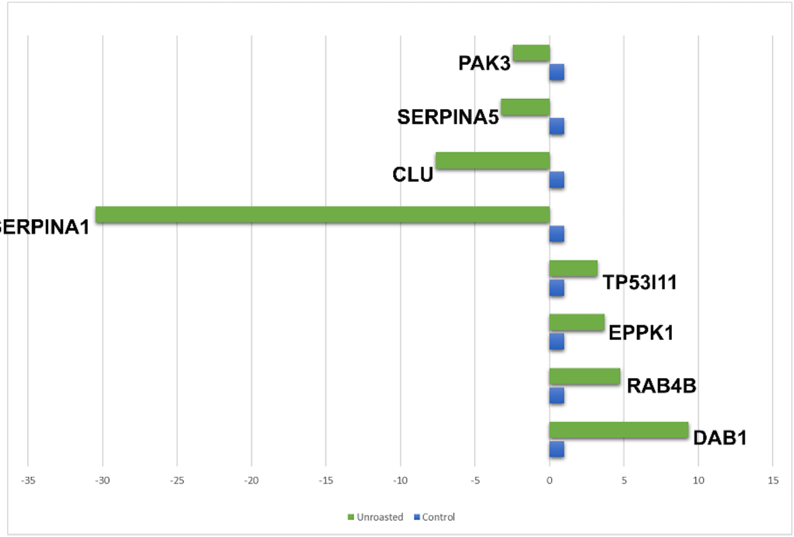


**Supplementary Fig. 1 Transcriptome profiling analysis of the RNA-sequencing data distinguishing the control and unroasted coffee EV-treated SK-MEL-28 cells. A)** The hierarchical clustering analysis representing the similarity of expression patterns between samples and genes. Experiment was performed with two biological replicates (SKMEL_UR_1 and SKMEL_UR_2 are RNA samples from unroasted coffee EV-treated SK-MEL-28 cells; SKMEL_CTRL_1 and SKMEL_CTRL_2 are RNA samples from control SK-MEL-28 cells). **B)** Graphical representation of selected genes showing decreased or increased expression following treatment.

| Gene_ID | Transcript_ID | Gene_Symbol | Description | gene_biotype | Protein_ID | HGNC | MIM | Ensembl |
|---|---|---|---|---|---|---|---|---|
| 1 | NM_130786 | A1BG | alpha-1-B glycoprotein | protein_coding | NP_570602.2 | 5 | 138670 | . |
| 2 | NM_000014,N | A2M | alpha-2-macroglobulin | protein_coding | NP_000005.3;NP_001334352.2;NP_001334353.2;NP_001334354.2;XP_0( | 7 | 103950 | . |
| 9 | NM_000662,N | NAT1 | N-acetyltransferase 1 | protein_coding | NP_000653.3;NP_001153642.1;NP_001153643.1;NP_001153644.1;NP_0( | 7645 | 108345 | . |
| 12 | NM_001085 | SERPINA3 | serpin family A member 3 | protein_coding | NP_001076.2 | 16 | 107280 | . |
| 14 | NM_001087,N | AAMP | angio associated migratory cell protein | protein_coding | NP_001078.2;NP_001289474.1;XP_024308480.1 | 18 | 603488 | . |
| 16 | NM_001605,XI | AARS1 | alanyl-tRNA synthetase 1 | protein_coding | NP_001596.2 | 20 | 601065 | . |
| 18 | NM_000663,N | ABAT | 4-aminobutyrate aminotransferase | protein_coding | NP_000654.2;NP_001120920.1;NP_065737.2;XP_011520702.1;XP_01152 | 23 | 137150 | . |
| 20 | NM_001606,N | ABCA2 | ATP binding cassette subfamily A member 2 | protein_coding | NP_001597.2;NP_997698.1;XP_006717059.1 | 32 | 600047 | . |
| 22 | NM_00127169 | ABCB7 | ATP binding cassette subfamily B member 7 | protein_coding | NP_001258625.1;NP_001258626.1;NP_001258627.1;NP_001258628.1;NI | 48 | 300135 | . |
| 23 | NM_00102509 | ABCF1 | ATP binding cassette subfamily F member 1 | protein_coding | NP_001020262.1;NP_001081.1 | 70 | 603429 | . |
| 25 | NM_005157,N | ABL1 | ABL proto-oncogene 1, non-receptor tyrosine kinase | protein_coding | NP_005148.2;NP_009297.2 | 76 | 189980 | . |
| 27 | NM_00113600 | ABL2 | ABL proto-oncogene 2, non-receptor tyrosine kinase | protein_coding | NP_001129472.1;NP_001129473.1;NP_001161708.1;NP_001161709.1;NI | 77 | 164690 | . |
| 29 | NM_001092,N | ABR | ABR activator of RhoGEF and GTPase | protein_coding | NP_001083.2;NP_001153218.1;NP_001243776.1;NP_001269078.1;NP_0( | 81 | 600365 | . |
| 30 | NM_00113041 | ACAA1 | acetyl-CoA acyltransferase 1 | protein_coding | NP_001123882.1;NP_001598.1;XP_006713185.1;XP_006713186.1;XP_01 | 82 | 604054 | . |
| 31 | NM_198834,N | ACACA | acetyl-CoA carboxylase alpha | protein_coding | NP_942131.1;NP_942133.1;NP_942134.1;NP_942135.1;NP_942136.1;XP_ | 84 | 200350 | . |
| 32 | NM_001093,XI | ACACB | acetyl-CoA carboxylase beta | protein_coding | NP_001084.3;XP_006719430.1;XP_011536561.1;XP_011536565.1;XP_01 | 85 | 601557 | . |
| 33 | NM_001608,XI | ACADL | acyl-CoA dehydrogenase long chain | protein_coding | NP_001599.1;XP_005246574.1;XP_016859444.1 | 88 | 609576 | . |
| 34 | NM_000016,N | ACADM | acyl-CoA dehydrogenase medium chain | protein_coding | NP_000007.1;NP_001120800.1;NP_001272971.1;NP_001272972.1;NP_0( | 89 | 607008 | . |
| 35 | NM_000017,N | ACADS | acyl-CoA dehydrogenase short chain | protein_coding | NP_000008.1;NP_001289483.1 | 90 | 606885 | . |
| 36 | NM_00133017 | ACADSB | acyl-CoA dehydrogenase short/branched chain | protein_coding | NP_001317103.1;NP_001600.1 | 91 | 600301 | . |
| 37 | NM_000018,N | ACADVL | acyl-CoA dehydrogenase very long chain | protein_coding | NP_000009.1;NP_001029031.1;NP_001257376.1;NP_001257377.1;XP_0( | 92 | 609575 | . |
| 38 | NM_000019,XI | ACAT1 | acetyl-CoA acetyltransferase 1 | protein_coding | NP_000010.1;XP_006718898.1;XP_016873170.1;XP_016873171.1;XP_01 | 93 | 607809 | . |
| 39 | NM_00130325 | ACAT2 | acetyl-CoA acetyltransferase 2 | protein_coding | NP_001290182.1;NP_005882.2 | 94 | 100678 | . |
| 41 | NM_001095,N | ASIC1 | acid sensing ion channel subunit 1 | protein_coding | NP_001086.2;NP_001243759.1;NP_064423.2;XP_011536652.1;XP_01153 | 100 | 602866 | . |
| 43 | NM_000665,N | ACHE | acetylcholinesterase (Cartwright blood group) | protein_coding | NP_000656.1;NP_001269378.1;NP_001289550.1;NP_001289551.1;NP_0( | 108 | 100740 | . |
| 47 | NM_001096,N | ACLY | ATP citrate lyase | protein_coding | NP_001087.2;NP_001290203.1;NP_001290204.1;NP_942127.1;XP_00525 | 115 | 108728 | . |
| 48 | NM_00127835 | ACO1 | aconitase 1 | protein_coding | NP_001265281.1;NP_001349769.1;NP_002188.1;XP_011516190.1 | 117 | 100880 | . |
| 50 | NM_001098,XI | ACO2 | aconitase 2 | protein_coding | NP_001089.1;XP_016884301.1;XP_024308018.1 | 118 | 100850 | . |
| 51 | NM_00118503 | ACOX1 | acyl-CoA oxidase 1 | protein_coding | NP_001171968.1;NP_004026.2;NP_009223.2;XP_011523170.1;XP_01152 | 119 | 609751 | . |
| 52 | NM_00104064 | ACP1 | acid phosphatase 1 | protein_coding | NP_001035739.1;NP_004291.1;NP_009030.1 | 122 | 171500 | . |
| 53 | NM_00130248 | ACP2 | acid phosphatase 2, lysosomal | protein_coding | NP_001289418.1;NP_001289419.1;NP_001289420.1;NP_001289421.1;NI | 123 | 171650 | . |
| 54 | NM_00111103 | ACP5 | acid phosphatase 5, tartrate resistant | protein_coding | NP_001104504.1;NP_001104505.1;NP_001104506.1;NP_001308952.1;NI | 124 | 171640 | . |
| 55 | NM_001099,N | ACP3 | acid phosphatase 3 | protein_coding | NP_001090.2;NP_001127666.1;NP_001278966.1;XP_011511248.1;XP_01 | 125 | 171790 | . |
| 59 | NM_00114194 | ACTA2 | actin alpha 2, smooth muscle | protein_coding | NP_001135417.1;NP_001307784.1;NP_001604.1 | 130 | 102620 | . |
| 60 | NM_001101 | ACTB | actin beta | protein_coding | NP_001092.1 | 132 | 102630 | . |
| 62 | gene-ACTBP2 | ACTBP2 | ACTB pseudogene 2 | pseudogene | . | 135 | . | . |
| 67 | gene-ACTBP7 | ACTBP7 | ACTB pseudogene 7 | pseudogene | . | 140 | . | . |
| 71 | NM_00119995 | ACTG1 | actin gamma 1 | protein_coding | NP_001186883.1;NP_001605.1 | 144 | 102560 | . |
| 72 | NM_00119989 | ACTG2 | actin gamma 2, smooth muscle | protein_coding | NP_001186822.1;NP_001606.1 | 145 | 102545 | . |
| 75 | gene-ACTG1P3 | ACTG1P3 | actin gamma 1 pseudogene 3 | pseudogene | . | 148 | . | . |
| 81 | NM_00132203 | ACTN4 | actinin alpha 4 | protein_coding | NP_001308962.1;NP_004915.2;XP_005259338.1;XP_006723469.1;XP_01 | 166 | 604638 | . |
| 86 | NM_004301,N | ACTL6A | actin like 6A | protein_coding | NP_004292.1;NP_817126.1;NP_829888.1 | 24124 | 604958 | . |
| 87 | NM_001102,N | ACTN1 | actinin alpha 1 | protein_coding | NP_001093.1;NP_001123476.1;NP_001123477.1;XP_011535567.1;XP_01 | 163 | 102575 | . |
| 88 | NM_001103,N | ACTN2 | actinin alpha 2 | protein_coding | NP_001094.1;NP_001265272.1;NP_001265273.1 | 164 | 102573 | . |
| 90 | NM_001105,N | ACVR1 | activin A receptor type 1 | protein_coding | NP_001096.1;NP_001104537.1;NP_001334592.1;NP_001334593.1;NP_0( | 171 | 102576 | . |
| 91 | NM_004302,N | ACVR1B | activin A receptor type 1B | protein_coding | NP_004293.1;NP_064732.3;NP_064733.3;XP_011537268.1;XP_01153726 | 172 | 601300 | . |
| 92 | NM_00127857 | ACVR2A | activin A receptor type 2A | protein_coding | NP_001265508.1;NP_001265509.1;NP_001607.1 | 173 | 102581 | . |
| 93 | NM_001106,XI | ACVR2B | activin A receptor type 2B | protein_coding | NP_001097.2;XP_005265640.1;XP_016863003.1;XP_016863004.1;XP_01( | 174 | 602730 | . |
| 95 | NM_000666,N | ACY1 | aminoacylase 1 | protein_coding | NP_000657.1;NP_001185824.1;NP_001185825.1;NP_001185826.1;NP_0( | 177 | 104620 | . |
| 97 | NM_001107,N | ACYP1 | acylphosphatase 1 | protein_coding | NP_001098.1;NP_001289545.1;NP_001289546.1 | 179 | 600875 | . |
| 98 | NM_00132058 | ACYP2 | acylphosphatase 2 | protein_coding | NP_001307515.1;NP_001307516.1;NP_001307517.1;NP_001307518.1;NI | 180 | 102595 | . |
| 100 | NM_000022,N | ADA | adenosine deaminase | protein_coding | NP_000013.2;NP_001308979.1;NP_001308980.1 | 186 | 608958 | . |
| 101 | NM_001109,N | ADAM8 | ADAM metallopeptidase domain 8 | protein_coding | NP_001100.3;NP_001157961.1;NP_001157962.1;XP_011537419.2;XP_01 | 215 | 602267 | . |
| 102 | NM_001110,N | ADAM10 | ADAM metallopeptidase domain 10 | protein_coding | NP_001101.1;NP_001307499.1;XP_024305586.1 | 188 | 602192 | . |
| 103 | NM_00102510 | ADAR | adenosine deaminase RNA specific | protein_coding | NP_001020278.1;NP_001102.3;NP_001180424.1;NP_001351974.1;NP_0( | 225 | 146920 | . |
| 104 | NM_001112,N | ADARB1 | adenosine deaminase RNA specific B1 | protein_coding | NP_001103.1;NP_001153702.1;NP_001333616.1;NP_001333617.1;NP_0! | 226 | 601218 | . |
| 107 | NM_00128176 | ADCY1 | adenylate cyclase 1 | protein_coding | NP_001268697.1;NP_066939.1;XP_005249641.1;XP_005249642.1 | 232 | 103072 | . |
| 108 | NM_020546,XI | ADCY2 | adenylate cyclase 2 | protein_coding | NP_065433.2;XP_011512244.1 | 233 | 103071 | . |
| 109 | NM_00132061 | ADCY3 | adenylate cyclase 3 | protein_coding | NP_001307542.1;NP_001364057.1;NP_001364058.1;NP_001364059.1;NI | 234 | 600291 | . |
| 112 | NM_015270,XI | ADCY6 | adenylate cyclase 6 | protein_coding | NP_056085.1;XP_006719273.1;XP_016874232.1 | 237 | 600294 | . |
| 113 | NM_001114,N | ADCY7 | adenylate cyclase 7 | protein_coding | NP_001105.1;NP_001272986.1;XP_011521137.1;XP_011521138.1;XP_01 | 238 | 600385 | . |
| 115 | NM_001116,XI | ADCY9 | adenylate cyclase 9 | protein_coding | NP_001107.2;XP_005255136.1;XP_011520655.1 | 240 | 603302 | . |
| 116 | NM_00109973 | ADCYAP1 | adenylate cyclase activating polypeptide 1 | protein_coding | NP_001093203.1;NP_001108.2;XP_005258138.2 | 241 | 102980 | . |
| 118 | NM_001119,N | ADD1 | adducin 1 | protein_coding | NP_001110.2;NP_001273574.1;NP_001341683.1;NP_001341684.1;NP_0( | 243 | 102680 | . |
| 119 | NM_00118505 | ADD2 | adducin 2 | protein_coding | NP_001171983.1;NP_001171984.1;NP_001608.1;NP_059516.2;NP_0595; | 244 | 102681 | . |
| 120 | NM_001121,N | ADD3 | adducin 3 | protein_coding | NP_001112.2;NP_001307520.1;NP_001307521.1;NP_001307522.1;NP_0( | 245 | 601568 | . |
| 123 | NM_001122,N | PLIN2 | perilipin 2 | protein_coding | NP_001113.2;XP_016869748.1 | 248 | 103195 | . |
| 128 | NM_000671 | ADH5 | alcohol dehydrogenase 5 (class III), chi polypeptide | protein_coding | NP_000662.3 | 253 | 103710 | . |
| 132 | NM_001123,N | ADK | adenosine kinase | protein_coding | NP_001114.2;NP_001189378.1;NP_001189379.1;NP_001356052.1;NP_0( | 257 | 102750 | . |
| 133 | NM_001124 | ADM | adrenomedullin | protein_coding | NP_001115.1 | 259 | 103275 | . |
| 134 | NM_000674,N | ADORA1 | adenosine A1 receptor | protein_coding | NP_000665.1;NP_001041695.1;NP_001351994.1;NP_001351995.1;XP_0( | 262 | 102775 | . |
| 136 | NM_000676,XI | ADORA2B | adenosine A2b receptor | protein_coding | NP_000667.1;XP_011521961.1;XP_011521963.1;XP_016879686.1 | 264 | 600446 | . |
| 141 | NM_001125,N | ADPRH | ADP-ribosylarginine hydrolase | protein_coding | NP_001116.1;NP_001278878.1;NP_001278879.1;NP_001358021.1 | 269 | 603081 | . |
| 142 | NM_001618 | PARP1 | poly(ADP-ribose) polymerase 1 | protein_coding | NP_001609.2 | 270 | 173870 | . |
| 143 | NM_006437,XI | PARP4 | poly(ADP-ribose) polymerase family member 4 | protein_coding | NP_006428.2;XP_011533233.1;XP_011533234.1 | 271 | 607519 | . |
| 150 | NM_000681 | ADRA2A | adrenoceptor alpha 2A | protein_coding | NP_000672.3 | 281 | 104210 | . |
| 151 | NM_000682 | ADRA2B | adrenoceptor alpha 2B | protein_coding | NP_000673.2 | 282 | 104260 | . |
| 152 | NM_000683 | ADRA2C | adrenoceptor alpha 2C | protein_coding | NP_000674.2 | 283 | 104250 | . |
| 153 | NM_000684 | ADRB1 | adrenoceptor beta 1 | protein_coding | NP_000675.1 | 285 | 109630 | . |
| 154 | NM_000024 | ADRB2 | adrenoceptor beta 2 | protein_coding | NP_000015.2 | 286 | 109690 | . |
| 156 | NM_001619,XI | GRK2 | G protein-coupled receptor kinase 2 | protein_coding | NP_001610.2;XP_011543075.1;XP_024304144.1 | 289 | 109635 | . |
| 157 | NM_00136277 | GRK3 | G protein-coupled receptor kinase 3 | protein_coding | NP_001349707.1;NP_005151.2;XP_011528277.1 | 290 | 109636 | . |
| 158 | NM_000026,N | ADSL | adenylosuccinate lyase | protein_coding | NP_000017.1;NP_001116850.1;NP_001304852.1;NP_001350769.1;XP_01 | 291 | 608222 | . |
| 159 | NM_001126,N | ADSS2 | adenylosuccinate synthase 2 | protein_coding | NP_001117.2;NP_001352002.1;XP_011542410.1 | 292 | 103060 | . |
| 160 | NM_014203,N | AP2A1 | adaptor related protein complex 2 subunit alpha 1 | protein_coding | NP_055018.2;NP_570603.2;XP_011524858.1;XP_011524859.1 | 561 | 601026 | . |
| 161 | NM_00124283 | AP2A2 | adaptor related protein complex 2 subunit alpha 2 | protein_coding | NP_001229766.1;NP_036437.1;XP_011518231.1;XP_011518232.1 | 562 | 607242 | . |
| 162 | NM_001127,N | AP1B1 | adaptor related protein complex 1 subunit beta 1 | protein_coding | NP_001118.3;NP_001159491.1;NP_001365491.1;NP_001365492.1;NP_0( | 554 | 600157 | . |
| 163 | NM_00103000 | AP2B1 | adaptor related protein complex 2 subunit beta 1 | protein_coding | NP_001025177.1;NP_001273.1;XP_005257994.1;XP_005257995.1;XP_00 | 563 | 601025 | . |
| 164 | NM_00103000 | AP1G1 | adaptor related protein complex 1 subunit gamma 1 | protein_coding | NP_001025178.1;NP_001119.3 | 555 | 603533 | . |
| 165 | NM_001129,XI | AEBP1 | AE binding protein 1 | protein_coding | NP_001120.3;XP_011513464.1 | 303 | 602981 | . |
| 166 | NM_001130,N | TLE5 | TLE family member 5, transcriptional modulator | protein_coding | NP_001121.2;NP_945320.1;NP_945321.1;XP_006722727.1 | 307 | 600188 | . |
| 172 | NR_003226,NF | AFG3L1P | AFG3 like matrix AAA peptidase subunit 1, pseudogene | transcribed_ps | . | 314 | 603020 | . |
| 175 | NM_000027,N | AGA | aspartylglucosaminidase | protein_coding | NP_000018.2;NP_001165459.1 | 318 | 613228 | . |
| 176 | NM_001135,N | ACAN | aggrecan | protein_coding | NP_001126.3;NP_001356197.1;NP_037359.3;XP_011519615.1;XP_01151 | 319 | 155760 | . |
| 177 | NM_001136,N | AGER | advanced glycosylation end-product specific receptor | protein_coding | NP_001127.1;NP_001193858.1;NP_001193861.1;NP_001193863.1;NP_0( | 320 | 600214 | . |
| 178 | NM_000028,N | AGL | amylo-alpha-1, 6-glucosidase, 4-alpha-glucanotransferase | protein_coding | NP_000019.2;NP_000633.2;NP_000634.2;NP_000635.2;NP_000637.2;XP_ | 321 | 610860 | . |
| 182 | NM_000214 | JAG1 | jagged canonical Notch ligand 1 | protein_coding | NP_000205.1 | 6188 | 601920 | . |
| 185 | NM_000685,N | AGTR1 | angiotensin II receptor type 1 | protein_coding | NP_000676.1;NP_001369665.1;NP_001369666.1;NP_004826.5;NP_0336; | 336 | 106165 | . |
| 189 | NM_000030 | AGXT | alanine--glyoxylate and serine--pyruvate aminotransferase | protein_coding | NP_000021.1 | 341 | 604285 | . |
| 191 | NM_000687,N | AHCY | adenosylhomocysteinase | protein_coding | NP_000678.1;NP_001155238.1;NP_001309013.1;NP_001309014.1;NP_0( | 343 | 180960 | . |
| 196 | NM_001621 | AHR | aryl hydrocarbon receptor | protein_coding | NP_001612.1 | 348 | 600253 | . |
| 202 | NM_00137124 | CRYBG1 | crystallin beta-gamma domain containing 1 | protein_coding | NP_001358171.1;NP_001615.2;XP_016865822.1 | 356 | 601797 | . |
| 203 | NM_000476,N | AK1 | adenylate kinase 1 | protein_coding | NP_000467.1;NP_001305050.1;NP_001305051.1;XP_016869917.1;XP_02 | 361 | 103000 | . |
| 204 | NM_00119919 | AK2 | adenylate kinase 2 | protein_coding | NP_001186128.1;NP_001306068.1;NP_001306069.1;NP_001306070.1;NI | 362 | 103020 | . |
| 205 | NM_00100535 | AK4 | adenylate kinase 4 | protein_coding | NP_001005353.1;NP_001317545.1;NP_037542.1;NP_982289.1;XP_01685 | 363 | 103030 | . |
| 206 | gene-AK4P1 | AK4P1 | adenylate kinase 4 pseudogene 1 | pseudogene | . | 364 | . | . |
| 207 | NM_00101443 | AKT1 | AKT serine/threonine kinase 1 | protein_coding | NP_001014431.1;NP_001014432.1;NP_001369359.1;NP_001369360.1;NI | 391 | 164730 | . |
| 208 | NM_00124302 | AKT2 | AKT serine/threonine kinase 2 | protein_coding | NP_001229956.1;NP_001229957.1;NP_001317440.1;NP_001617.1;XP_01 | 392 | 164731 | . |
| 210 | NM_000031,N | ALAD | aminolevulinate dehydratase | protein_coding | NP_000022.3;NP_001003945.1;NP_001304674.1;XP_011516666.1;XP_02 | 395 | 125270 | . |
| 211 | NM_000688,N | ALAS1 | 5'-aminolevulinate synthase 1 | protein_coding | NP_000679.1;NP_001291372.1;NP_001291373.1;NP_954635.1;XP_01153 | 396 | 125290 | . |
| 214 | NM_00124328 | ALCAM | activated leukocyte cell adhesion molecule | protein_coding | NP_001230209.1;NP_001230210.1;NP_001230212.1;NP_001618.2 | 400 | 601662 | . |
| 215 | NM_000033,XI | ABCD1 | ATP binding cassette subfamily D member 1 | protein_coding | NP_000024.2 | 61 | 300371 | . |
| 216 | NM_000689 | ALDH1A1 | aldehyde dehydrogenase 1 family member A1 | protein_coding | NP_000680.2 | 402 | 100640 | . |
| 219 | NM_000692,XI | ALDH1B1 | aldehyde dehydrogenase 1 family member B1 | protein_coding | NP_000683.3;XP_011516104.1 | 407 | 100670 | . |
| 220 | NM_000693,N | ALDH1A3 | aldehyde dehydrogenase 1 family member A3 | protein_coding | NP_000684.2;NP_001280744.1 | 409 | 600463 | . |
| 221 | NM_000694,N | ALDH3B1 | aldehyde dehydrogenase 3 family member B1 | protein_coding | NP_000685.1;NP_001025181.1;NP_001154945.1;NP_001276987.1;NP_0( | 410 | 600466 | . |
| 223 | NM_000696,N | ALDH9A1 | aldehyde dehydrogenase 9 family member A1 | protein_coding | NP_000687.3;NP_001352703.1 | 412 | 602733 | . |
| 224 | NM_000382,N | ALDH3A2 | aldehyde dehydrogenase 3 family member A2 | protein_coding | NP_000373.1;NP_001026976.1;NP_001356065.1;NP_001356066.1;NP_0( | 403 | 609523 | . |
| 226 | NM_00112761 | ALDOA | aldolase, fructose-bisphosphate A | protein_coding | NP_001121089.1;NP_001230106.1;NP_908930.1;NP_908932.1 | 414 | 103850 | . |
| 230 | NM_005165,XI | ALDOC | aldolase, fructose-bisphosphate C | protein_coding | NP_005156.1;XP_005258006.1;XP_011522858.1 | 418 | 103870 | . |
| 231 | NM_00134614 | AKR1B1 | aldo-keto reductase family 1 member B | protein_coding | NP_001333071.1;NP_001619.1;XP_005250291.1;XP_024302463.1 | 381 | 103880 | . |
| 238 | NM_00135376 | ALK | ALK receptor tyrosine kinase | protein_coding | NP_001340694.1;NP_004295.2;XP_024308546.1;XP_024308547.1 | 427 | 105590 | . |
| 262 | NM_00128721 | AMD1 | adenosylmethionine decarboxylase 1 | protein_coding | NP_001274143.1;NP_001274144.1;NP_001274145.1;NP_001625.2 | 457 | 180980 | . |
| 267 | NM_001144,N | AMFR | autocrine motility factor receptor | protein_coding | NP_001135.3;NP_001310440.1;NP_001310441.1;XP_005255947.1 | 463 | 603243 | . |
| 268 | NM_000479 | AMH | anti-Mullerian hormone | protein_coding | NP_000470.3 | 464 | 600957 | . |
| 271 | NM_00125736 | AMPD2 | adenosine monophosphate deaminase 2 | protein_coding | NP_001244290.1;NP_001295099.1;NP_001355738.1;NP_004028.4;NP_6: | 469 | 102771 | . |
| 272 | NM_000480,N | AMPD3 | adenosine monophosphate deaminase 3 | protein_coding | NP_000471.1;NP_001020560.1;NP_001020561.1;NP_001165901.1;NP_0( | 470 | 102772 | . |
| 273 | NM_001635,N | AMPH | amphiphysin | protein_coding | NP_001626.1;NP_647477.1;XP_006715752.1;XP_006715753.1;XP_00671 | 471 | 600418 | . |
| 274 | NM_00132063 | BIN1 | bridging integrator 1 | protein_coding | NP_001307561.1;NP_001307562.1;NP_001307563.1;NP_001307569.1;NI | 1052 | 601248 | . |
| 275 | NM_000481,N | AMT | aminomethyltransferase | protein_coding | NP_000472.2;NP_001158182.1;NP_001158183.1;NP_001158184.1 | 473 | 238310 | . |
| 280 | NM_020978 | AMY2B | amylase alpha 2B | protein_coding | NP_066188.1 | 478 | 104660 | . |
| 283 | NM_00109757 | ANG | angiogenin | protein_coding | NP_001091046.1;NP_001136.1 | 483 | 105850 | . |
| 284 | NM_001146,N | ANGPT1 | angiopoietin 1 | protein_coding | NP_001137.2;NP_001186788.1;NP_001300980.1 | 484 | 601667 | . |
| 287 | NM_00112749 | ANK2 | ankyrin 2 | protein_coding | NP_001120965.1;NP_001139.3;NP_001341154.1;NP_001341157.1;NP_0( | 493 | 106410 | . |
| 288 | NM_001149,N | ANK3 | ankyrin 3 | protein_coding | NP_001140.2;NP_001191332.1;NP_001191333.1;NP_001307803.1;NP_0( | 494 | 600465 | . |
| 291 | NM_001151 | SLC25A4 | solute carrier family 25 member 4 | protein_coding | NP_001142.2 | 10990 | 103220 | . |
| 292 | NM_001152 | SLC25A5 | solute carrier family 25 member 5 | protein_coding | NP_001143.2 | 10991 | 300150 | . |
| 293 | NM_001636,N | SLC25A6 | solute carrier family 25 member 6 | protein_coding | NP_001627.2 | 10992 | 403000 | . |
| 301 | NM_000700,XI | ANXA1 | annexin A1 | protein_coding | NP_000691.1;XP_011516911.1;XP_016870146.1 | 533 | 151690 | . |
| 302 | NM_00100285 | ANXA2 | annexin A2 | protein_coding | NP_001002857.1;NP_001002858.1;NP_001129487.1;NP_004030.1;XP_01 | 537 | 151740 | . |
| 304 | NR_003573 | ANXA2P2 | annexin A2 pseudogene 2 | transcribed_ps | . | 539 | . | . |
| 306 | NM_005139,XI | ANXA3 | annexin A3 | protein_coding | NP_005130.1 | 541 | 106490 | . |
| 307 | NM_001153,N | ANXA4 | annexin A4 | protein_coding | NP_001144.1;NP_001307627.1;NP_001307629.1;NP_001307631.1;NP_0( | 542 | 106491 | . |
| 308 | NM_001154,XI | ANXA5 | annexin A5 | protein_coding | NP_001145.1;XP_016863630.1 | 543 | 131230 | . |
| 309 | NM_001155,N | ANXA6 | annexin A6 | protein_coding | NP_001146.2;NP_001180473.1;NP_001350043.1 | 544 | 114070 | . |
| 310 | NM_001156,N | ANXA7 | annexin A7 | protein_coding | NP_001147.1;NP_001307808.1;NP_001307809.1;NP_004025.1;XP_01687 | 545 | 186360 | . |
| 311 | NM_001157,N | ANXA11 | annexin A11 | protein_coding | NP_001148.1;NP_001265336.1;NP_001265337.1;NP_001265338.1;NP_6( | 535 | 602572 | . |
| 314 | NM_001158,N | AOC2 | amine oxidase copper containing 2 | protein_coding | NP_001149.2;NP_033720.2 | 549 | 602268 | . |
| 317 | NM_001160,N | APAF1 | apoptotic peptidase activating factor 1 | protein_coding | NP_001151.1;NP_037361.1;NP_863651.1;NP_863658.1;NP_863659.1;XP_ | 576 | 602233 | . |
| 318 | NM_001161,N | NUDT2 | nudix hydrolase 2 | protein_coding | NP_001152.1;NP_001231319.1;NP_671701.1;NP_671702.1;XP_0243032! | 8049 | 602852 | . |
| 320 | NM_001163,XI | APBA1 | amyloid beta precursor protein binding family A member 1 | protein_coding | NP_001154.2;XP_005252025.1;XP_011516919.1;XP_016870159.1 | 578 | 602414 | . |
| 321 | NM_00113041 | APBA2 | amyloid beta precursor protein binding family A member 2 | protein_coding | NP_001123886.1;NP_001340717.1;NP_001340718.1;NP_001340719.1;NI | 579 | 602712 | . |
| 322 | NM_001164,N | APBB1 | amyloid beta precursor protein binding family B member 1 | protein_coding | NP_001155.1;NP_001244248.1;NP_001244249.1;NP_001244250.1;NP_0( | 581 | 602709 | . |
| 323 | NM_00116605 | APBB2 | amyloid beta precursor protein binding family B member 2 | protein_coding | NP_001159522.1;NP_001159523.1;NP_001159524.1;NP_001159525.1;NI | 582 | 602710 | . |
| 324 | NM_000038,N | APC | APC regulator of WNT signaling pathway | protein_coding | NP_000029.2;NP_001120982.1;NP_001120983.2;NP_001341824.1;NP_0( | 583 | 611731 | . |
| 327 | NM_001640,XI | APEH | acylaminoacyl-peptide hydrolase | protein_coding | NP_001631.3;XP_005265154.1;XP_005265155.1;XP_011531960.1;XP_01: | 586 | 102645 | . |
| 328 | NM_00124424 | APEX1 | apurinic/apyrimidinic endodeoxyribonuclease 1 | protein_coding | NP_001231178.1;NP_001632.2;NP_542379.1;NP_542380.1 | 587 | 107748 | . |
| 329 | NM_001166,N | BIRC2 | baculoviral IAP repeat containing 2 | protein_coding | NP_001157.1;NP_001243092.1;NP_001243095.1 | 590 | 601712 | . |
| 331 | NM_001167,N | XIAP | X-linked inhibitor of apoptosis | protein_coding | NP_001158.2;NP_001191330.1;NP_001365519.1;NP_001365520.1;NP_0( | 592 | 300079 | . |
| 332 | NM_00101227 | BIRC5 | baculoviral IAP repeat containing 5 | protein_coding | NP_001012270.1;NP_001012271.1;NP_001159.2 | 593 | 603352 | . |
| 333 | NM_00102480 | APLP1 | amyloid beta precursor like protein 1 | protein_coding | NP_001019978.1;NP_005157.1;XP_016882226.1;XP_016882227.1 | 597 | 104775 | . |

| | | | | | | | | | | | | | | | | | | | | | | | | | | | | | | | | | |
|---|---|---|---|---|---|---|---|---|---|---|---|---|---|---|---|---|---|---|---|---|---|---|---|---|---|---|---|---|---|---|---|---|---|
| 334 | NM_00114227 | APLP2 | amyloid beta precursor like protein 2 | protein_coding | NP_001135748.1;NP_001135749.1;NP_001135750.1;NP_001230228.1;NI | 598 | 104776 | . | . | 32.921241 | 30.643676 | 25.292129 | 27.135921 | 76.609078 | 70.834227 | 56.754810 | 62.043058 | -1.218182 | 3243.798863 | 0.11556799 | -2.46373873 | 0.01374963 | 0.388727919 | 11.798834 | 11.514074 | 11.773168 | 11.702868 | 11.549151 | 11.603955 | 4275 | 3683 | 2456 | 2836 |
| 341 | NM_00132106 | APOC1 | apolipoprotein C1 | protein_coding | NP_001307994.1;NP_001307995.1;NP_001366616.1;NP_001636.1 | 607 | 107710 | . | . | 1.307346 | 6.732400 | 1.424131 | 1.984752 | 3.042248 | 15.562244 | 3.195708 | 4.537899 | -2.375080 | 46.502034 | 0.625076566 | -1.99651751 | 0.045877621 | 0.653499062 | 6.033518 | 4.781347 | 5.383700 | 5.700692 | 5.391563 | 5.426662 | 26 | 117 | 20 | 30 |
| 347 | NM_001647 | APOD | apolipoprotein D | protein_coding | NP_001638.1 | 612 | 107740 | . | . | 17.502089 | 19.093161 | 33.194298 | 29.535128 | 40.728077 | 44.134697 | 74.487053 | 67.528564 | 1.690429 | 666.082509 | 0.159198669 | 4.75750874 | 1.95997E-06 | 0.000491881 | 8.951648 | 9.709153 | 9.141578 | 9.190453 | 9.566441 | 9.474446 | 548 | 555 | 774 | 735 |
| 348 | NM_000041,N | APOE | apolipoprotein E | protein_coding | NP_000032.1;NP_001289617.1;NP_001289618.1;NP_001289619.1;NP_0( | 613 | 107741 | . | . | 116.452404 | 118.845098 | 111.349620 | 130.382657 | 270.989561 | 274.715778 | 249.865336 | 298.104441 | 1.015536 | 4353.403292 | 0.116849499 | 0.190339929 | 0.849042767 | 0.998865404 | 12.076846 | 12.098925 | 12.075218 | 12.086445 | 12.030696 | 12.152906 | 4941 | 4693 | 3535 | 4414 |
| 351 | NM_000484,N | APP | amyloid beta precursor protein | protein_coding | NP_000475.1;NP_001129488.1;NP_001129601.1;NP_001129602.1;NP_0( | 620 | 104760 | . | . | 242.503231 | 231.262332 | 207.840189 | 214.960784 | 564.314974 | 534.573231 | 466.387377 | 491.482290 | -1.136371 | 24797.152663 | 0.099974622 | -1.84480768 | 0.065065519 | 0.731993537 | 14.687157 | 14.502726 | 14.674431 | 14.625711 | 14.530106 | 14.550397 | 31227 | 27668 | 19946 | 22023 |
| 353 | NM_000485,N | APRT | adenine phosphoribosyltransferase | protein_coding | NP_000476.1;NP_001025189.1 | 626 | 102600 | . | . | 58.108595 | 58.965207 | 51.224471 | 64.808775 | 135.221094 | 136.300722 | 114.946233 | 148.177559 | -1.034270 | 1588.211706 | 0.143209448 | -0.33945122 | 0.734269838 | 0.998865404 | 10.657516 | 10.608441 | 10.645903 | 10.643272 | 10.520691 | 10.707319 | 1863 | 1740 | 1194 | 1640 |
| 355 | NM_000043,N | FAS | Fas cell surface death receptor | protein_coding | NP_000034.1;NP_001307548.1;NP_690610.1;NP_690611.1;XP_0067178ε | 11920 | 134637 | . | . | 0.175177 | 0.193088 | 0.242768 | 0.328188 | 0.407644 | 0.446333 | 0.544764 | 0.750362 | -1.051069 | 31.682044 | 0.568554422 | -0.12638673 | 0.899425807 | 0.998865404 | 5.020416 | 4.949909 | 5.021695 | 4.953242 | 4.973240 | 4.990110 | 47 | 26 | 25 | 31 |
| 357 | NM_00132066 | SHROOM2 | shroom family member 2 | protein_coding | NP_001307592.1;NP_001307593.1;NP_001640.1;XP_005274557.2;XP_01 | 630 | 300103 | . | . | 4.477611 | 4.645992 | 5.501809 | 4.817654 | 10.419584 | 10.739419 | 12.345898 | 11.014991 | 1.114004 | 1131.783053 | 0.139373118 | 1.117533655 | 0.263766218 | 0.996308385 | 10.064233 | 10.220311 | 10.091270 | 10.111498 | 10.234273 | 10.127178 | 1216 | 1171 | 1111 | 1039 |
| 363 | NM_001652 | AQP6 | aquaporin 6 | protein_coding | NP_001643.2 | 639 | 601383 | . | . | 0.067804 | 0.098843 | 0.097448 | 0.071353 | 0.157782 | 0.228479 | 0.218670 | 0.163140 | 1.002094 | 7.217405 | 1.127432034 | 0.002677097 | 0.997863988 | NA | 2.849979 | 2.852980 | 2.843910 | 2.854434 | 2.852879 | 2.845617 | 7 | 9 | 7 | 6 |
| 367 | NM_000044,N | AR | androgen receptor | protein_coding | NP_000035.2;NP_001011645.1;NP_001334990.1;NP_001334992.1;NP_0( | 644 | 313700 | . | . | 0.482279 | 0.508121 | 0.428697 | 0.489638 | 1.122282 | 1.174545 | 0.961984 | 1.119500 | -1.045210 | 155.309176 | 0.264754453 | -0.2409491 | 0.809594569 | 0.998865404 | 7.310945 | 7.246330 | 7.281668 | 7.294264 | 7.246804 | 7.293491 | 180 | 174 | 124 | 151 |
| 369 | NM_00125619 | ARAF | A-Raf proto-oncogene, serine/threonine kinase | protein_coding | NP_001243125.1;NP_001243126.1;NP_001645.1 | 646 | 311010 | . | . | 13.911861 | 13.860257 | 16.173638 | 15.197699 | 32.373473 | 32.038603 | 36.293176 | 34.747733 | 1.101685 | 1052.462413 | 0.136670184 | 1.02225288 | 0.306661229 | 0.998865404 | 9.967869 | 10.107844 | 10.004363 | 9.999569 | 10.103699 | 10.042531 | 1157 | 1077 | 999 | 992 |
| 372 | NM_00114228 | ARCN1 | archain 1 | protein_coding | NP_001135753.1;NP_001646.2;XP_005271599.1 | 649 | 600820 | . | . | 20.899818 | 22.056364 | 19.052347 | 22.657333 | 48.634732 | 50.984277 | 42.752912 | 51.803300 | -1.044876 | 2624.401601 | 0.124741299 | -0.50770782 | 0.611658265 | 0.998865404 | 11.389235 | 11.325610 | 11.358203 | 11.390750 | 11.271840 | 11.401834 | 3027 | 2952 | 2052 | 2600 |
| 373 | NM_001656,N | TRIM23 | tripartite motif containing 23 | protein_coding | NP_001647.1;NP_150230.1;NP_150231.1;XP_016864933.1 | 660 | 601747 | . | . | 2.471650 | 2.720815 | 2.980762 | 2.211994 | 5.751631 | 6.289285 | 6.688743 | 5.057461 | -1.024111 | 310.703957 | 0.215308663 | -0.15963885 | 0.873165573 | 0.998865404 | 8.296014 | 8.262586 | 8.256644 | 8.308374 | 8.343483 | 8.190720 | 345 | 355 | 305 | 247 |
| 374 | NM_001657 | AREG | amphiregulin | protein_coding | NP_001648.1 | 651 | 104640 | . | . | 0.272267 | 0.283520 | 0.231462 | 0.397645 | 0.633577 | 0.655370 | 0.519394 | 0.909168 | 1.126986 | 11.902621 | 0.892593287 | 0.193223191 | 0.846784178 | NA | 3.486115 | 3.655341 | 3.568730 | 3.568010 | 3.559973 | 3.589165 | 13 | 12 | 8 | 15 |
| 375 | NM_00102422 | ARF1 | ADP ribosylation factor 1 | protein_coding | NP_001019397.1;NP_001019398.1;NP_001019399.1;NP_001649.1;XP_01 | 652 | 103180 | . | . | 124.255309 | 123.006358 | 111.461515 | 128.777534 | 289.147218 | 284.334701 | 250.116417 | 294.434532 | -1.064509 | 7116.016708 | 0.11173727 | -0.80714772 | 0.41958138 | 0.998865404 | 12.841295 | 12.751001 | 12.814127 | 12.830309 | 12.712003 | 12.823584 | 8370 | 7994 | 5597 | 6891 |
| 377 | NM_001659,XI | ARF3 | ADP ribosylation factor 3 | protein_coding | NP_001650.1;XP_005268913.1;XP_006719454.1;XP_024304740.1 | 654 | 103190 | . | . | 29.932405 | 32.640251 | 28.445426 | 32.398931 | 69.653939 | 75.449401 | 63.830714 | 74.076310 | -1.039414 | 3577.057542 | 0.118994955 | -0.46867421 | 0.639302524 | 0.998865404 | 11.832284 | 11.776288 | 11.782725 | 11.855390 | 11.736946 | 11.836122 | 4015 | 4107 | 2866 | 3487 |
| 378 | NM_001660 | ARF4 | ADP ribosylation factor 4 | protein_coding | NP_001651.1 | 655 | 601177 | . | . | 85.717049 | 74.237984 | 69.388321 | 71.024681 | 199.467117 | 171.604416 | 155.705383 | 162.389481 | -1.155601 | 3736.184376 | 0.119172953 | -1.75075771 | 0.079987654 | 0.779933816 | 11.967856 | 11.759314 | 11.985495 | 11.857658 | 11.798710 | 11.809186 | 4963 | 3994 | 2995 | 3271 |
| 379 | NM_001661,XI | ARL4D | ADP ribosylation factor like GTPase 4D | protein_coding | NP_001652.2;XP_011523084.1 | 656 | 600732 | . | . | 3.013136 | 3.344507 | 1.912839 | 3.093549 | 7.011692 | 7.730978 | 4.292356 | 7.073033 | -1.280082 | 140.241417 | 0.303818088 | -1.17253106 | 0.240983911 | 0.992733415 | 7.300128 | 6.941134 | 7.153978 | 7.191797 | 6.999163 | 7.156288 | 173 | 178 | 82 | 142 |
| 381 | NM_001662 | ARF5 | ADP ribosylation factor 5 | protein_coding | NP_001653.1 | 658 | 103188 | . | . | 57.189598 | 57.753071 | 46.861237 | 64.523499 | 133.082535 | 133.498810 | 105.155258 | 147.525299 | -1.046850 | 1819.083427 | 0.151148466 | -0.43701448 | 0.66210085 | 0.998865404 | 10.861875 | 10.795350 | 10.843775 | 10.844439 | 10.680088 | 10.921494 | 2142 | 2009 | 1308 | 1922 |
| 382 | NM_001663 | ARF6 | ADP ribosylation factor 6 | protein_coding | NP_001654.1 | 659 | 600464 | . | . | 19.037685 | 18.104128 | 18.642344 | 17.526918 | 44.301479 | 41.848503 | 41.832878 | 40.073219 | -1.041394 | 2213.572000 | 0.11962728 | -0.48915381 | 0.62473281 | 0.998865404 | 11.141012 | 11.082720 | 11.151413 | 11.103873 | 11.123822 | 11.066412 | 2677 | 2365 | 1954 | 1960 |
| 384 | NM_001172 | ARG2 | arginase 2 | protein_coding | NP_001163.1 | 664 | 107830 | . | . | 7.676244 | 6.829620 | 9.193566 | 8.569577 | 17.862934 | 15.786972 | 20.630096 | 19.593321 | 1.207591 | 480.642153 | 0.174709569 | 1.557626472 | 0.11932182 | 0.858706505 | 8.765945 | 9.038811 | 8.881603 | 8.804745 | 8.988083 | 8.935875 | 533 | 440 | 476 | 473 |
| 387 | NM_00131394 | RHOA | ras homolog family member A | protein_coding | NP_001300870.1;NP_001300872.1;NP_001300873.1;NP_001300874.1;NI | 667 | 165390 | . | . | 101.721685 | 105.572538 | 88.702622 | 111.606082 | 236.710530 | 244.035631 | 199.046116 | 255.174043 | -1.048218 | 5740.750877 | 0.125644934 | -0.54071866 | 0.588701514 | 0.998865404 | 12.520675 | 12.452568 | 12.492858 | 12.517199 | 12.369153 | 12.553449 | 6669 | 6432 | 4339 | 5832 |
| 388 | NM_004040 | RHOB | ras homolog family member B | protein_coding | NP_004031.1 | 668 | 165370 | . | . | 26.265705 | 26.203543 | 19.160570 | 22.689587 | 61.121380 | 60.570660 | 42.995766 | 51.877045 | -1.271414 | 1740.345257 | 0.13048943 | -2.6548799 | 0.007933665 | 0.280790707 | 10.928159 | 10.581367 | 10.855742 | 10.846740 | 10.601101 | 10.721061 | 2256 | 2091 | 1227 | 1550 |
| 389 | NM_00104267 | RHOC | ras homolog family member C | protein_coding | NP_001036143.1;NP_001036144.1;NP_786886.1 | 669 | 165380 | . | . | 245.866768 | 238.779115 | 190.523148 | 239.246152 | 572.142107 | 551.948643 | 427.528448 | 547.007893 | -1.139551 | 8157.053048 | 0.12426869 | -1.51659429 | 0.129369149 | 0.879150687 | 13.085040 | 12.896467 | 13.057598 | 13.032968 | 12.836761 | 13.023539 | 10149 | 9242 | 5894 | 7942 |
| 390 | NM_00125473 | RND3 | Rho family GTPase 3 | protein_coding | NP_001241667.1;NP_005159.1 | 671 | 602924 | . | . | 18.206609 | 16.944469 | 17.938179 | 15.885791 | 42.367527 | 39.167897 | 40.252757 | 36.320976 | -1.056333 | 1463.920643 | 0.13421785 | -0.58907875 | 0.555808445 | 0.998865404 | 10.554389 | 10.475784 | 10.568956 | 10.501088 | 10.542648 | 10.441848 | 1807 | 1552 | 1320 | 1247 |
| 391 | NM_001665,XI | RHOG | ras homolog family member G | protein_coding | NP_001656.2;XP_005252973.1;XP_016873208.1 | 672 | 179505 | . | . | 22.638675 | 25.307231 | 17.989820 | 23.153914 | 52.681125 | 58.498802 | 40.368635 | 52.938673 | -1.183871 | 906.851159 | 0.159570266 | -1.52605129 | 0.126997091 | 0.874064017 | 9.941747 | 9.697359 | 9.841748 | 9.918271 | 9.672152 | 9.838696 | 1074 | 1116 | 636 | 871 |
| 392 | NM_004308,XI | ARHGAP1 | Rho GTPase activating protein 1 | protein_coding | NP_004299.1;XP_011518397.1;XP_024304288.1 | 673 | 602732 | . | . | 7.064789 | 7.921517 | 7.594971 | 8.588674 | 16.440055 | 18.310943 | 17.042895 | 19.636985 | 1.068515 | 823.352701 | 0.150578445 | 0.634931473 | 0.525473109 | 0.998865404 | 9.637088 | 9.732082 | 9.622122 | 9.697420 | 9.667401 | 9.745599 | 868 | 905 | 699 | 841 |
| 393 | NM_00116474 | ARHGAP4 | Rho GTPase activating protein 4 | protein_coding | NP_001158213.1;NP_001657.3 | 674 | 300023 | . | . | 0.770302 | 0.637576 | 0.761456 | 0.495085 | 1.792525 | 1.473785 | 1.708685 | 1.131954 | -1.133524 | 70.729812 | 0.395923473 | -0.45669138 | 0.647892889 | 0.998865404 | 6.230303 | 6.052730 | 6.181332 | 6.130341 | 6.172797 | 6.077015 | 95 | 73 | 70 | 49 |
| 394 | NM_00103005 | ARHGAP5 | Rho GTPase activating protein 5 | protein_coding | NP_001025226.1;NP_001164.2;XP_005267692.1;XP_005267693.1;XP_01 | 675 | 602680 | . | . | 5.653218 | 5.009951 | 5.527045 | 4.176223 | 13.155270 | 11.580726 | 12.402529 | 9.548440 | -1.076741 | 1481.476071 | 0.166527223 | -0.6405662 | 0.521804563 | 0.998865404 | 10.584833 | 10.478860 | 10.612360 | 10.496764 | 10.624202 | 10.356811 | 1899 | 1535 | 1457 | 1105 |
| 396 | NM_00118507 | ARHGDIA | Rho GDP dissociation inhibitor alpha | protein_coding | NP_001172006.1;NP_001172007.1;NP_001288169.1;NP_001288170.1;NI | 678 | 601925 | . | . | 89.208371 | 101.269266 | 86.284195 | 103.354743 | 207.591534 | 234.088444 | 193.619231 | 236.308338 | -1.014699 | 5460.256625 | 0.123167092 | -0.17092599 | 0.86428196 | 0.998865404 | 12.425334 | 12.404094 | 12.372212 | 12.464979 | 12.333001 | 12.476620 | 5985 | 6262 | 4310 | 5516 |
| 397 | NM_001175,N | ARHGDIB | Rho GDP dissociation inhibitor beta | protein_coding | NP_001166.3;NP_001308349.1;NP_001308350.1;NP_001308351.1;NP_0( | 679 | 602843 | . | . | 1.648663 | 1.093706 | 0.373671 | 1.376651 | 3.836506 | 2.528150 | 0.838506 | 3.147549 | -1.517103 | 40.990525 | 0.588240484 | -1.02223313 | 0.306670575 | 0.998865404 | 5.628808 | 5.022362 | 5.416160 | 5.349231 | 5.243169 | 5.388142 | 67 | 44 | 12 | 48 |
| 400 | NM_001177,N | ARL1 | ADP ribosylation factor like GTPase 1 | protein_coding | NP_001168.1;NP_001287997.1 | 692 | 603425 | . | . | 18.417271 | 16.830804 | 15.769392 | 16.669215 | 42.857745 | 38.905158 | 35.386060 | 38.112181 | -1.101568 | 1686.568203 | 0.126260961 | -1.10531959 | 0.269021133 | 0.998865404 | 10.787946 | 10.648432 | 10.794611 | 10.716488 | 10.662986 | 10.697979 | 2136 | 1814 | 1364 | 1539 |
| 402 | NM_00119974 | ARL2 | ADP ribosylation factor like GTPase 2 | protein_coding | NP_001186674.1;NP_001658.2 | 693 | 601175 | . | . | 32.144929 | 31.078733 | 24.578222 | 32.835636 | 74.802572 | 71.839890 | 55.152816 | 75.074784 | -1.116453 | 869.483130 | 0.163016071 | -0.97489109 | 0.329614286 | 0.998865404 | 9.841615 | 9.682001 | 9.815817 | 9.784583 | 9.621213 | 9.812403 | 1079 | 969 | 617 | 875 |
| 403 | NM_004311,XI | ARL3 | ADP ribosylation factor like GTPase 3 | protein_coding | NP_004302.1;XP_016871749.1 | 694 | 604695 | . | . | 3.147728 | 3.117377 | 2.969311 | 3.056229 | 7.324895 | 7.205955 | 6.663047 | 6.987706 | -1.051533 | 369.910013 | 0.185403724 | -0.39100319 | 0.695794876 | 0.998865404 | 8.566874 | 8.494273 | 8.548882 | 8.542483 | 8.508344 | 8.524776 | 439 | 407 | 309 | 343 |
| 405 | NM_00119732 | ARNT | aryl hydrocarbon receptor nuclear translocator | protein_coding | NP_001184254.1;NP_001272964.1;NP_001272965.1;NP_001337153.1;NI | 700 | 126110 | . | . | 3.526671 | 3.909860 | 4.010908 | 4.285295 | 8.206708 | 9.037814 | 9.000363 | 9.797820 | 1.112518 | 566.680942 | 0.164542231 | 0.9348925 | 0.349843703 | 0.998865404 | 9.067752 | 9.220968 | 9.076319 | 9.141132 | 9.151708 | 9.205278 | 587 | 608 | 498 | 582 |
| 406 | NM_00103027 | ARNTL | aryl hydrocarbon receptor nuclear translocator like | protein_coding | NP_001025443.1;NP_001025444.1;NP_001169.3;NP_001284648.1;NP_0( | 701 | 602550 | . | . | 2.352613 | 2.325880 | 2.512463 | 2.549548 | 5.474631 | 5.376372 | 5.637895 | 5.829236 | 1.054458 | 221.579192 | 0.225901489 | 0.338647535 | 0.734875268 | 0.998865404 | 7.752955 | 7.829390 | 7.773752 | 7.782940 | 7.803900 | 7.802996 | 246 | 235 | 198 | 213 |
| 408 | NM_004041,N | ARRB1 | arrestin beta 1 | protein_coding | NP_004032.2;NP_064647.1;XP_011543336.1;XP_011543337.1;XP_01687 | 711 | 107940 | . | . | 4.339128 | 5.078943 | 5.265343 | 6.118518 | 10.097330 | 11.740204 | 11.815276 | 13.989262 | 1.193530 | 1190.064019 | 0.144842636 | 1.762155822 | 0.078042974 | 0.768898765 | 10.083900 | 10.338527 | 10.088252 | 10.197847 | 10.222024 | 10.329784 | 1157 | 1258 | 1047 | 1299 |
| 409 | NM_00125732 | ARRB2 | arrestin beta 2 | protein_coding | NP_001244257.1;NP_001244258.1;NP_001244259.1;NP_001244260.1;NI | 712 | 107941 | . | . | 11.417773 | 9.838987 | 10.799749 | 11.962530 | 26.569629 | 22.743262 | 24.234321 | 27.350905 | 1.049699 | 616.015714 | 0.164435852 | 0.425545987 | 0.670438705 | 0.998865404 | 9.231352 | 9.301443 | 9.299019 | 9.197075 | 9.253651 | 9.309094 | 745 | 599 | 526 | 616 |
| 410 | NM_000487,N | ARSA | arylsulfatase A | protein_coding | NP_000478.3;NP_001078894.2;NP_001078895.2;NP_001078896.2;NP_0( | 713 | 607574 | . | . | 20.870153 | 21.545129 | 24.318897 | 22.595282 | 48.565703 | 49.802533 | 54.570900 | 51.661424 | 1.101623 | 2009.377493 | 0.122903076 | 1.136097999 | 0.255915552 | 0.995760118 | 10.900914 | 11.040764 | 10.932543 | 10.931677 | 11.052845 | 10.963506 | 2203 | 2062 | 1934 | 1865 |
| 411 | NM_000046,N | ARSB | arylsulfatase B | protein_coding | NP_000037.2;NP_942002.1;XP_011541692.1;XP_011541693.1;XP_01154 | 714 | 611542 | . | . | 12.955976 | 13.348905 | 12.152599 | 10.953221 | 30.149087 | 30.856589 | 27.270082 | 25.043236 | -1.075942 | 1624.641071 | 0.125842244 | -0.83915185 | 0.401384103 | 0.998865404 | 10.717875 | 10.611993 | 10.669229 | 10.717442 | 10.608483 | 10.663781 | 1880 | 1873 | 1312 | 1520 |
| 412 | NM_000351,N | STS | steroid sulfatase | protein_coding | NP_000342.3;NP_001307679.1;NP_001307680.1;NP_001307681.2;NP_0( | 11425 | 300747 | . | . | 0.866952 | 0.798569 | 0.777739 | 0.695121 | 2.017431 | 1.845929 | 1.745224 | 1.589311 | -1.135047 | 163.426186 | 0.261498931 | -0.69886393 | 0.484637069 | 0.998865404 | 7.440419 | 7.258863 | 7.396704 | 7.357993 | 7.354289 | 7.297166 | 209 | 179 | 143 | 133 |
| 414 | NM_001669,N | ARSD | arylsulfatase D | protein_coding | NP_001660.2;NP_033667.2;XP_005274571.1;XP_005274572.1 | 717 | 300002 | . | . | 0.047018 | 0.159877 | 0.063149 | 0.062231 | 0.109412 | 0.369562 | 0.141704 | 0.142284 | -1.642521 | 14.663295 | 0.866889114 | -0.82584054 | 0.408894553 | NA | 4.189301 | 3.470304 | 3.839688 | 3.923932 | 3.853675 | 3.849764 | 10 | 30 | 10 | 10 |
| 415 | NM_000047,N | ARSL | arylsulfatase L | protein_coding | NP_000038.2;NP_001269557.1;NP_001269560.2;NP_001356008.1;NP_0( | 719 | 300180 | . | . | 0.948375 | 1.290117 | 0.588542 | 1.171159 | 2.206909 | 2.982163 | 1.320672 | 2.677717 | -1.482714 | 72.309164 | 0.399553219 | -1.42219015 | 0.154971057 | 0.923603412 | 6.433969 | 5.861992 | 6.179679 | 6.256703 | 6.076922 | 6.168599 | 86 | 106 | 39 | 67 |
| 419 | NM_00113001 | ART3 | ADP-ribosyltransferase 3 | protein_coding | NP_001123488.1;NP_001123489.1;NP_001170.2;NP_001364102.1;NP_0( | 725 | 603086 | . | . | 3.881837 | 5.294019 | 3.677305 | 3.847988 | 9.033195 | 12.237362 | 8.251769 | 8.797965 | -1.033545 | 334.688544 | 0.192229778 | -0.24762972 | 0.804420911 | 0.998865404 | 8.410299 | 8.362659 | 8.399347 | 8.392765 | 8.373346 | 8.381850 | 394 | 365 | 284 | 311 |
| 421 | NM_001670,XI | ARVCF | ARVCF delta catenin family member | protein_coding | NP_001661.1;XP_005261299.1;XP_005261300.1;XP_005261301.1;XP_00( | 728 | 602269 | . | . | 2.867844 | 2.530327 | 3.760093 | 2.888842 | 6.673596 | 5.848963 | 8.437538 | 6.604991 | 1.072776 | 304.301625 | 0.205996 | 0.491992405 | 0.622724713 | 0.998865404 | 8.197251 | 8.299649 | 8.261061 | 8.194349 | 8.301299 | 8.230075 | 359 | 297 | 292 | 276 |
| 427 | NM_00112750 | ASAH1 | N-acylsphingosine amidohydrolase 1 | protein_coding | NP_001120977.1;NP_001350672.1;NP_004306.3;NP_808592.2;XP_00527 | 735 | 613468 | . | . | 147.107053 | 132.605584 | 149.129209 | 121.047742 | 342.324165 | 306.523739 | 334.641633 | 276.761257 | -1.049520 | 11796.682357 | 0.125984991 | -0.55347296 | 0.579939594 | 0.998865404 | 13.560489 | 13.490858 | 13.594678 | 13.492014 | 13.589218 | 13.409619 | 14742 | 12258 | 11064 | 9657 |
| 432 | NM_00119721 | ASGR1 | asialoglycoprotein receptor 1 | protein_coding | NP_001184145.1;NP_001662.1;XP_011522163.1 | 742 | 108360 | . | . | 0.164029 | 0.662822 | 0.297930 | 0.138376 | 0.381703 | 1.532143 | 0.668547 | 0.316379 | -1.857025 | 14.417876 | 0.91286106 | -0.97823566 | 0.327957782 | NA | 4.227734 | 3.336262 | 3.806231 | 3.902533 | 3.834020 | 3.801451 | 9 | 32 | 12 | 6 |
| 435 | NM_000048,N | ASL | argininosuccinate lyase | protein_coding | NP_000039.2;NP_001020114.1;NP_001020115.1;NP_001020117.1 | 746 | 608310 | . | . | 10.143440 | 9.650453 | 9.573740 | 11.016025 | 23.604205 | 22.307454 | 21.483193 | 25.186832 | 1.041581 | 706.332718 | 0.154204714 | 0.381153417 | 0.703089413 | 0.998865404 | 9.434641 | 9.493173 | 9.469249 | 9.429318 | 9.438592 | 9.514862 | 820 | 725 | 592 | 713 |
| 436 | gene-ASLP1 | ASLP1 | argininosuccinate lyase pseudogene 1 | pseudogene | . | 747 | . | | . | 0.815254 | 0.820567 | 0.964722 | 0.930433 | 1.897129 | 1.896777 | 2.164807 | 2.127325 | 1.149232 | 34.696064 | 0.524141445 | 0.382854146 | 0.701827912 | 0.998865404 | 5.012757 | 5.213655 | 5.103361 | 5.104738 | 5.126253 | 5.119986 | 37 | 35 | 33 | 34 |
| 439 | NM_00137148 | GET3 | guided entry of tail-anchored proteins factor 3, ATPase | protein_coding | NP_001358417.1;NP_001358418.1;NP_004308.2 | 752 | 601913 | . | . | 47.308300 | 50.686584 | 44.411008 | 50.867659 | 110.088357 | 117.164309 | 99.657019 | 116.302865 | -1.042745 | 1920.223267 | 0.126525598 | -0.47726248 | 0.633175227 | 0.998865404 | 10.937103 | 10.876374 | 10.898782 | 10.946575 | 10.839110 | 10.937886 | 2190 | 2179 | 1532 | 1873 |
| 440 | NM_00117807 | ASNS | asparagine synthetase (glutamine-hydrolyzing) | protein_coding | NP_001171546.1;NP_001171547.1;NP_001171548.1;NP_001339425.1;NI | 753 | 108370 | . | . | 32.004148 | 28.774006 | 12.739026 | 24.017492 | 74.474964 | 66.512408 | 28.586006 | 54.913139 | -1.674997 | 1815.901523 | 0.439567594 | -1.69293365 | 0.090468097 | 0.805424371 | 11.151141 | 10.406850 | 11.030312 | 10.935367 | 10.329122 | 10.777729 | 2767 | 2315 | 826 | 1655 |
| 443 | NM_000049,N | ASPA | aspartoacylase | protein_coding | NP_000040.1;NP_001121557.1;XP_016880150.1;XP_024306532.1 | 756 | 608034 | . | . | 0.133093 | 0.160856 | 0.371103 | 0.182429 | 0.309712 | 0.371827 | 0.832744 | 0.417101 | 1.191647 | 28.083219 | 0.584419629 | 0.432834279 | 0.665135168 | 0.998865404 | 4.679488 | 4.932690 | 4.786807 | 4.805388 | 4.827112 | 4.805133 | 27 | 30 | 29 | 26 |
| 444 | NM_00116475 | ASPH | aspartate beta-hydroxylase | protein_coding | NP_001158222.1;NP_001158223.1;NP_001158224.1;NP_001158225.1;NI | 757 | 600582 | . | . | 49.227335 | 38.341073 | 37.657813 | 34.472628 | 114.554029 | 88.627114 | 84.503045 | 78.817569 | -1.204003 | 5310.066277 | 0.1290153 | -2.07602586 | 0.037891559 | 0.60652635 | 12.502149 | 12.234482 | 12.535410 | 12.347418 | 12.327348 | 12.251158 | 7413 | 5573 | 4384 | 4309 |
| 463 | NM_00116476 | ZFHX3 | zinc finger homeobox 3 | protein_coding | NP_001158238.1;NP_008816.3;XP_005256014.1;XP_016878718.1;XP_01 | 777 | 104155 | . | . | 1.231713 | 0.995359 | 1.065074 | 1.267127 | 2.866246 | 2.300817 | 2.389993 | 2.897136 | 1.093809 | 531.202201 | 0.167798228 | 0.770929465 | 0.440748741 | 0.998865404 | 8.987127 | 9.116221 | 9.041580 | 9.003077 | 9.042294 | 9.116788 | 602 | 531 | 455 | 550 |
| 466 | NM_005171,XI | ATF1 | activating transcription factor 1 | protein_coding | NP_005162.1;XP_011536688.1;XP_011536689.1;XP_011536690.1;XP_01( | 783 | 123803 | . | . | 3.407968 | 3.980685 | 4.071440 | 4.719247 | 7.930484 | 9.201531 | 9.136193 | 10.789996 | 1.155056 | 323.156664 | 0.199335698 | 1.043276735 | 0.296820143 | 0.998865404 | 8.228820 | 8.435933 | 8.263348 | 8.321190 | 8.343143 | 8.400193 | 327 | 341 | 286 | 341 |
| 467 | NM_00103028 | ATF3 | activating transcription factor 3 | protein_coding | NP_001025458.1;NP_001035709.1;NP_001193413.2;NP_001193415.1;NI | 785 | 603148 | . | . | 1.706436 | 1.900666 | 1.687302 | 2.640005 | 3.970949 | 4.393476 | 3.786255 | 6.036059 | 1.163764 | 127.193844 | 0.310975379 | 0.703586524 | 0.481690306 | 0.998865404 | 6.878630 | 7.095033 | 6.937562 | 6.975824 | 6.930887 | 7.084554 | 128 | 134 | 94 | 155 |
| 468 | NM_001675,N | ATF4 | activating transcription factor 4 | protein_coding | NP_001666.2;NP_877962.1;XP_016884296.1 | 786 | 604064 | . | . | 146.066035 | 168.337336 | 140.628366 | 184.251066 | 339.901688 | 389.119291 | 315.565963 | 421.268164 | 1.007761 | 9188.568974 | 0.117270743 | 0.095113465 | 0.924224703 | 0.998865404 | 13.160078 | 13.171150 | 13.169307 | 13.153310 | 13.087774 | 13.242210 | 10639 | 9784 | 7294 | 9431 |
| 471 | NM_004044,XI | ATIC | 5-aminoimidazole-4-carboxamide ribonucleotide formyltransferas | protein_coding | NP_004035.2;XP_016859676.1;XP_024308687.1 | 794 | 601731 | . | . | 90.755148 | 90.210768 | 90.060554 | 94.553859 | 211.190944 | 208.526231 | 202.093266 | 216.186166 | 1.006846 | 5692.628199 | 0.10550524 | 0.093292378 | 0.925671288 | 0.998865404 | 12.469970 | 12.479771 | 12.477102 | 12.466733 | 12.455749 | 12.499091 | 6573 | 6084 | 4839 | 5493 |
| 472 | NM_000051,N | ATM | ATM serine/threonine kinase | protein_coding | NP_000042.3;NP_001338763.1;NP_001338764.1;NP_001338765.1;NP_0( | 795 | 607585 | . | . | 14.596018 | 14.392986 | 18.638016 | 14.450090 | 33.965532 | 33.270034 | 41.823170 | 33.038411 | 1.178688 | 5713.631520 | 0.141164078 | 1.680188377 | 0.09292067 | 0.809520218 | 12.356654 | 12.593982 | 12.391565 | 12.415843 | 12.667156 | 12.400010 | 5951 | 5742 | 6213 | 4894 |
| 473 | NM_00104268 | RERE | arginine-glutamic acid dipeptide repeats | protein_coding | NP_001036146.1;NP_001036147.1;NP_036234.3 | 9965 | 605226 | . | . | 14.701251 | 15.843210 | 18.086759 | 17.714694 | 34.210415 | 36.622290 | 40.586162 | 40.502541 | 1.189911 | 3787.064142 | 0.116195245 | 2.158896918 | 0.030858165 | 0.558237552 | 11.756017 | 12.006824 | 11.760648 | 11.856763 | 11.977224 | 11.929694 | 3748 | 3952 | 3680 | 3749 |
| 475 | NM_004045 | ATOX1 | antioxidant 1 copper chaperone | protein_coding | NP_004036.1 | 798 | 602270 | . | . | 373.003387 | 380.648193 | 301.950531 | 432.879272 | 867.994202 | 879.885315 | 677.568237 | 989.726990 | -1.040783 | 5458.443036 | 0.14730943 | -0.3914809 | 0.695441805 | 0.998865404 | 12.442913 | 12.385054 | 12.422323 | 12.432150 | 12.237439 | 12.529148 | 6374 | 6043 | 3846 | 5883 |
| 476 | NM_000701,N | ATP1A1 | ATPase Na+/K+ transporting subunit alpha 1 | protein_coding | NP_000692.2;NP_001153705.1;NP_001153706.1;XP_016856849.1;XP_01 | 799 | 182310 | . | . | 754.265634 | 741.804341 | 689.458254 | 760.491838 | 1755.207147 | 1714.713835 | 1547.124361 | 1738.774182 | -1.045629 | 84441.663105 | 0.101333294 | -0.63523998 | 0.525271912 | 0.998865404 | 16.397496 | 16.333120 | 16.395162 | 16.373625 | 16.307606 | 16.382654 | 100634 | 91995 | 68670 | 80763 |
| 478 | NM_00125621 | ATP1A3 | ATPase Na+/K+ transporting subunit alpha 3 | protein_coding | NP_001243142.1;NP_001243143.1;NP_689509.1 | 801 | 182350 | . | . | 0.188600 | 0.201293 | 0.047430 | 0.083061 | 0.438878 | 0.465298 | 0.106433 | 0.189910 | -2.913000 | 15.057386 | 0.829319078 | -1.8599667 | 0.062890237 | NA | 4.487990 | 2.940155 | 3.948747 | 3.954514 | 3.880233 | 3.896210 | 25 | 25 | 5 | 9 |
| 481 | NM_001677 | ATP1B1 | ATPase Na+/K+ transporting subunit beta 1 | protein_coding | NP_001668.1 | 804 | 182330 | . | . | 28.357349 | 24.012211 | 29.284470 | 24.329594 | 65.988724 | 55.505302 | 65.713501 | 55.626724 | 1.009329 | 1830.141705 | 0.143939974 | 0.09307032 | 0.925847696 | 0.998865404 | 10.830764 | 10.844682 | 10.898456 | 10.760511 | 10.913341 | 10.759465 | 2280 | 1794 | 1755 | 1556 |
| 482 | NM_00130326 | ATP1B2 | ATPase Na+/K+ transporting subunit beta 2 | protein_coding | NP_001290192.1;NP_001669.3 | 805 | 182331 | . | . | 0.190177 | 0.125469 | 0.120783 | 0.073024 | 0.442551 | 0.290027 | 0.271033 | 0.166962 | -1.645403 | 13.981708 | 0.8373678 | -0.85797508 | 0.390906226 | NA | 4.118683 | 3.404839 | 3.840612 | 3.809887 | 3.804658 | 3.786675 | 24 | 15 | 11 | 8 |
| 483 | NM_001679 | ATP1B3 | ATPase Na+/K+ transporting subunit beta 3 | protein_coding | NP_001670.1 | 806 | 601867 | . | . | 192.111084 | 182.254868 | 195.826889 | 165.176453 | 447.050415 | 421.290253 | 439.429871 | 377.656342 | -1.051280 | 10582.512750 | 0.11795123 | -0.61167246 | 0.540754485 | 0.998865404 | 13.404971 | 13.332919 | 13.415169 | 13.362676 | 13.420344 | 13.267196 | 12872 | 11346 | 9780 | 8803 |
| 484 | gene-ATP1B3P | ATP1B3P1 | ATPase Na+/K+ transporting subunit beta 3 pseudogene 1 | pseudogene | . | 807 | . | | . | 0.069654 | 0.075734 | 0.110618 | 0.072717 | 0.162088 | 0.175063 | 0.248224 | 0.166258 | 1.393540 | 4.299322 | 1.44525516 | 0.331259241 | 0.740448676 | NA | 1.844134 | 2.324312 | 2.091499 | 2.092661 | 2.101856 | 2.095299 | 4 | 4 | 5 | 4 |
| 485 | gene-ATP1B1P | ATP1B1P1 | ATPase Na+/K+ transporting subunit beta 1 pseudogene 1 | pseudogene | . | 809 | . | | . | 0.037393 | 0.030218 | 0.033835 | 0.065911 | 0.087015 | 0.069851 | 0.075925 | 0.150697 | 1.634156 | 3.542205 | 1.593114532 | 0.444754905 | 0.656496886 | NA | 1.429096 | 2.134756 | 1.811221 | 1.812122 | 1.815310 | 1.823837 | 3 | 3 | 3 | 5 |
| 487 | NM_00128607 | ATP2A1 | ATPase sarcoplasmic/endoplasmic reticulum Ca2+ transporting 1 | protein_coding | NP_001273004.1;NP_004311.1;NP_775293.1 | 811 | 108730 | . | . | 0.568980 | 0.606026 | 0.533434 | 0.502169 | 1.324040 | 1.400856 | 1.197011 | 1.148151 | -1.102082 | 59.973765 | 0.406144661 | -0.34527564 | 0.729887156 | 0.998865404 | 5.974755 | 5.834349 | 5.903184 | 5.930557 | 5.901976 | 5.890653 | 68 | 72 | 51 | 52 |
| 488 | NM_001681,N | ATP2A2 | ATPase sarcoplasmic/endoplasmic reticulum Ca2+ transporting 2 | protein_coding | NP_001672.1;NP_733765.1;XP_005253945.1;XP_011536704.1 | 812 | 108740 | . | . | 47.459221 | 45.855906 | 43.688302 | 42.597210 | 110.439561 | 105.997989 | 98.035282 | 97.393460 | -1.095381 | 11421.960498 | 0.10193344 | -1.28940029 | 0.19725896 | 0.96621494 | 13.543727 | 13.412328 | 13.538297 | 13.495445 | 13.451986 | 13.426500 | 14093 | 12565 | 9623 | 10067 |
| 489 | NM_005173,N | ATP2A3 | ATPase sarcoplasmic/endoplasmic reticulum Ca2+ transporting 3 | protein_coding | NP_005164.2;NP_777613.1;NP_777614.1;NP_777615.1;NP_777616.1;NP_ | 813 | 601929 | . | . | 1.254740 | 1.252732 | 0.903618 | 1.070709 | 2.919831 | 2.895747 | 2.027692 | 2.448052 | -1.263534 | 168.652848 | 0.257827598 | -1.30887693 | 0.190576042 | 0.962151098 | 7.557263 | 7.218751 | 7.440757 | 7.448136 | 7.316953 | 7.377847 | 215 | 205 | 119 | 151 |
| 490 | NM_00100132 | ATP2B1 | ATPase plasma membrane Ca2+ transporting 1 | protein_coding | NP_001001323.1;NP_001353449.1;NP_001353450.1;NP_001353451.1;NI | 814 | 108731 | . | . | 6.743079 | 5.794425 | 6.362748 | 6.079058 | 15.691425 | 13.394073 | 14.277823 | 13.899042 | -1.025572 | 1369.579055 | 0.136685408 | -0.26651057 | 0.78984602 | 0.998865404 | 10.437429 | 10.401379 | 10.487420 | 10.365487 | 10.425685 | 10.392165 | 1723 | 1378 | 1202 | 1240 |
| 493 | NM_00100139 | ATP2B4 | ATPase plasma membrane Ca2+ transporting 4 | protein_coding | NP_001001396.1;NP_001352712.1;NP_001352713.1;NP_001675.3 | 817 | 108732 | . | . | 11.864787 | 11.873966 | 9.933400 | 11.171978 | 27.609845 | 27.447207 | 22.290262 | 25.543399 | -1.130470 | 2987.515837 | 0.115467853 | -1.53222326 | 0.125467351 | 0.873901718 | 11.630549 | 11.453483 | 11.599214 | 11.585967 | 11.452727 | 11.530433 | 3680 | 3394 | 2321 | 2756 |
| 498 | NM_00100193 | ATP5F1A | ATP synthase F1 subunit alpha | protein_coding | NP_001001935.1;NP_001001937.1;NP_001244263.1;NP_001244264.1;NI | 823 | 164360 | . | . | 182.304988 | 179.738216 | 152.946461 | 193.843320 | 424.231208 | 415.472939 | 343.207425 | 443.199738 | -1.061564 | 11223.318977 | 0.121552038 | -0.70909105 | 0.478267976 | 0.998865404 | 13.496701 | 13.410433 | 13.487835 | 13.467790 | 13.332133 | 13.513477 | 13465 | 12327 | 8460 | 11294 |
| 501 | NM_001182,N | ALDH7A1 | aldehyde dehydrogenase 7 family member A1 | protein_coding | NP_001173.2;NP_001188306.1;NP_001189333.2 | 877 | 107323 | . | . | 15.516194 | 14.147367 | 13.609134 | 16.082951 | 36.106820 | 32.702272 | 30.538502 | 36.771757 | -1.013342 | 2203.548352 | 0.129897716 | -0.14719863 | 0.882975246 | 0.998865404 | 11.115213 | 11.095948 | 11.149662 | 11.068624 | 11.035167 | 11.159646 | 2683 | 2273 | 1755 | 2212 |
| 506 | NM_001686 | ATP5F1B | ATP synthase F1 subunit beta | protein_coding | NP_001677.2 | 830 | 102910 | . | . | 326.987030 | 341.879730 | 272.244476 | 346.927063 | 760.912231 | 790.270325 | 610.908691 | 793.207520 | -1.095941 | 17644.498332 | 0.123692035 | -1.06854721 | 0.285273745 | 0.998865404 | 14.171533 | 14.039300 | 14.127750 | 14.158843 | 13.961126 | 14.158755 | 20866 | 20269 | 12949 | 17608 |
| 509 | NM_00100197 | ATP5F1C | ATP synthase F1 subunit gamma | protein_coding | NP_001001973.1;NP_001307815.1;NP_005165.1;XP_016871779.1 | 833 | 108729 | . | . | 119.897289 | 120.495544 | 108.341107 | 131.781359 | 279.005927 | 278.530862 | 243.114309 | 301.302421 | -1.015700 | 4149.994508 | 0.122027922 | -0.18416842 | 0.85388133 | 0.998865404 | 12.030177 | 12.007521 | 12.026093 | 12.022691 | 11.932873 | 12.084315 | 4826 | 4504 | 3254 | 4212 |
| 513 | NM_00100197 | ATP5F1D | ATP synthase F1 subunit delta | protein_coding | NP_001001975.1;NP_001678.1 | 837 | 603150 | . | . | 37.424402 | 36.266424 | 33.235976 | 38.703587 | 87.088123 | 83.831463 | 74.580567 | 88.491150 | -1.035303 | 953.711285 | 0.145417369 | -0.34420701 | 0.730690609 | 0.998865404 | 9.922397 | 9.871981 | 9.925503 | 9.893097 | 9.829818 | 9.935635 | 1142 | 1024 | 755 | 943 |
| 514 | NM_006886 | ATP5F1E | ATP synthase F1 subunit epsilon | protein_coding | NP_008817.1 | 838 | 606153 | . | . | 48.543991 | 43.385574 | 42.098427 | 48.679501 | 112.963860 | 100.287704 | 94.467644 | 111.299889 | -1.027434 | 5137.325853 | 0.120228349 | -0.32476372 | 0.745359903 | 0.998865404 | 12.346214 | 12.307124 | 12.386661 | 12.284986 | 12.256607 | 12.369135 | 6358 | 5279 | 4110 | 5071 |
| 515 | NM_001688 | ATP5PB | ATP synthase peripheral stalk-membrane subunit b | protein_coding | NP_001679.2 | 840 | 603270 | . | . | 76.888794 | 71.244781 | 68.708237 | 75.228035 | 178.923370 | 164.685501 | 154.179291 | 171.999969 | -1.043964 | 5978.973795 | 0.110789724 | -0.56026323 | 0.575299902 | 0.998865404 | 12.576394 | 12.514302 | 12.598395 | 12.526642 | 12.492718 | 12.560032 | 7331 | 6311 | 4883 | 5705 |
| 516 | NM_00100202 | ATP5MC1 | ATP synthase membrane subunit c locus 1 | protein_coding | NP_001002027.1;NP_005166.1 | 841 | 603192 | . | . | 108.816801 | 105.506768 | 95.659110 | 122.903483 | 253.221180 | 243.883609 | 214.656255 | 281.004208 | 1.002266 | 1961.820806 | 0.140679402 | 0.023208984 | 0.981483572 | 0.998865404 | 10.936511 | 10.939443 | 10.954959 | 10.915266 | 10.835854 | 11.029090 | 2313 | 2062 | 1500 | 2065 |
| 517 | NM_00100203 | ATP5MC2 | ATP synthase membrane subunit c locus 2 | protein_coding | NP_001002031.1;NP_001317198.1;NP_001356682.1;NP_001356683.1;NI | 842 | 603193 | . | . | 110.048278 | 118.655198 | 115.182723 | 125.940190 | 256.086871 | 274.276805 | 258.466694 | 287.947256 | 1.041690 | 2449.481870 | 0.117879136 | 0.499879139 | 0.617160182 | 0.998865404 | 11.228602 | 11.287322 | 11.220313 | 11.261806 | 11.242627 | 11.304735 | 2692 | 2656 | 2090 | 2433 |
| 518 | NM_00100225 | ATP5MC3 | ATP synthase membrane subunit c locus 3 | protein_coding | NP_001002258.1;NP_001177258.1;NP_001680.1 | 843 | 602736 | . | . | 95.759897 | 90.738052 | 78.794366 | 92.807766 | 222.837210 | 209.745057 | 176.812270 | 212.193925 | -1.103060 | 7297.688169 | 0.11524397 | -1.22792741 | 0.219474219 | 0.982008277 | 12.902280 | 12.760695 | 12.898755 | 12.845825 | 12.725209 | 12.850084 | 9088 | 8005 | 5586 | 6990 |
| 521 | NM_007100,N | ATP5ME | ATP synthase membrane subunit e | protein_coding | NP_009031.1 | 846 | 601519 | . | . | 89.561976 | 97.737252 | 87.569580 | 101.293713 | 208.414391 | 225.924022 | 196.503602 | 231.596057 | -1.003193 | 883.148151 | 0.147965342 | -0.03108547 | 0.975201377 | 0.998865404 | 9.789107 | 9.783911 | 9.759239 | 9.814178 | 9.736415 | 9.830664 | 980 | 991 | 716 | 881 |
| 522 | NM_00100369 | ATP5PF | ATP synthase peripheral stalk subunit F6 | protein_coding | NP_001003696.1;NP_001003697.1;NP_001003701.1;NP_001003703.1;NI | 847 | 603152 | . | . | 50.764751 | 45.090555 | 44.991828 | 52.026788 | 118.131657 | 104.228844 | 100.960354 | 118.953067 | 1.053618 | 1866.101910 | 0.130046412 | 0.5794246 | 0.562302709 | 0.998865404 | 10.827751 | 10.902894 | 10.867432 | 10.819931 | 10.822310 | 10.944382 | 2156 | 1902 | 1537 | 1933 |
| 523 | NM_001690 | ATP6V1A | ATPase H+ transporting V1 subunit A | protein_coding | NP_001681.2 | 851 | 607027 | . | . | 12.261069 | 11.420169 | 12.443395 | 10.881703 | 28.532013 | 26.398230 | 27.922617 | 24.879721 | -1.029217 | 1675.759526 | 0.132060796 | -0.31460784 | 0.753059426 | 0.998865404 | 10.731012 | 10.689894 | 10.751713 | 10.689041 | 10.753041 | 10.640850 | 2035 | 1761 | 1540 | 1437 |
| 525 | NM_001692,XI | ATP6V1B1 | ATPase H+ transporting V1 subunit B1 | protein_coding | NP_001683.2;XP_011531209.1 | 853 | 192132 | . | . | 0.177995 | 0.161150 | 0.120189 | 0.187285 | 0.414202 | 0.372505 | 0.269700 | 0.428204 | -1.114440 | 11.032238 | 0.922858815 | -0.16938498 | 0.86549384 | NA | 3.540965 | 3.381963 | 3.471776 | 3.467033 | 3.453848 | 3.474301 | 14 | 12 | 7 | 12 |
| 526 | NM_001693,XI | ATP6V1B2 | ATPase H+ transporting V1 subunit B2 | protein_coding | NP_001684.2 | 854 | 606939 | . | . | 66.210866 | 78.318425 | 70.396162 | 82.301832 | 154.075392 | 181.036537 | 157.966945 | 188.173359 | -1.001393 | 7363.538778 | 0.106966403 | -0.01877809 | 0.98501813 | 0.999197585 | 12.847230 | 12.845137 | 12.826442 | 12.866235 | 12.810684 | 12.878823 | 8286 | 8138 | 6141 | 7176 |
| 527 | NM_00119856 | ATP6V0C | ATPase H+ transporting V0 subunit c | protein_coding | NP_001185498.1;NP_001685.1 | 855 | 108745 | . | . | 118.530861 | 118.774084 | 126.290911 | 124.780476 | 275.826213 | 274.551603 | 283.393135 | 285.295749 | 1.043655 | 4151.880634 | 0.107086666 | 0.575654579 | 0.564848682 | 0.998865404 | 11.988380 | 12.050059 | 12.004348 | 11.998571 | 12.046075 | 12.027820 | 4695 | 4369 | 3728 | 3932 |
| 528 | NM_001695 | ATP6V1C1 | ATPase H+ transporting V1 subunit C1 | protein_coding | NP_001686.1 | 856 | 603097 | . | . | 24.312233 | 23.487499 | 26.628353 | 22.288912 | 56.575562 | 54.292408 | 59.753250 | 50.960949 | 1.009537 | 4245.640642 | 0.123340666 | 0.111025951 | 0.911595768 | 0.998865404 | 12.044798 | 12.058702 | 12.064348 | 12.028514 | 12.129761 | 11.973421 | 4970 | 4461 | 4058 | 3624 |
| 529 | NM_00103936 | ATP6V1E1 | ATPase H+ transporting V1 subunit E1 | protein_coding | NP_001034455.1;NP_001034456.1;NP_001687.1 | 857 | 108746 | . | . | 159.290075 | 155.847121 | 144.942857 | 175.433639 | 370.674512 | 360.247608 | 325.247552 | 401.108186 | 1.002195 | 6596.788699 | 0.118901168 | 0.026598704 | 0.978779807 | 0.998865404 | 12.686019 | 12.689076 | 12.698361 | 12.672765 | 12.608499 | 12.760786 | 7703 | 7003 | 5225 | 6749 |
| 533 | NM_00103945 | ATP6V0B | ATPase H+ transporting V0 subunit b | protein_coding | NP_001034546.1;NP_001281262.1;NP_004038.1 | 861 | 603717 | . | . | 131.979098 | 134.060542 | 125.315947 | 134.349360 | 307.120778 | 309.886935 | 281.205373 | 307.173845 | -1.038376 | 4150.951813 | 0.108063605 | -0.50274357 | 0.61514457 | 0.998865404 | 12.046171 | 11.991769 | 12.028142 | 12.041121 | 11.984355 | 12.021959 | 4832 | 4599 | 3461 | 3905 |
| 534 | NM_00120407 | ATP6V1G2 | ATPase H+ transporting V1 subunit G2 | protein_coding | NP_001191007.1;NP_569730.1;NP_612139.1 | 862 | 606853 | . | . | 0.392221 | 0.285457 | 0.155934 | 0.536316 | 0.912713 | 0.659846 | 0.349911 | 1.226224 | 1.035538 | 14.945762 | 0.853909792 | 0.059000302 | 0.952951868 | NA | 3.878555 | 3.924410 | 3.909496 | 3.885783 | 3.864476 | 3.934390 | 20 | 13 | 6 | 22 |
| 535 | NM_00113002 | ATP6V0A1 | ATPase H+ transporting V0 subunit a1 | protein_coding | NP_001123492.1;NP_001123493.1;NP_001365451.1;NP_001365452.1;NI | 865 | 192130 | . | . | 19.116685 | 21.149538 | 24.553944 | 21.340217 | 44.485319 | 48.888101 | 55.098338 | 48.791874 | 1.123668 | 2768.200098 | 0.12593182 | 1.335770824 | 0.181624192 | 0.953435029 | 11.348107 | 11.516450 | 11.350584 | 11.416554 | 11.542745 | 11.411188 | 2879 | 2928 | 2751 | 2527 |
| 537 | NM_001183,XI | ATP6AP1 | ATPase H+ transporting accessory protein 1 | protein_coding | NP_001174.2;XP_011529481.1 | 868 | 300197 | . | . | 68.715848 | 71.352549 | 63.065765 | 67.232416 | 159.904613 | 164.934612 | 141.517752 | 153.718918 | -1.094568 | 4350.196284 | 0.108968943 | -1.19631534 | 0.231573526 | 0.986726781 | 12.150624 | 12.020157 | 12.106003 | 12.140175 | 12.024261 | 12.071180 | 5129 | 5006 | 3509 | 4005 |
| 538 | NM_000052,N | ATP7A | ATPase copper transporting alpha | protein_coding | NP_000043.4;NP_001269153.1 | 869 | 300011 | . | . | 2.469423 | 2.131610 | 2.145449 | 1.718749 | 5.746449 | 4.927312 | 4.814325 | 3.929715 | -1.203360 | 558.222193 | 0.181139191 | -1.47438067 | 0.140379126 | 0.900674731 | 9.251504 | 8.985658 | 9.231411 | 9.124523 | 9.130546 | 8.981892 | 757 | 606 | 493 | 419 |
| 539 | NM_001697 | ATP5PO | ATP synthase peripheral stalk subunit OSCP | protein_coding | NP_001688.1 | 850 | 600828 | . | . | 121.415070 | 118.412498 | 99.523529 | 135.509048 | 282.537872 | 273.715790 | 223.327896 | 309.825348 | -1.035386 | 2795.523362 | 0.143984112 | -0.34843051 | 0.727516895 | 0.998865404 | 11.473916 | 11.423448 | 11.473930 | 11.446548 | 11.304388 | 11.545336 | 3330 | 3018 | 2035 | 2956 |
| 540 | NM_000053,N | ATP7B | ATPase copper transporting beta | protein_coding | NP_000044.2;NP_001005918.1;NP_001230111.1;NP_001317507.1;NP_0( | 870 | 606882 | . | . | 5.920226 | 4.354562 | 5.172969 | 4.758554 | 13.776611 | 10.065762 | 11.607992 | 10.879870 | -1.059647 | 1198.444804 | 0.131600317 | -0.63513177 | 0.525342479 | 0.998865404 | 10.268184 | 10.184498 | 10.259590 | 10.237060 | 10.184864 | 10.224349 | 1441 | 1311 | 985 | 1120 |
| 545 | NM_001184,N | ATR | ATR serine/threonine kinase | protein_coding | NP_001175.2;NP_001341508.1;XP_011511226.1;XP_011511227.1 | 882 | 601215 | . | . | 3.566071 | 3.636804 | 3.850987 | 3.334060 | 8.298394 | 8.406639 | 8.641503 | 7.622933 | -1.016146 | 915.077069 | 0.146525961 | -0.15770705 | 0.874687647 | 0.998865404 | 9.849024 | 9.826386 | 9.841341 | 9.844019 | 9.883943 | 9.775687 | 1060 | 997 | 849 | 787 |
| 546 | NM_000489,N | ATRX | ATRX chromatin remodeler | protein_coding | NP_000480.3;NP_612114.2;XP_005262210.2;XP_005262211.2;XP_00526 | 886 | 300032 | . | . | 2.086062 | 2.117952 | 2.234998 | 2.003280 | 4.854353 | 4.895741 | 5.015270 | 4.580260 | -1.008209 | 699.904356 | 0.154098049 | -0.07654026 | 0.938989284 | 0.998865404 | 9.456697 | 9.445309 | 9.448878 | 9.457609 | 9.487210 | 9.406699 | 804 | 763 | 642 | 615 |
| 547 | NM_00124400 | KIF1A | kinesin family member 1A | protein_coding | NP_001230937.1;NP_001307634.1;NP_001317218.1;NP_001317219.1;NI | 888 | 601255 | . | . | 0.029706 | 0.039472 | 0.040969 | 0.038765 | 0.069128 | 0.091241 | 0.091932 | 0.088631 | 1.177087 | 11.278790 | 0.908128456 | 0.259017763 | 0.795621535 | NA | 3.373794 | 3.607807 | 3.478693 | 3.494157 | 3.496517 | 3.492660 | 10 | 13 | 11 | 11 |
| 549 | NM_00130619 | AUH | AU RNA binding methylglutaconyl-CoA hydratase | protein_coding | NP_001293119.1;NP_001338360.1;NP_001338361.1;NP_001338362.1;NI | 890 | 600529 | . | . | 2.184749 | 2.659119 | 2.952270 | 2.815882 | 5.083999 | 6.146672 | 6.624807 | 6.438181 | 1.065170 | 279.605275 | 0.210944003 | 0.431788495 | 0.665895142 | 0.998865404 | 8.080847 | 8.172202 | 8.083744 | 8.133196 | 8.174979 | 8.107468 | 297 | 306 | 267 | 253 |
| 550 | NM_181575,N | AUP1 | AUP1 lipid droplet regulating VLDL assembly factor | protein_coding | NP_853553.1 | 891 | 602434 | . | . | 158.673614 | 158.453124 | 184.243809 | 174.654300 | 369.240020 | 366.271507 | 413.437760 | 399.326309 | 1.126197 | 8027.602056 | 0.104682398 | 1.637903342 | 0.101441859 | 0.819486355 | 12.882449 | 13.053963 | 12.923010 | 12.912953 | 13.049327 | 12.987486 | 8747 | 8099 | 7662 | 7685 |
| 558 | NM_00127859 | AXL | AXL receptor tyrosine kinase | protein_coding | NP_001265528.1;NP_001690.2;NP_068713.2 | 905 | 109135 | . | . | 0.189032 | 0.241694 | 0.156193 | 0.110501 | 0.439886 | 0.558686 | 0.350493 | 0.252648 | -1.618917 | 26.216948 | 0.614680265 | -1.13071568 | 0.258174782 | NA | 5.017958 | 4.324312 | 4.723966 | 4.754258 | 4.700860 | 4.676045 | 33 | 39 | 20 | 16 |
| 563 | NM_001185 | AZGP1 | alpha-2-glycoprotein 1, zinc-binding | protein_coding | NP_001176.1 | 910 | 194460 | . | . | 0.381549 | 2.053685 | 0.380137 | 0.451809 | 0.887879 | 4.747185 | 0.853017 | 1.033008 | -2.827478 | 30.693232 | 0.722225433 | -2.07624316 | 0.037871468 | 0.60652635 | 5.503930 | 4.000737 | 4.822316 | 5.066432 | 4.827665 | 4.835528 | 17 | 82 | 13 | 16 |
| 567 | NM_004048,XI | B2M | beta-2-microglobulin | protein_coding | NP_004039.1;XP_005254606.1 | 914 | 109700 | . | . | 431.372018 | 365.895155 | 388.054924 | 339.093778 | 1003.820404 | 845.782955 | 870.784017 | 775.297614 | -1.103455 | 11432.289789 | 0.123055675 | -1.15417429 | 0.248428722 | 0.992733415 | 13.550047 | 13.408117 | 13.587466 | 13.447432 | 13.498931 | 13.368847 | 14933 | 11891 | 10179 | 9408 |
| 570 | NM_00112761 | BAAT | bile acid-CoA:amino acid N-acyltransferase | protein_coding | NP_001121082.1;NP_001361644.1;NP_001692.1 | 932 | 602938 | . | . | 10.412432 | 11.189679 | 13.951100 | 13.425749 | 24.230157 | 25.865440 | 31.305865 | 30.696381 | 1.249670 | 1326.629349 | 0.129745159 | 2.478294127 | 0.013201227 | 0.380410307 | 10.203848 | 10.525374 | 10.257317 | 10.304600 | 10.469222 | 10.432440 | 1315 | 1313 | 1312 | 1349 |
| 571 | NM_001186,N | BACH1 | BTB domain and CNC homolog 1 | protein_coding | NP_001177.1;NP_996749.1 | 935 | 602751 | . | . | 3.076629 | 3.236738 | 3.362293 | 3.127096 | 7.159440 | 7.481865 | 7.544888 | 7.149735 | 1.033178 | 555.121900 | 0.163502327 | 0.288001892 | 0.77334529 | 0.998865404 | 9.092824 | 9.140110 | 9.089887 | 9.120512 | 9.153233 | 9.099280 | 614 | 603 | 511 | 507 |
| 572 | NM_004322,N | BAD | BCL2 associated agonist of cell death | protein_coding | NP_004313.1;NP_116784.1 | 936 | 603167 | . | . | 12.908535 | 10.879693 | 10.632145 | 13.469103 | 30.038693 | 25.148897 | 23.858225 | 30.795504 | 1.020856 | 379.434832 | 0.203821356 | 0.146107027 | 0.883836899 | 0.998865404 | 8.553015 | 8.582254 | 8.606203 | 8.509840 | 8.492809 | 8.643843 | 468 | 372 | 290 | 406 |
| 573 | NM_00117241 | BAG1 | BAG cochaperone 1 | protein_coding | NP_001165886.1;NP_001336215.1;NP_001336228.1;NP_004314.6 | 937 | 601497 | . | . | 1.920661 | 1.955522 | 3.249010 | 3.045540 | 4.469459 | 4.520275 | 7.290682 | 6.963264 | 1.596470 | 305.076401 | 0.200962341 | 3.3582676 | 0.000784326 | 0.057585413 | 7.876305 | 8.551468 | 8.099375 | 8.105837 | 8.376534 | 8.331320 | 269 | 255 | 339 | 338 |
| 575 | NM_001702,XI | ADGRB1 | adhesion G protein-coupled receptor B1 | protein_coding | NP_001693.2;XP_011515504.1;XP_016869180.1;XP_016869181.1;XP_01( | 943 | 602682 | . | . | 0.245220 | 0.189673 | 0.231234 | 0.210120 | 0.570637 | 0.438438 | 0.518883 | 0.480415 | 1.031906 | 43.093697 | 0.480539614 | 0.094293606 | 0.924875929 | 0.998865404 | 5.405581 | 5.452842 | 5.449431 | 5.400629 | 5.438580 | 5.422296 | 56 | 39 | 40 | 39 |
| 576 | NM_00129433 | ADGRB2 | adhesion G protein-coupled receptor B2 | protein_coding | NP_001281264.1;NP_001281265.1;NP_001351786.1;XP_011540150.1;XF | 944 | 602683 | . | . | 3.721849 | 3.478187 | 3.458675 | 3.862187 | 8.660897 | 8.039991 | 7.761170 | 8.830430 | -1.010744 | 598.597546 | 0.16440254 | -0.09378305 | 0.925281497 | 0.998865404 | 9.233163 | 9.217680 | 9.268186 | 9.187014 | 9.186791 | 9.254696 | 735 | 610 | 491 | 587 |
| 578 | NM_001188,XI | BAK1 | BCL2 antagonist/killer 1 | protein_coding | NP_001179.1;XP_011513081.1;XP_011513082.1 | 949 | 600516 | . | . | 9.400369 | 9.264774 | 7.619976 | 10.074188 | 21.875047 | 21.415939 | 17.099004 | 23.033435 | -1.070291 | 615.706048 | 0.175079878 | -0.55975912 | 0.575643748 | 0.998865404 | 9.314680 | 9.215823 | 9.295726 | 9.277561 | 9.148301 | 9.326247 | 743 | 678 | 447 | 633 |
| 579 | NM_001189 | NKX3-2 | NK3 homeobox 2 | protein_coding | NP_001180.1 | 951 | 602183 | . | . | 0.596485 | 0.406110 | 0.501214 | 0.447194 | 1.388045 | 0.938741 | 1.124710 | 1.022456 | -1.055446 | 34.732991 | 0.536424729 | -0.1451332 | 0.884605704 | 0.998865404 | 5.155481 | 5.080001 | 5.148925 | 5.093122 | 5.121056 | 5.106863 | 49 | 31 | 31 | 30 |
| 580 | NM_000465,N | BARD1 | BRCA1 associated RING domain 1 | protein_coding | NP_000456.2;NP_001269472.1;NP_001269474.1;NP_001269477.1;NP_0( | 952 | 601593 | . | . | 2.056221 | 2.075510 | 1.866726 | 2.015580 | 4.784912 | 4.797635 | 4.188879 | 4.608383 | -1.065593 | 339.732033 | 0.191809302 | -0.47785425 | 0.632753946 | 0.998865404 | 8.453480 | 8.361563 | 8.428149 | 8.424213 | 8.375034 | 8.405622 | 405 | 377 | 278 | 317 |
| 581 | NM_00129142 | BAX | BCL2 associated X, apoptosis regulator | protein_coding | NP_001278357.1;NP_001278358.1;NP_001278359.1;NP_001278360.1;NI | 959 | 600040 | . | . | 37.500244 | 36.209024 | 35.120501 | 37.297829 | 87.264613 | 83.698778 | 78.809390 | 85.277052 | 1.047861 | 1212.632741 | 0.130438452 | 0.517083487 | 0.60509788 | 0.998865404 | 10.209884 | 10.277185 | 10.219364 | 10.232370 | 10.246881 | 10.275031 | 1352 | 1289 | 1059 | 1185 |
| 582 | NM_024649 | BBS1 | Bardet-Biedl syndrome 1 | protein_coding | NP_078925.3 | 966 | 209901 | . | . | 2.647506 | 2.463603 | 2.784548 | 2.967172 | 6.160854 | 5.694727 | 6.248445 | 6.784088 | 1.109965 | 285.306006 | 0.206122716 | 0.730216752 | 0.465257704 | 0.998865404 | 8.079121 | 8.229685 | 8.147637 | 8.105788 | 8.167274 | 8.196275 | 324 | 280 | 254 | 289 |
| 583 | NM_00137745 | BBS2 | Bardet-Biedl syndrome 2 | protein_coding | NP_001364385.1;NP_114091.4 | 967 | 606151 | . | . | 8.980912 | 9.995966 | 9.703507 | 10.837470 | 20.898955 | 23.106120 | 21.774391 | 24.778586 | 1.075847 | 827.862780 | 0.147737074 | 0.713927151 | 0.475272227 | 0.998865404 | 9.639810 | 9.744777 | 9.635881 | 9.695255 | 9.685528 | 9.748916 | 880 | 897 | 713 | 840 |
| 585 | NM_00125267 | BBS4 | Bardet-Biedl syndrome 4 | protein_coding | NP_001239607.1;NP_001307594.1;NP_149017.2;XP_011520150.1;XP_01 | 969 | 600374 | . | . | 9.756324 | 10.307645 | 9.733726 | 12.527891 | 22.703365 | 23.826586 | 21.842200 | 28.643533 | 1.092213 | 846.268243 | 0.158530092 | 0.802716064 | 0.422138866 | 0.998865404 | 9.660344 | 9.786827 | 9.667034 | 9.713850 | 9.669405 | 9.831070 | 901 | 902 | 682 | 921 |
| 587 | NM_00116477 | BCAT2 | branched chain amino acid transaminase 2 | protein_coding | NP_001158245.1;NP_001181.2;NP_001271254.1;XP_024307398.1 | 977 | 113530 | . | . | 10.059815 | 10.016054 | 7.632291 | 10.394017 | 23.409604 | 23.152557 | 17.126637 | 23.764685 | -1.119888 | 458.171235 | 0.190457932 | -0.85768997 | 0.391063678 | 0.998865404 | 8.919647 | 8.755152 | 8.869157 | 8.874990 | 8.712452 | 8.884025 | 555 | 525 | 322 | 463 |
| 590 | NM_000055,N | BCHE | butyrylcholinesterase | protein_coding | NP_000046.1 | 983 | 177400 | . | . | 21.395535 | 19.252922 | 15.845315 | 17.825712 | 49.788286 | 44.503989 | 35.556429 | 40.756373 | -1.224029 | 1389.029533 | 0.136276841 | -2.14004267 | 0.032351319 | 0.56898561 | 10.578378 | 10.286620 | 10.556457 | 10.464122 | 10.313807 | 10.398315 | 1864 | 1553 | 1024 | 1238 |
| 593 | NM_000709,N | BCKDHA | branched chain keto acid dehydrogenase E1 subunit alpha | protein_coding | NP_000700.1;NP_001158255.1 | 986 | 608348 | . | . | 6.628291 | 6.149370 | 5.773074 | 7.465316 | 15.424306 | 14.214544 | 12.954610 | 17.068558 | 1.021655 | 353.809272 | 0.202341563 | 0.152749592 | 0.878595751 | 0.998865404 | 8.451679 | 8.481819 | 8.482689 | 8.434757 | 8.399494 | 8.538061 | 420 | 362 | 273 | 376 |
| 594 | NM_000056,N | BCKDHB | branched chain keto acid dehydrogenase E1 subunit beta | protein_coding | NP_000047.1;NP_001305904.1;NP_898871.1;XP_005248813.1;XP_01153 | 987 | 248611 | . | . | 2.948080 | 3.146757 | 2.987418 | 3.761910 | 6.860302 | 7.273872 | 6.703678 | 8.601160 | 1.030735 | 312.181579 | 0.201003796 | 0.217279021 | 0.827990905 | 0.998865404 | 8.264559 | 8.307604 | 8.278736 | 8.276034 | 8.256593 | 8.329692 | 355 | 331 | 257 | 317 |
| 595 | NM_053056 | CCND1 | cyclin D1 | protein_coding | NP_444284.1 | 1582 | 168461 | . | . | 102.393066 | 103.926613 | 89.885513 | 95.282997 | 238.272873 | 240.231033 | 201.700470 | 217.853241 | -1.129956 | 12925.520268 | 0.100849897 | -1.74781366 | 0.080496299 | 0.779933816 | 13.743390 | 13.567097 | 13.705230 | 13.710676 | 13.582821 | 13.623544 | 15742 | 14845 | 10300 | 11652 |
| 596 | NM_000633,N | BCL2 | BCL2 apoptosis regulator | protein_coding | NP_000624.2;NP_000648.2;XP_011524437.1;XP_016881406.1 | 990 | 151430 | . | . | 1.370951 | 1.684367 | 1.223813 | 1.303833 | 3.190261 | 3.893491 | 2.746202 | 2.981059 | -1.217669 | 269.250613 | 0.214746118 | -1.32306278 | 0.185814501 | 0.958175316 | 8.208212 | 7.923362 | 8.077360 | 8.162054 | 8.010175 | 8.028425 | 314 | 344 | 207 | 232 |
| 597 | NM_00111473 | BCL2A1 | BCL2 related protein A1 | protein_coding | NP_001108207.1;NP_004040.1 | 991 | 601056 | . | . | 81.185215 | 80.383705 | 79.305565 | 85.422177 | 188.921341 | 185.810531 | 177.959388 | 195.307663 | 1.006559 | 2040.690778 | 0.119062615 | 0.079216866 | 0.93686013 | 0.998865404 | 10.990177 | 10.999492 | 10.996811 | 10.987449 | 10.971864 | 11.022121 | 2356 | 2182 | 1724 | 1980 |
| 598 | NM_001191,N | BCL2L1 | BCL2 like 1 | protein_coding | NP_001182.1;NP_001304848.1;NP_001304849.1;NP_001304850.1;NP_0( | 992 | 600039 | . | . | 10.126341 | 9.784058 | 9.959983 | 11.748595 | 23.564414 | 22.612689 | 22.349913 | 26.861765 | 1.078039 | 812.960733 | 0.153722204 | 0.705229567 | 0.480667384 | 0.998865404 | 9.612033 | 9.720030 | 9.657090 | 9.619850 | 9.628192 | 9.751523 | 925 | 822 | 670 | 859 |
| 599 | NM_00119983 | BCL2L2 | BCL2 like 2 | protein_coding | NP_001186768.1;NP_004041.2 | 995 | 601931 | . | . | 13.159959 | 13.246549 | 13.068940 | 12.781168 | 30.623765 | 30.619989 | 29.326319 | 29.222624 | -1.034213 | 1443.981471 | 0.125436134 | -0.38692019 | 0.698815292 | 0.998865404 | 10.519853 | 10.471414 | 10.509278 | 10.508005 | 10.495356 | 10.470066 | 1692 | 1583 | 1255 | 1309 |
| 600 | gene-BAK1P1 | BAK1P1 | BCL2 antagonist/killer 1 pseudogene 1 | pseudogene | . | 996 | . | | . | 0.744620 | 0.791830 | 0.654911 | 0.673932 | 1.732760 | 1.830350 | 1.469601 | 1.540865 | -1.165745 | 47.923283 | 0.45016676 | -0.49148819 | 0.623081203 | 0.998865404 | 5.689216 | 5.467590 | 5.593449 | 5.602501 | 5.568753 | 5.572573 | 58 | 57 | 38 | 42 |
| 602 | NM_005178,XI | BCL3 | BCL3 transcription coactivator | protein_coding | NP_005169.2;XP_011525499.2;XP_011525500.2;XP_011525502.1;XP_01( | 998 | 109560 | . | . | 0.913438 | 1.024434 | 0.777697 | 0.777795 | 2.125607 | 2.368025 | 1.745128 | 1.778335 | -1.106424 | 70.031649 | 0.397320803 | -0.36722175 | 0.713453614 | 0.998865404 | 6.201746 | 6.054362 | 6.080275 | 6.192425 | 6.118588 | 6.108994 | 66 | 97 | 59 | 61 |
| 604 | NM_00113084 | BCL6 | BCL6 transcription repressor | protein_coding | NP_001124317.1;NP_001128210.1;NP_001697.2;XP_005247751.1;XP_01 | 1001 | 109565 | . | . | 5.238688 | 5.674471 | 6.112066 | 5.768890 | 12.190644 | 13.116794 | 13.715302 | 13.189884 | 1.019947 | 761.635860 | 0.154191793 | 0.184798403 | 0.85338716 | 0.998865404 | 9.558782 | 9.586996 | 9.509123 | 9.618073 | 9.589353 | 9.568085 | 801 | 877 | 679 | 710 |
| 605 | NM_00102480 | BCL7A | BAF chromatin remodeling complex subunit BCL7A | protein_coding | NP_001019979.1;NP_066273.1 | 1004 | 601406 | . | . | 2.942916 | 2.909312 | 2.520995 | 3.084735 | 6.848287 | 6.725004 | 5.657040 | 7.052879 | -1.052222 | 333.373836 | 0.199907942 | -0.36736379 | 0.713347679 | 0.998865404 | 8.417645 | 8.343394 | 8.400848 | 8.388450 | 8.313727 | 8.415944 | 398 | 365 | 256 | 333 |
| 606 | NR_027992 | NBEAP1 | neurobeachin pseudogene 1 | transcribed_ps | . | 1007 | 601889 | . | . | 0.673362 | 0.968379 | 0.469091 | 0.315591 | 1.566942 | 2.238450 | 1.052626 | 0.721561 | -2.158655 | 10.555687 | 0.965945878 | -1.14927006 | 0.250444641 | NA | 3.849979 | 2.742346 | 3.413471 | 3.433376 | 3.395462 | 3.383227 | 14 | 18 | 7 | 5 |
| 607 | NM_004326,XI | BCL9 | BCL9 transcription coactivator | protein_coding | NP_004317.2;XP_005273028.1;XP_005277474.1;XP_005277475.1;XP_01( | 1008 | 602597 | . | . | 4.916711 | 5.949146 | 4.906649 | 5.659770 | 11.441386 | 13.751718 | 11.010376 | 12.940393 | -1.043057 | 1027.120913 | 0.151372482 | -0.40177768 | 0.687847653 | 0.998865404 | 10.034813 | 9.973312 | 9.948974 | 10.083265 | 9.938385 | 10.033603 | 1099 | 1234 | 817 | 1004 |
| 610 | NM_001194 | HCN2 | hyperpolarization activated cyclic nucleotide gated potassium and | protein_coding | NP_001185.3 | 4846 | 602781 | . | . | 0.238178 | 0.193547 | 0.174548 | 0.184715 | 0.554251 | 0.447393 | 0.391680 | 0.422329 | -1.190199 | 21.796157 | 0.659996602 | -0.38061249 | 0.703490811 | NA | 4.565853 | 4.315285 | 4.467049 | 4.446714 | 4.439646 | 4.442442 | 30 | 23 | 17 | 19 |
| 613 | NM_004327,N | BCR | BCR activator of RhoGEF and GTPase | protein_coding | NP_004318.3;NP_067585.2 | 1014 | 151410 | . | . | 6.096874 | 5.947791 | 5.791866 | 6.527815 | 14.187676 | 13.748586 | 12.996781 | 14.925073 | 1.008631 | 1287.381911 | 0.133011954 | 0.093218493 | 0.925729983 | 0.998865404 | 10.324127 | 10.336296 | 10.339357 | 10.313741 | 10.291005 | 10.373491 | 1501 | 1360 | 1063 | 1278 |
| 617 | NM_00107986 | BCS1L | BCS1 homolog, ubiquinol-cytochrome c reductase complex chaper | protein_coding | NP_001073335.1;NP_001244271.1;NP_001244272.1;NP_001244273.1;NI | 1020 | 603647 | . | . | 26.272654 | 31.381088 | 31.387601 | 26.791620 | 61.137554 | 72.538790 | 70.432866 | 61.255856 | 1.060951 | 1832.753708 | 0.137224898 | 0.622028835 | 0.533922908 | 0.998865404 | 10.796389 | 10.881938 | 10.771605 | 10.855200 | 10.933248 | 10.783290 | 1939 | 2021 | 1796 | 1602 |
| 618 | NR_001568 | BCYRN1 | brain cytoplasmic RNA 1 | lncRNA | . | 1022 | 606089 | . | . | 14.742379 | 15.474198 | 14.505907 | 11.467568 | 34.306126 | 35.776392 | 32.550835 | 26.219233 | -1.173056 | 88.136541 | 0.344785461 | -0.66787007 | 0.504216537 | 0.998865404 | 6.571612 | 6.342650 | 6.476344 | 6.490010 | 6.469205 | 6.407033 | 107 | 105 | 79 | 67 |
| 619 | gene-BCYRN1P | BCYRN1P1 | brain cytoplasmic RNA 1, pseudogene 1 | pseudogene | . | 1023 | . | | . | 6.195616 | 7.671301 | 8.320897 | 4.882186 | 14.417455 | 17.732555 | 18.671852 | 11.162539 | -1.059688 | 41.989849 | 0.503417965 | -0.16614361 | 0.868043937 | 0.998865404 | 5.432022 | 5.350772 | 5.375485 | 5.407361 | 5.422558 | 5.340196 | 45 | 51 | 45 | 28 |
| 622 | NM_004051,N | BDH1 | 3-hydroxybutyrate dehydrogenase 1 | protein_coding | NP_004042.1;NP_976059.1;NP_976060.1;XP_005269409.1;XP_00526941 | 1027 | 603063 | . | . | 3.962255 | 3.987153 | 6.060117 | 4.863258 | 9.220327 | 9.216480 | 13.598728 | 11.119264 | 1.350296 | 509.697726 | 0.17075767 | 2.537371518 | 0.011168836 | 0.343348416 | 8.760284 | 9.194207 | 8.900421 | 8.870104 | 9.118252 | 9.033247 | 512 | 456 | 542 | 514 |
| 627 | NM_00114380 | BDNF | brain derived neurotrophic factor | protein_coding | NP_001137277.1;NP_001137278.1;NP_001137279.1;NP_001137280.1;NI | 1033 | 113505 | . | . | 0.504080 | 0.625282 | 1.105255 | 0.790122 | 1.173016 | 1.445369 | 2.480159 | 1.806522 | 1.609604 | 99.296429 | 0.342288699 | 2.006218643 | 0.044832926 | 0.649185583 | 6.249719 | 6.936619 | 6.502863 | 6.572179 | 6.736329 | 6.623297 | 76 | 93 | 124 | 96 |
| 631 | NM_00116170 | BFSP1 | beaded filament structural protein 1 | protein_coding | NP_001155177.1;NP_001186.1;NP_001265535.1;NP_001265536.1;NP_0( | 1040 | 603307 | . | . | 3.055604 | 2.973213 | 3.650655 | 3.010395 | 7.110516 | 6.872714 | 8.191964 | 6.882911 | 1.272390 | 211.719387 | 0.231526885 | 1.501080632 | 0.133334707 | 0.889091289 | 7.541868 | 7.889286 | 7.674038 | 7.652369 | 7.757265 | 7.791600 | 220 | 196 | 199 | 230 |
| 634 | NM_00102491 | CEACAM1 | CEA cell adhesion molecule 1 | protein_coding | NP_001020083.1;NP_001171742.1;NP_001171744.1;NP_001171745.1;NI | 1814 | 109770 | . | . | 164.730154 | 172.940809 | 171.940526 | 178.207289 | 383.333810 | 399.760398 | 385.829564 | 407.449793 | 1.023766 | 18476.760238 | 0.099721015 | 0.339803803 | 0.734004284 | 0.998865404 | 14.156391 | 14.190259 | 14.145312 | 14.180742 | 14.173100 | 14.193513 | 20592 | 20110 | 16063 | 17733 |
| 636 | NM_00100339 | BICD1 | BICD cargo adaptor 1 | protein_coding | NP_001003398.1;NP_001341115.1;NP_001341116.1;NP_001341117.1;NI | 1049 | 602204 | . | . | 4.125082 | 4.349457 | 3.537245 | 4.652556 | 9.599237 | 10.053966 | 7.937477 | 10.637515 | -1.007153 | 609.260745 | 0.166060777 | -0.06192566 | 0.950622036 | 0.998865404 | 9.256470 | 9.245340 | 9.218227 | 9.285469 | 9.193842 | 9.298421 | 669 | 693 | 486 | 614 |
| 637 | NM_001196,N | BID | BH3 interacting domain death agonist | protein_coding | NP_001187.1;NP_001231496.1;NP_001231498.1;NP_001231499.1;NP_0( | 1050 | 601997 | . | . | 10.842759 | 11.015936 | 10.685526 | 10.510331 | 25.231547 | 25.463828 | 23.978011 | 24.030621 | -1.046519 | 726.879950 | 0.148526947 | -0.44166189 | 0.658733891 | 0.998865404 | 9.537949 | 9.472455 | 9.519570 | 9.522776 | 9.499963 | 9.479417 | 854 | 804 | 627 | 656 |
| 638 | NM_001197 | BIK | BCL2 interacting killer | protein_coding | NP_001188.1 | 1051 | 603392 | . | . | 0.726668 | 0.813623 | 0.476695 | 0.863810 | 1.690986 | 1.880725 | 1.069689 | 1.974998 | -1.170733 | 22.037820 | 0.669335953 | -0.33975755 | 0.734039116 | NA | 4.572938 | 4.341619 | 4.466137 | 4.476448 | 4.434744 | 4.478385 | 26 | 27 | 13 | 24 |
| 641 | NM_000057,N | BLM | BLM RecQ like helicase | protein_coding | NP_000048.1;NP_001274175.1;NP_001274176.1;NP_001274177.1;XP_0( | 1058 | 604610 | . | . | 6.474780 | 6.208989 | 6.577946 | 5.155196 | 15.067081 | 14.352351 | 14.760718 | 11.786743 | -1.096443 | 979.446999 | 0.153145142 | -0.86735383 | 0.38574817 | 0.998865404 | 10.000360 | 9.868265 | 9.990317 | 9.942851 | 9.975771 | 9.817747 | 1217 | 1070 | 902 | 780 |
| 642 | NM_000386 | BLMH | bleomycin hydrolase | protein_coding | NP_000377.1 | 1059 | 602403 | . | . | 19.852558 | 19.958645 | 17.110601 | 19.594891 | 46.197712 | 46.135300 | 38.395695 | 44.801393 | -1.099545 | 1375.457922 | 0.132260228 | -1.03513005 | 0.300608182 | 0.998865404 | 10.492684 | 10.355446 | 10.462057 | 10.459270 | 10.339792 | 10.433934 | 1662 | 1552 | 1068 | 1305 |
| 644 | NM_000712,N | BLVRA | biliverdin reductase A | protein_coding | NP_000703.2;NP_001240752.1;XP_011513776.1;XP_016868009.1;XP_02 | 1062 | 109750 | . | . | 31.597611 | 28.924639 | 28.841738 | 37.114924 | 73.528939 | 66.860594 | 64.720025 | 84.858866 | 1.063228 | 1083.782189 | 0.152899309 | 0.578491469 | 0.562932355 | 0.998865404 | 10.037158 | 10.125217 | 10.088637 | 10.023784 | 10.012170 | 10.184842 | 1263 | 1084 | 859 | 1168 |
| 645 | NM_000713 | BLVRB | biliverdin reductase B | protein_coding | NP_000704.1 | 1063 | 600941 | . | . | 50.921249 | 50.889023 | 43.387154 | 55.991909 | 118.495842 | 117.632256 | 97.359512 | 128.018860 | -1.038964 | 1252.032672 | 0.148572928 | -0.3711722 | 0.71050928 | 0.998865404 | 10.317632 | 10.261944 | 10.306183 | 10.299417 | 10.178311 | 10.361024 | 1476 | 1371 | 938 | 1291 |
| 646 | NM_00130120 | BNC1 | basonuclin 1 | protein_coding | NP_001288135.1;NP_001708.3;XP_011520195.1;XP_011520196.1 | 1081 | 601930 | . | . | 0.126303 | 0.135139 | 0.128437 | 0.158578 | 0.293912 | 0.312380 | 0.288210 | 0.362571 | 1.090287 | 20.627587 | 0.675710688 | 0.184557375 | 0.853576222 | NA | 4.303566 | 4.426810 | 4.358406 | 4.364049 | 4.360668 | 4.376824 | 22 | 22 | 17 | 22 |
| 648 | NM_005180 | BMI1 | BMI1 proto-oncogene, polycomb ring finger | protein_coding | NP_005171.4 | 1066 | 164831 | . | . | 8.068342 | 5.777283 | 5.282320 | 5.297781 | 18.775364 | 13.354448 | 11.853374 | 12.112746 | -1.328149 | 675.012204 | 0.182758629 | -2.24020622 | 0.025077538 | 0.522089373 | 9.588741 | 9.179935 | 9.598066 | 9.353629 | 9.292345 | 9.287682 | 1037 | 690 | 506 | 542 |
| 649 | NM_001199,N | BMP1 | bone morphogenetic protein 1 | protein_coding | NP_001190.1;NP_006120.1 | 1067 | 112264 | . | . | 12.418698 | 13.557368 | 13.787746 | 13.934982 | 28.898818 | 31.338465 | 30.939302 | 31.860680 | 1.026225 | 1226.375172 | 0.133198214 | 0.280382655 | 0.779183943 | 0.998865404 | 10.241503 | 10.278627 | 10.212479 | 10.286137 | 10.266498 | 10.272332 | 1326 | 1370 | 1076 | 1169 |
| 650 | NM_001200 | BMP2 | bone morphogenetic protein 2 | protein_coding | NP_001191.1 | 1069 | 112261 | . | . | 2.604584 | 2.554896 | 2.781970 | 2.297361 | 6.060975 | 5.905756 | 6.242661 | 5.252642 | -1.030890 | 283.055471 | 0.212950624 | -0.20610885 | 0.83670589 | 0.998865404 | 8.166193 | 8.123371 | 8.158411 | 8.144675 | 8.186598 | 8.082150 | 335 | 306 | 267 | 235 |
| 652 | NM_001202,N | BMP4 | bone morphogenetic protein 4 | protein_coding | NP_001193.2;NP_001334841.1;NP_001334842.1;NP_001334843.1;NP_0( | 1071 | 112262 | . | . | 0.303898 | 0.385110 | 0.262491 | 0.412961 | 0.707183 | 0.890199 | 0.589022 | 0.944186 | -1.039908 | 20.234730 | 0.684228615 | -0.08251102 | 0.934240357 | NA | 4.367696 | 4.309235 | 4.338508 | 4.344295 | 4.327393 | 4.348629 | 23 | 23 | 15 | 21 |
| 654 | NM_001718 | BMP6 | bone morphogenetic protein 6 | protein_coding | NP_001709.1 | 1073 | 112266 | . | . | 0.203105 | 0.154007 | 0.098597 | 0.190998 | 0.472633 | 0.355994 | 0.221248 | 0.436694 | -1.220395 | 19.475250 | 0.714568938 | -0.4021275 | 0.687590197 | NA | 4.421266 | 4.131331 | 4.307623 | 4.282580 | 4.258372 | 4.298780 | 28 | 20 | 11 | 21 |
| 655 | NM_001719 | BMP7 | bone morphogenetic protein 7 | protein_coding | NP_001710.1 | 1074 | 112267 | . | . | 22.329985 | 21.664175 | 18.324961 | 22.798662 | 51.962791 | 50.077713 | 41.120682 | 52.126434 | -1.085239 | 2666.383383 | 0.130739669 | -0.9026534 | 0.366709877 | 0.998865404 | 11.438593 | 11.320321 | 11.427578 | 11.395751 | 11.258392 | 11.425379 | 3258 | 2937 | 1993 | 2646 |
| 656 | NM_001720,XI | BMP8B | bone morphogenetic protein 8b | protein_coding | NP_001711.2;XP_005271206.1;XP_011540324.1;XP_011540326.2;XP_01 | 1075 | 602284 | . | . | 12.213771 | 11.611852 | 13.292273 | 13.762815 | 28.421949 | 26.841318 | 29.827474 | 31.467043 | 1.095662 | 1560.130245 | 0.126485636 | 1.042041011 | 0.297392665 | 0.998865404 | 10.540042 | 10.671850 | 10.598121 | 10.542053 | 10.619802 | 10.662438 | 1777 | 1548 | 1380 | 1571 |
| 657 | NM_004329,XI | BMPR1A | bone morphogenetic protein receptor type 1A | protein_coding | NP_004320.2;XP_011538405.1;XP_011538406.1 | 1076 | 601299 | . | . | 5.312329 | 4.691079 | 6.051149 | 4.914228 | 12.362008 | 10.843639 | 13.578604 | 11.235797 | 1.083114 | 1049.666338 | 0.154454996 | 0.745750048 | 0.45581842 | 0.998865404 | 9.976582 | 10.092519 | 10.049199 | 9.954532 | 10.139481 | 9.978488 | 1235 | 1017 | 1052 | 913 |
| 658 | NM_001203,N | BMPR1B | bone morphogenetic protein receptor type 1B | protein_coding | NP_001194.1;NP_001243721.1;NP_001243722.1;NP_001243723.1;XP_01 | 1077 | 603248 | . | . | 1.586713 | 1.577149 | 1.688585 | 1.517148 | 3.692348 | 3.645651 | 3.789137 | 3.468778 | -1.004728 | 277.308797 | 0.208951772 | -0.03256904 | 0.974018262 | 0.998865404 | 8.118443 | 8.112249 | 8.119345 | 8.112753 | 8.141882 | 8.084668 | 322 | 298 | 254 | 245 |
| 659 | NM_001204,XI | BMPR2 | bone morphogenetic protein receptor type 2 | protein_coding | NP_001195.2;XP_011509989.1 | 1078 | 600799 | . | . | 3.904599 | 3.986484 | 3.039477 | 3.473381 | 9.086161 | 9.214936 | 6.820499 | 7.941473 | -1.227551 | 1355.468562 | 0.132534164 | -2.23175134 | 0.025631402 | 0.524911334 | 10.545069 | 10.248911 | 10.472659 | 10.482060 | 10.274263 | 10.365215 | 1710 | 1622 | 993 | 1211 |
| 661 | NM_001722,XI | POLR3D | RNA polymerase III subunit D | protein_coding | NP_001713.2 | 1080 | 187280 | . | . | 6.052985 | 6.637766 | 6.036420 | 6.204055 | 14.085544 | 15.343495 | 13.545550 | 14.184835 | -1.039571 | 1030.782303 | 0.13917258 | -0.40229592 | 0.687466254 | 0.998865404 | 10.037388 | 9.981112 | 9.983242 | 10.061985 | 9.988187 | 10.001195 | 1146 | 1194 | 871 | 956 |

| | | | | | | | | | | | | | | | | | | | | | | | | | | | | | | | | | |
|---|---|---|---|---|---|---|---|---|---|---|---|---|---|---|---|---|---|---|---|---|---|---|---|---|---|---|---|---|---|---|---|---|---|
| 662 | NM_001205,N | BNIP1 | BCL2 interacting protein 1 | protein_coding | NP_001196.2;NP_053581.2;NP_053582.2;NP_053583.2;XP_011532940.1 | 1082 | 603291 | . | . | 11.678319 | 11.471743 | 10.099430 | 12.275112 | 27.175928 | 26.517448 | 22.662827 | 28.065586 | -1.040374 | 422.748446 | 0.183177574 | -0.31173016 | 0.755245608 | 0.998865404 | 8.752273 | 8.694460 | 8.738202 | 8.732059 | 8.660024 | 8.759733 | 499 | 463 | 330 | 421 |
| 663 | NM_00132067 | BNIP2 | BCL2 interacting protein 2 | protein_coding | NP_001307603.2;NP_001307604.2;NP_001354986.1;NP_001354987.1;N | 1083 | 603292 | . | . | 6.939604 | 5.888359 | 6.636375 | 5.458796 | 16.148745 | 13.611207 | 14.891832 | 12.480889 | -1.074526 | 1190.140732 | 0.153432921 | -0.67586989 | 0.499123258 | 0.998865404 | 10.267484 | 10.164512 | 10.304430 | 10.173565 | 10.260400 | 10.108433 | 1541 | 1216 | 1100 | 966 |
| 664 | NM_004052 | BNIP3 | BCL2 interacting protein 3 | protein_coding | NP_004043.3 | 1084 | 603293 | . | . | 18.873722 | 18.120743 | 25.900953 | 26.824951 | 43.919930 | 41.886906 | 58.120991 | 61.332058 | 1.404415 | 1140.128944 | 0.132909909 | 3.686475445 | **0.000227381** | **0.022579461** | 9.889259 | 10.379291 | 10.029539 | 9.994545 | 10.255839 | 10.275277 | 1119 | 998 | 1144 | 1264 |
| 665 | NM_00133049 | BNIP3L | BCL2 interacting protein 3 like | protein_coding | NP_001317420.1;NP_004322.1 | 1085 | 605368 | . | . | 10.131133 | 8.199275 | 11.965656 | 9.261568 | 23.575564 | 18.952990 | 26.850588 | 21.175475 | 1.141750 | 1063.805613 | 0.168051073 | 1.138025088 | 0.255110007 | 0.995760118 | 9.955771 | 10.147876 | 10.076365 | 9.915966 | 10.192402 | 9.993287 | 1269 | 954 | 1117 | 923 |
| 666 | NM_032515,XI | BOK | BCL2 family apoptosis regulator BOK | protein_coding | NP_115904.1;XP_011509998.1;XP_011509999.1;XP_016860264.1;XP_01 | 1087 | 605404 | . | . | 7.186476 | 8.738226 | 6.851135 | 10.069761 | 16.723227 | 20.198801 | 15.373748 | 23.023313 | 1.042777 | 656.260803 | 0.191252829 | 0.315972091 | 0.752023692 | 0.998865404 | 9.328232 | 9.387412 | 9.266403 | 9.406299 | 9.244012 | 9.476769 | 664 | 765 | 483 | 735 |
| 667 | NM_00114476 | DST | dystonin | protein_coding | NP_001138241.1;NP_001138242.1;NP_001361651.1;NP_001361658.1;N | 1090 | 113810 | . | . | 8.648609 | 8.759191 | 10.350069 | 8.879378 | 20.125668 | 20.247261 | 23.225254 | 20.301641 | 1.084798 | 5101.704623 | 0.117992841 | 0.995195006 | 0.31964143 | 0.998865404 | 12.256790 | 12.374343 | 12.273259 | 12.288153 | 12.416005 | 12.277326 | 5585 | 5328 | 4996 | 4569 |
| 669 | NM_00129308 | BPGM | bisphosphoglycerate mutase | protein_coding | NP_001280014.1;NP_001715.1;NP_954655.1;XP_011514829.1 | 1093 | 613896 | . | . | 10.516875 | 10.597565 | 9.241326 | 10.438383 | 24.473199 | 24.496742 | 20.737267 | 23.866124 | -1.075077 | 553.283147 | 0.166497839 | -0.62727383 | 0.530479754 | 0.998865404 | 9.163432 | 9.058406 | 9.134580 | 9.135045 | 9.041807 | 9.131393 | 660 | 619 | 431 | 535 |
| 670 | NM_00130277 | BPHL | biphenyl hydrolase like | protein_coding | NP_001289706.1;NP_004323.2 | 1094 | 603156 | . | . | 5.085745 | 4.898644 | 6.074137 | 6.405916 | 11.834738 | 11.323435 | 13.630186 | 14.646366 | 1.238274 | 347.380108 | 0.19050162 | 1.618519596 | 0.105550667 | 0.833478683 | 8.277935 | 8.586355 | 8.389555 | 8.357607 | 8.485278 | 8.504161 | 368 | 325 | 329 | 366 |
| 672 | NM_007294,N | BRCA1 | BRCA1 DNA repair associated | protein_coding | NP_009225.1;NP_009228.2;NP_009229.2;NP_009230.2;NP_009231.2 | 1100 | 113705 | . | . | 9.811602 | 9.930461 | 10.601085 | 9.608918 | 22.832000 | 22.954704 | 23.788529 | 21.969649 | 1.059145 | 2053.136089 | 0.122974779 | 0.674120273 | 0.500234852 | 0.998865404 | 10.961549 | 11.044486 | 10.946278 | 11.012092 | 11.057294 | 10.992617 | 2201 | 2241 | 1911 | 1900 |
| 673 | NM_00135460 | BRAF | B-Raf proto-oncogene, serine/threonine kinase | protein_coding | NP_001341538.1;NP_001361173.1;NP_001361187.1;NP_001365396.1;N | 1097 | 164757 | . | . | 1.840680 | 1.833677 | 2.106391 | 1.794451 | 4.283340 | 4.238626 | 4.726680 | 4.102800 | 1.066304 | 555.161306 | 0.170357626 | 0.543674396 | 0.586665555 | 0.998865404 | 9.069372 | 9.162647 | 9.088778 | 9.098515 | 9.195091 | 9.072498 | 614 | 584 | 545 | 487 |
| 675 | NM_000059 | BRCA2 | BRCA2 DNA repair associated | protein_coding | NP_000050.3 | 1101 | 600185 | . | . | 1.582127 | 1.666579 | 2.303588 | 1.323283 | 3.681674 | 3.852372 | 5.169184 | 3.025529 | 1.101583 | 640.897464 | 0.218276787 | 0.639455631 | 0.522526568 | 0.998865404 | 9.251946 | 9.392530 | 9.264713 | 9.293524 | 9.517070 | 9.137320 | 687 | 672 | 745 | 457 |
| 677 | NM_00124469 | ZFP36L1 | ZFP36 ring finger protein like 1 | protein_coding | NP_001231627.1;NP_001231630.1;NP_004917.2 | 1107 | 601064 | . | . | 11.341627 | 13.056425 | 13.480710 | 12.440250 | 26.392433 | 30.180509 | 30.250322 | 28.443155 | 1.047480 | 1184.350755 | 0.139886886 | 0.478407721 | 0.632360039 | 0.998865404 | 10.176036 | 10.242949 | 10.140864 | 10.239038 | 10.260529 | 10.190199 | 1245 | 1330 | 1103 | 1083 |
| 678 | NM_006887 | ZFP36L2 | ZFP36 ring finger protein like 2 | protein_coding | NP_008818.3 | 1108 | 612053 | . | . | 8.078177 | 7.898101 | 9.984557 | 8.794314 | 18.798252 | 18.256815 | 22.405054 | 20.107155 | 1.158674 | 999.874425 | 0.142659174 | 1.489388069 | 0.136385208 | 0.89431118 | 9.855211 | 10.068146 | 9.918598 | 9.896483 | 10.065027 | 9.963803 | 1083 | 984 | 997 | 938 |
| 682 | NM_00132224 | BSG | basigin (Ok blood group) | protein_coding | NP_001309172.1;NP_001719.2;NP_940991.1;NP_940992.1;NP_940993.1 | 1116 | 109480 | . | . | 313.594545 | 329.354756 | 293.474802 | 321.126059 | 729.747358 | 761.318256 | 658.548906 | 734.216566 | -1.061852 | 15696.465528 | 0.103437185 | -0.83704978 | 0.40256459 | 0.998865404 | 13.980813 | 13.894191 | 13.946629 | 13.979395 | 13.876886 | 13.945289 | 18263 | 17777 | 12709 | 14846 |
| 684 | NM_004335 | BST2 | bone marrow stromal cell antigen 2 | protein_coding | NP_004326.1 | 1119 | 600534 | . | . | 0.647563 | 0.912964 | 0.521292 | 0.514200 | 1.506906 | 2.110357 | 1.169763 | 1.175656 | -1.490147 | 20.705329 | 0.684432212 | -0.84077718 | 0.400472771 | NA | 4.631445 | 4.055266 | 4.374781 | 4.408015 | 4.360048 | 4.354489 | 24 | 31 | 15 | 15 |
| 686 | NM_00128172 | BTD | biotinidase | protein_coding | NP_001268652.2;NP_001268653.2;NP_001268654.1;NP_001268655.2;N | 1122 | 609019 | . | . | 3.842793 | 3.900124 | 3.256849 | 4.362882 | 8.942337 | 9.015314 | 7.308275 | 9.975211 | 1.058393 | 996.731903 | 0.136917741 | 0.597993973 | 0.549843955 | 0.998865404 | 9.919522 | 10.001439 | 9.927593 | 9.951394 | 9.994735 | 9.967341 | 1097 | 1062 | 908 | 943 |
| 687 | NM_001206 | KLF9 | Kruppel like factor 9 | protein_coding | NP_001197.1 | 1123 | 602902 | . | . | 8.509641 | 8.024340 | 8.224055 | 8.545932 | 19.802284 | 18.548620 | 18.454542 | 19.539259 | 1.000424 | 1351.431406 | 0.12889215 | 0.004749033 | 0.996210834 | 0.999862634 | 10.399939 | 10.400606 | 10.425360 | 10.373933 | 10.389601 | 10.410911 | 1608 | 1409 | 1159 | 1285 |
| 688 | NM_00128681 | KLF5 | Kruppel like factor 5 | protein_coding | NP_001273747.1;NP_001721.2 | 6349 | 602903 | . | . | 0.672996 | 0.729483 | 0.973030 | 0.696839 | 1.566089 | 1.686231 | 2.183451 | 1.593239 | 1.194222 | 80.289912 | 0.3655788 | 0.700453408 | 0.483644192 | 0.998865404 | 6.192575 | 6.450230 | 6.289428 | 6.302933 | 6.388613 | 6.294970 | 82 | 81 | 89 | 68 |
| 689 | NM_00103763 | BTF3 | basic transcription factor 3 | protein_coding | NP_001032726.1;NP_001198.2 | 1125 | 602542 | . | . | 227.901045 | 217.052000 | 217.138697 | 230.366160 | 530.335102 | 501.725421 | 487.252932 | 526.704815 | -1.007438 | 7111.254585 | 0.105500921 | -0.10133837 | 0.919281857 | 0.998865404 | 12.801229 | 12.790528 | 12.822010 | 12.775177 | 12.771262 | 12.813479 | 8446 | 7489 | 6007 | 6808 |
| 694 | NM_001731 | BTG1 | BTG anti-proliferation factor 1 | protein_coding | NP_001722.1 | 1130 | 109580 | . | . | 4.293664 | 4.200676 | 3.745066 | 3.921119 | 9.991533 | 9.710051 | 8.403821 | 8.965174 | -1.124404 | 583.307807 | 0.159220878 | -1.06242812 | 0.288041373 | 0.998865404 | 9.270248 | 9.101019 | 9.236155 | 9.215127 | 9.137115 | 9.159725 | 722 | 656 | 469 | 524 |
| 699 | NM_00127861 | BUB1 | BUB1 mitotic checkpoint serine/threonine kinase | protein_coding | NP_001265545.1;NP_001265546.1;NP_004327.1 | 1148 | 602452 | . | . | 31.036429 | 30.204844 | 26.879760 | 28.093631 | 72.223048 | 69.819846 | 60.317401 | 64.232746 | -1.137177 | 3385.077915 | 0.110215417 | -1.68267824 | 0.092437396 | 0.809520218 | 11.814735 | 11.629254 | 11.787536 | 11.763420 | 11.656352 | 11.683125 | 4208 | 3831 | 2709 | 3019 |
| 701 | NM_001211 | BUB1B | BUB1 mitotic checkpoint serine/threonine kinase B | protein_coding | NP_001202.5 | 1149 | 602860 | . | . | 16.688629 | 17.086128 | 15.196034 | 17.900831 | 38.835125 | 39.495350 | 34.099461 | 40.928123 | -1.034724 | 1911.113005 | 0.128250048 | -0.38398355 | 0.700990639 | 0.998865404 | 10.924772 | 10.875197 | 10.907131 | 10.918807 | 10.823670 | 10.944476 | 2222 | 2113 | 1508 | 1896 |
| 705 | NM_004053 | BYSL | bystin like | protein_coding | NP_004044.3 | 1157 | 603871 | . | . | 12.465061 | 11.983323 | 10.659853 | 12.704459 | 29.006710 | 27.699987 | 23.920403 | 29.047235 | -1.060451 | 640.081968 | 0.162172561 | -0.52214308 | 0.601570713 | 0.998865404 | 9.364069 | 9.278900 | 9.357428 | 9.324500 | 9.245836 | 9.354179 | 777 | 694 | 496 | 630 |
| 706 | NM_000714,N | TSPO | translocator protein | protein_coding | NP_000705.2;NP_001243459.1;NP_001243460.1 | 1158 | 109610 | . | . | 102.674285 | 101.366460 | 83.530255 | 105.316889 | 238.927258 | 234.313087 | 187.439472 | 240.794553 | -1.087628 | 2599.404507 | 0.130229972 | -0.9305448 | 0.352089078 | 0.998865404 | 11.403420 | 11.281955 | 11.384310 | 11.367518 | 11.222392 | 11.386571 | 3151 | 2893 | 1944 | 2573 |
| 708 | NM_001212 | C1QBP | complement C1q binding protein | protein_coding | NP_001203.1 | 1243 | 601269 | . | . | 190.530289 | 198.605835 | 175.359863 | 212.030151 | 443.371796 | 459.086273 | 393.502441 | 484.781769 | -1.018948 | 7068.206773 | 0.118445295 | -0.22863565 | 0.819152112 | 0.998865404 | 12.800669 | 12.773460 | 12.779952 | 12.807972 | 12.699576 | 12.851104 | 8080 | 7826 | 5543 | 7152 |
| 715 | NM_00135434 | C1R | complement C1r | protein_coding | NP_001341275.1;NP_001724.4 | 1246 | 613785 | . | . | 1.836030 | 2.715065 | 1.740629 | 2.159949 | 4.272519 | 6.275994 | 3.905920 | 4.938468 | -1.174850 | 164.389057 | 0.284436113 | -0.81732246 | 0.413744172 | 0.998865404 | 7.473540 | 7.238868 | 7.302800 | 7.467268 | 7.283002 | 7.360468 | 166 | 228 | 118 | 156 |
| 716 | NM_00134685 | C1S | complement C1s | protein_coding | NP_001333779.1;NP_001725.1;NP_958850.1;XP_005253817.1 | 1247 | 120580 | . | . | 0.249176 | 1.289601 | 0.239759 | 0.099386 | 0.579844 | 2.980971 | 0.538012 | 0.227234 | **-4.622538** | 40.987122 | 0.698182147 | -3.16347996 | **0.001558951** | 0.092750856 | 6.073773 | 3.869520 | 5.191737 | 5.525450 | 5.188531 | 5.133621 | 26 | 121 | 18 | 8 |
| 718 | NM_000064 | C3 | complement C3 | protein_coding | NP_000055.2 | 1318 | 120700 | . | . | 0.119795 | 2.148226 | 0.133355 | 0.272231 | 0.278769 | 4.965719 | 0.299245 | 0.622424 | **-5.567042** | 109.464996 | 1.627304144 | -1.52209469 | 0.127985355 | 0.876241045 | 7.536094 | 5.058523 | 6.146139 | 7.107634 | 6.174562 | 6.267590 | 23 | 379 | 19 | 42 |
| 719 | NM_00132647 | C3AR1 | complement C3a receptor 1 | protein_coding | NP_001313404.1;NP_001313406.1;NP_004045.1 | 1319 | 605246 | . | . | 1.003230 | 1.135526 | 1.155402 | 0.781500 | 2.334557 | 2.624820 | 2.592689 | 1.786806 | -1.118815 | 109.787815 | 0.326228915 | -0.49649803 | 0.619543075 | 0.998865404 | 6.856450 | 6.696253 | 6.773748 | 6.811654 | 6.815468 | 6.694080 | 126 | 132 | 108 | 78 |
| 720 | NM_00125220 | C4A | complement C4A (Rodgers blood group) | protein_coding | NP_001239133.1;NP_009224.2 | 1323 | 120810 | . | . | 0.144864 | 0.610433 | 0.118280 | 0.140368 | 0.337105 | 1.411043 | 0.265415 | 0.320935 | **-2.913826** | 43.286613 | 0.621210257 | -2.48372384 | **0.013001659** | 0.377020806 | 6.011006 | 4.464871 | 5.304839 | 5.595916 | 5.291060 | 5.300970 | 29 | 112 | 18 | 22 |
| 721 | NM_00100202 | C4B | complement C4B (Chido blood group) | protein_coding | NP_001002029.3 | 1324 | 120820 | . | . | 0.122920 | 0.525129 | 0.092136 | 0.092999 | 0.286041 | 1.213860 | 0.206752 | 0.212631 | **-3.471027** | 35.926236 | 0.667597898 | -2.68928764 | **0.007160469** | 0.26784811 | 5.802262 | 4.004486 | 5.051628 | 5.312760 | 5.031951 | 5.029967 | 25 | 97 | 14 | 15 |
| 726 | NM_004055,XI | CAPN5 | calpain 5 | protein_coding | NP_004046.2;XP_011543527.1;XP_016873712.2 | 1482 | 602537 | . | . | 3.253616 | 2.974564 | 3.707211 | 3.507077 | 7.571297 | 6.875837 | 8.318873 | 8.018514 | 1.141439 | 457.780253 | 0.175310315 | 1.088665042 | 0.276301623 | 0.998865404 | 8.739610 | 8.931066 | 8.824459 | 8.763429 | 8.896121 | 8.856182 | 517 | 438 | 438 | 443 |
| 727 | NM_00131716 | C5 | complement C5 | protein_coding | NP_001304092.1;NP_001304093.1;NP_001726.2 | 1331 | 120900 | . | . | 1.032328 | 1.093523 | 1.150122 | 0.767094 | 2.402270 | 2.527727 | 2.580840 | 1.753870 | -1.124610 | 173.451808 | 0.275020408 | -0.61604245 | 0.537866504 | 0.998865404 | 7.519747 | 7.352171 | 7.447305 | 7.469286 | 7.487585 | 7.323927 | 206 | 203 | 171 | 122 |
| 733 | NM_000606,XI | C8G | complement C8 gamma chain | protein_coding | NP_000597.2 | 1354 | 120930 | . | . | 0.091966 | 0.342670 | 0.226307 | 0.404684 | 0.214009 | 0.792096 | 0.507825 | 0.925263 | 1.448060 | 11.127463 | 0.975115919 | 0.547752086 | 0.583862143 | NA | 3.189301 | 3.715150 | 3.430877 | 3.475114 | 3.454305 | 3.488242 | 5 | 15 | 8 | 16 |
| 734 | NM_00112611 | OSGIN2 | oxidative stress induced growth inhibitor family member 2 | protein_coding | NP_001119583.1;NP_004328.1;XP_011515589.1;XP_011515590.1 | 1355 | 604598 | . | . | 5.544997 | 5.309327 | 4.093900 | 4.228093 | 12.903436 | 12.272749 | 9.186595 | 9.667033 | -1.181424 | 688.871170 | 0.15125206 | -1.5902414 | 0.111780402 | 0.848562841 | 9.543385 | 9.302776 | 9.490502 | 9.474268 | 9.357587 | 9.379365 | 869 | 796 | 540 | 602 |
| 738 | NM_013265,N | VPS51 | VPS51 subunit of GARP complex | protein_coding | NP_037397.2 | 1172 | 615738 | . | . | 7.196923 | 6.976185 | 6.337340 | 8.018229 | 16.747536 | 16.125761 | 14.220806 | 18.332729 | -1.001235 | 591.810348 | 0.171265869 | -0.01039829 | 0.991703516 | 0.999576129 | 9.210228 | 9.207753 | 9.220731 | 9.195417 | 9.131487 | 9.277678 | 695 | 627 | 457 | 616 |
| 740 | NM_004927,N | MRPL49 | mitochondrial ribosomal protein L49 | protein_coding | NP_004918.1 | 1176 | 606866 | . | . | 26.648294 | 28.580922 | 28.332246 | 32.393193 | 62.011677 | 66.066075 | 63.576741 | 74.063191 | 1.086795 | 1831.086147 | 0.127068496 | 0.945003729 | 0.344656972 | 0.998865404 | 10.777372 | 10.897112 | 10.779713 | 10.827840 | 10.820345 | 10.916579 | 1959 | 1952 | 1558 | 1896 |
| 741 | NM_014205 | ZNHIT2 | zinc finger HIT-type containing 2 | protein_coding | NP_055020.1 | 1177 | 604575 | . | . | 5.187038 | 4.818378 | 5.806460 | 6.477581 | 12.070453 | 11.137897 | 13.029528 | 14.810219 | 1.209473 | 225.526579 | 0.227392091 | 1.206633137 | 0.227573459 | 0.982008277 | 7.673526 | 7.947768 | 7.785816 | 7.747500 | 7.832275 | 7.881931 | 245 | 211 | 204 | 243 |
| 744 | NM_00114539 | MPPED2 | metallophosphoesterase domain containing 2 | protein_coding | NP_001138871.1;NP_001364881.1;NP_001364882.1;NP_001364883.1;N | 1180 | 600911 | . | . | 0.654390 | 0.692277 | 0.718683 | 1.246096 | 1.522792 | 1.600229 | 1.612706 | 2.849052 | 1.392608 | 67.559268 | 0.396008518 | 1.206512117 | 0.22762009 | 0.982008277 | 5.820670 | 6.296445 | 6.024791 | 6.032164 | 6.057510 | 6.150864 | 64 | 62 | 56 | 87 |
| 745 | NM_00112739 | MYRF | myelin regulatory factor | protein_coding | NP_001120864.1;NP_037411.1;XP_005274279.1;XP_005274280.1;XP_00 | 1181 | 608329 | . | . | 0.149928 | 0.116860 | 0.105540 | 0.183613 | 0.348889 | 0.270126 | 0.236829 | 0.419808 | 1.056176 | 26.145843 | 0.622973342 | 0.126571907 | 0.899279233 | NA | 4.669607 | 4.746391 | 4.717359 | 4.690256 | 4.680199 | 4.738572 | 33 | 24 | 17 | 32 |
| 746 | NM_014206 | TMEM258 | transmembrane protein 258 | protein_coding | NP_055021.1 | 1164 | 617615 | . | . | 36.406544 | 36.057720 | 34.197113 | 35.967949 | 84.719528 | 83.349037 | 76.737328 | 82.236435 | -1.047524 | 642.556951 | 0.154355154 | -0.4339569 | 0.664319742 | 0.998865404 | 9.360837 | 9.293744 | 9.349530 | 9.337011 | 9.298821 | 9.324950 | 764 | 703 | 535 | 600 |
| 747 | NM_006133,XI | DAGLA | diacylglycerol lipase alpha | protein_coding | NP_006124.1;XP_016873727.1;XP_016873728.1;XP_016873729.1 | 1165 | 614015 | . | . | 0.857323 | 0.917983 | 0.905051 | 0.642762 | 1.995025 | 2.121960 | 2.030908 | 1.469600 | -1.136793 | 155.838469 | 0.281213929 | -0.65775526 | 0.510695406 | 0.998865404 | 7.372753 | 7.189229 | 7.281596 | 7.328989 | 7.318313 | 7.186907 | 180 | 189 | 149 | 113 |
| 752 | NM_005892,XI | FMNL1 | formin like 1 | protein_coding | NP_005883.3;XP_006722125.1;XP_006722126.1;XP_006722127.1;XP_00 | 1212 | 604656 | . | . | 0.054048 | 0.086057 | 0.090868 | 0.057124 | 0.125771 | 0.198926 | 0.203906 | 0.130607 | 1.109448 | 13.810556 | 0.839796483 | 0.178426594 | 0.858387966 | NA | 3.710171 | 3.861274 | 3.764806 | 3.788943 | 3.799476 | 3.766642 | 12 | 17 | 16 | 10 |
| 753 | NM_00100367 | LDLRAD4 | low density lipoprotein receptor class A domain containing 4 | protein_coding | NP_001003674.1;NP_001003675.1;NP_001263178.1;NP_001263180.1;N | 1224 | 606571 | . | . | 0.079973 | 0.081627 | 0.020391 | 0.064624 | 0.186098 | 0.188683 | 0.045756 | 0.147754 | -1.719150 | 23.727353 | 0.683432909 | -1.14377733 | 0.252715993 | NA | 4.909524 | 4.121007 | 4.595885 | 4.599894 | 4.511287 | 4.578933 | 34 | 33 | 8 | 24 |
| 754 | NM_00128682 | PTTG1IP | PTTG1 interacting protein | protein_coding | NP_001273751.1;NP_004330.1 | 13524 | 603784 | . | . | 178.821951 | 179.222916 | 182.477012 | 181.507718 | 416.126030 | 414.281780 | 409.473141 | 414.995829 | 1.002523 | 14627.008601 | 0.099132496 | 0.03667226 | 0.970746327 | 0.998865404 | 13.834524 | 13.838168 | 13.837823 | 13.832680 | 13.843961 | 13.830835 | 16872 | 15716 | 12833 | 13625 |
| 755 | NM_00127144 | CFAP410 | cilia and flagella associated protein 410 | protein_coding | NP_001258369.1;NP_001258370.1;NP_001258371.1;NP_004919.1;XP_0 | 1260 | 603191 | . | . | 2.162900 | 2.335161 | 3.151525 | 2.549279 | 5.033157 | 5.397829 | 7.071931 | 5.828625 | 1.253724 | 182.200906 | 0.253258961 | 1.288088378 | 0.197715201 | 0.96621494 | 7.336768 | 7.663541 | 7.431911 | 7.472535 | 7.594080 | 7.503629 | 177 | 183 | 194 | 171 |
| 757 | NM_006134,N | TMEM50B | transmembrane protein 50B | protein_coding | NP_006125.2;XP_011528048.1 | 1280 | 617894 | . | . | 5.818498 | 6.423664 | 7.416004 | 5.264516 | 13.539883 | 14.848588 | 16.641299 | 12.036690 | -1.045784 | 534.598760 | 0.189697974 | -0.34046351 | 0.733507497 | 0.998865404 | 9.093825 | 9.030097 | 9.043111 | 9.101806 | 9.141981 | 8.935845 | 601 | 616 | 528 | 411 |
| 767 | NM_00132183 | CA8 | carbonic anhydrase 8 | protein_coding | NP_001308766.1;NP_001308767.1;NP_001308768.1;NP_004047.3;XP_01 | 1382 | 114815 | . | . | 7.219035 | 7.165879 | 6.174282 | 5.565611 | 16.798993 | 16.564247 | 13.854911 | 12.725107 | -1.237818 | 1016.816767 | 0.144043614 | -2.13684584 | **0.032610533** | 0.570498328 | 10.135285 | 9.828080 | 10.084141 | 10.039466 | 9.956967 | 9.851625 | 1333 | 1178 | 849 | 789 |
| 770 | NM_001217,N | CA11 | carbonic anhydrase 11 | protein_coding | NP_001208.2 | 1370 | 604644 | . | . | 2.758123 | 2.378100 | 2.823552 | 2.790082 | 6.418265 | 5.497083 | 6.335967 | 6.379193 | 1.054345 | 173.181721 | 0.255880791 | 0.298367972 | 0.765422331 | 0.998865404 | 7.396951 | 7.474298 | 7.459367 | 7.386252 | 7.457496 | 7.433877 | 211 | 166 | 159 | 162 |
| 775 | NM_000719,N | CACNA1C | calcium voltage-gated channel subunit alpha1 C | protein_coding | NP_000710.5;NP_001123299.1;NP_001123301.1;NP_001123302.2;NP_0 | 1390 | 114205 | . | . | 0.008561 | 0.003924 | 0.026103 | 0.006720 | 0.019923 | 0.009071 | 0.058575 | 0.015365 | **2.215576** | 5.707638 | 1.332849102 | 0.861074041 | 0.389197265 | NA | 1.820514 | 2.978782 | 2.485389 | 2.469503 | 2.513035 | 2.481202 | 6 | 2 | 10 | 4 |
| 776 | NM_000720,N | CACNA1D | calcium voltage-gated channel subunit alpha1 D | protein_coding | NP_000711.1;NP_001122311.1;NP_001122312.1;XP_005265505.1;XP_01 | 1391 | 114206 | . | . | 0.483252 | 0.318806 | 0.334795 | 0.317303 | 1.124548 | 0.736936 | 0.751271 | 0.725476 | -1.275827 | 105.739679 | 0.331971847 | -1.05862208 | 0.289771926 | 0.998865404 | 6.888809 | 6.538755 | 6.824792 | 6.689052 | 6.683291 | 6.681680 | 163 | 103 | 81 | 87 |
| 777 | NM_000721,N | CACNA1E | calcium voltage-gated channel subunit alpha1 E | protein_coding | NP_000712.2;NP_001192222.1;NP_001192223.1;XP_016857732.1;XP_01 | 1392 | 601013 | . | . | 0.010723 | 0.030449 | 0.006914 | 0.009598 | 0.024953 | 0.070384 | 0.015515 | 0.021946 | **-2.191315** | 7.941747 | 1.133893674 | -0.99815063 | 0.318206322 | NA | 3.448488 | 2.312263 | 2.983616 | 3.027334 | 2.977789 | 2.980746 | 7 | 17 | 4 | 5 |
| 780 | NM_00120252 | DDR1 | discoidin domain receptor tyrosine kinase 1 | protein_coding | NP_001189450.1;NP_001189451.1;NP_001189452.1;NP_001284581.1;N | 2730 | 600408 | . | . | 0.420301 | 0.585455 | 0.438807 | 0.398240 | 0.978058 | 1.353306 | 0.984671 | 0.910529 | -1.210142 | 58.847957 | 0.420649776 | -0.65416982 | 0.513002401 | 0.998865404 | 6.010217 | 5.734467 | 5.858521 | 5.934121 | 5.868348 | 5.845898 | 62 | 81 | 49 | 47 |
| 782 | NM_000723,N | CACNB1 | calcium voltage-gated channel auxiliary subunit beta 1 | protein_coding | NP_000714.3;NP_954855.1;NP_954856.1;XP_005257702.1;XP_00525770 | 1401 | 114207 | . | . | 2.356133 | 2.492951 | 2.802883 | 2.188028 | 5.482822 | 5.762567 | 6.289589 | 5.002666 | 1.026437 | 247.427408 | 0.227774541 | 0.165274014 | 0.868728311 | 0.998865404 | 7.931424 | 7.970040 | 7.927178 | 7.956036 | 8.014176 | 7.891690 | 273 | 271 | 245 | 206 |
| 784 | NM_000725,N | CACNB3 | calcium voltage-gated channel auxiliary subunit beta 3 | protein_coding | NP_000716.2;NP_001193844.1;NP_001193845.1;NP_001193846.1;XP_01 | 1403 | 601958 | . | . | 4.088828 | 3.908748 | 3.661735 | 4.115880 | 9.514870 | 9.035246 | 8.216825 | 9.410470 | 1.008277 | 353.687813 | 0.188441358 | 0.06310608 | 0.949682033 | 0.998865404 | 8.460372 | 8.472269 | 8.471571 | 8.456244 | 8.463433 | 8.473816 | 411 | 375 | 306 | 336 |
| 785 | NM_000726,N | CACNB4 | calcium voltage-gated channel auxiliary subunit beta 4 | protein_coding | NP_000717.2;NP_001005746.1;NP_001005747.1;NP_001139270.1;NP_0 | 1404 | 601949 | . | . | 0.079583 | 0.097086 | 0.073388 | 0.052478 | 0.185193 | 0.224417 | 0.164681 | 0.119983 | -1.435266 | 23.685471 | 0.641879979 | -0.81217349 | 0.41669209 | NA | 4.802857 | 4.282298 | 4.570071 | 4.599915 | 4.559983 | 4.539932 | 28 | 34 | 19 | 16 |
| 788 | NM_000387 | SLC25A20 | solute carrier family 25 member 20 | protein_coding | NP_000378.1 | 1421 | 613698 | . | . | 7.090524 | 7.844774 | 8.318241 | 8.113832 | 16.499939 | 18.133545 | 18.665892 | 18.551315 | 1.084690 | 434.758287 | 0.177222522 | 0.661779149 | 0.508112771 | 0.998865404 | 8.704324 | 8.821439 | 8.709140 | 8.765450 | 8.797357 | 8.776900 | 458 | 471 | 400 | 417 |
| 790 | NM_00130607 | CAD | carbamoyl-phosphate synthetase 2, aspartate transcarbamylase, a | protein_coding | NP_001293008.1;NP_004332.2;XP_006712164.1;XP_024308899.1 | 1424 | 114010 | . | . | 21.590360 | 23.555518 | 23.695664 | 24.115670 | 50.241653 | 54.449631 | 53.172382 | 55.137612 | 1.047136 | 4781.283812 | 0.110884456 | 0.599257617 | 0.549001106 | 0.998865404 | 12.189640 | 12.255962 | 12.168975 | 12.236534 | 12.215521 | 12.267300 | 5161 | 5245 | 4122 | 4727 |
| 794 | NM_001740,N | CALB2 | calbindin 2 | protein_coding | NP_001731.2;NP_009019.1 | 1435 | 114051 | . | . | 1.056710 | 0.643901 | 0.804103 | 0.783916 | 2.459005 | 1.488405 | 1.804383 | 1.792330 | -1.085003 | 38.055722 | 0.5220383 | -0.2254617 | 0.821620117 | 0.998865404 | 5.306645 | 5.191126 | 5.293579 | 5.214915 | 5.243984 | 5.241844 | 57 | 32 | 32 | 34 |
| 800 | NM_004342,N | CALD1 | caldesmon 1 | protein_coding | NP_004333.1;NP_149129.2;NP_149130.1;NP_149131.1;NP_149347.2;XP_ | 1441 | 114213 | . | . | 54.727644 | 49.375480 | 48.306249 | 47.670195 | 127.353473 | 114.133630 | 108.397831 | 108.992233 | -1.096594 | 6683.337384 | 0.108121696 | -1.23037098 | 0.218558214 | 0.982008277 | 12.771301 | 12.638343 | 12.783660 | 12.702313 | 12.675982 | 12.655761 | 8434 | 7184 | 5611 | 5905 |
| 801 | NM_00136366 | CALM1 | calmodulin 1 | protein_coding | NP_001350598.1;NP_001350599.1;NP_008819.1 | 1442 | 114180 | . | . | 30.626210 | 30.362150 | 28.850649 | 28.950555 | 71.268449 | 70.183470 | 64.740019 | 66.192004 | -1.069293 | 3920.702008 | 0.107558618 | -0.89864623 | 0.368841128 | 0.998865404 | 11.984405 | 11.887770 | 11.969642 | 11.958449 | 11.911237 | 11.905839 | 4697 | 4343 | 3306 | 3541 |
| 805 | NM_00130562 | CALM2 | calmodulin 2 | protein_coding | NP_001292553.1;NP_001292554.1;NP_001292555.1;NP_001734.1 | 1445 | 114182 | . | . | 208.879108 | 191.861363 | 152.116879 | 159.819163 | 486.070297 | 443.496164 | 341.345871 | 365.407526 | -1.283553 | 6995.593110 | 0.108549977 | -3.31775899 | **0.000907427** | 0.064386964 | 12.941083 | 12.580963 | 12.910977 | 12.823904 | 12.645370 | 12.669143 | 9511 | 8059 | 5251 | 5827 |
| 808 | NM_00132992 | CALM3 | calmodulin 3 | protein_coding | NP_001316850.1;NP_001316851.1;NP_001316852.1;NP_001316853.1;N | 1449 | 114183 | . | . | 71.382433 | 75.452385 | 63.138470 | 78.404076 | 166.109850 | 174.411558 | 141.680893 | 179.261599 | -1.052027 | 4909.687317 | 0.124940066 | -0.58566 | 0.558104018 | 0.998865404 | 12.297634 | 12.224264 | 12.260905 | 12.300731 | 12.149399 | 12.320483 | 5660 | 5561 | 3734 | 4946 |
| 811 | NM_004343 | CALR | calreticulin | protein_coding | NP_004334.1 | 1455 | 109091 | . | . | 567.412537 | 573.509094 | 494.281372 | 518.887512 | 1320.392334 | 1325.692017 | 1109.153076 | 1186.374634 | -1.142022 | 31903.238288 | 0.098148621 | -1.95204858 | 0.050932437 | 0.672475152 | 15.054037 | 14.862437 | 15.015069 | 15.016932 | 14.885001 | 14.917693 | 39130 | 36747 | 25406 | 28462 |
| 813 | NM_00113067 | CALU | calumenin | protein_coding | NP_001124146.1;NP_001186600.1;NP_001186601.1;NP_001186602.1;N | 1458 | 603420 | . | . | 230.177017 | 232.463322 | 189.827632 | 181.357293 | 535.631414 | 537.349377 | 425.967697 | 414.651878 | -1.249829 | 34151.209814 | 0.099635017 | -3.22908922 | **0.001241851** | 0.080099418 | 15.211558 | 14.889842 | 15.162042 | 15.134405 | 14.978773 | 14.932338 | 44378 | 40299 | 27123 | 27682 |
| 814 | NM_00132337 | CAMK4 | calcium/calmodulin dependent protein kinase IV | protein_coding | NP_001310303.1;NP_001310304.1;NP_001310305.1;NP_001310306.1;N | 1464 | 114080 | . | . | 3.164227 | 2.731179 | 2.909130 | 2.809339 | 7.363287 | 6.313243 | 6.528004 | 6.423218 | -1.046845 | 1072.394814 | 0.141760087 | -0.46591585 | 0.641275735 | 0.998865404 | 10.099054 | 10.033441 | 10.134161 | 10.028467 | 10.067778 | 10.029665 | 1352 | 1100 | 935 | 957 |
| 817 | NM_001221,N | CAMK2D | calcium/calmodulin dependent protein kinase II delta | protein_coding | NP_001212.2;NP_001308495.1;NP_001308496.1;NP_001308497.1;NP_0 | 1462 | 607708 | . | . | 5.039686 | 4.927984 | 4.412442 | 4.628463 | 11.727551 | 11.391255 | 9.901388 | 10.582431 | -1.144829 | 804.308021 | 0.144486496 | -1.35051949 | 0.176849407 | 0.949538805 | 9.745813 | 9.550811 | 9.707987 | 9.686863 | 9.607301 | 9.597036 | 1003 | 913 | 657 | 698 |
| 818 | NM_00120449 | CAMK2G | calcium/calmodulin dependent protein kinase II gamma | protein_coding | NP_001191421.1;NP_001213.2;NP_001307827.1;NP_001354443.1;NP_0 | 1463 | 602123 | . | . | 4.200946 | 4.439458 | 3.958352 | 4.822302 | 9.775776 | 10.261999 | 8.882428 | 11.025618 | -1.004829 | 497.309279 | 0.175007422 | -0.03971535 | 0.968320062 | 0.998865404 | 8.961904 | 8.954085 | 8.936085 | 8.980999 | 8.902953 | 9.005604 | 554 | 557 | 396 | 503 |
| 819 | NM_001745 | CAMLG | calcium modulating ligand | protein_coding | NP_001736.1 | 1471 | 601118 | . | . | 12.734218 | 10.858578 | 11.654489 | 10.221251 | 29.633049 | 25.100088 | 26.152336 | 23.369677 | -1.093280 | 769.818860 | 0.160214449 | -0.80306644 | 0.421936336 | 0.998865404 | 9.650881 | 9.523039 | 9.675321 | 9.555866 | 9.602552 | 9.505314 | 1003 | 795 | 685 | 641 |
| 821 | NM_00102464 | CANX | calnexin | protein_coding | NP_001019820.1;NP_001350922.1;NP_001350923.1;NP_001350924.1;N | 1473 | 114217 | . | . | 377.305006 | 362.225129 | 346.164797 | 307.791725 | 878.004266 | 837.299624 | 776.783765 | 703.729219 | -1.142182 | 53135.880404 | 0.106455193 | -1.80162187 | 0.071604918 | 0.753985777 | 15.790102 | 15.598327 | 15.769223 | 15.733940 | 15.688586 | 15.582114 | 66562 | 59908 | 45821 | 43618 |
| 822 | NM_00125613 | CAPG | capping actin protein, gelsolin like | protein_coding | NP_001243068.1;NP_001243069.1;NP_001307661.1;NP_001307662.1;N | 1474 | 153615 | . | . | 778.354088 | 769.330951 | 680.578979 | 910.107257 | 1811.261928 | 1778.342976 | 1527.199489 | 2080.852200 | 1.019334 | 31493.267738 | 0.13036028 | 0.211926 | 0.832164766 | 0.998865404 | 14.928892 | 14.956488 | 14.939333 | 14.924806 | 14.821734 | 15.061524 | 36220 | 33374 | 23823 | 33931 |
| 823 | NM_00119886 | CAPN1 | calpain 1 | protein_coding | NP_001185797.1;NP_001185798.1;NP_005177.2;XP_006718761.1;XP_01 | 1476 | 114220 | . | . | 18.819339 | 19.346729 | 18.644207 | 22.678969 | 43.793373 | 44.720831 | 41.837060 | 51.852766 | 1.066120 | 1828.799034 | 0.1330583 | 0.694208469 | 0.487551482 | 0.998865404 | 10.789948 | 10.881947 | 10.802530 | 10.816980 | 10.783985 | 10.931087 | 2020 | 1928 | 1489 | 1933 |
| 824 | NM_00114606 | CAPN2 | calpain 2 | protein_coding | NP_001139540.1;NP_001739.3 | 1479 | 114230 | . | . | 57.714196 | 57.065950 | 51.982432 | 57.131527 | 134.303303 | 131.910498 | 116.647076 | 130.624458 | -1.065718 | 6494.053529 | 0.105592804 | -0.86961642 | 0.384510063 | 0.998865404 | 12.710110 | 12.618235 | 12.697575 | 12.684542 | 12.609019 | 12.664927 | 7775 | 7175 | 5285 | 6091 |
| 825 | NM_000070,N | CAPN3 | calpain 3 | protein_coding | NP_000061.1;NP_077320.1;NP_775110.1;NP_775111.1;NP_775112.1;NP_ | 1480 | 114240 | . | . | 80.714669 | 86.425710 | 74.184563 | 93.622630 | 187.826349 | 199.776905 | 166.468011 | 214.057024 | -1.021500 | 3762.648924 | 0.120075911 | -0.25557903 | 0.798275911 | 0.998865404 | 11.892907 | 11.861993 | 11.856107 | 11.914173 | 11.805930 | 11.926270 | 4226 | 4247 | 3001 | 3744 |
| 826 | NM_00100396 | CAPNS1 | calpain small subunit 1 | protein_coding | NP_001003962.1;NP_001289561.1;NP_001289562.1;NP_001740.1;XP_0 | 1481 | 114170 | . | . | 98.404322 | 103.875885 | 100.976632 | 111.967867 | 228.990889 | 240.113771 | 226.588622 | 256.001232 | 1.037315 | 4635.170575 | 0.112071954 | 0.471602408 | 0.637210607 | 0.998865404 | 12.151812 | 12.204520 | 12.136824 | 12.187650 | 12.153519 | 12.231143 | 5080 | 5060 | 3914 | 4630 |
| 828 | NM_004058,N | CAPS | calcyphosine | protein_coding | NP_004049.3;NP_542157.3 | 1487 | 114212 | . | . | 75.505680 | 73.841288 | 27.332032 | 47.752313 | 175.704831 | 170.687431 | 61.332286 | 109.179991 | **-2.018213** | 2673.100720 | 0.38656126 | -2.62074413 | **0.008773808** | 0.294943244 | 11.803724 | 10.790520 | 11.606475 | 11.580304 | 10.803406 | 11.207343 | 4178 | 3800 | 1127 | 2106 |
| 829 | NM_006135,XI | CAPZA1 | capping actin protein of muscle Z-line subunit alpha 1 | protein_coding | NP_006126.1;XP_011540527.1;XP_016857913.1 | 1488 | 601580 | . | . | 60.803167 | 54.859952 | 51.509813 | 48.475303 | 141.491461 | 126.811233 | 115.586530 | 110.833025 | -1.173087 | 3973.414496 | 0.115602768 | -1.99225822 | **0.046342734** | 0.654058028 | 12.066648 | 11.836511 | 12.062818 | 11.971722 | 11.914481 | 11.856572 | 5206 | 4379 | 3293 | 3307 |
| 830 | NM_006136 | CAPZA2 | capping actin protein of muscle Z-line subunit alpha 2 | protein_coding | NP_006127.1 | 1490 | 601571 | . | . | 11.177348 | 10.168977 | 10.070089 | 8.868339 | 26.010149 | 23.506046 | 22.596989 | 20.276403 | -1.143413 | 1591.440630 | 0.133945648 | -1.44346642 | 0.148889126 | 0.909660767 | 10.729336 | 10.536457 | 10.724931 | 10.643595 | 10.632012 | 10.527312 | 2055 | 1738 | 1380 | 1297 |
| 831 | NM_00104244 | CAST | calpastatin | protein_coding | NP_001035905.1;NP_001035906.1;NP_001035907.1;NP_001035908.1;N | 1515 | 114090 | . | . | 18.695396 | 19.878092 | 16.854868 | 21.918735 | 43.504951 | 45.949102 | 37.821833 | 50.114588 | -1.010375 | 2654.427277 | 0.137908232 | -0.10797281 | 0.914017264 | 0.998865404 | 11.381789 | 11.366541 | 11.354830 | 11.397610 | 11.260616 | 11.465140 | 2992 | 2955 | 2007 | 2787 |
| 832 | NM_00120654 | CAPZB | capping actin protein of muscle Z-line subunit beta | protein_coding | NP_001193469.1;NP_001193470.1;NP_001269091.1;NP_001300861.1;N | 1491 | 601572 | . | . | 149.868046 | 156.961721 | 128.418676 | 155.633304 | 348.749109 | 362.824060 | 288.167771 | 355.837089 | -1.098512 | 7530.144055 | 0.119631651 | -1.13306745 | 0.257185928 | 0.995760118 | 12.944708 | 12.809027 | 12.898632 | 12.932964 | 12.754172 | 12.912285 | 8896 | 8678 | 5654 | 7360 |
| 833 | NM_00101443 | CARS1 | cysteinyl-tRNA synthetase 1 | protein_coding | NP_001014437.1;NP_001181926.1;NP_001365065.1;NP_001365066.1;N | 1493 | 123859 | . | . | 33.353919 | 35.434419 | 26.016660 | 34.774629 | 77.615936 | 81.908245 | 58.380625 | 79.508052 | -1.120600 | 2699.400333 | 0.142099215 | -1.15603596 | 0.24766645 | 0.992733415 | 11.478403 | 11.313748 | 11.415163 | 11.465544 | 11.230135 | 11.453678 | 3185 | 3173 | 1906 | 2718 |
| 834 | NM_001223,N | CASP1 | caspase 1 | protein_coding | NP_001214.1;NP_001244047.1;NP_001244048.1;NP_150634.1;NP_1506 | 1499 | 147678 | . | . | 0.729584 | 0.852967 | 0.933170 | 1.184331 | 1.697770 | 1.971671 | 2.094007 | 2.707834 | 1.281510 | 52.146672 | 0.432206556 | 0.827948567 | 0.407699628 | 0.998865404 | 5.514721 | 5.872198 | 5.675988 | 5.673923 | 5.714169 | 5.730184 | 53 | 49 | 49 | 57 |
| 835 | NM_001224,N | CASP2 | caspase 2 | protein_coding | NP_001215.1;NP_116764.2;NP_116765.2 | 1503 | 600639 | . | . | 6.148025 | 6.962999 | 7.064054 | 7.047936 | 14.306705 | 16.095284 | 15.851533 | 16.114267 | 1.065595 | 866.464764 | 0.145570568 | 0.629651627 | 0.528922538 | 0.998865404 | 9.712566 | 9.803981 | 9.693558 | 9.774975 | 9.784739 | 9.776187 | 911 | 957 | 782 | 833 |
| 836 | NM_00135477 | CASP3 | caspase 3 | protein_coding | NP_001341706.1;NP_001341708.1;NP_001341709.1;NP_001341710.1;N | 1504 | 600636 | . | . | 7.697990 | 7.656472 | 6.918388 | 7.031285 | 17.913539 | 17.698278 | 15.524662 | 16.076198 | -1.114221 | 597.932421 | 0.157392243 | -0.99138163 | 0.321499267 | 0.998865404 | 9.299714 | 9.143750 | 9.262897 | 9.254702 | 9.190063 | 9.184294 | 730 | 676 | 494 | 528 |
| 837 | NM_001225,N | CASP4 | caspase 4 | protein_coding | NP_001216.1;NP_150649.1;XP_011541321.1;XP_016873886.1 | 1505 | 602664 | . | . | 18.448399 | 20.432032 | 19.624223 | 21.995329 | 42.930183 | 47.229556 | 44.036188 | 50.289707 | 1.055558 | 822.945140 | 0.148630873 | 0.524826459 | 0.599703836 | 0.998865404 | 9.645401 | 9.722864 | 9.635572 | 9.692620 | 9.665355 | 9.738643 | 885 | 899 | 697 | 833 |
| 839 | NM_001226,N | CASP6 | caspase 6 | protein_coding | NP_001217.2;NP_116787.1 | 1507 | 601532 | . | . | 3.915677 | 3.514473 | 3.408610 | 3.294503 | 9.111944 | 8.123862 | 7.648823 | 7.532491 | -1.121750 | 180.328306 | 0.248908633 | -0.66591095 | 0.505468018 | 0.998865404 | 7.574658 | 7.409587 | 7.544211 | 7.492953 | 7.478558 | 7.460884 | 232 | 194 | 151 | 156 |
| 840 | NM_001227,N | CASP7 | caspase 7 | protein_coding | NP_001218.1;NP_001253985.1;NP_001253986.1;NP_001253987.1;NP_0 | 1508 | 601761 | . | . | 6.538622 | 7.093840 | 7.142057 | 8.042573 | 15.215639 | 16.397735 | 16.026571 | 18.388388 | 1.098331 | 548.524166 | 0.164753658 | 0.821306688 | 0.411471599 | 0.998865404 | 9.030449 | 9.165227 | 9.043245 | 9.090631 | 9.102287 | 9.153003 | 580 | 585 | 480 | 559 |
| 841 | NM_00108012 | CASP8 | caspase 8 | protein_coding | NP_001073593.1;NP_001073594.1;NP_001219.2;NP_001358980.1;NP_2 | 1509 | 601763 | . | . | 12.571088 | 13.818572 | 16.087179 | 15.764315 | 29.253436 | 31.942247 | 36.099162 | 36.043236 | 1.172118 | 1198.568218 | 0.134314925 | 1.705826767 | 0.088040338 | 0.801685384 | 10.108165 | 10.336965 | 10.126578 | 10.198764 | 10.267672 | 10.297413 | 1209 | 1249 | 1103 | 1236 |
| 842 | NM_001229,N | CASP9 | caspase 9 | protein_coding | NP_001220.2;NP_001264983.1;NP_127463.2;XP_005246071.1;XP_01154 | 1511 | 602234 | . | . | 5.833416 | 5.748095 | 4.666953 | 6.395151 | 13.574600 | 13.286979 | 10.472509 | 14.621756 | -1.053144 | 482.457196 | 0.191978069 | -0.38912068 | 0.697186876 | 0.998865404 | 8.951631 | 8.875890 | 8.939017 | 8.916778 | 8.792971 | 8.988239 | 580 | 525 | 344 | 510 |
| 843 | NM_00120652 | CASP10 | caspase 10 | protein_coding | NP_001193453.1;NP_001193471.1;NP_001221.2;NP_001293012.1;NP_1 | 1500 | 601762 | . | . | 0.074218 | 0.082711 | 0.092145 | 0.023726 | 0.172709 | 0.191190 | 0.206770 | 0.054246 | -1.385165 | 12.791504 | 0.920353623 | -0.51073562 | 0.609536195 | NA | 3.889947 | 3.427357 | 3.678245 | 3.686412 | 3.691918 | 3.635750 | 16 | 17 | 15 | 4 |
| 845 | NM_001232 | CASQ2 | calsequestrin 2 | protein_coding | NP_001223.2 | 1513 | 114251 | . | . | 0.273878 | 0.240171 | 0.125404 | 0.355053 | 0.637326 | 0.555167 | 0.281402 | 0.811786 | -1.070226 | 20.297729 | 0.729692063 | -0.13418725 | 0.893254506 | NA | 4.393730 | 4.290932 | 4.351961 | 4.338444 | 4.300700 | 4.376777 | 26 | 21 | 9 | 27 |
| 847 | NM_001752 | CAT | catalase | protein_coding | NP_001743.1 | 1516 | 115500 | . | . | 9.167190 | 9.225447 | 8.945392 | 10.710208 | 21.332426 | 21.325033 | 20.073198 | 24.487619 | 1.054047 | 678.989480 | 0.159414345 | 0.476359618 | 0.633818199 | 0.998865404 | 9.369076 | 9.444431 | 9.388523 | 9.387569 | 9.364679 | 9.479768 | 762 | 713 | 555 | 708 |
| 857 | NM_00117289 | CAV1 | caveolin 1 | protein_coding | NP_001166366.1;NP_001166367.1;NP_001166368.1;NP_001744.2 | 1527 | 601047 | . | . | 29.187598 | 26.742186 | 25.366068 | 24.096374 | 67.920739 | 61.815765 | 56.920720 | 55.093491 | -1.150557 | 2102.795716 | 0.122478658 | -1.65198376 | 0.098537862 | 0.813553488 | 11.135595 | 10.933521 | 11.128762 | 11.051952 | 11.004036 | 10.953914 | 2713 | 2313 | 1755 | 1775 |
| 858 | NM_00120674 | CAV2 | caveolin 2 | protein_coding | NP_001193676.1;NP_001193677.1;NP_001224.1;NP_937855.1 | 1528 | 601048 | . | . | 5.336316 | 4.803570 | 3.988540 | 4.541278 | 12.417827 | 11.103666 | 8.950169 | 10.383091 | -1.203562 | 431.345013 | 0.182098717 | -1.46794235 | 0.142119882 | 0.903146012 | 8.880280 | 8.612731 | 8.840673 | 8.769504 | 8.658479 | 8.727285 | 573 | 480 | 320 | 389 |
| 860 | NM_00101505 | RUNX2 | RUNX family transcription factor 2 | protein_coding | NP_001015051.3;NP_001019801.3;NP_001265407.1;NP_001356334.1 | 10472 | 600211 | . | . | 0.290662 | 0.255452 | 0.325207 | 0.311235 | 0.676382 | 0.590489 | 0.729755 | 0.711602 | 1.157614 | 52.122252 | 0.434062661 | 0.486460567 | 0.626640657 | 0.998865404 | 5.593806 | 5.806050 | 5.700839 | 5.672008 | 5.719811 | 5.709086 | 60 | 48 | 50 | 51 |
| 861 | NM_00100189 | RUNX1 | RUNX family transcription factor 1 | protein_coding | NP_001001890.1;NP_001116079.1;NP_001745.2;XP_005261125.1;XP_00 | 10471 | 151385 | . | . | 1.682820 | 1.739896 | 1.615506 | 1.001042 | 3.915990 | 4.021848 | 3.625151 | 2.288765 | -1.256874 | 289.648651 | 0.228226294 | -1.44523104 | 0.148393002 | 0.909660767 | 8.332860 | 8.004857 | 8.250814 | 8.211798 | 8.197186 | 8.016160 | 384 | 336 | 264 | 197 |
| 864 | NM_00103168 | RUNX3 | RUNX family transcription factor 3 | protein_coding | NP_001026850.1;NP_001307601.1;NP_004341.1;XP_005246081.1;XP_01 | 10473 | 600210 | . | . | 21.585597 | 24.530139 | 20.621057 | 23.760616 | 50.230572 | 56.702517 | 46.273052 | 54.325826 | -1.060760 | 3014.658172 | 0.123816356 | -0.68729668 | 0.49189579 | 0.998865404 | 11.599837 | 11.514457 | 11.531027 | 11.627892 | 11.480358 | 11.581750 | 3364 | 3546 | 2385 | 2914 |
| 865 | NM_00136870 | CBFB | core-binding factor subunit beta | protein_coding | NP_001355636.1;NP_001355637.1;NP_001355638.1;NP_001355639.1;N | 1539 | 121360 | . | . | 11.181373 | 8.975777 | 9.286674 | 8.089099 | 26.019518 | 20.747913 | 20.839029 | 18.494762 | -1.175869 | 914.392487 | 0.161770865 | -1.44480729 | 0.148512023 | 0.909660767 | 9.948433 | 9.715516 | 9.971067 | 9.801780 | 9.824566 | 9.718915 | 1267 | 945 | 786 | 729 |
| 866 | NM_001756 | SERPINA6 | serpin family A member 6 | protein_coding | NP_001747.3 | 1540 | 122500 | . | . | 1.007830 | 0.850045 | 0.543699 | 0.543173 | 2.345262 | 1.964916 | 1.220044 | 1.241901 | -1.704886 | 34.411335 | 0.534836714 | -1.43908491 | 0.150126468 | 0.912861516 | 5.438724 | 4.669578 | 5.156653 | 5.128649 | 5.071544 | 5.072060 | 54 | 43 | 22 | 24 |
| 867 | NM_005188 | CBL | Cbl proto-oncogene | protein_coding | NP_005179.2 | 1541 | 165360 | . | . | 2.207392 | 2.304431 | 2.253560 | 2.499895 | 5.136692 | 5.326795 | 5.056923 | 5.715714 | 1.038989 | 806.134071 | 0.147108657 | 0.375095851 | 0.707589183 | 0.998865404 | 9.627233 | 9.681999 | 9.628628 | 9.652908 | 9.634221 | 9.700174 | 895 | 868 | 681 | 806 |
| 868 | NM_00132178 | CBLB | Cbl proto-oncogene B | protein_coding | NP_001308715.1;NP_001308717.1;NP_001308718.1;NP_001308719.1;N | 1542 | 604491 | . | . | 1.871020 | 1.988238 | 3.067008 | 2.484637 | 4.353941 | 4.595900 | 6.882277 | 5.680828 | 1.410588 | 484.221608 | 0.181829525 | 2.729461447 | **0.006343787** | 0.25089767 | 8.649825 | 9.146657 | 8.775497 | 8.813085 | 9.080567 | 8.935072 | 448 | 447 | 547 | 473 |
| 869 | NM_004352 | CBLN1 | cerebellin 1 precursor | protein_coding | NP_004343.1 | 1543 | 600432 | . | . | 0.271813 | 0.197041 | 0.060581 | 0.134194 | 0.632521 | 0.455469 | 0.135942 | 0.306818 | **-2.443647** | 13.183070 | 0.890855674 | -1.44696356 | 0.14790713 | NA | 4.227499 | 2.932304 | 3.767234 | 3.740983 | 3.693093 | 3.718857 | 25 | 17 | 4 | 10 |
| 871 | NM_00120701 | SERPINH1 | serpin family H member 1 | protein_coding | NP_001193943.1;NP_001226.2;XP_011543629.1;XP_024304524.1 | 1546 | 600943 | . | . | 147.761332 | 151.067959 | 122.693544 | 112.263331 | 343.846730 | 349.200346 | 275.320771 | 256.676768 | -1.288561 | 8597.770121 | 0.107177751 | -3.41265895 | **0.000643324** | 0.050453413 | 13.241036 | 12.875346 | 13.163696 | 13.171656 | 12.994181 | 12.908732 | 11097 | 10495 | 6870 | 6682 |
| 873 | NM_00128678 | CBR1 | carbonyl reductase 1 | protein_coding | NP_001273718.1;NP_001748.1 | 1548 | 114830 | . | . | 48.343240 | 49.117468 | 46.444533 | 56.366398 | 112.496705 | 113.537229 | 104.220185 | 128.875082 | 1.045108 | 1902.850682 | 0.131680003 | 0.48338297 | 0.628823838 | 0.998865404 | 10.861943 | 10.925256 | 10.873731 | 10.876729 | 10.835603 | 10.979459 | 2139 | 2012 | 1538 | 1988 |
| 874 | NM_001236,XI | CBR3 | carbonyl reductase 3 | protein_coding | NP_001227.1;XP_011528074.1 | 1549 | 603608 | . | . | 16.823678 | 16.378288 | 17.237633 | 23.385951 | 39.149390 | 37.859151 | 38.680745 | 53.469199 | 1.204589 | 572.244806 | 0.181666172 | 1.478211031 | 0.139351302 | 0.900674731 | 9.020412 | 9.288161 | 9.103566 | 9.080201 | 9.110094 | 9.308437 | 609 | 550 | 464 | 672 |
| 875 | NM_000071,N | CBS | cystathionine beta-synthase | protein_coding | NP_000062.1;NP_001171479.1;NP_001171480.1;NP_001307227.1;NP_0 | 1550 | 613381 | . | . | 3.438785 | 4.590498 | 2.135212 | 3.956147 | 8.002194 | 10.611141 | 4.791353 | 9.045261 | -1.327540 | 273.954731 | 0.276979224 | -1.47576085 | 0.140008105 | 0.900674731 | 8.288841 | 7.877529 | 8.089499 | 8.217260 | 7.846579 | 8.148171 | 319 | 376 | 140 | 290 |
| 883 | NM_00112267 | KYAT1 | kynurenine aminotransferase 1 | protein_coding | NP_001116143.1;NP_001116144.1;NP_001274319.1;NP_001339917.1;N | 1564 | 600547 | . | . | 3.505684 | 3.681701 | 4.252708 | 4.419982 | 8.157873 | 8.510420 | 9.542953 | 10.105760 | 1.248962 | 271.993855 | 0.209054991 | 1.534188319 | 0.12498333 | 0.873049043 | 7.918362 | 8.238747 | 8.006149 | 8.040499 | 8.134559 | 8.141934 | 269 | 270 | 261 | 285 |
| 889 | NM_00101340 | KRIT1 | KRIT1 ankyrin repeat containing | protein_coding | NP_001013424.1;NP_001337598.1;NP_001337599.1;NP_001337600.1;N | 1573 | 604214 | . | . | 8.675559 | 10.141786 | 10.927992 | 7.729246 | 20.188380 | 23.443194 | 24.522097 | 17.672001 | -1.061614 | 1345.790392 | 0.163910941 | -0.52625952 | 0.598707903 | 0.998865404 | 10.436509 | 10.350691 | 10.364730 | 10.458424 | 10.484883 | 10.233914 | 1499 | 1586 | 1323 | 1022 |
| 890 | NM_001237 | CCNA2 | cyclin A2 | protein_coding | NP_001228.2 | 1578 | 123835 | . | . | 16.884026 | 17.242851 | 11.986467 | 11.515625 | 39.289822 | 39.857624 | 26.897285 | 26.329109 | -1.472178 | 1235.738833 | 0.13066121 | -4.27021729 | **1.95283E-05** | **0.003369357** | 10.523265 | 9.965468 | 10.393177 | 10.402875 | 10.124675 | 10.088874 | 1684 | 1598 | 891 | 914 |
| 891 | NM_00135484 | CCNB1 | cyclin B1 | protein_coding | NP_001341773.1;NP_001341774.1;NP_114172.1 | 1579 | 123836 | . | . | 94.269228 | 99.028980 | 74.005400 | 87.312030 | 219.368365 | 228.909924 | 166.065971 | 199.628576 | -1.215318 | 5607.904140 | 0.116584632 | -2.41313224 | **0.01581608** | 0.422261805 | 12.587139 | 12.305644 | 12.511431 | 12.546289 | 12.299341 | 12.426858 | 6941 | 6775 | 4062 | 5113 |
| 892 | NM_00101339 | CCNC | cyclin C | protein_coding | NP_001013417.1;NP_001350466.1;NP_005181.2;XP_011534534.1;XP_01 | 1581 | 123838 | . | . | 18.166876 | 15.691369 | 15.701139 | 14.174053 | 42.275067 | 36.271300 | 35.232902 | 32.407287 | -1.143958 | 1076.296143 | 0.144464887 | -1.3431256 | 0.179231309 | 0.951511445 | 10.165372 | 9.971862 | 10.176459 | 10.057447 | 10.047675 | 9.989910 | 1426 | 1142 | 908 | 904 |
| 896 | NM_00113601 | CCND3 | cyclin D3 | protein_coding | NP_001129489.1;NP_001129597.1;NP_001129598.1;NP_001274356.1;N | 1585 | 123834 | . | . | 23.789033 | 23.593837 | 16.564056 | 24.328084 | 55.358059 | 54.538217 | 37.169261 | 55.623275 | -1.181615 | 1380.213251 | 0.162929423 | -1.47769544 | 0.139489316 | 0.900674731 | 10.546320 | 10.304947 | 10.498323 | 10.477186 | 10.220497 | 10.482458 | 1745 | 1592 | 908 | 1393 |
| 898 | NM_001238,N | CCNE1 | cyclin E1 | protein_coding | NP_001229.1;NP_001309188.1;NP_001309190.1;NP_001309191.1;XP_01 | 1589 | 123837 | . | . | 6.750660 | 6.624167 | 6.348488 | 7.330584 | 15.709065 | 15.312060 | 14.245823 | 16.760508 | 1.004248 | 404.994976 | 0.183000537 | 0.033415619 | 0.973343154 | 0.998865404 | 8.658934 | 8.664581 | 8.668648 | 8.651697 | 8.625347 | 8.698634 | 472 | 430 | 331 | 404 |
| 899 | NM_00132353 | CCNF | cyclin F | protein_coding | NP_001310467.1;NP_001752.2 | 1591 | 600227 | . | . | 7.373151 | 6.804831 | 5.866043 | 6.926872 | 17.157624 | 15.729671 | 13.163232 | 15.837469 | -1.122401 | 888.347079 | 0.150532398 | -1.1066622 | 0.268440004 | 0.998865404 | 9.876022 | 9.709112 | 9.866494 | 9.801249 | 9.695051 | 9.804694 | 1130 | 969 | 671 | 846 |
| 900 | NM_00136401 | CCNG1 | cyclin G1 | protein_coding | NP_001350944.1;NP_001350947.1;NP_001350948.1;NP_001350949.1;N | 1592 | 601578 | . | . | 52.005619 | 46.888608 | 51.431092 | 34.699546 | 121.019219 | 108.385111 | 115.409886 | 79.336390 | -1.158066 | 3313.682938 | 0.159022893 | -1.3313676 | 0.183068084 | 0.954800306 | 11.796028 | 11.584678 | 11.813769 | 11.675286 | 11.761521 | 11.470108 | 4433 | 3520 | 3148 | 2365 |
| 901 | NM_004354,XI | CCNG2 | cyclin G2 | protein_coding | NP_004345.1;XP_011530700.1;XP_011530701.1 | 1593 | 603203 | . | . | 1.778576 | 1.525018 | 2.476932 | 1.556136 | 4.138817 | 3.525146 | 5.558164 | 3.557923 | 1.180106 | 282.468139 | 0.243393421 | 0.98160529 | 0.32629434 | 0.998865404 | 8.016429 | 8.257408 | 8.151932 | 8.018920 | 8.273301 | 8.065746 | 337 | 243 | 317 | 232 |
| 902 | NM_00119918 | CCNH | cyclin H | protein_coding | NP_001186118.1;NP_001230.1;NP_001350468.1;NP_001351004.1;NP_0 | 1594 | 601953 | . | . | 27.482255 | 28.169633 | 24.757856 | 26.821122 | 63.952337 | 65.115363 | 55.555908 | 61.323309 | -1.024028 | 1455.293748 | 0.124871017 | -0.27432908 | 0.783831742 | 0.998865404 | 10.524145 | 10.489841 | 10.511337 | 10.521078 | 10.494580 | 10.501236 | 1685 | 1599 | 1245 | 1354 |
| 904 | NM_001240,N | CCNT1 | cyclin T1 | protein_coding | NP_001231.2;NP_001264771.1;XP_016875686.1 | 1599 | 143055 | . | . | 4.766126 | 5.076687 | 5.619513 | 4.328319 | 11.090972 | 11.734990 | 12.610024 | 9.896187 | 1.000192 | 1038.580527 | 0.157163884 | 0.001758454 | 0.998596958 | 0.999862634 | 10.019983 | 10.020812 | 9.998679 | 10.036557 | 10.111000 | 9.915892 | 1165 | 1149 | 1023 | 845 |
| 905 | NM_001241,N | CCNT2 | cyclin T2 | protein_coding | NP_001232.1;NP_001307677.1;NP_001307678.1;NP_490595.1;XP_0168 | 1600 | 603862 | . | . | 8.381796 | 8.494005 | 9.909387 | 6.775078 | 19.504785 | 19.634274 | 22.236378 | 15.490412 | 1.042263 | 1660.909610 | 0.172281027 | 0.346635571 | 0.728865119 | 0.998865404 | 10.667304 | 10.727582 | 10.692926 | 10.657544 | 10.860209 | 10.529032 | 1915 | 1715 | 1783 | 1257 |
| 908 | NM_00100918 | CCT6A | chaperonin containing TCP1 subunit 6A | protein_coding | NP_001009186.1;NP_001753.1 | 1620 | 104613 | . | . | 114.475283 | 109.211961 | 97.410361 | 98.543836 | 266.388696 | 252.448328 | 218.586028 | 225.308791 | -1.157524 | 8449.109327 | 0.103155748 | -2.04585795 | **0.040770349** | 0.62818612 | 13.146238 | 12.935220 | 13.126932 | 13.079386 | 12.978739 | 12.979897 | 10728 | 9512 | 6806 | 7348 |
| 928 | NM_00133031 | CD9 | CD9 molecule | protein_coding | NP_001317241.1;NP_001760.1;XP_005253871.1 | 1709 | 143030 | . | . | 160.157628 | 155.226665 | 145.869009 | 153.768340 | 372.693404 | 358.813379 | 327.325826 | 351.573003 | -1.067153 | 5878.514375 | 0.105590907 | -0.88802198 | 0.374528929 | 0.998865404 | 12.567365 | 12.473583 | 12.564625 | 12.530911 | 12.475372 | 12.510754 | 7128 | 6419 | 4841 | 5445 |
| 933 | NM_00118509 | CD22 | CD22 molecule | protein_coding | NP_001172028.1;NP_001172029.1;NP_001172030.1;NP_001265346.1;N | 1643 | 107266 | . | . | 5.250348 | 5.941160 | 3.691925 | 4.697950 | 12.217777 | 13.733256 | 8.284574 | 10.741304 | -1.339004 | 487.374006 | 0.18400831 | -2.28881236 | **0.022090257** | 0.491793249 | 9.124687 | 8.702274 | 8.975674 | 9.045500 | 8.757269 | 8.893587 | 609 | 634 | 320 | 436 |
| 948 | NM_000072,N | CD36 | CD36 molecule | protein_coding | NP_000063.2;NP_001001547.1;NP_001001548.1;NP_001120915.1;NP_0 | 1663 | 173510 | . | . | 0.577272 | 1.536687 | 0.478788 | 0.431836 | 1.343336 | 3.552122 | 1.074385 | 0.987342 | -1.417192 | 84.446323 | 0.35511375 | -1.41654572 | 0.156615789 | 0.925053243 | 6.629747 | 6.126516 | 6.404750 | 6.480821 | 6.362146 | 6.336647 | 100 | 120 | 64 | 62 |
| 949 | NM_00108295 | SCARB1 | scavenger receptor class B member 1 | protein_coding | NP_001076428.1;NP_001354910.1;NP_001354911.1;NP_001354912.1;N | 1664 | 601040 | . | . | 65.495393 | 68.847413 | 68.474100 | 71.583980 | 152.410467 | 159.143887 | 153.653888 | 163.668256 | 1.026677 | 7228.458163 | 0.10408451 | 0.364923604 | 0.715168434 | 0.998865404 | 12.800383 | 12.838312 | 12.791005 | 12.824976 | 12.813423 | 12.847081 | 8049 | 7852 | 6242 | 7002 |
| 950 | NM_00120425 | SCARB2 | scavenger receptor class B member 2 | protein_coding | NP_001191184.1;NP_005497.1 | 1665 | 602257 | . | . | 88.763063 | 82.573088 | 79.711358 | 74.905026 | 206.555285 | 190.871385 | 178.869986 | 171.261438 | -1.124128 | 11912.797764 | 0.10648445 | -1.58526592 | 0.112905928 | 0.850211417 | 13.622132 | 13.453386 | 13.622055 | 13.552366 | 13.518370 | 13.456519 | 15105 | 13056 | 10111 | 10131 |
| 953 | NM_00109817 | ENTPD1 | ectonucleoside triphosphate diphosphohydrolase 1 | protein_coding | NP_001091645.1;NP_001157650.1;NP_001157651.1;NP_001157653.1;N | 3363 | 601752 | . | . | 2.279824 | 2.270015 | 1.974147 | 1.895977 | 5.305250 | 5.247240 | 4.429926 | 4.334925 | -1.165539 | 773.671666 | 0.145900025 | -1.51471661 | 0.129844185 | 0.880707269 | 9.701857 | 9.480842 | 9.639406 | 9.654224 | 9.542715 | 9.536667 | 949 | 908 | 624 | 667 |
| 955 | NM_00111408 | ENTPD6 | ectonucleoside triphosphate diphosphohydrolase 6 | protein_coding | NP_001107561.1;NP_001238.3;NP_001304870.1;NP_001309307.1;NP_0 | 3368 | 603160 | . | . | 78.073310 | 84.350444 | 51.880483 | 55.397990 | 181.679803 | 194.979838 | 116.418301 | 126.660919 | -1.431302 | 6208.687849 | 0.107568435 | -4.80929736 | **1.51462E-06** | **0.000418125** | 12.835706 | 12.318295 | 12.739209 | 12.725233 | 12.394513 | 12.464758 | 8485 | 7825 | 4249 | 4995 |
| 957 | NM_001249,N | ENTPD5 | ectonucleoside triphosphate diphosphohydrolase 5 (inactive) | protein_coding | NP_001240.1;NP_001308913.1;NP_001308914.1;NP_001308915.1;NP_0 | 3367 | 603162 | . | . | 3.458355 | 2.945850 | 3.268764 | 2.624662 | 8.047730 | 6.809463 | 7.335011 | 6.000981 | -1.036561 | 469.760205 | 0.17363032 | -0.2983663 | 0.765423606 | 0.998865404 | 8.901246 | 8.849857 | 8.877742 | 8.895045 | 8.898437 | 8.828684 | 543 | 523 | 424 | 408 |
| 958 | NM_001250,N | CD40 | CD40 molecule | protein_coding | NP_001241.1;NP_001289682.1;NP_001309350.1;NP_001309351.1;NP_0 | 11919 | 109535 | . | . | 2.823187 | 2.994293 | 2.250395 | 2.824997 | 6.569675 | 6.921444 | 5.049821 | 6.459020 | -1.182968 | 142.711603 | 0.277515121 | -0.87350534 | 0.382387698 | 0.998865404 | 7.273635 | 7.030009 | 7.183580 | 7.193146 | 7.091627 | 7.157100 | 176 | 169 | 103 | 134 |
| 960 | NM_000610,N | CD44 | CD44 molecule (Indian blood group) | protein_coding | NP_000601.3;NP_001001389.1;NP_001001390.1;NP_001001391.1;NP_0 | 1681 | 107269 | . | . | 85.493853 | 88.379192 | 73.900383 | 70.459631 | 198.947723 | 204.292466 | 165.830309 | 161.097577 | -1.228665 | 10901.773121 | 0.101242192 | -2.9344691 | **0.003341188** | 0.163366554 | 13.553179 | 13.256105 | 13.486281 | 13.501345 | 13.334693 | 13.301263 | 13721 | 13083 | 8677 | 8989 |
| 961 | NM_00138230 | CD47 | CD47 molecule | protein_coding | NP_001369235.1;NP_001768.1;NP_942088.1;XP_005247966.1;XP_01686 | 1682 | 601028 | . | . | 6.784893 | 6.589383 | 7.361375 | 5.023791 | 15.788725 | 15.231653 | 16.518718 | 11.486302 | -1.076488 | 1118.983576 | 0.184966102 | -0.57487281 | 0.565377322 | 0.998865404 | 10.179752 | 10.074267 | 10.182612 | 10.112896 | 10.244739 | 9.915905 | 1397 | 1194 | 1145 | 787 |
| 965 | NM_00114482 | CD58 | CD58 molecule | protein_coding | NP_001138294.1;NP_001770.1;XP_016858358.1 | 1688 | 153420 | . | . | 27.450520 | 23.990711 | 26.860132 | 14.535593 | 63.878490 | 55.455603 | 60.273355 | 33.233907 | -1.231972 | 856.549917 | 0.456879993 | -0.65875003 | 0.510056302 | 0.998865404 | 9.884925 | 9.584221 | 9.843206 | 9.755324 | 9.844427 | 9.410704 | 1153 | 960 | 874 | 497 |
| 966 | NM_000611,N | CD59 | CD59 molecule (CD59 blood group) | protein_coding | NP_000602.1;NP_001120695.1;NP_001120697.1;NP_001120698.1;NP_0 | 1689 | 107271 | . | . | 45.687741 | 41.192187 | 41.115963 | 39.871245 | 106.317246 | 95.217590 | 92.263028 | 91.160869 | -1.086594 | 9919.345609 | 0.10843959 | -1.10488133 | 0.269211015 | 0.998865404 | 13.334662 | 13.214913 | 13.356927 | 13.260788 | 13.256878 | 13.221252 | 12561 | 10524 | 8442 | 8726 |
| 967 | NM_00125738 | CD63 | CD63 molecule | protein_coding | NP_001244318.1;NP_001244319.1;NP_001244320.1;NP_001244321.1;N | 1692 | 155740 | . | . | 1209.983672 | 1200.322639 | 1366.889783 | 1292.143406 | 2815.681740 | 2774.599608 | 3067.261102 | 2954.332338 | 1.092639 | 39007.910360 | 0.098513061 | 1.297455584 | 0.194474478 | 0.96621494 | 15.186151 | 15.313977 | 15.218463 | 15.205388 | 15.308795 | 15.267726 | 43256 | 39921 | 36259 | 37287 |
| 968 | NM_00104005 | CD68 | CD68 molecule | protein_coding | NP_001035148.1;NP_001242.2 | 1693 | 153634 | . | . | 193.307170 | 191.216876 | 171.834451 | 200.444704 | 449.833737 | 442.006422 | 385.591530 | 458.292952 | -1.047573 | 10016.539785 | 0.111793513 | -0.59976822 | 0.548660714 | 0.998865404 | 13.323265 | 13.256147 | 13.317209 | 13.300682 | 13.205802 | 13.329164 | 11916 | 10953 | 7890 | 9838 |
| 971 | NM_001782,XI | CD72 | CD72 molecule | protein_coding | NP_001773.1;XP_006716956.1 | 1696 | 107272 | . | . | 0.525995 | 0.441637 | 0.209908 | 0.294394 | 1.224012 | 1.020863 | 0.471028 | 0.673097 | -1.836163 | 17.945764 | 0.739578417 | -1.18539721 | 0.235860436 | NA | 4.539157 | 3.660102 | 4.203478 | 4.186649 | 4.140495 | 4.159457 | 29 | 23 | 9 | 14 |
| 972 | NM_00102515 | CD74 | CD74 molecule | protein_coding | NP_001020329.1;NP_001020330.1;NP_001351012.1;NP_001351013.1;N | 1697 | 142790 | . | . | 0.358731 | 0.706438 | 0.608557 | 0.220869 | 0.834782 | 1.632962 | 1.365585 | 0.504991 | -1.651892 | 34.145253 | 0.583263593 | -1.24149578 | 0.214422655 | 0.979353522 | 5.409647 | 4.688346 | 5.073387 | 5.161663 | 5.086601 | 5.015958 | 36 | 58 | 30 | 16 |
| 975 | NM_00129764 | CD81 | CD81 molecule | protein_coding | NP_001284578.1;NP_004347.1;XP_011518794.1 | 1701 | 186845 | . | . | 365.960352 | 368.274462 | 340.473003 | 347.982816 | 851.604767 | 851.282887 | 764.011504 | 795.621309 | -1.081848 | 16424.244289 | 0.098694892 | -1.14998485 | 0.25015011 | 0.993339639 | 14.059174 | 13.945673 | 14.037666 | 14.035101 | 13.964158 | 13.973556 | 19685 | 18392 | 13645 | 14874 |
| 976 | NM_00102516 | ADGRE5 | adhesion G protein-coupled receptor E5 | protein_coding | NP_001020331.1;NP_001775.2;NP_510966.1;XP_011526753.1 | 1711 | 601211 | . | . | 15.240397 | 15.602010 | 15.711538 | 14.702860 | 35.465030 | 36.064743 | 35.256237 | 33.616343 | -1.032026 | 1543.909562 | 0.126075086 | -0.36073033 | 0.718301048 | 0.998865404 | 10.614843 | 10.569546 | 10.598754 | 10.609712 | 10.610601 | 10.548210 | 1793 | 1704 | 1375 | 1367 |
| 977 | NM_00103949 | CD151 | CD151 molecule (Raph blood group) | protein_coding | NP_001034579.1;NP_004348.2;NP_620598.1;NP_620599.1;XP_0243045 | 1630 | 602243 | . | . | 185.848860 | 184.488222 | 157.171360 | 156.658519 | 432.477928 | 426.452797 | 352.687973 | 358.181086 | -1.194282 | 8220.762715 | 0.101670819 | -2.51933787 | **0.011757577** | 0.354930366 | 13.127444 | 12.871317 | 13.080978 | 13.070604 | 12.930658 | 12.919535 | 10366 | 9598 | 6559 | 6978 |
| 983 | NM_00117040 | CDK1 | cyclin dependent kinase 1 | protein_coding | NP_001163877.1;NP_001163878.1;NP_001307847.1;NP_001777.1;NP_2 | 1722 | 116940 | . | . | 35.228862 | 32.554092 | 26.425242 | 25.220274 | 81.979008 | 75.250244 | 59.297478 | 57.663160 | -1.331668 | 1760.337637 | 0.125089498 | -3.30350669 | **0.000954837** | 0.065898069 | 10.973393 | 10.560428 | 10.917821 | 10.846839 | 10.679412 | 10.636758 | 2416 | 2075 | 1350 | 1376 |
| 984 | NM_00129134 | CDK11B | cyclin dependent kinase 11B | protein_coding | NP_001278274.1;NP_001778.2;NP_277021.2;NP_277022.1;NP_277024.2 | 1729 | 176873 | . | . | 26.431375 | 29.064659 | 31.288914 | 29.658127 | 61.506897 | 67.184270 | 70.211416 | 67.809790 | 1.095316 | 2722.163873 | 0.117173421 | 1.120961803 | 0.262304122 | 0.995760118 | 11.343372 | 11.474716 | 11.338507 | 11.404265 | 11.470736 | 11.420390 | 2870 | 2918 | 2551 | 2586 |
| 987 | NM_00119928 | LRBA | LPS responsive beige-like anchor protein | protein_coding | NP_001186211.2;NP_001351834.1;NP_001354479.1;NP_006717.2;XP_0 | 1742 | 606453 | . | . | 4.569838 | 4.290018 | 4.519507 | 4.705799 | 10.634202 | 9.916571 | 10.141643 | 10.759249 | 1.031709 | 1433.087351 | 0.127544875 | 0.353096936 | 0.72401577 | 0.998865404 | 10.462203 | 10.507267 | 10.497588 | 10.446906 | 10.482074 | 10.510863 | 1678 | 1472 | 1242 | 1390 |
| 988 | NM_001253 | CDC5L | cell division cycle 5 like | protein_coding | NP_001244.1 | 1743 | 602868 | . | . | 6.153867 | 6.466687 | 6.676196 | 6.957363 | 14.320303 | 14.948037 | 14.981191 | 15.907183 | 1.065118 | 1276.115693 | 0.12935777 | 0.703576319 | 0.481696663 | 0.998865404 | 10.271409 | 10.362248 | 10.277267 | 10.308268 | 10.328832 | 10.352339 | 1394 | 1361 | 1127 | 1253 |
| 989 | NM_00101155 | SEPTIN7 | septin 7 | protein_coding | NP_001011553.2;NP_001229885.1;NP_001350644.1;NP_001362228.1;N | 1717 | 603151 | . | . | 44.953665 | 41.608290 | 31.203022 | 29.675560 | 104.609025 | 96.179428 | 70.018677 | 67.849637 | -1.120272 | 3867.141217 | 0.110097164 | -1.48821661 | 0.136693777 | 0.895483725 | 11.996623 | 11.832835 | 11.990420 | 11.933151 | 11.868118 | 11.868130 | 4866 | 4259 | 3172 | 3420 |
| 990 | NM_001254,XI | CDC6 | cell division cycle 6 | protein_coding | NP_001245.1;XP_011523843.1;XP_011523844.1 | 1744 | 602627 | . | . | 21.404974 | 20.417707 | 19.327238 | 24.763786 | 49.810257 | 47.196443 | 43.369760 | 56.619456 | -1.024330 | 2749.560497 | 0.120855623 | -0.28696079 | 0.774142342 | 0.998865404 | 11.442351 | 11.407408 | 11.408527 | 11.459217 | 11.362889 | 11.463731 | 3105 | 3096 | 2215 | 2705 |
| 991 | NM_001255 | CDC20 | cell division cycle 20 | protein_coding | NP_001246.2 | 1723 | 603618 | . | . | 78.730881 | 81.953217 | 62.664875 | 68.686577 | 183.209991 | 189.438522 | 140.618179 | 157.043701 | -1.240430 | 3752.911845 | 0.111886582 | -2.77817079 | **0.005466588** | 0.225239728 | 12.020925 | 11.709937 | 11.942819 | 11.969398 | 11.744436 | 11.810417 | 4710 | 4555 | 2794 | 3269 |
| 993 | NM_001789,N | CDC25A | cell division cycle 25A | protein_coding | NP_001780.2;NP_963861.1;XP_006713497.1;XP_006713498.1 | 1725 | 116947 | . | . | 6.938443 | 6.396503 | 6.034025 | 7.014672 | 16.146045 | 14.785802 | 13.540179 | 16.038216 | -1.034534 | 785.648695 | 0.153166949 | -0.31978553 | 0.749130921 | 0.998865404 | 9.642136 | 9.592925 | 9.661623 | 9.595336 | 9.558000 | 9.650159 | 963 | 822 | 627 | 772 |
| 994 | NM_00128751 | CDC25B | cell division cycle 25B | protein_coding | NP_001274445.1;NP_001274446.1;NP_001274447.1;NP_001274448.1;N | 1726 | 116949 | . | . | 47.612227 | 60.031727 | 51.548002 | 61.349894 | 110.795600 | 138.766019 | 115.672229 | 140.269243 | 1.037138 | 5606.360550 | 0.111086584 | 0.473570951 | 0.635805894 | 0.998865404 | 12.426365 | 12.478855 | 12.394657 | 12.477545 | 12.442106 | 12.492316 | 6025 | 6233 | 4816 | 5511 |
| 995 | NM_00128758 | CDC25C | cell division cycle 25C | protein_coding | NP_001274511.1;NP_001274512.1;NP_001305027.1;NP_001350955.1;N | 1727 | 157680 | . | . | 7.253404 | 7.047090 | 5.929027 | 8.035399 | 16.878971 | 16.289662 | 13.304567 | 18.371982 | -1.055889 | 440.240037 | 0.194600262 | -0.40317701 | 0.686818012 | 0.998865404 | 8.821404 | 8.741791 | 8.795489 | 8.797592 | 8.668253 | 8.849869 | 520 | 489 | 316 | 462 |
| 996 | NM_00111409 | CDC27 | cell division cycle 27 | protein_coding | NP_001107563.1;NP_001247.3;NP_001280018.1;NP_001280020.1;NP_0 | 1728 | 116946 | . | . | 17.203928 | 16.262722 | 16.646814 | 14.078002 | 40.034245 | 37.592004 | 37.354970 | 32.187678 | -1.114894 | 2799.696649 | 0.129221888 | -1.21423777 | 0.224656942 | 0.982008277 | 11.527225 | 11.370638 | 11.510122 | 11.474435 | 11.481433 | 11.320174 | 3472 | 3116 | 2533 | 2234 |
| 997 | NM_004359,XI | CDC34 | cell division cycle 34, ubiqiutin conjugating enzyme | protein_coding | NP_004350.1;XP_005259747.1;XP_006723015.1 | 1734 | 116948 | . | . | 21.742751 | 23.684846 | 21.972222 | 23.441156 | 50.596275 | 54.748580 | 49.305029 | 53.595418 | -1.012956 | 997.597764 | 0.139508229 | -0.13312422 | 0.894095141 | 0.998865404 | 9.971756 | 9.952811 | 9.938197 | 9.994626 | 9.934387 | 9.979634 | 1112 | 1125 | 835 | 958 |
| 998 | NM_00103980 | CDC42 | cell division cycle 42 | protein_coding | NP_001034891.1;NP_001782.1;NP_426359.1 | 1736 | 116952 | . | . | 17.034063 | 16.520027 | 16.172326 | 17.049042 | 39.638967 | 38.186783 | 36.290231 | 38.980609 | -1.031848 | 5092.196790 | 0.10983309 | -0.41181551 | 0.68047465 | 0.998865404 | 12.336545 | 12.291244 | 12.341043 | 12.312414 | 12.262026 | 12.337675 | 6057 | 5486 | 4161 | 4912 |
| 999 | NM_00131718 | CDH1 | cadherin 1 | protein_coding | NP_001304113.1;NP_001304114.1;NP_001304115.1;NP_004351.1 | 1748 | 192090 | . | . | 0.711516 | 0.513421 | 0.689691 | 1.108215 | 1.655729 | 1.186797 | 1.547648 | 2.533803 | 1.442143 | 114.312832 | 0.339437771 | 1.556144967 | 0.119673624 | 0.858998979 | 6.548978 | 7.076714 | 6.806239 | 6.706642 | 6.792804 | 6.958597 | 126 | 84 | 91 | 155 |
| 1000 | NM_00130817 | CDH2 | cadherin 2 | protein_coding | NP_001295105.1;NP_001783.2;XP_011524090.1;XP_016881003.1 | 1759 | 114020 | . | . | 18.907302 | 19.109368 | 16.490123 | 16.620875 | 43.998072 | 44.172160 | 37.003356 | 38.001655 | -1.161152 | 2223.947092 | 0.115647493 | -1.86391418 | 0.062333757 | 0.723723143 | 11.222664 | 11.007105 | 11.173535 | 11.178086 | 11.057279 | 11.055154 | 2744 | 2586 | 1792 | 1927 |
| 1005 | NM_00131721 | CDH7 | cadherin 7 | protein_coding | NP_001304143.1;NP_001349367.1;NP_004352.2;NP_387450.1 | 1766 | 605806 | . | . | 0.479000 | 0.563579 | 0.311511 | 0.404506 | 1.114655 | 1.302739 | 0.699022 | 0.924855 | -1.273577 | 111.937321 | 0.304633267 | -1.14526607 | 0.252098955 | 0.994172589 | 6.970402 | 6.621670 | 6.857164 | 6.833950 | 6.765064 | 6.768826 | 149 | 131 | 85 | 93 |
| 1010 | NM_00131722 | CDH12 | cadherin 12 | protein_coding | NP_001304156.1;NP_001304157.1;NP_001351033.1;NP_001351034.1;N | 1751 | 600562 | . | . | 0.172128 | 0.136718 | 0.096701 | 0.098345 | 0.400548 | 0.316031 | 0.216994 | 0.224855 | -1.583292 | 17.861095 | 0.732528142 | -0.90498569 | 0.365472967 | NA | 4.452135 | 3.790061 | 4.192302 | 4.171184 | 4.147423 | 4.147355 | 28 | 21 | 12 | 13 |
| 1017 | NM_00129023 | CDK2 | cyclin dependent kinase 2 | protein_coding | NP_001277159.1;NP_001789.2;NP_439892.2;XP_011536034.1 | 1771 | 116953 | . | . | 113.310822 | 117.147331 | 119.925231 | 137.360785 | 263.678944 | 270.791302 | 269.108735 | 314.059108 | 1.100119 | 8463.893151 | 0.110361691 | 1.247348944 | 0.212269578 | 0.975238634 | 12.976679 | 13.114254 | 12.993742 | 13.015507 | 13.029727 | 13.138841 | 9162 | 8810 | 7220 | 8842 |
| 1019 | NM_000075 | CDK4 | cyclin dependent kinase 4 | protein_coding | NP_000066.1 | 1773 | 123829 | . | . | 62.942696 | 63.410416 | 61.340981 | 70.762527 | 146.470245 | 146.576019 | 137.647369 | 161.790115 | 1.030845 | 3753.147048 | 0.1164052 | 0.376507176 | 0.706539882 | 0.998865404 | 11.851885 | 11.895554 | 11.861189 | 11.860149 | 11.829421 | 11.939153 | 4259 | 3986 | 3094 | 3808 |
| 1020 | NM_00116441 | CDK5 | cyclin dependent kinase 5 | protein_coding | NP_001157882.1;NP_004926.1 | 1774 | 123831 | . | . | 28.249740 | 28.846443 | 29.903148 | 32.980302 | 65.738303 | 66.679844 | 67.101799 | 75.405544 | 1.082035 | 1029.277324 | 0.137348657 | 0.828166987 | 0.407575937 | 0.998865404 | 9.949570 | 10.063032 | 9.970992 | 9.983417 | 10.005032 | 10.064445 | 1129 | 1076 | 893 | 1043 |
| 1021 | NM_00114530 | CDK6 | cyclin dependent kinase 6 | protein_coding | NP_001138778.1;NP_001250.1;XP_006715898.1 | 1777 | 603368 | . | . | 4.023730 | 4.387310 | 3.670670 | 3.196322 | 9.363385 | 10.141463 | 8.236876 | 7.308011 | -1.243985 | 1392.032452 | 0.136378466 | -2.30952557 | **0.020914435** | 0.478742856 | 10.591760 | 10.277067 | 10.489132 | 10.549439 | 10.406314 | 10.295980 | 1709 | 1729 | 1159 | 1077 |
| 1022 | NM_00132406 | CDK7 | cyclin dependent kinase 7 | protein_coding | NP_001310998.1;NP_001310999.1;NP_001311000.1;NP_001311001.1;N | 1778 | 601955 | . | . | 18.854725 | 18.956422 | 20.384916 | 18.561236 | 43.875721 | 43.818618 | 45.743162 | 42.438060 | 1.010415 | 842.451007 | 0.146913558 | 0.101744324 | 0.918959619 | 0.998865404 | 9.710755 | 9.726102 | 9.712681 | 9.715523 | 9.763588 | 9.677793 | 963 | 906 | 781 | 746 |

| | | | | | | | | | | | | | | | | | | | | | | | | | | | | | | | | | |
|---|---|---|---|---|---|---|---|---|---|---|---|---|---|---|---|---|---|---|---|---|---|---|---|---|---|---|---|---|---|---|---|---|---|
| 1024 | NM_001260,N | CDK8 | cyclin dependent kinase 8 | protein_coding | NP_001251.1;NP_001305297.1;NP_001333430.1;XP_011533167.1;XP_01 | 1779 | 603184 | . | . | 5.682059 | 6.074920 | 5.393975 | 4.785092 | 13.222384 | 14.042450 | 12.103925 | 10.940545 | -1.170178 | 524.808290 | 0.169511746 | -1.33753709 | 0.181047383 | 0.952877815 | 9.144375 | 8.918051 | 9.063163 | 9.102990 | 9.022122 | 8.941411 | 632 | 629 | 448 | 424 |
| 1025 | NM_001261 | CDK9 | cyclin dependent kinase 9 | protein_coding | NP_001252.1 | 1780 | 603251 | . | . | 11.756637 | 11.172870 | 11.457170 | 11.839036 | 27.358179 | 25.826588 | 25.709558 | 27.068550 | 1.002173 | 894.548452 | 0.141293345 | 0.022165695 | 0.982315783 | 0.998865404 | 9.803415 | 9.806615 | 9.825047 | 9.782786 | 9.797638 | 9.813732 | 1059 | 936 | 770 | 849 |
| 1026 | NM_000389,N | CDKN1A | cyclin dependent kinase inhibitor 1A | protein_coding | NP_000380.1;NP_001207706.1;NP_001207707.1;NP_001278478.1;NP_0 | 1784 | 116899 | . | . | 6.739563 | 6.951853 | 8.089043 | 10.138245 | 15.683237 | 16.069518 | 18.151576 | 23.179893 | 1.290396 | 506.773927 | 0.185220142 | 1.985819437 | 0.047053377 | 0.658032177 | 8.790033 | 9.157081 | 8.905285 | 8.881003 | 8.959537 | 9.142870 | 520 | 468 | 428 | 609 |
| 1027 | NM_004064 | CDKN1B | cyclin dependent kinase inhibitor 1B | protein_coding | NP_004055.1 | 1785 | 600778 | . | . | 8.948820 | 7.613939 | 7.889814 | 7.089571 | 20.824270 | 17.599960 | 17.704515 | 16.209459 | -1.120828 | 593.203856 | 0.169527462 | -0.97072555 | 0.331684966 | 0.998865404 | 9.291913 | 9.128214 | 9.303853 | 9.188337 | 9.209419 | 9.134029 | 783 | 619 | 515 | 494 |
| 1028 | NM_000076,N | CDKN1C | cyclin dependent kinase inhibitor 1C | protein_coding | NP_000067.1;NP_001116102.1;NP_001116103.1;NP_001349403.1;NP_0 | 1786 | 600856 | . | . | 0.626765 | 0.841964 | 0.827328 | 0.918225 | 1.458507 | 1.946238 | 1.856501 | 2.099412 | 1.151004 | 45.472323 | 0.465412213 | 0.43594097 | 0.662879559 | 0.998865404 | 5.402705 | 5.604106 | 5.471170 | 5.510139 | 5.509375 | 5.520471 | 43 | 51 | 41 | 47 |
| 1029 | NM_000077,N | CDKN2A | cyclin dependent kinase inhibitor 2A | protein_coding | NP_000068.1;NP_001182061.1;NP_001350692.1;NP_478102.2;NP_4781 | 1787 | 600160 | . | . | 50.343638 | 48.369255 | 48.853757 | 58.339555 | 117.151718 | 111.807699 | 109.626416 | 133.386461 | 1.071768 | 1707.141888 | 0.129758893 | 0.770601646 | 0.440943085 | 0.998865404 | 10.686619 | 10.786391 | 10.726801 | 10.690547 | 10.706193 | 10.816959 | 1942 | 1737 | 1428 | 1772 |
| 1030 | NM_004936,N | CDKN2B | cyclin dependent kinase inhibitor 2B | protein_coding | NP_004927.2;NP_511042.1 | 1788 | 600431 | . | . | 3.989088 | 4.341711 | 6.210071 | 5.354239 | 9.282770 | 10.036059 | 13.935220 | 12.241829 | 1.371491 | 598.317538 | 0.165867067 | 2.747653297 | 0.006002344 | 0.242252506 | 8.978810 | 9.434837 | 9.081399 | 9.129492 | 9.367883 | 9.259409 | 559 | 565 | 649 | 598 |
| 1031 | NM_001262,N | CDKN2C | cyclin dependent kinase inhibitor 2C | protein_coding | NP_001253.1;NP_523240.1 | 1789 | 603369 | . | . | 11.296940 | 11.691428 | 9.636251 | 12.284673 | 26.288443 | 27.025259 | 21.623469 | 28.087445 | -1.105688 | 695.542915 | 0.167357946 | -0.86607392 | 0.386449625 | 0.998865404 | 9.513070 | 9.367237 | 9.463791 | 9.484273 | 9.318310 | 9.484863 | 829 | 800 | 503 | 696 |
| 1032 | NM_001800,N | CDKN2D | cyclin dependent kinase inhibitor 2D | protein_coding | NP_001791.1;NP_524145.1 | 1790 | 600927 | . | . | 10.066392 | 9.073259 | 8.277066 | 10.823642 | 23.424906 | 20.973246 | 18.573497 | 24.746971 | -1.033987 | 414.569117 | 0.195014033 | -0.24725354 | 0.804712012 | 0.998865404 | 8.719732 | 8.670790 | 8.735400 | 8.671130 | 8.604957 | 8.756578 | 511 | 431 | 309 | 431 |
| 1033 | NM_00113085 | CDKN3 | cyclin dependent kinase inhibitor 3 | protein_coding | NP_001124323.1;NP_001317102.1;NP_005183.2;XP_024305226.1 | 1791 | 123832 | . | . | 31.577256 | 32.551173 | 24.680609 | 27.305719 | 73.481574 | 75.243491 | 55.382570 | 62.431287 | -1.236815 | 779.955099 | 0.148107774 | -2.07031322 | 0.038423023 | 0.611005738 | 9.752619 | 9.445569 | 9.663454 | 9.690096 | 9.499466 | 9.555360 | 975 | 948 | 581 | 681 |
| 1039 | NM_001802,XI | CDR2 | cerebellar degeneration related protein 2 | protein_coding | NP_001793.1;XP_024305911.1 | 1799 | 117340 | . | . | 9.869836 | 8.922304 | 6.866940 | 8.105277 | 22.967513 | 20.624306 | 15.409215 | 18.531752 | -1.266798 | 702.306736 | 0.160586689 | -2.12462322 | 0.033618065 | 0.575719787 | 9.616662 | 9.275082 | 9.566472 | 9.493241 | 9.313732 | 9.419764 | 950 | 804 | 495 | 628 |
| 1040 | NM_001263,XI | CDS1 | CDP-diacylglycerol synthase 1 | protein_coding | NP_001254.2;XP_005262744.1;XP_016863137.1;XP_016863138.1;XP_01 | 1800 | 603548 | . | . | 0.949714 | 0.789965 | 0.650287 | 0.647344 | 2.210024 | 1.826037 | 1.459224 | 1.480077 | -1.346879 | 105.214253 | 0.316527784 | -1.35728974 | 0.174689165 | 0.948677095 | 6.915734 | 6.486886 | 6.789863 | 6.733913 | 6.673045 | 6.665231 | 150 | 120 | 79 | 83 |
| 1043 | NM_001803 | CD52 | CD52 molecule | protein_coding | NP_001794.2 | 1804 | 114280 | . | . | 0.146037 | 0.561032 | 0.614256 | 1.047004 | 0.339834 | 1.296851 | 1.378373 | 2.393851 | 2.317123 | 9.036458 | 1.074049125 | 1.128751371 | 0.259002732 | NA | 2.452336 | 3.655341 | 3.122725 | 3.150035 | 3.153913 | 3.183639 | 3 | 9 | 8 | 15 |
| 1047 | NM_00113067 | CLGN | calmegin | protein_coding | NP_001124147.1;NP_004353.1 | 2060 | 601858 | . | . | 0.742265 | 0.541920 | 0.481153 | 0.728296 | 1.727281 | 1.252672 | 1.079693 | 1.665163 | -1.062749 | 54.968239 | 0.443368166 | -0.19803103 | 0.843020785 | 0.998865404 | 5.824308 | 5.735374 | 5.816225 | 5.753905 | 5.734627 | 5.810839 | 75 | 52 | 37 | 60 |
| 1050 | NM_00128582 | CEBPA | CCAAT enhancer binding protein alpha | protein_coding | NP_001272758.1;NP_001274353.1;NP_001274364.1;NP_004355.2 | 1833 | 116897 | . | . | 0.314322 | 0.203988 | 0.250386 | 0.325796 | 0.731439 | 0.471528 | 0.561859 | 0.744894 | 1.084627 | 22.756907 | 0.655052007 | 0.178914986 | 0.858004456 | NA | 4.448287 | 4.565787 | 4.520941 | 4.484619 | 4.498454 | 4.522764 | 30 | 19 | 18 | 25 |
| 1051 | NM_00128587 | CEBPB | CCAAT enhancer binding protein beta | protein_coding | NP_001272807.1;NP_001272808.1;NP_005185.2 | 1834 | 189965 | . | . | 18.388370 | 17.477811 | 18.560689 | 23.456703 | 42.790496 | 40.400748 | 41.649647 | 53.630964 | 1.150254 | 1122.476632 | 0.14965969 | 1.349411546 | 0.177204814 | 0.949886621 | 10.028191 | 10.229716 | 10.093675 | 10.057616 | 10.091371 | 10.261754 | 1232 | 1099 | 927 | 1252 |
| 1052 | NM_005195 | CEBPD | CCAAT enhancer binding protein delta | protein_coding | NP_005186.2 | 1835 | 116898 | . | . | 1.748109 | 1.577886 | 2.050283 | 1.844253 | 4.067922 | 3.647355 | 4.600777 | 4.216666 | 1.154062 | 70.883040 | 0.376302465 | 0.549347402 | 0.582767063 | 0.998865404 | 6.039657 | 6.247594 | 6.139342 | 6.113466 | 6.175498 | 6.146199 | 80 | 67 | 70 | 67 |
| 1054 | NM_00125229 | CEBPG | CCAAT enhancer binding protein gamma | protein_coding | NP_001239225.1;NP_001797.1;XP_024307095.1 | 1837 | 138972 | . | . | 4.839400 | 4.711051 | 3.861498 | 4.116569 | 11.261483 | 10.889805 | 8.665090 | 9.412046 | -1.221748 | 512.824698 | 0.166764151 | -1.73267045 | 0.083154259 | 0.784575614 | 9.139626 | 8.850562 | 9.077437 | 9.047489 | 8.919528 | 8.950582 | 664 | 595 | 391 | 444 |
| 1056 | NM_001807 | CEL | carboxyl ester lipase | protein_coding | NP_001798.3 | 1848 | 114840 | . | . | 0.063829 | 0.201863 | 0.140389 | 0.139697 | 0.148533 | 0.466614 | 0.315028 | 0.319402 | 1.060304 | 10.782932 | 0.970617375 | 0.087035948 | 0.930642938 | NA | 3.389932 | 3.470304 | 3.396205 | 3.444504 | 3.425262 | 3.421693 | 6 | 17 | 10 | 10 |
| 1058 | NM_00104242 | CENPA | centromere protein A | protein_coding | NP_001035891.1;NP_001800.1 | 1851 | 117139 | . | . | 16.400885 | 15.332250 | 13.926836 | 15.895523 | 38.165535 | 35.441187 | 31.251415 | 36.343226 | -1.076414 | 664.983099 | 0.158496621 | -0.67025447 | 0.502695589 | 0.998865404 | 9.429437 | 9.322946 | 9.428970 | 9.373048 | 9.309953 | 9.391061 | 826 | 714 | 523 | 637 |
| 1059 | NM_001810 | CENPB | centromere protein B | protein_coding | NP_001801.1 | 1852 | 117140 | . | . | 30.358509 | 29.942749 | 25.249256 | 30.954891 | 70.645493 | 69.214005 | 56.658600 | 70.774689 | -1.088297 | 2622.649825 | 0.128500653 | -0.94997156 | 0.342126702 | 0.998865404 | 11.416683 | 11.294343 | 11.398289 | 11.379978 | 11.239909 | 11.395133 | 3183 | 2917 | 1973 | 2582 |
| 1060 | NM_00136248 | CENPC | centromere protein C | protein_coding | NP_001349410.1;NP_001803.2;XP_011529844.1 | 1854 | 117141 | . | . | 1.800285 | 1.781677 | 1.848847 | 1.602153 | 4.189335 | 4.118424 | 4.148757 | 3.663133 | -1.116116 | 348.826025 | 0.205128172 | -0.77262721 | 0.439743035 | 0.998865404 | 8.522802 | 8.365648 | 8.487659 | 8.461226 | 8.484746 | 8.334471 | 433 | 388 | 326 | 267 |
| 1062 | NM_00128673 | CENPE | centromere protein E | protein_coding | NP_001273663.1;NP_001804.2;XP_011529846.1;XP_011529847.1;XP_01 | 1856 | 117143 | . | . | 6.500859 | 6.199966 | 4.970624 | 3.641243 | 15.127762 | 14.331497 | 11.153934 | 8.325268 | -1.492259 | 1375.346737 | 0.154719959 | -3.73253661 | 0.000189561 | 0.019822063 | 10.685325 | 10.108572 | 10.576000 | 10.534259 | 10.351468 | 10.137025 | 1945 | 1731 | 1102 | 882 |
| 1063 | NM_016343,XI | CENPF | centromere protein F | protein_coding | NP_057427.3;XP_011507384.1;XP_016855575.1 | 1857 | 600236 | . | . | 23.589055 | 23.527759 | 22.411333 | 21.354941 | 54.892700 | 54.385469 | 50.290385 | 48.825534 | -1.080345 | 7245.653454 | 0.10429583 | -1.06899262 | 0.285072994 | 0.998865404 | 12.877548 | 12.766100 | 12.850562 | 12.858690 | 12.813855 | 12.764067 | 8624 | 8159 | 6237 | 6337 |
| 1066 | NM_00102519 | CES1 | carboxylesterase 1 | protein_coding | NP_001020365.1;NP_001020366.1;NP_001257.4;XP_005255831.1 | 1863 | 114835 | . | . | 0.234563 | 1.702367 | 0.219544 | 0.279346 | 0.545838 | 3.935095 | 0.492650 | 0.638692 | -3.890933 | 37.819697 | 0.730969034 | -2.6815311 | 0.00732861 | 0.269034057 | 5.911840 | 3.947964 | 5.052845 | 5.401828 | 5.055143 | 5.067567 | 17 | 114 | 12 | 16 |
| 1069 | NM_004344 | CETN2 | centrin 2 | protein_coding | NP_004335.1 | 1867 | 300006 | . | . | 11.343726 | 10.982300 | 13.379864 | 11.785231 | 26.397316 | 25.386078 | 30.024029 | 26.945528 | 1.111297 | 523.225310 | 0.169555428 | 0.897900963 | 0.369238351 | 0.998865404 | 8.952810 | 9.105717 | 9.007283 | 8.982072 | 9.106223 | 9.017063 | 582 | 524 | 512 | 481 |
| 1070 | NM_00129776 | CETN3 | centrin 3 | protein_coding | NP_001284694.1;NP_001284697.1;NP_004356.2 | 1868 | 602907 | . | . | 7.238715 | 6.397656 | 5.498458 | 5.064683 | 16.844787 | 14.788468 | 12.338382 | 11.579795 | -1.339499 | 438.076947 | 0.180387695 | -2.33770865 | 0.019402367 | 0.462516353 | 8.970225 | 8.549255 | 8.896256 | 8.813825 | 8.699563 | 8.656277 | 614 | 507 | 338 | 338 |
| 1072 | NM_005507 | CFL1 | cofilin 1 | protein_coding | NP_005498.1 | 1874 | 601442 | . | . | 319.084076 | 313.743744 | 263.906616 | 300.286072 | 742.521729 | 725.232727 | 592.198730 | 686.568359 | -1.137779 | 11610.271187 | 0.108025825 | -1.72385386 | 0.084734229 | 0.788085288 | 13.593246 | 13.406974 | 13.566340 | 13.544335 | 13.393638 | 13.494064 | 14412 | 13166 | 8884 | 10788 |
| 1073 | NM_00124364 | CFL2 | cofilin 2 | protein_coding | NP_001230574.1;NP_068733.1;NP_619579.1;XP_011534665.1 | 1875 | 601443 | . | . | 6.350461 | 6.509047 | 6.725420 | 5.708126 | 14.777784 | 15.045952 | 15.091649 | 13.050952 | -1.058068 | 791.385921 | 0.15612005 | -0.52160008 | 0.601948808 | 0.998865404 | 9.668054 | 9.587291 | 9.651306 | 9.641750 | 9.672407 | 9.537085 | 943 | 872 | 734 | 651 |
| 1074 | gene-CFL1P5 | CFL1P5 | cofilin 1 pseudogene 5 | pseudogene | . | 1877 | . | . | . | 0.478350 | 0.664314 | 0.343620 | 0.599291 | 1.113140 | 1.535592 | 0.771074 | 1.370208 | -1.216100 | 18.861816 | 0.724581891 | -0.38955131 | 0.696868363 | NA | 4.373794 | 4.086744 | 4.232149 | 4.261163 | 4.212934 | 4.249039 | 20 | 26 | 11 | 20 |
| 1075 | NM_00111417 | CTSC | cathepsin C | protein_coding | NP_001107645.1;NP_001805.4;NP_680475.1 | 2528 | 602365 | . | . | 115.533014 | 111.609428 | 111.530006 | 100.026672 | 268.850071 | 257.990180 | 250.270118 | 228.699084 | -1.068374 | 7372.612355 | 0.115503307 | -0.82609997 | 0.408747384 | 0.998865404 | 12.894821 | 12.799526 | 12.893232 | 12.855446 | 12.882683 | 12.750131 | 8964 | 8036 | 6687 | 6158 |
| 1102 | NM_001268,N | RCBTB2 | RCC1 and BTB domain containing protein 2 | protein_coding | NP_001259.1;NP_001273759.1;NP_001273760.1;NP_001273761.1;NP_0 | 1914 | 603524 | . | . | 1.531985 | 1.553541 | 1.329256 | 1.205963 | 3.564991 | 3.591079 | 2.982810 | 2.757288 | -1.257183 | 133.411522 | 0.306749734 | -1.07642863 | 0.281735603 | 0.998865404 | 7.214397 | 6.886490 | 7.086349 | 7.104521 | 7.084572 | 6.934798 | 167 | 164 | 126 | 86 |
| 1104 | NM_00104819 | RCC1 | regulator of chromosome condensation 1 | protein_coding | NP_001041659.1;NP_001041660.1;NP_001041664.1;NP_001260.1;NP_0 | 1913 | 179710 | . | . | 29.702781 | 32.052843 | 27.607717 | 31.347518 | 69.119592 | 74.091585 | 61.950917 | 71.672377 | -1.060164 | 2267.122682 | 0.123438634 | -0.682827 | 0.494716166 | 0.998865404 | 11.188322 | 11.103732 | 11.144500 | 11.193337 | 11.072084 | 11.169981 | 2606 | 2594 | 1796 | 2190 |
| 1105 | NM_001270,N | CHD1 | chromodomain helicase DNA binding protein 1 | protein_coding | NP_001261.2;NP_001351042.1;NP_001363123.1;XP_016864480.1;XP_01 | 1915 | 602118 | . | . | 7.768626 | 7.794836 | 9.434784 | 7.778113 | 18.077912 | 18.018114 | 21.171379 | 17.783729 | 1.104313 | 2080.889283 | 0.131634199 | 1.087475182 | 0.276826861 | 0.998865404 | 10.949459 | 11.092943 | 10.986770 | 10.974366 | 11.134014 | 10.979924 | 2295 | 2117 | 2081 | 1852 |
| 1106 | NM_00104257 | CHD2 | chromodomain helicase DNA binding protein 2 | protein_coding | NP_001036037.1;NP_001262.3 | 1917 | 602119 | . | . | 6.772971 | 7.799016 | 7.090701 | 6.718908 | 15.760982 | 18.027777 | 15.911327 | 15.361984 | -1.035651 | 1974.480318 | 0.123576083 | -0.4089604 | 0.682568722 | 0.998865404 | 10.972321 | 10.921750 | 10.919598 | 11.000012 | 10.953672 | 10.911466 | 2197 | 2277 | 1732 | 1770 |
| 1107 | NM_00100527 | CHD3 | chromodomain helicase DNA binding protein 3 | protein_coding | NP_001005271.2;NP_001005273.1;NP_005843.2;XP_005256484.1;XP_00 | 1918 | 602120 | . | . | 16.181864 | 16.200958 | 16.281622 | 17.937341 | 37.655855 | 37.449242 | 36.535487 | 41.011598 | 1.036693 | 3777.980790 | 0.110947515 | 0.468593267 | 0.639360391 | 0.998865404 | 11.857220 | 11.909112 | 11.870025 | 11.866197 | 11.865299 | 11.929482 | 4282 | 3994 | 3215 | 3745 |
| 1108 | NM_001273,N | CHD4 | chromodomain helicase DNA binding protein 4 | protein_coding | NP_001264.2;NP_001284482.1;NP_001350535.1 | 1919 | 603277 | . | . | 38.164608 | 38.138904 | 34.504858 | 39.707068 | 88.810612 | 88.159786 | 77.427903 | 90.785495 | -1.043378 | 7542.907709 | 0.111225727 | -0.55078874 | 0.581778505 | 0.998865404 | 12.911253 | 12.849905 | 12.902158 | 12.894129 | 12.804883 | 12.916361 | 8912 | 8273 | 5996 | 7378 |
| 1111 | NM_00111412 | CHEK1 | checkpoint kinase 1 | protein_coding | NP_001107593.1;NP_001107594.1;NP_001231775.1;NP_001265.2;NP_0 | 1925 | 603078 | . | . | 6.193859 | 6.409689 | 5.683886 | 5.423456 | 14.413365 | 14.816285 | 12.754476 | 12.400086 | -1.048266 | 591.149796 | 0.163430138 | -0.41610884 | 0.67733036 | 0.998865404 | 9.240647 | 9.173327 | 9.247339 | 9.196373 | 9.229325 | 9.151563 | 723 | 628 | 532 | 509 |
| 1112 | NM_00108547 | FOXN3 | forkhead box N3 | protein_coding | NP_001078940.1;NP_005188.2 | 1928 | 602628 | . | . | 1.042178 | 0.752125 | 0.836882 | 1.010018 | 2.425193 | 1.738570 | 1.877940 | 2.309286 | 1.012060 | 227.449428 | 0.249440796 | 0.069334097 | 0.944723686 | 0.998865404 | 7.820612 | 7.838139 | 7.901007 | 7.738128 | 7.782208 | 7.872965 | 304 | 203 | 180 | 235 |
| 1119 | NM_001277,N | CHKA | choline kinase alpha | protein_coding | NP_001268.2;NP_001363148.1;NP_001363149.1;NP_001363150.1;NP_0 | 1937 | 118491 | . | . | 15.838054 | 14.646561 | 15.946517 | 13.505430 | 36.855803 | 33.856186 | 35.783521 | 30.878562 | 1.068253 | 1637.299849 | 0.152497102 | 0.624624937 | 0.532217249 | 0.998865404 | 10.628404 | 10.724211 | 10.675319 | 10.618871 | 10.808883 | 10.576383 | 1889 | 1646 | 1688 | 1352 |
| 1120 | NM_005198 | CHKB | choline kinase beta | protein_coding | NP_005189.2 | 1938 | 612395 | . | . | 13.753873 | 15.850337 | 17.098623 | 17.542070 | 32.005825 | 36.638767 | 38.368813 | 40.107857 | 1.154874 | 737.632760 | 0.153211984 | 1.35586949 | 0.175140696 | 0.949120257 | 9.419379 | 9.626697 | 9.426571 | 9.517576 | 9.567225 | 9.578628 | 736 | 788 | 682 | 747 |
| 1121 | NM_000390,N | CHM | CHM Rab escort protein | protein_coding | NP_000381.1;NP_001138886.1;NP_001307888.1;NP_001349446.1;NP_0 | 1940 | 300390 | . | . | 4.174783 | 4.064139 | 4.151809 | 4.012826 | 9.714893 | 9.394437 | 9.316540 | 9.174850 | 1.011828 | 640.655248 | 0.154242603 | 0.109981159 | 0.912424317 | 0.998865404 | 9.314812 | 9.331946 | 9.326352 | 9.312203 | 9.336300 | 9.318154 | 741 | 680 | 568 | 596 |
| 1122 | NM_00138185 | CHML | CHM like Rab escort protein | protein_coding | NP_001368782.1;NP_001368783.1;NP_001812.2 | 1941 | 118825 | . | . | 1.192630 | 2.271984 | 2.098720 | 0.756701 | 2.775299 | 5.251791 | 4.709470 | 1.730105 | -1.230455 | 378.887775 | 0.820209848 | -0.36477518 | 0.715279232 | 0.998865404 | 8.707409 | 8.408377 | 8.368527 | 8.759673 | 8.696539 | 8.150050 | 332 | 591 | 436 | 167 |
| 1123 | NM_00102520 | CHN1 | chimerin 1 | protein_coding | NP_001020372.2;NP_001193531.1;NP_001358442.1;NP_001358443.1;N | 1943 | 118423 | . | . | 5.024009 | 4.759854 | 4.127352 | 4.485213 | 11.691077 | 11.002614 | 9.261659 | 10.254906 | -1.174668 | 361.316477 | 0.188411803 | -1.23268651 | 0.217692746 | 0.982008277 | 8.608790 | 8.376074 | 8.524902 | 8.558023 | 8.434411 | 8.463975 | 437 | 433 | 281 | 320 |
| 1124 | NM_00103993 | CHN2 | chimerin 2 | protein_coding | NP_001035025.1;NP_001279998.1;NP_001279999.1;NP_001280000.1;N | 1944 | 602857 | . | . | 2.838588 | 3.101299 | 4.282819 | 4.757725 | 6.605511 | 7.168793 | 9.610519 | 10.877973 | 1.507778 | 375.289067 | 0.188601891 | 3.141135585 | 0.001682941 | 0.09884951 | 8.225888 | 8.817582 | 8.392174 | 8.432377 | 8.615410 | 8.684243 | 332 | 335 | 371 | 446 |
| 1128 | NM_000738,XI | CHRM1 | cholinergic receptor muscarinic 1 | protein_coding | NP_000729.2;XP_011543044.1 | 1950 | 118510 | . | . | 0.073751 | 0.173856 | 0.179356 | 0.241860 | 0.171622 | 0.401876 | 0.402469 | 0.552984 | 1.623547 | 14.247949 | 0.838480094 | 0.833829385 | 0.404377134 | NA | 3.444631 | 4.138173 | 3.785048 | 3.821300 | 3.821636 | 3.844301 | 8 | 16 | 13 | 19 |
| 1130 | NM_000081,N | LYST | lysosomal trafficking regulator | protein_coding | NP_000072.2;NP_001288294.1;XP_011542333.1;XP_011542334.1;XP_01 | 1968 | 606897 | . | . | 12.906194 | 12.986142 | 16.892900 | 13.947209 | 30.033235 | 30.018050 | 37.907179 | 31.888636 | 1.116397 | 4837.327581 | 0.132216655 | 1.201440542 | 0.229580357 | 0.98402759 | 12.158281 | 12.317326 | 12.203195 | 12.176577 | 12.383720 | 12.169518 | 5348 | 4853 | 4999 | 4180 |
| 1131 | NM_000740,N | CHRM3 | cholinergic receptor muscarinic 3 | protein_coding | NP_000731.1;NP_001334645.1;NP_001362907.1;NP_001362908.1;NP_0 | 1952 | 118494 | . | . | 0.660970 | 0.689611 | 0.954965 | 0.588230 | 1.538105 | 1.594071 | 2.142913 | 1.344920 | -1.041262 | 179.839406 | 0.247516356 | -0.23567339 | 0.81368612 | 0.998865404 | 7.519289 | 7.461258 | 7.492399 | 7.505833 | 7.497458 | 7.466020 | 208 | 201 | 159 | 159 |
| 1132 | NM_000741,N | CHRM4 | cholinergic receptor muscarinic 4 | protein_coding | NP_000732.2;NP_001353621.1 | 1953 | 118495 | . | . | 0.052901 | 0.090689 | 0.090464 | 0.079247 | 0.123102 | 0.209632 | 0.202998 | 0.181190 | 1.152399 | 7.786141 | 1.094683044 | 0.186940745 | 0.851707093 | NA | 2.855800 | 3.058876 | 2.942555 | 2.961494 | 2.960811 | 2.953263 | 6 | 10 | 8 | 7 |
| 1136 | NM_000743,N | CHRNA3 | cholinergic receptor nicotinic alpha 3 subunit | protein_coding | NP_000734.2;NP_001160166.1;XP_006720445.1 | 1957 | 118503 | . | . | 0.173516 | 0.337258 | 0.053758 | 0.131840 | 0.403778 | 0.779587 | 0.120632 | 0.301437 | -2.660640 | 18.035846 | 0.794279839 | -1.77742586 | 0.075498199 | NA | 4.714983 | 3.293999 | 4.171956 | 4.459050 | 4.119014 | 4.153330 | 21 | 37 | 5 | 13 |
| 1138 | NM_000745,N | CHRNA5 | cholinergic receptor nicotinic alpha 5 subunit | protein_coding | NP_000736.2;NP_001294874.1 | 1959 | 118505 | . | . | 5.278747 | 3.928054 | 3.838907 | 3.407895 | 12.283858 | 9.079874 | 8.614398 | 7.791748 | -1.278675 | 462.474749 | 0.197588794 | -1.79488971 | 0.072671274 | 0.758760658 | 9.019162 | 8.665704 | 9.015291 | 8.816092 | 8.807386 | 8.727873 | 688 | 475 | 377 | 355 |
| 1139 | NM_000746,N | CHRNA7 | cholinergic receptor nicotinic alpha 7 subunit | protein_coding | NP_000737.1;NP_001177384.1;XP_011519478.2;XP_011519479.1;XP_01 | 1960 | 118511 | . | . | 1.364974 | 1.370620 | 1.588548 | 1.411370 | 3.176349 | 3.168249 | 3.564657 | 3.226931 | 1.073529 | 267.737339 | 0.211925574 | 0.483002831 | 0.629079726 | 0.998865404 | 8.012268 | 8.115244 | 8.047637 | 8.041913 | 8.110505 | 8.051198 | 299 | 277 | 255 | 245 |
| 1140 | NM_000747 | CHRNB1 | cholinergic receptor nicotinic beta 1 subunit | protein_coding | NP_000738.2 | 1961 | 100710 | . | . | 1.069335 | 1.218516 | 1.448003 | 1.207298 | 2.488386 | 2.816655 | 3.249277 | 2.760344 | 1.145860 | 99.306648 | 0.326114028 | 0.602337674 | 0.546949388 | 0.998865404 | 6.531999 | 6.728924 | 6.590062 | 6.626549 | 6.678822 | 6.619802 | 100 | 106 | 101 | 90 |
| 1145 | NM_000080,XI | CHRNE | cholinergic receptor nicotinic epsilon subunit | protein_coding | NP_000071.1;XP_016879604.1 | 1966 | 100725 | . | . | 0.052491 | 0.087014 | 0.140747 | 0.034986 | 0.122150 | 0.201137 | 0.315832 | 0.079992 | 1.247069 | 6.613978 | 1.228266822 | 0.259341692 | 0.795371614 | NA | 2.555346 | 2.877719 | 2.699620 | 2.714124 | 2.731604 | 2.694982 | 5 | 8 | 10 | 3 |
| 1147 | NM_001278,N | CHUK | component of inhibitor of nuclear factor kappa B kinase complex | protein_coding | NP_001269.3;NP_001307857.1;XP_016871100.1;XP_016871101.1;XP_01 | 1974 | 600664 | . | . | 7.339602 | 7.239902 | 7.105460 | 6.041951 | 17.079555 | 16.735357 | 15.944444 | 13.814202 | -1.115344 | 783.884315 | 0.155403124 | -1.01341972 | 0.310859723 | 0.998865404 | 9.690774 | 9.533961 | 9.657642 | 9.643401 | 9.635465 | 9.508714 | 961 | 883 | 704 | 631 |
| 1152 | NM_00136253 | CKB | creatine kinase B | protein_coding | NP_001349460.1;NP_001814.2;XP_016876441.1 | 1991 | 123280 | . | . | 13.848555 | 12.326305 | 10.528751 | 14.037122 | 32.226156 | 28.492805 | 23.626212 | 32.094209 | -1.076563 | 562.582123 | 0.18296594 | -0.581706 | 0.56076473 | 0.998865404 | 9.188472 | 9.081381 | 9.197548 | 9.115661 | 9.016653 | 9.193808 | 713 | 591 | 406 | 578 |
| 1153 | NM_001280,N | CIRBP | cold inducible RNA binding protein | protein_coding | NP_001271.1;NP_001287744.1;NP_001287758.1;XP_006722700.1;XP_01 | 1982 | 602649 | . | . | 27.850229 | 23.487734 | 20.246541 | 23.418884 | 64.808629 | 54.292957 | 45.432651 | 53.544495 | -1.079218 | 1756.864884 | 0.122537842 | -0.89756916 | 0.369415289 | 0.998865404 | 10.832730 | 10.722749 | 10.831469 | 10.783838 | 10.733478 | 10.762484 | 2163 | 1909 | 1442 | 1614 |
| 1154 | NM_013324,N | CISH | cytokine inducible SH2 containing protein | protein_coding | NP_037456.5;NP_659508.1;XP_011531631.1 | 1984 | 602441 | . | . | 0.212624 | 0.137856 | 0.198941 | 0.152338 | 0.494784 | 0.318661 | 0.446417 | 0.348302 | -1.014866 | 27.425087 | 0.6059031 | -0.03513723 | 0.971970318 | 0.998865404 | 4.786188 | 4.768607 | 4.801931 | 4.749317 | 4.789301 | 4.759015 | 39 | 23 | 27 | 22 |
| 1155 | NM_001281,N | TBCB | tubulin folding cofactor B | protein_coding | NP_001272.2;NP_001287900.1 | 1989 | 601303 | . | . | 44.040705 | 45.963154 | 38.686794 | 45.758434 | 102.484526 | 106.245886 | 86.812042 | 104.621223 | -1.080760 | 1427.500558 | 0.133710107 | -0.83797964 | 0.402042137 | 0.998865404 | 10.534407 | 10.421954 | 10.499831 | 10.515988 | 10.390647 | 10.502533 | 1690 | 1617 | 1105 | 1381 |
| 1159 | NM_00137548 | CKMT1B | creatine kinase, mitochondrial 1B | protein_coding | NP_001362413.1;NP_066270.1;XP_005254207.1;XP_011519496.1;XP_01 | 1995 | 123290 | . | . | 1.285791 | 0.900061 | 1.479429 | 1.350809 | 2.992088 | 2.080531 | 3.319794 | 3.088463 | 1.280630 | 66.621900 | 0.400409828 | 0.891220943 | 0.372810651 | 0.998865404 | 5.867147 | 6.226395 | 6.060642 | 5.979996 | 6.092414 | 6.063319 | 80 | 51 | 69 | 66 |
| 1161 | NM_000082,N | ERCC8 | ERCC excision repair 8, CSA ubiquitin ligase complex subunit | protein_coding | NP_000073.1;NP_001007234.1;NP_001007235.1;NP_001277214.1 | 3439 | 609412 | . | . | 0.652212 | 0.637242 | 0.539614 | 0.481322 | 1.517724 | 1.473014 | 1.210878 | 1.100486 | -1.222117 | 135.307956 | 0.279298156 | -1.03610509 | 0.300153116 | 0.998865404 | 7.217479 | 6.928256 | 7.118070 | 7.114841 | 7.046909 | 7.039481 | 172 | 160 | 107 | 113 |
| 1163 | NM_001826,N | CKS1B | CDC28 protein kinase regulatory subunit 1B | protein_coding | NP_001817.1 | 19083 | 116900 | . | . | 58.340265 | 56.041191 | 44.302973 | 55.163055 | 135.760198 | 129.541722 | 99.414583 | 126.123766 | -1.163432 | 1341.218230 | 0.143121497 | -1.52588206 | 0.127039239 | 0.874064017 | 10.494587 | 10.275781 | 10.464919 | 10.420931 | 10.247138 | 10.402110 | 1708 | 1513 | 966 | 1283 |
| 1164 | NM_001827 | CKS2 | CDC28 protein kinase regulatory subunit 2 | protein_coding | NP_001818.1 | 2000 | 116901 | . | . | 111.402100 | 102.236694 | 104.802826 | 104.578789 | 259.237274 | 236.324722 | 235.174515 | 239.106964 | -1.034975 | 2030.172270 | 0.121421219 | -0.40845972 | 0.682936198 | 0.998865404 | 11.011891 | 10.962458 | 11.037756 | 10.962146 | 10.978301 | 10.968402 | 2490 | 2123 | 1746 | 1859 |
| 1173 | NM_00102520 | AP2M1 | adaptor related protein complex 2 subunit mu 1 | protein_coding | NP_001020376.1;NP_001298127.1;NP_004059.2 | 564 | 601024 | . | . | 129.450292 | 128.780133 | 96.315920 | 121.051005 | 301.236157 | 297.681052 | 216.130146 | 276.768741 | -1.205044 | 7161.893451 | 0.123824472 | -2.1731278 | 0.0297707 | 0.553061869 | 12.934463 | 12.665246 | 12.884653 | 12.872602 | 12.626232 | 12.807258 | 9077 | 8388 | 5035 | 6751 |
| 1174 | NM_001283 | AP1S1 | adaptor related protein complex 1 subunit sigma 1 | protein_coding | NP_001274.1 | 559 | 603531 | . | . | 58.666008 | 59.600853 | 48.790073 | 63.462414 | 136.518204 | 137.770050 | 109.483513 | 145.099274 | -1.068777 | 2201.133053 | 0.139307476 | -0.68883919 | 0.490924468 | 0.998865404 | 11.151399 | 11.055054 | 11.125548 | 11.131182 | 10.970257 | 11.169970 | 2610 | 2463 | 1618 | 2245 |
| 1175 | NM_00130107 | AP2S1 | adaptor related protein complex 2 subunit sigma 1 | protein_coding | NP_001288005.1;NP_001288007.1;NP_001288010.1;NP_004060.2;NP_0 | 565 | 602242 | . | . | 132.077502 | 128.319006 | 107.191294 | 137.639969 | 307.349781 | 296.615118 | 240.534163 | 314.697442 | -1.079240 | 3157.512264 | 0.134094976 | -0.82043218 | 0.411969777 | 0.998865404 | 11.678653 | 11.568386 | 11.667395 | 11.639159 | 11.486945 | 11.684987 | 3839 | 3477 | 2320 | 3193 |
| 1176 | NM_00100292 | AP3S1 | adaptor related protein complex 3 subunit sigma 1 | protein_coding | NP_001002924.2;NP_001275.1;NP_001305019.1;NP_001305020.1;NP_0 | 2013 | 601507 | . | . | 29.538871 | 24.825770 | 26.006360 | 23.423840 | 68.738167 | 57.385880 | 58.357518 | 53.555830 | -1.098337 | 1084.410272 | 0.148978706 | -0.90832234 | 0.363707942 | 0.998865404 | 10.148465 | 10.013783 | 10.180538 | 10.045103 | 10.088578 | 10.000828 | 1424 | 1115 | 953 | 911 |
| 1181 | NM_00117108 | CLCN2 | chloride voltage-gated channel 2 | protein_coding | NP_001164558.1;NP_001164559.1;NP_001164560.1;NP_004357.3;XP_0 | 2020 | 600570 | . | . | 8.342109 | 9.552933 | 8.386094 | 7.482146 | 19.412432 | 22.082032 | 18.818151 | 17.107041 | -1.055845 | 805.533408 | 0.150769228 | -0.51998601 | 0.603073327 | 0.998865404 | 9.692356 | 9.614187 | 9.642534 | 9.700629 | 9.675270 | 9.590335 | 914 | 929 | 723 | 690 |
| 1182 | NM_00124337 | CLCN3 | chloride voltage-gated channel 3 | protein_coding | NP_001230301.1;NP_001230303.1;NP_001820.2;NP_776297.2;XP_0052 | 2021 | 600580 | . | . | 19.103069 | 18.427885 | 21.962934 | 18.419462 | 44.453626 | 42.596878 | 49.284188 | 42.113913 | 1.063322 | 3674.887128 | 0.118194426 | 0.749430838 | 0.45359757 | 0.998865404 | 11.798408 | 11.887195 | 11.861082 | 11.770605 | 11.905965 | 11.827800 | 4327 | 3632 | 3449 | 3384 |
| 1183 | NM_00125694 | CLCN4 | chloride voltage-gated channel 4 | protein_coding | NP_001243873.1;NP_001821.2 | 2022 | 302910 | . | . | 1.268392 | 1.112311 | 1.363738 | 1.356578 | 2.951599 | 2.571157 | 3.060190 | 3.101654 | 1.125543 | 269.126861 | 0.212947564 | 0.80123336 | 0.422996562 | 0.998865404 | 7.983983 | 8.155224 | 8.073749 | 8.002890 | 8.104191 | 8.095594 | 312 | 254 | 250 | 265 |
| 1184 | NM_000084,N | CLCN5 | chloride voltage-gated channel 5 | protein_coding | NP_000075.1;NP_001121370.1;NP_001121371.1;NP_001259031.1;NP_0 | 2023 | 300008 | . | . | 4.174245 | 4.199473 | 5.627889 | 4.792302 | 9.713636 | 9.707271 | 12.628819 | 10.957029 | 1.220220 | 1444.887013 | 0.136361995 | 2.105731683 | 0.035227666 | 0.585116558 | 10.345832 | 10.633349 | 10.414875 | 10.411534 | 10.632464 | 10.496073 | 1502 | 1401 | 1501 | 1360 |
| 1185 | NM_00125695 | CLCN6 | chloride voltage-gated channel 6 | protein_coding | NP_001243888.2;NP_001277.2 | 2024 | 602726 | . | . | 10.839795 | 12.209664 | 17.107841 | 14.119857 | 25.224647 | 28.223185 | 38.389502 | 32.283376 | 1.334647 | 2367.516205 | 0.135174155 | 3.080901733 | 0.002063747 | 0.115796367 | 10.985876 | 11.402471 | 11.037534 | 11.118874 | 11.392623 | 11.223436 | 2214 | 2302 | 2594 | 2279 |
| 1186 | NM_00111433 | CLCN7 | chloride voltage-gated channel 7 | protein_coding | NP_001107803.1;NP_001278.1;XP_011520656.1 | 2025 | 602727 | . | . | 55.832902 | 55.840078 | 70.573262 | 64.456916 | 129.925454 | 129.076841 | 158.364350 | 147.373080 | 1.190271 | 8022.256886 | 0.1072144 | 2.343811653 | 0.019087808 | 0.461416831 | 12.838647 | 13.090003 | 12.894051 | 12.887905 | 13.080940 | 12.994892 | 8466 | 7875 | 7961 | 7766 |
| 1191 | NM_001831,N | CLU | clusterin | protein_coding | NP_001822.3 | 2095 | 185430 | . | . | 0.193284 | 1.837021 | 0.109183 | 0.130039 | 0.449780 | 4.246356 | 0.245004 | 0.297319 | -7.627712 | 49.965105 | 1.346186248 | -2.17744784 | 0.02944717 | 0.552254458 | 6.465245 | 3.532926 | 5.343821 | 5.848617 | 5.317110 | 5.325200 | 21 | 171 | 9 | 12 |
| 1192 | NM_00128759 | CLIC1 | chloride intracellular channel 1 | protein_coding | NP_001274522.1;NP_001274523.1;NP_001279.2 | 2062 | 602872 | . | . | 183.090974 | 204.317406 | 186.334838 | 213.667225 | 426.060215 | 472.288823 | 418.129965 | 488.524685 | 1.022772 | 7589.290344 | 0.114668011 | 0.283297202 | 0.776949022 | 0.998865404 | 12.873476 | 12.905841 | 12.837624 | 12.919527 | 12.843781 | 12.950284 | 8223 | 8489 | 6255 | 7646 |
| 1195 | NM_00116240 | CLK1 | CDC like kinase 1 | protein_coding | NP_001155879.1;NP_004062.2 | 2068 | 601951 | . | . | 13.910020 | 15.383971 | 20.066635 | 13.583677 | 32.369187 | 35.560737 | 45.028950 | 31.057465 | 1.142411 | 1179.560299 | 0.190777898 | 1.006833668 | 0.314014712 | 0.998865404 | 10.104485 | 10.297154 | 10.118324 | 10.167228 | 10.413230 | 10.039696 | 1226 | 1227 | 1359 | 894 |
| 1196 | NM_00129433 | CLK2 | CDC like kinase 2 | protein_coding | NP_001281267.1;NP_001281268.1;NP_001350633.1;NP_003984.2;XP_01 | 2069 | 602989 | . | . | 12.155701 | 13.639922 | 15.556417 | 13.689233 | 28.286816 | 31.529287 | 34.908149 | 31.298801 | 1.117283 | 917.142983 | 0.148126949 | 1.080115763 | 0.280090633 | 0.998865404 | 9.758718 | 9.918846 | 9.759549 | 9.834943 | 9.924138 | 9.829961 | 945 | 984 | 896 | 849 |
| 1198 | NM_00113002 | CLK3 | CDC like kinase 3 | protein_coding | NP_001123500.1;NP_003983.2 | 2071 | 602990 | . | . | 14.831594 | 15.693646 | 15.801842 | 16.083149 | 34.513729 | 36.276568 | 35.458879 | 36.772205 | 1.026881 | 1307.666713 | 0.12867957 | 0.297401973 | 0.766159637 | 0.998865404 | 10.333591 | 10.371716 | 10.323926 | 10.360675 | 10.356518 | 10.368627 | 1450 | 1426 | 1143 | 1252 |
| 1200 | NM_000391 | TPP1 | tripeptidyl peptidase 1 | protein_coding | NP_000382.3 | 2073 | 607998 | . | . | 48.970654 | 55.632942 | 57.977036 | 50.337208 | 113.956718 | 128.598308 | 130.098785 | 115.090042 | 1.021798 | 5790.596001 | 0.121130209 | 0.256828227 | 0.797331379 | 0.998865404 | 12.483833 | 12.514991 | 12.438022 | 12.537908 | 12.569126 | 12.441552 | 6204 | 6548 | 5475 | 5072 |
| 1201 | NM_000086,N | CLN3 | CLN3 lysosomal/endosomal transmembrane protein, battenin | protein_coding | NP_000077.1;NP_001035897.1;NP_001273033.1;NP_001273034.1;NP_0 | 2074 | 607042 | . | . | 19.429224 | 19.793623 | 23.359360 | 21.595021 | 45.212601 | 45.753846 | 52.417724 | 49.374448 | 1.114973 | 1213.897104 | 0.13177159 | 1.191518201 | 0.233450216 | 0.988893316 | 10.164667 | 10.321912 | 10.209203 | 10.195404 | 10.310653 | 10.257466 | 1334 | 1227 | 1152 | 1158 |
| 1203 | NM_00136662 | CLN5 | CLN5 intracellular trafficking protein | protein_coding | NP_001353553.1;NP_006484.2 | 2076 | 608102 | . | . | 6.185007 | 6.118427 | 5.657674 | 5.149306 | 14.392765 | 14.143018 | 12.695656 | 11.773275 | -1.154414 | 946.532705 | 0.141725442 | -1.46170569 | 0.143821877 | 0.906161673 | 9.986187 | 9.779429 | 9.943152 | 9.928858 | 9.868189 | 9.794325 | 1179 | 1084 | 804 | 781 |
| 1207 | NM_001293,N | CLNS1A | chloride nucleotide-sensitive channel 1A | protein_coding | NP_001284.1;NP_001298128.1;NP_001298129.1;NP_001298130.1;NP_0 | 2080 | 602158 | . | . | 18.150963 | 17.651967 | 17.242553 | 17.189496 | 42.238036 | 40.803313 | 38.691789 | 39.301735 | -1.047035 | 1589.917005 | 0.124255155 | -0.5336552 | 0.593580113 | 0.998865404 | 10.667431 | 10.601283 | 10.674345 | 10.629706 | 10.625771 | 10.607329 | 1926 | 1705 | 1367 | 1439 |
| 1209 | NM_00128217 | CLPTM1 | CLPTM1 regulator of GABA type A receptor forward trafficking | protein_coding | NP_001269104.1;NP_001269105.1;NP_001285.1 | 2087 | 604783 | . | . | 64.002128 | 67.424048 | 66.632790 | 67.397598 | 148.935583 | 155.853701 | 149.522034 | 154.096610 | 1.005818 | 5107.715404 | 0.105891616 | 0.079039961 | 0.937000838 | 0.998865404 | 12.314289 | 12.322624 | 12.297585 | 12.334030 | 12.320169 | 12.321457 | 5737 | 5615 | 4451 | 4806 |
| 1211 | NM_00107667 | CLTA | clathrin light chain A | protein_coding | NP_001070145.1;NP_001171689.1;NP_001171690.1;NP_001171691.1;N | 2090 | 118960 | . | . | 171.827688 | 179.338866 | 157.250289 | 186.944031 | 399.850099 | 414.549856 | 352.865095 | 427.425302 | -1.035258 | 5830.409671 | 0.117430807 | -0.42569768 | 0.670328153 | 0.998865404 | 12.534232 | 12.484095 | 12.509391 | 12.536428 | 12.422392 | 12.560534 | 6721 | 6503 | 4572 | 5813 |
| 1212 | NM_00136412 | CLTB | clathrin light chain B | protein_coding | NP_001351055.1;NP_001351056.1;NP_001825.1;NP_009028.1 | 2091 | 118970 | . | . | 38.470064 | 41.019168 | 30.200383 | 42.173807 | 89.521418 | 94.817644 | 67.768781 | 96.425391 | -1.082916 | 1358.289556 | 0.154167198 | -0.74543675 | 0.456007736 | 0.998865404 | 10.464205 | 10.348632 | 10.411999 | 10.457366 | 10.259443 | 10.476769 | 1579 | 1569 | 971 | 1398 |
| 1213 | NM_00128865 | CLTC | clathrin heavy chain | protein_coding | NP_001275582.1;NP_004850.1 | 2092 | 118955 | . | . | 57.184423 | 56.879299 | 53.806854 | 53.984284 | 133.070497 | 131.479050 | 120.741020 | 123.428656 | -1.072123 | 14470.551259 | 0.099141212 | -1.01341293 | 0.310862966 | 0.998865404 | 13.870190 | 13.769726 | 13.855961 | 13.843915 | 13.792425 | 13.788128 | 17363 | 16045 | 12176 | 13061 |
| 1230 | NM_001295 | CCR1 | C-C motif chemokine receptor 1 | protein_coding | NP_001286.1 | 1602 | 601159 | . | . | 5.319986 | 4.527916 | 4.321284 | 4.434137 | 12.379827 | 10.466481 | 9.696836 | 10.138127 | -1.139733 | 384.067736 | 0.18989324 | -0.99369467 | 0.320371538 | 0.998865404 | 8.676272 | 8.488025 | 8.672127 | 8.568876 | 8.541400 | 8.549136 | 511 | 404 | 310 | 339 |
| 1241 | NM_00114391 | LTB4R | leukotriene B4 receptor | protein_coding | NP_001137391.1;NP_858043.1 | 6713 | 601531 | . | . | 1.732527 | 1.568556 | 1.431465 | 1.856133 | 4.031661 | 3.625788 | 3.212165 | 4.243827 | 1.029058 | 206.173849 | 0.238485326 | 0.173277823 | 0.862433054 | 0.998865404 | 7.667169 | 7.707977 | 7.699703 | 7.660597 | 7.655590 | 7.729857 | 244 | 210 | 167 | 212 |
| 1244 | NM_000392,XI | ABCC2 | ATP binding cassette subfamily C member 2 | protein_coding | NP_000383.2;XP_006717693.1;XP_006717694.1;XP_011537593.1;XP_01 | 53 | 601107 | . | . | 38.419606 | 40.757681 | 57.833046 | 50.136736 | 89.404008 | 94.213205 | 129.775685 | 114.631697 | 1.318202 | 8279.229598 | 0.132283995 | 3.012999105 | 0.002586797 | 0.134524108 | 12.802231 | 13.200879 | 12.856901 | 12.910697 | 13.227486 | 12.997463 | 7958 | 7957 | 9267 | 7664 |
| 1263 | NM_004073,XI | PLK3 | polo like kinase 3 | protein_coding | NP_004064.2 | 2154 | 602913 | . | . | 5.311115 | 6.353255 | 5.857351 | 4.948994 | 12.359182 | 14.685835 | 13.143725 | 11.315287 | -1.085969 | 427.933393 | 0.190413308 | -0.62486999 | 0.532056389 | 0.998865404 | 8.799379 | 8.680665 | 8.709003 | 8.815542 | 8.765776 | 8.659062 | 469 | 522 | 389 | 350 |
| 1265 | NM_00130349 | CNN2 | calponin 2 | protein_coding | NP_001290428.1;NP_001290430.1;NP_004359.1;NP_958434.1 | 2156 | 602373 | . | . | 23.265812 | 24.937098 | 19.155201 | 23.997227 | 54.140500 | 57.643218 | 42.983712 | 54.866810 | -1.133864 | 1527.057509 | 0.14042644 | -1.29068991 | 0.196811224 | 0.96621494 | 10.664569 | 10.482786 | 10.598609 | 10.644971 | 10.442468 | 10.600621 | 1814 | 1803 | 1115 | 1481 |
| 1266 | NM_00128605 | CNN3 | calponin 3 | protein_coding | NP_001272984.1;NP_001272985.1;NP_001830.1;XP_016855734.1 | 2157 | 602374 | . | . | 50.694273 | 53.882809 | 44.863445 | 53.060570 | 117.967659 | 124.552530 | 100.672269 | 121.316683 | -1.083859 | 3143.740220 | 0.122439562 | -0.94884828 | 0.342697782 | 0.998865404 | 11.675311 | 11.558873 | 11.628378 | 11.670561 | 11.517604 | 11.645526 | 3670 | 3619 | 2417 | 3051 |
| 1267 | NM_00133021 | CNP | 2',3'-cyclic nucleotide 3' phosphodiesterase | protein_coding | NP_001317145.1;NP_149124.3;XP_011522642.1 | 2158 | 123830 | . | . | 39.408826 | 41.648216 | 36.668982 | 40.670787 | 91.705951 | 96.271714 | 82.284121 | 92.988924 | 1.022149 | 6110.522766 | 0.110517854 | 0.285975968 | 0.774896528 | 0.998865404 | 12.561250 | 12.592738 | 12.540133 | 12.593939 | 12.544198 | 12.625767 | 6737 | 6730 | 5112 | 6071 |
| 1270 | NM_000614 | CNTF | ciliary neurotrophic factor | protein_coding | NP_000605.1 | 2169 | 118945 | . | . | 0.475118 | 0.730839 | 1.085847 | 0.891232 | 1.105619 | 1.689368 | 2.436610 | 2.037697 | 1.616816 | 47.220211 | 0.471373119 | 1.470503074 | 0.141425554 | 0.903146012 | 5.174238 | 5.866231 | 5.469437 | 5.530185 | 5.605157 | 5.562860 | 33 | 47 | 56 | 49 |
| 1271 | NM_00120701 | CNTFR | ciliary neurotrophic factor receptor | protein_coding | NP_001193940.1;NP_001833.1;NP_671693.1;XP_016869749.1;XP_01686 | 2170 | 118946 | . | . | 0.314030 | 0.314830 | 0.159802 | 0.185724 | 0.730761 | 0.727746 | 0.358591 | 0.424636 | -1.809031 | 20.183786 | 0.691499102 | -1.23675825 | 0.216176846 | NA | 4.700498 | 3.844638 | 4.365753 | 4.364964 | 4.314258 | 4.318526 | 30 | 28 | 12 | 14 |
| 1272 | NM_00125606 | CNTN1 | contactin 1 | protein_coding | NP_001242992.1;NP_001242993.1;NP_001834.2;NP_778203.1;XP_0052 | 2171 | 600016 | . | . | 2.312397 | 1.824118 | 1.411227 | 2.020013 | 5.381045 | 4.216531 | 3.166752 | 4.618520 | -1.186232 | 333.857952 | 0.219954166 | -1.120171 | 0.262640901 | 0.995798899 | 8.501381 | 8.254228 | 8.484210 | 8.363873 | 8.243926 | 8.409935 | 460 | 351 | 224 | 331 |
| 1277 | NM_000088,XI | COL1A1 | collagen type I alpha 1 chain | protein_coding | NP_000079.2;XP_005257115.2;XP_005257116.2;XP_011522643.1 | 2197 | 120150 | . | . | 0.909312 | 0.982592 | 0.993231 | 0.981704 | 2.116006 | 2.271306 | 2.228781 | 2.244550 | 1.026382 | 178.707337 | 0.24792672 | 0.151526058 | 0.879560757 | 0.998865404 | 7.462639 | 7.500029 | 7.460099 | 7.490258 | 7.490615 | 7.482143 | 196 | 197 | 159 | 168 |
| 1280 | NM_001844,N | COL2A1 | collagen type II alpha 1 chain | protein_coding | NP_001835.3;NP_149162.2;XP_016874317.1;XP_016874318.1;XP_01687 | 2200 | 120140 | . | . | 0.139021 | 0.057901 | 0.142047 | 0.115802 | 0.323508 | 0.133842 | 0.318749 | 0.264767 | 1.282741 | 19.731676 | 0.728205976 | 0.493307961 | 0.621795004 | NA | 4.108966 | 4.473012 | 4.314072 | 4.251751 | 4.312044 | 4.296290 | 28 | 11 | 21 | 19 |
| 1281 | NM_000090 | COL3A1 | collagen type III alpha 1 chain | protein_coding | NP_000081.2 | 2201 | 120180 | . | . | 0.071638 | 0.032586 | 0.034951 | 0.040065 | 0.166705 | 0.075324 | 0.078429 | 0.091603 | -1.360397 | 8.470090 | 1.062585165 | -0.41787455 | 0.676038844 | NA | 3.286426 | 2.844638 | 3.108647 | 3.078678 | 3.080989 | 3.083349 | 15 | 7 | 6 | 7 |
| 1282 | NM_00130311 | COL4A1 | collagen type IV alpha 1 chain | protein_coding | NP_001290039.1;NP_001836.3;XP_011519350.1 | 2202 | 120130 | . | . | 5.184542 | 4.981925 | 4.084881 | 4.897747 | 12.064642 | 11.515944 | 9.166355 | 11.198114 | -1.162743 | 983.186464 | 0.146719395 | -1.48264436 | 0.138168924 | 0.899796454 | 10.046201 | 9.828215 | 10.008877 | 9.977342 | 9.820511 | 9.941831 | 1243 | 1117 | 723 | 925 |
| 1284 | NM_001846 | COL4A2 | collagen type IV alpha 2 chain | protein_coding | NP_001837.2 | 2203 | 120090 | . | . | 6.966110 | 7.185295 | 5.524251 | 6.146447 | 16.210424 | 16.609131 | 12.396258 | 14.053121 | -1.229373 | 1297.489957 | 0.131744601 | -2.26136165 | 0.023736872 | 0.510492798 | 10.482943 | 10.184685 | 10.405600 | 10.423423 | 10.221494 | 10.292601 | 1629 | 1562 | 963 | 1144 |
| 1286 | NM_000092,XI | COL4A4 | collagen type IV alpha 4 chain | protein_coding | NP_000083.3;XP_005246338.1;XP_005246339.1;XP_006712309.1;XP_01 | 2206 | 120131 | . | . | 0.172077 | 0.205817 | 0.180232 | 0.149266 | 0.400432 | 0.475754 | 0.404435 | 0.341278 | 1.030658 | 50.836533 | 0.444048662 | 0.098109 | 0.921845741 | 0.998865404 | 5.644780 | 5.690446 | 5.683893 | 5.641978 | 5.685551 | 5.651766 | 64 | 48 | 49 | 44 |
| 1287 | NM_000495,N | COL4A5 | collagen type IV alpha 5 chain | protein_coding | NP_000486.1;NP_203699.1;XP_011529151.2;XP_016884748.1;XP_01688 | 2207 | 303630 | . | . | 1.494346 | 1.680030 | 1.170793 | 1.495222 | 3.477406 | 3.883466 | 2.627225 | 3.418648 | -1.201297 | 297.333446 | 0.213389871 | -1.23995171 | 0.214993255 | 0.980036956 | 8.342828 | 8.076802 | 8.233154 | 8.289742 | 8.104532 | 8.219466 | 355 | 368 | 208 | 282 |
| 1290 | NM_000393,XI | COL5A2 | collagen type V alpha 2 chain | protein_coding | NP_000384.2;XP_011508875.1 | 2210 | 120190 | . | . | 0.637669 | 0.416610 | 0.512167 | 0.408838 | 1.483880 | 0.963014 | 1.149289 | 0.934760 | -1.162690 | 105.674342 | 0.341556448 | -0.63669352 | 0.524324494 | 0.998865404 | 6.826865 | 6.612114 | 6.813083 | 6.666338 | 6.728714 | 6.659594 | 159 | 96 | 95 | 81 |
| 1291 | NM_001848 | COL6A1 | collagen type VI alpha 1 chain | protein_coding | NP_001839.2 | 2211 | 120220 | . | . | 8.134918 | 7.655898 | 8.249998 | 7.805246 | 18.930290 | 17.696949 | 18.512756 | 17.845766 | 1.002417 | 1043.016069 | 0.13782795 | 0.025265853 | 0.97984291 | 0.998865404 | 10.024623 | 10.028465 | 10.050637 | 9.999304 | 10.051005 | 10.002795 | 1241 | 1085 | 938 | 947 |
| 1292 | NM_001849,N | COL6A2 | collagen type VI alpha 2 chain | protein_coding | NP_001840.3;NP_478054.2;NP_478055.2;XP_011527753.1 | 2212 | 120240 | . | . | 7.147990 | 8.056439 | 6.234270 | 6.893137 | 16.633665 | 18.622819 | 13.989522 | 15.760341 | -1.173481 | 741.875066 | 0.155478602 | -1.48441018 | 0.137700135 | 0.899514893 | 9.646136 | 9.414653 | 9.546590 | 9.626455 | 9.436661 | 9.514348 | 871 | 913 | 559 | 677 |
| 1293 | NM_004369,N | COL6A3 | collagen type VI alpha 3 chain | protein_coding | NP_004360.2;NP_476505.3;NP_476506.3;NP_476507.3;NP_476508.2;XP_ | 2213 | 120250 | . | . | 0.079894 | 0.071849 | 0.043635 | 0.057131 | 0.185917 | 0.166083 | 0.097916 | 0.130623 | -1.492441 | 21.291685 | 0.672444718 | -0.85906561 | 0.390304319 | NA | 4.672908 | 4.093800 | 4.441363 | 4.429169 | 4.391269 | 4.408593 | 31 | 26 | 13 | 18 |
| 1294 | NM_000094,XI | COL7A1 | collagen type VII alpha 1 chain | protein_coding | NP_000085.1;XP_016861177.1;XP_016861178.1;XP_016861179.1;XP_01 | 2214 | 120120 | . | . | 1.515023 | 1.838414 | 1.662352 | 1.178281 | 3.525518 | 4.249576 | 3.730269 | 2.694001 | -1.188314 | 444.765540 | 0.206371673 | -1.20615401 | 0.227758113 | 0.982008277 | 8.915615 | 8.667536 | 8.783355 | 8.898411 | 8.834165 | 8.629091 | 506 | 568 | 415 | 315 |
| 1297 | NM_00137728 | COL9A1 | collagen type IX alpha 1 chain | protein_coding | NP_001364218.1;NP_001364219.1;NP_001364220.1;NP_001842.3;NP_5 | 2217 | 120210 | . | . | 0.763747 | 0.501960 | 0.703772 | 0.511497 | 1.777269 | 1.160304 | 1.579244 | 1.169475 | 1.194952 | 67.981088 | 0.39202654 | 0.655447297 | 0.512179808 | 0.998865404 | 5.951979 | 6.210572 | 6.054951 | 6.063211 | 6.140043 | 6.059676 | 70 | 68 | 75 | 58 |
| 1298 | NM_001852,XI | COL9A2 | collagen type IX alpha 2 chain | protein_coding | NP_001843.1;XP_006710428.1;XP_011539017.1;XP_011539018.1;XP_01 | 2218 | 120260 | . | . | 1.526690 | 1.276279 | 1.015796 | 1.043688 | 3.552670 | 2.950175 | 2.279418 | 2.268326 | -1.098166 | 121.318207 | 0.297040951 | -0.45480741 | 0.649247797 | 0.998865404 | 6.988261 | 6.853917 | 6.970006 | 6.906705 | 6.908136 | 6.905237 | 159 | 125 | 101 | 108 |
| 1299 | NM_001853,XI | COL9A3 | collagen type IX alpha 3 chain | protein_coding | NP_001844.3;XP_011526847.1;XP_016883155.1 | 2219 | 120270 | . | . | 398.988107 | 398.782876 | 228.732445 | 289.722855 | 928.461718 | 921.804498 | 513.268981 | 662.416911 | -1.560841 | 25487.781713 | 0.122175875 | -5.25737002 | 1.4613E-07 | 6.30323E-05 | 14.923243 | 14.280872 | 14.802593 | 14.793891 | 14.315561 | 14.505217 | 35951 | 33363 | 15334 | 20791 |
| 1301 | NM_00119070 | COL11A1 | collagen type XI alpha 1 chain | protein_coding | NP_001177638.1;NP_001845.3;NP_542196.2;NP_542197.3;XP_0168558 | 2186 | 120280 | . | . | 28.853314 | 28.671273 | 21.550817 | 23.452635 | 67.142850 | 66.274932 | 48.359408 | 53.621660 | -1.286177 | 5798.368298 | 0.106566793 | -3.40715246 | 0.000656444 | 0.050903951 | 12.671611 | 12.308459 | 12.605807 | 12.590102 | 12.358123 | 12.413804 | 7581 | 6976 | 4261 | 4917 |
| 1302 | NM_00116377 | COL11A2 | collagen type XI alpha 2 chain | protein_coding | NP_001157243.1;NP_542410.2;NP_542411.2;NP_542412.2;XP_0115126 | 2187 | 120290 | . | . | 6.338087 | 6.238725 | 4.330448 | 4.567346 | 14.748987 | 14.421092 | 9.717399 | 10.442696 | -1.462526 | 972.517020 | 0.139168238 | -3.94100298 | 8.11416E-05 | 0.01086228 | 10.174032 | 9.625252 | 10.038475 | 10.054701 | 9.746573 | 9.793203 | 1316 | 1260 | 663 | 766 |
| 1303 | NM_004370,N | COL12A1 | collagen type XII alpha 1 chain | protein_coding | NP_004361.3;NP_542376.2;XP_011533736.1;XP_011533737.1;XP_01153 | 2188 | 120320 | . | . | 0.532044 | 0.484768 | 0.336109 | 0.437877 | 1.238089 | 1.120564 | 0.754219 | 1.001153 | -1.296811 | 160.148036 | 0.268316264 | -1.39748443 | 0.162267945 | 0.935295382 | 7.499094 | 7.122987 | 7.389099 | 7.356230 | 7.225174 | 7.309899 | 216 | 188 | 107 | 146 |
| 1306 | NM_001855,XI | COL15A1 | collagen type XV alpha 1 chain | protein_coding | NP_001846.3;XP_011516516.1 | 2192 | 120325 | . | . | 0.706145 | 0.854357 | 0.511459 | 0.807112 | 1.643228 | 1.974885 | 1.147699 | 1.845365 | -1.199862 | 121.483440 | 0.320128418 | -0.82113327 | 0.411570361 | 0.998865404 | 7.051403 | 6.785604 | 6.918493 | 6.984979 | 6.818084 | 6.956347 | 139 | 156 | 75 | 126 |
| 1307 | NM_001856,XI | COL16A1 | collagen type XVI alpha 1 chain | protein_coding | NP_001847.3;XP_005270538.1;XP_011539024.1;XP_011539025.1;XP_01 | 2193 | 120326 | . | . | 6.822279 | 6.947931 | 6.274412 | 6.400034 | 15.875724 | 16.060453 | 14.079597 | 14.632919 | -1.100513 | 878.110888 | 0.14236465 | -0.97057854 | 0.331758195 | 0.998865404 | 9.845578 | 9.707645 | 9.809213 | 9.813856 | 9.767737 | 9.717541 | 1056 | 996 | 753 | 756 |
| 1308 | NM_000494 | COL17A1 | collagen type XVII alpha 1 chain | protein_coding | NP_000485.3 | 2194 | 113811 | . | . | 0.098735 | 0.079511 | 0.068187 | 0.034827 | 0.229762 | 0.183794 | 0.153009 | 0.079627 | -1.742994 | 13.028729 | 0.872461715 | -0.91874219 | 0.358230439 | NA | 4.047242 | 3.251698 | 3.731443 | 3.716162 | 3.706595 | 3.679292 | 21 | 16 | 11 | 6 |
| 1310 | NM_001858,XI | COL19A1 | collagen type XIX alpha 1 chain | protein_coding | NP_001849.2;XP_011533739.1;XP_011533740.1;XP_016865742.1;XP_01 | 2196 | 120165 | . | . | 6.907850 | 7.476807 | 10.023040 | 7.468207 | 16.074850 | 17.282975 | 22.491413 | 17.075169 | 1.202443 | 2640.561083 | 0.142534724 | 1.865989958 | 0.06204277 | 0.723723143 | 11.227405 | 11.493591 | 11.251140 | 11.315865 | 11.540530 | 11.314641 | 2649 | 2692 | 2849 | 2335 |
| 1312 | NM_000754,N | COMT | catechol-O-methyltransferase | protein_coding | NP_000745.1;NP_001128633.1;NP_001128634.1;NP_001349757.1;NP_0 | 2228 | 116790 | . | . | 59.600182 | 56.372983 | 58.608963 | 59.273334 | 138.692082 | 130.308687 | 131.516812 | 135.521437 | 1.012936 | 4122.282670 | 0.109046318 | 0.170045315 | 0.864974499 | 0.998865404 | 11.999901 | 12.018495 | 12.029616 | 11.977174 | 12.012402 | 12.016449 | 4865 | 4280 | 3599 | 3899 |
| 1314 | NM_00109839 | COPA | COPI coat complex subunit alpha | protein_coding | NP_001091868.1;NP_004362.2 | 2230 | 601924 | . | . | 44.852788 | 46.301010 | 45.129306 | 51.303784 | 104.374285 | 107.026858 | 101.268849 | 117.300012 | 1.043154 | 7768.924023 | 0.109611197 | 0.556081498 | 0.578155143 | 0.998865404 | 12.892746 | 12.953610 | 12.895069 | 12.914606 | 12.889191 | 12.989716 | 8656 | 8301 | 6491 | 7876 |
| 1315 | NM_00114406 | COPB1 | COPI coat complex subunit beta 1 | protein_coding | NP_001137533.1;NP_001137534.1;NP_057535.1 | 2231 | 600959 | . | . | 37.720232 | 36.458177 | 33.214535 | 31.420286 | 87.776536 | 84.274712 | 74.532446 | 71.838749 | -1.162770 | 3635.588485 | 0.111506177 | -1.95115621 | 0.051038467 | 0.672475152 | 11.932596 | 11.715166 | 11.903894 | 11.869614 | 11.789397 | 11.734676 | 4593 | 4132 | 3022 | 3046 |
| 1316 | NM_00116012 | KLF6 | Kruppel like factor 6 | protein_coding | NP_001153596.1;NP_001153597.1;NP_001291.3 | 2235 | 602053 | . | . | 16.481777 | 16.692554 | 17.497196 | 15.873804 | 38.353772 | 38.585584 | 39.263196 | 36.293565 | -1.005973 | 2375.253852 | 0.119843192 | -0.07169061 | 0.942848132 | 0.998865404 | 11.218060 | 11.209660 | 11.213462 | 11.218191 | 11.254080 | 11.166261 | 2735 | 2578 | 2172 | 2099 |
| 1317 | NM_001859 | SLC31A1 | solute carrier family 31 member 1 | protein_coding | NP_001850.1 | 11016 | 603085 | . | . | 17.006739 | 17.239050 | 15.557119 | 17.096989 | 39.575382 | 39.848835 | 34.909725 | 39.090233 | -1.063183 | 2481.750425 | 0.116998119 | -0.75547714 | 0.449962746 | 0.998865404 | 11.320740 | 11.232186 | 11.299255 | 11.303312 | 11.217535 | 11.284811 | 2938 | 2767 | 2004 | 2350 |
| 1318 | NM_001860,XI | SLC31A2 | solute carrier family 31 member 2 | protein_coding | NP_001851.1;XP_011516518.1;XP_016869756.1 | 11017 | 603088 | . | . | 1.059672 | 0.832723 | 0.801002 | 0.784183 | 2.465900 | 1.924877 | 1.797425 | 1.792942 | -1.223873 | 53.555811 | 0.434524682 | -0.67074077 | 0.502385684 | 0.998865404 | 5.880698 | 5.590696 | 5.792211 | 5.733949 | 5.728961 | 5.719491 | 77 | 55 | 43 | 44 |
| 1325 | NM_001302 | CORT | cortistatin | protein_coding | NP_001293.3 | 2257 | 602784 | . | . | 0.011585 | 0.107846 | 0.046477 | 0.078175 | 0.026958 | 0.249292 | 0.104293 | 0.178737 | 1.209283 | 2.518843 | 1.905639286 | 0.143863333 | 0.88560839 | NA | 1.193905 | 1.459419 | 1.317363 | 1.330972 | 1.324370 | 1.328457 | 1 | 4 | 2 | 3 |
| 1326 | NM_00124413 | MAP3K8 | mitogen-activated protein kinase kinase kinase 8 | protein_coding | NP_001231063.1;NP_001307890.1;NP_005195.2;XP_016871197.1;XP_01 | 6860 | 191195 | . | . | 0.599867 | 0.764075 | 0.362623 | 0.355942 | 1.395915 | 1.766194 | 0.813715 | 0.813817 | -1.150300 | 76.419221 | 0.366669951 | -0.55093302 | 0.581679591 | 0.998865404 | 6.352958 | 6.151760 | 6.258263 | 6.287674 | 6.261017 | 6.213719 | 89 | 93 | 68 | 60 |
| 1327 | NM_00131878 | COX4I1 | cytochrome c oxidase subunit 4I1 | protein_coding | NP_001305715.1;NP_001305717.1;NP_001305723.1;NP_001305726.1;N | 2265 | 123864 | . | . | 194.531677 | 195.184378 | 171.529602 | 207.469698 | 452.683215 | 451.177443 | 384.907455 | 474.354821 | -1.077440 | 4659.358686 | 0.121736738 | -0.8839415 | 0.376727793 | 0.998865404 | 12.238774 | 12.131047 | 12.244487 | 12.185272 | 12.078689 | 12.222236 | 5761 | 5032 | 3563 | 4569 |
| 1329 | NM_001862 | COX5B | cytochrome c oxidase subunit 5B | protein_coding | NP_001853.2 | 2269 | 123866 | . | . | 269.218475 | 253.730988 | 230.155930 | 297.168060 | 626.482483 | 586.510559 | 516.463196 | 679.439514 | -1.006284 | 5642.652706 | 0.130966374 | -0.06900722 | 0.94498387 | 0.998865404 | 12.466733 | 12.457568 | 12.491042 | 12.433927 | 12.349564 | 12.555325 | 6739 | 5901 | 4294 | 5917 |
| 1337 | NM_004373 | COX6A1 | cytochrome c oxidase subunit 6A1 | protein_coding | NP_004364.2 | 2277 | 602072 | . | . | 412.679871 | 424.434875 | 384.717590 | 432.327240 | 960.323059 | 981.100281 | 863.295105 | 988.464844 | -1.039220 | 6917.953382 | 0.108869011 | -0.50979317 | 0.61019637 | 0.998865404 | 12.783659 | 12.728065 | 12.763536 | 12.780137 | 12.693509 | 12.783142 | 8040 | 7683 | 5586 | 6699 |
| 1340 | NM_001863 | COX6B1 | cytochrome c oxidase subunit 6B1 | protein_coding | NP_001854.1 | 2280 | 124089 | . | . | 216.778687 | 203.031006 | 169.157928 | 224.671112 | 504.452942 | 469.315247 | 379.585480 | 513.683838 | -1.081739 | 3092.521373 | 0.140264555 | -0.80813644 | 0.419012042 | 0.998865404 | 11.650246 | 11.536655 | 11.653825 | 11.592567 | 11.442393 | 11.664026 | 3838 | 3340 | 2232 | 3164 |
| 1345 | NM_004374,XI | COX6C | cytochrome c oxidase subunit 6C | protein_coding | NP_004365.1;XP_016868509.1 | 2285 | 124090 | . | . | 71.455632 | 70.188442 | 65.685584 | 73.695182 | 166.280178 | 162.243722 | 147.396537 | 168.495276 | -1.024946 | 1602.916927 | 0.126573069 | -0.28084611 | 0.778828436 | 0.998865404 | 10.664247 | 10.628499 | 10.664746 | 10.646802 | 10.597881 | 10.673857 | 1890 | 1731 | 1309 | 1557 |
| 1347 | NM_00136629 | COX7A2 | cytochrome c oxidase subunit 7A2 | protein_coding | NP_001353221.1;NP_001353222.1;NP_001856.2;XP_024302098.1 | 2288 | 123996 | . | . | 145.368823 | 143.153818 | 131.842945 | 160.804620 | 338.279244 | 330.906438 | 295.851742 | 367.660620 | -1.000936 | 2730.266280 | 0.126890231 | -0.01063372 | 0.991515678 | 0.999576129 | 11.415610 | 11.414041 | 11.429511 | 11.398729 | 11.336840 | 11.484654 | 3205 | 2892 | 2157 | 2791 |
| 1349 | NM_001866 | COX7B | cytochrome c oxidase subunit 7B | protein_coding | NP_001857.1 | 2291 | 300885 | . | . | 27.599688 | 25.805094 | 23.813765 | 27.549889 | 64.225609 | 59.649628 | 53.437401 | 62.989548 | -1.054673 | 1994.482899 | 0.127636997 | -0.60167672 | 0.547389349 | 0.998865404 | 10.999766 | 10.922805 | 11.011866 | 10.951236 | 10.884979 | 10.991542 | 2447 | 2126 | 1574 | 1943 |
| 1350 | NM_001867 | COX7C | cytochrome c oxidase subunit 7C | protein_coding | NP_001858.1 | 2292 | 603774 | . | . | 205.475510 | 186.655502 | 179.622299 | 202.855286 | 478.149933 | 431.462555 | 403.067261 | 463.804474 | -1.040223 | 3795.205075 | 0.118863638 | -0.47863639 | 0.632197323 | 0.998865404 | 11.918151 | 11.861211 | 11.947945 | 11.861041 | 11.825903 | 11.917605 | 4689 | 3958 | 3055 | 3682 |
| 1351 | NM_004074 | COX8A | cytochrome c oxidase subunit 8A | protein_coding | NP_004065.1 | 2294 | 123870 | . | . | 167.903320 | 174.142990 | 153.786438 | 185.817535 | 390.717926 | 402.539337 | 345.092316 | 424.849701 | -1.021357 | 2622.375084 | 0.126654636 | -0.24071313 | 0.80977746 | 0.998865404 | 11.371968 | 11.341185 | 11.352792 | 11.375654 | 11.272417 | 11.416614 | 3009 | 2900 | 2055 | 2649 |
| 1352 | NM_001303 | COX10 | cytochrome c oxidase assembly factor heme A:farnesyltransferase | protein_coding | NP_001294.2 | 2260 | 602125 | . | . | 4.978917 | 4.985906 | 4.941219 | 5.173423 | 11.586143 | 11.525146 | 11.087952 | 11.828417 | 1.000913 | 453.842991 | 0.172474402 | 0.007631074 | 0.993911343 | 0.999862634 | 8.825460 | 8.826639 | 8.827711 | 8.823856 | 8.815701 | 8.836829 | 524 | 488 | 388 | 433 |
| 1353 | NM_00116286 | COX11 | cytochrome c oxidase copper chaperone COX11 | protein_coding | NP_001156333.1;NP_001156334.1;NP_001308447.1;NP_004366.1;XP_01 | 2261 | 603648 | . | . | 9.395328 | 5.809213 | 8.681735 | 4.193111 | 21.863313 | 13.428256 | 19.481562 | 9.587049 | -1.161164 | 763.894353 | 0.187027806 | -1.15261884 | 0.249066868 | 0.992733415 | 9.680444 | 9.466058 | 9.700663 | 9.536894 | 9.625652 | 9.396937 | 1052 | 785 | 718 | 552 |
| 1355 | NM_00132097 | COX15 | cytochrome c oxidase assembly homolog COX15 | protein_coding | NP_001307903.1;NP_001307904.1;NP_001307905.1;NP_001358953.1;N | 2263 | 603646 | . | . | 7.777623 | 7.323160 | 7.940043 | 7.183698 | 18.098852 | 16.927813 | 17.817221 | 16.424672 | 1.013058 | 1127.650103 | 0.13920951 | 0.134455467 | 0.893042423 | 0.998865404 | 10.129483 | 10.148661 | 10.146762 | 10.119930 | 10.193181 | 10.090758 | 1313 | 1187 | 1055 | 991 |
| 1362 | NM_00119977 | CPD | carboxypeptidase D | protein_coding | NP_001186704.1;NP_001295.2 | 2301 | 603102 | . | . | 11.554337 | 10.862164 | 9.870712 | 8.616493 | 26.887413 | 25.108376 | 22.149591 | 19.700586 | -1.229391 | 2989.083360 | 0.122573207 | -2.43074123 | 0.015067972 | 0.411847949 | 11.686653 | 11.388996 | 11.652152 | 11.593366 | 11.511135 | 11.394549 | 3929 | 3432 | 2502 | 2331 |
| 1365 | NM_001306 | CLDN3 | claudin 3 | protein_coding | NP_001297.1 | 2045 | 602910 | . | . | 0.074338 | 0.138808 | 0.053462 | 0.043634 | 0.172988 | 0.320861 | 0.119967 | 0.099763 | -2.036202 | 3.366880 | 1.662005528 | -0.61725455 | 0.537066837 | NA | 2.175403 | 1.148375 | 1.761682 | 1.771387 | 1.756463 | 1.755602 | 4 | 6 | 2 | 2 |
| 1368 | NM_00100550 | CPM | carboxypeptidase M | protein_coding | NP_001005502.1;NP_001865.1;NP_938079.1 | 2311 | 114860 | . | . | 3.487554 | 3.221028 | 2.362619 | 2.745541 | 8.115679 | 7.445551 | 5.301649 | 6.278266 | -1.328476 | 609.247799 | 0.164196917 | -2.49561405 | 0.012573933 | 0.369272325 | 9.441442 | 9.031273 | 9.366475 | 9.305872 | 9.103111 | 9.190656 | 835 | 718 | 423 | 525 |
| 1369 | NM_001308 | CPN1 | carboxypeptidase N subunit 1 | protein_coding | NP_001299.1 | 2312 | 603103 | . | . | 10.146114 | 10.100796 | 12.623403 | 15.607617 | 23.610424 | 23.348442 | 28.326550 | 35.684963 | 1.375448 | 699.916513 | 0.161951158 | 2.83975507 | 0.004514818 | 0.199099166 | 9.203163 | 9.662506 | 9.330859 | 9.323005 | 9.470648 | 9.615376 | 683 | 632 | 634 | 836 |
| 1371 | NM_000097,XI | CPOX | coproporphyrinogen oxidase | protein_coding | NP_000088.3;XP_005247182.1 | 2321 | 612732 | . | . | 10.623402 | 9.698301 | 12.372274 | 12.223931 | 24.721090 | 22.418059 | 27.763025 | 27.948566 | 1.202220 | 973.976698 | 0.140668302 | 1.888848558 | 0.058912124 | 0.718513409 | 9.788607 | 10.054634 | 9.890676 | 9.818559 | 9.998575 | 9.980779 | 1070 | 906 | 936 | 985 |
| 1373 | NM_00112263 | CPS1 | carbamoyl-phosphate synthase 1 | protein_coding | NP_001116105.2;NP_001356185.1;NP_001356186.1;NP_001866.2 | 2323 | 608307 | . | . | 6.479328 | 7.030990 | 6.220799 | 7.488375 | 15.077660 | 16.252447 | 13.959293 | 17.121278 | 1.003436 | 1234.798374 | 0.141393451 | 0.035002119 | 0.972078052 | 0.998865404 | 10.267860 | 10.272256 | 10.241516 | 10.293491 | 10.203511 | 10.332921 | 1371 | 1376 | 983 | 1259 |
| 1374 | NM_00103184 | CPT1A | carnitine palmitoyltransferase 1A | protein_coding | NP_001027017.1;NP_001867.2;XP_005273819.1;XP_005273820.1;XP_01 | 2328 | 600528 | . | . | 6.966702 | 6.764905 | 9.274106 | 8.814462 | 16.211802 | 15.637381 | 20.810826 | 20.153220 | 1.289266 | 1275.167335 | 0.132529584 | 2.765797384 | 0.00567838 | 0.232577699 | 10.121406 | 10.488275 | 10.223188 | 10.197720 | 10.440402 | 10.364545 | 1305 | 1181 | 1310 | 1281 |
| 1375 | NM_00114513 | CPT1B | carnitine palmitoyltransferase 1B | protein_coding | NP_001138606.1;NP_001138607.1;NP_001138609.1;NP_004368.1;NP_6 | 2329 | 601987 | . | . | 3.550308 | 5.196625 | 4.988831 | 3.238698 | 8.261713 | 12.012231 | 11.194795 | 7.404897 | -1.076461 | 363.708766 | 0.24469226 | -0.43440964 | 0.663990999 | 0.998865404 | 8.558538 | 8.452805 | 8.402882 | 8.619679 | 8.590304 | 8.343812 | 355 | 481 | 371 | 257 |
| 1376 | NM_000098,N | CPT2 | carnitine palmitoyltransferase 2 | protein_coding | NP_000089.1;NP_001317518.1 | 2330 | 600650 | . | . | 8.044064 | 8.411822 | 7.960636 | 9.921833 | 18.718869 | 19.444305 | 17.863436 | 22.685092 | 1.072344 | 719.792617 | 0.161298886 | 0.624723804 | 0.532152347 | 0.998865404 | 9.440559 | 9.540582 | 9.453037 | 9.477365 | 9.439240 | 9.582509 | 786 | 763 | 581 | 770 |
| 1384 | NM_000755,N | CRAT | carnitine O-acetyltransferase | protein_coding | NP_000746.3;NP_001244292.2;NP_001333475.2;NP_001333476.2;NP_0 | 2342 | 600184 | . | . | 3.210234 | 3.437458 | 3.227321 | 3.526091 | 7.470345 | 7.945835 | 7.242015 | 8.061988 | -1.006056 | 293.061690 | 0.206291746 | -0.04222642 | 0.9663182 | 0.998865404 | 8.199843 | 8.190263 | 8.174422 | 8.217658 | 8.161440 | 8.223491 | 325 | 330 | 239 | 290 |
| 1385 | NM_00132079 | CREB1 | cAMP responsive element binding protein 1 | protein_coding | NP_001307722.1;NP_001358355.1;NP_001358356.1;NP_001358357.1;N | 2345 | 123810 | . | . | 2.558897 | 2.554467 | 2.848409 | 2.400357 | 5.954657 | 5.904759 | 6.391747 | 5.488132 | 1.014091 | 714.169816 | 0.154133826 | 0.130971419 | 0.895797917 | 0.998865404 | 9.469706 | 9.490466 | 9.494314 | 9.454383 | 9.523659 | 9.443445 | 840 | 743 | 662 | 635 |
| 1386 | NM_00125609 | ATF2 | activating transcription factor 2 | protein_coding | NP_001243019.1;NP_001243020.1;NP_001243021.1;NP_001243022.1;N | 784 | 123811 | . | . | 7.868150 | 7.632920 | 8.923507 | 8.089080 | 18.309509 | 17.643838 | 20.024092 | 18.494721 | 1.090504 | 1052.945706 | 0.139685672 | 0.894832344 | 0.370876717 | 0.998865404 | 9.976140 | 10.101565 | 10.020161 | 9.992260 | 10.113622 | 10.025454 | 1182 | 1066 | 1012 | 969 |
| 1387 | NM_00107984 | CREBBP | CREB binding protein | protein_coding | NP_001073315.1;NP_004371.2;XP_005255181.1;XP_005255182.1;XP_00 | 2348 | 600140 | . | . | 3.078414 | 3.677082 | 3.894230 | 3.952084 | 7.163598 | 8.499741 | 8.738540 | 9.035970 | 1.135332 | 1217.944671 | 0.139825734 | 1.309593042 | 0.190333544 | 0.961278413 | 10.155971 | 10.338711 | 10.135473 | 10.258922 | 10.287008 | 10.302195 | 1205 | 1333 | 1115 | 1226 |
| 1388 | NM_00113615 | ATF6B | activating transcription factor 6 beta | protein_coding | NP_001129625.1;NP_004372.3 | 2349 | 600984 | . | . | 25.698380 | 28.370618 | 30.275066 | 28.952977 | 59.801180 | 65.579951 | 67.936373 | 66.197537 | 1.080316 | 2316.372849 | 0.119981472 | 0.928922717 | 0.352929139 | 0.998865404 | 11.120863 | 11.232290 | 11.108336 | 11.180894 | 11.229140 | 11.185114 | 2448 | 2512 | 2149 | 2194 |
| 1389 | NM_001310 | CREBL2 | cAMP responsive element binding protein like 2 | protein_coding | NP_001301.1 | 2350 | 603476 | . | . | 8.365633 | 6.908775 | 7.715229 | 7.504443 | 19.467171 | 15.969943 | 17.312752 | 17.158016 | -1.018133 | 896.136206 | 0.153314229 | -0.16910233 | 0.865716149 | 0.998865404 | 9.820217 | 9.794819 | 9.882774 | 9.737539 | 9.813346 | 9.786324 | 1145 | 878 | 787 | 817 |
| 1390 | NM_00126756 | CREM | cAMP responsive element modulator | protein_coding | NP_001254491.1;NP_001254492.1;NP_001254493.1;NP_001254494.1;N | 2352 | 123812 | . | . | 6.207733 | 5.757402 | 6.731786 | 5.887545 | 14.445652 | 13.308489 | 15.105937 | 13.461175 | 1.048507 | 419.493582 | 0.183071996 | 0.373276842 | 0.70894241 | 0.998865404 | 8.677516 | 8.746665 | 8.711269 | 8.681871 | 8.770341 | 8.678342 | 483 | 431 | 402 | 372 |
| 1393 | NM_001882,XI | CRHBP | corticotropin releasing hormone binding protein | protein_coding | NP_001873.2 | 2356 | 122559 | . | . | 0.258214 | 0.275052 | 0.073682 | 0.251965 | 0.600876 | 0.635797 | 0.165341 | 0.576088 | -1.500379 | 11.580279 | 0.939699448 | -0.6228873 | 0.533358584 | NA | 3.799841 | 3.206844 | 3.545957 | 3.553853 | 3.509394 | 3.549249 | 15 | 16 | 4 | 13 |
| 1397 | NM_00127083 | CRIP2 | cysteine rich protein 2 | protein_coding | NP_001257766.1;NP_001257770.1;NP_001303.1;XP_024305248.1;XP_02 | 2361 | 601183 | . | . | 20.484366 | 19.051991 | 15.309461 | 20.014262 | 47.667961 | 44.039531 | 34.353984 | 45.760231 | -1.128540 | 710.462862 | 0.16989621 | -1.02684984 | 0.304491174 | 0.998865404 | 9.557525 | 9.382394 | 9.539495 | 9.481823 | 9.335394 | 9.512171 | 903 | 780 | 504 | 708 |
| 1398 | NM_005206,N | CRK | CRK proto-oncogene, adaptor protein | protein_coding | NP_005197.3;NP_058431.2 | 2362 | 164762 | . | . | 18.266121 | 19.453902 | 20.198255 | 19.794476 | 42.506012 | 44.968569 | 45.324297 | 45.257713 | 1.049541 | 2276.052736 | 0.116841845 | 0.597035434 | 0.550483724 | 0.998865404 | 11.117025 | 11.186769 | 11.111663 | 11.152687 | 11.183396 | 11.158837 | 2492 | 2458 | 2058 | 2152 |
| 1399 | NM_005207,N | CRKL | CRK like proto-oncogene, adaptor protein | protein_coding | NP_005198.1 | 2363 | 602007 | . | . | 15.857725 | 16.089762 | 16.177565 | 17.750431 | 36.901576 | 37.192208 | 36.301988 | 40.584256 | 1.047299 | 2659.769109 | 0.117094841 | 0.569398254 | 0.569085901 | 0.998865404 | 11.343459 | 11.409946 | 11.348982 | 11.366384 | 11.355449 | 11.433408 | 2960 | 2834 | 2253 | 2673 |
| 1407 | NM_004075,XI | CRY1 | cryptochrome circadian regulator 1 | protein_coding | NP_004066.1;XP_016874321.1;XP_024304612.1;XP_024304613.1 | 2384 | 601933 | . | . | 9.757893 | 9.261664 | 8.150010 | 9.407602 | 22.707019 | 21.408750 | 18.288386 | 21.509363 | -1.088643 | 851.501759 | 0.148763003 | -0.82366358 | 0.410130738 | 0.998865404 | 9.793988 | 9.671128 | 9.784044 | 9.741651 | 9.653991 | 9.748626 | 1052 | 930 | 661 | 816 |
| 1408 | NM_00112745 | CRY2 | cryptochrome circadian regulator 2 | protein_coding | NP_001120929.1;NP_066940.3 | 2385 | 603732 | . | . | 3.897144 | 4.385542 | 4.013317 | 3.699559 | 9.068813 | 10.137377 | 9.005766 | 8.458602 | -1.087848 | 515.022889 | 0.170464182 | -0.71262221 | 0.476079562 | 0.998865404 | 9.067909 | 8.946524 | 8.998787 | 9.068398 | 9.009171 | 8.950959 | 585 | 610 | 449 | 441 |
| 1410 | NM_00128980 | CRYAB | crystallin alpha B | protein_coding | NP_001276736.1;NP_001276737.1;NP_001317308.1;NP_001355174.1;N | 2389 | 123590 | . | . | 0.742167 | 0.829192 | 0.573346 | 0.448983 | 1.727054 | 1.916716 | 1.286572 | 1.026547 | -1.487374 | 17.295144 | 0.745394191 | -0.76840926 | 0.44224408 | NA | 4.369732 | 3.798723 | 4.123380 | 4.136750 | 4.110452 | 4.091679 | 22 | 24 | 14 | 11 |
| 1416 | NR_033733,NF | CRYBB2P1 | crystallin beta B2 pseudogene 1 | transcribed_ps | . | 2399 | . | . | . | 7.406206 | 6.742953 | 10.011542 | 7.974353 | 17.234545 | 15.586636 | 22.465608 | 18.232410 | 1.258084 | 363.987471 | 0.200837858 | 1.64922963 | 0.0991006 | 0.813553488 | 8.331947 | 8.664430 | 8.461359 | 8.402280 | 8.632036 | 8.492437 | 391 | 329 | 394 | 336 |

| | | | | | | | | | | | | | | | | | | | | | | | | | | | | | | | | | |
|---|---|---|---|---|---|---|---|---|---|---|---|---|---|---|---|---|---|---|---|---|---|---|---|---|---|---|---|---|---|---|---|---|---|
| 1427 | NM_017541 | CRYGS | crystallin gamma S | protein_coding | NP_060011.1 | 2417 | 123730 | . | . | 1.391495 | 1.053556 | 2.309914 | 1.281092 | 3.238065 | 2.435343 | 5.183380 | 2.929065 | 1.455297 | 40.056833 | 0.52187977 | 1.037238825 | 0.299624568 | 0.998865404 | 5.025590 | 5.571100 | 5.296803 | 5.257985 | 5.383444 | 5.284688 | 43 | 30 | 53 | 32 |
| 1428 | NM_00137625 | CRYM | crystallin mu | protein_coding | NP_001363185.1;NP_001879.1 | 2418 | 123740 | . | . | 0.465566 | 0.632616 | 0.472927 | 0.396379 | 1.083392 | 1.462320 | 1.061234 | 0.906274 | -1.259163 | 21.362780 | 0.670279114 | -0.49601046 | 0.619887031 | NA | 4.572938 | 4.242203 | 4.423627 | 4.433905 | 4.420333 | 4.395912 | 26 | 27 | 19 | 15 |
| 1429 | NM_00113004 | CRYZ | crystallin zeta | protein_coding | NP_001123514.1;NP_001123515.1;NP_001128231.1;NP_001880.2;XP_0( | 2419 | 123691 | . | . | 13.088208 | 11.893501 | 11.497390 | 12.656101 | 30.456800 | 27.492359 | 25.799809 | 28.936672 | -1.048602 | 771.362751 | 0.151589743 | -0.45166543 | 0.651510023 | 0.998865404 | 9.624933 | 9.556794 | 9.653648 | 9.558631 | 9.574202 | 9.573979 | 970 | 795 | 655 | 706 |
| 1431 | NM_004077 | CS | citrate synthase | protein_coding | NP_004068.2 | 2422 | 118950 | . | . | 45.892342 | 50.452675 | 43.075039 | 50.986248 | 106.793373 | 116.623604 | 96.659142 | 116.573990 | -1.038543 | 4322.167770 | 0.121353956 | -0.44960416 | 0.652995891 | 0.998865404 | 12.104668 | 12.049891 | 12.055501 | 12.127588 | 11.992998 | 12.124033 | 4853 | 4957 | 3396 | 4289 |
| 1432 | NM_001315,N | MAPK14 | mitogen-activated protein kinase 14 | protein_coding | NP_001306.1;NP_620581.1;NP_620582.1;NP_620583.1;XP_006715061.1 | 6876 | 600289 | . | . | 7.707428 | 7.988403 | 7.850584 | 8.209376 | 17.935499 | 18.465551 | 17.616487 | 18.769762 | -1.018018 | 997.000520 | 0.136286517 | -0.18904004 | 0.850061434 | 0.998865404 | 9.974324 | 9.948461 | 9.960132 | 9.975808 | 9.949638 | 9.960118 | 1146 | 1097 | 853 | 933 |
| 1434 | NM_00125613 | CSE1L | chromosome segregation 1 like | protein_coding | NP_001243064.1;NP_001307.2;NP_001349691.1 | 2431 | 601342 | . | . | 113.906098 | 107.869630 | 95.126186 | 81.377830 | 265.064171 | 249.345490 | 213.460413 | 186.060739 | -1.268865 | 11127.352255 | 0.116881824 | -2.9391921 | 0.00329069 | 0.163366554 | 13.603348 | 13.259906 | 13.560514 | 13.506961 | 13.400984 | 13.255466 | 14771 | 13017 | 9271 | 8397 |
| 1438 | NM_00116152 | CSF2RA | colony stimulating factor 2 receptor subunit alpha | protein_coding | NP_001155001.1;NP_001155002.1;NP_001155003.1;NP_001155004.1;NI | 2435 | 425000 | . | . | 2.713284 | 2.199950 | 2.421262 | 3.150038 | 6.313926 | 5.085274 | 5.433246 | 7.202186 | 1.086446 | 155.608867 | 0.277046477 | 0.431754892 | 0.665919567 | 0.998865404 | 7.220807 | 7.340281 | 7.307359 | 7.214920 | 7.251682 | 7.336667 | 192 | 142 | 126 | 168 |
| 1445 | NM_00112719 | CSK | C-terminal Src kinase | protein_coding | NP_001120662.1;NP_001341917.1;NP_004374.1 | 2444 | 124095 | . | . | 28.018565 | 30.342282 | 30.809144 | 32.880333 | 65.200355 | 70.137540 | 69.134825 | 75.176975 | 1.045853 | 2511.607905 | 0.116148585 | 0.556875515 | 0.577612485 | 0.998865404 | 11.261777 | 11.326293 | 11.262168 | 11.289238 | 11.284404 | 11.338819 | 2780 | 2694 | 2158 | 2488 |
| 1452 | NM_00102510 | CSNK1A1 | casein kinase 1 alpha 1 | protein_coding | NP_001020276.1;NP_001258670.1;NP_001258671.1;NP_001883.4 | 2451 | 600505 | . | . | 33.589085 | 32.630110 | 24.500738 | 25.617707 | 78.163182 | 75.425961 | 54.978946 | 58.571840 | -1.337993 | 4879.496029 | 0.106830324 | -3.93212516 | 8.41982E-05 | 0.011068455 | 12.447325 | 12.027234 | 12.378060 | 12.345662 | 12.103901 | 12.133160 | 6552 | 5913 | 3564 | 3984 |
| 1453 | NM_00136374 | CSNK1D | casein kinase 1 delta | protein_coding | NP_001350678.1;NP_001884.2;NP_620693.1;XP_005256393.1 | 2452 | 600864 | . | . | 25.304154 | 27.298152 | 26.702591 | 27.838772 | 58.883807 | 63.100903 | 59.919847 | 63.650049 | 1.022409 | 3150.925226 | 0.112624957 | 0.283888579 | 0.776495769 | 0.998865404 | 11.605538 | 11.637406 | 11.584187 | 11.639554 | 11.622259 | 11.638418 | 3467 | 3476 | 2743 | 3016 |
| 1454 | NM_001894,N | CSNK1E | casein kinase 1 epsilon | protein_coding | NP_001885.1;NP_689407.1 | 2453 | 600863 | . | . | 34.119529 | 35.505039 | 33.636608 | 37.636753 | 79.397549 | 82.071495 | 75.479584 | 86.051964 | 1.009393 | 3090.869077 | 0.115897824 | 0.116377081 | 0.907353698 | 0.998865404 | 11.587131 | 11.600432 | 11.576677 | 11.602452 | 11.554474 | 11.638276 | 3487 | 3372 | 2562 | 3060 |
| 1455 | NM_001319,XI | CSNK1G2 | casein kinase 1 gamma 2 | protein_coding | NP_001310.3;XP_005259555.1;XP_005259556.1;XP_005259557.1;XP_00! | 2455 | 602214 | . | . | 17.143109 | 17.947760 | 20.546513 | 21.659457 | 39.892720 | 41.487047 | 46.105780 | 49.521775 | 1.188350 | 1765.472612 | 0.121320503 | 2.052081992 | 0.040161695 | 0.626386296 | 10.656074 | 10.904889 | 10.698841 | 10.728070 | 10.835374 | 10.862643 | 1823 | 1774 | 1639 | 1828 |
| 1456 | NM_00103181 | CSNK1G3 | casein kinase 1 gamma 3 | protein_coding | NP_001026982.1;NP_001038188.1;NP_001257501.1;NP_001257502.1;NI | 2456 | 604253 | . | . | 6.095455 | 5.500399 | 5.919954 | 4.805696 | 14.184374 | 12.714421 | 13.284205 | 10.987653 | -1.090995 | 776.997078 | 0.161949017 | -0.77582425 | 0.437852766 | 0.998865404 | 9.662793 | 9.538042 | 9.665656 | 9.592564 | 9.640399 | 9.491997 | 981 | 830 | 716 | 622 |
| 1457 | NM_00136277 | CSNK2A1 | casein kinase 2 alpha 1 | protein_coding | NP_001349699.1;NP_001349700.1;NP_001886.1;NP_808227.1;NP_8082; | 2457 | 115440 | . | . | 7.655711 | 8.198931 | 7.225539 | 8.178527 | 17.815153 | 18.952195 | 16.213902 | 18.699230 | -1.077239 | 2776.997105 | 0.118398978 | -0.90658188 | 0.364627948 | 0.998865404 | 11.492077 | 11.384539 | 11.424530 | 11.510670 | 11.386913 | 11.428394 | 3143 | 3271 | 2265 | 2571 |
| 1459 | NM_001896,XI | CSNK2A2 | casein kinase 2 alpha 2 | protein_coding | NP_001887.1;XP_005255857.1;XP_005255858.1;XP_016878434.1 | 2459 | 115442 | . | . | 9.844855 | 10.519921 | 10.120384 | 11.014502 | 22.909383 | 24.317267 | 22.709848 | 25.183350 | 1.002629 | 581.614201 | 0.163291185 | 0.023199841 | 0.981490866 | 0.998865404 | 9.182335 | 9.185501 | 9.152457 | 9.211919 | 9.153614 | 9.213906 | 639 | 655 | 483 | 571 |
| 1460 | NM_00128238 | CSNK2B | casein kinase 2 beta | protein_coding | NP_001269314.1;NP_001311.3 | 2460 | 115441 | . | . | 65.419796 | 65.435931 | 68.351483 | 76.206429 | 152.234543 | 151.258085 | 153.378749 | 174.236944 | 1.089339 | 1997.126856 | 0.121868712 | 1.013004168 | 0.31105817 | 0.998865404 | 10.900760 | 11.024028 | 10.931331 | 10.925273 | 10.956179 | 11.034802 | 2210 | 2055 | 1722 | 2048 |
| 1462 | NM_00112633 | VCAN | versican | protein_coding | NP_001119808.1;NP_001157569.1;NP_001157570.1;NP_004376.2 | 2464 | 118661 | . | . | 1.282494 | 1.511558 | 1.541838 | 1.270987 | 2.984417 | 3.494034 | 3.459839 | 2.905961 | -1.081810 | 431.272538 | 0.185701674 | -0.6109082 | 0.541260359 | 0.998865404 | 8.807677 | 8.695037 | 8.764903 | 8.784047 | 8.787029 | 8.662958 | 508 | 491 | 398 | 348 |
| 1463 | NM_004386 | NCAN | neurocan | protein_coding | NP_004377.2 | 2465 | 600826 | . | . | 0.035513 | 0.059788 | 0.052051 | 0.049740 | 0.082639 | 0.138203 | 0.116800 | 0.113725 | 1.117220 | 10.507519 | 0.942859602 | 0.169605034 | 0.865320763 | NA | 3.312060 | 3.470304 | 3.376523 | 3.395983 | 3.393252 | 3.389705 | 9 | 13 | 10 | 10 |
| 1464 | NM_001897 | CSPG4 | chondroitin sulfate proteoglycan 4 | protein_coding | NP_001888.2 | 2466 | 601172 | . | . | 55.195221 | 55.934784 | 46.425392 | 56.353672 | 128.441544 | 129.295746 | 104.177238 | 128.845978 | -1.096881 | 13811.671795 | 0.116066389 | -1.14939856 | 0.250391674 | 0.993339639 | 13.818793 | 13.685321 | 13.788948 | 13.793002 | 13.630873 | 13.786676 | 16600 | 15629 | 10407 | 13480 |
| 1465 | NM_00119357 | CSRP1 | cysteine and glycine rich protein 1 | protein_coding | NP_001180499.1;NP_001180500.1;NP_001180501.1;NP_004069.1 | 2469 | 123876 | . | . | 121.484527 | 126.989892 | 113.946697 | 136.448240 | 282.699472 | 293.542826 | 255.693108 | 311.972695 | -1.007006 | 7071.472616 | 0.117013368 | -0.08608388 | 0.93139974 | 0.998865404 | 12.792885 | 12.782687 | 12.775100 | 12.804446 | 12.709955 | 12.852931 | 8030 | 7790 | 5610 | 7164 |
| 1466 | NM_00130096 | CSRP2 | cysteine and glycine rich protein 2 | protein_coding | NP_001287894.1;NP_001312.1 | 2470 | 601871 | . | . | 11.877719 | 11.029170 | 9.325021 | 10.433783 | 27.639938 | 25.494416 | 20.925077 | 23.855605 | -1.172289 | 302.390361 | 0.203383171 | -1.12756923 | 0.259501888 | 0.995760118 | 8.350498 | 8.120914 | 8.304163 | 8.258511 | 8.170531 | 8.221139 | 391 | 338 | 230 | 274 |
| 1468 | NM_00127088 | SLC25A10 | solute carrier family 25 member 10 | protein_coding | NP_001257817.1;NP_001257882.1;NP_036272.2;XP_016879709.1 | 10980 | 606794 | . | . | 8.274438 | 7.984791 | 6.229103 | 8.863659 | 19.254959 | 18.457201 | 13.977924 | 20.265702 | -1.067876 | 486.037171 | 0.19006688 | -0.49848099 | 0.618145063 | 0.998865404 | 8.971985 | 8.876273 | 8.959475 | 8.926547 | 8.804292 | 8.990532 | 593 | 528 | 347 | 507 |
| 1469 | NM_001898 | CST1 | cystatin SN | protein_coding | NP_001889.2 | 2473 | 123855 | . | . | 9.713654 | 8.911450 | 7.245879 | 9.076578 | 22.604071 | 20.599216 | 16.259542 | 20.752518 | -1.156034 | 207.411653 | 0.241720841 | -0.86539449 | 0.386822304 | 0.998865404 | 7.797555 | 7.587514 | 7.750594 | 7.705792 | 7.613562 | 7.708000 | 268 | 229 | 149 | 200 |
| 1471 | NM_000099,N | CST3 | cystatin C | protein_coding | NP_000090.1;NP_001275543.1 | 2475 | 604312 | . | . | 120.118688 | 122.681367 | 125.543248 | 132.961766 | 279.521098 | 283.583459 | 281.715425 | 304.001247 | 1.050301 | 3129.071908 | 0.111518913 | 0.634890282 | 0.525499976 | 0.998865404 | 11.575734 | 11.646438 | 11.581648 | 11.600170 | 11.612170 | 11.649752 | 3475 | 3331 | 2724 | 3074 |
| 1476 | NM_000100 | CSTB | cystatin B | protein_coding | NP_000091.1 | 2482 | 601145 | . | . | 303.831665 | 306.819611 | 314.176819 | 352.377502 | 707.028809 | 709.227234 | 705.003601 | 805.669312 | 1.076224 | 5846.408043 | 0.109571222 | 0.967207973 | 0.333440077 | 0.998865404 | 12.459420 | 12.565307 | 12.480738 | 12.481817 | 12.497837 | 12.586391 | 6481 | 6081 | 4995 | 5979 |
| 1477 | NM_00103352 | CSTF1 | cleavage stimulation factor subunit 1 | protein_coding | NP_001028693.1;NP_001028694.1;NP_001315.1;XP_011526902.1 | 2483 | 600369 | . | . | 17.270311 | 17.916980 | 16.365555 | 15.085053 | 40.188726 | 41.415903 | 36.723830 | 34.490180 | -1.079610 | 1810.904171 | 0.124303227 | -0.88903452 | 0.373984526 | 0.998865404 | 10.876577 | 10.766306 | 10.888561 | 10.813323 | 10.807433 | 10.775070 | 2267 | 1933 | 1546 | 1599 |
| 1478 | NM_00130620 | CSTF2 | cleavage stimulation factor subunit 2 | protein_coding | NP_001293135.1;NP_001293138.1;NP_001316.1 | 2484 | 300907 | . | . | 6.238139 | 6.406598 | 5.782360 | 6.174256 | 14.516405 | 14.809137 | 12.975447 | 14.116701 | -1.075195 | 648.773031 | 0.154606098 | -0.67654989 | 0.49869158 | 0.998865404 | 9.393074 | 9.288159 | 9.355742 | 9.375566 | 9.297474 | 9.335507 | 763 | 736 | 528 | 603 |
| 1479 | NM_00103350 | CSTF3 | cleavage stimulation factor subunit 3 | protein_coding | NP_001028677.1;NP_001028678.1;NP_001317.1 | 2485 | 600367 | . | . | 9.598624 | 8.490845 | 9.612958 | 8.696911 | 22.336394 | 19.626971 | 21.571198 | 19.884454 | -1.123453 | 661.628994 | 0.155811836 | -1.07783447 | 0.281107634 | 0.998865404 | 9.451259 | 9.283632 | 9.442720 | 9.371915 | 9.332864 | 9.325242 | 847 | 717 | 545 | 581 |
| 1486 | NM_004388 | CTBS | chitobiase | protein_coding | NP_004379.1 | 2496 | 600873 | . | . | 4.427432 | 3.798205 | 4.061301 | 3.657123 | 10.302816 | 8.779722 | 9.113441 | 8.361576 | -1.081304 | 816.985570 | 0.155688734 | -0.72434168 | 0.468855996 | 0.998865404 | 9.729106 | 9.617053 | 9.756513 | 9.640259 | 9.684044 | 9.604133 | 1056 | 842 | 722 | 694 |
| 1487 | NM_00101261 | CTBP1 | C-terminal binding protein 1 | protein_coding | NP_001012632.1;NP_001319.1;NP_001364115.1;NP_001364116.1;NP_0( | 2494 | 602618 | . | . | 14.934441 | 16.391792 | 20.617039 | 21.715688 | 34.753056 | 37.890360 | 46.264040 | 49.650338 | 1.013331 | 2517.803593 | 0.116792159 | 0.163585453 | 0.870057497 | 0.998865404 | 11.288286 | 11.307550 | 11.294276 | 11.290271 | 11.336569 | 11.268391 | 2887 | 2692 | 2297 | 2276 |
| 1488 | NM_00108391 | CTBP2 | C-terminal binding protein 2 | protein_coding | NP_001077383.1;NP_001277143.1;NP_001277144.1;NP_001307941.1;NI | 2495 | 602619 | . | . | 19.397090 | 20.382374 | 20.238821 | 25.449685 | 45.137825 | 47.114769 | 45.415326 | 58.187682 | -1.009287 | 2932.527113 | 0.112556509 | -0.11849238 | 0.905677529 | 0.998865404 | 11.524624 | 11.511202 | 11.523460 | 11.519825 | 11.491940 | 11.535717 | 3400 | 3172 | 2470 | 2810 |
| 1490 | NM_001901 | CCN2 | cellular communication network factor 2 | protein_coding | NP_001892.2 | 2500 | 121009 | . | . | 21.139303 | 21.014530 | 8.424598 | 11.602265 | 49.192024 | 48.576031 | 18.904551 | 26.527201 | -2.133570 | 1134.985630 | 0.159741858 | -6.84397462 | 7.70256E-12 | 1.18132E-08 | 10.594234 | 9.499982 | 10.362395 | 10.350046 | 9.691160 | 9.891215 | 1793 | 1656 | 533 | 783 |
| 1491 | NM_00119046 | CTH | cystathionine gamma-lyase | protein_coding | NP_001177392.1;NP_001893.2;NP_714964.2;XP_005270566.1;XP_01685 | 2501 | 607657 | . | . | 2.958465 | 2.837710 | 1.695145 | 2.863203 | 6.884471 | 6.559491 | 3.803857 | 6.546376 | -1.157516 | 238.096249 | 0.238014618 | -0.88663519 | 0.375275339 | 0.998865404 | 7.997619 | 7.785385 | 7.953981 | 7.902433 | 7.782377 | 7.926462 | 309 | 262 | 162 | 239 |
| 1493 | NM_00103763 | CTLA4 | cytotoxic T-lymphocyte associated protein 4 | protein_coding | NP_001032720.1;NP_005205.2 | 2505 | 123890 | . | . | 0.269261 | 0.544119 | 0.112588 | 0.287642 | 0.626580 | 1.257756 | 0.252643 | 0.657658 | -1.929061 | 19.243279 | 0.765516084 | -1.23824786 | 0.21562417 | NA | 4.666471 | 3.710567 | 4.251403 | 4.326785 | 4.216497 | 4.258891 | 19 | 37 | 7 | 17 |
| 1495 | NM_00129030 | CTNNA1 | catenin alpha 1 | protein_coding | NP_001277236.1;NP_001277238.1;NP_001277239.1;NP_001277241.1;NI | 2509 | 116805 | . | . | 96.482945 | 98.335886 | 83.505942 | 100.074976 | 224.519785 | 227.307782 | 187.384901 | 228.809548 | -1.079158 | 11046.569804 | 0.114881652 | -0.95669245 | 0.338722509 | 0.998865404 | 13.485255 | 13.375271 | 13.458526 | 13.465761 | 13.322188 | 13.466853 | 13149 | 12426 | 8453 | 10810 |
| 1496 | NM_00116488 | CTNNA2 | catenin alpha 2 | protein_coding | NP_001158355.1;NP_001269526.1;NP_001269527.1;NP_001269528.1;NI | 2510 | 114025 | . | . | 0.376606 | 0.326325 | 0.129990 | 0.194639 | 0.876378 | 0.754315 | 0.291695 | 0.445020 | -2.145916 | 33.745270 | 0.553597967 | -1.98988009 | 0.046604147 | 0.654401867 | 5.525622 | 4.421211 | 5.138658 | 5.111644 | 5.011644 | 5.044402 | 57 | 46 | 15 | 24 |
| 1497 | NM_00103168 | CTNS | cystinosin, lysosomal cystine transporter | protein_coding | NP_001026851.2;NP_001361421.1;NP_001361422.1;NP_001361423.1;NI | 2518 | 606272 | . | . | 11.398935 | 11.916268 | 14.297082 | 14.708692 | 26.525795 | 27.544988 | 32.082240 | 33.629674 | 1.263730 | 1332.782374 | 0.127786416 | 2.642600822 | 0.008227197 | 0.28822334 | 10.201539 | 10.539199 | 10.269810 | 10.296150 | 10.469160 | 10.453643 | 1332 | 1293 | 1307 | 1381 |
| 1499 | NM_00109820 | CTNNB1 | catenin beta 1 | protein_coding | NP_001091679.1;NP_001091680.1;NP_001317658.1;NP_001895.1;XP_0( | 2514 | 116806 | . | . | 75.841344 | 70.705125 | 65.474974 | 70.475149 | 176.485920 | 163.438058 | 146.923946 | 161.133050 | -1.085524 | 7442.203645 | 0.108459341 | -1.09157731 | 0.275018936 | 0.998865404 | 12.919504 | 12.801095 | 12.927979 | 12.860823 | 12.794008 | 12.856016 | 9272 | 8031 | 5977 | 6938 |
| 1500 | NM_00108545 | CTNND1 | catenin delta 1 | protein_coding | NP_001078927.1;NP_001078928.1;NP_001078929.1;NP_001078930.1;NI | 2515 | 601045 | . | . | 25.350733 | 27.412855 | 25.562321 | 27.823428 | 58.992195 | 63.366039 | 57.361107 | 63.614965 | -1.000920 | 5333.022147 | 0.110135732 | -0.01205067 | 0.990385188 | 0.999289967 | 12.381462 | 12.380013 | 12.353400 | 12.407542 | 12.345594 | 12.413303 | 5946 | 5943 | 4448 | 5199 |
| 1501 | NM_00128871 | CTNND2 | catenin delta 2 | protein_coding | NP_001275644.1;NP_001275645.1;NP_001275646.1;NP_001323.1;NP_0( | 2516 | 604275 | . | . | 0.114772 | 0.122232 | 0.156972 | 0.088641 | 0.267079 | 0.282546 | 0.352240 | 0.202668 | 1.018846 | 22.673925 | 0.658872119 | 0.040882084 | 0.967389901 | NA | 4.487990 | 4.517781 | 4.494592 | 4.500874 | 4.523024 | 4.475367 | 25 | 25 | 25 | 16 |
| 1503 | NM_00130123 | CTPS1 | CTP synthase 1 | protein_coding | NP_001288166.1;NP_001896.2;XP_024309320.1;XP_024309321.1;XP_02 | 2519 | 123860 | . | . | 46.180270 | 47.363718 | 53.454963 | 48.891505 | 107.463389 | 109.483356 | 119.951389 | 111.784618 | 1.083929 | 4460.025587 | 0.111962513 | 1.038470327 | 0.299051143 | 0.998865404 | 12.063481 | 12.179846 | 12.079107 | 12.096490 | 12.198086 | 12.110165 | 4877 | 4667 | 4246 | 4122 |
| 1506 | NM_001907 | CTRL | chymotrypsin like | protein_coding | NP_001898.1 | 2524 | 118888 | . | . | 0.961127 | 1.174797 | 1.219224 | 0.758842 | 2.236582 | 2.715596 | 2.735902 | 1.735001 | -1.092227 | 37.936367 | 0.521780694 | -0.24392111 | 0.80729192 | 0.998865404 | 5.306757 | 5.181547 | 5.232627 | 5.264074 | 5.267762 | 5.202055 | 41 | 47 | 39 | 26 |
| 1508 | NM_00131723 | CTSB | cathepsin B | protein_coding | NP_001304166.1;NP_001899.1;NP_680090.1;NP_680091.1;NP_680092.1 | 2527 | 116810 | . | . | 247.465053 | 247.800799 | 194.463567 | 238.741317 | 575.861355 | 572.802667 | 436.370606 | 545.853631 | -1.151076 | 27625.147705 | 0.11458061 | -1.77153392 | 0.076471957 | 0.767975096 | 14.851635 | 14.648618 | 14.812805 | 14.808056 | 14.608468 | 14.763820 | 34137 | 31816 | 20303 | 26270 |
| 1509 | NM_001909 | CTSD | cathepsin D | protein_coding | NP_001900.1 | 2529 | 116840 | . | . | 1635.461670 | 1630.445190 | 1466.380981 | 1615.814331 | 3805.786377 | 3768.847168 | 3290.516113 | 3694.367432 | -1.074749 | 101608.397655 | 0.100948747 | -1.03021796 | 0.302907702 | 0.998865404 | 16.683725 | 16.579721 | 16.667938 | 16.657802 | 16.562336 | 16.637129 | 121921 | 112930 | 81478 | 95810 |
| 1510 | NM_00131733 | CTSE | cathepsin E | protein_coding | NP_001304260.1;NP_001901.1;NP_683865.1;XP_011507546.1;XP_01150 | 2530 | 116890 | . | . | 0.337280 | 0.279944 | 0.144637 | 0.136805 | 0.784863 | 0.647102 | 0.324561 | 0.312789 | -2.286702 | 16.380744 | 0.777488811 | -1.53477215 | 0.124839806 | NA | 4.510114 | 3.318300 | 4.078688 | 4.057811 | 4.015267 | 4.011486 | 29 | 22 | 9 | 9 |
| 1511 | NM_001911,XI | CTSG | cathepsin G | protein_coding | NP_001902.1;XP_011534801.1 | 2532 | 116830 | . | . | 6.607525 | 5.942626 | 4.421860 | 6.276053 | 15.375982 | 13.736649 | 9.922526 | 14.349448 | -1.189957 | 166.285550 | 0.273092066 | -0.9187704 | 0.358215679 | 0.998865404 | 7.498164 | 7.245857 | 7.435677 | 7.385585 | 7.273767 | 7.401127 | 220 | 184 | 110 | 166 |
| 1512 | NM_00131913 | CTSH | cathepsin H | protein_coding | NP_001306066.1;NP_004381.2;XP_005254238.1;XP_016877440.1;XP_01 | 2535 | 116820 | . | . | 131.818641 | 108.465204 | 96.673697 | 111.809686 | 306.747382 | 250.722178 | 216.932971 | 255.639558 | -1.139375 | 6600.850766 | 0.120879715 | -1.55727331 | 0.119405609 | 0.858861713 | 12.779544 | 12.591185 | 12.756322 | 12.723400 | 12.546312 | 12.706914 | 8252 | 7441 | 4850 | 6341 |
| 1513 | NM_000396 | CTSK | cathepsin K | protein_coding | NP_000387.1 | 2536 | 601105 | . | . | 1.778491 | 2.105526 | 2.696369 | 2.564826 | 4.138621 | 4.867019 | 6.050574 | 5.864171 | 1.332922 | 116.367729 | 0.301680435 | 1.374274944 | 0.169356342 | 0.946181982 | 6.640659 | 7.054814 | 6.780465 | 6.828810 | 6.911277 | 6.890770 | 106 | 116 | 119 | 121 |
| 1514 | NM_00125797 | CTSL | cathepsin L | protein_coding | NP_001244900.1;NP_001244901.1;NP_001244902.1;NP_001369686.1;NI | 2537 | 116880 | . | . | 101.993065 | 102.001885 | 101.350003 | 87.987028 | 237.342044 | 235.781930 | 227.426477 | 201.171882 | -1.087924 | 4831.778973 | 0.116049056 | -1.04764613 | 0.294801673 | 0.998865404 | 12.297775 | 12.176348 | 12.274843 | 12.268925 | 12.260908 | 12.138407 | 5818 | 5414 | 4322 | 4019 |
| 1515 | NM_00120157 | CTSV | cathepsin V | protein_coding | NP_001188504.1;NP_001324.2 | 2538 | 603308 | . | . | 6.250299 | 5.953246 | 4.992931 | 5.707646 | 14.544703 | 13.761194 | 11.203993 | 13.049854 | -1.155889 | 778.514840 | 0.151244603 | -1.38188601 | 0.167006703 | 0.945199193 | 9.705452 | 9.496123 | 9.673541 | 9.631955 | 9.508283 | 9.592579 | 989 | 875 | 589 | 719 |
| 1519 | NM_001334 | CTSO | cathepsin O | protein_coding | NP_001325.1 | 2542 | 600550 | . | . | 0.585255 | 0.493608 | 0.646594 | 0.426239 | 1.361912 | 1.140998 | 1.450939 | 0.974544 | -1.024992 | 49.075651 | 0.461161278 | -0.07722286 | 0.938446253 | 0.998865404 | 5.633052 | 5.600637 | 5.631805 | 5.599813 | 5.647595 | 5.573878 | 62 | 49 | 51 | 36 |
| 1520 | NM_00119973 | CTSS | cathepsin S | protein_coding | NP_001186668.1;NP_004070.3 | 2545 | 116845 | . | . | 0.462045 | 0.387879 | 0.301698 | 0.228416 | 1.075197 | 0.896601 | 0.677001 | 0.522246 | -1.583622 | 43.064266 | 0.484917808 | -1.36771189 | 0.171402276 | 0.947127937 | 5.721207 | 5.060678 | 5.482550 | 5.447960 | 5.409372 | 5.373515 | 66 | 52 | 33 | 27 |
| 1522 | NM_001336 | CTSZ | cathepsin Z | protein_coding | NP_001327.2 | 2547 | 603169 | . | . | 286.791992 | 289.425964 | 267.303619 | 269.477844 | 667.376709 | 669.021118 | 599.821594 | 616.129028 | -1.088567 | 13023.115233 | 0.099369525 | -1.23207415 | 0.217921387 | 0.982008277 | 13.728704 | 13.606274 | 13.703777 | 13.704305 | 13.632336 | 13.630519 | 15627 | 14653 | 10856 | 11679 |
| 1523 | NM_00120254 | CUX1 | cut like homeobox 1 | protein_coding | NP_001189472.1;NP_001189473.1;NP_001189474.1;NP_001189475.1;NI | 2557 | 116896 | . | . | 2.951357 | 2.938014 | 2.659902 | 3.396793 | 6.867928 | 6.791351 | 5.968744 | 7.766362 | 1.013186 | 939.350150 | 0.149733217 | 0.126215528 | 0.89956132 | 0.998865404 | 9.866367 | 9.884614 | 9.847991 | 9.891068 | 9.811828 | 9.942176 | 1043 | 1037 | 748 | 966 |
| 1525 | NM_00120706 | CXADR | CXADR Ig-like cell adhesion molecule | protein_coding | NP_001193992.1;NP_001193993.1;NP_001193994.1;NP_001193995.1;NI | 2559 | 602621 | . | . | 2.326056 | 2.100563 | 2.011985 | 1.337241 | 5.412831 | 4.855547 | 4.514838 | 3.057444 | -1.321415 | 268.449831 | 0.250754035 | -1.60349834 | 0.108824675 | 0.841988225 | 8.254569 | 7.854849 | 8.196137 | 8.055664 | 8.081258 | 7.879488 | 396 | 288 | 244 | 171 |
| 1528 | NM_00119080 | CYB5A | cytochrome b5 type A | protein_coding | NP_001177736.1;NP_001905.1;NP_683725.1;XP_011524137.1 | 2570 | 613218 | . | . | 4.846745 | 5.896986 | 4.826647 | 5.750011 | 11.278574 | 13.631149 | 10.830856 | 13.146714 | -1.027588 | 502.435467 | 0.187246107 | -0.20968231 | 0.833915634 | 0.998865404 | 8.992917 | 8.952388 | 8.917240 | 9.037297 | 8.888010 | 9.030118 | 532 | 601 | 382 | 517 |
| 1534 | NM_00101791 | CYB561 | cytochrome b561 | protein_coding | NP_001017916.1;NP_001017917.1;NP_001317350.1;NP_001906.3 | 2571 | 600019 | . | . | 3.592922 | 3.195629 | 4.471786 | 4.209323 | 8.360878 | 7.386841 | 10.034558 | 9.624117 | 1.230031 | 357.081309 | 0.189961743 | 1.572391831 | 0.115859708 | 0.857396579 | 8.322719 | 8.622005 | 8.434341 | 8.394951 | 8.554043 | 8.511728 | 382 | 333 | 355 | 356 |
| 1535 | NM_000101,XI | CYBA | cytochrome b-245 alpha chain | protein_coding | NP_000092.2;XP_011521207.1 | 2577 | 608508 | . | . | 125.449397 | 122.029124 | 114.545776 | 126.429918 | 291.925885 | 282.075791 | 257.037398 | 289.066993 | -1.030968 | 2859.074618 | 0.113350905 | -0.3881747 | 0.697886752 | 0.998865404 | 11.503178 | 11.459151 | 11.510740 | 11.476124 | 11.450569 | 11.486523 | 3413 | 3066 | 2394 | 2698 |
| 1537 | NM_001916,XI | CYC1 | cytochrome c1 | protein_coding | NP_001907.3;XP_016868591.1;XP_024302840.1 | 2579 | 123980 | . | . | 89.987023 | 90.065566 | 76.266091 | 96.242580 | 209.403516 | 208.190591 | 171.138909 | 220.047231 | -1.070172 | 3387.425579 | 0.128381984 | -0.76211733 | 0.445989978 | 0.998865404 | 11.774179 | 11.676101 | 11.756929 | 11.747070 | 11.605394 | 11.779420 | 4057 | 3757 | 2543 | 3394 |
| 1540 | NM_00104235 | CYLD | CYLD lysine 63 deubiquitinase | protein_coding | NP_001035814.1;NP_001035877.1;NP_001365672.1;NP_001365673.1;NI | 2584 | 605018 | . | . | 1.403469 | 1.638908 | 1.713697 | 1.673552 | 3.265932 | 3.788413 | 3.845487 | 3.826380 | 1.108434 | 427.538514 | 0.179215452 | 0.828736429 | 0.407253567 | 0.998865404 | 8.663909 | 8.812108 | 8.671051 | 8.740600 | 8.773580 | 8.763926 | 440 | 463 | 394 | 418 |
| 1543 | NM_000499,N | CYP1A1 | cytochrome P450 family 1 subfamily A member 1 | protein_coding | NP_000490.1;NP_001306145.1;NP_001306146.1 | 2595 | 108330 | . | . | 0.743947 | 0.666414 | 1.054662 | 0.901209 | 1.731194 | 1.540447 | 2.366632 | 2.060506 | 1.372611 | 69.046357 | 0.382671844 | 1.194032906 | 0.232465096 | 0.988209339 | 5.862170 | 6.320558 | 6.074724 | 6.049793 | 6.156684 | 6.116302 | 71 | 59 | 75 | 69 |
| 1545 | NM_000104 | CYP1B1 | cytochrome P450 family 1 subfamily B member 1 | protein_coding | NP_000095.2 | 2597 | 601771 | . | . | 0.010566 | 0.017355 | 0.021630 | 0.022951 | 0.024587 | 0.040117 | 0.048537 | 0.052474 | 1.633271 | 3.577607 | 1.594525897 | 0.443871507 | 0.657135486 | NA | 1.444631 | 2.148375 | 1.817216 | 1.826716 | 1.830642 | 1.829233 | 2 | 4 | 4 | 4 |
| 1562 | NM_000772,N | CYP2C18 | cytochrome P450 family 2 subfamily C member 18 | protein_coding | NP_000763.1;NP_001122397.1 | 2620 | 601131 | . | . | 0.036745 | 0.075919 | 0.110331 | 0.060013 | 0.085508 | 0.175490 | 0.247580 | 0.137212 | 1.481702 | 5.604348 | 1.28298601 | 0.442136823 | 0.6583902 | NA | 2.175403 | 2.742346 | 2.464234 | 2.474203 | 2.485124 | 2.473179 | 4 | 6 | 7 | 5 |
| 1564 | NM_00134838 | CYP2D7 | cytochrome P450 family 2 subfamily D member 7 (gene/pseudoge | protein_coding | NP_001335315.1 | 2624 | . | | . | 0.465372 | 0.952437 | 0.683175 | 0.348628 | 1.082941 | 2.201601 | 1.533024 | 0.797096 | -1.407519 | 43.805578 | 0.535982939 | -0.92009279 | 0.357524274 | 0.998865404 | 5.677932 | 5.186506 | 5.404014 | 5.528909 | 5.456771 | 5.365003 | 40 | 73 | 42 | 23 |
| 1565 | NM_000106,N | CYP2D6 | cytochrome P450 family 2 subfamily D member 6 | protein_coding | NP_000097.3;NP_001020332.2 | 2625 | 124030 | . | . | 0.752809 | 0.745886 | 0.519392 | 0.697185 | 1.751817 | 1.724149 | 1.165500 | 1.594031 | -1.241227 | 33.965999 | 0.535298752 | -0.58241769 | 0.560285372 | 0.998865404 | 5.234218 | 4.920835 | 5.105825 | 5.101066 | 5.059459 | 5.089299 | 44 | 40 | 23 | 32 |
| 1568 | gene-CYP2D8P | CYP2D8P | ccytochrome P450 family 2 subfamily D member 8, pseudogene | pseudogene | . | 2628 | . | | . | 0.161113 | 0.549672 | 0.232209 | 0.326054 | 0.374916 | 1.270592 | 0.521070 | 0.745483 | -1.222727 | 15.316146 | 0.850511184 | -0.34109213 | 0.733034224 | NA | 4.077449 | 3.781347 | 3.894811 | 3.974902 | 3.912947 | 3.931940 | 9 | 28 | 10 | 15 |
| 1573 | NM_000775,N | CYP2J2 | cytochrome P450 family 2 subfamily J member 2 | protein_coding | NP_000766.2 | 2634 | 601258 | . | . | 5.137600 | 4.519278 | 4.178891 | 5.154376 | 11.955406 | 10.446517 | 9.377309 | 11.784870 | -1.052470 | 280.823928 | 0.215370332 | -0.34257075 | 0.731921407 | 0.998865404 | 8.170095 | 8.095998 | 8.184662 | 8.105956 | 8.075664 | 8.161125 | 356 | 288 | 218 | 278 |
| 1593 | NM_000784,XI | CYP27A1 | cytochrome P450 family 27 subfamily A member 1 | protein_coding | NP_000775.1;XP_016858977.1 | 2605 | 606530 | . | . | 72.695426 | 72.473999 | 73.171549 | 82.431128 | 169.165243 | 167.526894 | 164.194830 | 188.468990 | 1.057207 | 4546.603214 | 0.112325223 | 0.714518821 | 0.474906422 | 0.998865404 | 12.109946 | 12.190088 | 12.127173 | 12.125876 | 12.124889 | 12.218760 | 5094 | 4766 | 3836 | 4626 |
| 1594 | NM_000785 | CYP27B1 | cytochrome P450 family 27 subfamily B member 1 | protein_coding | NP_000776.1 | 2606 | 609506 | . | . | 0.409051 | 0.644977 | 0.682814 | 0.299309 | 0.951879 | 1.490893 | 1.532215 | 0.684335 | -1.090407 | 38.113380 | 0.558010859 | -0.22377051 | 0.822935879 | 0.998865404 | 5.312060 | 5.189802 | 5.213374 | 5.280748 | 5.287586 | 5.178864 | 36 | 52 | 44 | 21 |
| 1595 | NM_000786,N | CYP51A1 | cytochrome P450 family 51 subfamily A member 1 | protein_coding | NP_000777.1;NP_001139624.1 | 2649 | 601637 | . | . | 37.463502 | 36.019725 | 63.080194 | 44.446194 | 87.179103 | 83.261203 | 141.550128 | 101.620944 | 1.436528 | 4410.083785 | 0.14999213 | 3.484087617 | 0.000493818 | 0.040572423 | 11.821638 | 12.344437 | 11.951054 | 11.912913 | 12.374692 | 12.071565 | 4267 | 3813 | 5313 | 4023 |
| 1600 | NM_00135398 | DAB1 | DAB adaptor protein 1 | protein_coding | NP_001340909.1;NP_001340912.1;NP_001340914.1;NP_001340915.1;NI | 2661 | 603448 | . | . | 0.000231 | 0.000226 | 0.054409 | 0.088980 | 0.000539 | 0.000522 | 0.122093 | 0.203443 | 9.323510 | 4.636240 | 1.654324319 | 1.946941791 | 0.051541714 | NA | -0.155866 | 3.066070 | 2.146985 | 2.147558 | 2.190648 | 2.173403 | 1 | 1 | 9 | 6 |
| 1601 | NM_00124487 | DAB2 | DAB adaptor protein 2 | protein_coding | NP_001231800.1;NP_001334.2 | 2662 | 601236 | . | . | 11.142685 | 12.886761 | 11.378849 | 11.945929 | 25.929485 | 29.788326 | 25.533808 | 27.312946 | -1.052698 | 1631.575834 | 0.129662655 | -0.57141468 | 0.56771858 | 0.998865404 | 10.708755 | 10.634387 | 10.637552 | 10.742279 | 10.639751 | 10.662328 | 1799 | 1926 | 1362 | 1512 |
| 1603 | NM_001344 | DAD1 | defender against cell death 1 | protein_coding | NP_001335.1 | 2664 | 600243 | . | . | 122.809914 | 121.464516 | 131.728348 | 112.167694 | 285.783722 | 280.770691 | 295.594604 | 256.458099 | -1.015444 | 2601.727932 | 0.125662659 | -0.17595784 | 0.860327066 | 0.998865404 | 11.356128 | 11.334298 | 11.358369 | 11.342471 | 11.404900 | 11.266719 | 3048 | 2801 | 2437 | 2214 |
| 1604 | NM_000574,N | CD55 | CD55 molecule (Cromer blood group) | protein_coding | NP_000565.1;NP_001108224.1;NP_001287831.1;NP_001287832.1;NP_0( | 2665 | 125240 | . | . | 17.872805 | 16.824252 | 18.527777 | 15.931089 | 41.590754 | 38.890013 | 41.575793 | 36.424543 | -1.060030 | 1308.434852 | 0.131991169 | -0.63719825 | 0.523995716 | 0.998865404 | 10.394892 | 10.311145 | 10.396969 | 10.352659 | 10.363844 | 10.296771 | 1595 | 1411 | 1154 | 1138 |
| 1605 | NM_00116592 | DAG1 | dystroglycan 1 | protein_coding | NP_001159400.2;NP_001171105.1;NP_001171106.1;NP_001171107.1;NI | 2666 | 128239 | . | . | 69.029531 | 69.774378 | 59.416750 | 69.519192 | 160.634552 | 161.286601 | 133.329471 | 158.947374 | -1.090202 | 11631.849739 | 0.110153804 | -1.13110728 | 0.258009943 | 0.995760118 | 13.566779 | 13.442115 | 13.533684 | 13.548379 | 13.407336 | 13.523812 | 13853 | 13205 | 9014 | 11150 |
| 1606 | NM_001345,N | DGKA | diacylglycerol kinase alpha | protein_coding | NP_001336.2;NP_001337962.1;NP_001337963.1;NP_001337964.1;NP_0( | 2849 | 125855 | . | . | 2.178820 | 1.884263 | 2.212422 | 2.340784 | 5.070205 | 4.355558 | 4.964613 | 5.351924 | 1.097938 | 191.332795 | 0.24627282 | 0.547347751 | 0.584139844 | 0.998865404 | 7.510575 | 7.646123 | 7.597608 | 7.514511 | 7.597652 | 7.598857 | 230 | 178 | 174 | 188 |
| 1608 | NM_00108074 | DGKG | diacylglycerol kinase gamma | protein_coding | NP_001074213.1;NP_001074214.1;NP_001337.2 | 2853 | 601854 | . | . | 1.290330 | 1.253858 | 1.310047 | 1.388479 | 3.002651 | 2.898349 | 2.939708 | 3.174592 | 1.045821 | 237.405898 | 0.220265985 | 0.293447469 | 0.769180146 | 0.998865404 | 7.858589 | 7.923114 | 7.888326 | 7.871063 | 7.889775 | 7.913266 | 272 | 246 | 206 | 233 |
| 1609 | NM_001347,XI | DGKQ | diacylglycerol kinase theta | protein_coding | NP_001338.2;XP_011511713.1;XP_011511714.1;XP_011511716.1;XP_01: | 2856 | 601207 | . | . | 4.588809 | 4.239813 | 4.140747 | 4.043268 | 10.678346 | 9.800516 | 9.291718 | 9.244452 | -1.078302 | 588.169857 | 0.160945884 | -0.6757627 | 0.49919132 | 0.998865404 | 9.253228 | 9.144918 | 9.252069 | 9.195788 | 9.191318 | 9.157744 | 732 | 631 | 505 | 517 |
| 1611 | NM_00129196 | DAP | death associated protein | protein_coding | NP_001278892.1;NP_004385.1 | 2672 | 600954 | . | . | 15.860778 | 15.782183 | 14.466459 | 19.483078 | 36.908686 | 36.481223 | 32.462312 | 44.545744 | 1.058674 | 1170.275650 | 0.155429282 | 0.529232933 | 0.596643873 | 0.998865404 | 10.151212 | 10.232897 | 10.173294 | 10.163537 | 10.100632 | 10.310776 | 1318 | 1219 | 901 | 1285 |
| 1612 | NM_00128872 | DAPK1 | death associated protein kinase 1 | protein_coding | NP_001275658.1;NP_001275659.1;NP_001275660.1;NP_004929.2;XP_0( | 2674 | 600831 | . | . | 7.049138 | 7.102930 | 8.127365 | 8.307763 | 16.403635 | 16.418743 | 18.237572 | 18.994711 | 1.136327 | 1396.911292 | 0.126267967 | 1.460210574 | 0.14423221 | 0.906161673 | 10.352941 | 10.537227 | 10.395544 | 10.397577 | 10.481569 | 10.507150 | 1504 | 1413 | 1271 | 1416 |
| 1613 | NM_001348,N | DAPK3 | death associated protein kinase 3 | protein_coding | NP_001339.1;NP_001362587.1 | 2676 | 603289 | . | . | 14.144730 | 13.970091 | 11.279720 | 14.170566 | 32.915368 | 32.292490 | 25.311364 | 32.399313 | -1.119334 | 921.086797 | 0.154364861 | -1.05361451 | 0.292059443 | 0.998865404 | 9.926529 | 9.763240 | 9.889065 | 9.881811 | 9.721372 | 9.880023 | 1126 | 1045 | 671 | 906 |
| 1615 | NM_00129331 | DARS1 | aspartyl-tRNA synthetase 1 | protein_coding | NP_001280241.1;NP_001340.2 | 2678 | 603084 | . | . | 23.257577 | 24.513019 | 24.683804 | 24.389885 | 54.121340 | 56.662940 | 55.389740 | 55.764572 | 1.013028 | 2337.733881 | 0.115851325 | 0.161189221 | 0.87194437 | 0.998865404 | 11.181537 | 11.200193 | 11.167803 | 11.203011 | 11.204957 | 11.186891 | 2616 | 2561 | 2069 | 2181 |
| 1616 | NM_00114196 | DAXX | death domain associated protein | protein_coding | NP_001135441.1;NP_001135442.1;NP_001241646.1;NP_001341.1;XP_0( | 2681 | 603186 | . | . | 14.023877 | 14.300270 | 13.708650 | 16.734649 | 32.634139 | 33.055715 | 30.761811 | 38.261786 | 1.037644 | 1156.504266 | 0.138753363 | 0.384220164 | 0.700815274 | 0.998865404 | 10.148863 | 10.201762 | 10.157875 | 10.163955 | 10.132719 | 10.241330 | 1302 | 1230 | 950 | 1184 |
| 1617 | NM_004081,XI | DAZ1 | deleted in azoospermia 1 | protein_coding | NP_004072.3;XP_011529784.1;XP_011529785.1 | 2682 | 400003 | . | . | 0.589031 | 0.371182 | 0.568210 | 0.414300 | 1.370699 | 0.858004 | 1.275047 | 0.947248 | 1.009982 | 68.054097 | 0.412329667 | 0.034754346 | 0.972275625 | 0.998865404 | 6.079678 | 6.097487 | 6.135419 | 6.024900 | 6.119790 | 6.046754 | 96 | 56 | 69 | 54 |
| 1622 | NM_00107986 | DBI | diazepam binding inhibitor, acyl-CoA binding protein | protein_coding | NP_001073331.1;NP_001073332.1;NP_001171488.1;NP_001171512.1;NI | 2690 | 125950 | . | . | 415.795856 | 406.126598 | 432.343470 | 475.572061 | 967.574143 | 938.779733 | 970.166188 | 1087.339088 | 1.091159 | 7904.654030 | 0.105693971 | 1.190803226 | 0.233730842 | 0.988893316 | 12.884204 | 13.010026 | 12.921992 | 12.898097 | 12.949791 | 13.017317 | 8829 | 8042 | 6890 | 8039 |
| 1627 | NM_00136354 | DBN1 | drebrin 1 | protein_coding | NP_001350470.2;NP_001351080.2;NP_001351081.2;NP_004386.3;NP_5¿ | 2695 | 126660 | . | . | 52.278874 | 51.641379 | 38.710241 | 49.993300 | 121.655094 | 119.371356 | 86.864660 | 114.303749 | -1.188476 | 4336.056907 | 0.131149535 | -1.89945726 | 0.05750438 | 0.711571492 | 12.201451 | 11.952127 | 12.157162 | 12.139079 | 11.899567 | 12.099299 | 5481 | 5028 | 3026 | 4167 |
| 1628 | NM_001352,XI | DBP | D-box binding PAR bZIP transcription factor | protein_coding | NP_001343.2;XP_016881877.1 | 2697 | 124097 | . | . | 3.634185 | 3.082164 | 2.730136 | 4.116402 | 8.456898 | 7.124560 | 6.126347 | 9.411665 | 1.001842 | 224.597201 | 0.253129811 | 0.010489085 | 0.991631074 | 0.999576129 | 7.810441 | 7.811951 | 7.844572 | 7.767843 | 7.713374 | 7.894103 | 279 | 223 | 158 | 251 |
| 1629 | NM_001918,XI | DBT | dihydrolipoamide branched chain transacylase E2 | protein_coding | NP_001909.3;XP_005270602.1;XP_016855957.1;XP_016855958.1 | 2698 | 248610 | . | . | 0.935815 | 0.948061 | 1.418135 | 1.008996 | 2.177681 | 2.191487 | 3.182255 | 2.306949 | 1.126332 | 324.635302 | 0.194749625 | 0.881297642 | 0.378156742 | 0.998865404 | 8.254199 | 8.426039 | 8.306357 | 8.308340 | 8.385932 | 8.360220 | 351 | 330 | 306 | 315 |
| 1632 | NM_00117802 | ECI1 | enoyl-CoA delta isomerase 1 | protein_coding | NP_001171500.1;NP_001910.2 | 2703 | 600305 | . | . | 12.323274 | 12.070998 | 8.514182 | 13.579687 | 28.676765 | 27.902651 | 19.105576 | 31.048339 | -1.119761 | 545.218107 | 0.205918286 | -0.79250073 | 0.428068768 | 0.998865404 | 9.170548 | 9.006150 | 9.131646 | 9.110449 | 8.895659 | 9.179699 | 674 | 612 | 347 | 590 |
| 1633 | NM_000788 | DCK | deoxycytidine kinase | protein_coding | NP_000779.1 | 2704 | 125450 | . | . | 9.385838 | 8.943168 | 7.987092 | 6.749541 | 21.841230 | 20.672531 | 17.922804 | 15.432022 | -1.260484 | 646.788702 | 0.163315358 | -2.04498383 | 0.040856452 | 0.62818612 | 9.494124 | 9.160989 | 9.428653 | 9.388300 | 9.308590 | 9.193823 | 854 | 756 | 542 | 489 |
| 1634 | NM_001920,N | DCN | decorin | protein_coding | NP_001911.1;NP_598010.1;NP_598011.1;NP_598012.1;NP_598013.1;NP_ | 2705 | 125255 | . | . | 0.942251 | 0.721231 | 0.740045 | 0.846818 | 2.192658 | 1.667157 | 1.660639 | 1.936147 | -1.131492 | 117.240565 | 0.298501106 | -0.59706913 | 0.550461228 | 0.998865404 | 6.959700 | 6.781454 | 6.908876 | 6.880498 | 6.844549 | 6.861876 | 149 | 129 | 93 | 106 |
| 1635 | NM_00101273 | DCTD | dCMP deaminase | protein_coding | NP_001012750.1;NP_001338672.1;NP_001338673.1;NP_001338674.1;NI | 2710 | 607638 | . | . | 19.192609 | 18.044907 | 20.007059 | 22.388379 | 44.661989 | 41.711617 | 44.895259 | 51.188375 | 1.141432 | 1414.962091 | 0.130128624 | 1.4665891 | 0.142487867 | 0.903146012 | 10.368131 | 10.558677 | 10.404829 | 10.420729 | 10.472102 | 10.554817 | 1506 | 1441 | 1242 | 1489 |
| 1638 | NM_00112988 | DCT | dopachrome tautomerase | protein_coding | NP_001123361.1;NP_001309111.1;NP_001309112.1;NP_001309113.1;NI | 2709 | 191275 | . | . | 147.504560 | 133.784752 | 186.325668 | 168.391700 | 343.249176 | 309.249439 | 418.109393 | 385.007598 | 1.236121 | 24381.317437 | 0.107594393 | 2.842342145 | 0.00447834 | 0.198760547 | 14.412471 | 14.718326 | 14.518314 | 14.429467 | 14.692305 | 14.621467 | 26427 | 22296 | 24388 | 24250 |
| 1639 | NM_00113504 | DCTN1 | dynactin subunit 1 | protein_coding | NP_001128512.1;NP_001128513.1;NP_001177765.1;NP_001177766.1;NI | 2711 | 601143 | . | . | 38.152055 | 39.481776 | 36.302585 | 41.849843 | 88.781398 | 91.263891 | 81.461950 | 95.684689 | -1.008280 | 5038.313590 | 0.113995689 | -0.10435544 | 0.916887282 | 0.998865404 | 12.304732 | 12.292693 | 12.290123 | 12.313092 | 12.238442 | 12.348070 | 5746 | 5534 | 4077 | 5012 |
| 1641 | NM_000555,N | DCX | doublecortin | protein_coding | NP_000546.2;NP_001182482.1;NP_001356299.1;NP_001356300.1;NP_0( | 2714 | 300121 | . | . | 0.078954 | 0.051250 | 0.037335 | 0.081718 | 0.183730 | 0.118466 | 0.083778 | 0.186838 | -1.110002 | 19.026031 | 0.739032716 | -0.20372808 | 0.838565996 | NA | 4.324488 | 4.171250 | 4.273509 | 4.236286 | 4.219241 | 4.273317 | 28 | 17 | 10 | 23 |
| 1642 | NM_001923 | DDB1 | damage specific DNA binding protein 1 | protein_coding | NP_001914.3 | 2717 | 600045 | . | . | 46.556671 | 48.773331 | 41.149239 | 49.821495 | 108.339287 | 112.741737 | 92.337700 | 113.910927 | -1.062869 | 6159.207308 | 0.11951255 | -0.73602693 | 0.461714312 | 0.998865404 | 12.631914 | 12.543799 | 12.596504 | 12.628711 | 12.482800 | 12.634187 | 7170 | 6979 | 4724 | 6103 |
| 1643 | NM_000107,N | DDB2 | damage specific DNA binding protein 2 | protein_coding | NP_000098.1;NP_001287663.1 | 2718 | 600811 | . | . | 7.889833 | 7.716583 | 8.535647 | 8.739025 | 18.359966 | 17.837228 | 19.153746 | 19.980742 | 1.099788 | 453.578017 | 0.172449512 | 0.795746299 | 0.42617951 | 0.998865404 | 8.754980 | 8.892173 | 8.801025 | 8.787614 | 8.845076 | 8.860836 | 503 | 461 | 408 | 451 |
| 1645 | NM_001353,XI | AKR1C1 | aldo-keto reductase family 1 member C1 | protein_coding | NP_001344.2;XP_016871280.1 | 384 | 600449 | . | . | 0.355809 | 0.489965 | 0.577675 | 0.543935 | 0.827982 | 1.132576 | 1.296287 | 1.243642 | 1.306844 | 100.786705 | 0.330336836 | 1.168767509 | 0.242497314 | 0.992733415 | 6.449832 | 6.834876 | 6.558654 | 6.647440 | 6.697871 | 6.673709 | 85 | 109 | 103 | 103 |
| 1646 | NM_00113524 | AKR1C2 | aldo-keto reductase family 1 member C2 | protein_coding | NP_001128713.1;NP_001307956.1;NP_001345.1;NP_995317.1 | 385 | 600450 | . | . | 0.219987 | 0.283859 | 0.514090 | 0.586589 | 0.511919 | 0.656152 | 1.153602 | 1.341167 | 2.168866 | 45.600735 | 0.471913032 | 2.36683637 | 0.017940863 | 0.447807832 | 4.848520 | 5.963267 | 5.400379 | 5.431652 | 5.531066 | 5.558464 | 29 | 35 | 51 | 62 |
| 1647 | NM_00119974 | GADD45A | growth arrest and DNA damage inducible alpha | protein_coding | NP_001186670.1;NP_001186671.1;NP_001915.1 | 4095 | 126335 | . | . | 22.801853 | 21.786891 | 18.140523 | 21.266904 | 53.060845 | 50.361380 | 40.706806 | 48.624251 | -1.143484 | 881.972863 | 0.1483024 | -1.30433601 | 0.192119043 | 0.96570253 | 9.878249 | 9.684427 | 9.849434 | 9.810897 | 9.681521 | 9.784012 | 1112 | 989 | 663 | 828 |
| 1649 | NM_00119505 | DDIT3 | DNA damage inducible transcript 3 | protein_coding | NP_001181982.1;NP_001181983.1;NP_001181984.1;NP_001181985.1;NI | 2726 | 126337 | . | . | 21.075120 | 20.678908 | 22.444508 | 25.001708 | 49.042667 | 47.800220 | 50.364824 | 57.163434 | 1.106854 | 717.673450 | 0.151661012 | 0.965737617 | 0.334175491 | 0.998865404 | 9.412262 | 9.558406 | 9.453206 | 9.447353 | 9.485956 | 9.553521 | 788 | 732 | 624 | 741 |
| 1650 | NM_005216 | DDOST | dolichyl-diphosphooligosaccharide--protein glycosyltransferase no | protein_coding | NP_005207.3 | 2728 | 602202 | . | . | 426.095612 | 429.524658 | 375.922150 | 392.297394 | 991.541992 | 992.865479 | 843.558411 | 896.941406 | -1.129510 | 24555.245095 | 0.09832346 | -1.78693227 | 0.073948437 | 0.763883535 | 14.668924 | 14.493217 | 14.634177 | 14.633693 | 14.515182 | 14.542786 | 30003 | 28100 | 19729 | 21971 |
| 1652 | NM_00108439 | DDT | D-dopachrome tautomerase | protein_coding | NP_001077861.1;NP_001346.1;NP_001368781.1 | 2732 | 602750 | . | . | 120.485283 | 121.811941 | 118.784467 | 136.632572 | 280.374196 | 281.573756 | 266.548901 | 312.394160 | -1.020391 | 2482.231173 | 0.13632622 | -0.2136228 | 0.830841222 | 0.998865404 | 11.292074 | 11.262619 | 11.282905 | 11.284825 | 11.165140 | 11.360488 | 2884 | 2709 | 1878 | 2582 |
| 1653 | NM_004939 | DDX1 | DEAD-box helicase 1 | protein_coding | NP_004930.1 | 2734 | 601257 | . | . | 55.788269 | 53.552551 | 51.992699 | 55.536564 | 129.821594 | 123.789131 | 116.670113 | 126.977760 | -1.031442 | 4196.040533 | 0.109661572 | -0.4072738 | 0.683806906 | 0.998865404 | 12.056985 | 12.012295 | 12.067888 | 12.026533 | 11.998301 | 12.044525 | 5028 | 4484 | 3492 | 3981 |
| 1654 | NM_00119341 | DDX3X | DEAD-box helicase 3 X-linked | protein_coding | NP_001180345.1;NP_001180346.1;NP_001347.3;NP_001350748.1;XP_01 | 2745 | 300160 | . | . | 38.000387 | 39.524012 | 40.069113 | 35.874784 | 88.428461 | 91.361520 | 89.913930 | 82.023428 | -1.004853 | 6057.481061 | 0.1149059 | -0.06078187 | 0.951532931 | 0.998865404 | 12.567924 | 12.561072 | 12.578194 | 12.553262 | 12.622229 | 12.497568 | 7095 | 6455 | 5644 | 5246 |
| 1655 | NM_00132059 | DDX5 | DEAD-box helicase 5 | protein_coding | NP_001307524.1;NP_001307525.1;NP_001307526.1;NP_004387.1 | 2746 | 180630 | . | . | 137.783411 | 137.711149 | 141.414276 | 135.323593 | 320.627688 | 318.325496 | 317.329528 | 309.401312 | -1.007734 | 15907.667301 | 0.100659179 | -0.11041572 | 0.912079689 | 0.998865404 | 13.962969 | 13.951879 | 13.963621 | 13.957630 | 13.977653 | 13.929853 | 18452 | 17171 | 14164 | 14442 |
| 1656 | NM_00125719 | DDX6 | DEAD-box helicase 6 | protein_coding | NP_001244120.1;NP_004388.2;XP_005271474.1;XP_011540946.1;XP_01 | 2747 | 600326 | . | . | 7.079570 | 7.586561 | 7.809345 | 6.796276 | 16.474446 | 17.536673 | 17.523945 | 15.538877 | -1.015652 | 1352.242115 | 0.135428103 | -0.16544829 | 0.868591148 | 0.998865404 | 10.412111 | 10.390080 | 10.404271 | 10.408154 | 10.450911 | 10.334749 | 1565 | 1474 | 1256 | 1162 |
| 1657 | NM_00129032 | DMXL1 | Dmx like 1 | protein_coding | NP_001277250.1;NP_001277251.1;NP_001336168.1;NP_001336169.1;NI | 2937 | 605671 | . | . | 3.478442 | 3.530685 | 3.916438 | 2.964858 | 8.094476 | 8.161339 | 8.788374 | 6.778795 | -1.016577 | 1241.241687 | 0.14832884 | -0.15990981 | 0.872952123 | 0.998865404 | 10.289105 | 10.265939 | 10.282996 | 10.280603 | 10.355886 | 10.175234 | 1443 | 1348 | 1199 | 1016 |
| 1659 | NM_00130262 | DHX8 | DEAH-box helicase 8 | protein_coding | NP_001289552.1;NP_001309145.1;NP_001309146.1;NP_001309147.1;NI | 2749 | 600396 | . | . | 9.259522 | 10.193036 | 10.643661 | 10.982226 | 21.547287 | 23.561659 | 23.884068 | 25.109556 | 1.088057 | 1531.711104 | 0.126490575 | 0.962555716 | 0.33577053 | 0.998865404 | 10.518897 | 10.640403 | 10.519459 | 10.573675 | 10.593427 | 10.630599 | 1630 | 1639 | 1355 | 1532 |
| 1660 | NM_001357,N | DHX9 | DExH-box helicase 9 | protein_coding | NP_001348.2 | 2750 | 603115 | . | . | 45.885579 | 46.790609 | 43.210907 | 45.172592 | 106.777635 | 108.158583 | 96.964022 | 103.281757 | -1.062514 | 6363.795875 | 0.103787209 | -0.84290219 | 0.39928315 | 0.998865404 | 12.678769 | 12.591248 | 12.655191 | 12.666202 | 12.595876 | 12.623048 | 7501 | 7121 | 5279 | 5879 |
| 1662 | NM_004398 | DDX10 | DEAD-box helicase 10 | protein_coding | NP_004389.2 | 2735 | 601235 | . | . | 6.485099 | 6.494728 | 5.248382 | 6.406161 | 15.091092 | 15.012855 | 11.777216 | 14.646927 | -1.129214 | 617.742719 | 0.165366303 | -1.06018514 | 0.289060382 | 0.998865404 | 9.356194 | 9.180164 | 9.311695 | 9.307300 | 9.165334 | 9.287363 | 757 | 705 | 457 | 595 |
| 1663 | NM_00125714 | DDX11 | DEAD/H-box helicase 11 | protein_coding | NP_001244073.1;NP_001244074.1;NP_004390.3;NP_085911.2;NP_6896! | 2736 | 601150 | . | . | 28.521350 | 30.896720 | 34.494056 | 29.692406 | 66.370355 | 71.419151 | 77.403671 | 67.888156 | 1.069560 | 3861.889466 | 0.124029958 | 0.782210941 | 0.434090615 | 0.998865404 | 11.865695 | 11.962852 | 11.860189 | 11.909806 | 12.014202 | 11.862384 | 4168 | 4148 | 3787 | 3402 |
| 1665 | NM_001358,XI | DHX15 | DEAH-box helicase 15 | protein_coding | NP_001349.2 | 2738 | 603403 | . | . | 53.420318 | 58.392389 | 53.078096 | 55.288589 | 124.311271 | 134.976625 | 119.105707 | 126.410797 | -1.046139 | 5140.645987 | 0.108788806 | -0.59816963 | 0.549726751 | 0.998865404 | 12.359947 | 12.294785 | 12.311767 | 12.379403 | 12.295603 | 12.320878 | 5810 | 5900 | 4303 | 4782 |
| 1666 | NM_00133057 | DECR1 | 2,4-dienoyl-CoA reductase 1 | protein_coding | NP_001317504.1;NP_001350.1;XP_005250865.1;XP_011515182.1;XP_01 | 2753 | 222745 | . | . | 16.570947 | 15.870111 | 14.361261 | 17.072745 | 38.561276 | 36.684470 | 32.226252 | 39.034802 | 1.040184 | 1027.895677 | 0.143279212 | 0.396698005 | 0.691590171 | 0.998865404 | 9.976925 | 10.033477 | 10.011716 | 9.967919 | 9.964688 | 10.070519 | 1195 | 1055 | 846 | 1053 |
| 1676 | NM_004401,N | DFFA | DNA fragmentation factor subunit alpha | protein_coding | NP_004392.1;NP_998731.1 | 2772 | 601882 | . | . | 26.121756 | 28.410803 | 27.020209 | 27.444484 | 60.786398 | 65.672842 | 60.632563 | 62.748553 | 1.000196 | 4607.432305 | 0.109463272 | 0.002578639 | 0.997942546 | 0.999862634 | 12.169650 | 12.169844 | 12.135520 | 12.202330 | 12.160434 | 12.178671 | 5093 | 5170 | 3963 | 4367 |
| 1677 | NM_00128266 | DFFB | DNA fragmentation factor subunit beta | protein_coding | NP_001269598.1;NP_001307061.1;NP_001307065.1;NP_004393.1;XP_01 | 2773 | 601883 | . | . | 5.283261 | 5.112336 | 5.126062 | 4.369220 | 12.294365 | 11.817390 | 11.502737 | 9.989703 | 1.019431 | 403.679063 | 0.205323771 | 0.135224323 | 0.892434516 | 0.998865404 | 8.642521 | 8.671463 | 8.638890 | 8.654956 | 8.756278 | 8.550419 | 454 | 437 | 416 | 316 |
| 1678 | NM_00114595 | TIMM8A | translocase of inner mitochondrial membrane 8A | protein_coding | NP_001139423.1;NP_004076.1 | 11817 | 300356 | . | . | 4.934524 | 2.866903 | 4.652771 | 5.183598 | 11.482839 | 6.626974 | 10.440684 | 11.851680 | -1.007274 | 233.390904 | 0.221946314 | -0.04711335 | 0.96242289 | 0.998865404 | 7.871860 | 7.861329 | 7.854007 | 7.882165 | 7.871233 | 7.858052 | 262 | 260 | 205 | 215 |
| 1687 | NM_00112745 | GSDME | gasdermin E | protein_coding | NP_001120925.1;NP_001120926.1;NP_004394.1;XP_016867291.1;XP_02 | 2810 | 608798 | . | . | 6.604320 | 6.547050 | 5.181772 | 6.303187 | 15.368524 | 15.133801 | 11.627746 | 14.411487 | -1.194634 | 441.185040 | 0.184359175 | -1.39168121 | 0.164018955 | 0.940065727 | 8.908114 | 8.650918 | 8.862443 | 8.809562 | 8.670134 | 8.781894 | 576 | 497 | 316 | 413 |
| 1716 | NM_00131885 | DGUOK | deoxyguanosine kinase | protein_coding | NP_001305788.1;NP_001305789.1;NP_001305790.1;NP_001305791.1;NI | 2858 | 601465 | . | . | 52.997264 | 56.005803 | 53.106435 | 67.829659 | 123.326807 | 129.459926 | 119.169303 | 155.084453 | 1.115055 | 1704.538193 | 0.141964854 | 1.106716495 | 0.268416522 | 0.998865404 | 10.654697 | 10.811381 | 10.683783 | 10.694110 | 10.674401 | 10.865132 | 1844 | 1751 | 1375 | 1887 |
| 1717 | NM_00116381 | DHCR7 | 7-dehydrocholesterol reductase | protein_coding | NP_001157289.1;NP_001351.2;XP_011543079.1 | 2860 | 602858 | . | . | 49.629918 | 48.635586 | 84.115757 | 79.766895 | 115.490858 | 112.423334 | 188.753317 | 182.377538 | 1.652383 | 5459.584336 | 0.106690896 | 6.791092484 | 1.11288E-11 | 1.5361E-08 | 12.007242 | 12.731861 | 12.176984 | 12.156979 | 12.613766 | 12.561613 | 4800 | 4386 | 6092 | 6187 |
| 1718 | NM_014762 | DHCR24 | 24-dehydrocholesterol reductase | protein_coding | NP_055577.1 | 2859 | 606418 | . | . | 66.922745 | 67.206924 | 94.653465 | 98.746948 | 155.731979 | 155.351807 | 212.399628 | 225.773178 | 1.421783 | 10783.857083 | 0.100767197 | 5.038357354 | 4.69544E-07 | 0.000170556 | 13.120524 | 13.628212 | 13.229761 | 13.226240 | 13.513924 | 13.542141 | 10277 | 9589 | 10834 | 12061 |
| 1719 | NM_000791,N | DHFR | dihydrofolate reductase | protein_coding | NP_000782.1;NP_001277283.1;NP_001277286.1 | 2861 | 126060 | . | . | 25.493097 | 27.149441 | 23.984373 | 28.322701 | 59.323492 | 62.757147 | 53.820237 | 64.756493 | -1.020349 | 3201.841254 | 0.122218069 | -0.23779233 | 0.812042164 | 0.998865404 | 11.659270 | 11.629953 | 11.629490 | 11.674327 | 11.569658 | 11.697420 | 3623 | 3585 | 2541 | 3203 |
| 1723 | NM_001361,XI | DHODH | dihydroorotate dehydrogenase (quinone) | protein_coding | NP_001352.2;XP_005255884.1;XP_005255886.1;XP_016878479.1 | 2867 | 126064 | . | . | 2.260797 | 2.386182 | 1.829260 | 1.960182 | 5.260968 | 5.515768 | 4.104806 | 4.481723 | 1.004526 | 267.942121 | 0.211260388 | 0.030839647 | 0.975397422 | 0.998865404 | 8.062831 | 8.068718 | 8.052713 | 8.076004 | 8.043603 | 8.089215 | 301 | 295 | 223 | 263 |
| 1725 | NM_00120697 | DHPS | deoxyhypusine synthase | protein_coding | NP_001193903.1;NP_001356620.1;NP_001356621.1;NP_001356622.1;NI | 2869 | 600944 | . | . | 28.060339 | 27.724846 | 26.746707 | 29.863121 | 65.297571 | 64.087226 | 60.018835 | 68.278478 | 1.000206 | 1155.497506 | 0.13496607 | 0.002202363 | 0.99824277 | 0.999862634 | 10.174279 | 10.174318 | 10.179910 | 10.168076 | 10.136452 | 10.210380 | 1341 | 1237 | 955 | 1137 |
| 1727 | NM_000398,N | CYB5R3 | cytochrome b5 reductase 3 | protein_coding | NP_000389.1;NP_001123291.1;NP_001165131.1;NP_001165132.1;NP_0: | 2873 | 613213 | . | . | 53.295409 | 57.448610 | 44.154926 | 52.250385 | 124.020605 | 132.795024 | 99.082375 | 119.464294 | -1.165402 | 4707.615074 | 0.119475331 | -1.84830873 | 0.064557693 | 0.731823074 | 12.307065 | 12.086039 | 12.231825 | 12.288551 | 12.065584 | 12.195593 | 5640 | 5651 | 3482 | 4398 |
| 1728 | NM_000903,N | NQO1 | NAD(P)H quinone dehydrogenase 1 | protein_coding | NP_000894.1;NP_001020604.1;NP_001020605.1;NP_001273066.1 | 2874 | 125860 | . | . | 620.233706 | 622.378334 | 851.466397 | 906.426237 | 1443.309287 | 1438.655444 | 1910.665963 | 2072.435806 | 1.395036 | 58595.818868 | 0.098307568 | 4.885710465 | 1.03057E-06 | 0.000302657 | 15.578465 | 16.058763 | 15.679763 | 15.675757 | 15.942158 | 15.988719 | 56478 | 52673 | 57802 | 65672 |
| 1729 | NM_00107981 | DIAPH1 | diaphanous related formin 1 | protein_coding | NP_001073280.1;NP_001300936.1;NP_005210.3;XP_024310152.1;XP_02 | 2876 | 602121 | . | . | 24.947329 | 25.422056 | 24.045970 | 25.474349 | 58.053460 | 58.764221 | 53.958456 | 58.244073 | -1.031407 | 4465.041612 | 0.10740588 | -0.41537895 | 0.677864515 | 0.998865404 | 12.146626 | 12.101944 | 12.132999 | 12.141343 | 12.091073 | 12.131285 | 5195 | 4917 | 3730 | 4221 |
| 1730 | NM_006729,N | DIAPH2 | diaphanous related formin 2 | protein_coding | NP_006720.1;NP_009293.1 | 2877 | 300108 | . | . | 1.990476 | 2.322069 | 1.779264 | 1.866714 | 4.631918 | 5.367567 | 3.992617 | 4.268021 | -1.054969 | 417.916730 | 0.183230078 | -0.42132983 | 0.673514253 | 0.998865404 | 8.744794 | 8.668336 | 8.724757 | 8.719643 | 8.736198 | 8.641883 | 496 | 461 | 382 | 351 |
| 1731 | NM_00136597 | SEPTIN1 | septin 1 | protein_coding | NP_001352906.1;NP_443070.6;XP_016878480.1 | 2879 | 612897 | . | . | 0.702606 | 0.673239 | 0.559068 | 1.017432 | 1.634993 | 1.556224 | 1.254534 | 2.326237 | 1.120366 | 38.913019 | 0.518449387 | 0.316270334 | 0.751797327 | 0.998865404 | 5.199399 | 5.360470 | 5.271852 | 5.267143 | 5.243656 | 5.328072 | 43 | 39 | 26 | 49 |
| 1734 | NM_000793,N | DIO2 | iodothyronine deiodinase 2 | protein_coding | NP_000784.3;NP_001311391.2;NP_001353425.1;NP_054644.1 | 2884 | 601413 | . | . | 0.045367 | 0.042524 | 0.031782 | 0.028707 | 0.105572 | 0.098297 | 0.071318 | 0.065636 | -1.415770 | 8.030040 | 1.072292548 | -0.46777069 | 0.639948582 | NA | 3.234218 | 2.733338 | 3.018168 | 3.016764 | 3.006231 | 3.003904 | 11 | 10 | 6 | 6 |
| 1736 | NM_00114246 | DKC1 | dyskerin pseudouridine synthase 1 | protein_coding | NP_001135935.1;NP_001275676.1;NP_001354.1 | 2890 | 300126 | . | . | 26.593601 | 26.999221 | 26.615614 | 26.933108 | 61.884398 | 62.409912 | 59.724669 | 61.579352 | -1.000241 | 2124.142265 | 0.116824248 | -0.00296983 | 0.997630419 | 0.999862634 | 11.052811 | 11.052518 | 11.050363 | 11.055056 | 11.064151 | 11.040797 | 2439 | 2299 | 1874 | 1962 |
| 1737 | NM_00137203 | DLAT | dihydrolipoamide S-acetyltransferase | protein_coding | NP_001358960.1;NP_001358961.1;NP_001358962.1;NP_001358963.1;NI | 2896 | 608770 | . | . | 11.392584 | 10.741946 | 9.895365 | 10.259811 | 26.511007 | 24.830490 | 22.204910 | 23.457838 | -1.114989 | 1270.919106 | 0.130465327 | -1.20361191 | 0.228739615 | 0.982967904 | 10.388019 | 10.231026 | 10.375655 | 10.326641 | 10.258311 | 10.279662 | 1592 | 1400 | 1030 | 1143 |
| 1738 | NM_000108,N | DLD | dihydrolipoamide dehydrogenase | protein_coding | NP_000099.2;NP_001276679.1;NP_001276680.1;NP_001276681.1 | 2898 | 238331 | . | . | 30.199889 | 26.024282 | 25.195946 | 24.012254 | 70.276381 | 60.156298 | 56.538973 | 54.901171 | -1.158206 | 2906.306831 | 0.123886571 | -1.71036984 | 0.087197504 | 0.797077582 | 11.606923 | 11.395263 | 11.624177 | 11.493921 | 11.464638 | 11.417436 | 3873 | 3102 | 2411 | 2451 |
| 1739 | NM_00109842 | DLG1 | discs large MAGUK scaffold protein 1 | protein_coding | NP_001091894.1;NP_001191315.1;NP_001191316.1;NP_001191317.1;NI | 2900 | 601014 | . | . | 8.180514 | 8.014515 | 7.101586 | 6.720415 | 19.036390 | 18.525898 | 15.935751 | 15.365430 | -1.169002 | 1230.898574 | 0.130769768 | -1.72270455 | 0.084941967 | 0.788085288 | 10.373646 | 10.148577 | 10.327184 | 10.315638 | 10.224427 | 10.182572 | 1539 | 1421 | 1015 | 1034 |
| 1740 | NM_00114269 | DLG2 | discs large MAGUK scaffold protein 2 | protein_coding | NP_001136171.1;NP_001136172.1;NP_001136174.1;NP_001193698.1;NI | 2901 | 603583 | . | . | 0.138141 | 0.166113 | 0.279629 | 0.297365 | 0.321460 | 0.383976 | 0.627480 | 0.679891 | 1.840318 | 65.715684 | 0.392566386 | 2.241545544 | 0.024990759 | 0.521068648 | 5.532907 | 6.411637 | 5.930751 | 5.960446 | 6.074036 | 6.090114 | 49 | 54 | 72 | 82 |
| 1741 | NM_00116627 | DLG3 | discs large MAGUK scaffold protein 3 | protein_coding | NP_001159750.1;NP_065781.1;NP_066943.2;XP_005262305.1;XP_00672 | 2902 | 300189 | . | . | 0.685330 | 0.756115 | 0.432209 | 0.791520 | 1.594791 | 1.747794 | 0.969865 | 1.809716 | -1.114092 | 109.823477 | 0.326542503 | -0.4773313 | 0.633126224 | 0.998865404 | 6.856087 | 6.697650 | 6.778219 | 6.810811 | 6.695961 | 6.819562 | 127 | 131 | 72 | 117 |
| 1742 | NM_00112882 | DLG4 | discs large MAGUK scaffold protein 4 | protein_coding | NP_001122299.1;NP_001308003.1;NP_001308004.1;NP_001308005.1;NI | 2903 | 602887 | . | . | 1.084479 | 0.921357 | 1.139760 | 0.831492 | 2.523626 | 2.129756 | 2.557591 | 1.901109 | 1.028378 | 170.560365 | 0.271146549 | 0.148886412 | 0.881643264 | 0.998865404 | 7.392729 | 7.435235 | 7.443547 | 7.363407 | 7.476048 | 7.352163 | 212 | 164 | 173 | 138 |
| 1743 | NM_00124488 | DLST | dihydrolipoamide S-succinyltransferase | protein_coding | NP_001231812.1;NP_001924.2 | 2911 | 126063 | . | . | 34.709052 | 35.040608 | 33.468986 | 37.359821 | 80.769390 | 80.997934 | 75.103434 | 85.418792 | 1.004906 | 2964.310103 | 0.11615597 | 0.060786972 | 0.951528866 | 0.998865404 | 11.530033 | 11.536920 | 11.526564 | 11.535801 | 11.491953 | 11.576556 | 3386 | 3209 | 2450 | 2930 |
| 1745 | NM_00103849 | DLX1 | distal-less homeobox 1 | protein_coding | NP_001033582.1;NP_835221.2 | 2914 | 600029 | . | . | 12.815017 | 13.777147 | 10.170129 | 12.073624 | 29.821070 | 31.846488 | 22.821475 | 27.604906 | -1.213740 | 854.997092 | 0.151616953 | -1.84319076 | 0.065301164 | 0.731993537 | 9.873062 | 9.592910 | 9.785718 | 9.822159 | 9.605210 | 9.722931 | 1053 | 1037 | 614 | 786 |
| 1746 | NM_004405 | DLX2 | distal-less homeobox 2 | protein_coding | NP_004396.1 | 2915 | 126255 | . | . | 1.559287 | 1.855671 | 0.594043 | 0.942313 | 3.628525 | 4.289465 | 1.333015 | 2.154488 | -2.236690 | 95.224451 | 0.354228245 | -3.27857934 | 0.00104331 | 0.069569141 | 7.041288 | 5.876095 | 6.632998 | 6.694673 | 6.392415 | 6.478177 | 139 | 154 | 40 | 67 |
| 1748 | NM_001934,N | DLX4 | distal-less homeobox 4 | protein_coding | NP_001925.2;NP_612138.1;XP_016879780.1 | 2917 | 601911 | . | . | 1.245134 | 0.912365 | 0.958633 | 1.735052 | 2.897479 | 2.108973 | 2.151143 | 3.966992 | 1.212705 | 76.453409 | 0.403888079 | 0.688876178 | 0.490901192 | 0.998865404 | 6.111465 | 6.388293 | 6.258100 | 6.183157 | 6.190733 | 6.344526 | 92 | 63 | 52 | 101 |
| 1749 | NM_005221,XI | DLX5 | distal-less homeobox 5 | protein_coding | NP_005212.1;XP_005250242.1;XP_016867292.1 | 2918 | 600028 | . | . | 4.145698 | 3.847034 | 3.243819 | 3.187860 | 9.647210 | 8.892593 | 7.279035 | 7.288662 | -1.263355 | 164.143830 | 0.257400457 | -1.31025347 | 0.190110102 | 0.961278413 | 7.517454 | 7.180556 | 7.415414 | 7.394058 | 7.315646 | 7.304135 | 216 | 193 | 128 | 134 |
| 1750 | NM_005222 | DLX6 | distal-less homeobox 6 | protein_coding | NP_005213.3 | 2919 | 600030 | . | . | 1.838130 | 1.500586 | 1.853531 | 1.852969 | 4.277405 | 3.468671 | 4.159270 | 4.236594 | 1.091870 | 126.691571 | 0.292970719 | 0.432813731 | 0.665150097 | 0.998865404 | 6.919872 | 7.047653 | 7.003095 | 6.930244 | 7.001279 | 6.995551 | 154 | 117 | 116 | 123 |
| 1756 | NM_000109,N | DMD | dystrophin | protein_coding | NP_000100.3;NP_003997.2;NP_004000.1;NP_004001.1;NP_004002.3;NP_ | 2928 | 300377 | . | . | 0.310015 | 0.294531 | 0.280043 | 0.492268 | 0.721419 | 0.680822 | 0.628408 | 1.125511 | -1.321550 | 112.934159 | 0.303723498 | -1.32433233 | 0.185392698 | 0.958058933 | 7.006590 | 6.604106 | 6.871413 | 6.856563 | 6.764324 | 6.781499 | 151 | 136 | 82 | 94 |
| 1760 | NM_00108156 | DMPK | DM1 protein kinase | protein_coding | NP_001075029.1;NP_001075031.1;NP_001075032.1;NP_001275693.1;NI | 2933 | 605377 | . | . | 5.408084 | 6.118507 | 6.216045 | 4.883405 | 12.584835 | 14.143202 | 13.948624 | 11.165323 | -1.076931 | 524.704589 | 0.183102903 | -0.58396362 | 0.559244786 | 0.998865404 | 9.087548 | 8.981216 | 9.014059 | 9.096612 | 9.083935 | 8.927426 | 587 | 624 | 494 | 415 |
| 1761 | NM_00136376 | DMRT1 | doublesex and mab-3 related transcription factor 1 | protein_coding | NP_001350696.1;NP_068770.2;XP_006716795.1;XP_011516072.1;XP_01 | 2934 | 602424 | . | . | 0.073167 | 0.065648 | 0.054573 | 0.060065 | 0.170262 | 0.151747 | 0.122459 | 0.137332 | -1.209732 | 4.894846 | 1.359067692 | -0.20211421 | 0.839827441 | NA | 2.421266 | 2.148375 | 2.300838 | 2.294032 | 2.293956 | 2.292427 | 7 | 5 | 4 | 4 |
| 1762 | NM_004943 | DMWD | DM1 locus, WD repeat containing | protein_coding | NP_004934.1 | 2936 | 609857 | . | . | 6.705696 | 7.351583 | 8.004088 | 7.208206 | 15.604429 | 16.993515 | 17.960941 | 16.480705 | 1.067769 | 777.480226 | 0.151614798 | 0.623949315 | 0.532660872 | 0.998865404 | 9.554499 | 9.649269 | 9.548739 | 9.606634 | 9.663365 | 9.583273 | 830 | 845 | 738 | 710 |
| 1763 | NM_00108044 | DNA2 | DNA replication helicase/nuclease 2 | protein_coding | NP_001073918.2;XP_006717743.1;XP_011537719.1;XP_016871288.1 | 2939 | 601810 | . | . | 3.287502 | 3.077288 | 2.971956 | 2.202629 | 7.650153 | 7.113288 | 6.668985 | 5.036051 | -1.206889 | 386.930509 | 0.208240076 | -1.30278916 | 0.192646754 | 0.96621494 | 8.724482 | 8.454795 | 8.674726 | 8.613915 | 8.625579 | 8.434587 | 511 | 434 | 357 | 273 |
| 1767 | NM_001369,XI | DNAH5 | dynein axonemal heavy chain 5 | protein_coding | NP_001360.1;XP_005248319.2;XP_016864666.1;XP_016864667.1;XP_01( | 2950 | 603335 | . | . | 0.012996 | 0.023640 | 0.037070 | 0.030466 | 0.030242 | 0.054644 | 0.083186 | 0.069657 | 1.791448 | 15.137507 | 0.803606106 | 1.046689934 | 0.295242608 | NA | 3.440763 | 4.279207 | 3.869456 | 3.897591 | 3.926220 | 3.915876 | 9 | 15 | 18 | 17 |
| 1768 | NM_001370,XI | DNAH6 | dynein axonemal heavy chain 6 | protein_coding | NP_001361.1;XP_006712019.1;XP_011530951.1;XP_011530952.1;XP_01: | 2951 | 603336 | . | . | 0.041007 | 0.041536 | 0.045390 | 0.043145 | 0.095425 | 0.096012 | 0.101853 | 0.098647 | 1.044466 | 17.877480 | 0.722170694 | 0.086912094 | 0.930741386 | NA | 4.128334 | 4.191126 | 4.156925 | 4.158004 | 4.161334 | 4.160258 | 20 | 19 | 16 | 17 |
| 1770 | NM_001372,N | DNAH9 | dynein axonemal heavy chain 9 | protein_coding | NP_001363.2;NP_004653.2;XP_011522005.1;XP_016879781.1;XP_01687 | 2953 | 603330 | . | . | 2.555071 | 1.106005 | 0.902594 | 1.304377 | 5.945756 | 2.556582 | 2.025396 | 2.982304 | -1.001710 | 481.531383 | 0.198754113 | -0.01239828 | 0.990107854 | 0.999289967 | 8.913339 | 8.909630 | 8.900196 | 8.916591 | 8.786926 | 9.014769 | 548 | 527 | 342 | 533 |
| 1773 | NM_00135182 | DNASE1 | deoxyribonuclease 1 | protein_coding | NP_001338754.1;NP_005214.2;XP_006720917.1;XP_011520695.1;XP_01 | 2956 | 125505 | . | . | 2.338611 | 2.161482 | 2.604713 | 2.795839 | 5.442046 | 4.996361 | 5.844897 | 6.392355 | 1.150401 | 314.722398 | 0.20192077 | 1.001067594 | 0.316794131 | 0.998865404 | 8.193150 | 8.395624 | 8.234564 | 8.276134 | 8.369872 | 8.293193 | 324 | 328 | 313 | 294 |
| 1774 | NM_00100993 | DNASE1L1 | deoxyribonuclease 1 like 1 | protein_coding | NP_001009932.1;NP_001009933.1;NP_001009934.1;NP_001290549.1;NI | 2957 | 300081 | . | . | 5.625104 | 5.114174 | 4.967168 | 4.431768 | 13.089848 | 11.821642 | 11.146180 | 10.132710 | -1.132710 | 441.538367 | 0.177884194 | -1.01064891 | 0.3121845 | 0.998865404 | 8.873258 | 8.693965 | 8.859992 | 8.784853 | 8.755101 | 8.738847 | 572 | 476 | 365 | 383 |
| 1775 | NM_00130168 | DNASE1L2 | deoxyribonuclease 1 like 2 | protein_coding | NP_001288609.1;NP_001365.1 | 2958 | 602622 | . | . | 0.399786 | 0.697046 | 0.788559 | 0.629815 | 0.930319 | 1.611253 | 1.769504 | 1.439998 | 1.277449 | 26.760152 | 0.614981231 | 0.57443253 | 0.565675145 | NA | 4.555346 | 4.907274 | 4.691559 | 4.744360 | 4.758120 | 4.731894 | 20 | 32 | 29 | 25 |
| 1777 | NM_001375 | DNASE2 | deoxyribonuclease 2, lysosomal | protein_coding | NP_001366.1 | 2960 | 126350 | . | . | 34.676167 | 35.976250 | 29.244759 | 33.587444 | 80.692863 | 83.160713 | 65.624390 | 76.793701 | -1.139897 | 2015.572502 | 0.124767914 | -1.51404124 | 0.130015379 | 0.881002591 | 11.068480 | 10.879269 | 11.014997 | 11.037745 | 10.870977 | 10.971031 | 2438 | 2350 | 1533 | 1879 |
| 1778 | NM_001376 | DYNC1H1 | dynein cytoplasmic 1 heavy chain 1 | protein_coding | NP_001367.2 | 2961 | 600112 | . | . | 25.540390 | 26.385839 | 24.070177 | 27.780909 | 59.433537 | 60.992046 | 54.012779 | 63.517746 | -1.015771 | 16113.333047 | 0.108785616 | -0.207523 | 0.835601431 | 0.998865404 | 13.987235 | 13.964610 | 13.971730 | 13.992396 | 13.910113 | 14.024085 | 18475 | 17734 | 12978 | 15984 |
| 1780 | NM_00113555 | DYNC1I1 | dynein cytoplasmic 1 intermediate chain 1 | protein_coding | NP_001129028.1;NP_001129029.1;NP_001265350.1;NP_001265351.1;NI | 2963 | 603772 | . | . | 7.022710 | 5.722721 | 5.029011 | 4.811702 | 16.342134 | 13.228325 | 11.284953 | 11.001386 | -1.307167 | 528.881933 | 0.17650115 | -2.18946994 | 0.028562701 | 0.545298698 | 9.226819 | 8.841076 | 9.193201 | 9.051004 | 8.966907 | 8.938420 | 762 | 579 | 408 | 420 |
| 1781 | NM_00127178 | DYNC1I2 | dynein cytoplasmic 1 intermediate chain 2 | protein_coding | NP_001258714.1;NP_001258715.1;NP_001258716.1;NP_001258717.1;NI | 2964 | 603331 | . | . | 28.835792 | 31.211326 | 27.934364 | 29.514024 | 67.102074 | 72.146380 | 62.683904 | 67.480309 | -1.069140 | 3875.343831 | 0.1123714 | -0.8583159 | 0.390718051 | 0.998865404 | 11.967599 | 11.871001 | 11.916702 | 11.976328 | 11.864406 | 11.917943 | 4447 | 4476 | 3151 | 3626 |
| 1783 | NM_00128615 | DYNC1LI2 | dynein cytoplasmic 1 light intermediate chain 2 | protein_coding | NP_001273086.1;NP_001310884.1;NP_006132.1;XP_016878497.1 | 2966 | 611406 | . | . | 30.184997 | 29.431536 | 35.201236 | 28.446710 | 70.241724 | 68.032314 | 78.990557 | 65.040025 | 1.052285 | 4172.614066 | 0.127967287 | 0.574569226 | 0.56558267 | 0.998865404 | 11.989393 | 12.063136 | 12.016533 | 11.990332 | 12.133857 | 11.949924 | 4756 | 4318 | 4130 | 3571 |
| 1785 | NM_00100536 | DNM2 | dynamin 2 | protein_coding | NP_001005360.1;NP_001005361.1;NP_001005362.1;NP_001177645.1;NI | 2974 | 602378 | . | . | 20.953788 | 20.772093 | 17.924683 | 22.141555 | 48.760320 | 48.015625 | 40.222471 | 50.624034 | -1.055971 | 2318.795118 | 0.131004717 | -0.59974831 | 0.548673987 | 0.998865404 | 11.218056 | 11.139185 | 11.206687 | 11.192491 | 11.073072 | 11.232199 | 2767 | 2548 | 1766 | 2325 |
| 1786 | NM_00113082 | DNMT1 | DNA methyltransferase 1 | protein_coding | NP_001124295.1;NP_001305659.1;NP_001305660.1;NP_001370.1 | 2976 | 126375 | . | . | 30.174451 | 31.517680 | 29.732961 | 32.702837 | 70.217185 | 72.854530 | 66.719901 | 74.771152 | -1.002058 | 5066.847279 | 0.109460852 | -0.02709614 | 0.97838305 | 0.998865404 | 12.308412 | 12.305332 | 12.291788 | 12.322985 | 12.269700 | 12.340553 | 5732 | 5575 | 4215 | 4946 |
| 1787 | NM_00132100 | TRDMT1 | tRNA aspartic acid methyltransferase 1 | protein_coding | NP_001307935.1;NP_001307936.1;NP_001338148.1;NP_001338149.1;NI | 2977 | 602478 | . | . | 0.267855 | 0.289663 | 0.353193 | 0.183643 | 0.623310 | 0.669568 | 0.792554 | 0.419878 | -1.166558 | 76.828919 | 0.396535355 | -0.56049927 | 0.575138937 | 0.998865404 | 6.368661 | 6.150235 | 6.308292 | 6.245324 | 6.301938 | 6.169661 | 106 | 79 | 79 | 48 |

| | | | | | | | | | | |
|---|---|---|---|---|---|---|---|---|---|---|
| 1788 | NM_00132089 | DNMT3A | DNA methyltransferase 3 alpha | protein_coding | NP_001307821.1;NP_001307822.1;NP_001362748.1;NP_072046.2;NP_7 | 2978 | 602769 | . | . | [illegible] |
| 1789 | NM_00120705 | DNMT3B | DNA methyltransferase 3 beta | protein_coding | NP_001193984.1;NP_001193985.1;NP_008823.1;NP_787044.1;NP_7870 | 2979 | 602900 | . | . | [illegible] |
| 1793 | NM_00129022 | DOCK1 | dedicator of cytokinesis 1 | protein_coding | NP_001277152.2;NP_001364472.1;NP_001364473.1;NP_001364475.1;NI | 2987 | 601403 | . | . | [illegible] |
| 1794 | NM_004946,N | DOCK2 | dedicator of cytokinesis 2 | protein_coding | NP_004937.1;XP_005265887.1;XP_011532750.1;XP_011532751.1;XP_01 | 2988 | 603122 | . | . | [illegible] |
| 1795 | NM_004947,XI | DOCK3 | dedicator of cytokinesis 3 | protein_coding | NP_004938.1;XP_005264971.1;XP_005264972.1;XP_005264973.1;XP_00 | 2989 | 603123 | . | . | [illegible] |
| 1796 | NM_00119726 | DOK1 | docking protein 1 | protein_coding | NP_001184189.1;NP_001305795.1;NP_001305796.1;NP_001305797.1;NI | 2990 | 602919 | . | . | [illegible] |
| 1797 | NM_00137120 | DXO | decapping exoribonuclease | protein_coding | NP_001358134.1;NP_001358135.1;NP_005501.2;XP_006715068.1 | 2992 | 605996 | . | . | [illegible] |
| 1798 | NM_001382,XI | DPAGT1 | dolichyl-phosphate N-acetylglucosaminephosphotransferase 1 | protein_coding | NP_001373.2;XP_005271479.1;XP_011540950.1;XP_016872782.1;XP_01 | 2995 | 191350 | . | . | [illegible] |
| 1801 | NM_00134657 | DPH1 | diphthamide biosynthesis 1 | protein_coding | NP_001333503.1;NP_001333504.1;NP_001333505.1;NP_001374.4 | 3003 | 603527 | . | . | [illegible] |
| 1802 | NM_00103958 | DPH2 | diphthamide biosynthesis 2 | protein_coding | NP_001034678.1;NP_001306094.1;NP_001306095.1;NP_001306096.1;NI | 3004 | 603456 | . | . | [illegible] |
| 1806 | NM_000110,N | DPYD | dihydropyrimidine dehydrogenase | protein_coding | NP_000101.2;NP_001153773.1;XP_005270619.2;XP_006710460.1;XP_01 | 3012 | 612779 | . | . | [illegible] |
| 1808 | NM_00119729 | DPYSL2 | dihydropyrimidinase like 2 | protein_coding | NP_001184222.1;NP_001231533.1;NP_001377.1 | 3014 | 602463 | . | . | [illegible] |
| 1809 | NM_00119729 | DPYSL3 | dihydropyrimidinase like 3 | protein_coding | NP_001184223.1;NP_001378.1;XP_011535876.1 | 3015 | 601168 | . | . | [illegible] |
| 1810 | NM_001938 | DR1 | down-regulator of transcription 1 | protein_coding | NP_001929.1 | 3017 | 601482 | . | . | [illegible] |
| 1813 | NM_000795,N | DRD2 | dopamine receptor D2 | protein_coding | NP_000786.1;NP_057658.2;XP_016872785.1 | 3023 | 126450 | . | . | [illegible] |
| 1819 | NM_00133014 | DRG2 | developmentally regulated GTP binding protein 2 | protein_coding | NP_001317073.1;NP_001379.1;XP_005256556.1;XP_005256557.1;XP_01 | 3030 | 602986 | . | . | [illegible] |
| 1820 | NM_005224,XI | ARID3A | AT-rich interaction domain 3A | protein_coding | NP_005215.1;XP_005259570.1;XP_005259571.1;XP_016881934.1;XP_01 | 3031 | 603265 | . | . | [illegible] |
| 1821 | NM_00117118 | DRP2 | dystrophin related protein 2 | protein_coding | NP_001164655.1;NP_001930.2;XP_016884822.1;XP_016884823.1 | 3032 | 300052 | . | . | [illegible] |
| 1822 | NM_00100702 | ATN1 | atrophin 1 | protein_coding | NP_001007027.1;NP_001931.2 | 3033 | 607462 | . | . | [illegible] |
| 1827 | NM_00128538 | RCAN1 | regulator of calcineurin 1 | protein_coding | NP_001272318.1;NP_001272320.2;NP_001272321.1;NP_001272322.1;NI | 3040 | 602917 | . | . | [illegible] |
| 1829 | NM_001943,XI | DSG2 | desmoglein 2 | protein_coding | NP_001934.2;XP_024306863.1 | 3049 | 125671 | . | . | [illegible] |
| 1831 | NM_00101588 | TSC22D3 | TSC22 domain family member 3 | protein_coding | NP_001015881.1;NP_001305397.1;NP_001305399.1;NP_004080.2;NP_9 | 3051 | 300506 | . | . | [illegible] |
| 1832 | NM_00100884 | DSP | desmoplakin | protein_coding | NP_001008844.1;NP_001305963.1;NP_004406.2 | 3052 | 125647 | . | . | [illegible] |
| 1836 | NM_000112,XI | SLC26A2 | solute carrier family 26 member 2 | protein_coding | NP_000103.2;XP_016864680.1 | 10994 | 606718 | . | . | [illegible] |
| 1837 | NM_00112817 | DTNA | dystrobrevin alpha | protein_coding | NP_001121647.1;NP_001185867.1;NP_001185868.1;NP_001185869.1;NI | 3057 | 601239 | . | . | [illegible] |
| 1838 | NM_00125630 | DTNB | dystrobrevin beta | protein_coding | NP_001243232.1;NP_001243233.1;NP_001243237.1;NP_001307861.1;NI | 3058 | 602415 | . | . | [illegible] |
| 1839 | NM_001945 | HBEGF | heparin binding EGF like growth factor | protein_coding | NP_001936.1 | 3059 | 126150 | . | . | [illegible] |
| 1840 | NM_004416,XI | DTX1 | deltex E3 ubiquitin ligase 1 | protein_coding | NP_004407.2;XP_011536311.1 | 3060 | 602582 | . | . | [illegible] |
| 1841 | NM_00116503 | DTYMK | deoxythymidylate kinase | protein_coding | NP_001158503.1;NP_001307831.1;NP_001307832.1;NP_001307833.1;NI | 3061 | 188345 | . | . | [illegible] |
| 1843 | NM_004417 | DUSP1 | dual specificity phosphatase 1 | protein_coding | NP_004408.1 | 3064 | 600714 | . | . | [illegible] |
| 1844 | NM_004418,XI | DUSP2 | dual specificity phosphatase 2 | protein_coding | NP_004409.1;XP_016859035.1 | 3068 | 603068 | . | . | [illegible] |
| 1845 | NM_004090 | DUSP3 | dual specificity phosphatase 3 | protein_coding | NP_004081.1 | 3069 | 600183 | . | . | [illegible] |
| 1846 | NM_001394,N | DUSP4 | dual specificity phosphatase 4 | protein_coding | NP_001385.1;NP_476499.1;XP_011542730.1 | 3070 | 602747 | . | . | [illegible] |
| 1847 | NM_004419 | DUSP5 | dual specificity phosphatase 5 | protein_coding | NP_004410.3 | 3071 | 603069 | . | . | [illegible] |
| 1848 | NM_001946,N | DUSP6 | dual specificity phosphatase 6 | protein_coding | NP_001937.2;NP_073143.2 | 3072 | 602748 | . | . | [illegible] |
| 1849 | NM_001947 | DUSP7 | dual specificity phosphatase 7 | protein_coding | NP_001938.2 | 3073 | 602749 | . | . | [illegible] |
| 1850 | NM_004420,XI | DUSP8 | dual specificity phosphatase 8 | protein_coding | NP_004411.2;XP_011518234.1;XP_011518235.1 | 3074 | 602038 | . | . | [illegible] |
| 1852 | NM_00131850 | DUSP9 | dual specificity phosphatase 9 | protein_coding | NP_001305432.1;NP_001386.1;XP_011529425.1;XP_011529426.1 | 3076 | 300134 | . | . | [illegible] |
| 1854 | NM_00102524 | DUT | deoxyuridine triphosphatase | protein_coding | NP_001020419.1;NP_001020420.1;NP_001317215.1;NP_001939.1;XP_01 | 3078 | 601266 | . | . | [illegible] |
| 1855 | NM_00133031 | DVL1 | dishevelled segment polarity protein 1 | protein_coding | NP_001317240.1;NP_004412.2;XP_005244789.1;XP_005244790.1 | 3084 | 601365 | . | . | [illegible] |
| 1856 | NM_004422,XI | DVL2 | dishevelled segment polarity protein 2 | protein_coding | NP_004413.1;XP_005256559.1 | 3086 | 602151 | . | . | [illegible] |
| 1857 | NM_004423,XI | DVL3 | dishevelled segment polarity protein 3 | protein_coding | NP_004414.3;XP_005247229.1;XP_011510815.1 | 3087 | 601368 | . | . | [illegible] |
| 1859 | NM_00134772 | DYRK1A | dual specificity tyrosine phosphorylation regulated kinase 1A | protein_coding | NP_001334650.1;NP_001334651.1;NP_001334652.1;NP_001387.2;NP_5 | 3091 | 600855 | . | . | [illegible] |
| 1861 | NM_000113,XI | TOR1A | torsin family 1 member A | protein_coding | NP_000104.1 | 3098 | 605204 | . | . | [illegible] |
| 1869 | NM_005225 | E2F1 | E2F transcription factor 1 | protein_coding | NP_005216.1 | 3113 | 189971 | . | . | [illegible] |
| 1870 | NM_004091,XI | E2F2 | E2F transcription factor 2 | protein_coding | NP_004082.1;XP_005245805.1;XP_005245806.1;XP_011539170.1;XP_01 | 3114 | 600426 | . | . | [illegible] |
| 1871 | NM_00124307 | E2F3 | E2F transcription factor 3 | protein_coding | NP_001230005.1;NP_001940.1;XP_005248922.1;XP_005248923.1;XP_01 | 3115 | 600427 | . | . | [illegible] |
| 1874 | NM_001950,XI | E2F4 | E2F transcription factor 4 | protein_coding | NP_001941.2 | 3118 | 600659 | . | . | [illegible] |
| 1875 | NM_00108358 | E2F5 | E2F transcription factor 5 | protein_coding | NP_001077057.1;NP_001077058.1;NP_001942.2 | 3119 | 600967 | . | . | [illegible] |
| 1876 | NM_00127827 | E2F6 | E2F transcription factor 6 | protein_coding | NP_001265204.1;NP_001265205.1;NP_001265206.1;NP_001265207.1;NI | 3120 | 602944 | . | . | [illegible] |
| 1877 | NM_00128877 | E4F1 | E4F transcription factor 1 | protein_coding | NP_001275705.1;NP_001275707.1;NP_004415.4;XP_005255212.1;XP_00 | 3121 | 603022 | . | . | [illegible] |
| 1879 | NM_00129036 | EBF1 | EBF transcription factor 1 | protein_coding | NP_001277289.1;NP_001311030.1;NP_001311032.1;NP_001311035.1;NI | 3126 | 164343 | . | . | [illegible] |
| 1889 | NM_00111334 | ECE1 | endothelin converting enzyme 1 | protein_coding | NP_001106818.1;NP_001106819.1;NP_001106820.1;NP_001388.1;XP_00 | 3146 | 600423 | . | . | [illegible] |
| 1890 | NM_00111375 | TYMP | thymidine phosphorylase | protein_coding | NP_001107227.1;NP_001107228.1;NP_001244917.1;NP_001244918.1;NI | 3148 | 131222 | . | . | [illegible] |
| 1891 | NM_001398,XI | ECH1 | enoyl-CoA hydratase 1 | protein_coding | NP_001389.2;XP_016881937.1;XP_024307176.1;XP_024307177.1 | 3149 | 600696 | . | . | [illegible] |
| 1892 | NM_004092,XI | ECHS1 | enoyl-CoA hydratase, short chain 1 | protein_coding | NP_004083.3 | 3151 | 602292 | . | . | [illegible] |
| 1893 | NM_00120285 | ECM1 | extracellular matrix protein 1 | protein_coding | NP_001189787.1;NP_004416.2;NP_073155.2 | 3153 | 602201 | . | . | [illegible] |
| 1894 | NM_00125831 | ECT2 | epithelial cell transforming 2 | protein_coding | NP_001245244.1;NP_001245245.1;NP_001336023.1;NP_001336024.1;NI | 3155 | 600586 | . | . | [illegible] |
| 1896 | NM_00100560 | EDA | ectodysplasin A | protein_coding | NP_001005609.1;NP_001005610.2;NP_001005612.2;NP_001005613.1;NI | 3157 | 300451 | . | . | [illegible] |
| 1901 | NM_00132073 | S1PR1 | sphingosine-1-phosphate receptor 1 | protein_coding | NP_001307659.1;NP_001391.2 | 3165 | 601974 | . | . | [illegible] |
| 1902 | NM_00135139 | LPAR1 | lysophosphatidic acid receptor 1 | protein_coding | NP_001338326.1;NP_001338327.1;NP_001338328.1;NP_001338329.1;NI | 3166 | 602282 | . | . | [illegible] |
| 1903 | NM_005226 | S1PR3 | sphingosine-1-phosphate receptor 3 | protein_coding | NP_005217.2 | 3167 | 601965 | . | . | [illegible] |
| 1910 | NM_000115,N | EDNRB | endothelin receptor type B | protein_coding | NP_000106.1;NP_001116131.1;NP_001188326.1;NP_003982.1 | 3180 | 131244 | . | . | [illegible] |
| 1911 | NM_004426,XI | PHC1 | polyhomeotic homolog 1 | protein_coding | NP_004417.2;XP_005253391.1;XP_011518901.1;XP_011518902.1;XP_01 | 3182 | 602978 | . | . | [illegible] |
| 1912 | NM_00133048 | PHC2 | polyhomeotic homolog 2 | protein_coding | NP_001317417.1;NP_004418.2;NP_932157.1 | 3183 | 602979 | . | . | [illegible] |
| 1915 | NM_001402 | EEF1A1 | eukaryotic translation elongation factor 1 alpha 1 | protein_coding | NP_001393.1 | 3189 | 130590 | . | . | [illegible] |
| 1917 | NM_001958 | EEF1A2 | eukaryotic translation elongation factor 1 alpha 2 | protein_coding | NP_001949.1 | 3192 | 602959 | . | . | [illegible] |
| 1933 | NM_00103766 | EEF1B2 | eukaryotic translation elongation factor 1 beta 2 | protein_coding | NP_001032752.1;NP_001950.1;NP_066944.1 | 3208 | 600655 | . | . | [illegible] |
| 1936 | NM_00113005 | EEF1D | eukaryotic translation elongation factor 1 delta | protein_coding | NP_001123525.3;NP_001123527.1;NP_001123528.1;NP_001123529.1;NI | 3211 | 130592 | . | . | [illegible] |
| 1937 | NM_001404 | EEF1G | eukaryotic translation elongation factor 1 gamma | protein_coding | NP_001395.1 | 3213 | 130593 | . | . | [illegible] |
| 1938 | NM_001961 | EEF2 | eukaryotic translation elongation factor 2 | protein_coding | NP_001952.1 | 3214 | 130610 | . | . | [illegible] |
| 1939 | NM_00120147 | EIF2D | eukaryotic translation initiation factor 2D | protein_coding | NP_001188407.1;NP_008824.2;XP_005277566.1;XP_006711259.1;XP_01 | 6583 | 613709 | . | . | [illegible] |
| 1942 | NM_004428,N | EFNA1 | ephrin A1 | protein_coding | NP_004419.2;NP_872626.1 | 3221 | 191164 | . | . | [illegible] |
| 1944 | NM_004952 | EFNA3 | ephrin A3 | protein_coding | NP_004943.1 | 3223 | 601381 | . | . | [illegible] |
| 1945 | NM_005227,N | EFNA4 | ephrin A4 | protein_coding | NP_005218.1;NP_872631.1;NP_872632.2 | 3224 | 601380 | . | . | [illegible] |
| 1946 | NM_001962,XI | EFNA5 | ephrin A5 | protein_coding | NP_001953.1;XP_006714628.1;XP_011541552.1;XP_011541553.1;XP_01 | 3225 | 601535 | . | . | [illegible] |
| 1947 | NM_004429 | EFNB1 | ephrin B1 | protein_coding | NP_004420.1 | 3226 | 300035 | . | . | [illegible] |
| 1948 | NM_00137205 | EFNB2 | ephrin B2 | protein_coding | NP_001358985.1;NP_001358986.1;NP_001358987.1;NP_004084.1;XP_01 | 3227 | 600527 | . | . | [illegible] |
| 1949 | NM_001406 | EFNB3 | ephrin B3 | protein_coding | NP_001397.1 | 3228 | 602297 | . | . | [illegible] |
| 1950 | NM_00117813 | EGF | epidermal growth factor | protein_coding | NP_001171601.1;NP_001171602.1;NP_001343950.1;NP_001954.2;XP_00 | 3229 | 131530 | . | . | [illegible] |
| 1951 | NM_001407 | CELSR3 | cadherin EGF LAG seven-pass G-type receptor 3 | protein_coding | NP_001398.2 | 3230 | 604264 | . | . | [illegible] |
| 1952 | NM_001408 | CELSR2 | cadherin EGF LAG seven-pass G-type receptor 2 | protein_coding | NP_001399.1 | 3231 | 604265 | . | . | [illegible] |
| 1953 | NM_001409,XI | MEGF6 | multiple EGF like domains 6 | protein_coding | NP_001400.3;XP_006710469.1;XP_011539187.1;XP_011539188.1;XP_01 | 3232 | 604266 | . | . | [illegible] |
| 1954 | NM_00127193 | MEGF8 | multiple EGF like domains 8 | protein_coding | NP_001258867.1;NP_001401.2 | 3233 | 604267 | . | . | [illegible] |
| 1955 | NM_00108049 | MEGF9 | multiple EGF like domains 9 | protein_coding | NP_001073966.2 | 3234 | 604268 | . | . | [illegible] |
| 1956 | NM_00134689 | EGFR | epidermal growth factor receptor | protein_coding | NP_001333826.1;NP_001333827.1;NP_001333828.1;NP_001333829.1;NI | 3236 | 131550 | . | . | [illegible] |
| 1958 | NM_001964 | EGR1 | early growth response 1 | protein_coding | NP_001955.1 | 3238 | 128990 | . | . | [illegible] |
| 1959 | NM_000399,N | EGR2 | early growth response 2 | protein_coding | NP_000390.2;NP_001129649.1;NP_001129650.1;NP_001129651.1;NP_00 | 3239 | 129010 | . | . | [illegible] |
| 1960 | NM_00119988 | EGR3 | early growth response 3 | protein_coding | NP_001186809.1;NP_001186810.1;NP_004421.2;XP_005273482.1;XP_00 | 3240 | 602419 | . | . | [illegible] |
| 1962 | NM_00116641 | EHHADH | enoyl-CoA hydratase and 3-hydroxyacyl CoA dehydrogenase | protein_coding | NP_001159887.1;NP_001957.2;XP_006713588.1 | 3247 | 607037 | . | . | [illegible] |
| 1964 | NM_001412 | EIF1AX | eukaryotic translation initiation factor 1A X-linked | protein_coding | NP_001403.1 | 3250 | 300186 | . | . | [illegible] |
| 1965 | NM_004094 | EIF2S1 | eukaryotic translation initiation factor 2 subunit alpha | protein_coding | NP_004085.1 | 3265 | 603907 | . | . | [illegible] |
| 1967 | NM_001414 | EIF2B1 | eukaryotic translation initiation factor 2B subunit alpha | protein_coding | NP_001405.1 | 3257 | 606686 | . | . | [illegible] |
| 1968 | NM_001415 | EIF2S3 | eukaryotic translation initiation factor 2 subunit gamma | protein_coding | NP_001406.1 | 3267 | 300161 | . | . | [illegible] |
| 1969 | NM_00132909 | EPHA2 | EPH receptor A2 | protein_coding | NP_001316019.1;NP_004422.2;XP_016856026.1 | 3386 | 176946 | . | . | [illegible] |
| 1973 | NM_00120451 | EIF4A1 | eukaryotic translation initiation factor 4A1 | protein_coding | NP_001191439.1;NP_001407.1 | 3282 | 602641 | . | . | [illegible] |
| 1974 | NM_001967 | EIF4A2 | eukaryotic translation initiation factor 4A2 | protein_coding | NP_001958.2 | 3284 | 601102 | . | . | [illegible] |
| 1975 | NM_00130082 | EIF4B | eukaryotic translation initiation factor 4B | protein_coding | NP_001287750.1;NP_001317583.1;NP_001408.2 | 3285 | 603928 | . | . | [illegible] |
| 1977 | NM_00113067 | EIF4E | eukaryotic translation initiation factor 4E | protein_coding | NP_001124150.1;NP_001124151.1;NP_001317946.1;NP_001959.1 | 3287 | 133440 | . | . | [illegible] |
| 1978 | NM_004095 | EIF4EBP1 | eukaryotic translation initiation factor 4E binding protein 1 | protein_coding | NP_004086.1 | 3288 | 602223 | . | . | [illegible] |
| 1979 | NM_004096 | EIF4EBP2 | eukaryotic translation initiation factor 4E binding protein 2 | protein_coding | NP_004087.1 | 3289 | 602224 | . | . | [illegible] |
| 1981 | NM_00119494 | EIF4G1 | eukaryotic translation initiation factor 4 gamma 1 | protein_coding | NP_001181875.2;NP_001181876.1;NP_001278086.2;NP_004944.3;NP_8 | 3296 | 600495 | . | . | [illegible] |
| 1982 | NM_00104255 | EIF4G2 | eukaryotic translation initiation factor 4 gamma 2 | protein_coding | NP_001036024.3;NP_001166176.1;NP_001409.3 | 3297 | 602325 | . | . | [illegible] |
| 1983 | NM_001969,N | EIF5 | eukaryotic translation initiation factor 5 | protein_coding | NP_001960.2;NP_892116.2 | 3299 | 601710 | . | . | [illegible] |
| 1984 | NM_00114376 | EIF5A | eukaryotic translation initiation factor 5A | protein_coding | NP_001137232.1;NP_001137233.1;NP_001137234.1;NP_001357349.1;NI | 3300 | 600187 | . | . | [illegible] |
| 1992 | NM_030666,N | SERPINB1 | serpin family B member 1 | protein_coding | NP_109591.1;XP_011512635.1;XP_011512636.1;XP_011512637.1 | 3311 | 130135 | . | . | [illegible] |
| 1994 | NM_001419,XI | ELAVL1 | ELAV like RNA binding protein 1 | protein_coding | NP_001410.2 | 3312 | 603466 | . | . | [illegible] |
| 1997 | NM_00114535 | ELF1 | E74 like ETS transcription factor 1 | protein_coding | NP_001138825.1;NP_001357258.1;NP_001357259.1;NP_001357260.1;NI | 3316 | 189973 | . | . | [illegible] |
| 1998 | NM_00127645 | ELF2 | E74 like ETS transcription factor 2 | protein_coding | NP_001263386.1;NP_001263387.1;NP_001263388.1;NP_001317965.1;NI | 3317 | . | | . | [illegible] |
| 2000 | NM_00112719 | ELF4 | E74 like ETS transcription factor 4 | protein_coding | NP_001120669.1;NP_001412.1;XP_005262446.1;XP_011529609.1;XP_01 | 3319 | 300775 | . | . | [illegible] |
| 2002 | NM_00111412 | ELK1 | ETS transcription factor ELK1 | protein_coding | NP_001107595.1;NP_001244097.1;NP_005220.2 | 3321 | 311040 | . | . | [illegible] |
| 2004 | NM_00130351 | ELK3 | ETS transcription factor ELK3 | protein_coding | NP_001290440.1;NP_005221.2;XP_006719338.1 | 3325 | 600247 | . | . | [illegible] |
| 2005 | NM_001973,N | ELK4 | ETS transcription factor ELK4 | protein_coding | NP_001964.2;NP_068567.1 | 3326 | 600246 | . | . | [illegible] |
| 2009 | NM_00100870 | EML1 | EMAP like 1 | protein_coding | NP_001008707.1;NP_001362340.1;NP_001362341.1;NP_004425.2;XP_00 | 3330 | 602033 | . | . | [illegible] |
| 2010 | NM_000117,XI | EMD | emerin | protein_coding | NP_000108.1;XP_024308117.1 | 3331 | 300384 | . | . | [illegible] |
| 2011 | NM_00103946 | MARK2 | microtubule affinity regulating kinase 2 | protein_coding | NP_001034558.2;NP_001156768.1;NP_001156769.1;NP_004945.4;NP_0 | 3332 | 600526 | . | . | [illegible] |
| 2012 | NM_001423,XI | EMP1 | epithelial membrane protein 1 | protein_coding | NP_001414.1;XP_024304645.1 | 3333 | 602333 | . | . | [illegible] |
| 2013 | NM_001424,XI | EMP2 | epithelial membrane protein 2 | protein_coding | NP_001415.1;XP_006720927.1 | 3334 | 602334 | . | . | [illegible] |
| 2014 | NM_00131390 | EMP3 | epithelial membrane protein 3 | protein_coding | NP_001300834.1;NP_001416.1;XP_011524907.1 | 3335 | 602335 | . | . | [illegible] |
| 2015 | NM_00125625 | ADGRE1 | adhesion G protein-coupled receptor E1 | protein_coding | NP_001243181.1;NP_001243182.1;NP_001243183.1;NP_001243184.1;NI | 3336 | 600493 | . | . | [illegible] |
| 2017 | NM_00118474 | CTTN | cortactin | protein_coding | NP_001171669.1;NP_005222.2;NP_612632.1;XP_006718510.1;XP_00671 | 3338 | 164765 | . | . | [illegible] |
| 2018 | NM_00116592 | EMX2 | empty spiracles homeobox 2 | protein_coding | NP_001159396.1;NP_004089.1 | 3341 | 600035 | . | . | [illegible] |
| 2019 | NM_001426 | EN1 | engrailed homeobox 1 | protein_coding | NP_001417.3 | 3342 | 131290 | . | . | [illegible] |
| 2020 | NM_001427 | EN2 | engrailed homeobox 2 | protein_coding | NP_001418.2 | 3343 | 131310 | . | . | [illegible] |
| 2021 | NM_004435,XI | ENDOG | endonuclease G | protein_coding | NP_004426.2;XP_011516649.1 | 3346 | 600440 | . | . | [illegible] |
| 2022 | NM_000118,N | ENG | endoglin | protein_coding | NP_000109.1;NP_001108225.1;NP_001265067.1 | 3349 | 131195 | . | . | [illegible] |
| 2023 | NM_00120148 | ENO1 | enolase 1 | protein_coding | NP_001188412.1;NP_001340275.1;NP_001419.1 | 3350 | 172430 | . | . | [illegible] |
| 2026 | NM_001975 | ENO2 | enolase 2 | protein_coding | NP_001966.1 | 3353 | 131360 | . | . | [illegible] |
| 2027 | NM_00119350 | ENO3 | enolase 3 | protein_coding | NP_001180432.1;NP_001361452.1;NP_001361453.1;NP_001967.3;NP_4 | 3354 | 131370 | . | . | [illegible] |
| 2029 | NM_004436,N | ENSA | endosulfine alpha | protein_coding | NP_004427.1;NP_996925.1;NP_996926.1;NP_996927.1;NP_996928.1;NP | 3360 | 603061 | . | . | [illegible] |
| 2030 | NM_00107817 | SLC29A1 | solute carrier family 29 member 1 (Augustine blood group) | protein_coding | NP_001071643.1;NP_001071645.1;NP_001291391.1;NP_001291394.1;NI | 11003 | 602193 | . | . | [illegible] |
| 2033 | NM_00136284 | EP300 | E1A binding protein p300 | protein_coding | NP_001349772.1;NP_001420.2 | 3373 | 602700 | . | . | [illegible] |
| 2034 | NM_001430,XI | EPAS1 | endothelial PAS domain protein 1 | protein_coding | NP_001421.2;XP_011531000.1 | 3374 | 603349 | . | . | [illegible] |
| 2035 | NM_00116600 | EPB41 | erythrocyte membrane protein band 4.1 | protein_coding | NP_001159477.1;NP_001159478.1;NP_001159479.1;NP_001362942.1;NI | 3377 | 130500 | . | . | [illegible] |
| 2036 | NM_00125832 | EPB41L1 | erythrocyte membrane protein band 4.1 like 1 | protein_coding | NP_001245258.1;NP_001245259.1;NP_001245260.1;NP_036288.2;NP_8 | 3378 | 602879 | . | . | [illegible] |
| 2037 | NM_00113555 | EPB41L2 | erythrocyte membrane protein band 4.1 like 2 | protein_coding | NP_001129026.1;NP_001129027.1;NP_001186317.1;NP_001239589.1;NI | 3379 | 603237 | . | . | [illegible] |
| 2039 | NM_00111413 | DMTN | dematin actin binding protein | protein_coding | NP_001107607.1;NP_001107608.1;NP_001107609.1;NP_001107610.1;NI | 3382 | 125305 | . | . | [illegible] |
| 2040 | NM_00127052 | STOM | stomatin | protein_coding | NP_001257455.1;NP_001257456.1;NP_004090.4;NP_937837.1 | 3383 | 133090 | . | . | [illegible] |
| 2042 | NM_005233,N | EPHA3 | EPH receptor A3 | protein_coding | NP_005224.2;NP_872585.1;XP_005264772.1;XP_005264773.1 | 3387 | 179611 | . | . | [illegible] |
| 2043 | NM_00130453 | EPHA4 | EPH receptor A4 | protein_coding | NP_001291465.1;NP_001291466.1;NP_001350677.1;NP_004429.1 | 3388 | 602188 | . | . | [illegible] |
| 2044 | NM_00128176 | EPHA5 | EPH receptor A5 | protein_coding | NP_001268694.1;NP_001268695.1;NP_001268696.1;NP_001305690.1;NI | 3389 | 600004 | . | . | [illegible] |
| 2047 | NM_004441,XI | EPHB1 | EPH receptor B1 | protein_coding | NP_004432.1;XP_016861355.1;XP_016861356.1;XP_024309157.1;XP_02 | 3392 | 600600 | . | . | [illegible] |
| 2048 | NM_00130919 | EPHB2 | EPH receptor B2 | protein_coding | NP_001296121.1;NP_001296122.1;NP_004433.2;NP_059145.2;XP_00671 | 3393 | 600997 | . | . | [illegible] |
| 2049 | NM_004443 | EPHB3 | EPH receptor B3 | protein_coding | NP_004434.2 | 3394 | 601839 | . | . | [illegible] |
| 2050 | NM_004444,XI | EPHB4 | EPH receptor B4 | protein_coding | NP_004435.3;XP_016867305.1 | 3395 | 600011 | . | . | [illegible] |
| 2051 | NM_00128079 | EPHB6 | EPH receptor B6 | protein_coding | NP_001267723.2;NP_001267724.2;NP_004436.4;XP_011514181.1;XP_01 | 3396 | 602757 | . | . | [illegible] |
| 2052 | NM_000120,N | EPHX1 | epoxide hydrolase 1 | protein_coding | NP_000111.1;NP_001129490.1;NP_001278092.1;NP_001365355.1;NP_00 | 3401 | 132810 | . | . | [illegible] |
| 2053 | NM_00125648 | EPHX2 | epoxide hydrolase 2 | protein_coding | NP_001243411.1;NP_001243412.1;NP_001243413.1;NP_001970.2;XP_01 | 3402 | 132811 | . | . | [illegible] |
| 2054 | NM_00135104 | STX2 | syntaxin 2 | protein_coding | NP_001337978.2;NP_001337979.1;NP_001337980.1;NP_001337981.1;NI | 3403 | 132350 | . | . | [illegible] |
| 2055 | NM_018941,XI | CLN8 | CLN8 transmembrane ER and ERGIC protein | protein_coding | NP_061764.2;XP_005266078.1;XP_005266079.1;XP_005266080.1;XP_01 | 2079 | 607837 | . | . | [illegible] |
| 2057 | NM_000121,N | EPOR | erythropoietin receptor | protein_coding | NP_000112.1 | 3416 | 133171 | . | . | [illegible] |
| 2058 | NM_004446,XI | EPRS1 | glutamyl-prolyl-tRNA synthetase 1 | protein_coding | NP_004437.2;XP_016856103.1 | 3418 | 138295 | . | . | [illegible] |
| 2059 | NM_004447,XI | EPS8 | epidermal growth factor receptor pathway substrate 8 | protein_coding | NP_004438.3;XP_011518907.1;XP_024304646.1;XP_024304647.1;XP_02 | 3420 | 600206 | . | . | [illegible] |
| 2060 | NM_00115996 | EPS15 | epidermal growth factor receptor pathway substrate 15 | protein_coding | NP_001153441.1;NP_001972.1;XP_005270675.1;XP_016856104.1;XP_01 | 3419 | 600051 | . | . | [illegible] |
| 2063 | NM_005234 | NR2F6 | nuclear receptor subfamily 2 group F member 6 | protein_coding | NP_005225.2 | 7977 | 132880 | . | . | [illegible] |
| 2064 | NM_00100586 | ERBB2 | erb-b2 receptor tyrosine kinase 2 | protein_coding | NP_001005862.1;NP_001276865.1;NP_001276866.1;NP_001276867.1;NI | 3430 | 164870 | . | . | [illegible] |
| 2065 | NM_00100591 | ERBB3 | erb-b2 receptor tyrosine kinase 3 | protein_coding | NP_001005915.1;NP_001973.2 | 3431 | 190151 | . | . | [illegible] |
| 2066 | NM_00104259 | ERBB4 | erb-b2 receptor tyrosine kinase 4 | protein_coding | NP_001036064.1;NP_005226.1;XP_005246433.1;XP_005246434.1;XP_00 | 3432 | 600543 | . | . | [illegible] |
| 2067 | NM_00116604 | ERCC1 | ERCC excision repair 1, endonuclease non-catalytic subunit | protein_coding | NP_001159521.1;NP_001356337.1;NP_001356338.1;NP_001356339.1;NI | 3433 | 126380 | . | . | [illegible] |
| 2068 | NM_000400,N | ERCC2 | ERCC excision repair 2, TFIIH core complex helicase subunit | protein_coding | NP_000391.1;NP_001124339.1;XP_011524913.1;XP_016881956.1 | 3434 | 126340 | . | . | [illegible] |
| 2069 | NM_001432 | EREG | epiregulin | protein_coding | NP_001423.1 | 3443 | 602061 | . | . | [illegible] |
| 2070 | NM_00130101 | EYA4 | EYA transcriptional coactivator and phosphatase 4 | protein_coding | NP_001287941.1;NP_001287942.1;NP_001357387.1;NP_001357388.1;NI | 3522 | 603550 | . | . | [illegible] |

| | | | | | | | | | | | | | | | | | | | | | | | | | | | | | | | | | |
|---|---|---|---|---|---|---|---|---|---|---|---|---|---|---|---|---|---|---|---|---|---|---|---|---|---|---|---|---|---|---|---|---|---|
| 2071 | NM_000122,N | ERCC3 | ERCC excision repair 3, TFIIH core complex helicase subunit | protein_coding | NP_000113.1;NP_001290345.1;NP_001290347.1;XP_011509096.1;XP_01 | 3435 | 133510 | . | . | 26.019170 | 29.007470 | 29.124685 | 29.143062 | 60.547679 | 67.052063 | 65.354947 | 66.632143 | 1.048396 | 2405.696160 | 0.11882584 | 0.573815411 | 0.566092719 | 0.998865404 | 11.197802 | 11.265873 | 11.171928 | 11.251514 | 11.254572 | 11.246520 | 2571 | 2660 | 2152 | 2297 |
| 2072 | NM_005236,XI | ERCC4 | ERCC excision repair 4, endonuclease catalytic subunit | protein_coding | NP_005227.1;XP_011520726.1;XP_011520729.1;XP_016878532.1 | 3436 | 133520 | . | . | 1.817647 | 1.840698 | 1.911332 | 1.838889 | 4.229741 | 4.254856 | 4.288973 | 4.204402 | 1.017653 | 392.648543 | 0.182044985 | 0.138674892 | 0.88970706 | 0.998865404 | 8.604281 | 8.629796 | 8.610045 | 8.613070 | 8.639726 | 8.603830 | 447 | 421 | 355 | 360 |
| 2073 | NM_000123 | ERCC5 | ERCC excision repair 5, endonuclease | protein_coding | NP_000114.3 | 3437 | 133530 | . | . | 10.959213 | 10.743576 | 12.001072 | 12.319369 | 25.502539 | 24.834257 | 26.930058 | 28.166775 | 1.105372 | 1393.484054 | 0.1263362 | 1.144024518 | 0.252613469 | 0.994320829 | 10.370397 | 10.514945 | 10.415219 | 10.393656 | 10.474861 | 10.487603 | 1546 | 1408 | 1262 | 1383 |
| 2074 | NM_000124,N | ERCC6 | ERCC excision repair 6, chromatin remodeling factor | protein_coding | NP_000115.1;NP_001263987.1;NP_001263988.1;NP_001333369.1 | 3438 | 609413 | . | . | 2.642445 | 2.571640 | 2.988400 | 3.200146 | 6.149076 | 5.944460 | 6.705883 | 7.316752 | 1.086190 | 829.046329 | 0.144708689 | 0.824253692 | 0.409795425 | 0.998865404 | 9.634286 | 9.753855 | 9.669006 | 9.660006 | 9.749777 | 9.696112 | 921 | 852 | 779 | 779 |
| 2077 | NM_00130103 | ERF | ETS2 repressor factor | protein_coding | NP_001287964.1;NP_001295331.1;NP_001299585.1;NP_006485.2;XP_01 | 3444 | 611888 | . | . | 24.124835 | 24.508797 | 21.153527 | 21.646122 | 56.139480 | 56.653182 | 47.467904 | 49.491283 | -1.150625 | 1905.739926 | 0.118666167 | -1.70577454 | 0.088050065 | 0.801685384 | 10.993811 | 10.791364 | 10.946017 | 10.952212 | 10.835694 | 10.840723 | 2339 | 2209 | 1536 | 1667 |
| 2079 | NM_004450 | ERH | ERH mRNA splicing and mitosis factor | protein_coding | NP_004441.1 | 3447 | 601191 | . | . | 78.219620 | 73.064758 | 66.705093 | 65.077118 | 182.020264 | 168.892471 | 149.684311 | 148.791092 | -1.164102 | 1745.257950 | 0.123647839 | -1.77291481 | 0.076242824 | 0.767975096 | 10.874578 | 10.655567 | 10.856088 | 10.794418 | 10.719863 | 10.693082 | 2245 | 1948 | 1427 | 1486 |
| 2081 | NM_001433,XI | ERN1 | endoplasmic reticulum to nucleus signaling 1 | protein_coding | NP_001424.3;XP_016879836.1;XP_016879837.1 | 3449 | 604033 | . | . | 0.965098 | 1.207706 | 0.859528 | 1.053958 | 2.245823 | 2.791665 | 1.928755 | 2.409750 | 1.009467 | 321.673608 | 0.203466417 | 0.066812763 | 0.946730763 | 0.998865404 | 8.322981 | 8.335897 | 8.342802 | 8.308872 | 8.277218 | 8.382229 | 380 | 335 | 254 | 332 |
| 2086 | NM_00100725 | ERV3-1 | endogenous retrovirus group 3 member 1, envelope | protein_coding | NP_001007254.2 | 3454 | 131170 | . | . | 0.976073 | 1.536074 | 0.666657 | 1.266155 | 2.271362 | 3.550702 | 1.495958 | 2.894914 | -1.131724 | 37.301618 | 0.552449536 | -0.32314651 | 0.746584289 | 0.998865404 | 5.308769 | 5.127898 | 5.259838 | 5.195412 | 5.163253 | 5.255005 | 55 | 34 | 20 | 44 |
| 2091 | NM_001436,XI | FBL | fibrillarin | protein_coding | NP_001427.2;XP_005258708.1;XP_011524925.1 | 3599 | 134795 | . | . | 79.842222 | 79.317680 | 67.733093 | 77.179411 | 185.796129 | 183.346357 | 151.991104 | 176.461556 | 1.021158 | 2500.367289 | 0.120666726 | 0.250325733 | 0.802335458 | 0.998865404 | 11.272835 | 11.302857 | 11.287370 | 11.270465 | 11.244537 | 11.344630 | 2879 | 2642 | 2061 | 2515 |
| 2098 | NM_001984,XI | ESD | esterase D | protein_coding | NP_001975.1;XP_005266335.1;XP_011533256.1;XP_016875903.1 | 3465 | 133280 | . | . | 32.949022 | 29.379244 | 33.607373 | 34.034675 | 76.673726 | 67.911439 | 75.413973 | 77.816239 | 1.043446 | 1261.120859 | 0.130522311 | 0.47008212 | 0.638296348 | 0.998865404 | 10.269397 | 10.330929 | 10.308968 | 10.258170 | 10.320821 | 10.311433 | 1469 | 1287 | 1127 | 1200 |
| 2099 | NM_000125,N | ESR1 | estrogen receptor 1 | protein_coding | NP_000116.2;NP_001116212.1;NP_001116213.1;NP_001116214.1;NP_0 | 3467 | 133430 | . | . | 0.052003 | 0.054472 | 0.056851 | 0.026322 | 0.121013 | 0.125914 | 0.127572 | 0.060182 | -1.251143 | 10.098434 | 0.982120366 | -0.32913155 | 0.742056256 | NA | 3.486115 | 3.168566 | 3.340834 | 3.339988 | 3.346235 | 3.314282 | 13 | 12 | 11 | 5 |
| 2100 | NM_00104027 | ESR2 | estrogen receptor 2 | protein_coding | NP_001035365.1;NP_001201831.1;NP_001258805.1;NP_001258806.1;N | 3468 | 601663 | . | . | 0.157345 | 0.102922 | 0.160655 | 0.141397 | 0.366150 | 0.237908 | 0.360505 | 0.323286 | 1.143204 | 23.913554 | 0.635757486 | 0.303750044 | 0.761352615 | NA | 4.478589 | 4.674293 | 4.585428 | 4.553186 | 4.590959 | 4.574420 | 30 | 20 | 24 | 22 |
| 2101 | NM_00128245 | ESRRA | estrogen related receptor alpha | protein_coding | NP_001269379.1;NP_001269380.1;NP_004442.3;XP_016872802.1 | 3471 | 601998 | . | . | 11.729753 | 10.585679 | 9.804860 | 11.310639 | 27.295615 | 24.469269 | 22.001824 | 25.860434 | -1.075822 | 781.090624 | 0.153259371 | -0.6879767 | 0.491467456 | 0.998865404 | 9.661194 | 9.555565 | 9.669843 | 9.596800 | 9.539308 | 9.624285 | 980 | 829 | 614 | 749 |
| 2107 | NM_00125630 | ETF1 | eukaryotic translation termination factor 1 | protein_coding | NP_001243231.1;NP_001269114.1;NP_001278903.1;NP_001278904.1;N | 3477 | 600285 | . | . | 53.036732 | 50.226321 | 52.198997 | 52.724748 | 123.418663 | 116.100390 | 117.133034 | 120.548874 | 1.017551 | 6054.150514 | 0.105158349 | 0.23869245 | 0.811344071 | 0.998865404 | 12.551092 | 12.576217 | 12.578456 | 12.533527 | 12.566056 | 12.575725 | 7095 | 6303 | 5280 | 5758 |
| 2108 | NM_000126,N | ETFA | electron transfer flavoprotein subunit alpha | protein_coding | NP_000117.1;NP_001121188.1 | 3481 | 608053 | . | . | 42.577683 | 36.991449 | 30.170791 | 34.049722 | 99.080019 | 85.507396 | 67.702377 | 77.850648 | -1.074800 | 2100.332296 | 0.121792471 | -0.85447032 | 0.392844446 | 0.998865404 | 11.087558 | 10.983365 | 11.084746 | 11.043399 | 10.968567 | 11.043427 | 2570 | 2288 | 1677 | 1988 |
| 2109 | NM_00101476 | ETFB | electron transfer flavoprotein subunit beta | protein_coding | NP_001014763.1;NP_001976.1;XP_024307186.1 | 3482 | 130410 | . | . | 83.932799 | 74.079083 | 82.411039 | 96.960930 | 195.315044 | 171.237119 | 184.927982 | 221.689632 | 1.107584 | 2404.260617 | 0.125967993 | 1.170263531 | 0.241894933 | 0.992733415 | 11.155839 | 11.303157 | 11.224718 | 11.149457 | 11.213264 | 11.324041 | 2751 | 2346 | 2048 | 2530 |
| 2110 | NM_00128173 | ETFDH | electron transfer flavoprotein dehydrogenase | protein_coding | NP_001268666.1;NP_001268667.1;NP_004444.2;XP_024309703.1 | 3483 | 231675 | . | . | 5.299135 | 5.736945 | 6.108694 | 7.256921 | 12.331305 | 13.261206 | 13.707730 | 16.592087 | 1.075260 | 508.083378 | 0.167200687 | 0.626106119 | 0.531245339 | 0.998865404 | 8.935698 | 9.040251 | 8.942028 | 8.986703 | 9.017904 | 9.003563 | 544 | 547 | 463 | 488 |
| 2113 | NM_00114382 | ETS1 | ETS proto-oncogene 1, transcription factor | protein_coding | NP_001137292.1;NP_001155894.1;NP_001317380.1;NP_005229.1;XP_01 | 3488 | 164720 | . | . | 12.934447 | 12.735428 | 12.268144 | 13.314477 | 30.098991 | 29.438514 | 27.529359 | 30.441969 | -1.016945 | 2087.219170 | 0.119435085 | -0.20297412 | 0.839155259 | 0.998865404 | 11.039491 | 11.015139 | 11.044875 | 11.022854 | 10.991546 | 11.048630 | 2457 | 2240 | 1735 | 2010 |
| 2114 | NM_00125629 | ETS2 | ETS proto-oncogene 2, transcription factor | protein_coding | NP_001243224.1;NP_005230.1;XP_005260992.1;XP_016883779.1 | 3489 | 164740 | . | . | 3.562867 | 3.686537 | 4.527420 | 4.202101 | 8.290937 | 8.521598 | 10.159400 | 9.607604 | 1.215061 | 464.956901 | 0.173957436 | 1.615501737 | 0.106202084 | 0.833478683 | 8.713397 | 8.994808 | 8.789002 | 8.800114 | 8.950014 | 8.880049 | 479 | 457 | 468 | 452 |
| 2115 | NM_00116314 | ETV1 | ETS variant transcription factor 1 | protein_coding | NP_001156619.1;NP_001156620.1;NP_001156621.1;NP_001156622.1;N | 3490 | 600541 | . | . | 7.453696 | 6.919597 | 8.242702 | 7.527516 | 17.345055 | 15.994956 | 18.496385 | 17.210769 | 1.043219 | 1347.883047 | 0.129945726 | 0.469749693 | 0.63853386 | 0.998865404 | 10.365491 | 10.426816 | 10.406358 | 10.352308 | 10.432299 | 10.391450 | 1573 | 1373 | 1229 | 1256 |
| 2116 | NM_00130097 | ETV2 | ETS variant transcription factor 2 | protein_coding | NP_001287903.1;NP_001291478.1;NP_055024.2;XP_005258709.1;XP_01 | 3491 | 609358 | . | . | 0.317683 | 0.232508 | 0.359698 | 0.454976 | 0.739261 | 0.537452 | 0.807153 | 1.040249 | 1.457167 | 21.383729 | 0.670442875 | 0.810159652 | 0.417848424 | NA | 4.121102 | 4.664848 | 4.404157 | 4.381720 | 4.416852 | 4.436345 | 23 | 16 | 20 | 26 |
| 2117 | NM_00114531 | ETV3 | ETS variant transcription factor 3 | protein_coding | NP_001138784.1;NP_005231.1;XP_006711273.1 | 3492 | 164873 | . | . | 5.439619 | 6.580704 | 6.476509 | 6.268735 | 12.658217 | 15.211591 | 14.533098 | 14.332716 | 1.057245 | 977.082571 | 0.144604358 | 0.555376663 | 0.578637051 | 0.998865404 | 9.891710 | 9.971851 | 9.857445 | 9.961200 | 9.969291 | 9.932855 | 1016 | 1098 | 895 | 918 |
| 2118 | NM_00107967 | ETV4 | ETS variant transcription factor 4 | protein_coding | NP_001073143.1;NP_001248366.1;NP_001248367.1;NP_001248368.1;N | 3493 | 600711 | . | . | 39.112910 | 45.472937 | 56.635609 | 59.135529 | 91.017351 | 105.112732 | 127.088665 | 135.206357 | 1.324443 | 3434.591704 | 0.119690457 | 3.386950329 | 0.000706742 | 0.053599766 | 11.529162 | 11.934330 | 11.563187 | 11.669898 | 11.818359 | 11.879250 | 3177 | 3401 | 3281 | 3801 |
| 2119 | NM_004454 | ETV5 | ETS variant transcription factor 5 | protein_coding | NP_004445.1 | 3494 | 601600 | . | . | 45.118965 | 45.379700 | 50.568771 | 58.040501 | 104.993683 | 104.897209 | 113.474846 | 132.702713 | 1.183209 | 6325.370947 | 0.111664565 | 2.173519313 | 0.029741254 | 0.553061869 | 12.500534 | 12.743147 | 12.552399 | 12.550091 | 12.637083 | 12.746265 | 6682 | 6244 | 5582 | 6837 |
| 2120 | NM_001987,XI | ETV6 | ETS variant transcription factor 6 | protein_coding | NP_001978.1;XP_011518909.1;XP_011518910.1;XP_011518911.1;XP_01 | 3495 | 600618 | . | . | 2.775344 | 2.658795 | 2.895669 | 2.663932 | 6.458337 | 6.145922 | 6.497799 | 6.090764 | 1.001420 | 528.134273 | 0.168826026 | 0.012125565 | 0.990325436 | 0.999289967 | 9.043391 | 9.046130 | 9.061944 | 9.024767 | 9.084499 | 9.003051 | 625 | 553 | 489 | 464 |
| 2121 | NM_00130609 | EVC | EvC ciliary complex subunit 1 | protein_coding | NP_001293019.1;NP_001293021.1;NP_714928.1;XP_006713928.1;XP_00 | 3497 | 604831 | . | . | 7.218169 | 7.517097 | 7.879562 | 7.901977 | 16.796978 | 17.376107 | 17.681504 | 18.066935 | 1.121349 | 1885.429675 | 0.12133671 | 1.361791638 | 0.173263651 | 0.947527012 | 10.795700 | 10.960927 | 10.808892 | 10.856952 | 10.941297 | 10.905763 | 1984 | 1977 | 1772 | 1827 |
| 2122 | NM_00110507 | MECOM | MDS1 and EVI1 complex locus | protein_coding | NP_001098547.3;NP_001098548.2;NP_001157471.1;NP_001157472.1;N | 3498 | 165215 | . | . | 1.430074 | 1.427225 | 1.086914 | 1.433508 | 3.327840 | 3.299093 | 2.439005 | 3.277544 | -1.179959 | 223.325722 | 0.228600345 | -1.04433881 | 0.296328663 | 0.998865404 | 7.917841 | 7.678233 | 7.829668 | 7.852664 | 7.734625 | 7.789993 | 272 | 267 | 167 | 204 |
| 2123 | NM_00100392 | EVI2A | ecotropic viral integration site 2A | protein_coding | NP_001003927.1;NP_055025.2 | 3499 | 158380 | . | . | 8.593344 | 8.051012 | 7.337850 | 4.936398 | 19.997067 | 18.610275 | 16.465921 | 11.286486 | -1.374220 | 621.115677 | 0.192857533 | -2.3779911 | 0.017407246 | 0.442490276 | 9.489306 | 9.032050 | 9.394700 | 9.344008 | 9.274438 | 9.025526 | 857 | 748 | 545 | 394 |
| 2124 | NM_006495,XI | EVI2B | ecotropic viral integration site 2B | protein_coding | NP_006486.3;XP_005258003.1 | 3500 | 158381 | . | . | 1.753562 | 1.831938 | 1.790801 | 0.771141 | 4.080612 | 4.234608 | 4.018506 | 1.763121 | -1.395467 | 96.828543 | 0.397723107 | -1.20875006 | 0.226758882 | 0.982008277 | 6.816237 | 6.339233 | 6.617234 | 6.646823 | 6.636624 | 6.417414 | 124 | 127 | 100 | 44 |
| 2130 | NM_00116328 | EWSR1 | EWS RNA binding protein 1 | protein_coding | NP_001156757.1;NP_001156758.1;NP_001156759.1;NP_005234.1;NP_0 | 3508 | 133450 | . | . | 93.361847 | 93.074152 | 93.500460 | 95.315578 | 217.256859 | 215.145073 | 209.812325 | 217.927749 | 1.013046 | 6814.026034 | 0.103024552 | 0.181511329 | 0.855966242 | 0.998865404 | 12.724921 | 12.743602 | 12.717487 | 12.739917 | 12.738327 | 12.741091 | 7692 | 7402 | 5954 | 6440 |
| 2131 | NM_000127 | EXT1 | exostosin glycosyltransferase 1 | protein_coding | NP_000118.2 | 3512 | 608177 | . | . | 11.839344 | 11.859204 | 10.691922 | 11.684820 | 27.550640 | 27.413080 | 23.992363 | 26.715956 | -1.074053 | 2958.625150 | 0.114054381 | -0.90364979 | 0.366181131 | 0.998865404 | 11.581384 | 11.478194 | 11.562011 | 11.556192 | 11.468286 | 11.532099 | 3541 | 3295 | 2383 | 2780 |
| 2132 | NM_000401,N | EXT2 | exostosin glycosyltransferase 2 | protein_coding | NP_000392.3;NP_001171554.1;NP_997005.1;XP_011518252.1;XP_01151 | 3513 | 608210 | . | . | 20.733447 | 20.427466 | 15.706504 | 24.265300 | 48.247582 | 47.219003 | 35.244937 | 55.479726 | -1.189364 | 4656.065780 | 0.113708914 | -2.20026997 | 0.027787747 | 0.53804272 | 12.304622 | 12.054327 | 12.270641 | 12.234239 | 12.057976 | 12.155296 | 5950 | 5342 | 3479 | 4224 |
| 2134 | NM_004455,XI | EXTL1 | exostosin like glycosyltransferase 1 | protein_coding | NP_004446.2;XP_005245836.1;XP_016856139.1 | 3515 | 601738 | . | . | 23.454613 | 21.828017 | 23.080214 | 22.471131 | 54.579850 | 50.456440 | 51.791329 | 51.377570 | -1.004039 | 2738.665391 | 0.115701248 | -0.05026379 | 0.95991218 | 0.998865404 | 11.422075 | 11.416434 | 11.446432 | 11.394167 | 11.440430 | 11.393888 | 3261 | 2866 | 2445 | 2489 |
| 2135 | NM_00103302 | EXTL2 | exostosin like glycosyltransferase 2 | protein_coding | NP_001028197.1;NP_001248369.1;NP_001248370.1;NP_001248371.1;N | 3516 | 602411 | . | . | 13.387697 | 12.024559 | 11.003868 | 8.651316 | 31.153720 | 27.795304 | 24.692360 | 19.780206 | -1.308790 | 1034.749039 | 0.167675252 | -2.31538754 | 0.020591733 | 0.474989278 | 10.195753 | 9.808463 | 10.154906 | 10.046131 | 10.007511 | 9.788984 | 1446 | 1176 | 900 | 711 |
| 2137 | NM_001440,N | EXTL3 | exostosin like glycosyltransferase 3 | protein_coding | NP_001431.1;XP_011542742.1;XP_024302862.1;XP_024302863.1;XP_02 | 3518 | 605744 | . | . | 5.593673 | 6.366034 | 6.466588 | 6.004712 | 13.016706 | 14.715374 | 14.510833 | 13.729059 | -1.026710 | 1391.178298 | 0.130532184 | -0.29133223 | 0.770797241 | 0.998865404 | 10.461049 | 10.422882 | 10.473470 | 10.429834 | 10.396159 | 10.465948 | 1669 | 1478 | 1139 | 1346 |
| 2138 | NM_000503,N | EYA1 | EYA transcriptional coactivator and phosphatase 1 | protein_coding | NP_000494.2;NP_001275503.1;NP_001275504.1;NP_001357262.1;NP_0 | 3519 | 601653 | . | . | 5.707169 | 5.250141 | 6.600558 | 5.971846 | 13.280814 | 12.135935 | 14.811459 | 13.653915 | 1.131673 | 793.043079 | 0.149005499 | 1.197654335 | 0.231051614 | 0.986164394 | 9.539005 | 9.717960 | 9.605864 | 9.563224 | 9.707060 | 9.635707 | 883 | 778 | 769 | 750 |
| 2139 | NM_005244,N | EYA2 | EYA transcriptional coactivator and phosphatase 2 | protein_coding | NP_005235.3;NP_742108.2;XP_005260384.1;XP_016883210.1 | 3520 | 601654 | . | . | 0.390325 | 0.510655 | 0.072116 | 0.492395 | 0.908303 | 1.180402 | 0.161826 | 1.125803 | -1.559947 | 28.352585 | 0.72115172 | -0.88954496 | 0.373710271 | 0.998865404 | 5.113960 | 4.464338 | 4.831539 | 4.858636 | 4.726033 | 4.858945 | 36 | 41 | 5 | 36 |
| 2140 | NM_00128256 | EYA3 | EYA transcriptional coactivator and phosphatase 3 | protein_coding | NP_001269489.1;NP_001269490.1;NP_001269491.1;NP_001981.2;XP_0 | 3521 | 601655 | . | . | 3.966379 | 4.028691 | 4.787768 | 4.656887 | 9.229927 | 9.312499 | 10.743613 | 10.647421 | 1.184419 | 771.809206 | 0.147263446 | 1.658117085 | 0.097293836 | 0.813553488 | 9.464824 | 9.709053 | 9.512564 | 9.544162 | 9.665381 | 9.628023 | 795 | 780 | 747 | 764 |
| 2145 | NM_00132107 | EZH1 | enhancer of zeste 1 polycomb repressive complex 2 subunit | protein_coding | NP_001308008.1;NP_001308010.1;NP_001308011.1;NP_001982.2;XP_0 | 3526 | 601674 | . | . | 6.469971 | 6.888951 | 7.425541 | 6.878180 | 15.055890 | 15.924117 | 16.662697 | 15.726142 | 1.060542 | 991.627657 | 0.139245128 | 0.60901058 | 0.542517426 | 0.998865404 | 9.910568 | 9.995492 | 9.912379 | 9.949589 | 10.002162 | 9.945426 | 1080 | 1065 | 922 | 920 |
| 2146 | NM_00120324 | EZH2 | enhancer of zeste 2 polycomb repressive complex 2 subunit | protein_coding | NP_001190176.1;NP_001190177.1;NP_001190178.1;NP_004447.2;NP_6 | 3527 | 601573 | . | . | 18.875214 | 15.273699 | 17.004755 | 17.412619 | 43.923402 | 35.305844 | 38.158175 | 39.811885 | -1.119066 | 1711.695433 | 0.126728239 | -1.28065723 | 0.200314088 | 0.96621494 | 10.820080 | 10.657779 | 10.742857 | 10.822169 | 10.718609 | 10.671137 | 1978 | 2048 | 1448 | 1468 |
| 2149 | NM_00131131 | F2R | coagulation factor II thrombin receptor | protein_coding | NP_001298242.1;NP_001983.2 | 3537 | 187930 | . | . | 5.204727 | 4.710387 | 2.242304 | 2.965185 | 12.111613 | 10.888270 | 5.031665 | 6.779544 | -1.934344 | 440.507642 | 0.193391696 | -4.92184836 | 8.57306E-07 | 0.000262964 | 9.182232 | 8.229206 | 8.975002 | 8.902116 | 8.470835 | 8.607228 | 704 | 594 | 226 | 319 |
| 2150 | NM_005242,XI | F2RL1 | F2R like trypsin receptor 1 | protein_coding | NP_005233.4;XP_016864712.1 | 3538 | 600933 | . | . | 1.322107 | 1.381508 | 1.225394 | 1.263155 | 3.076598 | 3.193419 | 2.749748 | 2.888054 | -1.102737 | 116.319101 | 0.298432408 | -0.4727643 | 0.636381346 | 0.998865404 | 6.930908 | 6.789518 | 6.872743 | 6.885096 | 6.842333 | 6.850022 | 138 | 134 | 95 | 105 |
| 2152 | NM_00117809 | F3 | coagulation factor III, tissue factor | protein_coding | NP_001171567.1;NP_001984.1 | 3541 | 134390 | . | . | 5.362844 | 4.754937 | 1.935126 | 2.504910 | 12.479558 | 10.991250 | 4.342367 | 5.727179 | -2.315253 | 261.306513 | 0.228654342 | -5.29694707 | 1.17755E-07 | 5.70364E-05 | 8.512074 | 7.299500 | 8.221848 | 8.144018 | 7.722216 | 7.814652 | 447 | 369 | 120 | 166 |
| 2157 | NM_000132,N | F8 | coagulation factor VIII | protein_coding | NP_000123.1;NP_063916.1 | 3546 | 300841 | . | . | 0.942738 | 0.903710 | 0.883623 | 0.843410 | 2.193790 | 2.088967 | 1.982824 | 1.928355 | -1.058492 | 74.053610 | 0.366569716 | -0.22372207 | 0.822973569 | 0.998865404 | 6.250766 | 6.169074 | 6.225102 | 6.211364 | 6.205833 | 6.202474 | 90 | 80 | 63 | 67 |
| 2161 | NM_000505,XI | F12 | coagulation factor XII | protein_coding | NP_000496.2;XP_011532764.1 | 3530 | 610619 | . | . | 2.093900 | 1.940598 | 2.077865 | 1.711330 | 4.872591 | 4.485779 | 4.662667 | 3.912753 | -1.079518 | 124.693183 | 0.295385463 | -0.37370741 | 0.708622012 | 0.998865404 | 7.015651 | 6.906773 | 6.989392 | 6.959923 | 6.981840 | 6.913196 | 155 | 134 | 115 | 101 |
| 2166 | NM_001441 | FAAH | fatty acid amide hydrolase | protein_coding | NP_001432.2 | 3553 | 602935 | . | . | 0.289505 | 0.195502 | 0.271859 | 0.192409 | 0.673689 | 0.451912 | 0.610043 | 0.439919 | -1.031122 | 15.819936 | 0.780499119 | -0.05664896 | 0.954824836 | NA | 4.003609 | 3.963455 | 3.996938 | 3.969711 | 3.993041 | 3.969399 | 22 | 14 | 16 | 12 |
| 2167 | NM_001442 | FABP4 | fatty acid binding protein 4 | protein_coding | NP_001433.1 | 3559 | 600434 | . | . | 0.225295 | 0.204114 | 0.285793 | 0.124300 | 0.524272 | 0.471819 | 0.641311 | 0.284196 | -1.001197 | 6.725303 | 1.179688418 | -0.00146274 | 0.998832904 | NA | 2.747897 | 2.751299 | 2.746750 | 2.744601 | 2.756451 | 2.734670 | 8 | 7 | 8 | 4 |
| 2171 | NM_001444 | FABP5 | fatty acid binding protein 5 | protein_coding | NP_001435.1 | 3560 | 605168 | . | . | 106.906487 | 102.146088 | 90.674561 | 110.875099 | 248.775787 | 236.115265 | 203.471069 | 253.502747 | -1.052175 | 2162.093805 | 0.131743069 | -0.55694779 | 0.577563098 | 0.998865404 | 11.114561 | 11.040926 | 11.118723 | 11.075201 | 10.978098 | 11.129190 | 2622 | 2328 | 1658 | 2163 |
| 2173 | NM_00131903 | FABP7 | fatty acid binding protein 7 | protein_coding | NP_001305968.1;NP_001305970.1;NP_001305971.1;NP_001437.1 | 3562 | 602965 | . | . | 1.960004 | 2.163655 | 2.485751 | 1.611522 | 4.561011 | 5.001386 | 5.577954 | 3.684554 | 1.031924 | 66.515963 | 0.384971701 | 0.117766227 | 0.90625289 | 0.998865404 | 6.032674 | 6.078224 | 6.050117 | 6.052518 | 6.063328 | 6.053728 | 75 | 71 | 60 | 62 |
| 2175 | NM_000135,N | FANCA | FA complementation group A | protein_coding | NP_000126.2;NP_001018122.1;NP_001273096.1;NP_001338759.1 | 3582 | 607139 | . | . | 6.814306 | 8.012976 | 7.647957 | 6.354241 | 15.857169 | 18.522351 | 17.161796 | 14.528218 | -1.093281 | 990.958797 | 0.157357773 | -0.8176519 | 0.413555985 | 0.998865404 | 10.015490 | 9.887013 | 9.913238 | 10.046864 | 9.982708 | 9.846768 | 1085 | 1217 | 901 | 804 |
| 2176 | NM_000136,N | FANCC | FA complementation group C | protein_coding | NP_000127.2;NP_001230672.1;NP_001230673.1;XP_005251859.1;XP_00 | 3584 | 613899 | . | . | 2.452574 | 2.584874 | 2.290982 | 2.883514 | 5.707239 | 5.975052 | 5.140898 | 6.592812 | -1.023684 | 340.309283 | 0.200720397 | -0.16824284 | 0.866392241 | 0.998865404 | 8.427998 | 8.393198 | 8.411260 | 8.421558 | 8.341914 | 8.460820 | 393 | 375 | 261 | 349 |
| 2177 | NM_00101811 | FANCD2 | FA complementation group D2 | protein_coding | NP_001018125.1;NP_001306913.1;NP_001361182.1;NP_001361183.1;N | 3585 | 613984 | . | . | 9.737964 | 9.688115 | 9.655952 | 9.114580 | 22.660643 | 22.394514 | 21.667677 | 20.839406 | -1.032708 | 1525.680806 | 0.125157685 | -0.37098875 | 0.710645909 | 0.998865404 | 10.598172 | 10.551932 | 10.597056 | 10.577776 | 10.583929 | 10.540848 | 1807 | 1652 | 1342 | 1368 |
| 2178 | NM_021922,XI | FANCE | FA complementation group E | protein_coding | NP_068741.1;XP_005248942.1;XP_005248943.1;XP_005248944.1;XP_00 | 3586 | 613976 | . | . | 3.047279 | 3.759715 | 2.926668 | 2.919095 | 7.091144 | 8.690749 | 6.567360 | 6.674165 | -1.182945 | 254.364120 | 0.222507178 | -1.08932852 | 0.276009039 | 0.998865404 | 8.107160 | 7.864119 | 7.974676 | 8.082997 | 7.946991 | 7.944725 | 285 | 328 | 203 | 218 |
| 2180 | NM_00128670 | ACSL1 | acyl-CoA synthetase long chain family member 1 | protein_coding | NP_001273637.1;NP_001273639.1;NP_001273640.1;NP_001368806.1;N | 3569 | 152425 | . | . | 13.737223 | 13.925852 | 13.034321 | 13.366624 | 31.967085 | 32.190231 | 29.248637 | 30.561196 | -1.063499 | 1569.490130 | 0.123169296 | -0.72110751 | 0.47084337 | 0.998865404 | 10.659844 | 10.570947 | 10.635437 | 10.643745 | 10.583190 | 10.600283 | 1853 | 1755 | 1308 | 1442 |
| 2181 | NM_00135415 | ACSL3 | acyl-CoA synthetase long chain family member 3 | protein_coding | NP_001341087.1;NP_001341088.1;NP_004448.2;NP_976251.1 | 3570 | 602371 | . | . | 62.841915 | 56.317749 | 59.390014 | 52.233523 | 146.235722 | 130.181003 | 133.269471 | 119.425738 | -1.082293 | 9924.234679 | 0.117348621 | -0.97224191 | 0.330930213 | 0.998865404 | 13.332609 | 13.218621 | 13.359626 | 13.254681 | 13.298084 | 13.180800 | 12604 | 10452 | 8863 | 8318 |
| 2182 | NM_00131850 | ACSL4 | acyl-CoA synthetase long chain family member 4 | protein_coding | NP_001305438.1;NP_001305439.1;NP_004449.1;NP_075266.1;XP_0052 | 3571 | 300157 | . | . | 7.146186 | 6.384656 | 6.755485 | 7.091306 | 16.629468 | 14.758420 | 15.159114 | 16.213426 | 1.018264 | 971.867734 | 0.145156859 | 0.179884954 | 0.857242888 | 0.998865404 | 9.911274 | 9.937836 | 9.969928 | 9.861448 | 9.945783 | 9.914899 | 1191 | 962 | 871 | 900 |
| 2184 | NM_000137,N | FAH | fumarylacetoacetate hydrolase | protein_coding | NP_000128.1;NP_001361306.1;NP_001361309.1 | 3579 | 613871 | . | . | 30.881727 | 33.577761 | 32.356733 | 33.281264 | 71.863048 | 77.616502 | 72.607571 | 76.093652 | -1.014991 | 1677.718126 | 0.122924355 | -0.17463835 | 0.861363811 | 0.998865404 | 10.722919 | 10.701572 | 10.712210 | 10.723634 | 10.729142 | 10.683035 | 1932 | 1837 | 1491 | 1515 |
| 2185 | NM_004103,N | PTK2B | protein tyrosine kinase 2 beta | protein_coding | NP_004094.3;NP_775266.1;NP_775267.1;NP_775268.1;XP_005273504.1 | 9612 | 601212 | . | . | 1.032790 | 0.982171 | 0.829367 | 1.014521 | 2.403346 | 2.270332 | 1.861075 | 2.319582 | -1.054980 | 138.855950 | 0.280924318 | -0.27486461 | 0.783420254 | 0.998865404 | 7.156133 | 7.077692 | 7.122201 | 7.132587 | 7.071778 | 7.141874 | 162 | 156 | 106 | 139 |
| 2186 | NM_004459,N | BPTF | bromodomain PHD finger transcription factor | protein_coding | NP_004450.3;NP_872579.2;XP_005257207.1;XP_005257208.1;XP_00525 | 3581 | 601819 | . | . | 4.641674 | 5.023477 | 5.717248 | 5.003034 | 10.801367 | 11.611991 | 12.829339 | 11.438842 | 1.100272 | 1808.431282 | 0.131368739 | 1.049418612 | 0.293985496 | 0.998865404 | 10.749836 | 10.887908 | 10.755721 | 10.804493 | 10.917140 | 10.790483 | 1921 | 1916 | 1779 | 1635 |
| 2187 | NM_00101811 | FANCB | FA complementation group B | protein_coding | NP_001018123.1;NP_001311091.1;NP_689846.1;XP_011543772.1;XP_01 | 3583 | 300515 | . | . | 4.057309 | 3.737576 | 3.928829 | 2.636228 | 9.441525 | 8.639572 | 8.816183 | 6.027421 | -1.104178 | 407.660504 | 0.215497534 | -0.66345145 | 0.507041466 | 0.998865404 | 8.740319 | 8.598654 | 8.676426 | 8.712386 | 8.750910 | 8.505152 | 478 | 475 | 408 | 287 |
| 2188 | NM_022725 | FANCF | FA complementation group F | protein_coding | NP_073562.1 | 3587 | 613897 | . | . | 5.284045 | 4.604955 | 4.369959 | 4.886991 | 12.296188 | 10.644560 | 9.806063 | 11.173524 | -1.082895 | 491.223427 | 0.17602358 | -0.65271203 | 0.513941946 | 0.998865404 | 8.996549 | 8.881634 | 9.010313 | 8.916138 | 8.881873 | 8.945290 | 631 | 511 | 389 | 465 |
| 2189 | NM_004629 | FANCG | FA complementation group G | protein_coding | NP_004620.1 | 3588 | 602956 | . | . | 24.648563 | 26.738012 | 25.864941 | 25.801311 | 57.358219 | 61.806110 | 58.040180 | 58.991634 | -1.008376 | 2051.625004 | 0.119612829 | -0.10060162 | 0.919866714 | 0.998865404 | 11.008596 | 10.996481 | 10.976203 | 11.034368 | 11.004121 | 10.993903 | 2286 | 2304 | 1788 | 1903 |
| 2192 | NM_001996,N | FBLN1 | fibulin 1 | protein_coding | NP_001987.3;NP_006476.3;NP_006477.3;NP_006478.3 | 3600 | 135820 | . | . | 0.700656 | 0.501547 | 0.319659 | 0.600004 | 1.630456 | 1.159349 | 0.717306 | 1.371840 | -1.460709 | 42.420512 | 0.517838342 | -1.05567417 | 0.2911171 | 0.998865404 | 5.655579 | 5.105724 | 5.465746 | 5.406361 | 5.331041 | 5.418446 | 67 | 46 | 20 | 43 |
| 2193 | NM_004461 | FARSA | phenylalanyl-tRNA synthetase subunit alpha | protein_coding | NP_004452.1 | 3592 | 602918 | . | . | 37.275864 | 39.216679 | 38.604530 | 39.997028 | 86.742470 | 90.651108 | 86.627449 | 91.448463 | 1.013787 | 2187.610969 | 0.117021936 | 0.168814139 | 0.865942837 | 0.998865404 | 11.085287 | 11.104927 | 11.072488 | 11.106348 | 11.090286 | 11.110561 | 2449 | 2394 | 1891 | 2091 |
| 2194 | NM_004104,XI | FASN | fatty acid synthase | protein_coding | NP_004095.4;XP_011521840.1 | 3594 | 600212 | . | . | 50.778578 | 52.418059 | 67.524616 | 67.193882 | 118.163845 | 121.166690 | 151.523274 | 153.630826 | 1.287412 | 15671.500518 | 0.099239623 | 3.672668214 | 0.000240031 | 0.023665347 | 13.742140 | 14.106612 | 13.807973 | 13.827698 | 14.041153 | 14.027940 | 15593 | 14959 | 15456 | 16413 |
| 2195 | NM_005245,XI | FAT1 | FAT atypical cadherin 1 | protein_coding | NP_005236.2;XP_005262891.1;XP_005262892.1;XP_006714202.1 | 3595 | 600976 | . | . | 9.797820 | 10.993317 | 7.899519 | 8.801442 | 22.799932 | 25.411546 | 17.726292 | 20.123452 | -1.259758 | 4345.526453 | 0.116401982 | -2.86203971 | 0.004209242 | 0.192048182 | 12.242376 | 11.909044 | 12.127061 | 12.217510 | 11.941608 | 12.018910 | 5283 | 5506 | 3185 | 3777 |
| 2196 | NM_001447,XI | FAT2 | FAT atypical cadherin 2 | protein_coding | NP_001438.1;XP_006714824.1;XP_011535902.1;XP_011535905.1;XP_01 | 3596 | 604269 | . | . | 0.192759 | 0.185495 | 0.127429 | 0.170152 | 0.448559 | 0.428779 | 0.285947 | 0.389031 | -1.245416 | 79.175518 | 0.363174189 | -0.87183365 | 0.383299133 | 0.998865404 | 6.457409 | 6.139026 | 6.335617 | 6.336369 | 6.247236 | 6.309902 | 101 | 95 | 53 | 75 |
| 2197 | NM_001997 | FAU | FAU ubiquitin like and ribosomal protein S30 fusion | protein_coding | NP_001988.1 | 3597 | 134690 | . | . | 215.512650 | 207.057236 | 182.056732 | 224.294434 | 501.506805 | 478.622101 | 408.530060 | 512.822571 | -1.055045 | 3266.685009 | 0.127355524 | -0.6069948 | 0.543854361 | 0.998865404 | 11.711843 | 11.634339 | 11.713932 | 11.673868 | 11.565951 | 11.727669 | 3956 | 3532 | 2491 | 3275 |
| 2201 | NM_001999,XI | FBN2 | fibrillin 2 | protein_coding | NP_001990.2;XP_016864717.1 | 3604 | 612570 | . | . | 0.944480 | 0.917616 | 0.802007 | 0.888270 | 2.197845 | 2.121112 | 1.799681 | 2.030924 | -1.116146 | 302.795716 | 0.201886208 | -0.78522587 | 0.432321149 | 0.998865404 | 8.319459 | 8.160571 | 8.282288 | 8.260411 | 8.187421 | 8.236416 | 375 | 338 | 237 | 281 |
| 2202 | NM_00103934 | EFEMP1 | EGF containing fibulin extracellular matrix protein 1 | protein_coding | NP_001034437.1;NP_001034438.1;XP_005264262.1;XP_016859075.1;XP | 3218 | 601548 | . | . | 0.194375 | 0.180908 | 0.117415 | 0.183678 | 0.452318 | 0.418179 | 0.263475 | 0.419957 | -1.267573 | 16.818426 | 0.755994155 | -0.45247517 | 0.650926703 | NA | 4.234218 | 3.889137 | 4.086179 | 4.083559 | 4.053829 | 4.081625 | 22 | 20 | 10 | 17 |
| 2207 | NM_004106 | FCER1G | Fc fragment of IgE receptor Ig | protein_coding | NP_004097.1 | 3611 | 147139 | . | . | 10.829810 | 11.114873 | 6.655665 | 9.466369 | 25.201412 | 25.692526 | 14.935120 | 21.643728 | -1.374547 | 175.765804 | 0.266035331 | -1.7251683 | 0.084497147 | 0.788061307 | 7.669650 | 7.208709 | 7.513975 | 7.523209 | 7.319815 | 7.445514 | 232 | 222 | 107 | 162 |
| 2212 | NM_00113621 | FCGR2A | Fc fragment of IgG receptor IIa | protein_coding | NP_001129691.1;NP_001362225.1;NP_001362226.1;NP_067674.2;XP_01 | 3616 | 146790 | . | . | 2.023911 | 1.832707 | 1.237592 | 1.468195 | 4.709721 | 4.236385 | 2.777119 | 3.356851 | -1.380589 | 130.002023 | 0.291083506 | -1.59845647 | 0.109941426 | 0.843391194 | 7.236701 | 6.770595 | 7.098263 | 7.054772 | 6.932341 | 6.990635 | 183 | 154 | 86 | 112 |
| 2213 | NM_00100227 | FCGR2B | Fc fragment of IgG receptor IIb | protein_coding | NP_001002273.1;NP_001002274.1;NP_001002275.1;NP_001177757.1;N | 3618 | 604590 | . | . | 0.102112 | 0.219808 | 0.240367 | 0.235674 | 0.237619 | 0.508096 | 0.539376 | 0.538840 | 1.459507 | 14.471732 | 0.823567755 | 0.662339685 | 0.507753549 | NA | 3.558919 | 4.100822 | 3.812813 | 3.849119 | 3.855030 | 3.854502 | 9 | 17 | 15 | 16 |
| 2218 | NM_00107980 | FKTN | fukutin | protein_coding | NP_001073270.1;NP_001185892.1;NP_001338425.1;NP_001338426.1;N | 3622 | 607440 | . | . | 2.048890 | 1.833693 | 1.810618 | 1.712829 | 4.767852 | 4.238662 | 4.062972 | 3.916182 | 1.042238 | 396.106472 | 0.187824335 | 0.317766002 | 0.750662442 | 0.998865404 | 8.599792 | 8.659088 | 8.564561 | 8.664845 | 8.651166 | 8.629450 | 409 | 454 | 358 | 372 |
| 2222 | NM_00128774 | FDFT1 | farnesyl-diphosphate farnesyltransferase 1 | protein_coding | NP_001274671.1;NP_001274672.1;NP_001274673.1;NP_001274674.1;N | 3629 | 184420 | . | . | 88.924353 | 84.919354 | 113.742517 | 116.235263 | 206.930617 | 196.294885 | 255.234910 | 265.758125 | 1.324562 | 6926.944855 | 0.104515539 | 3.879949274 | 0.000104478 | 0.012762065 | 12.541060 | 12.946548 | 12.635597 | 12.619005 | 12.840676 | 12.888548 | 6932 | 6366 | 6675 | 7605 |
| 2224 | NM_00113582 | FDPS | farnesyl diphosphate synthase | protein_coding | NP_001129293.1;NP_001129294.1;NP_001229753.1;NP_001229754.1;N | 3631 | 134629 | . | . | 71.738709 | 71.062831 | 102.575974 | 109.892916 | 166.938921 | 164.264913 | 230.177513 | 251.257104 | 1.472177 | 3489.528803 | 0.11141194 | 5.007997857 | 5.49991E-07 | 0.000189788 | 11.463086 | 12.020950 | 11.587099 | 11.588134 | 11.875584 | 11.935712 | 3249 | 3048 | 3487 | 4033 |
| 2230 | NM_004109 | FDX1 | ferredoxin 1 | protein_coding | NP_004100.1 | 3638 | 103260 | . | . | 7.387494 | 6.856238 | 6.032613 | 7.950868 | 17.191002 | 15.848501 | 13.537009 | 18.178715 | -1.033193 | 691.477642 | 0.171293004 | -0.27502375 | 0.783297994 | 0.998865404 | 9.457217 | 9.409466 | 9.471111 | 9.413494 | 9.326293 | 9.506198 | 843 | 727 | 513 | 722 |
| 2231 | gene-FDX1P1 | FDX1P1 | ferredoxin 1 pseudogene 1 | pseudogene | . | 3640 | . | . | . | 0.018483 | 0.115365 | 0.081928 | 0.112607 | 0.043012 | 0.266671 | 0.183844 | 0.257463 | 1.344963 | 4.806000 | 1.414161368 | 0.302346321 | 0.762388071 | NA | 2.039857 | 2.459419 | 2.240110 | 2.262694 | 2.253922 | 2.262012 | 2 | 7 | 4 | 6 |
| 2232 | NM_00125801 | FDXR | ferredoxin reductase | protein_coding | NP_001244941.2;NP_001244942.2;NP_001244943.2;NP_001244944.1;N | 3642 | 103270 | . | . | 8.588474 | 9.044721 | 7.117415 | 8.851580 | 19.985736 | 20.907282 | 15.971273 | 20.238087 | -1.111257 | 487.841181 | 0.17868735 | -0.85172714 | 0.394365556 | 0.998865404 | 9.004816 | 8.851657 | 8.947001 | 8.976468 | 8.831431 | 8.955321 | 578 | 567 | 361 | 477 |
| 2235 | NM_000140,N | FECH | ferrochelatase | protein_coding | NP_000131.2;NP_001012533.1;NP_001358023.1;NP_001358024.1;NP_0 | 3647 | 612386 | . | . | 2.542554 | 2.502004 | 2.771841 | 3.520432 | 5.916627 | 5.783494 | 6.219929 | 8.049050 | 1.183738 | 660.075705 | 0.165014106 | 1.474718655 | 0.140288198 | 0.900674731 | 9.240036 | 9.482744 | 9.307371 | 9.300499 | 9.342792 | 9.488516 | 700 | 649 | 557 | 741 |
| 2237 | NM_004111 | FEN1 | flap structure-specific endonuclease 1 | protein_coding | NP_004102.1 | 3650 | 600393 | . | . | 39.838230 | 43.182667 | 40.861198 | 42.773914 | 92.705200 | 99.818672 | 91.691345 | 97.797478 | -1.006326 | 2616.928300 | 0.115957033 | -0.07845312 | 0.93746762 | 0.998865404 | 11.358276 | 11.349026 | 11.326302 | 11.384944 | 11.336385 | 11.365151 | 2914 | 2935 | 2228 | 2489 |
| 2239 | NM_001448 | GPC4 | glypican 4 | protein_coding | NP_001439.2 | 4452 | 300168 | . | . | 6.715619 | 5.585819 | 5.386289 | 5.301651 | 15.627521 | 12.911872 | 12.086677 | 12.121595 | -1.167649 | 889.139441 | 0.152746554 | -1.46390568 | 0.143219722 | 0.905131505 | 9.903525 | 9.680387 | 9.916994 | 9.774495 | 9.746337 | 9.727541 | 1209 | 934 | 723 | 759 |
| 2241 | NM_00130802 | FER | FER tyrosine kinase | protein_coding | NP_001294957.1;NP_001294960.1;NP_001294967.1;NP_005237.2;XP_01 | 3655 | 176942 | . | . | 1.955463 | 1.932360 | 1.861623 | 2.033464 | 4.550442 | 4.466735 | 4.177428 | 4.649275 | -1.082070 | 701.000966 | 0.152123681 | -0.74803543 | 0.454438786 | 0.998865404 | 9.508887 | 9.395428 | 9.478881 | 9.480323 | 9.454373 | 9.395830 | 838 | 787 | 611 | 604 |
| 2245 | NM_004463 | FGD1 | FYVE, RhoGEF and PH domain containing 1 | protein_coding | NP_004454.2 | 3663 | 300546 | . | . | 3.788401 | 4.636882 | 4.120998 | 4.875469 | 8.815765 | 10.718361 | 9.247401 | 11.147182 | 1.053545 | 589.010196 | 0.174557959 | 0.431099362 | 0.666396124 | 0.998865404 | 9.164554 | 9.238788 | 9.116663 | 9.244325 | 9.163163 | 9.267720 | 597 | 679 | 484 | 611 |
| 2246 | NM_000800,N | FGF1 | fibroblast growth factor 1 | protein_coding | NP_000791.1;NP_001138364.1;NP_001138406.1;NP_001138407.1;NP_0 | 3665 | 131220 | . | . | 1.775467 | 1.883652 | 2.489021 | 2.466667 | 4.131585 | 4.354145 | 5.585292 | 5.639741 | 1.344120 | 253.462574 | 0.214667054 | 1.987551071 | 0.046861362 | 0.656677546 | 7.756634 | 8.183206 | 7.889309 | 7.912585 | 8.055689 | 8.042676 | 243 | 239 | 256 | 269 |
| 2247 | NM_00136166 | FGF2 | fibroblast growth factor 2 | protein_coding | NP_001348594.1;NP_001997.5 | 3676 | 134920 | . | . | 1.048238 | 0.716522 | 0.883232 | 0.813913 | 2.439293 | 1.656272 | 1.981947 | 1.860915 | -1.053601 | 180.292374 | 0.271360965 | -0.27759585 | 0.781322619 | 0.998865404 | 7.530563 | 7.456886 | 7.583524 | 7.412485 | 7.496142 | 7.462073 | 254 | 161 | 159 | 158 |
| 2252 | NM_002009 | FGF7 | fibroblast growth factor 7 | protein_coding | NP_002000.1 | 3685 | 148180 | . | . | 0.464100 | 0.329255 | 0.341410 | 0.265155 | 1.079981 | 0.761088 | 0.766114 | 0.606246 | -1.324139 | 58.749133 | 0.427271087 | -0.94800472 | 0.343127051 | 0.998865404 | 6.063551 | 5.661523 | 5.943290 | 5.863072 | 5.870182 | 5.820238 | 90 | 60 | 50 | 41 |
| 2256 | NM_00130346 | FGF11 | fibroblast growth factor 11 | protein_coding | NP_001290389.1;NP_004103.1 | 3667 | 601514 | . | . | 0.267577 | 0.200206 | 0.137629 | 0.381481 | 0.622663 | 0.462785 | 0.308836 | 0.872210 | 1.073236 | 18.972166 | 0.74063168 | 0.137675619 | 0.890496789 | NA | 4.195961 | 4.293999 | 4.247336 | 4.232948 | 4.211785 | 4.278966 | 23 | 18 | 10 | 26 |
| 2257 | NM_00137729 | FGF12 | fibroblast growth factor 12 | protein_coding | NP_001364221.1;NP_001364222.1;NP_001364223.1;NP_004104.3;NP_0 | 3668 | 601513 | . | . | 0.098769 | 0.162736 | 0.035230 | 0.074343 | 0.229839 | 0.376170 | 0.079055 | 0.169977 | -1.869355 | 17.305253 | 0.776759305 | -1.16193117 | 0.245263426 | NA | 4.497331 | 3.587955 | 4.116009 | 4.160366 | 4.076881 | 4.110232 | 20 | 30 | 7 | 15 |
| 2258 | NM_00113949 | FGF13 | fibroblast growth factor 13 | protein_coding | NP_001132970.1;NP_001132972.1;NP_001132973.1;NP_001132974.1;N | 3670 | 300070 | . | . | 0.654557 | 0.543760 | 0.483058 | 0.536042 | 1.523182 | 1.256924 | 1.083966 | 1.225597 | -1.227295 | 333.754763 | 0.205013196 | -1.44128601 | 0.149503899 | 0.910705785 | 8.522892 | 8.227280 | 8.490677 | 8.377733 | 8.283496 | 8.358131 | 464 | 359 | 242 | 301 |
| 2259 | NM_00132193 | FGF14 | fibroblast growth factor 14 | protein_coding | NP_001308860.1;NP_001308861.1;NP_001308862.1;NP_001308863.1;N | 3671 | 601515 | . | . | 0.355881 | 0.459272 | 0.456835 | 0.441936 | 0.828148 | 1.061631 | 1.025124 | 1.010435 | -1.011056 | 93.946011 | 0.335779572 | -0.04724033 | 0.962321686 | 0.998865404 | 6.562390 | 6.545078 | 6.517982 | 6.588588 | 6.539336 | 6.561870 | 96 | 114 | 78 | 91 |
| 2260 | NM_00117406 | FGFR1 | fibroblast growth factor receptor 1 | protein_coding | NP_001167534.1;NP_001167535.1;NP_001167536.1;NP_001167537.1;N | 3688 | 136350 | . | . | 4.550251 | 4.142715 | 4.337842 | 4.549065 | 10.588613 | 9.576072 | 9.733992 | 10.400897 | 1.020065 | 739.968406 | 0.150192735 | 0.190826853 | 0.84866125 | 0.998865404 | 9.516937 | 9.545561 | 9.548077 | 9.499575 | 9.516503 | 9.558860 | 873 | 763 | 630 | 722 |
| 2262 | NM_004466,XI | GPC5 | glypican 5 | protein_coding | NP_004457.1;XP_011519356.1;XP_011519357.1;XP_011519358.1;XP_01 | 4453 | 602446 | . | . | 0.417420 | 0.533471 | 0.212022 | 0.267051 | 0.971354 | 1.233143 | 0.475772 | 0.610581 | -1.997491 | 42.188155 | 0.49428656 | -2.01945372 | 0.043440083 | 0.642590349 | 5.814036 | 4.813340 | 5.424048 | 5.473564 | 5.330975 | 5.352173 | 57 | 68 | 22 | 29 |
| 2264 | NM_00129198 | FGFR4 | fibroblast growth factor receptor 4 | protein_coding | NP_001278909.1;NP_001341913.1;NP_002002.3;NP_075252.2;NP_9988 | 3691 | 134935 | . | . | 12.231022 | 12.261181 | 10.663868 | 12.332915 | 28.462090 | 28.342272 | 23.929411 | 28.197747 | -1.063889 | 1121.641056 | 0.139301939 | -0.64139909 | 0.521263424 | 0.998865404 | 10.175541 | 10.085856 | 10.166272 | 10.140931 | 10.055513 | 10.156821 | 1354 | 1227 | 880 | 1089 |
| 2271 | NM_000143,XI | FH | fumarate hydratase | protein_coding | NP_000134.2;XP_011542434.1 | 3700 | 136850 | . | . | 42.331833 | 42.500894 | 38.722339 | 47.204819 | 98.507924 | 98.242728 | 86.891808 | 107.928200 | 1.000257 | 2391.723919 | 0.129933439 | 0.00284848 | 0.997727244 | 0.999862634 | 11.223798 | 11.223872 | 11.224139 | 11.221187 | 11.141257 | 11.298297 | 2759 | 2576 | 1877 | 2461 |
| 2272 | NM_00116624 | FHIT | fragile histidine triad diadenosine triphosphatase | protein_coding | NP_001159715.1;NP_001307828.1;NP_001307829.1;NP_001307830.1;N | 3701 | 601153 | . | . | 1.330227 | 1.762197 | 1.697089 | 4.077968 | 3.095495 | 4.073395 | 3.808217 | 9.323791 | 1.076953 | 79.554146 | 0.410311339 | 0.260667401 | 0.794349007 | 0.998865404 | 6.261164 | 6.364709 | 6.290227 | 6.298104 | 6.210453 | 6.407517 | 87 | 84 | 45 | 106 |
| 2273 | NM_00115969 | FHL1 | four and a half LIM domains 1 | protein_coding | NP_001153171.1;NP_001153172.1;NP_001153173.1;NP_001153174.1;N | 3702 | 300163 | . | . | 3.122963 | 3.417144 | 3.131924 | 3.399671 | 7.267265 | 7.898887 | 7.027945 | 7.772945 | -1.012965 | 246.479552 | 0.217588906 | -0.08541252 | 0.931933442 | 0.998865404 | 7.954846 | 7.935739 | 7.936651 | 7.960255 | 7.926102 | 7.957791 | 279 | 274 | 206 | 237 |
| 2274 | NM_00103949 | FHL2 | four and a half LIM domains 2 | protein_coding | NP_001034581.1;NP_001305823.1;NP_001305824.1;NP_001305825.1;N | 3703 | 602633 | . | . | 45.742657 | 48.838627 | 43.038803 | 41.338549 | 106.445046 | 112.892676 | 96.577828 | 94.515677 | -1.100984 | 2741.644017 | 0.113281071 | -1.22521412 | 0.220494556 | 0.982008277 | 11.488597 | 11.349713 | 11.440945 | 11.476095 | 11.371795 | 11.389015 | 3238 | 3167 | 2245 | 2473 |
| 2275 | NM_00124387 | FHL3 | four and a half LIM domains 3 | protein_coding | NP_001230807.1;NP_004459.2;XP_016856164.1;XP_024309867.1;XP_02 | 3704 | 602790 | . | . | 3.935748 | 4.899635 | 3.795032 | 5.517135 | 9.158647 | 11.325726 | 8.515940 | 12.614276 | 1.072065 | 219.731538 | 0.248551017 | 0.403910715 | 0.686278377 | 0.998865404 | 7.729554 | 7.827965 | 7.714789 | 7.801085 | 7.715260 | 7.862616 | 222 | 250 | 167 | 246 |
| 2280 | NM_000801,N | FKBP1A | FKBP prolyl isomerase 1A | protein_coding | NP_000792.1;NP_001186715.1;NP_463460.1 | 3711 | 186945 | . | . | 150.128311 | 142.396918 | 131.502009 | 133.339330 | 349.354747 | 329.156863 | 295.086727 | 304.864512 | -1.117929 | 6449.410021 | 0.105590716 | -1.52313171 | 0.127725757 | 0.875806568 | 12.733108 | 12.572317 | 12.728330 | 12.671028 | 12.605704 | 12.606337 | 8103 | 7100 | 5294 | 5712 |
| 2281 | NM_00132296 | FKBP1B | FKBP prolyl isomerase 1B | protein_coding | NP_001309892.1;NP_001309893.1;NP_004107.1;NP_473374.1;XP_0115 | 3712 | 600620 | . | . | 2.822281 | 3.163922 | 2.377284 | 4.433925 | 6.567571 | 7.313547 | 5.334553 | 10.137642 | 1.003549 | 111.980515 | 0.353157861 | 0.014470793 | 0.98845438 | 0.999197585 | 6.805789 | 6.808418 | 6.817442 | 6.780174 | 6.690242 | 6.899390 | 136 | 114 | 66 | 139 |
| 2286 | NM_00113520 | FKBP2 | FKBP prolyl isomerase 2 | protein_coding | NP_001128680.1;NP_001357289.1;NP_001357291.1;NP_001357292.1;N | 3718 | 186946 | . | . | 71.173858 | 58.968393 | 69.981443 | 70.657557 | 165.624495 | 136.308073 | 157.036328 | 161.550100 | 1.063128 | 1230.020446 | 0.143764914 | 0.614307162 | 0.539012371 | 0.998865404 | 10.219447 | 10.308124 | 10.312400 | 10.160867 | 10.286087 | 10.285428 | 1511 | 1157 | 1103 | 1188 |
| 2287 | NM_002013,XI | FKBP3 | FKBP prolyl isomerase 3 | protein_coding | NP_002004.1;XP_011534867.1;XP_016876581.1;XP_016876582.1 | 3719 | 186947 | . | . | 29.088209 | 28.882956 | 29.167079 | 29.443414 | 67.689457 | 66.764247 | 65.450083 | 67.318872 | 1.026878 | 1361.580064 | 0.128522832 | 0.29772458 | 0.765913382 | 0.998865404 | 10.391701 | 10.430175 | 10.404017 | 10.397057 | 10.447410 | 10.393655 | 1554 | 1443 | 1242 | 1248 |
| 2288 | NM_002014,XI | FKBP4 | FKBP prolyl isomerase 4 | protein_coding | NP_002005.1;XP_011519231.1 | 3720 | 600611 | . | . | 32.965572 | 32.676537 | 29.428853 | 33.491943 | 76.712244 | 75.533283 | 66.037495 | 76.575350 | -1.069466 | 3599.729734 | 0.115992336 | -0.83531463 | 0.403540582 | 0.998865404 | 11.861379 | 11.764335 | 11.845758 | 11.835012 | 11.732684 | 11.834555 | 4312 | 3989 | 2837 | 3464 |
| 2289 | NM_00114577 | FKBP5 | FKBP prolyl isomerase 5 | protein_coding | NP_001139247.1;NP_001139248.1;NP_001139249.1;NP_004108.1 | 3721 | 602623 | . | . | 3.259016 | 3.336128 | 2.301967 | 2.850639 | 7.583862 | 7.711609 | 5.165546 | 6.517646 | -1.310269 | 348.532540 | 0.197414264 | -1.97484947 | 0.048285233 | 0.660542298 | 8.627387 | 8.236518 | 8.513958 | 8.517190 | 8.311022 | 8.411716 | 454 | 428 | 237 | 310 |
| 2295 | NM_001452 | FOXF2 | forkhead box F2 | protein_coding | NP_001443.1 | 3810 | 603250 | . | . | 0.398400 | 0.398867 | 0.336454 | 0.446743 | 0.927093 | 0.921998 | 0.754993 | 1.021425 | -1.019785 | 30.607312 | 0.561079936 | -0.05037578 | 0.959822941 | 0.998865404 | 4.950620 | 4.920835 | 4.939076 | 4.936249 | 4.920038 | 4.950246 | 36 | 33 | 23 | 32 |
| 2296 | NM_001453 | FOXC1 | forkhead box C1 | protein_coding | NP_001444.2 | 3800 | 601090 | . | . | 1.089325 | 1.021326 | 1.105449 | 1.166495 | 2.534905 | 2.360841 | 2.480596 | 2.667052 | 1.063673 | 136.775399 | 0.27821189 | 0.320095054 | 0.748896275 | 0.998865404 | 7.050418 | 7.139536 | 7.096140 | 7.070356 | 7.098139 | 7.114053 | 158 | 138 | 120 | 135 |
| 2297 | NM_004472 | FOXD1 | forkhead box D1 | protein_coding | NP_004463.1 | 3802 | 601091 | . | . | 2.720074 | 2.568500 | 2.550027 | 3.064923 | 6.329723 | 5.937202 | 5.722188 | 7.007583 | 1.049713 | 213.841571 | 0.235078685 | 0.297749494 | 0.765894365 | 0.998865404 | 7.705301 | 7.774663 | 7.742679 | 7.712979 | 7.709464 | 7.789876 | 248 | 218 | 174 | 223 |
| 2303 | NM_005251 | FOXC2 | forkhead box C2 | protein_coding | NP_005242.1 | 3801 | 602402 | . | . | 0.047716 | 0.035960 | 0.032452 | 0.040704 | 0.111037 | 0.083123 | 0.072821 | 0.093065 | -1.155700 | 4.158556 | 1.470708199 | -0.14194993 | 0.887119558 | NA | 2.156660 | 1.947964 | 2.065350 | 2.058249 | 2.057237 | 2.060853 | 6 | 4 | 3 | 4 |
| 2305 | NM_00124308 | FOXM1 | forkhead box M1 | protein_coding | NP_001230017.1;NP_001230018.1;NP_068772.2;NP_973731.1;NP_9737 | 3818 | 602341 | . | . | 32.073741 | 34.012144 | 29.899733 | 34.933297 | 74.636916 | 78.620594 | 67.094131 | 79.870835 | -1.031995 | 3629.908581 | 0.118993668 | -0.38183366 | 0.702584748 | 0.998865404 | 11.848363 | 11.802710 | 11.816873 | 11.858667 | 11.751768 | 11.868670 | 4139 | 4079 | 2886 | 3587 |
| 2306 | NM_004474 | FOXD2 | forkhead box D2 | protein_coding | NP_004465.3 | 3803 | 602211 | . | . | 1.126252 | 0.983330 | 0.964692 | 0.976686 | 2.620835 | 2.273012 | 2.164741 | 2.233077 | -1.098425 | 84.177989 | 0.347985625 | -0.38920144 | 0.69712714 | 0.998865404 | 6.461182 | 6.326414 | 6.429901 | 6.387209 | 6.384223 | 6.382433 | 109 | 88 | 70 | 75 |
| 2308 | NM_002015,XI | FOXO1 | forkhead box O1 | protein_coding | NP_002006.2;XP_011533310.1;XP_011533312.1 | 3819 | 136533 | . | . | 0.775548 | 0.450803 | 0.774634 | 0.833897 | 1.804735 | 1.042050 | 1.738256 | 1.906607 | 1.113908 | 137.379532 | 0.285980437 | 0.544197494 | 0.586305576 | 0.998865404 | 7.021745 | 7.178069 | 7.122570 | 7.033382 | 7.106996 | 7.130780 | 168 | 123 | 122 | 140 |
| 2309 | NM_001455,N | FOXO3 | forkhead box O3 | protein_coding | NP_001446.1;NP_963853.1;XP_005266924.1;XP_005266925.1;XP_01153 | 3821 | 602681 | . | . | 2.964995 | 2.971494 | 3.082743 | 3.278116 | 6.899664 | 6.868737 | 6.917587 | 7.495021 | 1.111863 | 710.621749 | 0.149925201 | 1.020365199 | 0.307555292 | 0.998865404 | 9.394430 | 9.547393 | 9.446805 | 9.422224 | 9.496034 | 9.519043 | 789 | 713 | 640 | 713 |
| 2310 | NM_00136813 | FOXO3B | forkhead box O3B | protein_coding | NP_001355063.1;NP_001355064.1 | 3822 | . | . | . | 1.267360 | 1.449783 | 1.441060 | 1.348920 | 2.949198 | 3.351237 | 3.233696 | 3.084145 | 1.016608 | 255.039943 | 0.218001746 | 0.109002889 | 0.913200199 | 0.998865404 | 7.982705 | 8.006356 | 7.955152 | 8.022341 | 8.016156 | 7.978648 | 272 | 291 | 232 | 232 |
| 2314 | NM_00125626 | FLII | FLII actin remodeling protein | protein_coding | NP_001243193.1;NP_001243194.1;NP_002009.1;XP_005256612.1;XP_00 | 3750 | 600362 | . | . | 29.895735 | 31.785141 | 30.723978 | 32.840236 | 69.568597 | 73.472777 | 68.943715 | 75.085301 | 1.015837 | 4085.985094 | 0.109974474 | 0.206125961 | 0.83669252 | 0.998865404 | 11.985147 | 12.007701 | 11.967780 | 12.011254 | 11.980019 | 12.024983 | 4546 | 4489 | 3484 | 3965 |
| 2315 | NM_005511 | MLANA | melan-A | protein_coding | NP_005502.1 | 7124 | 605513 | . | . | 173.437088 | 163.407257 | 206.913254 | 192.396927 | 403.595215 | 377.723206 | 464.307343 | 439.892700 | 1.169052 | 13341.920341 | 0.105906095 | 2.12772285 | 0.033360069 | 0.575719787 | 13.586585 | 13.811978 | 13.661169 | 13.603733 | 13.801670 | 13.729905 | 14648 | 12822 | 13025 | 12924 |
| 2316 | NM_00111055 | FLNA | filamin A | protein_coding | NP_001104026.1;NP_001447.2 | 3754 | 300017 | . | . | 86.412282 | 92.428249 | 62.324272 | 76.594826 | 201.084938 | 213.652039 | 139.853859 | 175.124962 | -1.305704 | 21044.815573 | 0.11846735 | -3.24838758 | 0.001160611 | 0.075570712 | 14.540822 | 14.155939 | 14.437921 | 14.489646 | 14.148884 | 14.313612 | 26639 | 26476 | 14322 | 18786 |
| 2317 | NM_00116431 | FLNB | filamin B | protein_coding | NP_001157789.1;NP_001157790.1;NP_001157791.1;NP_001448.2 | 3755 | 603381 | . | . | 4.231395 | 4.301750 | 3.384685 | 3.661600 | 9.846632 | 9.943688 | 7.595136 | 8.371813 | -1.224999 | 1146.208496 | 0.133318507 | -2.19609597 | 0.028085075 | 0.539571956 | 10.301738 | 10.008717 | 10.230235 | 10.236883 | 10.057695 | 10.105608 | 1447 | 1368 | 866 | 998 |
| 2319 | NM_00133017 | FLOT2 | flotillin 2 | protein_coding | NP_001317099.1;NP_004466.2;XP_005258010.1;XP_016879883.1;XP_01 | 3758 | 131560 | . | . | 15.416875 | 15.730744 | 15.044522 | 17.859086 | 35.875706 | 36.362318 | 33.759473 | 40.832677 | 1.043644 | 1341.423455 | 0.136661333 | 0.45096995 | 0.652011207 | 0.998865404 | 10.358617 | 10.419832 | 10.367213 | 10.377186 | 10.340957 | 10.464693 | 1502 | 1426 | 1095 | 1388 |
| 2321 | NM_00115992 | FLT1 | fms related receptor tyrosine kinase 1 | protein_coding | NP_001153392.1;NP_001153502.1;NP_001153503.1;NP_002010.2;XP_01 | 3763 | 165070 | . | . | 0.468399 | 0.737079 | 0.858630 | 0.447876 | 1.089985 | 1.703790 | 1.926742 | 1.024016 | -1.046896 | 78.519323 | 0.413148083 | -0.16003565 | 0.872852993 | 0.998865404 | 6.326695 | 6.262543 | 6.239362 | 6.335318 | 6.356861 | 6.189171 | 76 | 102 | 89 | 48 |
| 2323 | NM_00120450 | FLT3LG | fms related receptor tyrosine kinase 3 ligand | protein_coding | NP_001191431.1;NP_001191432.1;NP_001265566.1;NP_001265567.1;N | 3766 | 600007 | . | . | 0.746806 | 0.700575 | 1.228031 | 1.386352 | 1.737848 | 1.619411 | 2.755666 | 3.169732 | 1.770848 | 38.506191 | 0.50189297 | 1.642662047 | 0.100452875 | 0.815618258 | 4.796818 | 5.621060 | 5.206658 | 5.207306 | 5.287294 | 5.294791 | 32 | 30 | 42 | 47 |
| 2329 | NM_002022,XI | FMO4 | flavin containing dimethylaniline monoxygenase 4 | protein_coding | NP_002013.1;XP_005245102.1;XP_005245103.1;XP_005245105.1;XP_00 | 3772 | 136131 | . | . | 1.434289 | 1.198097 | 1.089582 | 1.276913 | 3.337647 | 2.769456 | 2.444991 | 2.919510 | -1.110542 | 93.940603 | 0.331207847 | -0.456705 | 0.647883098 | 0.998865404 | 6.627152 | 6.476258 | 6.591700 | 6.547196 | 6.534120 | 6.544021 | 122 | 99 | 76 | 85 |
| 2331 | NM_002023,N | FMOD | fibromodulin | protein_coding | NP_002014.2 | 3774 | 600245 | . | . | 1.019943 | 0.625643 | 0.822951 | 1.054612 | 2.373451 | 1.446202 | 1.846679 | 2.411246 | 1.122628 | 81.235862 | 0.380892365 | 0.438129481 | 0.661292424 | 0.998865404 | 6.257813 | 6.425412 | 6.382234 | 6.259254 | 6.320972 | 6.384760 | 109 | 63 | 66 | 90 |
| 2332 | NM_00118507 | FMR1 | FMRP translational regulator 1 | protein_coding | NP_001172004.1;NP_001172005.1;NP_001172010.1;NP_001172011.1;N | 3775 | 309550 | . | . | 5.360014 | 4.779406 | 5.573816 | 4.423242 | 12.472972 | 11.047810 | 12.507482 | 10.113219 | -1.033816 | 689.921294 | 0.16987864 | -0.28243103 | 0.777613024 | 0.998865404 | 9.453581 | 9.406612 | 9.480194 | 9.396622 | 9.493879 | 9.333204 | 856 | 711 | 660 | 561 |
| 2334 | NM_00116912 | AFF2 | AF4/FMR2 family member 2 | protein_coding | NP_001162593.1;NP_001162594.1;NP_001162595.1;NP_001162596.1;N | 3776 | 300806 | . | . | 0.115243 | 0.122053 | 0.044481 | 0.100911 | 0.268176 | 0.282132 | 0.099814 | 0.230721 | -1.639236 | 41.850938 | 0.523763249 | -1.36134635 | 0.173404263 | 0.94763772 | 5.702115 | 4.983744 | 5.418766 | 5.431010 | 5.299423 | 5.392765 | 58 | 58 | 17 | 41 |
| 2335 | NM_00130612 | FN1 | fibronectin 1 | protein_coding | NP_001293058.2;NP_001293059.2;NP_001293060.2;NP_001293061.2;N | 3778 | 135600 | . | . | 714.456546 | 736.705632 | 717.006569 | 774.519619 | 1662.569776 | 1702.928032 | 1608.941924 | 1770.847051 | 1.013623 | 185300.535301 | 0.098789564 | 0.197598401 | 0.843359286 | 0.998865404 | 17.489716 | 17.509233 | 17.485444 | 17.501452 | 17.477132 | 17.532545 | 209970 | 200430 | 157048 | 180475 |
| 2339 | NM_002027,N | FNTA | farnesyltransferase, CAAX box, alpha | protein_coding | NP_002018.1 | 3782 | 134635 | . | . | 15.564592 | 17.551613 | 17.295476 | 16.604978 | 36.219444 | 40.571339 | 38.810548 | 37.965310 | 1.012574 | 868.653123 | 0.146153219 | 0.123347913 | 0.901831602 | 0.998865404 | 9.753648 | 9.771570 | 9.716240 | 9.797238 | 9.784070 | 9.748520 | 938 | 984 | 779 | 799 |
| 2342 | NM_002028 | FNTB | farnesyltransferase, CAAX box, beta | protein_coding | NP_002019.1 | 3785 | 134636 | . | . | 11.062338 | 12.875194 | 10.106055 | 12.525995 | 25.742514 | 29.761587 | 22.677696 | 28.639198 | -1.072226 | 983.277071 | 0.155908775 | -0.64531057 | 0.518725943 | 0.998865404 | 9.991269 | 9.889857 | 9.910210 | 10.016581 | 9.837550 | 9.984365 | 1088 | 1177 | 741 | 980 |
| 2346 | NM_00101498 | FOLH1 | folate hydrolase 1 | protein_coding | NP_001014986.1;NP_001180400.1;NP_001180401.1;NP_001180402.1;N | 3788 | 600934 | . | . | 0.890055 | 0.825475 | 0.743027 | 0.707855 | 2.071197 | 1.908121 | 1.667330 | 1.618425 | -1.193234 | 103.274571 | 0.316985975 | -0.80406524 | 0.421359297 | 0.998865404 | 6.811752 | 6.557766 | 6.735127 | 6.700468 | 6.674097 | 6.652209 | 136 | 115 | 85 | 85 |
| 2353 | NM_005252 | FOS | Fos proto-oncogene, AP-1 transcription factor subunit | protein_coding | NP_005243.1 | 3796 | 164810 | . | . | 6.117984 | 7.116478 | 5.655674 | 5.284133 | 14.236801 | 16.450060 | 12.691168 | 12.081542 | -1.225695 | 396.565101 | 0.187360694 | -1.56703261 | 0.117107068 | 0.857396579 | 8.770777 | 8.477135 | 8.642191 | 8.730836 | 8.589103 | 8.544783 | 467 | 505 | 322 | 321 |
| 2354 | NM_00111417 | FOSB | FosB proto-oncogene, AP-1 transcription factor subunit | protein_coding | NP_001107643.1;NP_006723.2;XP_005258748.1 | 3797 | 164772 | . | . | 1.146744 | 1.203665 | 1.378189 | 1.127903 | 2.668521 | 2.782325 | 3.092618 | 2.578817 | 1.044884 | 141.263849 | 0.278924376 | 0.227094797 | 0.82035003 | 0.998865404 | 7.109703 | 7.174076 | 7.129389 | 7.134372 | 7.184842 | 7.110668 | 158 | 150 | 138 | 122 |
| 2355 | NM_005253,XI | FOSL2 | FOS like 2, AP-1 transcription factor subunit | protein_coding | NP_005244.1;XP_005264288.1;XP_006712039.1;XP_006712040.1 | 3798 | 601575 | . | . | 5.087794 | 5.399746 | 4.214184 | 4.757861 | 11.839506 | 12.481758 | 9.456506 | 10.878285 | -1.186321 | 982.308032 | 0.139030027 | -1.77295654 | 0.076235909 | 0.767975096 | 10.058173 | 9.811346 | 9.988944 | 10.009157 | 9.846539 | 9.901266 | 1211 | 1166 | 751 | 875 |
| 2356 | NM_00101807 | FPGS | folylpolyglutamate synthase | protein_coding | NP_001018088.1;NP_001275732.1;NP_004948.4;XP_005251921.1;XP_01 | 3824 | 136510 | . | . | 11.788330 | 11.649101 | 10.044196 | 11.726144 | 27.431925 | 26.927419 | 22.538885 | 26.810434 | -1.089157 | 765.233808 | 0.148866885 | -0.82766561 | 0.4078599 | 0.998865404 | 9.640215 | 9.516654 | 9.609831 | 9.607984 | 9.516425 | 9.580787 | 920 | 860 | 607 | 719 |
| 2395 | NM_000144,N | FXN | frataxin | protein_coding | NP_000135.2;NP_852090.1 | 3951 | 606829 | . | . | 1.084206 | 0.984689 | 1.005468 | 1.220302 | 2.522991 | 2.276152 | 2.256241 | 2.790076 | 1.058196 | 234.395858 | 0.229049052 | 0.356286271 | 0.72162619 | 0.998865404 | 7.831692 | 7.912775 | 7.882386 | 7.831919 | 7.838379 | 7.929376 | 276 | 233 | 190 | 247 |
| 2444 | NM_002031,XI | FRK | fyn related Src family tyrosine kinase | protein_coding | NP_002022.1;XP_005266937.1;XP_005266938.1;XP_005266939.1;XP_01 | 3955 | 606573 | . | . | 0.074774 | 0.148190 | 0.146358 | 0.062964 | 0.174001 | 0.342548 | 0.328422 | 0.143959 | -1.170715 | 33.806251 | 0.541588322 | -0.41985782 | 0.674589324 | 0.998865404 | 5.188027 | 4.961528 | 5.071404 | 5.106035 | 5.085327 | 5.049971 | 37 | 44 | 31 | 25 |
| 2475 | NM_004958,XI | MTOR | mechanistic target of rapamycin kinase | protein_coding | NP_004949.1;XP_005263495.1;XP_011539468.1;XP_016856389.1;XP_01 | 3942 | 601231 | . | . | 20.209647 | 21.605871 | 20.256673 | 23.342635 | 47.028678 | 49.942936 | 45.455383 | 53.370159 | 1.030398 | 5785.058491 | 0.113881089 | 0.379353952 | 0.704425041 | 0.998865404 | 12.476424 | 12.519486 | 12.461756 | 12.507336 | 12.453665 | 12.563299 | 6384 | 6316 | 4774 | 5862 |
| 2483 | NM_004477,XI | FRG1 | FSHD region gene 1 | protein_coding | NP_004468.1;XP_016863447.1 | 3954 | 601278 | . | . | 22.221214 | 22.620119 | 17.761988 | 24.951820 | 51.709679 | 52.287424 | 39.857385 | 57.049370 | -1.051063 | 711.507093 | 0.165247902 | -0.43479513 | 0.663711143 | 0.998865404 | 9.510542 | 9.438016 | 9.504536 | 9.475975 | 9.372434 | 9.533013 | 857 | 771 | 532 | 727 |
| 2491 | NM_00131852 | CENPI | centromere protein I | protein_coding | NP_001305450.1;NP_001305452.1;NP_006724.2;XP_005262168.1;XP_01 | 3968 | 300065 | . | . | 4.390622 | 4.609557 | 4.093724 | 4.162136 | 10.217158 | 10.655199 | 9.186200 | 9.516229 | -1.124531 | 714.187788 | 0.150065937 | -1.12832957 | 0.259180761 | 0.995760118 | 9.562417 | 9.392928 | 9.500028 | 9.538207 | 9.436662 | 9.440159 | 847 | 838 | 583 | 632 |
| 2494 | NM_00127646 | NR5A2 | nuclear receptor subfamily 5 group A member 2 | protein_coding | NP_001263393.1;NP_003813.1;NP_995582.1;XP_005245119.1;XP_01150 | 7984 | 604453 | . | . | 0.044471 | 0.106706 | 0.043964 | 0.058697 | 0.103487 | 0.246656 | 0.098655 | 0.134204 | -1.439850 | 13.448634 | 0.872849095 | -0.60253133 | 0.546820516 | NA | 3.990716 | 3.459419 | 3.730236 | 3.786384 | 3.729964 | 3.745217 | 11 | 24 | 8 | 12 |
| 2495 | NM_002032 | FTH1 | ferritin heavy chain 1 | protein_coding | NP_002023.2 | 3976 | 134770 | . | . | 1553.516602 | 1587.349854 | 1991.578613 | 2348.401123 | 3615.097412 | 3669.230225 | 4469.044434 | 5369.340332 | 1.362145 | 69992.959008 | 0.109660918 | 4.065993324 | 4.78283E-05 | 0.00725165 | 15.854831 | 16.300700 | 15.940516 | 15.951477 | 16.142767 | 16.279770 | 67797 | 64362 | 64781 | 81516 |
| 2497 | gene-FTH1P2 | FTH1P2 | ferritin heavy chain 1 pseudogene 2 | pseudogene | . | 3989 | . | . | . | 10.107709 | 10.618726 | 16.750906 | 15.419298 | 23.521055 | 24.545660 | 37.588547 | 35.254395 | 1.529647 | 190.675703 | 0.242833504 | 2.525181759 | 0.011563846 | 0.352352697 | 7.235893 | 7.849336 | 7.454380 | 7.471357 | 7.671482 | 7.626610 | 170 | 166 | 210 | 206 |
| 2498 | NR_002201 | FTH1P3 | ferritin heavy chain 1 pseudogene 3 | transcribed_ps | . | 3990 | . | . | . | 0.299399 | 0.475878 | 0.899824 | 0.320157 | 0.696714 | 1.100012 | 2.019179 | 0.732000 | 1.523038 | 15.395041 | 0.837733811 | 0.724516303 | 0.468748822 | NA | 3.607672 | 4.217207 | 3.895134 | 3.919336 | 3.971877 | 3.896639 | 11 | 16 | 24 | 9 |
| 2500 | gene-FTH1P7 | FTH1P7 | ferritin heavy chain 1 pseudogene 7 | pseudogene | . | 3994 | . | . | . | 1.953602 | 1.609893 | 1.385950 | 3.371057 | 4.546113 | 3.721341 | 3.110033 | 7.707521 | 1.305248 | 60.207576 | 0.468207053 | 0.820842841 | 0.411735793 | 0.998865404 | 5.708479 | 6.090103 | 5.884655 | 5.847383 | 5.823788 | 6.014281 | 66 | 51 | 35 | 90 |
| 2501 | gene-FTH1P8 | FTH1P8 | ferritin heavy chain 1 pseudogene 8 | pseudogene | . | 3995 | . | . | . | 1.357684 | 1.733379 | 1.479544 | 1.824011 | 3.159387 | 4.006783 | 3.320054 | 4.170384 | 1.053525 | 45.718295 | 0.467259021 | 0.16099017 | 0.872101143 | 0.998865404 | 5.477744 | 5.550733 | 5.486493 | 5.529024 | 5.500950 | 5.532745 | 45 | 54 | 37 | 48 |
| 2502 | gene-FTH1P10 | FTH1P10 | ferritin heavy chain 1 pseudogene 10 | pseudogene | . | 3980 | . | . | . | 4.431739 | 4.357848 | 4.004667 | 6.964723 | 10.312839 | 10.073362 | 8.986356 | 15.924012 | 1.233172 | 141.492705 | 0.311076576 | 0.972024195 | 0.331038512 | 0.998865404 | 6.986638 | 7.286931 | 7.098041 | 7.088718 | 7.062244 | 7.271971 | 148 | 135 | 100 | 185 |
| 2503 | gene-FTH1P11 | FTH1P11 | ferritin heavy chain 1 pseudogene 11 | pseudogene | . | 3981 | . | . | . | 0.308763 | 0.462080 | 0.232562 | 0.845638 | 0.718505 | 1.068118 | 0.521863 | 1.933450 | 1.347050 | 13.721806 | 0.892169391 | 0.48175087 | 0.629982934 | NA | 3.551764 | 3.974217 | 3.748059 | 3.767926 | 3.737919 | 3.812604 | 11 | 15 | 6 | 23 |
| 2504 | gene-FTH1P12 | FTH1P12 | ferritin heavy chain 1 pseudogene 12 | pseudogene | . | 3982 | . | . | . | 0.172568 | 0.258678 | 0.216616 | 0.287452 | 0.401573 | 0.597946 | 0.486080 | 0.657225 | 1.225154 | 7.014448 | 1.143421424 | 0.256216632 | 0.797783566 | NA | 2.658167 | 2.947964 | 2.797164 | 2.807524 | 2.805172 | 2.812424 | 6 | 8 | 6 | 8 |
| 2507 | gene-FTH1P15 | FTH1P15 | ferritin heavy chain 1 pseudogene 15 | pseudogene | . | 3985 | . | . | . | 0.186953 | 0.273222 | 0.578652 | 0.519939 | 0.435047 | 0.631566 | 1.298479 | 1.188778 | 2.056873 | 6.193489 | 1.226833017 | 0.848079908 | 0.396393471 | NA | 2.019256 | 3.058876 | 2.599748 | 2.605427 | 2.625885 | 2.618445 | 4 | 5 | 8 | 7 |
| 2508 | gene-FTH1P16 | FTH1P16 | ferritin heavy chain 1 pseudogene 16 | pseudogene | . | 3986 | . | . | . | 0.520068 | 0.460429 | 0.553572 | 0.733089 | 1.210219 | 1.064303 | 1.242199 | 1.676121 | 1.268411 | 16.741528 | 0.750678131 | 0.456950349 | 0.647706734 | NA | 3.884262 | 4.226238 | 4.056126 | 4.048520 | 4.058402 | 4.080384 | 18 | 15 | 14 | 20 |
| 2509 | gene-FTH1P5 | FTH1P5 | ferritin heavy chain 1 pseudogene 5 | pseudogene | . | 3996 | . | . | . | 0.224675 | 0.152236 | 0.148239 | 0.348107 | 0.522827 | 0.351901 | 0.332643 | 0.795905 | 1.320495 | 6.711957 | 1.19080396 | 0.33681381 | 0.736257262 | NA | 2.533720 | 2.932304 | 2.743954 | 2.733182 | 2.733580 | 2.760325 | 8 | 5 | 4 | 10 |
| 2512 | NM_000146,XI | FTL | ferritin light chain | protein_coding | NP_000137.2;XP_024307215.1 | 3999 | 134790 | . | . | 435.918274 | 454.397133 | 525.270081 | 594.639468 | 1014.399719 | 1050.359261 | 1178.690796 | 1359.572446 | 1.240174 | 13627.808945 | 0.107238779 | 2.895808896 | 0.003781826 | 0.178158867 | 13.570684 | 13.881175 | 13.620404 | 13.648774 | 13.767950 | 13.867668 | 13774 | 13350 | 12371 | 14954 |
| 2517 | NM_000147,XI | FUCA1 | alpha-L-fucosidase 1 | protein_coding | NP_000138.2;XP_005245878.1;XP_011539469.1;XP_016856394.1 | 4006 | 612280 | . | . | 31.498190 | 29.612713 | 28.078305 | 27.589855 | 73.297578 | 68.451110 | 63.006907 | 63.080926 | -1.108134 | 1910.076055 | 0.120478315 | -1.22953313 | 0.218871984 | 0.982008277 | 10.971506 | 10.823530 | 10.961540 | 10.915096 | 10.868850 | 10.845889 | 2379 | 2104 | 1599 | 1674 |
| 2519 | NM_032020 | FUCA2 | alpha-L-fucosidase 2 | protein_coding | NP_114409.2 | 4008 | 136820 | . | . | 75.106743 | 73.638023 | 74.282135 | 74.344818 | 174.776505 | 170.217590 | 166.686951 | 169.980576 | -1.014853 | 5524.499073 | 0.104665796 | -0.20322535 | 0.838958902 | 0.998865404 | 12.442209 | 12.420969 | 12.449625 | 12.425768 | 12.429315 | 12.421457 | 6499 | 5920 | 4791 | 5117 |
| 2521 | NM_00117063 | FUS | FUS RNA binding protein | protein_coding | NP_001164105.1;NP_001164408.1;NP_004951.1;XP_005255290.1;XP_01 | 4010 | 137070 | . | . | 130.591069 | 137.078006 | 150.215653 | 138.472152 | 303.890783 | 316.861942 | 337.079556 | 316.600099 | 1.032526 | 8283.081094 | 0.105259059 | 0.438704957 | 0.660875335 | 0.998865404 | 12.992662 | 13.038871 | 12.986315 | 13.017399 | 13.059836 | 12.997546 | 9213 | 8956 | 7586 | 7601 |
| 2526 | NM_002033 | FUT4 | fucosyltransferase 4 | protein_coding | NP_002024.1 | 4015 | 104230 | . | . | 2.078120 | 2.019334 | 1.987436 | 1.774548 | 4.835869 | 4.667781 | 4.459748 | 4.057293 | -1.104775 | 370.559214 | 0.188796919 | -0.76141359 | 0.446410072 | 0.998865404 | 8.603308 | 8.460268 | 8.572320 | 8.550392 | 8.537707 | 8.468844 | 456 | 412 | 325 | 310 |
| 2530 | NM_00137153 | FUT8 | fucosyltransferase 8 | protein_coding | NP_001358462.1;NP_001358463.1;NP_001358465.1;NP_004471.4;NP_8 | 4019 | 602589 | . | . | 5.257843 | 4.754906 | 4.730517 | 4.436153 | 12.235218 | 10.991179 | 10.615145 | 10.142736 | -1.097211 | 628.372589 | 0.162122176 | -0.82555823 | 0.409054738 | 0.998865404 | 9.360504 | 9.227379 | 9.364240 | 9.284334 | 9.297482 | 9.227510 | 801 | 668 | 549 | 532 |
| 2531 | NM_002035,XI | KDSR | 3-ketodihydrosphingosine reductase | protein_coding | NP_002026.1;XP_005266734.1;XP_016881179.1 | 4021 | 136440 | . | . | 7.048113 | 7.154903 | 7.199510 | 6.216756 | 16.401249 | 16.538879 | 16.155493 | 14.213876 | -1.073973 | 1108.893741 | 0.141130819 | -0.72951849 | 0.465684561 | 0.998865404 | 10.165253 | 10.062737 | 10.137707 | 10.142466 | 10.143177 | 10.027820 | 1318 | 1243 | 1003 | 924 |
| 2533 | NM_00124309 | FYB1 | FYN binding protein 1 | protein_coding | NP_001230022.1;NP_001336262.1;NP_001456.3;NP_061064.2;NP_9553 | 4036 | 602731 | . | . | 0.805946 | 0.764374 | 0.660710 | 0.910701 | 1.875471 | 1.766886 | 1.482612 | 2.082210 | -1.047105 | 117.265372 | 0.306950479 | -0.21634359 | 0.828719934 | 0.998865404 | 6.907073 | 6.839400 | 6.893040 | 6.868743 | 6.820024 | 6.908844 | 143 | 125 | 86 | 122 |
| 2534 | NM_00137052 | FYN | FYN proto-oncogene, Src family tyrosine kinase | protein_coding | NP_001357458.1;NP_002028.1;NP_694592.1;NP_694593.1;XP_0052669 | 4037 | 137025 | . | . | 27.160649 | 27.949815 | 27.224243 | 30.541939 | 63.203945 | 64.607253 | 61.090413 | 69.830517 | 1.049481 | 2873.631029 | 0.121686388 | 0.572591347 | 0.56692142 | 0.998865404 | 11.453472 | 11.523008 | 11.490528 | 11.444820 | 11.448335 | 11.563034 | 3320 | 2941 | 2380 | 2952 |
| 2535 | NM_001466 | FZD2 | frizzled class receptor 2 | protein_coding | NP_001457.1 | 4040 | 600667 | . | . | 1.375416 | 1.688615 | 1.260377 | 1.371934 | 3.200649 | 3.903310 | 2.828249 | 3.136765 | -1.180557 | 168.164792 | 0.261406145 | -0.91607484 | 0.359627644 | 0.998865404 | 7.509032 | 7.268410 | 7.382006 | 7.468803 | 7.343237 | 7.372042 | 189 | 216 | 129 | 150 |
| 2537 | NM_002038,N | IFI6 | interferon alpha inducible protein 6 | protein_coding | NP_002029.3;NP_075010.1;NP_075011.1;XP_024301975.1 | 4054 | 147572 | . | . | 64.451378 | 58.002002 | 62.777351 | 62.117155 | 149.980999 | 134.074220 | 140.870551 | 142.023494 | 1.003491 | 1575.430321 | 0.128201788 | 0.039221311 | 0.968713943 | 0.998865404 | 10.618893 | 10.624163 | 10.662374 | 10.575259 | 10.630698 | 10.614296 | 1912 | 1602 | 1387 | 1464 |
| 2539 | NM_000402,N | G6PD | glucose-6-phosphate dehydrogenase | protein_coding | NP_000393.4;NP_001035810.1;NP_001346945.1 | 4057 | 305900 | . | . | 62.465096 | 64.909033 | 80.417950 | 78.278058 | 145.358840 | 150.040146 | 180.455541 | 178.973489 | 1.229392 | 4957.795849 | 0.106345591 | 2.801663951 | 0.00508398 | 0.215920545 | 12.118826 | 12.416784 | 12.170755 | 12.194349 | 12.371426 | 12.339177 | 5048 | 4868 | 4845 | 5029 |

| | | | | | | | | | | | | | | | | | | | | | | | | | | | | | | | | | |
|---|---|---|---|---|---|---|---|---|---|---|---|---|---|---|---|---|---|---|---|---|---|---|---|---|---|---|---|---|---|---|---|---|---|
| 2542 | NM_00116427 | SLC37A4 | solute carrier family 37 member 4 | protein_coding | NP_001157749.1;NP_001157750.1;NP_001157751.1;NP_001157752.1;NI | 4061 | 602671 | . | . | 7.947178 | 8.080958 | 6.416649 | 8.761933 | 18.493412 | 18.679500 | 14.398772 | 20.033118 | -1.078915 | 600.058464 | 0.169161269 | -0.64779143 | 0.517119862 | 0.998865404 | 9.283121 | 9.172684 | 9.242969 | 9.262490 | 9.131398 | 9.268368 | 707 | 682 | 449 | 598 |
| 2547 | NM_00128897 | XRCC6 | X-ray repair cross complementing 6 | protein_coding | NP_001275905.1;NP_001275906.1;NP_001275907.1;NP_001460.1 | 4055 | 152690 | . | . | 196.669631 | 204.340901 | 175.769743 | 210.590298 | 457.658339 | 472.343173 | 394.422218 | 481.489680 | -1.052561 | 13037.393370 | 0.114621689 | -0.64476588 | 0.519078915 | 0.998865404 | 13.706881 | 13.632906 | 13.677877 | 13.704043 | 13.572176 | 13.717045 | 15163 | 14648 | 10109 | 12920 |
| 2548 | NM_000152,N | GAA | glucosidase alpha, acid | protein_coding | NP_000143.2;NP_001073271.1;NP_001073272.1;XP_005257250.1;XP_00 | 4065 | 606800 | . | . | 13.055718 | 12.483081 | 11.806051 | 13.050639 | 30.381194 | 28.855197 | 26.492439 | 29.838735 | -1.069170 | 1321.182479 | 0.130175043 | -0.74124329 | 0.458545941 | 0.998865404 | 10.415138 | 10.318471 | 10.403128 | 10.381779 | 10.311031 | 10.371128 | 1594 | 1453 | 1066 | 1245 |
| 2549 | NM_002039,N | GAB1 | GRB2 associated binding protein 1 | protein_coding | NP_002030.2;NP_997006.1;XP_006714230.1;XP_006714231.1;XP_01686 | 4066 | 604439 | . | . | 1.133111 | 1.271445 | 1.169651 | 1.195015 | 2.636798 | 2.939001 | 2.624662 | 2.732259 | -1.035612 | 293.966414 | 0.205036159 | -0.24621958 | 0.805512257 | 0.998865404 | 8.224778 | 8.173786 | 8.175667 | 8.240227 | 8.185385 | 8.194093 | 324 | 342 | 249 | 273 |
| 2550 | NM_00131905 | GABBR1 | gamma-aminobutyric acid type B receptor subunit 1 | protein_coding | NP_001305982.1;NP_001461.1;NP_068703.1;NP_068704.2;XP_0052490 | 4070 | 603540 | . | . | 6.430799 | 6.499332 | 7.088747 | 5.909928 | 14.964734 | 15.023499 | 15.906943 | 13.512351 | -1.019664 | 827.921729 | 0.152290113 | -0.18447856 | 0.853638045 | 0.998865404 | 9.707063 | 9.679507 | 9.695097 | 9.703174 | 9.744878 | 9.622390 | 957 | 907 | 775 | 702 |
| 2551 | NM_00119729 | GABPA | GA binding protein transcription factor subunit alpha | protein_coding | NP_001184226.1;NP_002031.2;XP_005260995.1;XP_011527822.1;XP_01 | 4071 | 600609 | . | . | 2.980423 | 2.394442 | 2.423667 | 2.689052 | 6.935568 | 5.534860 | 5.438637 | 6.148200 | -1.068243 | 411.448080 | 0.189052135 | -0.50377906 | 0.614416643 | 0.998865404 | 8.731264 | 8.636307 | 8.762282 | 8.640296 | 8.643128 | 8.683677 | 538 | 413 | 334 | 386 |
| 2553 | NM_00132091 | GABPB1 | GA binding protein transcription factor subunit beta 1 | protein_coding | NP_001307839.1;NP_001307844.1;NP_002032.2;NP_005245.2;NP_0577 | 4074 | 600610 | . | . | 4.276487 | 4.556635 | 3.752164 | 3.422309 | 9.951561 | 10.532869 | 8.419744 | 7.824702 | -1.180143 | 369.167334 | 0.185852956 | -1.28575892 | 0.198527211 | 0.96621494 | 8.642732 | 8.403635 | 8.563475 | 8.584802 | 8.477891 | 8.479316 | 452 | 439 | 294 | 318 |
| 2556 | NM_000808,XI | GABRA3 | gamma-aminobutyric acid type A receptor subunit alpha3 | protein_coding | NP_000799.1;XP_006724874.1 | 4077 | 305660 | . | . | 0.751099 | 0.570292 | 0.618691 | 0.941982 | 1.747839 | 1.318256 | 1.388325 | 2.153729 | 1.166579 | 82.652051 | 0.373152988 | 0.595692642 | 0.551380575 | 0.998865404 | 6.254018 | 6.475451 | 6.376887 | 6.305706 | 6.326325 | 6.437378 | 100 | 71 | 62 | 100 |
| 2558 | NM_000810,N | GABRA5 | gamma-aminobutyric acid type A receptor subunit alpha5 | protein_coding | NP_000801.1;NP_001158509.1;XP_005268315.1;XP_006720522.1;XP_01 | 4079 | 137142 | . | . | 3.969102 | 3.407126 | 3.098857 | 3.226742 | 9.236260 | 7.875722 | 6.953744 | 7.377562 | -1.348722 | 341.539013 | 0.208970612 | -2.06533125 | **0.038891669** | 0.614916047 | 8.616019 | 8.183498 | 8.529916 | 8.451286 | 8.251992 | 8.389901 | 481 | 396 | 219 | 309 |
| 2562 | NM_000814,N | GABRB3 | gamma-aminobutyric acid type A receptor subunit beta3 | protein_coding | NP_000805.1;NP_001178249.1;NP_001178250.1;NP_001265560.1;NP_0 | 4083 | 137192 | . | . | 3.959872 | 4.107407 | 5.288048 | 4.571282 | 9.214785 | 9.494456 | 11.866224 | 10.451693 | 1.201086 | 798.358522 | 0.152591785 | 1.732328172 | 0.083215149 | 0.784575614 | 9.502455 | 9.767202 | 9.559340 | 9.581291 | 9.755332 | 9.640774 | 822 | 795 | 818 | 752 |
| 2564 | NM_004961,XI | GABRE | gamma-aminobutyric acid type A receptor subunit epsilon | protein_coding | NP_004952.2;XP_011529442.1;XP_016884876.1;XP_016884877.1;XP_01 | 4085 | 300093 | . | . | 0.079313 | 0.075958 | 0.067345 | 0.085893 | 0.184564 | 0.175580 | 0.151120 | 0.196386 | 1.028911 | 16.847445 | 0.747193187 | 0.055030823 | 0.956113908 | NA | 4.054853 | 4.093800 | 4.068616 | 4.077376 | 4.066134 | 4.083961 | 18 | 19 | 13 | 18 |
| 2567 | NM_00127087 | GABRG3 | gamma-aminobutyric acid type A receptor subunit gamma3 | protein_coding | NP_001257802.1;NP_150092.2;XP_011519732.1;XP_011519733.1;XP_01 | 4088 | 600233 | . | . | 1.121183 | 1.131958 | 0.208950 | 0.182540 | 2.609040 | 2.616573 | 0.468877 | 0.417357 | -1.017336 | 67.138986 | 0.385175476 | -0.06437587 | 0.948670939 | 0.998865404 | 6.080986 | 6.057072 | 6.070740 | 6.070413 | 6.082442 | 6.050521 | 78 | 73 | 62 | 58 |
| 2570 | NM_002043,XI | GABRR2 | gamma-aminobutyric acid type A receptor subunit rho2 | protein_coding | NP_002034.3;XP_011534015.1;XP_011534016.1;XP_011534017.1;XP_01 | 4091 | 137162 | . | . | 0.485657 | 0.672898 | 0.328336 | 0.559190 | 1.130146 | 1.555435 | 0.736778 | 1.278523 | -1.317015 | 74.386559 | 0.401017677 | -0.99065932 | 0.321851963 | 0.998865404 | 6.403699 | 6.002429 | 6.207543 | 6.284717 | 6.119550 | 6.234377 | 84 | 104 | 41 | 76 |
| 2571 | NM_000817,N | GAD1 | glutamate decarboxylase 1 | protein_coding | NP_000808.2;NP_038473.2;XP_005246501.1;XP_011509224.1;XP_01685 | 4092 | 605363 | . | . | 13.288187 | 14.904009 | 18.955845 | 12.710795 | 30.922157 | 34.451283 | 42.536368 | 29.061721 | 1.247989 | 964.031746 | 0.159672527 | 2.001628033 | **0.045324752** | 0.649654775 | 9.743912 | 10.064220 | 9.838555 | 9.806252 | 10.079231 | 9.879369 | 1010 | 904 | 1055 | 870 |
| 2580 | NM_00131813 | GAK | cyclin G associated kinase | protein_coding | NP_001305063.1;NP_005246.2;XP_005272325.1;XP_005272327.1;XP_01 | 4113 | 602052 | . | . | 16.843896 | 16.526435 | 17.582527 | 18.266536 | 39.196438 | 38.201593 | 39.454676 | 41.764262 | 1.050291 | 2420.867318 | 0.115054151 | 0.615264835 | 0.538379837 | 0.998865404 | 11.205459 | 11.276288 | 11.238283 | 11.203534 | 11.256183 | 11.265078 | 2778 | 2493 | 2145 | 2338 |
| 2582 | NM_000403,N | GALE | UDP-galactose-4-epimerase | protein_coding | NP_000394.2;NP_001008217.1;NP_001121093.1 | 4116 | 606953 | . | . | 81.854834 | 87.117427 | 80.533926 | 96.108530 | 190.479565 | 201.375835 | 180.715786 | 219.740742 | 1.032076 | 4184.104271 | 0.120490436 | 0.37803112 | 0.705407476 | 0.998865404 | 12.007854 | 12.053196 | 11.990803 | 12.041671 | 11.973588 | 12.107525 | 4597 | 4580 | 3399 | 4304 |
| 2583 | NM_00127646 | B4GALNT1 | beta-1,4-N-acetyl-galactosaminyltransferase 1 | protein_coding | NP_001263397.1;NP_001263398.1;NP_001469.1;XP_005268830.1;XP_01 | 4117 | 601873 | . | . | 0.288642 | 0.247027 | 0.166918 | 0.216458 | 0.671681 | 0.571013 | 0.374559 | 0.494904 | -1.354819 | 36.995829 | 0.514553833 | -0.85141828 | 0.394537043 | 0.998865404 | 5.412509 | 4.972679 | 5.234095 | 5.234106 | 5.175886 | 5.205057 | 49 | 46 | 24 | 33 |
| 2584 | NM_000154,N | GALK1 | galactokinase 1 | protein_coding | NP_000145.1;NP_001368914.1 | 4118 | 604313 | . | . | 11.209898 | 10.681009 | 11.077329 | 11.613800 | 26.085894 | 24.689631 | 24.857204 | 26.553573 | 1.023812 | 485.867228 | 0.169543041 | 0.200252688 | 0.841282962 | 0.998865404 | 8.907336 | 8.841301 | 8.934968 | 8.898212 | 8.919031 | 8.943933 | 569 | 503 | 419 | 470 |
| 2585 | NM_00100155 | GALK2 | galactokinase 2 | protein_coding | NP_001001556.1;NP_001275959.1;NP_001275960.1;NP_001338976.1;NI | 4119 | 137028 | . | . | 5.343071 | 6.132735 | 5.067567 | 6.299807 | 12.433541 | 14.176095 | 11.371471 | 14.403757 | 1.005907 | 686.679320 | 0.161006902 | 0.052775192 | 0.957911028 | 0.998865404 | 9.419625 | 9.427350 | 9.369892 | 9.468629 | 9.382869 | 9.464102 | 732 | 792 | 563 | 684 |
| 2588 | NM_000512,N | GALNS | galactosamine (N-acetyl)-6-sulfatase | protein_coding | NP_000503.1;NP_001310472.1;NP_001310473.1;XP_005256358.1;XP_01 | 4122 | 612222 | . | . | 18.777227 | 18.726137 | 20.238018 | 19.982775 | 43.695383 | 43.286300 | 45.413529 | 45.688237 | 1.013885 | 1607.857857 | 0.123579231 | 0.160986836 | 0.872103769 | 0.998865404 | 10.640863 | 10.660916 | 10.662380 | 10.628093 | 10.668309 | 10.643891 | 1879 | 1685 | 1430 | 1494 |
| 2589 | NM_020474 | GALNT1 | polypeptide N-acetylgalactosaminyltransferase 1 | protein_coding | NP_065207.2 | 4123 | 602273 | . | . | 21.632677 | 18.909649 | 15.360388 | 15.847763 | 50.340126 | 43.710503 | 34.468266 | 36.234028 | -1.317789 | 2147.404570 | 0.12555963 | -3.17076242 | **0.001520394** | 0.091626649 | 11.253697 | 10.855696 | 11.224931 | 11.107179 | 10.939159 | 10.955596 | 3012 | 2447 | 1594 | 1756 |
| 2590 | NM_00129186 | GALNT2 | polypeptide N-acetylgalactosaminyltransferase 2 | protein_coding | NP_001278795.1;NP_004472.1;XP_016856452.1;XP_016856453.1;XP_01 | 4124 | 602274 | . | . | 100.698745 | 99.945326 | 86.062999 | 91.016748 | 234.330130 | 231.028109 | 193.122879 | 208.098972 | -1.148019 | 12999.613147 | 0.10081566 | -1.97535175 | **0.048228245** | 0.660542298 | 13.762329 | 13.563165 | 13.729804 | 13.714902 | 13.585013 | 13.623102 | 16138 | 14865 | 10288 | 11603 |
| 2591 | NM_004482,XI | GALNT3 | polypeptide N-acetylgalactosaminyltransferase 3 | protein_coding | NP_004473.2;XP_005246506.1;XP_011509231.1;XP_016859259.1 | 4125 | 601756 | . | . | 1.926309 | 1.696861 | 1.942413 | 1.757611 | 4.482600 | 3.922368 | 4.358719 | 4.018570 | 1.047619 | 193.902664 | 0.254609655 | 0.263593594 | 0.792093095 | 0.998865404 | 7.565034 | 7.632553 | 7.538720 | 7.628870 | 7.656710 | 7.551494 | 196 | 225 | 193 | 164 |
| 2592 | NM_000155,N | GALT | galactose-1-phosphate uridylyltransferase | protein_coding | NP_000146.2;NP_001245261.1 | 4135 | 606999 | . | . | 16.996549 | 18.173586 | 22.623745 | 20.213080 | 39.551666 | 42.009057 | 50.767029 | 46.214804 | 1.198968 | 1014.746650 | 0.141817953 | 1.845980662 | 0.064895013 | 0.731823074 | 9.849936 | 10.111988 | 9.899208 | 9.930003 | 10.094939 | 9.998939 | 1040 | 1017 | 1024 | 971 |
| 2593 | NM_000156,N | GAMT | guanidinoacetate N-methyltransferase | protein_coding | NP_000147.1;NP_620279.1 | 4136 | 601240 | . | . | 8.709061 | 10.228105 | 6.819057 | 10.759935 | 20.266343 | 23.642721 | 15.301769 | 24.601311 | -1.115292 | 338.907106 | 0.224669354 | -0.70068232 | 0.483501295 | 0.998865404 | 8.482171 | 8.322930 | 8.393612 | 8.466278 | 8.255018 | 8.471631 | 386 | 410 | 223 | 360 |
| 2595 | NM_00130140 | GANC | glucosidase alpha, neutral C | protein_coding | NP_001288338.1;NP_001288339.1;NP_937784.2 | 4139 | 104180 | . | . | 6.962036 | 7.334313 | 8.586910 | 8.042999 | 16.200945 | 16.953594 | 19.268777 | 18.389362 | 1.135108 | 680.581716 | 0.151925862 | 1.203416952 | 0.228815011 | 0.98298587 | 9.316330 | 9.499132 | 9.349864 | 9.379478 | 9.460688 | 9.441679 | 718 | 703 | 638 | 669 |
| 2597 | NM_00125679 | GAPDH | glyceraldehyde-3-phosphate dehydrogenase | protein_coding | NP_001243728.1;NP_001276674.1;NP_001276675.1;NP_001344872.1;NI | 4141 | 138400 | . | . | 2270.185000 | 2368.703443 | 2192.895066 | 2580.043177 | 5282.813446 | 5475.363804 | 4920.792964 | 5898.962163 | 1.014326 | 94308.730770 | 0.109726799 | 0.187026442 | 0.851639901 | 0.998865404 | 16.514811 | 16.535323 | 16.503088 | 16.533035 | 16.462193 | 16.594529 | 105958 | 102788 | 76297 | 95830 |
| 2613 | gene-GAPDHP7 | GAPDHP71 | glyceraldehyde-3-phosphate dehydrogenase pseudogene 71 | pseudogene | . | 4149 | . | | . | 0.167537 | 0.297885 | 0.403918 | 0.153222 | 0.389865 | 0.688573 | 0.906381 | 0.350324 | 1.182543 | 10.373715 | 0.984952975 | 0.245588007 | 0.806001174 | NA | 3.247564 | 3.491830 | 3.346398 | 3.369964 | 3.386680 | 3.344753 | 8 | 13 | 14 | 6 |
| 2615 | NM_00112892 | LRRC32 | leucine rich repeat containing 32 | protein_coding | NP_001122394.1;NP_001357116.1;NP_001357117.1;NP_001357118.1;NI | 4161 | 137207 | . | . | 0.059544 | 0.075278 | 0.042093 | 0.050927 | 0.138562 | 0.174008 | 0.094455 | 0.116439 | -1.312890 | 8.311282 | 1.053456 | -0.37281699 | 0.709284654 | NA | 3.238681 | 2.844638 | 3.061144 | 3.067988 | 3.053309 | 3.055687 | 10 | 11 | 6 | 7 |
| 2617 | NM_00131677 | GARS1 | glycyl-tRNA synthetase 1 | protein_coding | NP_001303701.1;NP_002038.2;XP_006715749.1 | 4162 | 600287 | . | . | 67.466096 | 67.269471 | 54.101253 | 73.279113 | 156.996383 | 155.496409 | 121.401645 | 167.543982 | -1.073473 | 5212.577113 | 0.13796809 | -0.74137498 | 0.458466111 | 0.998865404 | 12.398107 | 12.295636 | 12.379565 | 12.369017 | 12.185493 | 12.429580 | 6254 | 5788 | 3734 | 5401 |
| 2618 | NM_000819,N | GART | phosphoribosylglycinamide formyltransferase, phosphoribosylglyc | protein_coding | NP_000810.1;NP_001129477.1;NP_001129478.1;NP_780294.1;XP_0052 | 4163 | 138440 | . | . | 28.881973 | 32.111232 | 28.685274 | 33.923517 | 67.209543 | 74.226556 | 64.368926 | 77.562090 | 1.028291 | 3035.638236 | 0.124731389 | 0.322684213 | 0.746934409 | 0.998865404 | 11.547661 | 11.587630 | 11.527391 | 11.582480 | 11.507603 | 11.643968 | 3331 | 3339 | 2454 | 3125 |
| 2619 | NM_002048 | GAS1 | growth arrest specific 1 | protein_coding | NP_002039.2 | 4165 | 139185 | . | . | 2.221892 | 1.876773 | 1.634266 | 1.191561 | 5.170433 | 4.338246 | 3.667247 | 2.724362 | -1.469236 | 170.279904 | 0.272647674 | -2.03583614 | **0.041766812** | 0.633524514 | 7.661801 | 7.109116 | 7.522279 | 7.438270 | 7.378319 | 7.263192 | 254 | 199 | 139 | 109 |
| 2620 | NM_00114383 | GAS2 | growth arrest specific 2 | protein_coding | NP_001137302.1;NP_001338153.1;NP_005247.1;NP_808221.1;XP_01151 | 4167 | 602835 | . | . | 0.618961 | 0.478637 | 0.702258 | 0.744056 | 1.440350 | 1.106391 | 1.575846 | 1.701196 | 1.251638 | 46.799014 | 0.458360942 | 0.706468149 | 0.479897054 | 0.998865404 | 5.376728 | 5.701810 | 5.540535 | 5.505678 | 5.565426 | 5.559653 | 53 | 40 | 46 | 48 |
| 2621 | NM_000820 | GAS6 | growth arrest specific 6 | protein_coding | NP_000811.1 | 4168 | 600441 | . | . | 2.766049 | 3.065487 | 2.568331 | 3.007694 | 6.436711 | 7.086011 | 5.763260 | 6.876736 | -1.059169 | 223.394993 | 0.230609325 | -0.35962654 | 0.719126432 | 0.998865404 | 7.844866 | 7.760816 | 7.793017 | 7.839883 | 7.755585 | 7.821988 | 252 | 260 | 175 | 218 |
| 2622 | NM_00128620 | GAS8 | growth arrest specific 8 | protein_coding | NP_001273134.1;NP_001273137.1;NP_001273138.1;NP_001472.1;XP_0 | 4166 | 605178 | . | . | 7.835865 | 8.557635 | 8.958197 | 8.076741 | 18.234384 | 19.781360 | 20.101936 | 18.466507 | 1.028854 | 828.852150 | 0.148952481 | 0.275508764 | 0.782925394 | 0.998865404 | 9.674217 | 9.715431 | 9.654967 | 9.712072 | 9.743099 | 9.663654 | 903 | 917 | 772 | 744 |
| 2624 | NM_00114566 | GATA2 | GATA binding protein 2 | protein_coding | NP_001139133.1;NP_001139134.1;NP_116027.2 | 4171 | 137295 | . | . | 1.093451 | 1.124826 | 0.651218 | 0.843023 | 2.544504 | 2.600085 | 1.461314 | 1.927471 | -1.471939 | 100.166663 | 0.327236471 | -1.70432644 | 0.088320117 | 0.802216049 | 6.899064 | 6.339539 | 6.692830 | 6.709715 | 6.553849 | 6.614379 | 134 | 132 | 62 | 85 |
| 2625 | NM_00100229 | GATA3 | GATA binding protein 3 | protein_coding | NP_001002295.1;NP_002042.1;XP_005252499.1;XP_005252500.1 | 4172 | 131320 | . | . | 0.244494 | 0.082705 | 0.137992 | 0.187432 | 0.568949 | 0.191177 | 0.309647 | 0.428540 | 1.060876 | 15.496196 | 0.819452932 | 0.104039794 | 0.917137769 | NA | 3.909377 | 3.996978 | 3.978304 | 3.918767 | 3.943745 | 3.962027 | 25 | 9 | 12 | 17 |
| 2626 | NM_00130809 | GATA4 | GATA binding protein 4 | protein_coding | NP_001295022.1;NP_001295023.1;NP_001361202.1;NP_001361203.1;NI | 4173 | 600576 | . | . | 7.446356 | 7.121757 | 6.873039 | 8.220085 | 17.327977 | 16.462261 | 15.422903 | 18.794246 | 1.022963 | 765.327957 | 0.149285136 | 0.21940301 | 0.826336126 | 0.998865404 | 9.563684 | 9.596004 | 9.559168 | 9.583951 | 9.553675 | 9.620024 | 856 | 831 | 641 | 760 |
| 2627 | NM_005257 | GATA6 | GATA binding protein 6 | protein_coding | NP_005248.2 | 4174 | 601656 | . | . | 0.905816 | 0.732771 | 0.690739 | 0.804548 | 2.107870 | 1.693832 | 1.549998 | 1.839503 | -1.111070 | 89.121257 | 0.344844683 | -0.4406324 | 0.659479145 | 0.998865404 | 6.551670 | 6.399726 | 6.525342 | 6.459038 | 6.443329 | 6.482382 | 120 | 90 | 68 | 85 |
| 2628 | NM_00132101 | GATM | glycine amidinotransferase | protein_coding | NP_001307944.1;NP_001473.1 | 4175 | 602360 | . | . | 0.056358 | 0.370589 | 0.174734 | 0.314139 | 0.131147 | 0.856632 | 0.392098 | 0.718242 | 1.170160 | 20.369927 | 0.824224711 | 0.275053411 | 0.783275204 | NA | 4.234452 | 4.453945 | 4.257387 | 4.378054 | 4.309001 | 4.357317 | 6 | 35 | 14 | 26 |
| 2629 | NM_000157,N | GBA | glucosylceramidase beta | protein_coding | NP_000148.2;NP_001005741.1;NP_001005742.1;NP_001165282.1;NP_0 | 4177 | 606463 | . | . | 82.582459 | 84.766442 | 81.955215 | 81.885695 | 192.172771 | 195.941426 | 183.905121 | 187.221912 | -1.034527 | 5937.153799 | 0.103882296 | -0.47141283 | 0.637345953 | 0.998865404 | 12.559832 | 12.510864 | 12.542412 | 12.556888 | 12.527160 | 12.515006 | 6893 | 6571 | 5112 | 5432 |
| 2630 | NR_002188 | GBAP1 | glucosylceramidase beta pseudogene 1 | transcribed_ps | . | 4178 | . | | . | 20.904819 | 21.863503 | 23.385870 | 22.093615 | 48.646374 | 50.538467 | 52.477215 | 50.514423 | 1.048559 | 1399.621914 | 0.128141803 | 0.533847503 | 0.593447045 | 0.998865404 | 10.416157 | 10.484672 | 10.418070 | 10.445698 | 10.493945 | 10.442135 | 1545 | 1501 | 1288 | 1298 |
| 2631 | NM_00120246 | NIPSNAP2 | nipsnap homolog 2 | protein_coding | NP_001189398.1;NP_001474.1 | 4179 | 603004 | . | . | 20.842807 | 17.343265 | 17.191734 | 20.337965 | 48.502066 | 40.089736 | 38.577753 | 46.500340 | -1.033278 | 1174.034932 | 0.151408559 | -0.31192883 | 0.755094609 | 0.998865404 | 10.220709 | 10.173423 | 10.278282 | 10.131784 | 10.122787 | 10.240105 | 1507 | 1163 | 925 | 1168 |
| 2632 | NM_000158 | GBE1 | 1,4-alpha-glucan branching enzyme 1 | protein_coding | NP_000149.4 | 4180 | 607839 | . | . | 6.652238 | 6.641238 | 6.726543 | 6.838696 | 15.480031 | 15.351519 | 15.094166 | 15.635868 | 1.006235 | 615.572123 | 0.155838913 | 0.057544186 | 0.954111709 | 0.998865404 | 9.261294 | 9.270260 | 9.266885 | 9.260472 | 9.265407 | 9.270254 | 710 | 659 | 535 | 581 |
| 2633 | NM_002053,XI | GBP1 | guanylate binding protein 1 | protein_coding | NP_002044.2;XP_024302119.1 | 4182 | 600411 | . | . | 5.655911 | 5.360162 | 3.922200 | 4.946165 | 13.161537 | 12.390254 | 8.801303 | 11.308818 | -1.257265 | 447.365167 | 0.185271263 | -1.78273278 | 0.074629812 | 0.764716896 | 8.961380 | 8.630284 | 8.890383 | 8.849520 | 8.664648 | 8.790703 | 592 | 521 | 307 | 412 |
| 2634 | NM_004120 | GBP2 | guanylate binding protein 2 | protein_coding | NP_004111.2 | 4183 | 600412 | . | . | 1.682250 | 1.805727 | 1.316670 | 1.771112 | 3.914666 | 4.174019 | 2.954570 | 4.049437 | -1.144626 | 209.674909 | 0.244878397 | -0.79580808 | 0.426143596 | 0.998865404 | 7.806936 | 7.610396 | 7.726644 | 7.755941 | 7.615595 | 7.738657 | 250 | 249 | 146 | 209 |
| 2635 | NM_00131917 | GBP3 | guanylate binding protein 3 | protein_coding | NP_001306108.1;NP_001306109.1;NP_001306110.1;NP_060754.2;XP_0 | 4184 | 600413 | . | . | 2.335855 | 2.507113 | 2.282740 | 2.309550 | 5.435630 | 5.795304 | 5.122403 | 5.280511 | -1.063150 | 216.832578 | 0.228599208 | -0.38586074 | 0.699599805 | 0.998865404 | 7.804136 | 7.715375 | 7.759148 | 7.790376 | 7.739362 | 7.752918 | 249 | 249 | 180 | 200 |
| 2637 | NM_00130168 | GBX2 | gastrulation brain homeobox 2 | protein_coding | NP_001288616.1;NP_001476.2 | 4186 | 601135 | . | . | 1.131359 | 0.739225 | 0.933996 | 0.871224 | 2.632719 | 1.708752 | 2.095861 | 1.991950 | -1.060759 | 62.761505 | 0.41416898 | -0.20546417 | 0.837209493 | 0.998865404 | 6.012689 | 5.929735 | 6.023727 | 5.926729 | 5.971321 | 5.955383 | 90 | 55 | 55 | 55 |
| 2639 | NM_000159,N | GCDH | glutaryl-CoA dehydrogenase | protein_coding | NP_000150.1;NP_039663.1 | 4189 | 608801 | . | . | 7.723232 | 8.534531 | 9.722412 | 8.282601 | 17.972275 | 19.727949 | 21.816811 | 18.937186 | 1.127612 | 511.294546 | 0.177640937 | 0.975396326 | 0.329363706 | 0.998865404 | 8.908580 | 9.082220 | 8.920398 | 8.989569 | 9.091771 | 8.969080 | 523 | 547 | 516 | 460 |
| 2643 | NM_000161,N | GCH1 | GTP cyclohydrolase 1 | protein_coding | NP_000152.1;NP_001019195.1;NP_001019241.1;NP_001019242.1;XP_01 | 4193 | 600225 | . | . | 1.069406 | 1.295956 | 1.393015 | 1.185328 | 2.488551 | 2.995662 | 3.125883 | 2.710112 | 1.060144 | 104.081687 | 0.317679652 | 0.265234823 | 0.79082857 | 0.998865404 | 6.658540 | 6.743359 | 6.675607 | 6.704307 | 6.734286 | 6.681691 | 111 | 114 | 101 | 92 |
| 2647 | NM_001487,N | BLOC1S1 | biogenesis of lysosomal organelles complex 1 subunit 1 | protein_coding | NP_001478.2 | 4200 | 601444 | . | . | 29.872892 | 31.768592 | 24.117726 | 31.619343 | 69.515449 | 73.434523 | 54.119473 | 72.293865 | -1.167848 | 593.815052 | 0.178988445 | -1.25065195 | 0.211061489 | 0.974817004 | 9.321951 | 9.097031 | 9.252788 | 9.269244 | 9.060970 | 9.246853 | 728 | 699 | 407 | 588 |
| 2648 | NM_00137622 | KAT2A | lysine acetyltransferase 2A | protein_coding | NP_001363156.1;NP_066564.2;XP_006721881.1;XP_016879937.1 | 4201 | 602301 | . | . | 35.915463 | 39.442315 | 48.635623 | 39.558213 | 83.576760 | 91.172682 | 109.136924 | 90.445155 | 1.153784 | 4114.724030 | 0.130079364 | 1.586514107 | 0.112622735 | 0.850211417 | 11.899638 | 12.106139 | 11.908163 | 11.975208 | 12.149727 | 11.964316 | 4220 | 4291 | 4255 | 3679 |
| 2649 | NM_00127854 | NR6A1 | nuclear receptor subfamily 6 group A member 1 | protein_coding | NP_001265475.1;NP_001480.3;NP_201591.2;XP_005251974.1;XP_00525 | 7985 | 602778 | . | . | 0.672136 | 0.558149 | 0.654769 | 0.657342 | 1.564088 | 1.290186 | 1.469281 | 1.502935 | 1.093462 | 156.656491 | 0.265158543 | 0.486133429 | 0.626872567 | 0.998865404 | 7.225030 | 7.354967 | 7.281830 | 7.260373 | 7.336374 | 7.276311 | 177 | 157 | 153 | 142 |
| 2650 | NM_00109763 | GCNT1 | glucosaminyl (N-acetyl) transferase 1 | protein_coding | NP_001091102.1;NP_001091103.1;NP_001091104.1;NP_001091105.1;NI | 4203 | 600391 | . | . | 1.115777 | 1.127094 | 0.750576 | 0.822640 | 2.596461 | 2.605330 | 1.684272 | 1.880868 | -1.429441 | 166.474952 | 0.257082732 | -2.00500239 | **0.044962792** | 0.649185583 | 7.614229 | 7.098193 | 7.445788 | 7.447714 | 7.285762 | 7.315866 | 225 | 212 | 114 | 134 |
| 2651 | NM_00137474 | GCNT2 | glucosaminyl (N-acetyl) transferase 2 (I blood group) | protein_coding | NP_001361676.1;NP_001482.1;NP_663624.1;NP_663630.2;XP_0052490 | 4204 | 600429 | . | . | 6.458874 | 6.574372 | 7.518653 | 8.505120 | 15.030065 | 15.196955 | 16.871638 | 19.445948 | 1.190874 | 998.956160 | 0.13788281 | 1.827791024 | 0.067580921 | 0.739745797 | 9.832901 | 10.084683 | 9.892788 | 9.898648 | 9.995994 | 10.050498 | 1046 | 988 | 910 | 1055 |
| 2653 | NM_004483,N | GCSH | glycine cleavage system protein H | protein_coding | NP_004474.2;XP_016878625.1;XP_016878626.1 | 4208 | 238330 | . | . | 16.164454 | 15.163384 | 14.530751 | 12.868827 | 37.615349 | 35.050843 | 32.606584 | 29.423046 | -1.180948 | 703.449301 | 0.15724564 | -1.52592741 | 0.127027943 | 0.874064017 | 9.572964 | 9.333735 | 9.539107 | 9.483925 | 9.442968 | 9.350587 | 911 | 790 | 600 | 563 |
| 2657 | NM_001492 | GDF1 | growth differentiation factor 1 | protein_coding | NP_001483.3 | 4214 | 602880 | . | . | 0.149864 | 0.141122 | 0.138674 | 0.119107 | 0.348741 | 0.326209 | 0.311180 | 0.272324 | -1.189711 | 11.543515 | 0.89677496 | -0.27945759 | 0.779893679 | NA | 3.648138 | 3.399154 | 3.538224 | 3.533737 | 3.531288 | 3.522997 | 15 | 13 | 10 | 9 |
| 2661 | NM_00128882 | GDF9 | growth differentiation factor 9 | protein_coding | NP_001275753.1;NP_001275754.1;NP_001275755.1;NP_001275756.1;NI | 4224 | 601918 | . | . | 0.442812 | 0.534262 | 0.499297 | 0.824375 | 1.030441 | 1.234970 | 1.120408 | 1.884833 | 1.471731 | 53.279413 | 0.433205636 | 1.286949061 | 0.198112044 | 0.96621494 | 5.431047 | 5.986776 | 5.673708 | 5.699015 | 5.739911 | 5.781354 | 46 | 50 | 50 | 65 |
| 2664 | NM_001493 | GDI1 | GDP dissociation inhibitor 1 | protein_coding | NP_001484.1 | 4226 | 300104 | . | . | 33.948277 | 38.520157 | 37.632015 | 37.654213 | 78.999031 | 89.041069 | 84.445152 | 86.091888 | 1.024889 | 2577.535324 | 0.119587876 | 0.296582831 | 0.766785021 | 0.998865404 | 11.313997 | 11.349339 | 11.272433 | 11.367875 | 11.345102 | 11.337141 | 2759 | 2909 | 2280 | 2434 |
| 2665 | NM_00111515 | GDI2 | GDP dissociation inhibitor 2 | protein_coding | NP_001108628.1;NP_001485.2;XP_016871560.1 | 4227 | 600767 | . | . | 67.848490 | 66.683318 | 59.047445 | 67.406959 | 157.886226 | 154.141483 | 132.500751 | 154.118015 | -1.065766 | 4717.386969 | 0.112727257 | -0.81516318 | 0.414978912 | 0.998865404 | 12.249038 | 12.157040 | 12.239741 | 12.219055 | 12.126920 | 12.223214 | 5674 | 5188 | 3741 | 4530 |
| 2669 | NM_005261,N | GEM | GTP binding protein overexpressed in skeletal muscle | protein_coding | NP_005252.1;NP_859053.1;XP_016868804.1 | 4234 | 600164 | . | . | 1.617644 | 1.720973 | 1.606390 | 1.556657 | 3.764323 | 3.978107 | 3.604693 | 3.559111 | -1.074854 | 110.891212 | 0.305372841 | -0.3410259 | 0.733084084 | 0.998865404 | 6.844134 | 6.739989 | 6.793973 | 6.815891 | 6.788450 | 6.774540 | 128 | 128 | 95 | 98 |
| 2671 | NM_005262 | GFER | growth factor, augmenter of liver regeneration | protein_coding | NP_005253.3 | 4236 | 600924 | . | . | 6.114977 | 6.448611 | 5.428823 | 6.587464 | 14.229803 | 14.906253 | 12.182122 | 15.061454 | -1.059564 | 453.349223 | 0.180735847 | -0.46183537 | 0.644199383 | 0.998865404 | 8.866051 | 8.781673 | 8.827076 | 8.855762 | 8.749412 | 8.858983 | 525 | 515 | 348 | 450 |
| 2672 | NM_00112721 | GFI1 | growth factor independent 1 transcriptional repressor | protein_coding | NP_001120687.1;NP_001120688.1;NP_005254.2;XP_005270806.1;XP_01 | 4237 | 600871 | . | . | 0.064621 | 0.115536 | 0.061636 | 0.043960 | 0.150376 | 0.267067 | 0.138309 | 0.100508 | -1.726663 | 12.141956 | 0.901177982 | -0.87439601 | 0.381902628 | NA | 3.942580 | 3.155137 | 3.600183 | 3.636296 | 3.596459 | 3.583666 | 13 | 21 | 9 | 7 |
| 2673 | NM_00124471 | GFPT1 | glutamine--fructose-6-phosphate transaminase 1 | protein_coding | NP_001231639.1;NP_002047.2;XP_016859290.1;XP_016859291.1 | 4241 | 138292 | . | . | 14.441566 | 13.984286 | 12.727909 | 12.134292 | 33.606120 | 32.325299 | 28.561057 | 27.743614 | -1.141302 | 3453.678760 | 0.111976628 | -1.70286284 | 0.088593739 | 0.802216049 | 11.846046 | 11.655503 | 11.825804 | 11.785248 | 11.720947 | 11.672480 | 4342 | 3876 | 2889 | 2934 |
| 2677 | NM_000821,N | GGCX | gamma-glutamyl carboxylase | protein_coding | NP_000812.2;NP_001135741.1;NP_001298241.1;XP_005264316.1;XP_01 | 4247 | 137167 | . | . | 24.055020 | 26.557461 | 23.452837 | 24.666763 | 55.977020 | 61.388760 | 52.627487 | 56.397619 | 1.004786 | 4007.805156 | 0.107198906 | 0.064254029 | 0.948767957 | 0.998865404 | 11.965146 | 11.972040 | 11.966079 | 11.967130 | 11.972832 | 11.968302 | 4601 | 4317 | 3503 | 3756 |
| 2678 | NM_00128883 | GGT1 | gamma-glutamyltransferase 1 | protein_coding | NP_001275762.1;NP_038265.2;NP_038347.2 | 4250 | 612346 | . | . | 0.568833 | 0.321244 | 0.691698 | 0.464559 | 1.323697 | 0.742570 | 1.552150 | 1.062158 | 1.234239 | 39.816243 | 0.521460166 | 0.582252773 | 0.560396432 | 0.998865404 | 5.153119 | 5.461051 | 5.323596 | 5.253585 | 5.353702 | 5.288078 | 51 | 29 | 46 | 33 |
| 2683 | NM_00137849 | B4GALT1 | beta-1,4-galactosyltransferase 1 | protein_coding | NP_001365424.1;NP_001365425.1;NP_001365426.1;NP_001488.2 | 924 | 137060 | . | . | 11.558740 | 11.027142 | 9.730650 | 9.885733 | 26.897661 | 25.489733 | 21.835297 | 22.602554 | -1.180479 | 1348.205090 | 0.128237258 | -1.86663554 | 0.061952502 | 0.723723143 | 10.511512 | 10.272225 | 10.478353 | 10.434523 | 10.328031 | 10.332505 | 1728 | 1531 | 1072 | 1163 |
| 2686 | NM_00135170 | GGT7 | gamma-glutamyltransferase 7 | protein_coding | NP_001338631.1;NP_821158.2;XP_011527084.1;XP_011527085.1;XP_01 | 4259 | 612342 | . | . | 22.595700 | 22.617069 | 25.767290 | 23.790027 | 52.581121 | 52.280368 | 57.821053 | 54.393072 | 1.131157 | 1908.002102 | 0.121193343 | 1.467065603 | 0.14235821 | 0.903146012 | 10.806135 | 10.984076 | 10.842002 | 10.850968 | 10.974344 | 10.912918 | 2050 | 1943 | 1829 | 1826 |
| 2690 | NM_000163,N | GHR | growth hormone receptor | protein_coding | NP_000154.1;NP_001229328.1;NP_001229329.1;NP_001229330.1;NP_0 | 4263 | 600946 | . | . | 5.155637 | 5.027704 | 5.673662 | 4.955962 | 11.997377 | 11.621763 | 12.731534 | 11.331218 | 1.015200 | 724.332516 | 0.152820382 | 0.142414317 | 0.886752754 | 0.998865404 | 9.489377 | 9.511555 | 9.491509 | 9.497644 | 9.545868 | 9.462682 | 824 | 779 | 673 | 643 |
| 2696 | NM_000164,N | GIPR | gastric inhibitory polypeptide receptor | protein_coding | NP_000155.1;NP_001295347.1;XP_011525011.1;XP_011525012.1;XP_01 | 4271 | 137241 | . | . | 0.534802 | 0.328262 | 0.710326 | 0.530506 | 1.244508 | 0.758794 | 1.593951 | 1.212938 | 1.386196 | 57.426370 | 0.440184208 | 1.070304677 | 0.28448219 | 0.998865404 | 5.586822 | 6.061578 | 5.837355 | 5.753794 | 5.897649 | 5.828070 | 68 | 40 | 67 | 53 |
| 2700 | NM_021954,XI | GJA3 | gap junction protein alpha 3 | protein_coding | NP_068773.2;XP_011533350.1 | 4277 | 121015 | . | . | 2.226940 | 1.863420 | 2.579938 | 1.682944 | 5.182180 | 4.307379 | 5.789305 | 3.847852 | 1.033809 | 338.909171 | 0.231726456 | 0.207007824 | 0.836003748 | 0.998865404 | 8.379622 | 8.429457 | 8.436119 | 8.337740 | 8.520110 | 8.279382 | 417 | 328 | 364 | 254 |
| 2702 | NM_005266,N | GJA5 | gap junction protein alpha 5 | protein_coding | NP_005257.2;NP_859054.1;XP_005273008.1;XP_016856533.1 | 4279 | 121013 | . | . | 0.065424 | 0.070529 | 0.021323 | 0.019512 | 0.152244 | 0.163030 | 0.047849 | 0.044613 | **-4.230687** | 7.209344 | 1.221784336 | -1.70315803 | 0.088538498 | NA | 3.544574 | 1.459419 | 2.883537 | 2.886640 | 2.843532 | 2.847093 | 13 | 13 | 2 | 3 |
| 2705 | NM_000166,N | GJB1 | gap junction protein beta 1 | protein_coding | NP_000157.1;NP_001091111.1;XP_011529209.1 | 4283 | 304040 | . | . | 128.210636 | 114.859003 | 142.716614 | 139.274620 | 298.351417 | 265.501732 | 320.251938 | 318.434867 | 1.176560 | 7397.234248 | 0.102462835 | 2.289362496 | **0.022058299** | 0.491793249 | 12.730712 | 12.965302 | 12.777861 | 12.782109 | 12.928253 | 12.906698 | 7807 | 7353 | 7042 | 7403 |
| 2706 | NM_004004,XI | GJB2 | gap junction protein beta 2 | protein_coding | NP_003995.2;XP_011533351.1 | 4284 | 121011 | . | . | 0.273991 | 0.198204 | 0.479505 | 0.311315 | 0.637588 | 0.458156 | 1.075995 | 0.711785 | 1.622608 | 23.429934 | 0.655586349 | 1.065175811 | 0.286796377 | NA | 4.156660 | 4.859205 | 4.522983 | 4.496841 | 4.582407 | 4.529935 | 24 | 16 | 31 | 21 |
| 2710 | NM_000167,N | GK | glycerol kinase | protein_coding | NP_000158.1;NP_001121599.1;NP_001191948.1;NP_976325.1;XP_00527 | 4289 | 300474 | . | . | 3.299107 | 3.503531 | 4.025247 | 4.336440 | 7.677153 | 8.098569 | 9.032536 | 9.914755 | 1.203336 | 524.524200 | 0.165666044 | 1.611913395 | 0.106980787 | 0.83501324 | 8.895387 | 9.162040 | 8.949227 | 8.991241 | 9.075471 | 9.102662 | 530 | 531 | 488 | 548 |
| 2713 | NR_026575 | GK3P | glycerol kinase 3 pseudogene | transcribed_ps | . | 4292 | 600149 | . | . | 0.036929 | 0.041721 | 0.024281 | 0.007379 | 0.085935 | 0.096440 | 0.054486 | 0.016872 | -1.881650 | 2.419730 | 1.944511059 | -0.46901154 | 0.639061387 | NA | 1.658167 | 0.751299 | 1.283501 | 1.288659 | 1.282086 | 1.276459 | 3 | 4 | 2 | 1 |
| 2717 | NM_000169,N | GLA | galactosidase alpha | protein_coding | NP_000160.1 | 4296 | 300644 | . | . | 50.847337 | 53.382251 | 62.492013 | 54.247143 | 118.323841 | 123.395468 | 140.230271 | 124.029648 | 1.105992 | 2272.358843 | 0.125732059 | 1.155957093 | 0.247698708 | 0.992733415 | 11.075371 | 11.220910 | 11.090997 | 11.122902 | 11.247701 | 11.124724 | 2435 | 2375 | 2233 | 2068 |
| 2719 | NM_00116461 | GPC3 | glypican 3 | protein_coding | NP_001158089.1;NP_001158090.1;NP_001158091.1;NP_004475.1;XP_01 | 4451 | 300037 | . | . | 2.642024 | 2.711907 | 1.510355 | 2.001304 | 6.148098 | 6.268694 | 3.389193 | 4.575743 | -1.517644 | 156.860138 | 0.269938464 | -2.22952061 | **0.025779285** | 0.525611611 | 7.563678 | 6.960374 | 7.368333 | 7.367034 | 7.164425 | 7.240331 | 218 | 204 | 96 | 130 |
| 2720 | NM_000404,N | GLB1 | galactosidase beta 1 | protein_coding | NP_000395.3;NP_001073279.1;NP_001129074.1;NP_001303969.1 | 4298 | 611458 | . | . | 28.957543 | 28.464803 | 25.688893 | 30.168545 | 67.385393 | 65.797661 | 57.645132 | 68.976795 | -1.043635 | 2228.022148 | 0.125309612 | -0.49171972 | 0.622917495 | 0.998865404 | 11.152147 | 11.090285 | 11.146614 | 11.129055 | 11.042167 | 11.161458 | 2649 | 2429 | 1754 | 2197 |
| 2729 | NM_00119711 | GCLC | glutamate-cysteine ligase catalytic subunit | protein_coding | NP_001184044.1;NP_001489.1;XP_016866238.1 | 4311 | 606857 | . | . | 5.132810 | 5.189637 | 4.917178 | 5.244266 | 11.944260 | 11.996077 | 11.034006 | 11.990392 | -1.029784 | 604.851458 | 0.157541158 | -0.26876789 | 0.788108306 | 0.998865404 | 9.261568 | 9.218992 | 9.249522 | 9.250923 | 9.212511 | 9.248482 | 706 | 663 | 504 | 574 |
| 2730 | NM_00130825 | GCLM | glutamate-cysteine ligase modifier subunit | protein_coding | NP_001295182.1;NP_002052.1;XP_011539563.1;XP_016856545.1;XP_01 | 4312 | 601176 | . | . | 8.866516 | 7.818825 | 11.881977 | 10.207546 | 20.632748 | 18.073565 | 26.662812 | 23.338341 | 1.307265 | 1511.566726 | 0.140151619 | 2.758093333 | **0.00581396** | 0.236029657 | 10.355346 | 10.742426 | 10.492693 | 10.397809 | 10.718533 | 10.584615 | 1604 | 1324 | 1615 | 1471 |
| 2731 | NM_000170 | GLDC | glycine decarboxylase | protein_coding | NP_000161.2 | 4313 | 238300 | . | . | 13.887553 | 15.346665 | 15.168716 | 17.321238 | 32.316902 | 35.474503 | 34.038162 | 39.602951 | 1.096194 | 1846.779468 | 0.128740995 | 1.029218678 | 0.303376927 | 0.998865404 | 10.783220 | 10.915348 | 10.776322 | 10.847584 | 10.835197 | 10.932389 | 1937 | 1988 | 1577 | 1921 |
| 2733 | NM_00100372 | GLE1 | GLE1 RNA export mediator | protein_coding | NP_001003722.1;NP_001490.1;XP_006717122.1;XP_006717123.1;XP_01 | 4315 | 603371 | . | . | 8.174976 | 8.794032 | 8.325631 | 8.841926 | 19.023506 | 20.327796 | 18.682474 | 20.216014 | 1.011417 | 887.320723 | 0.142852915 | 0.114650015 | 0.908722534 | 0.998865404 | 9.785277 | 9.801302 | 9.763802 | 9.813574 | 9.777574 | 9.816145 | 981 | 985 | 755 | 859 |
| 2734 | NM_00114566 | GLG1 | golgi glycoprotein 1 | protein_coding | NP_001139138.1;NP_001139139.1;NP_036333.2 | 4316 | 600753 | . | . | 16.863384 | 17.132521 | 13.892644 | 15.453134 | 39.241789 | 39.602586 | 31.174691 | 35.331756 | -1.174522 | 4364.953976 | 0.11296658 | -2.05435988 | **0.039940873** | 0.624353197 | 12.203192 | 11.970975 | 12.152142 | 12.157347 | 11.972805 | 12.066545 | 5413 | 5104 | 3291 | 3979 |
| 2735 | NM_00116004 | GLI1 | GLI family zinc finger 1 | protein_coding | NP_001153517.1;NP_001161081.1;NP_005260.1;XP_011536491.1;XP_01 | 4317 | 165220 | . | . | 0.467210 | 0.504653 | 0.297432 | 0.352064 | 1.087217 | 1.166528 | 0.667429 | 0.804953 | -1.521797 | 49.452067 | 0.447787277 | -1.35282091 | 0.176112851 | 0.949193943 | 5.899769 | 5.292767 | 5.658211 | 5.675090 | 5.582424 | 5.599721 | 66 | 67 | 32 | 39 |
| 2736 | NM_00137127 | GLI2 | GLI family zinc finger 2 | protein_coding | NP_001358200.1;NP_001361282.1;NP_001361283.1;NP_005261.2;XP_01 | 4318 | 165230 | . | . | 0.925939 | 0.883249 | 0.570825 | 0.759784 | 2.154697 | 2.041670 | 1.280915 | 1.737156 | -1.358180 | 175.347840 | 0.261755185 | -1.68735628 | 0.091534864 | 0.805877675 | 7.658726 | 7.215520 | 7.522476 | 7.504730 | 7.330640 | 7.436298 | 237 | 214 | 111 | 159 |
| 2737 | NM_000168,XI | GLI3 | GLI family zinc finger 3 | protein_coding | NP_000159.3;XP_011513576.1;XP_016867486.1 | 4319 | 165240 | . | . | 1.718474 | 1.903425 | 1.731982 | 1.830296 | 3.998960 | 4.399851 | 3.886517 | 4.184755 | -1.035539 | 469.482303 | 0.172975831 | -0.2912676 | 0.770846668 | 0.998865404 | 8.900121 | 8.849285 | 8.855099 | 8.915101 | 8.852880 | 8.874415 | 524 | 540 | 394 | 440 |
| 2738 | NM_138465 | GLI4 | GLI family zinc finger 4 | protein_coding | NP_612474.1 | 4320 | 165280 | . | . | 3.212816 | 3.706835 | 4.030957 | 4.900640 | 7.476354 | 8.568516 | 9.045351 | 11.204730 | 1.274351 | 164.934919 | 0.263656787 | 1.326583867 | 0.184646376 | 0.95742822 | 7.181119 | 7.529416 | 7.281904 | 7.333684 | 7.366716 | 7.446196 | 156 | 167 | 146 | 189 |
| 2739 | NM_006708 | GLO1 | glyoxalase I | protein_coding | NP_006699.2 | 4323 | 138750 | . | . | 51.474239 | 43.670147 | 45.518879 | 42.532555 | 119.782661 | 100.945503 | 102.143044 | 97.245644 | -1.095943 | 2878.624993 | 0.126958693 | -1.04107114 | 0.297842531 | 0.998865404 | 11.555611 | 11.423704 | 11.595104 | 11.452565 | 11.482643 | 11.419664 | 3765 | 2968 | 2482 | 2475 |
| 2743 | NM_000824,N | GLRB | glycine receptor beta | protein_coding | NP_000815.1;NP_001159532.1;NP_001159533.1;XP_016863523.1;XP_01 | 4329 | 138492 | . | . | 0.574529 | 0.704442 | 0.754987 | 0.768009 | 1.336952 | 1.628348 | 1.694169 | 1.755963 | 1.438421 | 76.752688 | 0.368857392 | 1.421920473 | 0.155049339 | 0.923671138 | 5.976092 | 6.500758 | 6.181998 | 6.222651 | 6.323198 | 6.264650 | 66 | 74 | 87 | 76 |
| 2744 | NM_00125631 | GLS | glutaminase | protein_coding | NP_001243239.1;NP_055720.3;XP_005246524.1;XP_006712498.1;XP_01 | 4331 | 138280 | . | . | 42.614321 | 35.095697 | 46.810503 | 30.265375 | 99.165276 | 81.125271 | 105.041409 | 69.198183 | -1.017211 | 5586.754813 | 0.346440955 | -0.07106114 | 0.943349099 | 0.998865404 | 12.460025 | 12.435460 | 12.530177 | 12.358233 | 12.601546 | 12.232241 | 7147 | 5462 | 5864 | 4064 |
| 2745 | NM_00111889 | GLRX | glutaredoxin | protein_coding | NP_001112362.1;NP_001230587.1;NP_001230588.1;NP_002055.1 | 4330 | 600443 | . | . | 38.626410 | 36.794790 | 34.659762 | 37.473609 | 89.885242 | 85.052809 | 77.775498 | 85.678956 | -1.040324 | 1119.585082 | 0.134378493 | -0.42441502 | 0.671263168 | 0.998865404 | 10.156998 | 10.099935 | 10.164401 | 10.121673 | 10.095638 | 10.131655 | 1352 | 1197 | 931 | 1054 |
| 2746 | NM_00131890 | GLUD1 | glutamate dehydrogenase 1 | protein_coding | NP_001305829.1;NP_001305830.1;NP_001305831.1;NP_001305833.1;NI | 4335 | 138130 | . | . | 22.005746 | 21.188555 | 20.674510 | 23.773158 | 51.208276 | 48.978301 | 46.392999 | 54.354501 | 1.014847 | 2255.079892 | 0.123712987 | 0.171866581 | 0.86354242 | 0.998865404 | 11.128381 | 11.149468 | 11.150774 | 11.113854 | 11.090368 | 11.195420 | 2637 | 2360 | 1846 | 2269 |
| 2747 | NM_012084 | GLUD2 | glutamate dehydrogenase 2 | protein_coding | NP_036216.2 | 4336 | 300144 | . | . | 1.860213 | 2.385044 | 1.812209 | 1.900540 | 4.328793 | 5.513136 | 4.066545 | 4.345358 | -1.155580 | 154.419369 | 0.272694715 | -0.7650194 | 0.44425999 | 0.998865404 | 7.371764 | 7.162040 | 7.246414 | 7.345999 | 7.232894 | 7.247675 | 168 | 200 | 122 | 137 |
| 2749 | NR_048575 | GLUD1P3 | glutamate dehydrogenase 1 pseudogene 3 | transcribed_ps | . | 4338 | . | | . | 0.905906 | 0.726941 | 1.204387 | 0.819049 | 2.108080 | 1.680358 | 2.702609 | 1.872658 | 1.245208 | 26.101888 | 0.614265493 | 0.515065399 | 0.606507321 | NA | 4.537347 | 4.857133 | 4.698605 | 4.674506 | 4.731692 | 4.688022 | 30 | 22 | 30 | 22 |
| 2752 | NM_00103304 | GLUL | glutamate-ammonia ligase | protein_coding | NP_001028216.1;NP_001028228.1;NP_002056.2;XP_006711341.1 | 4341 | 138290 | . | . | 4.850013 | 5.800651 | 7.510536 | 8.041173 | 11.286182 | 13.408466 | 16.853427 | 18.385189 | 1.461022 | 1471.326967 | 0.131030281 | 4.174440842 | **2.98719E-05** | **0.004794437** | 10.223868 | 10.770470 | 10.317453 | 10.393711 | 10.617437 | 10.678048 | 1306 | 1357 | 1462 | 1699 |
| 2760 | NM_000405,N | GM2A | GM2 ganglioside activator | protein_coding | NP_000396.2;NP_001161079.1 | 4367 | 613109 | . | . | 51.009730 | 48.463105 | 45.396551 | 49.837624 | 118.701743 | 112.024634 | 101.868542 | 113.947818 | -1.059089 | 5444.502651 | 0.109843942 | -0.75400895 | 0.450843848 | 0.998865404 | 12.451422 | 12.368557 | 12.459005 | 12.408248 | 12.350801 | 12.419545 | 6643 | 5862 | 4409 | 5162 |
| 2762 | NM_00125384 | GMDS | GDP-mannose 4,6-dehydratase | protein_coding | NP_001240775.1;NP_001491.1;XP_006715129.1;XP_011512802.1;XP_01 | 4369 | 602884 | . | . | 7.720825 | 6.979520 | 5.456433 | 7.628288 | 17.966678 | 16.133474 | 12.244078 | 17.441175 | -1.104708 | 379.110538 | 0.199620072 | -0.71969211 | 0.471714584 | 0.998865404 | 8.636955 | 8.492374 | 8.615163 | 8.571427 | 8.458239 | 8.606901 | 475 | 414 | 272 | 382 |
| 2764 | NM_004124 | GMFB | glia maturation factor beta | protein_coding | NP_004115.1 | 4373 | 601713 | . | . | 9.427774 | 6.159074 | 7.132363 | 4.571985 | 21.938818 | 14.236977 | 16.004816 | 10.453300 | -1.351204 | 870.500377 | 0.44242118 | -0.98152078 | 0.326335991 | 0.998865404 | 9.966425 | 9.532466 | 10.000714 | 9.676312 | 9.777032 | 9.469728 | 1398 | 849 | 788 | 539 |
| 2766 | NM_006877,XI | GMPR | guanosine monophosphate reductase | protein_coding | NP_006868.3;XP_011512810.1 | 4376 | 139265 | . | . | 23.381618 | 20.977780 | 22.593087 | 25.554556 | 54.409989 | 48.491082 | 50.698233 | 58.427448 | 1.069718 | 1087.736938 | 0.142195873 | 0.683777154 | 0.494115895 | 0.998865404 | 10.037710 | 10.134882 | 10.102803 | 10.016683 | 10.068756 | 10.150798 | 1281 | 1068 | 924 | 1112 |
| 2767 | NM_002067 | GNA11 | G protein subunit alpha 11 | protein_coding | NP_002058.2 | 4379 | 139313 | . | . | 10.287019 | 11.507619 | 10.473843 | 12.089916 | 23.938314 | 26.600376 | 23.502998 | 27.642157 | 1.021057 | 1447.404297 | 0.135707101 | 0.221527206 | 0.824681955 | 0.998865404 | 10.484385 | 10.513968 | 10.449575 | 10.529548 | 10.453757 | 10.556120 | 1564 | 1626 | 1187 | 1462 |
| 2768 | NM_00128244 | GNA12 | G protein subunit alpha 12 | protein_coding | NP_001269369.1;NP_001269370.1;NP_001280021.1;NP_031379.2 | 4380 | 604394 | . | . | 10.809107 | 11.894150 | 10.259206 | 12.859394 | 25.153238 | 27.493861 | 23.021360 | 29.401474 | 1.003806 | 1558.750693 | 0.142283157 | 0.038523086 | 0.969270626 | 0.998865404 | 10.603703 | 10.608641 | 10.568808 | 10.636408 | 10.519373 | 10.685790 | 1714 | 1752 | 1212 | 1621 |
| 2770 | NM_00125641 | GNAI1 | G protein subunit alpha i1 | protein_coding | NP_001243343.1;NP_002060.4 | 4384 | 139310 | . | . | 2.112304 | 1.754307 | 1.950261 | 1.741587 | 4.915418 | 4.055157 | 4.376329 | 3.981931 | -1.063132 | 594.289714 | 0.171940155 | -0.51367486 | 0.607479332 | 0.998865404 | 9.258062 | 9.170659 | 9.296412 | 9.166245 | 9.232404 | 9.152087 | 773 | 597 | 532 | 507 |
| 2771 | NM_00116642 | GNAI2 | G protein subunit alpha i2 | protein_coding | NP_001159897.1;NP_001269546.1;NP_001269547.1;NP_001269548.1;NI | 4385 | 139360 | . | . | 54.808981 | 57.870851 | 44.776029 | 51.868967 | 127.542753 | 133.771064 | 100.476113 | 118.592222 | -1.183541 | 3616.157461 | 0.117916202 | -2.06171299 | **0.039235069** | 0.616675358 | 11.936782 | 11.693454 | 11.865768 | 11.904556 | 11.690356 | 11.799001 | 4410 | 4328 | 2687 | 3313 |
| 2773 | NM_006496 | GNAI3 | G protein subunit alpha i3 | protein_coding | NP_006487.1 | 4387 | 139370 | . | . | 8.582144 | 8.115662 | 6.678659 | 7.434550 | 19.971001 | 18.759714 | 14.986717 | 16.998217 | -1.199632 | 2171.939546 | 0.122066975 | -2.15121082 | **0.031459563** | 0.561754658 | 11.210154 | 10.947439 | 11.178928 | 11.126119 | 10.968769 | 11.044366 | 2816 | 2474 | 1634 | 1941 |
| 2774 | NM_00114233 | GNAL | G protein subunit alpha L | protein_coding | NP_001135811.1;NP_001248372.1;NP_001248373.1;NP_001356316.1;NI | 4388 | 139312 | . | . | 0.841236 | 0.636877 | 0.820912 | 0.859694 | 1.957590 | 1.472168 | 1.842102 | 1.965589 | 1.134184 | 149.892151 | 0.285788291 | 0.635627077 | 0.525019517 | 0.998865404 | 7.133341 | 7.316417 | 7.264632 | 7.126665 | 7.248056 | 7.246085 | 191 | 124 | 139 | 149 |
| 2775 | NM_020988,N | GNAO1 | G protein subunit alpha o1 | protein_coding | NP_066268.1;NP_620073.2;XP_011521305.1 | 4389 | 139311 | . | . | 3.632216 | 3.207199 | 2.943521 | 4.052106 | 8.452314 | 7.413586 | 6.605175 | 9.264658 | 1.034033 | 627.612649 | 0.184515527 | 0.261671398 | 0.793574794 | 0.998865404 | 9.269764 | 9.317305 | 9.310743 | 9.246038 | 9.183793 | 9.406617 | 745 | 634 | 463 | 697 |
| 2776 | NM_002072,XI | GNAQ | G protein subunit alpha q | protein_coding | NP_002063.2;XP_016870117.1 | 4390 | 600998 | . | . | 5.306705 | 4.867132 | 5.282846 | 4.187540 | 12.348922 | 11.250594 | 11.854554 | 9.574311 | -1.090574 | 1053.143082 | 0.154949168 | -0.70721784 | 0.419502529 | 0.998865404 | 10.101306 | 9.976988 | 10.104676 | 10.033222 | 10.089061 | 9.914990 | 1325 | 1129 | 981 | 832 |
| 2778 | NM_000516,N | GNAS | GNAS complex locus | protein_coding | NP_000507.1;NP_001070956.1;NP_001070957.1;NP_001070958.1;NP_0 | 4392 | 139320 | . | . | 358.990112 | 382.240929 | 357.288101 | 401.914814 | 835.384735 | 883.567014 | 801.744153 | 918.930473 | 1.013725 | 21579.039902 | 0.105657562 | 0.186126923 | 0.852345229 | 0.998865404 | 14.387495 | 14.407124 | 14.369041 | 14.412601 | 14.357067 | 14.446521 | 24054 | 23712 | 17912 | 21425 |
| 2781 | NM_002073,XI | GNAZ | G protein subunit alpha z | protein_coding | NP_002064.1;XP_016884275.1 | 4395 | 139160 | . | . | 1.619476 | 1.774816 | 1.510425 | 2.373024 | 3.768588 | 4.102567 | 3.389348 | 5.425637 | 1.142537 | 352.378880 | 0.191044701 | 1.006261246 | 0.314289917 | 0.998865404 | 8.361995 | 8.553614 | 8.399370 | 8.440385 | 8.472783 | 8.518298 | 365 | 368 | 314 | 366 |
| 2782 | NM_00128253 | GNB1 | G protein subunit beta 1 | protein_coding | NP_001269467.1;NP_001269468.1;NP_002065.1;XP_016856548.1;XP_01 | 4396 | 139380 | . | . | 117.182237 | 121.208512 | 106.551212 | 119.703163 | 272.687873 | 280.178921 | 239.097821 | 273.687059 | -1.053092 | 11146.075001 | 0.108150984 | -0.69006256 | 0.490154849 | 0.998865404 | 13.481112 | 13.406417 | 13.459144 | 13.471880 | 13.368834 | 13.471269 | 13069 | 12430 | 8873 | 10792 |
| 2783 | NM_005273 | GNB2 | G protein subunit beta 2 | protein_coding | NP_005264.2 | 4398 | 139390 | . | . | 82.289314 | 86.638626 | 77.497025 | 94.545540 | 191.490616 | 200.269043 | 173.901077 | 216.167145 | 1.004023 | 4417.757199 | 0.123064193 | 0.047069 | 0.962458235 | 0.998865404 | 12.106301 | 12.111890 | 12.088256 | 12.124111 | 12.028220 | 12.185033 | 4968 | 4860 | 3487 | 4540 |
| 2784 | NM_00129757 | GNB3 | G protein subunit beta 3 | protein_coding | NP_001284500.1;NP_002066.1;XP_011519255.1 | 4400 | 139130 | . | . | 0.048222 | 0.111987 | 0.133815 | 0.037130 | 0.112214 | 0.258863 | 0.300278 | 0.084893 | -1.008502 | 4.531430 | 1.462690595 | -0.00835062 | 0.993337246 | NA | 2.184683 | 2.175234 | 2.161032 | 2.179398 | 2.180679 | 2.159355 | 3 | 7 | 6 | 2 |
| 2786 | NM_00109872 | GNG4 | G protein subunit gamma 4 | protein_coding | NP_001092191.1;NP_001092192.1;NP_004476.1;XP_006711824.1;XP_01 | 4407 | 604388 | . | . | 0.800066 | 1.062131 | 0.772749 | 0.861675 | 1.861786 | 2.455163 | 1.734027 | 1.970117 | -1.154508 | 133.968759 | 0.292292845 | -0.70914756 | 0.478232912 | 0.998865404 | 7.166385 | 6.957571 | 7.036008 | 7.140942 | 7.021405 | 7.053205 | 143 | 176 | 103 | 122 |
| 2787 | NM_005274 | GNG5 | G protein subunit gamma 5 | protein_coding | NP_005265.1 | 4408 | 600874 | . | . | 78.353172 | 72.905724 | 77.216171 | 71.867256 | 182.331070 | 168.524841 | 173.270859 | 164.315948 | -1.028877 | 1886.089055 | 0.124447053 | -0.33002933 | 0.741377799 | 0.998865404 | 10.901426 | 10.860650 | 10.923366 | 10.859436 | 10.901134 | 10.836877 | 2291 | 1981 | 1683 | 1672 |
| 2788 | NM_052847,XI | GNG7 | G protein subunit gamma 7 | protein_coding | NP_443079.1;XP_016882095.1 | 4410 | 604430 | . | . | 0.927015 | 0.665323 | 0.306667 | 0.668813 | 2.157203 | 1.537924 | 0.688152 | 1.529161 | -1.656344 | 85.743059 | 0.400716865 | -1.81675014 | 0.06925538 | 0.743915966 | 6.741623 | 6.010658 | 6.532925 | 6.421273 | 6.265391 | 6.412804 | 144 | 96 | 37 | 81 |
| 2790 | NM_00101799 | GNG10 | G protein subunit gamma 10 | protein_coding | NP_001017998.1;NP_001185593.1 | 4402 | 604389 | . | . | 13.761007 | 10.166084 | 11.998698 | 10.189434 | 32.022433 | 23.499358 | 26.924730 | 23.296929 | -1.093590 | 432.762725 | 0.203162287 | -0.63531605 | 0.52522231 | 0.998865404 | 8.819877 | 8.692162 | 8.871798 | 8.679852 | 8.776470 | 8.672947 | 600 | 413 | 391 | 354 |
| 2791 | NM_004126 | GNG11 | G protein subunit gamma 11 | protein_coding | NP_004117.1 | 4403 | 604390 | . | . | 14.163296 | 12.433914 | 14.033157 | 12.700476 | 32.958569 | 28.741550 | 31.490000 | 29.038130 | -1.009335 | 1254.937983 | 0.140171028 | -0.09563469 | 0.923810711 | 0.998865404 | 10.299827 | 10.286945 | 10.347171 | 10.241822 | 10.329478 | 10.246825 | 1552 | 1266 | 1146 | 1107 |
| 2794 | NM_005275,XI | GNL1 | G protein nucleolar 1 (putative) | protein_coding | NP_005266.2;XP_005249072.1 | 4413 | 143024 | . | . | 7.075333 | 6.857586 | 6.281596 | 7.549070 | 16.464591 | 15.851617 | 14.095719 | 17.260051 | -1.031366 | 1648.865159 | 0.135097801 | -0.32980888 | 0.741544374 | 0.998865404 | 10.709497 | 10.664670 | 10.724014 | 10.671921 | 10.603875 | 10.739819 | 1992 | 1747 | 1288 | 1655 |
| 2796 | NM_000825,N | GNRH1 | gonadotropin releasing hormone 1 | protein_coding | NP_000816.4;NP_001076580.1 | 4419 | 152760 | . | . | 0.395745 | 0.506593 | 0.810447 | 0.227161 | 0.920915 | 1.171014 | 1.818619 | 0.519377 | 1.101561 | 31.720847 | 0.625596219 | 0.22306533 | 0.823484661 | 0.998865404 | 4.913716 | 5.057426 | 4.949669 | 4.976462 | 5.040483 | 4.905513 | 31 | 36 | 44 | 15 |
| 2797 | NM_00131022 | GNRH2 | gonadotropin releasing hormone 2 | protein_coding | NP_001297149.1;NP_001492.1;NP_847901.1;NP_847902.1;XP_0168833 | 4420 | 602352 | . | . | 0.152530 | 0.082828 | 0.197925 | 0.071452 | 0.354945 | 0.191461 | 0.444138 | 0.163366 | 1.120918 | 12.747223 | 0.90541285 | 0.181884607 | 0.855673285 | NA | 3.583318 | 3.755755 | 3.675321 | 3.642269 | 3.690691 | 3.639106 | 18 | 9 | 17 | 7 |
| 2799 | NM_002076 | GNS | glucosamine (N-acetyl)-6-sulfatase | protein_coding | NP_002067.1 | 4422 | 607664 | . | . | 76.582100 | 76.980797 | 74.877457 | 74.104889 | 178.209686 | 177.944565 | 168.022858 | 169.432022 | -1.045186 | 11974.063639 | 0.099947498 | -0.63792822 | 0.523520406 | 0.998865404 | 13.579147 | 13.515400 | 13.567649 | 13.564789 | 13.537152 | 13.519668 | 14116 | 13184 | 10287 | 10865 |
| 2800 | NM_002077,XI | GOLGA1 | golgin A1 | protein_coding | NP_002068.2;XP_005251986.1;XP_006717125.1;XP_006717126.1 | 4424 | 602502 | . | . | 5.749204 | 5.155486 | 6.176866 | 5.996037 | 13.378636 | 11.917137 | 13.860708 | 13.709225 | 1.094921 | 867.610074 | 0.146686114 | 0.891879433 | 0.372457559 | 0.998865404 | 9.693786 | 9.825035 | 9.769729 | 9.681414 | 9.804452 | 9.778496 | 1013 | 838 | 803 | 835 |
| 2801 | NM_00136624 | GOLGA2 | golgin A2 | protein_coding | NP_001353173.1;NP_001353175.1;NP_004477.4;XP_005251989.1 | 4425 | 602580 | . | . | 15.778402 | 16.505883 | 14.916109 | 16.928491 | 36.716991 | 38.154090 | 33.471316 | 38.704982 | -1.025763 | 2116.646506 | 0.122659833 | -0.29917592 | 0.764805821 | 0.998865404 | 11.065931 | 11.028962 | 11.043178 | 11.071359 | 10.990898 | 11.080965 | 2425 | 2354 | 1714 | 2070 |
| 2802 | NM_00117255 | GOLGA3 | golgin A3 | protein_coding | NP_001166028.1;NP_005886.2;XP_005266221.1;XP_005266224.1;XP_00 | 4426 | 602581 | . | . | 8.576311 | 8.627630 | 10.330755 | 9.397920 | 19.957430 | 19.943156 | 23.181913 | 21.487232 | 1.124557 | 2551.551888 | 0.120456832 | 1.405959152 | 0.159736255 | 0.933525128 | 11.229901 | 11.399439 | 11.260926 | 11.272937 | 11.411993 | 11.309577 | 2745 | 2611 | 2496 | 2374 |
| 2803 | NM_00117271 | GOLGA4 | golgin A4 | protein_coding | NP_001166184.1;NP_002069.2;XP_005265126.1;XP_005265127.1;XP_00 | 4427 | 602509 | . | . | 10.603874 | 10.523089 | 12.155991 | 9.432302 | 24.675647 | 24.324590 | 27.277689 | 21.565837 | 1.007615 | 2583.453056 | 0.138648603 | 0.078939833 | 0.93708048 | 0.998865404 | 11.329430 | 11.340718 | 11.338725 | 11.320757 | 11.435583 | 11.225804 | 2996 | 2746 | 2547 | 2117 |
| 2804 | NM_00125648 | GOLGB1 | golgin B1 | protein_coding | NP_001243415.1;NP_001243416.1;NP_001243417.1;NP_001353211.1;NI | 4429 | 602500 | . | . | 7.239542 | 7.594408 | 10.060873 | 8.224527 | 16.846715 | 17.554811 | 22.576305 | 18.804398 | 1.211495 | 2908.919530 | 0.130215379 | 2.125620866 | **0.033534841** | 0.575719787 | 11.361132 | 11.638128 | 11.406282 | 11.435397 | 11.661456 | 11.486373 | 2974 | 2890 | 3063 | 2674 |
| 2805 | NM_002079 | GOT1 | glutamic-oxaloacetic transaminase 1 | protein_coding | NP_002070.1 | 4432 | 138180 | . | . | 33.647495 | 33.762264 | 30.464880 | 37.498981 | 78.299103 | 78.042984 | 68.362312 | 85.736961 | -1.005829 | 2085.395608 | 0.132214639 | -0.06342476 | 0.949428271 | 0.998865404 | 11.030457 | 11.021741 | 11.029489 | 11.025205 | 10.941188 | 11.098043 | 2415 | 2251 | 1630 | 2141 |
| 2806 | NM_00128622 | GOT2 | glutamic-oxaloacetic transaminase 2 | protein_coding | NP_001273149.1;NP_002071.2 | 4433 | 138150 | . | . | 39.723100 | 42.249917 | 37.662366 | 45.678788 | 92.437290 | 97.662572 | 84.513255 | 104.493323 | 1.002359 | 3114.281160 | 0.125702922 | 0.027044851 | 0.978423961 | 0.998865404 | 11.603121 | 11.606244 | 11.580666 | 11.624276 | 11.527826 | 11.675918 | 3487 | 3446 | 2466 | 3187 |
| 2817 | NM_002081,XI | GPC1 | glypican 1 | protein_coding | NP_002072.2;XP_011509278.1 | 4449 | 600395 | . | . | 21.884421 | 21.046173 | 21.351156 | 20.743336 | 50.925942 | 48.649175 | 47.911377 | 47.427172 | -1.034307 | 2463.662809 | 0.11561538 | -0.42091867 | 0.673814471 | 0.998865404 | 11.290649 | 11.242121 | 11.299034 | 11.260502 | 11.268163 | 11.237139 | 2953 | 2639 | 2147 | 2227 |
| 2819 | NM_00125719 | GPD1 | glycerol-3-phosphate dehydrogenase 1 | protein_coding | NP_001244128.1;NP_005267.2 | 4455 | 138420 | . | . | 0.066460 | 0.090559 | 0.040208 | 0.071196 | 0.154655 | 0.209331 | 0.090226 | 0.162781 | -1.307440 | 6.354924 | 1.204928465 | -0.3209689 | 0.748233958 | NA | 2.849979 | 2.459419 | 2.670961 | 2.681390 | 2.664583 | 2.672462 | 7 | 9 | 4 | 6 |
| 2820 | NM_000408,N | GPD2 | glycerol-3-phosphate dehydrogenase 2 | protein_coding | NP_000399.3;NP_001076581.2;XP_005246526.1;XP_011509279.1;XP_01 | 4456 | 138430 | . | . | 11.102759 | 11.381368 | 10.585313 | 9.357682 | 25.836577 | 26.308541 | 23.753134 | 21.395229 | -1.134995 | 1918.046859 | 0.12618627 | -1.44774594 | 0.147688118 | 0.90884489 | 10.993751 | 10.811331 | 10.945341 | 10.960396 | 10.903978 | 10.798818 | 2326 | 2221 | 1668 | 1571 |
| 2821 | NM_000175,N | GPI | glucose-6-phosphate isomerase | protein_coding | NP_000166.2;NP_001171651.1;NP_001276718.1;NP_001276719.1;NP_0 | 4458 | 172400 | . | . | 85.578388 | 92.699057 | 92.582395 | 111.592800 | 199.144415 | 214.278036 | 207.752206 | 255.143690 | 1.131020 | 12550.284398 | 0.116441554 | 1.525433916 | 0.127150905 | 0.874064017 | 13.523930 | 13.701476 | 13.531812 | 13.586340 | 13.587618 | 13.736564 | 13127 | 13119 | 10588 | 13563 |
| 2824 | NM_00100199 | GPM6B | glycoprotein M6B | protein_coding | NP_001001994.1;NP_001001995.1;NP_001001996.1;NP_001305658.1;NI | 4461 | 300051 | . | . | 22.212165 | 18.996757 | 20.927477 | 17.898507 | 51.688620 | 43.911856 | 46.960653 | 40.922809 | -1.094763 | 2722.079517 | 0.133859269 | -0.97579027 | 0.329168409 | 0.998865404 | 11.474157 | 11.343890 | 11.515389 | 11.368421 | 11.429020 | 11.309522 | 3568 | 2796 | 2428 | 2256 |
| 2827 | NM_005281 | GPR3 | G protein-coupled receptor 3 | protein_coding | NP_005272.1 | 4484 | 600241 | . | . | 4.209036 | 4.689110 | 4.116279 | 4.120118 | 9.794600 | 10.839087 | 9.236811 | 9.420160 | -1.094185 | 285.728569 | 0.206317279 | -0.62940362 | 0.529084847 | 0.998865404 | 8.222119 | 8.091948 | 8.154826 | 8.208026 | 8.136001 | 8.132626 | 327 | 338 | 238 | 255 |
| 2841 | NM_00109820 | GPR18 | G protein-coupled receptor 18 | protein_coding | NP_001091670.1;NP_005283.1;XP_006720009.1;XP_024305107.1 | 4472 | 602042 | . | . | 0.069695 | 0.083962 | 0.316080 | 0.185236 | 0.162184 | 0.194082 | 0.709275 | 0.423521 | 1.738272 | 9.244014 | 1.015252751 | 0.785670382 | 0.432060619 | NA | 2.754145 | 3.553545 | 3.176480 | 3.182910 | 3.215769 | 3.188182 | 7 | 8 | 13 | 8 |
| 2842 | NM_006143,XI | GPR19 | G protein-coupled receptor 19 | protein_coding | NP_006134.2;XP_011518925.1;XP_011518926.1;XP_011518927.1;XP_01 | 4473 | 602927 | . | . | 5.096572 | 6.151031 | 5.617887 | 4.431750 | 11.859932 | 14.218386 | 12.606377 | 10.132671 | -1.207532 | 389.297749 | 0.193837035 | -1.40355687 | 0.160450845 | 0.933525128 | 8.734369 | 8.462281 | 8.594458 | 8.713602 | 8.576164 | 8.512657 | 443 | 504 | 324 | 312 |
| 2850 | NM_018971 | GPR27 | G protein-coupled receptor 27 | protein_coding | NP_061844.1 | 4482 | 605187 | . | . | 0.189874 | 0.184015 | 0.119474 | 0.177805 | 0.441844 | 0.425359 | 0.268097 | 0.406531 | -1.273646 | 14.384542 | 0.811390184 | -0.43008175 | 0.667136177 | NA | 4.011454 | 3.660102 | 3.860432 | 3.857017 | 3.833934 | 3.853317 | 19 | 17 | 9 | 14 |
| 2852 | NM_00103996 | GPER1 | G protein-coupled estrogen receptor 1 | protein_coding | NP_001035055.1;NP_001091671.1;NP_001496.1 | 4485 | 601805 | . | . | 0.157549 | 0.115288 | 0.106491 | 0.178418 | 0.366622 | 0.266493 | 0.238962 | 0.407931 | -1.023045 | 12.791890 | 0.859434133 | -0.03824563 | 0.969491841 | NA | 3.694014 | 3.660102 | 3.687514 | 3.671096 | 3.668224 | 3.687804 | 17 | 12 | 9 | 14 |
| 2861 | NM_005302 | GPR37 | G protein-coupled receptor 37 | protein_coding | NP_005293.1 | 4494 | 602583 | . | . | 1.613836 | 1.753059 | 1.353874 | 1.642977 | 3.755464 | 4.052274 | 3.038054 | 3.756473 | -1.139698 | 263.244615 | 0.218477619 | -0.86348235 | 0.387872321 | 0.998865404 | 8.132071 | 7.942207 | 8.052419 | 8.091891 | 7.959483 | 8.049667 | 311 | 314 | 194 | 252 |
| 2868 | NM_00100405 | GRK4 | G protein-coupled receptor kinase 4 | protein_coding | NP_001004056.1;NP_001004057.1;NP_001337102.1;NP_005298.2;NP_8 | 4543 | 137026 | . | . | 0.504556 | 0.461651 | 0.427621 | 0.483827 | 1.174122 | 1.067128 | 0.959568 | 1.106213 | -1.006162 | 39.332278 | 0.494708757 | -0.01791495 | 0.985706706 | 0.999197585 | 5.302500 | 5.292767 | 5.296868 | 5.299913 | 5.288340 | 5.306496 | 45 | 43 | 32 | 39 |
| 2869 | NM_005308,XI | GRK5 | G protein-coupled receptor kinase 5 | protein_coding | NP_005299.1;XP_005269764.1;XP_005269765.1 | 4544 | 600870 | . | . | 8.404225 | 9.708732 | 7.205938 | 10.283218 | 19.556979 | 22.442167 | 16.169917 | 23.511356 | 1.145499 | 1211.420452 | 0.157083096 | 1.247594445 | 0.212179614 | 0.975238634 | 10.141440 | 10.336912 | 10.200846 | 10.170759 | 10.170370 | 10.393912 | 1327 | 1194 | 961 | 1389 |
| 2870 | NM_00100410 | GRK6 | G protein-coupled receptor kinase 6 | protein_coding | NP_001004105.1;NP_001004106.1;NP_001351093.1;NP_002073.2;XP_0 | 4545 | 600869 | . | . | 27.358989 | 28.324233 | 22.513336 | 27.819541 | 63.665487 | 65.472738 | 50.519272 | 63.606075 | -1.112287 | 2365.396796 | 0.130855781 | -1.17327181 | 0.240686823 | 0.992733415 | 11.282750 | 11.128882 | 11.242818 | 11.253486 | 11.079480 | 11.239033 | 2850 | 2706 | 1752 | 2310 |
| 2872 | NM_017572,N | MKNK2 | MAPK interacting serine/threonine kinase 2 | protein_coding | NP_060042.2;NP_951009.1;XP_024307239.1 | 7111 | 605069 | . | . | 11.499434 | 9.363090 | 9.300264 | 11.304446 | 26.759659 | 21.643203 | 20.869524 | 25.846275 | -1.128269 | 977.767536 | 0.141872439 | -1.22723395 | 0.219734672 | 0.982008277 | 10.017961 | 9.843461 | 9.978840 | 9.972353 | 9.842868 | 9.930091 | 1199 | 1114 | 750 | 914 |
| 2873 | NM_00132108 | GPS1 | G protein pathway suppressor 1 | protein_coding | NP_001308018.1;NP_001308019.1;NP_001308020.1;NP_001308021.1;NI | 4549 | 601934 | . | . | 39.829756 | 42.221750 | 39.762562 | 40.336449 | 92.685479 | 97.597471 | 89.226033 | 92.224505 | 1.001833 | 2623.436055 | 0.116239762 | 0.022732983 | 0.981863266 | 0.998865404 | 11.356012 | 11.358471 | 11.336128 | 11.376117 | 11.329244 | 11.385450 | 2944 | 2898 | 2204 | 2547 |
| 2874 | NM_004489 | GPS2 | G protein pathway suppressor 2 | protein_coding | NP_004480.1 | 4550 | 601935 | . | . | 52.610039 | 54.035057 | 55.879910 | 59.930149 | 122.425720 | 124.904449 | 125.392883 | 137.023178 | 1.070860 | 2036.961233 | 0.118994279 | 0.830033553 | 0.406519813 | 0.998865404 | 10.942052 | 11.040668 | 10.956943 | 10.971234 | 10.994282 | 11.042216 | 2245 | 2142 | 1777 | 2034 |
| 2875 | NM_00138266 | GPT | glutamic--pyruvic transaminase | protein_coding | NP_001369593.1;NP_001369594.1;NP_005300.1 | 4552 | 138200 | . | . | 0.119693 | 0.125669 | 0.090471 | 0.049859 | 0.278531 | 0.290490 | 0.203014 | 0.113996 | -1.609499 | 5.827842 | 1.266776229 | -0.54201471 | 0.587808376 | NA | 2.844134 | 2.161867 | 2.553993 | 2.556058 | 2.548410 | 2.536956 | 8 | 8 | 5 | 3 |
| 2876 | NM_000581,N | GPX1 | glutathione peroxidase 1 | protein_coding | NP_000572.2;NP_001316384.1;NP_001316431.1;NP_001316432.1;NP_9 | 4553 | 138320 | . | . | 115.680396 | 114.944947 | 77.311317 | 103.052916 | 269.193042 | 265.700375 | 173.484347 | 235.618256 | -1.296525 | 2905.405969 | 0.138880261 | -2.69764551 | **0.006983176** | 0.265533818 | 11.679866 | 11.304884 | 11.605861 | 11.592394 | 11.274282 | 11.488771 | 3808 | 3512 | 1903 | 2692 |
| 2879 | NM_00103984 | GPX4 | glutathione peroxidase 4 | protein_coding | NP_001034936.1;NP_001034937.1;NP_001354761.1;NP_002076.2 | 4556 | 138322 | . | . | 103.160989 | 103.546493 | 89.428950 | 118.130861 | 240.059839 | 239.352355 | 200.675975 | 270.092167 | -1.003041 | 2588.379065 | 0.141262253 | -0.03101339 | 0.975258864 | 0.998865404 | 11.340188 | 11.335475 | 11.337282 | 11.336381 | 11.216087 | 11.440278 | 2987 | 2796 | 1937 | 2757 |
| 2885 | NM_002086,N | GRB2 | growth factor receptor bound protein 2 | protein_coding | NP_002077.1;NP_987102.1 | 4566 | 108355 | . | . | 30.577712 | 30.585560 | 28.011167 | 33.643519 | 71.155594 | 70.699895 | 62.856243 | 76.921910 | -1.006015 | 3129.990218 | 0.123386016 | -0.07012102 | 0.94409734 | 0.998865404 | 11.616370 | 11.607501 | 11.617152 | 11.609979 | 11.535757 | 11.676594 | 3631 | 3373 | 2480 | 3177 |
| 2887 | NM_00100154 | GRB10 | growth factor receptor bound protein 10 | protein_coding | NP_001001549.1;NP_001001550.1;NP_001001555.1;NP_001337743.1;NI | 4564 | 601523 | . | . | 14.445390 | 15.067838 | 15.260850 | 17.732247 | 33.615017 | 34.829981 | 34.244897 | 40.542674 | 1.108042 | 2461.576736 | 0.1211716 | 1.221510524 | 0.221892783 | 0.982008277 | 11.189586 | 11.337366 | 11.213947 | 11.229724 | 11.251819 | 11.354883 | 2663 | 2545 | 2108 | 2579 |
| 2888 | NM_00130342 | GRB14 | growth factor receptor bound protein 14 | protein_coding | NP_001290351.1;NP_004481.2;XP_011509324.1;XP_016859388.1;XP_02 | 4565 | 601524 | . | . | 3.582587 | 3.181327 | 3.746718 | 3.414961 | 8.336825 | 7.353779 | 8.407527 | 7.807905 | -1.028217 | 243.969256 | 0.224782755 | -0.17859316 | 0.858257167 | 0.998865404 | 7.949922 | 7.910926 | 7.976735 | 7.894370 | 7.949563 | 7.895261 | 308 | 245 | 221 | 213 |
| 2889 | NM_00130427 | RAPGEF1 | Rap guanine nucleotide exchange factor 1 | protein_coding | NP_001291204.1;NP_001364864.1;NP_001364865.1;NP_001364866.1;NI | 4568 | 600303 | . | . | 6.820486 | 8.017176 | 7.925075 | 9.002850 | 15.871551 | 18.532066 | 17.783637 | 20.583947 | 1.116228 | 1524.220709 | 0.137580134 | 1.153013244 | 0.248904948 | 0.992733415 | 10.492620 | 10.650760 | 10.470286 | 10.583233 | 10.563373 | 10.660959 | 1537 | 1669 | 1311 | 1601 |
| 2894 | NM_017551,XI | GRID1 | glutamate ionotropic receptor delta type subunit 1 | protein_coding | NP_060021.1;XP_011538022.1 | 4575 | 610659 | . | . | 0.338973 | 0.293757 | 0.242645 | 0.245062 | 0.788804 | 0.679032 | 0.544489 | 0.560305 | -1.312284 | 54.370941 | 0.427547683 | -0.91704442 | 0.359119362 | 0.998865404 | 5.947216 | 5.555851 | 5.811486 | 5.773924 | 5.740324 | 5.739075 | 77 | 61 | 41 | 44 |
| 2896 | NM_002087,XI | GRN | granulin precursor | protein_coding | NP_002078.1;XP_005257310.1;XP_024306498.1 | 4601 | 138945 | . | . | 110.844322 | 112.612391 | 107.214328 | 112.713118 | 257.939284 | 260.308600 | 240.585867 | 257.705164 | -1.030450 | 7707.328262 | 0.103395969 | -0.41852929 | 0.67556018 | 0.998865404 | 12.933513 | 12.890192 | 12.916191 | 12.932828 | 12.878737 | 12.919061 | 8920 | 8524 | 6444 | 7287 |
| 2898 | NM_00116624 | GRIK2 | glutamate ionotropic receptor kainate type subunit 2 | protein_coding | NP_001159719.1;NP_068775.1;NP_786944.1;XP_005267003.1;XP_01153 | 4580 | 138244 | . | . | 1.837857 | 1.636025 | 1.006209 | 1.780805 | 4.276771 | 3.781744 | 2.257904 | 4.071599 | -1.102785 | 271.492665 | 0.208979312 | -0.67556611 | 0.499316166 | 0.998865404 | 8.153787 | 8.012283 | 8.103583 | 8.116259 | 8.047248 | 8.071455 | 324 | 311 | 219 | 248 |
| 2901 | NM_00130103 | GRIK5 | glutamate ionotropic receptor kainate type subunit 5 | protein_coding | NP_001287959.1;NP_002079.3;XP_005258878.1;XP_011525164.1;XP_01 | 4583 | 600283 | . | . | 0.030773 | 0.055271 | 0.083060 | 0.087199 | 0.071611 | 0.127760 | 0.186385 | 0.199372 | 1.915659 | 10.536116 | 0.954365355 | 0.982685302 | 0.325762347 | NA | 2.855800 | 3.790061 | 3.355283 | 3.374460 | 3.394195 | 3.394790 | 6 | 10 | 12 | 13 |
| 2903 | NM_000833,N | GRIN2A | glutamate ionotropic receptor NMDA type subunit 2A | protein_coding | NP_000824.1;NP_001127879.1;NP_001127880.1;XP_011520760.1;XP_01 | 4585 | 138253 | . | . | 0.020304 | 0.021014 | 0.011529 | 0.022470 | 0.047247 | 0.048575 | 0.025871 | 0.051375 | -1.199159 | 9.040899 | 1.021390568 | -0.25653559 | 0.797537304 | NA | 3.303566 | 3.037079 | 3.182884 | 3.185760 | 3.166273 | 3.187641 | 11 | 11 | 5 | 10 |
| 2904 | NM_000834,XI | GRIN2B | glutamate ionotropic receptor NMDA type subunit 2B | protein_coding | NP_000825.2;XP_005253408.1;XP_011518931.1;XP_016874708.1 | 4586 | 138252 | . | . | 0.014100 | 0.029768 | 0.012473 | 0.017001 | 0.032812 | 0.068810 | 0.027988 | 0.038870 | -1.429668 | 18.085138 | 0.75577039 | -0.68232426 | 0.495033939 | NA | 4.413600 | 3.893187 | 4.156434 | 4.220556 | 4.152471 | 4.170868 | 16 | 31 | 11 | 16 |
| 2907 | NM_000837,N | GRINA | glutamate ionotropic receptor NMDA type subunit associated prot | protein_coding | NP_000828.1;NP_001009184.1 | 4589 | 138251 | . | . | 67.751115 | 72.498379 | 73.763703 | 76.776386 | 157.659631 | 167.583254 | 165.523610 | 175.540093 | 1.061618 | 4246.837138 | 0.108962869 | 0.791693975 | 0.428539136 | 0.998865404 | 12.008442 | 12.094618 | 12.001680 | 12.050972 | 12.063649 | 12.088823 | 4603 | 4578 | 3746 | 4162 |
| 2908 | NM_000176,N | NR3C1 | nuclear receptor subfamily 3 group C member 1 | protein_coding | NP_000167.1;NP_001018084.1;NP_001018085.1;NP_001018086.1;NP_0 | 7978 | 138040 | . | . | 3.931657 | 4.199085 | 4.040849 | 3.822905 | 9.149127 | 9.706375 | 9.067549 | 8.740618 | -1.052155 | 836.187847 | 0.145960164 | -0.50251133 | 0.615307886 | 0.998865404 | 9.743883 | 9.670551 | 9.695386 | 9.754152 | 9.710013 | 9.667688 | 947 | 963 | 730 | 741 |
| 2909 | NM_004491,XI | ARHGAP35 | Rho GTPase activating protein 35 | protein_coding | NP_004482.4;XP_024307241.1 | 4591 | 605277 | . | . | 7.143497 | 7.852394 | 7.281850 | 8.827124 | 16.623212 | 18.151159 | 16.340260 | 20.182169 | 1.058277 | 2242.883022 | 0.132564908 | 0.616435299 | 0.537607262 | 0.998865404 | 11.089901 | 11.171230 | 11.071361 | 11.140750 | 11.073357 | 11.224993 | 2401 | 2454 | 1817 | 2365 |

| | | | | | | | | | | | | | | | | | | | | | | | | | | | | | | | | | |
|---|---|---|---|---|---|---|---|---|---|---|---|---|---|---|---|---|---|---|---|---|---|---|---|---|---|---|---|---|---|---|---|---|---|
| 2911 | NM_00127806 | GRM1 | glutamate metabotropic receptor 1 | protein_coding | NP_001264993.1;NP_001264994.1;NP_001264995.1;NP_001264996.1;XF | 4593 | 604473 | . | . | 0.045140 | 0.027944 | 0.012653 | 0.005854 | 0.105044 | 0.064595 | 0.028392 | 0.013384 | -2.728430 | 5.188723 | 1.378819973 | -1.05022503 | 0.293614666 | NA | 2.923304 | 1.481107 | 2.405745 | 2.395468 | 2.381899 | 2.375806 | 10 | 7 | 3 | 2 |
| 2919 | NM_001511,N | CXCL1 | C-X-C motif chemokine ligand 1 | protein_coding | NP_001502.1 | 4602 | 155730 | . | . | 9.529520 | 7.784448 | 4.513574 | 4.659313 | 22.175582 | 17.994103 | 10.128331 | 10.652969 | -1.909361 | 243.754099 | 0.229624864 | -4.06354102 | 4.83338E-05 | 0.00725165 | 8.321522 | 7.388863 | 8.113901 | 7.992840 | 7.738134 | 7.745843 | 407 | 309 | 144 | 159 |
| 2921 | NM_002090 | CXCL3 | C-X-C motif chemokine ligand 3 | protein_coding | NP_002081.2 | 4604 | 139111 | . | . | 0.214282 | 0.156306 | 0.519471 | 0.232699 | 0.498644 | 0.361308 | 1.165679 | 0.532038 | 2.005406 | 10.077343 | 0.992867334 | 1.011106349 | 0.311965532 | NA | 2.741623 | 3.751299 | 3.299496 | 3.289157 | 3.345377 | 3.303193 | 9 | 6 | 16 | 8 |
| 2923 | NM_005313 | PDIA3 | protein disulfide isomerase family A member 3 | protein_coding | NP_005304.3 | 4606 | 602046 | . | . | 179.061020 | 180.574356 | 174.037613 | 166.399353 | 416.682343 | 417.405670 | 390.535370 | 380.452332 | -1.071123 | 20069.595543 | 0.099987218 | -0.99136644 | 0.321506681 | 0.998865404 | 14.341426 | 14.242320 | 14.321539 | 14.321415 | 14.285669 | 14.238599 | 23905 | 22398 | 17317 | 17669 |
| 2925 | NM_005314 | GRPR | gastrin releasing peptide receptor | protein_coding | NP_005305.1 | 4609 | 305670 | . | . | 0.338425 | 0.222900 | 0.246951 | 0.186691 | 0.787530 | 0.515244 | 0.554152 | 0.426847 | -1.269239 | 19.536078 | 0.706624793 | -0.48677043 | 0.62642103 | NA | 4.448287 | 4.107809 | 4.318047 | 4.281541 | 4.289965 | 4.270519 | 30 | 19 | 17 | 14 |
| 2926 | NM_00109847 | GRSF1 | G-rich RNA sequence binding factor 1 | protein_coding | NP_001091947.1;NP_002083.4;XP_011530199.1 | 4610 | 604851 | . | . | 13.410622 | 13.574641 | 13.935657 | 13.828759 | 31.207068 | 31.378393 | 31.271214 | 31.617815 | 1.023219 | 2749.842606 | 0.114539748 | 0.289113263 | 0.772494704 | 0.998865404 | 11.408405 | 11.441670 | 11.432261 | 11.398455 | 11.457577 | 11.410331 | 3195 | 2872 | 2489 | 2532 |
| 2931 | NM_019884,XI | GSK3A | glycogen synthase kinase 3 alpha | protein_coding | NP_063937.2 | 4616 | 606784 | . | . | 19.306582 | 21.529739 | 19.268410 | 22.772179 | 44.927204 | 49.766956 | 43.237753 | 52.065884 | 1.014715 | 1448.493447 | 0.135266751 | 0.155798394 | 0.876191937 | 0.998865404 | 10.490000 | 10.510601 | 10.457409 | 10.529151 | 10.451151 | 10.556149 | 1579 | 1624 | 1182 | 1461 |
| 2932 | NM_00114615 | GSK3B | glycogen synthase kinase 3 beta | protein_coding | NP_001139628.1;NP_001341525.1;NP_002084.2;XP_006713673.1 | 4617 | 605004 | . | . | 6.491113 | 6.303902 | 6.600636 | 6.599021 | 15.105087 | 14.571754 | 14.811635 | 15.087879 | 1.020038 | 1566.325257 | 0.123573979 | 0.231621551 | 0.816831967 | 0.998865404 | 10.598736 | 10.627457 | 10.619283 | 10.591145 | 10.625464 | 10.616016 | 1818 | 1643 | 1384 | 1474 |
| 2934 | NM_000177,N | GSN | gelsolin | protein_coding | NP_000168.1;NP_001121134.1;NP_001121135.2;NP_001121136.1;NP_0( | 4620 | 137350 | . | . | 158.961700 | 153.815985 | 159.975022 | 185.396553 | 369.910425 | 355.552536 | 358.979313 | 423.887186 | 1.086982 | 13661.191707 | 0.110385868 | 1.090067048 | 0.27568361 | 0.998865404 | 13.676399 | 13.796686 | 13.714497 | 13.686087 | 13.709742 | 13.830391 | 15327 | 13891 | 11510 | 14273 |
| 2935 | NM_00113000 | GSPT1 | G1 to S phase transition 1 | protein_coding | NP_001123478.2;NP_001123479.1;NP_002085.3 | 4621 | 139259 | . | . | 18.988818 | 19.205347 | 18.027739 | 19.314464 | 44.187764 | 44.394019 | 40.453725 | 44.160227 | -1.037306 | 4202.370844 | 0.108537473 | -0.48685133 | 0.626363692 | 0.998865404 | 12.063205 | 12.010286 | 12.050839 | 12.053106 | 11.996919 | 12.045488 | 4921 | 4624 | 3482 | 3981 |
| 2936 | NM_000637,N | GSR | glutathione-disulfide reductase | protein_coding | NP_000628.2;NP_001182031.1;NP_001182032.1;NP_001182033.1 | 4623 | 138300 | . | . | 35.477863 | 37.248192 | 41.483312 | 44.059519 | 82.558448 | 86.100864 | 93.087354 | 100.736853 | 1.160618 | 3741.799207 | 0.110126266 | 1.951338732 | 0.051016764 | 0.672475152 | 11.758129 | 11.972917 | 11.787932 | 11.820854 | 11.906355 | 11.948338 | 3908 | 3813 | 3408 | 3863 |
| 2937 | NM_000178,N | GSS | glutathione synthetase | protein_coding | NP_000169.1;NP_001309423.1;NP_001309424.1 | 4624 | 601002 | . | . | 56.526145 | 55.398818 | 58.044199 | 65.020022 | 131.538654 | 128.056837 | 130.249488 | 148.660545 | 1.081065 | 3501.361498 | 0.114150898 | 0.985127547 | 0.324561424 | 0.998865404 | 11.716435 | 11.828779 | 11.750425 | 11.730095 | 11.761616 | 11.845767 | 3923 | 3586 | 3003 | 3583 |
| 2941 | NM_001512,XI | GSTA4 | glutathione S-transferase alpha 4 | protein_coding | NP_001503.1;XP_005249092.1;XP_011512836.1;XP_011512837.1 | 4629 | 605450 | . | . | 6.102407 | 6.550915 | 7.309804 | 5.669532 | 14.200552 | 15.142736 | 16.402988 | 12.962711 | 1.017463 | 251.761843 | 0.225428079 | 0.110796671 | 0.911777585 | 0.998865404 | 7.962906 | 7.988809 | 7.954509 | 7.983685 | 8.035016 | 7.917490 | 279 | 277 | 247 | 210 |
| 2944 | NM_000561,N | GSTM1 | glutathione S-transferase mu 1 | protein_coding | NP_000552.2;NP_666533.1;XP_005270839.1 | 4632 | 138350 | . | . | 3.283339 | 2.781150 | 3.654698 | 3.181444 | 7.640464 | 6.428751 | 8.201035 | 7.273993 | 1.109657 | 121.256200 | 0.299466867 | 0.501269224 | 0.616181662 | 0.998865404 | 6.844096 | 6.995750 | 6.930144 | 6.870513 | 6.961440 | 6.911100 | 144 | 113 | 119 | 111 |
| 2946 | NM_000848,N | GSTM2 | glutathione S-transferase mu 2 | protein_coding | NP_000839.1;NP_001135840.1 | 4634 | 138380 | . | . | 12.717294 | 11.396455 | 12.749224 | 12.142995 | 29.593664 | 26.343418 | 28.608889 | 27.763515 | 1.013954 | 446.621367 | 0.177582362 | 0.112578062 | 0.910365078 | 0.998865404 | 8.792566 | 8.813177 | 8.834068 | 8.760565 | 8.823392 | 8.789539 | 541 | 450 | 402 | 410 |
| 2947 | NM_000849,N | GSTM3 | glutathione S-transferase mu 3 | protein_coding | NP_000840.2 | 4635 | 138390 | . | . | 8.814006 | 8.135607 | 7.897939 | 9.444282 | 20.510553 | 18.805820 | 17.722748 | 21.593230 | 1.008536 | 1100.922793 | 0.145042528 | 0.084547441 | 0.932621185 | 0.998865404 | 10.098490 | 10.110480 | 10.133216 | 10.066394 | 10.042748 | 10.166678 | 1319 | 1130 | 881 | 1123 |
| 2948 | NM_000850,N | GSTM4 | glutathione S-transferase mu 4 | protein_coding | NP_000841.1;NP_671489.1;XP_016856574.1;XP_016856575.1;XP_01685 | 4636 | 138333 | . | . | 30.495163 | 31.433999 | 34.022750 | 37.319936 | 70.963495 | 72.661099 | 76.346064 | 85.327602 | 1.137127 | 1470.254456 | 0.126948526 | 1.460386641 | 0.144183842 | 0.906161673 | 10.426300 | 10.611463 | 10.459711 | 10.479489 | 10.539824 | 10.596730 | 1564 | 1504 | 1311 | 1520 |
| 2950 | NM_000852 | GSTP1 | glutathione S-transferase pi 1 | protein_coding | NP_000843.1 | 4638 | 134660 | . | . | 392.989166 | 406.286407 | 429.325836 | 514.336487 | 914.501953 | 939.149109 | 963.394714 | 1175.969238 | 1.163876 | 10050.261270 | 0.11540547 | 1.897117944 | 0.057812373 | 0.713122601 | 13.181372 | 13.400227 | 13.215381 | 13.236276 | 13.279102 | 13.426163 | 10564 | 10148 | 8602 | 10997 |
| 2953 | NR_126445 | GSTT2 | glutathione S-transferase theta 2 (gene/pseudogene) | transcribed_ps | . | 4642 | 600437 | . | . | 4.274727 | 4.339804 | 6.855057 | 6.415958 | 9.947465 | 10.031651 | 15.382547 | 14.669326 | 1.519590 | 189.264891 | 0.242728301 | 2.487069443 | 0.012880025 | 0.374881745 | 7.230892 | 7.834876 | 7.452090 | 7.457298 | 7.655093 | 7.619714 | 172 | 163 | 206 | 206 |
| 2954 | NM_00131266 | GSTZ1 | glutathione S-transferase zeta 1 | protein_coding | NP_001299589.1;NP_001350632.1;NP_665877.1;NP_665878.2;XP_01153 | 4643 | 603758 | . | . | 4.682814 | 4.911434 | 4.379072 | 5.694669 | 10.897098 | 11.353002 | 9.826511 | 13.020183 | 1.183727 | 205.453904 | 0.235804243 | 1.031940454 | 0.302100013 | 0.998865404 | 7.555946 | 7.799157 | 7.615187 | 7.663191 | 7.731614 | 7.701217 | 205 | 214 | 200 | 202 |
| 2956 | NM_000179,N | MSH6 | mutS homolog 6 | protein_coding | NP_000170.1;NP_001268421.1;NP_001268422.1;NP_001268423.1;XP_02 | 7329 | 600678 | . | . | 17.663444 | 18.227485 | 17.268480 | 18.565615 | 41.103561 | 42.133648 | 38.749969 | 42.448071 | -1.015885 | 2379.716086 | 0.11633804 | -0.19543702 | 0.845050841 | 0.998865404 | 11.227971 | 11.205086 | 11.212841 | 11.232710 | 11.185975 | 11.233618 | 2728 | 2620 | 1992 | 2279 |
| 2957 | NM_00127894 | GTF2A1 | general transcription factor IIA subunit 1 | protein_coding | NP_001265869.1;NP_056943.1;NP_963889.1 | 4646 | 600520 | . | . | 2.576580 | 2.721481 | 2.694883 | 2.090026 | 5.995806 | 6.290827 | 6.047238 | 4.778598 | -1.125009 | 499.728387 | 0.183906356 | -0.92404049 | 0.355465234 | 0.998865404 | 9.047045 | 8.878007 | 8.983786 | 9.013851 | 9.002465 | 8.839958 | 595 | 584 | 463 | 383 |
| 2958 | NM_00132092 | GTF2A2 | general transcription factor IIA subunit 2 | protein_coding | NP_001307858.1;NP_001307859.1;NP_004483.1 | 4647 | 600519 | . | . | 16.423769 | 14.315607 | 14.462568 | 13.060004 | 38.218783 | 33.091162 | 32.453581 | 29.860146 | -1.128396 | 712.654558 | 0.158876821 | -1.0969108 | 0.272680388 | 0.998865404 | 9.561225 | 9.387678 | 9.565606 | 9.464523 | 9.469033 | 9.396148 | 932 | 757 | 614 | 594 |
| 2959 | NM_001514,XI | GTF2B | general transcription factor IIB | protein_coding | NP_001505.1;XP_011539601.1 | 4648 | 189963 | . | . | 16.597872 | 15.528519 | 16.681223 | 16.736301 | 38.623930 | 35.894870 | 37.432180 | 38.265564 | -1.006943 | 802.856273 | 0.147952702 | -0.06746627 | 0.946210517 | 0.998865404 | 9.653750 | 9.644230 | 9.678554 | 9.622692 | 9.672272 | 9.619552 | 964 | 835 | 722 | 722 |
| 2960 | NM_005513 | GTF2E1 | general transcription factor IIE subunit 1 | protein_coding | NP_005504.2 | 4650 | 189962 | . | . | 4.376492 | 4.940302 | 4.381277 | 3.777505 | 10.184277 | 11.419729 | 9.831459 | 8.636816 | -1.157842 | 408.952556 | 0.187649628 | -1.12677154 | 0.259839086 | 0.995760118 | 8.777451 | 8.566418 | 8.681520 | 8.750845 | 8.674312 | 8.582188 | 477 | 500 | 356 | 327 |
| 2961 | NM_00134835 | GTF2E2 | general transcription factor IIE subunit 2 | protein_coding | NP_001335282.1;NP_002086.1;XP_011542811.1;XP_011542812.1;XP_01 | 4651 | 189964 | . | . | 14.084144 | 15.437823 | 13.270677 | 15.075551 | 32.774382 | 35.685223 | 29.779013 | 34.468456 | -1.040543 | 778.599960 | 0.151125831 | -0.37939461 | 0.704394853 | 0.998865404 | 9.633403 | 9.575493 | 9.599007 | 9.637066 | 9.545933 | 9.632584 | 890 | 880 | 622 | 760 |
| 2962 | NM_002096,XI | GTF2F1 | general transcription factor IIF subunit 1 | protein_coding | NP_002087.2;XP_024307243.1 | 4652 | 189968 | . | . | 33.039708 | 34.470023 | 31.388291 | 35.770260 | 76.884758 | 79.679002 | 70.434417 | 81.784450 | -1.017005 | 2561.994724 | 0.120667723 | -0.20159697 | 0.840231815 | 0.998865404 | 11.335287 | 11.310712 | 11.315063 | 11.343591 | 11.264466 | 11.364478 | 2923 | 2837 | 2072 | 2529 |
| 2963 | NM_004128,XI | GTF2F2 | general transcription factor IIF subunit 2 | protein_coding | NP_004119.1;XP_011533354.1;XP_011533355.1;XP_016876039.1;XP_01( | 4653 | 189969 | . | . | 15.269160 | 14.660244 | 15.522014 | 14.388894 | 35.531966 | 33.887811 | 34.830950 | 32.898493 | -1.002576 | 1031.013105 | 0.139357957 | -0.02663159 | 0.978753578 | 0.998865404 | 10.011490 | 10.008202 | 10.036397 | 9.983560 | 10.039556 | 9.976590 | 1231 | 1074 | 934 | 924 |
| 2965 | NM_00114230 | GTF2H1 | general transcription factor IIH subunit 1 | protein_coding | NP_001135779.1;NP_005307.1;XP_006718271.1;XP_024304225.1;XP_02 | 4655 | 189972 | . | . | 13.158733 | 14.994157 | 15.970272 | 14.840031 | 30.620913 | 34.659663 | 35.836828 | 33.929968 | 1.078659 | 1445.138313 | 0.134115931 | 0.814514695 | 0.415350157 | 0.998865404 | 10.441356 | 10.550562 | 10.415546 | 10.514227 | 10.556364 | 10.491320 | 1498 | 1597 | 1359 | 1347 |
| 2966 | NM_00136456 | GTF2H2 | general transcription factor IIH subunit 2 | protein_coding | NP_001351496.1;NP_001351497.1;NP_001351498.1;NP_001351499.1;NI | 4656 | 601748 | . | . | 12.209344 | 11.125975 | 11.510415 | 10.047021 | 28.411644 | 25.718191 | 25.829034 | 22.971316 | -1.087567 | 649.587147 | 0.162362071 | -0.74589334 | 0.455731846 | 0.998865404 | 9.402280 | 9.281971 | 9.407436 | 9.331478 | 9.358743 | 9.266415 | 822 | 690 | 579 | 543 |
| 2967 | NM_00127186 | GTF2H3 | general transcription factor IIH subunit 3 | protein_coding | NP_001258795.1;NP_001258796.1;NP_001258797.1;NP_001507.2;XP_01 | 4657 | 601750 | . | . | 7.636565 | 7.001063 | 7.424861 | 7.109429 | 17.770601 | 16.183269 | 16.661173 | 16.254865 | -1.019359 | 755.981981 | 0.150759483 | -0.18348146 | 0.854420267 | 0.998865404 | 9.575755 | 9.548533 | 9.604094 | 9.531548 | 9.571880 | 9.538269 | 924 | 781 | 667 | 685 |
| 2968 | NM_001517 | GTF2H4 | general transcription factor IIH subunit 4 | protein_coding | NP_001508.1 | 4658 | 601760 | . | . | 14.551026 | 14.799536 | 15.343001 | 16.692717 | 33.860836 | 34.209789 | 34.429253 | 38.165916 | 1.077035 | 818.535935 | 0.14508477 | 0.737946015 | 0.460547255 | 0.998865404 | 9.622520 | 9.729308 | 9.646687 | 9.652451 | 9.675475 | 9.727925 | 904 | 854 | 711 | 825 |
| 2969 | NM_00116363 | GTF2I | general transcription factor IIi | protein_coding | NP_001157108.1;NP_001267729.1;NP_001509.3;NP_127492.1;NP_1274 | 4659 | 601679 | . | . | 15.050068 | 15.449983 | 17.545397 | 19.801511 | 35.022128 | 35.713325 | 39.371363 | 45.273805 | 1.203029 | 2326.889356 | 0.119743483 | 2.227026852 | 0.025945479 | 0.527323808 | 11.044807 | 11.311279 | 11.098061 | 11.110902 | 11.209984 | 11.294441 | 2413 | 2298 | 2096 | 2505 |
| 2970 | NR_002206 | GTF2IP1 | general transcription factor IIi pseudogene 1 | transcribed_ps | . | 4660 | . | | . | 11.893728 | 11.963167 | 14.396729 | 14.416872 | 27.677193 | 27.653399 | 32.305847 | 32.962463 | 1.190931 | 1487.075947 | 0.124239358 | 2.029061658 | 0.042452011 | 0.636918588 | 10.406704 | 10.658820 | 10.467216 | 10.465515 | 10.604456 | 10.597743 | 1565 | 1463 | 1412 | 1509 |
| 2971 | NM_002097 | GTF3A | general transcription factor IIIA | protein_coding | NP_002088.2 | 4662 | 600860 | . | . | 39.113216 | 39.780228 | 41.238945 | 44.600563 | 91.018066 | 91.953781 | 92.538994 | 101.973892 | 1.072715 | 1850.732480 | 0.121237634 | 0.835278197 | 0.403561091 | 0.998865404 | 10.802444 | 10.903547 | 10.822023 | 10.828514 | 10.853189 | 10.907296 | 2048 | 1935 | 1609 | 1857 |
| 2972 | NM_00124278 | BRF1 | BRF1 RNA polymerase III transcription initiation factor subunit | protein_coding | NP_001229715.1;NP_001229716.1;NP_001229717.1;NP_001229718.1;NI | 11551 | 604902 | . | . | 6.630206 | 6.662567 | 6.474890 | 6.471234 | 15.428761 | 15.400819 | 14.529465 | 14.795706 | -1.059605 | 709.745957 | 0.149813026 | -0.55753738 | 0.57716033 | 0.998865404 | 9.512328 | 9.428780 | 9.478396 | 9.503138 | 9.457429 | 9.444484 | 826 | 802 | 605 | 640 |
| 2975 | NM_00128624 | GTF3C1 | general transcription factor IIIC subunit 1 | protein_coding | NP_001273171.1;NP_001511.2;XP_016878677.1 | 4664 | 603246 | . | . | 19.749244 | 19.428952 | 20.602313 | 22.729030 | 45.957304 | 44.910891 | 46.230996 | 51.967227 | 1.090056 | 4529.511981 | 0.110430641 | 1.126519938 | 0.259945506 | 0.995760118 | 12.081638 | 12.205964 | 12.117837 | 12.097642 | 12.141471 | 12.216536 | 5052 | 4620 | 3925 | 4627 |
| 2976 | NM_00103552 | GTF3C2 | general transcription factor IIIC subunit 2 | protein_coding | NP_001030598.1;NP_001305838.1;NP_001512.1;XP_005264329.1 | 4665 | 604883 | . | . | 17.392001 | 18.638339 | 21.423899 | 21.813359 | 40.471899 | 43.083352 | 48.074611 | 49.873654 | 1.183754 | 2457.466972 | 0.115051456 | 2.115304952 | 0.034403965 | 0.579826526 | 11.136209 | 11.379462 | 11.178696 | 11.202459 | 11.309926 | 11.343027 | 2553 | 2465 | 2270 | 2548 |
| 2977 | NM_000855,N | GUCY1A2 | guanylate cyclase 1 soluble subunit alpha 2 | protein_coding | NP_000846.1;NP_001243353.1 | 4684 | 601244 | . | . | 0.175692 | 0.164995 | 0.235136 | 0.144076 | 0.408842 | 0.381393 | 0.527637 | 0.329413 | 1.103835 | 91.463922 | 0.359750016 | 0.396174961 | 0.691975961 | 0.998865404 | 6.440662 | 6.585944 | 6.503829 | 6.481418 | 6.590413 | 6.446698 | 104 | 90 | 104 | 68 |
| 2979 | NM_002098 | GUCA1B | guanylate cyclase activator 1B | protein_coding | NP_002089.4 | 4679 | 602275 | . | . | 0.149691 | 0.043222 | 0.056783 | 0.150408 | 0.348337 | 0.099911 | 0.127419 | 0.343891 | 1.018185 | 7.565823 | 1.159648201 | 0.022420053 | 0.982112884 | NA | 2.906576 | 2.932304 | 2.937022 | 2.901935 | 2.906881 | 2.933444 | 13 | 4 | 4 | 10 |
| 2987 | NM_000858,N | GUK1 | guanylate kinase 1 | protein_coding | NP_000849.1;NP_001152862.1;NP_001152863.1;NP_001229768.1;NP_0( | 4693 | 139270 | . | . | 55.125365 | 57.722603 | 51.567384 | 59.046559 | 128.278990 | 133.428382 | 115.715721 | 135.002940 | -1.030504 | 1833.878473 | 0.128602079 | -0.33708818 | 0.736050429 | 0.998865404 | 10.862379 | 10.818654 | 10.824984 | 10.878056 | 10.773967 | 10.879392 | 2072 | 2076 | 1465 | 1807 |
| 2990 | NM_000181,N | GUSB | glucuronidase beta | protein_coding | NP_000172.2;NP_001271219.1;NP_001280033.1;NP_001280034.1;XP_0( | 4696 | 611499 | . | . | 27.896717 | 28.031450 | 24.418298 | 26.516878 | 64.916804 | 64.795948 | 54.793950 | 60.627691 | -1.124623 | 1781.303372 | 0.123423887 | -1.37284591 | 0.169800254 | 0.946389416 | 10.881050 | 10.711401 | 10.845433 | 10.840662 | 10.715607 | 10.784488 | 2178 | 2029 | 1393 | 1642 |
| 2992 | NM_00118472 | GYG1 | glycogenin 1 | protein_coding | NP_001171649.1;NP_001171650.1;NP_004121.2;XP_016861764.1;XP_01 | 4699 | 603942 | . | . | 13.425488 | 12.714408 | 13.677844 | 12.832192 | 31.241658 | 29.389923 | 30.692686 | 29.339283 | -1.224615 | 1054.098591 | 0.137058979 | -2.13286541 | 0.032935769 | 0.574731249 | 10.180509 | 9.888309 | 10.143016 | 10.079309 | 9.957885 | 9.966473 | 1392 | 1200 | 819 | 894 |
| 2995 | NM_00125658 | GYPC | glycophorin C (Gerbich blood group) | protein_coding | NP_001243513.1;NP_002092.1;NP_058131.1;XP_006712523.1 | 4704 | 110750 | . | . | 187.884775 | 186.946220 | 187.373673 | 229.321849 | 437.215623 | 432.134567 | 420.461067 | 524.317153 | 1.101409 | 6612.005381 | 0.118556013 | 1.175401662 | 0.239834059 | 0.992733415 | 12.619573 | 12.758813 | 12.651073 | 12.644272 | 12.649517 | 12.803240 | 7276 | 6763 | 5484 | 7083 |
| 2997 | NM_00116158 | GYS1 | glycogen synthase 1 | protein_coding | NP_001155059.1;NP_002094.2 | 4706 | 138570 | . | . | 12.618560 | 13.772557 | 14.006365 | 15.666021 | 29.363906 | 31.835880 | 31.429880 | 35.818496 | 1.106470 | 1553.966894 | 0.129529813 | 1.126881271 | 0.259792683 | 0.995760118 | 10.527103 | 10.672705 | 10.528252 | 10.591748 | 10.600892 | 10.675345 | 1629 | 1658 | 1352 | 1603 |
| 3005 | NM_005318 | H1-0 | H1.0 linker histone | protein_coding | NP_005309.1 | 4714 | 142708 | . | . | 22.490049 | 20.437412 | 19.548182 | 25.930609 | 52.335266 | 47.241997 | 43.865551 | 59.287258 | 1.043904 | 1517.443340 | 0.151704101 | 0.408617417 | 0.682820447 | 0.998865404 | 10.536273 | 10.597923 | 10.587426 | 10.506947 | 10.470122 | 10.681380 | 1799 | 1519 | 1165 | 1650 |
| 3006 | NM_005319 | H1-2 | H1.2 linker histone, cluster member | protein_coding | NP_005310.1 | 4716 | 142710 | . | . | 4.270192 | 3.865050 | 3.806617 | 6.413149 | 9.936913 | 8.934237 | 8.541940 | 14.662903 | 1.238294 | 105.108668 | 0.342896223 | 0.899263309 | 0.368512426 | 0.998865404 | 6.554358 | 6.860868 | 6.687607 | 6.656122 | 6.653196 | 6.818878 | 114 | 96 | 76 | 136 |
| 3012 | NM_021052 | H2AC8 | H2A clustered histone 8 | protein_coding | NP_066390.1 | 4724 | 602786 | . | . | 0.154645 | 0.245232 | 0.206832 | 0.161515 | 0.359864 | 0.566864 | 0.464126 | 0.369284 | -1.040207 | 3.976848 | 1.501826248 | -0.0378681 | 0.969792848 | NA | 2.019256 | 1.963455 | 1.987477 | 1.992960 | 1.992634 | 1.986359 | 4 | 5 | 4 | 3 |
| 3013 | NM_021065 | H2AC7 | H2A clustered histone 7 | protein_coding | NP_066409.1 | 4729 | 602792 | . | . | 0.186770 | 0.171646 | 0.603445 | 0.406388 | 0.434622 | 0.396768 | 1.354114 | 0.929159 | 2.670787 | 5.750943 | 1.293378869 | 1.095784972 | 0.273172887 | NA | 1.644780 | 3.066070 | 2.487473 | 2.484343 | 2.518527 | 2.501307 | 4 | 3 | 9 | 6 |
| 3014 | NM_002105 | H2AX | H2A.X variant histone | protein_coding | NP_002096.1 | 4739 | 601772 | . | . | 12.164454 | 14.446112 | 12.918591 | 15.155170 | 28.307184 | 33.392834 | 28.988943 | 34.650497 | 1.041502 | 678.417844 | 0.165236669 | 0.355039347 | 0.722560114 | 0.998865404 | 9.376851 | 9.434631 | 9.333091 | 9.444640 | 9.365812 | 9.467451 | 703 | 776 | 557 | 697 |
| 3015 | NM_002106 | H2AZ1 | H2A.Z variant histone 1 | protein_coding | NP_002097.1 | 4741 | 142763 | . | . | 410.295288 | 393.584442 | 321.983521 | 367.516449 | 954.773987 | 909.788025 | 722.521729 | 840.282715 | -1.182689 | 10077.166358 | 0.109837677 | -2.20389346 | 0.027531838 | 0.538018865 | 13.414790 | 13.172668 | 13.387141 | 13.343792 | 13.170191 | 13.272552 | 12890 | 11489 | 7540 | 9184 |
| 3017 | NM_021063,N | H2BC5 | H2B clustered histone 5 | protein_coding | NP_066407.1;NP_619790.1;XP_005249096.1 | 4747 | 602799 | . | . | 2.184553 | 1.592137 | 3.517960 | 2.537028 | 5.083546 | 3.680297 | 7.894201 | 5.800613 | 1.572705 | 60.894172 | 0.417621913 | 1.564210083 | 0.11776824 | 0.857396579 | 5.563187 | 6.219312 | 5.890064 | 5.833762 | 5.989501 | 5.927392 | 63 | 43 | 73 | 61 |
| 3020 | NM_00137904 | H3-3A | H3.3 histone A | protein_coding | NP_001365972.1;NP_001365974.1;NP_001365975.1;NP_001365976.1;NI | 4764 | 601128 | . | . | 85.523866 | 82.867411 | 78.504526 | 89.341890 | 199.017566 | 191.551750 | 176.161899 | 204.269615 | -1.002846 | 2838.871010 | 0.122571895 | -0.03345414 | 0.973312439 | 0.998865404 | 11.473238 | 11.468962 | 11.488955 | 11.453909 | 11.405965 | 11.528778 | 3344 | 3002 | 2279 | 2858 |
| 3021 | NM_005324 | H3-3B | H3.3 histone B | protein_coding | NP_005315.1 | 4765 | 601058 | . | . | 62.245689 | 61.757915 | 60.179367 | 57.729568 | 144.848282 | 142.756195 | 135.040756 | 131.991791 | -1.066446 | 5098.100254 | 0.106486424 | -0.87157908 | 0.383438045 | 0.998865404 | 12.361370 | 12.268628 | 12.348956 | 12.335029 | 12.309508 | 12.266380 | 6109 | 5631 | 4402 | 4506 |
| 3024 | NM_005325 | H1-1 | H1.1 linker histone, cluster member | protein_coding | NP_005316.1 | 4715 | 142709 | . | . | 0.118414 | 0.099862 | 0.061515 | 0.067630 | 0.275555 | 0.230835 | 0.138038 | 0.154628 | -1.412257 | 2.671821 | 1.820869166 | -0.27349696 | 0.784471225 | NA | 1.644780 | 1.148375 | 1.428682 | 1.425364 | 1.423205 | 1.422411 | 4 | 3 | 2 | 2 |
| 3028 | NM_00103781 | HSD17B10 | hydroxysteroid 17-beta dehydrogenase 10 | protein_coding | NP_001032900.1;NP_004484.1 | 4800 | 300256 | . | . | 59.579953 | 54.260735 | 51.570489 | 62.992360 | 138.645000 | 125.426120 | 115.722682 | 144.024537 | -1.007569 | 1696.060445 | 0.138400045 | -0.07860588 | 0.937346112 | 0.998865404 | 10.733513 | 10.722409 | 10.769149 | 10.688517 | 10.646944 | 10.795024 | 2061 | 1743 | 1330 | 1734 |
| 3029 | NM_00104042 | HAGH | hydroxyacylglutathione hydrolase | protein_coding | NP_001035517.1;NP_001273178.1;NP_001350841.1;NP_001350843.1;NI | 4805 | 138760 | . | . | 14.202358 | 13.182298 | 9.114675 | 14.313557 | 33.049472 | 30.471475 | 20.453063 | 32.726249 | -1.011556 | 761.030851 | 0.157636636 | -0.10515763 | 0.916250727 | 0.998865404 | 9.580420 | 9.563150 | 9.566437 | 9.583291 | 9.497129 | 9.631842 | 871 | 836 | 594 | 778 |
| 3030 | NM_000182 | HADHA | hydroxyacyl-CoA dehydrogenase trifunctional multienzyme compl | protein_coding | NP_000173.2 | 4801 | 600890 | . | . | 59.217892 | 59.460175 | 61.048573 | 72.155701 | 137.802460 | 137.444855 | 136.991226 | 164.975433 | 1.106413 | 5770.284437 | 0.116020222 | 1.257448933 | 0.20859112 | 0.970482251 | 12.419700 | 12.565472 | 12.451436 | 12.447759 | 12.465903 | 12.599081 | 6323 | 5899 | 4858 | 6128 |
| 3032 | NM_000183,N | HADHB | hydroxyacyl-CoA dehydrogenase trifunctional multienzyme compl | protein_coding | NP_000174.1;NP_001268441.1;NP_001268442.1;XP_011531105.1;XP_02 | 4803 | 143450 | . | . | 64.381620 | 63.975768 | 65.940468 | 71.332525 | 149.818675 | 147.882850 | 147.968499 | 163.093349 | 1.073860 | 4230.376990 | 0.109228054 | 0.941209379 | 0.346597571 | 0.998865404 | 11.994289 | 12.097024 | 12.021118 | 12.010882 | 12.045928 | 12.103728 | 4727 | 4375 | 3678 | 4249 |
| 3033 | NM_00118470 | HADH | hydroxyacyl-CoA dehydrogenase | protein_coding | NP_001171634.2;NP_001317956.1;NP_005318.6 | 4799 | 601609 | . | . | 14.252173 | 13.753660 | 12.071206 | 14.106776 | 33.165392 | 31.792205 | 27.087437 | 32.253465 | -1.087928 | 773.492291 | 0.15117522 | -0.80424969 | 0.421252788 | 0.998865404 | 9.654950 | 9.532958 | 9.633613 | 9.613995 | 9.518743 | 9.608685 | 941 | 858 | 602 | 740 |
| 3035 | NM_00125804 | HARS1 | histidyl-tRNA synthetase 1 | protein_coding | NP_001244969.1;NP_001244970.1;NP_001244971.1;NP_001276021.1;NI | 4816 | 142810 | . | . | 46.498382 | 49.337551 | 48.652798 | 54.797735 | 108.203642 | 114.045961 | 109.175467 | 125.288504 | 1.057511 | 3055.759257 | 0.11709547 | 0.688943325 | 0.490858934 | 0.998865404 | 11.536523 | 11.616986 | 11.530734 | 11.575851 | 11.555788 | 11.640935 | 3326 | 3294 | 2590 | 3097 |
| 3037 | NM_005328 | HAS2 | hyaluronan synthase 2 | protein_coding | NP_005319.1 | 4819 | 601636 | . | . | 8.977293 | 7.745065 | 9.143346 | 7.637866 | 20.890530 | 17.903065 | 20.517405 | 17.463072 | -1.010297 | 1107.220416 | 0.15219304 | -0.09710877 | 0.92264002 | 0.998865404 | 10.119735 | 10.105684 | 10.171241 | 10.054455 | 10.175007 | 10.034329 | 1381 | 1107 | 1049 | 935 |
| 3038 | NM_00119928 | HAS3 | hyaluronan synthase 3 | protein_coding | NP_001186209.1;NP_005320.2;NP_619515.1;XP_005255978.1;XP_01152 | 4820 | 602428 | . | . | 0.688374 | 0.600256 | 0.573444 | 0.700043 | 1.601875 | 1.387518 | 1.286793 | 1.600565 | -1.039730 | 85.533379 | 0.348191981 | -0.16142895 | 0.871755569 | 0.998865404 | 6.446409 | 6.389868 | 6.445237 | 6.402325 | 6.391581 | 6.434578 | 108 | 87 | 67 | 85 |
| 3039 | NM_000558 | HBA1 | hemoglobin subunit alpha 1 | protein_coding | NP_000549.1 | 4823 | 141800 | . | . | 0.127557 | 0.410598 | 0.285797 | 0.169886 | 0.296830 | 0.949117 | 0.641319 | 0.388425 | -1.119095 | 4.742289 | 1.416512308 | -0.11459994 | 0.908762225 | NA | 2.324708 | 2.161867 | 2.229917 | 2.252555 | 2.244563 | 2.233151 | 3 | 8 | 5 | 3 |
| 3040 | NM_000517 | HBA2 | hemoglobin subunit alpha 2 | protein_coding | NP_000508.1 | 4824 | 141850 | . | . | 0.128094 | 0.400091 | 0.286295 | 0.254974 | 0.298081 | 0.924828 | 0.642437 | 0.582967 | 1.109492 | 5.275623 | 1.336922816 | 0.112122971 | 0.910725904 | NA | 2.324708 | 2.470304 | 2.379620 | 2.402316 | 2.394457 | 2.392633 | 3 | 8 | 5 | 5 |
| 3049 | NM_005331 | HBQ1 | hemoglobin subunit theta 1 | protein_coding | NP_005322.1 | 4833 | 142240 | . | . | 0.944403 | 0.685432 | 1.589574 | 0.878003 | 2.197664 | 1.584407 | 3.566959 | 2.007449 | 1.444888 | 17.482707 | 0.753293062 | 0.704848447 | 0.480904556 | NA | 3.835322 | 4.370958 | 4.110639 | 4.090857 | 4.151572 | 4.104124 | 19 | 13 | 23 | 14 |
| 3052 | NM_00112260 | HCCS | holocytochrome c synthase | protein_coding | NP_001116080.1;NP_001165462.1;NP_005324.3;XP_024308136.1 | 4837 | 300056 | . | . | 11.360152 | 11.382698 | 10.371280 | 10.889969 | 26.435543 | 26.311612 | 23.272852 | 24.898621 | -1.087563 | 805.505236 | 0.145171937 | -0.83418057 | 0.404179241 | 0.998865404 | 9.713146 | 9.591803 | 9.680213 | 9.685348 | 9.602693 | 9.643869 | 963 | 909 | 651 | 745 |
| 3053 | NM_000185 | SERPIND1 | serpin family D member 1 | protein_coding | NP_000176.2 | 4838 | 142360 | . | . | 0.298831 | 0.489640 | 0.177860 | 0.191759 | 0.695393 | 1.131826 | 0.399112 | 0.438435 | -2.075552 | 20.412139 | 0.707792436 | -1.48842424 | 0.136639049 | NA | 4.784822 | 3.728813 | 4.353773 | 4.409947 | 4.319720 | 4.324270 | 24 | 37 | 11 | 13 |
| 3054 | NM_005334,XI | HCFC1 | host cell factor C1 | protein_coding | NP_005325.2;XP_006724878.1;XP_006724879.1;XP_011529449.1;XP_01: | 4839 | 300019 | . | . | 9.586032 | 10.632691 | 11.377634 | 10.923351 | 22.307090 | 24.577940 | 25.531097 | 24.974944 | 1.092671 | 2996.027182 | 0.116776674 | 1.094904689 | 0.273558395 | 0.998865404 | 11.483525 | 11.611316 | 11.467151 | 11.553003 | 11.597650 | 11.568993 | 3123 | 3253 | 2768 | 2882 |
| 3064 | NM_002111 | HTT | huntingtin | protein_coding | NP_002102.4 | 4851 | 613004 | . | . | 3.397529 | 3.563459 | 3.144198 | 3.473098 | 7.906188 | 8.237095 | 7.055489 | 7.940827 | -1.066442 | 1425.454958 | 0.129118256 | -0.71876413 | 0.472286265 | 0.998865404 | 10.523014 | 10.429897 | 10.485916 | 10.516463 | 10.417310 | 10.485101 | 1661 | 1619 | 1146 | 1351 |
| 3065 | NM_004964,XI | HDAC1 | histone deacetylase 1 | protein_coding | NP_004955.2;XP_011539611.1 | 4852 | 601241 | . | . | 50.396292 | 54.168973 | 51.865396 | 57.689346 | 117.274238 | 125.214000 | 116.384446 | 131.899837 | 1.036959 | 3578.418027 | 0.113820797 | 0.460016607 | 0.6455043 | 0.998865404 | 11.778775 | 11.830966 | 11.765310 | 11.813360 | 11.783621 | 11.854162 | 3928 | 3902 | 3033 | 3561 |
| 3066 | NM_001527,N | HDAC2 | histone deacetylase 2 | protein_coding | NP_001518.3;XP_016866288.1 | 4853 | 605164 | . | . | 8.776024 | 9.001740 | 8.275984 | 7.896144 | 20.422167 | 20.807926 | 18.571068 | 18.053594 | -1.100546 | 2527.356762 | 0.113951258 | -1.21296888 | 0.225141719 | 0.982008277 | 11.370846 | 11.232675 | 11.336247 | 11.345580 | 11.276184 | 11.250632 | 3032 | 2874 | 2126 | 2220 |
| 3068 | NM_00112605 | HDGF | heparin binding growth factor | protein_coding | NP_001119522.1;NP_001119523.1;NP_001306115.1;NP_001306116.1;NI | 4856 | 600339 | . | . | 74.657334 | 77.043111 | 68.271802 | 86.047218 | 173.730687 | 178.088608 | 153.199943 | 196.736723 | 1.002478 | 5445.744533 | 0.127452213 | 0.028013787 | 0.977651155 | 0.998865404 | 12.409214 | 12.412611 | 12.398408 | 12.418091 | 12.309171 | 12.502418 | 6190 | 5938 | 4197 | 5698 |
| 3069 | NM_00124390 | HDLBP | high density lipoprotein binding protein | protein_coding | NP_001230829.1;NP_001307894.1;NP_001307895.1;NP_001307896.1;NI | 4857 | 142695 | . | . | 106.886636 | 122.381607 | 111.372704 | 128.923591 | 248.729595 | 282.890562 | 249.917117 | 294.768467 | -1.049444 | 20956.409071 | 0.114012047 | -0.61068784 | 0.54140626 | 0.998865404 | 14.389521 | 14.319846 | 14.340420 | 14.407711 | 14.263300 | 14.398382 | 23752 | 24060 | 16384 | 20681 |
| 3070 | NM_00128906 | HELLS | helicase, lymphoid specific | protein_coding | NP_001275996.1;NP_001275997.1;NP_001275998.1;NP_001275999.1;NI | 4861 | 603946 | . | . | 4.875176 | 5.356200 | 4.751708 | 3.684398 | 11.344732 | 12.381099 | 10.662693 | 8.423939 | -1.096524 | 1055.128011 | 0.187446893 | -0.70920064 | 0.478199974 | 0.998865404 | 10.107722 | 9.975661 | 10.096292 | 10.042322 | 10.154583 | 9.821664 | 1316 | 1148 | 1074 | 730 |
| 3073 | NM_000520,N | HEXA | hexosaminidase subunit alpha | protein_coding | NP_000511.2;NP_001305754.1 | 4878 | 606869 | . | . | 64.906089 | 64.527451 | 65.100799 | 71.686896 | 151.039129 | 149.158087 | 146.084298 | 163.903557 | 1.040004 | 9789.433686 | 0.105496215 | 0.536403218 | 0.591679908 | 0.998865404 | 13.228461 | 13.285004 | 13.246124 | 13.233452 | 13.233307 | 13.311600 | 11134 | 10279 | 8282 | 9786 |
| 3074 | NM_000521,N | HEXB | hexosaminidase subunit beta | protein_coding | NP_000512.2;NP_001278933.1 | 4879 | 606873 | . | . | 141.943157 | 133.297421 | 119.737532 | 107.516268 | 330.307576 | 308.122949 | 268.687539 | 245.823184 | -1.228647 | 7107.609347 | 0.112463825 | -2.64147695 | 0.008254542 | 0.288449238 | 12.935995 | 12.639049 | 12.906020 | 12.844003 | 12.749728 | 12.650544 | 9348 | 8152 | 5878 | 5623 |
| 3077 | NM_000410,N | HFE | homeostatic iron regulator | protein_coding | NP_000401.1;NP_001287678.1;NP_620572.1;NP_620573.1;NP_620575.1 | 4886 | 613609 | . | . | 5.426444 | 5.496549 | 4.393230 | 3.864502 | 12.627559 | 12.705520 | 9.858280 | 8.835727 | -1.206177 | 461.664674 | 0.172657553 | -1.56634899 | 0.117266938 | 0.857396579 | 8.979451 | 8.709326 | 8.925395 | 8.885689 | 8.798242 | 8.781599 | 599 | 528 | 369 | 387 |
| 3084 | NM_00115999 | NRG1 | neuregulin 1 | protein_coding | NP_001153467.1;NP_001153468.1;NP_001153471.1;NP_001153473.1;NI | 7997 | 142445 | . | . | 1.018670 | 1.316491 | 0.942845 | 1.471086 | 2.370486 | 3.043128 | 2.115718 | 3.363462 | -1.205719 | 283.178745 | 0.21002634 | -1.28504899 | 0.198775165 | 0.96621494 | 8.274465 | 8.004017 | 8.150053 | 8.232334 | 8.091367 | 8.096383 | 330 | 359 | 222 | 242 |
| 3087 | NM_002729 | HHEX | hematopoietically expressed homeobox | protein_coding | NP_002720.1 | 4901 | 604420 | . | . | 5.638540 | 5.693495 | 6.040242 | 6.494031 | 13.121115 | 13.160769 | 13.554128 | 14.847829 | 1.092487 | 320.801365 | 0.196031351 | 0.650999962 | 0.515046512 | 0.998865404 | 8.260479 | 8.387787 | 8.299496 | 8.299984 | 8.330466 | 8.366030 | 353 | 331 | 282 | 324 |
| 3090 | NM_00109820 | HIC1 | HIC ZBTB transcriptional repressor 1 | protein_coding | NP_001091672.1;NP_006488.2 | 4909 | 603825 | . | . | 0.103868 | 0.133067 | 0.247616 | 0.169410 | 0.241705 | 0.307590 | 0.555644 | 0.387336 | 1.693943 | 34.564360 | 0.542672345 | 1.401186968 | 0.16115817 | 0.933525128 | 4.681128 | 5.442095 | 5.042148 | 5.068654 | 5.156467 | 5.094979 | 26 | 31 | 45 | 33 |
| 3091 | NM_00124308 | HIF1A | hypoxia inducible factor 1 subunit alpha | protein_coding | NP_001230013.1;NP_001521.1;NP_851397.1 | 4910 | 603348 | . | . | 81.888464 | 80.743851 | 78.236236 | 83.175447 | 190.557831 | 186.643026 | 175.559843 | 190.170777 | -1.022331 | 9946.401923 | 0.102443964 | -0.31101616 | 0.755788335 | 0.998865404 | 13.295813 | 13.263929 | 13.298639 | 13.279707 | 13.247713 | 13.292572 | 11708 | 10731 | 8328 | 9465 |
| 3092 | NM_00124319 | HIP1 | huntingtin interacting protein 1 | protein_coding | NP_001230127.1;NP_001369373.1;NP_001369374.1;NP_005329.3;XP_01 | 4913 | 601767 | . | . | 3.802209 | 4.110701 | 3.496238 | 4.276929 | 8.847896 | 9.502070 | 7.845459 | 9.778688 | -1.029886 | 983.017111 | 0.149462257 | -0.28425417 | 0.776215604 | 0.998865404 | 9.962489 | 9.919333 | 9.925527 | 9.975395 | 9.858726 | 9.994867 | 1110 | 1113 | 763 | 993 |
| 3093 | NM_00111111 | UBE2K | ubiquitin conjugating enzyme E2 K | protein_coding | NP_001104582.1;NP_001104583.1;NP_001299575.1;NP_001299576.1;NI | 4914 | 602846 | . | . | 8.286547 | 7.185781 | 7.751367 | 7.456877 | 19.283135 | 16.610256 | 17.393844 | 17.049263 | -1.030876 | 1230.994575 | 0.138835053 | -0.31599459 | 0.752006611 | 0.998865404 | 10.287177 | 10.243713 | 10.332369 | 10.216776 | 10.271460 | 10.234948 | 1548 | 1246 | 1080 | 1109 |
| 3094 | NM_005340,N | HINT1 | histidine triad nucleotide binding protein 1 | protein_coding | NP_005331.1 | 4912 | 601314 | . | . | 231.310631 | 223.367176 | 224.159731 | 243.195862 | 538.269355 | 516.323233 | 503.007926 | 556.038471 | 1.014790 | 6189.641641 | 0.106574499 | 0.198745643 | 0.842461718 | 0.998865404 | 12.585027 | 12.606176 | 12.606131 | 12.572087 | 12.573724 | 12.628759 | 7218 | 6496 | 5245 | 6036 |
| 3096 | NM_002114,XI | HIVEP1 | HIVEP zinc finger 1 | protein_coding | NP_002105.3;XP_011512848.1;XP_011512849.1;XP_011512850.1;XP_01: | 4920 | 194540 | . | . | 0.804454 | 1.111492 | 1.311180 | 1.252981 | 1.871997 | 2.569264 | 2.942250 | 2.864793 | 1.283302 | 320.384872 | 0.204969152 | 1.755681223 | 0.079142855 | 0.775642847 | 8.132836 | 8.492171 | 8.189909 | 8.298306 | 8.398670 | 8.366529 | 290 | 334 | 322 | 328 |
| 3097 | NM_006734,XI | HIVEP2 | HIVEP zinc finger 2 | protein_coding | NP_006725.3;XP_016866294.1;XP_024302184.1;XP_024302185.1;XP_02 | 4921 | 143054 | . | . | 1.534067 | 1.570754 | 1.802781 | 1.710839 | 3.569836 | 3.630867 | 4.045387 | 3.911630 | 1.134172 | 523.297148 | 0.165233307 | 1.099294051 | 0.271639826 | 0.998865404 | 8.937645 | 9.119595 | 8.993209 | 8.983607 | 9.089470 | 9.048525 | 569 | 525 | 499 | 505 |
| 3098 | NM_000188,N | HK1 | hexokinase 1 | protein_coding | NP_000179.2;NP_001309293.1;NP_001309294.1;NP_001309295.1;NP_0( | 4922 | 142600 | . | . | 46.045679 | 45.210918 | 46.133788 | 51.582639 | 107.150186 | 104.507064 | 103.522886 | 117.937564 | 1.055365 | 5300.594603 | 0.110257887 | 0.705090327 | 0.480754026 | 0.998865404 | 12.332583 | 12.410249 | 12.359711 | 12.337303 | 12.350423 | 12.435323 | 6019 | 5491 | 4495 | 5360 |
| 3099 | NM_000189,N | HK2 | hexokinase 2 | protein_coding | NP_000180.2;NP_001358454.1;XP_005264337.1;XP_011531109.1;XP_01 | 4923 | 601125 | . | . | 54.797374 | 54.216599 | 55.975556 | 53.632838 | 127.515747 | 125.324100 | 125.607525 | 122.625104 | -1.009720 | 9577.314487 | 0.102426171 | -0.13624684 | 0.891626151 | 0.998865404 | 13.232346 | 13.218431 | 13.237284 | 13.221481 | 13.242709 | 13.199219 | 11184 | 10288 | 8500 | 8707 |
| 3104 | NM_00127864 | ZBTB48 | zinc finger and BTB domain containing 48 | protein_coding | NP_001265576.1;NP_001265577.1;NP_005332.1;XP_016856599.1;XP_01 | 4930 | 165270 | . | . | 8.969170 | 9.853050 | 10.645117 | 9.959951 | 20.871626 | 22.775766 | 23.887341 | 22.772242 | 1.075360 | 687.155326 | 0.154690659 | 0.677605673 | 0.498021748 | 0.998865404 | 9.371110 | 9.475970 | 9.368713 | 9.426781 | 9.473849 | 9.421170 | 730 | 745 | 643 | 642 |
| 3105 | NM_002116 | HLA-A | major histocompatibility complex, class I, A | protein_coding | NP_002107.3 | 4931 | 142800 | . | . | 234.795776 | 228.752182 | 205.919922 | 223.732666 | 546.379456 | 528.770935 | 462.078339 | 511.538177 | -1.094265 | 10680.601343 | 0.103971083 | -1.24998318 | 0.211305693 | 0.974817004 | 13.446237 | 13.316246 | 13.434587 | 13.404756 | 13.311660 | 13.373210 | 13075 | 11835 | 8547 | 9910 |
| 3106 | NM_005514 | HLA-B | major histocompatibility complex, class I, B | protein_coding | NP_005505.2 | 4932 | 142830 | . | . | 105.081947 | 100.025742 | 90.631035 | 98.548683 | 244.529999 | 231.213989 | 203.373413 | 225.319839 | -1.099827 | 4718.228229 | 0.110192119 | -1.24579552 | 0.212839474 | 0.975238634 | 12.271056 | 12.133737 | 12.266371 | 12.217566 | 12.131982 | 12.192747 | 5856 | 5179 | 3764 | 4368 |
| 3107 | NM_002117 | HLA-C | major histocompatibility complex, class I, C | protein_coding | NP_002108.4 | 4933 | 142840 | . | . | 88.458427 | 85.046173 | 81.762848 | 79.791275 | 205.846390 | 196.588043 | 183.473450 | 182.433273 | -1.089203 | 4025.763878 | 0.109025476 | -1.13068035 | 0.258189656 | 0.995760118 | 12.035325 | 11.912140 | 12.029222 | 11.989146 | 11.952576 | 11.924294 | 4949 | 4421 | 3409 | 3551 |
| 3108 | NM_006120 | HLA-DMA | major histocompatibility complex, class II, DM alpha | protein_coding | NP_006111.2 | 4934 | 142855 | . | . | 2.882879 | 3.427251 | 2.355911 | 3.265975 | 6.708579 | 7.922243 | 5.286596 | 7.467264 | -1.140986 | 101.983084 | 0.330829088 | -0.57516276 | 0.565181229 | 0.998865404 | 6.765319 | 6.572624 | 6.664167 | 6.717839 | 6.605266 | 6.694069 | 115 | 127 | 70 | 103 |
| 3109 | NM_002118 | HLA-DMB | major histocompatibility complex, class II, DM beta | protein_coding | NP_002109.2 | 4935 | 142856 | . | . | 10.332094 | 8.549016 | 8.824969 | 8.659764 | 24.043209 | 19.761435 | 19.802975 | 19.799522 | -1.096719 | 382.423479 | 0.192577048 | -0.69163636 | 0.489165724 | 0.998865404 | 8.643740 | 8.511275 | 8.662361 | 8.543295 | 8.559144 | 8.542090 | 506 | 389 | 322 | 337 |
| 3112 | NM_002120 | HLA-DOB | major histocompatibility complex, class II, DO beta | protein_coding | NP_002111.1 | 4937 | 600629 | . | . | 1.330075 | 1.953282 | 2.622667 | 1.879520 | 3.095139 | 4.515100 | 5.885188 | 4.297299 | 1.356708 | 80.795355 | 0.380222363 | 1.157507573 | 0.247065047 | 0.992733415 | 6.099441 | 6.539528 | 6.233536 | 6.325055 | 6.409636 | 6.309380 | 64 | 88 | 95 | 72 |
| 3113 | NM_00124252 | HLA-DPA1 | major histocompatibility complex, class II, DP alpha 1 | protein_coding | NP_001229453.1;NP_001229454.1;NP_291032.2 | 4938 | 142880 | . | . | 18.737455 | 16.828873 | 13.602638 | 16.096552 | 43.602828 | 38.900691 | 30.523928 | 36.802853 | -1.221192 | 859.173579 | 0.152856926 | -1.88601049 | 0.059293541 | 0.718550266 | 9.883911 | 9.595290 | 9.850408 | 9.774083 | 9.613993 | 9.723526 | 1144 | 967 | 619 | 783 |
| 3115 | NM_002121 | HLA-DPB1 | major histocompatibility complex, class II, DP beta 1 | protein_coding | NP_002112.3 | 4940 | 142858 | . | . | 2.092170 | 1.645639 | 1.321788 | 1.489652 | 4.868567 | 3.803969 | 2.966054 | 3.405912 | -1.347833 | 204.197084 | 0.247363772 | -1.74092464 | 0.08169679 | 0.782498736 | 7.873060 | 7.442583 | 7.794913 | 7.672527 | 7.577870 | 7.621977 | 303 | 222 | 143 | 172 |
| 3119 | NM_00124396 | HLA-DQB1 | major histocompatibility complex, class II, DQ beta 1 | protein_coding | NP_001230890.1;NP_002114.3 | 4944 | 604305 | . | . | 4.763054 | 4.488039 | 3.440652 | 4.515620 | 11.083823 | 10.374304 | 7.720724 | 10.324428 | -1.177699 | 218.950701 | 0.239417064 | -0.9856041 | 0.324327424 | 0.998865404 | 7.888154 | 7.651038 | 7.827809 | 7.793968 | 7.674781 | 7.790015 | 282 | 247 | 152 | 213 |
| 3120 | NM_00119885 | HLA-DQB2 | major histocompatibility complex, class II, DQ beta 2 | protein_coding | NP_001185787.1;NP_001287719.1;XP_005249108.1;XP_011512862.1;XP | 4945 | 615161 | . | . | 3.973473 | 2.986970 | 2.981704 | 3.709220 | 9.246435 | 6.904515 | 6.690856 | 8.480691 | -1.058060 | 129.332792 | 0.301445187 | -0.27010499 | 0.787079486 | 0.998865404 | 7.055103 | 6.973635 | 7.074853 | 6.969128 | 6.967066 | 7.038149 | 175 | 123 | 98 | 130 |
| 3122 | NM_019111 | HLA-DRA | major histocompatibility complex, class II, DR alpha | protein_coding | NP_061984.2 | 4947 | 142860 | . | . | 1.892615 | 1.432629 | 1.366595 | 1.205027 | 4.404194 | 3.311585 | 3.066600 | 2.755152 | -1.312271 | 57.046940 | 0.42493812 | -0.92264287 | 0.356193349 | 0.998865404 | 6.015942 | 5.625935 | 5.893868 | 5.828547 | 5.819271 | 5.793425 | 85 | 60 | 46 | 43 |
| 3131 | NM_00133037 | HLF | HLF transcription factor, PAR bZIP family member | protein_coding | NP_001317304.1;NP_002117.1;XP_005257326.1;XP_011523007.1 | 4977 | 142385 | . | . | 0.127118 | 0.115890 | 0.056710 | 0.089871 | 0.295809 | 0.267885 | 0.127256 | 0.205478 | -1.596144 | 17.126878 | 0.752985101 | -0.89588877 | 0.370312175 | NA | 4.397726 | 3.719719 | 4.120485 | 4.122277 | 4.075845 | 4.099437 | 24 | 23 | 9 | 15 |
| 3133 | NM_005516,XI | HLA-E | major histocompatibility complex, class I, E | protein_coding | NP_005507.3;XP_016866296.1;XP_016866297.1;XP_016866298.1 | 4962 | 143010 | . | . | 33.716563 | 31.510649 | 31.349517 | 33.678999 | 78.459828 | 72.838267 | 70.347402 | 77.003033 | -1.019897 | 2595.352893 | 0.117178686 | -0.24256856 | 0.808339637 | 0.998865404 | 11.355865 | 11.327425 | 11.378666 | 11.319325 | 11.309111 | 11.357238 | 3128 | 2725 | 2168 | 2481 |
| 3134 | NM_00109847 | HLA-F | major histocompatibility complex, class I, F | protein_coding | NP_001091948.1;NP_001091949.1;NP_061823.2;XP_011512866.1;XP_01 | 4963 | 143110 | . | . | 44.732029 | 37.638019 | 27.320771 | 32.100074 | 104.093272 | 87.001969 | 61.307016 | 73.393004 | -1.427394 | 1675.046260 | 0.142769446 | -3.59589488 | 0.000323278 | 0.029164777 | 10.944112 | 10.430360 | 10.872550 | 10.787232 | 10.467149 | 10.631980 | 2387 | 2015 | 1067 | 1437 |
| 3136 | NR_001434 | HLA-H | major histocompatibility complex, class I, H (pseudogene) | transcribed_ps | . | 4965 | . | | . | 19.614929 | 17.511185 | 15.983996 | 18.620842 | 45.644745 | 40.477890 | 35.867626 | 42.574341 | -1.088513 | 953.296353 | 0.149236146 | -0.81989818 | 0.412274152 | 0.998865404 | 9.956754 | 9.834206 | 9.970166 | 9.879954 | 9.811943 | 9.914154 | 1214 | 1007 | 737 | 917 |
| 3139 | NR_027822 | HLA-L | major histocompatibility complex, class I, L (pseudogene) | transcribed_ps | . | 4970 | . | | . | 0.757326 | 0.695050 | 0.894784 | 0.551148 | 1.762330 | 1.606640 | 2.007869 | 1.260133 | -1.019177 | 104.759450 | 0.340220411 | -0.08055035 | 0.935799553 | 0.998865404 | 6.723199 | 6.698569 | 6.722252 | 6.693886 | 6.773436 | 6.625521 | 127 | 109 | 112 | 74 |
| 3140 | NM_00119499 | MR1 | major histocompatibility complex, class I-related | protein_coding | NP_001181928.1;NP_001181929.1;NP_001181964.1;NP_001297142.1;NI | 4975 | 600764 | . | . | 7.928476 | 7.443249 | 6.775739 | 8.163946 | 18.449891 | 17.205405 | 15.204561 | 18.665895 | -1.044831 | 1821.865517 | 0.13266069 | -0.47692594 | 0.633414857 | 0.998865404 | 10.862604 | 10.799099 | 10.875538 | 10.818449 | 10.744727 | 10.876525 | 2221 | 1937 | 1419 | 1811 |
| 3141 | NM_000411,N | HLCS | holocarboxylase synthetase | protein_coding | NP_000402.3;NP_001229713.1;NP_001229714.1;NP_001339443.1;NP_0( | 4976 | 609018 | . | . | 2.729676 | 2.360073 | 2.313053 | 2.567416 | 6.352066 | 5.455414 | 5.190423 | 5.870096 | 1.018939 | 549.396672 | 0.169026027 | 0.160139621 | 0.872771091 | 0.998865404 | 9.088366 | 9.114921 | 9.108480 | 9.081037 | 9.053002 | 9.158138 | 640 | 575 | 443 | 562 |
| 3142 | NM_021958 | HLX | H2.0 like homeobox | protein_coding | NP_068777.1 | 4978 | 142995 | . | . | 0.146531 | 0.187915 | 0.350473 | 0.137594 | 0.340984 | 0.434374 | 0.786452 | 0.314593 | 1.431656 | 14.805478 | 0.838936841 | 0.617072403 | 0.537186971 | NA | 3.604218 | 4.125132 | 3.849670 | 3.865664 | 3.912948 | 3.847032 | 12 | 15 | 22 | 9 |
| 3145 | NM_000190,N | HMBS | hydroxymethylbilane synthase | protein_coding | NP_000181.2;NP_001019553.1;NP_001245137.1;NP_001245138.1;XP_0( | 4982 | 609806 | . | . | 14.299343 | 13.531870 | 13.219791 | 15.747914 | 33.275161 | 31.279524 | 29.664827 | 36.005729 | 1.017766 | 672.176731 | 0.160920894 | 0.157877714 | 0.874553159 | 0.998865404 | 9.380140 | 9.405145 | 9.409311 | 9.361774 | 9.340219 | 9.452133 | 794 | 694 | 541 | 688 |
| 3146 | NM_00131389 | HMGB1 | high mobility group box 1 | protein_coding | NP_001300821.1;NP_001300822.1;NP_001350590.1;NP_001357268.1;NI | 4983 | 163905 | . | . | 40.623807 | 37.194285 | 35.993430 | 38.601680 | 94.533275 | 85.976254 | 80.768215 | 88.258158 | -1.144021 | 6034.912935 | 0.109877301 | -1.76664117 | 0.077288342 | 0.767975096 | 12.652870 | 12.458846 | 12.656020 | 12.567737 | 12.513291 | 12.485583 | 7799 | 6590 | 4977 | 5190 |
| 3148 | NM_00113068 | HMGB2 | high mobility group box 2 | protein_coding | NP_001124160.1;NP_001124161.1;NP_002120.1 | 5000 | 163906 | . | . | 37.670969 | 38.743943 | 29.611176 | 32.611153 | 87.661894 | 89.558360 | 66.446622 | 74.561526 | -1.242897 | 1563.414087 | 0.126164108 | -2.48649788 | 0.012900734 | 0.374881745 | 10.758946 | 10.444974 | 10.681004 | 10.696375 | 10.487828 | 10.551126 | 1976 | 1888 | 1161 | 1362 |
| 3149 | NM_00130122 | HMGB3 | high mobility group box 3 | protein_coding | NP_001288157.1;NP_001288158.1;NP_001288160.1;NP_005333.2;XP_02 | 5004 | 300193 | . | . | 4.939840 | 4.610866 | 2.844850 | 3.810229 | 11.495212 | 10.658223 | 6.383763 | 8.711635 | -1.459290 | 450.344825 | 0.191818174 | -2.84262148 | 0.004474418 | 0.198760547 | 9.062342 | 8.516015 | 8.942052 | 8.888959 | 8.603408 | 8.758917 | 640 | 554 | 274 | 391 |
| 3150 | NM_004965,XI | HMGN1 | high mobility group nucleosome binding domain 1 | protein_coding | NP_004956.5;XP_011527845.1;XP_011527846.1;XP_011527851.1;XP_01( | 4984 | 163920 | . | . | 59.684280 | 67.457451 | 64.015097 | 66.752437 | 138.887774 | 155.930930 | 143.648025 | 152.621516 | -1.064903 | 3507.261690 | 0.120379719 | -0.75363574 | 0.45106798 | 0.998865404 | 11.820673 | 11.730166 | 11.809035 | 11.792335 | 11.811498 | 11.682320 | 4207 | 3864 | 3187 | 2934 |
| 3151 | NM_005517 | HMGN2 | high mobility group nucleosomal binding domain 2 | protein_coding | NP_005508.1 | 4986 | 163910 | . | . | 72.354012 | 73.535187 | 63.433529 | 74.787888 | 168.370758 | 169.979874 | 142.343002 | 170.993622 | -1.070585 | 4293.235151 | 0.117864853 | -0.83485316 | 0.403800391 | 0.998865404 | 12.116294 | 12.017722 | 12.091769 | 12.098125 | 11.972435 | 12.099845 | 5093 | 4809 | 3328 | 4187 |
| 3155 | NM_000191,N | HMGCL | 3-hydroxy-3-methylglutaryl-CoA lyase | protein_coding | NP_000182.2;NP_001159531.1 | 5005 | 613898 | . | . | 12.813655 | 12.617168 | 12.927794 | 17.087349 | 29.817904 | 29.165147 | 29.009596 | 39.068192 | 1.152803 | 671.836823 | 0.16887031 | 1.214813333 | 0.224437294 | 0.982008277 | 9.286128 | 9.490570 | 9.345288 | 9.332216 | 9.345656 | 9.518106 | 726 | 667 | 548 | 758 |
| 3156 | NM_000859,N | HMGCR | 3-hydroxy-3-methylglutaryl-CoA reductase | protein_coding | NP_000850.1;NP_001124468.1;NP_001351116.1;XP_011541659.1;XP_01 | 5006 | 142910 | . | . | 21.671291 | 19.956013 | 32.372782 | 28.100764 | 50.429983 | 46.129220 | 72.643586 | 64.249056 | 1.433105 | 3598.004576 | 0.121864256 | 4.260020693 | 2.04408E-05 | 0.003483263 | 11.530045 | 12.049423 | 11.680065 | 11.607796 | 12.001322 | 11.878189 | 3560 | 3046 | 3952 | 3687 |
| 3157 | NM_00109827 | HMGCS1 | 3-hydroxy-3-methylglutaryl-CoA synthase 1 | protein_coding | NP_001091742.1;NP_001311148.1;NP_001311149.1;NP_001311151.1;NI | 5007 | 142940 | . | . | 13.286084 | 10.742952 | 28.365419 | 21.611232 | 30.917263 | 24.832819 | 63.651177 | 49.411517 | 2.056240 | 2963.475442 | 0.150090619 | 6.929205235 | 4.23211E-12 | 8.34513E-09 | 10.921061 | 11.961450 | 11.230399 | 11.090349 | 11.864139 | 11.621124 | 2441 | 1897 | 3950 | 3219 |
| 3159 | NM_00131907 | HMGA1 | high mobility group AT-hook 1 | protein_coding | NP_001306006.1;NP_001306007.1;NP_001306008.1;NP_001306009.1;NI | 5010 | 600701 | . | . | 75.611134 | 84.219725 | 77.820065 | 88.218185 | 175.950212 | 194.677656 | 174.625977 | 201.700407 | 1.031897 | 4862.747339 | 0.117506018 | 0.385507271 | 0.699861621 | 0.998865404 | 12.224819 | 12.269940 | 12.189476 | 12.275785 | 12.207967 | 12.308962 | 5227 | 5432 | 4036 | 4909 |
| 3161 | NM_00114255 | HMMR | hyaluronan mediated motility receptor | protein_coding | NP_001136028.1;NP_001136029.1;NP_036616.2;NP_036617.2 | 5012 | 600936 | . | . | 23.447558 | 21.822496 | 18.411067 | 15.419228 | 54.563428 | 50.443679 | 41.313903 | 35.254233 | -1.355142 | 1851.475037 | 0.134646702 | -3.2562527 | 0.001128932 | 0.074325569 | 11.056882 | 10.618926 | 10.993028 | 10.927572 | 10.788797 | 10.649203 | 2551 | 2207 | 1493 | 1339 |
| 3162 | NM_002133 | HMOX1 | heme oxygenase 1 | protein_coding | NP_002124.1 | 5013 | 141250 | . | . | 127.949089 | 131.572464 | 245.563812 | 223.862015 | 297.742798 | 304.135651 | 551.038086 | 511.833923 | 1.784183 | 8805.959231 | 0.107311839 | 7.783515918 | 7.05363E-15 | 2.43403E-11 | 12.626986 | 13.462288 | 12.801154 | 12.816901 | 13.344923 | 13.253647 | 7213 | 6892 | 10318 | 10038 |
| 3163 | NM_00112720 | HMOX2 | heme oxygenase 2 | protein_coding | NP_001120676.1;NP_001120677.1;NP_001120678.1;NP_001273196.1;NI | 5014 | 141251 | . | . | 28.824553 | 28.092780 | 26.513723 | 31.643757 | 67.075918 | 64.937718 | 59.496024 | 72.349696 | -1.028578 | 1530.215728 | 0.132154849 | -0.30760099 | 0.758385965 | 0.998865404 | 10.599809 | 10.558940 | 10.613557 | 10.565512 | 10.514466 | 10.618718 | 1842 | 1623 | 1223 | 1510 |
| 3164 | NM_00120223 | NR4A1 | nuclear receptor subfamily 4 group A member 1 | protein_coding | NP_001189162.1;NP_001189163.1;NP_002126.2;NP_775180.1;XP_0052( | 7980 | 139139 | . | . | 20.295320 | 20.204009 | 18.144023 | 21.480413 | 47.228043 | 46.702474 | 40.714662 | 49.112416 | -1.044450 | 1612.693939 | 0.132045475 | -0.4751679 | 0.634667307 | 0.998865404 | 10.686445 | 10.623379 | 10.679403 | 10.663298 | 10.574888 | 10.696317 | 1917 | 1760 | 1264 | 1595 |
| 3175 | NM_004498,N | ONECUT1 | one cut homeobox 1 | protein_coding | NP_004489.1;XP_011519789.1 | 8138 | 604164 | . | . | 0.174658 | 0.210733 | 0.347830 | 0.286208 | 0.406437 | 0.487118 | 0.780523 | 0.654380 | 1.463069 | 32.656979 | 0.543040492 | 1.010969648 | 0.312030957 | 0.998865404 | 4.729157 | 5.277658 | 4.984287 | 5.003221 | 5.048295 | 5.037616 | 28 | 31 | 35 | 35 |
| 3176 | NM_00102407 | HNMT | histamine N-methyltransferase | protein_coding | NP_001019245.1;NP_001019246.1;NP_008826.1;XP_011509366.1;XP_01 | 5028 | 605238 | . | . | 1.389178 | 1.883126 | 1.995581 | 2.386057 | 3.232673 | 4.352931 | 4.478027 | 5.455436 | 1.107391 | 87.072161 | 0.348745253 | 0.421982492 | 0.673037797 | 0.998865404 | 6.369732 | 6.514897 | 6.405734 | 6.447356 | 6.420564 | 6.488464 | 88 | 96 | 70 | 96 |
| 3177 | NM_00130086 | SLC29A2 | solute carrier family 29 member 2 | protein_coding | NP_001287797.1;NP_001287798.1;NP_001523.2;XP_011543276.1;XP_01 | 11004 | 602110 | . | . | 0.414277 | 0.381682 | 0.531140 | 0.741454 | 0.964038 | 0.882277 | 1.191863 | 1.695246 | 1.565034 | 51.591404 | 0.439517984 | 1.470234178 | 0.141498341 | 0.903146012 | 5.329726 | 5.976501 | 5.648910 | 5.617288 | 5.705933 | 5.728636 | 51 | 39 | 52 | 62 |
| 3178 | NM_002136,N | HNRNPA1 | heterogeneous nuclear ribonucleoprotein A1 | protein_coding | NP_002127.1;NP_112420.1 | 5031 | 164017 | . | . | 129.414924 | 128.077698 | 115.664276 | 132.241341 | 301.153845 | 296.057322 | 259.547289 | 302.354105 | -1.054504 | 13966.648332 | 0.108423634 | -0.70616084 | 0.480088121 | 0.998865404 | 13.807493 | 13.730887 | 13.804740 | 13.778167 | 13.691032 | 13.798270 | 16764 | 15232 | 11080 | 13547 |
| 3181 | NM_002137,N | HNRNPA2B1 | heterogeneous nuclear ribonucleoprotein A2/B1 | protein_coding | NP_002128.1;NP_112533.1;XP_005249786.1;XP_016867598.1;XP_01686 | 5033 | 600124 | . | . | 287.168185 | 302.446749 | 322.357529 | 293.687476 | 668.252137 | 699.119208 | 723.360931 | 671.481531 | 1.006185 | 28308.775607 | 0.104419746 | 0.085190496 | 0.932109946 | 0.998865404 | 14.784495 | 14.793415 | 14.794134 | 14.777654 | 14.836530 | 14.743900 | 32807 | 30161 | 26018 | 25193 |
| 3182 | NM_004499,N | HNRNPAB | heterogeneous nuclear ribonucleoprotein A/B | protein_coding | NP_004490.2;NP_112556.2 | 5034 | 602688 | . | . | 159.910313 | 159.842441 | 159.618687 | 178.574478 | 372.117875 | 369.482971 | 358.179734 | 408.289307 | 1.043902 | 8807.897966 | 0.106864069 | 0.580050492 | 0.561880568 | 0.998865404 | 13.073285 | 13.135215 | 13.090196 | 13.081239 | 13.078289 | 13.164395 | 9977 | 9251 | 7429 | 8860 |
| 3183 | NM_00107744 | HNRNPC | heterogeneous nuclear ribonucleoprotein C | protein_coding | NP_001070910.1;NP_001070911.1;NP_004491.2;NP_112604.2;XP_00672 | 5035 | 164020 | . | . | 138.255001 | 138.353009 | 132.065598 | 141.798321 | 321.725069 | 319.809176 | 296.351396 | 324.205010 | -1.003927 | 11481.205309 | 0.106478215 | -0.05310734 | 0.95764638 | 0.998865404 | 13.489834 | 13.484134 | 13.495683 | 13.480911 | 13.438427 | 13.529476 | 13361 | 12306 | 9433 | 11315 |
| 3184 | NM_00100381 | HNRNPD | heterogeneous nuclear ribonucleoprotein D | protein_coding | NP_001003810.1;NP_002129.2;NP_112737.1;NP_112738.1 | 5036 | 601324 | . | . | 46.006704 | 42.324168 | 45.559130 | 46.353924 | 107.059492 | 97.834211 | 102.233358 | 105.982744 | 1.018770 | 4006.264687 | 0.111160993 | 0.241341954 | 0.809290097 | 0.998865404 | 11.954541 | 11.981418 | 11.994434 | 11.924446 | 11.966648 | 11.984241 | 4763 | 4102 | 3479 | 3831 |
| 3185 | NM_00109820 | HNRNPF | heterogeneous nuclear ribonucleoprotein F | protein_coding | NP_001091674.1;NP_001091675.1;NP_001091676.1;NP_001091677.1;NI | 5039 | 601037 | . | . | 84.125959 | 82.406546 | 84.570332 | 98.002289 | 195.764561 | 190.486426 | 189.773376 | 224.070596 | 1.083215 | 7261.352708 | 0.111394965 | 1.035232924 | 0.300560146 | 0.998865404 | 12.767250 | 12.882495 | 12.800427 | 12.780859 | 12.801453 | 12.912257 | 8121 | 7435 | 6140 | 7539 |
| 3187 | NM_00125729 | HNRNPH1 | heterogeneous nuclear ribonucleoprotein H1 | protein_coding | NP_001244222.1;NP_001350501.1;NP_001351154.1;NP_001351155.1;NI | 5041 | 601035 | . | . | 139.414900 | 144.344506 | 162.467297 | 131.251191 | 324.424211 | 333.658785 | 364.571903 | 300.090263 | 1.016578 | 9879.027188 | 0.121420193 | 0.195361331 | 0.845110087 | 0.998865404 | 13.258204 | 13.282004 | 13.248740 | 13.274514 | 13.360903 | 13.182887 | 11111 | 10732 | 9564 | 8365 |
| 3188 | NM_00103239 | HNRNPH2 | heterogeneous nuclear ribonucleoprotein H2 | protein_coding | NP_001027565.1;NP_062543.1 | 5042 | 300610 | . | . | 16.755844 | 16.872335 | 16.521493 | 16.435791 | 38.991539 | 39.001162 | 37.073750 | 37.578483 | -1.036459 | 1197.581923 | 0.130212918 | -0.39675766 | 0.691546174 | 0.998865404 | 10.251494 | 10.199861 | 10.238937 | 10.239638 | 10.216199 | 10.208528 | 1403 | 1316 | 1028 | 1097 |
| 3189 | NM_00132243 | HNRNPH3 | heterogeneous nuclear ribonucleoprotein H3 | protein_coding | NP_001309363.1;NP_001309365.1;NP_001309366.1;NP_001309367.1;NI | 5043 | 602324 | . | . | 49.883160 | 50.225207 | 50.650175 | 47.159692 | 116.080168 | 116.079813 | 113.657525 | 107.825031 | -1.040454 | 3693.181059 | 0.112078618 | -0.51047328 | 0.60971993 | 0.998865404 | 11.878911 | 11.821821 | 11.867389 | 11.865761 | 11.869534 | 11.796886 | 4341 | 4060 | 3287 | 3244 |
| 3190 | NM_00131818 | HNRNPK | heterogeneous nuclear ribonucleoprotein K | protein_coding | NP_001305115.1;NP_001305116.1;NP_001305117.1;NP_002131.2;NP_1: | 5044 | 600712 | . | . | 164.935854 | 170.712532 | 159.351479 | 180.004513 | 383.812506 | 394.609666 | 357.580095 | 411.558930 | -1.003783 | 14736.208448 | 0.106178684 | -0.05130066 | 0.959085944 | 0.998865404 | 13.849822 | 13.844328 | 13.839312 | 13.857264 | 13.796674 | 13.891299 | 16823 | 16098 | 12084 | 14550 |
| 3191 | NM_00100533 | HNRNPL | heterogeneous nuclear ribonucleoprotein L | protein_coding | NP_001005335.1;NP_001524.2;XP_011525190.1;XP_024307247.1;XP_02 | 5045 | 603083 | . | . | 79.406817 | 82.788012 | 77.282113 | 85.750305 | 184.782924 | 191.368205 | 173.418827 | 196.057875 | -1.021729 | 5286.095134 | 0.110102989 | -0.28166331 | 0.778201687 | 0.998865404 | 12.383469 | 12.352337 | 12.359077 | 12.393952 | 12.318643 | 12.397043 | 6025 | 5885 | 4334 | 5132 |
| 3192 | NM_004501,N | HNRNPU | heterogeneous nuclear ribonucleoprotein U | protein_coding | NP_004492.2;NP_114032.2 | 5048 | 602869 | . | . | 57.600201 | 56.742245 | 53.122352 | 56.109678 | 134.038032 | 131.162243 | 119.205017 | 128.288128 | -1.062900 | 11860.482657 | 0.101161364 | -0.86994947 | 0.384328017 | 0.998865404 | 13.577214 | 13.489194 | 13.569026 | 13.549565 | 13.488851 | 13.525238 | 14233 | 13039 | 9785 | 11011 |
| 3198 | NM_005522,N | HOXA1 | homeobox A1 | protein_coding | NP_005513.2;NP_705873.3 | 5099 | 142955 | . | . | 1.852330 | 1.712870 | 1.744013 | 1.920527 | 4.310446 | 3.959375 | 3.913515 | 4.391059 | 1.022286 | 139.874901 | 0.276946942 | 0.11482076 | 0.908587194 | 0.998865404 | 7.112105 | 7.143708 | 7.139276 | 7.107556 | 7.113616 | 7.149611 | 166 | 143 | 117 | 139 |
| 3199 | NM_006735 | HOXA2 | homeobox A2 | protein_coding | NP_006726.1 | 5103 | 604685 | . | . | 0.292518 | 0.196535 | 0.197861 | 0.291040 | 0.680702 | 0.454299 | 0.443995 | 0.665428 | 1.028256 | 13.563161 | 0.836182947 | 0.048075471 | 0.961656095 | NA | 3.741623 | 3.781347 | 3.771328 | 3.751061 | 3.753277 | 3.772440 | 18 | 12 | 10 | 15 |
| 3200 | NM_030661,N | HOXA3 | homeobox A3 | protein_coding | NP_109377.1;NP_705895.1;XP_005249787.1;XP_005249788.1;XP_00524 | 5104 | 142954 | . | . | 2.414760 | 2.837144 | 3.397893 | 1.547335 | 5.619242 | 6.558187 | 7.624773 | 3.537798 | 1.014748 | 313.247673 | 0.2247993 | 0.09395646 | 0.925143743 | 0.998865404 | 8.280265 | 8.301973 | 8.220115 | 8.340321 | 8.370978 | 8.202118 | 319 | 372 | 317 | 250 |
| 3201 | NM_002141 | HOXA4 | homeobox A4 | protein_coding | NP_002132.3 | 5105 | 142953 | . | . | 3.081610 | 2.832139 | 2.080979 | 3.971363 | 7.171034 | 6.546617 | 4.669657 | 9.080050 | 1.007622 | 157.612296 | 0.312933512 | 0.035004114 | 0.97207646 | 0.998865404 | 7.295852 | 7.304607 | 7.308375 | 7.272118 | 7.163374 | 7.408814 | 189 | 162 | 95 | 194 |
| 3202 | NM_019102 | HOXA5 | homeobox A5 | protein_coding | NP_061975.2 | 5106 | 142952 | . | . | 2.889962 | 2.525076 | 2.843961 | 2.532679 | 6.725061 | 5.836826 | 6.381766 | 5.790669 | -1.021729 | 141.229259 | 0.278660883 | -0.11129032 | 0.911386127 | 0.998865404 | 7.156691 | 7.126946 | 7.172225 | 7.117413 | 7.159493 | 7.114350 | 176 | 143 | 129 | 123 |
| 3203 | NM_024014 | HOXA6 | homeobox A6 | protein_coding | NP_076919.1 | 5107 | 142951 | . | . | 0.861567 | 0.661175 | 0.818362 | 0.943034 | 2.004903 | 1.528336 | 1.836380 | 2.156136 | 1.121269 | 25.584070 | 0.611416314 | 0.270081062 | 0.787097891 | NA | 4.592063 | 4.757542 | 4.681062 | 4.657260 | 4.674229 | 4.688357 | 31 | 23 | 22 | 27 |
| 3206 | NM_018951,N | HOXA10 | homeobox A10 | protein_coding | NP_061824.3 | 5100 | 142957 | . | . | 0.060902 | 0.034798 | 0.050583 | 0.054827 | 0.141721 | 0.080436 | 0.113507 | 0.125356 | 1.241043 | 4.481030 | 1.423930064 | 0.218797845 | 0.826807526 | NA | 1.998355 | 2.312263 | 2.164367 | 2.152898 | 2.161422 | 2.164737 | 6 | 3 | 4 | 5 |
| 3207 | NM_005523 | HOXA11 | homeobox A11 | protein_coding | NP_005514.1 | 5101 | 142958 | . | . | 0.170365 | 0.148914 | 0.203467 | 0.140780 | 0.396445 | 0.344222 | 0.456573 | 0.321876 | 1.042677 | 14.180223 | 0.815729491 | 0.073912635 | 0.941079902 | NA | 3.793789 | 3.857133 | 3.825774 | 3.817757 | 3.835970 | 3.812568 | 17 | 14 | 15 | 11 |
| 3209 | NM_000522 | HOXA13 | homeobox A13 | protein_coding | NP_000513.2 | 5102 | 142959 | . | . | 0.666354 | 0.670632 | 0.780853 | 0.863706 | 1.550634 | 1.550196 | 1.752212 | 1.974762 | 1.206601 | 116.707375 | 0.29852886 | 0.907612486 | 0.364082995 | 0.998865404 | 6.725188 | 6.995658 | 6.831081 | 6.830492 | 6.877743 | 6.909572 | 122 | 114 | 106 | 125 |
| 3212 | NM_002145,XI | HOXB2 | homeobox B2 | protein_coding | NP_002136.1;XP_005257332.1 | 5113 | 142967 | . | . | 22.536870 | 21.328368 | 24.804040 | 21.549062 | 52.444224 | 49.301473 | 55.659546 | 49.269374 | 1.079081 | 1055.862436 | 0.146919316 | 0.747373753 | 0.454837985 | 0.998865404 | 9.987951 | 10.098355 | 10.032757 | 9.994276 | 10.137830 | 9.997642 | 1200 | 1067 | 1043 | 931 |
| 3213 | NM_00133032 | HOXB3 | homeobox B3 | protein_coding | NP_001317251.1;NP_001317252.1;NP_002137.4;XP_005257334.1;XP_00 | 5114 | 142966 | . | . | 1.210106 | 0.983752 | 0.785688 | 0.844778 | 2.815965 | 2.273986 | 1.763060 | 1.931483 | -1.012298 | 235.408253 | 0.22630091 | -0.07792148 | 0.937890515 | 0.998865404 | 7.888351 | 7.869631 | 7.867221 | 7.895554 | 7.833736 | 7.915486 | 265 | 263 | 186 | 238 |
| 3214 | NM_024015 | HOXB4 | homeobox B4 | protein_coding | NP_076920.1 | 5115 | 142965 | . | . | 3.461669 | 3.525667 | 3.831038 | 3.053659 | 8.055445 | 8.149737 | 8.596738 | 6.981829 | -1.027510 | 216.851844 | 0.237402901 | -0.16492249 | 0.869004991 | 0.998865404 | 7.779433 | 7.741449 | 7.763305 | 7.766995 | 7.805284 | 7.696878 | 252 | 238 | 208 | 177 |
| 3215 | NM_002147 | HOXB5 | homeobox B5 | protein_coding | NP_002138.1 | 5116 | 142960 | . | . | 1.572163 | 1.675107 | 2.372258 | 1.353161 | 3.658487 | 3.872084 | 5.323277 | 3.093842 | 1.135815 | 99.279296 | 0.354946672 | 0.517622253 | 0.604721853 | 0.998865404 | 6.537395 | 6.723470 | 6.594843 | 6.611433 | 6.723748 | 6.552723 | 104 | 103 | 117 | 72 |
| 3216 | NM_00136939 | HOXB6 | homeobox B6 | protein_coding | NP_001356326.1;NP_061825.2;XP_011523029.1 | 5117 | 142961 | . | . | 8.864501 | 9.685301 | 8.733301 | 8.915578 | 20.628059 | 22.388008 | 19.597274 | 20.384411 | -1.067800 | 528.083870 | 0.165671871 | -0.57126 | 0.56782341 | 0.998865404 | 9.091299 | 8.996387 | 9.037773 | 9.092106 | 9.021565 | 9.024600 | 602 | 613 | 443 | 480 |
| 3217 | NM_004502 | HOXB7 | homeobox B7 | protein_coding | NP_004493.3 | 5118 | 142962 | . | . | 16.884470 | 15.320175 | 17.262262 | 16.787294 | 39.290855 | 35.413273 | 38.736015 | 38.382149 | 1.043263 | 703.794500 | 0.153564713 | 0.397895967 | 0.690706873 | 0.998865404 | 9.427918 | 9.489447 | 9.479231 | 9.406753 | 9.485888 | 9.459656 | 835 | 704 | 637 | 661 |
| 3218 | NM_024016,XI | HOXB8 | homeobox B8 | protein_coding | NP_076921.1;XP_005257343.1;XP_016880053.1 | 5119 | 142963 | . | . | 0.856941 | 0.623919 | 0.958422 | 1.312829 | 1.994138 | 1.442217 | 2.150673 | 3.001627 | 1.224157 | 77.612033 | 0.365019466 | 0.799377353 | 0.424071638 | 0.998865404 | 6.125325 | 6.416432 | 6.260743 | 6.233172 | 6.268148 | 6.328896 | 85 | 71 | 66 | 89 |
| 3219 | NM_024017 | HOXB9 | homeobox B9 | protein_coding | NP_076922.1 | 5120 | 142964 | . | . | 4.844365 | 5.235707 | 3.993480 | 5.367140 | 11.273036 | 12.102571 | 8.961253 | 12.271330 | -1.090931 | 392.097453 | 0.199585181 | -0.62910196 | 0.52928231 | 0.998865404 | 8.677127 | 8.550220 | 8.617309 | 8.659233 | 8.502757 | 8.664685 | 455 | 457 | 280 | 401 |
| 3221 | NM_014620,N | HOXC4 | homeobox C4 | protein_coding | NP_055435.2;NP_705897.1 | 5126 | 142974 | . | . | 5.189236 | 5.605454 | 5.428238 | 6.449581 | 12.075566 | 12.957260 | 12.180808 | 14.746201 | 1.098114 | 303.571817 | 0.207135575 | 0.651881312 | 0.514477741 | 0.998865404 | 8.177362 | 8.311318 | 8.199782 | 8.236286 | 8.217382 | 8.317351 | 322 | 323 | 251 | 325 |
| 3222 | NM_018953,N | HOXC5 | homeobox C5 | protein_coding | NP_061826.1 | 5127 | 142973 | . | . | 0.066837 | 0.205611 | 0.332359 | 0.228478 | 0.155532 | 0.475280 | 0.745803 | 0.522388 | 1.992018 | 10.894415 | 0.97894509 | 1.015614593 | 0.309812953 | NA | 2.867373 | 3.857133 | 3.387066 | 3.423076 | 3.449009 | 3.425696 | 4 | 12 | 15 | 11 |
| 3223 | NM_004503,N | HOXC6 | homeobox C6 | protein_coding | NP_004494.1;NP_710160.1 | 5128 | 142972 | . | . | 4.456807 | 3.681934 | 5.772455 | 4.236328 | 10.371175 | 8.510955 | 12.953221 | 9.685861 | 1.229924 | 254.157768 | 0.232840265 | 1.282290158 | 0.199740879 | 0.96621494 | 7.781692 | 8.131882 | 7.952995 | 7.902961 | 8.108891 | 7.949856 | 276 | 233 | 281 | 223 |
| 3224 | NM_022658 | HOXC8 | homeobox C8 | protein_coding | NP_073149.1 | 5129 | 142970 | . | . | 1.463009 | 1.346030 | 1.990760 | 1.824319 | 3.404482 | 3.111409 | 4.467207 | 4.171090 | 1.341251 | 125.287743 | 0.291011487 | 1.455541895 | 0.145519301 | 0.906189003 | 6.741271 | 7.165812 | 6.923122 | 6.893332 | 7.028882 | 6.992507 | 129 | 110 | 131 | 128 |
| 3225 | NM_006897 | HOXC9 | homeobox C9 | protein_coding | NP_008828.1 | 5130 | 142971 | . | . | 1.482604 | 1.228437 | 1.745712 | 1.489667 | 3.450080 | 2.839587 | 3.917328 | 3.405946 | 1.170518 | 68.924608 | 0.384901979 | 0.590142431 | 0.555095164 | 0.998865404 | 5.987929 | 6.216890 | 6.104843 | 6.062218 | 6.140658 | 6.100648 | 80 | 62 | 70 | 64 |
| 3226 | NM_017409 | HOXC10 | homeobox C10 | protein_coding | NP_059105.2 | 5122 | 605560 | . | . | 7.368219 | 8.063487 | 7.371277 | 9.379370 | 17.146149 | 18.639112 | 16.540932 | 21.444817 | 1.069438 | 495.437830 | 0.181819591 | 0.532685369 | 0.594251396 | 0.998865404 | 8.903874 | 8.999657 | 8.902737 | 8.954354 | 8.895345 | 9.041742 | 529 | 538 | 394 | 535 |
| 3229 | NM_017410 | HOXC13 | homeobox C13 | protein_coding | NP_059106.2 | 5125 | 142976 | . | . | 2.016218 | 2.536438 | 2.410083 | 2.626910 | 4.691823 | 5.863091 | 5.408155 | 6.006117 | 1.091589 | 176.368094 | 0.257173464 | 0.491612448 | 0.622993342 | 0.998865404 | 7.398500 | 7.523677 | 7.393632 | 7.485856 | 7.461530 | 7.493985 | 173 | 202 | 154 | 179 |

| | | | | | | | | | | | | | | | | | | | | | | | | | | | | | | | | | |
|---|---|---|---|---|---|---|---|---|---|---|---|---|---|---|---|---|---|---|---|---|---|---|---|---|---|---|---|---|---|---|---|---|---|
| 3231 | NM_024501 | HOXD1 | homeobox D1 | protein_coding | NP_078777.1 | 5132 | 142987 | . | . | 0.417211 | 0.557419 | 0.902003 | 0.831111 | 0.970866 | 1.288499 | 2.024068 | 1.900236 | 1.743081 | 40.128284 | 0.497829781 | 1.610267671 | 0.107339434 | 0.835490835 | 4.871539 | 5.671937 | 5.247545 | 5.282242 | 5.356299 | 5.343045 | 29 | 36 | 46 | 46 |
| 3232 | NM_006898,XI | HOXD3 | homeobox D3 | protein_coding | NP_008829.3;XP_005246566.1;XP_005246568.1;XP_011509367.1;XP_01 | 5137 | 142980 | . | . | 0.670461 | 0.630294 | 0.598085 | 0.762146 | 1.560188 | 1.456953 | 1.342085 | 1.742557 | 1.164887 | 86.062821 | 0.350905588 | 0.627490579 | 0.53033771 | 0.998865404 | 6.314120 | 6.532277 | 6.379151 | 6.423636 | 6.409320 | 6.476886 | 84 | 93 | 71 | 97 |
| 3233 | NM_014621,XI | HOXD4 | homeobox D4 | protein_coding | NP_055436.2;XP_005246571.1 | 5138 | 142981 | . | . | 4.084466 | 4.652239 | 4.870628 | 5.768058 | 9.504722 | 10.753859 | 10.929549 | 13.187980 | 1.225842 | 548.790285 | 0.1681242 | 1.74735577 | 0.080575645 | 0.779933816 | 8.946161 | 9.239206 | 8.987852 | 9.065754 | 9.127747 | 9.190513 | 533 | 565 | 501 | 593 |
| 3234 | NM_00119974 | HOXD8 | homeobox D8 | protein_coding | NP_001186675.1;NP_001186676.1;NP_062458.1 | 5139 | 142985 | . | . | 5.848487 | 5.925137 | 5.799500 | 6.473201 | 13.609668 | 13.696216 | 13.013910 | 14.800207 | -1.007886 | 472.494533 | 0.177242354 | -0.06394001 | 0.949017997 | 0.998865404 | 8.890195 | 8.878088 | 8.879811 | 8.892146 | 8.826475 | 8.932661 | 541 | 517 | 374 | 479 |
| 3235 | NM_014213 | HOXD9 | homeobox D9 | protein_coding | NP_055028.3 | 5140 | 142982 | . | . | 0.350099 | 0.740908 | 0.762705 | 0.712694 | 0.814694 | 1.712641 | 1.711488 | 1.629491 | 1.345065 | 37.712158 | 0.541712617 | 0.78948942 | 0.429826011 | 0.998865404 | 5.008980 | 5.433778 | 5.152895 | 5.246770 | 5.250377 | 5.238578 | 24 | 47 | 39 | 39 |
| 3238 | NM_021193 | HOXD12 | homeobox D12 | protein_coding | NP_067016.3 | 5135 | 142988 | . | . | 0.252266 | 0.203290 | 0.189198 | 0.273545 | 0.587033 | 0.469914 | 0.424554 | 0.625427 | 1.026726 | 19.005916 | 0.70953255 | 0.053627939 | 0.957231596 | NA | 4.229742 | 4.266773 | 4.256570 | 4.238383 | 4.236514 | 4.262640 | 24 | 18 | 14 | 21 |
| 3239 | NM_000523,XI | HOXD13 | homeobox D13 | protein_coding | NP_000514.2;XP_011509370.1;XP_011509371.1 | 5136 | 142989 | . | . | 6.190140 | 5.253514 | 7.709719 | 7.593840 | 14.404710 | 12.143733 | 17.300384 | 17.362413 | 1.322830 | 527.810810 | 0.170410091 | 2.368567145 | 0.017857139 | 0.447807832 | 8.827644 | 9.231885 | 8.989530 | 8.894090 | 9.131680 | 9.113847 | 564 | 452 | 530 | 556 |
| 3241 | NM_00125835 | HPCAL1 | hippocalcin like 1 | protein_coding | NP_001245286.1;NP_001245287.1;NP_001245288.1;NP_002140.2;NP_6( | 5145 | 600207 | . | . | 19.040900 | 19.310659 | 19.988718 | 18.831955 | 44.308954 | 44.637459 | 44.854107 | 43.057029 | 1.016209 | 1107.578914 | 0.132698963 | 0.174807188 | 0.86123114 | 0.998865404 | 10.101510 | 10.124784 | 10.111165 | 10.103020 | 10.125253 | 10.112885 | 1272 | 1179 | 979 | 1038 |
| 3248 | NM_000860,N | HPGD | 15-hydroxyprostaglandin dehydrogenase | protein_coding | NP_000851.2;NP_001139288.1;NP_001243230.1;NP_001243234.1;NP_0( | 5154 | 601688 | . | . | 2.025490 | 1.639072 | 2.158673 | 1.843136 | 4.713399 | 3.788791 | 4.843999 | 4.214109 | 1.070016 | 157.784667 | 0.271739685 | 0.359287796 | 0.719379803 | 0.998865404 | 7.251289 | 7.350627 | 7.326386 | 7.242315 | 7.345888 | 7.277647 | 194 | 147 | 154 | 140 |
| 3251 | NM_000194 | HPRT1 | hypoxanthine phosphoribosyltransferase 1 | protein_coding | NP_000185.1 | 5157 | 308000 | . | . | 12.426476 | 10.676528 | 11.857558 | 9.988029 | 28.916920 | 24.679274 | 26.608017 | 22.836439 | -1.071121 | 489.132807 | 0.182399039 | -0.54343113 | 0.586833 | 0.998865404 | 8.982246 | 8.884255 | 9.003820 | 8.900713 | 8.964804 | 8.853283 | 629 | 502 | 448 | 403 |
| 3257 | NM_000195,N | HPS1 | HPS1 biogenesis of lysosomal organelles complex 3 subunit 1 | protein_coding | NP_000186.2;NP_001298274.1;NP_001309405.1;NP_001309406.1;NP_0( | 5163 | 604982 | . | . | 11.553215 | 11.963832 | 12.120544 | 13.225358 | 26.884812 | 27.654929 | 27.198146 | 30.238210 | 1.100001 | 1132.155776 | 0.133020061 | 1.033714611 | 0.301269601 | 0.998865404 | 10.074575 | 10.211873 | 10.095969 | 10.119477 | 10.161613 | 10.195747 | 1222 | 1182 | 1008 | 1137 |
| 3265 | NM_00113044 | HRAS | HRas proto-oncogene, GTPase | protein_coding | NP_001123914.1;NP_001304983.1;NP_005334.1;NP_789765.1 | 5173 | 190020 | . | . | 18.833348 | 21.837763 | 19.741574 | 21.827525 | 43.825977 | 50.478968 | 44.299520 | 49.906043 | 1.016093 | 721.473979 | 0.157368436 | 0.146356642 | 0.883639853 | 0.998865404 | 9.483575 | 9.505945 | 9.436300 | 9.537870 | 9.469071 | 9.528473 | 764 | 829 | 605 | 711 |
| 3267 | NM_00113518 | AGFG1 | ArfGAP with FG repeats 1 | protein_coding | NP_001128659.1;NP_001128660.1;NP_001128661.1;NP_004495.2 | 5175 | 600862 | . | . | 8.281769 | 8.059067 | 8.366677 | 7.600605 | 19.272018 | 18.628898 | 18.774581 | 17.377880 | -1.037698 | 2184.427390 | 0.121507865 | -0.43937222 | 0.66039185 | 0.998865404 | 11.119360 | 11.066230 | 11.121426 | 11.092887 | 11.119306 | 11.034401 | 2607 | 2358 | 1964 | 1903 |
| 3268 | NM_006076,XI | AGFG2 | ArfGAP with FG repeats 2 | protein_coding | NP_006067.3;XP_005250363.1;XP_016867601.1;XP_016867602.1;XP_01( | 5177 | 604019 | . | . | 2.841647 | 2.539950 | 2.716208 | 3.334709 | 6.612628 | 5.871208 | 6.095095 | 7.624416 | 1.098467 | 426.654356 | 0.188452572 | 0.718971562 | 0.472158442 | 0.998865404 | 8.667779 | 8.802906 | 8.742939 | 8.666945 | 8.701732 | 8.819903 | 498 | 411 | 351 | 459 |
| 3269 | NM_000861,N | HRH1 | histamine receptor H1 | protein_coding | NP_000852.1;NP_001091681.1;NP_001091682.1;NP_001091683.1 | 5182 | 600167 | . | . | 0.054274 | 0.078593 | 0.029888 | 0.071727 | 0.126297 | 0.181670 | 0.067068 | 0.163996 | -1.351421 | 8.999668 | 1.040033667 | -0.41775302 | 0.6761277 | NA | 3.373794 | 2.932304 | 3.172955 | 3.188192 | 3.155336 | 3.181712 | 10 | 13 | 4 | 10 |
| 3275 | NM_00124286 | PRMT2 | protein arginine methyltransferase 2 | protein_coding | NP_001229793.1;NP_001229794.1;NP_001229795.1;NP_001273605.1;NI | 5186 | 601961 | . | . | 27.282596 | 28.832168 | 23.846268 | 29.984085 | 63.487723 | 66.646845 | 53.510337 | 68.555049 | -1.015395 | 2642.036527 | 0.114493738 | -0.19251162 | 0.847341464 | 0.998865404 | 11.378476 | 11.356308 | 11.367514 | 11.379536 | 11.336768 | 11.384867 | 3043 | 2894 | 2212 | 2531 |
| 3276 | NM_00120704 | PRMT1 | protein arginine methyltransferase 1 | protein_coding | NP_001193971.1;NP_001527.3;NP_938074.2;XP_005258899.1;XP_01688 | 5187 | 602950 | . | . | 67.698688 | 73.469935 | 62.022140 | 71.459470 | 157.537630 | 169.829047 | 139.175890 | 163.383577 | -1.081241 | 2802.534149 | 0.122314855 | -0.92129012 | 0.356898976 | 0.998865404 | 11.507896 | 11.394926 | 11.448152 | 11.516518 | 11.365967 | 11.469768 | 3214 | 3273 | 2191 | 2687 |
| 3280 | NM_005524 | HES1 | hes family bHLH transcription factor 1 | protein_coding | NP_005515.1 | 5192 | 139605 | . | . | 12.969328 | 14.664146 | 20.924839 | 17.636106 | 30.180162 | 33.896832 | 46.954731 | 40.322861 | 1.376684 | 811.890569 | 0.157781934 | 2.923007698 | 0.003466679 | 0.166147835 | 9.416078 | 9.877471 | 9.497347 | 9.574400 | 9.827036 | 9.693713 | 742 | 779 | 892 | 802 |
| 3281 | NM_001537 | HSBP1 | heat shock factor binding protein 1 | protein_coding | NP_001528.1 | 5203 | 604553 | . | . | 9.942743 | 9.218182 | 9.033064 | 9.226134 | 23.137175 | 21.308241 | 20.269932 | 21.094460 | -1.064362 | 2521.764781 | 0.116896523 | -0.7698192 | 0.441407147 | 0.998865404 | 11.344470 | 11.254565 | 11.358477 | 11.289895 | 11.269978 | 11.278681 | 3120 | 2688 | 2113 | 2303 |
| 3292 | NM_000413,N | HSD17B1 | hydroxysteroid 17-beta dehydrogenase 1 | protein_coding | NP_000404.2;NP_001317148.1 | 5210 | 109684 | . | . | 5.539502 | 5.718474 | 4.630453 | 5.954813 | 12.890650 | 13.218507 | 10.390604 | 13.614974 | -1.089348 | 214.120272 | 0.24065398 | -0.51304134 | 0.607922407 | 0.998865404 | 7.803383 | 7.678469 | 7.753281 | 7.768993 | 7.661273 | 7.777742 | 253 | 245 | 155 | 217 |
| 3295 | NM_000414,N | HSD17B4 | hydroxysteroid 17-beta dehydrogenase 4 | protein_coding | NP_000405.1;NP_001186220.1;NP_001186221.1;NP_001278956.1;NP_0( | 5213 | 601860 | . | . | 30.671189 | 28.264808 | 30.139300 | 31.969361 | 71.373118 | 65.335376 | 67.631719 | 73.094146 | 1.037101 | 2498.350937 | 0.118291114 | 0.444297093 | 0.656827803 | 0.998865404 | 11.260226 | 11.312816 | 11.306768 | 11.235018 | 11.282070 | 11.319524 | 2950 | 2529 | 2161 | 2440 |
| 3297 | NM_005526,XI | HSF1 | heat shock transcription factor 1 | protein_coding | NP_005517.1;XP_005272372.1;XP_005272373.1;XP_005272374.1;XP_01 | 5224 | 140580 | . | . | 25.429498 | 25.785978 | 25.536764 | 28.376486 | 59.175485 | 59.605443 | 57.303759 | 64.879461 | 1.041896 | 1755.780503 | 0.123734553 | 0.478534928 | 0.63226952 | 0.998865404 | 10.748106 | 10.807085 | 10.753150 | 10.769352 | 10.758477 | 10.827226 | 1960 | 1875 | 1490 | 1753 |
| 3298 | NM_00113556 | HSF2 | heat shock transcription factor 2 | protein_coding | NP_001129036.1;NP_001230023.1;NP_004497.1 | 5225 | 140581 | . | . | 5.118304 | 4.115552 | 4.232351 | 4.431428 | 11.910506 | 9.513281 | 9.497271 | 10.131934 | -1.073841 | 365.759226 | 0.199381041 | -0.51549483 | 0.606207282 | 0.998865404 | 8.564965 | 8.462725 | 8.601738 | 8.460647 | 8.482008 | 8.503099 | 488 | 360 | 301 | 337 |
| 3299 | NM_00104066 | HSF4 | heat shock transcription factor 4 | protein_coding | NP_001035757.1;NP_001361603.1;NP_001361604.1;NP_001529.2 | 5227 | 602438 | . | . | 22.909073 | 23.825788 | 36.254310 | 29.687202 | 53.310353 | 55.074378 | 81.353625 | 67.876259 | 1.367626 | 1477.302083 | 0.139091201 | 3.247317109 | 0.001164985 | 0.075570712 | 10.285094 | 10.737138 | 10.391610 | 10.393661 | 10.710144 | 10.552219 | 1435 | 1348 | 1631 | 1442 |
| 3300 | NM_00103955 | DNAJB2 | DnaJ heat shock protein family (Hsp40) member B2 | protein_coding | NP_001034639.1;NP_006727.2 | 5228 | 604139 | . | . | 25.185982 | 26.759368 | 26.272831 | 27.353955 | 58.608818 | 61.855476 | 58.955477 | 62.541568 | 1.003333 | 2044.578827 | 0.119058976 | 0.04032474 | 0.96783423 | 0.998865404 | 10.995270 | 10.999002 | 10.978871 | 11.013172 | 10.979479 | 11.017666 | 2300 | 2250 | 1738 | 1966 |
| 3301 | NM_00131403 | DNAJA1 | DnaJ heat shock protein family (Hsp40) member A1 | protein_coding | NP_001300968.1;NP_001530.1 | 5229 | 602837 | . | . | 71.326258 | 70.167474 | 65.911850 | 69.077971 | 165.979139 | 162.195245 | 147.904265 | 157.938591 | -1.062863 | 4994.256166 | 0.106123312 | -0.82880193 | 0.407216496 | 0.998865404 | 12.329373 | 12.241394 | 12.321336 | 12.300600 | 12.244576 | 12.275171 | 5999 | 5485 | 4133 | 4623 |
| 3303 | NM_005345 | HSPA1A | heat shock protein family A (Hsp70) member 1A | protein_coding | NP_005336.3 | 5232 | 140550 | . | . | 58.491856 | 61.918392 | 43.448837 | 48.211391 | 136.112946 | 143.127121 | 97.497932 | 110.229622 | -1.332049 | 3967.392206 | 0.112982575 | -3.6611614 | 0.000251075 | 0.024405505 | 12.146098 | 11.732280 | 12.040109 | 12.081250 | 11.784888 | 11.860094 | 5093 | 5009 | 2820 | 3339 |
| 3304 | NM_005346 | HSPA1B | heat shock protein family A (Hsp70) member 1B | protein_coding | NP_005337.2 | 5233 | 603012 | . | . | 46.505089 | 50.035606 | 35.287876 | 38.577950 | 108.219254 | 115.659546 | 79.184975 | 88.203903 | -1.325207 | 3343.336092 | 0.114552003 | -3.5461468 | 0.000390908 | 0.033601598 | 11.896012 | 11.489607 | 11.783849 | 11.838329 | 11.548142 | 11.610787 | 4247 | 4245 | 2402 | 2802 |
| 3305 | NM_005527 | HSPA1L | heat shock protein family A (Hsp70) member 1 like | protein_coding | NP_005518.3 | 5234 | 140559 | . | . | 0.625310 | 0.508809 | 0.567175 | 0.522541 | 1.455122 | 1.176136 | 1.272725 | 1.194727 | -1.053608 | 48.361127 | 0.452086442 | -0.1666449 | 0.867649465 | 0.998865404 | 5.632204 | 5.558404 | 5.619839 | 5.581026 | 5.599720 | 5.582650 | 63 | 48 | 43 | 42 |
| 3306 | NM_021979 | HSPA2 | heat shock protein family A (Hsp70) member 2 | protein_coding | NP_068814.2 | 5235 | 140560 | . | . | 17.247656 | 17.377775 | 13.727034 | 16.077524 | 40.136002 | 40.169506 | 30.803064 | 36.759350 | -1.178222 | 1390.994012 | 0.134082502 | -1.76467114 | 0.077619051 | 0.767975096 | 10.555541 | 10.318538 | 10.501504 | 10.500468 | 10.319024 | 10.428534 | 1734 | 1623 | 1029 | 1285 |
| 3308 | NM_002154 | HSPA4 | heat shock protein family A (Hsp70) member 4 | protein_coding | NP_002145.3 | 5237 | 601113 | . | . | 63.119411 | 62.737999 | 60.230412 | 61.232529 | 146.881470 | 145.021698 | 135.155304 | 140.000900 | -1.050782 | 9196.921006 | 0.101014855 | -0.70744797 | 0.479288134 | 0.998865404 | 13.202223 | 13.130763 | 13.193915 | 13.181361 | 13.142855 | 13.148158 | 10932 | 10095 | 7775 | 8435 |
| 3309 | NM_005347 | HSPA5 | heat shock protein family A (Hsp70) member 5 | protein_coding | NP_005338.1 | 5238 | 138120 | . | . | 241.651932 | 252.753159 | 190.630463 | 214.488083 | 562.333923 | 584.250244 | 427.769226 | 490.401550 | -1.237727 | 27483.488743 | 0.105006499 | -2.93023349 | 0.003387074 | 0.164041341 | 14.891950 | 14.584227 | 14.814970 | 14.846849 | 14.600553 | 14.691695 | 34373 | 33403 | 20211 | 24267 |
| 3310 | NM_002155 | HSPA6 | heat shock protein family A (Hsp70) member 6 | protein_coding | NP_002146.2 | 5239 | 140555 | . | . | 1.154341 | 1.168160 | 0.599526 | 1.251440 | 2.686200 | 2.700253 | 1.345320 | 2.861269 | -1.257287 | 77.186350 | 0.405111973 | -0.81536436 | 0.414863777 | 0.998865404 | 6.427631 | 6.093628 | 6.295118 | 6.295471 | 6.155496 | 6.312197 | 99 | 93 | 39 | 86 |
| 3311 | NR_024151 | HSPA7 | heat shock protein family A (Hsp70) member 7 (pseudogene) | transcribed_ps | . | 5240 | 140556 | . | . | 0.135443 | 0.156422 | 0.179738 | 0.150535 | 0.315182 | 0.361577 | 0.403328 | 0.344180 | 1.099053 | 12.736794 | 0.854846186 | 0.159398084 | 0.873355249 | NA | 3.600756 | 3.737849 | 3.660837 | 3.668517 | 3.676785 | 3.663092 | 13 | 14 | 13 | 11 |
| 3312 | NM_006597,N | HSPA8 | heat shock protein family A (Hsp70) member 8 | protein_coding | NP_006588.1;NP_694881.1;XP_011541100.1 | 5241 | 600816 | . | . | 748.879879 | 798.192333 | 671.165868 | 747.432692 | 1742.674414 | 1845.057257 | 1506.076778 | 1708.916022 | -1.104768 | 52116.578008 | 0.104197393 | -1.37953215 | 0.167730741 | 0.945638095 | 15.739544 | 15.595785 | 15.685784 | 15.734993 | 15.581669 | 15.665579 | 61222 | 60701 | 40959 | 48696 |
| 3313 | NM_004134 | HSPA9 | heat shock protein family A (Hsp70) member 9 | protein_coding | NP_004125.3 | 5244 | 600548 | . | . | 112.161873 | 111.869720 | 94.593811 | 116.153633 | 261.005280 | 258.591858 | 212.265762 | 265.571472 | -1.078524 | 14912.330555 | 0.1177015 | -0.92656394 | 0.354152985 | 0.998865404 | 13.917745 | 13.808628 | 13.899495 | 13.889839 | 13.743451 | 13.909497 | 17920 | 16606 | 11264 | 14760 |
| 3314 | gene-HSPA8P1 | HSPA8P1 | heat shock protein family A (Hsp70) member 8 pseudogene 1 | pseudogene | | 5245 | . | | . | 0.028264 | 0.005549 | 0.005018 | 0.018759 | 0.065771 | 0.012826 | 0.011261 | 0.042889 | -1.076094 | 1.703845 | 2.268349467 | -0.04664372 | 0.962797187 | NA | 0.820514 | 0.715150 | 0.778739 | 0.770724 | 0.771939 | 0.776369 | 3 | 1 | 1 | 2 |
| 3315 | NM_001540 | HSPB1 | heat shock protein family B (small) member 1 | protein_coding | NP_001531.1 | 5246 | 602195 | . | . | 84.751358 | 96.428940 | 71.372078 | 92.231789 | 197.219910 | 222.899796 | 160.156876 | 210.877014 | -1.122959 | 2086.623601 | 0.143523878 | -1.16569556 | 0.243737556 | 0.992733415 | 11.108415 | 10.940618 | 11.019362 | 11.117057 | 10.875857 | 11.068503 | 2389 | 2526 | 1500 | 2068 |
| 3320 | NM_00101796 | HSP90AA1 | heat shock protein 90 alpha family class A member 1 | protein_coding | NP_001017963.2;NP_005339.3;XP_011535020.1 | 5253 | 140571 | . | . | 440.000924 | 451.068617 | 378.233683 | 415.702093 | 1023.900285 | 1042.665342 | 848.745434 | 950.453417 | -1.137853 | 42382.821895 | 0.101328717 | -1.83870797 | 0.065958151 | 0.73302766 | 15.461350 | 15.275022 | 15.417143 | 15.431378 | 15.276244 | 15.347827 | 51527 | 49079 | 33036 | 38727 |
| 3321 | NM_00100723 | IGSF3 | immunoglobulin superfamily member 3 | protein_coding | NP_001007238.1;NP_001533.2;XP_005270851.1;XP_006710656.1;XP_01 | 5950 | 603491 | . | . | 25.848099 | 25.410397 | 19.090189 | 22.352176 | 60.149587 | 58.737270 | 42.837829 | 51.105596 | -1.251478 | 5245.291610 | 0.115729839 | -2.79645497 | 0.005166661 | 0.218089973 | 12.509628 | 12.185860 | 12.452831 | 12.433089 | 12.192259 | 12.313928 | 6792 | 6220 | 3751 | 4692 |
| 3322 | gene-HSP90AA | HSP90AA3P | heat shock protein 90 alpha family class A member 3, pseudogene | pseudogene | . | 5254 | . | | . | 0.249378 | 0.157377 | 0.193434 | 0.185450 | 0.580312 | 0.363784 | 0.434061 | 0.424009 | -1.056472 | 18.218418 | 0.725978686 | -0.10917012 | 0.913067562 | NA | 4.225252 | 4.148375 | 4.206828 | 4.172879 | 4.188499 | 4.182898 | 26 | 16 | 16 | 16 |
| 3324 | gene-HSP90AA | HSP90AA2P | heat shock protein 90 alpha family class A member 2, pseudogene | pseudogene | . | 5256 | 140575 | . | . | 0.657528 | 0.477179 | 0.386236 | 0.458815 | 1.530094 | 1.103020 | 0.866702 | 1.049026 | -1.351026 | 45.399291 | 0.474572924 | -0.91462417 | 0.360388968 | 0.998865404 | 5.705258 | 5.271449 | 5.563475 | 5.498384 | 5.469248 | 5.491253 | 70 | 47 | 31 | 39 |
| 3326 | NM_00127196 | HSP90AB1 | heat shock protein 90 alpha family class B member 1 | protein_coding | NP_001258898.1;NP_001258899.1;NP_001258900.1;NP_001258901.1;NI | 5258 | 140572 | . | . | 490.294696 | 514.293237 | 445.944223 | 527.642143 | 1140.935935 | 1188.811866 | 1000.685922 | 1206.391117 | -1.046335 | 39379.142057 | 0.1112482 | -0.58738175 | 0.55694735 | 0.998865404 | 15.297459 | 15.232089 | 15.266497 | 15.300462 | 15.174103 | 15.310327 | 45464 | 44305 | 30824 | 38932 |
| 3327 | gene-HSP90AB | HSP90AB3P | heat shock protein 90 alpha family class B member 3, pseudogene | pseudogene | . | 5259 | . | | . | 0.886193 | 0.815779 | 0.643237 | 0.731997 | 2.062207 | 1.885709 | 1.443405 | 1.673623 | -1.244208 | 61.358312 | 0.402698864 | -0.78278858 | 0.433751277 | 0.998865404 | 6.088358 | 5.772795 | 5.975617 | 5.953082 | 5.907237 | 5.927457 | 82 | 70 | 45 | 54 |
| 3329 | NM_002156,N | HSPD1 | heat shock protein family D (Hsp60) member 1 | protein_coding | NP_002147.2;NP_955472.1 | 5261 | 118190 | . | . | 361.800421 | 355.841986 | 304.855558 | 311.442675 | 841.924419 | 822.544726 | 684.086979 | 712.076678 | -1.180662 | 23342.498740 | 0.098244115 | -2.43878616 | 0.014736686 | 0.406008931 | 14.625501 | 14.385906 | 14.588966 | 14.566837 | 14.430580 | 14.439629 | 29475 | 26928 | 18517 | 20178 |
| 3336 | NM_002157 | HSPE1 | heat shock protein family E (Hsp10) member 1 | protein_coding | NP_002148.1 | 5269 | 600141 | . | . | 330.801331 | 346.000854 | 297.669861 | 335.137512 | 769.788330 | 799.796448 | 667.962524 | 766.252075 | -1.084685 | 5681.784900 | 0.110821596 | -1.05824211 | 0.289945077 | 0.998865404 | 12.529634 | 12.412235 | 12.489530 | 12.520287 | 12.390435 | 12.480828 | 6685 | 6496 | 4483 | 5387 |
| 3337 | NM_00130091 | DNAJB1 | DnaJ heat shock protein family (Hsp40) member B1 | protein_coding | NP_001287843.1;NP_001300893.1;NP_006136.1;XP_006722796.1;XP_00 | 5270 | 604572 | . | . | 52.497850 | 57.269108 | 46.760264 | 56.440640 | 122.164653 | 132.380113 | 104.928677 | 129.044816 | -1.074652 | 3726.143203 | 0.125611199 | -0.82691331 | 0.408286196 | 0.998865404 | 11.914593 | 11.810464 | 11.857780 | 11.923985 | 11.753408 | 11.904555 | 4269 | 4331 | 2836 | 3677 |
| 3338 | NM_00130798 | DNAJC4 | DnaJ heat shock protein family (Hsp40) member C4 | protein_coding | NP_001294909.1;NP_001294910.1;NP_005519.2 | 5271 | 604189 | . | . | 7.983075 | 8.807758 | 8.117949 | 8.782140 | 18.576944 | 20.359529 | 18.216442 | 20.079321 | 1.000484 | 347.630334 | 0.193343643 | 0.003612962 | 0.99711728 | 0.999862634 | 8.441413 | 8.441407 | 8.408393 | 8.472174 | 8.421656 | 8.459874 | 378 | 396 | 292 | 337 |
| 3339 | NM_00129186 | HSPG2 | heparan sulfate proteoglycan 2 | protein_coding | NP_001278789.1;NP_005520.4;XP_011539620.1;XP_016856609.1;XP_01 | 5273 | 142461 | . | . | 21.519180 | 21.860853 | 14.008061 | 16.406941 | 50.076018 | 50.532338 | 31.433682 | 37.512517 | -1.446705 | 8253.482390 | 0.112796881 | -4.72327602 | 2.32076E-06 | 0.000561989 | 13.252763 | 12.719882 | 13.143848 | 13.149921 | 12.771197 | 12.892339 | 11200 | 10569 | 5433 | 6792 |
| 3340 | NM_00130106 | NDST1 | N-deacetylase and N-sulfotransferase 1 | protein_coding | NP_001287992.1;NP_001534.1 | 7680 | 600853 | . | . | 30.118821 | 29.589624 | 29.243773 | 32.592459 | 70.087736 | 68.397740 | 65.622175 | 74.518779 | 1.021504 | 7574.181466 | 0.10762397 | 0.285200035 | 0.775490894 | 0.998865404 | 12.871474 | 12.902112 | 12.888224 | 12.866642 | 12.853039 | 12.936260 | 8739 | 7983 | 6330 | 7528 |
| 3344 | NM_00137544 | FOXN2 | forkhead box N2 | protein_coding | NP_001362371.1;NP_001362372.1;NP_001362373.1;NP_001362374.1;NI | 5281 | 143089 | . | . | 1.973426 | 1.582381 | 2.174106 | 1.163538 | 4.592245 | 3.657743 | 4.878629 | 2.660291 | -1.121942 | 297.868474 | 0.265155868 | -0.62604027 | 0.531288526 | 0.998865404 | 8.298098 | 8.134320 | 8.283482 | 8.189111 | 8.320847 | 8.005893 | 393 | 311 | 318 | 184 |
| 3357 | NM_000867,N | HTR2B | 5-hydroxytryptamine receptor 2B | protein_coding | NP_000858.3;NP_001307687.1;XP_005246577.1;XP_006712545.1 | 5294 | 601122 | . | . | 0.407791 | 0.699699 | 0.516106 | 0.409259 | 0.948948 | 1.617386 | 1.158127 | 0.935722 | -1.068562 | 39.777223 | 0.517808534 | -0.18476104 | 0.853416469 | 0.998865404 | 5.360655 | 5.265518 | 5.277382 | 5.347909 | 5.331335 | 5.274037 | 37 | 54 | 40 | 29 |
| 3358 | NM_000868,N | HTR2C | 5-hydroxytryptamine receptor 2C | protein_coding | NP_000859.1;NP_001243689.1;NP_001243690.1 | 5295 | 312861 | . | . | 0.248156 | 0.235515 | 0.337522 | 0.234915 | 0.577468 | 0.544403 | 0.757390 | 0.537105 | 1.166663 | 39.842061 | 0.503636175 | 0.441564683 | 0.658804245 | 0.998865404 | 5.199399 | 5.424287 | 5.299819 | 5.295159 | 5.353453 | 5.287792 | 43 | 39 | 45 | 32 |
| 3362 | NM_000871 | HTR6 | 5-hydroxytryptamine receptor 6 | protein_coding | NP_000862.1 | 5301 | 601109 | . | . | 0.237235 | 0.191715 | 0.198029 | 0.342601 | 0.552054 | 0.443158 | 0.444371 | 0.783316 | 1.249590 | 29.127505 | 0.590473162 | 0.544401414 | 0.586165273 | 0.998865404 | 4.695638 | 5.015311 | 4.856676 | 4.833690 | 4.838891 | 4.903226 | 33 | 25 | 21 | 38 |
| 3363 | NM_000872,N | HTR7 | 5-hydroxytryptamine receptor 7 | protein_coding | NP_000863.1;NP_062873.1;NP_062874.1;XP_024303741.1 | 5302 | 182137 | . | . | 3.537806 | 2.927763 | 3.313950 | 2.656988 | 8.232622 | 6.767653 | 7.436410 | 6.074886 | -1.110734 | 347.084957 | 0.209534703 | -0.72309433 | 0.469621931 | 0.998865404 | 8.512177 | 8.362218 | 8.521858 | 8.405781 | 8.469117 | 8.339032 | 462 | 355 | 320 | 272 |
| 3364 | NM_00136368 | HUS1 | HUS1 checkpoint clamp component | protein_coding | NP_001350612.1;NP_004498.1 | 5309 | 603760 | . | . | 6.711316 | 7.647920 | 7.582539 | 7.912572 | 15.617507 | 17.678506 | 17.015000 | 18.091156 | 1.077483 | 599.898925 | 0.157271319 | 0.684580091 | 0.493608934 | 0.998865404 | 9.173813 | 9.281305 | 9.198741 | 9.206860 | 9.243438 | 9.261071 | 661 | 627 | 534 | 591 |
| 3371 | NM_002160,XI | TNC | tenascin C | protein_coding | NP_002151.2;XP_005252029.1;XP_005252030.1;XP_005252031.1;XP_00 | 5318 | 187380 | . | . | 332.922719 | 340.318535 | 289.170933 | 326.843676 | 774.724832 | 786.661584 | 648.891189 | 747.289225 | -1.105289 | 74540.010432 | 0.105363368 | -1.37071897 | 0.170462578 | 0.946389416 | 16.256138 | 16.111704 | 16.220357 | 16.233816 | 16.086136 | 16.191961 | 89430 | 85106 | 57787 | 70472 |
| 3376 | NM_00137429 | IARS1 | isoleucyl-tRNA synthetase 1 | protein_coding | NP_001361228.1;NP_001361229.1;NP_001361230.1;NP_001365498.1;NI | 5330 | 600709 | . | . | 39.163254 | 42.626572 | 35.044570 | 40.575686 | 91.134500 | 98.533229 | 78.639001 | 92.771488 | -1.100077 | 5568.861214 | 0.114671341 | -1.1999884 | 0.230143845 | 0.985321181 | 12.510402 | 12.372644 | 12.450565 | 12.511280 | 12.347367 | 12.452277 | 6478 | 6521 | 4321 | 5285 |
| 3382 | NM_00113602 | ICA1 | islet cell autoantigen 1 | protein_coding | NP_001129492.1;NP_001263407.1;NP_001337748.1;NP_001337749.1;NI | 5343 | 147625 | . | . | 1.713938 | 2.773452 | 1.646557 | 2.001818 | 3.988407 | 6.410959 | 3.694828 | 4.576918 | -1.243929 | 149.717286 | 0.301028607 | -1.04609373 | 0.295517758 | 0.998865404 | 7.376075 | 7.058700 | 7.168214 | 7.345747 | 7.132702 | 7.217788 | 151 | 217 | 102 | 140 |
| 3383 | NM_000201 | ICAM1 | intercellular adhesion molecule 1 | protein_coding | NP_000192.2 | 5344 | 147840 | . | . | 95.637016 | 105.351570 | 109.930321 | 103.591545 | 222.551270 | 243.524873 | 246.680435 | 236.849777 | 1.048008 | 9576.454804 | 0.107703725 | 0.628108305 | 0.529932993 | 0.998865404 | 13.191053 | 13.258705 | 13.166489 | 13.241316 | 13.273774 | 13.214366 | 10294 | 10536 | 8819 | 8869 |
| 3385 | NM_00132060 | ICAM3 | intercellular adhesion molecule 3 | protein_coding | NP_001307534.1;NP_001307535.1;NP_001307537.1;NP_002153.2 | 5346 | 146631 | . | . | 2.252877 | 2.463720 | 2.416069 | 2.357516 | 5.242537 | 5.695000 | 5.421589 | 5.390179 | 1.000769 | 152.237288 | 0.265933848 | 0.004171919 | 0.9966713 | 0.999862634 | 7.249690 | 7.250677 | 7.232998 | 7.265915 | 7.257351 | 7.242361 | 168 | 171 | 135 | 140 |
| 3396 | NM_00130326 | MRPL58 | mitochondrial ribosomal protein L58 | protein_coding | NP_001290194.1;NP_001536.1;XP_016880063.1 | 5359 | 603000 | . | . | 13.516283 | 14.404988 | 11.643710 | 14.222336 | 31.452947 | 33.297772 | 26.128146 | 32.517684 | -1.092922 | 375.806175 | 0.192525857 | -0.66583198 | 0.5055185 | 0.998865404 | 8.617005 | 8.487792 | 8.562596 | 8.594501 | 8.472714 | 8.578185 | 440 | 435 | 283 | 368 |
| 3397 | NM_002165,N | ID1 | inhibitor of DNA binding 1, HLH protein | protein_coding | NP_002156.2;NP_851998.1 | 5360 | 600349 | . | . | 25.121507 | 27.698630 | 1.579560 | 6.365143 | 58.458777 | 64.026628 | 3.544488 | 14.553145 | -6.465247 | 487.289353 | 0.876068393 | -3.0736248 | 0.002114752 | 0.116759697 | 9.721249 | 7.027808 | 9.132454 | 9.202509 | 7.897084 | 8.284212 | 928 | 952 | 47 | 194 |
| 3398 | NM_002166 | ID2 | inhibitor of DNA binding 2 | protein_coding | NP_002157.2 | 5361 | 600386 | . | . | 17.920570 | 15.776475 | 6.054732 | 8.351913 | 41.701900 | 36.468029 | 13.586643 | 19.095655 | -2.367529 | 488.523032 | 0.194768411 | -6.38390127 | 1.72632E-10 | 1.9857E-07 | 9.424391 | 8.179606 | 9.171488 | 9.072210 | 8.512132 | 8.666296 | 845 | 691 | 213 | 314 |
| 3399 | NM_002167 | ID3 | inhibitor of DNA binding 3, HLH protein | protein_coding | NP_002158.3 | 5362 | 600277 | . | . | 100.500473 | 97.360855 | 7.561135 | 23.767311 | 233.868744 | 225.053986 | 16.966969 | 54.341133 | -6.411592 | 1704.396301 | 0.738117238 | -3.63178419 | 0.000281468 | 0.026793856 | 11.525979 | 8.845038 | 11.049408 | 11.011796 | 9.323588 | 9.886508 | 3464 | 3118 | 195 | 652 |
| 3400 | NM_001546 | ID4 | inhibitor of DNA binding 4, HLH protein | protein_coding | NP_001537.1 | 5363 | 600581 | . | . | 2.968766 | 2.195060 | 2.003710 | 1.818221 | 6.908442 | 5.073978 | 4.496268 | 4.157146 | -1.371347 | 272.019799 | 0.22993726 | -1.98138058 | 0.04754861 | 0.659753773 | 8.296881 | 7.842648 | 8.242250 | 8.068340 | 8.021020 | 7.971396 | 418 | 287 | 210 | 204 |
| 3416 | NM_00116594 | IDE | insulin degrading enzyme | protein_coding | NP_001159418.1;NP_001309722.1;NP_001309723.1;NP_001309724.1;NI | 5381 | 146680 | . | . | 5.704890 | 4.909930 | 5.237943 | 4.968790 | 13.275515 | 11.349525 | 11.753794 | 11.360548 | -1.018145 | 949.760365 | 0.14351049 | -0.18077326 | 0.856545552 | 0.998865404 | 9.904105 | 9.878622 | 9.936872 | 9.856294 | 9.908322 | 9.860006 | 1164 | 977 | 846 | 853 |
| 3417 | NM_00128238 | IDH1 | isocitrate dehydrogenase (NADP(+)) 1 | protein_coding | NP_001269315.1;NP_001269316.1;NP_005887.2 | 5382 | 147700 | . | . | 76.248106 | 68.582344 | 73.630013 | 80.086875 | 177.432491 | 158.531153 | 165.223617 | 183.109136 | 1.046810 | 5403.418815 | 0.114107021 | 0.578400155 | 0.56299399 | 0.998865404 | 12.366275 | 12.432283 | 12.426929 | 12.329874 | 12.385512 | 12.449411 | 6432 | 5367 | 4624 | 5378 |
| 3418 | NM_00128991 | IDH2 | isocitrate dehydrogenase (NADP(+)) 2 | protein_coding | NP_001276839.1;NP_001277043.1;NP_002159.2 | 5383 | 147650 | . | . | 15.352675 | 17.520595 | 12.465479 | 17.110231 | 35.726308 | 40.499641 | 27.972173 | 39.120513 | -1.126654 | 1292.784033 | 0.160802541 | -1.06991324 | 0.284658364 | 0.998865404 | 10.420079 | 10.247281 | 10.328342 | 10.423227 | 10.165869 | 10.393619 | 1481 | 1569 | 896 | 1313 |
| 3419 | NM_005530,XI | IDH3A | isocitrate dehydrogenase (NAD(+)) 3 catalytic subunit alpha | protein_coding | NP_005521.1;XP_005254393.1;XP_024305679.1;XP_024305680.1 | 5384 | 601149 | . | . | 21.497752 | 18.924574 | 16.968929 | 16.287025 | 50.026157 | 43.744996 | 38.077782 | 37.238349 | -1.067019 | 2093.568492 | 0.122819581 | -0.76197789 | 0.446073198 | 0.998865404 | 11.077651 | 10.984337 | 11.093290 | 11.018626 | 11.033045 | 10.976224 | 2604 | 2224 | 1825 | 1831 |
| 3420 | NM_00125838 | IDH3B | isocitrate dehydrogenase (NAD(+)) 3 non-catalytic subunit beta | protein_coding | NP_001245313.1;NP_001317692.1;NP_008830.2;NP_777280.1 | 5385 | 604526 | . | . | 55.349569 | 57.176461 | 56.398930 | 63.048484 | 128.800723 | 132.165950 | 126.557560 | 144.152878 | 1.045357 | 2728.294813 | 0.117622796 | 0.544081347 | 0.586385495 | 0.998865404 | 11.381535 | 11.445328 | 11.382313 | 11.407751 | 11.388199 | 11.472291 | 3024 | 2924 | 2300 | 2749 |
| 3421 | NM_004135,N | IDH3G | isocitrate dehydrogenase (NAD(+)) 3 non-catalytic subunit gamma | protein_coding | NP_004126.1;NP_777358.1 | 5386 | 300089 | . | . | 28.901062 | 28.976545 | 27.159491 | 32.330058 | 67.253956 | 66.980583 | 60.945104 | 73.918839 | 1.015017 | 1254.712747 | 0.138630223 | 0.155113113 | 0.876732145 | 0.998865404 | 10.282544 | 10.303662 | 10.293465 | 10.279957 | 10.234075 | 10.357957 | 1447 | 1332 | 1009 | 1282 |
| 3422 | NM_00131795 | IDI1 | isopentenyl-diphosphate delta isomerase 1 | protein_coding | NP_001304884.1;NP_001304885.1;NP_001304886.1;NP_004499.2;XP_01 | 5387 | 604055 | . | . | 18.182786 | 15.489285 | 24.476590 | 20.173733 | 42.312091 | 35.804176 | 54.924761 | 46.124845 | 1.311872 | 1825.158805 | 0.143815967 | 2.723114769 | 0.006466959 | 0.253589295 | 10.624454 | 11.016593 | 10.771160 | 10.653140 | 11.009481 | 10.839573 | 1958 | 1572 | 1990 | 1739 |
| 3423 | NM_000202,N | IDS | iduronate 2-sulfatase | protein_coding | NP_000193.1;NP_001160022.1;NP_006114.1 | 5389 | 300823 | . | . | 15.783876 | 16.228246 | 19.055177 | 17.830257 | 36.729730 | 37.512319 | 42.759267 | 40.766765 | 1.070609 | 3667.183952 | 0.111712886 | 0.881118956 | 0.378253438 | 0.998865404 | 11.790377 | 11.888857 | 11.792849 | 11.829398 | 11.896285 | 11.838238 | 3988 | 3907 | 3413 | 3431 |
| 3425 | NM_000203,N | IDUA | alpha-L-iduronidase | protein_coding | NP_000194.2;NP_001350505.1;XP_011511763.1;XP_016863652.1 | 5391 | 252800 | . | . | 4.123232 | 4.504832 | 5.102343 | 3.795156 | 9.594932 | 10.413119 | 11.449513 | 8.677173 | 1.037401 | 319.422270 | 0.210962795 | 0.251107159 | 0.801731261 | 0.998865404 | 8.292008 | 8.346126 | 8.303097 | 8.308997 | 8.399490 | 8.247656 | 359 | 340 | 322 | 263 |
| 3428 | NM_00120656 | IFI16 | interferon gamma inducible protein 16 | protein_coding | NP_001193496.1;NP_001351796.1;NP_001363516.1;NP_001363517.1;NI | 5395 | 147586 | . | . | 87.837694 | 86.374421 | 104.483542 | 102.349125 | 204.401929 | 199.658352 | 234.458045 | 234.009113 | 1.166772 | 8296.233735 | 0.102765937 | 2.165329722 | 0.030362444 | 0.555089817 | 12.902672 | 13.125235 | 12.963400 | 12.934712 | 13.094176 | 13.065626 | 8969 | 8121 | 7899 | 8237 |
| 3430 | NM_00133023 | IFI35 | interferon induced protein 35 | protein_coding | NP_001317159.1;NP_005524.2;XP_016880073.1;XP_016880074.1 | 5399 | 600735 | . | . | 17.613775 | 16.534521 | 14.950387 | 20.808756 | 40.987978 | 38.220284 | 33.548234 | 47.576748 | 1.030380 | 646.756842 | 0.180518646 | 0.239183084 | 0.810963619 | 0.998865404 | 9.315722 | 9.358125 | 9.346786 | 9.300289 | 9.230594 | 9.445900 | 759 | 664 | 480 | 713 |
| 3431 | NM_00118501 | SP110 | SP110 nuclear body protein | protein_coding | NP_001171944.1;NP_001365371.1;NP_001365372.1;NP_001365373.1;NI | 5401 | 604457 | . | . | 7.031478 | 6.648468 | 7.978497 | 8.885646 | 16.362541 | 15.368235 | 17.903517 | 20.315975 | 1.193368 | 532.001894 | 0.171663808 | 1.48568882 | 0.137361449 | 0.898578239 | 8.922588 | 9.176803 | 8.968140 | 9.017327 | 9.052343 | 9.158469 | 537 | 544 | 463 | 586 |
| 3433 | NM_001547 | IFIT2 | interferon induced protein with tetratricopeptide repeats 2 | protein_coding | NP_001538.4 | 5409 | 147040 | . | . | 0.958033 | 0.814779 | 0.973902 | 1.089803 | 2.229383 | 1.883398 | 2.185408 | 2.491706 | 1.147752 | 101.805309 | 0.320759139 | 0.61981262 | 0.535381159 | 0.998865404 | 6.566740 | 6.765741 | 6.669179 | 6.620936 | 6.672051 | 6.703350 | 118 | 94 | 90 | 107 |
| 3434 | NM_00127092 | IFIT1 | interferon induced protein with tetratricopeptide repeats 1 | protein_coding | NP_001257856.1;NP_001257857.1;NP_001257858.1;NP_001257859.1;NI | 5407 | 147690 | . | . | 1.595904 | 1.359462 | 1.262480 | 1.379190 | 3.713733 | 3.142457 | 2.832969 | 3.153354 | -1.135494 | 188.798914 | 0.247275598 | -0.74135729 | 0.458476833 | 0.998865404 | 7.649374 | 7.466231 | 7.624924 | 7.549737 | 7.516772 | 7.547701 | 250 | 199 | 148 | 172 |
| 3437 | NM_00103168 | IFIT3 | interferon induced protein with tetratricopeptide repeats 3 | protein_coding | NP_001026853.1;NP_001276687.1;NP_001276688.1;NP_001540.2 | 5411 | 604650 | . | . | 4.060573 | 3.885542 | 3.741306 | 4.114339 | 9.449121 | 8.981606 | 8.395382 | 9.406948 | -1.025601 | 295.482131 | 0.203433986 | -0.17926772 | 0.85772749 | 0.998865404 | 8.225170 | 8.188450 | 8.226960 | 8.200356 | 8.177915 | 8.222035 | 353 | 315 | 243 | 285 |
| 3454 | NM_000629,XI | IFNAR1 | interferon alpha and beta receptor subunit 1 | protein_coding | NP_000620.2;XP_005261021.1;XP_011527854.1 | 5432 | 107450 | . | . | 8.371830 | 7.461481 | 7.590459 | 6.352915 | 19.481593 | 17.247549 | 17.032768 | 14.525187 | -1.198945 | 1379.349395 | 0.151746302 | -1.72501674 | 0.084524456 | 0.788061307 | 10.554411 | 10.293339 | 10.539924 | 10.443220 | 10.443461 | 10.255946 | 1838 | 1523 | 1228 | 1029 |
| 3455 | NM_000874,N | IFNAR2 | interferon alpha and beta receptor subunit 2 | protein_coding | NP_000865.2;NP_001276054.1;NP_001276055.1;NP_001276057.1;NP_9! | 5433 | 602376 | . | . | 7.337756 | 7.481492 | 6.526895 | 6.600463 | 17.075259 | 17.293806 | 14.646164 | 15.091175 | -1.171699 | 892.722029 | 0.140454739 | -1.62758659 | 0.103612575 | 0.826207034 | 9.911821 | 9.683271 | 9.849538 | 9.862971 | 9.751451 | 9.733560 | 1099 | 1049 | 724 | 761 |
| 3459 | NM_000416,N | IFNGR1 | interferon gamma receptor 1 | protein_coding | NP_000407.1;NP_001350455.1;NP_001350456.1;XP_006715533.1;XP_01 | 5439 | 107470 | . | . | 14.045862 | 14.587300 | 13.026529 | 11.613584 | 32.685299 | 33.719199 | 29.231155 | 26.553081 | -1.155830 | 891.838029 | 0.152024171 | -1.37431337 | 0.169344419 | 0.946181982 | 9.901022 | 9.692743 | 9.853322 | 9.845062 | 9.811591 | 9.674371 | 1107 | 1026 | 790 | 700 |
| 3460 | NM_00132912 | IFNGR2 | interferon gamma receptor 2 | protein_coding | NP_001316057.1;NP_005525.2;XP_011527855.1;XP_011527856.1 | 5440 | 147569 | . | . | 42.675661 | 42.034687 | 40.731129 | 50.188617 | 99.308022 | 97.165060 | 91.399466 | 114.750301 | 1.051672 | 2354.953110 | 0.131347517 | 0.553381414 | 0.580002263 | 0.998865404 | 11.164831 | 11.237226 | 11.184212 | 11.174930 | 11.134524 | 11.299033 | 2659 | 2463 | 1886 | 2493 |
| 3475 | NM_00100724 | IFRD1 | interferon related developmental regulator 1 | protein_coding | NP_001007246.1;NP_001184008.1;NP_001184009.1;NP_001541.2 | 5456 | 603502 | . | . | 18.957363 | 17.256013 | 18.363641 | 16.159005 | 44.114563 | 39.888042 | 41.207474 | 36.945643 | -1.059567 | 1840.383285 | 0.132080963 | -0.63200031 | 0.527386676 | 0.998865404 | 10.886716 | 10.803670 | 10.906192 | 10.826602 | 10.878269 | 10.761739 | 2289 | 1941 | 1670 | 1551 |
| 3476 | NM_00137019 | IGBP1 | immunoglobulin binding protein 1 | protein_coding | NP_001357121.1;NP_001357122.1;NP_001357123.1;NP_001542.1;XP_01 | 5461 | 300139 | . | . | 12.833167 | 13.192485 | 11.681797 | 13.040332 | 29.863310 | 30.495019 | 26.213613 | 29.815171 | -1.071374 | 714.112068 | 0.158998443 | -0.62555463 | 0.531607105 | 0.998865404 | 9.529149 | 9.429131 | 9.515202 | 9.490029 | 9.391578 | 9.514737 | 866 | 783 | 545 | 705 |
| 3480 | NM_000875,N | IGF1R | insulin like growth factor 1 receptor | protein_coding | NP_000866.1;NP_001278787.1;XP_011519818.1;XP_011519819.1;XP_01 | 5465 | 147370 | . | . | 18.487421 | 22.826502 | 19.659964 | 20.740612 | 43.020984 | 52.764482 | 44.116389 | 47.420947 | -1.063174 | 6662.854538 | 0.103843439 | -0.85106145 | 0.39473522 | 0.998865404 | 12.745441 | 12.657055 | 12.710298 | 12.743845 | 12.682406 | 12.668495 | 7751 | 7556 | 5663 | 6005 |
| 3481 | NM_000612,N | IGF2 | insulin like growth factor 2 | protein_coding | NP_000603.1;NP_001007140.2;NP_001121070.1;NP_001278790.1;NP_0( | 5466 | 147470 | . | . | 0.038885 | 0.272383 | 0.024609 | 0.021892 | 0.090488 | 0.629625 | 0.055222 | 0.050054 | -6.209356 | 16.001208 | 1.008611412 | -2.61195104 | 0.009002715 | NA | 4.784982 | 2.148375 | 3.913223 | 4.090864 | 3.905686 | 3.903112 | 8 | 52 | 4 | 4 |
| 3482 | NM_000876 | IGF2R | insulin like growth factor 2 receptor | protein_coding | NP_000867.3 | 5467 | 147280 | . | . | 35.081573 | 36.212280 | 32.476410 | 36.820919 | 81.636261 | 83.706299 | 72.876122 | 84.186661 | -1.043461 | 15395.096696 | 0.106664652 | -0.5754214 | 0.565006335 | 0.998865404 | 13.940569 | 13.879144 | 13.917752 | 13.937690 | 13.840838 | 13.939311 | 17895 | 17162 | 12348 | 14939 |
| 3485 | NM_000597,N | IGFBP2 | insulin like growth factor binding protein 2 | protein_coding | NP_000588.3;NP_001300919.1;NP_001300921.1;NP_001300922.1 | 5471 | 146731 | . | . | 20.417395 | 19.390232 | 15.420933 | 19.614565 | 47.512117 | 48.842932 | 34.604122 | 44.846372 | -1.152485 | 825.062841 | 0.159608402 | -1.28281206 | 0.199557927 | 0.96621494 | 9.787385 | 9.582034 | 9.756918 | 9.713188 | 9.552259 | 9.710611 | 1048 | 925 | 590 | 801 |
| 3486 | NM_000598,N | IGFBP3 | insulin like growth factor binding protein 3 | protein_coding | NP_000589.2;NP_001013416.1 | 5472 | 146732 | . | . | 31.063592 | 33.158646 | 22.292907 | 24.822697 | 72.286252 | 76.647696 | 50.024636 | 56.754147 | -1.381568 | 2270.154315 | 0.121024886 | -3.85298421 | 0.000116687 | 0.013534702 | 11.363087 | 10.896500 | 11.239903 | 11.286976 | 10.966457 | 11.041186 | 2947 | 2923 | 1577 | 1874 |
| 3488 | NM_000599 | IGFBP5 | insulin like growth factor binding protein 5 | protein_coding | NP_000590.1 | 5474 | 146734 | . | . | 0.312084 | 0.239846 | 0.179013 | 0.153174 | 0.726231 | 0.554414 | 0.401700 | 0.350214 | -1.665227 | 43.609812 | 0.483552184 | -1.52148843 | 0.128137312 | 0.876241045 | 5.767170 | 5.033773 | 5.514210 | 5.459988 | 5.416227 | 5.393093 | 71 | 51 | 31 | 28 |
| 3489 | NM_002178 | IGFBP6 | insulin like growth factor binding protein 6 | protein_coding | NP_002169.1 | 5475 | 146735 | . | . | 3.750710 | 3.219536 | 2.985159 | 2.709267 | 8.728057 | 7.442101 | 6.698611 | 6.194417 | -1.232102 | 94.007946 | 0.333249 | -0.90359376 | 0.366210849 | 0.998865404 | 6.696854 | 6.397016 | 6.610303 | 6.558970 | 6.537716 | 6.510354 | 129 | 103 | 77 | 75 |
| 3490 | NM_00125383 | IGFBP7 | insulin like growth factor binding protein 7 | protein_coding | NP_001240764.1;NP_001544.1 | 5476 | 602867 | . | . | 2.536007 | 2.049118 | 2.101742 | 2.064487 | 5.901392 | 4.736627 | 4.716249 | 4.720204 | -1.128902 | 96.083268 | 0.331612519 | -0.52748297 | 0.59785824 | 0.998865404 | 6.670433 | 6.496771 | 6.637765 | 6.569278 | 6.577025 | 6.559228 | 130 | 98 | 81 | 82 |
| 3491 | NM_001554 | CCN1 | cellular communication network factor 1 | protein_coding | NP_001545.2 | 2654 | 602369 | . | . | 20.297049 | 19.423403 | 7.814940 | 9.120841 | 47.232067 | 44.898064 | 17.536501 | 20.853718 | -2.376966 | 1008.559188 | 0.144825224 | -8.62502381 | 6.40793E-18 | 8.84487E-14 | 10.471649 | 9.221935 | 10.223180 | 10.179874 | 9.531170 | 9.618975 | 1678 | 1492 | 482 | 600 |
| 3493 | id-IGHA1 | IGHA1 | immunoglobulin heavy constant alpha 1 | C_region | . | 5478 | 146900 | . | IGHA1 | 0.101095 | 0.231738 | 0.061088 | 0.182082 | 0.235252 | 0.535673 | 0.137081 | 0.416309 | -1.348508 | 5.128606 | 1.379432903 | -0.3127114 | 0.754499933 | NA | 2.562483 | 2.121007 | 2.353321 | 2.375989 | 2.348694 | 2.366851 | 4 | 9 | 2 | 6 |
| 3503 | id-IGHG4 | IGHG4 | immunoglobulin heavy constant gamma 4 (G4m marker) | C_region | . | 5528 | 147130 | . | IGHG4 | 0.882975 | 1.268963 | 1.155549 | 1.060262 | 2.054719 | 2.933265 | 2.593019 | 2.424163 | 1.016724 | 34.089014 | 0.538639659 | 0.044423272 | 0.964567012 | 0.998865404 | 5.079808 | 5.102572 | 5.062257 | 5.111381 | 5.094344 | 5.084374 | 32 | 43 | 31 | 31 |
| 3508 | NM_002180,XI | IGHMBP2 | immunoglobulin mu DNA binding protein 2 | protein_coding | NP_002171.2;XP_005274031.1;XP_005274032.1;XP_005274033.1;XP_01 | 5542 | 600502 | . | . | 5.276844 | 5.423165 | 5.976888 | 5.098645 | 12.279434 | 12.535891 | 13.411963 | 11.657449 | 1.012012 | 606.116599 | 0.162784306 | 0.105824715 | 0.915721427 | 0.998865404 | 9.234575 | 9.252274 | 9.228173 | 9.248719 | 9.293748 | 9.197060 | 683 | 660 | 569 | 530 |
| 3516 | NM_00136357 | RBPJ | recombination signal binding protein for immunoglobulin kappa J | protein_coding | NP_001350506.1;NP_001361329.1;NP_001361330.1;NP_001361331.1;NI | 5724 | 147183 | . | . | 6.330380 | 6.569648 | 6.657688 | 4.976486 | 14.731057 | 15.186040 | 14.939660 | 11.378144 | -1.077750 | 1064.168878 | 0.157022767 | -0.68794026 | 0.491490401 | 0.998865404 | 10.108188 | 10.000839 | 10.080824 | 10.079508 | 10.120288 | 9.919745 | 1272 | 1190 | 1013 | 829 |
| 3550 | NM_006083,XI | IK | IK cytokine | protein_coding | NP_006074.2 | 5958 | 600549 | . | . | 53.069164 | 55.974863 | 55.278371 | 61.694916 | 123.494125 | 129.388400 | 124.043060 | 141.058097 | 1.056367 | 3355.403852 | 0.115282319 | 0.686236951 | 0.4925637 | 0.998865404 | 11.672268 | 11.751194 | 11.669049 | 11.708350 | 11.691846 | 11.774382 | 3668 | 3606 | 2848 | 3393 |
| 3551 | NM_00119072 | IKBKB | inhibitor of nuclear factor kappa B kinase subunit beta | protein_coding | NP_001177649.2;NP_001229707.1;NP_001547.1;XP_005273547.1;XP_00 | 5960 | 603258 | . | . | 9.445680 | 9.652753 | 12.001656 | 9.075964 | 21.980488 | 22.312763 | 26.931366 | 20.751114 | 1.079835 | 1199.380531 | 0.152170556 | 0.728203273 | 0.466489162 | 0.998865404 | 10.171314 | 10.282684 | 10.195322 | 10.196273 | 10.347817 | 10.150257 | 1327 | 1245 | 1224 | 1016 |
| 3553 | NM_000576,XI | IL1B | interleukin 1 beta | protein_coding | NP_000567.1;XP_016859477.1 | 5992 | 147720 | . | . | 6.795925 | 6.068405 | 6.087729 | 7.301169 | 15.814397 | 14.027390 | 13.660688 | 16.693256 | 1.026711 | 308.742079 | 0.206989305 | 0.183732463 | 0.854223341 | 0.998865404 | 8.251304 | 8.288967 | 8.294265 | 8.228593 | 8.229482 | 8.320900 | 372 | 309 | 249 | 318 |
| 3554 | NM_000877,N | IL1R1 | interleukin 1 receptor type 1 | protein_coding | NP_000868.1;NP_001276635.1;NP_001307907.1;NP_001307909.1;NP_0( | 5993 | 147810 | . | . | 0.233612 | 0.202053 | 0.310972 | 0.146711 | 0.543624 | 0.467056 | 0.697813 | 0.335438 | 1.019114 | 36.627061 | 0.551685684 | 0.049512555 | 0.960510832 | 0.998865404 | 5.178768 | 5.210731 | 5.196303 | 5.173289 | 5.238069 | 5.134188 | 45 | 36 | 44 | 22 |
| 3556 | NM_00116792 | IL1RAP | interleukin 1 receptor accessory protein | protein_coding | NP_001161400.1;NP_001161401.1;NP_001161402.1;NP_001161403.1;NI | 5995 | 602626 | . | . | 34.186429 | 31.716378 | 27.737711 | 25.973397 | 79.553222 | 73.313824 | 62.242614 | 59.385088 | -1.215970 | 3767.410679 | 0.119527505 | -2.36019375 | 0.018265393 | 0.455085227 | 12.013425 | 11.731543 | 11.999132 | 11.907024 | 11.837203 | 11.745575 | 5020 | 4218 | 3124 | 3008 |
| 3557 | NM_000577,N | IL1RN | interleukin 1 receptor antagonist | protein_coding | NP_000568.1;NP_001305843.1;NP_001366289.1;NP_776213.1;NP_7762 | 6000 | 147679 | . | . | 0.165824 | 0.045104 | 0.146442 | 0.073671 | 0.385879 | 0.104260 | 0.328611 | 0.168439 | -1.017597 | 7.887068 | 1.136210681 | -0.02214908 | 0.982329033 | NA | 2.987791 | 2.971139 | 2.995075 | 2.955959 | 2.985966 | 2.963974 | 14 | 4 | 9 | 5 |
| 3566 | NM_000418,N | IL4R | interleukin 4 receptor | protein_coding | NP_000409.1;NP_001244335.1;NP_001244336.1;NP_001244926.1;XP_0( | 6015 | 147781 | . | . | 0.372681 | 0.642475 | 0.139381 | 0.339313 | 0.867245 | 1.485110 | 0.312767 | 0.775800 | -1.753606 | 50.416939 | 0.493628066 | -1.64156871 | 0.100679416 | 0.816017607 | 6.006951 | 5.190803 | 5.679399 | 5.722785 | 5.542310 | 5.653063 | 67 | 76 | 19 | 48 |
| 3569 | NM_000600,N | IL6 | interleukin 6 | protein_coding | NP_000591.1;NP_001305024.1;NP_001358025.1;XP_005249802.1;XP_01 | 6018 | 147620 | . | . | 0.260694 | 0.284299 | 0.375825 | 0.212141 | 0.606645 | 0.657171 | 0.843341 | 0.485036 | -1.031565 | 14.124456 | 0.814734209 | -0.05503017 | 0.956114428 | NA | 3.841203 | 3.798723 | 3.822135 | 3.818258 | 3.827526 | 3.808930 | 17 | 15 | 14 | 11 |
| 3570 | NM_000565,N | IL6R | interleukin 6 receptor | protein_coding | NP_000556.1;NP_001193795.1;NP_001369698.1;NP_001369699.1;NP_0( | 6019 | 147880 | . | . | 5.793114 | 5.685419 | 5.020045 | 5.667451 | 13.480815 | 13.142100 | 11.264836 | 12.957955 | -1.071954 | 936.104832 | 0.14226664 | -0.70461237 | 0.481051496 | 0.998865404 | 9.919947 | 9.819353 | 9.897450 | 9.892738 | 9.805165 | 9.882623 | 1119 | 1042 | 743 | 893 |
| 3572 | NM_00119098 | IL6ST | interleukin 6 signal transducer | protein_coding | NP_001177910.1;NP_001351204.1;NP_001351205.1;NP_001351206.1;NI | 6021 | 600694 | . | . | 9.509461 | 8.148065 | 8.610541 | 6.529842 | 22.128907 | 18.834616 | 19.321806 | 14.929708 | -1.177305 | 2314.192965 | 0.152244144 | -1.54678223 | 0.121915767 | 0.864889544 | 11.289013 | 11.054015 | 11.299143 | 11.166411 | 11.212956 | 10.981477 | 3114 | 2482 | 2119 | 1702 |
| 3573 | gene-IL6STP1 | IL6STP1 | interleukin 6 signal transducer pseudogene 1 | pseudogene | . | 6022 | . | | . | 0.356943 | 0.133435 | 0.262484 | 0.223710 | 0.830620 | 0.308441 | 0.589007 | 0.511486 | -1.030020 | 48.392509 | 0.520014584 | -0.08205975 | 0.934599198 | 0.998865404 | 5.616002 | 5.577160 | 5.669032 | 5.500649 | 5.597402 | 5.568160 | 82 | 29 | 45 | 41 |
| 3574 | NM_000880,N | IL7 | interleukin 7 | protein_coding | NP_000871.1;NP_001186815.1;NP_001186816.1;NP_001186817.1;XP_01 | 6023 | 146660 | . | . | 0.010085 | 0.013965 | 0.012410 | 0.015829 | 0.023468 | 0.032282 | 0.027848 | 0.036191 | -1.017609 | 2.237600 | 1.972119472 | -0.01276997 | 0.989811317 | NA | 1.175403 | 1.148375 | 1.158018 | 1.163002 | 1.160711 | 1.159957 | 2 | 3 | 2 | 2 |
| 3575 | NM_002185,N | IL7R | interleukin 7 receptor | protein_coding | NP_002176.2;XP_005248356.1 | 6024 | 146661 | . | . | 0.775200 | 0.927209 | 0.290136 | 0.294635 | 1.803925 | 2.143285 | 0.651056 | 0.673649 | -2.934796 | 81.671043 | 0.368999857 | -4.20937846 | 2.56074E-05 | 0.004258544 | 6.928874 | 5.375024 | 6.418935 | 6.480893 | 6.185515 | 6.183886 | 128 | 143 | 36 | 39 |
| 3576 | NM_000584,N | CXCL8 | C-X-C motif chemokine ligand 8 | protein_coding | NP_000575.1;NP_001341769.1 | 6025 | 146930 | . | . | 14.858229 | 12.986673 | 12.465876 | 10.800849 | 34.575709 | 30.019278 | 27.973066 | 24.694858 | -1.215651 | 656.401396 | 0.165754066 | -1.69968487 | 0.089190217 | 0.802216049 | 9.492023 | 9.211202 | 9.469357 | 9.366270 | 9.336290 | 9.236421 | 890 | 720 | 555 | 513 |
| 3586 | NM_000572,N | IL10 | interleukin 10 | protein_coding | NP_000563.1;NP_001369553.1 | 5962 | 124092 | . | . | 0.360677 | 0.164491 | 0.260993 | 0.246383 | 0.839311 | 0.380229 | 0.585662 | 0.563325 | -1.153342 | 13.687410 | 0.842415233 | -0.24432231 | 0.806981209 | NA | 3.872825 | 3.669578 | 3.800712 | 3.760884 | 3.772179 | 3.772735 | 22 | 11 | 11 | 12 |
| 3588 | NM_000628 | IL10RB | interleukin 10 receptor subunit beta | protein_coding | NP_000619.3 | 5965 | 123889 | . | . | 19.341673 | 19.617527 | 17.657610 | 17.657427 | 45.008869 | 45.346794 | 39.623165 | 40.371613 | -1.118736 | 1123.429569 | 0.132236523 | -1.22408986 | 0.220918331 | 0.982008277 | 10.212349 | 10.050502 | 10.171907 | 10.176371 | 10.094465 | 10.086693 | 1362 | 1284 | 927 | 989 |
| 3589 | NM_000641,N | IL11 | interleukin 11 | protein_coding | NP_000632.1;NP_001254647.1 | 5966 | 147681 | . | . | 0.306445 | 0.274362 | 0.361719 | 0.230589 | 0.713109 | 0.634201 | 0.811687 | 0.527215 | 1.016406 | 22.125594 | 0.665109498 | 0.035297406 | 0.971842592 | NA | 4.454055 | 4.481107 | 4.470034 | 4.456851 | 4.486241 | 4.443175 | 27 | 22 | 24 | 16 |
| 3590 | NM_00114278 | IL11RA | interleukin 11 receptor subunit alpha | protein_coding | NP_001136256.1 | 5967 | 600939 | . | . | 1.975368 | 2.864210 | 2.734049 | 2.017121 | 4.596763 | 6.620750 | 6.135128 | 4.611907 | -1.043647 | 132.460186 | 0.309751952 | -0.19898022 | 0.842278216 | 0.998865404 | 7.079808 | 7.018368 | 6.982180 | 7.115459 | 7.085115 | 6.983490 | 128 | 172 | 129 | 104 |
| 3592 | NM_000882,N | IL12A | interleukin 12A | protein_coding | NP_000873.2;NP_001341511.1;NP_001341512.1 | 5969 | 161560 | . | . | 2.671566 | 2.890674 | 2.351291 | 2.455075 | 6.216845 | 6.681921 | 5.276228 | 5.613237 | -1.169227 | 116.979311 | 0.298572598 | -0.75544335 | 0.449983008 | 0.998865404 | 6.978727 | 6.752644 | 6.883739 | 6.909370 | 6.838450 | 6.849869 | 140 | 141 | 92 | 103 |
| 3595 | NM_00125821 | IL12RB2 | interleukin 12 receptor subunit beta 2 | protein_coding | NP_001245143.1;NP_001245144.1;NP_001245145.1;NP_001306162.1;NI | 5972 | 601642 | . | . | 7.255005 | 7.434199 | 6.895322 | 7.698105 | 16.882697 | 17.184484 | 15.472905 | 17.600802 | -1.011048 | 1197.595714 | 0.134919199 | -0.11748548 | 0.906475346 | 0.998865404 | 10.234024 | 10.217781 | 10.209525 | 10.249622 | 10.183474 | 10.257944 | 1350 | 1334 | 984 | 1172 |
| 3597 | NM_001560,XI | IL13RA1 | interleukin 13 receptor subunit alpha 1 | protein_coding | NP_001551.1;XP_011529638.1;XP_016884996.1 | 5974 | 300119 | . | . | 11.386887 | 11.435984 | 9.434120 | 8.124559 | 26.497752 | 26.434792 | 21.169891 | 18.575838 | -1.317940 | 1259.324370 | 0.138058604 | -2.88489652 | 0.003915423 | 0.182583072 | 10.483668 | 10.085865 | 10.394179 | 10.390897 | 10.240831 | 10.123829 | 1652 | 1542 | 1021 | 937 |
| 3598 | NM_000640 | IL13RA2 | interleukin 13 receptor subunit alpha 2 | protein_coding | NP_000631.1 | 5975 | 300130 | . | . | 0.916297 | 0.764470 | 0.502478 | 0.809642 | 2.132261 | 1.767107 | 1.127545 | 1.851150 | -1.275519 | 31.874705 | 0.562599987 | -0.62403928 | 0.532601791 | 0.998865404 | 5.160193 | 4.806913 | 5.025984 | 4.999011 | 4.957280 | 5.005590 | 45 | 35 | 19 | 32 |
| 3600 | NM_000585,N | IL15 | interleukin 15 | protein_coding | NP_000576.1;NP_751915.1 | 5977 | 600554 | . | . | 0.520531 | 0.662800 | 0.292285 | 0.411882 | 1.211299 | 1.532091 | 0.655880 | 0.941720 | -1.597680 | 35.019159 | 0.536591056 | -1.25976539 | 0.207754008 | 0.970482251 | 5.430072 | 4.750862 | 5.155304 | 5.172786 | 5.080719 | 5.120296 | 47 | 49 | 19 | 30 |
| 3601 | NM_00124353 | IL15RA | interleukin 15 receptor subunit alpha | protein_coding | NP_001230468.1;NP_001243694.1;NP_001338024.1;NP_001338025.1;NI | 5978 | 601070 | . | . | 0.150429 | 0.149338 | 0.184795 | 0.128459 | 0.350055 | 0.345201 | 0.414676 | 0.293706 | 1.077873 | 10.716304 | 0.929058349 | 0.116448879 | 0.907296799 | NA | 3.365657 | 3.475715 | 3.417624 | 3.416612 | 3.427222 | 3.414086 | 12 | 11 | 11 | 9 |
| 3603 | NM_00117212 | IL16 | interleukin 16 | protein_coding | NP_001165599.1;NP_001339613.1;NP_001339614.1;NP_001339615.1;NI | 5980 | 603035 | . | . | 14.793947 | 16.519675 | 19.077330 | 18.555561 | 34.426122 | 38.185970 | 42.808975 | 42.425089 | 1.178978 | 4913.834828 | 0.112019226 | 2.120501819 | 0.033963748 | 0.578053783 | 12.138946 | 12.376550 | 12.171538 | 12.206613 | 12.371341 | 12.280689 | 5083 | 4970 | 4871 | 4719 |
| 3607 | NM_004514,XI | FOXK2 | forkhead box K2 | protein_coding | NP_004505.2;XP_024306507.1 | 6036 | 147685 | . | . | 11.981715 | 12.255453 | 12.051657 | 13.770216 | 27.881941 | 28.329031 | 27.043571 | 31.483965 | 1.050216 | 2040.660065 | 0.123221881 | 0.573653655 | 0.566202196 | 0.998865404 | 10.959169 | 11.029612 | 10.970912 | 10.978166 | 10.965405 | 11.059383 | 2282 | 2158 | 1711 | 2075 |
| 3608 | NM_00126780 | ILF2 | interleukin enhancer binding factor 2 | protein_coding | NP_001254738.1;NP_004506.2 | 6037 | 603181 | . | . | 81.845844 | 84.743602 | 71.572192 | 80.634666 | 190.458665 | 195.888635 | 160.605927 | 184.361573 | -1.109526 | 4600.820210 | 0.112132144 | -1.337204 | 0.181156055 | 0.952877815 | 12.240767 | 12.090682 | 12.198106 | 12.220215 | 12.076868 | 12.166045 | 5500 | 5291 | 3588 | 4310 |
| 3609 | NM_00113767 | ILF3 | interleukin enhancer binding factor 3 | protein_coding | NP_001131145.1;NP_004507.2;NP_036350.2;NP_060090.2;NP_703194.1 | 6038 | 603182 | . | . | 48.252552 | 49.453216 | 50.651411 | 51.792766 | 112.285669 | 114.313323 | 113.660297 | 118.418005 | 1.016823 | 8056.356393 | 0.103207042 | 0.233210922 | 0.815597625 | 0.998865404 | 12.963824 | 12.987899 | 12.952397 | 12.984734 | 12.998110 | 12.967365 | 9024 | 8785 | 7192 | 7478 |
| 3611 | NM_00101479 | ILK | integrin linked kinase | protein_coding | NP_001014794.1;NP_001014795.1;NP_001265370.1;NP_001265371.1;NI | 6040 | 602366 | . | . | 32.023256 | 32.206682 | 29.442010 | 34.590348 | 74.519430 | 74.447192 | 66.067017 | 79.086721 | -1.025092 | 1773.545554 | 0.129602599 | -0.27586583 | 0.78265112 | 0.998865404 | 10.810351 | 10.774265 | 10.798933 | 10.804084 | 10.720053 | 10.840255 | 2061 | 1944 | 1406 | 1768 |
| 3612 | NM_00114487 | IMPA1 | inositol monophosphatase 1 | protein_coding | NP_001138350.1;NP_001138351.1;NP_005527.1 | 6050 | 602064 | . | . | 6.985237 | 6.238305 | 6.195717 | 5.392382 | 16.254933 | 14.420122 | 13.903011 | 12.329041 | -1.156258 | 653.114765 | 0.163562703 | -1.28063381 | 0.200322318 | 0.96621494 | 9.451733 | 9.243117 | 9.438467 | 9.354119 | 9.346736 | 9.249276 | 855 | 710 | 566 | 526 |
| 3613 | NM_014214,XI | IMPA2 | inositol monophosphatase 2 | protein_coding | NP_055029.1;XP_011523961.1;XP_011523963.1 | 6051 | 605922 | . | . | 9.057633 | 7.997616 | 6.942821 | 7.389284 | 21.077484 | 18.486848 | 15.579489 | 16.894718 | -1.205616 | 357.274393 | 0.192431702 | -1.40189948 | 0.160945265 | 0.933525128 | 8.609402 | 8.339798 | 8.570097 | 8.491661 | 8.412031 | 8.437515 | 479 | 394 | 275 | 311 |
| 3614 | NM_000883,N | IMPDH1 | inosine monophosphate dehydrogenase 1 | protein_coding | NP_000874.2;NP_001096075.1;NP_001136045.1;NP_001136046.1;NP_0( | 6052 | 146690 | . | . | 37.346095 | 39.755329 | 40.719431 | 41.978157 | 86.905897 | 91.896229 | 91.373221 | 95.978071 | 1.056955 | 3023.937608 | 0.112258468 | 0.711875639 | 0.476541789 | 0.998865404 | 11.521752 | 11.601570 | 11.517885 | 11.559644 | 11.574983 | 11.593424 | 3299 | 3254 | 2672 | 2946 |
| 3615 | NM_000884,XI | IMPDH2 | inosine monophosphate dehydrogenase 2 | protein_coding | NP_000875.2;XP_006713191.1;XP_016861838.1;XP_016861839.1 | 6053 | 146691 | . | . | 92.081212 | 98.118913 | 78.594150 | 98.613376 | 214.276776 | 226.806252 | 176.363002 | 225.467768 | -1.089092 | 4748.800345 | 0.12685313 | -0.97062059 | 0.331737251 | 0.998865404 | 12.273699 | 12.150360 | 12.223146 | 12.269519 | 12.082285 | 12.260377 | 5545 | 5490 | 3529 | 4719 |
| 3619 | NM_00104069 | INCENP | inner centromere protein | protein_coding | NP_001035784.1;NP_064623.2;XP_006718596.1;XP_011543297.1;XP_01 | 6058 | 604411 | . | . | 14.993888 | 16.678468 | 15.295282 | 17.469070 | 34.891392 | 38.553026 | 34.322168 | 39.940954 | 1.045651 | 2920.004391 | 0.111495348 | 0.577621112 | 0.563526027 | 0.998865404 | 11.479195 | 11.543596 | 11.501724 | 11.484565 | 11.523329 | 11.535878 | 3322 | 3048 | 2576 | 2820 |
| 3621 | NM_00126772 | ING1 | inhibitor of growth family member 1 | protein_coding | NP_001254657.1;NP_005528.4;NP_937860.1;NP_937861.1;NP_937862.1 | 6062 | 601566 | . | . | 3.665040 | 3.671265 | 3.763093 | 3.217758 | 8.528698 | 8.486293 | 8.444268 | 7.357023 | 1.000510 | 342.791259 | 0.194308036 | 0.00378823 | 0.996977437 | 0.999862634 | 8.421059 | 8.421314 | 8.431276 | 8.410158 | 8.384555 | 8.456154 | 402 | 363 | 279 | 342 |
| 3622 | NM_00129195 | ING2 | inhibitor of growth family member 2 | protein_coding | NP_001278888.1;NP_001555.1;XP_011530229.1 | 6063 | 604215 | . | . | 2.092400 | 2.025136 | 2.077010 | 2.025829 | 4.869104 | 4.681192 | 4.660748 | 4.631818 | -1.019741 | 157.092471 | 0.261401984 | -0.10788867 | 0.914084005 | 0.998865404 | 7.309355 | 7.281450 | 7.306984 | 7.291958 | 7.296686 | 7.286737 | 186 | 168 | 137 | 144 |
| 3623 | NM_002191 | INHA | inhibin subunit alpha | protein_coding | NP_002182.1 | 6065 | 147380 | . | . | 0.642630 | 0.720367 | 0.971704 | 0.505443 | 1.495426 | 1.665161 | 2.180476 | 1.155635 | 1.072282 | 30.276284 | 0.587740673 | 0.171307125 | 0.863982278 | 0.998865404 | 4.867222 | 4.971139 | 4.901975 | 4.913658 | 4.954918 | 4.876127 | 32 | 33 | 36 | 20 |
| 3624 | NM_002192,XI | INHBA | inhibin subunit beta A | protein_coding | NP_002183.1;XP_016867663.1;XP_016867664.1;XP_016867665.1 | 6066 | 147290 | . | . | 0.386870 | 0.270360 | 0.445003 | 0.457695 | 0.900263 | 0.624948 | 0.998575 | 1.046466 | 1.365522 | 77.124266 | 0.367059272 | 1.224470013 | 0.220774972 | 0.982008277 | 6.026236 | 6.476933 | 6.249770 | 6.189040 | 6.297337 | 6.301248 | 85 | 61 | 77 | 84 |
| 3628 | NM_00112892 | INPP1 | inositol polyphosphate-1-phosphatase | protein_coding | NP_001122400.1;NP_002185.1;XP_005246589.1;XP_024308643.1 | 6071 | 147263 | . | . | 11.673726 | 12.723366 | 14.393450 | 14.726211 | 27.165238 | 29.410631 | 32.298485 | 33.669732 | 1.160290 | 819.046635 | 0.146130905 | 1.467758481 | 0.142169839 | 0.903146012 | 9.566725 | 9.780934 | 9.588910 | 9.653216 | 9.723846 | 9.729725 | 833 | 856 | 762 | 828 |
| 3631 | NM_00113422 | INPP4A | inositol polyphosphate-4-phosphatase type I A | protein_coding | NP_001127696.1;NP_001127697.1;NP_001338353.1;NP_001338354.1;NI | 6074 | 600916 | . | . | 3.920895 | 4.854239 | 4.403205 | 5.020641 | 9.124081 | 11.220790 | 9.880666 | 11.479096 | 1.018555 | 1337.681225 | 0.135006559 | 0.196464303 | 0.844246771 | 0.998865404 | 10.372252 | 10.398665 | 10.325257 | 10.427618 | 10.411684 | 10.371298 | 1424 | 1526 | 1205 | 1232 |
| 3632 | NM_00132104 | INPP5A | inositol polyphosphate-5-phosphatase A | protein_coding | NP_001307971.1;NP_005530.3;XP_016871693.1;XP_016871694.1 | 6076 | 600106 | . | . | 3.138270 | 3.021705 | 2.701916 | 2.824987 | 7.302883 | 6.984806 | 6.063020 | 6.458998 | -1.104688 | 289.319907 | 0.203924676 | -0.70437073 | 0.481201932 | 0.998865404 | 8.246508 | 8.102966 | 8.217231 | 8.187977 | 8.145712 | 8.154368 | 359 | 319 | 237 | 260 |
| 3633 | NM_00129743 | INPP5B | inositol polyphosphate-5-phosphatase B | protein_coding | NP_001284363.1;NP_001337156.1;NP_001337157.1;NP_001352749.1;NI | 6077 | 147264 | . | . | 4.060085 | 3.442521 | 4.019685 | 3.993732 | 9.447988 | 7.957542 | 9.020056 | 9.131193 | 1.061881 | 534.013237 | 0.169221353 | 0.511883144 | 0.608732796 | 0.998865404 | 9.016527 | 9.103622 | 9.086948 | 8.990728 | 9.080940 | 9.076674 | 641 | 517 | 480 | 514 |
| 3636 | NM_001567,XI | INPPL1 | inositol polyphosphate phosphatase like 1 | protein_coding | NP_001558.3;XP_005274036.1;XP_011543301.1;XP_024304269.1;XP_02 | 6080 | 600829 | . | . | 36.162708 | 38.137660 | 42.837128 | 40.603594 | 84.152113 | 88.156910 | 96.125269 | 92.835291 | 1.111738 | 6055.518377 | 0.106542769 | 1.434324917 | 0.151479563 | 0.914154052 | 12.485593 | 12.638431 | 12.498817 | 12.535934 | 12.631748 | 12.581073 | 6457 | 6326 | 5710 | 5799 |
| 3638 | NM_00134659 | INSIG1 | insulin induced gene 1 | protein_coding | NP_001333519.1;NP_001333520.1;NP_001333521.1;NP_001333522.1;NI | 6083 | 602055 | . | . | 19.457814 | 16.600226 | 30.453451 | 24.237235 | 45.279134 | 38.372166 | 68.336665 | 55.415557 | 1.505115 | 2098.355269 | 0.145993488 | 4.040410884 | 5.33576E-05 | 0.007835058 | 10.709881 | 11.300230 | 10.907113 | 10.783597 | 11.261845 | 11.069199 | 2088 | 1658 | 2450 | 2087 |
| 3642 | NM_002196 | INSM1 | INSM transcriptional repressor 1 | protein_coding | NP_002187.1 | 6090 | 600010 | . | . | 0.384750 | 0.580907 | 0.478510 | 0.544133 | 0.895329 | 1.342793 | 1.073763 | 1.244095 | 1.046967 | 44.296115 | 0.480678429 | 0.137755804 | 0.890433415 | 0.998865404 | 5.436885 | 5.500628 | 5.427507 | 5.494436 | 5.459105 | 5.479301 | 40 | 56 | 37 | 45 |
| 3643 | NM_000208,N | INSR | insulin receptor | protein_coding | NP_000199.2;NP_001073285.1;XP_011526290.2;XP_011526291.2 | 6091 | 147670 | . | . | 0.930613 | 0.958582 | 0.744203 | 0.855989 | 2.165574 | 2.215807 | 1.669971 | 1.957116 | -1.197787 | 257.899747 | 0.215889553 | -1.20603831 | 0.227802718 | 0.982008277 | 8.135334 | 7.874198 | 8.050234 | 8.059078 | 7.932900 | 7.993189 | 321 | 306 | 191 | 234 |
| 3646 | NM_001568,XI | EIF3E | eukaryotic translation initiation factor 3 subunit E | protein_coding | NP_001559.1;XP_016868887.2 | 3277 | 602210 | . | . | 117.510075 | 111.074612 | 104.504066 | 93.869806 | 273.450788 | 256.753940 | 234.504097 | 214.622151 | -1.168445 | 6724.689876 | 0.112001979 | -2.00523344 | 0.044938097 | 0.649185583 | 12.823124 | 12.598658 | 12.804084 | 12.749098 | 12.693041 | 12.595632 | 8612 | 7569 | 5709 | 5475 |
| 3647 | gene-EIF3EP1 | EIF3EP1 | eukaryotic translation initiation factor 3 subunit E pseudogene 1 | pseudogene | . | 6102 | . | | . | 0.225985 | 0.397003 | 0.466530 | 0.409338 | 0.525877 | 0.917690 | 1.046880 | 0.935904 | 1.378135 | 17.718541 | 0.739433655 | 0.625772016 | 0.531464489 | NA | 3.898433 | 4.359293 | 4.108222 | 4.140822 | 4.154216 | 4.143482 | 13 | 20 | 19 | 18 |
| 3652 | NM_00114534 | IPP | intracisternal A particle-promoted polypeptide | protein_coding | NP_001138821.1;NP_005888.1;XP_005270896.1;XP_006710686.1;XP_01 | 6108 | 147485 | . | . | 3.115958 | 3.630603 | 2.317270 | 3.272831 | 7.250963 | 8.392301 | 5.199887 | 7.482940 | -1.289861 | 293.591760 | 0.212837036 | -1.72533733 | 0.084466699 | 0.788061307 | 8.370227 | 8.001629 | 8.234538 | 8.285020 | 8.070885 | 8.175859 | 364 | 373 | 199 | 266 |
| 3653 | NR_023915 | IPW | imprinted in Prader-Willi syndrome | lncRNA | . | 6109 | 601491 | . | . | 7.334908 | 6.302506 | 8.127082 | 7.074861 | 17.068632 | 14.568526 | 18.236937 | 16.175829 | 1.099231 | 1010.771189 | 0.151059936 | 0.903583351 | 0.366216373 | 0.998865404 | 9.911012 | 10.048209 | 9.999820 | 9.883729 | 10.066575 | 9.956655 | 1197 | 956 | 989 | 919 |
| 3654 | NM_00102524 | IRAK1 | interleukin 1 receptor associated kinase 1 | protein_coding | NP_001020413.1;NP_001020414.1;NP_001560.2;XP_005274725.1 | 6112 | 300283 | . | . | 25.552747 | 26.953317 | 24.591495 | 28.189211 | 59.462295 | 62.303800 | 55.182603 | 64.451282 | -1.010354 | 2926.178430 | 0.119944452 | -0.12389452 | 0.901398794 | 0.998865404 | 11.522334 | 11.507231 | 11.498695 | 11.537902 | 11.455777 | 11.561487 | 3306 | 3250 | 2367 | 2906 |
| 3655 | NM_000210,N | ITGA6 | integrin subunit alpha 6 | protein_coding | NP_000201.2;NP_001073286.1;NP_001303235.1;NP_001352458.1;NP_0( | 6142 | 147556 | . | . | 103.163963 | 95.118714 | 65.516123 | 64.567369 | 240.066794 | 219.871155 | 147.016271 | 147.625612 | -1.544148 | 14778.384212 | 0.105130661 | -5.96220971 | 2.48849E-09 | 1.90826E-06 | 14.130815 | 13.504041 | 14.051407 | 13.964777 | 13.649643 | 13.624075 | 21664 | 18413 | 10259 | 10723 |
| 3658 | NM_00132094 | IREB2 | iron responsive element binding protein 2 | protein_coding | NP_001307870.2;NP_001307871.2;NP_001307872.1;NP_001341923.1;NI | 6115 | 147582 | . | . | 8.566702 | 8.039424 | 9.392778 | 7.975287 | 19.935068 | 18.583489 | 21.077122 | 18.234545 | 1.029527 | 1669.907409 | 0.134842038 | 0.311340511 | 0.755541773 | 0.998865404 | 10.684180 | 10.726613 | 10.718590 | 10.666300 | 10.781754 | 10.645067 | 1958 | 1716 | 1602 | 1450 |
| 3659 | NM_00135492 | IRF1 | interferon regulatory factor 1 | protein_coding | NP_001341853.1;NP_001341854.1;NP_002189.1;XP_011541681.1 | 6116 | 147575 | . | . | 1.948431 | 2.214656 | 1.869553 | 2.236687 | 4.534080 | 5.119277 | 4.195223 | 5.113917 | 1.073881 | 217.487621 | 0.234996774 | 0.437596083 | 0.661679111 | 0.998865404 | 7.713117 | 7.814675 | 7.713270 | 7.777622 | 7.742934 | 7.814357 | 225 | 242 | 181 | 227 |
| 3660 | NM_002199,XI | IRF2 | interferon regulatory factor 2 | protein_coding | NP_002190.2;XP_024309802.1;XP_024309803.1;XP_024309804.1;XP_02 | 6117 | 147576 | . | . | 5.678645 | 5.413110 | 6.972761 | 4.839831 | 13.214438 | 12.512644 | 15.646671 | 11.065700 | 1.086334 | 463.979623 | 0.175904844 | 0.679164161 | 0.497033853 | 0.998865404 | 8.796629 | 8.916708 | 8.832401 | 8.827816 | 8.924978 | 8.836420 | 514 | 478 | 450 | 421 |
| 3661 | NM_00119712 | IRF3 | interferon regulatory factor 3 | protein_coding | NP_001184051.1;NP_001184052.1;NP_001184053.1;NP_001184054.1;NI | 6118 | 603734 | . | . | 23.201717 | 24.196507 | 24.929212 | 24.847070 | 53.991346 | 55.931309 | 55.940429 | 56.809873 | 1.089540 | 1223.875844 | 0.1297209 | 0.953729243 | 0.340220714 | 0.998865404 | 10.194068 | 10.317764 | 10.215347 | 10.231943 | 10.291795 | 10.284854 | 1334 | 1278 | 1115 | 1191 |
| 3662 | NM_00119528 | IRF4 | interferon regulatory factor 4 | protein_coding | NP_001182215.1;NP_002451.2;XP_006715153.1 | 6119 | 601900 | . | . | 13.161345 | 12.643533 | 16.315257 | 14.351044 | 30.626990 | 29.226093 | 36.610963 | 32.811956 | 1.171762 | 2336.151261 | 0.124204582 | 1.841147528 | 0.065599942 | 0.731993537 | 11.070897 | 11.299864 | 11.140563 | 11.102278 | 11.304555 | 11.190786 | 2537 | 2265 | 2345 | 2199 |
| 3663 | NM_00109862 | IRF5 | interferon regulatory factor 5 | protein_coding | NP_001092097.2;NP_001092099.1;NP_001092100.1;NP_001229381.1;NI | 6120 | 607218 | . | . | 0.112082 | 0.188095 | 0.189895 | 0.179498 | 0.260820 | 0.434791 | 0.426120 | 0.410402 | 1.266122 | 16.357707 | 0.764079465 | 0.445524237 | 0.655940784 | NA | 3.852893 | 4.191126 | 4.001950 | 4.030401 | 4.033365 | 4.032421 | 13 | 19 | 16 | 17 |
| 3665 | NM_001572,N | IRF7 | interferon regulatory factor 7 | protein_coding | NP_001563.2;NP_004020.1;NP_004022.2;XP_005252964.1;XP_00525296 | 6122 | 605047 | . | . | 2.503743 | 2.114998 | 2.150836 | 3.102889 | 5.826313 | 4.888911 | 4.826411 | 7.094384 | 1.129924 | 151.343520 | 0.287442543 | 0.613080461 | 0.539823139 | 0.998865404 | 7.151407 | 7.326641 | 7.245236 | 7.178495 | 7.187118 | 7.328425 | 178 | 140 | 115 | 177 |
| 3667 | NM_005544 | IRS1 | insulin receptor substrate 1 | protein_coding | NP_005535.1 | 6125 | 147545 | . | . | 1.661236 | 1.655408 | 1.295689 | 1.789282 | 3.865763 | 3.826549 | 2.907487 | 4.090981 | -1.088626 | 436.274916 | 0.19591404 | -0.62531455 | 0.531764634 | 0.998865404 | 8.829596 | 8.705936 | 8.796193 | 8.788611 | 8.644304 | 8.828268 | 527 | 488 | 307 | 452 |
| 3669 | NM_00130323 | ISG20 | interferon stimulated exonuclease gene 20 | protein_coding | NP_001290162.1;NP_001290163.1;NP_001290164.1;NP_001290165.1;NI | 6130 | 604533 | . | . | 8.537680 | 9.974445 | 7.068295 | 8.133772 | 19.867528 | 23.056372 | 15.861048 | 18.596905 | -1.201316 | 616.945686 | 0.15996056 | -1.65425821 | 0.09807506 | 0.813553488 | 9.395492 | 9.130341 | 9.323833 | 9.329878 | 9.165215 | 9.242059 | 772 | 730 | 458 | 557 |
| 3670 | NM_002202,XI | ISL1 | ISL LIM homeobox 1 | protein_coding | NP_002193.2;XP_011541682.1 | 6132 | 600366 | . | . | 0.995958 | 0.966149 | 0.805202 | 0.980082 | 2.317636 | 2.233298 | 1.806850 | 2.240841 | -1.026110 | 74.765705 | 0.372785112 | -0.0997502 | 0.920542646 | 0.998865404 | 6.243626 | 6.204721 | 6.226575 | 6.228026 | 6.185756 | 6.254756 | 87 | 82 | 55 | 79 |

| | | | | | | | | | | | | | | | | | | | | | | | | | | | | | | | | | |
|---|---|---|---|---|---|---|---|---|---|---|---|---|---|---|---|---|---|---|---|---|---|---|---|---|---|---|---|---|---|---|---|---|---|
| 3672 | NM_181501 | ITGA1 | integrin subunit alpha 1 | protein_coding | NP_852478.1 | 6134 | 192968 | . | . | 3.211967 | 2.918918 | 3.221674 | 2.736164 | 7.474378 | 6.747209 | 7.229342 | 6.255915 | -1.043448 | 1011.724959 | 0.148970913 | -0.41187938 | 0.680427836 | 0.998865404 | 10.012618 | 9.951947 | 10.034372 | 9.957560 | 10.026850 | 9.900558 | 1251 | 1057 | 936 | 848 |
| 3673 | NM_002203,N | ITGA2 | integrin subunit alpha 2 | protein_coding | NP_002194.2 | 6137 | 192974 | . | . | 3.377011 | 2.948237 | 2.256325 | 2.425214 | 7.858440 | 6.814982 | 5.063128 | 5.544965 | -1.369227 | 673.734574 | 0.159594171 | -2.84071717 | 0.004501221 | 0.199099166 | 9.604955 | 9.151657 | 9.542170 | 9.438662 | 9.258247 | 9.297898 | 961 | 780 | 479 | 550 |
| 3675 | NM_002204,XI | ITGA3 | integrin subunit alpha 3 | protein_coding | NP_002195.1;XP_005257365.1 | 6139 | 605025 | . | . | 10.492265 | 9.566371 | 6.941422 | 8.057231 | 24.415932 | 22.113096 | 15.576351 | 18.421899 | -1.368771 | 1324.646008 | 0.137285387 | -3.29882919 | 0.00097089 | 0.066342526 | 10.580238 | 10.127109 | 10.513751 | 10.440519 | 10.189096 | 10.286542 | 1844 | 1576 | 907 | 1119 |
| 3676 | NM_000885,N | ITGA4 | integrin subunit alpha 4 | protein_coding | NP_000876.3;NP_001303241.1 | 6140 | 192975 | . | . | 17.830048 | 16.907350 | 15.357547 | 13.611120 | 41.491255 | 39.082096 | 34.461889 | 31.120209 | -1.220032 | 3337.973366 | 0.119094532 | -2.40917219 | 0.015988753 | 0.425226882 | 11.840987 | 11.554308 | 11.802960 | 11.757275 | 11.669377 | 11.561643 | 4339 | 3851 | 2790 | 2631 |
| 3678 | NM_002205,XI | ITGA5 | integrin subunit alpha 5 | protein_coding | NP_002196.4;XP_024304738.1 | 6141 | 135620 | . | . | 4.665236 | 4.873885 | 3.162257 | 3.932736 | 10.856194 | 11.266203 | 7.096013 | 8.991732 | -1.360547 | 551.281709 | 0.172896546 | -2.5690882 | 0.01019665 | 0.325163413 | 9.312130 | 8.866956 | 9.183264 | 9.209177 | 8.932128 | 9.057092 | 718 | 699 | 364 | 483 |
| 3679 | NM_00114499 | ITGA7 | integrin subunit alpha 7 | protein_coding | NP_001138468.1;NP_001138469.1;NP_001354922.1;NP_001354923.1;NI | 6143 | 600536 | . | . | 6.019078 | 5.886591 | 6.192472 | 6.382826 | 14.006643 | 13.607116 | 13.895729 | 14.593572 | 1.042366 | 782.236021 | 0.146497582 | 0.408617201 | 0.682820606 | 0.998865404 | 9.581227 | 9.641073 | 9.613727 | 9.578868 | 9.610937 | 9.640007 | 904 | 806 | 680 | 764 |
| 3680 | NM_002207 | ITGA9 | integrin subunit alpha 9 | protein_coding | NP_002198.2 | 6145 | 603963 | . | . | 2.113211 | 2.260707 | 1.729925 | 2.273681 | 4.917530 | 5.225725 | 3.881901 | 5.198501 | -1.107185 | 513.095362 | 0.182666078 | -0.80418025 | 0.421292885 | 0.998865404 | 9.075199 | 8.927172 | 9.013969 | 9.052045 | 8.883579 | 9.045954 | 603 | 599 | 368 | 516 |
| 3682 | NM_002208,XI | ITGAE | integrin subunit alpha E | protein_coding | NP_002199.3;XP_011522127.1;XP_011522129.1;XP_011522130.1;XP_01( | 6147 | 604682 | . | . | 2.953479 | 2.790488 | 3.507021 | 4.164998 | 6.872868 | 6.450337 | 7.869654 | 9.522774 | 1.315514 | 454.162225 | 0.178594467 | 2.215224945 | 0.026744629 | 0.532994094 | 8.615923 | 9.011213 | 8.750311 | 8.713662 | 8.852811 | 8.947342 | 466 | 410 | 416 | 519 |
| 3685 | NM_00114499 | ITGAV | integrin subunit alpha V | protein_coding | NP_001138471.1;NP_001138472.2;NP_002201.2 | 6150 | 193210 | . | . | 26.162101 | 22.637811 | 23.935958 | 22.258845 | 60.880285 | 52.328312 | 53.711592 | 50.892203 | -1.071392 | 5208.381265 | 0.119145269 | -0.83499765 | 0.403719029 | 0.998865404 | 12.395428 | 12.296102 | 12.436442 | 12.307378 | 12.350003 | 12.281167 | 6677 | 5370 | 4554 | 4520 |
| 3687 | NM_000887,N | ITGAX | integrin subunit alpha X | protein_coding | NP_000878.2;NP_001273304.1;XP_011544154.1;XP_011544156.1;XP_02 | 6152 | 151510 | . | . | 0.648546 | 0.669032 | 0.741383 | 0.476190 | 1.509192 | 1.546497 | 1.663642 | 1.088752 | -1.118125 | 84.795346 | 0.356355737 | -0.45202351 | 0.651252046 | 0.998865404 | 6.483816 | 6.323562 | 6.390306 | 6.446627 | 6.422468 | 6.353294 | 93 | 106 | 79 | 65 |
| 3688 | NM_002211,N | ITGB1 | integrin subunit beta 1 | protein_coding | NP_002202.2;NP_391988.1;NP_596867.1 | 6153 | 135630 | . | . | 94.684304 | 86.663788 | 75.087642 | 70.385702 | 220.334262 | 200.327223 | 168.494489 | 160.928543 | -1.264076 | 9518.726336 | 0.108958345 | -3.10287088 | 0.001916532 | 0.108968719 | 13.375678 | 13.037675 | 13.349059 | 13.264759 | 13.138953 | 13.076696 | 12832 | 10913 | 7585 | 7589 |
| 3690 | NM_000212 | ITGB3 | integrin subunit beta 3 | protein_coding | NP_000203.2 | 6156 | 173470 | . | . | 90.293533 | 91.757141 | 86.598389 | 86.781334 | 210.116745 | 212.100739 | 194.324280 | 198.415207 | -1.064703 | 16448.497365 | 0.098684209 | -0.91656375 | 0.359371287 | 0.998865404 | 14.050184 | 13.959735 | 14.028733 | 14.035246 | 13.981621 | 13.974733 | 19460 | 18374 | 13911 | 14877 |
| 3692 | NM_00126781 | EIF6 | eukaryotic translation initiation factor 6 | protein_coding | NP_001254739.1;NP_002203.1;NP_852131.1;NP_852133.1 | 6159 | 602912 | . | . | 85.091806 | 76.176469 | 71.330871 | 83.184827 | 198.012131 | 176.085322 | 160.064406 | 190.192224 | -1.028479 | 2693.404936 | 0.121549496 | -0.33330287 | 0.738905676 | 0.998865404 | 11.415392 | 11.374753 | 11.433123 | 11.378120 | 11.331009 | 11.432729 | 3251 | 2848 | 2164 | 2646 |
| 3693 | NM_00135476 | ITGB5 | integrin subunit beta 5 | protein_coding | NP_001341693.1;NP_001341694.1;NP_001341695.1;NP_002204.2;XP_0( | 6160 | 147561 | . | . | 35.305232 | 38.730759 | 29.291486 | 29.819997 | 82.156722 | 89.527884 | 65.729240 | 68.179883 | -1.230508 | 4116.919752 | 0.107692023 | -2.77879596 | 0.005456078 | 0.225239728 | 12.149259 | 11.849943 | 12.081966 | 12.092802 | 11.900909 | 11.929144 | 5197 | 4933 | 3154 | 3521 |
| 3696 | NM_002214,XI | ITGB8 | integrin subunit beta 8 | protein_coding | NP_002205.1;XP_011513695.1;XP_011513696.1;XP_011513698.1;XP_01( | 6163 | 604160 | . | . | 6.117376 | 5.250091 | 5.990154 | 3.931647 | 14.235386 | 12.135821 | 13.441735 | 8.989243 | -1.138219 | 1454.812057 | 0.17677083 | -1.05661091 | 0.290689199 | 0.998865404 | 10.596641 | 10.410600 | 10.610161 | 10.482982 | 10.591471 | 10.282985 | 1927 | 1536 | 1423 | 1018 |
| 3700 | NM_00116644 | ITIH4 | inter-alpha-trypsin inhibitor heavy chain 4 | protein_coding | NP_001159921.1;NP_002209.2 | 6169 | 600564 | . | . | 0.263118 | 1.311373 | 0.197151 | 0.155052 | 0.612286 | 3.031297 | 0.442400 | 0.354508 | -4.367960 | 49.253549 | 0.633231176 | -3.35889925 | 0.000782536 | 0.057585413 | 6.324762 | 4.197690 | 5.440642 | 5.813901 | 5.416474 | 5.394149 | 32 | 143 | 18 | 15 |
| 3702 | NM_005546,XI | ITK | IL2 inducible T cell kinase | protein_coding | NP_005537.3;XP_016864932.1 | 6171 | 186973 | . | . | 0.071684 | 0.059950 | 0.046707 | 0.060286 | 0.166811 | 0.138578 | 0.104810 | 0.137837 | -1.274833 | 8.780484 | 1.025941535 | -0.34145081 | 0.732764232 | NA | 3.299300 | 2.947964 | 3.146958 | 3.141608 | 3.131268 | 3.138308 | 12 | 10 | 6 | 8 |
| 3703 | NM_00127850 | STT3A | STT3 oligosaccharyltransferase complex catalytic subunit A | protein_coding | NP_001265432.1;NP_001265433.1;NP_689926.1;XP_011541109.1 | 6172 | 601134 | . | . | 31.021339 | 34.673617 | 30.194838 | 29.155269 | 72.187930 | 80.149617 | 67.756341 | 66.660064 | -1.080602 | 4296.574241 | 0.109241933 | -1.02374054 | 0.305957841 | 0.998865404 | 12.123799 | 12.011977 | 12.076945 | 12.122995 | 12.054752 | 12.016383 | 4998 | 4948 | 3675 | 3782 |
| 3704 | NM_00126762 | ITPA | inosine triphosphatase | protein_coding | NP_001254552.1;NP_001311165.1;NP_001311166.1;NP_001311167.1;NI | 6176 | 147520 | . | . | 22.949027 | 22.576611 | 25.077585 | 29.747985 | 53.403324 | 52.186847 | 56.273372 | 68.015232 | 1.203471 | 846.748147 | 0.149569081 | 1.786474825 | 0.07402241 | 0.763883535 | 9.586240 | 9.853022 | 9.659523 | 9.648910 | 9.728944 | 9.840245 | 892 | 823 | 743 | 933 |
| 3705 | NM_00114259 | ITPK1 | inositol-tetrakisphosphate 1-kinase | protein_coding | NP_001136065.1;NP_001136066.1;NP_001350636.1;NP_055031.2;XP_01 | 6177 | 601838 | . | . | 25.619503 | 32.869401 | 23.877513 | 27.664940 | 59.617637 | 75.979090 | 53.580444 | 63.252603 | 1.046205 | 3137.346801 | 0.129670272 | 0.502544303 | 0.615284696 | 0.998865404 | 11.582522 | 11.647407 | 11.577957 | 11.611966 | 11.540981 | 11.715733 | 3458 | 3378 | 2494 | 3326 |
| 3706 | NM_002220,XI | ITPKA | inositol-trisphosphate 3-kinase A | protein_coding | NP_002211.1;XP_011519824.1 | 6178 | 147521 | . | . | 0.742073 | 0.495926 | 0.600749 | 0.446367 | 1.726834 | 1.146354 | 1.348064 | 1.020566 | -1.329786 | 31.625139 | 0.563770517 | -0.72936488 | 0.465778495 | 0.998865404 | 5.172951 | 4.764191 | 5.031782 | 4.972272 | 4.972961 | 4.960085 | 50 | 31 | 25 | 24 |
| 3707 | NM_002221,XI | ITPKB | inositol-trisphosphate 3-kinase B | protein_coding | NP_002212.3;XP_005273177.1;XP_016856700.1 | 6179 | 147522 | . | . | 13.621200 | 13.687017 | 14.533168 | 16.046957 | 31.697093 | 31.638156 | 32.612011 | 36.689459 | 1.099814 | 2743.756823 | 0.116502272 | 1.178168332 | 0.238729505 | 0.992733415 | 11.351759 | 11.488859 | 11.379591 | 11.383596 | 11.418677 | 11.497655 | 3002 | 2827 | 2379 | 2824 |
| 3708 | NM_00109995 | ITPR1 | inositol 1,4,5-trisphosphate receptor type 1 | protein_coding | NP_001093422.2;NP_001161744.1;NP_001365381.1;NP_002213.5;XP_0( | 6180 | 147265 | . | . | 3.183384 | 3.073036 | 2.838406 | 3.095408 | 7.407866 | 7.103459 | 6.369304 | 7.077283 | -1.068588 | 928.840294 | 0.140998299 | -0.67877087 | 0.497283051 | 0.998865404 | 9.906425 | 9.810556 | 9.897329 | 9.868132 | 9.808120 | 9.860770 | 1127 | 1015 | 752 | 873 |
| 3709 | NM_002223,XI | ITPR2 | inositol 1,4,5-trisphosphate receptor type 2 | protein_coding | NP_002214.2;XP_016874755.1;XP_016874756.1;XP_016874758.1 | 6181 | 600144 | . | . | 2.238175 | 2.867375 | 2.274736 | 2.144590 | 5.208325 | 6.628067 | 5.104442 | 4.903351 | -1.073990 | 893.439401 | 0.144107162 | -0.71461378 | 0.474847725 | 0.998865404 | 9.853786 | 9.750830 | 9.795695 | 9.859326 | 9.799905 | 9.752974 | 1019 | 1042 | 774 | 781 |
| 3710 | NM_002224,XI | ITPR3 | inositol 1,4,5-trisphosphate receptor type 3 | protein_coding | NP_002215.2;XP_011512879.1;XP_016866321.1 | 6182 | 147267 | . | . | 14.149482 | 14.977207 | 13.479282 | 14.031801 | 32.926426 | 34.620487 | 30.247118 | 32.082046 | -1.077555 | 3991.089471 | 0.109043378 | -0.98824325 | 0.323033531 | 0.998865404 | 12.015480 | 11.907639 | 11.970666 | 12.014282 | 11.920619 | 11.940733 | 4643 | 4584 | 3300 | 3646 |
| 3712 | NM_00115950 | IVD | isovaleryl-CoA dehydrogenase | protein_coding | NP_001152980.2;NP_001341526.1;NP_001341527.2;NP_001341528.2;NI | 6186 | 607036 | . | . | 7.214521 | 8.113539 | 6.924595 | 8.281687 | 16.788488 | 18.754809 | 15.538590 | 18.935092 | 1.006776 | 1012.125137 | 0.146237076 | 0.066619993 | 0.946884229 | 0.998865404 | 9.978606 | 9.987723 | 9.944479 | 10.013959 | 9.927620 | 10.038546 | 1107 | 1140 | 816 | 1024 |
| 3714 | NM_002226,N | JAG2 | jagged canonical Notch ligand 2 | protein_coding | NP_002217.3;NP_660142.1 | 6189 | 602570 | . | . | 0.148891 | 0.176519 | 0.116141 | 0.150738 | 0.346476 | 0.408031 | 0.260618 | 0.344644 | -1.230318 | 27.669378 | 0.590499527 | -0.50640484 | 0.612572479 | 0.998865404 | 4.932974 | 4.631770 | 4.796781 | 4.812220 | 4.769449 | 4.792890 | 33 | 35 | 19 | 26 |
| 3716 | NM_00132092 | JAK1 | Janus kinase 1 | protein_coding | NP_001307852.1;NP_001308781.1;NP_001308782.1;NP_001308783.1;NI | 6190 | 147795 | . | . | 14.917420 | 15.287912 | 14.283033 | 14.420968 | 34.713450 | 35.338694 | 32.050713 | 32.971832 | -1.067562 | 2337.533802 | 0.11481912 | -0.82146514 | 0.411381375 | 0.998865404 | 11.237171 | 11.142831 | 11.210236 | 11.222827 | 11.164290 | 11.163635 | 2758 | 2625 | 1967 | 2119 |
| 3717 | NM_00132219 | JAK2 | Janus kinase 2 | protein_coding | NP_001309123.1;NP_001309124.1;NP_001309125.1;NP_001309127.1;NI | 6192 | 147796 | . | . | 1.532174 | 1.277416 | 1.903829 | 1.715583 | 3.565433 | 2.952802 | 4.272136 | 3.922476 | 1.248271 | 312.441670 | 0.204257375 | 1.566312492 | 0.117275478 | 0.857396579 | 8.118141 | 8.438948 | 8.225583 | 8.212674 | 8.389934 | 8.287893 | 324 | 296 | 329 | 296 |
| 3720 | NM_00126704 | JARID2 | jumonji and AT-rich interaction domain containing 2 | protein_coding | NP_001253969.1;NP_004964.2;XP_011512881.1;XP_016866322.1;XP_01 | 6196 | 601594 | . | . | 5.115420 | 4.859217 | 5.771042 | 5.488239 | 11.903790 | 11.232299 | 12.950054 | 12.548206 | 1.127979 | 972.690141 | 0.137945572 | 1.259487329 | 0.207854364 | 0.970482251 | 9.836235 | 10.010197 | 9.890626 | 9.868320 | 9.983835 | 9.950688 | 1069 | 971 | 917 | 945 |
| 3725 | NM_002228 | JUN | Jun proto-oncogene, AP-1 transcription factor subunit | protein_coding | NP_002219.1 | 6204 | 165160 | . | . | 4.605468 | 4.901416 | 3.480160 | 3.633918 | 10.717112 | 11.329843 | 7.809378 | 8.308521 | -1.354883 | 422.664649 | 0.178134522 | -2.45976016 | 0.013902989 | 0.390841055 | 8.926024 | 8.487523 | 8.789480 | 8.823782 | 8.615143 | 8.632753 | 545 | 539 | 307 | 342 |
| 3726 | NM_002229 | JUNB | JunB proto-oncogene, AP-1 transcription factor subunit | protein_coding | NP_002220.1 | 6205 | 165161 | . | . | 6.411191 | 7.277942 | 5.100172 | 6.380012 | 14.919106 | 16.823288 | 11.444638 | 14.587139 | -1.207461 | 359.118757 | 0.199888228 | -1.360641 | 0.173627168 | 0.94763772 | 8.618504 | 8.345206 | 8.502783 | 8.573175 | 8.371866 | 8.485917 | 426 | 449 | 253 | 337 |
| 3727 | NM_00128696 | JUND | JunD proto-oncogene, AP-1 transcription factor subunit | protein_coding | NP_001273897.1;NP_005345.3 | 6206 | 165162 | . | . | 9.231969 | 8.805847 | 8.596323 | 10.922745 | 21.483171 | 20.355110 | 19.289900 | 24.973557 | 1.066323 | 564.444707 | 0.174415607 | 0.53117303 | 0.59529888 | 0.998865404 | 9.093955 | 9.185956 | 9.135679 | 9.098961 | 9.081886 | 9.231434 | 647 | 573 | 449 | 608 |
| 3728 | NM_00135277 | JUP | junction plakoglobin | protein_coding | NP_001339702.1;NP_001339703.1;NP_001339704.1;NP_001339705.1;NI | 6207 | 173325 | . | . | 0.465623 | 0.315262 | 0.334971 | 0.411273 | 1.083524 | 0.728742 | 0.751665 | 0.940328 | -1.046589 | 42.294637 | 0.489578979 | -0.13418762 | 0.893254214 | 0.998865404 | 5.434840 | 5.369220 | 5.433503 | 5.378055 | 5.381717 | 5.415599 | 58 | 39 | 32 | 43 |
| 3732 | NM_00102484 | CD82 | CD82 molecule | protein_coding | NP_001020015.1;NP_002222.1;XP_006718286.1;XP_011518369.1 | 6210 | 600623 | . | . | 37.616318 | 35.338171 | 24.931851 | 28.641132 | 87.534714 | 81.685772 | 55.946356 | 65.484549 | -1.375955 | 2188.234573 | 0.124512282 | -3.69789333 | 0.000217396 | 0.021989319 | 11.307555 | 10.846938 | 11.237678 | 11.179346 | 10.905903 | 10.999403 | 3021 | 2639 | 1504 | 1832 |
| 3735 | NM_00113008 | KARS1 | lysyl-tRNA synthetase 1 | protein_coding | NP_001123561.1;NP_001365077.1;NP_005539.1 | 6215 | 601421 | . | . | 51.897032 | 54.476041 | 47.422578 | 58.598419 | 120.766523 | 125.923821 | 106.414888 | 133.978315 | -1.011749 | 3373.664176 | 0.127170182 | -0.13251086 | 0.894580232 | 0.998865404 | 11.728634 | 11.711517 | 11.704342 | 11.742696 | 11.628830 | 11.792637 | 3817 | 3747 | 2626 | 3457 |
| 3737 | NM_00120426 | KCNA2 | potassium voltage-gated channel subfamily A member 2 | protein_coding | NP_001191198.1;NP_004965.1;XP_011539698.1;XP_011539699.1;XP_01 | 6220 | 176262 | . | . | 0.021002 | 0.020371 | 0.021791 | 0.027170 | 0.048872 | 0.047087 | 0.048899 | 0.062121 | 1.167474 | 9.222897 | 0.996892493 | 0.224086233 | 0.822690205 | NA | 3.089593 | 3.312263 | 3.200172 | 3.198634 | 3.202569 | 3.209011 | 10 | 9 | 8 | 10 |
| 3746 | NM_00111274 | KCNC1 | potassium voltage-gated channel subfamily C member 1 | protein_coding | NP_001106212.1;NP_004967.1 | 6233 | 176258 | . | . | 0.060136 | 0.062463 | 0.071016 | 0.101352 | 0.139937 | 0.144387 | 0.159358 | 0.231731 | 1.216272 | 13.447237 | 0.860058032 | 0.328426673 | 0.742589078 | NA | 3.604218 | 3.881002 | 3.730036 | 3.745882 | 3.725750 | 3.772823 | 12 | 15 | 8 | 19 |
| 3748 | NM_00137230 | KCNC3 | potassium voltage-gated channel subfamily C member 3 | protein_coding | NP_001359234.1;NP_004968.2 | 6235 | 176264 | . | . | 0.640520 | 0.577325 | 0.754459 | 0.704747 | 1.490516 | 1.334513 | 1.692984 | 1.611319 | 1.181823 | 141.143056 | 0.276010279 | 0.873208239 | 0.382549585 | 0.998865404 | 7.015001 | 7.256900 | 7.124642 | 7.085736 | 7.182473 | 7.153609 | 157 | 132 | 138 | 138 |
| 3749 | NM_00103957 | KCNC4 | potassium voltage-gated channel subfamily C member 4 | protein_coding | NP_001034663.1;NP_001364259.1;NP_001364260.1;NP_004969.2;XP_0( | 6236 | 176265 | . | . | 1.567994 | 1.914431 | 1.072126 | 1.408477 | 3.648786 | 4.425295 | 2.405819 | 3.220316 | -1.340103 | 203.012492 | 0.250088276 | -1.68877941 | 0.091261712 | 0.805877675 | 7.862027 | 7.437739 | 7.698147 | 7.752800 | 7.532751 | 7.650364 | 253 | 265 | 129 | 186 |
| 3750 | NM_004979,XI | KCND1 | potassium voltage-gated channel subfamily D member 1 | protein_coding | NP_004970.3;XP_011542212.1;XP_016884997.1;XP_016884998.1;XP_01( | 6237 | 300281 | . | . | 0.096781 | 0.196175 | 0.131246 | 0.145057 | 0.225213 | 0.453465 | 0.294512 | 0.331655 | -1.025153 | 19.633799 | 0.705942076 | -0.05076742 | 0.959510854 | NA | 4.314175 | 4.276108 | 4.270595 | 4.315274 | 4.291871 | 4.290404 | 17 | 27 | 17 | 18 |
| 3752 | NM_00137896 | KCND3 | potassium voltage-gated channel subfamily D member 3 | protein_coding | NP_001365898.1;NP_001365899.1;NP_004971.2;NP_751948.1;XP_00671 | 6239 | 605411 | . | . | 0.054361 | 0.081173 | 0.012869 | 0.032129 | 0.126501 | 0.187634 | 0.028877 | 0.073459 | -2.842102 | 11.566679 | 0.959835741 | -1.57001708 | 0.116411139 | NA | 4.099441 | 2.582949 | 3.549948 | 3.578057 | 3.504615 | 3.526299 | 16 | 22 | 3 | 8 |
| 3755 | NM_002237,XI | KCNG1 | potassium voltage-gated channel modifier subfamily G member 1 | protein_coding | NP_002228.2;XP_006723848.1;XP_011527102.1;XP_011527103.1;XP_01 | 6248 | 603788 | . | . | 2.953205 | 3.859878 | 2.109667 | 2.893640 | 6.872232 | 8.922280 | 4.734030 | 6.615963 | -1.419019 | 237.439934 | 0.24271387 | -2.08020171 | 0.037507035 | 0.605846634 | 8.122690 | 7.615861 | 7.891310 | 8.034556 | 7.744659 | 7.843703 | 278 | 341 | 151 | 205 |
| 3756 | NM_002238,N | KCNH1 | potassium voltage-gated channel subfamily H member 1 | protein_coding | NP_002229.1;NP_758872.1;XP_016856735.1 | 6250 | 603305 | . | . | 0.543288 | 0.607306 | 0.441566 | 0.403719 | 1.264254 | 1.403815 | 0.990862 | 0.923057 | -1.367176 | 125.969013 | 0.291568203 | -1.54748913 | 0.121745344 | 0.864429515 | 7.184851 | 6.733903 | 7.008425 | 7.047732 | 6.934377 | 6.904097 | 159 | 165 | 97 | 95 |
| 3759 | NM_000891 | KCNJ2 | potassium inwardly rectifying channel subfamily J member 2 | protein_coding | NP_000882.1 | 6263 | 600681 | . | . | 0.157855 | 0.124183 | 0.053254 | 0.051366 | 0.367336 | 0.287055 | 0.119499 | 0.117443 | -2.723080 | 16.492491 | 0.784812449 | -1.84150891 | 0.065547015 | NA | 4.592063 | 3.148375 | 4.095234 | 4.070834 | 4.019253 | 4.015686 | 31 | 23 | 8 | 8 |
| 3766 | NM_002241 | KCNJ10 | potassium inwardly rectifying channel subfamily J member 10 | protein_coding | NP_002232.2 | 6256 | 602208 | . | . | 1.434709 | 1.311605 | 1.217168 | 1.589794 | 3.338627 | 3.031834 | 2.731290 | 3.634876 | 1.007969 | 225.538714 | 0.236916607 | 0.048336 | 0.961448464 | 0.998865404 | 7.811939 | 7.822504 | 7.836446 | 7.790793 | 7.756264 | 7.875414 | 271 | 231 | 172 | 239 |
| 3768 | NM_021012,XI | KCNJ12 | potassium inwardly rectifying channel subfamily J member 12 | protein_coding | NP_066292.2;XP_005256682.1;XP_011522133.1 | 6258 | 602323 | . | . | 0.033734 | 0.046439 | 0.022300 | 0.025558 | 0.078500 | 0.107345 | 0.050040 | 0.058435 | -1.625532 | 5.821591 | 1.262241304 | -0.55529163 | 0.578695202 | NA | 2.849979 | 2.148375 | 2.549185 | 2.559564 | 2.542637 | 2.541023 | 7 | 9 | 4 | 4 |
| 3769 | NM_00117241 | KCNJ13 | potassium inwardly rectifying channel subfamily J member 13 | protein_coding | NP_001165887.1;NP_001165888.1;NP_002233.2 | 6259 | 603208 | . | . | 0.223915 | 0.333399 | 0.323449 | 0.285049 | 0.521059 | 0.770666 | 0.725811 | 0.651730 | 1.089714 | 32.995137 | 0.550184812 | 0.225287596 | 0.821755549 | 0.998865404 | 4.981427 | 5.104320 | 5.006649 | 5.059454 | 5.053898 | 5.035411 | 29 | 41 | 32 | 30 |
| 3770 | NM_013348 | KCNJ14 | potassium inwardly rectifying channel subfamily J member 14 | protein_coding | NP_037480.1 | 6260 | 603953 | . | . | 0.301433 | 0.255612 | 0.246300 | 0.285851 | 0.701448 | 0.590859 | 0.552690 | 0.653564 | -1.075195 | 33.221252 | 0.538886317 | -0.19410131 | 0.846096559 | 0.998865404 | 5.105434 | 5.000737 | 5.072107 | 5.048383 | 5.042487 | 5.058911 | 43 | 34 | 26 | 32 |
| 3776 | NM_00101742 | KCNK2 | potassium two pore domain channel subfamily K member 2 | protein_coding | NP_001017424.1;NP_001017425.2;NP_055032.1;XP_011507824.1;XP_01 | 6277 | 603219 | . | . | 1.667818 | 2.034767 | 1.658746 | 2.009364 | 3.881083 | 4.703455 | 3.722178 | 4.594172 | -1.034465 | 220.155407 | 0.237821087 | -0.20555118 | 0.837141515 | 0.998865404 | 7.807331 | 7.756986 | 7.739180 | 7.836175 | 7.737609 | 7.806702 | 232 | 266 | 174 | 218 |
| 3778 | NM_00101479 | KCNMA1 | potassium calcium-activated channel subfamily M alpha 1 | protein_coding | NP_001014797.1;NP_001154824.1;NP_001154825.1;NP_001258447.1;NI | 6284 | 600150 | . | . | 1.215365 | 0.851672 | 0.956646 | 1.209882 | 2.828204 | 1.968679 | 2.146683 | 2.766247 | -1.260402 | 146.233660 | 0.30056647 | -1.11085048 | 0.266632713 | 0.998274218 | 7.349556 | 7.015402 | 7.306295 | 7.149397 | 7.093209 | 7.189006 | 223 | 143 | 97 | 138 |
| 3780 | NM_002248,XI | KCNN1 | potassium calcium-activated channel subfamily N member 1 | protein_coding | NP_002239.2;XP_005259962.1;XP_005259963.1;XP_005259964.1;XP_01 | 6290 | 602982 | . | . | 0.237487 | 0.541536 | 0.216563 | 0.463378 | 0.552642 | 1.251786 | 0.485960 | 1.059458 | -1.032686 | 27.178766 | 0.672979087 | -0.06894954 | 0.945029787 | 0.998865404 | 4.790912 | 4.737408 | 4.711385 | 4.797791 | 4.704195 | 4.797939 | 20 | 41 | 13 | 36 |
| 3781 | NM_00127820 | KCNN2 | potassium calcium-activated channel subfamily N member 2 | protein_coding | NP_001265133.1;NP_001359162.1;NP_067627.3;NP_740721.1;XP_0115 | 6291 | 605879 | . | . | 6.504848 | 5.669647 | 3.933965 | 3.883079 | 15.137047 | 13.105644 | 8.827703 | 8.878197 | -1.550206 | 420.806191 | 0.185627159 | -3.40715396 | 0.000656441 | 0.050903951 | 8.998589 | 8.366816 | 8.883477 | 8.773386 | 8.580920 | 8.553833 | 638 | 506 | 294 | 302 |
| 3783 | NM_002250,XI | KCNN4 | potassium calcium-activated channel subfamily N member 4 | protein_coding | NP_002241.1;XP_005258939.1;XP_005258940.1 | 6293 | 602754 | . | . | 32.361390 | 31.590000 | 30.607832 | 32.955844 | 75.306284 | 73.021699 | 68.683086 | 75.349623 | -1.015829 | 2116.328549 | 0.120766825 | -0.18761106 | 0.851181555 | 0.998865404 | 11.058739 | 11.035867 | 11.048400 | 11.058341 | 11.001394 | 11.078690 | 2441 | 2316 | 1737 | 2064 |
| 3786 | NM_00120482 | KCNQ3 | potassium voltage-gated channel subfamily Q member 3 | protein_coding | NP_001191753.1;NP_004510.1;XP_005250971.1;XP_006716618.1;XP_01 | 6297 | 602232 | . | . | 0.179711 | 0.187125 | 0.155458 | 0.233331 | 0.418196 | 0.432547 | 0.348843 | 0.533485 | 1.142557 | 103.194131 | 0.327359687 | 0.587324859 | 0.556985549 | 0.998865404 | 6.590800 | 6.781347 | 6.664090 | 6.664254 | 6.651789 | 6.756801 | 111 | 104 | 80 | 120 |
| 3790 | NM_00128242 | KCNS3 | potassium voltage-gated channel modifier subfamily S member 3 | protein_coding | NP_001269357.1;NP_002243.3;XP_011531127.1 | 6302 | 603888 | . | . | 4.307430 | 3.714058 | 4.258837 | 5.027674 | 10.023566 | 8.585211 | 9.556705 | 11.495180 | 1.140700 | 316.372762 | 0.206195488 | 0.921064552 | 0.357016726 | 0.998865404 | 8.207456 | 8.397268 | 8.306771 | 8.223010 | 8.294504 | 8.379290 | 367 | 294 | 271 | 340 |
| 3791 | NM_002253 | KDR | kinase insert domain receptor | protein_coding | NP_002244.1 | 6307 | 191306 | . | . | 0.089932 | 0.139652 | 0.072885 | 0.162247 | 0.209276 | 0.322811 | 0.163553 | 0.370959 | 1.012199 | 21.746017 | 0.696720694 | 0.025106742 | 0.979969823 | NA | 4.436885 | 4.448451 | 4.419547 | 4.456557 | 4.406001 | 4.471756 | 20 | 28 | 12 | 28 |
| 3795 | NM_000221,N | KHK | ketohexokinase | protein_coding | NP_000212.1;NP_006479.1;XP_005264351.1;XP_005264353.1;XP_00526 | 6315 | 614058 | . | . | 3.574832 | 4.228264 | 3.982601 | 4.321337 | 8.318781 | 9.773819 | 8.936840 | 9.880223 | 1.042024 | 299.562687 | 0.206305273 | 0.287869116 | 0.773446929 | 0.998865404 | 8.197152 | 8.255682 | 8.172111 | 8.254842 | 8.218390 | 8.253923 | 312 | 341 | 257 | 296 |
| 3796 | NM_00109851 | KIF2A | kinesin family member 2A | protein_coding | NP_001091981.1;NP_001230881.2;NP_001230882.1;NP_004511.2 | 6318 | 602591 | . | . | 8.748063 | 8.386080 | 7.964441 | 7.546905 | 20.357104 | 19.384798 | 17.871975 | 17.255101 | -1.132454 | 1152.012699 | 0.136136091 | -1.3181837 | 0.18744216 | 0.960264078 | 10.256686 | 10.077645 | 10.232728 | 10.195496 | 10.158430 | 10.082967 | 1443 | 1288 | 988 | 961 |
| 3797 | NM_002254,XI | KIF3C | kinesin family member 3C | protein_coding | NP_002245.4;XP_005264356.2 | 6321 | 602845 | . | . | 21.550887 | 23.803692 | 27.197678 | 29.289366 | 50.149799 | 55.023304 | 61.030801 | 66.966652 | 1.228180 | 4281.768750 | 0.112551605 | 2.634537709 | 0.008425194 | 0.29000738 | 11.908206 | 12.204581 | 11.935802 | 12.006645 | 12.114619 | 12.170230 | 4234 | 4327 | 3970 | 4570 |
| 3799 | NM_004521,XI | KIF5B | kinesin family member 5B | protein_coding | NP_004512.1;XP_016871713.1 | 6324 | 602809 | . | . | 23.767882 | 22.083977 | 20.523879 | 20.318861 | 55.308842 | 51.048106 | 46.054988 | 46.456660 | -1.137668 | 3961.269482 | 0.111114962 | -1.67465963 | 0.094001033 | 0.810751527 | 12.041744 | 11.855761 | 12.036621 | 11.967626 | 11.904159 | 11.887324 | 5054 | 4363 | 3256 | 3439 |
| 3800 | NM_004522,N | KIF5C | kinesin family member 5C | protein_coding | NP_004513.1;XP_011509459.1;XP_016859551.1 | 6325 | 604593 | . | . | 1.665687 | 1.372787 | 1.304658 | 1.619153 | 3.876122 | 3.173258 | 2.927614 | 3.702002 | -1.058266 | 322.842440 | 0.211866593 | -0.38563027 | 0.699770512 | 0.998865404 | 8.375096 | 8.293113 | 8.404043 | 8.289615 | 8.260933 | 8.370517 | 421 | 322 | 245 | 324 |
| 3801 | NM_00113009 | KIFC3 | kinesin family member C3 | protein_coding | NP_001123571.1;NP_001123572.1;NP_001305639.1;NP_001305640.1;NI | 6326 | 604535 | . | . | 25.389368 | 27.482747 | 28.743421 | 28.552025 | 59.082101 | 63.527604 | 64.499402 | 65.280812 | 1.082571 | 3020.784086 | 0.114490436 | 0.999747021 | 0.31743295 | 0.998865404 | 11.502377 | 11.616770 | 11.491711 | 11.561147 | 11.600011 | 11.583718 | 3198 | 3264 | 2758 | 2915 |
| 3815 | NM_000222,N | KIT | KIT proto-oncogene, receptor tyrosine kinase | protein_coding | NP_000213.1;NP_001087241.1;XP_005265797.1;XP_005265798.1;XP_00 | 6342 | 164920 | . | . | 0.271662 | 0.289661 | 0.240814 | 0.198683 | 0.632169 | 0.669564 | 0.540380 | 0.454264 | -1.288701 | 40.663416 | 0.490685669 | -0.74572804 | 0.455831714 | 0.998865404 | 5.516559 | 5.151760 | 5.361018 | 5.372226 | 5.340921 | 5.313388 | 51 | 51 | 34 | 30 |
| 3831 | NM_00113010 | KLC1 | kinesin light chain 1 | protein_coding | NP_001123579.1;NP_005543.2;NP_891553.2 | 6387 | 600025 | . | . | 18.218896 | 17.358676 | 17.745324 | 20.107415 | 42.396119 | 40.125358 | 39.819995 | 45.973216 | 1.049199 | 1441.748563 | 0.130821873 | 0.529643985 | 0.596358792 | 0.998865404 | 10.458616 | 10.527763 | 10.498008 | 10.450098 | 10.470284 | 10.550593 | 1671 | 1471 | 1217 | 1456 |
| 3832 | NM_004523 | KIF11 | kinesin family member 11 | protein_coding | NP_004514.2 | 6388 | 148760 | . | . | 13.683925 | 12.788810 | 10.300295 | 10.318729 | 31.843058 | 29.561905 | 23.113562 | 23.592550 | -1.302248 | 1841.771080 | 0.1220913 | -3.1206505 | 0.001804521 | 0.104654615 | 11.024783 | 10.643908 | 10.971611 | 10.910133 | 10.737078 | 10.730738 | 2490 | 2163 | 1397 | 1494 |
| 3833 | NM_002263,XI | KIFC1 | kinesin family member C1 | protein_coding | NP_002254.2;XP_011512887.1;XP_011512889.1;XP_016866325.1;XP_01( | 6389 | 603763 | . | . | 24.348547 | 25.349078 | 19.286270 | 22.134695 | 56.660066 | 58.595532 | 43.277833 | 50.608352 | -1.192128 | 1709.489655 | 0.130208963 | -1.94716738 | 0.051514671 | 0.672667836 | 10.860734 | 10.606807 | 10.793117 | 10.814158 | 10.606836 | 10.721816 | 2113 | 2033 | 1254 | 1572 |
| 3835 | NM_00125626 | KIF22 | kinesin family member 22 | protein_coding | NP_001243198.1;NP_001243199.1;NP_015556.1;XP_024306038.1 | 6391 | 603213 | . | . | 32.856000 | 31.997101 | 30.023295 | 33.296244 | 76.457271 | 73.962730 | 67.371408 | 76.127913 | -1.035785 | 2099.367295 | 0.12063745 | -0.42047088 | 0.674141501 | 0.998865404 | 11.060942 | 11.010088 | 11.063705 | 11.034968 | 10.986113 | 11.055412 | 2504 | 2264 | 1715 | 2018 |
| 3836 | NM_002264,N | KPNA1 | karyopherin subunit alpha 1 | protein_coding | NP_002255.3;NP_005247494.1;XP_011511097.1;XP_011511099.1;XP_01( | 6394 | 600686 | . | . | 12.448234 | 12.803990 | 13.848853 | 12.221846 | 28.967553 | 29.596996 | 31.076425 | 27.943803 | 1.006414 | 2723.850074 | 0.118247581 | 0.077998945 | 0.937828892 | 0.998865404 | 11.406740 | 11.416108 | 11.396230 | 11.420339 | 11.457313 | 11.367960 | 3080 | 2973 | 2507 | 2421 |
| 3837 | NM_00127645 | KPNB1 | karyopherin subunit beta 1 | protein_coding | NP_001263382.1;NP_002256.2 | 6400 | 602738 | . | . | 71.143176 | 73.220287 | 69.580702 | 77.249038 | 165.553098 | 169.251970 | 156.137081 | 176.620757 | 1.002687 | 13733.773420 | 0.10460868 | 0.037001071 | 0.970484152 | 0.998865404 | 13.743526 | 13.747352 | 13.735343 | 13.752627 | 13.705138 | 13.785880 | 15634 | 14949 | 11395 | 13500 |
| 3838 | NM_00132061 | KPNA2 | karyopherin subunit alpha 2 | protein_coding | NP_001307540.1;NP_002257.1 | 6395 | 600685 | . | . | 172.515066 | 185.370391 | 141.563153 | 164.473881 | 401.449697 | 428.492015 | 317.663593 | 376.049994 | -1.186103 | 10239.891037 | 0.112706731 | -2.18469561 | 0.028911173 | 0.547408663 | 13.439824 | 13.193500 | 13.361879 | 13.416844 | 13.183677 | 13.301273 | 12389 | 12371 | 7576 | 9397 |
| 3839 | NM_002267,XI | KPNA3 | karyopherin subunit alpha 3 | protein_coding | NP_002258.2;XP_016876050.1 | 6396 | 601892 | . | . | 8.978411 | 8.543187 | 7.930337 | 7.123609 | 20.893133 | 19.747961 | 17.795446 | 16.287285 | -1.179967 | 1072.207870 | 0.139715854 | -1.70879967 | 0.087488064 | 0.798675762 | 10.180585 | 9.942327 | 10.145173 | 10.102211 | 10.043280 | 9.958166 | 1374 | 1217 | 906 | 868 |
| 3840 | NM_002268 | KPNA4 | karyopherin subunit alpha 4 | protein_coding | NP_002259.1 | 6397 | 602970 | . | . | 7.742051 | 7.638095 | 7.405953 | 6.982520 | 18.016069 | 17.655800 | 16.618746 | 15.964702 | -1.083954 | 2076.612444 | 0.119256646 | -0.97523423 | 0.329444085 | 0.998865404 | 11.076909 | 10.960788 | 11.060096 | 11.042148 | 11.013856 | 10.959370 | 2515 | 2305 | 1793 | 1804 |
| 3841 | NM_00136630 | KPNA5 | karyopherin subunit alpha 5 | protein_coding | NP_001353233.1;NP_001353234.1;NP_001353235.1;NP_001353236.1;NI | 6398 | 604545 | . | . | 0.666606 | 0.451532 | 0.785869 | 0.600501 | 1.551219 | 1.043735 | 1.763468 | 1.372973 | 1.139899 | 116.752560 | 0.308695512 | 0.611947787 | 0.540572303 | 0.998865404 | 6.768713 | 6.959598 | 6.872180 | 6.807581 | 6.921074 | 6.846345 | 138 | 106 | 120 | 104 |
| 3842 | NM_00136429 | TNPO1 | transportin 1 | protein_coding | NP_001351221.1;NP_001351222.1;NP_001351223.1;NP_001351224.1;NI | 6401 | 602901 | . | . | 19.852004 | 18.624427 | 19.861445 | 15.130517 | 46.196432 | 43.051200 | 44.568508 | 34.594125 | -1.150013 | 5650.492495 | 0.122234649 | -1.6496967 | 0.099004985 | 0.813553488 | 12.561381 | 12.359913 | 12.549187 | 12.487684 | 12.475622 | 12.321494 | 7211 | 6287 | 4995 | 4471 |
| 3843 | NM_002271,XI | IPO5 | importin 5 | protein_coding | NP_002262.4;XP_005254109.1;XP_005254110.1;XP_011519389.1;XP_01 | 6402 | 602008 | . | . | 19.819956 | 20.444378 | 17.394243 | 18.589420 | 46.121853 | 47.258091 | 39.032177 | 42.502499 | -1.098478 | 3620.568018 | 0.110277467 | -1.22877007 | 0.219158026 | 0.982008277 | 11.888213 | 11.752602 | 11.850150 | 11.868782 | 11.758705 | 11.804654 | 4316 | 4136 | 2916 | 3326 |
| 3844 | gene-KRASP1 | KRASP1 | KRAS proto-oncogene, GTPase pseudogene 1 | pseudogene | . | 6406 | . | | . | 0.047827 | 0.036319 | 0.021273 | 0.013311 | 0.111295 | 0.083954 | 0.047735 | 0.030434 | -1.950911 | 5.742883 | 1.283692548 | -0.75107372 | 0.452608293 | NA | 2.923304 | 1.963455 | 2.545734 | 2.535468 | 2.527055 | 2.520661 | 10 | 7 | 4 | 3 |
| 3845 | NM_00136978 | KRAS | KRAS proto-oncogene, GTPase | protein_coding | NP_001356715.1;NP_001356716.1;NP_004976.2;NP_203524.1 | 6407 | 190070 | . | . | 7.409905 | 6.935501 | 6.439995 | 5.256913 | 17.243151 | 16.031718 | 14.451163 | 12.019307 | -1.242299 | 1080.427254 | 0.1498183 | -2.08928359 | 0.036682202 | 0.596884093 | 10.225096 | 9.912809 | 10.179055 | 10.121329 | 10.063267 | 9.911244 | 1430 | 1243 | 927 | 808 |
| 3856 | NM_00125628 | KRT8 | keratin 8 | protein_coding | NP_001243211.1;NP_001243222.1;NP_002264.1 | 6446 | 148060 | . | . | 0.520377 | 0.509935 | 0.361172 | 0.538790 | 1.210940 | 1.178737 | 0.810459 | 1.231879 | -1.154932 | 32.577956 | 0.551798827 | -0.37660009 | 0.706470823 | 0.998865404 | 5.127131 | 4.916862 | 5.039912 | 5.034457 | 4.994541 | 5.041927 | 41 | 37 | 21 | 34 |
| 3858 | NM_000421,N | KRT10 | keratin 10 | protein_coding | NP_000412.4;NP_001366295.1 | 6413 | 148080 | . | . | 8.795501 | 9.039866 | 8.174811 | 10.294885 | 20.467491 | 20.896055 | 18.344038 | 23.538032 | 1.021273 | 607.168011 | 0.169502986 | 0.179161893 | 0.857810584 | 0.998865404 | 9.231102 | 9.260651 | 9.230793 | 9.244304 | 9.176030 | 9.321626 | 685 | 655 | 475 | 638 |
| 3866 | NM_002275,XI | KRT15 | keratin 15 | protein_coding | NP_002266.3;XP_011523086.1;XP_016880103.1 | 6421 | 148030 | . | . | 3.789212 | 4.410829 | 2.216757 | 2.753222 | 8.817651 | 10.195829 | 4.974338 | 6.294914 | -1.759284 | 212.583710 | 0.236493048 | -3.4461422 | 0.000568651 | 0.045370464 | 8.083004 | 7.266850 | 7.817999 | 7.862461 | 7.557419 | 7.606634 | 299 | 305 | 125 | 154 |
| 3868 | NM_005557 | KRT16 | keratin 16 | protein_coding | NP_005548.2 | 6423 | 148067 | . | . | 0.176961 | 0.313600 | 0.150184 | 0.216896 | 0.411795 | 0.724901 | 0.337009 | 0.495907 | -1.321573 | 11.504357 | 0.918107612 | -0.43813597 | 0.661287723 | NA | 3.713381 | 3.306201 | 3.516930 | 3.548900 | 3.511101 | 3.526883 | 11 | 18 | 7 | 11 |
| 3875 | NM_000224,N | KRT18 | keratin 18 | protein_coding | NP_000215.1;NP_954657.1 | 6430 | 148070 | . | . | 20.147448 | 20.539120 | 22.257769 | 20.893761 | 46.883939 | 47.477095 | 49.945789 | 47.771099 | 1.045509 | 924.152305 | 0.140503022 | 0.456967034 | 0.647694741 | 0.998865404 | 9.819419 | 9.883836 | 9.831456 | 9.838990 | 9.893092 | 9.841218 | 1035 | 980 | 851 | 854 |
| 3895 | NM_00107952 | KTN1 | kinectin 1 | protein_coding | NP_001072989.1;NP_001072990.1;NP_001257943.1;NP_004977.2;XP_0( | 6467 | 600381 | . | . | 32.622303 | 30.583472 | 27.434825 | 26.440854 | 75.913447 | 70.695062 | 61.562946 | 60.453870 | -1.178186 | 4338.724107 | 0.110779923 | -2.13546553 | 0.032723003 | 0.57174128 | 12.196434 | 11.960001 | 12.172113 | 12.121773 | 12.038925 | 11.981892 | 5565 | 4914 | 3585 | 3605 |
| 3897 | NM_000425,N | L1CAM | L1 cell adhesion molecule | protein_coding | NP_000416.1;NP_001137435.1;NP_001265045.1;NP_076493.1 | 6470 | 308840 | . | . | 151.705261 | 156.964543 | 140.983116 | 151.086399 | 353.024375 | 362.830551 | 316.362014 | 345.441124 | -1.064706 | 23497.293637 | 0.100399423 | -0.90094831 | 0.367615803 | 0.998865404 | 14.564736 | 14.474260 | 14.535401 | 14.557483 | 14.466466 | 14.517902 | 27549 | 26483 | 19198 | 21979 |
| 3898 | NM_005558 | LAD1 | ladinin 1 | protein_coding | NP_005549.2 | 6472 | 602314 | . | . | 0.293918 | 0.252942 | 0.141205 | 0.254908 | 0.683958 | 0.584686 | 0.316860 | 0.582816 | -1.365538 | 20.120752 | 0.70007771 | -0.64202742 | 0.520855377 | NA | 4.539157 | 4.086744 | 4.356880 | 4.340161 | 4.304169 | 4.340074 | 29 | 23 | 11 | 20 |
| 3899 | NM_00102510 | AFF3 | AF4/FMR2 family member 3 | protein_coding | NP_001020279.1;NP_002276.2;XP_005264000.2;XP_005264002.1;XP_01 | 6473 | 601464 | . | . | 1.333043 | 1.499847 | 0.848287 | 1.190335 | 3.102049 | 3.466963 | 1.903529 | 2.721559 | -1.198073 | 321.235310 | 0.197395125 | -1.32078315 | 0.186573679 | 0.95851159 | 8.452065 | 8.191126 | 8.348589 | 8.400729 | 8.279568 | 8.271541 | 385 | 395 | 256 | 272 |
| 3908 | NM_000426,N | LAMA2 | laminin subunit alpha 2 | protein_coding | NP_000417.3;NP_001073291.2;XP_005267038.1;XP_005267039.1;XP_01 | 6482 | 156225 | . | . | 0.015509 | 0.022913 | 0.061214 | 0.076501 | 0.036091 | 0.052965 | 0.137363 | 0.174910 | 3.530617 | 17.340671 | 0.774406745 | 2.350083507 | 0.0187692 | NA | 2.939842 | 4.755319 | 4.026343 | 4.042011 | 4.106887 | 4.131140 | 7 | 10 | 21 | 28 |
| 3909 | NM_000227,N | LAMA3 | laminin subunit alpha 3 | protein_coding | NP_000218.3;NP_001121189.2;NP_001121190.2;NP_001289925.1;NP_9: | 6483 | 600805 | . | . | 0.103641 | 0.179118 | 0.071452 | 0.093202 | 0.241176 | 0.414039 | 0.160336 | 0.213095 | -1.719906 | 37.514725 | 0.539107194 | -1.45115806 | 0.146735851 | 0.90884489 | 5.569492 | 4.783530 | 5.213011 | 5.310846 | 5.172355 | 5.198549 | 40 | 65 | 21 | 29 |
| 3910 | NM_00110520 | LAMA4 | laminin subunit alpha 4 | protein_coding | NP_001098676.2;NP_001098677.2;NP_001098678.1;NP_001098679.1;NI | 6484 | 600133 | . | . | 30.550774 | 31.087472 | 38.613777 | 41.892077 | 71.092910 | 71.860085 | 86.648197 | 95.781253 | 1.185291 | 6263.782585 | 0.104480771 | 2.347233019 | 0.018913422 | 0.458808379 | 12.485021 | 12.730211 | 12.526457 | 12.547131 | 12.662662 | 12.696972 | 6519 | 6263 | 5790 | 6498 |
| 3911 | NM_005560,XI | LAMA5 | laminin subunit alpha 5 | protein_coding | NP_005551.3;XP_006723859.1;XP_006723861.1;XP_011527120.1;XP_01 | 6485 | 601033 | . | . | 1.040247 | 0.869936 | 0.898382 | 0.948912 | 2.420698 | 2.010897 | 2.015944 | 2.169575 | -1.046122 | 334.852336 | 0.200783156 | -0.32398559 | 0.745948946 | 0.998865404 | 8.419328 | 8.354711 | 8.452451 | 8.342852 | 8.363046 | 8.384757 | 432 | 334 | 279 | 313 |
| 3912 | NM_002291,XI | LAMB1 | laminin subunit beta 1 | protein_coding | NP_002282.2;XP_016867690.1;XP_016867691.1 | 6486 | 150240 | . | . | 66.258834 | 67.418602 | 59.145353 | 60.963337 | 154.187041 | 155.841122 | 132.720452 | 139.385426 | -1.145189 | 11231.225302 | 0.100278172 | -1.95043093 | 0.051124778 | 0.672475152 | 13.549716 | 13.354114 | 13.505109 | 13.515590 | 13.386386 | 13.402905 | 13724 | 13018 | 9016 | 9913 |
| 3913 | NM_002292,XI | LAMB2 | laminin subunit beta 2 | protein_coding | NP_002283.3;XP_005265184.1 | 6487 | 150325 | . | . | 35.704468 | 38.422100 | 33.225053 | 34.306612 | 83.085763 | 88.814407 | 74.556062 | 78.437992 | -1.112549 | 6270.579022 | 0.105782426 | -1.45458162 | 0.145785123 | 0.906428851 | 12.689295 | 12.535367 | 12.630210 | 12.684212 | 12.558413 | 12.576844 | 7364 | 7353 | 5105 | 5627 |
| 3914 | NM_000228,N | LAMB3 | laminin subunit beta 3 | protein_coding | NP_000219.2;NP_001017402.1;NP_001121113.1;XP_005273181.1;XP_01 | 6490 | 150310 | . | . | 6.002087 | 6.170976 | 4.362891 | 4.804393 | 13.967103 | 14.264487 | 9.790198 | 10.984673 | -1.344523 | 711.971144 | 0.15155999 | -2.8179882 | 0.004832559 | 0.207154674 | 9.673645 | 9.246151 | 9.560132 | 9.575858 | 9.338487 | 9.389741 | 930 | 891 | 507 | 592 |
| 3915 | NM_002293 | LAMC1 | laminin subunit gamma 1 | protein_coding | NP_002284.3 | 6492 | 150290 | . | . | 78.351318 | 76.932335 | 80.215363 | 85.548149 | 182.326736 | 177.832535 | 180.000961 | 195.595688 | 1.052514 | 19820.004379 | 0.100388873 | 0.735538514 | 0.462011593 | 0.998865404 | 14.237283 | 14.311112 | 14.263576 | 14.240599 | 14.272539 | 14.319248 | 22537 | 20560 | 17198 | 19572 |
| 3916 | NM_005561,XI | LAMP1 | lysosomal associated membrane protein 1 | protein_coding | NP_005552.3;XP_011535796.1 | 6499 | 153330 | . | . | 125.080726 | 122.826607 | 114.023552 | 112.045471 | 291.067976 | 283.919220 | 255.865538 | 256.178650 | -1.111955 | 9972.937913 | 0.101347433 | -1.51062446 | 0.130884159 | 0.882587711 | 13.358309 | 13.205239 | 13.338606 | 13.315894 | 13.248408 | 13.225329 | 12252 | 11182 | 8325 | 8733 |
| 3918 | NM_005562,N | LAMC2 | laminin subunit gamma 2 | protein_coding | NP_005553.2;NP_061486.2;XP_016856762.1 | 6493 | 150292 | . | . | 0.129737 | 0.108839 | 0.041245 | 0.112262 | 0.301904 | 0.251587 | 0.092552 | 0.256673 | -1.535359 | 17.308165 | 0.773776409 | -0.79942534 | 0.424043822 | NA | 4.391727 | 3.768175 | 4.145350 | 4.123929 | 4.079743 | 4.126536 | 27 | 20 | 7 | 18 |
| 3920 | NM_00112260 | LAMP2 | lysosomal associated membrane protein 2 | protein_coding | NP_001116078.1;NP_002285.1;NP_054701.1 | 6501 | 309060 | . | . | 74.332625 | 66.003779 | 65.409180 | 50.524617 | 172.975071 | 152.570681 | 146.776307 | 115.518528 | -1.183000 | 9225.174957 | 0.128710991 | -1.88367966 | 0.059608319 | 0.719735192 | 13.287439 | 13.045119 | 13.282208 | 13.189302 | 13.185808 | 12.992655 | 12131 | 10209 | 8194 | 7014 |
| 3921 | NM_00130428 | RPSA | ribosomal protein SA | protein_coding | NP_001291217.1;NP_002286.2 | 6502 | 150370 | . | . | 337.022881 | 342.388340 | 311.494499 | 378.048827 | 784.266056 | 791.445992 | 698.984636 | 864.363786 | 1.000077 | 12156.720348 | 0.116429839 | 0.000956817 | 0.999236571 | 0.999862634 | 13.569447 | 13.569486 | 13.564943 | 13.571925 | 13.487283 | 13.643747 | 13941 | 13171 | 9605 | 12439 |
| 3924 | gene-RPSAP14 | RPSAP14 | ribosomal protein SA pseudogene 14 | pseudogene | . | 6506 | . | | . | 0.008209 | 0.000399 | 0.003555 | 0.003883 | 0.019104 | 0.000921 | 0.007977 | 0.008878 | 1.234655 | 1.002957 | 2.868339486 | 0.106022226 | 0.915564717 | NA | -0.155866 | 0.148375 | -0.001261 | -0.000972 | 0.000053 | -0.000315 | 1 | 1 | 1 | 1 |
| 3925 | NM_00114545 | STMN1 | stathmin 1 | protein_coding | NP_001138926.1;NP_005554.1;NP_981944.1;NP_981946.1 | 6510 | 151442 | . | . | 100.075480 | 100.355873 | 86.320901 | 98.971835 | 232.879750 | 231.977113 | 193.701596 | 226.287337 | -1.089700 | 4393.547945 | 0.113920817 | -1.08786999 | 0.276652507 | 0.998865404 | 12.161874 | 12.037801 | 12.135302 | 12.135765 | 12.011527 | 12.113757 | 5275 | 4946 | 3430 | 4186 |
| 3927 | NM_00127160 | LASP1 | LIM and SH3 protein 1 | protein_coding | NP_001258537.1;NP_006139.1 | 6513 | 602920 | . | . | 43.192052 | 44.221956 | 41.090029 | 45.519594 | 100.509682 | 102.221031 | 92.204835 | 104.075144 | -1.023775 | 5301.633138 | 0.109165301 | -0.31053051 | 0.756157563 | 0.998865404 | 12.389122 | 12.355120 | 12.375676 | 12.387850 | 12.322784 | 12.399842 | 6132 | 5830 | 4346 | 5138 |
| 3930 | NM_002296,N | LBR | lamin B receptor | protein_coding | NP_002287.2;NP_919424.1;XP_005273182.1;XP_011542487.1 | 6518 | 600024 | . | . | 16.412857 | 13.901830 | 14.308229 | 11.253335 | 38.193391 | 32.134704 | 32.107251 | 25.729415 | -1.205889 | 1633.875484 | 0.152649458 | -1.7693946 | 0.076828046 | 0.767975096 | 10.802532 | 10.533068 | 10.806894 | 10.667831 | 10.685010 | 10.493735 | 2235 | 1760 | 1450 | 1215 |
| 3931 | NM_000229 | LCAT | lecithin-cholesterol acyltransferase | protein_coding | NP_000220.1 | 6522 | 606967 | . | . | 0.699485 | 0.815553 | 0.540096 | 0.608758 | 1.627730 | 1.885188 | 1.211959 | 1.391852 | -1.327994 | 31.505881 | 0.554488858 | -0.73806372 | 0.460475731 | 0.998865404 | 5.168403 | 4.757542 | 4.986012 | 5.009626 | 4.955558 | 4.969132 | 38 | 42 | 22 | 27 |
| 3936 | NM_002298,XI | LCP1 | lymphocyte cytosolic protein 1 | protein_coding | NP_002289.2;XP_005266431.1 | 6528 | 153430 | . | . | 2.312585 | 2.560916 | 1.584555 | 2.168541 | 5.381478 | 5.919672 | 3.555696 | 4.958113 | -1.309382 | 246.145582 | 0.232351931 | -1.67369503 | 0.094190551 | 0.810751527 | 8.125507 | 7.734866 | 7.979950 | 8.027979 | 7.804867 | 7.932873 | 307 | 315 | 159 | 228 |
| 3937 | NM_005565 | LCP2 | lymphocyte cytosolic protein 2 | protein_coding | NP_005556.1 | 6529 | 601603 | . | . | 3.908290 | 3.403538 | 3.097731 | 3.420194 | 9.094752 | 7.867429 | 6.951217 | 7.819868 | -1.138442 | 456.609836 | 0.179525328 | -1.04197226 | 0.297424538 | 0.998865404 | 8.925305 | 8.738274 | 8.919076 | 8.824902 | 8.766513 | 8.818810 | 601 | 486 | 355 | 418 |
| 3939 | NM_00113523 | LDHA | lactate dehydrogenase A | protein_coding | NP_001128711.1;NP_001158886.1;NP_001158887.1;NP_001158888.1;NI | 6535 | 150000 | . | . | 316.860381 | 316.572988 | 325.947754 | 345.499107 | 737.347081 | 731.772668 | 731.417287 | 789.942569 | 1.044544 | 22695.144658 | 0.099387751 | 0.632611141 | 0.52698761 | 0.998865404 | 14.438323 | 14.501185 | 14.454923 | 14.446998 | 14.467091 | 14.509443 | 25682 | 23845 | 19671 | 22273 |
| 3945 | NM_00117409 | LDHB | lactate dehydrogenase B | protein_coding | NP_001167568.1;NP_001302466.1;NP_002291.1;XP_006719137.1 | 6541 | 150100 | . | . | 480.255154 | 469.026260 | 461.617770 | 515.298737 | 1117.573516 | 1084.175204 | 1035.856866 | 1178.169359 | 1.014466 | 19614.184801 | 0.104389491 | 0.198487516 | 0.842663652 | 0.998865404 | 14.249223 | 14.269922 | 14.267451 | 14.238519 | 14.222691 | 14.306457 | 22801 | 20659 | 16343 | 19421 |
| 3949 | NM_000527,N | LDLR | low density lipoprotein receptor | protein_coding | NP_000518.1;NP_001182727.1;NP_001182728.1;NP_001182729.1;NP_0( | 6547 | 606945 | . | . | 29.216352 | 31.615137 | 36.647902 | 37.595529 | 67.987650 | 73.079807 | 82.236822 | 85.957708 | 1.203231 | 5440.315354 | 0.107246019 | 2.488792911 | 0.01281776 | 0.374881745 | 12.269892 | 12.536735 | 12.296636 | 12.355576 | 12.474812 | 12.488761 | 5484 | 5519 | 5125 | 5616 |
| 3953 | NM_00100367 | LEPR | leptin receptor | protein_coding | NP_001003679.1;NP_001003680.1;NP_001185616.1;NP_001185617.1;NI | 6554 | 601007 | . | . | 0.201389 | 0.220912 | 0.265017 | 0.164135 | 0.468640 | 0.510649 | 0.594691 | 0.375276 | 1.010783 | 33.859853 | 0.547723267 | 0.028250475 | 0.97746238 | 0.998865404 | 5.072328 | 5.090622 | 5.073616 | 5.078874 | 5.105968 | 5.044265 | 38 | 37 | 37 | 24 |
| 3954 | NM_012318,XI | LETM1 | leucine zipper and EF-hand containing transmembrane protein 1 | protein_coding | NP_036450.1;XP_006713947.1 | 6556 | 604407 | . | . | 8.782510 | 9.154894 | 9.929182 | 9.558680 | 20.437264 | 21.161947 | 22.280797 | 21.854787 | 1.074591 | 1560.679417 | 0.124591814 | 0.833017608 | 0.404834799 | 0.998865404 | 10.555102 | 10.658947 | 10.565348 | 10.592518 | 10.654098 | 10.615267 | 1702 | 1652 | 1441 | 1478 |
| 3955 | NM_00104016 | LFNG | LFNG O-fucosylpeptide 3-beta-N-acetylglucosaminyltransferase | protein_coding | NP_001035257.1;NP_001035258.1;NP_001159827.1;NP_002295.1 | 6560 | 602576 | . | . | 0.834421 | 1.002978 | 0.431740 | 0.394922 | 1.941731 | 2.318428 | 0.968813 | 0.902943 | -2.329454 | 50.763065 | 0.452869963 | -2.69391253 | 0.007061869 | 0.266401081 | 6.150285 | 4.930721 | 5.710169 | 5.750666 | 5.588011 | 5.573355 | 75 | 83 | 28 | 27 |
| 3956 | NM_002305 | LGALS1 | galectin 1 | protein_coding | NP_002296.1 | 6561 | 150570 | . | . | 1450.549316 | 1436.577026 | 1014.435547 | 1187.814209 | 3375.488281 | 3320.712158 | 2276.364258 | 2715.796143 | -1.329882 | 20945.125657 | 0.110029462 | -3.73807457 | 0.000185435 | 0.019538615 | 14.545385 | 14.134047 | 14.472190 | 14.456133 | 14.155180 | 14.278940 | 27784 | 25566 | 14483 | 18097 |
| 3958 | NM_00135767 | LGALS3 | galectin 3 | protein_coding | NP_001344607.1;NP_002297.2 | 6563 | 153619 | . | . | 552.047509 | 545.125090 | 590.484134 | 717.016939 | 1284.637295 | 1260.080977 | 1325.029307 | 1639.374033 | 1.175019 | 18550.824502 | 0.112713655 | 2.064377617 | 0.038981927 | 0.615637916 | 14.058188 | 14.290834 | 14.111152 | 14.099025 | 14.169460 | 14.313066 | 19783 | 18275 | 15995 | 20337 |
| 3959 | NM_005567 | LGALS3BP | galectin 3 binding protein | protein_coding | NP_005558.1 | 6564 | 600626 | . | . | 292.530579 | 302.937347 | 272.109955 | 291.602997 | 680.730652 | 700.253235 | 610.606873 | 666.715576 | -1.071262 | 19881.156214 | 0.100703655 | -0.9861775 | 0.324046021 | 0.998865404 | 14.327927 | 14.228591 | 14.296411 | 14.319251 | 14.223183 | 14.273594 | 23368 | 22484 | 16201 | 18528 |
| 3964 | NM_006499,N | LGALS8 | galectin 8 | protein_coding | NP_006490.3;NP_963837.1;NP_963838.1;NP_963839.1;XP_011542490.1 | 6569 | 606099 | . | . | 0.317692 | 0.253041 | 0.454182 | 0.405064 | 0.739283 | 0.584916 | 1.019170 | 0.926130 | 1.302499 | 65.367985 | 0.40250555 | 0.947271991 | 0.3435002 | 0.998865404 | 5.825792 | 6.209760 | 6.014697 | 5.966151 | 6.086170 | 6.013311 | 73 | 54 | 74 | 59 |
| 3975 | NM_005568 | LHX1 | LIM homeobox 1 | protein_coding | NP_005559.2 | 6593 | 601999 | . | . | 0.992114 | 0.972613 | 0.975067 | 1.072909 | 2.308689 | 2.248239 | 2.188022 | 2.453078 | 1.030275 | 107.797504 | 0.309268666 | 0.139133692 | 0.889344505 | 0.998865404 | 6.730701 | 6.773344 | 6.751534 | 6.742582 | 6.742712 | 6.770290 | 124 | 113 | 91 | 107 |
| 3976 | NM_00125713 | LIF | LIF interleukin 6 family cytokine | protein_coding | NP_001244064.1;NP_002300.1;XP_024308007.1;XP_024308008.1 | 6596 | 159540 | . | . | 47.948917 | 52.865801 | 49.086881 | 45.660121 | 111.579093 | 122.201674 | 110.149534 | 104.396443 | -1.079003 | 5883.535394 | 0.111658222 | -0.9824544 | 0.325876039 | 0.998865404 | 12.576264 | 12.466587 | 12.514067 | 12.590798 | 12.523588 | 12.453642 | 6714 | 6888 | 5128 | 5084 |
| 3977 | NM_00112767 | LIFR | LIF receptor subunit alpha | protein_coding | NP_001121143.1;NP_001351226.1;NP_001351227.1;NP_002301.1;XP_01 | 6597 | 151443 | . | . | 0.844319 | 0.753875 | 0.694805 | 0.509991 | 1.964766 | 1.742615 | 1.559123 | 1.166034 | -1.344501 | 231.478626 | 0.241720648 | -1.76679299 | 0.077262904 | 0.767975096 | 8.051670 | 7.626603 | 7.949350 | 7.885450 | 7.843035 | 7.703691 | 324 | 269 | 199 | 156 |
| 3978 | NM_000234,N | LIG1 | DNA ligase 1 | protein_coding | NP_000225.1;NP_001275992.1;NP_001275993.1;NP_001307899.1;NP_0( | 6598 | 126391 | . | . | 15.841181 | 15.575543 | 14.541627 | 15.974529 | 36.863078 | 36.003567 | 32.630991 | 36.523861 | -1.031458 | 1615.504768 | 0.125884077 | -0.35496747 | 0.722613958 | 0.998865404 | 10.680045 | 10.635144 | 10.672826 | 10.666314 | 10.608407 | 10.681068 | 1897 | 1763 | 1318 | 1561 |
| 3980 | NM_002311,N | LIG3 | DNA ligase 3 | protein_coding | NP_002302.2;NP_039269.2;XP_005258027.1;XP_006721959.1;XP_01688 | 6600 | 600940 | . | . | 5.594977 | 6.248722 | 6.397957 | 5.373687 | 13.019740 | 14.444202 | 14.356827 | 12.286296 | 1.029238 | 1224.365328 | 0.132786287 | 0.313104563 | 0.754201222 | 0.998865404 | 10.237033 | 10.278309 | 10.238572 | 10.254524 | 10.233454 | 10.301831 | 1375 | 1316 | 1031 | 1217 |
| 3981 | NM_00109826 | LIG4 | DNA ligase 4 | protein_coding | NP_001091738.1;NP_001317524.1;NP_001339527.1;NP_001339528.1;NI | 6601 | 601837 | . | . | 3.493231 | 2.942588 | 3.684305 | 2.562879 | 8.128892 | 6.801925 | 8.267475 | 5.859719 | -1.037581 | 405.456048 | 0.209083427 | -0.25455673 | 0.79906548 | 0.998865404 | 8.688981 | 8.637361 | 8.718239 | 8.619060 | 8.743134 | 8.543074 | 514 | 409 | 405 | 310 |
| 3983 | NM_00100340 | ABLIM1 | actin binding LIM protein 1 | protein_coding | NP_001003407.1;NP_001309811.1;NP_001309812.1;NP_001309813.1;NI | 78 | 602330 | . | . | 0.372765 | 0.347827 | 0.286509 | 0.373628 | 0.867439 | 0.804017 | 0.642917 | 0.854256 | -1.096234 | 92.400943 | 0.339176264 | -0.39081562 | 0.69593353 | 0.998865404 | 6.595112 | 6.461466 | 6.558891 | 6.527730 | 6.481945 | 6.550618 | 117 | 99 | 67 | 93 |
| 3984 | NM_00120442 | LIMK1 | LIM domain kinase 1 | protein_coding | NP_001191355.1;NP_002305.1 | 6613 | 601329 | . | . | 44.571524 | 48.111233 | 38.461159 | 41.917385 | 103.719765 | 111.211262 | 86.305729 | 95.839127 | -1.169648 | 4379.140412 | 0.112448643 | -2.01047327 | 0.044381122 | 0.645513836 | 12.205116 | 11.978889 | 12.125010 | 12.189605 | 11.995068 | 12.058839 | 5229 | 5290 | 3373 | 3932 |
| 3985 | NM_00103180 | LIMK2 | LIM domain kinase 2 | protein_coding | NP_001026971.1;NP_005560.1;NP_057952.1 | 6614 | 601988 | . | . | 31.236233 | 31.043976 | 28.356624 | 35.006012 | 72.687995 | 71.759536 | 63.631448 | 80.037084 | -1.001796 | 3524.922278 | 0.125882799 | -0.02056054 | 0.983596216 | 0.998865404 | 11.784763 | 11.781988 | 11.798963 | 11.766751 | 11.698007 | 11.858617 | 4142 | 3733 | 2764 | 3623 |
| 3987 | NM_00119348 | LIMS1 | LIM zinc finger domain containing 1 | protein_coding | NP_001180411.1;NP_001180412.1;NP_001180413.1;NP_001180414.1;NI | 6616 | 602567 | . | . | 6.513583 | 5.447688 | 5.553375 | 5.110727 | 15.157377 | 12.592574 | 12.461613 | 11.685069 | -1.142603 | 768.361617 | 0.159593656 | -1.20508472 | 0.228170598 | 0.982008277 | 9.678259 | 9.486668 | 9.693771 | 9.560119 | 9.569564 | 9.501344 | 1031 | 802 | 655 | 639 |
| 3988 | NM_000235,N | LIPA | lipase A, lysosomal acid type | protein_coding | NP_000226.2;NP_001121077.1;NP_001275908.1;XP_024303791.1 | 6617 | 613497 | . | . | 44.862764 | 39.417038 | 41.024893 | 36.931067 | 104.397485 | 91.114251 | 92.058671 | 84.438492 | -1.090973 | 3207.232482 | 0.123359678 | -1.01828094 | 0.308544464 | 0.998865404 | 11.708427 | 11.583078 | 11.732719 | 11.626156 | 11.655694 | 11.561233 | 4101 | 3379 | 2823 | 2709 |
| 3991 | NM_005357,XI | LIPE | lipase E, hormone sensitive type | protein_coding | NP_005348.2;XP_005258994.1;XP_005258995.1;XP_005258996.2;XP_00! | 6621 | 151750 | . | . | 0.211239 | 0.235256 | 0.099742 | 0.186633 | 0.491561 | 0.543805 | 0.223819 | 0.426714 | -1.531914 | 22.226333 | 0.671326419 | -0.91659683 | 0.359353947 | NA | 4.751025 | 4.131331 | 4.494861 | 4.501921 | 4.439172 | 4.479215 | 30 | 30 | 11 | 21 |
| 3992 | NM_013402,XI | FADS1 | fatty acid desaturase 1 | protein_coding | NP_037534.5;XP_011543324.1 | 3574 | 606148 | . | . | 19.833618 | 21.286067 | 27.500211 | 27.749031 | 46.153641 | 49.203700 | 61.709679 | 63.444865 | 1.324596 | 3271.464387 | 0.111621004 | 3.633301782 | 0.000279817 | 0.026793856 | 11.458784 | 11.864261 | 11.523249 | 11.573228 | 11.779841 | 11.778916 | 3140 | 3132 | 3244 | 3493 |
| 3993 | NM_00101500 | LLGL2 | LLGL scribble cell polarity complex component 2 | protein_coding | NP_001015002.1;NP_001026973.1;NP_004515.2;XP_011523104.1;XP_01 | 6629 | 618483 | . | . | 1.563513 | 1.570471 | 1.477722 | 1.689265 | 3.638361 | 3.630211 | 3.315962 | 3.862302 | 1.022021 | 168.218571 | 0.253991777 | 0.123724828 | 0.901533153 | 0.998865404 | 7.378272 | 7.409940 | 7.385721 | 7.392624 | 7.409528 | 7.386754 | 190 | 181 | 152 | 155 |
| 3995 | NM_021727,XI | FADS3 | fatty acid desaturase 3 | protein_coding | NP_068373.1;XP_011543325.1;XP_016873212.1;XP_016873213.1 | 3576 | 606150 | . | . | 32.522211 | 33.279072 | 43.283894 | 40.130050 | 75.680523 | 76.926064 | 97.127806 | 91.752593 | 1.258823 | 2191.671413 | 0.121936718 | 2.723338195 | 0.006462586 | 0.253589295 | 10.922148 | 11.254397 | 10.990887 | 11.003858 | 11.227863 | 11.133369 | 2216 | 2111 | 2247 | 2158 |
| 3996 | NM_004140,XI | LLGL1 | LLGL scribble cell polarity complex component 1 | protein_coding | NP_004131.4;XP_011522151.1;XP_011522152.1;XP_011522153.1;XP_01 | 6628 | 600966 | . | . | 5.426413 | 6.266333 | 6.074547 | 6.299817 | 12.627483 | 14.484911 | 13.631106 | 14.403783 | 1.041357 | 790.526912 | 0.150999674 | 0.387187246 | 0.698617591 | 0.998865404 | 9.597362 | 9.655396 | 9.562163 | 9.658438 | 9.630466 | 9.649122 | 831 | 893 | 692 | 766 |
| 3998 | NM_005570 | LMAN1 | lectin, mannose binding 1 | protein_coding | NP_005561.1 | 6631 | 601567 | . | . | 39.451073 | 36.068619 | 31.816696 | 27.290529 | 91.804268 | 83.374229 | 71.395744 | 62.396553 | -1.295541 | 5063.935844 | 0.12167654 | -3.07006725 | 0.002140106 | 0.117381585 | 12.480674 | 12.107325 | 12.444956 | 12.360731 | 12.251527 | 12.118101 | 6904 | 5865 | 4150 | 3799 |
| 4000 | NM_00125737 | LMNA | lamin A/C | protein_coding | NP_001244303.1;NP_001269553.1;NP_001269554.1;NP_001269555.1;NI | 6636 | 150330 | . | . | 163.991417 | 165.644789 | 150.229404 | 184.260459 | 381.614757 | 382.895329 | 337.110431 | 421.289641 | 1.001629 | 13334.357783 | 0.117579608 | 0.019976216 | 0.984062345 | 0.99904476 | 13.701716 | 13.704005 | 13.709781 | 13.692270 | 13.617309 | 13.781065 | 15499 | 14230 | 10496 | 13706 |
| 4001 | NM_00119855 | LMNB1 | lamin B1 | protein_coding | NP_001185486.1;NP_005564.1 | 6637 | 150340 | . | . | 23.435577 | 25.198328 | 21.345015 | 23.539080 | 54.535554 | 58.247059 | 47.897595 | 53.819309 | -1.097911 | 2111.177530 | 0.121615533 | -1.10809301 | 0.267821648 | 0.998849558 | 11.109766 | 10.974740 | 11.058060 | 11.101012 | 10.968015 | 11.040903 | 2477 | 2448 | 1669 | 1974 |
| 4005 | NM_00114231 | LMO2 | LIM domain only 2 | protein_coding | NP_001135787.1;NP_001135788.1;NP_005565.2;XP_005252978.1;XP_01 | 6642 | 180385 | . | . | 0.909283 | 0.589560 | 0.474790 | 0.604027 | 2.115939 | 1.362796 | 1.065415 | 1.381036 | -1.552917 | 47.970058 | 0.468841434 | -1.35436141 | 0.175621102 | 0.949137853 | 5.866426 | 5.232645 | 5.664924 | 5.576310 | 5.541307 | 5.546669 | 81 | 50 | 32 | 36 |
| 4007 | NM_00130797 | PRICKLE3 | prickle planar cell polarity protein 3 | protein_coding | NP_001294908.1;NP_006141.2 | 6645 | 300111 | . | . | 2.783138 | 2.987380 | 3.381968 | 3.469463 | 6.476477 | 6.905462 | 7.589040 | 7.932514 | 1.179905 | 273.957282 | 0.208758784 | 1.143284686 | 0.252920411 | 0.994320829 | 7.973752 | 8.212035 | 8.030527 | 8.070709 | 8.132473 | 8.140533 | 278 | 282 | 256 | 280 |
| 4008 | NM_00130608 | LMO7 | LIM domain 7 | protein_coding | NP_001293009.1;NP_001317512.1;NP_001353561.1;NP_001353562.1;NI | 6646 | 604362 | . | . | 8.925257 | 7.597208 | 5.972687 | 6.166709 | 20.769441 | 17.561291 | 13.402534 | 14.099446 | -1.185713 | 1456.882715 | 0.125072443 | -1.96489866 | 0.049425968 | 0.664938243 | 10.626299 | 10.380590 | 10.579682 | 10.561501 | 10.440232 | 10.439098 | 1841 | 1686 | 1160 | 1249 |
| 4010 | NM_00117414 | LMX1B | LIM homeobox transcription factor 1 beta | protein_coding | NP_001167617.1;NP_001167618.1;NP_002307.2 | 6654 | 602575 | . | . | 0.056666 | 0.067354 | 0.060104 | 0.031200 | 0.131865 | 0.155692 | 0.134873 | 0.071334 | -1.371966 | 11.277270 | 0.925999498 | -0.4927056 | 0.62222062 | NA | 3.703730 | 3.251698 | 3.500599 | 3.508331 | 3.502407 | 3.475150 | 14 | 15 | 11 | 6 |
| 4012 | NM_005575,N | LNPEP | leucyl and cystinyl aminopeptidase | protein_coding | NP_005566.2;NP_787116.2;XP_024301813.1 | 6656 | 151300 | . | . | 3.892985 | 3.459918 | 3.615195 | 3.075659 | 9.059137 | 7.997756 | 8.112393 | 7.032131 | -1.114247 | 1330.151353 | 0.143053572 | -1.09098326 | 0.275280249 | 0.998865404 | 10.453014 | 10.297549 | 10.463197 | 10.366098 | 10.395686 | 10.267316 | 1715 | 1418 | 1188 | 1079 |
| 4013 | NM_00113014 | VWA5A | von Willebrand factor A domain containing 5A | protein_coding | NP_001123614.1;NP_055437.2;NP_938057.1;XP_011541130.1 | 6658 | 602929 | . | . | 3.064935 | 2.803319 | 3.224344 | 3.691230 | 7.132232 | 6.480000 | 7.235336 | 8.439559 | 1.118074 | 207.789585 | 0.233389041 | 0.689903644 | 0.490254784 | 0.998865404 | 7.616351 | 7.777131 | 7.681170 | 7.661183 | 7.703182 | 7.741386 | 231 | 207 | 183 | 214 |
| 4017 | NM_002318 | LOXL2 | lysyl oxidase like 2 | protein_coding | NP_002309.1 | 6666 | 606663 | . | . | 16.993401 | 17.391249 | 9.683474 | 9.312984 | 39.544342 | 40.200649 | 21.729433 | 21.293032 | -1.835004 | 1550.563775 | 0.125069797 | -7.00235743 | 2.51691E-12 | 5.79015E-09 | 10.970955 | 10.095326 | 10.778859 | 10.790906 | 10.334810 | 10.298773 | 2294 | 2182 | 975 | 1000 |
| 4023 | NM_000237 | LPL | lipoprotein lipase | protein_coding | NP_000228.1 | 6677 | 609708 | . | . | 0.303324 | 0.501405 | 0.318011 | 0.477512 | 0.705849 | 1.159020 | 0.713606 | 1.091774 | -1.022747 | 45.062919 | 0.494230675 | -0.06565661 | 0.94765122 | 0.998865404 | 5.512055 | 5.475451 | 5.447976 | 5.532030 | 5.453931 | 5.519282 | 40 | 61 | 31 | 50 |
| 4026 | NM_00116767 | LPP | LIM domain containing preferred translocation partner in lipoma | protein_coding | NP_001161143.1;NP_001161144.1;NP_001362384.1;NP_001362385.1;NI | 6679 | 600700 | . | . | 2.972414 | 2.898532 | 2.965721 | 2.625081 | 6.916932 | 6.700088 | 6.654992 | 6.001935 | -1.067979 | 1585.521140 | 0.139086074 | -0.68219487 | 0.49511574 | 0.998865404 | 10.677157 | 10.582783 | 10.679331 | 10.628739 | 10.678084 | 10.522764 | 1946 | 1710 | 1468 | 1293 |
| 4034 | NM_00128993 | LRCH4 | leucine rich repeats and calponin homology domain containing 4 | protein_coding | NP_001276863.1;NP_002310.2 | 6691 | . | | . | 14.649496 | 14.306598 | 16.121908 | 17.661239 | 34.089984 | 33.070342 | 36.177097 | 40.380323 | 1.151850 | 1537.513813 | 0.125451961 | 1.625746963 | 0.104003495 | 0.828395026 | 10.480861 | 10.684711 | 10.538824 | 10.518124 | 10.612168 | 10.663065 | 1668 | 1521 | 1385 | 1593 |
| 4035 | NM_002332,XI | LRP1 | LDL receptor related protein 1 | protein_coding | NP_002323.2;XP_016874792.1 | 6692 | 107770 | . | . | 4.627269 | 4.620924 | 3.256194 | 4.173056 | 10.767845 | 10.681474 | 7.306805 | 9.541196 | -1.250382 | 1948.036401 | 0.138910403 | -2.32069952 | 0.020303067 | 0.470997021 | 11.080194 | 10.757397 | 11.012275 | 11.004348 | 10.735692 | 10.918320 | 2505 | 2325 | 1327 | 1815 |
| 4036 | NM_004525,XI | LRP2 | LDL receptor related protein 2 | protein_coding | NP_004516.2;XP_011509485.1;XP_011509486.1 | 6694 | 600073 | . | . | 0.706741 | 0.577682 | 0.558387 | 0.669804 | 1.644614 | 1.335338 | 1.253003 | 1.531427 | -1.060389 | 306.878724 | 0.213630686 | -0.39597964 | 0.692120046 | 0.998865404 | 8.303258 | 8.218549 | 8.332650 | 8.215613 | 8.198108 | 8.287932 | 402 | 305 | 237 | 303 |
| 4037 | NM_002333,XI | LRP3 | LDL receptor related protein 3 | protein_coding | NP_002324.2;XP_005259002.1 | 6695 | 603159 | . | . | 0.299498 | 0.454009 | 0.259794 | 0.241248 | 0.696943 | 1.049463 | 0.582971 | 0.551586 | -1.414700 | 38.815050 | 0.506773958 | -0.98761153 | 0.323342938 | 0.998865404 | 5.507439 | 5.006356 | 5.279731 | 5.329025 | 5.257932 | 5.248058 | 45 | 56 | 29 | 29 |
| 4038 | NM_002334,XI | LRP4 | LDL receptor related protein 4 | protein_coding | NP_002325.2;XP_011518405.1;XP_011518406.1;XP_016873223.1 | 6696 | 604270 | . | . | 2.537437 | 3.090927 | 2.706375 | 3.602515 | 5.904719 | 7.144818 | 6.073025 | 8.236721 | 1.017193 | 761.742743 | 0.14952458 | 0.164479868 | 0.869353394 | 0.998865404 | 9.561037 | 9.585182 | 9.542561 | 9.590595 | 9.552286 | 9.604340 | 840 | 843 | 643 | 747 |
| 4040 | NM_002336,XI | LRP6 | LDL receptor related protein 6 | protein_coding | NP_002327.2;XP_006719141.1;XP_011518973.1 | 6698 | 603507 | . | . | 4.302852 | 4.287226 | 4.463819 | 4.015582 | 10.012915 | 9.910118 | 10.016681 | 9.181151 | -1.025484 | 1317.625313 | 0.131155763 | -0.27680858 | 0.781927093 | 0.998865404 | 10.381623 | 10.345601 | 10.375309 | 10.370469 | 10.390287 | 10.316042 | 1541 | 1435 | 1187 | 1160 |
| 4041 | NM_00129190 | LRP5 | LDL receptor related protein 5 | protein_coding | NP_001278831.1;NP_002326.2;XP_005274051.1;XP_011543331.1;XP_01 | 6697 | 603506 | . | . | 11.292039 | 10.647307 | 9.266896 | 10.117583 | 26.277035 | 24.611729 | 20.794643 | 23.132651 | -1.146856 | 1668.607035 | 0.125936721 | -1.56971393 | 0.116481679 | 0.857396579 | 10.799927 | 10.602158 | 10.782187 | 10.727788 | 10.618790 | 10.677160 | 2123 | 1858 | 1300 | 1513 |
| 4043 | NM_002337,N | LRPAP1 | LDL receptor related protein associated protein 1 | protein_coding | NP_002328.1 | 6701 | 104225 | . | . | 25.876570 | 25.215842 | 24.179399 | 26.524780 | 60.215838 | 58.287550 | 54.257868 | 60.645753 | -1.020766 | 8191.052884 | 0.106398587 | -0.27869296 | 0.780480468 | 0.998865404 | 13.014603 | 12.984911 | 13.024566 | 12.991688 | 12.954183 | 13.026156 | 9713 | 8757 | 6750 | 7923 |
| 4045 | NM_00131891 | LSAMP | limbic system associated membrane protein | protein_coding | NP_001305844.1;NP_002329.2;XP_011511142.1;XP_016861872.1;XP_01 | 6705 | 603241 | . | . | 1.297079 | 1.215290 | 1.253719 | 1.045313 | 3.018356 | 2.809198 | 2.813310 | 2.389985 | -1.104809 | 351.210317 | 0.198755098 | -0.72348854 | 0.469379794 | 0.998865404 | 8.525748 | 8.383111 | 8.502688 | 8.462132 | 8.481013 | 8.368072 | 439 | 384 | 320 | 281 |
| 4046 | NM_00101325 | LSP1 | lymphocyte specific protein 1 | protein_coding | NP_001013271.1;NP_001013272.1;NP_001013273.1;NP_001229861.1;NI | 6707 | 153432 | . | . | 0.017092 | 0.018251 | 0.018345 | 0.017106 | 0.039774 | 0.042187 | 0.041167 | 0.039111 | -1.620137 | 1.451754 | 2.440017378 | -0.28529139 | 0.775420906 | NA | 0.844134 | 0.148375 | 0.548030 | 0.548560 | 0.545314 | 0.544903 | 2 | 2 | 1 | 1 |
| 4047 | NM_00100143 | LSS | lanosterol synthase | protein_coding | NP_001001438.1;NP_001138908.1;NP_001138909.1;NP_002331.3 | 6708 | 600909 | . | . | 14.991713 | 14.504753 | 23.044403 | 22.074992 | 34.886331 | 33.528384 | 51.710977 | 50.471847 | 1.512856 | 2625.053186 | 0.114274354 | 5.226673896 | 1.72586E-07 | 7.0065E-05 | 11.028735 | 11.626123 | 11.175605 | 11.155640 | 11.520437 | 11.481182 | 2437 | 2225 | 2813 | 2894 |
| 4048 | NM_000895,N | LTA4H | leukotriene A4 hydrolase | protein_coding | NP_000886.1;NP_001243572.1;NP_001243573.1;XP_005268928.1;XP_01 | 6710 | 151570 | . | . | 40.952194 | 39.545467 | 43.190957 | 45.373077 | 95.297431 | 91.411118 | 96.919254 | 103.740145 | 1.081210 | 2839.435866 | 0.112932853 | 0.997472074 | 0.318535422 | 0.998865404 | 11.413973 | 11.526606 | 11.453359 | 11.423512 | 11.486856 | 11.517236 | 3200 | 2890 | 2518 | 2817 |
| 4052 | NM_000627,N | LTBP1 | latent transforming growth factor beta binding protein 1 | protein_coding | NP_000618.4;NP_001159736.2;NP_001159737.2;NP_001159738.2;NP_9! | 6714 | 150390 | . | . | 1.296874 | 1.409187 | 0.916159 | 1.333757 | 3.017876 | 3.257397 | 2.055834 | 3.049475 | -1.220517 | 246.305905 | 0.237525325 | -1.21036392 | 0.226139281 | 0.982008277 | 8.081779 | 7.792343 | 7.953366 | 8.023908 | 7.813828 | 7.961585 | 291 | 312 | 162 | 241 |
| 4053 | NM_000428,XI | LTBP2 | latent transforming growth factor beta binding protein 2 | protein_coding | NP_000419.1;XP_011535067.1 | 6715 | 602091 | . | . | 0.105204 | 0.094690 | 0.081634 | 0.139853 | 0.244815 | 0.218879 | 0.183184 | 0.319757 | 1.104353 | 28.215890 | 0.597811764 | 0.239542529 | 0.810684923 | 0.998865404 | 4.746331 | 4.887108 | 4.817669 | 4.802466 | 4.790424 | 4.851732 | 33 | 27 | 19 | 35 |
| 4054 | NM_00113014 | LTBP3 | latent transforming growth factor beta binding protein 3 | protein_coding | NP_001123616.1;NP_001157738.1;NP_066548.2;XP_011543334.1;XP_01 | 6716 | 602090 | . | . | 2.821039 | 2.782847 | 2.544174 | 2.603848 | 6.564673 | 6.432677 | 5.709053 | 5.953390 | -1.060336 | 389.679429 | 0.182155115 | -0.46400549 | 0.642643813 | 0.998865404 | 8.647835 | 8.563213 | 8.630143 | 8.616692 | 8.578584 | 8.599196 | 467 | 428 | 323 | 361 |
| 4055 | NM_00127098 | LTBR | lymphotoxin beta receptor | protein_coding | NP_001257916.1;NP_002333.1;XP_005253745.1;XP_006719046.1 | 6718 | 600979 | . | . | 79.423745 | 84.434705 | 94.428011 | 89.830823 | 184.822323 | 195.174591 | 211.893728 | 205.387499 | 1.109461 | 5756.635127 | 0.107315582 | 1.396438282 | 0.162582554 | 0.935295382 | 12.414125 | 12.564004 | 12.424230 | 12.466318 | 12.558436 | 12.506525 | 6126 | 6038 | 5427 | 5503 |
| 4059 | NM_00101325 | BCAM | basal cell adhesion molecule (Lutheran blood group) | protein_coding | NP_001013275.1;NP_005572.2 | 6722 | 612773 | . | . | 1.775935 | 2.342951 | 1.725708 | 1.518054 | 4.132674 | 5.415836 | 3.872439 | 3.470850 | -1.284972 | 163.155714 | 0.265169339 | -1.36417258 | 0.172513256 | 0.947431715 | 7.520000 | 7.157500 | 7.349710 | 7.442996 | 7.295626 | 7.295868 | 189 | 219 | 124 | 134 |
| 4060 | NM_002345 | LUM | lumican | protein_coding | NP_002336.1 | 6724 | 600616 | . | . | 26.060966 | 22.358501 | 23.482294 | 21.745802 | 60.644939 | 51.682678 | 52.693588 | 49.719193 | -1.086185 | 1949.328060 | 0.131829402 | -0.90472928 | 0.365608828 | 0.998865404 | 10.986987 | 10.868086 | 11.022554 | 10.892596 | 10.927746 | 10.859481 | 2526 | 2013 | 1696 | 1676 |

| | | | | | | | | | | | | | | | | | | | | | | | | | | | | | | | | | |
|---|---|---|---|---|---|---|---|---|---|---|---|---|---|---|---|---|---|---|---|---|---|---|---|---|---|---|---|---|---|---|---|---|---|
| 4061 | NM_00112721 | LY6E | lymphocyte antigen 6 family member E | protein_coding | NP_001120685.1;NP_002337.1 | 6727 | 601384 | . | . | 197.302749 | 203.863401 | 228.758847 | 240.527422 | 459.131649 | 471.239399 | 513.328220 | 549.937408 | 1.153506 | 7688.398019 | 0.103122851 | 1.997861145 | 0.045731716 | 0.653147492 | 12.801803 | 13.007786 | 12.835022 | 12.854595 | 12.948376 | 12.982885 | 8126 | 7795 | 7018 | 7877 |
| 4067 | NM_00111109 | LYN | LYN proto-oncogene, Src family tyrosine kinase | protein_coding | NP_001104567.1;NP_002341.1;XP_011515831.1;XP_016868904.1;XP_01 | 6735 | 165120 | . | . | 0.688427 | 0.681388 | 0.426987 | 0.638324 | 1.601998 | 1.575060 | 0.958146 | 1.459452 | -1.326321 | 113.812674 | 0.321246046 | -1.26827879 | 0.204698408 | 0.96993643 | 7.020808 | 6.611258 | 6.887645 | 6.860337 | 6.721096 | 6.837638 | 155 | 135 | 68 | 110 |
| 4069 | NM_000239 | LYZ | lysozyme | protein_coding | NP_000230.1 | 6740 | 153450 | . | . | 4.348027 | 3.385891 | 3.859193 | 4.577612 | 10.118038 | 7.826638 | 8.659918 | 10.466166 | 1.071689 | 188.238772 | 0.256527696 | 0.389377888 | 0.696996627 | 0.998865404 | 7.505516 | 7.605589 | 7.592588 | 7.481063 | 7.533258 | 7.603029 | 236 | 171 | 156 | 197 |
| 4071 | NM_014220,XI | TM4SF1 | transmembrane 4 L six family member 1 | protein_coding | NP_055035.1;XP_016861874.1 | 11853 | 191155 | . | . | 88.760712 | 76.799029 | 89.412147 | 70.813524 | 206.549809 | 177.524382 | 200.638276 | 161.906708 | -1.046861 | 3866.065195 | 0.140942875 | -0.46877343 | 0.639231593 | 0.998865404 | 11.949162 | 11.883389 | 11.997666 | 11.863095 | 11.992944 | 11.786059 | 4920 | 3923 | 3696 | 3099 |
| 4072 | NM_002354 | EPCAM | epithelial cell adhesion molecule | protein_coding | NP_002345.2 | 11529 | 185535 | . | . | 0.205007 | 0.131213 | 0.148888 | 0.149279 | 0.477059 | 0.303304 | 0.334101 | 0.341309 | -1.091740 | 8.106252 | 1.070230862 | -0.11831924 | 0.905814715 | NA | 3.079678 | 2.955730 | 3.031623 | 3.013456 | 3.020664 | 3.018077 | 12 | 7 | 7 | 7 |
| 4074 | NM_00120702 | M6PR | mannose-6-phosphate receptor, cation dependent | protein_coding | NP_001193953.1;NP_002346.1;XP_005253433.1 | 6752 | 154540 | . | . | 84.197242 | 85.343555 | 90.300251 | 85.613290 | 195.930426 | 197.275450 | 202.631147 | 195.744609 | 1.023906 | 6584.939484 | 0.105128813 | 0.324198709 | 0.7457876 | 0.998865404 | 12.667787 | 12.701920 | 12.672624 | 12.676855 | 12.721085 | 12.667641 | 7474 | 7039 | 5977 | 6049 |
| 4076 | NM_005898,N | CAPRIN1 | cell cycle associated protein 1 | protein_coding | NP_005889.3;NP_976240.1 | 6743 | 601178 | . | . | 26.861871 | 26.835626 | 25.743951 | 24.581469 | 62.508679 | 62.031752 | 57.768680 | 56.202605 | -1.080107 | 4313.567441 | 0.107494326 | -1.03423881 | 0.301024533 | 0.998865404 | 12.129151 | 12.018042 | 12.111348 | 12.100329 | 12.059087 | 12.024132 | 5192 | 4802 | 3683 | 3806 |
| 4077 | NM_00129157 | NBR1 | NBR1 autophagy cargo receptor | protein_coding | NP_001278500.1;NP_001278501.1;NP_005890.2;NP_114068.1;XP_0067 | 6746 | 166945 | . | . | 18.186311 | 18.002239 | 18.581371 | 21.169860 | 42.320294 | 41.612981 | 41.696058 | 48.402369 | 1.065368 | 2741.783305 | 0.120578558 | 0.757616715 | 0.448680471 | 0.998865404 | 11.374589 | 11.465769 | 11.404666 | 11.383188 | 11.389227 | 11.498722 | 3098 | 2827 | 2295 | 2829 |
| 4082 | NM_002356 | MARCKS | myristoylated alanine rich protein kinase C substrate | protein_coding | NP_002347.5 | 6759 | 177061 | . | . | 15.868873 | 11.316469 | 10.984134 | 9.931481 | 36.927521 | 26.158525 | 24.648077 | 22.707153 | -1.318972 | 1612.240316 | 0.163030578 | -2.44993056 | 0.014288377 | 0.399235756 | 10.840566 | 10.441648 | 10.878501 | 10.599706 | 10.573592 | 10.490660 | 2474 | 1639 | 1276 | 1232 |
| 4083 | gene-MARCKSI | MARCKSP1 | myristoylated alanine rich protein kinase C substrate pseudogene | pseudogene | . | 6760 | . | | . | 0.630661 | 0.589122 | 0.424408 | 0.497362 | 1.467573 | 1.361782 | 0.952359 | 1.137160 | -1.329226 | 46.287168 | 0.460671575 | -0.89127722 | 0.372780468 | 0.998865404 | 5.723597 | 5.312263 | 5.566644 | 5.552352 | 5.499643 | 5.521202 | 63 | 55 | 32 | 40 |
| 4084 | NM_00120251 | MXD1 | MAX dimerization protein 1 | protein_coding | NP_001189442.1;NP_001189443.1;NP_002348.1 | 6761 | 600021 | . | . | 0.968495 | 1.280163 | 1.354378 | 1.265332 | 2.253729 | 2.959154 | 3.039183 | 2.893031 | 1.152397 | 211.227886 | 0.243946383 | 0.838864762 | 0.401545204 | 0.998865404 | 7.617039 | 7.821067 | 7.622050 | 7.742664 | 7.769632 | 7.729604 | 196 | 240 | 205 | 203 |
| 4085 | NM_002358 | MAD2L1 | mitotic arrest deficient 2 like 1 | protein_coding | NP_002349.1 | 6763 | 601467 | . | . | 7.541132 | 6.679803 | 5.379804 | 5.647112 | 17.548525 | 15.440663 | 12.072124 | 12.911451 | -1.307698 | 1029.145375 | 0.141874182 | -2.72797746 | 0.006372396 | 0.2513091 | 10.187755 | 9.800844 | 10.145343 | 10.045906 | 9.886580 | 9.912744 | 1430 | 1177 | 761 | 852 |
| 4086 | NM_00100368 | SMAD1 | SMAD family member 1 | protein_coding | NP_001003688.1;NP_001341740.1;NP_001341741.1;NP_001341742.1;NI | 6767 | 601595 | . | . | 2.174507 | 1.979384 | 1.406310 | 1.881621 | 5.060168 | 4.575434 | 3.155720 | 4.302104 | -1.279811 | 174.552561 | 0.262675667 | -1.35501882 | 0.175411561 | 0.949120257 | 7.615065 | 7.257924 | 7.518981 | 7.472997 | 7.339261 | 7.443075 | 237 | 201 | 115 | 163 |
| 4087 | NM_00100365 | SMAD2 | SMAD family member 2 | protein_coding | NP_001003652.1;NP_001129409.1;NP_005892.1;XP_005258316.1;XP_00 | 6768 | 601366 | . | . | 1.568136 | 1.351532 | 1.973935 | 2.219915 | 3.649115 | 3.124126 | 4.429452 | 5.075574 | 1.094403 | 1188.395506 | 0.13403672 | 0.970958054 | 0.331569167 | 0.998865404 | 10.148081 | 10.278569 | 10.197478 | 10.160569 | 10.278728 | 10.214431 | 1339 | 1194 | 1126 | 1115 |
| 4088 | NM_00114510 | SMAD3 | SMAD family member 3 | protein_coding | NP_001138574.1;NP_001138575.1;NP_001138576.1;NP_005893.1;XP_01 | 6769 | 603109 | . | . | 8.438810 | 9.481321 | 8.391183 | 8.486729 | 19.637458 | 21.916498 | 18.829570 | 19.403896 | -1.073983 | 1750.906798 | 0.125323603 | -0.82163838 | 0.411282741 | 0.998865404 | 10.824533 | 10.721397 | 10.757837 | 10.842392 | 10.744825 | 10.744935 | 1977 | 2061 | 1468 | 1583 |
| 4089 | NM_005359 | SMAD4 | SMAD family member 4 | protein_coding | NP_005350.1 | 6770 | 600993 | . | . | 2.699221 | 2.702578 | 2.384626 | 2.220798 | 6.281199 | 6.247129 | 5.351033 | 5.077592 | -1.188850 | 684.701637 | 0.152950255 | -1.63168631 | 0.102745586 | 0.82309769 | 9.538565 | 9.289349 | 9.477540 | 9.472984 | 9.383554 | 9.330227 | 859 | 800 | 566 | 563 |
| 4090 | NM_00100141 | SMAD5 | SMAD family member 5 | protein_coding | NP_001001419.1;NP_001001420.1;NP_005894.3;XP_016864959.1;XP_02 | 6771 | 603110 | . | . | 5.518501 | 4.852433 | 5.426353 | 4.432989 | 12.841778 | 11.216613 | 12.176577 | 10.135503 | -1.065006 | 1104.421011 | 0.152372807 | -0.59631275 | 0.550966314 | 0.998865404 | 10.153432 | 10.063310 | 10.180339 | 10.077291 | 10.156315 | 10.005003 | 1401 | 1145 | 1026 | 900 |
| 4091 | NM_005585,N | SMAD6 | SMAD family member 6 | protein_coding | NP_005576.3;XP_011519863.1 | 6772 | 602931 | . | . | 0.746025 | 0.643436 | 0.232831 | 0.350771 | 1.736029 | 1.487333 | 0.522466 | 0.801997 | -2.303530 | 72.756028 | 0.389906403 | -3.08752685 | 0.002018295 | 0.11370829 | 6.665513 | 5.459419 | 6.291252 | 6.240446 | 6.045176 | 6.096194 | 126 | 101 | 32 | 48 |
| 4092 | NM_00119082 | SMAD7 | SMAD family member 7 | protein_coding | NP_001177750.1;NP_001177751.1;NP_001177752.1;NP_005895.1 | 6773 | 602932 | . | . | 3.558006 | 4.496111 | 3.569110 | 3.456695 | 8.279627 | 10.392964 | 8.008980 | 7.903324 | -1.206354 | 353.728476 | 0.198910782 | -1.36067586 | 0.173616147 | 0.94763772 | 8.595701 | 8.324577 | 8.451663 | 8.576452 | 8.417442 | 8.400655 | 399 | 461 | 283 | 296 |
| 4093 | NM_00112721 | SMAD9 | SMAD family member 9 | protein_coding | NP_001120689.1;NP_001365550.1;NP_005896.1;XP_005266458.1;XP_00 | 6774 | 603295 | . | . | 1.040575 | 0.798781 | 0.559336 | 0.543367 | 2.421460 | 1.846418 | 1.255133 | 1.242345 | -1.686418 | 124.835401 | 0.302043135 | -2.49620654 | 0.012552949 | 0.369272325 | 7.291832 | 6.538755 | 7.094579 | 6.986337 | 6.863256 | 6.860897 | 204 | 147 | 81 | 87 |
| 4094 | NM_00103180 | MAF | MAF bZIP transcription factor | protein_coding | NP_001026974.1;NP_005351.2;XP_016878722.1;XP_016878723.1;XP_01 | 6776 | 177075 | . | . | 3.820678 | 3.452544 | 2.867411 | 3.038920 | 8.890876 | 7.980709 | 6.434388 | 6.948131 | -1.288543 | 542.589001 | 0.178733962 | -2.04628444 | 0.040728393 | 0.6281612 | 9.254576 | 8.889867 | 9.225736 | 9.084566 | 9.035527 | 8.950508 | 776 | 591 | 441 | 414 |
| 4097 | NM_002359,N | MAFG | MAF bZIP transcription factor G | protein_coding | NP_002350.1;NP_116100.2 | 6781 | 602020 | . | . | 11.343398 | 11.648024 | 11.892630 | 12.201392 | 26.396552 | 26.924928 | 26.686715 | 27.897033 | 1.035223 | 1854.949632 | 0.119322515 | 0.418546655 | 0.675547482 | 0.998865404 | 10.832009 | 10.881889 | 10.837220 | 10.849281 | 10.864317 | 10.876791 | 2083 | 1982 | 1628 | 1783 |
| 4101 | NM_00128250 | MAGEA2 | MAGE family member A2 | protein_coding | NP_001269430.1;NP_001269431.1;NP_001269433.1;NP_001269434.1;NI | 6800 | 300173 | . | . | 5.722825 | 4.813034 | 5.219410 | 4.999957 | 13.317251 | 11.125545 | 11.712206 | 11.431809 | -1.027467 | 313.715606 | 0.203492233 | -0.19210285 | 0.847661643 | 0.998865404 | 8.312360 | 8.274013 | 8.345887 | 8.251879 | 8.294888 | 8.275585 | 397 | 314 | 274 | 285 |
| 4102 | NM_005362,XI | MAGEA3 | MAGE family member A3 | protein_coding | NP_005353.1;XP_005274733.1;XP_006724881.1;XP_011529462.1;XP_01 | 6801 | 300174 | . | . | 34.599655 | 34.676241 | 31.646464 | 36.777416 | 80.514823 | 80.155690 | 71.013750 | 84.087193 | -1.037365 | 1878.765534 | 0.126602099 | -0.41802738 | 0.675927096 | 0.998865404 | 10.901916 | 10.848732 | 10.895878 | 10.883031 | 10.805799 | 10.912219 | 2221 | 2048 | 1495 | 1846 |
| 4103 | NM_00101154 | MAGEA4 | MAGE family member A4 | protein_coding | NP_001011548.1;NP_001011549.1;NP_001011550.1;NP_002353.3;XP_00 | 6802 | 300175 | . | . | 113.645493 | 109.285850 | 89.857570 | 106.987886 | 264.457730 | 252.619134 | 201.637784 | 244.615083 | -1.148461 | 5637.323650 | 0.1180713 | -1.69136393 | 0.090767317 | 0.805424371 | 12.557246 | 12.357430 | 12.535482 | 12.494965 | 12.328720 | 12.464819 | 7105 | 6349 | 4189 | 5323 |
| 4105 | NM_005363,N | MAGEA6 | MAGE family member A6 | protein_coding | NP_005354.1;NP_787064.1 | 6804 | 300176 | . | . | 27.605247 | 25.478415 | 25.274928 | 26.425138 | 64.238541 | 58.894501 | 56.716198 | 60.417938 | -1.059704 | 1418.723516 | 0.127062561 | -0.65842757 | 0.510263426 | 0.998865404 | 10.511599 | 10.427943 | 10.512361 | 10.471697 | 10.435020 | 10.460247 | 1725 | 1534 | 1178 | 1313 |
| 4109 | NM_00101154 | MAGEA10 | MAGE family member A10 | protein_coding | NP_001011543.2;NP_001238757.2;NP_066386.3 | 6797 | 300343 | . | . | 11.981761 | 12.064813 | 10.861580 | 10.518419 | 27.882050 | 27.888358 | 24.373071 | 24.049116 | -1.139251 | 948.282540 | 0.138423606 | -1.35877165 | 0.174218958 | 0.948096927 | 9.980080 | 9.792151 | 9.937292 | 9.932011 | 9.853899 | 9.826139 | 1167 | 1086 | 786 | 815 |
| 4111 | NM_00116638 | MAGEA12 | MAGE family member A12 | protein_coding | NP_001159858.1;NP_001159859.1;NP_005358.2 | 6799 | 300177 | . | . | 18.196013 | 17.117907 | 14.189709 | 17.756964 | 42.342867 | 39.568805 | 31.857856 | 40.599187 | -1.121488 | 904.515470 | 0.155939034 | -1.06076102 | 0.28879852 | 0.998865404 | 9.901580 | 9.735655 | 9.886556 | 9.832087 | 9.696570 | 9.851391 | 1142 | 994 | 660 | 887 |
| 4116 | NM_002370 | MAGOH | mago homolog, exon junction complex subunit | protein_coding | NP_002361.1 | 6815 | 602603 | . | . | 34.371819 | 31.312025 | 28.460947 | 30.489988 | 79.984634 | 72.379143 | 63.865543 | 69.711731 | -1.129770 | 637.472579 | 0.158401805 | -1.11128007 | 0.266447814 | 0.998274218 | 9.401541 | 9.225534 | 9.390029 | 9.320244 | 9.254032 | 9.292930 | 818 | 693 | 505 | 578 |
| 4117 | NM_00124238 | MAK | male germ cell associated kinase | protein_coding | NP_001229314.1;NP_001229886.1;NP_001364191.1;NP_005897.1;XP_01 | 6816 | 154235 | . | . | 0.047117 | 0.060262 | 0.044344 | 0.040918 | 0.109644 | 0.139299 | 0.099507 | 0.093554 | -1.404538 | 7.308325 | 1.123358903 | -0.43627738 | 0.662635492 | NA | 3.094525 | 2.602870 | 2.877399 | 2.883995 | 2.868262 | 2.871008 | 9 | 10 | 5 | 6 |
| 4118 | NM_002371,N | MAL | mal, T cell differentiation protein | protein_coding | NP_002362.1;NP_071883.1;NP_071884.1;NP_071885.1 | 6817 | 188860 | . | . | 0.932600 | 0.816001 | 0.330170 | 0.408408 | 2.170198 | 1.886223 | 0.740892 | 0.933778 | -2.198633 | 21.157724 | 0.685770753 | -1.65741544 | 0.097435513 | NA | 4.862893 | 3.724273 | 4.447257 | 4.436043 | 4.368357 | 4.382144 | 35 | 30 | 10 | 14 |
| 4121 | NM_005907,XI | MAN1A1 | mannosidase alpha class 1A member 1 | protein_coding | NP_005898.2;XP_005267043.1;XP_011534135.1;XP_011534136.1 | 6821 | 604344 | . | . | 0.209264 | 0.117491 | 0.119824 | 0.122610 | 0.486967 | 0.271586 | 0.268881 | 0.280332 | -1.332155 | 22.571316 | 0.668300891 | -0.61912466 | 0.535834242 | NA | 4.687500 | 4.276108 | 4.542141 | 4.480722 | 4.485805 | 4.484097 | 38 | 20 | 17 | 18 |
| 4122 | NM_00132097 | MAN2A2 | mannosidase alpha class 2A member 2 | protein_coding | NP_001307906.1;NP_006113.2;XP_005254967.1;XP_011519867.1;XP_01 | 6825 | 600988 | . | . | 34.044048 | 35.022893 | 39.554628 | 37.347758 | 79.221895 | 80.956989 | 88.759432 | 85.391209 | 1.110093 | 7275.938343 | 0.105931668 | 1.422434299 | 0.154900212 | 0.923603412 | 12.751587 | 12.902313 | 12.774004 | 12.791545 | 12.904288 | 12.837138 | 7858 | 7519 | 6919 | 6895 |
| 4123 | NM_00125649 | MAN2C1 | mannosidase alpha class 2C member 1 | protein_coding | NP_001243423.1;NP_001243424.1;NP_001243425.1;NP_006706.2 | 6827 | 154580 | . | . | 13.554204 | 13.794096 | 16.503757 | 14.473111 | 31.541190 | 31.885672 | 37.033951 | 33.091047 | 1.103063 | 1468.548880 | 0.133926564 | 1.056665663 | 0.290664201 | 0.998865404 | 10.447513 | 10.589353 | 10.474176 | 10.485610 | 10.614482 | 10.493300 | 1596 | 1518 | 1444 | 1332 |
| 4124 | NM_002372,XI | MAN2A1 | mannosidase alpha class 2A member 1 | protein_coding | NP_002363.2;XP_011541697.1;XP_016864961.1;XP_024301816.1 | 6824 | 154582 | . | . | 4.926933 | 4.482836 | 4.386010 | 3.862983 | 11.465176 | 10.362275 | 9.842078 | 8.832254 | -1.203094 | 955.310736 | 0.157357083 | -1.69518775 | 0.09003981 | 0.804059937 | 10.026660 | 9.760758 | 10.003344 | 9.917136 | 9.903974 | 9.743566 | 1270 | 1061 | 840 | 721 |
| 4125 | NM_000528,N | MAN2B1 | mannosidase alpha class 2B member 1 | protein_coding | NP_000519.2;NP_001166969.1;XP_005259970.1;XP_024307286.1 | 6826 | 609458 | . | . | 18.995374 | 19.198933 | 16.103350 | 18.681421 | 44.203014 | 44.379190 | 36.135453 | 42.712850 | -1.114010 | 1812.800498 | 0.127621757 | -1.22049809 | 0.222276112 | 0.982008277 | 10.899931 | 10.743859 | 10.864590 | 10.864685 | 10.723864 | 10.832243 | 2200 | 2062 | 1387 | 1720 |
| 4126 | NM_005908,XI | MANBA | mannosidase beta | protein_coding | NP_005899.3;XP_016863692.1;XP_016863693.1;XP_016863694.1;XP_02 | 6831 | 609489 | . | . | 11.767466 | 12.168712 | 10.527142 | 11.650921 | 27.383376 | 28.128522 | 23.622600 | 26.638448 | -1.081320 | 1433.923984 | 0.127139252 | -0.88716499 | 0.374990078 | 0.998865404 | 10.541149 | 10.428144 | 10.510663 | 10.519487 | 10.428574 | 10.480566 | 1706 | 1617 | 1157 | 1336 |
| 4128 | NM_000240,N | MAOA | monoamine oxidase A | protein_coding | NP_000231.1;NP_001257387.1 | 6833 | 309850 | . | . | 2.388262 | 2.730078 | 2.188817 | 3.000330 | 5.557584 | 6.310698 | 4.911643 | 6.859897 | -1.014216 | 320.408625 | 0.216675837 | -0.09399066 | 0.925116574 | 0.998865404 | 8.334699 | 8.312756 | 8.298018 | 8.351379 | 8.227854 | 8.399158 | 354 | 365 | 233 | 345 |
| 4130 | NM_002373 | MAP1A | microtubule associated protein 1A | protein_coding | NP_002364.5 | 6835 | 600178 | . | . | 1.288140 | 1.402111 | 1.668199 | 1.465726 | 2.997555 | 3.241043 | 3.743389 | 3.351208 | 1.150046 | 466.209601 | 0.177025211 | 1.139337136 | 0.254562565 | 0.995760118 | 8.760304 | 8.962301 | 8.793364 | 8.840767 | 8.947495 | 8.859262 | 480 | 486 | 464 | 435 |
| 4131 | NM_00132425 | MAP1B | microtubule associated protein 1B | protein_coding | NP_001311184.1;NP_005900.2 | 6836 | 157129 | . | . | 1.697889 | 1.607290 | 1.942695 | 1.803054 | 3.951058 | 3.715323 | 4.359350 | 4.122469 | 1.117605 | 647.866234 | 0.156736043 | 1.023441688 | 0.306099057 | 0.998865404 | 9.256854 | 9.417766 | 9.320621 | 9.278095 | 9.403257 | 9.345424 | 727 | 639 | 620 | 614 |
| 4133 | NM_00103953 | MAP2 | microtubule associated protein 2 | protein_coding | NP_001034627.1;NP_001350839.1;NP_001350840.1;NP_001350842.1;NI | 6839 | 157130 | . | . | 2.501596 | 2.791701 | 2.584901 | 2.494099 | 5.821316 | 6.453142 | 5.800441 | 5.702457 | 1.030851 | 520.103362 | 0.167622805 | 0.261517246 | 0.793693652 | 0.998865404 | 9.000833 | 9.044151 | 8.982629 | 9.041219 | 9.013302 | 9.049745 | 563 | 578 | 446 | 509 |
| 4134 | NM_00113436 | MAP4 | microtubule associated protein 4 | protein_coding | NP_001127836.1;NP_002366.2;NP_112147.2;XP_011532005.1;XP_01153 | 6862 | 157132 | . | . | 21.704072 | 24.165140 | 19.562185 | 24.064413 | 50.506266 | 55.858801 | 43.896978 | 55.020427 | -1.058943 | 4427.138805 | 0.124429217 | -0.66403634 | 0.506667055 | 0.998865404 | 12.152988 | 12.070140 | 12.102551 | 12.165319 | 12.005722 | 12.161563 | 5048 | 5098 | 3388 | 4410 |
| 4137 | NM_00112306 | MAPT | microtubule associated protein tau | protein_coding | NP_001116538.2;NP_001116539.1;NP_001190180.1;NP_001190181.1;NI | 6893 | 157140 | . | . | 0.279550 | 0.329258 | 0.391877 | 0.554553 | 0.650523 | 0.761093 | 0.879361 | 1.267919 | 1.465064 | 76.506071 | 0.375590841 | 1.466925724 | 0.142396262 | 0.903146012 | 5.957404 | 6.505798 | 6.164749 | 6.224554 | 6.254335 | 6.326874 | 62 | 76 | 69 | 96 |
| 4139 | NM_00128612 | MARK1 | microtubule affinity regulating kinase 1 | protein_coding | NP_001273053.1;NP_001273055.1;NP_001273057.1;NP_001273058.1;NI | 6896 | 606511 | . | . | 0.072172 | 0.074889 | 0.099725 | 0.038723 | 0.167946 | 0.173109 | 0.223779 | 0.088536 | -1.670147 | 10.400213 | 0.955430225 | -0.77449404 | 0.438638692 | NA | 3.700498 | 2.963455 | 3.396658 | 3.396248 | 3.379570 | 3.367124 | 15 | 14 | 8 | 6 |
| 4140 | NM_00112891 | MARK3 | microtubule affinity regulating kinase 3 | protein_coding | NP_001122390.2;NP_001122391.2;NP_001122392.2;NP_001122393.2;NI | 6897 | 602678 | . | . | 12.915522 | 13.332478 | 14.789269 | 13.975308 | 30.054959 | 30.818626 | 33.186687 | 31.952878 | 1.098519 | 1471.601352 | 0.126113875 | 1.074895822 | 0.282421373 | 0.998865404 | 10.453720 | 10.589432 | 10.483949 | 10.486749 | 10.581974 | 10.533121 | 1612 | 1516 | 1382 | 1399 |
| 4141 | NM_004990,XI | MARS1 | methionyl-tRNA synthetase 1 | protein_coding | NP_004981.2;XP_006719461.1 | 6898 | 156560 | . | . | 54.723588 | 55.908878 | 41.219796 | 52.850795 | 127.344051 | 129.235882 | 92.496023 | 120.837069 | -1.195647 | 4463.314254 | 0.12980426 | -1.98600061 | 0.047033256 | 0.658032177 | 12.247146 | 11.989132 | 12.186391 | 12.198112 | 11.941337 | 12.135878 | 5559 | 5282 | 3116 | 4263 |
| 4144 | NM_005911 | MAT2A | methionine adenosyltransferase 2A | protein_coding | NP_005902.1 | 6904 | 601468 | . | . | 138.067764 | 147.587891 | 159.097549 | 138.465851 | 321.289368 | 341.156006 | 357.010284 | 316.585724 | 1.027726 | 12795.229598 | 0.112822609 | 0.349715026 | 0.726552576 | 0.998865404 | 13.623435 | 13.662931 | 13.605589 | 13.655205 | 13.716037 | 13.588255 | 14110 | 14013 | 12118 | 11255 |
| 4146 | NM_002379 | MATN1 | matrilin 1 | protein_coding | NP_002370.1 | 6907 | 115437 | . | . | 0.101409 | 0.228184 | 0.189227 | 0.147266 | 0.235983 | 0.527458 | 0.424620 | 0.336706 | 1.008665 | 20.490450 | 0.710548083 | 0.017517462 | 0.986023803 | NA | 4.351521 | 4.362218 | 4.316362 | 4.380632 | 4.359767 | 4.339702 | 15 | 30 | 20 | 17 |
| 4147 | NM_00131774 | MATN2 | matrilin 2 | protein_coding | NP_001304677.1;NP_002371.3;NP_085072.2;XP_005250977.1;XP_01686 | 6908 | 602108 | . | . | 5.061929 | 5.612062 | 3.536764 | 4.101385 | 11.779317 | 12.972534 | 7.936395 | 9.377328 | -1.438802 | 578.822909 | 0.165138456 | -3.17835124 | 0.001481152 | 0.090747667 | 9.415955 | 8.890401 | 9.243744 | 9.319352 | 9.016273 | 9.070434 | 744 | 777 | 394 | 465 |
| 4148 | NM_002381 | MATN3 | matrilin 3 | protein_coding | NP_002372.1 | 6909 | 602109 | . | . | 1.876894 | 2.260329 | 1.249782 | 1.284000 | 4.367609 | 5.224853 | 2.804476 | 2.935712 | -1.651275 | 133.518431 | 0.287797993 | -2.51419598 | 0.011930412 | 0.357990159 | 7.378046 | 6.653785 | 7.102703 | 7.175234 | 6.958611 | 6.961122 | 175 | 195 | 87 | 95 |
| 4149 | NM_00127106 | MAX | MYC associated factor X | protein_coding | NP_001257997.1;NP_001257998.1;NP_001307344.1;NP_002373.3;NP_6 | 6913 | 154950 | . | . | 18.491040 | 17.669992 | 17.721065 | 18.836324 | 43.029405 | 40.844975 | 39.765553 | 43.067011 | 1.009514 | 1110.598767 | 0.133646362 | 0.102214005 | 0.918586811 | 0.998865404 | 10.110300 | 10.123912 | 10.127015 | 10.099823 | 10.104877 | 10.135967 | 1296 | 1171 | 950 | 1068 |
| 4150 | NM_00104253 | MAZ | MYC associated zinc finger protein | protein_coding | NP_001036004.1;NP_001263204.1;NP_001263205.1;NP_002374.2;XP_0 | 6914 | 600999 | . | . | 51.939919 | 52.794538 | 52.162768 | 56.323301 | 120.866336 | 122.036943 | 117.051730 | 128.776535 | 1.023664 | 4348.776483 | 0.108689075 | 0.310447737 | 0.756220497 | 0.998865404 | 12.069469 | 12.103122 | 12.071194 | 12.081675 | 12.068339 | 12.122711 | 4918 | 4667 | 3701 | 4259 |
| 4151 | NM_00136284 | MB | myoglobin | protein_coding | NP_001349775.1;NP_001369738.1;NP_001369739.1;NP_001369740.1;NI | 6915 | 160000 | . | . | 0.084044 | 0.336472 | 0.196475 | 0.243882 | 0.195574 | 0.777771 | 0.440885 | 0.557607 | 1.082058 | 9.022858 | 1.073777664 | 0.106395022 | 0.915268943 | NA | 3.118936 | 3.226238 | 3.138293 | 3.186506 | 3.160205 | 3.171651 | 4 | 15 | 7 | 10 |
| 4152 | NM_00120413 | MBD1 | methyl-CpG binding domain protein 1 | protein_coding | NP_001191065.1;NP_001191066.1;NP_001191067.1;NP_001191068.1;NI | 6916 | 156535 | . | . | 13.822706 | 13.255272 | 13.857186 | 15.012121 | 32.166008 | 30.640152 | 31.095123 | 34.323441 | 1.028564 | 1601.314459 | 0.124065072 | 0.327507042 | 0.743284418 | 0.998865404 | 10.624564 | 10.665231 | 10.616959 | 10.650278 | 10.673109 | 10.637956 | 1779 | 1740 | 1444 | 1488 |
| 4154 | NM_00131405 | MBNL1 | muscleblind like splicing regulator 1 | protein_coding | NP_001300986.1;NP_001350799.1;NP_001363747.1;NP_001363748.1;NI | 6923 | 606516 | . | . | 14.091092 | 12.729929 | 13.754999 | 12.422481 | 32.790552 | 29.425793 | 30.865820 | 28.402527 | -1.042527 | 2299.844477 | 0.124428236 | -0.48288894 | 0.629173953 | 0.998865404 | 11.196898 | 11.137124 | 11.224435 | 11.140077 | 11.190353 | 11.107572 | 2844 | 2401 | 2060 | 2002 |
| 4155 | NM_00102508 | MBP | myelin basic protein | protein_coding | NP_001020252.1;NP_001020261.1;NP_001020263.1;NP_001020271.1;NI | 6925 | 159430 | . | . | 278.868041 | 285.959756 | 355.967881 | 391.014175 | 648.937395 | 661.008806 | 798.781585 | 894.007460 | 1.286244 | 23289.961594 | 0.100942931 | 3.597715333 | 0.000321025 | 0.029151999 | 14.314452 | 14.677597 | 14.384633 | 14.394033 | 14.574518 | 14.636708 | 23330 | 22107 | 21982 | 25437 |
| 4157 | NM_002386 | MC1R | melanocortin 1 receptor | protein_coding | NP_002377.4 | 6929 | 155555 | . | . | 41.354782 | 41.041470 | 40.331314 | 43.384354 | 96.234283 | 94.869194 | 90.502289 | 99.193176 | 1.001934 | 2731.242352 | 0.114184492 | 0.024410082 | 0.980525507 | 0.998865404 | 11.413992 | 11.416690 | 11.420908 | 11.407975 | 11.390321 | 11.441048 | 3167 | 2921 | 2303 | 2643 |
| 4160 | NM_005912 | MC4R | melanocortin 4 receptor | protein_coding | NP_005903.2 | 6932 | 155541 | . | . | 0.027866 | 0.067012 | 0.037174 | 0.102703 | 0.064846 | 0.154901 | 0.083417 | 0.234817 | 1.608698 | 3.535954 | 1.628658268 | 0.421139979 | 0.673652869 | NA | 1.444631 | 2.121007 | 1.803866 | 1.813339 | 1.807014 | 1.825463 | 2 | 4 | 2 | 6 |
| 4162 | NM_006500,XI | MCAM | melanoma cell adhesion molecule | protein_coding | NP_006491.2;XP_016873248.1;XP_016873249.1;XP_016873250.1;XP_01 | 6934 | 155735 | . | . | 118.147632 | 127.041770 | 97.304072 | 113.912560 | 274.934401 | 293.662717 | 218.347508 | 260.447528 | -1.174888 | 12059.912539 | 0.111791328 | -2.07997319 | 0.037527992 | 0.605846634 | 13.669550 | 13.436948 | 13.597385 | 13.646720 | 13.423733 | 13.541895 | 14577 | 14460 | 8968 | 11125 |
| 4163 | NM_00108537 | MCC | MCC regulator of WNT signaling pathway | protein_coding | NP_001078846.2;NP_002378.2;XP_005272048.1;XP_016864962.1;XP_01 | 6935 | 159350 | . | . | 7.345747 | 8.313813 | 7.205730 | 6.782462 | 17.093859 | 19.217752 | 16.169450 | 15.507295 | -1.179680 | 1958.179574 | 0.120557424 | -1.97744521 | 0.047991328 | 0.659785158 | 11.049528 | 10.811238 | 10.982902 | 11.011091 | 10.891439 | 10.840157 | 2398 | 2327 | 1614 | 1629 |
| 4166 | NM_021615,N | CHST6 | carbohydrate sulfotransferase 6 | protein_coding | NP_067628.1;XP_005256012.1;XP_011521387.1 | 6938 | 605294 | . | . | 3.617967 | 3.850939 | 3.528105 | 3.233834 | 8.419158 | 8.901616 | 7.916964 | 7.393778 | -1.118148 | 849.715273 | 0.142619721 | -1.12965108 | 0.258623278 | 0.995760118 | 9.809121 | 9.648057 | 9.758073 | 9.780500 | 9.701120 | 9.678398 | 1017 | 983 | 709 | 740 |
| 4170 | NM_00119732 | MCL1 | MCL1 apoptosis regulator, BCL2 family member | protein_coding | NP_001184249.1;NP_068779.1;NP_877495.1 | 6943 | 159552 | . | . | 25.598994 | 23.622468 | 23.033031 | 21.564381 | 59.569909 | 54.604393 | 51.685453 | 49.304397 | -1.117798 | 2882.829309 | 0.117796205 | -1.36388316 | 0.172604908 | 0.947431715 | 11.571262 | 11.410819 | 11.571774 | 11.500065 | 11.476479 | 11.414352 | 3656 | 3141 | 2459 | 2454 |
| 4171 | NM_004526,N | MCM2 | minichromosome maintenance complex component 2 | protein_coding | NP_004517.2;XP_024309299.1 | 6944 | 116945 | . | . | 18.767179 | 19.551970 | 15.612165 | 19.189997 | 43.671998 | 45.195255 | 35.033246 | 43.875645 | -1.116862 | 1956.410347 | 0.133358859 | -1.19564845 | 0.231833777 | 0.986726781 | 11.011709 | 10.851852 | 10.962528 | 10.987845 | 10.808169 | 10.961061 | 2340 | 2263 | 1450 | 1902 |
| 4172 | NM_00127047 | MCM3 | minichromosome maintenance complex component 3 | protein_coding | NP_001257401.2;NP_001353298.1;NP_001353299.1;NP_001353300.1;NI | 6945 | 602693 | . | . | 52.248261 | 55.470240 | 47.910598 | 56.426802 | 121.583855 | 128.221938 | 107.509994 | 129.013189 | -1.047632 | 5067.723044 | 0.117719338 | -0.57027352 | 0.568492196 | 0.998865404 | 12.340382 | 12.273077 | 12.302107 | 12.348510 | 12.220568 | 12.348499 | 5807 | 5750 | 3974 | 4996 |
| 4173 | NM_005914,N | MCM4 | minichromosome maintenance complex component 4 | protein_coding | NP_005905.2;NP_877423.1 | 6947 | 602638 | . | . | 44.554011 | 47.240479 | 41.510953 | 46.577121 | 103.679022 | 109.198471 | 93.149372 | 106.493054 | -1.055467 | 5697.714566 | 0.111153287 | -0.70067278 | 0.483507244 | 0.998865404 | 12.514645 | 12.436636 | 12.477018 | 12.518734 | 12.407139 | 12.496212 | 6572 | 6470 | 4564 | 5474 |
| 4174 | NM_006739,XI | MCM5 | minichromosome maintenance complex component 5 | protein_coding | NP_006730.2;XP_006724305.1 | 6948 | 602696 | . | . | 36.068117 | 37.147946 | 32.062355 | 39.017323 | 83.931990 | 85.869138 | 71.947004 | 89.208473 | -1.046730 | 3879.713250 | 0.124077891 | -0.53103563 | 0.595394088 | 0.998865404 | 11.954410 | 11.888301 | 11.930412 | 11.948196 | 11.819374 | 11.977379 | 4521 | 4328 | 2981 | 3894 |
| 4175 | NM_005915 | MCM6 | minichromosome maintenance complex component 6 | protein_coding | NP_005906.2 | 6949 | 601806 | . | . | 34.196793 | 36.671444 | 30.337811 | 41.235794 | 79.577347 | 84.767685 | 68.077171 | 94.280739 | -1.004414 | 4161.785362 | 0.140985431 | -0.04506528 | 0.964055279 | 0.998865404 | 12.026287 | 12.019679 | 11.996246 | 12.047333 | 11.888154 | 12.133944 | 4656 | 4639 | 3079 | 4466 |
| 4176 | NM_00127859 | MCM7 | minichromosome maintenance complex component 7 | protein_coding | NP_001265524.1;NP_005907.3;NP_877577.1;XP_005250405.1;XP_01686 | 6950 | 600592 | . | . | 81.006179 | 86.664920 | 70.490853 | 81.539156 | 188.504712 | 200.329840 | 158.179432 | 186.429589 | -1.101752 | 6400.509994 | 0.115378888 | -1.21165631 | 0.225643969 | 0.982008277 | 12.712244 | 12.572307 | 12.664450 | 12.700875 | 12.535867 | 12.662250 | 7562 | 7396 | 4896 | 6141 |
| 4179 | NM_002389,N | CD46 | CD46 molecule | protein_coding | NP_002380.3;NP_722548.1;NP_758860.1;NP_758861.1;NP_758862.1;NP_ | 6953 | 120920 | . | . | 35.075892 | 32.249724 | 35.422070 | 26.864612 | 81.623041 | 74.546687 | 79.486102 | 61.422744 | -1.098244 | 3412.333930 | 0.141802084 | -0.95343141 | 0.340371534 | 0.998865404 | 11.802405 | 11.667530 | 11.806499 | 11.734664 | 11.803695 | 11.571354 | 4290 | 3688 | 3230 | 2617 |
| 4185 | NM_00131893 | ADAM11 | ADAM metallopeptidase domain 11 | protein_coding | NP_001305862.1;NP_002381.2;XP_005257430.1;XP_024306522.1 | 189 | 155120 | . | . | 0.944373 | 0.816575 | 0.765807 | 0.885454 | 2.197595 | 1.887550 | 1.718448 | 2.024486 | -1.073164 | 132.163294 | 0.28804509 | -0.35365998 | 0.723593719 | 0.998865404 | 7.096306 | 6.994245 | 7.088254 | 7.028797 | 7.008569 | 7.057687 | 170 | 136 | 103 | 128 |
| 4189 | NM_012328 | DNAJB9 | DnaJ heat shock protein family (Hsp40) member B9 | protein_coding | NP_036460.1 | 6968 | 602634 | . | . | 11.575034 | 10.006384 | 11.035246 | 7.049590 | 26.935581 | 23.130205 | 24.762774 | 16.118050 | -1.209654 | 745.717965 | 0.199160921 | -1.37875399 | 0.167970616 | 0.945638095 | 9.672696 | 9.399347 | 9.652087 | 9.540927 | 9.606289 | 9.302375 | 1012 | 813 | 719 | 491 |
| 4190 | NM_00119911 | MDH1 | malate dehydrogenase 1 | protein_coding | NP_001186040.1;NP_001186041.1;NP_001303303.1;NP_005908.1 | 6970 | 154200 | . | . | 211.297439 | 196.437928 | 176.683871 | 202.477825 | 491.697871 | 454.075077 | 396.473488 | 462.941496 | -1.090086 | 7952.666740 | 0.113269736 | -1.09863809 | 0.271925957 | 0.998865404 | 13.018125 | 12.893636 | 13.026349 | 12.956675 | 12.864552 | 12.970502 | 9941 | 8587 | 6201 | 7583 |
| 4191 | NM_00128240 | MDH2 | malate dehydrogenase 2 | protein_coding | NP_001269332.1;NP_001269333.1;NP_005909.2 | 6971 | 154100 | . | . | 54.302660 | 58.685038 | 48.969178 | 59.942446 | 126.364523 | 135.653091 | 109.885420 | 137.051294 | -1.057671 | 3658.680634 | 0.126476098 | -0.63957223 | 0.522450741 | 0.998865404 | 11.877113 | 11.795961 | 11.829962 | 11.886603 | 11.729617 | 11.888698 | 4184 | 4197 | 2793 | 3656 |
| 4192 | NM_00101233 | MDK | midkine | protein_coding | NP_001012333.1;NP_001012334.1;NP_001257479.1;NP_001257480.1;NI | 6972 | 162096 | . | . | 0.399030 | 0.454190 | 0.633319 | 0.718921 | 0.928558 | 1.049880 | 1.421149 | 1.643728 | 1.579606 | 16.195560 | 0.764349862 | 0.862909384 | 0.388187291 | NA | 3.651489 | 4.309235 | 3.988981 | 3.992917 | 4.011618 | 4.033353 | 14 | 14 | 15 | 21 |
| 4193 | NM_00114533 | MDM2 | MDM2 proto-oncogene | protein_coding | NP_001138809.1;NP_001138811.1;NP_001138812.1;NP_001265391.1;NI | 6973 | 164785 | . | . | 8.807416 | 7.863330 | 10.877802 | 8.043296 | 20.495220 | 18.176443 | 24.409469 | 18.390042 | 1.158863 | 2149.825673 | 0.15029733 | 1.415262429 | 0.156991566 | 0.925698889 | 10.959487 | 11.172654 | 11.043867 | 10.963369 | 11.237026 | 10.993669 | 2410 | 2039 | 2307 | 1841 |
| 4194 | NM_00120417 | MDM4 | MDM4 regulator of p53 | protein_coding | NP_001191100.1;NP_001191101.1;NP_001265445.1;NP_001265446.1;NI | 6974 | 602704 | . | . | 1.626492 | 1.767448 | 2.419752 | 1.389452 | 3.784913 | 4.085537 | 5.429852 | 3.176817 | 1.128156 | 553.776617 | 0.227552672 | 0.764513491 | 0.4445613 | 0.998865404 | 9.023001 | 9.198015 | 9.036011 | 9.083531 | 9.312030 | 8.930325 | 577 | 582 | 656 | 394 |
| 4199 | NM_002395,XI | ME1 | malic enzyme 1 | protein_coding | NP_002386.1;XP_011534138.1 | 6983 | 154250 | . | . | 14.496499 | 14.301945 | 20.438052 | 17.009918 | 33.733946 | 33.059586 | 45.862390 | 38.891153 | 1.268863 | 1702.366440 | 0.135229677 | 2.540388533 | 0.011072938 | 0.341160196 | 10.551145 | 10.895059 | 10.639351 | 10.621589 | 10.890479 | 10.738097 | 1748 | 1600 | 1813 | 1616 |
| 4200 | NM_00116833 | ME2 | malic enzyme 2 | protein_coding | NP_001161807.1;NP_002387.1 | 6984 | 154270 | . | . | 7.617827 | 7.413071 | 6.052144 | 7.224887 | 17.726998 | 17.135645 | 13.580835 | 16.518843 | -1.157459 | 1972.075516 | 0.126333155 | -1.66988052 | 0.094943005 | 0.813553488 | 11.047251 | 10.836023 | 11.009662 | 10.990845 | 10.828116 | 10.937314 | 2465 | 2257 | 1478 | 1834 |
| 4201 | NM_00131894 | MEA1 | male-enhanced antigen 1 | protein_coding | NP_001305871.1;NP_001305872.1;NP_001350507.1;NP_055438.1;XP_01 | 6986 | 143170 | . | . | 32.494202 | 31.326054 | 32.722759 | 34.388078 | 75.615342 | 72.411580 | 73.428921 | 78.624257 | -1.012681 | 1622.883493 | 0.135308953 | -0.13435651 | 0.893120673 | 0.998865404 | 10.673588 | 10.655039 | 10.673636 | 10.662786 | 10.583700 | 10.728006 | 1894 | 1750 | 1272 | 1652 |
| 4204 | NM_00111079 | MECP2 | methyl-CpG binding protein 2 | protein_coding | NP_001104262.1;NP_001303266.1;NP_001356320.1;NP_001356321.1;NI | 6990 | 300005 | . | . | 3.129254 | 3.181207 | 3.351239 | 3.437717 | 7.281901 | 7.353500 | 7.520082 | 7.859932 | 1.060692 | 1066.683213 | 0.133889832 | 0.634888602 | 0.525501072 | 0.998865404 | 10.015826 | 10.100756 | 10.033915 | 10.039072 | 10.073109 | 10.086973 | 1188 | 1121 | 946 | 1039 |
| 4205 | NM_00113092 | MEF2A | myocyte enhancer factor 2A | protein_coding | NP_001124398.1;NP_001124399.1;NP_001124400.1;NP_001165365.1;NI | 6993 | 600660 | . | . | 8.667598 | 8.831022 | 9.398755 | 8.056107 | 20.169860 | 20.413304 | 21.090531 | 18.419328 | -1.006332 | 1470.403485 | 0.132347508 | -0.06880688 | 0.94514334 | 0.998865404 | 10.526384 | 10.517595 | 10.518065 | 10.529283 | 10.571911 | 10.463068 | 1686 | 1603 | 1365 | 1277 |
| 4207 | NM_005919,N | BORCS8-MEF2I | BORCS8-MEF2B readthrough | protein_coding | NP_005910.1 | 39979 | . | | . | 0.553765 | 0.662289 | 0.291612 | 0.406439 | 1.288635 | 1.530910 | 0.654369 | 0.929274 | -1.564188 | 26.133219 | 0.612950005 | -1.05296369 | 0.292357636 | NA | 4.995859 | 4.347534 | 4.725535 | 4.744736 | 4.675498 | 4.700458 | 34 | 37 | 15 | 22 |
| 4208 | NM_00113100 | MEF2C | myocyte enhancer factor 2C | protein_coding | NP_001124477.1;NP_001180276.1;NP_001180277.1;NP_001180278.1;NI | 6996 | 600662 | . | . | 2.723633 | 2.829254 | 2.189945 | 2.537855 | 6.338002 | 6.539948 | 4.914170 | 5.802503 | -1.178229 | 522.581703 | 0.168782906 | -1.40192126 | 0.160938762 | 0.933525128 | 9.143260 | 8.906024 | 9.075497 | 9.083680 | 8.933680 | 9.013553 | 647 | 614 | 390 | 479 |
| 4209 | NM_00127162 | MEF2D | myocyte enhancer factor 2D | protein_coding | NP_001258558.1;NP_005911.1;XP_005245226.1;XP_005245227.1;XP_00 | 6997 | 600663 | . | . | 4.531118 | 4.669261 | 4.545049 | 4.428853 | 10.544097 | 10.793203 | 10.198960 | 10.126048 | -1.037322 | 854.814108 | 0.142659036 | -0.37055562 | 0.710968545 | 0.998865404 | 9.765583 | 9.712869 | 9.746383 | 9.758275 | 9.744720 | 9.707425 | 994 | 947 | 749 | 766 |
| 4211 | NM_002398 | MEIS1 | Meis homeobox 1 | protein_coding | NP_002389.1 | 7000 | 601739 | . | . | 1.912289 | 1.818351 | 2.190412 | 2.028043 | 4.449976 | 4.203200 | 4.915220 | 4.636878 | 1.116791 | 283.073620 | 0.207019746 | 0.769776537 | 0.44143246 | 0.998865404 | 8.062814 | 8.222819 | 8.128619 | 8.098256 | 8.194238 | 8.148488 | 317 | 280 | 271 | 268 |
| 4212 | NM_00122048 | MEIS2 | Meis homeobox 2 | protein_coding | NP_001207411.1;NP_002390.1;NP_733774.1;NP_733775.1;NP_733776.1 | 7001 | 601740 | . | . | 3.324784 | 3.483810 | 3.893680 | 3.977375 | 7.736914 | 8.052978 | 8.737306 | 9.093794 | 1.020928 | 503.097198 | 0.168924321 | 0.176892068 | 0.859593169 | 0.998865404 | 8.959447 | 8.989780 | 8.995872 | 8.938573 | 8.993527 | 8.968054 | 599 | 513 | 451 | 467 |
| 4213 | NR_002211 | MEIS3P1 | Meis homeobox 3 pseudogene 1 | transcribed_ps | . | 7002 | . | | . | 1.608208 | 1.776349 | 1.428017 | 1.228601 | 3.742366 | 4.106111 | 3.204430 | 2.809051 | -1.284193 | 134.987591 | 0.284295643 | -1.26931953 | 0.204327122 | 0.969317911 | 7.245568 | 6.885374 | 7.102280 | 7.137364 | 7.055091 | 7.001797 | 167 | 171 | 111 | 102 |
| 4214 | NM_005921,XI | MAP3K1 | mitogen-activated protein kinase kinase kinase 1 | protein_coding | NP_005912.1;XP_016864973.1;XP_016864974.1 | 6848 | 600982 | . | . | 4.683726 | 4.638426 | 4.972021 | 4.212364 | 10.899224 | 10.721930 | 11.157070 | 9.631072 | -1.035600 | 1030.577645 | 0.145566393 | -0.34669563 | 0.728819994 | 0.998865404 | 10.033986 | 9.984057 | 10.025690 | 10.016092 | 10.058244 | 9.928243 | 1215 | 1124 | 959 | 865 |
| 4215 | NM_00133043 | MAP3K3 | mitogen-activated protein kinase kinase kinase 3 | protein_coding | NP_001317360.1;NP_001350697.1;NP_002392.2;NP_976226.1;XP_0052 | 6855 | 602539 | . | . | 6.760400 | 7.631373 | 7.470226 | 7.992160 | 15.731728 | 17.640259 | 16.762969 | 18.273122 | 1.064756 | 1100.033774 | 0.137853534 | 0.656656516 | 0.511401801 | 0.998865404 | 10.057558 | 10.147698 | 10.039473 | 10.116909 | 10.105191 | 10.145311 | 1163 | 1210 | 960 | 1092 |
| 4216 | NM_00129195 | MAP3K4 | mitogen-activated protein kinase kinase kinase 4 | protein_coding | NP_001278887.1;NP_001288001.1;NP_001350511.1;NP_005913.3;NP_0 | 6856 | 602425 | . | . | 2.292603 | 2.556749 | 2.383948 | 2.388518 | 5.334982 | 5.910041 | 5.349508 | 5.461063 | -1.020300 | 412.666403 | 0.181751456 | -0.15952407 | 0.873256 | 0.998865404 | 8.703427 | 8.674888 | 8.657004 | 8.730419 | 8.685278 | 8.679239 | 451 | 477 | 357 | 381 |
| 4217 | NM_005923,XI | MAP3K5 | mitogen-activated protein kinase kinase kinase 5 | protein_coding | NP_005914.1;XP_011534141.2;XP_016866359.1;XP_016866360.1;XP_01 | 6857 | 602448 | . | . | 0.661389 | 1.067784 | 0.962256 | 0.652009 | 1.539080 | 2.468231 | 2.159274 | 1.490742 | -1.044049 | 141.831788 | 0.314933628 | -0.19746636 | 0.843462603 | 0.998865404 | 7.178569 | 7.116845 | 7.064866 | 7.223193 | 7.199392 | 7.057216 | 133 | 188 | 144 | 105 |
| 4218 | NM_005370 | RAB8A | RAB8A, member RAS oncogene family | protein_coding | NP_005361.2 | 7007 | 165040 | . | . | 20.579754 | 22.278360 | 18.782335 | 21.764059 | 47.889931 | 51.497429 | 42.147018 | 49.760937 | -1.071575 | 1667.916966 | 0.130950082 | -0.76160651 | 0.446294889 | 0.998865404 | 10.753036 | 10.652890 | 10.701412 | 10.757232 | 10.621121 | 10.727202 | 1917 | 1928 | 1304 | 1613 |
| 4221 | NM_000244,N | MEN1 | menin 1 | protein_coding | NP_000235.2;NP_001357180.1;NP_001357188.2;NP_001357189.1;NP_0 | 7010 | 613733 | . | . | 12.781053 | 15.408329 | 16.324217 | 15.514079 | 29.742038 | 35.617039 | 36.631072 | 35.471097 | 1.111164 | 1362.372514 | 0.13323898 | 1.141341673 | 0.253727767 | 0.994807659 | 10.333960 | 10.485855 | 10.319557 | 10.416729 | 10.460854 | 10.438674 | 1391 | 1482 | 1265 | 1325 |
| 4232 | NM_00125390 | MEST | mesoderm specific transcript | protein_coding | NP_001240829.1;NP_001240830.1;NP_001240831.1;NP_002393.2;NP_8 | 7028 | 601029 | . | . | 0.191199 | 0.227529 | 0.363150 | 0.290960 | 0.444929 | 0.525944 | 0.814897 | 0.665246 | 1.651170 | 23.194071 | 0.642867992 | 1.125408429 | 0.260416001 | NA | 4.128334 | 4.852980 | 4.496147 | 4.497559 | 4.557468 | 4.531558 | 20 | 19 | 28 | 24 |
| 4233 | NM_000245,N | MET | MET proto-oncogene, receptor tyrosine kinase | protein_coding | NP_000236.2;NP_001120972.1;NP_001311330.1;NP_001311331.1;XP_0 | 7029 | 164860 | . | . | 7.156239 | 7.204010 | 5.634272 | 5.543706 | 16.652864 | 16.652395 | 12.643140 | 12.675027 | -1.307050 | 1331.145467 | 0.129082422 | -2.99277325 | 0.002764551 | 0.141329972 | 10.558616 | 10.172536 | 10.477879 | 10.464675 | 10.291088 | 10.243496 | 1751 | 1614 | 1036 | 1047 |
| 4234 | NM_005371,N | METTL1 | methyltransferase like 1 | protein_coding | NP_005362.3;NP_075422.3;XP_005268930.1;XP_016874794.2 | 7030 | 604466 | . | . | 8.805000 | 8.172353 | 7.566195 | 8.768318 | 20.489597 | 18.890760 | 16.978320 | 20.047719 | -1.035881 | 353.464232 | 0.192910014 | -0.26363605 | 0.792060378 | 0.998865404 | 8.490879 | 8.439505 | 8.487455 | 8.462914 | 8.415650 | 8.493065 | 423 | 380 | 281 | 348 |
| 4236 | NM_005926 | MFAP1 | microfibril associated protein 1 | protein_coding | NP_005917.2 | 7032 | 600215 | . | . | 13.073181 | 16.049641 | 16.408945 | 16.666428 | 30.421831 | 37.099461 | 36.821198 | 38.105804 | 1.120913 | 989.515134 | 0.149222234 | 1.103552295 | 0.269787395 | 0.998865404 | 9.866120 | 10.030364 | 9.831797 | 9.974318 | 9.986915 | 9.990962 | 969 | 1106 | 907 | 983 |
| 4237 | NM_00113524 | MFAP2 | microfibril associated protein 2 | protein_coding | NP_001128719.1;NP_001128720.1;NP_002394.1;NP_059453.1 | 7033 | 156790 | . | . | 7.021574 | 7.465493 | 6.582634 | 6.854976 | 16.339490 | 17.256822 | 14.771240 | 15.673089 | -1.092053 | 232.057254 | 0.22238386 | -0.57127518 | 0.567813125 | 0.998865404 | 7.920602 | 7.793263 | 7.867085 | 7.892016 | 7.832618 | 7.841540 | 272 | 268 | 191 | 210 |
| 4238 | NM_00113503 | MFAP3 | microfibril associated protein 3 | protein_coding | NP_001128509.1;NP_001229265.1;NP_005918.1 | 7034 | 600491 | . | . | 9.449574 | 9.677763 | 9.714873 | 7.780464 | 21.989548 | 22.370581 | 21.799893 | 17.789107 | -1.102338 | 1441.773070 | 0.144540957 | -0.9725018 | 0.330800969 | 0.998865404 | 10.561969 | 10.421889 | 10.521580 | 10.534727 | 10.539293 | 10.359503 | 1726 | 1645 | 1334 | 1134 |
| 4240 | NM_00111461 | MFGE8 | milk fat globule EGF and factor V/VIII domain containing | protein_coding | NP_001108086.1;NP_001297248.1;NP_001297249.1;NP_001297250.1;NI | 7036 | 602281 | . | . | 261.029791 | 270.227590 | 239.564166 | 250.735795 | 607.427071 | 624.643208 | 537.575040 | 573.277587 | -1.089087 | 15669.240988 | 0.099853859 | -1.23299279 | 0.217578453 | 0.982008277 | 13.995904 | 13.872764 | 13.959051 | 13.983058 | 13.884432 | 13.911242 | 18551 | 17874 | 12839 | 14285 |
| 4241 | NM_005929,N | MELTF | melanotransferrin | protein_coding | NP_005920.2;NP_201573.1;XP_006713706.2;XP_011511152.1;XP_01151 | 7037 | 155750 | . | . | 75.112637 | 71.880235 | 67.870733 | 71.799218 | 174.790195 | 166.154385 | 152.299961 | 164.160383 | -1.093117 | 5668.354020 | 0.10772782 | -1.19233401 | 0.233130303 | 0.988893316 | 12.531527 | 12.403041 | 12.524233 | 12.485046 | 12.401035 | 12.458188 | 6975 | 6241 | 4548 | 5252 |
| 4245 | NM_00111461 | MGAT1 | alpha-1,3-mannosyl-glycoprotein 2-beta-N-acetylglucosaminyltrar | protein_coding | NP_001108089.1;NP_001108090.1;NP_001108091.1;NP_001108092.1;NI | 7044 | 160995 | . | . | 60.774624 | 61.091471 | 64.806847 | 60.819117 | 141.425042 | 141.215673 | 145.424693 | 139.055687 | -1.029536 | 7516.878093 | 0.103743437 | -0.40479008 | 0.685631822 | 0.998865404 | 12.896768 | 12.854762 | 12.903752 | 12.872113 | 12.846288 | 12.880212 | 8945 | 8076 | 6312 | 7084 |
| 4247 | NM_002408 | MGAT2 | alpha-1,6-mannosyl-glycoprotein 2-beta-N-acetylglucosaminyltrar | protein_coding | NP_002399.1 | 7045 | 602616 | . | . | 13.184299 | 13.231422 | 11.730331 | 13.570178 | 30.680407 | 30.585028 | 26.322521 | 31.026598 | -1.059097 | 1081.022084 | 0.139821692 | -0.59242819 | 0.553563896 | 0.998865404 | 10.119196 | 10.035964 | 10.100752 | 10.096812 | 10.005220 | 10.104567 | 1284 | 1197 | 851 | 1051 |
| 4249 | NM_00137145 | MGAT5 | alpha-1,6-mannosylglycoprotein 6-beta-N-acetylglucosaminyltrans | protein_coding | NP_001358386.1;NP_002401.1;XP_005263726.1;XP_005263727.1;XP_00 | 7049 | 601774 | . | . | 34.856970 | 34.996550 | 35.891189 | 37.802569 | 81.113604 | 80.896091 | 80.538794 | 86.431083 | 1.045944 | 9601.881960 | 0.101583145 | 0.637955485 | 0.523502658 | 0.998865404 | 13.196343 | 13.261133 | 13.216481 | 13.202701 | 13.232318 | 13.263175 | 10896 | 10046 | 8383 | 9370 |
| 4253 | NM_00124798 | MIA2 | MIA SH3 domain ER export factor 2 | protein_coding | NP_001234917.1;NP_001234918.1;NP_001234919.1;NP_001316143.1;NI | 18432 | 602132 | . | . | 4.130978 | 3.571030 | 4.210900 | 4.341903 | 9.612956 | 8.254598 | 9.449139 | 9.927248 | 1.093220 | 460.864020 | 0.175169847 | 0.734051886 | 0.462917116 | 0.998865404 | 8.782261 | 8.911251 | 8.855606 | 8.780545 | 8.874503 | 8.873676 | 538 | 446 | 419 | 451 |
| 4254 | NM_000899,N | KITLG | KIT ligand | protein_coding | NP_000890.1;NP_003985.2 | 6343 | 184745 | . | . | 2.038048 | 1.777379 | 1.653854 | 1.249767 | 4.742620 | 4.108492 | 3.711201 | 2.857444 | -1.330629 | 285.721593 | 0.223713862 | -1.84212369 | 0.06545706 | 0.731993537 | 8.349080 | 7.938777 | 8.264700 | 8.184055 | 8.142792 | 8.002976 | 402 | 327 | 244 | 197 |
| 4255 | NM_002412,XI | MGMT | O-6-methylguanine-DNA methyltransferase | protein_coding | NP_002403.3;XP_005252739.1;XP_016871764.1 | 7059 | 156569 | . | . | 2.644442 | 2.416049 | 3.591739 | 2.497478 | 6.153726 | 5.584805 | 8.059759 | 5.710187 | -1.134215 | 336.709145 | 0.210264673 | -0.86411355 | 0.387525517 | 0.998865404 | 8.483767 | 8.301178 | 8.461101 | 8.393733 | 8.279338 | 8.429671 | 436 | 364 | 236 | 337 |
| 4256 | NM_000900,N | MGP | matrix Gla protein | protein_coding | NP_000891.2;NP_001177768.1 | 7060 | 154870 | . | . | 353.019966 | 324.521316 | 490.011776 | 513.946804 | 821.491997 | 750.145600 | 1099.572193 | 1175.078220 | 1.461272 | 21585.221217 | 0.103903627 | 5.26665562 | 1.38931E-07 | 6.18604E-05 | 14.098345 | 14.645582 | 14.250651 | 14.173448 | 14.521936 | 14.555851 | 21133 | 18049 | 21862 | 24488 |
| 4257 | NM_00126051 | MGST1 | microsomal glutathione S-transferase 1 | protein_coding | NP_001247440.1;NP_001247441.1;NP_001254527.1;NP_064696.1;NP_6 | 7061 | 138330 | . | . | 79.443384 | 78.197673 | 98.505634 | 82.755660 | 184.868026 | 180.757426 | 221.043805 | 189.210982 | 1.125190 | 2465.781894 | 0.127414219 | 1.335559216 | 0.181693388 | 0.953435029 | 11.180087 | 11.350540 | 11.224127 | 11.209693 | 11.382583 | 11.236931 | 2696 | 2481 | 2473 | 2230 |
| 4258 | NM_00120436 | MGST2 | microsomal glutathione S-transferase 2 | protein_coding | NP_001191295.1;NP_001191296.1;NP_001191297.1;NP_002404.1;XP_0 | 7063 | 601733 | . | . | 38.441091 | 35.519742 | 40.461021 | 36.129711 | 89.453997 | 82.105465 | 90.793354 | 82.606298 | 1.063891 | 931.326160 | 0.141689013 | 0.630605314 | 0.528298624 | 0.998865404 | 9.817644 | 9.907250 | 9.873738 | 9.804518 | 9.887220 | 9.882091 | 1089 | 927 | 838 | 897 |
| 4259 | NM_004528,XI | MGST3 | microsomal glutathione S-transferase 3 | protein_coding | NP_004519.1;XP_005245231.2 | 7064 | 604564 | . | . | 57.291312 | 57.668230 | 55.244997 | 65.655360 | 133.319241 | 133.302704 | 123.968166 | 150.113179 | 1.014418 | 1801.405557 | 0.131348037 | 0.157236381 | 0.875058557 | 0.998865404 | 10.804652 | 10.825090 | 10.834634 | 10.781523 | 10.756109 | 10.879417 | 2129 | 1865 | 1454 | 1834 |
| 4261 | NM_000246,N | CIITA | class II major histocompatibility complex transactivator | protein_coding | NP_000237.2;NP_001273331.1;NP_001273332.1;NP_001366259.1;NP_0 | 7067 | 600005 | . | . | 0.163661 | 0.151935 | 0.150906 | 0.284067 | 0.380845 | 0.351205 | 0.338626 | 0.649485 | 1.082665 | 33.804402 | 0.562258487 | 0.203798082 | 0.838511292 | 0.998865404 | 5.023141 | 5.133045 | 5.037429 | 5.097541 | 5.044728 | 5.111509 | 29 | 43 | 23 | 41 |
| 4267 | NM_00112289 | CD99 | CD99 molecule (Xg blood group) | protein_coding | NP_001116370.1;NP_001308296.1;NP_001308297.1;NP_001308298.1;NI | 7082 | 450000 | . | . | 221.398491 | 220.753443 | 234.714627 | 250.811297 | 515.203388 | 510.281451 | 526.692836 | 573.450207 | 1.082110 | 8311.573326 | 0.103061207 | 1.104657105 | 0.269308197 | 0.998865404 | 12.962874 | 13.076693 | 12.991732 | 12.980981 | 13.031096 | 13.075132 | 9252 | 8560 | 7319 | 8308 |
| 4277 | NM_00128916 | MICB | MHC class I polypeptide-related sequence B | protein_coding | NP_001276089.1;NP_001276090.1;NP_005922.2 | 7091 | 602436 | . | MICB | 9.077878 | 9.621695 | 7.959511 | 8.574099 | 21.124598 | 22.240980 | 17.860913 | 19.603659 | -1.147691 | 663.657019 | 0.154448411 | -1.28673727 | 0.198185878 | 0.96621494 | 9.470430 | 9.271292 | 9.401326 | 9.436772 | 9.304419 | 9.346599 | 796 | 785 | 520 | 598 |
| 4280 | gene-MICE | MICE | MHC class I polypeptide-related sequence E (pseudogene) | pseudogene | . | 7094 | . | | . | 3.341603 | 3.131321 | 3.015395 | 3.237804 | 7.776048 | 7.238189 | 6.766459 | 7.402855 | -1.053954 | 116.940306 | 0.29848646 | -0.25398962 | 0.799503581 | 0.998865404 | 6.907073 | 6.831186 | 6.891107 | 6.866819 | 6.851496 | 6.871140 | 143 | 125 | 96 | 110 |
| 4281 | NM_000381,N | MID1 | midline 1 | protein_coding | NP_000372.1;NP_001092094.1;NP_001180206.1;NP_001180207.1;NP_0 | 7095 | 300552 | . | . | 10.416753 | 10.905183 | 11.654061 | 11.139776 | 24.240212 | 25.207815 | 26.151376 | 25.469770 | 1.054124 | 2073.462058 | 0.118849737 | 0.6398364 | 0.522278965 | 0.998865404 | 10.979281 | 11.055368 | 10.979765 | 11.012328 | 11.058466 | 11.017518 | 2277 | 2223 | 1898 | 1944 |
| 4282 | NM_002415 | MIF | macrophage migration inhibitory factor | protein_coding | NP_002406.1 | 7097 | 153620 | . | . | 517.932251 | 517.997803 | 491.497650 | 606.626160 | 1205.249634 | 1197.375122 | 1102.906494 | 1386.978638 | 1.044898 | 9253.742886 | 0.119952439 | 0.528222201 | 0.597345122 | 0.998865404 | 13.143838 | 13.207111 | 13.159207 | 13.152037 | 13.104043 | 13.274880 | 10466 | 9725 | 7403 | 9750 |
| 4283 | NM_002416 | CXCL9 | C-X-C motif chemokine ligand 9 | protein_coding | NP_002407.1 | 7098 | 601704 | . | . | 0.071865 | 0.080648 | 0.078118 | 0.047848 | 0.167234 | 0.186421 | 0.175294 | 0.109400 | -1.288968 | 6.382002 | 1.202349992 | -0.30458424 | 0.760682822 | NA | 2.844134 | 2.481107 | 2.679081 | 2.681190 | 2.678689 | 2.666957 | 8 | 8 | 6 | 4 |
| 4285 | NM_005932,XI | MIPEP | mitochondrial intermediate peptidase | protein_coding | NP_005923.3;XP_011533399.1;XP_011533400.1 | 7104 | 602241 | . | . | 14.828429 | 14.037697 | 12.907173 | 17.479213 | 34.506362 | 32.448766 | 28.963322 | 39.964145 | 1.033458 | 1100.002195 | 0.159013216 | 0.298586991 | 0.765255192 | 0.998865404 | 10.079627 | 10.126573 | 10.110188 | 10.066075 | 10.000286 | 10.214055 | 1283 | 1133 | 833 | 1198 |
| 4286 | NM_000248,N | MITF | melanocyte inducing transcription factor | protein_coding | NP_000239.1;NP_001171896.1;NP_001171897.1;NP_001341533.1;NP_0 | 7105 | 156845 | . | . | 30.445104 | 29.926513 | 32.999982 | 32.467205 | 70.847008 | 69.176476 | 74.050998 | 74.232409 | 1.068209 | 4393.600947 | 0.107188431 | 0.888103334 | 0.37448517 | 0.998865404 | 12.052775 | 12.148029 | 12.087202 | 12.058507 | 12.137699 | 12.118079 | 4977 | 4505 | 3993 | 4205 |
| 4287 | NM_00112769 | ATXN3 | ataxin 3 | protein_coding | NP_001121168.1;NP_001121169.2;NP_001158246.1;NP_001158248.1;NI | 7106 | 607047 | . | . | 1.023604 | 1.287779 | 1.345281 | 1.141678 | 2.381970 | 2.976761 | 3.018772 | 2.610310 | 1.059649 | 259.071267 | 0.229139453 | 0.364787008 | 0.715270404 | 0.998865404 | 7.974773 | 8.058425 | 7.937767 | 8.055695 | 8.075333 | 7.977999 | 256 | 303 | 254 | 226 |
| 4288 | NM_00114596 | MKI67 | marker of proliferation Ki-67 | protein_coding | NP_001139438.1;NP_002408.3;XP_006717927.2;XP_011538120.1 | 7107 | 176741 | . | . | 22.536537 | 26.057512 | 23.752712 | 24.680130 | 52.443451 | 60.233102 | 53.300395 | 56.428182 | -1.027706 | 9206.204939 | 0.111285228 | -0.3542886 | 0.723122608 | 0.998865404 | 13.188006 | 13.148505 | 13.121082 | 13.234475 | 13.137501 | 13.174254 | 10033 | 10738 | 7725 | 8697 |
| 4289 | NM_00114535 | MKLN1 | muskelin 1 | protein_coding | NP_001138826.1;NP_001308245.1;NP_037387.2;XP_006716056.1;XP_01 | 7109 | 605623 | . | . | 4.937617 | 4.563708 | 5.455226 | 4.233848 | 11.490037 | 10.549219 | 12.241366 | 9.680190 | -1.037035 | 1632.028493 | 0.137610572 | -0.38125664 | 0.703012822 | 0.998865404 | 10.698199 | 10.646234 | 10.718210 | 10.650890 | 10.726398 | 10.582340 | 1995 | 1716 | 1523 | 1363 |
| 4291 | NM_00113015 | MLF1 | myeloid leukemia factor 1 | protein_coding | NP_001123628.1;NP_001123629.1;NP_001182361.2;NP_001182362.1;NI | 7125 | 601402 | . | . | 6.082000 | 5.356531 | 5.795565 | 6.329909 | 14.153064 | 12.381861 | 13.005082 | 14.472587 | 1.040068 | 402.114973 | 0.184478062 | 0.307230192 | 0.758668166 | 0.998865404 | 8.622818 | 8.679553 | 8.673809 | 8.602853 | 8.639444 | 8.684689 | 481 | 400 | 343 | 399 |
| 4292 | NM_000249,N | MLH1 | mutL homolog 1 | protein_coding | NP_000240.1;NP_001161089.1;NP_001161090.1;NP_001161091.1;NP_0 | 7127 | 120436 | . | . | 15.170103 | 16.694144 | 15.320240 | 17.715139 | 35.301458 | 38.589257 | 34.378172 | 40.503559 | 1.027718 | 1324.569425 | 0.14214089 | 0.277501097 | 0.781395364 | 0.998865404 | 10.351742 | 10.390611 | 10.320989 | 10.396707 | 10.309715 | 10.446183 | 1430 | 1480 | 1063 | 1371 |
| 4293 | NM_00128423 | MAP3K9 | mitogen-activated protein kinase kinase kinase 9 | protein_coding | NP_001271159.1;NP_001271160.1;NP_001271161.1;NP_149132.2;XP_0 | 6861 | 600136 | . | . | 2.004532 | 2.224515 | 2.097154 | 2.109494 | 4.664628 | 5.142068 | 4.705954 | 4.823108 | -1.013047 | 741.231415 | 0.149646647 | -0.12496381 | 0.900552205 | 0.998865404 | 9.543233 | 9.524265 | 9.509058 | 9.566323 | 9.524355 | 9.533650 | 824 | 838 | 636 | 695 |
| 4294 | NM_002446,XI | MAP3K10 | mitogen-activated protein kinase kinase kinase 10 | protein_coding | NP_002437.2;XP_006723286.1;XP_011525283.1;XP_011525284.1;XP_02 | 6849 | 600137 | . | . | 4.824245 | 4.614173 | 4.910921 | 5.274453 | 11.226218 | 10.665867 | 11.019966 | 12.059411 | 1.072478 | 536.793407 | 0.163825644 | 0.616192625 | 0.537765395 | 0.998865404 | 9.016950 | 9.117736 | 9.052637 | 9.036648 | 9.070646 | 9.108631 | 604 | 552 | 469 | 536 |
| 4296 | NM_002419,XI | MAP3K11 | mitogen-activated protein kinase kinase kinase 11 | protein_coding | NP_002410.1 | 6850 | 600050 | . | . | 15.947299 | 16.729397 | 19.795990 | 20.150199 | 37.110020 | 38.670754 | 44.421626 | 46.071034 | 1.207523 | 1999.832164 | 0.11826134 | 2.300417214 | 0.021424594 | 0.485588962 | 10.823278 | 11.095249 | 10.870579 | 10.900098 | 11.032418 | 11.038048 | 2047 | 1992 | 1896 | 2058 |
| 4297 | NM_00119710 | KMT2A | lysine methyltransferase 2A | protein_coding | NP_001184033.1;NP_005924.2;XP_006718902.2;XP_011541131.1;XP_01 | 7132 | 159555 | . | . | 1.606806 | 1.876450 | 1.878657 | 1.840868 | 3.739104 | 4.337499 | 4.215652 | 4.208927 | 1.023576 | 917.415982 | 0.14665128 | 0.229237798 | 0.818684096 | 0.998865404 | 9.824747 | 9.857927 | 9.777875 | 9.883703 | 9.836306 | 9.861285 | 968 | 1050 | 793 | 885 |
| 4298 | NM_005934,XI | MLLT1 | MLLT1 super elongation complex subunit | protein_coding | NP_005925.2;XP_011526323.1;XP_011526324.1;XP_011526325.1;XP_01 | 7134 | 159556 | . | . | 10.398320 | 11.326450 | 9.830606 | 11.547666 | 24.197319 | 26.181598 | 22.059593 | 26.402369 | -1.019940 | 1515.063751 | 0.133659918 | -0.21310795 | 0.831242767 | 0.998865404 | 10.579553 | 10.550628 | 10.548864 | 10.594847 | 10.497645 | 10.612522 | 1710 | 1700 | 1207 | 1511 |
| 4299 | NM_00116669 | AFF1 | AF4/FMR2 family member 1 | protein_coding | NP_001160165.1;NP_001300888.1;NP_001300889.1;NP_005926.1;XP_0 | 7135 | 159557 | . | . | 3.061226 | 3.219272 | 3.378850 | 3.171655 | 7.123598 | 7.441491 | 7.582040 | 7.251610 | 1.023852 | 933.436855 | 0.141530542 | 0.24028194 | 0.810111695 | 0.998865404 | 9.849211 | 9.883403 | 9.842093 | 9.872319 | 9.905399 | 9.842535 | 1040 | 1016 | 857 | 847 |
| 4300 | NM_00128669 | MLLT3 | MLLT3 super elongation complex subunit | protein_coding | NP_001273620.1;NP_004520.2;XP_016870215.1;XP_016870216.1 | 7136 | 159558 | . | . | 0.759010 | 0.733605 | 0.819504 | 0.914987 | 1.766248 | 1.695761 | 1.838944 | 2.092009 | 1.142292 | 169.377379 | 0.254656438 | 0.753686736 | 0.451037348 | 0.998865404 | 7.305134 | 7.496706 | 7.382210 | 7.365136 | 7.406703 | 7.450581 | 186 | 167 | 149 | 178 |
| 4301 | NM_00104000 | AFDN | afadin, adherens junction formation factor | protein_coding | NP_001035089.1;NP_001193937.1;NP_001278893.1;NP_001353248.1;NI | 7137 | 159559 | . | . | 5.203337 | 5.203150 | 5.055543 | 5.112509 | 12.108376 | 12.027313 | 11.344494 | 11.689145 | -1.038339 | 1246.926121 | 0.129018343 | -0.42069274 | 0.673979464 | 0.998865404 | 10.311037 | 10.256773 | 10.301523 | 10.295052 | 10.269231 | 10.270482 | 1469 | 1365 | 1063 | 1148 |
| 4302 | NM_005937 | MLLT6 | MLLT6, PHD finger containing | protein_coding | NP_005928.2 | 7138 | 600328 | . | . | 6.069268 | 6.867307 | 7.663952 | 7.402361 | 14.123435 | 15.874086 | 17.197687 | 16.924618 | 1.149219 | 1709.087097 | 0.127284232 | 1.576420104 | 0.114929018 | 0.855274326 | 10.635252 | 10.835804 | 10.635568 | 10.724674 | 10.807634 | 10.772094 | 1726 | 1815 | 1625 | 1675 |
| 4303 | NM_00117093 | FOXO4 | forkhead box O4 | protein_coding | NP_001164402.1;NP_005929.2;XP_011529259.1 | 7139 | 300033 | . | . | 1.791424 | 2.050054 | 2.228424 | 2.242131 | 4.168719 | 4.738793 | 5.000519 | 5.126367 | 1.155754 | 234.866123 | 0.224449368 | 0.930427782 | 0.35214964 | 0.998865404 | 7.767809 | 7.976071 | 7.802056 | 7.870172 | 7.906893 | 7.907013 | 233 | 252 | 219 | 236 |
| 4306 | NM_000901,N | NR3C2 | nuclear receptor subfamily 3 group C member 2 | protein_coding | NP_000892.2;NP_001159576.1;NP_001341748.1;XP_011530277.1;XP_01 | 7979 | 600983 | . | . | 0.021151 | 0.028382 | 0.109669 | 0.081460 | 0.049219 | 0.065605 | 0.246095 | 0.186249 | 3.599957 | 11.383889 | 0.948217006 | 1.948899622 | 0.051307413 | NA | 2.307819 | 4.155137 | 3.439014 | 3.445300 | 3.511999 | 3.487987 | 5 | 6 | 18 | 14 |
| 4308 | NM_00125202 | TRPM1 | transient receptor potential cation channel subfamily M member | protein_coding | NP_001238949.1;NP_001238953.1;NP_001238959.1;NP_002411.3 | 7146 | 603576 | . | . | 17.162736 | 17.043125 | 17.185538 | 21.745846 | 39.938394 | 39.395950 | 38.563850 | 49.719294 | 1.146052 | 912.800727 | 0.14556412 | 1.351107785 | 0.176660908 | 0.949538805 | 9.732724 | 9.929923 | 9.772301 | 9.790899 | 9.828747 | 9.928713 | 967 | 930 | 790 | 976 |
| 4311 | NM_000902,N | MME | membrane metalloendopeptidase | protein_coding | NP_000893.2;NP_001341571.1;NP_001341572.1;NP_001341573.1;NP_0 | 7154 | 120520 | . | . | 7.299933 | 7.325734 | 6.945205 | 6.425182 | 16.987241 | 16.933763 | 15.584842 | 14.690417 | -1.110513 | 1213.553615 | 0.13467964 | -1.12285958 | 0.261497135 | 0.995760118 | 10.318481 | 10.167621 | 10.288898 | 10.276710 | 10.245249 | 10.160999 | 1482 | 1367 | 1057 | 1017 |
| 4312 | NM_00114593 | MMP1 | matrix metallopeptidase 1 | protein_coding | NP_001139410.1;NP_002412.1 | 7155 | 120353 | . | . | 19.728291 | 18.328989 | 19.930364 | 15.956754 | 45.908546 | 42.368280 | 44.723163 | 36.483222 | -1.074407 | 1134.794900 | 0.151598314 | -0.68299265 | 0.494611487 | 0.998865404 | 10.198715 | 10.095885 | 10.204308 | 10.139615 | 10.201154 | 10.029830 | 1411 | 1214 | 1062 | 907 |
| 4313 | NM_00112789 | MMP2 | matrix metallopeptidase 2 | protein_coding | NP_001121363.1;NP_001289437.1;NP_001289438.1;NP_001289439.1;NI | 7166 | 120360 | . | . | 0.849860 | 1.043697 | 0.823030 | 0.818698 | 1.977660 | 2.412553 | 1.846855 | 1.871857 | -1.160865 | 97.506135 | 0.327990751 | -0.65611746 | 0.511748553 | 0.998865404 | 6.711333 | 6.495439 | 6.597811 | 6.659671 | 6.586664 | 6.582485 | 109 | 124 | 79 | 84 |
| 4314 | NM_002422 | MMP3 | matrix metallopeptidase 3 | protein_coding | NP_002413.1 | 7173 | 185250 | . | . | 1.066258 | 1.103442 | 1.638184 | 1.379162 | 2.481226 | 2.550655 | 3.676037 | 3.153289 | 1.370839 | 73.923162 | 0.370280052 | 1.228957962 | 0.219087568 | 0.982008277 | 5.962063 | 6.417976 | 6.156045 | 6.161370 | 6.260160 | 6.212250 | 71 | 68 | 81 | 73 |
| 4317 | NM_00130444 | MMP8 | matrix metallopeptidase 8 | protein_coding | NP_001291370.1;NP_001291371.1;NP_002415.1;XP_011541136.1;XP_01 | 7175 | 120355 | . | . | 60.280868 | 60.797161 | 43.813590 | 45.552868 | 140.276060 | 140.535366 | 98.316422 | 104.151222 | -1.372234 | 5035.039526 | 0.105483224 | -4.32795303 | 1.50502E-05 | 0.002769831 | 12.508084 | 12.051519 | 12.415348 | 12.416208 | 12.139554 | 12.162398 | 6704 | 6289 | 3639 | 4036 |
| 4323 | NM_004995 | MMP14 | matrix metallopeptidase 14 | protein_coding | NP_004986.1 | 7160 | 600754 | . | . | 177.271408 | 177.209412 | 112.466484 | 145.376495 | 412.517853 | 409.627472 | 252.371521 | 332.386108 | -1.394930 | 17664.844093 | 0.125612962 | -3.82279209 | 0.000131949 | 0.014928627 | 14.328829 | 13.848573 | 14.237261 | 14.229384 | 13.839657 | 14.044885 | 23801 | 22106 | 11255 | 15525 |
| 4324 | NM_002428 | MMP15 | matrix metallopeptidase 15 | protein_coding | NP_002419.1 | 7161 | 602261 | . | . | 20.760618 | 20.544634 | 13.170929 | 14.963106 | 48.310814 | 47.489841 | 29.555183 | 34.211361 | -1.488976 | 2199.828137 | 0.122079475 | -4.70447923 | 2.54515E-06 | 0.000605701 | 11.362044 | 10.787473 | 11.248594 | 11.232396 | 10.876970 | 10.965748 | 3060 | 2813 | 1447 | 1754 |
| 4325 | NM_005941,XI | MMP16 | matrix metallopeptidase 16 | protein_coding | NP_005932.2;XP_024302922.1 | 7162 | 602262 | . | . | 3.566428 | 3.426788 | 3.624322 | 2.815455 | 8.299223 | 7.921175 | 8.132872 | 6.437204 | -1.098601 | 1212.387884 | 0.153404243 | -0.88437647 | 0.376493022 | 0.998865404 | 10.309550 | 10.174564 | 10.294774 | 10.256243 | 10.299439 | 10.100966 | 1498 | 1335 | 1139 | 939 |
| 4326 | NM_016155,XI | MMP17 | matrix metallopeptidase 17 | protein_coding | NP_057239.4;XP_011536657.1;XP_011536658.1;XP_011536659.1;XP_01 | 7163 | 602285 | . | . | 93.299841 | 96.695125 | 103.568042 | 98.761115 | 217.112559 | 223.515120 | 232.403674 | 225.805543 | 1.044028 | 6952.624396 | 0.104529676 | 0.594668679 | 0.552064962 | 0.998865404 | 12.731914 | 12.794100 | 12.732869 | 12.756407 | 12.803259 | 12.758546 | 7724 | 7443 | 6339 | 6483 |
| 4327 | NM_00127210 | MMP19 | matrix metallopeptidase 19 | protein_coding | NP_001259030.1;NP_002420.1;XP_006719464.1;XP_011536661.1;XP_01 | 7165 | 601807 | . | . | 0.196116 | 0.099553 | 0.138336 | 0.386622 | 0.456371 | 0.230121 | 0.310422 | 0.883965 | 1.739219 | 21.882468 | 0.721347918 | 1.10687127 | 0.268349589 | NA | 3.998355 | 4.796139 | 4.427887 | 4.384813 | 4.408434 | 4.503247 | 24 | 12 | 14 | 37 |
| 4329 | NM_00127859 | ALDH6A1 | aldehyde dehydrogenase 6 family member A1 | protein_coding | NP_001265522.1;NP_001265523.1;NP_005580.1;XP_016876820.1 | 7179 | 603178 | . | . | 0.872599 | 0.829635 | 0.706986 | 0.971534 | 2.030573 | 1.917738 | 1.586456 | 2.221296 | -1.014506 | 145.341207 | 0.281450235 | -0.07382467 | 0.941149898 | 0.998865404 | 7.194269 | 7.172247 | 7.197389 | 7.172543 | 7.126202 | 7.230109 | 174 | 153 | 108 | 154 |
| 4330 | NM_002430 | MN1 | MN1 proto-oncogene, transcriptional regulator | protein_coding | NP_002421.3 | 7180 | 156100 | . | . | 0.161055 | 0.194096 | 0.112423 | 0.148121 | 0.374782 | 0.448661 | 0.252275 | 0.338662 | -1.377514 | 38.001282 | 0.512224762 | -0.90207803 | 0.367015419 | 0.998865404 | 5.461710 | 4.996978 | 5.257280 | 5.288980 | 5.209646 | 5.242849 | 46 | 52 | 24 | 34 |
| 4331 | NM_00117796 | MNAT1 | MNAT1 component of CDK activating kinase | protein_coding | NP_001171434.1;NP_002422.1;XP_005267744.1;XP_005267745.1;XP_01 | 7181 | 602659 | . | . | 5.552088 | 4.552736 | 3.959159 | 5.171267 | 12.919936 | 10.523851 | 8.884241 | 11.823486 | -1.134274 | 383.817068 | 0.19852582 | -0.91559457 | 0.359879579 | 0.998865404 | 8.672382 | 8.490435 | 8.677964 | 8.555668 | 8.495500 | 8.591824 | 517 | 396 | 286 | 366 |
| 4335 | NM_020310,XI | MNT | MAX network transcriptional repressor | protein_coding | NP_064706.1;XP_011522170.1;XP_011522171.1;XP_016880143.1;XP_02 | 7188 | 603039 | . | . | 5.518465 | 6.461781 | 7.225437 | 5.578044 | 12.841700 | 14.936697 | 16.213670 | 12.753533 | 1.016127 | 1126.606904 | 0.173145633 | 0.133298851 | 0.893957036 | 0.998865404 | 10.125967 | 10.149474 | 10.065183 | 10.184808 | 10.261303 | 9.999603 | 1184 | 1302 | 1160 | 879 |
| 4337 | NM_00107509 | MOCS1 | molybdenum cofactor synthesis 1 | protein_coding | NP_001068566.1;NP_001345458.1;NP_001345459.1;NP_001345460.1;NI | 7190 | 603707 | . | . | 3.171581 | 3.741369 | 3.017936 | 3.806114 | 7.380395 | 8.648345 | 6.772161 | 8.702228 | -1.027711 | 440.474703 | 0.190327776 | -0.20719586 | 0.835856901 | 0.998865404 | 8.803120 | 8.762423 | 8.743392 | 8.833558 | 8.703820 | 8.836923 | 477 | 517 | 336 | 452 |
| 4338 | NM_004531,N | MOCS2 | molybdenum cofactor synthesis 2 | protein_coding | NP_004522.1;NP_789776.1 | 7193 | 603708 | . | . | 16.514135 | 16.291194 | 15.163714 | 16.677368 | 38.429073 | 37.657824 | 34.026934 | 38.130817 | -1.031388 | 1940.183399 | 0.122453097 | -0.36411103 | 0.715775104 | 0.998865404 | 10.944190 | 10.899417 | 10.940668 | 10.927128 | 10.870252 | 10.947047 | 2288 | 2108 | 1580 | 1878 |
| 4343 | NM_00113007 | MOV10 | Mov10 RISC complex RNA helicase | protein_coding | NP_001123551.1;NP_001273001.1;NP_001308253.1;NP_001356436.1;NI | 7200 | 610742 | . | . | 24.906791 | 28.231290 | 31.544745 | 28.482215 | 57.959123 | 65.257891 | 70.785492 | 65.121197 | 1.100705 | 3674.513016 | 0.122159902 | 1.133169034 | 0.257143276 | 0.995760118 | 11.772461 | 11.910895 | 11.744038 | 11.855460 | 11.925745 | 11.833369 | 3755 | 4031 | 3534 | 3410 |
| 4345 | NM_00100419 | CD200 | CD200 molecule | protein_coding | NP_001004196.2;NP_001305755.1;NP_001305757.1;NP_001305759.1;NI | 7203 | 155970 | . | . | 16.620811 | 15.126315 | 15.054407 | 12.727569 | 38.677993 | 34.965155 | 33.781652 | 29.100073 | -1.158803 | 1042.106442 | 0.151627458 | -1.40235314 | 0.160809819 | 0.933525128 | 10.127349 | 9.915451 | 10.104470 | 10.045599 | 10.042333 | 9.885663 | 1338 | 1160 | 931 | 807 |
| 4350 | NM_00101505 | MPG | N-methylpurine DNA glycosylase | protein_coding | NP_001015052.1;NP_001015054.1;NP_002425.2;XP_024306050.1 | 7211 | 156565 | . | . | 54.897748 | 55.977594 | 47.281830 | 60.310348 | 127.749321 | 129.394723 | 106.099054 | 137.892446 | -1.049808 | 1798.005234 | 0.138531363 | -0.5062035 | 0.6127138 | 0.998865404 | 10.847033 | 10.776467 | 10.820294 | 10.838987 | 10.700200 | 10.874916 | 2096 | 2011 | 1355 | 1828 |
| 4351 | NM_00128915 | MPI | mannose phosphate isomerase | protein_coding | NP_001276084.1;NP_001276085.1;NP_001276086.1;NP_001317301.1;NI | 7216 | 154550 | . | . | 4.833329 | 4.724774 | 5.627824 | 6.477670 | 11.247357 | 10.921527 | 12.628671 | 14.810422 | 1.221046 | 930.189766 | 0.143179712 | 2.012281502 | 0.044190271 | 0.645513836 | 9.710359 | 9.998082 | 9.775811 | 9.790144 | 9.883689 | 9.969521 | 955 | 913 | 838 | 1014 |
| 4352 | NM_005373,XI | MPL | MPL proto-oncogene, thrombopoietin receptor | protein_coding | NP_005364.1;XP_016856809.1 | 7217 | 159530 | . | . | 0.036511 | 0.069295 | 0.067532 | 0.019079 | 0.084963 | 0.160179 | 0.151540 | 0.043822 | -1.250046 | 5.716517 | 1.318217212 | -0.24425475 | 0.807033529 | NA | 2.664814 | 2.348113 | 2.503529 | 2.521980 | 2.520079 | 2.493650 | 5 | 9 | 7 | 2 |
| 4354 | NM_00116646 | MPP1 | membrane palmitoylated protein 1 | protein_coding | NP_001159932.1;NP_001159933.1;NP_001159934.1;NP_002427.1;XP_01 | 7219 | 305360 | . | . | 6.005990 | 6.667535 | 5.432710 | 6.593996 | 13.976188 | 15.412308 | 12.190844 | 15.076389 | -1.071967 | 386.962475 | 0.191847667 | -0.52260051 | 0.601252288 | 0.998865404 | 8.645769 | 8.544403 | 8.585985 | 8.643838 | 8.520633 | 8.625307 | 439 | 453 | 295 | 382 |
| 4355 | NM_00127837 | MPP2 | membrane palmitoylated protein 2 | protein_coding | NP_001265299.1;NP_001265300.1;NP_001265301.1;NP_001265302.1;NI | 7220 | 600723 | . | . | 3.114600 | 3.558352 | 3.049576 | 3.550254 | 7.247803 | 8.225291 | 6.843161 | 8.117234 | -1.024327 | 437.544051 | 0.183207844 | -0.18927201 | 0.84987963 | 0.998865404 | 8.791003 | 8.755346 | 8.740131 | 8.817854 | 8.721115 | 8.806882 | 478 | 508 | 349 | 434 |
| 4356 | NM_00133023 | MPP3 | membrane palmitoylated protein 3 | protein_coding | NP_001317162.1;NP_001340009.1;NP_001923.2;XP_006721978.1;XP_00 | 7221 | 601114 | . | . | 1.039100 | 0.778036 | 1.195097 | 1.279891 | 2.418032 | 1.798467 | 2.681762 | 2.926320 | 1.439265 | 96.646054 | 0.325897219 | 1.611957935 | 0.106971094 | 0.83501324 | 6.307763 | 6.833823 | 6.542020 | 6.514874 | 6.642201 | 6.630966 | 96 | 81 | 101 | 105 |
| 4357 | NM_00101343 | MPST | mercaptopyruvate sulfurtransferase | protein_coding | NP_001013454.1;NP_001123989.1;NP_001356833.1;NP_001356834.1;NI | 7223 | 602496 | . | . | 19.341502 | 18.435946 | 16.222269 | 21.343084 | 45.008468 | 42.615515 | 36.402304 | 48.798427 | -1.003774 | 779.344910 | 0.169055365 | -0.0321434 | 0.974357693 | 0.998865404 | 9.609204 | 9.603026 | 9.616587 | 9.593516 | 9.495558 | 9.698999 | 914 | 829 | 578 | 835 |

| | | | | | | | | | | | | | | | | | | | | | | | | | | | | | | | | | |
|---|---|---|---|---|---|---|---|---|---|---|---|---|---|---|---|---|---|---|---|---|---|---|---|---|---|---|---|---|---|---|---|---|---|
| 4358 | NM_002437,XI | MPV17 | mitochondrial inner membrane protein MPV17 | protein_coding | NP_002428.1;XP_005264383.1;XP_006712084.1;XP_016859639.1;XP_01 | 7224 | 137960 | . | . | 28.002193 | 28.165394 | 25.990085 | 31.985090 | 65.162253 | 65.105570 | 58.320995 | 73.130110 | 1.123041 | 1370.687564 | 0.127600992 | 1.31198869 | 0.189523951 | 0.961057712 | 10.334561 | 10.501954 | 10.356205 | 10.391723 | 10.474342 | 10.451166 | 1452 | 1426 | 1280 | 1339 |
| 4359 | NM_000530,N | MPZ | myelin protein zero | protein_coding | NP_000521.2;NP_001302420.1;XP_016856810.1 | 7225 | 159440 | . | . | 292.230520 | 296.668526 | 295.125782 | 298.884272 | 680.032415 | 685.762604 | 662.253691 | 683.363378 | -1.005312 | 17972.895237 | 0.0983907 | -0.07768022 | 0.938082426 | 0.998865404 | 14.137353 | 14.129707 | 14.132927 | 14.138679 | 14.130177 | 14.132329 | 20681 | 19510 | 15568 | 16826 |
| 4361 | NM_00133034 | MRE11 | MRE11 homolog, double strand break repair nuclease | protein_coding | NP_001317276.1;NP_005581.2;NP_005582.1;XP_005274065.1;XP_00671 | 7230 | 600814 | . | . | 4.546184 | 4.365696 | 4.169504 | 4.012557 | 10.579158 | 10.091501 | 9.356246 | 9.174237 | -1.023340 | 813.643749 | 0.145341044 | -0.22901665 | 0.818855978 | 0.998865404 | 9.684640 | 9.651678 | 9.692942 | 9.659185 | 9.679404 | 9.640175 | 970 | 867 | 719 | 733 |
| 4363 | NM_004996,XI | ABCC1 | ATP binding cassette subfamily C member 1 | protein_coding | NP_004987.2;XP_011520799.1;XP_011520800.1;XP_016878726.1;XP_01 | 51 | 158343 | . | . | 9.024726 | 9.213892 | 9.286671 | 9.595679 | 21.000907 | 21.298326 | 20.839020 | 21.939377 | 1.019431 | 1872.242941 | 0.119181942 | 0.232960358 | 0.815792188 | 0.998865404 | 10.856638 | 10.884333 | 10.856850 | 10.868834 | 10.870459 | 10.885563 | 2119 | 2016 | 1628 | 1789 |
| 4430 | NM_00113015 | MYO1B | myosin IB | protein_coding | NP_001123630.1;NP_001155291.1;NP_001317166.1;NP_001317167.1;NI | 7596 | 606537 | . | . | 41.952011 | 41.320731 | 37.255742 | 39.932683 | 97.624053 | 95.514713 | 83.600812 | 91.301333 | -1.097713 | 6262.174258 | 0.105461213 | -1.27535408 | 0.202183943 | 0.967656221 | 12.678152 | 12.543609 | 12.659638 | 12.641099 | 12.546824 | 12.596234 | 7629 | 6995 | 5038 | 5763 |
| 4435 | NM_00114488 | CITED1 | Cbp/p300 interacting transactivator with Glu/Asp rich carboxy-teri | protein_coding | NP_001138357.1;NP_001138358.1;NP_001138359.1;NP_004134.2;XP_01 | 1986 | 300149 | . | . | 1.089469 | 0.570856 | 0.378631 | 0.728990 | 2.535238 | 1.319559 | 0.849637 | 1.666749 | -1.368159 | 22.713444 | 0.675659176 | -0.66932557 | 0.50328782 | NA | 4.714815 | 4.260516 | 4.547824 | 4.498201 | 4.471641 | 4.515564 | 37 | 22 | 12 | 23 |
| 4436 | NM_000251,N | MSH2 | mutS homolog 2 | protein_coding | NP_000242.1;NP_001245210.1;XP_005264389.2;XP_011531169.1 | 7325 | 609309 | . | . | 18.436016 | 18.936556 | 16.826361 | 16.311000 | 42.901366 | 43.772695 | 37.757866 | 37.293167 | -1.150501 | 1709.231728 | 0.121522852 | -1.66439474 | 0.096033571 | 0.813553488 | 10.836700 | 10.634485 | 10.784001 | 10.798955 | 10.693293 | 10.670245 | 2086 | 1992 | 1403 | 1468 |
| 4437 | NM_002439 | MSH3 | mutS homolog 3 | protein_coding | NP_002430.3 | 7326 | 600887 | . | . | 3.069368 | 2.996428 | 2.741067 | 2.697645 | 7.142546 | 6.926377 | 6.150875 | 6.167844 | -1.130347 | 398.636386 | 0.180807142 | -0.97764589 | 0.328249491 | 0.998865404 | 8.724492 | 8.547971 | 8.684362 | 8.664247 | 8.609903 | 8.593721 | 495 | 449 | 330 | 346 |
| 4439 | NM_002441,N | MSH5 | mutS homolog 5 | protein_coding | NP_002432.1;NP_079535.4;NP_751897.1;NP_751898.1 | 7328 | 603382 | . | . | 6.350992 | 6.312042 | 4.660441 | 1.169159 | 14.779021 | 14.590565 | 10.457893 | 2.673141 | -2.069245 | 408.329775 | 0.909655645 | -1.1532983 | 0.248787969 | 0.992733415 | 9.104702 | 8.055934 | 8.808013 | 8.806867 | 8.638089 | 8.017369 | 634 | 594 | 371 | 99 |
| 4440 | NM_002442,XI | MSI1 | musashi RNA binding protein 1 | protein_coding | NP_002433.1;XP_006719466.1;XP_006719467.1;XP_011536663.1;XP_01 | 7330 | 603328 | . | . | 0.678153 | 0.451695 | 0.842633 | 0.918836 | 1.578089 | 1.044115 | 1.890844 | 2.100810 | 1.518108 | 67.339981 | 0.399819124 | 1.506367189 | 0.131972939 | 0.885996989 | 5.740050 | 6.343985 | 6.049727 | 5.969756 | 6.096489 | 6.116924 | 74 | 46 | 68 | 79 |
| 4478 | NM_002444,XI | MSN | moesin | protein_coding | NP_002435.1;XP_005262326.1;XP_011529261.1;XP_016885034.1;XP_01 | 7373 | 309845 | . | . | 93.418306 | 101.006313 | 87.114411 | 106.541990 | 217.388236 | 233.480599 | 195.482207 | 243.595591 | -1.018464 | 11970.500955 | 0.119612554 | -0.22067364 | 0.825346561 | 0.998865404 | 13.560377 | 13.533893 | 13.523274 | 13.583340 | 13.453251 | 13.616427 | 13423 | 13492 | 9330 | 12180 |
| 4479 | gene-MSNP1 | MSNP1 | moesin pseudogene 1 | pseudogene | . | 7374 | . | | . | 0.016869 | 0.030087 | 0.021247 | 0.042720 | 0.039254 | 0.069548 | 0.047677 | 0.097673 | 1.218116 | 4.005577 | 1.504408192 | 0.18921186 | 0.849926773 | NA | 1.855800 | 2.134756 | 1.990769 | 2.000428 | 1.995011 | 2.003472 | 3 | 5 | 3 | 5 |
| 4482 | NM_00113567 | MSRA | methionine sulfoxide reductase A | protein_coding | NP_001129142.1;NP_001129143.1;NP_001186658.1;NP_036463.1;XP_01 | 7377 | 601250 | . | . | 3.407514 | 3.254651 | 3.745503 | 3.525279 | 7.929423 | 7.523274 | 8.404800 | 8.060133 | 1.080277 | 161.649738 | 0.259836472 | 0.428738561 | 0.668113495 | 0.998865404 | 7.279546 | 7.391728 | 7.335129 | 7.303392 | 7.367171 | 7.333971 | 186 | 161 | 152 | 151 |
| 4485 | NM_020998,N | MST1 | macrophage stimulating 1 | protein_coding | NP_066278.3;XP_006713229.1;XP_011532034.1;XP_011532040.1;XP_01 | 7380 | 142408 | . | . | 2.876365 | 2.858174 | 2.834194 | 2.851351 | 6.693423 | 6.606797 | 6.359847 | 6.519275 | -1.043867 | 201.296826 | 0.235586308 | -0.26291013 | 0.792619849 | 0.998865404 | 7.683725 | 7.621975 | 7.675366 | 7.650796 | 7.643034 | 7.644469 | 243 | 216 | 171 | 185 |
| 4487 | NM_002448 | MSX1 | msh homeobox 1 | protein_coding | NP_002439.2 | 7391 | 142983 | . | . | 2.710598 | 2.740114 | 2.155959 | 2.908044 | 6.307672 | 6.333896 | 4.837911 | 6.648896 | -1.089737 | 159.412575 | 0.270944635 | -0.45758367 | 0.647251576 | 0.998865404 | 7.378019 | 7.252494 | 7.332012 | 7.334229 | 7.243100 | 7.351017 | 191 | 180 | 114 | 163 |
| 4488 | NM_00136362 | MSX2 | msh homeobox 2 | protein_coding | NP_001350555.1;NP_002440.2 | 7392 | 123101 | . | . | 3.791066 | 4.907085 | 3.868319 | 4.243474 | 8.821966 | 11.342947 | 8.680396 | 9.702198 | -1.083809 | 288.543882 | 0.21796424 | -0.53270154 | 0.594240203 | 0.998865404 | 8.230096 | 8.112815 | 8.120202 | 8.258417 | 8.128087 | 8.169345 | 302 | 365 | 231 | 270 |
| 4494 | NM_00130127 | MT1F | metallothionein 1F | protein_coding | NP_001288201.1;NP_005940.1 | 7398 | 156352 | . | . | 0.506619 | 1.433299 | 0.815084 | 0.716834 | 1.178925 | 3.313135 | 1.829025 | 1.638955 | -1.269665 | 12.592695 | 0.90024569 | -0.38261518 | 0.702005111 | NA | 3.817846 | 3.470304 | 3.628342 | 3.684628 | 3.645671 | 3.641952 | 9 | 22 | 10 | 10 |
| 4501 | NM_005952 | MT1X | metallothionein 1X | protein_coding | NP_005943.1 | 7405 | 156359 | . | . | 17.685110 | 14.052329 | 12.251326 | 18.438810 | 41.153976 | 32.482590 | 27.491617 | 42.158150 | -1.052211 | 196.789978 | 0.269855071 | -0.27208654 | 0.785555478 | 0.998865404 | 7.657247 | 7.582819 | 7.681027 | 7.571504 | 7.515666 | 7.686873 | 260 | 192 | 134 | 215 |
| 4502 | NM_005953 | MT2A | metallothionein 2A | protein_coding | NP_005944.1 | 7406 | 156360 | . | . | 303.615173 | 319.387482 | 211.325989 | 303.759247 | 706.524963 | 738.278442 | 474.209381 | 694.509399 | -1.227069 | 3555.435429 | 0.150892259 | -1.95647367 | 0.050409381 | 0.670328218 | 11.936024 | 11.640488 | 11.853005 | 11.888425 | 11.550369 | 11.833729 | 4417 | 4317 | 2292 | 3515 |
| 4507 | NM_002451 | MTAP | methylthioadenosine phosphorylase | protein_coding | NP_002442.2 | 7413 | 156540 | . | . | 5.189012 | 5.455999 | 5.869699 | 4.732800 | 12.075045 | 12.611786 | 13.171436 | 10.820985 | -1.017854 | 998.961477 | 0.152606905 | -0.16729473 | 0.867138153 | 0.998865404 | 9.976734 | 9.951728 | 9.953386 | 9.984234 | 10.034877 | 9.870758 | 1136 | 1110 | 958 | 824 |
| 4508 | gene-ATP6 | ATP6 | ATP synthase F0 subunit 6 | protein_coding | YP_003024031.1 | 7414 | 516060 | . | . | 13108.467773 | 13425.564453 | 15892.435547 | 12843.672852 | 30503.945312 | 31033.794922 | 35662.164062 | 29365.533203 | 1.068623 | 293131.873293 | 0.120020134 | 0.797813147 | 0.424978931 | 0.998865404 | 18.112518 | 18.208274 | 18.123553 | 18.136803 | 18.280512 | 18.085976 | 323836 | 308156 | 292629 | 252372 |
| 4509 | gene-ATP8 | ATP8 | ATP synthase F0 subunit 8 | protein_coding | YP_003024030.1 | 7415 | 516070 | . | . | 9642.646484 | 10309.251953 | 11935.416016 | 10603.783203 | 22438.832031 | 23830.296875 | 26782.728516 | 24244.291016 | 1.114522 | 68479.467108 | 0.107681657 | 1.452661797 | 0.146317679 | 0.907376824 | 15.983049 | 16.139480 | 15.988725 | 16.038763 | 16.162515 | 16.050322 | 72409 | 71927 | 66802 | 63334 |
| 4511 | rna-TRNC | TRNC | tRNA-Cys | tRNA | . | 7477 | 590020 | . | . | 74.555031 | 60.069897 | 83.452980 | 67.835350 | 173.492615 | 138.854263 | 187.266083 | 155.097458 | 1.107139 | 147.411954 | 0.282072213 | 0.520563734 | 0.602670718 | 0.998865404 | 7.127401 | 7.276184 | 7.221112 | 7.136524 | 7.259247 | 7.176952 | 179 | 134 | 149 | 130 |
| 4512 | gene-COX1 | COX1 | cytochrome c oxidase subunit I | protein_coding | YP_003024028.1 | 7419 | 516030 | . | . | 21779.615234 | 21434.199219 | 24888.470703 | 19176.501953 | 50682.062500 | 49546.109375 | 55849.007812 | 43844.792969 | 1.006203 | 1048455.385423 | 0.128157871 | 0.069613407 | 0.944501365 | 0.998865404 | 19.995366 | 20.004288 | 20.005516 | 19.984211 | 20.111140 | 19.876103 | 1218318 | 1113994 | 1037675 | 853214 |
| 4513 | gene-COX2 | COX2 | cytochrome c oxidase subunit II | protein_coding | YP_003024029.1 | 7421 | 516040 | . | . | 24439.511719 | 23418.736328 | 27165.335938 | 24250.763672 | 56871.746094 | 54133.457031 | 60958.222656 | 55446.492188 | 1.059671 | 528815.047186 | 0.106174204 | 0.787533475 | 0.43096964 | 0.998865404 | 18.969989 | 19.053607 | 19.008084 | 18.963836 | 19.089010 | 18.981186 | 606422 | 539897 | 502401 | 478615 |
| 4514 | gene-COX3 | COX3 | cytochrome c oxidase subunit III | protein_coding | YP_003024032.1 | 7422 | 516050 | . | . | 16773.330078 | 16984.705078 | 19213.625000 | 16326.196289 | 39032.230469 | 39260.906250 | 43114.812500 | 37327.906250 | 1.038657 | 423135.627727 | 0.111877081 | 0.489105595 | 0.624766941 | 0.998865404 | 18.663140 | 18.717862 | 18.671456 | 18.674889 | 18.778637 | 18.628077 | 477048 | 448813 | 407291 | 369323 |
| 4515 | NM_00101802 | MTCP1 | mature T cell proliferation 1 | protein_coding | NP_001018025.1 | 7423 | 300116 | . | . | 1.645644 | 1.333219 | 1.545694 | 1.876108 | 3.829482 | 3.081795 | 3.468493 | 4.289499 | 1.132474 | 109.561166 | 0.316025462 | 0.567921851 | 0.570088034 | 0.998865404 | 6.683090 | 6.862520 | 6.785185 | 6.717879 | 6.761643 | 6.821985 | 131 | 99 | 92 | 119 |
| 4519 | gene-CYTB | CYTB | cytochrome b | protein_coding | YP_003024038.1 | 7427 | 516020 | . | . | 3679.064453 | 3934.259277 | 5182.543457 | 4305.634766 | 8561.333984 | 9094.215820 | 11629.477539 | 9844.322266 | 1.229748 | 151883.051514 | 0.116925677 | 2.551729409 | 0.010718973 | 0.333896806 | 17.055719 | 17.354085 | 17.089922 | 17.139812 | 17.377057 | 17.204838 | 152283 | 151301 | 159885 | 141752 |
| 4520 | NM_005955,XI | MTF1 | metal regulatory transcription factor 1 | protein_coding | NP_005946.2;XP_011539793.1;XP_011539795.1;XP_011539796.1 | 7428 | 600172 | . | . | 1.958689 | 1.897155 | 1.940960 | 1.728411 | 4.557950 | 4.385358 | 4.355458 | 3.951807 | -1.083851 | 460.171761 | 0.173308499 | -0.67028513 | 0.502676041 | 0.998865404 | 8.902722 | 8.787015 | 8.878311 | 8.862379 | 8.845391 | 8.795286 | 558 | 510 | 400 | 397 |
| 4521 | NM_00136755 | NUDT1 | nudix hydrolase 1 | protein_coding | NP_001354482.1;NP_001354483.1;NP_001354484.1;NP_002443.3;NP_9 | 8048 | 600312 | . | . | 41.735720 | 44.225677 | 38.163151 | 42.796214 | 97.120734 | 102.229631 | 85.636999 | 97.848452 | -1.078694 | 905.671561 | 0.145616177 | -0.75050128 | 0.45295285 | 0.998865404 | 9.876712 | 9.766887 | 9.827713 | 9.870336 | 9.749969 | 9.837140 | 1051 | 1044 | 711 | 867 |
| 4522 | NM_00136483 | MTHFD1 | methylenetetrahydrofolate dehydrogenase, cyclohydrolase and fo | protein_coding | NP_001351766.1;NP_005947.3 | 7432 | 172460 | . | . | 43.515490 | 46.254191 | 38.891839 | 47.972159 | 101.262336 | 106.918635 | 87.272158 | 109.682641 | -1.047879 | 4335.111609 | 0.125371657 | -0.53817983 | 0.590452904 | 0.998865404 | 12.115303 | 12.047610 | 12.077618 | 12.121523 | 11.974152 | 12.140663 | 4975 | 4913 | 3312 | 4367 |
| 4524 | NM_00133035 | MTHFR | methylenetetrahydrofolate reductase | protein_coding | NP_001317287.1;NP_005948.3;XP_005263517.1;XP_005263519.1;XP_00 | 7436 | 607093 | . | . | 7.318901 | 7.827306 | 8.713364 | 9.577804 | 17.031381 | 18.093170 | 19.552531 | 21.898511 | 1.228565 | 1575.590218 | 0.124013194 | 2.394696004 | 0.01663415 | 0.431581156 | 10.465651 | 10.762463 | 10.519271 | 10.551412 | 10.682230 | 10.708218 | 1594 | 1558 | 1486 | 1655 |
| 4528 | NM_00100536 | MTIF2 | mitochondrial translational initiation factor 2 | protein_coding | NP_001005369.1;NP_001307930.1;NP_001307931.1;NP_001307932.1;NI | 7441 | 603766 | . | . | 16.914099 | 18.128288 | 18.099239 | 19.326993 | 39.359806 | 41.904351 | 40.614166 | 44.188874 | 1.027570 | 1682.792247 | 0.121726301 | 0.32233888 | 0.747195981 | 0.998865404 | 10.696884 | 10.736132 | 10.696186 | 10.715314 | 10.735867 | 10.718200 | 1888 | 1813 | 1500 | 1581 |
| 4534 | NM_000252,N | MTM1 | myotubularin 1 | protein_coding | NP_000243.1;NP_001363835.1;NP_001363836.1;NP_001363837.1;XP_0 | 7448 | 300415 | . | . | 2.111648 | 1.995829 | 2.239770 | 1.950914 | 4.913892 | 4.613449 | 5.025977 | 4.460531 | -1.005556 | 222.495025 | 0.229985673 | -0.03475363 | 0.972276199 | 0.998865404 | 7.801122 | 7.794128 | 7.813282 | 7.782631 | 7.828882 | 7.761072 | 265 | 233 | 207 | 193 |
| 4535 | gene-ND1 | ND1 | NADH dehydrogenase subunit 1 | protein_coding | YP_003024026.1 | 7455 | 516000 | . | . | 2637.578857 | 2890.641846 | 3802.176758 | 3008.634033 | 6137.754395 | 6681.848633 | 8531.974609 | 6878.883789 | 1.215897 | 91825.830059 | 0.126080779 | 2.236831776 | 0.025297337 | 0.524911334 | 16.338718 | 16.620746 | 16.358032 | 16.428876 | 16.665306 | 16.450596 | 91473 | 93142 | 98281 | 82992 |
| 4536 | gene-ND2 | ND2 | NADH dehydrogenase subunit 2 | protein_coding | YP_003024027.1 | 7456 | 516001 | . | . | 4707.349609 | 5230.043945 | 6623.230957 | 5176.554688 | 10954.195312 | 12089.479492 | 14862.338867 | 11835.578125 | 1.171958 | 176338.871383 | 0.128714543 | 1.77852062 | 0.075318388 | 0.766002767 | 17.308993 | 17.537917 | 17.310775 | 17.393626 | 17.597189 | 17.372040 | 177939 | 183681 | 186602 | 155637 |
| 4537 | gene-ND3 | ND3 | NADH dehydrogenase subunit 3 | protein_coding | YP_003024033.1 | 7458 | 516002 | . | . | 3921.023682 | 4097.893066 | 4632.411133 | 3407.387939 | 9124.383789 | 9472.463867 | 10394.997070 | 7790.587891 | -1.010535 | 43288.228942 | 0.137128655 | -0.11025717 | 0.912205427 | 0.998865404 | 15.409216 | 15.394119 | 15.387624 | 15.418413 | 15.521856 | 15.248188 | 49216 | 47789 | 43338 | 34018 |
| 4538 | gene-ND4 | ND4 | NADH dehydrogenase subunit 4 | protein_coding | YP_003024035.1 | 7459 | 516003 | . | . | 11445.629883 | 12466.151367 | 13917.507812 | 11608.130859 | 26634.451172 | 28816.068359 | 31230.484375 | 26540.611328 | 1.053385 | 530524.505317 | 0.117128421 | 0.640602176 | 0.521781186 | 0.998865404 | 18.979055 | 19.054089 | 18.959031 | 19.025464 | 19.118036 | 18.951105 | 572157 | 578992 | 518549 | 461547 |
| 4539 | gene-ND4L | ND4L | NADH dehydrogenase subunit 4L | protein_coding | YP_003024034.1 | 7460 | 516004 | . | . | 9034.075195 | 9807.163086 | 10958.746094 | 7615.831055 | 21022.664062 | 22669.697266 | 24591.107422 | 17412.691406 | -1.027487 | 86582.083583 | 0.147763256 | -0.26474447 | 0.791206316 | 0.998865404 | 16.421203 | 16.382094 | 16.376625 | 16.440549 | 16.534357 | 16.211615 | 97335 | 98173 | 88003 | 65265 |
| 4540 | gene-ND5 | ND5 | NADH dehydrogenase subunit 5 | protein_coding | YP_003024036.1 | 7461 | 516005 | . | . | 2367.062500 | 2606.265137 | 2937.457520 | 2403.649170 | 5508.251953 | 6024.498047 | 6591.569336 | 5495.658203 | 1.059829 | 145546.701563 | 0.120909477 | 0.693344144 | 0.488093606 | 0.998865404 | 17.108596 | 17.192432 | 17.085274 | 17.160992 | 17.261912 | 17.078431 | 155595 | 159173 | 143916 | 125671 |
| 4541 | gene-ND6 | ND6 | NADH dehydrogenase subunit 6 | protein_coding | YP_003024037.1 | 7462 | 516006 | . | . | 745.466248 | 667.222717 | 855.597107 | 491.630951 | 1734.730713 | 1542.315063 | 1919.935181 | 1124.055786 | -1.062085 | 11296.084808 | 0.407990981 | -0.21299374 | 0.831331842 | 0.998865404 | 13.506321 | 13.419442 | 13.529107 | 13.426271 | 13.638408 | 13.161519 | 14198 | 11807 | 12146 | 7448 |
| 4542 | NM_00134835 | MYO1F | myosin IF | protein_coding | NP_001335284.1;NP_036467.2;XP_011526326.1;XP_011526327.1;XP_01 | 7600 | 601480 | . | . | 0.162844 | 0.180471 | 0.106088 | 0.175266 | 0.378946 | 0.417166 | 0.238059 | 0.400724 | -1.225801 | 20.768196 | 0.688017409 | -0.42691471 | 0.669441453 | NA | 4.517478 | 4.219802 | 4.383024 | 4.393105 | 4.350213 | 4.390064 | 25 | 26 | 12 | 22 |
| 4548 | NM_000254,N | MTR | 5-methyltetrahydrofolate-homocysteine methyltransferase | protein_coding | NP_000245.2;NP_001278868.1;NP_001278869.1;XP_005273198.1;XP_00 | 7468 | 156570 | . | . | 4.529304 | 4.278885 | 4.590921 | 4.700396 | 10.539875 | 9.890834 | 10.301896 | 10.746896 | 1.042154 | 1492.074179 | 0.126313323 | 0.471594593 | 0.637216187 | 0.998865404 | 10.512969 | 10.572621 | 10.553090 | 10.499594 | 10.551372 | 10.565793 | 1741 | 1522 | 1312 | 1441 |
| 4549 | rna-RNR1 | RNR1 | s-rRNA | rRNA | . | 7470 | 561000 | . | . | 1431.612305 | 1360.789917 | 1760.487427 | 1479.349121 | 3331.420410 | 3145.526855 | 3950.482422 | 3382.355957 | 1.144527 | 44808.976310 | 0.115810829 | 1.681633946 | 0.092639844 | 0.809520218 | 15.350829 | 15.545603 | 15.414252 | 15.363562 | 15.583291 | 15.422413 | 49545 | 43756 | 45411 | 40722 |
| 4550 | rna-RNR2 | RNR2 | l-rRNA | rRNA | . | 7471 | 561010 | . | . | 9177.684570 | 8736.606445 | 12066.220703 | 10054.453125 | 21356.849609 | 20195.056641 | 27076.251953 | 22988.310547 | 1.218138 | 485885.180235 | 0.116263199 | 2.448563064 | 0.014342732 | 0.399401052 | 18.740906 | 19.025585 | 18.821875 | 18.772499 | 19.050094 | 18.880477 | 519046 | 459071 | 508623 | 452282 |
| 4552 | NM_00136444 | MTRR | 5-methyltetrahydrofolate-homocysteine methyltransferase reduct | protein_coding | NP_001351369.1;NP_001351370.1;NP_001351371.1;NP_002445.2;NP_0 | 7473 | 602568 | . | . | 13.246848 | 12.913402 | 14.576945 | 14.751123 | 30.825956 | 29.849911 | 32.710244 | 33.726693 | 1.092922 | 1466.142116 | 0.124705694 | 1.027945513 | 0.303975459 | 0.998865404 | 10.452319 | 10.580455 | 10.481170 | 10.483432 | 10.543395 | 10.558077 | 1611 | 1514 | 1319 | 1449 |
| 4553 | rna-TRNA | TRNA | tRNA-Ala | tRNA | . | 7475 | 590000 | . | . | 19.113073 | 14.883671 | 26.917229 | 14.799884 | 44.476906 | 34.404270 | 60.401478 | 33.838173 | 1.219442 | 41.192496 | 0.514841596 | 0.555939258 | 0.57825238 | 0.998865404 | 5.211816 | 5.502214 | 5.355426 | 5.318442 | 5.416658 | 5.317801 | 48 | 35 | 51 | 30 |
| 4555 | rna-TRND | TRND | tRNA-Asp | tRNA | . | 7478 | 590015 | . | . | 56.898056 | 54.949776 | 69.273521 | 41.580788 | 132.404114 | 127.018867 | 155.447769 | 95.069534 | -1.021292 | 118.470894 | 0.327759895 | -0.09273735 | 0.926112225 | 0.998865404 | 6.902195 | 6.874449 | 6.893602 | 6.877010 | 6.959397 | 6.789826 | 141 | 126 | 128 | 82 |
| 4556 | rna-TRNE | TRNE | tRNA-Glu | tRNA | . | 7479 | 590025 | . | . | 129.551193 | 106.876343 | 163.413895 | 82.752037 | 301.470947 | 247.049438 | 366.696045 | 189.202698 | 1.025118 | 259.936331 | 0.285663056 | 0.12528618 | 0.900296993 | 0.998865404 | 8.002824 | 8.040953 | 8.050837 | 7.947745 | 8.169078 | 7.828826 | 325 | 249 | 305 | 165 |
| 4558 | rna-TRNF | TRNF | tRNA-Phe | tRNA | . | 7481 | 590070 | . | . | 54.751472 | 52.586582 | 88.915825 | 56.173866 | 127.408928 | 121.556236 | 199.524536 | 128.434875 | 1.332430 | 140.116899 | 0.303912449 | 1.362429485 | 0.173062384 | 0.947431715 | 6.907423 | 7.323636 | 7.070150 | 7.051722 | 7.255913 | 7.072664 | 142 | 126 | 171 | 116 |
| 4563 | rna-TRNG | TRNG | tRNA-Gly | tRNA | . | 7486 | 590035 | . | . | 3.853142 | 2.488292 | 3.274122 | 2.583661 | 8.966419 | 5.751799 | 7.347034 | 5.907234 | 1.011829 | 7.173678 | 1.134087183 | 0.014959477 | 0.98806451 | NA | 2.832372 | 2.852980 | 2.849158 | 2.834634 | 2.845910 | 2.838658 | 10 | 6 | 7 | 6 |
| 4564 | rna-TRNH | TRNH | tRNA-His | tRNA | . | 7487 | 590040 | . | . | 5.885814 | 6.641455 | 9.319135 | 9.070895 | 13.696531 | 15.352021 | 20.911869 | 20.739525 | 1.471343 | 17.205190 | 0.738365797 | 0.754549221 | 0.450519501 | NA | 3.799841 | 4.356363 | 4.076768 | 4.085080 | 4.109989 | 4.108478 | 15 | 16 | 18 | 19 |
| 4565 | rna-TRNI | TRNI | tRNA-Ile | tRNA | . | 7488 | 590045 | . | . | 81.948059 | 85.246346 | 123.027596 | 62.952465 | 190.696503 | 197.050735 | 276.070374 | 143.933319 | 1.096273 | 190.553850 | 0.301778145 | 0.439417153 | 0.6603593 | 0.998865404 | 7.505102 | 7.639862 | 7.532140 | 7.545575 | 7.716165 | 7.421482 | 206 | 199 | 230 | 126 |
| 4566 | rna-TRNK | TRNK | tRNA-Lys | tRNA | . | 7489 | 590060 | . | . | 47.287231 | 45.591034 | 56.413364 | 37.333897 | 110.039322 | 105.385719 | 126.589966 | 85.359520 | -1.006836 | 102.320967 | 0.335804018 | -0.02926997 | 0.97664928 | 0.998865404 | 6.680675 | 6.673231 | 6.681275 | 6.665882 | 6.732028 | 6.607138 | 121 | 108 | 107 | 76 |
| 4567 | rna-TRNL1 | TRNL1 | tRNA-Leu | tRNA | . | 7490 | 590050 | . | . | 755.749268 | 903.062866 | 1237.216064 | 895.318054 | 1758.659668 | 2087.470215 | 2776.276855 | 2047.038452 | 1.269013 | 2213.657485 | 0.159452833 | 2.155538999 | 0.03111968 | 0.560017015 | 10.930051 | 11.273937 | 10.935400 | 11.067825 | 11.322286 | 11.049516 | 2057 | 2283 | 2509 | 1938 |
| 4568 | rna-TRNL2 | TRNL2 | tRNA-Leu | tRNA | . | 7491 | 590055 | . | . | 1.045594 | 2.416252 | 1.980492 | 1.217914 | 2.433141 | 5.585275 | 4.444166 | 2.784616 | -1.046143 | 3.991423 | 1.512902382 | -0.04301703 | 0.965687959 | NA | 2.029593 | 1.963455 | 1.986518 | 2.000503 | 1.995847 | 1.989568 | 3 | 6 | 4 | 3 |
| 4569 | rna-TRNM | TRNM | tRNA-Met | tRNA | . | 7492 | 590065 | . | . | 77.319717 | 93.120865 | 130.555603 | 93.758629 | 179.926163 | 215.253052 | 292.962982 | 214.367966 | 1.299219 | 209.401591 | 0.25607692 | 1.474732606 | 0.140284446 | 0.900674731 | 7.508570 | 7.886948 | 7.596530 | 7.674559 | 7.836976 | 7.670293 | 191 | 214 | 241 | 184 |
| 4570 | rna-TRNN | TRNN | tRNA-Asn | tRNA | . | 7493 | 590010 | . | . | 20.249228 | 13.810588 | 20.907640 | 16.771675 | 47.120789 | 31.923792 | 46.916142 | 38.346439 | 1.103414 | 41.276032 | 0.498631835 | 0.284728611 | 0.775852069 | 0.998865404 | 5.292877 | 5.437942 | 5.385059 | 5.324131 | 5.388818 | 5.352079 | 54 | 34 | 42 | 36 |
| 4571 | rna-TRNP | TRNP | tRNA-Pro | tRNA | . | 7494 | 590075 | . | . | 46.398251 | 25.055717 | 42.348179 | 20.669285 | 107.970634 | 57.917412 | 95.028091 | 47.257870 | -1.155198 | 71.763342 | 0.449848734 | -0.46268978 | 0.643586747 | 0.998865404 | 6.263241 | 6.059954 | 6.244396 | 6.086324 | 6.208244 | 6.051639 | 115 | 58 | 78 | 41 |
| 4572 | rna-TRNQ | TRNQ | tRNA-Gln | tRNA | . | 7495 | 590030 | . | . | 75.824013 | 83.543709 | 125.468956 | 73.801117 | 176.445602 | 193.115036 | 281.548706 | 168.737488 | 1.234260 | 201.739896 | 0.279105015 | 1.087928429 | 0.276626704 | 0.998865404 | 7.495652 | 7.800934 | 7.574444 | 7.611657 | 7.806627 | 7.555510 | 199 | 203 | 245 | 154 |
| 4573 | rna-TRNR | TRNR | tRNA-Arg | tRNA | . | 7496 | 590005 | . | . | 4.089764 | 4.939631 | 6.675828 | 4.708970 | 9.517050 | 11.418179 | 14.980366 | 10.766500 | 1.238332 | 10.569577 | 0.939015971 | 0.328427036 | 0.742588804 | NA | 3.238681 | 3.548418 | 3.386169 | 3.393198 | 3.408664 | 3.390639 | 10 | 11 | 12 | 9 |
| 4574 | rna-TRNS1 | TRNS1 | tRNA-Ser | tRNA | . | 7497 | 590080 | . | . | 2688.740967 | 2673.339111 | 3060.582275 | 2483.595459 | 6256.811035 | 6179.542480 | 6867.857910 | 5678.445801 | 1.019967 | 5862.251078 | 0.125098455 | 0.22800217 | 0.819644553 | 0.998865404 | 12.502830 | 12.531506 | 12.513315 | 12.501291 | 12.612099 | 12.427724 | 6731 | 6218 | 5710 | 4945 |
| 4575 | rna-TRNS2 | TRNS2 | tRNA-Ser | tRNA | . | 7498 | 590085 | . | . | 2.553523 | 3.146682 | 4.816259 | 3.768751 | 5.942153 | 7.273695 | 10.807547 | 8.616801 | 1.424760 | 7.091082 | 1.136745144 | 0.449281975 | 0.653228259 | NA | 2.548173 | 3.058876 | 2.807869 | 2.813958 | 2.825926 | 2.818429 | 6 | 7 | 8 | 7 |
| 4576 | rna-TRNT | TRNT | tRNA-Thr | tRNA | . | 7499 | 590090 | . | . | 65.304916 | 34.859123 | 52.774982 | 24.747849 | 151.967224 | 80.578430 | 118.425529 | 56.583019 | -1.304436 | 92.269765 | 0.422029568 | -0.90853031 | 0.363598105 | 0.998865404 | 6.704813 | 6.325958 | 6.642359 | 6.442808 | 6.559765 | 6.371804 | 157 | 78 | 95 | 48 |
| 4577 | rna-TRNV | TRNV | tRNA-Val | tRNA | . | 7500 | 590105 | . | . | 164.415146 | 155.576096 | 223.314285 | 141.508957 | 382.600830 | 359.621094 | 501.110748 | 323.543427 | 1.124350 | 368.404779 | 0.227348411 | 0.74375272 | 0.457026089 | 0.998865404 | 8.437294 | 8.607958 | 8.499715 | 8.463145 | 8.676291 | 8.404968 | 412 | 362 | 417 | 282 |
| 4578 | rna-TRNW | TRNW | tRNA-Trp | tRNA | . | 7501 | 590095 | . | . | 15.476787 | 12.476019 | 21.626436 | 11.464995 | 36.015118 | 28.838875 | 48.529095 | 26.213348 | 1.157615 | 32.819288 | 0.568122192 | 0.371672654 | 0.710136588 | 0.998865404 | 4.924690 | 5.140213 | 5.031747 | 5.002936 | 5.077055 | 4.994228 | 39 | 29 | 40 | 23 |
| 4579 | rna-TRNY | TRNY | tRNA-Tyr | tRNA | . | 7502 | 590100 | . | . | 588.782837 | 554.647095 | 699.782715 | 527.277649 | 1370.121826 | 1282.091553 | 1570.292114 | 1205.557739 | 1.058707 | 1219.106156 | 0.15753823 | 0.522435974 | 0.601366814 | 0.998865404 | 10.209518 | 10.292505 | 10.251791 | 10.200460 | 10.374188 | 10.153293 | 1410 | 1234 | 1249 | 1005 |
| 4580 | NM_002455,N | MTX1 | metaxin 1 | protein_coding | NP_002446.3;NP_942584.2 | 7504 | 600605 | . | . | 13.498614 | 13.412924 | 11.666376 | 14.932720 | 31.411827 | 31.004573 | 26.179012 | 34.141891 | -1.025532 | 685.908179 | 0.166396421 | -0.21858685 | 0.826971894 | 0.998865404 | 9.440311 | 9.403194 | 9.434303 | 9.423855 | 9.328152 | 9.488900 | 806 | 744 | 519 | 710 |
| 4582 | NM_00101801 | MUC1 | mucin 1, cell surface associated | protein_coding | NP_001018016.1;NP_001018017.1;NP_001037855.1;NP_001037856.1;NI | 7508 | 158340 | . | . | 0.338120 | 0.240395 | 0.368862 | 0.366212 | 0.786821 | 0.555685 | 0.827716 | 0.837301 | 1.386147 | 39.492763 | 0.495090197 | 0.951503322 | 0.341348933 | 0.998865404 | 5.048513 | 5.520145 | 5.280509 | 5.260375 | 5.316035 | 5.324190 | 41 | 33 | 39 | 44 |
| 4585 | NM_004532,N | MUC4 | mucin 4, cell surface associated | protein_coding | NP_004523.3;NP_060876.5;NP_612154.2 | 7514 | 158372 | . | . | 0.034890 | 0.019285 | 0.023078 | 0.038562 | 0.081191 | 0.044577 | 0.051786 | 0.088167 | 1.123318 | 15.554077 | 0.801968643 | 0.209192532 | 0.834297942 | NA | 3.872825 | 4.040734 | 3.972572 | 3.932839 | 3.944582 | 3.976665 | 22 | 11 | 11 | 19 |
| 4591 | NM_00100520 | TRIM37 | tripartite motif containing 37 | protein_coding | NP_001005207.1;NP_001307916.1;NP_001307917.1;NP_001307918.1;NI | 7523 | 605073 | . | . | 9.300838 | 9.037788 | 10.095632 | 9.588724 | 21.643439 | 20.891252 | 22.654309 | 21.923479 | 1.076943 | 1310.186189 | 0.129601769 | 0.825159442 | 0.409281076 | 0.998865404 | 10.300968 | 10.408154 | 10.343408 | 10.308752 | 10.403402 | 10.361543 | 1486 | 1330 | 1214 | 1239 |
| 4594 | NM_000255,XI | MMUT | methylmalonyl-CoA mutase | protein_coding | NP_000246.2;XP_005249200.1 | 7526 | 609058 | . | . | 6.149379 | 5.319246 | 6.093377 | 6.394356 | 14.309856 | 12.295677 | 13.673362 | 14.619936 | 1.071341 | 688.572050 | 0.16085696 | 0.618048616 | 0.536543293 | 0.998865404 | 9.376798 | 9.476410 | 9.458656 | 9.342108 | 9.423977 | 9.473964 | 831 | 656 | 597 | 692 |
| 4595 | NM_00104817 | MUTYH | mutY DNA glycosylase | protein_coding | NP_001041636.2;NP_001041637.1;NP_001041638.1;NP_001041639.1;NI | 7527 | 604933 | . | . | 12.602829 | 12.669174 | 12.748655 | 12.868928 | 29.327303 | 29.285358 | 28.607613 | 29.423277 | 1.008763 | 749.438546 | 0.147731058 | 0.085201033 | 0.932101569 | 0.998865404 | 9.543278 | 9.556027 | 9.543040 | 9.549816 | 9.569795 | 9.535016 | 855 | 809 | 671 | 688 |
| 4597 | NM_002461,XI | MVD | mevalonate diphosphate decarboxylase | protein_coding | NP_002452.1;XP_011521388.1;XP_011521389.1;XP_011521390.1;XP_01 | 7529 | 603236 | . | . | 11.798441 | 11.685152 | 17.571160 | 18.417266 | 27.455458 | 27.010753 | 39.429177 | 42.108893 | 1.564121 | 873.045940 | 0.142025528 | 4.543919025 | 5.52179E-06 | 0.001190894 | 9.411343 | 10.056870 | 9.585665 | 9.581199 | 9.922058 | 9.883724 | 787 | 732 | 950 | 973 |
| 4598 | NM_000431,N | MVK | mevalonate kinase | protein_coding | NP_000422.1;NP_001107657.1;NP_001288111.1;XP_016874802.1;XP_01 | 7530 | 251170 | . | . | 7.906021 | 7.839271 | 12.153093 | 12.418463 | 18.397637 | 18.120825 | 27.271185 | 28.393337 | 1.642679 | 842.872805 | 0.143148882 | 5.002138382 | 5.66979E-07 | 0.000190878 | 9.317240 | 10.033178 | 9.511871 | 9.509907 | 9.846543 | 9.879086 | 736 | 687 | 892 | 1003 |
| 4601 | NM_00100854 | MXI1 | MAX interactor 1, dimerization protein | protein_coding | NP_001008541.1;NP_005953.4;NP_569157.2 | 7534 | 600020 | . | . | 32.085911 | 32.781519 | 53.972447 | 34.685797 | 74.665238 | 75.775945 | 121.112608 | 79.304952 | 1.345415 | 3951.788866 | 0.318308222 | 1.344770674 | 0.178699303 | 0.951511445 | 11.718414 | 12.146523 | 11.798010 | 11.813163 | 12.225239 | 11.843883 | 3844 | 3670 | 4814 | 3311 |
| 4602 | NM_00113017 | MYB | MYB proto-oncogene, transcription factor | protein_coding | NP_001123644.1;NP_001123645.1;NP_001155128.1;NP_001155129.1;NI | 7545 | 189990 | . | . | 0.543639 | 0.618826 | 0.777743 | 0.956392 | 1.265071 | 1.430446 | 1.745231 | 2.186678 | 1.476258 | 82.794236 | 0.354448035 | 1.585407963 | 0.112873673 | 0.850211417 | 6.064246 | 6.624599 | 6.293161 | 6.318513 | 6.378258 | 6.440716 | 73 | 76 | 77 | 102 |
| 4603 | NM_00108041 | MYBL1 | MYB proto-oncogene like 1 | protein_coding | NP_001073885.1;NP_001138227.1;NP_001281211.1;XP_011515835.1;XP | 7547 | 159405 | . | . | 4.905146 | 4.471219 | 4.210302 | 3.246404 | 11.414475 | 10.335422 | 9.447797 | 7.422516 | -1.258429 | 633.768519 | 0.174255716 | -1.9030881 | 0.057029051 | 0.708295137 | 9.463614 | 9.133124 | 9.419736 | 9.333108 | 9.298925 | 9.141502 | 863 | 715 | 548 | 462 |
| 4605 | NM_00127861 | MYBL2 | MYB proto-oncogene like 2 | protein_coding | NP_001265539.1;NP_002457.1 | 7548 | 601415 | . | . | 21.737993 | 21.791464 | 16.701603 | 21.454610 | 50.585202 | 50.371940 | 37.477913 | 49.053416 | -1.157432 | 1699.305906 | 0.141569169 | -1.48992427 | 0.136244151 | 0.894235858 | 10.832555 | 10.621168 | 10.786542 | 10.780996 | 10.573252 | 10.756839 | 2107 | 1961 | 1206 | 1653 |
| 4609 | NM_00135487 | MYC | MYC proto-oncogene, bHLH transcription factor | protein_coding | NP_001341799.1;NP_002458.2 | 7553 | 190080 | . | . | 18.337017 | 20.003835 | 27.717607 | 26.718444 | 42.670994 | 46.239762 | 62.197505 | 61.088544 | 1.391227 | 2840.213980 | 0.116041217 | 4.105074857 | 4.04184E-05 | 0.006268485 | 11.214050 | 11.690378 | 11.289645 | 11.351664 | 11.612850 | 11.566712 | 2628 | 2664 | 2953 | 3013 |
| 4610 | NM_00103308 | MYCL | MYCL proto-oncogene, bHLH transcription factor | protein_coding | NP_001028253.1;NP_001028254.2;NP_005367.2 | 7555 | 164850 | . | . | 0.122237 | 0.040786 | 0.038466 | 0.173737 | 0.284452 | 0.094278 | 0.086317 | 0.397229 | 1.022655 | 8.043350 | 1.143111618 | 0.028272811 | 0.977444566 | NA | 2.993083 | 3.022362 | 3.022553 | 2.991843 | 2.987535 | 3.028352 | 13 | 5 | 3 | 12 |
| 4615 | NM_00117256 | MYD88 | MYD88 innate immune signal transduction adaptor | protein_coding | NP_001166037.2;NP_001166038.2;NP_001166039.2;NP_001166040.2;NI | 7562 | 602170 | . | . | 7.819050 | 9.368850 | 8.860085 | 10.049412 | 18.195250 | 21.656520 | 19.881773 | 22.976787 | 1.086737 | 749.863812 | 0.159708519 | 0.751383229 | 0.452422053 | 0.998865404 | 9.489635 | 9.608871 | 9.457178 | 9.577400 | 9.536697 | 9.615449 | 756 | 843 | 641 | 773 |
| 4616 | NM_015675,XI | GADD45B | growth arrest and DNA damage inducible beta | protein_coding | NP_056490.2;XP_016882311.1 | 4096 | 604948 | . | . | 3.422855 | 3.334049 | 2.020383 | 1.774530 | 7.965123 | 7.706803 | 4.533682 | 4.057253 | -1.780679 | 116.519007 | 0.306032052 | -2.72006504 | 0.006526908 | 0.255215036 | 7.220836 | 6.390008 | 6.956607 | 6.936018 | 6.782728 | 6.730439 | 176 | 157 | 80 | 71 |
| 4621 | NM_002470,XI | MYH3 | myosin heavy chain 3 | protein_coding | NP_002461.2;XP_011522172.1;XP_011522173.1 | 7573 | 160720 | . | . | 0.338004 | 0.405428 | 0.511687 | 0.438295 | 0.786550 | 0.937166 | 1.148210 | 1.002110 | 1.265530 | 80.981891 | 0.357696427 | 0.949805066 | 0.342211311 | 0.998865404 | 6.159667 | 6.499430 | 6.278240 | 6.322424 | 6.384818 | 6.339990 | 75 | 84 | 85 | 78 |
| 4627 | NM_002473 | MYH9 | myosin heavy chain 9 | protein_coding | NP_002464.1 | 7579 | 160775 | . | . | 62.963001 | 65.408699 | 42.632896 | 51.588482 | 146.517487 | 151.195145 | 95.666985 | 117.950943 | -1.382029 | 12928.063037 | 0.116132954 | -4.01942792 | 5.83396E-05 | 0.008388145 | 13.872850 | 13.405979 | 13.767117 | 13.793082 | 13.425678 | 13.575536 | 17019 | 16427 | 8589 | 11092 |
| 4628 | NM_00125601 | MYH10 | myosin heavy chain 10 | protein_coding | NP_001242941.1;NP_001243024.1;NP_001362195.1;NP_005955.3;XP_01 | 7568 | 160776 | . | . | 56.326397 | 57.900101 | 57.518685 | 59.473923 | 131.073846 | 133.838679 | 129.070264 | 135.980063 | 1.011328 | 13904.959622 | 0.099611269 | 0.163139568 | 0.870408546 | 0.998865404 | 13.755171 | 13.771407 | 13.749896 | 13.766924 | 13.759765 | 13.776345 | 15763 | 15068 | 12042 | 13236 |
| 4636 | NM_00136365 | MYL5 | myosin light chain 5 | protein_coding | NP_001350579.1;NP_002468.1;XP_006713949.1;XP_016863734.1;XP_01 | 7586 | 160782 | . | . | 3.534911 | 3.766736 | 3.790516 | 3.424385 | 8.225881 | 8.706979 | 8.505808 | 7.829448 | -1.027633 | 172.027936 | 0.267537299 | -0.14698759 | 0.883141819 | 0.998865404 | 7.445495 | 7.407249 | 7.401357 | 7.455617 | 7.477838 | 7.353309 | 188 | 200 | 170 | 135 |
| 4637 | NM_021019,N | MYL6 | myosin light chain 6 | protein_coding | NP_066299.2;NP_524147.2 | 7587 | 609931 | . | . | 352.459533 | 361.486328 | 255.227372 | 319.406509 | 820.187775 | 835.591797 | 572.722824 | 730.285049 | -1.257437 | 6949.238869 | 0.123598475 | -2.67386499 | 0.007498263 | 0.2720434 | 12.918505 | 12.587871 | 12.842892 | 12.858138 | 12.563957 | 12.741898 | 8836 | 8428 | 4779 | 6391 |
| 4638 | NM_00132130 | MYLK | myosin light chain kinase | protein_coding | NP_001308238.1;NP_444253.3;NP_444254.3;NP_444255.3;NP_444256.3 | 7590 | 600922 | . | . | 0.490840 | 0.492551 | 0.250433 | 0.404563 | 1.142205 | 1.138555 | 0.561964 | 0.924984 | -1.414055 | 122.050371 | 0.314860324 | -1.58749135 | 0.112401405 | 0.850211417 | 7.160812 | 6.658318 | 6.979167 | 6.991690 | 6.800990 | 6.928246 | 162 | 157 | 69 | 115 |
| 4641 | NM_00108077 | MYO1C | myosin IC | protein_coding | NP_001074248.1;NP_001074419.1;NP_001350784.1;NP_203693.3;XP_02 | 7597 | 606538 | . | . | 30.103877 | 34.658124 | 30.186790 | 36.607165 | 70.052956 | 80.113809 | 67.738283 | 83.697930 | 1.015383 | 4863.646185 | 0.125887051 | 0.174950237 | 0.861118735 | 0.998865404 | 12.236876 | 12.258687 | 12.187244 | 12.290213 | 12.175649 | 12.324362 | 5216 | 5529 | 3883 | 5002 |
| 4642 | NM_00130327 | MYO1D | myosin ID | protein_coding | NP_001290208.1;NP_001290209.1;NP_056009.1;XP_016880174.1;XP_02 | 7598 | 606539 | . | . | 7.252398 | 8.975108 | 7.095293 | 9.122217 | 16.876629 | 20.746367 | 15.921630 | 20.856862 | -1.007267 | 1326.487139 | 0.152799491 | -0.06836684 | 0.945493616 | 0.998865404 | 10.378949 | 10.367820 | 10.323546 | 10.421858 | 10.269057 | 10.458571 | 1435 | 1529 | 1006 | 1393 |
| 4643 | NM_004998 | MYO1E | myosin IE | protein_coding | NP_004989.2 | 7599 | 601479 | . | . | 10.843672 | 10.387015 | 9.980665 | 11.277691 | 25.233669 | 24.010050 | 22.396324 | 25.785101 | -1.013184 | 2860.563107 | 0.118542487 | -0.15940691 | 0.873348292 | 0.998865404 | 11.491559 | 11.472545 | 11.507862 | 11.465722 | 11.429842 | 11.520728 | 3401 | 3027 | 2333 | 2813 |
| 4644 | NM_000259,N | MYO5A | myosin VA | protein_coding | NP_000250.3;NP_001135967.2;NP_001369276.1;NP_001369277.1;NP_0 | 7602 | 160777 | . | . | 12.881424 | 12.738881 | 16.867271 | 15.466353 | 29.975605 | 29.446496 | 37.849664 | 35.361979 | 1.287152 | 5331.204534 | 0.109517052 | 3.325348401 | 0.000883081 | 0.063156313 | 12.186647 | 12.550913 | 12.264221 | 12.264868 | 12.517023 | 12.433941 | 5366 | 5032 | 5471 | 5353 |
| 4645 | NM_00108046 | MYO5B | myosin VB | protein_coding | NP_001073936.1 | 7603 | 606540 | . | . | 0.166840 | 0.225966 | 0.142951 | 0.218174 | 0.388244 | 0.522330 | 0.320777 | 0.498830 | -1.090498 | 56.914042 | 0.435067085 | -0.28728105 | 0.773897131 | 0.998865404 | 5.893489 | 5.765080 | 5.807452 | 5.867012 | 5.781567 | 5.857081 | 59 | 73 | 38 | 61 |
| 4646 | NM_00130089 | MYO6 | myosin VI | protein_coding | NP_001287828.1;NP_001355065.1;NP_001355066.1;NP_001355067.1;NI | 7605 | 600970 | . | . | 2.387529 | 2.473377 | 1.714513 | 2.158252 | 5.555879 | 5.717319 | 3.847319 | 4.934587 | -1.242276 | 573.010822 | 0.171339139 | -1.82670015 | 0.067744866 | 0.740331407 | 9.310806 | 8.997001 | 9.234588 | 9.221753 | 9.017250 | 9.150055 | 738 | 679 | 397 | 530 |
| 4649 | NM_006901,XI | MYO9A | myosin IXA | protein_coding | NP_008832.2;XP_006720602.1;XP_011519915.1;XP_011519916.1;XP_01 | 7608 | 604875 | . | . | 1.571900 | 1.635770 | 1.557235 | 1.497049 | 3.657879 | 3.781153 | 3.494391 | 3.422824 | -1.064163 | 620.797533 | 0.156426787 | -0.57354936 | 0.566272789 | 0.998865404 | 9.322082 | 9.232485 | 9.288233 | 9.308518 | 9.273033 | 9.240463 | 726 | 701 | 536 | 550 |
| 4650 | NM_00113006 | MYO9B | myosin IXB | protein_coding | NP_001123537.1;NP_004136.2 | 7609 | 602129 | . | . | 19.699150 | 21.142030 | 22.490485 | 23.453144 | 45.840733 | 48.870747 | 50.467999 | 53.622830 | 1.109480 | 5143.143812 | 0.107405208 | 1.395491421 | 0.1628677 | 0.935523455 | 12.251590 | 12.401393 | 12.257176 | 12.308420 | 12.356675 | 12.383442 | 5442 | 5424 | 4630 | 5152 |
| 4651 | NM_012334,XI | MYO10 | myosin X | protein_coding | NP_036466.2;XP_005248363.1;XP_005248364.1;XP_006714538.1;XP_01 | 7593 | 601481 | . | . | 18.129168 | 21.414711 | 18.440096 | 17.686868 | 42.187319 | 49.501064 | 41.379042 | 40.438922 | -1.011853 | 5727.041972 | 0.104843461 | -0.16214405 | 0.871192419 | 0.998865404 | 12.492073 | 12.475025 | 12.474590 | 12.502173 | 12.467857 | 12.489121 | 6525 | 6318 | 4888 | 5405 |
| 4659 | NM_00114388 | PPP1R12A | protein phosphatase 1 regulatory subunit 12A | protein_coding | NP_001137357.1;NP_001137358.1;NP_001231919.1;NP_001231921.1;NI | 7618 | 602021 | . | . | 4.120995 | 4.263375 | 3.827893 | 3.786340 | 9.589726 | 9.854981 | 8.589680 | 8.657017 | -1.118103 | 693.037031 | 0.151083132 | -1.06599344 | 0.28642661 | 0.998865404 | 9.515072 | 9.354013 | 9.460092 | 9.487460 | 9.405659 | 9.389168 | 826 | 805 | 576 | 606 |
| 4660 | NM_00116785 | PPP1R12B | protein phosphatase 1 regulatory subunit 12B | protein_coding | NP_001161329.1;NP_001161330.1;NP_001184060.1;NP_001317958.1;NI | 7619 | 603768 | . | . | 1.429286 | 1.547772 | 1.846195 | 1.595013 | 3.326008 | 3.577746 | 4.142810 | 3.646805 | 1.040924 | 560.288657 | 0.185718388 | 0.311569533 | 0.755367691 | 0.998865404 | 9.100381 | 9.159075 | 9.090853 | 9.134425 | 9.236834 | 9.031300 | 611 | 612 | 575 | 452 |
| 4661 | NM_004535 | MYT1 | myelin transcription factor 1 | protein_coding | NP_004526.1 | 7622 | 600379 | . | . | 0.180546 | 0.137971 | 0.079932 | 0.133688 | 0.420138 | 0.318926 | 0.179365 | 0.305662 | -1.461534 | 23.661015 | 0.650492587 | -0.84164373 | 0.399987403 | NA | 4.813210 | 4.263648 | 4.604085 | 4.570302 | 4.533004 | 4.567685 | 37 | 26 | 13 | 22 |
| 4664 | NM_00132131 | NAB1 | NGFI-A binding protein 1 | protein_coding | NP_001308241.1;NP_001308242.1;NP_001308243.1;NP_001308244.1;NI | 7626 | 600800 | . | . | 11.259599 | 10.051524 | 12.345619 | 10.357919 | 26.201544 | 23.234541 | 27.703215 | 23.682152 | 1.063348 | 1433.643360 | 0.13980088 | 0.633857037 | 0.526174126 | 0.998865404 | 10.440208 | 10.529356 | 10.496984 | 10.421194 | 10.572325 | 10.437693 | 1679 | 1425 | 1397 | 1265 |
| 4665 | NM_00133030 | NAB2 | NGFI-A binding protein 2 | protein_coding | NP_001317234.1;NP_005958.1 | 7627 | 602381 | . | . | 15.940426 | 16.831170 | 15.678719 | 16.059505 | 37.094027 | 38.906004 | 35.182593 | 36.718146 | -1.035278 | 1215.339664 | 0.130770413 | -0.38249021 | 0.702097787 | 0.998865404 | 10.272018 | 10.221833 | 10.242991 | 10.276887 | 10.224921 | 10.242810 | 1392 | 1364 | 1026 | 1133 |
| 4666 | NM_00111320 | NACA | nascent polypeptide associated complex subunit alpha | protein_coding | NP_001106672.1;NP_001106673.1;NP_001106674.2;NP_001307122.1;NI | 7629 | 601234 | . | . | 255.064054 | 243.847619 | 231.152992 | 287.781791 | 593.544567 | 563.664691 | 518.700633 | 657.978876 | 1.038957 | 6837.174646 | 0.118319867 | 0.465992886 | 0.641220593 | 0.998865404 | 12.711404 | 12.766440 | 12.734445 | 12.709068 | 12.675952 | 12.826766 | 7839 | 7128 | 5521 | 7112 |
| 4668 | NM_000262,N | NAGA | alpha-N-acetylgalactosaminidase | protein_coding | NP_000253.1;NP_001349777.1;NP_001349779.1 | 7631 | 104170 | . | . | 22.108264 | 22.915757 | 18.701364 | 22.030142 | 51.446842 | 52.970800 | 41.965320 | 50.369301 | -1.122842 | 2444.226105 | 0.126127541 | -1.32528663 | 0.1850761 | 0.958025663 | 11.336465 | 11.169002 | 11.290702 | 11.308061 | 11.136752 | 11.270491 | 2946 | 2820 | 1834 | 2340 |
| 4669 | NM_000263,XI | NAGLU | N-acetyl-alpha-glucosaminidase | protein_coding | NP_000254.2;XP_006721983.1;XP_016880176.1;XP_024306539.1;XP_02 | 7632 | 609701 | . | . | 24.486535 | 22.508384 | 21.125763 | 22.610443 | 56.981174 | 52.029137 | 47.405600 | 51.696093 | -1.098389 | 1735.351599 | 0.128364202 | -1.05472115 | 0.291552874 | 0.998865404 | 10.827147 | 10.691699 | 10.835759 | 10.755136 | 10.686501 | 10.757623 | 2198 | 1861 | 1372 | 1622 |
| 4670 | NM_00129741 | HNRNPM | heterogeneous nuclear ribonucleoprotein M | protein_coding | NP_001284347.1;NP_005959.2;NP_112480.2;XP_005272535.1;XP_00527 | 5046 | 160994 | . | . | 69.148082 | 69.600084 | 62.633189 | 71.266489 | 160.910432 | 160.883713 | 140.547055 | 162.942365 | -1.057980 | 5282.012673 | 0.10992433 | -0.73971691 | 0.459471787 | 0.998865404 | 12.406981 | 12.325616 | 12.411343 | 12.367788 | 12.305932 | 12.377861 | 6415 | 5709 | 4271 | 5020 |
| 4671 | NM_00134687 | NAIP | NLR family apoptosis inhibitory protein | protein_coding | NP_001333799.1;NP_004527.2;NP_075043.1 | 7634 | 600355 | . | . | 0.483757 | 0.492031 | 0.804591 | 0.420229 | 1.125723 | 1.137352 | 1.805479 | 0.960804 | 1.241703 | 122.239296 | 0.34139959 | 0.914822777 | 0.360284675 | 0.998865404 | 6.767596 | 7.082394 | 6.877188 | 6.879950 | 7.062639 | 6.831182 | 125 | 118 | 155 | 87 |
| 4673 | NM_00130792 | NAP1L1 | nucleosome assembly protein 1 like 1 | protein_coding | NP_001294853.1;NP_001317160.1;NP_001317161.1;NP_004528.1;NP_6 | 7637 | 164060 | . | . | 26.657357 | 26.442685 | 22.734262 | 21.401578 | 62.032764 | 61.123452 | 51.015021 | 48.932170 | -1.138246 | 5237.868789 | 0.106481396 | -1.75441816 | 0.079358882 | 0.776573458 | 12.445129 | 12.258359 | 12.430523 | 12.381945 | 12.299183 | 12.296949 | 6605 | 5845 | 4266 | 4587 |
| 4675 | NM_004538 | NAP1L3 | nucleosome assembly protein 1 like 3 | protein_coding | NP_004529.2 | 7639 | 300117 | . | . | 0.020812 | 0.014084 | 0.040641 | 0.122671 | 0.048431 | 0.032555 | 0.091197 | 0.280473 | 3.945667 | 4.426742 | 1.536101912 | 1.289152248 | 0.197345162 | NA | 0.844134 | 2.819320 | 2.105415 | 2.106177 | 2.114027 | 2.146389 | 2 | 2 | 3 | 10 |
| 4676 | NM_00136937 | NAP1L4 | nucleosome assembly protein 1 like 4 | protein_coding | NP_001356304.1;NP_001356305.1;NP_001356306.1;NP_001356307.1;NI | 7640 | 601651 | . | . | 77.471919 | 80.901187 | 71.055831 | 77.854050 | 180.280341 | 187.006719 | 159.447231 | 178.004023 | -1.073707 | 6024.560821 | 0.107552584 | -0.953951 | 0.340108444 | 0.998865404 | 12.607074 | 12.504380 | 12.570468 | 12.600692 | 12.492553 | 12.557910 | 7056 | 6852 | 4855 | 5660 |
| 4677 | NM_004539,XI | NARS1 | asparaginyl-tRNA synthetase 1 | protein_coding | NP_004530.1;XP_005266757.1 | 7643 | 108410 | . | . | 73.054189 | 68.565758 | 63.262150 | 68.010622 | 170.000106 | 158.492820 | 141.958431 | 155.498195 | -1.095045 | 5824.373628 | 0.108654458 | -1.20556677 | 0.22798458 | 0.982008277 | 12.571902 | 12.440899 | 12.575170 | 12.513177 | 12.442849 | 12.493124 | 7267 | 6330 | 4689 | 5370 |
| 4678 | NM_00119519 | NASP | nuclear autoantigenic sperm protein | protein_coding | NP_001182122.1;NP_002473.2;NP_689511.2;XP_005270945.1;XP_00527 | 7644 | 603185 | . | . | 42.699071 | 42.527763 | 40.869781 | 38.469504 | 99.362493 | 98.304830 | 91.710607 | 87.955949 | -1.053502 | 3285.081969 | 0.110836754 | -0.67840781 | 0.497513158 | 0.998865404 | 11.718857 | 11.643589 | 11.704164 | 11.701326 | 11.639420 | 11.679963 | 3888 | 3631 | 2713 | 3074 |
| 4681 | NM_00120408 | NBL1 | NBL1, DAN family BMP antagonist | protein_coding | NP_001191013.1;NP_001191014.1;NP_001191015.1;NP_001265093.1;NI | 7650 | 600613 | . | . | 238.637493 | 240.002265 | 235.524175 | 271.073594 | 555.319280 | 554.775994 | 528.509415 | 619.777589 | 1.043667 | 15537.277031 | 0.108285366 | 0.569436409 | 0.569060014 | 0.998865404 | 13.892308 | 13.953926 | 13.905540 | 13.903098 | 13.883213 | 13.995678 | 17534 | 16384 | 12899 | 15848 |
| 4682 | NM_00127850 | NUBP1 | nucleotide binding protein 1 | protein_coding | NP_001265435.1;NP_001310523.1;NP_001310524.1;NP_001310525.1;NI | 8041 | 600280 | . | . | 18.371149 | 20.694179 | 18.592067 | 23.055330 | 42.750420 | 47.835522 | 41.720063 | 52.713271 | 1.041852 | 770.813206 | 0.158462443 | 0.373277734 | 0.708941746 | 0.998865404 | 9.560764 | 9.619121 | 9.541088 | 9.606575 | 9.536309 | 9.665400 | 829 | 853 | 621 | 805 |
| 4683 | NM_00102468 | NBN | nibrin | protein_coding | NP_001019859.1;NP_002476.2;XP_011515347.1;XP_011515348.1;XP_01 | 7652 | 602667 | . | . | 8.986163 | 8.930238 | 9.407842 | 7.791685 | 20.911173 | 20.642644 | 21.110927 | 17.814765 | -1.061070 | 1270.061907 | 0.142920666 | -0.59836871 | 0.54959394 | 0.998865404 | 10.352555 | 10.267559 | 10.336724 | 10.326012 | 10.360216 | 10.207303 | 1516 | 1401 | 1181 | 1038 |
| 4684 | NM_000615,N | NCAM1 | neural cell adhesion molecule 1 | protein_coding | NP_000606.3;NP_001070150.1;NP_001229536.1;NP_001229537.1;NP_8 | 7656 | 116930 | . | . | 6.980265 | 7.395986 | 5.859925 | 6.031322 | 16.243362 | 17.096155 | 13.149501 | 13.789900 | -1.224762 | 1156.082375 | 0.132799931 | -2.20257138 | 0.027624973 | 0.53804272 | 10.313971 | 10.021265 | 10.225868 | 10.265345 | 10.083567 | 10.105143 | 1428 | 1409 | 890 | 989 |
| 4685 | NM_00135259 | NCAM2 | neural cell adhesion molecule 2 | protein_coding | NP_001339520.1;NP_001339521.1;NP_001339522.1;NP_001339523.1;NI | 7657 | 602040 | . | . | 1.244005 | 0.994200 | 1.239006 | 0.931908 | 2.894850 | 2.298134 | 2.780295 | 2.130699 | -1.216269 | 179.042495 | 0.254742459 | -1.10881639 | 0.2675094 | 0.998546485 | 7.618899 | 7.335525 | 7.541923 | 7.506671 | 7.399650 | 7.480635 | 235 | 204 | 126 | 167 |
| 4686 | NM_00135150 | NCBP1 | nuclear cap binding protein subunit 1 | protein_coding | NP_001338433.1;NP_001338434.1;NP_001338435.1;NP_001338436.1;NI | 7658 | 600469 | . | . | 10.994644 | 10.952328 | 9.728074 | 10.059372 | 25.584985 | 25.316795 | 21.829518 | 22.999558 | -1.131693 | 1646.899758 | 0.121939878 | -1.46369689 | 0.143276785 | 0.905131505 | 10.772046 | 10.593508 | 10.728990 | 10.734884 | 10.629451 | 10.641428 | 2006 | 1894 | 1333 | 1460 |
| 4690 | NM_00119079 | NCK1 | NCK adaptor protein 1 | protein_coding | NP_001177725.1;NP_001278928.1;NP_006144.1;XP_024309306.1 | 7664 | 600508 | . | . | 3.388312 | 3.767943 | 3.537748 | 2.998737 | 7.884742 | 8.709768 | 7.938604 | 6.856258 | -1.049446 | 454.983811 | 0.179085595 | -0.38879327 | 0.69742908 | 0.998865404 | 8.863715 | 8.794805 | 8.841847 | 8.845098 | 8.864121 | 8.760901 | 535 | 504 | 419 | 381 |
| 4691 | NM_005381 | NCL | nucleolin | protein_coding | NP_005372.2 | 7667 | 164035 | . | . | 207.707184 | 211.720016 | 194.799026 | 206.196228 | 483.343170 | 489.400299 | 437.123383 | 471.443207 | -1.060740 | 25074.769665 | 0.099071404 | -0.85868237 | 0.390515781 | 0.998865404 | 14.655864 | 14.570779 | 14.634179 | 14.643349 | 14.567770 | 14.607771 | 29567 | 28002 | 20668 | 23347 |
| 4692 | NM_002487 | NDN | necdin, MAGE family member | protein_coding | NP_002478.1 | 7675 | 602117 | . | . | 29.183561 | 28.367401 | 29.430796 | 33.327663 | 67.911346 | 65.572517 | 66.041855 | 76.199745 | 1.075018 | 1785.420900 | 0.125942115 | 0.828636804 | 0.407309955 | 0.998865404 | 10.749019 | 10.853197 | 10.786576 | 10.757954 | 10.783105 | 10.873353 | 2018 | 1823 | 1517 | 1833 |
| 4694 | NM_004541 | NDUFA1 | NADH:ubiquinone oxidoreductase subunit A1 | protein_coding | NP_004532.1 | 7683 | 300078 | . | . | 95.836510 | 91.107307 | 81.891441 | 96.286583 | 223.015488 | 210.598602 | 183.762009 | 220.147858 | -1.063869 | 1197.651670 | 0.139567239 | -0.63997817 | 0.522186791 | 0.998865404 | 10.270108 | 10.180485 | 10.270595 | 10.225467 | 10.144034 | 10.256144 | 1464 | 1293 | 933 | 1170 |
| 4695 | NM_00118501 | NDUFA2 | NADH:ubiquinone oxidoreductase subunit A2 | protein_coding | NP_001171941.1;NP_002479.1 | 7685 | 602137 | . | . | 54.054703 | 55.009233 | 47.704068 | 62.367828 | 125.787519 | 127.156308 | 107.046546 | 142.596638 | 1.017919 | 1160.740280 | 0.145620876 | 0.175955379 | 0.860328998 | 0.998865404 | 10.168234 | 10.193316 | 10.167838 | 10.177999 | 10.105777 | 10.260565 | 1316 | 1250 | 913 | 1211 |
| 4696 | NM_004542,XI | NDUFA3 | NADH:ubiquinone oxidoreductase subunit A3 | protein_coding | NP_004533.1;XP_016882322.1 | 7686 | 603832 | . | . | 22.657663 | 22.797308 | 24.971159 | 25.069484 | 52.725313 | 52.697001 | 56.034555 | 57.318398 | 1.089500 | 697.368052 | 0.150022851 | 0.824312598 | 0.409761962 | 0.998865404 | 9.382593 | 9.506308 | 9.417168 | 9.413023 | 9.476686 | 9.471148 | 771 | 718 | 635 | 679 |
| 4697 | NM_002489 | NDUFA4 | NDUFA4 mitochondrial complex associated | protein_coding | NP_002480.1 | 7687 | 603833 | . | . | 55.298073 | 54.887386 | 53.347797 | 58.210701 | 128.680893 | 126.874649 | 119.710915 | 133.091843 | -1.001765 | 3514.308912 | 0.111835621 | -0.02275459 | 0.981846032 | 0.998865404 | 11.780343 | 11.777706 | 11.786233 | 11.772927 | 11.746014 | 11.809236 | 4083 | 3765 | 2936 | 3418 |
| 4698 | NM_00128241 | NDUFA5 | NADH:ubiquinone oxidoreductase subunit A5 | protein_coding | NP_001269348.1;NP_001269349.1;NP_001269350.1;NP_001269351.1;NI | 7688 | 601677 | . | . | 6.598011 | 5.749515 | 6.037606 | 5.450226 | 15.353840 | 13.290265 | 13.548213 | 12.461296 | -1.056045 | 996.569825 | 0.150984866 | -0.52105021 | 0.602331793 | 0.998865404 | 9.999320 | 9.921279 | 10.046913 | 9.906440 | 9.973787 | 9.903610 | 1290 | 1000 | 884 | 865 |
| 4700 | NM_002490 | NDUFA6 | NADH:ubiquinone oxidoreductase subunit A6 | protein_coding | NP_002481.3 | 7690 | 602138 | . | . | 54.585415 | 56.617043 | 50.133827 | 57.838387 | 127.022499 | 130.872833 | 112.498856 | 132.240601 | -1.044000 | 1830.304757 | 0.126999451 | -0.48915026 | 0.624735324 | 0.998865404 | 10.868754 | 10.806307 | 10.843020 | 10.865269 | 10.766962 | 10.870690 | 2123 | 2046 | 1454 | 1790 |
| 4701 | NM_005001,N | NDUFA7 | NADH:ubiquinone oxidoreductase subunit A7 | protein_coding | NP_004992.2 | 7691 | 602139 | . | . | 30.398650 | 33.650224 | 29.178190 | 37.849667 | 70.738911 | 77.784002 | 65.475014 | 86.538765 | 1.024299 | 614.907351 | 0.17128979 | 0.202212564 | 0.839750551 | 0.998865404 | 9.247284 | 9.280970 | 9.230480 | 9.277968 | 9.191035 | 9.343940 | 675 | 679 | 479 | 650 |
| 4702 | NM_00131819 | NDUFA8 | NADH:ubiquinone oxidoreductase subunit A8 | protein_coding | NP_001305124.1;NP_055037.1 | 7692 | 603359 | . | . | 58.266564 | 53.176903 | 44.684086 | 60.824916 | 135.588691 | 122.920789 | 100.269791 | 139.068961 | -1.071493 | 1359.661210 | 0.157090904 | -0.63417331 | 0.525967725 | 0.998865404 | 10.458194 | 10.358134 | 10.470265 | 10.392664 | 10.261376 | 10.485000 | 1701 | 1442 | 973 | 1412 |
| 4704 | NM_005002 | NDUFA9 | NADH:ubiquinone oxidoreductase subunit A9 | protein_coding | NP_004993.1 | 7693 | 603834 | . | . | 6.090350 | 5.994432 | 5.780161 | 6.786591 | 14.172495 | 13.856399 | 12.970513 | 15.516734 | 1.025046 | 1604.321220 | 0.131506768 | 0.271378614 | 0.786099849 | 0.998865404 | 10.629941 | 10.665336 | 10.646841 | 10.627820 | 10.596237 | 10.713499 | 1847 | 1689 | 1306 | 1637 |
| 4705 | NM_00132201 | NDUFA10 | NADH:ubiquinone oxidoreductase subunit A10 | protein_coding | NP_001308948.1;NP_001308949.1;NP_004535.1;XP_011509530.1 | 7684 | 603835 | . | . | 21.418701 | 21.039488 | 19.544346 | 22.040948 | 49.842199 | 48.633720 | 43.856944 | 50.394007 | -1.040217 | 3846.341292 | 0.109986535 | -0.51719622 | 0.605019189 | 0.998865404 | 11.937477 | 11.880503 | 11.925966 | 11.924648 | 11.864694 | 11.919806 | 4520 | 4229 | 3169 | 3653 |
| 4706 | NM_005003,XI | NDUFAB1 | NADH:ubiquinone oxidoreductase subunit AB1 | protein_coding | NP_004994.1;XP_011544158.2 | 7694 | 603836 | . | . | 88.980122 | 88.565250 | 69.408528 | 88.435842 | 207.060423 | 204.722549 | 155.750735 | 202.198052 | -1.140801 | 1740.109393 | 0.140054192 | -1.35695183 | 0.174796517 | 0.948677095 | 10.857062 | 10.666580 | 10.817660 | 10.808452 | 10.616445 | 10.794881 | 2148 | 1990 | 1250 | 1700 |
| 4707 | NM_004545 | NDUFB1 | NADH:ubiquinone oxidoreductase subunit B1 | protein_coding | NP_004536.3 | 7695 | 603837 | . | . | 105.338402 | 103.708466 | 95.169899 | 115.364052 | 245.126801 | 239.726761 | 213.558487 | 263.766205 | -1.007198 | 1036.272803 | 0.146010219 | -0.07086721 | 0.94350344 | 0.998865404 | 10.022584 | 10.011772 | 10.027782 | 10.010113 | 9.944565 | 10.078221 | 1212 | 1109 | 816 | 1056 |
| 4708 | NM_004546 | NDUFB2 | NADH:ubiquinone oxidoreductase subunit B2 | protein_coding | NP_004537.1 | 7697 | 603838 | . | . | 86.987854 | 88.478165 | 74.361626 | 101.830902 | 202.424316 | 204.521240 | 166.865326 | 232.824265 | -1.010193 | 1273.406964 | 0.157062482 | -0.09315247 | 0.925782437 | 0.998865404 | 10.322079 | 10.306836 | 10.312579 | 10.318832 | 10.186918 | 10.416049 | 1468 | 1387 | 935 | 1367 |
| 4709 | NM_00125710 | NDUFB3 | NADH:ubiquinone oxidoreductase subunit B3 | protein_coding | NP_001244031.1;NP_002482.1;XP_011509532.1;XP_016859675.1 | 7698 | 603839 | . | . | 108.467983 | 104.404799 | 100.540220 | 113.859952 | 252.409472 | 241.336377 | 225.609338 | 260.327266 | 1.008116 | 1768.995069 | 0.124594647 | 0.093598807 | 0.925427857 | 0.998865404 | 10.782937 | 10.794469 | 10.803398 | 10.766790 | 10.755144 | 10.826733 | 2076 | 1857 | 1474 | 1741 |
| 4710 | NM_00116833 | NDUFB4 | NADH:ubiquinone oxidoreductase subunit B4 | protein_coding | NP_001161803.1;NP_004538.2 | 7699 | 603840 | . | . | 113.013275 | 112.487595 | 102.179967 | 130.461256 | 262.986553 | 260.020131 | 229.288881 | 298.284175 | 1.021753 | 2369.595185 | 0.134587977 | 0.230674435 | 0.817567736 | 0.998865404 | 11.194970 | 11.225716 | 11.208951 | 11.191344 | 11.122413 | 11.304412 | 2729 | 2502 | 1848 | 2498 |
| 4711 | NM_00119995 | NDUFB5 | NADH:ubiquinone oxidoreductase subunit B5 | protein_coding | NP_001186886.1;NP_001186887.1;NP_002483.1 | 7700 | 603841 | . | . | 13.068385 | 11.327616 | 11.141652 | 11.041545 | 30.410668 | 26.184294 | 25.001541 | 25.245180 | -1.039537 | 1382.650649 | 0.132421987 | -0.4224432 | 0.672701553 | 0.998865404 | 10.460831 | 10.405073 | 10.494510 | 10.398179 | 10.411979 | 10.423664 | 1724 | 1426 | 1170 | 1281 |
| 4712 | NM_00119998 | NDUFB6 | NADH:ubiquinone oxidoreductase subunit B6 | protein_coding | NP_001186916.1;NP_002484.1;NP_877416.1 | 7701 | 603322 | . | . | 12.603490 | 11.578838 | 10.960307 | 9.616218 | 29.328837 | 26.765003 | 24.594612 | 21.986339 | -1.185555 | 457.001606 | 0.17821063 | -1.37793742 | 0.16822261 | 0.945638095 | 8.953226 | 8.708521 | 8.910416 | 8.860046 | 8.821003 | 8.738580 | 593 | 514 | 389 | 365 |
| 4713 | NM_004146,XI | NDUFB7 | NADH:ubiquinone oxidoreductase subunit B7 | protein_coding | NP_004137.2;XP_011526341.1 | 7702 | 603842 | . | . | 66.625955 | 66.404770 | 55.766977 | 66.869463 | 155.041327 | 153.497612 | 125.139479 | 152.889077 | -1.082744 | 1100.589205 | 0.140851049 | -0.81428062 | 0.41548421 | 0.998865404 | 10.160477 | 10.045347 | 10.134297 | 10.130629 | 10.016066 | 10.127400 | 1321 | 1232 | 849 | 1066 |
| 4714 | NM_00128436 | NDUFB8 | NADH:ubiquinone oxidoreductase subunit B8 | protein_coding | NP_001271296.1;NP_001271297.1;NP_004995.1 | 7703 | 602140 | . | . | 117.944778 | 116.764581 | 111.301961 | 127.744959 | 274.462329 | 269.906572 | 249.758358 | 292.073644 | 1.005560 | 2521.075225 | 0.120846343 | 0.066189531 | 0.947226931 | 0.998865404 | 11.295913 | 11.303723 | 11.305456 | 11.288744 | 11.249094 | 11.351453 | 2925 | 2685 | 2059 | 2520 |
| 4715 | NM_00127864 | NDUFB9 | NADH:ubiquinone oxidoreductase subunit B9 | protein_coding | NP_001265574.1;NP_001265575.1;NP_001298097.1;NP_004996.1 | 7704 | 601445 | . | . | 223.606525 | 217.565664 | 185.519863 | 212.799443 | 520.341581 | 502.912802 | 416.301174 | 486.540639 | -1.124021 | 4522.860555 | 0.114616151 | -1.47159978 | 0.141128986 | 0.902689247 | 12.224946 | 12.056160 | 12.204263 | 12.174314 | 12.038177 | 12.142847 | 5610 | 5074 | 3468 | 4246 |
| 4716 | NM_004548 | NDUFB10 | NADH:ubiquinone oxidoreductase subunit B10 | protein_coding | NP_004539.1 | 7696 | 603843 | . | . | 61.657711 | 59.648270 | 54.482315 | 61.200855 | 143.480026 | 137.879654 | 122.256737 | 139.928482 | -1.063659 | 1246.389625 | 0.133824926 | -0.66531451 | 0.505849351 | 0.998865404 | 10.327478 | 10.238221 | 10.321865 | 10.290368 | 10.219431 | 10.298082 | 1510 | 1358 | 995 | 1192 |
| 4717 | NM_00118498 | NDUFC1 | NADH:ubiquinone oxidoreductase subunit C1 | protein_coding | NP_001171915.1;NP_001171916.1;NP_001171917.1;NP_001171918.1;NI | 7705 | 603844 | . | . | 22.642788 | 21.430871 | 23.256984 | 20.474760 | 52.690698 | 49.538424 | 52.187999 | 46.813105 | -1.027945 | 694.890835 | 0.152005304 | -0.26159015 | 0.79363744 | 0.998865404 | 9.460246 | 9.420769 | 9.474319 | 9.424920 | 9.439058 | 9.423077 | 840 | 733 | 603 | 635 |
| 4718 | NM_00120405 | NDUFC2 | NADH:ubiquinone oxidoreductase subunit C2 | protein_coding | NP_001190983.1;NP_001190984.1;NP_004540.1 | 7706 | 603845 | . | . | 29.765546 | 26.834891 | 25.429189 | 25.556486 | 69.265649 | 62.030056 | 57.062368 | 58.431871 | -1.125432 | 1818.416686 | 0.125142669 | -1.36227545 | 0.173110973 | 0.947431715 | 10.911106 | 10.740805 | 10.917002 | 10.826836 | 10.781973 | 10.777755 | 2342 | 1961 | 1492 | 1600 |
| 4719 | NM_00119998 | NDUFS1 | NADH:ubiquinone oxidoreductase core subunit S1 | protein_coding | NP_001186910.1;NP_001186911.1;NP_001186912.1;NP_001186913.1;NI | 7707 | 157655 | . | . | 15.400153 | 15.352495 | 13.787362 | 17.753462 | 35.836789 | 35.487983 | 30.938442 | 40.591181 | 1.010999 | 5632.367237 | 0.128426666 | 0.122881939 | 0.902200588 | 0.998865404 | 12.451691 | 12.467318 | 12.457061 | 12.449368 | 12.357386 | 12.557604 | 6480 | 6017 | 4339 | 5940 |
| 4720 | NM_00116615 | NDUFS2 | NADH:ubiquinone oxidoreductase core subunit S2 | protein_coding | NP_001159631.1;NP_001364227.1;NP_001364228.1;NP_001364229.1;NI | 7708 | 602985 | . | . | 56.804483 | 56.831600 | 53.840861 | 63.840506 | 132.186367 | 131.368787 | 120.817338 | 145.963739 | 1.024057 | 2955.312961 | 0.121457922 | 0.282367272 | 0.777661907 | 0.998865404 | 11.511949 | 11.546040 | 11.522313 | 11.514940 | 11.473990 | 11.598234 | 3378 | 3137 | 2403 | 3016 |
| 4722 | NM_004551 | NDUFS3 | NADH:ubiquinone oxidoreductase core subunit S3 | protein_coding | NP_004542.1 | 7710 | 603846 | . | . | 41.868633 | 42.795818 | 41.153790 | 52.294674 | 97.430038 | 98.924446 | 92.347916 | 119.565567 | 1.088118 | 1239.500860 | 0.146922561 | 0.829248624 | 0.406963736 | 0.998865404 | 10.213618 | 10.334920 | 10.235834 | 10.246108 | 10.213863 | 10.388120 | 1358 | 1290 | 995 | 1349 |
| 4723 | NM_00116610 | NDUFV1 | NADH:ubiquinone oxidoreductase core subunit V1 | protein_coding | NP_001159574.1;NP_009034.2 | 7716 | 161015 | . | . | 68.923800 | 63.305023 | 62.456006 | 67.899857 | 160.388517 | 146.332399 | 140.149464 | 155.244965 | -1.029287 | 3190.402352 | 0.116827254 | -0.3564711 | 0.721487794 | 0.998865404 | 11.660202 | 11.618542 | 11.688693 | 11.611589 | 11.596489 | 11.656987 | 3901 | 3329 | 2633 | 3057 |
| 4724 | NM_00131805 | NDUFS4 | NADH:ubiquinone oxidoreductase subunit S4 | protein_coding | NP_001304980.1;NP_002486.1;XP_016864980.1 | 7711 | 602694 | . | . | 51.663699 | 51.367045 | 42.470010 | 47.346061 | 120.223550 | 118.737233 | 95.301465 | 108.251145 | -1.168654 | 1029.708363 | 0.138706045 | -1.62103473 | 0.10501019 | 0.832542015 | 10.116184 | 9.891085 | 10.075461 | 10.049352 | 9.912878 | 9.981529 | 1300 | 1177 | 786 | 933 |
| 4725 | NM_00118497 | NDUFS5 | NADH:ubiquinone oxidoreductase subunit S5 | protein_coding | NP_001171908.1;NP_004543.1 | 7712 | 603847 | . | . | 142.645786 | 149.104034 | 133.870758 | 153.101505 | 331.942596 | 344.660675 | 300.402092 | 350.048394 | -1.029386 | 2341.767237 | 0.122094558 | -0.34223116 | 0.73217694 | 0.998865404 | 11.214253 | 11.172204 | 11.192692 | 11.216221 | 11.129312 | 11.230441 | 2696 | 2601 | 1880 | 2300 |
| 4726 | NM_004553 | NDUFS6 | NADH:ubiquinone oxidoreductase subunit S6 | protein_coding | NP_004544.1 | 7713 | 603848 | . | . | 77.897766 | 75.761772 | 63.158123 | 79.674522 | 181.271301 | 175.126724 | 141.724991 | 182.166351 | -1.091124 | 1196.402568 | 0.147000919 | -0.85587592 | 0.392066416 | 0.998865404 | 10.286259 | 10.159952 | 10.268991 | 10.241204 | 10.104340 | 10.268058 | 1464 | 1323 | 885 | 1191 |
| 4728 | NM_002496 | NDUFS8 | NADH:ubiquinone oxidoreductase core subunit S8 | protein_coding | NP_002487.1 | 7715 | 602141 | . | . | 69.767471 | 76.509666 | 70.310440 | 85.076736 | 162.351776 | 176.855515 | 157.774582 | 194.517853 | 1.047841 | 1731.296535 | 0.135665172 | 0.496954062 | 0.619221447 | 0.998865404 | 10.723770 | 10.790728 | 10.704575 | 10.769982 | 10.701041 | 10.842732 | 1866 | 1901 | 1402 | 1810 |
| 4729 | NM_021074,XI | NDUFV2 | NADH:ubiquinone oxidoreductase core subunit V2 | protein_coding | NP_066552.2;XP_016881271.1 | 7717 | 600532 | . | . | 77.017492 | 69.138229 | 61.242130 | 59.461399 | 179.222861 | 159.816111 | 137.425562 | 135.951423 | 1.010763 | 2107.722015 | 0.127745692 | 0.12089925 | 0.903770836 | 0.998865404 | 11.033795 | 11.049102 | 11.070547 | 11.001124 | 10.988202 | 11.098656 | 2519 | 2164 | 1715 | 2124 |
| 4730 | gene-NDUFV2F | NDUFV2P1 | NADH:ubiquinone oxidoreductase core subunit V2 pseudogene 1 | pseudogene | . | 7718 | . | | . | 1.562580 | 2.312696 | 1.873857 | 2.041399 | 3.636188 | 5.345901 | 4.204881 | 4.667417 | 1.001636 | 49.170556 | 0.456943862 | 0.005160734 | 0.995882348 | 0.999862634 | 5.619607 | 5.619838 | 5.581038 | 5.651073 | 5.609603 | 5.624309 | 46 | 63 | 41 | 48 |
| 4731 | NM_00100150 | NDUFV3 | NADH:ubiquinone oxidoreductase subunit V3 | protein_coding | NP_001001503.1;NP_066553.3;XP_011527888.1;XP_016883848.1 | 7719 | 602184 | . | . | 4.292172 | 4.381065 | 3.950364 | 4.876776 | 9.988060 | 10.127030 | 8.864502 | 11.150169 | -1.007220 | 732.196923 | 0.159619349 | -0.06502299 | 0.948155696 | 0.998865404 | 9.521619 | 9.510536 | 9.511524 | 9.523647 | 9.441376 | 9.578900 | 839 | 800 | 571 | 752 |
| 4733 | NM_004147 | DRG1 | developmentally regulated GTP binding protein 1 | protein_coding | NP_004138.1 | 3029 | 603952 | . | . | 44.709850 | 45.022491 | 40.278225 | 48.194820 | 104.041649 | 104.071503 | 90.383171 | 110.191734 | -1.028568 | 2311.000105 | 0.126883455 | -0.32027205 | 0.748762109 | 0.998865404 | 11.194617 | 11.153696 | 11.185440 | 11.184122 | 11.092267 | 11.227366 | 2701 | 2527 | 1814 | 2316 |
| 4734 | NM_00128433 | NEDD4 | NEDD4 E3 ubiquitin protein ligase | protein_coding | NP_001271267.1;NP_001271268.1;NP_001271269.1;NP_001316141.1;NI | 7727 | 602278 | . | . | 4.164369 | 3.640223 | 3.088699 | 3.413768 | 9.690657 | 8.414536 | 6.930952 | 7.805177 | -1.214144 | 680.905465 | 0.152475781 | -1.83596064 | 0.066363478 | 0.733835894 | 9.544387 | 9.264700 | 9.492521 | 9.454247 | 9.350354 | 9.332645 | 883 | 784 | 542 | 569 |
| 4735 | NM_00100849 | SEPTIN2 | septin 2 | protein_coding | NP_001008491.1;NP_001008492.1;NP_001269901.1;NP_001269902.1;NI | 7729 | 601506 | . | . | 112.980748 | 108.170650 | 102.480417 | 105.278376 | 262.910840 | 250.041306 | 229.963075 | 240.706509 | -1.094026 | 11300.047024 | 0.101530503 | -1.27693279 | 0.201625977 | 0.967296129 | 13.527404 | 13.397767 | 13.522249 | 13.479592 | 13.418993 | 13.429830 | 13933 | 12425 | 9325 | 10183 |
| 4736 | NM_007104,XI | RPL10A | ribosomal protein L10a | protein_coding | NP_009035.3;XP_016866389.1 | 10299 | 615660 | . | . | 327.726801 | 323.983093 | 284.101504 | 346.050398 | 762.633736 | 748.901432 | 637.515519 | 791.203186 | -1.043641 | 7335.105477 | 0.120841133 | -0.50997638 | 0.610068013 | 0.998865404 | 12.871141 | 12.809402 | 12.863188 | 12.851341 | 12.733700 | 12.901580 | 8687 | 8028 | 5622 | 7398 |
| 4738 | NM_006156 | NEDD8 | NEDD8 ubiquitin like modifier | protein_coding | NP_006147.1 | 7732 | 603171 | . | . | 101.835350 | 99.258179 | 88.916290 | 95.222885 | 236.975021 | 229.439728 | 199.525589 | 217.715805 | -1.107547 | 1848.893599 | 0.121264363 | -1.21526197 | 0.224266193 | 0.982008277 | 10.924291 | 10.776836 | 10.905011 | 10.877568 | 10.787843 | 10.833252 | 2276 | 2061 | 1481 | 1693 |
| 4739 | NM_00114239 | NEDD9 | neural precursor cell expressed, developmentally down-regulated | protein_coding | NP_001135865.1;NP_001257962.1;NP_006394.1;NP_892011.2 | 7733 | 602265 | . | . | 3.639682 | 3.360655 | 3.734515 | 4.069408 | 8.469691 | 7.768304 | 8.380144 | 9.304218 | 1.103089 | 518.313170 | 0.168339227 | 0.840853514 | 0.400430004 | 0.998865404 | 8.945196 | 9.086696 | 9.011197 | 8.956903 | 9.021024 | 9.072625 | 592 | 509 | 454 | 530 |

| | | | | | | | | | | | | | | | | | | | | | | | | | | | | | | | | | |
|---|---|---|---|---|---|---|---|---|---|---|---|---|---|---|---|---|---|---|---|---|---|---|---|---|---|---|---|---|---|---|---|---|---|
| 4747 | NM_006158 | NEFL | neurofilament light | protein_coding | NP_006149.2 | 7739 | 162280 | . | . | 10.382795 | 10.234518 | 9.072164 | 10.418835 | 24.161194 | 23.657549 | 20.357672 | 23.821428 | -1.072343 | 1120.156142 | 0.138218086 | -0.7290432 | 0.465975237 | 0.998865404 | 10.179150 | 10.078047 | 10.163159 | 10.146155 | 10.054615 | 10.148330 | 1350 | 1237 | 880 | 1078 |
| 4750 | NM_00119939 | NEK1 | NIMA related kinase 1 | protein_coding | NP_001186326.1;NP_001186327.1;NP_001186328.1;NP_001186329.1;NI | 7744 | 604588 | . | . | 3.298041 | 3.318450 | 3.675295 | 3.248185 | 7.674680 | 7.670747 | 8.247257 | 7.426586 | 1.047436 | 618.349040 | 0.15998739 | 0.417922885 | 0.6760035 | 0.998865404 | 9.238318 | 9.305456 | 9.237761 | 9.273245 | 9.323423 | 9.248579 | 677 | 669 | 581 | 560 |
| 4751 | NM_00120418 | NEK2 | NIMA related kinase 2 | protein_coding | NP_001191111.1;NP_001191112.1;NP_002488.1;XP_005273204.1 | 7745 | 604043 | . | . | 17.116247 | 17.109322 | 14.406230 | 14.509065 | 39.830207 | 39.548963 | 32.327161 | 33.173253 | -1.202041 | 1044.937912 | 0.134632907 | -1.971922 | **0.048618512** | 0.661815897 | 10.155834 | 9.890375 | 10.096860 | 10.089356 | 9.958426 | 9.956971 | 1320 | 1225 | 826 | 889 |
| 4752 | NM_002498,N | NEK3 | NIMA related kinase 3 | protein_coding | NP_002489.1;NP_689933.1 | 7746 | 604044 | . | . | 0.609376 | 0.670349 | 0.766057 | 0.558974 | 1.418043 | 1.549542 | 1.719011 | 1.278028 | 1.011487 | 49.241391 | 0.452672159 | 0.036402156 | 0.970961695 | 0.998865404 | 5.612748 | 5.630795 | 5.610075 | 5.625578 | 5.648332 | 5.590704 | 54 | 55 | 50 | 39 |
| 4753 | NM_00114510 | NELL2 | neural EGFL like 2 | protein_coding | NP_001138579.1;NP_001138580.1;NP_001138581.1;NP_001138582.1;NI | 7751 | 602320 | . | . | 0.058995 | 0.049690 | 0.075133 | 0.110196 | 0.137285 | 0.114860 | 0.168597 | 0.251949 | 1.759362 | 8.626951 | 1.041729223 | 0.782403036 | 0.43397775 | NA | 2.644780 | 3.459419 | 3.089530 | 3.083200 | 3.099866 | 3.115718 | 8 | 6 | 8 | 12 |
| 4756 | NM_00117262 | NEO1 | neogenin 1 | protein_coding | NP_001166094.1;NP_001166095.1;NP_002490.2;XP_005254465.1;XP_01 | 7754 | 601907 | . | . | 7.072960 | 7.666450 | 6.708645 | 7.767281 | 16.459070 | 17.721342 | 15.054003 | 17.758966 | -1.045889 | 1606.872234 | 0.13176951 | -0.49123558 | 0.623259839 | 0.998865404 | 10.682243 | 10.617101 | 10.646150 | 10.686846 | 10.573642 | 10.686507 | 1843 | 1819 | 1266 | 1580 |
| 4758 | NM_000434 | NEU1 | neuraminidase 1 | protein_coding | NP_000425.1 | 7758 | 608272 | . | . | 38.336899 | 39.400978 | 37.481564 | 39.952507 | 89.211540 | 91.077126 | 84.107536 | 91.346657 | -1.017954 | 4052.426460 | 0.108786612 | -0.23598912 | 0.813441113 | 0.998865404 | 11.997393 | 11.971633 | 11.984049 | 11.999600 | 11.954657 | 11.998958 | 4664 | 4453 | 3399 | 3866 |
| 4763 | NM_000267,N | NF1 | neurofibromin 1 | protein_coding | NP_000258.1;NP_001035957.1;NP_001121619.1 | 7765 | 613113 | . | . | 4.206648 | 4.397065 | 4.119217 | 4.042276 | 9.789041 | 10.164014 | 9.243404 | 9.242184 | -1.073658 | 1611.036377 | 0.122965725 | -0.83385137 | 0.404364743 | 0.998865404 | 10.704140 | 10.601585 | 10.666829 | 10.694650 | 10.631687 | 10.619279 | 1887 | 1832 | 1362 | 1445 |
| 4771 | NM_000268,N | NF2 | neurofibromin 2 | protein_coding | NP_000259.1;NP_057502.2;NP_861546.1;NP_861966.1;NP_861967.1;NP_ | 7773 | 607379 | . | . | 7.728935 | 8.139646 | 7.307799 | 8.777630 | 17.985550 | 18.815160 | 16.398487 | 20.069011 | 1.005042 | 1480.779242 | 0.136353467 | 0.053216314 | 0.957559557 | 0.998865404 | 10.528737 | 10.535536 | 10.513701 | 10.544711 | 10.463591 | 10.598121 | 1667 | 1626 | 1178 | 1513 |
| 4772 | NM_00127866 | NFATC1 | nuclear factor of activated T cells 1 | protein_coding | NP_001265598.1;NP_001265599.1;NP_001265601.1;NP_001265602.1;NI | 7775 | 600489 | . | . | 3.768596 | 3.666061 | 3.884312 | 4.057946 | 8.769678 | 8.474265 | 8.716281 | 9.278013 | 1.042850 | 545.655002 | 0.162053885 | 0.373524854 | 0.708757851 | 0.998865404 | 9.061257 | 9.121798 | 9.085010 | 9.070594 | 9.100372 | 9.109682 | 622 | 570 | 481 | 526 |
| 4773 | NM_00113602 | NFATC2 | nuclear factor of activated T cells 2 | protein_coding | NP_001129493.1;NP_001245221.1;NP_001245223.1;NP_001245224.1;NI | 7776 | 600490 | . | . | 20.118395 | 21.842585 | 24.867058 | 26.753997 | 46.816329 | 50.490112 | 55.800958 | 61.169840 | 1.237878 | 5207.931399 | 0.108913991 | 2.826715186 | **0.004702814** | 0.203488824 | 12.184423 | 12.492184 | 12.225054 | 12.274405 | 12.400493 | 12.456460 | 5199 | 5173 | 4846 | 5578 |
| 4774 | NM_00113467 | NFIA | nuclear factor I A | protein_coding | NP_001128145.1;NP_001138983.1;NP_001138984.1;NP_005586.1;XP_01 | 7784 | 600727 | . | . | 0.663351 | 0.690479 | 0.661973 | 0.716319 | 1.543645 | 1.596071 | 1.485449 | 1.637778 | -1.059128 | 194.476461 | 0.239202227 | -0.34647512 | 0.728985676 | 0.998865404 | 7.644193 | 7.561526 | 7.609635 | 7.622871 | 7.603163 | 7.578167 | 227 | 219 | 169 | 172 |
| 4775 | NM_004555,N | NFATC3 | nuclear factor of activated T cells 3 | protein_coding | NP_004546.1;NP_775186.1;NP_775188.1 | 7777 | 602698 | . | . | 3.027934 | 3.104305 | 2.900754 | 3.408060 | 7.046126 | 7.175739 | 6.509209 | 7.792125 | 1.018217 | 617.599673 | 0.161833488 | 0.160935666 | 0.872144071 | 0.998865404 | 9.257755 | 9.283190 | 9.257300 | 9.270429 | 9.225703 | 9.323554 | 698 | 667 | 502 | 627 |
| 4778 | NM_00113602 | NFE2 | nuclear factor, erythroid 2 | protein_coding | NP_001129495.1;NP_001248390.1;NP_006154.1 | 7780 | 601490 | . | . | 0.348415 | 0.389465 | 0.107915 | 0.178824 | 0.810777 | 0.900267 | 0.242159 | 0.408859 | **-2.446837** | 14.244752 | 0.838614842 | -1.53934546 | 0.123719982 | NA | 4.339107 | 3.044381 | 3.857154 | 3.874088 | 3.808348 | 3.819476 | 21 | 24 | 6 | 9 |
| 4779 | NM_00133026 | NFE2L1 | nuclear factor, erythroid 2 like 1 | protein_coding | NP_001317190.1;NP_001317191.1;NP_003195.1;XP_005257467.1;XP_00 | 7781 | 163260 | . | . | 82.511708 | 81.630744 | 72.526790 | 80.602449 | 192.008135 | 188.693107 | 162.748009 | 184.287915 | -1.090616 | 11089.634805 | 0.103850398 | -1.20503749 | 0.228188828 | 0.982008277 | 13.498159 | 13.372975 | 13.476475 | 13.468875 | 13.363938 | 13.432125 | 13382 | 12430 | 8854 | 10346 |
| 4780 | NM_00114541 | NFE2L2 | nuclear factor, erythroid 2 like 2 | protein_coding | NP_001138884.1;NP_001138885.1;NP_001300829.1;NP_001300830.1;NI | 7782 | 600492 | . | . | 47.623115 | 49.948742 | 49.358747 | 53.424572 | 110.820948 | 115.458753 | 110.759599 | 122.148932 | 1.038768 | 4047.000689 | 0.10835123 | 0.506439819 | 0.612547928 | 0.998865404 | 11.954979 | 12.009775 | 11.956247 | 11.976757 | 11.980526 | 12.015367 | 4515 | 4337 | 3511 | 3947 |
| 4781 | NM_00119073 | NFIB | nuclear factor I B | protein_coding | NP_001177666.1;NP_001177667.1;NP_001269716.1;NP_001356387.1;NI | 7785 | 600728 | . | . | 1.013199 | 0.814795 | 0.871664 | 0.919703 | 2.357759 | 1.883437 | 1.955990 | 2.102795 | -1.019536 | 239.489131 | 0.226831131 | -0.12305573 | 0.902062965 | 0.998865404 | 7.917456 | 7.890046 | 7.956003 | 7.857073 | 7.888310 | 7.908236 | 306 | 235 | 202 | 227 |
| 4782 | NM_00124500 | NFIC | nuclear factor I C | protein_coding | NP_001231931.1;NP_001231933.1;NP_001231934.1;NP_005588.2;NP_9 | 7786 | 600729 | . | . | 4.737556 | 5.523418 | 4.734362 | 5.756477 | 11.024487 | 12.767630 | 10.623771 | 13.161500 | 1.010660 | 1261.104558 | 0.14858468 | 0.102956576 | 0.917997435 | 0.998865404 | 10.293133 | 10.307775 | 10.228662 | 10.356715 | 10.234411 | 10.366382 | 1323 | 1468 | 1006 | 1292 |
| 4783 | NM_00128999 | NFIL3 | nuclear factor, interleukin 3 regulated | protein_coding | NP_001276928.1;NP_001276929.1;NP_005375.2;XP_016870232.1;XP_01 | 7787 | 605327 | . | . | 3.309026 | 3.090520 | 3.391351 | 3.331905 | 7.700240 | 7.143876 | 7.610093 | 7.618004 | 1.046463 | 209.489582 | 0.231775962 | 0.282693858 | 0.777411525 | 0.998865404 | 7.677413 | 7.743304 | 7.711740 | 7.687204 | 7.727062 | 7.713955 | 242 | 215 | 189 | 198 |
| 4784 | NM_00127104 | NFIX | nuclear factor I X | protein_coding | NP_001257972.1;NP_001257973.1;NP_001352831.1;NP_001352911.1;NI | 7788 | 164005 | . | . | 4.179124 | 4.417582 | 4.019505 | 5.045212 | 9.724995 | 10.211439 | 9.019652 | 11.535276 | 1.068591 | 795.079443 | 0.158581734 | 0.603537639 | 0.546151084 | 0.998865404 | 9.586684 | 9.681664 | 9.595606 | 9.623241 | 9.577622 | 9.728996 | 868 | 846 | 638 | 852 |
| 4790 | NM_00116541 | NFKB1 | nuclear factor kappa B subunit 1 | protein_coding | NP_001158884.1;NP_001306155.1;NP_001369554.1;NP_001369555.1;NI | 7794 | 164011 | . | . | 6.975823 | 6.964558 | 6.742329 | 7.516459 | 16.233029 | 16.098886 | 15.129592 | 17.185488 | 1.014732 | 882.165841 | 0.14250569 | 0.148060417 | 0.882295088 | 0.998865404 | 9.774429 | 9.795308 | 9.785555 | 9.773368 | 9.762046 | 9.817036 | 1017 | 937 | 743 | 865 |
| 4791 | NM_00107749 | NFKB2 | nuclear factor kappa B subunit 2 | protein_coding | NP_001070962.1;NP_001248332.1;NP_001275653.1;NP_001309863.1;NI | 7795 | 164012 | . | . | 11.169054 | 11.774226 | 12.212631 | 12.272303 | 25.990848 | 27.216651 | 27.404790 | 28.059166 | 1.044287 | 1160.356221 | 0.133597419 | 0.467957381 | 0.639815068 | 0.998865404 | 10.148837 | 10.211193 | 10.133216 | 10.193024 | 10.197498 | 10.194391 | 1255 | 1274 | 1033 | 1109 |
| 4792 | NM_020529 | NFKBIA | NFKB inhibitor alpha | protein_coding | NP_065390.1 | 7797 | 164008 | . | . | 6.675690 | 6.210909 | 5.032866 | 6.115155 | 15.534604 | 14.356794 | 11.293606 | 13.981576 | -1.169978 | 292.665319 | 0.210324829 | -1.07681702 | 0.281562018 | 0.998865404 | 8.302220 | 8.075064 | 8.255228 | 8.210437 | 8.102331 | 8.194695 | 378 | 327 | 213 | 276 |
| 4793 | NM_00124311 | NFKBIB | NFKB inhibitor beta | protein_coding | NP_001230045.1;NP_001356628.1;NP_001356629.1;NP_002494.2;XP_00 | 7798 | 604495 | . | . | 13.850753 | 12.576781 | 14.108872 | 16.312090 | 32.231272 | 29.071792 | 31.659904 | 37.295655 | 1.124736 | 859.739870 | 0.150541146 | 1.126513458 | 0.259948247 | 0.995760118 | 9.660467 | 9.830064 | 9.748772 | 9.654721 | 9.749037 | 9.823319 | 994 | 815 | 751 | 897 |
| 4794 | NM_004556 | NFKBIE | NFKB inhibitor epsilon | protein_coding | NP_004547.3 | 7799 | 604548 | . | . | 5.976185 | 6.717490 | 7.356501 | 8.358169 | 13.906829 | 15.527780 | 16.507774 | 19.109959 | 1.221862 | 518.727518 | 0.17130155 | 1.687555603 | 0.091496568 | 0.805877675 | 8.867493 | 9.155794 | 8.914736 | 8.981377 | 9.037040 | 9.114245 | 509 | 531 | 467 | 566 |
| 4795 | NM_00114496 | NFKBIL1 | NFKB inhibitor like 1 | protein_coding | NP_001138433.1;NP_001138434.1;NP_001138435.1;NP_004998.3 | 7800 | 601022 | . | . | 8.372153 | 7.283841 | 6.948603 | 7.993636 | 19.482344 | 16.836924 | 15.592462 | 18.276502 | -1.062319 | 345.561358 | 0.198232595 | -0.43997456 | 0.659955532 | 0.998865404 | 8.475806 | 8.388469 | 8.490693 | 8.405776 | 8.379565 | 8.449148 | 440 | 356 | 273 | 334 |
| 4796 | NM_013432,XI | TONSL | tonsoku like, DNA repair protein | protein_coding | NP_038460.4;XP_011515350.1;XP_011515351.1;XP_011515352.1 | 7801 | 604546 | . | . | 20.648097 | 23.009011 | 23.962150 | 20.960441 | 48.048968 | 53.186358 | 53.770369 | 47.923563 | 1.020717 | 3099.667074 | 0.12274447 | 0.241007353 | 0.809549417 | 0.998865404 | 11.582979 | 11.612663 | 11.551802 | 11.623925 | 11.661476 | 11.545917 | 3377 | 3456 | 2915 | 2729 |
| 4798 | NM_00114383 | NFRKB | nuclear factor related to kappaB binding protein | protein_coding | NP_001137307.1;NP_006156.2;XP_005271624.1;XP_005271625.1;XP_01 | 7802 | 164013 | . | . | 3.340817 | 3.135582 | 3.444773 | 3.364522 | 7.774216 | 7.248040 | 7.729969 | 7.692578 | 1.040876 | 531.896979 | 0.165263067 | 0.349732684 | 0.726539322 | 0.998865404 | 9.025624 | 9.083796 | 9.066742 | 9.015536 | 9.077026 | 9.057583 | 624 | 540 | 479 | 501 |
| 4799 | NM_00131875 | NFX1 | nuclear transcription factor, X-box binding 1 | protein_coding | NP_001305687.1;NP_002495.2;NP_667345.1 | 7803 | 603255 | . | . | 6.318153 | 7.325475 | 7.412321 | 7.012617 | 14.702604 | 16.933168 | 16.633031 | 16.033515 | 1.059464 | 921.908300 | 0.148305844 | 0.561908331 | 0.574178477 | 0.998865404 | 9.806298 | 9.889462 | 9.767690 | 9.881104 | 9.889633 | 9.845292 | 950 | 1042 | 850 | 862 |
| 4800 | NM_002505,N | NFYA | nuclear transcription factor Y subunit alpha | protein_coding | NP_002496.1;NP_068351.1 | 7804 | 189903 | . | . | 4.755970 | 5.175022 | 5.612759 | 5.027781 | 11.067339 | 11.962295 | 12.594330 | 11.495425 | 1.058743 | 991.594557 | 0.142948015 | 0.576101992 | 0.564546245 | 0.998865404 | 9.911743 | 9.994290 | 9.904578 | 9.957446 | 10.016197 | 9.928301 | 1069 | 1077 | 940 | 899 |
| 4801 | NM_006166,XI | NFYB | nuclear transcription factor Y subunit beta | protein_coding | NP_006157.1;XP_005268965.1;XP_005268966.1;XP_005268967.1;XP_02 | 7805 | 189904 | . | . | 3.946258 | 3.818519 | 3.446094 | 3.920287 | 9.183104 | 8.826677 | 7.732936 | 8.963270 | -1.072601 | 395.598288 | 0.183303755 | -0.55162083 | 0.581208165 | 0.998865404 | 8.677722 | 8.576280 | 8.660566 | 8.636159 | 8.578971 | 8.634165 | 481 | 433 | 316 | 375 |
| 4802 | NM_00114258 | NFYC | nuclear transcription factor Y subunit gamma | protein_coding | NP_001136059.1;NP_001136060.1;NP_001136061.1;NP_001136062.1;NI | 7806 | 605344 | . | . | 16.737240 | 18.102139 | 17.653314 | 19.405364 | 38.948242 | 41.843899 | 39.613525 | 44.368055 | 1.055192 | 1200.067487 | 0.133530006 | 0.580438704 | 0.561618811 | 0.998865404 | 10.189791 | 10.266976 | 10.179150 | 10.235955 | 10.226525 | 10.269507 | 1294 | 1308 | 1041 | 1188 |
| 4804 | NM_002507 | NGFR | nerve growth factor receptor | protein_coding | NP_002498.1 | 7809 | 162010 | . | . | 4.186462 | 4.379371 | 3.062391 | 2.659153 | 9.742068 | 10.123111 | 6.871917 | 6.079836 | -1.516584 | 380.460458 | 0.18951914 | -3.17026074 | **0.001523022** | 0.091626649 | 8.840671 | 8.240576 | 8.664973 | 8.688340 | 8.474678 | 8.394202 | 518 | 504 | 283 | 262 |
| 4807 | NM_005598 | NHLH1 | nescient helix-loop-helix 1 | protein_coding | NP_005589.1 | 7817 | 162360 | . | . | 0.060734 | 0.083980 | 0.050696 | 0.047103 | 0.141331 | 0.194123 | 0.113760 | 0.107696 | -1.422917 | 5.372794 | 1.306590236 | -0.38945022 | 0.696943125 | NA | 2.658167 | 2.148375 | 2.429451 | 2.439652 | 2.426831 | 2.425258 | 6 | 8 | 4 | 4 |
| 4809 | NM_00100379 | SNU13 | small nuclear ribonucleoprotein 13 | protein_coding | NP_001003796.1;NP_004999.1;XP_016884300.1 | 7819 | 601304 | . | . | 56.535341 | 59.190204 | 49.618839 | 62.852059 | 131.560057 | 136.820803 | 111.343246 | 143.703769 | -1.050567 | 2675.550382 | 0.129507947 | -0.54952454 | 0.582645527 | 0.998865404 | 11.420917 | 11.349438 | 11.392980 | 11.415328 | 11.279649 | 11.442024 | 3114 | 2999 | 2041 | 2692 |
| 4810 | NM_00113602 | NHS | NHS actin remodeling regulator | protein_coding | NP_001129496.1;NP_001278796.1;NP_001278797.1;NP_938011.1;XP_01 | 7820 | 300457 | . | . | 0.032260 | 0.026439 | 0.014657 | 0.032099 | 0.075071 | 0.061116 | 0.032890 | 0.073390 | -1.256288 | 8.023367 | 1.087071759 | -0.30280193 | 0.762040815 | NA | 3.161369 | 2.827809 | 3.016846 | 3.011217 | 2.994879 | 3.016703 | 11 | 9 | 4 | 9 |
| 4811 | NM_002508,XI | NID1 | nidogen 1 | protein_coding | NP_002499.2;XP_011542497.1 | 7821 | 131390 | . | . | 0.467922 | 0.408328 | 0.348925 | 0.473776 | 1.088874 | 0.943868 | 0.782977 | 1.083232 | -1.076754 | 77.174325 | 0.370775629 | -0.28774584 | 0.773541294 | 0.998865404 | 6.322968 | 6.215115 | 6.299190 | 6.259484 | 6.223912 | 6.296066 | 99 | 80 | 55 | 80 |
| 4814 | NM_004148,XI | NINJ1 | ninjurin 1 | protein_coding | NP_004139.2;XP_011517018.1 | 7824 | 602062 | . | . | 13.902843 | 16.968265 | 11.219300 | 15.110894 | 32.352487 | 39.222901 | 25.175785 | 34.549263 | -1.187818 | 551.825541 | 0.18944781 | -1.31072398 | 0.189951032 | 0.961278413 | 9.227509 | 8.977839 | 9.092403 | 9.220055 | 8.953164 | 9.131218 | 625 | 708 | 376 | 540 |
| 4817 | NM_00118509 | NIT1 | nitrilase 1 | protein_coding | NP_001172021.1;NP_001172022.1;NP_001172023.1;NP_005591.1;XP_00 | 7828 | 604618 | . | . | 13.656698 | 13.302928 | 13.360970 | 13.634308 | 31.779699 | 30.750314 | 29.981631 | 31.173224 | -1.038673 | 667.199604 | 0.152232676 | -0.35958782 | 0.719155392 | 0.998865404 | 9.409068 | 9.354362 | 9.402909 | 9.386754 | 9.366534 | 9.371629 | 792 | 725 | 567 | 616 |
| 4820 | NM_00134912 | NKTR | natural killer cell triggering receptor | protein_coding | NP_001336053.1;NP_001336054.1;NP_001336055.1;NP_005376.2;XP_00 | 7833 | 161565 | . | . | 6.841179 | 6.634051 | 8.131746 | 6.674419 | 15.919703 | 15.334903 | 18.247406 | 15.260264 | 1.075469 | 1688.224002 | 0.144407372 | 0.726877863 | 0.467300779 | 0.998865404 | 10.667617 | 10.773309 | 10.695480 | 10.683864 | 10.842771 | 10.640956 | 1887 | 1742 | 1716 | 1431 |
| 4824 | NM_00125633 | NKX3-1 | NK3 homeobox 1 | protein_coding | NP_001243268.1;NP_006158.2 | 7838 | 602041 | . | . | 0.759402 | 0.824633 | 0.566257 | 0.534184 | 1.767159 | 1.906177 | 1.270664 | 1.221349 | -1.391670 | 67.597480 | 0.387624795 | -1.23010012 | 0.218659617 | 0.982008277 | 6.297753 | 5.820801 | 6.093822 | 6.137956 | 6.047744 | 6.032247 | 83 | 92 | 51 | 51 |
| 4825 | NM_006168 | NKX6-1 | NK6 homeobox 1 | protein_coding | NP_006159.2 | 7839 | 602563 | . | . | 2.653006 | 2.882553 | 2.118031 | 3.271845 | 6.173653 | 6.663151 | 4.752799 | 7.480684 | -1.041750 | 95.367998 | 0.346674899 | -0.17021478 | 0.864841232 | 0.998865404 | 6.605901 | 6.544308 | 6.567995 | 6.590612 | 6.505028 | 6.625869 | 108 | 109 | 64 | 106 |
| 4828 | NM_021077,N | NMB | neuromedin B | protein_coding | NP_066563.2;NP_995580.1;XP_016877728.1 | 7842 | 162340 | . | . | 2.967553 | 2.949690 | 3.887772 | 2.997532 | 6.905619 | 6.818344 | 8.724048 | 6.853502 | 1.151018 | 68.015778 | 0.386709376 | 0.524709974 | 0.599784822 | 0.998865404 | 5.982023 | 6.186344 | 6.067175 | 6.066022 | 6.128111 | 6.069874 | 73 | 68 | 71 | 60 |
| 4830 | NM_000269,N | NME1 | NME/NM23 nucleoside diphosphate kinase 1 | protein_coding | NP_000260.1;NP_937818.1 | 7849 | 156490 | . | . | 142.353695 | 143.571991 | 130.003289 | 153.611061 | 331.262879 | 331.873078 | 291.723587 | 351.213425 | -1.012336 | 4250.222108 | 0.119865966 | -0.14757071 | 0.882681569 | 0.998865404 | 12.062230 | 12.044360 | 12.052278 | 12.062907 | 11.975251 | 12.114702 | 4893 | 4644 | 3365 | 4292 |
| 4831 | NM_00101813 | NME2 | NME/NM23 nucleoside diphosphate kinase 2 | protein_coding | NP_001018147.1;NP_001018148.1;NP_001018149.1;NP_001185611.1;NI | 7850 | 156491 | . | . | 425.905985 | 410.983180 | 398.637492 | 473.695942 | 991.100714 | 950.006017 | 894.531042 | 1083.049506 | 1.027264 | 9184.142239 | 0.114922511 | 0.337678624 | 0.735605385 | 0.998865404 | 13.145428 | 13.184170 | 13.171151 | 13.133597 | 13.104706 | 13.241590 | 10664 | 9561 | 7445 | 9425 |
| 4832 | NM_002513,XI | NME3 | NME/NM23 nucleoside diphosphate kinase 3 | protein_coding | NP_002504.2;XP_005255389.1;XP_011520805.1;XP_011520806.1 | 7851 | 601817 | . | . | 14.341942 | 13.390545 | 15.166566 | 16.648553 | 33.374286 | 30.952841 | 34.033335 | 38.064936 | 1.130044 | 475.884694 | 0.173208131 | 1.0183032 | 0.308533888 | 0.998865404 | 8.803646 | 8.979910 | 8.877706 | 8.828987 | 8.900460 | 8.958698 | 535 | 463 | 419 | 495 |
| 4833 | NM_00128643 | NME4 | NME/NM23 nucleoside diphosphate kinase 4 | protein_coding | NP_001273362.1;NP_001273364.1;NP_001273365.1;NP_001273367.1;NI | 7852 | 601818 | . | . | 72.404060 | 76.774522 | 64.964416 | 80.243521 | 168.487216 | 177.467744 | 145.778261 | 183.467280 | -1.029294 | 2408.370468 | 0.135122426 | -0.30827846 | 0.757870451 | 0.998865404 | 11.254702 | 11.212675 | 11.224510 | 11.262883 | 11.125315 | 11.307555 | 2747 | 2699 | 1830 | 2477 |
| 4835 | NM_000904,N | NQO2 | N-ribosyldihydronicotinamide:quinone reductase 2 | protein_coding | NP_000895.2;NP_001277150.1;NP_001277151.1;NP_001305869.1 | 7856 | 160998 | . | . | 38.739272 | 40.963946 | 43.698978 | 51.410643 | 90.147886 | 94.689994 | 98.059228 | 117.544322 | 1.167403 | 1505.508876 | 0.134223784 | 1.66366357 | 0.09617968 | 0.813553488 | 10.440290 | 10.663180 | 10.475062 | 10.508395 | 10.547523 | 10.671613 | 1565 | 1532 | 1299 | 1640 |
| 4836 | NM_021079,XI | NMT1 | N-myristoyltransferase 1 | protein_coding | NP_066565.1 | 7857 | 160993 | . | . | 24.862472 | 25.842571 | 24.279439 | 28.937410 | 57.855999 | 59.736265 | 54.482355 | 66.161952 | 1.033506 | 3979.280290 | 0.119867551 | 0.396661271 | 0.691617263 | 0.998865404 | 11.934422 | 11.981773 | 11.931105 | 11.956072 | 11.901738 | 12.035785 | 4439 | 4287 | 3234 | 4097 |
| 4837 | NM_00137204 | NNMT | nicotinamide N-methyltransferase | protein_coding | NP_001358974.1;NP_001358975.1;NP_001358976.1;NP_006160.1 | 7861 | 600008 | . | . | 6.446452 | 5.450430 | 3.952299 | 6.000217 | 15.001157 | 12.598915 | 8.868843 | 13.718784 | -1.212698 | 344.116236 | 0.225683087 | -1.2327943 | 0.217652517 | 0.982008277 | 8.559671 | 8.280333 | 8.524678 | 8.421632 | 8.254376 | 8.465561 | 472 | 372 | 217 | 349 |
| 4839 | NM_00103371 | NOP2 | NOP2 nucleolar protein | protein_coding | NP_001028886.1;NP_001245237.1;NP_001245238.1;NP_001245239.1;NI | 7867 | 164031 | . | . | 29.095559 | 29.747710 | 32.402759 | 31.540971 | 67.706559 | 68.763161 | 72.710848 | 72.114680 | 1.074596 | 2568.337889 | 0.113874519 | 0.911483857 | 0.362040487 | 0.998865404 | 11.273770 | 11.377601 | 11.288914 | 11.304208 | 11.368908 | 11.340603 | 2824 | 2697 | 2354 | 2451 |
| 4841 | NM_00114540 | NONO | non-POU domain containing octamer binding | protein_coding | NP_001138880.1;NP_001138881.1;NP_001138882.1;NP_031389.3 | 7871 | 300084 | . | . | 103.926637 | 113.051064 | 106.279826 | 117.670395 | 241.841545 | 261.322586 | 238.488852 | 269.039366 | 1.017209 | 9159.069990 | 0.108466159 | 0.226949275 | 0.820463186 | 0.998865404 | 13.148666 | 13.173200 | 13.120583 | 13.184970 | 13.128941 | 13.205164 | 10061 | 10170 | 7673 | 9047 |
| 4846 | NM_000603,N | NOS3 | nitric oxide synthase 3 | protein_coding | NP_000594.2;NP_001153581.1;NP_001153582.1;NP_001153583.1 | 7876 | 163729 | . | . | 0.049435 | 0.081430 | 0.048539 | 0.061394 | 0.115038 | 0.188229 | 0.108919 | 0.140369 | -1.167858 | 8.375414 | 1.056054559 | -0.21198226 | 0.832120873 | NA | 3.175403 | 2.947964 | 3.060668 | 3.079821 | 3.060937 | 3.068030 | 8 | 12 | 6 | 8 |
| 4848 | NM_00119930 | CNOT2 | CCR4-NOT transcription complex subunit 2 | protein_coding | NP_001186231.1;NP_001186232.1;NP_055330.1;XP_006719492.1;XP_00 | 7878 | 604909 | . | . | 10.938033 | 10.823185 | 11.943225 | 10.177944 | 25.453248 | 25.018277 | 26.800256 | 23.270660 | 1.020674 | 1013.747308 | 0.139941627 | 0.210965225 | 0.832914407 | 0.998865404 | 9.970499 | 10.000312 | 9.968839 | 9.985734 | 10.032255 | 9.950995 | 1142 | 1095 | 940 | 907 |
| 4849 | NM_014516,XI | CNOT3 | CCR4-NOT transcription complex subunit 3 | protein_coding | NP_055331.1;XP_005278336.1;XP_005278337.1;XP_005278338.1;XP_00 | 7879 | 604910 | . | . | 14.930193 | 15.715456 | 15.684424 | 15.403645 | 34.743173 | 36.326979 | 35.195395 | 35.218605 | 1.008776 | 1387.682431 | 0.128470762 | 0.098126516 | 0.921831833 | 0.998865404 | 10.432121 | 10.444774 | 10.413924 | 10.455338 | 10.462766 | 10.419883 | 1548 | 1531 | 1246 | 1270 |
| 4850 | NM_00100822 | CNOT4 | CCR4-NOT transcription complex subunit 4 | protein_coding | NP_001008226.1;NP_001177776.1;NP_001177777.1;NP_001177778.1;NI | 7880 | 604911 | . | . | 5.673752 | 6.174753 | 6.637159 | 7.239441 | 13.203056 | 14.273218 | 14.893591 | 16.552122 | 1.141713 | 699.344578 | 0.153589665 | 1.244876128 | 0.213177287 | 0.97614175 | 9.351356 | 9.542065 | 9.373008 | 9.429059 | 9.466776 | 9.517005 | 721 | 734 | 625 | 724 |
| 4851 | NM_017617,XI | NOTCH1 | notch receptor 1 | protein_coding | NP_060087.3;XP_011517019.2 | 7881 | 190198 | . | . | 0.794974 | 0.894994 | 0.601633 | 0.521528 | 1.849937 | 2.068818 | 1.350047 | 1.192411 | -1.524746 | 210.128822 | 0.23855784 | -2.55103247 | **0.010740432** | 0.333896806 | 7.987319 | 7.379361 | 7.771246 | 7.827928 | 7.638114 | 7.575114 | 276 | 289 | 156 | 144 |
| 4853 | NM_00120000 | NOTCH2 | notch receptor 2 | protein_coding | NP_001186930.1;NP_077719.2 | 7882 | 600275 | . | . | 32.908736 | 33.702222 | 29.779278 | 29.000879 | 76.579981 | 77.904205 | 66.823833 | 66.307063 | -1.149812 | 11144.764839 | 0.100992883 | -1.99417817 | **0.046132587** | 0.654058028 | 13.541258 | 13.339877 | 13.493768 | 13.507617 | 13.397297 | 13.365730 | 13617 | 12967 | 9185 | 9538 |
| 4854 | NM_000435,XI | NOTCH3 | notch receptor 3 | protein_coding | NP_000426.2;XP_005259981.1 | 7883 | 600276 | . | . | 0.187907 | 0.179063 | 0.095807 | 0.196200 | 0.437268 | 0.413912 | 0.214988 | 0.448588 | -1.265349 | 45.251690 | 0.497402952 | -0.68261739 | 0.494848643 | 0.998865404 | 5.661407 | 5.318005 | 5.525343 | 5.513770 | 5.427906 | 5.528684 | 60 | 53 | 23 | 50 |
| 4855 | NM_004557,N | NOTCH4 | notch receptor 4 | protein_coding | NP_004548.3 | 7884 | 164951 | . | . | 0.101422 | 0.102216 | 0.113879 | 0.089067 | 0.236014 | 0.236277 | 0.255542 | 0.203641 | -1.014884 | 21.825598 | 0.65868349 | -0.03235876 | 0.97418595 | NA | 4.457887 | 4.437942 | 4.446513 | 4.448790 | 4.456389 | 4.436086 | 25 | 24 | 21 | 18 |
| 4856 | NM_002514 | CCN3 | cellular communication network factor 3 | protein_coding | NP_002505.1 | 7885 | 164958 | . | . | 4.182945 | 4.741254 | 3.738711 | 3.224257 | 9.733886 | 10.959621 | 8.389560 | 7.371881 | -1.298068 | 305.402948 | 0.207121199 | -1.81712672 | 0.069197712 | 0.743875399 | 8.430316 | 8.054406 | 8.284818 | 8.352027 | 8.216607 | 8.138743 | 374 | 394 | 249 | 230 |
| 4857 | NM_00136639 | NOVA1 | NOVA alternative splicing regulator 1 | protein_coding | NP_001353319.1;NP_001353320.1;NP_001353321.1;NP_001353322.1;NI | 7886 | 602157 | . | . | 0.226205 | 0.120095 | 0.575106 | 0.240868 | 0.526389 | 0.277607 | 1.290522 | 0.550715 | **2.056376** | 31.129882 | 0.59887538 | 1.736761948 | 0.082429198 | 0.78412713 | 4.349460 | 5.387998 | 4.854646 | 4.912227 | 5.008151 | 4.933451 | 16 | 29 | 45 | 30 |
| 4860 | NM_000270 | PNP | purine nucleoside phosphorylase | protein_coding | NP_000261.2 | 7892 | 164050 | . | . | 5.296929 | 4.741475 | 5.591940 | 6.931152 | 12.326172 | 10.960132 | 12.548152 | 15.847256 | 1.229976 | 259.900941 | 0.22229669 | 1.343386288 | 0.179146924 | 0.951511445 | 7.865040 | 8.163216 | 7.990175 | 7.934106 | 8.012338 | 8.119869 | 284 | 237 | 224 | 296 |
| 4861 | NM_00132108 | NPAS1 | neuronal PAS domain protein 1 | protein_coding | NP_001308015.1;NP_002508.2;XP_016882333.1;XP_024307292.1;XP_02 | 7894 | 603346 | . | . | 22.695773 | 20.419444 | 10.792282 | 13.691739 | 52.813998 | 47.200460 | 24.217566 | 31.304533 | -1.788207 | 987.105411 | 0.164160239 | -5.107897 | **3.25764E-07** | **0.000124903** | 10.306509 | 9.467236 | 10.159985 | 10.068824 | 9.594018 | 9.783165 | 1542 | 1288 | 527 | 759 |
| 4862 | NM_002518,XI | NPAS2 | neuronal PAS domain protein 2 | protein_coding | NP_002509.2;XP_005264010.1;XP_005264016.1;XP_005264017.1;XP_00 | 7895 | 603347 | . | . | 29.432884 | 32.912608 | 37.621474 | 35.372721 | 68.491530 | 76.078969 | 84.421497 | 80.875519 | 1.152640 | 4251.156634 | 0.117342847 | 1.746521137 | 0.080720437 | 0.780241035 | 11.947536 | 12.152472 | 11.940773 | 12.038460 | 12.146174 | 12.070378 | 4278 | 4515 | 4137 | 4076 |
| 4863 | NM_00132130 | NPAT | nuclear protein, coactivator of histone transcription | protein_coding | NP_001308236.1;NP_002510.2;XP_011541156.1;XP_016873291.1;XP_01 | 7896 | 601448 | . | . | 6.158887 | 6.348928 | 8.753669 | 6.588963 | 14.331985 | 14.675832 | 19.642976 | 15.064881 | 1.181579 | 1367.594205 | 0.153579516 | 1.567368474 | 0.117028588 | 0.857396579 | 10.291774 | 10.533003 | 10.346235 | 10.346792 | 10.580794 | 10.355618 | 1443 | 1353 | 1474 | 1189 |
| 4864 | NM_000271,XI | NPC1 | NPC intracellular cholesterol transporter 1 | protein_coding | NP_000262.2;XP_005258334.1;XP_005258335.1;XP_005258336.1;XP_00 | 7897 | 607623 | . | . | 29.353492 | 29.914764 | 37.102067 | 35.213706 | 68.306784 | 69.149309 | 83.255962 | 80.511954 | 1.201410 | 4924.936574 | 0.10761507 | 2.459957348 | **0.013895353** | 0.390841055 | 12.127428 | 12.392218 | 12.181482 | 12.186332 | 12.365488 | 12.308632 | 5137 | 4842 | 4831 | 4871 |
| 4867 | NM_000272,N | NPHP1 | nephrocystin 1 | protein_coding | NP_000263.2;NP_001121650.1;NP_001121651.1;NP_001361185.1;NP_00 | 7905 | 607100 | . | . | 3.424278 | 3.157746 | 3.570641 | 3.260275 | 7.968438 | 7.299269 | 8.012415 | 7.454232 | -1.034036 | 263.211301 | 0.212357614 | -0.22738276 | 0.820126125 | 0.998865404 | 8.063758 | 8.016002 | 8.053325 | 8.043478 | 8.053207 | 8.009152 | 311 | 286 | 235 | 232 |
| 4869 | NM_00103773 | NPM1 | nucleophosmin 1 | protein_coding | NP_001032827.1;NP_001341935.1;NP_001341936.1;NP_001341938.1;NI | 7910 | 164040 | . | . | 774.161711 | 724.256235 | 684.644832 | 706.716542 | 1801.506089 | 1674.150703 | 1536.323157 | 1615.823456 | -1.074430 | 33248.978063 | 0.100418168 | -1.03140562 | 0.302350645 | 0.998865404 | 15.071879 | 14.968307 | 15.076184 | 15.025328 | 14.970793 | 15.007122 | 40821 | 36076 | 27285 | 30689 |
| 4880 | NM_024409,XI | NPPC | natriuretic peptide C | protein_coding | NP_077720.1;XP_011509547.1 | 7941 | 600296 | . | . | 3.711176 | 3.545375 | 3.866540 | 3.793281 | 8.636060 | 8.195293 | 8.676405 | 8.672887 | 1.037110 | 145.812447 | 0.270646084 | 0.194235237 | 0.845991693 | 0.998865404 | 7.161400 | 7.214060 | 7.168683 | 7.191331 | 7.205401 | 7.182518 | 160 | 159 | 133 | 135 |
| 4882 | NM_00137892 | NPR2 | natriuretic peptide receptor 2 | protein_coding | NP_001365852.1;NP_003986.2;XP_024303324.1;XP_024303325.1;XP_02 | 7944 | 108961 | . | . | 7.039013 | 7.310071 | 7.696219 | 7.328772 | 16.380071 | 16.897557 | 17.270093 | 16.756366 | 1.015114 | 964.417076 | 0.137738667 | 0.157121204 | 0.875149327 | 0.998865404 | 9.902644 | 9.924302 | 9.897105 | 9.918560 | 9.928283 | 9.909271 | 1086 | 1048 | 856 | 899 |
| 4884 | NM_002522 | NPTX1 | neuronal pentraxin 1 | protein_coding | NP_002513.2 | 7952 | 602367 | . | . | 0.120935 | 0.100349 | 0.244729 | 0.112173 | 0.281420 | 0.231962 | 0.549165 | 0.256469 | 1.571123 | 24.762460 | 0.65558755 | 0.994217049 | 0.320117209 | NA | 4.264940 | 4.921224 | 4.593103 | 4.579211 | 4.675198 | 4.586953 | 24 | 19 | 36 | 18 |
| 4885 | NM_002523 | NPTX2 | neuronal pentraxin 2 | protein_coding | NP_002514.1 | 7953 | 600750 | . | . | 23.010345 | 22.570757 | 17.641247 | 20.130178 | 53.546021 | 52.173313 | 39.586449 | 46.025265 | -1.223594 | 1762.526623 | 0.126636709 | -2.29889941 | **0.021510652** | 0.485943585 | 10.921775 | 10.630396 | 10.868343 | 10.845710 | 10.651337 | 10.743677 | 2265 | 2064 | 1295 | 1576 |
| 4891 | NM_000617,N | SLC11A2 | solute carrier family 11 member 2 | protein_coding | NP_000608.1;NP_001167596.1;NP_001167597.1;NP_001167598.1;NP_00 | 10908 | 600523 | . | . | 14.725849 | 14.700042 | 17.523293 | 14.654219 | 34.267656 | 33.979798 | 39.321763 | 33.505132 | 1.083481 | 1908.993660 | 0.134025005 | 0.863071859 | 0.388097959 | 0.998865404 | 10.839410 | 10.955451 | 10.870793 | 10.858032 | 11.004198 | 10.845810 | 2127 | 1961 | 1898 | 1677 |
| 4893 | NM_002524 | NRAS | NRAS proto-oncogene, GTPase | protein_coding | NP_002515.1 | 7989 | 164790 | . | . | 20.202280 | 17.563131 | 17.122435 | 17.743740 | 47.011536 | 40.597965 | 38.422249 | 40.568951 | -1.098925 | 2448.322296 | 0.124068546 | -1.09691607 | 0.272678084 | 0.998865404 | 11.323962 | 11.187991 | 11.352750 | 11.231330 | 11.207574 | 11.227768 | 3171 | 2561 | 2003 | 2215 |
| 4897 | NM_00103713 | NRCAM | neuronal cell adhesion molecule | protein_coding | NP_001032209.1;NP_001180511.1;NP_001180512.1;NP_001180513.1;NI | 7994 | 601581 | . | . | 8.121927 | 7.538165 | 6.535867 | 6.219597 | 18.900059 | 17.424806 | 14.666292 | 14.220366 | -1.235421 | 1412.358916 | 0.129618033 | -2.35308751 | **0.018618252** | 0.458012736 | 10.608207 | 10.303517 | 10.568174 | 10.508853 | 10.400717 | 10.353253 | 1865 | 1621 | 1134 | 1147 |
| 4898 | NM_00110166 | NRDC | nardilysin convertase | protein_coding | NP_001095132.1;NP_001229290.1;NP_002516.2 | 7995 | 602651 | . | . | 33.759469 | 35.303465 | 34.450271 | 33.216956 | 78.559668 | 81.605537 | 77.305408 | 75.946621 | -1.031488 | 3892.625046 | 0.109864973 | -0.40710606 | 0.683930095 | 0.998865404 | 11.948696 | 11.904013 | 11.924241 | 11.953750 | 11.936559 | 11.889773 | 4471 | 4341 | 3427 | 3491 |
| 4899 | NM_00104011 | NRF1 | nuclear respiratory factor 1 | protein_coding | NP_001035199.1;NP_001280092.1;NP_001280093.1;NP_005002.3 | 7996 | 600879 | . | . | 2.462835 | 2.630727 | 2.587541 | 2.238306 | 5.731121 | 6.081044 | 5.806366 | 5.117621 | -1.075464 | 272.838176 | 0.212083789 | -0.49489764 | 0.620672379 | 0.998865404 | 8.143193 | 8.038719 | 8.094257 | 8.125358 | 8.105568 | 8.038012 | 316 | 314 | 244 | 230 |
| 4900 | NM_00112618 | NRGN | neurogranin | protein_coding | NP_001119653.1;NP_006167.1 | 8000 | 602350 | . | . | 5.223546 | 5.548899 | 5.143662 | 7.021254 | 12.155406 | 12.826529 | 11.542228 | 16.053261 | 1.115188 | 217.233807 | 0.242881686 | 0.647588117 | 0.517251385 | 0.998865404 | 7.683133 | 7.838875 | 7.720315 | 7.746493 | 7.712272 | 7.852090 | 230 | 228 | 170 | 246 |
| 4901 | NM_00135476 | NRL | neural retina leucine zipper | protein_coding | NP_001341697.1;NP_001341698.1;NP_001341699.1;NP_006168.1;XP_00 | 8002 | 162080 | . | . | 0.110721 | 0.062856 | 0.050861 | 0.122482 | 0.257652 | 0.145295 | 0.114131 | 0.280042 | 1.012062 | 12.489399 | 0.902737115 | 0.019161754 | 0.984712068 | NA | 3.634657 | 3.650563 | 3.660269 | 3.623455 | 3.623248 | 3.661788 | 19 | 9 | 7 | 16 |
| 4902 | NM_004558,XI | NRTN | neurturin | protein_coding | NP_004549.1;XP_011526343.1 | 8007 | 602018 | . | . | 0.907144 | 0.964578 | 0.642642 | 1.171759 | 2.110962 | 2.229665 | 1.442070 | 2.679087 | -1.067113 | 33.471240 | 0.552477472 | -0.1696238 | 0.865306004 | 0.998865404 | 5.112745 | 5.015311 | 5.058781 | 5.078727 | 5.027842 | 5.091553 | 37 | 40 | 21 | 38 |
| 4904 | NM_004559,N | YBX1 | Y-box binding protein 1 | protein_coding | NP_004550.2 | 8014 | 154030 | . | . | 245.645480 | 253.392073 | 234.146972 | 283.610487 | 571.627123 | 585.727194 | 525.419023 | 648.441639 | 1.022641 | 23576.817241 | 0.114844977 | 0.281241837 | 0.778524918 | 0.998865404 | 14.508861 | 14.541121 | 14.504671 | 14.523956 | 14.452499 | 14.608855 | 26544 | 25438 | 18857 | 24371 |
| 4905 | NM_006178,N | NSF | N-ethylmaleimide sensitive factor, vesicle fusing ATPase | protein_coding | NP_006169.2 | 8016 | 601633 | . | . | 9.696447 | 9.403460 | 9.659952 | 10.346784 | 22.564031 | 21.736519 | 21.676650 | 23.656692 | 1.032901 | 1213.622874 | 0.131693145 | 0.354630567 | 0.722866374 | 0.998865404 | 10.221608 | 10.268225 | 10.247312 | 10.217649 | 10.234620 | 10.278793 | 1402 | 1263 | 1041 | 1190 |
| 4907 | NM_00120481 | NT5E | 5'-nucleotidase ecto | protein_coding | NP_001191742.1;NP_002517.1 | 8021 | 129190 | . | . | 25.271841 | 25.217464 | 20.180098 | 17.577945 | 58.808607 | 58.291300 | 45.283556 | 40.189880 | -1.356984 | 2470.558033 | 0.12214008 | -3.60572362 | **0.000311284** | **0.028454684** | 11.473963 | 11.033842 | 11.385497 | 11.375396 | 11.186317 | 11.079471 | 3295 | 3051 | 1948 | 1831 |
| 4913 | NM_00131819 | NTHL1 | nth like DNA glycosylase 1 | protein_coding | NP_001305122.2;NP_001305123.1;NP_002519.2 | 8028 | 602656 | . | . | 9.596641 | 10.713328 | 9.242519 | 11.512595 | 22.331776 | 24.764337 | 20.739946 | 26.322180 | 1.011804 | 327.778226 | 0.205737392 | 0.082290691 | 0.934415557 | 0.998865404 | 8.348734 | 8.364376 | 8.324011 | 8.378851 | 8.295611 | 8.417046 | 357 | 369 | 255 | 343 |
| 4914 | NM_00100779 | NTRK1 | neurotrophic receptor tyrosine kinase 1 | protein_coding | NP_001007793.1;NP_001012331.1;NP_002520.2 | 8031 | 191315 | . | . | 0.063941 | 0.112192 | 0.159163 | 0.104429 | 0.148793 | 0.259336 | 0.357157 | 0.238764 | 1.372391 | 9.651579 | 0.9949227 | 0.459021862 | 0.646218467 | NA | 3.024434 | 3.481107 | 3.242295 | 3.261510 | 3.276770 | 3.254074 | 7 | 11 | 12 | 8 |
| 4915 | NM_00100709 | NTRK2 | neurotrophic receptor tyrosine kinase 2 | protein_coding | NP_001007098.1;NP_001018074.1;NP_001018075.1;NP_001018076.1;NI | 8032 | 600456 | . | . | 0.213889 | 0.176826 | 0.127162 | 0.162686 | 0.497730 | 0.408742 | 0.285347 | 0.371962 | -1.398957 | 45.562668 | 0.473855225 | -1.02215035 | 0.306709747 | 0.998865404 | 5.732243 | 5.246648 | 5.561908 | 5.517561 | 5.461286 | 5.504287 | 68 | 51 | 28 | 41 |
| 4916 | NM_00100715 | NTRK3 | neurotrophic receptor tyrosine kinase 3 | protein_coding | NP_001007157.1;NP_001012338.1;NP_001230030.1;NP_001307063.1;NI | 8033 | 191316 | . | . | 0.063737 | 0.095199 | 0.185926 | 0.039355 | 0.148317 | 0.220055 | 0.417212 | 0.089981 | 1.076552 | 15.088927 | 0.829678009 | 0.128263881 | 0.897940148 | NA | 3.861598 | 3.967302 | 3.875754 | 3.928283 | 3.923568 | 3.890546 | 10 | 22 | 17 | 11 |
| 4919 | NM_00108359 | ROR1 | receptor tyrosine kinase like orphan receptor 1 | protein_coding | NP_001077061.1;NP_005003.2;XP_011539828.1;XP_016856865.1;XP_01 | 10256 | 602336 | . | . | 2.203661 | 1.970873 | 1.587175 | 2.132289 | 5.128010 | 4.555762 | 3.561576 | 4.875226 | -1.207707 | 304.764844 | 0.227292218 | -1.19788645 | 0.230961227 | 0.986164394 | 8.381754 | 8.108424 | 8.343629 | 8.248543 | 8.102030 | 8.280458 | 416 | 330 | 198 | 304 |
| 4920 | NM_00131820 | ROR2 | receptor tyrosine kinase like orphan receptor 2 | protein_coding | NP_001305133.1;NP_004551.2;XP_005252065.1;XP_005252066.1;XP_00 | 10257 | 602337 | . | . | 0.156159 | 0.191256 | 0.075614 | 0.192721 | 0.363389 | 0.442097 | 0.169676 | 0.440635 | -1.280374 | 28.001539 | 0.633004779 | -0.56328948 | 0.57323778 | 0.998865404 | 4.977427 | 4.614701 | 4.805401 | 4.835825 | 4.748716 | 4.834821 | 32 | 38 | 12 | 33 |
| 4921 | NM_00101479 | DDR2 | discoidin domain receptor tyrosine kinase 2 | protein_coding | NP_001014796.1;NP_001341911.1;NP_001341912.1;NP_006173.2;XP_01 | 2731 | 191311 | . | . | 0.231067 | 0.380712 | 0.293853 | 0.353258 | 0.537703 | 0.880033 | 0.659396 | 0.807683 | -1.119708 | 95.687566 | 0.364842119 | -0.44710521 | 0.654799095 | 0.998865404 | 6.661079 | 6.494643 | 6.513036 | 6.661897 | 6.507938 | 6.603287 | 89 | 135 | 65 | 99 |
| 4922 | NM_006183 | NTS | neurotensin | protein_coding | NP_006174.1 | 8038 | 162650 | . | . | 0.528239 | 0.551238 | 0.435048 | 0.157585 | 1.229234 | 1.274212 | 0.976234 | 0.360300 | -1.830492 | 16.683525 | 0.795515813 | -1.09643501 | 0.272888451 | NA | 4.429096 | 3.563745 | 4.079886 | 4.085609 | 4.063945 | 4.017226 | 24 | 24 | 15 | 6 |
| 4924 | NM_006184,XI | NUCB1 | nucleobindin 1 | protein_coding | NP_006175.2;XP_016882334.1 | 8043 | 601323 | . | . | 88.524293 | 92.868823 | 79.739426 | 88.068057 | 205.999647 | 214.670446 | 178.932968 | 201.357155 | -1.087992 | 6592.310120 | 0.107533371 | -1.13144671 | 0.25786712 | 0.995760118 | 12.746165 | 12.624403 | 12.702803 | 12.738765 | 12.614834 | 12.683396 | 7744 | 7570 | 5266 | 6162 |
| 4925 | NM_00133022 | NUCB2 | nucleobindin 2 | protein_coding | NP_001317156.1;NP_001339590.1;NP_001339591.1;NP_001339592.1;NI | 8044 | 608020 | . | . | 37.597681 | 34.598127 | 31.260181 | 25.304915 | 87.491348 | 79.975117 | 70.146946 | 57.856679 | -1.293998 | 2030.730106 | 0.138619754 | -2.68241549 | **0.007309262** | 0.269034057 | 11.161546 | 10.790188 | 11.102266 | 11.056540 | 10.960785 | 10.781011 | 2711 | 2403 | 1719 | 1467 |
| 4926 | NM_00128656 | NUMA1 | nuclear mitotic apparatus protein 1 | protein_coding | NP_001273490.1;NP_006176.2;XP_006718627.1;XP_011543356.1;XP_01 | 8059 | 164009 | . | . | 18.742251 | 20.272692 | 20.734087 | 22.411681 | 43.613981 | 46.861236 | 46.526689 | 51.241649 | 1.083880 | 4559.208733 | 0.110189946 | 1.054584023 | 0.291615614 | 0.998865404 | 12.095359 | 12.211442 | 12.091788 | 12.146980 | 12.160881 | 12.212213 | 4871 | 4879 | 3998 | 4582 |
| 4927 | NM_00132065 | NUP88 | nucleoporin 88 | protein_coding | NP_001307582.1;NP_002523.2;XP_011522195.1;XP_016880183.1 | 8067 | 602552 | . | . | 16.087089 | 17.426956 | 14.216782 | 16.196666 | 37.435318 | 40.283189 | 31.902050 | 37.031749 | -1.128241 | 1778.168656 | 0.132465872 | -1.3141166 | 0.188806969 | 0.960264078 | 10.880798 | 10.706280 | 10.806103 | 10.874724 | 10.684154 | 10.804246 | 2078 | 2122 | 1340 | 1688 |
| 4928 | NM_00136512 | NUP98 | nucleoporin 98 and 96 precursor | protein_coding | NP_001352054.1;NP_001352055.1;NP_001352056.1;NP_001352057.1;NI | 8068 | 601021 | . | . | 23.763376 | 25.210858 | 23.727861 | 26.332270 | 55.298352 | 58.276031 | 53.244631 | 60.205606 | 1.005828 | 4773.880517 | 0.112467428 | 0.074545701 | 0.940576179 | 0.998865404 | 12.216822 | 12.225059 | 12.193837 | 12.241860 | 12.179463 | 12.264624 | 5324 | 5284 | 3951 | 4717 |
| 4929 | NM_006186,N | NR4A2 | nuclear receptor subfamily 4 group A member 2 | protein_coding | NP_006177.1;NP_775265.1;XP_005246678.1;XP_006712616.1;XP_01150 | 7981 | 601828 | . | . | 1.429849 | 1.513192 | 1.551983 | 1.637848 | 3.327316 | 3.497809 | 3.482609 | 3.744747 | 1.057435 | 161.986890 | 0.25803527 | 0.312242311 | 0.754856378 | 0.998865404 | 7.299061 | 7.379291 | 7.317694 | 7.337183 | 7.345043 | 7.355146 | 177 | 174 | 143 | 158 |
| 4931 | NM_00124314 | NVL | nuclear VCP like | protein_coding | NP_001230075.1;NP_001230076.1;NP_002524.2;NP_996671.1;XP_0115 | 8070 | 602426 | . | . | 7.745398 | 7.786436 | 10.795063 | 8.475113 | 18.023863 | 17.998699 | 24.223808 | 19.377338 | 1.182092 | 817.372256 | 0.156892864 | 1.538263591 | 0.123984178 | 0.871805202 | 9.548992 | 9.790603 | 9.572417 | 9.645931 | 9.791828 | 9.660755 | 817 | 851 | 840 | 755 |
| 4935 | NM_000273,XI | GPR143 | G protein-coupled receptor 143 | protein_coding | NP_000264.2;XP_005274598.1;XP_024308155.1;XP_024308156.1 | 20145 | 300808 | . | . | 59.728916 | 56.473133 | 77.815922 | 68.574950 | 138.991640 | 130.540175 | 174.616676 | 156.788474 | 1.251484 | 3318.591399 | 0.120046198 | 2.695965371 | **0.007018496** | 0.266143692 | 11.525350 | 11.849222 | 11.626153 | 11.564663 | 11.833563 | 11.724384 | 3524 | 3059 | 3417 | 3234 |
| 4938 | NM_00103240 | OAS1 | 2'-5'-oligoadenylate synthetase 1 | protein_coding | NP_001027581.1;NP_001307080.1;NP_002525.2;NP_058132.2;XP_00671 | 8086 | 164350 | . | . | 6.139542 | 6.652743 | 5.169289 | 6.313920 | 14.286967 | 15.378115 | 11.599735 | 14.436025 | -1.188141 | 334.389906 | 0.21158143 | -1.17546408 | 0.2398091 | 0.992733415 | 8.505114 | 8.254818 | 8.397913 | 8.460332 | 8.253058 | 8.407295 | 396 | 413 | 227 | 328 |
| 4939 | NM_00103273 | OAS2 | 2'-5'-oligoadenylate synthetase 2 | protein_coding | NP_001027903.1;NP_002526.2;NP_058197.2 | 8087 | 603350 | . | . | 0.220401 | 0.359030 | 0.202072 | 0.328938 | 0.512882 | 0.829913 | 0.453445 | 0.752078 | -1.245796 | 31.364953 | 0.60705652 | -0.52230447 | 0.601458359 | 0.998865404 | 5.123643 | 4.800458 | 4.935914 | 5.021167 | 4.914003 | 4.988795 | 28 | 49 | 16 | 35 |
| 4940 | NM_006187,XI | OAS3 | 2'-5'-oligoadenylate synthetase 3 | protein_coding | NP_006178.2;XP_005253946.1;XP_016874852.1 | 8088 | 603351 | . | . | 3.474887 | 3.880551 | 3.167311 | 3.976436 | 8.086207 | 8.970067 | 7.107353 | 9.091649 | -1.042042 | 746.459698 | 0.163080701 | -0.36431908 | 0.715619754 | 0.998865404 | 9.573760 | 9.513450 | 9.520354 | 9.591821 | 9.452082 | 9.598364 | 833 | 864 | 568 | 758 |
| 4942 | NM_000274,N | OAT | ornithine aminotransferase | protein_coding | NP_000265.1;NP_001165285.1;NP_001309894.1;NP_001309895.1;NP_00 | 8091 | 613349 | . | . | 51.362459 | 46.836641 | 51.809032 | 77.487104 | 119.522554 | 108.264999 | 116.257960 | 177.165057 | 1.028807 | 4998.580586 | 0.10891868 | 0.376172091 | 0.706788963 | 0.998865404 | 12.266659 | 12.307655 | 12.307105 | 12.242005 | 12.284586 | 12.312915 | 5895 | 5110 | 4334 | 4833 |
| 4943 | NM_00134826 | TBC1D25 | TBC1 domain family member 25 | protein_coding | NP_001335191.1;NP_001335192.1;NP_001335193.1;NP_001335194.1;NI | 8092 | 311240 | . | . | 2.956286 | 3.081647 | 3.099218 | 3.250962 | 6.879399 | 7.123364 | 6.954557 | 7.432940 | 1.058809 | 369.464567 | 0.185943256 | 0.443375132 | 0.65749442 | 0.998865404 | 8.487639 | 8.569777 | 8.497384 | 8.526311 | 8.537100 | 8.552760 | 403 | 397 | 326 | 361 |
| 4946 | NM_00130102 | OAZ1 | ornithine decarboxylase antizyme 1 | protein_coding | NP_001287949.1;NP_004143.1 | 8095 | 601579 | . | . | 223.008421 | 227.976570 | 214.957256 | 264.840013 | 518.949794 | 526.978029 | 482.357872 | 605.525206 | 1.048822 | 8555.896225 | 0.120072473 | 0.572737453 | 0.566822474 | 0.998865404 | 13.027961 | 13.096628 | 13.035113 | 13.046673 | 12.993268 | 13.162586 | 9552 | 9074 | 6863 | 9025 |
| 4947 | NM_00130130 | OAZ2 | ornithine decarboxylase antizyme 2 | protein_coding | NP_001288231.1;NP_002528.1 | 8096 | 604152 | . | . | 25.157524 | 25.613753 | 23.304136 | 29.226620 | 58.542592 | 59.207340 | 52.293807 | 66.823193 | 1.020232 | 1556.008787 | 0.139894289 | 0.206568775 | 0.836346648 | 0.998865404 | 10.589339 | 10.617790 | 10.590942 | 10.598009 | 10.523200 | 10.689848 | 1766 | 1670 | 1220 | 1631 |
| 4948 | NM_000275,N | OCA2 | OCA2 melanosomal transmembrane protein | protein_coding | NP_000266.2;NP_001287913.1;XP_011519942.1;XP_016877744.1;XP_01 | 8101 | 611409 | . | . | 8.310234 | 7.514088 | 9.101281 | 7.482713 | 19.338261 | 17.369150 | 20.423012 | 17.108333 | 1.015994 | 799.400469 | 0.160777599 | 0.142378159 | 0.886781313 | 0.998865404 | 9.630848 | 9.654604 | 9.678406 | 9.589968 | 9.714524 | 9.573681 | 970 | 802 | 771 | 680 |
| 4952 | NM_000276,N | OCRL | OCRL inositol polyphosphate-5-phosphatase | protein_coding | NP_000267.2;NP_001305713.1;NP_001578.2 | 8108 | 300535 | . | . | 4.658015 | 4.701512 | 4.833651 | 4.718691 | 10.839394 | 10.867756 | 10.846571 | 10.788725 | 1.005903 | 761.311727 | 0.146596785 | 0.057918759 | 0.953813341 | 0.998865404 | 9.568033 | 9.576642 | 9.569354 | 9.571007 | 9.585774 | 9.562746 | 873 | 820 | 675 | 704 |
| 4953 | NM_00128718 | ODC1 | ornithine decarboxylase 1 | protein_coding | NP_001274117.1;NP_001274118.1;NP_001274119.1;NP_002530.1 | 8109 | 165640 | . | . | 22.550393 | 24.289879 | 21.391883 | 23.967592 | 52.475688 | 56.147140 | 48.002767 | 54.799055 | -1.062153 | 1791.783121 | 0.124847793 | -0.69678457 | 0.485937623 | 0.998865404 | 10.850161 | 10.762879 | 10.815341 | 10.844921 | 10.742222 | 10.821609 | 2086 | 2029 | 1434 | 1712 |
| 4957 | NM_00124235 | ODF2 | outer dense fiber of sperm tails 2 | protein_coding | NP_001229281.1;NP_001229282.1;NP_001229283.1;NP_001338506.1;NI | 8114 | 602015 | . | . | 11.394093 | 12.605802 | 11.425167 | 12.329886 | 26.514523 | 29.138875 | 25.637741 | 28.190819 | -1.035616 | 1423.810308 | 0.13073099 | -0.38620788 | 0.699342716 | 0.998865404 | 10.500736 | 10.449899 | 10.452477 | 10.523358 | 10.434049 | 10.488306 | 1592 | 1635 | 1173 | 1358 |
| 4967 | NM_00100394 | OGDH | oxoglutarate dehydrogenase | protein_coding | NP_001003941.1;NP_001158508.1;NP_001350452.1;NP_002532.2;XP_00 | 8124 | 613022 | . | . | 19.467762 | 20.618286 | 20.786063 | 24.008760 | 45.302284 | 47.660094 | 46.643322 | 54.893179 | 1.112290 | 2727.241145 | 0.120175356 | 1.277575336 | 0.201399201 | 0.967111885 | 11.334545 | 11.487838 | 11.348929 | 11.386198 | 11.402103 | 11.503754 | 2905 | 2851 | 2343 | 2859 |
| 4968 | NM_00135464 | OGG1 | 8-oxoguanine DNA glycosylase | protein_coding | NP_001341577.1;NP_001341578.1;NP_001341579.1;NP_001341580.1;NI | 8125 | 601982 | . | . | 6.722780 | 6.284018 | 6.431560 | 7.366924 | 15.644186 | 14.525788 | 14.432236 | 16.843598 | 1.061198 | 376.901506 | 0.189766575 | 0.451575734 | 0.651574654 | 0.998865404 | 8.514709 | 8.600115 | 8.565079 | 8.513212 | 8.534333 | 8.612203 | 440 | 377 | 315 | 388 |
| 4974 | NM_002544 | OMG | oligodendrocyte myelin glycoprotein | protein_coding | NP_002535.3 | 8135 | 164345 | . | . | 0.870303 | 0.703361 | 1.079987 | 0.795124 | 2.025232 | 1.625850 | 2.423460 | 1.817956 | 1.156487 | 48.220809 | 0.459980183 | 0.455995781 | 0.648393012 | 0.998865404 | 5.481415 | 5.693933 | 5.591265 | 5.554092 | 5.627322 | 5.569684 | 57 | 43 | 52 | 41 |
| 4976 | NM_00135466 | OPA1 | OPA1 mitochondrial dynamin like GTPase | protein_coding | NP_001341592.1;NP_001341593.1;NP_056375.2;NP_570844.1;NP_5708 | 8140 | 605290 | . | . | 20.948492 | 20.150633 | 16.845826 | 17.955825 | 48.748001 | 46.579087 | 37.801544 | 41.053858 | -1.195576 | 3761.280057 | 0.110296093 | -2.3364896 | **0.019465739** | 0.462516353 | 12.000128 | 11.742392 | 11.965840 | 11.926449 | 11.778992 | 11.817774 | 4827 | 4317 | 2908 | 3289 |
| 4983 | NM_002547,XI | OPHN1 | oligophrenin 1 | protein_coding | NP_002538.1;XP_005262327.1;XP_006724716.1;XP_011529263.1;XP_01 | 8148 | 300127 | . | . | 2.022833 | 1.956974 | 1.946406 | 1.760274 | 4.707215 | 4.523632 | 4.367680 | 4.024656 | -1.069338 | 444.638246 | 0.176949048 | -0.54658486 | 0.58466397 | 0.998865404 | 8.843758 | 8.747617 | 8.819887 | 8.812803 | 8.810494 | 8.738984 | 532 | 493 | 396 | 379 |
| 4985 | NM_000911 | OPRD1 | opioid receptor delta 1 | protein_coding | NP_000902.3 | 8153 | 165195 | . | . | 3.978545 | 4.116440 | 3.005725 | 2.999154 | 9.258239 | 9.515335 | 6.744760 | 6.857210 | -1.365860 | 1024.704096 | 0.13577074 | -3.31301119 | **0.000922973** | 0.064386964 | 10.208439 | 9.758624 | 10.094103 | 10.113492 | 9.879672 | 9.870470 | 1345 | 1293 | 758 | 807 |
| 4987 | NM_000913,N | OPRL1 | opioid related nociceptin receptor 1 | protein_coding | NP_000904.1;NP_001186948.1;NP_001305782.1;NP_001305783.1;NP_00 | 8155 | 602548 | . | . | 0.265562 | 0.377429 | 0.452153 | 0.464080 | 0.617975 | 0.872444 | 1.014619 | 1.061063 | 1.440645 | 41.741473 | 0.4864228 | 1.082833703 | 0.278882243 | 0.998865404 | 5.096985 | 5.622280 | 5.323604 | 5.362214 | 5.401589 | 5.400705 | 34 | 42 | 43 | 46 |
| 4992 | NM_00137063 | OR1F1 | olfactory receptor family 1 subfamily F member 1 | protein_coding | NP_001357568.1;NP_001357569.1;NP_001357570.1;NP_036492.1;XP_01 | 8194 | 603232 | . | . | 0.089195 | 0.070662 | 0.048658 | 0.078116 | 0.207560 | 0.163338 | 0.109188 | 0.178601 | -1.255462 | 6.809550 | 1.167886968 | -0.28103624 | 0.778682606 | NA | 2.923304 | 2.592943 | 2.782132 | 2.771917 | 2.763761 | 2.776320 | 10 | 7 | 4 | 7 |
| 4998 | NM_00119081 | ORC1 | origin recognition complex subunit 1 | protein_coding | NP_001177747.1;NP_001177748.1;NP_004144.2;XP_011539829.1;XP_01 | 8487 | 601902 | . | . | 8.894129 | 9.653908 | 8.783044 | 10.001008 | 20.697007 | 22.315444 | 19.708897 | 22.866115 | -1.000855 | 913.276575 | 0.14544298 | -0.00847917 | 0.993234686 | 0.999862634 | 9.835794 | 9.834022 | 9.808723 | 9.860134 | 9.790846 | 9.875372 | 1015 | 1021 | 747 | 906 |
| 4999 | NM_006190,N | ORC2 | origin recognition complex subunit 2 | protein_coding | NP_006181.1;XP_006712618.1;XP_011509554.1;XP_011509555.1 | 8488 | 601182 | . | . | 8.788169 | 9.553969 | 9.486422 | 8.066829 | 20.450431 | 22.084431 | 21.287258 | 18.443847 | -1.023015 | 1221.534865 | 0.143353434 | -0.22899639 | 0.818871725 | 0.998865404 | 10.270637 | 10.238139 | 10.232727 | 10.289768 | 10.311385 | 10.173119 | 1369 | 1383 | 1147 | 1028 |
| 5000 | NM_00119087 | ORC4 | origin recognition complex subunit 4 | protein_coding | NP_001177808.1;NP_001177810.1;NP_001177811.1;NP_001361199.1;NI | 8490 | 603056 | . | . | 3.127115 | 3.017026 | 3.071686 | 2.185595 | 7.276923 | 6.973992 | 6.892774 | 4.997102 | -1.181167 | 584.706064 | 0.187534399 | -1.28090061 | 0.200228578 | 0.96621494 | 9.306124 | 9.067124 | 9.255215 | 9.224855 | 9.234964 | 9.014137 | 745 | 668 | 547 | 416 |
| 5001 | NM_002553,N | ORC5 | origin recognition complex subunit 5 | protein_coding | NP_002544.1;NP_859531.1;XP_011514575.1 | 8491 | 602331 | . | . | 21.895026 | 20.856748 | 22.302287 | 16.804560 | 50.950627 | 48.211307 | 50.045687 | 38.421627 | -1.108014 | 1237.158130 | 0.156914683 | -0.94303399 | 0.345663517 | 0.998865404 | 10.344571 | 10.197301 | 10.329789 | 10.284750 | 10.333314 | 10.115497 | 1541 | 1362 | 1170 | 940 |
| 5002 | NM_00131550 | SLC22A18 | solute carrier family 22 member 18 | protein_coding | NP_001302430.1;NP_001302431.1;NP_002546.3;NP_899056.2;XP_01151 | 10964 | 602631 | . | . | 24.949402 | 23.757656 | 20.129205 | 23.399496 | 58.058284 | 54.916885 | 45.169355 | 53.500164 | -1.126010 | 1107.654457 | 0.141563186 | -1.20948892 | 0.226475064 | 0.982008277 | 10.196513 | 10.024975 | 10.178581 | 10.131508 | 10.011690 | 10.119366 | 1393 | 1227 | 839 | 1049 |
| 5003 | NM_00130286 | SLC22A18AS | solute carrier family 22 member 18 antisense | protein_coding | NP_001289791.1;NP_009036.2;XP_016873321.1;XP_016873322.1;XP_01 | 10965 | 603240 | . | . | 5.071214 | 5.936979 | 3.733200 | 7.047919 | 11.800926 | 13.723591 | 8.377195 | 16.114231 | -1.016377 | 387.649213 | 0.198009738 | -0.11835447 | 0.905786798 | 0.998865404 | 8.610694 | 8.586420 | 8.623878 | 8.579178 | 8.514078 | 8.664569 | 467 | 406 | 291 | 407 |
| 5007 | NM_002556 | OSBP | oxysterol binding protein | protein_coding | NP_002547.1 | 8503 | 167040 | . | . | 12.862035 | 12.847975 | 11.693397 | 13.010180 | 29.930483 | 29.698668 | 26.239643 | 29.746233 | -1.055213 | 1851.167758 | 0.123192815 | -0.62937477 | 0.529103732 | 0.998865404 | 10.892565 | 10.814828 | 10.878922 | 10.870897 | 10.794093 | 10.869377 | 2200 | 2041 | 1491 | 1770 |
| 5013 | NM_00119977 | OTX1 | orthodenticle homeobox 1 | protein_coding | NP_001186699.1;NP_055377.1 | 8521 | 600036 | . | . | 0.984790 | 1.099566 | 1.044149 | 1.656706 | 2.291647 | 2.541697 | 2.343041 | 3.787862 | 1.252588 | 107.184417 | 0.334816171 | 0.970417689 | 0.331838335 | 0.998865404 | 6.573821 | 6.896118 | 6.676559 | 6.719103 | 6.693305 | 6.841136 | 102 | 110 | 81 | 136 |
| 5016 | NM_002557 | OVGP1 | oviductal glycoprotein 1 | protein_coding | NP_002548.3 | 8524 | 603578 | . | . | 0.105891 | 0.288005 | 0.235452 | 0.277201 | 0.246413 | 0.665736 | 0.528347 | 0.633788 | 1.256085 | 15.876003 | 0.808451636 | 0.406869128 | 0.684104111 | NA | 3.817846 | 4.141582 | 3.941491 | 3.998095 | 3.979251 | 3.992416 | 9 | 22 | 14 | 18 |
| 5018 | NM_005015 | OXA1L | OXA1L mitochondrial inner membrane protein | protein_coding | NP_005006.4 | 8526 | 601066 | . | . | 39.902283 | 41.656654 | 44.989365 | 44.006855 | 92.854248 | 96.291229 | 100.954826 | 100.616447 | 1.076267 | 3726.383286 | 0.109193738 | 0.97107555 | 0.331510658 | 0.998865404 | 11.809564 | 11.915608 | 11.817961 | 11.846322 | 11.906415 | 11.879437 | 4062 | 3940 | 3414 | 3563 |
| 5019 | NM_000436,N | OXCT1 | 3-oxoacid CoA-transferase 1 | protein_coding | NP_000427.1;NP_001351228.1;NP_001351229.1;NP_001351230.1;NP_00 | 8527 | 601424 | . | . | 16.277078 | 16.068157 | 13.158471 | 14.839937 | 37.877430 | 37.142263 | 29.527228 | 33.929751 | -1.157626 | 1508.497377 | 0.131245032 | -1.60896796 | 0.107623347 | 0.835804188 | 10.660769 | 10.449280 | 10.619630 | 10.605227 | 10.446426 | 10.549368 | 1881 | 1731 | 1133 | 1400 |
| 5021 | NM_000916,N | OXTR | oxytocin receptor | protein_coding | NP_000907.2;NP_001341582.1;NP_001341583.1;NP_001341584.1;NP_00 | 8529 | 167055 | . | . | 0.211267 | 0.140243 | 0.094548 | 0.078055 | 0.491627 | 0.324179 | 0.212163 | 0.178463 | **-2.007374** | 18.363758 | 0.740241479 | -1.35808264 | 0.174437464 | NA | 4.614376 | 3.612728 | 4.251155 | 4.207900 | 4.184209 | 4.170321 | 34 | 21 | 12 | 10 |
| 5025 | NM_00125679 | P2RX4 | purinergic receptor P2X 4 | protein_coding | NP_001243725.1;NP_001248326.1;NP_001248327.1;NP_002551.2;XP_01 | 8535 | 600846 | . | . | 9.878877 | 10.429248 | 9.243630 | 10.591044 | 22.988553 | 24.107674 | 20.742440 | 24.215161 | -1.036342 | 552.626140 | 0.166431806 | -0.30943403 | 0.756991388 | 0.998865404 | 9.136007 | 9.083841 | 9.106196 | 9.136377 | 9.055208 | 9.139120 | 633 | 621 | 441 | 542 |
| 5027 | NM_002562,N | P2RX7 | purinergic receptor P2X 7 | protein_coding | NP_002553.3;XP_011536721.1;XP_011536722.1;XP_016874853.1;XP_01 | 8537 | 602566 | . | . | 21.697748 | 20.475113 | 30.036961 | 25.827898 | 50.491554 | 47.329144 | 67.402074 | 59.052421 | 1.315026 | 3969.873185 | 0.123746771 | 3.192741857 | **0.001409289** | 0.087623478 | 11.743727 | 12.139033 | 11.853622 | 11.803722 | 12.130139 | 11.976554 | 4071 | 3586 | 4276 | 3847 |
| 5028 | NM_002563 | P2RY1 | purinergic receptor P2Y1 | protein_coding | NP_002554.1 | 8539 | 601167 | . | . | 1.097434 | 1.121109 | 0.938707 | 0.677474 | 2.553774 | 2.591493 | 2.106432 | 1.548964 | -1.389052 | 189.437992 | 0.258550612 | -1.83368771 | 0.066700364 | 0.734768658 | 7.782472 | 7.310216 | 7.626677 | 7.631952 | 7.549410 | 7.418132 | 252 | 239 | 161 | 124 |
| 5033 | NM_000917,N | P4HA1 | prolyl 4-hydroxylase subunit alpha 1 | protein_coding | NP_000908.2;NP_001017962.1;NP_001136067.1;NP_001136068.1 | 8546 | 176710 | . | . | 42.737394 | 38.296989 | 41.151563 | 34.788302 | 99.451670 | 88.525209 | 92.342917 | 79.539318 | -1.082254 | 3489.991641 | 0.128330446 | -0.88863631 | 0.374198572 | 0.998865404 | 11.824763 | 11.711012 | 11.847967 | 11.748241 | 11.804309 | 11.659002 | 4427 | 3680 | 3174 | 2864 |
| 5034 | NM_000918,XI | P4HB | prolyl 4-hydroxylase subunit beta | protein_coding | NP_000909.2;XP_024306545.1 | 8548 | 176790 | . | . | 573.009883 | 572.034414 | 524.646573 | 568.304472 | 1333.417460 | 1322.283186 | 1177.291672 | 1299.360512 | -1.060963 | 42901.145920 | 0.099754383 | -0.85584724 | 0.392082282 | 0.998865404 | 15.430789 | 15.345405 | 15.417622 | 15.409649 | 15.333055 | 15.391134 | 51095 | 47443 | 34980 | 40348 |
| 5036 | NM_006191 | PA2G4 | proliferation-associated 2G4 | protein_coding | NP_006182.2 | 8550 | 602145 | . | . | 88.340965 | 94.881973 | 79.395020 | 94.405716 | 205.573059 | 219.323914 | 178.160126 | 215.847443 | -1.069160 | 6634.405067 | 0.117614059 | -0.82028573 | 0.412053241 | 0.998865404 | 12.743253 | 12.646632 | 12.695415 | 12.748880 | 12.595225 | 12.731941 | 7647 | 7631 | 5123 | 6500 |
| 5037 | NM_002567 | PEBP1 | phosphatidylethanolamine binding protein 1 | protein_coding | NP_002558.1 | 8630 | 604591 | . | . | 119.901718 | 121.844978 | 120.079910 | 135.679626 | 279.016235 | 281.650116 | 269.455841 | 310.215363 | 1.043194 | 5543.249294 | 0.110747281 | 0.550870506 | 0.581722447 | 0.998865404 | 12.405744 | 12.466646 | 12.414980 | 12.421230 | 12.405389 | 12.499653 | 6225 | 5877 | 4647 | 5603 |
| 5042 | NM_030979 | PABPC3 | poly(A) binding protein cytoplasmic 3 | protein_coding | NP_112241.2 | 8556 | 604680 | . | . | 0.274166 | 0.308566 | 0.241938 | 0.232080 | 0.637997 | 0.713263 | 0.542902 | 0.530623 | -1.253532 | 20.584836 | 0.677237504 | -0.48136519 | 0.630256974 | NA | 4.517478 | 4.191126 | 4.371148 | 4.381225 | 4.357324 | 4.356024 | 25 | 26 | 16 | 17 |
| 5045 | NM_00128982 | FURIN | furin, paired basic amino acid cleaving enzyme | protein_coding | NP_001276752.1;NP_001276753.1;NP_001369548.1;NP_001369549.1;NI | 8568 | 136950 | . | . | 30.880335 | 31.050124 | 26.051900 | 28.347170 | 71.859806 | 71.773747 | 58.459707 | 64.812441 | -1.155138 | 3800.406483 | 0.110704489 | -1.87946962 | 0.060180398 | 0.719735192 | 11.992272 | 11.784102 | 11.950196 | 11.947035 | 11.798297 | 11.859216 | 4700 | 4388 | 2952 | 3430 |
| 5046 | NM_00129130 | PCSK6 | proprotein convertase subtilisin/kexin type 6 | protein_coding | NP_001278238.1;NP_002561.1;NP_612192.1;NP_612195.1;NP_612196.1 | 8569 | 167405 | . | . | 0.842499 | 0.959118 | 0.687021 | 0.930165 | 1.960532 | 2.217045 | 1.541655 | 2.126712 | -1.039279 | 111.857049 | 0.311960816 | -0.17817467 | 0.858585801 | 0.998865404 | 6.833925 | 6.776529 | 6.792090 | 6.829868 | 6.760165 | 6.835448 | 124 | 130 | 84 | 115 |
| 5047 | NM_00101804 | PAEP | progestagen associated endometrial protein | protein_coding | NP_001018058.1;NP_001018059.1;NP_002562.2;XP_011517047.1;XP_01 | 8573 | 173310 | . | . | 1.938999 | 1.925769 | 1.129902 | 1.345894 | 4.512133 | 4.451503 | 2.535465 | 3.077226 | -1.785040 | 46.848398 | 0.463729391 | -1.80268095 | 0.071438336 | 0.753295146 | 5.908824 | 5.071091 | 5.598360 | 5.598900 | 5.486239 | 5.513752 | 69 | 65 | 26 | 35 |
| 5048 | NM_000430,XI | PAFAH1B1 | platelet activating factor acetylhydrolase 1b regulatory subunit 1 | protein_coding | NP_000421.1;XP_011522203.1;XP_011522204.1;XP_011522205.1;XP_01 | 8574 | 601545 | . | . | 20.828606 | 20.457785 | 20.026586 | 20.609262 | 48.469023 | 47.289088 | 44.939082 | 47.120628 | -1.045231 | 3499.992497 | 0.109132042 | -0.5848174 | 0.558670501 | 0.998865404 | 11.804671 | 11.740896 | 11.799478 | 11.782660 | 11.761719 | 11.747534 | 4161 | 3821 | 3002 | 3181 |
| 5049 | NM_00118474 | PAFAH1B2 | platelet activating factor acetylhydrolase 1b catalytic subunit 2 | protein_coding | NP_001171675.1;NP_001171676.1;NP_001171677.1;NP_001296360.1;NI | 8575 | 602508 | . | . | 7.644926 | 6.767575 | 6.478608 | 6.896287 | 17.790058 | 15.643555 | 14.537806 | 15.767539 | -1.085479 | 900.428409 | 0.146152901 | -0.8096453 | 0.418144064 | 0.998865404 | 9.872366 | 9.754148 | 9.888546 | 9.795152 | 9.765924 | 9.801508 | 1148 | 947 | 732 | 830 |
| 5050 | NM_00114593 | PAFAH1B3 | platelet activating factor acetylhydrolase 1b catalytic subunit 3 | protein_coding | NP_001139411.1;NP_001139412.1;NP_002564.1;XP_016882335.1;XP_01 | 8576 | 603074 | . | . | 27.576066 | 27.888253 | 26.086429 | 30.685018 | 64.170650 | 64.464949 | 58.537189 | 70.157650 | 1.000734 | 917.489635 | 0.146399553 | 0.0072283 | 0.994232701 | 0.999862634 | 9.841258 | 9.841838 | 9.839877 | 9.841615 | 9.785628 | 9.893802 | 1055 | 991 | 738 | 925 |
| 5051 | NM_000437,XI | PAFAH2 | platelet activating factor acetylhydrolase 2 | protein_coding | NP_000428.2;XP_006710732.1;XP_006710733.1;XP_011539830.1;XP_01 | 8579 | 602344 | . | . | 4.474060 | 5.165571 | 4.832723 | 5.144904 | 10.411322 | 11.940452 | 10.844490 | 11.763211 | 1.022139 | 538.020525 | 0.167616907 | 0.188475786 | 0.850503695 | 0.998865404 | 9.055951 | 9.086917 | 9.026176 | 9.100022 | 9.054509 | 9.100516 | 578 | 607 | 456 | 528 |
| 5052 | NM_00120243 | PRDX1 | peroxiredoxin 1 | protein_coding | NP_001189360.1;NP_002565.1;NP_859047.1;NP_859048.1 | 9352 | 176763 | . | . | 498.165176 | 472.321463 | 554.532902 | 643.136693 | 1159.250828 | 1091.792250 | 1244.355873 | 1470.455646 | 1.214542 | 16021.756438 | 0.110919917 | 2.528058333 | **0.011469529** | 0.350252022 | 13.820751 | 14.101137 | 13.902440 | 13.852856 | 13.982497 | 14.103472 | 17151 | 15155 | 14223 | 17617 |
| 5054 | NM_000602 | SERPINE1 | serpin family E member 1 | protein_coding | NP_000593.1 | 8583 | 173360 | . | . | 0.464311 | 0.728714 | 0.377207 | 0.509604 | 1.080471 | 1.684455 | 0.846442 | 1.165148 | -1.349931 | 51.816118 | 0.458055679 | -0.94504978 | 0.344633464 | 0.998865404 | 5.897022 | 5.460784 | 5.673457 | 5.765875 | 5.641609 | 5.686109 | 54 | 78 | 33 | 47 |
| 5055 | NM_00114381 | SERPINB2 | serpin family B member 2 | protein_coding | NP_001137290.1;NP_002566.1;XP_024306960.1 | 8584 | 173390 | . | . | 2.661482 | 3.083520 | 3.873572 | 3.679983 | 6.193377 | 7.127695 | 8.692185 | 8.413846 | 1.297386 | 198.674339 | 0.240030248 | 1.564834938 | 0.117621617 | 0.857396579 | 7.434430 | 7.809752 | 7.538327 | 7.594826 | 7.697843 | 7.669724 | 186 | 199 | 201 | 204 |
| 5058 | NM_00112862 | PAK1 | p21 (RAC1) activated kinase 1 | protein_coding | NP_001122092.1;NP_001363197.1;NP_001363198.1;NP_001363199.1;NI | 8590 | 602590 | . | . | 10.838254 | 10.658328 | 8.779160 | 11.171898 | 25.221062 | 24.637199 | 19.700180 | 25.543216 | -1.087941 | 1034.091248 | 0.14674295 | -0.82866571 | 0.407293593 | 0.998865404 | 10.073908 | 9.951805 | 10.052013 | 10.035176 | 9.910792 | 10.047468 | 1255 | 1150 | 780 | 1016 |
| 5062 | NM_002577,XI | PAK2 | p21 (RAC1) activated kinase 2 | protein_coding | NP_002568.2;XP_011511172.1;XP_016861990.1 | 8591 | 605022 | . | . | 9.438832 | 9.957839 | 10.154799 | 9.059042 | 21.964547 | 23.017991 | 22.787071 | 20.712425 | -1.021928 | 1844.107419 | 0.126117563 | -0.24812665 | 0.804036416 | 0.998865404 | 10.864169 | 10.833078 | 10.840883 | 10.872164 | 10.886361 | 10.790630 | 2104 | 2051 | 1683 | 1606 |
| 5063 | NM_00112816 | PAK3 | p21 (RAC1) activated kinase 3 | protein_coding | NP_001121638.1;NP_001121639.1;NP_001121640.1;NP_001121644.1;NI | 8592 | 300142 | . | . | 0.072673 | 0.074595 | 0.047310 | 0.009616 | 0.169113 | 0.172430 | 0.106161 | 0.021986 | **-2.446462** | 14.572247 | 0.883394331 | -1.46106546 | 0.143997479 | NA | 4.367696 | 3.087437 | 3.890928 | 3.896334 | 3.863821 | 3.817443 | 23 | 23 | 12 | 3 |
| 5064 | NM_00104013 | PALM | paralemmin | protein_coding | NP_001035224.1;NP_002570.2;XP_005259622.1;XP_005259623.1;XP_01 | 8594 | 608134 | . | . | 0.135366 | 0.164305 | 0.151443 | 0.309077 | 0.315003 | 0.379797 | 0.339833 | 0.706669 | 1.482018 | 18.370680 | 0.730883285 | 0.77654339 | 0.437428212 | NA | 3.889947 | 4.453945 | 4.169603 | 4.178142 | 4.180172 | 4.228318 | 16 | 17 | 14 | 26 |
| 5066 | NM_000919,N | PAM | peptidylglycine alpha-amidating monooxygenase | protein_coding | NP_000910.2;NP_001170777.1;NP_001306872.1;NP_001351511.1;NP_00 | 8596 | 170270 | . | . | 22.633118 | 18.905233 | 18.709697 | 19.447387 | 52.668198 | 43.700299 | 41.984015 | 44.464143 | -1.031803 | 2739.700512 | 0.120186677 | -0.37581067 | 0.707057655 | 0.998865404 | 11.442220 | 11.397031 | 11.472892 | 11.389257 | 11.372078 | 11.439751 | 3368 | 2849 | 2247 | 2634 |
| 5073 | NM_00113447 | PARN | poly(A)-specific ribonuclease | protein_coding | NP_001127949.1;NP_001229921.1;NP_002573.1;XP_011520812.1;XP_01 | 8609 | 604212 | . | . | 18.041343 | 18.174893 | 17.428445 | 18.108692 | 41.982947 | 42.012079 | 39.108928 | 41.403370 | -1.029440 | 1700.447126 | 0.121458541 | -0.34464391 | 0.73036209 | 0.998865404 | 10.752516 | 10.710576 | 10.742806 | 10.743187 | 10.707807 | 10.732545 | 1986 | 1862 | 1434 | 1596 |
| 5074 | NM_00135473 | PAWR | pro-apoptotic WT1 regulator | protein_coding | NP_001341661.1;NP_001341662.1;NP_002574.2;XP_006719498.1;XP_01 | 8614 | 601936 | . | . | 0.957965 | 1.136458 | 0.968974 | 1.077671 | 2.229227 | 2.626974 | 2.174349 | 2.463967 | -1.130536 | 229.685596 | 0.225407494 | -0.78527275 | 0.432293671 | 0.998865404 | 7.929201 | 7.752420 | 7.853009 | 7.890938 | 7.833981 | 7.792145 | 270 | 273 | 196 | 193 |
| 5077 | NM_000438,N | PAX3 | paired box 3 | protein_coding | NP_000429.2;NP_001120838.1;NP_039230.1;NP_852122.1;NP_852123.1 | 8617 | 606597 | . | . | 9.657128 | 10.139543 | 11.081005 | 10.702563 | 22.472534 | 23.438008 | 24.865456 | 24.470139 | 1.039112 | 993.782605 | 0.136318402 | 0.406042793 | 0.684711158 | 0.998865404 | 9.928890 | 9.984154 | 9.940195 | 9.944679 | 9.962346 | 9.978580 | 1119 | 1055 | 871 | 960 |
| 5082 | NM_005388 | PDCL | phosducin like | protein_coding | NP_005379.3 | 8770 | 604421 | . | . | 4.031905 | 3.867326 | 4.659904 | 4.555026 | 9.382408 | 8.939496 | 10.456688 | 10.414525 | 1.150096 | 402.648968 | 0.180411396 | 1.118303132 | 0.26343755 | 0.996308385 | 8.548800 | 8.750886 | 8.623038 | 8.594160 | 8.702035 | 8.681669 | 442 | 394 | 381 | 397 |
| 5083 | NM_00137207 | PAX9 | paired box 9 | protein_coding | NP_001359005.1;NP_006185.1 | 8623 | 167416 | . | . | 2.974066 | 3.110264 | 3.188825 | 2.528220 | 6.920774 | 7.189515 | 7.155631 | 5.780475 | -1.078633 | 383.510980 | 0.195997364 | -0.55717238 | 0.577409657 | 0.998865404 | 8.636237 | 8.527980 | 8.592335 | 8.613640 | 8.626087 | 8.486844 | 450 | 437 | 360 | 304 |
| 5088 | gene-PBX2P1 | PBX2P1 | PBX homeobox 2 pseudogene 1 | pseudogene | . | 8635 | . | . | . | 0.224535 | 0.167723 | 0.220950 | 0.269881 | 0.522503 | 0.387699 | 0.495805 | 0.617050 | 1.209246 | 22.680562 | 0.64963922 | 0.421939318 | 0.67306931 | NA | 4.359524 | 4.634191 | 4.503258 | 4.478057 | 4.501214 | 4.516234 | 27 | 19 | 20 | 25 |
| 5089 | NM_002586 | PBX2 | PBX homeobox 2 | protein_coding | NP_002577.2 | 8633 | 176311 | . | . | 12.807211 | 13.247252 | 14.574241 | 14.602098 | 29.802908 | 30.621616 | 32.704174 | 33.385963 | 1.104479 | 1388.887172 | 0.126247597 | 1.135594676 | 0.256126238 | 0.995760118 | 10.366266 | 10.509602 | 10.390551 | 10.409493 | 10.479335 | 10.472975 | 1501 | 1442 | 1273 | 1361 |
| 5090 | NM_00113477 | PBX3 | PBX homeobox 3 | protein_coding | NP_001128250.1;NP_001317711.1;NP_006186.1;XP_006717193.1;XP_00 | 8634 | 176312 | . | . | 6.203739 | 5.616411 | 5.441078 | 5.149877 | 14.436357 | 12.982585 | 12.209622 | 11.774583 | -1.140989 | 480.993826 | 0.176362122 | -1.07894321 | 0.280613045 | 0.998865404 | 9.001529 | 8.812000 | 8.996147 | 8.899763 | 8.889949 | 8.842649 | 634 | 512 | 406 | 405 |

| | | | | | | | | | | | | | | | | | | | | | | | | | | | | | | | | | |
|---|---|---|---|---|---|---|---|---|---|---|---|---|---|---|---|---|---|---|---|---|---|---|---|---|---|---|---|---|---|---|---|---|---|
| 5091 | NM_000920,N | PC | pyruvate carboxylase | protein_coding | NP_000911.2;NP_001035806.1;NP_071504.2;XP_005274088.1;XP_00527 | 8636 | 608786 | . | . | 13.328550 | 13.241326 | 15.271863 | 17.007817 | 31.016082 | 30.607918 | 34.269617 | 38.886354 | 1.199462 | 1970.098187 | 0.122281957 | 2.145753971 | 0.031892613 | 0.564468799 | 10.806994 | 11.069211 | 10.871962 | 10.861457 | 10.970161 | 11.050280 | 2077 | 1920 | 1776 | 2114 |
| 5092 | NM_000281,N | PCBD1 | pterin-4 alpha-carbinolamine dehydratase 1 | protein_coding | NP_000272.1;NP_001276726.1;NP_001309933.1 | 8646 | 126090 | . | . | 27.010865 | 29.684293 | 29.169576 | 29.562115 | 62.855391 | 68.616569 | 65.455682 | 67.590264 | 1.031138 | 702.582420 | 0.152251323 | 0.290555398 | 0.771391376 | 0.998865404 | 9.434363 | 9.478349 | 9.414166 | 9.475846 | 9.466152 | 9.466796 | 761 | 780 | 620 | 669 |
| 5093 | NM_006196 | PCBP1 | poly(rC) binding protein 1 | protein_coding | NP_006187.2 | 8647 | 601209 | . | . | 141.773285 | 145.521133 | 134.792633 | 161.649338 | 329.912262 | 336.378601 | 302.470764 | 369.592010 | 1.017251 | 7849.246047 | 0.116430312 | 0.211933702 | 0.832158757 | 0.998865404 | 12.926002 | 12.950570 | 12.922724 | 12.937545 | 12.869384 | 13.014722 | 8883 | 8471 | 6295 | 8056 |
| 5094 | NM_00109862 | PCBP2 | poly(rC) binding protein 2 | protein_coding | NP_001092090.1;NP_001122383.1;NP_001122384.1;NP_001122385.1;NI | 8648 | 601210 | . | . | 64.219294 | 66.876319 | 64.073881 | 68.873407 | 149.440937 | 154.587612 | 143.779929 | 157.470864 | 1.002227 | 6425.796764 | 0.105716526 | 0.030358026 | 0.97578152 | 0.998865404 | 12.648090 | 12.651227 | 12.635119 | 12.661893 | 12.624201 | 12.676024 | 7274 | 7036 | 5422 | 6216 |
| 5095 | NM_000282,N | PCCA | propionyl-CoA carboxylase subunit alpha | protein_coding | NP_000273.2;NP_001121164.1;NP_001171475.1;NP_001339534.1;NP_0( | 8653 | 232000 | . | . | 3.492268 | 3.384000 | 4.343813 | 3.861119 | 8.126651 | 7.822266 | 9.747390 | 8.827989 | 1.190120 | 297.988213 | 0.204006916 | 1.23087519 | 0.218369548 | 0.982008277 | 8.087756 | 8.339499 | 8.171901 | 8.163138 | 8.294085 | 8.226528 | 316 | 291 | 299 | 285 |
| 5096 | NM_000532,N | PCCB | propionyl-CoA carboxylase subunit beta | protein_coding | NP_000523.2;NP_001171485.1;XP_011511175.1 | 8654 | 232050 | . | . | 19.654088 | 19.060771 | 15.813206 | 20.763471 | 45.735870 | 44.059825 | 35.484377 | 47.473205 | -1.082003 | 1050.213833 | 0.15341367 | -0.74115922 | 0.458596907 | 0.998865404 | 10.092465 | 9.978208 | 10.079890 | 10.046622 | 9.913174 | 10.089360 | 1285 | 1152 | 771 | 1060 |
| 5097 | NM_00127861 | PCDH1 | protocadherin 1 | protein_coding | NP_001265542.1;NP_001265544.1;NP_002578.2;NP_115796.2;XP_0052€ | 8655 | 603626 | . | . | 8.337395 | 8.922686 | 4.967632 | 6.563241 | 19.401461 | 20.625188 | 11.147222 | 15.006070 | -1.524028 | 874.885676 | 0.164432538 | -3.69689529 | 0.000218252 | 0.021989319 | 10.045502 | 9.436629 | 9.888472 | 9.913031 | 9.500006 | 9.690507 | 1197 | 1159 | 516 | 743 |
| 5098 | NM_002588,N | PCDHGC3 | protocadherin gamma subfamily C, 3 | protein_coding | NP_002579.2;NP_115778.1;NP_115779.1 | 8716 | 603627 | . | . | 31.213334 | 33.791561 | 34.369373 | 34.381020 | 72.634709 | 78.110708 | 77.123875 | 78.608115 | 1.005009 | 4535.885561 | 0.107447192 | 0.067086242 | 0.946513046 | 0.998865404 | 12.143589 | 12.150739 | 12.123458 | 12.166101 | 12.143835 | 12.154424 | 5077 | 5007 | 3929 | 4290 |
| 5099 | NM_00117352 | PCDH7 | protocadherin 7 | protein_coding | NP_001166994.1;NP_002580.2;NP_115832.1;NP_115833.2;XP_00524822 | 8659 | 602988 | . | . | 9.256349 | 9.509150 | 9.505828 | 9.050947 | 21.539904 | 21.980827 | 21.330803 | 20.693916 | -1.027728 | 1872.700527 | 0.120033243 | -0.32873412 | 0.74235666 | 0.998865404 | 10.890458 | 10.851082 | 10.875054 | 10.887992 | 10.878274 | 10.841327 | 2168 | 2065 | 1644 | 1691 |
| 5101 | NM_00131837 | PCDH9 | protocadherin 9 | protein_coding | NP_001305301.1;NP_001305302.1;NP_001305303.1;NP_065136.1;NP_9ℇ | 8661 | 603581 | . | . | 0.242550 | 0.296930 | 0.340622 | 0.319123 | 0.564426 | 0.686367 | 0.764344 | 0.729636 | 1.176994 | 185.839443 | 0.249175744 | 0.943539699 | 0.345404919 | 0.998865404 | 7.414950 | 7.651214 | 7.508862 | 7.485603 | 7.608965 | 7.525987 | 202 | 179 | 191 | 171 |
| 5104 | NM_000624 | SERPINA5 | serpin family A member 5 | protein_coding | NP_000615.3 | 8723 | 601841 | . | . | 0.311390 | 0.431995 | 0.132116 | 0.079836 | 0.724618 | 0.998576 | 0.296465 | 0.182534 | -3.238001 | 17.709646 | 0.779280571 | -2.17521586 | 0.029613944 | NA | 4.757258 | 3.066070 | 4.168226 | 4.205359 | 4.116001 | 4.098018 | 26 | 34 | 9 | 6 |
| 5106 | NM_00101807 | PCK2 | phosphoenolpyruvate carboxykinase 2, mitochondrial | protein_coding | NP_001018083.2;NP_001278485.1;NP_001294983.1;NP_004554.3;XP_0( | 8725 | 614095 | . | . | 12.264795 | 12.539884 | 9.025128 | 13.154943 | 28.540681 | 28.986502 | 20.252126 | 30.077214 | -1.131435 | 799.486057 | 0.182566048 | -0.97583228 | 0.329147585 | 0.998865404 | 9.729726 | 9.550574 | 9.670817 | 9.690920 | 9.451175 | 9.718578 | 964 | 929 | 526 | 839 |
| 5108 | NM_00131550 | PCM1 | pericentriolar material 1 | protein_coding | NP_001302436.2;NP_001302437.1;NP_001339561.2;NP_001339562.2;NI | 8727 | 600299 | . | . | 9.990440 | 10.306221 | 10.918547 | 10.714030 | 23.248162 | 23.823285 | 24.500899 | 24.496361 | 1.065012 | 2757.203649 | 0.112103626 | 0.810588271 | 0.417602155 | 0.998865404 | 11.382829 | 11.473719 | 11.401139 | 11.404276 | 11.459826 | 11.448047 | 3069 | 2887 | 2491 | 2647 |
| 5110 | NM_00125204 | PCMT1 | protein-L-isoaspartate (D-aspartate) O-methyltransferase | protein_coding | NP_001238978.1;NP_001238979.1;NP_001238980.1;NP_001238981.1;NI | 8728 | 176851 | . | . | 64.063815 | 55.762921 | 55.804934 | 45.628856 | 149.079127 | 128.898495 | 125.224650 | 104.324957 | -1.145772 | 2447.962279 | 0.126305491 | -1.55432653 | 0.120106548 | 0.859972379 | 11.352030 | 11.156050 | 11.349038 | 11.267140 | 11.252475 | 11.142988 | 3160 | 2680 | 2121 | 1992 |
| 5111 | NM_002592,N | PCNA | proliferating cell nuclear antigen | protein_coding | NP_002583.1;NP_872590.1 | 8729 | 176740 | . | . | 119.103021 | 117.758837 | 105.566369 | 118.699096 | 277.157612 | 272.204857 | 236.887872 | 271.391365 | -1.068287 | 4601.573097 | 0.112095357 | -0.85016542 | 0.395233124 | 0.998865404 | 12.214825 | 12.119421 | 12.203776 | 12.185262 | 12.093380 | 12.183421 | 5534 | 5073 | 3659 | 4398 |
| 5116 | NM_00131552 | PCNT | pericentrin | protein_coding | NP_001302458.1;NP_006022.3;XP_005261181.1;XP_011527896.1;XP_01 | 16068 | 605925 | . | . | 4.028323 | 4.083319 | 4.702046 | 4.345386 | 9.374072 | 9.438776 | 10.551255 | 9.935209 | 1.087585 | 1406.681318 | 0.129454864 | 0.935681074 | 0.349437418 | 0.998865404 | 10.396110 | 10.517497 | 10.426409 | 10.422222 | 10.525929 | 10.450357 | 1556 | 1450 | 1337 | 1307 |
| 5118 | NM_002593,XI | PCOLCE | procollagen C-endopeptidase enhancer | protein_coding | NP_002584.2;XP_024302553.1 | 8738 | 600270 | . | . | 2.849128 | 3.553091 | 3.641973 | 3.792187 | 6.630038 | 8.213128 | 8.172482 | 8.670386 | 1.149151 | 161.420935 | 0.265713326 | 0.754829935 | 0.45035103 | 0.998865404 | 7.231481 | 7.430994 | 7.257619 | 7.344759 | 7.352464 | 7.364206 | 154 | 180 | 148 | 164 |
| 5119 | NM_00108331 | CHMP1A | charged multivesicular body protein 1A | protein_coding | NP_001076783.1;NP_002759.2;XP_011521400.1 | 8740 | 164010 | . | . | 16.417920 | 17.503808 | 14.917028 | 18.766516 | 38.205176 | 40.460839 | 33.473378 | 42.907409 | -1.030380 | 1217.623193 | 0.147071161 | -0.29357843 | 0.76908006 | 0.998865404 | 10.271579 | 10.227793 | 10.241190 | 10.277850 | 10.150369 | 10.316783 | 1389 | 1366 | 927 | 1249 |
| 5125 | NM_00119048 | PCSK5 | proprotein convertase subtilisin/kexin type 5 | protein_coding | NP_001177411.1;NP_001358972.1;NP_006191.2;XP_011517071.1;XP_01 | 8747 | 600488 | . | . | 0.100687 | 0.062756 | 0.022846 | 0.098797 | 0.234303 | 0.145064 | 0.051266 | 0.225888 | -1.307340 | 18.829458 | 0.76981617 | -0.50224276 | 0.615496765 | NA | 4.417335 | 4.026055 | 4.267398 | 4.226880 | 4.191118 | 4.259767 | 30 | 18 | 7 | 23 |
| 5126 | NM_00120152 | PCSK2 | proprotein convertase subtilisin/kexin type 2 | protein_coding | NP_001188457.1;NP_001188458.1;NP_002585.2 | 8744 | 162151 | . | . | 0.203346 | 0.291628 | 0.353389 | 0.375913 | 0.473193 | 0.674110 | 0.792995 | 0.859481 | 1.452938 | 45.312830 | 0.472451306 | 1.140800558 | 0.253952929 | 0.994807659 | 5.208527 | 5.745491 | 5.430708 | 5.483993 | 5.518210 | 5.524166 | 35 | 47 | 46 | 51 |
| 5127 | NM_00117046 | CDK16 | cyclin dependent kinase 16 | protein_coding | NP_001163931.1;NP_006192.1;NP_148978.2;XP_011542222.1;XP_01154 | 8749 | 311550 | . | . | 23.860042 | 25.314578 | 24.522182 | 26.883766 | 55.523305 | 58.515785 | 55.027066 | 61.466536 | 1.022589 | 2528.922062 | 0.116689461 | 0.276177326 | 0.782411872 | 0.998865404 | 11.288196 | 11.320240 | 11.271507 | 11.317959 | 11.288107 | 11.337288 | 2797 | 2776 | 2156 | 2470 |
| 5128 | NM_00117046 | CDK17 | cyclin dependent kinase 17 | protein_coding | NP_001163935.1;NP_002586.2;XP_016874894.1;XP_016874895.1;XP_01 | 8750 | 603440 | . | . | 4.700878 | 4.177488 | 4.462723 | 4.793079 | 10.939136 | 9.656449 | 10.014221 | 10.958803 | 1.000749 | 609.902666 | 0.159348416 | 0.006775253 | 0.994594171 | 0.999862634 | 9.251787 | 9.253083 | 9.284817 | 9.217902 | 9.246702 | 9.258182 | 737 | 625 | 526 | 577 |
| 5129 | NM_002596,N | CDK18 | cyclin dependent kinase 18 | protein_coding | NP_002587.2;NP_997667.1;NP_997668.1;XP_011507904.1;XP_01685691 | 8751 | 169190 | . | . | 3.930738 | 3.943251 | 4.438870 | 4.686070 | 9.146990 | 9.115003 | 9.960696 | 10.714146 | 1.183233 | 438.302347 | 0.176723712 | 1.373521602 | 0.16959025 | 0.946181982 | 8.649500 | 8.891895 | 8.725963 | 8.714379 | 8.791792 | 8.853006 | 467 | 429 | 393 | 467 |
| 5130 | NM_00131267 | PCYT1A | phosphate cytidylyltransferase 1, choline, alpha | protein_coding | NP_001299602.1;NP_005008.2 | 8754 | 123695 | . | . | 15.747516 | 16.676067 | 18.206424 | 19.041107 | 36.645119 | 38.547475 | 40.854685 | 43.535227 | 1.133888 | 3010.020635 | 0.112459442 | 1.611939881 | 0.106975023 | 0.83501324 | 11.462132 | 11.643300 | 11.481965 | 11.521121 | 11.590710 | 11.617853 | 3170 | 3118 | 2736 | 3048 |
| 5134 | NM_00119946 | PDCD2 | programmed cell death 2 | protein_coding | NP_001186390.1;NP_001186391.1;NP_001186392.1;NP_001186393.1;NI | 8762 | 600866 | . | . | 14.452347 | 15.262230 | 14.409460 | 15.387217 | 33.631206 | 35.279327 | 32.334413 | 35.181041 | 1.007124 | 1168.320809 | 0.133880253 | 0.076490922 | 0.939028536 | 0.998865404 | 10.185190 | 10.195234 | 10.197070 | 10.177587 | 10.160248 | 10.224023 | 1358 | 1240 | 976 | 1146 |
| 5137 | NM_00119105 | PDE1C | phosphodiesterase 1C | protein_coding | NP_001177985.1;NP_001177986.1;NP_001177987.2;NP_001177988.1;NI | 8776 | 602987 | . | . | 1.352581 | 1.155870 | 0.994264 | 1.177714 | 3.147511 | 2.671840 | 2.231100 | 2.692702 | -1.351543 | 263.608731 | 0.217042485 | -2.00240577 | 0.045241108 | 0.649595749 | 8.242794 | 7.809269 | 8.120556 | 8.103647 | 7.999195 | 7.917713 | 354 | 322 | 212 | 192 |
| 5139 | NM_000921,N | PDE3A | phosphodiesterase 3A | protein_coding | NP_000912.3;NP_001231612.1;NP_001365336.1;NP_001365337.1;NP_0( | 8778 | 123805 | . | . | 0.136325 | 0.165176 | 0.177517 | 0.107178 | 0.317234 | 0.381811 | 0.398342 | 0.245048 | -1.216390 | 54.923191 | 0.436513585 | -0.64741715 | 0.517362 | 0.998865404 | 5.913018 | 5.632007 | 5.776509 | 5.816558 | 5.790292 | 5.724477 | 63 | 71 | 51 | 38 |
| 5140 | NM_000922,N | PDE3B | phosphodiesterase 3B | protein_coding | NP_000913.2;NP_001350498.1;NP_001350499.1;XP_006718312.1;XP_01 | 8779 | 602047 | . | . | 5.580981 | 4.904387 | 7.148609 | 5.812724 | 12.987172 | 11.336712 | 16.042177 | 13.290100 | 1.210148 | 1124.550581 | 0.160661461 | 1.712812911 | 0.086746964 | 0.795065303 | 9.990565 | 10.266525 | 10.111460 | 9.996202 | 10.292933 | 10.093338 | 1264 | 1011 | 1211 | 1004 |
| 5141 | NM_00111130 | PDE4A | phosphodiesterase 4A | protein_coding | NP_001104777.1;NP_001104778.1;NP_001104779.1;NP_001230050.1;NI | 8780 | 600126 | . | . | 0.456165 | 0.569608 | 0.568292 | 0.600011 | 1.061514 | 1.316673 | 1.275230 | 1.371855 | 1.126771 | 85.100709 | 0.347979074 | 0.494840711 | 0.620712565 | 0.998865404 | 6.323023 | 6.494107 | 6.364233 | 6.417712 | 6.418873 | 6.429019 | 83 | 95 | 77 | 86 |
| 5142 | NM_00103733 | PDE4B | phosphodiesterase 4B | protein_coding | NP_001032416.1;NP_001032417.1;NP_001032418.1;NP_001284369.1;NI | 8781 | 600127 | . | . | 11.655748 | 11.336146 | 14.702225 | 13.360464 | 27.123407 | 26.204011 | 32.991365 | 30.547113 | 1.170871 | 1568.106809 | 0.134504777 | 1.692004396 | 0.090645135 | 0.805424371 | 10.496288 | 10.724326 | 10.585579 | 10.510550 | 10.726154 | 10.613643 | 1745 | 1482 | 1576 | 1473 |
| 5143 | NM_000923,N | PDE4C | phosphodiesterase 4C | protein_coding | NP_000914.2;NP_001092288.1;NP_001092289.1;NP_001317101.1;NP_0( | 8782 | 600128 | . | . | 0.564837 | 0.415493 | 0.541960 | 0.464606 | 1.314400 | 0.960431 | 1.216142 | 1.062267 | 1.021965 | 83.956467 | 0.35895139 | 0.087326515 | 0.930411978 | 0.998865404 | 6.374702 | 6.408242 | 6.426626 | 6.343702 | 6.413963 | 6.369379 | 110 | 76 | 80 | 73 |
| 5144 | NM_00110463 | PDE4D | phosphodiesterase 4D | protein_coding | NP_001098101.1;NP_001159371.1;NP_001184147.1;NP_001184148.1;NI | 8783 | 600129 | . | . | 2.214406 | 1.949778 | 2.445620 | 2.487841 | 5.153012 | 4.506998 | 5.487899 | 5.688154 | 1.010537 | 477.308097 | 0.184530183 | 0.081950926 | 0.934685738 | 0.998865404 | 8.890610 | 8.906898 | 8.935531 | 8.849156 | 8.962983 | 8.832900 | 584 | 477 | 461 | 403 |
| 5145 | NM_000440,XI | PDE6A | phosphodiesterase 6A | protein_coding | NP_000431.2;XP_011535952.1;XP_011535953.1;XP_011535954.1;XP_01: | 8785 | 180071 | . | . | 0.061447 | 0.055004 | 0.071875 | 0.038356 | 0.142990 | 0.127145 | 0.161285 | 0.087697 | 1.016567 | 9.965860 | 0.970189028 | 0.02443351 | 0.980506819 | NA | 3.303566 | 3.330299 | 3.311702 | 3.314650 | 3.325075 | 3.302579 | 11 | 11 | 11 | 7 |
| 5147 | NM_00129101 | PDE6D | phosphodiesterase 6D | protein_coding | NP_001277947.1;NP_002592.1;XP_011509644.1 | 8788 | 602676 | . | . | 21.916667 | 23.311446 | 17.833188 | 23.759567 | 51.000984 | 53.885456 | 40.017155 | 54.323424 | -1.102098 | 774.576819 | 0.166074555 | -0.84451337 | 0.398382596 | 0.998865404 | 9.666110 | 9.524969 | 9.614431 | 9.642645 | 9.464265 | 9.648043 | 917 | 894 | 555 | 783 |
| 5150 | NM_00124231 | PDE7A | phosphodiesterase 7A | protein_coding | NP_001229247.1;NP_002594.1;XP_011515842.1;XP_016869027.1 | 8791 | 171885 | . | . | 1.501851 | 1.419492 | 1.512294 | 1.426867 | 3.494869 | 3.281220 | 3.393544 | 3.262363 | -1.007434 | 314.301867 | 0.198964269 | -0.05370633 | 0.957169139 | 0.998865404 | 8.301034 | 8.290962 | 8.309491 | 8.285641 | 8.318449 | 8.268293 | 371 | 333 | 285 | 280 |
| 5151 | NM_00124313 | PDE8A | phosphodiesterase 8A | protein_coding | NP_001230066.1;NP_002596.1;NP_775656.1;XP_011519970.1;XP_01687 | 8793 | 602972 | . | . | 10.182109 | 10.592526 | 10.661929 | 10.691368 | 23.694187 | 24.485102 | 23.925057 | 24.444542 | 1.027972 | 1302.305251 | 0.127974469 | 0.311002383 | 0.75579881 | 0.998865404 | 10.326804 | 10.366625 | 10.330012 | 10.342261 | 10.364316 | 10.349900 | 1467 | 1397 | 1159 | 1226 |
| 5154 | NM_002607,N | PDGFA | platelet derived growth factor subunit A | protein_coding | NP_002598.4;NP_148983.1;XP_011513717.1;XP_011513718.1;XP_01151 | 8799 | 173430 | . | . | 3.702564 | 3.438761 | 3.670401 | 3.541748 | 8.616018 | 7.948849 | 8.236275 | 8.097787 | 1.044463 | 297.563084 | 0.201366308 | 0.31167439 | 0.755287993 | 0.998865404 | 8.185270 | 8.248149 | 8.197970 | 8.211393 | 8.239252 | 8.216655 | 331 | 318 | 270 | 279 |
| 5156 | NM_00134782 | PDGFRA | platelet derived growth factor receptor alpha | protein_coding | NP_001334756.1;NP_001334757.1;NP_001334758.1;NP_001334759.1;NI | 8803 | 173490 | . | . | 0.210989 | 0.105382 | 0.271657 | 0.180988 | 0.490981 | 0.243596 | 0.609590 | 0.413807 | 1.510166 | 34.762457 | 0.54894124 | 1.083371772 | 0.27864344 | 0.998865404 | 4.789233 | 5.387998 | 5.093314 | 5.056008 | 5.167527 | 5.093209 | 37 | 25 | 45 | 30 |
| 5157 | NM_00137207 | PDGFRL | platelet derived growth factor receptor like | protein_coding | NP_001359002.1;NP_006198.1 | 8805 | 604584 | . | . | 14.225165 | 12.737312 | 10.363634 | 12.824380 | 33.102543 | 29.442869 | 23.255696 | 29.321419 | -1.176610 | 590.513237 | 0.172146326 | -1.3630044 | 0.172881125 | 0.947431715 | 9.318596 | 9.083487 | 9.295991 | 9.214344 | 9.083420 | 9.209505 | 779 | 648 | 425 | 559 |
| 5160 | NM_000284,N | PDHA1 | pyruvate dehydrogenase E1 subunit alpha 1 | protein_coding | NP_000275.1;NP_001166925.1;NP_001166926.1;NP_001166927.1;XP_01 | 8806 | 300502 | . | . | 30.219550 | 29.262248 | 25.683560 | 30.798683 | 70.322131 | 67.640998 | 57.633166 | 70.417527 | -1.072195 | 2926.388267 | 0.125523655 | -0.80118086 | 0.423026949 | 0.998865404 | 11.564437 | 11.463612 | 11.540338 | 11.543335 | 11.408283 | 11.556040 | 3481 | 3274 | 2232 | 2889 |
| 5162 | NM_000925,N | PDHB | pyruvate dehydrogenase E1 subunit beta | protein_coding | NP_000916.2;NP_001166939.1;NP_001302465.1 | 8808 | 179060 | . | . | 31.909633 | 29.851435 | 33.771240 | 33.550515 | 74.255030 | 69.002934 | 75.781688 | 76.709272 | 1.073313 | 1516.660969 | 0.126069989 | 0.80963188 | 0.41815178 | 0.998865404 | 10.514654 | 10.616900 | 10.565944 | 10.509842 | 10.598979 | 10.587169 | 1746 | 1521 | 1376 | 1461 |
| 5163 | NM_00127854 | PDK1 | pyruvate dehydrogenase kinase 1 | protein_coding | NP_001265478.1;NP_002601.1;XP_006712657.1;XP_006712658.1;XP_01 | 8809 | 602524 | . | . | 6.513146 | 5.487269 | 8.390189 | 5.159737 | 15.156356 | 12.684071 | 18.827339 | 11.797127 | 1.168110 | 978.615038 | 0.148820421 | 1.506352483 | 0.131976712 | 0.885996989 | 9.817891 | 10.042565 | 9.870487 | 9.875186 | 10.055277 | 9.912736 | 1036 | 977 | 1005 | 894 |
| 5164 | NM_00119989 | PDK2 | pyruvate dehydrogenase kinase 2 | protein_coding | NP_001186827.1;NP_001186828.1;NP_001186829.1;NP_002602.2 | 8810 | 602525 | . | . | 3.955099 | 3.628488 | 3.666407 | 3.825325 | 9.203678 | 8.387413 | 8.227310 | 8.746153 | -1.031954 | 377.587319 | 0.184372233 | -0.24612873 | 0.805582582 | 0.998865404 | 8.583190 | 8.537785 | 8.581133 | 8.558636 | 8.542423 | 8.560724 | 450 | 406 | 318 | 354 |
| 5165 | NM_00114238 | PDK3 | pyruvate dehydrogenase kinase 3 | protein_coding | NP_001135858.1;NP_005382.1 | 8811 | 300906 | . | . | 2.144348 | 1.773957 | 2.732836 | 2.082815 | 4.989987 | 4.100581 | 6.132405 | 4.762107 | 1.135521 | 795.039196 | 0.171046462 | 1.071956954 | 0.283739367 | 0.998865404 | 9.539767 | 9.724113 | 9.656229 | 9.507526 | 9.750114 | 9.591811 | 949 | 717 | 817 | 705 |
| 5166 | NM_002612 | PDK4 | pyruvate dehydrogenase kinase 4 | protein_coding | NP_002603.1 | 8812 | 602527 | . | . | 0.026528 | 0.030347 | 0.081555 | 0.161804 | 0.061731 | 0.070147 | 0.183007 | 0.369946 | 3.815559 | 8.628471 | 1.100740324 | 1.755086542 | 0.079244506 | NA | 1.844134 | 3.772579 | 3.052438 | 3.053919 | 3.079366 | 3.118369 | 4 | 4 | 8 | 17 |
| 5167 | NM_006208 | ENPP1 | ectonucleotide pyrophosphatase/phosphodiesterase 1 | protein_coding | NP_006199.2 | 3356 | 173335 | . | . | 17.654037 | 17.354391 | 13.379108 | 13.805901 | 41.081669 | 40.115456 | 30.022329 | 31.565552 | -1.306044 | 3606.159944 | 0.109075215 | -3.53154262 | 0.000413143 | 0.034985378 | 11.996036 | 11.610821 | 11.927257 | 11.905391 | 11.687654 | 11.704422 | 4764 | 4351 | 2691 | 2963 |
| 5168 | NM_00104009 | ENPP2 | ectonucleotide pyrophosphatase/phosphodiesterase 2 | protein_coding | NP_001035181.1;NP_001124335.1;NP_001317529.1;NP_006200.3;XP_0( | 3357 | 601060 | . | . | 0.626108 | 0.814685 | 0.651200 | 0.526443 | 1.456978 | 1.883181 | 1.461273 | 1.203649 | -1.215220 | 68.917239 | 0.389692974 | -0.72163916 | 0.470516355 | 0.998865404 | 6.240351 | 5.959598 | 6.097922 | 6.157218 | 6.105391 | 6.058691 | 77 | 91 | 60 | 52 |
| 5169 | NM_005021,N | ENPP3 | ectonucleotide pyrophosphatase/phosphodiesterase 3 | protein_coding | NP_005012.2;XP_011534199.1;XP_016866421.1 | 3358 | 602182 | . | . | 0.184728 | 0.167344 | 0.132686 | 0.221892 | 0.429869 | 0.386823 | 0.297744 | 0.507329 | 1.013183 | 17.996708 | 0.732478186 | 0.025795014 | 0.979420839 | NA | 4.161369 | 4.177906 | 4.174775 | 4.164132 | 4.152422 | 4.188248 | 22 | 18 | 12 | 21 |
| 5170 | NM_00126181 | PDPK1 | 3-phosphoinositide dependent protein kinase 1 | protein_coding | NP_001248745.1;NP_002604.1;NP_112558.2;XP_011520823.1;XP_01152 | 8816 | 605213 | . | . | 4.619430 | 4.688138 | 5.001488 | 4.712254 | 10.749605 | 10.836839 | 11.223195 | 10.774005 | 1.027268 | 1067.738108 | 0.135574291 | 0.286285179 | 0.774659709 | 0.998865404 | 10.040708 | 10.079713 | 10.045761 | 10.054169 | 10.095510 | 10.043718 | 1206 | 1143 | 974 | 979 |
| 5172 | NM_000441,XI | SLC26A4 | solute carrier family 26 member 4 | protein_coding | NP_000432.1;XP_005250482.1;XP_006716088.1;XP_016867807.1 | 8818 | 605646 | . | . | 0.260343 | 0.218376 | 0.181507 | 0.172839 | 0.605829 | 0.504787 | 0.407298 | 0.395175 | -1.243373 | 29.817872 | 0.570762921 | -0.55059562 | 0.58191091 | 0.998865404 | 5.045969 | 4.733338 | 4.931054 | 4.895716 | 4.891519 | 4.883294 | 43 | 31 | 24 | 24 |
| 5184 | NM_000285,N | PEPD | peptidase D | protein_coding | NP_000276.2;NP_001159528.1;NP_001159529.1 | 8840 | 613230 | . | . | 39.565295 | 36.794837 | 36.240888 | 42.835505 | 92.070056 | 85.052911 | 81.323499 | 97.938294 | 1.020220 | 2304.997053 | 0.127836479 | 0.22591686 | 0.821266085 | 0.998865404 | 11.156115 | 11.184840 | 11.194130 | 11.127917 | 11.113581 | 11.237821 | 2736 | 2361 | 1868 | 2351 |
| 5187 | NM_002616,XI | PER1 | period circadian regulator 1 | protein_coding | NP_002607.2;XP_005256746.1;XP_005256747.1;XP_024306571.1 | 8845 | 602260 | . | . | 3.427717 | 3.397661 | 3.539110 | 3.555191 | 7.976437 | 7.853845 | 7.941661 | 8.128519 | 1.045247 | 481.598054 | 0.171290224 | 0.37272615 | 0.709352272 | 0.998865404 | 8.879307 | 8.943354 | 8.926434 | 8.866936 | 8.917392 | 8.932391 | 568 | 484 | 423 | 467 |
| 5188 | NM_00136334 | GATB | glutamyl-tRNA amidotransferase subunit B | protein_coding | NP_001350270.1;NP_004555.1 | 8849 | 603645 | . | . | 9.525788 | 8.782501 | 8.551703 | 10.966191 | 22.166899 | 20.301144 | 19.189776 | 25.072892 | 1.042976 | 694.496688 | 0.168606731 | 0.360048145 | 0.71881113 | 0.998865404 | 9.409427 | 9.469594 | 9.453365 | 9.392154 | 9.365395 | 9.531817 | 818 | 701 | 542 | 745 |
| 5189 | NM_000466,N | PEX1 | peroxisomal biogenesis factor 1 | protein_coding | NP_000457.1;NP_001269606.1;NP_001269607.1;XP_016867808.1 | 8850 | 602136 | . | . | 7.639481 | 7.967812 | 8.038481 | 7.942400 | 17.777385 | 18.417952 | 18.038117 | 18.159355 | -1.004146 | 1049.766359 | 0.134331447 | -0.04443503 | 0.964557636 | 0.998865404 | 10.038842 | 10.032856 | 10.033520 | 10.041233 | 10.035159 | 10.033488 | 1205 | 1141 | 912 | 981 |
| 5190 | NM_000287,N | PEX6 | peroxisomal biogenesis factor 6 | protein_coding | NP_000278.3;NP_001303242.1;XP_011512963.1 | 8859 | 601498 | . | . | 7.199021 | 6.447552 | 7.009055 | 7.982666 | 16.752419 | 14.903804 | 15.728116 | 18.251415 | 1.087356 | 784.716687 | 0.149867178 | 0.806210347 | 0.420121573 | 0.998865404 | 9.554302 | 9.675221 | 9.622341 | 9.546271 | 9.623759 | 9.663648 | 913 | 767 | 691 | 788 |
| 5191 | NM_000288,XI | PEX7 | peroxisomal biogenesis factor 7 | protein_coding | NP_000279.1;XP_005267076.1;XP_006715565.1;XP_016866423.1 | 8860 | 601757 | . | . | 3.828289 | 3.645856 | 2.751936 | 3.391327 | 8.908585 | 8.427561 | 6.175265 | 7.753863 | -1.221304 | 160.487317 | 0.265212242 | -1.0875137 | 0.276809847 | 0.998865404 | 7.463805 | 7.174329 | 7.373555 | 7.357354 | 7.247206 | 7.321397 | 207 | 187 | 113 | 149 |
| 5192 | NM_00137442 | PEX10 | peroxisomal biogenesis factor 10 | protein_coding | NP_001361354.1;NP_001361355.1;NP_001361356.1;NP_002608.1;NP_7: | 8851 | 602859 | . | . | 10.135707 | 11.693745 | 11.040973 | 12.429845 | 23.586204 | 27.030615 | 24.775621 | 28.419363 | 1.032229 | 994.246448 | 0.144115261 | 0.317544888 | 0.750830185 | 0.998865404 | 9.934642 | 9.979922 | 9.892997 | 9.994359 | 9.945104 | 9.990428 | 1049 | 1129 | 851 | 976 |
| 5193 | NM_000286 | PEX12 | peroxisomal biogenesis factor 12 | protein_coding | NP_000277.1 | 8854 | 601758 | . | . | 2.425299 | 2.406290 | 2.958302 | 2.782378 | 5.643771 | 5.562248 | 6.638344 | 6.361577 | 1.171899 | 216.142028 | 0.229092497 | 0.998933448 | 0.317826932 | 0.998865404 | 7.636649 | 7.865923 | 7.718373 | 7.711335 | 7.805676 | 7.772155 | 231 | 213 | 210 | 211 |
| 5194 | NM_002618 | PEX13 | peroxisomal biogenesis factor 13 | protein_coding | NP_002609.1 | 8855 | 601789 | . | . | 2.934617 | 3.320068 | 3.097503 | 2.977558 | 6.828975 | 7.674486 | 6.950707 | 6.807834 | -1.043419 | 430.113296 | 0.179400357 | -0.3417944 | 0.732505628 | 0.998865404 | 8.778974 | 8.717517 | 8.725126 | 8.795445 | 8.749591 | 8.720347 | 477 | 501 | 375 | 385 |
| 5195 | NM_004565,XI | PEX14 | peroxisomal biogenesis factor 14 | protein_coding | NP_004556.1;XP_005263527.1;XP_011539879.1;XP_011539880.1;XP_01: | 8856 | 601791 | . | . | 16.983519 | 14.848067 | 15.352524 | 16.153001 | 39.521346 | 34.321970 | 34.450618 | 36.931918 | -1.042165 | 1022.842223 | 0.138549363 | -0.43005457 | 0.667155943 | 0.998865404 | 10.027784 | 9.968339 | 10.047187 | 9.978015 | 9.975251 | 9.990442 | 1258 | 1074 | 862 | 949 |
| 5198 | NM_012393,XI | PFAS | phosphoribosylformylglycinamidine synthase | protein_coding | NP_036525.1;XP_006721609.1;XP_016880249.1;XP_024306572.1 | 8863 | 602133 | . | . | 16.797123 | 17.239886 | 18.247304 | 16.931012 | 39.087597 | 39.850770 | 40.946421 | 38.710744 | 1.015519 | 2800.152440 | 0.115482218 | 0.192383007 | 0.847442202 | 0.998865404 | 11.440075 | 11.462418 | 11.438706 | 11.450613 | 11.491897 | 11.421424 | 3176 | 3020 | 2560 | 2531 |
| 5199 | NM_00114525 | CFP | complement factor properdin | protein_coding | NP_001138724.1;NP_002612.1;XP_016885064.1 | 8864 | 300383 | . | . | 0.127650 | 0.236412 | 0.101661 | 0.126434 | 0.297046 | 0.546477 | 0.228124 | 0.289075 | -1.534931 | 11.202709 | 0.932407597 | -0.66298677 | 0.507339026 | NA | 3.763465 | 3.141582 | 3.480703 | 3.516686 | 3.474783 | 3.481295 | 11 | 19 | 7 | 9 |
| 5201 | NM_002622,XI | PFDN1 | prefoldin subunit 1 | protein_coding | NP_002613.2;XP_005268522.1 | 8866 | 604897 | . | . | 35.353852 | 36.801896 | 34.677272 | 41.169564 | 82.269870 | 85.069231 | 77.814795 | 94.129313 | 1.036179 | 1536.452197 | 0.133783654 | 0.383253564 | 0.701531766 | 0.998865404 | 10.559731 | 10.610595 | 10.558527 | 10.582623 | 10.533484 | 10.658822 | 1711 | 1654 | 1250 | 1584 |
| 5202 | NM_012394,XI | PFDN2 | prefoldin subunit 2 | protein_coding | NP_036526.2;XP_011507926.1 | 8867 | 613466 | . | . | 39.172713 | 39.388020 | 31.763316 | 42.409974 | 91.156518 | 91.047169 | 71.275959 | 96.965368 | -1.072733 | 713.999646 | 0.16943276 | -0.59782805 | 0.549954673 | 0.998865404 | 9.529946 | 9.427806 | 9.502405 | 9.501089 | 9.356046 | 9.541794 | 852 | 797 | 517 | 734 |
| 5203 | NM_002623,XI | PFDN4 | prefoldin subunit 4 | protein_coding | NP_002614.2;XP_016883368.1 | 8868 | 604898 | . | . | 28.025452 | 24.165884 | 22.364335 | 14.656744 | 65.216371 | 55.860521 | 50.184919 | 33.510904 | -1.429290 | 858.049735 | 0.190929171 | -2.6988991 | 0.006956926 | 0.265533818 | 9.979188 | 9.465088 | 9.908394 | 9.791352 | 9.730342 | 9.443343 | 1252 | 1005 | 745 | 522 |
| 5204 | NM_002624,N | PFDN5 | prefoldin subunit 5 | protein_coding | NP_002615.2;NP_665904.1 | 8869 | 604899 | . | . | 153.225484 | 158.299145 | 143.311853 | 170.823867 | 356.561983 | 365.915552 | 321.587644 | 390.568472 | -1.005687 | 2899.159498 | 0.123712645 | -0.06613642 | 0.947269214 | 0.998865404 | 11.505633 | 11.497193 | 11.492862 | 11.512947 | 11.428367 | 11.563661 | 3307 | 3176 | 2305 | 2935 |
| 5207 | NM_00127180 | PFKFB1 | 6-phosphofructo-2-kinase/fructose-2,6-biphosphatase 1 | protein_coding | NP_001258733.1;NP_001258734.1;NP_002616.2;XP_016885065.1;XP_01 | 8872 | 311790 | . | . | 0.185595 | 0.152441 | 0.155754 | 0.036227 | 0.431887 | 0.352376 | 0.349507 | 0.082829 | -1.938854 | 13.217110 | 0.900015272 | -1.06131992 | 0.288544535 | NA | 4.121102 | 3.175234 | 3.757093 | 3.734210 | 3.729526 | 3.687927 | 23 | 16 | 12 | 4 |
| 5208 | NM_00101805 | PFKFB2 | 6-phosphofructo-2-kinase/fructose-2,6-biphosphatase 2 | protein_coding | NP_001018063.1;NP_006203.2;XP_005273219.1;XP_024303422.1;XP_02 | 8873 | 171835 | . | . | 5.096977 | 4.658922 | 5.200324 | 5.654988 | 11.860877 | 10.769309 | 11.669376 | 12.929457 | 1.067313 | 1054.703331 | 0.136342006 | 0.689323575 | 0.490619666 | 0.998865404 | 9.994886 | 10.088828 | 10.037512 | 9.997476 | 10.045164 | 10.085997 | 1207 | 1071 | 910 | 1049 |
| 5209 | NM_00114544 | PFKFB3 | 6-phosphofructo-2-kinase/fructose-2,6-biphosphatase 3 | protein_coding | NP_001138915.1;NP_001269559.1;NP_001300992.1;NP_001309945.1;NI | 8874 | 605319 | . | . | 28.881508 | 29.726755 | 20.605127 | 21.866514 | 67.208457 | 68.714728 | 46.237307 | 49.995187 | -1.425214 | 3428.779259 | 0.111102962 | -4.60094149 | 4.20586E-06 | 0.000921483 | 11.976596 | 11.465296 | 11.862056 | 11.880904 | 11.557112 | 11.602441 | 4589 | 4397 | 2388 | 2727 |
| 5210 | NM_00131713 | PFKFB4 | 6-phosphofructo-2-kinase/fructose-2,6-biphosphatase 4 | protein_coding | NP_001304063.1;NP_001304064.1;NP_001304065.1;NP_001304066.1;NI | 8875 | 605320 | . | . | 20.447820 | 21.836319 | 20.690861 | 21.148489 | 47.582913 | 50.475626 | 46.429691 | 48.353506 | -1.036547 | 2543.962710 | 0.115587159 | -0.44802586 | 0.654134533 | 0.998865404 | 11.338571 | 11.286686 | 11.298891 | 11.354567 | 11.295656 | 11.300386 | 2880 | 2890 | 2166 | 2349 |
| 5211 | NM_00100202 | PFKL | phosphofructokinase, liver type | protein_coding | NP_001002021.2;NP_002617.3;XP_005261192.1;XP_005261194.1;XP_00 | 8876 | 171860 | . | . | 19.718383 | 21.185522 | 20.326565 | 21.695295 | 45.885484 | 48.971285 | 45.612224 | 49.603715 | 1.012816 | 1881.520952 | 0.122038233 | 0.150547353 | 0.880332794 | 0.998865404 | 10.868581 | 10.886729 | 10.846393 | 10.897810 | 10.860379 | 10.903969 | 2083 | 2083 | 1601 | 1824 |
| 5213 | NM_000289,N | PFKM | phosphofructokinase, muscle | protein_coding | NP_000280.1;NP_001160158.1;NP_001160159.1;NP_001160160.1;NP_0( | 8877 | 610681 | . | . | 44.398471 | 43.510542 | 44.535810 | 48.330296 | 103.317067 | 100.576577 | 99.937069 | 110.501479 | 1.027692 | 4562.783789 | 0.106868306 | 0.368749171 | 0.712314694 | 0.998865404 | 12.135891 | 12.175238 | 12.137300 | 12.150922 | 12.150172 | 12.183352 | 5140 | 4896 | 3942 | 4422 |
| 5214 | NM_00124233 | PFKP | phosphofructokinase, platelet | protein_coding | NP_001229268.1;NP_001309996.1;NP_001309997.1;NP_001309998.1;NI | 8878 | 171840 | . | . | 59.467747 | 63.980322 | 57.618375 | 67.802657 | 138.383896 | 147.893374 | 129.293955 | 155.022720 | 1.001501 | 5100.515296 | 0.118164652 | 0.018312435 | 0.985389607 | 0.999197585 | 12.315432 | 12.317422 | 12.287324 | 12.341944 | 12.249941 | 12.378425 | 5678 | 5679 | 4098 | 5152 |
| 5216 | NM_00137599 | PFN1 | profilin 1 | protein_coding | NP_001362920.1;NP_005013.1 | 8881 | 176610 | . | . | 522.969445 | 588.995266 | 459.987817 | 565.127364 | 1216.971484 | 1361.489033 | 1032.199285 | 1292.096573 | -1.100648 | 13656.074791 | 0.122313873 | -1.13113508 | 0.257998242 | 0.995760118 | 13.804814 | 13.666376 | 13.727181 | 13.821699 | 13.607674 | 13.772381 | 15578 | 16286 | 10212 | 13368 |
| 5217 | NM_002628,N | PFN2 | profilin 2 | protein_coding | NP_002619.1;NP_444252.1 | 8882 | 176590 | . | . | 90.736271 | 84.225986 | 69.084450 | 66.239320 | 211.147026 | 194.692146 | 155.023498 | 151.448333 | -1.307041 | 4491.018856 | 0.110958741 | -3.48151391 | 0.000498588 | 0.040721944 | 12.313031 | 11.926857 | 12.265610 | 12.201132 | 12.034826 | 11.986221 | 6080 | 5284 | 3487 | 3541 |
| 5218 | NM_00128713 | CDK14 | cyclin dependent kinase 14 | protein_coding | NP_001274064.1;NP_001274065.1;NP_001274066.1;NP_036527.1 | 8883 | 610679 | . | . | 1.931154 | 1.814151 | 1.450387 | 1.778809 | 4.493874 | 4.193492 | 3.254626 | 4.067035 | -1.177156 | 279.823451 | 0.213835945 | -1.10040398 | 0.271156144 | 0.998865404 | 8.241544 | 8.005564 | 8.191857 | 8.145943 | 8.037717 | 8.128112 | 363 | 313 | 203 | 263 |
| 5223 | NM_00131707 | PGAM1 | phosphoglycerate mutase 1 | protein_coding | NP_001304008.1;NP_002620.1 | 8888 | 172250 | . | . | 136.099508 | 138.476301 | 124.626616 | 156.224423 | 316.709117 | 320.094180 | 279.658512 | 357.188592 | 1.008273 | 7792.911307 | 0.123079003 | 0.096574579 | 0.923064245 | 0.998865404 | 12.922050 | 12.933820 | 12.919372 | 12.926742 | 12.834681 | 13.017246 | 8898 | 8411 | 6073 | 8124 |
| 5226 | NM_00130445 | PGD | phosphogluconate dehydrogenase | protein_coding | NP_001291380.1;NP_001291381.1;NP_002622.2 | 8891 | 172200 | . | . | 107.010572 | 117.644684 | 152.003035 | 160.135826 | 249.017982 | 271.940939 | 341.090399 | 366.131536 | 1.370233 | 9469.376225 | 0.105965709 | 4.288377898 | 1.79983E-05 | 0.003144685 | 12.964062 | 13.418419 | 13.023564 | 13.095058 | 13.309577 | 13.346055 | 8804 | 8994 | 9322 | 10478 |
| 5228 | NM_00120701 | PGF | placental growth factor | protein_coding | NP_001193941.1;NP_001280572.1;NP_002623.2 | 8893 | 601121 | . | . | 4.488444 | 4.179464 | 3.821779 | 5.137379 | 10.444793 | 9.661019 | 8.575962 | 11.746005 | 1.021037 | 260.516961 | 0.227099301 | 0.132254797 | 0.89478276 | 0.998865404 | 8.010638 | 8.039682 | 8.038449 | 7.997018 | 7.955800 | 8.096181 | 309 | 267 | 197 | 281 |
| 5229 | NM_005023,XI | PGGT1B | protein geranylgeranyltransferase type I subunit beta | protein_coding | NP_005014.2;XP_005272077.2;XP_011541792.1;XP_011541794.1 | 8895 | 602031 | . | . | 1.568516 | 1.295718 | 1.530384 | 1.357753 | 3.650000 | 2.995109 | 3.434139 | 3.104340 | -1.018066 | 432.906979 | 0.192091251 | -0.13447284 | 0.893028683 | 0.998865404 | 8.770149 | 8.745573 | 8.820753 | 8.697542 | 8.803293 | 8.694463 | 553 | 424 | 407 | 366 |
| 5230 | NM_000291 | PGK1 | phosphoglycerate kinase 1 | protein_coding | NP_000282.1 | 8896 | 311800 | . | . | 77.565529 | 75.916245 | 73.946671 | 88.289688 | 180.498169 | 175.483795 | 165.934189 | 201.863876 | 1.041937 | 11827.976756 | 0.114461271 | 0.517798289 | 0.604598663 | 0.998865404 | 13.500007 | 13.559217 | 13.523975 | 13.498334 | 13.472289 | 13.615427 | 13541 | 12313 | 9622 | 12259 |
| 5231 | gene-PGK1P1 | PGK1P1 | phosphoglycerate kinase 1 pseudogene 1 | pseudogene | . | 8897 | . | | . | 0.799100 | 0.886403 | 0.728344 | 0.934163 | 1.859538 | 2.048961 | 1.634383 | 2.135854 | -1.039638 | 46.663082 | 0.460582436 | -0.1217609 | 0.903088383 | 0.998865404 | 5.572938 | 5.514897 | 5.538480 | 5.557107 | 5.520747 | 5.560939 | 52 | 54 | 35 | 48 |
| 5236 | NM_00117281 | PGM1 | phosphoglucomutase 1 | protein_coding | NP_001166289.1;NP_001166290.1;NP_002624.2 | 8905 | 171900 | . | . | 46.786485 | 48.787488 | 48.084973 | 53.016974 | 108.874068 | 112.774465 | 107.901285 | 121.217010 | 1.040956 | 3589.473600 | 0.112725102 | 0.513717095 | 0.607449798 | 0.998865404 | 11.780387 | 11.838148 | 11.778879 | 11.805931 | 11.789363 | 11.860533 | 3984 | 3858 | 3047 | 3580 |
| 5238 | NM_00119991 | PGM3 | phosphoglucomutase 3 | protein_coding | NP_001186846.1;NP_001186847.1;NP_001186848.1;NP_001354215.1;NI | 8907 | 172100 | . | . | 6.130225 | 6.417479 | 6.211935 | 5.592235 | 14.265285 | 14.834288 | 13.939400 | 12.785981 | -1.071387 | 1121.731167 | 0.140418442 | -0.708449 | 0.478666474 | 0.998865404 | 10.180158 | 10.081167 | 10.166100 | 10.145638 | 10.158702 | 10.047644 | 1354 | 1235 | 1013 | 939 |
| 5239 | NM_021965,XI | PGM5 | phosphoglucomutase 5 | protein_coding | NP_068800.2;XP_011517084.1;XP_011517085.1;XP_024303341.1 | 8908 | 600981 | . | . | 0.612383 | 0.438563 | 0.669375 | 0.887230 | 1.425039 | 1.013757 | 1.502057 | 2.028545 | 1.450908 | 74.874550 | 0.383252896 | 1.401048702 | 0.16119951 | 0.933525128 | 5.932902 | 6.470171 | 6.202157 | 6.129760 | 6.221821 | 6.295758 | 82 | 55 | 67 | 94 |
| 5245 | NM_00128149 | PHB | prohibitin | protein_coding | NP_001268425.1;NP_001268426.1;NP_001268644.1;NP_002625.1;XP_01 | 8912 | 176705 | . | . | 121.564398 | 120.349211 | 107.920998 | 120.653513 | 282.885365 | 278.192599 | 242.171599 | 275.859934 | -1.070089 | 6743.075555 | 0.108081408 | -0.90422732 | 0.365874875 | 0.998865404 | 12.767262 | 12.669463 | 12.754860 | 12.738876 | 12.647921 | 12.729712 | 8103 | 7452 | 5389 | 6405 |
| 5250 | NM_002635,N | SLC25A3 | solute carrier family 25 member 3 | protein_coding | NP_002626.1;NP_005879.1;NP_998776.1 | 10989 | 600370 | . | . | 52.982660 | 54.778233 | 50.998218 | 62.074616 | 123.292830 | 126.622338 | 114.438521 | 141.926237 | 1.047524 | 9926.838780 | 0.117670706 | 0.569242963 | 0.569191268 | 0.998865404 | 13.243282 | 13.310180 | 13.252460 | 13.259414 | 13.212173 | 13.372590 | 11120 | 10506 | 8006 | 10413 |
| 5251 | NM_000444,N | PHEX | phosphate regulating endopeptidase homolog X-linked | protein_coding | NP_000435.3;NP_001269683.1;XP_011543835.1;XP_011543838.1;XP_01 | 8918 | 300550 | . | . | 0.167151 | 0.129236 | 0.203382 | 0.130089 | 0.388967 | 0.298736 | 0.456383 | 0.297434 | 1.120425 | 32.707251 | 0.558451837 | 0.293752234 | 0.768947237 | 0.998865404 | 4.945169 | 5.113028 | 5.033864 | 5.001507 | 5.062821 | 5.001432 | 40 | 29 | 37 | 25 |
| 5252 | NM_002636,N | PHF1 | PHD finger protein 1 | protein_coding | NP_002627.2;NP_077084.2;XP_011512964.1;XP_011512965.1;XP_01151 | 8919 | 602881 | . | . | 16.483040 | 15.888849 | 18.367366 | 18.503922 | 38.356709 | 36.727785 | 41.215837 | 42.307015 | 1.122123 | 1210.932926 | 0.130359575 | 1.275173346 | 0.202247893 | 0.967656221 | 10.156342 | 10.322673 | 10.209819 | 10.182846 | 10.285415 | 10.280951 | 1338 | 1209 | 1117 | 1197 |
| 5253 | NM_005392,XI | PHF2 | PHD finger protein 2 | protein_coding | NP_005383.3;XP_005252108.1;XP_006717206.1 | 8920 | 604351 | . | . | 2.866170 | 3.037443 | 2.699093 | 3.148513 | 6.669697 | 7.021186 | 6.056687 | 7.198702 | -1.019914 | 491.200911 | 0.173099393 | -0.16433907 | 0.869464228 | 0.998865404 | 8.954676 | 8.925515 | 8.933290 | 8.958566 | 8.888530 | 8.976436 | 560 | 546 | 393 | 488 |
| 5255 | NM_00112267 | PHKA1 | phosphorylase kinase regulatory subunit alpha 1 | protein_coding | NP_001116142.1;NP_001165907.1;NP_002628.2;XP_006724724.1 | 8925 | 311870 | . | . | 2.220134 | 2.283388 | 2.048563 | 2.478392 | 5.166342 | 5.278153 | 4.596915 | 5.666548 | 1.007058 | 432.775384 | 0.181349277 | 0.055947894 | 0.955383316 | 0.998865404 | 8.752760 | 8.762174 | 8.751394 | 8.758087 | 8.708814 | 8.806617 | 494 | 468 | 347 | 440 |
| 5256 | NM_000292,XI | PHKA2 | phosphorylase kinase regulatory subunit alpha 2 | protein_coding | NP_000283.1;XP_005274605.1;XP_005274607.1;XP_006724559.1;XP_00 | 8926 | 300798 | . | . | 4.063263 | 4.869965 | 4.236179 | 5.009246 | 9.455381 | 11.257141 | 9.505864 | 11.453048 | -1.061221 | 816.508971 | 0.151373496 | -0.5663166 | 0.571178577 | 0.998865404 | 9.715850 | 9.629509 | 9.648508 | 9.736436 | 9.616505 | 9.684817 | 909 | 963 | 655 | 779 |
| 5257 | NM_000293,N | PHKB | phosphorylase kinase regulatory subunit beta | protein_coding | NP_000284.1;NP_001027005.1;NP_001350766.1 | 8927 | 172490 | . | . | 3.785978 | 3.931949 | 3.851945 | 4.181519 | 8.810124 | 9.088878 | 8.643655 | 9.560547 | 1.035038 | 681.887133 | 0.153165048 | 0.324377773 | 0.745652045 | 0.998865404 | 9.388534 | 9.437824 | 9.387122 | 9.415094 | 9.399280 | 9.449343 | 756 | 738 | 581 | 674 |
| 5260 | NM_00125845 | PHKG1 | phosphorylase kinase catalytic subunit gamma 1 | protein_coding | NP_001245388.1;NP_001245389.1;NP_006204.1;XP_005271829.1;XP_01 | 8930 | 172470 | . | . | 0.273192 | 0.353825 | 0.293273 | 0.230052 | 0.635730 | 0.817881 | 0.658097 | 0.525987 | -1.184019 | 23.664512 | 0.638385352 | -0.38173168 | 0.702660401 | NA | 4.681128 | 4.437942 | 4.560121 | 4.586028 | 4.566745 | 4.546340 | 26 | 31 | 21 | 18 |
| 5261 | NM_000294,N | PHKG2 | phosphorylase kinase catalytic subunit gamma 2 | protein_coding | NP_000285.1;NP_001165903.1 | 8931 | 172471 | . | . | 5.779845 | 6.449119 | 6.760563 | 7.189038 | 13.449938 | 14.907427 | 15.170507 | 16.436881 | 1.117231 | 1048.976115 | 0.137974526 | 1.159112319 | 0.246410405 | 0.992733415 | 9.952786 | 10.112337 | 9.965675 | 10.017098 | 10.046153 | 10.099838 | 1101 | 1107 | 928 | 1075 |
| 5264 | NM_00103753 | PHYH | phytanoyl-CoA 2-hydroxylase | protein_coding | NP_001032626.1;NP_001310009.1;NP_001310011.1;NP_001310012.1;NI | 8940 | 602026 | . | . | 7.973936 | 8.102036 | 7.685627 | 7.540471 | 18.555674 | 18.728220 | 17.246326 | 17.240390 | -1.067098 | 378.769240 | 0.183802629 | -0.5097494 | 0.61022704 | 0.998865404 | 8.611243 | 8.517588 | 8.578377 | 8.589673 | 8.552159 | 8.540169 | 446 | 426 | 322 | 340 |
| 5265 | NM_000295,N | SERPINA1 | serpin family A member 1 | protein_coding | NP_000286.3;NP_001002235.1;NP_001002236.1;NP_001121172.1;NP_0( | 8941 | 107400 | . | . | 0.240723 | 7.690151 | 0.131025 | 0.085415 | 0.560171 | 17.776127 | 0.294016 | 0.195292 | -30.447369 | 192.931749 | 1.796306608 | -2.7435437 | 0.006077996 | 0.243173429 | 8.545307 | 6.371632 | 6.407605 | 8.014577 | 6.392434 | 6.354811 | 28 | 780 | 13 | 9 |
| 5266 | NM_002638 | PI3 | peptidase inhibitor 3 | protein_coding | NP_002629.1 | 8947 | 182257 | . | . | 1.066461 | 0.974035 | 3.116225 | 1.804144 | 2.481699 | 2.251526 | 6.992719 | 4.124962 | 2.309909 | 30.424003 | 0.589852897 | 2.047690317 | 0.040590353 | 0.62818612 | 4.198253 | 5.409088 | 4.844693 | 4.838788 | 4.983192 | 4.895732 | 22 | 19 | 47 | 29 |
| 5269 | NM_00119529 | SERPINB6 | serpin family B member 6 | protein_coding | NP_001182220.2;NP_001258751.1;NP_001258752.1;NP_001258753.1;NI | 8950 | 173321 | . | . | 30.618880 | 29.582330 | 34.448581 | 35.540952 | 71.251391 | 68.380885 | 77.301617 | 81.260165 | 1.174833 | 1538.135231 | 0.127302829 | 1.826008626 | 0.067848963 | 0.740331407 | 10.466140 | 10.698451 | 10.533450 | 10.507228 | 10.608542 | 10.680684 | 1657 | 1500 | 1379 | 1629 |
| 5270 | NM_00113652 | SERPINE2 | serpin family E member 2 | protein_coding | NP_001130000.1;NP_001130002.1;NP_006207.1;XP_005246698.1;XP_01 | 8951 | 177010 | . | . | 1081.242334 | 1026.699412 | 948.378040 | 823.992960 | 2516.095259 | 2373.261608 | 2128.132840 | 1883.961994 | -1.203803 | 66483.636916 | 0.110443657 | -2.42294349 | 0.015395321 | 0.417236976 | 16.148308 | 15.880725 | 16.122722 | 16.066614 | 15.995223 | 15.869896 | 86304 | 75871 | 56318 | 52413 |
| 5271 | NM_00103184 | SERPINB8 | serpin family B member 8 | protein_coding | NP_001027018.1;NP_001263419.1;NP_001335296.1;NP_001335297.1;NI | 8952 | 601697 | . | . | 3.080591 | 3.446779 | 3.094725 | 2.968167 | 7.168659 | 7.967385 | 6.944474 | 6.786360 | -1.090154 | 329.268151 | 0.195643118 | -0.63652545 | 0.524434 | 0.998865404 | 8.424123 | 8.299421 | 8.359469 | 8.412928 | 8.346597 | 8.330507 | 377 | 388 | 278 | 291 |
| 5274 | NM_00112275 | SERPINI1 | serpin family I member 1 | protein_coding | NP_001116224.1;NP_005016.1;XP_016862107.1 | 8943 | 602445 | . | . | 3.406967 | 2.442192 | 2.663225 | 2.778811 | 7.928151 | 5.645237 | 5.976199 | 6.353419 | -1.103419 | 143.610770 | 0.292804305 | -0.48489941 | 0.627747703 | 0.998865404 | 7.234719 | 7.093886 | 7.254034 | 7.104286 | 7.139197 | 7.149898 | 206 | 132 | 117 | 130 |
| 5277 | NM_002641,N | PIGA | phosphatidylinositol glycan anchor biosynthesis class A | protein_coding | NP_002632.1;NP_065206.3;XP_011543841.1 | 8957 | 311770 | . | . | 3.579373 | 3.692760 | 4.276898 | 3.725452 | 8.329346 | 8.535980 | 9.597237 | 8.517807 | 1.065581 | 441.152205 | 0.181751554 | 0.504208784 | 0.614114668 | 0.998865404 | 8.738154 | 8.830630 | 8.773776 | 8.753304 | 8.855667 | 8.746677 | 500 | 453 | 431 | 389 |
| 5278 | gene-PIGAP1 | PIGAP1 | phosphatidylinositol glycan anchor biosynthesis class A pseudoger | pseudogene | . | 8958 | . | | . | 0.208028 | 0.207096 | 0.270890 | 0.038298 | 0.484089 | 0.478712 | 0.607870 | 0.087564 | -1.327222 | 20.483131 | 0.814789291 | -0.50124541 | 0.61619842 | NA | 4.542770 | 4.142249 | 4.356579 | 4.355183 | 4.386056 | 4.273282 | 27 | 25 | 27 | 4 |
| 5279 | NM_002642,N | PIGC | phosphatidylinositol glycan anchor biosynthesis class C | protein_coding | NP_002633.1;NP_714969.1 | 8960 | 601730 | . | . | 13.130630 | 12.265687 | 12.867672 | 12.582515 | 30.555516 | 28.352684 | 28.874686 | 28.768428 | -1.008568 | 578.799532 | 0.16088487 | -0.07649991 | 0.939021387 | 0.998865404 | 9.182884 | 9.170931 | 9.204744 | 9.153653 | 9.184062 | 9.163819 | 695 | 603 | 509 | 532 |
| 5281 | NM_002643,N | PIGF | phosphatidylinositol glycan anchor biosynthesis class F | protein_coding | NP_002634.1;NP_775097.1;XP_005264426.1;XP_011531210.1 | 8962 | 600153 | . | . | 11.011927 | 9.167027 | 8.685308 | 6.984994 | 25.625204 | 21.189996 | 19.489583 | 15.970361 | -1.301649 | 336.518840 | 0.214138425 | -1.77614329 | 0.075709303 | 0.767975096 | 8.571467 | 8.192850 | 8.511886 | 8.405125 | 8.383230 | 8.236940 | 477 | 374 | 290 | 236 |
| 5283 | NM_00136369 | PIGH | phosphatidylinositol glycan anchor biosynthesis class H | protein_coding | NP_001350623.1;NP_004560.1;XP_011535140.1;XP_016876860.1 | 8964 | 600154 | . | . | 19.767350 | 19.525983 | 22.872066 | 18.922444 | 45.999437 | 45.135179 | 51.324252 | 43.263921 | 1.049699 | 891.240114 | 0.151813976 | 0.460929182 | 0.644849271 | 0.998865404 | 9.763945 | 9.834533 | 9.788097 | 9.772587 | 9.882241 | 9.743956 | 1012 | 928 | 870 | 774 |
| 5286 | NM_00132137 | PIK3C2A | phosphatidylinositol-4-phosphate 3-kinase catalytic subunit type 2 | protein_coding | NP_001308307.1;NP_001308309.1;NP_002636.2;XP_011518488.1;XP_01 | 8971 | 603601 | . | . | 5.292668 | 5.519486 | 5.577135 | 4.343463 | 12.316257 | 12.758537 | 12.514927 | 9.930812 | -1.024979 | 1276.184359 | 0.155722199 | -0.22857369 | 0.819200275 | 0.998865404 | 10.334954 | 10.300077 | 10.372136 | 10.273681 | 10.400516 | 10.200897 | 1583 | 1304 | 1241 | 1026 |
| 5287 | NM_00137733 | PIK3C2B | phosphatidylinositol-4-phosphate 3-kinase catalytic subunit type 2 | protein_coding | NP_001364263.1;NP_001364264.1;NP_002637.3;XP_005245315.1;XP_01 | 8972 | 602838 | . | . | 1.743883 | 1.895790 | 1.452789 | 2.295464 | 4.058088 | 4.382202 | 3.260016 | 5.248307 | 1.002908 | 449.251541 | 0.19150605 | 0.021875106 | 0.982547582 | 0.998865404 | 8.809711 | 8.813046 | 8.828996 | 8.787499 | 8.722114 | 8.891117 | 534 | 468 | 337 | 480 |
| 5289 | NM_00130802 | PIK3C3 | phosphatidylinositol 3-kinase catalytic subunit type 3 | protein_coding | NP_001294949.1;NP_002638.2;XP_011524333.1 | 8974 | 602609 | . | . | 3.282270 | 3.318306 | 3.667417 | 3.095559 | 7.637978 | 7.670413 | 8.229578 | 7.077626 | -1.049324 | 899.679585 | 0.143761268 | -0.4831655 | 0.628978229 | 0.998865404 | 9.847492 | 9.778211 | 9.808604 | 9.850923 | 9.827896 | 9.761801 | 1030 | 1023 | 799 | 785 |
| 5290 | NM_006218,XI | PIK3CA | phosphatidylinositol-4,5-bisphosphate 3-kinase catalytic subunit a | protein_coding | NP_006209.2;XP_006713721.1;XP_011511196.1 | 8975 | 171834 | . | . | 1.797626 | 1.877576 | 1.970977 | 1.454497 | 4.183150 | 4.340102 | 4.422816 | 3.325534 | -1.086817 | 512.658055 | 0.188527695 | -0.63709071 | 0.524065759 | 0.998865404 | 9.060171 | 8.941078 | 9.013389 | 9.035950 | 9.064677 | 8.869494 | 604 | 586 | 494 | 389 |
| 5291 | NM_00125604 | PIK3CB | phosphatidylinositol-4,5-bisphosphate 3-kinase catalytic subunit b | protein_coding | NP_001242974.1;NP_006210.1;XP_006713722.1;XP_011511197.1;XP_01 | 8976 | 602925 | . | . | 8.835272 | 7.600980 | 7.741332 | 8.274334 | 20.560041 | 17.570007 | 17.371325 | 18.918282 | -1.061661 | 1467.025226 | 0.131470818 | -0.65659816 | 0.511439329 | 0.998865404 | 10.561088 | 10.474983 | 10.590429 | 10.488776 | 10.494428 | 10.495165 | 1853 | 1524 | 1237 | 1335 |
| 5292 | NM_00124318 | PIM1 | Pim-1 proto-oncogene, serine/threonine kinase | protein_coding | NP_001230115.1;NP_002639.1 | 8986 | 164960 | . | . | 1.855284 | 1.757650 | 1.497794 | 1.939015 | 4.317322 | 4.062885 | 3.361008 | 4.433328 | -1.065957 | 149.688697 | 0.276254395 | -0.33356687 | 0.738706424 | 0.998865404 | 7.271732 | 7.178402 | 7.249228 | 7.226217 | 7.164401 | 7.259207 | 183 | 162 | 110 | 153 |
| 5293 | NM_00135023 | PIK3CD | phosphatidylinositol-4,5-bisphosphate 3-kinase catalytic subunit d | protein_coding | NP_001337163.1;NP_001337164.1;NP_005017.3;XP_006710750.1;XP_00 | 8977 | 602839 | . | . | 24.777232 | 27.001671 | 28.118119 | 30.796242 | 57.657641 | 62.415579 | 63.096243 | 70.411955 | 1.117139 | 4583.022161 | 0.111629097 | 1.431599049 | 0.152258602 | 0.916288189 | 12.080041 | 12.239710 | 12.088388 | 12.138356 | 12.168795 | 12.242027 | 4835 | 4813 | 4023 | 4731 |
| 5295 | NM_00124246 | PIK3R1 | phosphoinositide-3-kinase regulatory subunit 1 | protein_coding | NP_001229395.1;NP_852556.2;NP_852664.1;NP_852665.1;XP_0052485S | 8979 | 171833 | . | . | 7.876424 | 7.791782 | 7.839010 | 6.923798 | 18.328762 | 18.011052 | 17.590511 | 15.830441 | -1.073225 | 1645.146143 | 0.129047438 | -0.79004354 | 0.429502342 | 0.998865404 | 10.733921 | 10.632289 | 10.711985 | 10.708772 | 10.705964 | 10.602086 | 1965 | 1834 | 1473 | 1388 |
| 5296 | NM_005027,N | PIK3R2 | phosphoinositide-3-kinase regulatory subunit 2 | protein_coding | NP_005018.2 | 8980 | 603157 | . | . | 9.516247 | 10.306788 | 9.546886 | 13.341154 | 22.144696 | 23.824602 | 21.422933 | 30.502967 | 1.137113 | 1319.602375 | 0.15945861 | 1.162527797 | 0.24502114 | 0.992733415 | 10.270569 | 10.455297 | 10.283692 | 10.338132 | 10.280605 | 10.523266 | 1371 | 1381 | 1030 | 1522 |
| 5297 | NM_00136286 | PI4KA | phosphatidylinositol 4-kinase alpha | protein_coding | NP_001349791.1;NP_001349792.1;NP_477352.3;XP_005261692.1;XP_01 | 8983 | 600286 | . | . | 16.663950 | 18.030222 | 19.807823 | 19.590380 | 38.777697 | 41.677667 | 44.448176 | 44.791077 | 1.138213 | 3795.910310 | 0.10893413 | 1.714528619 | 0.086431686 | 0.795065303 | 11.793840 | 11.980582 | 11.815744 | 11.851997 | 11.946809 | 11.935523 | 3998 | 3916 | 3538 | 3763 |
| 5298 | NM_00119877 | PI4KB | phosphatidylinositol 4-kinase beta | protein_coding | NP_001185702.1;NP_001185703.1;NP_001185704.1;NP_001317650.1;NI | 8984 | 602758 | . | . | 12.376957 | 12.762952 | 11.975804 | 13.494566 | 28.801687 | 29.502135 | 26.873358 | 30.853720 | 1.011446 | 1570.813792 | 0.129568588 | 0.126717873 | 0.899163699 | 0.998865404 | 10.609242 | 10.625306 | 10.596849 | 10.627744 | 10.572257 | 10.667778 | 1763 | 1719 | 1289 | 1572 |
| 5300 | NM_006221,N | PIN1 | peptidylprolyl cis/trans isomerase, NIMA-interacting 1 | protein_coding | NP_006212.1;XP_011526370.1 | 8988 | 601052 | . | . | 31.986948 | 33.756795 | 28.152236 | 35.415650 | 74.434946 | 78.030352 | 63.172800 | 80.973681 | -1.047711 | 1013.552169 | 0.152403342 | -0.44120735 | 0.659062892 | 0.998865404 | 10.018776 | 9.950834 | 9.980962 | 10.020175 | 9.881416 | 10.044757 | 1163 | 1149 | 764 | 1032 |
| 5303 | NM_00117074 | PIN4 | peptidylprolyl cis/trans isomerase, NIMA-interacting 4 | protein_coding | NP_001164218.1;NP_006214.3 | 8992 | 300252 | . | . | 6.831752 | 7.721123 | 5.616606 | 7.571426 | 15.897769 | 17.847719 | 12.603500 | 17.311165 | -1.116906 | 269.304367 | 0.227121347 | -0.70230204 | 0.482490823 | 0.998865404 | 8.151453 | 7.990232 | 8.066953 | 8.130229 | 7.967687 | 8.111252 | 308 | 325 | 189 | 273 |
| 5305 | NM_00133006 | PIP4K2A | phosphatidylinositol-5-phosphate 4-kinase type 2 alpha | protein_coding | NP_001316991.1;NP_005019.2;XP_006717513.1;XP_016871819.1;XP_01 | 8997 | 603140 | . | . | 9.740121 | 9.364378 | 7.547526 | 8.228906 | 22.665661 | 21.646180 | 16.936430 | 18.814415 | -1.220760 | 1035.178347 | 0.138147115 | -2.08313478 | 0.03723894 | 0.602769983 | 10.152454 | 9.864532 | 10.105926 | 10.062342 | 9.910423 | 9.963954 | 1348 | 1193 | 780 | 907 |
| 5306 | NM_006224,XI | PITPNA | phosphatidylinositol transfer protein alpha | protein_coding | NP_006215.1;XP_011522241.1 | 9001 | 600174 | . | . | 16.594003 | 16.706500 | 17.422789 | 18.150080 | 38.614926 | 38.617824 | 39.096230 | 41.498001 | 1.054538 | 2080.570850 | 0.117356767 | 0.652809931 | 0.513878821 | 0.998865404 | 10.983984 | 11.060528 | 11.002715 | 10.999481 | 11.029416 | 11.057211 | 2337 | 2181 | 1825 | 2037 |
| 5308 | NM_000325,N | PITX2 | paired like homeodomain 2 | protein_coding | NP_000316.2;NP_001191326.1;NP_001191327.1;NP_001191328.1;NP_7( | 9005 | 601542 | . | . | 1.544294 | 1.678221 | 1.799591 | 1.642482 | 3.593635 | 3.879283 | 4.038229 | 3.755343 | 1.013971 | 136.022760 | 0.290530607 | 0.068893596 | 0.945074316 | 0.998865404 | 7.077942 | 7.097401 | 7.029031 | 7.132498 | 7.105070 | 7.068947 | 134 | 166 | 125 | 122 |
| 5310 | NM_000296,N | PKD1 | polycystin 1, transient receptor potential channel interacting | protein_coding | NP_000287.4;NP_001009944.3;XP_005255427.1;XP_011520830.1;XP_01 | 9008 | 601313 | . | . | 4.790956 | 4.863916 | 4.773039 | 5.038120 | 11.148752 | 11.243160 | 10.710558 | 11.519064 | -1.052419 | 2001.112115 | 0.12099343 | -0.60919457 | 0.542395478 | 0.998865404 | 11.002924 | 10.929310 | 10.967853 | 11.004457 | 10.974760 | 10.915934 | 2309 | 2265 | 1759 | 1760 |
| 5311 | NM_000297,N | PKD2 | polycystin 2, transient receptor potential cation channel | protein_coding | NP_000288.1;XP_011530330.1;XP_011530331.1;XP_011530332.1 | 9009 | 173910 | . | . | 8.158282 | 8.090024 | 8.748886 | 6.651973 | 18.984658 | 18.700454 | 19.632248 | 15.208946 | -1.056817 | 1243.685372 | 0.156000299 | -0.51105901 | 0.609309738 | 0.998865404 | 10.319393 | 10.240336 | 10.314028 | 10.282088 | 10.359469 | 10.142150 | 1502 | 1350 | 1204 | 970 |
| 5314 | NM_138694,N | PKHD1 | PKHD1 ciliary IPT domain containing fibrocystin/polyductin | protein_coding | NP_619639.3;NP_733842.2;XP_011512982.1;XP_011512984.1;XP_01151 | 9016 | 606702 | . | . | 0.045282 | 0.026222 | 0.011933 | 0.023186 | 0.105374 | 0.060613 | 0.026777 | 0.053013 | -2.289628 | 14.353159 | 0.87260246 | -1.36959619 | 0.170812988 | NA | 4.322387 | 3.121007 | 3.897849 | 3.852444 | 3.808690 | 3.843694 | 29 | 16 | 4 | 12 |
| 5315 | NM_00120679 | PKM | pyruvate kinase M1/2 | protein_coding | NP_001193725.1;NP_001193726.1;NP_001193727.1;NP_001193728.1;NI | 9021 | 179050 | . | . | 957.734939 | 989.045784 | 997.527101 | 1151.326548 | 2228.688656 | 2286.223763 | 2238.421847 | 2632.371455 | 1.088783 | 74137.701528 | 0.106763184 | 1.149429433 | 0.250378949 | 0.993339639 | 16.115263 | 16.237969 | 16.129460 | 16.149317 | 16.153625 | 16.269345 | 80797 | 77482 | 62754 | 77261 |
| 5316 | NM_00128625 | PKNOX1 | PBX/knotted 1 homeobox 1 | protein_coding | NP_001273187.1;NP_001307623.1;NP_004562.2 | 9022 | 602100 | . | . | 1.803352 | 1.979116 | 1.934551 | 1.967554 | 4.196475 | 4.574816 | 4.341077 | 4.498580 | 1.019960 | 317.810840 | 0.197527103 | 0.144344816 | 0.885228191 | 0.998865404 | 8.297872 | 8.326039 | 8.282402 | 8.329333 | 8.314831 | 8.319027 | 347 | 354 | 278 | 302 |
| 5318 | NM_00100524 | PKP2 | plakophilin 2 | protein_coding | NP_001005242.2;NP_004563.2 | 9024 | 602861 | . | . | 0.120726 | 0.145295 | 0.124692 | 0.215123 | 0.280933 | 0.335857 | 0.279806 | 0.491853 | 1.251201 | 21.218380 | 0.682788685 | 0.473518759 | 0.635843121 | NA | 4.238681 | 4.558154 | 4.384609 | 4.397965 | 4.385235 | 4.436808 | 20 | 22 | 15 | 28 |
| 5321 | NM_00131119 | PLA2G4A | phospholipase A2 group IVA | protein_coding | NP_001298122.2;NP_077734.2;XP_005245324.2;XP_011507944.1 | 9035 | 600522 | . | . | 9.090710 | 9.567497 | 10.219114 | 8.712041 | 21.154455 | 22.115698 | 22.931396 | 19.919047 | 1.001451 | 831.591253 | 0.153292093 | 0.013650612 | 0.989108726 | 0.999197585 | 9.698446 | 9.701015 | 9.683110 | 9.712530 | 9.760838 | 9.633359 | 937 | 915 | 789 | 710 |
| 5324 | NM_00111463 | PLAG1 | PLAG1 zinc finger | protein_coding | NP_001108106.1;NP_001108107.1;NP_002646.2;XP_011515846.1;XP_01 | 9045 | 603026 | . | . | 0.074813 | 0.083213 | 0.094914 | 0.088707 | 0.174092 | 0.192351 | 0.212985 | 0.202819 | 1.143601 | 19.736667 | 0.689277503 | 0.280849448 | 0.778825875 | NA | 4.202828 | 4.396303 | 4.289058 | 4.298326 | 4.308135 | 4.301726 | 20 | 21 | 19 | 19 |
| 5325 | NM_00108095 | PLAGL1 | PLAG1 like zinc finger 1 | protein_coding | NP_001074420.1;NP_001074421.1;NP_001074422.1;NP_001074423.1;NI | 9046 | 603044 | . | . | 2.745359 | 2.795049 | 3.493362 | 3.063054 | 6.388564 | 6.460878 | 7.839005 | 7.003302 | 1.168153 | 321.912201 | 0.201008913 | 1.11551703 | 0.264628925 | 0.996598404 | 8.213688 | 8.438601 | 8.278306 | 8.287510 | 8.414945 | 8.320353 | 339 | 323 | 327 | 298 |
| 5326 | NM_002657,XI | PLAGL2 | PLAG1 like zinc finger 2 | protein_coding | NP_002648.1;XP_005260493.1;XP_011527166.1 | 9047 | 604866 | . | . | 2.904692 | 3.403510 | 3.730414 | 3.746981 | 6.759337 | 7.867367 | 8.370940 | 8.567026 | 1.172511 | 608.194445 | 0.163002694 | 1.408578295 | 0.158959901 | 0.931687264 | 9.129251 | 9.358434 | 9.140204 | 9.239211 | 9.299264 | 9.294282 | 596 | 650 | 573 | 613 |
| 5327 | NM_000930,N | PLAT | plasminogen activator, tissue type | protein_coding | NP_000921.1;NP_001306118.1;NP_127509.1 | 9051 | 173370 | . | . | 213.363739 | 216.959884 | 139.122453 | 150.378681 | 496.506198 | 501.512504 | 312.186777 | 343.822983 | -1.505580 | 17133.611379 | 0.101274265 | -5.82891578 | 5.57886E-09 | 4.0529E-06 | 14.329729 | 13.739379 | 14.212756 | 14.217468 | 13.836107 | 13.891258 | 23647 | 22278 | 11496 | 13249 |
| 5329 | NM_00100537 | PLAUR | plasminogen activator, urokinase receptor | protein_coding | NP_001005376.1;NP_001005377.1;NP_001287966.1;NP_002650.1;XP_0( | 9053 | 173391 | . | . | 23.488349 | 24.632586 | 16.652640 | 19.672321 | 54.658354 | 56.939327 | 37.368042 | 44.978426 | -1.329650 | 892.373151 | 0.147458315 | -2.78754407 | 0.005310923 | 0.222141426 | 9.992684 | 9.581060 | 9.888965 | 9.899159 | 9.637027 | 9.739256 | 1165 | 1107 | 616 | 772 |
| 5331 | NM_000932,N | PLCB3 | phospholipase C beta 3 | protein_coding | NP_000923.1;NP_001171812.1;NP_001303243.1;XP_011543403.1;XP_01 | 9056 | 600230 | . | . | 6.919449 | 8.256571 | 8.199178 | 7.802884 | 16.101841 | 19.085433 | 18.398720 | 17.840364 | 1.038063 | 1010.391644 | 0.144963478 | 0.371775228 | 0.710060209 | 0.998865404 | 9.953643 | 10.007257 | 9.904308 | 10.023775 | 10.001320 | 9.984858 | 1049 | 1157 | 905 | 954 |
| 5332 | NM_000933,N | PLCB4 | phospholipase C beta 4 | protein_coding | NP_000924.3;NP_001166117.1;NP_001364063.1;NP_001364064.1;NP_0( | 9059 | 600810 | . | . | 6.600872 | 7.341048 | 8.720671 | 5.623248 | 15.360504 | 16.969165 | 19.568930 | 12.856888 | 1.017665 | 1237.578969 | 0.164041018 | 0.15399965 | 0.877610012 | 0.998865404 | 10.260311 | 10.286183 | 10.256611 | 10.268699 | 10.396970 | 10.138867 | 1401 | 1334 | 1271 | 971 |
| 5333 | NM_00113096 | PLCD1 | phospholipase C delta 1 | protein_coding | NP_001124436.1;NP_006216.2;XP_016862111.1 | 9060 | 602142 | . | . | 1.256512 | 1.165960 | 1.099673 | 1.567166 | 2.923954 | 2.695168 | 2.467634 | 3.583141 | 1.109435 | 110.471231 | 0.315141178 | 0.475421509 | 0.634486567 | 0.998865404 | 6.711291 | 6.859935 | 6.779005 | 6.756180 | 6.754386 | 6.846330 | 125 | 109 | 87 | 124 |
| 5334 | NM_006226,XI | PLCL1 | phospholipase C like 1 (inactive) | protein_coding | NP_006217.3;XP_005246700.1;XP_005246701.1;XP_011509653.1;XP_01( | 9063 | 600597 | . | . | 1.668303 | 1.510987 | 2.269439 | 2.317985 | 3.882209 | 3.492714 | 5.092555 | 5.299797 | 1.393086 | 369.755280 | 0.197488307 | 2.421834563 | 0.015442379 | 0.417236976 | 8.272154 | 8.749409 | 8.405640 | 8.433559 | 8.539131 | 8.678481 | 347 | 342 | 332 | 449 |
| 5335 | NM_002660,N | PLCG1 | phospholipase C gamma 1 | protein_coding | NP_002651.2;NP_877963.1;XP_005260495.1;XP_011527167.1;XP_01152 | 9065 | 172420 | . | . | 25.578054 | 28.659646 | 37.904660 | 33.607104 | 59.521185 | 66.248058 | 85.056958 | 76.838649 | 1.309613 | 5292.061232 | 0.113781874 | 3.420059394 | 0.000626075 | 0.049381191 | 12.161927 | 12.551131 | 12.223941 | 12.265788 | 12.523411 | 12.416427 | 5142 | 5071 | 5544 | 5276 |
| 5336 | NM_002661 | PLCG2 | phospholipase C gamma 2 | protein_coding | NP_002652.2 | 9066 | 600220 | . | . | 1.262203 | 1.168722 | 0.962365 | 1.288368 | 2.937198 | 2.701553 | 2.159520 | 2.945701 | -1.094030 | 316.536468 | 0.213276709 | -0.60790342 | 0.543251531 | 0.998865404 | 8.370073 | 8.239426 | 8.352493 | 8.304786 | 8.202484 | 8.351160 | 397 | 342 | 226 | 323 |
| 5337 | NM_00113008 | PLD1 | phospholipase D1 | protein_coding | NP_001123553.1;NP_002653.1;XP_005247590.1;XP_005247591.1;XP_01 | 9067 | 602382 | . | . | 3.169874 | 3.250579 | 2.688362 | 3.072603 | 7.376427 | 7.513862 | 6.032605 | 7.025143 | -1.119544 | 568.269896 | 0.163885675 | -0.99405582 | 0.320195693 | 0.998865404 | 9.229849 | 9.066388 | 9.182031 | 9.190423 | 9.073049 | 9.149864 | 687 | 652 | 438 | 533 |
| 5338 | NM_00124310 | PLD2 | phospholipase D2 | protein_coding | NP_001230037.1;NP_002654.3;XP_005256753.1;XP_016880253.1 | 9068 | 602384 | . | . | 6.550403 | 6.803332 | 8.164941 | 7.637155 | 15.243057 | 15.726210 | 18.321888 | 17.461449 | 1.186292 | 769.264967 | 0.147068648 | 1.67581307 | 0.093774813 | 0.810751527 | 9.458778 | 9.705371 | 9.515200 | 9.531045 | 9.662928 | 9.621845 | 801 | 768 | 747 | 760 |
| 5339 | NM_000445,N | PLEC | plectin | protein_coding | NP_000436.2;NP_958780.1;NP_958781.1;NP_958782.1;NP_958783.1;NP | 9069 | 601282 | . | . | 41.470111 | 43.525279 | 42.253524 | 36.372540 | 96.502648 | 100.610635 | 94.815680 | 83.161490 | -1.098812 | 16685.778159 | 0.114104153 | -1.19140255 | 0.233495592 | 0.988893316 | 14.092686 | 13.956777 | 14.035464 | 14.092864 | 14.049642 | 13.913148 | 19451 | 19477 | 14933 | 13714 |
| 5342 | NM_002665 | PLGLB2 | plasminogen like B2 | protein_coding | NP_002656.1 | 9073 | . | | . | 0.028868 | 0.043945 | 0.106011 | 0.041754 | 0.067176 | 0.101582 | 0.237885 | 0.095466 | 1.799479 | 5.680982 | 1.296005817 | 0.653993247 | 0.513116157 | NA | 2.019256 | 2.869520 | 2.475517 | 2.481160 | 2.506490 | 2.479661 | 4 | 5 | 9 | 4 |
| 5343 | NM_00103239 | PLGLB1 | plasminogen like B1 | protein_coding | NP_001027564.1 | 9072 | 173340 | . | . | 0.041540 | 0.055202 | 0.111805 | 0.025307 | 0.096666 | 0.127602 | 0.250886 | 0.057861 | 1.353401 | 6.382292 | 1.245468513 | 0.350542139 | 0.725931871 | NA | 2.436885 | 2.877719 | 2.647804 | 2.657995 | 2.679735 | 2.643104 | 5 | 7 | 10 | 3 |
| 5346 | NM_00114531 | PLIN1 | perilipin 1 | protein_coding | NP_001138783.1;NP_002657.3;XP_005254991.1 | 9076 | 170290 | . | . | 0.328060 | 0.311248 | 0.215240 | 0.255923 | 0.763409 | 0.719463 | 0.482993 | 0.585137 | -1.360785 | 26.033553 | 0.607841593 | -0.73117528 | 0.464672096 | NA | 4.908124 | 4.462148 | 4.724439 | 4.721149 | 4.680100 | 4.700048 | 35 | 32 | 17 | 23 |
| 5347 | NM_005030 | PLK1 | polo like kinase 1 | protein_coding | NP_005021.2 | 9077 | 602098 | . | . | 46.074902 | 50.009769 | 35.300831 | 39.679577 | 107.218193 | 115.599823 | 79.214043 | 90.722641 | -1.299168 | 2880.100833 | 0.119234179 | -3.16677836 | 0.001541377 | 0.092102308 | 11.668491 | 11.290647 | 11.557593 | 11.618898 | 11.332182 | 11.416594 | 3611 | 3641 | 2062 | 2474 |
| 5349 | NM_00113600 | FXYD3 | FXYD domain containing ion transport regulator 3 | protein_coding | NP_001129479.1;NP_001129480.1;NP_001129481.1;NP_001129482.1;NI | 4027 | 604996 | . | . | 254.361841 | 263.814050 | 304.259863 | 324.355930 | 591.910480 | 609.817990 | 682.750311 | 741.601290 | 1.192546 | 11824.973508 | 0.103851049 | 2.446240612 | 0.014435463 | 0.400910849 | 13.396967 | 13.650966 | 13.439547 | 13.459194 | 13.564507 | 13.634124 | 12276 | 11775 | 10737 | 12543 |
| 5351 | NM_000302,N | PLOD1 | procollagen-lysine,2-oxoglutarate 5-dioxygenase 1 | protein_coding | NP_000293.2;NP_001303249.1 | 9081 | 153454 | . | . | 243.630456 | 254.778096 | 194.357047 | 216.696907 | 566.938092 | 588.931000 | 436.131567 | 495.451727 | -1.229909 | 21127.473254 | 0.104528931 | -2.85615812 | 0.004288015 | 0.193526306 | 14.508415 | 14.209828 | 14.433030 | 14.465020 | 14.228362 | 14.312077 | 26346 | 25608 | 15662 | 18644 |
| 5352 | NM_000935,N | PLOD2 | procollagen-lysine,2-oxoglutarate 5-dioxygenase 2 | protein_coding | NP_000926.2;NP_891988.1;XP_005247592.1;XP_016862114.1;XP_02430 | 9082 | 601865 | . | . | 47.325830 | 39.019356 | 42.157849 | 31.642373 | 110.129156 | 90.194991 | 94.600990 | 72.346529 | -1.182840 | 4716.703516 | 0.150330851 | -1.61147493 | 0.107076247 | 0.83501324 | 12.319490 | 12.077501 | 12.349451 | 12.174708 | 12.240105 | 11.996319 | 6489 | 4950 | 4313 | 3454 |
| 5354 | NM_000533,N | PLP1 | proteolipid protein 1 | protein_coding | NP_000524.3;NP_001122306.1;NP_001291933.1;NP_955772.1 | 9086 | 300401 | . | . | 104.121588 | 97.632308 | 97.100345 | 90.767183 | 242.295231 | 225.681453 | 217.890354 | 207.528373 | -1.088820 | 8882.247782 | 0.107365554 | -1.14343093 | 0.252859718 | 0.994320829 | 13.176749 | 13.054062 | 13.182844 | 13.119098 | 13.110657 | 13.046722 | 11059 | 9620 | 7676 | 7671 |
| 5355 | NM_002668 | PLP2 | proteolipid protein 2 | protein_coding | NP_002659.1 | 9087 | 300112 | . | . | 112.459061 | 118.474106 | 124.683487 | 122.849777 | 261.696869 | 273.858185 | 279.786133 | 280.881409 | 1.057217 | 4109.430681 | 0.108289139 | 0.741266254 | 0.458532018 | 0.998865404 | 11.964034 | 12.044296 | 11.962820 | 11.998928 | 12.037651 | 12.016726 | 4500 | 4405 | 3719 | 3910 |
| 5356 | NM_00120156 | PLRG1 | pleiotropic regulator 1 | protein_coding | NP_001188493.1;NP_002660.1 | 9089 | 605961 | . | . | 15.459785 | 16.118562 | 13.284772 | 14.959593 | 35.929013 | 37.258777 | 29.810644 | 34.203332 | -1.133060 | 1546.900768 | 0.128443957 | -1.40313261 | 0.160577298 | 0.933525128 | 10.682621 | 10.502063 | 10.627814 | 10.654895 | 10.502625 | 10.584699 | 1860 | 1804 | 1193 | 1433 |
| 5357 | NM_00114531 | PLS1 | plastin 1 | protein_coding | NP_001138791.1;NP_001165783.1;NP_002661.2;XP_006713723.1;XP_01 | 9090 | 602734 | . | . | 4.591438 | 3.267449 | 3.454269 | 3.337245 | 10.684465 | 7.552856 | 7.751279 | 7.630211 | -1.168524 | 424.981747 | 0.204083693 | -1.1009576 | 0.270915112 | 0.998865404 | 8.838788 | 8.615062 | 8.875934 | 8.658229 | 8.690811 | 8.666516 | 618 | 409 | 348 | 360 |
| 5358 | NM_00113602 | PLS3 | plastin 3 | protein_coding | NP_001129497.1;NP_001165806.1;NP_001269266.1;NP_001269267.1;NI | 9091 | 300131 | . | . | 33.599381 | 32.775182 | 26.927833 | 29.095354 | 78.187135 | 75.761298 | 60.425278 | 66.523064 | -1.200566 | 3138.518144 | 0.113032554 | -2.33308456 | 0.019643707 | 0.464284407 | 11.741744 | 11.477950 | 11.700100 | 11.671823 | 11.508940 | 11.562810 | 4010 | 3633 | 2397 | 2764 |
| 5359 | NM_00136387 | PLSCR1 | phospholipid scramblase 1 | protein_coding | NP_001350801.1;NP_001350803.1;NP_066928.1;XP_005247595.1;XP_01 | 9092 | 604170 | . | . | 15.244574 | 14.501720 | 15.546934 | 15.986748 | 35.474749 | 33.521376 | 34.886869 | 36.551799 | 1.032744 | 904.001635 | 0.139968041 | 0.332096423 | 0.739816451 | 0.998865404 | 9.796818 | 9.843173 | 9.808153 | 9.808501 | 9.818108 | 9.844740 | 1024 | 960 | 784 | 877 |
| 5360 | NM_00124292 | PLTP | phospholipid transfer protein | protein_coding | NP_001229849.1;NP_001229850.1;NP_006218.1;NP_872617.1 | 9093 | 172425 | . | . | 1.494606 | 1.628278 | 2.010756 | 1.442616 | 3.478011 | 3.763836 | 4.512077 | 3.298370 | 1.086433 | 98.854680 | 0.333499809 | 0.358617631 | 0.719881155 | 0.998865404 | 6.565455 | 6.686482 | 6.598550 | 6.620996 | 6.683070 | 6.584420 | 105 | 106 | 104 | 81 |
| 5361 | NM_032242,XI | PLXNA1 | plexin A1 | protein_coding | NP_115618.3;XP_011511210.1 | 9099 | 601055 | . | . | 14.843937 | 14.735972 | 14.179801 | 14.539136 | 34.542452 | 34.062860 | 31.819064 | 33.242005 | -1.040357 | 4141.175712 | 0.1071612 | -0.53264001 | 0.5942828 | 0.998865404 | 12.044079 | 11.987006 | 12.040981 | 12.023050 | 11.994523 | 12.003819 | 4915 | 4508 | 3510 | 3827 |
| 5362 | NM_025179,XI | PLXNA2 | plexin A2 | protein_coding | NP_079455.3;XP_005273221.1;XP_005273222.1 | 9100 | 601054 | . | . | 0.874500 | 0.727034 | 0.637474 | 0.775339 | 2.034997 | 1.680571 | 1.430474 | 1.772720 | -1.027094 | 288.409603 | 0.217868401 | -0.17702712 | 0.859487087 | 0.998865404 | 8.191250 | 8.152440 | 8.228127 | 8.124306 | 8.113034 | 8.211371 | 369 | 285 | 224 | 292 |
| 5364 | NM_00113008 | PLXNB1 | plexin B1 | protein_coding | NP_001123554.1;NP_002664.2;XP_011532135.1;XP_011532136.1;XP_01 | 9103 | 601053 | . | . | 0.329052 | 0.562679 | 0.440590 | 0.423973 | 0.765718 | 1.300661 | 0.988674 | 0.969364 | -1.048714 | 148.200933 | 0.300314628 | -0.22849787 | 0.819259209 | 0.998865404 | 7.245903 | 7.176074 | 7.116178 | 7.303117 | 7.208185 | 7.182745 | 134 | 202 | 130 | 131 |
| 5365 | NM_00116325 | PLXNB3 | plexin B3 | protein_coding | NP_001156729.1;NP_005384.2 | 9105 | 300214 | . | . | 10.506478 | 11.789411 | 14.450067 | 11.233402 | 24.449007 | 27.251749 | 32.425530 | 25.683839 | 1.136555 | 2305.400728 | 0.143494423 | 1.286927383 | 0.1981196 | 0.96621494 | 11.075400 | 11.260284 | 11.071460 | 11.155224 | 11.317797 | 11.105748 | 2354 | 2451 | 2410 | 2001 |

| | | | | | | | | | | |
|---|---|---|---|---|---|---|---|---|---|---|
| 5366 | NM_00138261 | PMAIP1 | phorbol-12-myristate-13-acetate-induced protein 1 | protein_coding | NP_001369544.1;NP_001369545.1;NP_001369546.1;NP_001369547.1;NI | 9108 | 604959 | . | . | [illegible] |
| 5371 | NM_002675,N | PML | PML nuclear body scaffold | protein_coding | NP_002666.1;NP_150241.2;NP_150242.1;NP_150243.2;NP_150247.2;NP_ | 9113 | 102578 | . | . | [illegible] |
| 5372 | NM_002676,XI | PMM1 | phosphomannomutase 1 | protein_coding | NP_002667.2;XP_005261695.1;XP_011528531.1;XP_011528532.1;XP_01 | 9114 | 601786 | . | . | [illegible] |
| 5373 | NM_000303,XI | PMM2 | phosphomannomutase 2 | protein_coding | NP_000294.1;XP_005255431.1 | 9115 | 601785 | . | . | [illegible] |
| 5375 | NM_00134838 | PMP2 | peripheral myelin protein 2 | protein_coding | NP_001335310.1;NP_002668.1 | 9117 | 170715 | . | . | [illegible] |
| 5376 | NM_000304,N | PMP22 | peripheral myelin protein 22 | protein_coding | NP_000295.1;NP_001268384.1;NP_001268385.1;NP_001317072.1;NP_69 | 9118 | 601097 | . | . | [illegible] |
| 5378 | NM_000534,N | PMS1 | PMS1 homolog 1, mismatch repair system component | protein_coding | NP_000525.1;NP_001121615.1;NP_001121616.1;NP_001276337.1;NP_0( | 9121 | 600258 | . | . | [illegible] |
| 5379 | NR_003613 | PMS2P1 | PMS1 homolog 2, mismatch repair system component pseudogen | transcribed_ps | . | 9123 | 605038 | . | . | [illegible] |
| 5380 | NR_003614 | PMS2P2 | PMS1 homolog 2, mismatch repair system component pseudogen | transcribed_ps | . | 9127 | . | . | . | [illegible] |
| 5382 | NR_022007,NF | PMS2P4 | PMS1 homolog 2, mismatch repair system component pseudogen | transcribed_ps | . | 9129 | . | . | . | [illegible] |
| 5383 | NR_027775,NF | PMS2P5 | PMS1 homolog 2, mismatch repair system component pseudogen | transcribed_ps | . | 9130 | . | . | . | [illegible] |
| 5387 | NR_028059 | PMS2P3 | PMS1 homolog 2, mismatch repair system component pseudogen | transcribed_ps | . | 9128 | . | . | . | [illegible] |
| 5393 | NM_00103419 | EXOSC9 | exosome component 9 | protein_coding | NP_001029366.1;NP_005024.2;XP_011530337.1 | 9137 | 606180 | . | . | [illegible] |
| 5394 | NM_00100199 | EXOSC10 | exosome component 10 | protein_coding | NP_001001998.1;NP_002676.1;XP_005263532.2;XP_016856984.1;XP_01 | 9138 | 605960 | . | . | [illegible] |
| 5395 | NM_000535,N | PMS2 | PMS1 homolog 2, mismatch repair system component | protein_coding | NP_000526.2;NP_001308932.1;NP_001308933.1;NP_001308934.1;NP_0( | 9122 | 600259 | . | . | [illegible] |
| 5411 | NM_002687 | PNN | pinin, desmosome associated protein | protein_coding | NP_002678.3 | 9162 | 603154 | . | . | [illegible] |
| 5412 | NM_007106 | UBL3 | ubiquitin like 3 | protein_coding | NP_009037.1 | 12504 | 604711 | . | . | [illegible] |
| 5413 | NM_00100993 | SEPTIN5 | septin 5 | protein_coding | NP_001009939.1;NP_002679.2 | 9164 | 602724 | . | . | [illegible] |
| 5414 | NM_00103870 | SEPTIN4 | septin 4 | protein_coding | NP_001033793.3;NP_001185642.1;NP_001243711.1;NP_001243751.1;NI | 9165 | 603696 | . | . | [illegible] |
| 5420 | NM_00101811 | PODXL | podocalyxin like | protein_coding | NP_001018121.1;NP_005388.2 | 9171 | 602632 | . | . | [illegible] |
| 5422 | NM_00133036 | POLA1 | DNA polymerase alpha 1, catalytic subunit | protein_coding | NP_001317289.1;NP_001365232.1;NP_058633.2;XP_006724562.1;XP_01 | 9173 | 312040 | . | . | [illegible] |
| 5423 | NM_002690,XI | POLB | DNA polymerase beta | protein_coding | NP_002681.1;XP_005273592.1;XP_005273593.1;XP_005273594.1;XP_00 | 9174 | 174760 | . | . | [illegible] |
| 5424 | NM_00125684 | POLD1 | DNA polymerase delta 1, catalytic subunit | protein_coding | NP_001243778.1;NP_001295561.1;NP_002682.2;XP_005259065.1;XP_01 | 9175 | 174761 | . | . | [illegible] |
| 5425 | NM_00112721 | POLD2 | DNA polymerase delta 2, accessory subunit | protein_coding | NP_001120690.1;NP_001243808.1;NP_006221.2;XP_024302570.1 | 9176 | 600815 | . | . | [illegible] |
| 5426 | NM_006231,XI | POLE | DNA polymerase epsilon, catalytic subunit | protein_coding | NP_006222.2;XP_011533097.1;XP_011533099.1;XP_011533101.1;XP_01 | 9177 | 174762 | . | . | [illegible] |
| 5427 | NM_00119733 | POLE2 | DNA polymerase epsilon 2, accessory subunit | protein_coding | NP_001184259.1;NP_001184260.1;NP_001335313.1;NP_001335314.1;NI | 9178 | 602670 | . | . | [illegible] |
| 5428 | NM_00112613 | POLG | DNA polymerase gamma, catalytic subunit | protein_coding | NP_001119603.1;NP_002684.1 | 9179 | 174763 | . | . | [illegible] |
| 5429 | NM_00129196 | POLH | DNA polymerase eta | protein_coding | NP_001278898.1;NP_001278899.1;NP_006493.1;XP_005249243.1;XP_01 | 9181 | 603968 | . | . | [illegible] |
| 5430 | NM_000937 | POLR2A | RNA polymerase II subunit A | protein_coding | NP_000928.1 | 9187 | 180660 | . | . | [illegible] |
| 5431 | NM_000938,N | POLR2B | RNA polymerase II subunit B | protein_coding | NP_000929.1;NP_001290197.1;NP_001290198.1 | 9188 | 180661 | . | . | [illegible] |
| 5432 | NM_032940 | POLR2C | RNA polymerase II subunit C | protein_coding | NP_116558.1 | 9189 | 180663 | . | . | [illegible] |
| 5433 | NM_004805 | POLR2D | RNA polymerase II subunit D | protein_coding | NP_004796.1 | 9191 | 606017 | . | . | [illegible] |
| 5434 | NM_00131632 | POLR2E | RNA polymerase II subunit E | protein_coding | NP_001303252.1;NP_001303253.1;NP_002686.3;XP_011526372.1;XP_01 | 9192 | 180664 | . | . | [illegible] |
| 5435 | NM_00130112 | POLR2F | RNA polymerase II subunit F | protein_coding | NP_001288058.1;NP_001288059.1;NP_001288060.1;NP_001350754.1;NI | 9193 | 604414 | . | . | [illegible] |
| 5436 | NM_002696 | POLR2G | RNA polymerase II subunit G | protein_coding | NP_002687.1 | 9194 | 602013 | . | . | [illegible] |
| 5437 | NM_00127869 | POLR2H | RNA polymerase II subunit H | protein_coding | NP_001265627.1;NP_001265628.1;NP_001265629.1;NP_001265643.1;NI | 9195 | 606023 | . | . | [illegible] |
| 5438 | NM_006233 | POLR2I | RNA polymerase II subunit I | protein_coding | NP_006224.1 | 9196 | 180662 | . | . | [illegible] |
| 5439 | NM_00137110 | POLR2J | RNA polymerase II subunit J | protein_coding | NP_001358029.1;NP_006225.1;XP_005250509.1;XP_016867842.1 | 9197 | 604150 | . | . | [illegible] |
| 5440 | NM_005034 | POLR2K | RNA polymerase II subunit K | protein_coding | NP_005025.1 | 9198 | 606033 | . | . | [illegible] |
| 5441 | NM_021128 | POLR2L | RNA polymerase II subunit L | protein_coding | NP_066951.1 | 9199 | 601189 | . | . | [illegible] |
| 5442 | NM_005035,XI | POLRMT | RNA polymerase mitochondrial | protein_coding | NP_005026.3;XP_005259637.3 | 9200 | 601778 | . | . | [illegible] |
| 5445 | NM_000305,N | PON2 | paraoxonase 2 | protein_coding | NP_000296.2;NP_001018171.1;XP_005250510.1;XP_016867846.1;XP_01 | 9205 | 602447 | . | . | [illegible] |
| 5447 | NM_000941,N | POR | cytochrome p450 oxidoreductase | protein_coding | NP_000932.3;NP_001354491.1;NP_001369584.1;NP_001369586.1;NP_0( | 9208 | 124015 | . | . | [illegible] |
| 5451 | NM_00119878 | POU2F1 | POU class 2 homeobox 1 | protein_coding | NP_001185712.1;NP_001185715.1;NP_001352777.1;NP_001352778.1;NI | 9212 | 164175 | . | . | [illegible] |
| 5452 | NM_00120702 | POU2F2 | POU class 2 homeobox 2 | protein_coding | NP_001193954.1;NP_001193955.1;NP_001234923.1;NP_002689.1;XP_0( | 9213 | 164176 | . | . | [illegible] |
| 5453 | NM_002699 | POU3F1 | POU class 3 homeobox 1 | protein_coding | NP_002690.3 | 9214 | 602479 | . | . | [illegible] |
| 5454 | NM_005604 | POU3F2 | POU class 3 homeobox 2 | protein_coding | NP_005595.2 | 9215 | 600494 | . | . | [illegible] |
| 5455 | NM_006236 | POU3F3 | POU class 3 homeobox 3 | protein_coding | NP_006227.1 | 9216 | 602480 | . | . | [illegible] |
| 5457 | NM_006237 | POU4F1 | POU class 4 homeobox 1 | protein_coding | NP_006228.3 | 9218 | 601632 | . | . | [illegible] |
| 5463 | NM_00133042 | POU6F1 | POU class 6 homeobox 1 | protein_coding | NP_001317351.1;NP_001338017.1;NP_001354937.1;XP_016874997.1;XP | 9224 | 618043 | . | . | [illegible] |
| 5464 | NM_021129,XI | PPA1 | inorganic pyrophosphatase 1 | protein_coding | NP_066952.1 | 9226 | 179030 | . | . | [illegible] |
| 5465 | NM_00100192 | PPARA | peroxisome proliferator activated receptor alpha | protein_coding | NP_001001928.1;NP_001349801.1;NP_001349802.1;NP_005027.2;XP_0( | 9232 | 170998 | . | . | [illegible] |
| 5467 | NM_00117181 | PPARD | peroxisome proliferator activated receptor delta | protein_coding | NP_001165289.1;NP_001165290.1;NP_001165291.1;NP_006229.1;NP_8( | 9235 | 600409 | . | . | [illegible] |
| 5468 | NM_00133061 | PPARG | peroxisome proliferator activated receptor gamma | protein_coding | NP_001317544.2;NP_001341595.2;NP_001341596.2;NP_001341597.1;NI | 9236 | 601487 | . | . | [illegible] |
| 5469 | NM_004774,XI | MED1 | mediator complex subunit 1 | protein_coding | NP_004765.2;XP_005257522.1;XP_006722020.1;XP_016880268.1 | 9234 | 604311 | . | . | [illegible] |
| 5471 | NM_002703,N | PPAT | phosphoribosyl pyrophosphate amidotransferase | protein_coding | NP_002694.3 | 9238 | 172450 | . | . | [illegible] |
| 5475 | NM_00137798 | PPEF1 | protein phosphatase with EF-hand domain 1 | protein_coding | NP_001364915.1;NP_001364922.1;NP_001364923.1;NP_001364924.1;NI | 9243 | 300109 | . | . | [illegible] |
| 5476 | NM_000308,N | CTSA | cathepsin A | protein_coding | NP_000299.3;NP_001121167.1;NP_001161066.1 | 9251 | 613111 | . | . | [illegible] |
| 5478 | NM_00130098 | PPIA | peptidylprolyl isomerase A | protein_coding | NP_001287910.1;NP_066953.1;XP_024302576.1;XP_024302577.1 | 9253 | 123840 | . | . | [illegible] |
| 5479 | NM_000942 | PPIB | peptidylprolyl isomerase B | protein_coding | NP_000933.1 | 9255 | 123841 | . | . | [illegible] |
| 5480 | NM_000943 | PPIC | peptidylprolyl isomerase C | protein_coding | NP_000934.1 | 9256 | 123842 | . | . | [illegible] |
| 5481 | NM_005038 | PPID | peptidylprolyl isomerase D | protein_coding | NP_005029.1 | 9257 | 601753 | . | . | [illegible] |
| 5483 | gene-PPIAP11 | PPIAP11 | peptidylprolyl isomerase A pseudogene 11 | pseudogene | . | 9263 | . | . | . | [illegible] |
| 5494 | NM_021003,N | PPM1A | protein phosphatase, Mg2+/Mn2+ dependent 1A | protein_coding | NP_066283.1;NP_808820.1;NP_808821.2;XP_005267836.1;XP_00526783 | 9275 | 606108 | . | . | [illegible] |
| 5495 | NM_00103355 | PPM1B | protein phosphatase, Mg2+/Mn2+ dependent 1B | protein_coding | NP_001028729.1;NP_002697.1;NP_808907.1;NP_808908.1;XP_01153123 | 9276 | 603770 | . | . | [illegible] |
| 5496 | NM_177983 | PPM1G | protein phosphatase, Mg2+/Mn2+ dependent 1G | protein_coding | NP_817092.1 | 9278 | 605119 | . | . | [illegible] |
| 5498 | NM_000309,N | PPOX | protoporphyrinogen oxidase | protein_coding | NP_000300.1;NP_001116236.1;NP_001337057.1;NP_001337058.1;NP_0( | 9280 | 600923 | . | . | [illegible] |
| 5499 | NM_00100870 | PPP1CA | protein phosphatase 1 catalytic subunit alpha | protein_coding | NP_001008709.1;NP_002699.1;NP_996756.1 | 9281 | 176875 | . | . | [illegible] |
| 5500 | NM_002709,N | PPP1CB | protein phosphatase 1 catalytic subunit beta | protein_coding | NP_002700.1;NP_996759.1 | 9282 | 600590 | . | . | [illegible] |
| 5501 | NM_00124497 | PPP1CC | protein phosphatase 1 catalytic subunit gamma | protein_coding | NP_001231903.1;NP_002701.1;XP_011536806.1;XP_011536807.1 | 9283 | 176914 | . | . | [illegible] |
| 5504 | NM_00129150 | PPP1R2 | protein phosphatase 1 regulatory inhibitor subunit 2 | protein_coding | NP_001278433.1;NP_001278434.1;NP_001303254.1;NP_006232.1;XP_0( | 9288 | 601792 | . | . | [illegible] |
| 5507 | NM_005398 | PPP1R3C | protein phosphatase 1 regulatory subunit 3C | protein_coding | NP_005389.1 | 9293 | 602999 | . | . | [illegible] |
| 5509 | NM_006242 | PPP1R3D | protein phosphatase 1 regulatory subunit 3D | protein_coding | NP_006233.1 | 9294 | 603326 | . | . | [illegible] |
| 5510 | NM_00128240 | PPP1R7 | protein phosphatase 1 regulatory subunit 7 | protein_coding | NP_001269338.1;NP_001269339.1;NP_001269340.1;NP_001269341.1;NI | 9295 | 602877 | . | . | [illegible] |
| 5511 | NM_002713,N | PPP1R8 | protein phosphatase 1 regulatory subunit 8 | protein_coding | NP_002704.1;NP_054829.2;NP_612568.1;XP_016857083.1 | 9296 | 602636 | . | . | [illegible] |
| 5512 | gene-PPP1R8P | PPP1R8P1 | protein phosphatase 1 regulatory subunit 8 pseudogene 1 | pseudogene | . | 9297 | . | . | . | [illegible] |
| 5514 | NM_00137619 | PPP1R10 | protein phosphatase 1 regulatory subunit 10 | protein_coding | NP_001363124.1;NP_002705.2;XP_006715193.1;XP_011513024.1;XP_01 | 9284 | 603771 | . | . | [illegible] |
| 5515 | NM_00135501 | PPP2CA | protein phosphatase 2 catalytic subunit alpha | protein_coding | NP_001341948.1;NP_002706.1 | 9299 | 176915 | . | . | [illegible] |
| 5516 | NM_00100955 | PPP2CB | protein phosphatase 2 catalytic subunit beta | protein_coding | NP_001009552.1 | 9300 | 176916 | . | . | [illegible] |
| 5517 | gene-PPP2CBP | PPP2CBP1 | protein phosphatase 2 catalytic subunit beta pseudogene 1 | pseudogene | . | 9301 | . | . | . | [illegible] |
| 5518 | NM_00136365 | PPP2R1A | protein phosphatase 2 scaffold subunit Aalpha | protein_coding | NP_001350585.1;NP_055040.2 | 9302 | 605983 | . | . | [illegible] |
| 5519 | NM_00117756 | PPP2R1B | protein phosphatase 2 scaffold subunit Abeta | protein_coding | NP_001171033.1;NP_001171034.1;NP_002707.3;NP_859050.1;NP_8590! | 9303 | 603113 | . | . | [illegible] |
| 5520 | NM_00117759 | PPP2R2A | protein phosphatase 2 regulatory subunit Balpha | protein_coding | NP_001171062.1;NP_002708.1;XP_005273616.1;XP_005273617.1;XP_01 | 9304 | 604941 | . | . | [illegible] |
| 5521 | NM_00127189 | PPP2R2B | protein phosphatase 2 regulatory subunit Bbeta | protein_coding | NP_001258828.1;NP_001258829.1;NP_001258877.1;NP_858060.2;NP_8! | 9305 | 604325 | . | . | [illegible] |
| 5523 | NM_00119044 | PPP2R3A | protein phosphatase 2 regulatory subunit B''alpha | protein_coding | NP_001177376.1;NP_002709.2;NP_871626.1;XP_006713749.1;XP_01151 | 9307 | 604944 | . | . | [illegible] |
| 5524 | NM_00119339 | PTPA | protein phosphatase 2 phosphatase activator | protein_coding | NP_001180326.1;NP_001258761.1;NP_066954.2;NP_821067.1;NP_8210( | 9308 | 600756 | . | . | [illegible] |
| 5525 | NM_00119975 | PPP2R5A | protein phosphatase 2 regulatory subunit B'alpha | protein_coding | NP_001186685.1;NP_006234.1 | 9309 | 601643 | . | . | [illegible] |
| 5526 | NM_006244,XI | PPP2R5B | protein phosphatase 2 regulatory subunit B'beta | protein_coding | NP_006235.1;XP_011543434.1;XP_011543435.1;XP_011543436.1 | 9310 | 601644 | . | . | [illegible] |
| 5527 | NM_00116172 | PPP2R5C | protein phosphatase 2 regulatory subunit B'gamma | protein_coding | NP_001155197.1;NP_001155198.1;NP_001339841.1;NP_001339842.1;NI | 9311 | 601645 | . | . | [illegible] |
| 5528 | NM_00127047 | PPP2R5D | protein phosphatase 2 regulatory subunit B'delta | protein_coding | NP_001257405.1;NP_006236.1;NP_851307.1;NP_851308.1 | 9312 | 601646 | . | . | [illegible] |
| 5529 | NM_00128217 | PPP2R5E | protein phosphatase 2 regulatory subunit B'epsilon | protein_coding | NP_001269108.1;NP_001269109.1;NP_001269110.1;NP_001269111.1;NI | 9313 | 601647 | . | . | [illegible] |
| 5530 | NM_000944,N | PPP3CA | protein phosphatase 3 catalytic subunit alpha | protein_coding | NP_000935.1;NP_001124163.1;NP_001124164.1;XP_016863854.1;XP_02 | 9314 | 114105 | . | . | [illegible] |
| 5531 | NM_00130350 | PPP4C | protein phosphatase 4 catalytic subunit | protein_coding | NP_001290432.1;NP_001290433.1;NP_001290435.1;NP_001290436.1;NI | 9319 | 602035 | . | . | [illegible] |
| 5532 | NM_00114235 | PPP3CB | protein phosphatase 3 catalytic subunit beta | protein_coding | NP_001135825.1;NP_001135826.1;NP_001276897.1;NP_001276898.1;NI | 9315 | 114106 | . | . | [illegible] |
| 5533 | NM_00124397 | PPP3CC | protein phosphatase 3 catalytic subunit gamma | protein_coding | NP_001230903.1;NP_001230904.1;NP_005596.2;XP_016869100.1 | 9316 | 114107 | . | . | [illegible] |
| 5534 | NM_000945 | PPP3R1 | protein phosphatase 3 regulatory subunit B, alpha | protein_coding | NP_000936.1 | 9317 | 601302 | . | . | [illegible] |
| 5536 | NM_00120428 | PPP5C | protein phosphatase 5 catalytic subunit | protein_coding | NP_001191213.1;NP_006238.1;XP_016882424.1 | 9322 | 600658 | . | . | [illegible] |
| 5537 | NM_00112335 | PPP6C | protein phosphatase 6 catalytic subunit | protein_coding | NP_001116827.1;NP_001116841.1;NP_002712.1;XP_011517149.1;XP_02 | 9323 | 612725 | . | . | [illegible] |
| 5538 | NM_000310,N | PPT1 | palmitoyl-protein thioesterase 1 | protein_coding | NP_000301.1;NP_001136076.1;NP_001350624.1 | 9325 | 600722 | . | . | [illegible] |
| 5546 | NM_005973,XI | PRCC | proline rich mitotic checkpoint control factor | protein_coding | NP_005964.3;XP_005245370.1;XP_005245371.1;XP_011508028.1 | 9343 | 179755 | . | . | [illegible] |
| 5547 | NM_00131921 | PRCP | prolylcarboxypeptidase | protein_coding | NP_001306143.1;NP_005031.1;NP_955450.2;XP_005274150.1 | 9344 | 176785 | . | . | [illegible] |
| 5549 | NM_002725,N | PRELP | proline and arginine rich end leucine rich repeat protein | protein_coding | NP_002716.1;NP_958505.1 | 9357 | 601914 | . | . | [illegible] |
| 5550 | NM_002726,XI | PREP | prolyl endopeptidase | protein_coding | NP_002717.3;XP_005267101.1;XP_011534227.1 | 9358 | 600400 | . | . | [illegible] |
| 5552 | NM_00132105 | SRGN | serglycin | protein_coding | NP_001307982.1;NP_001307983.1;NP_002718.2;XP_016871881.1;XP_02 | 9361 | 177040 | . | . | [illegible] |
| 5553 | NM_00124324 | PRG2 | proteoglycan 2, pro eosinophil major basic protein | protein_coding | NP_001230174.1;NP_001289855.1;NP_001289856.1;NP_002719.3 | 9362 | 605601 | . | . | [illegible] |
| 5554 | NM_00129131 | PRH1 | proline rich protein HaeIII subfamily 1 | protein_coding | NP_001278243.1;NP_001278244.1 | 9366 | 168730 | . | . | [illegible] |
| 5557 | NM_000946 | PRIM1 | DNA primase subunit 1 | protein_coding | NP_000937.1 | 9369 | 176635 | . | . | [illegible] |
| 5558 | NM_000947,N | PRIM2 | DNA primase subunit 2 | protein_coding | NP_000938.2;NP_001269416.1;NP_001269417.1;XP_011513033.1;XP_01 | 9370 | 176636 | . | . | [illegible] |
| 5562 | NM_00135502 | PRKAA1 | protein kinase AMP-activated catalytic subunit alpha 1 | protein_coding | NP_001341957.1;NP_001341958.1;NP_001341963.1;NP_001341964.1;NI | 9376 | 602739 | . | . | [illegible] |
| 5563 | NM_006252,XI | PRKAA2 | protein kinase AMP-activated catalytic subunit alpha 2 | protein_coding | NP_006243.2;XP_016857181.1;XP_016857182.1;XP_016857183.1 | 9377 | 600497 | . | . | [illegible] |
| 5564 | NM_006253,XI | PRKAB1 | protein kinase AMP-activated non-catalytic subunit beta 1 | protein_coding | NP_006244.2;XP_005253966.1 | 9378 | 602740 | . | . | [illegible] |
| 5565 | NM_005399,N | PRKAB2 | protein kinase AMP-activated non-catalytic subunit beta 2 | protein_coding | NP_005390.1;XP_011508031.1;XP_016857186.1 | 9379 | 602741 | . | . | [illegible] |
| 5566 | NM_00130434 | PRKACA | protein kinase cAMP-activated catalytic subunit alpha | protein_coding | NP_001291278.1;NP_002721.1;NP_997401.1;XP_016882437.1 | 9380 | 601639 | . | . | [illegible] |
| 5567 | NM_00124285 | PRKACB | protein kinase cAMP-activated catalytic subunit beta | protein_coding | NP_001229786.1;NP_001229787.1;NP_001229788.1;NP_001229789.1;NI | 9381 | 176892 | . | . | [illegible] |
| 5571 | NM_00120670 | PRKAG1 | protein kinase AMP-activated non-catalytic subunit gamma 1 | protein_coding | NP_001193638.1;NP_001193639.1;NP_002724.1;XP_005269076.1;XP_00 | 9385 | 602742 | . | . | [illegible] |
| 5573 | NM_00127628 | PRKAR1A | protein kinase cAMP-dependent type I regulatory subunit alpha | protein_coding | NP_001263218.1;NP_001263219.1;NP_001265362.1;NP_001356318.1;NI | 9388 | 188830 | . | . | [illegible] |
| 5575 | NM_00116475 | PRKAR1B | protein kinase cAMP-dependent type I regulatory subunit beta | protein_coding | NP_001158230.1;NP_001158231.1;NP_001158232.1;NP_001158233.1;NI | 9390 | 176911 | . | . | [illegible] |
| 5576 | NM_00132198 | PRKAR2A | protein kinase cAMP-dependent type II regulatory subunit alpha | protein_coding | NP_001308911.1;NP_001308912.1;NP_001308918.1;NP_004148.1;XP_0( | 9391 | 176910 | . | . | [illegible] |
| 5577 | NM_002736,XI | PRKAR2B | protein kinase cAMP-dependent type II regulatory subunit beta | protein_coding | NP_002727.2;XP_011514700.1;XP_024302604.1 | 9392 | 176912 | . | . | [illegible] |
| 5578 | NM_002737,XI | PRKCA | protein kinase C alpha | protein_coding | NP_002728.2;XP_016880325.1;XP_016880326.1;XP_016880330.1;XP_02 | 9393 | 176960 | . | . | [illegible] |
| 5579 | NM_002738,N | PRKCB | protein kinase C beta | protein_coding | NP_002729.2;NP_997700.1 | 9395 | 176970 | . | . | [illegible] |
| 5580 | NM_00131632 | PRKCD | protein kinase C delta | protein_coding | NP_001303256.1;NP_001341605.1;NP_001341607.1;NP_001341608.1;NI | 9399 | 176977 | . | . | [illegible] |
| 5581 | NM_005400,XI | PRKCE | protein kinase C epsilon | protein_coding | NP_005391.1;XP_005264485.1;XP_005264488.1;XP_006712113.1;XP_01 | 9401 | 176975 | . | . | [illegible] |
| 5583 | NM_006255,XI | PRKCH | protein kinase C eta | protein_coding | NP_006246.2;XP_011535256.1;XP_011535257.1;XP_011535258.1;XP_01 | 9403 | 605437 | . | . | [illegible] |
| 5584 | NM_002740,XI | PRKCI | protein kinase C iota | protein_coding | NP_002731.4;XP_016862349.1 | 9404 | 600539 | . | . | [illegible] |
| 5585 | NM_002741,N | PKN1 | protein kinase N1 | protein_coding | NP_002732.3;NP_998725.1 | 9405 | 601032 | . | . | [illegible] |
| 5586 | NM_00132070 | PKN2 | protein kinase N2 | protein_coding | NP_001307636.1;NP_001307637.1;NP_001307638.1;NP_006247.1;XP_01 | 9406 | 602549 | . | . | [illegible] |
| 5587 | NM_00133006 | PRKD1 | protein kinase D1 | protein_coding | NP_001316998.1;NP_001335319.1;NP_002733.2;XP_011535267.1;XP_01 | 9407 | 605435 | . | . | [illegible] |
| 5589 | NM_00100132 | PRKCSH | protein kinase C substrate 80K-H | protein_coding | NP_001001329.1;NP_001276031.1;NP_001276032.1;NP_001276033.1;NI | 9411 | 177060 | . | . | [illegible] |
| 5590 | NM_00103358 | PRKCZ | protein kinase C zeta | protein_coding | NP_001028753.1;NP_001028754.1;NP_001229803.1;NP_001337732.1;NI | 9412 | 176982 | . | . | [illegible] |
| 5591 | NM_00108164 | PRKDC | protein kinase, DNA-activated, catalytic subunit | protein_coding | NP_001075109.1;NP_008835.5 | 9413 | 600899 | . | . | [illegible] |
| 5592 | NM_00109851 | PRKG1 | protein kinase cGMP-dependent 1 | protein_coding | NP_001091982.1;NP_001361710.1;NP_001361711.1;NP_006249.1;XP_01 | 9414 | 176894 | . | . | [illegible] |
| 5593 | NM_00128248 | PRKG2 | protein kinase cGMP-dependent 2 | protein_coding | NP_001269409.1;NP_001269410.1;NP_001269411.1;NP_001269412.1;NI | 9416 | 601591 | . | . | [illegible] |
| 5594 | NM_002745,N | MAPK1 | mitogen-activated protein kinase 1 | protein_coding | NP_002736.3;NP_620407.1 | 6871 | 176948 | . | . | [illegible] |
| 5595 | NM_00104005 | MAPK3 | mitogen-activated protein kinase 3 | protein_coding | NP_001035145.1;NP_001103361.1;NP_002737.2 | 6877 | 601795 | . | . | [illegible] |
| 5596 | NM_00129203 | MAPK4 | mitogen-activated protein kinase 4 | protein_coding | NP_001278968.1;NP_001278969.1;NP_002738.2;XP_005258356.1;XP_01 | 6878 | 176949 | . | . | [illegible] |
| 5597 | NM_002748,XI | MAPK6 | mitogen-activated protein kinase 6 | protein_coding | NP_002739.1;XP_005254594.1;XP_005254595.1;XP_011520084.1 | 6879 | 602904 | . | . | [illegible] |
| 5598 | NM_002749,N | MAPK7 | mitogen-activated protein kinase 7 | protein_coding | NP_002740.2;NP_620601.1;NP_620602.2;NP_620603.2;XP_006721620.1 | 6880 | 602521 | . | . | [illegible] |
| 5599 | NM_00127854 | MAPK8 | mitogen-activated protein kinase 8 | protein_coding | NP_001265476.1;NP_001265477.1;NP_001310231.1;NP_001310249.1;NI | 6881 | 601158 | . | . | [illegible] |
| 5600 | NM_002751,N | MAPK11 | mitogen-activated protein kinase 11 | protein_coding | NP_002742.3 | 6873 | 602898 | . | . | [illegible] |
| 5601 | NM_00113504 | MAPK9 | mitogen-activated protein kinase 9 | protein_coding | NP_001128516.1;NP_001295173.1;NP_001351536.1;NP_001351537.1;NI | 6886 | 602896 | . | . | [illegible] |
| 5603 | NM_002754,N | MAPK13 | mitogen-activated protein kinase 13 | protein_coding | NP_002745.1;XP_024302259.1 | 6875 | 602899 | . | . | [illegible] |
| 5604 | NM_002755,XI | MAP2K1 | mitogen-activated protein kinase kinase 1 | protein_coding | NP_002746.1;XP_011520085.1;XP_016877900.1;XP_016877901.1;XP_01( | 6840 | 176872 | . | . | [illegible] |
| 5605 | NM_030662,XI | MAP2K2 | mitogen-activated protein kinase kinase 2 | protein_coding | NP_109587.1;XP_006722862.1;XP_016882478.1;XP_016882479.1;XP_01( | 6842 | 601263 | . | . | [illegible] |
| 5606 | NM_00131633 | MAP2K3 | mitogen-activated protein kinase kinase 3 | protein_coding | NP_001303261.1;NP_002747.2;NP_659731.1;XP_005256780.1;XP_01152 | 6843 | 602315 | . | . | [illegible] |
| 5607 | NM_00120680 | MAP2K5 | mitogen-activated protein kinase kinase 5 | protein_coding | NP_001193733.1;NP_002748.1;NP_660143.1;XP_011520089.1;XP_01152 | 6845 | 602520 | . | . | [illegible] |
| 5608 | NM_00133045 | MAP2K6 | mitogen-activated protein kinase kinase 6 | protein_coding | NP_001317379.1;NP_002749.2;XP_005257573.1;XP_006722038.1;XP_01 | 6846 | 601254 | . | . | [illegible] |
| 5609 | NM_00129755 | MAP2K7 | mitogen-activated protein kinase kinase 7 | protein_coding | NP_001284484.1;NP_001284485.1;NP_660186.1;XP_006722863.1 | 6847 | 603014 | . | . | [illegible] |
| 5610 | NM_00113565 | EIF2AK2 | eukaryotic translation initiation factor 2 alpha kinase 2 | protein_coding | NP_001129123.1;NP_001129124.1;NP_002750.1;XP_011531289.1;XP_01 | 9437 | 176871 | . | . | [illegible] |
| 5611 | NM_006260,XI | DNAJC3 | DnaJ heat shock protein family (Hsp40) member C3 | protein_coding | NP_006251.1;XP_011519406.1;XP_011519407.1;XP_016876163.1;XP_01( | 9439 | 601184 | . | . | [illegible] |
| 5612 | NM_004705,N | THAP12 | THAP domain containing 12 | protein_coding | NP_004696.2 | 9440 | 607374 | . | . | [illegible] |
| 5613 | NM_005044 | PRKX | protein kinase X-linked | protein_coding | NP_005035.1 | 9441 | 300083 | . | . | [illegible] |
| 5621 | NM_000311,N | PRNP | prion protein | protein_coding | NP_000302.1;NP_001073590.1;NP_001073591.1;NP_001073592.1;NP_0( | 9449 | 176640 | . | . | [illegible] |
| 5625 | NM_00119522 | PRODH | proline dehydrogenase 1 | protein_coding | NP_001182155.2;NP_057419.5 | 9453 | 606810 | . | . | [illegible] |
| 5626 | NM_006261 | PROP1 | PROP paired-like homeobox 1 | protein_coding | NP_006252.4 | 9455 | 601538 | . | . | [illegible] |
| 5627 | NM_000313,N | PROS1 | protein S | protein_coding | NP_000304.2;NP_001301006.1 | 9456 | 176880 | . | . | [illegible] |
| 5629 | NM_00127061 | PROX1 | prospero homeobox 1 | protein_coding | NP_001257545.1;NP_002754.2;XP_005273251.1;XP_005273252.1;XP_01 | 9459 | 601546 | . | . | [illegible] |
| 5630 | NM_006262,XI | PRPH | peripherin | protein_coding | NP_006253.2;XP_005269082.1 | 9461 | 170710 | . | . | [illegible] |
| 5631 | NM_00120440 | PRPS1 | phosphoribosyl pyrophosphate synthetase 1 | protein_coding | NP_001191331.1;NP_002755.1 | 9462 | 311850 | . | . | [illegible] |
| 5634 | NM_00103909 | PRPS2 | phosphoribosyl pyrophosphate synthetase 2 | protein_coding | NP_001034180.1;NP_002756.1 | 9465 | 311860 | . | . | [illegible] |
| 5635 | NM_00133050 | PRPSAP1 | phosphoribosyl pyrophosphate synthetase associated protein 1 | protein_coding | NP_001317432.1;NP_001353165.1;NP_002757.2 | 9466 | 601249 | . | . | [illegible] |
| 5636 | NM_00124393 | PRPSAP2 | phosphoribosyl pyrophosphate synthetase associated protein 2 | protein_coding | NP_001230865.1;NP_001230869.1;NP_001230870.1;NP_001230871.1;NI | 9467 | 603762 | . | . | [illegible] |

| | | | | | | | | | | |
|---|---|---|---|---|---|---|---|---|---|---|
| 5638 | NM_000950,N | PRRG1 | proline rich and Gla domain 1 | protein_coding | NP_000941.1;NP_001135867.1;NP_001166957.1;NP_001166960.1;NP_0( | 9469 | 300935 | . | . | [illegible] |
| 5639 | NM_000951,N | PRRG2 | proline rich and Gla domain 2 | protein_coding | NP_000942.1;NP_001303264.1;XP_006723349.1;XP_011525426.1 | 9470 | 604429 | . | . | [illegible] |
| 5641 | NM_00100853 | LGMN | legumain | protein_coding | NP_001008530.1;NP_001350625.1;NP_001350628.1;NP_005597.3;XP_0( | 9472 | 602620 | . | . | [illegible] |
| 5653 | NM_00101296 | KLK6 | kallikrein related peptidase 6 | protein_coding | NP_001012982.1;NP_001012983.1;NP_001306877.1;NP_001306878.1;NI | 6367 | 602652 | . | . | [illegible] |
| 5654 | NM_002775 | HTRA1 | HtrA serine peptidase 1 | protein_coding | NP_002766.1 | 9476 | 602194 | . | . | [illegible] |
| 5660 | NM_00104246 | PSAP | prosaposin | protein_coding | NP_001035930.1;NP_001035931.1;NP_002769.1 | 9498 | 176801 | . | . | [illegible] |
| 5663 | NM_000021,N | PSEN1 | presenilin 1 | protein_coding | NP_000012.1;NP_015557.2;XP_005267921.1;XP_005267923.1;XP_01153 | 9508 | 104311 | . | . | [illegible] |
| 5664 | NM_000447,N | PSEN2 | presenilin 2 | protein_coding | NP_000438.2;NP_036618.2;XP_005273256.1;XP_016857324.1;XP_01685 | 9509 | 600759 | . | . | [illegible] |
| 5681 | NM_006742 | PSKH1 | protein serine kinase H1 | protein_coding | NP_006733.1 | 9529 | 177015 | . | . | [illegible] |
| 5682 | NM_00114393 | PSMA1 | proteasome 20S subunit alpha 1 | protein_coding | NP_001137409.1;NP_002777.1;NP_683877.1 | 9530 | 602854 | . | . | [illegible] |
| 5683 | NM_002787 | PSMA2 | proteasome 20S subunit alpha 2 | protein_coding | NP_002778.1 | 9531 | 176842 | . | . | [illegible] |
| 5684 | NM_002788,N | PSMA3 | proteasome 20S subunit alpha 3 | protein_coding | NP_002779.1;NP_687033.1 | 9532 | 176843 | . | . | [illegible] |
| 5685 | NM_00110266 | PSMA4 | proteasome 20S subunit alpha 4 | protein_coding | NP_001096137.1;NP_001096138.1;NP_001317602.1;NP_001317604.1;NI | 9533 | 176846 | . | . | [illegible] |
| 5686 | NM_00119977 | PSMA5 | proteasome 20S subunit alpha 5 | protein_coding | NP_001186701.1;NP_001186702.1;NP_001186703.1;NP_002781.2 | 9534 | 176844 | . | . | [illegible] |
| 5687 | NM_00128223 | PSMA6 | proteasome 20S subunit alpha 6 | protein_coding | NP_001269161.1;NP_001269162.1;NP_001269163.1;NP_002782.1;XP_01 | 9535 | 602855 | . | . | [illegible] |
| 5688 | NM_002792 | PSMA7 | proteasome 20S subunit alpha 7 | protein_coding | NP_002783.1 | 9536 | 606607 | . | . | [illegible] |
| 5689 | NM_002793 | PSMB1 | proteasome 20S subunit beta 1 | protein_coding | NP_002784.1 | 9537 | 602017 | . | . | [illegible] |
| 5690 | NM_00119977 | PSMB2 | proteasome 20S subunit beta 2 | protein_coding | NP_001186708.1;NP_001186709.1;NP_002785.1 | 9539 | 602175 | . | . | [illegible] |
| 5691 | NM_002795,N | PSMB3 | proteasome 20S subunit beta 3 | protein_coding | NP_002786.2 | 9540 | 602176 | . | . | [illegible] |
| 5692 | NM_002796 | PSMB4 | proteasome 20S subunit beta 4 | protein_coding | NP_002787.2 | 9541 | 602177 | . | . | [illegible] |
| 5693 | NM_00113072 | PSMB5 | proteasome 20S subunit beta 5 | protein_coding | NP_001124197.1;NP_001138404.1;NP_002788.1 | 9542 | 600306 | . | . | [illegible] |
| 5694 | NM_00127048 | PSMB6 | proteasome 20S subunit beta 6 | protein_coding | NP_001257410.1;NP_002789.1 | 9543 | 600307 | . | . | [illegible] |
| 5695 | NM_002799 | PSMB7 | proteasome 20S subunit beta 7 | protein_coding | NP_002790.1 | 9544 | 604030 | . | . | [illegible] |
| 5696 | NM_004159,N | PSMB8 | proteasome 20S subunit beta 8 | protein_coding | NP_004150.1;NP_683720.2 | 9545 | 177046 | . | . | [illegible] |
| 5698 | NM_002800 | PSMB9 | proteasome 20S subunit beta 9 | protein_coding | NP_002791.1 | 9546 | 177045 | . | . | [illegible] |
| 5699 | NM_002801 | PSMB10 | proteasome 20S subunit beta 10 | protein_coding | NP_002792.1 | 9538 | 176847 | . | . | [illegible] |
| 5700 | NM_00133021 | PSMC1 | proteasome 26S subunit, ATPase 1 | protein_coding | NP_001317141.1;NP_002793.2;XP_016876959.1 | 9547 | 602706 | . | . | [illegible] |
| 5701 | NM_00120445 | PSMC2 | proteasome 26S subunit, ATPase 2 | protein_coding | NP_001191382.1;NP_002794.1 | 9548 | 154365 | . | . | [illegible] |
| 5702 | NM_002804,XI | PSMC3 | proteasome 26S subunit, ATPase 3 | protein_coding | NP_002795.2;XP_016873515.1 | 9549 | 186852 | . | . | [illegible] |
| 5704 | NM_006503,N | PSMC4 | proteasome 26S subunit, ATPase 4 | protein_coding | NP_006494.1;NP_694546.1 | 9551 | 602707 | . | . | [illegible] |
| 5705 | NM_00119916 | PSMC5 | proteasome 26S subunit, ATPase 5 | protein_coding | NP_001186092.1;NP_002796.4;XP_024306608.1;XP_024306609.1 | 9552 | 601681 | . | . | [illegible] |
| 5706 | NM_00136641 | PSMC6 | proteasome 26S subunit, ATPase 6 | protein_coding | NP_001353343.1;NP_002797.4 | 9553 | 602708 | . | . | [illegible] |
| 5707 | NM_00119103 | PSMD1 | proteasome 26S subunit, non-ATPase 1 | protein_coding | NP_001177966.1;NP_002798.2;XP_016860006.1 | 9554 | 617842 | . | . | [illegible] |
| 5708 | NM_00127870 | PSMD2 | proteasome 26S subunit, non-ATPase 2 | protein_coding | NP_001265637.1;NP_001265638.1;NP_002799.3 | 9559 | 606223 | . | . | [illegible] |
| 5709 | NM_002809 | PSMD3 | proteasome 26S subunit, non-ATPase 3 | protein_coding | NP_002800.2 | 9560 | 617676 | . | . | [illegible] |
| 5710 | NM_00133069 | PSMD4 | proteasome 26S subunit, non-ATPase 4 | protein_coding | NP_001317621.1;NP_002801.1 | 9561 | 601648 | . | . | [illegible] |
| 5711 | NM_00127042 | PSMD5 | proteasome 26S subunit, non-ATPase 5 | protein_coding | NP_001257356.1;NP_005038.1 | 9563 | 604452 | . | . | [illegible] |
| 5713 | NM_002811 | PSMD7 | proteasome 26S subunit, non-ATPase 7 | protein_coding | NP_002802.2 | 9565 | 157970 | . | . | [illegible] |
| 5714 | NM_002812 | PSMD8 | proteasome 26S subunit, non-ATPase 8 | protein_coding | NP_002803.2 | 9566 | 617844 | . | . | [illegible] |
| 5715 | NM_00126140 | PSMD9 | proteasome 26S subunit, non-ATPase 9 | protein_coding | NP_001248329.1;NP_002804.2 | 9567 | 603146 | . | . | [illegible] |
| 5716 | NM_002814,N | PSMD10 | proteasome 26S subunit, non-ATPase 10 | protein_coding | NP_002805.1;NP_736606.1 | 9555 | 300880 | . | . | [illegible] |
| 5717 | NM_00127048 | PSMD11 | proteasome 26S subunit, non-ATPase 11 | protein_coding | NP_001257411.1;NP_002806.2 | 9556 | 604449 | . | . | [illegible] |
| 5718 | NM_00131634 | PSMD12 | proteasome 26S subunit, non-ATPase 12 | protein_coding | NP_001303270.1;NP_002807.1;NP_777360.1;XP_024306610.1;XP_02430 | 9557 | 604450 | . | . | [illegible] |
| 5719 | NM_002817,N | PSMD13 | proteasome 26S subunit, non-ATPase 13 | protein_coding | NP_002808.3;NP_787128.2;XP_011518537.1 | 9558 | 603481 | . | . | [illegible] |
| 5720 | NM_00128152 | PSME1 | proteasome activator subunit 1 | protein_coding | NP_001268457.1;NP_001268458.1;NP_006254.1;NP_788955.1 | 9568 | 600654 | . | . | [illegible] |
| 5721 | NM_002818,XI | PSME2 | proteasome activator subunit 2 | protein_coding | NP_002809.2;XP_006720276.1 | 9569 | 602161 | . | . | [illegible] |
| 5723 | NM_00137050 | PSPH | phosphoserine phosphatase | protein_coding | NP_001357432.1;NP_001357433.1;NP_001357434.1;NP_001357435.1;NI | 9577 | 172480 | . | . | [illegible] |
| 5724 | NM_000952,N | PTAFR | platelet activating factor receptor | protein_coding | NP_000943.1;NP_001158193.1;NP_001158194.1;NP_001158195.1 | 9582 | 173393 | . | . | [illegible] |
| 5725 | NM_002819,N | PTBP1 | polypyrimidine tract binding protein 1 | protein_coding | NP_002810.1;NP_114367.1;NP_114368.1;XP_005259654.1;XP_00525965 | 9583 | 600693 | . | . | [illegible] |
| 5727 | NM_000264,N | PTCH1 | patched 1 | protein_coding | NP_000255.2;NP_001077071.1;NP_001077072.1;NP_001077073.1;NP_0( | 9585 | 601309 | . | . | [illegible] |
| 5728 | NM_000314,N | PTEN | phosphatase and tensin homolog | protein_coding | NP_000305.3;NP_001291646.4;NP_001291647.1 | 9588 | 601728 | . | . | [illegible] |
| 5734 | NM_000958,XI | PTGER4 | prostaglandin E receptor 4 | protein_coding | NP_000949.1;XP_016865145.1;XP_016865146.1;XP_016865147.1;XP_01( | 9596 | 601586 | . | . | [illegible] |
| 5738 | NM_020440,XI | PTGFRN | prostaglandin F2 receptor inhibitor | protein_coding | NP_065173.2;XP_016857363.1 | 9601 | 601204 | . | . | [illegible] |
| 5743 | NM_000963 | PTGS2 | prostaglandin-endoperoxide synthase 2 | protein_coding | NP_000954.1 | 9605 | 600262 | . | . | [illegible] |
| 5744 | NM_002820,N | PTHLH | parathyroid hormone like hormone | protein_coding | NP_002811.1;NP_945315.1;NP_945316.1;NP_945317.1;XP_011519076.1 | 9607 | 168470 | . | . | [illegible] |
| 5746 | NM_00130951 | PTH2R | parathyroid hormone 2 receptor | protein_coding | NP_001296445.1;NP_001358834.1;NP_001358835.1;NP_001358836.1;NI | 9609 | 601469 | . | . | [illegible] |
| 5747 | NM_00119964 | PTK2 | protein tyrosine kinase 2 | protein_coding | NP_001186578.1;NP_001303271.1;NP_001339623.1;NP_001339624.1;NI | 9611 | 600758 | . | . | [illegible] |
| 5753 | NM_00125635 | PTK6 | protein tyrosine kinase 6 | protein_coding | NP_001243287.1;NP_005966.1;XP_016883471.1 | 9617 | 602004 | . | . | [illegible] |
| 5754 | NM_00127039 | PTK7 | protein tyrosine kinase 7 (inactive) | protein_coding | NP_001257327.1;NP_002812.2;NP_690619.1;NP_690620.1;NP_690621.1 | 9618 | 601890 | . | . | [illegible] |
| 5756 | NM_00124239 | TWF1 | twinfilin actin binding protein 1 | protein_coding | NP_001229326.1;NP_002813.3 | 9620 | 610932 | . | . | [illegible] |
| 5757 | NM_00109928 | PTMA | prothymosin alpha | protein_coding | NP_001092755.1;NP_002814.3 | 9623 | 188390 | . | . | [illegible] |
| 5763 | NM_00133033 | PTMS | parathymosin | protein_coding | NP_001317262.1;NP_002815.3 | 9629 | 168440 | . | . | [illegible] |
| 5767 | gene-PTP4A2P | PTP4A2P1 | PTP4A2 pseudogene 1 | pseudogene | . | 9637 | | . | . | [illegible] |
| 5768 | NM_00100412 | QSOX1 | quiescin sulfhydryl oxidase 1 | protein_coding | NP_001004128.1;NP_002817.2 | 9756 | 603120 | . | . | [illegible] |
| 5770 | NM_00127861 | PTPN1 | protein tyrosine phosphatase non-receptor type 1 | protein_coding | NP_001265547.1;NP_002818.1 | 9642 | 176885 | . | . | [illegible] |
| 5771 | NM_00120701 | PTPN2 | protein tyrosine phosphatase non-receptor type 2 | protein_coding | NP_001193942.1;NP_001295216.1;NP_002819.2;NP_536347.1;NP_5363 | 9650 | 176887 | . | . | [illegible] |
| 5774 | NM_00114536 | PTPN3 | protein tyrosine phosphatase non-receptor type 3 | protein_coding | NP_001138840.1;NP_001138841.1;NP_001138842.1;NP_001138843.1;NI | 9655 | 176877 | . | . | [illegible] |
| 5775 | NM_002830 | PTPN4 | protein tyrosine phosphatase non-receptor type 4 | protein_coding | NP_002821.1 | 9656 | 176878 | . | . | [illegible] |
| 5780 | NM_002833 | PTPN9 | protein tyrosine phosphatase non-receptor type 9 | protein_coding | NP_002824.1 | 9661 | 600768 | . | . | [illegible] |
| 5781 | NM_00133043 | PTPN11 | protein tyrosine phosphatase non-receptor type 11 | protein_coding | NP_001317366.1;NP_001361554.1;NP_002825.3;NP_542168.1;XP_01153 | 9644 | 176876 | . | . | [illegible] |
| 5782 | NM_00113100 | PTPN12 | protein tyrosine phosphatase non-receptor type 12 | protein_coding | NP_001124480.1;NP_001124481.1;NP_002826.3;XP_005250575.1;XP_00 | 9645 | 600079 | . | . | [illegible] |
| 5783 | NM_006264,N | PTPN13 | protein tyrosine phosphatase non-receptor type 13 | protein_coding | NP_006255.1;NP_542414.1;NP_542415.1;NP_542416.1;XP_011530467.1 | 9646 | 600267 | . | . | [illegible] |
| 5784 | NM_005401,XI | PTPN14 | protein tyrosine phosphatase non-receptor type 14 | protein_coding | NP_005392.2;XP_016857430.1;XP_024304527.1 | 9647 | 603155 | . | . | [illegible] |
| 5786 | NM_002836,N | PTPRA | protein tyrosine phosphatase receptor type A | protein_coding | NP_002827.1;NP_543030.1;NP_543031.1 | 9664 | 176884 | . | . | [illegible] |
| 5789 | NM_00104071 | PTPRD | protein tyrosine phosphatase receptor type D | protein_coding | NP_001035802.1;NP_001164496.1;NP_001364875.1;NP_001364876.1;NI | 9668 | 601598 | . | . | [illegible] |
| 5791 | NM_00131667 | PTPRE | protein tyrosine phosphatase receptor type E | protein_coding | NP_001303605.1;NP_001303606.1;NP_001310283.1;NP_001310284.1;NI | 9669 | 600926 | . | . | [illegible] |
| 5792 | NM_00132913 | PTPRF | protein tyrosine phosphatase receptor type F | protein_coding | NP_001316066.1;NP_001316067.1;NP_001316068.1;NP_001316069.1;NI | 9670 | 179590 | . | . | [illegible] |
| 5793 | NM_00137547 | PTPRG | protein tyrosine phosphatase receptor type G | protein_coding | NP_001362400.1;NP_002832.3;XP_016862450.1;XP_016862451.1;XP_01 | 9671 | 176886 | . | . | [illegible] |
| 5794 | NM_00116144 | PTPRH | protein tyrosine phosphatase receptor type H | protein_coding | NP_001154912.2;NP_002833.4;XP_011525485.1;XP_011525490.1;XP_01 | 9672 | 602510 | . | . | [illegible] |
| 5795 | NM_00109850 | PTPRJ | protein tyrosine phosphatase receptor type J | protein_coding | NP_001091973.1;NP_002834.3;XP_016873572.1;XP_016873573.1;XP_01 | 9673 | 600925 | . | . | [illegible] |
| 5796 | NM_00113564 | PTPRK | protein tyrosine phosphatase receptor type K | protein_coding | NP_001129120.1;NP_001278910.1;NP_001278911.1;NP_001278912.1;NI | 9674 | 602545 | . | . | [illegible] |
| 5797 | NM_00110524 | PTPRM | protein tyrosine phosphatase receptor type M | protein_coding | NP_001098714.1;NP_001365071.1;NP_001365072.1;NP_001365073.1;NI | 9675 | 176888 | . | . | [illegible] |
| 5798 | NM_00119976 | PTPRN | protein tyrosine phosphatase receptor type N | protein_coding | NP_001186692.1;NP_001186693.1;NP_002837.1;XP_011509868.1;XP_01 | 9676 | 601773 | . | . | [illegible] |
| 5800 | NM_002848,N | PTPRO | protein tyrosine phosphatase receptor type O | protein_coding | NP_002839.1;NP_109592.1;NP_109593.1;NP_109594.1;NP_109595.1;NP_ | 9678 | 600579 | . | . | [illegible] |
| 5802 | NM_002850,N | PTPRS | protein tyrosine phosphatase receptor type S | protein_coding | NP_002841.3;NP_570923.2;NP_570924.2;NP_570925.2;XP_005259657.1 | 9681 | 601576 | . | . | [illegible] |
| 5803 | NM_00120683 | PTPRZ1 | protein tyrosine phosphatase receptor type Z1 | protein_coding | NP_001193767.1;NP_001193768.1;NP_001356324.1;NP_001356325.1;NI | 9685 | 176891 | . | . | [illegible] |
| 5805 | NM_000317 | PTS | 6-pyruvoyltetrahydropterin synthase | protein_coding | NP_000308.1 | 9689 | 612719 | . | . | [illegible] |
| 5806 | NM_002852 | PTX3 | pentraxin 3 | protein_coding | NP_002843.2 | 9692 | 602492 | . | . | [illegible] |
| 5810 | NM_002853,N | RAD1 | RAD1 checkpoint DNA exonuclease | protein_coding | NP_002844.1 | 9806 | 603153 | . | . | [illegible] |
| 5813 | NM_005859 | PURA | purine rich element binding protein A | protein_coding | NP_005850.1 | 9701 | 600473 | . | . | [illegible] |
| 5814 | NM_033224 | PURB | purine rich element binding protein B | protein_coding | NP_150093.1 | 9702 | 608887 | . | . | [illegible] |
| 5817 | NM_00113576 | PVR | PVR cell adhesion molecule | protein_coding | NP_001129240.1;NP_001129241.1;NP_001129242.2;NP_006496.4 | 9705 | 173850 | . | . | [illegible] |
| 5818 | NM_002855,N | NECTIN1 | nectin cell adhesion molecule 1 | protein_coding | NP_002846.3;NP_976030.1;NP_976031.1 | 9706 | 600644 | . | . | [illegible] |
| 5819 | NM_00104272 | NECTIN2 | nectin cell adhesion molecule 2 | protein_coding | NP_001036189.1;NP_002847.1 | 9707 | 600798 | . | . | [illegible] |
| 5820 | NR_003367 | PVT1 | Pvt1 oncogene | lncRNA | . | 9709 | 165140 | . | . | [illegible] |
| 5822 | NM_005049,XI | PWP2 | PWP2 small subunit processome component | protein_coding | NP_005040.2;XP_011527969.1 | 9711 | 601475 | . | . | [illegible] |
| 5824 | NM_00119364 | PEX19 | peroxisomal biogenesis factor 19 | protein_coding | NP_001180573.1;NP_002848.1 | 9713 | 600279 | . | . | [illegible] |
| 5825 | NM_00112267 | ABCD3 | ATP binding cassette subfamily D member 3 | protein_coding | NP_001116146.1;NP_002849.1;XP_006710865.2;XP_011540179.1 | 67 | 170995 | . | . | [illegible] |
| 5826 | NM_00135359 | ABCD4 | ATP binding cassette subfamily D member 4 | protein_coding | NP_001340520.1;NP_001340521.1;NP_001340522.1;NP_001340523.1;NI | 68 | 603214 | . | . | [illegible] |
| 5827 | NM_018663 | PXMP2 | peroxisomal membrane protein 2 | protein_coding | NP_061133.1 | 9716 | 617399 | . | . | [illegible] |
| 5828 | NM_000318,N | PEX2 | peroxisomal biogenesis factor 2 | protein_coding | NP_000309.2;NP_001073336.1;NP_001165557.1;NP_001165558.1 | 9717 | 170993 | . | . | [illegible] |
| 5829 | NM_00108085 | PXN | paxillin | protein_coding | NP_001074324.1;NP_001230685.1;NP_002850.2;NP_079433.3;XP_0052 | 9718 | 602505 | . | . | [illegible] |
| 5830 | NM_000319,N | PEX5 | peroxisomal biogenesis factor 5 | protein_coding | NP_000310.2;NP_001124495.1;NP_001124496.1;NP_001124497.1;NP_0( | 9719 | 600414 | . | . | [illegible] |
| 5831 | NM_00128227 | PYCR1 | pyrroline-5-carboxylate reductase 1 | protein_coding | NP_001269208.1;NP_001269209.1;NP_001269210.1;NP_001317452.1;NI | 9721 | 179035 | . | . | [illegible] |
| 5832 | NM_00101742 | ALDH18A1 | aldehyde dehydrogenase 18 family member A1 | protein_coding | NP_001017423.1;NP_001310341.1;NP_001310342.1;NP_001310343.1;NI | 9722 | 138250 | . | . | [illegible] |
| 5833 | NM_00118491 | PCYT2 | phosphate cytidylyltransferase 2, ethanolamine | protein_coding | NP_001171846.1;NP_001243362.1;NP_001243363.1;NP_001243364.1;NI | 8756 | 602679 | . | . | [illegible] |
| 5834 | NM_002862,XI | PYGB | glycogen phosphorylase B | protein_coding | NP_002853.2 | 9723 | 138550 | . | . | [illegible] |
| 5836 | NM_00116394 | PYGL | glycogen phosphorylase L | protein_coding | NP_001157412.1;NP_002854.3 | 9725 | 613741 | . | . | [illegible] |
| 5859 | NM_00127207 | QARS1 | glutaminyl-tRNA synthetase 1 | protein_coding | NP_001259002.1;NP_005042.1;XP_016862454.2 | 9751 | 603727 | . | . | [illegible] |
| 5860 | NM_000320,N | QDPR | quinoid dihydropteridine reductase | protein_coding | NP_000311.2;NP_001293069.1 | 9752 | 612676 | . | . | [illegible] |
| 5861 | NM_004161,N | RAB1A | RAB1A, member RAS oncogene family | protein_coding | NP_004152.1;NP_056358.1;XP_005264525.1 | 9758 | 179508 | . | . | [illegible] |
| 5862 | NM_00124264 | RAB2A | RAB2A, member RAS oncogene family | protein_coding | NP_001229573.1;NP_002856.1 | 9763 | 179509 | . | . | [illegible] |
| 5863 | NM_00124373 | RGL2 | ral guanine nucleotide dissociation stimulator like 2 | protein_coding | NP_001230667.1;NP_004752.1;XP_011513083.1;XP_011513087.1;XP_02 | 9769 | 602306 | . | . | [illegible] |
| 5864 | NM_002866,XI | RAB3A | RAB3A, member RAS oncogene family | protein_coding | NP_002857.1;XP_011526466.1 | 9777 | 179490 | . | . | [illegible] |
| 5866 | NM_00127168 | RAB3IL1 | RAB3A interacting protein like 1 | protein_coding | NP_001258615.1;NP_037533.2;XP_005274197.1;XP_006718710.1;XP_01 | 9780 | | . | . | [illegible] |
| 5867 | NM_00127199 | RAB4A | RAB4A, member RAS oncogene family | protein_coding | NP_001258927.1;NP_004569.2 | 9781 | 179511 | . | . | [illegible] |
| 5868 | NM_00129204 | RAB5A | RAB5A, member RAS oncogene family | protein_coding | NP_001278977.1;NP_004153.2 | 9783 | 179512 | . | . | [illegible] |
| 5869 | NM_00125203 | RAB5B | RAB5B, member RAS oncogene family | protein_coding | NP_001238965.1;NP_001238966.1;NP_002859.1;XP_005269108.1;XP_00 | 9784 | 179514 | . | . | [illegible] |
| 5870 | NM_00124371 | RAB6A | RAB6A, member RAS oncogene family | protein_coding | NP_001230647.1;NP_001230648.1;NP_002860.2;NP_942599.1 | 9786 | 179513 | . | . | [illegible] |
| 5871 | NM_00130799 | MAP4K2 | mitogen-activated protein kinase kinase kinase kinase 2 | protein_coding | NP_001294919.1;NP_004570.2;XP_011543506.1;XP_016873582.1;XP_01 | 6864 | 603166 | . | . | [illegible] |
| 5872 | NM_00127203 | RAB13 | RAB13, member RAS oncogene family | protein_coding | NP_001258967.1;NP_002861.1;XP_016857448.1 | 9762 | 602672 | . | . | [illegible] |
| 5873 | NM_004580,N | RAB27A | RAB27A, member RAS oncogene family | protein_coding | NP_004571.2;NP_899057.1;NP_899058.1;NP_899059.1;XP_005254633.1 | 9766 | 603868 | . | . | [illegible] |
| 5874 | NM_00137532 | RAB27B | RAB27B, member RAS oncogene family | protein_coding | NP_001362256.1;NP_004154.2;XP_006722581.1;XP_011524433.1;XP_01 | 9767 | 603869 | . | . | [illegible] |
| 5875 | NM_004581,N | RABGGTA | Rab geranylgeranyltransferase subunit alpha | protein_coding | NP_004572.3;NP_878256.1 | 9795 | 601905 | . | . | [illegible] |
| 5876 | NM_004582,N | RABGGTB | Rab geranylgeranyltransferase subunit beta | protein_coding | NP_004573.2 | 9796 | 179080 | . | . | [illegible] |
| 5877 | NM_002871 | RABIF | RAB interacting factor | protein_coding | NP_002862.2 | 9797 | 603417 | . | . | [illegible] |
| 5878 | NM_00125203 | RAB5C | RAB5C, member RAS oncogene family | protein_coding | NP_001238968.1;NP_004574.2;NP_958842.1 | 9785 | 604037 | . | . | [illegible] |
| 5879 | NM_006908,N | RAC1 | Rac family small GTPase 1 | protein_coding | NP_008839.2;NP_061485.1 | 9801 | 602048 | . | . | [illegible] |
| 5881 | NM_00131630 | RAC3 | Rac family small GTPase 3 | protein_coding | NP_001303236.1;NP_005043.1 | 9803 | 602050 | . | . | [illegible] |
| 5883 | NM_00124322 | RAD9A | RAD9 checkpoint clamp component A | protein_coding | NP_001230153.1;NP_004575.1;XP_006718715.1;XP_016873586.1;XP_01 | 9827 | 603761 | . | . | [illegible] |
| 5884 | NM_00127862 | RAD17 | RAD17 checkpoint clamp loader component | protein_coding | NP_001265551.1;NP_002864.1;NP_579916.1;NP_579917.1;NP_579918.1 | 9807 | 603139 | . | . | [illegible] |
| 5885 | NM_006265 | RAD21 | RAD21 cohesin complex component | protein_coding | NP_006256.1 | 9811 | 606462 | . | . | [illegible] |
| 5886 | NM_00127036 | RAD23A | RAD23 homolog A, nucleotide excision repair protein | protein_coding | NP_001257291.1;NP_001257292.1;NP_005044.1;XP_024307399.1 | 9812 | 600061 | . | . | [illegible] |
| 5887 | NM_00124471 | RAD23B | RAD23 homolog B, nucleotide excision repair protein | protein_coding | NP_001231642.1;NP_001231653.1;NP_002865.1 | 9813 | 600062 | . | . | [illegible] |
| 5888 | NM_00116426 | RAD51 | RAD51 recombinase | protein_coding | NP_001157741.1;NP_001157742.1;NP_002866.2;NP_597994.3;XP_0067 | 9817 | 179617 | . | . | [illegible] |
| 5889 | NM_002876,N | RAD51C | RAD51 paralog C | protein_coding | NP_002867.1;NP_478123.1;XP_006722064.1;XP_006722065.1;XP_0067 | 9820 | 602774 | . | . | [illegible] |
| 5890 | NM_00132180 | RAD51B | RAD51 paralog B | protein_coding | NP_001308738.1;NP_001308739.1;NP_001308741.1;NP_001308743.1;NI | 9822 | 602948 | . | . | [illegible] |
| 5891 | NM_00127201 | MOK | MOK protein kinase | protein_coding | NP_001258940.1;NP_001317163.1;NP_001340756.1;NP_001340757.1;NI | 9833 | 605762 | . | . | [illegible] |
| 5892 | NM_00114257 | RAD51D | RAD51 paralog D | protein_coding | NP_001136043.1;NP_002869.3;NP_598332.1 | 9823 | 602954 | . | . | [illegible] |
| 5893 | NM_00129741 | RAD52 | RAD52 homolog, DNA repair protein | protein_coding | NP_001284348.1;NP_001284349.1;NP_001284350.1;NP_001284351.1;NI | 9824 | 600392 | . | . | [illegible] |
| 5894 | NM_00135468 | RAF1 | Raf-1 proto-oncogene, serine/threonine kinase | protein_coding | NP_001341618.1;NP_001341619.1;NP_001341620.1;NP_001341621.1;NI | 9829 | 164760 | . | . | [illegible] |
| 5898 | NM_005402,XI | RALA | RAS like proto-oncogene A | protein_coding | NP_005393.2;XP_006715825.1;XP_011513768.1;XP_011513769.1 | 9839 | 179550 | . | . | [illegible] |
| 5899 | NM_00136940 | RALB | RAS like proto-oncogene B | protein_coding | NP_001356329.1;NP_002872.1;XP_005263785.1;XP_011509876.1 | 9840 | 179551 | . | . | [illegible] |
| 5900 | NM_00104236 | RALGDS | ral guanine nucleotide dissociation stimulator | protein_coding | NP_001035827.1;NP_001258703.1;NP_001258704.1;NP_001258705.1;NI | 9842 | 601619 | . | . | [illegible] |
| 5901 | NM_00130079 | RAN | RAN, member RAS oncogene family | protein_coding | NP_001287725.1;NP_001287726.1;NP_006316.1;XP_016875261.1;XP_01 | 9846 | 601179 | . | . | [illegible] |
| 5902 | NM_00127863 | RANBP1 | RAN binding protein 1 | protein_coding | NP_001265568.1;NP_001265569.1;NP_001265570.1;NP_002873.1;XP_01 | 9847 | 601180 | . | . | [illegible] |
| 5903 | NM_006267,XI | RANBP2 | RAN binding protein 2 | protein_coding | NP_006258.3;XP_005264059.1;XP_005264060.1;XP_005264061.1;XP_00 | 9848 | 601181 | . | . | [illegible] |
| 5905 | NM_00127865 | RANGAP1 | Ran GTPase activating protein 1 | protein_coding | NP_001265580.1;NP_001304859.1;NP_002874.1;XP_005261752.1;XP_00 | 9854 | 602362 | . | . | [illegible] |
| 5906 | NM_00101093 | RAP1A | RAP1A, member of RAS oncogene family | protein_coding | NP_001010935.1;NP_001278825.1;NP_001357145.1;NP_001357146.1;NI | 9855 | 179520 | . | . | [illegible] |
| 5908 | NM_00101094 | RAP1B | RAP1B, member of RAS oncogene family | protein_coding | NP_001010942.1;NP_001238846.1;NP_001238847.1;NP_001238850.1;NI | 9857 | 179530 | . | . | [illegible] |
| 5909 | NM_00114565 | RAP1GAP | RAP1 GTPase activating protein | protein_coding | NP_001139129.1;NP_001139130.1;NP_001317312.1;NP_001337453.1;NI | 9858 | 600278 | . | . | [illegible] |
| 5910 | NM_00110042 | RAP1GDS1 | Rap1 GTPase-GDP dissociation stimulator 1 | protein_coding | NP_001093896.1;NP_001093897.1;NP_001093898.1;NP_001093899.1;NI | 9859 | 179502 | . | . | [illegible] |
| 5911 | NM_021033 | RAP2A | RAP2A, member of RAS oncogene family | protein_coding | NP_066361.1 | 9861 | 179540 | . | . | [illegible] |
| 5912 | NM_002886 | RAP2B | RAP2B, member of RAS oncogene family | protein_coding | NP_002877.2 | 9862 | 179541 | . | . | [illegible] |
| 5914 | NM_000964,N | RARA | retinoic acid receptor alpha | protein_coding | NP_000955.1;NP_001019980.1;NP_001138773.1;NP_001138774.1;XP_0( | 9864 | 180240 | . | . | [illegible] |
| 5916 | NM_000966,N | RARG | retinoic acid receptor gamma | protein_coding | NP_000957.1;NP_001036193.1;NP_001230659.1;NP_001230660.1;NP_0( | 9866 | 180190 | . | . | [illegible] |
| 5917 | NM_002887 | RARS1 | arginyl-tRNA synthetase 1 | protein_coding | NP_002878.2 | 9870 | 107820 | . | . | [illegible] |
| 5921 | NM_002890,N | RASA1 | RAS p21 protein activator 1 | protein_coding | NP_002881.1;NP_072179.1;XP_011541827.1;XP_011541829.1 | 9871 | 139150 | . | . | [illegible] |
| 5922 | NM_00130324 | RASA2 | RAS p21 protein activator 2 | protein_coding | NP_001290174.1;NP_001290175.1;NP_006497.2;XP_011511361.1;XP_01 | 9872 | 601589 | . | . | [illegible] |
| 5923 | NM_00114564 | RASGRF1 | Ras protein specific guanine nucleotide releasing factor 1 | protein_coding | NP_001139120.1;NP_002882.3;NP_722522.1;XP_011520168.1;XP_01152 | 9875 | 606600 | . | . | [illegible] |
| 5925 | NM_000321,XI | RB1 | RB transcriptional corepressor 1 | protein_coding | NP_000312.2;XP_011533473.1 | 9884 | 614041 | . | . | [illegible] |
| 5926 | NM_002892,N | ARID4A | AT-rich interaction domain 4A | protein_coding | NP_002883.3;NP_075376.2;NP_075377.2;XP_005268021.1;XP_00526802 | 9885 | 180201 | . | . | [illegible] |

| | | | | | | | | | | | | | | | | | | | | | | | | | | | | | | | | | |
|---|---|---|---|---|---|---|---|---|---|---|---|---|---|---|---|---|---|---|---|---|---|---|---|---|---|---|---|---|---|---|---|---|---|
| 5927 | NM_00104260 | KDM5A | lysine demethylase 5A | protein_coding | NP_001036068.1 | 9886 | 180202 | . | . | 3.412886 | 3.656064 | 3.403214 | 3.848501 | 7.941926 | 8.451156 | 7.636715 | 8.799141 | 1.012074 | 1193.513488 | 0.136318839 | 0.127021826 | 0.898923124 | 0.998865404 | 10.212534 | 10.229415 | 10.193007 | 10.238545 | 10.181678 | 10.266393 | 1325 | 1319 | 985 | 1189 |
| 5928 | NM_00113525 | RBBP4 | RB binding protein 4, chromatin remodeling factor | protein_coding | NP_001128727.1;NP_001128728.1;NP_005601.1 | 9887 | 602923 | . | . | 20.594329 | 21.625117 | 22.524558 | 21.561092 | 47.923852 | 49.987426 | 50.544456 | 49.296873 | 1.029898 | 5299.562430 | 0.107027281 | 0.39710733 | 0.691288315 | 0.998865404 | 12.350238 | 12.392764 | 12.341941 | 12.375776 | 12.406223 | 12.360706 | 5891 | 5748 | 4803 | 4905 |
| 5929 | NM_00119327 | RBBP5 | RB binding protein 5, histone lysine methyltransferase complex su | protein_coding | NP_001180201.1;NP_001180202.1;NP_005048.2;XP_011508135.1 | 9888 | 600697 | . | . | 6.568100 | 5.725778 | 5.917450 | 6.384525 | 15.284238 | 13.235391 | 13.278587 | 14.597460 | -1.014579 | 831.772598 | 0.150694869 | -0.13856716 | 0.889792199 | 0.998865404 | 9.710383 | 9.689633 | 9.754683 | 9.651771 | 9.672369 | 9.715735 | 1034 | 839 | 696 | 798 |
| 5930 | NM_006910,N | RBBP6 | RB binding protein 6, ubiquitin ligase | protein_coding | NP_008841.2;NP_061173.1;NP_116015.2 | 9889 | 600938 | . | . | 11.149117 | 11.548876 | 12.460335 | 10.735895 | 25.944454 | 26.695741 | 27.960631 | 24.546350 | 1.011024 | 2378.210525 | 0.126480472 | 0.125055127 | 0.900479908 | 0.998865404 | 11.207609 | 11.223668 | 11.200040 | 11.220317 | 11.284128 | 11.149767 | 2689 | 2584 | 2253 | 2055 |
| 5931 | NM_00119871 | RBBP7 | RB binding protein 7, chromatin remodeling factor | protein_coding | NP_001185648.1;NP_002884.1 | 9890 | 300825 | . | . | 52.717951 | 50.367271 | 50.371642 | 52.358763 | 122.676833 | 116.426210 | 113.032505 | 119.712082 | -1.018048 | 3658.700133 | 0.110185149 | -0.23420661 | 0.81482458 | 0.998865404 | 11.849955 | 11.824161 | 11.866442 | 11.821816 | 11.817748 | 11.841310 | 4365 | 3876 | 3108 | 3448 |
| 5932 | NM_002894,N | RBBP8 | RB binding protein 8, endonuclease | protein_coding | NP_002885.1;NP_976036.1;NP_976037.1;XP_005258382.1;XP_00525838 | 9891 | 604124 | . | . | 10.978978 | 10.785023 | 10.940579 | 11.491012 | 25.548532 | 24.930065 | 24.550342 | 26.272834 | 1.013622 | 1128.786931 | 0.132815113 | 0.146973048 | 0.883153293 | 0.998865404 | 10.130789 | 10.150261 | 10.146350 | 10.124369 | 10.131687 | 10.159121 | 1310 | 1192 | 970 | 1085 |
| 5933 | NM_00132328 | RBL1 | RB transcriptional corepressor like 1 | protein_coding | NP_001310210.1;NP_001310211.1;NP_002886.2;NP_899662.1;XP_00672 | 9893 | 116957 | . | . | 5.023517 | 5.360456 | 4.700853 | 4.541502 | 11.689933 | 12.390933 | 10.548576 | 10.383604 | -1.145572 | 864.427713 | 0.143054341 | -1.37058157 | 0.170505433 | 0.946389416 | 9.850301 | 9.654247 | 9.780637 | 9.823070 | 9.719226 | 9.690640 | 1032 | 1025 | 715 | 740 |
| 5934 | NM_00132360 | RBL2 | RB transcriptional corepressor like 2 | protein_coding | NP_001310537.1;NP_001310538.1;NP_001310539.1;NP_001310540.1;NI | 9894 | 180203 | . | . | 7.676386 | 7.597333 | 8.421393 | 8.241399 | 17.863266 | 17.561575 | 18.897362 | 18.842980 | 1.065386 | 1190.742476 | 0.130877306 | 0.698182029 | 0.485063363 | 0.998865404 | 10.171165 | 10.262676 | 10.203889 | 10.182147 | 10.249622 | 10.231719 | 1347 | 1226 | 1081 | 1138 |
| 5935 | NM_006743,XI | RBM3 | RNA binding motif protein 3 | protein_coding | NP_006734.1;XP_024308181.1;XP_024308182.1 | 9900 | 300027 | . | . | 34.145170 | 34.501053 | 35.062393 | 29.802000 | 79.457211 | 79.750731 | 78.678990 | 68.138734 | -1.016413 | 2847.979361 | 0.113820465 | -0.20635309 | 0.836515112 | 0.998865404 | 11.487350 | 11.464002 | 11.494229 | 11.469948 | 11.493613 | 11.443674 | 3355 | 3052 | 2532 | 2567 |
| 5936 | NM_00119884 | RBM4 | RNA binding motif protein 4 | protein_coding | NP_001185772.1;NP_001185773.1;NP_002887.2 | 9901 | 602571 | . | . | 33.079360 | 37.016763 | 37.629140 | 35.475777 | 76.977030 | 85.565905 | 84.438707 | 81.111153 | 1.020642 | 1788.884760 | 0.127619151 | 0.230972255 | 0.817336357 | 0.998865404 | 10.790008 | 10.819531 | 10.756393 | 10.834258 | 10.848278 | 10.774696 | 1938 | 2005 | 1645 | 1615 |
| 5937 | NM_002897,N | RBMS1 | RNA binding motif single stranded interacting protein 1 | protein_coding | NP_002888.1;NP_058520.1;XP_005246794.1;XP_005246795.1;XP_00524 | 9907 | 602310 | . | . | 13.074623 | 12.719013 | 11.429888 | 11.164255 | 30.425185 | 29.400572 | 25.648338 | 25.525737 | -1.150522 | 1597.195612 | 0.124119976 | -1.62978646 | 0.103146635 | 0.824396646 | 10.738838 | 10.536741 | 10.707795 | 10.678540 | 10.602682 | 10.565772 | 2004 | 1810 | 1323 | 1359 |
| 5939 | NM_002898,XI | RBMS2 | RNA binding motif single stranded interacting protein 2 | protein_coding | NP_002889.1;XP_005269116.1;XP_005269117.1;XP_005269118.1;XP_00 | 9909 | 602387 | . | . | 3.392961 | 3.073145 | 3.160783 | 2.962852 | 7.895557 | 7.103710 | 7.092704 | 6.774205 | -1.085472 | 805.094843 | 0.150108381 | -0.78824634 | 0.430552626 | 0.998865404 | 9.710689 | 9.592939 | 9.715244 | 9.644993 | 9.656033 | 9.588941 | 1012 | 860 | 704 | 689 |
| 5954 | NM_002901 | RCN1 | reticulocalbin 1 | protein_coding | NP_002892.1 | 9934 | 602735 | . | . | 81.826042 | 72.906456 | 68.962517 | 62.013695 | 190.412567 | 168.526550 | 154.749893 | 141.786942 | -1.198128 | 5276.362262 | 0.118505981 | -2.20058449 | **0.027765452** | 0.53804272 | 12.489755 | 12.229148 | 12.486740 | 12.381276 | 12.330935 | 12.235276 | 7033 | 5822 | 4418 | 4239 |
| 5955 | NM_00127183 | RCN2 | reticulocalbin 2 | protein_coding | NP_001258766.1;NP_002893.1 | 9935 | 602584 | . | . | 26.125312 | 22.651220 | 20.146057 | 20.112591 | 60.794672 | 52.359312 | 45.207165 | 45.985048 | -1.228460 | 4327.962895 | 0.117635446 | -2.52347996 | **0.011619968** | 0.353282869 | 12.220225 | 11.923488 | 12.220480 | 12.092341 | 11.991823 | 11.982496 | 5908 | 4761 | 3397 | 3621 |
| 5959 | NM_00119977 | RDH5 | retinol dehydrogenase 5 | protein_coding | NP_001186700.1;NP_002896.2 | 9940 | 601617 | . | . | 2.415388 | 2.452660 | 0.794977 | 1.320809 | 5.620706 | 5.669433 | 1.783905 | 3.019874 | **-2.252243** | 71.478339 | 0.399117337 | -2.9348827 | **0.003336738** | 0.163366554 | 6.630552 | 5.455315 | 6.238435 | 6.237927 | 6.011802 | 6.087306 | 114 | 107 | 29 | 51 |
| 5962 | NM_00126049 | RDX | radixin | protein_coding | NP_001247421.1;NP_001247422.1;NP_001247423.1;NP_001247424.1;NI | 9944 | 179410 | . | . | 30.537822 | 30.055343 | 28.604805 | 29.533926 | 71.062775 | 69.474266 | 64.188354 | 67.525812 | -1.066759 | 3970.782329 | 0.108392662 | -0.86015351 | 0.389704425 | 0.998865404 | 12.001091 | 11.907817 | 11.989346 | 11.973420 | 11.910852 | 11.944488 | 4765 | 4381 | 3273 | 3676 |
| 5965 | NM_002907,N | RECQL | RecQ like helicase | protein_coding | NP_002898.2;NP_116559.1;XP_005253518.1;XP_005253519.1;XP_00525 | 9948 | 600537 | . | . | 25.453818 | 23.959320 | 24.921401 | 22.889897 | 59.232081 | 55.383041 | 55.922900 | 52.335030 | -1.044230 | 2636.870397 | 0.117503139 | -0.53138273 | 0.59515359 | 0.998865404 | 11.395389 | 11.333162 | 11.404550 | 11.358102 | 11.379617 | 11.312438 | 3190 | 2824 | 2337 | 2318 |
| 5966 | NM_00129174 | REL | REL proto-oncogene, NF-kB subunit | protein_coding | NP_001278675.1;NP_002899.1;XP_011531312.1;XP_016860116.1 | 9954 | 164910 | . | . | 1.188997 | 1.355237 | 1.588726 | 1.235430 | 2.766847 | 3.132693 | 3.565057 | 2.824664 | 1.068213 | 452.286022 | 0.192571479 | 0.494361549 | 0.621050864 | 0.998865404 | 8.772297 | 8.868290 | 8.771343 | 8.821410 | 8.921831 | 8.745620 | 483 | 491 | 464 | 376 |
| 5970 | NM_00114513 | RELA | RELA proto-oncogene, NF-kB subunit | protein_coding | NP_001138610.1;NP_001230913.1;NP_001230914.1;NP_068810.3;XP_01 | 9955 | 164014 | . | . | 12.948834 | 13.506999 | 13.489916 | 16.074467 | 30.132466 | 31.222030 | 30.270985 | 36.752361 | 1.103356 | 1089.627512 | 0.143462623 | 0.989099049 | 0.322614683 | 0.998865404 | 10.017202 | 10.158574 | 10.036269 | 10.065040 | 10.058271 | 10.185087 | 1170 | 1140 | 910 | 1162 |
| 5971 | NM_006509,XI | RELB | RELB proto-oncogene, NF-kB subunit | protein_coding | NP_006500.2;XP_005259184.1;XP_005259185.1 | 9956 | 604758 | . | . | 1.967025 | 2.069411 | 1.394113 | 2.143795 | 4.577349 | 4.783536 | 3.128348 | 4.901533 | -1.154782 | 133.836118 | 0.301803561 | -0.68793325 | 0.49149482 | 0.998865404 | 7.165476 | 6.955541 | 7.080673 | 7.095916 | 6.964159 | 7.102742 | 162 | 158 | 86 | 140 |
| 5973 | NM_002910,XI | RENBP | renin binding protein | protein_coding | NP_002901.2;XP_016885187.1 | 9959 | 312420 | . | . | 0.612212 | 0.830072 | 0.732644 | 1.167969 | 1.424642 | 1.918748 | 1.644032 | 2.670422 | 1.309940 | 35.830167 | 0.537975546 | 0.724012482 | 0.469058072 | 0.998865404 | 4.957404 | 5.343100 | 5.117623 | 5.152143 | 5.136136 | 5.208800 | 31 | 38 | 27 | 47 |
| 5976 | NM_00129754 | UPF1 | UPF1 RNA helicase and ATPase | protein_coding | NP_001284478.1;NP_002902.2;XP_016882594.1;XP_016882595.1 | 9962 | 601430 | . | . | 19.171136 | 20.019854 | 18.977592 | 20.426839 | 44.612020 | 46.276787 | 42.585162 | 46.703538 | -1.008221 | 3273.651950 | 0.112423776 | -0.10506266 | 0.916326085 | 0.998865404 | 11.682646 | 11.670700 | 11.668719 | 11.690989 | 11.644127 | 11.701346 | 3734 | 3595 | 2736 | 3163 |
| 5977 | NM_00133030 | DPF2 | double PHD fingers 2 | protein_coding | NP_001317237.1;NP_006259.1;XP_016873590.1;XP_016873591.1;XP_02 | 9964 | 601671 | . | . | 8.928413 | 9.746323 | 9.077199 | 9.891084 | 20.776784 | 22.529060 | 20.368975 | 22.614787 | 1.031961 | 1027.910330 | 0.138715216 | 0.327203536 | 0.743513947 | 0.998865404 | 9.982813 | 10.027833 | 9.983201 | 10.003639 | 9.979517 | 10.052196 | 1149 | 1107 | 863 | 1027 |
| 5978 | NM_00119350 | REST | RE1 silencing transcription factor | protein_coding | NP_001180437.1;NP_001350382.1;NP_005603.3;XP_016864016.1 | 9966 | 600571 | . | . | 2.381431 | 2.153554 | 2.040353 | 2.006677 | 5.541689 | 4.978036 | 4.578495 | 4.588027 | -1.137713 | 488.154550 | 0.172070283 | -1.08174564 | 0.279365569 | 0.998865404 | 9.021042 | 8.835377 | 9.007372 | 8.931642 | 8.898077 | 8.880088 | 633 | 528 | 403 | 422 |
| 5979 | NM_00135521 | RET | ret proto-oncogene | protein_coding | NP_001342145.1;NP_065681.1;NP_066124.1 | 9967 | 164761 | . | . | 0.067054 | 0.073374 | 0.026403 | 0.056325 | 0.156039 | 0.169607 | 0.059248 | 0.128781 | -1.525791 | 10.387290 | 0.961410403 | -0.63402393 | 0.526065203 | NA | 3.651489 | 3.037079 | 3.391943 | 3.395472 | 3.363909 | 3.385087 | 14 | 14 | 5 | 10 |
| 5980 | NM_00128643 | REV3L | REV3 like, DNA directed polymerase zeta catalytic subunit | protein_coding | NP_001273360.1;NP_001273361.1;NP_001359007.1;NP_002903.3;XP_01 | 9968 | 602776 | . | . | 5.574341 | 5.572501 | 6.456320 | 5.591283 | 12.971717 | 12.881089 | 14.487800 | 12.783804 | 1.053421 | 1926.408539 | 0.128282226 | 0.585290527 | 0.558352383 | 0.998865404 | 10.873494 | 10.948916 | 10.903433 | 10.875875 | 10.989552 | 10.869132 | 2198 | 1989 | 1849 | 1713 |
| 5981 | NM_00120474 | RFC1 | replication factor C subunit 1 | protein_coding | NP_001191676.1;NP_001350424.1;NP_001350425.1;NP_002904.3;XP_01 | 9969 | 102579 | . | . | 11.592992 | 10.820100 | 10.295664 | 11.827500 | 26.977370 | 25.011145 | 23.103175 | 27.042175 | -1.021503 | 1685.210093 | 0.130330974 | -0.2355028 | 0.813818509 | 0.998865404 | 10.734057 | 10.703204 | 10.757127 | 10.694745 | 10.658726 | 10.758310 | 2039 | 1765 | 1357 | 1663 |
| 5982 | NM_00127879 | RFC2 | replication factor C subunit 2 | protein_coding | NP_001265720.1;NP_001265721.1;NP_001265722.1;NP_002905.2;NP_8 | 9970 | 600404 | . | . | 23.217030 | 22.003920 | 22.221419 | 24.084938 | 54.026982 | 50.863043 | 49.864219 | 55.067352 | -1.001537 | 1217.101979 | 0.132378898 | -0.01673619 | 0.986647074 | 0.999197585 | 10.250466 | 10.248001 | 10.248779 | 10.250584 | 10.216389 | 10.279527 | 1401 | 1316 | 1013 | 1188 |
| 5983 | NM_002915,N | RFC3 | replication factor C subunit 3 | protein_coding | NP_002906.1;NP_853536.2;XP_011533474.1;XP_011533475.1;XP_01153 | 9971 | 600405 | . | . | 16.473076 | 17.338550 | 15.808594 | 15.565581 | 38.333523 | 40.078828 | 35.474024 | 35.588855 | -1.098110 | 1236.107338 | 0.130713158 | -1.03296984 | 0.301618013 | 0.998865404 | 10.337430 | 10.202597 | 10.306464 | 10.305185 | 10.257733 | 10.212291 | 1491 | 1395 | 1056 | 1071 |
| 5984 | NM_002916,N | RFC4 | replication factor C subunit 4 | protein_coding | NP_002907.1;NP_853551.1 | 9972 | 102577 | . | . | 40.676621 | 39.980235 | 45.179042 | 38.505741 | 94.656167 | 92.416105 | 101.380461 | 88.038798 | 1.024596 | 1768.150547 | 0.132016076 | 0.265540068 | 0.790593447 | 0.998865404 | 10.770201 | 10.805633 | 10.786613 | 10.767864 | 10.861872 | 10.726717 | 2035 | 1862 | 1690 | 1534 |
| 5985 | NM_00113011 | RFC5 | replication factor C subunit 5 | protein_coding | NP_001123584.1;NP_001123585.1;NP_001193730.1;NP_001333744.1;NI | 9973 | 600407 | . | . | 16.802482 | 18.540120 | 15.601898 | 16.232656 | 39.100064 | 42.856318 | 35.010206 | 37.114041 | -1.071140 | 1369.149408 | 0.140013278 | -0.7081307 | 0.478864097 | 0.998865404 | 10.467518 | 10.368926 | 10.478892 | 10.407019 | 10.453727 | 10.324571 | 1706 | 1457 | 1248 | 1134 |
| 5986 | NM_002917,XI | RFNG | RFNG O-fucosylpeptide 3-beta-N-acetylglucosaminyltransferase | protein_coding | NP_002908.1;XP_011521889.1 | 9974 | 602578 | . | . | 15.213147 | 14.056218 | 15.774180 | 16.899295 | 35.401619 | 32.491578 | 35.396805 | 38.638229 | 1.100282 | 898.473915 | 0.143341674 | 0.961848928 | 0.336125497 | 0.998865404 | 9.740736 | 9.878636 | 9.804974 | 9.744478 | 9.822754 | 9.864817 | 1027 | 884 | 795 | 908 |
| 5987 | NM_006510 | TRIM27 | tripartite motif containing 27 | protein_coding | NP_006501.1 | 9975 | 602165 | . | . | 16.043879 | 15.460519 | 17.047184 | 18.801352 | 37.334763 | 35.737682 | 38.253384 | 42.987057 | 1.122018 | 1553.755518 | 0.127170579 | 1.306084761 | 0.191523735 | 0.964114555 | 10.516171 | 10.682146 | 10.571995 | 10.536878 | 10.608775 | 10.677816 | 1725 | 1544 | 1366 | 1608 |
| 5989 | NM_002918,XI | RFX1 | regulatory factor X1 | protein_coding | NP_002909.4;XP_011526467.1;XP_011526468.1;XP_011526469.1;XP_01 | 9982 | 600006 | . | . | 2.190403 | 2.796807 | 2.362624 | 2.550361 | 5.097158 | 6.464943 | 5.301659 | 5.831098 | -1.029273 | 338.805001 | 0.204292214 | -0.20375222 | 0.838547135 | 0.998865404 | 8.425482 | 8.382825 | 8.342114 | 8.474801 | 8.374559 | 8.413589 | 350 | 414 | 280 | 324 |
| 5990 | NM_000635,N | RFX2 | regulatory factor X2 | protein_coding | NP_000626.2;NP_602309.1;XP_011526474.1;XP_016882596.1 | 9983 | 142765 | . | . | 0.810366 | 0.711646 | 0.670634 | 0.612300 | 1.885753 | 1.644999 | 1.504884 | 1.399953 | -1.194463 | 87.564175 | 0.342822837 | -0.74779856 | 0.454581669 | 0.998865404 | 6.574217 | 6.319053 | 6.497195 | 6.456924 | 6.441134 | 6.414485 | 117 | 96 | 73 | 71 |
| 5991 | NM_00128211 | RFX3 | regulatory factor X3 | protein_coding | NP_001269045.1;NP_001269046.1;NP_001364928.1;NP_002910.1;NP_6 | 9984 | 601337 | . | . | 0.470072 | 0.602319 | 0.546313 | 0.403629 | 1.093877 | 1.392288 | 1.225909 | 0.922850 | -1.040472 | 126.525456 | 0.300684876 | -0.19036007 | 0.849026984 | 0.998865404 | 7.011162 | 6.954857 | 6.958786 | 7.015170 | 7.020353 | 6.924204 | 137 | 150 | 123 | 100 |
| 5993 | NM_000449,N | RFX5 | regulatory factor X5 | protein_coding | NP_000440.1;NP_001020774.1;NP_001366341.1;NP_001366342.1;NP_0 | 9986 | 601863 | . | . | 15.212665 | 16.000418 | 18.033055 | 19.103977 | 35.400495 | 36.985682 | 40.465650 | 43.678973 | 1.174693 | 1914.047023 | 0.120369281 | 1.929757867 | 0.053636846 | 0.686359128 | 10.781694 | 11.013802 | 10.818482 | 10.850845 | 10.941666 | 10.981982 | 1985 | 1939 | 1753 | 1987 |
| 5994 | NM_000538 | RFXAP | regulatory factor X associated protein | protein_coding | NP_000529.1 | 9988 | 601861 | . | . | 2.037158 | 1.722158 | 1.573038 | 1.173916 | 4.740551 | 3.980846 | 3.529851 | 2.684020 | -1.378761 | 117.700384 | 0.312696009 | -1.48186126 | 0.138377216 | 0.900064053 | 7.091104 | 6.630188 | 6.961309 | 6.893330 | 6.863710 | 6.774753 | 171 | 134 | 99 | 79 |
| 5995 | NM_00101272 | RGR | retinal G protein coupled receptor | protein_coding | NP_001012738.1;NP_001012740.1;NP_002912.2;XP_024303886.1 | 9990 | 600342 | . | . | 0.113154 | 0.203267 | 0.222118 | 0.293787 | 0.263314 | 0.469860 | 0.498427 | 0.671709 | 1.301324 | 16.638949 | 0.764552409 | 0.496996551 | 0.619191483 | NA | 3.855800 | 4.232645 | 4.019967 | 4.056483 | 4.055454 | 4.058985 | 12 | 20 | 16 | 18 |
| 5996 | NM_002922 | RGS1 | regulator of G protein signaling 1 | protein_coding | NP_002913.3 | 9991 | 600323 | . | . | 0.309066 | 0.199676 | 0.545485 | 0.533749 | 0.719209 | 0.461561 | 1.224053 | 1.220353 | 1.956423 | 17.140504 | 0.75191494 | 1.287669865 | 0.197860908 | NA | 3.533720 | 4.504604 | 4.067742 | 4.047049 | 4.106435 | 4.104446 | 16 | 10 | 20 | 21 |
| 5997 | NM_002923 | RGS2 | regulator of G protein signaling 2 | protein_coding | NP_002914.1 | 9998 | 600861 | . | . | 4.012570 | 4.146153 | 2.511408 | 2.123003 | 9.337415 | 9.584019 | 5.635526 | 4.853994 | -1.783194 | 135.142210 | 0.286142976 | -2.9162479 | **0.003542689** | 0.168619779 | 7.435620 | 6.602372 | 7.162469 | 7.171795 | 6.985620 | 6.932711 | 197 | 189 | 92 | 83 |
| 5998 | NM_00127626 | RGS3 | regulator of G protein signaling 3 | protein_coding | NP_001263189.1;NP_001263190.1;NP_001263191.1;NP_001269851.1;NI | 9999 | 602189 | . | . | 4.894039 | 5.160344 | 3.267345 | 4.537592 | 11.388629 | 11.928366 | 7.331829 | 10.374664 | -1.323189 | 241.702576 | 0.23610979 | -1.71114913 | 0.087053588 | 0.796289376 | 8.105771 | 7.699964 | 7.967602 | 7.991310 | 7.768398 | 7.910801 | 310 | 304 | 152 | 226 |
| 6001 | NM_00100533 | RGS10 | regulator of G protein signaling 10 | protein_coding | NP_001005339.1;NP_002916.1 | 9992 | 602856 | . | . | 46.432007 | 43.735969 | 39.836269 | 47.669292 | 108.049185 | 101.097650 | 89.391434 | 108.990166 | -1.045650 | 1278.103464 | 0.140114935 | -0.45961997 | 0.64578902 | 0.998865404 | 10.351780 | 10.287073 | 10.361493 | 10.309253 | 10.236470 | 10.363121 | 1556 | 1362 | 995 | 1270 |
| 6002 | NM_002926,N | RGS12 | regulator of G protein signaling 12 | protein_coding | NP_002917.1;NP_937870.1;NP_937872.1;XP_006713968.1;XP_00671396 | 9994 | 602512 | . | . | 17.103985 | 19.855484 | 30.595407 | 21.998117 | 39.801675 | 45.896839 | 68.655202 | 50.296085 | 1.485602 | 3414.062748 | 0.120134671 | 4.753392821 | **2.00031E-06** | **0.000493041** | 11.423575 | 11.994796 | 11.554693 | 11.546848 | 11.939637 | 11.809269 | 3179 | 2949 | 3820 | 3534 |
| 6004 | NM_002928,XI | RGS16 | regulator of G protein signaling 16 | protein_coding | NP_002919.3;XP_024304564.1 | 9997 | 602514 | . | . | 0.239835 | 0.310592 | 0.261713 | 0.355683 | 0.558105 | 0.717946 | 0.587275 | 0.813226 | 1.121142 | 22.424716 | 0.655245717 | 0.251765936 | 0.801221989 | NA | 4.403699 | 4.565787 | 4.465639 | 4.491064 | 4.476937 | 4.501306 | 21 | 26 | 18 | 25 |
| 6006 | NM_00133043 | RHCE | Rh blood group CcEe antigens | protein_coding | NP_001317359.1;NP_065231.3;NP_619522.3;NP_619523.3;NP_619524.5 | 10008 | 111700 | . | . | 0.348644 | 0.285400 | 0.147672 | 0.147704 | 0.811311 | 0.659715 | 0.331371 | 0.337708 | -1.979096 | 25.545892 | 0.623784393 | -1.57881747 | 0.114377931 | NA | 5.084644 | 4.100822 | 4.730679 | 4.695309 | 4.644669 | 4.643215 | 44 | 32 | 15 | 16 |
| 6008 | gene-RHEBP1 | RHEBP1 | RHEB pseudogene 1 | pseudogene | . | 10010 | . | . | . | 1.207153 | 1.373265 | 1.303377 | 0.758910 | 2.809095 | 3.174364 | 2.924741 | 1.735157 | -1.259644 | 49.205444 | 0.468745779 | -0.7104405 | 0.477431018 | 0.998865404 | 5.776407 | 5.446236 | 5.625218 | 5.651541 | 5.638802 | 5.552370 | 59 | 63 | 48 | 30 |
| 6009 | NM_005614,XI | RHEB | Ras homolog, mTORC1 binding | protein_coding | NP_005605.1;XP_011514759.1;XP_024302622.1 | 10011 | 601293 | . | . | 33.872376 | 31.939141 | 29.155251 | 30.702873 | 78.822406 | 73.828757 | 65.423533 | 70.198473 | -1.100910 | 2021.693347 | 0.119816742 | -1.15757442 | 0.247037754 | 0.992733415 | 11.049020 | 10.910346 | 11.044126 | 10.991769 | 10.929381 | 10.954231 | 2519 | 2212 | 1647 | 1833 |
| 6015 | NM_002931 | RING1 | ring finger protein 1 | protein_coding | NP_002922.2 | 10018 | 602045 | . | . | 17.066097 | 15.565559 | 18.297245 | 16.868773 | 39.713509 | 35.980488 | 41.058487 | 38.568443 | 1.062754 | 919.759195 | 0.145097248 | 0.605162078 | 0.54507131 | 0.998865404 | 9.800273 | 9.888600 | 9.857092 | 9.785192 | 9.898531 | 9.832178 | 1078 | 914 | 862 | 848 |
| 6016 | NM_00125682 | RIT1 | Ras like without CAAX 1 | protein_coding | NP_001243749.1;NP_001243750.1;NP_008843.1 | 10023 | 609591 | . | . | 4.753978 | 4.808137 | 5.824943 | 5.186616 | 11.062702 | 11.114222 | 13.071004 | 11.858582 | 1.136167 | 541.636120 | 0.166156031 | 1.108448792 | 0.267668043 | 0.998546485 | 8.985901 | 9.170555 | 9.037880 | 9.035497 | 9.157855 | 9.079524 | 585 | 546 | 531 | 508 |
| 6017 | NM_000326,XI | RLBP1 | retinaldehyde binding protein 1 | protein_coding | NP_000317.1;XP_011520172.1;XP_016877949.1 | 10024 | 180090 | . | . | 24.818688 | 23.332560 | 28.365831 | 36.323273 | 57.754108 | 53.934259 | 63.652103 | 83.048854 | 1.322231 | 1664.096855 | 0.137271722 | 2.935594846 | **0.003329088** | 0.163366554 | 10.485179 | 10.887860 | 10.592333 | 10.565454 | 10.713215 | 10.874665 | 1680 | 1519 | 1483 | 1954 |
| 6018 | NM_012421,XI | RLF | RLF zinc finger | protein_coding | NP_036553.2;XP_016857505.1;XP_016857506.1;XP_016857507.1 | 10025 | 180610 | . | . | 3.240989 | 3.022317 | 4.125432 | 2.810753 | 7.541914 | 6.986222 | 9.257350 | 6.426454 | 1.088759 | 640.456665 | 0.193126197 | 0.635259016 | 0.525259498 | 0.998865404 | 9.259717 | 9.383541 | 9.312754 | 9.261322 | 9.471709 | 9.200189 | 733 | 636 | 696 | 502 |
| 6023 | NR_003051 | RMRP | RNA component of mitochondrial RNA processing endoribonuclea | RNase_MRP_R | . | 10031 | 157660 | . | . | 1.219023 | 0.306910 | 0.949621 | 0.627387 | 2.836718 | 0.709436 | 2.130922 | 1.434446 | 1.008382 | 6.646173 | 1.225460943 | 0.009826202 | 0.992159952 | NA | 2.722634 | 2.742346 | 2.747365 | 2.711764 | 2.735899 | 2.723931 | 12 | 3 | 7 | 5 |
| 6029 | NR_002715 | RN7SL1 | RNA component of signal recognition particle 7SL1 | scRNA | . | 10038 | 612177 | . | . | 17.915339 | 18.603657 | 17.120512 | 18.016090 | 41.689732 | 43.003189 | 38.417931 | 41.191650 | -1.052859 | 167.455178 | 0.254459263 | -0.29204012 | 0.770255948 | 0.998865404 | 7.424478 | 7.349819 | 7.392596 | 7.404586 | 7.369337 | 7.385280 | 195 | 188 | 139 | 156 |
| 6030 | gene-RN7SL4P | RN7SL4P | RNA, 7SL, cytoplasmic 4, pseudogene | pseudogene | . | 10039 | . | . | . | 2.762673 | 2.890857 | 1.563655 | 1.926772 | 6.428853 | 6.682346 | 3.508797 | 4.405334 | -1.596488 | 20.051213 | 0.692094246 | -0.97515832 | 0.329481732 | NA | 4.624641 | 3.947964 | 4.349308 | 4.352023 | 4.305127 | 4.318421 | 28 | 27 | 12 | 16 |
| 6038 | NM_00128219 | RNASE4 | ribonuclease A family member 4 | protein_coding | NP_001269121.1;NP_001269122.1;NP_002928.1;NP_919412.1 | 10047 | 601030 | . | . | 1.894456 | 1.585477 | 1.774082 | 1.347473 | 4.408475 | 3.664902 | 3.980989 | 3.080837 | -1.144916 | 107.289214 | 0.322016484 | -0.6063089 | 0.544309649 | 0.998865404 | 6.838633 | 6.645640 | 6.793639 | 6.735714 | 6.762309 | 6.680088 | 143 | 113 | 99 | 81 |
| 6041 | NM_021133,XI | RNASEL | ribonuclease L | protein_coding | NP_066956.1 | 10050 | 180435 | . | . | 2.163142 | 2.174491 | 2.186406 | 2.429763 | 5.033719 | 5.026431 | 4.906232 | 5.555367 | 1.053379 | 291.744182 | 0.205200946 | 0.365611989 | 0.714654628 | 0.998865404 | 8.150846 | 8.225313 | 8.175027 | 8.172750 | 8.169743 | 8.232189 | 328 | 306 | 245 | 297 |
| 6043 | NR_002586 | SNORA63 | small nucleolar RNA, H/ACA box 63 | snoRNA | . | 10106 | 180647 | . | . | 12.142390 | 11.941752 | 12.227856 | 8.271004 | 28.255842 | 27.603897 | 27.438955 | 18.910667 | -1.187592 | 47.506585 | 0.463418255 | -0.5352387 | 0.592484811 | 0.998865404 | 5.687585 | 5.442095 | 5.586664 | 5.582190 | 5.584358 | 5.522691 | 60 | 55 | 45 | 33 |
| 6044 | NR_002324 | SNORA62 | small nucleolar RNA, H/ACA box 62 | snoRNA | . | 10107 | 180646 | . | . | 5.128974 | 5.716810 | 2.705962 | 3.362350 | 11.935333 | 13.214662 | 6.072100 | 7.687614 | -1.776141 | 20.696715 | 0.683925338 | -1.2117492 | 0.225608399 | NA | 4.727571 | 3.897226 | 4.395574 | 4.406231 | 4.347920 | 4.356655 | 29 | 30 | 12 | 15 |
| 6045 | NM_007212,XI | RNF2 | ring finger protein 2 | protein_coding | NP_009143.1;XP_005245470.1;XP_011508153.1;XP_011508154.1 | 10061 | 608985 | . | . | 4.391236 | 3.987529 | 3.728666 | 3.560967 | 10.218587 | 9.217348 | 8.367020 | 8.141726 | -1.166534 | 405.381387 | 0.181574691 | -1.22389646 | 0.220991289 | 0.982008277 | 8.769784 | 8.547971 | 8.734584 | 8.679118 | 8.623661 | 8.607419 | 525 | 450 | 330 | 346 |
| 6046 | NM_00111318 | BRD2 | bromodomain containing 2 | protein_coding | NP_001106653.1;NP_001186384.1;NP_001186385.1;NP_001278915.1;NI | 1103 | 601540 | . | . | 28.849609 | 31.852962 | 34.739259 | 34.824883 | 67.134224 | 73.629549 | 77.953891 | 79.622955 | 1.097358 | 4487.226343 | 0.110283055 | 1.215370598 | 0.224224777 | 0.982008277 | 12.063098 | 12.197012 | 12.062357 | 12.119551 | 12.144357 | 12.192508 | 4757 | 4777 | 3966 | 4528 |
| 6047 | NM_00118500 | RNF4 | ring finger protein 4 | protein_coding | NP_001171938.1;NP_001171939.1;NP_002929.1 | 10067 | 602850 | . | . | 16.786195 | 18.545288 | 16.017293 | 20.254684 | 39.062168 | 42.868267 | 35.942343 | 46.309933 | 1.012545 | 1628.444825 | 0.142939921 | 0.125831108 | 0.899865617 | 0.998865404 | 10.660528 | 10.677977 | 10.626540 | 10.697610 | 10.581176 | 10.755878 | 1779 | 1826 | 1265 | 1708 |
| 6048 | NM_006913 | RNF5 | ring finger protein 5 | protein_coding | NP_008844.1 | 10068 | 602677 | . | . | 30.423851 | 27.286234 | 26.655588 | 36.076805 | 70.797554 | 63.073353 | 59.814365 | 82.485329 | 1.071451 | 1047.723955 | 0.163352232 | 0.609517234 | 0.542181653 | 0.998865404 | 9.982625 | 10.081759 | 10.045877 | 9.960113 | 9.941225 | 10.157025 | 1233 | 1028 | 806 | 1163 |
| 6049 | NM_005977,N | RNF6 | ring finger protein 6 | protein_coding | NP_005968.1;NP_898864.1;NP_898865.1;NP_898866.1;XP_005266542.1 | 10069 | 604242 | . | . | 8.818452 | 8.695531 | 8.961859 | 9.594043 | 20.520902 | 20.100111 | 20.110151 | 21.935637 | 1.065380 | 963.489789 | 0.138332845 | 0.660493783 | 0.508937007 | 0.998865404 | 9.865821 | 9.956990 | 9.885972 | 9.890624 | 9.913545 | 9.954762 | 1071 | 1010 | 840 | 958 |
| 6050 | NM_002939,N | RNH1 | ribonuclease/angiogenin inhibitor 1 | protein_coding | NP_002930.2;NP_976317.1;NP_976318.1;NP_976319.1;NP_976320.1;NP | 10074 | 173320 | . | . | 85.381512 | 87.704765 | 77.733945 | 92.765050 | 198.686289 | 202.733501 | 174.432726 | 212.096267 | -1.034944 | 4955.060829 | 0.117456834 | -0.42188045 | 0.673112279 | 0.998865404 | 12.299331 | 12.249615 | 12.276981 | 12.299275 | 12.191529 | 12.323023 | 5727 | 5511 | 3901 | 4925 |
| 6051 | NM_00131918 | RNPEP | arginyl aminopeptidase | protein_coding | NP_001306111.1;NP_001306112.1;NP_001306113.1;NP_064601.3;XP_00 | 10078 | 602675 | . | . | 27.925580 | 26.475273 | 27.141523 | 33.034249 | 64.983979 | 61.198771 | 60.904791 | 75.528886 | 1.091335 | 2107.495208 | 0.13111548 | 0.961704241 | 0.336198192 | 0.998865404 | 10.977008 | 11.102875 | 11.026133 | 10.981758 | 10.998163 | 11.144443 | 2386 | 2114 | 1739 | 2249 |
| 6059 | NM_00104087 | ABCE1 | ATP binding cassette subfamily E member 1 | protein_coding | NP_001035809.1;NP_002931.2 | 69 | 601213 | . | . | 20.034836 | 19.149990 | 18.720673 | 13.062364 | 46.621887 | 44.266064 | 42.008644 | 29.865543 | -1.249212 | 2152.094957 | 0.159273242 | -2.0155195 | **0.043850245** | 0.644584591 | 11.222899 | 10.902388 | 11.171519 | 11.127404 | 11.105252 | 10.818413 | 2826 | 2510 | 1969 | 1466 |
| 6060 | NR_004421 | RNU1-4 | RNA, U1 small nuclear 4 | snRNA | . | 10128 | 180680 | . | . | 0.565499 | 0.317301 | 0.914767 | 0.500021 | 1.315940 | 0.733456 | 2.052711 | 1.143238 | 1.671915 | 3.569283 | 1.602038051 | 0.462848637 | 0.643472866 | NA | 1.413393 | 2.161867 | 1.821784 | 1.814090 | 1.831907 | 1.820551 | 4 | 2 | 5 | 3 |
| 6079 | NR_000005 | SNORD15A | small nucleolar RNA, C/D box 15A | snoRNA | . | 10114 | 600455 | . | . | 0.916531 | 0.974558 | 0.925359 | 0.746567 | 2.132806 | 2.252734 | 2.076478 | 1.706936 | -1.012468 | 4.460625 | 1.415462877 | -0.01262893 | 0.989923838 | NA | 2.166062 | 2.148375 | 2.156438 | 2.157811 | 2.157616 | 2.156126 | 5 | 5 | 4 | 4 |
| 6080 | NR_002907 | SNORA73A | small nucleolar RNA, H/ACA box 73A | snoRNA | . | 10115 | 180645 | . | . | 5.445432 | 9.225662 | 10.744955 | 5.845026 | 12.671745 | 21.325533 | 24.111366 | 13.363957 | 1.120666 | 50.831578 | 0.490189657 | 0.335292434 | 0.737404501 | 0.998865404 | 5.582624 | 5.747948 | 5.593954 | 5.688221 | 5.720305 | 5.603118 | 41 | 65 | 61 | 35 |
| 6082 | NR_002908 | SNORD20 | small nucleolar RNA, C/D box 20 | snoRNA | . | 10143 | 604012 | . | . | 0.603860 | 1.769884 | 1.080245 | 1.938755 | 1.405206 | 4.091166 | 2.424038 | 4.432733 | 1.386013 | 3.788467 | 1.561599097 | 0.301576318 | 0.762975063 | NA | 1.671431 | 2.134756 | 1.903225 | 1.917078 | 1.911590 | 1.920087 | 2 | 5 | 3 | 5 |
| 6083 | NR_000006 | SNORD21 | small nucleolar RNA, C/D box 21 | snoRNA | . | 10144 | 603635 | . | . | 1.855930 | 0.782320 | 1.730705 | 1.018591 | 4.318825 | 1.808369 | 3.883651 | 2.328888 | 1.005086 | 4.452300 | 1.443259733 | 0.005071028 | 0.995953923 | NA | 2.147197 | 2.161867 | 2.160441 | 2.144875 | 2.158443 | 2.147069 | 7 | 3 | 5 | 3 |
| 6091 | NM_00114584 | ROBO1 | roundabout guidance receptor 1 | protein_coding | NP_001139317.1;NP_002932.1;NP_598334.2;XP_006713340.1;XP_01153 | 10249 | 602430 | . | . | 4.831101 | 5.096581 | 3.943794 | 4.265789 | 11.242172 | 11.780977 | 8.849761 | 9.753222 | -1.224748 | 1059.218337 | 0.136806092 | -2.13795648 | **0.032520275** | 0.569641321 | 10.187781 | 9.894954 | 10.104572 | 10.132824 | 9.941241 | 9.996020 | 1318 | 1282 | 796 | 927 |
| 6092 | NM_00112892 | ROBO2 | roundabout guidance receptor 2 | protein_coding | NP_001122401.1;NP_001276968.1;NP_001276969.1;NP_001276994.1;NI | 10250 | 602431 | . | . | 0.329542 | 0.334218 | 0.325708 | 0.307067 | 0.766855 | 0.772558 | 0.730881 | 0.702073 | -1.062165 | 91.448157 | 0.333540339 | -0.26086018 | 0.794200337 | 0.998865404 | 6.557487 | 6.470981 | 6.519682 | 6.527625 | 6.520131 | 6.492398 | 107 | 103 | 81 | 79 |
| 6093 | NM_005406 | ROCK1 | Rho associated coiled-coil containing protein kinase 1 | protein_coding | NP_005397.1 | 10251 | 601702 | . | . | 4.405443 | 4.466899 | 4.133727 | 3.934968 | 10.251647 | 10.325437 | 9.275965 | 8.996836 | -1.115100 | 1247.252984 | 0.130393368 | -1.20537295 | 0.22805936 | 0.982008277 | 10.360905 | 10.203903 | 10.322130 | 10.326383 | 10.263774 | 10.219703 | 1510 | 1423 | 1056 | 1073 |
| 6094 | NM_000327 | ROM1 | retinal outer segment membrane protein 1 | protein_coding | NP_000318.2 | 10254 | 180721 | . | . | 0.878886 | 0.886029 | 1.218819 | 1.046862 | 2.045203 | 2.048095 | 2.734994 | 2.393527 | 1.266156 | 43.189212 | 0.473788279 | 0.718581038 | 0.4723991 | 0.998865404 | 5.251870 | 5.593187 | 5.407146 | 5.406550 | 5.458006 | 5.432921 | 44 | 41 | 45 | 42 |
| 6095 | NM_002943,N | RORA | RAR related orphan receptor A | protein_coding | NP_002934.1;NP_599022.1;NP_599023.1;NP_599024.1;XP_005254641.1 | 10258 | 600825 | . | . | 0.075596 | 0.071278 | 0.083550 | 0.052201 | 0.175914 | 0.164762 | 0.187484 | 0.119350 | -1.110466 | 24.706627 | 0.630481685 | -0.23976198 | 0.810514786 | NA | 4.698880 | 4.550984 | 4.635084 | 4.626979 | 4.641689 | 4.598476 | 31 | 27 | 25 | 17 |
| 6096 | NM_00136502 | RORB | RAR related orphan receptor B | protein_coding | NP_001351952.1;NP_008845.2 | 10259 | 601972 | . | . | 0.069451 | 0.093483 | 0.109003 | 0.061029 | 0.161614 | 0.216091 | 0.244599 | 0.139535 | 1.019052 | 25.480683 | 0.634619322 | 0.042903944 | 0.965778105 | NA | 4.656500 | 4.686013 | 4.648543 | 4.678353 | 4.695685 | 4.634398 | 25 | 31 | 29 | 17 |
| 6100 | NM_203288,XI | RP9 | RP9 pre-mRNA splicing factor | protein_coding | NP_976033.1;XP_011513770.1 | 10288 | 607331 | . | . | 11.031884 | 12.095403 | 9.523652 | 10.710577 | 25.671648 | 27.959064 | 21.370801 | 24.488462 | -1.158613 | 385.839679 | 0.185854267 | -1.14282309 | 0.253112051 | 0.994507726 | 8.694473 | 8.481381 | 8.611516 | 8.654824 | 8.520174 | 8.573271 | 460 | 463 | 296 | 351 |
| 6102 | NM_006915 | RP2 | RP2 activator of ARL3 GTPase | protein_coding | NP_008846.2 | 10274 | 300757 | . | . | 2.869979 | 3.424647 | 2.843572 | 3.460823 | 6.678559 | 7.916225 | 6.380894 | 7.912761 | -1.012178 | 363.568727 | 0.200084639 | -0.08727907 | 0.930449687 | 0.998865404 | 8.515434 | 8.496673 | 8.457806 | 8.555661 | 8.446756 | 8.552056 | 386 | 428 | 285 | 370 |
| 6103 | NM_000328,N | RPGR | retinitis pigmentosa GTPase regulator | protein_coding | NP_000319.1;NP_001030025.1;NP_001354174.1;NP_001354175.1;NP_0 | 10295 | 312610 | . | . | 2.089282 | 2.082353 | 2.294015 | 2.417067 | 4.861847 | 4.813454 | 5.147705 | 5.526336 | 1.175846 | 209.735962 | 0.23148797 | 1.009551265 | 0.312710328 | 0.998865404 | 7.591019 | 7.824413 | 7.669201 | 7.673581 | 7.733474 | 7.758447 | 221 | 209 | 192 | 218 |
| 6117 | NM_00135512 | RPA1 | replication protein A1 | protein_coding | NP_001342049.1;NP_001342050.1;NP_002936.1;XP_024306631.1 | 10289 | 179835 | . | . | 22.186034 | 22.589880 | 19.603606 | 23.075964 | 51.627809 | 52.217521 | 43.989923 | 52.760450 | -1.062787 | 3298.826638 | 0.120384496 | -0.72977046 | 0.465530504 | 0.998865404 | 11.731101 | 11.643030 | 11.706048 | 11.717229 | 11.597038 | 11.722168 | 3888 | 3693 | 2568 | 3228 |
| 6118 | NM_00128607 | RPA2 | replication protein A2 | protein_coding | NP_001273005.1;NP_001284487.1;NP_001342057.1;NP_001342058.1;NI | 10290 | 179836 | . | . | 30.797437 | 34.754026 | 31.411067 | 36.353388 | 71.666908 | 80.335486 | 70.485530 | 83.117698 | 1.014821 | 1654.456576 | 0.133950917 | 0.15845415 | 0.874098947 | 0.998865404 | 10.681719 | 10.702490 | 10.640085 | 10.728894 | 10.642047 | 10.748740 | 1784 | 1873 | 1350 | 1669 |
| 6119 | NM_002947 | RPA3 | replication protein A3 | protein_coding | NP_002938.1 | 10291 | 179837 | . | . | 15.530837 | 14.871638 | 12.633723 | 16.925070 | 36.140900 | 34.376457 | 28.349709 | 38.697163 | -1.043132 | 943.845158 | 0.161874507 | -0.37635044 | 0.706656383 | 0.998865404 | 9.912850 | 9.851306 | 9.914473 | 9.876057 | 9.759013 | 9.960367 | 1139 | 1013 | 691 | 987 |
| 6120 | NM_00127828 | RPE | ribulose-5-phosphate-3-epimerase | protein_coding | NP_001265211.1;NP_001265212.1;NP_001265214.1;NP_001265215.1;NI | 10293 | 180480 | . | . | 21.172911 | 20.365445 | 22.756201 | 20.043167 | 49.270236 | 47.075643 | 51.064254 | 45.826322 | -1.092355 | 2015.472774 | 0.132446085 | -0.96221663 | 0.335940797 | 0.998865404 | 11.039051 | 10.911956 | 11.079000 | 10.935801 | 10.968217 | 10.910454 | 2638 | 2069 | 1737 | 1740 |
| 6122 | NM_000967,N | RPL3 | ribosomal protein L3 | protein_coding | NP_000958.1;NP_001029025.1 | 10332 | 604163 | . | . | 492.736845 | 503.251224 | 434.815621 | 556.153587 | 1146.618866 | 1163.287773 | 975.713734 | 1271.579121 | -1.019822 | 19965.373139 | 0.123429988 | -0.22941784 | 0.818544172 | 0.998865404 | 14.299326 | 14.270959 | 14.287093 | 14.296814 | 14.169671 | 14.370666 | 23085 | 21878 | 15186 | 20695 |
| 6124 | NM_000968 | RPL4 | ribosomal protein L4 | protein_coding | NP_000959.2 | 10353 | 180479 | . | . | 251.248627 | 255.616714 | 238.076721 | 299.381714 | 584.665894 | 590.869507 | 534.237305 | 684.500793 | 1.045130 | 22282.533222 | 0.120694383 | 0.527639228 | 0.597749761 | 0.998865404 | 14.411454 | 14.475095 | 14.419128 | 14.426604 | 14.362076 | 14.551057 | 24983 | 23616 | 17645 | 23678 |
| 6125 | NM_000969,N | RPL5 | ribosomal protein L5 | protein_coding | NP_000960.2 | 10360 | 603634 | . | . | 299.238911 | 288.217463 | 268.506008 | 321.646196 | 696.341202 | 666.227597 | 602.519644 | 735.405871 | -1.009838 | 9423.262886 | 0.116247626 | -0.12149622 | 0.903298012 | 0.998865404 | 13.209089 | 13.194898 | 13.224615 | 13.185523 | 13.121949 | 13.266751 | 11154 | 9984 | 7461 | 9538 |
| 6126 | gene-RPL9P7 | RPL9P7 | ribosomal protein L9 pseudogene 7 | pseudogene | . | 30335 | . | . | . | 7.585473 | 7.069444 | 6.506910 | 9.754967 | 17.651707 | 16.341337 | 14.601317 | 22.303572 | 1.092524 | 169.491946 | 0.275958563 | 0.462621587 | 0.643635633 | 0.998865404 | 7.340611 | 7.466778 | 7.397975 | 7.365656 | 7.333660 | 7.497200 | 194 | 168 | 124 | 198 |
| 6128 | NM_000970,N | RPL6 | ribosomal protein L6 | protein_coding | NP_000961.2;NP_001019833.1;NP_001307066.1;NP_001307067.1;NP_0 | 10362 | 603703 | . | . | 484.620599 | 491.215711 | 472.407791 | 600.951310 | 1127.732070 | 1135.467122 | 1060.069334 | 1374.003751 | 1.059779 | 16342.711523 | 0.115735049 | 0.723749958 | 0.469219257 | 0.998865404 | 13.953897 | 14.037608 | 13.968652 | 13.971146 | 13.937401 | 14.095992 | 18246 | 17148 | 13282 | 17212 |
| 6129 | NM_000971,N | RPL7 | ribosomal protein L7 | protein_coding | NP_000962.2;NP_001350666.1 | 10363 | 604166 | . | . | 345.148434 | 340.504427 | 310.187304 | 347.483996 | 803.174598 | 787.091228 | 696.051296 | 794.480877 | -1.055896 | 13001.366911 | 0.1053987 | -0.74448518 | 0.456583003 | 0.998865404 | 13.705094 | 13.626590 | 13.699719 | 13.678022 | 13.599324 | 13.683657 | 15572 | 14229 | 10454 | 12444 |
| 6130 | NM_000972 | RPL7A | ribosomal protein L7a | protein_coding | NP_000963.1 | 10364 | 185640 | . | . | 559.305481 | 579.946960 | 509.587799 | 634.334961 | 1301.526855 | 1340.573364 | 1143.500244 | 1450.331543 | -1.010342 | 15767.626429 | 0.119959771 | -0.12374425 | 0.901517773 | 0.998865404 | 13.952112 | 13.937205 | 13.935779 | 13.959609 | 13.845655 | 14.024423 | 17997 | 17339 | 12222 | 16235 |
| 6132 | NM_000973,N | RPL8 | ribosomal protein L8 | protein_coding | NP_000964.1;NP_001304700.1;NP_001304711.1;NP_150644.1 | 10368 | 604177 | . | . | 670.768352 | 696.571125 | 646.534078 | 757.201728 | 1560.905481 | 1610.155310 | 1450.803820 | 1731.251789 | 1.014253 | 18646.163470 | 0.1113124 | 0.183423929 | 0.854465402 | 0.998865404 | 14.176370 | 14.196741 | 14.167989 | 14.191374 | 14.123644 | 14.255889 | 21029 | 20250 | 15075 | 18956 |
| 6133 | NM_000661,N | RPL9 | ribosomal protein L9 | protein_coding | NP_000652.2;NP_001020092.1 | 10369 | 603686 | . | . | 294.857294 | 308.744496 | 277.504713 | 353.167112 | 686.145085 | 713.676731 | 622.712558 | 807.474575 | 1.029963 | 7043.064588 | 0.12616308 | 0.337603318 | 0.735662142 | 0.998865404 | 12.760605 | 12.803058 | 12.751373 | 12.783874 | 12.691154 | 12.885126 | 7837 | 7632 | 5506 | 7464 |
| 6134 | NM_00125657 | RPL10 | ribosomal protein L10 | protein_coding | NP_001243506.2;NP_001243509.2;NP_001290553.1;NP_001290554.1;NI | 10298 | 312173 | . | . | 134.994562 | 134.460878 | 118.933012 | 152.068594 | 314.137888 | 310.812352 | 266.882235 | 347.686766 | -1.006237 | 9252.685206 | 0.125461225 | -0.07149719 | 0.94300206 | 0.998865404 | 13.180182 | 13.171116 | 13.182644 | 13.170651 | 13.066878 | 13.266183 | 10763 | 9945 | 7084 | 9657 |
| 6135 | NM_000975,N | RPL11 | ribosomal protein L11 | protein_coding | NP_000966.2;NP_001186731.1 | 10301 | 604175 | . | . | 615.025162 | 617.544205 | 598.577469 | 661.325821 | 1431.188873 | 1427.481018 | 1343.190521 | 1512.042938 | 1.000261 | 17939.511507 | 0.106710937 | 0.003532066 | 0.997181825 | 0.999862634 | 14.130681 | 14.131025 | 14.137622 | 14.122945 | 14.077798 | 14.180678 | 20831 | 19190 | 14668 | 17827 |
| 6136 | NM_000976 | RPL12 | ribosomal protein L12 | protein_coding | NP_000967.1 | 10302 | 180475 | . | . | 617.901794 | 614.129272 | 553.133728 | 730.021973 | 1437.882935 | 1419.587402 | 1241.216064 | 1669.108521 | 1.026354 | 12413.573133 | 0.130002467 | 0.288678548 | 0.77282738 | 0.998865404 | 13.580782 | 13.618236 | 13.592493 | 13.579964 | 13.487764 | 13.716570 | 14212 | 13124 | 9482 | 13355 |
| 6137 | NM_000977,N | RPL13 | ribosomal protein L13 | protein_coding | NP_000968.2;NP_001230060.1;NP_150254.1 | 10303 | 113703 | . | . | 137.148463 | 133.360912 | 121.522072 | 162.173999 | 319.150135 | 308.269726 | 272.691972 | 370.791585 | 1.063884 | 10524.167623 | 0.126436655 | 0.706602762 | 0.479813372 | 0.998865404 | 13.316095 | 13.405361 | 13.351419 | 13.313470 | 13.277873 | 13.482890 | 12006 | 10759 | 8311 | 11383 |
| 6138 | NM_00125337 | RPL15 | ribosomal protein L15 | protein_coding | NP_001240308.1;NP_001240309.1;NP_001240311.1;NP_001240312.1;NI | 10306 | 604174 | . | . | 174.513121 | 175.640434 | 159.238226 | 195.046186 | 406.099210 | 406.000722 | 357.325949 | 445.949912 | 1.013542 | 12102.582239 | 0.121536846 | 0.159675197 | 0.87313694 | 0.998865404 | 13.553329 | 13.572661 | 13.555506 | 13.556199 | 13.470910 | 13.655468 | 13837 | 12977 | 9454 | 12652 |
| 6139 | NM_000985,N | RPL17 | ribosomal protein L17 | protein_coding | NP_000976.1;NP_001030178.1;NP_001186269.1;NP_001186270.1;NP_0 | 10307 | 603661 | . | . | 459.617038 | 452.462336 | 413.628658 | 490.355009 | 1069.547730 | 1045.886941 | 928.170721 | 1121.138453 | -1.023036 | 10858.984080 | 0.113252148 | -0.29011725 | 0.771726537 | 0.998865404 | 13.422969 | 13.390046 | 13.422278 | 13.408693 | 13.326204 | 13.461437 | 12747 | 11757 | 8595 | 10862 |
| 6141 | NM_000979,N | RPL18 | ribosomal protein L18 | protein_coding | NP_000970.1;NP_001257419.1 | 10310 | 604179 | . | . | 403.248588 | 410.092837 | 387.550489 | 460.491786 | 938.376017 | 947.947948 | 869.652053 | 1052.859668 | 1.027444 | 8310.027478 | 0.114705254 | 0.340520922 | 0.73346427 | 0.998865404 | 13.001024 | 13.039988 | 13.004673 | 13.011806 | 12.959912 | 13.097891 | 9400 | 8883 | 6731 | 8535 |
| 6142 | NM_000980 | RPL18A | ribosomal protein L18a | protein_coding | NP_000971.1 | 10311 | 604178 | . | . | 446.574432 | 446.178864 | 431.523865 | 544.085144 | 1039.197021 | 1031.362427 | 968.326965 | 1243.985962 | 1.077159 | 9220.348526 | 0.123338267 | 0.869405635 | 0.384625302 | 0.998865404 | 13.116042 | 13.223181 | 13.140598 | 13.132671 | 13.100782 | 13.290333 | 10271 | 9535 | 7398 | 9954 |
| 6143 | NM_000981,N | RPL19 | ribosomal protein L19 | protein_coding | NP_000972.1;NP_001317129.1 | 10312 | 180466 | . | . | 801.649933 | 823.294937 | 726.939392 | 884.137970 | 1865.472290 | 1903.083008 | 1631.230896 | 2021.476379 | -1.020490 | 21080.808882 | 0.116312832 | -0.25158251 | 0.80136378 | 0.998865404 | 14.378222 | 14.348914 | 14.362228 | 14.380154 | 14.269047 | 14.432025 | 24266 | 23217 | 16428 | 21414 |
| 6144 | NM_000982 | RPL21 | ribosomal protein L21 | protein_coding | NP_000973.2 | 10313 | 603636 | . | . | 371.598389 | 360.696472 | 338.070251 | 411.649384 | 864.724792 | 833.765991 | 758.619812 | 941.187317 | 1.009155 | 6566.202152 | 0.120014833 | 0.109553685 | 0.912763343 | 0.998865404 | 12.674307 | 12.687351 | 12.691973 | 12.659603 | 12.601907 | 12.759222 | 7671 | 6918 | 5202 | 6759 |
| 6146 | NM_000983 | RPL22 | ribosomal protein L22 | protein_coding | NP_000974.1 | 10315 | 180474 | . | . | 161.467621 | 154.637589 | 148.472656 | 169.861496 | 375.741791 | 357.451721 | 333.168335 | 388.368134 | -1.007333 | 10182.537907 | 0.110167276 | -0.09568501 | 0.923770743 | 0.998865404 | 13.319093 | 13.308507 | 13.338272 | 13.294032 | 13.255848 | 13.361724 | 12073 | 10742 | 8274 | 10102 |
| 6147 | NM_000984 | RPL23A | ribosomal protein L23a | protein_coding | NP_000975.2 | 10317 | 602326 | . | . | 290.161743 | 293.315002 | 289.865387 | 328.817596 | 675.218262 | 678.010742 | 650.449524 | 751.802368 | 1.045428 | 9079.000705 | 0.108325532 | 0.591678601 | 0.554065829 | 0.998865404 | 13.115949 | 13.179976 | 13.127767 | 13.129667 | 13.115849 | 13.214766 | 10211 | 9590 | 7603 | 9204 |
| 6148 | gene-RPL23AP | RPL23AP1 | ribosomal protein L23a pseudogene 1 | pseudogene | . | 10318 | . | . | . | 0.012374 | 0.013017 | 0.029030 | 0.133350 | 0.028794 | 0.030090 | 0.065142 | 0.304888 | **2.432439** | 1.536291 | 2.42076221 | 0.52975204 | 0.596283861 | NA | -0.155866 | 1.121007 | 0.602056 | 0.602388 | 0.603562 | 0.612940 | 1 | 1 | 1 | 3 |
| 6150 | NM_021134,XI | MRPL23 | mitochondrial ribosomal protein L23 | protein_coding | NP_066957.3;XP_006718334.1;XP_011518575.1;XP_011518577.1;XP_01 | 10322 | 600789 | . | . | 59.209969 | 62.247857 | 53.574130 | 69.936968 | 137.784032 | 143.888702 | 120.218793 | 159.902549 | -1.002723 | 1290.067519 | 0.151028389 | -0.02597373 | 0.979278293 | 0.998865404 | 10.335499 | 10.330959 | 10.315314 | 10.348982 | 10.226311 | 10.424280 | 1455 | 1425 | 974 | 1365 |
| 6152 | NM_000986 | RPL24 | ribosomal protein L24 | protein_coding | NP_000977.1 | 10325 | 604180 | . | . | 238.855469 | 235.183945 | 220.920837 | 268.914917 | 555.826477 | 543.638245 | 495.739960 | 614.842041 | 1.018636 | 4203.211334 | 0.122640456 | 0.217206485 | 0.828047431 | 0.998865404 | 12.023979 | 12.050452 | 12.038427 | 12.018507 | 11.963456 | 12.118359 | 4853 | 4440 | 3346 | 4346 |
| 6154 | NM_000987,N | RPL26 | ribosomal protein L26 | protein_coding | NP_000978.1;NP_001302459.1;NP_001302460.1 | 10327 | 603704 | . | . | 737.586410 | 693.962133 | 633.860660 | 775.609761 | 1716.393955 | 1604.124483 | 1422.364753 | 1773.339561 | -1.029745 | 12257.914810 | 0.119152456 | -0.3549012 | 0.722663606 | 0.998865404 | 13.602440 | 13.560102 | 13.622820 | 13.561563 | 13.484285 | 13.644589 | 14833 | 12942 | 9518 | 12385 |
| 6155 | NM_000988,N | RPL27 | ribosomal protein L27 | protein_coding | NP_000979.1;NP_001336850.1;NP_001336851.1 | 10328 | 607526 | . | . | 651.303155 | 647.926566 | 596.599230 | 721.291553 | 1515.609228 | 1497.711230 | 1338.751510 | 1649.147415 | 1.000563 | 10695.168599 | 0.11940924 | 0.006802404 | 0.994572509 | 0.999862634 | 13.384303 | 13.385040 | 13.393309 | 13.373088 | 13.295602 | 13.464863 | 12458 | 11401 | 8381 | 11023 |
| 6156 | NM_000989 | RPL30 | ribosomal protein L30 | protein_coding | NP_000980.1 | 10333 | 180467 | . | . | 573.521179 | 538.795959 | 473.529724 | 581.850708 | 1334.607422 | 1245.451050 | 1062.587036 | 1330.332520 | -1.069380 | 8408.016496 | 0.121538515 | -0.79625013 | 0.425886667 | 0.998865404 | 13.085163 | 12.988311 | 13.094084 | 13.033303 | 12.921312 | 13.086310 | 10362 | 9044 | 6377 | 8361 |
| 6157 | NM_000990 | RPL27A | ribosomal protein L27a | protein_coding | NP_000981.1 | 10329 | 603637 | . | . | 78.458885 | 77.152328 | 75.923882 | 92.318077 | 182.577057 | 178.341064 | 170.370987 | 211.074310 | 1.065711 | 11432.904312 | 0.117162141 | 0.783664484 | 0.433237008 | 0.998865404 | 13.434299 | 13.526051 | 13.462765 | 13.441206 | 13.423818 | 13.582879 | 12908 | 11793 | 9310 | 12081 |
| 6158 | NM_000991,N | RPL28 | ribosomal protein L28 | protein_coding | NP_000982.2;NP_001129606.1;NP_001129607.1;NP_001129608.1;NP_0 | 10330 | 603638 | . | . | 60.719980 | 59.187722 | 60.781079 | 71.045601 | 141.297885 | 136.815078 | 136.390980 | 162.437321 | 1.036548 | 5940.744998 | 0.135223211 | 0.382968558 | 0.701743077 | 0.998865404 | 12.510384 | 12.562010 | 12.516807 | 12.520643 | 12.423276 | 12.660904 | 6703 | 6310 | 4522 | 6463 |
| 6159 | NM_000992,XI | RPL29 | ribosomal protein L29 | protein_coding | NP_000983.1;XP_024309466.1 | 10331 | 601832 | . | . | 252.339494 | 266.727719 | 243.149072 | 299.316955 | 587.204343 | 616.553087 | 545.619509 | 684.352651 | 1.030423 | 6235.758220 | 0.122125973 | 0.354033118 | 0.723314064 | 0.998865404 | 12.584646 | 12.627731 | 12.572501 | 12.611833 | 12.531050 | 12.697406 | 6908 | 6783 | 4963 | 6516 |
| 6160 | NM_000993,N | RPL31 | ribosomal protein L31 | protein_coding | NP_000984.1;NP_001092047.1;NP_001093163.1 | 10334 | 617415 | . | . | 322.846104 | 298.657472 | 264.780078 | 318.448358 | 751.276136 | 690.360129 | 594.158836 | 728.094331 | -1.027445 | 10943.363423 | 0.123176746 | -0.31711956 | 0.751152883 | 0.998865404 | 13.437201 | 13.398071 | 13.450357 | 13.404616 | 13.307713 | 13.493142 | 13113 | 11649 | 8368 | 11219 |
| 6161 | NM_000994,N | RPL32 | ribosomal protein L32 | protein_coding | NP_000985.1;NP_001007074.1;NP_001007075.1 | 10336 | . | . | . | 102.663700 | 106.098295 | 97.228664 | 116.464109 | 238.902666 | 245.250951 | 218.178303 | 266.281327 | 1.010135 | 6858.718301 | 0.117064414 | 0.124278951 | 0.901094413 | 0.998865404 | 12.736493 | 12.750918 | 12.728064 | 12.748985 | 12.672671 | 12.816415 | 7761 | 7455 | 5485 | 7011 |
| 6164 | NM_000995,N | RPL34 | ribosomal protein L34 | protein_coding | NP_000986.2;NP_001306161.1;NP_001306163.1;NP_001306164.1;NP_0 | 10340 | 616862 | . | . | 118.374466 | 169.300468 | 103.932727 | 198.402493 | 275.462264 | 391.345620 | 233.222030 | 453.623747 | -1.011045 | 3354.571033 | 0.128555212 | -0.12326803 | 0.901894858 | 0.998865404 | 11.719916 | 11.703864 | 11.731507 | 11.698454 | 11.616206 | 11.788590 | 3962 | 3567 | 2597 | 3455 |
| 6165 | NM_000996,N | RPL35A | ribosomal protein L35a | protein_coding | NP_000987.2;NP_001303240.1 | 10345 | 180468 | . | . | 142.425843 | 150.619942 | 138.358498 | 173.580726 | 331.430786 | 348.164749 | 310.472419 | 396.871689 | 1.040389 | 5963.629696 | 0.128321513 | 0.44515666 | 0.656206545 | 0.998865404 | 12.513216 | 12.570172 | 12.504241 | 12.542072 | 12.452275 | 12.651401 | 6580 | 6450 | 4667 | 6371 |
| 6166 | NM_001001 | RPL36AL | ribosomal protein L36a like | protein_coding | NP_000992.1 | 10346 | 180469 | . | . | 82.398613 | 78.598770 | 75.664505 | 88.691872 | 191.744949 | 181.684586 | 169.788940 | 202.783417 | 1.006259 | 1712.667455 | 0.130880228 | 0.068774478 | 0.945169134 | 0.998865404 | 10.737637 | 10.746409 | 10.760580 | 10.716745 | 10.684015 | 10.800118 | 2021 | 1791 | 1383 | 1730 |
| 6167 | NM_000997,N | RPL37 | ribosomal protein L37 | protein_coding | NP_000988.1 | 10347 | 604181 | . | . | 80.247648 | 78.155369 | 68.008257 | 89.615566 | 186.739574 | 180.659629 | 152.608558 | 204.895332 | -1.052750 | 10905.501155 | 0.109630684 | -0.67648051 | 0.498735616 | 0.998865404 | 13.449392 | 13.375191 | 13.459454 | 13.407538 | 13.338604 | 13.438796 | 13270 | 11705 | 8695 | 10548 |
| 6168 | NM_000998 | RPL37A | ribosomal protein L37a | protein_coding | NP_000989.1 | 10348 | 613314 | . | . | 144.990326 | 145.487457 | 129.952225 | 169.995880 | 337.398468 | 336.300751 | 291.609070 | 388.675354 | 1.017630 | 13752.206692 | 0.128174363 | 0.196713074 | 0.844052078 | 0.998865404 | 13.734749 | 13.759893 | 13.740045 | 13.735631 | 13.636421 | 13.857298 | 15738 | 14672 | 10513 | 14676 |
| 6169 | NM_000999,N | RPL38 | ribosomal protein L38 | protein_coding | NP_000990.1;NP_001030335.1 | 10349 | 604182 | . | . | 147.927759 | 146.561123 | 133.500557 | 160.769869 | 344.233985 | 338.782593 | 299.571377 | 367.581176 | -1.014639 | 5217.316969 | 0.119387956 | -0.17562129 | 0.860591475 | 0.998865404 | 12.359606 | 12.338502 | 12.361935 | 12.346544 | 12.265960 | 12.412862 | 6107 | 5619 | 4109 | 5281 |
| 6170 | NM_001000 | RPL39 | ribosomal protein L39 | protein_coding | NP_000991.1 | 10350 | 300899 | . | . | 365.822205 | 349.471008 | 326.331116 | 392.433197 | 851.283264 | 807.817810 | 732.277527 | 897.251892 | -1.009463 | 4355.550421 | 0.121587814 | -0.11175149 | 0.91102045 | 0.998865404 | 12.095483 | 12.081763 | 12.114015 | 12.068686 | 12.008823 | 12.153510 | 5176 | 4594 | 3442 | 4417 |
| 6171 | NM_00103526 | RPL41 | ribosomal protein L41 | protein_coding | NP_001030344.1;NP_066927.1 | 10354 | 613315 | . | . | 804.142570 | 770.860321 | 684.578872 | 822.421676 | 1871.272767 | 1781.877991 | 1536.175232 | 1880.369323 | -1.068400 | 13789.020610 | 0.11458028 | -0.83305467 | 0.404813898 | 0.998865404 | 13.798194 | 13.702690 | 13.793384 | 13.762503 | 13.646725 | 13.790929 | 16697 | 15096 | 10614 | 13556 |
| 6172 | gene-RPL41P5 | RPL41P5 | ribosomal protein L41 pseudogene 5 | pseudogene | . | 10355 | . | . | . | 50.521320 | 46.515724 | 41.373444 | 53.122559 | 117.565186 | 107.523178 | 92.840805 | 121.458427 | -1.041819 | 632.876150 | 0.171823633 | -0.34398304 | 0.730859038 | 0.998865404 | 9.335316 | 9.275626 | 9.348749 | 9.286065 | 9.208409 | 9.365980 | 778 | 665 | 475 | 650 |
| 6173 | NM_021029 | RPL36A | ribosomal protein L36a | protein_coding | NP_066357.3 | 10359 | 300902 | . | . | 128.566040 | 132.472000 | 121.321465 | 150.850082 | 299.178436 | 306.214935 | 272.241852 | 344.900787 | 1.027996 | 3159.350502 | 0.128379964 | 0.310285236 | 0.756344058 | 0.998865404 | 11.605491 | 11.645063 | 11.603959 | 11.621355 | 11.546593 | 11.716896 | 3550 | 3398 | 2497 | 3313 |
| 6174 | gene-RPL7P1 | RPL7P1 | ribosomal protein L7 pseudogene 1 | pseudogene | . | 10367 | . | . | . | 8.390825 | 7.495327 | 6.625065 | 8.157690 | 19.525795 | 17.325787 | 14.866452 | 18.651590 | -1.088698 | 198.686291 | 0.245405592 | -0.49959421 | 0.617360836 | 0.998865404 | 7.694644 | 7.571423 | 7.680824 | 7.624138 | 7.570888 | 7.656530 | 253 | 210 | 149 | 196 |
| 6175 | NM_001002,N | RPLP0 | ribosomal protein lateral stalk subunit P0 | protein_coding | NP_000993.1;NP_444505.1 | 10371 | 180510 | . | . | 819.573550 | 838.055847 | 703.402909 | 896.431900 | 1907.181358 | 1937.203522 | 1578.415832 | 2049.585022 | -1.035876 | 28775.286264 | 0.12483661 | -0.40734483 | 0.683752099 | 0.998865404 | 14.837763 | 14.786873 | 14.807592 | 14.843624 | 14.685898 | 14.894261 | 33017 | 32255 | 21615 | 29699 |
| 6176 | NM_001003,N | RPLP1 | ribosomal protein lateral stalk subunit P1 | protein_coding | NP_000994.1;NP_998890.1 | 10372 | 180520 | . | . | 330.949873 | 334.837165 | 300.418227 | 361.624198 | 770.133990 | 773.991144 | 674.129758 | 826.810741 | -1.020165 | 12128.367665 | 0.115206945 | -0.25000223 | 0.802585621 | 0.998865404 | 13.580461 | 13.551590 | 13.572316 | 13.575036 | 13.479092 | 13.628637 | 14083 | 13239 | 9528 | 12241 |
| 6181 | NM_001004 | RPLP2 | ribosomal protein lateral stalk subunit P2 | protein_coding | NP_000995.1 | 10377 | 180530 | . | . | 359.442261 | 369.545868 | 330.102417 | 431.559174 | 836.436951 | 854.221863 | 740.740234 | 986.708740 | 1.029775 | 5361.202373 | 0.131884099 | 0.320956867 | 0.748243078 | 0.998865404 | 12.367112 | 12.409262 | 12.365850 | 12.381889 | 12.284621 | 12.501018 | 6025 | 5755 | 4124 | 5753 |
| 6182 | NM_002949 | MRPL12 | mitochondrial ribosomal protein L12 | protein_coding | NP_002940.2 | 10378 | 602375 | . | . | 72.676979 | 74.251579 | 65.578636 | 83.420006 | 169.122299 | 171.635864 | 147.156540 | 190.729919 | -1.000295 | 2325.317504 | 0.135299848 | -0.00314179 | 0.997493219 | 0.999862634 | 11.183599 | 11.182825 | 11.176604 | 11.187014 | 11.085158 | 11.269542 | 2661 | 2526 | 1790 | 2429 |
| 6183 | NM_021107,N | MRPS12 | mitochondrial ribosomal protein S12 | protein_coding | NP_066930.1;NP_203526.1;NP_203527.1 | 10380 | 603021 | . | . | 36.210694 | 35.087200 | 30.208131 | 39.055038 | 84.263774 | 81.105638 | 67.786165 | 89.294705 | -1.036949 | 1091.795351 | 0.153592646 | -0.34080219 | 0.733252503 | 0.998865404 | 10.118704 | 10.065784 | 10.113912 | 10.094253 | 9.979385 | 10.166181 | 1297 | 1184 | 814 | 1132 |
| 6184 | NM_002950 | RPN1 | ribophorin I | protein_coding | NP_002941.1 | 10381 | 180470 | . | . | 128.742432 | 124.874466 | 105.555695 | 117.161331 | 299.588928 | 288.652924 | 236.863907 | 267.875458 | -1.155094 | 8476.937429 | 0.107101516 | -1.94217505 | 0.052115912 | 0.675217033 | 13.149609 | 12.941551 | 13.123937 | 13.090306 | 12.944571 | 13.023424 | 10668 | 9614 | 6519 | 7722 |
| 6185 | NM_00113577 | RPN2 | ribophorin II | protein_coding | NP_001129243.1;NP_001311228.1;NP_001311230.1;NP_001311231.1;NI | 10382 | 180490 | . | . | 329.685550 | 341.843983 | 292.796893 | 306.508309 | 767.191831 | 790.187675 | 657.027761 | 700.794877 | -1.120834 | 22373.644479 | 0.098487241 | -1.67100329 | 0.094721028 | 0.81257573 | 14.529458 | 14.364876 | 14.484054 | 14.509039 | 14.392735 | 14.404164 | 26839 | 25885 | 18224 | 19913 |
| 6187 | NM_002952 | RPS2 | ribosomal protein S2 | protein_coding | NP_002943.2 | 10404 | 603624 | . | . | 843.055725 | 853.360046 | 755.993103 | 956.884277 | 1961.825562 | 1972.579956 | 1696.426514 | 2187.802246 | -1.004849 | 25083.773437 | 0.121564832 | -0.05741054 | 0.954218164 | 0.998865404 | 14.617971 | 14.610954 | 14.613483 | 14.616575 | 14.509286 | 14.703301 | 28902 | 27181 | 19317 | 26092 |
| 6188 | NM_001005,N | RPS3 | ribosomal protein S3 | protein_coding | NP_000996.2;NP_001243731.1;NP_001247435.1;NP_001247436.1 | 10420 | 600454 | . | . | 239.601589 | 239.456178 | 221.860768 | 268.749969 | 557.562733 | 553.513692 | 497.849099 | 614.464827 | 1.010576 | 15489.825616 | 0.115545359 | 0.131357738 | 0.895492319 | 0.998865404 | 13.911451 | 13.926576 | 13.916996 | 13.910023 | 13.841681 | 13.997466 | 17815 | 16559 | 12313 | 15921 |
| 6189 | NM_001006,N | RPS3A | ribosomal protein S3A | protein_coding | NP_000997.1;NP_001254628.1 | 10421 | 180478 | . | . | 451.599872 | 465.226726 | 429.050177 | 525.342430 | 1050.891524 | 1075.392376 | 962.776128 | 1201.133059 | 1.025122 | 12821.728576 | 0.116664905 | 0.306828801 | 0.758973685 | 0.998865404 | 13.628330 | 13.664055 | 13.625000 | 13.644023 | 13.573598 | 13.731878 | 14419 | 13816 | 10248 | 13290 |
| 6191 | NM_001007 | RPS4X | ribosomal protein S4 X-linked | protein_coding | NP_000998.1 | 10424 | 312760 | . | . | 227.211868 | 229.138870 | 216.289246 | 265.397064 | 528.731323 | 529.664795 | 485.346802 | 606.798828 | 1.040472 | 10764.025064 | 0.118476642 | 0.483120855 | 0.629009927 | 0.998865404 | 13.365067 | 13.422227 | 13.375682 | 13.375564 | 13.322995 | 13.489253 | 12150 | 11384 | 8621 | 11288 |
| 6192 | NM_001008 | RPS4Y1 | ribosomal protein S4 Y-linked 1 | protein_coding | NP_000999.1 | 10425 | 470000 | . | . | 64.933411 | 67.230087 | 64.600540 | 67.913872 | 151.102722 | 155.405365 | 144.961731 | 155.277008 | -1.011463 | 2450.978576 | 0.114982584 | -0.1430071 | 0.886284576 | 0.998865404 | 11.267396 | 11.250841 | 11.252956 | 11.274329 | 11.238346 | 11.270313 | 2801 | 2695 | 2077 | 2330 |
| 6193 | NM_001009 | RPS5 | ribosomal protein S5 | protein_coding | NP_001000.2 | 10426 | 603630 | . | . | 385.506653 | 391.191345 | 341.570343 | 406.744537 | 897.089844 | 904.256348 | 766.473938 | 929.973022 | -1.052865 | 8800.529017 | 0.114783677 | -0.64748777 | 0.517316304 | 0.998865404 | 13.140102 | 13.065688 | 13.121493 | 13.126746 | 13.008438 | 13.147426 | 10363 | 9771 | 6844 | 8697 |
| 6194 | NM_001010 | RPS6 | ribosomal protein S6 | protein_coding | NP_001001.2 | 10429 | 180460 | . | . | 247.190842 | 258.625610 | 217.510345 | 290.354156 | 575.223267 | 597.824707 | 488.086914 | 663.860291 | -1.010508 | 10803.975034 | 0.132655844 | -0.11368415 | 0.909488175 | 0.998865404 | 13.406845 | 13.391664 | 13.385591 | 13.416927 | 13.267033 | 13.504205 | 12277 | 11934 | 8052 | 11470 |
| 6195 | NM_00100666 | RPS6KA1 | ribosomal protein S6 kinase A1 | protein_coding | NP_001006666.1;NP_001317370.1;NP_002944.2;XP_024304639.1 | 10430 | 601684 | . | . | 12.141242 | 13.078607 | 12.017751 | 13.313595 | 28.253171 | 30.231787 | 26.967483 | 30.439953 | -1.009456 | 1257.321693 | 0.133684533 | -0.10156429 | 0.919102523 | 0.998865404 | 10.303103 | 10.289139 | 10.273563 | 10.324585 | 10.256914 | 10.326207 | 1406 | 1409 | 1038 | 1227 |
| 6196 | NM_00100693 | RPS6KA2 | ribosomal protein S6 kinase A2 | protein_coding | NP_001006933.3;NP_001305865.2;NP_001305866.1;NP_001305867.1;NI | 10431 | 601685 | . | . | 3.697492 | 3.257214 | 3.351921 | 4.149938 | 8.604217 | 7.529195 | 7.521613 | 9.488343 | 1.063907 | 657.743004 | 0.169440331 | 0.527454348 | 0.597878111 | 0.998865404 | 9.316171 | 9.405216 | 9.382966 | 9.290994 | 9.307408 | 9.447719 | 784 | 641 | 529 | 701 |
| 6197 | NM_004586,XI | RPS6KA3 | ribosomal protein S6 kinase A3 | protein_coding | NP_004577.1;XP_005274630.1;XP_005274634.1;XP_006724570.1;XP_01 | 10432 | 300075 | . | . | 5.777908 | 5.047025 | 4.733800 | 4.132179 | 13.445432 | 11.666425 | 10.622507 | 9.447737 | -1.244538 | 1181.435362 | 0.143072518 | -2.20594807 | **0.027387636** | 0.538018865 | 10.355250 | 10.040239 | 10.331024 | 10.229447 | 10.165156 | 10.067673 | 1607 | 1321 | 977 | 921 |
| 6198 | NM_00127204 | RPS6KB1 | ribosomal protein S6 kinase B1 | protein_coding | NP_001258971.1;NP_001258972.1;NP_001258973.1;NP_001258989.1;NI | 10436 | 608938 | . | . | 3.704003 | 3.391679 | 4.267215 | 3.320759 | 8.619361 | 7.840020 | 9.575506 | 7.592524 | 1.050152 | 626.774063 | 0.174697778 | 0.404112945 | 0.686129666 | 0.998865404 | 9.255555 | 9.327160 | 9.296807 | 9.247017 | 9.390054 | 9.212795 | 730 | 635 | 632 | 523 |
| 6199 | NM_003952,XI | RPS6KB2 | ribosomal protein S6 kinase B2 | protein_coding | NP_003943.2;XP_005274221.1;XP_005274222.1;XP_006718718.1;XP_00 | 10437 | 608939 | . | . | 13.160203 | 14.530556 | 13.864615 | 16.399997 | 30.624337 | 33.588032 | 31.111791 | 37.496644 | 1.095900 | 797.206197 | 0.155390381 | 0.850219261 | 0.395203195 | 0.998865404 | 9.571618 | 9.703010 | 9.574829 | 9.632694 | 9.605991 | 9.727521 | 840 | 855 | 663 | 848 |
| 6201 | NM_001011 | RPS7 | ribosomal protein S7 | protein_coding | NP_001002.1 | 10440 | 603658 | . | . | 659.688354 | 646.821533 | 577.926575 | 754.844055 | 1535.121948 | 1495.156738 | 1296.850586 | 1725.861084 | 1.005315 | 15041.196050 | 0.127970363 | 0.059758073 | 0.952348319 | 0.998865404 | 13.872833 | 13.880420 | 13.884398 | 13.860178 | 13.760998 | 13.981072 | 17518 | 15959 | 11439 | 15944 |
| 6202 | NM_001012 | RPS8 | ribosomal protein S8 | protein_coding | NP_001003.1 | 10441 | 600357 | . | . | 575.423279 | 591.316467 | 531.412109 | 672.737549 | 1339.033691 | 1366.854614 | 1192.473389 | 1538.134521 | 1.017253 | 14359.710330 | 0.122742 | 0.201060393 | 0.840651352 | 0.998865404 | 13.797381 | 13.821992 | 13.792803 | 13.808792 | 13.714243 | 13.907656 | 16241 | 15506 | 11179 | 15102 |
| 6203 | NM_001013,N | RPS9 | ribosomal protein S9 | protein_coding | NP_001004.2;NP_001308630.1;NP_001308631.1;NP_001308633.1;NP_0 | 10442 | 603631 | . | . | 299.342300 | 296.779639 | 267.629228 | 333.559281 | 696.581825 | 686.019423 | 600.552262 | 762.643635 | -1.010210 | 6794.788339 | 0.122258378 | -0.11987037 | 0.904585839 | 0.998865404 | 12.737582 | 12.722806 | 12.739030 | 12.727436 | 12.634121 | 12.807805 | 7918 | 7319 | 5275 | 6988 |
| 6204 | NM_001014,N | RPS10 | ribosomal protein S10 | protein_coding | NP_001005.1;NP_001190174.1;NP_001191020.1 | 10383 | 603632 | . | . | 470.433134 | 453.378376 | 385.946144 | 528.823444 | 1094.717193 | 1048.004364 | 866.051987 | 1209.092026 | -1.024804 | 8853.343927 | 0.133464355 | -0.26485221 | 0.791123313 | 0.998865404 | 13.129620 | 13.094176 | 13.141784 | 13.097742 | 12.975080 | 13.209827 | 10586 | 9421 | 6557 | 9327 |
| 6205 | NM_001015 | RPS11 | ribosomal protein S11 | protein_coding | NP_001006.1 | 10384 | 180471 | . | . | 708.501953 | 716.160522 | 652.061462 | 825.978943 | 1648.713135 | 1655.437134 | 1463.207031 | 1888.502441 | 1.022563 | 12948.702850 | 0.122981016 | 0.261745084 | 0.79351798 | 0.998865404 | 13.644372 | 13.676490 | 13.648413 | 13.650306 | 13.567020 | 13.760784 | 14728 | 13832 | 10103 | 13657 |
| 6206 | NM_001016 | RPS12 | ribosomal protein S12 | protein_coding | NP_001007.2 | 10385 | 603660 | . | . | 633.149231 | 635.430969 | 596.528748 | 726.615784 | 1473.364258 | 1468.827148 | 1338.593262 | 1661.320679 | 1.028083 | 10149.153442 | 0.117573456 | 0.339845461 | 0.733972911 | 0.998865404 | 13.288996 | 13.328872 | 13.298230 | 13.293967 | 13.236745 | 13.396465 | 11554 | 10773 | 8113 | 10546 |
| 6207 | NM_001017 | RPS13 | ribosomal protein S13 | protein_coding | NP_001008.1 | 10386 | 180476 | . | . | 271.197754 | 254.853149 | 242.143951 | 269.817413 | 631.088318 | 589.104492 | 543.364014 | 616.905396 | -1.042277 | 4260.835501 | 0.113883843 | -0.52455362 | 0.599893536 | 0.998865404 | 12.086509 | 12.026713 | 12.102662 | 12.043522 | 11.997807 | 12.078727 | 5185 | 4527 | 3451 | 4103 |
| 6208 | NM_00102507 | RPS14 | ribosomal protein S14 | protein_coding | NP_001020241.1;NP_001020242.1;NP_005608.1 | 10387 | 130620 | . | . | 144.089824 | 145.581566 | 130.015209 | 160.488236 | 335.302933 | 336.518317 | 291.750359 | 366.937286 | -1.025005 | 10123.479345 | 0.114530825 | -0.31110606 | 0.755719995 | 0.998865404 | 13.323162 | 13.287453 | 13.317132 | 13.312973 | 13.220770 | 13.362300 | 11830 | 11032 | 7972 | 10152 |
| 6209 | NM_001018,N | RPS15 | ribosomal protein S15 | protein_coding | NP_001009.1;NP_001295155.1 | 10388 | 180535 | . | . | 371.216896 | 377.699995 | 351.443405 | 429.659853 | 863.837017 | 873.070484 | 788.628838 | 982.366253 | 1.027738 | 6040.441100 | 0.120599339 | 0.32730161 | 0.743439775 | 0.998865404 | 12.540637 | 12.579971 | 12.544312 | 12.551105 | 12.486988 | 12.648159 | 6833 | 6455 | 4817 | 6287 |
| 6210 | NM_001019,N | RPS15A | ribosomal protein S15a | protein_coding | NP_001010.2;NP_001025180.1 | 10389 | 603674 | . | . | 126.295482 | 121.847721 | 108.352218 | 135.389946 | 293.894776 | 281.656487 | 243.139237 | 309.553016 | -1.032455 | 8339.827622 | 0.123117482 | -0.37426796 | 0.708204969 | 0.998865404 | 13.048703 | 13.002531 | 13.055866 | 13.019483 | 12.916279 | 13.097126 | 9965 | 8948 | 6376 | 8513 |
| 6217 | NM_001020,N | RPS16 | ribosomal protein S16 | protein_coding | NP_001011.1;NP_001308040.1;NP_001350789.1 | 10396 | 603675 | . | . | 399.734302 | 395.513214 | 366.206266 | 432.201306 | 930.198144 | 914.246404 | 821.756244 | 988.176884 | -1.008770 | 7847.835311 | 0.113919402 | -0.11057832 | 0.911950737 | 0.998865404 | 12.944407 | 12.931723 | 12.948786 | 12.933326 | 12.866697 | 12.996521 | 9159 | 8428 | 6274 | 7886 |
| 6218 | NM_001021,N | RPS17 | ribosomal protein S17 | protein_coding | NP_001012.1 | 10397 | 180472 | . | . | 710.285574 | 686.826731 | 626.455691 | 757.795826 | 1652.863663 | 1587.630720 | 1405.748486 | 1732.610083 | -1.023531 | 10733.288377 | 0.117404157 | -0.28580951 | 0.775024027 | 0.998865404 | 13.406518 | 13.372895 | 13.413829 | 13.383196 | 13.297557 | 13.454002 | 12727 | 11507 | 8379 | 10857 |
| 6222 | NM_022551 | RPS18 | ribosomal protein S18 | protein_coding | NP_072045.1 | 10401 | 180473 | . | . | 821.944519 | 800.709778 | 684.945068 | 873.770264 | 1912.698730 | 1850.876343 | 1536.996826 | 1997.771973 | -1.056278 | 13601.429615 | 0.124425536 | -0.63483399 | 0.525536692 | 0.998865404 | 13.770456 | 13.691403 | 13.767685 | 13.737801 | 13.601847 | 13.800429 | 16370 | 14817 | 10168 | 13842 |
| 6223 | NM_001022,N | RPS19 | ribosomal protein S19 | protein_coding | NP_001013.1;NP_001308412.1;NP_001308413.1;NP_001308414.1;XP_01 | 10402 | 603474 | . | . | 146.448364 | 143.550220 | 137.477915 | 153.382247 | 340.791349 | 331.822782 | 308.496443 | 350.690281 | 1.003329 | 8770.087830 | 0.11025183 | 0.04349061 | 0.96531045 | 0.998865404 | 13.096007 | 13.100741 | 13.109952 | 13.082802 | 13.044111 | 13.151618 | 10243 | 9297 | 7156 | 8756 |
| 6224 | NM_001023,N | RPS20 | ribosomal protein S20 | protein_coding | NP_001014.1;NP_001139699.1 | 10405 | 603682 | . | . | 650.310137 | 623.495847 | 592.048881 | 700.985159 | 1513.298332 | 1441.238441 | 1328.540471 | 1602.719221 | -1.001799 | 10413.653032 | 0.114269322 | -0.02268881 | 0.981898505 | 0.998865404 | 13.347513 | 13.344863 | 13.365929 | 13.325987 | 13.274847 | 13.409927 | 12283 | 10984 | 8331 | 10526 |

| | | | | | | | | | |
|---|---|---|---|---|---|---|---|---|---|
| 6225 | gene-RPS20P3 | RPS20P3 | ribosomal protein S20 pseudogene 3 | pseudogene | . | 10406 | 180463 | . | . |
| 6227 | NM_001024,XI | RPS21 | ribosomal protein S21 | protein_coding | NP_001015.1;XP_024307727.1;XP_024307728.1 | 10409 | 180477 | . | . |
| 6228 | NM_001025 | RPS23 | ribosomal protein S23 | protein_coding | NP_001016.1 | 10410 | 603683 | . | . |
| 6229 | NM_001026,N | RPS24 | ribosomal protein S24 | protein_coding | NP_001017.1;NP_001135754.1;NP_001135755.1;NP_001135756.1;NP_0( | 10411 | 602412 | . | . |
| 6230 | NM_001028 | RPS25 | ribosomal protein S25 | protein_coding | NP_001019.1 | 10413 | 180465 | . | . |
| 6231 | NM_001029 | RPS26 | ribosomal protein S26 | protein_coding | NP_001020.2 | 10414 | 603701 | . | . |
| 6232 | NM_001030,N | RPS27 | ribosomal protein S27 | protein_coding | NP_001021.1;NP_001336875.1;NP_001336876.1 | 10416 | 603702 | . | . |
| 6233 | NM_00113559 | RPS27A | ribosomal protein S27a | protein_coding | NP_001129064.1;NP_001170884.1;NP_002945.1 | 10417 | 191343 | . | . |
| 6234 | NM_001031 | RPS28 | ribosomal protein S28 | protein_coding | NP_001022.1 | 10418 | 603685 | . | . |
| 6235 | NM_00103000 | RPS29 | ribosomal protein S29 | protein_coding | NP_001023.1;NP_001025172.1;NP_001338304.1 | 10419 | 603633 | . | . |
| 6237 | NM_006270 | RRAS | RAS related | protein_coding | NP_006261.1 | 10447 | 165090 | . | . |
| 6238 | NM_00104257 | RRBP1 | ribosome binding protein 1 | protein_coding | NP_001036041.2;NP_001352542.1;NP_004578.3 | 10448 | 601418 | . | . |
| 6239 | NM_00100369 | RREB1 | ras responsive element binding protein 1 | protein_coding | NP_001003698.1;NP_001003699.1;NP_001003700.1;NP_001161816.1 | 10449 | 602209 | . | . |
| 6240 | NM_001033,N | RRM1 | ribonucleotide reductase catalytic subunit M1 | protein_coding | NP_001024.1;NP_001304993.1;NP_001304994.1;NP_001317122.1 | 10451 | 180410 | . | . |
| 6241 | NM_001034,N | RRM2 | ribonucleotide reductase regulatory subunit M2 | protein_coding | NP_001025.1;NP_001159403.1 | 10452 | 180390 | . | . |
| 6242 | NM_00101505 | RTKN | rhotekin | protein_coding | NP_001015055.1;NP_001015056.1;NP_149035.1;XP_005264535.1;XP_00 | 10466 | 602288 | . | . |
| 6248 | NM_006511 | RSC1A1 | regulator of solute carriers 1 | protein_coding | NP_006502.1 | 10458 | 601966 | . | . |
| 6249 | NM_00124799 | CLIP1 | CAP-Gly domain containing linker protein 1 | protein_coding | NP_001234926.1;NP_002947.1;NP_937883.1;XP_005253650.1;XP_00671 | 10461 | 179838 | . | . |
| 6251 | NM_012425,N | RSU1 | Ras suppressor protein 1 | protein_coding | NP_036557.1;NP_689937.2;XP_005252609.1 | 10464 | 179555 | . | . |
| 6252 | NM_00136370 | RTN1 | reticulon 1 | protein_coding | NP_001350631.1;NP_066959.1;NP_996734.1;XP_011535365.1 | 10467 | 600865 | . | . |
| 6253 | NM_005619,N | RTN2 | reticulon 2 | protein_coding | NP_005610.1;NP_996783.1;NP_996784.1 | 10468 | 603183 | . | . |
| 6256 | NM_00129192 | RXRA | retinoid X receptor alpha | protein_coding | NP_001278849.1;NP_001278850.1;NP_002948.1 | 10477 | 180245 | . | . |
| 6257 | NM_00127040 | RXRB | retinoid X receptor beta | protein_coding | NP_001257330.1;NP_001278918.1;NP_068811.1;XP_005249335.1;XP_01 | 10478 | 180246 | . | . |
| 6258 | NM_00125657 | RXRG | retinoid X receptor gamma | protein_coding | NP_001243499.1;NP_001243500.1;NP_008848.1 | 10479 | 180247 | . | . |
| 6259 | NM_00100586 | RYK | receptor like tyrosine kinase | protein_coding | NP_001005861.1;NP_002949.2 | 10481 | 600524 | . | . |
| 6262 | NM_001035,XI | RYR2 | ryanodine receptor 2 | protein_coding | NP_001026.2;XP_006711865.1;XP_006711866.1;XP_006711867.1;XP_00 | 10484 | 180902 | . | . |
| 6271 | NM_006271 | S100A1 | S100 calcium binding protein A1 | protein_coding | NP_006262.1 | 10486 | 176940 | . | . |
| 6272 | NM_00120522 | SORT1 | sortilin 1 | protein_coding | NP_001192157.1;NP_002950.3;XP_005271157.1;XP_005271158.1;XP_00 | 11186 | 602458 | . | . |
| 6273 | NM_00136640 | S100A2 | S100 calcium binding protein A2 | protein_coding | NP_001353335.1;NP_001353336.1;NP_005969.2 | 10492 | 176993 | . | . |
| 6274 | NM_002960 | S100A3 | S100 calcium binding protein A3 | protein_coding | NP_002951.1 | 10493 | 176992 | . | . |
| 6275 | NM_002961,N | S100A4 | S100 calcium binding protein A4 | protein_coding | NP_002952.1;NP_062427.1 | 10494 | 114210 | . | . |
| 6276 | NM_002962,XI | S100A5 | S100 calcium binding protein A5 | protein_coding | NP_002953.2;XP_011508162.1;XP_016857518.1;XP_016857520.1;XP_01 | 10495 | 176991 | . | . |
| 6277 | NM_014624,XI | S100A6 | S100 calcium binding protein A6 | protein_coding | NP_055439.1;XP_016857522.1 | 10496 | 114110 | . | . |
| 6281 | NM_002966 | S100A10 | S100 calcium binding protein A10 | protein_coding | NP_002957.1 | 10487 | 114085 | . | . |
| 6282 | NM_005620 | S100A11 | S100 calcium binding protein A11 | protein_coding | NP_005611.1 | 10488 | 603114 | . | . |
| 6284 | NM_00102421 | S100A13 | S100 calcium binding protein A13 | protein_coding | NP_001019381.1;NP_001019382.1;NP_001019383.1;NP_001019384.1;NI | 10490 | 601989 | . | . |
| 6285 | NM_006272,XI | S100B | S100 calcium binding protein B | protein_coding | NP_006263.1;XP_016883913.1 | 10500 | 176990 | . | . |
| 6288 | NM_000331,N | SAA1 | serum amyloid A1 | protein_coding | NP_000322.2;NP_001171477.1;NP_954630.2 | 10513 | 104750 | . | . |
| 6289 | NM_00112738 | SAA2 | serum amyloid A2 | protein_coding | NP_001120852.1;NP_110381.2 | 10514 | 104751 | . | . |
| 6291 | NM_006512 | SAA4 | serum amyloid A4, constitutive | protein_coding | NP_006503.2 | 10516 | 104752 | . | . |
| 6293 | NM_00128917 | VPS52 | VPS52 subunit of GARP complex | protein_coding | NP_001276103.1;NP_001276104.1;NP_001276105.1;NP_072047.4;XP_01 | 10518 | 603443 | . | . |
| 6294 | NM_00120133 | SAFB | scaffold attachment factor B | protein_coding | NP_001188267.1;NP_001188268.1;NP_001188269.1;NP_001307500.1;NI | 10520 | 602895 | . | . |
| 6296 | NM_005622,N | ACSM3 | acyl-CoA synthetase medium chain family member 3 | protein_coding | NP_005613.2;NP_973729.1;XP_011544213.1;XP_016879012.1;XP_02430 | 10522 | 145505 | . | . |
| 6297 | NM_00129144 | SALL2 | spalt like transcription factor 2 | protein_coding | NP_001278375.1;NP_001278376.1;NP_001351493.1;NP_005398.2;XP_01 | 10526 | 602219 | . | . |
| 6299 | NM_00112789 | SALL1 | spalt like transcription factor 1 | protein_coding | NP_001121364.1;NP_002959.2 | 10524 | 602218 | . | . |
| 6300 | NM_00130325 | MAPK12 | mitogen-activated protein kinase 12 | protein_coding | NP_001290181.1;NP_002960.2 | 6874 | 602399 | . | . |
| 6301 | NM_00133066 | SARS1 | seryl-tRNA synthetase 1 | protein_coding | NP_001317598.1;NP_006504.2 | 10537 | 607529 | . | . |
| 6302 | NM_00133016 | TSPAN31 | tetraspanin 31 | protein_coding | NP_001317097.1;NP_001317098.1;NP_005972.1;XP_024304891.1 | 10539 | 181035 | . | . |
| 6303 | NM_002970,N | SAT1 | spermidine/spermine N1-acetyltransferase 1 | protein_coding | NP_002961.1;XP_024308189.1 | 10540 | 313020 | . | . |
| 6304 | NM_00113101 | SATB1 | SATB homeobox 1 | protein_coding | NP_001124482.1;NP_001182399.1;NP_001309800.1;NP_001309801.1;NI | 10541 | 602075 | . | . |
| 6305 | NM_00136581 | SBF1 | SET binding factor 1 | protein_coding | NP_001352748.1;NP_002963.2;XP_005261992.1;XP_011529011.1;XP_01 | 10542 | 603560 | . | . |
| 6307 | NM_00101736 | MSMO1 | methylsterol monooxygenase 1 | protein_coding | NP_001017369.1;NP_006736.1;XP_005263233.1 | 10545 | 607545 | . | . |
| 6309 | NM_00102495 | SC5D | sterol-C5-desaturase | protein_coding | NP_001020127.1;NP_008849.2 | 10547 | 602286 | . | . |
| 6310 | NM_000332,N | ATXN1 | ataxin 1 | protein_coding | NP_000323.2;NP_001121636.1;NP_001344786.1 | 10548 | 601556 | . | . |
| 6311 | NM_00131012 | ATXN2 | ataxin 2 | protein_coding | NP_001297050.1;NP_001297052.1;NP_001359503.1;NP_002964.4 | 10555 | 601517 | . | . |
| 6314 | NM_000333,N | ATXN7 | ataxin 7 | protein_coding | NP_000324.1;NP_001121621.2;NP_001170858.1;NP_001364334.1;NP_0( | 10560 | 607640 | . | . |
| 6319 | NM_005063 | SCD | stearoyl-CoA desaturase | protein_coding | NP_005054.3 | 10571 | 604031 | . | . |
| 6320 | NM_002975 | CLEC11A | C-type lectin domain containing 11A | protein_coding | NP_002966.1 | 10576 | 604713 | . | . |
| 6322 | NM_00103753 | SCML1 | Scm polycomb group protein like 1 | protein_coding | NP_001032624.1;NP_001032625.1;NP_001032629.1;NP_006737.2;XP_0( | 10580 | 300227 | . | . |
| 6324 | NM_001037,N | SCN1B | sodium voltage-gated channel beta subunit 1 | protein_coding | NP_001028.1;NP_001308534.1;NP_950238.1 | 10586 | 600235 | . | . |
| 6326 | NM_00104014 | SCN2A | sodium voltage-gated channel alpha subunit 2 | protein_coding | NP_001035232.1;NP_001035233.1;NP_001358175.1;NP_001358176.1;NI | 10588 | 182390 | . | . |
| 6330 | NM_00114234 | SCN4B | sodium voltage-gated channel beta subunit 4 | protein_coding | NP_001135820.1;NP_001135821.1;NP_777594.1 | 10592 | 608256 | . | . |
| 6331 | NM_000335,N | SCN5A | sodium voltage-gated channel alpha subunit 5 | protein_coding | NP_000326.2;NP_001092874.1;NP_001092875.1;NP_001153632.1;NP_0( | 10593 | 600163 | . | . |
| 6334 | NM_00117798 | SCN8A | sodium voltage-gated channel alpha subunit 8 | protein_coding | NP_001171455.1;NP_001317189.1;NP_001356717.1;NP_055006.1 | 10596 | 600702 | . | . |
| 6335 | NM_00136553 | SCN9A | sodium voltage-gated channel alpha subunit 9 | protein_coding | NP_001352465.1;NP_002968.1;XP_011509918.1;XP_011509919.1;XP_01 | 10597 | 603415 | . | . |
| 6339 | NM_00113041 | SCNN1D | sodium channel epithelial 1 subunit delta | protein_coding | NP_001123885.2 | 10601 | 601328 | . | . |
| 6341 | NM_004589 | SCO1 | synthesis of cytochrome C oxidase 1 | protein_coding | NP_004580.1 | 10603 | 603644 | . | . |
| 6342 | NM_00100709 | SCP2 | sterol carrier protein 2 | protein_coding | NP_001007099.1;NP_001007100.1;NP_001007101.1;NP_001007251.1;NI | 10606 | 184755 | . | . |
| 6347 | NM_002982 | CCL2 | C-C motif chemokine ligand 2 | protein_coding | NP_002973.1 | 10618 | 158105 | . | . |
| 6348 | NM_002983,N | CCL3 | C-C motif chemokine ligand 3 | protein_coding | NP_002974.1 | 10627 | 182283 | . | . |
| 6352 | NM_00127873 | CCL5 | C-C motif chemokine ligand 5 | protein_coding | NP_001265665.1;NP_002976.2 | 10632 | 187011 | . | . |
| 6361 | NM_002987,XI | CCL17 | C-C motif chemokine ligand 17 | protein_coding | NP_002978.1;XP_011521558.1;XP_016879019.1 | 10615 | 601520 | . | . |
| 6362 | NM_002988 | CCL18 | C-C motif chemokine ligand 18 | protein_coding | NP_002979.1 | 10616 | 603757 | . | . |
| 6364 | NM_00113004 | CCL20 | C-C motif chemokine ligand 20 | protein_coding | NP_001123518.1;NP_004582.1 | 10619 | 601960 | . | . |
| 6376 | NM_00130439 | CX3CL1 | C-X3-C motif chemokine ligand 1 | protein_coding | NP_001291321.1;NP_002987.1 | 10647 | 601880 | . | . |
| 6382 | NM_00100694 | SDC1 | syndecan 1 | protein_coding | NP_001006947.1;NP_002988.4;XP_005262677.1;XP_005262678.3;XP_00 | 10658 | 186355 | . | . |
| 6383 | NM_002998,XI | SDC2 | syndecan 2 | protein_coding | NP_002989.2;XP_011515514.1;XP_024302996.1 | 10659 | 142460 | . | . |
| 6385 | NM_002999,XI | SDC4 | syndecan 4 | protein_coding | NP_002990.2;XP_011527279.1 | 10661 | 600017 | . | . |
| 6386 | NM_00100706 | SDCBP | syndecan binding protein | protein_coding | NP_001007068.1;NP_001007069.1;NP_001007070.1;NP_001007071.1;NI | 10662 | 602217 | . | . |
| 6388 | NM_006923,N | SDF2 | stromal cell derived factor 2 | protein_coding | NP_008854.2;XP_011523408.1 | 10675 | 602934 | . | . |
| 6389 | NM_00129433 | SDHA | succinate dehydrogenase complex flavoprotein subunit A | protein_coding | NP_001281261.1;NP_001317687.1;NP_004159.2;XP_011512374.1;XP_01 | 10680 | 600857 | . | . |
| 6390 | NM_003000 | SDHB | succinate dehydrogenase complex iron sulfur subunit B | protein_coding | NP_002991.2 | 10681 | 185470 | . | . |
| 6391 | NM_00103551 | SDHC | succinate dehydrogenase complex subunit C | protein_coding | NP_001030588.1;NP_001030589.1;NP_001030590.1;NP_001265101.1;NI | 10682 | 602413 | . | . |
| 6392 | NM_00127650 | SDHD | succinate dehydrogenase complex subunit D | protein_coding | NP_001263432.1;NP_001263433.1;NP_001263435.1;NP_002993.1 | 10683 | 602690 | . | . |
| 6396 | NM_00113602 | SEC13 | SEC13 homolog, nuclear pore and COPII coat complex component | protein_coding | NP_001129498.1;NP_001129704.1;NP_001265875.1;NP_109598.2;NP_8! | 10697 | 600152 | . | . |
| 6397 | NM_00103957 | SEC14L1 | SEC14 like lipid binding 1 | protein_coding | NP_001034662.3;NP_001137470.2;NP_001137471.2;NP_001137473.2;NI | 10698 | 601504 | . | . |
| 6399 | NM_00101165 | TRAPPC2 | trafficking protein particle complex 2 | protein_coding | NP_001011658.1;NP_001122307.2;NP_055378.1;XP_011543867.1;XP_01 | 23068 | 300202 | . | . |
| 6400 | NM_00124498 | SEL1L | SEL1L adaptor subunit of ERAD E3 ubiquitin ligase | protein_coding | NP_001231913.1;NP_005056.3;XP_005268045.1;XP_005268046.1 | 10717 | 602329 | . | . |
| 6402 | NM_000655,N | SELL | selectin L | protein_coding | NP_000646.3 | 10720 | 153240 | . | . |
| 6404 | NM_00120660 | SELPLG | selectin P ligand | protein_coding | NP_001193538.1;NP_002997.2 | 10722 | 600738 | . | . |
| 6415 | NM_003009 | SELENOW | selenoprotein W | protein_coding | NP_003000.1 | 10752 | 603235 | . | . |
| 6416 | NM_00128143 | MAP2K4 | mitogen-activated protein kinase kinase 4 | protein_coding | NP_001268364.1;NP_003001.1;XP_005256810.1;XP_005256812.1;XP_01 | 6844 | 601335 | . | . |
| 6418 | NM_00112282 | SET | SET nuclear proto-oncogene | protein_coding | NP_001116293.1;NP_001234929.1;NP_001234930.1;NP_001361255.1;NI | 10760 | 600960 | . | . |
| 6419 | NM_00124372 | SETMAR | SET domain and mariner transposase fusion gene | protein_coding | NP_001230652.1;NP_001263254.1;NP_001307605.1;NP_001307606.1;NI | 10762 | 609834 | . | . |
| 6421 | NM_005066,N | SFPQ | splicing factor proline and glutamine rich | protein_coding | NP_005057.1;XP_005271169.1;XP_005271170.1;XP_005271172.1;XP_01 | 10774 | 605199 | . | . |
| 6422 | NM_003012 | SFRP1 | secreted frizzled related protein 1 | protein_coding | NP_003003.3 | 10776 | 604156 | . | . |
| 6424 | NM_003014 | SFRP4 | secreted frizzled related protein 4 | protein_coding | NP_003005.2 | 10778 | 606570 | . | . |
| 6426 | NM_00107816 | SRSF1 | serine and arginine rich splicing factor 1 | protein_coding | NP_001071634.1;NP_008855.1 | 10780 | 600812 | . | . |
| 6427 | NM_00119542 | SRSF2 | serine and arginine rich splicing factor 2 | protein_coding | NP_001182356.1;NP_003007.2;XP_016880431.1 | 10783 | 600813 | . | . |
| 6428 | NM_003017,N | SRSF3 | serine and arginine rich splicing factor 3 | protein_coding | NP_003008.1 | 10785 | 603364 | . | . |
| 6429 | NM_005626,XI | SRSF4 | serine and arginine rich splicing factor 4 | protein_coding | NP_005617.2;XP_011540253.1 | 10786 | 601940 | . | . |
| 6430 | NM_00103946 | SRSF5 | serine and arginine rich splicing factor 5 | protein_coding | NP_001034554.1;NP_001307143.1;NP_008856.2;XP_005268056.1;XP_00 | 10787 | 600914 | . | . |
| 6431 | NM_006275,N | SRSF6 | serine and arginine rich splicing factor 6 | protein_coding | NP_006266.2 | 10788 | 601944 | . | . |
| 6432 | NM_00103168 | SRSF7 | serine and arginine rich splicing factor 7 | protein_coding | NP_001026854.1;NP_001182375.1;NP_001350731.1;XP_005264542.1;XP | 10789 | 600572 | . | . |
| 6433 | NM_00126141 | SFSWAP | splicing factor SWAP | protein_coding | NP_001248340.1;NP_004583.2;XP_011536955.1;XP_011536956.1;XP_01 | 10790 | 601945 | . | . |
| 6434 | NM_00124387 | TRA2B | transformer 2 beta homolog | protein_coding | NP_001230808.1;NP_004584.1;XP_005247760.1;XP_024309474.1 | 10781 | 602719 | . | . |
| 6440 | NM_00117235 | SFTPC | surfactant protein C | protein_coding | NP_001165828.1;NP_001165881.1;NP_001304707.1;NP_001304708.1;NI | 10802 | 178620 | . | . |
| 6443 | NM_000232 | SGCB | sarcoglycan beta | protein_coding | NP_000223.1 | 10806 | 600900 | . | . |
| 6444 | NM_000337,N | SGCD | sarcoglycan delta | protein_coding | NP_000328.2;NP_001121681.1;NP_758447.1;XP_005266023.1;XP_00526 | 10807 | 601411 | . | . |
| 6446 | NM_00114367 | SGK1 | serum/glucocorticoid regulated kinase 1 | protein_coding | NP_001137148.1;NP_001137149.1;NP_001137150.1;NP_001278924.1;NI | 10810 | 602958 | . | . |
| 6447 | NM_00114475 | SCG5 | secretogranin V | protein_coding | NP_001138229.1;NP_003011.1 | 10816 | 173120 | . | . |
| 6448 | NM_000199,N | SGSH | N-sulfoglucosamine sulfohydrolase | protein_coding | NP_000190.1;NP_001339850.1;NP_001339851.1;XP_005257640.1;XP_01 | 10818 | 605270 | . | . |
| 6449 | NM_003021,XI | SGTA | small glutamine rich tetratricopeptide repeat containing alpha | protein_coding | NP_003012.1;XP_011526480.1 | 10819 | 603419 | . | . |
| 6450 | NM_00100171 | SH3BGR | SH3 domain binding glutamate rich protein | protein_coding | NP_001001713.1;NP_001304669.1;NP_001304670.1;NP_001304671.1;NI | 10822 | 602230 | . | . |
| 6451 | NM_003022,XI | SH3BGRL | SH3 domain binding glutamate rich protein like | protein_coding | NP_003013.1;XP_011529315.1;XP_011529316.1 | 10823 | 300190 | . | . |
| 6452 | NM_00112268 | SH3BP2 | SH3 domain binding protein 2 | protein_coding | NP_001116153.1;NP_001139327.1;NP_001139328.1;NP_003014.3 | 10825 | 602104 | . | . |
| 6453 | NM_00100113 | ITSN1 | intersectin 1 | protein_coding | NP_001001132.1;NP_001317937.1;NP_001317938.1;NP_001317939.1;NI | 6183 | 602442 | . | . |
| 6455 | NM_00119994 | SH3GL1 | SH3 domain containing GRB2 like 1, endophilin A2 | protein_coding | NP_001186872.1;NP_001186873.1;NP_003016.1;XP_011526485.1;XP_01 | 10830 | 601768 | . | . |
| 6456 | NM_003026,XI | SH3GL2 | SH3 domain containing GRB2 like 2, endophilin A1 | protein_coding | NP_003017.1;XP_011516307.1 | 10831 | 604465 | . | . |
| 6458 | NR_033412 | SH3GL1P1 | SH3 domain containing GRB2 like 1, endophilin A2 pseudogene 1 | transcribed_ps | . | 10835 | . | . | . |
| 6459 | NR_033420 | SH3GL1P2 | SH3 domain containing GRB2 like 1, endophilin A2 pseudogene 2 | transcribed_ps | . | 10836 | . | . | . |
| 6461 | NM_003028 | SHB | SH2 domain containing adaptor protein B | protein_coding | NP_003019.2 | 10838 | 600314 | . | . |
| 6462 | NM_001040,N | SHBG | sex hormone binding globulin | protein_coding | NP_001031.2;NP_001139751.1;NP_001139752.1;NP_001139753.1;NP_0( | 10839 | 182205 | . | . |
| 6464 | NM_00113004 | SHC1 | SHC adaptor protein 1 | protein_coding | NP_001123512.1;NP_001123513.1;NP_001189788.1;NP_003020.2;NP_8! | 10840 | 600560 | . | . |
| 6468 | NM_00132354 | FBXW4 | F-box and WD repeat domain containing 4 | protein_coding | NP_001310470.1;NP_071322.2;XP_005270110.2;XP_016872059.1 | 10847 | 608071 | . | . |
| 6470 | NM_00128178 | SHMT1 | serine hydroxymethyltransferase 1 | protein_coding | NP_001268715.1;NP_004160.3;NP_683718.1;XP_005256824.1;XP_01152 | 10850 | 182144 | . | . |
| 6472 | NM_00116635 | SHMT2 | serine hydroxymethyltransferase 2 | protein_coding | NP_001159828.1;NP_001159829.1;NP_001159830.1;NP_001159831.1;NI | 10852 | 138450 | . | . |
| 6474 | NM_00116367 | SHOX2 | short stature homeobox 2 | protein_coding | NP_001157150.1;NP_003021.3;NP_006875.2;XP_006713790.1;XP_00671 | 10854 | 602504 | . | . |
| 6477 | NM_00100661 | SIAH1 | siah E3 ubiquitin protein ligase 1 | protein_coding | NP_001006611.1;NP_003022.3;XP_006721309.1;XP_011521581.1;XP_02 | 10857 | 602212 | . | . |
| 6478 | NM_005067 | SIAH2 | siah E3 ubiquitin protein ligase 2 | protein_coding | NP_005058.3 | 10858 | 602213 | . | . |
| 6480 | NM_00135391 | ST6GAL1 | ST6 beta-galactoside alpha-2,6-sialyltransferase 1 | protein_coding | NP_001340845.1;NP_003023.1;NP_775323.1;NP_775324.1 | 10860 | 109675 | . | . |
| 6482 | NM_003033,N | ST3GAL1 | ST3 beta-galactoside alpha-2,3-sialyltransferase 1 | protein_coding | NP_003024.1;NP_775479.1;XP_005251082.1;XP_006716680.1;XP_01151 | 10862 | 607187 | . | . |
| 6483 | NM_006927 | ST3GAL2 | ST3 beta-galactoside alpha-2,3-sialyltransferase 2 | protein_coding | NP_008858.1 | 10863 | 607188 | . | . |
| 6484 | NM_00125475 | ST3GAL4 | ST3 beta-galactoside alpha-2,3-sialyltransferase 4 | protein_coding | NP_001241686.1;NP_001241687.1;NP_001241688.1;NP_001335325.1;NI | 10864 | 104240 | . | . |
| 6487 | NM_00127045 | ST3GAL3 | ST3 beta-galactoside alpha-2,3-sialyltransferase 3 | protein_coding | NP_001257388.1;NP_001257389.1;NP_001257390.1;NP_001257391.1;NI | 10866 | 606494 | . | . |
| 6489 | NM_00130445 | ST8SIA1 | ST8 alpha-N-acetyl-neuraminide alpha-2,8-sialyltransferase 1 | protein_coding | NP_001291379.1;NP_003025.1 | 10869 | 601123 | . | . |
| 6490 | NM_00120005 | PMEL | premelanosome protein | protein_coding | NP_001186982.1;NP_001186983.1;NP_001307050.1;NP_001307051.1;NI | 10880 | 155550 | . | . |
| 6491 | NM_00104816 | STIL | STIL centriolar assembly protein | protein_coding | NP_001041631.1;NP_001269865.1;NP_001269866.1;NP_001269867.1;NI | 10879 | 181590 | . | . |
| 6494 | NM_006747,N | SIPA1 | signal-induced proliferation-associated 1 | protein_coding | NP_006738.3;NP_694985.29;XP_005274246.1 | 10885 | 602180 | . | . |
| 6495 | NM_005982,XI | SIX1 | SIX homeobox 1 | protein_coding | NP_005973.1;XP_016877091.1 | 10887 | 601205 | . | . |
| 6497 | NM_003036,XI | SKI | SKI proto-oncogene | protein_coding | NP_003027.1;XP_005244832.1;XP_005244833.1;XP_016857617.1 | 10896 | 164780 | . | . |
| 6498 | NM_00114509 | SKIL | SKI like proto-oncogene | protein_coding | NP_001138569.1;NP_001138570.1;NP_001234937.1;NP_005405.2;XP_0( | 10897 | 165340 | . | . |
| 6499 | NM_006929,XI | SKIV2L | Ski2 like RNA helicase | protein_coding | NP_008860.4;XP_011513117.1 | 10898 | 600478 | . | . |
| 6500 | NM_006930,N | SKP1 | S-phase kinase associated protein 1 | protein_coding | NP_008861.2;NP_733779.1 | 10899 | 601434 | . | . |
| 6502 | NM_00124312 | SKP2 | S-phase kinase associated protein 2 | protein_coding | NP_001230049.1;NP_005974.2;NP_116026.1;XP_011512384.1;XP_01151 | 10901 | 601436 | . | . |
| 6505 | NM_004170,XI | SLC1A1 | solute carrier family 1 member 1 | protein_coding | NP_004161.4;XP_011516309.1;XP_011516310.1;XP_011516311.1;XP_01 | 10939 | 133550 | . | . |
| 6507 | NM_00116669 | SLC1A3 | solute carrier family 1 member 3 | protein_coding | NP_001160167.1;NP_001160168.1;NP_001276868.1;NP_001276869.1;NI | 10941 | 600111 | . | . |
| 6508 | NM_00132655 | SLC4A3 | solute carrier family 4 member 3 | protein_coding | NP_001313488.1;NP_005061.3;NP_963868.2;XP_005246847.1;XP_01150 | 11029 | 106195 | . | . |
| 6509 | NM_00119349 | SLC1A4 | solute carrier family 1 member 4 | protein_coding | NP_001180422.1;NP_001335335.1;NP_001335336.1;NP_003029.2 | 10942 | 600229 | . | . |
| 6510 | NM_00114514 | SLC1A5 | solute carrier family 1 member 5 | protein_coding | NP_001138616.1;NP_001138617.1;NP_005619.1 | 10943 | 109190 | . | . |
| 6513 | NM_006516 | SLC2A1 | solute carrier family 2 member 1 | protein_coding | NP_006507.2 | 11005 | 138140 | . | . |
| 6515 | NM_006931 | SLC2A3 | solute carrier family 2 member 3 | protein_coding | NP_008862.1 | 11007 | 138170 | . | . |
| 6517 | NM_001042 | SLC2A4 | solute carrier family 2 member 4 | protein_coding | NP_001033.1 | 11009 | 138190 | . | . |
| 6520 | NM_00101266 | SLC3A2 | solute carrier family 3 member 2 | protein_coding | NP_001012680.1;NP_001012682.1;NP_001013269.1;NP_002385.3 | 11026 | 158070 | . | . |
| 6522 | NM_00119969 | SLC4A2 | solute carrier family 4 member 2 | protein_coding | NP_001186621.1;NP_001186622.1;NP_001186623.1;NP_003031.3;XP_0( | 11028 | 109280 | . | . |
| 6525 | NM_00120701 | SMTN | smoothelin | protein_coding | NP_001193946.1;NP_001193947.1;NP_001369567.1;NP_001369568.1;NI | 11126 | 602127 | . | . |
| 6526 | NM_006933 | SLC5A3 | solute carrier family 5 member 3 | protein_coding | NP_008864.4 | 11038 | 600444 | . | . |
| 6527 | NM_014227,XI | SLC5A4 | solute carrier family 5 member 4 | protein_coding | NP_055042.1;XP_006724371.1;XP_011528644.1;XP_011528645.1;XP_01 | 11039 | 618633 | . | . |
| 6533 | NM_00113436 | SLC6A6 | solute carrier family 6 member 6 | protein_coding | NP_001127839.2;NP_001127840.1;NP_003034.2;XP_006713370.1;XP_01 | 11052 | 186854 | . | . |
| 6535 | NM_00114280 | SLC6A8 | solute carrier family 6 member 8 | protein_coding | NP_001136277.1;NP_001136278.1;NP_005620.1 | 11055 | 300036 | . | . |
| 6536 | NM_00102484 | SLC6A9 | solute carrier family 6 member 9 | protein_coding | NP_001020016.1;NP_001248309.1;NP_001315555.1;NP_001315556.1;NI | 11056 | 601019 | . | . |
| 6539 | NM_00112284 | SLC6A12 | solute carrier family 6 member 12 | protein_coding | NP_001116319.1;NP_001116320.1;NP_001193860.1;NP_003035.3;XP_0( | 11045 | 603080 | . | . |

| | | | | | | | | | | |
|---|---|---|---|---|---|---|---|---|---|---|
| 6541 | NM_003045,XI | SLC7A1 | solute carrier family 7 member 1 | protein_coding | NP_003036.1;XP_005266564.1;XP_016876202.1 | 11057 | 104615 | . | . | [illegible] |
| 6542 | NM_00100853 | SLC7A2 | solute carrier family 7 member 2 | protein_coding | NP_001008539.3;NP_001158243.1;NP_001357266.1;NP_001357267.1;NI | 11060 | 601872 | . | . | [illegible] |
| 6545 | NM_004173 | SLC7A4 | solute carrier family 7 member 4 | protein_coding | NP_004164.2 | 11062 | 603752 | . | . | [illegible] |
| 6546 | NM_00111280 | SLC8A1 | solute carrier family 8 member A1 | protein_coding | NP_001106271.1;NP_001106272.1;NP_001106273.1;NP_001239553.1;NI | 11068 | 182305 | . | . | [illegible] |
| 6548 | NM_003047,N | SLC9A1 | solute carrier family 9 member A1 | protein_coding | NP_003038.2;XP_011540323.1 | 11071 | 107310 | . | . | [illegible] |
| 6549 | NM_003048 | SLC9A2 | solute carrier family 9 member A2 | protein_coding | NP_003039.2 | 11072 | 600530 | . | . | [illegible] |
| 6550 | NM_00128435 | SLC9A3 | solute carrier family 9 member A3 | protein_coding | NP_001271280.1;NP_004165.2 | 11073 | 182307 | . | . | [illegible] |
| 6553 | NM_00132397 | SLC9A5 | solute carrier family 9 member A5 | protein_coding | NP_001310900.1;NP_001310901.1;NP_001310902.1;NP_001310903.1;NI | 11078 | 600477 | . | . | [illegible] |
| 6556 | NM_000578,XI | SLC11A1 | solute carrier family 11 member 1 | protein_coding | NP_000569.3;XP_005246850.1;XP_005246851.1;XP_006712772.1;XP_00 | 10907 | 600266 | . | . | [illegible] |
| 6558 | NM_001046,N | SLC12A2 | solute carrier family 12 member 2 | protein_coding | NP_001037.1;NP_001243390.1;XP_011541890.1;XP_016865260.1 | 10911 | 600840 | . | . | [illegible] |
| 6560 | NM_00114596 | SLC12A4 | solute carrier family 12 member 4 | protein_coding | NP_001139433.1;NP_001139434.1;NP_001139435.1;NP_001139436.1;NI | 10913 | 604119 | . | . | [illegible] |
| 6565 | NM_00114599 | SLC15A2 | solute carrier family 15 member 2 | protein_coding | NP_001139470.1;NP_066568.3;XP_005247779.1;XP_006713799.1;XP_01 | 10921 | 602339 | . | . | [illegible] |
| 6566 | NM_00116649 | SLC16A1 | solute carrier family 16 member 1 | protein_coding | NP_001159968.1;NP_003042.3 | 10922 | 600682 | . | . | [illegible] |
| 6567 | NM_006517 | SLC16A2 | solute carrier family 16 member 2 | protein_coding | NP_006508.2 | 10923 | 300095 | . | . | [illegible] |
| 6569 | NM_00116757 | SLC34A1 | solute carrier family 34 member 1 | protein_coding | NP_001161051.1;NP_003043.3;XP_016865262.1;XP_016865263.1;XP_01 | 11019 | 182309 | . | . | [illegible] |
| 6570 | NM_00113569 | SLC18A1 | solute carrier family 18 member A1 | protein_coding | NP_001129163.1;NP_001135796.1;NP_001135797.1;NP_003044.1;XP_01 | 10934 | 193002 | . | . | [illegible] |
| 6573 | NM_00120520 | SLC19A1 | solute carrier family 19 member 1 | protein_coding | NP_001192135.1;NP_001192136.1;NP_001339439.1;NP_001339440.1;NI | 10937 | 600424 | . | . | [illegible] |
| 6574 | NM_005415,XI | SLC20A1 | solute carrier family 20 member 1 | protein_coding | NP_005406.3;XP_016860259.1;XP_016860260.1;XP_024308820.1 | 10946 | 137570 | . | . | [illegible] |
| 6575 | NM_00125718 | SLC20A2 | solute carrier family 20 member 2 | protein_coding | NP_001244109.1;NP_001244110.1;NP_006740.1;XP_005273670.1;XP_00 | 10947 | 158378 | . | . | [illegible] |
| 6576 | NM_00125653 | SLC25A1 | solute carrier family 25 member 1 | protein_coding | NP_001243463.1;NP_001274316.1;NP_005975.1 | 10979 | 190315 | . | . | [illegible] |
| 6580 | NM_003057,N | SLC22A1 | solute carrier family 22 member 1 | protein_coding | NP_003048.1;NP_694857.1;XP_005267159.1;XP_005267160.1;XP_00526 | 10963 | 602607 | . | . | [illegible] |
| 6583 | NM_003059,XI | SLC22A4 | solute carrier family 22 member 4 | protein_coding | NP_003050.2;XP_006714738.1;XP_011541891.1;XP_016865265.1 | 10968 | 604190 | . | . | [illegible] |
| 6584 | NM_00130812 | SLC22A5 | solute carrier family 22 member 5 | protein_coding | NP_001295051.1;NP_003051.1;XP_011541892.1;XP_016865267.1 | 10969 | 603377 | . | . | [illegible] |
| 6586 | NM_00127194 | SLIT3 | slit guidance ligand 3 | protein_coding | NP_001258875.2;NP_003053.2;XP_016865268.1 | 11087 | 603745 | . | . | [illegible] |
| 6591 | NM_003068 | SNAI2 | snail family transcriptional repressor 2 | protein_coding | NP_003059.1 | 11094 | 602150 | . | . | [illegible] |
| 6594 | NM_00128287 | SMARCA1 | SWI/SNF related, matrix associated, actin dependent regulator of | protein_coding | NP_001269803.1;NP_001269804.1;NP_001365190.1;NP_001365191.1;NI | 11097 | 300012 | . | . | [illegible] |
| 6595 | NM_00128939 | SMARCA2 | SWI/SNF related, matrix associated, actin dependent regulator of | protein_coding | NP_001276325.1;NP_001276326.1;NP_001276327.1;NP_001276328.1;NI | 11098 | 600014 | . | . | [illegible] |
| 6596 | NM_00131893 | HLTF | helicase like transcription factor | protein_coding | NP_001305863.1;NP_001305864.1;NP_003062.2;NP_620636.1;XP_01151 | 11099 | 603257 | . | . | [illegible] |
| 6597 | NM_00112884 | SMARCA4 | SWI/SNF related, matrix associated, actin dependent regulator of | protein_coding | NP_001122316.1;NP_001122317.1;NP_001122318.1;NP_001122319.1;NI | 11100 | 603254 | . | . | [illegible] |
| 6598 | NM_00100746 | SMARCB1 | SWI/SNF related, matrix associated, actin dependent regulator of | protein_coding | NP_001007469.1;NP_001304875.1;NP_001349806.1;NP_003064.2 | 11103 | 601607 | . | . | [illegible] |
| 6599 | NM_003074 | SMARCC1 | SWI/SNF related, matrix associated, actin dependent regulator of | protein_coding | NP_003065.3 | 11104 | 601732 | . | . | [illegible] |
| 6601 | NM_00113042 | SMARCC2 | SWI/SNF related, matrix associated, actin dependent regulator of | protein_coding | NP_001123892.1;NP_001317217.1;NP_003066.2;NP_620706.1;XP_00526 | 11105 | 601734 | . | . | [illegible] |
| 6602 | NM_003076,N | SMARCD1 | SWI/SNF related, matrix associated, actin dependent regulator of | protein_coding | NP_003067.3;NP_620710.2;XP_005269164.2;XP_011536997.1 | 11106 | 601735 | . | . | [illegible] |
| 6603 | NM_00109842 | SMARCD2 | SWI/SNF related, matrix associated, actin dependent regulator of | protein_coding | NP_001091896.1;NP_001317368.1;NP_001317369.1 | 11107 | 601736 | . | . | [illegible] |
| 6604 | NM_00100380 | SMARCD3 | SWI/SNF related, matrix associated, actin dependent regulator of | protein_coding | NP_001003801.1;NP_001003802.1;NP_003069.2;XP_011514823.1;XP_02 | 11108 | 601737 | . | . | [illegible] |
| 6605 | NM_003079 | SMARCE1 | SWI/SNF related, matrix associated, actin dependent regulator of | protein_coding | NP_003070.3 | 11109 | 603111 | . | . | [illegible] |
| 6606 | NM_000344,N | SMN1 | survival of motor neuron 1, telomeric | protein_coding | NP_000335.1;NP_001284644.1;NP_075012.1;XP_011541898.1;XP_01154 | 11117 | 600354 | . | . | [illegible] |
| 6607 | NM_017411,N | SMN2 | survival of motor neuron 2, centromeric | protein_coding | NP_059107.1;NP_075013.1;NP_075014.1;NP_075015.1;XP_011541901.1 | 11118 | 601627 | . | . | [illegible] |
| 6608 | NM_005631,XI | SMO | smoothened, frizzled class receptor | protein_coding | NP_005622.1;XP_024302659.1 | 11119 | 601500 | . | . | [illegible] |
| 6609 | NM_000543,N | SMPD1 | sphingomyelin phosphodiesterase 1 | protein_coding | NP_000534.3;NP_001007594.2;NP_001305016.1;NP_001305017.1;NP_00 | 11120 | 607608 | . | . | [illegible] |
| 6610 | NM_003080,XI | SMPD2 | sphingomyelin phosphodiesterase 2 | protein_coding | NP_003071.2;XP_005267166.1;XP_011534381.1;XP_024302294.1;XP_02 | 11121 | 603498 | . | . | [illegible] |
| 6611 | NM_00125842 | SMS | spermine synthase | protein_coding | NP_001245352.1;NP_004586.2;XP_005274639.1;XP_011543870.1;XP_01 | 11123 | 300105 | . | . | [illegible] |
| 6612 | NM_00128641 | SUMO3 | small ubiquitin like modifier 3 | protein_coding | NP_001273345.1;NP_008867.2 | 11124 | 602231 | . | . | [illegible] |
| 6613 | NM_00100584 | SUMO2 | small ubiquitin like modifier 2 | protein_coding | NP_001005849.1;NP_008868.3 | 11125 | 603042 | . | . | [illegible] |
| 6615 | NM_005985 | SNAI1 | snail family transcriptional repressor 1 | protein_coding | NP_005976.2 | 11128 | 604238 | . | . | [illegible] |
| 6616 | NM_00132290 | SNAP25 | synaptosome associated protein 25 | protein_coding | NP_001309831.1;NP_001309832.1;NP_001309833.1;NP_001309834.1;NI | 11132 | 600322 | . | . | [illegible] |
| 6617 | NM_003082 | SNAPC1 | small nuclear RNA activating complex polypeptide 1 | protein_coding | NP_003073.1 | 11134 | 600591 | . | . | [illegible] |
| 6618 | NM_003083,N | SNAPC2 | small nuclear RNA activating complex polypeptide 2 | protein_coding | NP_003074.1 | 11135 | 605076 | . | . | [illegible] |
| 6619 | NM_00103969 | SNAPC3 | small nuclear RNA activating complex polypeptide 3 | protein_coding | NP_001034786.1;NP_001356576.1;NP_001356577.1;NP_001356578.1;NI | 11136 | 602348 | . | . | [illegible] |
| 6620 | NM_00100150 | SNCB | synuclein beta | protein_coding | NP_001001502.1;NP_001304963.1;NP_001304964.1;NP_001304965.1;NI | 11140 | 602569 | . | . | [illegible] |
| 6621 | NM_003086,XI | SNAPC4 | small nuclear RNA activating complex polypeptide 4 | protein_coding | NP_003077.2;XP_005266153.1;XP_006717304.1;XP_006717305.1;XP_00 | 11137 | 602777 | . | . | [illegible] |
| 6622 | NM_000345,N | SNCA | synuclein alpha | protein_coding | NP_000336.1;NP_001139526.1;NP_001139527.1;NP_001362214.1;NP_00 | 11138 | 163890 | . | . | [illegible] |
| 6624 | NM_003088 | FSCN1 | fascin actin-bundling protein 1 | protein_coding | NP_003079.1 | 11148 | 602689 | . | . | [illegible] |
| 6625 | NM_00130106 | SNRNP70 | small nuclear ribonucleoprotein U1 subunit 70 | protein_coding | NP_001287998.1;NP_003080.2;XP_011525542.2;XP_011525543.1 | 11150 | 180740 | . | . | [illegible] |
| 6626 | NM_004596 | SNRPA | small nuclear ribonucleoprotein polypeptide A | protein_coding | NP_004587.1 | 11151 | 182285 | . | . | [illegible] |
| 6627 | NM_003090,N | SNRPA1 | small nuclear ribonucleoprotein polypeptide A' | protein_coding | NP_003081.2 | 11152 | 603521 | . | . | [illegible] |
| 6628 | NM_003091,N | SNRPB | small nuclear ribonucleoprotein polypeptides B and B1 | protein_coding | NP_003082.1;NP_937859.1 | 11153 | 182282 | . | . | [illegible] |
| 6629 | NM_003092,N | SNRPB2 | small nuclear ribonucleoprotein polypeptide B2 | protein_coding | NP_003083.1;NP_937863.1 | 11155 | 603520 | . | . | [illegible] |
| 6631 | NM_003093,N | SNRPC | small nuclear ribonucleoprotein polypeptide C | protein_coding | NP_003084.1 | 11157 | 603522 | . | . | [illegible] |
| 6632 | NM_00129191 | SNRPD1 | small nuclear ribonucleoprotein D1 polypeptide | protein_coding | NP_001278845.1;NP_008869.1 | 11158 | 601063 | . | . | [illegible] |
| 6633 | NM_00136975 | SNRPD2 | small nuclear ribonucleoprotein D2 polypeptide | protein_coding | NP_001356680.1;NP_001356681.1;NP_004588.1;NP_808210.2 | 11159 | 601061 | . | . | [illegible] |
| 6634 | NM_00127865 | SNRPD3 | small nuclear ribonucleoprotein D3 polypeptide | protein_coding | NP_001265585.1;NP_004166.1 | 11160 | 601062 | . | . | [illegible] |
| 6635 | NM_00130446 | SNRPE | small nuclear ribonucleoprotein polypeptide E | protein_coding | NP_001291393.1;NP_001315566.1;NP_001315567.1;NP_003085.1 | 11161 | 128260 | . | . | [illegible] |
| 6636 | NM_003095 | SNRPF | small nuclear ribonucleoprotein polypeptide F | protein_coding | NP_003086.1 | 11162 | 603541 | . | . | [illegible] |
| 6637 | NM_00131716 | SNRPG | small nuclear ribonucleoprotein polypeptide G | protein_coding | NP_001304094.1;NP_001304095.1;NP_001304096.1;NP_001304097.1;NI | 11163 | 603542 | . | . | [illegible] |
| 6638 | NM_00134945 | SNRPN | small nuclear ribonucleoprotein polypeptide N | protein_coding | NP_001336383.1;NP_001336384.1;NP_001336385.1;NP_001336386.1;NI | 11164 | 182279 | . | . | [illegible] |
| 6640 | NM_003098,XI | SNTA1 | syntrophin alpha 1 | protein_coding | NP_003089.1;XP_005260574.1;XP_011527309.1;XP_011527310.1;XP_02 | 11167 | 601017 | . | . | [illegible] |
| 6641 | NM_021021,XI | SNTB1 | syntrophin beta 1 | protein_coding | NP_066301.1;XP_011515541.1 | 11168 | 600026 | . | . | [illegible] |
| 6642 | NM_00124293 | SNX1 | sorting nexin 1 | protein_coding | NP_001229862.1;NP_003090.2;NP_683758.1;XP_005254668.1 | 11172 | 601272 | . | . | [illegible] |
| 6643 | NM_00127819 | SNX2 | sorting nexin 2 | protein_coding | NP_001265128.1;NP_003091.2 | 11173 | 605929 | . | . | [illegible] |
| 6645 | NM_006750 | SNTB2 | syntrophin beta 2 | protein_coding | NP_006741.1 | 11169 | 600027 | . | . | [illegible] |
| 6646 | NM_00125251 | SOAT1 | sterol O-acyltransferase 1 | protein_coding | NP_001239440.1;NP_001239441.1;NP_003092.4;XP_011508213.1;XP_01 | 11177 | 102642 | . | . | [illegible] |
| 6647 | NM_000454 | SOD1 | superoxide dismutase 1 | protein_coding | NP_000445.1 | 11179 | 147450 | . | . | [illegible] |
| 6648 | NM_000636,N | SOD2 | superoxide dismutase 2 | protein_coding | NP_000627.2;NP_001019636.1;NP_001019637.1;NP_001309743.1;NP_00 | 11180 | 147460 | . | . | [illegible] |
| 6650 | NM_005632,XI | CAPN15 | calpain 15 | protein_coding | NP_005623.1;XP_011520922.1;XP_011520923.1;XP_011520924.1;XP_01 | 11182 | 603267 | . | . | [illegible] |
| 6651 | NM_00129141 | SON | SON DNA and RNA binding protein | protein_coding | NP_001278340.2;NP_001278341.1;NP_115571.3;NP_620305.3 | 11183 | 182465 | . | . | [illegible] |
| 6652 | NM_003104,N | SORD | sorbitol dehydrogenase | protein_coding | NP_003095.2 | 11184 | 182500 | . | . | [illegible] |
| 6653 | NM_003105 | SORL1 | sortilin related receptor 1 | protein_coding | NP_003096.2 | 11185 | 602005 | . | . | [illegible] |
| 6654 | NM_00138239 | SOS1 | SOS Ras/Rac guanine nucleotide exchange factor 1 | protein_coding | NP_001369323.1;NP_001369324.1;NP_005624.2;XP_011531366.1 | 11187 | 182530 | . | . | [illegible] |
| 6655 | NM_006939 | SOS2 | SOS Ras/Rho guanine nucleotide exchange factor 2 | protein_coding | NP_008870.2 | 11188 | 601247 | . | . | [illegible] |
| 6657 | NM_003106 | SOX2 | SRY-box transcription factor 2 | protein_coding | NP_003097.1 | 11195 | 184429 | . | . | [illegible] |
| 6659 | NM_003107 | SOX4 | SRY-box transcription factor 4 | protein_coding | NP_003098.1 | 11200 | 184430 | . | . | [illegible] |
| 6660 | NM_00126141 | SOX5 | SRY-box transcription factor 5 | protein_coding | NP_001248343.1;NP_001248344.1;NP_001317714.1;NP_008871.3;NP_69 | 11201 | 604975 | . | . | [illegible] |
| 6662 | NM_000346 | SOX9 | SRY-box transcription factor 9 | protein_coding | NP_000337.1 | 11204 | 608160 | . | . | [illegible] |
| 6663 | NM_006941 | SOX10 | SRY-box transcription factor 10 | protein_coding | NP_008872.1 | 11190 | 602229 | . | . | [illegible] |
| 6666 | NM_006943 | SOX12 | SRY-box transcription factor 12 | protein_coding | NP_008874.2 | 11198 | 601947 | . | . | [illegible] |
| 6667 | NM_00125182 | SP1 | Sp1 transcription factor | protein_coding | NP_001238754.1;NP_003100.1;NP_612482.2 | 11205 | 189906 | . | . | [illegible] |
| 6668 | NM_003110,XI | SP2 | Sp2 transcription factor | protein_coding | NP_003101.3;XP_006722086.1;XP_006722088.1;XP_006722089.1;XP_00 | 11207 | 601801 | . | . | [illegible] |
| 6670 | NM_00101737 | SP3 | Sp3 transcription factor | protein_coding | NP_001017371.3;NP_001166183.1;NP_003102.1 | 11208 | 601804 | . | . | [illegible] |
| 6671 | NM_00132654 | SP4 | Sp4 transcription factor | protein_coding | NP_001313471.1;NP_001313472.1;NP_003103.2;XP_005249886.1;XP_01 | 11209 | 600540 | . | . | [illegible] |
| 6672 | NM_00108039 | SP100 | SP100 nuclear antigen | protein_coding | NP_001073860.1;NP_001193630.1;NP_001193631.1;NP_001193632.1;NI | 11206 | 604585 | . | . | [illegible] |
| 6674 | NM_00137432 | SPAG1 | sperm associated antigen 1 | protein_coding | NP_001361250.1;NP_003105.2;NP_757367.1;XP_011515542.1;XP_01151 | 11212 | 603395 | . | . | [illegible] |
| 6675 | NM_00132411 | UAP1 | UDP-N-acetylglucosamine pyrophosphorylase 1 | protein_coding | NP_001311042.1;NP_001311043.1;NP_001311044.1;NP_001311045.1;NI | 12457 | 602862 | . | . | [illegible] |
| 6676 | NM_00131793 | SPAG4 | sperm associated antigen 4 | protein_coding | NP_001304860.1;NP_003107.1;XP_005260577.1;XP_011527311.1 | 11214 | 603038 | . | . | [illegible] |
| 6678 | NM_00130944 | SPARC | secreted protein acidic and cysteine rich | protein_coding | NP_001296372.1;NP_001296373.1;NP_003109.1 | 11219 | 182120 | . | . | [illegible] |
| 6683 | NM_00136382 | SPAST | spastin | protein_coding | NP_001350752.1;NP_001350804.1;NP_001364888.1;NP_055761.2;NP_9 | 11233 | 604277 | . | . | [illegible] |
| 6687 | NM_00136385 | SPG7 | SPG7 matrix AAA peptidase subunit, paraplegin | protein_coding | NP_001350779.1;NP_003110.1;NP_955399.1;XP_005256378.1;XP_01687 | 11237 | 602783 | . | . | [illegible] |
| 6690 | NM_00135496 | SPINK1 | serine peptidase inhibitor Kazal type 1 | protein_coding | NP_001341895.1;NP_001366539.1;NP_003113.2 | 11244 | 167790 | . | . | [illegible] |
| 6691 | NM_00127171 | SPINK2 | serine peptidase inhibitor Kazal type 2 | protein_coding | NP_001258647.1;NP_001258648.1;NP_001258649.1;NP_001258650.1;NI | 11245 | 605753 | . | . | [illegible] |
| 6695 | NM_004598 | SPOCK1 | SPARC (osteonectin), cwcv and kazal like domains proteoglycan 1 | protein_coding | NP_004589.1 | 11251 | 602264 | . | . | [illegible] |
| 6696 | NM_000582,N | SPP1 | secreted phosphoprotein 1 | protein_coding | NP_000573.1;NP_001035147.1;NP_001035149.1;NP_001238758.1;NP_00 | 11255 | 166490 | . | . | [illegible] |
| 6697 | NM_003124 | SPR | sepiapterin reductase | protein_coding | NP_003115.1 | 11257 | 182125 | . | . | [illegible] |
| 6703 | NM_00138224 | SPRR2D | small proline rich protein 2D | protein_coding | NP_001369177.1;NP_008876.3 | 11264 | 617587 | . | . | [illegible] |
| 6709 | NM_00113043 | SPTAN1 | spectrin alpha, non-erythrocytic 1 | protein_coding | NP_001123910.1;NP_001182461.1;NP_001350688.1;NP_001350694.1;NI | 11273 | 182810 | . | . | [illegible] |
| 6710 | NM_00102485 | SPTB | spectrin beta, erythrocytic | protein_coding | NP_001020029.1;NP_001342365.1;NP_001342366.1;XP_011535407.1;XP | 11274 | 182870 | . | . | [illegible] |
| 6711 | NM_003128,N | SPTBN1 | spectrin beta, non-erythrocytic 1 | protein_coding | NP_003119.2;NP_842565.2;XP_005264574.1;XP_005264575.1;XP_00671 | 11275 | 182790 | . | . | [illegible] |
| 6712 | NM_006946,XI | SPTBN2 | spectrin beta, non-erythrocytic 2 | protein_coding | NP_008877.2;XP_005274250.1;XP_006718732.1;XP_006718734.1;XP_01 | 11276 | 604985 | . | . | [illegible] |
| 6713 | NM_003129,XI | SQLE | squalene epoxidase | protein_coding | NP_003120.2;XP_011515548.1 | 11279 | 602019 | . | . | [illegible] |
| 6714 | NM_005417,N | SRC | SRC proto-oncogene, non-receptor tyrosine kinase | protein_coding | NP_005408.1;NP_938033.1;XP_011527315.1;XP_016883513.1;XP_01688 | 11283 | 190090 | . | . | [illegible] |
| 6715 | NM_001047,N | SRD5A1 | steroid 5 alpha-reductase 1 | protein_coding | NP_001038.1;NP_001311251.1;NP_001311252.1 | 11284 | 184753 | . | . | [illegible] |
| 6717 | NM_00125689 | SRI | sorcin | protein_coding | NP_001243820.1;NP_001243821.1;NP_003121.1;NP_944490.1;XP_01151 | 11292 | 182520 | . | . | [illegible] |
| 6720 | NM_00100529 | SREBF1 | sterol regulatory element binding transcription factor 1 | protein_coding | NP_001005291.1;NP_001308025.1;NP_004167.3;XP_016880459.1;XP_01 | 11289 | 184756 | . | . | [illegible] |
| 6721 | NM_004599,N | SREBF2 | sterol regulatory element binding transcription factor 2 | protein_coding | NP_004590.2;XP_006724373.1;XP_011528649.1;XP_016884410.1;XP_01 | 11290 | 600481 | . | . | [illegible] |
| 6722 | NM_00129200 | SRF | serum response factor | protein_coding | NP_001278930.1;NP_003122.1 | 11291 | 600589 | . | . | [illegible] |
| 6723 | NM_003132,XI | SRM | spermidine synthase | protein_coding | NP_003123.2;XP_016857668.1 | 11296 | 182891 | . | . | [illegible] |
| 6726 | NM_00113044 | SRP9 | signal recognition particle 9 | protein_coding | NP_001123912.1;NP_003124.1 | 11304 | 600707 | . | . | [illegible] |
| 6727 | NM_00130943 | SRP14 | signal recognition particle 14 | protein_coding | NP_001296363.1;NP_003125.3 | 11299 | 600708 | . | . | [illegible] |
| 6728 | NM_00120419 | SRP19 | signal recognition particle 19 | protein_coding | NP_001191122.1;NP_001191123.1;NP_001191125.1;NP_001191128.1;NI | 11300 | 182175 | . | . | [illegible] |
| 6729 | NM_00114628 | SRP54 | signal recognition particle 54 | protein_coding | NP_001139754.1;NP_003127.1;XP_005268081.1;XP_011535408.1;XP_01 | 11301 | 604857 | . | . | [illegible] |
| 6730 | NM_00126050 | SRP68 | signal recognition particle 68 | protein_coding | NP_001247431.1;NP_001247432.1;NP_055045.2 | 11302 | 604858 | . | . | [illegible] |
| 6731 | NM_00126772 | SRP72 | signal recognition particle 72 | protein_coding | NP_001254651.1;NP_008878.3;XP_024309960.1 | 11303 | 602122 | . | . | [illegible] |
| 6732 | NM_003137,N | SRPK1 | SRSF protein kinase 1 | protein_coding | NP_003128.3 | 11305 | 601939 | . | . | [illegible] |
| 6733 | NM_00127827 | SRPK2 | SRSF protein kinase 2 | protein_coding | NP_001265202.1;NP_001337667.1;NP_001337668.1;NP_001337669.1;NI | 11306 | 602980 | . | . | [illegible] |
| 6734 | NM_00117784 | SRPRA | SRP receptor subunit alpha | protein_coding | NP_001171313.1;NP_003130.2;XP_016873668.1 | 11307 | 182180 | . | . | [illegible] |
| 6737 | NM_003141 | TRIM21 | tripartite motif containing 21 | protein_coding | NP_003132.2 | 11312 | 109092 | . | . | [illegible] |
| 6738 | NM_00104236 | RO60 | Ro60, Y RNA binding protein | protein_coding | NP_001035828.1;NP_001035829.2;NP_001166995.1;NP_001166996.1;NI | 11313 | 600063 | . | . | [illegible] |
| 6741 | NM_00129414 | SSB | small RNA binding exonuclease protection factor La | protein_coding | NP_001281074.1;NP_003133.1 | 11316 | 109090 | . | . | [illegible] |
| 6742 | NM_00125651 | SSBP1 | single stranded DNA binding protein 1 | protein_coding | NP_001243439.1;NP_001243440.1;NP_001243441.1;NP_001243442.1;NI | 11317 | 600439 | . | . | [illegible] |
| 6744 | NM_00113044 | ITPRID2 | ITPR interacting domain containing 2 | protein_coding | NP_001123917.1;NP_001274432.1;NP_001274433.1;NP_001274434.1;NI | 11319 | 118990 | . | . | [illegible] |
| 6745 | NM_00129200 | SSR1 | signal sequence receptor subunit 1 | protein_coding | NP_001278937.1;NP_003135.2 | 11323 | 600868 | . | . | [illegible] |
| 6746 | NM_003145 | SSR2 | signal sequence receptor subunit 2 | protein_coding | NP_003136.1 | 11324 | 600867 | . | . | [illegible] |
| 6747 | NM_00130819 | SSR3 | signal sequence receptor subunit 3 | protein_coding | NP_001295126.1;NP_001295133.1;NP_001295134.1;NP_009038.1 | 11325 | 606213 | . | . | [illegible] |
| 6748 | NM_00120452 | SSR4 | signal sequence receptor subunit 4 | protein_coding | NP_001191455.1;NP_001191456.1;NP_006271.1;XP_016885245.1;XP_01 | 11326 | 300090 | . | . | [illegible] |
| 6749 | NM_003146,XI | SSRP1 | structure specific recognition protein 1 | protein_coding | NP_003137.1;XP_016873669.1;XP_024304432.1;XP_024304433.1;XP_02 | 11327 | 604328 | . | . | [illegible] |
| 6752 | NM_001050 | SSTR2 | somatostatin receptor 2 | protein_coding | NP_001041.1 | 11331 | 182452 | . | . | [illegible] |
| 6756 | NM_00127869 | SSX1 | SSX family member 1 | protein_coding | NP_001265620.1;NP_005626.1 | 11335 | 312820 | . | . | [illegible] |
| 6760 | NM_00100755 | SS18 | SS18 subunit of BAF chromatin remodeling complex | protein_coding | NP_001007560.1;NP_001295130.1;NP_005628.2;XP_006722590.1;XP_01 | 11340 | 600192 | . | . | [illegible] |
| 6764 | NM_00137649 | DENND2B | DENN domain containing 2B | protein_coding | NP_001363424.1;NP_001363425.1;NP_001363426.1;NP_001363427.1;NI | 11350 | 140750 | . | . | [illegible] |
| 6767 | NM_00127858 | ST13 | ST13 Hsp70 interacting protein | protein_coding | NP_001265518.1;NP_003923.2 | 11343 | 606796 | . | . | [illegible] |
| 6769 | NM_00129204 | STAC | SH3 and cysteine rich domain | protein_coding | NP_001278978.1;NP_003140.1;XP_011532339.1;XP_011532340.1;XP_01 | 11353 | 602317 | . | . | [illegible] |
| 6772 | NM_007315,N | STAT1 | signal transducer and activator of transcription 1 | protein_coding | NP_009330.1;NP_644671.1;XP_006712781.1;XP_016860272.1 | 11362 | 600555 | . | . | [illegible] |
| 6773 | NM_005419,N | STAT2 | signal transducer and activator of transcription 2 | protein_coding | NP_005410.1;NP_938146.1;XP_011536999.1;XP_011537000.1;XP_01153 | 11363 | 600556 | . | . | [illegible] |
| 6774 | NM_00136951 | STAT3 | signal transducer and activator of transcription 3 | protein_coding | NP_001356441.1;NP_001356442.1;NP_001356443.1;NP_001356445.1;NI | 11364 | 102582 | . | . | [illegible] |
| 6776 | NM_00128871 | STAT5A | signal transducer and activator of transcription 5A | protein_coding | NP_001275647.1;NP_001275648.1;NP_001275649.1;NP_003143.2;XP_00 | 11366 | 601511 | . | . | [illegible] |
| 6777 | NM_012448,XI | STAT5B | signal transducer and activator of transcription 5B | protein_coding | NP_036580.2;XP_005257683.1;XP_016880466.1;XP_024306665.1;XP_02 | 11367 | 604260 | . | . | [illegible] |
| 6778 | NM_00117807 | STAT6 | signal transducer and activator of transcription 6 | protein_coding | NP_001171549.1;NP_001171550.1;NP_001171551.1;NP_001171552.1;NI | 11368 | 601512 | . | . | [illegible] |
| 6780 | NM_00103732 | STAU1 | staufen double-stranded RNA binding protein 1 | protein_coding | NP_001032405.1;NP_001306063.1;NP_001306064.1;NP_001309856.1;NI | 11370 | 601716 | . | . | [illegible] |
| 6781 | NM_003155 | STC1 | stanniocalcin 1 | protein_coding | NP_003146.1 | 11373 | 601185 | . | . | [illegible] |
| 6782 | NM_006948 | HSPA13 | heat shock protein family A (Hsp70) member 13 | protein_coding | NP_008879.3 | 11375 | 601100 | . | . | [illegible] |
| 6785 | NM_022726 | ELOVL4 | ELOVL fatty acid elongase 4 | protein_coding | NP_073563.1 | 14415 | 605512 | . | . | [illegible] |
| 6786 | NM_00127796 | STIM1 | stromal interaction molecule 1 | protein_coding | NP_001264890.1;NP_001264891.1;NP_001369495.1;NP_001369496.1;NI | 11386 | 605921 | . | . | [illegible] |
| 6787 | NM_00119353 | NEK4 | NIMA related kinase 4 | protein_coding | NP_001180462.1;NP_001335341.1;NP_001335342.1;NP_001335343.1;NI | 11399 | 601959 | . | . | [illegible] |
| 6788 | NM_00125631 | STK3 | serine/threonine kinase 3 | protein_coding | NP_001243241.1;NP_001243242.1;NP_006272.2;XP_011515550.2;XP_01 | 11406 | 605030 | . | . | [illegible] |
| 6789 | NM_00135238 | STK4 | serine/threonine kinase 4 | protein_coding | NP_001339314.1;NP_006273.1;XP_005260587.1;XP_005260588.1;XP_00 | 11408 | 604965 | . | . | [illegible] |
| 6790 | NM_00132330 | AURKA | aurora kinase A | protein_coding | NP_001310232.1;NP_001310233.1;NP_001310234.1;NP_003591.2;NP_9 | 11393 | 603072 | . | . | [illegible] |
| 6791 | NR_001587 | AURKAP1 | aurora kinase A pseudogene 1 | transcribed_ps | . | 18611 | . | . | . | [illegible] |
| 6792 | NM_00103734 | CDKL5 | cyclin dependent kinase like 5 | protein_coding | NP_001032420.1;NP_001310218.1;NP_003150.1 | 11411 | 300203 | . | . | [illegible] |
| 6793 | NM_005990,XI | STK10 | serine/threonine kinase 10 | protein_coding | NP_005981.3;XP_016865277.1 | 11388 | 603919 | . | . | [illegible] |
| 6794 | NM_000455 | STK11 | serine/threonine kinase 11 | protein_coding | NP_000446.1 | 11389 | 602216 | . | . | [illegible] |
| 6801 | NM_003162,XI | STRN | striatin | protein_coding | NP_003153.2;XP_005264576.1;XP_011531375.1 | 11424 | 614765 | . | . | [illegible] |
| 6804 | NM_00116590 | STX1A | syntaxin 1A | protein_coding | NP_001159375.1;NP_004594.1;XP_016868056.1 | 11433 | 186590 | . | . | [illegible] |
| 6809 | NM_00117804 | STX3 | syntaxin 3 | protein_coding | NP_001171511.1;NP_004168.1;XP_005274252.1;XP_005274255.1;XP_00 | 11438 | 600876 | . | . | [illegible] |

| | | | | | | | | | | |
|---|---|---|---|---|---|---|---|---|---|---|
| 6810 | NM_00127209 | STX4 | syntaxin 4 | protein_coding | NP_001259024.1;NP_001259025.1;NP_004595.2;XP_005255578.1 | 11439 | 186591 | . | . | [illegible] |
| 6811 | NM_00124466 | STX5 | syntaxin 5 | protein_coding | NP_001231595.1;NP_001317223.1;NP_003155.2;XP_011543526.1 | 11440 | 603189 | . | . | [illegible] |
| 6812 | NM_00103222 | STXBP1 | syntaxin binding protein 1 | protein_coding | NP_001027392.1;NP_001361235.1;NP_001361236.1;NP_001361237.1;NI | 11444 | 602926 | . | . | [illegible] |
| 6813 | NM_00112739 | STXBP2 | syntaxin binding protein 2 | protein_coding | NP_001120868.1;NP_001258963.1;NP_008880.2;XP_011526512.1;XP_01 | 11445 | 601717 | . | . | [illegible] |
| 6814 | NM_007269 | STXBP3 | syntaxin binding protein 3 | protein_coding | NP_009200.2 | 11446 | 608339 | . | . | [illegible] |
| 6815 | NM_00113070 | STYX | serine/threonine/tyrosine interacting protein | protein_coding | NP_001124173.1;NP_660294.1;XP_011535410.1;XP_011535411.1 | 11447 | 615814 | . | . | [illegible] |
| 6817 | NM_001055,N | SULT1A1 | sulfotransferase family 1A member 1 | protein_coding | NP_001046.2;NP_803565.1;NP_803566.1;NP_803878.1;NP_803880.1;XP_ | 11453 | 171150 | . | . | [illegible] |
| 6818 | NM_177552 | SULT1A3 | sulfotransferase family 1A member 3 | protein_coding | NP_808220.1 | 11455 | 600641 | . | . | [illegible] |
| 6819 | NM_001056,N | SULT1C2 | sulfotransferase family 1C member 2 | protein_coding | NP_001047.1;NP_789795.1;XP_005264070.1 | 11456 | 602385 | . | . | [illegible] |
| 6820 | NM_004605,N | SULT2B1 | sulfotransferase family 2B member 1 | protein_coding | NP_004596.2;NP_814444.1 | 11459 | 604125 | . | . | [illegible] |
| 6821 | NM_000456,N | SUOX | sulfite oxidase | protein_coding | NP_000447.2;NP_001027558.1;NP_001027559.1;XP_016875394.1;XP_01 | 11460 | 606887 | . | . | [illegible] |
| 6822 | NM_003167 | SULT2A1 | sulfotransferase family 2A member 1 | protein_coding | NP_003158.2 | 11458 | 125263 | . | . | [illegible] |
| 6827 | NM_003168,N | SUPT4H1 | SPT4 homolog, DSIF elongation factor subunit | protein_coding | NP_003159.1 | 11467 | 603555 | . | . | [illegible] |
| 6829 | NM_00111102 | SUPT5H | SPT5 homolog, DSIF elongation factor subunit | protein_coding | NP_001104490.1;NP_001124296.1;NP_001124297.1;NP_001306919.1;NI | 11469 | 602102 | . | . | [illegible] |
| 6830 | NM_00132075 | SUPT6H | SPT6 homolog, histone chaperone and transcription elongation fac | protein_coding | NP_001307684.1;NP_003161.2;XP_016880467.1;XP_016880468.1;XP_01 | 11470 | 601333 | . | . | [illegible] |
| 6832 | NM_00130168 | SUPV3L1 | Suv3 like RNA helicase | protein_coding | NP_001288612.1;NP_001310513.1;NP_001310514.1;NP_001310515.1;NI | 11471 | 605122 | . | . | [illegible] |
| 6834 | NM_00128078 | SURF1 | SURF1 cytochrome c oxidase assembly factor | protein_coding | NP_001267716.1;NP_003163.1;XP_011517244.1 | 11474 | 185620 | . | . | [illegible] |
| 6835 | NM_00127892 | SURF2 | surfeit 2 | protein_coding | NP_001265857.1;NP_059973.4 | 11475 | 185630 | . | . | [illegible] |
| 6836 | NM_00128078 | SURF4 | surfeit 4 | protein_coding | NP_001267717.1;NP_001267718.1;NP_001267719.1;NP_001267720.1;NI | 11476 | 185660 | . | . | [illegible] |
| 6837 | NM_133640,N | MED22 | mediator complex subunit 22 | protein_coding | NP_598395.1;NP_852468.1 | 11477 | 185641 | . | . | [illegible] |
| 6838 | NM_00127894 | SURF6 | surfeit 6 | protein_coding | NP_001265871.1;NP_006744.2 | 11478 | 185642 | . | . | [illegible] |
| 6839 | NM_00128216 | SUV39H1 | suppressor of variegation 3-9 homolog 1 | protein_coding | NP_001269095.1;NP_003164.1 | 11479 | 300254 | . | . | [illegible] |
| 6840 | NM_00132359 | SVIL | supervillin | protein_coding | NP_001310528.1;NP_001310529.1;NP_003165.2;NP_068506.2;XP_0052! | 11480 | 604126 | . | . | [illegible] |
| 6843 | NM_00129743 | VAMP1 | vesicle associated membrane protein 1 | protein_coding | NP_001284367.1;NP_055046.1;NP_058439.1;NP_954740.1 | 12642 | 185880 | . | . | [illegible] |
| 6844 | NM_00133012 | VAMP2 | vesicle associated membrane protein 2 | protein_coding | NP_001317054.1;NP_055047.2 | 12643 | 185881 | . | . | [illegible] |
| 6845 | NM_00114514 | VAMP7 | vesicle associated membrane protein 7 | protein_coding | NP_001138621.1;NP_001172112.1;NP_005629.1;XP_011529490.1;XP_01 | 11486 | 300053 | . | . | [illegible] |
| 6850 | NM_00113505 | SYK | spleen associated tyrosine kinase | protein_coding | NP_001128524.1;NP_001167638.1;NP_001167639.1;NP_003168.2;XP_0C | 11491 | 600085 | . | . | [illegible] |
| 6853 | NM_006950,N | SYN1 | synapsin I | protein_coding | NP_008881.2;NP_598006.1 | 11494 | 313440 | . | . | [illegible] |
| 6855 | NM_003179 | SYP | synaptophysin | protein_coding | NP_003170.1 | 11506 | 313475 | . | . | [illegible] |
| 6856 | NM_00138191 | SYPL1 | synaptophysin like 1 | protein_coding | NP_001368839.1;NP_001368840.1;NP_001368841.1;NP_001368842.1;NI | 11507 | 616665 | . | . | [illegible] |
| 6861 | NM_00129777 | SYT5 | synaptotagmin 5 | protein_coding | NP_001284703.1;NP_003171.2;XP_006723402.1;XP_006723403.1;XP_00 | 11513 | 600782 | . | . | [illegible] |
| 6865 | NM_001057 | TACR2 | tachykinin receptor 2 | protein_coding | NP_001048.2 | 11527 | 162321 | . | . | [illegible] |
| 6867 | NM_00112282 | TACC1 | transforming acidic coiled-coil containing protein 1 | protein_coding | NP_001116296.1;NP_001139688.1;NP_001317450.1;NP_001339707.1;NI | 11522 | 605301 | . | . | [illegible] |
| 6868 | NM_00138277 | ADAM17 | ADAM metallopeptidase domain 17 | protein_coding | NP_001369706.1;NP_001369707.1;NP_003174.3;XP_011508677.1;XP_01 | 195 | 603639 | . | . | [illegible] |
| 6871 | NM_00116610 | TADA2A | transcriptional adaptor 2A | protein_coding | NP_001159577.2;NP_001278847.1;NP_001479.4;NP_597683.3;XP_00672 | 11531 | 602276 | . | . | [illegible] |
| 6872 | NM_00128607 | TAF1 | TATA-box binding protein associated factor 1 | protein_coding | NP_001273003.2;NP_004597.3;NP_620278.2;XP_005262352.1;XP_00526 | 11535 | 313650 | . | . | [illegible] |
| 6873 | NM_003184,XI | TAF2 | TATA-box binding protein associated factor 2 | protein_coding | NP_003175.2;XP_006716684.1;XP_011515561.1;XP_016869268.1;XP_01 | 11536 | 604912 | . | . | [illegible] |
| 6874 | NM_003185 | TAF4 | TATA-box binding protein associated factor 4 | protein_coding | NP_003176.2 | 11537 | 601796 | . | . | [illegible] |
| 6875 | NM_00129372 | TAF4B | TATA-box binding protein associated factor 4b | protein_coding | NP_001280654.1;NP_005631.1;XP_005258396.1;XP_011524455.1;XP_01 | 11538 | 601689 | . | . | [illegible] |
| 6876 | NM_00100152 | TAGLN | transgelin | protein_coding | NP_001001522.1;NP_003177.2 | 11553 | 600818 | . | . | [illegible] |
| 6877 | NM_006951,N | TAF5 | TATA-box binding protein associated factor 5 | protein_coding | NP_008882.2;NP_620640.1 | 11539 | 601787 | . | . | [illegible] |
| 6878 | NM_00119041 | TAF6 | TATA-box binding protein associated factor 6 | protein_coding | NP_001177344.1;NP_001351927.1;NP_001351928.1;NP_001351929.1;NI | 11540 | 602955 | . | . | [illegible] |
| 6879 | NM_005642 | TAF7 | TATA-box binding protein associated factor 7 | protein_coding | NP_005633.2 | 11541 | 600573 | . | . | [illegible] |
| 6880 | NM_00101589 | TAF9 | TATA-box binding protein associated factor 9 | protein_coding | NP_001015892.1;NP_003178.1 | 11542 | 600822 | . | . | [illegible] |
| 6881 | NM_006284 | TAF10 | TATA-box binding protein associated factor 10 | protein_coding | NP_006275.1 | 11543 | 600475 | . | . | [illegible] |
| 6882 | NM_00127048 | TAF11 | TATA-box binding protein associated factor 11 | protein_coding | NP_001257417.1;NP_005634.1;XP_011513129.1 | 11544 | 600772 | . | . | [illegible] |
| 6883 | NM_00113521 | TAF12 | TATA-box binding protein associated factor 12 | protein_coding | NP_001128690.1;NP_005635.1;XP_016857675.1;XP_024305128.1;XP_02 | 11545 | 600773 | . | . | [illegible] |
| 6884 | NM_005645 | TAF13 | TATA-box binding protein associated factor 13 | protein_coding | NP_005636.1 | 11546 | 600774 | . | . | [illegible] |
| 6885 | NM_003188,N | MAP3K7 | mitogen-activated protein kinase kinase kinase 7 | protein_coding | NP_003179.1;NP_663304.1;NP_663305.1;NP_663306.1;XP_006715616.1 | 6859 | 602614 | . | . | [illegible] |
| 6888 | NM_006755 | TALDO1 | transaldolase 1 | protein_coding | NP_006746.1 | 11559 | 602063 | . | . | [illegible] |
| 6890 | NM_000593,N | TAP1 | transporter 1, ATP binding cassette subfamily B member | protein_coding | NP_000584.3;NP_001278951.1 | 43 | 170260 | . | . | [illegible] |
| 6891 | NM_00129004 | TAP2 | transporter 2, ATP binding cassette subfamily B member | protein_coding | NP_001276972.1;NP_061313.2 | 44 | 170261 | . | . | [illegible] |
| 6892 | NM_003190,N | TAPBP | TAP binding protein | protein_coding | NP_003181.3;NP_757345.2;NP_757346.2;XP_011513130.1;XP_01686671 | 11566 | 601962 | . | . | [illegible] |
| 6894 | NM_005646,XI | TARBP1 | TAR (HIV-1) RNA binding protein 1 | protein_coding | NP_005637.3;XP_005273291.1;XP_011542565.1;XP_016857683.1;XP_01 | 11568 | 605052 | . | . | [illegible] |
| 6895 | NM_004178,N | TARBP2 | TARBP2 subunit of RISC loading complex | protein_coding | NP_004169.3;NP_599150.1;NP_599151.2;XP_005269171.1;XP_00526917 | 11569 | 605053 | . | . | [illegible] |
| 6897 | NM_00125843 | TARS1 | threonyl-tRNA synthetase 1 | protein_coding | NP_001245366.1;NP_001245367.1;NP_689508.3 | 11572 | 187790 | . | . | [illegible] |
| 6901 | NM_000116,N | TAZ | tafazzin | protein_coding | NP_000107.1;NP_001290394.1;NP_851828.1;NP_851829.1;NP_851830.1 | 11577 | 300394 | . | . | [illegible] |
| 6902 | NM_00129773 | TBCA | tubulin folding cofactor A | protein_coding | NP_001284667.1;NP_001284669.1;NP_004598.1 | 11579 | 610058 | . | . | [illegible] |
| 6903 | NM_003192 | TBCC | tubulin folding cofactor C | protein_coding | NP_003183.2 | 11580 | 602971 | . | . | [illegible] |
| 6904 | NM_005993,XI | TBCD | tubulin folding cofactor D | protein_coding | NP_005984.3;XP_005256453.1;XP_005256456.1;XP_005256461.1;XP_00 | 11581 | 604649 | . | . | [illegible] |
| 6905 | NM_00107951 | TBCE | tubulin folding cofactor E | protein_coding | NP_001072983.1;NP_001274730.1;NP_001274731.1;NP_003184.1 | 11582 | 604934 | . | . | [illegible] |
| 6907 | NM_00113946 | TBL1X | transducin beta like 1 X-linked | protein_coding | NP_001132938.1;NP_001132939.1;NP_001132940.1;NP_005638.1 | 11585 | 300196 | . | . | [illegible] |
| 6908 | NM_00117208 | TBP | TATA-box binding protein | protein_coding | NP_001165556.1;NP_003185.1 | 11588 | 600075 | . | . | [illegible] |
| 6909 | NM_005994 | TBX2 | T-box transcription factor 2 | protein_coding | NP_005985.3 | 11597 | 600747 | . | . | [illegible] |
| 6911 | NM_004608,XI | TBX6 | T-box transcription factor 6 | protein_coding | NP_004599.2;XP_005255580.1;XP_011544228.1;XP_016879103.1 | 11605 | 602427 | . | . | [illegible] |
| 6916 | NM_001061,N | TBXAS1 | thromboxane A synthase 1 | protein_coding | NP_001052.3;NP_001124438.2;NP_001159725.2;NP_001159726.1;NP_0C | 11609 | 274180 | . | . | [illegible] |
| 6917 | NM_006756,N | TCEA1 | transcription elongation factor A1 | protein_coding | NP_006747.1;NP_958845.1;XP_006716530.1;XP_011515880.1 | 11612 | 601425 | . | . | [illegible] |
| 6919 | NM_003195,N | TCEA2 | transcription elongation factor A2 | protein_coding | NP_003186.1;NP_942016.1;XP_005260286.1;XP_011527324.1;XP_02430 | 11614 | 604784 | . | . | [illegible] |
| 6920 | NM_003196,XI | TCEA3 | transcription elongation factor A3 | protein_coding | NP_003187.1;XP_006710927.1;XP_011540355.1;XP_016857689.1;XP_01 | 11615 | 604128 | . | . | [illegible] |
| 6921 | NM_00120485 | ELOC | elongin C | protein_coding | NP_001191786.1;NP_001191787.1;NP_001191788.1;NP_001191789.1;NI | 11617 | 600788 | . | . | [illegible] |
| 6922 | gene-SKP1P1 | SKP1P1 | S-phase kinase associated protein 1 pseudogene 1 | pseudogene | . | 33696 | . | . | . | [illegible] |
| 6923 | NM_007108,N | ELOB | elongin B | protein_coding | NP_009039.1;NP_996896.1 | 11619 | 600787 | . | . | [illegible] |
| 6924 | NM_003198 | ELOA | elongin A | protein_coding | NP_003189.2 | 11620 | 600786 | . | . | [illegible] |
| 6925 | NM_00108396 | TCF4 | transcription factor 4 | protein_coding | NP_001077431.1;NP_001230155.2;NP_001230156.1;NP_001230157.1;NI | 11634 | 602272 | . | . | [illegible] |
| 6926 | NM_005996,N | TBX3 | T-box transcription factor 3 | protein_coding | NP_005987.3;NP_057653.3 | 11602 | 601621 | . | . | [illegible] |
| 6929 | NM_00113613 | TCF3 | transcription factor 3 | protein_coding | NP_001129611.1;NP_001338707.1;NP_001338708.1;NP_003191.1;XP_0C | 11633 | 147141 | . | . | [illegible] |
| 6932 | NM_00113485 | TCF7 | transcription factor 7 | protein_coding | NP_001128323.2;NP_001333354.1;NP_001333379.1;NP_001353431.1;NI | 11639 | 189908 | . | . | [illegible] |
| 6934 | NM_00114627 | TCF7L2 | transcription factor 7 like 2 | protein_coding | NP_001139746.1;NP_001139755.1;NP_001139756.1;NP_001139757.1;NI | 11641 | 602228 | . | . | [illegible] |
| 6935 | NM_00112812 | ZEB1 | zinc finger E-box binding homeobox 1 | protein_coding | NP_001121600.1;NP_001167564.1;NP_001167565.1;NP_001167566.1;NI | 11642 | 189909 | . | . | [illegible] |
| 6936 | NM_00120133 | GCFC2 | GC-rich sequence DNA-binding factor 2 | protein_coding | NP_001188263.1;NP_001188264.1;NP_003194.3;XP_005264577.1;XP_01 | 1317 | 189901 | . | . | [illegible] |
| 6938 | NM_00130621 | TCF12 | transcription factor 12 | protein_coding | NP_001293148.1;NP_001293149.1;NP_001309080.1;NP_001309081.1;NI | 11623 | 600480 | . | . | [illegible] |
| 6940 | NM_00132433 | ZNF354A | zinc finger protein 354A | protein_coding | NP_001311268.1;NP_005640.2;XP_011532947.1;XP_016865280.1 | 11628 | 602444 | . | . | [illegible] |
| 6941 | NM_00107751 | TCF19 | transcription factor 19 | protein_coding | NP_001070979.1;NP_001305837.1;NP_009040.2 | 11629 | 600912 | . | . | [illegible] |
| 6942 | NM_00137841 | TCF20 | transcription factor 20 | protein_coding | NP_001365347.1;NP_005641.1;NP_852469.1;XP_005261779.1;XP_01152 | 11631 | 603107 | . | . | [illegible] |
| 6944 | NM_00127108 | VPS72 | vacuolar protein sorting 72 homolog | protein_coding | NP_001258016.1;NP_001258017.1;NP_005988.1;XP_016857694.1 | 11644 | 600607 | . | . | [illegible] |
| 6945 | NM_170607,N | MLX | MAX dimerization protein MLX | protein_coding | NP_733752.1;NP_937847.1;NP_937848.1 | 11645 | 602976 | . | . | [illegible] |
| 6947 | NM_001062 | TCN1 | transcobalamin 1 | protein_coding | NP_001053.2 | 11652 | 189905 | . | . | [illegible] |
| 6949 | NM_000356,N | TCOF1 | treacle ribosome biogenesis factor 1 | protein_coding | NP_000347.2;NP_001008657.1;NP_001128715.1;NP_001128716.1;NP_0C | 11654 | 606847 | . | . | [illegible] |
| 6950 | NM_00100889 | TCP1 | t-complex 1 | protein_coding | NP_001008897.1;NP_110379.2 | 11655 | 186980 | . | . | [illegible] |
| 6988 | NM_022171 | TCTA | T cell leukemia translocation altered | protein_coding | NP_071503.1 | 11692 | 600690 | . | . | [illegible] |
| 6990 | NM_006520 | DYNLT3 | dynein light chain Tctex-type 3 | protein_coding | NP_006511.1 | 11694 | 300302 | . | . | [illegible] |
| 6991 | NM_174910,N | TCTE3 | t-complex-associated-testis-expressed 3 | protein_coding | NP_777570.1;XP_006715617.1;XP_011534395.1 | 11695 | 186977 | . | . | [illegible] |
| 6992 | NM_021959,XI | PPP1R11 | protein phosphatase 1 regulatory inhibitor subunit 11 | protein_coding | NP_068778.1;XP_006715237.1 | 9285 | 606670 | . | . | [illegible] |
| 6993 | NM_00129160 | DYNLT1 | dynein light chain Tctex-type 1 | protein_coding | NP_001278531.1;NP_001278532.1;NP_006510.1 | 11697 | 601554 | . | . | [illegible] |
| 6996 | NM_00136361 | TDG | thymine DNA glycosylase | protein_coding | NP_001350541.1;NP_003202.3;XP_005269182.1;XP_011537017.1 | 11700 | 601423 | . | . | [illegible] |
| 6999 | NM_005651 | TDO2 | tryptophan 2,3-dioxygenase | protein_coding | NP_005642.1 | 11708 | 191070 | . | . | [illegible] |
| 7001 | NM_005809 | PRDX2 | peroxiredoxin 2 | protein_coding | NP_005800.3 | 9353 | 600538 | . | . | [illegible] |
| 7003 | NM_021961 | TEAD1 | TEA domain transcription factor 1 | protein_coding | NP_068780.2 | 11714 | 189967 | . | . | [illegible] |
| 7004 | NM_003213,N | TEAD4 | TEA domain transcription factor 4 | protein_coding | NP_003204.2;NP_958849.1;NP_958851.1 | 11717 | 601714 | . | . | [illegible] |
| 7005 | NM_003214 | TEAD3 | TEA domain transcription factor 3 | protein_coding | NP_003205.2 | 11716 | 603170 | . | . | [illegible] |
| 7006 | NM_003215,XI | TEC | tec protein tyrosine kinase | protein_coding | NP_003206.2;XP_011512039.1;XP_011512043.1;XP_016864058.1;XP_02 | 11719 | 600583 | . | . | [illegible] |
| 7008 | NM_00114539 | TEF | TEF transcription factor, PAR bZIP family member | protein_coding | NP_001138870.1;NP_003207.1 | 11722 | 188595 | . | . | [illegible] |
| 7009 | NM_00109857 | TMBIM6 | transmembrane BAX inhibitor motif containing 6 | protein_coding | NP_001092046.1;NP_003208.2;XP_005269183.1;XP_024304937.1;XP_02 | 11723 | 600748 | . | . | [illegible] |
| 7011 | NM_00131903 | TEP1 | telomerase associated protein 1 | protein_coding | NP_001305964.1;NP_009041.2;XP_005268084.1;XP_011535412.1 | 11726 | 601686 | . | . | [illegible] |
| 7013 | NM_003218,N | TERF1 | telomeric repeat binding factor 1 | protein_coding | NP_003209.2;NP_059523.2;XP_005251348.1;XP_005251349.1;XP_01151 | 11728 | 600951 | . | . | [illegible] |
| 7014 | NM_005652,XI | TERF2 | telomeric repeat binding factor 2 | protein_coding | NP_005643.2;XP_005256178.1;XP_005256179.1;XP_005256180.1;XP_00! | 11729 | 602027 | . | . | [illegible] |
| 7015 | NM_00119337 | TERT | telomerase reverse transcriptase | protein_coding | NP_001180305.1;NP_937983.2 | 11730 | 187270 | . | . | [illegible] |
| 7016 | NM_00131823 | TESK1 | testis associated actin remodelling kinase 1 | protein_coding | NP_001305159.1;NP_006276.2 | 11731 | 601782 | . | . | [illegible] |
| 7018 | NM_001063,N | TF | transferrin | protein_coding | NP_001054.2;NP_001341632.2;NP_001341633.2 | 11740 | 190000 | . | . | [illegible] |
| 7019 | NM_00127078 | TFAM | transcription factor A, mitochondrial | protein_coding | NP_001257711.1;NP_003192.1;XP_011538423.1 | 11741 | 600438 | . | . | [illegible] |
| 7020 | NM_00103228 | TFAP2A | transcription factor AP-2 alpha | protein_coding | NP_001027451.1;NP_001035890.1;NP_001358995.1;XP_006715238.1;XP | 11742 | 107580 | . | . | [illegible] |
| 7022 | NM_003222 | TFAP2C | transcription factor AP-2 gamma | protein_coding | NP_003213.1 | 11744 | 601602 | . | . | [illegible] |
| 7023 | NM_003223,XI | TFAP4 | transcription factor AP-4 | protein_coding | NP_003214.1;XP_011520935.1;XP_011520937.1 | 11745 | 600743 | . | . | [illegible] |
| 7024 | NM_00117345 | TFCP2 | transcription factor CP2 | protein_coding | NP_001166923.1;NP_001166924.1;NP_005644.2 | 11748 | 189889 | . | . | [illegible] |
| 7025 | NM_005654,XI | NR2F1 | nuclear receptor subfamily 2 group F member 1 | protein_coding | NP_005645.1;XP_016865286.1 | 7975 | 132890 | . | . | [illegible] |
| 7026 | NM_00114515 | NR2F2 | nuclear receptor subfamily 2 group F member 2 | protein_coding | NP_001138627.1;NP_001138628.1;NP_001138629.1;NP_066285.1 | 7976 | 107773 | . | . | [illegible] |
| 7027 | NM_007111,N | TFDP1 | transcription factor Dp-1 | protein_coding | NP_009042.1;XP_005268384.1;XP_005268385.1;XP_011535821.1;XP_01 | 11749 | 189902 | . | . | [illegible] |
| 7029 | NM_00117813 | TFDP2 | transcription factor Dp-2 | protein_coding | NP_001171609.1;NP_001171610.1;NP_001171611.1;NP_001171612.1;NI | 11751 | 602160 | . | . | [illegible] |
| 7030 | NM_00128214 | TFE3 | transcription factor binding to IGHM enhancer 3 | protein_coding | NP_001269071.1;NP_006512.2;XP_024308200.1 | 11752 | 314310 | . | . | [illegible] |
| 7035 | NM_00103228 | TFPI | tissue factor pathway inhibitor | protein_coding | NP_001027452.1;NP_001305870.1;NP_001316168.1;NP_001316169.1;NI | 11760 | 152310 | . | . | [illegible] |
| 7036 | NM_00120685 | TFR2 | transferrin receptor 2 | protein_coding | NP_001193784.1;NP_003218.2 | 11762 | 604720 | . | . | [illegible] |
| 7037 | NM_00112814 | TFRC | transferrin receptor | protein_coding | NP_001121620.1;NP_001300894.1;NP_001300895.1;NP_003225.2;XP_02 | 11763 | 190010 | . | . | [illegible] |
| 7039 | NM_00109969 | TGFA | transforming growth factor alpha | protein_coding | NP_001093161.1;NP_001295087.1;NP_001295088.1;NP_003227.1 | 11765 | 190170 | . | . | [illegible] |
| 7040 | NM_000660,XI | TGFB1 | transforming growth factor beta 1 | protein_coding | NP_000651.3;XP_011525544.1 | 11766 | 190180 | . | . | [illegible] |
| 7041 | NM_00104245 | TGFB1I1 | transforming growth factor beta 1 induced transcript 1 | protein_coding | NP_001035919.1;NP_001158191.1;NP_057011.2;XP_024306180.1 | 11767 | 602353 | . | . | [illegible] |
| 7042 | NM_00113559 | TGFB2 | transforming growth factor beta 2 | protein_coding | NP_001129071.1;NP_003229.1 | 11768 | 190220 | . | . | [illegible] |
| 7043 | NM_00132993 | TGFB3 | transforming growth factor beta 3 | protein_coding | NP_001316867.1;NP_001316868.1;NP_003230.1 | 11769 | 190230 | . | . | [illegible] |
| 7045 | NM_000358 | TGFBI | transforming growth factor beta induced | protein_coding | NP_000349.1 | 11771 | 601692 | . | . | [illegible] |
| 7046 | NM_00113091 | TGFBR1 | transforming growth factor beta receptor 1 | protein_coding | NP_001124388.1;NP_001293139.1;NP_004603.1;XP_011517250.1;XP_01 | 11772 | 190181 | . | . | [illegible] |
| 7048 | NM_00102484 | TGFBR2 | transforming growth factor beta receptor 2 | protein_coding | NP_001020018.1;NP_003233.4;XP_011532345.1;XP_011532347.1;XP_01 | 11773 | 190182 | . | . | [illegible] |
| 7049 | NM_00119568 | TGFBR3 | transforming growth factor beta receptor 3 | protein_coding | NP_001182612.1;NP_001182613.1;NP_003234.2;XP_006710930.1 | 11774 | 600742 | . | . | [illegible] |
| 7050 | NM_00127868 | TGIF1 | TGFB induced factor homeobox 1 | protein_coding | NP_001265611.1;NP_001265613.1;NP_001265615.1;NP_001361325.1;NI | 11776 | 602630 | . | . | [illegible] |
| 7052 | NM_00132331 | TGM2 | transglutaminase 2 | protein_coding | NP_001310245.1;NP_001310246.1;NP_001310247.1;NP_004604.2;NP_9 | 11778 | 190196 | . | . | [illegible] |
| 7057 | NM_003246,XI | THBS1 | thrombospondin 1 | protein_coding | NP_003237.2;XP_011520273.1 | 11785 | 188060 | . | . | [illegible] |
| 7058 | NM_00138193 | THBS2 | thrombospondin 2 | protein_coding | NP_001368868.1;NP_001368869.1;NP_001368870.1;NP_001368871.1;NI | 11786 | 188061 | . | . | [illegible] |
| 7059 | NM_00125260 | THBS3 | thrombospondin 3 | protein_coding | NP_001239536.1;NP_001239537.1;NP_009043.1;XP_006711561.1;XP_01 | 11787 | 188062 | . | . | [illegible] |
| 7060 | NM_00130621 | THBS4 | thrombospondin 4 | protein_coding | NP_001293141.1;NP_001293142.1;NP_001293143.1;NP_003239.2;XP_01 | 11788 | 600715 | . | . | [illegible] |
| 7062 | NM_007113 | TCHH | trichohyalin | protein_coding | NP_009044.2 | 11791 | 190370 | . | . | [illegible] |
| 7064 | NM_003249,XI | THOP1 | thimet oligopeptidase 1 | protein_coding | NP_003240.1;XP_011526530.1 | 11793 | 601117 | . | . | [illegible] |
| 7067 | NM_00119091 | THRA | thyroid hormone receptor alpha | protein_coding | NP_001177847.1;NP_001177848.1;NP_003241.2;NP_955366.1 | 11796 | 190120 | . | . | [illegible] |
| 7068 | NM_000461,N | THRB | thyroid hormone receptor beta | protein_coding | NP_000452.2;NP_001121648.1;NP_001121649.1;NP_001239563.1;NP_0C | 11799 | 190160 | . | . | [illegible] |
| 7071 | NM_00103228 | KLF10 | Kruppel like factor 10 | protein_coding | NP_001027453.1;NP_005646.1 | 11810 | 601878 | . | . | [illegible] |
| 7072 | NM_00135150 | TIA1 | TIA1 cytotoxic granule associated RNA binding protein | protein_coding | NP_001338437.1;NP_001338438.1;NP_001338439.1;NP_001338440.1;NI | 11802 | 603518 | . | . | [illegible] |
| 7073 | NM_00103392 | TIAL1 | TIA1 cytotoxic granule associated RNA binding protein like 1 | protein_coding | NP_001029097.1;NP_001310893.1;NP_001310894.1;NP_001310896.1;NI | 11804 | 603413 | . | . | [illegible] |
| 7074 | NM_00135368 | TIAM1 | TIAM Rac1 associated GEF 1 | protein_coding | NP_001340613.1;NP_001340614.1;NP_001340615.1;NP_001340616.1;NI | 11805 | 600687 | . | . | [illegible] |
| 7076 | NM_003254,XI | TIMP1 | TIMP metallopeptidase inhibitor 1 | protein_coding | NP_003245.1;XP_016885255.1 | 11820 | 305370 | . | . | [illegible] |
| 7077 | NM_003255 | TIMP2 | TIMP metallopeptidase inhibitor 2 | protein_coding | NP_003246.1 | 11821 | 188825 | . | . | [illegible] |
| 7078 | NM_000362 | TIMP3 | TIMP metallopeptidase inhibitor 3 | protein_coding | NP_000353.1 | 11822 | 188826 | . | . | [illegible] |
| 7082 | NM_00130102 | TJP1 | tight junction protein 1 | protein_coding | NP_001287954.2;NP_001287955.1;NP_001317168.1;NP_001341941.1;NI | 11827 | 601009 | . | . | [illegible] |
| 7083 | NM_00134666 | TK1 | thymidine kinase 1 | protein_coding | NP_001333592.1;NP_001350777.1;NP_003249.3 | 11830 | 188300 | . | . | [illegible] |
| 7084 | NM_00117264 | TK2 | thymidine kinase 2 | protein_coding | NP_001166114.1;NP_001166115.1;NP_001166116.1;NP_001258863.1;NI | 11831 | 188250 | . | . | [illegible] |
| 7086 | NM_001064,N | TKT | transketolase | protein_coding | NP_001055.1;NP_001128527.1;NP_001244957.1;XP_011532356.1;XP_01 | 11834 | 606781 | . | . | [illegible] |
| 7087 | NM_003259,XI | ICAM5 | intercellular adhesion molecule 5 | protein_coding | NP_003250.3;XP_011526531.1;XP_016882674.1 | 5348 | 601852 | . | . | [illegible] |
| 7088 | NM_00130310 | TLE1 | TLE family member 1, transcriptional corepressor | protein_coding | NP_001290032.1;NP_001290033.1;NP_005068.2;XP_005252208.1;XP_00 | 11837 | 600189 | . | . | [illegible] |
| 7089 | NM_00114476 | TLE2 | TLE family member 2, transcriptional corepressor | protein_coding | NP_001138233.1;NP_001138234.1;NP_001287775.1;NP_003251.2;XP_0C | 11838 | 601041 | . | . | [illegible] |
| 7090 | NM_00110519 | TLE3 | TLE family member 3, transcriptional corepressor | protein_coding | NP_001098662.1;NP_001269908.1;NP_001269909.1;NP_001269910.1;NI | 11839 | 600190 | . | . | [illegible] |
| 7091 | NM_00128274 | TLE4 | TLE family member 4, transcriptional corepressor | protein_coding | NP_001269677.1;NP_001269678.1;NP_001269682.1;NP_001269689.1;NI | 11840 | 605132 | . | . | [illegible] |
| 7093 | NM_012465 | TLL2 | tolloid like 2 | protein_coding | NP_036597.1 | 11844 | 606743 | . | . | [illegible] |
| 7094 | NM_006289 | TLN1 | talin 1 | protein_coding | NP_006280.3 | 11845 | 186745 | . | . | [illegible] |
| 7095 | NM_003262,XI | SEC62 | SEC62 homolog, preprotein translocation factor | protein_coding | NP_003253.1;XP_011511416.1 | 11846 | 602173 | . | . | [illegible] |
| 7096 | NM_003263,XI | TLR1 | toll like receptor 1 | protein_coding | NP_003254.2;XP_005262719.1;XP_011512044.1;XP_011512047.1;XP_01 | 11847 | 601194 | . | . | [illegible] |
| 7098 | NM_003265 | TLR3 | toll like receptor 3 | protein_coding | NP_003256.1 | 11849 | 603029 | . | . | [illegible] |

| | | | | | | | | | | | | | | | | | | | | | | | | | | | | | | | | | |
|---|---|---|---|---|---|---|---|---|---|---|---|---|---|---|---|---|---|---|---|---|---|---|---|---|---|---|---|---|---|---|---|---|---|
| 7099 | NM_003266,N | TLR4 | toll like receptor 4 | protein_coding | NP_003257.1;NP_612564.1;NP_612567.1 | 11850 | 603030 | . | . | 0.313505 | 0.389297 | 0.290324 | 0.311030 | 0.729538 | 0.899879 | 0.651478 | 0.711132 | -1.191186 | 130.289965 | 0.290730805 | -0.86815087 | 0.385311745 | 0.998865404 | 7.146759 | 6.893286 | 7.014329 | 7.095123 | 6.985405 | 7.000225 | 146 | 169 | 101 | 114 |
| 7100 | NM_003268,XI | TLR5 | toll like receptor 5 | protein_coding | NP_003259.2;XP_005273298.2;XP_005273299.2;XP_005273300.2;XP_00( | 11851 | 603031 | . | . | 0.109817 | 0.105550 | 0.073333 | 0.032913 | 0.255548 | 0.243984 | 0.164557 | 0.075251 | -1.935027 | 13.940898 | 0.846837974 | -1.12459956 | 0.260758759 | NA | 4.198253 | 3.251698 | 3.829145 | 3.822311 | 3.800790 | 3.773445 | 22 | 19 | 11 | 6 |
| 7102 | NM_004615 | TSPAN7 | tetraspanin 7 | protein_coding | NP_004606.2 | 11854 | 300096 | . | . | 15.564658 | 14.256558 | 12.198658 | 14.077113 | 36.219601 | 32.954670 | 27.373436 | 32.185646 | -1.151385 | 761.898653 | 0.154532444 | -1.31603937 | 0.188160829 | 0.960264078 | 9.671682 | 9.468049 | 9.653967 | 9.585215 | 9.475760 | 9.565594 | 984 | 838 | 575 | 708 |
| 7105 | NM_00127874 | TSPAN6 | tetraspanin 6 | protein_coding | NP_001265669.1;NP_001265670.1;NP_001265671.1;NP_001265672.1;NI | 11858 | 300191 | . | . | 6.946479 | 6.070321 | 6.865301 | 5.831612 | 16.164742 | 14.031821 | 15.405534 | 13.333288 | -1.034148 | 760.302423 | 0.161233976 | -0.30045194 | 0.763832449 | 0.998865404 | 9.594024 | 9.546443 | 9.631503 | 9.526594 | 9.613751 | 9.497306 | 956 | 772 | 705 | 642 |
| 7106 | NM_00102523 | TSPAN4 | tetraspanin 4 | protein_coding | NP_001020405.1;NP_001020406.1;NP_001020407.1;NP_001020408.1;NI | 11859 | 602644 | . | . | 68.335991 | 69.424746 | 66.817208 | 64.980463 | 159.020664 | 160.478414 | 149.935873 | 148.570103 | -1.070370 | 2954.153506 | 0.111907219 | -0.87670264 | 0.380648172 | 0.998865404 | 11.576715 | 11.478678 | 11.555141 | 11.556149 | 11.517091 | 11.482893 | 3515 | 3298 | 2534 | 2619 |
| 7107 | NM_003272,XI | GPR137B | G protein-coupled receptor 137B | protein_coding | NP_003263.1;XP_016857698.1;XP_016857699.1 | 11862 | 604658 | . | . | 29.339452 | 31.492856 | 26.630992 | 30.590911 | 68.274108 | 72.797143 | 59.759176 | 69.942483 | -1.038988 | 2392.518470 | 0.115192968 | -0.47901754 | 0.631926152 | 0.998865404 | 11.251672 | 11.196428 | 11.222145 | 11.256604 | 11.207248 | 11.210102 | 2748 | 2687 | 2037 | 2204 |
| 7108 | NM_00127723 | TM7SF2 | transmembrane 7 superfamily member 2 | protein_coding | NP_001264162.1;NP_003264.2 | 11863 | 603414 | . | . | 1.542816 | 1.591202 | 2.683697 | 2.423104 | 3.590197 | 3.678133 | 6.022137 | 5.540138 | 1.652245 | 108.688105 | 0.313651276 | 2.30965954 | 0.02090701 | 0.478742856 | 6.355988 | 7.081777 | 6.674981 | 6.648991 | 6.852890 | 6.790088 | 99 | 84 | 128 | 116 |
| 7109 | NM_00135170 | TRAPPC10 | trafficking protein particle complex 10 | protein_coding | NP_001338638.1;NP_003265.3;XP_005261225.1;XP_011528016.1;XP_01 | 11868 | 602103 | . | . | 3.916002 | 4.492956 | 4.835250 | 4.639241 | 9.112694 | 10.385670 | 10.850162 | 10.607078 | 1.093183 | 880.366973 | 0.145931526 | 0.880787497 | 0.378432848 | 0.998865404 | 9.716494 | 9.844586 | 9.705582 | 9.789123 | 9.794134 | 9.829589 | 912 | 961 | 780 | 883 |
| 7110 | NM_00136387 | TMF1 | TATA element modulatory factor 1 | protein_coding | NP_001350808.1;NP_009045.2 | 11870 | 601126 | . | . | 20.590214 | 20.994811 | 21.358144 | 20.299761 | 47.914277 | 48.530449 | 47.927055 | 46.412995 | -1.012265 | 4461.426349 | 0.108158943 | -0.16259918 | 0.870834032 | 0.998865404 | 12.132024 | 12.114502 | 12.123604 | 12.132807 | 12.143218 | 12.092319 | 5140 | 4870 | 3974 | 4030 |
| 7111 | NM_00116611 | TMOD1 | tropomodulin 1 | protein_coding | NP_001159588.1;NP_003266.1;XP_024303428.1 | 11871 | 190930 | . | . | 0.737349 | 0.663306 | 1.045490 | 1.033046 | 1.715841 | 1.533262 | 2.346049 | 2.361936 | 1.400653 | 88.111252 | 0.339920959 | 1.43003698 | 0.152706406 | 0.916437617 | 6.197681 | 6.684032 | 6.416909 | 6.390815 | 6.489333 | 6.506656 | 89 | 75 | 87 | 99 |
| 7112 | NM_00103228 | TMPO | thymopoietin | protein_coding | NP_001027454.1;NP_001027455.1;NP_001294904.1;NP_003267.1;XP_0( | 11875 | 188380 | . | . | 36.795862 | 35.043747 | 30.960571 | 33.937940 | 85.625481 | 81.005190 | 69.474620 | 77.595069 | -1.109773 | 4105.579617 | 0.112487141 | -1.33584054 | 0.181601398 | 0.953435029 | 12.076573 | 11.926253 | 12.070617 | 12.018420 | 11.923454 | 11.992015 | 5128 | 4516 | 3243 | 3801 |
| 7113 | NM_00113509 | TMPRSS2 | transmembrane serine protease 2 | protein_coding | NP_001128571.1;NP_001369649.1;NP_005647.3 | 11876 | 602060 | . | . | 0.118745 | 0.028694 | 0.076952 | 0.039984 | 0.276324 | 0.066328 | 0.172678 | 0.091418 | -1.250559 | 7.550018 | 1.171193747 | -0.27542224 | 0.78299186 | NA | 3.064677 | 2.751299 | 2.940491 | 2.897480 | 2.922410 | 2.900647 | 15 | 4 | 8 | 4 |
| 7114 | NM_021109 | TMSB4X | thymosin beta 4 X-linked | protein_coding | NP_066932.1 | 11881 | 300159 | . | . | 241.402985 | 222.010605 | 167.971069 | 221.640472 | 561.754700 | 513.187439 | 376.922211 | 506.754578 | -1.207188 | 4134.287074 | 0.137571415 | -1.97461141 | 0.048312264 | 0.660542298 | 12.142948 | 11.871112 | 12.122788 | 12.044541 | 11.813562 | 12.030626 | 5448 | 4655 | 2825 | 3978 |
| 7115 | gene-TMSB4XF | TMSB4XP1 | TMSB4X pseudogene 1 | pseudogene | . | 11883 | . | . | . | 0.975100 | 0.541401 | 0.270152 | 0.212316 | 2.269097 | 1.251473 | 0.606213 | 0.485435 | -3.027807 | 10.060800 | 1.017863317 | -1.57022392 | 0.116363027 | NA | 3.917750 | 2.324312 | 3.381292 | 3.345418 | 3.322078 | 3.315061 | 22 | 12 | 5 | 4 |
| 7117 | gene-TMSB4XF | TMSB4XP8 | TMSB4X pseudogene 8 | pseudogene | . | 11885 | . | . | . | 13.055855 | 10.615866 | 8.416709 | 8.638067 | 30.381512 | 24.539049 | 18.886850 | 19.749914 | -1.407618 | 196.419303 | 0.247209368 | -1.99529458 | 0.046010761 | 0.654054105 | 7.843180 | 7.350555 | 7.738975 | 7.631654 | 7.532703 | 7.536966 | 293 | 221 | 141 | 154 |
| 7118 | gene-TMSB4XF | TMSB4XP4 | TMSB4X pseudogene 4 | pseudogene | . | 11886 | . | . | . | 0.205828 | 0.257766 | 0.396498 | 0.048858 | 0.478970 | 0.595839 | 0.889730 | 0.111708 | -1.092737 | 4.754791 | 1.459837931 | -0.08764422 | 0.930159453 | NA | 2.307819 | 2.188478 | 2.238788 | 2.244519 | 2.255193 | 2.223968 | 5 | 6 | 7 | 1 |
| 7120 | gene-TMSB4XF | TMSB4XP6 | TMSB4X pseudogene 6 | pseudogene | . | 11888 | . | . | . | 0.177525 | 0.423507 | 0.212701 | 0.242635 | 0.413108 | 0.978956 | 0.477296 | 0.554755 | -1.185486 | 5.436926 | 1.314045782 | -0.18681164 | 0.851808321 | NA | 2.562483 | 2.312263 | 2.432718 | 2.455461 | 2.438363 | 2.441579 | 4 | 9 | 4 | 5 |
| 7126 | NM_021137,XI | TNFAIP1 | TNF alpha induced protein 1 | protein_coding | NP_066960.1;XP_016880482.1 | 11894 | 191161 | . | . | 8.605332 | 9.254991 | 8.638389 | 9.909454 | 20.024959 | 21.393329 | 19.384294 | 22.656790 | 1.029413 | 1016.702286 | 0.142325868 | 0.293842834 | 0.768878003 | 0.998865404 | 9.968880 | 10.010187 | 9.954504 | 10.001610 | 9.951889 | 10.044917 | 1117 | 1116 | 840 | 1028 |
| 7127 | NM_00137122 | TNFAIP2 | TNF alpha induced protein 2 | protein_coding | NP_001358149.1;NP_001358150.1;NP_006282.2;XP_006720306.1;XP_01 | 11895 | 603300 | . | . | 0.056320 | 0.042200 | 0.022045 | 0.040533 | 0.131059 | 0.097548 | 0.049469 | 0.092674 | -1.536554 | 6.255390 | 1.226486953 | -0.50526298 | 0.613374144 | NA | 2.923304 | 2.300113 | 2.665639 | 2.655403 | 2.642070 | 2.654984 | 10 | 7 | 3 | 6 |
| 7128 | NM_00127050 | TNFAIP3 | TNF alpha induced protein 3 | protein_coding | NP_001257436.1;NP_001257437.1;NP_006281.1;XP_005267176.1;XP_00 | 11896 | 191163 | . | . | 1.941603 | 1.514067 | 1.840624 | 1.794144 | 4.518188 | 3.499833 | 4.130310 | 4.102099 | 1.042672 | 259.257079 | 0.221850515 | 0.271736561 | 0.785824587 | 0.998865404 | 7.987267 | 8.048562 | 8.059552 | 7.948953 | 8.039615 | 8.014347 | 324 | 244 | 236 | 242 |
| 7130 | NM_007115 | TNFAIP6 | TNF alpha induced protein 6 | protein_coding | NP_009046.2 | 11898 | 600410 | . | . | 6.072232 | 5.751392 | 4.937313 | 4.776847 | 14.130334 | 13.294600 | 11.079187 | 10.921693 | -1.236426 | 239.008555 | 0.22071724 | -1.38718567 | 0.165385153 | 0.94058973 | 8.045684 | 7.739989 | 7.967110 | 7.933498 | 7.857305 | 7.836742 | 314 | 276 | 190 | 196 |
| 7132 | NM_001065,N | TNFRSF1A | TNF receptor superfamily member 1A | protein_coding | NP_001056.1;NP_001333020.1;NP_001333021.1 | 11916 | 191190 | . | . | 34.242676 | 36.829461 | 31.723642 | 34.834701 | 79.684113 | 85.132949 | 71.186934 | 79.645404 | -1.082966 | 2326.139152 | 0.119784066 | -0.95995966 | 0.337075519 | 0.998865404 | 11.240186 | 11.124957 | 11.188299 | 11.240431 | 11.117296 | 11.182769 | 2696 | 2694 | 1862 | 2180 |
| 7138 | NM_00112613 | TNNT1 | troponin T1, slow skeletal type | protein_coding | NP_001119604.1;NP_001119605.1;NP_001278703.1;NP_003274.3;XP_01 | 11948 | 191041 | . | . | 19.706997 | 19.641990 | 18.227860 | 25.050378 | 45.858991 | 45.403335 | 40.902784 | 57.274715 | 1.090509 | 721.155676 | 0.173570357 | 0.720178383 | 0.471415172 | 0.998865404 | 9.430717 | 9.554943 | 9.469138 | 9.451394 | 9.408547 | 9.620687 | 805 | 735 | 555 | 812 |
| 7145 | NM_00130802 | TNS1 | tensin 1 | protein_coding | NP_001294951.1;NP_001294952.1;NP_072174.3;XP_016860300.1;XP_01 | 11973 | 600076 | . | . | 3.412594 | 3.671708 | 4.715468 | 5.555310 | 7.941245 | 8.487315 | 10.581370 | 12.701557 | 1.431736 | 1418.512008 | 0.134278008 | 3.855923922 | 0.000115293 | 0.01348639 | 10.188434 | 10.705762 | 10.280490 | 10.346012 | 10.539044 | 10.636617 | 1284 | 1315 | 1364 | 1661 |
| 7150 | NM_003286 | TOP1 | DNA topoisomerase I | protein_coding | NP_003277.1 | 11986 | 126420 | . | . | 19.042965 | 19.928621 | 19.193270 | 21.274925 | 44.313766 | 46.065903 | 43.069141 | 48.642590 | 1.023974 | 2310.075249 | 0.119458666 | 0.286113618 | 0.774791102 | 0.998865404 | 11.156647 | 11.190602 | 11.148833 | 11.178627 | 11.144728 | 11.219496 | 2580 | 2509 | 1938 | 2293 |
| 7151 | NR_002719 | TOP1P1 | DNA topoisomerase I pseudogene 1 | transcribed_ps | . | 11987 | . | . | . | 0.055203 | 0.021479 | 0.043552 | 0.007763 | 0.128460 | 0.049651 | 0.097729 | 0.017750 | -1.382723 | 2.678072 | 1.861946172 | -0.25108785 | 0.801746191 | NA | 1.631267 | 1.175234 | 1.432721 | 1.420893 | 1.428210 | 1.417447 | 5 | 2 | 3 | 1 |
| 7153 | NM_001067,XI | TOP2A | DNA topoisomerase II alpha | protein_coding | NP_001058.2;XP_005257689.1;XP_011523467.1 | 11989 | 126430 | . | . | 45.771118 | 45.814121 | 36.189022 | 38.695834 | 106.511265 | 105.901395 | 81.207111 | 88.473427 | -1.241320 | 7379.117875 | 0.104193308 | -2.99323393 | 0.002760381 | 0.141329972 | 12.996780 | 12.684858 | 12.937077 | 12.930616 | 12.725956 | 12.775095 | 9447 | 8787 | 5556 | 6356 |
| 7155 | NM_001068,N | TOP2B | DNA topoisomerase II beta | protein_coding | NP_001059.2;NP_001317629.1;XP_011532359.1 | 11990 | 126431 | . | . | 12.399720 | 11.690996 | 11.021927 | 9.967162 | 28.854658 | 27.024260 | 24.732887 | 22.788730 | -1.135524 | 1935.993213 | 0.125071454 | -1.46603085 | 0.142639883 | 0.903146012 | 11.007476 | 10.824438 | 10.999287 | 10.932919 | 10.902554 | 10.828208 | 2468 | 2130 | 1652 | 1619 |
| 7156 | NM_00132075 | TOP3A | DNA topoisomerase III alpha | protein_coding | NP_001307688.1;NP_004609.1;XP_011522303.1;XP_024306671.1 | 11992 | 601243 | . | . | 7.729579 | 7.888470 | 8.635419 | 9.267917 | 17.987047 | 18.234551 | 19.377628 | 21.189992 | 1.088610 | 1822.544979 | 0.120377912 | 1.017524205 | 0.308904123 | 0.998865404 | 10.769247 | 10.891635 | 10.797593 | 10.796612 | 10.845982 | 10.881632 | 2011 | 1882 | 1615 | 1821 |
| 7157 | NM_000546,N | TP53 | tumor protein p53 | protein_coding | NP_000537.3;NP_001119584.1;NP_001119585.1;NP_001119586.1;NP_0( | 11998 | 191170 | . | . | 6.046663 | 6.341264 | 7.662938 | 7.843639 | 14.070833 | 14.658118 | 17.195412 | 17.933548 | 1.234425 | 555.282178 | 0.161498198 | 1.881380268 | 0.059920209 | 0.719735192 | 8.957284 | 9.260924 | 9.029560 | 9.054900 | 9.173526 | 9.185192 | 561 | 547 | 529 | 580 |
| 7158 | NM_00114197 | TP53BP1 | tumor protein p53 binding protein 1 | protein_coding | NP_001135451.1;NP_001135452.1;NP_001341930.1;NP_005648.1;XP_01 | 11999 | 605230 | . | . | 8.103520 | 8.189339 | 8.594034 | 9.742706 | 18.857225 | 18.930027 | 19.284763 | 22.275542 | 1.101079 | 2754.557658 | 0.114574192 | 1.212467961 | 0.2253333 | 0.982008277 | 11.356528 | 11.495344 | 11.392082 | 11.381625 | 11.435481 | 11.494020 | 3039 | 2811 | 2421 | 2803 |
| 7159 | NM_00103168 | TP53BP2 | tumor protein p53 binding protein 2 | protein_coding | NP_001026855.2;NP_005417.1;XP_011542570.1;XP_011542571.1 | 12000 | 602143 | . | . | 8.868448 | 8.943977 | 9.012536 | 10.046778 | 20.637242 | 20.674403 | 20.223870 | 22.970765 | 1.065151 | 1338.411584 | 0.130268158 | 0.699004502 | 0.484549218 | 0.998865404 | 10.340170 | 10.431012 | 10.366078 | 10.356621 | 10.374016 | 10.443683 | 1502 | 1390 | 1146 | 1353 |
| 7161 | NM_00112624 | TP73 | tumor protein p73 | protein_coding | NP_001119712.1;NP_001119713.1;NP_001119714.1;NP_001191113.1;NI | 12003 | 601990 | . | . | 2.454920 | 2.905205 | 2.697593 | 2.867396 | 5.712700 | 6.715511 | 6.053320 | 6.555962 | 1.027113 | 440.400030 | 0.180515776 | 0.213803253 | 0.830700488 | 0.998865404 | 8.763588 | 8.801504 | 8.725110 | 8.819545 | 8.775388 | 8.803727 | 462 | 505 | 379 | 428 |
| 7162 | NM_00116639 | TPBG | trophoblast glycoprotein | protein_coding | NP_001159864.1;NP_001363851.1;NP_006661.1 | 12004 | 190920 | . | . | 4.727597 | 4.314017 | 4.905861 | 4.890390 | 11.001313 | 9.972044 | 11.008610 | 11.181298 | 1.069660 | 429.745828 | 0.179196604 | 0.542152973 | 0.587713129 | 0.998865404 | 8.697766 | 8.795266 | 8.761401 | 8.688518 | 8.762823 | 8.770302 | 507 | 421 | 384 | 419 |
| 7163 | NM_00102525 | TPD52 | tumor protein D52 | protein_coding | NP_001020423.1;NP_001020424.1;NP_001274069.1;NP_001274071.1;NI | 12005 | 604068 | . | . | 19.117609 | 14.874106 | 15.340707 | 14.568816 | 44.487466 | 34.382166 | 34.424100 | 33.309867 | -1.151912 | 2021.153796 | 0.143083495 | -1.42595661 | 0.153880867 | 0.920284923 | 11.079210 | 10.875535 | 11.131192 | 10.923183 | 10.944284 | 10.895136 | 2810 | 2036 | 1685 | 1706 |
| 7164 | NM_00100339 | TPD52L1 | TPD52 like 1 | protein_coding | NP_001003395.1;NP_001003396.1;NP_001003397.1;NP_001278955.1;NI | 12006 | 604069 | . | . | 22.431018 | 20.579523 | 21.655520 | 24.120919 | 52.197902 | 47.570485 | 48.594358 | 55.149619 | 1.089457 | 1218.038109 | 0.134325066 | 0.920224365 | 0.357455524 | 0.998865404 | 10.187113 | 10.310919 | 10.256651 | 10.174658 | 10.276528 | 10.285709 | 1416 | 1189 | 1098 | 1198 |
| 7165 | NM_00124389 | TPD52L2 | TPD52 like 2 | protein_coding | NP_001230820.1;NP_001230821.1;NP_001230823.1;NP_001230824.1;NI | 12007 | 603747 | . | . | 41.819548 | 44.057563 | 38.990216 | 45.258729 | 97.315805 | 101.841032 | 87.492921 | 103.478708 | -1.035227 | 3019.982316 | 0.120902111 | -0.41312378 | 0.679515925 | 0.998865404 | 11.585203 | 11.535010 | 11.557593 | 11.589643 | 11.484779 | 11.602502 | 3469 | 3380 | 2394 | 2983 |
| 7166 | NM_004179 | TPH1 | tryptophan hydroxylase 1 | protein_coding | NP_004170.1 | 12008 | 191060 | . | . | 0.020416 | 0.079313 | 0.075566 | 0.055034 | 0.047509 | 0.183336 | 0.169568 | 0.125828 | 1.294098 | 8.888633 | 1.06931756 | 0.347836139 | 0.727963248 | NA | 2.956191 | 3.324312 | 3.113114 | 3.153097 | 3.149931 | 3.137109 | 4 | 13 | 10 | 8 |
| 7167 | NM_000365,N | TPI1 | triosephosphate isomerase 1 | protein_coding | NP_000356.1;NP_001152759.1;NP_001244955.1 | 12009 | 190450 | . | . | 280.939669 | 239.024456 | 225.679979 | 277.158165 | 653.758188 | 552.515754 | 506.419312 | 633.689208 | 1.039609 | 12689.783789 | 0.114344165 | 0.490111605 | 0.624054926 | 0.998865404 | 13.603119 | 13.659098 | 13.614687 | 13.612627 | 13.571228 | 13.717366 | 14346 | 13411 | 10293 | 13158 |
| 7168 | NM_000366,N | TPM1 | tropomyosin 1 | protein_coding | NP_000357.3;NP_001018004.1;NP_001018005.1;NP_001018006.1;NP_0( | 12010 | 191010 | . | . | 17.294872 | 18.844790 | 10.592074 | 11.432065 | 40.245879 | 43.560577 | 23.768308 | 26.138060 | -1.653578 | 784.500836 | 0.148451883 | -4.88771983 | 1.02011E-06 | 0.000302657 | 9.933309 | 9.207649 | 9.731169 | 9.792317 | 9.418796 | 9.403397 | 1080 | 1098 | 520 | 548 |
| 7169 | NM_00130122 | TPM2 | tropomyosin 2 | protein_coding | NP_001288155.1;NP_001288156.1;NP_003280.2;NP_998839.1;XP_01687 | 12011 | 190990 | . | . | 39.704113 | 42.713203 | 50.527231 | 39.237569 | 92.393102 | 98.733483 | 113.381644 | 89.712032 | 1.065158 | 1678.262758 | 0.146955865 | 0.619694053 | 0.535459232 | 0.998865404 | 10.666288 | 10.757768 | 10.673010 | 10.696567 | 10.835447 | 10.622285 | 1843 | 1780 | 1709 | 1404 |
| 7170 | NM_00104335 | TPM3 | tropomyosin 3 | protein_coding | NP_001036816.1;NP_001036817.1;NP_001036818.1;NP_001265117.1;NI | 12012 | 191030 | . | . | 66.197832 | 65.795764 | 55.566401 | 64.568671 | 154.045064 | 152.089852 | 124.689397 | 147.628606 | -1.113345 | 6298.317443 | 0.112581052 | -1.37590412 | 0.168851321 | 0.945638095 | 12.696173 | 12.541167 | 12.668157 | 12.659587 | 12.515528 | 12.627967 | 7680 | 7125 | 4834 | 5964 |
| 7171 | NM_00114516 | TPM4 | tropomyosin 4 | protein_coding | NP_001138632.1;NP_001354765.1;NP_001354766.1;NP_001354767.1;NI | 12013 | 600317 | . | . | 88.743753 | 96.151347 | 62.181680 | 91.233921 | 206.510367 | 222.258140 | 139.533887 | 208.595516 | -1.216248 | 7437.840804 | 0.119970115 | -2.354236 | 0.018560823 | 0.457491143 | 12.995042 | 12.712481 | 12.938633 | 12.935232 | 12.687027 | 12.848715 | 9419 | 8792 | 5270 | 6903 |
| 7172 | NM_000367,N | TPMT | thiopurine S-methyltransferase | protein_coding | NP_000358.1;NP_001333746.1;NP_001333747.1 | 12014 | 187680 | . | . | 9.647449 | 9.308924 | 9.042340 | 8.654686 | 22.450007 | 21.517994 | 20.290747 | 19.787909 | -1.095403 | 918.603532 | 0.141118048 | -0.93156833 | 0.351559658 | 0.998865404 | 9.907379 | 9.776239 | 9.895780 | 9.854219 | 9.828429 | 9.790234 | 1137 | 1007 | 783 | 800 |
| 7174 | NM_00133058 | TPP2 | tripeptidyl peptidase 2 | protein_coding | NP_001317517.1;NP_001354876.1;NP_003282.2;XP_005254127.1 | 12016 | 190470 | . | . | 9.030938 | 9.813256 | 10.173913 | 8.734602 | 21.015364 | 22.683781 | 22.829970 | 19.970631 | -1.017458 | 1552.457350 | 0.134471302 | -0.18568033 | 0.852695456 | 0.998865404 | 10.612638 | 10.587932 | 10.581958 | 10.629491 | 10.652909 | 10.528646 | 1748 | 1741 | 1446 | 1327 |
| 7175 | NM_003292 | TPR | translocated promoter region, nuclear basket protein | protein_coding | NP_003283.2 | 12017 | 189940 | . | . | 15.305874 | 15.573758 | 16.189642 | 14.434197 | 35.617397 | 35.999443 | 36.329090 | 33.002079 | -1.022216 | 4617.194606 | 0.113652465 | -0.27891525 | 0.780309869 | 0.998865404 | 12.188501 | 12.156927 | 12.177680 | 12.185096 | 12.213787 | 12.109793 | 5351 | 5059 | 4219 | 4014 |
| 7178 | NM_00128627 | TPT1 | tumor protein, translationally-controlled 1 | protein_coding | NP_001273201.1;NP_001273202.1;NP_003286.1 | 12022 | 600763 | . | . | 146.104621 | 148.526397 | 144.410878 | 164.347317 | 339.991459 | 343.325443 | 324.053818 | 375.760597 | 1.032139 | 21988.686072 | 0.105813801 | 0.431300011 | 0.666250243 | 0.998865404 | 14.401490 | 14.447097 | 14.406821 | 14.413753 | 14.385844 | 14.486724 | 24819 | 23446 | 18289 | 22163 |
| 7179 | NM_00129022 | TPTE | transmembrane phosphatase with tensin homology | protein_coding | NP_001277153.2;NP_954868.2;NP_954869.2;NP_954870.3 | 12023 | 604336 | . | . | 0.773386 | 0.607435 | 1.374150 | 0.725735 | 1.799702 | 1.404112 | 3.083552 | 1.659306 | 1.490858 | 57.932820 | 0.452466665 | 1.273337111 | 0.202898443 | 0.968254058 | 5.537347 | 6.117362 | 5.814288 | 5.771104 | 5.945267 | 5.800014 | 60 | 44 | 79 | 45 |
| 7181 | NM_00103228 | NR2C1 | nuclear receptor subfamily 2 group C member 1 | protein_coding | NP_001027458.1;NP_001120834.1;NP_003288.2;XP_005269190.1;XP_00 | 7971 | 601529 | . | . | 7.721536 | 6.424065 | 7.894594 | 5.690060 | 17.968334 | 14.849514 | 17.715237 | 13.009646 | -1.061492 | 778.583885 | 0.171863716 | -0.50094389 | 0.616410612 | 0.998865404 | 9.646623 | 9.561540 | 9.660136 | 9.582770 | 9.676857 | 9.474092 | 973 | 818 | 753 | 605 |
| 7182 | NM_00129169 | NR2C2 | nuclear receptor subfamily 2 group C member 2 | protein_coding | NP_001278623.1;NP_003289.2;XP_011532360.1;XP_011532361.1;XP_01 | 7972 | 601426 | . | . | 3.726781 | 3.239651 | 3.382028 | 3.319208 | 8.672372 | 7.488598 | 7.589172 | 7.588973 | -1.057819 | 902.365346 | 0.150810555 | -0.5377169 | 0.590772508 | 0.998865404 | 9.857305 | 9.776705 | 9.900099 | 9.768732 | 9.809358 | 9.782747 | 1164 | 911 | 777 | 807 |
| 7184 | NM_003299 | HSP90B1 | heat shock protein 90 beta family member 1 | protein_coding | NP_003290.1 | 12028 | 191175 | . | . | 579.286133 | 575.118103 | 519.466309 | 508.687256 | 1348.022705 | 1329.411255 | 1165.667358 | 1163.052856 | -1.138512 | 47309.955481 | 0.097498481 | -1.91951691 | 0.054918948 | 0.695455271 | 15.620396 | 15.433252 | 15.590199 | 15.576424 | 15.484480 | 15.457858 | 58463 | 53927 | 39075 | 40834 |
| 7185 | NM_00119094 | TRAF1 | TNF receptor associated factor 1 | protein_coding | NP_001177874.1;NP_001177876.1;NP_005649.1 | 12031 | 601711 | . | . | 1.066314 | 1.022376 | 0.811870 | 0.865309 | 2.481355 | 2.363267 | 1.821812 | 1.978426 | -1.254389 | 128.398160 | 0.286568565 | -1.14103601 | 0.253854937 | 0.994807659 | 7.158753 | 6.831714 | 7.054773 | 7.033450 | 6.957538 | 6.970340 | 169 | 150 | 97 | 109 |
| 7186 | NM_021138,XI | TRAF2 | TNF receptor associated factor 2 | protein_coding | NP_066961.2;XP_011517276.1;XP_011517278.1;XP_011517279.1 | 12032 | 601895 | . | . | 10.011679 | 10.013193 | 10.890733 | 9.830186 | 23.297589 | 23.145942 | 24.438490 | 22.475552 | 1.009465 | 762.540697 | 0.152565741 | 0.089079239 | 0.929018936 | 0.998865404 | 9.567589 | 9.581717 | 9.576301 | 9.564677 | 9.625815 | 9.526216 | 881 | 812 | 714 | 667 |
| 7187 | NM_00119942 | TRAF3 | TNF receptor associated factor 3 | protein_coding | NP_001186356.1;NP_003291.2;NP_663777.1;NP_663778.1;XP_01153541 | 12033 | 601896 | . | . | 3.956309 | 3.906699 | 3.954686 | 4.717331 | 9.206493 | 9.030509 | 8.874202 | 10.785616 | 1.089626 | 1023.186047 | 0.142041472 | 0.871808479 | 0.383312868 | 0.998865404 | 9.935766 | 10.059296 | 9.979823 | 9.951135 | 9.977123 | 10.077663 | 1150 | 1036 | 865 | 1068 |
| 7188 | NM_00103391 | TRAF5 | TNF receptor associated factor 5 | protein_coding | NP_001029082.1;NP_001306136.1;NP_004610.1;NP_665702.1;XP_0115( | 12035 | 602356 | . | . | 5.897498 | 5.563511 | 6.057526 | 5.606837 | 13.723720 | 12.860306 | 13.592914 | 12.819366 | 1.025809 | 658.119943 | 0.154599759 | 0.237787579 | 0.812045853 | 0.998865404 | 9.343513 | 9.380661 | 9.366542 | 9.339447 | 9.393898 | 9.346455 | 763 | 687 | 600 | 603 |
| 7189 | NM_004620,N | TRAF6 | TNF receptor associated factor 6 | protein_coding | NP_004611.1;NP_665802.1;XP_016873709.1 | 12036 | 602355 | . | . | 0.908039 | 0.920636 | 0.853255 | 0.992948 | 2.113046 | 2.128092 | 1.914679 | 2.270257 | 1.013561 | 221.608704 | 0.228576484 | 0.085017629 | 0.932247375 | 0.998865404 | 7.782472 | 7.801209 | 7.785483 | 7.791246 | 7.765339 | 7.823183 | 252 | 239 | 182 | 222 |
| 7190 | NR_073383 | HSP90B2P | heat shock protein 90 beta family member 2, pseudogene | transcribed_ps | . | 12099 | . | . | . | 0.458885 | 0.309928 | 0.203584 | 0.135162 | 1.067844 | 0.716413 | 0.456836 | 0.309032 | -2.225565 | 24.239214 | 0.658644283 | -1.75234477 | 0.079714543 | NA | 5.062293 | 3.913269 | 4.664707 | 4.610648 | 4.575883 | 4.547100 | 46 | 29 | 16 | 11 |
| 7203 | NM_00100880 | CCT3 | chaperonin containing TCP1 subunit 3 | protein_coding | NP_001008800.1;NP_005989.3 | 1616 | 600114 | . | . | 135.206842 | 137.120471 | 117.951142 | 140.331336 | 314.631862 | 316.960111 | 264.678934 | 320.850918 | -1.069623 | 8181.808402 | 0.11489975 | -0.84510773 | 0.398050692 | 0.998865404 | 13.045986 | 12.948785 | 13.023486 | 13.027251 | 12.897193 | 13.034589 | 9717 | 9146 | 6316 | 8015 |
| 7204 | NM_007118,N | TRIO | trio Rho guanine nucleotide exchange factor | protein_coding | NP_009049.2;XP_011512409.1;XP_011512410.1;XP_011512411.1;XP_01: | 12303 | 601893 | . | . | 6.790540 | 6.842571 | 6.298797 | 6.972507 | 15.801865 | 15.816909 | 14.134317 | 15.941808 | 1.011984 | 2442.485127 | 0.114209624 | 0.150482958 | 0.880383595 | 0.998865404 | 11.245536 | 11.262681 | 11.250792 | 11.247780 | 11.249185 | 11.268494 | 2801 | 2615 | 2111 | 2331 |
| 7205 | NM_003302 | TRIP6 | thyroid hormone receptor interactor 6 | protein_coding | NP_003293.2 | 12311 | 602933 | . | . | 29.768810 | 32.252186 | 28.382017 | 30.924946 | 69.273247 | 74.552376 | 63.688423 | 70.706215 | -1.060231 | 1600.304554 | 0.126922323 | -0.66480664 | 0.506174177 | 0.998865404 | 10.685851 | 10.601168 | 10.637805 | 10.693946 | 10.591339 | 10.649172 | 1829 | 1841 | 1300 | 1511 |
| 7220 | NM_00125184 | TRPC1 | transient receptor potential cation channel subfamily C member 1 | protein_coding | NP_001238774.1;NP_003295.1;XP_005247795.1;XP_005247796.1;XP_01 | 12333 | 602343 | . | . | 0.906007 | 1.387106 | 1.586516 | 0.879572 | 2.108317 | 3.206358 | 3.560094 | 2.011036 | -1.070999 | 177.695384 | 0.295381307 | -0.33501481 | 0.737613912 | 0.998865404 | 7.520893 | 7.424006 | 7.456525 | 7.500168 | 7.560943 | 7.323807 | 201 | 208 | 192 | 115 |
| 7222 | NM_00113069 | TRPC3 | transient receptor potential cation channel subfamily C member 3 | protein_coding | NP_001124170.1;NP_001353408.1;NP_003296.1;XP_011530519.1;XP_01 | 12335 | 602345 | . | . | 0.022904 | 0.014721 | 0.016346 | 0.016338 | 0.053299 | 0.034028 | 0.036679 | 0.037355 | -1.209732 | 4.894846 | 1.359067692 | -0.20211421 | 0.839827441 | NA | 2.421266 | 2.148375 | 2.300838 | 2.294032 | 2.293956 | 2.292427 | 7 | 5 | 4 | 4 |
| 7226 | NM_00132035 | TRPM2 | transient receptor potential cation channel subfamily M member : | protein_coding | NP_001307279.1;NP_001307280.1;NP_001307281.1;NP_003298.1;XP_0( | 12339 | 603749 | . | . | 5.728061 | 3.991089 | 4.861160 | 7.999730 | 13.329434 | 9.225581 | 10.908302 | 18.290435 | 1.068600 | 757.027711 | 0.157223654 | 0.608826605 | 0.542639377 | 0.998865404 | 9.515940 | 9.610903 | 9.498855 | 9.578136 | 9.531377 | 9.636705 | 795 | 835 | 629 | 788 |
| 7227 | NM_00128290 | TRPS1 | transcriptional repressor GATA binding 1 | protein_coding | NP_001269831.1;NP_001269832.1;NP_001317528.1;NP_054831.2 | 12340 | 604386 | . | . | 0.622967 | 0.521788 | 0.615844 | 0.636280 | 1.449669 | 1.206136 | 1.381937 | 1.454778 | 1.099899 | 182.680708 | 0.24892148 | 0.551863616 | 0.581041798 | 0.998865404 | 7.442572 | 7.580494 | 7.523266 | 7.456652 | 7.527743 | 7.535562 | 216 | 173 | 165 | 181 |
| 7247 | NM_00126140 | TSN | translin | protein_coding | NP_001248330.1;NP_004613.1 | 12379 | 600575 | . | . | 43.147092 | 40.816524 | 43.728774 | 40.602066 | 100.405070 | 94.349220 | 98.126104 | 92.831798 | -1.009772 | 4330.117075 | 0.112085654 | -0.1251692 | 0.900389601 | 0.998865404 | 12.087115 | 12.073233 | 12.110149 | 12.056872 | 12.109991 | 12.040480 | 5170 | 4545 | 3904 | 3871 |
| 7248 | NM_000368,N | TSC1 | TSC complex subunit 1 | protein_coding | NP_000359.1;NP_001155898.1;NP_001155899.1;NP_001349106.1;XP_0( | 12362 | 605284 | . | . | 2.804648 | 3.239457 | 3.417301 | 3.351205 | 6.526533 | 7.488152 | 7.668323 | 7.662132 | 1.101591 | 862.550876 | 0.147915568 | 0.943703918 | 0.345320971 | 0.998865404 | 9.681112 | 9.820456 | 9.665466 | 9.763497 | 9.795436 | 9.774002 | 880 | 947 | 798 | 835 |
| 7249 | NM_000548,N | TSC2 | TSC complex subunit 2 | protein_coding | NP_000539.2;NP_001070651.1;NP_001107854.1;NP_001305756.1;NP_0( | 12363 | 191092 | . | . | 7.867604 | 8.232616 | 8.878105 | 8.026290 | 18.308237 | 19.030058 | 19.922213 | 18.351162 | 1.024320 | 1438.233142 | 0.130554694 | 0.265531949 | 0.790599701 | 0.998865404 | 10.472547 | 10.507406 | 10.463041 | 10.497016 | 10.539916 | 10.455783 | 1600 | 1567 | 1335 | 1290 |
| 7251 | NM_006292,XI | TSG101 | tumor susceptibility 101 | protein_coding | NP_006283.1;XP_005253165.1 | 15971 | 601387 | . | . | 26.453675 | 29.014932 | 25.705819 | 27.151474 | 61.558790 | 67.069314 | 57.683113 | 62.078616 | -1.063746 | 1306.488623 | 0.129954886 | -0.68603502 | 0.492691025 | 0.998865404 | 10.395471 | 10.306103 | 10.347359 | 10.400851 | 10.317204 | 10.337508 | 1497 | 1504 | 1086 | 1203 |
| 7257 | NM_005999 | TSNAX | translin associated factor X | protein_coding | NP_005990.1 | 12380 | 602964 | . | . | 16.246273 | 13.055734 | 13.473065 | 14.917642 | 37.805748 | 30.178915 | 30.233171 | 34.107414 | -1.047712 | 1184.322558 | 0.150532148 | -0.44670095 | 0.655090998 | 0.998865404 | 10.243004 | 10.175909 | 10.307722 | 10.135914 | 10.155923 | 10.223798 | 1553 | 1160 | 960 | 1134 |
| 7259 | NM_003309 | TSPYL1 | TSPY like 1 | protein_coding | NP_003300.1 | 12382 | 604714 | . | . | 6.280746 | 5.336377 | 5.829360 | 6.886117 | 14.615554 | 12.335275 | 13.080914 | 15.744289 | 1.078504 | 961.304217 | 0.154258941 | 0.706805233 | 0.479687522 | 0.998865404 | 9.853338 | 9.962303 | 9.938759 | 9.815181 | 9.875146 | 9.989438 | 1156 | 913 | 800 | 1008 |
| 7260 | NM_00133053 | EIPR1 | EARP complex and GARP complex interacting protein 1 | protein_coding | NP_001317459.1;NP_001317460.1;NP_003301.1;XP_006711956.1;XP_01 | 12383 | 608998 | . | . | 14.854532 | 13.941065 | 13.038057 | 15.153552 | 34.567108 | 32.225398 | 29.257023 | 34.646794 | -1.037070 | 741.036069 | 0.154548713 | -0.33978789 | 0.73401627 | 0.998865404 | 9.559577 | 9.506739 | 9.568778 | 9.521676 | 9.471698 | 9.566355 | 898 | 787 | 589 | 729 |
| 7262 | NM_003311 | PHLDA2 | pleckstrin homology like domain family A member 2 | protein_coding | NP_003302.1 | 12385 | 602131 | . | . | 26.738029 | 26.108582 | 21.431753 | 23.514954 | 62.220493 | 60.351154 | 48.092232 | 53.764145 | -1.191418 | 701.733804 | 0.152737466 | -1.65922797 | 0.097069864 | 0.813553488 | 9.575711 | 9.322777 | 9.523425 | 9.500106 | 9.364154 | 9.418228 | 893 | 810 | 534 | 625 |
| 7263 | NM_00127048 | TST | thiosulfate sulfurtransferase | protein_coding | NP_001257412.1;NP_003303.2 | 12388 | 180370 | . | . | 6.691295 | 7.084888 | 6.876263 | 9.479882 | 15.570919 | 16.377037 | 15.430134 | 21.674625 | 1.171063 | 264.318887 | 0.228326819 | 0.997772317 | 0.318389777 | 0.998865404 | 7.928511 | 8.154889 | 7.987994 | 8.011427 | 7.996716 | 8.157754 | 274 | 269 | 210 | 308 |
| 7264 | NM_00131778 | GFUS | GDP-L-fucose synthase | protein_coding | NP_001304712.1;NP_003304.1;XP_005251108.2;XP_011515571.1 | 12390 | 137020 | . | . | 25.844992 | 26.036240 | 29.435688 | 28.746104 | 60.142360 | 60.183935 | 66.052835 | 65.724554 | 1.103789 | 1154.510295 | 0.13162988 | 1.082312085 | 0.279113878 | 0.998865404 | 10.100028 | 10.242583 | 10.134004 | 10.135000 | 10.220093 | 10.196566 | 1263 | 1185 | 1070 | 1118 |
| 7265 | NM_00128250 | TTC1 | tetratricopeptide repeat domain 1 | protein_coding | NP_001269429.1;NP_003305.1 | 12391 | 601963 | . | . | 31.239872 | 32.639980 | 26.935987 | 35.159344 | 72.696463 | 75.448767 | 60.443574 | 80.387664 | -1.041847 | 1419.994437 | 0.147890742 | -0.39991145 | 0.689221742 | 0.998865404 | 10.501217 | 10.441504 | 10.471446 | 10.499224 | 10.351687 | 10.546605 | 1639 | 1592 | 1055 | 1470 |
| 7266 | NM_00114476 | DNAJC7 | DnaJ heat shock protein family (Hsp40) member C7 | protein_coding | NP_001138238.1;NP_003306.3;XP_011523469.1;XP_011523470.1;XP_01 | 12392 | 601964 | . | . | 34.690014 | 37.395499 | 34.488518 | 35.493595 | 80.725088 | 86.441365 | 77.391233 | 81.151889 | -1.033257 | 2046.756771 | 0.118400186 | -0.39864634 | 0.690153815 | 0.998865404 | 11.022595 | 10.975265 | 10.994645 | 11.029049 | 10.974257 | 10.997407 | 2344 | 2293 | 1725 | 1915 |
| 7267 | NM_00100189 | TTC3 | tetratricopeptide repeat domain 3 | protein_coding | NP_001001894.1;NP_001307632.1;NP_001307633.1;NP_001317610.1;NI | 12393 | 602259 | . | . | 13.867507 | 14.736568 | 17.604508 | 15.001342 | 32.270256 | 34.064237 | 39.504004 | 34.298785 | 1.076352 | 3693.495149 | 0.115734131 | 0.917182482 | 0.359047025 | 0.998865404 | 11.796642 | 11.902942 | 11.818662 | 11.819247 | 11.930510 | 11.826582 | 4095 | 3840 | 3538 | 3366 |
| 7268 | NM_00129133 | TTC4 | tetratricopeptide repeat domain 4 | protein_coding | NP_001278262.1;NP_004614.3 | 12394 | 606753 | . | . | 11.396208 | 11.702611 | 11.738813 | 13.399165 | 26.519441 | 27.051107 | 26.341558 | 30.635597 | 1.074607 | 883.777319 | 0.14534376 | 0.714228547 | 0.475085767 | 0.998865404 | 9.734931 | 9.838296 | 9.752220 | 9.768619 | 9.767411 | 9.854732 | 970 | 930 | 748 | 910 |
| 7270 | NM_00120529 | TTF1 | transcription termination factor 1 | protein_coding | NP_001192225.1;NP_031370.2;XP_006717336.2 | 12397 | 600777 | . | . | 5.119865 | 5.494845 | 4.144528 | 5.163989 | 11.914138 | 12.701580 | 9.300201 | 11.806849 | -1.112815 | 474.360905 | 0.178792441 | -0.86252604 | 0.38839811 | 0.998865404 | 8.965331 | 8.810182 | 8.905431 | 8.937798 | 8.796870 | 8.909631 | 561 | 553 | 354 | 460 |
| 7272 | NM_00116669 | TTK | TTK protein kinase | protein_coding | NP_001160163.1;NP_003309.2;XP_011534401.1;XP_011534402.1;XP_01 | 12401 | 604092 | . | . | 10.190487 | 9.851283 | 8.696224 | 6.984743 | 23.713687 | 22.771682 | 19.514075 | 15.969785 | -1.320129 | 843.204104 | 0.159077915 | -2.51875633 | 0.011777013 | 0.354930366 | 9.905864 | 9.505992 | 9.810375 | 9.805587 | 9.688950 | 9.526072 | 1108 | 1032 | 708 | 600 |
| 7273 | NM_00125685 | TTN | titin | protein_coding | NP_001243779.1;NP_001254479.2;NP_003310.4;NP_596869.4;NP_5968( | 12403 | 188840 | . | . | 0.044840 | 0.071874 | 0.085595 | 0.047408 | 0.104343 | 0.166141 | 0.192074 | 0.108392 | 1.457703 | 105.422843 | 0.320748206 | 1.695089514 | 0.090058441 | 0.804059937 | 6.422299 | 6.966728 | 6.631538 | 6.659140 | 6.805022 | 6.720683 | 94 | 97 | 122 | 103 |
| 7274 | NM_000370,XI | TTPA | alpha tocopherol transfer protein | protein_coding | NP_000361.1;XP_006716531.1 | 12404 | 600415 | . | . | 0.232608 | 0.282600 | 0.311945 | 0.273289 | 0.541289 | 0.653243 | 0.699996 | 0.624843 | 1.108381 | 23.229737 | 0.639729153 | 0.232057639 | 0.816493246 | NA | 4.461710 | 4.610270 | 4.520287 | 4.538123 | 4.547339 | 4.531018 | 23 | 26 | 23 | 21 |
| 7275 | NM_003320,N | TUB | TUB bipartite transcription factor | protein_coding | NP_003311.2;NP_813977.1;XP_005253166.1;XP_011518646.1 | 12406 | 601197 | . | . | 6.072034 | 6.356448 | 6.277891 | 6.114164 | 14.129874 | 14.693213 | 14.087404 | 13.979312 | -1.016759 | 1245.735714 | 0.130066271 | -0.18434492 | 0.853742873 | 0.998865404 | 10.294709 | 10.270756 | 10.274474 | 10.303226 | 10.289417 | 10.263209 | 1419 | 1381 | 1093 | 1138 |
| 7280 | NM_001069,N | TUBB2A | tubulin beta 2A class IIa | protein_coding | NP_001060.1;NP_001297244.1 | 12412 | 615101 | . | . | 90.750062 | 90.669123 | 74.941868 | 82.231134 | 211.179131 | 209.585723 | 168.167377 | 188.011717 | -1.168998 | 4275.702908 | 0.109934793 | -2.04914696 | 0.040447747 | 0.628009277 | 12.170235 | 11.944865 | 12.127133 | 12.119508 | 11.960121 | 12.025600 | 5331 | 4951 | 3291 | 3844 |
| 7283 | NM_001070,XI | TUBG1 | tubulin gamma 1 | protein_coding | NP_001061.2;XP_024306672.1 | 12417 | 191135 | . | . | 70.329838 | 77.327183 | 65.340628 | 74.042421 | 163.660420 | 178.745257 | 146.622480 | 169.289198 | -1.074288 | 3654.531215 | 0.117605136 | -0.87904904 | 0.379374689 | 0.998865404 | 11.886338 | 11.782745 | 11.826466 | 11.899893 | 11.759312 | 11.847967 | 4167 | 4265 | 2898 | 3482 |
| 7284 | NM_00136536 | TUFM | Tu translation elongation factor, mitochondrial | protein_coding | NP_001352289.1;NP_003312.3 | 12420 | 602389 | . | . | 133.829178 | 133.947921 | 129.450780 | 135.939097 | 311.426005 | 309.626603 | 290.483791 | 310.808572 | -1.023634 | 8338.837436 | 0.102353752 | -0.32924504 | 0.741970484 | 0.998865404 | 13.042394 | 13.008670 | 13.039002 | 13.031862 | 12.998605 | 13.032301 | 9755 | 9065 | 7024 | 7880 |
| 7286 | NM_00112633 | TUFT1 | tuftelin 1 | protein_coding | NP_001119809.1;NP_001288246.1;NP_064512.1;XP_011508263.1;XP_01 | 12422 | 600087 | . | . | 1.903834 | 2.287259 | 2.162661 | 2.261980 | 4.430301 | 5.287100 | 4.852952 | 5.171748 | 1.055408 | 207.344265 | 0.236986632 | 0.328294513 | 0.742688993 | 0.998865404 | 7.656895 | 7.733847 | 7.644857 | 7.717605 | 7.696242 | 7.716794 | 214 | 235 | 181 | 204 |
| 7289 | NM_00116040 | TULP3 | TUB like protein 3 | protein_coding | NP_001153880.1;NP_003315.2 | 12425 | 604730 | . | . | 11.011661 | 11.897676 | 11.037743 | 12.299760 | 25.624590 | 27.502010 | 24.768375 | 28.121943 | 1.005760 | 1099.119366 | 0.138638896 | 0.059769236 | 0.952339428 | 0.998865404 | 10.098208 | 10.106046 | 10.080993 | 10.117846 | 10.059207 | 10.146369 | 1231 | 1212 | 902 | 1094 |
| 7290 | NM_003325 | HIRA | histone cell cycle regulator | protein_coding | NP_003316.3 | 4916 | 600237 | . | . | 10.114735 | 10.233225 | 9.643631 | 10.602941 | 23.537405 | 23.654558 | 21.640028 | 24.242363 | -1.019384 | 1281.505250 | 0.130962504 | -0.21149578 | 0.832500426 | 0.998865404 | 10.337528 | 10.309584 | 10.329791 | 10.331750 | 10.283814 | 10.347454 | 1488 | 1398 | 1057 | 1240 |
| 7291 | NM_000474,N | TWIST1 | twist family bHLH transcription factor 1 | protein_coding | NP_000465.1 | 12428 | 601622 | . | . | 6.133001 | 6.549146 | 6.167801 | 6.587079 | 14.271745 | 15.138644 | 13.840366 | 15.060573 | -1.025326 | 306.544624 | 0.201424433 | -0.17913599 | 0.857830924 | 0.998865404 | 8.278220 | 8.241452 | 8.255078 | 8.278079 | 8.223964 | 8.281239 | 350 | 342 | 249 | 299 |
| 7292 | NM_00129756 | TNFSF4 | TNF superfamily member 4 | protein_coding | NP_001284491.1;NP_003317.1;XP_011508266.2;XP_016857717.1;XP_01 | 11934 | 603594 | . | . | 0.218462 | 0.193170 | 0.251700 | 0.258296 | 0.508368 | 0.446523 | 0.564807 | 0.590564 | 1.168496 | 70.371763 | 0.376268588 | 0.597054857 | 0.550470756 | 0.998865404 | 6.019836 | 6.245220 | 6.127811 | 6.101657 | 6.155823 | 6.148317 | 79 | 66 | 67 | 70 |
| 7294 | NM_003328,XI | TXK | TXK tyrosine kinase | protein_coding | NP_003319.2;XP_011512049.1;XP_011512050.1;XP_016864070.1;XP_01( | 12434 | 600058 | . | . | 0.115036 | 0.184096 | 0.110328 | 0.128867 | 0.267694 | 0.425545 | 0.247573 | 0.294639 | -1.180038 | 28.302668 | 0.589459987 | -0.40517253 | 0.685350697 | 0.998865404 | 4.938471 | 4.697185 | 4.809715 | 4.854852 | 4.806446 | 4.820621 | 29 | 39 | 21 | 26 |
| 7295 | NM_00124493 | TXN | thioredoxin | protein_coding | NP_001231867.1;NP_003320.2 | 12435 | 187700 | . | . | 190.215424 | 181.057541 | 223.101678 | 217.704312 | 442.639069 | 418.522644 | 500.633724 | 497.755071 | 1.170691 | 4659.212164 | 0.108128668 | 2.102687805 | 0.035493069 | 0.586017737 | 12.067669 | 12.295117 | 12.139700 | 12.092098 | 12.262325 | 12.233051 | 5086 | 4498 | 4446 | 4630 |
| 7296 | NM_00109377 | TXNRD1 | thioredoxin reductase 1 | protein_coding | NP_001087240.1;NP_001248374.1;NP_001248375.1;NP_003321.3;NP_8( | 12437 | 601112 | . | . | 126.069184 | 131.465240 | 148.179164 | 157.939745 | 293.368174 | 303.887787 | 332.509725 | 361.110469 | 1.174912 | 16283.246365 | 0.100993832 | 2.302640508 | 0.021299078 | 0.484334718 | 13.870159 | 14.102684 | 13.903304 | 13.932440 | 14.033643 | 14.078157 | 16945 | 16436 | 14906 | 16916 |
| 7297 | NM_003331,XI | TYK2 | tyrosine kinase 2 | protein_coding | NP_003322.3;XP_011526547.1;XP_011526548.1;XP_011526549.1;XP_01: | 12440 | 176941 | . | . | 17.681546 | 18.914817 | 21.717735 | 19.062350 | 41.145686 | 43.722452 | 48.733968 | 43.583796 | 1.099652 | 2613.161611 | 0.125311736 | 1.09365061 | 0.274108243 | 0.998865404 | 11.281349 | 11.418552 | 11.284479 | 11.336199 | 11.449949 | 11.321586 | 2774 | 2772 | 2567 | 2365 |
| 7298 | NM_001071,N | TYMS | thymidylate synthetase | protein_coding | NP_001062.1;NP_001341796.1;NP_001341797.1;XP_024307010.1 | 12441 | 188350 | . | . | 115.865907 | 118.280055 | 90.025402 | 107.217013 | 269.624726 | 273.409651 | 202.014398 | 245.138946 | -1.202443 | 5378.240349 | 0.11801737 | -2.25364066 | 0.024218783 | 0.514295174 | 12.519854 | 12.253727 | 12.461280 | 12.467950 | 12.238032 | 12.376962 | 6736 | 6362 | 3889 | 4964 |
| 7299 | NM_000372,XI | TYR | tyrosinase | protein_coding | NP_000363.1;XP_011541272.1 | 12442 | 606933 | . | . | 104.090143 | 106.969473 | 113.858043 | 106.523588 | 242.222037 | 247.264723 | 255.494129 | 243.553528 | 1.030405 | 6891.582419 | 0.106090986 | 0.40730406 | 0.683784681 | 0.998865404 | 12.728826 | 12.772088 | 12.729942 | 12.745189 | 12.795186 | 12.729636 | 7746 | 7391 | 6318 | 6304 |
| 7300 | gene-TYRL | TYRL | tyrosinase like (pseudogene) | pseudogene | . | 12443 | 191270 | . | . | 3.125313 | 3.317991 | 4.473367 | 3.213604 | 7.272732 | 7.669684 | 10.038105 | 7.347523 | 1.172067 | 89.231989 | 0.349361634 | 0.655638856 | 0.512056519 | 0.998865404 | 6.359578 | 6.590194 | 6.442434 | 6.457397 | 6.543385 | 6.443861 | 92 | 91 | 98 | 75 |
| 7301 | NM_00133026 | TYRO3 | TYRO3 protein tyrosine kinase | protein_coding | NP_001317193.1;NP_006284.2;XP_016878032.1 | 12446 | 600341 | . | . | 8.617129 | 8.197445 | 7.923896 | 8.832346 | 20.052414 | 18.948760 | 17.780992 | 20.194109 | -1.003182 | 2005.319353 | 0.122985464 | -0.03726811 | 0.970271236 | 0.998865404 | 10.971987 | 10.967241 | 10.985317 | 10.955606 | 10.925270 | 11.008998 | 2356 | 2127 | 1649 | 1976 |
| 7306 | NM_000550,XI | TYRP1 | tyrosinase related protein 1 | protein_coding | NP_000541.1 | 12450 | 115501 | . | . | 16.705351 | 14.564483 | 19.895872 | 20.393760 | 38.874043 | 33.666456 | 44.645757 | 46.627910 | 1.282742 | 1600.532903 | 0.129034135 | 2.783999541 | 0.005369311 | 0.223905136 | 10.453461 | 10.812884 | 10.584377 | 10.489458 | 10.723352 | 10.740191 | 1716 | 1418 | 1548 | 1704 |
| 7307 | NM_00102520 | U2AF1 | U2 small nuclear RNA auxiliary factor 1 | protein_coding | NP_001020374.1;NP_001020375.1;NP_006749.1;XP_011528045.1;XP_01 | 12453 | 191317 | . | . | 22.552987 | 20.998410 | 19.831654 | 21.733999 | 52.481728 | 48.538763 | 44.501660 | 49.692204 | -1.064581 | 652.775004 | 0.155173704 | -0.58183458 | 0.560678108 | 0.998865404 | 9.394924 | 9.304545 | 9.390493 | 9.351744 | 9.308426 | 9.349099 | 797 | 706 | 533 | 611 |
| 7310 | gene-ZRSR2P1 | ZRSR2P1 | ZRSR2 pseudogene 1 | pseudogene | . | 12456 | 601079 | . | . | 0.117709 | 0.193120 | 0.076686 | 0.103107 | 0.273913 | 0.446405 | 0.172081 | 0.235743 | -1.541851 | 6.319944 | 1.214227649 | -0.51445308 | 0.606935259 | NA | 2.939842 | 2.312263 | 2.664480 | 2.679125 | 2.658099 | 2.661217 | 7 | 10 | 4 | 5 |
| 7311 | NM_00103393 | UBA52 | ubiquitin A-52 residue ribosomal protein fusion product 1 | protein_coding | NP_001029102.1;NP_001307946.1;NP_001307947.1;NP_001307948.1;NI | 12458 | 191321 | . | . | 128.820788 | 113.240914 | 102.731852 | 141.116386 | 299.771265 | 261.761452 | 230.527304 | 322.645848 | -1.024766 | 8691.256677 | 0.119029632 | -0.29651802 | 0.766834506 | 0.998865404 | 13.102934 | 13.067548 | 13.102749 | 13.086349 | 12.989258 | 13.151826 | 10228 | 9402 | 6751 | 8818 |
| 7314 | NM_00128171 | UBB | ubiquitin B | protein_coding | NP_001268645.1;NP_001268646.1;NP_001268647.1;NP_001268648.1;NI | 12463 | 191339 | . | . | 612.908595 | 636.656933 | 527.953596 | 649.955426 | 1426.263536 | 1471.661047 | 1184.712581 | 1486.045954 | -1.075934 | 17760.249897 | 0.117700001 | -0.89710525 | 0.369662755 | 0.998865404 | 14.168220 | 14.062575 | 14.133060 | 14.158527 | 13.996130 | 14.163016 | 20887 | 20158 | 13428 | 17605 |
| 7315 | gene-UBBP1 | UBBP1 | ubiquitin B pseudogene 1 | pseudogene | . | 12464 | . | . | . | 0.024116 | 0.077525 | 0.150480 | 0.089034 | 0.056119 | 0.179202 | 0.337673 | 0.203566 | 2.054720 | 2.076297 | 2.080611993 | 0.499344301 | 0.617536851 | NA | 0.444631 | 1.481107 | 1.028646 | 1.033470 | 1.040700 | 1.034794 | 1 | 2 | 3 | 2 |
| 7316 | NM_021009 | UBC | ubiquitin C | protein_coding | NP_066289.3 | 12468 | 191340 | . | . | 425.878448 | 433.646210 | 452.512268 | 491.238495 | 991.036682 | 1002.392639 | 1015.424377 | 1123.158325 | 1.082607 | 30785.256529 | 0.100538935 | 1.138958723 | 0.254720371 | 0.995760118 | 14.851569 | 14.966064 | 14.870510 | 14.878666 | 14.911386 | 14.973773 | 33881 | 32053 | 26832 | 31084 |
| 7317 | NM_003334,N | UBA1 | ubiquitin like modifier activating enzyme 1 | protein_coding | NP_003325.2;NP_695012.1;XP_005272706.1;XP_011542256.1;XP_01688 | 12469 | 314370 | . | . | 62.345872 | 62.857128 | 58.892185 | 67.235994 | 145.081400 | 145.297063 | 132.152358 | 153.727109 | -1.003441 | 7202.293944 | 0.108702547 | -0.04558401 | 0.963641818 | 0.998865404 | 12.816756 | 12.811721 | 12.815380 | 12.815343 | 12.765387 | 12.857405 | 8308 | 7785 | 5912 | 7107 |
| 7318 | NM_003335,XI | UBA7 | ubiquitin like modifier activating enzyme 7 | protein_coding | NP_003326.2;XP_005265487.1;XP_006713384.1;XP_011532372.1;XP_01: | 12471 | 191325 | . | . | 0.223553 | 0.119152 | 0.271089 | 0.130737 | 0.520217 | 0.275425 | 0.608315 | 0.298915 | 1.037896 | 20.874301 | 0.714696103 | 0.075083873 | 0.940147979 | NA | 4.353365 | 4.413324 | 4.398456 | 4.349722 | 4.407674 | 4.346513 | 30 | 16 | 25 | 13 |
| 7319 | NM_00128216 | UBE2A | ubiquitin conjugating enzyme E2 A | protein_coding | NP_001269090.1;NP_003327.2;NP_861427.1 | 12472 | 312180 | . | . | 16.530594 | 14.689287 | 16.458854 | 14.711865 | 38.467370 | 33.954944 | 36.933191 | 33.636929 | -1.016089 | 863.061525 | 0.151214761 | -0.15227821 | 0.878967514 | 0.998865404 | 9.764456 | 9.742096 | 9.802620 | 9.711248 | 9.789028 | 9.702610 | 1065 | 879 | 790 | 754 |
| 7320 | NM_003337 | UBE2B | ubiquitin conjugating enzyme E2 B | protein_coding | NP_003328.1 | 12473 | 179095 | . | . | 10.634159 | 9.573592 | 9.167833 | 8.513153 | 24.746122 | 22.129789 | 20.572351 | 19.464314 | -1.159442 | 662.321211 | 0.158470488 | -1.34681746 | 0.17803903 | 0.951511445 | 9.473874 | 9.261060 | 9.457691 | 9.378590 | 9.346128 | 9.290551 | 865 | 724 | 556 | 551 |
| 7321 | NM_00120488 | UBE2D1 | ubiquitin conjugating enzyme E2 D1 | protein_coding | NP_001191809.1;NP_003329.1 | 12474 | 602961 | . | . | 2.743200 | 2.344006 | 2.021834 | 2.196740 | 6.383539 | 5.418274 | 4.536936 | 5.022584 | -1.222890 | 191.006107 | 0.246519194 | -1.17757218 | 0.238967206 | 0.992733415 | 7.715257 | 7.425131 | 7.658459 | 7.580082 | 7.516710 | 7.546042 | 262 | 208 | 144 | 167 |
| 7322 | NM_003339,N | UBE2D2 | ubiquitin conjugating enzyme E2 D2 | protein_coding | NP_003330.1;NP_862821.1;XP_016865309.1 | 12475 | 602962 | . | . | 22.199391 | 22.555879 | 21.609345 | 20.740517 | 51.658890 | 52.138924 | 48.490745 | 47.420729 | -1.069738 | 1747.209161 | 0.121722686 | -0.79900526 | 0.424287363 | 0.998865404 | 10.818585 | 10.721453 | 10.794550 | 10.798775 | 10.764655 | 10.722531 | 2074 | 1954 | 1508 | 1540 |
| 7323 | NM_00130079 | UBE2D3 | ubiquitin conjugating enzyme E2 D3 | protein_coding | NP_001287724.1;NP_003331.1;NP_871615.1;NP_871616.1;NP_871617.1 | 12476 | 602963 | . | . | 35.093005 | 31.957045 | 32.791422 | 24.909039 | 81.662863 | 73.870142 | 73.583000 | 56.951558 | -1.178496 | 3757.719098 | 0.139648014 | -1.69674575 | 0.089744739 | 0.803337638 | 11.989116 | 11.752474 | 11.977675 | 11.895212 | 11.906410 | 11.684418 | 4910 | 4172 | 3408 | 2795 |
| 7324 | NM_00120247 | UBE2E1 | ubiquitin conjugating enzyme E2 E1 | protein_coding | NP_001189405.1;NP_003332.1;NP_872607.1;XP_005265488.1 | 12477 | 602916 | . | . | 16.890941 | 15.795131 | 16.606514 | 15.643195 | 39.305910 | 36.511156 | 37.264541 | 35.766306 | -1.024840 | 889.636676 | 0.144071033 | -0.24570547 | 0.805910237 | 0.998865404 | 9.814449 | 9.779484 | 9.833856 | 9.776245 | 9.813528 | 9.761721 | 1078 | 933 | 792 | 794 |
| 7325 | NM_00137022 | UBE2E2 | ubiquitin conjugating enzyme E2 E2 | protein_coding | NP_001357154.1;NP_001357155.1;NP_001357156.1;NP_001357157.1;NI | 12478 | 602163 | . | . | 3.279584 | 2.804407 | 2.929834 | 3.341374 | 7.631727 | 6.482512 | 6.574463 | 7.639654 | 1.071899 | 272.717855 | 0.21174853 | 0.473057884 | 0.636171883 | 0.998865404 | 8.040456 | 8.140346 | 8.091832 | 8.051766 | 8.076901 | 8.137942 | 315 | 273 | 231 | 280 |
| 7326 | NM_003342 | UBE2G1 | ubiquitin conjugating enzyme E2 G1 | protein_coding | NP_003333.1 | 12482 | 601569 | . | . | 7.749019 | 7.595997 | 7.845559 | 6.220721 | 18.032286 | 17.558489 | 17.605206 | 14.222939 | -1.105865 | 964.931286 | 0.155305314 | -0.93477227 | 0.349905673 | 0.998865404 | 9.984702 | 9.840249 | 9.958348 | 9.937486 | 9.957459 | 9.784953 | 1183 | 1078 | 893 | 756 |
| 7327 | NM_00120248 | UBE2G2 | ubiquitin conjugating enzyme E2 G2 | protein_coding | NP_001189418.1;NP_003334.2;NP_872630.1;XP_024307900.1 | 12483 | 603124 | . | . | 16.365236 | 15.830987 | 16.101384 | 16.685683 | 38.082576 | 36.594034 | 36.131042 | 38.149829 | 1.023970 | 1685.434933 | 0.124638522 | 0.274185966 | 0.783941712 | 0.998865404 | 10.701665 | 10.735942 | 10.742702 | 10.674742 | 10.720405 | 10.735237 | 2001 | 1719 | 1469 | 1614 |
| 7328 | NM_00120249 | UBE2H | ubiquitin conjugating enzyme E2 H | protein_coding | NP_001189427.1;NP_003335.1;NP_874356.1 | 12484 | 601082 | . | . | 11.931956 | 11.810796 | 10.698459 | 11.951840 | 27.766153 | 27.301186 | 24.007033 | 27.326463 | -1.062368 | 1864.888050 | 0.12342797 | -0.70715792 | 0.479468344 | 0.998865404 | 10.907951 | 10.820470 | 10.895160 | 10.881056 | 10.800484 | 10.878585 | 2232 | 2055 | 1494 | 1780 |
| 7329 | NM_003345,N | UBE2I | ubiquitin conjugating enzyme E2 I | protein_coding | NP_003336.1;NP_919235.1;NP_919236.1;NP_919237.1;XP_005255597.1 | 12485 | 601661 | . | . | 20.454719 | 22.399967 | 21.590767 | 22.144727 | 47.598972 | 51.778532 | 48.449050 | 50.631286 | 1.001201 | 1921.481893 | 0.119341019 | 0.014505876 | 0.988426391 | 0.999197585 | 10.907198 | 10.908809 | 10.892904 | 10.921861 | 10.897546 | 10.919119 | 2171 | 2110 | 1649 | 1827 |
| 7332 | NM_00125635 | UBE2L3 | ubiquitin conjugating enzyme E2 L3 | protein_coding | NP_001243284.1;NP_001243285.1;NP_003338.1;XP_016884424.1 | 12488 | 603721 | . | . | 40.682369 | 41.400584 | 42.051816 | 50.926811 | 94.669548 | 95.699309 | 94.363057 | 116.438091 | 1.102212 | 3906.871407 | 0.121487683 | 1.155687724 | 0.247808911 | 0.992733415 | 11.859991 | 12.000200 | 11.882097 | 11.896067 | 11.894445 | 12.039894 | 4244 | 4045 | 3253 | 4174 |
| 7333 | gene-UBE2L4 | UBE2L4 | ubiquitin conjugating enzyme E2 L4 (pseudogene) | pseudogene | . | 12489 | . | . | . | 0.071059 | 0.050118 | 0.012459 | 0.054483 | 0.165357 | 0.115851 | 0.027958 | 0.124569 | -1.089958 | 1.718421 | 2.237283933 | -0.0555461 | 0.955703407 | NA | 0.844134 | 0.715150 | 0.784731 | 0.785283 | 0.782101 | 0.786536 | 2 | 2 | 1 | 2 |
| 7334 | NM_003348,XI | UBE2N | ubiquitin conjugating enzyme E2 N | protein_coding | NP_003339.1 | 12492 | 603679 | . | . | 9.130821 | 8.373664 | 7.840674 | 6.889787 | 21.247795 | 19.356102 | 17.594244 | 15.752678 | -1.201157 | 1165.602905 | 0.144272706 | -1.83281261 | 0.066830444 | 0.735028378 | 10.312734 | 10.048946 | 10.283696 | 10.215743 | 10.175202 | 10.047197 | 1529 | 1312 | 1002 | 906 |
| 7335 | NM_00103228 | UBE2V1 | ubiquitin conjugating enzyme E2 V1 | protein_coding | NP_001027459.1;NP_001244322.1;NP_001244323.1;NP_001244324.1;NI | 12494 | 602995 | . | . | 59.679322 | 60.562791 | 61.349056 | 65.371269 | 138.876233 | 139.993602 | 137.665506 | 149.463641 | 1.043061 | 4026.632531 | 0.108296073 | 0.561646928 | 0.5743566 | 0.998865404 | 11.944660 | 12.005416 | 11.953311 | 11.961711 | 11.973391 | 12.011222 | 4516 | 4275 | 3494 | 3942 |
| 7336 | NM_003350,XI | UBE2V2 | ubiquitin conjugating enzyme E2 V2 | protein_coding | NP_003341.1;XP_011515885.1;XP_016869297.1 | 12495 | 603001 | . | . | 7.981061 | 6.823584 | 7.453916 | 6.052431 | 18.572256 | 15.773020 | 16.726370 | 13.838164 | -1.114157 | 954.015869 | 0.158594639 | -0.98334336 | 0.325438477 | 0.998865404 | 9.973340 | 9.818232 | 9.990812 | 9.873315 | 9.927207 | 9.777990 | 1249 | 999 | 867 | 758 |
| 7337 | NM_000462,N | UBE3A | ubiquitin protein ligase E3A | protein_coding | NP_000453.2;NP_001341434.1;NP_001341435.1;NP_001341436.1;NP_0( | 12496 | 601623 | . | . | 6.969741 | 6.217876 | 6.095380 | 5.757807 | 16.218873 | 14.372900 | 13.677856 | 13.164544 | -1.117949 | 1657.016813 | 0.128064533 | -1.25603668 | 0.209102673 | 0.971472297 | 10.772432 | 10.611846 | 10.781754 | 10.687955 | 10.662565 | 10.635150 | 2133 | 1776 | 1385 | 1441 |
| 7341 | NM_00100578 | SUMO1 | small ubiquitin like modifier 1 | protein_coding | NP_001005781.1;NP_001005782.1;NP_001358321.1;NP_001358322.1;NI | 12502 | 601912 | . | . | 56.687977 | 54.734721 | 53.116414 | 52.498104 | 131.915258 | 126.521751 | 119.191692 | 120.030671 | -1.073447 | 2559.719857 | 0.114502967 | -0.89299319 | 0.371860821 | 0.998865404 | 11.371940 | 11.269793 | 11.368371 | 11.330753 | 11.301679 | 11.283064 | 3122 | 2794 | 2172 | 2288 |
| 7342 | NM_00112816 | UBP1 | upstream binding protein 1 | protein_coding | NP_001121632.1;NP_001121633.1;NP_055332.3 | 12507 | 609784 | . | . | 11.650273 | 11.026203 | 11.595491 | 11.063279 | 27.110665 | 25.487565 | 26.019946 | 25.294874 | -1.014215 | 1425.045917 | 0.126247187 | -0.16130223 | 0.871855364 | 0.998865404 | 10.486868 | 10.466647 | 10.497708 | 10.466310 | 10.480301 | 10.462287 | 1686 | 1517 | 1245 | 1311 |
| 7343 | NM_00107668 | UBTF | upstream binding transcription factor | protein_coding | NP_001070151.1;NP_001070152.1;NP_055048.1;XP_006722122.1;XP_00 | 12511 | 600673 | . | . | 18.775635 | 19.613740 | 19.245556 | 22.235533 | 43.691675 | 45.338041 | 43.186472 | 50.838901 | 1.004162 | 2788.893568 | 0.121101774 | 0.049479911 | 0.960536847 | 0.998865404 | 11.442602 | 11.448347 | 11.429656 | 11.456613 | 11.388375 | 11.501536 | 3152 | 3053 | 2261 | 2802 |
| 7347 | NM_00127095 | UCHL3 | ubiquitin C-terminal hydrolase L3 | protein_coding | NP_001257881.1;NP_005993.1;XP_011533514.1;XP_011533515.1;XP_01 | 12515 | 603090 | . | . | 16.440208 | 15.693295 | 13.788598 | 13.953245 | 38.257037 | 36.275751 | 30.941213 | 31.902434 | -1.172939 | 472.871846 | 0.170493515 | -1.34977658 | 0.177087661 | 0.949886621 | 8.995669 | 8.765795 | 8.939903 | 8.926040 | 8.845640 | 8.821656 | 594 | 545 | 386 | 400 |
| 7349 | NM_003353 | UCN | urocortin | protein_coding | NP_003344.1 | 12516 | 600945 | . | . | 0.373864 | 1.016509 | 1.162118 | 0.508455 | 0.869996 | 2.349706 | 2.607759 | 1.162522 | 1.170333 | 19.262770 | 0.776345351 | 0.292291785 | 0.77006354 | NA | 4.149817 | 4.376754 | 4.205489 | 4.278387 | 4.288055 | 4.221858 | 11 | 28 | 25 | 12 |
| 7351 | NM_00138194 | UCP2 | uncoupling protein 2 | protein_coding | NP_001368872.1;NP_001368873.1;NP_001368874.1;NP_001368876.1;NI | 12518 | 601693 | . | . | 16.648664 | 18.012591 | 15.566548 | 18.788894 | 38.742127 | 41.636911 | 34.930884 | 42.958575 | -1.015261 | 889.399302 | 0.151811809 | -0.14393466 | 0.885552067 | 0.998865404 | 9.807917 | 9.785370 | 9.774801 | 9.826911 | 9.724484 | 9.851788 | 995 | 1002 | 699 | 901 |
| 7352 | NM_003356,N | UCP3 | uncoupling protein 3 | protein_coding | NP_003347.1;NP_073714.1 | 12519 | 602044 | . | . | 0.238246 | 0.265408 | 0.144254 | 0.100516 | 0.554407 | 0.613504 | 0.323703 | 0.229819 | -2.119862 | 13.878840 | 0.840059549 | -1.29034976 | 0.196929247 | NA | 4.236451 | 3.155137 | 3.820673 | 3.825690 | 3.786422 | 3.773521 | 21 | 21 | 9 | 7 |
| 7353 | NM_00103524 | UFD1 | ubiquitin recognition factor in ER associated degradation 1 | protein_coding | NP_001030324.2;NP_001349839.1;NP_005650.2 | 12520 | 601754 | . | . | 44.937608 | 45.198742 | 44.344672 | 52.467535 | 104.571653 | 104.478915 | 99.508154 | 119.960777 | 1.061544 | 2611.999628 | 0.12373694 | 0.696349262 | 0.48621013 | 0.998865404 | 11.307334 | 11.393264 | 11.326779 | 11.324114 | 11.307631 | 11.435973 | 2923 | 2730 | 2155 | 2720 |
| 7355 | NM_00103228 | SLC35A2 | solute carrier family 35 member A2 | protein_coding | NP_001027460.1;NP_001035963.1;NP_001269576.1;NP_001269577.1;NI | 11022 | 314375 | . | . | 17.924344 | 17.511618 | 18.277146 | 17.566114 | 41.710680 | 40.478887 | 41.013382 | 40.162830 | -1.032881 | 1195.026997 | 0.130405782 | -0.35791695 | 0.720405466 | 0.998865404 | 10.245969 | 10.199309 | 10.229319 | 10.240468 | 10.216158 | 10.204999 | 1388 | 1320 | 1030 | 1094 |
| 7357 | NM_003358,XI | UGCG | UDP-glucose ceramide glucosyltransferase | protein_coding | NP_003349.1;XP_005252243.1;XP_016870596.1 | 12524 | 602874 | . | . | 17.175011 | 15.810297 | 19.400314 | 13.102324 | 39.966959 | 36.546213 | 43.533741 | 29.956903 | -1.031512 | 2014.719882 | 0.170291561 | -0.26284911 | 0.792666884 | 0.998865404 | 10.998314 | 10.954074 | 11.021030 | 10.943069 | 11.110648 | 10.779074 | 2470 | 2100 | 2083 | 1474 |
| 7358 | NM_00118470 | UGDH | UDP-glucose 6-dehydrogenase | protein_coding | NP_001171629.1;NP_001171630.1;NP_003350.1;XP_005262724.1 | 12525 | 603370 | . | . | 11.726805 | 12.401422 | 11.977948 | 12.259268 | 27.288755 | 28.666442 | 26.878172 | 28.029360 | -1.010996 | 1138.633555 | 0.132579456 | -0.11900292 | 0.905273043 | 0.998865404 | 10.161032 | 10.145099 | 10.140855 | 10.173160 | 10.141602 | 10.156179 | 1290 | 1262 | 975 | 1072 |
| 7360 | NM_00100152 | UGP2 | UDP-glucose pyrophosphorylase 2 | protein_coding | NP_001001521.1;NP_001364453.1;NP_001364454.1;NP_001364455.1;NI | 12527 | 191760 | . | . | 50.851455 | 46.961071 | 45.634067 | 44.443260 | 118.333429 | 108.552625 | 102.401521 | 101.614240 | -1.100447 | 3131.869180 | 0.114629359 | -1.20465794 | 0.228335378 | 0.982008277 | 11.680135 | 11.542186 | 11.686638 | 11.613124 | 11.584792 | 11.559318 | 3946 | 3384 | 2634 | 2752 |
| 7364 | NM_001074,N | UGT2B7 | UDP glucuronosyltransferase family 2 member B7 | protein_coding | NP_001065.2;NP_001317648.1;NP_001336497.1 | 12554 | 600068 | . | . | 8.788722 | 9.864448 | 9.399541 | 9.924453 | 20.451719 | 22.802114 | 21.092298 | 22.691080 | 1.019914 | 472.074172 | 0.17325996 | 0.16418898 | 0.869582376 | 0.998865404 | 8.868833 | 8.896771 | 8.842650 | 8.908636 | 8.875555 | 8.901023 | 510 | 531 | 406 | 456 |
| 7368 | NM_00112817 | UGT8 | UDP glycosyltransferase 8 | protein_coding | NP_001121646.2;NP_001309041.1;NP_001309042.1;NP_001309043.1;NI | 12555 | 601291 | . | . | 2.906871 | 2.798163 | 2.303361 | 2.251933 | 6.764409 | 6.468077 | 5.168675 | 5.148778 | -1.317195 | 334.488802 | 0.195459804 | -2.03350576 | 0.042001452 | 0.633602228 | 8.570606 | 8.173827 | 8.477409 | 8.435501 | 8.325771 | 8.280323 | 453 | 396 | 262 | 259 |
| 7371 | NM_00136356 | UCK2 | uridine-cytidine kinase 2 | protein_coding | NP_001350497.1;NP_036606.2 | 12562 | 609329 | . | . | 10.281935 | 10.790993 | 10.588887 | 12.283951 | 23.926485 | 24.943864 | 23.761157 | 28.085796 | 1.067096 | 1650.437143 | 0.130183391 | 0.719678867 | 0.471722741 | 0.998865404 | 10.641216 | 10.734540 | 10.646227 | 10.677549 | 10.656182 | 10.766078 | 1802 | 1758 | 1378 | 1709 |
| 7372 | NM_000373,N | UMPS | uridine monophosphate synthetase | protein_coding | NP_000364.1 | 12563 | 613891 | . | . | 4.895537 | 5.353114 | 5.134199 | 5.234203 | 11.392118 | 12.373966 | 11.520993 | 11.967382 | -1.002598 | 1076.669748 | 0.135431668 | -0.0276373 | 0.977951435 | 0.998865404 | 10.074310 | 10.070407 | 10.046159 | 10.099465 | 10.071195 | 10.071175 | 1197 | 1205 | 935 | 1008 |
| 7373 | NM_021110,XI | COL14A1 | collagen type XIV alpha 1 chain | protein_coding | NP_066933.1;XP_005251116.1;XP_006716714.1;XP_016869298.1 | 2191 | 120324 | . | . | 0.062465 | 0.096321 | 0.093972 | 0.065835 | 0.145359 | 0.222651 | 0.210871 | 0.150523 | 1.008473 | 22.192221 | 0.665648597 | 0.018286311 | 0.985410448 | NA | 4.465522 | 4.478414 | 4.450367 | 4.483615 | 4.484772 | 4.450753 | 21 | 28 | 23 | 17 |
| 7374 | NM_003362,N | UNG | uracil DNA glycosylase | protein_coding | NP_003353.1;NP_550433.1 | 12572 | 191525 | . | . | 20.111398 | 18.920273 | 18.926117 | 20.861343 | 46.800051 | 43.735059 | 42.469657 | 47.696986 | 1.007087 | 1264.592706 | 0.133348021 | 0.076406738 | 0.939095509 | 0.998865404 | 10.299407 | 10.309490 | 10.327253 | 10.274989 | 10.273014 | 10.339445 | 1501 | 1313 | 1055 | 1242 |
| 7375 | NM_00125187 | USP4 | ubiquitin specific peptidase 4 | protein_coding | NP_001238806.1;NP_003354.2;NP_955475.1 | 12627 | 603486 | . | . | 9.694430 | 10.909019 | 10.750369 | 11.781393 | 22.559338 | 25.216686 | 24.123517 | 26.936755 | 1.073687 | 1354.914508 | 0.133923625 | 0.765913427 | 0.443727816 | 0.998865404 | 10.352000 | 10.454164 | 10.332658 | 10.416521 | 10.397672 | 10.460641 | 1422 | 1488 | 1170 | 1369 |
| 7376 | NM_00125664 | NR1H2 | nuclear receptor subfamily 1 group H member 2 | protein_coding | NP_001243576.2;NP_009052.4 | 7965 | 600380 | . | . | 13.964051 | 15.676892 | 14.011944 | 17.406765 | 32.494921 | 36.237836 | 31.442400 | 39.798497 | 1.048712 | 1137.759775 | 0.147404423 | 0.465512211 | 0.641564694 | 0.998865404 | 10.117597 | 10.185563 | 10.094300 | 10.169267 | 10.093683 | 10.236976 | 1215 | 1259 | 916 | 1196 |
| 7378 | NM_00128742 | UPP1 | uridine phosphorylase 1 | protein_coding | NP_001274355.1;NP_001274357.1;NP_001274358.1;NP_001274359.1;NI | 12576 | 191730 | . | . | 60.203177 | 67.943634 | 60.566157 | 65.772160 | 140.095265 | 157.054753 | 135.908703 | 150.380235 | -1.021043 | 3190.458077 | 0.116253721 | -0.25843552 | 0.796070806 | 0.998865404 | 11.654591 | 11.624346 | 11.615413 | 11.678991 | 11.595628 | 11.664294 | 3569 | 3613 | 2630 | 3084 |
| 7381 | NM_00119997 | UQCRB | ubiquinol-cytochrome c reductase binding protein | protein_coding | NP_001186904.1;NP_001241681.1;NP_006285.1 | 12582 | 191330 | . | . | 15.322032 | 13.494529 | 13.532661 | 15.553258 | 35.654997 | 31.193210 | 30.366898 | 35.560676 | -1.004593 | 3828.454035 | 0.123400517 | -0.0535705 | 0.957277363 | 0.998865404 | 11.905865 | 11.899219 | 11.958135 | 11.845630 | 11.845343 | 11.951168 | 4718 | 3860 | 3109 | 3810 |
| 7384 | NM_003365,XI | UQCRC1 | ubiquinol-cytochrome c reductase core protein 1 | protein_coding | NP_003356.2 | 12585 | 191328 | . | . | 88.870040 | 84.857192 | 72.054100 | 89.937627 | 206.804245 | 196.151198 | 161.687296 | 205.631708 | -1.088979 | 4208.382456 | 0.125686934 | -0.97842895 | 0.327862215 | 0.998865404 | 12.099305 | 11.976168 | 12.095037 | 12.049001 | 11.915170 | 12.080795 | 5191 | 4605 | 3152 | 4156 |
| 7385 | NM_003366 | UQCRC2 | ubiquinol-cytochrome c reductase core protein 2 | protein_coding | NP_003357.2 | 12586 | 191329 | . | . | 69.930168 | 72.557472 | 64.408897 | 75.844696 | 162.730377 | 167.719849 | 144.531693 | 173.409897 | -1.030337 | 4214.382428 | 0.118181242 | -0.36483206 | 0.715236774 | 0.998865404 | 12.062594 | 12.019292 | 12.040881 | 12.064486 | 11.962513 | 12.089230 | 4856 | 4681 | 3334 | 4189 |
| 7386 | NM_006003 | UQCRFS1 | ubiquinol-cytochrome c reductase, Rieske iron-sulfur polypeptide : | protein_coding | NP_005994.2 | 12587 | 191327 | . | . | 32.966839 | 30.778292 | 27.854361 | 33.054260 | 76.715195 | 71.145401 | 62.504379 | 75.574631 | -1.061620 | 2996.703637 | 0.124706004 | -0.69176602 | 0.489084278 | 0.998865404 | 11.591722 | 11.505305 | 11.603585 | 11.539989 | 11.456068 | 11.586438 | 3691 | 3202 | 2325 | 2944 |
| 7388 | NM_00129756 | UQCRH | ubiquinol-cytochrome c reductase hinge protein | protein_coding | NP_001284494.1;NP_001284495.1;NP_005995.2 | 12590 | 613844 | . | . | 275.067272 | 270.834717 | 253.745996 | 269.787748 | 640.092823 | 626.046610 | 569.398685 | 616.837620 | -1.057058 | 4544.947270 | 0.107562755 | -0.74426006 | 0.456719158 | 0.998865404 | 12.189544 | 12.109440 | 12.180073 | 12.165299 | 12.104521 | 12.147854 | 5426 | 4996 | 3742 | 4251 |
| 7389 | NM_000374,N | UROD | uroporphyrinogen decarboxylase | protein_coding | NP_000365.3 | 12591 | 613521 | . | . | 67.272801 | 65.034837 | 64.006165 | 73.558697 | 156.546580 | 150.330937 | 143.627977 | 168.183218 | 1.023708 | 2546.192883 | 0.121387012 | 0.278488641 | 0.780637283 | 0.998865404 | 11.297205 | 11.330851 | 11.321002 | 11.286799 | 11.269930 | 11.373349 | 2959 | 2658 | 2097 | 2569 |
| 7390 | NM_000375,N | UROS | uroporphyrinogen III synthase | protein_coding | NP_000366.1;NP_001310965.1;NP_001310966.1;NP_001310967.1;NP_0( | 12592 | 606938 | . | . | 15.959228 | 17.421723 | 15.615708 | 21.858394 | 37.137785 | 40.271089 | 35.041199 | 49.976623 | 1.087626 | 989.269489 | 0.168530471 | 0.719057892 | 0.47210525 | 0.998865404 | 9.888741 | 10.009186 | 9.909173 | 9.920708 | 9.844572 | 10.093082 | 1083 | 1031 | 746 | 1128 |
| 7391 | NM_00127637 | USF1 | upstream transcription factor 1 | protein_coding | NP_001263302.1;NP_009053.1;NP_996888.1 | 12593 | 191523 | . | . | 10.553853 | 10.549590 | 13.486619 | 11.964778 | 24.559249 | 24.385851 | 30.263581 | 27.356043 | 1.191285 | 651.875777 | 0.157735638 | 1.600895434 | 0.109400081 | 0.843131948 | 9.216426 | 9.469405 | 9.285964 | 9.282296 | 9.444917 | 9.358522 | 687 | 640 | 655 | 623 |
| 7392 | NM_00132115 | USF2 | upstream transcription factor 2, c-fos interacting | protein_coding | NP_001308079.1;NP_003358.1;NP_997174.1;XP_005259254.1;XP_01152 | 12594 | 600390 | . | . | 56.296596 | 62.096684 | 57.657615 | 66.707995 | 131.004492 | 143.539267 | 129.382006 | 152.519905 | 1.040204 | 3166.464476 | 0.118269447 | 0.480824659 | 0.630641123 | 0.998865404 | 11.600058 | 11.656701 | 11.584939 | 11.637869 | 11.595546 | 11.690219 | 3459 | 3458 | 2646 | 3201 |
| 7398 | NM_00101741 | USP1 | ubiquitin specific peptidase 1 | protein_coding | NP_001017415.1;NP_001017416.1;NP_003359.3 | 12607 | 603478 | . | . | 14.903981 | 13.323112 | 11.988838 | 9.956935 | 34.682178 | 30.796972 | 26.902605 | 22.765348 | -1.305180 | 1409.325458 | 0.144242686 | -2.66390454 | 0.007723948 | 0.276918574 | 10.639882 | 10.256270 | 10.599153 | 10.503172 | 10.418404 | 10.272923 | 1948 | 1618 | 1168 | 1034 |
| 7399 | NM_007123,N | USH2A | usherin | protein_coding | NP_009054.6;NP_996816.3 | 12601 | 608400 | . | . | 0.013561 | 0.011572 | 0.010306 | 0.011904 | 0.031558 | 0.026748 | 0.023126 | 0.027218 | -1.120695 | 7.616223 | 1.095567932 | -0.15005327 | 0.880722588 | NA | 3.008843 | 2.844638 | 2.936803 | 2.930901 | 2.928723 | 2.931170 | 10 | 8 | 6 | 7 |
| 7402 | NM_00137532 | UTRN | utrophin | protein_coding | NP_001362252.1;NP_009055.2;XP_005267184.1;XP_005267187.1;XP_00 | 12635 | 128240 | . | . | 2.332196 | 2.148449 | 1.941031 | 2.098770 | 5.427115 | 4.966232 | 4.355617 | 4.798589 | -1.160554 | 846.630258 | 0.148755668 | -1.44406861 | 0.148719676 | 0.909660767 | 9.829078 | 9.614097 | 9.809902 | 9.740549 | 9.632390 | 9.706089 | 1097 | 935 | 645 | 774 |
| 7403 | NM_00129141 | KDM6A | lysine demethylase 6A | protein_coding | NP_001278344.1;NP_001278345.1;NP_001278346.1;NP_001278347.1;NI | 12637 | 300128 | . | . | 2.215739 | 2.265159 | 2.468277 | 2.157822 | 5.156115 | 5.236016 | 5.538745 | 4.933603 | 1.046672 | 418.976446 | 0.17961783 | 0.366387028 | 0.714076298 | 0.998865404 | 8.677219 | 8.743471 | 8.694177 | 8.697746 | 8.754220 | 8.691864 | 470 | 443 | 392 | 381 |
| 7404 | NM_00125824 | UTY | ubiquitously transcribed tetratricopeptide repeat containing, Y-linl | protein_coding | NP_001245178.1;NP_001245179.1;NP_001245180.1;NP_001245181.1;NI | 12638 | 400009 | . | . | 0.038300 | 0.032724 | 0.025596 | 0.045702 | 0.089125 | 0.075643 | 0.057437 | 0.104492 | 1.043263 | 9.131687 | 1.011859993 | 0.06038678 | 0.951847588 | NA | 3.161369 | 3.219802 | 3.193312 | 3.187747 | 3.181705 | 3.202743 | 11 | 9 | 6 | 11 |
| 7405 | NM_003369,XI | UVRAG | UV radiation resistance associated | protein_coding | NP_003360.2;XP_011543528.1;XP_011543529.1;XP_011543530.1;XP_01( | 12640 | 602493 | . | . | 3.872917 | 4.084483 | 4.263242 | 4.604887 | 9.012436 | 9.441464 | 9.566594 | 10.528530 | 1.114880 | 643.368080 | 0.155821418 | 1.006851607 | 0.314006091 | 0.998865404 | 9.249137 | 9.405623 | 9.270875 | 9.310332 | 9.343132 | 9.384421 | 680 | 676 | 572 | 655 |
| 7407 | NM_006295,XI | VARS1 | valyl-tRNA synthetase 1 | protein_coding | NP_006286.1;XP_005249419.1;XP_016866735.1;XP_024302305.1 | 12651 | 192150 | . | . | 25.655127 | 27.863844 | 26.099680 | 29.162856 | 59.700532 | 64.408526 | 58.566924 | 66.677403 | 1.008254 | 3447.830240 | 0.114450852 | 0.103619878 | 0.917471012 | 0.998865404 | 11.745620 | 11.757302 | 11.723067 | 11.771894 | 11.717914 | 11.789773 | 3837 | 3815 | 2879 | 3387 |

| | | | | | | | | | |
|---|---|---|---|---|---|---|---|---|---|
| 7408 | NM_003370,XI | VASP | vasodilator stimulated phosphoprotein | protein_coding | NP_003361.1;XP_005259256.1;XP_005259257.1;XP_016882689.1 | 12652 | 601703 | . | . |
| 7410 | NM_00113439 | VAV2 | vav guanine nucleotide exchange factor 2 | protein_coding | NP_001127870.1;NP_003362.2;XP_005272270.1;XP_016870597.1;XP_01 | 12658 | 600428 | . | . |
| 7411 | NM_00130354 | VBP1 | VHL binding protein 1 | protein_coding | NP_001290472.1;NP_001290473.1;NP_001290474.1;NP_003363.1 | 12662 | 300133 | . | . |
| 7414 | NM_003373,N | VCL | vinculin | protein_coding | NP_003364.1;NP_054706.1 | 12665 | 193065 | . | . |
| 7415 | NM_00135492 | VCP | valosin containing protein | protein_coding | NP_001341856.1;NP_001341857.1;NP_009057.1 | 12666 | 601023 | . | . |
| 7416 | NM_003374,N | VDAC1 | voltage dependent anion channel 1 | protein_coding | NP_003365.1;XP_005272132.1;XP_016865310.1;XP_016865311.1;XP_01 | 12669 | 604492 | . | . |
| 7417 | NM_00118478 | VDAC2 | voltage dependent anion channel 2 | protein_coding | NP_001171712.1;NP_001171752.1;NP_001311016.1;NP_001311017.1;NI | 12672 | 193245 | . | . |
| 7419 | NM_00113569 | VDAC3 | voltage dependent anion channel 3 | protein_coding | NP_001129166.1;NP_005653.3;XP_006716457.1 | 12674 | 610029 | . | . |
| 7421 | NM_000376,N | VDR | vitamin D receptor | protein_coding | NP_000367.1;NP_001017535.1;NP_001017536.1;NP_001351014.1;NP_0( | 12679 | 601769 | . | . |
| 7422 | NM_00102536 | VEGFA | vascular endothelial growth factor A | protein_coding | NP_001020537.2;NP_001020538.2;NP_001020539.2;NP_001020540.2;NI | 12680 | 192240 | . | . |
| 7423 | NM_00124373 | VEGFB | vascular endothelial growth factor B | protein_coding | NP_001230662.1;NP_003368.1 | 12681 | 601398 | . | . |
| 7424 | NM_005429 | VEGFC | vascular endothelial growth factor C | protein_coding | NP_005420.1 | 12682 | 601528 | . | . |
| 7425 | NM_003378,XI | VGF | VGF nerve growth factor inducible | protein_coding | NP_003369.2;XP_005250618.1;XP_011514851.1 | 12684 | 602186 | . | . |
| 7428 | NM_000551,N | VHL | von Hippel-Lindau tumor suppressor | protein_coding | NP_000542.1;NP_001341652.1;NP_937799.1 | 12687 | 608537 | . | . |
| 7430 | NM_00111107 | EZR | ezrin | protein_coding | NP_001104547.1;NP_003370.2;XP_011534412.1 | 12691 | 123900 | . | . |
| 7431 | NM_003380,XI | VIM | vimentin | protein_coding | NP_003371.2;XP_006717563.1 | 12692 | 193060 | . | . |
| 7436 | NM_00101805 | VLDLR | very low density lipoprotein receptor | protein_coding | NP_001018066.1;NP_001309154.1;NP_001309155.1;NP_003374.3 | 12698 | 192977 | . | . |
| 7439 | NM_00113944 | BEST1 | bestrophin 1 | protein_coding | NP_001132915.1;NP_001287715.1;NP_001287716.1;NP_001350520.1;NI | 12703 | 607854 | . | . |
| 7442 | NM_018727,N | TRPV1 | transient receptor potential cation channel subfamily V member 1 | protein_coding | NP_061197.4;NP_542435.2;NP_542436.2;NP_542437.2 | 12716 | 602076 | . | . |
| 7443 | NM_003384,XI | VRK1 | VRK serine/threonine kinase 1 | protein_coding | NP_003375.1;XP_006720310.1;XP_011535434.1;XP_016877113.1;XP_01 | 12718 | 602168 | . | . |
| 7444 | NM_00113048 | VRK2 | VRK serine/threonine kinase 2 | protein_coding | NP_001123952.1;NP_001123953.1;NP_001123954.1;NP_001123955.1;NI | 12719 | 602169 | . | . |
| 7448 | NM_000638 | VTN | vitronectin | protein_coding | NP_000629.3 | 12724 | 193190 | . | . |
| 7450 | NM_000552 | VWF | von Willebrand factor | protein_coding | NP_000543.3 | 12726 | 613160 | . | . |
| 7453 | NM_004184,N | WARS1 | tryptophanyl-tRNA synthetase 1 | protein_coding | NP_004175.2;NP_776049.1;NP_998810.1;NP_998811.1;XP_005268101.1 | 12729 | 191050 | . | . |
| 7456 | NM_00107726 | WIPF1 | WAS/WASL interacting protein family member 1 | protein_coding | NP_001070737.1;NP_001362761.1;NP_001362762.1;NP_001362763.1;NI | 12736 | 602357 | . | . |
| 7458 | NM_022170,N | EIF4H | eukaryotic translation initiation factor 4H | protein_coding | NP_071496.1;NP_114381.1 | 12741 | 603431 | . | . |
| 7461 | NM_003388,N | CLIP2 | CAP-Gly domain containing linker protein 2 | protein_coding | NP_003379.4;NP_115797.2 | 2586 | 603432 | . | . |
| 7462 | NM_014146,N | LAT2 | linker for activation of T cells family member 2 | protein_coding | NP_054865.2;NP_115852.1;NP_115853.2;XP_011514860.1 | 12749 | 605719 | . | . |
| 7464 | NM_003389,N | CORO2A | coronin 2A | protein_coding | NP_003380.3;NP_438171.1;XP_011517288.1 | 2255 | 602159 | . | . |
| 7465 | NM_00114397 | WEE1 | WEE1 G2 checkpoint kinase | protein_coding | NP_001137448.1;NP_003381.1 | 12761 | 193525 | . | . |
| 7466 | NM_00114585 | WFS1 | wolframin ER transmembrane glycoprotein | protein_coding | NP_001139325.1;NP_005996.2;XP_016864075.1 | 12762 | 606201 | . | . |
| 7468 | NM_00104242 | NSD2 | nuclear receptor binding SET domain protein 2 | protein_coding | NP_001035889.1;NP_015627.1;NP_579877.1;NP_579878.1;NP_579889.1 | 12766 | 602952 | . | . |
| 7469 | NM_005663,XI | NELFA | negative elongation factor complex member A | protein_coding | NP_005654.4;XP_016864078.1 | 12768 | 606026 | . | . |
| 7474 | NM_00125610 | WNT5A | Wnt family member 5A | protein_coding | NP_001243034.1;NP_001364200.1;NP_001364201.1;NP_003383.4;XP_01 | 12784 | 164975 | . | . |
| 7480 | NM_003394,XI | WNT10B | Wnt family member 10B | protein_coding | NP_003385.2;XP_011537024.1;XP_011537026.1;XP_016875408.1;XP_02 | 12775 | 601906 | . | . |
| 7482 | NM_00129188 | WNT2B | Wnt family member 2B | protein_coding | NP_001278809.1;NP_004176.2;NP_078613.1 | 12781 | 601968 | . | . |
| 7483 | NM_003395,XI | WNT9A | Wnt family member 9A | protein_coding | NP_003386.1;XP_011542573.1 | 12778 | 602863 | . | . |
| 7485 | NM_00114621 | GET1 | guided entry of tail-anchored proteins factor 1 | protein_coding | NP_001139690.1;NP_001337222.1;NP_001337223.1;NP_001337224.1;NI | 12790 | 602915 | . | . |
| 7486 | NM_000553,XI | WRN | WRN RecQ like helicase | protein_coding | NP_000544.2;XP_011542941.1;XP_011542942.1;XP_024303033.1 | 12791 | 604611 | . | . |
| 7494 | NM_00107953 | XBP1 | X-box binding protein 1 | protein_coding | NP_001073007.1;NP_005071.2 | 12801 | 194355 | . | . |
| 7507 | NM_000380,N | XPA | XPA, DNA damage recognition and repair factor | protein_coding | NP_000371.1;NP_001341904.1;XP_006717341.1 | 12814 | 611153 | . | . |
| 7508 | NM_00135472 | XPC | XPC complex subunit, DNA damage recognition and repair factor | protein_coding | NP_001341655.1;NP_001341656.1;NP_001341658.1;NP_001341659.1;NI | 12816 | 613208 | . | . |
| 7511 | NM_00116760 | XPNPEP1 | X-prolyl aminopeptidase 1 | protein_coding | NP_001161076.1;NP_001311057.1;NP_001311060.1;NP_001311061.1;NI | 12822 | 602443 | . | . |
| 7514 | NM_003400,XI | XPO1 | exportin 1 | protein_coding | NP_003391.1;XP_005264601.1;XP_005264603.1;XP_006712157.1;XP_01 | 12825 | 602559 | . | . |
| 7515 | NM_006297 | XRCC1 | X-ray repair cross complementing 1 | protein_coding | NP_006288.2 | 12828 | 194360 | . | . |
| 7516 | NM_005431 | XRCC2 | X-ray repair cross complementing 2 | protein_coding | NP_005422.1 | 12829 | 600375 | . | . |
| 7517 | NM_00110011 | XRCC3 | X-ray repair cross complementing 3 | protein_coding | NP_001093588.1;NP_001093589.1;NP_001358158.1;NP_001358160.1;NI | 12830 | 600675 | . | . |
| 7518 | NM_00131801 | XRCC4 | X-ray repair cross complementing 4 | protein_coding | NP_001304941.1;NP_001304942.1;NP_003392.1;NP_071801.1;NP_0720 | 12831 | 194363 | . | . |
| 7520 | NM_021141 | XRCC5 | X-ray repair cross complementing 5 | protein_coding | NP_066964.1 | 12833 | 194364 | . | . |
| 7525 | NM_005433,XI | YES1 | YES proto-oncogene 1, Src family tyrosine kinase | protein_coding | NP_005424.1;XP_005258196.1;XP_016881449.1;XP_024307011.1;XP_02 | 12841 | 164880 | . | . |
| 7528 | NM_003403 | YY1 | YY1 transcription factor | protein_coding | NP_003394.1 | 12856 | 600013 | . | . |
| 7529 | NM_003404,N | YWHAB | tyrosine 3-monooxygenase/tryptophan 5-monooxygenase activati | protein_coding | NP_003395.1;NP_647539.1;XP_016883528.1;XP_016883529.1 | 12849 | 601289 | . | . |
| 7531 | NM_006761,N | YWHAE | tyrosine 3-monooxygenase/tryptophan 5-monooxygenase activati | protein_coding | NP_006752.1;XP_005256841.1;XP_016880494.1 | 12851 | 605066 | . | . |
| 7532 | NM_012479 | YWHAG | tyrosine 3-monooxygenase/tryptophan 5-monooxygenase activati | protein_coding | NP_036611.2 | 12852 | 605356 | . | . |
| 7533 | NM_003405 | YWHAH | tyrosine 3-monooxygenase/tryptophan 5-monooxygenase activati | protein_coding | NP_003396.1 | 12853 | 113508 | . | . |
| 7534 | NM_00113569 | YWHAZ | tyrosine 3-monooxygenase/tryptophan 5-monooxygenase activati | protein_coding | NP_001129171.1;NP_001129172.1;NP_001129173.1;NP_001129174.1;NI | 12855 | 601288 | . | . |
| 7536 | NM_00117803 | SF1 | splicing factor 1 | protein_coding | NP_001171501.1;NP_001171502.1;NP_001333292.1;NP_001333293.1;NI | 12950 | 601516 | . | . |
| 7538 | NM_003407 | ZFP36 | ZFP36 ring finger protein | protein_coding | NP_003398.3 | 12862 | 190700 | . | . |
| 7539 | NM_00128251 | ZFP37 | ZFP37 zinc finger protein | protein_coding | NP_001269444.1;NP_001269447.1;NP_003399.1 | 12863 | 602951 | . | . |
| 7541 | NM_00114382 | ZBTB14 | zinc finger and BTB domain containing 14 | protein_coding | NP_001137295.1;NP_001230631.1;NP_001230633.1;NP_003400.2;XP_02 | 12860 | 602126 | . | . |
| 7542 | NM_006782 | ZFPL1 | zinc finger protein like 1 | protein_coding | NP_006773.2 | 12868 | . | . | . |
| 7543 | NM_00117808 | ZFX | zinc finger protein X-linked | protein_coding | NP_001171555.1;NP_001171556.1;NP_001171557.1;NP_001171566.1;NI | 12869 | 314980 | . | . |
| 7544 | NM_00114527 | ZFY | zinc finger protein Y-linked | protein_coding | NP_001138747.1;NP_001138748.1;NP_001356631.1;NP_001356632.1;NI | 12870 | 490000 | . | . |
| 7545 | NM_003412 | ZIC1 | Zic family member 1 | protein_coding | NP_003403.2 | 12872 | 600470 | . | . |
| 7546 | NM_007129 | ZIC2 | Zic family member 2 | protein_coding | NP_009060.2 | 12873 | 603073 | . | . |
| 7549 | NM_00101739 | ZNF2 | zinc finger protein 2 | protein_coding | NP_001017396.1;NP_001269327.1;NP_001278533.1;NP_001278534.1;NI | 12991 | 194500 | . | . |
| 7551 | NM_00127828 | ZNF3 | zinc finger protein 3 | protein_coding | NP_001265213.1;NP_001265216.1;NP_001265219.1;NP_001265220.1;NI | 13089 | 194510 | . | . |
| 7553 | NM_00128279 | ZNF7 | zinc finger protein 7 | protein_coding | NP_001269724.1;NP_001269725.1;NP_001269726.1;NP_001317552.1;NI | 13139 | 194531 | . | . |
| 7554 | NM_021089 | ZNF8 | zinc finger protein 8 | protein_coding | NP_066575.2 | 13154 | 194532 | . | . |
| 7555 | NM_00112719 | CNBP | CCHC-type zinc finger nucleic acid binding protein | protein_coding | NP_001120664.1;NP_001120665.1;NP_001120666.1;NP_001120667.1;NI | 13164 | 116955 | . | . |
| 7556 | NM_015394 | ZNF10 | zinc finger protein 10 | protein_coding | NP_056209.2 | 12879 | 194538 | . | . |
| 7559 | NM_006956,N | ZNF12 | zinc finger protein 12 | protein_coding | NP_008887.2;NP_057349.2 | 12902 | 194536 | . | . |
| 7561 | NM_021030 | ZNF14 | zinc finger protein 14 | protein_coding | NP_066358.2 | 12924 | 194556 | . | . |
| 7562 | NM_00129756 | ZNF708 | zinc finger protein 708 | protein_coding | NP_001284489.1;NP_001284490.1;NP_067092.2;XP_016882690.1;XP_01 | 12945 | . | . | . |
| 7564 | NM_00102997 | ZNF16 | zinc finger protein 16 | protein_coding | NP_001025147.2;NP_008889.2;XP_005272399.1;XP_011515600.1;XP_02 | 12947 | 601262 | . | . |
| 7565 | NM_00133061 | ZNF17 | zinc finger protein 17 | protein_coding | NP_001317546.1;NP_008890.2 | 12958 | . | . | . |
| 7566 | NM_00130328 | ZNF18 | zinc finger protein 18 | protein_coding | NP_001290210.1;NP_001290211.1;NP_653281.2;XP_011522304.1;XP_01 | 12969 | 194524 | . | . |
| 7567 | NM_006961 | ZNF19 | zinc finger protein 19 | protein_coding | NP_008892.2 | 12981 | 194525 | . | . |
| 7568 | NM_00120325 | ZNF20 | zinc finger protein 20 | protein_coding | NP_001190179.1;NP_066966.2 | 12992 | 194557 | . | . |
| 7569 | NM_00100708 | ZNF182 | zinc finger protein 182 | protein_coding | NP_001007089.1;NP_001171570.1;NP_008893.1 | 13001 | 314993 | . | . |
| 7570 | NM_006963 | ZNF22 | zinc finger protein 22 | protein_coding | NP_008894.2 | 13012 | 194529 | . | . |
| 7571 | NM_00130449 | ZNF23 | zinc finger protein 23 | protein_coding | NP_001291421.1;NP_001291422.1;NP_001291423.1;NP_001368894.1;NI | 13023 | 194527 | . | . |
| 7572 | NM_00130812 | ZNF24 | zinc finger protein 24 | protein_coding | NP_001295052.1;NP_001362744.1;NP_008896.2 | 13032 | 194534 | . | . |
| 7574 | NM_00125627 | ZNF26 | zinc finger protein 26 | protein_coding | NP_001243208.1;NP_001243209.1;NP_001317442.1;NP_001317443.1;NI | 13053 | 194537 | . | . |
| 7576 | NM_00136976 | ZNF28 | zinc finger protein 28 | protein_coding | NP_001356690.1;NP_001356691.1;NP_001356692.1;NP_001356693.1;NI | 13073 | . | . | . |
| 7579 | NM_00137737 | ZSCAN20 | zinc finger and SCAN domain containing 20 | protein_coding | NP_001364305.1;NP_001364306.1;NP_001364307.1;NP_001364308.1;NI | 13093 | 611315 | . | . |
| 7580 | NM_00100536 | ZNF32 | zinc finger protein 32 | protein_coding | NP_001005368.1;NP_001311093.1;NP_001311094.1;NP_001311095.1;NI | 13095 | 194539 | . | . |
| 7581 | NM_00127817 | ZNF33A | zinc finger protein 33A | protein_coding | NP_001265099.1;NP_001265100.1;NP_001265102.1;NP_001265103.1;NI | 13096 | 194521 | . | . |
| 7582 | NM_00130503 | ZNF33B | zinc finger protein 33B | protein_coding | NP_001291962.1;NP_001291964.1;NP_001291965.1;NP_001291966.1;NI | 13097 | 194522 | . | . |
| 7584 | NM_003420 | ZNF35 | zinc finger protein 35 | protein_coding | NP_003411.3 | 13099 | 194533 | . | . |
| 7586 | NM_00128705 | ZKSCAN1 | zinc finger with KRAB and SCAN domains 1 | protein_coding | NP_001273983.1;NP_001273984.1;NP_001333508.1;NP_001333509.1;NI | 13101 | 601260 | . | . |
| 7587 | NM_00100709 | ZNF37A | zinc finger protein 37A | protein_coding | NP_001007095.1;NP_001171572.1;NP_001311174.1;NP_001311175.1;NI | 13102 | 616085 | . | . |
| 7589 | NM_00136277 | ZSCAN21 | zinc finger and SCAN domain containing 21 | protein_coding | NP_001349708.1;NP_001349709.1;NP_001349710.1;NP_666019.1;XP_0C | 13104 | 601261 | . | . |
| 7592 | NM_00132413 | ZNF41 | zinc finger protein 41 | protein_coding | NP_001311068.1;NP_001311069.1;NP_001311070.1;NP_001311071.1;NI | 13107 | 314995 | . | . |
| 7593 | NM_00126703 | MZF1 | myeloid zinc finger 1 | protein_coding | NP_001253962.1;NP_003413.2;NP_932172.1;XP_005259261.1;XP_01152 | 13108 | 194550 | . | . |
| 7594 | NM_00125664 | ZNF43 | zinc finger protein 43 | protein_coding | NP_001243577.1;NP_001243578.1;NP_001243579.1;NP_001243580.1;NI | 13109 | 603972 | . | . |
| 7596 | NM_003425,XI | ZNF45 | zinc finger protein 45 | protein_coding | NP_003416.1;XP_011525569.1;XP_011525571.1;XP_011525573.1;XP_01 | 13111 | 194554 | . | . |
| 7597 | NM_00130450 | ZBTB25 | zinc finger and BTB domain containing 25 | protein_coding | NP_001291436.1;NP_001291437.1;NP_001341611.1;NP_001341612.1;NI | 13112 | 194541 | . | . |
| 7608 | NM_00135519 | ZNF56 | zinc finger protein 56 | protein_coding | NP_001342123.1 | 13124 | . | . | . |
| 7617 | NM_00135519 | ZNF66 | zinc finger protein 66 | protein_coding | NP_001342126.2 | 13135 | . | . | . |
| 7621 | NM_021916,XI | ZNF70 | zinc finger protein 70 | protein_coding | NP_068735.1;XP_024308037.1;XP_024308038.1;XP_024308039.1 | 13140 | 194544 | . | . |
| 7625 | NM_00125652 | ZNF74 | zinc finger protein 74 | protein_coding | NP_001243452.1;NP_001243453.1;NP_001243454.1;NP_003417.2 | 13144 | 194548 | . | . |
| 7626 | NM_00118506 | ZNF75D | zinc finger protein 75D | protein_coding | NP_001171992.1;NP_009062.2;XP_005262528.1;XP_011529697.1;XP_01 | 13145 | 314997 | . | . |
| 7627 | NM_00130210 | ZNF75A | zinc finger protein 75a | protein_coding | NP_001289038.1;NP_001310970.1;NP_001311014.1;NP_001339424.1;NI | 13146 | 601473 | . | . |
| 7629 | NM_00129203 | ZNF76 | zinc finger protein 76 | protein_coding | NP_001278961.1;NP_003418.2;XP_011513148.1;XP_011513149.1;XP_01 | 13149 | 194549 | . | . |
| 7633 | NM_00128669 | ZNF79 | zinc finger protein 79 | protein_coding | NP_001273625.1;NP_001273626.1;NP_001273627.1;NP_001309189.1;NI | 13153 | 194552 | . | . |
| 7637 | NM_00112737 | ZNF84 | zinc finger protein 84 | protein_coding | NP_001120844.1;NP_001276900.1;NP_001276901.1;NP_003419.3;XP_0C | 13159 | 618554 | . | . |
| 7638 | NM_00129758 | ZNF221 | zinc finger protein 221 | protein_coding | NP_001284517.1;NP_001284518.1;NP_037491.2;XP_016882721.1;XP_01 | 13014 | . | . | . |
| 7639 | NM_00125617 | ZNF85 | zinc finger protein 85 | protein_coding | NP_001243100.1;NP_001243101.1;NP_001243102.1;NP_003420.2;XP_01 | 13160 | 603899 | . | . |
| 7643 | NM_007138,XI | ZNF90 | zinc finger protein 90 | protein_coding | NP_009069.1 | 13165 | 603973 | . | . |
| 7644 | NM_00130095 | ZNF91 | zinc finger protein 91 | protein_coding | NP_001287880.1;NP_003421.2;XP_006722943.1;XP_011526567.1;XP_01 | 13166 | 603971 | . | . |
| 7652 | NM_00108040 | ZNF99 | zinc finger protein 99 | protein_coding | NP_001073878.2 | 13175 | 603981 | . | . |
| 7673 | NM_00112999 | ZNF222 | zinc finger protein 222 | protein_coding | NP_001123468.1;NP_037492.2;XP_005259265.1;XP_005259266.1;XP_01 | 13015 | 617357 | . | . |
| 7675 | NM_00100872 | ZNF121 | zinc finger protein 121 | protein_coding | NP_001008727.1;NP_001295198.1;XP_016882728.1 | 12904 | 194628 | . | . |
| 7678 | NM_00124374 | ZNF124 | zinc finger protein 124 | protein_coding | NP_001230669.1;NP_001284496.1;NP_001284497.1;NP_001284498.1;NI | 12907 | 194631 | . | . |
| 7681 | NM_005664 | MKRN3 | makorin ring finger protein 3 | protein_coding | NP_005655.1 | 7114 | 603856 | . | . |
| 7690 | NM_00129754 | ZNF131 | zinc finger protein 131 | protein_coding | NP_001284477.1;NP_001317633.1;NP_001317634.1;NP_001317635.1;NI | 12915 | 604073 | . | . |
| 7692 | NM_00108333 | ZNF133 | zinc finger protein 133 | protein_coding | NP_001076799.2;NP_001269924.1;NP_001269925.2;NP_001269926.1;NI | 12917 | 604075 | . | . |
| 7693 | NM_003435 | ZNF134 | zinc finger protein 134 | protein_coding | NP_003426.3 | 12918 | 604076 | . | . |
| 7695 | NM_00134801 | ZNF136 | zinc finger protein 136 | protein_coding | NP_001334942.1;NP_001334943.1;NP_003428.1 | 12920 | 604078 | . | . |
| 7697 | NM_00116018 | ZNF138 | zinc finger protein 138 | protein_coding | NP_001153655.1;NP_001258566.1;NP_001258567.2;NP_001258568.2;NI | 12922 | 604080 | . | . |
| 7699 | NM_00130077 | ZNF140 | zinc finger protein 140 | protein_coding | NP_001287705.1;NP_001287706.1;NP_001287707.1;NP_003431.2;XP_0C | 12925 | 604082 | . | . |
| 7700 | NM_00134827 | ZNF141 | zinc finger protein 141 | protein_coding | NP_001335206.1;NP_001335207.1;NP_001335208.1;NP_001335209.1;NI | 12926 | 194648 | . | . |
| 7701 | NM_00110553 | ZNF142 | zinc finger protein 142 | protein_coding | NP_001099007.1;NP_001353216.1;NP_001353217.1;NP_001353218.1;NI | 12927 | 604083 | . | . |
| 7702 | NM_00128265 | ZNF143 | zinc finger protein 143 | protein_coding | NP_001269585.1;NP_001269586.1;NP_003433.3;XP_005253178.1;XP_0C | 12928 | 603433 | . | . |
| 7703 | NM_00136961 | PCGF2 | polycomb group ring finger 2 | protein_coding | NP_001356543.1;NP_001356544.1;NP_009075.1;XP_005257697.1;XP_01 | 12929 | 600346 | . | . |
| 7705 | NM_00109963 | ZNF146 | zinc finger protein 146 | protein_coding | NP_001093108.1;NP_001093109.1;NP_009076.2;XP_005259271.1;XP_01 | 12931 | 601505 | . | . |
| 7706 | NM_005082 | TRIM25 | tripartite motif containing 25 | protein_coding | NP_005073.2 | 12932 | 600453 | . | . |
| 7707 | NM_00134842 | ZNF148 | zinc finger protein 148 | protein_coding | NP_001335353.1;NP_001335354.1;NP_001335355.1;NP_001335356.1;NI | 12933 | 601897 | . | . |
| 7709 | NM_00124288 | ZBTB17 | zinc finger and BTB domain containing 17 | protein_coding | NP_001229813.1;NP_001274532.1;NP_001274533.1;NP_001311066.1;NI | 12936 | 604084 | . | . |
| 7710 | NM_00108538 | ZNF154 | zinc finger protein 154 | protein_coding | NP_001078853.1 | 12939 | 604085 | . | . |
| 7711 | NM_00126048 | ZNF155 | zinc finger protein 155 | protein_coding | NP_001247415.1;NP_001247416.1;NP_001247417.1;NP_003436.3;NP_9: | 12940 | 604086 | . | . |
| 7716 | NM_00133039 | VEZF1 | vascular endothelial zinc finger 1 | protein_coding | NP_001317322.1;NP_009077.2;XP_005257700.1;XP_005257701.1;XP_01 | 12949 | 606747 | . | . |
| 7718 | NM_00137649 | ZNF165 | zinc finger protein 165 | protein_coding | NP_001363420.1;NP_001363421.1;NP_001363422.1;NP_001363423.1;NI | 12953 | 600834 | . | . |
| 7726 | NM_00124278 | TRIM26 | tripartite motif containing 26 | protein_coding | NP_001229712.1;NP_003440.1;XP_005249431.1;XP_005249432.1;XP_0C | 12962 | 600830 | . | . |
| 7727 | NM_00103229 | ZNF174 | zinc finger protein 174 | protein_coding | NP_001027463.1;NP_001334797.1;NP_001334798.1;NP_001334799.1;NI | 12963 | 603900 | . | . |
| 7728 | NM_007147,N | ZNF175 | zinc finger protein 175 | protein_coding | NP_009078.1;XP_016882738.1 | 12964 | 601139 | . | . |
| 7733 | NM_00127850 | ZNF180 | zinc finger protein 180 | protein_coding | NP_001265437.2;NP_001265438.2;NP_001275688.2;NP_001275689.1;NI | 12970 | 606740 | . | . |
| 7737 | NM_006978 | RNF113A | ring finger protein 113A | protein_coding | NP_008909.1 | 12974 | 300951 | . | . |
| 7738 | NM_00131889 | ZNF184 | zinc finger protein 184 | protein_coding | NP_001305820.1;NP_001305821.1;NP_001305822.1;NP_001334761.1;NI | 12975 | 602277 | . | . |
| 7739 | NM_00117810 | ZNF185 | zinc finger protein 185 with LIM domain | protein_coding | NP_001171577.1;NP_001171578.1;NP_001171579.1;NP_001171580.1;NI | 12976 | 300381 | . | . |
| 7741 | NM_00102356 | ZSCAN26 | zinc finger and SCAN domain containing 26 | protein_coding | NP_001018854.2;NP_001104509.1;NP_001274350.1;NP_001274351.1;NI | 12978 | 616474 | . | . |
| 7743 | NM_00127823 | ZNF189 | zinc finger protein 189 | protein_coding | NP_001265160.1;NP_001265161.1;NP_001265169.1;NP_003443.2;NP_9: | 12980 | 603132 | . | . |
| 7745 | NM_00127811 | ZKSCAN8 | zinc finger with KRAB and SCAN domains 8 | protein_coding | NP_001265048.1;NP_001265050.1;NP_001265051.1;NP_006289.2;XP_01 | 12983 | 602240 | . | . |
| 7746 | NM_00119947 | ZSCAN9 | zinc finger and SCAN domain containing 9 | protein_coding | NP_001186408.1;NP_001186409.1;NP_006290.1;XP_011513177.1;XP_01 | 12984 | 602246 | . | . |
| 7748 | NM_00113051 | ZNF195 | zinc finger protein 195 | protein_coding | NP_001123991.1;NP_001123992.1;NP_001229770.1;NP_001229771.1;NI | 12986 | 602187 | . | . |
| 7750 | NM_00119096 | ZMYM2 | zinc finger MYM-type containing 2 | protein_coding | NP_001177893.1;NP_001177894.1;NP_001340086.1;NP_001340088.1;NI | 12989 | 602221 | . | . |
| 7752 | NM_00114544 | ZNF200 | zinc finger protein 200 | protein_coding | NP_001138918.1;NP_001138919.1;NP_001138920.1;NP_003445.2;NP_9: | 12993 | 603231 | . | . |
| 7753 | NM_00130177 | ZNF202 | zinc finger protein 202 | protein_coding | NP_001288708.1;NP_001288709.1;NP_001288748.1;NP_003446.2;XP_0C | 12994 | 603430 | . | . |
| 7754 | NR_002722,NF | ZNF204P | zinc finger protein 204, pseudogene | transcribed_ps | . | 12995 | 603282 | . | . |
| 7755 | NM_00104242 | ZNF205 | zinc finger protein 205 | protein_coding | NP_001035893.1;NP_001265087.1;NP_003447.2;XP_005255615.1 | 12996 | 603436 | . | . |
| 7756 | NM_00103229 | ZNF207 | zinc finger protein 207 | protein_coding | NP_001027464.1;NP_001091977.1;NP_003448.1 | 12998 | 603428 | . | . |
| 7757 | NM_00132997 | ZNF208 | zinc finger protein 208 | protein_coding | NP_001316900.1;NP_001316901.1;NP_001316902.1;NP_001316903.1;NI | 12999 | 603977 | . | . |
| 7760 | NM_00113465 | ZNF213 | zinc finger protein 213 | protein_coding | NP_001128127.1;NP_004211.1;XP_011520954.1 | 13005 | 608387 | . | . |
| 7762 | NM_00135485 | ZNF215 | zinc finger protein 215 | protein_coding | NP_001341782.1;NP_001341783.1;NP_001341784.1;NP_001341785.1;NI | 13007 | 605016 | . | . |
| 7763 | NM_00110242 | ZFAND5 | zinc finger AN1-type containing 5 | protein_coding | NP_001095890.1;NP_001095891.1;NP_001265172.1;NP_001265173.1;NI | 13008 | 604761 | . | . |
| 7764 | NM_006526,XI | ZNF217 | zinc finger protein 217 | protein_coding | NP_006517.1;XP_005260602.1;XP_006723938.1;XP_011527338.1;XP_01 | 13009 | 602967 | . | . |
| 7766 | NM_013361,XI | ZNF223 | zinc finger protein 223 | protein_coding | NP_037493.3;XP_016882747.1;XP_016882748.1;XP_016882749.1 | 13016 | . | . | . |
| 7767 | NM_00132164 | ZNF224 | zinc finger protein 224 | protein_coding | NP_001308574.1;NP_037530.2;XP_016882750.1 | 13017 | 194555 | . | . |
| 7768 | NM_00132168 | ZNF225 | zinc finger protein 225 | protein_coding | NP_001308614.1;NP_037494.2;XP_005259280.1;XP_011525587.1;XP_01 | 13018 | . | . | . |
| 7769 | NM_00103237 | ZNF226 | zinc finger protein 226 | protein_coding | NP_001027544.1;NP_001027545.1;NP_001027546.1;NP_001139692.1;NI | 13019 | . | . | . |
| 7770 | NM_00128916 | ZNF227 | zinc finger protein 227 | protein_coding | NP_001276095.1;NP_001276096.1;NP_001276097.1;NP_001276098.1;NI | 13020 | . | . | . |

| | | | | | | | | | | | | | | | | | | | | | | | | | | | | | | | | | |
|---|---|---|---|---|---|---|---|---|---|---|---|---|---|---|---|---|---|---|---|---|---|---|---|---|---|---|---|---|---|---|---|---|---|
| 7771 | NM_00108333 | ZNF112 | zinc finger protein 112 | protein_coding | NP_001076804.1;NP_001335210.1;NP_001335211.1;NP_001335212.1;NI | 12892 | 603994 | . | . | 1.372097 | 1.516203 | 1.879085 | 1.311817 | 3.192927 | 3.504772 | 4.216613 | 2.999313 | 1.081269 | 175.301294 | 0.271544497 | 0.415129195 | 0.678047329 | 0.998865404 | 7.395505 | 7.509624 | 7.411504 | 7.450109 | 7.540691 | 7.381277 | 185 | 190 | 188 | 139 |
| 7772 | NM_00127851 | ZNF229 | zinc finger protein 229 | protein_coding | NP_001265439.2;NP_055333.3;XP_006723435.1;XP_011525594.1;XP_01 | 13022 | . | . | . | 0.977931 | 0.906634 | 1.285461 | 1.129478 | 2.275685 | 2.095726 | 2.884537 | 2.582418 | 1.224731 | 178.377295 | 0.252199379 | 1.15965621 | 0.246188806 | 0.992733415 | 7.324543 | 7.618124 | 7.445325 | 7.412895 | 7.549224 | 7.481075 | 192 | 166 | 184 | 170 |
| 7773 | NM_006300 | ZNF230 | zinc finger protein 230 | protein_coding | NP_006291.2 | 13024 | . | . | . | 0.894075 | 1.080329 | 1.227308 | 1.042136 | 2.080548 | 2.497229 | 2.754042 | 2.382721 | 1.134689 | 136.299037 | 0.286439071 | 0.636426074 | 0.524498751 | 0.998865404 | 6.996518 | 7.178980 | 7.030086 | 7.094105 | 7.141337 | 7.075817 | 134 | 150 | 137 | 124 |
| 7775 | NM_00132095 | ZNF232 | zinc finger protein 232 | protein_coding | NP_001307881.1;NP_001307882.1;NP_001307883.1;NP_001307884.1;NI | 13026 | 616463 | . | . | 6.696119 | 8.171670 | 9.866178 | 8.703316 | 15.582141 | 18.889180 | 22.139418 | 19.899102 | 1.222384 | 464.503964 | 0.181864772 | 1.592929584 | 0.111175989 | 0.848126978 | 8.707436 | 8.997139 | 8.740637 | 8.836920 | 8.957707 | 8.869652 | 443 | 487 | 475 | 446 |
| 7776 | NM_00130608 | ZNF236 | zinc finger protein 236 | protein_coding | NP_001293018.1;NP_031371.3;XP_011524467.1;XP_011524468.1;XP_01 | 13028 | 604760 | . | . | 1.474724 | 1.077924 | 1.519139 | 1.146637 | 3.431741 | 2.491668 | 3.408903 | 2.621651 | 1.150638 | 343.884527 | 0.196426331 | 1.030585801 | 0.302735097 | 0.998865404 | 8.320643 | 8.523773 | 8.378625 | 8.385305 | 8.508648 | 8.411703 | 366 | 347 | 347 | 316 |
| 7779 | NM_021194 | SLC30A1 | solute carrier family 30 member 1 | protein_coding | NP_067017.2 | 11012 | 609521 | . | . | 9.627101 | 7.361314 | 6.877287 | 4.181761 | 22.402658 | 17.016008 | 15.432433 | 9.561100 | -1.556783 | 1290.947109 | 0.407071409 | -1.56868803 | 0.116720649 | 0.857396579 | 10.618362 | 9.980024 | 10.573082 | 10.350762 | 10.294567 | 9.937586 | 2059 | 1463 | 1096 | 712 |
| 7781 | NM_00131894 | SLC30A3 | solute carrier family 30 member 3 | protein_coding | NP_001305878.1;NP_001305879.1;NP_001305880.1;NP_003450.2;XP_0C | 11014 | 602878 | . | . | 1.914620 | 1.413402 | 1.493401 | 1.361020 | 4.455399 | 3.267141 | 3.351149 | 3.111812 | -1.159844 | 131.519774 | 0.288830886 | -0.74067852 | 0.458888387 | 0.998865404 | 7.141674 | 6.928749 | 7.103408 | 7.025718 | 7.018622 | 7.005164 | 179 | 137 | 108 | 112 |
| 7782 | NM_00132103 | SLC30A4 | solute carrier family 30 member 4 | protein_coding | NP_001307965.1;NP_037441.2;XP_011520299.1;XP_016878049.1;XP_01 | 11015 | 602095 | . | . | 1.382486 | 1.543102 | 1.405222 | 1.273896 | 3.217101 | 3.566948 | 3.153279 | 2.912612 | -1.035653 | 218.829354 | 0.250531796 | -0.20173527 | 0.840123691 | 0.998865404 | 7.797952 | 7.748957 | 7.765764 | 7.789229 | 7.843866 | 7.671091 | 250 | 246 | 222 | 164 |
| 7784 | NM_00111035 | ZP3 | zona pellucida glycoprotein 3 | protein_coding | NP_001103824.1;NP_009086.4;XP_016868075.1 | 13189 | 182889 | . | . | 0.329442 | 0.249076 | 0.298877 | 0.213506 | 0.766626 | 0.575750 | 0.670672 | 0.488155 | -1.189069 | 12.749007 | 0.857453388 | -0.2913662 | 0.770771259 | NA | 3.790753 | 3.543272 | 3.687881 | 3.671737 | 3.674464 | 3.665855 | 18 | 13 | 11 | 10 |
| 7786 | NM_00119351 | MAP3K12 | mitogen-activated protein kinase kinase kinase 12 | protein_coding | NP_001180440.1;NP_006292.3;XP_005269195.1;XP_005269197.1;XP_00 | 6851 | 600447 | . | . | 5.756077 | 6.461574 | 8.125296 | 8.611230 | 13.394624 | 14.936219 | 18.232929 | 19.688554 | 1.320819 | 986.093550 | 0.140184811 | 2.863599792 | 0.004188569 | 0.192048182 | 9.731162 | 10.132217 | 9.800779 | 9.863945 | 10.016363 | 10.055314 | 935 | 958 | 951 | 1079 |
| 7789 | NM_007156 | ZXDA | zinc finger X-linked duplicated A | protein_coding | NP_009087.1 | 13198 | 300235 | . | . | 0.185162 | 0.277302 | 0.229252 | 0.189078 | 0.430879 | 0.640996 | 0.514436 | 0.432304 | -1.100200 | 36.172116 | 0.529455021 | -0.26020371 | 0.794706645 | 0.998865404 | 5.244239 | 5.106066 | 5.150361 | 5.210290 | 5.180766 | 5.153554 | 35 | 49 | 33 | 29 |
| 7791 | NM_00101097 | ZYX | zyxin | protein_coding | NP_001010972.1;NP_001349712.1;NP_003452.1 | 13200 | 602002 | . | . | 33.847856 | 36.611999 | 27.917455 | 33.674875 | 78.765348 | 84.630280 | 62.645957 | 76.993600 | -1.160275 | 2289.098166 | 0.130193408 | -1.64729318 | 0.0994978 | 0.813730268 | 11.263993 | 11.049142 | 11.186573 | 11.245312 | 11.023933 | 11.164873 | 2730 | 2749 | 1678 | 2164 |
| 7798 | NM_00114254 | LUZP1 | leucine zipper protein 1 | protein_coding | NP_001136018.1;NP_361013.3;XP_011540392.1;XP_011540393.1;XP_01 | 14985 | 601422 | . | . | 6.632963 | 7.621108 | 7.197977 | 7.784640 | 15.435175 | 17.616532 | 16.152050 | 17.798655 | 1.035466 | 1924.270336 | 0.127306954 | 0.394956656 | 0.692874876 | 0.998865404 | 10.884898 | 10.934861 | 10.843106 | 10.945242 | 10.895874 | 10.949315 | 2037 | 2172 | 1645 | 1897 |
| 7799 | NM_00100725 | PRDM2 | PR/SET domain 2 | protein_coding | NP_001007258.1;NP_001129082.1;NP_036363.2;NP_056950.2;XP_01685 | 9347 | 601196 | . | . | 9.947446 | 10.214972 | 10.071586 | 9.672057 | 23.148111 | 23.612369 | 22.600350 | 22.114008 | 1.014069 | 2034.683376 | 0.121854922 | 0.165411123 | 0.868620399 | 0.998865404 | 10.980466 | 11.000641 | 10.953142 | 11.015148 | 11.021423 | 10.969387 | 2236 | 2265 | 1840 | 1858 |
| 7803 | NM_003463,XI | PTP4A1 | protein tyrosine phosphatase 4A1 | protein_coding | NP_003454.1;XP_011534413.1;XP_011534414.1;XP_016866759.1;XP_01 | 9634 | 601585 | . | . | 19.322552 | 17.778966 | 20.231868 | 16.974207 | 44.964376 | 41.096876 | 45.399725 | 38.809502 | -1.006309 | 2940.433671 | 0.129444411 | -0.07009718 | 0.944116312 | 0.998865404 | 11.526183 | 11.517431 | 11.561950 | 11.482393 | 11.590995 | 11.439247 | 3561 | 3028 | 2786 | 2493 |
| 7804 | NM_00101805 | LRP8 | LDL receptor related protein 8 | protein_coding | NP_001018064.1;NP_004622.2;NP_059992.3;NP_150643.2;XP_00527123 | 6700 | 602600 | . | . | 17.465616 | 17.405571 | 22.900131 | 17.912069 | 40.643204 | 40.233760 | 51.387230 | 40.953815 | 1.224882 | 3249.008607 | 0.143006803 | 2.04635271 | 0.040721681 | 0.62818612 | 11.511903 | 11.804822 | 11.585251 | 11.561202 | 11.857959 | 11.607075 | 3413 | 3104 | 3575 | 2854 |
| 7805 | NM_006762,XI | LAPTM5 | lysosomal protein transmembrane 5 | protein_coding | NP_006753.1;XP_011540400.1 | 29612 | 601476 | . | . | 1.295576 | 1.398088 | 0.514206 | 0.543322 | 3.014858 | 3.231745 | 1.153863 | 1.242242 | -2.545042 | 64.703384 | 0.40366849 | -3.33860524 | 0.000842001 | 0.060848921 | 6.537847 | 5.189480 | 6.086064 | 6.107482 | 5.896764 | 5.901329 | 103 | 104 | 31 | 35 |
| 7812 | NM_00100755 | CSDE1 | cold shock domain containing E1 | protein_coding | NP_001007554.1;NP_001123995.1;NP_001229820.1;NP_001229821.1;NI | 29905 | 191510 | . | . | 108.591638 | 110.280193 | 105.395283 | 116.137654 | 252.697200 | 254.917608 | 236.503947 | 265.534947 | 1.000735 | 13779.925021 | 0.103659205 | 0.01022973 | 0.991837999 | 0.999576129 | 13.749769 | 13.750791 | 13.747433 | 13.752057 | 13.712454 | 13.786922 | 15819 | 14904 | 11466 | 13485 |
| 7813 | NM_00130824 | EVI5 | ecotropic viral integration site 5 | protein_coding | NP_001295177.1;NP_001337126.1;NP_001337127.1;NP_001364139.1;NI | 3501 | 602942 | . | . | 1.794455 | 1.640572 | 1.872111 | 1.882479 | 4.175772 | 3.792256 | 4.200962 | 4.304064 | 1.060249 | 433.699485 | 0.179482024 | 0.470255977 | 0.638172146 | 0.998865404 | 8.717547 | 8.802312 | 8.781120 | 8.700365 | 8.772104 | 8.782342 | 517 | 424 | 385 | 422 |
| 7818 | NM_00119984 | DAP3 | death associated protein 3 | protein_coding | NP_001186778.1;NP_001186779.1;NP_001186780.1;NP_004623.1;NP_3 | 2673 | 602074 | . | . | 31.339946 | 33.936443 | 34.998365 | 40.612315 | 72.929343 | 78.445609 | 78.535317 | 92.855238 | 1.019404 | 2002.179074 | 0.14036332 | 0.19753034 | 0.843412541 | 0.998865404 | 10.953647 | 10.980935 | 10.937193 | 10.978087 | 10.871559 | 11.065325 | 2225 | 2195 | 1544 | 2125 |
| 7832 | NM_006763 | BTG2 | BTG anti-proliferation factor 2 | protein_coding | NP_006754.1 | 1131 | 601597 | . | . | 1.861511 | 1.902583 | 2.514987 | 2.988194 | 4.331812 | 4.397906 | 5.643558 | 6.832150 | 1.443078 | 197.117695 | 0.242111796 | 2.185557305 | 0.028848009 | 0.547128317 | 7.334685 | 7.863036 | 7.523970 | 7.528499 | 7.649256 | 7.729124 | 185 | 175 | 186 | 236 |
| 7837 | NM_012293,XI | PXDN | peroxidasin | protein_coding | NP_036425.1;XP_005264764.1;XP_011508698.1 | 14966 | 605158 | . | . | 0.269288 | 0.247648 | 0.341905 | 0.283846 | 0.626643 | 0.572450 | 0.767224 | 0.648981 | 1.199707 | 61.085445 | 0.405559169 | 0.64770263 | 0.517177303 | 0.998865404 | 5.794547 | 6.058876 | 5.917366 | 5.897841 | 5.969967 | 5.923201 | 67 | 57 | 64 | 56 |
| 7840 | NM_00137845 | ALMS1 | ALMS1 centrosome and basal body associated protein | protein_coding | NP_001365383.1;NP_055935.4 | 428 | 606844 | . | . | 1.410245 | 1.563865 | 1.543160 | 1.638482 | 3.281698 | 3.614944 | 3.462807 | 3.746194 | 1.055720 | 620.183742 | 0.158846577 | 0.492467613 | 0.622388812 | 0.998865404 | 9.237148 | 9.314908 | 9.225408 | 9.288630 | 9.276712 | 9.310303 | 662 | 682 | 540 | 612 |
| 7841 | NM_00114615 | MOGS | mannosyl-oligosaccharide glucosidase | protein_coding | NP_001139630.1;NP_006293.2 | 24862 | 601336 | . | . | 40.219127 | 38.851863 | 35.645891 | 36.779466 | 93.591557 | 89.807823 | 79.988351 | 84.091882 | -1.106955 | 3376.877517 | 0.110432329 | -1.32748358 | 0.184348769 | 0.956844295 | 11.792908 | 11.646318 | 11.779234 | 11.744109 | 11.669072 | 11.687339 | 4172 | 3748 | 2756 | 3039 |
| 7844 | NM_00119895 | RNF103 | ring finger protein 103 | protein_coding | NP_001185880.1;NP_001185881.1;NP_005658.1 | 12859 | 602507 | . | . | 7.309425 | 6.138030 | 6.479808 | 5.589751 | 17.009332 | 14.188329 | 14.540500 | 12.780299 | -1.117656 | 667.069181 | 0.171560206 | -0.93538938 | 0.34958767 | 0.998865404 | 9.459218 | 9.299763 | 9.481236 | 9.346339 | 9.399498 | 9.278581 | 889 | 686 | 598 | 537 |
| 7846 | NM_00127039 | TUBA1A | tubulin alpha 1a | protein_coding | NP_001257328.1;NP_001257329.1;NP_006000.2 | 20766 | 602529 | . | . | 111.237526 | 109.226985 | 103.079931 | 128.180643 | 258.854290 | 252.483078 | 231.308375 | 293.069793 | 1.019443 | 6019.683638 | 0.121691043 | 0.22829723 | 0.819415177 | 0.998865404 | 12.541579 | 12.569232 | 12.553086 | 12.539149 | 12.475667 | 12.642252 | 6922 | 6380 | 4764 | 6259 |
| 7849 | NM_003466,N | PAX8 | paired box 8 | protein_coding | NP_003457.1;NP_039246.1;NP_039247.1;NP_054698.1 | 8622 | 167415 | . | . | 0.326692 | 0.324762 | 0.375303 | 0.392314 | 0.760227 | 0.750701 | 0.842170 | 0.896980 | 1.155573 | 45.405696 | 0.460775015 | 0.452734296 | 0.65074008 | 0.998865404 | 5.396728 | 5.605340 | 5.491049 | 5.487886 | 5.512705 | 5.515722 | 49 | 45 | 42 | 46 |
| 7850 | NM_00126141 | IL1R2 | interleukin 1 receptor type 2 | protein_coding | NP_001248348.1;NP_004624.1;XP_006712797.1;XP_006712799.1;XP_01 | 5994 | 147811 | . | . | 0.197954 | 0.398086 | 0.047423 | 0.475913 | 0.460648 | 0.920195 | 0.106417 | 1.088117 | -1.493141 | 16.296461 | 0.819757462 | -0.70551424 | 0.480490275 | NA | 4.288580 | 3.705970 | 4.068455 | 4.019265 | 3.990167 | 4.041996 | 29 | 15 | 6 | 18 |
| 7852 | NM_00100854 | CXCR4 | C-X-C motif chemokine receptor 4 | protein_coding | NP_001008540.1;NP_001334985.1;NP_001334988.1;NP_001334989.1;NI | 2561 | 162643 | . | . | 0.101969 | 0.031290 | 0.185712 | 0.080194 | 0.237287 | 0.072329 | 0.416732 | 0.183354 | 2.106936 | 6.191415 | 1.278806159 | 0.840742206 | 0.400492371 | NA | 1.987791 | 3.073227 | 2.607701 | 2.587738 | 2.631339 | 2.604339 | 7 | 2 | 10 | 5 |
| 7855 | NM_003468,XI | FZD5 | frizzled class receptor 5 | protein_coding | NP_003459.2;XP_024308898.1 | 4043 | 601723 | . | . | 0.974191 | 1.005200 | 0.870633 | 0.834773 | 2.266981 | 2.323566 | 1.953674 | 1.908610 | -1.190739 | 217.429341 | 0.230058469 | -1.09475074 | 0.273625852 | 0.998865404 | 7.884796 | 7.633040 | 7.827580 | 7.780422 | 7.713492 | 7.731507 | 285 | 243 | 169 | 190 |
| 7857 | NM_003469 | SCG2 | secretogranin II | protein_coding | NP_003460.2 | 10575 | 118930 | . | . | 4.409953 | 4.154352 | 4.101600 | 3.582836 | 10.262141 | 9.602970 | 9.203870 | 8.191728 | -1.130944 | 308.501381 | 0.203578692 | -0.87203341 | 0.383190149 | 0.998865404 | 8.354717 | 8.178150 | 8.320514 | 8.281268 | 8.271449 | 8.196100 | 390 | 341 | 270 | 252 |
| 7862 | NM_00100369 | BRPF1 | bromodomain and PHD finger containing 1 | protein_coding | NP_001003694.1;NP_001305978.1;NP_001305979.1;NP_004625.2;XP_0C | 14255 | 602410 | . | . | 3.141614 | 3.655461 | 3.601058 | 3.717591 | 7.310665 | 8.449761 | 8.080666 | 8.499831 | 1.058459 | 525.228010 | 0.168700898 | 0.485860125 | 0.627066342 | 0.998865404 | 8.995448 | 9.077000 | 8.974161 | 9.058655 | 9.052757 | 9.054609 | 549 | 587 | 469 | 507 |
| 7866 | NM_006764 | IFRD2 | interferon related developmental regulator 2 | protein_coding | NP_006755.4 | 5457 | 602725 | . | . | 18.675114 | 18.712290 | 20.040703 | 19.728168 | 43.457752 | 43.254295 | 44.970757 | 45.106110 | 1.048830 | 1267.497196 | 0.128759754 | 0.53418206 | 0.59321558 | 0.998865404 | 10.272922 | 10.341790 | 10.291501 | 10.286955 | 10.335067 | 10.315597 | 1429 | 1331 | 1143 | 1201 |
| 7867 | NM_00124392 | MAPKAPK3 | MAPK activated protein kinase 3 | protein_coding | NP_001230854.1;NP_001230855.1;NP_004626.1 | 6888 | 602130 | . | . | 8.893253 | 9.602770 | 9.668443 | 11.589274 | 20.694966 | 22.197232 | 21.695705 | 26.497498 | 1.131187 | 788.423497 | 0.155269514 | 1.1453484 | 0.252064864 | 0.994172589 | 9.531538 | 9.708682 | 9.552917 | 9.598971 | 9.599351 | 9.722466 | 824 | 825 | 665 | 852 |
| 7869 | NM_00100591 | SEMA3B | semaphorin 3B | protein_coding | NP_001005914.1;NP_001276989.1;NP_001276990.1;NP_001276991.1;NI | 10724 | 601281 | . | . | 57.170296 | 62.613447 | 38.544330 | 46.477677 | 133.037629 | 144.733776 | 86.492364 | 106.265688 | -1.419213 | 4648.867369 | 0.112984925 | -4.4704268 | 7.80637E-06 | 0.001608228 | 12.413271 | 11.908049 | 12.322148 | 12.299275 | 11.970782 | 12.064539 | 6364 | 5808 | 3147 | 3813 |
| 7871 | NM_00130442 | SLMAP | sarcolemma associated protein | protein_coding | NP_001291349.1;NP_001291350.1;NP_001291351.1;NP_001291352.1;NI | 16643 | 602701 | . | . | 3.448792 | 3.572657 | 3.589865 | 3.470658 | 8.025481 | 8.258361 | 8.055549 | 7.935247 | -1.002992 | 660.737041 | 0.153907057 | -0.02800275 | 0.977659959 | 0.998865404 | 9.370029 | 9.365832 | 9.355243 | 9.382019 | 9.385654 | 9.347432 | 747 | 728 | 590 | 601 |
| 7873 | NM_006010 | MANF | mesencephalic astrocyte derived neurotrophic factor | protein_coding | NP_006001.5 | 15461 | 601916 | . | . | 103.726944 | 104.878883 | 102.263817 | 101.170959 | 241.376877 | 242.432236 | 229.477066 | 231.315384 | -1.039788 | 2917.866947 | 0.111400283 | -0.50529055 | 0.613354781 | 0.998865404 | 11.538552 | 11.482296 | 11.525384 | 11.527471 | 11.503092 | 11.486022 | 3421 | 3214 | 2514 | 2654 |
| 7874 | NM_00128645 | USP7 | ubiquitin specific peptidase 7 | protein_coding | NP_001273386.2;NP_001273387.1;NP_001308787.1;NP_003461.2;XP_01 | 12630 | 602519 | . | . | 12.537311 | 12.864626 | 12.478799 | 11.951535 | 29.174835 | 29.737162 | 28.002063 | 27.325767 | -1.062139 | 2246.588459 | 0.116006082 | -0.74972499 | 0.453420353 | 0.998865404 | 11.176325 | 11.089407 | 11.152414 | 11.161990 | 11.122344 | 11.095264 | 2649 | 2512 | 1927 | 2008 |
| 7879 | NM_004637 | RAB7A | RAB7A, member RAS oncogene family | protein_coding | NP_004628.4 | 9788 | 602298 | . | . | 131.386826 | 143.999237 | 143.815872 | 159.912231 | 305.742554 | 332.860687 | 322.718658 | 365.620300 | 1.087621 | 9849.605443 | 0.109310274 | 1.108554001 | 0.267622631 | 0.998546485 | 13.204032 | 13.325128 | 13.192823 | 13.263056 | 13.258126 | 13.340269 | 10415 | 10605 | 8497 | 10082 |
| 7881 | NM_00130821 | KCNAB1 | potassium voltage-gated channel subfamily A member regulatory | protein_coding | NP_001295146.1;NP_001295151.1;NP_003462.2;NP_751891.1;NP_7518 | 6228 | 601141 | . | . | 0.567198 | 0.568583 | 0.314697 | 0.387754 | 1.319894 | 1.314305 | 0.706172 | 0.886555 | -1.630298 | 100.843263 | 0.324992585 | -2.1696977 | 0.030029755 | 0.553375672 | 6.966407 | 6.259886 | 6.726227 | 6.720487 | 6.551736 | 6.599702 | 145 | 134 | 60 | 79 |
| 7884 | NM_00130607 | SLBP | stem-loop binding protein | protein_coding | NP_001293003.1;NP_001293004.1;NP_006518.1 | 10904 | 602422 | . | . | 23.590333 | 17.331713 | 16.737172 | 21.327156 | 54.895672 | 40.063033 | 37.557732 | 48.762013 | -1.104951 | 1283.656112 | 0.140878975 | -1.02202636 | 0.306768423 | 0.998865404 | 10.396274 | 10.252217 | 10.415245 | 10.305559 | 10.239704 | 10.330487 | 1662 | 1351 | 996 | 1213 |
| 7903 | NM_005668,N | ST8SIA4 | ST8 alpha-N-acetyl-neuraminide alpha-2,8-sialyltransferase 4 | protein_coding | NP_005659.1;NP_778222.1;XP_005272135.1;XP_011541932.1 | 10871 | 602547 | . | . | 0.958502 | 0.688382 | 0.441672 | 0.831312 | 2.230474 | 1.591224 | 0.991099 | 1.900694 | -1.158378 | 76.837437 | 0.376623085 | -0.56317838 | 0.573313421 | 0.998865404 | 6.365604 | 6.154128 | 6.329759 | 6.231065 | 6.229233 | 6.258867 | 112 | 73 | 58 | 71 |
| 7905 | NM_005669,XI | REEP5 | receptor accessory protein 5 | protein_coding | NP_005660.4;XP_016865333.1 | 30077 | 125265 | . | . | 29.803752 | 28.130739 | 28.120759 | 25.978000 | 69.354555 | 65.025458 | 63.102169 | 59.395607 | -1.086171 | 2620.221845 | 0.118728776 | -1.00440548 | 0.315183211 | 0.998865404 | 11.413754 | 11.294738 | 11.414596 | 11.360049 | 11.356647 | 11.283543 | 3246 | 2846 | 2284 | 2248 |
| 7913 | NM_00113470 | DEK | DEK proto-oncogene | protein_coding | NP_001128181.1;NP_003463.1;XP_016866762.1;XP_024302311.1;XP_02 | 2768 | 125264 | . | . | 22.875631 | 20.299947 | 22.296819 | 18.203341 | 53.232533 | 46.924240 | 50.033416 | 41.619774 | -1.104529 | 2001.644064 | 0.139660404 | -1.02699664 | 0.304422046 | 0.998865404 | 11.036674 | 10.893726 | 11.056302 | 10.947525 | 11.000989 | 10.841767 | 2582 | 2114 | 1822 | 1603 |
| 7915 | NM_001080,N | ALDH5A1 | aldehyde dehydrogenase 5 family member A1 | protein_coding | NP_001071.1;NP_001355883.1;NP_733936.1 | 408 | 610045 | . | . | 0.088109 | 0.119378 | 0.125225 | 0.084956 | 0.205032 | 0.275947 | 0.281002 | 0.194242 | 1.009109 | 17.197841 | 0.747931068 | 0.017491611 | 0.986044426 | NA | 4.096985 | 4.111291 | 4.089585 | 4.110354 | 4.114894 | 4.086268 | 17 | 21 | 18 | 13 |
| 7916 | NM_004638,N | PRRC2A | proline rich coiled-coil 2A | protein_coding | NP_004629.3;NP_542417.2;XP_016866763.1 | 13918 | 142580 | . | . | 21.686944 | 25.486172 | 23.801187 | 26.133522 | 50.466412 | 58.912427 | 53.409176 | 59.751195 | 1.044104 | 5200.568762 | 0.119302478 | 0.521911707 | 0.601731807 | 0.998865404 | 12.313068 | 12.375171 | 12.258608 | 12.387639 | 12.325954 | 12.395421 | 5408 | 5914 | 4429 | 5186 |
| 7917 | NM_00109853 | BAG6 | BAG cochaperone 6 | protein_coding | NP_001092004.1;NP_001186626.1;NP_001186627.1;NP_542433.1;NP_5 | 13919 | 142590 | . | . | 43.574768 | 47.085963 | 43.737638 | 50.974523 | 101.400279 | 108.841306 | 98.145981 | 116.547195 | 1.025605 | 5406.404240 | 0.11689772 | 0.312031484 | 0.755016596 | 0.998865404 | 12.382182 | 12.418497 | 12.361750 | 12.415526 | 12.346685 | 12.469943 | 5950 | 5945 | 4412 | 5508 |
| 7918 | NM_00119923 | GPANK1 | G-patch domain and ankyrin repeats 1 | protein_coding | NP_001186166.1;NP_001186167.1;NP_001186168.1;NP_001186169.1;NI | 13920 | 142610 | . | . | 8.086132 | 8.232506 | 8.525463 | 9.593001 | 18.816765 | 19.029806 | 19.130894 | 21.933256 | 1.090990 | 540.350411 | 0.165991607 | 0.756894758 | 0.449112916 | 0.998865404 | 9.013806 | 9.138982 | 9.047700 | 9.048314 | 9.065857 | 9.141257 | 595 | 558 | 462 | 559 |
| 7919 | NM_004640,N | DDX39B | DExD-box helicase 39B | protein_coding | NP_004631.1;NP_542165.1 | 13917 | 142560 | . | . | 98.122495 | 104.496782 | 99.381282 | 102.623174 | 228.335076 | 241.549003 | 223.008718 | 234.635708 | -1.016001 | 5218.861848 | 0.106819417 | -0.21439498 | 0.830239057 | 0.998865404 | 12.360960 | 12.337987 | 12.332564 | 12.378992 | 12.330350 | 12.354907 | 5890 | 5833 | 4436 | 4925 |
| 7920 | NM_00117751 | ABHD16A | abhydrolase domain containing 16A, phospholipase | protein_coding | NP_001170986.1;NP_066983.1 | 13921 | 142620 | . | . | 14.256945 | 15.465767 | 14.504428 | 14.585570 | 33.176500 | 35.749813 | 32.547514 | 33.348174 | -1.041169 | 902.401771 | 0.1417912 | -0.41048896 | 0.681447297 | 0.998865404 | 9.846543 | 9.788118 | 9.803253 | 9.859743 | 9.799238 | 9.806276 | 1019 | 1032 | 765 | 833 |
| 7922 | NM_00107751 | SLC39A7 | solute carrier family 39 member 7 | protein_coding | NP_001070984.1;NP_001275706.1;NP_008910.2 | 4927 | 601416 | . | . | 77.378357 | 73.939812 | 65.874535 | 68.145485 | 180.062615 | 170.915192 | 147.820549 | 155.806557 | -1.144398 | 5041.189949 | 0.107182109 | -1.81549956 | 0.069447174 | 0.744724233 | 12.393566 | 12.198974 | 12.375380 | 12.330701 | 12.225955 | 12.254756 | 6359 | 5653 | 4017 | 4485 |
| 7923 | NM_014234 | HSD17B8 | hydroxysteroid 17-beta dehydrogenase 8 | protein_coding | NP_055049.1 | 3554 | 601417 | . | . | 1.336998 | 1.095649 | 1.257846 | 1.419984 | 3.111249 | 2.532642 | 2.822570 | 3.246626 | 1.090077 | 39.701813 | 0.495199228 | 0.251272863 | 0.801603154 | 0.998865404 | 5.247449 | 5.372125 | 5.317643 | 5.288239 | 5.307498 | 5.325014 | 48 | 37 | 34 | 41 |
| 7936 | NM_002904,XI | NELFE | negative elongation factor complex member E | protein_coding | NP_002895.3;XP_006715268.1;XP_011513215.1;XP_016866788.1 | 13974 | 154040 | . | . | 37.024351 | 38.307640 | 32.487400 | 42.830548 | 86.157188 | 88.549825 | 72.900786 | 97.926955 | -1.008634 | 1616.690102 | 0.144055908 | -0.08609205 | 0.931393243 | 0.998865404 | 10.665258 | 10.652368 | 10.652487 | 10.668311 | 10.552505 | 10.745615 | 1851 | 1770 | 1226 | 1696 |
| 7942 | NM_00116782 | TFEB | transcription factor EB | protein_coding | NP_001161299.2;NP_001258872.1;NP_001258873.1;NP_001258874.1;NI | 11753 | 600744 | . | . | 0.399256 | 0.406918 | 0.482014 | 0.573965 | 0.929086 | 0.940610 | 1.081625 | 1.312303 | 1.301975 | 35.609331 | 0.520198857 | 0.731837832 | 0.464267555 | 0.998865404 | 4.951979 | 5.331500 | 5.127125 | 5.131971 | 5.153070 | 5.179649 | 35 | 34 | 32 | 41 |
| 7957 | NM_00101804 | EPM2A | EPM2A glucan phosphatase, laforin | protein_coding | NP_001018051.1;NP_001346986.1;NP_001346993.1;NP_001347000.1;NI | 3413 | 607566 | . | . | 1.306719 | 1.270329 | 1.832431 | 1.658355 | 3.040790 | 2.936421 | 4.111920 | 3.791632 | 1.334589 | 151.311604 | 0.267534303 | 1.556418249 | 0.119608669 | 0.858998979 | 7.017924 | 7.434820 | 7.172270 | 7.178754 | 7.310902 | 7.264168 | 148 | 141 | 159 | 153 |
| 7965 | NM_00132660 | AIMP2 | aminoacyl tRNA synthetase complex interacting multifunctional p | protein_coding | NP_001313535.1;NP_001313536.1;NP_001313538.1;NP_001313539.1;NI | 20609 | 600859 | . | . | 21.379600 | 23.839387 | 22.843493 | 25.065671 | 49.751209 | 55.105809 | 51.260139 | 57.309681 | 1.048751 | 869.523166 | 0.145925137 | 0.470596707 | 0.63792876 | 0.998865404 | 9.729588 | 9.797768 | 9.714581 | 9.776891 | 9.749064 | 9.810287 | 935 | 956 | 741 | 870 |
| 7975 | NM_002360,XI | MAFK | MAF bZIP transcription factor K | protein_coding | NP_002351.1;XP_005249908.2;XP_006715836.1 | 6782 | 600197 | . | . | 5.066017 | 6.089343 | 5.816541 | 4.676785 | 11.788832 | 14.075788 | 13.052148 | 10.692914 | -1.073259 | 562.163660 | 0.18238717 | -0.55924341 | 0.575995609 | 0.998865404 | 9.184794 | 9.083107 | 9.087717 | 9.217416 | 9.177320 | 9.034617 | 605 | 689 | 524 | 452 |
| 7976 | NM_017412,N | FZD3 | frizzled class receptor 3 | protein_coding | NP_059108.1;NP_665873.1;XP_016869330.1;XP_016869331.1;XP_01686 | 4041 | 606143 | . | . | 1.508746 | 0.657340 | 0.757686 | 0.652302 | 3.510914 | 1.519471 | 1.700225 | 1.491410 | -1.015435 | 303.554234 | 0.220110956 | -0.10039488 | 0.920030834 | 0.998865404 | 8.256069 | 8.235479 | 8.320188 | 8.167810 | 8.280354 | 8.195376 | 401 | 284 | 283 | 260 |
| 7978 | NM_00130113 | MTERF1 | mitochondrial transcription termination factor 1 | protein_coding | NP_001288063.1;NP_001288064.1;NP_008911.1;XP_005250650.1;XP_00 | 21463 | 602318 | . | . | 3.876746 | 4.274461 | 4.030087 | 3.909024 | 9.021347 | 9.880610 | 9.043398 | 8.937521 | -1.145501 | 437.590327 | 0.199438135 | -0.98265603 | 0.325776759 | 0.998865404 | 8.867673 | 8.672613 | 8.773014 | 8.845470 | 8.816412 | 8.629185 | 507 | 533 | 411 | 322 |
| 7979 | NM_006304,N | SEM1 | SEM1 26S proteasome complex subunit | protein_coding | NP_006295.1;XP_024302702.1 | 10845 | 601285 | . | . | 111.783647 | 103.488718 | 95.013452 | 106.300362 | 260.125123 | 239.218804 | 213.207429 | 243.043132 | -1.082815 | 1629.687579 | 0.128956299 | -0.89012363 | 0.373399505 | 0.998865404 | 10.726679 | 10.611794 | 10.733556 | 10.665774 | 10.599814 | 10.675231 | 2035 | 1750 | 1294 | 1539 |
| 7980 | NM_00127100 | TFPI2 | tissue factor pathway inhibitor 2 | protein_coding | NP_001257932.1;NP_001257933.1;NP_006519.1 | 11761 | 600033 | . | . | 0.089287 | 0.072807 | 0.083126 | 0.130292 | 0.207774 | 0.168297 | 0.186532 | 0.297896 | 1.228822 | 6.964470 | 1.152038105 | 0.258043411 | 0.796373405 | NA | 2.644780 | 2.940155 | 2.797386 | 2.790955 | 2.792152 | 2.809134 | 8 | 6 | 5 | 9 |
| 7982 | NM_00136959 | ST7 | suppression of tumorigenicity 7 | protein_coding | NP_001356527.1;NP_001356528.1;NP_001356529.1;NP_001356530.1;NI | 11351 | 600833 | . | . | 27.327442 | 28.859025 | 32.727519 | 29.758965 | 63.592076 | 66.708928 | 73.439604 | 68.040342 | 1.080577 | 2248.508801 | 0.121491358 | 0.920245251 | 0.357444612 | 0.998865404 | 11.077691 | 11.189643 | 11.087426 | 11.116759 | 11.208340 | 11.118562 | 2443 | 2375 | 2144 | 2068 |
| 7984 | NM_005435,XI | ARHGEF5 | Rho guanine nucleotide exchange factor 5 | protein_coding | NP_005426.2;XP_016868112.1 | 13209 | 600888 | . | . | 1.109045 | 1.079787 | 1.284457 | 1.456018 | 2.580795 | 2.495975 | 2.882284 | 3.329011 | 1.232790 | 210.044114 | 0.233582678 | 1.292589693 | 0.196153006 | 0.96621494 | 7.555923 | 7.857448 | 7.669065 | 7.651839 | 7.728551 | 7.784786 | 221 | 199 | 190 | 230 |
| 7988 | NM_012256 | ZNF212 | zinc finger protein 212 | protein_coding | NP_036388.2 | 13004 | 602386 | . | . | 2.925102 | 2.764892 | 2.579860 | 3.876642 | 6.806832 | 6.391171 | 5.789132 | 8.863481 | 1.118104 | 265.482826 | 0.238290632 | 0.675837694 | 0.499120841 | 0.998865404 | 7.970402 | 8.130129 | 8.033734 | 8.001730 | 7.967681 | 8.172129 | 298 | 262 | 196 | 314 |
| 7991 | NM_00135642 | TUSC3 | tumor suppressor candidate 3 | protein_coding | NP_001343358.1;NP_006756.2;NP_839952.1;XP_011542953.1;XP_01686 | 30242 | 601385 | . | . | 34.333467 | 33.306359 | 29.442107 | 30.960991 | 79.895391 | 76.989140 | 66.067232 | 70.788626 | -1.115563 | 3708.142552 | 0.109863371 | -1.43607742 | 0.150980312 | 0.913627824 | 11.933171 | 11.775485 | 11.921241 | 11.878139 | 11.820133 | 11.798942 | 4619 | 4111 | 3088 | 3244 |
| 7993 | NM_00128218 | UBXN8 | UBX domain protein 8 | protein_coding | NP_001269118.1;NP_001269128.1;NP_005662.2;XP_011542957.1;XP_01 | 30307 | 602155 | . | . | 15.400245 | 13.931178 | 17.163631 | 13.416527 | 35.837004 | 32.202541 | 38.514694 | 30.675293 | 1.042983 | 701.143505 | 0.170000902 | 0.357151617 | 0.720978305 | 0.998865404 | 9.422389 | 9.484083 | 9.470218 | 9.400850 | 9.548319 | 9.374894 | 830 | 703 | 701 | 587 |
| 7994 | NM_00130587 | KAT6A | lysine acetyltransferase 6A | protein_coding | NP_001292807.1;NP_006757.2;XP_011542958.1;XP_011542959.1;XP_01 | 13013 | 601408 | . | . | 3.272029 | 3.247515 | 2.945649 | 3.450199 | 7.506777 | 7.614146 | 6.609952 | 7.888469 | -1.036449 | 926.724048 | 0.145878579 | -0.35405204 | 0.723299885 | 0.998865404 | 9.881801 | 9.829721 | 9.874881 | 9.861882 | 9.788180 | 9.893829 | 1096 | 1009 | 733 | 916 |
| 8000 | NM_005672,N | PSCA | prostate stem cell antigen | protein_coding | NP_005663.2 | 9500 | 602470 | . | . | 17.049973 | 15.849345 | 15.574954 | 17.246667 | 39.675989 | 36.636471 | 34.949743 | 39.432458 | 1.003702 | 524.005225 | 0.166940342 | 0.031935012 | 0.974523878 | 0.998865404 | 9.030826 | 9.036044 | 9.054014 | 9.009590 | 9.009671 | 9.058524 | 623 | 545 | 439 | 511 |
| 8013 | NM_006981,N | NR4A3 | nuclear receptor subfamily 4 group A member 3 | protein_coding | NP_008912.2;NP_775291.1;NP_775292.1;XP_016870651.1 | 7982 | 600542 | . | . | 1.311590 | 1.197115 | 1.219332 | 1.151945 | 3.052123 | 2.767183 | 2.736146 | 2.633784 | -1.096424 | 212.778058 | 0.234779135 | -0.56566283 | 0.571623005 | 0.998865404 | 7.797555 | 7.665852 | 7.775847 | 7.731262 | 7.742214 | 7.679217 | 268 | 229 | 189 | 177 |
| 8019 | NM_007371,XI | BRD3 | bromodomain containing 3 | protein_coding | NP_031397.1;XP_006717354.1;XP_011517354.1;XP_016870654.1;XP_01 | 1104 | 601541 | . | . | 2.114016 | 2.059244 | 2.003562 | 2.250177 | 4.760035 | 4.919402 | 4.495937 | 5.144760 | 1.008899 | 354.394500 | 0.189099948 | 0.06759151 | 0.946110814 | 0.998865404 | 8.462984 | 8.475414 | 8.464714 | 8.468472 | 8.452189 | 8.490628 | 405 | 382 | 299 | 345 |
| 8021 | NM_00131832 | NUP214 | nucleoporin 214 | protein_coding | NP_001305253.1;NP_001305254.1;NP_005076.3 | 8064 | 114350 | . | . | 5.989707 | 6.224157 | 6.178613 | 6.970482 | 13.938296 | 14.387419 | 13.864627 | 15.937178 | 1.061111 | 1493.818579 | 0.129411714 | 0.661268505 | 0.508440136 | 0.998865404 | 10.501533 | 10.586786 | 10.509188 | 10.532559 | 10.522556 | 10.608955 | 1644 | 1588 | 1263 | 1522 |
| 8027 | NM_00132428 | STAM | signal transducing adaptor molecule | protein_coding | NP_001311211.1;NP_001311212.1;NP_001311213.1;NP_001311214.1;NI | 11357 | 601899 | . | . | 9.472050 | 8.758553 | 9.931008 | 9.034477 | 22.041848 | 20.245786 | 22.284895 | 20.656258 | 1.019975 | 1110.620745 | 0.139806417 | 0.204098529 | 0.838276504 | 0.998865404 | 10.102561 | 10.131594 | 10.142508 | 10.074223 | 10.163760 | 10.082217 | 1324 | 1132 | 1029 | 994 |
| 8028 | NM_00119562 | MLLT10 | MLLT10 histone lysine methyltransferase DOT1L cofactor | protein_coding | NP_001182555.1;NP_001182556.1;NP_001182557.1;NP_001182559.1;NI | 16063 | 602409 | . | . | 5.715130 | 5.582561 | 7.456232 | 6.319376 | 13.299344 | 12.904341 | 16.731572 | 14.448503 | 1.232133 | 873.059750 | 0.148571675 | 2.027018865 | 0.042660483 | 0.63734958 | 9.611297 | 9.912847 | 9.680370 | 9.695513 | 9.891939 | 9.780268 | 891 | 853 | 902 | 835 |
| 8029 | NM_001081,XI | CUBN | cubilin | protein_coding | NP_001072.2;XP_011518010.1;XP_011518011.1;XP_011518012.1;XP_01 | 2548 | 602997 | . | . | 0.582295 | 0.749598 | 0.538878 | 0.590827 | 1.355024 | 1.732729 | 1.209229 | 1.350857 | -1.192408 | 226.656524 | 0.23605797 | -1.07548738 | 0.282156578 | 0.998865404 | 7.946264 | 7.691205 | 7.800910 | 7.922012 | 7.762422 | 7.795575 | 250 | 298 | 173 | 201 |
| 8030 | NM_005436 | CCDC6 | coiled-coil domain containing 6 | protein_coding | NP_005427.2 | 18782 | 601985 | . | . | 4.167450 | 4.075021 | 3.806703 | 3.889007 | 9.697827 | 9.419594 | 8.542130 | 8.891754 | -1.085833 | 711.442680 | 0.149555311 | -0.79437425 | 0.426977589 | 0.998865404 | 9.532764 | 9.414000 | 9.513224 | 9.491606 | 9.442185 | 9.449355 | 866 | 787 | 590 | 643 |
| 8031 | NM_00114526 | NCOA4 | nuclear receptor coactivator 4 | protein_coding | NP_001138732.1;NP_001138733.1;NP_001138734.1;NP_001138735.1;NI | 7671 | 601984 | . | . | 33.151384 | 31.500860 | 28.970930 | 30.702346 | 77.144627 | 72.815649 | 65.009926 | 70.197263 | -1.097603 | 3439.253661 | 0.111879798 | -1.20089994 | 0.229790017 | 0.984376273 | 11.813501 | 11.679131 | 11.809014 | 11.760223 | 11.689079 | 11.727034 | 4266 | 3770 | 2785 | 3146 |
| 8034 | NM_00132431 | SLC25A16 | solute carrier family 25 member 16 | protein_coding | NP_001311241.1;NP_001311242.1;NP_001311243.1;NP_001311244.1;NI | 10986 | 139080 | . | . | 4.202202 | 4.617120 | 6.725386 | 4.124189 | 9.778699 | 10.672679 | 15.091574 | 9.429470 | 1.198892 | 610.057206 | 0.19857887 | 1.317870471 | 0.187547012 | 0.960264078 | 9.115429 | 9.378221 | 9.194475 | 9.167524 | 9.437541 | 9.146486 | 652 | 586 | 701 | 492 |
| 8036 | NM_00126903 | SHOC2 | SHOC2 leucine rich repeat scaffold protein | protein_coding | NP_001255968.1;NP_001311265.1;NP_001311266.1;NP_031399.2 | 15454 | 602775 | . | . | 4.629344 | 4.931104 | 4.116313 | 3.834721 | 10.772672 | 11.398467 | 9.236885 | 8.767632 | -1.230428 | 528.335578 | 0.166119419 | -1.80087661 | 0.07172233 | 0.753985777 | 9.187067 | 8.888093 | 9.089526 | 9.125648 | 8.996922 | 8.951911 | 653 | 646 | 427 | 428 |
| 8038 | NM_00128897 | ADAM12 | ADAM metallopeptidase domain 12 | protein_coding | NP_001275902.1;NP_001275903.1;NP_001275904.1;NP_003465.3;NP_0 | 190 | 602714 | . | . | 0.094822 | 0.083602 | 0.185696 | 0.068801 | 0.220655 | 0.193249 | 0.416695 | 0.157306 | 1.276948 | 26.024313 | 0.648604082 | 0.543782712 | 0.586591007 | NA | 4.511958 | 4.869520 | 4.679327 | 4.666558 | 4.746424 | 4.649353 | 28 | 23 | 36 | 16 |
| 8045 | NM_00114399 | RASSF7 | Ras association domain family member 7 | protein_coding | NP_001137465.1;NP_001137466.1;NP_003466.1 | 1166 | 143023 | . | . | 5.719424 | 6.697164 | 7.133512 | 6.992852 | 13.309333 | 15.480795 | 16.007395 | 15.988326 | 1.131438 | 363.379670 | 0.190640928 | 0.934515721 | 0.350037931 | 0.998865404 | 8.413756 | 8.591444 | 8.427417 | 8.504583 | 8.538599 | 8.536852 | 366 | 393 | 336 | 361 |
| 8050 | NM_00113502 | PDHX | pyruvate dehydrogenase complex component X | protein_coding | NP_001128496.1;NP_001159630.1;NP_003468.2;XP_011518692.1 | 21350 | 608769 | . | . | 11.022395 | 9.248448 | 9.465174 | 9.866615 | 25.649564 | 21.378205 | 21.239574 | 22.558843 | -1.061604 | 771.772907 | 0.156681637 | -0.55045033 | 0.582010533 | 0.998865404 | 9.634364 | 9.548421 | 9.674900 | 9.544388 | 9.558091 | 9.581607 | 999 | 779 | 640 | 714 |
| 8061 | NM_00130084 | FOSL1 | FOS like 1, AP-1 transcription factor subunit | protein_coding | NP_001287773.1;NP_001287784.1;NP_001287785.1;NP_001287786.1;NI | 13718 | 136515 | . | . | 30.226438 | 36.203942 | 26.512805 | 27.898306 | 70.338167 | 83.687034 | 59.493967 | 63.786161 | -1.235612 | 1580.059380 | 0.135883403 | -2.24623664 | 0.024688854 | 0.517665397 | 10.770472 | 10.464905 | 10.627887 | 10.769686 | 10.521408 | 10.551909 | 1833 | 2052 | 1204 | 1352 |
| 8065 | NM_003478,XI | CUL5 | cullin 5 | protein_coding | NP_003469.2;XP_005271739.1;XP_011541315.1;XP_016873852.1;XP_01 | 2556 | 601741 | . | . | 3.670348 | 3.131155 | 3.775703 | 2.331954 | 8.541050 | 7.237807 | 8.472568 | 5.331737 | -1.155543 | 601.114285 | 0.207021809 | -1.00748476 | 0.313701878 | 0.998865404 | 9.331352 | 9.124210 | 9.329250 | 9.208289 | 9.308495 | 9.020349 | 807 | 634 | 593 | 407 |
| 8073 | NM_00119510 | PTP4A2 | protein tyrosine phosphatase 4A2 | protein_coding | NP_001182029.1;NP_001182030.1;NP_001356787.1;NP_001356788.1;NI | 9635 | 601584 | . | . | 29.713999 | 32.672564 | 34.914465 | 35.615187 | 69.145694 | 75.524097 | 78.347047 | 81.429890 | 1.015626 | 3557.286244 | 0.1113865 | 0.200824034 | 0.84083617 | 0.998865404 | 11.785314 | 11.807722 | 11.818506 | 11.760587 | 11.793348 | 11.812092 | 4207 | 3675 | 3082 | 3399 |
| 8078 | NM_00109853 | USP5 | ubiquitin specific peptidase 5 | protein_coding | NP_001092006.1;NP_001369517.1;NP_001369518.1;NP_001369519.1;NI | 12628 | 601447 | . | . | 25.063526 | 26.797909 | 24.443374 | 29.773767 | 58.323859 | 61.944573 | 54.850217 | 68.074186 | 1.028415 | 2567.140152 | 0.12915727 | 0.31297119 | 0.75430255 | 0.998865404 | 11.305761 | 11.345853 | 11.288625 | 11.337488 | 11.257262 | 11.409017 | 2827 | 2814 | 2052 | 2668 |
| 8079 | NM_00138222 | MLF2 | myeloid leukemia factor 2 | protein_coding | NP_001369154.1;NP_001369155.1;NP_005430.1 | 7126 | 601401 | . | . | 76.792601 | 80.293756 | 73.252618 | 83.768921 | 178.699545 | 185.602621 | 164.376746 | 191.527679 | -1.013846 | 3866.614724 | 0.114753457 | -0.17288025 | 0.862745556 | 0.998865404 | 11.926825 | 11.906816 | 11.909052 | 11.935075 | 11.861539 | 11.957603 | 4413 | 4267 | 3145 | 3809 |
| 8082 | NM_00113582 | SSPN | sarcospan | protein_coding | NP_001129295.1;NP_005077.2;XP_011519155.1;XP_011519156.1;XP_01 | 11322 | 601599 | . | . | 0.051299 | 0.081588 | 0.153809 | 0.064571 | 0.119374 | 0.188595 | 0.345143 | 0.147634 | 1.581383 | 12.828294 | 0.889250143 | 0.743533279 | 0.457158881 | NA | 3.312060 | 3.974966 | 3.638656 | 3.658334 | 3.699529 | 3.647696 | 9 | 13 | 19 | 9 |
| 8085 | NM_003482,XI | KMT2D | lysine methyltransferase 2D | protein_coding | NP_003473.3;XP_006719677.1;XP_006719679.1;XP_011537072.1;XP_01 | 7133 | 602113 | . | . | 2.567775 | 2.689439 | 3.146173 | 3.010380 | 5.975318 | 6.216758 | 7.059922 | 6.882876 | 1.154438 | 1842.074784 | 0.121010593 | 1.71217396 | 0.086864615 | 0.795087717 | 10.739778 | 10.947018 | 10.773287 | 10.801004 | 10.920797 | 10.879678 | 1934 | 1878 | 1761 | 1803 |
| 8086 | NM_00117346 | AAAS | aladin WD repeat nucleoporin | protein_coding | NP_001166937.1;NP_056480.1;XP_011537080.1;XP_011537082.1 | 13666 | 605378 | . | . | 20.055467 | 20.947538 | 21.998878 | 23.809503 | 46.669895 | 48.421172 | 49.364848 | 54.437600 | 1.099601 | 1200.352684 | 0.134448222 | 1.018834856 | 0.308281375 | 0.998865404 | 10.159316 | 10.295936 | 10.175897 | 10.207462 | 10.224736 | 10.300130 | 1289 | 1260 | 1039 | 1237 |
| 8087 | NM_00101343 | FXR1 | FMR1 autosomal homolog 1 | protein_coding | NP_001013456.1;NP_001013457.1;NP_001350811.1;NP_005078.2;XP_0C | 4023 | 600819 | . | . | 42.634920 | 28.719313 | 25.537963 | 24.181785 | 99.213207 | 66.385979 | 57.306447 | 55.288778 | -1.186823 | 3764.640997 | 0.112791304 | -2.19081715 | 0.028465027 | 0.544186654 | 11.996606 | 11.749411 | 11.962339 | 11.926995 | 11.766444 | 11.838473 | 4804 | 4317 | 2861 | 3371 |
| 8089 | NM_00130095 | YEATS4 | YEATS domain containing 4 | protein_coding | NP_001287879.1;NP_006521.1 | 24859 | 602116 | . | . | 6.920693 | 7.335326 | 6.376418 | 6.329764 | 16.104735 | 16.955933 | 14.308497 | 14.472253 | -1.116159 | 321.712176 | 0.197046953 | -0.80459142 | 0.421055496 | 0.998865404 | 8.406748 | 8.248149 | 8.335900 | 8.381803 | 8.309925 | 8.287160 | 375 | 381 | 270 | 279 |
| 8091 | NM_00130091 | HMGA2 | high mobility group AT-hook 2 | protein_coding | NP_001287847.1;NP_001287848.1;NP_001317119.1;NP_003474.1;NP_0 | 5009 | 600698 | . | . | 16.720778 | 16.370898 | 14.887265 | 12.974283 | 38.909937 | 37.842066 | 33.406591 | 29.664155 | -1.199377 | 1692.366490 | 0.134310704 | -1.95282153 | 0.050840747 | 0.672475152 | 10.849810 | 10.587979 | 10.805617 | 10.775414 | 10.715652 | 10.576718 | 2165 | 1954 | 1460 | 1312 |
| 8092 | NM_006982,XI | ALX1 | ALX homeobox 1 | protein_coding | NP_008913.2;XP_011537084.1 | 1494 | 601527 | . | . | 5.612910 | 6.217730 | 7.835582 | 7.251874 | 13.061472 | 14.372562 | 17.582819 | 16.580548 | 1.256855 | 279.686906 | 0.209253025 | 1.576169317 | 0.114986788 | 0.855244326 | 7.953337 | 8.283188 | 8.036567 | 8.083881 | 8.203768 | 8.155499 | 272 | 280 | 283 | 279 |
| 8099 | NM_00127043 | CDK2AP1 | cyclin dependent kinase 2 associated protein 1 | protein_coding | NP_001257362.1;NP_001257363.1;NP_004633.1 | 14002 | 602198 | . | . | 122.566874 | 98.364170 | 97.066856 | 76.101880 | 285.218153 | 227.373196 | 217.815207 | 173.997895 | -1.303630 | 4036.161227 | 0.14760009 | -2.59169452 | 0.009550453 | 0.310907798 | 12.157262 | 11.775030 | 12.170134 | 11.971001 | 11.949544 | 11.751397 | 5872 | 4355 | 3420 | 2884 |
| 8100 | NM_00131849 | IFT88 | intraflagellar transport 88 | protein_coding | NP_001305420.1;NP_001305422.1;NP_001340494.1;NP_001340495.1;NI | 20606 | 600595 | . | . | 5.405271 | 5.716468 | 5.210084 | 4.842475 | 12.578291 | 13.213870 | 11.691276 | 11.071745 | -1.096825 | 499.082458 | 0.168752487 | -0.7901106 | 0.429463177 | 0.998865404 | 9.028123 | 8.895079 | 8.982796 | 9.000299 | 8.958271 | 8.907421 | 593 | 571 | 431 | 428 |
| 8106 | NM_00136055 | PABPN1 | poly(A) binding protein nuclear 1 | protein_coding | NP_001347480.1;NP_001347481.1;NP_004634.1 | 8565 | 602279 | . | . | 34.575686 | 36.622199 | 39.249937 | 35.255377 | 80.459046 | 84.653851 | 88.075718 | 80.607224 | 1.024833 | 3080.926535 | 0.114209133 | 0.309862963 | 0.756665169 | 0.998865404 | 11.571293 | 11.606786 | 11.569078 | 11.588241 | 11.632979 | 11.563508 | 3463 | 3322 | 2827 | 2800 |
| 8110 | NM_00128054 | DPF3 | double PHD fingers 3 | protein_coding | NP_001267471.1;NP_001267472.1;NP_001267473.1;NP_036206.3 | 17427 | 601672 | . | . | 0.119075 | 0.105609 | 0.184069 | 0.132019 | 0.277093 | 0.244121 | 0.413046 | 0.301846 | 1.375921 | 19.693230 | 0.696268802 | 0.661235033 | 0.508461597 | NA | 4.049783 | 4.512525 | 4.280366 | 4.273408 | 4.318009 | 4.288561 | 20 | 17 | 23 | 18 |
| 8120 | NM_00127851 | AP3B2 | adaptor related protein complex 3 subunit beta 2 | protein_coding | NP_001265440.1;NP_001265441.1;NP_001335369.1;NP_001335370.1;NI | 567 | 602166 | . | . | 1.071058 | 0.830528 | 1.108250 | 1.140847 | 2.492395 | 1.919802 | 2.486880 | 2.608410 | 1.078286 | 118.472053 | 0.301068078 | 0.361181807 | 0.717963542 | 0.998865404 | 6.832411 | 6.942303 | 6.905722 | 6.840341 | 6.908790 | 6.889426 | 144 | 111 | 110 | 112 |
| 8123 | NR_022008 | PWAR5 | Prader Willi/Angelman region RNA 5 | lncRNA | . | 30090 | 600162 | . | . | 2.025734 | 1.803176 | 2.355042 | 0.799037 | 4.713966 | 4.168123 | 5.284646 | 1.826904 | -1.227325 | 183.950197 | 0.738236753 | -0.40030132 | 0.688934593 | 0.998865404 | 7.663047 | 7.368264 | 7.570954 | 7.511843 | 7.635162 | 7.226725 | 248 | 205 | 215 | 78 |
| 8125 | NM_006305 | ANP32A | acidic nuclear phosphoprotein 32 family member A | protein_coding | NP_006296.1 | 13233 | 600832 | . | . | 29.226982 | 29.482273 | 26.722326 | 29.845589 | 68.012390 | 68.149590 | 59.964127 | 68.238388 | -1.052633 | 2191.105758 | 0.120377603 | -0.61474845 | 0.538720859 | 0.998865404 | 11.134068 | 11.059865 | 11.117492 | 11.117405 | 11.035588 | 11.115474 | 2588 | 2425 | 1763 | 2102 |
| 8128 | NM_00133041 | ST8SIA2 | ST8 alpha-N-acetyl-neuraminide alpha-2,8-sialyltransferase 2 | protein_coding | NP_001317345.1;NP_006002.1;XP_016878131.1 | 10870 | 602546 | . | . | 0.198692 | 0.148982 | 0.133393 | 0.108738 | 0.462363 | 0.344379 | 0.299331 | 0.248616 | -1.464303 | 26.674908 | 0.60852969 | -0.90416993 | 0.365905305 | NA | 4.985276 | 4.437942 | 4.780258 | 4.740564 | 4.729707 | 4.709158 | 42 | 29 | 21 | 18 |
| 8131 | NM_00103947 | NPRL3 | NPR3 like, GATOR1 complex subunit | protein_coding | NP_001034565.1;NP_001070818.1;NP_001230176.1;NP_001230177.1;NI | 14124 | 600928 | . | . | 8.116505 | 8.348577 | 8.161060 | 8.430512 | 18.887442 | 19.298110 | 18.313183 | 19.275364 | -1.009082 | 776.769939 | 0.145794133 | -0.08946491 | 0.92871244 | 0.998865404 | 9.607928 | 9.594729 | 9.597815 | 9.611216 | 9.588402 | 9.607812 | 890 | 850 | 663 | 735 |
| 8139 | NM_00137748 | GAN | gigaxonin | protein_coding | NP_001364415.1;NP_071324.1 | 4137 | 605379 | . | . | 0.621659 | 0.516180 | 0.408033 | 0.515466 | 1.446627 | 1.193174 | 0.915614 | 1.178551 | -1.248988 | 244.214247 | 0.232927569 | -1.37708103 | 0.168487197 | 0.945638095 | 8.083731 | 7.762423 | 8.031296 | 7.931307 | 7.821909 | 7.922142 | 342 | 265 | 168 | 226 |
| 8140 | NM_003486,XI | SLC7A5 | solute carrier family 7 member 5 | protein_coding | NP_003477.4;XP_006721349.2;XP_016879224.1;XP_016879225.1;XP_01 | 11063 | 600182 | . | . | 106.106979 | 110.342961 | 78.056825 | 83.182939 | 246.915299 | 255.062706 | 175.157238 | 190.187911 | -1.362005 | 13299.514685 | 0.101484973 | -4.39209502 | 1.12264E-05 | 0.00224576 | 13.904821 | 13.459053 | 13.805229 | 13.828720 | 13.530302 | 13.574538 | 17425 | 16772 | 9532 | 10838 |
| 8148 | NM_003487,N | TAF15 | TATA-box binding protein associated factor 15 | protein_coding | NP_003478.1;NP_631961.1;XP_024306727.1 | 11547 | 601574 | . | . | 39.492359 | 38.443324 | 40.191210 | 39.125603 | 91.900346 | 88.863464 | 90.187908 | 89.456040 | 1.002823 | 2644.289762 | 0.113959665 | 0.035687927 | 0.971531198 | 0.998865404 | 11.366573 | 11.370753 | 11.381620 | 11.352996 | 11.384838 | 11.354417 | 3094 | 2799 | 2346 | 2436 |
| 8161 | NM_004645 | COIL | coilin | protein_coding | NP_004636.1 | 2184 | 600272 | . | . | 7.296549 | 6.926702 | 7.270518 | 8.592696 | 16.979370 | 16.011383 | 16.314831 | 19.646179 | 1.099920 | 617.351891 | 0.163732415 | 0.839165521 | 0.401376433 | 0.998865404 | 9.199820 | 9.336827 | 9.256050 | 9.215397 | 9.244797 | 9.351417 | 698 | 615 | 518 | 654 |
| 8165 | NM_00124290 | AKAP1 | A-kinase anchoring protein 1 | protein_coding | NP_001229831.1;NP_001229832.1;NP_001357352.1;NP_001357353.1;NI | 367 | 602449 | . | . | 13.832469 | 13.703991 | 13.744538 | 13.127880 | 32.188726 | 31.677388 | 30.842347 | 30.015340 | -1.039058 | 1661.945914 | 0.122908397 | -0.44973798 | 0.652899381 | 0.998865404 | 10.725950 | 10.670839 | 10.720904 | 10.705756 | 10.703759 | 10.662818 | 1969 | 1810 | 1455 | 1488 |
| 8175 | NM_007165 | SF3A2 | splicing factor 3a subunit 2 | protein_coding | NP_009096.2 | 10766 | 600796 | . | . | 18.182102 | 21.757681 | 19.726362 | 22.762066 | 42.310497 | 50.293850 | 44.265381 | 52.042759 | 1.049809 | 1039.245129 | 0.150159307 | 0.467019896 | 0.640485643 | 0.998865404 | 9.986175 | 10.055629 | 9.936591 | 10.062641 | 9.986877 | 10.085199 | 1068 | 1188 | 864 | 1064 |
| 8178 | NM_006532,XI | ELL | elongation factor for RNA polymerase II | protein_coding | NP_006523.1;XP_011526632.2;XP_016882824.1;XP_016882825.1;XP_01 | 23114 | 600284 | . | . | 3.570637 | 3.285600 | 3.098799 | 3.805263 | 8.309020 | 7.594810 | 6.953615 | 8.700282 | -1.007384 | 421.685361 | 0.189554976 | -0.05599244 | 0.95534783 | 0.998865404 | 8.725564 | 8.714460 | 8.756113 | 8.684616 | 8.653679 | 8.775630 | 516 | 430 | 328 | 434 |
| 8187 | NM_00109928 | ZNF239 | zinc finger protein 239 | protein_coding | NP_001092752.1;NP_001092753.1;NP_001092754.1;NP_001311276.1;NI | 13031 | 601069 | . | . | 1.167458 | 1.111479 | 1.404285 | 1.169233 | 2.716723 | 2.569236 | 3.151173 | 2.673314 | 1.166451 | 151.950769 | 0.283071418 | 0.784698963 | 0.432630091 | 0.998865404 | 7.131637 | 7.354672 | 7.179676 | 7.238676 | 7.332367 | 7.203536 | 149 | 163 | 165 | 129 |
| 8189 | NM_004819,XI | SYMPK | symplekin | protein_coding | NP_004810.2;XP_005259343.1;XP_011525656.1 | 22935 | 602388 | . | . | 15.434529 | 14.863299 | 14.442212 | 14.550533 | 35.916783 | 34.357183 | 32.407908 | 33.268065 | -1.057569 | 1892.005033 | 0.119517672 | -0.67565045 | 0.499262601 | 0.998865404 | 10.925472 | 10.844801 | 10.923521 | 10.891088 | 10.866034 | 10.860377 | 2285 | 2056 | 1605 | 1718 |
| 8190 | NM_00120255 | MIA | MIA SH3 domain containing | protein_coding | NP_001189482.1;NP_006524.1 | 7076 | 601340 | . | . | 194.398633 | 174.821689 | 237.709202 | 294.902024 | 452.373596 | 404.108169 | 533.412583 | 674.258438 | 1.425073 | 3602.861577 | 0.131025684 | 3.900269127 | 9.60858E-05 | 0.012046912 | 11.536946 | 12.047902 | 11.692504 | 11.602926 | 11.850269 | 12.024543 | 3614 | 3026 | 3307 | 4374 |
| 8192 | NM_006012 | CLPP | caseinolytic mitochondrial matrix peptidase proteolytic subunit | protein_coding | NP_006003.1 | 2084 | 601119 | . | . | 11.339995 | 9.974442 | 9.490536 | 12.453835 | 26.388636 | 23.056370 | 21.296488 | 28.474213 | 1.014905 | 812.429750 | 0.16800458 | 0.127044398 | 0.898905258 | 0.998865404 | 9.655606 | 9.676517 | 9.705592 | 9.608896 | 9.572746 | 9.756496 | 992 | 811 | 619 | 867 |
| 8193 | NM_00113515 | DPF1 | double PHD fingers 1 | protein_coding | NP_001128627.1;NP_001128628.1;NP_001276907.1;NP_001350508.1;NI | 20225 | 601670 | . | . | 3.158421 | 3.238080 | 3.397310 | 2.640895 | 7.349778 | 7.484967 | 7.623467 | 6.038092 | -1.066811 | 228.598869 | 0.234461876 | -0.39795247 | 0.69066522 | 0.998865404 | 7.881984 | 7.789896 | 7.845439 | 7.854888 | 7.877115 | 7.757856 | 269 | 257 | 217 | 181 |
| 8195 | NM_018848,N | MKKS | McKusick-Kaufman syndrome | protein_coding | NP_061336.1;NP_740754.1 | 7108 | 604896 | . | . | 3.886640 | 4.007374 | 4.375773 | 4.376322 | 9.044371 | 9.263225 | 9.819108 | 10.005939 | 1.091311 | 863.321941 | 0.141369544 | 0.891725934 | 0.372539849 | 0.998865404 | 9.689364 | 9.815394 | 9.713909 | 9.728481 | 9.785908 | 9.781434 | 941 | 900 | 786 | 842 |
| 8202 | NM_00117408 | NCOA3 | nuclear receptor coactivator 3 | protein_coding | NP_001167558.1;NP_001167559.1;NP_006525.2;NP_858045.1 | 7670 | 601937 | . | . | 2.452797 | 2.852137 | 3.481921 | 3.370731 | 5.707756 | 6.592840 | 7.813329 | 7.706776 | 1.274473 | 753.689913 | 0.153538076 | 2.278921088 | 0.022671757 | 0.497517101 | 9.372443 | 9.722080 | 9.415206 | 9.511303 | 9.648311 | 9.618136 | 709 | 766 | 750 | 775 |
| 8204 | NM_003489,XI | NRIP1 | nuclear receptor interacting protein 1 | protein_coding | NP_003480.2;XP_005261120.1;XP_005261122.1;XP_011528049.1;XP_01 | 8001 | 602490 | . | . | 1.678509 | 1.535882 | 1.498193 | 1.432039 | 3.905958 | 3.550258 | 3.361902 | 3.274187 | -1.059723 | 374.606175 | 0.190407965 | -0.43951298 | 0.660289882 | 0.998865404 | 8.590025 | 8.507249 | 8.595894 | 8.533522 | 8.566305 | 8.495778 | 467 | 394 | 336 | 320 |
| 8208 | NM_005441,XI | CHAF1B | chromatin assembly factor 1 subunit B | protein_coding | NP_005432.1;XP_011528057.1;XP_016883966.1;XP_016883967.1 | 1911 | 601245 | . | . | 5.346170 | 5.721484 | 4.479899 | 5.518701 | 12.440758 | 13.225466 | 10.052762 | 12.617853 | -1.076670 | 702.247743 | 0.162422804 | -0.65616091 | 0.511720595 | 0.998865404 | 9.508558 | 9.401114 | 9.443471 | 9.514718 | 9.364961 | 9.488763 | 796 | 826 | 534 | 692 |
| 8209 | NM_00132038 | GATD3A | glutamine amidotransferase like class 1 domain containing 3A | protein_coding | NP_001307312.1;NP_001307313.1;NP_004640.4;NP_937798.4;XP_0168 | 1273 | 601659 | . | . | 14.066936 | 12.981461 | 12.426394 | 15.648749 | 32.734339 | 30.007234 | 27.884469 | 35.779005 | 1.051544 | 639.735065 | 0.164927943 | 0.439640374 | 0.660197594 | 0.998865404 | 9.284847 | 9.356915 | 9.327612 | 9.277574 | 9.272875 | 9.396726 | 745 | 648 | 518 | 671 |
| 8214 | NM_005675,XI | DGCR6 | DiGeorge syndrome critical region gene 6 | protein_coding | NP_005666.2;XP_011528696.1 | 2846 | 601279 | . | . | 12.592741 | 11.859430 | 11.679833 | 15.036427 | 29.303826 | 27.413604 | 26.209208 | 34.379004 | 1.079529 | 450.150573 | 0.186223743 | 0.592846828 | 0.553283666 | 0.998865404 | 8.758353 | 8.868088 | 8.809243 | 8.766744 | 8.761920 | 8.903651 | 516 | 451 | 360 | 488 |
| 8216 | NM_006767 | LZTR1 | leucine zipper like transcription regulator 1 | protein_coding | NP_006758.2 | 6742 | 600574 | . | . | 22.215643 | 22.925014 | 26.422852 | 25.836964 | 51.696716 | 52.992199 | 59.292114 | 59.073143 | 1.142011 | 3247.095346 | 0.110511026 | 1.733552978 | 0.08299743 | 0.78429781 | 11.565957 | 11.757555 | 11.598333 | 11.616983 | 11.730113 | 11.703081 | 3451 | 3309 | 3060 | 3193 |
| 8218 | NM_001835,N | CLTCL1 | clathrin heavy chain like 1 | protein_coding | NP_001826.3;NP_009029.3;XP_011528703.1;XP_016884442.1;XP_01688 | 2093 | 601273 | . | . | 1.225017 | 1.294375 | 1.437517 | 1.483315 | 2.850664 | 2.992006 | 3.225746 | 3.391422 | 1.197841 | 229.941150 | 0.224715803 | 1.158959244 | 0.246472798 | 0.992733415 | 7.709325 | 7.969227 | 7.771318 | 7.821749 | 7.879051 | 7.888223 | 228 | 238 | 216 | 237 |
| 8220 | NM_022719,N | ESS2 | ess-2 splicing factor homolog | protein_coding | NP_073210.1;XP_005261339.1;XP_006724392.1;XP_006724393.1;XP_0 | 16817 | 601755 | . | . | 7.163925 | 8.081610 | 7.517316 | 8.869478 | 16.670749 | 18.681002 | 16.868638 | 20.279007 | -1.010886 | 868.693483 | 0.145901021 | -0.10706134 | 0.914740316 | 0.998865404 | 9.770329 | 9.755038 | 9.755491 | 9.776389 | 9.799701 | 9.715239 | 991 | 956 | 796 | 762 |
| 8224 | NM_00113577 | SYN3 | synapsin III | protein_coding | NP_001129246.1;NP_001356836.1;NP_001356837.1;NP_001356838.1;NI | 11496 | 602705 | . | . | 0.478789 | 0.372075 | 0.262032 | 0.279032 | 1.114162 | 0.860067 | 0.587993 | 0.637973 | -1.238070 | 75.537697 | 0.366385048 | -0.84090022 | 0.400403835 | 0.998865404 | 6.385307 | 6.076444 | 6.245223 | 6.286597 | 6.210222 | 6.215041 | 89 | 97 | 58 | 64 |
| 8225 | NM_012227,N | GTPBP6 | GTP binding protein 6 (putative) | protein_coding | NP_036359.3;XP_006724510.1;XP_011544486.1 | 30189 | 300124 | . | . | 11.653744 | 11.468116 | 11.480604 | 12.005948 | 27.118740 | 26.509062 | 25.762140 | 27.450172 | 1.003070 | 721.883024 | 0.148676983 | 0.029742526 | 0.976272396 | 0.998865404 | 9.493436 | 9.497803 | 9.497919 | 9.491193 | 9.489323 | 9.503955 | 834 | 774 | 622 | 685 |
| 8226 | NM_00113556 | PUDP | pseudouridine 5'-phosphatase | protein_coding | NP_001129037.1;NP_001171606.1;NP_001171607.1;NP_036212.3 | 16818 | 306480 | . | . | 4.652458 | 4.069982 | 4.612422 | 4.496993 | 10.826462 | 9.407944 | 10.350141 | 10.281841 | 1.021223 | 292.533300 | 0.206771021 | 0.146525995 | 0.88350617 | 0.998865404 | 8.176896 | 8.207852 | 8.223526 | 8.147208 | 8.205283 | 8.189337 | 357 | 290 | 261 | 273 |
| 8227 | NM_005088,N | AKAP17A | A-kinase anchoring protein 17A | protein_coding | NP_005079.2 | 18783 | 465000 | . | . | 10.785261 | 11.858953 | 11.046784 | 11.334752 | 25.097745 | 27.412500 | 24.788663 | 25.915561 | -1.029930 | 1129.801005 | 0.134667566 | -0.31593722 | 0.752050157 | 0.998865404 | 10.163080 | 10.120309 | 10.121253 | 10.183024 | 10.124499 | 10.136536 | 1266 | 1288 | 960 | 1052 |
| 8228 | NM_00114238 | PNPLA4 | patatin like phospholipase domain containing 4 | protein_coding | NP_001135861.1;NP_001166143.1;NP_004641.1;XP_011543890.1 | 24887 | 300102 | . | . | 3.035195 | 3.078241 | 3.261617 | 3.951016 | 7.063023 | 7.115493 | 7.318974 | 9.033527 | 1.160041 | 345.664488 | 0.196804014 | 1.088268972 | 0.276476384 | 0.998865404 | 8.322601 | 8.535974 | 8.384668 | 8.389853 | 8.418053 | 8.521626 | 367 | 347 | 294 | 379 |
| 8233 | NM_005089,XI | ZRSR2 | zinc finger CCCH-type, RNA binding motif and serine/arginine rich | protein_coding | NP_005080.1;XP_005274654.2;XP_011543891.2;XP_016885370.1;XP_01 | 23019 | 300028 | . | . | 3.453308 | 4.976796 | 4.528848 | 4.890750 | 8.035993 | 11.504088 | 10.162605 | 11.182116 | 1.102942 | 219.983889 | 0.251926046 | 0.561105437 | 0.574725662 | 0.998865404 | 7.709568 | 7.849546 | 7.658360 | 7.837358 | 7.791759 | 7.804269 | 195 | 269 | 198 | 219 |
| 8237 | NM_00137107 | USP11 | ubiquitin specific peptidase 11 | protein_coding | NP_001358001.1;XP_005272731.1;XP_011542290.1 | 12609 | 300050 | . | . | 13.219469 | 14.533403 | 13.841903 | 16.276537 | 30.762247 | 33.594611 | 31.060829 | 37.214367 | 1.071907 | 1439.604674 | 0.136281673 | 0.735090106 | 0.462284619 | 0.998865404 | 10.440747 | 10.540445 | 10.430801 | 10.494420 | 10.456014 | 10.573423 | 1534 | 1562 | 1197 | 1502 |
| 8239 | NM_00103959 | USP9X | ubiquitin specific peptidase 9 X-linked | protein_coding | NP_001034679.2;NP_001034680.2;XP_005272732.1;XP_005272733.1 | 12632 | 300072 | . | . | 7.083277 | 6.597012 | 6.346967 | 6.533242 | 16.483077 | 15.249288 | 14.242410 | 14.937480 | -1.022453 | 2418.531331 | 0.117861925 | -0.27180151 | 0.785774642 | 0.998865404 | 11.255775 | 11.223880 | 11.283437 | 11.212467 | 11.236546 | 11.224852 | 2939 | 2523 | 2095 | 2226 |
| 8241 | NM_00120446 | RBM10 | RNA binding motif protein 10 | protein_coding | NP_001191395.1;NP_001191396.1;NP_001191397.1;NP_005667.2;NP_6 | 9896 | 300080 | . | . | 13.034422 | 13.565377 | 12.996139 | 12.876859 | 30.331635 | 31.356976 | 29.162961 | 29.441409 | -1.075783 | 1345.749226 | 0.128128983 | -0.8225062 | 0.410788858 | 0.998865404 | 10.446019 | 10.340437 | 10.409753 | 10.433067 | 10.347911 | 10.383148 | 1582 | 1528 | 1101 | 1244 |
| 8242 | NM_00114670 | KDM5C | lysine demethylase 5C | protein_coding | NP_001140174.1;NP_001269551.1;NP_001340907.1;NP_001340908.1;NI | 11114 | 314690 | . | . | 12.283959 | 12.708151 | 14.724553 | 14.938108 | 28.585278 | 29.375462 | 33.041472 | 34.154209 | 1.173225 | 2459.811337 | 0.113978205 | 2.022134833 | 0.043162424 | 0.639862814 | 11.144516 | 11.374956 | 11.189107 | 11.203004 | 11.321848 | 11.327952 | 2584 | 2464 | 2301 | 2499 |
| 8243 | NM_00128146 | SMC1A | structural maintenance of chromosomes 1A | protein_coding | NP_001268392.1;NP_006297.2 | 11111 | 300040 | . | . | 9.993295 | 10.471390 | 10.381451 | 11.929908 | 23.254807 | 24.205090 | 23.295676 | 27.276319 | 1.075110 | 3234.612184 | 0.118247012 | 0.88361561 | 0.376903749 | 0.998865404 | 11.606297 | 11.710573 | 11.612690 | 11.643685 | 11.633163 | 11.739516 | 3522 | 3428 | 2727 | 3344 |
| 8260 | NM_00125611 | NAA10 | N-alpha-acetyltransferase 10, NatA catalytic subunit | protein_coding | NP_001243048.1;NP_001243049.1;NP_003482.1 | 18704 | 300013 | . | . | 66.451124 | 64.721440 | 57.811286 | 65.207710 | 154.634492 | 149.606499 | 129.726855 | 149.089681 | -1.081942 | 3172.788761 | 0.116439544 | -0.97580693 | 0.329160149 | 0.998865404 | 11.687292 | 11.573538 | 11.676911 | 11.648262 | 11.552200 | 11.641898 | 3863 | 3497 | 2504 | 3015 |
| 8263 | NM_012151 | F8A1 | coagulation factor VIII associated 1 | protein_coding | NP_036283.2 | 3547 | 305423 | . | . | 6.095185 | 6.495374 | 6.250041 | 7.431362 | 14.183745 | 15.014347 | 14.024910 | 16.990925 | 1.071169 | 348.401138 | 0.195600157 | 0.507088531 | 0.612092702 | 0.998865404 | 8.394631 | 8.492907 | 8.407997 | 8.438766 | 8.415205 | 8.507457 | 377 | 373 | 288 | 365 |
| 8266 | NM_014235 | UBL4A | ubiquitin like 4A | protein_coding | NP_055050.1 | 12505 | 312070 | . | . | 13.983749 | 13.675907 | 11.822871 | 14.401072 | 32.540760 | 31.612471 | 26.530180 | 32.926338 | -1.070104 | 976.768793 | 0.148370181 | -0.65883588 | 0.510001165 | 0.998865404 | 9.980159 | 9.881915 | 9.966766 | 9.943803 | 9.836447 | 9.970652 | 1181 | 1073 | 744 | 967 |
| 8269 | NM_003492,XI | TMEM187 | transmembrane protein 187 | protein_coding | NP_003483.1;XP_011529501.1 | 13705 | 300059 | . | . | 3.062889 | 3.579601 | 2.724444 | 3.658768 | 7.127467 | 8.274409 | 6.113574 | 8.365339 | -1.062087 | 148.225967 | 0.282559368 | -0.30755184 | 0.758423368 | 0.998865404 | 7.255453 | 7.166484 | 7.191862 | 7.250647 | 7.143423 | 7.250387 | 163 | 177 | 107 | 154 |
| 8270 | NM_006014 | LAGE3 | L antigen family member 3 | protein_coding | NP_006005.2 | 26058 | 300060 | . | . | 13.380498 | 14.872561 | 12.033426 | 14.240591 | 31.136969 | 34.378590 | 27.002661 | 32.559422 | -1.090174 | 403.206132 | 0.187503042 | -0.6643035 | 0.506496084 | 0.998865404 | 8.716789 | 8.591227 | 8.650018 | 8.709155 | 8.581991 | 8.672901 | 461 | 476 | 309 | 390 |
| 8273 | NM_00114239 | SLC10A3 | solute carrier family 10 member 3 | protein_coding | NP_001135863.1;NP_001135864.1;NP_062822.1;XP_005277970.1;XP_00 | 22979 | 312090 | . | . | 5.780737 | 6.129760 | 6.368241 | 5.149849 | 13.452012 | 14.169212 | 14.290151 | 11.774515 | -1.017797 | 392.093549 | 0.197950714 | -0.12856712 | 0.897700184 | 0.998865404 | 8.627292 | 8.602712 | 8.595931 | 8.638825 | 8.682064 | 8.526650 | 439 | 442 | 383 | 316 |
| 8284 | NM_00114670 | KDM5D | lysine demethylase 5D | protein_coding | NP_001140177.1;NP_001140178.1;NP_004644.2;XP_005262617.1;XP_00 | 11115 | 426000 | . | . | 0.101477 | 0.127230 | 0.108075 | 0.101811 | 0.236141 | 0.294098 | 0.242517 | 0.232780 | -1.081991 | 24.232826 | 0.626814461 | -0.18137526 | 0.856073034 | NA | 4.654832 | 4.540693 | 4.591786 | 4.613882 | 4.598570 | 4.591384 | 26 | 30 | 21 | 21 |
| 8287 | NM_004654,XI | USP9Y | ubiquitin specific peptidase 9 Y-linked | protein_coding | NP_004645.2;XP_016885567.1 | 12633 | 400005 | . | . | 0.038427 | 0.035651 | 0.017915 | 0.031593 | 0.089421 | 0.082409 | 0.040201 | 0.072233 | -1.469904 | 10.155603 | 0.970542385 | -0.57258894 | 0.566923052 | NA | 3.597285 | 3.037079 | 3.360343 | 3.359713 | 3.333224 | 3.353432 | 14 | 13 | 5 | 10 |
| 8289 | NM_006015,N | ARID1A | AT-rich interaction domain 1A | protein_coding | NP_006006.3;NP_624361.1 | 11110 | 603024 | . | . | 5.798398 | 6.406852 | 5.495028 | 6.459182 | 13.493110 | 14.809725 | 12.330684 | 14.768153 | -1.032644 | 1615.440041 | 0.1345352 | -0.34447217 | 0.730491222 | 0.998865404 | 10.680927 | 10.634116 | 10.630653 | 10.706481 | 10.584169 | 10.700694 | 1799 | 1857 | 1278 | 1602 |
| 8291 | NM_00113045 | DYSF | dysferlin | protein_coding | NP_001123927.1;NP_001124448.1;NP_001124449.1;NP_001124450.1;NI | 3097 | 603009 | . | . | 0.081135 | 0.064419 | 0.025484 | 0.016301 | 0.188804 | 0.148908 | 0.057186 | 0.037270 | -3.212653 | 11.723835 | 0.94419116 | -1.78328855 | 0.074539344 | NA | 4.159017 | 2.481107 | 3.596711 | 3.577899 | 3.539569 | 3.527492 | 23 | 17 | 6 | 4 |
| 8293 | NM_021967,N | SERF1A | small EDRK-rich factor 1A | protein_coding | NP_068802.1;NP_075257.1 | 10755 | 603011 | . | . | 11.580974 | 11.176514 | 12.808144 | 13.068994 | 26.949401 | 25.835012 | 28.741103 | 29.880700 | 1.178005 | 417.680558 | 0.177976089 | 1.327964525 | 0.184189827 | 0.956498185 | 8.583081 | 8.819737 | 8.665269 | 8.640575 | 8.762382 | 8.740558 | 451 | 405 | 400 | 416 |
| 8295 | NM_00124458 | TRRAP | transformation/transcription domain associated protein | protein_coding | NP_001231509.1;NP_001362453.1;NP_003487.1 | 12347 | 603015 | . | . | 5.879489 | 6.453046 | 6.537080 | 6.602636 | 13.681811 | 14.916504 | 14.669017 | 15.096141 | 1.051910 | 2503.742250 | 0.11708926 | 0.623554882 | 0.532919949 | 0.998865404 | 11.252964 | 11.325857 | 11.233385 | 11.302944 | 11.308441 | 11.310804 | 2689 | 2747 | 2229 | 2410 |
| 8301 | NM_00100866 | PICALM | phosphatidylinositol binding clathrin assembly protein | protein_coding | NP_001008660.1;NP_001193875.1;NP_001193876.1;NP_009097.2;XP_0C | 15514 | 603025 | . | . | 29.763405 | 27.749393 | 27.196353 | 22.975245 | 69.260662 | 64.143968 | 61.027829 | 52.530166 | -1.164471 | 3235.192790 | 0.125742367 | -1.74701785 | 0.080634242 | 0.779954058 | 11.765164 | 11.545776 | 11.741314 | 11.693751 | 11.664325 | 11.516057 | 4122 | 3649 | 2838 | 2546 |
| 8303 | NM_003498,XI | SNN | stannin | protein_coding | NP_003489.1;XP_016879230.1;XP_016879231.1 | 11149 | 603032 | . | . | 7.876359 | 7.856812 | 7.261084 | 8.574680 | 18.328613 | 18.161373 | 16.293661 | 19.604988 | -1.008081 | 803.444517 | 0.151534016 | -0.07662538 | 0.93892157 | 0.998865404 | 9.656092 | 9.643992 | 9.656921 | 9.647789 | 9.590972 | 9.699284 | 935 | 865 | 642 | 808 |
| 8310 | NM_00110166 | ACOX3 | acyl-CoA oxidase 3, pristanoyl | protein_coding | NP_001095137.1;NP_001362712.1;NP_001362713.1;NP_001362714.1;NI | 121 | 603402 | . | . | 3.626767 | 3.476307 | 3.709013 | 3.819680 | 8.439635 | 8.035641 | 8.322914 | 8.733245 | 1.057301 | 336.484036 | 0.19195254 | 0.418778244 | 0.675378206 | 0.998865404 | 8.353574 | 8.434091 | 8.387392 | 8.368627 | 8.409674 | 8.408917 | 383 | 347 | 301 | 324 |
| 8312 | NM_003502,N | AXIN1 | axin 1 | protein_coding | NP_003493.1;NP_851393.1;XP_011520984.1;XP_011520985.1;XP_01152 | 903 | 603816 | . | . | 6.405272 | 6.398192 | 6.248243 | 5.938796 | 14.905333 | 14.789712 | 14.020878 | 13.578351 | -1.067100 | 707.761484 | 0.150882346 | -0.62098297 | 0.534610829 | 0.998865404 | 9.513064 | 9.419663 | 9.492061 | 9.485922 | 9.467425 | 9.420835 | 845 | 785 | 616 | 620 |
| 8313 | NM_00136381 | AXIN2 | axin 2 | protein_coding | NP_001350742.1;NP_004646.3;XP_011523621.1;XP_011523622.1;XP_01 | 904 | 604025 | . | . | 0.785624 | 0.908016 | 0.672980 | 1.081431 | 1.828180 | 2.098918 | 1.510147 | 2.472563 | 1.045550 | 116.659125 | 0.322371076 | 0.199339863 | 0.841996899 | 0.998865404 | 6.835031 | 6.896623 | 6.826943 | 6.880804 | 6.796696 | 6.936886 | 121 | 133 | 82 | 135 |
| 8314 | NM_004656,XI | BAP1 | BRCA1 associated protein 1 | protein_coding | NP_004647.1;XP_011532451.1;XP_011532452.1;XP_011532453.1;XP_01 | 950 | 603089 | . | . | 15.637085 | 16.268828 | 15.820198 | 17.006942 | 36.388140 | 37.606122 | 35.500064 | 38.884354 | 1.015459 | 1817.918624 | 0.121724279 | 0.181822915 | 0.855721701 | 0.998865404 | 10.817062 | 10.838998 | 10.809100 | 10.834469 | 10.808933 | 10.858231 | 2044 | 1978 | 1543 | 1771 |
| 8315 | NM_006768,XI | BRAP | BRCA1 associated protein | protein_coding | NP_006759.3;XP_005254001.1;XP_011537091.1;XP_016875481.1;XP_01 | 1099 | 604986 | . | . | 4.751422 | 4.722903 | 5.595165 | 5.187851 | 11.056757 | 10.917201 | 12.555389 | 11.861403 | 1.129112 | 625.566450 | 0.157812536 | 1.110103015 | 0.266954638 | 0.998546485 | 9.198626 | 9.374081 | 9.236087 | 9.254957 | 9.358227 | 9.294821 | 667 | 643 | 604 | 593 |
| 8317 | NM_00113441 | CDC7 | cell division cycle 7 | protein_coding | NP_001127891.1;NP_001127892.1;NP_003494.1;XP_005271298.1;XP_00 | 1745 | 603311 | . | . | 9.754863 | 9.410136 | 9.843909 | 8.875429 | 22.699968 | 21.751954 | 22.089446 | 20.292615 | -1.042167 | 953.567377 | 0.143277634 | -0.41587921 | 0.677498393 | 0.998865404 | 9.926438 | 9.867339 | 9.926790 | 9.895933 | 9.924784 | 9.836486 | 1144 | 1028 | 862 | 822 |
| 8318 | NM_00117801 | CDC45 | cell division cycle 45 | protein_coding | NP_001171481.1;NP_001171482.1;NP_001356220.1;NP_003495.1;XP_01 | 1739 | 603465 | . | . | 32.230097 | 33.228554 | 29.574916 | 29.897593 | 75.000769 | 76.809294 | 66.365255 | 68.357289 | -1.106853 | 1825.499943 | 0.11972824 | -1.22329858 | 0.221216942 | 0.982008277 | 10.905476 | 10.758957 | 10.864614 | 10.880968 | 10.789321 | 10.796327 | 2186 | 2091 | 1500 | 1632 |
| 8320 | NM_00127818 | EOMES | eomesodermin | protein_coding | NP_001265111.1;NP_001265112.1;NP_005433.2;XP_005265567.1 | 3372 | 604615 | . | . | 0.055050 | 0.074979 | 0.170437 | 0.063088 | 0.128104 | 0.173317 | 0.382455 | 0.144243 | 1.762867 | 10.576118 | 0.97661434 | 0.837509082 | 0.402306474 | NA | 2.934350 | 3.755755 | 3.363211 | 3.369913 | 3.418098 | 3.366363 | 8 | 9 | 17 | 7 |
| 8321 | NM_003505 | FZD1 | frizzled class receptor 1 | protein_coding | NP_003496.1 | 4038 | 603408 | . | . | 4.807131 | 4.910432 | 3.564590 | 4.310086 | 11.186392 | 11.350683 | 7.998837 | 9.854500 | -1.250796 | 945.521211 | 0.148318313 | -2.1767123 | 0.02950204 | 0.552532775 | 10.037651 | 9.714188 | 9.955316 | 9.964190 | 9.732964 | 9.857761 | 1203 | 1141 | 665 | 858 |
| 8322 | NM_012193 | FZD4 | frizzled class receptor 4 | protein_coding | NP_036325.2 | 4042 | 604579 | . | . | 0.699930 | 0.870969 | 0.746988 | 0.754813 | 1.628766 | 2.013284 | 1.676220 | 1.725789 | -1.057953 | 177.150068 | 0.255433387 | -0.31818735 | 0.750342826 | 0.998865404 | 7.509234 | 7.427227 | 7.433079 | 7.524085 | 7.456598 | 7.454640 | 188 | 217 | 150 | 161 |
| 8323 | NM_00116461 | FZD6 | frizzled class receptor 6 | protein_coding | NP_001158087.1;NP_001158088.1;NP_001304725.1;NP_003497.2 | 4044 | 603409 | . | . | 19.191631 | 18.260092 | 16.942365 | 16.063213 | 44.659713 | 42.209016 | 38.018173 | 36.726628 | -1.158263 | 1992.219022 | 0.121550603 | -1.74382918 | 0.081188881 | 0.782373954 | 11.062148 | 10.850412 | 11.045131 | 10.984722 | 10.919822 | 10.877590 | 2554 | 2221 | 1649 | 1684 |
| 8324 | NM_003507 | FZD7 | frizzled class receptor 7 | protein_coding | NP_003498.1 | 4045 | 603410 | . | . | 10.707522 | 10.288578 | 9.827936 | 10.314143 | 24.916845 | 23.782509 | 22.053606 | 23.582066 | -1.056746 | 1234.590289 | 0.130793953 | -0.60881211 | 0.542648988 | 0.998865404 | 10.309090 | 10.229444 | 10.308531 | 10.271762 | 10.234312 | 10.262782 | 1496 | 1336 | 1024 | 1147 |
| 8325 | NM_031866 | FZD8 | frizzled class receptor 8 | protein_coding | NP_114072.1 | 4046 | 606146 | . | . | 2.476100 | 2.269315 | 2.592234 | 2.503733 | 5.761986 | 5.245623 | 5.816898 | 5.724487 | 1.059774 | 244.840733 | 0.219123917 | 0.382234618 | 0.702287346 | 0.998865404 | 7.892903 | 7.977264 | 7.943868 | 7.895978 | 7.959881 | 7.937983 | 287 | 244 | 224 | 231 |
| 8327 | gene-GABPAP | GABPAP | GABPA pseudogene | pseudogene | . | 4072 | . | . | . | 0.077047 | 0.108803 | 0.112291 | 0.101654 | 0.179292 | 0.251504 | 0.251978 | 0.232420 | 1.116445 | 15.255637 | 0.785838913 | 0.202219417 | 0.839745194 | NA | 3.849979 | 4.008225 | 3.913115 | 3.933442 | 3.935455 | 3.925759 | 14 | 18 | 15 | 14 |

| | | | | | | | | | | | | | | | | | | | | | | | | | | | | | | | | | |
|---|---|---|---|---|---|---|---|---|---|---|---|---|---|---|---|---|---|---|---|---|---|---|---|---|---|---|---|---|---|---|---|---|---|
| 8329 | NM_003509 | H2AC13 | H2A clustered histone 13 | protein_coding | NP_003500.1 | 4725 | 602787 | . | . | 0.322830 | 0.307406 | 0.537811 | 0.485945 | 0.751239 | 0.710583 | 1.206832 | 1.111056 | 1.546454 | 6.859396 | 1.155404864 | 0.544366743 | 0.586189127 | NA | 2.429096 | 3.058876 | 2.759559 | 2.761315 | 2.777550 | 2.770069 | 6 | 6 | 8 | 7 |
| 8330 | NM_003510 | H2AC15 | H2A clustered histone 15 | protein_coding | NP_003501.1 | 4726 | 602788 | . | . | 0.226158 | 0.119635 | 0.090808 | 0.170879 | 0.526280 | 0.276541 | 0.203770 | 0.390696 | -1.127667 | 2.923912 | 1.754679452 | -0.09878791 | 0.921306672 | NA | 1.631267 | 1.459419 | 1.559607 | 1.547801 | 1.550083 | 1.554103 | 5 | 2 | 2 | 3 |
| 8331 | NM_021066 | H2AC14 | H2A clustered histone 14 | protein_coding | NP_066544.1 | 4727 | 602791 | . | . | 0.036983 | 0.375836 | 0.074622 | 0.072810 | 0.086061 | 0.868761 | 0.167449 | 0.166471 | -2.223658 | 2.659741 | 1.952295605 | -0.59055332 | 0.554819751 | NA | 1.878854 | 0.715150 | 1.406556 | 1.432977 | 1.408442 | 1.412766 | 1 | 7 | 1 | 2 |
| 8334 | NM_003512 | H2AC6 | H2A clustered histone 6 | protein_coding | NP_003503.1 | 4733 | 602794 | . | . | 3.226789 | 2.003092 | 4.061898 | 2.018524 | 7.508872 | 4.630237 | 9.114781 | 4.615115 | 1.138243 | 46.238247 | 0.508469341 | 0.367394041 | 0.713325116 | 0.998865404 | 5.431920 | 5.623738 | 5.544462 | 5.468163 | 5.586628 | 5.468554 | 61 | 36 | 57 | 31 |
| 8336 | NM_003514 | H2AC17 | H2A clustered histone 17 | protein_coding | NP_003505.1 | 4735 | 602796 | . | . | 0.279096 | 0.462652 | 0.121059 | 0.424172 | 0.649467 | 1.069440 | 0.271652 | 0.969819 | -1.342855 | 5.114030 | 1.366749727 | -0.31117908 | 0.755664482 | NA | 2.555346 | 2.121007 | 2.355548 | 2.369830 | 2.346796 | 2.364945 | 5 | 8 | 2 | 6 |
| 8337 | NM_003516 | H2AC18 | H2A clustered histone 18 | protein_coding | NP_003507.1 | 4736 | 142720 | . | . | 0.952357 | 1.218244 | 0.855693 | 1.144130 | 2.216173 | 2.816027 | 1.920150 | 2.615918 | -1.078248 | 17.759615 | 0.73072912 | -0.14873998 | 0.881758814 | NA | 4.205110 | 4.093800 | 4.145794 | 4.162793 | 4.139719 | 4.157442 | 19 | 22 | 13 | 18 |
| 8338 | NM_003517 | H2AC20 | H2A clustered histone 20 | protein_coding | NP_003508.1 | 4738 | 602797 | . | . | 0.481774 | 0.487761 | 0.250795 | 0.466669 | 1.121107 | 1.127481 | 0.562776 | 1.066983 | -1.335426 | 7.055812 | 1.146863109 | -0.36386236 | 0.715960792 | NA | 3.014059 | 2.592943 | 2.828337 | 2.830686 | 2.814086 | 2.826619 | 9 | 9 | 4 | 7 |
| 8341 | NM_003520 | H2BC15 | H2B clustered histone 15 | protein_coding | NP_003511.1 | 4749 | 602801 | . | . | 0.600750 | 0.237390 | 0.278355 | 0.582456 | 1.397970 | 0.548738 | 0.624620 | 1.331716 | 1.090421 | 7.867893 | 1.111448865 | 0.112362756 | 0.910535784 | NA | 2.912173 | 3.037079 | 2.987463 | 2.960674 | 2.966415 | 2.987940 | 12 | 5 | 5 | 10 |
| 8345 | NM_003524 | H2BC9 | H2B clustered histone 9 | protein_coding | NP_003515.1 | 4755 | 602806 | . | . | 0.905043 | 1.350518 | 0.749014 | 0.588244 | 2.106073 | 3.121782 | 1.680766 | 1.344951 | -1.712416 | 13.579134 | 0.847009146 | -0.91620388 | 0.359559972 | NA | 4.099441 | 3.324312 | 3.768047 | 3.796418 | 3.757858 | 3.744753 | 16 | 22 | 10 | 8 |
| 8347 | NM_00138198 | H2BC4 | H2B clustered histone 4 | protein_coding | NP_001368918.1;NP_003517.2 | 4757 | 602847 | . | . | 0.281932 | 0.464902 | 0.369119 | 0.833061 | 0.656068 | 1.074642 | 0.828292 | 1.904695 | 1.260218 | 29.560755 | 0.595704784 | 0.56013258 | 0.575389009 | 0.998865404 | 4.711777 | 5.040734 | 4.836025 | 4.888802 | 4.860122 | 4.919876 | 23 | 35 | 22 | 38 |
| 8349 | NM_003528 | H2BC21 | H2B clustered histone 21 | protein_coding | NP_003519.1 | 4760 | 601831 | . | . | 0.435455 | 0.397179 | 0.507401 | 0.693049 | 1.013322 | 0.918097 | 1.138592 | 1.584574 | 1.417176 | 35.678870 | 0.52555631 | 0.957117117 | 0.338508145 | 0.998865404 | 4.884262 | 5.386280 | 5.130558 | 5.116548 | 5.148523 | 5.195438 | 36 | 30 | 31 | 45 |
| 8351 | NM_00137693 | H3C4 | H3 clustered histone 4 | protein_coding | NP_001363866.1;NP_003521.2 | 4767 | 602811 | . | . | 0.305487 | 0.172867 | 0.440307 | 0.201929 | 0.710881 | 0.399589 | 0.988038 | 0.461686 | 2.113246 | 6.247645 | 1.292524875 | 0.835156514 | 0.403629591 | NA | 1.998355 | 3.087437 | 2.609223 | 2.597792 | 2.646932 | 2.600299 | 6 | 3 | 12 | 3 |
| 8352 | NM_003531 | H3C3 | H3 clustered histone 3 | protein_coding | NP_003522.1 | 4768 | 602812 | . | . | 0.253275 | 0.111821 | 0.174804 | 0.261240 | 0.589382 | 0.258479 | 0.392255 | 0.597295 | 1.242873 | 3.478073 | 1.613783807 | 0.194375115 | 0.845882173 | NA | 1.631267 | 1.947964 | 1.800000 | 1.788194 | 1.795747 | 1.799443 | 5 | 2 | 3 | 4 |
| 8353 | NM_00138199 | H3C6 | H3 clustered histone 6 | protein_coding | NP_001368928.1;NP_003523.1 | 4769 | 602813 | . | . | 0.555772 | 0.316136 | 1.036703 | 0.804068 | 1.293304 | 0.730762 | 2.326331 | 1.838406 | 2.054253 | 57.807849 | 0.443766645 | 2.340449903 | 0.019260522 | 0.462516353 | 5.239678 | 6.282298 | 5.786134 | 5.707036 | 5.923261 | 5.860903 | 55 | 30 | 76 | 64 |
| 8355 | NM_003534 | H3C8 | H3 clustered histone 8 | protein_coding | NP_003525.1 | 4772 | 602815 | . | . | 0.604190 | 0.228017 | 0.834251 | 0.363205 | 1.405975 | 0.527071 | 1.872036 | 0.830426 | 1.514357 | 8.364884 | 1.097920998 | 0.545308561 | 0.585541317 | NA | 2.728992 | 3.336262 | 3.055400 | 3.028163 | 3.073183 | 3.041076 | 11 | 4 | 12 | 6 |
| 8357 | NM_003536 | H3C10 | H3 clustered histone 10 | protein_coding | NP_003527.1 | 4775 | 602818 | . | . | 0.263945 | 0.364476 | 0.793400 | 0.642510 | 0.614212 | 0.842502 | 1.780367 | 1.469022 | 2.356020 | 8.304767 | 1.070509737 | 1.154918763 | 0.248123695 | NA | 2.307819 | 3.543272 | 3.013469 | 3.019483 | 3.051013 | 3.042711 | 5 | 6 | 11 | 10 |
| 8358 | NM_003537 | H3C2 | H3 clustered histone 2 | protein_coding | NP_003528.1 | 4776 | 602819 | . | . | 0.277738 | 0.236845 | 0.217131 | 0.067823 | 0.646308 | 0.547477 | 0.487236 | 0.155069 | -1.468409 | 4.144402 | 1.491696646 | -0.37155921 | 0.710221065 | NA | 2.299300 | 1.751299 | 2.060770 | 2.058005 | 2.057729 | 2.046623 | 6 | 5 | 4 | 2 |
| 8362 | NM_003541 | H4C12 | H4 clustered histone 12 | protein_coding | NP_003532.1 | 4784 | 602825 | . | . | 0.901306 | 0.971219 | 0.883764 | 0.719092 | 2.097377 | 2.245015 | 1.983140 | 1.644118 | -1.106769 | 11.109294 | 0.91253313 | -0.16038283 | 0.872579511 | NA | 3.544574 | 3.399154 | 3.475399 | 3.478800 | 3.476238 | 3.467977 | 13 | 13 | 10 | 9 |
| 8363 | NM_021968 | H4C11 | H4 clustered histone 11 | protein_coding | NP_068803.1 | 4785 | 602826 | . | . | 0.798446 | 0.689837 | 0.636352 | 0.497512 | 1.858016 | 1.594590 | 1.427955 | 1.137503 | -1.364666 | 8.534644 | 1.042111769 | -0.43042204 | 0.666888669 | NA | 3.299300 | 2.852980 | 3.106769 | 3.101400 | 3.095974 | 3.088620 | 12 | 10 | 7 | 6 |
| 8364 | NM_003542 | H4C3 | H4 clustered histone 3 | protein_coding | NP_003533.1 | 4787 | 602827 | . | . | 0.128042 | 0.146516 | 0.305606 | 0.222831 | 0.297958 | 0.338677 | 0.685771 | 0.509476 | 2.170210 | 2.847568 | 1.783980699 | 0.6265958 | 0.530924222 | NA | 0.844134 | 1.963455 | 1.485636 | 1.486278 | 1.498845 | 1.492632 | 2 | 2 | 4 | 3 |
| 8367 | NM_003545 | H4C5 | H4 clustered histone 5 | protein_coding | NP_003536.1 | 4790 | 602830 | . | . | 0.129841 | 0.144026 | 0.533279 | 0.231538 | 0.302145 | 0.332923 | 1.196662 | 0.529384 | 2.806931 | 3.422555 | 1.670243255 | 0.891483094 | 0.372670057 | NA | 0.844134 | 2.336262 | 1.739091 | 1.739779 | 1.762747 | 1.746248 | 2 | 2 | 6 | 3 |
| 8370 | NM_003548 | H4C14 | H4 clustered histone 14 | protein_coding | NP_003539.1 | 4794 | 142750 | . | . | 0.745046 | 0.700639 | 0.430452 | 0.488285 | 1.733753 | 1.619559 | 0.965922 | 1.116405 | -1.548230 | 7.742546 | 1.094712193 | -0.57605988 | 0.564574711 | NA | 3.234218 | 2.602870 | 2.968719 | 2.967296 | 2.951701 | 2.954402 | 11 | 10 | 5 | 6 |
| 8372 | NM_00120002 | HYAL3 | hyaluronidase 3 | protein_coding | NP_001186958.1;NP_001186959.1;NP_001186960.1;NP_001186961.1;NI | 5322 | 604038 | . | . | 2.404534 | 2.154821 | 2.366173 | 2.376591 | 5.595451 | 4.980966 | 5.309623 | 5.433793 | 1.077828 | 128.629049 | 0.285156888 | 0.379186056 | 0.704549706 | 0.998865404 | 6.951979 | 7.060139 | 6.985799 | 6.998657 | 7.024891 | 7.012970 | 140 | 136 | 118 | 123 |
| 8379 | NM_00101383 | MAD1L1 | mitotic arrest deficient 1 like 1 | protein_coding | NP_001013858.1;NP_001013859.1;NP_001291452.1;NP_001291453.1;NI | 6762 | 602686 | . | . | 29.762126 | 29.757818 | 31.624055 | 34.052651 | 69.257696 | 68.786527 | 70.963466 | 77.857341 | 1.071187 | 1839.560603 | 0.124555178 | 0.796513685 | 0.425733525 | 0.998865404 | 10.794767 | 10.893824 | 10.832687 | 10.800914 | 10.829245 | 10.911369 | 2087 | 1878 | 1569 | 1876 |
| 8382 | NM_003551,XI | NME5 | NME/NM23 family member 5 | protein_coding | NP_003542.1;XP_005272156.1;XP_016865434.1;XP_024301995.1;XP_02 | 7853 | 603575 | . | . | 1.084896 | 0.427260 | 0.580500 | 0.650849 | 2.524597 | 0.987631 | 1.302625 | 1.488088 | -1.279254 | 26.404480 | 0.653854918 | -0.54339658 | 0.586856781 | NA | 4.888379 | 4.535520 | 4.785464 | 4.677467 | 4.700816 | 4.711481 | 49 | 18 | 19 | 23 |
| 8387 | NM_003553 | OR1E1 | olfactory receptor family 1 subfamily E member 1 | protein_coding | NP_003544.2 | 8189 | . | . | . | 0.129540 | 0.101440 | 0.113190 | 0.106067 | 0.301444 | 0.234484 | 0.253994 | 0.242510 | 1.033245 | 5.449006 | 1.287464181 | 0.036647048 | 0.97076643 | NA | 2.421266 | 2.470304 | 2.448596 | 2.441817 | 2.446940 | 2.445101 | 7 | 5 | 5 | 5 |
| 8388 | NM_003554 | OR1E2 | olfactory receptor family 1 subfamily E member 2 | protein_coding | NP_003545.1 | 8190 | . | . | . | 0.023961 | 0.088953 | 0.152200 | 0.016945 | 0.055759 | 0.205619 | 0.341533 | 0.038744 | 1.867360 | 2.616304 | 1.928728158 | 0.467147446 | 0.640394391 | NA | 0.867373 | 1.769039 | 1.352782 | 1.362089 | 1.375300 | 1.354143 | 1 | 3 | 5 | 1 |
| 8394 | NM_00113563 | PIP5K1A | phosphatidylinositol-4-phosphate 5-kinase type 1 alpha | protein_coding | NP_001129108.1;NP_001129109.1;NP_001129110.1;NP_001317618.1;NI | 8994 | 603275 | . | . | 14.850838 | 16.566606 | 15.577172 | 16.974585 | 34.558510 | 38.294458 | 34.954717 | 38.810370 | 1.015987 | 1869.115446 | 0.125512416 | 0.182307524 | 0.855341394 | 0.998865404 | 10.856809 | 10.879383 | 10.822372 | 10.898987 | 10.842953 | 10.903762 | 2032 | 2098 | 1575 | 1834 |
| 8396 | NM_003559,XI | PIP4K2B | phosphatidylinositol-5-phosphate 4-kinase type 2 beta | protein_coding | NP_003550.1;XP_011523628.1;XP_011523629.1;XP_011523632.1;XP_01 | 8998 | 603261 | . | . | 13.574991 | 13.974483 | 15.616912 | 15.882248 | 31.589563 | 32.302642 | 35.043899 | 36.312872 | 1.124318 | 2498.575030 | 0.113835274 | 1.485046067 | 0.137531621 | 0.898839475 | 11.199905 | 11.368927 | 11.227710 | 11.247070 | 11.333518 | 11.330645 | 2676 | 2569 | 2304 | 2475 |
| 8398 | NM_00100442 | PLA2G6 | phospholipase A2 group VI | protein_coding | NP_001004426.1;NP_001186491.1;NP_001336793.1;NP_001336794.1;NI | 9039 | 603604 | . | . | 78.011576 | 80.166982 | 75.946068 | 74.155383 | 181.536137 | 185.309571 | 170.420771 | 169.547460 | -1.063813 | 7390.337794 | 0.102850177 | -0.86771239 | 0.385551802 | 0.998865404 | 12.895371 | 12.806098 | 12.862665 | 12.891221 | 12.819274 | 12.830000 | 8626 | 8359 | 6188 | 6757 |
| 8402 | NM_00116541 | SLC25A11 | solute carrier family 25 member 11 | protein_coding | NP_001158889.1;NP_001158890.1;NP_003553.2;XP_024306762.1 | 10981 | 604165 | . | . | 44.630790 | 42.811565 | 33.251106 | 46.657409 | 103.857683 | 98.960854 | 74.614517 | 106.676617 | -1.104967 | 2266.575292 | 0.146772964 | -0.98113293 | 0.326527192 | 0.998865404 | 11.216688 | 11.072298 | 11.189831 | 11.175190 | 10.976160 | 11.214539 | 2765 | 2545 | 1592 | 2321 |
| 8405 | NM_00100722 | SPOP | speckle type BTB/POZ protein | protein_coding | NP_001007227.1;NP_001007228.1;NP_001007229.1;NP_001007230.1;NI | 11254 | 602650 | . | . | 7.874271 | 7.917049 | 8.547715 | 8.706658 | 18.323755 | 18.300615 | 19.180824 | 19.906742 | 1.056332 | 739.518370 | 0.147994762 | 0.534233779 | 0.593179801 | 0.998865404 | 9.490306 | 9.569491 | 9.507173 | 9.514228 | 9.562666 | 9.534740 | 824 | 780 | 674 | 698 |
| 8406 | NM_00117075 | SRPX | sushi repeat containing protein X-linked | protein_coding | NP_001164221.1;NP_001164222.1;NP_001164223.1;NP_006298.1;XP_01 | 11309 | 300187 | . | . | 128.725267 | 131.011887 | 105.093644 | 116.088200 | 299.549006 | 302.839850 | 235.827096 | 265.421859 | -1.167935 | 6776.087302 | 0.107517218 | -2.08301259 | 0.037250076 | 0.602769983 | 12.833917 | 12.609866 | 12.775790 | 12.800287 | 12.619073 | 12.693398 | 8287 | 7991 | 5193 | 6122 |
| 8407 | NM_00127722 | TAGLN2 | transgelin 2 | protein_coding | NP_001264152.1;NP_001264153.1;NP_003555.1 | 11554 | 604634 | . | . | 69.205771 | 70.641451 | 53.313791 | 64.080919 | 161.044678 | 163.290873 | 119.634601 | 146.513397 | -1.208160 | 2758.456645 | 0.124953543 | -2.18329939 | 0.02901377 | 0.548598713 | 11.559742 | 11.286644 | 11.495415 | 11.506368 | 11.276043 | 11.413341 | 3453 | 3279 | 1985 | 2544 |
| 8408 | NM_003565,XI | ULK1 | unc-51 like autophagy activating kinase 1 | protein_coding | NP_003556.2;XP_011537100.1;XP_011537101.1 | 12558 | 603168 | . | . | 4.137984 | 4.123000 | 5.375383 | 4.133241 | 9.629259 | 9.530498 | 12.062203 | 9.450164 | 1.141469 | 721.840025 | 0.168280597 | 1.134362853 | 0.256642374 | 0.995760118 | 9.396547 | 9.588166 | 9.439952 | 9.449001 | 9.625192 | 9.438665 | 771 | 732 | 758 | 626 |
| 8409 | NM_004182,N | UXT | ubiquitously expressed prefoldin like chaperone | protein_coding | NP_004173.1;NP_705582.1 | 12641 | 300234 | . | . | 32.720133 | 33.262804 | 30.675570 | 37.849960 | 76.141096 | 76.888464 | 68.835088 | 86.539431 | 1.046085 | 675.812053 | 0.162036026 | 0.40114279 | 0.688314998 | 0.998865404 | 9.367937 | 9.432302 | 9.386563 | 9.380941 | 9.346427 | 9.478024 | 764 | 710 | 543 | 710 |
| 8411 | NM_003566,XI | EEA1 | early endosome antigen 1 | protein_coding | NP_003557.3;XP_011537116.1;XP_011537117.1;XP_016875507.1 | 3185 | 605070 | . | . | 1.508956 | 1.345074 | 1.439505 | 1.135624 | 3.511403 | 3.109198 | 3.230208 | 2.596470 | -1.125087 | 416.829696 | 0.194282435 | -0.87520494 | 0.381462406 | 0.998865404 | 8.785199 | 8.616500 | 8.772824 | 8.696827 | 8.733690 | 8.591046 | 539 | 447 | 383 | 323 |
| 8412 | NM_00126140 | BCAR3 | BCAR3 adaptor protein, NSP family member | protein_coding | NP_001248337.1;NP_001248338.1;NP_001248339.1;NP_001295180.1;NI | 973 | 604704 | . | . | 12.538485 | 12.669392 | 15.409975 | 15.600999 | 29.177572 | 29.285867 | 34.579540 | 35.669826 | 1.211353 | 1413.298426 | 0.125519277 | 2.203801514 | 0.027538306 | 0.538018865 | 10.319923 | 10.596539 | 10.384844 | 10.386886 | 10.533264 | 10.532796 | 1470 | 1381 | 1347 | 1451 |
| 8417 | NM_00132657 | STX7 | syntaxin 7 | protein_coding | NP_001313507.1;NP_001313508.1;NP_001313509.1;NP_003560.2 | 11442 | 603217 | . | . | 6.518949 | 5.542410 | 7.515905 | 6.121045 | 15.169863 | 12.811527 | 16.865474 | 13.995042 | 1.112688 | 2536.388021 | 0.123666279 | 1.245682929 | 0.212880823 | 0.975238634 | 11.229338 | 11.383656 | 11.315754 | 11.207006 | 11.377587 | 11.320288 | 2952 | 2415 | 2405 | 2417 |
| 8420 | NR_002909,NF | SNHG3 | small nucleolar RNA host gene 3 | lncRNA | . | 10118 | 603238 | . | . | 33.923788 | 31.915493 | 40.677274 | 26.791176 | 78.942051 | 73.774097 | 91.278623 | 61.254840 | 1.012450 | 2331.753964 | 0.16832808 | 0.106050221 | 0.915542506 | 0.998865404 | 11.178025 | 11.196316 | 11.204370 | 11.147510 | 11.342667 | 11.002065 | 2762 | 2412 | 2468 | 1739 |
| 8425 | NM_00104254 | LTBP4 | latent transforming growth factor beta binding protein 4 | protein_coding | NP_001036009.1;NP_001036010.1;NP_003564.2 | 6717 | 604710 | . | . | 3.314161 | 3.646152 | 3.180718 | 3.429757 | 7.712189 | 8.428242 | 7.137439 | 7.841732 | -1.066773 | 544.408477 | 0.165422378 | -0.56372688 | 0.572940024 | 0.998865404 | 9.134677 | 9.040891 | 9.079836 | 9.137380 | 9.046043 | 9.087603 | 619 | 633 | 443 | 510 |
| 8427 | NM_00130348 | ZNF282 | zinc finger protein 282 | protein_coding | NP_001290410.1;NP_003566.1;XP_006716214.1 | 13076 | 603397 | . | . | 5.671080 | 6.838423 | 7.106832 | 7.363864 | 13.196837 | 15.807319 | 15.947525 | 16.836600 | 1.139562 | 763.387277 | 0.157948887 | 1.19328989 | 0.232755861 | 0.988836302 | 9.479258 | 9.667170 | 9.457461 | 9.591663 | 9.607771 | 9.628048 | 742 | 845 | 697 | 773 |
| 8428 | NM_00103229 | STK24 | serine/threonine kinase 24 | protein_coding | NP_001027467.2;NP_001273578.1;NP_003567.2;XP_005254136.1;XP_01 | 11403 | 604984 | . | . | 10.706998 | 14.475675 | 14.569967 | 12.409046 | 24.915626 | 33.461171 | 32.694586 | 28.371812 | 1.077844 | 2835.377559 | 0.128835317 | 0.83942982 | 0.401228156 | 0.998865404 | 11.414228 | 11.522395 | 11.502425 | 11.366394 | 11.450525 | 11.542971 | 3406 | 2698 | 2414 | 2913 |
| 8434 | NM_00131634 | RECK | reversion inducing cysteine rich protein with kazal motifs | protein_coding | NP_001303274.1;NP_001303275.1;NP_001303276.1;NP_001303277.1;NI | 11345 | 605227 | . | . | 7.665623 | 7.898778 | 6.517543 | 7.837374 | 17.838224 | 18.258378 | 14.625178 | 17.919223 | -1.125152 | 721.307081 | 0.160158838 | -1.06219726 | 0.288146143 | 0.998865404 | 9.577358 | 9.406528 | 9.532568 | 9.533553 | 9.381727 | 9.516667 | 879 | 825 | 531 | 700 |
| 8438 | NM_00114254 | RAD54L | RAD54 like | protein_coding | NP_001136020.1;NP_001357695.1;NP_003570.2;XP_011540601.1;XP_01 | 9826 | 603615 | . | . | 10.128378 | 10.523619 | 10.840066 | 11.220595 | 23.569149 | 24.325816 | 24.324794 | 25.654558 | 1.047558 | 1006.055051 | 0.135893798 | 0.493253618 | 0.621833397 | 0.998865404 | 9.940626 | 10.007584 | 9.957288 | 9.956790 | 9.982694 | 9.999416 | 1132 | 1060 | 885 | 976 |
| 8439 | NM_00114477 | NSMAF | neutral sphingomyelinase activation associated factor | protein_coding | NP_001138244.1;NP_003571.2 | 8017 | 603043 | . | . | 8.644514 | 8.463049 | 8.744268 | 8.261299 | 20.116138 | 19.562716 | 19.621886 | 18.888477 | -1.018428 | 947.208621 | 0.139956983 | -0.18823101 | 0.850695565 | 0.998865404 | 9.900494 | 9.874466 | 9.906129 | 9.881564 | 9.905854 | 9.855027 | 1119 | 1014 | 845 | 849 |
| 8440 | NM_00100472 | NCK2 | NCK adaptor protein 2 | protein_coding | NP_001004720.1;NP_001004722.1;NP_003572.2;XP_006712860.1;XP_01 | 7665 | 604930 | . | . | 42.497846 | 43.402573 | 30.814571 | 42.325249 | 98.894230 | 100.326988 | 69.147001 | 96.771654 | -1.170248 | 3079.504703 | 0.137669653 | -1.64752493 | 0.099450197 | 0.813730268 | 11.697591 | 11.470442 | 11.628025 | 11.666588 | 11.407331 | 11.618943 | 3736 | 3667 | 2144 | 3009 |
| 8443 | NM_00131635 | GNPAT | glyceronephosphate O-acyltransferase | protein_coding | NP_001303279.1;NP_055051.1;XP_005273370.1;XP_011542605.1;XP_01 | 4416 | 602744 | . | . | 14.154575 | 13.727426 | 13.732631 | 15.021413 | 32.938280 | 31.731560 | 30.815626 | 34.344679 | 1.040191 | 1150.960030 | 0.133954064 | 0.424383114 | 0.671286434 | 0.998865404 | 10.140040 | 10.196649 | 10.154385 | 10.152484 | 10.151139 | 10.213562 | 1301 | 1216 | 978 | 1146 |
| 8444 | NM_00100402 | DYRK3 | dual specificity tyrosine phosphorylation regulated kinase 3 | protein_coding | NP_001004023.1;NP_003573.2;XP_005273372.1;XP_011508363.1 | 3094 | 603497 | . | . | 3.645295 | 3.293163 | 3.245880 | 3.253196 | 8.482753 | 7.612294 | 7.283662 | 7.438045 | -1.080089 | 852.944012 | 0.145655466 | -0.76310497 | 0.445400796 | 0.998865404 | 9.790683 | 9.679801 | 9.800788 | 9.723702 | 9.709890 | 9.706274 | 1074 | 905 | 715 | 767 |
| 8445 | NM_003583,N | DYRK2 | dual specificity tyrosine phosphorylation regulated kinase 2 | protein_coding | NP_003574.1;NP_006473.2;XP_016875521.1 | 3093 | 603496 | . | . | 2.796121 | 2.952317 | 2.747854 | 2.315333 | 6.506690 | 6.824415 | 6.166107 | 5.293734 | -1.146422 | 746.504879 | 0.157255572 | -1.2536164 | 0.209981467 | 0.972608791 | 9.638966 | 9.442356 | 9.571298 | 9.605517 | 9.552416 | 9.431286 | 896 | 881 | 659 | 594 |
| 8446 | NM_003584 | DUSP11 | dual specificity phosphatase 11 | protein_coding | NP_003575.2 | 3066 | 603092 | . | . | 10.315172 | 9.652747 | 9.700878 | 10.364305 | 24.003830 | 22.312756 | 21.768490 | 23.696751 | -1.009693 | 511.581904 | 0.167784478 | -0.08294693 | 0.933893741 | 0.998865404 | 9.005772 | 8.991837 | 9.025510 | 8.977482 | 8.977331 | 9.013598 | 614 | 534 | 430 | 491 |
| 8448 | NM_00128206 | DOC2A | double C2 domain alpha | protein_coding | NP_001268991.1;NP_001268992.1;NP_001268997.1;NP_003577.2;XP_01 | 2985 | 604567 | . | . | 2.037471 | 2.289831 | 2.397603 | 1.945420 | 4.741279 | 5.293047 | 5.380150 | 4.447974 | 1.002455 | 156.047601 | 0.264551828 | 0.013372811 | 0.989330359 | 0.999248276 | 7.284141 | 7.287542 | 7.260727 | 7.307834 | 7.297391 | 7.273508 | 169 | 178 | 140 | 142 |
| 8449 | NM_00116423 | DHX16 | DEAH-box helicase 16 | protein_coding | NP_001157711.1;NP_001350444.1;NP_003578.2;XP_011513240.1;XP_01 | 2739 | 603405 | . | . | 19.667552 | 20.419677 | 17.617538 | 19.885201 | 45.767199 | 47.200995 | 39.533242 | 45.465150 | -1.076613 | 2076.700457 | 0.123386357 | -0.86314368 | 0.388058474 | 0.998865404 | 11.072482 | 10.965697 | 11.035508 | 11.061301 | 10.941735 | 11.035208 | 2440 | 2361 | 1636 | 1986 |
| 8450 | NM_00107987 | CUL4B | cullin 4B | protein_coding | NP_001073341.1;NP_001317553.1;NP_001356074.1;NP_003579.3 | 2555 | 300304 | . | . | 7.941574 | 6.985670 | 6.969938 | 5.310307 | 18.480367 | 16.147687 | 15.640341 | 12.141386 | -1.197175 | 1179.225296 | 0.162364674 | -1.59907768 | 0.109803344 | 0.843391194 | 10.327232 | 10.068424 | 10.323647 | 10.200647 | 10.227655 | 10.019989 | 1597 | 1276 | 1066 | 864 |
| 8451 | NM_00100889 | CUL4A | cullin 4A | protein_coding | NP_001008895.1;NP_001265442.1;NP_001265443.1;NP_001341867.1;NI | 2554 | 603137 | . | . | 10.212881 | 10.273221 | 10.286701 | 12.188512 | 23.765791 | 23.747015 | 23.083056 | 27.867588 | 1.058539 | 1637.051763 | 0.124281243 | 0.660392207 | 0.509002172 | 0.998865404 | 10.635307 | 10.717297 | 10.668944 | 10.638861 | 10.672879 | 10.723200 | 1867 | 1683 | 1417 | 1629 |
| 8452 | NM_00125719 | CUL3 | cullin 3 | protein_coding | NP_001244126.1;NP_001244127.1;NP_003581.1;XP_006712863.2;XP_01 | 2553 | 603136 | . | . | 12.189110 | 11.934250 | 12.026416 | 9.279590 | 28.364561 | 27.586550 | 26.986931 | 21.216682 | -1.180129 | 2237.838972 | 0.130693842 | -1.82828072 | 0.067507431 | 0.739527835 | 11.242287 | 11.003635 | 11.173090 | 11.205559 | 11.128820 | 10.979244 | 2734 | 2666 | 1955 | 1742 |
| 8453 | NM_00119877 | CUL2 | cullin 2 | protein_coding | NP_001185706.1;NP_001185707.1;NP_001185708.1;NP_001311304.1;NI | 2552 | 603135 | . | . | 7.559622 | 8.084445 | 7.041095 | 6.052467 | 17.591554 | 18.687558 | 15.800014 | 13.838248 | -1.214760 | 945.196780 | 0.147780213 | -1.89924729 | 0.05753197 | 0.711571492 | 10.017811 | 9.737540 | 9.925256 | 9.973758 | 9.863753 | 9.750637 | 1155 | 1155 | 802 | 736 |
| 8454 | NM_00137066 | CUL1 | cullin 1 | protein_coding | NP_001357589.1;NP_001357590.1;NP_001357591.1;NP_001357592.1;NI | 2551 | 603134 | . | . | 29.919850 | 32.179092 | 29.864998 | 32.526531 | 69.624718 | 74.383408 | 67.016187 | 74.368046 | -1.009032 | 3054.913661 | 0.115263273 | -0.11254205 | 0.910393629 | 0.998865404 | 11.583479 | 11.570323 | 11.553582 | 11.606414 | 11.542077 | 11.602840 | 3420 | 3418 | 2546 | 2957 |
| 8455 | NM_00120704 | ATRN | attractin | protein_coding | NP_001193976.1;NP_001310261.1;NP_647537.1;NP_647538.1 | 885 | 603130 | . | . | 14.746145 | 14.914611 | 13.736179 | 14.887284 | 34.314882 | 34.475789 | 30.823584 | 34.038006 | -1.080716 | 3133.868638 | 0.112991716 | -0.99111295 | 0.321630428 | 0.998865404 | 11.668705 | 11.556575 | 11.624408 | 11.664435 | 11.557702 | 11.603703 | 3658 | 3598 | 2545 | 2904 |
| 8458 | NM_003594,XI | TTF2 | transcription termination factor 2 | protein_coding | NP_003585.3;XP_005271334.1;XP_011540605.1;XP_016858039.1;XP_01 | 12398 | 604718 | . | . | 7.165841 | 8.101970 | 8.602893 | 9.043118 | 16.675209 | 18.728066 | 19.304640 | 20.676016 | 1.142648 | 2430.386779 | 0.122440672 | 1.571213857 | 0.116132982 | 0.857396579 | 11.147704 | 11.339844 | 11.137173 | 11.240103 | 11.278386 | 11.316128 | 2444 | 2606 | 2202 | 2486 |
| 8459 | NM_00100856 | TPST2 | tyrosylprotein sulfotransferase 2 | protein_coding | NP_001008566.1;NP_001349851.1;NP_001349852.1;NP_003586.3;XP_02 | 12021 | 603126 | . | . | 2.609366 | 2.494520 | 2.326360 | 2.690964 | 6.072100 | 5.766194 | 5.220284 | 6.152569 | -1.031697 | 445.851179 | 0.178335259 | -0.25244323 | 0.80069849 | 0.998865404 | 8.822964 | 8.777515 | 8.825738 | 8.793405 | 8.749146 | 8.830200 | 535 | 476 | 356 | 439 |
| 8460 | NM_003596,XI | TPST1 | tyrosylprotein sulfotransferase 1 | protein_coding | NP_003587.1;XP_005250699.1;XP_011514936.1;XP_011514937.1;XP_01 | 12020 | 603125 | . | . | 15.903234 | 16.152088 | 11.241452 | 12.593638 | 37.007482 | 37.336273 | 25.225493 | 28.793859 | -1.367516 | 1116.628010 | 0.137301981 | -3.28879527 | 0.001006172 | 0.068323392 | 10.333294 | 9.881316 | 10.223751 | 10.234569 | 9.955153 | 10.036067 | 1475 | 1402 | 773 | 935 |
| 8462 | NM_00117771 | KLF11 | Kruppel like factor 11 | protein_coding | NP_001171187.1;NP_001171189.1;NP_003588.1 | 11811 | 603301 | . | . | 2.721395 | 2.749193 | 2.667209 | 3.003156 | 6.332798 | 6.354882 | 5.985141 | 6.866358 | 1.022103 | 350.790913 | 0.191256217 | 0.164915664 | 0.869010363 | 0.998865404 | 8.438870 | 8.469898 | 8.447044 | 8.448758 | 8.429834 | 8.489775 | 399 | 375 | 292 | 350 |
| 8463 | NM_00125665 | TEAD2 | TEA domain transcription factor 2 | protein_coding | NP_001243587.1;NP_001243588.1;NP_001243589.1;NP_001243590.1;NI | 11715 | 601729 | . | . | 1.684175 | 1.861729 | 1.773703 | 1.755100 | 3.919142 | 4.303466 | 3.980140 | 4.012828 | -1.026642 | 130.149509 | 0.28362351 | -0.13374285 | 0.893605919 | 0.998865404 | 7.042850 | 7.004954 | 7.024461 | 7.033301 | 7.023588 | 7.015322 | 150 | 144 | 113 | 119 |
| 8464 | NM_00126182 | SUPT3H | SPT3 homolog, SAGA and STAGA complex component | protein_coding | NP_001248752.1;NP_001337253.1;NP_001337254.1;NP_001337255.1;NI | 11466 | 602947 | . | . | 1.448481 | 1.761097 | 1.253064 | 1.000526 | 3.370675 | 4.070857 | 2.811839 | 2.287578 | -1.043473 | 139.228884 | 0.281326837 | -0.2182261 | 0.82725295 | 0.998865404 | 7.151111 | 7.090890 | 7.164026 | 7.092012 | 7.126416 | 7.099112 | 179 | 139 | 123 | 123 |
| 8467 | NM_003601,XI | SMARCA5 | SWI/SNF related, matrix associated, actin dependent regulator of | protein_coding | NP_003592.3 | 11101 | 603375 | . | . | 12.224050 | 11.927106 | 11.430156 | 11.411357 | 28.445866 | 27.570038 | 25.648939 | 26.090712 | -1.109646 | 2462.612826 | 0.114322243 | -1.31295345 | 0.189198636 | 0.960264078 | 11.339090 | 11.188954 | 11.313114 | 11.299965 | 11.211542 | 11.233522 | 3007 | 2773 | 2002 | 2219 |
| 8470 | NM_00114567 | SORBS2 | sorbin and SH3 domain containing 2 | protein_coding | NP_001139142.1;NP_001139143.1;NP_001139144.1;NP_001139145.1;NI | 24098 | 616349 | . | . | 1.102824 | 1.198376 | 0.682355 | 0.788747 | 2.566317 | 2.770098 | 1.531184 | 1.803378 | -1.586815 | 259.986838 | 0.217217223 | -3.06667163 | 0.002164565 | 0.117381585 | 8.317642 | 7.650499 | 8.107330 | 8.138672 | 7.861981 | 7.919492 | 357 | 354 | 163 | 201 |
| 8473 | NM_181672,N | OGT | O-linked N-acetylglucosamine (GlcNAc) transferase | protein_coding | NP_858058.1;NP_858059.1;XP_016885397.1;XP_024308235.1 | 8127 | 300255 | . | . | 14.971688 | 14.787666 | 16.005932 | 13.582028 | 34.839731 | 34.182354 | 35.916847 | 31.053693 | -1.065663 | 3113.060929 | 0.125121936 | -0.73329245 | 0.463380081 | 0.998865404 | 11.649174 | 11.557610 | 11.606809 | 11.649849 | 11.646011 | 11.501938 | 3602 | 3556 | 2853 | 2576 |
| 8476 | NM_00136601 | CDC42BPA | CDC42 binding protein kinase alpha | protein_coding | NP_001352939.1;NP_001352940.1;NP_001352948.1;NP_003598.2;NP_0 | 1737 | 603412 | . | . | 5.161867 | 5.015407 | 5.701911 | 5.442429 | 12.011876 | 11.593339 | 12.794925 | 12.443469 | 1.079448 | 1788.642017 | 0.121291594 | 0.909328335 | 0.363176838 | 0.998865404 | 10.748365 | 10.858820 | 10.786092 | 10.760725 | 10.851677 | 10.815360 | 2013 | 1826 | 1652 | 1701 |
| 8479 | NM_00119732 | HIRIP3 | HIRA interacting protein 3 | protein_coding | NP_001184252.1;NP_003600.2 | 4917 | 603365 | . | . | 11.634019 | 11.662198 | 11.134708 | 10.758766 | 27.072841 | 26.957693 | 24.985964 | 24.598639 | -1.082904 | 867.514224 | 0.141897601 | -0.80977742 | 0.418068112 | 0.998865404 | 9.816964 | 9.702243 | 9.789588 | 9.787588 | 9.748673 | 9.714403 | 1040 | 972 | 742 | 762 |
| 8480 | NM_00101588 | RAE1 | ribonucleic acid export 1 | protein_coding | NP_001015885.1;NP_003601.1;XP_005260639.2;XP_005260640.1;XP_01 | 9828 | 603343 | . | . | 39.420490 | 38.459748 | 33.876030 | 41.395879 | 91.733101 | 88.901432 | 76.016833 | 94.646754 | -1.019080 | 2324.974420 | 0.120608783 | -0.22608333 | 0.821136612 | 0.998865404 | 11.196688 | 11.169179 | 11.178930 | 11.201450 | 11.129830 | 11.218209 | 2665 | 2568 | 1892 | 2278 |
| 8481 | NM_00133020 | OFD1 | OFD1 centriole and centriolar satellite protein | protein_coding | NP_001317138.1;NP_001317139.1;NP_003602.1;XP_005274663.1;XP_01 | 2567 | 300170 | . | . | 5.256013 | 5.305861 | 6.110035 | 4.513787 | 12.230960 | 12.264734 | 13.710743 | 10.320235 | -1.027070 | 728.791725 | 0.176051996 | -0.21888069 | 0.826742992 | 0.998865404 | 9.528016 | 9.490465 | 9.531309 | 9.498512 | 9.601497 | 9.379965 | 870 | 778 | 726 | 566 |
| 8482 | NM_00114602 | SEMA7A | semaphorin 7A (John Milton Hagen blood group) | protein_coding | NP_001139501.1;NP_001139502.1;NP_003603.1 | 10741 | 607961 | . | . | 10.214247 | 9.691121 | 5.974580 | 7.245886 | 23.768973 | 22.401459 | 13.406786 | 16.566856 | -1.523806 | 871.524908 | 0.1523683 | -3.98822842 | 6.65686E-05 | 0.009472636 | 10.039705 | 9.431449 | 9.918225 | 9.873914 | 9.534186 | 9.653744 | 1247 | 1103 | 546 | 706 |
| 8487 | NM_00100918 | GEMIN2 | gem nuclear organelle associated protein 2 | protein_coding | NP_001009182.1;NP_001009183.1;NP_003607.1;XP_016877198.1;XP_02 | 10884 | 602595 | . | . | 5.890849 | 5.488536 | 5.805828 | 5.180562 | 13.708249 | 12.686996 | 13.028113 | 11.844738 | -1.032708 | 234.428744 | 0.225509836 | -0.20590052 | 0.836868619 | 0.998865404 | 7.895536 | 7.850118 | 7.904832 | 7.854867 | 7.893532 | 7.834923 | 288 | 244 | 213 | 203 |
| 8490 | NM_00119530 | RGS5 | regulator of G protein signaling 5 | protein_coding | NP_001182232.1;NP_001241677.1;NP_001241678.1;NP_003608.1 | 10001 | 603276 | . | . | 1.295274 | 1.393116 | 1.206015 | 1.551212 | 3.014154 | 3.220250 | 2.706263 | 3.546660 | -1.008857 | 77.997516 | 0.366067475 | -0.03475389 | 0.97227599 | 0.998865404 | 6.292397 | 6.278281 | 6.294073 | 6.277998 | 6.247807 | 6.318522 | 93 | 82 | 58 | 83 |
| 8491 | NM_00127042 | MAP4K3 | mitogen-activated protein kinase kinase kinase kinase 3 | protein_coding | NP_001257354.1;NP_003609.2;XP_024308951.1 | 6865 | 604921 | . | . | 5.733900 | 6.021372 | 4.715868 | 5.119605 | 13.343021 | 13.918671 | 10.582273 | 11.705371 | -1.194292 | 726.607703 | 0.150414653 | -1.70299513 | 0.08856898 | 0.802216049 | 9.627605 | 9.371072 | 9.545136 | 9.580773 | 9.419799 | 9.460678 | 888 | 875 | 558 | 640 |
| 8492 | NM_003619,XI | PRSS12 | serine protease 12 | protein_coding | NP_003610.2;XP_005263375.1;XP_011530689.1 | 9477 | 606709 | . | . | 1.668284 | 1.809377 | 1.556678 | 1.664788 | 3.882164 | 4.182455 | 3.493141 | 3.806339 | -1.163391 | 222.838057 | 0.22977533 | -0.95021574 | 0.342002644 | 0.998865404 | 7.905356 | 7.686019 | 7.812795 | 7.856116 | 7.735092 | 7.790113 | 264 | 270 | 168 | 205 |
| 8493 | NM_003620,XI | PPM1D | protein phosphatase, Mg2+/Mn2+ dependent 1D | protein_coding | NP_003611.1 | 9277 | 605100 | . | . | 1.932017 | 1.807631 | 1.839218 | 2.159982 | 4.495882 | 4.178421 | 4.127151 | 4.938541 | 1.048815 | 288.940468 | 0.208809067 | 0.329299584 | 0.741929259 | 0.998865404 | 8.140066 | [illegible] | 8.180531 | 8.140958 | 8.145736 | 8.224933 | 337 | 293 | 238 | 298 |
| 8495 | NM_00125656 | PPFIBP2 | PPFIA binding protein 2 | protein_coding | NP_001243497.2;NP_001243498.2;NP_001338782.2;NP_001338783.2;NI | 9250 | 603142 | . | . | 10.268599 | 11.072442 | 9.177422 | 10.318914 | 23.895449 | 25.594444 | 20.593867 | 23.592965 | -1.126049 | 1092.212333 | 0.138830323 | -1.23366227 | 0.217328779 | 0.982008277 | 10.176352 | 10.004615 | 10.107356 | 10.162339 | 10.008652 | 10.083877 | 1284 | 1294 | 847 | 1013 |
| 8496 | NM_00119891 | PPFIBP1 | PPFIA binding protein 1 | protein_coding | NP_001185844.1;NP_001185845.2;NP_003613.4;NP_803193.3;XP_00671 | 9249 | 603141 | . | . | 35.347320 | 36.288050 | 39.277632 | 38.230981 | 82.254665 | 83.881451 | 88.137862 | 87.410598 | 1.079182 | 6284.826075 | 0.103455929 | 1.06265803 | 0.287937061 | 0.998865404 | 12.561651 | 12.671570 | 12.575507 | 12.593656 | 12.646262 | 12.651449 | 6886 | 6594 | 5656 | 6135 |
| 8497 | NM_00130433 | PPFIA4 | PTPRF interacting protein alpha 4 | protein_coding | NP_001291260.1;NP_001291261.1;XP_006711649.1;XP_011508372.1;XP | 9248 | 603145 | . | . | 1.464295 | 1.564520 | 3.454923 | 2.027206 | 3.407474 | 3.616460 | 7.752749 | 4.634965 | 1.592653 | 369.210715 | 0.232434124 | 2.888696306 | 0.003868425 | 0.181618599 | 8.153523 | 8.825522 | 8.303674 | 8.413077 | 8.759286 | 8.478849 | 294 | 339 | 482 | 331 |
| 8498 | NM_00130086 | RANBP3 | RAN binding protein 3 | protein_coding | NP_001287794.1;NP_003615.2;NP_015559.2;NP_015561.1;XP_0067229 | 9850 | 603327 | . | . | 11.362439 | 12.004661 | 11.918259 | 13.618431 | 26.440865 | 27.749308 | 26.744227 | 31.136922 | 1.063175 | 1161.028659 | 0.138409953 | 0.638534203 | 0.523125997 | 0.998865404 | 10.136568 | 10.224469 | 10.130686 | 10.182765 | 10.155136 | 10.248348 | 1251 | 1257 | 976 | 1191 |
| 8500 | NM_00137800 | PPFIA1 | PTPRF interacting protein alpha 1 | protein_coding | NP_001364935.1;NP_003617.1;NP_803172.1;XP_006718779.1;XP_01154 | 9245 | 611054 | . | . | 19.713638 | 20.009440 | 17.667911 | 18.809530 | 45.874449 | 46.252718 | 39.646278 | 43.005760 | -1.102913 | 3072.106904 | 0.112291571 | -1.25850532 | 0.208209064 | 0.970482251 | 11.654000 | 11.512561 | 11.612845 | 11.634556 | 11.521518 | 11.565127 | 3662 | 3523 | 2472 | 2813 |
| 8501 | NM_00119881 | SLC43A1 | solute carrier family 43 member 1 | protein_coding | NP_001185739.1;NP_003618.1;XP_005274415.1;XP_011543622.1;XP_01 | 9225 | 603733 | . | . | 4.663790 | 4.353143 | 5.174250 | 4.639050 | 10.852832 | 10.062483 | 11.610866 | 10.606635 | 1.082498 | 364.235414 | 0.190414142 | 0.600608151 | 0.548101008 | 0.998865404 | 8.450009 | 8.565149 | 8.505429 | 8.462092 | 8.561975 | 8.496470 | 418 | 363 | 348 | 335 |
| 8502 | NM_00100547 | PKP4 | plakophilin 4 | protein_coding | NP_001005476.1;NP_001291898.1;NP_001291899.1;NP_001291900.1;NI | 9026 | 604276 | . | . | 16.319296 | 16.378900 | 17.065137 | 17.303498 | 37.975673 | 37.860555 | 38.293676 | 39.562394 | 1.033697 | 2337.248349 | 0.114770068 | 0.41660224 | 0.676969371 | 0.998865404 | 11.166486 | 11.214308 | 11.182282 | 11.171803 | 11.201712 | 11.205726 | 2664 | 2464 | 2061 | 2233 |
| 8503 | NM_00111417 | PIK3R3 | phosphoinositide-3-kinase regulatory subunit 3 | protein_coding | NP_001107644.1;NP_001290357.1;NP_001290358.1;NP_001315577.1;NI | 8981 | 606076 | . | . | 29.038164 | 29.594885 | 34.908545 | 33.925446 | 67.572994 | 68.409898 | 78.333765 | 77.566501 | 1.155318 | 5530.987775 | 0.105175166 | 1.980409002 | 0.047657589 | 0.659753773 | 12.325404 | 12.533725 | 12.365310 | 12.373702 | 12.508080 | 12.472993 | 5880 | 5566 | 5263 | 5444 |
| 8504 | NM_003630 | PEX3 | peroxisomal biogenesis factor 3 | protein_coding | NP_003621.1 | 8858 | 603164 | . | . | 8.530380 | 8.800934 | 7.904928 | 7.711874 | 19.850544 | 20.343752 | 17.738430 | 17.632282 | -1.125185 | 706.063265 | 0.150132214 | -1.1334119 | 0.257041321 | 0.995760118 | 9.546188 | 9.376109 | 9.494839 | 9.510889 | 9.433677 | 9.409924 | 851 | 816 | 588 | 612 |
| 8505 | NM_00130348 | PARG | poly(ADP-ribose) glycohydrolase | protein_coding | NP_001290415.1;NP_001290416.1;NP_001290418.1;NP_001311310.1;NI | 8605 | 603501 | . | . | 4.105507 | 3.985401 | 4.277789 | 4.249503 | 9.553683 | 9.212432 | 9.599236 | 9.715985 | 1.017176 | 543.927031 | 0.163246149 | 0.150502184 | 0.880368428 | 0.998865404 | 9.074793 | 9.099639 | 9.101262 | 9.061321 | 9.099225 | 9.085901 | 640 | 564 | 482 | 509 |
| 8506 | NM_003632,XI | CNTNAP1 | contactin associated protein 1 | protein_coding | NP_003623.1;XP_005257805.1;XP_016880727.1;XP_024306779.1 | 8011 | 602346 | . | . | 0.230268 | 0.147201 | 0.284102 | 0.262254 | 0.535841 | 0.340261 | 0.637517 | 0.599612 | 1.358942 | 52.112011 | 0.451169136 | 0.980750092 | 0.326715995 | 0.998865404 | 5.463517 | 5.909275 | 5.693152 | 5.631724 | 5.751476 | 5.689795 | 60 | 39 | 60 | 48 |
| 8507 | NM_00125657 | ENC1 | ectodermal-neural cortex 1 | protein_coding | NP_001243503.1;NP_001243504.1;NP_001243505.1;NP_003624.1;XP_01 | 3345 | 605173 | . | . | 4.734798 | 5.311374 | 4.904914 | 4.247104 | 11.018071 | 12.277479 | 11.006483 | 9.710499 | -1.106460 | 829.330092 | 0.153232176 | -0.95248411 | 0.340851519 | 0.998865404 | 9.766796 | 9.621134 | 9.692768 | 9.765113 | 9.709578 | 9.603356 | 953 | 987 | 737 | 682 |
| 8508 | NM_00120250 | NIPSNAP1 | nipsnap homolog 1 | protein_coding | NP_001189431.1;NP_003625.2 | 7827 | 603249 | . | . | 19.206661 | 19.376391 | 16.070344 | 24.078892 | 44.694688 | 44.789397 | 36.061388 | 55.053530 | 1.024539 | 1223.249950 | 0.171599611 | 0.203817765 | 0.83849591 | 0.998865404 | 10.239243 | 10.273560 | 10.242355 | 10.243237 | 10.104842 | 10.396335 | 1391 | 1305 | 869 | 1384 |
| 8509 | NM_00133010 | NDST2 | N-deacetylase and N-sulfotransferase 2 | protein_coding | NP_001317036.1;NP_003626.1;XP_005270313.1;XP_011538612.1 | 7681 | 603268 | . | . | 3.911604 | 3.621375 | 4.184558 | 3.440714 | 9.102464 | 8.370969 | 9.390028 | 7.866782 | 1.070937 | 412.042478 | 0.191256411 | 0.516969056 | 0.60517776 | 0.998865404 | 8.635803 | 8.735764 | 8.678471 | 8.647387 | 8.772379 | 8.631411 | 470 | 418 | 414 | 353 |
| 8510 | NM_006983,XI | MMP23B | matrix metallopeptidase 23B | protein_coding | NP_008914.1;XP_016858104.1;XP_016858106.1 | 7171 | 603321 | . | . | 0.052368 | 0.146486 | 0.093933 | 0.145530 | 0.121863 | 0.338609 | 0.210782 | 0.332737 | 1.029669 | 9.204566 | 1.038418342 | 0.040620013 | 0.967598829 | NA | 3.184683 | 3.219802 | 3.178967 | 3.214730 | 3.187676 | 3.208735 | 6 | 14 | 6 | 11 |
| 8512 | NR_002724 | MBL1P | mannose binding lectin 1, pseudogene | transcribed_ps | . | 6921 | . | . | . | 0.450788 | 0.428051 | 1.022180 | 0.443516 | 1.049003 | 0.989460 | 2.293742 | 1.014046 | 1.639517 | 26.642474 | 0.639245617 | 1.115801245 | 0.264507221 | NA | 4.332860 | 5.050189 | 4.687390 | 4.681724 | 4.787302 | 4.686096 | 24 | 21 | 40 | 19 |
| 8514 | NM_00119986 | KCNAB2 | potassium voltage-gated channel subfamily A regulatory beta sub | protein_coding | NP_001186789.1;NP_001186790.1;NP_001186791.1;NP_001186792.1;NI | 6229 | 601142 | . | . | 70.517349 | 71.767114 | 58.343941 | 77.577018 | 164.096773 | 165.892900 | 130.922118 | 177.370640 | -1.054646 | 8339.669739 | 0.132925444 | -0.57746231 | 0.563627191 | 0.998865404 | 13.063704 | 12.986821 | 13.040954 | 13.050450 | 12.876807 | 13.110827 | 9804 | 9290 | 6086 | 8659 |
| 8515 | NM_00130304 | ITGA10 | integrin subunit alpha 10 | protein_coding | NP_001289969.1;NP_001289970.1;NP_003628.2;XP_005277493.1;XP_01 | 6135 | 604042 | . | . | 3.735488 | 3.681304 | 3.550721 | 3.893040 | 8.692637 | 8.509499 | 7.967714 | 8.900972 | 1.033483 | 573.634448 | 0.160847901 | 0.295399885 | 0.767688421 | 0.998865404 | 9.140148 | 9.187440 | 9.160154 | 9.145440 | 9.150814 | 9.197439 | 657 | 602 | 489 | 566 |
| 8517 | NM_00109985 | IKBKG | inhibitor of nuclear factor kappa B kinase regulatory subunit gamn | protein_coding | NP_001093326.2;NP_001093327.1;NP_001138727.1;NP_001308325.1;NI | 5961 | 300248 | . | . | 9.044231 | 9.042062 | 9.672031 | 9.674052 | 21.046297 | 20.901131 | 21.703759 | 22.118573 | 1.035848 | 581.832947 | 0.158563263 | 0.320456257 | 0.748622485 | 0.998865404 | 9.158834 | 9.209641 | 9.173345 | 9.171742 | 9.194614 | 9.196879 | 659 | 616 | 514 | 556 |
| 8518 | NM_00131836 | ELP1 | elongator complex protein 1 | protein_coding | NP_001305289.1;NP_001317678.1;NP_003631.2;XP_011517438.1;XP_01 | 5959 | 603722 | . | . | 8.370201 | 8.851478 | 8.578796 | 8.873749 | 19.477801 | 20.460587 | 19.250570 | 20.288770 | 1.014559 | 1558.217661 | 0.123572326 | 0.168748088 | 0.865994793 | 0.998865404 | 10.595267 | 10.616020 | 10.588422 | 10.611433 | 10.604473 | 10.617830 | 1755 | 1694 | 1353 | 1484 |
| 8519 | NM_003641 | IFITM1 | interferon induced transmembrane protein 1 | protein_coding | NP_003632.4 | 5412 | 604456 | . | . | 0.188355 | 0.289359 | 0.145816 | 0.283298 | 0.438310 | 0.668867 | 0.327207 | 0.647727 | -1.094900 | 5.169837 | 1.335249377 | -0.09795823 | 0.92196546 | NA | 2.436885 | 2.300113 | 2.367924 | 2.377962 | 2.364269 | 2.377353 | 5 | 7 | 3 | 6 |
| 8520 | NM_003642,N | HAT1 | histone acetyltransferase 1 | protein_coding | NP_003633.2;XP_006712871.1;XP_016860625.1;XP_016860626.1;XP_02 | 4821 | 603053 | . | . | 65.370807 | 60.350556 | 55.905336 | 50.822576 | 152.120560 | 139.503022 | 125.449946 | 116.199788 | -1.203705 | 3118.778573 | 0.125467551 | -2.13187976 | 0.033016733 | 0.575268292 | 11.734180 | 11.466998 | 11.707400 | 11.645444 | 11.591805 | 11.453685 | 4068 | 3540 | 2673 | 2426 |
| 8522 | NM_00113083 | GAS7 | growth arrest specific 7 | protein_coding | NP_001124303.1;NP_003635.2;NP_958836.1;NP_958839.1;XP_0052568 | 4169 | 603127 | . | . | 32.850691 | 33.991568 | 34.614229 | 38.877938 | 76.444908 | 78.573033 | 77.673334 | 88.889780 | 1.082420 | 8768.262285 | 0.107494004 | 1.062948387 | 0.287805358 | 0.998865404 | 13.039852 | 13.154040 | 13.052931 | 13.073188 | 13.084799 | 13.174112 | 9581 | 9196 | 7513 | 8991 |
| 8525 | NM_00110554 | DGKZ | diacylglycerol kinase zeta | protein_coding | NP_001099010.1;NP_001186195.1;NP_001186196.1;NP_001186197.1;NI | 2857 | 601441 | . | . | 6.038027 | 6.415129 | 5.998264 | 5.516940 | 14.050736 | 14.828860 | 13.459931 | 12.613829 | -1.092339 | 670.910294 | 0.155245743 | -0.82076506 | 0.411780103 | 0.998865404 | 9.452179 | 9.324970 | 9.399410 | 9.438179 | 9.388800 | 9.328269 | 784 | 777 | 583 | 574 |
| 8526 | NM_003647,XI | DGKE | diacylglycerol kinase epsilon | protein_coding | NP_003638.1;XP_011523696.1;XP_011523697.1;XP_011523698.1;XP_01 | 2852 | 601440 | . | . | 0.154144 | 0.178021 | 0.245745 | 0.153390 | 0.358698 | 0.411503 | 0.551446 | 0.350708 | 1.316849 | 50.070990 | 0.463540391 | 0.856644824 | 0.391641205 | 0.998865404 | 5.432996 | 5.831393 | 5.587273 | 5.626604 | 5.703576 | 5.610174 | 44 | 52 | 61 | 41 |
| 8527 | NM_00137725 | DGKD | diacylglycerol kinase delta | protein_coding | NP_001364188.1;NP_003639.2;NP_690618.2;XP_005246164.1;XP_01151 | 2851 | 601826 | . | . | 5.683529 | 6.248361 | 6.473828 | 6.535738 | 13.225803 | 14.443367 | 14.527084 | 14.943186 | 1.081080 | 1226.627584 | 0.132260995 | 0.850387094 | 0.395109909 | 0.998865404 | 10.203249 | 10.315530 | 10.197537 | 10.261138 | 10.287077 | 10.290100 | 1300 | 1326 | 1106 | 1197 |
| 8530 | NM_003650 | CST7 | cystatin F | protein_coding | NP_003641.3 | 2479 | 603253 | . | . | 1.557023 | 2.531384 | 1.929970 | 2.411240 | 3.623257 | 5.851409 | 4.330797 | 5.513014 | 1.041379 | 58.958012 | 0.431952005 | 0.135422162 | 0.892278103 | 0.998865404 | 5.853620 | 5.909079 | 5.820983 | 5.920351 | 5.859180 | 5.901312 | 51 | 77 | 47 | 62 |
| 8531 | NM_00114542 | YBX3 | Y-box binding protein 3 | protein_coding | NP_001138898.1;NP_003642.3;XP_011519173.1;XP_016875611.1;XP_01 | 2428 | 603437 | . | . | 47.296093 | 50.623589 | 52.113631 | 58.000313 | 110.059954 | 117.018688 | 116.941474 | 132.610822 | 1.100780 | 2965.658890 | 0.118846471 | 1.165585609 | 0.243782029 | 0.992733415 | 11.463333 | 11.601628 | 11.468159 | 11.517255 | 11.522857 | 11.617744 | 3153 | 3139 | 2547 | 3081 |
| 8533 | NM_00119912 | COPS3 | COP9 signalosome subunit 3 | protein_coding | NP_001186054.1;NP_001303283.1;NP_001303284.1;NP_001303285.1;NI | 2239 | 604665 | . | . | 35.133145 | 37.442170 | 32.005603 | 34.824794 | 81.756282 | 86.549251 | 71.819639 | 79.622754 | -1.099264 | 1958.149579 | 0.121917545 | -1.11992154 | 0.262747199 | 0.995798899 | 11.002049 | 10.865261 | 10.951755 | 10.990910 | 10.864505 | 10.927263 | 2304 | 2267 | 1557 | 1819 |
| 8534 | NM_003654,XI | CHST1 | carbohydrate sulfotransferase 1 | protein_coding | NP_003645.1;XP_006718419.1;XP_016873948.1 | 1969 | 603797 | . | . | 0.754365 | 0.721907 | 0.755525 | 0.739346 | 1.755439 | 1.668721 | 1.695377 | 1.690428 | -1.001857 | 90.872194 | 0.333444488 | -0.00802667 | 0.99359571 | 0.999862634 | 6.506929 | 6.504604 | 6.512342 | 6.499643 | 6.509189 | 6.501777 | 107 | 96 | 80 | 84 |
| 8535 | NM_003655,XI | CBX4 | chromobox 4 | protein_coding | NP_003646.2;XP_011523701.1 | 1554 | 603079 | . | . | 22.052534 | 20.270348 | 20.575144 | 23.232902 | 51.317155 | 46.855821 | 46.170027 | 53.119271 | 1.033669 | 1911.665202 | 0.124328977 | 0.384255881 | 0.700788805 | 0.998865404 | 10.876581 | 10.924253 | 10.912218 | 10.861030 | 10.873995 | 10.950786 | 2235 | 1963 | 1608 | 1910 |
| 8539 | NM_00114293 | API5 | apoptosis inhibitor 5 | protein_coding | NP_001136402.1;NP_001136403.1;NP_001230676.1;NP_006586.1;XP_0 | 594 | 609774 | . | . | 17.418397 | 15.862835 | 16.565979 | 15.653866 | 40.533326 | 36.667653 | 37.173575 | 35.790705 | -1.025388 | 2021.823531 | 0.126247937 | -0.28650332 | 0.774492652 | 0.998865404 | 10.999251 | 10.963409 | 11.034132 | 10.945236 | 11.006523 | 10.933860 | 2488 | 2086 | 1816 | 1786 |
| 8540 | NM_003659,XI | AGPS | alkylglycerone phosphate synthase | protein_coding | NP_003650.1;XP_011510343.1;XP_011510344.1;XP_011510345.1 | 327 | 603051 | . | . | 8.810350 | 8.761216 | 9.353129 | 8.238146 | 20.502049 | 20.251945 | 20.988150 | 18.835543 | -1.013119 | 2095.035396 | 0.124641766 | -0.15086189 | 0.880084663 | 0.998865404 | 11.041992 | 11.023466 | 11.042265 | 11.032295 | 11.079573 | 10.971596 | 2443 | 2261 | 1933 | 1819 |
| 8541 | NM_003660,N | PPFIA3 | PTPRF interacting protein alpha 3 | protein_coding | NP_003651.1;XP_016882896.1 | 9247 | 603144 | . | . | 3.593320 | 3.587072 | 3.594692 | 3.849709 | 8.361803 | 8.291677 | 8.066385 | 8.801902 | 1.027612 | 536.118605 | 0.163800921 | 0.239897118 | 0.810410015 | 0.998865404 | 9.046718 | 9.085833 | 9.063743 | 9.050954 | 9.056290 | 9.093196 | 615 | 565 | 459 | 524 |
| 8542 | NM_00113654 | APOL1 | apolipoprotein L1 | protein_coding | NP_001130012.1;NP_001130013.1;NP_001349856.1;NP_003652.2;NP_6 | 618 | 603743 | . | . | 0.517456 | 0.624351 | 0.403811 | 0.404536 | 1.204141 | 1.443215 | 0.906139 | 0.924923 | -1.403657 | 44.598743 | 0.468463482 | -1.0442444 | 0.296372329 | 0.998865404 | 5.702923 | 5.213655 | 5.499167 | 5.518227 | 5.455684 | 5.448789 | 57 | 59 | 33 | 34 |
| 8543 | NM_00136949 | LMO4 | LIM domain only 4 | protein_coding | NP_001356420.1;NP_006760.1 | 6644 | 603129 | . | . | 4.677438 | 4.051419 | 2.586285 | 3.808087 | 10.884592 | 9.365038 | 5.803548 | 8.706738 | -1.384370 | 587.943266 | 0.194579997 | -2.41149966 | 0.015887067 | 0.423338192 | 9.415546 | 8.945382 | 9.344161 | 9.237341 | 8.958227 | 9.184447 | 846 | 681 | 350 | 547 |
| 8544 | NM_00101810 | PIR | pirin | protein_coding | NP_001018119.1;NP_003653.1 | 30048 | 300931 | . | . | 88.588557 | 87.354267 | 86.780106 | 109.915949 | 206.149214 | 201.923307 | 194.732050 | 251.309757 | 1.108557 | 3750.250645 | 0.128989678 | 1.152672313 | 0.24904491 | 0.992733415 | 11.796613 | 11.945110 | 11.839869 | 11.813990 | 11.815856 | 12.000426 | 4161 | 3778 | 3051 | 4104 |
| 8545 | NM_00100839 | CGGBP1 | CGG triplet repeat binding protein 1 | protein_coding | NP_001008391.1;NP_001182237.1;NP_003654.3;XP_024309565.1 | 1888 | 603363 | . | . | 16.170661 | 13.894353 | 16.139253 | 11.849291 | 37.629792 | 32.117418 | 36.216017 | 27.091995 | -1.089563 | 2019.077909 | 0.157996 | -0.78324316 | 0.433484335 | 0.998865404 | 11.039768 | 10.916564 | 11.071179 | 10.942111 | 11.058947 | 10.806592 | 2618 | 2090 | 1949 | 1523 |
| 8546 | NM_00127176 | AP3B1 | adaptor related protein complex 3 subunit beta 1 | protein_coding | NP_001258698.1;NP_003655.3;XP_005248675.1;XP_005248676.1;XP_01 | 566 | 603401 | . | . | 13.796894 | 14.084378 | 11.952943 | 12.419051 | 32.105936 | 32.556668 | 26.822057 | 28.394683 | -1.132260 | 1717.455318 | 0.122098818 | -1.46770519 | 0.142184321 | 0.903146012 | 10.832860 | 10.653695 | 10.778123 | 10.807063 | 10.710299 | 10.680403 | 2062 | 2004 | 1428 | 1481 |
| 8548 | NM_00132097 | BLZF1 | basic leucine zipper nuclear factor 1 | protein_coding | NP_001307901.1;NP_001307902.1;NP_003657.1 | 1065 | 608692 | . | . | 5.906378 | 6.510008 | 7.178642 | 6.562774 | 13.744384 | 15.048175 | 16.108665 | 15.005004 | 1.082342 | 500.092822 | 0.169993528 | 0.671534311 | 0.501880209 | 0.998865404 | 8.907792 | 9.022051 | 8.914981 | 8.963009 | 9.015721 | 8.962444 | 532 | 538 | 471 | 467 |
| 8550 | NM_00137147 | MAPKAPK5 | MAPK activated protein kinase 5 | protein_coding | NP_001358408.1;NP_001358409.1;NP_001358410.1;NP_001358411.1;NI | 6889 | 606723 | . | . | 3.496138 | 2.704567 | 3.264625 | 2.782875 | 8.135658 | 6.251732 | 7.325724 | 6.362714 | 1.117846 | 962.018545 | 0.143178393 | 1.122528688 | 0.261637717 | 0.995760118 | 9.827169 | 9.988183 | 9.853473 | 9.879739 | 9.993390 | 9.900339 | 1027 | 998 | 939 | 892 |
| 8552 | NR_024616 | INE1 | inactivation escape 1 | lncRNA | . | 6060 | 300164 | . | . | 0.462488 | 0.769045 | 0.645408 | 0.726555 | 1.076229 | 1.777681 | 1.448276 | 1.661182 | 1.088621 | 21.205070 | 0.678227074 | 0.180620648 | 0.856665348 | NA | 4.345328 | 4.464871 | 4.377734 | 4.418611 | 4.400269 | 4.411682 | 18 | 27 | 18 | 22 |
| 8553 | NM_003670,XI | BHLHE40 | basic helix-loop-helix family member e40 | protein_coding | NP_003661.1;XP_024309566.1 | 1046 | 604256 | . | . | 25.927032 | 26.970484 | 31.802675 | 31.360724 | 60.333273 | 62.343478 | 71.364281 | 71.702565 | 1.177796 | 2849.483659 | 0.112330837 | 2.10174022 | 0.035576038 | 0.58668585 | 11.353616 | 11.589705 | 11.392999 | 11.418251 | 11.549449 | 11.528638 | 2966 | 2868 | 2714 | 2853 |
| 8554 | NM_00132068 | PIAS1 | protein inhibitor of activated STAT 1 | protein_coding | NP_001307616.1;NP_057250.1;XP_011520428.1;XP_011520429.1;XP_01 | 2752 | 603566 | . | . | 2.283480 | 2.643059 | 2.381818 | 2.456782 | 5.313750 | 6.109552 | 5.344731 | 5.617140 | -1.039573 | 595.170348 | 0.161511865 | -0.3466691 | 0.728839927 | 0.998865404 | 9.245090 | 9.188676 | 9.190693 | 9.267009 | 9.195488 | 9.211982 | 659 | 692 | 501 | 554 |
| 8555 | NM_00107718 | CDC14B | cell division cycle 14B | protein_coding | NP_001070649.1;NP_001338496.1;NP_001338497.1;NP_001338498.1;NI | 1719 | 603505 | . | . | 2.199324 | 2.026019 | 2.003803 | 3.159986 | 5.117917 | 4.683227 | 4.496478 | 7.224931 | 1.040165 | 340.209636 | 0.199944699 | 0.284138636 | 0.77630414 | 0.998865404 | 8.381893 | 8.438119 | 8.419472 | 8.375840 | 8.365186 | 8.471964 | 399 | 346 | 273 | 356 |
| 8556 | NM_00131921 | CDC14A | cell division cycle 14A | protein_coding | NP_001306139.1;NP_001306140.1;NP_001306141.1;NP_003663.2;NP_2 | 1718 | 603504 | . | . | 0.917618 | 1.034660 | 0.966530 | 1.230355 | 2.135335 | 2.391661 | 2.168864 | 2.813059 | 1.148931 | 128.243475 | 0.297683448 | 0.672836651 | 0.501051218 | 0.998865404 | 6.900121 | 7.098545 | 6.956719 | 6.989842 | 6.968507 | 7.074310 | 131 | 135 | 102 | 147 |
| 8558 | NM_00109853 | CDK10 | cyclin dependent kinase 10 | protein_coding | NP_001092003.2;NP_001153839.1;NP_443713.2;NP_443714.3;XP_00672 | 1770 | 603464 | . | . | 24.209352 | 22.931864 | 30.884448 | 24.564657 | 56.336158 | 53.008029 | 69.303801 | 56.164172 | 1.166928 | 1421.072376 | 0.14229398 | 1.565179583 | 0.117540807 | 0.857396579 | 10.356861 | 10.580045 | 10.415584 | 10.400703 | 10.608014 | 10.438958 | 1525 | 1401 | 1475 | 1280 |
| 8559 | NM_003675,XI | PRPF18 | pre-mRNA processing factor 18 | protein_coding | NP_003666.1;XP_011518056.1;XP_016872349.1 | 17351 | 604993 | . | . | 3.294362 | 2.622420 | 3.138856 | 4.238984 | 7.666116 | 6.061842 | 7.043499 | 9.691933 | -1.033222 | 445.980576 | 0.175418149 | -0.26878545 | 0.788094794 | 0.998865404 | 8.823976 | 8.777321 | 8.832077 | 8.788696 | 8.808684 | 8.772324 | 540 | 472 | 393 | 399 |
| 8560 | NM_00132154 | DEGS1 | delta 4-desaturase, sphingolipid 1 | protein_coding | NP_001308470.1;NP_001308471.1;NP_003667.1;XP_016858137.1 | 13709 | 615843 | . | . | 72.137749 | 69.664093 | 66.229649 | 56.968226 | 167.867514 | 161.031672 | 148.617413 | 130.251068 | -1.165991 | 4203.329912 | 0.119771997 | -1.84982012 | 0.06433948 | 0.731823074 | 12.143749 | 11.922403 | 12.116163 | 12.077421 | 12.033214 | 11.901273 | 5330 | 4771 | 3647 | 3346 |
| 8562 | NM_003677 | DENR | density regulated re-initiation and release factor | protein_coding | NP_003668.2 | 2769 | 604550 | . | . | 22.976686 | 21.354996 | 20.417793 | 19.733152 | 53.467693 | 49.363033 | 45.816936 | 45.117508 | -1.119662 | 1790.713269 | 0.123702837 | -1.31818312 | 0.187442353 | 0.960264078 | 10.885439 | 10.722606 | 10.881025 | 10.815801 | 10.777174 | 10.742719 | 2267 | 1958 | 1502 | 1549 |
| 8563 | NM_00100287 | THOC5 | THO complex 5 | protein_coding | NP_001002877.1;NP_001002878.1;NP_001002879.1;NP_003669.4;XP_0 | 19074 | 612733 | . | . | 34.304206 | 32.887253 | 37.932797 | 39.376531 | 79.827289 | 76.020356 | 85.120097 | 90.029753 | 1.112668 | 3192.581071 | 0.110918847 | 1.388606342 | 0.164952488 | 0.940065727 | 11.561486 | 11.715425 | 11.585235 | 11.604538 | 11.666982 | 11.697702 | 3439 | 3300 | 2870 | 3211 |
| 8565 | NM_003680,XI | YARS1 | tyrosyl-tRNA synthetase 1 | protein_coding | NP_003671.1;XP_011540649.1;XP_011540650.1;XP_016858140.1 | 12840 | 603623 | . | . | 50.016241 | 48.941193 | 36.073429 | 46.610401 | 116.389852 | 113.129761 | 80.947725 | 106.569137 | -1.174463 | 3998.498523 | 0.134072628 | -1.73041103 | 0.083556865 | 0.785005898 | 12.076707 | 11.844440 | 12.006936 | 12.045592 | 11.783522 | 11.991697 | 4860 | 4768 | 2791 | 3886 |
| 8566 | NM_00133103 | PDXK | pyridoxal kinase | protein_coding | NP_001317959.1;NP_003672.1;XP_005261252.2;XP_005261253.1;XP_00 | 8819 | 179020 | . | . | 6.105281 | 6.379622 | 5.690403 | 6.639878 | 14.207234 | 14.746781 | 12.769098 | 15.181290 | -1.040481 | 1310.082015 | 0.13225814 | -0.43287127 | 0.665108297 | 0.998865404 | 10.383941 | 10.326368 | 10.366353 | 10.373943 | 10.297479 | 10.380555 | 1531 | 1449 | 1055 | 1270 |
| 8567 | NM_00113594 | MADD | MAP kinase activating death domain | protein_coding | NP_001129415.1;NP_001129416.1;NP_001363500.1;NP_001363501.1;NI | 6766 | 603584 | . | . | 5.605442 | 5.683504 | 5.877559 | 6.482805 | 13.044092 | 13.137666 | 13.189072 | 14.822166 | 1.084110 | 1083.014285 | 0.135727322 | 0.858421127 | 0.390659966 | 0.998865404 | 10.021545 | 10.137787 | 10.045402 | 10.053921 | 10.078720 | 10.139349 | 1190 | 1128 | 940 | 1099 |
| 8568 | NM_003683,XI | RRP1 | ribosomal RNA processing 1 | protein_coding | NP_003674.1;XP_016883974.1;XP_016883975.1;XP_024307902.1 | 18785 | 610653 | . | . | 19.207453 | 18.538769 | 19.144133 | 18.308837 | 44.696531 | 42.853197 | 42.958879 | 41.860979 | 1.027951 | 1317.499542 | 0.127837399 | 0.311109721 | 0.755717211 | 0.998865404 | 10.343617 | 10.383284 | 10.350245 | 10.355587 | 10.361706 | 10.385893 | 1491 | 1407 | 1143 | 1272 |
| 8569 | NM_00113555 | MKNK1 | MAPK interacting serine/threonine kinase 1 | protein_coding | NP_001129025.2;NP_001364266.1;NP_001364267.1;NP_001364270.1;NI | 7110 | 606724 | . | . | 9.215703 | 7.531492 | 8.445198 | 8.700281 | 21.445324 | 17.409383 | 18.950782 | 19.892155 | 1.014408 | 856.577099 | 0.143907056 | 0.143413869 | 0.88596333 | 0.998865404 | 9.731922 | 9.752880 | 9.762859 | 9.710464 | 9.761170 | 9.733244 | 1015 | 884 | 765 | 793 |
| 8570 | NM_00136629 | KHSRP | KH-type splicing regulatory protein | protein_coding | NP_001353228.1;NP_001353229.1;NP_003676.2 | 6316 | 603445 | . | . | 53.502976 | 57.251834 | 55.245086 | 61.283607 | 124.503623 | 132.340179 | 123.968374 | 140.117691 | 1.057317 | 6807.525598 | 0.109104632 | 0.736975237 | 0.461137408 | 0.998865404 | 12.692203 | 12.772509 | 12.676625 | 12.739503 | 12.721202 | 12.788850 | 7336 | 7408 | 5842 | 6821 |
| 8572 | NM_00113102 | PDLIM4 | PDZ and LIM domain 4 | protein_coding | NP_001124499.1;NP_003678.2;XP_016865491.1 | 16501 | 603422 | . | . | 40.414189 | 39.342171 | 33.497139 | 38.841354 | 94.045484 | 90.941181 | 75.166607 | 88.806135 | -1.116640 | 2669.727660 | 0.121498906 | -1.3100003 | 0.190195734 | 0.961278413 | 11.459955 | 11.300600 | 11.439337 | 11.410764 | 11.279094 | 11.388986 | 3300 | 2987 | 2043 | 2527 |
| 8573 | NM_00112605 | CASK | calcium/calmodulin dependent serine protein kinase | protein_coding | NP_001119526.1;NP_001119527.1;NP_001354650.1;NP_003679.2;XP_0 | 1497 | 300172 | . | . | 2.486404 | 2.270642 | 2.334094 | 2.702689 | 5.785963 | 5.248690 | 5.237638 | 6.179377 | 1.069797 | 654.278633 | 0.159166054 | 0.611540628 | 0.54084173 | 0.998865404 | 9.304403 | 9.401487 | 9.354179 | 9.304381 | 9.331311 | 9.417302 | 755 | 657 | 551 | 674 |
| 8574 | NM_00132097 | AKR7A2 | aldo-keto reductase family 7 member A2 | protein_coding | NP_001307908.1;NP_003680.2 | 389 | 603418 | . | . | 35.515735 | 35.578369 | 30.781795 | 36.110507 | 82.646576 | 82.240997 | 69.073456 | 82.562385 | -1.064887 | 2305.014700 | 0.130834431 | -0.6932427 | 0.488157258 | 0.998865404 | 11.215357 | 11.124328 | 11.191657 | 11.196900 | 11.061605 | 11.219593 | 2729 | 2574 | 1749 | 2300 |
| 8575 | NM_00113951 | PRKRA | protein activator of interferon induced protein kinase EIF2AK2 | protein_coding | NP_001132989.1;NP_001132990.1;NP_001303291.1;NP_003681.1;XP_01 | 9438 | 603424 | . | . | 19.951994 | 19.570815 | 20.124011 | 19.163568 | 46.429113 | 45.238816 | 45.157699 | 43.815226 | -1.031439 | 1126.500286 | 0.135124478 | -0.33049997 | 0.741022213 | 0.998865404 | 10.159636 | 10.115287 | 10.155010 | 10.142525 | 10.158619 | 10.091914 | 1328 | 1224 | 1008 | 993 |
| 8576 | NM_00100891 | STK16 | serine/threonine kinase 16 | protein_coding | NP_001008910.1;NP_001317142.1;NP_001317143.1;NP_001317144.1;XI | 11394 | 604719 | . | . | 7.886661 | 8.622658 | 8.032926 | 9.570526 | 18.352584 | 19.931658 | 18.025655 | 21.881870 | 1.017968 | 714.851507 | 0.154631966 | 0.166151385 | 0.868037816 | 0.998865404 | 9.468900 | 9.493991 | 9.452512 | 9.496605 | 9.444339 | 9.527800 | 790 | 789 | 589 | 717 |
| 8577 | NM_003692 | TMEFF1 | transmembrane protein with EGF like and two follistatin like doma | protein_coding | NP_003683.2 | 11866 | 603421 | . | . | 1.390912 | 1.050095 | 0.854228 | 0.935102 | 3.236710 | 2.427342 | 1.916863 | 2.137999 | -1.379796 | 85.455784 | 0.354513903 | -1.31011843 | 0.190155773 | 0.961278413 | 6.630507 | 6.166566 | 6.501778 | 6.414592 | 6.362396 | 6.380354 | 130 | 92 | 60 | 70 |
| 8578 | NM_003693,N | SCARF1 | scavenger receptor class F member 1 | protein_coding | NP_003684.2;NP_663325.1;XP_016880745.1 | 16820 | 607873 | . | . | 0.064117 | 0.062905 | 0.074082 | 0.043970 | 0.149203 | 0.145409 | 0.166239 | 0.100532 | -1.140434 | 7.153273 | 1.134076775 | -0.16716898 | 0.867237088 | NA | 2.928838 | 2.742346 | 2.842494 | 2.840596 | 2.843290 | 2.831276 | 9 | 8 | 7 | 5 |
| 8601 | NM_00128667 | RGS20 | regulator of G protein signaling 20 | protein_coding | NP_001273602.1;NP_001273603.1;NP_001273604.1;NP_003693.2;NP_7 | 14600 | 607193 | . | . | 8.068178 | 7.789300 | 9.851863 | 11.074932 | 18.774981 | 18.005317 | 22.107294 | 25.321515 | 1.334221 | 347.938534 | 0.191661092 | 2.170484213 | 0.029970183 | 0.553375672 | 8.220005 | 8.635552 | 8.348491 | 8.353146 | 8.483859 | 8.542262 | 342 | 323 | 329 | 391 |
| 8602 | NM_00129197 | NOP14 | NOP14 nucleolar protein | protein_coding | NP_001278907.1;NP_001278908.1;NP_003694.1 | 16821 | 611526 | . | . | 12.653810 | 12.089350 | 12.908305 | 12.813880 | 29.445939 | 27.945076 | 28.965861 | 29.297414 | 1.035904 | 1362.430261 | 0.127053667 | 0.400545133 | 0.688755048 | 0.998865404 | 10.386276 | 10.437208 | 10.411412 | 10.384814 | 10.420923 | 10.429412 | 1568 | 1419 | 1199 | 1307 |
| 8603 | NM_00125666 | FAM193A | family with sequence similarity 193 member A | protein_coding | NP_001243595.1;NP_001243596.1;NP_001243597.1;NP_001353245.1;NI | 16822 | . | . | . | 3.348074 | 4.082263 | 4.127351 | 3.925137 | 7.791108 | 9.436335 | 9.261654 | 8.974360 | 1.040526 | 613.214640 | 0.163949182 | 0.34958159 | 0.726652729 | 0.998865404 | 9.231109 | 9.288811 | 9.223053 | 9.267443 | 9.322470 | 9.219096 | 669 | 670 | 586 | 541 |
| 8604 | NM_003705,N | SLC25A12 | solute carrier family 25 member 12 | protein_coding | NP_003696.2;XP_011510372.1 | 10982 | 603667 | . | . | 8.764163 | 8.782173 | 7.717091 | 8.635696 | 20.394569 | 20.300386 | 17.316929 | 19.744490 | -1.092073 | 1024.333299 | 0.142232107 | -0.89339497 | 0.371645695 | 0.998865404 | 10.062779 | 9.935347 | 10.044620 | 10.018516 | 9.915107 | 10.015965 | 1253 | 1134 | 792 | 982 |
| 8605 | NM_00115932 | PLA2G4C | phospholipase A2 group IVC | protein_coding | NP_001152794.1;NP_001152795.1;NP_003697.2;XP_011525733.1;XP_01 | 9037 | 603602 | . | . | 0.109253 | 0.094696 | 0.066088 | 0.078288 | 0.254236 | 0.218893 | 0.148299 | 0.178996 | -1.193033 | 9.097129 | 1.007723542 | -0.25268251 | 0.800513565 | NA | 3.307819 | 3.051647 | 3.185822 | 3.196921 | 3.183221 | 3.185256 | 10 | 12 | 7 | 8 |
| 8607 | NM_00131908 | RUVBL1 | RuvB like AAA ATPase 1 | protein_coding | NP_001306013.1;NP_001306015.1;NP_003698.1;XP_005247898.1;XP_01 | 10474 | 603449 | . | . | 26.460760 | 25.767383 | 22.159987 | 26.092129 | 61.575276 | 59.562461 | 49.726368 | 59.656552 | -1.097679 | 1486.337921 | 0.133792088 | -1.00496279 | 0.314914767 | 0.998865404 | 10.603362 | 10.468584 | 10.585262 | 10.558422 | 10.439432 | 10.556939 | 1822 | 1650 | 1137 | 1431 |
| 8609 | NM_00127094 | KLF7 | Kruppel like factor 7 | protein_coding | NP_001257871.1;NP_001257872.1;NP_001257873.1;NP_003700.1;XP_0 | 6350 | 604865 | . | . | 1.402457 | 1.741003 | 1.620106 | 1.615362 | 3.263571 | 4.024407 | 3.635473 | 3.693331 | 1.019227 | 381.630273 | 0.189734672 | 0.144813128 | 0.884858417 | 0.998865404 | 8.562490 | 8.589448 | 8.518990 | 8.617252 | 8.582131 | 8.578564 | 399 | 442 | 336 | 360 |
| 8611 | NM_003711,N | PLPP1 | phospholipid phosphatase 1 | protein_coding | NP_003702.2;NP_795714.1;XP_006714787.1 | 9228 | 607124 | . | . | 8.621151 | 6.378083 | 7.549389 | 5.894729 | 20.061773 | 14.743226 | 16.940609 | 13.477596 | -1.127561 | 350.489602 | 0.217743759 | -0.79545667 | 0.426347908 | 0.998865404 | 8.536441 | 8.364919 | 8.562452 | 8.395261 | 8.479686 | 8.344116 | 488 | 344 | 322 | 271 |
| 8612 | NM_003712,N | PLPP2 | phospholipid phosphatase 2 | protein_coding | NP_003703.1;NP_803545.1;NP_808211.1;XP_011526698.1 | 9230 | 607126 | . | . | 28.784609 | 25.682677 | 26.082663 | 29.494937 | 66.982970 | 59.366658 | 58.528740 | 67.436668 | 1.005768 | 1098.030231 | 0.142264989 | 0.058327625 | 0.953487663 | 0.998865404 | 10.096579 | 10.104813 | 10.141929 | 10.051737 | 10.060612 | 10.141929 | 1337 | 1110 | 905 | 1089 |
| 8613 | NM_003713 | PLPP3 | phospholipid phosphatase 3 | protein_coding | NP_003704.3 | 9229 | 607125 | . | . | 8.714650 | 8.508352 | 7.050481 | 7.584360 | 20.279350 | 19.667439 | 15.821074 | 17.340738 | -1.193241 | 812.858780 | 0.145412836 | -1.75283769 | 0.079629872 | 0.778116089 | 9.788760 | 9.533704 | 9.737837 | 9.714056 | 9.579287 | 9.622528 | 1035 | 939 | 624 | 717 |
| 8615 | NM_00129004 | USO1 | USO1 vesicle transport factor | protein_coding | NP_001276978.1;NP_003706.2;XP_006714459.1;XP_006714460.1 | 30904 | 603344 | . | . | 21.123128 | 19.540578 | 18.553994 | 17.546512 | 49.154392 | 45.168919 | 41.634628 | 40.118017 | -1.142359 | 2468.173399 | 0.119329918 | -1.60911925 | 0.107590268 | 0.835804188 | 11.361938 | 11.170149 | 11.355078 | 11.284026 | 11.238803 | 11.186353 | 3162 | 2717 | 2070 | 2089 |
| 8618 | NM_003716,N | CADPS | calcium dependent secretion activator | protein_coding | NP_003707.2;NP_899630.1;NP_899631.1;XP_006713441.2;XP_01153247 | 1426 | 604667 | . | . | 6.667863 | 7.018796 | 10.485735 | 8.821499 | 15.516389 | 16.224258 | 23.529684 | 20.169307 | 1.375035 | 1271.140640 | 0.141504809 | 3.24702926 | 0.001166164 | 0.075570712 | 10.063809 | 10.523537 | 10.153164 | 10.197699 | 10.485139 | 10.343998 | 1195 | 1190 | 1395 | 1256 |
| 8620 | NM_00132029 | NPFF | neuropeptide FF-amide peptide precursor | protein_coding | NP_001307225.1;NP_003708.1 | 7901 | 604643 | . | . | 0.447907 | 0.401787 | 0.767895 | 0.532639 | 1.042298 | 0.928750 | 1.723133 | 1.217816 | 1.476862 | 11.663322 | 0.895393777 | 0.628254502 | 0.529837232 | NA | 3.234218 | 3.798723 | 3.523706 | 3.522546 | 3.552216 | 3.533739 | 11 | 10 | 14 | 11 |
| 8621 | NM_003718,N | CDK13 | cyclin dependent kinase 13 | protein_coding | NP_003709.3;NP_112557.2;XP_011513899.1;XP_016868239.1;XP_01686 | 1733 | 603309 | . | . | 4.549882 | 5.082073 | 4.949328 | 4.593692 | 10.587763 | 11.747440 | 11.106146 | 10.502930 | 1.032727 | 1143.840874 | 0.133206373 | 0.348777175 | 0.727256601 | 0.998865404 | 10.136269 | 10.182699 | 10.125715 | 10.168320 | 10.183283 | 10.159161 | 1259 | 1249 | 1027 | 1072 |
| 8623 | NM_00117347 | ASMTL | acetylserotonin O-methyltransferase like | protein_coding | NP_001166944.1;NP_001166945.1;NP_004183.2;XP_005274491.1;XP_01 | 751 | 400011 | . | . | 1.253471 | 1.364045 | 0.667327 | 0.883538 | 2.916879 | 3.153051 | 1.497462 | 2.020107 | -1.609620 | 71.928549 | 0.379985537 | -1.80722661 | 0.070726954 | 0.751131579 | 6.472070 | 5.783530 | 6.222415 | 6.223046 | 6.085345 | 6.131956 | 102 | 96 | 42 | 58 |
| 8624 | NM_00126182 | PSMG1 | proteasome assembly chaperone 1 | protein_coding | NP_001248753.1;NP_001307724.1;NP_003711.1;NP_982257.1;XP_0243 | 3043 | 605296 | . | . | 21.472106 | 24.754244 | 25.213069 | 22.026572 | 49.966474 | 57.220546 | 56.577395 | 50.361138 | -1.047589 | 1122.414908 | 0.14281644 | -0.46964144 | 0.638611214 | 0.998865404 | 10.165285 | 10.098728 | 10.154574 | 10.142291 | 10.176614 | 10.047195 | 1333 | 1229 | 1037 | 938 |
| 8625 | NM_00127872 | RFXANK | regulatory factor X associated ankyrin containing protein | protein_coding | NP_001265656.1;NP_001265657.1;NP_001357162.1;NP_001357163.1;NI | 9987 | 603200 | . | . | 23.425326 | 25.877506 | 22.822066 | 27.376434 | 54.511697 | 59.817013 | 51.212056 | 62.592966 | 1.003342 | 893.747814 | 0.148044829 | 0.032517609 | 0.974059274 | 0.998865404 | 9.801608 | 9.805837 | 9.783709 | 9.819759 | 9.749033 | 9.856414 | 1002 | 987 | 720 | 902 |
| 8629 | NM_00107752 | JRK | Jrk helix-turn-helix protein | protein_coding | NP_001070995.2;NP_001266281.1;NP_003715.3;XP_006716740.1;XP_00 | 6199 | 603210 | . | . | 3.894985 | 3.726266 | 7.479848 | 4.318931 | 9.063789 | 8.613430 | 16.784562 | 9.874720 | 1.064850 | 1240.537032 | 0.138541585 | 0.654315268 | 0.512908711 | 0.998865404 | 10.230588 | 10.321479 | 10.225733 | 10.276347 | 10.352909 | 10.241930 | 1337 | 1340 | 1194 | 1112 |
| 8630 | NM_003725,XI | HSD17B6 | hydroxysteroid 17-beta dehydrogenase 6 | protein_coding | NP_003716.2;XP_005269264.1;XP_005269265.1;XP_005269266.1;XP_00 | 23316 | 606623 | . | . | 0.484744 | 1.089481 | 0.396700 | 0.620379 | 1.128019 | 2.518383 | 0.890183 | 1.418425 | -1.548994 | 36.528285 | 0.56286235 | -1.12164466 | 0.262013554 | 0.995760118 | 5.474063 | 4.838350 | 5.150318 | 5.273483 | 5.134505 | 5.169256 | 33 | 65 | 21 | 31 |
| 8633 | NM_003728,XI | UNC5C | unc-5 netrin receptor C | protein_coding | NP_003719.3;XP_005263378.1;XP_016864278.1;XP_016864279.1 | 12569 | 603610 | . | . | 0.020787 | 0.014656 | 0.015258 | 0.013181 | 0.048371 | 0.033877 | 0.034238 | 0.030138 | -1.154122 | 8.352514 | 1.047924673 | -0.1973383 | 0.84356281 | NA | 3.161269 | 2.955730 | 3.070438 | 3.064820 | 3.063569 | 3.060959 | 11 | 9 | 7 | 7 |

| | | | | | | | | | | | | | | | | | | | | | | | | | | | | | | | | | |
|---|---|---|---|---|---|---|---|---|---|---|---|---|---|---|---|---|---|---|---|---|---|---|---|---|---|---|---|---|---|---|---|---|---|
| 8634 | NM_00113084 | RTCA | RNA 3'-terminal phosphate cyclase | protein_coding | NP_001124313.1;NP_003720.1;XP_005271355.1;XP_005271356.1;XP_01 | 17981 | 611286 | . | . | 9.651593 | 9.140361 | 8.107639 | 7.455866 | 22.459651 | 21.128354 | 18.193307 | 17.046954 | -1.227744 | 681.551455 | 0.154970263 | -1.91011033 | 0.05611901 | 0.703410744 | 9.552864 | 9.257391 | 9.499331 | 9.456273 | 9.368473 | 9.307095 | 891 | 786 | 557 | 547 |
| 8635 | NM_003730,X | RNASET2 | ribonuclease T2 | protein_coding | NP_003721.2;XP_016866886.1;XP_016866887.1;XP_016866888.1;XP_02 | 21686 | 612944 | . | . | 3.590309 | 1.723693 | 2.269452 | 2.081564 | 8.354796 | 3.984393 | 5.092584 | 4.759248 | 1.099028 | 370.996802 | 0.187019459 | 0.728420385 | 0.466356289 | 0.998865404 | 8.465829 | 8.601508 | 8.491753 | 8.520405 | 8.536526 | 8.584466 | 397 | 391 | 324 | 379 |
| 8636 | NM_003731 | SSNA1 | SS nuclear autoantigen 1 | protein_coding | NP_003722.2 | 11321 | 610882 | . | . | 27.031227 | 29.682699 | 26.236305 | 31.498327 | 62.902779 | 68.612892 | 58.873508 | 72.017181 | 1.004427 | 769.636459 | 0.156870158 | 0.040621142 | 0.967597929 | 0.998865404 | 9.585223 | 9.590838 | 9.555232 | 9.614903 | 9.526825 | 9.646485 | 847 | 864 | 613 | 785 |
| 8637 | NM_003732 | EIF4EBP3 | eukaryotic translation initiation factor 4E binding protein 3 | protein_coding | NP_003723.1 | 3290 | 603483 | . | . | 1.658303 | 1.034942 | 1.737798 | 1.391969 | 3.858940 | 2.392314 | 3.899569 | 3.182572 | 1.134315 | 31.864754 | 0.56394232 | 0.32241023 | 0.747141935 | 0.998865404 | 4.898285 | 5.083550 | 5.008024 | 4.952980 | 5.012421 | 4.981183 | 42 | 25 | 33 | 28 |
| 8639 | NM_00127773 | AOC3 | amine oxidase copper containing 3 | protein_coding | NP_001264660.1;NP_001264661.1;NP_003725.1;XP_011523721.1;XP_01 | 550 | 603735 | . | . | 0.220601 | 0.223455 | 0.175378 | 0.176866 | 0.513347 | 0.516527 | 0.393542 | 0.404384 | -1.228310 | 26.490964 | 0.600453772 | -0.49408447 | 0.621246525 | NA | 4.868663 | 4.570852 | 4.731373 | 4.750291 | 4.715633 | 4.721618 | 31 | 34 | 20 | 23 |
| 8641 | NM_003736,N | PCDHGB4 | protocadherin gamma subfamily B, 4 | protein_coding | NP_003727.1;NP_115269.1 | 8711 | 603058 | . | . | 0.097156 | 0.072699 | 0.010176 | 0.041345 | 0.226087 | 0.168048 | 0.022834 | 0.094532 | -2.446415 | 6.598823 | 1.228381452 | -1.05070725 | 0.293393063 | NA | 3.229742 | 1.932304 | 2.754096 | 2.744424 | 2.717728 | 2.730880 | 12 | 9 | 2 | 5 |
| 8643 | NM_00116629 | PTCH2 | patched 2 | protein_coding | NP_001159764.1;NP_003729.3 | 9586 | 603673 | . | . | 0.086397 | 0.071940 | 0.090093 | 0.090960 | 0.201050 | 0.166294 | 0.202166 | 0.207969 | 1.137921 | 11.950526 | 0.879521041 | 0.211933624 | 0.832158818 | NA | 3.482357 | 3.669578 | 3.577730 | 3.568838 | 3.579745 | 3.580460 | 14 | 11 | 11 | 12 |
| 8644 | NM_00125390 | AKR1C3 | aldo-keto reductase family 1 member C3 | protein_coding | NP_001240837.1;NP_001240838.1;NP_003730.4 | 386 | 603966 | . | . | 3.302677 | 4.292591 | 6.018687 | 4.609648 | 7.685466 | 9.922516 | 13.505758 | 10.539412 | 1.383878 | 169.568953 | 0.273607653 | 1.713098003 | 0.086694511 | 0.795065303 | 7.152356 | 7.621179 | 7.273346 | 7.370332 | 7.514275 | 7.396088 | 143 | 173 | 194 | 160 |
| 8645 | NM_003740,X | KCNK5 | potassium two pore domain channel subfamily K member 5 | protein_coding | NP_003731.1;XP_005249513.1;XP_006715298.1 | 6280 | 603493 | . | . | 2.157754 | 2.157166 | 2.733371 | 2.691945 | 5.021183 | 4.986386 | 6.133605 | 6.154812 | 1.230460 | 285.011632 | 0.204415014 | 1.463680229 | 0.14328134 | 0.905131505 | 7.997472 | 8.296786 | 8.093984 | 8.095256 | 8.210438 | 8.196771 | 294 | 276 | 277 | 291 |
| 8646 | NM_00130447 | CHRD | chordin | protein_coding | NP_001291401.1;NP_001291402.1;NP_001291403.1;NP_003732.2;XP_01 | 1949 | 603475 | . | . | 0.280448 | 0.145445 | 0.218004 | 0.102345 | 0.652613 | 0.336201 | 0.489195 | 0.234000 | -1.262796 | 24.693865 | 0.677470302 | -0.49688024 | 0.619273507 | NA | 4.781608 | 4.451739 | 4.669470 | 4.594119 | 4.640978 | 4.575323 | 42 | 20 | 26 | 13 |
| 8648 | NM_00136295 | NCOA1 | nuclear receptor coactivator 1 | protein_coding | NP_001349879.1;NP_001349881.1;NP_001349883.1;NP_001349884.1;N | 7668 | 602691 | . | . | 3.126052 | 3.390208 | 3.326500 | 3.473907 | 7.274449 | 7.836619 | 7.464567 | 7.942674 | 1.040116 | 762.610138 | 0.147975086 | 0.383474059 | 0.701368301 | 0.998865404 | 9.546307 | 9.602745 | 9.537619 | 9.583028 | 9.575326 | 9.600461 | 833 | 833 | 664 | 742 |
| 8649 | NM_00124373 | LAMTOR3 | late endosomal/lysosomal adaptor, MAPK and MTOR activator 3 | protein_coding | NP_001230665.1;NP_068805.1 | 15606 | 603296 | . | . | 3.845790 | 3.465773 | 4.222311 | 3.348124 | 8.949311 | 8.011290 | 9.474743 | 7.655089 | 1.024412 | 485.326925 | 0.185012122 | 0.188072146 | 0.850820095 | 0.998865404 | 8.904715 | 8.940687 | 8.947572 | 8.876385 | 8.999835 | 8.850687 | 583 | 488 | 478 | 406 |
| 8650 | NM_00100574 | NUMB | NUMB endocytic adaptor protein | protein_coding | NP_001005743.1;NP_001005744.1;NP_001005745.1;NP_001307043.1;N | 8060 | 603728 | . | . | 11.402631 | 11.347321 | 12.707787 | 14.267789 | 26.534390 | 26.229842 | 28.515903 | 32.621605 | 1.170746 | 1362.994509 | 0.131199381 | 1.733452792 | 0.083015221 | 0.78429781 | 10.294501 | 10.521692 | 10.353868 | 10.342709 | 10.425597 | 10.510148 | 1457 | 1345 | 1209 | 1453 |
| 8651 | NM_003745 | SOCS1 | suppressor of cytokine signaling 1 | protein_coding | NP_003736.1 | 19383 | 603597 | . | . | 0.313107 | 0.416961 | 0.926002 | 0.766709 | 0.728614 | 0.963823 | 2.077921 | 1.752989 | 2.236386 | 23.322203 | 0.65252433 | 1.779503459 | 0.075157258 | NA | 3.849979 | 5.010091 | 4.468204 | 4.489186 | 4.567144 | 4.540638 | 14 | 18 | 31 | 27 |
| 8653 | NM_00112266 | DDX3Y | DEAD-box helicase 3 Y-linked | protein_coding | NP_001116137.1;NP_001289481.1;NP_001311124.1;NP_004651.2 | 2699 | 400010 | . | . | 0.239623 | 0.323694 | 0.551940 | 0.308943 | 0.557613 | 0.748234 | 1.238539 | 0.706361 | 1.514473 | 53.830885 | 0.46135164 | 1.297960742 | 0.194300825 | 0.96621494 | 5.419405 | 6.019385 | 5.662487 | 5.712609 | 5.828526 | 5.704965 | 42 | 53 | 72 | 44 |
| 8654 | NM_001083,N | PDE5A | phosphodiesterase 5A | protein_coding | NP_001074.2;NP_236914.2;NP_246273.2;XP_016864280.1 | 8784 | 603310 | . | . | 0.508542 | 0.409416 | 0.369436 | 0.416284 | 1.183398 | 0.946384 | 0.829005 | 0.951785 | -1.163487 | 95.594601 | 0.331284777 | -0.65941615 | 0.509628575 | 0.998865404 | 6.683909 | 6.465551 | 6.629843 | 6.572774 | 6.543320 | 6.570463 | 129 | 101 | 73 | 87 |
| 8655 | NM_00103749 | DYNLL1 | dynein light chain LC8-type 1 | protein_coding | NP_001032583.1;NP_001032584.1;NP_003737.1 | 15476 | 601562 | . | . | 257.590646 | 255.092675 | 229.403172 | 252.888527 | 599.423989 | 589.658133 | 514.774067 | 578.199603 | -1.077638 | 5154.617421 | 0.108902246 | -0.99054978 | 0.321905469 | 0.998865404 | 12.384616 | 12.276664 | 12.369613 | 12.354352 | 12.262901 | 12.334482 | 6213 | 5718 | 4129 | 4852 |
| 8658 | NM_003747,X | TNKS | tankyrase | protein_coding | NP_003738.2;XP_006716326.1;XP_011542147.1;XP_011542148.1 | 11941 | 603303 | . | . | 3.844734 | 3.523032 | 4.043733 | 3.610543 | 8.946855 | 8.143644 | 9.074019 | 8.255077 | 1.022579 | 1125.483845 | 0.141450301 | 0.227726601 | 0.819858791 | 0.998865404 | 10.119857 | 10.152617 | 10.163940 | 10.088522 | 10.190190 | 10.095057 | 1346 | 1140 | 1053 | 999 |
| 8659 | NM_00116150 | ALDH4A1 | aldehyde dehydrogenase 4 family member A1 | protein_coding | NP_001154976.1;NP_001306147.1;NP_003739.2;NP_733844.1 | 406 | 606811 | . | . | 5.588068 | 6.299894 | 6.064120 | 7.315322 | 13.003664 | 14.562485 | 13.607709 | 16.725616 | 1.109551 | 614.584528 | 0.168021406 | 0.89259662 | 0.37207323 | 0.998865404 | 9.187003 | 9.336083 | 9.192584 | 9.257975 | 9.230446 | 9.355785 | 638 | 660 | 510 | 662 |
| 8660 | NM_003749 | IRS2 | insulin receptor substrate 2 | protein_coding | NP_003740.2 | 6126 | 600797 | . | . | 7.329010 | 7.074543 | 7.389873 | 7.053100 | 17.054909 | 16.353125 | 16.582661 | 16.126076 | -1.011401 | 1832.823712 | 0.121670579 | -0.13442681 | 0.893065084 | 0.998865404 | 10.847907 | 10.831753 | 10.861398 | 10.826541 | 10.857382 | 10.812637 | 2169 | 1945 | 1630 | 1660 |
| 8661 | NM_003750 | EIF3A | eukaryotic translation initiation factor 3 subunit A | protein_coding | NP_003741.1 | 3271 | 602039 | . | . | 40.967003 | 40.227180 | 35.881023 | 41.101269 | 95.331902 | 92.986938 | 80.515976 | 93.973160 | -1.069835 | 8204.267552 | 0.110296914 | -0.88296968 | 0.377252654 | 0.998865404 | 13.050065 | 12.952607 | 13.041100 | 13.018175 | 12.918046 | 13.023942 | 9897 | 9029 | 6461 | 7898 |
| 8662 | NM_00103728 | EIF3B | eukaryotic translation initiation factor 3 subunit B | protein_coding | NP_001032360.1;NP_001349720.1;NP_001349721.1;NP_001349722.1;N | 3280 | 603917 | . | . | 90.633091 | 97.698197 | 97.316744 | 106.647863 | 210.906919 | 225.833769 | 218.375960 | 243.837658 | 1.066233 | 9376.262294 | 0.106807743 | 0.866259728 | 0.386347746 | 0.998865404 | 13.147832 | 13.240282 | 13.138904 | 13.193500 | 13.185884 | 13.255154 | 10114 | 10109 | 8074 | 9439 |
| 8663 | NM_00103780 | EIF3C | eukaryotic translation initiation factor 3 subunit C | protein_coding | NP_001032897.1;NP_001186071.1;NP_001254503.1;NP_001273407.1;N | 3279 | 603916 | . | . | 65.376062 | 69.063608 | 57.479010 | 70.185108 | 152.132781 | 159.643623 | 128.981231 | 160.469917 | -1.077649 | 6110.972215 | 0.119487293 | -0.90292439 | 0.366566026 | 0.998865404 | 12.630194 | 12.522157 | 12.594186 | 12.619547 | 12.465044 | 12.617065 | 7191 | 6943 | 4651 | 6015 |
| 8664 | NM_003753,N | EIF3D | eukaryotic translation initiation factor 3 subunit D | protein_coding | NP_003744.1 | 3278 | 603915 | . | . | 120.102119 | 122.842034 | 114.484431 | 134.985749 | 279.482562 | 283.954866 | 256.899739 | 308.628875 | 1.011692 | 7211.492118 | 0.114423332 | 0.146560601 | 0.883478854 | 0.998865404 | 12.807727 | 12.824389 | 12.804580 | 12.816263 | 12.752835 | 12.883482 | 8199 | 7790 | 5821 | 7324 |
| 8665 | NM_003754 | EIF3F | eukaryotic translation initiation factor 3 subunit F | protein_coding | NP_003745.1 | 3275 | 603914 | . | . | 12.134439 | 12.687084 | 11.619882 | 13.299274 | 28.237341 | 29.326763 | 26.074677 | 30.407207 | -1.010150 | 2680.892349 | 0.120104296 | -0.12130456 | 0.903449805 | 0.998865404 | 11.395884 | 11.381073 | 11.377387 | 11.406710 | 11.332235 | 11.433027 | 3047 | 2960 | 2175 | 2656 |
| 8666 | NM_003755 | EIF3G | eukaryotic translation initiation factor 3 subunit G | protein_coding | NP_003746.2 | 3274 | 603913 | . | . | 99.704773 | 100.779106 | 98.669281 | 110.145050 | 232.017090 | 232.955414 | 221.410995 | 251.833557 | 1.026982 | 3515.426420 | 0.113661521 | 0.337942116 | 0.735406809 | 0.998865404 | 11.760220 | 11.798494 | 11.767226 | 11.768999 | 11.747978 | 11.830391 | 3990 | 3747 | 2943 | 3506 |
| 8667 | NM_003756 | EIF3H | eukaryotic translation initiation factor 3 subunit H | protein_coding | NP_003747.1 | 3273 | 603912 | . | . | 34.802406 | 34.231480 | 32.787907 | 35.584805 | 80.986633 | 79.127594 | 73.575111 | 81.360435 | -1.024112 | 4239.018467 | 0.109624168 | -0.31355972 | 0.75385546 | 0.998865404 | 12.066632 | 12.032191 | 12.069570 | 12.048757 | 12.009320 | 12.068574 | 5001 | 4571 | 3512 | 4067 |
| 8668 | NM_003757,X | EIF3I | eukaryotic translation initiation factor 3 subunit I | protein_coding | NP_003748.1;XP_024306286.1 | 3272 | 603911 | . | . | 105.908917 | 106.857246 | 94.571149 | 109.687348 | 246.454394 | 247.005308 | 212.214915 | 250.787081 | -1.054773 | 6214.047521 | 0.113116086 | -0.68012306 | 0.496426547 | 0.998865404 | 12.639325 | 12.562281 | 12.623517 | 12.622216 | 12.518083 | 12.634356 | 7352 | 6879 | 4895 | 6063 |
| 8669 | NM_00128433 | EIF3J | eukaryotic translation initiation factor 3 subunit J | protein_coding | NP_001271264.1;NP_001271265.1;NP_003749.2 | 3270 | 603910 | . | . | 20.055973 | 17.636430 | 18.670583 | 14.088584 | 46.671078 | 40.767401 | 41.896242 | 32.211871 | -1.166420 | 1358.723855 | 0.158542266 | -1.40081046 | 0.16127076 | 0.933525128 | 10.514473 | 10.293118 | 10.513145 | 10.405558 | 10.446101 | 10.228899 | 1800 | 1470 | 1249 | 1006 |
| 8671 | NM_00109848 | SLC4A4 | solute carrier family 4 member 4 | protein_coding | NP_001091954.1;NP_001128214.1;NP_003750.1;XP_011530692.1;XP_01 | 11030 | 603345 | . | . | 0.980544 | 0.459256 | 0.588932 | 0.771227 | 2.281766 | 1.061590 | 1.321547 | 1.763320 | -1.147363 | 175.554139 | 0.300581755 | -0.65979345 | 0.509386387 | 0.998865404 | 7.551399 | 7.353354 | 7.587816 | 7.350586 | 7.362366 | 7.470361 | 273 | 149 | 123 | 174 |
| 8672 | NM_00119880 | EIF4G3 | eukaryotic translation initiation factor 4 gamma 3 | protein_coding | NP_001185730.1;NP_001185731.1;NP_001185732.1;NP_003751.2;XP_01 | 3298 | 603929 | . | . | 20.659703 | 20.770639 | 22.218828 | 23.343106 | 48.075980 | 48.012259 | 49.858412 | 53.371239 | 1.093080 | 4322.040821 | 0.107374264 | 1.195810309 | 0.231770592 | 0.986726781 | 12.011896 | 12.140245 | 12.039435 | 12.038954 | 12.095249 | 12.130917 | 4757 | 4455 | 3842 | 4322 |
| 8673 | NM_003761,X | VAMP8 | vesicle associated membrane protein 8 | protein_coding | NP_003752.2;XP_016860659.1 | 12647 | 603177 | . | . | 7.732338 | 9.509278 | 8.816698 | 11.567920 | 17.993467 | 21.981125 | 19.784415 | 26.448673 | 1.155914 | 202.491129 | 0.250916031 | 0.83308433 | 0.40479717 | 0.998865404 | 7.554405 | 7.761593 | 7.580357 | 7.664326 | 7.628453 | 7.746014 | 196 | 222 | 165 | 229 |
| 8674 | NM_00118512 | VAMP4 | vesicle associated membrane protein 4 | protein_coding | NP_001172056.1;NP_003753.2;XP_016858202.1 | 12645 | 606909 | . | . | 2.008512 | 1.144105 | 2.056691 | 0.990964 | 4.673890 | 2.644651 | 4.615154 | 2.265721 | -1.124526 | 187.304952 | 0.27114155 | -0.6244622 | 0.532324088 | 0.998865404 | 7.630295 | 7.463370 | 7.633792 | 7.501477 | 7.583255 | 7.449427 | 261 | 183 | 178 | 139 |
| 8675 | NM_00100143 | STX16 | syntaxin 16 | protein_coding | NP_001001433.1;NP_001128244.1;NP_001128245.1;NP_001191797.1;N | 11431 | 603666 | . | . | 28.741442 | 28.912127 | 34.007553 | 27.146978 | 66.882516 | 66.831678 | 76.311961 | 62.068339 | 1.049289 | 4492.125942 | 0.129186887 | 0.53729701 | 0.59106247 | 0.998865404 | 12.097961 | 12.167565 | 12.113131 | 12.108818 | 12.244469 | 12.048626 | 5061 | 4718 | 4467 | 3810 |
| 8676 | NM_003764,X | STX11 | syntaxin 11 | protein_coding | NP_003755.2;XP_011534515.1;XP_011534516.1;XP_011534519.1;XP_01 | 11429 | 605014 | . | . | 0.188892 | 0.158512 | 0.115046 | 0.173816 | 0.439557 | 0.366407 | 0.258161 | 0.397409 | -1.259826 | 25.997306 | 0.630059796 | -0.52887766 | 0.596890318 | NA | 4.858551 | 4.522505 | 4.731653 | 4.698325 | 4.662414 | 4.717941 | 38 | 27 | 14 | 28 |
| 8677 | NM_00127160 | STX10 | syntaxin 10 | protein_coding | NP_001258538.1;NP_001258539.1;NP_001258540.1;NP_003756.1;XP_01 | 11428 | 603765 | . | . | 22.852544 | 20.237438 | 23.809260 | 20.386840 | 53.178814 | 46.779749 | 53.427288 | 46.612087 | -1.000028 | 893.719908 | 0.154812235 | -0.0002585 | 0.99979375 | 0.999938637 | 9.803311 | 9.804047 | 9.846121 | 9.755940 | 9.864380 | 9.736008 | 1093 | 904 | 847 | 763 |
| 8678 | NM_00131399 | BECN1 | beclin 1 | protein_coding | NP_001300927.1;NP_001300928.1;NP_001300929.1;NP_003757.1;XP_0 | 1034 | 604378 | . | . | 45.246590 | 43.099615 | 45.761798 | 48.620610 | 105.290682 | 99.626687 | 102.688149 | 111.165246 | 1.057692 | 2924.632893 | 0.114401809 | 0.707323066 | 0.479365734 | 0.998865404 | 11.472999 | 11.553946 | 11.519516 | 11.460531 | 11.521211 | 11.551053 | 3392 | 2956 | 2567 | 2870 |
| 8682 | NM_00129757 | PEA15 | proliferation and apoptosis adaptor protein 15 | protein_coding | NP_001284505.1;NP_001284506.1;NP_001284507.1;NP_003759.1;XP_0 | 8822 | 603434 | . | . | 84.918889 | 91.461525 | 72.513277 | 85.514858 | 197.609734 | 211.417418 | 162.717695 | 195.519554 | -1.130787 | 6338.074302 | 0.117528462 | -1.50879703 | 0.131350659 | 0.884838043 | 12.715840 | 12.538362 | 12.650695 | 12.705907 | 12.506342 | 12.639209 | 7496 | 7494 | 4761 | 6021 |
| 8683 | NM_003769 | SRSF9 | serine and arginine rich splicing factor 9 | protein_coding | NP_003760.1 | 10791 | 601943 | . | . | 77.440331 | 77.048866 | 73.943741 | 82.392372 | 180.206833 | 178.101898 | 165.927612 | 188.380356 | -1.002201 | 2788.777314 | 0.116465664 | -0.02723945 | 0.978268749 | 0.998865404 | 11.447075 | 11.443757 | 11.451655 | 11.440455 | 11.403771 | 11.483147 | 3237 | 2992 | 2304 | 2739 |
| 8690 | NM_00126183 | JRKL | JRK like | protein_coding | NP_001248762.1;NP_003763.2 | 6200 | 603211 | . | . | 3.841054 | 3.933933 | 4.338202 | 3.074015 | 8.938290 | 9.093464 | 9.734800 | 7.028371 | -1.048694 | 406.113770 | 0.205724667 | -0.33342268 | 0.738815242 | 0.998865404 | 8.698992 | 8.631704 | 8.676397 | 8.675139 | 8.746485 | 8.537924 | 479 | 448 | 406 | 306 |
| 8692 | NM_003773,N | HYAL2 | hyaluronidase 2 | protein_coding | NP_003764.3;NP_149348.2;XP_005265581.1;XP_005265582.1 | 5321 | 603551 | . | . | 14.017191 | 14.419607 | 12.008729 | 12.469121 | 32.618580 | 33.331566 | 26.947242 | 28.509160 | -1.161289 | 962.262252 | 0.139108258 | -1.55078967 | 0.120952097 | 0.862345966 | 10.014254 | 9.798241 | 9.954885 | 9.969589 | 9.829175 | 9.876983 | 1179 | 1127 | 748 | 863 |
| 8693 | NM_003774 | GALNT4 | polypeptide N-acetylgalactosaminyltransferase 4 | protein_coding | NP_003765.2 | 4126 | 603565 | . | . | 0.817931 | 0.780416 | 0.470275 | 0.579157 | 1.903359 | 1.803966 | 1.055283 | 1.324173 | -1.541720 | 111.474187 | 0.31051238 | -2.01132284 | 0.044291368 | 0.645513836 | 7.079743 | 6.454079 | 6.875655 | 6.854492 | 6.699989 | 6.750399 | 160 | 142 | 69 | 90 |
| 8694 | NM_012079,X | DGAT1 | diacylglycerol O-acyltransferase 1 | protein_coding | NP_036211.2;XP_011515658.1 | 2843 | 604900 | . | . | 11.600223 | 11.983162 | 12.190032 | 13.000543 | 26.994197 | 27.699613 | 27.354080 | 29.724197 | 1.037841 | 1367.159211 | 0.126618618 | 0.423201087 | 0.672148554 | 0.998865404 | 10.389963 | 10.443472 | 10.394991 | 10.409921 | 10.425830 | 10.435865 | 1530 | 1462 | 1203 | 1314 |
| 8697 | NM_004661 | CDC23 | cell division cycle 23 | protein_coding | NP_004652.2 | 1724 | 603462 | . | . | 14.170791 | 13.026859 | 13.837467 | 13.585734 | 32.976013 | 30.112169 | 31.050879 | 31.062166 | -1.005595 | 1333.528661 | 0.130864209 | -0.0615108 | 0.950952412 | 0.998865404 | 10.384924 | 10.377131 | 10.417700 | 10.346761 | 10.389465 | 10.367773 | 1611 | 1376 | 1173 | 1229 |
| 8702 | NM_003778,N | B4GALT4 | beta-1,4-galactosyltransferase 4 | protein_coding | NP_003769.1;NP_997708.1;XP_005247912.1;XP_006713861.1;XP_00671 | 927 | 604015 | . | . | 20.309149 | 19.951578 | 16.730819 | 16.381881 | 47.260224 | 46.118970 | 37.543477 | 37.455229 | -1.218921 | 1313.884253 | 0.127876661 | -2.23343749 | 0.025520108 | 0.524911334 | 10.495345 | 10.209793 | 10.439374 | 10.420621 | 10.280726 | 10.278830 | 1682 | 1539 | 1031 | 1109 |
| 8703 | NM_00119987 | B4GALT3 | beta-1,4-galactosyltransferase 3 | protein_coding | NP_001186802.1;NP_001186803.1;NP_003770.1;XP_005245623.1;XP_01 | 926 | 604014 | . | . | 18.034624 | 17.642501 | 18.715441 | 21.239440 | 41.967311 | 40.781432 | 41.996906 | 48.561459 | 1.110645 | 1231.697891 | 0.135786275 | 1.114974021 | 0.264861555 | 0.996598404 | 10.188919 | 10.339993 | 10.227885 | 10.221402 | 10.252324 | 10.352781 | 1350 | 1254 | 1053 | 1295 |
| 8704 | NM_00100541 | B4GALT2 | beta-1,4-galactosyltransferase 2 | protein_coding | NP_001005417.1;NP_003771.1;NP_085076.2;XP_016858205.1;XP_01685 | 925 | 604013 | . | . | 12.935803 | 12.662991 | 11.960694 | 13.982115 | 30.102148 | 29.271068 | 26.839450 | 31.968446 | -1.018307 | 869.245117 | 0.149650544 | -0.17489047 | 0.861165697 | 0.998865404 | 9.776856 | 9.750260 | 9.783581 | 9.755199 | 9.697929 | 9.811681 | 1030 | 928 | 689 | 872 |
| 8706 | NM_00103862 | B3GALNT1 | beta-1,3-N-acetylgalactosaminyltransferase 1 (globoside blood gr | protein_coding | NP_001033717.1;NP_001336059.1;NP_001336060.1;NP_001336061.1;N | 918 | 603094 | . | . | 4.736719 | 3.553469 | 3.729502 | 3.205429 | 11.022539 | 8.214008 | 8.368890 | 7.328829 | -1.172214 | 421.250419 | 0.199316005 | -1.15011397 | 0.250096933 | 0.993339639 | 8.827995 | 8.600082 | 8.840931 | 8.675503 | 8.713873 | 8.615526 | 592 | 426 | 366 | 333 |
| 8707 | NM_003783 | B3GALT2 | beta-1,3-galactosyltransferase 2 | protein_coding | NP_003774.1 | 917 | 603018 | . | . | 0.042540 | 0.077579 | 0.159768 | 0.026113 | 0.098991 | 0.179327 | 0.358514 | 0.059704 | 1.487706 | 9.019422 | 1.11379667 | 0.514536621 | 0.606876864 | NA | 2.855800 | 3.432932 | 3.128848 | 3.147951 | 3.186724 | 3.120631 | 6 | 10 | 16 | 3 |
| 8708 | NM_020981,X | B3GALT1 | beta-1,3-galactosyltransferase 1 | protein_coding | NP_066191.1;XP_005246988.1;XP_006712882.1;XP_011510387.1 | 916 | 603093 | . | . | 0.680237 | 0.686444 | 0.946659 | 1.006571 | 1.582939 | 1.586748 | 2.124274 | 2.301407 | 1.394921 | 53.143246 | 0.42865562 | 1.120208763 | 0.26262481 | 0.995798899 | 5.472070 | 5.951852 | 5.690157 | 5.691323 | 5.747295 | 5.762401 | 51 | 48 | 52 | 60 |
| 8712 | NM_003785,X | PAGE1 | PAGE family member 1 | protein_coding | NP_003776.2;XP_011542300.1 | 4107 | 300288 | . | . | 100.203321 | 101.551221 | 98.147550 | 115.143250 | 233.177254 | 234.740198 | 220.240253 | 263.261374 | 1.040267 | 2142.611406 | 0.125734801 | 0.452965925 | 0.650573278 | 0.998865404 | 11.036536 | 11.093216 | 11.046923 | 11.050063 | 11.018354 | 11.138077 | 2414 | 2271 | 1758 | 2201 |
| 8717 | NM_00132355 | TRADD | TNFRSF1A associated via death domain | protein_coding | NP_001310481.1;NP_003780.1;XP_005256270.1;XP_016879304.1 | 12030 | 603500 | . | . | 4.261472 | 4.655805 | 4.505968 | 4.813150 | 9.916621 | 10.762100 | 10.111260 | 11.004696 | 1.092486 | 247.684065 | 0.231263354 | 0.551814611 | 0.581075376 | 0.998865404 | 7.887855 | 8.014099 | 7.857655 | 7.990790 | 7.945747 | 7.994275 | 236 | 290 | 214 | 254 |
| 8718 | NM_00103966 | TNFRSF25 | TNF receptor superfamily member 25 | protein_coding | NP_001034753.1;NP_003781.1;NP_683866.1;NP_683867.1;NP_683868.1 | 11910 | 603366 | . | . | 1.146446 | 1.177802 | 1.256969 | 1.233678 | 2.667828 | 2.722543 | 2.820600 | 2.820657 | 1.158568 | 62.285626 | 0.399167028 | 0.531964767 | 0.594750398 | 0.998865404 | 5.850632 | 6.063200 | 5.955969 | 5.927281 | 5.965394 | 5.982380 | 71 | 58 | 56 | 65 |
| 8720 | NM_003791,X | MBTPS1 | membrane bound transcription factor peptidase, site 1 | protein_coding | NP_003782.1;XP_016879305.1;XP_016879306.1;XP_024306245.1 | 15456 | 603355 | . | . | 24.311268 | 23.569126 | 22.795931 | 24.552348 | 56.573320 | 54.481092 | 51.153413 | 56.136025 | -1.017898 | 3255.387400 | 0.113665605 | -0.2251776 | 0.821841116 | 0.998865404 | 11.681389 | 11.655724 | 11.693411 | 11.657609 | 11.628573 | 11.692570 | 3864 | 3467 | 2696 | 3143 |
| 8721 | NM_00128129 | EDF1 | endothelial differentiation related factor 1 | protein_coding | NP_001268226.1;NP_001268227.1;NP_001268228.1;NP_003783.1;NP_6 | 3164 | 605107 | . | . | 117.314799 | 113.148767 | 111.822794 | 133.720088 | 272.996347 | 261.548452 | 250.927113 | 305.735081 | 1.050313 | 2542.734922 | 0.12597418 | 0.562172363 | 0.573998589 | 0.998865404 | 11.276423 | 11.347044 | 11.309448 | 11.272333 | 11.261293 | 11.395778 | 2922 | 2615 | 2078 | 2644 |
| 8722 | NM_003793,X | CTSF | cathepsin F | protein_coding | NP_003784.2;XP_011543630.1 | 2531 | 603539 | . | . | 9.722362 | 8.701498 | 9.323822 | 10.362350 | 22.624334 | 20.113903 | 20.922386 | 23.692282 | 1.054148 | 607.095146 | 0.163110218 | 0.46641576 | 0.640917933 | 0.998865404 | 9.207251 | 9.283305 | 9.265683 | 9.187013 | 9.229824 | 9.293534 | 721 | 600 | 516 | 612 |
| 8723 | NM_003794,N | SNX4 | sorting nexin 4 | protein_coding | NP_003785.1;XP_016862903.1 | 11175 | 605931 | . | . | 12.599513 | 11.087013 | 11.377404 | 11.247951 | 29.319585 | 25.628125 | 25.530565 | 25.717103 | -1.056043 | 915.466630 | 0.145511536 | -0.5406342 | 0.588759739 | 0.998865404 | 9.876992 | 9.798672 | 9.905623 | 9.806525 | 9.823095 | 9.812844 | 1156 | 946 | 780 | 829 |
| 8724 | NM_00130092 | SNX3 | sorting nexin 3 | protein_coding | NP_001287857.1;NP_001287858.1;NP_003786.1;NP_690040.1 | 11174 | 605930 | . | . | 54.190193 | 49.439205 | 48.419926 | 38.256629 | 126.102809 | 114.280943 | 108.652911 | 87.469240 | -1.213322 | 2272.250231 | 0.140871681 | -1.98025909 | 0.047674424 | 0.659753773 | 11.282447 | 11.003945 | 11.260488 | 11.178593 | 11.156893 | 10.964603 | 3011 | 2554 | 2004 | 1690 |
| 8725 | NM_00125264 | URI1 | URI1 prefoldin like chaperone | protein_coding | NP_001239570.1;NP_003787.2;XP_005259419.1;XP_005259420.1;XP_01 | 13236 | 603494 | . | . | 15.970457 | 16.457335 | 14.412355 | 16.140994 | 37.163912 | 38.041865 | 32.340907 | 36.904467 | -1.055488 | 1600.391791 | 0.124307317 | -0.62675797 | 0.530817897 | 0.998865404 | 10.682687 | 10.604677 | 10.676568 | 10.653000 | 10.599780 | 10.645322 | 1921 | 1747 | 1314 | 1503 |
| 8726 | NM_00130800 | EED | embryonic ectoderm development | protein_coding | NP_001294936.1;NP_001317263.1;NP_003788.2;XP_011543632.1;XP_01 | 3188 | 605984 | . | . | 8.856725 | 9.582602 | 9.748139 | 6.824422 | 20.609966 | 22.150613 | 21.874537 | 15.603232 | -1.061504 | 703.786384 | 0.154902737 | -0.55589426 | 0.578283145 | 0.998865404 | 9.501185 | 9.415531 | 9.473244 | 9.483973 | 9.481434 | 9.392022 | 828 | 788 | 633 | 598 |
| 8727 | NM_00128697 | CTNNAL1 | catenin alpha like 1 | protein_coding | NP_001273903.1;NP_003789.1;XP_005252348.1;XP_016870739.1;XP_01 | 2512 | 604785 | . | . | 31.700994 | 30.950233 | 26.267295 | 25.903179 | 73.769518 | 71.542849 | 58.943047 | 59.224236 | -1.217078 | 2203.836911 | 0.116759587 | -2.42739024 | 0.015207888 | 0.414850745 | 11.240516 | 10.957201 | 11.193889 | 11.164425 | 11.032084 | 11.011929 | 2836 | 2564 | 1751 | 1840 |
| 8728 | NM_033274,X | ADAM19 | ADAM metallopeptidase domain 19 | protein_coding | NP_150377.1;XP_016865498.1 | 197 | 603640 | . | . | 18.241393 | 18.551576 | 16.925561 | 19.050163 | 42.448473 | 42.882797 | 37.980469 | 43.555933 | -1.038292 | 3669.096556 | 0.114311881 | -0.47424646 | 0.635324163 | 0.998865404 | 11.868136 | 11.813770 | 11.852477 | 11.859998 | 11.778524 | 11.869704 | 4285 | 4052 | 2957 | 3562 |
| 8729 | NM_00119937 | GBF1 | golgi brefeldin A resistant guanine nucleotide exchange factor 1 | protein_coding | NP_001186307.1;NP_001186308.1;NP_001364066.1;NP_001364067.1;N | 4181 | 603698 | . | . | 12.198060 | 13.506990 | 12.957810 | 13.804411 | 28.385383 | 31.222018 | 29.076953 | 31.562144 | 1.029404 | 2599.726010 | 0.117676017 | 0.355296726 | 0.722367308 | 0.998865404 | 11.323184 | 11.364803 | 11.295844 | 11.366871 | 11.335619 | 11.374763 | 2821 | 2886 | 2238 | 2532 |
| 8731 | NM_00130826 | RNMT | RNA guanine-7 methyltransferase | protein_coding | NP_001295192.1;NP_001365061.1;NP_001365063.1;NP_001365064.1;N | 10075 | 603514 | . | . | 10.156494 | 11.015598 | 12.440494 | 10.020986 | 23.634579 | 25.463044 | 27.916108 | 22.911795 | 1.075247 | 1847.611076 | 0.137472567 | 0.761373318 | 0.446434119 | 0.998865404 | 10.797969 | 10.903011 | 10.819068 | 10.821469 | 10.961401 | 10.786542 | 2047 | 1924 | 1848 | 1598 |
| 8732 | NM_00128642 | RNGTT | RNA guanylyltransferase and 5'-phosphatase | protein_coding | NP_001273355.1;NP_001273357.1;NP_003791.3;XP_016866890.1;XP_01 | 10073 | 603512 | . | . | 3.972996 | 3.329500 | 2.965491 | 2.914318 | 9.245325 | 7.696289 | 6.654476 | 6.663243 | -1.252500 | 493.771780 | 0.175799188 | -1.84762045 | 0.06465727 | 0.731823074 | 9.100724 | 8.776499 | 9.068340 | 8.955219 | 8.880582 | 8.862330 | 686 | 542 | 387 | 405 |
| 8733 | NM_003801 | GPAA1 | glycosylphosphatidylinositol anchor attachment 1 | protein_coding | NP_003792.1 | 4446 | 603048 | . | . | 66.044960 | 65.884628 | 57.438496 | 57.442387 | 153.689331 | 152.295288 | 128.890305 | 131.335205 | -1.164649 | 3950.675206 | 0.107500089 | -2.04553295 | 0.040802344 | 0.62818612 | 12.053634 | 11.833764 | 12.012398 | 12.003128 | 11.885625 | 11.877439 | 4922 | 4562 | 3190 | 3405 |
| 8737 | NM_00131706 | RIPK1 | receptor interacting serine/threonine kinase 1 | protein_coding | NP_001303990.1;NP_001341859.1;NP_001341860.1;NP_001341861.1;N | 10019 | 603453 | . | . | 4.864787 | 5.205587 | 5.602778 | 6.036456 | 11.320560 | 12.032945 | 12.572471 | 13.801639 | 1.115475 | 670.445135 | 0.153494613 | 1.027121802 | 0.304363114 | 0.998865404 | 9.308182 | 9.465483 | 9.334923 | 9.364439 | 9.402712 | 9.444891 | 714 | 699 | 596 | 683 |
| 8738 | NM_00132009 | CRADD | CASP2 and RIPK1 domain containing adaptor with death domain | protein_coding | NP_001307028.1;NP_001307029.1;NP_001307030.1;NP_001317055.1;N | 2340 | 603454 | . | . | 2.865121 | 1.980271 | 2.825965 | 2.440148 | 6.667255 | 4.577486 | 6.341386 | 5.579105 | 1.112839 | 93.288773 | 0.351806288 | 0.43843744 | 0.661069211 | 0.998865404 | 6.463041 | 6.619958 | 6.575194 | 6.465929 | 6.585538 | 6.519857 | 121 | 77 | 95 | 82 |
| 8739 | NM_003806,N | HRK | harakiri, BCL2 interacting protein | protein_coding | NP_003797.1;XP_016875635.1 | 5185 | 603447 | . | . | 0.844215 | 1.221974 | 1.369749 | 1.309063 | 1.964523 | 2.824648 | 3.073678 | 2.993018 | 1.275827 | 216.446589 | 0.250187523 | 1.404675747 | 0.160117721 | 0.933525128 | 7.572101 | 7.922420 | 7.606793 | 7.766288 | 7.815119 | 7.792131 | 180 | 242 | 216 | 222 |
| 8740 | NM_00137688 | TNFSF14 | TNF superfamily member 14 | protein_coding | NP_001363816.1;NP_003798.2;NP_742011.2;XP_016882907.1 | 11930 | 604520 | . | . | 0.055866 | 0.047586 | 0.053801 | 0.112322 | 0.130002 | 0.109998 | 0.120728 | 0.256811 | 1.563428 | 10.303718 | 0.970736239 | 0.664148511 | 0.506595264 | NA | 3.008843 | 3.650563 | 3.348189 | 3.342461 | 3.345950 | 3.384920 | 10 | 8 | 7 | 16 |
| 8741 | NM_00119862 | TNFSF13 | TNF superfamily member 13 | protein_coding | NP_001185551.1;NP_001185552.1;NP_001185553.1;NP_003799.1;NP_7 | 11928 | 604472 | . | . | 0.079025 | 0.134617 | 0.125011 | 0.184688 | 0.183895 | 0.311173 | 0.280521 | 0.422267 | 1.422434 | 9.834939 | 0.984107589 | 0.51657111 | 0.605455588 | NA | 3.024434 | 3.527725 | 3.271746 | 3.290944 | 3.286551 | 3.306880 | 7 | 11 | 8 | 13 |
| 8742 | NM_003809,N | TNFSF12 | TNF superfamily member 12 | protein_coding | NP_003800.1 | 11927 | 602695 | . | . | 3.754044 | 4.250616 | 4.575722 | 4.351510 | 8.735814 | 9.825490 | 10.267788 | 9.949209 | 1.083547 | 192.288017 | 0.242127428 | 0.478104484 | 0.632575841 | 0.998865404 | 7.528154 | 7.643780 | 7.543166 | 7.591019 | 7.615763 | 7.589809 | 201 | 210 | 179 | 182 |
| 8744 | NM_003811 | TNFSF9 | TNF superfamily member 9 | protein_coding | NP_003802.1 | 11939 | 606182 | . | . | 5.528437 | 5.523578 | 4.687007 | 4.630953 | 12.864901 | 12.767999 | 10.517509 | 10.588123 | -1.200457 | 260.075223 | 0.212253042 | -1.24183544 | 0.214297283 | 0.979127905 | 8.148492 | 7.885070 | 8.069719 | 8.065295 | 7.980973 | 7.969250 | 328 | 305 | 208 | 219 |
| 8745 | NM_003812,X | ADAM23 | ADAM metallopeptidase domain 23 | protein_coding | NP_003803.1;XP_005246989.1;XP_011510388.1 | 202 | 603710 | . | . | 28.151101 | 27.645846 | 30.396499 | 27.899621 | 65.508768 | 63.904609 | 68.208861 | 63.789164 | 1.030887 | 5604.221511 | 0.10942311 | 0.401070805 | 0.688367995 | 0.998865404 | 12.430135 | 12.474127 | 12.449805 | 12.428114 | 12.504552 | 12.423071 | 6437 | 5878 | 5187 | 5076 |
| 8747 | NM_003813 | ADAM21 | ADAM metallopeptidase domain 21 | protein_coding | NP_003804.2 | 200 | 603713 | . | . | 0.208178 | 0.200314 | 0.222311 | 0.304641 | 0.484439 | 0.463034 | 0.498860 | 0.696526 | 1.293721 | 19.512462 | 0.698655266 | 0.531773657 | 0.59488277 | NA | 4.089593 | 4.459419 | 4.271804 | 4.268926 | 4.276684 | 4.306449 | 20 | 18 | 16 | 24 |
| 8751 | NM_00126146 | ADAM15 | ADAM metallopeptidase domain 15 | protein_coding | NP_001248393.1;NP_001248394.1;NP_001248395.1;NP_003806.3;NP_9 | 193 | 605548 | . | . | 35.425700 | 34.663684 | 31.986066 | 36.708347 | 82.437058 | 80.126658 | 71.775801 | 83.929273 | -1.030596 | 2981.104048 | 0.119767525 | -0.36302545 | 0.716585875 | 0.998865404 | 11.563286 | 11.519646 | 11.568173 | 11.538275 | 11.472613 | 11.581759 | 3548 | 3206 | 2382 | 2937 |
| 8754 | NM_003816,N | ADAM9 | ADAM metallopeptidase domain 9 | protein_coding | NP_003807.1;XP_011542984.1;XP_016869431.1 | 216 | 602713 | . | . | 35.020106 | 33.545740 | 30.689307 | 31.045305 | 81.493227 | 77.542482 | 68.865914 | 70.981403 | -1.125395 | 4181.811847 | 0.10818514 | -1.57536317 | 0.115172641 | 0.855571358 | 12.112592 | 11.942205 | 12.098134 | 12.055346 | 11.979366 | 11.978336 | 5229 | 4657 | 3424 | 3687 |
| 8760 | NM_003818,X | CDS2 | CDP-diacylglycerol synthase 2 | protein_coding | NP_003809.1;XP_006723723.1;XP_011527692.1;XP_024307784.1 | 1801 | 603549 | . | . | 5.901233 | 5.956361 | 5.831968 | 5.669717 | 13.732413 | 13.768396 | 13.086768 | 12.963135 | -1.044973 | 1657.862545 | 0.122247794 | -0.51915726 | 0.60365108 | 0.998865404 | 10.726447 | 10.663075 | 10.711269 | 10.712798 | 10.692463 | 10.662756 | 1949 | 1830 | 1437 | 1491 |
| 8761 | NM_00113565 | PABPC4 | poly(A) binding protein cytoplasmic 4 | protein_coding | NP_001129125.1;NP_001129126.1;NP_003810.1 | 8557 | 603407 | . | . | 57.424619 | 60.933431 | 58.948679 | 63.618573 | 133.629451 | 140.850356 | 132.279124 | 145.456305 | 1.021919 | 5843.187367 | 0.10734798 | 0.291394615 | 0.770749532 | 0.998865404 | 12.496861 | 12.528050 | 12.481765 | 12.524114 | 12.494260 | 12.547765 | 6488 | 6394 | 4973 | 5713 |
| 8763 | NM_00114240 | CD164 | CD164 molecule | protein_coding | NP_001135873.1;NP_001135874.1;NP_001135875.1;NP_001135876.1;N | 1632 | 603356 | . | . | 76.929751 | 56.113789 | 64.211420 | 38.910866 | 179.018681 | 129.709541 | 144.088556 | 88.965072 | -1.307263 | 5569.761434 | 0.406518683 | -0.95087652 | 0.341667064 | 0.998865404 | 12.623744 | 12.237247 | 12.669515 | 12.379635 | 12.497708 | 12.076979 | 8479 | 5675 | 5259 | 3383 |
| 8764 | NM_00129760 | TNFRSF14 | TNF receptor superfamily member 14 | protein_coding | NP_001284534.1;NP_003811.2;XP_006711082.1;XP_011540685.1;XP_01 | 11912 | 602746 | . | . | 258.690220 | 271.773169 | 352.957117 | 305.717374 | 601.982770 | 628.215863 | 792.025550 | 698.986471 | 1.225626 | 22931.600961 | 0.112851547 | 2.600932843 | 0.009297065 | 0.307003327 | 14.330822 | 14.624370 | 14.381850 | 14.398657 | 14.636380 | 14.489453 | 23493 | 22456 | 23803 | 21694 |
| 8766 | NM_00120683 | RAB11A | RAB11A, member RAS oncogene family | protein_coding | NP_001193765.1;NP_004654.1 | 9760 | 605570 | . | . | 13.262619 | 12.018879 | 13.650044 | 12.476824 | 30.862658 | 27.782179 | 30.630305 | 28.526776 | 1.019024 | 1960.601317 | 0.127120392 | 0.21387953 | 0.830641003 | 0.998865404 | 10.923247 | 10.950782 | 10.970469 | 10.885324 | 10.983075 | 10.903049 | 2356 | 1983 | 1808 | 1762 |
| 8767 | NM_00137536 | RIPK2 | receptor interacting serine/threonine kinase 2 | protein_coding | NP_001362289.1;NP_003812.1;XP_011515659.1 | 10020 | 603455 | . | . | 6.861515 | 6.761764 | 5.855529 | 6.120248 | 15.967028 | 15.630121 | 13.139637 | 13.993220 | -1.153557 | 490.302990 | 0.168019721 | -1.22657547 | 0.219982194 | 0.982008277 | 9.036890 | 8.830816 | 8.987739 | 8.974274 | 8.888276 | 8.893995 | 611 | 561 | 394 | 429 |
| 8771 | NM_003823 | TNFRSF6B | TNF receptor superfamily member 6b | protein_coding | NP_003814.1 | 11921 | 603361 | . | . | 0.634840 | 0.093132 | 0.131145 | 0.093582 | 1.477299 | 0.215278 | 0.294286 | 0.213964 | -2.723383 | 9.292863 | 1.145269519 | -1.26206108 | 0.206926808 | NA | 3.763141 | 2.324312 | 3.278671 | 3.189383 | 3.199832 | 3.192850 | 27 | 4 | 5 | 4 |
| 8772 | NM_003824 | FADD | Fas associated via death domain | protein_coding | NP_003815.1 | 3573 | 602457 | . | . | 18.253292 | 16.046371 | 14.679332 | 18.768366 | 42.476162 | 37.091908 | 32.939999 | 42.911633 | -1.040587 | 901.217443 | 0.161553328 | -0.35528738 | 0.722374312 | 0.998865404 | 9.844329 | 9.786555 | 9.876715 | 9.776811 | 9.712105 | 9.879491 | 1131 | 924 | 678 | 925 |
| 8773 | NM_003825,N | SNAP23 | synaptosome associated protein 23 | protein_coding | NP_003816.2;NP_570710.1;XP_006720788.1;XP_016878181.1;XP_01687 | 11131 | 602534 | . | . | 16.956807 | 16.974882 | 15.431741 | 15.901600 | 39.459187 | 39.238201 | 34.628380 | 36.357119 | -1.107110 | 1181.932365 | 0.130831375 | -1.12204128 | 0.261844889 | 0.995760118 | 10.278481 | 10.131648 | 10.249286 | 10.238729 | 10.160521 | 10.174794 | 1440 | 1331 | 966 | 1062 |
| 8774 | NM_003826,X | NAPG | NSF attachment protein gamma | protein_coding | NP_003817.1;XP_011524056.1;XP_011524058.1;XP_016881552.1 | 7642 | 603216 | . | . | 7.092781 | 6.738691 | 6.779709 | 5.956720 | 16.505193 | 15.576787 | 15.213473 | 13.619333 | -1.096734 | 737.401895 | 0.154787798 | -0.86062003 | 0.389447349 | 0.998865404 | 9.591062 | 9.458509 | 9.578093 | 9.535067 | 9.539876 | 9.443653 | 915 | 807 | 655 | 613 |
| 8775 | NM_003827,N | NAPA | NSF attachment protein alpha | protein_coding | NP_003818.2;XP_011525738.1;XP_011525739.1 | 7641 | 603215 | . | . | 41.501725 | 43.663510 | 39.955867 | 50.350815 | 96.576222 | 100.930150 | 89.659801 | 115.121152 | 1.032583 | 2291.094299 | 0.135679457 | 0.34093684 | 0.73315113 | 0.998865404 | 11.138695 | 11.184583 | 11.131871 | 11.162318 | 11.077080 | 11.260489 | 2547 | 2479 | 1794 | 2430 |
| 8776 | NM_00130614 | MTMR1 | myotubularin related protein 1 | protein_coding | NP_001293073.1;NP_001293074.1;NP_001340919.1;NP_001340920.1;N | 7449 | 300171 | . | . | 3.259595 | 3.138468 | 3.968107 | 3.966113 | 7.585210 | 7.254710 | 8.904317 | 9.068042 | 1.252003 | 466.034607 | 0.170802042 | 1.898328123 | 0.057652871 | 0.712428456 | 8.693051 | 9.017349 | 8.784161 | 8.790967 | 8.934655 | 8.919690 | 474 | 449 | 456 | 480 |
| 8777 | NM_00126140 | MPDZ | multiple PDZ domain crumbs cell polarity complex component | protein_coding | NP_001248335.1;NP_001248336.1;NP_001317566.1;NP_001362342.1;N | 7208 | 603785 | . | . | 6.026331 | 6.478568 | 6.882181 | 6.162701 | 14.023517 | 14.975498 | 15.443419 | 14.090283 | 1.023284 | 1545.906366 | 0.130701129 | 0.254065363 | 0.799445061 | 0.998865404 | 10.577419 | 10.610862 | 10.568730 | 10.600009 | 10.651975 | 10.550325 | 1724 | 1682 | 1449 | 1370 |
| 8780 | NM_00134819 | RIOK3 | RIO kinase 3 | protein_coding | NP_001335122.1;NP_003822.2;XP_011524544.1;XP_011524545.1 | 11451 | 603579 | . | . | 8.372218 | 8.159926 | 7.477435 | 7.673277 | 19.482499 | 18.862033 | 16.779148 | 17.544037 | -1.103881 | 898.046075 | 0.140167861 | -1.01724256 | 0.309038056 | 0.998865404 | 9.880160 | 9.737612 | 9.855998 | 9.834468 | 9.771066 | 9.777339 | 1101 | 1002 | 739 | 804 |
| 8787 | NM_00108195 | RGS9 | regulator of G protein signaling 9 | protein_coding | NP_001075424.1;NP_001159405.1;NP_003826.2 | 10004 | 604067 | . | . | 0.339195 | 0.467475 | 0.128695 | 0.160285 | 0.789320 | 1.080590 | 0.288786 | 0.366472 | -2.221326 | 29.266603 | 0.589876341 | -1.95197027 | 0.050941734 | 0.672475152 | 5.335987 | 4.181222 | 4.912240 | 4.921975 | 4.814974 | 4.840282 | 45 | 45 | 13 | 20 |
| 8790 | NM_00119932 | FPGT | fucose-1-phosphate guanylyltransferase | protein_coding | NP_001186257.2;NP_001186258.2;NP_003829.3 | 3825 | 603609 | . | . | 2.823978 | 2.417957 | 2.617881 | 1.694257 | 6.571513 | 5.589216 | 5.874449 | 3.873717 | -1.231773 | 350.962645 | 0.231767558 | -1.29757914 | 0.194431994 | 0.96621494 | 8.596828 | 8.298081 | 8.554960 | 8.451735 | 8.500186 | 8.257873 | 484 | 382 | 335 | 229 |
| 8792 | NM_00127094 | TNFRSF11A | TNF receptor superfamily member 11a | protein_coding | NP_001257878.1;NP_001257879.1;NP_001257880.1;NP_001265197.1;N | 11908 | 603499 | . | . | 0.688024 | 0.667451 | 0.638098 | 0.696347 | 1.601060 | 1.542845 | 1.431873 | 1.592113 | -1.067260 | 112.564024 | 0.320487516 | -0.29302898 | 0.7695 | 0.998865404 | 6.861447 | 6.766185 | 6.849493 | 6.797742 | 6.745502 | 6.856485 | 144 | 116 | 78 | 120 |
| 8795 | NM_003842,N | TNFRSF10B | TNF receptor superfamily member 10b | protein_coding | NP_003833.4;NP_671716.2 | 11905 | 603612 | . | . | 29.377423 | 28.527976 | 30.114229 | 28.554799 | 68.362470 | 65.943695 | 67.575454 | 65.287158 | 1.001645 | 3592.109865 | 0.111222397 | 0.021319655 | 0.982990665 | 0.998865404 | 11.809367 | 11.811863 | 11.824295 | 11.794966 | 11.837353 | 11.784248 | 4206 | 3804 | 3227 | 3262 |
| 8796 | NM_00116070 | SCEL | sciellin | protein_coding | NP_001154178.1;NP_003834.3;NP_659001.2;XP_005266635.1;XP_00671 | 10573 | 604112 | . | . | 0.051869 | 0.041387 | 0.261741 | 0.080038 | 0.120701 | 0.095668 | 0.587339 | 0.182997 | 3.251324 | 11.423891 | 0.987104598 | 1.723249453 | 0.084843424 | NA | 2.421266 | 4.128572 | 3.443427 | 3.437007 | 3.531272 | 3.455648 | 7 | 5 | 23 | 8 |
| 8798 | NM_00128228 | DYRK4 | dual specificity tyrosine phosphorylation regulated kinase 4 | protein_coding | NP_001269214.1;NP_001269215.1;NP_001358230.1;NP_003836.1 | 3095 | 609181 | . | . | 7.460045 | 7.827105 | 6.615898 | 7.870423 | 17.359833 | 18.092707 | 14.845886 | 17.994787 | -1.041324 | 239.281943 | 0.227186363 | -0.25714087 | 0.797070032 | 0.998865404 | 7.932113 | 7.872404 | 7.889754 | 7.933297 | 7.844332 | 7.937133 | 269 | 275 | 184 | 241 |
| 8799 | NM_00118479 | PEX11B | peroxisomal biogenesis factor 11 beta | protein_coding | NP_001171724.1;NP_003837.1 | 8853 | 603867 | . | . | 5.992900 | 7.100013 | 7.449873 | 7.730783 | 13.945724 | 16.411999 | 16.717299 | 17.675515 | 1.127910 | 363.809713 | 0.191606753 | 0.906296525 | 0.364778924 | 0.998865404 | 8.417917 | 8.590977 | 8.425357 | 8.511420 | 8.534329 | 8.542671 | 364 | 397 | 333 | 364 |
| 8800 | NM_00127157 | PEX11A | peroxisomal biogenesis factor 11 alpha | protein_coding | NP_001258501.1;NP_001258502.1;NP_003838.1 | 8852 | 603866 | . | . | 4.679406 | 5.106692 | 4.663401 | 5.206726 | 10.889169 | 11.804349 | 10.464538 | 11.904560 | -1.005237 | 412.672293 | 0.181728592 | -0.0414662 | 0.96692424 | 0.998865404 | 8.692960 | 8.684734 | 8.667567 | 8.712151 | 8.656843 | 8.715986 | 459 | 463 | 340 | 405 |
| 8801 | NM_00117759 | SUCLG2 | succinate-CoA ligase GDP-forming subunit beta | protein_coding | NP_001171070.1;NP_003839.2;XP_016862909.1 | 11450 | 603922 | . | . | 8.574255 | 8.112689 | 8.120581 | 8.854776 | 19.952642 | 18.752844 | 18.222348 | 20.245393 | -1.030000 | 657.063984 | 0.153074172 | -0.2785859 | 0.780562638 | 0.998865404 | 9.381024 | 9.338442 | 9.380790 | 9.358923 | 9.347621 | 9.352190 | 780 | 708 | 561 | 609 |
| 8802 | NM_003849 | SUCLG1 | succinate-CoA ligase GDP/ADP-forming subunit alpha | protein_coding | NP_003840.2 | 11449 | 611224 | . | . | 48.626720 | 52.587025 | 43.538349 | 52.939316 | 113.156372 | 121.557266 | 97.698799 | 121.039452 | -1.063698 | 1955.494217 | 0.133257926 | -0.66854574 | 0.503785296 | 0.998865404 | 10.977380 | 10.887866 | 10.928160 | 10.983724 | 10.832192 | 10.977074 | 2241 | 2251 | 1496 | 1940 |
| 8803 | NM_003850 | SUCLA2 | succinate-CoA ligase ADP-forming subunit beta | protein_coding | NP_003841.1 | 11448 | 603921 | . | . | 8.964441 | 9.604841 | 8.018942 | 9.593684 | 20.860622 | 22.202021 | 17.994272 | 21.934818 | -1.068770 | 595.300400 | 0.166294158 | -0.57699506 | 0.563942793 | 0.998865404 | 9.265044 | 9.168282 | 9.217938 | 9.258738 | 9.137866 | 9.247781 | 687 | 684 | 458 | 585 |
| 8804 | NM_003851 | CREG1 | cellular repressor of E1A stimulated genes 1 | protein_coding | NP_003842.1 | 2351 | 618055 | . | . | 27.875681 | 26.653830 | 29.837261 | 25.806612 | 64.867859 | 61.611523 | 66.953957 | 59.003754 | 1.006471 | 1690.364454 | 0.131760325 | 0.070629355 | 0.943692747 | 0.998865404 | 10.718254 | 10.727967 | 10.740474 | 10.698337 | 10.783506 | 10.662543 | 1992 | 1769 | 1589 | 1467 |
| 8805 | NM_003852,N | TRIM24 | tripartite motif containing 24 | protein_coding | NP_003843.3;NP_056989.2;XP_024302749.1 | 11812 | 603406 | . | . | 3.016869 | 3.422878 | 2.351178 | 2.632819 | 7.020377 | 7.912135 | 5.275977 | 6.019628 | -1.093951 | 635.113185 | 0.154854794 | -0.83657697 | 0.402830392 | 0.998865404 | 9.374115 | 9.244725 | 9.352904 | 9.327825 | 9.284535 | 9.275747 | 778 | 703 | 531 | 565 |
| 8809 | NM_00128239 | IL18R1 | interleukin 18 receptor 1 | protein_coding | NP_001269328.1;NP_001358347.1;NP_001358348.1;NP_001358349.1;N | 5988 | 604494 | . | . | 0.055520 | 0.156256 | 0.094436 | 0.045653 | 0.129198 | 0.361193 | 0.211913 | 0.104380 | -1.238705 | 10.655090 | 0.969219832 | -0.31864076 | 0.749998938 | NA | 3.558919 | 3.251698 | 3.396015 | 3.431990 | 3.417653 | 3.390395 | 9 | 17 | 11 | 6 |
| 8811 | NM_003857,X | GALR2 | galanin receptor 2 | protein_coding | NP_003848.1;XP_011523729.1 | 4133 | 603691 | . | . | 0.097398 | 0.046851 | 0.253930 | 0.140376 | 0.226648 | 0.108299 | 0.569812 | 0.320954 | 2.207272 | 9.263997 | 1.027452178 | 1.111744536 | 0.266248005 | NA | 2.526439 | 3.674293 | 3.185591 | 3.166496 | 3.211818 | 3.203224 | 9 | 4 | 12 | 11 |
| 8812 | NM_00109940 | CCNK | cyclin K | protein_coding | NP_001092872.1;XP_005268211.1;XP_011535577.1 | 1596 | 603544 | . | . | 6.620568 | 7.473060 | 7.470729 | 8.634134 | 15.406333 | 17.274315 | 16.764099 | 19.740921 | 1.099923 | 712.984262 | 0.159774249 | 0.859975636 | 0.389802473 | 0.998865404 | 9.407801 | 9.544419 | 9.398567 | 9.484476 | 9.456877 | 9.556330 | 733 | 779 | 603 | 750 |
| 8813 | NM_00131703 | DPM1 | dolichyl-phosphate mannosyltransferase subunit 1, catalytic | protein_coding | NP_001303963.1;NP_001303964.1;NP_001303965.1;NP_003850.1 | 3005 | 603503 | . | . | 62.000267 | 60.176789 | 61.687363 | 58.814590 | 144.277176 | 139.101340 | 138.424661 | 134.472576 | -1.030984 | 2022.789555 | 0.119338919 | -0.36888497 | 0.712213468 | 0.998865404 | 11.003888 | 10.960040 | 11.007592 | 10.980247 | 10.991830 | 10.947307 | 2405 | 2178 | 1781 | 1815 |
| 8814 | NM_00128223 | CDKL1 | cyclin dependent kinase like 1 | protein_coding | NP_001269165.1;NP_001353993.1;NP_001353994.1;NP_004127.2;XP_0 | 1781 | 603441 | . | . | 0.388911 | 0.355010 | 0.367013 | 0.400637 | 0.905011 | 0.820620 | 0.823566 | 0.916010 | -1.043088 | 61.367602 | 0.405615322 | -0.15004477 | 0.880729296 | 0.998865404 | 5.970066 | 5.908079 | 5.953244 | 5.935645 | 5.910318 | 5.959782 | 75 | 65 | 46 | 63 |
| 8815 | NM_00114398 | BANF1 | BAF nuclear assembly factor 1 | protein_coding | NP_001137457.1;NP_003851.1;XP_016874003.1;XP_016874004.1 | 17397 | 603811 | . | . | 67.254297 | 74.579805 | 62.986981 | 75.544408 | 156.503516 | 172.394564 | 141.340967 | 172.723311 | -1.037903 | 1833.085802 | 0.12983058 | -0.41339842 | 0.679314732 | 0.998865404 | 10.866840 | 10.812771 | 10.821909 | 10.884939 | 10.768282 | 10.877663 | 2065 | 2095 | 1455 | 1804 |
| 8816 | NM_00128420 | DCAF5 | DDB1 and CUL4 associated factor 5 | protein_coding | NP_001271135.1;NP_001271136.1;NP_001271137.1;NP_003852.1;XP_0 | 20224 | 603812 | . | . | 6.672102 | 6.745463 | 7.498533 | 7.493342 | 15.526257 | 15.592441 | 16.826491 | 17.132638 | 1.125610 | 1298.136595 | 0.128306476 | 1.330478313 | 0.183360727 | 0.954800306 | 10.254375 | 10.425034 | 10.282437 | 10.307575 | 10.388376 | 10.381468 | 1383 | 1340 | 1201 | 1283 |
| 8817 | NM_003862 | FGF18 | fibroblast growth factor 18 | protein_coding | NP_003853.1 | 3674 | 603726 | . | . | 0.112150 | 0.253031 | 0.415302 | 0.292478 | 0.260978 | 0.584891 | 0.931927 | 0.668717 | 1.822860 | 17.270034 | 0.772859337 | 1.120777525 | 0.262382573 | NA | 3.614556 | 4.478414 | 4.045552 | 4.086352 | 4.126620 | 4.092600 | 9 | 18 | 23 | 17 |
| 8818 | NM_00137843 | DPM2 | dolichyl-phosphate mannosyltransferase subunit 2, regulatory | protein_coding | NP_001365365.1;NP_001365366.1;NP_003854.1 | 3006 | 603564 | . | . | 26.171395 | 26.394976 | 35.361418 | 26.955632 | 60.901913 | 61.013169 | 79.350002 | 61.630846 | 1.138060 | 819.021338 | 0.15978469 | 1.16767256 | 0.242938869 | 0.992733415 | 9.581128 | 9.768318 | 9.620886 | 9.633684 | 9.799845 | 9.631468 | 874 | 834 | 847 | 722 |
| 8819 | NM_003864 | SAP30 | Sin3A associated protein 30 | protein_coding | NP_003855.1 | 10532 | 603378 | . | . | 9.119424 | 8.870542 | 8.880061 | 8.223533 | 21.221275 | 20.504654 | 19.926600 | 18.802130 | -1.064382 | 300.302518 | 0.202089534 | -0.44542639 | 0.656011463 | 0.998865404 | 8.274301 | 8.184859 | 8.256793 | 8.236639 | 8.235822 | 8.190360 | 363 | 328 | 264 | 261 |
| 8820 | NM_00137605 | HESX1 | HESX homeobox 1 | protein_coding | NP_001362987.1;NP_001362988.1;NP_001362989.1;NP_001362990.1;N | 4877 | 601802 | . | . | 0.613294 | 0.418862 | 0.533899 | 0.125424 | 1.427161 | 0.968219 | 1.198053 | 0.286768 | -1.638054 | 22.339665 | 0.743356102 | -0.95779502 | 0.338166129 | NA | 4.790753 | 4.087437 | 4.513896 | 4.483369 | 4.496977 | 4.407682 | 36 | 26 | 24 | 6 |
| 8821 | NM_00110166 | INPP4B | inositol polyphosphate-4-phosphatase type II B | protein_coding | NP_001095139.1;NP_001317969.1;NP_003857.2;XP_005263380.1;XP_00 | 6075 | 607494 | . | . | 15.517488 | 15.526588 | 17.710727 | 15.603116 | 36.109832 | 35.890404 | 39.742358 | 35.674674 | 1.107297 | 3088.353769 | 0.111144862 | 1.322975417 | 0.185843553 | 0.958175316 | 11.517223 | 11.664276 | 11.538070 | 11.560276 | 11.644191 | 11.621037 | 3329 | 3206 | 2862 | 3000 |
| 8824 | NM_00136540 | CES2 | carboxylesterase 2 | protein_coding | NP_001352334.1;NP_001352335.1;NP_001352336.1;NP_001352337.1;N | 1864 | 605278 | . | . | 4.348624 | 4.492490 | 4.567099 | 5.114703 | 10.119428 | 10.384597 | 10.248438 | 11.694159 | 1.070726 | 542.544971 | 0.165961392 | 0.594053378 | 0.552476412 | 0.998865404 | 9.033709 | 9.131821 | 9.059735 | 9.066091 | 9.065197 | 9.142289 | 603 | 566 | 459 | 557 |
| 8826 | NM_003870,X | IQGAP1 | IQ motif containing GTPase activating protein 1 | protein_coding | NP_003861.1 | 6110 | 603379 | . | . | 65.406662 | 64.150314 | 56.843723 | 65.024124 | 152.203986 | 148.286324 | 127.555669 | 148.669937 | -1.077756 | 14092.744364 | 0.108248877 | -0.99798486 | 0.318286702 | 0.998865404 | 13.835690 | 13.727617 | 13.825471 | 13.801291 | 13.694585 | 13.801183 | 17071 | 15555 | 11058 | 13513 |
| 8828 | NM_003872,N | NRP2 | neuropilin 2 | protein_coding | NP_003863.2;NP_061004.3;NP_957716.1;NP_957718.1;NP_957719.1;NP | 8005 | 602070 | . | . | 93.789453 | 98.779710 | 87.465250 | 92.464965 | 218.251905 | 228.333718 | 196.269479 | 211.410173 | -1.085804 | 19083.802486 | 0.100603972 | -1.18050442 | 0.237799652 | 0.992733415 | 14.278233 | 14.159445 | 14.236436 | 14.271905 | 14.163533 | 14.203217 | 22410 | 21879 | 15542 | 17554 |
| 8829 | NM_00102462 | NRP1 | neuropilin 1 | protein_coding | NP_001019799.2;NP_001019800.2;NP_001231901.2;NP_001231902.2;N | 8004 | 602069 | . | . | 8.553049 | 7.671379 | 6.648967 | 6.796272 | 19.903297 | 17.732732 | 14.920094 | 15.538871 | -1.260299 | 1037.087917 | 0.13632431 | -2.44832623 | 0.014352164 | 0.399401052 | 10.175501 | 9.841902 | 10.119235 | 10.074159 | 9.931700 | 9.922608 | 1371 | 1211 | 803 | 855 |
| 8831 | NM_00113006 | SYNGAP1 | synaptic Ras GTPase activating protein 1 | protein_coding | NP_001123538.1;NP_006763.2 | 11497 | 603384 | . | . | 3.071090 | 3.491705 | 4.296860 | 2.893062 | 7.146554 | 8.071234 | 9.642029 | 6.614643 | 1.084972 | 641.727405 | 0.196250359 | 0.599529666 | 0.548819734 | 0.998865404 | 9.265393 | 9.383811 | 9.249636 | 9.330444 | 9.475781 | 9.197395 | 666 | 704 | 699 | 499 |
| 8833 | NM_003875,X | GMPS | guanine monophosphate synthase | protein_coding | NP_003866.1;XP_011511565.1;XP_016862911.1 | 4378 | 600358 | . | . | 8.426230 | 8.977804 | 7.902270 | 8.290214 | 19.608185 | 20.752598 | 17.732464 | 18.954589 | -1.086638 | 2255.967905 | 0.117162422 | -1.02312497 | 0.30624876 | 0.998865404 | 11.198292 | 11.078274 | 11.151746 | 11.191690 | 11.090308 | 11.119929 | 2639 | 2598 | 1845 | 2065 |
| 8834 | NM_003876,N | TMEM11 | transmembrane protein 11 | protein_coding | NP_003867.1 | 16823 | 618817 | . | . | 26.232260 | 24.953567 | 20.996489 | 26.520533 | 61.043552 | 57.681292 | 47.115515 | 60.636048 | -1.054658 | 832.153697 | 0.155127969 | -0.49491404 | 0.620660801 | 0.998865404 | 9.738892 | 9.661482 | 9.719257 | 9.717613 | 9.607063 | 9.748951 | 985 | 921 | 633 | 836 |
| 8835 | NM_00127046 | SOCS2 | suppressor of cytokine signaling 2 | protein_coding | NP_001257396.1;NP_001257397.1;NP_001257398.1;NP_001257399.1;N | 19382 | 605117 | . | . | 0.805416 | 0.641906 | 0.521018 | 0.583647 | 1.874238 | 1.483795 | 1.169149 | 1.334439 | -1.384432 | 50.778758 | 0.444832751 | -1.05498983 | 0.291429978 | 0.998865404 | 5.881411 | 5.413048 | 5.723134 | 5.671237 | 5.637254 | 5.637152 | 76 | 56 | 37 | 40 |
| 8836 | NM_003878,X | GGH | gamma-glutamyl hydrolase | protein_coding | NP_003869.1;XP_011515925.1 | 4248 | 601509 | . | . | 110.921319 | 98.297590 | 90.916239 | 86.379314 | 258.118464 | 227.219303 | 204.013406 | 197.496048 | -1.199659 | 3802.270643 | 0.116960586 | -2.24541256 | 0.02474166 | 0.517665397 | 12.017912 | 11.755459 | 12.013699 | 11.909157 | 11.835282 | 11.788476 | 5071 | 4198 | 3093 | 3148 |
| 8837 | NM_00112718 | CFLAR | CASP8 and FADD like apoptosis regulator | protein_coding | NP_001120655.1;NP_001120656.1;NP_001189444.1;NP_001189445.1;N | 1876 | 603599 | . | . | 5.489331 | 5.217802 | 5.450665 | 6.166583 | 12.773896 | 12.061178 | 12.231131 | 14.099158 | 1.155870 | 1110.147950 | 0.135023653 | 1.54772353 | 0.121688874 | 0.864429515 | 10.008138 | 10.217356 | 10.059219 | 10.059094 | 10.197716 | 10.135410 | 1186 | 1111 | 1078 | 1070 |
| 8841 | NM_00135503 | HDAC3 | histone deacetylase 3 | protein_coding | NP_001341968.1;NP_001341969.1;NP_001341970.1;NP_003874.2 | 4854 | 605166 | . | . | 25.056868 | 26.685637 | 24.569736 | 27.758348 | 58.308359 | 61.685046 | 55.133773 | 63.466166 | -1.005497 | 1554.798268 | 0.13014969 | -0.0607638 | 0.951547321 | 0.998865404 | 10.606648 | 10.598363 | 10.583400 | 10.624581 | 10.551627 | 10.645851 | 1748 | 1727 | 1266 | 1542 |
| 8842 | NM_00114584 | PROM1 | prominin 1 | protein_coding | NP_001139319.1;NP_001139320.1;NP_001139321.1;NP_001139322.1;N | 9454 | 604365 | . | . | 0.286670 | 0.342749 | 0.241470 | 0.298308 | 0.667091 | 0.792279 | 0.541851 | 0.682046 | -1.133649 | 40.625356 | 0.493656556 | -0.36659942 | 0.713917844 | 0.998865404 | 5.432996 | 5.249810 | 5.335020 | 5.373804 | 5.321573 | 5.348333 | 44 | 52 | 30 | 39 |
| 8844 | NM_00136781 | KSR1 | kinase suppressor of ras 1 | protein_coding | NP_001354739.1;NP_055053.1;XP_006722217.1;XP_011523731.1;XP_01 | 6465 | 601132 | . | . | 1.017634 | 0.949145 | 0.879221 | 0.956289 | 2.368077 | 2.193991 | 1.972946 | 2.186443 | -1.072750 | 183.825777 | 0.246442661 | -0.41110308 | 0.680996952 | 0.998865404 | 7.572031 | 7.470576 | 7.555707 | 7.519286 | 7.490346 | 7.523931 | 228 | 197 | 148 | 173 |
| 8846 | NM_006020,X | ALKBH1 | alkB homolog 1, histone H2A dioxygenase | protein_coding | NP_006011.2 | 17911 | 605345 | . | . | 3.801396 | 3.392344 | 3.386327 | 3.281049 | 8.846005 | 7.841554 | 7.598819 | 7.501731 | -1.091664 | 280.826525 | 0.209062419 | -0.60522107 | 0.545032118 | 0.998865404 | 8.195115 | 8.069210 | 8.188434 | 8.122530 | 8.119901 | 8.100415 | 358 | 297 | 238 | 247 |
| 8847 | NR_152566,N | DLEU2 | deleted in lymphocytic leukemia 2 | lncRNA | . | 13748 | 605766 | . | . | 3.306078 | 3.830565 | 3.372565 | 3.416312 | 7.693381 | 8.854522 | 7.567943 | 7.810994 | -1.079672 | 157.274473 | 0.265697152 | -0.41623579 | 0.677237473 | 0.998865404 | 7.350871 | 7.241331 | 7.342583 | 7.280633 | 7.297376 | 7.265557 | 202 | 163 | 137 | 136 |
| 8848 | NM_00124379 | TSC22D1 | TSC22 domain family member 1 | protein_coding | NP_001230726.1;NP_001230727.1;NP_001230728.1;NP_006013.1;NP_9 | 16826 | 607715 | . | . | 15.106635 | 14.204835 | 12.147943 | 13.027773 | 35.153761 | 32.835111 | 27.259632 | 29.786451 | -1.177833 | 1627.570739 | 0.126540744 | -1.86607898 | 0.062030317 | 0.723723143 | 10.781796 | 10.545553 | 10.753898 | 10.702880 | 10.569879 | 10.631455 | 2092 | 1839 | 1247 | 1458 |
| 8850 | NM_003884,X | KAT2B | lysine acetyltransferase 2B | protein_coding | NP_003875.3;XP_005265585.1;XP_016862912.1;XP_016862913.1 | 8638 | 602303 | . | . | 13.354299 | 13.926419 | 17.304040 | 15.256936 | 31.076005 | 32.191543 | 38.829764 | 34.883174 | 1.176769 | 2054.307638 | 0.124958676 | 1.879272613 | 0.060207279 | 0.719735192 | 10.882140 | 11.117172 | 10.924681 | 10.945145 | 11.117086 | 11.009491 | 2145 | 2063 | 2060 | 1944 |
| 8851 | NM_003885,X | CDK5R1 | cyclin dependent kinase 5 regulatory subunit 1 | protein_coding | NP_003876.1;XP_011523740.1;XP_016880770.1 | 1775 | 603460 | . | . | 9.846008 | 10.760715 | 8.514307 | 9.612320 | 22.912068 | 24.873872 | 19.105858 | 21.977425 | -1.018416 | 1260.943778 | 0.136676019 | -0.19262091 | 0.847255869 | 0.998865404 | 10.313320 | 10.287138 | 10.271982 | 10.339709 | 10.335675 | 10.247728 | 1400 | 1434 | 1150 | 1103 |
| 8853 | NM_00113519 | ASAP2 | ArfGAP with SH3 domain, ankyrin repeat and PH domain 2 | protein_coding | NP_001128663.1;NP_003878.1;XP_006711961.1;XP_006711964.1;XP_00 | 2721 | 603817 | . | . | 11.718467 | 12.336238 | 11.154537 | 12.028605 | 27.269357 | 28.515767 | 25.030460 | 27.501978 | -1.046408 | 2040.213288 | 0.120564916 | -0.5428263 | 0.587249405 | 0.998865404 | 11.026965 | 10.961296 | 10.994432 | 11.029607 | 10.946079 | 11.005057 | 2350 | 2301 | 1669 | 1939 |
| 8854 | NM_00120689 | ALDH1A2 | aldehyde dehydrogenase 1 family member A2 | protein_coding | NP_001193826.1;NP_003879.2;NP_733797.1;NP_733798.1 | 15472 | 603687 | . | . | 0.105042 | 0.149260 | 0.106886 | 0.125572 | 0.244436 | 0.345021 | 0.239850 | 0.287105 | -1.108723 | 14.554460 | 0.804773052 | -0.1850196 | 0.853213663 | NA | 3.937099 | 3.785711 | 3.857282 | 3.877638 | 3.856169 | 3.865809 | 15 | 19 | 11 | 14 |
| 8859 | NM_004197,N | STK19 | serine/threonine kinase 19 | protein_coding | NP_004188.1;NP_115830.1 | 11398 | 604977 | . | . | 3.314290 | 3.597081 | 3.394109 | 3.153195 | 7.712488 | 8.314815 | 7.616281 | 7.209406 | -1.108313 | 186.126285 | 0.244035344 | -0.60796409 | 0.543211292 | 0.998865404 | 7.612533 | 7.463917 | 7.545580 | 7.579059 | 7.519130 | 7.516293 | 217 | 219 | 154 | 165 |
| 8861 | NM_00111340 | LDB1 | LIM domain binding 1 | protein_coding | NP_001106878.1;NP_001308541.1;NP_003884.1;XP_016872356.1;XP_01 | 6532 | 603451 | . | . | 20.955334 | 21.857605 | 22.076930 | 20.324101 | 48.763921 | 50.524835 | 49.539992 | 46.468641 | -1.021047 | 2134.989100 | 0.120487327 | -0.24939913 | 0.80305206 | 0.998865404 | 11.074907 | 11.044964 | 11.049409 | 11.086186 | 11.084467 | 11.017165 | 2427 | 2381 | 1915 | 1897 |
| 8863 | NM_00128986 | PER3 | period circadian regulator 3 | protein_coding | NP_001276790.1;NP_001276791.1;NP_001276792.1;NP_001276793.1;N | 8847 | 603427 | . | . | 8.539019 | 8.929109 | 8.664582 | 9.405526 | 19.870648 | 20.640036 | 19.443071 | 21.504618 | 1.017675 | 1611.563795 | 0.123579034 | 0.204545826 | 0.837926985 | 0.998865404 | 10.641469 | 10.666910 | 10.654232 | 10.640118 | 10.681197 | 10.640199 | 1856 | 1708 | 1451 | 1484 |
| 8864 | NM_022817,X | PER2 | period circadian regulator 2 | protein_coding | NP_073728.1;XP_005246168.1;XP_006712887.1 | 8846 | 603426 | . | . | 0.622351 | 0.925901 | 0.771226 | 0.858071 | 1.448236 | 2.140261 | 1.730609 | 1.961878 | 1.040861 | 158.244068 | 0.282983853 | 0.204172172 | 0.838218957 | 0.998865404 | 7.277735 | 7.333737 | 7.214052 | 7.366335 | 7.290960 | 7.326850 | 145 | 199 | 134 | 158 |
| 8867 | NM_00116030 | SYNJ1 | synaptojanin 1 | protein_coding | NP_001153774.1;NP_001153778.1;NP_003886.3;NP_982271.3;XP_01688 | 11503 | 604297 | . | . | 1.605547 | 2.129037 | 2.423211 | 1.977821 | 3.736175 | 4.921360 | 5.437613 | 4.522051 | 1.090176 | 508.127286 | 0.169133301 | 0.736463603 | 0.46144861 | 0.998865404 | 8.925141 | 9.050241 | 8.990014 | 8.928659 | 9.032756 | 8.996703 | 587 | 499 | 474 | 483 |
| 8869 | NM_00104243 | ST3GAL5 | ST3 beta-galactoside alpha-2,3-sialyltransferase 5 | protein_coding | NP_001035902.1;NP_001341152.1;NP_001341153.1;NP_001341155.1;N | 10872 | 604402 | . | . | 24.817813 | 25.147873 | 26.490981 | 29.030773 | 57.752074 | 58.130435 | 59.444994 | 66.375413 | 1.195348 | 2726.066221 | 0.125580953 | 2.049919774 | 0.04037226 | 0.627543139 | 11.278005 | 11.535712 | 11.352088 | 11.316057 | 11.547024 | 11.407081 | 2924 | 2619 | 2800 | 2548 |
| 8870 | NM_003897 | IER3 | immediate early response 3 | protein_coding | NP_003888.2 | 5392 | 602996 | . | . | 137.058014 | 128.998581 | 187.516556 | 168.248184 | 318.939636 | 298.186005 | 420.781708 | 384.679474 | 1.318693 | 5986.447144 | 0.113229492 | 3.524778936 | 0.000423836 | 0.03567203 | 12.334098 | 12.733338 | 12.447226 | 12.390989 | 12.700983 | 12.599550 | 6151 | 5379 | 6272 | 6006 |
| 8871 | NM_00117808 | SYNJ2 | synaptojanin 2 | protein_coding | NP_001171559.1;NP_003889.1;XP_005267255.1;XP_005267256.1;XP_00 | 11504 | 609410 | . | . | 1.762469 | 2.146639 | 2.109539 | 1.771885 | 4.101337 | 4.962055 | 4.733744 | 4.051204 | 1.002616 | 454.157731 | 0.186397505 | 0.020222025 | 0.983866258 | 0.999015705 | 8.825025 | 8.829071 | 8.773718 | 8.871958 | 8.880687 | 8.767529 | 481 | 528 | 432 | 387 |
| 8872 | NM_006023,X | CDC123 | cell division cycle 123 | protein_coding | NP_006014.2;XP_005252695.1 | 16827 | 617708 | . | . | 48.102457 | 50.687809 | 49.024057 | 50.036574 | 111.936399 | 117.167151 | 110.008559 | 114.402678 | -1.011976 | 2033.122328 | 0.1181746 | -0.1453373 | 0.884444565 | 0.998865404 | 10.998091 | 10.980820 | 10.977375 | 11.010694 | 10.979061 | 10.990230 | 2306 | 2253 | 1745 | 1908 |
| 8874 | NM_00111351 | ARHGEF7 | Rho guanine nucleotide exchange factor 7 | protein_coding | NP_001106983.1;NP_001106984.1;NP_001106985.1;NP_001307780.1;N | 15607 | 605477 | . | . | 8.194345 | 8.208205 | 9.618030 | 9.139846 | 19.068578 | 18.973638 | 21.582576 | 20.897171 | 1.128992 | 1359.570777 | 0.129123004 | 1.355568423 | 0.175236524 | 0.949120257 | 10.318646 | 10.493906 | 10.363274 | 10.356748 | 10.483159 | 10.421553 | 1477 | 1372 | 1304 | 1298 |
| 8877 | NM_00114260 | SPHK1 | sphingosine kinase 1 | protein_coding | NP_001136073.1;NP_001136074.1;NP_001342068.1;NP_068807.2;NP_8 | 11240 | 603730 | . | . | 19.011770 | 20.236040 | 19.579260 | 20.350137 | 44.241170 | 46.776514 | 43.935289 | 46.528167 | 1.001285 | 1190.195686 | 0.133710011 | 0.013852622 | 0.98894756 | 0.999197585 | 10.216225 | 10.217741 | 10.195280 | 10.236885 | 10.188176 | 10.245325 | 1332 | 1319 | 996 | 1159 |
| 8878 | NM_00114229 | SQSTM1 | sequestosome 1 | protein_coding | NP_001135770.1;NP_001135771.1;NP_003891.1 | 11280 | 601530 | . | . | 507.615256 | 539.542209 | 760.346665 | 712.082235 | 1181.241540 | 1247.175931 | 1706.195875 | 1628.091466 | 1.387265 | 55678.829512 | 0.101749286 | 4.641244757 | 3.46317E-06 | 0.000796701 | 15.509481 | 15.981727 | 15.584595 | 15.628729 | 15.922863 | 15.856028 | 52305 | 51653 | 58391 | 58382 |
| 8879 | NM_003901,X | SGPL1 | sphingosine-1-phosphate lyase 1 | protein_coding | NP_003892.2;XP_005270320.1;XP_006718116.1;XP_011538618.1;XP_01 | 10817 | 603729 | . | . | 5.209870 | 4.796336 | 4.477620 | 4.435078 | 12.123584 | 11.086946 | 11.169633 | 10.140278 | -1.058755 | 852.974420 | 0.146689959 | -0.56151412 | 0.574447111 | 0.998865404 | 9.776707 | 9.694849 | 9.783477 | 9.727652 | 9.748362 | 9.681305 | 1049 | 910 | 755 | 740 |
| 8880 | NM_00130343 | FUBP1 | far upstream element binding protein 1 | protein_coding | NP_001290362.1;NP_001362984.1;NP_001362985.1;NP_001362986.1;N | 4004 | 603444 | . | . | 16.258529 | 15.873622 | 15.705607 | 16.033845 | 37.834269 | 36.692589 | 35.242928 | 36.659485 | -1.081785 | 2901.309600 | 0.115295956 | -0.98367399 | 0.325275834 | 0.998865404 | 11.558169 | 11.444573 | 11.514114 | 11.552644 | 11.437371 | 11.500407 | 3391 | 3330 | 2329 | 2715 |
| 8881 | NM_00107864 | CDC16 | cell division cycle 16 | protein_coding | NP_001072113.1;NP_001305446.1;NP_001305447.1;NP_001317030.1;N | 1720 | 603461 | . | . | 49.718498 | 49.344631 | 51.209593 | 48.521413 | 115.696983 | 114.062327 | 114.912841 | 110.938438 | 1.012237 | 3421.850904 | 0.113880658 | 0.154088866 | 0.877539667 | 0.998865404 | 11.731687 | 11.749382 | 11.739352 | 11.730964 | 11.784932 | 11.703958 | 3936 | 3651 | 3141 | 3069 |
| 8882 | NM_00131708 | ZPR1 | ZPR1 zinc finger | protein_coding | NP_001304015.1;NP_003895.1 | 13051 | 603901 | . | . | 3.657491 | 4.274443 | 4.005521 | 4.234773 | 8.511132 | 9.880565 | 8.988273 | 9.682306 | 1.008227 | 768.582427 | 0.156151457 | 0.07569921 | 0.939658405 | 0.998865404 | 9.580443 | 9.591647 | 9.520894 | 9.639914 | 9.564651 | 9.610248 | 807 | 896 | 649 | 747 |
| 8883 | NM_00101815 | NAE1 | NEDD8 activating enzyme E1 subunit 1 | protein_coding | NP_001018169.1;NP_001018170.1;NP_001273429.1;NP_003896.1;XP_0 | 621 | 603385 | . | . | 24.306278 | 23.469540 | 22.828151 | 14.571703 | 56.561708 | 54.250892 | 51.225709 | 33.316466 | -1.289731 | 1221.847266 | 0.182465639 | -2.01172346 | 0.044249096 | 0.645513836 | 10.426379 | 10.060136 | 10.356550 | 10.316434 | 10.295709 | 9.967816 | 1625 | 1447 | 1136 | 777 |
| 8884 | NM_021095,N | SLC5A6 | solute carrier family 5 member 6 | protein_coding | NP_066918.2;XP_006712191.1;XP_006712192.1;XP_006712193.1;XP_01 | 11041 | 604024 | . | . | 82.435909 | 84.904172 | 110.599932 | 95.841887 | 191.831748 | 196.259802 | 248.183042 | 219.131083 | 1.211590 | 8931.918962 | 0.112725335 | 2.456423699 | 0.01403276 | 0.393687366 | 12.979634 | 13.256602 | 13.029094 | 13.044551 | 13.262347 | 13.133925 | 9215 | 8796 | 9142 | 8494 |
| 8886 | NM_006773 | DDX18 | DEAD-box helicase 18 | protein_coding | NP_006764.3 | 2741 | 606355 | . | . | 27.781862 | 27.779026 | 28.236126 | 26.197229 | 64.649536 | 64.212463 | 63.361046 | 59.896851 | -1.034889 | 3215.964562 | 0.113483733 | -0.43597692 | 0.662853475 | 0.998865404 | 11.675493 | 11.626156 | 11.668337 | 11.661077 | 11.670891 | 11.601178 | 3783 | 3514 | 2866 | 2837 |
| 8887 | NM_00107986 | TAX1BP1 | Tax1 binding protein 1 | protein_coding | NP_001073333.1;NP_001193830.1;NP_001193831.1;NP_001349723.1;N | 11575 | 605326 | . | . | 54.993072 | 55.755567 | 52.421918 | 52.876438 | 127.971135 | 128.881492 | 117.633265 | 120.895691 | -1.068158 | 5681.578634 | 0.103974059 | -0.91489196 | 0.360248352 | 0.998865404 | 12.518858 | 12.423726 | 12.495950 | 12.502420 | 12.443114 | 12.444414 | 6732 | 6357 | 4774 | 5155 |
| 8888 | NM_003906,X | MCM3AP | minichromosome maintenance complex component 3 associated | protein_coding | NP_003897.2;XP_005261260.1;XP_005261261.1;XP_005261262.1 | 6946 | 603294 | . | . | 12.472044 | 13.703520 | 14.260540 | 14.671409 | 29.022957 | 31.676298 | 32.000236 | 33.544431 | 1.088970 | 2876.136703 | 0.114939883 | 1.069814823 | 0.284702669 | 0.998865404 | 11.427189 | 11.550030 | 11.419254 | 11.487222 | 11.517424 | 11.529102 | 3038 | 3096 | 2589 | 2831 |
| 8890 | NM_00103411 | EIF2B4 | eukaryotic translation initiation factor 2B subunit delta | protein_coding | NP_001029288.1;NP_001305894.1;NP_001305895.1;NP_001305896.1;N | 3260 | 606687 | . | . | 25.877712 | 27.064257 | 26.846736 | 27.432616 | 60.218499 | 62.560242 | 60.243301 | 62.721418 | -1.005710 | 1447.174208 | 0.126844972 | -0.06475985 | 0.948365208 | 0.998865404 | 10.503235 | 10.494799 | 10.483272 | 10.518670 | 10.477075 | 10.515886 | 1633 | 1602 | 1223 | 1387 |
| 8891 | NM_00116658 | EIF2B3 | eukaryotic translation initiation factor 2B subunit gamma | protein_coding | NP_001160060.1;NP_001248347.1;NP_065098.1;XP_016858234.1;XP_01 | 3259 | 606273 | . | . | 12.073994 | 12.752962 | 9.795769 | 12.048455 | 28.096681 | 29.479045 | 21.981424 | 27.547360 | -1.123880 | 672.383367 | 0.162029288 | -1.03986321 | 0.298403457 | 0.998865404 | 9.475240 | 9.306085 | 9.436465 | 9.425389 | 9.286186 | 9.412480 | 826 | 762 | 498 | 650 |
| 8892 | NM_014239 | EIF2B2 | eukaryotic translation initiation factor 2B subunit beta | protein_coding | NP_055054.1 | 3258 | 606454 | . | . | 9.087371 | 9.544890 | 10.107347 | 10.323998 | 21.146685 | 22.063442 | 22.680595 | 23.604595 | 1.082045 | 1309.478606 | 0.12830281 | 0.886658981 | 0.375262529 | 0.998865404 | 10.296836 | 10.410480 | 10.308193 | 10.339013 | 10.379377 | 10.387903 | 1419 | 1385 | 1177 | 1283 |
| 8893 | NM_003907,X | EIF2B5 | eukaryotic translation initiation factor 2B subunit epsilon | protein_coding | NP_003898.2;XP_011511567.1;XP_011511568.1 | 3261 | 603945 | . | . | 21.255319 | 21.716178 | 21.896395 | 22.573179 | 49.461999 | 50.197922 | 49.134874 | 51.610887 | 1.015846 | 1769.544602 | 0.120441586 | 0.188318919 | 0.850626655 | 0.998865404 | 10.777817 | 10.800419 | 10.776019 | 10.789764 | 10.787192 | 10.803230 | 2004 | 1911 | 1535 | 1689 |
| 8894 | NM_00131636 | EIF2S2 | eukaryotic translation initiation factor 2 subunit beta | protein_coding | NP_001303292.1;NP_001303293.1;NP_003899.2;XP_016883607.1 | 3266 | 603908 | . | . | 82.972894 | 78.855337 | 67.363138 | 74.965098 | 193.081339 | 182.277638 | 151.160937 | 171.398791 | -1.146654 | 5903.809659 | 0.108103629 | -1.82630208 | 0.067804773 | 0.740331407 | 12.622798 | 12.425310 | 12.597793 | 12.566229 | 12.432169 | 12.500549 | 7397 | 6680 | 4586 | 5369 |

| | | | | | | | | | | | | | | | | | | | | | | | | | | | | | | | | | |
|---|---|---|---|---|---|---|---|---|---|---|---|---|---|---|---|---|---|---|---|---|---|---|---|---|---|---|---|---|---|---|---|---|---|
| 8895 | NM_003909,X | CPNE3 | copine 3 | protein_coding | NP_003900.1;XP_005251150.1;XP_016869434.1;XP_024303090.1 | 2316 | 604207 | . | . | 9.867248 | 8.725644 | 8.715040 | 8.521901 | 22.961490 | 20.169717 | 19.556299 | 19.484312 | -1.094468 | 1363.144925 | 0.133548873 | -0.9751423 | 0.329489677 | 0.998865404 | 10.476219 | 10.346304 | 10.494614 | 10.394467 | 10.389442 | 10.364119 | 1746 | 1438 | 1151 | 1200 |
| 8896 | NM_00137040 | BUD31 | BUD31 homolog | protein_coding | NP_001357336.1;NP_001357339.1;NP_003901.2;XP_005250727.1;XP_00 | 29629 | 603477 | . | . | 29.614501 | 27.626597 | 25.240834 | 31.271081 | 68.914164 | 63.860118 | 56.639697 | 71.497614 | -1.039248 | 882.847542 | 0.161975332 | -0.3428937 | 0.73167843 | 0.998865404 | 9.813800 | 9.757696 | 9.825238 | 9.769632 | 9.673364 | 9.858358 | 1076 | 934 | 655 | 917 |
| 8897 | NM_021090,N | MTMR3 | myotubularin related protein 3 | protein_coding | NP_066576.1;NP_694690.1;NP_694691.1 | 7451 | 603558 | . | . | 2.892953 | 2.925118 | 3.240400 | 3.173858 | 6.732021 | 6.761541 | 7.271364 | 7.256650 | 1.086304 | 854.982935 | 0.141952566 | 0.841324581 | 0.400166123 | 0.998865404 | 9.678748 | 9.798280 | 9.707620 | 9.710319 | 9.779777 | 9.756240 | 942 | 886 | 787 | 821 |
| 8898 | NM_00124357 | MTMR2 | myotubularin related protein 2 | protein_coding | NP_001230500.1;NP_057240.3;NP_958435.1;NP_958438.1;XP_00527443 | 7450 | 603557 | . | . | 8.704920 | 8.790172 | 8.752536 | 8.360616 | 20.256706 | 20.318878 | 19.640438 | 19.115554 | -1.039112 | 1236.522121 | 0.129937414 | -0.4259845 | 0.670119143 | 0.998865404 | 10.299417 | 10.244199 | 10.287388 | 10.285347 | 10.273177 | 10.241464 | 1453 | 1358 | 1077 | 1113 |
| 8899 | NM_003913,N | PRPF4B | pre-mRNA processing factor 4B | protein_coding | NP_003904.3;XP_011513272.1;XP_016866899.1;XP_016866900.1;XP_01 | 17346 | 602338 | . | . | 13.460341 | 12.699103 | 14.927599 | 11.043349 | 31.322773 | 29.354551 | 33.497096 | 25.249300 | -1.049369 | 2389.858761 | 0.158330349 | -0.43909283 | 0.660594272 | 0.998865404 | 11.256831 | 11.187761 | 11.276893 | 11.196511 | 11.332797 | 11.044479 | 2957 | 2510 | 2391 | 1796 |
| 8900 | NM_00111104 | CCNA1 | cyclin A1 | protein_coding | NP_001104515.1;NP_001104516.1;NP_001104517.1;NP_003905.1;XP_01 | 1577 | 604036 | . | . | 9.045935 | 10.662826 | 7.784354 | 8.970299 | 21.050263 | 24.647598 | 17.467864 | 20.509521 | -1.155006 | 515.002794 | 0.17671358 | -1.17648546 | 0.239400939 | 0.992733415 | 9.109118 | 8.900198 | 9.005122 | 9.096176 | 8.908727 | 9.007768 | 592 | 637 | 382 | 484 |
| 8904 | NM_00119886 | CPNE1 | copine 1 | protein_coding | NP_001185792.1;NP_003906.2;NP_690902.1;NP_690903.1;NP_690904.1 | 2314 | 604205 | . | . | 86.671834 | 90.333171 | 91.409555 | 96.630571 | 201.688920 | 208.809171 | 205.120396 | 220.934314 | 1.045980 | 5583.628420 | 0.105808833 | 0.612945486 | 0.539912386 | 0.998865404 | 12.414230 | 12.479017 | 12.413048 | 12.442143 | 12.447026 | 12.483526 | 6176 | 5992 | 4857 | 5468 |
| 8905 | NM_00127207 | AP1S2 | adaptor related protein complex 1 subunit sigma 2 | protein_coding | NP_001259000.1;NP_001355923.1;NP_001355936.1;NP_001355937.1;N | 560 | 300629 | . | . | 37.044584 | 29.390642 | 31.828903 | 20.671052 | 86.204274 | 67.937785 | 71.423130 | 47.261910 | -1.273278 | 2126.139114 | 0.345132715 | -1.00989529 | 0.312545459 | 0.998865404 | 11.217723 | 10.869337 | 11.229456 | 11.030923 | 11.102212 | 10.750810 | 3072 | 2261 | 1991 | 1361 |
| 8906 | NM_00128247 | AP1G2 | adaptor related protein complex 1 subunit gamma 2 | protein_coding | NP_001269403.1;NP_001269404.1;NP_001341602.1;NP_001341603.1;N | 556 | 603534 | . | . | 21.852303 | 24.505887 | 29.907698 | 24.927128 | 50.851201 | 56.646445 | 67.112014 | 56.992911 | 1.185166 | 2391.478493 | 0.13641031 | 1.796703224 | 0.072382747 | 0.758556671 | 11.095851 | 11.341112 | 11.108410 | 11.187976 | 11.370977 | 11.193348 | 2394 | 2480 | 2500 | 2169 |
| 8907 | NM_00113052 | AP1M1 | adaptor related protein complex 1 subunit mu 1 | protein_coding | NP_001123996.1;NP_115882.1 | 13667 | 603535 | . | . | 5.108956 | 5.308971 | 4.699637 | 5.476595 | 11.888749 | 12.271922 | 10.545850 | 12.521584 | -1.038054 | 2063.118631 | 0.125922313 | -0.42789783 | 0.668725507 | 0.998865404 | 11.037455 | 10.983258 | 11.012722 | 11.036745 | 10.936972 | 11.049794 | 2384 | 2302 | 1635 | 2033 |
| 8908 | NM_00107985 | GYG2 | glycogenin 2 | protein_coding | NP_001073324.1;NP_001171631.1;NP_001171632.1;NP_001171633.1;N | 4700 | 300198 | . | . | 15.373360 | 15.851434 | 15.449236 | 19.901612 | 35.774439 | 36.641302 | 34.667640 | 45.502668 | 1.116150 | 1675.510060 | 0.142739352 | 1.110631069 | 0.266727184 | 0.998274218 | 10.629185 | 10.787256 | 10.651907 | 10.675531 | 10.649327 | 10.841008 | 1796 | 1735 | 1351 | 1857 |
| 8910 | NM_00109940 | SGCE | sarcoglycan epsilon | protein_coding | NP_001092870.1;NP_001092871.1;NP_001288068.1;NP_001333642.1;N | 10808 | 604149 | . | . | 45.041852 | 47.352033 | 42.461833 | 40.138771 | 104.814243 | 109.456339 | 95.283120 | 91.772533 | -1.125635 | 2601.995733 | 0.115510773 | -1.47812379 | 0.139374648 | 0.900674731 | 11.428213 | 11.257548 | 11.375910 | 11.406523 | 11.320325 | 11.269965 | 3114 | 3029 | 2196 | 2223 |
| 8913 | NM_00125632 | CACNA1G | calcium voltage-gated channel subunit alpha1 G | protein_coding | NP_001243253.1;NP_001243254.1;NP_001243255.1;NP_001243256.1;N | 1394 | 604065 | . | . | 0.050640 | 0.027770 | 0.048535 | 0.105950 | 0.117841 | 0.064192 | 0.108912 | 0.242243 | 1.742046 | 17.666875 | 0.763750689 | 1.048487116 | 0.294414235 | NA | 3.687500 | 4.488632 | 4.123370 | 4.091048 | 4.121997 | 4.174574 | 19 | 10 | 14 | 27 |
| 8914 | NM_00133029 | TIMELESS | timeless circadian regulator | protein_coding | NP_001317224.1;NP_003911.2 | 11813 | 603887 | . | . | 15.343856 | 16.521246 | 17.660624 | 18.168107 | 35.705781 | 38.189597 | 39.629927 | 41.539216 | 1.107216 | 2715.974445 | 0.114424848 | 1.284129031 | 0.199096812 | 0.96621494 | 11.331974 | 11.478800 | 11.337604 | 11.389987 | 11.440595 | 11.453108 | 2872 | 2872 | 2463 | 2696 |
| 8915 | NM_00132071 | BCL10 | BCL10 immune signaling adaptor | protein_coding | NP_001307644.1;NP_003912.1;XP_011540699.1;XP_011540700.1;XP_01 | 989 | 603517 | . | . | 2.959822 | 2.795014 | 2.755212 | 3.247378 | 6.887628 | 6.460799 | 6.182616 | 7.424744 | 1.021671 | 258.894253 | 0.216607387 | 0.142793429 | 0.886453327 | 0.998865404 | 8.000889 | 8.031388 | 8.027422 | 7.992751 | 7.986133 | 8.054789 | 305 | 267 | 212 | 262 |
| 8916 | NM_00127160 | HERC3 | HECT and RLD domain containing E3 ubiquitin protein ligase 3 | protein_coding | NP_001258531.1;NP_001305434.1;NP_001362406.1;NP_001362407.1;N | 4876 | 605200 | . | . | 3.302639 | 3.578759 | 4.140604 | 3.333318 | 7.685375 | 8.272459 | 9.291395 | 7.621233 | 1.110206 | 564.481299 | 0.174340766 | 0.86513264 | 0.386965992 | 0.998865404 | 9.063067 | 9.214524 | 9.084740 | 9.120387 | 9.243447 | 9.093606 | 599 | 593 | 575 | 494 |
| 8924 | NM_004667,X | HERC2 | HECT and RLD domain containing E3 ubiquitin protein ligase 2 | protein_coding | NP_004658.3;XP_005268333.1;XP_006720789.1;XP_006720790.1;XP_01 | 4868 | 605837 | . | . | 6.667951 | 6.900900 | 6.951656 | 7.092329 | 15.516597 | 15.951738 | 15.599313 | 16.215770 | 1.076789 | 2865.446736 | 0.112359738 | 0.949944654 | 0.342140376 | 0.998865404 | 11.430213 | 11.536903 | 11.436066 | 11.470563 | 11.515691 | 11.511822 | 3110 | 3041 | 2590 | 2779 |
| 8925 | NM_003922,X | HERC1 | HECT and RLD domain containing E3 ubiquitin protein ligase family | protein_coding | NP_003913.3;XP_016878188.1;XP_016878189.1;XP_016878190.1;XP_01 | 4867 | 605109 | . | . | 3.295841 | 3.258130 | 3.325127 | 3.702624 | 7.669558 | 7.531311 | 7.461490 | 8.465609 | 1.063094 | 1613.375497 | 0.126355938 | 0.698578007 | 0.484815792 | 0.998865404 | 10.611157 | 10.699232 | 10.638825 | 10.623441 | 10.639693 | 10.716391 | 1819 | 1671 | 1375 | 1635 |
| 8926 | NM_005678,N | SNURF | SNRPN upstream reading frame | protein_coding | NP_005669.2;NP_073715.1 | 11171 | . | . | . | 3.783467 | 3.266790 | 2.087414 | 3.134411 | 8.804283 | 7.551332 | 4.684096 | 7.166460 | -1.361159 | 149.296083 | 0.289813008 | -1.5349038 | 0.124807461 | 0.873049043 | 7.428094 | 6.981555 | 7.305665 | 7.245838 | 7.090061 | 7.220005 | 212 | 173 | 89 | 141 |
| 8927 | NM_003458 | BSN | bassoon presynaptic cytomatrix protein | protein_coding | NP_003449.2 | 1117 | 604020 | . | . | 0.052481 | 0.058122 | 0.050235 | 0.055862 | 0.122125 | 0.134353 | 0.112727 | 0.127722 | -1.066063 | 27.402579 | 0.588725455 | -0.15676765 | 0.875427968 | 0.998865404 | 4.822157 | 4.728813 | 4.775074 | 4.786444 | 4.768771 | 4.778442 | 31 | 32 | 22 | 26 |
| 8928 | NM_003923 | FOXH1 | forkhead box H1 | protein_coding | NP_003914.1 | 3814 | 603621 | . | . | 0.334122 | 0.293018 | 0.297144 | 0.418641 | 0.777516 | 0.677324 | 0.666782 | 0.957173 | 1.149209 | 21.615547 | 0.665473046 | 0.30150079 | 0.763032648 | NA | 4.330772 | 4.530328 | 4.432206 | 4.418494 | 4.423356 | 4.452490 | 25 | 20 | 17 | 25 |
| 8930 | NM_00127627 | MBD4 | methyl-CpG binding domain 4, DNA glycosylase | protein_coding | NP_001263199.1;NP_001263200.1;NP_001263201.1;NP_001263202.1;N | 6919 | 603574 | . | . | 14.924364 | 13.736310 | 14.258320 | 13.024698 | 34.729608 | 31.752098 | 31.995256 | 29.779421 | -1.085004 | 1139.444918 | 0.139010545 | -0.8467032 | 0.397160575 | 0.998865404 | 10.211526 | 10.094325 | 10.211535 | 10.154066 | 10.165937 | 10.076975 | 1417 | 1231 | 1008 | 963 |
| 8932 | NM_003927,N | MBD2 | methyl-CpG binding domain protein 2 | protein_coding | NP_003918.1;NP_056647.1 | 6917 | 603547 | . | . | 5.360203 | 5.387057 | 5.132795 | 6.210603 | 12.473413 | 12.452424 | 11.517843 | 14.199806 | 1.032480 | 867.717422 | 0.150601529 | 0.306195642 | 0.75945569 | 0.998865404 | 9.738110 | 9.783693 | 9.749811 | 9.747484 | 9.707341 | 9.831674 | 985 | 920 | 700 | 898 |
| 8933 | NM_00107817 | RTL8C | retrotransposon Gag like 8C | protein_coding | NP_001071639.1 | 2569 | 300213 | . | . | 21.866068 | 24.057487 | 22.957504 | 25.003628 | 50.883236 | 55.609962 | 51.515976 | 57.167820 | 1.029790 | 871.505768 | 0.1453142 | 0.291442892 | 0.770712614 | 0.998865404 | 9.746269 | 9.788159 | 9.724674 | 9.786984 | 9.749955 | 9.803716 | 946 | 967 | 740 | 860 |
| 8934 | NM_00113566 | RAB29 | RAB29, member RAS oncogene family | protein_coding | NP_001129134.1;NP_001129135.1;NP_001129136.1;NP_003920.1;XP_00 | 9789 | 603949 | . | . | 17.769133 | 18.493974 | 19.673072 | 18.989722 | 41.349499 | 42.749649 | 44.145802 | 43.417744 | 1.037961 | 1883.811669 | 0.119794739 | 0.448705987 | 0.653643765 | 0.998865404 | 10.852285 | 10.906092 | 10.855127 | 10.873490 | 10.909967 | 10.877419 | 2104 | 2018 | 1703 | 1762 |
| 8935 | NM_00130346 | SKAP2 | src kinase associated phosphoprotein 2 | protein_coding | NP_001290397.1;NP_003921.2;XP_016868260.1 | 15687 | 605215 | . | . | 11.007364 | 11.562012 | 12.299622 | 13.065715 | 25.614589 | 26.726107 | 27.599998 | 29.873206 | 1.108306 | 1435.669641 | 0.1268199 | 1.169815396 | 0.242075267 | 0.992733415 | 10.411534 | 10.559680 | 10.430376 | 10.461848 | 10.505775 | 10.544274 | 1536 | 1500 | 1280 | 1450 |
| 8936 | NM_00102493 | WASF1 | WASP family member 1 | protein_coding | NP_001020105.1;NP_001020106.1;NP_001020107.1;NP_003922.1 | 12732 | 605035 | . | . | 2.835767 | 2.783972 | 3.447781 | 3.234566 | 6.598946 | 6.435274 | 7.736721 | 7.395452 | 1.179809 | 292.225998 | 0.203146752 | 1.174292022 | 0.240278078 | 0.992733415 | 8.066568 | 8.305437 | 8.143224 | 8.142510 | 8.248058 | 8.213111 | 309 | 289 | 284 | 287 |
| 8938 | NM_00119909 | BAIAP3 | BAI1 associated protein 3 | protein_coding | NP_001186025.1;NP_001186026.1;NP_001186027.1;NP_001186028.1;N | 948 | 604009 | . | . | 0.174502 | 0.104710 | 0.087859 | 0.106132 | 0.406073 | 0.242042 | 0.197153 | 0.242658 | -1.319983 | 19.186104 | 0.711548078 | -0.56288433 | 0.57351365 | NA | 4.448287 | 4.048019 | 4.295463 | 4.258970 | 4.248341 | 4.261417 | 30 | 19 | 13 | 17 |
| 8939 | NM_003934,X | FUBP3 | far upstream element binding protein 3 | protein_coding | NP_003925.1;XP_005272289.1;XP_006717376.1;XP_011517474.1;XP_01 | 4005 | 603536 | . | . | 13.856821 | 13.091193 | 13.581920 | 16.032396 | 32.245393 | 30.260882 | 30.477432 | 36.656167 | 1.067799 | 1401.865931 | 0.136017072 | 0.695799814 | 0.486554206 | 0.998865404 | 10.405120 | 10.499599 | 10.460424 | 10.391399 | 10.422065 | 10.528718 | 1632 | 1397 | 1172 | 1451 |
| 8940 | NM_00128211 | TOP3B | DNA topoisomerase III beta | protein_coding | NP_001269041.1;NP_001269042.1;NP_001336774.1;NP_001336776.1;N | 11993 | 603582 | . | . | 12.982064 | 13.123499 | 13.253937 | 12.763974 | 30.209798 | 30.335555 | 29.741446 | 29.183309 | -1.031580 | 1331.423764 | 0.127828654 | -0.35090357 | 0.725660692 | 0.998865404 | 10.400972 | 10.356188 | 10.383590 | 10.397290 | 10.380769 | 10.352618 | 1543 | 1472 | 1161 | 1206 |
| 8941 | NM_003936 | CDK5R2 | cyclin dependent kinase 5 regulatory subunit 2 | protein_coding | NP_003927.1 | 1776 | 603764 | . | . | 0.297045 | 0.275825 | 0.294271 | 0.336528 | 0.691236 | 0.637581 | 0.660336 | 0.769431 | 1.087532 | 23.838993 | 0.629461157 | 0.19231968 | 0.847491803 | NA | 4.513800 | 4.634191 | 4.572438 | 4.567326 | 4.570737 | 4.585641 | 27 | 24 | 20 | 25 |
| 8942 | NM_00103299 | KYNU | kynureninase | protein_coding | NP_001028170.1;NP_001186170.1;NP_003928.1;XP_011510404.1;XP_01 | 6469 | 605197 | . | . | 1.415235 | 0.427817 | 1.215398 | 0.458000 | 3.293311 | 0.988920 | 2.727316 | 1.047166 | 1.109874 | 208.449560 | 0.259608564 | 0.579318379 | 0.562374366 | 0.998865404 | 7.625236 | 7.777839 | 7.722264 | 7.619865 | 7.798697 | 7.635799 | 253 | 189 | 224 | 170 |
| 8943 | NM_00126182 | AP3D1 | adaptor related protein complex 3 subunit delta 1 | protein_coding | NP_001248755.1;NP_001361728.1;NP_003929.4;XP_016882911.1 | 568 | 607246 | . | . | 29.459937 | 30.906206 | 27.954900 | 30.909405 | 68.554482 | 71.441079 | 62.729984 | 70.670683 | -1.040792 | 4671.098513 | 0.110493356 | -0.52203987 | 0.601642568 | 0.998865404 | 12.218160 | 12.160353 | 12.191716 | 12.219678 | 12.134207 | 12.209161 | 5395 | 5227 | 3801 | 4485 |
| 8944 | NR_000007 | SNORD73A | small nucleolar RNA, C/D box 73A | snoRNA | . | 10235 | 603568 | . | . | 0.002776 | 0.001428 | 0.003140 | 0.001286 | 0.006460 | 0.003300 | 0.007045 | 0.002940 | 1.234655 | 1.002957 | 2.868339486 | 0.106022226 | 0.915564717 | NA | -0.155866 | 0.148375 | -0.001261 | -0.000972 | 0.000053 | -0.000315 | 1 | 1 | 1 | 1 |
| 8945 | NM_00125685 | BTRC | beta-transducin repeat containing E3 ubiquitin protein ligase | protein_coding | NP_001243785.1;NP_003930.1;NP_378663.1;XP_006718117.1;XP_01153 | 1144 | 603482 | . | . | 2.821588 | 2.900542 | 3.173008 | 3.167470 | 6.565949 | 6.704730 | 7.120138 | 7.242048 | 1.118957 | 556.037215 | 0.161048055 | 1.006874656 | 0.313995013 | 0.998865404 | 9.035617 | 9.197897 | 9.087398 | 9.073300 | 9.157773 | 9.149413 | 611 | 560 | 514 | 547 |
| 8969 | NM_021064 | H2AC11 | H2A clustered histone 11 | protein_coding | NP_066408.1 | 4737 | 615012 | . | . | 0.239714 | 0.206762 | 0.706372 | 1.103903 | 0.557825 | 0.477939 | 1.585079 | 2.523942 | **3.550009** | 9.153902 | 1.052788806 | 1.736172368 | 0.082533361 | NA | 2.008843 | 3.836248 | 3.141645 | 3.138996 | 3.174530 | 3.198953 | 5 | 4 | 10 | 16 |
| 8970 | NM_021058 | H2BC11 | H2B clustered histone 11 | protein_coding | NP_066402.2 | 4761 | 615044 | . | . | 0.653328 | 0.386780 | 0.849854 | 0.262361 | 1.520322 | 0.894059 | 1.907047 | 0.599857 | 1.062301 | 8.743719 | 1.075850534 | 0.081044852 | 0.935406283 | NA | 3.079678 | 3.175234 | 3.127519 | 3.109336 | 3.141369 | 3.099746 | 12 | 7 | 12 | 4 |
| 8971 | NM_006026 | H1-10 | H1.10 linker histone | protein_coding | NP_006017.1 | 4722 | 602785 | . | . | 17.153416 | 18.615509 | 14.338017 | 22.565229 | 39.916702 | 43.030582 | 32.174095 | 51.592712 | 1.017126 | 858.030586 | 0.190001115 | 0.128937448 | 0.897407144 | 0.998865404 | 9.733075 | 9.756600 | 9.706168 | 9.758849 | 9.575844 | 9.889939 | 944 | 952 | 588 | 988 |
| 8972 | NM_00136569 | MGAM | maltase-glucoamylase | protein_coding | NP_001352622.1;NP_004659.2;XP_011514972.1;XP_011514973.1;XP_01 | 7043 | 154360 | . | . | 0.207228 | 0.175687 | 0.171109 | 0.190913 | 0.482228 | 0.406107 | 0.383964 | 0.436501 | -1.027817 | 53.761807 | 0.431266133 | -0.09178527 | 0.926868649 | 0.998865404 | 5.767942 | 5.728813 | 5.774513 | 5.727753 | 5.737344 | 5.754658 | 70 | 52 | 44 | 52 |
| 8973 | NM_00119927 | CHRNA6 | cholinergic receptor nicotinic alpha 6 subunit | protein_coding | NP_001186208.1;NP_004189.1 | 15963 | 606888 | . | . | 3.715726 | 3.861263 | 2.835795 | 2.914267 | 8.646647 | 8.925484 | 6.563442 | 6.663126 | -1.330019 | 249.451210 | 0.216228079 | -1.90283704 | 0.057061812 | 0.708295137 | 8.153787 | 7.742181 | 8.022327 | 8.034762 | 7.884865 | 7.889892 | 324 | 311 | 185 | 202 |
| 8974 | NM_00101797 | P4HA2 | prolyl 4-hydroxylase subunit alpha 2 | protein_coding | NP_001017974.1;NP_001136070.1;NP_001136071.1;NP_001352606.1;N | 8547 | 600608 | . | . | 24.629490 | 22.965262 | 20.680181 | 21.006557 | 57.313837 | 53.085232 | 46.405726 | 48.028999 | -1.173175 | 3342.361697 | 0.111457374 | -2.06732522 | **0.03870352** | 0.61335056 | 11.817256 | 11.586859 | 11.793001 | 11.743663 | 11.628456 | 11.646374 | 4278 | 3779 | 2645 | 2916 |
| 8975 | NM_003940,X | USP13 | ubiquitin specific peptidase 13 | protein_coding | NP_003931.2;XP_011511571.1;XP_016862914.1;XP_016862915.1;XP_01 | 12611 | 603591 | . | . | 3.801383 | 3.670888 | 2.934979 | 3.385296 | 8.845972 | 8.485422 | 6.586006 | 7.740077 | -1.167592 | 808.610205 | 0.148305039 | -1.50727266 | 0.131740782 | 0.885996989 | 9.766920 | 9.543001 | 9.719212 | 9.703436 | 9.561377 | 9.640776 | 1014 | 930 | 611 | 740 |
| 8976 | NM_003941 | WASL | WASP like actin nucleation promoting factor | protein_coding | NP_003932.3 | 12735 | 605056 | . | . | 8.187676 | 7.822762 | 8.555024 | 7.162971 | 19.053059 | 18.082663 | 19.197226 | 16.377281 | -1.032731 | 1078.947965 | 0.1462126 | -0.3177882 | 0.750645601 | 0.998865404 | 10.098153 | 10.052302 | 10.105834 | 10.065674 | 10.129049 | 9.990793 | 1296 | 1151 | 1010 | 902 |
| 8985 | NM_001084 | PLOD3 | procollagen-lysine,2-oxoglutarate 5-dioxygenase 3 | protein_coding | NP_001075.1 | 9083 | 603066 | . | . | 176.601852 | 185.078079 | 161.389175 | 161.935684 | 410.959778 | 427.816254 | 362.152618 | 370.246735 | -1.134212 | 15055.299196 | 0.099829206 | -1.82000627 | 0.06875805 | 0.741130004 | 13.965974 | 13.784277 | 13.912825 | 13.946004 | 13.824497 | 13.818757 | 18073 | 17598 | 12310 | 13182 |
| 8986 | NM_00100694 | RPS6KA4 | ribosomal protein S6 kinase A4 | protein_coding | NP_001006945.1;NP_001287731.1;NP_001305290.1;NP_003933.1;XP_00 | 10433 | 603606 | . | . | 9.121904 | 10.077813 | 9.042742 | 10.517399 | 21.227046 | 23.295315 | 20.291650 | 24.046784 | 1.001564 | 938.898091 | 0.147036542 | 0.015329893 | 0.987768994 | 0.999197585 | 9.874004 | 9.875645 | 9.840299 | 9.905747 | 9.823874 | 9.922752 | 1032 | 1058 | 761 | 941 |
| 8987 | NM_003943 | STBD1 | starch binding domain 1 | protein_coding | NP_003934.1 | 24854 | 607406 | . | . | 5.587494 | 5.496194 | 5.906853 | 5.495395 | 13.002328 | 12.704698 | 13.254807 | 12.564569 | 1.014163 | 517.278560 | 0.166782046 | 0.121651086 | 0.903175353 | 0.998865404 | 9.004398 | 9.025123 | 9.018104 | 9.001517 | 9.045206 | 8.991969 | 599 | 547 | 472 | 468 |
| 8989 | NM_007332,X | TRPA1 | transient receptor potential cation channel subfamily A member 1 | protein_coding | NP_015628.2;XP_011515926.1;XP_011515927.1;XP_016869435.1 | 497 | 604775 | . | . | 4.478871 | 3.996569 | 3.665606 | 3.110052 | 10.422517 | 9.238248 | 8.225513 | 7.110764 | -1.269266 | 617.747507 | 0.168299384 | -2.04394228 | **0.04095925** | 0.628504734 | 9.432214 | 9.089190 | 9.385196 | 9.299672 | 9.238692 | 9.127199 | 844 | 700 | 515 | 466 |
| 8992 | NM_003945 | ATP6V0E1 | ATPase H+ transporting V0 subunit e1 | protein_coding | NP_003936.1 | 863 | 603931 | . | . | 206.036087 | 208.992035 | 238.216293 | 205.717041 | 479.454407 | 483.094452 | 534.550476 | 470.347595 | 1.055203 | 9492.434756 | 0.113429291 | 0.683425256 | 0.494338166 | 0.998865404 | 13.173243 | 13.250839 | 13.186004 | 13.190881 | 13.299118 | 13.165020 | 10606 | 9996 | 9140 | 8423 |
| 8994 | NM_014240,X | LIMD1 | LIM domains containing 1 | protein_coding | NP_055055.1;XP_011532509.1 | 6612 | 604543 | . | . | 13.394384 | 14.576766 | 18.991082 | 16.956577 | 31.169279 | 33.694849 | 42.615435 | 38.769200 | 1.274752 | 5016.895872 | 0.118796902 | 2.94803014 | **0.003198059** | 0.159937703 | 12.106838 | 12.457112 | 12.145928 | 12.215350 | 12.445635 | 12.319493 | 4864 | 4961 | 5250 | 4883 |
| 8996 | NM_00118505 | NOL3 | nucleolar protein 3 | protein_coding | NP_001171986.1;NP_001263236.1;NP_001263238.1;NP_001263240.1;N | 7869 | 605235 | . | . | 6.922460 | 7.074270 | 8.683783 | 8.699767 | 16.108852 | 16.352491 | 19.486157 | 19.890986 | 1.215407 | 384.913769 | 0.18252274 | 1.54194137 | 0.123087842 | 0.867712706 | 8.440795 | 8.722279 | 8.524262 | 8.527750 | 8.644515 | 8.635919 | 399 | 376 | 370 | 393 |
| 8997 | NM_00102466 | KALRN | kalirin RhoGEF kinase | protein_coding | NP_001019831.2;NP_001309917.1;NP_001309918.1;NP_001309919.1;N | 4814 | 604605 | . | . | 2.131518 | 1.330496 | 1.349485 | 1.484265 | 4.960132 | 3.075503 | 3.028207 | 3.393597 | 1.114878 | 298.846882 | 0.202016161 | 0.776597762 | 0.437396122 | 0.998865404 | 8.142434 | 8.299802 | 8.210509 | 8.172933 | 8.261927 | 8.238473 | 337 | 294 | 280 | 289 |
| 8999 | NM_00133072 | CDKL2 | cyclin dependent kinase like 2 | protein_coding | NP_001317653.1;NP_003939.1;XP_006714469.1;XP_016864298.1;XP_01 | 1782 | 603442 | . | . | 0.430440 | 0.462510 | 0.630212 | 0.479690 | 1.001651 | 1.069114 | 1.414179 | 1.096753 | 1.140441 | 74.448989 | 0.367590678 | 0.515767341 | 0.606016915 | 0.998865404 | 6.119956 | 6.310141 | 6.191686 | 6.204479 | 6.249134 | 6.212213 | 78 | 77 | 74 | 69 |
| 9013 | NM_00124315 | TAF1C | TATA-box binding protein associated factor, RNA polymerase I sub | protein_coding | NP_001230085.2;NP_001230086.1;NP_001230087.1;NP_001230088.1;N | 11534 | 604905 | . | . | 18.942942 | 18.028145 | 24.624264 | 20.418575 | 44.081006 | 41.672864 | 55.256134 | 46.684645 | 1.203741 | 2432.430600 | 0.130828073 | 2.044859274 | **0.040868734** | 0.62818612 | 11.108046 | 11.375904 | 11.189395 | 11.143737 | 11.393234 | 11.233777 | 2615 | 2313 | 2535 | 2250 |
| 9014 | NM_00131897 | TAF1B | TATA-box binding protein associated factor, RNA polymerase I sub | protein_coding | NP_001305905.1;NP_001305906.1;NP_005671.3;XP_011508712.1;XP_01 | 11533 | 604904 | . | . | 8.662528 | 7.510841 | 7.971229 | 7.922029 | 20.158059 | 17.361646 | 17.887205 | 18.112780 | -1.042686 | 565.215096 | 0.166542109 | -0.36210011 | 0.717277221 | 0.998865404 | 9.172268 | 9.112424 | 9.204124 | 9.104759 | 9.132037 | 9.124756 | 714 | 576 | 484 | 516 |
| 9015 | NM_00120153 | TAF1A | TATA-box binding protein associated factor, RNA polymerase I sub | protein_coding | NP_001188465.1;NP_005672.1;NP_647603.1;XP_005273400.1;XP_00671 | 11532 | 604903 | . | . | 3.844898 | 4.256840 | 3.741784 | 3.014146 | 8.947236 | 9.839877 | 8.396458 | 6.891486 | -1.226043 | 256.231051 | 0.23042864 | -1.27592556 | 0.201981836 | 0.967656221 | 8.140395 | 7.847353 | 8.010864 | 8.082789 | 8.008554 | 7.877111 | 303 | 325 | 228 | 186 |
| 9016 | NM_00128219 | SLC25A14 | solute carrier family 25 member 14 | protein_coding | NP_001269124.1;NP_001269125.1;NP_001269126.1;NP_001269127.1;X | 10984 | 300242 | . | . | 7.971624 | 7.775171 | 7.194707 | 7.196393 | 18.550297 | 17.972655 | 16.144713 | 16.453696 | -1.100569 | 345.860587 | 0.190563027 | -0.72547633 | 0.468159861 | 0.998865404 | 8.501405 | 8.363389 | 8.475737 | 8.445679 | 8.409904 | 8.403494 | 428 | 381 | 288 | 307 |
| 9019 | NM_00114619 | MPZL1 | myelin protein zero like 1 | protein_coding | NP_001139663.1;NP_003944.1;NP_078845.3 | 7226 | 604376 | . | . | 23.644190 | 23.522370 | 26.804111 | 24.124420 | 55.021003 | 54.373012 | 60.147651 | 55.157617 | 1.063539 | 3792.444946 | 0.114768459 | 0.774361524 | 0.43871703 | 0.998865404 | 11.843712 | 11.932739 | 11.867692 | 11.856907 | 11.961277 | 11.863142 | 4260 | 3940 | 3600 | 3449 |
| 9020 | NM_003954,X | MAP3K14 | mitogen-activated protein kinase kinase kinase 14 | protein_coding | NP_003945.2;XP_011523743.1 | 6853 | 604655 | . | . | 2.965214 | 3.440386 | 3.333462 | 3.120159 | 6.900174 | 7.952608 | 7.480192 | 7.133875 | 1.007137 | 450.914420 | 0.178170958 | 0.057584013 | 0.954079984 | 0.998865404 | 8.811698 | 8.821705 | 8.769030 | 8.856377 | 8.831131 | 8.804626 | 481 | 519 | 402 | 415 |
| 9021 | NM_00137893 | SOCS3 | suppressor of cytokine signaling 3 | protein_coding | NP_001365861.1;NP_001365862.1;NP_003946.3 | 19391 | 604176 | . | . | 2.129910 | 2.241505 | 0.762639 | 1.262749 | 4.956388 | 5.181339 | 1.711340 | 2.887126 | **-2.112212** | 134.217844 | 0.311323254 | -3.46506197 | **0.000530109** | **0.043041758** | 7.510393 | 6.428069 | 7.149868 | 7.195571 | 6.843827 | 6.969028 | 199 | 207 | 57 | 100 |
| 9024 | NM_00125662 | BRSK2 | BR serine/threonine kinase 2 | protein_coding | NP_001243556.1;NP_001243558.1;NP_001243559.1;NP_001269147.1;N | 11405 | 609236 | . | . | 3.665256 | 3.285534 | 3.866395 | 3.579835 | 8.529205 | 7.594659 | 8.676080 | 8.184861 | 1.052118 | 474.789540 | 0.171283982 | 0.427924965 | 0.66870575 | 0.998865404 | 8.853807 | 8.927540 | 8.893238 | 8.855253 | 8.923190 | 8.889036 | 549 | 484 | 435 | 444 |
| 9025 | NM_003958,N | RNF8 | ring finger protein 8 | protein_coding | NP_003949.1;NP_898901.1;XP_006715304.1;XP_006715305.1;XP_01686 | 10071 | 611685 | . | . | 5.406642 | 5.265960 | 5.847381 | 5.686956 | 12.581476 | 12.172502 | 13.121353 | 13.002546 | 1.085849 | 569.216734 | 0.165972708 | 0.715918465 | 0.474041696 | 0.998865404 | 9.092532 | 9.210717 | 9.078827 | 9.167210 | 9.156704 | 9.198513 | 586 | 629 | 499 | 573 |
| 9026 | NM_00130309 | HIP1R | huntingtin interacting protein 1 related | protein_coding | NP_001290026.1;NP_001290028.1;NP_003950.1;XP_011537265.1;XP_02 | 18415 | 605613 | . | . | 13.925521 | 12.956365 | 14.451118 | 14.561974 | 32.405261 | 29.949216 | 32.427890 | 33.294221 | 1.074717 | 1903.719073 | 0.122013247 | 0.852011446 | 0.394207739 | 0.998865404 | 10.841631 | 10.945702 | 10.898653 | 10.830214 | 10.917331 | 10.926788 | 2205 | 1894 | 1705 | 1860 |
| 9028 | NM_00127872 | RHBDL1 | rhomboid like 1 | protein_coding | NP_001265649.1;NP_001265650.1;NP_001305662.1;XP_016879338.1;X | 10007 | 603264 | . | . | 1.046256 | 0.936072 | 0.483963 | 0.256317 | 2.434679 | 2.163773 | 1.085999 | 0.586039 | **-2.787311** | 32.837024 | 0.592233795 | -2.4971121 | **0.012520937** | 0.369272325 | 5.592935 | 4.121684 | 5.109436 | 5.072344 | 4.995391 | 4.938155 | 61 | 47 | 21 | 10 |
| 9031 | NM_00137040 | BAZ1B | bromodomain adjacent to zinc finger domain 1B | protein_coding | NP_001357331.1;NP_115784.1 | 961 | 605681 | . | . | 14.231793 | 15.169690 | 14.391977 | 15.950506 | 33.117970 | 35.065419 | 32.295178 | 36.468935 | 1.018828 | 2969.289695 | 0.11547957 | 0.233031968 | 0.815736582 | 0.998865404 | 11.522472 | 11.549208 | 11.517438 | 11.538391 | 11.504153 | 11.580702 | 3344 | 3215 | 2484 | 2941 |
| 9037 | NM_003966,X | SEMA5A | semaphorin 5A | protein_coding | NP_003957.2;XP_006714569.1;XP_006714570.1;XP_011512457.1;XP_01 | 10736 | 609297 | . | . | 0.953825 | 1.210303 | 1.324672 | 1.153125 | 2.219590 | 2.797670 | 2.972524 | 2.636482 | 1.212557 | 482.163682 | 0.177546036 | 1.566090105 | 0.117327524 | 0.857396579 | 8.767413 | 9.045924 | 8.826764 | 8.865379 | 9.022362 | 8.909101 | 486 | 485 | 500 | 452 |
| 9039 | NM_00136386 | UBA3 | ubiquitin like modifier activating enzyme 3 | protein_coding | NP_001350790.1;NP_003959.3;NP_937838.1;XP_011532512.1 | 12470 | 603172 | . | . | 68.764075 | 62.110100 | 67.599987 | 64.769129 | 160.016826 | 143.570279 | 151.692407 | 148.086921 | -1.002749 | 4320.461702 | 0.114150664 | -0.03470026 | 0.972318754 | 0.998865404 | 12.078875 | 12.075062 | 12.122701 | 12.030462 | 12.098167 | 12.052094 | 5257 | 4410 | 3856 | 3933 |
| 9040 | NM_003969 | UBE2M | ubiquitin conjugating enzyme E2 M | protein_coding | NP_003960.1 | 12491 | 603173 | . | . | 58.032864 | 60.995823 | 59.588104 | 66.013519 | 135.044846 | 140.994583 | 133.713974 | 150.932053 | 1.040821 | 2207.696092 | 0.12028829 | 0.479867386 | 0.631321693 | 0.998865404 | 11.079295 | 11.136784 | 11.075098 | 11.108155 | 11.086119 | 11.159977 | 2440 | 2383 | 1868 | 2208 |
| 9043 | NM_00113052 | SPAG9 | sperm associated antigen 9 | protein_coding | NP_001123999.1;NP_001124000.1;NP_001238900.1;NP_003962.3;XP_00 | 14524 | 605430 | . | . | 15.003303 | 14.656056 | 13.377199 | 14.506457 | 34.913300 | 33.878134 | 30.018049 | 33.167287 | -1.115390 | 3584.106767 | 0.109312272 | -1.44127139 | 0.149508028 | 0.910705785 | 11.884034 | 11.726462 | 11.861078 | 11.840415 | 11.748035 | 11.773214 | 4406 | 4028 | 2901 | 3226 |
| 9044 | NM_003972,N | BTAF1 | B-TFIID TATA-box binding protein associated factor 1 | protein_coding | NP_003963.1;XP_011538628.1;XP_011538629.1;XP_011538630.1;XP_01 | 17307 | 605191 | . | . | 5.801180 | 5.941398 | 5.978542 | 4.767596 | 13.499588 | 13.733809 | 13.415675 | 10.900544 | -1.105255 | 1455.465012 | 0.141364491 | -1.02132738 | 0.307099359 | 0.998865404 | 10.577410 | 10.433533 | 10.548712 | 10.537481 | 10.544731 | 10.381087 | 1772 | 1637 | 1332 | 1157 |
| 9045 | NM_00103499 | RPL14 | ribosomal protein L14 | protein_coding | NP_001030168.1;NP_003964.3 | 10305 | 617414 | . | . | 26.888437 | 29.233787 | 27.593275 | 32.684664 | 62.570499 | 67.575209 | 61.918508 | 74.729601 | 1.060083 | 6426.518130 | 0.118063074 | 0.712986807 | 0.475853921 | 0.998865404 | 12.607194 | 12.691226 | 12.591234 | 12.655268 | 12.603401 | 12.738097 | 6911 | 6989 | 5295 | 6692 |
| 9049 | NM_00130295 | AIP | aryl hydrocarbon receptor interacting protein | protein_coding | NP_001289888.1;NP_001289889.1;NP_003968.3;XP_024304529.1 | 358 | 605555 | . | . | 23.239812 | 23.739898 | 21.244537 | 25.266971 | 54.080002 | 54.875840 | 47.672128 | 57.769930 | -1.023374 | 901.536556 | 0.148439837 | -0.22455749 | 0.822323541 | 0.998865404 | 9.833086 | 9.799200 | 9.818172 | 9.829630 | 9.746587 | 9.864267 | 1042 | 992 | 711 | 904 |
| 9052 | NM_003979 | GPRC5A | G protein-coupled receptor class C group 5 member A | protein_coding | NP_003970.1 | 9836 | 604138 | . | . | 0.296028 | 0.210119 | 0.199115 | 0.289223 | 0.688869 | 0.485701 | 0.446809 | 0.661274 | -1.051028 | 51.320539 | 0.455645313 | -0.15758091 | 0.87478705 | 0.998865404 | 5.717216 | 5.644805 | 5.718500 | 5.650783 | 5.644249 | 5.706452 | 71 | 47 | 36 | 55 |
| 9053 | NM_00119860 | MAP7 | microtubule associated protein 7 | protein_coding | NP_001185537.1;NP_001185538.1;NP_001185540.1;NP_001185543.1;N | 6869 | 604108 | . | . | 0.207312 | 0.182353 | 0.186698 | 0.125301 | 0.482423 | 0.421518 | 0.418943 | 0.286488 | -1.211848 | 30.718251 | 0.560669828 | -0.49442439 | 0.621006492 | 0.998865404 | 5.072328 | 4.796563 | 4.951586 | 4.956671 | 4.943446 | 4.917345 | 38 | 37 | 27 | 23 |
| 9054 | NM_00119898 | NFS1 | NFS1 cysteine desulfurase | protein_coding | NP_001185918.1;NP_066923.3 | 15910 | 603485 | . | . | 13.076012 | 12.852900 | 11.679554 | 13.762017 | 30.428416 | 29.710052 | 26.208584 | 31.465215 | -1.034543 | 1205.880083 | 0.13862345 | -0.35343388 | 0.723763189 | 0.998865404 | 10.260338 | 10.210981 | 10.258545 | 10.237344 | 10.163714 | 10.277757 | 1432 | 1305 | 952 | 1196 |
| 9055 | NM_00126758 | PRC1 | protein regulator of cytokinesis 1 | protein_coding | NP_001254509.2;NP_003972.2;NP_955445.2;XP_005255044.1;XP_00672 | 9341 | 603484 | . | . | 79.454949 | 80.472923 | 64.425762 | 75.559791 | 184.894946 | 186.016762 | 144.559549 | 172.781346 | -1.154439 | 7046.772174 | 0.113820202 | -1.82033988 | 0.068707261 | 0.741130004 | 12.882678 | 12.675375 | 12.837703 | 12.842066 | 12.654037 | 12.778887 | 8674 | 8170 | 5263 | 6591 |
| 9057 | NM_00107678 | SLC7A6 | solute carrier family 7 member 6 | protein_coding | NP_001070253.1;NP_003974.3;XP_011521735.1;XP_011521736.1;XP_01 | 11064 | 605641 | . | . | 11.992578 | 12.649433 | 12.399369 | 12.289893 | 27.907224 | 29.239726 | 27.823826 | 28.099383 | 1.013794 | 2360.631850 | 0.116145788 | 0.170169284 | 0.864877008 | 0.998865404 | 11.195053 | 11.214794 | 11.177977 | 11.220146 | 11.220436 | 11.200055 | 2629 | 2596 | 2093 | 2200 |
| 9060 | NM_00101588 | PAPSS2 | 3'-phosphoadenosine 5'-phosphosulfate synthase 2 | protein_coding | NP_001015880.1;NP_004661.2 | 8604 | 603005 | . | . | 1.719672 | 1.755364 | 1.103226 | 1.547122 | 4.001748 | 4.057601 | 2.475607 | 3.537310 | -1.326451 | 176.067319 | 0.264929849 | -1.53841277 | 0.123947723 | 0.871805202 | 7.650233 | 7.240778 | 7.512570 | 7.517373 | 7.331804 | 7.455867 | 230 | 218 | 110 | 165 |
| 9061 | NM_005443,X | PAPSS1 | 3'-phosphoadenosine 5'-phosphosulfate synthase 1 | protein_coding | NP_005434.4;XP_011530702.1;XP_011530703.1 | 8603 | 603262 | . | . | 28.842601 | 29.014729 | 29.118410 | 29.736221 | 67.117918 | 67.068838 | 65.340866 | 67.988341 | 1.003942 | 2287.714580 | 0.115109156 | 0.04930359 | 0.960677359 | 0.998865404 | 11.156860 | 11.162517 | 11.158990 | 11.157223 | 11.157122 | 11.165381 | 2632 | 2461 | 1983 | 2160 |
| 9063 | NM_00132404 | PIAS2 | protein inhibitor of activated STAT 2 | protein_coding | NP_001310975.1;NP_001310976.1;NP_001310977.1;NP_001310978.1;N | 17311 | 603567 | . | . | 2.849156 | 1.964520 | 2.251744 | 1.658691 | 6.630102 | 4.541074 | 5.052846 | 3.792398 | -1.050853 | 595.814643 | 0.176509262 | -0.40542118 | 0.685167946 | 0.998865404 | 9.253579 | 9.182997 | 9.240333 | 9.224657 | 9.284817 | 9.107142 | 711 | 651 | 574 | 471 |
| 9064 | NM_00129760 | MAP3K6 | mitogen-activated protein kinase kinase kinase 6 | protein_coding | NP_001284538.1;NP_004663.3;XP_016858260.1;XP_016858261.1 | 6858 | 604468 | . | . | 6.145586 | 7.152009 | 8.263495 | 7.864583 | 14.301032 | 16.532188 | 18.543043 | 17.981435 | 1.140277 | 1036.227104 | 0.142954631 | 1.324783018 | 0.185243129 | 0.958025663 | 9.919410 | 10.108639 | 9.912394 | 10.015630 | 10.083408 | 10.040123 | 1036 | 1119 | 988 | 1005 |
| 9066 | NM_00125206 | SYT7 | synaptotagmin 7 | protein_coding | NP_001238994.1;NP_001287702.1;NP_001352738.1;NP_001357139.1;N | 11514 | 604146 | . | . | 0.048810 | 0.049081 | 0.052370 | 0.052302 | 0.113584 | 0.113454 | 0.117517 | 0.119582 | 1.087590 | 11.698435 | 0.887527918 | 0.136485937 | 0.891437141 | NA | 3.486115 | 3.607807 | 3.544462 | 3.543726 | 3.550374 | 3.546483 | 13 | 12 | 11 | 11 |
| 9068 | NM_00137676 | ANGPTL1 | angiopoietin like 1 | protein_coding | NP_001363692.1;NP_004664.1 | 489 | 603874 | . | . | 0.017795 | 0.115247 | 0.087851 | 0.081355 | 0.041409 | 0.266400 | 0.197136 | 0.186008 | 1.285718 | 8.882382 | 1.093594776 | 0.331543803 | 0.74023376 | NA | 2.961600 | 3.318300 | 3.107141 | 3.155462 | 3.143170 | 3.140011 | 3 | 14 | 9 | 9 |
| 9069 | NM_00118507 | CLDN12 | claudin 12 | protein_coding | NP_001172001.1;NP_001172002.1;NP_036261.1 | 2034 | 611232 | . | . | 20.764853 | 17.503304 | 18.821111 | 12.010421 | 48.320663 | 40.459671 | 42.234027 | 27.460399 | -1.254136 | 1887.034654 | 0.351666745 | -0.92898551 | 0.352896596 | 0.998865404 | 11.035951 | 10.709423 | 11.028011 | 10.882016 | 10.939160 | 10.586932 | 2634 | 2063 | 1786 | 1214 |
| 9070 | NM_00110521 | ASH2L | ASH2 like, histone lysine methyltransferase complex subunit | protein_coding | NP_001098684.1;NP_001248761.1;NP_001269201.1;NP_004665.2;XP_00 | 744 | 604782 | . | . | 29.335409 | 27.478486 | 21.029907 | 28.198513 | 68.264705 | 63.517755 | 47.190500 | 64.472550 | -1.137413 | 2374.161233 | 0.138609097 | -1.34014641 | 0.180197749 | 0.952541638 | 11.303254 | 11.117153 | 11.271242 | 11.250873 | 11.052170 | 11.252584 | 2946 | 2693 | 1688 | 2345 |
| 9076 | NM_021101 | CLDN1 | claudin 1 | protein_coding | NP_066924.1 | 2032 | 603718 | . | . | 0.331620 | 0.255737 | 0.121459 | 0.147274 | 0.771693 | 0.591148 | 0.272551 | 0.336725 | **-2.156465** | 23.519151 | 0.653518507 | -1.69646051 | 0.089798703 | NA | 5.006229 | 3.897226 | 4.613574 | 4.577409 | 4.519649 | 4.528161 | 42 | 30 | 12 | 15 |
| 9079 | NM_00113083 | LDB2 | LIM domain binding 2 | protein_coding | NP_001124306.1;NP_001281.1;NP_001291363.1;NP_001291364.1;XP_00 | 6533 | 603450 | . | . | 0.365751 | 0.206239 | 0.136719 | 0.379660 | 0.851118 | 0.476730 | 0.306794 | 0.868047 | -1.137489 | 25.028486 | 0.682272246 | -0.27240173 | 0.785313145 | NA | 4.736899 | 4.547915 | 4.679490 | 4.618882 | 4.592031 | 4.678849 | 39 | 21 | 11 | 32 |
| 9086 | NM_00127861 | EIF1AY | eukaryotic translation initiation factor 1A Y-linked | protein_coding | NP_001265541.1;NP_004672.2 | 3252 | 400014 | . | . | 8.254487 | 9.239994 | 9.191096 | 9.106852 | 19.208527 | 21.358661 | 20.624554 | 20.821738 | 1.031664 | 374.511167 | 0.187110783 | 0.240355277 | 0.810054846 | 0.998865404 | 8.526342 | 8.571041 | 8.507540 | 8.569575 | 8.562806 | 8.551354 | 402 | 419 | 334 | 353 |
| 9088 | NM_00125845 | PKMYT1 | protein kinase, membrane associated tyrosine/threonine 1 | protein_coding | NP_001245379.1;NP_001245380.1;NP_004194.3;NP_872629.1;XP_01152 | 29650 | 602474 | . | . | 27.535214 | 27.429385 | 25.541640 | 25.816818 | 64.075576 | 63.404248 | 57.314703 | 59.027085 | -1.088459 | 1703.805760 | 0.12110095 | -1.00979189 | 0.312595005 | 0.998865404 | 10.794372 | 10.672130 | 10.771177 | 10.762585 | 10.704468 | 10.696632 | 2056 | 1906 | 1426 | 1522 |
| 9091 | NM_004204,N | PIGQ | phosphatidylinositol glycan anchor biosynthesis class Q | protein_coding | NP_004195.2;NP_683721.1 | 14135 | 605754 | . | . | 10.816494 | 10.075790 | 9.256459 | 10.076118 | 25.170423 | 23.290638 | 20.771226 | 23.037847 | -1.096664 | 925.753954 | 0.142896491 | -0.93159325 | 0.351546773 | 0.998865404 | 9.919547 | 9.786350 | 9.915993 | 9.856718 | 9.793087 | 9.846339 | 1160 | 1003 | 739 | 859 |
| 9092 | NM_005146,X | SART1 | spliceosome associated factor 1, recruiter of U4/U6.U5 tri-snRNP | protein_coding | NP_005137.1;XP_011543646.1;XP_011543647.1 | 10538 | 605941 | . | . | 14.700338 | 18.517340 | 15.018332 | 17.613729 | 34.208291 | 42.803662 | 33.700701 | 40.271703 | -1.051752 | 1580.717685 | 0.128217239 | -0.56774471 | 0.570208328 | 0.998865404 | 10.662462 | 10.589340 | 10.610212 | 10.679347 | 10.577097 | 10.634217 | 1784 | 1826 | 1290 | 1498 |
| 9093 | NM_00113511 | DNAJA3 | DnaJ heat shock protein family (Hsp40) member A3 | protein_coding | NP_001128582.1;NP_001273445.1;NP_005138.3 | 11808 | 608382 | . | . | 26.738957 | 28.988607 | 28.307731 | 29.559134 | 62.222649 | 67.008468 | 63.521723 | 67.583448 | 1.023391 | 2316.678093 | 0.117720957 | 0.283362114 | 0.776899267 | 0.998865404 | 11.161146 | 11.194346 | 11.138925 | 11.196810 | 11.174592 | 11.198864 | 2542 | 2560 | 2007 | 2230 |
| 9094 | NM_00133016 | UNC119 | unc-119 lipid binding chaperone | protein_coding | NP_001317095.1;NP_005139.1;NP_473376.1;XP_011523761.1 | 12565 | 604011 | . | . | 15.029446 | 14.502546 | 12.635261 | 16.479659 | 34.974142 | 33.523286 | 28.353161 | 37.678784 | -1.024971 | 673.861030 | 0.168798209 | -0.21080048 | 0.833042962 | 0.998865404 | 9.414340 | 9.378046 | 9.417334 | 9.388726 | 9.299124 | 9.466382 | 802 | 721 | 507 | 701 |
| 9095 | NM_005149 | TBX19 | T-box transcription factor 19 | protein_coding | NP_005140.1 | 11596 | 604614 | . | . | 0.580058 | 0.630322 | 0.576020 | 0.605701 | 1.349819 | 1.457018 | 1.292573 | 1.384863 | -1.030656 | 56.151416 | 0.417612025 | -0.1043159 | 0.916918663 | 0.998865404 | 5.833188 | 5.788975 | 5.806103 | 5.822951 | 5.804082 | 5.812811 | 63 | 64 | 47 | 53 |
| 9096 | NM_00108050 | TBX18 | T-box transcription factor 18 | protein_coding | NP_001073977.1;XP_006715665.2;XP_011534549.2;XP_011534550.2;XP_ | 11595 | 604613 | . | . | 3.062252 | 2.756266 | 3.666995 | 2.346657 | 7.125986 | 6.371233 | 8.228628 | 5.365351 | 1.022375 | 589.258936 | 0.206718014 | 0.154433736 | 0.877267755 | 0.998865404 | 9.186034 | 9.219290 | 9.230866 | 9.143056 | 9.341632 | 9.041867 | 715 | 587 | 635 | 433 |
| 9097 | NM_00103733 | USP14 | ubiquitin specific peptidase 14 | protein_coding | NP_001032411.1;NP_005142.1 | 12612 | 607274 | . | . | 14.315731 | 14.530525 | 14.481323 | 13.068810 | 33.313293 | 33.587959 | 32.495668 | 29.880284 | -1.062478 | 2184.359214 | 0.121449681 | -0.71990936 | 0.471580805 | 0.998865404 | 11.135945 | 11.048726 | 11.113807 | 11.118931 | 11.111353 | 11.022620 | 2583 | 2436 | 1945 | 1875 |
| 9099 | NM_00124375 | USP2 | ubiquitin specific peptidase 2 | protein_coding | NP_001230688.1;NP_004196.4;NP_741994.1;XP_005271778.1;XP_00527 | 12618 | 604725 | . | . | 0.808686 | 0.568473 | 0.744298 | 0.575971 | 1.881848 | 1.314051 | 1.670182 | 1.316889 | -1.053742 | 76.983804 | 0.377697776 | -0.19995092 | 0.841518968 | 0.998865404 | 6.302445 | 6.229602 | 6.313732 | 6.225418 | 6.285950 | 6.227595 | 106 | 71 | 73 | 62 |
| 9100 | NM_00127207 | USP10 | ubiquitin specific peptidase 10 | protein_coding | NP_001259004.1;NP_005144.2;XP_006721395.1;XP_011521745.1;XP_01 | 12608 | 609818 | . | . | 15.839731 | 16.928243 | 16.934613 | 18.976951 | 36.859703 | 39.130391 | 38.000781 | 43.388545 | 1.076243 | 1781.406433 | 0.12695499 | 0.834967468 | 0.403736025 | 0.998865404 | 10.745001 | 10.850667 | 10.742833 | 10.793686 | 10.781002 | 10.869608 | 1912 | 1912 | 1516 | 1828 |
| 9101 | NM_00112861 | USP8 | ubiquitin specific peptidase 8 | protein_coding | NP_001122082.1;NP_001269978.1;NP_005145.3;XP_006720824.1;XP_00 | 12631 | 603158 | . | . | 4.991192 | 5.996255 | 5.327666 | 5.532302 | 11.614710 | 13.860614 | 11.955129 | 12.648952 | -1.095467 | 1285.166309 | 0.12986011 | -1.01298674 | 0.311066496 | 0.998865404 | 10.391920 | 10.260570 | 10.361212 | 10.361317 | 10.316604 | 10.267087 | 1547 | 1450 | 1102 | 1112 |
| 9103 | NR_047648 | FCGR2C | Fc fragment of IgG receptor IIc (gene/pseudogene) | transcribed_ps | . | 15626 | 612169 | . | . | 0.286825 | 0.245526 | 0.182847 | 0.206120 | 0.667454 | 0.567544 | 0.410304 | 0.471268 | -1.358494 | 18.247701 | 0.720172565 | -0.613753 | 0.539378561 | NA | 4.393730 | 3.951852 | 4.213095 | 4.199450 | 4.180555 | 4.184726 | 26 | 21 | 13 | 15 |
| 9107 | NM_004685,X | MTMR6 | myotubularin related protein 6 | protein_coding | NP_004676.3;XP_011533608.1;XP_011533609.1;XP_016876334.1;XP_01 | 7453 | 603561 | . | . | 5.879882 | 4.696590 | 4.021944 | 4.328169 | 13.682726 | 10.856379 | 9.025124 | 9.895844 | -1.297007 | 714.568859 | 0.160404978 | -2.33899495 | **0.019335695** | 0.462516353 | 9.656325 | 9.281219 | 9.623345 | 9.497453 | 9.357732 | 9.408223 | 1010 | 795 | 520 | 606 |
| 9110 | NM_00137806 | MTMR4 | myotubularin related protein 4 | protein_coding | NP_001364995.1;NP_001364996.1;NP_004678.3;XP_005257843.1;XP_00 | 7452 | 603559 | . | . | 6.623780 | 7.175977 | 7.513514 | 7.472160 | 15.413809 | 16.587591 | 16.860107 | 17.084206 | 1.067695 | 1325.220460 | 0.128419542 | 0.735861057 | 0.461815259 | 0.998865404 | 10.324009 | 10.418478 | 10.328047 | 10.364074 | 10.405600 | 10.386511 | 1441 | 1416 | 1205 | 1267 |
| 9111 | NM_004688,X | NMI | N-myc and STAT interactor | protein_coding | NP_004679.2;XP_005246998.1;XP_016860736.1 | 7854 | 603525 | . | . | 4.947066 | 4.806246 | 6.649076 | 4.883128 | 11.512025 | 11.109855 | 14.920333 | 11.164691 | 1.177833 | 210.953274 | 0.248706323 | 0.949452396 | 0.342390573 | 0.998865404 | 7.597068 | 7.834716 | 7.679254 | 7.671279 | 7.826357 | 7.669971 | 225 | 207 | 231 | 179 |
| 9112 | NM_00120325 | MTA1 | metastasis associated 1 | protein_coding | NP_001190187.1;NP_004680.2;XP_011535607.1;XP_011535608.1;XP_01 | 7410 | 603526 | . | . | 13.573108 | 14.771441 | 15.000975 | 15.138794 | 31.585184 | 34.144850 | 33.661754 | 34.613055 | 1.051282 | 1319.922730 | 0.129931084 | 0.555293098 | 0.578694199 | 0.998865404 | 10.329771 | 10.401806 | 10.316394 | 10.375804 | 10.390727 | 10.378361 | 1424 | 1443 | 1186 | 1258 |
| 9113 | NM_00127051 | LATS1 | large tumor suppressor kinase 1 | protein_coding | NP_001257448.1;NP_001337268.1;NP_001337269.1;NP_001337321.1;N | 6514 | 603473 | . | . | 2.375358 | 2.155749 | 2.136366 | 3.481634 | 5.527558 | 4.983111 | 4.793943 | 7.960345 | -1.046780 | 479.914373 | 0.178576493 | -0.36935445 | 0.711863542 | 0.998865404 | 8.938895 | 8.873633 | 8.977834 | 8.858353 | 8.903538 | 8.878936 | 618 | 480 | 416 | 431 |
| 9114 | NM_004691,X | ATP6V0D1 | ATPase H+ transporting V0 subunit d1 | protein_coding | NP_004682.2;XP_011521746.1;XP_011521747.1;XP_011521748.1 | 13724 | 607028 | . | . | 44.158239 | 45.860388 | 44.997743 | 48.358597 | 102.758041 | 106.008341 | 100.973628 | 110.566184 | 1.022251 | 2337.345253 | 0.116903672 | 0.271586175 | 0.785940231 | 0.998865404 | 11.174774 | 11.206363 | 11.169754 | 11.193307 | 11.173951 | 11.223920 | 2623 | 2531 | 1991 | 2284 |
| 9117 | NM_00120157 | SEC22C | SEC22 homolog C, vesicle trafficking protein | protein_coding | NP_001188501.1;NP_001188513.1;NP_004197.1;NP_116752.1;XP_00526 | 16828 | 604028 | . | . | 6.619466 | 6.099159 | 5.967651 | 7.648946 | 15.403771 | 14.098482 | 13.391240 | 17.488410 | 1.024736 | 1115.967145 | 0.136947864 | 0.257415942 | 0.796857697 | 0.998865404 | 10.106438 | 10.141507 | 10.133489 | 10.095231 | 10.094679 | 10.169311 | 1302 | 1159 | 933 | 1112 |
| 9120 | NM_00117416 | SLC16A6 | solute carrier family 16 member 6 | protein_coding | NP_001167637.1;NP_004685.2;XP_005257846.2;XP_011523763.1;XP_01 | 10927 | 603880 | . | . | 54.756917 | 51.344193 | 50.047315 | 39.734316 | 127.421595 | 118.684407 | 112.304722 | 90.847793 | -1.199377 | 5722.667604 | 0.130946378 | -2.00299735 | **0.045177573** | 0.649595749 | 12.607571 | 12.345487 | 12.586019 | 12.517243 | 12.494413 | 12.294557 | 7475 | 6464 | 5071 | 4291 |
| 9121 | NM_00127176 | SLC16A5 | solute carrier family 16 member 5 | protein_coding | NP_001258694.1;NP_001356597.1;NP_004686.1;XP_005257847.1;XP_01 | 10926 | 603879 | . | . | 4.021308 | 4.186563 | 4.255320 | 4.780154 | 9.357750 | 9.677427 | 9.548815 | 10.929253 | 1.072085 | 461.201244 | 0.174042075 | 0.576981921 | 0.563951669 | 0.998865404 | 8.798415 | 8.898360 | 8.824483 | 8.828622 | 8.836043 | 8.902105 | 511 | 482 | 393 | 471 |
| 9122 | NM_00120154 | SLC16A4 | solute carrier family 16 member 4 | protein_coding | NP_001188475.1;NP_001188476.1;NP_001188477.1;NP_001188478.1;N | 10925 | 603878 | . | . | 13.496735 | 13.623528 | 16.854394 | 15.245857 | 31.407454 | 31.491396 | 37.820770 | 34.857840 | 1.146024 | 1193.929412 | 0.13497044 | 1.456891538 | 0.145146322 | 0.906189003 | 10.119642 | 10.316642 | 10.180042 | 10.153639 | 10.309243 | 10.228189 | 1304 | 1179 | 1168 | 1132 |
| 9123 | NM_00104242 | SLC16A3 | solute carrier family 16 member 3 | protein_coding | NP_001035887.1;NP_001035888.1;NP_001193879.1;NP_001193880.1;N | 10924 | 603877 | . | . | 19.955658 | 19.228982 | 16.913589 | 16.650374 | 46.437638 | 44.448654 | 37.953604 | 38.069100 | -1.185863 | 1209.148541 | 0.132605891 | -1.8546467 | 0.063646695 | 0.729663894 | 10.357425 | 10.111674 | 10.330809 | 10.268928 | 10.175926 | 10.168294 | 1571 | 1359 | 967 | 1032 |
| 9124 | NM_020992,X | PDLIM1 | PDZ and LIM domain 1 | protein_coding | NP_066272.1;XP_011538632.1 | 2067 | 605900 | . | . | 13.083649 | 14.131355 | 12.735033 | 14.617702 | 30.446189 | 32.665264 | 28.577044 | 33.421640 | -1.007607 | 607.513625 | 0.162310737 | -0.06735561 | 0.946298605 | 0.998865404 | 9.252567 | 9.240955 | 9.225299 | 9.271839 | 9.201180 | 9.284393 | 678 | 681 | 493 | 603 |
| 9125 | NM_00127163 | CNOT9 | CCR4-NOT transcription complex subunit 9 | protein_coding | NP_001258563.1;NP_001258564.1;NP_005435.1;XP_011510440.1;XP_01 | 10445 | 612054 | . | . | 22.542271 | 23.837163 | 21.763960 | 23.092010 | 52.456789 | 55.100668 | 48.837697 | 52.797137 | -1.007339 | 2608.117983 | 0.116089481 | -0.09087408 | 0.92759265 | 0.998865404 | 11.354147 | 11.343420 | 11.329413 | 11.373303 | 11.320410 | 11.370180 | 2933 | 2901 | 2190 | 2511 |
| 9126 | NM_005445 | SMC3 | structural maintenance of chromosomes 3 | protein_coding | NP_005436.1 | 2468 | 606062 | . | . | 12.555983 | 12.499607 | 12.655454 | 11.612998 | 29.218287 | 28.893402 | 28.398472 | 26.551741 | -1.046851 | 2122.346866 | 0.120623575 | -0.54762245 | 0.58395117 | 0.998865404 | 11.083990 | 11.018149 | 11.074426 | 11.063854 | 11.070039 | 10.993891 | 2516 | 2327 | 1890 | 1851 |
| 9127 | NM_00115955 | P2RX6 | purinergic receptor P2X 6 | protein_coding | NP_001153026.1;NP_001336803.1;NP_001336804.1;NP_001336805.1;N | 8538 | 608077 | . | . | 4.515879 | 3.664337 | 4.446519 | 4.128277 | 10.508636 | 8.470280 | 9.977856 | 9.438817 | 1.093298 | 407.123949 | 0.185523494 | 0.69364197 | 0.487906766 | 0.998865404 | 8.603080 | 8.732659 | 8.674292 | 8.605137 | 8.728939 | 8.657145 | 474 | 395 | 392 | 375 |
| 9128 | NM_00124492 | PRPF4 | pre-mRNA processing factor 4 | protein_coding | NP_001231855.1;NP_001309195.1;NP_001309196.1;NP_004688.2 | 17349 | 607795 | . | . | 13.435624 | 13.893722 | 12.911400 | 14.665417 | 31.265252 | 32.115961 | 28.972807 | 33.530737 | -1.003384 | 1239.691404 | 0.134391075 | -0.03626951 | 0.97106746 | 0.998865404 | 10.278386 | 10.273140 | 10.266552 | 10.286797 | 10.229741 | 10.316531 | 1411 | 1358 | 1014 | 1227 |
| 9129 | NM_00135052 | PRPF3 | pre-mRNA processing factor 3 | protein_coding | NP_001337458.1;NP_004689.1;XP_011508432.1;XP_011508433.1;XP_01 | 17348 | 607301 | . | . | 12.085360 | 12.611957 | 12.960733 | 12.620786 | 28.123127 | 29.153102 | 29.083510 | 28.855926 | -1.020159 | 1089.914415 | 0.137961259 | -0.20871461 | 0.834671034 | 0.998865404 | 10.104129 | 10.075730 | 10.106055 | 10.087611 | 10.123419 | 10.039234 | 1283 | 1173 | 992 | 954 |
| 9130 | NM_004699 | FAM50A | family with sequence similarity 50 member A | protein_coding | NP_004690.1 | 18786 | 300453 | . | . | 42.496071 | 42.875050 | 38.497391 | 40.021648 | 98.890114 | 99.107597 | 86.387032 | 91.504745 | -1.102917 | 1707.455839 | 0.121310659 | -1.16497476 | 0.244029211 | 0.992733415 | 10.806600 | 10.665202 | 10.774736 | 10.774976 | 10.687172 | 10.709007 | 2062 | 1933 | 1392 | 1544 |
| 9131 | NM_00113084 | AIFM1 | apoptosis inducing factor mitochondria associated 1 | protein_coding | NP_001124318.2;NP_001124319.1;NP_004199.1;NP_665811.1 | 8768 | 310490 | . | . | 25.099572 | 25.330278 | 22.207172 | 27.291579 | 58.407736 | 58.552072 | 49.832252 | 62.398944 | -1.034531 | 1728.794950 | 0.13545547 | -0.36157265 | 0.717671408 | 0.998865404 | 10.780026 | 10.730654 | 10.767048 | 10.768387 | 10.661179 | 10.814986 | 2023 | 1899 | 1332 | 1750 |
| 9133 | NM_004701 | CCNB2 | cyclin B2 | protein_coding | NP_004692.1 | 1580 | 602755 | . | . | 25.950628 | 25.525637 | 21.569212 | 24.467810 | 60.388172 | 59.003654 | 48.400688 | 55.942734 | -1.134050 | 1130.773042 | 0.137025766 | -1.3244573 | 0.185351216 | 0.958058933 | 10.231130 | 10.049335 | 10.197362 | 10.178551 | 10.050940 | 10.135466 | 1401 | 1281 | 868 | 1051 |
| 9134 | NM_057749,X | CCNE2 | cyclin E2 | protein_coding | NP_477097.1;XP_011515668.1;XP_016869447.1;XP_016869448.1 | 1590 | 603775 | . | . | 4.998114 | 4.904497 | 4.459305 | 3.362172 | 11.630817 | 11.336965 | 10.006552 | 7.687207 | -1.266754 | 363.040090 | 0.204020057 | -1.67207069 | 0.094510384 | 0.81228321 | 8.663887 | 8.324125 | 8.571788 | 8.558485 | 8.504995 | 8.348162 | 472 | 433 | 319 | 257 |
| 9135 | NM_00108358 | RABEP1 | rabaptin, RAB GTPase binding effector protein 1 | protein_coding | NP_001077054.1;NP_001278510.1;NP_001278511.1;NP_004694.2;XP_01 | 17677 | 603616 | . | . | 13.971640 | 14.448299 | 14.706001 | 15.772733 | 32.512581 | 33.397890 | 32.999841 | 36.062481 | 1.074831 | 2473.068461 | 0.117737746 | 0.884251293 | 0.376560575 | 0.998865404 | 11.219182 | 11.323120 | 11.238891 | 11.244876 | 11.261842 | 11.337024 | 2735 | 2582 | 2125 | 2513 |
| 9136 | NM_004704 | RRP9 | ribosomal RNA processing 9, U3 small nucleolar RNA binding prot | protein_coding | NP_004695.1 | 16829 | . | . | . | 14.333296 | 14.044264 | 13.614019 | 13.454154 | 33.354168 | 32.463947 | 30.549465 | 30.761324 | -1.062733 | 666.283298 | 0.152539844 | -0.57544748 | 0.564988699 | 0.998865404 | 9.423131 | 9.335523 | 9.410446 | 9.390297 | 9.366529 | 9.351689 | 802 | 730 | 568 | 599 |
| 9138 | NM_004706,N | ARHGEF1 | Rho guanine nucleotide exchange factor 1 | protein_coding | NP_004697.2;NP_945328.1;NP_945353.1 | 681 | 601855 | . | . | 22.776386 | 22.583190 | 24.089293 | 24.287666 | 53.001583 | 52.202057 | 54.055681 | 55.530863 | 1.052026 | 2356.367186 | 0.114662032 | 0.638143062 | 0.523380557 | 0.998865404 | 11.165285 | 11.238484 | 11.187972 | 11.174960 | 11.223061 | 11.221580 | 2666 | 2458 | 2103 | 2263 |
| 9139 | NM_00103299 | CBFA2T2 | CBFA2/RUNX1 partner transcriptional co-repressor 2 | protein_coding | NP_001028171.1;NP_001034798.1;NP_005084.1;XP_011527403.1;XP_01 | 1536 | 603672 | . | . | 2.718065 | 2.779826 | 3.073545 | 2.825922 | 6.325047 | 6.425693 | 6.896946 | 6.461135 | 1.044217 | 650.935736 | 0.155214037 | 0.40216388 | 0.687563423 | 0.998865404 | 9.314687 | 9.377375 | 9.325170 | 9.336759 | 9.387583 | 9.332264 | 727 | 693 | 602 | 598 |
| 9140 | NM_00127778 | ATG12 | autophagy related 12 | protein_coding | NP_001264712.1;NP_004698.3 | 588 | 609608 | . | . | 7.996795 | 7.783754 | 9.429554 | 7.194199 | 18.608871 | 17.992496 | 21.159648 | 16.448680 | 1.040111 | 1025.889598 | 0.158650151 | 0.357623395 | 0.720625168 | 0.998865404 | 9.973650 | 10.031097 | 9.997927 | 9.972274 | 10.111017 | 9.907550 | 1178 | 1066 | 1035 | 846 |
| 9141 | NM_004708,X | PDCD5 | programmed cell death 5 | protein_coding | NP_004699.1;XP_005259449.1 | 8764 | 604583 | . | . | 73.769844 | 78.791327 | 56.744558 | 56.332830 | 171.665467 | 182.129669 | 127.333142 | 128.798325 | -1.304861 | 1387.221430 | 0.127441222 | -3.01233804 | **0.002592437** | 0.134524108 | 10.617190 | 10.233314 | 10.513367 | 10.547954 | 10.338770 | 10.315634 | 1769 | 1732 | 1063 | 1111 |
| 9143 | NM_004209 | SYNGR3 | synaptogyrin 3 | protein_coding | NP_004200.2 | 11501 | 603927 | . | . | 0.270733 | 0.212415 | 0.060730 | 0.367023 | 0.630007 | 0.491008 | 0.136275 | 0.839155 | -1.109083 | 14.834147 | 0.885722913 | -0.16863894 | 0.866080651 | NA | 3.966708 | 3.810781 | 3.900344 | 3.884913 | 3.845382 | 3.925184 | 20 | 15 | 4 | 22 |
| 9144 | NM_00132052 | SYNGR2 | synaptogyrin 2 | protein_coding | NP_001307452.1;NP_001350707.1;NP_004701.1 | 11499 | 603926 | . | . | 28.040554 | 26.890006 | 28.153256 | 25.824359 | 65.251527 | 62.157452 | 63.175089 | 59.044327 | -1.034162 | 1301.026038 | 0.132016841 | -0.36709033 | 0.713551643 | 0.998865404 | 10.369288 | 10.321179 | 10.378583 | 10.336402 | 10.365336 | 10.298174 | 1565 | 1388 | 1162 | 1146 |
| 9145 | NM_004711,N | SYNGR1 | synaptogyrin 1 | protein_coding | NP_004702.2;NP_663783.1;NP_663791.1 | 11498 | 603925 | . | . | 51.254897 | 51.487201 | 54.017029 | 56.627302 | 119.272257 | 119.014973 | 121.212645 | 129.471598 | 1.031680 | 2939.569449 | 0.112014365 | 0.401697036 | 0.687907006 | 0.998865404 | 11.498743 | 11.543685 | 11.513521 | 11.503333 | 11.516720 | 11.550777 | 3353 | 3103 | 2542 | 2857 |
| 9146 | NM_004712 | HGS | hepatocyte growth factor-regulated tyrosine kinase substrate | protein_coding | NP_004703.1 | 4897 | 604375 | . | . | 38.356018 | 40.108906 | 49.127003 | 40.936600 | 89.256027 | 92.713531 | 110.239571 | 93.596680 | 1.132493 | 3796.516456 | 0.124728564 | 1.439147198 | 0.150108822 | 0.912861516 | 11.797833 | 11.977498 | 11.820450 | 11.849970 | 12.015715 | 11.854859 | 4026 | 3911 | 3843 | 3418 |
| 9147 | NM_00130173 | NEMF | nuclear export mediator factor | protein_coding | NP_001288661.1;NP_004704.3;XP_011535619.1;XP_011535620.1;XP_01 | 10663 | 608378 | . | . | 7.189641 | 7.552401 | 8.381061 | 6.094264 | 16.730587 | 17.457715 | 18.806856 | 13.933811 | -1.079412 | 1251.689881 | 0.149040344 | -0.73970485 | 0.459479102 | 0.998865404 | 10.343494 | 10.233742 | 10.298845 | 10.332730 | 10.344996 | 10.163470 | 1463 | 1433 | 1174 | 992 |
| 9149 | NM_004714,N | DYRK1B | dual specificity tyrosine phosphorylation regulated kinase 1B | protein_coding | NP_004705.1;NP_006474.1;NP_006475.1;XP_005259455.1 | 3092 | 604556 | . | . | 1.994823 | 1.960616 | 2.483878 | 2.098108 | 4.642035 | 4.532052 | 5.573750 | 4.797075 | 1.172866 | 167.564928 | 0.258007146 | 0.891594804 | 0.372610156 | 0.998865404 | 7.268491 | 7.499430 | 7.354130 | 7.347837 | 7.450402 | 7.382188 | 179 | 165 | 170 | 156 |
| 9150 | NM_00120250 | CTDP1 | CTD phosphatase subunit 1 | protein_coding | NP_001189433.1;NP_001305440.1;NP_004706.3;NP_430255.2;XP_01152 | 2498 | 604927 | . | . | 7.090190 | 6.336299 | 6.137514 | 6.455740 | 16.499166 | 14.646638 | 13.772402 | 14.760281 | -1.007040 | 824.520406 | 0.151514329 | -0.06679569 | 0.946744356 | 0.998865404 | 9.692527 | 9.682278 | 9.728325 | 9.648603 | 9.641515 | 9.725244 | 1006 | 843 | 672 | 816 |
| 9156 | NM_00131922 | EXO1 | exonuclease 1 | protein_coding | NP_001306153.1;NP_003677.4;NP_006018.4;NP_569082.2;XP_0067119 | 3511 | 606063 | . | . | 10.897016 | 11.753901 | 9.230736 | 11.037611 | 25.357808 | 27.169669 | 20.713503 | 25.236185 | -1.138241 | 1137.922313 | 0.142802982 | -1.30813466 | 0.19082764 | 0.963069074 | 10.242843 | 10.055449 | 10.173440 | 10.221758 | 10.038192 | 10.160232 | 1351 | 1349 | 848 | 1081 |
| 9158 | NM_004214,N | FIBP | FGF1 intracellular binding protein | protein_coding | NP_004205.2;NP_942600.1;XP_024304531.1 | 3705 | 608296 | . | . | 31.625293 | 36.148724 | 28.584604 | 35.228796 | 73.593360 | 83.559396 | 64.143020 | 80.546457 | -1.069128 | 1369.273434 | 0.144393436 | -0.66786074 | 0.50422249 | 0.998865404 | 10.466902 | 10.369856 | 10.397379 | 10.486434 | 10.314776 | 10.462970 | 1536 | 1615 | 1038 | 1362 |
| 9159 | NM_004716,X | PCSK7 | proprotein convertase subtilisin/kexin type 7 | protein_coding | NP_004707.2;XP_006719003.1;XP_024304532.1;XP_024304533.1 | 8748 | 604872 | . | . | 7.178135 | 6.992279 | 6.721248 | 7.902302 | 16.703818 | 16.162967 | 15.082286 | 18.067675 | 1.014338 | 932.792618 | 0.145496794 | 0.141163987 | 0.887740396 | 0.998865404 | 9.855302 | 9.875453 | 9.869430 | 9.849821 | 9.816497 | 9.920604 | 1081 | 986 | 758 | 944 |
| 9162 | NM_00132170 | DGKI | diacylglycerol kinase iota | protein_coding | NP_001308637.1;NP_001308638.1;NP_001308639.1;NP_004708.1;XP_01 | 2855 | 604072 | . | . | 0.545263 | 0.518405 | 0.570438 | 0.400895 | 1.268851 | 1.198316 | 1.280045 | 0.916598 | -1.108636 | 210.117834 | 0.251712777 | -0.59109404 | 0.55445741 | 0.998865404 | 7.786629 | 7.639744 | 7.749762 | 7.720255 | 7.763022 | 7.606108 | 262 | 231 | 204 | 154 |
| 9166 | NM_00127893 | EBAG9 | estrogen receptor binding site associated antigen 9 | protein_coding | NP_001265867.1;NP_004206.1;NP_936056.1;XP_016869449.1 | 3123 | 605772 | . | . | 5.562517 | 5.363568 | 5.493656 | 4.945439 | 12.944205 | 12.398126 | 12.327606 | 11.307159 | -1.056927 | 385.138638 | 0.183510003 | -0.43526591 | 0.663369429 | 0.998865404 | 8.628439 | 8.548934 | 8.610969 | 8.599586 | 8.590980 | 8.554496 | 460 | 423 | 336 | 340 |
| 9167 | NM_00131903 | COX7A2L | cytochrome c oxidase subunit 7A2 like | protein_coding | NP_001305965.1;NP_001305966.1;NP_001305967.1;NP_001305969.1;N | 2289 | 605771 | . | . | 38.170345 | 29.045458 | 23.875032 | 32.778990 | 88.823969 | 67.139883 | 53.574883 | 74.945267 | -1.159388 | 1903.343859 | 0.133175499 | -1.60212087 | 0.109128884 | 0.842810112 | 10.997148 | 10.783598 | 10.991180 | 10.905102 | 10.767917 | 10.892237 | 2479 | 2088 | 1410 | 1785 |
| 9168 | NM_021103 | TMSB10 | thymosin beta 10 | protein_coding | NP_066926.1 | 11879 | 188399 | . | . | 580.797241 | 543.381714 | 488.070953 | 616.631714 | 1351.539062 | 1256.051270 | 1095.217041 | 1409.855103 | -1.032600 | 8001.731102 | 0.126179072 | -0.36679266 | 0.713773687 | 0.998865404 | 12.989094 | 12.942727 | 13.009395 | 12.945482 | 12.851882 | 13.040811 | 9713 | 8444 | 6084 | 8203 |
| 9169 | NM_004719,X | SCAF11 | SR-related CTD associated factor 11 | protein_coding | NP_004710.2;XP_005269287.2;XP_011537286.1;XP_011537287.1;XP_01 | 10784 | 603668 | . | . | 6.153663 | 5.603771 | 5.845542 | 5.383766 | 14.319828 | 12.953369 | 13.117228 | 12.309342 | -1.044822 | 1411.392505 | 0.128970033 | -0.49047898 | 0.623795003 | 0.998865404 | 10.494040 | 10.431081 | 10.495137 | 10.463085 | 10.476575 | 10.413860 | 1695 | 1524 | 1250 | 1241 |
| 9170 | NM_004720,X | LPAR2 | lysophosphatidic acid receptor 2 | protein_coding | NP_004711.2;XP_016882957.1;XP_016882958.1;XP_024307542.1 | 3168 | 605110 | . | . | 4.242136 | 4.880611 | 7.474982 | 4.985433 | 9.871626 | 11.281751 | 16.773641 | 11.398599 | 1.302123 | 353.216753 | 0.203363726 | 1.872828207 | 0.061092112 | 0.722512417 | 8.261179 | 8.642515 | 8.344195 | 8.410011 | 8.595186 | 8.456285 | 332 | 351 | 388 | 331 |
| 9172 | NM_003970,X | MYOM2 | myomesin 2 | protein_coding | NP_003961.3;XP_006716300.1 | 7614 | 603509 | . | . | 0.805122 | 0.801254 | 0.854017 | 0.894963 | 1.873554 | 1.852135 | 1.916390 | 2.046226 | 1.075087 | 130.220436 | 0.283493347 | 0.36845048 | 0.712537363 | 0.998865404 | 6.971743 | 7.075998 | 7.011769 | 7.009583 | 7.029375 | 7.043871 | 145 | 135 | 115 | 129 |
| 9173 | NM_00128240 | IL1RL1 | interleukin 1 receptor like 1 | protein_coding | NP_001269337.1;NP_003847.2;NP_057316.3;XP_006712902.1;XP_01151 | 5998 | 601203 | . | . | 0.524505 | 0.675073 | 1.174799 | 0.920081 | 1.220544 | 1.560462 | 2.636214 | 2.103656 | 1.735664 | 100.356291 | 0.335400719 | 2.371753249 | **0.017703909** | 0.446233408 | 6.197167 | 6.992543 | 6.506109 | 6.569505 | 6.749080 | 6.658351 | 74 | 89 | 125 | 104 |
| 9175 | NM_00124231 | MAP3K13 | mitogen-activated protein kinase kinase kinase 13 | protein_coding | NP_001229243.1;NP_001229246.1;NP_004712.1;XP_011511612.1;XP_01 | 6852 | 604915 | . | . | 8.214489 | 9.146076 | 7.894713 | 7.942902 | 19.115453 | 21.141563 | 17.715505 | 18.160502 | -1.095276 | 461.849607 | 0.175437549 | -0.74838367 | 0.45422877 | 0.998865404 | 8.915518 | 8.784046 | 8.840038 | 8.915506 | 8.834128 | 8.809790 | 523 | 552 | 391 | 405 |
| 9179 | NM_00136367 | AP4M1 | adaptor related protein complex 4 subunit mu 1 | protein_coding | NP_001350600.1;NP_004713.2;XP_005250746.1;XP_005250747.1;XP_00 | 574 | 602296 | . | . | 7.449733 | 8.127492 | 7.370943 | 9.089788 | 17.335835 | 18.787061 | 16.540183 | 20.782719 | -1.117177 | 644.767011 | 0.155138885 | -1.03041916 | 0.302813282 | 0.998865404 | 9.410484 | 9.250342 | 9.370294 | 9.367704 | 9.270976 | 9.316926 | 785 | 733 | 511 | 591 |
| 9180 | NM_00116835 | OSMR | oncostatin M receptor | protein_coding | NP_001161827.1;NP_001310433.1;NP_001310434.1;NP_001310435.1;N | 8507 | 601743 | . | . | 4.745181 | 5.541032 | 5.301713 | 4.918414 | 11.042230 | 12.808345 | 11.896890 | 11.245368 | -1.049623 | 679.287514 | 0.160326125 | -0.43580532 | 0.662977982 | 0.998865404 | 9.442441 | 9.372468 | 9.368061 | 9.475533 | 9.416239 | 9.363326 | 739 | 809 | 599 | 597 |
| 9181 | NM_00116238 | ARHGEF2 | Rho/Rac guanine nucleotide exchange factor 2 | protein_coding | NP_001155855.1;NP_001155856.1;NP_001337039.1;NP_001337040.1;N | 682 | 607560 | . | . | 37.762390 | 40.978511 | 44.041954 | 50.841191 | 87.874629 | 94.723665 | 98.828864 | 116.242338 | 1.187442 | 5614.320564 | 0.115201106 | 2.151516665 | **0.031435442** | 0.561754658 | 12.325730 | 12.573441 | 12.349450 | 12.405007 | 12.463453 | 12.577318 | 5713 | 5725 | 4947 | 6095 |
| 9183 | NM_004724,X | ZW10 | zw10 kinetochore protein | protein_coding | NP_004715.1;XP_016874047.1;XP_016874048.1 | 13194 | 603954 | . | . | 8.089058 | 7.828986 | 7.442874 | 7.468378 | 18.823571 | 18.097053 | 16.701594 | 17.075559 | -1.082859 | 691.598734 | 0.151013989 | -0.76049422 | 0.446959224 | 0.998865404 | 9.490008 | 9.375295 | 9.474823 | 9.446110 | 9.407907 | 9.404303 | 845 | 760 | 579 | 621 |
| 9184 | NM_00100779 | BUB3 | BUB3 mitotic checkpoint protein | protein_coding | NP_001007794.1;NP_004716.1 | 1151 | 603719 | . | . | 24.654647 | 28.019259 | 23.767243 | 31.024651 | 57.372376 | 64.767773 | 53.333002 | 70.934181 | -1.114651 | 3216.123507 | 0.117392099 | -1.33392779 | 0.182227524 | 0.953843954 | 11.727175 | 11.570804 | 11.720087 | 11.666314 | 11.647388 | 11.559816 | 4030 | 3540 | 2788 | 2698 |
| 9185 | NM_00108097 | REPS2 | RALBP1 associated Eps domain containing 2 | protein_coding | NP_001074444.1;NP_004717.2;XP_005274682.1;XP_005274683.1;XP_01 | 9963 | 300317 | . | . | 0.428249 | 0.290639 | 0.387493 | 0.331063 | 0.996554 | 0.671825 | 0.869522 | 0.756937 | 1.082015 | 86.988106 | 0.349788784 | 0.325112993 | 0.745095556 | 0.998865404 | 6.383744 | 6.499430 | 6.461952 | 6.394244 | 6.472802 | 6.427853 | 108 | 79 | 85 | 78 |
| 9187 | NM_00125474 | SLC24A1 | solute carrier family 24 member 1 | protein_coding | NP_001241669.1;NP_001287960.1;NP_001287961.1;NP_001287962.1;N | 10975 | 603617 | . | . | 1.875503 | 2.054291 | 2.117144 | 1.851435 | 4.364374 | 4.748585 | 4.750809 | 4.233084 | -1.027897 | 382.247465 | 0.183809117 | -0.21595869 | 0.829019952 | 0.998865404 | 8.597925 | 8.558532 | 8.602055 | 8.570495 | 8.577637 | 8.563022 | 458 | 407 | 332 | 349 |
| 9188 | NM_00125691 | DDX21 | DExD-box helicase 21 | protein_coding | NP_001243839.1;NP_004719.2;XP_011538638.1;XP_016872399.1 | 2744 | 606357 | . | . | 28.196233 | 28.473092 | 27.984710 | 27.886111 | 65.613793 | 65.816828 | 62.796882 | 63.758275 | -1.029264 | 4088.977031 | 0.107045872 | -0.38874594 | 0.697464098 | 0.998865404 | 12.018174 | 11.976575 | 12.009178 | 12.009617 | 11.990292 | 11.980581 | 4775 | 4477 | 3525 | 3756 |
| 9189 | NM_00117113 | ZBED1 | zinc finger BED-type containing 1 | protein_coding | NP_001164606.1;NP_001164607.1;NP_004720.1 | 447 | 300178 | . | . | 4.434354 | 4.432371 | 3.882187 | 4.825635 | 10.318927 | 10.245624 | 8.711515 | 11.033242 | -1.026702 | 621.394598 | 0.166312192 | -0.22859134 | 0.819186557 | 0.998865404 | 9.298586 | 9.259886 | 9.292765 | 9.281761 | 9.201379 | 9.333343 | 731 | 674 | 480 | 632 |
| 9191 | NM_00103971 | DEDD | death effector domain containing | protein_coding | NP_001034800.1;NP_001034801.1;NP_001317694.1;NP_127491.1;XP_00 | 2755 | 606841 | . | . | 7.544368 | 8.007488 | 6.715167 | 7.314474 | 17.556056 | 18.509669 | 15.068641 | 16.723672 | -1.079806 | 596.293127 | 0.160357828 | -0.69077683 | 0.4897058 | 0.998865404 | 9.274322 | 9.163298 | 9.210921 | 9.277275 | 9.195614 | 9.191793 | 678 | 701 | 500 | 536 |
| 9194 | NM_00127062 | SLC16A7 | solute carrier family 16 member 7 | protein_coding | NP_001257551.1;NP_001257552.1;NP_004722.2;XP_005269288.1;XP_01 | 10928 | 603654 | . | . | 0.211782 | 0.136920 | 0.152725 | 0.158664 | 0.492825 | 0.316497 | 0.342711 | 0.362765 | -1.107221 | 65.221755 | 0.413160491 | -0.35565624 | 0.722098019 | 0.998865404 | 6.098085 | 5.952823 | 6.093189 | 5.976400 | 6.010042 | 6.016693 | 98 | 56 | 53 | 59 |
| 9196 | NM_004732,X | KCNAB3 | potassium voltage-gated channel subfamily A regulatory beta sub | protein_coding | NP_004723.2;XP_011522370.1;XP_016880793.1;XP_016880794.1;XP_01 | 6230 | 604111 | . | . | 0.764375 | 1.117815 | 1.053021 | 0.924629 | 1.778733 | 2.583878 | 2.362949 | 2.114056 | 1.015889 | 77.680937 | 0.373479107 | 0.060895059 | 0.951442785 | 0.998865404 | 6.268275 | 6.290616 | 6.228955 | 6.315756 | 6.299143 | 6.255046 | 74 | 97 | 74 | 67 |
| 9197 | NM_00119099 | SLC33A1 | solute carrier family 33 member 1 | protein_coding | NP_001177921.1;NP_001350812.1;NP_004724.1;XP_011511613.1;XP_01 | 95 | 603690 | . | . | 8.437847 | 6.764504 | 9.137867 | 7.263035 | 19.635222 | 15.636452 | 20.505107 | 16.606068 | -1.215923 | 1154.229806 | 0.148965308 | -1.89341038 | 0.058303316 | 0.715342823 | 10.306552 | 10.025180 | 10.257832 | 10.220990 | 10.172816 | 10.007598 | 1493 | 1334 | 1009 | 866 |
| 9200 | NM_014241,X | HACD1 | 3-hydroxyacyl-CoA dehydratase 1 | protein_coding | NP_055056.3;XP_005252698.1 | 9639 | 610467 | . | . | 14.022524 | 12.521635 | 10.043493 | 9.606969 | 32.630989 | 28.944320 | 22.537305 | 21.965193 | -1.164356 | 563.866377 | 0.175849429 | -1.24840894 | 0.211881338 | 0.975238634 | 9.244298 | 9.025860 | 9.219009 | 9.149782 | 9.154165 | 9.012419 | 734 | 621 | 504 | 434 |
| 9201 | NM_00119541 | DCLK1 | doublecortin like kinase 1 | protein_coding | NP_001182344.1;NP_001182345.1;NP_001182359.1;NP_001317000.1;N | 2700 | 604742 | . | . | 0.613969 | 0.519208 | 0.500649 | 0.518639 | 1.428729 | 1.200173 | 1.123443 | 1.185807 | -1.150468 | 115.350153 | 0.303523089 | -0.66624654 | 0.505253533 | 0.998865404 | 6.947216 | 6.745491 | 6.902843 | 6.843159 | 6.822565 | 6.830966 | 154 | 122 | 92 | 102 |
| 9202 | NM_00135013 | ZMYM4 | zinc finger MYM-type containing 4 | protein_coding | NP_001337067.1;NP_001337068.1;NP_001337069.1;NP_001362582.1;N | 13055 | 613568 | . | . | 5.702687 | 6.165368 | 6.424890 | 5.992622 | 13.270387 | 14.251525 | 14.417268 | 13.701417 | 1.029616 | 1398.334396 | 0.130226858 | 0.323330858 | 0.746444687 | 0.998865404 | 10.428248 | 10.470431 | 10.410953 | 10.463763 | 10.490252 | 10.429023 | 1532 | 1538 | 1283 | 1277 |
| 9203 | NM_00117116 | ZMYM3 | zinc finger MYM-type containing 3 | protein_coding | NP_001164633.1;NP_001164634.1;NP_005087.1;NP_963893.1;XP_00526 | 13054 | 300061 | . | . | 5.757679 | 5.998760 | 6.907245 | 7.283280 | 13.398356 | 13.866399 | 15.499657 | 16.652352 | 1.198759 | 1092.126708 | 0.133562004 | 1.958206665 | 0.050205767 | 0.668264424 | 9.956315 | 10.217710 | 10.011172 | 10.030340 | 10.144558 | 10.165986 | 1129 | 1086 | 1021 | 1132 |

| | | | | | | | | | | | | | | | | | | | | | | | | | | | | | | | | | |
|---|---|---|---|---|---|---|---|---|---|---|---|---|---|---|---|---|---|---|---|---|---|---|---|---|---|---|---|---|---|---|---|---|---|
| 9204 | NM_007167 | ZMYM6 | zinc finger MYM-type containing 6 | protein_coding | NP_009098.3 | 13050 | 613567 | . | . | 3.343163 | 3.467241 | 3.797182 | 2.517695 | 7.779679 | 8.014682 | 8.520766 | 5.756412 | -1.093185 | 524.511089 | 0.202072356 | -0.63609788 | 0.524712625 | 0.998865404 | 9.097099 | 8.969751 | 9.048437 | 9.066271 | 9.122363 | 8.861558 | 622 | 599 | 526 | 373 |
| 9205 | NM_00103964 | ZMYM5 | zinc finger MYM-type containing 5 | protein_coding | NP_001034738.1;NP_001034739.1;NP_001136156.1;XP_005266650.1;XP | 13029 | 616443 | . | . | 3.794243 | 2.468919 | 3.044787 | 3.425524 | 8.829357 | 5.707013 | 6.832415 | 7.832051 | -1.133154 | 271.139572 | 0.234348869 | -0.76955091 | 0.441566336 | 0.998865404 | 8.169626 | 7.990608 | 8.202995 | 8.009260 | 8.063770 | 8.028447 | 391 | 255 | 229 | 230 |
| 9208 | NM_00113755 | LRRFIP1 | LRR binding FLII interacting protein 1 | protein_coding | NP_001131022.1;NP_001131023.1;NP_001131024.1;NP_001131025.1;NI | 6702 | 603256 | . | . | 12.555578 | 15.324779 | 13.457167 | 17.400460 | 29.217353 | 35.423919 | 30.197496 | 39.784080 | 1.038818 | 3444.829841 | 0.134468526 | 0.408597008 | 0.682835427 | 0.998865404 | 11.722630 | 11.777286 | 11.704185 | 11.759819 | 11.657902 | 11.859618 | 3760 | 3770 | 2681 | 3691 |
| 9209 | NM_00113436 | LRRFIP2 | LRR binding FLII interacting protein 2 | protein_coding | NP_001127841.1;NP_001269620.1;NP_001335226.1;NP_001335227.1;NI | 6703 | 614043 | . | . | 14.478996 | 15.654865 | 13.992751 | 14.846630 | 33.693222 | 36.186921 | 31.399323 | 33.945064 | -1.025382 | 1309.047215 | 0.131216264 | -0.27558646 | 0.782865709 | 0.998865404 | 10.372429 | 10.335943 | 10.341079 | 10.385628 | 10.315175 | 10.372867 | 1482 | 1472 | 1081 | 1258 |
| 9212 | NM_00125683 | AURKB | aurora kinase B | protein_coding | NP_001243763.1;NP_001271455.1;NP_001300879.1;NP_001300880.1;NI | 11390 | 604970 | . | . | 55.154009 | 55.148546 | 51.930665 | 51.463092 | 128.345644 | 127.478343 | 116.530908 | 117.664246 | -1.094137 | 1997.444465 | 0.117856431 | -1.10128495 | 0.270772659 | 0.998865404 | 11.027360 | 10.897603 | 10.995541 | 11.001780 | 10.935320 | 10.919080 | 2394 | 2261 | 1676 | 1770 |
| 9213 | NM_00113566 | XPR1 | xenotropic and polytropic retrovirus receptor 1 | protein_coding | NP_001129141.1;NP_001315591.1;NP_004727.2 | 12827 | 605237 | . | . | 4.317678 | 3.838487 | 3.867983 | 3.449173 | 10.047414 | 8.872833 | 8.679641 | 7.886124 | -1.128857 | 1011.810896 | 0.146816973 | -1.19102687 | 0.233643035 | 0.988893316 | 10.067210 | 9.892981 | 10.071645 | 9.974984 | 9.981032 | 9.888574 | 1315 | 1083 | 880 | 834 |
| 9214 | NM_00114247 | FCMR | Fc fragment of IgM receptor | protein_coding | NP_001135945.1;NP_001180267.1;NP_005440.1;XP_005273408.1;XP_00 | 14315 | 606015 | . | . | 4.001534 | 4.916003 | 4.822156 | 3.977775 | 9.311734 | 11.363562 | 10.820778 | 9.094710 | -1.042518 | 407.556495 | 0.193819699 | -0.30993697 | 0.756608886 | 0.998865404 | 8.700530 | 8.640559 | 8.619497 | 8.738884 | 8.702360 | 8.607693 | 432 | 493 | 375 | 344 |
| 9215 | NM_00136294 | LARGE1 | LARGE xylosyl- and glucuronyltransferase 1 | protein_coding | NP_001349878.1;NP_001349880.1;NP_001349882.1;NP_001365553.1;NI | 6511 | 603590 | . | . | 5.638213 | 5.908351 | 5.549059 | 7.174128 | 13.120355 | 13.657418 | 12.451926 | 16.402787 | 1.063921 | 863.943782 | 0.153867076 | 0.580962674 | 0.56126561 | 0.998865404 | 9.709767 | 9.798458 | 9.708957 | 9.752254 | 9.703523 | 9.841799 | 935 | 931 | 700 | 915 |
| 9217 | NM_00119567 | VAPB | VAMP associated protein B and C | protein_coding | NP_001182606.1;NP_004729.1 | 12649 | 605704 | . | . | 9.387842 | 10.097092 | 10.440387 | 10.590591 | 21.845894 | 23.339880 | 23.427923 | 24.214128 | 1.067045 | 2436.255921 | 0.11573459 | 0.808934267 | 0.418552957 | 0.998865404 | 11.202908 | 11.296475 | 11.199715 | 11.246695 | 11.281617 | 11.269950 | 2635 | 2618 | 2203 | 2341 |
| 9218 | NM_003574,N | VAPA | VAMP associated protein A | protein_coding | NP_003565.4;NP_919415.2;XP_011524071.1;XP_016881568.1;XP_02430 | 12648 | 605703 | . | . | 8.789608 | 9.379336 | 8.348131 | 8.304924 | 20.453779 | 21.680756 | 18.732964 | 18.988222 | -1.100630 | 1806.290940 | 0.120695463 | -1.14610423 | 0.251752026 | 0.994172589 | 10.886364 | 10.747946 | 10.837101 | 10.873366 | 10.777642 | 10.782156 | 2130 | 2089 | 1491 | 1617 |
| 9219 | NM_00133029 | MTA2 | metastasis associated 1 family member 2 | protein_coding | NP_001317221.1;NP_004730.2 | 7411 | 603947 | . | . | 21.557646 | 24.279262 | 23.671725 | 25.930553 | 50.165526 | 56.122602 | 53.118663 | 59.287131 | 1.069074 | 2282.319328 | 0.123923819 | 0.777584038 | 0.436814274 | 0.998865404 | 11.107449 | 11.203522 | 11.080303 | 11.173882 | 11.148990 | 11.213410 | 2392 | 2520 | 1969 | 2299 |
| 9220 | NM_004740 | TIAF1 | TGFB1-induced anti-apoptotic factor 1 | protein_coding | NP_004731.2 | 11803 | 609517 | . | . | 0.399033 | 0.657093 | 1.127148 | 0.579105 | 0.928566 | 1.518899 | 2.529288 | 1.324055 | 1.573234 | 45.352603 | 0.515058289 | 1.269240721 | 0.204355221 | 0.969317911 | 5.139116 | 5.793528 | 5.402941 | 5.471226 | 5.577521 | 5.447482 | 31 | 47 | 64 | 35 |
| 9221 | NM_00128438 | NOLC1 | nucleolar and coiled-body phosphoprotein 1 | protein_coding | NP_001271317.1;NP_001271318.1;NP_004732.2;XP_005270330.1 | 15608 | 602394 | . | . | 36.919357 | 38.568811 | 35.038461 | 40.896534 | 85.912860 | 89.153539 | 78.625290 | 93.505062 | -1.000558 | 4459.337387 | 0.113901791 | -0.0070605 | 0.994366582 | 0.999862634 | 12.123096 | 12.122131 | 12.101475 | 12.142938 | 12.073202 | 12.168374 | 5009 | 4933 | 3655 | 4418 |
| 9223 | NM_00103305 | MAGI1 | membrane associated guanylate kinase, WW and PDZ domain cor | protein_coding | NP_001028229.1;NP_001352832.1;NP_001352833.1;NP_001352834.1;NI | 946 | 602625 | . | . | 6.230262 | 6.839093 | 6.575543 | 6.626953 | 14.498076 | 15.808870 | 14.755326 | 15.151742 | 1.002011 | 1589.387089 | 0.125585932 | 0.023074492 | 0.981590853 | 0.998865404 | 10.632882 | 10.635627 | 10.600132 | 10.665334 | 10.633959 | 10.635555 | 1752 | 1785 | 1382 | 1493 |
| 9227 | NM_00130164 | LRAT | lecithin retinol acyltransferase | protein_coding | NP_001288574.1;NP_004735.2 | 6685 | 604863 | . | . | 1.172751 | 1.317233 | 0.846993 | 1.292594 | 2.729037 | 3.044844 | 1.900628 | 2.955364 | -1.185554 | 119.877086 | 0.301873776 | -0.81345519 | 0.41595713 | 0.998865404 | 7.023720 | 6.776529 | 6.924831 | 6.943301 | 6.837997 | 6.912710 | 146 | 144 | 84 | 115 |
| 9230 | NM_004218 | RAB11B | RAB11B, member RAS oncogene family | protein_coding | NP_004209.2 | 9761 | 604198 | . | . | 20.014338 | 20.716471 | 21.603088 | 21.183498 | 46.574181 | 47.887047 | 48.476704 | 48.433552 | 1.035625 | 1034.447638 | 0.135358767 | 0.373091748 | 0.70908016 | 0.998865404 | 9.989156 | 10.039691 | 9.991720 | 10.010991 | 10.038147 | 10.016118 | 1155 | 1111 | 929 | 972 |
| 9231 | NM_004747,XI | DLG5 | discs large MAGUK scaffold protein 5 | protein_coding | NP_004738.3;XP_005270333.1;XP_006718119.1;XP_006718120.1;XP_01 | 2904 | 604090 | . | . | 3.355443 | 4.073393 | 3.435091 | 3.598053 | 7.808252 | 9.415830 | 7.708245 | 8.226522 | -1.042724 | 811.767359 | 0.156475444 | -0.38572922 | 0.699697223 | 0.998865404 | 9.695143 | 9.634055 | 9.617008 | 9.736379 | 9.607449 | 9.688258 | 875 | 969 | 651 | 788 |
| 9232 | NM_00128238 | PTTG1 | PTTG1 regulator of sister chromatid separation, securin | protein_coding | NP_001269311.1;NP_001269312.1;NP_004210.1;XP_024302028.1 | 9690 | 604147 | . | . | 182.945651 | 191.971808 | 150.580816 | 218.394402 | 425.722083 | 443.751447 | 337.899002 | 499.332891 | -1.011200 | 4628.540970 | 0.125550788 | -0.12798246 | 0.898162851 | 0.998865404 | 12.184437 | 12.168201 | 12.189050 | 12.170212 | 12.078643 | 12.254489 | 5420 | 4966 | 3581 | 4769 |
| 9236 | NM_00120445 | CCPG1 | cell cycle progression 1 | protein_coding | NP_001191379.1;NP_001191380.1;NP_004739.3;NP_065790.2 | 24227 | 611326 | . | . | 27.078003 | 22.587899 | 23.006653 | 18.929300 | 63.011623 | 52.212942 | 51.626263 | 43.279596 | -1.209625 | 4323.016915 | 0.143260644 | -1.91650285 | 0.055301121 | 0.697001222 | 12.208453 | 11.934177 | 12.217202 | 12.075703 | 12.094678 | 11.874380 | 5903 | 4682 | 3860 | 3176 |
| 9238 | NM_00126183 | TBRG4 | transforming growth factor beta regulator 4 | protein_coding | NP_001248763.1;NP_004740.2;NP_112162.1;NP_954573.1 | 17443 | 611325 | . | . | 57.344460 | 58.077537 | 57.117585 | 70.131847 | 133.442909 | 134.248828 | 128.170222 | 160.348141 | 1.087725 | 4154.402414 | 0.123916942 | 0.978993157 | 0.32758336 | 0.998865404 | 11.958591 | 12.079718 | 11.981065 | 11.985427 | 11.968007 | 12.131709 | 4571 | 4306 | 3397 | 4454 |
| 9240 | NM_006029 | PNMA1 | PNMA family member 1 | protein_coding | NP_006020.4 | 9158 | 604010 | . | . | 9.388424 | 9.087490 | 9.623817 | 10.256821 | 21.847248 | 21.006140 | 21.595568 | 23.451002 | 1.061796 | 777.751648 | 0.146853161 | 0.589067228 | 0.555816176 | 0.998865404 | 9.559304 | 9.645733 | 9.595921 | 9.566642 | 9.604425 | 9.642062 | 887 | 797 | 678 | 771 |
| 9241 | NM_005450 | NOG | noggin | protein_coding | NP_005441.1 | 7866 | 602991 | . | . | 0.297328 | 0.134056 | 0.077142 | 0.139732 | 0.691894 | 0.309877 | 0.173104 | 0.319482 | **-2.020429** | 9.927805 | 1.018512839 | -0.99621847 | 0.319144007 | NA | 3.732160 | 2.715150 | 3.357700 | 3.313029 | 3.297220 | 3.314144 | 21 | 9 | 4 | 8 |
| 9242 | NM_005098 | MSC | musculin | protein_coding | NP_005089.2 | 7321 | 603628 | . | . | 31.748419 | 32.864548 | 47.483582 | 48.103622 | 73.879883 | 75.967880 | 106.551781 | 109.983223 | 1.459004 | 2564.726059 | 0.11352543 | 4.800547429 | **1.58232E-06** | **0.000418567** | 11.026542 | 11.571480 | 11.141007 | 11.161282 | 11.455488 | 11.457922 | 2371 | 2280 | 2643 | 2857 |
| 9244 | NM_004750 | CRLF1 | cytokine receptor like factor 1 | protein_coding | NP_004741.1 | 2364 | 604237 | . | . | 1.324052 | 1.037682 | 1.255566 | 1.160594 | 3.081123 | 2.398648 | 2.817453 | 2.653561 | 1.017758 | 65.353094 | 0.395497152 | 0.06420816 | 0.94880448 | 0.998865404 | 6.016592 | 6.043649 | 6.055602 | 5.997303 | 6.041083 | 6.020552 | 84 | 61 | 60 | 59 |
| 9245 | NM_004751 | GCNT3 | glucosaminyl (N-acetyl) transferase 3, mucin type | protein_coding | NP_004742.1 | 4205 | 606836 | . | . | 0.144818 | 0.213084 | 0.142496 | 0.083038 | 0.336998 | 0.492553 | 0.319757 | 0.189857 | -1.565584 | 22.935449 | 0.672318919 | -0.96189652 | 0.336101586 | NA | 4.805868 | 4.161867 | 4.516365 | 4.561182 | 4.518150 | 4.475426 | 26 | 36 | 20 | 12 |
| 9246 | NM_004223,N | UBE2L6 | ubiquitin conjugating enzyme E2 L6 | protein_coding | NP_004214.1;NP_937826.1 | 12490 | 603890 | . | . | 14.186806 | 13.555702 | 15.657624 | 18.417683 | 33.013275 | 31.334614 | 35.135258 | 42.109845 | 1.216528 | 589.129752 | 0.166150823 | 1.701887615 | 0.088776438 | 0.802216049 | 9.054354 | 9.336734 | 9.149913 | 9.112806 | 9.204611 | 9.314390 | 630 | 557 | 517 | 655 |
| 9249 | NM_00131922 | DHRS3 | dehydrogenase/reductase 3 | protein_coding | NP_001306154.1;NP_001311299.1;NP_004744.2;XP_006711099.1;XP_01 | 17693 | 612830 | . | . | 6.280275 | 6.157838 | 4.754883 | 4.780529 | 14.614458 | 14.234118 | 10.669821 | 10.930112 | -1.357831 | 336.645277 | 0.193442771 | -2.28131311 | **0.022529926** | 0.49659767 | 8.599021 | 8.157500 | 8.453590 | 8.492030 | 8.302650 | 8.303118 | 432 | 432 | 248 | 268 |
| 9252 | NM_00132222 | RPS6KA5 | ribosomal protein S6 kinase A5 | protein_coding | NP_001309156.1;NP_001309157.1;NP_001309158.1;NP_001309159.1;NI | 10434 | 603607 | . | . | 2.102060 | 1.088745 | 0.810798 | 1.055979 | 4.891577 | 2.516684 | 1.819410 | 2.414373 | -1.127216 | 693.539205 | 0.150788491 | -1.14573482 | 0.251904892 | 0.994172589 | 9.521670 | 9.348823 | 9.483987 | 9.471225 | 9.385087 | 9.406055 | 854 | 786 | 558 | 621 |
| 9253 | NM_00128997 | NUMBL | NUMB like endocytic adaptor protein | protein_coding | NP_001276908.1;NP_001276909.1;NP_004747.1 | 8061 | 604018 | . | . | 4.816406 | 5.762589 | 5.871261 | 6.720408 | 11.207974 | 13.320481 | 13.174940 | 15.365414 | 1.175462 | 642.357695 | 0.166412474 | 1.401502654 | 0.161063815 | 0.933525128 | 9.206377 | 9.438742 | 9.209310 | 9.322687 | 9.332599 | 9.418415 | 622 | 692 | 566 | 691 |
| 9255 | NM_00114241 | AIMP1 | aminoacyl tRNA synthetase complex interacting multifunctional p | protein_coding | NP_001135887.1;NP_001135888.2;NP_004748.2;XP_016864324.1;XP_01 | 10648 | 603605 | . | . | 12.564072 | 12.327832 | 12.587018 | 12.763550 | 29.237111 | 28.496338 | 28.244904 | 29.182343 | 1.003832 | 1223.817655 | 0.129728105 | 0.042537157 | 0.966070491 | 0.998865404 | 10.254368 | 10.259972 | 10.260255 | 10.251084 | 10.266122 | 10.251010 | 1415 | 1310 | 1077 | 1138 |
| 9258 | NM_004225,XI | MFHAS1 | malignant fibrous histiocytoma amplified sequence 1 | protein_coding | NP_004216.2;XP_011542154.1;XP_024303098.1 | 16982 | 605352 | . | . | 2.534304 | 2.636805 | 2.631370 | 2.267374 | 5.897429 | 6.095093 | 5.904718 | 5.184081 | -1.065916 | 496.245085 | 0.172890906 | -0.53267068 | 0.594261566 | 0.998865404 | 9.000012 | 8.908351 | 8.955829 | 8.991997 | 8.976251 | 8.889486 | 573 | 568 | 447 | 419 |
| 9260 | NM_005451,N | PDLIM7 | PDZ and LIM domain 7 | protein_coding | NP_005442.2;NP_976227.1;NP_998801.1;XP_011533000.1;XP_01153300 | 22958 | 605903 | . | . | 43.788978 | 46.967623 | 38.220309 | 39.144324 | 101.898749 | 108.567769 | 85.765268 | 89.498846 | -1.185387 | 2043.027020 | 0.119675953 | -2.05018378 | **0.0403465** | 0.627543139 | 11.113997 | 10.868562 | 11.030846 | 11.088832 | 10.928862 | 10.920905 | 2461 | 2477 | 1634 | 1744 |
| 9261 | NM_004759,N | MAPKAPK2 | MAPK activated protein kinase 2 | protein_coding | NP_004750.1;NP_116584.2;XP_005273410.1;XP_016858299.1 | 6887 | 602006 | . | . | 15.485284 | 15.880606 | 17.307442 | 18.820335 | 36.034896 | 36.708733 | 38.837401 | 43.030458 | 1.138536 | 1682.393306 | 0.124270321 | 1.506229316 | 0.132008316 | 0.885996989 | 10.619789 | 10.806756 | 10.657272 | 10.668353 | 10.730772 | 10.796653 | 1799 | 1710 | 1493 | 1749 |
| 9262 | NM_004226,XI | STK17B | serine/threonine kinase 17b | protein_coding | NP_004217.1;XP_011510470.1;XP_011510471.1;XP_011510472.1;XP_01 | 11396 | 604727 | . | . | 6.287900 | 5.872996 | 5.127173 | 4.290886 | 14.632202 | 13.575694 | 11.505227 | 9.810602 | -1.310858 | 887.061782 | 0.152582319 | -2.55935262 | **0.010486731** | 0.33047568 | 9.974641 | 9.584898 | 9.908215 | 9.851789 | 9.747484 | 9.621535 | 1202 | 1045 | 729 | 654 |
| 9263 | NM_004760,XI | STK17A | serine/threonine kinase 17a | protein_coding | NP_004751.2;XP_016868281.1 | 11395 | 604726 | . | . | 23.305034 | 20.388699 | 16.849746 | 18.115191 | 54.231768 | 47.129391 | 37.810342 | 41.418235 | -1.264309 | 2351.896448 | 0.126561856 | -2.67339122 | **0.007508862** | 0.2720434 | 11.358861 | 11.020571 | 11.347990 | 11.219449 | 11.075086 | 11.118615 | 3259 | 2614 | 1756 | 2002 |
| 9265 | NM_00136758 | CYTH3 | cytohesin 3 | protein_coding | NP_001354509.1;NP_001354510.1;NP_001354511.1;NP_004218.1 | 9504 | 605081 | . | . | 31.884122 | 33.915053 | 35.467504 | 38.666366 | 74.195661 | 78.396163 | 79.588054 | 88.406059 | 1.110867 | 4959.089691 | 0.110127782 | 1.37736866 | 0.168398297 | 0.945638095 | 12.198087 | 12.349653 | 12.210443 | 12.249002 | 12.280972 | 12.353465 | 5285 | 5188 | 4344 | 5103 |
| 9266 | NM_004228,N | CYTH2 | cytohesin 2 | protein_coding | NP_004219.3;NP_059431.1;XP_006723535.1;XP_006723536.1 | 9502 | 602488 | . | . | 9.280491 | 9.656427 | 10.016542 | 9.061049 | 21.596082 | 22.321262 | 22.476830 | 20.717010 | -1.007305 | 1313.435848 | 0.132660594 | -0.0791523 | 0.936911483 | 0.998865404 | 10.364239 | 10.354003 | 10.352905 | 10.369792 | 10.398254 | 10.311793 | 1501 | 1438 | 1203 | 1157 |
| 9267 | NM_00129201 | CYTH1 | cytohesin 1 | protein_coding | NP_001278947.1;NP_001278948.1;NP_001351966.1;NP_001351967.1;NI | 9501 | 182115 | . | . | 5.695975 | 5.370871 | 5.048788 | 5.249526 | 13.254771 | 12.415007 | 11.329333 | 12.002423 | -1.063218 | 578.991623 | 0.159683467 | -0.553827 | 0.57969725 | 0.998865404 | 9.220836 | 9.132612 | 9.213720 | 9.180676 | 9.162279 | 9.151826 | 704 | 628 | 492 | 522 |
| 9270 | NM_00131906 | ITGB1BP1 | integrin subunit beta 1 binding protein 1 | protein_coding | NP_001305995.1;NP_001305996.1;NP_001305997.1;NP_001305998.1;NI | 23927 | 607153 | . | . | 25.683596 | 25.761928 | 26.165234 | 30.917757 | 59.766787 | 59.549849 | 58.714022 | 70.689776 | 1.093588 | 2726.868668 | 0.119743553 | 1.077886148 | 0.281084568 | 0.998865404 | 11.347146 | 11.476035 | 11.379824 | 11.370062 | 11.393947 | 11.498667 | 3018 | 2794 | 2319 | 2841 |
| 9274 | NM_00128652 | BCL7C | BAF chromatin remodeling complex subunit BCL7C | protein_coding | NP_001273455.1;NP_004756.2;XP_011544282.1;XP_016879374.1;XP_01 | 1006 | 605847 | . | . | 13.779328 | 13.192926 | 12.046335 | 15.932609 | 32.065060 | 30.496039 | 27.031627 | 36.428018 | 1.003634 | 383.080434 | 0.201105938 | 0.026022874 | 0.979239094 | 0.998865404 | 8.579370 | 8.583634 | 8.595063 | 8.561424 | 8.494092 | 8.659916 | 453 | 401 | 286 | 411 |
| 9275 | NM_00119724 | BCL7B | BAF chromatin remodeling complex subunit BCL7B | protein_coding | NP_001184173.1;NP_001287990.1;NP_001698.2 | 1005 | 605846 | . | . | 11.902069 | 12.524000 | 11.322155 | 13.794821 | 27.696603 | 28.949787 | 25.406588 | 31.540223 | 1.009672 | 643.869445 | 0.16399389 | 0.084675912 | 0.932519047 | 0.998865404 | 9.324039 | 9.337180 | 9.317161 | 9.335594 | 9.266917 | 9.394573 | 728 | 701 | 509 | 664 |
| 9276 | NM_004766,N | COPB2 | COPI coat complex subunit beta 2 | protein_coding | NP_004757.1;XP_016863002.1;XP_024309599.1 | 2232 | 606990 | . | . | 56.439928 | 59.015222 | 48.118848 | 54.090276 | 131.338030 | 136.416321 | 107.977293 | 123.670994 | -1.146091 | 5923.931718 | 0.107186003 | -1.83533414 | 0.066456196 | 0.733835894 | 12.627357 | 12.430620 | 12.612389 | 12.560634 | 12.450444 | 12.493454 | 7507 | 6620 | 4676 | 5310 |
| 9277 | NM_00116426 | WDR46 | WD repeat domain 46 | protein_coding | NP_001157739.1;NP_005443.3;XP_016866973.1;XP_016866974.1;XP_01 | 13923 | 611440 | . | . | 32.519939 | 32.358023 | 34.036547 | 36.021102 | 75.675240 | 74.797018 | 76.377027 | 82.357971 | 1.028064 | 2426.523672 | 0.115870192 | 0.344606439 | 0.730390264 | 0.998865404 | 11.224644 | 11.264432 | 11.215400 | 11.250851 | 11.237965 | 11.272832 | 2695 | 2639 | 2093 | 2356 |
| 9278 | NM_00114533 | ZBTB22 | zinc finger and BTB domain containing 22 | protein_coding | NP_001138810.1;NP_005444.4 | 13085 | 611439 | . | . | 3.098319 | 3.708866 | 3.176092 | 3.916126 | 7.209915 | 8.573210 | 7.127058 | 8.953758 | 1.027643 | 288.381135 | 0.217728703 | 0.1806808 | 0.85661813 | 0.998865404 | 8.152784 | 8.190634 | 8.114689 | 8.207955 | 8.121148 | 8.229255 | 299 | 334 | 228 | 302 |
| 9282 | NM_004229,XI | MED14 | mediator complex subunit 14 | protein_coding | NP_004220.2;XP_005272758.1;XP_011542302.1;XP_016885451.1 | 2370 | 300182 | . | . | 5.076611 | 5.553567 | 5.640197 | 5.583513 | 11.813483 | 12.837319 | 12.656439 | 12.766038 | -1.005288 | 1327.968615 | 0.130252457 | -0.05841187 | 0.953420561 | 0.998865404 | 10.378911 | 10.371089 | 10.343691 | 10.408868 | 10.366330 | 10.379043 | 1468 | 1498 | 1142 | 1252 |
| 9284 | NM_006985 | NPIPA1 | nuclear pore complex interacting protein family member A1 | protein_coding | NP_008916.2 | 7909 | 606406 | . | . | 2.424063 | 2.451068 | 4.212340 | 2.965644 | 5.640894 | 5.665753 | 9.452371 | 6.780592 | 1.448105 | 102.143579 | 0.332585557 | 1.606103048 | 0.10825127 | 0.838332591 | 6.381780 | 6.917652 | 6.600261 | 6.601007 | 6.773979 | 6.651572 | 96 | 90 | 124 | 93 |
| 9289 | NM_00114577 | ADGRG1 | adhesion G protein-coupled receptor G1 | protein_coding | NP_001139242.1;NP_001139243.1;NP_001139244.1;NP_001139245.1;NI | 4512 | 604110 | . | . | 129.494945 | 137.759945 | 141.549941 | 142.189441 | 301.340059 | 318.438291 | 317.633963 | 325.099271 | 1.029629 | 17460.703501 | 0.100430574 | 0.41944064 | 0.674894128 | 0.998865404 | 14.070609 | 14.112732 | 14.058106 | 14.099582 | 14.116614 | 14.091494 | 19333 | 19015 | 15630 | 16366 |
| 9290 | NM_005683,XI | GPR55 | G protein-coupled receptor 55 | protein_coding | NP_005674.2;XP_005247009.1;XP_011510477.1;XP_011510478.1 | 4511 | 604107 | . | . | 3.818319 | 4.449610 | 3.584498 | 3.889974 | 8.885384 | 10.285473 | 8.043510 | 8.893963 | -1.109503 | 516.752557 | 0.170860612 | -0.87739976 | 0.380269544 | 0.998865404 | 9.086628 | 8.936107 | 8.999092 | 9.089298 | 8.962650 | 8.994388 | 583 | 627 | 415 | 471 |
| 9294 | NM_004230 | S1PR2 | sphingosine-1-phosphate receptor 2 | protein_coding | NP_004221.3 | 3169 | 605111 | . | . | 2.280842 | 2.280626 | 2.336388 | 2.161713 | 5.307614 | 5.271769 | 5.242787 | 4.942499 | -1.027549 | 257.882234 | 0.213996678 | -0.18321516 | 0.854629203 | 0.998865404 | 8.029797 | 7.991080 | 8.019070 | 8.015369 | 8.025229 | 7.981386 | 302 | 281 | 231 | 228 |
| 9295 | NM_00119098 | SRSF11 | serine and arginine rich splicing factor 11 | protein_coding | NP_001177916.1;NP_001337534.1;NP_001337535.1;NP_001337536.1;NI | 10782 | 602010 | . | . | 44.145167 | 41.924700 | 47.471936 | 33.947558 | 102.727614 | 96.910818 | 106.525646 | 77.617060 | -1.004742 | 6073.126218 | 0.156406851 | -0.04364127 | 0.965190352 | 0.998865404 | 12.571541 | 12.564899 | 12.588888 | 12.540963 | 12.715783 | 12.379400 | 7208 | 6382 | 6311 | 4557 |
| 9296 | NM_00119890 | ATP6V1F | ATPase H+ transporting V1 subunit F | protein_coding | NP_001185838.1;NP_004222.2 | 16832 | 607160 | . | . | 139.016216 | 152.553539 | 131.349075 | 158.558692 | 323.496455 | 352.634365 | 294.743499 | 362.525611 | -1.019684 | 3078.569226 | 0.127232259 | -0.22103325 | 0.825066542 | 0.998865404 | 11.602186 | 11.573763 | 11.559627 | 11.628941 | 11.502866 | 11.649498 | 3429 | 3496 | 2413 | 3114 |
| 9297 | NR_002559 | SNORD29 | small nucleolar RNA, C/D box 29 | snoRNA | . | 10151 | 603228 | . | . | 24.563780 | 23.807848 | 26.692047 | 21.591578 | 57.160923 | 55.032906 | 59.896183 | 49.366577 | -1.018448 | 49.317313 | 0.446933866 | -0.05900751 | 0.95294613 | 0.998865404 | 5.636438 | 5.611499 | 5.627160 | 5.622395 | 5.640008 | 5.602515 | 58 | 53 | 47 | 41 |
| 9298 | NR_002560 | SNORD31 | small nucleolar RNA, C/D box 31 | snoRNA | . | 10158 | 603230 | . | . | 10.997509 | 9.529813 | 13.096488 | 10.147933 | 25.591654 | 22.028587 | 29.388136 | 23.202044 | 1.118234 | 23.696865 | 0.637033424 | 0.253082477 | 0.80020448 | NA | 4.482357 | 4.646240 | 4.565567 | 4.548748 | 4.583019 | 4.553141 | 28 | 22 | 25 | 20 |
| 9299 | NR_002561 | SNORD30 | small nucleolar RNA, C/D box 30 | snoRNA | . | 10157 | 603229 | . | . | 10.761274 | 7.822320 | 9.876606 | 7.328543 | 25.041927 | 18.081644 | 22.162817 | 16.755842 | -1.110324 | 19.943511 | 0.696665976 | -0.21671901 | 0.828427331 | NA | 4.389723 | 4.242203 | 4.336874 | 4.307777 | 4.325671 | 4.301330 | 28 | 19 | 19 | 15 |
| 9300 | NR_002562 | SNORD28 | small nucleolar RNA, C/D box 28 | snoRNA | . | 10150 | 603227 | . | . | 10.036579 | 8.127268 | 11.063660 | 3.861496 | 23.355532 | 18.786543 | 24.826532 | 8.828853 | -1.219991 | 19.956857 | 0.736145615 | -0.38969316 | 0.696763455 | NA | 4.452135 | 4.171904 | 4.333117 | 4.312061 | 4.340391 | 4.270064 | 28 | 21 | 23 | 9 |
| 9301 | NR_002563 | SNORD27 | small nucleolar RNA, C/D box 27 | snoRNA | . | 10149 | 603226 | . | . | 45.162479 | 56.156487 | 46.108753 | 46.228916 | 105.094948 | 129.808228 | 103.466705 | 105.696915 | -1.107940 | 109.101148 | 0.313113994 | -0.47228587 | 0.636722757 | 0.998865404 | 6.841974 | 6.693239 | 6.748389 | 6.820717 | 6.752344 | 6.752022 | 118 | 137 | 90 | 97 |
| 9302 | NR_002564 | SNORD26 | small nucleolar RNA, C/D box 26 | snoRNA | . | 10148 | 603225 | . | . | 7.671177 | 11.440844 | 12.918995 | 6.844548 | 17.851145 | 26.446022 | 28.989851 | 15.649245 | 1.041449 | 23.040548 | 0.671528605 | 0.087252231 | 0.930471023 | NA | 4.495467 | 4.556102 | 4.496867 | 4.534068 | 4.549674 | 4.488413 | 21 | 29 | 27 | 15 |
| 9303 | NR_002565 | SNORD25 | small nucleolar RNA, C/D box 25 | snoRNA | . | 10147 | 603224 | . | . | 0.001554 | 0.001965 | 0.003003 | 0.002769 | 0.003617 | 0.004541 | 0.006739 | 0.006331 | 1.234655 | 1.002957 | 2.868339486 | 0.106022226 | 0.915564717 | NA | -0.155866 | 0.148375 | -0.001261 | -0.000972 | 0.000053 | -0.000315 | 1 | 1 | 1 | 1 |
| 9304 | NR_000008 | SNORD22 | small nucleolar RNA, C/D box 22 | snoRNA | . | 10145 | 603223 | . | . | 19.096498 | 21.931499 | 37.961143 | 21.081474 | 44.438335 | 50.695644 | 85.183708 | 48.200283 | 1.414523 | 98.791733 | 0.362946911 | 1.378480871 | 0.16805487 | 0.945638095 | 6.353472 | 6.855678 | 6.533718 | 6.569856 | 6.749189 | 6.554802 | 88 | 94 | 130 | 77 |
| 9306 | NM_004232,XI | SOCS6 | suppressor of cytokine signaling 6 | protein_coding | NP_004223.2;XP_005266840.1;XP_016881575.1;XP_016881576.1 | 16833 | 605118 | . | . | 2.901257 | 3.029844 | 2.785000 | 2.705154 | 6.751346 | 7.003619 | 6.249461 | 6.185013 | -1.088161 | 513.505574 | 0.166207859 | -0.73337389 | 0.463330419 | 0.998865404 | 9.063844 | 8.942058 | 9.018722 | 9.042414 | 8.991741 | 8.961582 | 605 | 588 | 438 | 450 |
| 9308 | NM_00104028 | CD83 | CD83 molecule | protein_coding | NP_001035370.1;NP_001238830.1;NP_004224.1 | 1703 | 604534 | . | . | 7.839651 | 8.788374 | 6.054474 | 7.388288 | 18.243190 | 20.314720 | 13.586064 | 16.892442 | -1.255466 | 546.844159 | 0.173999641 | -1.8863432 | 0.059248722 | 0.718550266 | 9.250243 | 8.920984 | 9.126227 | 9.198098 | 8.956063 | 9.071097 | 664 | 692 | 382 | 497 |
| 9310 | NM_004234,XI | ZNF235 | zinc finger protein 235 | protein_coding | NP_004225.3;XP_005259469.1 | 12866 | 604749 | . | . | 1.171325 | 1.528870 | 1.827611 | 1.886443 | 2.725723 | 3.534051 | 4.101108 | 4.313127 | 1.399947 | 174.340876 | 0.258253263 | 1.879442905 | 0.060184043 | 0.719735192 | 7.183146 | 7.667864 | 7.317853 | 7.403111 | 7.518106 | 7.487101 | 149 | 174 | 183 | 184 |
| 9311 | NM_004769,N | ASIC3 | acid sensing ion channel subunit 3 | protein_coding | NP_004760.1;NP_064717.1;NP_064718.1 | 101 | 611741 | . | . | 0.820236 | 1.000676 | 0.882525 | 0.446880 | 1.908722 | 2.313107 | 1.980361 | 1.021738 | -1.346880 | 56.435905 | 0.452040199 | -0.95040615 | 0.341905918 | 0.998865404 | 6.016010 | 5.589693 | 5.825968 | 5.859884 | 5.836630 | 5.726289 | 69 | 75 | 55 | 31 |
| 9314 | NM_00131405 | KLF4 | Kruppel like factor 4 | protein_coding | NP_001300981.1;NP_004226.3 | 6348 | 602253 | . | . | 1.130342 | 1.293690 | 1.142067 | 1.321525 | 2.630352 | 2.990422 | 2.562765 | 3.021509 | 1.028992 | 114.332744 | 0.304503116 | 0.135404607 | 0.892291982 | 0.998865404 | 6.816984 | 6.856930 | 6.810923 | 6.851346 | 6.818308 | 6.862755 | 122 | 129 | 94 | 116 |
| 9315 | NM_00114247 | NREP | neuronal regeneration related protein | protein_coding | NP_001135946.1;NP_001135947.1;NP_001135948.1;NP_001135949.1;NI | 16834 | 607332 | . | . | 6.230018 | 5.666837 | 3.509197 | 4.151701 | 14.497510 | 13.099149 | 7.874538 | 9.492371 | -1.398605 | 352.796890 | 0.195462324 | -2.4761219 | **0.013281822** | 0.381141357 | 8.684737 | 8.200148 | 8.571995 | 8.524401 | 8.317759 | 8.397075 | 492 | 427 | 236 | 297 |
| 9317 | NM_00100148 | PTER | phosphotriesterase related | protein_coding | NP_001001484.1;NP_001248765.1;NP_001248766.1;NP_001248767.1;NI | 9590 | 604446 | . | . | 3.112036 | 2.909239 | 2.431925 | 2.519257 | 7.241834 | 6.724839 | 5.457169 | 5.759982 | -1.215927 | 320.020649 | 0.196403333 | -1.43611091 | 0.150970786 | 0.913627824 | 8.456092 | 8.174203 | 8.388545 | 8.357900 | 8.266021 | 8.266316 | 415 | 369 | 251 | 271 |
| 9318 | NM_00114388 | COPS2 | COP9 signalosome subunit 2 | protein_coding | NP_001137359.1;NP_004227.1 | 30747 | 604508 | . | . | 8.583564 | 7.226776 | 7.517971 | 5.441428 | 19.974305 | 16.705017 | 16.870107 | 12.441177 | -1.236100 | 1478.138962 | 0.165959375 | -1.84259382 | 0.065388337 | 0.731993537 | 10.674048 | 10.368988 | 10.670650 | 10.527008 | 10.554217 | 10.306546 | 2051 | 1604 | 1339 | 1036 |
| 9319 | NM_00116626 | TRIP13 | thyroid hormone receptor interactor 13 | protein_coding | NP_001159732.1;NP_004228.1;XP_011512465.1 | 12307 | 604507 | . | . | 12.683598 | 11.891380 | 12.091575 | 12.738228 | 29.515252 | 27.487458 | 27.133143 | 29.124446 | 1.000337 | 927.016861 | 0.141566386 | 0.003432262 | 0.997261456 | 0.999862634 | 9.856195 | 9.856708 | 9.884176 | 9.827285 | 9.839021 | 9.873445 | 1109 | 961 | 787 | 890 |
| 9320 | NM_00128421 | TRIP12 | thyroid hormone receptor interactor 12 | protein_coding | NP_001271143.1;NP_001271144.1;NP_001271145.1;NP_001335244.1;NI | 12306 | 604506 | . | . | 30.222424 | 30.047088 | 32.029924 | 32.099169 | 70.328825 | 69.455195 | 71.874222 | 73.390935 | 1.058901 | 9071.759384 | 0.100968248 | 0.817764678 | 0.41349157 | 0.998865404 | 13.105294 | 13.187858 | 13.123627 | 13.120996 | 13.167475 | 13.174505 | 10163 | 9494 | 8081 | 8784 |
| 9321 | NM_00132185 | TRIP11 | thyroid hormone receptor interactor 11 | protein_coding | NP_001308780.1;NP_004230.2;XP_016877276.1;XP_016877277.1 | 12305 | 604505 | . | . | 3.291594 | 3.071685 | 3.222694 | 2.756292 | 7.659674 | 7.100337 | 7.231632 | 6.301936 | -1.152458 | 929.041240 | 0.159283674 | -1.28521607 | 0.198716788 | 0.96621494 | 9.957939 | 9.754062 | 9.932721 | 9.879735 | 9.889804 | 9.710399 | 1186 | 1035 | 846 | 707 |
| 9322 | NM_00128896 | TRIP10 | thyroid hormone receptor interactor 10 | protein_coding | NP_001275891.1;NP_001275892.1;NP_004231.1;XP_005259740.1;XP_00 | 12304 | 604504 | . | . | 13.484790 | 14.112089 | 14.452577 | 14.324312 | 31.379660 | 32.620727 | 32.431164 | 32.750839 | 1.023003 | 885.443743 | 0.140408128 | 0.233677957 | 0.815235002 | 0.998865404 | 9.773767 | 9.806561 | 9.775472 | 9.788326 | 9.801248 | 9.795237 | 999 | 953 | 782 | 836 |
| 9324 | NM_00120136 | HMGN3 | high mobility group nucleosomal binding domain 3 | protein_coding | NP_001188291.1;NP_001188292.1;NP_001305813.1;NP_001305814.1;NI | 12312 | 604502 | . | . | 34.820795 | 32.293193 | 29.862333 | 33.931938 | 81.029421 | 74.647162 | 67.010203 | 77.581343 | -1.044243 | 932.390549 | 0.143808485 | -0.43430755 | 0.664065119 | 0.998865404 | 9.895713 | 9.833191 | 9.913556 | 9.845269 | 9.818205 | 9.878154 | 1148 | 980 | 760 | 891 |
| 9325 | NM_00132192 | TRIP4 | thyroid hormone receptor interactor 4 | protein_coding | NP_001308853.1;NP_057297.2 | 12310 | 604501 | . | . | 6.756558 | 7.507337 | 6.144869 | 7.334264 | 15.722787 | 17.353546 | 13.788907 | 16.768924 | -1.069222 | 433.870771 | 0.184059281 | -0.52462001 | 0.599847371 | 0.998865404 | 8.809084 | 8.711509 | 8.751850 | 8.808363 | 8.686818 | 8.789723 | 493 | 506 | 333 | 427 |
| 9326 | NM_00128143 | ZNHIT3 | zinc finger HIT-type containing 3 | protein_coding | NP_001268361.1;NP_001268362.1;NP_001268363.1;NP_004764.1 | 12309 | 604500 | . | . | 26.391443 | 27.876611 | 25.030372 | 28.942909 | 61.413976 | 64.438038 | 56.167428 | 66.174523 | -1.021999 | 800.446085 | 0.150192629 | -0.20902644 | 0.834427599 | 0.998865404 | 9.660546 | 9.628598 | 9.637401 | 9.666248 | 9.589361 | 9.681446 | 913 | 891 | 643 | 791 |
| 9328 | NM_00112282 | GTF3C5 | general transcription factor IIIC subunit 5 | protein_coding | NP_001116295.1;NP_001273638.1;NP_036219.2;XP_016870804.1;XP_01 | 4668 | 604890 | . | . | 17.898572 | 15.225204 | 16.850681 | 18.670601 | 41.650716 | 35.193739 | 37.812441 | 42.688112 | 1.002509 | 1443.009993 | 0.135664701 | 0.026651401 | 0.978737776 | 0.998865404 | 10.493164 | 10.496565 | 10.523086 | 10.462294 | 10.436406 | 10.550428 | 1725 | 1494 | 1163 | 1455 |
| 9329 | NM_012204,N | GTF3C4 | general transcription factor IIIC subunit 4 | protein_coding | NP_036336.2;XP_016870807.1 | 4667 | 604892 | . | . | 3.618729 | 4.177702 | 3.692840 | 3.740392 | 8.420931 | 9.656946 | 8.286625 | 8.551962 | -1.052874 | 1017.004013 | 0.139633135 | -0.53234739 | 0.594485414 | 0.998865404 | 10.026895 | 9.952361 | 9.967897 | 10.047469 | 9.974334 | 9.966738 | 1137 | 1186 | 866 | 924 |
| 9330 | NM_00120677 | GTF3C3 | general transcription factor IIIC subunit 3 | protein_coding | NP_001193703.1;NP_036218.1;XP_005247022.1 | 4666 | 604888 | . | . | 16.998027 | 16.153616 | 15.017517 | 15.257733 | 39.555106 | 37.339808 | 33.698875 | 34.884994 | -1.115513 | 2090.476955 | 0.118382429 | -1.3321842 | 0.182799669 | 0.954698793 | 11.106285 | 10.948644 | 11.094723 | 11.047895 | 10.981497 | 10.987467 | 2612 | 2310 | 1712 | 1860 |
| 9331 | NM_00133057 | B4GALT6 | beta-1,4-galactosyltransferase 6 | protein_coding | NP_001317499.1;NP_001365038.1;NP_001365039.1;NP_004766.2;XP_0( | 929 | 604017 | . | . | 8.813841 | 7.624396 | 7.737875 | 7.195166 | 20.510169 | 17.624133 | 17.363566 | 16.450892 | -1.124117 | 1190.817557 | 0.138501313 | -1.21870129 | 0.22295759 | 0.982008277 | 10.299463 | 10.131101 | 10.308895 | 10.207531 | 10.195524 | 10.147285 | 1547 | 1270 | 1007 | 1017 |
| 9334 | NM_004776 | B4GALT5 | beta-1,4-galactosyltransferase 5 | protein_coding | NP_004767.1 | 928 | 604016 | . | . | 35.956421 | 35.528614 | 37.280964 | 37.750282 | 83.672066 | 82.125992 | 83.657410 | 86.311539 | 1.034972 | 5387.789479 | 0.104601795 | 0.474106491 | 0.63542397 | 0.998865404 | 12.370462 | 12.420067 | 12.389410 | 12.372202 | 12.408690 | 12.410705 | 6160 | 5655 | 4760 | 5144 |
| 9337 | NM_00130107 | CNOT8 | CCR4-NOT transcription complex subunit 8 | protein_coding | NP_001288002.1;NP_001288003.1;NP_001288004.1;NP_001288006.1;NI | 9207 | 603731 | . | . | 13.644778 | 14.310563 | 15.325331 | 13.211914 | 31.751961 | 33.079509 | 34.389596 | 30.207469 | 1.030312 | 1215.426095 | 0.14110228 | 0.305323005 | 0.760120156 | 0.998865404 | 10.225318 | 10.268846 | 10.233270 | 10.235956 | 10.322694 | 10.187374 | 1376 | 1294 | 1169 | 1053 |
| 9338 | NM_00100663 | TCEAL1 | transcription elongation factor A like 1 | protein_coding | NP_001006640.1;NP_001006641.1;NP_004771.2 | 11616 | 300237 | . | . | 5.725847 | 4.232446 | 5.673263 | 4.848128 | 13.324283 | 9.783491 | 12.730638 | 11.084670 | 1.029186 | 192.506970 | 0.25873272 | 0.160410461 | 0.872557749 | 0.998865404 | 7.566939 | 7.610270 | 7.643399 | 7.509835 | 7.627561 | 7.554463 | 251 | 174 | 184 | 168 |
| 9341 | NM_004781,XI | VAMP3 | vesicle associated membrane protein 3 | protein_coding | NP_004772.1;XP_016858330.1 | 12644 | 603657 | . | . | 59.219873 | 54.073748 | 54.173111 | 55.152962 | 137.807078 | 124.993893 | 121.562890 | 126.100703 | -1.048976 | 3764.528363 | 0.113525466 | -0.60762635 | 0.543435323 | 0.998865404 | 11.912289 | 11.843395 | 11.940165 | 11.852797 | 11.855969 | 11.859772 | 4671 | 3941 | 3188 | 3453 |
| 9342 | NM_004782 | SNAP29 | synaptosome associated protein 29 | protein_coding | NP_004773.1 | 11133 | 604202 | . | . | 9.970596 | 10.677734 | 10.764317 | 11.773112 | 23.201988 | 24.682060 | 24.154816 | 26.917824 | 1.076518 | 1432.101095 | 0.129128297 | 0.823775672 | 0.410067033 | 0.998865404 | 10.429913 | 10.535973 | 10.431327 | 10.477037 | 10.479903 | 10.541171 | 1541 | 1533 | 1240 | 1447 |
| 9343 | NM_00114260 | EFTUD2 | elongation factor Tu GTP binding domain containing 2 | protein_coding | NP_001136077.1;NP_001245282.1;NP_001245283.1;NP_004238.3 | 30858 | 603892 | . | . | 32.092744 | 32.251137 | 30.434032 | 37.217953 | 74.681139 | 74.549948 | 68.293087 | 85.094428 | -1.000911 | 4192.243670 | 0.112422572 | -0.01169082 | 0.990672285 | 0.999408937 | 12.034228 | 12.032786 | 12.031908 | 12.035262 | 11.987914 | 12.075768 | 4820 | 4535 | 3451 | 4136 |
| 9344 | NM_00125204 | TAOK2 | TAO kinase 2 | protein_coding | NP_001238972.1;NP_004774.1;NP_057235.2;XP_011544284.1;XP_01154 | 16835 | 613199 | . | . | 16.763712 | 16.121297 | 17.383804 | 16.150517 | 39.009846 | 37.265100 | 39.008752 | 36.926238 | 1.003385 | 2554.486567 | 0.115458447 | 0.042228402 | 0.966316621 | 0.998865404 | 11.316302 | 11.321329 | 11.330608 | 11.303732 | 11.344028 | 11.295505 | 2985 | 2706 | 2292 | 2327 |
| 9348 | NM_004784,N | NDST3 | N-deacetylase and N-sulfotransferase 3 | protein_coding | NP_004775.1;XP_006714479.1;XP_011530717.1;XP_011530718.1;XP_01 | 7682 | 603950 | . | . | 0.069768 | 0.100790 | 0.093701 | 0.054068 | 0.162351 | 0.232983 | 0.210263 | 0.123619 | -1.211250 | 16.054540 | 0.781645135 | -0.35373664 | 0.723536259 | NA | 4.135531 | 3.861274 | 3.997473 | 4.022158 | 4.012978 | 3.980091 | 17 | 22 | 16 | 10 |
| 9349 | NM_000978 | RPL23 | ribosomal protein L23 | protein_coding | NP_000969.1 | 10316 | 603662 | . | . | 135.482635 | 132.939331 | 126.704300 | 148.002335 | 315.273651 | 307.295227 | 284.320770 | 338.389740 | 1.008849 | 11443.719114 | 0.111450299 | 0.114039519 | 0.909206466 | 0.998865404 | 13.475927 | 13.488582 | 13.489532 | 13.466032 | 13.421547 | 13.545430 | 13300 | 12125 | 9271 | 11556 |
| 9351 | NM_00113001 | SLC9A3R2 | SLC9A3 regulator 2 | protein_coding | NP_001123484.1;NP_001239002.1;NP_001239004.1;NP_001239005.1;NI | 11076 | 606553 | . | . | 1.741701 | 1.487528 | 1.694754 | 2.076579 | 4.053010 | 3.438486 | 3.802978 | 4.747852 | 1.141707 | 122.516300 | 0.302214631 | 0.632637143 | 0.526970626 | 0.998865404 | 6.838265 | 7.029089 | 6.940756 | 6.878413 | 6.915676 | 6.994546 | 144 | 112 | 101 | 136 |
| 9352 | NM_004786,N | TXNL1 | thioredoxin like 1 | protein_coding | NP_004777.1;XP_011524566.1;XP_016881583.1;XP_024307057.1;XP_02 | 12436 | 603049 | . | . | 6.173269 | 7.509102 | 6.596513 | 6.264615 | 14.365452 | 17.357626 | 14.802383 | 14.323298 | -1.100299 | 1210.955555 | 0.134792688 | -1.02301903 | 0.306298847 | 0.998865404 | 10.309210 | 10.171359 | 10.242446 | 10.309355 | 10.230997 | 10.177400 | 1397 | 1429 | 1039 | 1042 |
| 9353 | NM_00128913 | SLIT2 | slit guidance ligand 2 | protein_coding | NP_001276064.1;NP_001276065.1;NP_004778.1;XP_005248268.1;XP_00 | 11086 | 603746 | . | . | 0.013672 | 0.031798 | 0.026108 | 0.008084 | 0.031815 | 0.073502 | 0.058584 | 0.018483 | -1.409027 | 5.429023 | 1.343463721 | -0.36822639 | 0.712704433 | NA | 2.664814 | 2.175234 | 2.433425 | 2.452032 | 2.445009 | 2.423690 | 5 | 9 | 6 | 2 |
| 9354 | NM_00120407 | UBE4A | ubiquitination factor E4A | protein_coding | NP_001191006.1;NP_004779.2 | 12499 | 603753 | . | . | 6.798660 | 6.375721 | 7.404984 | 5.701942 | 15.820761 | 14.737765 | 16.616570 | 13.036811 | -1.018577 | 1247.565313 | 0.153701226 | -0.17277118 | 0.862831289 | 0.998865404 | 10.297781 | 10.271902 | 10.316515 | 10.261273 | 10.371557 | 10.169564 | 1502 | 1309 | 1219 | 1004 |
| 9360 | NM_004792,XI | PPIG | peptidylprolyl isomerase G | protein_coding | NP_004783.2;XP_005247023.1;XP_005247024.1;XP_016860791.1 | 14650 | 606093 | . | . | 9.149638 | 9.947863 | 8.919032 | 8.685833 | 21.291582 | 22.994929 | 20.014051 | 19.859125 | -1.099686 | 1802.445792 | 0.121883917 | -1.12477304 | 0.260685222 | 0.995760118 | 10.882677 | 10.745546 | 10.826265 | 10.876876 | 10.786997 | 10.767115 | 2105 | 2102 | 1512 | 1589 |
| 9361 | NM_00127647 | LONP1 | lon peptidase 1, mitochondrial | protein_coding | NP_001263408.1;NP_001263409.1;NP_004784.2;XP_011526743.1 | 9479 | 605490 | . | . | 38.485149 | 38.539767 | 34.433653 | 40.579285 | 89.556516 | 89.086406 | 77.268119 | 92.779720 | -1.040995 | 3647.016533 | 0.119000213 | -0.48708462 | 0.626198364 | 0.998865404 | 11.861278 | 11.803138 | 11.852325 | 11.844207 | 11.751044 | 11.875221 | 4305 | 3995 | 2873 | 3603 |
| 9363 | NM_004794,XI | RAB33A | RAB33A, member RAS oncogene family | protein_coding | NP_004785.1;XP_016885452.1 | 9773 | 300333 | . | . | 0.815904 | 1.001260 | 1.651284 | 1.256374 | 1.898643 | 2.314456 | 3.705434 | 2.872549 | 1.543961 | 34.326133 | 0.534436827 | 1.172517841 | 0.240989213 | 0.992733415 | 4.754145 | 5.380804 | 5.046388 | 5.069235 | 5.132179 | 5.099992 | 28 | 32 | 40 | 35 |
| 9364 | NM_00101797 | RAB28 | RAB28, member RAS oncogene family | protein_coding | NP_001017979.1;NP_001153073.1;NP_004240.2 | 9768 | 612994 | . | . | 8.024971 | 7.991498 | 7.171151 | 8.562770 | 18.674436 | 18.472703 | 16.091854 | 19.577758 | -1.032220 | 431.807966 | 0.181002281 | -0.25276323 | 0.800451183 | 0.998865404 | 8.777240 | 8.730880 | 8.770589 | 8.756394 | 8.696964 | 8.789196 | 511 | 468 | 341 | 429 |
| 9367 | NM_00119532 | RAB9A | RAB9A, member RAS oncogene family | protein_coding | NP_001182257.1;NP_004242.1 | 9792 | 300284 | . | . | 11.854110 | 11.688954 | 13.921874 | 11.320490 | 27.584999 | 27.019543 | 31.240281 | 25.882954 | 1.027727 | 640.161253 | 0.168259166 | 0.234503437 | 0.814594165 | 0.998865404 | 9.301946 | 9.342354 | 9.355697 | 9.264372 | 9.389245 | 9.266391 | 776 | 635 | 616 | 553 |
| 9368 | NM_004252,XI | SLC9A3R1 | SLC9A3 regulator 1 | protein_coding | NP_004243.1 | 11075 | 604990 | . | . | 8.321684 | 9.900505 | 7.304277 | 10.795927 | 19.364899 | 22.885464 | 16.390585 | 24.683604 | -1.048317 | 575.987619 | 0.195498059 | -0.3482146 | 0.727679025 | 0.998865404 | 9.204176 | 9.134777 | 9.138614 | 9.221640 | 9.020230 | 9.264648 | 637 | 676 | 397 | 626 |
| 9369 | NM_00110525 | NRXN3 | neurexin 3 | protein_coding | NP_001098720.1;NP_001258949.1;NP_001317124.1;NP_001353354.1;NI | 8010 | 600567 | . | . | 1.443146 | 1.301479 | 1.370352 | 1.229234 | 3.358259 | 3.008426 | 3.075032 | 2.810497 | -1.083936 | 383.611545 | 0.191639384 | -0.60676317 | 0.544008092 | 0.998865404 | 8.639982 | 8.524721 | 8.650109 | 8.559262 | 8.593290 | 8.522783 | 494 | 398 | 340 | 324 |
| 9371 | NM_004798,XI | KIF3B | kinesin family member 3B | protein_coding | NP_004789.1 | 6320 | 603754 | . | . | 7.256392 | 7.845329 | 7.912008 | 8.323339 | 16.885923 | 18.134830 | 17.754315 | 19.030327 | 1.060703 | 1494.476360 | 0.126602507 | 0.671558477 | 0.50186482 | 0.998865404 | 10.502403 | 10.587200 | 10.495628 | 10.547541 | 10.552281 | 10.582201 | 1614 | 1618 | 1312 | 1470 |
| 9372 | NM_004799,N | ZFYVE9 | zinc finger FYVE-type containing 9 | protein_coding | NP_004790.2;NP_015563.2;XP_011540739.1;XP_016858334.1 | 6775 | 603755 | . | . | 3.292775 | 3.206510 | 3.378875 | 3.905384 | 7.662423 | 7.411991 | 7.582099 | 8.929194 | 1.037470 | 582.984602 | 0.163207282 | 0.325167838 | 0.745054049 | 0.998865404 | 9.160650 | 9.213494 | 9.197231 | 9.151258 | 9.161976 | 9.234131 | 682 | 596 | 488 | 587 |
| 9373 | NM_00103168 | PLAA | phospholipase A2 activating protein | protein_coding | NP_001026859.1;NP_001308475.1;XP_011516374.1;XP_016870808.1 | 9043 | 603873 | . | . | 4.163133 | 3.978257 | 3.795848 | 3.563492 | 9.687783 | 9.195918 | 8.517770 | 8.147500 | -1.135698 | 561.456782 | 0.163037709 | -1.12599639 | 0.260167048 | 0.995760118 | 9.221675 | 9.038583 | 9.192868 | 9.152635 | 9.113046 | 9.066617 | 708 | 625 | 474 | 475 |
| 9374 | NM_00120410 | PPT2 | palmitoyl-protein thioesterase 2 | protein_coding | NP_001191032.1;NP_005146.4;NP_619731.2 | 9326 | 603298 | . | . | 42.733528 | 43.652556 | 43.255473 | 43.618598 | 99.442681 | 100.904830 | 97.064027 | 99.728740 | -1.015869 | 2468.562761 | 0.113772444 | -0.1996473 | 0.841756432 | 0.998865404 | 11.280767 | 11.258055 | 11.277814 | 11.273785 | 11.262061 | 11.264111 | 2872 | 2678 | 2127 | 2299 |
| 9375 | NM_004800 | TM9SF2 | transmembrane 9 superfamily member 2 | protein_coding | NP_004791.1 | 11865 | 604678 | . | . | 60.297985 | 56.671310 | 52.958378 | 43.233639 | 140.315887 | 130.998260 | 118.837051 | 98.848579 | -1.232763 | 5678.623311 | 0.126021589 | -2.39558081 | **0.016594056** | 0.431581156 | 12.614303 | 12.312603 | 12.581314 | 12.520709 | 12.459495 | 12.284196 | 7475 | 6527 | 4893 | 4263 |
| 9377 | NM_004255 | COX5A | cytochrome c oxidase subunit 5A | protein_coding | NP_004246.2 | 2267 | 603773 | . | . | 78.167122 | 76.550026 | 65.914024 | 87.141945 | 181.898117 | 176.948822 | 147.909164 | 199.239700 | -1.025610 | 2811.456845 | 0.139147379 | -0.26218591 | 0.793178117 | 0.998865404 | 11.475369 | 11.438601 | 11.477255 | 11.453074 | 11.329920 | 11.547661 | 3327 | 3027 | 2091 | 2950 |
| 9379 | NM_00137626 | NRXN2 | neurexin 2 | protein_coding | NP_001363191.1;NP_001363192.1;NP_001363194.1;NP_001363195.1;NI | 8009 | 600566 | . | . | 0.074406 | 0.035542 | 0.040693 | 0.069177 | 0.173147 | 0.082156 | 0.091315 | 0.158165 | 1.044775 | 11.809338 | 0.907015912 | 0.069670901 | 0.944455603 | NA | 3.530084 | 3.592943 | 3.575161 | 3.546043 | 3.550618 | 3.575228 | 17 | 9 | 8 | 14 |
| 9380 | NM_012203,XI | GRHPR | glyoxylate and hydroxypyruvate reductase | protein_coding | NP_036335.1;XP_005251688.1;XP_016870809.1;XP_016870810.1;XP_01 | 4570 | 604296 | . | . | 38.655381 | 39.430525 | 35.042066 | 41.993019 | 89.952658 | 91.145419 | 78.633378 | 96.012046 | -1.010934 | 1726.765249 | 0.133131342 | -0.11784222 | 0.906192679 | 0.998865404 | 10.761889 | 10.745779 | 10.737596 | 10.776552 | 10.679248 | 10.813547 | 1950 | 1920 | 1365 | 1748 |
| 9382 | NM_018714 | COG1 | component of oligomeric golgi complex 1 | protein_coding | NP_061184.1 | 6545 | 606973 | . | . | 7.030278 | 7.497575 | 8.437279 | 9.429765 | 16.359747 | 17.330978 | 18.933008 | 21.560038 | 1.213589 | 760.231870 | 0.150571834 | 1.854795273 | 0.063625467 | 0.729663894 | 9.424178 | 9.702966 | 9.477734 | 9.515956 | 9.594955 | 9.667876 | 769 | 762 | 688 | 821 |
| 9391 | NM_004804,XI | CIAO1 | cytosolic iron-sulfur assembly component 1 | protein_coding | NP_004795.1;XP_024309015.1 | 14280 | 604333 | . | . | 37.039264 | 38.737723 | 42.142543 | 43.794693 | 86.191887 | 89.543984 | 94.566651 | 100.131361 | 1.119262 | 5035.956761 | 0.105982626 | 1.533716568 | 0.125099396 | 0.873049916 | 12.214518 | 12.377010 | 12.234488 | 12.263161 | 12.332242 | 12.354019 | 5378 | 5217 | 4566 | 5051 |
| 9392 | NM_00114262 | TGFBRAP1 | transforming growth factor beta receptor associated protein 1 | protein_coding | NP_001136093.1;NP_001315575.1;NP_004248.2;XP_016860828.1 | 16836 | 606237 | . | . | 17.069928 | 18.985179 | 16.466727 | 18.894279 | 39.722424 | 43.885089 | 36.950855 | 43.199529 | -1.015794 | 3114.012136 | 0.121143265 | -0.18662168 | 0.851957271 | 0.998865404 | 11.615946 | 11.593081 | 11.573679 | 11.645644 | 11.544216 | 11.647898 | 3456 | 3535 | 2516 | 3080 |
| 9394 | NM_004807 | HS6ST1 | heparan sulfate 6-O-sulfotransferase 1 | protein_coding | NP_004798.3 | 5201 | 604846 | . | . | 6.160269 | 6.164473 | 5.627390 | 6.600759 | 14.335199 | 14.249454 | 12.627702 | 15.091850 | -1.021799 | 807.897925 | 0.150815733 | -0.20628953 | 0.836564758 | 0.998865404 | 9.673743 | 9.642142 | 9.667849 | 9.662477 | 9.596184 | 9.700744 | 944 | 878 | 643 | 805 |
| 9397 | NM_00130829 | NMT2 | N-myristoyltransferase 2 | protein_coding | NP_001295224.1;NP_004799.1;XP_005252699.1;XP_006717602.1;XP_01 | 7858 | 603801 | . | . | 7.232862 | 7.094428 | 10.720588 | 8.345819 | 16.831167 | 16.399087 | 24.056688 | 19.081724 | 1.087412 | 963.974653 | 0.137451231 | 0.879578991 | 0.379087424 | 0.998865404 | 9.851101 | 9.972067 | 9.890906 | 9.871277 | 9.941815 | 9.942561 | 1078 | 983 | 873 | 942 |
| 9398 | NM_00125610 | CD101 | CD101 molecule | protein_coding | NP_001243035.1;NP_001243038.1;NP_001243040.1;NP_004249.2;XP_01 | 5949 | 604516 | . | . | 0.156341 | 0.245663 | 0.255831 | 0.208774 | 0.363812 | 0.567860 | 0.574079 | 0.477338 | 1.125851 | 44.069318 | 0.479869806 | 0.356378838 | 0.721556875 | 0.998865404 | 5.373794 | 5.544561 | 5.419080 | 5.472078 | 5.484119 | 5.446974 | 40 | 52 | 45 | 39 |
| 9399 | NM_00125667 | STOML1 | stomatin like 1 | protein_coding | NP_001243601.1;NP_001243602.1;NP_001243603.1;NP_001243604.1;NI | 14560 | 608326 | . | . | 3.106999 | 2.400620 | 2.332415 | 3.270600 | 7.230111 | 5.549140 | 5.233869 | 7.477838 | -1.228431 | 352.036993 | 0.202505439 | -1.46572563 | 0.142723049 | 0.90325825 | 8.600602 | 8.303272 | 8.558629 | 8.467017 | 8.341507 | 8.449312 | 481 | 387 | 247 | 326 |
| 9400 | NM_00100371 | RECQL5 | RecQ like helicase 5 | protein_coding | NP_001003715.1;NP_001003716.1;NP_004250.4;XP_005257875.1;XP_00 | 9950 | 603781 | . | . | 12.951982 | 11.715571 | 15.644305 | 12.652217 | 30.139792 | 27.081067 | 35.105366 | 28.927791 | 1.137449 | 1468.660071 | 0.1395558 | 1.331384397 | 0.183062561 | 0.954800306 | 10.424128 | 10.610431 | 10.491722 | 10.440888 | 10.637963 | 10.488902 | 1635 | 1433 | 1489 | 1326 |
| 9401 | NM_004260,XI | RECQL4 | RecQ like helicase 4 | protein_coding | NP_004251.4;XP_011515686.1;XP_016869480.1;XP_016869481.1;XP_01 | 9949 | 603780 | . | . | 24.993501 | 26.340659 | 29.719920 | 26.805627 | 58.160901 | 60.887615 | 66.690641 | 61.287882 | 1.083672 | 3220.097929 | 0.118477057 | 0.978489831 | 0.327832118 | 0.998865404 | 11.593688 | 11.709756 | 11.604422 | 11.631879 | 11.737166 | 11.628539 | 3499 | 3391 | 3104 | 2934 |
| 9402 | NM_00129182 | GRAP2 | GRB2 related adaptor protein 2 | protein_coding | NP_001278753.1;NP_001278754.1;NP_001278755.1;NP_001278757.1;NI | 4563 | 604518 | . | . | 0.255203 | 0.176450 | 0.086327 | 0.212900 | 0.593868 | 0.407873 | 0.193715 | 0.486771 | -1.368081 | 22.000054 | 0.690802664 | -0.65453321 | 0.512768336 | NA | 4.669607 | 4.213338 | 4.490802 | 4.463490 | 4.419212 | 4.477178 | 33 | 24 | 10 | 24 |
| 9403 | NM_004261,N | SELENOF | selenoprotein F | protein_coding | NP_004252.2;NP_976086.1 | 17705 | 606254 | . | . | 75.426748 | 61.363033 | 65.349788 | 49.883402 | 175.521156 | 141.843407 | 146.643036 | 114.052472 | -1.200665 | 3082.035945 | 0.152410935 | -1.73106653 | 0.083439899 | 0.785005898 | 11.715386 | 11.451940 | 11.743260 | 11.560960 | 11.612118 | 11.388379 | 4293 | 3234 | 2774 | 2262 |
| 9404 | NM_00114399 | LPXN | leupaxin | protein_coding | NP_001137467.1;NP_001294880.1;NP_004802.1;XP_006718813.1;XP_01 | 14061 | 605390 | . | . | 35.753020 | 33.093001 | 36.666561 | 28.954998 | 83.198748 | 76.495950 | 82.278700 | 66.202163 | -1.064350 | 1946.772030 | 0.142186941 | -0.63277879 | 0.526878107 | 0.998865404 | 10.970921 | 10.881428 | 10.976296 | 10.920938 | 10.992859 | 10.797171 | 2391 | 2091 | 1844 | 1549 |
| 9406 | NM_005455,N | ZRANB2 | zinc finger RANBP2-type containing 2 | protein_coding | NP_005446.2;NP_976225.1 | 13058 | 604347 | . | . | 30.026859 | 26.390415 | 30.236210 | 24.660828 | 69.873734 | 61.002633 | 67.849181 | 56.384053 | -1.042595 | 2495.795283 | 0.138219309 | -0.43538228 | 0.663284967 | 0.998865404 | 11.314859 | 11.255090 | 11.354608 | 11.242050 | 11.348587 | 11.176773 | 3136 | 2559 | 2352 | 2047 |
| 9409 | NM_004813,N | PEX16 | peroxisomal biogenesis factor 16 | protein_coding | NP_004804.2;NP_476515.2 | 8857 | 603360 | . | . | 4.229938 | 4.811388 | 3.417725 | 4.847839 | 9.843239 | 11.121738 | 7.669275 | 11.084009 | -1.119780 | 223.497094 | 0.243581132 | -0.67006504 | 0.502816332 | 0.998865404 | 7.884299 | 7.719204 | 7.798979 | 7.857355 | 7.702289 | 7.842278 | 256 | 270 | 155 | 228 |
| 9410 | NM_004814 | SNRNP40 | small nuclear ribonucleoprotein U5 subunit 40 | protein_coding | NP_004805.2 | 30857 | 607797 | . | . | 20.607128 | 20.392534 | 18.914856 | 20.841175 | 47.953632 | 47.138256 | 42.444386 | 47.650871 | -1.046139 | 1016.342657 | 0.138236185 | -0.47074503 | 0.637822822 | 0.998865404 | 10.021460 | 9.956143 | 10.012418 | 9.998477 | 9.940035 | 10.003539 | 1208 | 1111 | 826 | 972 |
| 9411 | NM_00132866 | ARHGAP29 | Rho GTPase activating protein 29 | protein_coding | NP_001315593.1;NP_001315594.1;NP_001315595.1;NP_001315596.1;NI | 30207 | 610496 | . | . | 0.902452 | 0.744016 | 0.825432 | 0.640920 | 2.100044 | 1.719826 | 1.852245 | 1.465386 | -1.137083 | 218.511127 | 0.24547901 | -0.75500414 | 0.450246496 | 0.998865404 | 7.860362 | 7.676938 | 7.846025 | 7.748494 | 7.794770 | 7.678203 | 294 | 226 | 201 | 167 |
| 9412 | NM_00127181 | MED21 | mediator complex subunit 21 | protein_coding | NP_001258740.1;NP_004255.2 | 11473 | 603800 | . | . | 7.184781 | 6.419219 | 7.461161 | 4.907381 | 16.719281 | 14.838313 | 16.742628 | 11.220146 | -1.113712 | 542.052228 | 0.204411686 | -0.7601129 | 0.447187104 | 0.998865404 | 9.157207 | 9.003264 | 9.149638 | 9.069328 | 9.165398 | 8.898409 | 697 | 579 | 540 | 380 |
| 9413 | NM_00112760 | FAM189A2 | family with sequence similarity 189 member A2 | protein_coding | NP_001121080.1;NP_001334924.1;NP_004807.3;XP_016870813.1;XP_01 | 24820 | 607710 | . | . | 4.351505 | 4.402648 | 4.006605 | 3.615419 | 10.126130 | 10.176916 | 8.990704 | 8.266224 | -1.169406 | 323.586527 | 0.197790736 | -1.14148711 | 0.253667272 | 0.994807659 | 8.446194 | 8.221047 | 8.380029 | 8.378698 | 8.324316 | 8.260938 | 402 | 376 | 275 | 263 |
| 9414 | NM_00117041 | TJP2 | tight junction protein 2 | protein_coding | NP_001163885.1;NP_001163886.1;NP_001163887.1;NP_001356799.1;NI | 11828 | 607709 | . | . | 9.011336 | 9.730968 | 5.404258 | 7.123698 | 20.969751 | 22.493570 | 12.126997 | 16.287488 | -1.515555 | 1094.182336 | 0.154789649 | -3.87523421 | **0.000106522** | **0.012897595** | 10.364944 | 9.764250 | 10.197799 | 10.254899 | 9.826825 | 10.002980 | 1463 | 1475 | 660 | 919 |
| 9415 | NM_00128150 | FADS2 | fatty acid desaturase 2 | protein_coding | NP_001268430.1;NP_001268431.1;NP_004256.1 | 3575 | 606149 | . | . | 3.219671 | 3.425468 | 4.602878 | 3.817028 | 7.492308 | 7.918121 | 10.328725 | 8.727182 | 1.237434 | 356.464217 | 0.197018934 | 1.560011021 | 0.118757277 | 0.858706505 | 8.315413 | 8.623407 | 8.395405 | 8.422227 | 8.587836 | 8.470346 | 358 | 352 | 377 | 333 |
| 9416 | NM_004818 | DDX23 | DEAD-box helicase 23 | protein_coding | NP_004809.2 | 17347 | 612172 | . | . | 24.377314 | 26.366915 | 24.901569 | 28.108435 | 56.727009 | 60.948303 | 55.878399 | 64.266602 | 1.030254 | 2630.518491 | 0.120537106 | 0.356733069 | 0.721291646 | 0.998865404 | 11.339603 | 11.382343 | 11.319743 | 11.376307 | 11.326575 | 11.416421 | 2880 | 2894 | 2193 | 2641 |
| 9419 | NM_014171 | CRIPT | CXXC repeat containing interactor of PDZ3 domain | protein_coding | NP_054890.1 | 14312 | 604594 | . | . | 2.691475 | 2.332933 | 2.883466 | 1.887636 | 6.263173 | 5.392680 | 6.470414 | 4.315856 | -1.068833 | 483.288382 | 0.211687384 | -0.45366876 | 0.650067253 | 0.998865404 | 8.963208 | 8.868726 | 8.978256 | 8.881363 | 9.012713 | 8.748699 | 618 | 498 | 493 | 345 |
| 9420 | NM_00132411 | CYP7B1 | cytochrome P450 family 7 subfamily B member 1 | protein_coding | NP_001311041.1;NP_004811.1;XP_016869491.1 | 2652 | 603711 | . | . | 6.020855 | 5.870915 | 4.387645 | 4.269313 | 14.010777 | 13.570884 | 9.845751 | 9.761277 | -1.160769 | 1276.451990 | 0.143549576 | -1.49830209 | 0.134054782 | 0.891755273 | 10.421172 | 10.206712 | 10.405748 | 10.333766 | 10.324812 | 10.185527 | 1652 | 1411 | 1124 | 1004 |
| 9422 | NM_003417,XI | ZNF264 | zinc finger protein 264 | protein_coding | NP_003408.1;XP_011525824.1 | 13057 | 604668 | . | . | 0.608255 | 0.515759 | 0.724116 | 0.627378 | 1.415433 | 1.192200 | 1.624895 | 1.434426 | 1.016148 | 216.764934 | 0.237627346 | 0.097253979 | 0.922524702 | 0.998865404 | 7.748464 | 7.771420 | 7.795047 | 7.713405 | 7.725200 | 7.798549 | 269 | 212 | 175 | 221 |
| 9423 | NM_004822,XI | NTN1 | netrin 1 | protein_coding | NP_004813.2;XP_006721658.1 | 8029 | 601614 | . | . | 1.072982 | 0.788028 | 0.582366 | 0.684565 | 2.496872 | 1.821562 | 1.306813 | 1.565177 | -1.480737 | 146.470088 | 0.289127505 | -1.95870485 | 0.050147362 | 0.668264424 | 7.449971 | 6.883749 | 7.326839 | 7.189526 | 7.086670 | 7.134954 | 233 | 159 | 95 | 119 |
| 9425 | NM_00114397 | CDYL | chromodomain Y like | protein_coding | NP_001137442.1;NP_001137443.1;NP_001355054.1;NP_001355055.1;NI | 1811 | 603778 | . | . | 5.691317 | 5.989542 | 5.805763 | 6.647210 | 13.243929 | 13.845098 | 13.027965 | 15.198055 | 1.041295 | 652.454866 | 0.156877549 | 0.372132565 | 0.709794151 | 0.998865404 | 9.320518 | 9.378370 | 9.318477 | 9.352018 | 9.327212 | 9.396841 | 718 | 707 | 549 | 656 |
| 9429 | NM_00125738 | ABCG2 | ATP binding cassette subfamily G member 2 (Junior blood group) | protein_coding | NP_001244315.1;NP_001335914.1;NP_001335915.1;NP_001335916.1;NI | 74 | 603756 | . | . | 0.158365 | 0.142094 | 0.133564 | 0.142468 | 0.368518 | 0.328457 | 0.299714 | 0.325736 | -1.097740 | 22.637811 | 0.647247886 | -0.20786008 | 0.835338219 | NA | 4.565853 | 4.432387 | 4.516819 | 4.496527 | 4.498995 | 4.496796 | 30 | 23 | 19 | 20 |
| 9435 | NM_004267 | CHST2 | carbohydrate sulfotransferase 2 | protein_coding | NP_004258.2 | 1970 | 603798 | . | . | 6.896084 | 6.590589 | 4.796736 | 5.358390 | 16.047470 | 15.234443 | 10.763737 | 12.251323 | -1.346536 | 764.131526 | 0.150443217 | -2.85325363 | **0.004327407** | 0.193932454 | 9.776520 | 9.346979 | 9.691189 | 9.651542 | 9.430794 | 9.496185 | 1037 | 921 | 538 | 641 |
| 9439 | NM_00127052 | MED23 | mediator complex subunit 23 | protein_coding | NP_001257450.1;NP_001257451.1;NP_001363446.1;NP_001363447.1;NI | 2372 | 605042 | . | . | 5.306184 | 5.573388 | 5.829588 | 4.860185 | 12.347711 | 12.883137 | 13.081427 | 11.112234 | -1.026486 | 862.348967 | 0.157049864 | -0.24013981 | 0.810221874 | 0.998865404 | 9.770623 | 9.733393 | 9.732887 | 9.785674 | 9.814771 | 9.661602 | 969 | 977 | 820 | 712 |
| 9440 | NM_004268 | MED17 | mediator complex subunit 17 | protein_coding | NP_004259.3 | 2375 | 603810 | . | . | 4.716146 | 4.696209 | 5.340271 | 4.800756 | 10.974665 | 10.855497 | 11.983415 | 10.976359 | 1.062113 | 775.150179 | 0.149837012 | 0.580213682 | 0.561770528 | 0.998865404 | 9.553984 | 9.641358 | 9.580286 | 9.571566 | 9.658556 | 9.576407 | 871 | 806 | 735 | 705 |
| 9441 | NM_004831 | MED26 | mediator complex subunit 26 | protein_coding | NP_004822.2 | 2376 | 605043 | . | . | 2.522352 | 2.463645 | 2.651889 | 2.986863 | 5.869617 | 5.694827 | 5.950762 | 6.829108 | 1.115084 | 262.959546 | 0.21330345 | 0.736752969 | 0.461272589 | 0.998865404 | 7.958197 | 8.114942 | 8.016795 | 8.000715 | 8.035688 | 8.092103 | 291 | 264 | 228 | 274 |
| 9442 | NM_00125388 | MED27 | mediator complex subunit 27 | protein_coding | NP_001240810.1;NP_001240811.1;NP_004260.2;XP_005272293.1;XP_01 | 2377 | 605044 | . | . | 17.129001 | 16.064341 | 13.502352 | 17.404709 | 39.859889 | 37.133443 | 30.298887 | 39.793802 | -1.061740 | 810.034998 | 0.152700914 | -0.56601602 | 0.571382893 | 0.998865404 | 9.704600 | 9.617774 | 9.703708 | 9.660053 | 9.582913 | 9.693424 | 990 | 873 | 629 | 795 |
| 9443 | NM_00110081 | MED7 | mediator complex subunit 7 | protein_coding | NP_001094286.1;NP_004261.1 | 2378 | 605045 | . | . | 5.912066 | 5.433781 | 5.834704 | 5.671312 | 13.757620 | 12.560431 | 13.092904 | 12.966781 | -1.030038 | 315.496381 | 0.199567553 | -0.21394982 | 0.830586187 | 0.998865404 | 8.322325 | 8.280329 | 8.333745 | 8.284075 | 8.314111 | 8.271695 | 385 | 330 | 281 | 280 |
| 9444 | NM_00130108 | QKI | QKI, KH domain containing RNA binding | protein_coding | NP_001288014.1;NP_006766.1;NP_996735.1;NP_996736.1;NP_996737.1 | 21100 | 609590 | . | . | 8.162571 | 8.300517 | 7.955402 | 7.368179 | 18.994636 | 19.187015 | 17.851690 | 16.846468 | -1.107576 | 2187.549720 | 0.121995056 | -1.20829716 | 0.226932982 | 0.982008277 | 11.166800 | 11.019650 | 11.172360 | 11.094620 | 11.078338 | 11.026067 | 2774 | 2362 | 1865 | 1882 |
| 9445 | NM_021999 | ITM2B | integral membrane protein 2B | protein_coding | NP_068839.1 | 6174 | 603904 | . | . | 9.550525 | 8.232698 | 8.224256 | 6.862237 | 22.224463 | 19.030252 | 18.454992 | 15.689689 | -1.195422 | 2585.675454 | 0.136682946 | -1.88406633 | 0.059556004 | 0.719735192 | 11.459165 | 11.202045 | 11.465285 | 11.335458 | 11.330942 | 11.179260 | 3496 | 2800 | 2244 | 1998 |
| 9446 | NM_00119100 | GSTO1 | glutathione S-transferase omega 1 | protein_coding | NP_001177931.1;NP_001177932.1;NP_004823.1 | 13312 | 605482 | . | . | 191.620996 | 197.669481 | 187.908758 | 245.504066 | 445.909917 | 456.921875 | 421.661815 | 561.315878 | 1.097077 | 5237.137811 | 0.132013183 | 1.012514352 | 0.311292188 | 0.998865404 | 12.286277 | 12.419762 | 12.301788 | 12.322860 | 12.275959 | 12.492789 | 5679 | 5458 | 4156 | 5793 |
| 9448 | NM_00124255 | MAP4K4 | mitogen-activated protein kinase kinase kinase kinase 4 | protein_coding | NP_001229488.1;NP_001229489.1;NP_004825.3;NP_663719.2;NP_6637 | 6866 | 604666 | . | . | 27.294474 | 28.371500 | 24.612525 | 27.341119 | 63.515365 | 65.581984 | 55.229789 | 62.512227 | -1.076002 | 6188.313244 | 0.110097056 | -0.95988902 | 0.337111075 | 0.998865404 | 12.647257 | 12.541465 | 12.612048 | 12.637998 | 12.516764 | 12.607836 | 7274 | 7028 | 4902 | 5893 |
| 9451 | NM_00131391 | EIF2AK3 | eukaryotic translation initiation factor 2 alpha kinase 3 | protein_coding | NP_001300844.1;NP_004827.4;XP_016860865.1 | 3255 | 604032 | . | . | 7.060039 | 7.033916 | 7.120467 | 6.636320 | 16.428999 | 16.259212 | 15.978121 | 15.173157 | -1.038092 | 984.846233 | 0.138753886 | -0.38870435 | 0.697494864 | 0.998865404 | 9.970327 | 9.916684 | 9.961474 | 9.952647 | 9.958317 | 9.900556 | 1162 | 1076 | 874 | 870 |
| 9452 | NM_00117158 | ITM2A | integral membrane protein 2A | protein_coding | NP_001165052.1;NP_004858.1 | 6173 | 300222 | . | . | 1.613888 | 1.782687 | 1.617693 | 1.202273 | 3.755584 | 4.120759 | 3.630057 | 2.748853 | -1.217984 | 78.942795 | 0.365909524 | -0.77750269 | 0.436862247 | 0.998865404 | 6.437319 | 6.154294 | 6.313223 | 6.340182 | 6.309878 | 6.240447 | 95 | 98 | 71 | 57 |
| 9453 | NM_00103727 | GGPS1 | geranylgeranyl diphosphate synthase 1 | protein_coding | NP_001032354.1;NP_001032355.1;NP_001358406.1;NP_001358407.1;NI | 4249 | 606982 | . | . | 5.538595 | 5.277470 | 6.853830 | 6.500640 | 12.888539 | 12.199109 | 15.379795 | 14.862938 | 1.203369 | 633.201106 | 0.158409993 | 1.685999996 | 0.0917958 | 0.806781001 | 9.166544 | 9.434108 | 9.273185 | 9.203545 | 9.378622 | 9.348078 | 697 | 587 | 615 | 634 |
| 9454 | NM_00114572 | HOMER3 | homer scaffold protein 3 | protein_coding | NP_001139193.1;NP_001139194.1;NP_001139196.1;NP_004829.3;XP_0( | 17514 | 604800 | . | . | 15.510055 | 16.858499 | 17.018428 | 20.411296 | 36.092534 | 38.969179 | 38.188856 | 46.668004 | 1.132964 | 861.816477 | 0.152450087 | 1.181384986 | 0.23744982 | 0.992733415 | 9.658748 | 9.838152 | 9.667108 | 9.738118 | 9.733228 | 9.848192 | 884 | 916 | 733 | 926 |
| 9455 | NM_004839,N | HOMER2 | homer scaffold protein 2 | protein_coding | NP_004830.2;NP_955362.1;XP_005272505.1;XP_005272506.1;XP_00672 | 17513 | 604799 | . | . | 5.152952 | 4.040503 | 4.665358 | 6.310436 | 11.991132 | 9.339803 | 10.468928 | 14.428061 | -1.034386 | 477.876789 | 0.174086075 | -0.28017416 | 0.779343889 | 0.998865404 | 8.924986 | 8.875581 | 8.907743 | 8.913799 | 8.845644 | 8.931516 | 557 | 527 | 380 | 471 |
| 9456 | NM_00127707 | HOMER1 | homer scaffold protein 1 | protein_coding | NP_001264006.1;NP_001264007.1;NP_004263.1;XP_016865548.1 | 17512 | 604798 | . | . | 2.208434 | 1.627467 | 2.002702 | 2.088625 | 5.139116 | 3.761962 | 4.494005 | 4.775392 | 1.028856 | 354.436883 | 0.208676785 | 0.196672386 | 0.844083921 | 0.998865404 | 8.448299 | 8.490167 | 8.542719 | 8.366216 | 8.469696 | 8.479550 | 464 | 319 | 310 | 340 |
| 9462 | NM_004841,N | RASAL2 | RAS protein activator like 2 | protein_coding | NP_004832.1;NP_733793.2;XP_005245679.1;XP_011508468.1;XP_01150 | 9874 | 606136 | . | . | 3.060067 | 3.612770 | 3.223035 | 3.578770 | 7.120902 | 8.351080 | 7.232397 | 8.182436 | 1.003447 | 1060.244904 | 0.147780615 | 0.033591307 | 0.973203054 | 0.998865404 | 10.048011 | 10.052349 | 9.984293 | 10.107392 | 10.006598 | 10.091740 | 1118 | 1237 | 870 | 1053 |
| 9463 | NM_00103958 | PICK1 | protein interacting with PRKCA 1 | protein_coding | NP_001034672.1;NP_001034673.1;NP_036539.1;XP_006724442.1;XP_00 | 9394 | 605926 | . | . | 18.787753 | 21.028465 | 22.150983 | 22.843765 | 43.719874 | 48.608236 | 49.706163 | 52.229550 | 1.107260 | 1493.913901 | 0.125278593 | 1.173336153 | 0.24066103 | 0.992733415 | 10.469602 | 10.616426 | 10.490261 | 10.517273 | 10.566817 | 10.597885 | 1604 | 1557 | 1338 | 1501 |
| 9464 | NM_021973 | HAND2 | heart and neural crest derivatives expressed 2 | protein_coding | NP_068808.1 | 4808 | 602407 | . | . | 0.167882 | 0.120778 | 0.092205 | 0.255505 | 0.390668 | 0.279185 | 0.206905 | 0.584182 | 1.179567 | 14.083571 | 0.858258956 | 0.277605016 | 0.781315581 | NA | 3.694014 | 3.928362 | 3.814441 | 3.798108 | 3.785677 | 3.846434 | 17 | 12 | 7 | 21 |
| 9465 | NM_00137657 | AKAP7 | A-kinase anchoring protein 7 | protein_coding | NP_001363499.1;NP_004833.1;NP_057461.2;NP_619539.1;XP_0052672 | 377 | 604693 | . | . | 1.504702 | 0.788454 | 0.953773 | 0.951403 | 3.501507 | 1.822546 | 2.140239 | 2.175270 | -1.139174 | 77.124412 | 0.373169242 | -0.50376019 | 0.614429907 | 0.998865404 | 6.359012 | 6.173244 | 6.325646 | 6.241718 | 6.271139 | 6.231030 | 109 | 75 | 68 | 62 |
| 9466 | NM_004843 | IL27RA | interleukin 27 receptor subunit alpha | protein_coding | NP_004834.1 | 17290 | 605350 | . | . | 5.685798 | 6.412061 | 6.666113 | 7.542504 | 13.231085 | 14.821765 | 14.958563 | 17.245039 | 1.159059 | 604.449416 | 0.163905515 | 1.299246918 | 0.193859205 | 0.96621494 | 9.129469 | 9.341689 | 9.149531 | 9.222263 | 9.244880 | 9.322370 | 609 | 638 | 532 | 643 |
| 9467 | NM_00101800 | SH3BP5 | SH3 domain binding protein 5 | protein_coding | NP_001018009.2;NP_004835.2;XP_011532553.1;XP_016863010.1;XP_01 | 10827 | 605612 | . | . | 24.823812 | 27.242370 | 23.077194 | 26.754713 | 57.766028 | 62.971960 | 51.784551 | 61.171473 | -1.045924 | 2503.302902 | 0.125281032 | -0.51705756 | 0.605115977 | 0.998865404 | 11.321790 | 11.256710 | 11.290800 | 11.322558 | 11.203855 | 11.332873 | 2890 | 2816 | 1957 | 2478 |
| 9468 | NM_00116326 | PCYT1B | phosphate cytidylyltransferase 1, choline, beta | protein_coding | NP_001156736.1;NP_001156737.1;NP_004836.2;XP_016885466.1 | 8755 | 300948 | . | . | 0.076834 | 0.094184 | 0.044011 | 0.033400 | 0.178797 | 0.217710 | 0.098759 | 0.076366 | **-2.178607** | 11.473686 | 0.920971441 | -1.21980575 | 0.222538522 | NA | 3.974755 | 2.852980 | 3.542863 | 3.551071 | 3.513430 | 3.505851 | 17 | 18 | 7 | 6 |
| 9469 | NM_004273,XI | CHST3 | carbohydrate sulfotransferase 3 | protein_coding | NP_004264.2;XP_006718138.1;XP_011538671.1 | 1971 | 603799 | . | . | 5.725408 | 6.296324 | 4.919190 | 5.760002 | 13.323260 | 14.554235 | 11.038518 | 13.169560 | -1.156662 | 1246.489721 | 0.139837204 | -1.50151584 | 0.133222193 | 0.888770389 | 10.385080 | 10.174557 | 10.296159 | 10.373251 | 10.171841 | 10.276028 | 1463 | 1515 | 934 | 1160 |
| 9470 | NM_00127633 | EIF4E2 | eukaryotic translation initiation factor 4E family member 2 | protein_coding | NP_001263265.1;NP_001263266.1;NP_001269887.1;NP_001317130.1;NI | 3293 | 605895 | . | . | 31.700085 | 32.455408 | 30.455424 | 33.660645 | 73.767399 | 75.022128 | 68.341091 | 76.961063 | 1.031338 | 1751.434059 | 0.125141875 | 0.355736015 | 0.722038271 | 0.998865404 | 10.752007 | 10.796295 | 10.763782 | 10.759692 | 10.743126 | 10.826926 | 1991 | 1856 | 1464 | 1756 |
| 9472 | NM_004274,XI | AKAP6 | A-kinase anchoring protein 6 | protein_coding | NP_004265.3;XP_011535680.1;XP_011535681.1;XP_011535685.1;XP_01 | 376 | 604691 | . | . | 0.172626 | 0.213722 | 0.220076 | 0.122037 | 0.401709 | 0.494026 | 0.493844 | 0.279024 | 1.027286 | 57.978407 | 0.419755362 | 0.092524108 | 0.926281638 | 0.998865404 | 5.838342 | 5.876296 | 5.819714 | 5.882727 | 5.864947 | 5.849582 | 56 | 71 | 53 | 53 |
| 9473 | NM_00103947 | THEMIS2 | thymocyte selection associated family member 2 | protein_coding | NP_001034566.1;NP_001099026.1;NP_001273042.1;NP_001273044.1;NI | 16839 | 617856 | . | . | 0.591233 | 0.748089 | 0.711583 | 0.894409 | 1.375823 | 1.729240 | 1.596772 | 2.044960 | 1.141811 | 61.633035 | 0.405526155 | 0.471790296 | 0.637076478 | 0.998865404 | 5.847790 | 6.037258 | 5.903763 | 5.950415 | 5.939797 | 5.972203 | 59 | 69 | 53 | 66 |
| 9474 | NM_00128610 | ATG5 | autophagy related 5 | protein_coding | NP_001273035.1;NP_001273036.1;NP_001273037.1;NP_001273040.1;NI | 589 | 604261 | . | . | 6.420991 | 5.905664 | 5.744130 | 5.446170 | 14.941909 | 13.651206 | 12.889662 | 12.452024 | -1.163408 | 566.492042 | 0.169354845 | -1.2893482 | 0.197277061 | 0.96621494 | 9.250552 | 9.033085 | 9.228070 | 9.155298 | 9.140334 | 9.044461 | 739 | 622 | 490 | 454 |
| 9475 | NM_00132164 | ROCK2 | Rho associated coiled-coil containing protein kinase 2 | protein_coding | NP_001308572.1;NP_004841.2;XP_005246247.1;XP_011508719.1;XP_01 | 10252 | 604002 | . | . | 11.329613 | 11.049248 | 11.364276 | 10.367292 | 26.364474 | 25.540834 | 25.501107 | 23.703580 | -1.038065 | 2841.967736 | 0.116641033 | -0.46207365 | 0.644028502 | 0.998865404 | 11.499273 | 11.445576 | 11.502652 | 11.471847 | 11.495266 | 11.417267 | 3396 | 3065 | 2542 | 2489 |

| | | | | | | | | |
|---|---|---|---|---|---|---|---|---|
| 9477 | NM_00130545 | MED20 | mediator complex subunit 20 | protein_coding | NP_001292384.1;NP_001292385.1;NP_001292386.1;NP_004266.2 | 16840 | 612915 | [illegible] |
| 9478 | NM_00103367 | CABP1 | calcium binding protein 1 | protein_coding | NP_001028849.1;NP_004267.2;NP_112482.1;XP_016875724.1;XP_01687 | 1384 | 605563 | [illegible] |
| 9479 | NM_005456 | MAPK8IP1 | mitogen-activated protein kinase 8 interacting protein 1 | protein_coding | NP_005447.1 | 6882 | 604641 | [illegible] |
| 9480 | NM_004852 | ONECUT2 | one cut homeobox 2 | protein_coding | NP_004843.2 | 8139 | 604894 | [illegible] |
| 9481 | NM_00120405 | SLC25A27 | solute carrier family 25 member 27 | protein_coding | NP_001190980.1;NP_001190981.1;NP_004268.3;XP_005249540.1 | 21065 | 613725 | [illegible] |
| 9482 | NM_004853,N | STX8 | syntaxin 8 | protein_coding | NP_004844.1;XP_011522381.1 | 11443 | 604203 | [illegible] |
| 9486 | NM_004854,XI | CHST10 | carbohydrate sulfotransferase 10 | protein_coding | NP_004845.1;XP_011510509.1;XP_011510510.1;XP_011510512.1;XP_01 | 19650 | 606376 | [illegible] |
| 9487 | NM_004278,XI | PIGL | phosphatidylinositol glycan anchor biosynthesis class L | protein_coding | NP_004269.1;XP_011522382.1;XP_016880838.1;XP_016880839.1;XP_01 | 8966 | 605947 | [illegible] |
| 9488 | NM_004855,XI | PIGB | phosphatidylinositol glycan anchor biosynthesis class B | protein_coding | NP_004846.4;XP_005254852.1;XP_011520537.1;XP_011520538.1;XP_01 | 8959 | 604122 | [illegible] |
| 9489 | NM_024419,N | PGS1 | phosphatidylglycerophosphate synthase 1 | protein_coding | NP_077733.3;XP_011523789.1;XP_011523790.1;XP_011523791.1;XP_01 | 30029 | 614942 | [illegible] |
| 9491 | NM_00132340 | PSMF1 | proteasome inhibitor subunit 1 | protein_coding | NP_001310336.1;NP_001310337.1;NP_001310338.1;NP_001310339.1;NI | 9571 | 617858 | [illegible] |
| 9493 | NM_00128130 | KIF23 | kinesin family member 23 | protein_coding | NP_001268230.1;NP_001354733.1;NP_001354734.1;NP_004847.2;NP_6 | 6392 | 605064 | [illegible] |
| 9495 | NM_004857 | AKAP5 | A-kinase anchoring protein 5 | protein_coding | NP_004848.3 | 375 | 604688 | [illegible] |
| 9497 | NM_00125837 | SLC4A7 | solute carrier family 4 member 7 | protein_coding | NP_001245308.1;NP_001245309.1;NP_001308032.1;NP_001308033.1;NI | 11033 | 603353 | [illegible] |
| 9498 | NM_00103996 | SLC4A8 | solute carrier family 4 member 8 | protein_coding | NP_001035049.1;NP_001245330.1;NP_001245331.1;NP_001245332.1;NI | 11034 | 605024 | [illegible] |
| 9500 | NM_00100533 | MAGED1 | MAGE family member D1 | protein_coding | NP_001005332.1;NP_001005333.1;NP_008917.3;XP_011529137.1;XP_01 | 6813 | 300224 | [illegible] |
| 9508 | NM_014243,XI | ADAMTS3 | ADAM metallopeptidase with thrombospondin type 1 motif 3 | protein_coding | NP_055058.2;XP_011530723.1;XP_011530724.1 | 219 | 605011 | [illegible] |
| 9510 | NM_006988 | ADAMTS1 | ADAM metallopeptidase with thrombospondin type 1 motif 1 | protein_coding | NP_008919.3 | 217 | 605174 | [illegible] |
| 9512 | NM_004279,XI | PMPCB | peptidase, mitochondrial processing subunit beta | protein_coding | NP_004270.2;XP_005250774.1;XP_006716244.1 | 9119 | 603131 | [illegible] |
| 9513 | NM_004860,XI | FXR2 | FMR1 autosomal homolog 2 | protein_coding | NP_004851.2 | 4024 | 605339 | [illegible] |
| 9516 | NM_00113647 | LITAF | lipopolysaccharide induced TNF factor | protein_coding | NP_001129944.1;NP_001129945.1;NP_004853.2;XP_006721045.1;XP_00 | 16841 | 603795 | [illegible] |
| 9517 | NM_004863,XI | SPTLC2 | serine palmitoyltransferase long chain base subunit 2 | protein_coding | NP_004854.1;XP_011535686.1 | 11278 | 605713 | [illegible] |
| 9518 | NM_004864,XI | GDF15 | growth differentiation factor 15 | protein_coding | NP_004855.2;XP_024307557.1 | 30142 | 605312 | [illegible] |
| 9519 | NM_00125367 | TBPL1 | TATA-box binding protein like 1 | protein_coding | NP_001240605.1;NP_004856.1;XP_016867002.1;XP_024302360.1 | 11589 | 605521 | [illegible] |
| 9520 | NM_00133025 | NPEPPS | aminopeptidase puromycin sensitive | protein_coding | NP_001317186.1;NP_006301.3;XP_016880861.1;XP_016880862.1;XP_02 | 7900 | 606793 | [illegible] |
| 9521 | NM_00113565 | EEF1E1 | eukaryotic translation elongation factor 1 epsilon 1 | protein_coding | NP_001129122.1;NP_004271.1 | 3212 | 609206 | [illegible] |
| 9522 | NM_00129022 | SCAMP1 | secretory carrier membrane protein 1 | protein_coding | NP_001277158.1;NP_004857.4;XP_011542029.1 | 10563 | 606911 | [illegible] |
| 9524 | NM_00132117 | TECR | trans-2,3-enoyl-CoA reductase | protein_coding | NP_001308099.1;NP_612510.1;XP_006723008.1;XP_011526744.1;XP_02 | 4551 | 610057 | [illegible] |
| 9525 | NM_004869,XI | VPS4B | vacuolar protein sorting 4 homolog B | protein_coding | NP_004860.2;XP_006722645.1;XP_011524570.1 | 10895 | 609983 | [illegible] |
| 9526 | NM_00133007 | MPDU1 | mannose-P-dolichol utilization defect 1 | protein_coding | NP_001317002.1;NP_004861.2;XP_006721660.1;XP_006721661.1;XP_01 | 7207 | 604041 | [illegible] |
| 9527 | NM_00100702 | GOSR1 | golgi SNAP receptor complex member 1 | protein_coding | NP_001007025.1;NP_001007026.1;NP_004862.1;XP_005258127.1;XP_00 | 4430 | 604026 | [illegible] |
| 9528 | NM_00130504 | TMEM59 | transmembrane protein 59 | protein_coding | NP_001291972.1;NP_001291978.1;NP_001291979.1;NP_001291980.1;NI | 1239 | 617084 | [illegible] |
| 9529 | NM_00101504 | BAG5 | BAG cochaperone 5 | protein_coding | NP_001015048.1;NP_001015049.1;NP_004864.1 | 941 | 603885 | [illegible] |
| 9530 | NM_00120487 | BAG4 | BAG cochaperone 4 | protein_coding | NP_001191807.1;NP_004865.1 | 940 | 603884 | [illegible] |
| 9531 | NM_004281,XI | BAG3 | BAG cochaperone 3 | protein_coding | NP_004272.2;XP_005270344.1 | 939 | 603883 | [illegible] |
| 9532 | NM_004282,XI | BAG2 | BAG cochaperone 2 | protein_coding | NP_004273.1;XP_005249547.1;XP_011513300.1;XP_011513301.1;XP_02 | 938 | 603882 | [illegible] |
| 9533 | NM_00131887 | POLR1C | RNA polymerase I and III subunit C | protein_coding | NP_001305805.1;NP_001350587.1;NP_976035.1 | 20194 | 610060 | [illegible] |
| 9534 | NM_00127866 | ZNF254 | zinc finger protein 254 | protein_coding | NP_001265590.1;NP_001265591.1;NP_001265592.1;NP_001265593.1;NI | 13047 | 604768 | [illegible] |
| 9536 | NM_004878 | PTGES | prostaglandin E synthase | protein_coding | NP_004869.1 | 9599 | 605172 | [illegible] |
| 9537 | NM_00125832 | TP53I11 | tumor protein p53 inducible protein 11 | protein_coding | NP_001245249.1;NP_001245250.1;NP_001245251.1;NP_001245252.1;NI | 16842 | 617867 | [illegible] |
| 9538 | NM_00129013 | EI24 | EI24 autophagy associated transmembrane protein | protein_coding | NP_001277064.1;NP_001317348.1;NP_004870.3;XP_011541371.1;XP_01 | 13276 | 605170 | [illegible] |
| 9540 | NM_00120680 | TP53I3 | tumor protein p53 inducible protein 3 | protein_coding | NP_001193731.1;NP_004872.2;NP_671713.1;XP_005264707.1;XP_00671 | 19373 | 605171 | [illegible] |
| 9541 | NM_004882 | CIR1 | corepressor interacting with RBPJ, CIR1 | protein_coding | NP_004873.3 | 24217 | 605228 | [illegible] |
| 9542 | NM_00118493 | NRG2 | neuregulin 2 | protein_coding | NP_001171864.1;NP_004874.1;NP_053584.1;NP_053585.1;NP_053586.1 | 7998 | 603818 | [illegible] |
| 9545 | NM_004283 | RAB3D | RAB3D, member RAS oncogene family | protein_coding | NP_004274.1 | 9779 | 604350 | [illegible] |
| 9546 | NM_004886,XI | APBA3 | amyloid beta precursor protein binding family A member 3 | protein_coding | NP_004877.1;XP_006723013.1;XP_006723014.1 | 580 | 604262 | [illegible] |
| 9550 | NM_004888 | ATP6V1G1 | ATPase H+ transporting V1 subunit G1 | protein_coding | NP_004879.1 | 864 | 607296 | [illegible] |
| 9551 | NM_00100371 | ATP5MF | ATP synthase membrane subunit f | protein_coding | NP_001003713.1;NP_001003714.1;NP_001034267.1;NP_004880.1 | 848 | . | [illegible] |
| 9552 | NM_004890,XI | SPAG7 | sperm associated antigen 7 | protein_coding | NP_004881.2;XP_006721663.1 | 11216 | 610056 | [illegible] |
| 9553 | NM_004891,N | MRPL33 | mitochondrial ribosomal protein L33 | protein_coding | NP_004882.1;NP_663303.1 | 14487 | 610059 | [illegible] |
| 9554 | NM_004892,XI | SEC22B | SEC22 homolog B, vesicle trafficking protein | protein_coding | NP_004883.3;XP_016858345.1 | 10700 | 604029 | [illegible] |
| 9555 | NM_00104015 | MACROH2A1 | macroH2A.1 histone | protein_coding | NP_001035248.1;NP_004884.1;NP_613075.1;NP_613258.2;XP_0052721 | 4740 | 610054 | [illegible] |
| 9556 | NM_00112739 | ATP5MPL | ATP synthase membrane subunit 6.8PL | protein_coding | NP_001120865.1;NP_004885.1 | 1188 | 604573 | [illegible] |
| 9557 | NM_00125633 | CHD1L | chromodomain helicase DNA binding protein 1 like | protein_coding | NP_001243265.1;NP_001243266.1;NP_001243267.1;NP_001335380.1;NI | 1916 | 613039 | [illegible] |
| 9559 | NM_00103526 | VPS26A | VPS26, retromer complex component A | protein_coding | NP_001030337.1;NP_001305873.1;NP_001305874.1;NP_001305875.1;NI | 12711 | 605506 | [illegible] |
| 9562 | NM_00117811 | MINPP1 | multiple inositol-polyphosphate phosphatase 1 | protein_coding | NP_001171588.1;NP_001171589.1;NP_004888.2;XP_006718141.1;XP_01 | 7102 | 605391 | [illegible] |
| 9563 | NM_00128258 | H6PD | hexose-6-phosphate dehydrogenase/glucose 1-dehydrogenase | protein_coding | NP_001269516.1;NP_004276.2;XP_005263597.1;XP_006711115.1;XP_01 | 4795 | 138090 | [illegible] |
| 9564 | NM_00117071 | BCAR1 | BCAR1 scaffold protein, Cas family member | protein_coding | NP_001164185.1;NP_001164186.1;NP_001164187.1;NP_001164188.1;NI | 971 | 602941 | [illegible] |
| 9567 | NM_004286,XI | GTPBP1 | GTP binding protein 1 | protein_coding | NP_004277.2;XP_011528839.1;XP_016884588.1;XP_016884589.1;XP_01 | 4669 | 602245 | [illegible] |
| 9568 | NM_005458,XI | GABBR2 | gamma-aminobutyric acid type B receptor subunit 2 | protein_coding | NP_005449.5;XP_005252373.1;XP_016870820.1;XP_016870821.1 | 4507 | 607340 | [illegible] |
| 9569 | NM_00119920 | GTF2IRD1 | GTF2I repeat domain containing 1 | protein_coding | NP_001186136.1;NP_005676.3;NP_057412.1;XP_006716245.1;XP_00671 | 4661 | 604318 | [illegible] |
| 9570 | NM_00101251 | GOSR2 | golgi SNAP receptor complex member 2 | protein_coding | NP_001012529.1;NP_001308062.1;NP_001308063.1;NP_001317181.1;NI | 4431 | 604027 | [illegible] |
| 9572 | NM_021724 | NR1D1 | nuclear receptor subfamily 1 group D member 1 | protein_coding | NP_068370.1 | 7962 | 602408 | [illegible] |
| 9575 | NM_00126784 | CLOCK | clock circadian regulator | protein_coding | NP_001254772.1;NP_004889.1;XP_005265844.1;XP_011532711.1;XP_01 | 2082 | 601851 | [illegible] |
| 9577 | NM_00126184 | BABAM2 | BRISC and BRCA1 A complex member 2 | protein_coding | NP_001248769.1;NP_001316041.1;NP_001316042.1;NP_001316043.1;NI | 1106 | 610497 | [illegible] |
| 9578 | NM_006035,XI | CDC42BPB | CDC42 binding protein kinase beta | protein_coding | NP_006026.3;XP_005268284.1;XP_005268285.1;XP_005268286.1;XP_00 | 1738 | 614062 | [illegible] |
| 9580 | NM_005686,XI | SOX13 | SRY-box transcription factor 13 | protein_coding | NP_005677.2;XP_005245680.1 | 11192 | 604748 | [illegible] |
| 9581 | NM_00104238 | PREPL | prolyl endopeptidase like | protein_coding | NP_001035844.1;NP_001035845.1;NP_001165074.1;NP_001165077.1;NI | 30228 | 609557 | [illegible] |
| 9582 | NM_00127041 | APOBEC3B | apolipoprotein B mRNA editing enzyme catalytic subunit 3B | protein_coding | NP_001257340.2;NP_004891.5 | 17352 | 607110 | [illegible] |
| 9583 | NM_00112893 | ENTPD4 | ectonucleoside triphosphate diphosphohydrolase 4 | protein_coding | NP_001122402.1;NP_004892.1 | 14573 | 607577 | [illegible] |
| 9584 | NM_00124259 | RBM39 | RNA binding motif protein 39 | protein_coding | NP_001229528.1;NP_001229529.1;NP_001310351.1;NP_001310352.1;NI | 15923 | 604739 | [illegible] |
| 9585 | NM_00128425 | KIF20B | kinesin family member 20B | protein_coding | NP_001271188.1;NP_001369435.1;NP_057279.2 | 7212 | 605498 | [illegible] |
| 9586 | NM_00101166 | CREB5 | cAMP responsive element binding protein 5 | protein_coding | NP_001011666.1;NP_004895.2;NP_878901.2;NP_878902.2;XP_0052499 | 16844 | 618262 | [illegible] |
| 9587 | NM_00100369 | MAD2L1BP | MAD2L1 binding protein | protein_coding | NP_001003690.1;NP_055443.1 | 21059 | 618136 | [illegible] |
| 9588 | NM_004905 | PRDX6 | peroxiredoxin 6 | protein_coding | NP_004896.1 | 16753 | 602316 | [illegible] |
| 9589 | NM_00127053 | WTAP | WT1 associated protein | protein_coding | NP_001257460.1;NP_001257461.1;NP_001257462.1;NP_004897.2;NP_6 | 16846 | 605442 | [illegible] |
| 9590 | NM_00137034 | AKAP12 | A-kinase anchoring protein 12 | protein_coding | NP_001357275.1;NP_005091.2;NP_653080.1;XP_016867006.1;XP_01686 | 370 | 604698 | [illegible] |
| 9592 | NM_004907 | IER2 | immediate early response 2 | protein_coding | NP_004898.2 | 28871 | . | [illegible] |
| 9600 | NM_00113084 | PITPNM1 | phosphatidylinositol transfer protein membrane associated 1 | protein_coding | NP_001124320.1;NP_004901.2;XP_011543698.1;XP_016874075.1 | 9003 | 608794 | [illegible] |
| 9601 | NM_00137124 | PDIA4 | protein disulfide isomerase family A member 4 | protein_coding | NP_001358173.1;NP_001358174.1;NP_004902.1 | 30167 | . | [illegible] |
| 9603 | NM_004289 | NFE2L3 | nuclear factor, erythroid 2 like 3 | protein_coding | NP_004280.5 | 7783 | 604135 | [illegible] |
| 9604 | NM_00120136 | RNF14 | ring finger protein 14 | protein_coding | NP_001188294.1;NP_004281.1;NP_899645.1;NP_899646.1;NP_899647.1 | 10058 | 605675 | [illegible] |
| 9605 | NM_004913,XI | VPS9D1 | VPS9 domain containing 1 | protein_coding | NP_004904.2;XP_005256386.1;XP_005256387.1;XP_005256388.1;XP_00 | 13526 | . | [illegible] |
| 9609 | NM_00134987 | RAB36 | RAB36, member RAS oncogene family | protein_coding | NP_001336806.1;NP_001336807.1;NP_004905.2;XP_006724444.1;XP_01 | 9775 | 605662 | [illegible] |
| 9610 | NM_00136355 | RIN1 | Ras and Rab interactor 1 | protein_coding | NP_001350488.1;NP_001350489.1;NP_004283.2;XP_016874076.1 | 18749 | 605965 | [illegible] |
| 9611 | NM_00119043 | NCOR1 | nuclear receptor corepressor 1 | protein_coding | NP_001177367.1;NP_001177369.1;NP_006302.2;XP_005256923.1;XP_00 | 7672 | 600849 | [illegible] |
| 9612 | NM_00107726 | NCOR2 | nuclear receptor corepressor 2 | protein_coding | NP_001070729.2;NP_001193583.1;NP_006303.4 | 7673 | 600848 | [illegible] |
| 9616 | NM_00120137 | RNF7 | ring finger protein 7 | protein_coding | NP_001188299.1;NP_055060.1;NP_899060.1 | 10070 | 603863 | [illegible] |
| 9617 | NM_00135407 | MTRF1 | mitochondrial translation release factor 1 | protein_coding | NP_001341002.1;NP_001341003.1;NP_001341005.1;NP_004285.2;XP_00 | 7469 | 604601 | [illegible] |
| 9618 | NM_004295,XI | TRAF4 | TNF receptor associated factor 4 | protein_coding | NP_004286.2;XP_011523806.1;XP_011523807.1;XP_011523808.1;XP_01 | 12034 | 602464 | [illegible] |
| 9620 | NM_00137832 | CELSR1 | cadherin EGF LAG seven-pass G-type receptor 1 | protein_coding | NP_001365257.1;NP_055061.1;XP_011528855.1;XP_011528856.1;XP_01 | 1850 | 604523 | [illegible] |
| 9625 | NM_00108039 | AATK | apoptosis associated tyrosine kinase | protein_coding | NP_001073864.2;NP_004911.2 | 21 | 605276 | [illegible] |
| 9631 | NM_00127831 | NUP155 | nucleoporin 155 | protein_coding | NP_001265241.1;NP_004289.1;NP_705618.1;XP_011512467.1 | 8063 | 606694 | [illegible] |
| 9632 | NM_004922,N | SEC24C | SEC24 homolog C, COPII coat complex component | protein_coding | NP_004913.2;NP_940999.1;XP_011538682.1;XP_011538683.1;XP_01687 | 10705 | 607185 | [illegible] |
| 9633 | NM_00103965 | TESMIN | testis expressed metallothionein like protein | protein_coding | NP_001034745.1;NP_004914.2;XP_011543704.1;XP_011543705.1;XP_01 | 7446 | 604374 | [illegible] |
| 9635 | NM_006536,XI | CLCA2 | chloride channel accessory 2 | protein_coding | NP_006527.1;XP_011540750.1 | 2016 | 604003 | [illegible] |
| 9636 | NM_005101 | ISG15 | ISG15 ubiquitin like modifier | protein_coding | NP_005092.1 | 4053 | 147571 | [illegible] |
| 9637 | NM_00104254 | FEZ2 | fasciculation and elongation protein zeta 2 | protein_coding | NP_001036013.1;NP_005093.2;XP_006712216.1 | 3660 | 604826 | [illegible] |
| 9639 | NM_00130815 | ARHGEF10 | Rho guanine nucleotide exchange factor 10 | protein_coding | NP_001295081.1;NP_001295082.1;NP_055444.2;XP_005266098.1;XP_01 | 14103 | 608136 | [illegible] |
| 9640 | NM_014630,XI | ZNF592 | zinc finger protein 592 | protein_coding | NP_055445.2;XP_005255053.1;XP_011520548.1;XP_011520549.1;XP_01 | 28986 | 613624 | [illegible] |
| 9641 | NM_00119332 | IKBKE | inhibitor of nuclear factor kappa B kinase subunit epsilon | protein_coding | NP_001180250.1;NP_001180251.1;NP_054721.1;XP_005273413.1;XP_01 | 14552 | 605048 | [illegible] |
| 9643 | NM_00114241 | MORF4L2 | mortality factor 4 like 2 | protein_coding | NP_001135890.1;NP_001135891.1;NP_001135892.1;NP_001135893.1;NI | 16849 | 300409 | [illegible] |
| 9644 | NM_00136507 | SH3PXD2A | SH3 and PX domains 2A | protein_coding | NP_001352008.1;NP_055446.2 | 23664 | . | [illegible] |
| 9645 | NM_00128266 | MICAL2 | microtubule associated monooxygenase, calponin and LIM domai | protein_coding | NP_001269592.1;NP_001269593.1;NP_001269594.1;NP_001269595.1;NI | 24693 | 608881 | [illegible] |
| 9646 | NM_00134627 | CTR9 | CTR9 homolog, Paf1/RNA polymerase II complex component | protein_coding | NP_001333208.1;NP_055448.1 | 16850 | 609366 | [illegible] |
| 9647 | NM_014634,XI | PPM1F | protein phosphatase, Mg2+/Mn2+ dependent 1F | protein_coding | NP_055449.1;XP_011528858.1;XP_016884597.1 | 19388 | . | [illegible] |
| 9648 | NM_181453,N | GCC2 | GRIP and coiled-coil domain containing 2 | protein_coding | NP_852118.2;XP_006712933.1;XP_006712934.1;XP_006712935.1;XP_01 | 23218 | 612711 | [illegible] |
| 9649 | NM_00119072 | RALGPS1 | Ral GEF with PH domain and SH3 binding motif 1 | protein_coding | NP_001177657.1;NP_001177658.1;NP_001177659.1;NP_001309249.1;NI | 16851 | 614444 | [illegible] |
| 9650 | NM_00114583 | MTFR1 | mitochondrial fission regulator 1 | protein_coding | NP_001139310.1;NP_001139311.1;NP_055452.3;XP_006716547.2;XP_01 | 29510 | . | [illegible] |
| 9652 | NM_014639,XI | TTC37 | tetratricopeptide repeat domain 37 | protein_coding | NP_055454.1 | 23639 | 614589 | [illegible] |
| 9653 | NM_00113449 | HS2ST1 | heparan sulfate 2-O-sulfotransferase 1 | protein_coding | NP_001127964.1;NP_036394.1 | 5193 | 604844 | [illegible] |
| 9654 | NM_014640,XI | TTLL4 | tubulin tyrosine ligase like 4 | protein_coding | NP_055455.3;XP_006712938.1;XP_016860876.1;XP_016860877.1;XP_01 | 28976 | . | [illegible] |
| 9655 | NM_014011,N | SOCS5 | suppressor of cytokine signaling 5 | protein_coding | NP_054730.1;NP_659198.1 | 16852 | 607094 | [illegible] |
| 9656 | NM_014641,XI | MDC1 | mediator of DNA damage checkpoint 1 | protein_coding | NP_055456.2;XP_005249549.1;XP_005249550.1;XP_005249551.1;XP_00 | 21163 | 607593 | [illegible] |
| 9657 | NM_00102357 | IQCB1 | IQ motif containing B1 | protein_coding | NP_001018864.2;NP_001018865.2;NP_001306036.1;XP_005247968.1;XP | 28949 | 609237 | [illegible] |
| 9658 | NM_014643,XI | ZNF516 | zinc finger protein 516 | protein_coding | NP_055458.1;XP_011524571.1;XP_011524572.1;XP_011524573.1;XP_01 | 28990 | 615114 | [illegible] |
| 9659 | NM_00100281 | PDE4DIP | phosphodiesterase 4D interacting protein | protein_coding | NP_001002810.1;NP_001002811.2;NP_001002812.2;NP_001182189.1;NI | 15580 | 608117 | [illegible] |
| 9662 | NM_025009,XI | CEP135 | centrosomal protein 135 | protein_coding | NP_079285.2;XP_005265845.1;XP_006714118.1;XP_011532714.1 | 29086 | 611423 | [illegible] |
| 9663 | NM_00137580 | LPIN2 | lipin 2 | protein_coding | NP_001362737.1;NP_001362738.1;NP_055461.1;XP_005258234.1;XP_01 | 14450 | 605519 | [illegible] |
| 9665 | NM_00118499 | MARF1 | meiosis regulator and mRNA stability factor 1 | protein_coding | NP_001171927.1;NP_001171928.1;NP_055462.2;XP_005255761.1;XP_00 | 29562 | 614593 | [illegible] |
| 9666 | NM_014648,XI | DZIP3 | DAZ interacting zinc finger protein 3 | protein_coding | NP_055463.1;XP_005247971.1;XP_005247972.1;XP_005247973.1;XP_00 | 30938 | 608672 | [illegible] |
| 9667 | NM_014649,XI | SAFB2 | scaffold attachment factor B2 | protein_coding | NP_055464.1;XP_011526751.1;XP_011526752.1;XP_016883009.1;XP_01 | 21605 | 608066 | [illegible] |
| 9668 | NM_00132228 | ZNF432 | zinc finger protein 432 | protein_coding | NP_001309213.1;NP_001309214.1;NP_055465.1;XP_024307574.1 | 20810 | . | [illegible] |
| 9669 | NM_015904 | EIF5B | eukaryotic translation initiation factor 5B | protein_coding | NP_056988.3 | 30793 | 606086 | [illegible] |
| 9670 | NM_014652,XI | IPO13 | importin 13 | protein_coding | NP_055467.3;XP_024306837.1;XP_024306838.1 | 16853 | 610411 | [illegible] |
| 9672 | NM_014654,XI | SDC3 | syndecan 3 | protein_coding | NP_055469.3;XP_011540765.1;XP_011540766.1;XP_011540768.1 | 10660 | 186357 | [illegible] |
| 9673 | NM_00128618 | SLC25A44 | solute carrier family 25 member 44 | protein_coding | NP_001273113.1;NP_001364314.1;NP_001364315.1;NP_001364316.1;NI | 29036 | 610824 | [illegible] |
| 9674 | NM_00116289 | KIAA0040 | KIAA0040 | protein_coding | NP_001156365.1;NP_001156366.1;NP_001156367.1;NP_001306159.1;NI | 28950 | 616696 | [illegible] |
| 9675 | NM_00130345 | TTI1 | TELO2 interacting protein 1 | protein_coding | NP_001290386.1;NP_055472.1;XP_011527416.1;XP_016883637.1 | 29029 | 614425 | [illegible] |
| 9677 | NM_00113085 | PPIP5K1 | diphosphoinositol pentakisphosphate kinase 1 | protein_coding | NP_001124330.1;NP_001124331.1;NP_001177143.1;NP_001341311.1;NI | 29023 | 610979 | [illegible] |
| 9678 | NM_00100715 | PHF14 | PHD finger protein 14 | protein_coding | NP_001007158.1;NP_055475.2 | 22203 | . | [illegible] |
| 9679 | NM_014661 | FAM53B | family with sequence similarity 53 member B | protein_coding | NP_055476.3 | 28968 | 617289 | [illegible] |
| 9681 | NM_00100718 | DEPDC5 | DEP domain containing 5, GATOR1 subcomplex subunit | protein_coding | NP_001007189.1;NP_001129501.1;NP_001229825.1;NP_001229826.1;NI | 18423 | 614191 | [illegible] |
| 9682 | NM_014663,XI | KDM4A | lysine demethylase 4A | protein_coding | NP_055478.2;XP_005271411.1;XP_005271412.1 | 22978 | 609764 | [illegible] |
| 9683 | NM_153029,XI | N4BP1 | NEDD4 binding protein 1 | protein_coding | NP_694574.3;XP_011521784.1 | 29850 | . | [illegible] |
| 9684 | NM_00127203 | LRRC14 | leucine rich repeat containing 14 | protein_coding | NP_001258965.1;NP_055480.1;XP_005272415.1;XP_005272416.1;XP_00 | 20419 | . | [illegible] |
| 9685 | NM_00119555 | CLINT1 | clathrin interactor 1 | protein_coding | NP_001182484.1;NP_001182485.1;NP_055481.1;XP_011533003.1;XP_01 | 23186 | 607265 | [illegible] |
| 9686 | NM_00112821 | VGLL4 | vestigial like family member 4 | protein_coding | NP_001121691.1;NP_001121692.1;NP_001121693.1;NP_001271319.1;NI | 28966 | 618692 | [illegible] |
| 9687 | NM_014668,N | GREB1 | growth regulating estrogen receptor binding 1 | protein_coding | NP_055483.2;NP_149081.1;NP_683701.2;XP_005246249.1;XP_00524625 | 24885 | 611736 | [illegible] |
| 9688 | NM_00124279 | NUP93 | nucleoporin 93 | protein_coding | NP_001229724.1;NP_001229725.1;NP_055484.3;XP_005256320.1 | 28958 | 614351 | [illegible] |
| 9689 | NM_00120706 | BZW1 | basic leucine zipper and W2 domains 1 | protein_coding | NP_001193996.1;NP_001193997.1;NP_001193998.1;NP_001308617.1;NI | 18380 | . | [illegible] |
| 9690 | NM_014671,XI | UBE3C | ubiquitin protein ligase E3C | protein_coding | NP_055486.2;XP_005249621.1;XP_016868307.1 | 16803 | 614454 | [illegible] |
| 9692 | NM_00125667 | PRORP | protein only RNase P catalytic subunit | protein_coding | NP_001243607.1;NP_001243608.1;NP_001243609.1;NP_001243610.1;NI | 19958 | 609947 | [illegible] |
| 9693 | NM_00135172 | RAPGEF2 | Rap guanine nucleotide exchange factor 2 | protein_coding | NP_001338653.1;NP_001338654.1;NP_001338655.1;NP_001338656.1;NI | 16854 | 609530 | [illegible] |
| 9694 | NM_00132949 | EMC2 | ER membrane protein complex subunit 2 | protein_coding | NP_001316422.1;NP_001316423.1;NP_001316424.1;NP_055488.1 | 28963 | 607722 | [illegible] |
| 9695 | NM_014674,XI | EDEM1 | ER degradation enhancing alpha-mannosidase like protein 1 | protein_coding | NP_055489.1;XP_011532573.1;XP_011532574.1 | 18967 | 607673 | [illegible] |
| 9696 | NM_014675,XI | CROCC | ciliary rootlet coiled-coil, rootletin | protein_coding | NP_055490.4;XP_006711121.1;XP_011540770.2;XP_011540772.2;XP_01 | 21299 | 615776 | [illegible] |
| 9697 | NM_012288,XI | TRAM2 | translocation associated membrane protein 2 | protein_coding | NP_036420.1;XP_011513307.1 | 16855 | 608485 | [illegible] |
| 9698 | NM_00102065 | PUM1 | pumilio RNA binding family member 1 | protein_coding | NP_001018494.1;NP_055491.1 | 14957 | 607204 | [illegible] |
| 9699 | NM_00110011 | RIMS2 | regulating synaptic membrane exocytosis 2 | protein_coding | NP_001093587.1;NP_001269810.1;NP_001269811.1;NP_001335413.1;NI | 17283 | 606630 | [illegible] |
| 9700 | NM_012291,XI | ESPL1 | extra spindle pole bodies like 1, separase | protein_coding | NP_036423.4;XP_006719768.1;XP_011537326.1;XP_011537327.1;XP_01 | 16856 | 604143 | [illegible] |
| 9701 | NM_00124289 | PPP6R2 | protein phosphatase 6 regulatory subunit 2 | protein_coding | NP_001229827.1;NP_001229828.1;NP_001229829.1;NP_001338570.1;NI | 19253 | 610877 | [illegible] |
| 9702 | NM_00124377 | CEP57 | centrosomal protein 57 | protein_coding | NP_001230705.1;NP_001230706.1;NP_001350533.1;NP_055494.2;XP_00 | 30794 | 607951 | [illegible] |
| 9703 | NM_00132156 | KIAA0100 | KIAA0100 | protein_coding | NP_001308489.1;NP_001308490.1;NP_001350755.1;NP_001350756.1;NI | 28960 | 610664 | [illegible] |
| 9704 | NM_014681,XI | DHX34 | DExH-box helicase 34 | protein_coding | NP_055496.2;XP_005259500.1;XP_011525852.1;XP_011525853.1;XP_01 | 16719 | 615475 | [illegible] |
| 9706 | NM_00114261 | ULK2 | unc-51 like autophagy activating kinase 2 | protein_coding | NP_001136082.1;NP_055498.3;XP_011522389.1;XP_016880913.1;XP_01 | 13480 | 608650 | [illegible] |
| 9708 | NM_014004,N | PCDHGA8 | protocadherin gamma subfamily A, 8 | protein_coding | NP_054723.1;NP_114477.1 | 8706 | 606295 | [illegible] |
| 9709 | NM_00101098 | HERPUD1 | homocysteine inducible ER protein with ubiquitin like domain 1 | protein_coding | NP_001010989.1;NP_001259032.1;NP_055500.1 | 13744 | 608070 | [illegible] |
| 9710 | NM_014686 | GARRE1 | granule associated Rac and RHOG effector 1 | protein_coding | NP_055501.2 | 29016 | . | [illegible] |
| 9711 | NM_00114564 | RUBCN | rubicon autophagy regulator | protein_coding | NP_001139114.1;NP_001333802.1;NP_055502.1;XP_005269431.1;XP_00 | 28991 | 613516 | [illegible] |
| 9712 | NM_00108049 | USP6NL | USP6 N-terminal like | protein_coding | NP_001073960.1;NP_055503.1;XP_006717605.1;XP_011518064.1;XP_01 | 16858 | 605405 | [illegible] |
| 9716 | NM_014691 | AQR | aquarius intron-binding spliceosomal factor | protein_coding | NP_055506.1 | 29513 | 610548 | [illegible] |

| | | | | | | | | | | |
|---|---|---|---|---|---|---|---|---|---|---|
| 9720 | NM_00138200 | CCDC144A | coiled-coil domain containing 144A | protein_coding | NP_001368929.1;NP_055510.1;XP_016880918.1;XP_016880919.1;XP_02 | 29072 | . | . | . | [illegible] |
| 9722 | NM_00112606 | NOS1AP | nitric oxide synthase 1 adaptor protein | protein_coding | NP_001119532.2;NP_001158229.1;NP_055512.1 | 16859 | 605551 | . | . | [illegible] |
| 9724 | NM_021645 | UTP14C | UTP14C small subunit processome component | protein_coding | NP_067677.4 | 20321 | 608969 | . | . | [illegible] |
| 9725 | NM_014698,XI | TMEM63A | transmembrane protein 63A | protein_coding | NP_055513.2;XP_006711904.1;XP_011542630.1;XP_011542631.1;XP_01 | 29118 | 618685 | . | . | [illegible] |
| 9726 | NM_014699,XI | ZNF646 | zinc finger protein 646 | protein_coding | NP_055514.3;XP_005255767.1;XP_005255768.1;XP_005255769.1;XP_01 | 29004 | . | . | . | [illegible] |
| 9727 | NM_00114227 | RAB11FIP3 | RAB11 family interacting protein 3 | protein_coding | NP_001135744.2;NP_001357330.1;NP_055515.1;XP_005255771.1;XP_00 | 17224 | 608738 | . | . | [illegible] |
| 9728 | NM_00119348 | SECISBP2L | SECIS binding protein 2 like | protein_coding | NP_001180418.1;NP_055516.2 | 28997 | 615756 | . | . | [illegible] |
| 9730 | NM_00117190 | DCAF1 | DDB1 and CUL4 associated factor 1 | protein_coding | NP_001165375.1;NP_001336097.1;NP_001336098.1;NP_001336099.1;NI | 30911 | 617259 | . | . | [illegible] |
| 9731 | NM_014704,XI | CEP104 | centrosomal protein 104 | protein_coding | NP_055519.1;XP_005244872.1;XP_011540776.1;XP_016858407.1;XP_01 | 24866 | 616690 | . | . | [illegible] |
| 9732 | NM_00136354 | DOCK4 | dedicator of cytokinesis 4 | protein_coding | NP_001350469.1;NP_055520.3;XP_006716252.1;XP_016868308.1;XP_01 | 19192 | 607679 | . | . | [illegible] |
| 9733 | NM_014706,XI | SART3 | spliceosome associated factor 3, U4/U6 recycling protein | protein_coding | NP_055521.1;XP_005269298.1;XP_024305052.1 | 16860 | 611684 | . | . | [illegible] |
| 9734 | NM_00120414 | HDAC9 | histone deacetylase 9 | protein_coding | NP_001191073.1;NP_001191074.1;NP_001191075.1;NP_001191076.1;NI | 14065 | 606543 | . | . | [illegible] |
| 9735 | NM_014708,XI | KNTC1 | kinetochore associated 1 | protein_coding | NP_055523.1;XP_006719769.1;XP_011537329.1;XP_011537331.1;XP_01 | 17255 | 607363 | . | . | [illegible] |
| 9736 | NM_014709 | USP34 | ubiquitin specific peptidase 34 | protein_coding | NP_055524.3 | 20066 | 615295 | . | . | [illegible] |
| 9737 | NM_00109941 | GPRASP1 | G protein-coupled receptor associated sorting protein 1 | protein_coding | NP_001092880.1;NP_001092881.1;NP_001171656.1;NP_055525.3 | 24834 | 300417 | . | . | [illegible] |
| 9738 | NM_00119902 | CCP110 | centriolar coiled-coil protein 110 | protein_coding | NP_001185951.2;NP_001310498.1;NP_001310499.1;NP_001310500.1;NI | 24342 | 609544 | . | . | [illegible] |
| 9739 | NM_014712,XI | SETD1A | SET domain containing 1A, histone lysine methyltransferase | protein_coding | NP_055527.1;XP_005255780.1;XP_006721169.1;XP_016879398.1;XP_02 | 29010 | 611052 | . | . | [illegible] |
| 9741 | NM_014713 | LAPTM4A | lysosomal protein transmembrane 4 alpha | protein_coding | NP_055528.1 | 6924 | 618837 | . | . | [illegible] |
| 9742 | NM_014714,XI | IFT140 | intraflagellar transport 140 | protein_coding | NP_055529.2;XP_005255782.1;XP_005255783.1;XP_006721053.1;XP_00 | 29077 | 614620 | . | . | [illegible] |
| 9743 | NM_00114268 | ARHGAP32 | Rho GTPase activating protein 32 | protein_coding | NP_001136157.1;NP_001364953.1;NP_001364954.1;NP_055530.2;XP_01 | 17399 | 608541 | . | . | [illegible] |
| 9744 | NM_014716,XI | ACAP1 | ArfGAP with coiled-coil, ankyrin repeat and PH domains 1 | protein_coding | NP_055531.1;XP_024306815.1 | 16467 | 607763 | . | . | [illegible] |
| 9745 | NM_00135226 | ZNF536 | zinc finger protein 536 | protein_coding | NP_001339189.1;NP_001363039.1;NP_001363040.1;NP_055532.1;XP_01 | 29025 | 618037 | . | . | [illegible] |
| 9746 | NM_014718,XI | CLSTN3 | calsyntenin 3 | protein_coding | NP_055533.2;XP_006719226.1 | 18371 | 611324 | . | . | [illegible] |
| 9747 | NM_00120693 | TCAF1 | TRPM8 channel associated factor 1 | protein_coding | NP_001193867.2;NP_001193870.1;NP_055534.2;XP_005250131.1;XP_00 | 22201 | 616251 | . | . | [illegible] |
| 9748 | NM_00130474 | SLK | STE20 like kinase | protein_coding | NP_001291672.1;NP_055535.2;XP_011538703.1 | 11088 | 616563 | . | . | [illegible] |
| 9749 | NM_00110016 | PHACTR2 | phosphatase and actin regulator 2 | protein_coding | NP_001093634.1;NP_001093635.1;NP_001093636.1;NP_055536.2 | 20956 | 608724 | . | . | [illegible] |
| 9750 | NM_00128644 | RIPOR2 | RHO family interacting cell polarization regulator 2 | protein_coding | NP_001273374.1;NP_001273375.1;NP_001273376.1;NP_001332960.1;NI | 13872 | 611410 | . | . | [illegible] |
| 9751 | NM_00131823 | SNPH | syntaphilin | protein_coding | NP_001305163.1;NP_055538.2;XP_005260946.1;XP_011527704.1;XP_01 | 15931 | 604942 | . | . | [illegible] |
| 9753 | NM_00116339 | ZSCAN12 | zinc finger and SCAN domain containing 12 | protein_coding | NP_001156863.1;NP_001355053.1;XP_011513316.1;XP_011513317.1;XP | 13172 | 603978 | . | . | [illegible] |
| 9755 | NM_014726,XI | TBKBP1 | TBK1 binding protein 1 | protein_coding | NP_055541.1;XP_005257916.1;XP_005257917.1;XP_005257918.1;XP_00 | 30140 | 608476 | . | . | [illegible] |
| 9757 | NM_014727,XI | KMT2B | lysine methyltransferase 2B | protein_coding | NP_055542.1;XP_011525863.2;XP_011525864.1;XP_016883033.1;XP_01 | 15840 | 606834 | . | . | [illegible] |
| 9759 | NM_00137841 | HDAC4 | histone deacetylase 4 | protein_coding | NP_001365343.1;NP_001365344.1;NP_001365345.1;NP_001365346.1;NI | 14063 | 605314 | . | . | [illegible] |
| 9760 | NM_014729,XI | TOX | thymocyte selection associated high mobility group box | protein_coding | NP_055544.1;XP_016869574.1 | 18988 | 606863 | . | . | [illegible] |
| 9761 | NM_00130362 | MLEC | malectin | protein_coding | NP_001290556.1;NP_001290557.1;NP_055545.1;XP_011537333.1;XP_01 | 28973 | 613802 | . | . | [illegible] |
| 9762 | NM_00128253 | LZTS3 | leucine zipper tumor suppressor family member 3 | protein_coding | NP_001269462.1;NP_001352547.1;NP_001354538.1;XP_005260950.1;XP | 30139 | 610484 | . | . | [illegible] |
| 9764 | NM_00128656 | KIAA0513 | KIAA0513 | protein_coding | NP_001273494.1;NP_001273495.1;NP_001284695.1;NP_055547.1;XP_00 | 29058 | 611675 | . | . | [illegible] |
| 9765 | NM_00110525 | ZFYVE16 | zinc finger FYVE-type containing 16 | protein_coding | NP_001098721.2;NP_001271165.2;NP_001271166.2;NP_001336363.2;NI | 20756 | 608880 | . | . | [illegible] |
| 9766 | NM_014734 | SUSD6 | sushi domain containing 6 | protein_coding | NP_055549.1 | 19956 | 616761 | . | . | [illegible] |
| 9767 | NM_00107744 | JADE3 | jade family PHD finger 3 | protein_coding | NP_001070913.1;NP_055550.1 | 22982 | 300618 | . | . | [illegible] |
| 9768 | NM_00102998 | PCLAF | PCNA clamp associated factor | protein_coding | NP_001025160.1;NP_055551.1 | 28961 | 610696 | . | . | [illegible] |
| 9770 | NM_014737,N | RASSF2 | Ras association domain family member 2 | protein_coding | NP_055552.1;NP_739580.1;XP_005260952.1;XP_011527712.1;XP_01152 | 9883 | 609492 | . | . | [illegible] |
| 9771 | NM_00136760 | RAPGEF5 | Rap guanine nucleotide exchange factor 5 | protein_coding | NP_001354529.1;NP_001354530.1;NP_001354531.1;NP_001354532.1;NI | 16862 | 609527 | . | . | [illegible] |
| 9772 | NM_00132114 | TMEM94 | transmembrane protein 94 | protein_coding | NP_001308077.1;NP_001308078.1;NP_001338131.1;NP_001338132.1;NI | 28983 | 618163 | . | . | [illegible] |
| 9774 | NM_00107744 | BCLAF1 | BCL2 associated transcription factor 1 | protein_coding | NP_001070908.1;NP_001070909.1;NP_001287967.1;NP_001350588.1;NI | 16863 | 612588 | . | . | [illegible] |
| 9775 | NM_014740,XI | EIF4A3 | eukaryotic translation initiation factor 4A3 | protein_coding | NP_055555.1;XP_011523824.1 | 18683 | 608546 | . | . | [illegible] |
| 9776 | NM_00114267 | ATG13 | autophagy related 13 | protein_coding | NP_001136145.1;NP_001192048.1;NP_001192049.1;NP_001192050.1;NI | 29091 | 615088 | . | . | [illegible] |
| 9777 | NM_00136373 | TM9SF4 | transmembrane 9 superfamily member 4 | protein_coding | NP_001350660.1;NP_055557.2;XP_016883643.1;XP_016883644.1;XP_01 | 30797 | 617727 | . | . | [illegible] |
| 9778 | NM_00110059 | KIAA0232 | KIAA0232 | protein_coding | NP_001094060.1;NP_055558.2;XP_005248098.1;XP_011511900.1;XP_01 | 28992 | . | . | . | [illegible] |
| 9779 | NM_00113438 | TBC1D5 | TBC1 domain family member 5 | protein_coding | NP_001127853.1;NP_001336002.1;NP_001336003.1;NP_001336004.1;NI | 19166 | 615740 | . | . | [illegible] |
| 9780 | NM_00114286 | PIEZO1 | piezo type mechanosensitive ion channel component 1 | protein_coding | NP_001136336.2 | 28993 | 611184 | . | . | [illegible] |
| 9781 | NM_00134918 | RNF144A | ring finger protein 144A | protein_coding | NP_001336110.1;NP_001336111.1;NP_001336112.1;NP_001336113.1;NI | 20457 | . | . | . | [illegible] |
| 9782 | NM_00119495 | MATR3 | matrin 3 | protein_coding | NP_001181883.1;NP_001181884.1;NP_001181885.1;NP_001269207.1;NI | 6912 | 164015 | . | . | [illegible] |
| 9783 | NM_014747,XI | RIMS3 | regulating synaptic membrane exocytosis 3 | protein_coding | NP_055562.2;XP_011540781.1 | 21292 | 611600 | . | . | [illegible] |
| 9784 | NM_00126705 | SNX17 | sorting nexin 17 | protein_coding | NP_001253988.1;NP_001253989.1;NP_001253990.1;NP_055563.1;XP_01 | 14979 | 605963 | . | . | [illegible] |
| 9785 | NM_014003,XI | DHX38 | DEAH-box helicase 38 | protein_coding | NP_054722.2;XP_005256326.1;XP_011521786.1;XP_011521787.1;XP_01 | 17211 | 605584 | . | . | [illegible] |
| 9786 | NM_00124418 | KIAA0586 | KIAA0586 | protein_coding | NP_001231118.1;NP_001231119.1;NP_001231120.1;NP_001231121.1;NI | 19960 | 610178 | . | . | [illegible] |
| 9787 | NM_00114601 | DLGAP5 | DLG associated protein 5 | protein_coding | NP_001139487.1;NP_055565.3;XP_016877329.1 | 16864 | 617859 | . | . | [illegible] |
| 9788 | NM_00128297 | MTSS1 | MTSS I-BAR domain containing 1 | protein_coding | NP_001269900.1;NP_001269903.1;NP_001350223.1;NP_001350224.1;NI | 20443 | 608486 | . | . | [illegible] |
| 9789 | NM_014752 | SPCS2 | signal peptidase complex subunit 2 | protein_coding | NP_055567.2 | 28962 | . | . | . | [illegible] |
| 9790 | NM_014753,XI | BMS1 | BMS1 ribosome biogenesis factor | protein_coding | NP_055568.3;XP_005271903.1;XP_005271905.1;XP_011538704.1;XP_01 | 23505 | 611448 | . | . | [illegible] |
| 9791 | NM_00129022 | PTDSS1 | phosphatidylserine synthase 1 | protein_coding | NP_001277154.1;NP_055569.1 | 9587 | 612792 | . | . | [illegible] |
| 9792 | NM_014755,XI | SERTAD2 | SERTA domain containing 2 | protein_coding | NP_055570.1;XP_005264726.1;XP_011531507.1 | 30784 | 617851 | . | . | [illegible] |
| 9793 | NM_00100893 | CKAP5 | cytoskeleton associated protein 5 | protein_coding | NP_001008938.1;NP_055571.2 | 28959 | 611142 | . | . | [illegible] |
| 9794 | NM_014757 | MAML1 | mastermind like transcriptional coactivator 1 | protein_coding | NP_055572.1 | 13632 | 605424 | . | . | [illegible] |
| 9796 | NM_00109933 | PHYHIP | phytanoyl-CoA 2-hydroxylase interacting protein | protein_coding | NP_001092805.1;NP_001350240.1;NP_001350241.1;NP_055574.3;XP_01 | 16865 | 608511 | . | . | [illegible] |
| 9797 | NM_014760 | TATDN2 | TatD DNase domain containing 2 | protein_coding | NP_055575.3 | 28988 | . | . | . | [illegible] |
| 9798 | NM_00127097 | IST1 | IST1 factor associated with ESCRT-III | protein_coding | NP_001257904.1;NP_001257905.1;NP_001257906.1;NP_001257907.1;NI | 28977 | 616434 | . | . | [illegible] |
| 9801 | NM_014763 | MRPL19 | mitochondrial ribosomal protein L19 | protein_coding | NP_055578.2 | 14052 | 611832 | . | . | [illegible] |
| 9802 | NM_00113626 | DAZAP2 | DAZ associated protein 2 | protein_coding | NP_001129736.1;NP_001129738.1;NP_001129739.1;NP_001129740.1;NI | 2684 | 607431 | . | . | [illegible] |
| 9804 | NM_014765 | TOMM20 | translocase of outer mitochondrial membrane 20 | protein_coding | NP_055580.1 | 20947 | 601848 | . | . | [illegible] |
| 9805 | NM_00114551 | SCRN1 | secernin 1 | protein_coding | NP_001138985.1;NP_001138986.1;NP_001138987.1;NP_055581.3;XP_02 | 22192 | 614965 | . | . | [illegible] |
| 9806 | NM_00113443 | SPOCK2 | SPARC (osteonectin), cwcv and kazal like domains proteoglycan 2 | protein_coding | NP_001127906.1;NP_001231879.1;NP_055582.1;XP_011538706.1;XP_01 | 13564 | 607988 | . | . | [illegible] |
| 9807 | NM_00100611 | IP6K1 | inositol hexakisphosphate kinase 1 | protein_coding | NP_001006115.1;NP_001229758.1;NP_695005.1 | 18360 | 606991 | . | . | [illegible] |
| 9810 | NM_00120703 | RNF40 | ring finger protein 40 | protein_coding | NP_001193962.1;NP_001193963.1;NP_001273501.1;NP_055586.1;XP_01 | 16867 | 607700 | . | . | [illegible] |
| 9811 | NM_00114239 | CTIF | cap binding complex dependent translation initiation factor | protein_coding | NP_001135869.1;NP_055587.1;XP_005258449.1;XP_006722646.1;XP_00 | 23925 | 613178 | . | . | [illegible] |
| 9812 | NM_00114260 | DELE1 | DAP3 binding cell death enhancer 1 | protein_coding | NP_001136075.1;NP_055588.3;XP_005268604.1;XP_005268605.1;XP_00 | 28969 | 615741 | . | . | [illegible] |
| 9813 | NM_014774 | EFCAB14 | EF-hand calcium binding domain 14 | protein_coding | NP_055589.1 | 29051 | . | . | . | [illegible] |
| 9814 | NM_00100746 | SFI1 | SFI1 centrin binding protein | protein_coding | NP_001007468.1;NP_001245254.1;NP_001245255.1;NP_001245256.1;NI | 29064 | 612765 | . | . | [illegible] |
| 9815 | NM_00113521 | GIT2 | GIT ArfGAP 2 | protein_coding | NP_001128685.1;NP_001128686.1;NP_001317082.1;NP_001317083.1;NI | 4273 | 608564 | . | . | [illegible] |
| 9816 | NM_00131402 | URB2 | URB2 ribosome biogenesis homolog | protein_coding | NP_001300950.1;NP_055592.2;XP_005273417.1 | 28967 | . | . | . | [illegible] |
| 9817 | NM_012289,N | KEAP1 | kelch like ECH associated protein 1 | protein_coding | NP_036421.2;NP_987096.1;XP_005260230.1;XP_005260231.1;XP_01152 | 23177 | 606016 | . | . | [illegible] |
| 9818 | NM_00100856 | NUP58 | nucleoporin 58 | protein_coding | NP_001008564.1;NP_054808.1;XP_005266657.1;XP_011533630.1;XP_01 | 20261 | 607615 | . | . | [illegible] |
| 9819 | NM_00130326 | TSC22D2 | TSC22 domain family member 2 | protein_coding | NP_001290193.1;NP_055594.1;XP_011511639.1 | 29095 | 617724 | . | . | [illegible] |
| 9820 | NM_00116837 | CUL7 | cullin 7 | protein_coding | NP_001161842.2;NP_001361801.1;NP_001361802.1;NP_001361803.1;NI | 21024 | 609577 | . | . | [illegible] |
| 9821 | NM_00108361 | RB1CC1 | RB1 inducible coiled-coil 1 | protein_coding | NP_001077086.1;NP_055596.3;XP_011515945.1;XP_011515946.1;XP_01 | 15574 | 606837 | . | . | [illegible] |
| 9823 | NM_00128223 | ARMCX2 | armadillo repeat containing X-linked 2 | protein_coding | NP_001269160.1;NP_055597.1;NP_808818.1;XP_005278166.1;XP_00527 | 16869 | 300363 | . | . | [illegible] |
| 9824 | NM_00128647 | ARHGAP11A | Rho GTPase activating protein 11A | protein_coding | NP_001273408.1;NP_001273409.1;NP_055598.1;NP_955389.1 | 15783 | 610589 | . | . | [illegible] |
| 9825 | NM_00113577 | SPATA2 | spermatogenesis associated 2 | protein_coding | NP_001129245.1;NP_006029.1;XP_006723957.1 | 14681 | 607662 | . | . | [illegible] |
| 9826 | NM_00137741 | ARHGEF11 | Rho guanine nucleotide exchange factor 11 | protein_coding | NP_001364347.1;NP_001364348.1;NP_055599.1;NP_937879.1;XP_00524 | 14580 | 605708 | . | . | [illegible] |
| 9827 | NM_00108049 | RGP1 | RGP1 homolog, RAB6A GEF complex partner 1 | protein_coding | NP_001073965.2 | 21965 | 615742 | . | . | [illegible] |
| 9828 | NM_014786,XI | ARHGEF17 | Rho guanine nucleotide exchange factor 17 | protein_coding | NP_055601.2;XP_016874112.1;XP_016874113.1 | 21726 | 617043 | . | . | [illegible] |
| 9829 | NM_00125686 | DNAJC6 | DnaJ heat shock protein family (Hsp40) member C6 | protein_coding | NP_001243793.1;NP_001243794.1;NP_055602.1 | 15469 | 608375 | . | . | [illegible] |
| 9830 | NM_014788,N | TRIM14 | tripartite motif containing 14 | protein_coding | NP_055603.2;NP_150088.1;XP_005252377.1;XP_006717401.1;XP_01151 | 16283 | 606556 | . | . | [illegible] |
| 9831 | NM_00108248 | ZNF623 | zinc finger protein 623 | protein_coding | NP_001075949.1;NP_001248772.1;NP_055604.3;XP_006716771.1 | 29084 | . | . | . | [illegible] |
| 9832 | NM_00127093 | JAKMIP2 | janus kinase and microtubule interacting protein 2 | protein_coding | NP_001257863.1;NP_001257870.1;NP_001269211.1;NP_055605.2;XP_01 | 29067 | 611197 | . | . | [illegible] |
| 9833 | NM_00125668 | MELK | maternal embryonic leucine zipper kinase | protein_coding | NP_001243614.1;NP_001243616.1;NP_001243617.1;NP_001243618.1;NI | 16870 | 607025 | . | . | [illegible] |
| 9836 | NM_014793 | LCMT2 | leucine carboxyl methyltransferase 2 | protein_coding | NP_055608.2 | 17558 | 611246 | . | . | [illegible] |
| 9837 | NM_021067,N | GINS1 | GINS complex subunit 1 | protein_coding | NP_066545.3;XP_016883651.1 | 28980 | 610608 | . | . | [illegible] |
| 9839 | NM_00117165 | ZEB2 | zinc finger E-box binding homeobox 2 | protein_coding | NP_001165124.1;NP_055610.1 | 14881 | 605802 | . | . | [illegible] |
| 9841 | NM_00116431 | ZBTB24 | zinc finger and BTB domain containing 24 | protein_coding | NP_001157785.1;NP_055612.2 | 21143 | 614064 | . | . | [illegible] |
| 9842 | NM_00135282 | PLEKHM1 | pleckstrin homology and RUN domain containing M1 | protein_coding | NP_001339754.1;NP_055613.1;XP_006722264.1;XP_011523825.1;XP_01 | 29017 | 611466 | . | . | [illegible] |
| 9844 | NM_00103945 | ELMO1 | engulfment and cell motility 1 | protein_coding | NP_001034548.1;NP_001193409.1;NP_001193411.1;NP_055615.8;NP_5 | 16286 | 606420 | . | . | [illegible] |
| 9846 | NM_012296,N | GAB2 | GRB2 associated binding protein 2 | protein_coding | NP_036428.1;NP_536739.1;XP_006718816.1;XP_011543710.1;XP_02430 | 14458 | 606203 | . | . | [illegible] |
| 9847 | NM_00128617 | C2CD5 | C2 calcium dependent domain containing 5 | protein_coding | NP_001273102.1;NP_001273103.1;NP_001273104.1;NP_001273105.1;NI | 29062 | 618044 | . | . | [illegible] |
| 9848 | NM_00100955 | MFAP3L | microfibril associated protein 3 like | protein_coding | NP_001009554.1;NP_001288576.1;NP_001288577.1;NP_067679.6;XP_00 | 29083 | 616523 | . | . | [illegible] |
| 9849 | NM_00127852 | ZNF518A | zinc finger protein 518A | protein_coding | NP_001265453.1;NP_001265454.1;NP_001265455.1;NP_001317661.1;NI | 29009 | 617733 | . | . | [illegible] |
| 9851 | NM_00135122 | KIAA0753 | KIAA0753 | protein_coding | NP_001338154.1;NP_055619.2;XP_006721675.1;XP_011522392.1;XP_01 | 29110 | 617112 | . | . | [illegible] |
| 9852 | NM_014805 | EPM2AIP1 | EPM2A interacting protein 1 | protein_coding | NP_055620.1 | 19735 | 607911 | . | . | [illegible] |
| 9853 | NM_00113599 | RUSC2 | RUN and SH3 domain containing 2 | protein_coding | NP_001129471.1;NP_001317669.1;NP_055621.2 | 23625 | 611053 | . | . | [illegible] |
| 9854 | NM_00129047 | C2CD2L | C2CD2 like | protein_coding | NP_001277403.1;NP_001369540.1;NP_001369541.1;NP_001369542.1;NI | 29000 | 617582 | . | . | [illegible] |
| 9855 | NM_00128298 | FARP2 | FERM, ARH/RhoGEF and pleckstrin domain protein 2 | protein_coding | NP_001269912.1;NP_001269913.1;NP_055623.1;XP_005247107.1;XP_01 | 16460 | 617586 | . | . | [illegible] |
| 9856 | NM_00116837 | KIAA0319 | KIAA0319 | protein_coding | NP_001161846.1;NP_001161847.1;NP_001161848.1;NP_001161849.1;NI | 21580 | 609269 | . | . | [illegible] |
| 9857 | NM_014810,XI | CEP350 | centrosomal protein 350 | protein_coding | NP_055625.4;XP_011508493.1;XP_011508500.1;XP_016858418.1;XP_01 | 24238 | 617870 | . | . | [illegible] |
| 9858 | NM_014811,XI | PPP1R26 | protein phosphatase 1 regulatory subunit 26 | protein_coding | NP_055626.3;XP_005263468.1;XP_016870844.1;XP_016870845.1;XP_01 | 29089 | 614056 | . | . | [illegible] |
| 9859 | NM_00104240 | CEP170 | centrosomal protein 170 | protein_coding | NP_001035863.1;NP_001035864.1;NP_055627.2;XP_006711906.1;XP_01 | 28920 | 613023 | . | . | [illegible] |
| 9860 | NM_00131268 | LRIG2 | leucine rich repeats and immunoglobulin like domains 2 | protein_coding | NP_001299615.1;NP_055628.1;XP_005271426.1;XP_016858441.1;XP_02 | 20889 | 608869 | . | . | [illegible] |
| 9861 | NM_00127177 | PSMD6 | proteasome 26S subunit, non-ATPase 6 | protein_coding | NP_001258708.1;NP_001258709.1;NP_001258710.1;NP_055629.1;XP_00 | 9564 | 617857 | . | . | [illegible] |
| 9862 | NM_00107951 | MED24 | mediator complex subunit 24 | protein_coding | NP_001072986.1;NP_001254726.1;NP_001317140.1;NP_055630.2 | 22963 | 607000 | . | . | [illegible] |
| 9863 | NM_00130112 | MAGI2 | membrane associated guanylate kinase, WW and PDZ domain cor | protein_coding | NP_001288057.1;NP_036433.2;XP_011515020.1;XP_011515021.1;XP_01 | 18957 | 606382 | . | . | [illegible] |
| 9866 | NM_014818,XI | TRIM66 | tripartite motif containing 66 | protein_coding | NP_055633.1;XP_006718460.1;XP_006718461.1;XP_011518806.1;XP_01 | 29005 | 612000 | . | . | [illegible] |
| 9867 | NM_014819,XI | PJA2 | praja ring finger ubiquitin ligase 2 | protein_coding | NP_055634.3;XP_016865588.1 | 17481 | . | . | . | [illegible] |
| 9868 | NM_014820 | TOMM70 | translocase of outer mitochondrial membrane 70 | protein_coding | NP_055635.3 | 11985 | 606081 | . | . | [illegible] |
| 9869 | NM_00114541 | SETDB1 | SET domain bifurcated histone lysine methyltransferase 1 | protein_coding | NP_001138887.1;NP_001230420.1;NP_001353346.1;NP_001353347.1;NI | 10761 | 604396 | . | . | [illegible] |
| 9870 | NM_00103947 | AREL1 | apoptosis resistant E3 ubiquitin protein ligase 1 | protein_coding | NP_001034568.1;XP_005268301.1;XP_006720407.1;XP_011535717.1;XP | 20363 | 615380 | . | . | [illegible] |
| 9871 | NM_00131806 | SEC24D | SEC24 homolog D, COPII coat complex component | protein_coding | NP_001304995.1;NP_055637.2;XP_005263436.1;XP_016864364.1;XP_02 | 10706 | 607186 | . | . | [illegible] |
| 9873 | NM_014824,XI | FCHSD2 | FCH and double SH3 domains 2 | protein_coding | NP_055639.2;XP_011543711.1;XP_011543712.1;XP_016874121.1 | 29114 | 617556 | . | . | [illegible] |
| 9874 | NM_00113655 | TLK1 | tousled like kinase 1 | protein_coding | NP_001130026.1;NP_001130027.1;NP_036422.3;XP_011510540.1;XP_01 | 11841 | 608438 | . | . | [illegible] |
| 9875 | NM_014825 | URB1 | URB1 ribosome biogenesis homolog | protein_coding | NP_055640.2 | 17344 | 608865 | . | . | [illegible] |
| 9877 | NM_00131923 | ZC3H11A | zinc finger CCCH-type containing 11A | protein_coding | NP_001306167.1;NP_001306168.1;NP_001337190.1;NP_001337191.1;NI | 29093 | 613513 | . | . | [illegible] |
| 9878 | NM_00130352 | TOX4 | TOX high mobility group box family member 4 | protein_coding | NP_001290452.1;NP_055643.1 | 20161 | 614032 | . | . | [illegible] |
| 9879 | NM_00130086 | DDX46 | DEAD-box helicase 46 | protein_coding | NP_001287789.1;NP_055644.2;XP_011542060.1 | 18681 | 617848 | . | . | [illegible] |
| 9880 | NM_014830 | ZBTB39 | zinc finger and BTB domain containing 39 | protein_coding | NP_055645.1 | 29014 | . | . | . | [illegible] |
| 9881 | NM_00132999 | TRANK1 | tetratricopeptide repeat and ankyrin repeat containing 1 | protein_coding | NP_001316927.1;NP_055646.2;XP_011532591.1;XP_016863059.1;XP_01 | 29011 | . | . | . | [illegible] |
| 9882 | NM_00128665 | TBC1D4 | TBC1 domain family member 4 | protein_coding | NP_001273587.1;NP_001273588.1;NP_055647.2;XP_005266660.1;XP_00 | 19165 | 612465 | . | . | [illegible] |
| 9883 | NM_00125719 | POM121 | POM121 transmembrane nucleoporin | protein_coding | NP_001244119.1;NP_001354539.1;NP_742017.1;XP_005250783.1;XP_00 | 19702 | 615753 | . | . | [illegible] |
| 9884 | NM_014834 | LRRC37A | leucine rich repeat containing 37A | protein_coding | NP_055649.4 | 29069 | 616555 | . | . | [illegible] |
| 9885 | NM_00127864 | OSBPL2 | oxysterol binding protein like 2 | protein_coding | NP_001265578.1;NP_001350807.1;NP_055650.1;NP_653081.1 | 15761 | 606731 | . | . | [illegible] |
| 9886 | NM_00124235 | RHOBTB1 | Rho related BTB domain containing 1 | protein_coding | NP_001229288.1;NP_001337831.1;NP_001337832.1;NP_001337833.1;NI | 18738 | 607351 | . | . | [illegible] |
| 9887 | NM_00117406 | SMG7 | SMG7 nonsense mediated mRNA decay factor | protein_coding | NP_001167532.1;NP_001317936.1;NP_001337148.1;NP_001337149.1;NI | 16792 | 610964 | . | . | [illegible] |
| 9889 | NM_014838,XI | ZBED4 | zinc finger BED-type containing 4 | protein_coding | NP_055653.2;XP_024308080.1 | 20721 | 612552 | . | . | [illegible] |
| 9891 | NM_014840 | NUAK1 | NUAK family kinase 1 | protein_coding | NP_055655.1 | 14311 | 608130 | . | . | [illegible] |
| 9894 | NM_00135184 | TELO2 | telomere maintenance 2 | protein_coding | NP_001338775.1;NP_057195.2;XP_011521075.1;XP_011521076.1;XP_01 | 29099 | 611140 | . | . | [illegible] |
| 9895 | NM_00117263 | TECPR2 | tectonin beta-propeller repeat containing 2 | protein_coding | NP_001166102.1;NP_055659.2 | 19957 | 615000 | . | . | [illegible] |
| 9896 | NM_014845,XI | FIG4 | FIG4 phosphoinositide 5-phosphatase | protein_coding | NP_055660.1;XP_011534583.1;XP_016867080.1;XP_016867081.1;XP_01 | 16873 | 609390 | . | . | [illegible] |
| 9897 | NM_00133060 | WASHC5 | WASH complex subunit 5 | protein_coding | NP_001317538.1;NP_055661.3;XP_011515711.1;XP_016869602.1 | 28984 | 610657 | . | . | [illegible] |
| 9898 | NM_00112732 | UBAP2L | ubiquitin associated protein 2 like | protein_coding | NP_001120792.1;NP_001274744.1;NP_001274745.1;NP_001317659.1;NI | 29877 | 616472 | . | . | [illegible] |
| 9900 | NM_00127871 | SV2A | synaptic vesicle glycoprotein 2A | protein_coding | NP_001265648.1;NP_001315603.1;NP_001315604.1;NP_055664.3 | 20566 | 185860 | . | . | [illegible] |
| 9901 | NM_00103311 | SRGAP3 | SLIT-ROBO Rho GTPase activating protein 3 | protein_coding | NP_001028289.1;NP_055665.1;XP_011532597.1;XP_011532598.1;XP_01 | 19744 | 606525 | . | . | [illegible] |
| 9902 | NM_006039,XI | MRC2 | mannose receptor C type 2 | protein_coding | NP_006030.2;XP_011523845.1 | 16875 | 612264 | . | . | [illegible] |
| 9903 | NM_00132430 | KLHL21 | kelch like family member 21 | protein_coding | NP_001311238.1;NP_055666.2 | 29041 | 616262 | . | . | [illegible] |
| 9904 | NM_00114669 | RBM19 | RNA binding motif protein 19 | protein_coding | NP_001140170.1;NP_001140171.1;NP_057280.2;XP_016875769.1;XP_01 | 29098 | 616444 | . | . | [illegible] |
| 9905 | NM_00109850 | SGSM2 | small G protein signaling modulator 2 | protein_coding | NP_001091979.1;NP_001333629.1;NP_055668.2;XP_011522403.1;XP_01 | 29026 | 611418 | . | . | [illegible] |
| 9906 | NM_00119978 | SLC35E2A | solute carrier family 35 member E2A | protein_coding | NP_001186716.1;NP_878258.1 | 20863 | . | . | . | [illegible] |
| 9907 | NM_00136485 | AP5Z1 | adaptor related protein complex 5 subunit zeta 1 | protein_coding | NP_001351787.1;NP_055670.1 | 22197 | 613653 | . | . | [illegible] |
| 9908 | NM_012297,N | G3BP2 | G3BP stress granule assembly factor 2 | protein_coding | NP_036429.2;NP_987100.1;NP_987101.1;XP_005263439.1;XP_00526344 | 30291 | . | . | . | [illegible] |
| 9909 | NM_00136746 | DENND4B | DENN domain containing 4B | protein_coding | NP_001354395.1;NP_055671.2;XP_005245736.1;XP_011508521.1;XP_01 | 29044 | . | . | . | [illegible] |
| 9910 | NM_00103523 | RABGAP1L | RAB GTPase activating protein 1 like | protein_coding | NP_001030307.1;NP_001230692.1;NP_001230693.1;NP_001230694.1;NI | 24663 | 609238 | . | . | [illegible] |
| 9911 | NM_00124292 | TMCC2 | transmembrane and coiled-coil domain family 2 | protein_coding | NP_001229854.1;NP_001284540.1;NP_001284542.1;NP_001317963.1;NI | 24239 | . | . | . | [illegible] |
| 9912 | NM_00132116 | ARHGAP44 | Rho GTPase activating protein 44 | protein_coding | NP_001308093.1;NP_001308095.1;NP_001308096.1;NP_001308097.1;NI | 29096 | 617716 | . | . | [illegible] |
| 9913 | NM_00128272 | SUPT7L | SPT7 like, STAGA complex subunit gamma | protein_coding | NP_001269658.1;NP_001269659.1;NP_001269660.1;NP_001269661.1;NI | 30632 | 612762 | . | . | [illegible] |
| 9915 | NM_014862 | ARNT2 | aryl hydrocarbon receptor nuclear translocator 2 | protein_coding | NP_055677.3 | 16876 | 606036 | . | . | [illegible] |
| 9917 | NM_00132431 | FAM20B | FAM20B glycosaminoglycan xylosylkinase | protein_coding | NP_001311239.1;NP_001311240.1;NP_055679.1;XP_016858490.1;XP_01 | 23017 | 611063 | . | . | [illegible] |

| | | | | | | | | | |
|---|---|---|---|---|---|---|---|---|---|
| 9918 | NM_014865 | NCAPD2 | non-SMC condensin I complex subunit D2 | protein_coding | NP_055680.3 | 24305 | 615638 | . | . |
| 9919 | NM_00127641 | SEC16A | SEC16 homolog A, endoplasmic reticulum export factor | protein_coding | NP_001263347.1;NP_055681.1;XP_005266191.1;XP_011517548.1;XP_01 | 29006 | 612854 | . | . |
| 9920 | NM_014867,XI | KBTBD11 | kelch repeat and BTB domain containing 11 | protein_coding | NP_055682.1;XP_011533073.1;XP_011533074.1;XP_016869603.1;XP_01 | 29104 | 618794 | . | . |
| 9921 | NM_00133047 | RNF10 | ring finger protein 10 | protein_coding | NP_001317403.1;NP_055683.3;XP_005254070.1;XP_016875771.1;XP_01 | 10055 | 615998 | . | . |
| 9922 | NM_00113438 | IQSEC1 | IQ motif and Sec7 domain ArfGEF 1 | protein_coding | NP_001127854.1;NP_001317548.1;NP_001363867.1;NP_055684.3;XP_01 | 29112 | 610166 | . | . |
| 9923 | NM_00108362 | ZBTB40 | zinc finger and BTB domain containing 40 | protein_coding | NP_001077090.1;NP_001317327.1;NP_055685.3;XP_011540801.1;XP_01 | 29045 | 612106 | . | . |
| 9924 | NM_00112746 | PAN2 | poly(A) specific ribonuclease subunit PAN2 | protein_coding | NP_001120932.2;NP_001159751.2;NP_055686.4 | 20074 | 617447 | . | . |
| 9925 | NM_014872,XI | ZBTB5 | zinc finger and BTB domain containing 5 | protein_coding | NP_055687.1;XP_005251691.1 | 23836 | 616590 | . | . |
| 9926 | NM_00132080 | LPGAT1 | lysophosphatidylglycerol acyltransferase 1 | protein_coding | NP_001307737.1;NP_001362770.1;NP_001362771.1;NP_001362773.1;NI | 28985 | 610473 | . | . |
| 9927 | NM_00112766 | MFN2 | mitofusin 2 | protein_coding | NP_001121132.1;NP_055689.1;XP_005263600.1;XP_005263602.1;XP_00 | 16877 | 608507 | . | . |
| 9928 | NM_00130579 | KIF14 | kinesin family member 14 | protein_coding | NP_001292721.1;NP_055690.1;XP_011508533.1;XP_011508534.1;XP_01 | 19181 | 611279 | . | . |
| 9929 | NM_00136023 | JOSD1 | Josephin domain containing 1 | protein_coding | NP_001347164.1;NP_001347165.1;NP_055691.1;XP_005261935.1 | 28953 | 615323 | . | . |
| 9931 | NM_00133044 | HELZ | helicase with zinc finger | protein_coding | NP_001317376.2;NP_055692.3;XP_005257945.1;XP_006722277.1;XP_00 | 16878 | 606699 | . | . |
| 9933 | NM_014878 | PUM3 | pumilio RNA binding family member 3 | protein_coding | NP_055693.4 | 29676 | 609960 | . | . |
| 9935 | NM_005461 | MAFB | MAF bZIP transcription factor B | protein_coding | NP_005452.2 | 6408 | 608968 | . | . |
| 9936 | NM_00119876 | CD302 | CD302 molecule | protein_coding | NP_001185692.1;NP_001185693.1;NP_055695.2 | 30843 | 612246 | . | . |
| 9937 | NM_00127181 | DCLRE1A | DNA cross-link repair 1A | protein_coding | NP_001258745.1;NP_055696.3;XP_006718153.1;XP_011538731.1 | 17660 | 609682 | . | . |
| 9939 | NM_005105 | RBM8A | RNA binding motif protein 8A | protein_coding | NP_005096.1 | 9905 | 605313 | . | . |
| 9941 | NM_00114546 | EXOG | exo/endonuclease G | protein_coding | NP_001138936.1;NP_005098.2;XP_005265692.1;XP_006713505.1;XP_01 | 3347 | 604051 | . | . |
| 9942 | NM_00134917 | XYLB | xylulokinase | protein_coding | NP_001336107.1;NP_001336108.1;NP_001336109.1;NP_005099.2;XP_01 | 12839 | 604049 | . | . |
| 9943 | NM_005109,XI | OXSR1 | oxidative stress responsive kinase 1 | protein_coding | NP_005100.1;XP_005265695.1;XP_011532633.1;XP_016863090.1;XP_02 | 8508 | 604046 | . | . |
| 9945 | NM_005110 | GFPT2 | glutamine-fructose-6-phosphate transaminase 2 | protein_coding | NP_005101.1 | 4242 | 603865 | . | . |
| 9946 | NM_145858 | CRYZL1 | crystallin zeta like 1 | protein_coding | NP_665857.2 | 2420 | 603920 | . | . |
| 9947 | NM_005462,XI | MAGEC1 | MAGE family member C1 | protein_coding | NP_005453.2;XP_011529720.1 | 6812 | 300223 | . | . |
| 9948 | NM_005112,N | WDR1 | WD repeat domain 1 | protein_coding | NP_005103.2;NP_059830.1;XP_016864369.1 | 12754 | 604734 | . | . |
| 9949 | NM_00102558 | AMMECR1 | AMMECR nuclear protein 1 | protein_coding | NP_001020751.1;NP_001165160.1;NP_056180.1 | 467 | 300195 | . | . |
| 9950 | NM_005113,XI | GOLGA5 | golgin A5 | protein_coding | NP_005104.4;XP_011535722.1 | 4428 | 606918 | . | . |
| 9953 | NM_006041,N | HS3ST3B1 | heparan sulfate-glucosamine 3-sulfotransferase 3B1 | protein_coding | NP_006032.1;XP_016880968.1 | 5198 | 604058 | . | . |
| 9955 | NM_006042,XI | HS3ST3A1 | heparan sulfate-glucosamine 3-sulfotransferase 3A1 | protein_coding | NP_006033.1;XP_011522416.1;XP_016880969.1 | 5196 | 604057 | . | . |
| 9957 | NM_005114,XI | HS3ST1 | heparan sulfate-glucosamine 3-sulfotransferase 1 | protein_coding | NP_005105.1;XP_011512215.1 | 5194 | 603244 | . | . |
| 9958 | NM_00125207 | USP15 | ubiquitin specific peptidase 15 | protein_coding | NP_001239007.1;NP_001239008.1;NP_001338088.1;NP_001338089.1;NI | 12613 | 604731 | . | . |
| 9960 | NM_00125670 | USP3 | ubiquitin specific peptidase 3 | protein_coding | NP_001243631.1;NP_006528.2;XP_016878252.1;XP_016878253.1;XP_01 | 12626 | 604728 | . | . |
| 9961 | NM_00129320 | MVP | major vault protein | protein_coding | NP_001280133.1;NP_001280134.1;NP_005106.2;NP_059447.2 | 7531 | 605088 | . | . |
| 9962 | NM_005116,N | SLC23A2 | solute carrier family 23 member 2 | protein_coding | NP_005107.4;NP_976072.1 | 10973 | 603791 | . | . |
| 9967 | NM_00132147 | THRAP3 | thyroid hormone receptor associated protein 3 | protein_coding | NP_001308400.1;NP_001308402.1;NP_005110.2;XP_024307069.1;XP_02 | 22964 | 603809 | . | . |
| 9968 | NM_005120 | MED12 | mediator complex subunit 12 | protein_coding | NP_005111.2 | 11957 | 300188 | . | . |
| 9969 | NM_005121,XI | MED13 | mediator complex subunit 13 | protein_coding | NP_005112.2;XP_011523853.1;XP_011523854.1;XP_011523855.1 | 22474 | 603808 | . | . |
| 9972 | NM_00127820 | NUP153 | nucleoporin 153 | protein_coding | NP_001265138.1;NP_001265139.1;NP_005115.2;XP_005249564.1;XP_00 | 8062 | 603948 | . | . |
| 9973 | NM_005125,XI | CCS | copper chaperone for superoxide dismutase | protein_coding | NP_005116.1 | 1613 | 603864 | . | . |
| 9975 | NM_00114542 | NR1D2 | nuclear receptor subfamily 1 group D member 2 | protein_coding | NP_001138897.1;NP_005117.3;XP_006713514.1 | 7963 | 602304 | . | . |
| 9976 | NM_005127 | CLEC2B | C-type lectin domain family 2 member B | protein_coding | NP_005118.2 | 2053 | 603242 | . | . |
| 9978 | NM_014248 | RBX1 | ring-box 1 | protein_coding | NP_055063.1 | 9928 | 603814 | . | . |
| 9980 | NM_00132071 | DOP1B | DOP1 leucine zipper like protein B | protein_coding | NP_001307643.1;NP_005119.2;XP_016883998.1 | 1291 | 604803 | . | . |
| 9984 | NM_005131,XI | THOC1 | THO complex 1 | protein_coding | NP_005122.2;XP_011524074.1;XP_011524075.1;XP_011524076.1;XP_01 | 19070 | 606930 | . | . |
| 9985 | NM_00104820 | REC8 | REC8 meiotic recombination protein | protein_coding | NP_001041670.1;NP_005123.2;XP_011535723.1;XP_016877330.1;XP_02 | 16879 | 608193 | . | . |
| 9986 | NM_00103227 | RCE1 | Ras converting CAAX endopeptidase 1 | protein_coding | NP_001027450.1;NP_005124.1 | 13721 | 605385 | . | . |
| 9987 | NM_00120700 | HNRNPDL | heterogeneous nuclear ribonucleoprotein D like | protein_coding | NP_001193929.1;NP_112740.1 | 5037 | 607137 | . | . |
| 9988 | NM_00114232 | DMTF1 | cyclin D binding myb like transcription factor 1 | protein_coding | NP_001135798.1;NP_001135799.1;NP_066968.3;XP_011515037.1;XP_01 | 14603 | 608491 | . | . |
| 9989 | NM_00104238 | PPP4R1 | protein phosphatase 4 regulatory subunit 1 | protein_coding | NP_001035847.1;NP_001369491.1;NP_005125.1;XP_011524077.1;XP_01 | 9320 | 604908 | . | . |
| 9990 | NM_00104249 | SLC12A6 | solute carrier family 12 member 6 | protein_coding | NP_001035959.1;NP_001035960.1;NP_001035961.1;NP_001035962.1;NI | 10914 | 604878 | . | . |
| 9991 | NM_00116378 | PTBP3 | polypyrimidine tract binding protein 3 | protein_coding | NP_001157260.1;NP_001157262.1;NP_001231825.1;NP_001231826.1;NI | 10253 | 607527 | . | . |
| 9993 | NM_00117353 | DGCR2 | DiGeorge syndrome critical region gene 2 | protein_coding | NP_001167004.1;NP_001167005.1;NP_001171710.1;NP_005128.1 | 2845 | 600594 | . | . |
| 9994 | NM_00113766 | CASP8AP2 | caspase 8 associated protein 2 | protein_coding | NP_001131139.1;NP_001131140.1;NP_036247.1 | 1510 | 606880 | . | . |
| 9997 | NM_00116910 | SCO2 | synthesis of cytochrome C oxidase 2 | protein_coding | NP_001162580.1;NP_001162581.1;NP_001162582.1;NP_005129.2 | 10604 | 604272 | . | . |
| 10000 | NM_00120672 | AKT3 | AKT serine/threonine kinase 3 | protein_coding | NP_001193658.1;NP_001357003.1;NP_005456.1;NP_859029.1;XP_0115 | 393 | 611223 | . | . |
| 10001 | NM_00128420 | MED6 | mediator complex subunit 6 | protein_coding | NP_001271138.1;NP_001271139.1;NP_001271140.1;NP_005457.2 | 19970 | 602984 | . | . |
| 10005 | NM_005469,XI | ACOT8 | acyl-CoA thioesterase 8 | protein_coding | NP_005460.2;XP_005260296.1;XP_011526782.1 | 15919 | 608123 | . | . |
| 10006 | NM_00101275 | ABI1 | abl interactor 1 | protein_coding | NP_001012768.1;NP_001012769.1;NP_001012770.1;NP_001171587.1;NI | 11320 | 603050 | . | . |
| 10007 | NM_005471,XI | GNPDA1 | glucosamine-6-phosphate deaminase 1 | protein_coding | NP_005462.1;XP_005268405.1;XP_006714810.1;XP_011535839.1;XP_02 | 4417 | 601798 | . | . |
| 10009 | NM_00118474 | ZBTB33 | zinc finger and BTB domain containing 33 | protein_coding | NP_001171671.1;NP_006768.1 | 16682 | 300329 | . | . |
| 10010 | NM_00119913 | TANK | TRAF family member associated NFKB activator | protein_coding | NP_001186064.1;NP_004171.2;NP_597841.1;XP_005246264.2;XP_01685 | 11562 | 603893 | . | . |
| 10011 | NM_00103523 | SRA1 | steroid receptor RNA activator 1 | protein_coding | NP_001030312.2;NP_001240693.1 | 11281 | 603819 | . | . |
| 10013 | NM_00132122 | HDAC6 | histone deacetylase 6 | protein_coding | NP_001308154.1;NP_001308155.1;NP_001308156.1;NP_001308157.1;NI | 14064 | 300272 | . | . |
| 10014 | NM_00101505 | HDAC5 | histone deacetylase 5 | protein_coding | NP_001015053.1;NP_001369322.1;NP_005465.2;XP_005256962.1;XP_01 | 14068 | 605315 | . | . |
| 10015 | NM_00116242 | PDCD6IP | programmed cell death 6 interacting protein | protein_coding | NP_001155901.1;NP_001243121.1;NP_037506.2;XP_011531554.1;XP_01 | 8766 | 608074 | . | . |
| 10016 | NM_00126755 | PDCD6 | programmed cell death 6 | protein_coding | NP_001254485.1;NP_001254486.1;NP_001254487.1;NP_001254488.1;NI | 8765 | 601057 | . | . |
| 10018 | NM_00120410 | BCL2L11 | BCL2 like 11 | protein_coding | NP_001191035.1;NP_001191036.1;NP_001191037.1;NP_001191038.1;NI | 994 | 603827 | . | . |
| 10019 | NM_00129142 | SH2B3 | SH2B adaptor protein 3 | protein_coding | NP_001278353.1;NP_005466.1;XP_005253875.1;XP_005253876.1;XP_00 | 29605 | 605093 | . | . |
| 10020 | NM_00112822 | GNE | glucosamine (UDP-N-acetyl)-2-epimerase/N-acetylmannosamine k | protein_coding | NP_001121699.1;NP_001177312.1;NP_001177313.1;NP_001177317.2;NI | 23657 | 603824 | . | . |
| 10023 | NM_005479 | FRAT1 | FRAT regulator of WNT signaling pathway 1 | protein_coding | NP_005470.2 | 3944 | 602503 | . | . |
| 10024 | NM_00110062 | TROAP | trophinin associated protein | protein_coding | NP_001094090.1;NP_001265253.1;NP_005471.3;XP_006719244.1;XP_01 | 12327 | 603872 | . | . |
| 10025 | NM_005481,XI | MED16 | mediator complex subunit 16 | protein_coding | NP_005472.2;XP_016881609.1;XP_016881610.1 | 17556 | 604062 | . | . |
| 10026 | NM_005482 | PIGK | phosphatidylinositol glycan anchor biosynthesis class K | protein_coding | NP_005473.1 | 8965 | 605087 | . | . |
| 10036 | NM_005483,XI | CHAF1A | chromatin assembly factor 1 subunit A | protein_coding | NP_005474.2;XP_011525907.1;XP_011525908.1;XP_011525909.1 | 1910 | 601246 | . | . |
| 10038 | NM_00104261 | PARP2 | poly(ADP-ribose) polymerase 2 | protein_coding | NP_001036083.1;NP_005475.2;XP_005267304.1;XP_016876401.1 | 272 | 607725 | . | . |
| 10039 | NM_00100393 | PARP3 | poly(ADP-ribose) polymerase family member 3 | protein_coding | NP_001003931.3;NP_001357168.1;NP_001357169.1;NP_005476.4;XP_01 | 273 | 607726 | . | . |
| 10040 | NM_00132117 | TOM1L1 | target of myb1 like 1 membrane trafficking protein | protein_coding | NP_001308102.1;NP_001308103.1;NP_001308104.1;NP_001308105.1;NI | 11983 | 604701 | . | . |
| 10042 | NM_00100368 | HMGXB4 | HMG-box containing 4 | protein_coding | NP_001003681.1;NP_001349901.1;XP_006724164.1;XP_006724165.1 | 5003 | 604702 | . | . |
| 10043 | NM_00113572 | TOM1 | target of myb1 membrane trafficking protein | protein_coding | NP_001129201.1;NP_001129202.1;NP_001129204.1;NP_005479.1;XP_01 | 11982 | 604700 | . | . |
| 10044 | NM_00114253 | SH2D3C | SH2 domain containing 3C | protein_coding | NP_001136003.1;NP_001136004.1;NP_001136005.1;NP_001136006.1;NI | 16884 | 604722 | . | . |
| 10046 | NM_00117746 | MAMLD1 | mastermind like domain containing 1 | protein_coding | NP_001170936.1;NP_001170937.1;NP_005482.2;XP_006724865.1;XP_00 | 2568 | 300120 | . | . |
| 10048 | NM_005493,XI | RANBP9 | RAN binding protein 9 | protein_coding | NP_005484.2;XP_006715008.1;XP_011512507.1;XP_016865638.1 | 13727 | 603854 | . | . |
| 10049 | NM_00136367 | DNAJB6 | DnaJ heat shock protein family (Hsp40) member B6 | protein_coding | NP_001350605.1;NP_005485.1;NP_490647.1;XP_005249572.1;XP_00524 | 14888 | 611332 | . | . |
| 10051 | NM_00100280 | SMC4 | structural maintenance of chromosomes 4 | protein_coding | NP_001002800.1;NP_001275682.1;NP_005487.3;XP_006713522.1;XP_01 | 14013 | 605575 | . | . |
| 10052 | NM_00108038 | GJC1 | gap junction protein gamma 1 | protein_coding | NP_001073852.1;NP_005488.2;XP_005256977.1;XP_024306293.1;XP_02 | 4280 | 608655 | . | . |
| 10054 | NM_005499,XI | UBA2 | ubiquitin like modifier activating enzyme 2 | protein_coding | NP_005490.1;XP_005258461.2;XP_006723025.1;XP_011524606.1;XP_01 | 30661 | 613295 | . | . |
| 10055 | NM_00114571 | SAE1 | SUMO1 activating enzyme subunit 1 | protein_coding | NP_001139185.1;NP_001139186.1;NP_005491.1 | 30660 | 613294 | . | . |
| 10056 | NM_005687,N | FARSB | phenylalanyl-tRNA synthetase subunit beta | protein_coding | NP_005678.3;XP_006712232.1;XP_011508768.1;XP_016858599.1;XP_02 | 17800 | 609690 | . | . |
| 10057 | NM_00102358 | ABCC5 | ATP binding cassette subfamily C member 5 | protein_coding | NP_001018881.1;NP_001306961.1;NP_005679.2;XP_005247115.1;XP_00 | 56 | 605251 | . | . |
| 10058 | NM_00134982 | ABCB6 | ATP binding cassette subfamily B member 6 (Langereis blood grou | protein_coding | NP_001336757.1;NP_005680.1 | 47 | 605452 | . | . |
| 10059 | NM_00127846 | DNM1L | dynamin 1 like | protein_coding | NP_001265392.1;NP_001265393.1;NP_001265394.1;NP_001265395.1;NI | 2973 | 603850 | . | . |
| 10060 | NM_00137727 | ABCC9 | ATP binding cassette subfamily C member 9 | protein_coding | NP_001364202.1;NP_001364203.1;NP_005682.2;NP_064693.2;XP_00525 | 60 | 601439 | . | . |
| 10061 | NM_007189 | ABCF2 | ATP binding cassette subfamily F member 2 | protein_coding | NP_009120.1 | 71 | 612510 | . | . |
| 10062 | NM_00113010 | NR1H3 | nuclear receptor subfamily 1 group H member 3 | protein_coding | NP_001123573.1;NP_001123574.1;NP_001238863.1;NP_001238864.1;NI | 7966 | 602423 | . | . |
| 10063 | NM_00138200 | COX17 | cytochrome c oxidase copper chaperone COX17 | protein_coding | NP_001368931.1;NP_001368932.1;NP_005685.1 | 2264 | 604813 | . | . |
| 10064 | gene-VDAC1P2 | VDAC1P2 | voltage dependent anion channel 1 pseudogene 2 | pseudogene | . | 12670 | . | . | . |
| 10066 | NM_00132077 | SCAMP2 | secretory carrier membrane protein 2 | protein_coding | NP_001307707.1;NP_005688.2;XP_016877345.1 | 10564 | 606912 | . | . |
| 10067 | NM_005698,N | SCAMP3 | secretory carrier membrane protein 3 | protein_coding | NP_005689.2;NP_443069.1 | 10565 | 606913 | . | . |
| 10068 | NM_00103965 | IL18BP | interleukin 18 binding protein | protein_coding | NP_001034748.1;NP_001034749.1;NP_001138527.1;NP_001138529.1;NI | 5987 | 604113 | . | . |
| 10069 | NM_00132072 | RWDD2B | RWD domain containing 2B | protein_coding | NP_001307653.1;NP_058636.1 | 1302 | 617843 | . | . |
| 10072 | NM_00125667 | DPP3 | dipeptidyl peptidase 3 | protein_coding | NP_001243599.1;NP_005691.2;NP_569710.2 | 3008 | 606818 | . | . |
| 10073 | NM_00104258 | SNUPN | snurportin 1 | protein_coding | NP_001036046.1;NP_001036053.1;NP_005692.1 | 14245 | 607902 | . | . |
| 10075 | NM_031407,XI | HUWE1 | HECT, UBA and WWE domain containing E3 ubiquitin protein ligas | protein_coding | NP_113584.3;XP_005262022.1;XP_011529049.1;XP_011529053.1;XP_01 | 30892 | 300697 | . | . |
| 10076 | NM_00119500 | PTPRU | protein tyrosine phosphatase receptor type U | protein_coding | NP_001181930.1;NP_005695.3;NP_573438.3;NP_573439.2;XP_0067103 | 9683 | 602454 | . | . |
| 10078 | NM_00129765 | TSSC4 | tumor suppressing subtransferable candidate 4 | protein_coding | NP_001284587.1;NP_001284588.1;NP_001284589.1;NP_001284590.1;NI | 12386 | 603852 | . | . |
| 10079 | NM_006045,XI | ATP9A | ATPase phospholipid transporting 9A (putative) | protein_coding | NP_006036.1;XP_011526783.1 | 13540 | 609126 | . | . |
| 10081 | NM_005707 | PDCD7 | programmed cell death 7 | protein_coding | NP_005698.1 | 8767 | 608138 | . | . |
| 10082 | NM_005708,XI | GPC6 | glypican 6 | protein_coding | NP_005699.1;XP_011519346.1;XP_016875787.1;XP_016875788.1;XP_01 | 4454 | 604404 | . | . |
| 10084 | NM_00103238 | PQBP1 | polyglutamine binding protein 1 | protein_coding | NP_001027553.1;NP_001027554.1;NP_001027555.1;NP_001027556.1;NI | 9330 | 300463 | . | . |
| 10085 | NM_00127864 | EDIL3 | EGF like repeats and discoidin domains 3 | protein_coding | NP_001265571.1;NP_005702.3 | 3173 | 606018 | . | . |
| 10087 | NM_00113010 | CERT1 | ceramide transporter 1 | protein_coding | NP_001123577.1;NP_001365931.1;NP_001365932.1;NP_001365933.1;NI | 2205 | 604677 | . | . |
| 10090 | NM_005715,XI | UST | uronyl 2-sulfotransferase | protein_coding | NP_005706.1;XP_011533680.1;XP_016865641.1 | 17223 | 610752 | . | . |
| 10092 | NM_00127043 | ARPC5 | actin related protein 2/3 complex subunit 5 | protein_coding | NP_001257368.1;NP_005708.1 | 708 | 604227 | . | . |
| 10093 | NM_00102495 | ARPC4 | actin related protein 2/3 complex subunit 4 | protein_coding | NP_001020130.1;NP_001020131.1;NP_001185709.1;NP_005709.1 | 707 | 604226 | . | . |
| 10094 | NM_00127855 | ARPC3 | actin related protein 2/3 complex subunit 3 | protein_coding | NP_001265485.1;NP_001274151.1 | 706 | 604225 | . | . |
| 10095 | NM_005720,XI | ARPC1B | actin related protein 2/3 complex subunit 1B | protein_coding | NP_005711.1;XP_006715888.1;XP_006715889.1;XP_024302396.1;XP_02 | 704 | 604223 | . | . |
| 10096 | NM_00127714 | ACTR3 | actin related protein 3 | protein_coding | NP_001264069.1;NP_005712.1 | 170 | 604222 | . | . |
| 10097 | NM_00100538 | ACTR2 | actin related protein 2 | protein_coding | NP_001005386.1;NP_005713.1 | 169 | 604221 | . | . |
| 10098 | NM_005723,XI | TSPAN5 | tetraspanin 5 | protein_coding | NP_005714.2;XP_005262737.1 | 17753 | 613136 | . | . |
| 10099 | NM_00116841 | TSPAN3 | tetraspanin 3 | protein_coding | NP_001161884.1;NP_005715.1;NP_944492.1;XP_016877346.1 | 17752 | 613134 | . | . |
| 10101 | NM_00128450 | NUBP2 | nucleotide binding protein 2 | protein_coding | NP_001271430.1;NP_001271431.1;NP_036357.1;XP_005255084.1;XP_00 | 8042 | 610779 | . | . |
| 10102 | NM_00117269 | TSFM | Ts translation elongation factor, mitochondrial | protein_coding | NP_001166166.1;NP_001166167.1;NP_001166168.1;NP_005717.3 | 12367 | 604723 | . | . |
| 10103 | NM_005727,XI | TSPAN1 | tetraspanin 1 | protein_coding | NP_005718.2;XP_011538762.1 | 20657 | 613170 | . | . |
| 10105 | NM_005729,XI | PPIF | peptidylprolyl isomerase F | protein_coding | NP_005720.1;XP_005269436.1 | 9259 | 604486 | . | . |
| 10106 | NM_005730,XI | CTDSP2 | CTD small phosphatase 2 | protein_coding | NP_005721.3;XP_005268613.1 | 17077 | 608711 | . | . |
| 10109 | NM_005731,N | ARPC2 | actin related protein 2/3 complex subunit 2 | protein_coding | NP_005722.1;NP_690601.1;XP_016858602.1;XP_016858603.1;XP_01685 | 705 | 604224 | . | . |
| 10111 | NM_005732 | RAD50 | RAD50 double strand break repair protein | protein_coding | NP_005723.2 | 9816 | 604040 | . | . |
| 10112 | NM_005733 | KIF20A | kinesin family member 20A | protein_coding | NP_005724.1 | 9787 | 605664 | . | . |
| 10113 | NM_00133048 | PREB | prolactin regulatory element binding | protein_coding | NP_001317413.1;NP_001317414.1;NP_001317415.1;NP_001317416.1;NI | 9356 | 606395 | . | . |
| 10114 | NM_00104820 | HIPK3 | homeodomain interacting protein kinase 3 | protein_coding | NP_001041665.1;NP_001265091.1;NP_001265092.1;NP_005725.3;XP_0C | 4915 | 604424 | . | . |
| 10116 | NM_015322 | FEM1B | fem-1 homolog B | protein_coding | NP_056137.1 | 3649 | 613539 | . | . |
| 10120 | NM_005735,XI | ACTR1B | actin related protein 1B | protein_coding | NP_005726.1;XP_005263911.1;XP_016858605.1 | 168 | 605144 | . | . |
| 10121 | NM_005736 | ACTR1A | actin related protein 1A | protein_coding | NP_005727.1 | 167 | 605143 | . | . |
| 10123 | NM_00128243 | ARL4C | ADP ribosylation factor like GTPase 4C | protein_coding | NP_001269360.1;NP_005728.2 | 698 | 604787 | . | . |
| 10124 | NM_00103716 | ARL4A | ADP ribosylation factor like GTPase 4A | protein_coding | NP_001032241.1;NP_001182325.1;NP_005729.1;NP_997625.1 | 695 | 604786 | . | . |
| 10126 | NM_005740 | DNAL4 | dynein axonemal light chain 4 | protein_coding | NP_005731.1 | 2955 | 610565 | . | . |
| 10127 | NM_005741,XI | ZNF263 | zinc finger protein 263 | protein_coding | NP_005732.2;XP_006720894.1;XP_006720895.1;XP_006720896.1;XP_00 | 13056 | 604191 | . | . |
| 10128 | NM_133259,XI | LRPPRC | leucine rich pentatricopeptide repeat containing | protein_coding | NP_573566.2;XP_006711978.1;XP_006711979.1;XP_011530775.1;XP_01 | 15714 | 607544 | . | . |
| 10129 | NM_023037,XI | FRY | FRY microtubule binding protein | protein_coding | NP_075463.2;XP_006719812.1;XP_016875794.1;XP_016875795.1;XP_01 | 20367 | 614818 | . | . |
| 10130 | NM_00128270 | PDIA6 | protein disulfide isomerase family A member 6 | protein_coding | NP_001269633.1;NP_001269634.1;NP_001269635.1;NP_001269636.1;NI | 30168 | 611099 | . | . |
| 10131 | NM_00127204 | TRAP1 | TNF receptor associated protein 1 | protein_coding | NP_001258978.1;NP_057376.2;XP_011520647.1 | 16264 | 606219 | . | . |
| 10133 | NM_00100821 | OPTN | optineurin | protein_coding | NP_001008212.1;NP_001008213.1;NP_001008214.1;NP_068815.2 | 17142 | 602432 | . | . |
| 10134 | NM_00113944 | BCAP31 | B cell receptor associated protein 31 | protein_coding | NP_001132913.1;NP_001132929.1;NP_001243376.1;NP_005736.3 | 16695 | 300398 | . | . |
| 10135 | NM_005746 | NAMPT | nicotinamide phosphoribosyltransferase | protein_coding | NP_005737.1 | 30092 | 608764 | . | . |
| 10137 | NM_00119883 | RBM12 | RNA binding motif protein 12 | protein_coding | NP_001185767.1;NP_001185769.1;NP_006038.2;NP_690051.1 | 9898 | 607179 | . | . |
| 10138 | NM_00119097 | YAF2 | YY1 associated factor 2 | protein_coding | NP_001177906.1;NP_001177908.1;NP_001177909.1;NP_001307009.1;NI | 17363 | 607534 | . | . |
| 10139 | NM_00113475 | ARFRP1 | ADP ribosylation factor related protein 1 | protein_coding | NP_001128230.1;NP_001254473.1;NP_001254474.1;NP_001254475.1;NI | 662 | 604699 | . | . |
| 10140 | NM_00124387 | TOB1 | transducer of ERBB2, 1 | protein_coding | NP_001230806.1;NP_001230814.1;NP_005740.1 | 11979 | 605523 | . | . |
| 10142 | NM_00137927 | AKAP9 | A-kinase anchoring protein 9 | protein_coding | NP_001366206.1;NP_005742.4;NP_671714.1;XP_016867131.1;XP_01686 | 379 | 604001 | . | . |
| 10144 | NM_00101504 | FAM13A | family with sequence similarity 13 member A | protein_coding | NP_001015045.1;NP_001252507.1;NP_001252508.1;NP_001252509.1;NI | 19367 | 613299 | . | . |
| 10146 | NM_005754,N | G3BP1 | G3BP stress granule assembly factor 1 | protein_coding | NP_005745.1;NP_938405.1 | 30292 | 608431 | . | . |
| 10147 | NM_00101739 | SUGP2 | SURP and G-patch domain containing 2 | protein_coding | NP_001017392.2;NP_001308626.1;NP_001308627.1;NP_001308628.1;NI | 18641 | 607993 | . | . |
| 10149 | NM_00107985 | ADGRG2 | adhesion G protein-coupled receptor G2 | protein_coding | NP_001073327.1;NP_001073328.1;NP_001073329.1;NP_001171762.1;NI | 4516 | 300572 | . | . |
| 10150 | NM_00130607 | MBNL2 | muscleblind like splicing regulator 2 | protein_coding | NP_001292999.1;NP_001369578.1;NP_001369579.1;NP_001369580.1;NI | 16746 | 607327 | . | . |
| 10152 | NM_00128292 | ABI2 | abl interactor 2 | protein_coding | NP_001269854.1;NP_001269855.1;NP_001269856.1;NP_001269861.1;NI | 24011 | 606442 | . | . |
| 10153 | NM_005760 | CEBPZ | CCAAT enhancer binding protein zeta | protein_coding | NP_005751.2 | 24218 | 612828 | . | . |
| 10154 | NM_005761,N | PLXNC1 | plexin C1 | protein_coding | NP_005752.1;XP_006719249.1;XP_011536032.1;XP_011536033.1;XP_01 | 9106 | 604259 | . | . |
| 10155 | NM_005762,XI | TRIM28 | tripartite motif containing 28 | protein_coding | NP_005753.1;XP_024307077.1 | 16384 | 601742 | . | . |
| 10156 | NM_00107987 | RASA4 | RAS p21 protein activator 4 | protein_coding | NP_001073346.2;NP_008920.5;XP_011514025.1;XP_016867150.1;XP_01 | 23181 | 607943 | . | . |
| 10157 | NM_005763,XI | AASS | aminoadipate-semialdehyde synthase | protein_coding | NP_005754.2;XP_011514027.1 | 17366 | 605113 | . | . |
| 10159 | NM_005765 | ATP6AP2 | ATPase H+ transporting accessory protein 2 | protein_coding | NP_005756.2 | 18305 | 300556 | . | . |
| 10160 | NM_00100171 | FARP1 | FERM, ARH/RhoGEF and pleckstrin domain protein 1 | protein_coding | NP_001001715.2;NP_001273768.1;NP_005757.1;XP_011519348.1;XP_01 | 3591 | 602654 | . | . |

| | | | | | | | | | | |
|---|---|---|---|---|---|---|---|---|---|---|
| 10161 | NM_00116249 | LPAR6 | lysophosphatidic acid receptor 6 | protein_coding | NP_001155969.1;NP_001155970.1;NP_001364245.1;NP_001364246.1;NI | 15520 | 609239 | . | . | [illegible] |
| 10162 | NM_005768,XI | LPCAT3 | lysophosphatidylcholine acyltransferase 3 | protein_coding | NP_005759.4;XP_024304561.1 | 30244 | 611950 | . | . | [illegible] |
| 10163 | NM_00120140 | WASF2 | WASP family member 2 | protein_coding | NP_001188333.1;NP_008921.1 | 12733 | 605875 | . | . | [illegible] |
| 10165 | NM_00116021 | SLC25A13 | solute carrier family 25 member 13 | protein_coding | NP_001153682.1;NP_055066.1;XP_006715894.1;XP_011514029.1;XP_01 | 10983 | 603859 | . | . | [illegible] |
| 10166 | NM_014252 | SLC25A15 | solute carrier family 25 member 15 | protein_coding | NP_055067.1 | 10985 | 603861 | . | . | [illegible] |
| 10168 | NM_00102485 | ZNF197 | zinc finger protein 197 | protein_coding | NP_001020026.1;NP_001310222.1;NP_001310223.1;NP_001310224.1;NI | 12988 | 618359 | . | . | [illegible] |
| 10169 | NM_00101810 | SERF2 | small EDRK-rich factor 2 | protein_coding | NP_001018118.1;NP_001186804.1;NP_001186805.1;NP_001186806.1;NI | 10757 | 605054 | . | . | [illegible] |
| 10171 | NM_00128669 | RCL1 | RNA terminal phosphate cyclase like 1 | protein_coding | NP_001273628.1;NP_001273629.1;NP_001273630.1;NP_005763.3;XP_0C | 17687 | 611405 | . | . | [illegible] |
| 10172 | NM_00137540 | ZNF256 | zinc finger protein 256 | protein_coding | NP_001362332.1;NP_005764.2 | 13049 | 606956 | . | . | [illegible] |
| 10174 | NM_00101800 | SORBS3 | sorbin and SH3 domain containing 3 | protein_coding | NP_001018003.1;NP_005766.3;XP_005273428.1;XP_006716329.1;XP_01 | 30907 | 610795 | . | . | [illegible] |
| 10175 | NM_005776 | CNIH1 | cornichon family AMPA receptor auxiliary protein 1 | protein_coding | NP_005767.1 | 19431 | 611287 | . | . | [illegible] |
| 10178 | NM_00116327 | TENM1 | teneurin transmembrane protein 1 | protein_coding | NP_001156750.1;NP_001156751.1;NP_055068.2;XP_011529532.1;XP_01 | 8117 | 300588 | . | . | [illegible] |
| 10179 | NM_00128604 | RBM7 | RNA binding motif protein 7 | protein_coding | NP_001272974.1;NP_001272975.1;NP_001272976.1;NP_001272977.1;NI | 9904 | 612413 | . | . | [illegible] |
| 10180 | NM_00116758 | RBM6 | RNA binding motif protein 6 | protein_coding | NP_001161054.1;NP_001336119.1;NP_001336120.1;NP_001336121.1;NI | 9903 | 606886 | . | . | [illegible] |
| 10181 | NM_005778,N | RBM5 | RNA binding motif protein 5 | protein_coding | NP_005769.1;XP_006712980.1;XP_011531563.1;XP_011531564.1;XP_01 | 9902 | 606884 | . | . | [illegible] |
| 10184 | NM_005779,XI | LHFPL2 | LHFPL tetraspan subfamily member 2 | protein_coding | NP_005770.1;XP_006714578.1;XP_024310089.1;XP_024310090.1;XP_02 | 6588 | 609718 | . | . | [illegible] |
| 10186 | NM_005780,XI | LHFPL6 | LHFPL tetraspan subfamily member 6 | protein_coding | NP_005771.1;XP_011533163.1 | 6586 | 606710 | . | . | [illegible] |
| 10188 | NM_00101093 | TNK2 | tyrosine kinase non receptor 2 | protein_coding | NP_001010938.2;NP_001294975.1;NP_001369200.1;NP_001369201.1;NI | 19297 | 606994 | . | . | [illegible] |
| 10189 | NM_005782,N | ALYREF | Aly/REF export factor | protein_coding | NP_005773.3 | 19071 | 604171 | . | . | [illegible] |
| 10190 | NM_005783,XI | TXNDC9 | thioredoxin domain containing 9 | protein_coding | NP_005774.2;XP_016858636.1 | 24110 | 612564 | . | . | [illegible] |
| 10193 | NM_00124282 | RNF41 | ring finger protein 41 | protein_coding | NP_001229755.1;NP_005776.1;NP_919339.1;NP_919340.1;XP_00526861 | 18401 | . | . | . | [illegible] |
| 10194 | NM_00130821 | TSHZ1 | teashirt zinc finger homeobox 1 | protein_coding | NP_001295139.1;NP_005777.3;XP_005266698.1 | 10669 | 614427 | . | . | [illegible] |
| 10195 | NM_00100694 | ALG3 | ALG3 alpha-1,3- mannosyltransferase | protein_coding | NP_001006942.1;NP_005778.1;XP_011510624.1;XP_011510625.1;XP_02 | 23056 | 608750 | . | . | [illegible] |
| 10196 | NM_00114516 | PRMT3 | protein arginine methyltransferase 3 | protein_coding | NP_001138638.1;NP_001138639.1;NP_005779.1;XP_011518138.1;XP_01 | 30163 | 603190 | . | . | [illegible] |
| 10197 | NM_00126704 | PSME3 | proteasome activator subunit 3 | protein_coding | NP_001253974.1;NP_001317158.1;NP_005780.2;NP_789839.1 | 9570 | 605129 | . | . | [illegible] |
| 10198 | NM_022782,N | MPHOSPH9 | M-phase phosphoprotein 9 | protein_coding | NP_073619.3;XP_011536042.1;XP_011536043.1;XP_016874162.1;XP_01 | 7215 | 605501 | . | . | [illegible] |
| 10199 | NM_005791 | MPHOSPH10 | M-phase phosphoprotein 10 | protein_coding | NP_005782.1 | 7213 | 605503 | . | . | [illegible] |
| 10200 | NM_005792,XI | MPHOSPH6 | M-phase phosphoprotein 6 | protein_coding | NP_005783.2;XP_011521110.1 | 7214 | 605500 | . | . | [illegible] |
| 10201 | NM_00130842 | NME6 | NME/NM23 nucleoside diphosphate kinase 6 | protein_coding | NP_001295355.1;NP_001295356.1;NP_001295357.1;NP_001295359.1;NI | 20567 | 608294 | . | . | [illegible] |
| 10204 | NM_00132203 | NUTF2 | nuclear transport factor 2 | protein_coding | NP_001308967.1;NP_001308968.1;NP_001308969.1;NP_001308970.1;NI | 13722 | 605813 | . | . | [illegible] |
| 10206 | NM_00100727 | TRIM13 | tripartite motif containing 13 | protein_coding | NP_001007279.1;NP_005789.2;NP_434698.1;NP_998755.1 | 9976 | 605661 | . | . | [illegible] |
| 10207 | NM_00135014 | PATJ | PATJ crumbs cell polarity complex component | protein_coding | NP_001337074.2;NP_795352.3;XP_005270404.1;XP_006710341.1;XP_01 | 28881 | 603199 | . | . | [illegible] |
| 10208 | NM_00132153 | USPL1 | ubiquitin specific peptidase like 1 | protein_coding | NP_001308461.1;NP_001308462.1;NP_001308463.1;NP_005791.3;XP_0C | 20294 | 617470 | . | . | [illegible] |
| 10209 | NM_005801 | EIF1 | eukaryotic translation initiation factor 1 | protein_coding | NP_005792.1 | 3249 | . | . | . | [illegible] |
| 10210 | NM_00119562 | TOPORS | TOP1 binding arginine/serine rich protein, E3 ubiquitin ligase | protein_coding | NP_001182551.1;NP_005793.2 | 21653 | 609507 | . | . | [illegible] |
| 10211 | NM_00131887 | FLOT1 | flotillin 1 | protein_coding | NP_001305804.1;NP_005794.1;XP_005248837.1;XP_006715010.1;XP_01 | 3757 | 606998 | . | . | [illegible] |
| 10212 | NM_005804,N | DDX39A | DExD-box helicase 39A | protein_coding | NP_005795.2;XP_006722669.1;XP_011525922.1;XP_011525923.1;XP_02 | 17821 | . | . | . | [illegible] |
| 10213 | NM_005805 | PSMD14 | proteasome 26S subunit, non-ATPase 14 | protein_coding | NP_005796.1 | 16889 | 607173 | . | . | [illegible] |
| 10215 | NM_005806,XI | OLIG2 | oligodendrocyte transcription factor 2 | protein_coding | NP_005797.1;XP_005260965.1 | 9398 | 606386 | . | . | [illegible] |
| 10217 | NM_00100839 | CTDSPL | CTD small phosphatase like | protein_coding | NP_001008393.1;NP_005799.2;XP_016861008.1;XP_016861009.1 | 16890 | 608592 | . | . | [illegible] |
| 10219 | NM_00132909 | KLRG1 | killer cell lectin like receptor G1 | protein_coding | NP_001316028.1;NP_001316030.1;NP_001316031.1;NP_001316032.1;NI | 6380 | 604874 | . | . | [illegible] |
| 10220 | NM_005811,XI | GDF11 | growth differentiation factor 11 | protein_coding | NP_005802.1;XP_006719257.1 | 4216 | 603936 | . | . | [illegible] |
| 10221 | NM_00128298 | TRIB1 | tribbles pseudokinase 1 | protein_coding | NP_001269914.1;NP_079471.1 | 16891 | 609461 | . | . | [illegible] |
| 10224 | NM_005815 | ZNF443 | zinc finger protein 443 | protein_coding | NP_005806.3 | 20878 | 606697 | . | . | [illegible] |
| 10225 | NM_00131888 | CD96 | CD96 molecule | protein_coding | NP_001305818.1;NP_005807.1;NP_937839.1;XP_005247120.1;XP_00671 | 16892 | 606037 | . | . | [illegible] |
| 10226 | NM_00116418 | PLIN3 | perilipin 3 | protein_coding | NP_001157661.1;NP_001157666.1;NP_005808.3 | 16893 | 602702 | . | . | [illegible] |
| 10227 | NM_001120,N | MFSD10 | major facilitator superfamily domain containing 10 | protein_coding | NP_001111.3;NP_001139541.1;NP_001350608.1;XP_005247984.1;XP_02 | 16894 | 610977 | . | . | [illegible] |
| 10228 | NM_00128621 | STX6 | syntaxin 6 | protein_coding | NP_001273139.1;NP_005810.1;XP_016855496.1 | 11441 | 603944 | . | . | [illegible] |
| 10229 | NM_00119098 | COQ7 | coenzyme Q7, hydroxylase | protein_coding | NP_001177912.1;NP_001357418.1;NP_001357419.1;NP_001357420.1;NI | 2244 | 601683 | . | . | [illegible] |
| 10230 | NR_003108,NF | NBR2 | neighbor of BRCA1 lncRNA 2 | lncRNA | . | 20691 | 618708 | . | . | [illegible] |
| 10233 | NM_00113521 | LRRC23 | leucine rich repeat containing 23 | protein_coding | NP_001128689.1;NP_008923.1;NP_964013.1 | 19138 | . | . | . | [illegible] |
| 10234 | NM_00103169 | LRRC17 | leucine rich repeat containing 17 | protein_coding | NP_001026862.1;NP_005815.2;XP_005250165.1 | 16895 | 618749 | . | . | [illegible] |
| 10235 | NM_00109867 | RASGRP2 | RAS guanyl releasing protein 2 | protein_coding | NP_001092140.1;NP_001092141.1;NP_001305327.1;NP_722541.1;XP_01 | 9879 | 605577 | . | . | [illegible] |
| 10236 | NM_00110239 | HNRNPR | heterogeneous nuclear ribonucleoprotein R | protein_coding | NP_001095867.1;NP_001095868.1;NP_001095869.1;NP_001284549.1;NI | 5047 | 607201 | . | . | [illegible] |
| 10237 | NM_00127878 | SLC35B1 | solute carrier family 35 member B1 | protein_coding | NP_001265713.1;NP_005818.3;XP_005256986.1;XP_006721695.1;XP_01 | 20798 | 610790 | . | . | [illegible] |
| 10238 | NM_005828,N | DCAF7 | DDB1 and CUL4 associated factor 7 | protein_coding | NP_005819.3 | 30915 | 605973 | . | . | [illegible] |
| 10239 | NM_005829,N | AP3S2 | adaptor related protein complex 3 subunit sigma 2 | protein_coding | NP_005820.1 | 571 | 602416 | . | . | [illegible] |
| 10240 | NM_005830,XI | MRPS31 | mitochondrial ribosomal protein S31 | protein_coding | NP_005821.2 | 16632 | 611992 | . | . | [illegible] |
| 10241 | NM_00126139 | CALCOCO2 | calcium binding and coiled-coil domain 2 | protein_coding | NP_001248319.1;NP_001248320.1;NP_001248322.1;NP_001248324.1;NI | 29912 | 604587 | . | . | [illegible] |
| 10243 | NM_00102421 | GPHN | gephyrin | protein_coding | NP_001019389.1;NP_001364443.1;NP_001364444.1;NP_001364445.1;NI | 15465 | 603930 | . | . | [illegible] |
| 10244 | NM_00117415 | RABEPK | Rab9 effector protein with kelch motifs | protein_coding | NP_001167623.1;NP_001167624.1;NP_005824.2;XP_005251697.1;XP_00 | 16896 | 605962 | . | . | [illegible] |
| 10245 | NM_00116794 | TIMM17B | translocase of inner mitochondrial membrane 17B | protein_coding | NP_001161419.1;NP_005825.1 | 17310 | 300249 | . | . | [illegible] |
| 10247 | NM_005836 | RIDA | reactive intermediate imine deaminase A homolog | protein_coding | NP_005827.1 | 16897 | 602487 | . | . | [illegible] |
| 10248 | NM_005837 | POP7 | POP7 homolog, ribonuclease P/MRP subunit | protein_coding | NP_005828.2 | 19949 | 606113 | . | . | [illegible] |
| 10250 | NM_00130344 | SRRM1 | serine and arginine repetitive matrix 1 | protein_coding | NP_001290377.1;NP_001290378.1;NP_001353494.1;NP_001353495.1;NI | 16638 | 605975 | . | . | [illegible] |
| 10251 | NM_00130499 | SPRY3 | sprouty RTK signaling antagonist 3 | protein_coding | NP_001291919.1;NP_005831.1 | 11271 | 300531 | . | . | [illegible] |
| 10252 | NM_00125803 | SPRY1 | sprouty RTK signaling antagonist 1 | protein_coding | NP_001244967.1;NP_001244968.1;NP_001362339.1;NP_005832.1;NP_9 | 11269 | 602465 | . | . | [illegible] |
| 10253 | NM_00131853 | SPRY2 | sprouty RTK signaling antagonist 2 | protein_coding | NP_001305465.1;NP_001305466.1;NP_001305467.1;NP_005833.1 | 11270 | 602466 | . | . | [illegible] |
| 10254 | NM_005843,XI | STAM2 | signal transducing adaptor molecule 2 | protein_coding | NP_005834.4 | 11358 | 606244 | . | . | [illegible] |
| 10257 | NM_00110551 | ABCC4 | ATP binding cassette subfamily C member 4 | protein_coding | NP_001098985.1;NP_001288758.1;NP_001288759.1;NP_005836.2;XP_0C | 55 | 605250 | . | . | [illegible] |
| 10260 | NM_00114482 | DENND4A | DENN domain containing 4A | protein_coding | NP_001138295.1;NP_001307764.1;NP_001363848.1;NP_001363849.1;NI | 24321 | 600382 | . | . | [illegible] |
| 10262 | NM_005850 | SF3B4 | splicing factor 3b subunit 4 | protein_coding | NP_005841.1 | 10771 | 605593 | . | . | [illegible] |
| 10263 | NM_00127184 | CDK2AP2 | cyclin dependent kinase 2 associated protein 2 | protein_coding | NP_001258778.1;NP_005842.1 | 30833 | . | . | . | [illegible] |
| 10265 | NM_00125219 | IRX5 | iroquois homeobox 5 | protein_coding | NP_001239126.1;NP_005844.4;XP_011521111.1 | 14361 | 606195 | . | . | [illegible] |
| 10266 | NM_005854 | RAMP2 | receptor activity modifying protein 2 | protein_coding | NP_005845.2 | 9844 | 605154 | . | . | [illegible] |
| 10269 | NM_005857,XI | ZMPSTE24 | zinc metallopeptidase STE24 | protein_coding | NP_005848.2 | 12877 | 606480 | . | . | [illegible] |
| 10270 | NM_005858 | AKAP8 | A-kinase anchoring protein 8 | protein_coding | NP_005849.1 | 378 | 604692 | . | . | [illegible] |
| 10272 | NM_005860 | FSTL3 | follistatin like 3 | protein_coding | NP_005851.1 | 3973 | 605343 | . | . | [illegible] |
| 10273 | NM_00129319 | STUB1 | STIP1 homology and U-box containing protein 1 | protein_coding | NP_001280126.1;NP_005852.2 | 11427 | 607207 | . | . | [illegible] |
| 10274 | NM_005862,XI | STAG1 | stromal antigen 1 | protein_coding | NP_005853.2;XP_011510631.1;XP_011510633.1;XP_016861012.1;XP_01 | 11354 | 604358 | . | . | [illegible] |
| 10276 | NM_00104716 | NET1 | neuroepithelial cell transforming 1 | protein_coding | NP_001040625.1;NP_005854.2;XP_024303523.1 | 14592 | 606450 | . | . | [illegible] |
| 10277 | NM_00110556 | UBE4B | ubiquitination factor E4B | protein_coding | NP_001099032.1;NP_006039.2;XP_005263479.1;XP_011538790.1;XP_01 | 12500 | 613565 | . | . | [illegible] |
| 10279 | NM_005865,XI | PRSS16 | serine protease 16 | protein_coding | NP_005856.1;XP_016865650.1;XP_016865651.1;XP_016865652.1;XP_01 | 9480 | 607169 | . | . | [illegible] |
| 10280 | NM_00128220 | SIGMAR1 | sigma non-opioid intracellular receptor 1 | protein_coding | NP_001269134.1;NP_001269135.1;NP_001269136.1;NP_001269137.1;NI | 8157 | 601978 | . | . | [illegible] |
| 10281 | NR_147130 | DSCR4 | Down syndrome critical region 4 | lncRNA | . | 3045 | 604829 | . | . | [illegible] |
| 10282 | NM_00131773 | BET1 | Bet1 golgi vesicular membrane trafficking protein | protein_coding | NP_001304668.1;NP_005859.1 | 14562 | 605456 | . | . | [illegible] |
| 10283 | NM_00129764 | CWC27 | CWC27 spliceosome associated cyclophilin | protein_coding | NP_001284573.1;NP_001284574.1;NP_001304929.1;NP_001351407.1;NI | 10664 | 617170 | . | . | [illegible] |
| 10284 | NM_00136664 | SAP18 | Sin3A associated protein 18 | protein_coding | NP_001353572.1;NP_005861.2 | 10530 | 602949 | . | . | [illegible] |
| 10285 | NM_005871,XI | SMNDC1 | survival motor neuron domain containing 1 | protein_coding | NP_005862.1;XP_005269439.1 | 16900 | 603519 | . | . | [illegible] |
| 10286 | NM_005872 | BCAS2 | BCAS2 pre-mRNA processing factor | protein_coding | NP_005863.1 | 975 | 605783 | . | . | [illegible] |
| 10287 | NM_00103946 | RGS19 | regulator of G protein signaling 19 | protein_coding | NP_001034556.1;NP_005864.1;XP_005260239.1;XP_005260240.1;XP_01 | 13735 | 605071 | . | . | [illegible] |
| 10289 | NM_005875 | EIF1B | eukaryotic translation initiation factor 1B | protein_coding | NP_005866.1 | 30792 | . | . | . | [illegible] |
| 10290 | NM_00117347 | SPEG | striated muscle enriched protein kinase | protein_coding | NP_001166947.1;NP_005867.3;XP_005246294.1;XP_005246296.1;XP_00 | 16901 | 615950 | . | . | [illegible] |
| 10291 | NM_005877 | SF3A1 | splicing factor 3a subunit 1 | protein_coding | NP_005868.1 | 10765 | 605595 | . | . | [illegible] |
| 10293 | NM_005879,XI | TRAIP | TRAF interacting protein | protein_coding | NP_005870.2;XP_011531566.1;XP_016861015.1 | 30764 | 605958 | . | . | [illegible] |
| 10294 | NM_005880 | DNAJA2 | DnaJ heat shock protein family (Hsp40) member A2 | protein_coding | NP_005871.1 | 14884 | 611322 | . | . | [illegible] |
| 10295 | NM_00112295 | BCKDK | branched chain keto acid dehydrogenase kinase | protein_coding | NP_001116429.1;NP_001258855.1;NP_005872.2;XP_016878348.1 | 16902 | 614901 | . | . | [illegible] |
| 10296 | NM_00101740 | MAEA | macrophage erythroblast attacher, E3 ubiquitin ligase | protein_coding | NP_001017405.1;NP_001284359.1;NP_001284360.1;NP_001284361.1;NI | 13731 | 606801 | . | . | [illegible] |
| 10297 | NM_00135127 | APC2 | APC regulator of WNT signaling pathway 2 | protein_coding | NP_001338202.1;NP_005874.1;XP_005259532.1;XP_006722670.1;XP_00 | 24036 | 612034 | . | . | [illegible] |
| 10298 | NM_00101483 | PAK4 | p21 (RAC1) activated kinase 4 | protein_coding | NP_001014831.1;NP_001014832.1;NP_001014834.1;NP_001014835.1;NI | 16059 | 605451 | . | . | [illegible] |
| 10299 | NM_00127066 | MARCHF6 | membrane associated ring-CH-type finger 6 | protein_coding | NP_001257589.1;NP_001257590.1;NP_005876.2;XP_011512234.1;XP_01 | 30550 | 613297 | . | . | [illegible] |
| 10300 | NM_005886 | KATNB1 | katanin regulatory subunit B1 | protein_coding | NP_005877.2 | 6217 | 602703 | . | . | [illegible] |
| 10301 | NR_002605,NF | DLEU1 | deleted in lymphocytic leukemia 1 | lncRNA | . | 13747 | 605765 | . | . | [illegible] |
| 10302 | NM_00132961 | SNAPC5 | small nuclear RNA activating complex polypeptide 5 | protein_coding | NP_001316542.1;NP_001316543.1;NP_001316544.1;NP_006040.1 | 15484 | 605979 | . | . | [illegible] |
| 10307 | NM_006051,N | APBB3 | amyloid beta precursor protein binding family B member 3 | protein_coding | NP_006042.3;NP_573418.2;NP_573419.2;NP_573420.2 | 20708 | 602711 | . | . | [illegible] |
| 10308 | NM_00126558 | ZNF267 | zinc finger protein 267 | protein_coding | NP_001252517.2;NP_003405.4 | 13060 | 604752 | . | . | [illegible] |
| 10309 | NM_021147,N | CCNO | cyclin O | protein_coding | NP_066970.3 | 18576 | 607752 | . | . | [illegible] |
| 10311 | NM_00133101 | VPS26C | VPS26 endosomal protein sorting factor C | protein_coding | NP_001317947.1;NP_001317950.1;NP_001317951.1;NP_006043.1 | 3044 | 605298 | . | . | [illegible] |
| 10312 | NM_00135105 | TCIRG1 | T cell immune regulator 1, ATPase H+ transporting V0 subunit a3 | protein_coding | NP_001337988.1;NP_006010.2;NP_006044.1;XP_024304088.1;XP_02430 | 11647 | 604592 | . | . | [illegible] |
| 10313 | NM_00126558 | RTN3 | reticulon 3 | protein_coding | NP_001252518.1;NP_001252519.1;NP_001252520.1;NP_006045.1;NP_9 | 10469 | 604249 | . | . | [illegible] |
| 10314 | NM_00113657 | LANCL1 | LanC like 1 | protein_coding | NP_001130046.1;NP_001130047.1;NP_006046.1;XP_005246300.1 | 6508 | 604155 | . | . | [illegible] |
| 10317 | NM_00127865 | B3GALT5 | beta-1,3-galactosyltransferase 5 | protein_coding | NP_001265579.1;NP_001343265.1;NP_001343267.1;NP_001343268.1;NI | 920 | 604066 | . | . | [illegible] |
| 10318 | NM_00125238 | TNIP1 | TNFAIP3 interacting protein 1 | protein_coding | NP_001239314.1;NP_001239315.1;NP_001239319.1;NP_001239320.1;NI | 16903 | 607714 | . | . | [illegible] |
| 10319 | NM_006059,XI | LAMC3 | laminin subunit gamma 3 | protein_coding | NP_006050.3;XP_006716984.1;XP_011516423.1 | 6494 | 604349 | . | . | [illegible] |
| 10322 | NM_006062 | SMYD5 | SMYD family member 5 | protein_coding | NP_006053.2 | 16258 | . | . | . | [illegible] |
| 10324 | NM_006063 | KLHL41 | kelch like family member 41 | protein_coding | NP_006054.2 | 16905 | 607701 | . | . | [illegible] |
| 10325 | NM_00135401 | RRAGB | Ras related GTP binding B | protein_coding | NP_001340940.1;NP_001340942.1;NP_006055.3;NP_057740.2;XP_0168 | 19901 | 300725 | . | . | [illegible] |
| 10326 | NM_00108391 | SIRPB1 | signal regulatory protein beta 1 | protein_coding | NP_001077379.1;NP_001129316.1;NP_001316086.1;NP_001317568.1;NI | 15928 | 603889 | . | . | [illegible] |
| 10327 | NM_00120241 | AKR1A1 | aldo-keto reductase family 1 member A1 | protein_coding | NP_001189342.1;NP_001189343.1;NP_006057.1;NP_697021.1;XP_0115 | 380 | 103830 | . | . | [illegible] |
| 10328 | NM_00114228 | EMC8 | ER membrane protein complex subunit 8 | protein_coding | NP_001135760.1;NP_006058.1;XP_011521115.1;XP_016878356.1 | 7864 | 604886 | . | . | [illegible] |
| 10329 | NM_00127823 | RXYLT1 | ribitol xylosyltransferase 1 | protein_coding | NP_001265166.1;NP_055069.1;XP_005268619.1;XP_005268620.1;XP_01 | 13530 | 605862 | . | . | [illegible] |
| 10330 | NM_00119099 | CNPY2 | canopy FGF signaling regulator 2 | protein_coding | NP_001177920.1;NP_055070.1 | 13529 | 605861 | . | . | [illegible] |
| 10333 | NM_006068,XI | TLR6 | toll like receptor 6 | protein_coding | NP_006059.2;XP_005262694.1;XP_011511915.1;XP_011511916.1;XP_02 | 16711 | 605403 | . | . | [illegible] |
| 10336 | NM_00131783 | PCGF3 | polycomb group ring finger 3 | protein_coding | NP_001304765.1;NP_006306.2 | 10066 | 617543 | . | . | [illegible] |
| 10342 | NM_00100756 | TFG | trafficking from ER to golgi regulator | protein_coding | NP_001007566.1;NP_001182407.1;NP_001182408.1;NP_006061.2;XP_0C | 11758 | 602498 | . | . | [illegible] |
| 10347 | NM_019112,XI | ABCA7 | ATP binding cassette subfamily A member 7 | protein_coding | NP_061985.2;XP_006722679.1;XP_006722680.1;XP_011525935.1;XP_01 | 37 | 605414 | . | . | [illegible] |
| 10349 | NM_00137732 | ABCA10 | ATP binding cassette subfamily A member 10 | protein_coding | NP_001364250.1;NP_525021.3 | 30 | 612508 | . | . | [illegible] |
| 10352 | NM_00137822 | WARS2 | tryptophanyl tRNA synthetase 2, mitochondrial | protein_coding | NP_001365155.1;NP_001365156.1;NP_001365157.1;NP_001365158.1;NI | 12730 | 604733 | . | . | [illegible] |
| 10354 | gene-HMGB1P | HMGB1P5 | high mobility group box 1 pseudogene 5 | pseudogene | . | 4997 | . | . | . | [illegible] |
| 10356 | gene-HMGB1P | HMGB1P3 | high mobility group box 1 pseudogene 3 | pseudogene | . | 4995 | . | . | . | [illegible] |
| 10357 | gene-HMGB1P | HMGB1P1 | high mobility group box 1 pseudogene 1 | pseudogene | . | 4993 | . | . | . | [illegible] |
| 10360 | NM_006993 | NPM3 | nucleophosmin/nucleoplasmin 3 | protein_coding | NP_008924.1 | 7931 | 606456 | . | . | [illegible] |
| 10362 | NM_006339,XI | HMG20B | high mobility group 20B | protein_coding | NP_006330.2;XP_016881633.1;XP_024307094.1 | 5002 | 605535 | . | . | [illegible] |
| 10363 | NM_00130450 | HMG20A | high mobility group 20A | protein_coding | NP_001291433.1;NP_001291434.1;NP_060670.1;XP_011519460.1 | 5001 | 605534 | . | . | [illegible] |
| 10365 | NM_016270 | KLF2 | Kruppel like factor 2 | protein_coding | NP_057354.1 | 6347 | 602016 | . | . | [illegible] |
| 10367 | NM_00119551 | MICU1 | mitochondrial calcium uptake 1 | protein_coding | NP_001182447.1;NP_001182448.1;NP_001350442.1;NP_006068.2;XP_0C | 1530 | 605084 | . | . | [illegible] |
| 10370 | NM_00116838 | CITED2 | Cbp/p300 interacting transactivator with Glu/Asp rich carboxy-ter | protein_coding | NP_001161860.1;NP_001161861.2;NP_006070.2 | 1987 | 602937 | . | . | [illegible] |
| 10371 | NM_006080,XI | SEMA3A | semaphorin 3A | protein_coding | NP_006071.1;XP_005250167.1;XP_005250168.1;XP_006715902.1;XP_01 | 10723 | 603961 | . | . | [illegible] |
| 10376 | NM_006082 | TUBA1B | tubulin alpha 1b | protein_coding | NP_006073.2 | 18809 | 602530 | . | . | [illegible] |
| 10379 | NM_006084 | IRF9 | interferon regulatory factor 9 | protein_coding | NP_006075.3 | 6131 | 147574 | . | . | [illegible] |
| 10380 | NM_00128614 | BPNT1 | 3'(2'), 5'-bisphosphate nucleotidase 1 | protein_coding | NP_001273078.1;NP_001273079.1;NP_001273080.1;NP_006076.4;XP_0C | 1096 | 604053 | . | . | [illegible] |
| 10381 | NM_00119718 | TUBB3 | tubulin beta 3 class III | protein_coding | NP_001184110.1;NP_006077.2 | 20772 | 602661 | . | . | [illegible] |
| 10382 | NM_00128912 | TUBB4A | tubulin beta 4A class IVa | protein_coding | NP_001276052.1;NP_001276056.1;NP_001276058.1;NP_001276059.1;NI | 20774 | 602662 | . | . | [illegible] |
| 10383 | NM_006088 | TUBB4B | tubulin beta 4B class IVb | protein_coding | NP_006079.1 | 20771 | 602660 | . | . | [illegible] |
| 10384 | NM_00124280 | BTN3A3 | butyrophilin subfamily 3 member A3 | protein_coding | NP_001229732.1;NP_008925.1;NP_932078.2 | 1140 | 613595 | . | . | [illegible] |
| 10385 | NM_00119723 | BTN2A2 | butyrophilin subfamily 2 member A2 | protein_coding | NP_001184166.1;NP_001184167.1;NP_001184168.1;NP_001184169.1;NI | 1137 | 613591 | . | . | [illegible] |
| 10388 | NM_014258,XI | SYCP2 | synaptonemal complex protein 2 | protein_coding | NP_055073.2;XP_011526789.1;XP_011526790.1;XP_011526791.1;XP_01 | 11490 | 604105 | . | . | [illegible] |
| 10389 | NM_006089,N | SCML2 | Scm polycomb group protein like 2 | protein_coding | NP_006080.1;XP_006724521.1;XP_016884708.1;XP_016884709.1;XP_01 | 10581 | 300208 | . | . | [illegible] |
| 10390 | NM_00100779 | CEPT1 | choline/ethanolamine phosphotransferase 1 | protein_coding | NP_001007795.1;NP_001317672.1;NP_006081.1;XP_011538798.1;XP_02 | 24289 | 616751 | . | . | [illegible] |
| 10391 | NM_00119045 | CORO2B | coronin 2B | protein_coding | NP_001177385.1;NP_001177386.1;NP_001310943.1;NP_001310944.1;NI | 2256 | 605002 | . | . | [illegible] |
| 10392 | NM_00135484 | NOD1 | nucleotide binding oligomerization domain containing 1 | protein_coding | NP_001341778.1;NP_006083.1;XP_005249625.1;XP_005249629.1;XP_00 | 16390 | 605980 | . | . | [illegible] |
| 10393 | NM_00125670 | ANAPC10 | anaphase promoting complex subunit 10 | protein_coding | NP_001243635.1;NP_001243636.1;NP_001243637.1;NP_001243638.1;NI | 24077 | 613745 | . | . | [illegible] |
| 10395 | NM_00116427 | DLC1 | DLC1 Rho GTPase activating protein | protein_coding | NP_001157743.1;NP_001303597.1;NP_001335010.1;NP_001335011.1;NI | 2897 | 604258 | . | . | [illegible] |
| 10396 | NM_00110552 | ATP8A1 | ATPase phospholipid transporting 8A1 | protein_coding | NP_001098999.1;NP_006086.1;XP_005248100.1;XP_011511917.1;XP_01 | 13531 | 609542 | . | . | [illegible] |
| 10397 | NM_00113524 | NDRG1 | N-myc downstream regulated 1 | protein_coding | NP_001128714.1;NP_001245361.1;NP_001245362.1;NP_001361773.1;NI | 7679 | 605262 | . | . | [illegible] |
| 10398 | NM_006097,N | MYL9 | myosin light chain 9 | protein_coding | NP_006088.2;NP_852667.1 | 15754 | 609905 | . | . | [illegible] |
| 10399 | NM_006098 | RACK1 | receptor for activated C kinase 1 | protein_coding | NP_006089.1 | 4399 | 176981 | . | . | [illegible] |
| 10400 | NM_00126755 | PEMT | phosphatidylethanolamine N-methyltransferase | protein_coding | NP_001254480.1;NP_001254481.1;NP_009100.2;NP_680477.1;NP_6804 | 8830 | 602391 | . | . | [illegible] |
| 10401 | NM_006099 | PIAS3 | protein inhibitor of activated STAT 3 | protein_coding | NP_006090.2 | 16861 | 605987 | . | . | [illegible] |
| 10402 | NM_00127114 | ST3GAL6 | ST3 beta-galactoside alpha-2,3-sialyltransferase 6 | protein_coding | NP_001258071.1;NP_001258074.1;NP_001258075.1;NP_001258076.1;NI | 18080 | 607156 | . | . | [illegible] |
| 10403 | NM_006101 | NDC80 | NDC80 kinetochore complex component | protein_coding | NP_006092.1 | 16909 | 607272 | . | . | [illegible] |
| 10404 | NM_016134 | CPQ | carboxypeptidase Q | protein_coding | NP_057218.1 | 16910 | 618754 | . | . | [illegible] |

| | | | | | | | | | | |
|---|---|---|---|---|---|---|---|---|---|---|
| 10409 | NM_00127160 | BASP1 | brain abundant membrane attached signal protein 1 | protein_coding | NP_001258535.1;NP_006308.3 | 957 | 605940 | . | . | [illegible] |
| 10410 | NM_021034,N | IFITM3 | interferon induced transmembrane protein 3 | protein_coding | NP_066362.2 | 5414 | 605579 | . | . | [illegible] |
| 10411 | NM_00109853 | RAPGEF3 | Rap guanine nucleotide exchange factor 3 | protein_coding | NP_001092001.2;NP_001092002.1;NP_006096.2;XP_005268628.1;XP_01 | 16629 | 606057 | . | . | [illegible] |
| 10412 | NM_00127166 | NSA2 | NSA2 ribosome biogenesis factor | protein_coding | NP_001258594.1;NP_001351435.1;NP_055701.1 | 30728 | 612497 | . | . | [illegible] |
| 10413 | NM_00113014 | YAP1 | Yes1 associated transcriptional regulator | protein_coding | NP_001123617.1;NP_001181973.1;NP_001181974.1;NP_001269026.1;NI | 16262 | 606608 | . | . | [illegible] |
| 10417 | NM_00112832 | SPON2 | spondin 2 | protein_coding | NP_001121797.1;NP_001185950.1;NP_036577.2 | 11253 | 605918 | . | . | [illegible] |
| 10419 | NM_00103961 | PRMT5 | protein arginine methyltransferase 5 | protein_coding | NP_001034708.1;NP_001269882.1;NP_001269883.1;NP_001269884.1;NI | 10894 | 604045 | . | . | [illegible] |
| 10420 | NM_00132080 | TESK2 | testis associated actin remodelling kinase 2 | protein_coding | NP_001307729.1;NP_009101.2 | 11732 | 604746 | . | . | [illegible] |
| 10421 | NM_00124364 | CD2BP2 | CD2 cytoplasmic tail binding protein 2 | protein_coding | NP_001230575.1;NP_006101.1 | 1656 | 604470 | . | . | [illegible] |
| 10422 | NM_016172 | UBAC1 | UBA domain containing 1 | protein_coding | NP_057256.2 | 30221 | 608129 | . | . | [illegible] |
| 10423 | NM_00128658 | CDIPT | CDP-diacylglycerol--inositol 3-phosphatidyltransferase | protein_coding | NP_001273514.1;NP_001273515.1;NP_006310.1;XP_005255095.1 | 1769 | 605893 | . | . | [illegible] |
| 10424 | NM_006320,XI | PGRMC2 | progesterone receptor membrane component 2 | protein_coding | NP_006311.3;XP_011529835.1 | 16089 | 607735 | . | . | [illegible] |
| 10425 | NM_00131733 | ARIH2 | ariadne RBR E3 ubiquitin protein ligase 2 | protein_coding | NP_001304262.1;NP_001304263.1;NP_001336138.1;NP_001336139.1;NI | 690 | 605615 | . | . | [illegible] |
| 10426 | NM_00128627 | TUBGCP3 | tubulin gamma complex associated protein 3 | protein_coding | NP_001273206.1;NP_001273207.1;NP_001273208.1;NP_006313.1;XP_0C | 18598 | 617818 | . | . | [illegible] |
| 10427 | NM_00104273 | SEC24B | SEC24 homolog B, COPII coat complex component | protein_coding | NP_001036199.1;NP_001287742.1;NP_001305014.1;NP_001305015.1;NI | 10704 | 607184 | . | . | [illegible] |
| 10428 | NM_006324,XI | CFDP1 | craniofacial development protein 1 | protein_coding | NP_006315.1;XP_011521116.1;XP_011521117.1 | 1873 | 608108 | . | . | [illegible] |
| 10430 | NM_00124259 | TMEM147 | transmembrane protein 147 | protein_coding | NP_001229526.1;NP_001229527.1;NP_116024.1 | 30414 | 613585 | . | . | [illegible] |
| 10432 | NM_00119883 | RBM14 | RNA binding motif protein 14 | protein_coding | NP_001185765.1;NP_001185766.1;NP_006319.1 | 14219 | 612409 | . | . | [illegible] |
| 10434 | NM_00127935 | LYPLA1 | lysophospholipase 1 | protein_coding | NP_001266285.1;NP_001266286.1;NP_001266287.1;NP_001266288.1;NI | 6737 | 605599 | . | . | [illegible] |
| 10435 | NM_006779 | CDC42EP2 | CDC42 effector protein 2 | protein_coding | NP_006770.1 | 16263 | 606132 | . | . | [illegible] |
| 10436 | NM_00132004 | EMG1 | EMG1 N1-specific pseudouridine methyltransferase | protein_coding | NP_001306978.1;NP_006322.4 | 16912 | 611531 | . | . | [illegible] |
| 10437 | NM_006332 | IFI30 | IFI30 lysosomal thiol reductase | protein_coding | NP_006323.2 | 5398 | 604664 | . | . | [illegible] |
| 10438 | NM_00119026 | C1D | C1D nuclear receptor corepressor | protein_coding | NP_001177192.1;NP_001177194.1;NP_006324.1;NP_775269.1 | 29911 | 606997 | . | . | [illegible] |
| 10439 | NM_00128261 | OLFM1 | olfactomedin 1 | protein_coding | NP_001269540.1;NP_001269541.1;NP_006325.1;NP_055094.1 | 17187 | 605366 | . | . | [illegible] |
| 10440 | NM_006335 | TIMM17A | translocase of inner mitochondrial membrane 17A | protein_coding | NP_006326.1 | 17315 | 605057 | . | . | [illegible] |
| 10443 | NM_00127843 | N4BP2L2 | NEDD4 binding protein 2 like 2 | protein_coding | NP_001265361.1;NP_001307765.1;NP_055702.1;NP_149102.3;XP_0115: | 26916 | 615788 | . | . | [illegible] |
| 10444 | NM_00137595 | ZER1 | zyg-11 related cell cycle regulator | protein_coding | NP_001362883.1;NP_001362884.1;NP_001362885.1;NP_001362886.1;NI | 30960 | 617764 | . | . | [illegible] |
| 10445 | NM_00101230 | MCRS1 | microspherule protein 1 | protein_coding | NP_001012300.1;NP_001265270.1;NP_006328.2;XP_005268629.1;XP_01 | 6960 | 609504 | . | . | [illegible] |
| 10447 | NM_00104002 | FAM3C | FAM3 metabolism regulating signaling molecule C | protein_coding | NP_001035109.1;NP_055703.1;XP_011514038.1;XP_011514039.1 | 18664 | 608618 | . | . | [illegible] |
| 10449 | NM_006111 | ACAA2 | acetyl-CoA acyltransferase 2 | protein_coding | NP_006102.2 | 83 | 604770 | . | . | [illegible] |
| 10450 | NM_00119500 | PPIE | peptidylprolyl isomerase E | protein_coding | NP_001181936.1;NP_001306222.1;NP_006103.1;NP_982281.1;XP_00671 | 9258 | 602435 | . | . | [illegible] |
| 10451 | NM_00107987 | VAV3 | vav guanine nucleotide exchange factor 3 | protein_coding | NP_001073343.1;NP_006104.4;XP_005270417.1;XP_005270418.1;XP_01 | 12659 | 605541 | . | . | [illegible] |
| 10452 | NM_00112891 | TOMM40 | translocase of outer mitochondrial membrane 40 | protein_coding | NP_001122388.1;NP_001122389.1;NP_006105.1;XP_005258468.1 | 18001 | 608061 | . | . | [illegible] |
| 10454 | NM_006116,N | TAB1 | TGF-beta activated kinase 1 (MAP3K7) binding protein 1 | protein_coding | NP_006107.1;NP_705717.1 | 18157 | 602615 | . | . | [illegible] |
| 10455 | NM_00116601 | ECI2 | enoyl-CoA delta isomerase 2 | protein_coding | NP_001159482.1;NP_006108.2;NP_996667.2;XP_006715020.1;XP_01686 | 14601 | 608024 | . | . | [illegible] |
| 10456 | NM_00101883 | HAX1 | HCLS1 associated protein X-1 | protein_coding | NP_001018238.1;NP_006109.2 | 16915 | 605998 | . | . | [illegible] |
| 10457 | NM_00100534 | GPNMB | glycoprotein nmb | protein_coding | NP_001005340.1;NP_002501.1;XP_005249635.1;XP_016867165.1;XP_01 | 4462 | 604368 | . | . | [illegible] |
| 10458 | NM_00114488 | BAIAP2 | BAR/IMD domain containing adaptor protein 2 | protein_coding | NP_001138360.1;NP_006331.1;NP_059344.1;NP_059345.1;XP_0052570C | 947 | 605475 | . | . | [illegible] |
| 10459 | NM_00112732 | MAD2L2 | mitotic arrest deficient 2 like 2 | protein_coding | NP_001120797.1;NP_006332.3;XP_024306175.1 | 6764 | 604094 | . | . | [illegible] |
| 10460 | NM_006342,XI | TACC3 | transforming acidic coiled-coil containing protein 3 | protein_coding | NP_006333.1;XP_005247986.1;XP_005247987.1;XP_011511688.1;XP_01( | 11524 | 605303 | . | . | [illegible] |
| 10461 | NM_006343 | MERTK | MER proto-oncogene, tyrosine kinase | protein_coding | NP_006334.2 | 7027 | 604705 | . | . | [illegible] |
| 10463 | NM_006345,XI | SLC30A9 | solute carrier family 30 member 9 | protein_coding | NP_006336.3;XP_016863143.1 | 1329 | 604604 | . | . | [illegible] |
| 10464 | NM_00134965 | PIBF1 | progesterone immunomodulatory binding factor 1 | protein_coding | NP_001336584.1;NP_006337.2;XP_011533184.1;XP_011533186.1;XP_01 | 23352 | 607532 | . | . | [illegible] |
| 10465 | NM_00133051 | PPIH | peptidylprolyl isomerase H | protein_coding | NP_001317439.1;NP_006338.1;XP_005270419.1;XP_005270423.1;XP_01 | 14651 | 606095 | . | . | [illegible] |
| 10466 | NM_00116152 | COG5 | component of oligomeric golgi complex 5 | protein_coding | NP_001154992.2;NP_001366440.1;NP_001366441.1;NP_001366442.1;NI | 14857 | 606821 | . | . | [illegible] |
| 10467 | NM_006349,XI | ZNHIT1 | zinc finger HIT-type containing 1 | protein_coding | NP_006340.1;XP_011514041.1 | 21688 | 618617 | . | . | [illegible] |
| 10468 | NM_006350,N | FST | follistatin | protein_coding | NP_006341.1;NP_037541.1;XP_005248457.1;XP_005248458.1;XP_00524 | 3971 | 136470 | . | . | [illegible] |
| 10469 | NM_006351 | TIMM44 | translocase of inner mitochondrial membrane 44 | protein_coding | NP_006342.2 | 17316 | 605058 | . | . | [illegible] |
| 10471 | NM_00118518 | PFDN6 | prefoldin subunit 6 | protein_coding | NP_001172110.1;NP_001252524.1;NP_001252525.1;NP_055075.1 | 4926 | 605660 | . | . | [illegible] |
| 10472 | NM_00127819 | ZBTB18 | zinc finger and BTB domain containing 18 | protein_coding | NP_001265125.1;NP_006343.2;NP_991331.1;XP_005273063.1;XP_01685 | 13030 | 608433 | . | . | [illegible] |
| 10473 | NM_006353,XI | HMGN4 | high mobility group nucleosomal binding domain 4 | protein_coding | NP_006344.1;XP_016865661.1;XP_016865662.1 | 4989 | . | . | . | [illegible] |
| 10474 | NM_00127827 | TADA3 | transcriptional adaptor 3 | protein_coding | NP_001265199.1;NP_006345.1;NP_597814.1 | 19422 | 602945 | . | . | [illegible] |
| 10475 | NM_006355,XI | TRIM38 | tripartite motif containing 38 | protein_coding | NP_006346.1;XP_005248856.1;XP_005248857.1;XP_016865664.1;XP_02 | 10059 | . | . | . | [illegible] |
| 10476 | NM_00100378 | ATP5PD | ATP synthase peripheral stalk subunit d | protein_coding | NP_001003785.1;NP_006347.1 | 845 | 618121 | . | . | [illegible] |
| 10477 | NM_00127855 | UBE2E3 | ubiquitin conjugating enzyme E2 E3 | protein_coding | NP_001265483.1;NP_001265484.1;NP_006348.1;NP_872619.1;XP_00524 | 12479 | 604151 | . | . | [illegible] |
| 10478 | NM_00128272 | SLC25A17 | solute carrier family 25 member 17 | protein_coding | NP_001269655.1;NP_001269656.1;NP_006349.1 | 10987 | 606795 | . | . | [illegible] |
| 10479 | NM_00104253 | SLC9A6 | solute carrier family 9 member A6 | protein_coding | NP_001036002.1;NP_001171122.1;NP_001317581.1;NP_001366039.1;NI | 11079 | 300231 | . | . | [illegible] |
| 10480 | NM_00130792 | EIF3M | eukaryotic translation initiation factor 3 subunit M | protein_coding | NP_001294858.1;NP_006351.2;XP_011518142.1 | 24460 | 609641 | . | . | [illegible] |
| 10481 | NM_006361 | HOXB13 | homeobox B13 | protein_coding | NP_006352.2 | 5112 | 604607 | . | . | [illegible] |
| 10482 | NM_00108149 | NXF1 | nuclear RNA export factor 1 | protein_coding | NP_001074960.1;NP_006353.2 | 8071 | 602647 | . | . | [illegible] |
| 10483 | NM_00117274 | SEC23B | SEC23 homolog B, COPII coat complex component | protein_coding | NP_001166216.1;NP_001166217.1;NP_006354.2;NP_116780.1;NP_11678 | 10702 | 610512 | . | . | [illegible] |
| 10484 | NM_006364,XI | SEC23A | Sec23 homolog A, COPII coat complex component | protein_coding | NP_006355.2;XP_005267319.1;XP_011534657.1;XP_016876417.1 | 10701 | 610511 | . | . | [illegible] |
| 10486 | NM_00136353 | CAP2 | cyclase associated actin cytoskeleton regulatory protein 2 | protein_coding | NP_001350462.1;NP_001350463.1;NP_006357.1 | 20039 | 618385 | . | . | [illegible] |
| 10487 | NM_00110553 | CAP1 | cyclase associated actin cytoskeleton regulatory protein 1 | protein_coding | NP_001099000.2;NP_001317431.2;NP_001337404.2;NP_001337405.2;NI | 20040 | 617801 | . | . | [illegible] |
| 10488 | NM_006368 | CREB3 | cAMP responsive element binding protein 3 | protein_coding | NP_006359.3 | 2347 | 606443 | . | . | [illegible] |
| 10489 | NM_006369,XI | LRRC41 | leucine rich repeat containing 41 | protein_coding | NP_006360.3 | 16917 | 618753 | . | . | [illegible] |
| 10490 | NM_006370 | VTI1B | vesicle transport through interaction with t-SNAREs 1B | protein_coding | NP_006361.1 | 17793 | 603207 | . | . | [illegible] |
| 10491 | NM_006371 | CRTAP | cartilage associated protein | protein_coding | NP_006362.1 | 2379 | 605497 | . | . | [illegible] |
| 10492 | NM_00115967 | SYNCRIP | synaptotagmin binding cytoplasmic RNA interacting protein | protein_coding | NP_001153145.1;NP_001153146.1;NP_001153147.1;NP_001153148.1;NI | 16918 | 616686 | . | . | [illegible] |
| 10493 | NM_006373 | VAT1 | vesicle amine transport 1 | protein_coding | NP_006364.2 | 16919 | 604631 | . | . | [illegible] |
| 10494 | NM_00127197 | STK25 | serine/threonine kinase 25 | protein_coding | NP_001258906.1;NP_001258907.1;NP_001258908.1;NP_001258909.1;NI | 11404 | 602255 | . | . | [illegible] |
| 10495 | NM_00128173 | ENOX2 | ecto-NOX disulfide-thiol exchanger 2 | protein_coding | NP_001268665.1;NP_001369445.1;NP_001369446.1;NP_001369447.1;NI | 2259 | 300282 | . | . | [illegible] |
| 10497 | NM_00133065 | UNC13B | unc-13 homolog B | protein_coding | NP_001317582.1;NP_001358115.1;NP_001358116.1;NP_001358117.1;NI | 12566 | 605836 | . | . | [illegible] |
| 10498 | NM_00137008 | CARM1 | coactivator associated arginine methyltransferase 1 | protein_coding | NP_001357017.1;NP_001357018.1;NP_954592.1;XP_011525940.1 | 23393 | 603934 | . | . | [illegible] |
| 10499 | NM_00132170 | NCOA2 | nuclear receptor coactivator 2 | protein_coding | NP_001308632.1;NP_001308636.1;NP_001308640.1;NP_001308641.1;NI | 7669 | 601993 | . | . | [illegible] |
| 10500 | NM_00117806 | SEMA6C | semaphorin 6C | protein_coding | NP_001171532.1;NP_001171533.1;NP_112175.2;XP_005244892.1;XP_01 | 10740 | 609294 | . | . | [illegible] |
| 10501 | NM_032108,XI | SEMA6B | semaphorin 6B | protein_coding | NP_115484.2;XP_011525941.1;XP_011525942.1;XP_011525943.1 | 10739 | 608873 | . | . | [illegible] |
| 10505 | NM_00127166 | SEMA4F | ssemaphorin 4F | protein_coding | NP_001258590.1;NP_001258591.1;NP_004254.2;XP_011530777.1;XP_01 | 10734 | 603706 | . | . | [illegible] |
| 10507 | NM_00114228 | SEMA4D | semaphorin 4D | protein_coding | NP_001135759.1;NP_001358123.1;NP_001358124.1;NP_001358125.1;NI | 10732 | 601866 | . | . | [illegible] |
| 10509 | NM_00132402 | SEMA4B | semaphorin 4B | protein_coding | NP_001310958.1;NP_001310959.1;NP_001310960.1;NP_001310961.1;NI | 10730 | 617029 | . | . | [illegible] |
| 10512 | NM_00135012 | SEMA3C | semaphorin 3C | protein_coding | NP_001337049.1;NP_001337050.1;NP_006370.1 | 10725 | 602645 | . | . | [illegible] |
| 10513 | NM_00128247 | APPBP2 | amyloid beta precursor protein binding protein 2 | protein_coding | NP_001269405.1;NP_006371.2;XP_011522500.1 | 622 | 605324 | . | . | [illegible] |
| 10514 | NM_00110553 | MYBBP1A | MYB binding protein 1a | protein_coding | NP_001099008.1;NP_055335.2;XP_011521918.1;XP_024306304.1 | 7546 | 604885 | . | . | [illegible] |
| 10516 | NM_006329,XI | FBLN5 | fibulin 5 | protein_coding | NP_006320.2;XP_005267324.1;XP_011534658.1;XP_011534659.1;XP_01 | 3602 | 604580 | . | . | [illegible] |
| 10517 | NM_00126758 | FBXW10 | F-box and WD repeat domain containing 10 | protein_coding | NP_001254514.1;NP_001254515.1;XP_011521927.1;XP_011521928.1;XP | 1211 | 611679 | . | . | [illegible] |
| 10518 | NM_00127188 | CIB2 | calcium and integrin binding family member 2 | protein_coding | NP_001258817.1;NP_001258818.1;NP_001288153.1;NP_006374.1;XP_0C | 24579 | 605564 | . | . | [illegible] |
| 10519 | NM_00127776 | CIB1 | calcium and integrin binding 1 | protein_coding | NP_001264693.1;NP_006375.2;XP_006720438.1 | 16920 | 602293 | . | . | [illegible] |
| 10520 | NM_00126559 | ZNF211 | zinc finger protein 211 | protein_coding | NP_001252526.1;NP_001252527.1;NP_001252528.1;NP_001252529.1;NI | 13003 | 601856 | . | . | [illegible] |
| 10521 | NM_00109850 | DDX17 | DEAD-box helicase 17 | protein_coding | NP_001091974.1;NP_006377.2 | 2740 | 608469 | . | . | [illegible] |
| 10522 | NM_00129363 | DEAF1 | DEAF1 transcription factor | protein_coding | NP_001280563.1;NP_001354319.1;NP_066288.2;XP_011518144.1;XP_02 | 14677 | 602635 | . | . | [illegible] |
| 10523 | NM_006387 | CHERP | calcium homeostasis endoplasmic reticulum protein | protein_coding | NP_006378.3 | 16930 | 618539 | . | . | [illegible] |
| 10524 | NM_00120683 | KAT5 | lysine acetyltransferase 5 | protein_coding | NP_001193762.1;NP_006379.2;NP_874368.1;NP_874369.1;XP_0067184{ | 5275 | 601409 | . | . | [illegible] |
| 10525 | NM_00113099 | HYOU1 | hypoxia up-regulated 1 | protein_coding | NP_001124463.1;NP_006380.1;XP_005271449.1;XP_005271450.1;XP_00 | 16931 | 601746 | . | . | [illegible] |
| 10526 | NM_00119099 | IPO8 | importin 8 | protein_coding | NP_001177924.1;NP_006381.2;XP_016874180.1;XP_016874181.1;XP_01 | 9853 | 605600 | . | . | [illegible] |
| 10527 | NM_006391 | IPO7 | importin 7 | protein_coding | NP_006382.1 | 9852 | 605586 | . | . | [illegible] |
| 10528 | NM_006392,N | NOP56 | NOP56 ribonucleoprotein | protein_coding | NP_006383.2 | 15911 | 614154 | . | . | [illegible] |
| 10531 | NM_00124230 | PITRM1 | pitrilysin metallopeptidase 1 | protein_coding | NP_001229236.1;NP_001229238.1;NP_001334654.1;NP_001334655.1;NI | 17663 | 618211 | . | . | [illegible] |
| 10533 | NM_00113603 | ATG7 | autophagy related 7 | protein_coding | NP_001129503.2;NP_001138384.1;NP_001336161.1;NP_001336162.1;NI | 16935 | 608760 | . | . | [illegible] |
| 10534 | NM_00130302 | ZNRD2 | zinc ribbon domain containing 2 | protein_coding | NP_001289953.1;NP_006387.1 | 11328 | 606044 | . | . | [illegible] |
| 10535 | NM_006397 | RNASEH2A | ribonuclease H2 subunit A | protein_coding | NP_006388.2 | 18518 | 606034 | . | . | [illegible] |
| 10536 | NM_014262 | P3H3 | prolyl 3-hydroxylase 3 | protein_coding | NP_055077.2 | 19318 | 610342 | . | . | [illegible] |
| 10538 | NM_006399 | BATF | basic leucine zipper ATF-like transcription factor | protein_coding | NP_006390.1 | 958 | 612476 | . | . | [illegible] |
| 10539 | NM_00119986 | GLRX3 | glutaredoxin 3 | protein_coding | NP_001186797.1;NP_001308909.1;NP_006532.2;XP_016870963.1 | 15987 | 612754 | . | . | [illegible] |
| 10540 | NM_00126141 | DCTN2 | dynactin subunit 2 | protein_coding | NP_001248341.1;NP_001248342.1;NP_001334994.1;NP_001334995.1;NI | 2712 | 607376 | . | . | [illegible] |
| 10541 | NM_006401 | ANP32B | acidic nuclear phosphoprotein 32 family member B | protein_coding | NP_006392.1 | 16677 | . | . | . | [illegible] |
| 10542 | NM_00138229 | LAMTOR5 | late endosomal/lysosomal adaptor, MAPK and MTOR activator 5 | protein_coding | NP_001369222.1;NP_006393.2 | 17955 | 608521 | . | . | [illegible] |
| 10544 | NM_006404,XI | PROCR | protein C receptor | protein_coding | NP_006395.2;XP_011526798.1 | 9452 | 600646 | . | . | [illegible] |
| 10548 | NM_00101484 | TM9SF1 | transmembrane 9 superfamily member 1 | protein_coding | NP_001014842.1;NP_001275935.1;NP_006396.2 | 11864 | . | . | . | [illegible] |
| 10549 | NM_006406 | PRDX4 | peroxiredoxin 4 | protein_coding | NP_006397.1 | 17169 | 300927 | . | . | [illegible] |
| 10550 | NM_006407 | ARL6IP5 | ADP ribosylation factor like GTPase 6 interacting protein 5 | protein_coding | NP_006398.1 | 16937 | 605709 | . | . | [illegible] |
| 10552 | NM_00119099 | ARPC1A | actin related protein 2/3 complex subunit 1A | protein_coding | NP_001177925.1;NP_006400.2 | 703 | 604220 | . | . | [illegible] |
| 10553 | NM_00109852 | HTATIP2 | HIV-1 Tat interactive protein 2 | protein_coding | NP_001091990.1;NP_001091991.1;NP_001091992.1;NP_001091993.1;NI | 16637 | 605628 | . | . | [illegible] |
| 10554 | NM_00137143 | AGPAT1 | 1-acylglycerol-3-phosphate O-acyltransferase 1 | protein_coding | NP_001358366.1;NP_001358367.1;NP_001358368.1;NP_006402.1;NP_1: | 324 | 603099 | . | . | [illegible] |
| 10555 | NM_00101272 | AGPAT2 | 1-acylglycerol-3-phosphate O-acyltransferase 2 | protein_coding | NP_001012745.1;NP_006403.2 | 325 | 603100 | . | . | [illegible] |
| 10556 | NM_00110454 | RPP30 | ribonuclease P/MRP subunit p30 | protein_coding | NP_001098016.1;NP_006404.1 | 17688 | 606115 | . | . | [illegible] |
| 10557 | NM_00109759 | RPP38 | ribonuclease P/MRP subunit p38 | protein_coding | NP_001091059.1;NP_001252530.1;NP_006405.2;NP_892117.1;XP_00671 | 30329 | 606116 | . | . | [illegible] |
| 10558 | NM_00128130 | SPTLC1 | serine palmitoyltransferase long chain base subunit 1 | protein_coding | NP_001268232.1;NP_001355201.1;NP_001355202.1;NP_006406.1;NP_8 | 11277 | 605712 | . | . | [illegible] |
| 10559 | NM_00116839 | SLC35A1 | solute carrier family 35 member A1 | protein_coding | NP_001161870.1;NP_006407.1 | 11021 | 605634 | . | . | [illegible] |
| 10560 | NM_00131966 | SLC19A2 | solute carrier family 19 member 2 | protein_coding | NP_001306596.1;NP_008927.1 | 10938 | 603941 | . | . | [illegible] |
| 10561 | NM_006417,N | IFI44 | interferon induced protein 44 | protein_coding | NP_006408.3;XP_005270437.1;XP_011538818.1;XP_016855572.1 | 16938 | 610468 | . | . | [illegible] |
| 10564 | NM_006420,XI | ARFGEF2 | ADP ribosylation factor guanine nucleotide exchange factor 2 | protein_coding | NP_006411.2;XP_005260309.1 | 15853 | 605371 | . | . | [illegible] |
| 10565 | NM_006421,XI | ARFGEF1 | ADP ribosylation factor guanine nucleotide exchange factor 1 | protein_coding | NP_006412.2;XP_005251191.1;XP_005251192.1;XP_005251193.1 | 15772 | 604141 | . | . | [illegible] |
| 10566 | NM_00127830 | AKAP3 | A-kinase anchoring protein 3 | protein_coding | NP_001265238.2;NP_006413.3;XP_011519210.1;XP_011519211.1;XP_01 | 373 | 604689 | . | . | [illegible] |
| 10567 | NM_006423 | RABAC1 | Rab acceptor 1 | protein_coding | NP_006414.2 | 9794 | 604925 | . | . | [illegible] |
| 10569 | NM_00136451 | SLU7 | SLU7 homolog, splicing factor | protein_coding | NP_001351446.1;NP_001351447.1;NP_001351449.1;NP_001351450.1;NI | 16939 | 605974 | . | . | [illegible] |
| 10570 | NM_006426,XI | DPYSL4 | dihydropyrimidinase like 4 | protein_coding | NP_006417.2;XP_005252715.1;XP_011537427.1;XP_016870971.1;XP_01( | 3016 | 608407 | . | . | [illegible] |
| 10572 | NM_006427,N | SIVA1 | SIVA1 apoptosis inducing factor | protein_coding | NP_006418.2;NP_068355.1;XP_011534662.1 | 17712 | 605567 | . | . | [illegible] |
| 10573 | NM_006428,XI | MRPL28 | mitochondrial ribosomal protein L28 | protein_coding | NP_006419.2;XP_005255098.1;XP_011520653.1 | 14484 | 604853 | . | . | [illegible] |
| 10574 | NM_00100957 | CCT7 | chaperonin containing TCP1 subunit 7 | protein_coding | NP_001009570.1;NP_001159756.1;NP_001159757.1;NP_006420.1;XP_01 | 1622 | 605140 | . | . | [illegible] |
| 10575 | NM_00125672 | CCT4 | chaperonin containing TCP1 subunit 4 | protein_coding | NP_001243650.1;NP_006421.2 | 1617 | 605142 | . | . | [illegible] |
| 10576 | NM_00119884 | CCT2 | chaperonin containing TCP1 subunit 2 | protein_coding | NP_001185771.1;NP_006422.1 | 1615 | 605139 | . | . | [illegible] |
| 10577 | NM_00136368 | NPC2 | NPC intracellular cholesterol transporter 2 | protein_coding | NP_001350617.1;NP_001362369.1;NP_006423.1 | 14537 | 601015 | . | . | [illegible] |
| 10579 | NM_00129187 | TACC2 | transforming acidic coiled-coil containing protein 2 | protein_coding | NP_001278805.2;NP_001278806.2;NP_001278807.1;NP_001278808.1;NI | 11523 | 605302 | . | . | [illegible] |
| 10580 | NM_00103495 | SORBS1 | sorbin and SH3 domain containing 1 | protein_coding | NP_001030126.2;NP_001030127.2;NP_001030128.2;NP_001030129.2;NI | 14565 | 605264 | . | . | [illegible] |
| 10585 | NM_00107736 | POMT1 | protein O-mannosyltransferase 1 | protein_coding | NP_001070833.1;NP_001070834.1;NP_001129585.1;NP_001129586.1;NI | 9202 | 607423 | . | . | [illegible] |
| 10587 | NM_00128251 | TXNRD2 | thioredoxin reductase 2 | protein_coding | NP_001269441.1;NP_001339229.1;NP_001339230.1;NP_001339231.1;NI | 18155 | 606448 | . | . | [illegible] |
| 10588 | NM_00119975 | MTHFS | methenyltetrahydrofolate synthetase | protein_coding | NP_001186687.1;NP_006432.1 | 7437 | 604197 | . | . | [illegible] |
| 10589 | NM_006442 | DRAP1 | DR1 associated protein 1 | protein_coding | NP_006433.2 | 3019 | 602289 | . | . | [illegible] |
| 10590 | NM_006998,XI | SCGN | secretagogin, EF-hand calcium binding protein | protein_coding | NP_008929.2;XP_016865670.1 | 16941 | 609202 | . | . | [illegible] |
| 10591 | NM_006443,N | DNPH1 | 2'-deoxynucleoside 5'-phosphate N-hydrolase 1 | protein_coding | NP_006434.1;NP_954653.1 | 21218 | 618762 | . | . | [illegible] |
| 10592 | NM_00104255 | SMC2 | structural maintenance of chromosomes 2 | protein_coding | NP_001036015.1;NP_001036016.1;NP_001252531.1;NP_006435.2;XP_0C | 14011 | 605576 | . | . | [illegible] |
| 10594 | NM_006445,XI | PRPF8 | pre-mRNA processing factor 8 | protein_coding | NP_006436.3;XP_024306305.1 | 17340 | 607300 | . | . | [illegible] |
| 10597 | NM_00135520 | TRAPPC2B | trafficking protein particle complex 2B | protein_coding | NP_001342133.1 | 10710 | . | . | . | [illegible] |
| 10598 | NM_00132144 | AHSA1 | activator of HSP90 ATPase activity 1 | protein_coding | NP_001308370.1;NP_036243.1 | 1189 | 608466 | . | . | [illegible] |
| 10600 | NM_00100199 | USP16 | ubiquitin specific peptidase 16 | protein_coding | NP_001001992.1;NP_001027582.1;NP_006438.1;XP_016883746.1;XP_01 | 12614 | 604735 | . | . | [illegible] |
| 10602 | NM_00127043 | CDC42EP3 | CDC42 effector protein 3 | protein_coding | NP_001257365.1;NP_001257366.1;NP_001257367.1;NP_001358498.1;NI | 16943 | 606133 | . | . | [illegible] |
| 10603 | NM_00135922 | SH2B2 | SH2B adaptor protein 2 | protein_coding | NP_001346157.1;NP_001346158.1;NP_001346159.1;NP_001346160.1;NI | 17381 | 605300 | . | . | [illegible] |
| 10605 | NM_006451,N | PAIP1 | poly(A) binding protein interacting protein 1 | protein_coding | NP_006442.2;NP_877590.1;NP_899152.1;XP_005248287.1;XP_01686444 | 16945 | 605184 | . | . | [illegible] |
| 10606 | NM_00107952 | PAICS | phosphoribosylaminoimidazole carboxylase and phosphoribosylar | protein_coding | NP_001072992.1;NP_001072993.1;NP_006443.1;XP_011532668.1 | 8587 | 172439 | . | . | [illegible] |
| 10607 | NM_006453 | TBL3 | transducin beta like 3 | protein_coding | NP_006444.2 | 11587 | 605915 | . | . | [illegible] |
| 10608 | NM_006454 | MXD4 | MAX dimerization protein 4 | protein_coding | NP_006445.1 | 13906 | . | . | . | [illegible] |
| 10609 | NM_006455,XI | P3H4 | prolyl 3-hydroxylase family member 4 (inactive) | protein_coding | NP_006446.1;XP_006721703.1;XP_024306306.1 | 16946 | 617419 | . | . | [illegible] |
| 10610 | NM_006456,XI | ST6GALNAC2 | ST6 N-acetylgalactosaminide alpha-2,6-sialyltransferase 2 | protein_coding | NP_006447.2;XP_005257011.1;XP_024306307.1 | 10867 | 610137 | . | . | [illegible] |
| 10611 | NM_00101151 | PDLIM5 | PDZ and LIM domain 5 | protein_coding | NP_001011513.4;NP_001011515.1;NP_001011516.1;NP_001243354.1;NI | 17468 | 605904 | . | . | [illegible] |
| 10612 | NM_00124800 | TRIM3 | tripartite motif containing 3 | protein_coding | NP_001234935.1;NP_001234936.1;NP_006449.2;NP_150594.2;XP_01151 | 10064 | 605493 | . | . | [illegible] |
| 10613 | NM_00110062 | ERLIN1 | ER lipid raft associated 1 | protein_coding | NP_001094096.1;NP_001334785.1;NP_001334786.1;NP_001334787.1;NI | 16947 | 611604 | . | . | [illegible] |
| 10614 | NM_006460 | HEXIM1 | HEXIM P-TEFb complex subunit 1 | protein_coding | NP_006451.1 | 24953 | 607328 | . | . | [illegible] |
| 10615 | NM_006461 | SPAG5 | sperm associated antigen 5 | protein_coding | NP_006452.3 | 13452 | 615562 | . | . | [illegible] |
| 10616 | NM_00132395 | RBCK1 | RANBP2-type and C3HC4-type zinc finger containing 1 | protein_coding | NP_001310885.1;NP_001310887.1;NP_001310889.1;NP_006453.1;NP_1: | 15864 | 610924 | . | . | [illegible] |
| 10617 | NM_00135396 | STAMBP | STAM binding protein | protein_coding | NP_001340896.1;NP_001340897.1;NP_001340898.1;NP_001340899.1;NI | 16950 | 606247 | . | . | [illegible] |
| 10618 | NM_00120684 | TGOLN2 | trans-golgi network protein 2 | protein_coding | NP_001193773.1;NP_001355024.1;NP_001355025.1;NP_006455.2 | 15450 | 603062 | . | . | [illegible] |
| 10620 | NM_00130793 | ARID3B | AT-rich interaction domain 3B | protein_coding | NP_001294868.1;NP_006456.1 | 14350 | 612457 | . | . | [illegible] |

| | | | | | | | | | | | | | | | | | | | | | | | | | | | | | | | | | |
|---|---|---|---|---|---|---|---|---|---|---|---|---|---|---|---|---|---|---|---|---|---|---|---|---|---|---|---|---|---|---|---|---|---|
| 10621 | NM_00128252 | POLR3F | RNA polymerase III subunit F | protein_coding | NP_001269455.1;NP_006457.2;XP_011527443.1;XP_016883085.1 | 15763 | 617455 | . | . | 4.026242 | 4.742930 | 4.731946 | 4.371263 | 9.369230 | 10.963493 | 10.618348 | 9.994376 | 1.025239 | 302.188225 | 0.206259015 | 0.174347039 | 0.861592734 | 0.998865404 | 8.221289 | 8.257096 | 8.188222 | 8.272283 | 8.266338 | 8.222218 | 317 | 347 | 277 | 275 |
| 10622 | NM_00137035 | POLR3G | RNA polymerase III subunit G | protein_coding | NP_001357280.1;NP_001357281.1;NP_001357282.1;NP_001357283.1;NI | 30075 | 617456 | . | . | 28.576752 | 24.910548 | 23.949492 | 23.399172 | 66.499276 | 57.581847 | 53.741967 | 53.499426 | -1.145584 | 2626.132656 | 0.122426625 | -1.60163565 | 0.109236202 | 0.842810112 | 11.453340 | 11.257465 | 11.467121 | 11.351039 | 11.314797 | 11.286130 | 3456 | 2813 | 2167 | 2254 |
| 10623 | NM_00130345 | POLR3C | RNA polymerase III subunit C | protein_coding | NP_001290385.1;NP_006459.3;XP_005272984.1;XP_005272985.1;XP_01 | 30076 | 617454 | . | . | 6.109036 | 6.524814 | 6.461297 | 7.414691 | 14.215982 | 15.082397 | 14.498963 | 16.952810 | 1.064891 | 696.868517 | 0.157740524 | 0.575029604 | 0.565271276 | 0.998865404 | 9.399034 | 9.489047 | 9.394627 | 9.448189 | 9.415104 | 9.512089 | 747 | 757 | 581 | 721 |
| 10625 | NM_006469,XI | IVNS1ABP | influenza virus NS1A binding protein | protein_coding | NP_006460.2;XP_005244900.1;XP_011507381.1;XP_011507382.1;XP_01( | 16951 | 609209 | . | . | 58.619301 | 58.329843 | 52.951161 | 53.231088 | 136.409531 | 134.832041 | 118.820878 | 121.706568 | -1.141845 | 7731.870196 | 0.101859417 | -1.87873511 | 0.060280671 | 0.719735192 | 13.009112 | 12.817751 | 12.969822 | 12.970660 | 12.861052 | 12.854701 | 9488 | 8900 | 6302 | 6743 |
| 10626 | NM_00134811 | TRIM16 | tripartite motif containing 16 | protein_coding | NP_001335048.1;NP_001335049.1;NP_001335050.1;NP_001335051.1;NI | 17241 | 609505 | . | . | 21.019628 | 21.898820 | 32.118454 | 28.962432 | 48.913534 | 50.620108 | 72.072879 | 66.219157 | 1.356808 | 2614.446752 | 0.114598517 | 3.841382535 | 0.000122343 | 0.013956234 | 11.115462 | 11.555667 | 11.196364 | 11.233238 | 11.477413 | 11.446755 | 2495 | 2450 | 2666 | 2770 |
| 10627 | NM_00130304 | MYL12A | myosin light chain 12A | protein_coding | NP_001289976.1;NP_001289977.1;NP_001289978.1;NP_006462.1 | 16701 | . | . | . | 106.996325 | 100.874234 | 91.636448 | 93.087566 | 248.984850 | 233.175306 | 205.629530 | 212.833665 | -1.142269 | 2959.318726 | 0.113469449 | -1.69122008 | 0.090794778 | 0.805424371 | 11.623779 | 11.431948 | 11.611284 | 11.553630 | 11.473078 | 11.475373 | 3760 | 3287 | 2399 | 2594 |
| 10628 | NM_00131397 | TXNIP | thioredoxin interacting protein | protein_coding | NP_001300901.1;NP_006463.3;XP_016855574.1 | 16952 | 606599 | . | . | 11.068362 | 9.989903 | 7.826089 | 8.893734 | 25.756532 | 23.092106 | 17.561515 | 20.334467 | -1.276441 | 855.459011 | 0.150473297 | -2.34013203 | 0.019276924 | 0.462516353 | 9.905977 | 9.553678 | 9.862737 | 9.776070 | 9.606551 | 9.684524 | 1169 | 975 | 616 | 745 |
| 10629 | NM_006473,XI | TAF6L | TATA-box binding protein associated factor 6 like | protein_coding | NP_006464.1;XP_005273771.1;XP_016872589.1 | 17305 | 602946 | . | . | 6.358053 | 6.919614 | 7.909198 | 7.698425 | 14.795451 | 15.994997 | 17.748013 | 17.601535 | 1.152346 | 441.570349 | 0.175860994 | 1.163270538 | 0.244719755 | 0.992733415 | 8.680697 | 8.885070 | 8.713256 | 8.765561 | 8.833703 | 8.819208 | 452 | 462 | 416 | 438 |
| 10632 | NM_006476,N | ATP5MG | ATP synthase membrane subunit g | protein_coding | NP_006467.4 | 14247 | 617473 | . | . | 49.031301 | 46.972934 | 40.017989 | 53.970324 | 114.097851 | 108.580049 | 89.799213 | 123.396721 | -1.037151 | 1677.181372 | 0.149591004 | -0.35179849 | 0.724989392 | 0.998865404 | 10.738102 | 10.685057 | 10.743928 | 10.702822 | 10.578034 | 10.799850 | 2018 | 1795 | 1228 | 1763 |
| 10633 | NM_006477,XI | RASL10A | RAS like family 10 member A | protein_coding | NP_006468.1;XP_011528123.1;XP_011528124.1;XP_011528125.1 | 16954 | 602220 | . | . | 0.237288 | 0.383459 | 0.205103 | 0.435095 | 0.552179 | 0.886381 | 0.460246 | 0.994792 | 1.295301 | 15.951082 | 0.802217976 | 0.465318839 | 0.641703146 | NA | 3.799841 | 4.167911 | 3.974811 | 3.983024 | 3.964331 | 4.028243 | 15 | 16 | 9 | 24 |
| 10634 | NM_00127873 | GAS2L1 | growth arrest specific 2 like 1 | protein_coding | NP_001265659.1;NP_001349914.1;NP_006469.2;NP_689422.1;XP_01152 | 16955 | 602128 | . | . | 11.614933 | 11.602587 | 10.162808 | 11.740080 | 27.028425 | 26.819900 | 22.805044 | 26.842301 | -1.072312 | 918.788837 | 0.144957148 | -0.69486174 | 0.487141953 | 0.998865404 | 9.893319 | 9.792084 | 9.854521 | 9.881355 | 9.770307 | 9.862466 | 1075 | 1045 | 721 | 885 |
| 10635 | NM_00113086 | RAD51AP1 | RAD51 associated protein 1 | protein_coding | NP_001124334.1;NP_006470.1 | 16956 | 603070 | . | . | 8.867163 | 7.733689 | 8.080999 | 7.801421 | 20.634254 | 17.876768 | 18.133527 | 17.837022 | -1.059556 | 540.064860 | 0.168599724 | -0.49502128 | 0.620585101 | 0.998865404 | 9.117822 | 9.034966 | 9.139491 | 9.048257 | 9.074029 | 9.040917 | 684 | 558 | 468 | 479 |
| 10636 | NM_00136661 | RGS14 | regulator of G protein signaling 14 | protein_coding | NP_001353546.1;NP_001353547.1;NP_006471.2;XP_024310096.1 | 9996 | 602513 | . | . | 0.159082 | 0.113901 | 0.197867 | 0.155970 | 0.370189 | 0.263287 | 0.444009 | 0.356608 | 1.149211 | 12.461381 | 0.870380423 | 0.230523572 | 0.817684949 | NA | 3.533720 | 3.737849 | 3.642204 | 3.621227 | 3.646185 | 3.632517 | 16 | 10 | 13 | 11 |
| 10640 | NM_006544,XI | EXOC5 | exocyst complex component 5 | protein_coding | NP_006535.1;XP_005267329.1 | 10696 | 604469 | . | . | 3.991411 | 3.438543 | 4.201314 | 2.918158 | 9.288176 | 7.948345 | 9.427627 | 6.672022 | -1.106507 | 1139.621221 | 0.158211892 | -0.92288941 | 0.356064842 | 0.998865404 | 10.225124 | 10.079900 | 10.232967 | 10.141506 | 10.206611 | 10.009542 | 1461 | 1214 | 1067 | 879 |
| 10641 | NM_006545,XI | NPRL2 | NPR2 like, GATOR1 complex subunit | protein_coding | NP_006536.3;XP_005264865.1;XP_011531590.1;XP_016861044.1;XP_01( | 24969 | 607072 | . | . | 10.715375 | 11.159201 | 12.020478 | 11.275399 | 24.935123 | 25.794996 | 26.973602 | 25.779861 | 1.044671 | 536.221733 | 0.16454108 | 0.383179906 | 0.701586376 | 0.998865404 | 9.034752 | 9.097928 | 9.036787 | 9.066622 | 9.100817 | 9.058854 | 590 | 579 | 492 | 497 |
| 10642 | NM_00116042 | IGF2BP1 | insulin like growth factor 2 mRNA binding protein 1 | protein_coding | NP_001153895.1;NP_006537.3;XP_005257012.2;XP_011522503.1;XP_01 | 28866 | 608288 | . | . | 4.074693 | 4.671560 | 4.668517 | 4.429643 | 9.481978 | 10.798519 | 10.476020 | 10.127852 | 1.030485 | 1220.522230 | 0.137309149 | 0.315522365 | 0.752365073 | 0.998865404 | 10.231501 | 10.274740 | 10.191156 | 10.288601 | 10.287002 | 10.239763 | 1295 | 1381 | 1111 | 1125 |
| 10643 | NM_006547,XI | IGF2BP3 | insulin like growth factor 2 mRNA binding protein 3 | protein_coding | NP_006538.2;XP_006715702.1;XP_011513391.1;XP_011513392.1;XP_01: | 28868 | 608259 | . | . | 25.267987 | 26.805655 | 33.326439 | 27.728964 | 58.799639 | 61.962471 | 74.783570 | 63.398983 | 1.190874 | 3726.432813 | 0.126160422 | 1.997621536 | 0.045757707 | 0.653147492 | 11.732007 | 11.984149 | 11.753686 | 11.816086 | 12.006224 | 11.847541 | 3767 | 3811 | 3855 | 3440 |
| 10644 | NM_00100722 | IGF2BP2 | insulin like growth factor 2 mRNA binding protein 2 | protein_coding | NP_001007226.1;NP_001278798.1;NP_001278801.1;NP_001278802.1;NI | 28867 | 608289 | . | . | 30.220750 | 32.122700 | 31.323935 | 33.036727 | 70.324928 | 74.253058 | 70.290002 | 75.534556 | 1.046564 | 3700.690225 | 0.110243019 | 0.595593269 | 0.551446974 | 0.998865404 | 11.820420 | 11.885992 | 11.808971 | 11.859049 | 11.863495 | 11.880238 | 4038 | 4021 | 3258 | 3584 |
| 10645 | NM_00127048 | CAMKK2 | calcium/calmodulin dependent protein kinase kinase 2 | protein_coding | NP_001257414.1;NP_001257415.1;NP_006540.3;NP_705719.2;NP_7057: | 1470 | 615002 | . | . | 3.548521 | 3.900441 | 3.386064 | 3.967079 | 8.257560 | 9.016044 | 7.598230 | 9.070255 | -1.078913 | 561.161179 | 0.166117608 | -0.65964593 | 0.509481073 | 0.998865404 | 9.186266 | 9.076177 | 9.165519 | 9.147163 | 9.060275 | 9.151060 | 680 | 620 | 436 | 542 |
| 10650 | NM_00114240 | PRELID3A | PRELI domain containing 3A | protein_coding | NP_001135877.1;NP_001135878.1;NP_006544.2;XP_024306843.1;XP_02 | 24639 | 616545 | . | . | 1.880023 | 1.690993 | 2.905206 | 2.201074 | 4.374893 | 3.908806 | 6.519197 | 5.032494 | 1.361864 | 120.038679 | 0.306123125 | 1.455567328 | 0.145512266 | 0.906189003 | 6.666341 | 7.113811 | 6.860522 | 6.823415 | 6.996946 | 6.895610 | 124 | 103 | 137 | 112 |
| 10651 | NM_00100663 | MTX2 | metaxin 2 | protein_coding | NP_001006636.1;NP_001306026.1;NP_001306027.1;NP_006545.1;XP_01 | 7506 | 608555 | . | . | 37.661419 | 36.994303 | 33.939653 | 36.187718 | 87.639666 | 85.513991 | 76.159597 | 82.738915 | -1.084008 | 1555.465450 | 0.124985588 | -0.93111116 | 0.351796066 | 0.998865404 | 10.660171 | 10.543741 | 10.650629 | 10.616299 | 10.552003 | 10.589724 | 1904 | 1708 | 1266 | 1434 |
| 10652 | NM_00136367 | YKT6 | YKT6 v-SNARE homolog | protein_coding | NP_001350607.1;NP_006546.1;XP_005249639.1 | 16959 | 606209 | . | . | 35.740161 | 35.010509 | 34.958012 | 39.361068 | 83.168821 | 80.928362 | 78.444766 | 89.994409 | 1.033967 | 3128.449221 | 0.11601492 | 0.41537915 | 0.677864368 | 0.998865404 | 11.586999 | 11.635065 | 11.608415 | 11.585499 | 11.579823 | 11.667013 | 3592 | 3273 | 2619 | 3140 |
| 10654 | NM_00132301 | PMVK | phosphomevalonate kinase | protein_coding | NP_001309940.1;NP_001309941.1;NP_001335625.1;NP_006547.1 | 9141 | 607622 | . | . | 18.622716 | 18.063519 | 17.596656 | 19.009097 | 43.335821 | 41.754636 | 39.486386 | 43.462040 | -1.011550 | 579.396774 | 0.160506225 | -0.10322231 | 0.917786533 | 0.998865404 | 9.186691 | 9.170077 | 9.200766 | 9.163164 | 9.157603 | 9.191261 | 690 | 611 | 488 | 554 |
| 10656 | NM_006558,XI | KHDRBS3 | KH RNA binding domain containing, signal transduction associatec | protein_coding | NP_006549.1;XP_005250814.1;XP_005250815.1;XP_011515098.1;XP_01: | 18117 | 610421 | . | . | 11.270231 | 10.731197 | 12.700735 | 13.375586 | 26.226290 | 24.805644 | 28.500083 | 30.581686 | 1.136819 | 724.049445 | 0.149720803 | 1.235648955 | 0.216589079 | 0.982008277 | 9.404447 | 9.589511 | 9.472517 | 9.433141 | 9.531088 | 9.552108 | 803 | 710 | 660 | 733 |
| 10657 | NM_00127187 | KHDRBS1 | KH RNA binding domain containing, signal transduction associatec | protein_coding | NP_001258807.1;NP_006550.1 | 18116 | 602489 | . | . | 43.287878 | 42.876558 | 41.567735 | 39.037417 | 100.732669 | 99.111086 | 93.276788 | 89.254411 | -1.091849 | 3492.791107 | 0.110862854 | -1.14351259 | 0.252825832 | 0.994320829 | 11.832107 | 11.705443 | 11.812848 | 11.797496 | 11.757263 | 11.707819 | 4237 | 3898 | 2992 | 3036 |
| 10658 | NM_00102559 | CELF1 | CUGBP Elav-like family member 1 | protein_coding | NP_001020767.1;NP_001166110.1;NP_001166111.1;NP_001317201.1;NI | 2549 | 601074 | . | . | 6.823172 | 6.937901 | 7.409948 | 6.328976 | 15.877788 | 16.037274 | 16.627705 | 14.470437 | -1.023887 | 1605.214992 | 0.133340145 | -0.25540963 | 0.798406735 | 0.998865404 | 10.665276 | 10.631630 | 10.668402 | 10.644878 | 10.700752 | 10.572151 | 1898 | 1726 | 1494 | 1364 |
| 10659 | NM_00102507 | CELF2 | CUGBP Elav-like family member 2 | protein_coding | NP_001020247.1;NP_001020248.1;NP_001077060.1;NP_001313246.1;NI | 2550 | 602538 | . | . | 5.768696 | 5.745130 | 7.471474 | 6.380367 | 13.423993 | 13.280125 | 16.765771 | 14.587949 | 1.109832 | 1499.420545 | 0.129534133 | 1.160633511 | 0.245790972 | 0.992733415 | 10.472886 | 10.623560 | 10.522268 | 10.492860 | 10.630703 | 10.544145 | 1668 | 1504 | 1448 | 1397 |
| 10664 | NM_00119102 | CTCF | CCCTC-binding factor | protein_coding | NP_001177951.1;NP_001350845.1;NP_006556.1;XP_005255832.1;XP_01 | 13723 | 604167 | . | . | 9.605449 | 9.935858 | 10.747003 | 10.723242 | 22.352274 | 22.967183 | 24.115959 | 24.517420 | 1.084663 | 1216.969809 | 0.130084273 | 0.901310054 | 0.36742349 | 0.998865404 | 10.189283 | 10.306492 | 10.205859 | 10.228663 | 10.282472 | 10.274601 | 1324 | 1279 | 1107 | 1181 |
| 10666 | NM_00130361 | CD226 | CD226 molecule | protein_coding | NP_001290547.1;NP_001290548.1;NP_006557.2;XP_005266699.1;XP_00 | 16961 | 605397 | . | . | 0.025204 | 0.049139 | 0.070236 | 0.037362 | 0.058652 | 0.113588 | 0.157607 | 0.085425 | 1.423260 | 18.103917 | 0.757568472 | 0.672149253 | 0.501488684 | NA | 3.901250 | 4.410501 | 4.127114 | 4.167808 | 4.200457 | 4.148340 | 12 | 21 | 24 | 14 |
| 10667 | NM_00131887 | FARS2 | phenylalanyl-tRNA synthetase 2, mitochondrial | protein_coding | NP_001305801.1;NP_001361804.1;NP_001361805.1;NP_001361806.1;NI | 21062 | 611592 | . | . | 4.848562 | 4.277647 | 3.517967 | 4.981935 | 11.282801 | 9.887973 | 7.894218 | 11.390598 | -1.071303 | 239.962099 | 0.238477863 | -0.41667108 | 0.676919012 | 0.998865404 | 7.955997 | 7.855582 | 7.952970 | 7.888900 | 7.810648 | 7.959051 | 304 | 251 | 170 | 251 |
| 10668 | NM_006568,XI | CGRRF1 | cell growth regulator with ring finger domain 1 | protein_coding | NP_006559.1;XP_011534664.1;XP_016876420.1;XP_016876421.1 | 15528 | 606138 | . | . | 3.900920 | 3.464557 | 2.907556 | 2.856225 | 9.077603 | 8.008478 | 6.524470 | 6.530420 | -1.294435 | 203.810799 | 0.238190395 | -1.56313137 | 0.118021698 | 0.857396579 | 7.845010 | 7.473284 | 7.759861 | 7.691582 | 7.613620 | 7.605027 | 283 | 231 | 156 | 165 |
| 10669 | NM_00116623 | CGREF1 | cell growth regulator with EF-hand domain 1 | protein_coding | NP_001159711.1;NP_001159712.1;NP_001288253.1;NP_006560.3 | 16962 | 606137 | . | . | 12.471406 | 14.876137 | 12.273568 | 13.575345 | 29.021475 | 34.386856 | 27.541532 | 31.038413 | -1.072345 | 794.769603 | 0.155935296 | -0.64622674 | 0.518132523 | 0.998865404 | 9.684218 | 9.582786 | 9.594994 | 9.716491 | 9.577820 | 9.637642 | 867 | 963 | 638 | 750 |
| 10670 | NM_006570 | RRAGA | Ras related GTP binding A | protein_coding | NP_006561.1 | 16963 | 612194 | . | . | 17.074482 | 17.012350 | 18.256561 | 17.859287 | 39.733021 | 39.324810 | 40.967194 | 40.833138 | 1.044889 | 874.466102 | 0.14116076 | 0.448772273 | 0.653595942 | 0.998865404 | 9.740161 | 9.803658 | 9.760529 | 9.751538 | 9.799320 | 9.775650 | 991 | 917 | 790 | 824 |
| 10671 | NM_006571 | DCTN6 | dynactin subunit 6 | protein_coding | NP_006562.1 | 16964 | 612963 | . | . | 16.047249 | 14.648953 | 15.098516 | 16.275661 | 37.342606 | 33.861710 | 33.880634 | 37.212368 | 1.007436 | 509.924145 | 0.169303721 | 0.063133444 | 0.949660243 | 0.998865404 | 8.988766 | 8.999492 | 9.022734 | 8.959143 | 8.975939 | 9.015991 | 614 | 521 | 431 | 495 |
| 10672 | NM_00128242 | GNA13 | G protein subunit alpha 13 | protein_coding | NP_001269354.1;NP_006563.2;XP_011522504.1 | 4381 | 604406 | . | . | 8.572031 | 7.564489 | 8.616758 | 5.984478 | 19.947471 | 17.485657 | 19.335752 | 13.682797 | -1.113005 | 1473.958568 | 0.169755156 | -0.9098969 | 0.362876886 | 0.998865404 | 10.600310 | 10.446556 | 10.609966 | 10.508461 | 10.611348 | 10.324478 | 1903 | 1567 | 1441 | 1063 |
| 10675 | NM_00120694 | CSPG5 | chondroitin sulfate proteoglycan 5 | protein_coding | NP_001193871.1;NP_001193872.1;NP_001193873.1;NP_001193874.1;NI | 2467 | 606775 | . | . | 1.223335 | 0.861414 | 1.186241 | 1.304715 | 2.846749 | 1.991195 | 2.661890 | 2.983076 | 1.176662 | 78.145528 | 0.371082326 | 0.632474511 | 0.527076859 | 0.998865404 | 6.165415 | 6.401149 | 6.304095 | 6.218285 | 6.291110 | 6.316976 | 97 | 64 | 70 | 83 |
| 10677 | NM_006576,XI | AVIL | advillin | protein_coding | NP_006567.3;XP_016874199.1;XP_016874200.1;XP_016874201.1;XP_01( | 14188 | 613397 | . | . | 0.075528 | 0.194365 | 0.141365 | 0.091408 | 0.175756 | 0.449283 | 0.317219 | 0.208994 | -1.190381 | 17.130375 | 0.786152902 | -0.31981487 | 0.749108677 | NA | 4.218727 | 3.967302 | 4.064912 | 4.129137 | 4.101207 | 4.068129 | 13 | 28 | 17 | 11 |
| 10678 | NM_00131907 | B3GNT2 | UDP-GlcNAc:betaGal beta-1,3-N-acetylglucosaminyltransferase 2 | protein_coding | NP_001306004.1;NP_006568.2 | 15629 | 605581 | . | . | 11.084867 | 9.456284 | 8.954880 | 9.896524 | 25.794940 | 21.858622 | 20.094492 | 22.627229 | -1.102337 | 847.225137 | 0.153366724 | -0.91653036 | 0.359388793 | 0.998865404 | 9.795147 | 9.654636 | 9.817512 | 9.697996 | 9.657876 | 9.721281 | 1107 | 880 | 669 | 790 |
| 10681 | NM_00137934 | GNB5 | G protein subunit beta 5 | protein_coding | NP_001366272.1;NP_006569.1;NP_057278.2;XP_011519464.1;XP_01687 | 4401 | 604447 | . | . | 2.256394 | 2.251043 | 1.796399 | 2.613499 | 5.250721 | 5.203390 | 4.031068 | 5.975456 | 1.031743 | 450.817430 | 0.187588188 | 0.24033609 | 0.810069719 | 0.998865404 | 8.794083 | 8.838376 | 8.821516 | 8.788065 | 8.745340 | 8.897136 | 525 | 466 | 349 | 482 |
| 10682 | NM_006579 | EBP | EBP cholestenol delta-isomerase | protein_coding | NP_006570.1 | 3133 | 300205 | . | . | 38.629303 | 37.071438 | 47.703449 | 45.134975 | 89.891968 | 85.692291 | 107.045158 | 103.195755 | 1.209682 | 1476.508228 | 0.127190401 | 2.159189876 | 0.030835439 | 0.558237552 | 10.383994 | 10.658879 | 10.467577 | 10.430226 | 10.621084 | 10.569774 | 1577 | 1406 | 1452 | 1466 |
| 10683 | NM_016941,N | DLL3 | delta like canonical Notch ligand 3 | protein_coding | NP_058637.1;NP_982353.1 | 2909 | 602768 | . | . | 2.434039 | 2.584000 | 3.052545 | 2.889162 | 5.664109 | 5.973030 | 6.849822 | 6.605725 | 1.141341 | 178.322566 | 0.24938635 | 0.764798528 | 0.444391524 | 0.998865404 | 7.379788 | 7.570598 | 7.429696 | 7.463675 | 7.520501 | 7.485488 | 184 | 187 | 172 | 171 |
| 10687 | NM_007257,XI | PNMA2 | PNMA family member 2 | protein_coding | NP_009188.1;XP_011542667.1 | 9159 | 603970 | . | . | 2.994392 | 3.356900 | 3.230244 | 3.688459 | 6.968074 | 7.759624 | 7.248575 | 8.433221 | 1.075439 | 489.201000 | 0.174586115 | 0.601000025 | 0.54783997 | 0.998865404 | 8.881289 | 8.985400 | 8.875656 | 8.942648 | 8.916076 | 8.993131 | 514 | 536 | 414 | 504 |
| 10691 | NM_00131967 | GMEB1 | glucocorticoid modulatory element binding protein 1 | protein_coding | NP_001306603.1;NP_006573.2;NP_077808.1;XP_011538820.1;XP_01153 | 4370 | 604409 | . | . | 1.139285 | 1.583162 | 1.093881 | 1.333387 | 2.651163 | 3.659553 | 2.454637 | 3.048628 | -1.032021 | 241.126005 | 0.229419555 | -0.19820715 | 0.842882997 | 0.998865404 | 7.936909 | 7.889997 | 7.874256 | 7.963725 | 7.863323 | 7.943819 | 257 | 288 | 189 | 241 |
| 10693 | NM_00119352 | CCT6B | chaperonin containing TCP1 subunit 6B | protein_coding | NP_001180458.1;NP_001180459.1;NP_006575.2;XP_011522505.1;XP_01 | 1621 | 610730 | . | . | 0.616348 | 0.465206 | 0.431808 | 0.413183 | 1.434267 | 1.075344 | 0.968965 | 0.944694 | -1.272568 | 28.295735 | 0.586841016 | -0.59256683 | 0.55347108 | 0.998865404 | 4.985276 | 4.639023 | 4.858573 | 4.818963 | 4.813033 | 4.809689 | 42 | 29 | 22 | 23 |
| 10694 | NM_00128290 | CCT8 | chaperonin containing TCP1 subunit 8 | protein_coding | NP_001269836.1;NP_001269837.1;NP_001269838.1;NP_006576.2 | 1623 | 617786 | . | . | 106.389345 | 105.488224 | 94.867367 | 94.107179 | 247.572382 | 243.840750 | 212.879629 | 215.164872 | -1.138013 | 5801.554354 | 0.104173614 | -1.79044146 | 0.073382971 | 0.762854575 | 12.592456 | 12.405970 | 12.562205 | 12.546051 | 12.452816 | 12.438138 | 7179 | 6601 | 4761 | 5043 |
| 10695 | NM_00131884 | CNPY3 | canopy FGF signaling regulator 3 | protein_coding | NP_001305771.1;NP_001305774.1;NP_001305776.1;NP_001305777.1;NI | 11968 | 610774 | . | . | 46.232178 | 41.360055 | 47.486202 | 42.020578 | 107.584170 | 95.605618 | 106.557669 | 96.075055 | 1.027743 | 2135.584317 | 0.119097085 | 0.33149601 | 0.740269854 | 0.998865404 | 11.040496 | 11.080063 | 11.079188 | 11.018219 | 11.066127 | 11.075947 | 2518 | 2186 | 1871 | 2042 |
| 10712 | NM_00126760 | FAM189B | family with sequence similarity 189 member B | protein_coding | NP_001254537.1;NP_006580.2;NP_937995.1;XP_005244902.1 | 1233 | . | . | . | 8.141997 | 8.299783 | 9.891230 | 10.009008 | 18.946761 | 19.185321 | 22.195634 | 22.884409 | 1.188760 | 897.008203 | 0.140497469 | 1.775526949 | 0.07581092 | 0.767975096 | 9.678897 | 9.928293 | 9.750621 | 9.734176 | 9.852405 | 9.880796 | 955 | 874 | 831 | 931 |
| 10713 | NM_00125672 | USP39 | ubiquitin specific peptidase 39 | protein_coding | NP_001243654.1;NP_001243655.1;NP_001243656.1;NP_001243657.1;NI | 20071 | 611594 | . | . | 28.272762 | 29.261009 | 26.784764 | 31.103011 | 65.791878 | 67.638135 | 60.104232 | 71.113338 | -1.005004 | 1955.380427 | 0.127217003 | -0.05660222 | 0.954862069 | 0.998865404 | 10.936992 | 10.929466 | 10.923845 | 10.945318 | 10.871132 | 10.986755 | 2227 | 2144 | 1572 | 1962 |
| 10714 | NM_00136359 | POLD3 | DNA polymerase delta 3, accessory subunit | protein_coding | NP_001350526.1;NP_006582.1;XP_005273773.1;XP_011543036.1 | 20932 | 611415 | . | . | 6.200050 | 6.758391 | 6.334190 | 5.696088 | 14.427771 | 15.622323 | 14.213738 | 13.023430 | -1.066041 | 714.047647 | 0.155647982 | -0.59276988 | 0.55333517 | 0.998865404 | 9.525087 | 9.433203 | 9.482198 | 9.519123 | 9.503034 | 9.408426 | 826 | 816 | 643 | 603 |
| 10715 | NM_00129026 | CERS1 | ceramide synthase 1 | protein_coding | NP_001277194.1;NP_067090.1;NP_937850.1 | 14253 | 606919 | . | . | 7.702782 | 7.297123 | 9.188807 | 8.101673 | 17.924690 | 16.867627 | 20.619418 | 18.523511 | 1.127867 | 540.987994 | 0.168096678 | 1.032718615 | 0.3017356 | 0.998865404 | 8.989677 | 9.163968 | 9.056329 | 9.018052 | 9.158097 | 9.070849 | 603 | 532 | 532 | 502 |
| 10717 | NM_00125385 | AP4B1 | adaptor related protein complex 4 subunit beta 1 | protein_coding | NP_001240781.1;NP_001240782.1;NP_001295241.1;NP_006585.2;XP_01 | 572 | 607245 | . | . | 7.817411 | 7.014001 | 8.139390 | 8.215194 | 18.191432 | 16.213180 | 18.264558 | 18.783066 | -1.014536 | 634.329425 | 0.155184429 | -0.13416209 | 0.893274399 | 0.998865404 | 9.319538 | 9.298562 | 9.296478 | 9.331000 | 9.303435 | 9.304854 | 717 | 707 | 547 | 591 |
| 10718 | NM_00101084 | NRG3 | neuregulin 3 | protein_coding | NP_001010848.2;NP_001159444.1;NP_001159445.1;NP_001357010.1;NI | 7999 | 605533 | . | . | 0.986618 | 1.209691 | 1.774186 | 2.332914 | 2.295899 | 2.796255 | 3.981224 | 5.333933 | 1.296088 | 201.848366 | 0.23520776 | 1.590778365 | 0.111659464 | 0.848562841 | 7.458126 | 7.831978 | 7.582859 | 7.597995 | 7.699878 | 7.715654 | 199 | 193 | 195 | 217 |
| 10721 | NM_199420 | POLQ | DNA polymerase theta | protein_coding | NP_955452.3 | 9186 | 604419 | . | . | 4.375060 | 4.531475 | 5.285069 | 3.956424 | 10.180944 | 10.474706 | 11.859541 | 9.045894 | 1.023843 | 1238.388459 | 0.157221543 | 0.216219188 | 0.828816898 | 0.998865404 | 10.256868 | 10.291421 | 10.251931 | 10.272257 | 10.386867 | 10.160932 | 1390 | 1338 | 1252 | 1000 |
| 10723 | NM_006598,XI | SLC12A7 | solute carrier family 12 member 7 | protein_coding | NP_006589.2;XP_005248288.1;XP_011512241.1;XP_011512242.1;XP_01: | 10915 | 604879 | . | . | 4.236119 | 4.649188 | 6.178891 | 5.066369 | 9.857625 | 10.746807 | 13.865253 | 11.583650 | 1.248425 | 831.865308 | 0.158130938 | 2.024326377 | 0.042936581 | 0.639324307 | 9.531089 | 9.851562 | 9.582869 | 9.641321 | 9.843469 | 9.691359 | 816 | 832 | 887 | 776 |
| 10724 | NM_00114243 | OGA | O-GlcNAcase | protein_coding | NP_001135906.1;NP_036347.1;XP_016871074.1;XP_016871075.1;XP_02 | 7056 | 604039 | . | . | 9.879478 | 10.573614 | 11.111680 | 9.436505 | 22.989948 | 24.441381 | 24.934290 | 21.575450 | -1.008144 | 1642.634414 | 0.134625824 | -0.08692106 | 0.930734259 | 0.998865404 | 10.687497 | 10.676072 | 10.660866 | 10.706227 | 10.742890 | 10.607878 | 1844 | 1831 | 1546 | 1401 |
| 10725 | NM_00111317 | NFAT5 | nuclear factor of activated T cells 5 | protein_coding | NP_001106649.1;NP_001354638.1;NP_006590.1;NP_619727.2;NP_6197: | 7774 | 604708 | . | . | 3.282977 | 3.229947 | 3.837573 | 3.433554 | 7.639620 | 7.466166 | 8.611404 | 7.850412 | 1.000007 | 1337.754784 | 0.164091668 | 5.98821E-05 | 0.999952221 | 0.999952221 | 10.385280 | 10.385916 | 10.387388 | 10.375505 | 10.507373 | 10.238266 | 1552 | 1432 | 1368 | 1034 |
| 10726 | NM_006600,XI | NUDC | nuclear distribution C, dynein complex regulator | protein_coding | NP_006591.1;XP_011538832.1;XP_016855583.1;XP_016855584.1;XP_01( | 8045 | 610325 | . | . | 83.234201 | 85.224799 | 83.719735 | 91.159585 | 193.689406 | 197.000929 | 187.864669 | 208.425560 | 1.003951 | 4523.335025 | 0.10979945 | 0.051815212 | 0.958675934 | 0.998865404 | 12.140375 | 12.145962 | 12.139004 | 12.143653 | 12.108420 | 12.179508 | 5184 | 4885 | 3773 | 4430 |
| 10728 | NM_00128260 | PTGES3 | prostaglandin E synthase 3 | protein_coding | NP_001269530.1;NP_001269531.1;NP_001269532.1;NP_001269533.1;NI | 16049 | 607061 | . | . | 83.220071 | 74.106489 | 74.826144 | 47.040664 | 193.656529 | 171.300479 | 167.907691 | 107.552890 | -1.314272 | 4266.249081 | 0.332002145 | -1.18753565 | 0.235016402 | 0.990818043 | 12.242424 | 11.848238 | 12.210080 | 12.097830 | 12.093910 | 11.729230 | 5945 | 4886 | 3920 | 2687 |
| 10730 | NM_00125386 | YME1L1 | YME1 like 1 ATPase | protein_coding | NP_001240795.1;NP_055078.1;NP_647473.1;XP_011517602.1 | 12843 | 607472 | . | . | 24.755547 | 24.399196 | 22.301958 | 23.123837 | 57.607178 | 56.399834 | 50.044946 | 52.869907 | -1.101370 | 3105.356059 | 0.110990206 | -1.25506151 | 0.209456432 | 0.971807438 | 11.668528 | 11.529195 | 11.646479 | 11.631123 | 11.547085 | 11.572263 | 3783 | 3480 | 2530 | 2814 |
| 10732 | NM_00130172 | TCFL5 | transcription factor like 5 | protein_coding | NP_001288655.1;NP_006593.2;XP_005260242.1;XP_011526799.1;XP_02 | 11646 | 604745 | . | . | 20.050435 | 19.985861 | 23.876036 | 23.090819 | 46.658183 | 46.198210 | 53.577134 | 52.794411 | 1.147408 | 1778.961463 | 0.120701229 | 1.643550218 | 0.100269144 | 0.815618258 | 10.694186 | 10.892634 | 10.736228 | 10.743448 | 10.863056 | 10.832537 | 1899 | 1796 | 1685 | 1748 |
| 10733 | NM_00119079 | PLK4 | polo like kinase 4 | protein_coding | NP_001177728.1;NP_001177730.1;NP_055079.3;XP_005262758.1;XP_01 | 11397 | 605031 | . | . | 5.630888 | 5.209259 | 5.332375 | 4.602001 | 13.103308 | 12.041436 | 11.965692 | 10.521929 | -1.108491 | 618.844614 | 0.164313339 | -0.90435509 | 0.365807143 | 0.998865404 | 9.345434 | 9.197650 | 9.334806 | 9.276288 | 9.286520 | 9.185625 | 781 | 672 | 550 | 508 |
| 10734 | NM_00128271 | STAG3 | stromal antigen 3 | protein_coding | NP_001269645.1;NP_001269646.1;NP_001269647.1;NP_001362367.1;NI | 11356 | 608489 | . | . | 22.077331 | 25.656305 | 23.812632 | 22.996717 | 51.374854 | 59.305699 | 53.434860 | 52.579264 | -1.031843 | 3089.085328 | 0.121060292 | -0.37356339 | 0.708729172 | 0.998865404 | 11.615436 | 11.570136 | 11.543492 | 11.662966 | 11.597275 | 11.560901 | 3351 | 3631 | 2703 | 2787 |
| 10735 | NM_00104274 | STAG2 | stromal antigen 2 | protein_coding | NP_001036214.1;NP_001036215.1;NP_001036216.1;NP_001269347.1;NI | 11355 | 300826 | . | . | 10.646789 | 9.640682 | 10.431054 | 7.549734 | 24.775518 | 22.284865 | 23.406980 | 17.261567 | -1.094508 | 1627.517644 | 0.135851994 | -0.95900502 | 0.337556216 | 0.998865404 | 10.731895 | 10.602102 | 10.736710 | 10.665037 | 10.693825 | 10.564604 | 2047 | 1752 | 1465 | 1335 |
| 10741 | NM_006606,XI | RBBP9 | RB binding protein 9, serine hydrolase | protein_coding | NP_006597.2;XP_005260709.1 | 9892 | 602908 | . | . | 4.089420 | 3.385632 | 3.806366 | 3.517857 | 9.516247 | 7.826039 | 8.541375 | 8.043162 | -1.035355 | 443.794502 | 0.185150287 | -0.27072945 | 0.786599124 | 0.998865404 | 8.818137 | 8.768940 | 8.862603 | 8.739625 | 8.808040 | 8.754916 | 571 | 439 | 396 | 391 |
| 10743 | NM_030665,XI | RAI1 | retinoic acid induced 1 | protein_coding | NP_109590.3;XP_016879514.1;XP_016879515.1;XP_016879516.1;XP_01( | 9834 | 607642 | . | . | 14.309774 | 15.652193 | 16.122947 | 16.803665 | 33.299430 | 36.180744 | 36.179428 | 38.419585 | 1.086886 | 3718.232626 | 0.111271919 | 1.080239853 | 0.280035386 | 0.998865404 | 11.799106 | 11.919199 | 11.793499 | 11.854958 | 11.882892 | 11.904101 | 3950 | 3989 | 3325 | 3677 |
| 10745 | NM_00132304 | PHTF1 | putative homeodomain transcription factor 1 | protein_coding | NP_001309970.1;NP_001309971.1;NP_001309972.1;NP_001309973.1;NI | 8939 | 604950 | . | . | 10.518443 | 9.409665 | 12.842238 | 12.247357 | 24.476851 | 21.750864 | 28.817613 | 28.002123 | 1.225971 | 1169.010191 | 0.139679201 | 2.104283708 | 0.035353706 | 0.585116558 | 10.036393 | 10.330761 | 10.160146 | 10.052809 | 10.280367 | 10.240721 | 1298 | 1050 | 1149 | 1176 |
| 10746 | NM_00137191 | MAP3K2 | mitogen-activated protein kinase kinase kinase 2 | protein_coding | NP_001358839.1;NP_001358840.1;NP_006600.3 | 6854 | 609487 | . | . | 3.100632 | 3.040782 | 3.033353 | 2.677426 | 7.215300 | 7.028903 | 6.806757 | 6.121615 | -1.093335 | 1044.396814 | 0.14071342 | -0.91487656 | 0.360256438 | 0.998865404 | 10.091167 | 9.962891 | 10.068852 | 10.050912 | 10.040983 | 9.945292 | 1271 | 1163 | 925 | 874 |
| 10748 | NR_028045 | KLRA1P | killer cell lectin like receptor A1, pseudogene | transcribed_ps | . | 6372 | 604274 | . | . | 0.520645 | 0.473363 | 0.636346 | 0.312371 | 1.211563 | 1.094200 | 1.427941 | 0.714200 | -1.077174 | 33.717435 | 0.564895011 | -0.18986057 | 0.849418393 | 0.998865404 | 5.125927 | 5.023086 | 5.081277 | 5.068564 | 5.107721 | 5.020185 | 42 | 36 | 38 | 20 |
| 10749 | NM_006612,XI | KIF1C | kinesin family member 1C | protein_coding | NP_006603.2;XP_005256481.1 | 6317 | 603060 | . | . | 16.068392 | 15.495400 | 13.995714 | 15.971668 | 37.391808 | 35.818312 | 31.405975 | 36.517320 | -1.155358 | 3500.819771 | 0.122486988 | -1.70091859 | 0.088958276 | 0.802216049 | 11.873988 | 11.665436 | 11.835138 | 11.823469 | 11.637128 | 11.778358 | 4352 | 4022 | 2579 | 3310 |
| 10752 | NM_00125338 | CHL1 | cell adhesion molecule L1 like | protein_coding | NP_001240316.1;NP_001240317.1;NP_006605.2;XP_006713001.1;XP_00 | 1939 | 607416 | . | . | 7.757266 | 6.824811 | 6.230861 | 6.719498 | 18.051473 | 15.775853 | 13.981870 | 15.363331 | -1.146273 | 1500.143965 | 0.129699687 | -1.51851149 | 0.128885505 | 0.87852179 | 10.645999 | 10.449056 | 10.639994 | 10.560533 | 10.472576 | 10.518025 | 1938 | 1642 | 1179 | 1350 |
| 10755 | NM_005716,N | GIPC1 | GIPC PDZ domain containing family member 1 | protein_coding | NP_005707.1;NP_974197.1;NP_974198.1;NP_974199.1;NP_974223.1;XP_ | 1226 | 605072 | . | . | 52.073255 | 50.553146 | 44.942518 | 57.622865 | 121.176610 | 116.855865 | 100.849695 | 131.747841 | -1.013463 | 3032.983487 | 0.133685382 | -0.14431797 | 0.88524939 | 0.998865404 | 11.576258 | 11.556720 | 11.585135 | 11.555275 | 11.457668 | 11.651769 | 3580 | 3235 | 2309 | 3159 |
| 10758 | NM_00116428 | TRAF3IP2 | TRAF3 interacting protein 2 | protein_coding | NP_001157753.1;NP_001157755.1;NP_671733.2;NP_679211.2 | 1343 | 607043 | . | . | 9.724588 | 10.030019 | 10.799835 | 11.995566 | 22.629516 | 23.184837 | 24.234515 | 27.426436 | 1.141108 | 1950.710614 | 0.122524424 | 1.554266955 | 0.120120752 | 0.859972379 | 10.831552 | 11.021752 | 10.864240 | 10.884739 | 10.939256 | 11.017529 | 2071 | 1992 | 1720 | 2044 |
| 10761 | NM_00131688 | PLAC1 | placenta enriched 1 | protein_coding | NP_001303816.1;NP_001303817.1;NP_001303818.1;NP_068568.1;XP_01 | 9044 | 300296 | . | . | 1.553666 | 1.158636 | 1.261343 | 1.002259 | 3.615445 | 2.678237 | 2.830415 | 2.291547 | -1.174889 | 43.443935 | 0.478415371 | -0.48602973 | 0.626946085 | 0.998865404 | 5.551670 | 5.321310 | 5.474924 | 5.434857 | 5.442545 | 5.414364 | 60 | 45 | 38 | 34 |
| 10762 | NM_007172,N | NUP50 | nucleoporin 50 | protein_coding | NP_009103.2;NP_705931.1;XP_005261369.1;XP_005261371.1;XP_00672 | 8065 | 604646 | . | . | 34.597483 | 35.252269 | 42.463437 | 39.517822 | 80.509768 | 81.487194 | 95.286724 | 90.352803 | 1.163637 | 6074.932432 | 0.108133397 | 2.0219568 | 0.043180815 | 0.639862814 | 12.455158 | 12.673867 | 12.496521 | 12.505551 | 12.667306 | 12.588707 | 6431 | 6092 | 5944 | 5844 |
| 10763 | NM_006617 | NES | nestin | protein_coding | NP_006608.1 | 7756 | 600915 | . | . | 1.299706 | 1.360207 | 1.554970 | 1.304533 | 3.024469 | 3.144180 | 3.489306 | 2.982659 | 1.060841 | 239.972742 | 0.225335483 | 0.378140368 | 0.705326322 | 0.998865404 | 7.863093 | 7.949080 | 7.880549 | 7.899245 | 7.963984 | 7.871926 | 263 | 256 | 235 | 210 |
| 10765 | NM_00131404 | KDM5B | lysine demethylase 5B | protein_coding | NP_001300971.1;NP_001334520.1;NP_006609.3;XP_011507392.1;XP_01 | 18039 | 605393 | . | . | 9.374376 | 8.805276 | 8.788019 | 9.155261 | 21.814557 | 20.353790 | 19.720058 | 20.932416 | 1.021522 | 2551.158099 | 0.113784371 | 0.269985943 | 0.787171068 | 0.998865404 | 11.301526 | 11.332184 | 11.294907 | 11.321319 | 11.321415 | 11.329387 | 2859 | 2768 | 2231 | 2429 |
| 10766 | NM_016272,XI | TOB2 | transducer of ERBB2, 2 | protein_coding | NP_057356.1;XP_005261372.1;XP_006724168.1;XP_016884028.1 | 11980 | 607396 | . | . | 7.200059 | 6.883870 | 5.516940 | 6.956020 | 16.754833 | 15.912373 | 12.379852 | 15.904113 | -1.109086 | 958.692516 | 0.142238109 | -1.05015005 | 0.293649132 | 0.998865404 | 9.977797 | 9.828175 | 9.959602 | 9.922596 | 9.826884 | 9.903403 | 1190 | 1061 | 748 | 898 |
| 10767 | NM_00114515 | HBS1L | HBS1 like translational GTPase | protein_coding | NP_001138630.1;NP_001138679.1;NP_001350615.1;NP_006611.1;XP_01 | 4834 | 612450 | . | . | 10.885549 | 9.073387 | 7.876402 | 9.164726 | 25.331119 | 20.973545 | 17.674420 | 20.954055 | -1.011596 | 1359.647192 | 0.129098219 | -0.12884718 | 0.897478573 | 0.998865404 | 10.417440 | 10.400544 | 10.401645 | 10.424834 | 10.375945 | 10.432100 | 1551 | 1498 | 1132 | 1314 |
| 10768 | NM_00124267 | AHCYL1 | adenosylhomocysteinase like 1 | protein_coding | NP_001229602.1;NP_001229603.1;NP_001229604.1;NP_001229605.1;NI | 344 | 607826 | . | . | 37.734618 | 38.208038 | 34.299912 | 36.523314 | 87.810012 | 88.319593 | 76.968007 | 83.506217 | -1.078336 | 4498.239747 | 0.107270561 | -1.01432686 | 0.31042681 | 0.998865404 | 12.188556 | 12.079681 | 12.161069 | 12.170435 | 12.083367 | 12.122011 | 5345 | 5065 | 3676 | 4153 |
| 10769 | NM_00125222 | PLK2 | polo like kinase 2 | protein_coding | NP_001239155.1;NP_006613.2 | 19699 | 607023 | . | . | 15.569380 | 16.194793 | 18.753428 | 16.380186 | 36.230588 | 37.434989 | 42.082146 | 37.451347 | 1.090279 | 1452.004002 | 0.133331505 | 0.935244373 | 0.349662375 | 0.998865404 | 10.439988 | 10.564966 | 10.456653 | 10.480218 | 10.590043 | 10.477218 | 1575 | 1522 | 1413 | 1317 |
| 10771 | NM_00120246 | ZMYND11 | zinc finger MYND-type containing 11 | protein_coding | NP_001189393.1;NP_001189394.1;NP_001189395.1;NP_001189396.1;NI | 16966 | 608668 | . | . | 7.311488 | 6.760007 | 6.526116 | 6.392480 | 17.014129 | 15.626061 | 14.644415 | 14.615645 | -1.118374 | 859.003688 | 0.143107454 | -1.12784634 | 0.259384821 | 0.995760118 | 9.824852 | 9.663690 | 9.807710 | 9.761982 | 9.713990 | 9.696613 | 1077 | 948 | 714 | 751 |
| 10772 | NM_00119100 | SRSF10 | serine and arginine rich splicing factor 10 | protein_coding | NP_001177934.1;NP_001177935.1;NP_001177936.1;NP_001177938.1;NI | 16713 | 605221 | . | . | 36.692079 | 34.550058 | 46.347890 | 29.714219 | 85.383974 | 79.864010 | 104.003316 | 67.938032 | 1.062802 | 6629.742288 | 0.326881133 | 0.268824065 | 0.788065074 | 0.998865404 | 12.650110 | 12.738025 | 12.691509 | 12.631544 | 12.887512 | 12.504000 | 7665 | 6689 | 7265 | 4977 |
| 10773 | NM_006626 | ZBTB6 | zinc finger and BTB domain containing 6 | protein_coding | NP_006617.1 | 16764 | 605976 | . | . | 1.528187 | 1.346952 | 1.338062 | 1.064336 | 3.556153 | 3.113540 | 3.002571 | 2.433478 | -1.215964 | 169.263069 | 0.26478632 | -1.06538993 | 0.286699513 | 0.998865404 | 7.536465 | 7.256191 | 7.470033 | 7.410559 | 7.405552 | 7.312055 | 228 | 187 | 149 | 126 |
| 10775 | NM_006627,N | POP4 | POP4 homolog, ribonuclease P/MRP subunit | protein_coding | NP_006618.1 | 30081 | 606114 | . | . | 11.823938 | 13.092759 | 12.217844 | 14.898334 | 27.514789 | 30.264499 | 27.416489 | 34.063270 | 1.073399 | 1034.508165 | 0.14921326 | 0.684835568 | 0.493447688 | 0.998865404 | 9.963079 | 10.064594 | 9.951929 | 10.020030 | 9.966647 | 10.105764 | 1096 | 1127 | 844 | 1098 |
| 10776 | NM_00130619 | ARPP19 | cAMP regulated phosphoprotein 19 | protein_coding | NP_001293120.1;NP_001293124.1;NP_001293125.1;NP_001317238.1;NI | 16967 | 605487 | . | . | 14.761549 | 11.926913 | 14.789990 | 12.119702 | 34.350729 | 27.569596 | 33.188307 | 27.710261 | -1.007592 | 2244.760210 | 0.147390832 | -0.07403211 | 0.940984837 | 0.998865404 | 11.137561 | 11.127112 | 11.216984 | 11.040354 | 11.206488 | 11.039793 | 2878 | 2165 | 2147 | 1879 |
| 10777 | NM_00102506 | ARPP21 | cAMP regulated phosphoprotein 21 | protein_coding | NP_001020239.1;NP_001020240.1;NP_001254545.1;NP_001254546.1;NI | 16968 | 605488 | . | . | 0.005856 | 0.019764 | 0.012585 | 0.031267 | 0.013627 | 0.045685 | 0.028241 | 0.071489 | 1.607411 | 8.875287 | 1.078375534 | 0.634972218 | 0.525446534 | NA | 2.772726 | 3.448451 | 3.110040 | 3.141582 | 3.126931 | 3.161987 | 4 | 11 | 6 | 14 |
| 10778 | NR_024565,NF | ZNF271P | zinc finger protein 271, pseudogene | transcribed_ps | . | 13065 | 604754 | . | . | 3.487007 | 3.446201 | 3.121212 | 3.922789 | 8.114410 | 7.966047 | 7.003909 | 8.968991 | 1.000852 | 557.636539 | 0.173276254 | 0.007089846 | 0.994343169 | 0.999862634 | 9.122945 | 9.123418 | 9.127849 | 9.115584 | 9.048910 | 9.190467 | 648 | 596 | 432 | 580 |
| 10780 | NM_00114482 | ZNF234 | zinc finger protein 234 | protein_coding | NP_001138296.1;NP_006621.1;XP_006723037.1;XP_016881638.1;XP_01 | 13027 | 604750 | . | . | 2.558547 | 2.902868 | 3.552612 | 2.183096 | 5.953845 | 6.710109 | 7.971960 | 4.991388 | 1.073211 | 366.932179 | 0.225599194 | 0.45183517 | 0.651387731 | 0.998865404 | 8.466768 | 8.570121 | 8.487526 | 8.494369 | 8.655419 | 8.391424 | 405 | 384 | 405 | 276 |
| 10781 | NM_00127131 | ZNF266 | zinc finger protein 266 | protein_coding | NP_001258243.1;NP_001357303.1;NP_001357304.1;NP_001357305.1;NI | 13059 | 604751 | . | . | 7.267628 | 7.938840 | 8.732632 | 8.302147 | 16.912072 | 18.350985 | 19.595770 | 18.981871 | 1.102451 | 974.213270 | 0.139743754 | 1.006951272 | 0.313958191 | 0.998865404 | 9.855992 | 9.996763 | 9.867940 | 9.912856 | 9.985916 | 9.937081 | 1034 | 1031 | 918 | 926 |
| 10782 | NM_00127873 | ZNF274 | zinc finger protein 274 | protein_coding | NP_001265663.1;NP_057408.2;NP_057409.1;NP_598009.1;XP_01152462 | 13068 | 605467 | . | . | 4.034074 | 3.655563 | 5.489358 | 5.054955 | 9.387458 | 8.450000 | 12.317962 | 11.557554 | 1.379984 | 402.984393 | 0.180922178 | 2.56823771 | 0.010221703 | 0.325163413 | 8.403398 | 8.868446 | 8.562746 | 8.528503 | 8.748526 | 8.725719 | 402 | 354 | 414 | 430 |
| 10783 | NM_00114500 | NEK6 | NIMA related kinase 6 | protein_coding | NP_001138473.1;NP_001159639.1;NP_001159640.1;NP_001159641.1;NI | 7749 | 604884 | . | . | 11.589759 | 12.690208 | 12.298761 | 13.215857 | 26.969842 | 29.333983 | 27.598064 | 30.216487 | 1.024691 | 1248.309238 | 0.130203749 | 0.270259531 | 0.786960597 | 0.998865404 | 10.268160 | 10.303147 | 10.262390 | 10.290098 | 10.278284 | 10.310880 | 1394 | 1355 | 1075 | 1210 |
| 10785 | NM_00126047 | WDR4 | WD repeat domain 4 | protein_coding | NP_001247403.1;NP_001247404.1;NP_001247405.1;NP_001247406.1;NI | 12756 | 605924 | . | . | 8.152002 | 11.193351 | 9.530481 | 11.042261 | 18.970048 | 25.873934 | 21.386127 | 25.246819 | 1.004323 | 543.980524 | 0.16959397 | 0.036697182 | 0.970726456 | 0.998865404 | 9.084632 | 9.090185 | 9.033601 | 9.134200 | 9.068660 | 9.106561 | 577 | 631 | 460 | 526 |
| 10787 | NM_013436,N | NCKAP1 | NCK associated protein 1 | protein_coding | NP_038464.1;NP_995314.1;XP_006712263.1;XP_006712264.1 | 7666 | 604891 | . | . | 34.742732 | 32.532264 | 16.008636 | 17.180345 | 80.847765 | 75.199782 | 35.922916 | 39.280815 | -1.196213 | 6470.820505 | 0.129868179 | -1.9902815 | 0.046559935 | 0.654401867 | 12.783107 | 12.524814 | 12.770024 | 12.685821 | 12.669467 | 12.476663 | 8514 | 7233 | 5718 | 4885 |
| 10788 | NM_00128546 | IQGAP2 | IQ motif containing GTPase activating protein 2 | protein_coding | NP_001272389.1;NP_001272390.2;NP_001272391.2;NP_006624.3;XP_0C | 6111 | 605401 | . | . | 0.184163 | 0.297857 | 0.085368 | 0.117748 | 0.428554 | 0.688510 | 0.191562 | 0.269217 | -2.309762 | 31.031668 | 0.587297198 | -2.05644506 | 0.039739636 | 0.621911794 | 5.437855 | 4.226238 | 4.962607 | 5.036446 | 4.890960 | 4.911585 | 39 | 57 | 14 | 20 |
| 10789 | gene-NUDCP2 | NUDCP2 | nuclear distribution C pseudogene 2 | pseudogene | . | 8047 | . | . | . | 0.342046 | 0.379066 | 0.484761 | 0.475334 | 0.795955 | 0.876229 | 1.087789 | 1.086794 | 1.337582 | 12.589645 | 0.857717303 | 0.489237714 | 0.624673412 | NA | 3.429096 | 3.848815 | 3.637499 | 3.640821 | 3.657414 | 3.652965 | 12 | 12 | 13 | 13 |
| 10790 | gene-NUDCP1 | NUDCP1 | nuclear distribution C pseudogene 1 | pseudogene | . | 8046 | . | . | . | 0.021791 | 0.046908 | 0.089158 | 0.042045 | 0.050709 | 0.108430 | 0.200068 | 0.096131 | 1.488926 | 2.812587 | 1.788565834 | 0.321079512 | 0.748150136 | NA | 1.175403 | 1.751299 | 1.472828 | 1.477882 | 1.485999 | 1.474918 | 2 | 3 | 4 | 2 |
| 10791 | NM_006634 | VAMP5 | vesicle associated membrane protein 5 | protein_coding | NP_006625.1 | 12646 | 607029 | . | . | 1.953110 | 3.629230 | 1.432559 | 4.054280 | 4.544967 | 8.389129 | 3.214620 | 9.269629 | -1.020679 | 57.245619 | 0.51581273 | -0.05724759 | 0.954347968 | 0.998865404 | 5.856526 | 5.821447 | 5.758438 | 5.884917 | 5.720735 | 5.910060 | 47 | 81 | 26 | 78 |
| 10793 | NM_021148,N | ZNF273 | zinc finger protein 273 | protein_coding | NP_066971.2;XP_024302403.1;XP_024302404.1;XP_024302405.1;XP_02· | 13067 | 604756 | . | . | 0.556311 | 0.378640 | 0.405448 | 0.285290 | 1.294560 | 0.875241 | 0.909812 | 0.652280 | -1.296719 | 62.179205 | 0.429689953 | -0.87240922 | 0.382985172 | 0.998865404 | 6.131874 | 5.761088 | 6.010082 | 5.953542 | 5.976873 | 5.871639 | 90 | 67 | 60 | 37 |
| 10794 | NM_00133062 | ZNF460 | zinc finger protein 460 | protein_coding | NP_001317551.1;NP_006626.3 | 21628 | 604755 | . | . | 0.473971 | 0.599575 | 0.594686 | 0.429240 | 1.102949 | 1.385943 | 1.334460 | 0.981406 | -1.062622 | 57.240597 | 0.422092647 | -0.20760376 | 0.835538367 | 0.998865404 | 5.882199 | 5.794399 | 5.814799 | 5.871157 | 5.846044 | 5.815581 | 59 | 72 | 52 | 48 |
| 10795 | NM_00116588 | ZNF268 | zinc finger protein 268 | protein_coding | NP_001159353.1;NP_001159354.1;NP_001159355.1;NP_001159356.2;NI | 13061 | 604753 | . | . | 0.761563 | 0.745875 | 0.885894 | 0.759888 | 1.772191 | 1.724123 | 1.987921 | 1.737391 | 1.078076 | 331.605123 | 0.198814682 | 0.545528816 | 0.585389868 | 0.998865404 | 8.317628 | 8.426946 | 8.358493 | 8.341663 | 8.437459 | 8.344760 | 373 | 339 | 324 | 296 |
| 10797 | NM_006636,N | MTHFD2 | methylenetetrahydrofolate dehydrogenase (NADP+ dependent) 2, | protein_coding | NP_006627.2;XP_006711987.1 | 7434 | 604887 | . | . | 32.021075 | 27.372286 | 21.910229 | 26.615305 | 74.514357 | 63.272266 | 49.165916 | 60.852731 | -1.209185 | 3158.869610 | 0.129385685 | -2.11796837 | 0.034177747 | 0.578889643 | 11.755806 | 11.481567 | 11.727690 | 11.665002 | 11.456771 | 11.618787 | 4131 | 3592 | 2236 | 2951 |
| 10799 | NM_00128613 | RPP40 | ribonuclease P/MRP subunit p40 | protein_coding | NP_001273061.1;NP_001273062.1;NP_006629.2;XP_011512555.1;XP_02 | 20992 | 606117 | . | . | 3.427808 | 3.399358 | 3.184103 | 2.455712 | 7.976652 | 7.857773 | 7.145035 | 5.614693 | -1.232654 | 145.867592 | 0.281355439 | -1.07255203 | 0.283472159 | 0.998865404 | 7.330799 | 7.030649 | 7.222610 | 7.228385 | 7.195182 | 7.093730 | 184 | 175 | 130 | 105 |
| 10801 | NM_00111349 | SEPTIN9 | septin 9 | protein_coding | NP_001106963.1;NP_001106964.1;NP_001106965.1;NP_001106966.1;NI | 7323 | 604061 | . | . | 110.887034 | 120.200759 | 119.825717 | 140.586326 | 258.038686 | 277.849436 | 268.885432 | 321.433935 | 1.144243 | 14280.494991 | 0.117814422 | 1.650002223 | 0.098942481 | 0.813553488 | 13.701328 | 13.895650 | 13.703302 | 13.775640 | 13.776789 | 13.928823 | 14684 | 14986 | 12093 | 15539 |
| 10802 | NM_00125223 | SEC24A | SEC24 homolog A, COPII coat complex component | protein_coding | NP_001239160.1;NP_068817.1;XP_006714586.1;XP_016864450.1;XP_01 | 10703 | 607183 | . | . | 9.314969 | 9.162635 | 8.993746 | 8.190504 | 21.676316 | 21.179840 | 20.181705 | 18.726611 | -1.099708 | 1662.116910 | 0.124705761 | -1.09955565 | 0.271525775 | 0.998865404 | 10.765619 | 10.628748 | 10.758559 | 10.710032 | 10.683263 | 10.637183 | 2066 | 1821 | 1418 | 1440 |
| 10806 | NM_00135024 | SDCCAG8 | SHH signaling and ciliogenesis regulator SDCCAG8 | protein_coding | NP_001337175.1;NP_001337176.1;NP_001337177.1;NP_001337178.1;NI | 10671 | 613524 | . | . | 6.376378 | 6.445648 | 7.093122 | 6.562589 | 14.838095 | 14.899402 | 15.916758 | 15.004587 | 1.161324 | 734.335479 | 0.150181238 | 1.436733616 | 0.150793699 | 0.913627824 | 9.408586 | 9.623973 | 9.452085 | 9.477623 | 9.540117 | 9.596237 | 768 | 747 | 659 | 769 |
| 10807 | NM_00103970 | ENTR1 | endosome associated trafficking regulator 1 | protein_coding | NP_001034796.1;NP_001034797.1;NP_006634.3;XP_005266107.1;XP_00 | 10667 | 618289 | . | . | 16.285783 | 17.399661 | 18.538316 | 17.454990 | 37.897683 | 40.220097 | 41.599443 | 39.908761 | 1.049430 | 1256.259934 | 0.132436689 | 0.525575183 | 0.599183403 | 0.998865404 | 10.259598 | 10.329397 | 10.264205 | 10.287319 | 10.347134 | 10.276430 | 1390 | 1343 | 1171 | 1150 |
| 10808 | NM_00128650 | HSPH1 | heat shock protein family H (Hsp110) member 1 | protein_coding | NP_001273432.1;NP_001273433.1;NP_001273434.1;NP_001336633.1;NI | 16969 | 610703 | . | . | 34.832617 | 33.943370 | 33.489445 | 30.283213 | 81.056934 | 78.461615 | 75.149340 | 69.238967 | -1.113346 | 5221.061943 | 0.113218544 | -1.36817195 | 0.17125826 | 0.947127937 | 12.425434 | 12.270673 | 12.404016 | 12.381816 | 12.354427 | 12.248886 | 6417 | 5858 | 4570 | 4338 |
| 10809 | NM_006645 | STARD10 | StAR related lipid transfer domain containing 10 | protein_coding | NP_006636.2 | 10666 | 617382 | . | . | 9.113530 | 9.393121 | 9.388916 | 10.704042 | 21.207560 | 21.712618 | 21.068457 | 24.473520 | 1.069971 | 597.738676 | 0.161428108 | 0.604428568 | 0.545558747 | 0.998865404 | 9.174032 | 9.271079 | 9.192614 | 9.206884 | 9.203620 | 9.283825 | 658 | 630 | 505 | 614 |
| 10810 | NM_00129196 | WASF3 | WASP family member 3 | protein_coding | NP_001278894.1;NP_001278895.1;NP_006637.2;XP_011533191.1;XP_02 | 12734 | 605068 | . | . | 4.131492 | 4.650216 | 4.401174 | 4.287621 | 9.614153 | 10.749180 | 9.876107 | 9.803137 | -1.023898 | 659.552194 | 0.156489064 | -0.21772503 | 0.827643358 | 0.998865404 | 9.382361 | 9.348122 | 9.335254 | 9.409194 | 9.367778 | 9.345788 | 727 | 759 | 576 | 601 |
| 10811 | NM_00125606 | NOXA1 | NADPH oxidase activator 1 | protein_coding | NP_001242996.1;NP_001242997.1;NP_006638.1;XP_011516460.1;XP_01 | 10668 | 611255 | . | . | 0.412948 | 0.224910 | 0.051339 | 0.280209 | 0.960947 | 0.519889 | 0.115203 | 0.640665 | -1.752089 | 15.686493 | 0.856510249 | -0.94461918 | 0.344853331 | NA | 4.322387 | 3.506731 | 4.017073 | 3.971994 | 3.928772 | 3.986068 | 29 | 16 | 4 | 17 |
| 10813 | NM_00116622 | UTP14A | UTP14A small subunit processome component | protein_coding | NP_001159693.1;NP_006640.2;XP_016884726.1;XP_016884727.1 | 10665 | 300508 | . | . | 8.900973 | 9.670949 | 9.568937 | 9.934648 | 20.712932 | 22.354833 | 21.472417 | 22.714393 | 1.033864 | 747.587452 | 0.148947579 | 0.322570355 | 0.747020648 | 0.998865404 | 9.522025 | 9.569777 | 9.508578 | 9.558948 | 9.547784 | 9.566467 | 816 | 822 | 652 | 722 |
| 10817 | NM_006653,XI | FRS3 | fibroblast growth factor receptor substrate 3 | protein_coding | NP_006644.1;XP_011512556.1;XP_011512557.1;XP_016865682.1 | 16970 | 607744 | . | . | 1.388212 | 0.836748 | 0.811176 | 1.109531 | 3.230425 | 1.934180 | 1.820255 | 2.536811 | 1.105878 | 107.476936 | 0.319279999 | 0.454748203 | 0.649290398 | 0.998865404 | 6.674144 | 6.818036 | 6.737827 | 6.719763 | 6.713069 | 6.806719 | 121 | 107 | 84 | 121 |
| 10818 | NM_00104255 | FRS2 | fibroblast growth factor receptor substrate 2 | protein_coding | NP_001036020.1;NP_001265280.1;NP_001265282.1;NP_001265283.1;NI | 16971 | 607743 | . | . | 1.426972 | 1.340208 | 1.013370 | 1.065136 | 3.320623 | 3.097951 | 2.273974 | 2.435307 | -1.353916 | 259.675803 | 0.213892754 | -2.04372694 | 0.04098053 | 0.628504734 | 8.222653 | 7.785492 | 8.108768 | 8.073270 | 7.929903 | 7.947547 | 355 | 312 | 188 | 211 |
| 10825 | NM_00136786 | NEU3 | neuraminidase 3 | protein_coding | NP_001354789.1;NP_001354790.1;NP_001354791.1;NP_001354792.1;NI | 7760 | 604617 | . | . | 3.921638 | 4.388766 | 4.105779 | 4.007817 | 9.125811 | 10.144831 | 9.213251 | 9.163400 | 1.004895 | 735.700067 | 0.15322831 | 0.045979559 | 0.963326542 | 0.998865404 | 9.519473 | 9.526467 | 9.480110 | 9.559515 | 9.547044 | 9.500332 | 797 | 837 | 663 | 668 |
| 10826 | NM_032385,XI | FAXDC2 | fatty acid hydroxylase domain containing 2 | protein_coding | NP_115761.2;XP_006714816.1;XP_016864454.1 | 1334 | . | . | . | 6.115352 | 6.202793 | 7.419934 | 7.008740 | 14.230674 | 14.338033 | 16.650118 | 16.024651 | 1.220875 | 702.738993 | 0.151883266 | 1.895640328 | 0.058007621 | 0.713843573 | 9.305746 | 9.593608 | 9.358302 | 9.405781 | 9.535072 | 9.503561 | 703 | 707 | 687 | 708 |
| 10827 | NM_00131799 | FAM114A2 | family with sequence similarity 114 member A2 | protein_coding | NP_001304922.1;NP_001304923.1;NP_001304924.1;NP_061161.2;XP_0C | 1333 | . | . | . | 5.257348 | 5.441485 | 5.089240 | 5.543292 | 12.234066 | 12.578240 | 11.420110 | 12.674078 | -1.064195 | 671.416726 | 0.153652522 | -0.58419519 | 0.559088997 | 0.998865404 | 9.435182 | 9.345556 | 9.437505 | 9.385609 | 9.364044 | 9.375112 | 827 | 719 | 561 | 615 |
| 10838 | NM_00108048 | ZNF275 | zinc finger protein 275 | protein_coding | NP_001073954.3;NP_001354686.1 | 13069 | . | . | . | 1.178342 | 1.355444 | 1.240750 | 1.118313 | 2.742050 | 3.133170 | 2.784207 | 2.556890 | -1.087329 | 245.386432 | 0.222643073 | -0.5425186 | 0.587461296 | 0.998865404 | 7.997982 | 7.877319 | 7.924336 | 7.991518 | 7.943816 | 7.889668 | 275 | 294 | 216 | 208 |
| 10844 | NM_00125661 | TUBGCP2 | tubulin gamma complex associated protein 2 | protein_coding | NP_001243546.1;NP_001243547.1;NP_006650.1 | 18599 | 617817 | . | . | 13.726444 | 15.318783 | 14.587758 | 15.238859 | 31.941998 | 35.410054 | 32.734507 | 34.841842 | 1.012925 | 1806.386171 | 0.124701432 | 0.148573598 | 0.881890107 | 0.998865404 | 10.809715 | 10.828009 | 10.772264 | 10.853056 | 10.810573 | 10.836227 | 1961 | 2036 | 1555 | 1732 |
| 10845 | NM_006660,N | CLPX | caseinolytic mitochondrial matrix peptidase chaperone subunit X | protein_coding | NP_006651.2;XP_011519466.1 | 2088 | 615611 | . | . | 6.492209 | 6.903298 | 7.002495 | 6.936333 | 15.107639 | 15.957281 | 15.713397 | 15.859101 | 1.016336 | 974.189016 | 0.137524509 | 0.169982994 | 0.865023511 | 0.998865404 | 9.916410 | 9.939613 | 9.916934 | 9.927131 | 9.918384 | 9.949016 | 1105 | 1050 | 836 | 940 |
| 10846 | NM_00113069 | PDE10A | phosphodiesterase 10A | protein_coding | NP_001124162.1;NP_006652.1;XP_006715384.1;XP_011533689.2;XP_01 | 8772 | 610652 | . | . | 0.195220 | 0.139645 | 0.118679 | 0.091573 | 0.454284 | 0.322795 | 0.266313 | 0.209372 | -1.543872 | 43.806385 | 0.483687952 | -1.29536649 | 0.195193832 | 0.96621494 | 5.731451 | 5.107809 | 5.513787 | 5.462399 | 5.435209 | 5.399083 | 69 | 50 | 34 | 28 |
| 10847 | NM_006662 | SRCAP | Snf2 related CREBBP activator protein | protein_coding | NP_006653.2 | 16974 | 611421 | . | . | 6.654795 | 7.716440 | 8.833774 | 8.220605 | 15.485982 | 17.836897 | 19.822733 | 18.795437 | 1.170768 | 2868.152561 | 0.123156902 | 1.846875349 | 0.064765208 | 0.731823074 | 11.367723 | 11.595136 | 11.359449 | 11.471546 | 11.579082 | 11.511012 | 2831 | 3050 | 2801 | 2781 |
| 10848 | NM_00114250 | PPP1R13L | protein phosphatase 1 regulatory subunit 13 like | protein_coding | NP_001135974.1;NP_006654.2;XP_016881666.1;XP_016881667.1;XP_01 | 18838 | 607463 | . | . | 0.909483 | 0.893975 | 0.622936 | 0.845563 | 2.116403 | 2.066463 | 1.397850 | 1.933279 | -1.231632 | 82.351521 | 0.357910492 | -0.83979376 | 0.401024034 | 0.998865404 | 6.506929 | 6.204721 | 6.398454 | 6.385405 | 6.301617 | 6.369965 | 107 | 96 | 55 | 79 |
| 10849 | NM_00129759 | POLR1G | RNA polymerase I subunit G | protein_coding | NP_001284519.1;NP_036231.1 | 24219 | 107325 | . | . | 2.715568 | 2.455015 | 2.961078 | 2.485361 | 6.319238 | 5.674878 | 6.644572 | 5.682482 | 1.039059 | 234.197373 | 0.229065735 | 0.241319871 | 0.809307211 | 0.998865404 | 7.843016 | 7.899591 | 7.884959 | 7.834928 | 7.923056 | 7.832852 | 278 | 235 | 227 | 203 |
| 10850 | NM_006664 | CCL27 | C-C motif chemokine ligand 27 | protein_coding | NP_006655.1 | 10626 | 604833 | . | . | 0.793267 | 0.400129 | 0.639286 | 0.526645 | 1.845964 | 0.924916 | 1.434540 | 1.204112 | -1.014208 | 8.379170 | 1.063978213 | -0.01912928 | 0.984737976 | NA | 3.074695 | 3.058876 | 3.079825 | 3.053382 | 3.069920 | 3.062340 | 13 | 6 | 8 | 7 |
| 10855 | NM_00109854 | HPSE | heparanase | protein_coding | NP_001092010.1;NP_001159970.1;NP_001186759.1;NP_006656.2 | 5164 | 604724 | . | . | 3.395580 | 3.295394 | 2.852899 | 2.843416 | 7.901653 | 7.617449 | 6.401823 | 6.501132 | -1.190996 | 442.635142 | 0.17439317 | -1.44597902 | 0.148183086 | 0.909660767 | 8.910467 | 8.658493 | 8.853636 | 8.828321 | 8.738875 | 8.729803 | 565 | 509 | 354 | 376 |
| 10856 | NM_00132119 | RUVBL2 | RuvB like AAA ATPase 2 | protein_coding | NP_001308119.1;NP_001308120.1;NP_006657.1;XP_011524632.1 | 10475 | 604788 | . | . | 77.499146 | 77.504218 | 61.940631 | 67.940245 | 180.343715 | 179.154477 | 138.992987 | 155.337298 | -1.139064 | 3587.881260 | 0.119303831 | -1.57454373 | 0.115361801 | 0.856096201 | 11.899875 | 11.711835 | 11.863674 | 11.855741 | 11.688346 | 11.812222 | 4421 | 4104 | 2695 | 3384 |
| 10857 | NM_00128262 | PGRMC1 | progesterone receptor membrane component 1 | protein_coding | NP_001269550.1;NP_006658.1 | 16090 | 300435 | . | . | 45.455368 | 43.569171 | 47.546734 | 41.794621 | 105.776514 | 100.712091 | 106.693490 | 95.558431 | -1.012385 | 2608.290787 | 0.12212953 | -0.1453991 | 0.884395773 | 0.998865404 | 11.357604 | 11.340121 | 11.373779 | 11.332114 | 11.396233 | 11.287471 | 3099 | 2759 | 2406 | 2267 |
| 10865 | NM_00131908 | ARID5A | AT-rich interaction domain 5A | protein_coding | NP_001306014.1;NP_001306016.1;NP_001306021.1;NP_001306022.1;NI | 17361 | 611583 | . | . | 2.290098 | 2.572872 | 3.084601 | 3.227207 | 5.329152 | 5.947310 | 6.921752 | 7.378626 | 1.268191 | 193.441020 | 0.240341342 | 1.426189816 | 0.153813558 | 0.920281122 | 7.414483 | 7.756763 | 7.517513 | 7.554098 | 7.635857 | 7.646710 | 188 | 192 | 186 | 205 |
| 10867 | NM_00116832 | TSPAN9 | tetraspanin 9 | protein_coding | NP_001161792.1;NP_006666.1;XP_011519214.1;XP_011519215.1 | 21640 | 613137 | . | . | 5.421160 | 5.601657 | 5.541385 | 5.844811 | 12.615264 | 12.948484 | 12.434706 | 13.363464 | 1.017498 | 753.237021 | 0.147524054 | 0.169636578 | 0.865295955 | 0.998865404 | 9.544511 | 9.569302 | 9.541876 | 9.559605 | 9.547282 | 9.578096 | 849 | 816 | 646 | 728 |
| 10868 | NM_00100856 | USP20 | ubiquitin specific peptidase 20 | protein_coding | NP_001008563.2;NP_001103773.2;NP_006667.3;XP_005251722.1;XP_01 | 12619 | 615143 | . | . | 4.360041 | 4.776864 | 5.160959 | 5.471159 | 10.145993 | 11.041939 | 11.581043 | 12.509154 | 1.145212 | 675.951368 | 0.153973167 | 1.270449286 | 0.203924637 | 0.969101675 | 9.299862 | 9.495090 | 9.323559 | 9.378748 | 9.429849 | 9.457690 | 696 | 708 | 615 | 690 |
| 10869 | NM_00119916 | USP19 | ubiquitin specific peptidase 19 | protein_coding | NP_001186089.1;NP_001186090.1;NP_001186091.1;NP_001338027.1;NI | 12617 | 614471 | . | . | 10.432646 | 11.706868 | 12.142722 | 11.746614 | 24.277197 | 27.060945 | 27.247917 | 26.857239 | 1.084273 | 1708.638511 | 0.124833355 | 0.935069148 | 0.349752665 | 0.998865404 | 10.679118 | 10.795787 | 10.671178 | 10.738823 | 10.784666 | 10.752941 | 1806 | 1846 | 1577 | 1633 |
| 10870 | NM_00100746 | HCST | hematopoietic cell signal transducer | protein_coding | NP_001007470.1;NP_055081.1;XP_016881682.1 | 16977 | 604089 | . | . | 0.713524 | 0.707169 | 0.532396 | 0.564822 | 1.660398 | 1.634652 | 1.194682 | 1.291398 | -1.276886 | 13.185301 | 0.842306213 | -0.41864788 | 0.67547349 | NA | 3.887108 | 3.532926 | 3.732367 | 3.732548 | 3.713240 | 3.723599 | 17 | 16 | 9 | 12 |
| 10873 | NM_00101481 | ME3 | malic enzyme 3 | protein_coding | NP_001014811.1;NP_001155058.1;NP_001338863.1;NP_006671.2;XP_01 | 6985 | 604626 | . | . | 0.256815 | 0.272269 | 0.182409 | 0.256663 | 0.597621 | 0.629360 | 0.409322 | 0.586831 | -1.229050 | 37.411165 | 0.515848803 | -0.57680406 | 0.564071825 | 0.998865404 | 5.367696 | 5.067511 | 5.237349 | 5.247689 | 5.189525 | 5.235078 | 46 | 46 | 24 | 37 |
| 10882 | NM_006688 | C1QL1 | complement C1q like 1 | protein_coding | NP_006679.1 | 24182 | 611586 | . | . | 2.197210 | 2.633614 | 3.035712 | 2.829408 | 5.112998 | 6.087717 | 6.812049 | 6.469106 | 1.199612 | 127.554343 | 0.291081702 | 0.902041623 | 0.367034758 | 0.998865404 | 6.857901 | 7.120135 | 6.927033 | 6.986220 | 7.036578 | 7.006277 | 122 | 136 | 126 | 125 |
| 10884 | NM_016640 | MRPS30 | mitochondrial ribosomal protein S30 | protein_coding | NP_057724.2 | 8769 | 611991 | . | . | 35.532600 | 35.242958 | 29.407316 | 28.058352 | 82.685822 | 81.465675 | 65.989174 | 64.152084 | -1.248619 | 1647.905612 | 0.123368358 | -2.59655844 | 0.009416289 | 0.309459602 | 10.837630 | 10.517481 | 10.773389 | 10.759721 | 10.614945 | 10.571579 | 2125 | 1958 | 1311 | 1335 |
| 10885 | NM_006784 | WDR3 | WD repeat domain 3 | protein_coding | NP_006775.1 | 12755 | 604737 | . | . | 13.927275 | 14.349171 | 12.772209 | 13.927895 | 32.409344 | 33.168747 | 28.660467 | 31.844477 | -1.074020 | 4284.180529 | 0.109811589 | -0.93815491 | 0.348164806 | 0.998865404 | 12.115449 | 12.012316 | 12.084710 | 12.102565 | 12.002164 | 12.065276 | 5055 | 4839 | 3455 | 4021 |
| 10890 | NM_016131,XI | RAB10 | RAB10, member RAS oncogene family | protein_coding | NP_057215.3;XP_024308333.1 | 9759 | 612672 | . | . | 26.986178 | 25.117831 | 26.023739 | 24.494544 | 62.797945 | 58.060993 | 58.396519 | 56.003861 | -1.044436 | 2845.497176 | 0.117061365 | -0.53581718 | 0.592084906 | 0.998865404 | 11.505386 | 11.442867 | 11.524225 | 11.457824 | 11.484343 | 11.427299 | 3483 | 3010 | 2506 | 2518 |
| 10891 | NM_00133075 | PPARGC1A | PPARG coactivator 1 alpha | protein_coding | NP_001317680.1;NP_001317681.1;NP_001317682.1;NP_001341754.1;NI | 9237 | 604517 | . | . | 1.818041 | 1.640870 | 1.363438 | 1.354278 | 4.230654 | 3.792942 | 3.059512 | 3.096395 | -1.295221 | 304.219602 | 0.202088637 | -1.84670351 | 0.064790123 | 0.731823074 | 8.423363 | 8.050558 | 8.342463 | 8.284632 | 8.178251 | 8.171349 | 415 | 352 | 232 | 247 |
| 10892 | NM_006785,N | MALT1 | MALT1 paracaspase | protein_coding | NP_006776.1;NP_776216.1;XP_011524096.1 | 6819 | 604860 | . | . | 2.386009 | 2.493537 | 2.917095 | 2.251661 | 5.552342 | 5.763923 | 6.545877 | 5.148155 | 1.049139 | 709.939446 | 0.17176207 | 0.402913431 | 0.68701191 | 0.998865404 | 9.436166 | 9.506091 | 9.435304 | 9.467587 | 9.578395 | 9.381790 | 779 | 765 | 722 | 585 |
| 10893 | NM_006690,XI | MMP24 | matrix metallopeptidase 24 | protein_coding | NP_006681.1;XP_011526802.1;XP_016883086.1;XP_016883087.1 | 7172 | 604871 | . | . | 0.085638 | 0.104186 | 0.080921 | 0.118827 | 0.199284 | 0.240830 | 0.181584 | 0.271683 | 1.061072 | 13.888131 | 0.825195606 | 0.103638944 | 0.917455881 | NA | 3.754145 | 3.836248 | 3.786709 | 3.798742 | 3.784454 | 3.808135 | 14 | 16 | 10 | 16 |
| 10897 | NM_00130086 | YIF1A | Yip1 interacting factor homolog A, membrane trafficking protein | protein_coding | NP_001287790.1;NP_065203.2 | 16688 | 611484 | . | . | 34.261520 | 36.331686 | 32.764604 | 33.586490 | 79.727962 | 83.982319 | 73.522819 | 76.791520 | -1.062550 | 1134.680414 | 0.132775398 | -0.65923664 | 0.509743823 | 0.998865404 | 10.191252 | 10.103556 | 10.153815 | 10.186609 | 10.116761 | 10.133204 | 1317 | 1289 | 946 | 1043 |
| 10898 | NM_00108155 | CPSF4 | cleavage and polyadenylation specific factor 4 | protein_coding | NP_001075028.1;NP_001305089.1;NP_001305090.1;NP_001305091.1;NI | 2327 | 603052 | . | . | 11.374854 | 10.982180 | 12.017703 | 13.471980 | 26.469755 | 25.385801 | 26.967373 | 30.802077 | 1.108935 | 757.230091 | 0.148315255 | 1.005794573 | 0.314514399 | 0.998865404 | 9.488212 | 9.637123 | 9.527481 | 9.525897 | 9.571415 | 9.625429 | 828 | 774 | 666 | 775 |
| 10899 | NM_006694 | JTB | jumping translocation breakpoint | protein_coding | NP_006685.1 | 6201 | 604671 | . | . | 36.079906 | 34.626957 | 33.910225 | 42.360821 | 83.959427 | 80.041763 | 76.093567 | 96.852982 | 1.063301 | 1456.934911 | 0.141602486 | 0.625338761 | 0.531748745 | 0.998865404 | 10.463951 | 10.552143 | 10.502068 | 10.463773 | 10.445242 | 10.608870 | 1667 | 1486 | 1168 | 1556 |
| 10901 | NM_00128298 | DHRS4 | dehydrogenase/reductase 4 | protein_coding | NP_001269916.1;NP_001269917.1;NP_001269918.1;NP_001269919.1;NI | 16985 | 611596 | . | . | 8.654913 | 7.231773 | 6.268312 | 7.074490 | 20.140340 | 16.716566 | 14.065912 | 16.174979 | -1.231150 | 261.577146 | 0.224384098 | -1.33702481 | 0.181214538 | 0.952877815 | 8.173419 | 7.873173 | 8.132063 | 8.025818 | 7.934858 | 8.013061 | 365 | 281 | 187 | 238 |
| 10902 | NM_00116432 | BRD8 | bromodomain containing 8 | protein_coding | NP_001157798.1;NP_001287890.1;NP_001287891.1;NP_001287895.1;NI | 19874 | 602848 | . | . | 23.382970 | 24.698991 | 29.741531 | 26.432767 | 54.413130 | 57.092825 | 66.739135 | 60.435379 | 1.148842 | 2563.982382 | 0.12247239 | 1.634493305 | 0.102155311 | 0.821228746 | 11.220538 | 11.420856 | 11.242616 | 11.285598 | 11.430308 | 11.320275 | 2674 | 2644 | 2544 | 2398 |
| 10903 | NM_00114586 | MTMR11 | myotubularin related protein 11 | protein_coding | NP_001139334.1;NP_870988.2;XP_006711200.1;XP_011507401.1;XP_02 | 24307 | . | . | . | 1.196783 | 1.349132 | 1.197170 | 1.169098 | 2.784961 | 3.118578 | 2.686413 | 2.673002 | -1.024390 | 136.290530 | 0.28119242 | -0.12363641 | 0.901603163 | 0.998865404 | 7.107907 | 7.072965 | 7.068186 | 7.119662 | 7.097363 | 7.073232 | 148 | 159 | 121 | 122 |
| 10904 | NM_00116782 | BLCAP | BLCAP apoptosis inducing factor | protein_coding | NP_001161292.1;NP_001161293.1;NP_001161294.1;NP_001161295.1;NI | 1055 | 613110 | . | . | 51.921326 | 54.090782 | 54.342174 | 52.773434 | 120.823062 | 125.033259 | 121.942261 | 120.660199 | -1.005383 | 3396.384105 | 0.110667371 | -0.06998786 | 0.944203319 | 0.998865404 | 11.733646 | 11.725911 | 11.714839 | 11.748659 | 11.742008 | 11.712729 | 3841 | 3750 | 2998 | 3119 |
| 10905 | NM_006699,XI | MAN1A2 | mannosidase alpha class 1A member 2 | protein_coding | NP_006690.1;XP_006710365.1;XP_011538838.1;XP_016855604.1 | 6822 | 604345 | . | . | 10.514038 | 10.157726 | 10.996355 | 9.757536 | 24.466600 | 23.480035 | 24.675502 | 22.309447 | 1.013802 | 2678.594790 | 0.120729156 | 0.163803784 | 0.869885613 | 0.998865404 | 11.377208 | 11.397243 | 11.405140 | 11.356311 | 11.440088 | 11.342056 | 3155 | 2784 | 2487 | 2376 |
| 10906 | NM_00114390 | TRAFD1 | TRAF-type zinc finger domain containing 1 | protein_coding | NP_001137378.1;NP_006691.1 | 24808 | 613197 | . | . | 9.737829 | 10.796723 | 9.905947 | 10.734263 | 22.660329 | 24.957108 | 22.228657 | 24.542618 | -1.012358 | 881.785026 | 0.149070676 | -0.1188715 | 0.905377158 | 0.998865404 | 9.793407 | 9.775101 | 9.738119 | 9.835472 | 9.746763 | 9.810218 | 953 | 1022 | 728 | 858 |
| 10907 | NM_00130347 | TXNL4A | thioredoxin like 4A | protein_coding | NP_001290400.1;NP_001292486.1;NP_001292492.1;NP_001292493.1;NI | 30551 | 611595 | . | . | 19.005022 | 18.538774 | 18.257449 | 21.151724 | 44.225464 | 42.853205 | 40.969186 | 48.360908 | 1.031976 | 2103.187981 | 0.12523306 | 0.362599649 | 0.716903972 | 0.998865404 | 11.015576 | 11.060794 | 11.042357 | 11.007486 | 10.994629 | 11.102780 | 2436 | 2185 | 1732 | 2138 |
| 10908 | NM_00116611 | PNPLA6 | patatin like phospholipase domain containing 6 | protein_coding | NP_001159583.1;NP_001159584.1;NP_001159585.1;NP_001159586.1;NI | 16268 | 603197 | . | . | 15.737942 | 14.865559 | 17.265634 | 15.831490 | 36.622839 | 34.362409 | 38.743585 | 36.196822 | 1.071673 | 2180.940086 | 0.121485378 | 0.822033166 | 0.411058021 | 0.998865404 | 11.039803 | 11.139929 | 11.087966 | 11.034968 | 11.152581 | 11.080707 | 2505 | 2196 | 2050 | 2021 |
| 10910 | NM_00113091 | SUGT1 | SGT1 homolog, MIS12 kinetochore complex assembly cochaperon | protein_coding | NP_001124384.1;NP_001307760.1;NP_006695.1;XP_011533193.1;XP_01 | 16987 | 604098 | . | . | 2.425939 | 2.388568 | 2.355378 | 1.792128 | 5.645258 | 5.521285 | 5.285401 | 4.097484 | -1.180525 | 996.839644 | 0.159878239 | -1.4975692 | 0.134245214 | 0.891755273 | 10.075612 | 9.836966 | 10.026324 | 10.006269 | 9.990468 | 9.789703 | 1259 | 1149 | 907 | 737 |
| 10914 | NM_00125200 | PAPOLA | poly(A) polymerase alpha | protein_coding | NP_001238935.1;NP_001238936.1;NP_001280556.1;NP_001280557.1;NI | 14981 | 605553 | . | . | 28.117873 | 28.626386 | 26.361594 | 23.512502 | 65.431449 | 66.171168 | 59.154654 | 53.758546 | -1.151442 | 3732.299279 | 0.115116597 | -1.76726442 | 0.077183955 | 0.767975096 | 11.963913 | 11.760630 | 11.919317 | 11.922552 | 11.852317 | 11.754089 | 4591 | 4319 | 3199 | 3057 |
| 10915 | NM_00104000 | TCERG1 | transcription elongation regulator 1 | protein_coding | NP_001035095.1;NP_001369477.1;NP_006697.2;XP_016864466.1;XP_01 | 15630 | 605409 | . | . | 19.804333 | 20.005863 | 20.139602 | 17.533535 | 46.085499 | 46.244448 | 45.192682 | 40.088342 | -1.060213 | 2691.139021 | 0.133797867 | -0.63045762 | 0.528395223 | 0.998865404 | 11.435424 | 11.351354 | 11.405258 | 11.425820 | 11.457193 | 11.270708 | 3149 | 3026 | 2534 | 2167 |
| 10916 | NM_014599,N | MAGED2 | MAGE family member D2 | protein_coding | NP_055414.2;NP_803182.1;NP_957516.1 | 16353 | 300470 | . | . | 143.069169 | 136.585102 | 142.869156 | 168.499595 | 332.927845 | 315.722588 | 320.594267 | 385.254315 | 1.096888 | 9468.830923 | 0.114376996 | 1.166458792 | 0.243429 | 0.992733415 | 13.140749 | 13.274111 | 13.190415 | 13.144035 | 13.178051 | 13.310186 | 10685 | 9478 | 7952 | 10001 |
| 10919 | NM_00128941 | EHMT2 | euchromatic histone lysine methyltransferase 2 | protein_coding | NP_001276342.1;NP_001305762.1;NP_001350618.1;NP_006700.3;NP_0: | 14129 | 604599 | . | . | 9.085082 | 10.551940 | 10.459357 | 10.436254 | 21.141356 | 24.391283 | 23.470496 | 23.861261 | 1.052645 | 1267.112063 | 0.135621842 | 0.545778642 | 0.585218106 | 0.998865404 | 10.269970 | 10.343744 | 10.234021 | 10.336999 | 10.328242 | 10.322954 | 1325 | 1423 | 1134 | 1214 |
| 10920 | NM_006710,N | COPS8 | COP9 signalosome subunit 8 | protein_coding | NP_006701.1;NP_937832.1 | 24335 | 616011 | . | . | 33.234015 | 33.178945 | 34.410864 | 37.155409 | 77.336918 | 76.694612 | 77.216983 | 84.951428 | 1.031915 | 3604.858635 | 0.110376556 | 0.410629714 | 0.681344072 | 0.998865404 | 11.792925 | 11.838174 | 11.806743 | 11.798223 | 11.802913 | 11.853272 | 4107 | 3809 | 3087 | 3537 |
| 10921 | NM_00128662 | RNPS1 | RNA binding protein with serine rich domain 1 | protein_coding | NP_001273554.1;NP_001273555.1;NP_001273556.1;NP_006702.1;NP_5· | 10080 | 606447 | . | . | 35.933507 | 38.719166 | 36.880382 | 42.845227 | 83.618746 | 89.501090 | 82.758502 | 97.960520 | 1.053831 | 2358.333476 | 0.12399753 | 0.610038494 | 0.541836309 | 0.998865404 | 11.165388 | 11.240732 | 11.156903 | 11.205293 | 11.167728 | 11.276395 | 2565 | 2553 | 1963 | 2421 |
| 10922 | NM_00125846 | FASTK | Fas activated serine/threonine kinase | protein_coding | NP_001245390.1;NP_006703.1;NP_148936.2;XP_005249989.1;XP_00524 | 24676 | 606965 | . | . | 47.615938 | 50.489430 | 50.246667 | 52.580572 | 110.804244 | 116.708579 | 112.752061 | 120.219225 | 1.047825 | 2899.246517 | 0.112845829 | 0.597254102 | 0.550337743 | 0.998865404 | 11.467423 | 11.534717 | 11.461642 | 11.501879 | 11.509201 | 11.530755 | 3180 | 3131 | 2546 | 2818 |
| 10923 | NM_006713,XI | SUB1 | SUB1 regulator of transcription | protein_coding | NP_006704.3;XP_011512246.1;XP_016864475.1;XP_016864476.1 | 19985 | 600503 | . | . | 13.199486 | 12.455991 | 11.877721 | 12.562780 | 30.715744 | 28.792580 | 26.653262 | 28.723306 | -1.064080 | 1347.813323 | 0.129414492 | -0.69239619 | 0.488688557 | 0.998865404 | 10.440513 | 10.350905 | 10.444763 | 10.393759 | 10.355040 | 10.389048 | 1653 | 1450 | 1110 | 1252 |
| 10924 | NM_00128613 | SMPDL3A | sphingomyelin phosphodiesterase acid like 3A | protein_coding | NP_001273067.1;NP_006705.1;XP_005266854.1;XP_005266855.1;XP_00 | 17389 | 610728 | . | . | 12.132227 | 10.521509 | 14.774312 | 12.010915 | 28.232190 | 24.320941 | 33.153124 | 27.461529 | 1.177335 | 699.519487 | 0.168229231 | 1.400026693 | 0.161505325 | 0.933525128 | 9.327137 | 9.563624 | 9.434365 | 9.339206 | 9.572169 | 9.422959 | 792 | 644 | 728 | 634 |
| 10926 | NM_00131806 | DBF4 | DBF4 zinc finger | protein_coding | NP_001304989.1;NP_001304990.1;NP_001304991.1;NP_006707.1 | 17364 | 604281 | . | . | 11.336113 | 9.578324 | 9.663587 | 7.613127 | 26.379601 | 22.140724 | 21.684806 | 17.406513 | -1.227335 | 1086.202279 | 0.160711256 | -1.83887943 | 0.065932922 | 0.73302766 | 10.224906 | 9.930227 | 10.218408 | 10.083672 | 10.085669 | 9.909777 | 1499 | 1178 | 952 | 803 |

| | | | | | | | | | | |
|---|---|---|---|---|---|---|---|---|---|---|
| 10927 | NM_006717 | SPIN1 | spindlin 1 | protein_coding | NP_006708.2 | 11243 | 609936 | . | . | [illegible] |
| 10928 | NM_006788 | RALBP1 | ralA binding protein 1 | protein_coding | NP_006779.1 | 9841 | 605801 | . | . | [illegible] |
| 10929 | NM_032102,N | SRSF8 | serine and arginine rich splicing factor 8 | protein_coding | NP_115285.1;XP_016872631.1 | 16988 | 603269 | . | . | [illegible] |
| 10933 | NM_00126560 | MORF4L1 | mortality factor 4 like 1 | protein_coding | NP_001252532.1;NP_001252533.1;NP_001252534.1;NP_006782.1;NP_9 | 16989 | 607303 | . | . | [illegible] |
| 10935 | NM_00130227 | PRDX3 | peroxiredoxin 3 | protein_coding | NP_001289201.1;NP_006784.1 | 9354 | 604769 | . | . | [illegible] |
| 10936 | NM_006794 | GPR75 | G protein-coupled receptor 75 | protein_coding | NP_006785.1 | 4526 | 606704 | . | . | [illegible] |
| 10938 | NM_00128244 | EHD1 | EH domain containing 1 | protein_coding | NP_001269373.1;NP_001269374.1;NP_006786.2 | 3242 | 605888 | . | . | [illegible] |
| 10939 | NM_006796,XI | AFG3L2 | AFG3 like matrix AAA peptidase subunit 2 | protein_coding | NP_006787.2;XP_011523903.1 | 315 | 604581 | . | . | [illegible] |
| 10940 | NM_00114586 | POP1 | POP1 homolog, ribonuclease P/MRP subunit | protein_coding | NP_001139332.1;NP_001139333.1;NP_055844.2;XP_011515103.1 | 30129 | 602486 | . | . | [illegible] |
| 10943 | NM_00119327 | MSL3 | MSL complex subunit 3 | protein_coding | NP_001180199.1;NP_001269103.1;NP_006791.2;NP_523352.1;NP_5233 | 7370 | 300609 | . | . | [illegible] |
| 10944 | NM_014267,XI | C11orf58 | chromosome 11 open reading frame 58 | protein_coding | NP_055082.1;XP_016872632.1 | 16990 | . | | . | [illegible] |
| 10945 | NM_006801 | KDELR1 | KDEL endoplasmic reticulum protein retention receptor 1 | protein_coding | NP_006792.1 | 6304 | 131235 | . | . | [illegible] |
| 10946 | NM_00132083 | SF3A3 | splicing factor 3a subunit 3 | protein_coding | NP_001307759.1;NP_006793.1 | 10767 | 605596 | . | . | [illegible] |
| 10947 | NM_00113429 | AP3M2 | adaptor related protein complex 3 subunit mu 2 | protein_coding | NP_001127768.1;NP_006794.1;XP_016868466.1 | 570 | 610469 | . | . | [illegible] |
| 10948 | NM_00116593 | STARD3 | StAR related lipid transfer domain containing 3 | protein_coding | NP_001159409.1;NP_001159410.1;NP_006795.3;XP_016879530.1;XP_02 | 17579 | 607048 | . | . | [illegible] |
| 10949 | NM_006805 | HNRNPA0 | heterogeneous nuclear ribonucleoprotein A0 | protein_coding | NP_006796.1 | 5030 | 609409 | . | . | [illegible] |
| 10950 | NM_00113091 | BTG3 | BTG anti-proliferation factor 3 | protein_coding | NP_001124386.1;NP_006797.3;XP_011527743.1;XP_016883754.1;XP_01 | 1132 | 605674 | . | . | [illegible] |
| 10951 | NM_00112722 | CBX1 | chromobox 1 | protein_coding | NP_001120700.1;NP_006798.1 | 1551 | 604511 | . | . | [illegible] |
| 10952 | NM_006808 | SEC61B | SEC61 translocon subunit beta | protein_coding | NP_006799.1 | 16993 | 609214 | . | . | [illegible] |
| 10953 | NM_006809,XI | TOMM34 | translocase of outer mitochondrial membrane 34 | protein_coding | NP_006800.2;XP_005260311.1;XP_011526803.1;XP_016883089.1 | 15746 | 616049 | . | . | [illegible] |
| 10954 | NM_006810,N | PDIA5 | protein disulfide isomerase family A member 5 | protein_coding | NP_006801.1 | 24811 | 616942 | . | . | [illegible] |
| 10955 | NM_006811,N | SERINC3 | serine incorporator 3 | protein_coding | NP_006802.1;NP_945179.1 | 11699 | 607165 | . | . | [illegible] |
| 10956 | NM_00101795 | OS9 | OS9 endoplasmic reticulum lectin | protein_coding | NP_001017956.1;NP_001017957.1;NP_001017958.1;NP_001248349.1;NI | 16994 | 609677 | . | . | [illegible] |
| 10957 | NM_006813,XI | PNRC1 | proline rich nuclear receptor coactivator 1 | protein_coding | NP_006804.1 | 17278 | 606714 | . | . | [illegible] |
| 10959 | NM_00132144 | TMED2 | transmembrane p24 trafficking protein 2 | protein_coding | NP_001308374.1;NP_006806.1 | 16996 | . | | . | [illegible] |
| 10960 | NM_006816 | LMAN2 | lectin, mannose binding 2 | protein_coding | NP_006807.1 | 16986 | 609551 | . | . | [illegible] |
| 10961 | NM_00103402 | ERP29 | endoplasmic reticulum protein 29 | protein_coding | NP_001029197.1;NP_006808.1;XP_016874209.1 | 13799 | 602287 | . | . | [illegible] |
| 10962 | NM_006818 | MLLT11 | MLLT11 transcription factor 7 cofactor | protein_coding | NP_006809.1 | 16997 | 604684 | . | . | [illegible] |
| 10963 | NM_00128265 | STIP1 | stress induced phosphoprotein 1 | protein_coding | NP_001269581.1;NP_001269582.1;NP_006810.1 | 11387 | 605063 | . | . | [illegible] |
| 10965 | NM_00136417 | ACOT2 | acyl-CoA thioesterase 2 | protein_coding | NP_001351106.1;NP_001351107.1;NP_006812.3 | 18431 | 609972 | . | . | [illegible] |
| 10966 | NM_006822,XI | RAB40B | RAB40B, member RAS oncogene family | protein_coding | NP_006813.1;XP_005256391.1;XP_006722334.1;XP_011521830.1;XP_01 | 18284 | . | | . | [illegible] |
| 10969 | NM_00115993 | EBNA1BP2 | EBNA1 binding protein 2 | protein_coding | NP_001153408.1;NP_006815.2 | 15531 | 614443 | . | . | [illegible] |
| 10970 | NM_006825 | CKAP4 | cytoskeleton associated protein 4 | protein_coding | NP_006816.2 | 16991 | 618595 | . | . | [illegible] |
| 10971 | NM_006826 | YWHAQ | tyrosine 3-monooxygenase/tryptophan 5-monooxygenase activati | protein_coding | NP_006817.1 | 12854 | 609009 | . | . | [illegible] |
| 10972 | NM_006827 | TMED10 | transmembrane p24 trafficking protein 10 | protein_coding | NP_006818.3 | 16998 | 605406 | . | . | [illegible] |
| 10973 | NM_00128427 | ASCC3 | activating signal cointegrator 1 complex subunit 3 | protein_coding | NP_001271200.1;NP_006819.2;NP_071374.1;XP_011533696.1;XP_01153 | 18697 | 614217 | . | . | [illegible] |
| 10975 | NM_006830 | UQCR11 | ubiquinol-cytochrome c reductase, complex III subunit XI | protein_coding | NP_006821.1 | 30862 | 609711 | . | . | [illegible] |
| 10978 | NM_00114259 | CLP1 | cleavage factor polyribonucleotide kinase subunit 1 | protein_coding | NP_001136069.1;NP_006822.1 | 16999 | 608757 | . | . | [illegible] |
| 10979 | NM_00113499 | FERMT2 | fermitin family member 2 | protein_coding | NP_001128471.1;NP_001128472.1;NP_006823.1;XP_005267342.1;XP_00 | 15767 | 607746 | . | . | [illegible] |
| 10980 | NM_006833 | COPS6 | COP9 signalosome subunit 6 | protein_coding | NP_006824.2 | 21749 | 614729 | . | . | [illegible] |
| 10981 | NM_006834 | RAB32 | RAB32, member RAS oncogene family | protein_coding | NP_006825.1 | 9772 | 612906 | . | . | [illegible] |
| 10982 | NM_00114382 | MAPRE2 | microtubule associated protein RP/EB family member 2 | protein_coding | NP_001137298.1;NP_001137299.1;NP_001243349.1;NP_055083.1 | 6891 | 605789 | . | . | [illegible] |
| 10983 | NM_00134813 | CCNI | cyclin I | protein_coding | NP_001335061.1;NP_001335062.1;NP_001335063.1;NP_001335064.1;NI | 1595 | 618783 | . | . | [illegible] |
| 10984 | NR_002728 | KCNQ1OT1 | KCNQ1 opposite strand/antisense transcript 1 | lncRNA | . | 6295 | 604115 | . | . | [illegible] |
| 10985 | NM_006836 | GCN1 | GCN1 activator of EIF2AK4 | protein_coding | NP_006827.1 | 4199 | 605614 | . | . | [illegible] |
| 10987 | NM_006837 | COPS5 | COP9 signalosome subunit 5 | protein_coding | NP_006828.2 | 2240 | 604850 | . | . | [illegible] |
| 10988 | NM_00131718 | METAP2 | methionyl aminopeptidase 2 | protein_coding | NP_001304111.1;NP_001304112.1;NP_001317175.1;NP_006829.1 | 16672 | 601870 | . | . | [illegible] |
| 10989 | NM_00110016 | IMMT | inner membrane mitochondrial protein | protein_coding | NP_001093639.1;NP_001093640.1;NP_006830.2;XP_005264167.1;XP_00 | 6047 | 600378 | . | . | [illegible] |
| 10992 | NM_006842,XI | SF3B2 | splicing factor 3b subunit 2 | protein_coding | NP_006833.2;XP_005273783.1;XP_011543042.1;XP_016872633.1 | 10769 | 605591 | . | . | [illegible] |
| 10994 | NM_006844,XI | ILVBL | ilvB acetolactate synthase like | protein_coding | NP_006835.2;XP_005259774.1;XP_011525953.1;XP_024307098.1 | 6041 | 605770 | . | . | [illegible] |
| 10998 | NM_00132119 | SLC27A5 | solute carrier family 27 member 5 | protein_coding | NP_001308125.1;NP_036386.1;XP_011524666.1;XP_016881703.1 | 10999 | 603314 | . | . | [illegible] |
| 10999 | NM_005094,XI | SLC27A4 | solute carrier family 27 member 4 | protein_coding | NP_005085.2;XP_016869711.1;XP_024303159.1 | 10998 | 604194 | . | . | [illegible] |
| 11000 | NM_00131792 | SLC27A3 | solute carrier family 27 member 3 | protein_coding | NP_001304858.3;NP_077306.3 | 10997 | 604193 | . | . | [illegible] |
| 11004 | NM_00129765 | KIF2C | kinesin family member 2C | protein_coding | NP_001284584.1;NP_001284585.1;NP_001284586.1;NP_006836.2;XP_01 | 6393 | 604538 | . | . | [illegible] |
| 11007 | NM_006848 | CCDC85B | coiled-coil domain containing 85B | protein_coding | NP_006839.2 | 24926 | 605360 | . | . | [illegible] |
| 11009 | NM_00118515 | IL24 | interleukin 24 | protein_coding | NP_001172085.1;NP_001172086.1;NP_001172087.1;NP_006841.1;XP_01 | 11346 | 604136 | . | . | [illegible] |
| 11011 | NM_00128433 | TLK2 | tousled like kinase 2 | protein_coding | NP_001271262.1;NP_001271292.1;NP_001317347.1;NP_001362198.1;NI | 11842 | 608439 | . | . | [illegible] |
| 11013 | NM_021992 | TMSB15A | thymosin beta 15a | protein_coding | NP_068832.1 | 30744 | 300939 | . | . | [illegible] |
| 11014 | NM_00110060 | KDELR2 | KDEL endoplasmic reticulum protein retention receptor 2 | protein_coding | NP_001094073.1;NP_006845.1 | 6305 | 609024 | . | . | [illegible] |
| 11015 | NM_006855,N | KDELR3 | KDEL endoplasmic reticulum protein retention receptor 3 | protein_coding | NP_006846.1;NP_057839.1 | 6306 | . | | . | [illegible] |
| 11016 | NM_00113006 | ATF7 | activating transcription factor 7 | protein_coding | NP_001123532.1;NP_001193611.1;NP_001193612.1;NP_001353484.1;NI | 792 | 606371 | . | . | [illegible] |
| 11017 | NM_006857,N | SNRNP27 | small nuclear ribonucleoprotein U4/U6.U5 subunit 27 | protein_coding | NP_006848.1 | 30240 | . | | . | [illegible] |
| 11018 | NM_006858,N | TMED1 | transmembrane p24 trafficking protein 1 | protein_coding | NP_006849.1;XP_006722694.1 | 17291 | 605395 | . | . | [illegible] |
| 11019 | NM_00127859 | LIAS | lipoic acid synthetase | protein_coding | NP_001265519.1;NP_001265520.1;NP_001265521.1;NP_001350629.1;NI | 16429 | 607031 | . | . | [illegible] |
| 11020 | NM_00117770 | IFT27 | intraflagellar transport 27 | protein_coding | NP_001171172.1;NP_001349932.1;NP_006851.1;XP_016884029.1 | 18626 | 615870 | . | . | [illegible] |
| 11021 | NM_00116760 | RAB35 | RAB35, member RAS oncogene family | protein_coding | NP_001161078.1;NP_006852.1;XP_011536081.1;XP_024304569.1 | 9774 | 604199 | . | . | [illegible] |
| 11022 | NM_00108396 | TDRKH | tudor and KH domain containing | protein_coding | NP_001077432.1;NP_001077433.1;NP_001077434.1;NP_006853.2;XP_00 | 11713 | 609501 | . | . | [illegible] |
| 11023 | NM_00111270 | VAX1 | ventral anterior homeobox 1 | protein_coding | NP_001106175.1;NP_954582.1 | 12660 | 604294 | . | . | [illegible] |
| 11030 | NM_00100871 | RBPMS | RNA binding protein, mRNA processing factor | protein_coding | NP_001008710.1;NP_001008711.1;NP_001008712.1;NP_006858.1;XP_01 | 19097 | 601558 | . | . | [illegible] |
| 11031 | NM_006868,XI | RAB31 | RAB31, member RAS oncogene family | protein_coding | NP_006859.2;XP_016881017.1;XP_016881018.1 | 9771 | 605694 | . | . | [illegible] |
| 11033 | NM_00128430 | ADAP1 | ArfGAP with dual PH domains 1 | protein_coding | NP_001271237.1;NP_001271238.1;NP_001271239.1;NP_001271240.1;NI | 16486 | 608114 | . | . | [illegible] |
| 11034 | NM_00101154 | DSTN | destrin, actin depolymerizing factor | protein_coding | NP_001011546.1;NP_006861.1;XP_011527446.1 | 15750 | 609114 | . | . | [illegible] |
| 11037 | NM_00119859 | STON1 | stonin 1 | protein_coding | NP_001185524.1;NP_006864.2 | 17003 | 605357 | . | . | [illegible] |
| 11039 | NR_024054,NF | GUSBP14 | GUSB pseudogene 14 | transcribed_ps | . | 54496 | . | | . | [illegible] |
| 11040 | NM_006875 | PIM2 | Pim-2 proto-oncogene, serine/threonine kinase | protein_coding | NP_006866.2 | 8987 | 300295 | . | . | [illegible] |
| 11041 | NM_006876 | B4GAT1 | beta-1,4-glucuronyltransferase 1 | protein_coding | NP_006867.1 | 15685 | 605517 | . | . | [illegible] |
| 11042 | NR_034021 | GUSBP15 | GUSB pseudogene 15 | transcribed_ps | . | 54497 | . | | . | [illegible] |
| 11044 | NM_00117180 | TENT4A | terminal nucleotidyltransferase 4A | protein_coding | NP_001165276.2;NP_008930.2;XP_011512247.1;XP_016864477.1;XP_01 | 16705 | 605198 | . | . | [illegible] |
| 11045 | NM_00128144 | UPK1A | uroplakin 1A | protein_coding | NP_001268372.1;NP_008931.1 | 12577 | 611557 | . | . | [illegible] |
| 11046 | NM_00128699 | SLC35D2 | solute carrier family 35 member D2 | protein_coding | NP_001273919.1;NP_008932.2;XP_005251730.1;XP_005251731.1;XP_00 | 20799 | 609182 | . | . | [illegible] |
| 11047 | NM_00128143 | ADRM1 | adhesion regulating molecule 1 | protein_coding | NP_001268366.1;NP_001268367.1;NP_008933.2;NP_783163.1;XP_01152 | 15759 | 610650 | . | . | [illegible] |
| 11051 | NM_007006 | NUDT21 | nudix hydrolase 21 | protein_coding | NP_008937.1 | 13870 | 604978 | . | . | [illegible] |
| 11052 | NM_00130094 | CPSF6 | cleavage and polyadenylation specific factor 6 | protein_coding | NP_001287876.1;NP_008938.2;XP_005268645.1;XP_005268647.1 | 13871 | 604979 | . | . | [illegible] |
| 11054 | NM_007346 | OGFR | opioid growth factor receptor | protein_coding | NP_031372.2 | 15768 | 606459 | . | . | [illegible] |
| 11056 | NM_00129147 | DDX52 | DExD-box helicase 52 | protein_coding | NP_001278405.1;NP_008941.3;XP_011522534.1;XP_011522535.1 | 20038 | 612500 | . | . | [illegible] |
| 11057 | NM_007011,N | ABHD2 | abhydrolase domain containing 2, acylglycerol lipase | protein_coding | NP_008942.3;NP_690888.1;XP_005254889.1;XP_005254890.1;XP_01151 | 18717 | 612196 | . | . | [illegible] |
| 11059 | NM_007013,XI | WWP1 | WW domain containing E3 ubiquitin protein ligase 1 | protein_coding | NP_008944.1;XP_005250817.1;XP_005250818.1;XP_016868482.1;XP_01 | 17004 | 602307 | . | . | [illegible] |
| 11060 | NM_00127045 | WWP2 | WW domain containing E3 ubiquitin protein ligase 2 | protein_coding | NP_001257382.1;NP_001257383.1;NP_001257384.1;NP_008945.2;NP_9 | 16804 | 602308 | . | . | [illegible] |
| 11062 | NM_00127041 | DUS4L | dihydrouridine synthase 4 like | protein_coding | NP_001257348.1;NP_853559.1 | 21517 | . | | . | [illegible] |
| 11064 | NM_00133076 | CNTRL | centriolin | protein_coding | NP_001317691.1;NP_001356821.1;NP_001356822.1;NP_001356823.1;NI | 1858 | 605496 | . | . | [illegible] |
| 11065 | NM_00128174 | UBE2C | ubiquitin conjugating enzyme E2 C | protein_coding | NP_001268670.1;NP_001268671.1;NP_008950.1;NP_861515.1;NP_8615 | 15937 | 605574 | . | . | [illegible] |
| 11066 | NM_022717,N | SNRNP35 | small nuclear ribonucleoprotein U11/U12 subunit 35 | protein_coding | NP_073208.1;NP_851030.1;XP_006719267.1;XP_016874213.1 | 30852 | . | | . | [illegible] |
| 11068 | NM_00129128 | CYB561D2 | cytochrome b561 family member D2 | protein_coding | NP_001278213.1;NP_008953.1 | 30253 | 607068 | . | . | [illegible] |
| 11069 | NM_00110039 | RAPGEF4 | Rap guanine nucleotide exchange factor 4 | protein_coding | NP_001093867.1;NP_001269828.1;NP_001269829.1;NP_001269830.1;NI | 16626 | 606058 | . | . | [illegible] |
| 11070 | NM_007024 | TMEM115 | transmembrane protein 115 | protein_coding | NP_008955.1 | 30055 | 607069 | . | . | [illegible] |
| 11072 | NM_007026,XI | DUSP14 | dual specificity phosphatase 14 | protein_coding | NP_008957.1;XP_005257034.1;XP_011522536.1 | 17007 | 606618 | . | . | [illegible] |
| 11073 | NM_00136388 | TOPBP1 | DNA topoisomerase II binding protein 1 | protein_coding | NP_001350818.1;NP_008958.2;XP_011510659.1;XP_016861125.1;XP_01 | 17008 | 607760 | . | . | [illegible] |
| 11076 | NM_007030,XI | TPPP | tubulin polymerization promoting protein | protein_coding | NP_008961.1;XP_005248294.2;XP_016864482.1;XP_024310114.1 | 24164 | 608773 | . | . | [illegible] |
| 11077 | NM_007031,XI | HSF2BP | heat shock transcription factor 2 binding protein | protein_coding | NP_008962.1;XP_011527745.2;XP_011527747.2;XP_011527748.1;XP_01 | 5226 | 604554 | . | . | [illegible] |
| 11078 | NM_00103914 | TRIOBP | TRIO and F-actin binding protein | protein_coding | NP_001034230.1;NP_008963.3;NP_619538.2 | 17009 | 609761 | . | . | [illegible] |
| 11079 | NM_007033 | RER1 | retention in endoplasmic reticulum sorting receptor 1 | protein_coding | NP_008964.3 | 30309 | . | | . | [illegible] |
| 11080 | NM_00131709 | DNAJB4 | DnaJ heat shock protein family (Hsp40) member B4 | protein_coding | NP_001304028.1;NP_001304029.1;NP_001304030.1;NP_001304031.1;NI | 14886 | 611327 | . | . | [illegible] |
| 11083 | NM_00119336 | DIDO1 | death inducer-obliterator 1 | protein_coding | NP_001180298.1;NP_001180299.1;NP_071388.2;NP_149072.2;NP_5429 | 2680 | 604140 | . | . | [illegible] |
| 11091 | NM_017588,N | WDR5 | WD repeat domain 5 | protein_coding | NP_060058.1;NP_438172.1;XP_005272220.1;XP_024303161.1;XP_02430 | 12757 | 609012 | . | . | [illegible] |
| 11092 | NM_00131689 | SPACA9 | sperm acrosome associated 9 | protein_coding | NP_001303826.1;NP_001303827.1;NP_001303829.1;NP_061829.3;XP_01 | 1367 | 618552 | . | . | [illegible] |
| 11093 | NM_139025,N | ADAMTS13 | ADAM metallopeptidase with thrombospondin type 1 motif 13 | protein_coding | NP_620594.1;NP_620595.1;NP_620596.2;XP_011516478.1;XP_01151648 | 1366 | 604134 | . | . | [illegible] |
| 11094 | NM_00113577 | CACFD1 | calcium channel flower domain containing 1 | protein_coding | NP_001129247.1;NP_001229298.1;NP_001229299.1;NP_060056.1;XP_01 | 1365 | 613104 | . | . | [illegible] |
| 11097 | NM_00137044 | NUP42 | nucleoporin 42 | protein_coding | NP_001357372.1;NP_001357373.1;NP_001357374.1;NP_001357375.1;NI | 17010 | . | | . | [illegible] |
| 11098 | NM_00129317 | PRSS23 | serine protease 23 | protein_coding | NP_001280108.1;NP_001280109.1;NP_009104.3 | 14370 | 618376 | . | . | [illegible] |
| 11099 | NM_007039,XI | PTPN21 | protein tyrosine phosphatase non-receptor type 21 | protein_coding | NP_008970.2;XP_005267344.1;XP_006720074.1;XP_011534669.1;XP_01 | 9651 | 603271 | . | . | [illegible] |
| 11100 | NM_00130101 | HNRNPUL1 | heterogeneous nuclear ribonucleoprotein U like 1 | protein_coding | NP_001287945.1;NP_001308137.1;NP_001308140.1;NP_008971.2;NP_6 | 17011 | 605800 | . | . | [illegible] |
| 11101 | NM_00100197 | ATE1 | arginyltransferase 1 | protein_coding | NP_001001976.1;NP_001275663.1;NP_001275664.1;NP_001275665.1;NI | 782 | 607103 | . | . | [illegible] |
| 11102 | NM_00109878 | RPP14 | ribonuclease P/MRP subunit p14 | protein_coding | NP_001092253.1;NP_008973.1 | 30327 | 606112 | . | . | [illegible] |
| 11103 | NM_007043 | KRR1 | KRR1 small subunit processome component homolog | protein_coding | NP_008974.5 | 5176 | 612817 | . | . | [illegible] |
| 11104 | NM_00120407 | KATNA1 | katanin catalytic subunit A1 | protein_coding | NP_001191005.1;NP_008975.1;XP_005266861.1;XP_005266862.1;XP_01 | 6216 | 606696 | . | . | [illegible] |
| 11105 | NM_00109817 | PRDM7 | PR/SET domain 7 | protein_coding | NP_001091643.1;XP_005256331.1;XP_011521131.1;XP_011521133.1;XP | 9351 | 609759 | . | . | [illegible] |
| 11107 | NM_00130082 | PRDM5 | PR/SET domain 5 | protein_coding | NP_001287752.1;NP_001287753.1;NP_001366033.1;NP_001366035.1;NI | 9349 | 614161 | . | . | [illegible] |
| 11108 | NM_012406,XI | PRDM4 | PR/SET domain 4 | protein_coding | NP_036538.3;XP_005268650.1;XP_024304575.1 | 9348 | 605780 | . | . | [illegible] |
| 11112 | NM_152740,XI | HIBADH | 3-hydroxyisobutyrate dehydrogenase | protein_coding | NP_689953.1;XP_016867208.1 | 4907 | 608475 | . | . | [illegible] |
| 11113 | NM_00120699 | CIT | citron rho-interacting serine/threonine kinase | protein_coding | NP_001193928.1;NP_009105.1;XP_006719269.1;XP_011536085.1;XP_01 | 1985 | 605629 | . | . | [illegible] |
| 11116 | NM_00127869 | CEP43 | centrosomal protein 43 | protein_coding | NP_001265619.1;NP_008976.1;NP_919410.1 | 17012 | 605392 | . | . | [illegible] |
| 11117 | NM_007046 | EMILIN1 | elastin microfibril interfacer 1 | protein_coding | NP_008977.1 | 19880 | 130660 | . | . | [illegible] |
| 11118 | NM_00119724 | BTN3A2 | butyrophilin subfamily 3 member A2 | protein_coding | NP_001184175.1;NP_001184176.1;NP_001184177.1;NP_001184178.1;NI | 1139 | 613594 | . | . | [illegible] |
| 11119 | NM_00114500 | BTN3A1 | butyrophilin subfamily 3 member A1 | protein_coding | NP_001138480.1;NP_001138481.1;NP_008979.3;NP_919423.1;XP_0052 | 1138 | 613593 | . | . | [illegible] |
| 11120 | NM_00119723 | BTN2A1 | butyrophilin subfamily 2 member A1 | protein_coding | NP_001184162.1;NP_001184163.1;NP_008980.1;NP_510961.1 | 1136 | 613590 | . | . | [illegible] |
| 11123 | NM_00125197 | RCAN3 | RCAN family member 3 | protein_coding | NP_001238906.1;NP_001238907.1;NP_001238908.1;NP_001238909.1;NI | 3042 | 605860 | . | . | [illegible] |
| 11124 | NM_007051,XI | FAF1 | Fas associated factor 1 | protein_coding | NP_008982.1;XP_016855625.1;XP_024308502.1;XP_024308504.1 | 3578 | 604460 | . | . | [illegible] |
| 11127 | NM_00130079 | KIF3A | kinesin family member 3A | protein_coding | NP_001287720.1;NP_001287721.1;NP_008985.3;XP_006714589.1;XP_01 | 6319 | 604683 | . | . | [illegible] |
| 11128 | NM_007055 | POLR3A | RNA polymerase III subunit A | protein_coding | NP_008986.2 | 30074 | 614258 | . | . | [illegible] |
| 11129 | NM_00127843 | CLASRP | CLK4 associating serine/arginine rich protein | protein_coding | NP_001265368.1;NP_008987.2;XP_011524698.1;XP_011524699.1 | 17731 | 618532 | . | . | [illegible] |
| 11130 | NM_00100541 | ZWINT | ZW10 interacting kinetochore protein | protein_coding | NP_001005413.1;NP_008988.2;NP_127490.1;XP_016871094.1;XP_02430 | 13195 | 609177 | . | . | [illegible] |
| 11132 | NM_023083,N | CAPN10 | calpain 10 | protein_coding | NP_075571.2;NP_075573.3 | 1477 | 605286 | . | . | [illegible] |
| 11133 | NM_00129129 | KPTN | kaptin, actin binding protein | protein_coding | NP_001278225.1;NP_008990.2;XP_016881715.1;XP_016881716.1;XP_01 | 6404 | 615620 | . | . | [illegible] |
| 11137 | NM_00131796 | PWP1 | PWP1 homolog, endonuclein | protein_coding | NP_001304891.1;NP_001304892.1;NP_008993.1 | 17015 | . | | . | [illegible] |
| 11138 | NM_00110242 | TBC1D8 | TBC1 domain family member 8 | protein_coding | NP_001095896.1;NP_001317277.1;XP_005263919.1;XP_005263921.1;XP | 17791 | . | | . | [illegible] |
| 11140 | NM_007065,XI | CDC37 | cell division cycle 37, HSP90 cochaperone | protein_coding | NP_008996.1;XP_011525954.1 | 1735 | 605065 | . | . | [illegible] |
| 11141 | NM_014271,XI | IL1RAPL1 | interleukin 1 receptor accessory protein like 1 | protein_coding | NP_055086.1;XP_016884729.1;XP_016884730.1 | 5996 | 300206 | . | . | [illegible] |
| 11142 | NM_00128144 | PKIG | cAMP-dependent protein kinase inhibitor gamma | protein_coding | NP_001268373.1;NP_001268374.1;NP_008997.1;NP_861520.1;NP_8615 | 9019 | 604932 | . | . | [illegible] |
| 11143 | NM_00119915 | KAT7 | lysine acetyltransferase 7 | protein_coding | NP_001186084.1;NP_001186085.1;NP_001186086.1;NP_001186087.1;NI | 17016 | 609880 | . | . | [illegible] |
| 11144 | NM_00127820 | DMC1 | DNA meiotic recombinase 1 | protein_coding | NP_001265137.1;NP_001349946.1;NP_008999.2;XP_011528136.1;XP_01 | 2927 | 602721 | . | . | [illegible] |
| 11145 | NM_00112820 | PLAAT3 | phospholipase A and acyltransferase 3 | protein_coding | NP_001121675.1;NP_009000.2;XP_006718489.1;XP_011543043.1 | 17825 | 613867 | . | . | [illegible] |
| 11146 | NM_00131968 | GLMN | glomulin, FKBP associated protein | protein_coding | NP_001306612.1;NP_444504.1;XP_005270458.1;XP_006710372.1;XP_01 | 14373 | 601749 | . | . | [illegible] |
| 11147 | NR_027404,NF | HHLA3 | HERV-H LTR-associating 3 | lncRNA | . | 4906 | 604372 | . | . | [illegible] |
| 11149 | NM_00119956 | BVES | blood vessel epicardial substance | protein_coding | NP_001186492.1;NP_009004.2;NP_671488.1;XP_011533700.1 | 1152 | 604577 | . | . | [illegible] |
| 11151 | NM_00119333 | CORO1A | coronin 1A | protein_coding | NP_001180262.1;NP_009005.1;XP_011544016.1;XP_016878374.1;XP_01 | 2252 | 605000 | . | . | [illegible] |
| 11152 | NM_00102989 | WDR45 | WD repeat domain 45 | protein_coding | NP_001025067.1;NP_009006.2 | 28912 | 300526 | . | . | [illegible] |
| 11153 | NM_007076 | FICD | FIC domain protein adenylyltransferase | protein_coding | NP_009007.2 | 18416 | . | | . | [illegible] |
| 11154 | NM_00112812 | AP4S1 | adaptor related protein complex 4 subunit sigma 1 | protein_coding | NP_001121598.1;NP_001241655.1;NP_001241656.1;NP_001241657.1;NI | 575 | 607243 | . | . | [illegible] |
| 11155 | NM_00108011 | LDB3 | LIM domain binding 3 | protein_coding | NP_001073583.1;NP_001073584.1;NP_001073585.1;NP_001165081.1;NI | 15710 | 605906 | . | . | [illegible] |
| 11156 | NM_007079,N | PTP4A3 | protein tyrosine phosphatase 4A3 | protein_coding | NP_009010.2;NP_116000.1;XP_005250821.1;XP_006716563.1;XP_01151 | 9636 | 606449 | . | . | [illegible] |
| 11157 | NM_007080,XI | LSM6 | LSM6 homolog, U6 small nuclear RNA and mRNA degradation ass | protein_coding | NP_009011.1;XP_016863161.1;XP_016863162.1 | 17017 | 607286 | . | . | [illegible] |
| 11158 | NM_00100378 | RABL2B | RAB, member of RAS oncogene family like 2B | protein_coding | NP_001003789.1;NP_001124391.1;NP_001124392.1;NP_001124393.1;NI | 9800 | 605413 | . | . | [illegible] |
| 11159 | NM_00130615 | RABL2A | RAB, member of RAS oncogene family like 2A | protein_coding | NP_001293087.1;NP_001293088.1;NP_001293089.1;NP_001293090.1;NI | 9799 | 605412 | . | . | [illegible] |
| 11160 | NM_00100379 | ERLIN2 | ER lipid raft associated 2 | protein_coding | NP_001003790.1;NP_001003791.1;NP_001349807.1;NP_001349809.1;NI | 1356 | 611605 | . | . | [illegible] |
| 11161 | NM_007176 | ERG28 | ergosterol biosynthesis 28 homolog | protein_coding | NP_009107.1 | 1187 | 604576 | . | . | [illegible] |
| 11162 | NM_007083,N | NUDT6 | nudix hydrolase 6 | protein_coding | NP_009014.2;NP_932158.1 | 8053 | 606261 | . | . | [illegible] |
| 11163 | NM_00130102 | NUDT4 | nudix hydrolase 4 | protein_coding | NP_001287951.1;NP_001287952.1;NP_001287953.1;NP_061967.3;NP_9 | 8051 | 609229 | . | . | [illegible] |
| 11164 | NM_00132164 | NUDT5 | nudix hydrolase 5 | protein_coding | NP_001308576.1;NP_001308577.1;NP_054861.2 | 8052 | 609230 | . | . | [illegible] |
| 11165 | NM_006703 | NUDT3 | nudix hydrolase 3 | protein_coding | NP_006694.1 | 8050 | 609228 | . | . | [illegible] |
| 11167 | NM_007085,XI | FSTL1 | follistatin like 1 | protein_coding | NP_009016.1;XP_024309095.1 | 3972 | 605547 | . | . | [illegible] |
| 11168 | NM_00112821 | PSIP1 | PC4 and SFRS1 interacting protein 1 | protein_coding | NP_001121689.1;NP_001304827.1;NP_001304829.1;NP_066967.3;NP_1 | 9527 | 603620 | . | . | [illegible] |
| 11169 | NM_00100839 | WDHD1 | WD repeat and HMG-box DNA binding protein 1 | protein_coding | NP_001008397.1;NP_009017.1;XP_006720075.1;XP_011534675.1;XP_01 | 23170 | 608126 | . | . | [illegible] |

| | | | | | | | | | | | | | | | | | | | | | | | | | | | | | | | | | |
|---|---|---|---|---|---|---|---|---|---|---|---|---|---|---|---|---|---|---|---|---|---|---|---|---|---|---|---|---|---|---|---|---|---|
| 11171 | NM_007178,XI | STRAP | serine/threonine kinase receptor associated protein | protein_coding | NP_009109.3;XP_005253351.1 | 30796 | 605986 | . | . | 68.898382 | 65.020791 | 61.081731 | 63.502858 | 160.329367 | 150.298480 | 137.065636 | 145.191747 | -1.089867 | 3766.459224 | 0.110686095 | -1.12165541 | 0.262008981 | 0.995760118 | 11.939723 | 11.815595 | 11.941206 | 11.884874 | 11.831140 | 11.853273 | 4676 | 4096 | 3092 | 3425 |
| 11173 | NM_014272,XI | ADAMTS7 | ADAM metallopeptidase with thrombospondin type 1 motif 7 | protein_coding | NP_055087.2;XP_005254194.1;XP_011519468.1 | 223 | 605009 | . | . | 0.088011 | 0.121807 | 0.056025 | 0.060919 | 0.204806 | 0.281561 | 0.125719 | 0.139284 | -1.751734 | 15.206345 | 0.797535591 | -1.01410324 | 0.310533492 | NA | 4.275837 | 3.464871 | 3.937345 | 3.962073 | 3.911163 | 3.916716 | 19 | 24 | 9 | 11 |
| 11174 | NM_197941,N | ADAMTS6 | ADAM metallopeptidase with thrombospondin type 1 motif 6 | protein_coding | NP_922932.2;XP_011541415.1;XP_011541416.1;XP_011541417.1;XP_01 | 222 | 605008 | . | . | 0.194930 | 0.197519 | 0.178599 | 0.105406 | 0.453609 | 0.456572 | 0.400772 | 0.240998 | -1.344903 | 39.896745 | 0.509757567 | -0.83863711 | 0.401672979 | 0.998865404 | 5.514721 | 5.090622 | 5.341207 | 5.338521 | 5.326946 | 5.261087 | 53 | 49 | 37 | 24 |
| 11176 | NM_00130090 | BAZ2A | bromodomain adjacent to zinc finger domain 2A | protein_coding | NP_001287834.1;NP_001338085.1;NP_038477.2;XP_011536103.1;XP_01 | 962 | 605682 | . | . | 10.589385 | 10.952285 | 11.334259 | 11.847869 | 24.641937 | 25.316698 | 25.433748 | 27.088748 | 1.065628 | 2985.828317 | 0.112030206 | 0.81856583 | 0.413034169 | 0.998865404 | 11.497373 | 11.589004 | 11.518932 | 11.515534 | 11.549065 | 11.588866 | 3336 | 3113 | 2614 | 2958 |
| 11177 | NM_013448,N | BAZ1A | bromodomain adjacent to zinc finger domain 1A | protein_coding | NP_038476.2;NP_872589.1;XP_011534676.1;XP_011534678.3;XP_01687 | 960 | 605680 | . | . | 6.189541 | 6.218881 | 6.301001 | 5.597949 | 14.403314 | 14.375219 | 14.139263 | 12.799045 | -1.039591 | 1070.141807 | 0.139433595 | -0.40173616 | 0.687878208 | 0.998865404 | 10.091132 | 10.035504 | 10.077252 | 10.077229 | 10.094487 | 10.000595 | 1256 | 1177 | 971 | 921 |
| 11178 | NM_00136288 | LZTS1 | leucine zipper tumor suppressor 1 | protein_coding | NP_001349813.1;NP_066300.1;XP_011542686.1;XP_011542687.1;XP_01 | 13861 | 606551 | . | . | 33.229608 | 33.621059 | 35.296646 | 39.569344 | 77.326665 | 77.716587 | 79.204651 | 90.470595 | 1.103093 | 6297.536202 | 0.109041957 | 1.298162617 | 0.194231461 | 0.96621494 | 12.548105 | 12.689572 | 12.574086 | 12.580725 | 12.614961 | 12.703409 | 6869 | 6488 | 5446 | 6515 |
| 11179 | NM_021994,XI | ZNF277 | zinc finger protein 277 | protein_coding | NP_068834.2;XP_005250186.1;XP_011514070.1;XP_016867209.1 | 13070 | 605465 | . | . | 4.320626 | 3.598063 | 4.665495 | 2.965988 | 10.054275 | 8.317085 | 10.469237 | 6.781378 | -1.095299 | 297.977594 | 0.226558969 | -0.57964655 | 0.562152994 | 0.998865404 | 8.282357 | 8.152858 | 8.276282 | 8.200456 | 8.276007 | 8.096973 | 383 | 313 | 290 | 221 |
| 11180 | NM_00132054 | WDR6 | WD repeat domain 6 | protein_coding | NP_001307475.1;NP_001307476.1;NP_060501.4 | 12758 | 606031 | . | . | 24.777111 | 26.226899 | 27.412789 | 25.393917 | 57.657357 | 60.624653 | 61.513503 | 58.060178 | 1.023164 | 3314.446035 | 0.114723107 | 0.287969648 | 0.773369973 | 0.998865404 | 11.677902 | 11.711012 | 11.663150 | 11.705736 | 11.739762 | 11.665945 | 3675 | 3627 | 3046 | 3002 |
| 11182 | NM_00114509 | SLC2A6 | solute carrier family 2 member 6 | protein_coding | NP_001138571.1;NP_060055.2;XP_011516491.2;XP_016869725.1;XP_01 | 11011 | 606813 | . | . | 6.699815 | 7.103515 | 8.988500 | 9.290203 | 15.590745 | 16.420093 | 20.169933 | 21.240943 | 1.289436 | 602.134917 | 0.158680809 | 2.311181411 | 0.020822837 | 0.478232305 | 9.039131 | 9.405551 | 9.116158 | 9.164529 | 9.302342 | 9.316177 | 583 | 589 | 584 | 642 |
| 11183 | NM_006575,N | MAP4K5 | mitogen-activated protein kinase kinase kinase kinase 5 | protein_coding | NP_006566.2;NP_942089.1;XP_006720076.1;XP_011534679.1;XP_01153 | 6867 | 604923 | . | . | 7.477996 | 6.721740 | 8.530904 | 5.838888 | 17.401603 | 15.537601 | 19.143101 | 13.349926 | 1.001284 | 975.702625 | 0.179717843 | 0.010298343 | 0.991783256 | 0.999576129 | 9.928915 | 9.931679 | 9.962682 | 9.884697 | 10.057881 | 9.773522 | 1182 | 996 | 1014 | 738 |
| 11184 | NM_00104260 | MAP4K1 | mitogen-activated protein kinase kinase kinase kinase 1 | protein_coding | NP_001036065.1;NP_009112.1;XP_011524705.1;XP_011524706.1;XP_01 | 6863 | 601983 | . | . | 0.547889 | 0.385239 | 0.390185 | 0.477460 | 1.274959 | 0.890496 | 0.875564 | 1.091656 | -1.074722 | 40.022056 | 0.502843746 | -0.20675238 | 0.836203246 | 0.998865404 | 5.373688 | 5.269892 | 5.356497 | 5.300116 | 5.302040 | 5.332868 | 56 | 37 | 30 | 40 |
| 11186 | NM_00120695 | RASSF1 | Ras association domain family member 1 | protein_coding | NP_001193886.1;NP_009113.3;NP_733830.1;NP_733831.1;NP_733832.1 | 9882 | 605082 | . | . | 8.454003 | 7.291563 | 6.144811 | 6.463240 | 19.672811 | 16.854776 | 13.788778 | 14.777430 | -1.277198 | 395.190852 | 0.187044735 | -1.88715602 | 0.059139344 | 0.718513409 | 8.792092 | 8.439192 | 8.736221 | 8.648675 | 8.531127 | 8.566655 | 546 | 445 | 292 | 336 |
| 11188 | NM_00127629 | NISCH | nischarin | protein_coding | NP_001263222.2;NP_001263223.2;NP_009115.3;XP_005264896.1;XP_00 | 18006 | 615507 | . | . | 10.047203 | 10.343041 | 10.878224 | 9.976193 | 23.380257 | 23.908404 | 24.410420 | 22.809384 | 1.017706 | 1506.706727 | 0.129189618 | 0.196002701 | 0.844608052 | 0.998865404 | 10.544342 | 10.569910 | 10.542757 | 10.556794 | 10.606461 | 10.518560 | 1704 | 1626 | 1397 | 1344 |
| 11190 | NM_00131821 | CEP250 | centrosomal protein 250 | protein_coding | NP_001305148.1;NP_009117.2;XP_005260319.1;XP_005260320.1;XP_00 | 1859 | 609689 | . | . | 7.294292 | 6.822864 | 7.775066 | 7.790541 | 16.974119 | 15.771358 | 17.447024 | 17.812146 | 1.163267 | 2042.858933 | 0.121911329 | 1.789679112 | 0.073505512 | 0.762854575 | 10.883039 | 11.101450 | 10.941389 | 10.923155 | 11.092699 | 11.011286 | 2200 | 2014 | 2007 | 1956 |
| 11191 | NR_023917 | PTENP1 | phosphatase and tensin homolog pseudogene 1 | transcribed_ps | . | 9589 | 613531 | . | . | 0.290058 | 0.246661 | 0.143688 | 0.238174 | 0.674978 | 0.570167 | 0.322432 | 0.544556 | -1.418318 | 28.564186 | 0.592605285 | -0.85078748 | 0.39488742 | 0.998865404 | 5.067319 | 4.560703 | 4.872065 | 4.847818 | 4.798939 | 4.840473 | 42 | 33 | 16 | 27 |
| 11193 | NM_007187,XI | WBP4 | WW domain binding protein 4 | protein_coding | NP_009118.1;XP_005266302.1;XP_016875869.1 | 12739 | 604981 | . | . | 3.490514 | 3.788944 | 4.102617 | 3.640422 | 8.122571 | 8.758314 | 9.206153 | 8.323394 | 1.046943 | 279.785042 | 0.210026432 | 0.315118268 | 0.752671859 | 0.998865404 | 8.094509 | 8.161073 | 8.096628 | 8.132395 | 8.171884 | 8.104374 | 304 | 305 | 265 | 251 |
| 11194 | NM_00128229 | ABCB8 | ATP binding cassette subfamily B member 8 | protein_coding | NP_001269220.1;NP_001269221.1;NP_001269222.1;NP_009119.2 | 49 | 605464 | . | . | 11.073180 | 11.730115 | 11.242797 | 11.875242 | 25.767744 | 27.114684 | 25.228511 | 27.151329 | 1.000479 | 1660.012599 | 0.123717732 | 0.005579481 | 0.995548241 | 0.999862634 | 10.696737 | 10.697220 | 10.676326 | 10.716633 | 10.678048 | 10.715561 | 1862 | 1837 | 1409 | 1594 |
| 11196 | NM_007190,N | SEC23IP | SEC23 interacting protein | protein_coding | NP_009121.1;XP_005269526.1;XP_005269527.1;XP_016871099.1 | 17018 | 617852 | . | . | 8.858972 | 8.655648 | 8.302148 | 10.471882 | 20.615189 | 20.007918 | 18.629778 | 23.942714 | 1.029165 | 1668.928730 | 0.122540674 | 0.338453078 | 0.735021781 | 0.998865404 | 10.683887 | 10.725230 | 10.691514 | 10.695009 | 10.695788 | 10.735197 | 1890 | 1779 | 1435 | 1627 |
| 11198 | NM_007192,XI | SUPT16H | SPT16 homolog, facilitates chromatin remodeling subunit | protein_coding | NP_009123.1;XP_011534683.1 | 11465 | 605012 | . | . | 29.788962 | 31.271408 | 29.291854 | 31.598812 | 69.320138 | 72.285260 | 65.730067 | 72.246929 | -1.018425 | 4229.297945 | 0.109732332 | -0.24003774 | 0.810301002 | 0.998865404 | 12.059369 | 12.032914 | 12.037081 | 12.069882 | 12.010708 | 12.065330 | 4818 | 4696 | 3524 | 4058 |
| 11200 | NM_00100573 | CHEK2 | checkpoint kinase 2 | protein_coding | NP_001005735.1;NP_001244316.1;NP_001336885.1;NP_009125.1;NP_6 | 16627 | 604373 | . | . | 16.024595 | 16.732524 | 15.624529 | 14.955512 | 37.289889 | 38.677974 | 35.060991 | 34.194002 | -1.085614 | 917.492399 | 0.140041159 | -0.84626267 | 0.397406228 | 0.998865404 | 9.899588 | 9.781084 | 9.854500 | 9.886258 | 9.822235 | 9.800023 | 1076 | 1053 | 777 | 812 |
| 11201 | NM_00135161 | POLI | DNA polymerase iota | protein_coding | NP_001338539.1;NP_001338540.1;NP_001338541.1;NP_001338542.1;NI | 9182 | 605252 | . | . | 2.670067 | 2.149420 | 2.563001 | 2.228771 | 6.213357 | 4.968482 | 5.751301 | 5.095821 | -1.036727 | 414.852269 | 0.193873502 | -0.26839946 | 0.788391855 | 0.998865404 | 8.721616 | 8.670845 | 8.732081 | 8.676995 | 8.755309 | 8.606102 | 508 | 435 | 399 | 334 |
| 11209 | NR_027504 | MST1P2 | macrophage stimulating 1 pseudogene 2 | transcribed_ps | . | 7383 | . | | . | 4.005468 | 3.725338 | 4.459789 | 4.296391 | 9.320889 | 8.611284 | 10.007637 | 9.823188 | 1.115577 | 338.681558 | 0.19331273 | 0.81624487 | 0.414360099 | 0.998865404 | 8.322456 | 8.480774 | 8.393544 | 8.347533 | 8.445535 | 8.418847 | 384 | 331 | 318 | 327 |
| 11212 | NM_00134934 | PLPBP | pyridoxal phosphate binding protein | protein_coding | NP_001336275.1;NP_001336276.1;NP_001336277.1;NP_001336278.1;NI | 9457 | 604436 | . | . | 7.326683 | 7.980526 | 8.766932 | 8.374394 | 17.049492 | 18.447344 | 19.672744 | 19.147057 | 1.107194 | 615.137584 | 0.157720372 | 0.931446727 | 0.351622529 | 0.998865404 | 9.189431 | 9.336360 | 9.208625 | 9.248573 | 9.314344 | 9.278825 | 652 | 649 | 577 | 590 |
| 11214 | NM_00127054 | AKAP13 | A-kinase anchoring protein 13 | protein_coding | NP_001257475.1;NP_006729.4;NP_009131.2 | 371 | 604686 | . | . | 2.188639 | 2.317582 | 2.121935 | 2.478627 | 5.093055 | 5.357196 | 4.761559 | 5.667086 | -1.061283 | 912.617497 | 0.14955237 | -0.57377345 | 0.566121114 | 0.998865404 | 9.876443 | 9.789995 | 9.837033 | 9.871250 | 9.747463 | 9.871137 | 1057 | 1038 | 703 | 902 |
| 11215 | NM_016248,XI | AKAP11 | A-kinase anchoring protein 11 | protein_coding | NP_057332.1;XP_005266304.1;XP_005266305.1;XP_005266306.1;XP_00 | 369 | 604696 | . | . | 4.436998 | 4.452023 | 4.556614 | 4.245321 | 10.325079 | 10.291047 | 10.224911 | 9.706422 | -1.013163 | 1346.923959 | 0.12917675 | -0.14604632 | 0.883884825 | 0.998865404 | 10.404730 | 10.386116 | 10.410001 | 10.390303 | 10.417544 | 10.362323 | 1581 | 1444 | 1206 | 1209 |
| 11216 | NM_00133015 | AKAP10 | A-kinase anchoring protein 10 | protein_coding | NP_001317081.1;NP_009133.2 | 368 | 604694 | . | . | 7.228808 | 7.926378 | 7.244977 | 6.837446 | 16.821733 | 18.322179 | 16.257516 | 15.633009 | -1.019454 | 867.915973 | 0.146154236 | -0.19018269 | 0.849165975 | 0.998865404 | 9.775308 | 9.747380 | 9.725410 | 9.808444 | 9.768651 | 9.739028 | 951 | 1000 | 763 | 789 |
| 11218 | NM_007204 | DDX20 | DEAD-box helicase 20 | protein_coding | NP_009135.4 | 2743 | 606168 | . | . | 8.824690 | 8.889118 | 8.078644 | 7.299212 | 20.535416 | 20.547594 | 18.128242 | 16.688782 | -1.167492 | 928.975341 | 0.142752153 | -1.56503997 | 0.117573538 | 0.857396579 | 9.966701 | 9.743682 | 9.913086 | 9.911937 | 9.839109 | 9.760545 | 1153 | 1079 | 787 | 759 |
| 11219 | NM_080701 | TREX2 | three prime repair exonuclease 2 | protein_coding | NP_542432.2 | 12270 | 300370 | . | . | 0.092209 | 0.040260 | 0.187907 | 0.134523 | 0.214575 | 0.093063 | 0.421657 | 0.307570 | 2.148831 | 4.894714 | 1.395550027 | 0.790764719 | 0.429081301 | NA | 1.631267 | 2.742346 | 2.270001 | 2.258225 | 2.286525 | 2.274617 | 5 | 2 | 7 | 5 |
| 11221 | NM_007207,N | DUSP10 | dual specificity phosphatase 10 | protein_coding | NP_009138.1;XP_016855636.1 | 3065 | 608867 | . | . | 11.489019 | 9.932893 | 10.068904 | 10.043880 | 26.735420 | 22.960327 | 22.594331 | 22.964140 | -1.083466 | 861.633835 | 0.14735179 | -0.78488148 | 0.43252306 | 0.998865404 | 9.807465 | 9.692091 | 9.825503 | 9.729282 | 9.719093 | 9.723508 | 1100 | 903 | 717 | 778 |
| 11222 | NM_007208 | MRPL3 | mitochondrial ribosomal protein L3 | protein_coding | NP_009139.1 | 10379 | 607118 | . | . | 47.324902 | 48.714844 | 43.825752 | 44.426098 | 110.126991 | 112.606544 | 98.343719 | 101.574997 | -1.103567 | 2452.552129 | 0.114106986 | -1.24596804 | 0.212776127 | 0.975238634 | 11.329421 | 11.187213 | 11.290259 | 11.306837 | 11.217650 | 11.220471 | 2933 | 2805 | 2024 | 2190 |
| 11223 | NM_00127173 | MST1L | macrophage stimulating 1 like | protein_coding | NP_001258662.1 | 7390 | . | | . | 1.129012 | 1.189217 | 1.200430 | 1.240810 | 2.627258 | 2.748929 | 2.693730 | 2.836964 | 1.038438 | 173.741264 | 0.250405118 | 0.217306798 | 0.82796926 | 0.998865404 | 7.413496 | 7.467590 | 7.422934 | 7.441551 | 7.442854 | 7.453373 | 192 | 188 | 152 | 168 |
| 11224 | NM_007209 | RPL35 | ribosomal protein L35 | protein_coding | NP_009140.1 | 10344 | 618315 | . | . | 564.302124 | 592.616882 | 522.987122 | 693.544006 | 1313.154175 | 1369.860474 | 1173.567871 | 1585.706055 | 1.036451 | 8350.744850 | 0.132521281 | 0.389762548 | 0.69671214 | 0.998865404 | 13.001696 | 13.053222 | 12.992711 | 13.027024 | 12.918232 | 13.149635 | 9253 | 9029 | 6392 | 9046 |
| 11226 | NM_007210,XI | GALNT6 | polypeptide N-acetylgalactosaminyltransferase 6 | protein_coding | NP_009141.2;XP_005268664.1;XP_006719277.1;XP_011536121.1;XP_01 | 4128 | 605148 | . | . | 0.177428 | 0.169490 | 0.219283 | 0.153241 | 0.412883 | 0.391784 | 0.492065 | 0.350366 | 1.053584 | 30.380937 | 0.567493601 | 0.132697861 | 0.894432335 | 0.998865404 | 4.885686 | 4.963455 | 4.921927 | 4.915048 | 4.946592 | 4.902656 | 35 | 31 | 32 | 24 |
| 11228 | NM_00116474 | RASSF8 | Ras association domain family member 8 | protein_coding | NP_001158218.1;NP_001158219.1;NP_001158220.1;NP_009142.2;XP_0 | 13232 | 608231 | . | . | 13.565258 | 12.460985 | 12.152983 | 10.953487 | 31.566914 | 28.804125 | 27.270942 | 25.043845 | -1.154250 | 2157.016892 | 0.123134419 | -1.68072888 | 0.092815589 | 0.809520218 | 11.174452 | 10.967798 | 11.156342 | 11.100828 | 11.055254 | 10.971840 | 2752 | 2409 | 1834 | 1778 |
| 11230 | NM_007213 | PRAF2 | PRA1 domain family member 2 | protein_coding | NP_009144.1 | 28911 | 300840 | . | . | 14.086916 | 14.274476 | 13.479083 | 14.396460 | 32.780830 | 32.996090 | 30.246674 | 32.915794 | -1.030968 | 552.433123 | 0.162099489 | -0.27143846 | 0.786053825 | 0.998865404 | 9.131614 | 9.087359 | 9.117720 | 9.121506 | 9.081925 | 9.117212 | 644 | 607 | 460 | 524 |
| 11231 | NM_007214,XI | SEC63 | SEC63 homolog, protein translocation regulator | protein_coding | NP_009145.1;XP_016865707.1 | 21082 | 608648 | . | . | 19.476045 | 20.538166 | 18.569415 | 16.217694 | 45.321559 | 47.474891 | 41.669229 | 37.079828 | -1.185696 | 3003.654262 | 0.13973684 | -1.75854655 | 0.078654559 | 0.773268428 | 11.669974 | 11.424612 | 11.663776 | 11.565488 | 11.572564 | 11.370672 | 3973 | 3309 | 2690 | 2254 |
| 11232 | NM_007215 | POLG2 | DNA polymerase gamma 2, accessory subunit | protein_coding | NP_009146.2 | 9180 | 604983 | . | . | 6.226947 | 5.653888 | 6.405107 | 6.448778 | 14.490363 | 13.069215 | 14.372874 | 14.744365 | 1.065690 | 305.134731 | 0.201349071 | 0.4558642 | 0.648487635 | 0.998865404 | 8.206475 | 8.298658 | 8.262809 | 8.206365 | 8.269722 | 8.269133 | 358 | 302 | 274 | 295 |
| 11234 | NM_007216,N | HPS5 | HPS5 biogenesis of lysosomal organelles complex 2 subunit 2 | protein_coding | NP_009147.3;NP_852608.1;NP_852609.1;XP_011518164.1;XP_01151816 | 17022 | 607521 | . | . | 12.037587 | 12.278835 | 13.107853 | 13.029477 | 28.011965 | 28.383077 | 29.413644 | 29.790352 | 1.052382 | 1872.535065 | 0.118984942 | 0.619056521 | 0.535879125 | 0.998865404 | 10.833496 | 10.907119 | 10.848944 | 10.851368 | 10.885290 | 10.895647 | 2098 | 1972 | 1659 | 1812 |
| 11235 | NM_007217,N | PDCD10 | programmed cell death 10 | protein_coding | NP_009148.2;NP_665858.1;NP_665859.1;XP_005247143.1;XP_00524714 | 8761 | 609118 | . | . | 27.371970 | 24.325877 | 23.627292 | 16.953711 | 63.695702 | 56.230354 | 53.018952 | 38.762648 | -1.130465 | 1077.393711 | 0.141979833 | -1.24606765 | 0.212739561 | 0.975238634 | 10.158819 | 9.982454 | 10.153410 | 10.077819 | 10.067931 | 9.981503 | 1382 | 1172 | 932 | 892 |
| 11236 | NM_007218 | RNF139 | ring finger protein 139 | protein_coding | NP_009149.2 | 17023 | 603046 | . | . | 6.678995 | 6.413228 | 6.104005 | 3.573753 | 15.542295 | 14.824463 | 13.697211 | 8.170961 | -1.372608 | 568.601234 | 0.218207605 | -2.09396539 | 0.036263064 | 0.593055768 | 9.361087 | 8.905668 | 9.253900 | 9.218995 | 9.180858 | 8.851280 | 776 | 692 | 528 | 330 |
| 11237 | NM_00113433 | RNF24 | ring finger protein 24 | protein_coding | NP_001127809.1;NP_001127810.1;NP_001308678.1;NP_009150.1;XP_01 | 13779 | 612489 | . | . | 2.905630 | 2.416857 | 2.666076 | 2.642878 | 6.761520 | 5.586674 | 5.982595 | 6.042629 | 1.025728 | 556.836166 | 0.167121295 | 0.219292444 | 0.826422248 | 0.998865404 | 9.102553 | 9.139429 | 9.157456 | 9.064487 | 9.113381 | 9.143679 | 678 | 551 | 479 | 541 |
| 11243 | NM_00119965 | PMF1 | polyamine modulated factor 1 | protein_coding | NP_001186582.1;NP_001186583.1;NP_009152.2 | 9112 | 609176 | . | . | 17.713898 | 19.729945 | 19.614794 | 21.478545 | 41.220968 | 45.606654 | 44.015029 | 49.108136 | 1.083663 | 684.294088 | 0.155952786 | 0.743279699 | 0.45731236 | 0.998865404 | 9.359648 | 9.474992 | 9.355246 | 9.422569 | 9.415008 | 9.472583 | 719 | 744 | 593 | 695 |
| 11244 | NM_00101792 | ZHX1 | zinc fingers and homeoboxes 1 | protein_coding | NP_001017926.1;NP_009153.3 | 12871 | 604764 | . | . | 3.546225 | 3.409382 | 3.778496 | 3.562731 | 8.252210 | 7.880941 | 8.478835 | 8.145761 | 1.041536 | 569.641108 | 0.162853564 | 0.360528046 | 0.718452287 | 0.998865404 | 9.123985 | 9.183226 | 9.163424 | 9.115476 | 9.191445 | 9.140976 | 666 | 580 | 525 | 524 |
| 11245 | NM_00127185 | GPR176 | G protein-coupled receptor 176 | protein_coding | NP_001258783.1;NP_001258784.1;NP_009154.1;XP_011519469.1;XP_01 | 32370 | 612183 | . | . | 2.580875 | 3.028349 | 2.208166 | 2.468632 | 6.005803 | 7.000165 | 4.955062 | 5.644234 | -1.132780 | 374.031855 | 0.187513163 | -0.95922583 | 0.337444993 | 0.998865404 | 8.634450 | 8.453942 | 8.573639 | 8.590811 | 8.478390 | 8.540227 | 451 | 435 | 288 | 347 |
| 11247 | NM_007224,XI | NXPH4 | neurexophilin 4 | protein_coding | NP_009155.1;XP_016874236.1 | 8078 | 604637 | . | . | 3.660074 | 3.563193 | 3.339478 | 3.683316 | 8.517143 | 8.236480 | 7.493693 | 8.421464 | -1.030962 | 194.524427 | 0.242183013 | -0.18164189 | 0.855863772 | 0.998865404 | 7.625897 | 7.581374 | 7.623667 | 7.597393 | 7.566440 | 7.626672 | 234 | 207 | 155 | 192 |
| 11250 | NM_007227 | GPR45 | G protein-coupled receptor 45 | protein_coding | NP_009158.3 | 4503 | 604838 | . | . | 0.252046 | 0.105923 | 0.226716 | 0.369002 | 0.586521 | 0.244845 | 0.508744 | 0.843680 | 1.613153 | 13.324670 | 0.870188936 | 0.792796855 | 0.427896189 | NA | 3.349244 | 4.040734 | 3.727872 | 3.694167 | 3.722388 | 3.754819 | 16 | 7 | 11 | 19 |
| 11252 | NM_00118497 | PACSIN2 | protein kinase C and casein kinase substrate in neurons 2 | protein_coding | NP_001171899.1;NP_001171900.1;NP_001336897.1;NP_001336898.1;NI | 8571 | 604960 | . | . | 87.795404 | 88.502682 | 70.686937 | 87.030640 | 204.303501 | 204.577899 | 158.619440 | 198.985218 | -1.133407 | 8461.548468 | 0.119816921 | -1.50784997 | 0.131592929 | 0.885605656 | 13.134263 | 12.953490 | 13.096938 | 13.096091 | 12.903959 | 13.069520 | 10358 | 9697 | 6219 | 8168 |
| 11253 | NM_016219,N | MAN1B1 | mannosidase alpha class 1B member 1 | protein_coding | NP_057303.2;XP_006717008.1;XP_016869728.1;XP_024303171.1 | 6823 | 604346 | . | . | 42.426615 | 40.675382 | 33.281667 | 40.108007 | 98.728478 | 94.022967 | 74.683093 | 91.702192 | -1.145878 | 3359.603657 | 0.120468351 | -1.63074936 | 0.102943216 | 0.823247517 | 11.809040 | 11.612416 | 11.784220 | 11.749305 | 11.592062 | 11.713934 | 4218 | 3791 | 2518 | 3156 |
| 11257 | NR_015381 | TP53TG1 | TP53 target 1 | lncRNA | . | 17026 | 616403 | . | . | 6.278948 | 4.400558 | 4.147686 | 5.257813 | 14.611370 | 10.172087 | 9.307287 | 12.021363 | -1.153136 | 111.530136 | 0.326642412 | -0.62932066 | 0.529139149 | 0.998865404 | 6.900399 | 6.694866 | 6.882345 | 6.758166 | 6.740993 | 6.809050 | 162 | 106 | 80 | 108 |
| 11258 | NM_00128142 | DCTN3 | dynactin subunit 3 | protein_coding | NP_001268354.1;NP_001268355.1;NP_001268356.1;NP_009165.1;NP_0 | 2713 | 607387 | . | . | 51.559511 | 52.701906 | 41.310035 | 54.404698 | 119.981099 | 121.822821 | 92.698518 | 124.389880 | -1.097517 | 1515.816677 | 0.137854361 | -0.97380702 | 0.330152366 | 0.998865404 | 10.631660 | 10.496956 | 10.594388 | 10.605388 | 10.450570 | 10.599531 | 1814 | 1724 | 1134 | 1487 |
| 11260 | NM_007235,XI | XPOT | exportin for tRNA | protein_coding | NP_009166.2;XP_016874237.1 | 12826 | 603180 | . | . | 17.903626 | 17.151668 | 15.069447 | 15.354631 | 41.662475 | 39.646846 | 33.815405 | 35.106540 | -1.168521 | 3245.473108 | 0.11113478 | -2.02172731 | 0.043204531 | 0.639862814 | 11.772168 | 11.547522 | 11.745503 | 11.703440 | 11.593627 | 11.600303 | 4129 | 3679 | 2592 | 2818 |
| 11261 | NM_007236,XI | CHP1 | calcineurin like EF-hand protein 1 | protein_coding | NP_009167.1;XP_016877368.1 | 17433 | 606988 | . | . | 19.587039 | 19.373448 | 21.251461 | 23.367248 | 45.579847 | 44.782593 | 47.687658 | 53.426434 | 1.129634 | 2034.597437 | 0.120646944 | 1.457603932 | 0.144949745 | 0.906189003 | 10.899968 | 11.075737 | 10.955744 | 10.923107 | 11.005185 | 11.066958 | 2246 | 2019 | 1805 | 2101 |
| 11262 | NM_00100517 | SP140 | SP140 nuclear body protein | protein_coding | NP_001005176.1;NP_001265380.1;NP_001265381.1;NP_001265382.1;NI | 17133 | 608602 | . | . | 0.548691 | 0.477057 | 0.580201 | 0.617637 | 1.276827 | 1.102738 | 1.301955 | 1.412154 | -1.124713 | 28.825603 | 0.574110526 | -0.29533747 | 0.767736096 | 0.998865404 | 4.931597 | 4.761978 | 4.853551 | 4.861648 | 4.845800 | 4.841949 | 34 | 34 | 24 | 25 |
| 11264 | NM_007238,N | PXMP4 | peroxisomal membrane protein 4 | protein_coding | NP_009169.3;NP_899634.1 | 15920 | 616397 | . | . | 1.029411 | 0.985157 | 0.873309 | 1.176411 | 2.395482 | 2.277236 | 1.959679 | 2.689724 | 1.000458 | 180.059460 | 0.258373792 | 0.002556233 | 0.997960424 | 0.999862634 | 7.492619 | 7.492040 | 7.500549 | 7.481166 | 7.431260 | 7.547459 | 212 | 190 | 135 | 192 |
| 11266 | NM_007240,XI | DUSP12 | dual specificity phosphatase 12 | protein_coding | NP_009171.1;XP_005244919.1 | 3067 | 604835 | . | . | 12.621817 | 12.054184 | 10.347464 | 10.188014 | 29.371484 | 27.863785 | 23.219407 | 23.293683 | -1.198874 | 458.924833 | 0.172592533 | -1.51617255 | 0.129475728 | 0.879150687 | 8.966902 | 8.705501 | 8.910877 | 8.879790 | 8.791828 | 8.775276 | 590 | 527 | 368 | 386 |
| 11267 | NM_00131719 | SNF8 | SNF8 subunit of ESCRT-II | protein_coding | NP_001304121.1;NP_001304122.1;NP_001304123.1;NP_009172.2;XP_01 | 17028 | 610904 | . | . | 31.547329 | 36.626452 | 28.616276 | 47.242646 | 73.411929 | 84.663690 | 64.214092 | 108.014695 | -1.038543 | 2045.148682 | 0.131773564 | -0.41405201 | 0.678836016 | 0.998865404 | 11.025203 | 10.970253 | 10.992417 | 11.031022 | 10.905670 | 11.052400 | 2342 | 2303 | 1583 | 2055 |
| 11269 | NM_00101444 | DDX19B | DEAD-box helicase 19B | protein_coding | NP_001014449.1;NP_001014451.1;NP_001244101.1;NP_001244102.1;NI | 2742 | 605812 | . | . | 6.170965 | 6.571292 | 5.846490 | 6.969388 | 14.360088 | 15.189833 | 13.119356 | 15.934678 | -1.005663 | 653.579205 | 0.161258872 | -0.05051976 | 0.9597082 | 0.998865404 | 9.356650 | 9.347773 | 9.334402 | 9.372151 | 9.293962 | 9.402099 | 734 | 727 | 521 | 660 |
| 11270 | NM_00127070 | NRM | nurim | protein_coding | NP_001257636.1;NP_001257637.1;NP_001257638.1;NP_001257639.1;NI | 8003 | . | | . | 21.519441 | 21.155672 | 20.684999 | 21.930464 | 50.076625 | 48.902278 | 46.416535 | 50.141400 | -1.032695 | 970.211283 | 0.138658526 | -0.33474123 | 0.737820299 | 0.998865404 | 9.945248 | 9.898687 | 9.943984 | 9.923488 | 9.887375 | 9.932701 | 1151 | 1049 | 804 | 923 |
| 11272 | NM_00109853 | PRR4 | proline rich 4 | protein_coding | NP_001092008.2;NP_009175.2 | 18020 | 605359 | . | . | 14.790433 | 13.432278 | 16.603962 | 16.513009 | 34.417946 | 31.049311 | 37.258808 | 37.755033 | 1.156600 | 268.835348 | 0.211549786 | 0.992153472 | 0.321122666 | 0.998865404 | 7.961540 | 8.171953 | 8.057129 | 8.001497 | 8.109074 | 8.100130 | 303 | 254 | 253 | 268 |
| 11273 | NM_00130823 | ATXN2L | ataxin 2 like | protein_coding | NP_001295159.1;NP_009176.2;NP_059867.3;NP_663760.1;NP_680780.1 | 31326 | 607931 | . | . | 30.781368 | 36.228160 | 34.981558 | 39.326022 | 71.629513 | 83.743013 | 78.497602 | 89.914275 | 1.076729 | 4595.499776 | 0.120262846 | 0.886848249 | 0.375160607 | 0.998865404 | 12.111785 | 12.218263 | 12.065653 | 12.196926 | 12.162784 | 12.226293 | 4694 | 5153 | 3987 | 4636 |
| 11274 | NM_017414,XI | USP18 | ubiquitin specific peptidase 18 | protein_coding | NP_059110.2;XP_006724137.1 | 12616 | 607057 | . | . | 3.988676 | 4.009332 | 4.063051 | 3.739833 | 9.281813 | 9.267751 | 9.117369 | 8.550684 | -1.025536 | 241.253129 | 0.219121669 | -0.16601901 | 0.86814199 | 0.998865404 | 7.932268 | 7.896316 | 7.925412 | 7.915616 | 7.923975 | 7.892144 | 284 | 261 | 214 | 216 |
| 11275 | NM_00116152 | KLHL2 | kelch like family member 2 | protein_coding | NP_001154993.1;NP_001154994.1;NP_001317952.1;NP_001317953.1;NI | 6353 | 605774 | . | . | 3.552233 | 3.717664 | 3.346928 | 3.734581 | 8.266190 | 8.593549 | 7.510411 | 8.538674 | -1.025727 | 347.585346 | 0.191072538 | -0.19179243 | 0.847904798 | 0.998865404 | 8.459668 | 8.422540 | 8.442228 | 8.454653 | 8.410638 | 8.456815 | 401 | 384 | 286 | 335 |
| 11276 | NM_00116354 | SYNRG | synergin gamma | protein_coding | NP_001157016.1;NP_001157017.1;NP_001157018.1;NP_001157019.1;NI | 557 | 607291 | . | . | 5.045410 | 5.265059 | 6.472249 | 5.886322 | 11.740878 | 12.170417 | 14.523536 | 13.458381 | 1.157988 | 1353.293673 | 0.127886611 | 1.654749216 | 0.097975379 | 0.813553488 | 10.292508 | 10.504248 | 10.339119 | 10.346666 | 10.475416 | 10.433885 | 1437 | 1360 | 1297 | 1325 |
| 11277 | NM_007248,N | TREX1 | three prime repair exonuclease 1 | protein_coding | NP_009179.2;NP_338599.1 | 12269 | 606609 | . | . | 2.024756 | 1.692752 | 1.368142 | 2.017530 | 4.711690 | 3.912873 | 3.070071 | 4.612843 | -1.114220 | 61.036092 | 0.416506127 | -0.37462765 | 0.707937408 | 0.998865404 | 6.008190 | 5.850695 | 5.964539 | 5.921825 | 5.882720 | 5.956225 | 81 | 63 | 41 | 64 |
| 11278 | NM_007249,XI | KLF12 | Kruppel like factor 12 | protein_coding | NP_009180.3;XP_005266308.1;XP_011533209.1;XP_011533210.1;XP_01 | 6346 | 607531 | . | . | 0.807942 | 0.915546 | 0.956887 | 0.947488 | 1.880112 | 2.116328 | 2.147230 | 2.166319 | 1.052133 | 296.274967 | 0.203911497 | 0.35955594 | 0.719179238 | 0.998865404 | 8.173901 | 8.246764 | 8.165840 | 8.225515 | 8.220841 | 8.225581 | 314 | 329 | 263 | 286 |
| 11279 | NM_00115929 | KLF8 | Kruppel like factor 8 | protein_coding | NP_001152768.1;NP_001311028.1;NP_001311029.1;NP_001311031.1;NI | 6351 | 300286 | . | . | 0.137283 | 0.144427 | 0.107966 | 0.349127 | 0.319464 | 0.333848 | 0.242272 | 0.798236 | 1.019650 | 44.895395 | 0.465059406 | 0.06036777 | 0.951862728 | 0.998865404 | 5.474910 | 5.501955 | 5.475275 | 5.496917 | 5.483462 | 5.495420 | 48 | 51 | 38 | 44 |
| 11282 | NM_014275,N | MGAT4B | alpha-1,3-mannosyl-glycoprotein 4-beta-N-acetylglucosaminyltrar | protein_coding | NP_055090.1;NP_463459.1;XP_024310115.1;XP_024310116.1;XP_02431 | 7048 | 604561 | . | . | 96.930238 | 100.689792 | 113.691415 | 112.189000 | 225.560636 | 232.748971 | 255.120226 | 256.506817 | 1.135350 | 8292.933918 | 0.102448435 | 1.787603079 | 0.073840068 | 0.763883535 | 12.923194 | 13.106332 | 12.947521 | 12.974914 | 13.080286 | 13.057629 | 8798 | 8518 | 7770 | 8158 |
| 11284 | NM_007254 | PNKP | polynucleotide kinase 3'-phosphatase | protein_coding | NP_009185.2 | 9154 | 605610 | . | . | 20.206030 | 22.181732 | 23.254232 | 22.323538 | 47.020267 | 51.274063 | 52.181824 | 51.040115 | 1.060607 | 1185.960818 | 0.134232032 | 0.632414751 | 0.527115897 | 0.998865404 | 10.168790 | 10.253644 | 10.156377 | 10.220167 | 10.252167 | 10.213331 | 1269 | 1295 | 1089 | 1115 |
| 11285 | NM_007255,XI | B4GALT7 | beta-1,4-galactosyltransferase 7 | protein_coding | NP_009186.1;XP_006714879.1;XP_016864488.1 | 930 | 604327 | . | . | 24.259627 | 24.172194 | 22.057641 | 24.731490 | 56.453156 | 55.875115 | 49.496706 | 56.545613 | -1.049174 | 1263.193081 | 0.132656163 | -0.5220519 | 0.601634192 | 0.998865404 | 10.337203 | 10.267679 | 10.324956 | 10.316284 | 10.245021 | 10.321914 | 1498 | 1388 | 1017 | 1215 |
| 11309 | NM_00114521 | SLCO2B1 | solute carrier organic anion transporter family member 2B1 | protein_coding | NP_001138683.1;NP_001138684.1;NP_009187.1;XP_016872646.1 | 10962 | 604988 | . | . | 0.615055 | 0.672274 | 0.635340 | 0.452607 | 1.431258 | 1.553992 | 1.425684 | 1.034832 | -1.150498 | 80.198133 | 0.366338886 | -0.55210851 | 0.580874012 | 0.998865404 | 6.422299 | 6.221730 | 6.328185 | 6.355105 | 6.346053 | 6.262169 | 94 | 97 | 76 | 58 |
| 11311 | NM_00127935 | VPS45 | vacuolar protein sorting 45 homolog | protein_coding | NP_001266282.1;NP_001266283.1;NP_009190.2;XP_024308559.1 | 14579 | 610035 | . | . | 13.163946 | 12.346883 | 13.032116 | 12.664350 | 30.633043 | 28.540374 | 29.243692 | 28.955532 | -1.005742 | 932.816654 | 0.140812118 | -0.05866552 | 0.953218524 | 0.998865404 | 9.869430 | 9.861459 | 9.894591 | 9.839241 | 9.874815 | 9.851510 | 1118 | 971 | 822 | 858 |
| 11313 | NM_007260,XI | LYPLA2 | lysophospholipase 2 | protein_coding | NP_009191.1;XP_005245785.1;XP_016855654.1;XP_024308564.1;XP_02 | 6738 | 616143 | . | . | 41.605399 | 38.318200 | 42.697762 | 45.552896 | 96.817473 | 88.574237 | 95.812541 | 104.151285 | 1.088143 | 2490.361018 | 0.119822249 | 1.017069845 | 0.309120204 | 0.998865404 | 11.220000 | 11.341712 | 11.257327 | 11.235584 | 11.267521 | 11.359874 | 2784 | 2539 | 2129 | 2571 |
| 11315 | NM_00112337 | PARK7 | Parkinsonism associated deglycase | protein_coding | NP_001116849.1;NP_009193.2;XP_005263481.1 | 16369 | 602533 | . | . | 277.623483 | 288.426772 | 249.827306 | 276.980470 | 646.041278 | 666.711407 | 560.605240 | 633.282921 | -1.040900 | 8241.665109 | 0.117315823 | -0.49295418 | 0.622044964 | 0.998865404 | 13.037400 | 12.979459 | 13.012734 | 13.036289 | 12.912605 | 13.062899 | 9547 | 9197 | 6400 | 8243 |
| 11316 | NM_00133046 | COPE | COPI coat complex subunit epsilon | protein_coding | NP_001317398.1;NP_009194.2;NP_955474.1;NP_955476.1;XP_0115259 | 2234 | 606942 | . | . | 82.209442 | 78.724833 | 70.879231 | 81.125913 | 191.304755 | 181.975973 | 159.050932 | 185.484756 | -1.071149 | 2730.624639 | 0.120391739 | -0.82364305 | 0.410142406 | 0.998865404 | 11.463819 | 11.364503 | 11.460540 | 11.422904 | 11.333780 | 11.435593 | 3327 | 2978 | 2148 | 2628 |
| 11319 | NM_00113575 | ECD | ecdysoneless cell cycle regulator | protein_coding | NP_001129224.1;NP_001129225.1;NP_009196.1 | 17029 | 616464 | . | . | 14.315281 | 15.136912 | 13.780619 | 15.814916 | 33.312245 | 34.989650 | 30.923312 | 36.158926 | 1.003299 | 1367.620053 | 0.133055336 | 0.0357115 | 0.9715124 | 0.998865404 | 10.415271 | 10.419630 | 10.400805 | 10.430487 | 10.368951 | 10.465083 | 1542 | 1502 | 1116 | 1365 |
| 11320 | NM_00116015 | MGAT4A | alpha-1,3-mannosyl-glycoprotein 4-beta-N-acetylglucosaminyltrar | protein_coding | NP_001153626.1;NP_036346.1 | 7047 | 604623 | . | . | 1.242754 | 0.982131 | 1.642631 | 1.209078 | 2.891939 | 2.270239 | 3.686015 | 2.764414 | 1.029791 | 293.379324 | 0.213759427 | 0.198129212 | 0.84294397 | 0.998865404 | 8.175151 | 8.217781 | 8.243652 | 8.127280 | 8.176004 | 8.228933 | 369 | 278 | 246 | 293 |
| 11321 | NM_00114504 | GPN1 | GPN-loop GTPase 1 | protein_coding | NP_001138519.1;NP_001138520.1;NP_001138521.1;NP_009197.3 | 17030 | 611479 | . | . | 12.857313 | 13.176785 | 11.979424 | 13.071518 | 29.919496 | 30.458726 | 26.881481 | 29.886475 | -1.006635 | 950.865517 | 0.140036163 | -0.06812748 | 0.945684158 | 0.998865404 | 9.897958 | 9.888221 | 9.904937 | 9.885775 | 9.862030 | 9.918272 | 1113 | 1016 | 792 | 923 |
| 11322 | NM_00112719 | TMC6 | transmembrane channel like 6 | protein_coding | NP_001120670.1;NP_001308114.1;NP_001361522.1;NP_001361523.1;NI | 18021 | 605828 | . | . | 26.774926 | 30.011936 | 23.380708 | 30.049426 | 62.306361 | 69.373933 | 52.465622 | 68.704439 | -1.161891 | 3290.399591 | 0.112943381 | -1.91666265 | 0.055280804 | 0.697001222 | 11.788249 | 11.571729 | 11.764754 | 11.719518 | 11.594299 | 11.643543 | 4183 | 3713 | 2566 | 2941 |
| 11325 | NM_007372,N | DDX42 | DEAD-box helicase 42 | protein_coding | NP_031398.2;NP_987095.1;XP_006721720.1;XP_011522561.1;XP_01687 | 18676 | 613369 | . | . | 28.664866 | 29.326244 | 34.659269 | 32.462168 | 66.704324 | 67.788918 | 77.774396 | 74.220890 | 1.101508 | 3879.445729 | 0.10969889 | 1.271483536 | 0.203556681 | 0.968525635 | 11.850182 | 11.989715 | 11.870417 | 11.889483 | 11.984665 | 11.934906 | 4202 | 4030 | 3642 | 3699 |
| 11328 | NM_00128434 | FKBP9 | FKBP prolyl isomerase 9 | protein_coding | NP_001271270.1;NP_001271272.1;NP_009201.2;XP_011513417.1 | 3725 | 616257 | . | . | 40.876699 | 45.164790 | 38.730876 | 43.596872 | 95.121759 | 104.400427 | 86.910966 | 99.679066 | -1.052086 | 4445.255334 | 0.115120837 | -0.63631525 | 0.524570968 | 0.998865404 | 12.154298 | 12.080869 | 12.111200 | 12.164842 | 12.045693 | 12.143915 | 5081 | 5076 | 3544 | 4302 |
| 11329 | NM_00130510 | STK38 | serine/threonine kinase 38 | protein_coding | NP_001292031.1;NP_009202.1;XP_006715051.1;XP_006715052.1;XP_01 | 17847 | 606964 | . | . | 9.154750 | 9.839641 | 10.343732 | 9.876968 | 21.303478 | 22.744771 | 23.211034 | 22.582516 | 1.047510 | 1072.345844 | 0.135938457 | 0.492609525 | 0.622288516 | 0.998865404 | 10.032683 | 10.099649 | 10.025046 | 10.071816 | 10.100665 | 10.065149 | 1168 | 1166 | 977 | 1004 |
| 11331 | NM_00114483 | PHB2 | prohibitin 2 | protein_coding | NP_001138303.1;NP_001254629.1 | 30306 | 610704 | . | . | 80.228285 | 88.955618 | 74.762746 | 92.691520 | 186.694520 | 205.624898 | 167.765427 | 211.928160 | -1.024711 | 3612.799333 | 0.129611352 | -0.27170782 | 0.785846689 | 0.998865404 | 11.836540 | 11.801044 | 11.787231 | 11.866148 | 11.719761 | 11.887752 | 4011 | 4134 | 2787 | 3686 |
| 11332 | NM_007274,N | ACOT7 | acyl-CoA thioesterase 7 | protein_coding | NP_009205.3;NP_863654.1;NP_863655.1;NP_863656.1 | 24157 | 602587 | . | . | 177.188219 | 183.722949 | 155.488583 | 189.700825 | 412.324252 | 424.683834 | 348.911877 | 433.728394 | -1.084148 | 9201.034846 | 0.116622089 | -0.99948674 | 0.317558958 | 0.998865404 | 13.224733 | 13.108070 | 13.189473 | 13.210628 | 13.054018 | 13.203169 | 10887 | 10457 | 7001 | 9007 |
| 11333 | NM_014891 | PDAP1 | PDGFA associated protein 1 | protein_coding | NP_055706.1 | 14634 | 607075 | . | . | 22.709227 | 22.182314 | 19.495079 | 22.764427 | 52.845306 | 51.275410 | 43.746395 | 52.048157 | -1.077375 | 1768.007708 | 0.129042783 | -0.83321268 | 0.404724795 | 0.998865404 | 10.840797 | 10.733008 | 10.828810 | 10.803059 | 10.699611 | 10.812004 | 2146 | 1947 | 1373 | 1711 |
| 11334 | NM_007275 | TUSC2 | tumor suppressor 2, mitochondrial calcium regulator | protein_coding | NP_009206.1 | 17034 | 607052 | . | . | 11.199728 | 12.751726 | 9.736077 | 11.769949 | 26.062229 | 29.476185 | 21.847473 | 26.910591 | -1.128734 | 596.642049 | 0.16986759 | -1.02848266 | 0.303722844 | 0.998865404 | 9.305886 | 9.130213 | 9.216785 | 9.298575 | 9.119331 | 9.233567 | 685 | 724 | 444 | 572 |
| 11335 | NM_007276,N | CBX3 | chromobox 3 | protein_coding | NP_009207.2;NP_057671.2;XP_005249668.1 | 1553 | 604477 | . | . | 79.957973 | 69.017738 | 79.016498 | 68.459866 | 186.065487 | 159.537588 | 177.310726 | 156.525357 | -1.052919 | 5122.032325 | 0.12624806 | -0.58927663 | 0.555675723 | 0.998865404 | 12.359118 | 12.284930 | 12.399216 | 12.281893 | 12.368052 | 12.225135 | 6470 | 5275 | 4711 | 4278 |
| 11336 | NM_007277 | EXOC3 | exocyst complex component 3 | protein_coding | NP_009208.2 | 30378 | 608186 | . | . | 9.346885 | 8.856119 | 10.722635 | 9.317157 | 21.750582 | 20.471315 | 24.061283 | 21.302570 | 1.086052 | 834.975351 | 0.150910286 | 0.789161999 | 0.430017329 | 0.998865404 | 9.644492 | 9.764205 | 9.695553 | 9.652131 | 9.785346 | 9.677703 | 950 | 837 | 813 | 754 |
| 11337 | NM_007278 | GABARAP | GABA type A receptor-associated protein | protein_coding | NP_009209.1 | 4067 | 605125 | . | . | 62.723923 | 66.155342 | 64.961555 | 81.258034 | 145.961151 | 152.921051 | 145.771835 | 185.786835 | 1.118382 | 2824.309509 | 0.131120258 | 1.231028339 | 0.218312266 | 0.982008277 | 11.380879 | 11.541990 | 11.396511 | 11.432701 | 11.414280 | 11.589997 | 3002 | 2942 | 2317 | 3093 |
| 11338 | NM_00101247 | U2AF2 | U2 small nuclear RNA auxiliary factor 2 | protein_coding | NP_001012496.1;NP_009210.1;XP_006723057.1;XP_011524712.1 | 23156 | 191318 | . | . | 85.344617 | 91.278874 | 98.389096 | 101.187425 | 198.600432 | 210.995208 | 220.782276 | 231.353019 | 1.126597 | 7139.669404 | 0.104298024 | 1.648851437 | 0.099178074 | 0.813553488 | 12.713122 | 12.885042 | 12.727230 | 12.770312 | 12.837742 | 12.861653 | 7533 | 7432 | 6486 | 7191 |
| 11339 | NM_00131786 | OIP5 | Opa interacting protein 5 | protein_coding | NP_001304789.1;NP_009211.1 | 20300 | 606020 | . | . | 8.966822 | 9.121513 | 8.096057 | 7.806398 | 20.866164 | 21.084785 | 18.167318 | 17.848398 | -1.152359 | 318.259920 | 0.196460246 | -1.04138067 | 0.29769891 | 0.998865404 | 8.412633 | 8.208259 | 8.349992 | 8.353809 | 8.286171 | 8.262132 | 391 | 369 | 263 | 271 |
| 11340 | NM_181503,XI | EXOSC8 | exosome component 8 | protein_coding | NP_852480.1;XP_006719826.1 | 17035 | 606019 | . | . | 19.947147 | 20.796996 | 21.857159 | 20.425570 | 46.417830 | 48.073189 | 49.046832 | 46.700639 | 1.007822 | 768.971744 | 0.148730833 | 0.075581356 | 0.93975217 | 0.998865404 | 9.581079 | 9.592472 | 9.565151 | 9.601771 | 9.616433 | 9.561053 | 859 | 848 | 698 | 695 |
| 11341 | NM_00132959 | SCRG1 | stimulator of chondrogenesis 1 | protein_coding | NP_001316526.1;NP_009212.1;XP_024309650.1 | 17036 | 603163 | . | . | 0.363953 | 0.386437 | 0.301095 | 0.287039 | 0.846934 | 0.893266 | 0.675649 | 0.656280 | -1.282040 | 42.340183 | 0.478613378 | -0.74891689 | 0.453907298 | 0.998865404 | 5.572054 | 5.213655 | 5.420086 | 5.431647 | 5.389358 | 5.382599 | 53 | 53 | 33 | 34 |
| 11342 | NM_00137828 | RNF13 | ring finger protein 13 | protein_coding | NP_001365214.1;NP_001365215.1;NP_001365216.1;NP_001365217.1;NI | 10057 | 609247 | . | . | 19.672570 | 17.435175 | 18.938439 | 15.305916 | 45.778880 | 40.302189 | 42.497311 | 34.995156 | -1.097596 | 1350.531372 | 0.146416965 | -0.91756435 | 0.358846989 | 0.998865404 | 10.464617 | 10.330909 | 10.484497 | 10.376748 | 10.431820 | 10.285107 | 1740 | 1419 | 1229 | 1090 |
| 11343 | NM_00100379 | MGLL | monoglyceride lipase | protein_coding | NP_001003794.1;NP_001243514.1;NP_009214.1;XP_011510679.1;XP_01 | 17038 | 609699 | . | . | 13.769707 | 14.064742 | 10.134748 | 13.108026 | 32.042675 | 32.511279 | 22.742080 | 29.969945 | -1.219615 | 1761.054859 | 0.138549022 | -2.0673224 | 0.038703785 | 0.61335056 | 10.918547 | 10.631663 | 10.851796 | 10.855869 | 10.608981 | 10.779440 | 2223 | 2094 | 1226 | 1653 |
| 11344 | NM_007284 | TWF2 | twinfilin actin binding protein 2 | protein_coding | NP_009215.1 | 9621 | 607433 | . | . | 22.720819 | 23.472139 | 17.415213 | 23.796947 | 52.872280 | 54.256905 | 39.079231 | 54.408894 | -1.136729 | 1099.065935 | 0.16017373 | -1.15429646 | 0.248378646 | 0.992733415 | 10.191892 | 10.006266 | 10.136509 | 10.154292 | 9.934626 | 10.153650 | 1331 | 1277 | 760 | 1109 |
| 11345 | NM_007285 | GABARAPL2 | GABA type A receptor associated protein like 2 | protein_coding | NP_009216.1 | 13291 | 607452 | . | . | 33.979195 | 29.054012 | 29.434444 | 32.098087 | 79.070984 | 67.159653 | 66.050034 | 73.388466 | -1.038631 | 945.673462 | 0.148983329 | -0.36704294 | 0.713586989 | 0.998865404 | 9.912226 | 9.857655 | 9.956750 | 9.835587 | 9.842668 | 9.897506 | 1201 | 954 | 776 | 903 |
| 11346 | NM_00110997 | SYNPO | synaptopodin | protein_coding | NP_001103444.1;NP_001159680.1;NP_001159681.1;NP_009217.3;XP_0 | 30672 | 608155 | . | . | 0.070644 | 0.064968 | 0.108124 | 0.081868 | 0.164391 | 0.150178 | 0.242628 | 0.187183 | 1.318472 | 16.623551 | 0.753882646 | 0.529083587 | 0.596747466 | NA | 3.841203 | 4.242203 | 4.039525 | 4.035806 | 4.069407 | 4.045205 | 17 | 15 | 19 | 15 |
| 22794 | NM_007359,XI | CASC3 | CASC3 exon junction complex subunit | protein_coding | NP_031385.2;XP_005257220.1 | 17040 | 606504 | . | . | 14.101599 | 15.483041 | 17.236857 | 18.284125 | 32.814999 | 35.789742 | 38.679005 | 41.804481 | 1.182906 | 2009.561416 | 0.122328762 | 1.981018321 | 0.047589219 | 0.659753773 | 10.846545 | 11.088640 | 10.864447 | 10.937042 | 11.015055 | 11.054237 | 2022 | 2079 | 1848 | 2091 |
| 22796 | NM_00114503 | COG2 | component of oligomeric golgi complex 2 | protein_coding | NP_001138508.1;NP_031383.1 | 6546 | 606974 | . | . | 8.277011 | 8.028680 | 7.828798 | 9.096072 | 19.260944 | 18.558654 | 17.567596 | 20.797086 | 1.023414 | 759.619307 | 0.152653335 | 0.218729815 | 0.826860522 | 0.998865404 | 9.552534 | 9.585543 | 9.573534 | 9.547157 | 9.527227 | 9.623477 | 881 | 795 | 622 | 770 |
| 22800 | NM_00110266 | RRAS2 | RAS related 2 | protein_coding | NP_001096139.1;NP_001170785.1;NP_001170786.1;NP_036382.2;XP_01 | 17271 | 600098 | . | . | 4.690748 | 3.839636 | 4.259241 | 4.201656 | 10.915562 | 8.875492 | 9.557613 | 9.606587 | -1.026340 | 310.120506 | 0.20818613 | -0.18016607 | 0.857022195 | 0.998865404 | 8.294901 | 8.258236 | 8.341030 | 8.220428 | 8.276010 | 8.260881 | 401 | 302 | 270 | 283 |
| 22803 | NM_00131796 | XRN2 | 5'-3' exoribonuclease 2 | protein_coding | NP_001304889.1;NP_036387.2;XP_016883211.1;XP_016883212.1;XP_01 | 12836 | 608851 | . | . | 30.558256 | 31.714458 | 28.833636 | 30.796176 | 71.110321 | 73.309392 | 64.701839 | 70.411800 | -1.059080 | 3237.803434 | 0.111877862 | -0.74019325 | 0.459182742 | 0.998865404 | 11.701670 | 11.618737 | 11.671632 | 11.695943 | 11.613000 | 11.659840 | 3779 | 3647 | 2656 | 3033 |
| 22808 | NM_00108504 | MRAS | muscle RAS oncogene homolog | protein_coding | NP_001078518.1;NP_001239019.1;NP_001239020.1;NP_001239021.1;NI | 7227 | 608435 | . | . | 6.962964 | 6.778147 | 6.083026 | 10.058593 | 16.203103 | 15.667992 | 13.650135 | 22.997779 | 1.129765 | 981.352733 | 0.193673306 | 0.908866443 | 0.363420629 | 0.998865404 | 9.848345 | 10.023592 | 9.882656 | 9.886114 | 9.777224 | 10.138528 | 1059 | 997 | 687 | 1211 |
| 22809 | NM_00119364 | ATF5 | activating transcription factor 5 | protein_coding | NP_001180575.1;NP_001277675.1;NP_036200.2;XP_011524931.1 | 790 | 606398 | . | . | 11.545834 | 14.415501 | 14.666591 | 12.734801 | 26.867629 | 33.322080 | 32.911404 | 29.116607 | 1.020653 | 901.648060 | 0.146892283 | 0.200771921 | 0.840876919 | 0.998865404 | 9.801571 | 9.831119 | 9.767559 | 9.847127 | 9.857565 | 9.786915 | 971 | 1016 | 831 | 812 |
| 22818 | NM_00127173 | COPZ1 | COPI coat complex subunit zeta 1 | protein_coding | NP_001258663.1;NP_001258664.1;NP_001258665.1;NP_057141.1 | 2243 | 615472 | . | . | 34.922833 | 36.048639 | 34.925954 | 39.592629 | 81.266874 | 83.328040 | 78.372830 | 90.523838 | 1.044068 | 2151.844473 | 0.121389651 | 0.512525748 | 0.608283105 | 0.998865404 | 11.040035 | 11.102015 | 11.044694 | 11.062219 | 11.044446 | 11.129571 | 2398 | 2297 | 1810 | 2170 |
| 22820 | NM_016128,XI | COPG1 | COPI coat complex subunit gamma 1 | protein_coding | NP_057212.1;XP_011510851.1 | 2236 | 615525 | . | . | 58.570930 | 65.409524 | 59.198088 | 69.751470 | 136.296969 | 151.197044 | 132.838807 | 159.478452 | 1.026041 | 6037.450363 | 0.119610265 | 0.310081923 | 0.756498658 | 0.998865404 | 12.541142 | 12.578069 | 12.505255 | 12.588639 | 12.501822 | 12.632739 | 6523 | 6750 | 4905 | 6177 |
| 22821 | NM_00132082 | RASA3 | RAS p21 protein activator 3 | protein_coding | NP_001307750.1;NP_001307751.1;NP_031394.2;XP_011533143.1;XP_01 | 20331 | 605182 | . | . | 11.114955 | 11.132525 | 10.686464 | 13.044174 | 25.864960 | 25.733329 | 23.980117 | 29.823954 | 1.065717 | 1526.013674 | 0.134389902 | 0.683273217 | 0.494434215 | 0.998865404 | 10.529098 | 10.620557 | 10.549657 | 10.549251 | 10.531040 | 10.662068 | 1702 | 1594 | 1254 | 1600 |
| 22822 | NM_007350 | PHLDA1 | pleckstrin homology like domain family A member 1 | protein_coding | NP_031376.3 | 8933 | 605335 | . | . | 22.903946 | 22.354570 | 22.600153 | 22.520193 | 53.298428 | 51.673592 | 50.714088 | 51.489750 | -1.017116 | 4162.859807 | 0.107465779 | -0.22783492 | 0.819774581 | 0.998865404 | 12.035526 | 12.011089 | 12.043859 | 12.016636 | 12.021942 | 12.010529 | 4913 | 4456 | 3614 | 3843 |
| 22823 | NM_00116439 | MTF2 | metal response element binding transcription factor 2 | protein_coding | NP_001157863.1;NP_001157864.1;NP_001157865.1;NP_031384.1 | 29535 | 609882 | . | . | 4.156438 | 4.024263 | 4.020342 | 4.075452 | 9.672200 | 9.302264 | 9.021529 | 9.318038 | -1.024302 | 515.785067 | 0.165368306 | -0.20947826 | 0.834074904 | 0.998865404 | 9.027793 | 8.993253 | 9.031150 | 9.005357 | 9.001398 | 9.004629 | 613 | 552 | 442 | 479 |
| 22824 | NM_00131738 | HSPA4L | heat shock protein family A (Hsp70) member 4 like | protein_coding | NP_001304310.1;NP_001304311.1;NP_001304312.1;NP_055093.2;XP_01 | 17041 | . | | . | 1.543542 | 1.473466 | 0.936294 | 0.954760 | 3.591885 | 3.405983 | 2.101017 | 2.182946 | -1.454761 | 375.651621 | 0.193891041 | -2.78910169 | 0.005285447 | 0.221747783 | 8.798196 | 8.258040 | 8.706889 | 8.587210 | 8.437939 | 8.426032 | 561 | 435 | 269 | 284 |
| 22826 | NM_014280,N | DNAJC8 | DnaJ heat shock protein family (Hsp40) member C8 | protein_coding | NP_055095.2 | 15470 | . | | . | 65.538005 | 63.798313 | 56.208761 | 66.862428 | 152.509633 | 147.472665 | 126.130835 | 152.872998 | -1.066033 | 3473.657369 | 0.121792582 | -0.75745021 | 0.448780187 | 0.998865404 | 11.807713 | 11.715286 | 11.802480 | 11.771845 | 11.664815 | 11.799753 | 4204 | 3797 | 2683 | 3412 |
| 22827 | NM_00113603 | PUF60 | poly(U) binding splicing factor 60 | protein_coding | NP_001129505.1;NP_001258025.1;NP_001258026.1;NP_001258027.1;NI | 17042 | 604819 | . | . | 56.092383 | 60.996817 | 56.163065 | 61.803333 | 130.529282 | 140.996873 | 126.028284 | 141.305979 | -1.012756 | 3599.076866 | 0.112554498 | -0.16246896 | 0.870936571 | 0.998865404 | 11.822601 | 11.804163 | 11.795587 | 11.840756 | 11.777888 | 11.837052 | 4057 | 4015 | 2998 | 3473 |
| 22828 | NM_00128618 | SCAF8 | SR-related CTD associated factor 8 | protein_coding | NP_001273117.1;NP_001273118.1;NP_001273123.1;NP_001273128.1;NI | 20959 | 616024 | . | . | 6.011694 | 6.383619 | 6.054975 | 6.084258 | 13.989457 | 14.756021 | 13.587188 | 13.910933 | -1.033253 | 979.577417 | 0.137585358 | -0.34301734 | 0.731585414 | 0.998865404 | 9.959469 | 9.912175 | 9.929393 | 9.965952 | 9.925782 | 9.922052 | 1118 | 1101 | 840 | 901 |
| 22832 | NM_00128620 | CEP162 | centrosomal protein 162 | protein_coding | NP_001273135.1;NP_055710.2;XP_005248731.1;XP_006715443.1;XP_01 | 21107 | 610201 | . | . | 1.387130 | 1.454628 | 1.435372 | 1.060412 | 3.227911 | 3.362438 | 3.220934 | 2.424505 | -1.083175 | 214.098245 | 0.238389329 | -0.48351881 | 0.628727406 | 0.998865404 | 7.798141 | 7.683854 | 7.745287 | 7.772585 | 7.771576 | 7.669660 | 249 | 247 | 199 | 171 |
| 22834 | NM_00114536 | ZNF652 | zinc finger protein 652 | protein_coding | NP_001138837.1;NP_055712.1;XP_024306421.1;XP_024306422.1;XP_02 | 29147 | 613907 | . | . | 1.192550 | 1.158506 | 1.325596 | 1.534404 | 2.775115 | 2.677938 | 2.974600 | 3.508232 | 1.094456 | 418.751609 | 0.18867642 | 0.690146941 | 0.490101786 | 0.998865404 | 8.642804 | 8.774111 | 8.728631 | 8.628226 | 8.776905 | 8.689726 | 499 | 395 | 408 | 381 |
| 22835 | NM_00132066 | ZFP30 | ZFP30 zinc finger protein | protein_coding | NP_001307595.1;NP_001307596.1;NP_001307597.1;NP_001307598.1;NI | 29555 | 617317 | . | . | 1.049826 | 0.800632 | 1.045119 | 0.690271 | 2.442989 | 1.850696 | 2.345216 | 1.578223 | -1.054129 | 171.107471 | 0.286221132 | -0.26570651 | 0.790465252 | 0.998865404 | 7.454989 | 7.381596 | 7.477495 | 7.366568 | 7.482293 | 7.313782 | 228 | 165 | 175 | 124 |
| 22836 | NM_014899,XI | RHOBTB3 | Rho related BTB domain containing 3 | protein_coding | NP_055714.3;XP_011541581.1;XP_016864725.1;XP_016864726.1 | 18757 | 607353 | . | . | 13.788124 | 13.520025 | 11.116264 | 10.621983 | 32.085532 | 31.252145 | 24.944572 | 24.285899 | -1.268971 | 2031.075343 | 0.11907439 | -2.88609079 | 0.003900596 | 0.182508233 | 11.149575 | 10.806084 | 11.084212 | 11.065878 | 10.907482 | 10.863614 | 2645 | 2424 | 1601 | 1631 |
| 22837 | NM_00127845 | COBLL1 | cordon-bleu WH2 repeat protein like 1 | protein_coding | NP_001265387.1;NP_001265389.1;NP_001265390.1;NP_001352599.1;NI | 23571 | 610318 | . | . | 1.594327 | 1.758832 | 1.653916 | 1.842267 | 3.710066 | 4.065618 | 3.711340 | 4.212125 | 1.029580 | 483.566829 | 0.172001659 | 0.244505567 | 0.806839291 | 0.998865404 | 8.896702 | 8.938143 | 8.887183 | 8.928125 | 8.895383 | 8.955792 | 531 | 531 | 406 | 482 |
| 22838 | NM_014901,XI | RNF44 | ring finger protein 44 | protein_coding | NP_055716.1;XP_005265897.1;XP_005265898.1;XP_005265899.1;XP_00 | 19180 | . | | . | 6.904533 | 7.150414 | 7.265058 | 7.147269 | 16.067132 | 16.528503 | 16.302583 | 16.341384 | 1.017943 | 900.629032 | 0.140344958 | 0.182816882 | 0.854941702 | 0.998865404 | 9.801886 | 9.827578 | 9.796987 | 9.819242 | 9.832608 | 9.809248 | 1012 | 978 | 803 | 838 |
| 22839 | NM_00104248 | DLGAP4 | DLG associated protein 4 | protein_coding | NP_001035951.1;NP_001352550.1;NP_055717.2;NP_892118.1;XP_0052 | 24476 | 616191 | . | . | 24.153714 | 25.981857 | 22.138538 | 23.587722 | 56.206686 | 60.058228 | 49.678236 | 53.930521 | -1.112499 | 2676.615390 | 0.116862393 | -1.31610792 | 0.188137825 | 0.960264078 | 11.461142 | 11.307138 | 11.400714 | 11.454048 | 11.312685 | 11.368872 | 3141 | 3141 | 2126 | 2459 |
| 22841 | NM_00133016 | RAB11FIP2 | RAB11 family interacting protein 2 | protein_coding | NP_001317096.1;NP_055719.1;XP_016871415.1 | 29152 | 608599 | . | . | 2.323036 | 2.454881 | 2.555391 | 2.179152 | 5.405801 | 5.674568 | 5.734224 | 4.982374 | -1.021411 | 475.126387 | 0.176997972 | -0.17268009 | 0.862902898 | 0.998865404 | 8.907091 | 8.877088 | 8.882939 | 8.912191 | 8.934867 | 8.831593 | 540 | 530 | 443 | 404 |
| 22843 | NM_014906,N | PPM1E | protein phosphatase, Mg2+/Mn2+ dependent 1E | protein_coding | NP_055721.3;XP_016879848.1;XP_024306425.1 | 19322 | . | | . | 1.277952 | 1.084011 | 0.987535 | 1.302597 | 2.973847 | 2.505740 | 2.216000 | 2.978232 | -1.046385 | 237.292249 | 0.237123521 | -0.27586576 | 0.78265117 | 0.998865404 | 7.923149 | 7.857138 | 7.944482 | 7.853017 | 7.815233 | 7.936328 | 305 | 238 | 176 | 245 |
| 22844 | NM_00137122 | FRMPD1 | FERM and PDZ domain containing 1 | protein_coding | NP_001358152.1;NP_001358153.1;NP_001358154.1;NP_055722.2;XP_01 | 29159 | 616919 | . | . | 0.054814 | 0.042752 | 0.039207 | 0.072214 | 0.127553 | 0.098824 | 0.087979 | 0.165109 | 1.117841 | 10.386736 | 0.955404461 | 0.168217052 | 0.866412527 | NA | 3.295021 | 3.453945 | 3.379879 | 3.366408 | 3.365671 | 3.390831 | 13 | 9 | 7 | 13 |
| 22845 | NM_014908 | DOLK | dolichol kinase | protein_coding | NP_055723.1 | 23406 | 610746 | . | . | 10.360583 | 11.973063 | 9.324468 | 10.570409 | 24.109503 | 27.676273 | 20.923834 | 24.167984 | -1.137607 | 682.426753 | 0.160500198 | -1.15889148 | 0.246500421 | 0.992733415 | 9.504889 | 9.318133 | 9.407685 | 9.502547 | 9.329021 | 9.405533 | 780 | 837 | 523 | 633 |
| 22846 | NM_014909,XI | VASH1 | vasohibin 1 | protein_coding | NP_055724.1;XP_016876576.1;XP_016876577.1;XP_016876578.1;XP_01 | 19964 | 609011 | . | . | 1.990975 | 1.688864 | 2.130030 | 1.740614 | 4.633082 | 3.903882 | 4.779724 | 3.979708 | 1.117711 | 400.094854 | 0.187430117 | 0.856571769 | 0.391681593 | 0.998865404 | 8.561330 | 8.722564 | 8.658066 | 8.555814 | 8.681955 | 8.666891 | 473 | 372 | 372 | 391 |
| 22847 | NM_00113615 | ZNF507 | zinc finger protein 507 | protein_coding | NP_001129628.1;NP_055725.2 | 23783 | . | | . | 2.942760 | 2.939805 | 3.244355 | 3.144219 | 6.847924 | 6.795491 | 7.280237 | 7.188884 | 1.072766 | 737.761366 | 0.148139208 | 0.684054018 | 0.493941056 | 0.998865404 | 9.475358 | 9.576878 | 9.505243 | 9.497990 | 9.564868 | 9.535778 | 824 | 764 | 678 | 701 |
| 22848 | NM_00137157 | AAK1 | AP2 associated kinase 1 | protein_coding | NP_001358504.1;NP_001358506.1;NP_055726.4 | 19679 | 616405 | . | . | 2.324821 | 3.506083 | 2.454088 | 2.282926 | 5.409956 | 8.104468 | 5.506902 | 5.219639 | 1.014664 | 1539.406785 | 0.126488348 | 0.166036932 | 0.868127886 | 0.998865404 | 10.577490 | 10.598749 | 10.593305 | 10.570862 | 10.624877 | 10.561166 | 1785 | 1625 | 1404 | 1394 |
| 22849 | NM_00117813 | CPEB3 | cytoplasmic polyadenylation element binding protein 3 | protein_coding | NP_001171608.1;NP_055727.3;XP_005269687.1;XP_006717777.1;XP_00 | 21746 | 610606 | . | . | 0.217010 | 0.147100 | 0.132179 | 0.142393 | 0.504991 | 0.340029 | 0.296607 | 0.325565 | -1.108593 | 33.321630 | 0.536794311 | -0.27707155 | 0.781725169 | 0.998865404 | 5.130737 | 4.982216 | 5.057594 | 5.073969 | 5.059021 | 5.045282 | 38 | 40 | 29 | 28 |
| 22850 | NM_014913,XI | ADNP2 | ADNP homeobox 2 | protein_coding | NP_055728.1;XP_005266713.1;XP_011524185.1 | 23803 | 617422 | . | . | 4.311646 | 3.883026 | 4.220745 | 4.454518 | 10.033378 | 8.975787 | 9.471231 | 10.184725 | 1.044701 | 678.466018 | 0.156136706 | 0.404069565 | 0.686161565 | 0.998865404 | 9.374165 | 9.437407 | 9.429613 | 9.349563 | 9.404328 | 9.435935 | 809 | 674 | 589 | 665 |
| 22852 | NM_00125605 | ANKRD26 | ankyrin repeat domain 26 | protein_coding | NP_001242982.1;NP_055730.2;XP_006717486.1;XP_006717488.1;XP_01 | 29186 | 610855 | . | . | 2.127749 | 1.910154 | 2.097583 | 1.696090 | 4.951363 | 4.415406 | 4.706915 | 3.877909 | -1.051948 | 436.285239 | 0.18539745 | -0.39408993 | 0.693514648 | 0.998865404 | 8.804671 | 8.732687 | 8.812866 | 8.752223 | 8.807594 | 8.693826 | 540 | 459 | 405 | 361 |
| 22853 | NM_014916,XI | LMTK2 | lemur tyrosine kinase 2 | protein_coding | NP_055731.2;XP_011514283.1 | 17880 | 610989 | . | . | 3.069533 | 2.876190 | 2.964957 | 3.098893 | 7.142929 | 6.648442 | 6.653279 | 7.085250 | 1.006330 | 839.589182 | 0.144543662 | 0.062985418 | 0.949778116 | 0.998865404 | 9.708947 | 9.718118 | 9.736945 | 9.684609 | 9.703535 | 9.727563 | 999 | 870 | 720 | 803 |
| 22856 | NM_014918,XI | CHSY1 | chondroitin sulfate synthase 1 | protein_coding | NP_055733.2;XP_006720498.1;XP_011519666.1;XP_016877500.1;XP_02 | 17198 | 608183 | . | . | 14.699117 | 12.683572 | 14.208917 | 12.105020 | 34.205447 | 29.318646 | 31.884397 | 27.676685 | -1.061202 | 1896.140118 | 0.136334837 | -0.62859538 | 0.529613985 | 0.998865404 | 10.930837 | 10.845604 | 10.963814 | 10.854147 | 10.926245 | 10.796855 | 2402 | 1962 | 1732 | 1583 |
| 22858 | NM_00137539 | CILK1 | ciliogenesis associated kinase 1 | protein_coding | NP_001362326.1;NP_001362327.1;NP_001362328.1;NP_001362329.1;NI | 21219 | 612325 | . | . | 2.959901 | 2.856061 | 4.289942 | 3.488964 | 6.887810 | 6.601914 | 9.626506 | 7.977102 | 1.319649 | 654.349688 | 0.168120977 | 2.380158956 | 0.017305171 | 0.44170732 | 9.139552 | 9.540514 | 9.262834 | 9.232090 | 9.512140 | 9.352558 | 665 | 594 | 721 | 619 |
| 22859 | NM_00100870 | ADGRL1 | adhesion G protein-coupled receptor L1 | protein_coding | NP_001008701.1;NP_055736.2;XP_005259875.1;XP_011526098.1;XP_01 | 20973 | 616416 | . | . | 2.524301 | 2.061760 | 2.672106 | 2.800008 | 5.874152 | 4.765852 | 5.996127 | 6.401886 | 1.174483 | 616.182095 | 0.16729441 | 1.386928354 | 0.165463611 | 0.940648361 | 9.146326 | 9.378749 | 9.271096 | 9.141881 | 9.300621 | 9.331277 | 717 | 551 | 566 | 638 |
| 22861 | NM_00103305 | NLRP1 | NLR family pyrin domain containing 1 | protein_coding | NP_001028225.1;NP_055737.1;NP_127497.1;NP_127499.1;NP_127500.1 | 14374 | 606636 | . | . | 3.004171 | 3.084488 | 4.844907 | 3.419044 | 6.990830 | 7.129929 | 10.871830 | 7.817241 | 1.330917 | 602.732173 | 0.187395527 | 2.200804124 | 0.027749893 | 0.53804272 | 9.013969 | 9.427277 | 9.122232 | 9.127521 | 9.429054 | 9.186711 | 593 | 560 | 703 | 533 |
| 22862 | NM_00107967 | FNDC3A | fibronectin type III domain containing 3A | protein_coding | NP_001073141.1;NP_001265367.1;NP_055738.3;XP_011533299.1;XP_01 | 20296 | 615794 | . | . | 7.424575 | 7.831988 | 7.978658 | 7.506637 | 17.277289 | 18.103994 | 17.903879 | 17.163037 | -1.013274 | 1443.435464 | 0.127960076 | -0.14867138 | 0.881812948 | 0.998865404 | 10.504732 | 10.485788 | 10.479538 | 10.520219 | 10.515472 | 10.463802 | 1629 | 1609 | 1289 | 1299 |
| 22863 | NM_014924,XI | ATG14 | autophagy related 14 | protein_coding | NP_055739.2;XP_011534865.1 | 19962 | 613515 | . | . | 3.148652 | 3.147549 | 4.386528 | 2.816606 | 7.327042 | 7.275699 | 9.843243 | 6.439836 | 1.126109 | 493.842080 | 0.208632778 | 0.821285396 | 0.411483724 | 0.998865404 | 8.859041 | 9.031613 | 8.906373 | 8.896449 | 9.107641 | 8.826585 | 539 | 497 | 554 | 384 |
| 22864 | NM_00133012 | R3HDM2 | R3H domain containing 2 | protein_coding | NP_001317050.1;NP_001317051.1;NP_001317052.1;NP_001338133.1;NI | 29167 | . | | . | 4.523825 | 4.989060 | 5.100786 | 5.156647 | 10.527125 | 11.532435 | 11.446020 | 11.790062 | 1.065459 | 688.917998 | 0.156302576 | 0.585244623 | 0.558383245 | 0.998865404 | 9.381800 | 9.473131 | 9.361299 | 9.446947 | 9.467058 | 9.429383 | 720 | 765 | 635 | 648 |
| 22866 | NM_00116864 | CNKSR2 | connector enhancer of kinase suppressor of Ras 2 | protein_coding | NP_001162118.1;NP_001162119.1;NP_001162120.1;NP_001317699.1;NI | 19701 | 300724 | . | . | 0.253176 | 0.286855 | 0.215125 | 0.299567 | 0.589150 | 0.663077 | 0.482734 | 0.684924 | 1.039463 | 56.735536 | 0.423017725 | 0.131999235 | 0.894984897 | 0.998865404 | 5.799086 | 5.852776 | 5.810767 | 5.830771 | 5.799037 | 5.857544 | 61 | 63 | 43 | 62 |
| 22868 | NM_00113619 | FASTKD2 | FAST kinase domains 2 | protein_coding | NP_001129665.1;NP_001129666.1;NP_055744.2 | 29160 | 612322 | . | . | 9.009193 | 8.342430 | 9.086614 | 6.655241 | 20.964763 | 19.283901 | 20.390099 | 15.216417 | -1.115562 | 1751.513973 | 0.155228384 | -1.01637487 | 0.309450906 | 0.998865404 | 10.850844 | 10.693650 | 10.845120 | 10.778698 | 10.842791 | 10.595662 | 2215 | 1910 | 1669 | 1306 |
| 22869 | NM_00131405 | ZNF510 | zinc finger protein 510 | protein_coding | NP_001300988.1;NP_001300989.1;NP_055745.1;XP_005251865.1;XP_01 | 29161 | . | | . | 1.121386 | 1.233405 | 1.312981 | 1.320420 | 2.609511 | 2.851072 | 2.946291 | 3.018984 | 1.103509 | 256.835795 | 0.214312266 | 0.663044765 | 0.507301885 | 0.998865404 | 7.932113 | 8.073813 | 7.955860 | 7.999382 | 8.024556 | 8.030456 | 269 | 275 | 233 | 254 |
| 22870 | NM_014931 | PPP6R1 | protein phosphatase 6 regulatory subunit 1 | protein_coding | NP_055746.3 | 29195 | 610875 | . | . | 31.650631 | 33.377178 | 32.020390 | 34.075188 | 73.652321 | 77.152847 | 71.852821 | 77.908875 | 1.002513 | 4068.351373 | 0.109509528 | 0.033064615 | 0.973623061 | 0.998865404 | 11.988470 | 11.991985 | 11.970431 | 12.007368 | 11.969310 | 12.012447 | 4575 | 4482 | 3450 | 3918 |
| 22871 | NM_00136592 | NLGN1 | neuroligin 1 | protein_coding | NP_001352852.1;NP_001352853.1;NP_001352854.1;NP_001352855.1;NI | 14291 | 600568 | . | . | 1.432454 | 1.313570 | 1.641859 | 1.168351 | 3.333379 | 3.036371 | 3.684286 | 2.671298 | 1.011194 | 707.726629 | 0.182239591 | 0.088124921 | 0.929777389 | 0.998865404 | 9.458466 | 9.475580 | 9.482319 | 9.434645 | 9.580849 | 9.338013 | 838 | 733 | 728 | 550 |
| 22872 | NM_00107720 | SEC31A | SEC31 homolog A, COPII coat complex component | protein_coding | NP_001070674.1;NP_001070675.1;NP_001070676.1;NP_001177978.1;NI | 17052 | 610257 | . | . | 21.297293 | 22.449108 | 21.535040 | 22.080582 | 49.559675 | 51.892123 | 48.324005 | 50.484626 | -1.025665 | 2885.910846 | 0.112973554 | -0.32361266 | 0.746231298 | 0.998865404 | 11.513017 | 11.476372 | 11.482032 | 11.527455 | 11.479474 | 11.489080 | 3272 | 3241 | 2463 | 2687 |
| 22873 | NM_014934,N | DZIP1 | DAZ interacting zinc finger protein 1 | protein_coding | NP_055749.1;NP_945319.1;XP_005254089.1;XP_006719979.1;XP_00671 | 20908 | 608671 | . | . | 4.753410 | 5.482050 | 5.744180 | 5.049604 | 11.061380 | 12.672004 | 12.889771 | 11.545318 | 1.140738 | 832.787583 | 0.148530448 | 1.278982797 | 0.200903114 | 0.96631375 | 9.603476 | 9.793857 | 9.649373 | 9.653834 | 9.792777 | 9.695492 | 893 | 842 | 824 | 776 |
| 22874 | NM_014935,XI | PLEKHA6 | pleckstrin homology domain containing A6 | protein_coding | NP_055750.2;XP_005245023.2;XP_005245024.2;XP_005245025.2;XP_00 | 17053 | 607771 | . | . | 0.082697 | 0.055115 | 0.048942 | 0.052086 | 0.192439 | 0.127399 | 0.109825 | 0.119088 | -1.280973 | 21.429956 | 0.672999829 | -0.53081729 | 0.595545397 | NA | 4.588571 | 4.232645 | 4.455394 | 4.416083 | 4.412196 | 4.415286 | 33 | 21 | 16 | 18 |
| 22875 | NM_014936 | ENPP4 | ectonucleotide pyrophosphatase/phosphodiesterase 4 | protein_coding | NP_055751.1 | 3359 | 617000 | . | . | 2.685642 | 2.184262 | 1.459771 | 1.238904 | 6.249600 | 5.049019 | 3.275684 | 2.832606 | -1.824694 | 275.019903 | 0.224323254 | -3.86787443 | 0.000109788 | 0.012952189 | 8.472388 | 7.606326 | 8.289891 | 8.161220 | 7.957372 | 7.881949 | 453 | 342 | 184 | 167 |
| 22876 | NM_00124319 | INPP5F | inositol polyphosphate-5-phosphatase F | protein_coding | NP_001230123.1;NP_001230124.1;NP_055752.1;XP_006717783.1;XP_01 | 17054 | 609389 | . | . | 26.785491 | 26.537521 | 29.412292 | 30.299674 | 62.330935 | 61.342673 | 66.000330 | 69.276602 | 1.108864 | 4476.053304 | 0.106289038 | 1.402611677 | 0.160732668 | 0.933525128 | 12.051529 | 12.200643 | 12.090015 | 12.076530 | 12.175276 | 12.163523 | 4928 | 4543 | 4122 | 4382 |
| 22877 | NM_014938,XI | MLXIP | MLX interacting protein | protein_coding | NP_055753.3;XP_006719353.1;XP_006719354.1;XP_006719355.1;XP_00 | 17055 | 608090 | . | . | 18.742831 | 20.502030 | 22.953106 | 23.776732 | 43.615336 | 47.391359 | 51.506104 | 54.362668 | 1.151091 | 5474.822237 | 0.109052019 | 1.861517346 | 0.062671153 | 0.723723143 | 12.313559 | 12.516503 | 12.316629 | 12.394399 | 12.479006 | 12.469748 | 5587 | 5750 | 5123 | 5463 |
| 22878 | NM_014939,XI | TRAPPC8 | trafficking protein particle complex 8 | protein_coding | NP_055754.3;XP_005258290.1;XP_005258291.1;XP_005258292.1;XP_00 | 29169 | 614136 | . | . | 5.878074 | 5.573283 | 6.190945 | 4.676161 | 13.678523 | 12.882892 | 13.892300 | 10.691487 | -1.017735 | 1024.401892 | 0.162128312 | -0.15643402 | 0.875690922 | 0.998865404 | 10.012834 | 9.988193 | 10.009149 | 9.998735 | 10.095657 | 9.874846 | 1198 | 1107 | 1016 | 809 |
| 22879 | NM_00128663 | MON1B | MON1 homolog B, secretory trafficking associated | protein_coding | NP_001273568.1;NP_001273569.1;NP_055755.1 | 25020 | 608954 | . | . | 4.974990 | 5.142181 | 5.594802 | 5.641846 | 11.577006 | 11.886380 | 12.554573 | 12.899411 | 1.095430 | 967.033053 | 0.137281092 | 0.957869117 | 0.338128761 | 0.998865404 | 9.850205 | 9.981647 | 9.874027 | 9.891558 | 9.949833 | 9.948610 | 1050 | 1008 | 880 | 947 |
| 22880 | NM_00130325 | MORC2 | MORC family CW-type zinc finger 2 | protein_coding | NP_001290185.1;NP_001290186.1;NP_055756.1;XP_011528306.1;XP_01 | 23573 | 616661 | . | . | 4.306708 | 4.558691 | 5.011731 | 5.219168 | 10.021885 | 10.537618 | 11.246177 | 11.933008 | 1.180450 | 818.905702 | 0.149656016 | 1.599243096 | 0.109766598 | 0.843391194 | 9.553173 | 9.792056 | 9.565543 | 9.660569 | 9.720175 | 9.742774 | 806 | 866 | 759 | 844 |
| 22881 | NM_00124280 | ANKRD6 | ankyrin repeat domain 6 | protein_coding | NP_001229738.1;NP_001229740.1;NP_001229742.1;NP_001229743.1;NI | 17280 | 610583 | . | . | 3.041629 | 2.756984 | 3.152169 | 3.105905 | 7.077998 | 6.372893 | 7.073376 | 7.101281 | 1.077319 | 505.463210 | 0.168733582 | 0.636779283 | 0.524268622 | 0.998865404 | 8.926515 | 9.034393 | 8.986060 | 8.925900 | 9.013622 | 8.994026 | 587 | 500 | 463 | 484 |
| 22882 | NM_00136279 | ZHX2 | zinc fingers and homeoboxes 2 | protein_coding | NP_001349726.1;NP_055758.1;XP_005250893.1;XP_011515234.1 | 18513 | 609185 | . | . | 0.973056 | 0.926896 | 0.691141 | 0.788889 | 2.264340 | 2.142562 | 1.550900 | 1.803702 | -1.305648 | 115.865974 | 0.301708217 | -1.27529194 | 0.202205929 | 0.967656221 | 7.036170 | 6.650798 | 6.909391 | 6.890137 | 6.793116 | 6.829040 | 155 | 138 | 82 | 100 |
| 22883 | NM_00100956 | CLSTN1 | calsyntenin 1 | protein_coding | NP_001009566.1;NP_001289812.1;NP_055759.3 | 17447 | 611321 | . | . | 70.597764 | 74.284370 | 69.296770 | 68.882489 | 164.283893 | 171.711645 | 155.499939 | 157.491626 | -1.035798 | 10985.316572 | 0.101141507 | -0.50170608 | 0.615874285 | 0.998865404 | 13.448451 | 13.397680 | 13.422929 | 13.453008 | 13.397696 | 13.418284 | 12646 | 12274 | 9271 | 10240 |
| 22884 | NM_014023 | WDR37 | WD repeat domain 37 | protein_coding | NP_054742.2 | 31406 | 618586 | . | . | 2.456113 | 2.731014 | 2.935304 | 2.542209 | 5.715477 | 6.312860 | 6.586737 | 5.812459 | 1.041885 | 426.854936 | 0.18384881 | 0.321984711 | 0.747464277 | 0.998865404 | 8.707526 | 8.767064 | 8.693512 | 8.752452 | 8.793841 | 8.700644 | 458 | 473 | 408 | 377 |
| 22885 | NM_00130101 | ABLIM3 | actin binding LIM protein family member 3 | protein_coding | NP_001287944.1;NP_001287947.1;NP_001287957.1;NP_001332787.1;NI | 29132 | 611305 | . | . | 11.423357 | 11.570622 | 10.244203 | 11.426112 | 26.582622 | 26.746010 | 22.987693 | 26.124448 | -1.073039 | 1419.639749 | 0.128706136 | -0.79018935 | 0.429417196 | 0.998865404 | 10.521390 | 10.419427 | 10.494772 | 10.500638 | 10.408603 | 10.477206 | 1686 | 1592 | 1137 | 1342 |
| 22887 | NM_00119885 | FOXJ3 | forkhead box J3 | protein_coding | NP_001185779.1;NP_001185780.1;NP_001185781.1;NP_055762.3;XP_0 | 29178 | 616035 | . | . | 9.520748 | 9.771583 | 10.934915 | 9.361854 | 22.155174 | 22.587450 | 24.537635 | 21.404767 | 1.029967 | 1614.486793 | 0.132306509 | 0.321966972 | 0.747477716 | 0.998865404 | 10.635236 | 10.678165 | 10.640225 | 10.648239 | 10.728011 | 10.602902 | 1822 | 1725 | 1539 | 1413 |
| 22888 | NM_00126758 | UBOX5 | U-box domain containing 5 | protein_coding | NP_001254513.1;NP_055763.1;NP_955447.1 | 17777 | . | | . | 1.222011 | 1.399509 | 1.087074 | 1.327065 | 2.843671 | 3.235027 | 2.439362 | 3.034175 | -1.104624 | 168.977127 | 0.260881364 | -0.55026887 | 0.582134972 | 0.998865404 | 7.471384 | 7.326340 | 7.392009 | 7.448366 | 7.343209 | 7.414169 | 191 | 204 | 127 | 164 |
| 22889 | NM_014949 | KHDC4 | KH domain containing 4, pre-mRNA splicing factor | protein_coding | NP_055764.2 | 29145 | . | | . | 17.785166 | 18.229679 | 22.750443 | 18.307882 | 41.386814 | 42.138721 | 51.051338 | 41.858799 | 1.124726 | 1777.713848 | 0.139219705 | 1.218029853 | 0.223212628 | 0.982008277 | 10.708337 | 10.878276 | 10.739288 | 10.751951 | 10.924486 | 10.744086 | 1911 | 1820 | 1822 | 1565 |
| 22890 | NM_00112332 | ZBTB1 | zinc finger and BTB domain containing 1 | protein_coding | NP_001116801.1;NP_055765.2;XP_005267467.1;XP_011534868.1;XP_01 | 20259 | 616578 | . | . | 8.741334 | 7.447935 | 7.688060 | 5.463109 | 20.341444 | 17.216237 | 17.251782 | 12.490750 | -1.252634 | 894.466046 | 0.1804372 | -1.80098832 | 0.071704721 | 0.753985777 | 9.957756 | 9.633870 | 9.934341 | 9.812818 | 9.830930 | 9.580761 | 1238 | 986 | 815 | 611 |
| 22891 | NM_014951,N | ZNF365 | zinc finger protein 365 | protein_coding | NP_055766.2;NP_955522.1;XP_016871426.1 | 18194 | 607818 | . | . | 6.252375 | 6.501490 | 5.043083 | 5.551654 | 14.549533 | 15.028486 | 11.316530 | 12.693197 | -1.235982 | 673.345518 | 0.153383657 | -1.99276298 | 0.046287407 | 0.654058028 | 9.540091 | 9.234123 | 9.441670 | 9.483038 | 9.306808 | 9.330409 | 832 | 827 | 514 | 575 |
| 22893 | NM_00130113 | BAHD1 | bromo adjacent homology domain containing 1 | protein_coding | NP_001288061.1;NP_055767.3;XP_011519668.2;XP_011519669.2;XP_01 | 29153 | 613880 | . | . | 4.302514 | 4.136384 | 3.576153 | 4.721420 | 10.012129 | 9.561438 | 8.024783 | 10.794966 | -1.017786 | 600.141203 | 0.170047237 | -0.14956961 | 0.881104185 | 0.998865404 | 9.242253 | 9.215944 | 9.220933 | 9.246596 | 9.148040 | 9.290884 | 684 | 666 | 461 | 618 |

| | | | | | | | | | | | | | | | | | | | | | | | | | | | | | | | | | |
|---|---|---|---|---|---|---|---|---|---|---|---|---|---|---|---|---|---|---|---|---|---|---|---|---|---|---|---|---|---|---|---|---|---|
| 22894 | NM_00112822 | DIS3 | DIS3 homolog, exosome endoribonuclease and 3'-5' exoribonucle | protein_coding | NP_001121698.1;NP_001309277.1;NP_001309278.1;NP_055768.3 | 20604 | 607533 | . | . | 2.270023 | 2.040854 | 2.214518 | 1.587880 | 5.282438 | 4.717523 | 4.969314 | 3.630496 | -1.150386 | 670.301429 | 0.183060511 | -1.10410174 | 0.269549005 | 0.998865404 | 9.485633 | 9.284709 | 9.469369 | 9.390226 | 9.440364 | 9.218372 | 872 | 730 | 633 | 487 |
| 22895 | NM_00114385 | RPH3A | rabphilin 3A | protein_coding | NP_001137326.1;NP_001334881.1;NP_001334882.1;NP_001334883.1;NI | 17056 | 612159 | . | . | 8.837760 | 10.869467 | 13.078897 | 13.456074 | 20.565831 | 25.125262 | 29.348666 | 30.765714 | 1.329186 | 1701.962469 | 0.135394129 | 3.032205908 | 0.002427735 | 0.128390924 | 10.513354 | 10.923553 | 10.532986 | 10.679455 | 10.824674 | 10.838274 | 1521 | 1729 | 1675 | 1836 |
| 22897 | NM_00127193 | CEP164 | centrosomal protein 164 | protein_coding | NP_001258862.1;NP_055771.4;XP_005271510.1;XP_005271513.1;XP_00 | 29182 | 614848 | . | . | 4.033816 | 4.316551 | 4.140470 | 4.344063 | 9.386860 | 9.977900 | 9.291095 | 9.932184 | 1.052968 | 816.680158 | 0.145086782 | 0.513217219 | 0.607799383 | 0.998865404 | 9.635814 | 9.710474 | 9.648308 | 9.660800 | 9.715626 | 9.666094 | 908 | 866 | 754 | 758 |
| 22898 | NM_00135289 | DENND3 | DENN domain containing 3 | protein_coding | NP_001339819.2;NP_001339820.2;NP_001349727.1;NP_055772.3;XP_00 | 29134 | 617503 | . | . | 0.601973 | 0.500274 | 0.631376 | 1.183219 | 1.400815 | 1.156405 | 1.416791 | 2.705290 | 1.175112 | 114.817903 | 0.306867785 | 0.758627003 | 0.448075715 | 0.998865404 | 6.722403 | 6.954667 | 6.831396 | 6.792585 | 6.830358 | 6.900184 | 129 | 107 | 97 | 128 |
| 22900 | NM_00118490 | CARD8 | caspase recruitment domain family member 8 | protein_coding | NP_001171829.1;NP_001171830.1;NP_001171831.1;NP_001171832.1;NI | 17057 | 609051 | . | . | 2.407558 | 2.912952 | 2.850569 | 2.997769 | 5.602490 | 6.733423 | 6.396591 | 6.854047 | 1.038180 | 388.014566 | 0.186180894 | 0.290345028 | 0.771552295 | 0.998865404 | 8.572607 | 8.626818 | 8.561351 | 8.613538 | 8.638844 | 8.579962 | 419 | 429 | 361 | 352 |
| 22901 | NM_00126772 | ARSG | arylsulfatase G | protein_coding | NP_001254656.1;NP_001339828.1;NP_001339829.1;NP_001339830.1;NI | 24102 | 610008 | . | . | 3.461408 | 4.404125 | 3.995015 | 4.564721 | 8.054836 | 10.180336 | 8.964697 | 10.436690 | 1.082704 | 427.099958 | 0.185573826 | 0.617754645 | 0.536737085 | 0.998865404 | 8.680492 | 8.794131 | 8.688322 | 8.735437 | 8.701444 | 8.815676 | 454 | 460 | 350 | 455 |
| 22902 | NM_00103744 | RUFY3 | RUN and FYVE domain containing 3 | protein_coding | NP_001032519.1;NP_001124181.1;NP_001278922.1;NP_001278923.1;NI | 30285 | 611194 | . | . | 6.208569 | 5.879343 | 7.600508 | 5.679189 | 14.447596 | 13.590362 | 17.055322 | 12.984792 | 1.060178 | 885.856168 | 0.171388171 | 0.491907571 | 0.622784686 | 0.998865404 | 9.747702 | 9.832898 | 9.792582 | 9.737531 | 9.918777 | 9.682973 | 1028 | 892 | 922 | 716 |
| 22903 | NM_00128255 | BTBD3 | BTB domain containing 3 | protein_coding | NP_001269479.1;NP_001269480.1;NP_001269481.1;NP_001269483.1;NI | 15854 | 615566 | . | . | 8.874600 | 9.314987 | 8.679218 | 9.284433 | 20.651557 | 21.532011 | 19.475914 | 21.227753 | -1.015491 | 1327.407394 | 0.12948624 | -0.17127679 | 0.864006131 | 0.998865404 | 10.385570 | 10.363134 | 10.365433 | 10.394647 | 10.343364 | 10.392732 | 1511 | 1471 | 1108 | 1275 |
| 22904 | NM_00110012 | SBNO2 | strawberry notch homolog 2 | protein_coding | NP_001093592.1;NP_055778.2;XP_011526106.1 | 29158 | 615729 | . | . | 10.212635 | 11.225410 | 12.552263 | 11.255271 | 23.765220 | 25.948038 | 28.166912 | 25.733840 | 1.095898 | 1726.222525 | 0.129366422 | 1.021231203 | 0.307144912 | 0.998865404 | 10.685778 | 10.817999 | 10.681713 | 10.748001 | 10.833110 | 10.739262 | 1816 | 1853 | 1663 | 1592 |
| 22905 | NM_00110266 | EPN2 | epsin 2 | protein_coding | NP_001096134.1;NP_055779.2;NP_683723.2 | 18639 | 607263 | . | . | 3.173227 | 3.611706 | 3.871639 | 4.251363 | 7.384230 | 8.348620 | 8.687847 | 9.720236 | 1.166538 | 538.054302 | 0.167453226 | 1.327133757 | 0.18446444 | 0.956844295 | 8.956540 | 9.178172 | 8.979313 | 9.054929 | 9.097586 | 9.136506 | 538 | 568 | 489 | 559 |
| 22906 | NM_00104264 | TRAK1 | trafficking kinesin protein 1 | protein_coding | NP_001036111.1;NP_001252537.1;NP_001252538.1;NP_001252539.1;NI | 29947 | 608112 | . | . | 7.327244 | 6.900351 | 10.462553 | 9.195796 | 17.050799 | 15.950466 | 23.477665 | 21.025090 | 1.335604 | 1278.948652 | 0.133658932 | 3.123567808 | 0.001786727 | 0.10405988 | 10.096745 | 10.514591 | 10.212399 | 10.184677 | 10.460753 | 10.373615 | 1285 | 1159 | 1343 | 1295 |
| 22907 | NM_00133099 | DHX30 | DExH-box helicase 30 | protein_coding | NP_001317919.1;NP_055781.2;NP_619520.1;XP_006713096.1;XP_01153 | 16716 | 616423 | . | . | 15.710423 | 18.246331 | 17.553380 | 16.860213 | 36.558797 | 42.177209 | 39.389276 | 38.548873 | 1.001135 | 2172.197775 | 0.118653993 | 0.013796612 | 0.988992246 | 0.999197585 | 11.084195 | 11.085684 | 11.055061 | 11.112779 | 11.076315 | 11.093961 | 2410 | 2427 | 1869 | 2060 |
| 22908 | NM_00131907 | SACM1L | SAC1 like phosphatidylinositide phosphatase | protein_coding | NP_001306000.1;NP_001306001.1;NP_001306002.1;NP_054735.3;XP_01 | 17059 | 606569 | . | . | 11.685410 | 10.849902 | 12.764447 | 11.140052 | 27.192430 | 25.080034 | 28.643047 | 25.470404 | 1.045969 | 1294.138111 | 0.138220552 | 0.469104936 | 0.638994631 | 0.998865404 | 10.304732 | 10.370080 | 10.349829 | 10.287836 | 10.407835 | 10.296369 | 1516 | 1309 | 1235 | 1150 |
| 22909 | NM_00114609 | FAN1 | FANCD2 and FANCI associated nuclease 1 | protein_coding | NP_001139566.1;NP_001139567.1;NP_001139568.1;NP_055782.3;XP_0C | 29170 | 613534 | . | . | 11.162088 | 10.458007 | 13.000310 | 10.394222 | 25.974633 | 24.174151 | 29.172320 | 23.765152 | 1.074851 | 2015.826880 | 0.145354796 | 0.716427217 | 0.473727594 | 0.998865404 | 10.923925 | 11.028492 | 10.958934 | 10.930754 | 11.108203 | 10.884725 | 2277 | 2059 | 2069 | 1686 |
| 22911 | NM_00114255 | WDR47 | WD repeat domain 47 | protein_coding | NP_001136022.1;NP_001136023.1;NP_055784.3;XP_006710523.1;XP_01 | 29141 | 615734 | . | . | 4.822691 | 4.831293 | 4.751418 | 4.667055 | 11.222602 | 11.167752 | 10.662044 | 10.670665 | -1.045608 | 633.169641 | 0.154909865 | -0.41535444 | 0.677882453 | 0.998865404 | 9.338171 | 9.274013 | 9.325627 | 9.316974 | 9.303233 | 9.279423 | 750 | 694 | 548 | 570 |
| 22913 | NM_007367,N | RALY | RALY heterogeneous nuclear ribonucleoprotein | protein_coding | NP_031393.2;NP_057951.1;XP_005260391.1;XP_005260393.1;XP_01152 | 15921 | 614663 | . | . | 72.542704 | 77.237035 | 63.278885 | 70.768066 | 168.809861 | 178.536861 | 141.995977 | 161.802771 | -1.042328 | 5461.483223 | 0.122154705 | -0.48962405 | 0.624399953 | 0.998865404 | 12.444750 | 12.384781 | 12.426593 | 12.435659 | 12.310549 | 12.475370 | 6385 | 6048 | 4193 | 5507 |
| 22916 | NM_00104254 | NCBP2 | nuclear cap binding protein subunit 2 | protein_coding | NP_001036005.1;NP_001294965.1;NP_031388.2;XP_011510858.1;XP_01 | 7659 | 605133 | . | . | 32.365571 | 30.820109 | 34.818856 | 28.302765 | 75.316017 | 71.242059 | 78.132509 | 64.710913 | 1.004864 | 2172.437902 | 0.139855395 | 0.050049082 | 0.960083276 | 0.998865404 | 11.081380 | 11.088809 | 11.110133 | 11.051084 | 11.177258 | 10.983744 | 2587 | 2252 | 2123 | 1795 |
| 22919 | NM_012325,XI | MAPRE1 | microtubule associated protein RP/EB family member 1 | protein_coding | NP_036457.1;XP_011526998.1 | 6890 | 603108 | . | . | 53.877005 | 55.105556 | 49.533528 | 55.054615 | 125.373997 | 127.378949 | 111.151803 | 125.875847 | -1.055760 | 4267.342985 | 0.111051938 | -0.70491059 | 0.480865877 | 0.998865404 | 12.097792 | 12.019388 | 12.075726 | 12.086423 | 11.996924 | 12.073429 | 5015 | 4760 | 3443 | 4072 |
| 22920 | NM_00120451 | KIFAP3 | kinesin associated protein 3 | protein_coding | NP_001191443.1;NP_001191445.1;NP_001191446.1;NP_001362759.1;NI | 17060 | 601836 | . | . | 10.197895 | 10.570606 | 9.982041 | 9.057222 | 23.730922 | 24.434430 | 22.399412 | 20.708263 | -1.104716 | 916.132250 | 0.143578566 | -1.00067307 | 0.316984893 | 0.998865404 | 9.909305 | 9.765960 | 9.864423 | 9.883317 | 9.841504 | 9.761100 | 1093 | 1051 | 800 | 770 |
| 22921 | NM_012228,XI | MSRB2 | methionine sulfoxide reductase B2 | protein_coding | NP_036360.3;XP_011517728.1 | 17061 | 613782 | . | . | 3.041676 | 2.973760 | 2.448224 | 3.524301 | 7.078106 | 6.873979 | 5.493743 | 8.057895 | -1.090994 | 153.271102 | 0.293123388 | -0.42863799 | 0.668186697 | 0.998865404 | 7.322470 | 7.194581 | 7.273082 | 7.274286 | 7.148537 | 7.322624 | 184 | 173 | 98 | 169 |
| 22924 | NM_00130305 | MAPRE3 | microtubule associated protein RP/EB family member 3 | protein_coding | NP_001289979.1;NP_036458.2;XP_006712030.1;XP_011531002.1;XP_01 | 6892 | 605788 | . | . | 11.463178 | 10.771731 | 12.588093 | 12.647006 | 26.675286 | 24.899341 | 28.247316 | 28.915877 | 1.121448 | 693.057086 | 0.152011996 | 1.087823689 | 0.27667295 | 0.998865404 | 9.351651 | 9.517260 | 9.419845 | 9.370137 | 9.476849 | 9.470983 | 780 | 679 | 640 | 684 |
| 22926 | NM_007348,XI | ATF6 | activating transcription factor 6 | protein_coding | NP_031374.2;XP_006711287.1;XP_011507610.1;XP_011507611.1;XP_01 | 791 | 605537 | . | . | 9.026510 | 8.937811 | 7.158373 | 8.139843 | 21.005062 | 20.660146 | 16.063180 | 18.610781 | -1.176443 | 1901.518487 | 0.125780565 | -1.86381322 | 0.062347938 | 0.723723143 | 11.005517 | 10.770821 | 10.960848 | 10.945943 | 10.773545 | 10.873633 | 2389 | 2198 | 1421 | 1744 |
| 22927 | NM_014282,XI | HABP4 | hyaluronan binding protein 4 | protein_coding | NP_055097.2;XP_005251869.1;XP_016869973.1 | 17062 | 617369 | . | . | 9.503179 | 9.419208 | 7.915023 | 10.373868 | 22.114290 | 21.772923 | 17.761084 | 23.718619 | -1.048607 | 769.082311 | 0.164597744 | -0.41600861 | 0.6774037 | 0.998865404 | 9.621191 | 9.551967 | 9.606639 | 9.596930 | 9.475729 | 9.653357 | 913 | 844 | 569 | 794 |
| 22928 | NM_012248 | SEPHS2 | selenophosphate synthetase 2 | protein_coding | NP_036380.2 | 19686 | 606218 | . | . | 61.056614 | 60.954639 | 60.563587 | 68.646080 | 142.081253 | 140.899384 | 135.902924 | 156.951096 | 1.044000 | 4389.434107 | 0.113001601 | 0.549745622 | 0.58249386 | 0.998865404 | 12.068484 | 12.130489 | 12.085430 | 12.076970 | 12.067869 | 12.163967 | 4971 | 4611 | 3675 | 4445 |
| 22929 | NM_00119560 | SEPHS1 | selenophosphate synthetase 1 | protein_coding | NP_001182531.1;NP_001182533.1;NP_001362698.1;NP_036379.2;XP_01 | 19685 | 600902 | . | . | 9.605342 | 10.767973 | 8.474023 | 10.404703 | 22.352024 | 24.890649 | 19.015465 | 23.789119 | -1.092923 | 996.177119 | 0.151989984 | -0.84342568 | 0.398990413 | 0.998865404 | 10.023288 | 9.894349 | 9.948015 | 10.031677 | 9.853852 | 9.992313 | 1131 | 1186 | 748 | 978 |
| 22930 | NM_00117243 | RAB3GAP1 | RAB3 GTPase activating protein catalytic subunit 1 | protein_coding | NP_001165906.1;NP_036365.1;XP_011509125.1;XP_011509127.1 | 17063 | 602536 | . | . | 12.662264 | 12.138357 | 13.639936 | 13.176813 | 29.465609 | 28.058353 | 30.607623 | 30.127217 | 1.037582 | 1855.199762 | 0.121766086 | 0.437111856 | 0.66203023 | 0.998865404 | 10.830393 | 10.883830 | 10.863248 | 10.820929 | 10.893331 | 10.849203 | 2153 | 1912 | 1689 | 1722 |
| 22931 | NM_00125641 | RAB18 | RAB18, member RAS oncogene family | protein_coding | NP_001243339.1;NP_001243340.1;NP_001243341.1;NP_001243344.1;NI | 14244 | 602207 | . | . | 8.523447 | 6.662277 | 6.708710 | 4.966548 | 19.834410 | 15.400148 | 15.054152 | 11.355423 | -1.314180 | 1021.716155 | 0.177565323 | -2.21982145 | 0.02643089 | 0.532178003 | 10.180015 | 9.786837 | 10.174116 | 9.983915 | 9.986295 | 9.767557 | 1501 | 1097 | 887 | 700 |
| 22932 | NM_012230,N | POMZP3 | POM121 and ZP3 fusion | protein_coding | NP_036362.3;NP_694537.1 | 9203 | 600587 | . | . | 1.478371 | 1.908254 | 1.941325 | 1.222935 | 3.440230 | 4.411015 | 4.356277 | 2.796096 | -1.084003 | 79.647047 | 0.382568177 | -0.3041793 | 0.760991297 | 0.998865404 | 6.371764 | 6.257054 | 6.287007 | 6.352456 | 6.355692 | 6.239248 | 84 | 100 | 82 | 55 |
| 22933 | NM_00119328 | SIRT2 | sirtuin 2 | protein_coding | NP_001180215.1;NP_036369.2;NP_085096.1;XP_006723174.1;XP_01152 | 10886 | 604480 | . | . | 16.326820 | 19.213528 | 19.757540 | 22.545147 | 37.993183 | 44.412930 | 44.335349 | 51.546801 | 1.179687 | 1152.695655 | 0.143290633 | 1.663784022 | 0.096155598 | 0.813553488 | 10.046982 | 10.284819 | 10.047986 | 10.156974 | 10.184440 | 10.268684 | 1128 | 1226 | 1025 | 1234 |
| 22934 | NM_144563 | RPIA | ribose 5-phosphate isomerase A | protein_coding | NP_653164.2 | 10297 | 180430 | . | . | 7.170947 | 7.415209 | 6.648888 | 8.378126 | 16.687088 | 17.140589 | 14.919911 | 19.155592 | 1.015759 | 418.451315 | 0.188619498 | 0.119592986 | 0.904805576 | 0.998865404 | 8.698091 | 8.719660 | 8.694968 | 8.710791 | 8.643643 | 8.776347 | 472 | 454 | 326 | 439 |
| 22936 | NM_012081,XI | ELL2 | elongation factor for RNA polymerase II 2 | protein_coding | NP_036213.2;XP_006714638.1;XP_016864728.1;XP_016864729.1;XP_01 | 17064 | 601874 | . | . | 2.259095 | 2.188232 | 1.852020 | 1.778835 | 5.257006 | 5.058199 | 4.155879 | 4.067094 | -1.251219 | 361.586812 | 0.188140161 | -1.71858037 | 0.085690809 | 0.791792328 | 8.650641 | 8.327724 | 8.570739 | 8.546288 | 8.445286 | 8.415474 | 472 | 425 | 287 | 293 |
| 22937 | NM_00132004 | SCAP | SREBF chaperone | protein_coding | NP_001306973.1;NP_036367.2;XP_005265024.1;XP_005265025.1;XP_00 | 30634 | 601510 | . | . | 12.262927 | 11.846016 | 12.954393 | 13.448284 | 28.536336 | 27.382598 | 29.069285 | 30.747902 | 1.069886 | 1638.873596 | 0.122583162 | 0.795025652 | 0.426598581 | 0.998865404 | 10.628938 | 10.726394 | 10.663830 | 10.638709 | 10.694245 | 10.713733 | 1853 | 1681 | 1455 | 1608 |
| 22938 | NM_00131884 | SNW1 | SNW domain containing 1 | protein_coding | NP_001305773.1;NP_036377.1;XP_005267470.1 | 16696 | 603055 | . | . | 23.320738 | 24.387899 | 22.647215 | 25.219547 | 54.268321 | 56.373717 | 50.819690 | 57.661495 | -1.011481 | 1590.819834 | 0.126543507 | -0.13014118 | 0.896454728 | 0.998865404 | 10.643907 | 10.627154 | 10.626365 | 10.653022 | 10.594536 | 10.665837 | 1811 | 1756 | 1312 | 1551 |
| 22943 | NM_012242 | DKK1 | dickkopf WNT signaling pathway inhibitor 1 | protein_coding | NP_036374.1 | 2891 | 605189 | . | . | 9.351323 | 7.456862 | 4.961447 | 5.328118 | 21.760912 | 17.236872 | 11.133344 | 12.182109 | -1.655051 | 382.268571 | 0.198490838 | -3.6620118 | 0.000250242 | 0.024405505 | 8.896479 | 8.169868 | 8.773793 | 8.621293 | 8.393664 | 8.422732 | 613 | 454 | 243 | 278 |
| 22944 | NM_012311,N | KIN | Kin17 DNA and RNA binding protein | protein_coding | NP_036443.1;XP_006717497.1;XP_006717498.1;XP_011517729.1;XP_01 | 6327 | 601720 | . | . | 7.903086 | 7.385056 | 10.919461 | 6.045173 | 18.390808 | 17.070888 | 24.502952 | 13.821567 | 1.048445 | 919.565417 | 0.160143437 | 0.426189481 | 0.669969782 | 0.998865404 | 9.809904 | 9.878889 | 9.834767 | 9.814632 | 9.950801 | 9.758516 | 1048 | 955 | 927 | 766 |
| 22948 | NM_00130615 | CCT5 | chaperonin containing TCP1 subunit 5 | protein_coding | NP_001293082.1;NP_001293083.1;NP_001293084.1;NP_001293085.1;NI | 1618 | 610150 | . | . | 70.490491 | 74.551827 | 62.545460 | 65.060392 | 164.034268 | 172.329891 | 140.350191 | 148.752850 | -1.152180 | 7010.036815 | 0.104239965 | -1.96052982 | 0.049933895 | 0.667071261 | 12.873797 | 12.669381 | 12.811400 | 12.852333 | 12.700609 | 12.723995 | 8436 | 8293 | 5585 | 6193 |
| 22949 | NM_00114610 | PTGR1 | prostaglandin reductase 1 | protein_coding | NP_001139580.1;NP_001139581.1;NP_036344.2;XP_011516696.1;XP_01 | 18429 | 601274 | . | . | 17.948765 | 14.866654 | 16.991748 | 18.896413 | 41.767516 | 34.364937 | 38.128993 | 43.204400 | 1.096853 | 791.675027 | 0.156188772 | 0.853902018 | 0.393159281 | 0.998865404 | 9.560445 | 9.693994 | 9.653369 | 9.530447 | 9.630551 | 9.686474 | 946 | 743 | 692 | 807 |
| 22950 | NM_018158 | SLC4A1AP | solute carrier family 4 member 1 adaptor protein | protein_coding | NP_060628.2 | 13813 | 602655 | . | . | 12.282852 | 13.422256 | 12.629777 | 13.997451 | 28.582701 | 31.026146 | 28.340853 | 32.003510 | 1.021945 | 1210.646674 | 0.135424461 | 0.231258296 | 0.817114143 | 0.998865404 | 10.226028 | 10.256931 | 10.202301 | 10.262585 | 10.213816 | 10.283201 | 1324 | 1344 | 1015 | 1200 |
| 22954 | NM_00109967 | TRIM32 | tripartite motif containing 32 | protein_coding | NP_001093149.1;NP_001365977.1;NP_001365978.1;NP_001365979.1;NI | 16380 | 602290 | . | . | 5.379887 | 5.480049 | 5.044284 | 5.255022 | 12.519220 | 12.667380 | 11.319228 | 12.014983 | -1.068282 | 615.460467 | 0.156129437 | -0.61034586 | 0.541632721 | 0.998865404 | 9.312453 | 9.217013 | 9.284509 | 9.290234 | 9.233786 | 9.252653 | 729 | 689 | 510 | 566 |
| 22955 | NM_00103169 | SCMH1 | Scm polycomb group protein homolog 1 | protein_coding | NP_001026864.1;NP_001165689.1;NP_001165690.1;NP_001165691.1;NI | 19003 | 616396 | . | . | 5.644759 | 6.234356 | 6.748871 | 6.349481 | 13.135586 | 14.410991 | 15.144275 | 14.517333 | 1.101154 | 642.153846 | 0.155905748 | 0.891671121 | 0.372569237 | 0.998865404 | 9.255576 | 9.394625 | 9.271490 | 9.312980 | 9.376536 | 9.337850 | 682 | 680 | 602 | 613 |
| 22969 | gene-RPL41P3 | RPL41P3 | ribosomal protein L41 pseudogene 3 | pseudogene | . | 10358 | . | | . | 13.976716 | 0.560303 | 8.967670 | 0.479932 | 32.524391 | 1.295166 | 20.123192 | 1.097308 | -1.555512 | 79.744081 | 2.172043729 | -0.29345135 | 0.769177182 | 0.998865404 | 6.600872 | 5.964021 | 6.493137 | 5.867995 | 6.289001 | 5.873148 | 215 | 8 | 103 | 6 |
| 22970 | gene-RPL41P2 | RPL41P2 | ribosomal protein L41 pseudogene 2 | pseudogene | . | 10357 | . | | . | 1.536339 | 0.623940 | 0.819633 | 1.655174 | 3.575124 | 1.442265 | 1.839233 | 3.784359 | 1.161350 | 15.770766 | 0.825684166 | 0.261362434 | 0.793813023 | NA | 3.867072 | 4.083203 | 3.996489 | 3.940752 | 3.956079 | 4.001867 | 24 | 9 | 10 | 21 |
| 22971 | gene-RPL41P1 | RPL41P1 | ribosomal protein L41 pseudogene 1 | pseudogene | . | 10356 | . | | . | 97.850845 | 95.625481 | 79.056534 | 98.068336 | 227.702942 | 221.042572 | 177.400574 | 224.221588 | -1.107923 | 1221.501773 | 0.144143427 | -1.02577122 | 0.304999436 | 0.998865404 | 10.326699 | 10.178371 | 10.302877 | 10.278712 | 10.135142 | 10.286191 | 1502 | 1364 | 905 | 1197 |
| 22974 | NM_012112,XI | TPX2 | TPX2 microtubule nucleation factor | protein_coding | NP_036244.2;XP_011526999.1;XP_011527001.1;XP_011527002.1 | 1249 | 605917 | . | . | 42.452315 | 43.483200 | 34.892606 | 40.715172 | 98.788282 | 100.513364 | 78.297996 | 93.090399 | -1.151012 | 4357.376971 | 0.116866526 | -1.73618849 | 0.082530512 | 0.78412713 | 12.187214 | 11.984134 | 12.136404 | 12.152423 | 11.965417 | 12.085587 | 5319 | 5080 | 3265 | 4076 |
| 22976 | NM_007349,XI | PAXIP1 | PAX interacting protein 1 | protein_coding | NP_031375.3;XP_005249596.1;XP_011514284.1;XP_011514285.1;XP_01 | 8624 | 608254 | . | . | 3.819771 | 4.792494 | 4.602875 | 5.328855 | 8.888763 | 11.078065 | 10.328717 | 12.183795 | 1.139586 | 592.390717 | 0.171673066 | 1.098072523 | 0.272172822 | 0.998865404 | 9.113580 | 9.301139 | 9.103142 | 9.219580 | 9.200785 | 9.295676 | 581 | 651 | 510 | 633 |
| 22977 | NM_012067,XI | AKR7A3 | aldo-keto reductase family 7 member A3 | protein_coding | NP_036199.2;XP_011539348.1;XP_016856203.1 | 390 | 608477 | . | . | 0.187076 | 0.469014 | 0.323432 | 0.796853 | 0.435333 | 1.084146 | 0.725771 | 1.821909 | 1.299826 | 21.390793 | 0.720793519 | 0.524864463 | 0.599677415 | NA | 4.220984 | 4.592943 | 4.356782 | 4.429446 | 4.393704 | 4.440770 | 12 | 29 | 16 | 28 |
| 22978 | NM_00113437 | NT5C2 | 5'-nucleotidase, cytosolic II | protein_coding | NP_001127845.1;NP_001338098.1;NP_001338099.1;NP_001338100.1;NI | 8022 | 600417 | . | . | 7.368505 | 7.603370 | 6.121129 | 7.049473 | 17.146807 | 17.575538 | 13.735635 | 16.117778 | -1.148181 | 856.303908 | 0.14597145 | -1.36567857 | 0.172039873 | 0.947431715 | 9.838441 | 9.638602 | 9.765838 | 9.811831 | 9.656693 | 9.722932 | 1021 | 1019 | 660 | 783 |
| 22979 | NM_00131909 | EFR3B | EFR3 homolog B | protein_coding | NP_001306028.1;NP_055786.1;XP_011531003.1 | 29155 | 616797 | . | . | 3.327787 | 3.789159 | 3.065191 | 3.722498 | 7.743896 | 8.758812 | 6.878199 | 8.511050 | -1.035210 | 789.814944 | 0.157376836 | -0.31722424 | 0.751073459 | 0.998865404 | 9.650506 | 9.599790 | 9.603277 | 9.668521 | 9.546156 | 9.673617 | 883 | 907 | 613 | 794 |
| 22980 | NM_014972,XI | TCF25 | transcription factor 25 | protein_coding | NP_055787.1;XP_005256354.1;XP_005256355.1;XP_005256356.1;XP_01 | 29181 | 612326 | . | . | 30.695290 | 30.964718 | 28.407388 | 31.362227 | 71.429201 | 71.576331 | 63.745355 | 71.706005 | -1.024550 | 2202.218067 | 0.118741018 | -0.29467515 | 0.768242045 | 0.998865404 | 11.122219 | 11.087050 | 11.109327 | 11.119181 | 11.062410 | 11.126159 | 2551 | 2420 | 1816 | 2121 |
| 22981 | NM_00131822 | NINL | ninein like | protein_coding | NP_001305155.1;NP_079452.3;XP_011527488.1;XP_011527489.1;XP_01 | 29163 | 609580 | . | . | 1.760307 | 2.124740 | 2.208305 | 1.906863 | 4.096306 | 4.911431 | 4.955372 | 4.359816 | 1.084460 | 306.026985 | 0.209343235 | 0.558778554 | 0.576312861 | 0.998865404 | 8.197869 | 8.314793 | 8.181593 | 8.280725 | 8.311287 | 8.240006 | 307 | 346 | 295 | 279 |
| 22982 | NM_014974,XI | DIP2C | disco interacting protein 2 homolog C | protein_coding | NP_055789.1;XP_005252483.1;XP_005252484.1;XP_005252485.1;XP_00 | 29150 | 611380 | . | . | 5.971789 | 6.071745 | 5.940458 | 6.789703 | 13.896600 | 14.035114 | 13.330218 | 15.523850 | 1.034764 | 1519.925151 | 0.12849815 | 0.383679213 | 0.701216221 | 0.998865404 | 10.545069 | 10.594084 | 10.551324 | 10.560737 | 10.539317 | 10.623745 | 1710 | 1622 | 1271 | 1528 |
| 22983 | NM_014975 | MAST1 | microtubule associated serine/threonine kinase 1 | protein_coding | NP_055790.1 | 19034 | 612256 | . | . | 0.378689 | 0.359039 | 0.381502 | 0.548547 | 0.881226 | 0.829936 | 0.856080 | 1.254188 | 1.248674 | 63.411404 | 0.404914131 | 0.79127178 | 0.428785413 | 0.998865404 | 5.818436 | 6.137320 | 5.961984 | 5.949391 | 5.964605 | 6.043790 | 67 | 59 | 51 | 77 |
| 22984 | NM_014976,XI | PDCD11 | programmed cell death 11 | protein_coding | NP_055791.1;XP_005269704.1;XP_011537840.1;XP_011537841.1;XP_01 | 13408 | 612333 | . | . | 10.641874 | 11.680477 | 11.247175 | 11.193314 | 24.764075 | 26.999945 | 25.238334 | 25.592186 | -1.008146 | 2253.890941 | 0.118812638 | -0.09851562 | 0.921522865 | 0.998865404 | 11.144084 | 11.132296 | 11.106700 | 11.174611 | 11.141645 | 11.127660 | 2496 | 2545 | 1969 | 2086 |
| 22985 | NM_00116481 | ACIN1 | apoptotic chromatin condensation inducer 1 | protein_coding | NP_001158286.1;NP_001158287.1;NP_001158288.1;NP_001158289.1;NI | 17066 | 604562 | . | . | 42.291703 | 42.481914 | 50.811996 | 42.998838 | 98.414531 | 98.198847 | 114.020636 | 98.311734 | 1.170405 | 5750.878274 | 0.114074569 | 1.989997665 | 0.046591193 | 0.654401867 | 12.371560 | 12.598646 | 12.401421 | 12.436255 | 12.607556 | 12.491971 | 5974 | 5838 | 5759 | 5421 |
| 22990 | NM_00130816 | PCNX1 | pecanex 1 | protein_coding | NP_001295089.1;NP_055797.2;XP_005267477.1;XP_005267478.1;XP_00 | 19740 | 617655 | . | . | 3.348345 | 3.320734 | 3.086552 | 2.994323 | 7.791734 | 7.676030 | 6.926133 | 6.846161 | -1.110777 | 1274.197725 | 0.129368399 | -1.17160493 | 0.241355708 | 0.992733415 | 10.389085 | 10.237692 | 10.360788 | 10.346879 | 10.291301 | 10.257086 | 1558 | 1434 | 1074 | 1106 |
| 22992 | NM_00125640 | KDM2A | lysine demethylase 2A | protein_coding | NP_001243334.1;NP_036440.1;XP_006718543.1;XP_011543162.1;XP_01 | 13606 | 605657 | . | . | 7.404922 | 7.781664 | 7.917464 | 8.248770 | 17.231557 | 17.987665 | 17.766560 | 18.859832 | 1.040619 | 1591.258392 | 0.127912166 | 0.449068756 | 0.653382058 | 0.998865404 | 10.606977 | 10.664357 | 10.579223 | 10.658708 | 10.669658 | 10.631317 | 1704 | 1769 | 1446 | 1484 |
| 22993 | NM_00136650 | HMGXB3 | HMG-box containing 3 | protein_coding | NP_001353430.1;NP_055798.3 | 28982 | . | | . | 27.289770 | 28.155207 | 29.266764 | 31.268536 | 63.504418 | 65.082025 | 65.673772 | 71.491787 | 1.076023 | 4735.738480 | 0.107359853 | 0.984622381 | 0.324809592 | 0.998865404 | 12.155592 | 12.261223 | 12.168535 | 12.187075 | 12.216156 | 12.261372 | 5195 | 4978 | 4154 | 4726 |
| 22994 | NM_00100981 | CEP131 | centrosomal protein 131 | protein_coding | NP_001009811.2;NP_001306157.1;NP_001306158.1;NP_055799.2;XP_01 | 29511 | 613479 | . | . | 9.216792 | 9.924811 | 11.131008 | 10.671770 | 21.447852 | 22.941642 | 24.977660 | 24.399734 | 1.127983 | 1164.616268 | 0.13347959 | 1.301666079 | 0.19303056 | 0.96621494 | 10.096150 | 10.269899 | 10.114648 | 10.160946 | 10.247275 | 10.208797 | 1221 | 1218 | 1101 | 1128 |
| 22995 | NM_00119499 | CEP152 | centrosomal protein 152 | protein_coding | NP_001181927.1;NP_055800.2;XP_006720500.1;XP_011519675.1;XP_01 | 29298 | 613529 | . | . | 2.253342 | 3.093872 | 2.629557 | 2.602503 | 5.243619 | 7.151622 | 5.900647 | 5.950314 | -1.102174 | 495.412641 | 0.170220833 | -0.82453129 | 0.409637742 | 0.998865404 | 9.020995 | 8.880562 | 8.951267 | 9.012316 | 8.934489 | 8.906830 | 570 | 587 | 419 | 432 |
| 22996 | NM_00108049 | TTC39A | tetratricopeptide repeat domain 39A | protein_coding | NP_001073963.1;NP_001138304.1;NP_001284591.1;NP_001284592.1;NI | 18657 | . | | . | 12.054136 | 14.475739 | 13.756034 | 15.109312 | 28.050470 | 33.461315 | 30.868144 | 34.545644 | 1.071318 | 746.257221 | 0.159149623 | 0.624488058 | 0.53230711 | 0.998865404 | 9.493350 | 9.592021 | 9.453802 | 9.577433 | 9.533314 | 9.596251 | 756 | 847 | 641 | 756 |
| 22997 | NM_00127728 | IGSF9B | immunoglobulin superfamily member 9B | protein_coding | NP_001264214.1 | 32326 | 613773 | . | . | 0.094370 | 0.110175 | 0.105658 | 0.116907 | 0.219603 | 0.254674 | 0.237093 | 0.267294 | 1.080585 | 57.478789 | 0.414587471 | 0.269695406 | 0.787394595 | 0.998865404 | 5.788552 | 5.899241 | 5.819939 | 5.849874 | 5.843277 | 5.858617 | 59 | 64 | 50 | 58 |
| 22998 | NM_00111271 | LIMCH1 | LIM and calponin homology domains 1 | protein_coding | NP_001106188.1;NP_001106189.1;NP_001106190.1;NP_001106191.1;NI | 29191 | 617750 | . | . | 3.348853 | 3.748384 | 2.720597 | 2.484774 | 7.792917 | 8.664560 | 6.104939 | 5.681139 | -1.387532 | 583.136019 | 0.165800644 | -2.84993185 | 0.00437286 | 0.19533522 | 9.404597 | 8.932297 | 9.244117 | 9.321581 | 9.101729 | 9.034102 | 737 | 772 | 448 | 433 |
| 23001 | NM_014991,XI | WDFY3 | WD repeat and FYVE domain containing 3 | protein_coding | NP_055806.2;XP_005262915.1;XP_011530059.1;XP_011530061.1;XP_01 | 20751 | 617485 | . | . | 1.682980 | 1.718807 | 1.607877 | 1.783430 | 3.916366 | 3.973099 | 3.608030 | 4.077600 | 1.021752 | 744.271326 | 0.148696461 | 0.20877936 | 0.834620484 | 0.998865404 | 9.524167 | 9.555037 | 9.540297 | 9.523632 | 9.524236 | 9.569099 | 858 | 785 | 633 | 728 |
| 23002 | NM_00127052 | DAAM1 | dishevelled associated activator of morphogenesis 1 | protein_coding | NP_001257449.1;NP_055807.1;XP_005267487.1;XP_005267488.1 | 18142 | 606626 | . | . | 3.475861 | 3.816164 | 3.778352 | 3.548722 | 8.088470 | 8.821233 | 8.478514 | 8.113732 | -1.008061 | 672.072065 | 0.155463529 | -0.07450701 | 0.940606965 | 0.998865404 | 9.398231 | 9.386690 | 9.365185 | 9.423421 | 9.413264 | 9.364685 | 744 | 759 | 603 | 605 |
| 23005 | NM_00112860 | MAPKBP1 | mitogen-activated protein kinase binding protein 1 | protein_coding | NP_001122080.1;NP_001252540.1;NP_055809.2 | 29536 | 616786 | . | . | 6.229631 | 6.573994 | 6.210972 | 6.224121 | 14.496608 | 15.196082 | 13.937240 | 14.230713 | -1.044016 | 1412.415947 | 0.12644827 | -0.49145696 | 0.623103285 | 0.998865404 | 10.494719 | 10.432509 | 10.462676 | 10.497540 | 10.449890 | 10.444451 | 1624 | 1592 | 1206 | 1291 |
| 23007 | NM_00113096 | PLCH1 | phospholipase C eta 1 | protein_coding | NP_001124432.1;NP_001124433.1;NP_001336179.1;NP_001336180.1;NI | 29185 | 612835 | . | . | 0.026225 | 0.030574 | 0.010197 | 0.027284 | 0.061026 | 0.070674 | 0.022882 | 0.062382 | -1.364794 | 5.835745 | 1.261735045 | -0.35560817 | 0.722134027 | NA | 2.754145 | 2.300113 | 2.552246 | 2.558382 | 2.540640 | 2.553622 | 7 | 8 | 3 | 6 |
| 23008 | NM_014997,XI | KLHDC10 | kelch domain containing 10 | protein_coding | NP_055812.1;XP_005250290.1 | 22194 | 615152 | . | . | 5.234459 | 5.488006 | 5.505116 | 5.556133 | 12.180802 | 12.685772 | 12.353319 | 12.703439 | 1.019094 | 1082.775381 | 0.133779259 | 0.20397457 | 0.838373371 | 0.998865404 | 10.066846 | 10.094062 | 10.060006 | 10.086535 | 10.088557 | 10.086158 | 1213 | 1178 | 952 | 1023 |
| 23011 | NM_014999 | RAB21 | RAB21, member RAS oncogene family | protein_coding | NP_055814.1 | 18263 | 612398 | . | . | 1.526757 | 1.566427 | 1.572809 | 1.442833 | 3.552826 | 3.620866 | 3.529339 | 3.298866 | -1.040089 | 740.716126 | 0.15041332 | -0.37701019 | 0.706166036 | 0.998865404 | 9.560697 | 9.504306 | 9.539846 | 9.552077 | 9.551382 | 9.485082 | 862 | 822 | 662 | 648 |
| 23012 | NM_015000,XI | STK38L | serine/threonine kinase 38 like | protein_coding | NP_055815.1;XP_005253399.1;XP_006719121.1;XP_006719122.1;XP_01 | 17848 | 615836 | . | . | 5.445555 | 5.263915 | 4.326027 | 5.172788 | 12.672032 | 12.167778 | 9.707481 | 11.826966 | -1.148643 | 771.840583 | 0.154025536 | -1.29803258 | 0.19427614 | 0.96621494 | 9.688900 | 9.488462 | 9.650053 | 9.626307 | 9.484524 | 9.594715 | 966 | 876 | 574 | 728 |
| 23013 | NM_015001 | SPEN | spen family transcriptional repressor | protein_coding | NP_055816.2 | 17575 | 613484 | . | . | 9.106928 | 11.733664 | 16.831013 | 11.962402 | 21.192194 | 27.122887 | 37.768303 | 27.350613 | 1.364259 | 4784.048616 | 0.162135095 | 2.763856088 | 0.005712273 | 0.233204322 | 11.982599 | 12.430767 | 11.969743 | 12.168594 | 12.473729 | 12.172262 | 4092 | 4899 | 5637 | 4275 |
| 23014 | NM_015002,N | FBXO21 | F-box protein 21 | protein_coding | NP_055817.1;NP_296373.1;XP_016874526.1;XP_016874527.1;XP_01687 | 13592 | 609095 | . | . | 3.751954 | 3.407443 | 3.177278 | 3.715969 | 8.730954 | 7.876460 | 7.129720 | 8.496122 | -1.034720 | 649.719545 | 0.160822255 | -0.30618071 | 0.759467061 | 0.998865404 | 9.368242 | 9.318679 | 9.381274 | 9.327268 | 9.283157 | 9.377621 | 791 | 685 | 516 | 641 |
| 23015 | NM_00136807 | GOLGA8A | golgin A8 family member A | protein_coding | NP_001355000.1;NP_001355001.1;NP_851422.1 | 31972 | 616180 | . | . | 21.065125 | 22.046022 | 31.298684 | 20.316048 | 49.019412 | 50.960365 | 70.233338 | 46.450227 | 1.153178 | 4746.245165 | 0.162252889 | 1.267253371 | 0.205064708 | 0.96993643 | 12.106009 | 12.311800 | 12.120784 | 12.166779 | 12.427632 | 12.067969 | 4935 | 4889 | 5350 | 3765 |
| 23016 | NM_015004,N | EXOSC7 | exosome component 7 | protein_coding | NP_055819.2;XP_016861419.1 | 28112 | 606488 | . | . | 14.242649 | 11.610286 | 10.942596 | 15.194989 | 33.143229 | 26.837699 | 24.554868 | 34.741539 | -1.016168 | 505.329372 | 0.18270174 | -0.12664961 | 0.899217727 | 0.998865404 | 8.993047 | 8.969013 | 8.992497 | 8.976528 | 8.888793 | 9.052885 | 594 | 543 | 379 | 531 |
| 23019 | NM_00126561 | CNOT1 | CCR4-NOT transcription complex subunit 1 | protein_coding | NP_001252541.1;NP_057368.3;NP_996882.1 | 7877 | 604917 | . | . | 13.632604 | 14.626487 | 14.314062 | 15.062501 | 31.723629 | 33.809786 | 32.120339 | 34.438617 | 1.031088 | 3730.485487 | 0.111807034 | 0.395027806 | 0.692822367 | 0.998865404 | 11.842964 | 11.886996 | 11.823812 | 11.879682 | 11.855139 | 11.899163 | 4087 | 4098 | 3206 | 3645 |
| 23020 | NM_014014 | SNRNP200 | small nuclear ribonucleoprotein U5 subunit 200 | protein_coding | NP_054733.2 | 30859 | 601664 | . | . | 54.749653 | 59.198032 | 54.339062 | 62.660763 | 127.404701 | 136.838898 | 121.935272 | 143.266388 | 1.012406 | 12935.257606 | 0.111308678 | 0.159806233 | 0.873033714 | 0.998865404 | 13.650135 | 13.667853 | 13.622908 | 13.682355 | 13.605432 | 13.718616 | 14289 | 14354 | 10570 | 13007 |
| 23022 | NM_00116610 | PALLD | palladin, cytoskeletal associated protein | protein_coding | NP_001159580.1;NP_001159581.1;NP_001159582.1;NP_001354496.1;NI | 17068 | 608092 | . | . | 15.192628 | 14.594914 | 11.378496 | 11.296962 | 35.353866 | 33.736795 | 25.533014 | 25.829161 | -1.297655 | 1734.711631 | 0.122600184 | -3.06612365 | 0.002168536 | 0.117381585 | 10.936189 | 10.560365 | 10.879168 | 10.827503 | 10.647852 | 10.651180 | 2328 | 2047 | 1310 | 1419 |
| 23023 | NM_00101739 | TMCC1 | transmembrane and coiled-coil domain family 1 | protein_coding | NP_001017395.2;NP_001121696.1;NP_001336192.1;NP_001336193.1;NI | 29116 | 616242 | . | . | 1.242672 | 1.339426 | 1.047809 | 1.230498 | 2.891748 | 3.096145 | 2.351255 | 2.813383 | -1.133692 | 239.788371 | 0.222529576 | -0.81350594 | 0.415928042 | 0.998865404 | 7.993716 | 7.811788 | 7.920951 | 7.950390 | 7.844307 | 7.903297 | 285 | 283 | 183 | 224 |
| 23024 | NM_00130313 | PDZRN3 | PDZ domain containing ring finger 3 | protein_coding | NP_001290068.1;NP_001290069.1;NP_001290070.1;NP_001290071.1;NI | 17704 | 609729 | . | . | 5.707761 | 5.903399 | 4.132176 | 5.375220 | 13.282194 | 13.645970 | 9.272484 | 12.289800 | -1.237869 | 700.359580 | 0.16720819 | -1.84116867 | 0.065596846 | 0.731993537 | 9.598099 | 9.289314 | 9.509985 | 9.528047 | 9.286852 | 9.451363 | 881 | 847 | 476 | 660 |
| 23025 | NM_00108042 | UNC13A | unc-13 homolog A | protein_coding | NP_001073890.2;XP_011526112.1;XP_011526113.1;XP_016881991.1 | 23150 | 609894 | . | . | 0.204397 | 0.153590 | 0.297241 | 0.264612 | 0.475641 | 0.355029 | 0.666999 | 0.605003 | 1.544402 | 72.151292 | 0.382152477 | 1.640831806 | 0.100832334 | 0.816490589 | 5.824308 | 6.453528 | 6.134659 | 6.073098 | 6.226460 | 6.192338 | 75 | 52 | 81 | 77 |
| 23026 | NM_00119895 | MYO16 | myosin XVI | protein_coding | NP_001185879.1;NP_055826.1;XP_011519364.1 | 29822 | 615479 | . | . | 0.040945 | 0.036642 | 0.102567 | 0.060107 | 0.095282 | 0.084700 | 0.230157 | 0.137426 | 2.055481 | 13.689931 | 0.84523028 | 1.229813999 | 0.218766765 | NA | 3.161369 | 4.204225 | 3.733890 | 3.728606 | 3.791657 | 3.753663 | 11 | 9 | 20 | 13 |
| 23028 | NM_00100999 | KDM1A | lysine demethylase 1A | protein_coding | NP_001009999.1;NP_001350583.1;NP_055828.2;XP_005245843.1;XP_00 | 29079 | 609132 | . | . | 34.826100 | 36.325209 | 32.132044 | 36.084670 | 81.041765 | 83.967349 | 72.103374 | 82.503312 | -1.039011 | 3409.710818 | 0.116669019 | -0.47323157 | 0.636047974 | 0.998865404 | 11.762872 | 11.707463 | 11.735676 | 11.765304 | 11.668468 | 11.767169 | 3930 | 3817 | 2733 | 3324 |
| 23029 | NM_00134673 | RBM34 | RNA binding motif protein 34 | protein_coding | NP_001333667.1;NP_055829.2;XP_011542435.1 | 28965 | . | | . | 15.077908 | 16.713963 | 16.605408 | 16.248648 | 35.086912 | 38.635075 | 37.262053 | 37.150600 | 1.025818 | 902.556203 | 0.143130505 | 0.256932448 | 0.797230921 | 0.998865404 | 9.799265 | 9.836244 | 9.792495 | 9.823408 | 9.859520 | 9.792359 | 1004 | 982 | 832 | 817 |
| 23030 | NM_00137009 | KDM4B | lysine demethylase 4B | protein_coding | NP_001357022.1;NP_001357023.1;NP_055830.1;XP_005259578.2;XP_01 | 29136 | 609765 | . | . | 2.565977 | 2.486140 | 3.172622 | 3.037756 | 5.971133 | 5.746822 | 7.119273 | 6.945469 | 1.203224 | 485.584422 | 0.169336988 | 1.576176259 | 0.114985188 | 0.855274326 | 8.783789 | 9.051012 | 8.877596 | 8.842960 | 8.986034 | 8.967472 | 522 | 462 | 468 | 490 |
| 23031 | NM_015016,XI | MAST3 | microtubule associated serine/threonine kinase 3 | protein_coding | NP_055831.1;XP_005259880.1;XP_005259881.1;XP_005259882.1;XP_00 | 19036 | 612258 | . | . | 1.948741 | 2.208587 | 2.231195 | 2.227876 | 4.534798 | 5.105250 | 5.006736 | 5.093773 | 1.044993 | 409.352693 | 0.179857456 | 0.353019671 | 0.724073694 | 0.998865404 | 8.645244 | 8.708465 | 8.640972 | 8.685249 | 8.688312 | 8.690920 | 444 | 448 | 363 | 393 |
| 23032 | NM_00137743 | USP33 | ubiquitin specific peptidase 33 | protein_coding | NP_001364359.1;NP_001364360.1;NP_001364361.1;NP_001364362.1;NI | 20059 | 615146 | . | . | 11.587808 | 10.500000 | 10.757525 | 10.472056 | 26.965301 | 24.271222 | 24.139574 | 23.943109 | -1.057862 | 1456.152574 | 0.130248458 | -0.62305341 | 0.533249426 | 0.998865404 | 10.547814 | 10.466944 | 10.571014 | 10.484900 | 10.498717 | 10.472355 | 1819 | 1526 | 1252 | 1304 |
| 23033 | NM_00119994 | DOP1A | DOP1 leucine zipper like protein A | protein_coding | NP_001186871.1;NP_055833.2;XP_011533921.1;XP_016866048.1;XP_01 | 21194 | 616823 | . | . | 1.274043 | 1.445477 | 1.269402 | 1.176045 | 2.964750 | 3.341287 | 2.848502 | 2.688890 | -1.124259 | 378.567433 | 0.189268914 | -0.89277695 | 0.371976632 | 0.998865404 | 8.646399 | 8.477472 | 8.559214 | 8.633337 | 8.554018 | 8.502700 | 433 | 459 | 324 | 319 |
| 23034 | NM_00116157 | SAMD4A | sterile alpha motif domain containing 4A | protein_coding | NP_001155048.2;NP_001155049.1;NP_056404.4;XP_024305282.1;XP_02 | 23023 | 610747 | . | . | 6.390017 | 6.632738 | 7.359571 | 7.404605 | 14.869832 | 15.331871 | 16.514664 | 16.929748 | 1.109778 | 1527.199213 | 0.124043942 | 1.211431728 | 0.225729986 | 0.982008277 | 10.499571 | 10.649861 | 10.525774 | 10.543344 | 10.625118 | 10.605237 | 1648 | 1580 | 1415 | 1487 |
| 23035 | NM_00128900 | PHLPP2 | PH domain and leucine rich repeat protein phosphatase 2 | protein_coding | NP_001275932.1;NP_055835.2 | 29149 | 611066 | . | . | 1.452372 | 1.651494 | 1.811815 | 1.891966 | 3.379729 | 3.817502 | 4.065660 | 4.325756 | 1.178391 | 441.212986 | 0.177680015 | 1.332835134 | 0.182585913 | 0.954272382 | 8.662364 | 8.898636 | 8.694747 | 8.765520 | 8.820253 | 8.841774 | 439 | 463 | 408 | 455 |
| 23036 | NM_00135144 | ZNF292 | zinc finger protein 292 | protein_coding | NP_001338373.1;NP_055836.1;XP_005248754.1;XP_011533927.1;XP_01 | 18410 | 616213 | . | . | 1.888525 | 1.611991 | 2.215726 | 1.224908 | 4.394674 | 3.726191 | 4.972025 | 2.800607 | 1.019400 | 654.667747 | 0.22196576 | 0.12488628 | 0.90061358 | 0.998865404 | 9.340098 | 9.368995 | 9.356496 | 9.316824 | 9.528113 | 9.134102 | 768 | 679 | 740 | 442 |
| 23038 | NM_00127625 | WDTC1 | WD and tetratricopeptide repeats 1 | protein_coding | NP_001263181.1;NP_055838.2;XP_011539359.1;XP_011539360.1;XP_01 | 29175 | . | | . | 5.208293 | 5.213452 | 4.679454 | 6.496918 | 12.119911 | 12.051129 | 10.500560 | 14.854430 | 1.053983 | 789.324062 | 0.171318685 | 0.44274856 | 0.657947614 | 0.998865404 | 9.586459 | 9.661513 | 9.604832 | 9.600533 | 9.523703 | 9.743623 | 888 | 827 | 595 | 877 |
| 23039 | NM_00110016 | XPO7 | exportin 7 | protein_coding | NP_001093631.1;NP_001349731.1;NP_055839.3 | 14108 | 606140 | . | . | 15.018352 | 16.603814 | 14.826325 | 16.060542 | 34.948320 | 38.380457 | 33.269841 | 36.720521 | -1.037581 | 2373.366907 | 0.12050469 | -0.44167427 | 0.658724933 | 0.998865404 | 11.239206 | 11.185738 | 11.189570 | 11.263232 | 11.168600 | 11.225165 | 2657 | 2727 | 1954 | 2261 |
| 23041 | NM_00127846 | MON2 | MON2 homolog, regulator of endosome-to-Golgi trafficking | protein_coding | NP_001265398.1;NP_001265399.1;NP_001265400.1;NP_001265401.1;NI | 29177 | 616822 | . | . | 1.560478 | 1.458401 | 1.694409 | 1.401547 | 3.631295 | 3.371159 | 3.802205 | 3.204470 | -1.033369 | 618.708066 | 0.179723076 | -0.26349358 | 0.792170169 | 0.998865404 | 9.296053 | 9.249806 | 9.307813 | 9.253216 | 9.352886 | 9.155879 | 753 | 651 | 608 | 486 |
| 23042 | NM_00128544 | PDXDC1 | pyridoxal dependent decarboxylase domain containing 1 | protein_coding | NP_001272373.1;NP_001272374.1;NP_001272376.1;NP_001272377.1;NI | 28995 | 614244 | . | . | 56.020657 | 53.499258 | 64.572126 | 66.958180 | 130.362370 | 123.665929 | 144.897985 | 153.091920 | 1.154679 | 7525.447165 | 0.10221468 | 2.029958141 | 0.042360794 | 0.636241616 | 12.770099 | 12.977571 | 12.807141 | 12.819927 | 12.929430 | 12.941115 | 7982 | 7595 | 6966 | 7613 |
| 23043 | NM_00116156 | TNIK | TRAF2 and NCK interacting kinase | protein_coding | NP_001155032.1;NP_001155033.1;NP_001155034.1;NP_001155035.1;NI | 30765 | 610005 | . | . | 4.496276 | 4.555842 | 4.716654 | 4.511917 | 10.463020 | 10.531032 | 10.584033 | 10.315965 | -1.014077 | 1156.754890 | 0.1322394 | -0.15250513 | 0.878788544 | 0.998865404 | 10.185832 | 10.165833 | 10.198145 | 10.163579 | 10.178826 | 10.162308 | 1372 | 1228 | 1010 | 1065 |
| 23046 | NM_00125210 | KIF21B | kinesin family member 21B | protein_coding | NP_001239029.1;NP_001239031.1;NP_001239032.1;NP_060066.2;XP_01 | 29442 | 608322 | . | . | 0.104591 | 0.117591 | 0.093184 | 0.175504 | 0.243389 | 0.271818 | 0.209102 | 0.401270 | 1.175480 | 38.059153 | 0.524672668 | 0.444563148 | 0.656635485 | 0.998865404 | 5.130737 | 5.360470 | 5.224012 | 5.240655 | 5.213257 | 5.297845 | 38 | 40 | 26 | 49 |
| 23047 | NM_015032,XI | PDS5B | PDS5 cohesin associated factor B | protein_coding | NP_055847.1;XP_005266355.1;XP_011533301.1;XP_011533302.1;XP_01 | 20418 | 605333 | . | . | 4.187013 | 4.281710 | 4.037185 | 4.092961 | 9.743348 | 9.897366 | 9.059325 | 9.358069 | -1.045820 | 909.467137 | 0.141652345 | -0.45629008 | 0.648181397 | 0.998865404 | 9.860807 | 9.796226 | 9.822856 | 9.865983 | 9.832326 | 9.791981 | 1039 | 1033 | 795 | 810 |
| 23048 | NM_00136375 | FNBP1 | formin binding protein 1 | protein_coding | NP_001350684.1;NP_055848.1;XP_005251872.1;XP_005251877.1;XP_00 | 17069 | 606191 | . | . | 4.044496 | 4.179411 | 4.048853 | 4.574924 | 9.411704 | 9.660899 | 9.085510 | 10.460019 | 1.091225 | 598.627549 | 0.16025442 | 0.785926906 | 0.43191031 | 0.998865404 | 9.161392 | 9.286909 | 9.191641 | 9.198428 | 9.217458 | 9.287074 | 656 | 621 | 515 | 616 |
| 23049 | NM_015092,XI | SMG1 | SMG1 nonsense mediated mRNA decay associated PI3K related ki | protein_coding | NP_055907.3;XP_005255239.1;XP_005255240.1;XP_005255241.1;XP_01 | 30045 | 607032 | . | . | 6.436572 | 6.126585 | 7.014338 | 5.981103 | 14.978164 | 14.161875 | 15.739972 | 13.675082 | 1.021092 | 3198.925318 | 0.124164037 | 0.242526505 | 0.808372222 | 0.998865404 | 11.628103 | 11.658481 | 11.657726 | 11.608982 | 11.718174 | 11.578874 | 3753 | 3314 | 3049 | 2774 |
| 23051 | NM_015035,XI | ZHX3 | zinc fingers and homeoboxes 3 | protein_coding | NP_055850.1;XP_005260398.1;XP_005260400.1;XP_006723811.1;XP_00 | 15935 | 609598 | . | . | 4.320922 | 4.191779 | 4.424682 | 5.301431 | 10.054964 | 9.689485 | 9.928858 | 12.121095 | 1.086807 | 1126.841392 | 0.144244707 | 0.832581759 | 0.40508065 | 0.998865404 | 10.077053 | 10.196608 | 10.091802 | 10.117850 | 10.094782 | 10.234222 | 1222 | 1186 | 926 | 1202 |
| 23052 | NM_015036 | ENDOD1 | endonuclease domain containing 1 | protein_coding | NP_055851.1 | 29129 | . | | . | 17.977936 | 16.081583 | 15.721218 | 16.141233 | 41.835396 | 37.173298 | 35.277962 | 36.905014 | -1.084049 | 2389.323366 | 0.121104225 | -0.96140266 | 0.336349746 | 0.998865404 | 11.279372 | 11.163057 | 11.301445 | 11.203518 | 11.182265 | 11.195008 | 3034 | 2521 | 1978 | 2167 |
| 23053 | NM_00124248 | ZSWIM8 | zinc finger SWIM-type containing 8 | protein_coding | NP_001229416.1;NP_001229417.1;NP_001354728.1;NP_055852.2;XP_0C | 23528 | . | | . | 18.444925 | 20.004233 | 23.843911 | 21.429061 | 42.922095 | 46.240681 | 53.505041 | 48.995001 | 1.162627 | 3883.955490 | 0.116365165 | 1.868161101 | 0.061739619 | 0.723723143 | 11.810488 | 12.027947 | 11.829385 | 11.883568 | 12.029730 | 11.931415 | 3999 | 4004 | 3844 | 3686 |
| 23054 | NM_00124253 | NCOA6 | nuclear receptor coactivator 6 | protein_coding | NP_001229468.1;NP_001305169.1;NP_054790.2;XP_006723818.1;XP_01 | 15936 | 605299 | . | . | 3.863313 | 4.242940 | 4.958523 | 5.181711 | 8.990088 | 9.807746 | 11.126783 | 11.847365 | 1.210588 | 956.929975 | 0.141432836 | 1.949392955 | 0.051248515 | 0.672475152 | 9.757995 | 10.033419 | 9.788497 | 9.864320 | 9.964031 | 9.968064 | 944 | 984 | 909 | 985 |
| 23057 | NM_015039,N | NMNAT2 | nicotinamide nucleotide adenylyltransferase 2 | protein_coding | NP_055854.1;NP_733820.1;XP_024310045.1 | 16789 | 608701 | . | . | 0.156593 | 0.122591 | 0.189668 | 0.139822 | 0.364399 | 0.283374 | 0.425610 | 0.319686 | 1.176227 | 26.746968 | 0.604560158 | 0.387333766 | 0.698509131 | NA | 4.617806 | 4.855058 | 4.740581 | 4.713158 | 4.762014 | 4.727142 | 32 | 23 | 29 | 23 |
| 23059 | NM_00133045 | CLUAP1 | clusterin associated protein 1 | protein_coding | NP_001317383.1;NP_055856.1;NP_079069.1;XP_006720930.1;XP_01687 | 19009 | 616787 | . | . | 2.287050 | 2.040658 | 1.987528 | 2.406452 | 5.322061 | 4.717073 | 4.459956 | 5.502068 | -1.073441 | 268.334661 | 0.216497856 | -0.47226047 | 0.63674088 | 0.998865404 | 8.118276 | 8.015679 | 8.116586 | 8.053568 | 8.010760 | 8.086123 | 339 | 282 | 208 | 261 |
| 23060 | NM_015042,XI | ZNF609 | zinc finger protein 609 | protein_coding | NP_055857.1;XP_016877510.1 | 29003 | 617474 | . | . | 3.403499 | 3.419186 | 2.944318 | 3.968485 | 7.920082 | 7.903602 | 6.606965 | 9.074369 | -1.000728 | 963.250169 | 0.16172648 | -0.00649395 | 0.994818616 | 0.999862634 | 9.912641 | 9.910892 | 9.911160 | 9.908306 | 9.799296 | 10.008180 | 1112 | 1038 | 717 | 1032 |
| 23061 | NM_015043,N | TBC1D9B | TBC1 domain family member 9B | protein_coding | NP_055858.2;NP_942568.2 | 29097 | 618039 | . | . | 27.370573 | 28.754586 | 29.454257 | 31.847547 | 63.692443 | 66.467509 | 66.094496 | 72.815630 | 1.077396 | 4718.292733 | 0.108882991 | 0.987749959 | 0.32327512 | 0.998865404 | 12.149328 | 12.256771 | 12.155106 | 12.188397 | 12.205124 | 12.262265 | 5126 | 5000 | 4111 | 4744 |
| 23062 | NM_015044,XI | GGA2 | golgi associated, gamma adaptin ear containing, ARF binding prot | protein_coding | NP_055859.1;XP_016878564.1;XP_024305968.1 | 16064 | 606005 | . | . | 14.237112 | 15.977463 | 16.586164 | 17.190161 | 33.130347 | 36.932622 | 37.218871 | 39.303258 | 1.078188 | 2919.415270 | 0.115993227 | 0.936334135 | 0.349101178 | 0.998865404 | 11.456190 | 11.564697 | 11.438484 | 11.518985 | 11.544634 | 11.537254 | 3075 | 3182 | 2646 | 2827 |
| 23063 | NM_00131832 | WAPL | WAPL cohesin release factor | protein_coding | NP_001305257.1;NP_055860.1;XP_011537850.1;XP_016871468.1;XP_01 | 23293 | 610754 | . | . | 7.270676 | 7.380293 | 7.724319 | 6.512840 | 16.919163 | 17.059877 | 17.333145 | 14.890836 | -1.036012 | 1408.217148 | 0.136696261 | -0.37338509 | 0.708861855 | 0.998865404 | 10.484748 | 10.434116 | 10.470686 | 10.473468 | 10.508843 | 10.377037 | 1647 | 1549 | 1307 | 1185 |
| 23064 | NM_00135152 | SETX | senataxin | protein_coding | NP_001338456.1;NP_001338457.1;NP_055861.3;XP_005272229.1;XP_00 | 445 | 608465 | . | . | 4.943270 | 4.249211 | 5.297594 | 4.377393 | 11.503191 | 9.822242 | 11.887646 | 10.008389 | 1.120073 | 1488.109429 | 0.147484468 | 1.109216459 | 0.267336814 | 0.998546485 | 10.454913 | 10.618956 | 10.485257 | 10.496895 | 10.675512 | 10.472152 | 1604 | 1526 | 1545 | 1283 |
| 23065 | NM_00127142 | EMC1 | ER membrane protein complex subunit 1 | protein_coding | NP_001258356.1;NP_001258357.1;NP_001258358.1;NP_001362749.1;NI | 28957 | 616846 | . | . | 49.825608 | 49.784547 | 43.062618 | 42.909832 | 115.946231 | 115.079217 | 96.631266 | 98.108232 | -1.174921 | 9602.592866 | 0.100832715 | -2.30643059 | 0.021086584 | 0.4816697 | 13.340803 | 13.108253 | 13.298041 | 13.289870 | 13.162656 | 13.151012 | 12003 | 11142 | 7732 | 8221 |
| 23067 | NM_00135334 | SETD1B | SET domain containing 1B, histone lysine methyltransferase | protein_coding | NP_001340274.1;XP_006719359.1;XP_024304666.1 | 29187 | 611055 | . | . | 2.146321 | 2.603215 | 2.642910 | 2.413426 | 4.994577 | 6.017449 | 5.930614 | 5.518012 | 1.050495 | 653.688509 | 0.165543589 | 0.429310713 | 0.667697122 | 0.998865404 | 9.316549 | 9.387498 | 9.269813 | 9.393785 | 9.400872 | 9.331978 | 667 | 751 | 612 | 596 |
| 23070 | NM_015050,XI | CMTR1 | cap methyltransferase 1 | protein_coding | NP_055865.1;XP_005249012.1 | 21077 | 616189 | . | . | 12.611043 | 12.819291 | 13.605263 | 13.794568 | 29.346416 | 29.632365 | 30.529816 | 31.539641 | 1.058145 | 1673.851473 | 0.121628166 | 0.670381653 | 0.502614527 | 0.998865404 | 10.667648 | 10.749114 | 10.678353 | 10.694198 | 10.724841 | 10.736047 | 1854 | 1773 | 1486 | 1625 |
| 23071 | NM_015051 | ERP44 | endoplasmic reticulum protein 44 | protein_coding | NP_055866.1 | 18311 | 609170 | . | . | 16.891855 | 17.284323 | 15.361935 | 12.666296 | 39.308044 | 39.953487 | 34.471737 | 28.959982 | -1.235680 | 2331.990034 | 0.131293455 | -2.32536039 | 0.020052697 | 0.468337364 | 11.331788 | 11.026817 | 11.259308 | 11.270888 | 11.171544 | 11.012411 | 2947 | 2801 | 1998 | 1758 |
| 23074 | NM_00100694 | UHRF1BP1L | UHRF1 binding protein 1 like | protein_coding | NP_001006948.1;NP_055869.1;XP_005268795.1;XP_005268796.1;XP_01 | 29102 | . | | . | 3.984579 | 3.468479 | | 2.255107 | 9.272280 | 8.017546 | 7.579244 | 5.156033 | -1.267595 | 529.463167 | 0.179700705 | -1.90368424 | 0.056951321 | 0.708295137 | 9.208800 | 8.867879 | 9.165518 | 9.066811 | 9.026206 | 8.900374 | 731 | 592 | 448 | 393 |
| 23075 | NM_00129771 | SWAP70 | switching B cell complex subunit SWAP70 | protein_coding | NP_001284643.1;NP_055870.2;XP_005252887.1 | 17070 | 604762 | . | . | 10.462553 | 10.251584 | 11.056811 | 11.247104 | 24.346791 | 23.696996 | 24.811163 | 25.715165 | 1.056689 | 1620.046967 | 0.122761037 | 0.648018798 | 0.516972792 | 0.998865404 | 10.621453 | 10.701088 | 10.654187 | 10.625119 | 10.685076 | 10.680414 | 1848 | 1668 | 1452 | 1556 |
| 23076 | NM_015056 | RRP1B | ribosomal RNA processing 1B | protein_coding | NP_055871.1 | 23818 | 610654 | . | . | 9.701194 | 10.150948 | 10.767618 | 11.374414 | 22.575077 | 23.464369 | 24.162222 | 26.006247 | 1.099861 | 1660.600715 | 0.122986267 | 1.116556077 | 0.264184181 | 0.996308385 | 10.627286 | 10.764435 | 10.644530 | 10.672927 | 10.715396 | 10.750474 | 1788 | 1738 | 1479 | 1667 |
| 23077 | NM_015057,XI | MYCBP2 | MYC binding protein 2 | protein_coding | NP_055872.4;XP_005266356.1;XP_006719842.1;XP_006719843.1;XP_00 | 23386 | 610392 | . | . | 2.748032 | 2.998737 | 2.909270 | 2.929002 | 6.394784 | 6.931715 | 6.528318 | 6.696815 | 1.044314 | 1365.535232 | 0.126431081 | 0.494783574 | 0.6207529 | 0.998865404 | 10.383626 | 10.446197 | 10.399631 | 10.396989 | 10.432587 | 10.430351 | 1541 | 1439 | 1215 | 1306 |
| 23078 | NM_00100981 | VWA8 | von Willebrand factor A domain containing 8 | protein_coding | NP_001009814.1;NP_055873.1 | 29071 | 617509 | . | . | 2.235931 | 2.207540 | 2.581589 | 2.240214 | 5.203103 | 5.102826 | 5.793010 | 5.121984 | -1.000564 | 460.334859 | 0.171815402 | -0.00473124 | 0.996225029 | 0.999862634 | 8.846848 | 8.846232 | 8.846371 | 8.846802 | 8.859676 | 8.832703 | 530 | 497 | 409 | 422 |
| 23080 | NM_015060,XI | AVL9 | AVL9 cell migration associated | protein_coding | NP_055875.1;XP_005249725.1;XP_005249726.1;XP_005249727.1;XP_00 | 28994 | 612927 | . | . | 7.559463 | 7.425271 | 7.743112 | 7.373070 | 17.591182 | 17.163846 | 17.375321 | 16.857651 | -1.016429 | 1545.998864 | 0.124870031 | -0.18827109 | 0.850664148 | 0.998865404 | 10.605941 | 10.582612 | 10.605415 | 10.595496 | 10.611067 | 10.564195 | 1806 | 1671 | 1374 | 1394 |
| 23081 | NM_00114669 | KDM4C | lysine demethylase 4C | protein_coding | NP_001140167.1;NP_001140168.1;NP_001291268.1;NP_001291269.1;NI | 17071 | 605469 | . | . | 1.718229 | 2.237844 | 2.184010 | 1.900393 | 3.998391 | 5.172879 | 4.900853 | 4.345022 | 1.042595 | 300.807210 | 0.208820737 | 0.288184265 | 0.773205693 | 0.998865404 | 8.202700 | 8.262079 | 8.163630 | 8.272620 | 8.235896 | 8.247914 | 305 | 350 | 264 | 291 |
| 23082 | NM_00128872 | PPRC1 | PPARG related coactivator 1 | protein_coding | NP_001275656.1;NP_001275657.1;NP_055877.3;XP_005269713.1;XP_00 | 30025 | 617462 | . | . | 8.100236 | 8.766198 | 9.768287 | 8.765116 | 18.849581 | 20.263451 | 21.919755 | 20.040395 | 1.100363 | 1461.424379 | 0.131580407 | 1.048634736 | 0.294346262 | 0.998865404 | 10.442425 | 10.580587 | 10.452798 | 10.495118 | 10.593806 | 10.499972 | 1558 | 1543 | 1412 | 1349 |
| 23085 | NM_00130124 | ERC1 | ELKS/RAB6-interacting/CAST family member 1 | protein_coding | NP_001288177.1;NP_829883.1;NP_829884.1;XP_011519243.1;XP_01687 | 17072 | 607127 | . | . | 4.266075 | 4.350056 | 4.401713 | 5.017290 | 9.927330 | 10.055351 | 9.877318 | 11.471439 | 1.035215 | 1271.230000 | 0.136306769 | 0.366309851 | 0.714133879 | 0.998865404 | 10.287037 | 10.336557 | 10.290861 | 10.305298 | 10.267735 | 10.377748 | 1425 | 1361 | 1043 | 1300 |
| 23086 | NM_00114476 | EXPH5 | exophilin 5 | protein_coding | NP_001138235.1;NP_001138236.1;NP_001138237.1;NP_001294948.1;NI | 30578 | 612878 | . | . | 0.067860 | 0.046451 | 0.053777 | 0.074489 | 0.157913 | 0.107373 | 0.120674 | 0.170309 | 1.080145 | 22.252303 | 0.662255318 | 0.167948678 | 0.866623654 | NA | 4.419302 | 4.530328 | 4.487243 | 4.454589 | 4.463622 | 4.492670 | 29 | 19 | 17 | 25 |
| 23087 | NM_00130449 | TRIM35 | tripartite motif containing 35 | protein_coding | NP_001291424.1;NP_001349742.1;NP_741983.2;XP_011542755.1 | 16285 | 617007 | . | . | 3.081473 | 3.018854 | 3.055864 | 3.157405 | 7.170715 | 6.978214 | 6.857269 | 7.219030 | 1.007143 | 402.885995 | 0.180027245 | 0.057037392 | 0.954515409 | 0.998865404 | 8.649081 | 8.659356 | 8.663320 | 8.640673 | 8.648547 | 8.663987 | 471 | 425 | 347 | 384 |
| 23089 | NM_00104015 | PEG10 | paternally expressed 10 | protein_coding | NP_001035242.1;NP_001165908.1;NP_001165909.1;NP_001171890.1;NI | 14005 | 609810 | . | . | 101.343012 | 105.931123 | 90.755530 | 106.719165 | 235.829346 | 244.864539 | 203.652776 | 244.000695 | -1.063530 | 20760.528579 | 0.111036634 | -0.80028491 | 0.423545744 | 0.998865404 | 14.385323 | 14.296418 | 14.351941 | 14.381421 | 14.246761 | 14.376666 | 24222 | 23485 | 16171 | 20293 |
| 23091 | NM_00107678 | ZC3H13 | zinc finger CCCH-type containing 13 | protein_coding | NP_001070256.1;NP_001317493.1;NP_001317494.1;NP_001317495.1;NI | 20368 | 616453 | . | . | 10.248616 | 9.872668 | 11.142485 | 10.897740 | 23.848948 | 22.821113 | 25.003415 | 24.916387 | 1.085571 | 2648.210948 | 0.114361231 | 1.035794642 | 0.300297958 | 0.998865404 | 11.310292 | 11.428876 | 11.352362 | 11.319964 | 11.420670 | 11.384907 | 2983 | 2685 | 2450 | 2528 |
| 23092 | NM_00113560 | ARHGAP26 | Rho GTPase activating protein 26 | protein_coding | NP_001129080.1;NP_001336476.1;NP_055886.1;XP_005268455.2;XP_00 | 17073 | 605370 | . | . | 1.471417 | 1.773217 | 1.836891 | 1.353312 | 3.424050 | 4.098870 | 4.121930 | 3.094184 | -1.099164 | 287.328370 | 0.213430898 | -0.63911333 | 0.522749209 | 0.998865404 | 8.233226 | 8.096657 | 8.234469 | 8.144442 | 8.103418 | 8.176254 | 375 | 298 | 221 | 275 |
| 23093 | NM_015072 | TTLL5 | tubulin tyrosine ligase like 5 | protein_coding | NP_055887.3 | 19963 | 612268 | . | . | 4.239168 | 4.701851 | 4.435796 | 5.484167 | 9.864717 | 10.868539 | 9.953797 | 12.538897 | 1.094332 | 692.844954 | 0.164518337 | 0.790495288 | 0.429238573 | 0.998865404 | 9.370354 | 9.499533 | 9.369968 | 9.434513 | 9.392096 | 9.531884 | 726 | 748 | 566 | 747 |
| 23094 | NM_015073,XI | SIPA1L3 | signal induced proliferation associated 1 like 3 | protein_coding | NP_055888.1;XP_005258728.1;XP_011524959.1;XP_016882007.1 | 23801 | 616655 | . | . | 1.745719 | 1.828938 | 1.765697 | 2.186150 | 4.062358 | 4.227673 | 3.962173 | 4.998372 | 1.141399 | 482.295397 | 0.173355116 | 1.100651033 | 0.271048565 | 0.998865404 | 8.815584 | 9.005703 | 8.850072 | 8.887880 | 8.912596 | 8.990009 | 503 | 501 | 420 | 511 |
| 23095 | NM_00136595 | KIF1B | kinesin family member 1B | protein_coding | NP_001352880.1;NP_001352881.1;NP_001352882.1;NP_055889.2;NP_9 | 16636 | 605995 | . | . | 10.651431 | 10.797604 | 9.588954 | 10.099816 | 24.786316 | 24.959144 | 21.517335 | 23.092029 | -1.095473 | 2859.394215 | 0.112673763 | -1.16756188 | 0.242983534 | 0.992733415 | 11.545806 | 11.414181 | 11.522412 | 11.512639 | 11.423995 | 11.462049 | 3463 | 3207 | 2317 | 2619 |
| 23096 | NM_00111112 | IQSEC2 | IQ motif and Sec7 domain ArfGEF 2 | protein_coding | NP_001104595.1;NP_001230126.1;NP_055890.1;XP_006724642.1;XP_00 | 29059 | 300522 | . | . | 5.425762 | 4.598967 | 4.252087 | 4.603099 | 12.625975 | 10.630719 | 9.541558 | 10.524437 | -1.041676 | 332.017078 | 0.198083832 | -0.29738551 | 0.766172204 | 0.998865404 | 8.404379 | 8.345243 | 8.414730 | 8.356821 | 8.330458 | 8.395309 | 410 | 347 | 266 | 323 |
| 23097 | NM_00130096 | CDK19 | cyclin dependent kinase 19 | protein_coding | NP_001287889.1;NP_001287892.1;NP_001287893.1;NP_055891.1;XP_0C | 19338 | 614720 | . | . | 3.285573 | 3.239328 | 3.966875 | 3.313709 | 7.645668 | 7.487852 | 8.901551 | 7.576401 | 1.103750 | 686.974571 | 0.158375842 | 0.899215553 | 0.368537858 | 0.998865404 | 9.350827 | 9.493855 | 9.395833 | 9.379487 | 9.509072 | 9.398752 | 761 | 696 | 677 | 622 |
| 23098 | NM_015077 | SARM1 | sterile alpha and TIR motif containing 1 | protein_coding | NP_055892.2 | 17074 | 607732 | . | . | 2.614955 | 2.973413 | 3.404413 | 3.471848 | 6.085107 | 6.873178 | 7.639403 | 7.937968 | 1.214154 | 996.955234 | 0.141126026 | 1.983699434 | 0.047289357 | 0.659753773 | 9.814805 | 10.094436 | 9.840614 | 9.926470 | 10.022612 | 10.030244 | 975 | 1030 | 946 | 1030 |
| 23099 | NM_00113577 | ZBTB43 | zinc finger and BTB domain containing 43 | protein_coding | NP_001129248.1;NP_054726.1;XP_005251892.1;XP_011516711.1;XP_01 | 17908 | 618676 | . | . | 1.680225 | 2.087391 | 2.042935 | 1.683387 | 3.909953 | 4.825099 | 4.584288 | 3.848864 | -1.006385 | 350.702143 | 0.209636264 | -0.04379809 | 0.96506535 | 0.998865404 | 8.458510 | 8.449681 | 8.390760 | 8.510478 | 8.513320 | 8.379905 | 363 | 419 | 340 | 289 |
| 23101 | NM_015078,XI | MCF2L2 | MCF.2 cell line derived transforming sequence-like 2 | protein_coding | NP_055893.3;XP_011510887.1;XP_011510888.1;XP_016861432.2;XP_01 | 30319 | . | | . | 0.492198 | 0.320219 | 0.336489 | 0.375185 | 1.145365 | 0.740202 | 0.755070 | 0.857815 | -1.167221 | 68.245334 | 0.399374243 | -0.55856765 | 0.57645682 | 0.998865404 | 6.199339 | 5.977456 | 6.160327 | 6.056119 | 6.066461 | 6.080259 | 102 | 63 | 53 | 61 |
| 23102 | NM_015079,N | TBC1D2B | TBC1 domain family member 2B | protein_coding | NP_055894.6;NP_653173.1;XP_011519689.1;XP_016877511.1 | 29183 | . | | . | 5.086046 | 5.252496 | 4.842817 | 5.279260 | 11.835434 | 12.141379 | 10.867145 | 12.070400 | -1.039449 | 957.338135 | 0.140154899 | -0.39826755 | 0.690432974 | 0.998865404 | 9.930692 | 9.874531 | 9.907036 | 9.926379 | 9.857847 | 9.918277 | 1109 | 1067 | 782 | 917 |
| 23107 | NM_00128674 | MRPS27 | mitochondrial ribosomal protein S27 | protein_coding | NP_001273677.1;NP_001273680.1;NP_055899.2 | 14512 | 611989 | . | . | 20.931857 | 22.422244 | 22.313448 | 24.192034 | 48.709289 | 51.830019 | 50.070732 | 55.312211 | 1.057182 | 1938.550091 | 0.1220753 | 0.657169498 | 0.511071936 | 0.998865404 | 10.880218 | 10.960198 | 10.873550 | 10.921853 | 10.913325 | 10.969801 | 2104 | 2096 | 1671 | 1934 |
| 23108 | NM_00110039 | RAP1GAP2 | RAP1 GTPase activating protein 2 | protein_coding | NP_001093868.1;NP_001316987.1;NP_055900.4;XP_005256599.1;XP_01 | 29176 | 618714 | . | . | 1.740291 | 2.014535 | 2.037159 | 2.147560 | 4.049729 | 4.656689 | 4.571327 | 4.910141 | 1.094780 | 378.776988 | 0.188944762 | 0.691425672 | 0.489298076 | 0.998865404 | 8.498782 | 8.628704 | 8.494709 | 8.576866 | 8.573369 | 8.605072 | 387 | 418 | 335 | 381 |
| 23109 | NM_015086,XI | DDN | dendrin | protein_coding | NP_055901.2;XP_011536357.1 | 24458 | 610588 | . | . | 0.097480 | 0.089525 | 0.047726 | 0.038092 | 0.226839 | 0.206940 | 0.107096 | 0.087093 | -2.425980 | 9.488176 | 1.014772777 | -1.25995468 | 0.207685712 | NA | 3.747897 | 2.470304 | 3.276744 | 3.272364 | 3.240664 | 3.238465 | 16 | 14 | 5 | 5 |
| 23111 | NM_00114229 | SPART | spartin | protein_coding | NP_001135766.1;NP_001135767.1;NP_001135768.1;NP_055902.1;XP_0C | 18514 | 607111 | . | . | 13.238026 | 13.002239 | 14.741527 | 15.999367 | 30.805430 | 30.055255 | 33.079560 | 36.580652 | 1.155528 | 2196.545785 | 0.118187294 | 1.764592739 | 0.077632237 | 0.767975096 | 10.993027 | 11.201491 | 11.049914 | 11.029718 | 11.127878 | 11.182564 | 2377 | 2171 | 1979 | 2282 |
| 23112 | NM_00102484 | TNRC6B | trinucleotide repeat containing adaptor 6B | protein_coding | NP_001020014.1;NP_001155973.1;NP_055903.2 | 29190 | 610740 | . | . | 2.052767 | 2.288645 | 2.231069 | 2.411325 | 4.776873 | 5.290306 | 5.006452 | 5.513209 | 1.057180 | 1258.944556 | 0.1343747 | 0.596993324 | 0.550511838 | 0.998865404 | 10.257529 | 10.337365 | 10.237529 | 10.313095 | 10.291221 | 10.344006 | 1339 | 1387 | 1086 | 1255 |
| 23113 | NM_015089,XI | CUL9 | cullin 9 | protein_coding | NP_055904.1;XP_011512724.1;XP_011512725.1;XP_011512726.1;XP_01 | 15982 | 607489 | . | . | 3.528464 | 3.382951 | 4.049544 | 4.054598 | 8.210882 | 7.819842 | 9.087060 | 9.270359 | 1.090905 | 866.306579 | 0.146530436 | 0.856653332 | 0.391636502 | 0.998865404 | 9.694356 | 9.820362 | 9.737448 | 9.713734 | 9.831840 | 9.742332 | 970 | 879 | 835 | 795 |
| 23114 | NM_00100538 | NFASC | neurofascin | protein_coding | NP_001005388.2;NP_001005389.2;NP_001153803.1;NP_001153804.1;NI | 29866 | 609145 | . | . | 0.262702 | 0.161431 | 0.315577 | 0.282330 | 0.611318 | 0.373154 | 0.708148 | 0.645513 | 1.258304 | 72.633813 | 0.378828818 | 0.875012933 | 0.381566869 | 0.998865404 | 6.006229 | 6.339685 | 6.177531 | 6.116175 | 6.220169 | 6.186685 | 84 | 60 | 75 | 71 |
| 23116 | NM_00130812 | TOGARAM1 | TOG array regulator of axonemal microtubules 1 | protein_coding | NP_001295049.1;NP_055906.2;XP_011534873.1;XP_016876587.1;XP_01 | 19959 | 617618 | . | . | 1.795467 | 1.442149 | 2.120716 | 1.620932 | 4.178125 | 3.333594 | 4.758825 | 3.706067 | 1.093498 | 310.512379 | 0.20806736 | 0.619751296 | 0.535421538 | 0.998865404 | 8.211987 | 8.342094 | 8.265733 | 8.235685 | 8.357953 | 8.237791 | 351 | 311 | 313 | 271 |
| 23117 | NM_130464 | NPIPB3 | nuclear pore complex interacting protein family member B3 | protein_coding | NP_569731.2 | 28989 | . | | . | 7.510056 | 8.141432 | 12.231874 | 8.382972 | 17.476208 | 18.819284 | 27.447973 | 19.166670 | 1.300116 | 1036.235799 | 0.17665267 | 2.143414397 | 0.032079838 | 0.566237858 | 9.815122 | 10.194307 | 9.879108 | 9.930668 | 10.235185 | 9.941726 | 1000 | 1007 | 1214 | 888 |
| 23118 | NM_00129203 | TAB2 | TGF-beta activated kinase 1 (MAP3K7) binding protein 2 | protein_coding | NP_001278963.1;NP_001278964.1;NP_001356435.1;NP_055908.1;XP_01 | 17075 | 605101 | . | . | 7.889918 | 7.426271 | 7.486493 | 8.008673 | 18.360164 | 17.166161 | 16.799474 | 18.310879 | -1.022813 | 1076.230827 | 0.136066965 | -0.23916864 | 0.81097482 | 0.998865404 | 10.087979 | 10.055380 | 10.101627 | 10.057861 | 10.041340 | 10.084706 | 1290 | 1140 | 898 | 1027 |
| 23119 | NM_015094,XI | HIC2 | HIC ZBTB transcriptional repressor 2 | protein_coding | NP_055909.2;XP_011528309.1;XP_011528310.1;XP_011528311.1;XP_01 | 18595 | 607712 | . | . | 1.279305 | 1.372790 | 1.210242 | 1.207677 | 2.976994 | 3.173264 | 2.715748 | 2.761211 | -1.107933 | 273.611881 | 0.20841351 | -0.70950669 | 0.478010102 | 0.998865404 | 8.168095 | 8.020084 | 8.107274 | 8.136934 | 8.072074 | 8.066240 | 322 | 319 | 227 | 242 |
| 23120 | NM_00136665 | ATP10B | ATPase phospholipid transporting 10B (putative) | protein_coding | NP_001353581.1;NP_001353582.1;NP_001353583.1;NP_001353584.1;NI | 13543 | . | | . | 7.548150 | 8.674976 | 8.673263 | 9.198155 | 17.564856 | 20.052601 | 19.462550 | 21.030491 | 1.081531 | 1892.702513 | 0.124172624 | 0.910630413 | 0.362490141 | 0.998865404 | 10.828719 | 10.941540 | 10.812801 | 10.893385 | 10.897351 | 10.933900 | 1987 | 2063 | 1671 | 1886 |
| 23122 | NM_00120704 | CLASP2 | cytoplasmic linker associated protein 2 | protein_coding | NP_001193973.1;NP_001352556.1;NP_001352557.1;NP_001352558.1;NI | 17078 | 605853 | . | . | 4.564771 | 4.360150 | 4.730615 | 4.625674 | 10.622412 | 10.078681 | 10.615361 | 10.576055 | 1.043640 | 1012.071785 | 0.136931057 | 0.450040675 | 0.652681112 | 0.998865404 | 9.951838 | 10.013690 | 9.986344 | 9.947069 | 10.009722 | 9.986784 | 1171 | 1040 | 913 | 954 |
| 23125 | NM_00117116 | CAMTA2 | calmodulin binding transcription activator 2 | protein_coding | NP_001164637.1;NP_001164638.1;NP_001164639.1;NP_055914.2;XP_0C | 18807 | 611508 | . | . | 8.138981 | 8.270576 | 9.014112 | 9.106367 | 18.939741 | 19.117811 | 20.227401 | 20.820628 | 1.053173 | 1241.226002 | 0.129280068 | 0.578147278 | 0.56316469 | 0.998865404 | 10.239727 | 10.314407 | 10.257863 | 10.257117 | 10.288144 | 10.305011 | 1393 | 1304 | 1095 | 1207 |
| 23126 | NM_00119493 | POGZ | pogo transposable element derived with ZNF domain | protein_coding | NP_001181866.1;NP_001181867.1;NP_055915.2;NP_665739.3;NP_9970! | 18801 | 614787 | . | . | 6.143312 | 6.535220 | 7.222323 | 6.099689 | 14.295738 | 15.106451 | 16.206683 | 13.946212 | 1.113717 | 1439.637157 | 0.129992866 | 1.195312821 | 0.231964832 | 0.986726781 | 10.411661 | 10.567132 | 10.421981 | 10.472980 | 10.564561 | 10.495513 | 1516 | 1519 | 1378 | 1359 |
| 23127 | NM_00130342 | COLGALT2 | collagen beta(1-O)galactosyltransferase 2 | protein_coding | NP_001290349.1;NP_001290350.1;NP_055916.1 | 16790 | 617533 | . | . | 2.402620 | 2.804721 | 1.940093 | 2.589437 | 5.590996 | 6.483239 | 4.353512 | 5.920440 | -1.150625 | 390.336551 | 0.202127123 | -1.00143898 | 0.316614628 | 0.998865404 | 8.706933 | 8.503017 | 8.603341 | 8.687584 | 8.482520 | 8.638468 | 448 | 482 | 272 | 387 |
| 23129 | NM_015103,XI | PLXND1 | plexin D1 | protein_coding | NP_055918.3;XP_011510890.1;XP_011510891.1;XP_011510892.1;XP_01 | 9107 | 604282 | . | . | 5.721366 | 6.251594 | 3.643003 | 4.549935 | 13.313852 | 14.450841 | 8.174794 | 10.402884 | -1.449838 | 1063.655481 | 0.149926611 | -3.57436262 | 0.000351082 | 0.031064013 | 10.298333 | 9.761676 | 10.150232 | 10.197178 | 9.822779 | 9.971358 | 1406 | 1400 | 674 | 901 |
| 23130 | NM_00136797 | ATG2A | autophagy related 2A | protein_coding | NP_001354900.1;NP_001354901.1;NP_055919.2;XP_011543165.1;XP_01 | 29028 | 616225 | . | . | 3.534371 | 4.044277 | 4.539175 | 4.368407 | 8.224627 | 9.348531 | 10.185776 | 9.987845 | 1.159413 | 811.403984 | 0.149363532 | 1.428696229 | 0.153091562 | 0.91755225 | 9.553727 | 9.766954 | 9.564184 | 9.650144 | 9.729810 | 9.695124 | 812 | 861 | 776 | 797 |
| 23131 | NM_00100290 | GPATCH8 | G-patch domain containing 8 | protein_coding | NP_001002909.1;NP_001291868.1;NP_001291869.1;NP_001291870.1;NI | 29066 | 614396 | . | . | 7.543906 | 8.318837 | 8.846303 | 9.575121 | 17.554980 | 19.229364 | 19.850848 | 21.892377 | 1.139649 | 1861.768169 | 0.122899351 | 1.534503252 | 0.124905894 | 0.873049043 | 10.765209 | 10.953563 | 10.776327 | 10.838728 | 10.891203 | 10.930721 | 1924 | 1953 | 1682 | 1905 |
| 23132 | NM_00132225 | RAD54L2 | RAD54 like 2 | protein_coding | NP_001309182.1;NP_001309185.1;NP_001309186.1;NP_055921.2 | 29123 | . | | . | 1.956046 | 2.178203 | 2.279754 | 2.104950 | 4.551799 | 5.035015 | 5.115703 | 4.812718 | 1.046783 | 662.728431 | 0.157782761 | 0.418059705 | 0.675903465 | 0.998865404 | 9.338899 | 9.404894 | 9.319788 | 9.390171 | 9.415592 | 9.356789 | 707 | 735 | 615 | 608 |
| 23133 | NM_00118489 | PHF8 | PHD finger protein 8 | protein_coding | NP_001171825.1;NP_001171826.1;NP_001171827.1;NP_055922.1;XP_0C | 20672 | 300560 | . | . | 4.316725 | 4.787363 | 5.237665 | 5.205490 | 10.045198 | 11.066205 | 11.753168 | 11.901737 | 1.075043 | 892.522666 | 0.145816576 | 0.715928931 | 0.474035233 | 0.998865404 | 9.748705 | 9.852904 | 9.727907 | 9.818484 | 9.838541 | 9.813694 | 928 | 987 | 818 | 852 |
| 23135 | NM_00108042 | KDM6B | lysine demethylase 6B | protein_coding | NP_001073893.1;NP_001335645.1;XP_005256606.1;XP_005256607.1;XP | 29012 | 611577 | . | . | 3.122287 | 3.171646 | 3.631207 | 3.463139 | 7.265693 | 7.331401 | 8.148323 | 7.918058 | 1.125312 | 719.935079 | 0.152768722 | 1.114921132 | 0.26488422 | 0.996598404 | 9.403908 | 9.574498 | 9.433975 | 9.462154 | 9.567029 | 9.491162 | 764 | 746 | 699 | 676 |
| 23136 | NM_00128153 | EPB41L3 | erythrocyte membrane protein band 4.1 like 3 | protein_coding | NP_001268462.1;NP_001268463.1;NP_001268464.1;NP_001317486.1;NI | 3380 | 605331 | . | . | 16.236570 | 17.363281 | 11.291228 | 13.898540 | 37.783168 | 40.135992 | 25.337188 | 31.777362 | -1.374056 | 1818.931136 | 0.138246798 | -3.31610715 | 0.000912808 | 0.064386964 | 11.040177 | 10.581226 | 10.922476 | 10.955932 | 10.602218 | 10.769117 | 2375 | 2319 | 1185 | 1595 |
| 23137 | NM_015110,XI | SMC5 | structural maintenance of chromosomes 5 | protein_coding | NP_055925.2;XP_005251894.1;XP_005251895.1;XP_005251896.1;XP_01 | 20465 | 609386 | . | . | 6.571896 | 6.757695 | 7.246820 | 5.294414 | 15.293075 | 15.620714 | 16.261655 | 12.105050 | -1.033916 | 1144.574687 | 0.165372414 | -0.2909704 | 0.771073959 | 0.998865404 | 10.184066 | 10.136738 | 10.205415 | 10.131649 | 10.258413 | 10.016138 | 1405 | 1194 | 1138 | 884 |
| 23138 | NM_015111,XI | N4BP3 | NEDD4 binding protein 3 | protein_coding | NP_055926.1;XP_006714897.1;XP_011532775.1;XP_011532776.1 | 29852 | . | | . | 0.708897 | 0.608710 | 0.674638 | 0.721406 | 1.649632 | 1.407060 | 1.513869 | 1.649411 | -1.017067 | 130.353822 | 0.288584386 | -0.08460133 | 0.932578342 | 0.998865404 | 7.038377 | 7.014099 | 7.059280 | 6.997975 | 7.006041 | 7.039729 | 164 | 130 | 107 | 127 |
| 23139 | NM_00131924 | MAST2 | microtubule associated serine/threonine kinase 2 | protein_coding | NP_001306174.1;NP_001311249.1;NP_001311250.1;NP_055927.2;XP_0C | 19035 | 612257 | . | . | 6.454479 | 6.818764 | 5.839978 | 6.931341 | 15.019837 | 15.761874 | 13.104742 | 15.847687 | -1.047511 | 1170.172882 | 0.139787187 | -0.47904892 | 0.631903829 | 0.998865404 | 10.225849 | 10.158374 | 10.190324 | 10.227578 | 10.116782 | 10.228499 | 1345 | 1324 | 919 | 1152 |

| | | | | | | | | |
|---|---|---|---|---|---|---|---|---|
| 23140 | NM_015113,XI | ZZEF1 | zinc finger ZZ-type and EF-hand domain containing 1 | protein_coding | NP_055928.3;XP_005256618.1;XP_011522061.1;XP_016879871.1;XP_01 | 29027 | . | [illegible] |
| 23141 | NM_015114,XI | ANKLE2 | ankyrin repeat and LEM domain containing 2 | protein_coding | NP_055929.1;XP_005266216.1;XP_005266217.1;XP_006719798.1;XP_01 | 29101 | 616062 | [illegible] |
| 23142 | NM_00104040 | DCUN1D4 | defective in cullin neddylation 1 domain containing 4 | protein_coding | NP_001035492.1;NP_001274684.1;NP_001274686.1;NP_055930.2;XP_00 | 28998 | 612977 | [illegible] |
| 23143 | NM_00116421 | LRCH1 | leucine rich repeats and calponin homology domain containing 1 | protein_coding | NP_001157683.2;NP_001157685.2;NP_055931.2;XP_016875972.1;XP_01 | 20309 | 610368 | [illegible] |
| 23144 | NM_015117,XI | ZC3H3 | zinc finger CCCH-type containing 3 | protein_coding | NP_055932.2;XP_006716599.2;XP_011515245.1;XP_011515246.1;XP_01 | 28972 | 618640 | [illegible] |
| 23145 | NR_163594 | SSPOP | SCO-spondin, pseudogene | transcribed_ps | . | 21998 | 617356 | [illegible] |
| 23149 | NM_00116135 | FCHO1 | FCH and mu domain containing endocytic adaptor 1 | protein_coding | NP_001154829.1;NP_001154830.1;NP_001154831.1;NP_055937.1;XP_00 | 29002 | 613437 | [illegible] |
| 23150 | NM_015123,XI | FRMD4B | FERM domain containing 4B | protein_coding | NP_055938.2;XP_005264779.1;XP_005264780.1;XP_016861478.1;XP_01 | 24886 | 617467 | [illegible] |
| 23151 | NM_00136666 | GRAMD4 | GRAM domain containing 4 | protein_coding | NP_001353589.1;NP_055939.1;XP_005261453.1;XP_006724232.1;XP_00 | 29113 | 613691 | [illegible] |
| 23152 | NM_00130481 | CIC | capicua transcriptional repressor | protein_coding | NP_001291744.1;NP_001366409.1;NP_001366411.1;NP_001366413.1;NI | 14214 | 612082 | [illegible] |
| 23154 | NM_00101483 | NCDN | neurochondrin | protein_coding | NP_001014839.1;NP_001014841.1;NP_055099.1 | 17597 | 608458 | [illegible] |
| 23155 | NM_00104821 | CLCC1 | chloride channel CLIC like 1 | protein_coding | NP_001041675.1;NP_001265131.1;NP_001265132.1;NP_001364387.1;NI | 29675 | 617539 | [illegible] |
| 23157 | NM_015129,N | SEPTIN6 | septin 6 | protein_coding | NP_055944.2;NP_665798.1;NP_665799.1;NP_665801.1;XP_005262457.1 | 15848 | 300683 | [illegible] |
| 23158 | NM_015130 | TBC1D9 | TBC1 domain family member 9 | protein_coding | NP_055945.2 | 21710 | 618035 | [illegible] |
| 23160 | NM_015131 | WDR43 | WD repeat domain 43 | protein_coding | NP_055946.1 | 28945 | 616195 | [illegible] |
| 23161 | NM_00135086 | SNX13 | sorting nexin 13 | protein_coding | NP_001337791.1;NP_001337792.1;NP_001337793.1;NP_001337795.1;NI | 21335 | 606589 | [illegible] |
| 23162 | NM_00104043 | MAPK8IP3 | mitogen-activated protein kinase 8 interacting protein 3 | protein_coding | NP_001035529.1;NP_001305781.1;NP_055948.2;XP_005255247.1;XP_01 | 6884 | 605431 | [illegible] |
| 23163 | NM_00117270 | GGA3 | golgi associated, gamma adaptin ear containing, ARF binding prot | protein_coding | NP_001166174.1;NP_001166175.1;NP_001278570.1;NP_001278571.1;NI | 17079 | 606006 | [illegible] |
| 23164 | NM_00136471 | MPRIP | myosin phosphatase Rho interacting protein | protein_coding | NP_001351645.2;NP_055949.2;NP_958431.2;XP_005256620.1;XP_00525 | 30321 | 612935 | [illegible] |
| 23165 | NM_00132943 | NUP205 | nucleoporin 205 | protein_coding | NP_001316363.2;NP_055950.2 | 18658 | 614352 | [illegible] |
| 23167 | NM_00132355 | EFR3A | EFR3 homolog A | protein_coding | NP_001310482.1;NP_001310483.1;NP_001310484.1;NP_001310485.1;NI | 28970 | 611798 | [illegible] |
| 23168 | NM_015138 | RTF1 | RTF1 homolog, Paf1/RNA polymerase II complex component | protein_coding | NP_055953.3 | 28996 | [illegible] | [illegible] |
| 23169 | NM_015139,XI | SLC35D1 | solute carrier family 35 member D1 | protein_coding | NP_055954.1;XP_006710541.1;XP_011539372.1 | 20800 | [illegible] | [illegible] |
| 23170 | NM_015140 | TTLL12 | tubulin tyrosine ligase like 12 | protein_coding | NP_055955.1 | 28974 | . | [illegible] |
| 23171 | NM_015141,XI | GPD1L | glycerol-3-phosphate dehydrogenase 1 like | protein_coding | NP_055956.1;XP_006713131.1 | 28956 | [illegible] | [illegible] |
| 23172 | NM_032182,XI | ABRAXAS2 | abraxas 2, BRISC complex subunit | protein_coding | NP_115558.3;XP_011537856.1;XP_016871475.1 | 28975 | [illegible] | [illegible] |
| 23173 | NM_015143,XI | METAP1 | methionyl aminopeptidase 1 | protein_coding | NP_055958.2;XP_011530079.1;XP_011530080.1;XP_011530081.1;XP_01 | 15789 | [illegible] | [illegible] |
| 23174 | NM_015144,XI | ZCCHC14 | zinc finger CCHC-type containing 14 | protein_coding | NP_055959.2;XP_005255915.2;XP_011521266.1;XP_016878571.1 | 24134 | . | [illegible] |
| 23175 | NM_00126142 | LPIN1 | lipin 1 | protein_coding | NP_001248356.1;NP_001248357.1;NP_001336128.1;NP_001336129.1;NI | 13345 | [illegible] | [illegible] |
| 23176 | NM_00109881 | SEPTIN8 | septin 8 | protein_coding | NP_001092281.1;NP_001092282.1;NP_001092283.1;NP_001287727.1;NI | 16511 | [illegible] | [illegible] |
| 23177 | NM_00131910 | CEP68 | centrosomal protein 68 | protein_coding | NP_001306029.1;NP_001306030.1;NP_055962.2;XP_005264276.1;XP_01 | 29076 | [illegible] | [illegible] |
| 23178 | NM_00125211 | PASK | PAS domain containing serine/threonine kinase | protein_coding | NP_001239048.1;NP_001239049.1;NP_001239051.1;NP_001239053.1;NI | 17270 | [illegible] | [illegible] |
| 23179 | NM_00129766 | RGL1 | ral guanine nucleotide dissociation stimulator like 1 | protein_coding | NP_001284598.1;NP_001284599.1;NP_001284600.1;NP_001284601.1;NI | 30281 | [illegible] | [illegible] |
| 23180 | NM_015150,XI | RFTN1 | raftlin, lipid raft linker 1 | protein_coding | NP_055965.1;XP_005265042.1;XP_005265043.1;XP_006713132.1;XP_01 | 30278 | [illegible] | [illegible] |
| 23181 | NM_00114611 | DIP2A | disco interacting protein 2 homolog A | protein_coding | NP_001139587.1;NP_001139588.1;NP_001340871.1;NP_001340872.1;NI | 17217 | [illegible] | [illegible] |
| 23184 | NM_015154,N | MESD | mesoderm development LRP chaperone | protein_coding | NP_055969.1 | 13520 | [illegible] | [illegible] |
| 23185 | NM_00135127 | LARP4B | La ribonucleoprotein 4B | protein_coding | NP_001338206.1;NP_055970.1;XP_005252492.1;XP_011517737.1;XP_01 | 28987 | [illegible] | [illegible] |
| 23186 | NM_015156,XI | RCOR1 | REST corepressor 1 | protein_coding | NP_055971.2 | 17441 | [illegible] | [illegible] |
| 23187 | NM_00114475 | PHLDB1 | pleckstrin homology like domain family B member 1 | protein_coding | NP_001138230.1;NP_001138231.1;NP_055972.1;XP_005271525.1;XP_00 | 23697 | [illegible] | [illegible] |
| 23189 | NM_00125687 | KANK1 | KN motif and ankyrin repeat domains 1 | protein_coding | NP_001243805.1;NP_001243806.1;NP_001341260.1;NP_001341261.1;NI | 19309 | [illegible] | [illegible] |
| 23190 | NM_014607 | UBXN4 | UBX domain protein 4 | protein_coding | NP_055422.1 | 14860 | [illegible] | [illegible] |
| 23191 | NM_00103302 | CYFIP1 | cytoplasmic FMR1 interacting protein 1 | protein_coding | NP_001028200.1;NP_001274739.1;NP_001311048.1;NP_001311049.1;NI | 13759 | [illegible] | [illegible] |
| 23192 | NM_013325,N | ATG4B | autophagy related 4B cysteine peptidase | protein_coding | NP_037457.3;NP_847896.1;XP_005247049.2;XP_005247050.2;XP_00524 | 20790 | [illegible] | [illegible] |
| 23193 | NM_00127819 | GANAB | glucosidase II alpha subunit | protein_coding | NP_001265121.1;NP_001265122.1;NP_001265123.1;NP_001316151.1;NI | 4138 | [illegible] | [illegible] |
| 23194 | NM_00127831 | FBXL7 | F-box and leucine rich repeat protein 7 | protein_coding | NP_001265246.1;NP_036436.1;XP_011512300.1;XP_016864751.1 | 13604 | [illegible] | [illegible] |
| 23195 | NM_014611,XI | MDN1 | midasin AAA ATPase 1 | protein_coding | NP_055426.1;XP_005248757.1;XP_006715468.1;XP_011533937.1;XP_01 | 18302 | [illegible] | [illegible] |
| 23196 | NM_00128672 | FAM120A | family with sequence similarity 120A | protein_coding | NP_001273651.1;NP_001273652.1;NP_001273653.1;NP_055427.2;XP_00 | 13247 | [illegible] | [illegible] |
| 23197 | NM_014613,XI | FAF2 | Fas associated factor family member 2 | protein_coding | NP_055428.1;XP_011532777.1 | 24666 | [illegible] | [illegible] |
| 23198 | NM_014614,XI | PSME4 | proteasome activator subunit 4 | protein_coding | NP_055429.2;XP_006712032.1;XP_011531007.1;XP_011531008.1;XP_01 | 20635 | [illegible] | [illegible] |
| 23199 | NM_00113447 | GSE1 | Gse1 coiled-coil protein | protein_coding | NP_001127945.1;NP_001265113.1;NP_055430.1;XP_005255916.3;XP_00 | 28979 | [illegible] | [illegible] |
| 23200 | NM_014616,XI | ATP11B | ATPase phospholipid transporting 11B (putative) | protein_coding | NP_055431.1;XP_005247298.1;XP_005247300.1;XP_011510895.1;XP_01 | 13553 | [illegible] | [illegible] |
| 23201 | NM_00128605 | FAM168A | family with sequence similarity 168 member A | protein_coding | NP_001272979.1;NP_001272980.1;NP_055974.1;XP_016872904.1;XP_01 | 28999 | [illegible] | [illegible] |
| 23203 | NM_00128294 | PMPCA | peptidase, mitochondrial processing subunit alpha | protein_coding | NP_001269873.1;NP_001269875.1;NP_055975.1;XP_005266116.1;XP_01 | 18667 | [illegible] | [illegible] |
| 23204 | NM_00131385 | ARL6IP1 | ADP ribosylation factor like GTPase 6 interacting protein 1 | protein_coding | NP_001300787.1;NP_055976.1 | 697 | [illegible] | [illegible] |
| 23205 | NM_00119937 | ACSBG1 | acyl-CoA synthetase bubblegum family member 1 | protein_coding | NP_001186306.1;NP_055977.3;XP_011519692.1;XP_011519693.2;XP_01 | 29567 | [illegible] | [illegible] |
| 23207 | NM_015164,XI | PLEKHM2 | pleckstrin homology and RUN domain containing M2 | protein_coding | NP_055979.2;XP_005245847.1;XP_005245848.1;XP_016856246.1;XP_01 | 29131 | [illegible] | [illegible] |
| 23208 | NM_152280,XI | SYT11 | synaptotagmin 11 | protein_coding | NP_689493.3;XP_005245071.1;XP_016856248.1 | 19239 | [illegible] | [illegible] |
| 23209 | NM_00137647 | MLC1 | modulator of VRAC current 1 | protein_coding | NP_001363401.1;NP_001363402.1;NP_001363403.1;NP_001363404.1;NI | 17082 | [illegible] | [illegible] |
| 23210 | NM_00108146 | JMJD6 | jumonji domain containing 6, arginine demethylase and lysine hyd | protein_coding | NP_001074930.1;NP_055982.2 | 19355 | [illegible] | [illegible] |
| 23211 | NM_015168,XI | ZC3H4 | zinc finger CCCH-type containing 4 | protein_coding | NP_055983.1;XP_005258733.1;XP_005258734.1;XP_005258735.1;XP_00 | 17808 | . | [illegible] |
| 23212 | NM_015169 | RRS1 | ribosome biogenesis regulator 1 homolog | protein_coding | NP_055984.1 | 17083 | [illegible] | [illegible] |
| 23213 | NM_00112820 | SULF1 | sulfatase 1 | protein_coding | NP_001121676.1;NP_001121677.1;NP_001121678.1;NP_055985.2;XP_00 | 20391 | [illegible] | [illegible] |
| 23214 | NM_00127094 | XPO6 | exportin 6 | protein_coding | NP_001257869.1;NP_055986.1;XP_005255255.1;XP_005255256.1;XP_01 | 19733 | [illegible] | [illegible] |
| 23215 | NM_015172,XI | PRRC2C | proline rich coiled-coil 2C | protein_coding | NP_055987.2;XP_005245072.1;XP_005245073.1;XP_005245074.1;XP_00 | 24903 | [illegible] | [illegible] |
| 23216 | NM_00125391 | TBC1D1 | TBC1 domain family member 1 | protein_coding | NP_001240841.1;NP_001240842.1;NP_001240843.1;NP_001240844.1;NI | 11578 | [illegible] | [illegible] |
| 23217 | NM_00114564 | ZFR2 | zinc finger RNA binding protein 2 | protein_coding | NP_001139112.1;NP_055989.1;XP_011526133.1 | 29189 | . | [illegible] |
| 23218 | NM_00136511 | NBEAL2 | neurobeachin like 2 | protein_coding | NP_001352045.1;NP_055990.1;XP_006713135.1;XP_011531835.1;XP_01 | 31928 | [illegible] | [illegible] |
| 23219 | NM_00113611 | FBXO28 | F-box protein 28 | protein_coding | NP_001129587.1;NP_055991.1 | 29046 | [illegible] | [illegible] |
| 23220 | NM_00130072 | DTX4 | deltex E3 ubiquitin ligase 4 | protein_coding | NP_001287656.1;NP_055992.1;XP_006718546.1 | 29151 | [illegible] | [illegible] |
| 23221 | NM_00116003 | RHOBTB2 | Rho related BTB domain containing 2 | protein_coding | NP_001153508.1;NP_001153509.1;NP_001361720.1;NP_055993.2;XP_01 | 18756 | [illegible] | [illegible] |
| 23223 | NM_00114511 | RRP12 | ribosomal RNA processing 12 homolog | protein_coding | NP_001138586.1;NP_001271266.1;NP_055994.2;XP_011537858.1;XP_02 | 29100 | [illegible] | [illegible] |
| 23224 | NM_015180,N | SYNE2 | spectrin repeat containing nuclear envelope protein 2 | protein_coding | NP_055995.4;NP_878914.1;NP_878917.1;NP_878918.2;XP_005267511.1 | 17084 | [illegible] | [illegible] |
| 23225 | NM_024923,XI | NUP210 | nucleoporin 210 | protein_coding | NP_079199.2;XP_005265051.1;XP_011531840.1;XP_016861510.1;XP_02 | 30052 | [illegible] | [illegible] |
| 23228 | NM_00114438 | PLCL2 | phospholipase C like 2 | protein_coding | NP_001137854.1;NP_055999.2;XP_006713136.1;XP_016861511.1;XP_01 | 9064 | [illegible] | [illegible] |
| 23229 | NM_00117347 | ARHGEF9 | Cdc42 guanine nucleotide exchange factor 9 | protein_coding | NP_001166950.1;NP_001166951.1;NP_001317424.1;NP_001340850.1;NI | 14561 | [illegible] | [illegible] |
| 23230 | NM_00101803 | VPS13A | vacuolar protein sorting 13 homolog A | protein_coding | NP_001018047.1;NP_001018048.1;NP_056001.1;NP_150648.2 | 1908 | [illegible] | [illegible] |
| 23231 | NM_00129759 | SEL1L3 | SEL1L family member 3 | protein_coding | NP_001284521.1;NP_001284523.1;NP_056002.2;XP_011512121.2;XP_02 | 29108 | . | [illegible] |
| 23232 | NM_015188,XI | TBC1D12 | TBC1 domain family member 12 | protein_coding | NP_056003.1;XP_006717795.2;XP_006717796.1;XP_011537860.1;XP_01 | 29082 | . | [illegible] |
| 23233 | NM_00132172 | EXOC6B | exocyst complex component 6B | protein_coding | NP_001308658.1;NP_001308659.1;NP_001308660.1;NP_001308662.1;NI | 17085 | [illegible] | [illegible] |
| 23234 | NM_015190 | DNAJC9 | DnaJ heat shock protein family (Hsp40) member C9 | protein_coding | NP_056005.1 | 19123 | [illegible] | [illegible] |
| 23235 | NM_015191,XI | SIK2 | salt inducible kinase 2 | protein_coding | NP_056006.1;XP_016872906.1;XP_016872907.1 | 21680 | [illegible] | [illegible] |
| 23236 | NM_015192,N | PLCB1 | phospholipase C beta 1 | protein_coding | NP_056007.1;NP_877398.1 | 15917 | [illegible] | [illegible] |
| 23237 | NM_015193 | ARC | activity regulated cytoskeleton associated protein | protein_coding | NP_056008.1 | 648 | [illegible] | [illegible] |
| 23239 | NM_194449,XI | PHLPP1 | PH domain and leucine rich repeat protein phosphatase 1 | protein_coding | NP_919431.2;XP_011524188.1;XP_024306873.1 | 20610 | [illegible] | [illegible] |
| 23240 | NM_00113100 | TMEM131L | transmembrane 131 like | protein_coding | NP_001124479.1;NP_056011.3;XP_011530082.1;XP_011530083.1;XP_01 | 29146 | [illegible] | [illegible] |
| 23241 | NM_00110091 | PACS2 | phosphofurin acidic cluster sorting protein 2 | protein_coding | NP_001094383.2;NP_001230056.1;NP_056012.2;XP_006720153.1;XP_00 | 23794 | [illegible] | [illegible] |
| 23242 | NM_00128743 | COBL | cordon-bleu WH2 repeat protein | protein_coding | NP_001274365.1;NP_001274367.1;NP_001333370.1;NP_001333371.1;NI | 22199 | [illegible] | [illegible] |
| 23243 | NM_00119509 | ANKRD28 | ankyrin repeat domain 28 | protein_coding | NP_001182027.1;NP_001182028.1;NP_001336206.1;NP_001336207.1;NI | 29024 | [illegible] | [illegible] |
| 23244 | NM_00110039 | PDS5A | PDS5 cohesin associated factor A | protein_coding | NP_001093869.1;NP_001093870.1;XP_011511974.1;XP_016863417.1 | 29088 | [illegible] | [illegible] |
| 23245 | NM_00118473 | ASTN2 | astrotactin 2 | protein_coding | NP_001171663.1;NP_001171664.1;NP_001351997.1;NP_001351998.1;NI | 17021 | [illegible] | [illegible] |
| 23246 | NM_015201 | BOP1 | BOP1 ribosomal biogenesis factor | protein_coding | NP_056016.1 | 15519 | [illegible] | [illegible] |
| 23247 | NM_015202,XI | KATNIP | katanin interacting protein | protein_coding | NP_056017.4;XP_005255258.1;XP_005255259.1;XP_005255260.1;XP_00 | 29068 | [illegible] | [illegible] |
| 23248 | NM_00129767 | RPRD2 | regulation of nuclear pre-mRNA domain containing 2 | protein_coding | NP_001284602.1;NP_001284603.1;NP_056018.2;XP_005245090.1;XP_00 | 29039 | [illegible] | [illegible] |
| 23250 | NM_015205,N | ATP11A | ATPase phospholipid transporting 11A | protein_coding | NP_056020.2;NP_115565.3;XP_005268356.1;XP_005268357.1;XP_00526 | 13552 | [illegible] | [illegible] |
| 23251 | NM_015206,XI | MINAR1 | membrane integral NOTCH2 associated receptor 1 | protein_coding | NP_056021.1;XP_011519694.1;XP_011519695.1;XP_011519697.1;XP_01 | 29172 | [illegible] | [illegible] |
| 23252 | NM_015207,XI | OTUD3 | OTU deubiquitinase 3 | protein_coding | NP_056022.1;XP_005245849.1;XP_005245850.1;XP_005245851.1;XP_02 | 29038 | [illegible] | [illegible] |
| 23253 | NM_00108362 | ANKRD12 | ankyrin repeat domain 12 | protein_coding | NP_001077094.1;NP_001190985.1;NP_056023.3;XP_005258149.1;XP_00 | 29135 | [illegible] | [illegible] |
| 23254 | NM_00101799 | KAZN | kazrin, periplakin interacting protein | protein_coding | NP_001017999.1;NP_001018000.1;NP_001018001.1;NP_001357158.1;NI | 29173 | [illegible] | [illegible] |
| 23255 | NM_00137820 | MTCL1 | microtubule crosslinking factor 1 | protein_coding | NP_001365134.1;NP_001365135.1;NP_001365136.1;NP_056025.2;XP_01 | 29121 | [illegible] | [illegible] |
| 23256 | NM_00125737 | SCFD1 | sec1 family domain containing 1 | protein_coding | NP_001244305.1;NP_001269960.1;NP_001269961.1;NP_001269962.1;NI | 20726 | [illegible] | [illegible] |
| 23258 | NM_00124325 | DENND5A | DENN domain containing 5A | protein_coding | NP_001230183.1;NP_001335678.1;NP_001335679.1;NP_056028.2 | 19344 | [illegible] | [illegible] |
| 23259 | NM_00116423 | DDHD2 | DDHD domain containing 2 | protein_coding | NP_001157704.1;NP_001157706.1;NP_001349840.1;NP_001349841.1;NI | 29106 | [illegible] | [illegible] |
| 23261 | NM_00119556 | CAMTA1 | calmodulin binding transcription activator 1 | protein_coding | NP_001182492.1;NP_001229630.1;NP_001336537.1;NP_001336538.1;NI | 18806 | [illegible] | [illegible] |
| 23262 | NM_00127627 | PPIP5K2 | diphosphoinositol pentakisphosphate kinase 2 | protein_coding | NP_001263206.1;NP_001268400.1;NP_001332800.1;NP_001332801.1;NI | 29035 | [illegible] | [illegible] |
| 23264 | NM_017590,XI | ZC3H7B | zinc finger CCCH-type containing 7B | protein_coding | NP_060060.3;XP_011528318.1;XP_024307950.1 | 30869 | [illegible] | [illegible] |
| 23265 | NM_00101383 | EXOC7 | exocyst complex component 7 | protein_coding | NP_001013861.1;NP_001138769.1;NP_001138770.1;NP_001138771.1;NI | 23214 | [illegible] | [illegible] |
| 23266 | NM_00129770 | ADGRL2 | adhesion G protein-coupled receptor L2 | protein_coding | NP_001284633.1;NP_001284634.1;NP_001284635.1;NP_001317574.1;NI | 18582 | [illegible] | [illegible] |
| 23268 | NM_00131832 | DNMBP | dynamin binding protein | protein_coding | NP_001305255.1;NP_001305256.1;NP_056036.1;XP_006717798.1;XP_01 | 30373 | [illegible] | [illegible] |
| 23269 | NM_00108054 | MGA | MAX dimerization protein MGA | protein_coding | NP_001074010.2;NP_001157745.1;XP_005254300.1;XP_005254301.1;XP | 14010 | [illegible] | [illegible] |
| 23270 | NM_021648 | TSPYL4 | TSPY like 4 | protein_coding | NP_067680.3 | 21559 | . | [illegible] |
| 23271 | NM_00129770 | CAMSAP2 | calmodulin regulated spectrin associated protein family member 2 | protein_coding | NP_001284636.1;NP_001284637.1;NP_982284.1;XP_005245098.1;XP_01 | 29188 | [illegible] | [illegible] |
| 23272 | NM_00111273 | TASOR | transcription activation suppressor | protein_coding | NP_001106207.1;NP_001350869.1;NP_001352564.1;NP_001352565.1;NI | 30314 | [illegible] | [illegible] |
| 23274 | NM_00124340 | CLEC16A | C-type lectin domain containing 16A | protein_coding | NP_001230332.1;NP_056041.1;XP_005255267.1;XP_005255268.1;XP_00 | 29013 | [illegible] | [illegible] |
| 23275 | NM_015227,N | POFUT2 | protein O-fucosyltransferase 2 | protein_coding | NP_056042.1;NP_598368.2;XP_006724044.1;XP_011527806.1;XP_01152 | 14683 | [illegible] | [illegible] |
| 23276 | NM_025010,XI | KLHL18 | kelch like family member 18 | protein_coding | NP_079286.2;XP_005265058.2;XP_005265059.2;XP_005265060.2;XP_01 | 29120 | . | [illegible] |
| 23277 | NM_00136666 | CLUH | clustered mitochondria homolog | protein_coding | NP_001353590.1;NP_001353591.1;NP_056044.4;XP_011522070.1;XP_02 | 29094 | [illegible] | [illegible] |
| 23279 | NM_00131839 | NUP160 | nucleoporin 160 | protein_coding | NP_001305328.1;NP_056046.1 | 18017 | [illegible] | [illegible] |
| 23281 | NM_00103360 | MTUS2 | microtubule associated scaffold protein 2 | protein_coding | NP_001028774.3;NP_001353579.1;NP_001353580.1;NP_056048.1;XP_01 | 20595 | . | [illegible] |
| 23283 | NM_015235 | CSTF2T | cleavage stimulation factor subunit 2 tau variant | protein_coding | NP_056050.1 | 17086 | [illegible] | [illegible] |
| 23284 | NM_00132224 | ADGRL3 | adhesion G protein-coupled receptor L3 | protein_coding | NP_001309175.1;NP_001309331.1;NP_001358271.1;NP_001358272.1;NI | 20974 | [illegible] | [illegible] |
| 23286 | NM_00116166 | WWC1 | WW and C2 domain containing 1 | protein_coding | NP_001155133.1;NP_001155134.1;NP_056053.1;XP_005265907.1;XP_00 | 29435 | [illegible] | [illegible] |
| 23287 | NM_00128671 | AGTPBP1 | ATP/GTP binding protein 1 | protein_coding | NP_001273644.1;NP_001273646.1;NP_001317630.1;NP_056054.2;XP_00 | 17258 | [illegible] | [illegible] |
| 23288 | NM_00128749 | IQCE | IQ motif containing E | protein_coding | NP_001274428.1;NP_001274429.1;NP_001274430.1;NP_001274431.1;NI | 29171 | [illegible] | [illegible] |
| 23291 | NM_00137897 | FBXW11 | F-box and WD repeat domain containing 11 | protein_coding | NP_001365903.1;NP_001365904.1;NP_001365905.1;NP_001365906.1;NI | 13607 | [illegible] | [illegible] |
| 23293 | NM_00125682 | SMG6 | SMG6 nonsense mediated mRNA decay factor | protein_coding | NP_001243756.1;NP_001243757.1;NP_001269255.1;NP_060045.4;XP_00 | 17809 | [illegible] | [illegible] |
| 23294 | NM_015245,XI | ANKS1A | ankyrin repeat and sterile alpha motif domain containing 1A | protein_coding | NP_056060.2;XP_005249021.1;XP_006715099.1;XP_011512733.1;XP_01 | 20961 | [illegible] | [illegible] |
| 23295 | NM_00114228 | MGRN1 | mahogunin ring finger 1 | protein_coding | NP_001135761.2;NP_001135762.1;NP_001135763.2;NP_056061.1 | 20254 | [illegible] | [illegible] |
| 23299 | NM_00100380 | BICD2 | BICD cargo adaptor 2 | protein_coding | NP_001003800.1;NP_056065.1;XP_016870040.1 | 17208 | [illegible] | [illegible] |
| 23300 | NM_00130072 | ATMIN | ATM interactor | protein_coding | NP_001287657.1;NP_056066.2 | 29034 | [illegible] | [illegible] |
| 23301 | NM_00114261 | EHBP1 | EH domain binding protein 1 | protein_coding | NP_001136086.1;NP_001136087.1;NP_001136088.1;NP_001341141.1;NI | 29144 | [illegible] | [illegible] |
| 23303 | NM_015254,XI | KIF13B | kinesin family member 13B | protein_coding | NP_056069.2;XP_005273515.1;XP_011542759.1;XP_011542760.1;XP_01 | 14405 | [illegible] | [illegible] |
| 23304 | NM_00118480 | UBR2 | ubiquitin protein ligase E3 component n-recognin 2 | protein_coding | NP_001171730.1;NP_001350634.1;NP_056070.1;XP_005249023.1;XP_01 | 21289 | [illegible] | [illegible] |
| 23305 | NM_00100918 | ACSL6 | acyl-CoA synthetase long chain family member 6 | protein_coding | NP_001009185.1;NP_001192176.1;NP_001192177.1;NP_001192179.1;NI | 16496 | [illegible] | [illegible] |
| 23306 | NM_00113096 | NEMP1 | nuclear envelope integral membrane protein 1 | protein_coding | NP_001124435.1;NP_056072.2;XP_011536358.1;XP_011536359.1;XP_01 | 29001 | [illegible] | [illegible] |
| 23307 | NM_015258,XI | FKBP15 | FKBP prolyl isomerase 15 | protein_coding | NP_056073.1;XP_006717081.1;XP_006717082.1;XP_016870041.1 | 23397 | [illegible] | [illegible] |
| 23308 | NM_00128305 | ICOSLG | inducible T cell costimulator ligand | protein_coding | NP_001269979.1;NP_001269980.1;NP_001269981.1;NP_001352688.1;NI | 17087 | [illegible] | [illegible] |
| 23309 | NM_00129759 | SIN3B | SIN3 transcription regulator family member B | protein_coding | NP_001284524.1;NP_001284526.1;NP_056075.1;XP_006722767.1 | 19354 | [illegible] | [illegible] |
| 23310 | NM_00137206 | NCAPD3 | non-SMC condensin II complex subunit D3 | protein_coding | NP_001358994.1;NP_001358997.1;NP_001358998.1;NP_001358999.1;NI | 28952 | [illegible] | [illegible] |
| 23312 | NM_00117411 | DMXL2 | Dmx like 2 | protein_coding | NP_001167587.1;NP_001167588.1;NP_001365386.1;NP_001365387.1;NI | 2938 | 612186 | [illegible] |
| 23313 | NM_00100988 | KIAA0930 | KIAA0930 | protein_coding | NP_001009880.1;NP_056079.1;XP_011528319.1;XP_016884161.1 | 1314 | . | [illegible] |
| 23314 | NM_00117250 | SATB2 | SATB homeobox 2 | protein_coding | NP_001165980.1;NP_001165988.1;NP_056080.1;XP_005246453.1;XP_01 | 21637 | 608148 | [illegible] |
| 23315 | NM_00126049 | SLC9A8 | solute carrier family 9 member A8 | protein_coding | NP_001247420.1;NP_056081.1;XP_006723819.1;XP_011527038.1;XP_01 | 20728 | 612730 | [illegible] |
| 23317 | NM_00132912 | DNAJC13 | DnaJ heat shock protein family (Hsp40) member C13 | protein_coding | NP_001316055.1;NP_056083.3;XP_016861525.1 | 30343 | 614334 | [illegible] |
| 23318 | NM_00100988 | TUT4 | terminal uridylyl transferase 4 | protein_coding | NP_001009881.1;NP_056084.1;XP_005270733.1;XP_005270735.1;XP_00 | 28981 | 613692 | [illegible] |
| 23321 | NM_00113006 | TRIM2 | tripartite motif containing 2 | protein_coding | NP_001123539.1;NP_001289621.1;NP_001289622.1;NP_001289623.1;NI | 15974 | 614141 | [illegible] |
| 23322 | NM_00112789 | RPGRIP1L | RPGRIP1 like | protein_coding | NP_001121369.1;NP_001295263.1;NP_001315351.1;NP_001315352.1;NI | 29168 | 610937 | [illegible] |
| 23324 | NM_00129203 | MAN2B2 | mannosidase alpha class 2B member 2 | protein_coding | NP_001278967.1;NP_056089.1 | 29623 | . | [illegible] |
| 23325 | NM_00129364 | WASHC4 | WASH complex subunit 4 | protein_coding | NP_001280569.1;NP_056090.1;XP_011536375.1;XP_011536376.1;XP_01 | 29174 | 615748 | [illegible] |
| 23326 | NM_015276,XI | USP22 | ubiquitin specific peptidase 22 | protein_coding | NP_056091.1;XP_005256632.1 | 12621 | 612116 | [illegible] |
| 23327 | NM_00114496 | NEDD4L | NEDD4 like E3 ubiquitin protein ligase | protein_coding | NP_001138436.1;NP_001138437.1;NP_001138438.1;NP_001138439.1;NI | 7728 | 606384 | [illegible] |
| 23328 | NM_00134650 | SASH1 | SAM and SH3 domain containing 1 | protein_coding | NP_001333434.1;NP_001333435.1;NP_001333436.1;NP_001333437.1;NI | 19182 | 607955 | [illegible] |
| 23329 | NM_00133018 | TBC1D30 | TBC1 domain family member 30 | protein_coding | NP_001317115.1;NP_001317116.1;NP_001317117.1;NP_001351767.1;NI | 29164 | 615077 | [illegible] |
| 23331 | NM_00114541 | TTC28 | tetratricopeptide repeat domain 28 | protein_coding | NP_001138890.1;XP_005261462.1;XP_006724234.1;XP_011528320.1;XP_ | 29179 | 615098 | [illegible] |
| 23332 | NM_00114227 | CLASP1 | cytoplasmic linker associated protein 1 | protein_coding | NP_001135745.1;NP_001135746.1;NP_001193980.1;NP_001364932.1;NI | 17088 | 605852 | [illegible] |
| 23333 | NM_00136667 | DPY19L1 | dpy-19 like C-mannosyltransferase 1 | protein_coding | NP_001353602.1;NP_056098.1;XP_011513548.1 | 22205 | 613892 | [illegible] |
| 23334 | NM_00136599 | SZT2 | SZT2 subunit of KICSTOR complex | protein_coding | NP_001352928.1;NP_056099.3;XP_005270743.1;XP_011539408.1;XP_01 | 29040 | 615463 | [illegible] |
| 23335 | NM_00138248 | WDR7 | WD repeat domain 7 | protein_coding | NP_001369414.1;NP_001369416.1;NP_056100.2;NP_443066.2;XP_01152 | 13490 | 613473 | [illegible] |
| 23336 | NM_015286,N | SYNM | synemin | protein_coding | NP_056101.5;NP_663780.2;XP_016877524.1 | 24466 | 606087 | [illegible] |
| 23338 | NM_00128998 | JADE2 | jade family PHD finger 2 | protein_coding | NP_001276913.1;NP_001276914.1;NP_001295072.1;NP_056103.4;XP_00 | 22984 | 610515 | [illegible] |
| 23339 | NM_00130113 | VPS39 | VPS39 subunit of HOPS complex | protein_coding | NP_001288067.1;NP_056104.2;XP_011519705.1;XP_011519706.1;XP_01 | 20593 | 612188 | [illegible] |
| 23341 | NM_00128781 | DNAJC16 | DnaJ heat shock protein family (Hsp40) member C16 | protein_coding | NP_001274740.1;NP_056106.1 | 29157 | . | [illegible] |
| 23344 | NM_00118479 | ESYT1 | extended synaptotagmin 1 | protein_coding | NP_001171725.1;NP_056107.1 | 29534 | 616670 | [illegible] |

| | | | | | | | | | | | | | | | | | | | | | | | | | | | | | | | | | |
|---|---|---|---|---|---|---|---|---|---|---|---|---|---|---|---|---|---|---|---|---|---|---|---|---|---|---|---|---|---|---|---|---|---|
| 23345 | NM_00134770 | SYNE1 | spectrin repeat containing nuclear envelope protein 1 | protein_coding | NP_001334630.1;NP_001334631.1;NP_149062.1;NP_892006.3;XP_00671 | 17089 | 608441 | . | . | 0.788731 | 0.885276 | 0.783188 | 1.440686 | 1.835409 | 2.046359 | 1.757450 | 3.293955 | 1.036088 | 745.013388 | 0.162488127 | 0.314773341 | 0.752933754 | 0.998865404 | 9.515767 | 9.566040 | 9.489212 | 9.563493 | 9.478021 | 9.620059 | 798 | 832 | 592 | 783 |
| 23347 | NM_015295,XI | SMCHD1 | structural maintenance of chromosomes flexible hinge domain co | protein_coding | NP_056110.2;XP_011523944.1;XP_011523945.1;XP_016881173.1 | 29090 | 614982 | . | . | 4.766254 | 4.799393 | 4.844109 | 3.718476 | 11.091270 | 11.094014 | 10.870040 | 8.501855 | -1.151669 | 1233.173995 | 0.154054732 | -1.32242807 | 0.186025648 | 0.958175316 | 10.366135 | 10.163045 | 10.317323 | 10.315960 | 10.308330 | 10.101605 | 1521 | 1423 | 1136 | 925 |
| 23348 | NM_00113004 | DOCK9 | dedicator of cytokinesis 9 | protein_coding | NP_001123520.1;NP_001123521.1;NP_001123522.1;NP_001305778.1;NI | 14132 | 607325 | . | . | 4.773734 | 4.326094 | 3.702864 | 4.585296 | 11.108677 | 9.999956 | 8.309117 | 10.483730 | -1.102470 | 1028.565213 | 0.150592292 | -0.93457392 | 0.350007924 | 0.998865404 | 10.075210 | 9.934180 | 10.084288 | 9.993942 | 9.900068 | 10.032141 | 1317 | 1094 | 773 | 1001 |
| 23349 | NM_00130433 | PHF24 | PHD finger protein 24 | protein_coding | NP_001291262.1;NP_001334911.1;NP_001334912.1;NP_056112.1;XP_01 | 29180 | . | | . | 0.032707 | 0.069309 | 0.027790 | 0.099782 | 0.076112 | 0.160211 | 0.062361 | 0.228141 | 1.202208 | 11.948000 | 0.959373558 | 0.276937604 | 0.781828018 | NA | 3.444631 | 3.701358 | 3.545905 | 3.581906 | 3.542309 | 3.604498 | 8 | 16 | 5 | 19 |
| 23350 | NM_00108041 | U2SURP | U2 snRNP associated SURP domain containing | protein_coding | NP_001073884.1;NP_001307148.1;NP_001307149.1;NP_001307151.1;XF | 30855 | 617849 | . | . | 17.710273 | 17.339182 | 16.844597 | 15.757258 | 41.212534 | 40.080297 | 37.798789 | 36.027096 | -1.084172 | 3870.210577 | 0.110247566 | -1.05756401 | 0.290254258 | 0.998865404 | 11.975262 | 11.858781 | 11.959372 | 11.941786 | 11.911385 | 11.855263 | 4685 | 4299 | 3340 | 3363 |
| 23351 | NM_00129025 | KHNYN | KH and NYN domain containing | protein_coding | NP_001277185.1;NP_001277186.1;NP_056114.1;XP_005267531.1;XP_01 | 20166 | . | | . | 5.964848 | 5.692949 | 7.027529 | 6.822083 | 13.880445 | 13.159507 | 15.769572 | 15.597881 | 1.154955 | 1307.539033 | 0.12893381 | 1.61196501 | 0.106969554 | 0.83501324 | 10.244891 | 10.452895 | 10.314568 | 10.274222 | 10.411192 | 10.397199 | 1435 | 1274 | 1230 | 1302 |
| 23352 | NM_020765,XI | UBR4 | ubiquitin protein ligase E3 component n-recognin 4 | protein_coding | NP_065816.2;XP_011539410.2;XP_011539411.2;XP_011539412.2;XP_01 | 30313 | 609890 | . | . | 19.532132 | 19.821914 | 23.473632 | 23.093784 | 45.452072 | 45.819241 | 52.674148 | 52.801196 | 1.130957 | 11283.083219 | 0.101979594 | 1.740972396 | 0.081688417 | 0.782498736 | 13.370391 | 13.547900 | 13.384782 | 13.428697 | 13.500542 | 13.523302 | 11877 | 11724 | 10277 | 11376 |
| 23353 | NM_00113096 | SUN1 | Sad1 and UNC84 domain containing 1 | protein_coding | NP_001124437.1;NP_001165415.1;NP_001165416.1;NP_001165417.1;NI | 18587 | 607723 | . | . | 29.278702 | 27.962180 | 26.072270 | 24.979406 | 68.132738 | 64.635830 | 58.505420 | 57.112449 | -1.139820 | 3608.671200 | 0.11015473 | -1.71400486 | 0.086527834 | 0.795065303 | 11.908523 | 11.719815 | 11.886817 | 11.850402 | 11.775973 | 11.745502 | 4523 | 4058 | 2988 | 3103 |
| 23354 | NM_015302,XI | HAUS5 | HAUS augmin like complex subunit 5 | protein_coding | NP_056117.1;XP_011524986.1 | 29130 | 613432 | . | . | 10.543577 | 10.931434 | 11.373357 | 10.484887 | 24.535337 | 25.268497 | 25.521485 | 23.972447 | 1.009579 | 1448.608163 | 0.128418812 | 0.107096884 | 0.914712119 | 0.998865404 | 10.493472 | 10.507398 | 10.484701 | 10.507949 | 10.537359 | 10.469083 | 1635 | 1579 | 1322 | 1304 |
| 23355 | NM_00100992 | VPS8 | VPS8 subunit of CORVET complex | protein_coding | NP_001009921.1;NP_001336221.1;NP_001336222.1;NP_001336223.1;NI | 29122 | 618366 | . | . | 5.286852 | 5.105823 | 6.424618 | 5.631607 | 12.302723 | 11.802342 | 14.416660 | 12.876001 | 1.201397 | 846.936756 | 0.149675152 | 1.768582055 | 0.076963648 | 0.767975096 | 9.587498 | 9.852565 | 9.641404 | 9.668170 | 9.838664 | 9.729914 | 869 | 846 | 864 | 802 |
| 23357 | NM_00137074 | ANGEL1 | angel homolog 1 | protein_coding | NP_001357675.1;NP_001357676.1;NP_001357677.1;NP_056120.2 | 19961 | . | | . | 4.419660 | 4.643436 | 5.194290 | 5.447562 | 10.284731 | 10.733510 | 11.655836 | 12.455204 | 1.184962 | 803.587320 | 0.146151364 | 1.675254183 | 0.093884371 | 0.810751527 | 9.522877 | 9.767397 | 9.561100 | 9.610387 | 9.692329 | 9.718932 | 817 | 822 | 744 | 832 |
| 23358 | NM_015306,XI | USP24 | ubiquitin specific peptidase 24 | protein_coding | NP_056121.2;XP_005270747.1;XP_016856320.1;XP_016856321.1;XP_01 | 12623 | 610569 | . | . | 5.902776 | 6.041546 | 6.419305 | 6.299359 | 13.736006 | 13.965306 | 14.404735 | 14.402733 | 1.064576 | 2043.411186 | 0.117641296 | 0.767411883 | 0.442836666 | 0.998865404 | 10.950875 | 11.041238 | 10.975533 | 10.966738 | 11.033304 | 11.008520 | 2292 | 2124 | 1861 | 1945 |
| 23360 | NM_00131833 | FNBP4 | formin binding protein 4 | protein_coding | NP_001305268.1;NP_056123.2;XP_024304176.1;XP_024304177.1 | 19752 | 615265 | . | . | 9.714149 | 10.603443 | 12.511095 | 9.481001 | 22.605223 | 24.510331 | 28.074538 | 21.677184 | 1.059070 | 1318.406772 | 0.152555045 | 0.542740671 | 0.587308367 | 0.998865404 | 10.322385 | 10.405576 | 10.306295 | 10.371613 | 10.479526 | 10.277171 | 1411 | 1441 | 1335 | 1104 |
| 23361 | NM_00108041 | ZNF629 | zinc finger protein 629 | protein_coding | NP_001073886.1;NP_001332899.1 | 29008 | . | | . | 0.209819 | 0.174415 | 0.207262 | 0.191366 | 0.488258 | 0.403168 | 0.465090 | 0.437536 | 1.030362 | 37.673842 | 0.507764745 | 0.08498231 | 0.932275454 | 0.998865404 | 5.212950 | 5.257685 | 5.247074 | 5.217240 | 5.244179 | 5.229076 | 47 | 36 | 35 | 34 |
| 23362 | NM_00136281 | PSD3 | pleckstrin and Sec7 domain containing 3 | protein_coding | NP_001349748.1;NP_056125.3;NP_996792.1;XP_011542769.1;XP_01154 | 19093 | 614440 | . | . | 1.151687 | 1.470495 | 1.200317 | 1.128154 | 2.680022 | 3.399114 | 2.693478 | 2.579387 | -1.005678 | 462.275412 | 0.172346109 | -0.04739645 | 0.962197256 | 0.998865404 | 8.856522 | 8.848685 | 8.855600 | 8.852661 | 8.872086 | 8.828834 | 535 | 499 | 415 | 417 |
| 23363 | NM_00117340 | OBSL1 | obscurin like cytoskeletal adaptor 1 | protein_coding | NP_001166879.1;NP_001166902.1;NP_056126.1;XP_011509159.1;XP_01 | 29092 | 610991 | . | . | 14.679576 | 13.993278 | 15.237034 | 17.540450 | 34.159980 | 32.346092 | 34.191463 | 40.104155 | 1.122166 | 1807.513054 | 0.130744395 | 1.271841459 | 0.203429455 | 0.968254058 | 10.734353 | 10.900450 | 10.797177 | 10.745316 | 10.799981 | 10.921095 | 2027 | 1777 | 1536 | 1928 |
| 23365 | NM_00119866 | ARHGEF12 | Rho guanine nucleotide exchange factor 12 | protein_coding | NP_001185594.1;NP_001288013.1;NP_056128.1;XP_006718868.1;XP_01 | 14193 | 604763 | . | . | 3.980224 | 3.722503 | 3.799962 | 4.293881 | 9.262143 | 8.604732 | 8.527004 | 9.817447 | 1.040930 | 1252.927708 | 0.134957145 | 0.428826067 | 0.668049807 | 0.998865404 | 10.261931 | 10.319667 | 10.302127 | 10.247521 | 10.264911 | 10.344500 | 1465 | 1277 | 1053 | 1261 |
| 23366 | NM_00110042 | KIAA0895 | KIAA0895 | protein_coding | NP_001093895.1;NP_001186635.1;NP_001186636.1;NP_001186637.1;NI | 22206 | . | | . | 1.865160 | 1.490621 | 1.810826 | 1.987683 | 4.340303 | 3.445639 | 4.063440 | 4.544602 | 1.102902 | 238.628740 | 0.22977687 | 0.614961599 | 0.538580081 | 0.998865404 | 7.825977 | 7.967787 | 7.925121 | 7.815282 | 7.901886 | 7.937196 | 291 | 217 | 210 | 243 |
| 23367 | NM_00136771 | LARP1 | La ribonucleoprotein 1, translational regulator | protein_coding | NP_001354642.1;NP_001354643.1;NP_001354644.1;NP_001354645.1;NI | 29531 | 612059 | . | . | 30.222169 | 33.102886 | 33.204029 | 35.504644 | 70.328226 | 76.518796 | 74.508882 | 81.177149 | 1.053285 | 7196.878451 | 0.10885333 | 0.688045859 | 0.491423903 | 0.998865404 | 12.775264 | 12.850077 | 12.746736 | 12.832298 | 12.816347 | 12.851973 | 7663 | 7948 | 6286 | 7067 |
| 23368 | NM_015316,XI | PPP1R13B | protein phosphatase 1 regulatory subunit 13B | protein_coding | NP_056131.2;XP_005267544.1;XP_011534895.1;XP_016876605.1;XP_01 | 14950 | 606455 | . | . | 3.594864 | 4.112898 | 4.776891 | 4.803295 | 8.365396 | 9.507148 | 10.719203 | 10.982162 | 1.227944 | 740.116953 | 0.152830602 | 1.938389927 | 0.052575668 | 0.678753563 | 9.376092 | 9.671982 | 9.408353 | 9.497218 | 9.597766 | 9.594778 | 715 | 764 | 711 | 763 |
| 23369 | NM_00128275 | PUM2 | pumilio RNA binding family member 2 | protein_coding | NP_001269681.1;NP_001269719.1;NP_001269720.1;NP_001339846.1;NI | 14958 | 607205 | . | . | 11.035881 | 11.053591 | 11.652553 | 11.107476 | 25.680948 | 25.550875 | 26.147987 | 25.395928 | 1.017844 | 2198.037072 | 0.117373367 | 0.217396098 | 0.827899672 | 0.998865404 | 11.089119 | 11.114767 | 11.098672 | 11.090707 | 11.131742 | 11.085558 | 2521 | 2339 | 1984 | 2019 |
| 23370 | NM_00113095 | ARHGEF18 | Rho/Rac guanine nucleotide exchange factor 18 | protein_coding | NP_001124427.2;NP_001354752.1;NP_001354753.1;NP_056133.2;XP_00 | 17090 | 616432 | . | . | 4.626935 | 5.082641 | 4.842397 | 4.613131 | 10.767066 | 11.748752 | 10.866200 | 10.547374 | -1.039860 | 803.967732 | 0.147056104 | -0.3834513 | 0.70138517 | 0.998865404 | 9.678913 | 9.622524 | 9.636140 | 9.691839 | 9.655519 | 9.617715 | 907 | 919 | 704 | 719 |
| 23371 | NM_015319,N | TNS2 | tensin 2 | protein_coding | NP_056134.2;NP_736610.2;NP_938072.1;XP_006719365.1;XP_00671936 | 19737 | 607717 | . | . | 5.831967 | 7.049638 | 6.713380 | 6.801720 | 13.571228 | 16.295555 | 15.064633 | 15.551324 | 1.015342 | 1006.897227 | 0.145923064 | 0.150532724 | 0.880344335 | 0.998865404 | 9.964890 | 9.986431 | 9.904599 | 10.029762 | 9.970857 | 9.989335 | 1053 | 1170 | 871 | 963 |
| 23373 | NM_00109848 | CRTC1 | CREB regulated transcription coactivator 1 | protein_coding | NP_001091952.1;NP_056136.2;XP_005259890.1;XP_005259891.1;XP_00 | 16062 | 607536 | . | . | 2.580063 | 2.672077 | 3.518049 | 2.841756 | 6.003913 | 6.176627 | 7.894397 | 6.497337 | 1.137614 | 470.113702 | 0.175817398 | 1.057978308 | 0.290065332 | 0.998865404 | 8.780676 | 8.967041 | 8.814224 | 8.851720 | 8.956024 | 8.869288 | 491 | 489 | 465 | 437 |
| 23376 | NM_015323 | UFL1 | UFM1 specific ligase 1 | protein_coding | NP_056138.1 | 23039 | 613372 | . | . | 7.274422 | 7.667494 | 7.054298 | 6.354388 | 16.927877 | 17.723755 | 15.829638 | 14.528551 | -1.129837 | 934.202401 | 0.143520351 | -1.22710398 | 0.219783514 | 0.982008277 | 9.952824 | 9.777006 | 9.933301 | 9.925891 | 9.861561 | 9.779554 | 1116 | 1093 | 807 | 775 |
| 23378 | NM_015324,XI | RRP8 | ribosomal RNA processing 8 | protein_coding | NP_056139.1;XP_011518257.1;XP_016872912.1 | 29030 | 615818 | . | . | 2.236445 | 2.636465 | 2.796121 | 2.286007 | 5.204301 | 6.094309 | 6.274414 | 5.226683 | -1.024184 | 464.150644 | 0.184468633 | -0.18688662 | 0.851749526 | 0.998865404 | 8.875437 | 8.841259 | 8.815307 | 8.910191 | 8.903206 | 8.791936 | 500 | 545 | 435 | 391 |
| 23379 | NM_015325,XI | ICE1 | interactor of little elongation complex ELL subunit 1 | protein_coding | NP_056140.1;XP_011512301.1;XP_016864774.1 | 29154 | 617958 | . | . | 4.073329 | 4.598429 | 5.298780 | 4.705973 | 9.478804 | 10.629476 | 11.890308 | 10.759645 | 1.127795 | 1152.232660 | 0.142097595 | 1.2210293 | 0.222074926 | 0.982008277 | 10.080813 | 10.254402 | 10.077122 | 10.164909 | 10.258951 | 10.162414 | 1173 | 1238 | 1130 | 1072 |
| 23380 | NM_00117063 | SRGAP2 | SLIT-ROBO Rho GTPase activating protein 2 | protein_coding | NP_001164108.1;NP_001287881.1;NP_001364373.1;NP_001364374.1;NI | 19751 | 606524 | . | . | 12.933200 | 15.244942 | 15.890884 | 15.454399 | 30.096090 | 35.239369 | 35.658681 | 35.334643 | 1.101035 | 2871.795431 | 0.119640785 | 1.160641032 | 0.245787911 | 0.992733415 | 11.416692 | 11.555448 | 11.392667 | 11.497624 | 11.536724 | 11.513062 | 2944 | 3141 | 2655 | 2781 |
| 23381 | NM_00132361 | SMG5 | SMG5 nonsense mediated mRNA decay factor | protein_coding | NP_001310543.1;NP_001310544.1;NP_001310545.1;NP_001310546.1;NI | 24644 | 610962 | . | . | 26.962963 | 28.995403 | 30.607033 | 33.106043 | 62.743921 | 67.024169 | 68.681294 | 75.693032 | 1.120880 | 4248.389733 | 0.111325287 | 1.478834205 | 0.139184632 | 0.900674731 | 11.968109 | 12.132605 | 11.975663 | 12.029697 | 12.068225 | 12.126606 | 4462 | 4465 | 3769 | 4356 |
| 23382 | NM_00113072 | AHCYL2 | adenosylhomocysteinase like 2 | protein_coding | NP_001124192.1;NP_001124194.2;NP_001124195.1;NP_056143.1;XP_00 | 22204 | 616520 | . | . | 4.680646 | 5.546983 | 5.915694 | 5.750159 | 10.892054 | 12.822098 | 13.274645 | 13.147053 | 1.128845 | 859.068219 | 0.149280901 | 1.171263669 | 0.24149281 | 0.992733415 | 9.656687 | 9.831291 | 9.646074 | 9.751645 | 9.801029 | 9.772280 | 860 | 936 | 808 | 837 |
| 23383 | NM_015329,XI | MAU2 | MAU2 sister chromatid cohesion factor | protein_coding | NP_056144.3;XP_005259894.2;XP_005259895.3;XP_006722774.2;XP_01 | 29140 | 614560 | . | . | 6.601618 | 6.908257 | 7.532257 | 7.179809 | 15.362238 | 15.968747 | 16.902167 | 16.415781 | 1.064717 | 1054.172469 | 0.136853844 | 0.661066668 | 0.50856956 | 0.998865404 | 9.995883 | 10.086485 | 10.002085 | 10.033109 | 10.091525 | 10.035834 | 1151 | 1125 | 981 | 981 |
| 23384 | NM_00114546 | SPECC1L | sperm antigen with calponin homology and coiled-coil domains 1 l | protein_coding | NP_001138940.4;NP_001241661.3;NP_001241662.1;NP_056145.5 | 29022 | 614140 | . | . | 9.747391 | 9.848981 | 10.136078 | 10.830579 | 22.682579 | 22.766360 | 22.745065 | 24.762835 | 1.105349 | 2065.525622 | 0.122644653 | 1.17821243 | 0.238711928 | 0.992733415 | 10.938366 | 11.082616 | 10.930288 | 11.009113 | 11.028228 | 11.071682 | 2147 | 2223 | 1835 | 2088 |
| 23385 | NM_00129018 | NCSTN | nicastrin | protein_coding | NP_001277113.1;NP_001277115.1;NP_001336658.1;NP_056146.1;XP_00 | 17091 | 605254 | . | . | 42.077031 | 44.645367 | 40.462447 | 43.509497 | 97.914984 | 103.199772 | 90.796567 | 99.479292 | -1.049606 | 3719.028866 | 0.112563671 | -0.62051789 | 0.534916879 | 0.998865404 | 11.895287 | 11.825285 | 11.856942 | 11.902937 | 11.808245 | 11.871023 | 4265 | 4224 | 3035 | 3532 |
| 23386 | NM_015332,XI | NUDCD3 | NudC domain containing 3 | protein_coding | NP_056147.2;XP_011513549.1;XP_016867397.1;XP_016867398.1;XP_01 | 22208 | 610296 | . | . | 7.783738 | 10.686475 | 8.996325 | 11.146822 | 18.113078 | 24.702264 | 20.187492 | 25.485881 | 1.010372 | 2030.848018 | 0.121753754 | 0.122263349 | 0.902690458 | 0.998865404 | 10.980509 | 10.995186 | 10.945573 | 11.019977 | 10.977610 | 11.005358 | 2218 | 2282 | 1744 | 1947 |
| 23387 | NM_00128174 | SIK3 | SIK family kinase 3 | protein_coding | NP_001268677.1;NP_001268678.1;NP_001353615.1;NP_001353616.1;NI | 29165 | 614776 | . | . | 1.836949 | 2.039218 | 2.093778 | 2.516686 | 4.274655 | 4.713742 | 4.698374 | 5.754103 | 1.055873 | 386.039095 | 0.183435046 | 0.427594813 | 0.668946143 | 0.998865404 | 8.553050 | 8.631101 | 8.558747 | 8.592690 | 8.596024 | 8.619250 | 420 | 417 | 338 | 379 |
| 23389 | NM_015335,XI | MED13L | mediator complex subunit 13L | protein_coding | NP_056150.1;XP_011536382.1;XP_011536383.1;XP_011536384.1;XP_01 | 22962 | 608771 | . | . | 3.934047 | 3.823085 | 3.349329 | 3.881733 | 9.154689 | 8.837233 | 7.515798 | 8.875119 | -1.088424 | 1150.009939 | 0.138674662 | -0.88149339 | 0.37805083 | 0.998865404 | 10.227403 | 10.104857 | 10.215186 | 10.180381 | 10.083637 | 10.182902 | 1412 | 1264 | 893 | 1102 |
| 23390 | NM_00135962 | ZDHHC17 | zinc finger DHHC-type palmitoyltransferase 17 | protein_coding | NP_001346555.1;NP_056151.2 | 18412 | 607799 | . | . | 2.558458 | 1.881018 | 2.770220 | 1.528374 | 5.953636 | 4.348058 | 6.216294 | 3.494446 | -1.039231 | 317.969837 | 0.264790009 | -0.20966063 | 0.833932557 | 0.998865404 | 8.339156 | 8.285844 | 8.394706 | 8.219036 | 8.431826 | 8.123669 | 431 | 295 | 348 | 210 |
| 23392 | NM_00136375 | ECPAS | Ecm29 proteasome adaptor and scaffold | protein_coding | NP_001350685.1;NP_001351858.1;NP_001351859.1;NP_001351860.1;XF | 29020 | 616694 | . | . | 10.278202 | 11.026309 | 11.233956 | 11.441240 | 23.917800 | 25.487808 | 25.208670 | 26.159036 | 1.065625 | 2398.347059 | 0.115989666 | 0.790586344 | 0.429185418 | 0.998865404 | 11.181305 | 11.272891 | 11.178082 | 11.224318 | 11.245536 | 11.259681 | 2597 | 2578 | 2133 | 2340 |
| 23394 | NM_00128253 | ADNP | activity dependent neuroprotector homeobox | protein_coding | NP_001269460.1;NP_001269461.1;NP_001334440.1;NP_056154.1;NP_8 | 15766 | 611386 | . | . | 18.804792 | 19.516646 | 18.858070 | 18.994276 | 43.759524 | 45.113602 | 42.316966 | 43.428151 | -1.029449 | 3906.243980 | 0.108064592 | -0.38747859 | 0.698401929 | 0.998865404 | 11.952358 | 11.910471 | 11.930383 | 11.956376 | 11.923957 | 11.914822 | 4492 | 4343 | 3366 | 3589 |
| 23395 | NM_00136826 | LARS2 | leucyl-tRNA synthetase 2, mitochondrial | protein_coding | NP_001355192.1;NP_056155.1;XP_011531856.1;XP_016861531.1 | 17095 | 604544 | . | . | 5.894633 | 7.037804 | 6.113711 | 7.274567 | 13.717053 | 16.268202 | 13.718991 | 16.632435 | 1.019316 | 961.663029 | 0.151649898 | 0.182008554 | 0.855786013 | 0.998865404 | 9.895882 | 9.922768 | 9.846399 | 9.952924 | 9.858102 | 9.968464 | 1017 | 1103 | 780 | 979 |
| 23396 | NM_00119573 | PIP5K1C | phosphatidylinositol-4-phosphate 5-kinase type 1 gamma | protein_coding | NP_001182662.1;NP_001287778.1;NP_036530.1;XP_011526147.1;XP_01 | 8996 | 606102 | . | . | 6.786552 | 7.173162 | 7.435462 | 8.767451 | 15.792586 | 16.581084 | 16.684962 | 20.045739 | 1.145082 | 1162.606849 | 0.140371962 | 1.392378152 | 0.163807917 | 0.939750905 | 10.082384 | 10.277331 | 10.107080 | 10.150284 | 10.171365 | 10.285970 | 1212 | 1204 | 998 | 1251 |
| 23397 | NM_00128171 | NCAPH | non-SMC condensin I complex subunit H | protein_coding | NP_001268639.1;NP_001268640.1;NP_001268641.1;NP_056156.2;XP_00 | 1112 | 602332 | . | . | 17.099681 | 19.588946 | 16.040317 | 19.718535 | 39.791661 | 45.280726 | 35.994004 | 45.084787 | -1.107790 | 2849.141915 | 0.120759372 | -1.22296208 | 0.221344017 | 0.982008277 | 11.548390 | 11.400442 | 11.478648 | 11.552260 | 11.387205 | 11.475992 | 3295 | 3376 | 2221 | 2674 |
| 23398 | NM_00127892 | PPWD1 | peptidylprolyl isomerase domain and WD repeat containing 1 | protein_coding | NP_001265855.1;NP_001265856.1;NP_001265858.1;NP_056157.1;XP_00 | 28954 | 618274 | . | . | 9.723009 | 8.857967 | 9.589230 | 8.450646 | 22.625839 | 20.475589 | 21.517952 | 19.321399 | -1.040639 | 600.949362 | 0.164692692 | -0.34894683 | 0.727129226 | 0.998865404 | 9.259168 | 9.202474 | 9.276567 | 9.208960 | 9.259274 | 9.173061 | 741 | 628 | 546 | 516 |
| 23399 | NM_00114377 | CTDNEP1 | CTD nuclear envelope phosphatase 1 | protein_coding | NP_001137247.1;NP_056158.2 | 19085 | 610684 | . | . | 44.865536 | 44.339731 | 42.495573 | 48.373431 | 104.403950 | 102.493275 | 95.358831 | 110.600105 | -1.000717 | 2500.479162 | 0.120361435 | -0.00858981 | 0.993146411 | 0.999862634 | 11.288602 | 11.287375 | 11.293360 | 11.282388 | 11.236882 | 11.335243 | 2900 | 2681 | 2041 | 2486 |
| 23400 | NM_00114197 | ATP13A2 | ATPase cation transporting 13A2 | protein_coding | NP_001135445.1;NP_001135446.1;NP_071372.1;XP_005245867.1;XP_00 | 30213 | 610513 | . | . | 53.080332 | 55.657634 | 46.340237 | 48.840631 | 123.520112 | 128.655113 | 103.986151 | 111.668300 | -1.158295 | 6330.457127 | 0.104780156 | -2.02330659 | 0.043041548 | 0.639862814 | 12.730232 | 12.518170 | 12.670768 | 12.702645 | 12.547128 | 12.578640 | 7678 | 7469 | 5004 | 5604 |
| 23401 | NM_012083 | FRAT2 | FRAT regulator of WNT signaling pathway 2 | protein_coding | NP_036215.1 | 16048 | 605006 | . | . | 3.136077 | 3.075039 | 2.972442 | 3.790087 | 7.297781 | 7.108090 | 6.670074 | 8.665584 | 1.073775 | 226.196582 | 0.233612993 | 0.439582083 | 0.66023982 | 0.998865404 | 7.769697 | 7.871378 | 7.809229 | 7.795301 | 7.778785 | 7.890616 | 255 | 232 | 180 | 245 |
| 23403 | NM_00108046 | FBXO46 | F-box protein 46 | protein_coding | NP_001073938.1;NP_001316561.1;NP_001316562.1;NP_001316563.1 | 25069 | 609117 | . | . | 4.272250 | 4.382087 | 5.858263 | 5.167326 | 9.941700 | 10.129389 | 13.145770 | 11.814477 | 1.143117 | 491.138354 | 0.174871744 | 1.103511554 | 0.269805077 | 0.998865404 | 8.839937 | 9.033543 | 8.891837 | 8.895902 | 9.030889 | 8.923058 | 526 | 496 | 494 | 450 |
| 23404 | NM_00128270 | EXOSC2 | exosome component 2 | protein_coding | NP_001269637.1;NP_001269638.1;NP_055100.2;XP_005272233.1;XP_00 | 17097 | 602238 | . | . | 17.524430 | 17.066957 | 17.985895 | 18.835892 | 40.780069 | 39.451037 | 40.359826 | 43.066027 | 1.049655 | 1221.540445 | 0.130498754 | 0.535757684 | 0.592126031 | 0.998865404 | 10.219021 | 10.289100 | 10.239765 | 10.231512 | 10.290101 | 10.254059 | 1380 | 1279 | 1114 | 1145 |
| 23405 | NM_00119557 | DICER1 | dicer 1, ribonuclease III | protein_coding | NP_001182502.1;NP_001258211.1;NP_001278557.1;NP_085124.2;NP_8 | 17098 | 606241 | . | . | 5.825949 | 5.855452 | 6.059981 | 5.200601 | 13.557221 | 13.535141 | 13.598420 | 11.890559 | -1.040410 | 1942.827566 | 0.125705413 | -0.45465621 | 0.649356586 | 0.998865404 | 10.952100 | 10.895224 | 10.939131 | 10.938572 | 10.958944 | 10.853437 | 2282 | 2137 | 1767 | 1666 |
| 23406 | NM_021149 | COTL1 | coactosin like F-actin binding protein 1 | protein_coding | NP_066972.1 | 18304 | 606748 | . | . | 128.317963 | 137.180374 | 97.593185 | 120.193748 | 298.601166 | 317.098602 | 218.996292 | 274.808716 | -1.236467 | 6936.234243 | 0.122135536 | -2.50724328 | 0.012167693 | 0.364318163 | 12.905009 | 12.598630 | 12.816553 | 12.866679 | 12.577148 | 12.741070 | 8575 | 8517 | 4861 | 6389 |
| 23408 | NM_00119326 | SIRT5 | sirtuin 5 | protein_coding | NP_001180196.1;NP_001229756.1;NP_001363727.1;NP_001363728.1;NI | 14933 | 604483 | . | . | 2.834660 | 3.291878 | 3.500843 | 3.322841 | 6.596374 | 7.609325 | 7.855793 | 7.597285 | 1.139375 | 331.040175 | 0.203551852 | 0.924789388 | 0.355075472 | 0.998865404 | 8.274072 | 8.461566 | 8.267689 | 8.389132 | 8.403535 | 8.402146 | 317 | 371 | 307 | 330 |
| 23409 | NM_012240,XI | SIRT4 | sirtuin 4 | protein_coding | NP_036372.1;XP_005253922.1;XP_006719371.1;XP_006719372.1;XP_02 | 14932 | 604482 | . | . | 0.376690 | 0.340369 | 0.588317 | 0.569351 | 0.876572 | 0.786779 | 1.320166 | 1.301753 | 1.829244 | 19.617693 | 0.698542258 | 1.247237231 | 0.212310525 | NA | 3.793789 | 4.664848 | 4.257160 | 4.250016 | 4.293099 | 4.312804 | 17 | 14 | 20 | 26 |
| 23410 | NM_00101752 | SIRT3 | sirtuin 3 | protein_coding | NP_001017524.1;NP_001357239.1;NP_001357241.1;NP_001357243.1;NI | 14931 | 604481 | . | . | 4.407091 | 5.527011 | 5.732824 | 5.681709 | 10.255486 | 12.775933 | 12.864292 | 12.990550 | 1.168502 | 502.575405 | 0.175390318 | 1.280917299 | 0.200222716 | 0.96621494 | 8.856732 | 9.080957 | 8.860410 | 8.974401 | 9.028686 | 9.006730 | 485 | 546 | 479 | 499 |
| 23411 | NM_00114249 | SIRT1 | sirtuin 1 | protein_coding | NP_001135970.1;NP_001300978.1;NP_036370.2 | 14929 | 604479 | . | . | 2.554139 | 2.639070 | 3.085321 | 2.637835 | 5.943586 | 6.100328 | 6.923370 | 6.031096 | 1.096158 | 347.091344 | 0.196447642 | 0.674252869 | 0.500150563 | 0.998865404 | 8.371103 | 8.504173 | 8.398051 | 8.422482 | 8.510680 | 8.411853 | 373 | 365 | 343 | 311 |
| 23412 | NM_012071 | COMMD3 | COMM domain containing 3 | protein_coding | NP_036203.1 | 23332 | 616700 | . | . | 18.386185 | 18.364819 | 15.667899 | 21.456141 | 42.785404 | 42.451099 | 35.158310 | 49.056915 | -1.003480 | 533.213879 | 0.185200914 | -0.02705889 | 0.978412765 | 0.998865404 | 9.061572 | 9.055563 | 9.060224 | 9.054432 | 8.953687 | 9.147037 | 618 | 574 | 393 | 574 |
| 23413 | NM_00112882 | NCS1 | neuronal calcium sensor 1 | protein_coding | NP_001122298.1;NP_055101.2 | 3953 | 603315 | . | . | 6.133906 | 7.203836 | 5.710574 | 7.023256 | 14.273851 | 16.651991 | 12.814361 | 16.057840 | -1.061550 | 1048.831568 | 0.153889334 | -0.55996196 | 0.575505385 | 0.998865404 | 10.077381 | 9.990444 | 9.996420 | 10.109604 | 9.936371 | 10.079190 | 1150 | 1254 | 798 | 1047 |
| 23414 | NM_00136283 | ZFPM2 | zinc finger protein, FOG family member 2 | protein_coding | NP_001349765.1;NP_001349766.1;NP_036214.2;XP_011515249.1 | 16700 | 603693 | . | . | 0.010343 | 0.015660 | 0.020226 | 0.018112 | 0.024069 | 0.036199 | 0.045387 | 0.041412 | 1.763961 | 5.668480 | 1.315071284 | 0.622642016 | 0.533519795 | NA | 2.039857 | 2.852980 | 2.466179 | 2.488897 | 2.495514 | 2.488353 | 2 | 7 | 7 | 6 |
| 23415 | NM_012285,XI | KCNH4 | potassium voltage-gated channel subfamily H member 4 | protein_coding | NP_036417.1;XP_016879889.1;XP_016879890.1;XP_016879891.1;XP_01 | 6253 | 604528 | . | . | 0.099666 | 0.085926 | 0.065076 | 0.211868 | 0.231927 | 0.198622 | 0.146029 | 0.484411 | 1.427934 | 15.718974 | 0.825006653 | 0.622939294 | 0.533324415 | NA | 3.697260 | 4.206844 | 3.955113 | 3.946928 | 3.935603 | 4.013540 | 16 | 13 | 8 | 26 |
| 23416 | NM_00131403 | KCNH3 | potassium voltage-gated channel subfamily H member 3 | protein_coding | NP_001300959.1;NP_036416.1;XP_011536387.1;XP_011536388.1;XP_01 | 6252 | 604527 | . | . | 0.477590 | 0.444671 | 0.548973 | 0.569071 | 1.111373 | 1.027877 | 1.231880 | 1.301111 | 1.122050 | 68.830161 | 0.383989408 | 0.432658671 | 0.665262759 | 0.998865404 | 6.019836 | 6.185357 | 6.102019 | 6.075794 | 6.091808 | 6.139853 | 79 | 66 | 57 | 75 |
| 23417 | NM_012213 | MLYCD | malonyl-CoA decarboxylase | protein_coding | NP_036345.2 | 7150 | 606761 | . | . | 0.461603 | 0.577589 | 0.444280 | 0.490425 | 1.074169 | 1.335124 | 0.996952 | 1.121299 | -1.127722 | 201.097027 | 0.24506164 | -0.70762467 | 0.479178368 | 0.998865404 | 7.736431 | 7.561782 | 7.624748 | 7.727108 | 7.603616 | 7.641603 | 219 | 255 | 157 | 185 |
| 23420 | NM_014287 | NOMO1 | NODAL modulator 1 | protein_coding | NP_055102.3 | 30060 | 609157 | . | . | 83.421829 | 83.381706 | 80.849815 | 85.804176 | 194.126022 | 192.740555 | 181.424637 | 196.181046 | -1.015092 | 11198.233572 | 0.101477397 | -0.21295877 | 0.83135912 | 0.998865404 | 13.461760 | 13.440126 | 13.461143 | 13.453411 | 13.423293 | 13.465246 | 13050 | 12119 | 9427 | 10676 |
| 23421 | NM_00120673 | ITGB3BP | integrin subunit beta 3 binding protein | protein_coding | NP_001193668.1;NP_001334074.1;NP_001334076.1;NP_001334077.1;NI | 6157 | 605494 | . | . | 16.787951 | 12.856668 | 13.301531 | 8.659207 | 39.066250 | 29.718764 | 29.848251 | 19.798251 | -1.360540 | 420.114857 | 0.219993927 | -2.01905207 | 0.043481807 | 0.642590349 | 8.918916 | 8.476592 | 8.870405 | 8.708523 | 8.711070 | 8.492215 | 627 | 457 | 370 | 269 |
| 23423 | NM_00130120 | TMED3 | transmembrane p24 trafficking protein 3 | protein_coding | NP_001288132.1;NP_001317305.1;NP_001350664.1;NP_031390.1 | 28889 | . | | . | 125.287082 | 126.949194 | 126.183387 | 130.479457 | 291.548202 | 293.448744 | 283.151867 | 298.325790 | 1.002951 | 5694.642459 | 0.104213118 | 0.04079873 | 0.967456352 | 0.998865404 | 12.473276 | 12.477500 | 12.471088 | 12.477181 | 12.467376 | 12.485753 | 6524 | 6158 | 4905 | 5405 |
| 23424 | NM_00130288 | TDRD7 | tudor domain containing 7 | protein_coding | NP_001289813.1;NP_055105.2;XP_006717089.1 | 30831 | 611258 | . | . | 17.774702 | 18.516372 | 21.495413 | 22.487096 | 41.362464 | 42.801420 | 48.235080 | 51.414072 | 1.186460 | 2289.985478 | 0.115823894 | 2.129638482 | 0.033201471 | 0.575540992 | 11.032578 | 11.279145 | 11.075133 | 11.100828 | 11.215385 | 11.235209 | 2373 | 2297 | 2135 | 2358 |
| 23426 | NM_00117807 | GRIP1 | glutamate receptor interacting protein 1 | protein_coding | NP_001171545.1;NP_001353651.1;NP_001353652.1;NP_001353653.1;NI | 18708 | 604597 | . | . | 0.559541 | 0.586370 | 0.436706 | 0.464946 | 1.302074 | 1.355422 | 0.979956 | 1.063043 | -1.296835 | 83.832040 | 0.347501593 | -1.07911854 | 0.280534886 | 0.998865404 | 6.565011 | 6.189480 | 6.416512 | 6.432811 | 6.349554 | 6.360021 | 106 | 105 | 62 | 70 |
| 23428 | NM_00126703 | SLC7A8 | solute carrier family 7 member 8 | protein_coding | NP_001253965.1;NP_001253966.1;NP_036376.2;NP_877392.1 | 11066 | 604235 | . | . | 64.888188 | 66.930985 | 43.026716 | 51.441568 | 150.997493 | 154.713958 | 96.550704 | 117.615029 | -1.433121 | 7371.713127 | 0.12071362 | -4.30076355 | 1.70211E-05 | 0.003012073 | 13.084197 | 12.564890 | 12.970813 | 12.989258 | 12.591319 | 12.753250 | 9894 | 9470 | 4748 | 6242 |
| 23429 | NM_012234 | RYBP | RING1 and YY1 binding protein | protein_coding | NP_036366.3 | 10480 | 607535 | . | . | 24.676350 | 20.671911 | 25.871330 | 20.749012 | 57.422882 | 47.784054 | 58.054516 | 47.440147 | 1.013900 | 3188.346119 | 0.142274276 | 0.13997705 | 0.888678123 | 0.998865404 | 11.628426 | 11.648688 | 11.703451 | 11.551120 | 11.731881 | 11.542521 | 3983 | 3100 | 3113 | 2664 |
| 23431 | NM_00125212 | AP4E1 | adaptor related protein complex 4 subunit epsilon 1 | protein_coding | NP_001239056.1;NP_031373.2;XP_005254321.1;XP_006720510.1;XP_01 | 573 | 607244 | . | . | 2.217406 | 1.770981 | 2.492443 | 1.739375 | 5.159997 | 4.093701 | 5.592969 | 3.976875 | 1.050584 | 426.480294 | 0.21387279 | 0.332871045 | 0.739231627 | 0.998865404 | 8.699455 | 8.772296 | 8.787140 | 8.637221 | 8.844606 | 8.634303 | 538 | 393 | 446 | 339 |
| 23432 | NM_00126760 | GPR161 | G protein-coupled receptor 161 | protein_coding | NP_001254538.1;NP_001254539.1;NP_001254540.1;NP_001254541.1;NI | 23694 | 612250 | . | . | 2.435207 | 2.920916 | 3.114242 | 3.290019 | 5.666828 | 6.751828 | 6.988269 | 7.522237 | 1.188727 | 739.896718 | 0.15755786 | 1.583019611 | 0.113416995 | 0.850211417 | 9.401399 | 9.650243 | 9.402924 | 9.524909 | 9.573555 | 9.597703 | 709 | 795 | 687 | 766 |
| 23433 | NM_012249,XI | RHOQ | ras homolog family member Q | protein_coding | NP_036381.2;XP_005264286.1;XP_011531028.1;XP_011531030.1;XP_01 | 17736 | 605857 | . | . | 39.881309 | 33.942772 | 36.888096 | 27.615293 | 92.805447 | 78.460241 | 82.775814 | 63.139083 | -1.163358 | 4560.970346 | 0.134624456 | -1.62151482 | 0.104907274 | 0.832204085 | 12.260030 | 12.041989 | 12.271448 | 12.150649 | 12.175646 | 11.989101 | 6050 | 4916 | 4082 | 3506 |
| 23435 | NM_007375,XI | TARDBP | TAR DNA binding protein | protein_coding | NP_031401.1;XP_016856352.1;XP_016856353.1;XP_016856354.1;XP_01 | 11571 | 605078 | . | . | 42.420553 | 40.658229 | 38.131151 | 39.075281 | 98.714366 | 93.983322 | 85.565203 | 89.340988 | -1.019556 | 5267.947000 | 0.116657947 | -0.23951728 | 0.810704502 | 0.998865404 | 12.376850 | 12.349066 | 12.386606 | 12.353740 | 12.415887 | 12.288412 | 6244 | 5627 | 4882 | 4519 |
| 23438 | NM_00127873 | HARS2 | histidyl-tRNA synthetase 2, mitochondrial | protein_coding | NP_001265660.1;NP_001265661.1;NP_001350464.1;NP_001350465.1;NI | 4817 | 600783 | . | . | 18.684536 | 19.801523 | 19.825731 | 20.652271 | 43.479683 | 45.772108 | 44.488367 | 47.218961 | 1.041431 | 1514.154739 | 0.125435483 | 0.466906078 | 0.640567076 | 0.998865404 | 10.534807 | 10.593197 | 10.526126 | 10.569780 | 10.568877 | 10.590064 | 1660 | 1646 | 1325 | 1468 |
| 23443 | NM_00127168 | SLC35A3 | solute carrier family 35 member A3 | protein_coding | NP_001258613.1;NP_001258614.1;NP_036375.1;XP_005270748.1;XP_01 | 11023 | 605632 | . | . | 0.743210 | 0.478727 | 0.897237 | 0.469723 | 1.729481 | 1.106598 | 2.013374 | 1.073968 | -1.157345 | 213.324091 | 0.307157919 | -0.68635578 | 0.492488784 | 0.998865404 | 7.837162 | 7.629153 | 7.811335 | 7.696329 | 7.832904 | 7.516599 | 294 | 218 | 233 | 120 |
| 23446 | NM_00128673 | SLC44A1 | solute carrier family 44 member 1 | protein_coding | NP_001273659.1;NP_001317660.1;NP_536856.2;XP_005251912.1;XP_00 | 18798 | 606105 | . | . | 17.052020 | 14.556513 | 18.702077 | 13.296339 | 39.680750 | 33.648031 | 41.966917 | 30.400498 | 1.005217 | 4715.763006 | 0.148658925 | 0.050493917 | 0.959728797 | 0.998865404 | 12.199389 | 12.207151 | 12.278981 | 12.108468 | 12.314274 | 12.075237 | 5966 | 4559 | 4697 | 3802 |
| 23450 | NM_012426 | SF3B3 | splicing factor 3b subunit 3 | protein_coding | NP_036558.3 | 10770 | 605592 | . | . | 15.538988 | 17.139000 | 16.028387 | 17.239225 | 36.159863 | 39.617565 | 35.967239 | 39.415440 | 1.004058 | 4976.863876 | 0.111293291 | 0.052495298 | 0.958134042 | 0.998865404 | 12.278161 | 12.283876 | 12.240884 | 12.315723 | 12.255869 | 12.308106 | 5464 | 5599 | 4201 | 4821 |
| 23451 | NM_00100552 | SF3B1 | splicing factor 3b subunit 1 | protein_coding | NP_001005526.1;NP_001295753.1;NP_036565.2 | 10768 | 605590 | . | . | 76.482359 | 71.641353 | 65.901593 | 66.647013 | 177.977585 | 165.602204 | 147.881263 | 152.380472 | -1.091425 | 11258.399365 | 0.105376713 | -1.19773381 | 0.231020664 | 0.986164394 | 13.520413 | 13.394259 | 13.510241 | 13.478842 | 13.457438 | 13.381075 | 13777 | 12448 | 9784 | 9639 |
| 23452 | NM_012098,XI | ANGPTL2 | angiopoietin like 2 | protein_coding | NP_036230.1;XP_006717093.1 | 490 | 605001 | . | . | 2.914946 | 2.516381 | 2.419171 | 2.990156 | 6.783197 | 5.816728 | 5.428550 | 6.836638 | -1.019148 | 321.159418 | 0.208524119 | -0.13122611 | 0.895596439 | 0.998865404 | 8.340986 | 8.313171 | 8.371597 | 8.288812 | 8.262929 | 8.375087 | 401 | 324 | 248 | 329 |
| 23456 | NM_012089,XI | ABCB10 | ATP binding cassette subfamily B member 10 | protein_coding | NP_036221.2;XP_011542437.1;XP_011542438.1 | 41 | 605454 | . | . | 4.838358 | 4.441956 | 4.129594 | 4.850060 | 11.259058 | 10.267780 | 9.266692 | 11.089086 | -1.009326 | 558.006079 | 0.162399107 | -0.08246403 | 0.934277719 | 0.998865404 | 9.130939 | 9.117303 | 9.134682 | 9.119534 | 9.096744 | 9.144747 | 653 | 598 | 465 | 540 |
| 23457 | NM_00124301 | ABCB9 | ATP binding cassette subfamily B member 9 | protein_coding | NP_001229942.1;NP_001229943.1;NP_062570.1;NP_062571.1;NP_9822 | 50 | 605453 | . | . | 6.493412 | 6.579949 | 8.128987 | 7.150022 | 15.110435 | 15.209841 | 18.241209 | 16.347676 | 1.140574 | 752.876013 | 0.154689041 | 1.226719825 | 0.219927914 | 0.982008277 | 9.457935 | 9.648326 | 9.524477 | 9.488692 | 9.651102 | 9.543285 | 831 | 739 | 751 | 695 |
| 23461 | NM_018672,N | ABCA5 | ATP binding cassette subfamily A member 5 | protein_coding | NP_061142.2;NP_758424.1 | 35 | 612503 | . | . | 3.181754 | 2.857788 | 4.674253 | 2.156491 | 7.404072 | 6.605906 | 10.488889 | 4.930561 | 1.116418 | 927.698962 | 0.550191331 | 0.288767849 | 0.772759037 | 0.998865404 | 9.775796 | 9.934848 | 9.837186 | 9.753561 | 10.119312 | 9.556613 | 1068 | 891 | 1168 | 576 |
| 23462 | NM_00104070 | HEY1 | hes related family bHLH transcription factor with YRPW motif 1 | protein_coding | NP_001035798.1;NP_001269780.1;NP_036390.3 | 4880 | 602953 | . | . | 21.710669 | 21.335192 | 24.972782 | 21.985633 | 50.521621 | 49.317255 | 56.038200 | 50.267535 | 1.074056 | 1552.809148 | 0.131107628 | 0.786146971 | 0.431781389 | 0.998865404 | 10.548023 | 10.651453 | 10.582016 | 10.560556 | 10.679794 | 10.571143 | 1748 | 1593 | 1496 | 1403 |
| 23463 | NM_012405,XI | ICMT | isoprenylcysteine carboxyl methyltransferase | protein_coding | NP_036537.1;XP_011539442.1 | 5350 | 605851 | . | . | 54.575034 | 56.479224 | 51.310887 | 53.392033 | 126.998354 | 130.554255 | 115.140141 | 122.074540 | -1.076594 | 8029.703280 | 0.102648321 | -1.03727039 | 0.299609861 | 0.998865404 | 13.023405 | 12.916892 | 12.991122 | 13.012006 | 12.924392 | 12.953391 | 9470 | 9094 | 6609 | 7375 |
| 23464 | NM_00117169 | GCAT | glycine C-acetyltransferase | protein_coding | NP_001165161.1;NP_055106.1;XP_005261466.1;XP_005261467.1;XP_00 | 4188 | 607422 | . | . | 16.120574 | 16.672762 | 17.032164 | 21.194429 | 37.513234 | 38.539836 | 38.219682 | 48.458541 | 1.163411 | 895.949811 | 0.154599518 | 1.412432305 | 0.15782271 | 0.92817506 | 9.694315 | 9.912027 | 9.736808 | 9.759176 | 9.775100 | 9.932596 | 939 | 908 | 748 | 1000 |
| 23466 | NM_00130349 | CBX6 | chromobox 6 | protein_coding | NP_001290423.1;NP_055107.3 | 1556 | 617438 | . | . | 13.322102 | 14.542038 | 13.330334 | 15.575114 | 31.001078 | 33.614573 | 29.912883 | 35.610649 | 1.023043 | 2667.469176 | 0.124552847 | 0.263876962 | 0.791874728 | 0.998865404 | 11.364882 | 11.397446 | 11.337758 | 11.403453 | 11.328991 | 11.446648 | 2914 | 2961 | 2176 | 2712 |
| 23467 | NM_014293 | NPTXR | neuronal pentraxin receptor | protein_coding | NP_055108.2 | 7954 | 609474 | . | . | 1.072342 | 1.015350 | 1.122862 | 1.072041 | 2.495384 | 2.347027 | 2.519669 | 2.451093 | 1.039533 | 195.061867 | 0.239321527 | 0.233726791 | 0.815197087 | 0.998865404 | 7.579260 | 7.635762 | 7.612036 | 7.584542 | 7.628582 | 7.602683 | 227 | 200 | 178 | 181 |
| 23468 | NM_00112732 | CBX5 | chromobox 5 | protein_coding | NP_001120793.1;NP_001120794.1;NP_036249.1 | 1555 | 604478 | . | . | 2.121725 | 2.279725 | 2.168222 | 2.132889 | 4.937342 | 5.269685 | 4.865429 | 4.876598 | -1.037523 | 783.486415 | 0.146668785 | -0.3623387 | 0.717098943 | 0.998865404 | 9.640121 | 9.586918 | 9.603982 | 9.648481 | 9.610180 | 9.590909 | 890 | 888 | 678 | 711 |
| 23469 | NM_00129025 | PHF3 | PHD finger protein 3 | protein_coding | NP_001277188.1;NP_001277189.1;NP_001357277.1;NP_001357278.1;NI | 8921 | 607789 | . | . | 1.802430 | 1.998314 | 2.223908 | 2.280805 | 4.194325 | 4.619190 | 4.990388 | 5.214789 | 1.085036 | 1149.718080 | 0.134649326 | 0.874443065 | 0.381877014 | 0.998865404 | 10.106988 | 10.224739 | 10.107907 | 10.161627 | 10.217877 | 10.173910 | 1224 | 1233 | 1071 | 1089 |
| 23471 | NM_00131780 | TRAM1 | translocation associated membrane protein 1 | protein_coding | NP_001304733.1;NP_001304734.1;NP_055109.1;XP_016868755.1;XP_01 | 20568 | 605190 | . | . | 33.725164 | 26.077868 | 28.062988 | 17.935028 | 78.479845 | 60.280161 | 62.972526 | 41.006308 | -1.315789 | 2543.629366 | 0.367111673 | -1.07849522 | 0.280812813 | 0.998865404 | 11.497044 | 11.101239 | 11.507475 | 11.291857 | 11.347379 | 10.986215 | 3756 | 2718 | 2345 | 1591 |
| 23473 | NM_00137608 | CAPN7 | calpain 7 | protein_coding | NP_001363015.1;NP_001363016.1;NP_001363018.1;NP_001363019.1;NI | 1484 | 606400 | . | . | 5.545110 | 5.397313 | 5.281649 | 5.029093 | 12.903697 | 12.476132 | 11.851866 | 11.498426 | -1.064537 | 656.221465 | 0.155537356 | -0.58009284 | 0.561852011 | 0.998865404 | 9.402231 | 9.312451 | 9.389099 | 9.368062 | 9.366253 | 9.305681 | 791 | 719 | 578 | 569 |
| 23474 | NM_00132086 | ETHE1 | ETHE1 persulfide dioxygenase | protein_coding | NP_001307796.1;NP_001307797.1;NP_001307798.1;NP_055112.2;XP_00 | 23287 | 608451 | . | . | 10.818409 | 11.786191 | 10.275870 | 9.950478 | 25.174886 | 27.244307 | 23.058752 | 22.750587 | -1.075657 | 291.834604 | 0.203471361 | -0.51711594 | 0.605075225 | 0.998865404 | 8.240739 | 8.135351 | 8.190395 | 8.225762 | 8.172703 | 8.166244 | 337 | 337 | 246 | 262 |
| 23475 | NM_00131824 | QPRT | quinolinate phosphoribosyltransferase | protein_coding | NP_001305178.1;NP_001305179.2;NP_055113.3;XP_005255280.2;XP_01 | 9755 | 606248 | . | . | 57.397642 | 56.458653 | 63.548980 | 75.539401 | 133.566662 | 130.506713 | 142.602071 | 172.711862 | 1.207504 | 4348.492194 | 0.119561779 | 2.27520669 | 0.022893534 | 0.499999124 | 11.943955 | 12.215852 | 12.012591 | 11.990552 | 12.086383 | 12.226121 | 4598 | 4194 | 3796 | 4827 |
| 23476 | NM_00133038 | BRD4 | bromodomain containing 4 | protein_coding | NP_001317313.1;NP_001366220.1;NP_001366221.1;NP_055114.1;NP_4 | 13575 | 608749 | . | . | 7.263254 | 8.879823 | 9.334220 | 9.183110 | 16.901892 | 20.526110 | 20.945721 | 20.996091 | 1.139108 | 1558.853373 | 0.138802463 | 1.353751271 | 0.175815745 | 0.949193943 | 10.509347 | 10.697089 | 10.475228 | 10.622955 | 10.680367 | 10.624505 | 1517 | 1724 | 1495 | 1501 |
| 23478 | NM_00127191 | SEC11A | SEC11 homolog A, signal peptidase complex subunit | protein_coding | NP_001258847.1;NP_001258848.1;NP_001258849.1;NP_001258850.1;NI | 17718 | 618258 | . | . | 118.289185 | 112.061689 | 120.359810 | 108.711725 | 275.263803 | 259.035633 | 270.083921 | 248.556446 | -1.020885 | 4052.509922 | 0.116042549 | -0.25697227 | 0.797200179 | 0.998865404 | 11.999339 | 11.969709 | 12.020010 | 11.964285 | 12.023007 | 11.925588 | 4872 | 4270 | 3693 | 3539 |
| 23479 | NM_00130114 | ISCU | iron-sulfur cluster assembly enzyme | protein_coding | NP_001288069.1;NP_001288070.1;NP_001306971.1;NP_055116.1;NP_9 | 29882 | 611911 | . | . | 27.605373 | 28.608615 | 29.567467 | 26.911219 | 64.238839 | 66.130098 | 66.348537 | 61.529310 | 1.016498 | 1051.955104 | 0.136468979 | 0.172988903 | 0.86266015 | 0.998865404 | 10.026923 | 10.050694 | 10.021714 | 10.043081 | 10.071154 | 10.017310 | 1184 | 1142 | 956 | 958 |
| 23480 | NM_00101245 | SEC61G | SEC61 translocon subunit gamma | protein_coding | NP_001012474.1;NP_055117.1 | 18277 | 609215 | . | . | 59.032610 | 50.169280 | 55.570983 | 48.457763 | 137.371311 | 115.968543 | 124.699674 | 110.792913 | -1.060907 | 712.868695 | 0.163666906 | -0.52117167 | 0.602247185 | 0.998865404 | 9.519087 | 9.434663 | 9.552426 | 9.436144 | 9.505239 | 9.403108 | 915 | 726 | 647 | 600 |
| 23481 | NM_00124322 | PES1 | pescadillo ribosomal biogenesis factor 1 | protein_coding | NP_001230154.1;NP_001269256.1;NP_001269257.1;NP_055118.1;XP_00 | 8848 | 605819 | . | . | 48.609312 | 48.114643 | 44.673934 | 50.621643 | 113.115879 | 111.219161 | 100.247012 | 115.740367 | -1.026031 | 3432.805706 | 0.115045959 | -0.32225295 | 0.747261075 | 0.998865404 | 11.763657 | 11.726448 | 11.762819 | 11.747856 | 11.688163 | 11.778174 | 4040 | 3718 | 2785 | 3351 |
| 23483 | NM_00130443 | TGDS | TDP-glucose 4,6-dehydratase | protein_coding | NP_001291359.1;NP_055120.1;XP_011519367.1;XP_011519368.1 | 20324 | 616146 | . | . | 12.883778 | 11.119522 | 13.817231 | 9.248423 | 29.981083 | 25.703272 | 31.005464 | 21.145424 | -1.068038 | 669.847358 | 0.19671689 | -0.4827393 | 0.629280858 | 0.998865404 | 9.433784 | 9.340072 | 9.449312 | 9.355777 | 9.489865 | 9.208665 | 850 | 696 | 681 | 481 |
| 23484 | NM_00112820 | LEPROTL1 | leptin receptor overlapping transcript like 1 | protein_coding | NP_001121680.1;NP_056159.2 | 6555 | 607338 | . | . | 14.456694 | 12.478727 | 13.053824 | 10.051586 | 33.641318 | 28.845138 | 29.292406 | 22.981758 | -1.177859 | 1225.446405 | 0.160250285 | -1.47373475 | 0.140553023 | 0.900674731 | 10.371984 | 10.136610 | 10.377961 | 10.246820 | 10.285264 | 10.087481 | 1655 | 1309 | 1110 | 914 |
| 23492 | NM_00134674 | CBX7 | chromobox 7 | protein_coding | NP_001333672.1;NP_001333673.1;NP_783640.1 | 1557 | 608457 | . | . | 4.344595 | 4.021525 | 3.155814 | 3.577435 | 10.110049 | 9.295935 | 7.081556 | 8.179379 | -1.256124 | 484.065402 | 0.173738461 | -1.8935268 | 0.058287847 | 0.715342823 | 9.074325 | 8.745043 | 9.012372 | 8.958395 | 8.808789 | 8.875536 | 646 | 558 | 352 | 425 |
| 23493 | NM_012259,XI | HEY2 | hes related family bHLH transcription factor with YRPW motif 2 | protein_coding | NP_036391.1;XP_016866116.1;XP_016866117.1;XP_016866118.1 | 4881 | 604674 | . | . | 1.228613 | 1.293379 | 0.973665 | 0.877063 | 2.859032 | 2.989705 | 2.184876 | 2.005301 | -1.305151 | 97.347362 | 0.3318816 | -1.157693 | 0.246989342 | 0.992733415 | 6.783598 | 6.401288 | 6.653821 | 6.631734 | 6.597609 | 6.528354 | 131 | 115 | 83 | 69 |
| 23499 | NM_012090 | MACF1 | microtubule actin crosslinking factor 1 | protein_coding | NP_036222.3 | 13664 | 608271 | . | . | 10.981183 | 11.370513 | 12.474551 | 12.338735 | 25.553663 | 26.283449 | 27.992531 | 28.211048 | 1.094856 | 6495.751683 | 0.103591239 | 1.262084451 | 0.2069184 | 0.970482251 | 12.598429 | 12.729171 | 12.614648 | 12.636726 | 12.711172 | 12.693171 | 7047 | 6780 | 5967 | 6298 |
| 23500 | NM_00120142 | DAAM2 | dishevelled associated activator of morphogenesis 2 | protein_coding | NP_001188356.1;NP_056160.2;XP_006715102.3;XP_006715103.3;XP_00 | 18143 | 606627 | . | . | 24.481229 | 26.094112 | 23.365628 | 26.569979 | 56.968826 | 60.317706 | 52.431794 | 60.749102 | -1.042250 | 5008.719369 | 0.115226531 | -0.51811839 | 0.604375671 | 0.998865404 | 12.319848 | 12.259983 | 12.265321 | 12.346531 | 12.227126 | 12.315081 | 5603 | 5783 | 4040 | 4841 |
| 23503 | NM_015346,XI | ZFYVE26 | zinc finger FYVE-type containing 26 | protein_coding | NP_056161.2;XP_011534911.1;XP_016876613.1;XP_016876614.1;XP_01 | 20761 | 612012 | . | . | 4.485424 | 4.361028 | 6.721876 | 6.082962 | 10.437766 | 10.080709 | 15.083692 | 13.907969 | 1.414430 | 1616.936176 | 0.126166256 | 3.964776098 | 7.34649E-05 | 0.010039957 | 10.387212 | 10.887693 | 10.519970 | 10.493213 | 10.811902 | 10.740070 | 1570 | 1419 | 1721 | 1697 |
| 23505 | NM_015348,XI | TMEM131 | transmembrane protein 131 | protein_coding | NP_056163.1;XP_005263966.1;XP_005263967.1;XP_005263968.1;XP_00 | 30366 | 615659 | . | . | 16.996535 | 18.653494 | 18.289097 | 16.591549 | 39.551634 | 43.118387 | 41.040205 | 37.934607 | -1.030736 | 3695.526145 | 0.117893928 | -0.37045423 | 0.711044076 | 0.998865404 | 11.873202 | 11.829597 | 11.827384 | 11.897992 | 11.884576 | 11.789868 | 4136 | 4220 | 3347 | 3216 |
| 23506 | NM_00131881 | BICRAL | BRD4 interacting chromatin remodeling complex associated prote | protein_coding | NP_001305748.1;NP_056164.1;XP_024302157.1;XP_024302158.1 | 21111 | 618502 | . | . | 0.768681 | 0.822745 | 0.843619 | 0.963757 | 1.788752 | 1.901812 | 1.893056 | 2.203516 | 1.171150 | 175.090268 | 0.255784818 | 0.891085911 | 0.372883083 | 0.998865404 | 7.334191 | 7.560828 | 7.380705 | 7.446296 | 7.453094 | 7.507545 | 171 | 188 | 154 | 188 |
| 23507 | NM_00113447 | LRRC8B | leucine rich repeat containing 8 VRAC subunit B | protein_coding | NP_001127948.1;NP_001356746.1;NP_001356748.1;NP_056165.1;XP_00 | 30692 | 612888 | . | . | 1.116477 | 0.895388 | 0.901308 | 0.895940 | 2.598089 | 2.069728 | 2.022509 | 2.048463 | -1.151580 | 234.800463 | 0.227587245 | -0.89466553 | 0.370965912 | 0.998865404 | 7.973047 | 7.770427 | 7.950706 | 7.863572 | 7.857057 | 7.822026 | 314 | 248 | 197 | 197 |
| 23508 | NM_015351 | TTC9 | tetratricopeptide repeat domain 9 | protein_coding | NP_056166.1 | 20267 | 610488 | . | . | 1.028286 | 1.006466 | 0.623822 | 0.780120 | 2.392864 | 2.326490 | 1.399839 | 1.783652 | -1.470938 | 136.940945 | 0.284858841 | -1.95442923 | 0.050650477 | 0.672238973 | 7.349610 | 6.791583 | 7.170020 | 7.155192 | 6.987400 | 7.054222 | 191 | 173 | 86 | 115 |
| 23509 | NM_015352,N | POFUT1 | protein O-fucosyltransferase 1 | protein_coding | NP_056167.1;NP_758436.1 | 14988 | 607491 | . | . | 22.012327 | 21.974501 | 18.966554 | 21.951835 | 51.223587 | 50.795045 | 42.560398 | 50.190266 | -1.092882 | 3430.161889 | 0.116549472 | -1.09942679 | 0.271581952 | 0.998865404 | 11.806784 | 11.678486 | 11.785371 | 11.772955 | 11.653992 | 11.755420 | 4156 | 3837 | 2673 | 3264 |
| 23510 | NM_015353,N | KCTD2 | potassium channel tetramerization domain containing 2 | protein_coding | NP_056168.1 | 21294 | 613422 | . | . | 6.096575 | 6.187486 | 6.065128 | 7.922115 | 14.186983 | 14.302651 | 13.609971 | 18.112973 | 1.121954 | 748.564009 | 0.165718969 | 1.0017804 | 0.316449665 | 0.998865404 | 9.462984 | 9.628250 | 9.501666 | 9.506271 | 9.489267 | 9.670691 | 810 | 764 | 600 | 837 |
| 23511 | NM_015354 | NUP188 | nucleoporin 188 | protein_coding | NP_056169.1 | 17859 | 615587 | . | . | 12.344665 | 13.771816 | 12.454831 | 13.488096 | 28.726543 | 31.834169 | 27.948280 | 30.838930 | -1.020472 | 2306.585423 | 0.121709929 | -0.24021353 | 0.810164725 | 0.998865404 | 11.186214 | 11.156721 | 11.137759 | 11.219093 | 11.134561 | 11.190392 | 2548 | 2641 | 1916 | 2215 |
| 23512 | NM_00132120 | SUZ12 | SUZ12 polycomb repressive complex 2 subunit | protein_coding | NP_001308136.1;NP_056170.2;XP_006721857.1;XP_016879898.1 | 17101 | 606245 | . | . | 10.098448 | 8.207013 | 10.045088 | 6.611684 | 23.499507 | 18.970879 | 22.540887 | 15.116830 | -1.098818 | 1207.097044 | 0.187562591 | -0.72483434 | 0.468553669 | 0.998865404 | 10.303287 | 10.168204 | 10.349601 | 10.172041 | 10.337757 | 10.025968 | 1623 | 1206 | 1209 | 854 |
| 23513 | NM_015356,N | SCRIB | scribble planar cell polarity protein | protein_coding | NP_056171.3;NP_874365.3 | 30377 | 607733 | . | . | 14.065214 | 14.227895 | 13.899243 | 13.486691 | 32.730330 | 32.888416 | 31.189500 | 30.835721 | -1.048303 | 2412.690234 | 0.1149442 | -0.59207625 | 0.553799533 | 0.998865404 | 11.270017 | 11.202036 | 11.253525 | 11.256715 | 11.233178 | 11.200810 | 2838 | 2670 | 2090 | 2164 |
| 23514 | NM_00108039 | SPIDR | scaffold protein involved in DNA repair | protein_coding | NP_001073863.1;NP_001269845.1;NP_001269848.1;NP_001339860.1;NI | 28971 | 615384 | . | . | 8.371665 | 8.565465 | 7.960351 | 8.632436 | 19.481212 | 19.799455 | 17.862800 | 19.737035 | -1.024963 | 916.513579 | 0.142133633 | -0.25026842 | 0.80237978 | 0.998865404 | 9.857882 | 9.821918 | 9.829117 | 9.867968 | 9.802173 | 9.858657 | 1040 | 1028 | 756 | 882 |
| 23515 | NM_00132044 | MORC3 | MORC family CW-type zinc finger 3 | protein_coding | NP_001307374.1;NP_001307375.1;NP_056173.1;XP_011527820.1;XP_01 | 23572 | 610078 | . | . | 5.076790 | 5.590980 | 6.267631 | 5.184505 | 11.813897 | 12.923800 | 14.064382 | 11.853756 | 1.064388 | 730.461732 | 0.163004798 | 0.552277456 | 0.580758273 | 0.998865404 | 9.466721 | 9.557191 | 9.457585 | 9.518262 | 9.602441 | 9.455794 | 779 | 797 | 724 | 632 |
| 23516 | NM_00112843 | SLC39A14 | solute carrier family 39 member 14 | protein_coding | NP_001121903.1;NP_001128625.1;NP_001128626.1;NP_001338584.1;NI | 20858 | 608736 | . | . | 62.743318 | 58.279413 | 61.632352 | 62.074033 | 146.006282 | 134.715463 | 138.301207 | 141.924904 | 1.008481 | 9065.611714 | 0.105335378 | 0.115669653 | 0.907914358 | 0.998865404 | 13.140064 | 13.152288 | 13.178783 | 13.104578 | 13.150545 | 13.148551 | 10848 | 9316 | 7926 | 8525 |
| 23517 | NM_015360 | MTREX | Mtr4 exosome RNA helicase | protein_coding | NP_056175.3 | 18734 | 618122 | . | . | 18.336445 | 17.926426 | 16.896671 | 14.464647 | 42.669659 | 41.437737 | 37.915638 | 33.071697 | -1.172236 | 2088.173142 | 0.12834384 | -1.78632041 | 0.074047393 | 0.763883535 | 11.137943 | 10.909038 | 11.099713 | 11.074474 | 11.023127 | 10.893977 | 2635 | 2394 | 1810 | 1654 |
| 23518 | NM_00128279 | R3HDM1 | R3H domain containing 1 | protein_coding | NP_001269727.1;NP_001269728.1;NP_001269729.1;NP_001341128.1;NI | 9757 | . | | . | 8.827702 | 9.513336 | 8.623948 | 10.671088 | 20.542422 | 21.990500 | 19.351888 | 24.398176 | 1.029628 | 1332.251137 | 0.142547742 | 0.295506879 | 0.767606698 | 0.998865404 | 10.358723 | 10.400278 | 10.335874 | 10.397424 | 10.312964 | 10.460193 | 1451 | 1474 | 1062 | 1389 |
| 23521 | NM_00127049 | RPL13A | ribosomal protein L13a | protein_coding | NP_001257420.1;NP_036555.1 | 10304 | . | | . | 742.047277 | 731.522802 | 711.695106 | 871.032046 | 1726.774446 | 1690.947838 | 1597.023135 | 1991.511140 | 1.058633 | 26819.998386 | 0.116283743 | 0.706916649 | 0.479618277 | 0.998865404 | 14.669350 | 14.751523 | 14.694496 | 14.675074 | 14.647620 | 14.813647 | 30343 | 27798 | 21693 | 28332 |
| 23522 | NM_00125646 | KAT6B | lysine acetyltransferase 6B | protein_coding | NP_001243397.1;NP_001243398.1;NP_001357061.1;NP_001357062.1;NI | 17582 | 605880 | . | . | 1.197724 | 1.048429 | 1.050120 | 1.460654 | 2.787153 | 2.423491 | 2.356438 | 3.339611 | 1.175271 | 293.052594 | 0.217533988 | 1.071067308 | 0.284139167 | 0.998865404 | 8.074497 | 8.306238 | 8.138788 | 8.153764 | 8.153012 | 8.305715 | 306 | 295 | 237 | 338 |
| 23523 | NM_00119928 | CABIN1 | calcineurin binding protein 1 | protein_coding | NP_001186210.1;NP_001188358.1;NP_036427.1;XP_005261472.1;XP_00 | 24187 | 604251 | . | . | 10.041011 | 11.148036 | 12.735121 | 12.519857 | 23.365838 | 25.769182 | 28.577245 | 28.625163 | 1.175988 | 2779.211588 | 0.120132766 | 1.946789368 | 0.051559992 | 0.672667836 | 11.318848 | 11.552613 | 11.316667 | 11.420663 | 11.515014 | 11.488611 | 2751 | 2935 | 2654 | 2771 |
| 23524 | NM_016333,XI | SRRM2 | serine/arginine repetitive matrix 2 | protein_coding | NP_057417.3;XP_005255283.1;XP_006720937.1 | 16639 | 606032 | . | . | 36.675946 | 42.072155 | 46.706330 | 40.503918 | 85.346437 | 97.251671 | 104.807654 | 92.607396 | 1.093675 | 14255.096805 | 0.11828308 | 1.092159526 | 0.274762989 | 0.998865404 | 13.733145 | 13.862344 | 13.701162 | 13.812713 | 13.896957 | 13.769213 | 14652 | 15657 | 13909 | 12929 |
| 23526 | NM_00125832 | ARHGAP45 | Rho GTPase activating protein 45 | protein_coding | NP_001245257.1;NP_001269263.1;NP_001269264.1;NP_001308161.1;NI | 17102 | 601155 | . | . | 0.176282 | 0.131535 | 0.127795 | 0.206048 | 0.410215 | 0.304050 | 0.286769 | 0.471106 | 1.029410 | 27.603848 | 0.603093597 | 0.069337676 | 0.944720836 | 0.998865404 | 4.766397 | 4.806913 | 4.801131 | 4.767200 | 4.762858 | 4.811738 | 36 | 25 | 19 | 32 |
| 23527 | NM_012287,XI | ACAP2 | ArfGAP with coiled-coil, ankyrin repeat and PH domains 2 | protein_coding | NP_036419.3;XP_006713620.1;XP_011510904.1;XP_011510905.1;XP_01 | 16469 | 607766 | . | . | 6.604965 | 5.866656 | 6.548756 | 5.478714 | 15.370024 | 13.561038 | 14.695218 | 12.526428 | -1.073630 | 1334.904757 | 0.142233136 | -0.72062574 | 0.471139815 | 0.998865404 | 10.432576 | 10.330666 | 10.455389 | 10.357621 | 10.413243 | 10.290390 | 1692 | 1397 | 1211 | 1109 |
| 23528 | NM_00128129 | ZNF281 | zinc finger protein 281 | protein_coding | NP_001268222.1;NP_001268223.1;NP_036614.1 | 13075 | 618703 | . | . | 4.909878 | 4.962842 | 4.745916 | 4.417631 | 11.425488 | 11.471829 | 10.649700 | 10.100388 | -1.090547 | 721.192925 | 0.150742725 | -0.82957561 | 0.40677877 | 0.998865404 | 9.555264 | 9.430523 | 9.521467 | 9.525151 | 9.491364 | 9.434892 | 864 | 814 | 625 | 620 |
| 23529 | NM_00116621 | CLCF1 | cardiotrophin like cytokine factor 1 | protein_coding | NP_001159684.1;NP_037378.1 | 17412 | 607672 | . | . | 0.126159 | 0.112098 | 0.097109 | 0.065854 | 0.293577 | 0.259120 | 0.217909 | 0.150566 | -1.431024 | 6.079933 | 1.230813055 | -0.42008645 | 0.6744223 | NA | 2.838265 | 2.324312 | 2.617902 | 2.611665 | 2.608349 | 2.601673 | 9 | 7 | 5 | 4 |
| 23530 | NM_00133102 | NNT | nicotinamide nucleotide transhydrogenase | protein_coding | NP_001317955.1;NP_036475.3;NP_892022.2;XP_005248331.1;XP_00524 | 7863 | 607878 | . | . | 15.717751 | 15.606352 | 13.602767 | 15.230382 | 36.575855 | 36.074777 | 30.524216 | 34.822457 | -1.068267 | 2855.728826 | 0.118143046 | -0.80641766 | 0.420002067 | 0.998865404 | 11.526578 | 11.431130 | 11.511686 | 11.499592 | 11.402754 | 11.498505 | 3422 | 3160 | 2259 | 2742 |
| 23531 | NM_012329,XI | MMD | monocyte to macrophage differentiation associated | protein_coding | NP_036461.2;XP_006721858.1 | 7153 | 604467 | . | . | 3.671163 | 2.959757 | 3.108895 | 2.497167 | 8.542947 | 6.841609 | 6.976269 | 5.709477 | -1.184676 | 249.329724 | 0.234144921 | -1.04419354 | 0.296395856 | 0.998865404 | 8.078172 | 7.835454 | 8.057255 | 7.939275 | 7.967634 | 7.860978 | 346 | 259 | 221 | 190 |
| 23532 | NM_00129171 | PRAME | preferentially expressed antigen in melanoma | protein_coding | NP_001278644.1;NP_001278645.1;NP_001278646.1;NP_001278648.1;NI | 9336 | 606021 | . | . | 201.606306 | 205.234514 | 204.486113 | 225.897333 | 469.146174 | 474.408789 | 458.860889 | 516.487402 | 1.032089 | 14927.751613 | 0.103202256 | 0.441532491 | 0.658827545 | 0.998865404 | 13.842763 | 13.888296 | 13.849830 | 13.853766 | 13.841788 | 13.914648 | 16879 | 15890 | 12627 | 14818 |
| 23534 | NM_00119102 | TNPO3 | transportin 3 | protein_coding | NP_001177957.2;NP_001369145.1;NP_001369146.1;NP_001369147.1;NI | 17103 | 610032 | . | . | 22.736348 | 24.516398 | 23.550210 | 25.114440 | 52.908414 | 56.670756 | 52.845987 | 57.421177 | 1.014467 | 3268.588686 | 0.113967622 | 0.181825362 | 0.85571978 | 0.998865404 | 11.664134 | 11.684697 | 11.637262 | 11.698374 | 11.656617 | 11.702876 | 3598 | 3632 | 2782 | 3173 |
| 23536 | NM_00132444 | ADAT1 | adenosine deaminase tRNA specific 1 | protein_coding | NP_001311373.1;NP_001311374.1;NP_001311375.1;NP_001311377.1;NI | 228 | 604230 | . | . | 6.084539 | 6.712543 | 7.989845 | 7.869102 | 14.158971 | 15.516343 | 17.928983 | 17.991771 | 1.236998 | 1225.446097 | 0.132063685 | 2.32344592 | 0.02015521 | 0.469936417 | 10.097482 | 10.404407 | 10.156020 | 10.188199 | 10.358295 | 10.305674 | 1234 | 1208 | 1219 | 1227 |
| 23541 | NM_00120420 | SEC14L2 | SEC14 like lipid binding 2 | protein_coding | NP_001191133.1;NP_001278861.1;NP_036561.1;NP_203740.1 | 10699 | 607558 | . | . | 10.145909 | 11.224654 | 10.071664 | 11.184042 | 23.609943 | 25.946291 | 22.600523 | 25.570984 | -1.032077 | 1186.737852 | 0.141420995 | -0.3220966 | 0.747379518 | 0.998865404 | 10.235660 | 10.189542 | 10.185219 | 10.260903 | 10.144503 | 10.252392 | 1319 | 1366 | 942 | 1174 |
| 23542 | NM_012324,XI | MAPK8IP2 | mitogen-activated protein kinase 8 interacting protein 2 | protein_coding | NP_036456.1;XP_011528981.1;XP_011528982.1;XP_011528983.1 | 6883 | 607755 | . | . | 2.029983 | 1.629602 | 2.757218 | 2.514873 | 4.723855 | 3.766898 | 6.187118 | 5.749960 | 1.414002 | 396.450310 | 0.194607892 | 2.568162178 | 0.010223931 | 0.325163413 | 8.358897 | 8.859829 | 8.572778 | 8.447456 | 8.747344 | 8.685769 | 421 | 314 | 424 | 414 |
| 23543 | NM_00103169 | RBFOX2 | RNA binding fox-1 homolog 2 | protein_coding | NP_001026865.1;NP_001076045.1;NP_001076046.1;NP_001076047.2;NI | 9906 | 612149 | . | . | 11.938585 | 12.691602 | 13.998890 | 14.479047 | 27.781568 | 29.337201 | 31.413103 | 33.104615 | 1.128021 | 3302.085159 | 0.113650917 | 1.529195179 | 0.126216065 | 0.874064017 | 11.599734 | 11.773368 | 11.612673 | 11.661256 | 11.707070 | 11.764393 | 3467 | 3449 | 2939 | 3395 |
| 23545 | NM_012463,XI | ATP6V0A2 | ATPase H+ transporting V0 subunit a2 | protein_coding | NP_036595.2;XP_024304678.1;XP_024304679.1;XP_024304680.1 | 18481 | 611716 | . | . | 3.086426 | 2.709586 | 3.341358 | 2.511906 | 7.182241 | 6.263329 | 7.497910 | 5.743173 | 1.014721 | 583.726466 | 0.183097547 | 0.115148015 | 0.908327803 | 0.998865404 | 9.177985 | 9.200226 | 9.212131 | 9.148534 | 9.284299 | 9.087541 | 699 | 595 | 588 | 469 |
| 23548 | NM_012382,XI | TTC33 | tetratricopeptide repeat domain 33 | protein_coding | NP_036514.1;XP_011512305.1 | 29959 | . | | . | 1.402960 | 1.360141 | 1.225190 | 1.310708 | 3.264745 | 3.144027 | 2.749290 | 2.996778 | -1.104706 | 227.485563 | 0.224396492 | -0.64021455 | 0.522033125 | 0.998865404 | 7.899750 | 7.755929 | 7.862659 | 7.844390 | 7.793827 | 7.817865 | 280 | 253 | 183 | 208 |
| 23549 | NM_00131911 | DNPEP | aspartyl aminopeptidase | protein_coding | NP_001306045.1;NP_001306046.1;NP_001306047.1;NP_001306048.1;NI | 2981 | 611367 | . | . | 8.500296 | 9.225228 | 9.536839 | 10.242669 | 19.780539 | 21.324530 | 21.400391 | 23.418645 | 1.098165 | 1124.828252 | 0.135558463 | 0.996582975 | 0.318966973 | 0.998865404 | 10.066541 | 10.201292 | 10.071154 | 10.126014 | 10.144965 | 10.191740 | 1189 | 1200 | 992 | 1138 |
| 23550 | NM_012455,XI | PSD4 | pleckstrin and Sec7 domain containing 4 | protein_coding | NP_036587.2;XP_005263691.1;XP_006712455.1;XP_006712456.1;XP_00 | 19096 | 614442 | . | . | 0.051953 | 0.052629 | 0.041971 | 0.065924 | 0.120898 | 0.121653 | 0.094183 | 0.150727 | -1.001915 | 19.245927 | 0.707248398 | -0.00390326 | 0.996885653 | NA | 4.269309 | 4.263648 | 4.266086 | 4.267637 | 4.248932 | 4.284212 | 22 | 21 | 13 | 22 |
| 23551 | NM_00136672 | RASD2 | RASD family member 2 | protein_coding | NP_001353654.1;NP_001363444.1;NP_001363445.1;NP_055125.2;XP_01 | 18229 | 612842 | . | . | 0.303397 | 0.315232 | 0.083778 | 0.240110 | 0.706017 | 0.728672 | 0.187995 | 0.548983 | -2.077846 | 31.274591 | 0.58029721 | -1.81818623 | 0.069035674 | 0.743155658 | 5.401710 | 4.341619 | 4.994948 | 5.026118 | 4.901750 | 4.944183 | 44 | 50 | 13 | 24 |
| 23552 | NM_00103980 | CDK20 | cyclin dependent kinase 20 | protein_coding | NP_001034892.1;NP_001164110.1;NP_001164111.1;NP_036251.2;NP_8 | 21420 | 610076 | . | . | 4.350705 | 5.226904 | 7.241162 | 5.236847 | 10.124271 | 12.082224 | 16.248959 | 11.973429 | 1.277482 | 379.450280 | 0.209409983 | 1.68713346 | 0.091577692 | 0.805877675 | 8.380054 | 8.733844 | 8.433468 | 8.531986 | 8.714712 | 8.531060 | 350 | 391 | 426 | 339 |
| 23555 | NM_00135126 | TSPAN15 | tetraspanin 15 | protein_coding | NP_001338192.1;NP_036471.1;XP_005269723.1;XP_011537864.1;XP_01 | 23298 | 613140 | . | . | 2.306583 | 2.161160 | 3.004075 | 2.670124 | 5.367514 | 4.995617 | 6.741058 | 6.104920 | 1.212200 | 124.553404 | 0.292419809 | 0.949413219 | 0.34241049 | 0.998865404 | 6.814744 | 7.093092 | 6.916868 | 6.925586 | 7.016444 | 6.959737 | 128 | 123 | 128 | 118 |
| 23556 | NM_012327,N | PIGN | phosphatidylinositol glycan anchor biosynthesis class N | protein_coding | NP_036459.1;NP_789744.1;XP_011524191.1;XP_011524192.1;XP_01152 | 8967 | 606097 | . | . | 4.240219 | 4.764323 | 4.272091 | 3.349693 | 9.867167 | 11.012945 | 9.586450 | 7.658674 | -1.209591 | 839.971589 | 0.172372064 | -1.59260035 | 0.111249875 | 0.848126978 | 9.844462 | 9.570992 | 9.832270 | 9.713298 | 9.734237 | 9.534682 | 1144 | 912 | 757 | 610 |
| 23557 | NM_012437,N | SNAPIN | SNAP associated protein | protein_coding | NP_036569.1 | 17145 | 607007 | . | . | 12.229672 | 14.368872 | 11.757739 | 15.709159 | 28.458948 | 33.214293 | 26.384026 | 35.917123 | 1.018500 | 422.993795 | 0.198568942 | 0.133181725 | 0.894049663 | 0.998865404 | 8.711925 | 8.736952 | 8.668640 | 8.761405 | 8.640842 | 8.806320 | 446 | 487 | 320 | 455 |
| 23558 | NM_00133049 | WBP2 | WW domain binding protein 2 | protein_coding | NP_001317428.1;NP_001335099.1;NP_036610.2 | 12738 | 606962 | . | . | 40.185371 | 40.674562 | 44.965571 | 47.532868 | 93.513003 | 94.021071 | 100.901439 | 108.678254 | 1.128280 | 2510.447585 | 0.114481865 | 1.5209897 | 0.128262422 | 0.87643872 | 11.204086 | 11.378126 | 11.241315 | 11.244073 | 11.321439 | 11.358594 | 2712 | 2550 | 2262 | 2552 |
| 23559 | NM_012477 | WBP1 | WW domain binding protein 1 | protein_coding | NP_036609.1 | 12737 | 606961 | . | . | 20.798315 | 21.463209 | 22.099510 | 24.636288 | 48.398537 | 49.613174 | 49.590660 | 56.327946 | 1.090869 | 814.783385 | 0.14677225 | 0.854917925 | 0.392596586 | 0.998865404 | 9.606375 | 9.731460 | 9.629672 | 9.646294 | 9.663832 | 9.733895 | 887 | 851 | 703 | 836 |
| 23560 | NM_012341,XI | GTPBP4 | GTP binding protein 4 | protein_coding | NP_036473.2;XP_024303689.1 | 21535 | . | | . | 17.425589 | 17.047158 | 15.060809 | 15.850267 | 40.550064 | 39.405272 | 33.796016 | 36.239753 | -1.141998 | 2353.015336 | 0.115684675 | -1.65587806 | 0.097746515 | 0.813553488 | 11.292918 | 11.101315 | 11.262588 | 11.239988 | 11.129437 | 11.159503 | 2929 | 2670 | 1874 | 2099 |
| 23562 | NM_00114607 | CLDN14 | claudin 14 | protein_coding | NP_001139549.1;NP_001139550.1;NP_001139551.1;NP_036262.1;NP_6 | 2035 | 605608 | . | . | 2.752189 | 2.959351 | 3.129442 | 3.342392 | 6.404456 | 6.840671 | 7.022377 | 7.641982 | 1.071806 | 136.634893 | 0.280288992 | 0.35692979 | 0.721144367 | 0.998865404 | 7.043806 | 7.142858 | 7.067902 | 7.091867 | 7.081306 | 7.129114 | 147 | 147 | 115 | 141 |
| 23564 | NM_00130300 | DDAH2 | dimethylarginine dimethylaminohydrolase 2 | protein_coding | NP_001289936.1;NP_001289937.1;NP_039268.1;XP_011512750.1 | 2716 | 604744 | . | . | 6.774140 | 6.674064 | 5.071956 | 7.679256 | 15.763702 | 15.427398 | 11.381323 | 17.557708 | -1.073890 | 274.240404 | 0.232527644 | -0.44229604 | 0.658274997 | 0.998865404 | 8.150550 | 8.046156 | 8.119416 | 8.110155 | 7.979247 | 8.167384 | 330 | 304 | 187 | 294 |
| 23566 | NM_012152,XI | LPAR3 | lysophosphatidic acid receptor 3 | protein_coding | NP_036284.1;XP_024301895.1;XP_024301896.1 | 14298 | 605106 | . | . | 10.893440 | 11.428270 | 10.512886 | 9.484740 | 25.349482 | 26.416958 | 23.590610 | 21.685733 | -1.124294 | 1163.883295 | 0.136643436 | -1.23693944 | 0.216109567 | 0.982008277 | 10.266635 | 10.097895 | 10.211685 | 10.241185 | 10.181228 | 10.094725 | 1391 | 1355 | 1009 | 967 |
| 23567 | NM_00130821 | ZNF346 | zinc finger protein 346 | protein_coding | NP_001295142.1;NP_001295143.1;NP_001295144.1;NP_001295145.1;NI | 16403 | 605308 | . | . | 4.428579 | 5.318836 | 4.942476 | 5.116137 | 10.305488 | 12.294727 | 11.090773 | 11.697440 | -1.042526 | 586.778520 | 0.1704276 | -0.35254083 | 0.724432703 | 0.998865404 | 9.226745 | 9.165959 | 9.138769 | 9.274688 | 9.164937 | 9.197462 | 620 | 712 | 487 | 552 |
| 23568 | NM_012106 | ARL2BP | ADP ribosylation factor like GTPase 2 binding protein | protein_coding | NP_036238.1 | 17146 | 615407 | . | . | 17.389814 | 17.632786 | 16.457294 | 18.299740 | 40.466812 | 40.758972 | 36.929695 | 41.840179 | -1.021894 | 1070.638211 | 0.136924559 | -0.22819429 | 0.819495203 | 0.998865404 | 10.079946 | 10.048392 | 10.070129 | 10.074067 | 10.020411 | 10.090348 | 1243 | 1171 | 877 | 1040 |
| 23576 | NM_00113444 | DDAH1 | dimethylarginine dimethylaminohydrolase 1 | protein_coding | NP_001127917.1;NP_001317584.1;NP_036269.1;XP_005270764.1;XP_00 | 2715 | 604743 | . | . | 1.473508 | 1.828838 | 1.384418 | 1.212280 | 3.428914 | 4.227442 | 3.106594 | 2.771734 | -1.286967 | 181.459126 | 0.256680921 | -1.41800795 | 0.156188443 | 0.925053243 | 7.673981 | 7.310141 | 7.504053 | 7.599531 | 7.473146 | 7.415731 | 211 | 243 | 148 | 138 |
| 23580 | NM_012121,XI | CDC42EP4 | CDC42 effector protein 4 | protein_coding | NP_036253.2;XP_005257239.1 | 17147 | 605468 | . | . | 7.001658 | 6.391769 | 5.180062 | 6.394822 | 16.293147 | 14.774861 | 11.623907 | 14.621002 | -1.147709 | 628.317967 | 0.166937668 | -1.19060435 | 0.233808943 | 0.988893316 | 9.391566 | 9.192258 | 9.365099 | 9.312050 | 9.181012 | 9.307889 | 803 | 697 | 459 | 602 |
| 23582 | NM_012142,N | CCNDBP1 | cyclin D1 binding protein 1 | protein_coding | NP_036274.3;XP_006720511.1;XP_024305650.1 | 1587 | 607089 | . | . | 5.954450 | 5.594210 | 8.472885 | 6.868163 | 13.856252 | 12.931268 | 19.012909 | 15.703238 | 1.052536 | 565.489346 | 0.168605702 | 0.43812231 | 0.661297622 | 0.998865404 | 9.105586 | 9.180162 | 9.176852 | 9.070423 | 9.179951 | 9.136983 | 686 | 546 | 521 | 526 |
| 23583 | NM_00124378 | SMUG1 | single-strand-selective monofunctional uracil-DNA glycosylase 1 | protein_coding | NP_001230716.1;NP_001230717.1;NP_001230718.1;NP_001230719.1;NI | 17148 | 607753 | . | . | 10.242379 | 11.760341 | 10.193471 | 14.125511 | 23.834438 | 27.184554 | 22.873854 | 32.296298 | 1.082249 | 746.365159 | 0.163646087 | 0.696827234 | 0.485910921 | 0.998865404 | 9.485939 | 9.599310 | 9.521319 | 9.505789 | 9.481120 | 9.648974 | 835 | 765 | 594 | 814 |
| 23585 | NM_014313,XI | TMEM50A | transmembrane protein 50A | protein_coding | NP_055128.1;XP_005245874.1;XP_011539461.1 | 30590 | 605348 | . | . | 33.144406 | 32.590941 | 34.759962 | 29.715199 | 77.128388 | 75.335420 | 78.000349 | 67.940264 | -1.034182 | 2300.465111 | 0.1268009 | -0.3824148 | 0.702153713 | 0.998865404 | 11.191598 | 11.143419 | 11.189958 | 11.170240 | 11.217411 | 11.084803 | 2726 | 2493 | 2130 | 1945 |
| 23586 | NM_014314 | DDX58 | DExD/H-box helicase 58 | protein_coding | NP_055129.2 | 19102 | 609631 | . | . | 0.503165 | 0.488082 | 0.709661 | 0.603206 | 1.170885 | 1.128223 | 1.592458 | 1.379158 | 1.299144 | 83.481167 | 0.349901216 | 1.079052033 | 0.280564532 | 0.998865404 | 6.181668 | 6.560316 | 6.344023 | 6.336003 | 6.434407 | 6.385928 | 85 | 77 | 89 | 81 |
| 23587 | NM_015362,N | ELP5 | elongator acetyltransferase complex subunit 5 | protein_coding | NP_056177.3;NP_981958.1;NP_981959.1;NP_981960.1;XP_011522081.1 | 30617 | 615019 | . | . | 21.604192 | 22.663524 | 21.755492 | 23.655695 | 50.273840 | 52.387748 | 48.818692 | 54.085943 | 1.027407 | 1180.002332 | 0.133365165 | 0.292491219 | 0.769911073 | 0.998865404 | 10.185093 | 10.223796 | 10.178554 | 10.209418 | 10.185839 | 10.241977 | 1313 | 1282 | 1001 | 1163 |
| 23588 | NM_014315,XI | KLHDC2 | kelch domain containing 2 | protein_coding | NP_055130.1;XP_006720157.1;XP_011534912.1 | 20231 | 611280 | . | . | 5.463786 | 6.122415 | 6.240998 | 5.904793 | 12.714456 | 14.152239 | 14.004618 | 13.500605 | 1.099216 | 739.709064 | 0.148046307 | 0.921839774 | 0.356612158 | 0.998865404 | 9.460903 | 9.597494 | 9.511130 | 9.481539 | 9.561389 | 9.562921 | 829 | 744 | 673 | 727 |
| 23589 | NM_00104247 | CARHSP1 | calcium regulated heat stable protein 1 | protein_coding | NP_001035941.1;NP_001265189.1;NP_001265190.1;NP_001265191.1;NI | 17150 | 616885 | . | . | 26.207116 | 25.826291 | 19.300315 | 25.358083 | 60.985032 | 59.698634 | 43.309352 | 57.978244 | -1.190271 | 2074.610717 | 0.14053795 | -1.78805982 | 0.073766358 | 0.763838735 | 11.138980 | 10.887308 | 11.096774 | 11.067659 | 10.840434 | 11.036868 | 2643 | 2390 | 1440 | 1999 |
| 23590 | NM_00132197 | PDSS1 | decaprenyl diphosphate synthase subunit 1 | protein_coding | NP_001308907.1;NP_001308908.1;NP_055132.2;XP_011517739.1;XP_01 | 17759 | 607429 | . | . | 9.529335 | 8.481490 | 6.780942 | 7.275064 | 22.175154 | 19.605343 | 15.216238 | 16.633571 | -1.268417 | 458.652509 | 0.176736111 | -1.94090991 | 0.052269206 | 0.676168554 | 9.002677 | 8.659476 | 8.941211 | 8.876450 | 8.735371 | 8.789741 | 620 | 526 | 335 | 397 |
| 23592 | NM_00116761 | LEMD3 | LEM domain containing 3 | protein_coding | NP_001161086.1;NP_055134.2 | 28887 | 607844 | . | . | 4.055670 | 3.810264 | 3.887093 | 2.868829 | 9.437711 | 8.807595 | 8.722525 | 6.559236 | -1.180443 | 545.310426 | 0.186534738 | -1.28302169 | 0.199484476 | 0.96621494 | 9.205066 | 8.966992 | 9.163027 | 9.114545 | 9.124826 | 8.929314 | 704 | 614 | 503 | 396 |
| 23593 | NM_00132638 | HEBP2 | heme binding protein 2 | protein_coding | NP_001313309.1;NP_001313310.1;NP_055135.1 | 15716 | 605825 | . | . | 2.201691 | 2.006505 | 2.034442 | 2.337729 | 5.123427 | 4.638127 | 4.565229 | 5.344941 | 1.019053 | 667.770292 | 0.16005535 | 0.170125663 | 0.864911312 | 0.998865404 | 9.369628 | 9.396661 | 9.410557 | 9.340234 | 9.343712 | 9.431943 | 801 | 677 | 548 | 673 |

| | | | | | | | | | | |
|---|---|---|---|---|---|---|---|---|---|---|
| 23594 | NM_014321,N | ORC6 | origin recognition complex subunit 6 | protein_coding | NP_055136.1;XP_011521280.1 | 17151 | 607213 | . | . | [illegible] |
| 23595 | NM_00119725 | ORC3 | origin recognition complex subunit 3 | protein_coding | NP_001184188.1;NP_036513.2;NP_862820.1;XP_005248761.1;XP_00524 | 8489 | 604972 | . | . | [illegible] |
| 23596 | NM_00138185 | OPN3 | opsin 3 | protein_coding | NP_001368784.1;NP_001368785.1;NP_055137.2 | 14007 | 606695 | . | . | [illegible] |
| 23597 | NM_00103358 | ACOT9 | acyl-CoA thioesterase 9 | protein_coding | NP_001028755.2;NP_001032248.1;NP_001317188.1 | 17152 | 300862 | . | . | [illegible] |
| 23598 | NM_014323,N | PATZ1 | POZ/BTB and AT hook containing zinc finger 1 | protein_coding | NP_055138.2;NP_114439.1;NP_114440.1;NP_114441.1 | 13071 | 605165 | . | . | [illegible] |
| 23600 | NM_00116759 | AMACR | alpha-methylacyl-CoA racemase | protein_coding | NP_001161067.1;NP_055139.4;NP_976316.1 | 451 | 604489 | . | . | [illegible] |
| 23603 | NM_00110523 | CORO1C | coronin 1C | protein_coding | NP_001098707.1;NP_001263400.1;NP_055140.1;XP_011536427.1;XP_01 | 2254 | 605269 | . | . | [illegible] |
| 23604 | NM_00136373 | DAPK2 | death associated protein kinase 2 | protein_coding | NP_001350659.1;NP_055141.2;XP_011519715.1;XP_011519716.1;XP_01 | 2675 | 616567 | . | . | [illegible] |
| 23607 | NM_012120,XI | CD2AP | CD2 associated protein | protein_coding | NP_036252.1;XP_005249033.1;XP_011512751.1;XP_016866130.1 | 14258 | 604241 | . | . | [illegible] |
| 23608 | NM_00114512 | MKRN1 | makorin ring finger protein 1 | protein_coding | NP_001138597.1;NP_001278592.1;NP_038474.2;XP_011514298.1;XP_01 | 7112 | 607754 | . | . | [illegible] |
| 23609 | NM_00127170 | MKRN2 | makorin ring finger protein 2 | protein_coding | NP_001258636.1;NP_054879.3 | 7113 | 608426 | . | . | [illegible] |
| 23612 | NM_012396,N | PHLDA3 | pleckstrin homology like domain family A member 3 | protein_coding | NP_036528.1 | 8934 | 607054 | . | . | [illegible] |
| 23613 | NM_00128176 | ZMYND8 | zinc finger MYND-type containing 8 | protein_coding | NP_001268698.1;NP_001268700.1;NP_001268701.1;NP_001268702.1;NI | 9397 | 615713 | . | . | [illegible] |
| 23615 | NR_003064 | PYY2 | peptide YY 2 (pseudogene) | transcribed_ps | . | 9749 | 606637 | . | . | [illegible] |
| 23616 | NM_00135005 | SH3BP1 | SH3 domain binding protein 1 | protein_coding | NP_001336984.1;NP_061830.3 | 10824 | 617368 | . | . | [illegible] |
| 23621 | NM_00120704 | BACE1 | beta-secretase 1 | protein_coding | NP_001193977.1;NP_001193978.1;NP_036236.1;NP_620427.1;NP_6204 | 933 | 604252 | . | . | [illegible] |
| 23623 | NM_00110520 | RUSC1 | RUN and SH3 domain containing 1 | protein_coding | NP_001098673.1;NP_001098674.1;NP_001098675.1;NP_001265156.1;NI | 17153 | 617318 | . | . | [illegible] |
| 23625 | NM_00109878 | FAM89B | family with sequence similarity 89 member B | protein_coding | NP_001092254.1;NP_001092255.1;NP_690045.1 | 16708 | 616128 | . | . | [illegible] |
| 23632 | NM_012113,XI | CA14 | carbonic anhydrase 14 | protein_coding | NP_036245.1;XP_005245116.1;XP_005245117.1;XP_006711322.1;XP_00 | 1372 | 604832 | . | . | [illegible] |
| 23633 | NM_012316,XI | KPNA6 | karyopherin subunit alpha 6 | protein_coding | NP_036448.1;XP_005270768.1;XP_024301933.1;XP_024301934.1;XP_02 | 6399 | 610563 | . | . | [illegible] |
| 23635 | NM_00125673 | SSBP2 | single stranded DNA binding protein 2 | protein_coding | NP_001243661.1;NP_001243662.1;NP_001243663.1;NP_001243664.1;NI | 15831 | 607389 | . | . | [illegible] |
| 23636 | NM_00119335 | NUP62 | nucleoporin 62 | protein_coding | NP_001180286.1;NP_036478.2;NP_057637.2;NP_714940.1;NP_714941.1 | 8066 | 605815 | . | . | [illegible] |
| 23637 | NM_012197,XI | RABGAP1 | RAB GTPase activating protein 1 | protein_coding | NP_036329.3;XP_011516742.1;XP_011516743.1;XP_011516744.1;XP_01 | 17155 | 615882 | . | . | [illegible] |
| 23639 | NM_00132196 | LRRC6 | leucine rich repeat containing 6 | protein_coding | NP_001308890.1;NP_001308891.1;NP_001308892.1;NP_001308893.1;NI | 16725 | 614930 | . | . | [illegible] |
| 23640 | NM_00113010 | HSPBP1 | HSPA (Hsp70) binding protein 1 | protein_coding | NP_001123578.1;NP_001284529.1;NP_036399.3;XP_005258757.1;XP_00 | 24989 | 612939 | . | . | [illegible] |
| 23641 | NM_012317 | LDOC1 | LDOC1 regulator of NFKB signaling | protein_coding | NP_036449.1 | 6548 | 300402 | . | . | [illegible] |
| 23642 | NR_003098,NF | SNHG1 | small nucleolar RNA host gene 1 | lncRNA | . | 32688 | 603222 | . | . | [illegible] |
| 23643 | NM_00119579 | LY96 | lymphocyte antigen 96 | protein_coding | NP_001182726.1;NP_056179.4;XP_011515810.1;XP_016868788.1;XP_01 | 17156 | 605243 | . | . | [illegible] |
| 23644 | NM_014329 | EDC4 | enhancer of mRNA decapping 4 | protein_coding | NP_055144.3 | 17157 | 606030 | . | . | [illegible] |
| 23645 | NM_014330 | PPP1R15A | protein phosphatase 1 regulatory subunit 15A | protein_coding | NP_055145.3 | 14375 | 611048 | . | . | [illegible] |
| 23646 | NM_00103169 | PLD3 | phospholipase D family member 3 | protein_coding | NP_001026866.1;NP_001278240.1;NP_036400.2;XP_005258761.1;XP_00 | 17158 | 615698 | . | . | [illegible] |
| 23647 | NM_00124285 | ARFIP2 | ADP ribosylation factor interacting protein 2 | protein_coding | NP_001229783.1;NP_001229784.1;NP_001229785.1;NP_001357335.1;NI | 17160 | 601638 | . | . | [illegible] |
| 23648 | NM_00100995 | SSBP3 | single stranded DNA binding protein 3 | protein_coding | NP_001009955.1;NP_060540.2;NP_663768.1;XP_006710608.1;XP_01685 | 15674 | 607390 | . | . | [illegible] |
| 23649 | NM_002689,XI | POLA2 | DNA polymerase alpha 2, accessory subunit | protein_coding | NP_002680.2;XP_011543179.1;XP_011543180.1;XP_011543181.1;XP_01 | 30073 | . | . | . | [illegible] |
| 23654 | NM_00137686 | PLXNB2 | plexin B2 | protein_coding | NP_001363793.1;NP_001363794.1;NP_001363795.1;NP_001363796.1;NI | 9104 | 604293 | . | . | [illegible] |
| 23657 | NM_014331,XI | SLC7A11 | solute carrier family 7 member 11 | protein_coding | NP_055146.1;XP_011530104.1 | 11059 | 607933 | . | . | [illegible] |
| 23658 | NM_00113071 | LSM5 | LSM5 homolog, U6 small nuclear RNA and mRNA degradation ass | protein_coding | NP_001124182.1;NP_001132971.1;NP_036454.1 | 17162 | 607285 | . | . | [illegible] |
| 23659 | NM_00136355 | PLA2G15 | phospholipase A2 group XV | protein_coding | NP_001350480.1;NP_036452.1;XP_011521281.1;XP_011521282.2 | 17163 | 609362 | . | . | [illegible] |
| 23660 | NM_00131808 | ZKSCAN5 | zinc finger with KRAB and SCAN domains 5 | protein_coding | NP_001305011.1;NP_001305012.1;NP_001305013.1;NP_055384.1;NP_6 | 12867 | 611272 | . | . | [illegible] |
| 23666 | NR_144546 | UBBP4 | ubiquitin B pseudogene 4 | transcribed_ps | . | 12467 | . | . | . | [illegible] |
| 23670 | NM_00113582 | CEMIP2 | cell migration inducing hyaluronidase 2 | protein_coding | NP_001129292.1;NP_001336713.1;NP_037522.1;XP_005251926.1 | 11869 | 605835 | . | . | [illegible] |
| 23671 | NM_00130513 | TMEFF2 | transmembrane protein with EGF like and two follistatin like doma | protein_coding | NP_001292063.1;NP_001292074.1;NP_057276.2;XP_011509192.1;XP_01 | 11867 | 605734 | . | . | [illegible] |
| 23673 | NM_177424 | STX12 | syntaxin 12 | protein_coding | NP_803173.1 | 11430 | 606892 | . | . | [illegible] |
| 23676 | NM_014332,N | SMPX | small muscle protein X-linked | protein_coding | NP_055147.1 | 11122 | 300226 | . | . | [illegible] |
| 23677 | NM_00137130 | SH3BP4 | SH3 domain binding protein 4 | protein_coding | NP_001358231.1;NP_001358232.1;NP_001358233.1;NP_001358234.1;NI | 10826 | 605611 | . | . | [illegible] |
| 23678 | NM_00103357 | SGK3 | serum/glucocorticoid regulated kinase family member 3 | protein_coding | NP_001028750.1;NP_037389.4;NP_733827.2 | 10812 | 607591 | . | . | [illegible] |
| 23682 | NM_022337,XI | RAB38 | RAB38, member RAS oncogene family | protein_coding | NP_071732.1;XP_016872944.1;XP_016872945.1 | 9776 | 606281 | . | . | [illegible] |
| 23683 | NM_005813,XI | PRKD3 | protein kinase D3 | protein_coding | NP_005804.1;XP_005264294.1;XP_005264295.1;XP_016859230.1;XP_01 | 9408 | 607077 | . | . | [illegible] |
| 23705 | NM_00109851 | CADM1 | cell adhesion molecule 1 | protein_coding | NP_001091987.1;NP_001287972.1;NP_001287973.1;NP_001287974.1;NI | 5951 | 605686 | . | . | [illegible] |
| 23708 | NM_018094 | GSPT2 | G1 to S phase transition 2 | protein_coding | NP_060564.2 | 4622 | 300418 | . | . | [illegible] |
| 23710 | NM_00136359 | GABARAPL1 | GABA type A receptor associated protein like 1 | protein_coding | NP_001350527.1;NP_113600.1 | 4068 | 607420 | . | . | [illegible] |
| 23729 | NM_013276 | SHPK | sedoheptulokinase | protein_coding | NP_037408.2 | 1492 | 605060 | . | . | [illegible] |
| 23731 | NM_032012,XI | TMEM245 | transmembrane protein 245 | protein_coding | NP_114401.2;XP_011516748.1;XP_011516749.1;XP_011516751.1;XP_01 | 1363 | . | . | . | [illegible] |
| 23741 | NM_014335 | EID1 | EP300 interacting inhibitor of differentiation 1 | protein_coding | NP_055150.1 | 1191 | 605894 | . | . | [illegible] |
| 23753 | NM_022044 | SDF2L1 | stromal cell derived factor 2 like 1 | protein_coding | NP_071327.2 | 10676 | 607551 | . | . | [illegible] |
| 23755 | gene-SIRPAP1 | SIRPAP1 | signal regulatory protein alpha pseudogene 1 | pseudogene | . | 9663 | . | . | . | [illegible] |
| 23759 | NM_00131799 | PPIL2 | peptidylprolyl isomerase like 2 | protein_coding | NP_001304925.1;NP_055152.1;NP_680480.1;NP_680481.1;XP_0052615C | 9261 | 607588 | . | . | [illegible] |
| 23760 | NM_00128427 | PITPNB | phosphatidylinositol transfer protein beta | protein_coding | NP_001271206.1;NP_001271207.1;NP_036531.1;XP_011528354.1;XP_01 | 9002 | 606876 | . | . | [illegible] |
| 23761 | NM_00132641 | PISD | phosphatidylserine decarboxylase | protein_coding | NP_001313340.1;NP_001313341.1;NP_001313342.1;NP_001313343.1;NI | 8999 | 612770 | . | . | [illegible] |
| 23762 | NM_00128273 | OSBP2 | oxysterol binding protein 2 | protein_coding | NP_001269667.1;NP_001269668.1;NP_001269669.1;NP_001269670.1;NI | 8504 | 606729 | . | . | [illegible] |
| 23764 | NM_00116157 | MAFF | MAF bZIP transcription factor F | protein_coding | NP_001155044.1;NP_001155045.1;NP_001155046.1;NP_036455.1 | 6780 | 604877 | . | . | [illegible] |
| 23765 | NM_00128990 | IL17RA | interleukin 17 receptor A | protein_coding | NP_001276834.1;NP_055154.3 | 5985 | 605461 | . | . | [illegible] |
| 23767 | NM_013281,N | FLRT3 | fibronectin leucine rich transmembrane protein 3 | protein_coding | NP_037413.1;NP_938205.1;XP_005260739.1;XP_011527506.1;XP_01152 | 3762 | 604808 | . | . | [illegible] |
| 23768 | NM_00134614 | FLRT2 | fibronectin leucine rich transmembrane protein 2 | protein_coding | NP_001333072.1;NP_001333073.1;NP_001333074.1;NP_001333075.1;NI | 3761 | 604807 | . | . | [illegible] |
| 23770 | NM_00130837 | FKBP8 | FKBP prolyl isomerase 8 | protein_coding | NP_001295302.1;NP_036313.3 | 3724 | 604840 | . | . | [illegible] |
| 23774 | NM_00130480 | BRD1 | bromodomain containing 1 | protein_coding | NP_001291737.1;NP_001291738.1;NP_001336869.1;NP_001336870.1;NI | 1102 | 604589 | . | . | [illegible] |
| 23780 | NM_030882,N | APOL2 | apolipoprotein L2 | protein_coding | NP_112092.2;NP_663612.2;XP_011528376.1;XP_011528377.1;XP_01152 | 619 | 607252 | . | . | [illegible] |
| 23786 | NM_00127072 | BCL2L13 | BCL2 like 13 | protein_coding | NP_001257655.1;NP_001257656.1;NP_001257657.1;NP_001257658.1;NI | 17164 | . | . | . | [illegible] |
| 23787 | NM_00127164 | MTCH1 | mitochondrial carrier 1 | protein_coding | NP_001258570.1;NP_055156.1;XP_005249035.1;XP_005249036.1;XP_01 | 17586 | 610449 | . | . | [illegible] |
| 23788 | NM_00131723 | MTCH2 | mitochondrial carrier 2 | protein_coding | NP_001304160.1;NP_001304161.1;NP_001304162.1;NP_055157.1;XP_00 | 17587 | 613221 | . | . | [illegible] |
| 24137 | NM_012310 | KIF4A | kinesin family member 4A | protein_coding | NP_036442.3 | 13339 | 300521 | . | . | [illegible] |
| 24138 | NM_012420 | IFIT5 | interferon induced protein with tetratricopeptide repeats 5 | protein_coding | NP_036552.1 | 13328 | 616135 | . | . | [illegible] |
| 24139 | NM_00119326 | EML2 | EMAP like 2 | protein_coding | NP_001180197.1;NP_001180198.1;NP_001338980.1;NP_001338981.1;NI | 18035 | 617494 | . | . | [illegible] |
| 24140 | NM_00128215 | FTSJ1 | FtsJ RNA 2'-O-methyltransferase 1 | protein_coding | NP_001269086.1;NP_036412.1;NP_803188.1;XP_005272652.1;XP_02430 | 13254 | 300499 | . | . | [illegible] |
| 24142 | NM_00120001 | NAA80 | N-alpha-acetyltransferase 80, NatH catalytic subunit | protein_coding | NP_001186945.1;NP_001186947.1;NP_036323.2 | 30252 | 607073 | . | . | [illegible] |
| 24144 | NM_00100869 | TFIP11 | tuftelin interacting protein 11 | protein_coding | NP_001008697.1;NP_001333786.1;NP_001333787.1;NP_001333788.1;NI | 17165 | 612747 | . | . | [illegible] |
| 24145 | NM_015368,XI | PANX1 | pannexin 1 | protein_coding | NP_056183.2;XP_011541036.1;XP_016872953.1 | 8599 | 608420 | . | . | [illegible] |
| 24146 | NM_00118508 | CLDN15 | claudin 15 | protein_coding | NP_001172009.1;NP_055158.1 | 2036 | 615778 | . | . | [illegible] |
| 24147 | NM_014344 | FJX1 | four-jointed box kinase 1 | protein_coding | NP_055159.2 | 17166 | 612206 | . | . | [illegible] |
| 24148 | NM_012469,XI | PRPF6 | pre-mRNA processing factor 6 | protein_coding | NP_036601.2;XP_006723832.1 | 15860 | 613979 | . | . | [illegible] |
| 24149 | NM_014345 | ZNF318 | zinc finger protein 318 | protein_coding | NP_055160.2 | 13578 | 617512 | . | . | [illegible] |
| 24150 | NM_016212,N | TP53TG3 | TP53 target 3 | protein_coding | NP_057296.1;XP_006721089.1;XP_011544081.1;XP_016878592.1 | 30759 | 617482 | . | . | [illegible] |
| 25758 | NM_012194,XI | KIAA1549L | KIAA1549 like | protein_coding | NP_036326.3;XP_005252904.2;XP_005252905.1;XP_011518271.1;XP_01 | 24836 | 612297 | . | . | [illegible] |
| 25759 | NM_012435,XI | SHC2 | SHC adaptor protein 2 | protein_coding | NP_036567.2;XP_011526195.1;XP_011526196.1;XP_011526197.1;XP_01 | 29869 | 605217 | . | . | [illegible] |
| 25764 | NM_00119988 | HYPK | huntingtin interacting protein K | protein_coding | NP_001186814.1;NP_057484.4 | 18418 | 612784 | . | . | [illegible] |
| 25766 | NM_00103169 | PRPF40B | pre-mRNA processing factor 40 homolog B | protein_coding | NP_001026868.2;NP_001350536.2;NP_001365959.1;NP_001365960.1;NI | 25031 | . | . | . | [illegible] |
| 25771 | NM_00128430 | TBC1D22A | TBC1 domain family member 22A | protein_coding | NP_001271232.1;NP_001271233.1;NP_001271234.1;NP_055161.1;XP_00 | 1309 | 616879 | . | . | [illegible] |
| 25776 | NM_00100288 | CBY1 | chibby family member 1, beta catenin antagonist | protein_coding | NP_001002880.2;NP_056188.1 | 1307 | 607757 | . | . | [illegible] |
| 25777 | NM_00119957 | SUN2 | Sad1 and UNC84 domain containing 2 | protein_coding | NP_001186508.1;NP_001186509.1;NP_056189.1;XP_011528406.1;XP_01 | 14210 | 613569 | . | . | [illegible] |
| 25778 | NM_015375,N | DSTYK | dual serine/threonine and tyrosine protein kinase | protein_coding | NP_056190.1;NP_955749.1;XP_011507694.1;XP_011507695.1;XP_01150 | 29043 | 612666 | . | . | [illegible] |
| 25780 | NM_00113948 | RASGRP3 | RAS guanyl releasing protein 3 | protein_coding | NP_001132960.1;NP_001336904.1;NP_001336905.1;NP_001336906.1;NI | 14545 | 609531 | . | . | [illegible] |
| 25782 | NM_012414 | RAB3GAP2 | RAB3 GTPase activating non-catalytic protein subunit 2 | protein_coding | NP_036546.2 | 17168 | 609275 | . | . | [illegible] |
| 25786 | NR_024157 | DGCR11 | DiGeorge syndrome critical region gene 11 | lncRNA | . | 17226 | . | . | . | [illegible] |
| 25788 | NM_00120526 | RAD54B | RAD54 homolog B | protein_coding | NP_001192191.1;NP_001192192.1;NP_036547.1 | 17228 | 604289 | . | . | [illegible] |
| 25790 | NM_012337 | CFAP45 | cilia and flagella associated protein 45 | protein_coding | NP_036469.2 | 17229 | 605152 | . | . | [illegible] |
| 25792 | NM_00113101 | CIZ1 | CDKN1A interacting zinc finger protein 1 | protein_coding | NP_001124487.1;NP_001124488.1;NP_001124489.1;NP_001124490.1;NI | 16744 | 611420 | . | . | [illegible] |
| 25793 | NM_00103302 | FBXO7 | F-box protein 7 | protein_coding | NP_001028196.1;NP_001244919.1;NP_036311.3;XP_011528408.1;XP_02 | 13586 | 605648 | . | . | [illegible] |
| 25794 | NM_00107718 | FSCN2 | fascin actin-bundling protein 2, retinal | protein_coding | NP_001070650.1;NP_036550.1;XP_011522889.1;XP_011522892.1 | 3960 | 607643 | . | . | [illegible] |
| 25796 | NM_012088,XI | PGLS | 6-phosphogluconolactonase | protein_coding | NP_036220.1;XP_011526199.1 | 8903 | 604951 | . | . | [illegible] |
| 25797 | NM_012413 | QPCT | glutaminyl-peptide cyclotransferase | protein_coding | NP_036545.1 | 9753 | 607065 | . | . | [illegible] |
| 25798 | NM_00115949 | BRI3 | brain protein I3 | protein_coding | NP_001152963.1;NP_056194.1;XP_016867420.1;XP_016867421.1;XP_01 | 1109 | 615628 | . | . | [illegible] |
| 25799 | NM_014347,XI | ZNF324 | zinc finger protein 324 | protein_coding | NP_055162.1;XP_005258770.1 | 14096 | 617477 | . | . | [illegible] |
| 25800 | NM_00109940 | SLC39A6 | solute carrier family 39 member 6 | protein_coding | NP_001092876.1;NP_036451.4;XP_011524202.1;XP_011524203.1 | 18607 | 608731 | . | . | [illegible] |
| 25801 | NM_00133026 | GCA | grancalcin | protein_coding | NP_001317194.1;NP_001317195.1;NP_001317196.1;NP_001317197.1;NI | 15990 | 607030 | . | . | [illegible] |
| 25804 | NM_00125212 | LSM4 | LSM4 homolog, U6 small nuclear RNA and mRNA degradation ass | protein_coding | NP_001239058.1;NP_036453.1 | 17259 | 607284 | . | . | [illegible] |
| 25805 | NM_012342 | BAMBI | BMP and activin membrane bound inhibitor | protein_coding | NP_036474.1 | 30251 | 604444 | . | . | [illegible] |
| 25806 | NM_012476,XI | VAX2 | ventral anterior homeobox 2 | protein_coding | NP_036608.1;XP_006712045.1;XP_011531052.1;XP_011531053.1 | 12661 | 604295 | . | . | [illegible] |
| 25807 | NM_00132953 | RHBDD3 | rhomboid domain containing 3 | protein_coding | NP_001316465.1;NP_036397.1;XP_006724287.1;XP_016884239.1;XP_02 | 1308 | . | . | . | [illegible] |
| 25809 | NM_012263,N | TTLL1 | tubulin tyrosine ligase like 1 | protein_coding | NP_036395.1;XP_006724288.1;XP_011528410.1;XP_011528412.1;XP_01 | 1312 | 608955 | . | . | [illegible] |
| 25813 | NM_015380 | SAMM50 | SAMM50 sorting and assembly machinery component | protein_coding | NP_056195.3 | 24276 | 612058 | . | . | [illegible] |
| 25814 | NM_00116762 | ATXN10 | ataxin 10 | protein_coding | NP_001161093.1;NP_037368.1 | 10549 | 611150 | . | . | [illegible] |
| 25816 | NM_00107765 | TNFAIP8 | TNF alpha induced protein 8 | protein_coding | NP_001071122.1;NP_001273742.1;NP_001273743.1;NP_001273744.1;NI | 17260 | 612111 | . | . | [illegible] |
| 25817 | NM_00108296 | TAFA5 | TAFA chemokine like family member 5 | protein_coding | NP_001076436.1;NP_056196.2 | 21592 | 617499 | . | . | [illegible] |
| 25819 | NM_012118 | NOCT | nocturnin | protein_coding | NP_036250.2 | 14254 | 608468 | . | . | [illegible] |
| 25820 | NM_005744 | ARIH1 | ariadne RBR E3 ubiquitin protein ligase 1 | protein_coding | NP_005735.2 | 689 | 605624 | . | . | [illegible] |
| 25821 | NM_00112322 | MTO1 | mitochondrial tRNA translation optimization 1 | protein_coding | NP_001116698.1;NP_036255.2;NP_598400.1 | 19261 | 614667 | . | . | [illegible] |
| 25822 | NM_00113500 | DNAJB5 | DnaJ heat shock protein family (Hsp40) member B5 | protein_coding | NP_001128476.3;NP_001128477.1;NP_001336652.1;NP_001336653.1;NI | 14887 | 611328 | . | . | [illegible] |
| 25824 | NM_00135851 | PRDX5 | peroxiredoxin 5 | protein_coding | NP_001345440.2;NP_001345445.1;NP_036226.2;NP_857634.2;NP_8576 | 9355 | 606583 | . | . | [illegible] |
| 25825 | NM_012105,N | BACE2 | beta-secretase 2 | protein_coding | NP_036237.2;NP_620476.1;NP_620477.1;XP_016883803.1 | 934 | 605668 | . | . | [illegible] |
| 25827 | NM_00134931 | FBXL2 | F-box and leucine rich repeat protein 2 | protein_coding | NP_001336245.1;NP_001336248.1;NP_001336249.1;NP_001336250.1;NI | 13598 | 605652 | . | . | [illegible] |
| 25828 | NM_012473,XI | TXN2 | thioredoxin 2 | protein_coding | NP_036605.2;XP_006724289.1;XP_011528413.1 | 17772 | 609063 | . | . | [illegible] |
| 25829 | NM_00119507 | TMEM184B | transmembrane protein 184B | protein_coding | NP_001182000.1;NP_001182001.1;NP_036396.2;XP_011528414.2;XP_01 | 1310 | . | . | . | [illegible] |
| 25830 | NM_014351,XI | SULT4A1 | sulfotransferase family 4A member 1 | protein_coding | NP_055166.1;XP_011528422.1;XP_011528423.1;XP_024307980.1 | 14903 | 608359 | . | . | [illegible] |
| 25831 | NM_015382,XI | HECTD1 | HECT domain E3 ubiquitin protein ligase 1 | protein_coding | NP_056197.3;XP_005267559.2;XP_011534921.1;XP_011534923.1;XP_01 | 20157 | 618649 | . | . | [illegible] |
| 25832 | NM_015383 | NBPF14 | NBPF member 14 | protein_coding | NP_056198.2 | 25232 | 614003 | . | . | [illegible] |
| 25836 | NM_015384,N | NIPBL | NIPBL cohesin loading factor | protein_coding | NP_056199.2;NP_597677.2;XP_005248337.1;XP_005248339.3;XP_00671 | 28862 | 608667 | . | . | [illegible] |
| 25837 | NM_00130805 | RAB26 | RAB26, member RAS oncogene family | protein_coding | NP_001294982.1;NP_055168.2;XP_011520750.1;XP_011520752.1 | 14259 | 605455 | . | . | [illegible] |
| 25839 | NM_00119513 | COG4 | component of oligomeric golgi complex 4 | protein_coding | NP_001182068.2;NP_001352355.1;NP_056201.2;XP_024305992.1 | 18620 | 606976 | . | . | [illegible] |
| 25840 | NM_014033 | METTL7A | methyltransferase like 7A | protein_coding | NP_054752.3 | 24550 | 618338 | . | . | [illegible] |
| 25841 | NM_145804 | ABTB2 | ankyrin repeat and BTB domain containing 2 | protein_coding | NP_665803.2 | 23842 | . | . | . | [illegible] |
| 25842 | NM_014034 | ASF1A | anti-silencing function 1A histone chaperone | protein_coding | NP_054753.1 | 20995 | 609189 | . | . | [illegible] |
| 25843 | NM_00110081 | MOB4 | MOB family member 4, phocein | protein_coding | NP_001094289.1;NP_001191023.1;NP_056202.2;NP_955776.1 | 17261 | 609361 | . | . | [illegible] |
| 25844 | NM_015388 | YIPF3 | Yip1 domain family member 3 | protein_coding | NP_056203.2 | 21023 | 609775 | . | . | [illegible] |
| 25847 | NM_00124237 | ANAPC13 | anaphase promoting complex subunit 13 | protein_coding | NP_001229303.1;NP_001229304.1;NP_056206.1 | 24540 | 614484 | . | . | [illegible] |
| 25849 | NM_015393,XI | PARM1 | prostate androgen-regulated mucin-like protein 1 | protein_coding | NP_056208.2;XP_011530135.1 | 24536 | 617688 | . | . | [illegible] |
| 25850 | NM_00124247 | ZNF345 | zinc finger protein 345 | protein_coding | NP_001229401.1;NP_001229403.1;NP_001229404.1;NP_001229405.1;NI | 16367 | . | . | . | [illegible] |
| 25851 | NM_015395,XI | TECPR1 | tectonin beta-propeller repeat containing 1 | protein_coding | NP_056210.1;XP_005250310.1;XP_016867426.1;XP_016867427.1 | 22214 | 614781 | . | . | [illegible] |
| 25852 | NM_00126704 | ARMC8 | armadillo repeat containing 8 | protein_coding | NP_001253970.1;NP_001253971.1;NP_001269271.1;NP_001350870.1;NI | 24999 | 618521 | . | . | [illegible] |
| 25853 | NM_015397,XI | DCAF12 | DDB1 and CUL4 associated factor 12 | protein_coding | NP_056212.1;XP_011516145.1 | 19911 | . | . | . | [illegible] |
| 25854 | NM_00100665 | FAM149A | family with sequence similarity 149 member A | protein_coding | NP_001006656.1;NP_001337107.1;NP_001337108.1;NP_001354697.1;NI | 24527 | . | . | . | [illegible] |
| 25855 | NM_00102495 | BRMS1 | BRMS1 transcriptional repressor and anoikis regulator | protein_coding | NP_001020128.1;NP_056214.1;XP_024304193.1;XP_024304194.1 | 17262 | 606259 | . | . | [illegible] |
| 25861 | NM_00108388 | WHRN | whirlin | protein_coding | NP_001077354.2;NP_001166896.1;NP_001333819.1;NP_056219.3;XP_01 | 16361 | 607928 | . | . | [illegible] |
| 25862 | NM_00128655 | USP49 | ubiquitin specific peptidase 49 | protein_coding | NP_001273483.1;NP_061031.2 | 20078 | . | . | . | [illegible] |
| 25864 | NM_015407 | ABHD14A | abhydrolase domain containing 14A | protein_coding | NP_056222.2 | 24538 | 618771 | . | . | [illegible] |
| 25865 | NM_00107988 | PRKD2 | protein kinase D2 | protein_coding | NP_001073349.1;NP_001073350.1;NP_001073351.1;NP_057541.2;XP_00 | 17293 | 607074 | . | . | [illegible] |
| 25870 | NM_00104246 | SUMF2 | sulfatase modifying factor 2 | protein_coding | NP_001035933.3;NP_001035934.3;NP_001035935.3;NP_001123541.2;NI | 20415 | 607940 | . | . | [illegible] |
| 25871 | NM_00131910 | NEPRO | nucleolus and neural progenitor protein | protein_coding | NP_001306038.1;NP_001306039.1;NP_001306040.1;NP_001306041.1;NI | 24496 | 617089 | . | . | [illegible] |
| 25873 | NM_015414,N | RPL36 | ribosomal protein L36 | protein_coding | NP_056229.2;NP_378669.1 | 13631 | 617893 | . | . | [illegible] |
| 25874 | NM_00114367 | MPC2 | mitochondrial pyruvate carrier 2 | protein_coding | NP_001137146.1;NP_056230.1;XP_006711329.1 | 24515 | 614737 | . | . | [illegible] |
| 25875 | NM_00124368 | LETMD1 | LETM1 domain containing 1 | protein_coding | NP_001230618.1;NP_001287694.1;NP_001338241.1;NP_001338242.1;NI | 24241 | . | . | . | [illegible] |
| 25876 | NM_015417 | SPEF1 | sperm flagellar 1 | protein_coding | NP_056232.2 | 15874 | 610674 | . | . | [illegible] |
| 25879 | NM_015420 | DCAF13 | DDB1 and CUL4 associated factor 13 | protein_coding | NP_056235.5 | 24535 | 616196 | . | . | [illegible] |
| 25880 | NM_015421 | TMEM186 | transmembrane protein 186 | protein_coding | NP_056236.2 | 24530 | . | . | . | [illegible] |
| 25885 | NM_015425 | POLR1A | RNA polymerase I subunit A | protein_coding | NP_056240.2 | 17264 | 616404 | . | . | [illegible] |
| 25886 | NM_00116158 | POC1A | POC1 centriolar protein A | protein_coding | NP_001155052.1;NP_001155053.1;NP_056241.3;XP_011531862.1;XP_01 | 24488 | 614783 | . | . | [illegible] |
| 25888 | NM_00100665 | ZNF473 | zinc finger protein 473 | protein_coding | NP_001006657.1;NP_001295353.1;NP_056243.1 | 23239 | 617908 | . | . | [illegible] |
| 25893 | NM_015431 | TRIM58 | tripartite motif containing 58 | protein_coding | NP_056246.3 | 24150 | . | . | . | [illegible] |
| 25894 | NM_00112972 | PLEKHG4 | pleckstrin homology and RhoGEF domain containing G4 | protein_coding | NP_001123199.1;NP_001123200.1;NP_001123201.1;NP_001123203.1;XF | 24501 | 609526 | . | . | [illegible] |
| 25895 | NM_015433,N | EEF1AKMT3 | EEF1A lysine methyltransferase 3 | protein_coding | NP_056248.2;NP_996797.1 | 24936 | 615258 | . | . | [illegible] |
| 25896 | NM_00119980 | INTS7 | integrator complex subunit 7 | protein_coding | NP_001186738.1;NP_001186740.1;NP_001186741.1;NP_056249.1;XP_01 | 24484 | 611350 | . | . | [illegible] |
| 25897 | NM_00128053 | RNF19A | ring finger protein 19A, RBR E3 ubiquitin protein ligase | protein_coding | NP_001267468.1;NP_001340766.1;NP_001340767.1;NP_056250.3;NP_9 | 13432 | 607119 | . | . | [illegible] |
| 25898 | NM_00100992 | RCHY1 | ring finger and CHY zinc finger domain containing 1 | protein_coding | NP_001009922.1;NP_001265465.1;NP_001265466.1;NP_001265467.1;NI | 17479 | 607680 | . | . | [illegible] |

| | | | | | | | | | | | | | | | | | | | | | | | | | | | | | | | | | |
|---|---|---|---|---|---|---|---|---|---|---|---|---|---|---|---|---|---|---|---|---|---|---|---|---|---|---|---|---|---|---|---|---|---|
| 25900 | NM_00103967 | IFFO1 | intermediate filament family orphan 1 | protein_coding | NP_001034759.1;NP_001180386.1;NP_001317253.1;NP_001317254.1;NI | 24970 | 610495 | . | . | 14.539794 | 15.111488 | 18.692476 | 15.994592 | 33.834695 | 34.930885 | 41.945376 | 36.569733 | 1.136696 | 1504.305275 | 0.133877004 | 1.380721686 | 0.167364551 | 0.945199193 | 10.459342 | 10.644485 | 10.491320 | 10.512313 | 10.661139 | 10.537241 | 1599 | 1540 | 1502 | 1382 |
| 25901 | NM_00137907 | CCDC28A | coiled-coil domain containing 28A | protein_coding | NP_001366000.1;NP_056254.2 | 21098 | 615353 | . | . | 4.942960 | 5.281569 | 5.382944 | 6.057964 | 11.502471 | 12.208585 | 12.079170 | 13.850816 | 1.101883 | 210.755964 | 0.233127309 | 0.600405959 | 0.548235717 | 0.998865404 | 7.648160 | 7.787344 | 7.681041 | 7.708825 | 7.714764 | 7.764254 | 224 | 223 | 182 | 218 |
| 25902 | NM_00124276 | MTHFD1L | methylenetetrahydrofolate dehydrogenase (NADP+ dependent) 1 | protein_coding | NP_001229696.1;NP_001229697.1;NP_001229698.1;NP_001337415.1;NI | 21055 | 611427 | . | . | 10.154647 | 10.169489 | 6.872132 | 10.124607 | 23.630284 | 23.507226 | 15.420869 | 23.148715 | -1.207281 | 864.404765 | 0.161933318 | -1.6782295 | 0.093302305 | 0.809520218 | 9.885379 | 9.612901 | 9.832164 | 9.802230 | 9.587150 | 9.769647 | 1111 | 1000 | 591 | 831 |
| 25903 | NM_00129771 | OLFML2B | olfactomedin like 2B | protein_coding | NP_001284642.1;NP_001334629.1;NP_056256.1;XP_011507700.1;XP_01 | 24558 | . | . | . | 0.245518 | 0.184145 | 0.069575 | 0.221357 | 0.571330 | 0.425660 | 0.156124 | 0.506105 | -1.521344 | 30.989034 | 0.604048751 | -1.00214887 | 0.316271697 | 0.998865404 | 5.226376 | 4.617152 | 5.007407 | 4.952327 | 4.889995 | 4.966315 | 51 | 33 | 13 | 32 |
| 25904 | NM_00125674 | CNOT10 | CCR4-NOT transcription complex subunit 10 | protein_coding | NP_001243670.1;NP_001243671.1;NP_056257.1;XP_006713147.1;XP_00 | 23817 | . | . | . | 6.278369 | 6.305635 | 7.019477 | 5.868218 | 14.610023 | 14.575760 | 15.751502 | 13.416982 | 1.003677 | 541.268819 | 0.170052718 | 0.031136404 | 0.975160758 | 0.998865404 | 9.077201 | 9.083196 | 9.081013 | 9.074666 | 9.133916 | 9.023856 | 625 | 580 | 512 | 465 |
| 25906 | NM_00127848 | ANAPC15 | anaphase promoting complex subunit 15 | protein_coding | NP_001265414.1;NP_001265415.1;NP_001265416.1;NP_001265417.1;NI | 24531 | 614717 | . | . | 15.442898 | 18.725616 | 17.509540 | 16.538496 | 35.936261 | 43.285099 | 39.290905 | 37.813307 | 1.041451 | 457.500472 | 0.180070153 | 0.325402493 | 0.744876469 | 0.998865404 | 8.808260 | 8.866413 | 8.766491 | 8.877440 | 8.856285 | 8.840680 | 470 | 527 | 411 | 432 |
| 25907 | NM_015444 | TMEM158 | transmembrane protein 158 | protein_coding | NP_056259.2 | 30293 | . | . | . | 6.125066 | 5.103821 | 5.284153 | 5.139603 | 14.253281 | 11.797711 | 11.857487 | 11.751092 | -1.091416 | 307.981738 | 0.206508427 | -0.6111202 | 0.541120006 | 0.998865404 | 8.328039 | 8.202638 | 8.340845 | 8.234950 | 8.251861 | 8.231900 | 405 | 314 | 261 | 271 |
| 25909 | NM_00132334 | AHCTF1 | AT-hook containing transcription factor 1 | protein_coding | NP_001310271.1;NP_001310272.1;NP_056261.4;XP_006711821.1;XP_01 | 24618 | 610853 | . | . | 8.449212 | 8.586124 | 8.771464 | 7.284105 | 19.661662 | 19.847210 | 19.682912 | 16.654243 | -1.077121 | 2291.102215 | 0.131141352 | -0.81728538 | 0.413765362 | 0.998865404 | 11.214267 | 11.107407 | 11.187072 | 11.192278 | 11.206562 | 11.047479 | 2727 | 2572 | 2110 | 1862 |
| 25911 | NM_00132974 | DPCD | deleted in primary ciliary dyskinesia homolog (mouse) | protein_coding | NP_001316671.1;NP_001316672.1;NP_001316673.1;NP_001316674.1;NI | 24542 | 616467 | . | . | 13.933701 | 15.367070 | 16.218394 | 15.841537 | 32.424297 | 35.521673 | 36.393605 | 36.219790 | 1.059785 | 406.941059 | 0.181755093 | 0.460907368 | 0.644865065 | 0.998865404 | 8.626239 | 8.709901 | 8.620879 | 8.678547 | 8.699182 | 8.670118 | 433 | 447 | 373 | 383 |
| 25912 | NM_00109861 | C1orf43 | chromosome 1 open reading frame 43 | protein_coding | NP_001092086.1;NP_001284646.1;NP_001284647.1;NP_001284649.1;NI | 29876 | 617428 | . | . | 99.252837 | 96.776278 | 100.231941 | 111.681698 | 230.965444 | 223.702702 | 224.917564 | 255.346927 | 1.068561 | 5222.096692 | 0.110095965 | 0.868957629 | 0.384870307 | 0.998865404 | 12.301822 | 12.397421 | 12.334596 | 12.308504 | 12.335689 | 12.417123 | 5905 | 5363 | 4465 | 5302 |
| 25913 | NM_00104259 | POT1 | protection of telomeres 1 | protein_coding | NP_001036059.1;NP_056265.2 | 17284 | 606478 | . | . | 6.065337 | 5.521508 | 5.335504 | 4.415325 | 14.114292 | 12.763213 | 11.972715 | 10.095118 | -1.210408 | 659.018707 | 0.165887909 | -1.66071712 | 0.096770274 | 0.813553488 | 9.494928 | 9.220381 | 9.458098 | 9.388191 | 9.356027 | 9.229250 | 872 | 740 | 569 | 505 |
| 25914 | NM_00131852 | RTTN | rotatin | protein_coding | NP_001305449.1;NP_775901.3;XP_006722497.1;XP_006722498.1;XP_01 | 18654 | 610436 | . | . | 3.596285 | 3.128310 | 2.883280 | 2.867484 | 8.368706 | 7.231231 | 6.469997 | 6.556162 | -1.116492 | 738.601243 | 0.148038949 | -1.07386089 | 0.282885036 | 0.998865404 | 9.605884 | 9.447050 | 9.577435 | 9.554007 | 9.493986 | 9.484787 | 912 | 827 | 611 | 650 |
| 25915 | NM_199069,N | NDUFAF3 | NADH:ubiquinone oxidoreductase complex assembly factor 3 | protein_coding | NP_951032.1;NP_951033.1;NP_951047.1;NP_951056.1 | 29918 | 612911 | . | . | 23.687398 | 24.816843 | 24.743220 | 28.963393 | 55.121551 | 57.365240 | 55.523073 | 66.221358 | 1.092745 | 720.400862 | 0.155604446 | 0.822319473 | 0.410895097 | 0.998865404 | 9.427577 | 9.554926 | 9.446193 | 9.473855 | 9.468635 | 9.571316 | 777 | 758 | 606 | 757 |
| 25917 | NM_00111409 | THUMPD3 | THUMP domain containing 3 | protein_coding | NP_001107564.1;NP_056268.2;XP_005265078.1;XP_005265079.1;XP_00 | 24493 | . | . | . | 9.606132 | 8.806453 | 10.077627 | 9.512842 | 22.353862 | 20.356511 | 22.613905 | 21.749988 | 1.014718 | 1085.916649 | 0.137741437 | 0.15303515 | 0.878370556 | 0.998865404 | 10.073935 | 10.095381 | 10.114586 | 10.041656 | 10.108742 | 10.070337 | 1302 | 1106 | 976 | 999 |
| 25920 | NM_015456 | NELFB | negative elongation factor complex member B | protein_coding | NP_056271.3 | 24324 | 611180 | . | . | 14.463724 | 12.096530 | 12.275836 | 13.786490 | 33.657684 | 27.961670 | 27.546619 | 31.521172 | -1.033605 | 1041.679892 | 0.149918423 | -0.31807143 | 0.750430751 | 0.998865404 | 10.048310 | 10.000689 | 10.102688 | 9.963629 | 9.971966 | 10.048887 | 1333 | 1036 | 844 | 1011 |
| 25921 | NM_015457,XI | ZDHHC5 | zinc finger DHHC-type palmitoyltransferase 5 | protein_coding | NP_056272.2;XP_011543201.1;XP_011543202.1;XP_011543203.1;XP_01 | 18472 | 614586 | . | . | 17.810856 | 20.095177 | 17.596994 | 19.731634 | 41.446597 | 46.450901 | 39.487146 | 45.114039 | -1.030793 | 2599.152506 | 0.123377535 | -0.35464565 | 0.722855074 | 0.998865404 | 11.365679 | 11.321636 | 11.307465 | 11.401089 | 11.289045 | 11.370685 | 2864 | 3011 | 2113 | 2521 |
| 25923 | NM_00129004 | ATL3 | atlastin GTPase 3 | protein_coding | NP_001276977.1;NP_056274.3;XP_006718556.1;XP_024304196.1 | 24526 | 609369 | . | . | 9.324996 | 8.505788 | 7.649511 | 7.711471 | 21.699648 | 19.661510 | 17.165281 | 17.631365 | -1.171444 | 1796.253849 | 0.124609612 | -1.83202148 | 0.066948221 | 0.735737493 | 10.920341 | 10.692200 | 10.910323 | 10.827080 | 10.744224 | 10.745956 | 2347 | 1983 | 1437 | 1553 |
| 25924 | NM_00128442 | MYRIP | myosin VIIA and Rab interacting protein | protein_coding | NP_001271352.1;NP_001271353.1;NP_001271354.1;NP_001271355.1;NI | 19156 | 611790 | . | . | 0.057621 | 0.158836 | 0.033709 | 0.037744 | 0.134086 | 0.367157 | 0.075643 | 0.086296 | -2.914081 | 11.912898 | 0.965277274 | -1.59854679 | 0.109921341 | NA | 4.149817 | 2.602870 | 3.560895 | 3.633028 | 3.545173 | 3.547499 | 11 | 28 | 5 | 6 |
| 25925 | NM_00130822 | ZNF521 | zinc finger protein 521 | protein_coding | NP_001295154.1;NP_056276.1;XP_011524211.1;XP_011524212.1;XP_01 | 24605 | 610974 | . | . | 1.956313 | 1.479119 | 2.000491 | 2.181270 | 4.552420 | 3.419049 | 4.489043 | 4.987215 | 1.198142 | 292.354738 | 0.220716857 | 1.181600867 | 0.237364109 | 0.992733415 | 8.054945 | 8.316380 | 8.209181 | 8.058578 | 8.208297 | 8.257062 | 351 | 245 | 265 | 312 |
| 25926 | NM_00130327 | NOL11 | nucleolar protein 11 | protein_coding | NP_001290201.1;NP_056277.2 | 24557 | 615366 | . | . | 18.470240 | 16.156410 | 16.165689 | 15.240712 | 42.981008 | 37.346268 | 36.275341 | 34.846078 | -1.118261 | 1463.452902 | 0.134473769 | -1.19916831 | 0.230462507 | 0.985768201 | 10.593349 | 10.432491 | 10.610061 | 10.498847 | 10.496295 | 10.442883 | 1905 | 1549 | 1244 | 1250 |
| 25927 | NM_00111110 | CNRIP1 | cannabinoid receptor interacting protein 1 | protein_coding | NP_001104571.1;NP_001358533.1;NP_001358534.1;NP_001358535.1;NI | 24546 | 618538 | . | . | 22.136892 | 22.151393 | 18.082694 | 22.009719 | 51.513457 | 51.203936 | 40.577041 | 50.322608 | -1.170327 | 1146.777691 | 0.137412094 | -1.65132425 | 0.098672382 | 0.813553488 | 10.272551 | 10.045244 | 10.218877 | 10.219089 | 10.053749 | 10.146811 | 1424 | 1335 | 860 | 1054 |
| 25929 | NM_00125215 | GEMIN5 | gem nuclear organelle associated protein 5 | protein_coding | NP_001239085.1;NP_056280.2 | 20043 | 607005 | . | . | 7.815711 | 7.823725 | 7.015512 | 8.355775 | 18.187480 | 18.084890 | 15.742606 | 19.104487 | -1.031268 | 1305.685126 | 0.13754693 | -0.32293625 | 0.74674352 | 0.998865404 | 10.372829 | 10.328006 | 10.364669 | 10.358454 | 10.274125 | 10.398171 | 1533 | 1425 | 1026 | 1304 |
| 25930 | NM_00130448 | PTPN23 | protein tyrosine phosphatase non-receptor type 23 | protein_coding | NP_001291411.1;NP_056281.1;XP_005265088.2 | 14406 | 606584 | . | . | 8.801153 | 10.682671 | 10.877724 | 11.341565 | 20.480646 | 24.693471 | 24.409298 | 25.931143 | 1.119868 | 1694.668812 | 0.134811466 | 1.211535164 | 0.225690366 | 0.982008277 | 10.642991 | 10.805984 | 10.605099 | 10.748694 | 10.756226 | 10.779519 | 1671 | 1885 | 1534 | 1703 |
| 25932 | NM_013943 | CLIC4 | chloride intracellular channel 4 | protein_coding | NP_039234.1 | 13518 | 606536 | . | . | 78.067986 | 65.121666 | 66.545692 | 64.267967 | 181.667419 | 150.531647 | 149.326599 | 146.941071 | -1.110326 | 9081.275830 | 0.119340928 | -1.2651408 | 0.205820858 | 0.970176829 | 13.222157 | 13.071258 | 13.268761 | 13.105713 | 13.120240 | 13.082232 | 12045 | 9335 | 7653 | 7887 |
| 25934 | NM_00132957 | NIPSNAP3A | nipsnap homolog 3A | protein_coding | NP_001316499.1;NP_056284.1 | 23619 | 608871 | . | . | 4.733079 | 5.153127 | 4.535375 | 4.624859 | 11.014068 | 11.911682 | 10.177252 | 10.574192 | -1.093140 | 243.224851 | 0.218935959 | -0.58682844 | 0.557318936 | 0.998865404 | 7.989123 | 7.860295 | 7.927621 | 7.967816 | 7.901596 | 7.906402 | 281 | 285 | 201 | 219 |
| 25936 | NM_00104254 | NSL1 | NSL1 component of MIS12 kinetochore complex | protein_coding | NP_001036014.1;NP_001284665.1;NP_001284666.1;NP_001284668.1;NI | 24548 | 609174 | . | . | 1.646083 | 1.679960 | 1.552798 | 1.576176 | 3.830503 | 3.883303 | 3.484435 | 3.603737 | -1.030404 | 644.172686 | 0.155349896 | -0.27814288 | 0.780902678 | 0.998865404 | 9.352674 | 9.309612 | 9.365641 | 9.316355 | 9.316555 | 9.325695 | 780 | 680 | 548 | 599 |
| 25937 | NM_00116827 | WWTR1 | WW domain containing transcription regulator 1 | protein_coding | NP_001161750.1;NP_001161752.1;NP_001335291.1;NP_056287.1;XP_01 | 24042 | 607392 | . | . | 37.977098 | 37.735968 | 35.514462 | 38.585739 | 88.374273 | 87.228385 | 79.693429 | 88.221710 | -1.035873 | 5874.717607 | 0.106692407 | -0.47657888 | 0.633662023 | 0.998865404 | 12.545533 | 12.494626 | 12.540834 | 12.528797 | 12.474407 | 12.535006 | 6933 | 6405 | 4837 | 5606 |
| 25938 | NM_015473 | HEATR5A | HEAT repeat containing 5A | protein_coding | NP_056288.2 | 20276 | . | . | . | 4.389711 | 4.611728 | 4.041033 | 4.634048 | 10.215039 | 10.660215 | 9.067962 | 10.595201 | -1.051927 | 1075.770497 | 0.13975358 | -0.52259785 | 0.601254134 | 0.998865404 | 10.107437 | 10.033936 | 10.073710 | 10.104672 | 10.004302 | 10.096926 | 1244 | 1215 | 854 | 1045 |
| 25939 | NM_00136372 | SAMHD1 | SAM and HD domain containing deoxynucleoside triphosphate trip | protein_coding | NP_001350658.1;NP_001350662.1;NP_056289.2 | 15925 | 606754 | . | . | 2.055151 | 2.104167 | 2.143272 | 2.325480 | 4.782422 | 4.863876 | 4.809441 | 5.316933 | 1.025738 | 310.715997 | 0.198119612 | 0.185052577 | 0.853187797 | 0.998865404 | 8.261164 | 8.297512 | 8.264027 | 8.280417 | 8.274868 | 8.297092 | 348 | 336 | 268 | 301 |
| 25940 | NM_00130453 | FAM98A | family with sequence similarity 98 member A | protein_coding | NP_001291467.1;NP_056290.3 | 24520 | . | . | . | 23.953996 | 23.040852 | 22.986645 | 20.011471 | 55.741935 | 53.259965 | 51.581370 | 45.753848 | -1.108865 | 1929.522575 | 0.128353294 | -1.16151055 | 0.245434333 | 0.992733415 | 10.986470 | 10.837755 | 10.971848 | 10.933429 | 10.927681 | 10.811181 | 2392 | 2137 | 1710 | 1588 |
| 25941 | NM_00127194 | TPGS2 | tubulin polyglutamylase complex subunit 2 | protein_coding | NP_001258878.1;NP_001258879.1;NP_001258880.1;NP_001258881.1;NI | 24561 | . | . | . | 17.385394 | 19.313643 | 16.369193 | 17.791311 | 40.456524 | 44.644348 | 36.731994 | 40.677721 | 1.001447 | 1800.599962 | 0.128889897 | 0.016184326 | 0.98708734 | 0.999197585 | 10.813408 | 10.815115 | 10.776585 | 10.848324 | 10.766553 | 10.859519 | 1978 | 2030 | 1474 | 1789 |
| 25942 | NM_00114535 | SIN3A | SIN3 transcription regulator family member A | protein_coding | NP_001138829.1;NP_001138830.1;NP_056292.1;XP_006720528.1;XP_00 | 19353 | 607776 | . | . | 5.114675 | 5.700945 | 5.511140 | 5.719799 | 11.902058 | 13.177987 | 12.366838 | 13.077641 | 1.036373 | 1150.953694 | 0.135621191 | 0.3800583 | 0.703902139 | 0.998865404 | 10.142792 | 10.193982 | 10.120350 | 10.187908 | 10.160026 | 10.202117 | 1243 | 1275 | 990 | 1129 |
| 25943 | NM_00100998 | C20orf194 | chromosome 20 open reading frame 194 | protein_coding | NP_001009984.1;XP_005260741.1;XP_005260744.1;XP_011527510.1;XP_ | 17721 | 614146 | . | . | 3.738385 | 3.589873 | 3.958450 | 4.104738 | 8.699376 | 8.298153 | 8.882648 | 9.384993 | 1.080238 | 797.553280 | 0.146188963 | 0.761679903 | 0.446251073 | 0.998865404 | 9.582532 | 9.694182 | 9.633803 | 9.587676 | 9.677775 | 9.653003 | 912 | 800 | 733 | 763 |
| 25945 | NM_00124328 | NECTIN3 | nectin cell adhesion molecule 3 | protein_coding | NP_001230215.1;NP_001230217.1;NP_056295.1;XP_011510965.1;XP_01 | 17664 | 607147 | . | . | 2.973475 | 2.517268 | 2.551783 | 1.661005 | 6.919399 | 5.818780 | 5.726127 | 3.797690 | -1.301387 | 582.980800 | 0.191096822 | -1.98878098 | 0.046725383 | 0.655437462 | 9.364247 | 8.985586 | 9.305297 | 9.218853 | 9.193633 | 8.974081 | 806 | 667 | 518 | 392 |
| 25946 | NM_00113096 | ZNF385A | zinc finger protein 385A | protein_coding | NP_001124439.1;NP_001124440.1;NP_001276930.1;NP_001276931.1;NI | 17521 | 609124 | . | . | 1.066289 | 1.193277 | 0.877089 | 1.347784 | 2.481299 | 2.758312 | 1.968163 | 3.081548 | -1.029971 | 93.734537 | 0.348045354 | -0.12240917 | 0.902574974 | 0.998865404 | 6.572938 | 6.527725 | 6.537127 | 6.568004 | 6.485011 | 6.599798 | 104 | 108 | 64 | 104 |
| 25948 | NM_015483,XI | KBTBD2 | kelch repeat and BTB domain containing 2 | protein_coding | NP_056298.2;XP_005249753.1;XP_005249754.1;XP_011513558.1;XP_01 | 21751 | . | . | . | 16.318529 | 16.816988 | 18.548200 | 16.401908 | 37.973887 | 38.873222 | 41.621621 | 37.501014 | 1.013085 | 1822.437284 | 0.128135866 | 0.14637422 | 0.883625977 | 0.998865404 | 10.822096 | 10.841148 | 10.824706 | 10.826724 | 10.891644 | 10.777297 | 2082 | 1956 | 1711 | 1595 |
| 25949 | NM_015484,N | SYF2 | SYF2 pre-mRNA splicing factor | protein_coding | NP_056299.1;NP_997053.1 | 19824 | 607090 | . | . | 12.522009 | 11.598635 | 12.898209 | 10.847804 | 29.139228 | 26.810763 | 28.943204 | 24.802214 | -1.029625 | 654.586595 | 0.164237658 | -0.2564551 | 0.797599444 | 0.998865404 | 9.374926 | 9.333660 | 9.391600 | 9.333380 | 9.402402 | 9.280637 | 797 | 686 | 612 | 550 |
| 25950 | NM_00112814 | RWDD3 | RWD domain containing 3 | protein_coding | NP_001121614.1;NP_001186611.1;NP_001265176.1;NP_001265177.1;NI | 21393 | 615875 | . | . | 5.529229 | 4.110814 | 5.211667 | 3.006057 | 12.866743 | 9.502332 | 11.694826 | 6.872992 | -1.206793 | 177.957713 | 0.298471096 | -0.90855725 | 0.36358388 | 0.998865404 | 7.603263 | 7.335073 | 7.576199 | 7.428679 | 7.521289 | 7.318484 | 260 | 176 | 176 | 113 |
| 25953 | NM_00107739 | PNKD | PNKD metallo-beta-lactamase domain containing | protein_coding | NP_001070867.1;NP_056303.3;NP_072094.1;XP_016859260.1;XP_01685 | 9153 | 609023 | . | . | 73.203286 | 73.970211 | 64.809997 | 77.426362 | 170.347051 | 170.985459 | 145.431755 | 177.026174 | -1.002015 | 2220.807255 | 0.122172049 | -0.02376701 | 0.981038454 | 0.998865404 | 11.118425 | 11.115311 | 11.122621 | 11.111944 | 11.065917 | 11.163007 | 2577 | 2383 | 1812 | 2206 |
| 25956 | NM_015490 | SEC31B | SEC31 homolog B, COPII coat complex component | protein_coding | NP_056305.1 | 23197 | 610258 | . | . | 0.856723 | 0.727102 | 0.972067 | 0.749918 | 1.993629 | 1.680728 | 2.181291 | 1.714597 | 1.069509 | 119.900809 | 0.306924113 | 0.315871929 | 0.75209972 | 0.998865404 | 6.855323 | 6.954372 | 6.919674 | 6.860266 | 6.957434 | 6.867338 | 145 | 114 | 122 | 101 |
| 25957 | NM_00132240 | PNISR | PNN interacting serine and arginine rich protein | protein_coding | NP_001309334.1;NP_001309335.1;NP_001309337.1;NP_001309339.1;NI | 21222 | 616653 | . | . | 14.631935 | 16.234518 | 19.540273 | 14.057011 | 34.049114 | 37.526809 | 43.847809 | 32.139691 | 1.075124 | 2670.100198 | 0.148569457 | 0.703391756 | 0.481811643 | 0.998865404 | 11.329349 | 11.434106 | 11.315453 | 11.381507 | 11.527744 | 11.272393 | 2842 | 2890 | 2780 | 2191 |
| 25959 | NM_00113619 | KANK2 | KN motif and ankyrin repeat domains 2 | protein_coding | NP_001129663.1;NP_001316380.1;NP_001366477.1;NP_001366478.1;NI | 29300 | 614610 | . | . | 7.040463 | 7.637175 | 6.404798 | 7.256787 | 16.383446 | 17.653676 | 14.372183 | 16.591789 | -1.106845 | 1157.318191 | 0.142034513 | -1.03111445 | 0.302487151 | 0.998865404 | 10.248211 | 10.101184 | 10.187696 | 10.237101 | 10.074629 | 10.194919 | 1355 | 1355 | 877 | 1114 |
| 25960 | NM_032777,XI | ADGRA2 | adhesion G protein-coupled receptor A2 | protein_coding | NP_116166.9;XP_011542783.1;XP_011542784.1;XP_011542785.1 | 17849 | 606823 | . | . | 1.157043 | 1.223467 | 1.009290 | 1.576043 | 2.692486 | 2.828098 | 2.264818 | 3.603436 | 1.074119 | 267.798139 | 0.242552749 | 0.425281384 | 0.670631563 | 0.998865404 | 8.013408 | 8.114815 | 8.031111 | 8.050257 | 7.959008 | 8.183534 | 292 | 284 | 189 | 316 |
| 25961 | NM_00128301 | NUDT13 | nudix hydrolase 13 | protein_coding | NP_001269943.1;NP_001269944.1;NP_001269945.1;NP_001269946.1;NI | 18827 | 609233 | . | . | 2.146658 | 2.539597 | 3.206345 | 3.046542 | 4.995362 | 5.870392 | 7.194944 | 6.965555 | 1.278089 | 180.670801 | 0.2499632 | 1.416159693 | 0.156728754 | 0.925124297 | 7.309648 | 7.663186 | 7.405588 | 7.466692 | 7.551019 | 7.532381 | 169 | 184 | 180 | 186 |
| 25962 | NM_015496,N | VIRMA | vir like m6A methyltransferase associated | protein_coding | NP_056311.2;NP_892121.1;XP_016868795.1;XP_016868796.1;XP_01686 | 24500 | 616447 | . | . | 10.399628 | 11.955292 | 10.589163 | 10.711822 | 24.200358 | 27.635192 | 23.761775 | 24.491309 | -1.074990 | 1996.603419 | 0.119433953 | -0.87347958 | 0.382401731 | 0.998865404 | 11.014604 | 10.910171 | 10.966371 | 11.015684 | 10.930872 | 10.936827 | 2309 | 2301 | 1667 | 1811 |
| 25963 | NM_00111050 | TMEM87A | transmembrane protein 87A | protein_coding | NP_001103973.1;NP_001273416.1;NP_056312.2;XP_005254344.1;XP_00 | 24522 | . | . | . | 32.774640 | 30.446369 | 36.057191 | 32.435574 | 76.267939 | 70.378151 | 80.911286 | 74.160085 | 1.026049 | 2844.754042 | 0.130330546 | 0.284655976 | 0.775907722 | 0.998865404 | 11.455255 | 11.492679 | 11.493431 | 11.429521 | 11.560889 | 11.398864 | 3360 | 2911 | 2754 | 2434 |
| 25966 | NM_015500,N | C2CD2 | C2 calcium dependent domain containing 2 | protein_coding | NP_056315.1;NP_950251.1;XP_005261166.1;XP_011527824.1;XP_01152 | 1266 | 617581 | . | . | 4.564134 | 4.514379 | 5.903497 | 5.744313 | 10.620927 | 10.435187 | 13.247277 | 13.133688 | 1.250125 | 1036.792276 | 0.135782893 | 2.371963617 | 0.017693833 | 0.446233408 | 9.847797 | 10.170065 | 9.934991 | 9.921790 | 10.111570 | 10.073865 | 1071 | 985 | 1027 | 1053 |
| 25970 | NM_00114579 | SH2B1 | SH2B adaptor protein 1 | protein_coding | NP_001139267.1;NP_001139268.1;NP_001139269.1;NP_001139284.1;NI | 30417 | 608937 | . | . | 12.197508 | 12.017248 | 14.307905 | 11.869555 | 28.384103 | 27.778406 | 32.106524 | 27.138327 | 1.056498 | 1475.765675 | 0.129245543 | 0.613483786 | 0.5395565 | 0.998865404 | 10.486968 | 10.566434 | 10.489295 | 10.520541 | 10.586304 | 10.507184 | 1619 | 1580 | 1386 | 1349 |
| 25972 | NM_00133035 | UNC50 | unc-50 inner nuclear membrane RNA binding protein | protein_coding | NP_001317282.1;NP_001317283.1;NP_054763.2;XP_006712466.1 | 16046 | 617826 | . | . | 37.750935 | 35.964026 | 40.215698 | 33.468061 | 87.847979 | 83.132458 | 90.242859 | 76.520743 | -1.026101 | 1337.525445 | 0.140465579 | -0.26463644 | 0.791289543 | 0.998865404 | 10.403554 | 10.366915 | 10.414949 | 10.372273 | 10.442451 | 10.301384 | 1603 | 1421 | 1255 | 1123 |
| 25973 | NM_152268 | PARS2 | prolyl-tRNA synthetase 2, mitochondrial | protein_coding | NP_689481.2 | 30563 | 612036 | . | . | 2.336594 | 2.693434 | 2.617692 | 2.829032 | 5.437351 | 6.225995 | 5.874025 | 6.468245 | 1.070289 | 192.481063 | 0.243266347 | 0.402854674 | 0.687055138 | 0.998865404 | 7.539204 | 7.636307 | 7.542183 | 7.600701 | 7.586209 | 7.617289 | 200 | 214 | 167 | 193 |
| 25974 | NM_00133054 | MMACHC | metabolism of cobalamin associated C | protein_coding | NP_001317469.1;NP_056321.2;XP_005270781.1 | 24525 | 609831 | . | . | 2.436483 | 2.859543 | 2.830123 | 2.820302 | 5.669798 | 6.609961 | 6.350713 | 6.448286 | 1.126843 | 414.353346 | 0.190941934 | 0.902299117 | 0.366897994 | 0.998865404 | 8.606355 | 8.777979 | 8.582541 | 8.722088 | 8.731429 | 8.721197 | 396 | 470 | 385 | 408 |
| 25976 | NM_00118471 | TIPARP | TCDD inducible poly(ADP-ribose) polymerase | protein_coding | NP_001171646.1;NP_001171647.1;NP_056323.2 | 23696 | 612480 | . | . | 8.508720 | 7.105209 | 6.830007 | 7.342821 | 19.800141 | 16.424006 | 15.326338 | 16.788487 | -1.118976 | 897.688123 | 0.152450738 | -1.0638116 | 0.287414054 | 0.998865404 | 9.888797 | 9.726799 | 9.914324 | 9.777489 | 9.745329 | 9.788817 | 1193 | 928 | 714 | 819 |
| 25977 | NM_015509,N | NECAP1 | NECAP endocytosis associated 1 | protein_coding | NP_056324.2 | 24539 | 611623 | . | . | 9.117445 | 9.094691 | 9.468954 | 10.191936 | 21.216669 | 21.022785 | 21.248059 | 23.302651 | 1.065039 | 777.030615 | 0.146694837 | 0.61969323 | 0.535459774 | 0.998865404 | 9.555761 | 9.646468 | 9.582432 | 9.575449 | 9.600625 | 9.645038 | 871 | 808 | 675 | 775 |
| 25978 | NM_00124464 | CHMP2B | charged multivesicular body protein 2B | protein_coding | NP_001231573.1;NP_054762.2;XP_011531878.1 | 24537 | 609512 | . | . | 14.507042 | 11.503064 | 12.194158 | 11.121689 | 33.758482 | 26.589851 | 27.363338 | 25.428422 | -1.128878 | 990.697542 | 0.157428978 | -1.1109106 | 0.266606834 | 0.998274218 | 10.036777 | 9.862569 | 10.077450 | 9.900851 | 9.941336 | 9.866075 | 1352 | 1000 | 852 | 827 |
| 25979 | NM_00133015 | DHRS7B | dehydrogenase/reductase 7B | protein_coding | NP_001317088.1;NP_056325.2;XP_005266652.2;XP_011522088.1;XP_01 | 24547 | 616160 | . | . | 8.848940 | 6.998721 | 7.907855 | 9.235432 | 20.591846 | 16.177858 | 17.744996 | 21.115719 | 1.121653 | 485.994653 | 0.179414992 | 0.923148142 | 0.355930009 | 0.998865404 | 8.839450 | 9.005375 | 8.944188 | 8.823019 | 8.933184 | 8.981651 | 579 | 446 | 430 | 500 |
| 25980 | NM_00127187 | AAR2 | AAR2 splicing factor | protein_coding | NP_001258803.1;NP_056326.2;XP_006723833.1;XP_011527064.1;XP_01 | 15886 | 617365 | . | . | 13.847572 | 13.621023 | 12.551080 | 15.159685 | 32.223868 | 31.485603 | 28.164257 | 34.660821 | 1.033621 | 1126.363311 | 0.150855207 | 0.316247297 | 0.751814811 | 0.998865404 | 10.113704 | 10.160824 | 10.119861 | 10.126683 | 10.054047 | 10.233707 | 1270 | 1201 | 876 | 1202 |
| 25981 | NM_015512,XI | DNAH1 | dynein axonemal heavy chain 1 | protein_coding | NP_056327.4;XP_016861618.1;XP_016861619.1;XP_016861620.1;XP_01 | 2940 | 603332 | . | . | 0.103130 | 0.133682 | 0.138250 | 0.109728 | 0.239990 | 0.309013 | 0.310230 | 0.250880 | 1.035184 | 62.296904 | 0.408476207 | 0.122129162 | 0.902796727 | 0.998865404 | 5.935725 | 5.986014 | 5.926478 | 5.979686 | 5.987745 | 5.933509 | 62 | 74 | 62 | 52 |
| 25983 | NM_00104263 | NGDN | neuroguidin | protein_coding | NP_001036100.1;NP_056329.1 | 20271 | 610777 | . | . | 17.012729 | 18.208043 | 15.688128 | 17.817150 | 39.589319 | 42.088710 | 35.203706 | 40.736793 | -1.048182 | 626.408942 | 0.15832954 | -0.42878695 | 0.668078275 | 0.998865404 | 9.324760 | 9.256351 | 9.286574 | 9.325931 | 9.244851 | 9.303873 | 717 | 712 | 508 | 599 |
| 25987 | NM_00125821 | TSKU | tsukushi, small leucine rich proteoglycan | protein_coding | NP_001245139.1;NP_001305406.1;NP_001305407.1;NP_001305408.1;NI | 28850 | 608015 | . | . | 12.242714 | 12.887835 | 12.097078 | 12.988440 | 28.489298 | 29.790806 | 27.145494 | 29.696526 | -1.006064 | 1032.499696 | 0.137397889 | -0.06347726 | 0.949386466 | 0.998865404 | 10.016440 | 10.007397 | 9.991650 | 10.035882 | 9.987933 | 10.030533 | 1157 | 1151 | 869 | 993 |
| 25988 | NM_00124325 | HINFP | histone H4 transcription factor | protein_coding | NP_001230188.1;NP_001338886.1;NP_001338887.1;NP_001338888.1;NI | 17850 | 607099 | . | . | 1.619527 | 1.791668 | 2.261797 | 2.146712 | 3.768707 | 4.141519 | 5.075408 | 4.908202 | 1.249724 | 196.127743 | 0.243044886 | 1.323249943 | 0.185752272 | 0.958175316 | 7.445978 | 7.767451 | 7.526422 | 7.589480 | 7.681033 | 7.634426 | 186 | 202 | 199 | 194 |
| 25989 | NM_00109943 | ULK3 | unc-51 like kinase 3 | protein_coding | NP_001092906.3;NP_001271293.2;NP_001271294.1;XP_005254346.1;XF | 19703 | 613472 | . | . | 28.327737 | 32.032710 | 34.826900 | 33.351056 | 65.919813 | 74.045045 | 78.150559 | 76.253230 | 1.134871 | 2680.081186 | 0.11875926 | 1.536960921 | 0.124302879 | 0.87182553 | 11.293938 | 11.476418 | 11.291204 | 11.374703 | 11.457587 | 11.415050 | 2741 | 2850 | 2542 | 2602 |
| 25994 | NM_00109966 | HIGD1A | HIG1 hypoxia inducible domain family member 1A | protein_coding | NP_001093138.1;NP_001093139.1;NP_054775.2 | 29527 | 618623 | . | . | 18.447102 | 16.721510 | 15.443659 | 14.780999 | 42.927162 | 38.652516 | 34.655120 | 33.795000 | -1.178404 | 1392.910866 | 0.13188496 | -1.79576144 | 0.072532466 | 0.758556671 | 10.557299 | 10.320794 | 10.545124 | 10.459156 | 10.396650 | 10.357762 | 1830 | 1537 | 1141 | 1168 |
| 25996 | NM_015523 | REXO2 | RNA exonuclease 2 | protein_coding | NP_056338.2 | 17851 | 607149 | . | . | 55.850628 | 54.519337 | 55.930641 | 60.061047 | 129.966721 | 126.023880 | 125.506744 | 137.322464 | 1.036215 | 1904.116709 | 0.120699792 | 0.425220468 | 0.670675965 | 0.998865404 | 10.869057 | 10.920300 | 10.893370 | 10.867440 | 10.884230 | 10.932411 | 2189 | 1985 | 1634 | 1872 |
| 25998 | NM_00130090 | IBTK | inhibitor of Bruton tyrosine kinase | protein_coding | NP_001287835.1;NP_056340.2;XP_006715516.1;XP_006715517.1 | 17853 | 606457 | . | . | 9.007514 | 7.790525 | 8.692391 | 7.091670 | 20.960855 | 18.008148 | 19.505474 | 16.214262 | -1.076279 | 1529.923617 | 0.147381433 | -0.71957457 | 0.471786972 | 0.998865404 | 10.631001 | 10.525560 | 10.663309 | 10.542897 | 10.626944 | 10.462630 | 1966 | 1580 | 1419 | 1234 |
| 25999 | NM_00119957 | CLIP3 | CAP-Gly domain containing linker protein 3 | protein_coding | NP_001186499.1;NP_056341.1 | 24314 | 607382 | . | . | 3.501945 | 4.028935 | 4.323661 | 4.839787 | 8.149170 | 9.313062 | 9.702169 | 11.065599 | 1.200831 | 436.406789 | 0.181280808 | 1.45648967 | 0.145257303 | 0.906189003 | 8.631942 | 8.895131 | 8.668675 | 8.746352 | 8.787198 | 8.851221 | 427 | 456 | 393 | 469 |
| 26000 | NM_015527,XI | TBC1D10B | TBC1 domain family member 10B | protein_coding | NP_056342.3;XP_011544091.1;XP_011544092.1 | 24510 | 613620 | . | . | 8.519313 | 9.577771 | 9.779896 | 11.430871 | 19.824792 | 22.139446 | 21.945803 | 26.135326 | 1.136971 | 1071.334267 | 0.144364489 | 1.28283138 | 0.199551158 | 0.96621494 | 9.969934 | 10.154550 | 9.963075 | 10.062984 | 10.066436 | 10.150160 | 1076 | 1156 | 936 | 1128 |
| 26001 | NM_00132035 | RNF167 | ring finger protein 167 | protein_coding | NP_001307285.1;NP_001307286.1;NP_001307287.1;NP_001307288.1;NI | 24544 | 610431 | . | . | 53.921226 | 60.125885 | 61.933917 | 60.436550 | 125.476905 | 138.983682 | 138.977913 | 138.181000 | 1.036328 | 3413.642509 | 0.11024324 | 0.466968126 | 0.640522682 | 0.998865404 | 11.711161 | 11.762574 | 11.703058 | 11.740846 | 11.746726 | 11.756214 | 3772 | 3701 | 3004 | 3276 |
| 26002 | NM_015529,XI | MOXD1 | monooxygenase DBH like 1 | protein_coding | NP_056344.2;XP_016866203.1 | 21063 | 609000 | . | . | 18.853114 | 19.555965 | 14.335707 | 13.665075 | 43.871972 | 45.204490 | 32.168912 | 31.243567 | -1.389964 | 1546.299122 | 0.125115952 | -3.79686061 | 0.00014654 | 0.016053122 | 10.812609 | 10.337689 | 10.697467 | 10.718309 | 10.473953 | 10.431936 | 2044 | 1966 | 1157 | 1179 |
| 26003 | NM_00120142 | GORASP2 | golgi reassembly stacking protein 2 | protein_coding | NP_001188357.1;NP_056345.3 | 17500 | 608693 | . | . | 52.112564 | 55.901612 | 45.542179 | 52.041962 | 121.268082 | 129.219085 | 102.195335 | 118.987770 | -1.122445 | 3919.395987 | 0.116631244 | -1.42881886 | 0.153056304 | 0.91755225 | 12.017428 | 11.850582 | 11.955639 | 12.007596 | 11.834557 | 11.935195 | 4626 | 4612 | 3014 | 3675 |
| 26005 | NM_00128657 | C2CD3 | C2 domain containing 3 centriole elongation regulator | protein_coding | NP_001273506.1;NP_056346.3;XP_011543206.1;XP_011543207.1;XP_01 | 24564 | 615944 | . | . | 3.150204 | 3.743094 | 3.293480 | 3.628846 | 7.330653 | 8.652327 | 7.390476 | 8.296927 | -1.003533 | 828.930917 | 0.157640487 | -0.03227433 | 0.974253283 | 0.998865404 | 9.698016 | 9.692194 | 9.625113 | 9.760255 | 9.652869 | 9.730128 | 866 | 981 | 680 | 818 |
| 26007 | NM_00135197 | TKFC | triokinase and FMN cyclase | protein_coding | NP_001338905.1;NP_001338906.1;NP_001338907.1;NP_001338908.1;NI | 24552 | 615844 | . | . | 4.276469 | 4.445757 | 3.955005 | 3.814497 | 9.951521 | 10.276566 | 8.874917 | 8.721395 | -1.127560 | 601.046697 | 0.158510596 | -1.09269917 | 0.274525905 | 0.998865404 | 9.315281 | 9.142197 | 9.255307 | 9.284067 | 9.208074 | 9.172048 | 718 | 702 | 505 | 515 |
| 26009 | NM_00130823 | ZZZ3 | zinc finger ZZ-type containing 3 | protein_coding | NP_001295166.1;NP_001363075.1;NP_001363076.1;NP_001363077.1;NI | 24523 | . | . | . | 5.537104 | 5.723521 | 7.252850 | 5.571464 | 12.885070 | 13.230177 | 16.275184 | 12.738492 | 1.133148 | 1216.223733 | 0.151927761 | 1.186988267 | 0.235232249 | 0.990818043 | 10.154920 | 10.335800 | 10.196318 | 10.196177 | 10.385462 | 10.186224 | 1313 | 1230 | 1270 | 1054 |
| 26010 | NM_00110042 | SPATS2L | spermatogenesis associated serine rich 2 like | protein_coding | NP_001093892.1;NP_001093893.1;NP_001093894.1;NP_001269664.1;NI | 24574 | 613817 | . | . | 34.676544 | 33.671179 | 23.701189 | 27.759025 | 80.693744 | 77.832439 | 53.184786 | 63.467709 | -1.195444 | 2675.437357 | 0.126795262 | -2.03119799 | 0.042234914 | 0.63521717 | 11.508731 | 11.250881 | 11.446839 | 11.458772 | 11.232306 | 11.378075 | 3331 | 3167 | 1925 | 2494 |
| 26012 | NM_00113096 | NSMF | NMDA receptor synaptonuclear signaling and neuronal migration | protein_coding | NP_001124441.1;NP_001124442.1;NP_001124443.1;NP_001171535.1;NI | 29843 | 608137 | . | . | 9.311089 | 8.647588 | 8.676884 | 10.456948 | 21.667286 | 19.989290 | 19.470678 | 23.908570 | 1.046286 | 971.258513 | 0.145615948 | 0.448284849 | 0.653947634 | 0.998865404 | 9.890939 | 9.955757 | 9.908609 | 9.903684 | 9.878368 | 9.996308 | 1097 | 1021 | 794 | 1006 |
| 26013 | NM_00137730 | L3MBTL1 | L3MBTL histone methyl-lysine binding protein 1 | protein_coding | NP_001364232.1;NP_001364235.1;NP_001364236.1;NP_001364237.1;NI | 15905 | 608802 | . | . | 2.490553 | 2.364865 | 3.484397 | 2.362487 | 5.795621 | 5.466491 | 7.818885 | 5.401543 | 1.185970 | 310.099457 | 0.215095704 | 1.14398905 | 0.252628178 | 0.994320829 | 8.147603 | 8.394980 | 8.234754 | 8.210871 | 8.393603 | 8.231709 | 334 | 299 | 335 | 270 |
| 26015 | NM_015540,XI | RPAP1 | RNA polymerase II associated protein 1 | protein_coding | NP_056355.2;XP_005254354.1 | 24567 | 611475 | . | . | 5.920640 | 6.466711 | 7.033303 | 7.350459 | 13.777574 | 14.948093 | 15.782528 | 16.805950 | 1.138694 | 977.953108 | 0.140441112 | 1.33422995 | 0.182128507 | 0.953687323 | 9.837077 | 10.024108 | 9.848344 | 9.917138 | 9.963589 | 9.992312 | 1003 | 1034 | 888 | 995 |
| 26017 | NM_014077,XI | FAM32A | family with sequence similarity 32 member A | protein_coding | NP_054796.1;XP_011526200.1 | 24563 | 614554 | . | . | 35.354395 | 35.415616 | 33.737305 | 41.804082 | 82.271133 | 81.864783 | 75.705530 | 95.580072 | 1.053282 | 1685.099237 | 0.137635731 | 0.544126074 | 0.586354719 | 0.998865404 | 10.680886 | 10.755388 | 10.698942 | 10.694251 | 10.651521 | 10.816126 | 1896 | 1766 | 1346 | 1790 |
| 26018 | NM_00137734 | LRIG1 | leucine rich repeats and immunoglobulin like domains 1 | protein_coding | NP_001364273.1;NP_001364274.1;NP_001364275.1;NP_001364276.1;NI | 17360 | 608868 | . | . | 5.500138 | 6.188698 | 6.418671 | 7.004005 | 12.799045 | 14.305452 | 14.403314 | 16.013825 | 1.123101 | 1030.373943 | 0.140774507 | 1.189755998 | 0.234142309 | 0.989548772 | 9.922983 | 10.090086 | 9.917769 | 10.007517 | 10.035590 | 10.062192 | 1049 | 1112 | 931 | 1040 |
| 26019 | NM_015542,N | UPF2 | UPF2 regulator of nonsense mediated mRNA decay | protein_coding | NP_056357.1;NP_542166.1;XP_011517751.1 | 17854 | 605529 | . | . | 5.457080 | 5.777429 | 6.073996 | 5.788125 | 12.698849 | 13.354786 | 13.629870 | 13.233859 | 1.037150 | 933.958749 | 0.13958523 | 0.377004201 | 0.706170483 | 0.998865404 | 9.840626 | 9.893323 | 9.840531 | 9.866364 | 9.897357 | 9.862307 | 1037 | 1007 | 847 | 870 |
| 26020 | NM_00132922 | LRP10 | LDL receptor related protein 10 | protein_coding | NP_001316155.1;NP_054764.2;XP_005267567.1 | 14553 | 609921 | . | . | 15.308169 | 15.049735 | 15.971804 | 16.780700 | 35.622740 | 34.788137 | 35.840263 | 38.367078 | 1.066760 | 3374.408359 | 0.110916831 | 0.840592778 | 0.400576106 | 0.998865404 | 11.673060 | 11.766272 | 11.707955 | 11.677980 | 11.727748 | 11.764410 | 3829 | 3459 | 2962 | 3338 |
| 26022 | NM_00103350 | TMEM98 | transmembrane protein 98 | protein_coding | NP_001028676.1;NP_001288675.1;NP_056359.2 | 24529 | 615949 | . | . | 17.886315 | 21.177108 | 24.816991 | 33.922337 | 41.622190 | 48.951835 | 55.688609 | 77.559393 | 1.171507 | 2142.139280 | 0.117106341 | 1.950069582 | 0.051167826 | 0.672475152 | 10.946184 | 11.174468 | 10.998554 | 10.997187 | 11.106729 | 11.141355 | 2274 | 2127 | 1967 | 2213 |
| 26024 | NM_015545 | PTCD1 | pentatricopeptide repeat domain 1 | protein_coding | NP_056360.2 | 22198 | 614774 | . | . | 1.571178 | 3.614154 | 2.984288 | 1.976765 | 3.656196 | 8.354280 | 6.696656 | 4.519639 | -1.057389 | 432.026186 | 0.639671623 | -0.125855 | 0.899846703 | 0.998865404 | 8.794685 | 8.714140 | 8.465629 | 8.964007 | 8.828660 | 8.577409 | 312 | 666 | 441 | 312 |
| 26027 | NM_015547,N | ACOT11 | acyl-CoA thioesterase 11 | protein_coding | NP_056362.1;NP_671517.1 | 18156 | 606803 | . | . | 0.805695 | 1.138359 | 0.782514 | 0.798197 | 1.874885 | 2.631369 | 1.755938 | 1.824983 | -1.170141 | 89.445684 | 0.352398073 | -0.64325615 | 0.520057907 | 0.998865404 | 6.592590 | 6.364265 | 6.445572 | 6.555934 | 6.451999 | 6.463642 | 91 | 123 | 70 | 79 |
| 26030 | NM_00130814 | PLEKHG3 | pleckstrin homology and RhoGEF domain containing G3 | protein_coding | NP_001295076.1;XP_011534927.1;XP_011534929.1;XP_011534930.1;XP_ | 20364 | 617940 | . | . | 2.427978 | 3.035762 | 2.459355 | 3.590743 | 5.650003 | 7.017298 | 5.518721 | 8.209804 | -1.020109 | 815.212478 | 0.150137728 | -0.19131004 | 0.848282692 | 0.998865404 | 9.685581 | 9.656335 | 9.631159 | 9.721818 | 9.640321 | 9.685712 | 888 | 945 | 679 | 781 |
| 26031 | NM_015550,N | OSBPL3 | oxysterol binding protein like 3 | protein_coding | NP_056365.1;NP_663160.1;NP_663161.1;NP_663162.1;XP_005249755.1 | 16370 | 606732 | . | . | 11.062433 | 10.499151 | 12.481828 | 11.844710 | 25.742736 | 24.269253 | 28.008859 | 27.081526 | 1.109914 | 2437.228578 | 0.11822613 | 1.272545918 | 0.20317922 | 0.968254058 | 11.173731 | 11.324389 | 11.227120 | 11.186309 | 11.323503 | 11.257755 | 2728 | 2429 | 2320 | 2307 |
| 26032 | NM_015551,XI | SUSD5 | sushi domain containing 5 | protein_coding | NP_056366.1;XP_005265091.1;XP_016861626.1 | 29061 | . | . | . | 7.543612 | 7.551369 | 6.262965 | 7.117402 | 17.554295 | 17.455327 | 14.053913 | 16.273093 | -1.163868 | 1029.491639 | 0.140105781 | -1.56259106 | 0.118148813 | 0.857722481 | 10.113169 | 9.893944 | 10.075270 | 10.045966 | 9.907293 | 9.989098 | 1300 | 1172 | 780 | 943 |
| 26033 | NM_00127628 | ATRNL1 | attractin like 1 | protein_coding | NP_001263211.1;NP_997186.1;XP_011537889.1;XP_011537890.1;XP_01 | 29063 | 612869 | . | . | 0.945068 | 1.547033 | 2.085186 | 2.057570 | 2.199211 | 3.576035 | 4.679095 | 4.704386 | -1.003928 | 216.128569 | 0.235361745 | -0.02402852 | 0.980829857 | 0.998865404 | 7.757954 | 7.753534 | 7.790542 | 7.719131 | 7.783658 | 7.722917 | 268 | 216 | 200 | 189 |
| 26035 | NM_00132409 | GLCE | glucuronic acid epimerase | protein_coding | NP_001311020.1;NP_001311021.1;NP_001311022.1;NP_001311023.1;NI | 17855 | 612134 | . | . | 7.701229 | 6.793709 | 7.487449 | 6.451964 | 17.921078 | 15.703964 | 16.801620 | 14.751650 | -1.036523 | 1168.075756 | 0.145693788 | -0.35521117 | 0.722431396 | 0.998865404 | 10.215248 | 10.164136 | 10.248354 | 10.153059 | 10.234637 | 10.111893 | 1455 | 1202 | 1080 | 987 |
| 26036 | NM_00103162 | ZNF451 | zinc finger protein 451 | protein_coding | NP_001026794.1;NP_001244202.1;NP_056370.2;XP_005249051.1;XP_01 | 21091 | 615708 | . | . | 3.437215 | 3.536564 | 3.507342 | 3.031214 | 7.998542 | 8.174925 | 7.870376 | 6.930510 | -1.023641 | 599.523228 | 0.168085973 | -0.20055473 | 0.841046762 | 0.998865404 | 9.244035 | 9.211121 | 9.241436 | 9.226447 | 9.282396 | 9.150628 | 706 | 647 | 567 | 500 |
| 26037 | NM_00128424 | SIPA1L1 | signal induced proliferation associated 1 like 1 | protein_coding | NP_001271174.1;NP_001271175.1;NP_001271176.1;NP_001341214.1;NI | 20284 | 617504 | . | . | 1.801168 | 1.770693 | 1.787600 | 1.905318 | 4.191390 | 4.093036 | 4.011320 | 4.356279 | 1.021929 | 453.187736 | 0.173926558 | 0.17993578 | 0.857202986 | 0.998865404 | 8.808418 | 8.839346 | 8.816712 | 8.817334 | 8.804844 | 8.855182 | 516 | 484 | 382 | 447 |
| 26038 | NM_015557 | CHD5 | chromodomain helicase DNA binding protein 5 | protein_coding | NP_056372.1 | 16816 | 610771 | . | . | 0.080632 | 0.114058 | 0.088330 | 0.054241 | 0.187635 | 0.263650 | 0.198211 | 0.124015 | -1.354092 | 26.266794 | 0.622548148 | -0.70247701 | 0.482381736 | NA | 4.916503 | 4.481107 | 4.708994 | 4.750314 | 4.718860 | 4.675684 | 29 | 38 | 24 | 16 |
| 26039 | NM_00130177 | SS18L1 | SS18L1 subunit of BAF chromatin remodeling complex | protein_coding | NP_001288707.1;NP_945173.1;XP_005260446.1;XP_005260448.1;XP_01 | 15592 | 606472 | . | . | 27.568608 | 28.829328 | 39.793418 | 27.286961 | 64.153281 | 66.640278 | 89.295275 | 62.388392 | 1.187768 | 4374.378265 | 0.153300029 | 1.619395824 | 0.105362125 | 0.832940096 | 11.965295 | 12.213746 | 12.005287 | 12.022947 | 12.303941 | 11.987335 | 4555 | 4361 | 4897 | 3627 |
| 26040 | NM_00113011 | SETBP1 | SET binding protein 1 | protein_coding | NP_001123582.1;NP_001366070.1;NP_001366071.1;NP_056374.2;XP_02 | 15573 | 611060 | . | . | 0.632489 | 0.739939 | 0.898923 | 0.972235 | 1.471828 | 1.710403 | 2.017159 | 2.222897 | 1.244649 | 252.845816 | 0.222400459 | 1.419684081 | 0.155699675 | 0.924751554 | 7.815884 | 8.131155 | 7.871122 | 7.965805 | 8.046441 | 8.011742 | 234 | 267 | 252 | 254 |
| 26043 | NM_015562,XI | UBXN7 | UBX domain protein 7 | protein_coding | NP_056377.1;XP_011510972.1;XP_011510973.1 | 29119 | 616379 | . | . | 2.947275 | 3.026126 | 3.101753 | 2.831858 | 6.858432 | 6.995024 | 6.960246 | 6.474708 | -1.019588 | 976.326354 | 0.140509719 | -0.19918091 | 0.842121234 | 0.998865404 | 9.945001 | 9.917306 | 9.931361 | 9.944322 | 9.960326 | 9.885915 | 1125 | 1073 | 884 | 860 |
| 26045 | NM_015564 | LRRTM2 | leucine rich repeat transmembrane neuronal 2 | protein_coding | NP_056379.1 | 19409 | 610868 | . | . | 0.074129 | 0.054551 | 0.098310 | 0.056539 | 0.172501 | 0.126098 | 0.220604 | 0.129269 | 1.163411 | 14.025171 | 0.832795731 | 0.262202101 | 0.793165633 | NA | 3.694014 | 3.917252 | 3.806372 | 3.789987 | 3.826047 | 3.788456 | 17 | 12 | 17 | 10 |
| 26046 | NM_00132076 | LTN1 | listerin E3 ubiquitin protein ligase 1 | protein_coding | NP_001307695.2;NP_056380.3;XP_006724050.1;XP_016883805.1;XP_01 | 13082 | 613083 | . | . | 3.641709 | 2.885645 | 3.643788 | 2.222447 | 8.474407 | 6.670300 | 8.176555 | 5.081362 | -1.127864 | 741.266050 | 0.212453286 | -0.81708758 | 0.41387838 | 0.998865404 | 9.617437 | 9.445116 | 9.648639 | 9.473597 | 9.635791 | 9.298737 | 1015 | 744 | 755 | 493 |
| 26048 | NM_00130345 | ZNF500 | zinc finger protein 500 | protein_coding | NP_001290379.1;NP_067678.1;XP_005255300.1;XP_011520755.1;XP_01 | 23716 | . | . | . | 1.312645 | 1.183031 | 2.094849 | 1.402959 | 3.054582 | 2.734628 | 4.700779 | 3.207697 | 1.247831 | 229.302863 | 0.234867484 | 1.360012228 | 0.173826054 | 0.947973539 | 7.671899 | 7.992543 | 7.783172 | 7.775039 | 7.948215 | 7.818947 | 237 | 218 | 250 | 208 |
| 26049 | NM_00137604 | FAM169A | family with sequence similarity 169 member A | protein_coding | NP_001362978.1;NP_001362979.1;NP_001362980.1;NP_001362981.1;NI | 29138 | 615769 | . | . | 0.317644 | 0.507472 | 0.291247 | 0.589838 | 0.739171 | 1.173044 | 0.653550 | 1.348597 | 1.042302 | 85.956848 | 0.385998533 | 0.154854097 | 0.876936342 | 0.998865404 | 6.397279 | 6.453259 | 6.364767 | 6.456582 | 6.344034 | 6.496633 | 81 | 106 | 55 | 105 |
| 26050 | NM_015567,XI | SLITRK5 | SLIT and NTRK like family member 5 | protein_coding | NP_056382.1;XP_005254095.1;XP_005254096.1 | 20295 | 609680 | . | . | 0.061837 | 0.024832 | 0.038324 | 0.053145 | 0.143898 | 0.057399 | 0.085998 | 0.121510 | 1.023488 | 7.622053 | 1.117015762 | 0.029985083 | 0.97607895 | NA | 2.912173 | 2.947964 | 2.943573 | 2.916760 | 2.927455 | 2.934623 | 12 | 5 | 6 | 8 |
| 26053 | NM_00112723 | AUTS2 | activator of transcription and developmental regulator AUTS2 | protein_coding | NP_001120703.1;NP_001120704.1;NP_056385.1;XP_005250314.1;XP_01 | 14262 | 607270 | . | . | 2.748733 | 2.364262 | 1.935351 | 2.078614 | 6.396417 | 5.465095 | 4.342872 | 4.752504 | -1.238683 | 492.289809 | 0.171973091 | -1.79566763 | 0.072547392 | 0.758556671 | 9.089585 | 8.780634 | 9.036228 | 8.975779 | 8.846143 | 8.896971 | 656 | 561 | 366 | 430 |
| 26054 | NM_00110040 | SENP6 | SUMO specific peptidase 6 | protein_coding | NP_001093879.1;NP_001291721.1;NP_056386.2;XP_006715520.1;XP_01 | 20944 | 605003 | . | . | 5.591309 | 5.285410 | 7.378188 | 4.575701 | 13.011205 | 12.217461 | 16.556443 | 10.461795 | 1.036908 | 1064.961681 | 0.197827627 | 0.264308864 | 0.791541931 | 0.998865404 | 10.029789 | 10.082894 | 10.056059 | 10.011380 | 10.235726 | 9.850148 | 1240 | 1094 | 1185 | 756 |
| 26056 | NM_00137127 | RAB11FIP5 | RAB11 family interacting protein 5 | protein_coding | NP_001358201.1;NP_056285.1 | 24845 | 605536 | . | . | 5.402906 | 5.676593 | 5.616356 | 6.671934 | 12.572786 | 13.121701 | 12.602941 | 15.254585 | 1.069611 | 948.820635 | 0.145300241 | 0.668179951 | 0.504018731 | 0.998865404 | 9.840845 | 9.937519 | 9.869640 | 9.858696 | 9.856848 | 9.966196 | 1064 | 982 | 789 | 988 |
| 26057 | NM_00128677 | ANKRD17 | ankyrin repeat domain 17 | protein_coding | NP_001273700.1;NP_056389.1;NP_115593.3;NP_942592.1;XP_00526572 | 23575 | 615929 | . | . | 10.372154 | 10.402780 | 9.031175 | 10.542996 | 24.136427 | 24.046494 | 20.265697 | 24.105307 | -1.003307 | 2898.381837 | 0.113440247 | -0.04198485 | 0.966510779 | 0.998865404 | 11.503475 | 11.498585 | 11.494894 | 11.509622 | 11.473280 | 11.525146 | 3313 | 3161 | 2436 | 2799 |
| 26058 | NM_00110314 | GIGYF2 | GRB10 interacting GYF protein 2 | protein_coding | NP_001096616.1;NP_001096617.1;NP_001096618.1;NP_056390.2 | 11960 | 612003 | . | . | 12.500390 | 12.289622 | 13.565321 | 12.396619 | 29.088922 | 28.408013 | 30.440192 | 28.343393 | 1.054490 | 2863.313899 | 0.114415177 | 0.669014324 | 0.503486341 | 0.998865404 | 11.444620 | 11.521302 | 11.465044 | 11.457204 | 11.535627 | 11.472279 | 3225 | 2993 | 2655 | 2649 |
| 26060 | NM_012096,XI | APPL1 | adaptor protein, phosphotyrosine interacting with PH domain and | protein_coding | NP_036228.1;XP_011531885.1 | 24035 | 604299 | . | . | 3.811626 | 3.664082 | 3.776064 | 3.552356 | 8.869809 | 8.469689 | 8.473377 | 8.122040 | -1.033828 | 702.059852 | 0.152173565 | -0.3154034 | 0.752455385 | 0.998865404 | 9.479044 | 9.431464 | 9.482727 | 9.450120 | 9.469049 | 9.417933 | 841 | 752 | 623 | 623 |
| 26061 | NM_00128441 | HACL1 | 2-hydroxyacyl-CoA lyase 1 | protein_coding | NP_001271342.1;NP_001271344.1;NP_001271345.1;NP_036392.2 | 17856 | 604300 | . | . | 9.264163 | 8.584439 | 8.087996 | 9.555452 | 21.558087 | 19.843317 | 18.149228 | 21.847406 | -1.025733 | 553.340199 | 0.170086028 | -0.21551367 | 0.829366858 | 0.998865404 | 9.130406 | 9.093403 | 9.150403 | 9.087794 | 9.051682 | 9.151836 | 676 | 576 | 438 | 552 |
| 26063 | NM_020664 | DECR2 | 2,4-dienoyl-CoA reductase 2 | protein_coding | NP_065715.1 | 2754 | 615839 | . | . | 6.702908 | 6.017940 | 5.875127 | 7.044486 | 15.597942 | 13.910737 | 13.183616 | 16.106380 | 1.001751 | 310.326539 | 0.206586896 | 0.01222083 | 0.990249431 | 0.999289967 | 8.276580 | 8.278706 | 8.308002 | 8.243564 | 8.230182 | 8.322127 | 378 | 315 | 247 | 316 |
| 26064 | NM_00114552 | RAI14 | retinoic acid induced 14 | protein_coding | NP_001138992.1;NP_001138993.1;NP_001138994.1;NP_001138995.1;NI | 14873 | 606586 | . | . | 8.054104 | 8.659446 | 8.117678 | 8.536517 | 18.742233 | 20.016695 | 18.215837 | 19.517731 | -1.017378 | 1286.294414 | 0.130502357 | -0.19046525 | 0.848944568 | 0.998865404 | 10.341503 | 10.316398 | 10.310033 | 10.360145 | 10.306606 | 10.337632 | 1445 | 1446 | 1086 | 1220 |
| 26065 | NM_00111409 | LSM14A | LSM14A mRNA processing body assembly factor | protein_coding | NP_001107565.1;NP_056393.2;XP_005258776.1;XP_005258777.1;XP_00 | 24489 | 610677 | . | . | 18.009206 | 18.412326 | 17.945972 | 17.435854 | 41.908161 | 42.560916 | 40.270236 | 39.865005 | -1.044764 | 2019.358082 | 0.118603513 | -0.53266949 | 0.594262386 | 0.998865404 | 11.010875 | 10.947797 | 10.993049 | 11.000503 | 10.981759 | 10.941856 | 2365 | 2237 | 1761 | 1805 |
| 26071 | NM_00107817 | RTL8A | retrotransposon Gag like 8A | protein_coding | NP_001071640.1;NP_001127793.1 | 24514 | . | . | . | 17.438883 | 16.988966 | 21.067299 | 22.501472 | 40.581001 | 39.270756 | 47.274406 | 51.446943 | 1.160354 | 570.096579 | 0.161216958 | 1.330910495 | 0.183218463 | 0.954800306 | 9.043869 | 9.258296 | 9.114671 | 9.091148 | 9.178018 | 9.222191 | 619 | 559 | 515 | 593 |
| 26073 | NM_00129014 | POLDIP2 | DNA polymerase delta interacting protein 2 | protein_coding | NP_001277074.1;NP_056399.1 | 23781 | 611519 | . | . | 36.921025 | 40.668969 | 36.880197 | 42.325152 | 85.916741 | 94.008144 | 82.758094 | 96.771439 | 1.001913 | 3298.495456 | 0.120030535 | 0.022972909 | 0.981671883 | 0.998865404 | 11.686332 | 11.688851 | 11.658278 | 11.713323 | 11.630698 | 11.741612 | 3668 | 3675 | 2677 | 3304 |
| 26074 | NM_00116781 | CFAP61 | cilia and flagella associated protein 61 | protein_coding | NP_001161288.1;NP_056400.3;XP_005260745.1;XP_005260746.1;XP_00 | 15872 | . | . | . | 0.993252 | 1.416950 | 1.010303 | 1.157699 | 2.311338 | 3.275346 | 2.267091 | 2.646941 | -1.098965 | 99.705676 | 0.331114547 | -0.41117388 | 0.680945042 | 0.998865404 | 6.706954 | 6.568955 | 6.608158 | 6.694852 | 6.604200 | 6.642854 | 104 | 128 | 77 | 95 |
| 26082 | NR_026771 | LINC02249 | long intergenic non-protein coding RNA 2249 | lncRNA | . | 32351 | . | . | . | 0.917756 | 0.984568 | 1.387247 | 0.956811 | 2.135657 | 2.275872 | 3.112942 | 2.187636 | 1.214286 | 141.194783 | 0.293895706 | 0.95308905 | 0.340544954 | 0.998865404 | 6.993881 | 7.275485 | 7.083414 | 7.107121 | 7.239497 | 7.091965 | 142 | 142 | 160 | 118 |
| 26083 | XM_01152459 | TBC1D29P | TBC1 domain family member 29, pseudogene, transcript variant X | protein_coding | XP_011522898.1 | 24509 | . | . | . | 0.048452 | 0.050834 | 0.061182 | 0.033444 | 0.112749 | 0.117505 | 0.137292 | 0.076466 | -1.070014 | 3.478495 | 1.609933276 | -0.06064235 | 0.951644048 | NA | 1.844134 | 1.751299 | 1.796180 | 1.797278 | 1.801166 | 1.790082 | 4 | 4 | 4 | 2 |
| 26084 | NM_00125196 | ARHGEF26 | Rho guanine nucleotide exchange factor 26 | protein_coding | NP_001238891.1;NP_001238892.1;NP_056410.3;XP_011510974.1;XP_01 | 24490 | 617552 | . | . | 2.252350 | 2.639228 | 2.775569 | 2.525201 | 5.241313 | 6.100692 | 6.228294 | 5.773571 | 1.064382 | 408.670493 | 0.186635969 | 0.482308423 | 0.629586865 | 0.998865404 | 8.629099 | 8.719087 | 8.608654 | 8.697443 | 8.722803 | 8.658840 | 422 | 459 | 386 | 374 |
| 26086 | NM_00114563 | GPSM1 | G protein signaling modulator 1 | protein_coding | NP_001139110.2;NP_001139111.1;NP_001186932.1;NP_056412.5;XP_01 | 17858 | 609491 | . | . | 6.161314 | 6.531001 | 6.268239 | 5.487126 | 14.337633 | 15.096705 | 14.065744 | 12.545662 | -1.081011 | 654.919657 | 0.158203173 | -0.71036205 | 0.477479649 | 0.998865404 | 9.410126 | 9.298046 | 9.358747 | 9.401668 | 9.366631 | 9.287389 | 759 | 757 | 580 | 555 |
| 26088 | NM_00100156 | GGA1 | golgi associated, gamma adaptin ear containing, ARF binding prot | protein_coding | NP_001001560.1;NP_001166158.1;NP_001166159.1;NP_001350700.1;NI | 17842 | 606004 | . | . | 33.909719 | 34.747036 | 33.596400 | 30.886386 | 78.909313 | 80.319328 | 75.389343 | 70.618047 | -1.056253 | 3384.930163 | 0.109798844 | -0.71908734 | 0.472087105 | 0.998865404 | 11.763830 | 11.684912 | 11.741561 | 11.752421 | 11.715041 | 11.688753 | 3978 | 3777 | 2909 | 3037 |
| 26090 | NM_00104247 | ABHD12 | abhydrolase domain containing 12, lysophospholipase | protein_coding | NP_001035937.1;NP_056415.1;XP_011527516.1;XP_016883285.1;XP_01 | 15868 | 613599 | . | . | 50.948257 | 53.634169 | 45.332897 | 48.875247 | 118.558690 | 123.977777 | 101.725704 | 111.747447 | -1.137633 | 3022.502177 | 0.1130276 | -1.64592796 | 0.099778591 | 0.814937213 | 11.651613 | 11.465441 | 11.600549 | 11.623188 | 11.480915 | 11.531282 | 3654 | 3519 | 2382 | 2734 |
| 26091 | NM_00127818 | HERC4 | HECT and RLD domain containing E3 ubiquitin protein ligase 4 | protein_coding | NP_001265114.1;NP_001265115.1;NP_001265116.1;NP_056416.2;NP_0 | 24521 | 609248 | . | . | 7.345684 | 7.701523 | 9.058865 | 7.507832 | 17.093710 | 17.802418 | 20.327834 | 17.165766 | 1.083496 | 1080.614831 | 0.146197809 | 0.791355483 | 0.42873658 | 0.998865404 | 10.018411 | 10.134527 | 10.031554 | 10.059090 | 10.174064 | 10.030987 | 1172 | 1140 | 1071 | 953 |
| 26092 | NM_00126757 | TOR1AIP1 | torsin 1A interacting protein 1 | protein_coding | NP_001254507.1;NP_056417.2 | 29456 | 614512 | . | . | 28.472466 | 28.538364 | 28.815712 | 26.600521 | 66.256592 | 65.967705 | 64.661629 | 60.818929 | -1.042901 | 3334.790887 | 0.113419808 | -0.53432211 | 0.593118696 | 0.998865404 | 11.733295 | 11.672832 | 11.722811 | 11.717489 | 11.721859 | 11.648134 | 3933 | 3662 | 2967 | 2923 |
| 26093 | NM_015603,XI | CCDC9 | coiled-coil domain containing 9 | protein_coding | NP_056418.1;XP_016882067.1 | 24560 | . | . | . | 7.358358 | 8.011000 | 6.389338 | 7.698012 | 17.123201 | 18.517785 | 14.337490 | 17.600588 | -1.107227 | 477.343832 | 0.177390835 | -0.82840432 | 0.407441563 | 0.998865404 | 8.970922 | 8.823061 | 8.907240 | 8.951169 | 8.813654 | 8.914913 | 558 | 560 | 361 | 460 |
| 26094 | NM_00116350 | DCAF4 | DDB1 and CUL4 associated factor 4 | protein_coding | NP_001156980.1;NP_001156981.1;NP_001339376.1;NP_001339377.1;NI | 20229 | 616372 | . | . | 4.108134 | 4.000877 | 4.377279 | 4.505991 | 9.559796 | 9.248206 | 9.822488 | 10.302411 | 1.050183 | 337.262823 | 0.191888459 | 0.368136634 | 0.712771356 | 0.998865404 | 8.361854 | 8.432735 | 8.387956 | 8.378615 | 8.420998 | 8.400487 | 382 | 352 | 306 | 318 |
| 26095 | NM_00104235 | PTPN20 | protein tyrosine phosphatase non-receptor type 20 | protein_coding | NP_001035816.1;NP_001035817.1;NP_001035818.1;NP_001035819.1;NI | 23423 | 610631 | . | . | 1.761472 | 1.896446 | 2.086455 | 1.591804 | 4.099019 | 4.383717 | 4.681941 | 3.639471 | 1.012144 | 155.396203 | 0.276729018 | 0.062931364 | 0.94982116 | 0.998865404 | 7.270399 | 7.289155 | 7.260870 | 7.286956 | 7.337579 | 7.216462 | 172 | 172 | 157 | 124 |
| 26097 | NM_00120661 | CHTOP | chromatin target of PRMT1 | protein_coding | NP_001193541.1;NP_001231593.1;NP_001304006.1;NP_056422.2;XP_01 | 24511 | 614206 | . | . | 22.002400 | 21.226050 | 20.584150 | 22.391394 | 51.200493 | 49.064966 | 46.190236 | 51.195261 | -1.013372 | 1405.380864 | 0.129810564 | -0.14762516 | 0.882638601 | 0.998865404 | 10.466375 | 10.447051 | 10.478458 | 10.444480 | 10.415608 | 10.485924 | 1665 | 1493 | 1158 | 1369 |
| 26098 | NM_00120243 | EDRF1 | erythroid differentiation regulatory factor 1 | protein_coding | NP_001189367.1;NP_056423.2;XP_011537924.1;XP_016871555.1 | 24640 | . | . | . | 5.341537 | 5.821137 | 6.379459 | 3.935399 | 12.429977 | 13.455817 | 14.315323 | 8.997821 | -1.106881 | 815.802000 | 0.186779308 | -0.78434872 | 0.432835522 | 0.998865404 | 9.743049 | 9.597428 | 9.684704 | 9.717958 | 9.764095 | 9.474781 | 964 | 946 | 813 | 576 |
| 26099 | NM_00111460 | SZRD1 | SUZ RNA binding domain containing 1 | protein_coding | NP_001108072.1;NP_001258798.1 | 30232 | . | . | . | 42.264940 | 45.894733 | 39.770446 | 49.264091 | 98.352253 | 106.087719 | 89.243729 | 112.636487 | -1.004032 | 4747.611175 | 0.125971475 | -0.0460836 | 0.96324362 | 0.998865404 | 12.215991 | 12.209975 | 12.181771 | 12.243898 | 12.121744 | 12.291038 | 5275 | 5324 | 3703 | 4891 |
| 26100 | NM_00103351 | WIPI2 | WD repeat domain, phosphoinositide interacting 2 | protein_coding | NP_001028690.1;NP_001028691.1;NP_001028692.1;NP_001265228.1;NI | 32225 | 609225 | . | . | 11.642093 | 12.150906 | 13.213047 | 14.511059 | 27.091630 | 28.087365 | 29.649696 | 33.177814 | 1.149147 | 1766.044198 | 0.123796566 | 1.620106673 | 0.105209364 | 0.832886861 | 10.682673 | 10.882991 | 10.714225 | 10.742674 | 10.803546 | 10.870789 | 1858 | 1806 | 1573 | 1845 |
| 26108 | NM_00133032 | PYGO1 | pygopus family PHD finger 1 | protein_coding | NP_001317255.1;NP_001354735.1;NP_056432.1 | 30256 | 606902 | . | . | 0.732435 | 0.829684 | 1.073180 | 0.857871 | 1.704406 | 1.917854 | 2.408186 | 1.961418 | 1.272413 | 236.708777 | 0.231388884 | 1.502088824 | 0.133074171 | 0.888450802 | 7.702317 | 8.050647 | 7.801196 | 7.835709 | 7.994977 | 7.873523 | 231 | 233 | 258 | 219 |
| 26115 | NM_025185,XI | TANC2 | tetratricopeptide repeat, ankyrin repeat and coiled-coil containing | protein_coding | NP_079461.2;XP_006721869.1;XP_006721870.1;XP_006721873.1;XP_00 | 30212 | 615047 | . | . | 2.719872 | 3.261502 | 3.325429 | 3.761868 | 6.329255 | 7.539111 | 7.462165 | 8.601065 | 1.163348 | 1241.660180 | 0.143799442 | 1.517965612 | 0.129023068 | 0.878737612 | 10.165096 | 10.382807 | 10.151666 | 10.276927 | 10.284994 | 10.373912 | 1211 | 1343 | 1094 | 1324 |
| 26118 | NM_00134835 | WSB1 | WD repeat and SOCS box containing 1 | protein_coding | NP_001335279.1;NP_056441.6;NP_599027.1;XP_005258020.1;XP_01687 | 19221 | 610091 | . | . | 21.266943 | 21.027837 | 23.642064 | 17.132634 | 49.489049 | 48.606790 | 53.052107 | 39.171731 | 1.033609 | 3274.295754 | 0.154637947 | 0.308399543 | 0.757778329 | 0.998865404 | 11.652771 | 11.700768 | 11.672995 | 11.644468 | 11.826836 | 11.523076 | 3772 | 3414 | 3424 | 2550 |
| 26119 | NM_015627,XI | LDLRAP1 | low density lipoprotein receptor adaptor protein 1 | protein_coding | NP_056442.2;XP_006710622.1;XP_006710623.1;XP_006710624.1;XP_01 | 18640 | 605747 | . | . | 12.756830 | 13.093049 | 11.984909 | 13.416367 | 29.685669 | 30.265167 | 26.893787 | 30.674932 | -1.031237 | 1209.674705 | 0.13169022 | -0.33697251 | 0.736137623 | 0.998865404 | 10.262520 | 10.217943 | 10.252716 | 10.251021 | 10.202616 | 10.254029 | 1416 | 1324 | 1000 | 1155 |
| 26121 | NM_015629,XI | PRPF31 | pre-mRNA processing factor 31 | protein_coding | NP_056444.3;XP_006723200.1 | 15446 | 606419 | . | . | 34.290626 | 33.631473 | 37.265808 | 39.428817 | 79.795695 | 77.740658 | 83.623393 | 90.149304 | 1.116364 | 2147.516432 | 0.117566164 | 1.350788312 | 0.176763253 | 0.949538805 | 10.986908 | 11.145636 | 11.028610 | 11.016452 | 11.089292 | 11.131185 | 2355 | 2173 | 1919 | 2179 |
| 26122 | NM_015630,XI | EPC2 | enhancer of polycomb homolog 2 | protein_coding | NP_056445.3;XP_011509243.1;XP_011509245.1 | 24543 | 611000 | . | . | 2.754558 | 3.197870 | 3.697841 | 3.647854 | 6.409972 | 7.392019 | 8.297845 | 8.340386 | 1.211533 | 429.572228 | 0.176097733 | 1.572043454 | 0.115940474 | 0.857396579 | 8.601708 | 8.878546 | 8.672208 | 8.694373 | 8.809618 | 8.789510 | 439 | 427 | 416 | 434 |
| 26123 | NM_00114397 | TCTN3 | tectonic family member 3 | protein_coding | NP_001137445.1;NP_056446.4;XP_005269747.1;XP_011537929.1;XP_01 | 24519 | 613847 | . | . | 30.983646 | 32.634782 | 29.549381 | 31.910276 | 72.100221 | 75.436758 | 66.307952 | 72.959048 | -1.049204 | 2451.122376 | 0.116742635 | -0.59357011 | 0.552799677 | 0.998865404 | 11.293545 | 11.224072 | 11.260527 | 11.295593 | 11.213207 | 11.264905 | 2828 | 2767 | 2013 | 2315 |
| 26127 | NM_00117188 | FGFR1OP2 | FGFR1 oncogene partner 2 | protein_coding | NP_001165358.1;NP_001165359.1;NP_056448.1 | 23098 | 608858 | . | . | 17.257487 | 15.592476 | 14.684462 | 15.105183 | 40.158884 | 36.042706 | 32.951503 | 34.536205 | -1.056097 | 952.338857 | 0.142347256 | -0.55317234 | 0.580145407 | 0.998865404 | 9.934090 | 9.855502 | 9.954687 | 9.872870 | 9.864618 | 9.885198 | 1189 | 997 | 794 | 881 |
| 26128 | NM_015634,XI | KIFBP | kinesin family binding protein | protein_coding | NP_056449.1;XP_016871556.1 | 23419 | 609367 | . | . | 12.845928 | 12.630399 | 11.696612 | 12.360843 | 29.893003 | 29.195732 | 26.246859 | 28.261598 | -1.074269 | 957.805047 | 0.13827589 | -0.74745813 | 0.45478707 | 0.998865404 | 9.954371 | 9.850953 | 9.939895 | 9.918397 | 9.863615 | 9.890366 | 1159 | 1055 | 788 | 882 |
| 26130 | NM_00128267 | GAPVD1 | GTPase activating protein and VPS9 domains 1 | protein_coding | NP_001269608.1;NP_001269609.1;NP_001269610.1;NP_001317706.1;NI | 23375 | 611714 | . | . | 4.846816 | 5.381382 | 5.573304 | 5.568518 | 11.278740 | 12.439305 | 12.506328 | 12.731753 | 1.061870 | 1426.197249 | 0.130003281 | 0.666188988 | 0.50529031 | 0.998865404 | 10.434031 | 10.520586 | 10.416636 | 10.489746 | 10.519458 | 10.480099 | 1517 | 1564 | 1310 | 1342 |
| 26133 | NM_015638,N | TRPC4AP | transient receptor potential cation channel subfamily C member 4 | protein_coding | NP_056453.1;NP_955400.1;XP_016883288.1;XP_016883289.1;XP_01688 | 16181 | 608430 | . | . | 30.932022 | 30.620436 | 32.494415 | 34.287299 | 71.980088 | 70.780507 | 72.916525 | 78.393834 | 1.049024 | 3307.641018 | 0.112254352 | 0.615100738 | 0.538488195 | 0.998865404 | 11.656712 | 11.725638 | 11.644427 | 11.697670 | 11.696014 | 11.725146 | 3597 | 3597 | 2893 | 3231 |
| 26134 | NR_104269 | RALGAPA1P1 | RALGAPA1 pseudogene 1 | transcribed_ps | . | 23203 | . | . | . | 0.133134 | 0.186106 | 0.117852 | 0.067887 | 0.309808 | 0.430192 | 0.264457 | 0.155215 | -1.760796 | 30.053873 | 0.592903065 | -1.37666334 | 0.168616359 | 0.945638095 | 5.259576 | 4.446236 | 4.915595 | 4.964378 | 4.897377 | 4.849665 | 37 | 48 | 24 | 15 |
| 26135 | NM_00101806 | SERBP1 | SERPINE1 mRNA binding protein 1 | protein_coding | NP_001018077.1;NP_001018078.1;NP_001018079.1;NP_056455.3 | 17860 | 607378 | . | . | 40.655431 | 38.754653 | 37.766537 | 40.142023 | 94.606861 | 89.583119 | 84.747015 | 91.779962 | -1.033580 | 8201.910120 | 0.104915284 | -0.45417604 | 0.649702128 | 0.998865404 | 13.025377 | 12.977718 | 13.039441 | 12.990871 | 12.965168 | 13.009170 | 9874 | 8740 | 6832 | 7759 |
| 26136 | NM_015641,N | TES | testin LIM domain protein | protein_coding | NP_056456.1;NP_690042.1;XP_005250315.1 | 14620 | 606085 | . | . | 17.915507 | 16.741381 | 16.874545 | 17.221505 | 41.690123 | 38.698452 | 37.865992 | 39.374925 | -1.031327 | 1464.958923 | 0.127082602 | -0.35018196 | 0.726202147 | 0.998865404 | 10.538666 | 10.494282 | 10.557411 | 10.498317 | 10.501327 | 10.507779 | 1778 | 1544 | 1249 | 1358 |
| 26137 | NM_00116434 | ZBTB20 | zinc finger and BTB domain containing 20 | protein_coding | NP_001157814.1;NP_001157815.1;NP_001157816.1;NP_001157817.1;NI | 13503 | 606025 | . | . | 0.127329 | 0.129502 | 0.136103 | 0.088719 | 0.296300 | 0.299349 | 0.305409 | 0.202847 | -1.161554 | 104.797322 | 0.332778489 | -0.64925072 | 0.51617663 | 0.998865404 | 6.814370 | 6.600637 | 6.729937 | 6.732778 | 6.743861 | 6.621222 | 129 | 122 | 102 | 72 |
| 26140 | NM_00102593 | TTLL3 | tubulin tyrosine ligase like 3 | protein_coding | NP_001021100.3;NP_001352980.1 | 24483 | . | . | . | 1.111148 | 1.057564 | 1.439573 | 1.053131 | 2.585686 | 2.444605 | 3.230361 | 2.407859 | 1.135228 | 140.215538 | 0.288362508 | 0.63455395 | 0.52571937 | 0.998865404 | 7.036170 | 7.220924 | 7.111141 | 7.092620 | 7.207716 | 7.087779 | 155 | 138 | 150 | 118 |
| 26145 | NM_015649 | IRF2BP1 | interferon regulatory factor 2 binding protein 1 | protein_coding | NP_056464.1 | 21728 | 615331 | . | . | 7.584118 | 8.520566 | 7.693491 | 9.434656 | 17.648554 | 19.695671 | 17.263971 | 21.571222 | 1.049333 | 656.007373 | 0.166303553 | 0.417748287 | 0.676131164 | 0.998865404 | 9.322878 | 9.391444 | 9.302571 | 9.375029 | 9.304574 | 9.435028 | 698 | 728 | 528 | 690 |
| 26146 | NM_00113949 | TRAF3IP1 | TRAF3 interacting protein 1 | protein_coding | NP_001132962.1;NP_056465.2;XP_006712477.1;XP_011509246.1;XP_01 | 17861 | 607380 | . | . | 4.872805 | 4.321169 | 5.187700 | 5.062774 | 11.339218 | 9.988578 | 11.641049 | 11.575428 | 1.091813 | 605.230440 | 0.161290424 | 0.7856975 | 0.432044728 | 0.998865404 | 9.176322 | 9.303555 | 9.248860 | 9.170154 | 9.282701 | 9.254743 | 706 | 587 | 561 | 580 |
| 26147 | NM_00100993 | PHF19 | PHD finger protein 19 | protein_coding | NP_001009936.1;NP_001273769.1;NP_001273771.1;NP_001273772.1;NI | 24566 | 609740 | . | . | 39.420564 | 51.526868 | 40.236536 | 46.560056 | 91.733261 | 119.106660 | 90.289615 | 106.454045 | 1.001709 | 3117.853402 | 0.118581219 | 0.02077483 | 0.983425276 | 0.998865404 | 11.605220 | 11.607454 | 11.567783 | 11.641191 | 11.565439 | 11.645892 | 3427 | 3512 | 2580 | 3069 |
| 26149 | NM_00131791 | ZNF658 | zinc finger protein 658 | protein_coding | NP_001304845.1;NP_149350.3;XP_005272572.1;XP_011543981.1;XP_01 | 25226 | 616290 | . | . | 0.854505 | 1.020118 | 0.773470 | 1.010423 | 1.988466 | 2.358049 | 1.735643 | 2.310209 | -1.072065 | 137.434065 | 0.288584967 | -0.34787879 | 0.727931214 | 0.998865404 | 7.152652 | 7.050741 | 7.061043 | 7.163659 | 7.071439 | 7.103965 | 142 | 174 | 110 | 130 |
| 26150 | NM_015653,XI | RIBC2 | RIB43A domain with coiled-coils 2 | protein_coding | NP_056468.3;XP_005261581.1;XP_011528428.1;XP_016884255.1 | 13241 | . | . | . | 5.160773 | 6.107928 | 4.587612 | 6.990188 | 12.009329 | 14.118749 | 10.294466 | 15.982235 | 1.007907 | 252.956069 | 0.245321187 | 0.046317931 | 0.963056848 | 0.998865404 | 7.978077 | 7.987394 | 7.936262 | 8.014464 | 7.876472 | 8.073884 | 268 | 293 | 177 | 285 |
| 26151 | NM_00130507 | NAT9 | N-acetyltransferase 9 (putative) | protein_coding | NP_001292006.1;NP_001292007.1;NP_001292008.1;NP_001292009.1;NI | 23133 | . | . | . | 22.720329 | 18.160767 | 21.028552 | 21.843894 | 52.871144 | 41.979419 | 47.187460 | 49.943469 | 1.024812 | 1315.674730 | 0.137331359 | 0.257477089 | 0.7968105 | 0.998865404 | 10.343655 | 10.379299 | 10.410974 | 10.287931 | 10.369228 | 10.370865 | 1617 | 1289 | 1157 | 1250 |
| 26152 | NM_00129026 | ZNF337 | zinc finger protein 337 | protein_coding | NP_001277190.1;NP_056470.1;XP_005260759.1;XP_006723621.1;XP_01 | 15809 | . | . | . | 2.451866 | 2.489053 | 2.261080 | 3.619272 | 5.705594 | 5.753559 | 5.073797 | 8.275035 | 1.156836 | 427.514368 | 0.181965832 | 1.155076071 | 0.248059276 | 0.992733415 | 8.631177 | 8.840868 | 8.704036 | 8.679720 | 8.731202 | 8.826938 | 466 | 419 | 368 | 463 |
| 26153 | NM_015656,XI | KIF26A | kinesin family member 26A | protein_coding | NP_056471.1;XP_011534943.1 | 20226 | 613231 | . | . | 0.290946 | 0.352838 | 0.232164 | 0.332630 | 0.677043 | 0.815602 | 0.520970 | 0.760520 | -1.156132 | 64.094103 | 0.40580134 | -0.51578341 | 0.606005693 | 0.998865404 | 6.104275 | 5.892176 | 5.995889 | 6.039578 | 5.950761 | 6.019370 | 72 | 81 | 43 | 65 |
| 26154 | NM_015657,N | ABCA12 | ATP binding cassette subfamily A member 12 | protein_coding | NP_056472.2;NP_775099.2;XP_011509253.1 | 14637 | 607800 | . | . | 0.248963 | 0.297108 | 0.134674 | 0.319889 | 0.579347 | 0.686779 | 0.302204 | 0.731389 | -1.205878 | 73.495182 | 0.428932159 | -0.6296653 | 0.528913591 | 0.998865404 | 6.330304 | 6.055815 | 6.193328 | 6.237745 | 6.076990 | 6.254493 | 85 | 94 | 35 | 87 |
| 26155 | NM_015658 | NOC2L | NOC2 like nucleolar associated transcriptional repressor | protein_coding | NP_056473.3 | 24517 | 610770 | . | . | 57.038273 | 56.487469 | 54.013237 | 55.934826 | 132.730408 | 130.573318 | 121.204147 | 127.888336 | -1.047150 | 4799.371750 | 0.105902554 | -0.62763572 | 0.530242602 | 0.998865404 | 12.261488 | 12.195005 | 12.255186 | 12.239935 | 12.198775 | 12.219273 | 5705 | 5250 | 4027 | 4450 |
| 26156 | NM_015659 | RSL1D1 | ribosomal L1 domain containing 1 | protein_coding | NP_056474.2 | 24534 | 615874 | . | . | 23.687992 | 23.628349 | 22.170395 | 24.577389 | 55.122932 | 54.617992 | 49.749725 | 56.193279 | -1.026659 | 3985.204886 | 0.111733114 | -0.33970796 | 0.734076463 | 0.998865404 | 11.979345 | 11.941280 | 11.975620 | 11.966460 | 11.910104 | 11.986839 | 4675 | 4333 | 3261 | 3858 |
| 26157 | NM_015660 | GIMAP2 | GTPase, IMAP family member 2 | protein_coding | NP_056475.1 | 21789 | 608085 | . | . | 1.268192 | 1.037411 | 0.848340 | 0.885559 | 2.951136 | 2.398022 | 1.903651 | 2.024726 | -1.342905 | 46.003852 | 0.463993534 | -0.91673107 | 0.359283581 | 0.998865404 | 5.720410 | 5.295830 | 5.571350 | 5.530105 | 5.499863 | 5.500788 | 67 | 51 | 34 | 37 |
| 26160 | NM_015662,XI | IFT172 | intraflagellar transport 172 | protein_coding | NP_056477.1;XP_005264311.1;XP_006712049.1;XP_006712050.1;XP_01 | 30391 | 607386 | . | . | 7.340497 | 7.983604 | 8.559741 | 7.184391 | 17.081634 | 18.454459 | 19.207809 | 16.426257 | 1.026433 | 1274.584025 | 0.140244976 | 0.268380804 | 0.788406214 | 0.998865404 | 10.296706 | 10.334666 | 10.282649 | 10.326004 | 10.387979 | 10.256715 | 1407 | 1396 | 1220 | 1106 |
| 26164 | NM_015666,XI | MTG2 | mitochondrial ribosome associated GTPase 2 | protein_coding | NP_056481.1;XP_005260450.1;XP_006723837.1;XP_011527078.1;XP_01 | 16239 | 610919 | . | . | 11.801685 | 12.492945 | 14.769123 | 13.583699 | 27.463005 | 28.878004 | 33.141486 | 31.057511 | 1.074366 | 1655.016013 | 0.124268389 | 0.832761871 | 0.404979043 | 0.998865404 | 10.639907 | 10.743493 | 10.649313 | 10.677531 | 10.746635 | 10.691730 | 1804 | 1753 | 1543 | 1551 |
| 26167 | NM_015669 | PCDHB5 | protocadherin beta 5 | protein_coding | NP_056484.2 | 8690 | 606331 | . | . | 3.056468 | 3.324231 | 2.605322 | 3.598594 | 7.112527 | 7.684110 | 5.846266 | 8.227757 | -1.042025 | 334.973355 | 0.212522066 | -0.27945518 | 0.779895528 | 0.998865404 | 8.418027 | 8.357136 | 8.375350 | 8.418357 | 8.285230 | 8.455393 | 379 | 383 | 241 | 355 |
| 26168 | NM_015670 | SENP3 | SUMO specific peptidase 3 | protein_coding | NP_056485.2 | 17862 | 612844 | . | . | 34.633820 | 36.053310 | 34.475723 | 38.839542 | 80.594315 | 83.338844 | 77.362526 | 88.802002 | 1.022567 | 2878.431879 | 0.117444319 | 0.274136825 | 0.783979475 | 0.998865404 | 11.474983 | 11.506974 | 11.468576 | 11.494319 | 11.454226 | 11.543145 | 3226 | 3120 | 2393 | 2877 |
| 26173 | NM_00108045 | INTS1 | integrator complex subunit 1 | protein_coding | NP_001073922.2;XP_011513562.1;XP_011513564.1;XP_016867448.1;XP_ | 24555 | 611345 | . | . | 24.807323 | 26.797498 | 26.242050 | 26.008594 | 57.727664 | 61.943615 | 58.886398 | 59.465563 | 1.001105 | 5625.455333 | 0.107185635 | 0.01487026 | 0.988135686 | 0.999197585 | 12.456974 | 12.458534 | 12.425700 | 12.487596 | 12.464960 | 12.451046 | 6236 | 6290 | 4935 | 5233 |
| 26175 | NM_00109862 | TMEM251 | transmembrane protein 251 | protein_coding | NP_001092091.2;NP_056491.1 | 20218 | . | . | . | 14.307965 | 13.258912 | 15.229190 | 12.469596 | 33.295221 | 30.648569 | 34.173863 | 28.510249 | -1.006077 | 1016.156382 | 0.151672441 | -0.05762971 | 0.954043585 | 0.998865404 | 9.992915 | 9.984887 | 10.020170 | 9.957620 | 10.058618 | 9.905931 | 1223 | 1053 | 973 | 851 |
| 26190 | NM_00137588 | FBXW2 | F-box and WD repeat domain containing 2 | protein_coding | NP_001362817.1;NP_001362818.1;NP_001362819.1;NP_036296.2;XP_00 | 13608 | 609071 | . | . | 5.157644 | 5.600227 | 5.765338 | 5.684789 | 12.002046 | 12.945180 | 12.937249 | 12.997596 | 1.053650 | 1448.462155 | 0.127360326 | 0.591989698 | 0.553857487 | 0.998865404 | 10.462135 | 10.537494 | 10.452994 | 10.505312 | 10.533649 | 10.505553 | 1569 | 1574 | 1316 | 1368 |

| | | | | | | | | | | |
|---|---|---|---|---|---|---|---|---|---|---|
| 26191 | NM_00119343 | PTPN22 | protein tyrosine phosphatase non-receptor type 22 | protein_coding | NP_001180360.1;NP_001295226.1;NP_036543.4;NP_057051.3;XP_01153 | 9652 | 600716 | . | . | [illegible] |
| 26205 | NM_012384,XI | GMEB2 | glucocorticoid modulatory element binding protein 2 | protein_coding | NP_036516.1;XP_005260259.1;XP_006723839.1;XP_011527081.1 | 4371 | 607451 | . | . | [illegible] |
| 26206 | NM_00103959 | SPAG8 | sperm associated antigen 8 | protein_coding | NP_001034681.1;NP_001353689.1;NP_758516.1;XP_024303278.1;XP_02 | 14105 | 605731 | . | . | [illegible] |
| 26207 | NM_012417,N | PITPNC1 | phosphatidylinositol transfer protein cytoplasmic 1 | protein_coding | NP_036549.2;NP_858057.1;XP_005257273.1;XP_016879931.1;XP_01687 | 21045 | 605134 | . | . | [illegible] |
| 26220 | NR_002733,NF | DGCR5 | DiGeorge syndrome critical region gene 5 | lncRNA | . | 16757 | 618040 | . | . | [illegible] |
| 26224 | NM_012158,XI | FBXL3 | F-box and leucine rich repeat protein 3 | protein_coding | NP_036290.1;XP_005266393.1;XP_005266394.1;XP_016876027.1 | 13599 | 605653 | . | . | [illegible] |
| 26225 | NM_00103717 | ARL5A | ADP ribosylation factor like GTPase 5A | protein_coding | NP_001032251.1;NP_036229.1;NP_817114.2 | 696 | 608960 | . | . | [illegible] |
| 26226 | NR_033408 | FBXW4P1 | F-box and WD repeat domain containing 4 pseudogene 1 | transcribed_ps | . | 13609 | . | | . | [illegible] |
| 26227 | NM_006623,XI | PHGDH | phosphoglycerate dehydrogenase | protein_coding | NP_006614.2;XP_011539528.1;XP_011539529.1;XP_011539530.1;XP_01 | 8923 | 606879 | . | . | [illegible] |
| 26229 | NM_00128872 | B3GAT3 | beta-1,3-glucuronyltransferase 3 | protein_coding | NP_001275650.1;NP_001275651.1;NP_001275652.1;NP_036332.2 | 923 | 606374 | . | . | [illegible] |
| 26230 | NM_00101092 | TIAM2 | TIAM Rac1 associated GEF 2 | protein_coding | NP_001010927.2;NP_036586.2 | 11806 | 604709 | . | . | [illegible] |
| 26231 | NM_00100405 | LRRC29 | leucine rich repeat containing 29 | protein_coding | NP_001004055.1;NP_001352087.1;NP_001352088.1;NP_036295.1;XP_01 | 13605 | . | | . | [illegible] |
| 26232 | NM_012168 | FBXO2 | F-box protein 2 | protein_coding | NP_036300.2 | 13581 | 607112 | . | . | [illegible] |
| 26233 | NM_012162,N | FBXL6 | F-box and leucine rich repeat protein 6 | protein_coding | NP_036294.2;NP_078831.4 | 13603 | 609076 | . | . | [illegible] |
| 26234 | NM_00119353 | FBXL5 | F-box and leucine rich repeat protein 5 | protein_coding | NP_001180463.1;NP_001180464.1;NP_036293.1;XP_006714022.1;XP_01 | 13602 | 605655 | . | . | [illegible] |
| 26235 | NM_00127871 | FBXL4 | F-box and leucine rich repeat protein 4 | protein_coding | NP_001265645.1;NP_036292.2;XP_005266987.1;XP_011534050.1;XP_01 | 13601 | 605654 | . | . | [illegible] |
| 26240 | NM_012135,XI | FAM50B | family with sequence similarity 50 member B | protein_coding | NP_036267.1;XP_016866218.1 | 18789 | 614686 | . | . | [illegible] |
| 26249 | NM_00125719 | KLHL3 | kelch like family member 3 | protein_coding | NP_001244123.1;NP_001244124.1;NP_059111.2 | 6354 | 605775 | . | . | [illegible] |
| 26257 | NM_014360 | NKX2-8 | NK2 homeobox 8 | protein_coding | NP_055175.2 | 16364 | 603245 | . | . | [illegible] |
| 26258 | NM_00131125 | BLOC1S6 | biogenesis of lysosomal organelles complex 1 subunit 6 | protein_coding | NP_001298184.1;NP_001298185.1;NP_036520.1 | 8549 | 604310 | . | . | [illegible] |
| 26259 | NM_012174,N | FBXW8 | F-box and WD repeat domain containing 8 | protein_coding | NP_036306.1;NP_699179.2;XP_005253924.1;XP_005253925.1;XP_01687 | 13597 | 609073 | . | . | [illegible] |
| 26260 | NM_012173,N | FBXO25 | F-box protein 25 | protein_coding | NP_036305.2;NP_904356.1;NP_904357.1;XP_011533050.1;XP_01153305 | 13596 | 609098 | . | . | [illegible] |
| 26261 | NM_00116349 | FBXO24 | F-box protein 24 | protein_coding | NP_001156971.1;NP_036304.2;NP_277041.1;XP_005250316.1;XP_01151 | 13595 | 609097 | . | . | [illegible] |
| 26262 | NM_00100661 | TSPAN17 | tetraspanin 17 | protein_coding | NP_001006617.2;NP_001353420.1;NP_001353421.1;NP_036303.1;NP_5 | 13594 | . | | . | [illegible] |
| 26263 | NM_012170,N | FBXO22 | F-box protein 22 | protein_coding | NP_036302.1;NP_671717.1;XP_011519749.1 | 13593 | 609096 | . | . | [illegible] |
| 26267 | NM_012166,XI | FBXO10 | F-box protein 10 | protein_coding | NP_036298.2;XP_005251496.1;XP_006716817.1;XP_016870108.1 | 13589 | 609092 | . | . | [illegible] |
| 26268 | NM_012347,N | FBXO9 | F-box protein 9 | protein_coding | NP_036479.1;NP_258441.1;NP_258442.2;XP_005249052.1;XP_00524905 | 13588 | 609091 | . | . | [illegible] |
| 26269 | NM_012180,XI | FBXO8 | F-box protein 8 | protein_coding | NP_036312.2;XP_016863509.1 | 13587 | 605649 | . | . | [illegible] |
| 26270 | NM_018438,XI | FBXO6 | F-box protein 6 | protein_coding | NP_060908.1;XP_005263505.1;XP_005263506.1;XP_005263508.1;XP_00 | 13585 | 605647 | . | . | [illegible] |
| 26271 | NM_00114252 | FBXO5 | F-box protein 5 | protein_coding | NP_001135994.1;NP_036309.1 | 13584 | 606013 | . | . | [illegible] |
| 26272 | NM_00129743 | FBXO4 | F-box protein 4 | protein_coding | NP_001284366.1;NP_036308.1;NP_277019.1;XP_006714533.1;XP_01151 | 13583 | 609090 | . | . | [illegible] |
| 26273 | NM_012175,N | FBXO3 | F-box protein 3 | protein_coding | NP_036307.2;NP_208385.1;XP_011518282.1;XP_011518283.1;XP_01151 | 13582 | 609089 | . | . | [illegible] |
| 26275 | NM_014362,N | HIBCH | 3-hydroxyisobutyryl-CoA hydrolase | protein_coding | NP_055177.2;NP_932164.1;XP_011509255.1;XP_011509256.1 | 4908 | 610690 | . | . | [illegible] |
| 26276 | NM_00128914 | VPS33B | VPS33B late endosome and lysosome associated | protein_coding | NP_001276077.1;NP_001276078.1;NP_061138.3;XP_005254944.1;XP_01 | 12712 | 608552 | . | . | [illegible] |
| 26277 | NM_00109927 | TINF2 | TERF1 interacting nuclear factor 2 | protein_coding | NP_001092744.1;NP_001350597.1;NP_036593.2;XP_011534944.1;XP_01 | 11824 | 604319 | . | . | [illegible] |
| 26278 | NM_00127805 | SACS | sacsin molecular chaperone | protein_coding | NP_001264984.1;NP_055178.3;XP_005266395.1;XP_011533341.1;XP_01 | 10519 | 604490 | . | . | [illegible] |
| 26280 | NM_017416,XI | IL1RAPL2 | interleukin 1 receptor accessory protein like 2 | protein_coding | NP_059112.1;XP_011529207.1 | 5997 | 300277 | . | . | [illegible] |
| 26284 | NM_00131798 | ERAL1 | Era like 12S mitochondrial rRNA chaperone 1 | protein_coding | NP_001304914.1;NP_001304915.1;NP_005693.1 | 3424 | 607435 | . | . | [illegible] |
| 26286 | NM_00114229 | ARFGAP3 | ADP ribosylation factor GTPase activating protein 3 | protein_coding | NP_001135765.1;NP_055385.3;XP_005261582.1 | 661 | 612439 | . | . | [illegible] |
| 26289 | NM_012093,N | AK5 | adenylate kinase 5 | protein_coding | NP_036225.2;NP_777283.1;XP_005270796.1;XP_006710635.1;XP_01685 | 365 | 608009 | . | . | [illegible] |
| 26292 | NM_012333,N | MYCBP | MYC binding protein | protein_coding | NP_036465.2 | 7554 | 606535 | . | . | [illegible] |
| 26297 | NM_012139,N | SERGEF | secretion regulating guanine nucleotide exchange factor | protein_coding | NP_036271.1 | 17499 | 606051 | . | . | [illegible] |
| 26330 | NM_014364 | GAPDHS | glyceraldehyde-3-phosphate dehydrogenase, spermatogenic | protein_coding | NP_055179.1 | 24864 | 609169 | . | . | [illegible] |
| 26353 | NM_014365 | HSPB8 | heat shock protein family B (small) member 8 | protein_coding | NP_055180.1 | 30171 | 608014 | . | . | [illegible] |
| 26354 | NM_014366,N | GNL3 | G protein nucleolar 3 | protein_coding | NP_055181.3;NP_996561.1;NP_996562.1 | 29931 | 608011 | . | . | [illegible] |
| 26355 | NM_014367 | FAM162A | family with sequence similarity 162 member A | protein_coding | NP_055182.3 | 17865 | 608017 | . | . | [illegible] |
| 26468 | NM_00124233 | LHX6 | LIM homeobox 6 | protein_coding | NP_001229262.1;NP_001229263.1;NP_001229264.1;NP_001335119.1;NI | 21735 | 608215 | . | . | [illegible] |
| 26469 | NM_00114237 | PTPN18 | protein tyrosine phosphatase non-receptor type 18 | protein_coding | NP_001135842.1;NP_055184.2;XP_006712479.1;XP_006712480.1;XP_01 | 9649 | 606587 | . | . | [illegible] |
| 26470 | NM_00111409 | SEZ6L2 | seizure related 6 homolog like 2 | protein_coding | NP_001107571.1;NP_001107572.1;NP_001230261.1;NP_001230262.1;NI | 30844 | 616667 | . | . | [illegible] |
| 26471 | NM_00104248 | NUPR1 | nuclear protein 1, transcriptional regulator | protein_coding | NP_001035948.1;NP_036517.1 | 29990 | 614812 | . | . | [illegible] |
| 26472 | NM_138689 | PPP1R14B | protein phosphatase 1 regulatory inhibitor subunit 14B | protein_coding | NP_619634.1 | 9057 | 601140 | . | . | [illegible] |
| 26502 | NM_00103861 | NARF | nuclear prelamin A recognition factor | protein_coding | NP_001033707.1;NP_001077077.1;NP_036468.1;NP_114174.1;XP_0052 | 29916 | 605349 | . | . | [illegible] |
| 26503 | NM_00138262 | SLC17A5 | solute carrier family 17 member 5 | protein_coding | NP_001369558.1;NP_001369559.1;NP_001369560.1;NP_001369561.1;NI | 10933 | 604322 | . | . | [illegible] |
| 26504 | NM_020184,XI | CNNM4 | cyclin and CBS domain divalent metal cation transport mediator 4 | protein_coding | NP_064569.3;XP_005263971.1;XP_005263972.1;XP_011509257.1;XP_01 | 105 | 607805 | . | . | [illegible] |
| 26505 | NM_017623,N | CNNM3 | cyclin and CBS domain divalent metal cation transport mediator 3 | protein_coding | NP_060093.3;NP_951060.1;XP_011509259.1;XP_016859289.1 | 104 | 607804 | . | . | [illegible] |
| 26509 | NM_013451,N | MYOF | myoferlin | protein_coding | NP_038479.1;NP_579899.1;XP_005269750.1;XP_005269751.1;XP_01153 | 3656 | 604603 | . | . | [illegible] |
| 26511 | NM_012110,XI | CHIC2 | cysteine rich hydrophobic domain 2 | protein_coding | NP_036242.1;XP_006714100.1;XP_011532684.1 | 1935 | 604332 | . | . | [illegible] |
| 26512 | NM_00103993 | INTS6 | integrator complex subunit 6 | protein_coding | NP_001035026.1;NP_001035027.1;NP_001293020.1;NP_036273.1;XP_01 | 14879 | 604331 | . | . | [illegible] |
| 26515 | NM_012192 | TIMM10B | translocase of inner mitochondrial membrane 10B | protein_coding | NP_036324.1 | 4022 | 607388 | . | . | [illegible] |
| 26517 | NM_012458 | TIMM13 | translocase of inner mitochondrial membrane 13 | protein_coding | NP_036590.1 | 11816 | 607383 | . | . | [illegible] |
| 26519 | NM_012456,XI | TIMM10 | translocase of inner mitochondrial membrane 10 | protein_coding | NP_036588.1;XP_024304204.1 | 11814 | 602251 | . | . | [illegible] |
| 26520 | NM_00130448 | TIMM9 | translocase of inner mitochondrial membrane 9 | protein_coding | NP_001291414.1;NP_001291415.1;NP_001291416.1;NP_001291417.1;NI | 11819 | 607384 | . | . | [illegible] |
| 26521 | NM_012459,N | TIMM8B | translocase of inner mitochondrial membrane 8 homolog B | protein_coding | NP_036591.3 | 11818 | 606659 | . | . | [illegible] |
| 26523 | NM_00131712 | AGO1 | argonaute RISC component 1 | protein_coding | NP_001304051.1;NP_001304052.1;NP_036331.1;XP_011539538.1;XP_01 | 3262 | 606228 | . | . | [illegible] |
| 26524 | NM_014572,XI | LATS2 | large tumor suppressor kinase 2 | protein_coding | NP_055387.2;XP_005266399.1;XP_011533344.1;XP_016876030.1;XP_01 | 6515 | 604861 | . | . | [illegible] |
| 26528 | NM_00135203 | DAZAP1 | DAZ associated protein 1 | protein_coding | NP_001338962.1;NP_001338963.1;NP_001338964.1;NP_061832.2;NP_7 | 2683 | 607430 | . | . | [illegible] |
| 26548 | NM_00130327 | ITGB1BP2 | integrin subunit beta 1 binding protein 2 | protein_coding | NP_001290206.1;NP_036410.1 | 6154 | 300332 | . | . | [illegible] |
| 26574 | NM_012138,XI | AATF | apoptosis antagonizing transcription factor | protein_coding | NP_036270.1;XP_011522913.1 | 19235 | 608463 | . | . | [illegible] |
| 26575 | NM_012419,XI | RGS17 | regulator of G protein signaling 17 | protein_coding | NP_036551.3;XP_016866222.1 | 14088 | 607191 | . | . | [illegible] |
| 26577 | NM_013363 | PCOLCE2 | procollagen C-endopeptidase enhancer 2 | protein_coding | NP_037495.1 | 8739 | 607064 | . | . | [illegible] |
| 26578 | NM_012383,XI | OSTF1 | osteoclast stimulating factor 1 | protein_coding | NP_036515.4;XP_006717116.1;XP_006717118.1;XP_011516826.1;XP_01 | 8510 | 610180 | . | . | [illegible] |
| 26579 | NM_00129329 | MYEOV | myeloma overexpressed | protein_coding | NP_001280220.1;NP_001280223.1;NP_001280225.1;NP_001287852.1;NI | 7563 | 605625 | . | . | [illegible] |
| 26580 | NM_00112295 | BSCL2 | BSCL2 lipid droplet biogenesis associated, seipin | protein_coding | NP_001116427.1;NP_001124174.2;NP_116056.3 | 15832 | 606158 | . | . | [illegible] |
| 26586 | NM_00109852 | CKAP2 | cytoskeleton associated protein 2 | protein_coding | NP_001091995.1;NP_001273615.1;NP_001273616.1;NP_060674.3;XP_00 | 1990 | 611569 | . | . | [illegible] |
| 26589 | NM_022163 | MRPL46 | mitochondrial ribosomal protein L46 | protein_coding | NP_071446.2 | 1192 | 611851 | . | . | [illegible] |
| 26608 | NM_00136266 | TBL2 | transducin beta like 2 | protein_coding | NP_001349589.1;NP_001349590.1;NP_001349591.1;NP_001349592.1;NI | 11586 | 605842 | . | . | [illegible] |
| 26609 | NM_013452,XI | VCX | variable charge X-linked | protein_coding | NP_038480.2;XP_011543792.1 | 12667 | 300229 | . | . | [illegible] |
| 26610 | NM_00128872 | ELP4 | elongator acetyltransferase complex subunit 4 | protein_coding | NP_001275654.1;NP_001275655.1;NP_061913.3 | 1171 | 606985 | . | . | [illegible] |
| 26683 | NM_00100522 | OR4F3 | olfactory receptor family 4 subfamily F member 3 | protein_coding | NP_001005224.1;XP_016864827.1 | 8300 | . | | . | [illegible] |
| 26740 | NM_054107,XI | OR1J2 | olfactory receptor family 1 subfamily J member 2 | protein_coding | NP_473448.1;XP_024303284.1;XP_024303285.1 | 8209 | . | | . | [illegible] |
| 26747 | NM_012345,XI | NUFIP1 | nuclear FMR1 interacting protein 1 | protein_coding | NP_036477.2 | 8057 | 604354 | . | . | [illegible] |
| 26750 | NM_00113613 | RPS6KC1 | ribosomal protein S6 kinase C1 | protein_coding | NP_001129610.1;NP_001274147.1;NP_001274148.1;NP_001274149.1;NI | 10439 | 617517 | . | . | [illegible] |
| 26751 | NM_00115959 | SH3YL1 | SH3 and SYLF domain containing 1 | protein_coding | NP_001153069.1;NP_001269611.1;NP_001269616.1;NP_056492.2 | 29546 | 617314 | . | . | [illegible] |
| 26765 | NR_002433 | SNORD12C | small nucleolar RNA, C/D box 12C | snoRNA | . | 10105 | . | | . | [illegible] |
| 26768 | NR_004404 | SNORA73B | small nucleolar RNA, H/ACA box 73B | snoRNA | . | 10116 | 603239 | . | . | [illegible] |
| 26769 | NR_003938 | SNORD81 | small nucleolar RNA, C/D box 81 | snoRNA | . | 10101 | . | | . | [illegible] |
| 26770 | NR_003939 | SNORD79 | small nucleolar RNA, C/D box 79 | snoRNA | . | 10100 | . | | . | [illegible] |
| 26771 | NR_002574 | SNORD102 | small nucleolar RNA, C/D box 102 | snoRNA | . | 10099 | . | | . | [illegible] |
| 26772 | NR_000009 | SNORD4B | small nucleolar RNA, C/D box 4B | snoRNA | . | 10098 | . | | . | [illegible] |
| 26773 | NR_000010 | SNORD4A | small nucleolar RNA, C/D box 4A | snoRNA | . | 10097 | . | | . | [illegible] |
| 26774 | NR_003940 | SNORD80 | small nucleolar RNA, C/D box 80 | snoRNA | . | 10096 | . | | . | [illegible] |
| 26775 | NR_002581 | SNORA72 | small nucleolar RNA, H/ACA box 72 | snoRNA | . | 10234 | . | | . | [illegible] |
| 26776 | NR_002910 | SNORA71B | small nucleolar RNA, H/ACA box 71B | snoRNA | . | 10233 | . | | . | [illegible] |
| 26778 | NR_000011 | SNORA70 | small nucleolar RNA, H/ACA box 70 | snoRNA | . | 10231 | . | | . | [illegible] |
| 26779 | NR_002584 | SNORA69 | small nucleolar RNA, H/ACA box 69 | snoRNA | . | 10226 | . | | . | [illegible] |
| 26780 | NR_000012 | SNORA68 | small nucleolar RNA, H/ACA box 68 | snoRNA | . | 10225 | . | | . | [illegible] |
| 26781 | NR_002912 | SNORA67 | small nucleolar RNA, H/ACA box 67 | snoRNA | . | 10224 | . | | . | [illegible] |
| 26782 | NR_002444 | SNORA66 | small nucleolar RNA, H/ACA box 66 | snoRNA | . | 10223 | . | | . | [illegible] |
| 26783 | NR_002449 | SNORA65 | small nucleolar RNA, H/ACA box 65 | snoRNA | . | 10222 | . | | . | [illegible] |
| 26784 | NR_002326 | SNORA64 | small nucleolar RNA, H/ACA box 64 | snoRNA | . | 10221 | . | | . | [illegible] |
| 26785 | NR_002913 | SNORD63 | small nucleolar RNA, C/D box 63 | snoRNA | . | 10220 | . | | . | [illegible] |
| 26788 | NR_002736 | SNORD60 | small nucleolar RNA, C/D box 60 | snoRNA | . | 10217 | . | | . | [illegible] |
| 26789 | NR_002737 | SNORD59A | small nucleolar RNA, C/D box 59A | snoRNA | . | 10210 | . | | . | [illegible] |
| 26790 | NR_002572 | SNORD58B | small nucleolar RNA, C/D box 58B | snoRNA | . | 10209 | . | | . | [illegible] |
| 26791 | NR_002571 | SNORD58A | small nucleolar RNA, C/D box 58A | snoRNA | . | 10208 | . | | . | [illegible] |
| 26792 | NR_002738 | SNORD57 | small nucleolar RNA, C/D box 57 | snoRNA | . | 10207 | . | | . | [illegible] |
| 26793 | NR_002739 | SNORD56 | small nucleolar RNA, C/D box 56 | snoRNA | . | 10206 | . | | . | [illegible] |
| 26795 | NR_002437 | SNORD54 | small nucleolar RNA, C/D box 54 | snoRNA | . | 10204 | . | | . | [illegible] |
| 26797 | NR_002742 | SNORD52 | small nucleolar RNA, C/D box 52 | snoRNA | . | 10202 | . | | . | [illegible] |
| 26798 | NR_002589 | SNORD51 | small nucleolar RNA, C/D box 51 | snoRNA | . | 10201 | . | | . | [illegible] |
| 26800 | NR_002744 | SNORD49A | small nucleolar RNA, C/D box 49A | snoRNA | . | 10189 | . | | . | [illegible] |
| 26801 | NR_002745 | SNORD48 | small nucleolar RNA, C/D box 48 | snoRNA | . | 10188 | . | | . | [illegible] |
| 26802 | NR_002746 | SNORD47 | small nucleolar RNA, C/D box 47 | snoRNA | . | 10187 | . | | . | [illegible] |
| 26804 | NR_002748 | SNORD45B | small nucleolar RNA, C/D box 45B | snoRNA | . | 10185 | . | | . | [illegible] |
| 26805 | NR_002749 | SNORD45A | small nucleolar RNA, C/D box 45A | snoRNA | . | 10184 | . | | . | [illegible] |
| 26806 | NR_002750 | SNORD44 | small nucleolar RNA, C/D box 44 | snoRNA | . | 10183 | . | | . | [illegible] |
| 26807 | NR_002439 | SNORD43 | small nucleolar RNA, C/D box 43 | snoRNA | . | 10182 | 611068 | . | . | [illegible] |
| 26808 | NR_000013 | SNORD42B | small nucleolar RNA, C/D box 42B | snoRNA | . | 10181 | . | | . | [illegible] |
| 26809 | NR_000014 | SNORD42A | small nucleolar RNA, C/D box 42A | snoRNA | . | 10180 | . | | . | [illegible] |
| 26811 | NR_000015 | SNORD55 | small nucleolar RNA, C/D box 55 | snoRNA | . | 10205 | . | | . | [illegible] |
| 26812 | NR_002602 | SNORD37 | small nucleolar RNA, C/D box 37 | snoRNA | . | 10166 | . | | . | [illegible] |
| 26813 | NR_000016 | SNORD36C | small nucleolar RNA, C/D box 36C | snoRNA | . | 10165 | . | | . | [illegible] |
| 26814 | NR_000017 | SNORD36B | small nucleolar RNA, C/D box 36B | snoRNA | . | 10164 | . | | . | [illegible] |
| 26815 | NR_002448 | SNORD36A | small nucleolar RNA, C/D box 36A | snoRNA | . | 10163 | . | | . | [illegible] |
| 26816 | NR_000018 | SNORD35A | small nucleolar RNA, C/D box 35A | snoRNA | . | 10162 | . | | . | [illegible] |
| 26817 | NR_000019 | SNORD34 | small nucleolar RNA, C/D box 34 | snoRNA | . | 10161 | . | | . | [illegible] |
| 26818 | NR_000020 | SNORD33 | small nucleolar RNA, C/D box 33 | snoRNA | . | 10160 | . | | . | [illegible] |
| 26819 | NR_000021 | SNORD32A | small nucleolar RNA, C/D box 32A | snoRNA | . | 10159 | . | | . | [illegible] |
| 26820 | NR_002447 | SNORD24 | small nucleolar RNA, C/D box 24 | snoRNA | . | 10146 | . | | . | [illegible] |
| 26821 | NR_002915 | SNORA74A | small nucleolar RNA, H/ACA box 74A | snoRNA | . | 10119 | . | | . | [illegible] |
| 26822 | NR_000022 | SNORD14A | small nucleolar RNA, C/D box 14A | snoRNA | . | 10113 | . | | . | [illegible] |
| 26827 | NR_004394 | RNU6-1 | RNA, U6 small nuclear 1 | snRNA | . | 10227 | 180692 | . | . | [illegible] |
| 26851 | NR_003271 | SNORD3B-1 | small nucleolar RNA, C/D box 3B-1 | snoRNA | . | 10168 | . | | . | [illegible] |
| 26863 | NR_004400 | RNVU1-18 | RNA, variant U1 small nuclear 18 | snRNA | . | 37496 | . | | . | [illegible] |
| 26864 | NR_004426 | RNVU1-7 | RNA, variant U1 small nuclear 7 | snRNA | . | 37500 | . | | . | [illegible] |
| 26869 | NR_004408 | RNU1-3 | RNA, U1 small nuclear 3 | snRNA | . | 10130 | . | | . | [illegible] |
| 26870 | NR_004427 | RNU1-2 | RNA, U1 small nuclear 2 | snRNA | . | 10123 | . | | . | [illegible] |
| 26871 | NR_004430 | RNU1-1 | RNA, U1 small nuclear 1 | snRNA | . | 10120 | . | | . | [illegible] |
| 26872 | NM_012449 | STEAP1 | STEAP family member 1 | protein_coding | NP_036581.1 | 11378 | 604415 | . | . | [illegible] |
| 26953 | NM_00124320 | RANBP6 | RAN binding protein 6 | protein_coding | NP_001230131.1;NP_001230132.1;NP_036548.1;XP_016870113.1 | 9851 | . | | . | [illegible] |
| 26958 | NM_00129003 | COPG2 | COPI coat complex subunit gamma 2 | protein_coding | NP_001276962.1;NP_036265.3 | 2237 | 604355 | . | . | [illegible] |
| 26959 | NM_00124426 | HBP1 | HMG-box transcription factor 1 | protein_coding | NP_001231191.1;NP_036389.2;XP_005250323.1;XP_005250324.1;XP_01 | 23200 | 616714 | . | . | [illegible] |
| 26960 | NM_00120419 | NBEA | neurobeachin | protein_coding | NP_001191126.1;NP_001366174.1;NP_056493.3;XP_005266403.1;XP_00 | 7648 | 604889 | . | . | [illegible] |
| 26973 | NM_00114407 | CHORDC1 | cysteine and histidine rich domain containing 1 | protein_coding | NP_001137545.1;NP_036256.2;XP_016873029.1;XP_016873030.1 | 14525 | 604353 | . | . | [illegible] |
| 26974 | NM_00129148 | ZNF285 | zinc finger protein 285 | protein_coding | NP_001278417.1;NP_001278418.1;NP_001278419.1;NP_001278420.1;NI | 13079 | . | | . | [illegible] |
| 26984 | NM_012430,XI | SEC22A | SEC22 homolog A, vesicle trafficking protein | protein_coding | NP_036562.2;XP_011510975.1;XP_011510977.1;XP_011510978.1 | 20260 | 612442 | . | . | [illegible] |
| 26985 | NM_00132026 | AP3M1 | adaptor related protein complex 3 subunit mu 1 | protein_coding | NP_001307192.1;NP_001307193.1;NP_001307194.1;NP_036227.1;NP_9 | 569 | 610366 | . | . | [illegible] |
| 26986 | NM_002568,XI | PABPC1 | poly(A) binding protein cytoplasmic 1 | protein_coding | NP_002559.2;XP_005250918.1 | 8554 | 604679 | . | . | [illegible] |
| 26993 | NM_00129147 | AKAP8L | A-kinase anchoring protein 8 like | protein_coding | NP_001278407.1;NP_055186.3 | 29857 | 609475 | . | . | [illegible] |
| 26994 | NM_014372 | RNF11 | ring finger protein 11 | protein_coding | NP_055187.1 | 10056 | 612598 | . | . | [illegible] |
| 26995 | NM_00132986 | TRUB2 | TruB pseudouridine synthase family member 2 | protein_coding | NP_001316790.1;NP_001316791.1;NP_001316792.1;NP_056494.1 | 17170 | 610727 | . | . | [illegible] |
| 26996 | NM_014373,XI | GPR160 | G protein-coupled receptor 160 | protein_coding | NP_055188.1;XP_005247403.1;XP_005247404.1;XP_016861650.1;XP_02 | 23693 | . | | . | [illegible] |
| 26999 | NM_00103733 | CYFIP2 | cytoplasmic FMR1 interacting protein 2 | protein_coding | NP_001032410.1;NP_001278650.1;NP_001278651.1;NP_055191.2;XP_01 | 13760 | 606323 | . | . | [illegible] |
| 27000 | NM_00112988 | DNAJC2 | DnaJ heat shock protein family (Hsp40) member C2 | protein_coding | NP_001123359.1;NP_001349596.1;NP_001349597.1;NP_055192.1;XP_01 | 13192 | 605502 | . | . | [illegible] |
| 27005 | NM_00101444 | USP21 | ubiquitin specific peptidase 21 | protein_coding | NP_001014443.1;NP_001306776.1;NP_001306777.1;NP_036607.3;XP_01 | 12620 | 604729 | . | . | [illegible] |
| 27006 | NM_00130081 | FGF22 | fibroblast growth factor 22 | protein_coding | NP_001287741.1;NP_065688.1 | 3679 | 605831 | . | . | [illegible] |
| 27010 | NM_00104248 | TPK1 | thiamin pyrophosphokinase 1 | protein_coding | NP_001035947.1;NP_001337808.1;NP_001337809.1;NP_001337810.1;NI | 17358 | 606370 | . | . | [illegible] |
| 27013 | NM_00132138 | CNPPD1 | cyclin Pas1/PHO80 domain containing 1 | protein_coding | NP_001308318.1;NP_001308319.2;NP_001308320.2;NP_056495.4;XP_02 | 25220 | . | | . | [illegible] |
| 27018 | NM_00128267 | BEX3 | brain expressed X-linked 3 | protein_coding | NP_001269603.1;NP_055195.1;NP_996798.1;NP_996800.1 | 13388 | 300361 | . | . | [illegible] |
| 27019 | NM_00128142 | DNAI1 | dynein axonemal intermediate chain 1 | protein_coding | NP_001268357.1;NP_036276.1 | 2954 | 604366 | . | . | [illegible] |
| 27020 | NM_00116136 | NPTN | neuroplastin | protein_coding | NP_001154835.1;NP_001154836.1;NP_036560.1;NP_059429.1;XP_01151 | 17867 | 612820 | . | . | [illegible] |
| 27022 | NM_012183 | FOXD3 | forkhead box D3 | protein_coding | NP_036315.1 | 3804 | 611539 | . | . | [illegible] |

| | | | | | | | | | | | | | | | | | | | | | | | | | | | | | | | | | |
|---|---|---|---|---|---|---|---|---|---|---|---|---|---|---|---|---|---|---|---|---|---|---|---|---|---|---|---|---|---|---|---|---|---|
| 27030 | NM_00104010 | MLH3 | mutL homolog 3 | protein_coding | NP_001035197.1;NP_055196.2;XP_005267589.1;XP_005267590.1;XP_00 | 7128 | 604395 | . | . | 0.716004 | 0.753260 | 0.921955 | 0.941917 | 1.666170 | 1.741194 | 2.068842 | 2.153582 | 1.215698 | 185.555217 | 0.244076995 | 1.15449089 | 0.248298971 | 0.992733415 | 7.387767 | 7.669873 | 7.503412 | 7.472823 | 7.572732 | 7.573607 | 200 | 174 | 177 | 191 |
| 27031 | NM_153240 | NPHP3 | nephrocystin 3 | protein_coding | NP_694972.3 | 7907 | 608002 | . | . | 3.401215 | 3.618442 | 3.877917 | 2.963653 | 7.914768 | 8.364191 | 8.701932 | 6.776040 | -1.039985 | 578.001296 | 0.179037768 | -0.31592309 | 0.752060883 | 0.998865404 | 9.202535 | 9.146784 | 9.166790 | 9.202979 | 9.245407 | 9.065471 | 660 | 653 | 561 | 458 |
| 27032 | NM_00100148 | ATP2C1 | ATPase secretory pathway Ca2+ transporting 1 | protein_coding | NP_001001485.1;NP_001001486.1;NP_001001487.1;NP_001186108.1;NI | 13211 | 604384 | . | . | 25.561165 | 24.997670 | 24.591641 | 19.895886 | 59.481886 | 57.783234 | 55.182929 | 45.489578 | -1.097078 | 3092.825727 | 0.132175829 | -1.01127535 | 0.311884659 | 0.998865404 | 11.659849 | 11.526490 | 11.646414 | 11.612951 | 11.642580 | 11.457073 | 3801 | 3421 | 2858 | 2450 |
| 27034 | NM_014384,XI | ACAD8 | acyl-CoA dehydrogenase family member 8 | protein_coding | NP_055199.1;XP_005271562.1;XP_011541052.1;XP_016873031.1;XP_01 | 87 | 604773 | . | . | 7.596445 | 7.976415 | 9.094989 | 7.658567 | 17.677241 | 18.437837 | 20.408892 | 17.510399 | 1.025916 | 705.081562 | 0.153365006 | 0.240680477 | 0.809802771 | 0.998865404 | 9.442832 | 9.480218 | 9.461201 | 9.442956 | 9.506653 | 9.431644 | 812 | 741 | 655 | 633 |
| 27037 | NM_00125799 | TRMT2A | tRNA methyltransferase 2 homolog A | protein_coding | NP_001244923.1;NP_001317968.1;NP_073564.3;NP_892029.2;XP_01152 | 24974 | 611151 | . | . | 26.374420 | 27.812359 | 29.108336 | 28.312691 | 61.374358 | 64.289513 | 65.318263 | 64.733605 | 1.049307 | 2448.638547 | 0.115365752 | 0.601888648 | 0.547248261 | 0.998865404 | 11.222617 | 11.292075 | 11.222286 | 11.253134 | 11.292671 | 11.260432 | 2699 | 2628 | 2224 | 2304 |
| 27040 | NM_00101498 | LAT | linker for activation of T cells | protein_coding | NP_001014987.1;NP_001014988.1;NP_001014989.2;NP_055202.1 | 18874 | 602354 | . | . | 2.340187 | 2.513933 | 3.190384 | 1.988565 | 5.445713 | 5.811071 | 7.159130 | 4.546618 | 1.042423 | 131.141610 | 0.310002781 | 0.193357765 | 0.84667879 | 0.998865404 | 7.003678 | 7.065621 | 7.011400 | 7.029937 | 7.114563 | 6.950202 | 144 | 142 | 143 | 97 |
| 27042 | NM_014388,XI | UTP25 | UTP25 small subunit processor component | protein_coding | NP_055203.4;XP_006711338.1;XP_011507720.1 | 28440 | . | . | . | 2.045582 | 1.940020 | 2.577368 | 2.651636 | 4.760153 | 4.484444 | 5.783540 | 6.062651 | 1.293420 | 608.628395 | 0.157600544 | 2.355264189 | **0.018509541** | 0.457491143 | 9.051837 | 9.423170 | 9.174138 | 9.134970 | 9.319979 | 9.331948 | 630 | 555 | 592 | 649 |
| 27043 | NM_00127824 | PELP1 | proline, glutamate and leucine rich protein 1 | protein_coding | NP_001265170.1;NP_055204.4 | 30134 | 609455 | . | . | 10.765207 | 11.588585 | 12.066827 | 12.231328 | 25.051078 | 26.787535 | 27.077610 | 27.965478 | 1.072238 | 1846.928962 | 0.121112061 | 0.83084109 | 0.406063406 | 0.998865404 | 10.799784 | 10.900291 | 10.796658 | 10.847665 | 10.876006 | 10.878909 | 1986 | 1986 | 1659 | 1795 |
| 27044 | NM_014390,XI | SND1 | staphylococcal nuclease and tudor domain containing 1 | protein_coding | NP_055205.2;XP_016867476.1 | 30646 | 602181 | . | . | 86.045710 | 85.717128 | 70.743291 | 86.058601 | 200.231902 | 198.138972 | 158.745891 | 196.762750 | -1.113383 | 8848.305621 | 0.117960204 | -1.31358323 | 0.188986495 | 0.960264078 | 13.186627 | 13.031581 | 13.159328 | 13.149020 | 12.981610 | 13.138054 | 10800 | 10000 | 6606 | 8578 |
| 27065 | NM_00104010 | NSG1 | neuronal vesicle trafficking associated 1 | protein_coding | NP_001035190.1;NP_001274692.1;NP_001274693.1;NP_001369156.1;NI | 18790 | 607645 | . | . | 1.158033 | 1.477432 | 1.697787 | 2.005459 | 2.694791 | 3.415150 | 3.809785 | 4.585243 | 1.408965 | 113.749997 | 0.311284441 | 1.589015929 | 0.112056797 | 0.849366267 | 6.562390 | 7.055178 | 6.724190 | 6.795154 | 6.845467 | 6.900371 | 96 | 114 | 107 | 134 |
| 27067 | NM_00116438 | STAU2 | staufen double-stranded RNA binding protein 2 | protein_coding | NP_001157852.1;NP_001157853.1;NP_001157854.1;NP_001157855.1;NI | 11371 | 605920 | . | . | 3.440853 | 3.259938 | 3.145933 | 2.892505 | 8.007004 | 7.535492 | 7.059382 | 6.613369 | -1.095423 | 366.599306 | 0.189262711 | -0.69473811 | 0.487219437 | 0.998865404 | 8.581995 | 8.451160 | 8.548578 | 8.538381 | 8.524893 | 8.456003 | 445 | 410 | 323 | 308 |
| 27068 | NM_006903,N | PPA2 | inorganic pyrophosphatase 2 | protein_coding | NP_008834.3;NP_789842.2;NP_789843.2;NP_789845.1 | 28883 | 609988 | . | . | 6.190088 | 6.063779 | 6.192189 | 5.064983 | 14.404591 | 14.016695 | 13.895093 | 11.580482 | -1.106313 | 304.710450 | 0.208019133 | -0.700701 | 0.483489631 | 0.998865404 | 8.321813 | 8.177153 | 8.283778 | 8.269802 | 8.275616 | 8.166222 | 373 | 341 | 278 | 243 |
| 27069 | NM_014394 | GHITM | growth hormone inducible transmembrane protein | protein_coding | NP_055209.2 | 17281 | . | . | . | 62.284958 | 60.410774 | 62.073380 | 63.129566 | 144.939651 | 139.642212 | 139.290863 | 144.338242 | 1.006024 | 4599.082050 | 0.106646897 | 0.081249294 | 0.935243698 | 0.998865404 | 12.162779 | 12.171469 | 12.180752 | 12.148121 | 12.166811 | 12.172359 | 5383 | 4851 | 3998 | 4339 |
| 27072 | NM_014396,N | VPS41 | VPS41 subunit of HOPS complex | protein_coding | NP_055211.2;NP_542198.2;XP_016867477.1 | 12713 | 605485 | . | . | 7.854935 | 8.295750 | 8.688736 | 8.919612 | 18.278759 | 19.175998 | 19.497271 | 20.393633 | 1.095434 | 1512.519423 | 0.125433823 | 1.048380907 | 0.294463147 | 0.998865404 | 10.495561 | 10.626925 | 10.501993 | 10.549527 | 10.593562 | 10.599435 | 1611 | 1606 | 1370 | 1488 |
| 27074 | NM_014398,XI | LAMP3 | lysosomal associated membrane protein 3 | protein_coding | NP_055213.2;XP_005247417.1;XP_006713649.1;XP_011510990.1;XP_02 | 14582 | 605883 | . | . | 0.696216 | 0.653803 | 0.551177 | 0.741297 | 1.620123 | 1.511294 | 1.236826 | 1.694886 | -1.044781 | 67.006641 | 0.39181793 | -0.16130185 | 0.871855669 | 0.998865404 | 6.098149 | 6.033593 | 6.080978 | 6.061969 | 6.029917 | 6.091925 | 82 | 71 | 49 | 70 |
| 27075 | NM_014399 | TSPAN13 | tetraspanin 13 | protein_coding | NP_055214.1 | 21643 | 613139 | . | . | 25.966612 | 21.375542 | 17.672663 | 19.016926 | 60.425377 | 49.410522 | 39.656944 | 43.479939 | -1.309725 | 1227.416595 | 0.145701174 | -2.671662 | **0.007547663** | 0.272411452 | 10.442895 | 10.053764 | 10.429211 | 10.270269 | 10.126498 | 10.171968 | 1765 | 1350 | 895 | 1028 |
| 27077 | NM_00124347 | B9D1 | B9 domain containing 1 | protein_coding | NP_001230402.1;NP_001308143.1;NP_001308144.1;NP_001308145.1;NI | 24123 | 614144 | . | . | 10.960524 | 12.048383 | 10.861342 | 13.352972 | 25.505591 | 27.850371 | 24.372539 | 30.529985 | 1.078211 | 537.586518 | 0.170661277 | 0.636576911 | 0.524400469 | 0.998865404 | 9.015287 | 9.123394 | 8.987608 | 9.097431 | 9.089812 | 9.095275 | 545 | 606 | 483 | 525 |
| 27079 | NM_00128640 | RPUSD2 | RNA pseudouridine synthase domain containing 2 | protein_coding | NP_001273336.1;NP_689473.1 | 24180 | . | . | . | 2.957656 | 2.678071 | 2.742437 | 3.000502 | 6.882585 | 6.190481 | 6.153949 | 6.860290 | 1.029327 | 207.531379 | 0.235111148 | 0.177367478 | 0.859219751 | 0.998865404 | 7.676180 | 7.717890 | 7.714168 | 7.665258 | 7.685446 | 7.720808 | 248 | 209 | 176 | 205 |
| 27085 | NM_022045,XI | MTBP | MDM2 binding protein | protein_coding | NP_071328.2;XP_011515264.1;XP_011515265.1 | 7417 | 605927 | . | . | 3.146150 | 3.031988 | 2.962661 | 2.224604 | 7.321222 | 7.008578 | 6.648124 | 5.086292 | -1.198350 | 272.563228 | 0.222975886 | -1.1707501 | 0.241699242 | 0.992733415 | 8.214382 | 7.954857 | 8.145952 | 8.120152 | 8.106239 | 7.968740 | 350 | 313 | 246 | 200 |
| 27086 | NM_00101250 | FOXP1 | forkhead box P1 | protein_coding | NP_001012523.1;NP_001231737.1;NP_001231739.1;NP_001231741.1;NI | 3823 | 605515 | . | . | 1.060328 | 1.916636 | 1.715563 | 1.182889 | 2.467427 | 4.430393 | 3.849672 | 2.704536 | -1.000075 | 252.777500 | 0.217423487 | -0.00049526 | 0.999604836 | 0.999862634 | 7.981707 | 7.981741 | 7.957935 | 8.003520 | 8.003665 | 7.957738 | 278 | 285 | 230 | 226 |
| 27087 | NM_00136797 | B3GAT1 | beta-1,3-glucuronyltransferase 1 | protein_coding | NP_001354902.1;NP_061114.2;NP_473366.1;XP_011541053.1;XP_01154 | 921 | 151290 | . | . | 3.744596 | 3.435197 | 4.061695 | 2.796592 | 8.713829 | 7.940613 | 9.114328 | 6.394076 | -1.044082 | 427.906196 | 0.20405338 | -0.30499207 | 0.760372195 | 0.998865404 | 8.771218 | 8.710443 | 8.777708 | 8.721420 | 8.823421 | 8.613710 | 526 | 450 | 428 | 324 |
| 27089 | NM_014402 | UQCRQ | ubiquinol-cytochrome c reductase complex III subunit VII | protein_coding | NP_055217.2 | 29594 | 612080 | . | . | 52.459816 | 49.869350 | 45.609432 | 59.841602 | 122.076141 | 115.275238 | 102.346245 | 136.820709 | 1.015764 | 2544.878343 | 0.13998197 | 0.161202521 | 0.871933896 | 0.998865404 | 11.302190 | 11.324486 | 11.328637 | 11.280610 | 11.206603 | 11.417525 | 2994 | 2644 | 1940 | 2717 |
| 27090 | NM_175039,N | ST6GALNAC4 | ST6 N-acetylgalactosaminide alpha-2,6-sialyltransferase 4 | protein_coding | NP_778204.1;NP_778205.1;XP_011516829.1;XP_011516831.1;XP_02430 | 17846 | 606378 | . | . | 6.166068 | 6.563100 | 7.387566 | 7.951365 | 14.348697 | 15.170900 | 16.577482 | 18.179853 | 1.198647 | 377.830692 | 0.185037929 | 1.412717404 | 0.157738832 | 0.928077196 | 8.425212 | 8.686193 | 8.486990 | 8.521008 | 8.595552 | 8.623308 | 384 | 382 | 350 | 395 |
| 27094 | NM_00116367 | KCNMB3 | potassium calcium-activated channel subfamily M regulatory beta | protein_coding | NP_001157149.1;NP_055222.3;NP_741979.1;NP_741980.1;NP_741981.1 | 6287 | 605222 | . | . | 0.416496 | 0.440067 | 0.522539 | 0.525963 | 0.969202 | 1.017235 | 1.172561 | 1.202551 | 1.232336 | 31.050015 | 0.553398379 | 0.54462601 | 0.586010762 | 0.998865404 | 4.798330 | 5.099069 | 4.934911 | 4.942661 | 4.961117 | 4.968546 | 31 | 31 | 29 | 33 |
| 27095 | NM_00127089 | TRAPPC3 | trafficking protein particle complex 3 | protein_coding | NP_001257823.1;NP_001257824.1;NP_001257825.1;NP_001257826.1;NI | 19942 | 610955 | . | . | 37.584654 | 37.033600 | 37.682827 | 40.677531 | 87.461036 | 85.604827 | 84.559163 | 93.004340 | 1.034879 | 1570.856449 | 0.125467978 | 0.394217411 | 0.693420532 | 0.998865404 | 10.592446 | 10.641803 | 10.617989 | 10.589109 | 10.602798 | 10.656963 | 1811 | 1635 | 1341 | 1550 |
| 27097 | NM_00102524 | TAF5L | TATA-box binding protein associated factor 5 like | protein_coding | NP_001020418.1;NP_055224.1;XP_005273156.1;XP_005273157.1;XP_01 | 17304 | . | . | . | 7.187419 | 7.002390 | 7.432879 | 7.773742 | 16.725423 | 16.186339 | 16.679168 | 17.773738 | 1.065080 | 797.569030 | 0.14487957 | 0.627847933 | 0.530103563 | 0.998865404 | 9.593230 | 9.684265 | 9.630213 | 9.602556 | 9.657486 | 9.664363 | 907 | 817 | 712 | 775 |
| 27099 | NR_027330 | SND1-IT1 | SND1 intronic transcript 1 | lncRNA | . | 24158 | . | . | . | 0.010101 | 0.027329 | 0.014111 | 0.058971 | 0.023506 | 0.063172 | 0.031665 | 0.134831 | 1.793279 | 2.532997 | 1.944663636 | 0.433288302 | 0.664805335 | NA | 0.867373 | 1.696731 | 1.321350 | 1.330587 | 1.323204 | 1.342131 | 1 | 3 | 1 | 5 |
| 27101 | NM_00100721 | CACYBP | calcyclin binding protein | protein_coding | NP_001007215.1;NP_055227.1;XP_016856535.1 | 30423 | 606186 | . | . | 41.131867 | 38.895059 | 33.978624 | 35.023575 | 95.715550 | 89.907674 | 76.247053 | 80.077239 | -1.175712 | 3178.096456 | 0.111486002 | -2.09474221 | **0.036193914** | 0.593055768 | 11.746007 | 11.512445 | 11.711873 | 11.681122 | 11.548632 | 11.579152 | 4028 | 3638 | 2492 | 2791 |
| 27102 | NM_00113433 | EIF2AK1 | eukaryotic translation initiation factor 2 alpha kinase 1 | protein_coding | NP_001127807.1;NP_055228.2;XP_024302486.1 | 24921 | 613635 | . | . | 23.252923 | 23.691043 | 24.715341 | 26.091158 | 54.110503 | 54.762908 | 55.460514 | 59.654335 | 1.021914 | 3249.494479 | 0.111975163 | 0.279293906 | 0.780019283 | 0.998865404 | 11.650341 | 11.681490 | 11.643197 | 11.670421 | 11.650420 | 11.698327 | 3640 | 3526 | 2773 | 3169 |
| 27106 | NM_00102560 | ARRDC2 | arrestin domain containing 2 | protein_coding | NP_001020775.1;NP_001273755.1;NP_056498.1 | 25225 | . | . | . | 3.495993 | 3.391733 | 3.798952 | 4.296713 | 8.135320 | 7.840145 | 8.524737 | 9.823926 | 1.156516 | 296.084883 | 0.20413702 | 1.027667313 | 0.304106349 | 0.998865404 | 8.101353 | 8.310787 | 8.178651 | 8.156561 | 8.214770 | 8.275041 | 323 | 290 | 261 | 314 |
| 27107 | NM_014415,XI | ZBTB11 | zinc finger and BTB domain containing 11 | protein_coding | NP_055230.2;XP_011510991.1;XP_011510992.1 | 16740 | 618181 | . | . | 2.724970 | 2.721188 | 2.774392 | 2.574772 | 6.341117 | 6.290148 | 6.225655 | 5.886911 | -1.030784 | 482.261262 | 0.170616921 | -0.25637758 | 0.797659302 | 0.998865404 | 8.935174 | 8.891843 | 8.925557 | 8.920225 | 8.930505 | 8.876707 | 566 | 526 | 431 | 426 |
| 27109 | NM_00100380 | DMAC2L | distal membrane arm assembly complex 2 like | protein_coding | NP_001003803.2;NP_001003805.2;NP_001357534.1;NP_001369436.1;NI | 18799 | 618579 | . | . | 4.626184 | 3.599570 | 3.625920 | 3.510248 | 10.765319 | 8.320567 | 8.136456 | 8.025764 | -1.154479 | 338.954387 | 0.202502339 | -1.02340605 | 0.3061159 | 0.998865404 | 8.504603 | 8.297894 | 8.505104 | 8.376624 | 8.355236 | 8.369329 | 464 | 349 | 270 | 299 |
| 27112 | NM_015686,XI | FAM155B | family with sequence similarity 155 member B | protein_coding | NP_056501.2;XP_011529210.1;XP_011529211.1 | 30701 | . | . | . | 0.709319 | 0.642839 | 1.069141 | 0.740350 | 1.650613 | 1.485952 | 2.399121 | 1.692721 | 1.261672 | 114.598684 | 0.309311328 | 1.084140347 | 0.278302576 | 0.998865404 | 6.662238 | 6.999042 | 6.796705 | 6.789740 | 6.915959 | 6.822782 | 118 | 108 | 126 | 104 |
| 27113 | NM_00112724 | BBC3 | BCL2 binding component 3 | protein_coding | NP_001120712.1;NP_001120713.1;NP_001120714.1;NP_055232.1;XP_00 | 17868 | 605854 | . | . | 2.448787 | 2.193535 | 1.492717 | 2.703657 | 5.698429 | 5.070455 | 3.349615 | 6.181592 | -1.080523 | 125.995262 | 0.330056071 | -0.33851703 | 0.734973592 | 0.998865404 | 7.032962 | 6.919249 | 7.019676 | 6.950220 | 6.859932 | 7.047817 | 165 | 128 | 76 | 145 |
| 27115 | NM_018945 | PDE7B | phosphodiesterase 7B | protein_coding | NP_061818.1 | 8792 | 604645 | . | . | 0.326985 | 0.270573 | 0.255717 | 0.247336 | 0.760907 | 0.625440 | 0.573822 | 0.565505 | -1.195739 | 46.804142 | 0.45838493 | -0.56263363 | 0.573684385 | 0.998865404 | 5.671258 | 5.414458 | 5.583169 | 5.548146 | 5.538611 | 5.530537 | 64 | 50 | 38 | 39 |
| 27124 | NM_00100283 | INPP5J | inositol polyphosphate-5-phosphatase J | protein_coding | NP_001002837.1;NP_001271214.1;NP_001271215.1;NP_001271216.1;NI | 8956 | 606481 | . | . | 0.156549 | 0.101598 | 0.121957 | 0.167127 | 0.364295 | 0.234849 | 0.273669 | 0.382116 | 1.085496 | 15.337389 | 0.788010989 | 0.150193811 | 0.880611709 | NA | 3.878555 | 3.996978 | 3.947199 | 3.923474 | 3.931674 | 3.949984 | 20 | 13 | 12 | 17 |
| 27125 | NM_014423,XI | AFF4 | AF4/FMR2 family member 4 | protein_coding | NP_055238.1;XP_005272020.1;XP_005272021.1;XP_006714650.1 | 17869 | 604417 | . | . | 11.977777 | 11.949982 | 13.001960 | 11.239274 | 27.872780 | 27.622923 | 29.176024 | 25.697264 | -1.002120 | 3557.780175 | 0.119940552 | -0.02547859 | 0.979673225 | 0.998865404 | 11.798189 | 11.795333 | 11.801283 | 11.792330 | 11.857507 | 11.728766 | 4123 | 3822 | 3332 | 3072 |
| 27127 | NM_00129150 | SMC1B | structural maintenance of chromosomes 1B | protein_coding | NP_001278430.1;NP_683515.4;XP_011528446.1;XP_011528447.1 | 11112 | 608685 | . | . | 1.078999 | 0.942064 | 0.694479 | 1.037704 | 2.510875 | 2.177624 | 1.558391 | 2.372587 | -1.162614 | 120.158033 | 0.311647968 | -0.69749108 | 0.485495517 | 0.998865404 | 7.014025 | 6.795269 | 6.960346 | 6.905302 | 6.825611 | 6.934782 | 160 | 129 | 80 | 122 |
| 27130 | NM_00131838 | INVS | inversin | protein_coding | NP_001305310.1;NP_001305311.1;NP_055240.2 | 17870 | 243305 | . | . | 3.780104 | 4.074527 | 4.007426 | 5.153592 | 8.796458 | 9.418453 | 8.992547 | 11.783076 | 1.138234 | 654.745200 | 0.170028614 | 1.098624923 | 0.271931702 | 0.998865404 | 9.258853 | 9.444742 | 9.284127 | 9.328596 | 9.307641 | 9.473897 | 682 | 683 | 533 | 732 |
| 27131 | NM_00128245 | SNX5 | sorting nexin 5 | protein_coding | NP_001269383.1;NP_055241.1;NP_689413.1 | 14969 | 605937 | . | . | 30.805301 | 31.629904 | 29.370007 | 27.178478 | 71.685203 | 73.113945 | 65.905449 | 62.140359 | -1.137964 | 1993.351249 | 0.119433125 | -1.56116402 | 0.118485056 | 0.858706505 | 11.051140 | 10.864808 | 11.004993 | 11.015069 | 10.930999 | 10.883342 | 2428 | 2304 | 1671 | 1695 |
| 27132 | NM_014427,N | CPNE7 | copine 7 | protein_coding | NP_055242.1;NP_705900.1;XP_011521302.1;XP_011521303.1;XP_01687 | 2320 | 605689 | . | . | 0.080261 | 0.056585 | 0.211296 | 0.080745 | 0.186770 | 0.130800 | 0.474140 | 0.184613 | 1.881186 | 11.584904 | 0.930494046 | 0.979740136 | 0.327214409 | NA | 3.003609 | 3.921224 | 3.503034 | 3.489019 | 3.550747 | 3.503679 | 11 | 7 | 18 | 9 |
| 27143 | NM_014431,XI | PALD1 | phosphatase domain containing paladin 1 | protein_coding | NP_055246.2;XP_011537939.1;XP_011537940.1;XP_016871561.1 | 23530 | 614656 | . | . | 0.381230 | 0.413821 | 0.408406 | 0.396310 | 0.887138 | 0.956565 | 0.916450 | 0.906116 | -1.041320 | 69.033286 | 0.379564991 | -0.15389636 | 0.877691454 | 0.998865404 | 6.137859 | 6.080001 | 6.110714 | 6.117205 | 6.116659 | 6.092026 | 80 | 77 | 62 | 60 |
| 27148 | NM_00124331 | STK36 | serine/threonine kinase 36 | protein_coding | NP_001230242.1;NP_001356352.1;NP_056505.2;XP_011509261.1;XP_01 | 17209 | 607652 | . | . | 15.521627 | 16.984047 | 22.870982 | 18.900822 | 36.119465 | 39.259383 | 51.321821 | 43.214479 | 1.272242 | 2681.739474 | 0.133147904 | 2.608923865 | **0.009082745** | 0.30269315 | 11.204733 | 11.552291 | 11.254268 | 11.306798 | 11.562657 | 11.382523 | 2628 | 2631 | 2894 | 2511 |
| 27152 | NM_015693,XI | INTU | inturned planar cell polarity protein | protein_coding | NP_056508.2;XP_011530146.1;XP_011530147.1;XP_011530148.1;XP_01 | 29239 | 610621 | . | . | 0.831577 | 1.490919 | 0.991256 | 1.444484 | 1.935114 | 3.446325 | 2.224348 | 3.302641 | 1.083068 | 304.655976 | 0.199863776 | 0.57601109 | 0.564607686 | 0.998865404 | 8.192287 | 8.307488 | 8.217048 | 8.238855 | 8.283816 | 8.258282 | 330 | 322 | 282 | 290 |
| 27153 | NM_015694,XI | ZNF777 | zinc finger protein 777 | protein_coding | NP_056509.2;XP_005250037.1;XP_011514357.1 | 22213 | . | . | . | 2.540082 | 3.098997 | 3.295200 | 3.285520 | 5.910876 | 7.163470 | 7.394333 | 7.511952 | 1.147887 | 299.383394 | 0.208296119 | 0.955278987 | 0.339436631 | 0.998865404 | 8.123278 | 8.321612 | 8.132321 | 8.234397 | 8.261984 | 8.255221 | 290 | 330 | 280 | 298 |
| 27154 | NM_015695,XI | BRPF3 | bromodomain and PHD finger containing 3 | protein_coding | NP_056510.2;XP_005249067.1;XP_005249068.1;XP_011512791.1;XP_01 | 14256 | 616856 | . | . | 4.272132 | 4.333658 | 4.667369 | 4.551635 | 9.941426 | 10.017445 | 10.473439 | 10.406774 | 1.054440 | 818.079082 | 0.143477363 | 0.533023023 | 0.594017643 | 0.998865404 | 9.637336 | 9.713843 | 9.652048 | 9.661399 | 9.700786 | 9.687729 | 911 | 865 | 737 | 780 |
| 27158 | NM_00114402 | NDOR1 | NADPH dependent diflavin oxidoreductase 1 | protein_coding | NP_001137498.1;NP_001137499.1;NP_001137500.1;NP_055249.1;XP_01 | 29838 | 606073 | . | . | 7.889656 | 8.970856 | 9.280817 | 7.749312 | 18.359553 | 20.736538 | 20.825885 | 17.717881 | -1.014419 | 1335.210678 | 0.145932976 | -0.14152382 | 0.887456142 | 0.998865404 | 10.393028 | 10.372603 | 10.335471 | 10.434980 | 10.446880 | 10.298944 | 1448 | 1545 | 1265 | 1122 |
| 27161 | NM_00116462 | AGO2 | argonaute RISC catalytic component 2 | protein_coding | NP_001158095.1;NP_036286.2;XP_011515267.1;XP_011515268.1;XP_01 | 3263 | 606229 | . | . | 2.344398 | 2.614737 | 2.680700 | 2.626363 | 5.455509 | 6.044084 | 6.015414 | 6.004868 | 1.054888 | 1178.237132 | 0.136933743 | 0.562972759 | 0.573453434 | 0.998865404 | 10.163414 | 10.240388 | 10.135137 | 10.225374 | 10.239099 | 10.203221 | 1241 | 1312 | 1077 | 1107 |
| 27163 | NM_00104240 | NAAA | N-acylethanolamine acid amidase | protein_coding | NP_001035861.1;NP_001350648.1;NP_055250.2;XP_005262977.1;XP_00 | 736 | 607469 | . | . | 10.150620 | 9.691811 | 10.011098 | 9.607240 | 23.620907 | 22.403052 | 22.464617 | 21.965813 | -1.043803 | 608.158870 | 0.156745236 | -0.39458768 | 0.693147207 | 0.998865404 | 9.278816 | 9.217134 | 9.269807 | 9.255138 | 9.242199 | 9.225748 | 723 | 663 | 524 | 551 |
| 27166 | NM_00127182 | PRELID1 | PRELI domain containing 1 | protein_coding | NP_001258757.1;NP_037369.1 | 30255 | 605733 | . | . | 180.293346 | 187.745578 | 165.740992 | 204.505765 | 419.550013 | 433.982313 | 371.917948 | 467.578114 | -1.009096 | 7036.079588 | 0.121431725 | -0.10758039 | 0.914328556 | 0.998865404 | 12.787140 | 12.773942 | 12.769871 | 12.796323 | 12.687702 | 12.856347 | 8012 | 7746 | 5484 | 7220 |
| 27173 | NM_00127195 | SLC39A1 | solute carrier family 39 member 1 | protein_coding | NP_001258886.1;NP_001258887.1;NP_001258888.1;NP_001258889.1;NI | 12876 | 604740 | . | . | 81.960141 | 83.046461 | 76.760317 | 80.398932 | 190.724615 | 191.965624 | 172.247921 | 183.822601 | -1.050130 | 5523.673266 | 0.105932287 | -0.66615489 | 0.505312103 | 0.998865404 | 12.466298 | 12.395662 | 12.438506 | 12.464443 | 12.390710 | 12.429816 | 6416 | 6200 | 4576 | 5170 |
| 27175 | NM_00132050 | TUBG2 | tubulin gamma 2 | protein_coding | NP_001307438.1;NP_057521.1;XP_011522916.1;XP_016879942.1;XP_01 | 12419 | 605785 | . | . | 16.426293 | 16.192171 | 18.259780 | 17.472820 | 38.224655 | 37.428926 | 40.974414 | 39.949526 | 1.139565 | 1658.480142 | 0.133576406 | 1.411052007 | 0.15822928 | 0.929304099 | 10.598213 | 10.786913 | 10.615042 | 10.666127 | 10.805277 | 10.676822 | 1726 | 1728 | 1661 | 1522 |
| 27183 | NM_013245 | VPS4A | vacuolar protein sorting 4 homolog A | protein_coding | NP_037377.1 | 13488 | 609982 | . | . | 13.033557 | 13.611834 | 12.200280 | 14.578896 | 30.329622 | 31.464363 | 27.377075 | 33.332912 | -1.008859 | 1708.479390 | 0.132394704 | -0.09611552 | 0.923428826 | 0.998865404 | 10.745045 | 10.731920 | 10.727457 | 10.754711 | 10.666568 | 10.797524 | 1942 | 1884 | 1355 | 1728 |
| 27185 | NM_00101295 | DISC1 | DISC1 scaffold protein | protein_coding | NP_001012975.1;NP_001012976.1;NP_001012977.1;NP_001158009.1;NI | 2888 | 605210 | . | . | 1.002136 | 0.793637 | 1.011653 | 1.285534 | 2.332010 | 1.834527 | 2.270120 | 2.939221 | 1.798907 | 174.063490 | 0.273667078 | 3.095441591 | **0.001965201** | 0.111170762 | 6.957636 | 7.806266 | 7.353173 | 7.217115 | 7.514868 | 7.549108 | 171 | 108 | 188 | 217 |
| 27190 | NM_00131798 | IL17B | interleukin 17B | protein_coding | NP_001304916.1;NP_055258.1;XP_011535924.1;XP_016864835.1;XP_01 | 5982 | 604627 | . | . | 40.999172 | 36.386812 | 19.447514 | 23.098017 | 95.406750 | 84.109744 | 43.639657 | 52.810867 | -1.841878 | 668.131458 | 0.164586446 | -5.35388948 | **8.60834E-08** | **4.57004E-05** | 9.758489 | 8.876835 | 9.594831 | 9.499069 | 9.089946 | 9.181512 | 1061 | 875 | 377 | 475 |
| 27229 | NM_00128641 | TUBGCP4 | tubulin gamma complex associated protein 4 | protein_coding | NP_001273343.1;NP_055259.2;XP_011519756.1;XP_011519757.1;XP_01 | 16691 | 609610 | . | . | 2.142826 | 2.279275 | 2.436938 | 2.126263 | 4.986446 | 5.268644 | 5.468416 | 4.861448 | 1.078636 | 451.890374 | 0.175412035 | 0.622580745 | 0.533560068 | 0.998865404 | 8.764052 | 8.873530 | 8.780526 | 8.809026 | 8.871648 | 8.810534 | 489 | 480 | 428 | 418 |
| 27230 | NM_014445 | SERP1 | stress associated endoplasmic reticulum protein 1 | protein_coding | NP_055260.1 | 10759 | 617674 | . | . | 24.576651 | 22.431900 | 21.527164 | 17.957409 | 57.190876 | 51.852337 | 48.306332 | 41.057480 | -1.207482 | 2037.817123 | 0.135200838 | -2.01183405 | **0.044237434** | 0.645513836 | 11.122201 | 10.850658 | 11.101646 | 11.020100 | 10.983423 | 10.834779 | 2695 | 2285 | 1759 | 1566 |
| 27235 | NM_00135892 | COQ2 | coenzyme Q2, polyprenyltransferase | protein_coding | NP_001345850.1;NP_056512.5 | 25223 | 609825 | . | . | 10.512619 | 10.888531 | 8.400873 | 8.529977 | 24.463294 | 25.169328 | 18.851316 | 19.502779 | -1.201874 | 534.237770 | 0.166500428 | -1.59330066 | 0.111092759 | 0.848126978 | 9.187758 | 8.922767 | 9.152324 | 9.081185 | 8.999150 | 8.998845 | 707 | 596 | 422 | 455 |
| 27236 | NM_00102559 | ARFIP1 | ADP ribosylation factor interacting protein 1 | protein_coding | NP_001020764.1;NP_001020766.1;NP_001274360.1;NP_001274361.1;NI | 21496 | 605928 | . | . | 6.119015 | 5.521367 | 5.578130 | 5.846265 | 14.239200 | 12.762886 | 12.517162 | 13.366793 | 1.012040 | 580.737807 | 0.163001379 | 0.10592533 | 0.915641597 | 0.998865404 | 9.172924 | 9.190509 | 9.216701 | 9.136332 | 9.182564 | 9.187949 | 705 | 585 | 506 | 550 |
| 27237 | NM_014448,XI | ARHGEF16 | Rho guanine nucleotide exchange factor 16 | protein_coding | NP_055263.2;XP_016856538.1;XP_016856539.1;XP_016856540.1;XP_02 | 15515 | 618871 | . | . | 0.477724 | 0.657951 | 1.023547 | 0.973572 | 1.111683 | 1.520881 | 2.296812 | 2.225957 | 1.533460 | 72.803727 | 0.371793098 | 1.658962715 | 0.097123306 | 0.813553488 | 5.844866 | 6.461599 | 6.108424 | 6.129284 | 6.234741 | 6.211077 | 63 | 65 | 80 | 79 |
| 27238 | NM_015698,XI | GPKOW | G-patch domain and KOW motifs | protein_coding | NP_056513.2;XP_016884904.1 | 30677 | 301003 | . | . | 9.143818 | 8.963920 | 8.680771 | 11.044821 | 21.278042 | 20.720502 | 19.479401 | 25.252668 | 1.071614 | 489.586955 | 0.180858941 | 0.551731549 | 0.581132292 | 0.998865404 | 8.885072 | 8.984072 | 8.918894 | 8.903816 | 8.879637 | 9.024936 | 551 | 504 | 390 | 529 |
| 27241 | NM_00103360 | BBS9 | Bardet-Biedl syndrome 9 | protein_coding | NP_001028776.1;NP_001028777.1;NP_001334965.1;NP_001334966.1;NI | 30000 | 607968 | . | . | 2.913771 | 2.393078 | 3.157461 | 2.998844 | 6.780467 | 5.531706 | 7.085250 | 6.856496 | 1.144523 | 357.036457 | 0.198705374 | 0.98007875 | 0.327047248 | 0.998865404 | 8.378738 | 8.574483 | 8.494840 | 8.375739 | 8.534340 | 8.494134 | 424 | 321 | 343 | 345 |
| 27242 | NM_014452,XI | TNFRSF21 | TNF receptor superfamily member 21 | protein_coding | NP_055267.1;XP_016866233.1 | 13469 | 605732 | . | . | 10.647327 | 11.130463 | 9.405485 | 10.926584 | 24.776767 | 25.728564 | 21.105637 | 24.982337 | -1.089073 | 1166.768972 | 0.137388308 | -0.89600486 | 0.370250165 | 0.998865404 | 10.248757 | 10.125205 | 10.202738 | 10.235271 | 10.107555 | 10.200449 | 1371 | 1341 | 910 | 1113 |
| 27243 | NM_014453,N | CHMP2A | charged multivesicular body protein 2A | protein_coding | NP_055268.1;NP_940818.1;XP_005258803.1;XP_005258804.1 | 30216 | 610893 | . | . | 60.242179 | 60.314411 | 57.499109 | 68.252757 | 140.186023 | 139.419461 | 129.026328 | 156.051822 | 1.015794 | 1823.759415 | 0.131548301 | 0.171862066 | 0.86354597 | 0.998865404 | 10.821518 | 10.843796 | 10.828343 | 10.822916 | 10.766937 | 10.904212 | 2091 | 1946 | 1459 | 1873 |
| 27244 | NM_00119993 | SESN1 | sestrin 1 | protein_coding | NP_001186862.1;NP_001186863.1;NP_055269.1 | 21595 | 606103 | . | . | 2.639228 | 2.222025 | 2.254078 | 2.595563 | 6.141591 | 5.136311 | 5.058085 | 5.934445 | 1.065045 | 269.638986 | 0.216771945 | 0.419398399 | 0.674924994 | 0.998865404 | 8.028170 | 8.120135 | 8.102273 | 8.008718 | 8.110384 | 8.067509 | 328 | 256 | 252 | 250 |
| 27245 | NM_00102988 | AHDC1 | AT-hook DNA binding motif containing 1 | protein_coding | NP_001025053.1;NP_001358857.1;XP_005245905.1;XP_005245906.1;XP | 25230 | 615790 | . | . | 2.884667 | 3.063657 | 3.697429 | 3.421970 | 6.712735 | 7.081775 | 8.296923 | 7.823928 | 1.156590 | 674.807121 | 0.15553193 | 1.349421342 | 0.17720167 | 0.949886621 | 9.289552 | 9.499481 | 9.317190 | 9.372252 | 9.471778 | 9.415687 | 691 | 703 | 655 | 651 |
| 27246 | NM_014455,XI | RNF115 | ring finger protein 115 | protein_coding | NP_055270.1;XP_005273009.1 | 18154 | . | . | . | 2.651642 | 3.054136 | 3.115087 | 4.222175 | 6.170481 | 7.059773 | 6.990164 | 9.653501 | 1.100977 | 752.500635 | 0.147517369 | 0.940796472 | 0.346809169 | 0.998865404 | 9.484608 | 9.623165 | 9.507869 | 9.532927 | 9.575224 | 9.599492 | 810 | 787 | 674 | 752 |
| 27247 | NM_00100275 | NFU1 | NFU1 iron-sulfur cluster scaffold | protein_coding | NP_001002755.1;NP_001002756.1;NP_001361213.1;NP_056515.2 | 16287 | 608100 | . | . | 33.586052 | 34.246134 | 29.180449 | 37.203711 | 78.156122 | 79.161471 | 65.480080 | 85.061856 | -1.041015 | 971.676877 | 0.155017217 | -0.37409032 | 0.708337118 | 0.998865404 | 9.953366 | 9.894703 | 9.933102 | 9.941609 | 9.816954 | 9.991359 | 1135 | 1076 | 728 | 1000 |
| 27248 | NM_00112739 | ERLEC1 | endoplasmic reticulum lectin 1 | protein_coding | NP_001120869.1;NP_001120870.1;NP_056516.2;XP_011531068.1;XP_02 | 25222 | 611229 | . | . | 38.582252 | 34.522943 | 33.365388 | 27.915162 | 89.782482 | 79.801327 | 74.870961 | 63.824703 | -1.200420 | 2575.722306 | 0.131895233 | -1.99809673 | **0.045706174** | 0.653147492 | 11.456349 | 11.193190 | 11.447305 | 11.347819 | 11.319453 | 11.177508 | 3430 | 2850 | 2218 | 1999 |
| 27249 | NM_015702 | MMADHC | metabolism of cobalamin associated D | protein_coding | NP_056517.1 | 25221 | 611935 | . | . | 61.073677 | 57.165901 | 50.688084 | 49.106876 | 142.120956 | 132.141541 | 113.742592 | 112.277046 | -1.201818 | 2366.071254 | 0.118017958 | -2.24726983 | **0.024622788** | 0.517303408 | 11.334720 | 11.069680 | 11.307562 | 11.246295 | 11.146034 | 11.113202 | 3085 | 2683 | 1908 | 1973 |
| 27250 | NM_00118642 | PDCD4 | programmed cell death 4 | protein_coding | NP_001186421.1;NP_055271.2;NP_663314.1 | 8763 | 608610 | . | . | 7.673845 | 7.465668 | 7.022895 | 7.522757 | 17.857351 | 17.257226 | 15.759175 | 17.199891 | -1.050157 | 809.095620 | 0.145578461 | -0.48500096 | 0.627675667 | 0.998865404 | 9.695112 | 9.624354 | 9.689581 | 9.664800 | 9.619756 | 9.665047 | 971 | 879 | 664 | 764 |
| 27252 | NM_014458,XI | KLHL20 | kelch like family member 20 | protein_coding | NP_055273.2;XP_005245150.1;XP_011507722.1;XP_016856541.1;XP_01 | 25056 | 617679 | . | . | 5.601728 | 5.441902 | 6.521625 | 6.304774 | 13.035450 | 12.579201 | 14.634336 | 14.415116 | 1.122358 | 665.839428 | 0.152490147 | 1.092089523 | 0.274793754 | 0.998865404 | 9.293269 | 9.459979 | 9.345463 | 9.329761 | 9.425582 | 9.406494 | 731 | 669 | 621 | 651 |
| 27255 | NM_00128908 | CNTN6 | contactin 6 | protein_coding | NP_001276009.1;NP_001276010.1;NP_001336279.1;NP_001336280.1;NI | 2176 | 607220 | . | . | 0.121189 | 0.177536 | 0.124463 | 0.120201 | 0.282013 | 0.410383 | 0.279291 | 0.274825 | -1.234026 | 19.176810 | 0.707734254 | -0.42865339 | 0.668175483 | NA | 4.405684 | 4.100822 | 4.254183 | 4.287152 | 4.254233 | 4.253299 | 20 | 27 | 15 | 16 |
| 27257 | NM_014462,N | LSM1 | LSM1 homolog, mRNA degradation associated | protein_coding | NP_055277.1 | 20472 | 607281 | . | . | 24.705298 | 25.273283 | 26.781413 | 26.484131 | 57.490249 | 58.421255 | 60.096715 | 60.552817 | 1.051456 | 738.279260 | 0.147791025 | 0.489799068 | 0.62427609 | 0.998865404 | 9.491353 | 9.563784 | 9.504314 | 9.515709 | 9.553146 | 9.536535 | 822 | 783 | 666 | 701 |
| 27258 | NM_014463 | LSM3 | LSM3 homolog, U6 small nuclear RNA and mRNA degradation ass | protein_coding | NP_055278.1 | 17874 | 607283 | . | . | 10.827369 | 10.538684 | 10.655222 | 10.724147 | 25.195732 | 24.360640 | 23.910009 | 24.519487 | -1.013727 | 1118.846414 | 0.132749585 | -0.10816419 | 0.882213189 | 0.998865404 | 10.137553 | 10.117974 | 10.146010 | 10.119513 | 10.124007 | 10.121431 | 1320 | 1194 | 968 | 1040 |
| 27286 | NM_014467 | SRPX2 | sushi repeat containing protein X-linked 2 | protein_coding | NP_055282.1 | 30668 | 300642 | . | . | 0.142522 | 0.111950 | 0.130097 | 0.116352 | 0.331653 | 0.258778 | 0.291933 | 0.266025 | -1.046555 | 26.655890 | 0.599848936 | -0.1094402 | 0.912853347 | NA | 4.767942 | 4.704117 | 4.752349 | 4.725872 | 4.740364 | 4.728016 | 35 | 26 | 24 | 23 |
| 27289 | NM_014470 | RND1 | Rho family GTPase 1 | protein_coding | NP_055285.1 | 18314 | 609038 | . | . | 0.135392 | 0.256082 | 0.251655 | 0.430537 | 0.315063 | 0.591945 | 0.564706 | 0.984372 | 1.673411 | 14.479213 | 0.829608102 | 0.895352618 | 0.370598621 | NA | 3.440763 | 4.177906 | 3.811231 | 3.839283 | 3.838960 | 3.875362 | 9 | 15 | 12 | 21 |
| 27291 | NM_00125661 | R3HCC1L | R3H domain and coiled-coil containing 1 like | protein_coding | NP_001243548.2;NP_001243549.1;NP_001243550.2;NP_001337939.2;NI | 23512 | . | . | . | 5.357137 | 5.451638 | 6.056127 | 6.668082 | 12.466281 | 12.601707 | 13.589775 | 15.245776 | 1.104415 | 637.438442 | 0.158266686 | 0.90531982 | 0.365295979 | 0.998865404 | 9.243016 | 9.385740 | 9.256853 | 9.304151 | 9.314465 | 9.379367 | 673 | 677 | 554 | 657 |
| 27292 | NM_00134807 | DIMT1 | DIMT1 rRNA methyltransferase and ribosome maturation factor | protein_coding | NP_001335005.1;NP_001335006.1;NP_055288.1 | 30217 | 612499 | . | . | 6.388788 | 6.153657 | 7.318042 | 6.706432 | 14.866971 | 14.224452 | 16.421474 | 15.333462 | 1.089684 | 582.790474 | 0.160806104 | 0.770556059 | 0.440970115 | 0.998865404 | 9.123314 | 9.247674 | 9.173824 | 9.139693 | 9.237599 | 9.189011 | 659 | 586 | 548 | 549 |
| 27293 | NM_00100956 | SMPDL3B | sphingomyelin phosphodiesterase acid like 3B | protein_coding | NP_001009568.1;NP_001291508.1;NP_055289.2;XP_011539561.1;XP_01 | 21416 | 617737 | . | . | 1.595205 | 1.193646 | 1.367393 | 0.973256 | 3.712106 | 2.759164 | 3.068389 | 2.225233 | -1.221831 | 74.753562 | 0.383031079 | -0.7546234 | 0.45047498 | 0.998865404 | 6.360036 | 6.073945 | 6.282060 | 6.210106 | 6.232791 | 6.159736 | 107 | 77 | 68 | 53 |
| 27294 | NM_014475,XI | DHDH | dihydrodiol dehydrogenase | protein_coding | NP_055290.1;XP_005258805.1;XP_016882087.1 | 17887 | 606377 | . | . | 0.596586 | 0.773656 | 0.995991 | 0.910148 | 1.388281 | 1.788340 | 2.234973 | 2.080946 | 1.191888 | 30.581236 | 0.561915385 | 0.450689345 | 0.652213463 | 0.998865404 | 4.802857 | 5.055266 | 4.903161 | 4.933429 | 4.944774 | 4.935234 | 28 | 34 | 30 | 30 |
| 27295 | NM_00111410 | PDLIM3 | PDZ and LIM domain 3 | protein_coding | NP_001107579.1;NP_001244891.1;NP_001244892.1;NP_055291.2 | 20767 | 605889 | . | . | 1.966980 | 2.177163 | 1.119841 | 1.669274 | 4.577242 | 5.032612 | 2.512891 | 3.816596 | -1.457529 | 139.943535 | 0.29370896 | -1.85055572 | 0.064233495 | 0.731823074 | 7.376022 | 6.830027 | 7.173606 | 7.209549 | 6.992395 | 7.108097 | 183 | 187 | 81 | 126 |
| 27297 | NM_00104064 | CRCP | CGRP receptor component | protein_coding | NP_001035737.1;NP_001035738.1;NP_001135886.1;NP_055293.1 | 17888 | 606121 | . | . | 9.103540 | 9.028199 | 9.555343 | 8.954832 | 21.184312 | 20.869087 | 21.441912 | 20.474158 | 1.002115 | 789.893075 | 0.146629615 | 0.020790586 | 0.983412708 | 0.998865404 | 9.623842 | 9.627183 | 9.626424 | 9.622533 | 9.652497 | 9.598899 | 911 | 849 | 714 | 713 |
| 27300 | NM_00132076 | ZNF544 | zinc finger protein 544 | protein_coding | NP_001307696.1;NP_001307698.1;NP_001307699.1;NP_001307700.1;NI | 16759 | . | . | . | 6.384845 | 6.620047 | 7.361414 | 7.266083 | 14.857803 | 15.302532 | 16.518800 | 16.613033 | 1.147076 | 736.546850 | 0.154734687 | 1.279356792 | 0.200771442 | 0.966265067 | 9.422176 | 9.620295 | 9.441567 | 9.503948 | 9.610437 | 9.524812 | 754 | 774 | 726 | 693 |
| 27301 | NM_00127174 | APEX2 | apurinic/apyrimidinic endodeoxyribonuclease 2 | protein_coding | NP_001258677.1;NP_055296.2 | 17889 | 300773 | . | . | 7.357498 | 8.141606 | 8.093823 | 8.809879 | 17.121200 | 18.819689 | 18.162304 | 20.142740 | 1.075851 | 815.855506 | 0.147827959 | 0.713521136 | 0.475523339 | 0.998865404 | 9.618713 | 9.723717 | 9.612400 | 9.676630 | 9.670812 | 9.721768 | 864 | 887 | 709 | 821 |
| 27303 | NM_00100379 | RBMS3 | RNA binding motif single stranded interacting protein 3 | protein_coding | NP_001003792.1;NP_001003793.1;NP_001171182.1;NP_001171183.1;NI | 13427 | 605786 | . | . | 0.497386 | 0.590601 | 0.351740 | 0.228326 | 1.157442 | 1.365201 | 0.789294 | 0.522040 | -1.454816 | 66.697945 | 0.389395832 | -1.38891349 | 0.164859059 | 0.940065727 | 6.304630 | 5.764191 | 6.086482 | 6.115349 | 6.027357 | 6.004507 | 86 | 90 | 50 | 48 |
| 27304 | NM_014484 | MOCS3 | molybdenum cofactor synthesis 3 | protein_coding | NP_055299.1 | 15765 | 609277 | . | . | 3.410782 | 3.413714 | 3.684411 | 3.992460 | 7.937029 | 7.890954 | 8.267712 | 9.128284 | 1.108974 | 577.582577 | 0.160282018 | 0.931017773 | 0.351844371 | 0.998865404 | 9.097474 | 9.246448 | 9.140296 | 9.135841 | 9.182793 | 9.228702 | 633 | 589 | 510 | 589 |
| 27309 | NM_00129200 | ZNF330 | zinc finger protein 330 | protein_coding | NP_001278931.1;NP_055302.1;XP_016863522.1;XP_024309754.1 | 15462 | 609550 | . | . | 15.241158 | 15.345798 | 14.789881 | 14.194830 | 35.466801 | 35.472499 | 33.188062 | 32.454790 | -1.064974 | 872.048983 | 0.14167435 | -0.64103063 | 0.521502785 | 0.998865404 | 9.812884 | 9.722222 | 9.787838 | 9.792958 | 9.762093 | 9.728384 | 1032 | 974 | 752 | 773 |
| 27314 | NM_00128605 | RAB30 | RAB30, member RAS oncogene family | protein_coding | NP_001272988.1;NP_001272989.1;NP_001272990.1;NP_055303.2 | 9770 | 605693 | . | . | 0.833908 | 0.850140 | 1.016152 | 0.923239 | 1.940538 | 1.965136 | 2.280215 | 2.110876 | 1.138508 | 279.579869 | 0.207884107 | 0.900235254 | 0.367995069 | 0.998865404 | 8.030280 | 8.217860 | 8.086481 | 8.093625 | 8.184408 | 8.130718 | 299 | 284 | 272 | 265 |
| 27315 | NM_00114543 | PGAP2 | post-GPI attachment to proteins 2 | protein_coding | NP_001138910.1;NP_001243164.1;NP_001243165.1;NP_001243166.1;NI | 17893 | 615187 | . | . | 12.426859 | 11.622227 | 13.238812 | 11.256224 | 28.917811 | 26.865302 | 29.707512 | 25.736020 | -1.010238 | 696.508583 | 0.153611509 | -0.0956628 | 0.923788388 | 0.998865404 | 9.451091 | 9.436868 | 9.457298 | 9.436656 | 9.474944 | 9.404164 | 818 | 744 | 634 | 616 |
| 27316 | NM_00116480 | RBMX | RNA binding motif protein X-linked | protein_coding | NP_001158275.1;NP_002130.2 | 9910 | 300199 | . | . | 67.817418 | 68.708844 | 59.490020 | 61.491518 | 157.813925 | 158.823570 | 133.493883 | 140.593035 | -1.141926 | 4034.851059 | 0.107419482 | -1.78244006 | 0.074677497 | 0.764716896 | 12.070876 | 11.879373 | 12.030065 | 12.031721 | 11.911555 | 11.930003 | 4949 | 4647 | 3239 | 3572 |
| 27319 | NM_152414 | BHLHE22 | basic helix-loop-helix family member e22 | protein_coding | NP_689627.1 | 11963 | 613483 | . | . | 0.214380 | 0.073102 | 0.145666 | 0.056998 | 0.498871 | 0.168978 | 0.326870 | 0.130318 | -1.456915 | 13.067466 | 0.917043888 | -0.59202957 | 0.553830788 | NA | 3.950477 | 3.416142 | 3.747595 | 3.684219 | 3.713407 | 3.676424 | 26 | 9 | 13 | 6 |
| 27324 | NM_00108043 | TOX3 | TOX high mobility group box family member 3 | protein_coding | NP_001073899.2;NP_001139660.1;XP_005255949.1;XP_011521304.1;XP | 11972 | 611416 | . | . | 0.481413 | 0.371653 | 0.834619 | 0.687998 | 1.120268 | 0.859093 | 1.872860 | 1.573028 | 1.611925 | 87.525892 | 0.348128976 | 1.978531892 | **0.047868734** | 0.659753773 | 6.065436 | 6.755984 | 6.396137 | 6.336596 | 6.511967 | 6.485575 | 87 | 63 | 98 | 97 |
| 27327 | NM_00133052 | TNRC6A | trinucleotide repeat containing adaptor 6A | protein_coding | NP_001317449.1;NP_001338779.1;NP_055309.2;XP_005255314.1;XP_01 | 11969 | 610739 | . | . | 7.022424 | 8.020852 | 8.347528 | 8.173239 | 16.341470 | 18.540559 | 18.731613 | 18.687142 | 1.083333 | 2042.792189 | 0.123725293 | 0.93333344 | 0.350647835 | 0.998865404 | 10.937496 | 11.052852 | 10.914792 | 11.008952 | 11.037673 | 11.015483 | 2124 | 2242 | 1873 | 1964 |
| 27332 | NM_00101497 | ZNF638 | zinc finger protein 638 | protein_coding | NP_001014972.1;NP_001239541.1;NP_001239542.1;NP_055312.2 | 17894 | 614349 | . | . | 13.530170 | 13.502017 | 12.706013 | 12.820491 | 31.485261 | 31.210512 | 28.511926 | 29.312529 | -1.074479 | 2663.370094 | 0.112606964 | -0.92034405 | 0.357392997 | 0.998865404 | 11.429918 | 11.326297 | 11.411692 | 11.403376 | 11.350036 | 11.348471 | 3193 | 2962 | 2235 | 2405 |
| 27333 | NM_00130815 | GOLIM4 | golgi integral membrane protein 4 | protein_coding | NP_001295084.1;NP_055313.1;XP_005247421.1;XP_005247422.1;XP_01 | 15448 | 606805 | . | . | 16.518735 | 15.994035 | 11.147645 | 13.105292 | 38.439779 | 36.970933 | 25.014991 | 29.963694 | -1.368612 | 1879.015302 | 0.129017881 | -3.50892301 | **0.000449925** | **0.037638281** | 11.084563 | 10.631542 | 11.002138 | 10.969605 | 10.678719 | 10.794452 | 2550 | 2297 | 1274 | 1601 |
| 27335 | NM_00130099 | EIF3K | eukaryotic translation initiation factor 3 subunit K | protein_coding | NP_001287921.1;NP_001295322.1;NP_037366.1 | 24656 | 609596 | . | . | 150.461458 | 156.951871 | 132.974058 | 155.089896 | 350.130015 | 362.801277 | 298.389921 | 354.594615 | -1.083681 | 3542.064225 | 0.119007014 | -0.97423254 | 0.329941092 | 0.998865404 | 11.847284 | 11.731128 | 11.807703 | 11.836240 | 11.694707 | 11.813232 | 4170 | 4044 | 2742 | 3418 |
| 27336 | NM_00116328 | HTATSF1 | HIV-1 Tat specific factor 1 | protein_coding | NP_001156752.1;NP_055315.2;XP_005262461.1 | 5276 | 300346 | . | . | 14.361523 | 14.532868 | 12.404921 | 15.344432 | 33.419853 | 33.593374 | 27.836280 | 35.083223 | -1.081066 | 1239.193649 | 0.143698007 | -0.78257838 | 0.433874741 | 0.998865404 | 10.330570 | 10.217590 | 10.301014 | 10.304836 | 10.164380 | 10.317654 | 1479 | 1393 | 929 | 1231 |
| 27338 | NM_014501,XI | UBE2S | ubiquitin conjugating enzyme E2 S | protein_coding | NP_055316.2;XP_011525054.1 | 17895 | 610309 | . | . | 81.517101 | 85.192243 | 63.222628 | 81.796411 | 189.693642 | 196.925680 | 141.869748 | 187.017766 | -1.165348 | 6165.467080 | 0.129900687 | -1.69945924 | 0.089232689 | 0.802216049 | 12.696237 | 12.475302 | 12.634564 | 12.663866 | 12.411798 | 12.618315 | 7511 | 7284 | 4338 | 6000 |
| 27339 | NM_014502 | PRPF19 | pre-mRNA processing factor 19 | protein_coding | NP_055317.1 | 17896 | 608330 | . | . | 34.894218 | 37.896904 | 31.687822 | 39.222600 | 81.200287 | 87.600395 | 71.106560 | 89.676552 | -1.041161 | 2615.428241 | 0.131641853 | -0.44205598 | 0.658448694 | 0.998865404 | 11.381808 | 11.323261 | 11.337036 | 11.397452 | 11.249991 | 11.413236 | 2959 | 2986 | 2003 | 2645 |
| 27340 | NM_014503 | UTP20 | UTP20 small subunit processome component | protein_coding | NP_055318.2 | 17897 | 612822 | . | . | 7.268662 | 7.347690 | 7.453973 | 6.995930 | 16.914476 | 16.984514 | 16.726498 | 15.995362 | -1.025644 | 2045.098577 | 0.119622362 | -0.30537669 | 0.760079276 | 0.998865404 | 11.016027 | 10.979653 | 11.006902 | 11.008628 | 11.016594 | 10.957908 | 2382 | 2237 | 1821 | 1823 |
| 27341 | NM_015703 | RRP7A | ribosomal RNA processing 7 homolog A | protein_coding | NP_056518.2 | 24286 | . | . | . | 11.743219 | 12.220537 | 10.644731 | 11.403780 | 27.326956 | 28.248320 | 23.886467 | 26.073389 | -1.102042 | 1942.886943 | 0.120200458 | -1.16621618 | 0.243527053 | 0.992733415 | 10.992462 | 10.852098 | 10.948551 | 10.973787 | 10.860973 | 10.906938 | 2309 | 2233 | 1560 | 1784 |
| 27342 | NM_00128706 | RABGEF1 | RAB guanine nucleotide exchange factor 1 | protein_coding | NP_001273989.1;NP_001273990.1;NP_001273991.1;NP_001354651.1;NI | 17676 | 609700 | . | . | 7.414586 | 7.172857 | 7.027231 | 7.703504 | 17.254047 | 16.580384 | 15.768898 | 17.613144 | 1.025929 | 742.092771 | 0.148030324 | 0.249483716 | 0.802986635 | 0.998865404 | 9.516886 | 9.553790 | 9.538169 | 9.514400 | 9.532838 | 9.555313 | 858 | 777 | 643 | 716 |
| 27343 | NM_00117408 | POLL | DNA polymerase lambda | protein_coding | NP_001167555.1;NP_001167556.1;NP_001295311.1;NP_037406.1;XP_00 | 9184 | 606343 | . | . | 5.072191 | 5.515190 | 5.806278 | 5.208007 | 11.803195 | 12.748608 | 13.029121 | 11.907487 | 1.055297 | 436.635001 | 0.176809617 | 0.439171101 | 0.660537563 | 0.998865404 | 8.730810 | 8.808706 | 8.739947 | 8.765428 | 8.811986 | 8.758585 | 479 | 468 | 407 | 402 |
| 27344 | NM_013271 | PCSK1N | proprotein convertase subtilisin/kexin type 1 inhibitor | protein_coding | NP_037403.1 | 17301 | 300399 | . | . | 3.512687 | 2.844424 | 4.207016 | 3.793649 | 8.174167 | 6.575015 | 9.440420 | 8.673728 | 1.239861 | 115.148504 | 0.306928784 | 1.010587308 | 0.312213993 | 0.998865404 | 6.683090 | 6.994808 | 6.837861 | 6.770709 | 6.895401 | 6.857257 | 131 | 99 | 117 | 113 |
| 27345 | NM_014505,XI | KCNMB4 | potassium calcium-activated channel subfamily M regulatory beta | protein_coding | NP_055320.4;XP_011536490.1 | 6289 | 605223 | . | . | 0.506323 | 0.436916 | 0.383492 | 0.429160 | 1.178236 | 1.009952 | 0.860544 | 0.981224 | -1.151425 | 63.124505 | 0.396990885 | -0.51240529 | 0.608367392 | 0.998865404 | 6.077877 | 5.875273 | 6.012751 | 5.980701 | 5.969727 | 5.961826 | 83 | 68 | 52 | 54 |
| 27346 | NM_014573,XI | TMEM97 | transmembrane protein 97 | protein_coding | NP_055388.2;XP_005258022.1 | 28106 | 612912 | . | . | 30.835490 | 30.844754 | 54.892114 | 43.954983 | 71.755456 | 71.299028 | 123.176311 | 100.497841 | 1.585038 | 3129.099139 | 0.131846986 | 5.040065634 | **4.65372E-07** | **0.000170556** | 11.241211 | 11.905949 | 11.392806 | 11.388199 | 11.860655 | 11.665559 | 2796 | 2604 | 3717 | 3188 |
| 27347 | NM_013233,XI | STK39 | serine/threonine kinase 39 | protein_coding | NP_037365.2;XP_005246522.1;XP_016859302.1;XP_016859303.1;XP_01 | 17717 | 607648 | . | . | 5.501739 | 5.337827 | 5.217881 | 5.871536 | 12.802772 | 12.338623 | 11.708774 | 13.424568 | -1.013672 | 524.491341 | 0.167370029 | -0.11705286 | 0.906818169 | 0.998865404 | 9.044658 | 9.024825 | 9.056843 | 9.020780 | 8.998953 | 9.060528 | 625 | 554 | 431 | 512 |
| 27348 | NM_00131789 | TOR1B | torsin family 1 member B | protein_coding | NP_001304822.1;NP_001304823.1;NP_055321.1;XP_005251984.1 | 11995 | 608050 | . | . | 15.038452 | 14.516158 | 13.838764 | 14.429188 | 34.995096 | 33.554748 | 31.053783 | 32.990626 | -1.049567 | 1174.770630 | 0.133499768 | -0.52280743 | 0.601108268 | 0.998865404 | 10.232615 | 10.162869 | 10.244308 | 10.186761 | 10.166319 | 10.192871 | 1438 | 1249 | 979 | 1094 |
| 27349 | NM_014507,N | MCAT | malonyl-CoA-acyl carrier protein transacylase | protein_coding | NP_055322.1;NP_775738.3;XP_011528458.1 | 29622 | 614479 | . | . | 8.921544 | 9.190632 | 9.218070 | 10.412965 | 20.760800 | 21.244559 | 20.685082 | 23.808007 | 1.056170 | 572.748219 | 0.161776163 | 0.487349154 | 0.62601092 | 0.998865404 | 9.121972 | 9.200475 | 9.146523 | 9.139602 | 9.147989 | 9.208803 | 645 | 598 | 488 | 577 |
| 27350 | NM_014508,XI | APOBEC3C | apolipoprotein B mRNA editing enzyme catalytic subunit 3C | protein_coding | NP_055323.2;XP_024307986.1 | 17353 | 607750 | . | . | 16.073001 | 18.168965 | 15.768986 | 18.457636 | 37.402538 | 41.998381 | 35.385148 | 42.201188 | -1.016017 | 1413.177947 | 0.138203652 | -0.16587983 | 0.868251522 | 0.998865404 | 10.476404 | 10.452956 | 10.425918 | 10.512017 | 10.399645 | 10.512305 | 1549 | 1623 | 1129 | 1411 |
| 27351 | NM_015704,XI | DESI1 | desumoylating isopeptidase 1 | protein_coding | NP_056519.1;XP_005261628.1 | 24577 | 614637 | . | . | 16.980178 | 18.325414 | 18.247401 | 19.889783 | 39.513571 | 42.360015 | 40.946641 | 45.475626 | 1.057009 | 1980.991945 | 0.124268015 | 0.643670888 | 0.519788872 | 0.998865404 | 10.911614 | 10.991300 | 10.896429 | 10.960723 | 10.935733 | 11.008903 | 2128 | 2163 | 1689 | 1996 |
| 27352 | NM_00130184 | SGSM3 | small G protein signaling modulator 3 | protein_coding | NP_001288778.1;NP_001336968.1;NP_001336969.1;NP_001336970.1;NI | 25228 | 610440 | . | . | 30.106862 | 29.583476 | 35.136828 | 32.234108 | 70.059910 | 68.383530 | 78.846029 | 73.699454 | 1.113525 | 2919.306507 | 0.117210607 | 1.323553748 | 0.185651295 | 0.958175316 | 11.431654 | 11.586987 | 11.479699 | 11.450626 | 11.596229 | 11.508182 | 3238 | 2927 | 2818 | 2730 |
| 27429 | NM_00132172 | HTRA2 | HtrA serine peptidase 2 | protein_coding | NP_001308656.1;NP_001308657.1;NP_037379.1;NP_659540.1 | 14348 | 606441 | . | . | 14.504344 | 14.153983 | 15.387274 | 14.440927 | 33.752206 | 32.717565 | 34.528598 | 33.017461 | 1.102764 | 959.351647 | 0.138549489 | 1.018585109 | 0.308399977 | 0.998865404 | 9.833532 | 9.974841 | 9.888567 | 9.848375 | 9.944805 | 9.934962 | 1080 | 957 | 881 | 937 |
| 27430 | NM_013283,N | MAT2B | methionine adenosyltransferase 2B | protein_coding | NP_037415.1;NP_877725.1 | 6905 | 605527 | . | . | 20.435837 | 17.472850 | 17.672570 | 17.651460 | 47.555031 | 40.389276 | 39.656737 | 40.357968 | -1.072728 | 1229.175126 | 0.136773312 | -0.74052313 | 0.458982634 | 0.998865404 | 10.313087 | 10.212095 | 10.340349 | 10.234692 | 10.240727 | 10.231228 | 1566 | 1278 | 1038 | 1105 |
| 27433 | NM_00108534 | TOR2A | torsin family 2 member A | protein_coding | NP_001078816.2;NP_001127902.1;NP_001127903.1;NP_001238947.2;NI | 11996 | 608052 | . | . | 4.934903 | 5.091848 | 4.290164 | 4.926320 | 11.483720 | 11.770034 | 9.627004 | 11.263445 | -1.046604 | 243.355473 | 0.21789121 | -0.30160019 | 0.76295686 | 0.998865404 | 7.959549 | 7.893539 | 7.931993 | 7.944712 | 7.907098 | 7.924574 | 283 | 272 | 203 | 227 |
| 27434 | NM_00128433 | POLM | DNA polymerase mu | protein_coding | NP_001271259.1;NP_001271260.1;NP_001349612.1;NP_037416.1 | 9185 | 606344 | . | . | 7.211002 | 8.672181 | 10.316139 | 6.635753 | 16.780299 | 20.046136 | 23.149119 | 15.171860 | 1.072470 | 719.630251 | 0.201701949 | 0.500429568 | 0.616772638 | 0.998865404 | 9.439441 | 9.540996 | 9.391751 | 9.520374 | 9.650074 | 9.339721 | 725 | 819 | 791 | 544 |
| 27436 | NM_00114507 | EML4 | EMAP like 4 | protein_coding | NP_001138548.2;NP_061936.3;XP_005264324.1;XP_005264325.1;XP_00 | 1316 | 607442 | . | . | 14.166973 | 13.554956 | 14.237520 | 12.126403 | 32.967126 | 31.332886 | 31.948583 | 27.725580 | -1.068952 | 2339.972848 | 0.127825138 | -0.75256799 | 0.451709559 | 0.998865404 | 11.239408 | 11.143552 | 11.237201 | 11.197446 | 11.229834 | 11.093614 | 2852 | 2545 | 2135 | 1940 |
| 27439 | NM_00116307 | TMEM121B | transmembrane protein 121B | protein_coding | NP_001156551.1;NP_114096.1;XP_011544426.1;XP_011544427.1 | 1844 | . | . | . | 0.023337 | 0.050700 | 0.041941 | 0.068100 | 0.054307 | 0.117195 | 0.094113 | 0.155703 | 1.390734 | 7.562358 | 1.119379548 | 0.4250982 | 0.67076509 | NA | 2.664814 | 3.134756 | 2.895322 | 2.914130 | 2.907641 | 2.924258 | 5 | 9 | 6 | 10 |
| 27440 | NM_017829,N | HDHD5 | haloacid dehalogenase like hydrolase domain containing 5 | protein_coding | NP_060299.4;NP_149061.1;XP_005261293.1;XP_011544428.1;XP_01154 | 1843 | . | . | . | 19.459719 | 20.697624 | 21.610694 | 22.532749 | 45.283565 | 47.843486 | 48.493767 | 51.518460 | 1.009859 | 1396.785998 | 0.126327268 | 0.112045262 | 0.910787518 | 0.998865404 | 10.440856 | 10.454901 | 10.440939 | 10.447237 | 10.438618 | 10.464373 | 1594 | 1506 | 1200 | 1338 |
| 27443 | NM_00129004 | CECR2 | CECR2 histone acetyl-lysine reader | protein_coding | NP_001276975.1;NP_001276976.1;XP_006724142.1;XP_011544430.1;XP | 1840 | 607576 | . | . | 0.088352 | 0.161598 | 0.145893 | 0.208559 | 0.205599 | 0.373542 | 0.327379 | 0.476845 | 1.487497 | 50.376749 | 0.44729057 | 1.280793878 | 0.200266075 | 0.96621494 | 5.341184 | 5.912076 | 5.581675 | 5.627960 | 5.668541 | 5.688221 | 40 | 50 | 50 | 59 |
| 27445 | NM_014510,N | PCLO | piccolo presynaptic cytomatrix protein | protein_coding | NP_055325.2;NP_149015.2;XP_016867495.1;XP_016867496.1 | 13406 | 604918 | . | . | 0.438921 | 0.757009 | 0.797557 | 0.676594 | 1.021387 | 1.749860 | 1.789696 | 1.546952 | 1.310168 | 334.260430 | 0.199615003 | 1.952518666 | 0.050876657 | 0.672475152 | 8.177377 | 8.566166 | 8.254535 | 8.342092 | 8.425220 | 8.474208 | 306 | 338 | 316 | 370 |
| 28227 | NM_013239,N | PPP2R3B | protein phosphatase 2 regulatory subunit B''beta | protein_coding | NP_037371.2;XP_011544479.1 | 13417 | 300339 | . | . | 5.776480 | 6.184787 | 6.228677 | 7.295800 | 13.442103 | 14.296410 | 13.976969 | 16.680974 | 1.119636 | 480.661066 | 0.175449581 | 0.929211226 | 0.352779631 | 0.998865404 | 8.825469 | 8.987723 | 8.853499 | 8.887737 | 8.892025 | 8.988674 | 508 | 503 | 408 | 512 |
| 28231 | NM_016354,XI | SLCO4A1 | solute carrier organic anion transporter family member 4A1 | protein_coding | NP_057438.3;XP_005260260.1;XP_011527094.1;XP_016883315.1;XP_01 | 10953 | 612436 | . | . | 7.242500 | 7.024488 | 11.315809 | 9.495745 | 16.853600 | 16.237420 | 25.392343 | 21.710885 | 1.468281 | 1019.461168 | 0.148707965 | 3.726281398 | **0.000194325** | **0.020167475** | 9.689873 | 10.244356 | 9.816000 | 9.845765 | 10.184820 | 10.035732 | 931 | 910 | 1157 | 1027 |
| 28232 | NM_00114504 | SLCO3A1 | solute carrier organic anion transporter family member 3A1 | protein_coding | NP_001138516.1;NP_037404.2;XP_005254946.1;XP_005254948.1;XP_01 | 10952 | 612435 | . | . | 6.896224 | 6.460018 | 6.595211 | 7.108818 | 16.047796 | 14.932620 | 14.799464 | 16.253469 | 1.028667 | 954.601585 | 0.140812943 | 0.289578209 | 0.77213894 | 0.998865404 | 9.878199 | 9.919022 | 9.918038 | 9.857166 | 9.891316 | 9.925635 | 1129 | 973 | 822 | 929 |
| 28511 | NM_00100134 | NKIRAS2 | NFKB inhibitor interacting Ras like 2 | protein_coding | NP_001001349.1;NP_001138399.1;NP_001138400.1;NP_001138401.1;NI | 17898 | 604497 | . | . | 12.327716 | 12.603026 | 12.069342 | 14.352630 | 28.687103 | 29.132463 | 27.083255 | 32.815581 | 1.047249 | 975.084456 | 0.145829579 | 0.456726441 | 0.647867683 | 0.998865404 | 9.895947 | 9.962062 | 9.910029 | 9.912826 | 9.882959 | 10.003493 | 1095 | 1030 | 796 | 1012 |
| 28512 | NM_00137735 | NKIRAS1 | NFKB inhibitor interacting Ras like 1 | protein_coding | NP_001364280.1;NP_001364281.1;NP_001364282.1;NP_001364283.1;NI | 17899 | 604496 | . | . | 5.123382 | 4.369624 | 5.282344 | 4.988323 | 11.922323 | 10.100578 | 11.853425 | 11.405208 | 1.066187 | 634.344801 | 0.163482281 | 0.56557229 | 0.571684569 | 0.998865404 | 9.261814 | 9.354931 | 9.340239 | 9.228220 | 9.348461 | 9.309200 | 766 | 607 | 586 | 596 |
| 28513 | NM_00127102 | CDH19 | cadherin 19 | protein_coding | NP_001257957.1;NP_066976.1;XP_011524233.1;XP_011524234.1;XP_01 | 1758 | 603016 | . | . | 1.427104 | 1.992762 | 1.208111 | 2.525283 | 3.320929 | 4.606358 | 2.710966 | 5.773761 | -1.076387 | 239.671694 | 0.219165599 | -0.48454916 | 0.627996186 | 0.998865404 | 7.957030 | 7.850848 | 7.915758 | 7.930220 | 7.892364 | 7.881545 | 282 | 272 | 203 | 214 |
| 28514 | NM_005618,XI | DLL1 | delta like canonical Notch ligand 1 | protein_coding | NP_005609.3;XP_005266991.1 | 2908 | 606582 | . | . | 0.378424 | 0.304131 | 0.368508 | 0.226461 | 0.880607 | 0.703013 | 0.826921 | 0.517777 | -1.292470 | 35.672791 | 0.52955194 | -0.69895136 | 0.484582429 | 0.998865404 | 5.328680 | 4.961528 | 5.195130 | 5.155606 | 5.157348 | 5.121948 | 52 | 38 | 31 | 25 |
| 28951 | NM_021643,N | TRIB2 | tribbles pseudokinase 2 | protein_coding | NP_067675.1 | 30809 | 609462 | . | . | 58.104035 | 56.033595 | 82.807877 | 69.636314 | 135.210492 | 129.524165 | 185.818466 | 159.215149 | 1.317933 | 9144.304968 | 0.11827364 | 3.367422614 | **0.000758743** | 0.056765469 | 12.945764 | 13.344138 | 13.046845 | 13.010125 | 13.340293 | 13.180721 | 9286 | 8325 | 9870 | 8857 |
| 28952 | NM_014008,XI | CCDC22 | coiled-coil domain containing 22 | protein_coding | NP_054727.1;XP_005272656.1 | 28909 | 300859 | . | . | 5.213744 | 4.853464 | 5.506340 | 5.479980 | 12.132597 | 11.218999 | 12.356067 | 12.529326 | 1.077796 | 378.216213 | 0.185195795 | 0.583620983 | 0.55947534 | 0.998865404 | 8.507841 | 8.616257 | 8.562927 | 8.516492 | 8.586651 | 8.580767 | 436 | 377 | 343 | 366 |
| 28955 | NM_014015 | DEXI | Dexi homolog | protein_coding | NP_054734.2 | 13267 | 617901 | . | . | 10.585234 | 9.339586 | 9.818840 | 10.800493 | 24.632273 | 21.588873 | 22.033194 | 24.694044 | 1.019987 | 495.655491 | 0.17372942 | 0.164337628 | 0.869465362 | 0.998865404 | 8.938819 | 8.967427 | 8.987896 | 8.902700 | 8.931881 | 8.984845 | 603 | 494 | 417 | 489 |
| 28956 | NM_00114526 | LAMTOR2 | late endosomal/lysosomal adaptor, MAPK and MTOR activator 2 | protein_coding | NP_001138736.1;NP_054736.1 | 29796 | 610389 | . | . | 26.434430 | 23.384051 | 26.722415 | 31.489544 | 61.514005 | 54.053283 | 59.964324 | 71.997097 | 1.152467 | 519.491117 | 0.174629295 | 1.17234301 | 0.24105937 | 0.992733415 | 8.915046 | 9.119618 | 9.011489 | 8.929903 | 9.012894 | 9.109599 | 592 | 487 | 448 | 560 |
| 28957 | NM_014018 | MRPS28 | mitochondrial ribosomal protein S28 | protein_coding | NP_054737.1 | 14513 | 611990 | . | . | 25.071802 | 26.686689 | 22.575777 | 26.791336 | 58.343113 | 61.687481 | 50.659386 | 61.255211 | -1.063056 | 660.343493 | 0.160660314 | -0.54909311 | 0.582941553 | 0.998865404 | 9.410861 | 9.321914 | 9.368530 | 9.405077 | 9.290575 | 9.397273 | 763 | 754 | 512 | 648 |
| 28958 | NM_00104043 | COA3 | cytochrome c oxidase assembly factor 3 | protein_coding | NP_001035521.1 | 24990 | 614775 | . | . | 44.054893 | 51.827717 | 45.061840 | 56.044838 | 102.517555 | 119.802086 | 101.117470 | 128.139862 | 1.040431 | 1196.464960 | 0.151955706 | 0.376298688 | 0.706694855 | 0.998865404 | 10.196049 | 10.252523 | 10.148057 | 10.263517 | 10.155973 | 10.312037 | 1247 | 1363 | 951 | 1262 |
| 28960 | NM_00135023 | DCPS | decapping enzyme, scavenger | protein_coding | NP_001337165.1;NP_054745.1 | 29812 | 610534 | . | . | 3.031992 | 2.984153 | 2.808121 | 2.949452 | 7.055568 | 6.898002 | 6.301344 | 6.743573 | -1.060401 | 498.499329 | 0.167343766 | -0.50560703 | 0.613132547 | 0.998865404 | 9.003177 | 8.918475 | 8.987765 | 8.972103 | 8.931354 | 8.954159 | 598 | 547 | 413 | 462 |
| 28962 | NM_014028,XI | OSTM1 | osteoclastogenesis associated transmembrane protein 1 | protein_coding | NP_054747.2 | 21652 | 607649 | . | . | 41.553357 | 35.346480 | 37.465226 | 31.252415 | 96.696362 | 81.704969 | 84.070880 | 71.454936 | -1.146385 | 4969.370757 | 0.132839196 | -1.48368717 | 0.137891931 | 0.899796454 | 12.373923 | 12.177062 | 12.396692 | 12.261474 | 12.298580 | 12.128327 | 6598 | 5272 | 4441 | 3895 |
| 28964 | NM_00108545 | GIT1 | GIT ArfGAP 1 | protein_coding | NP_001078923.1;NP_054749.2;XP_011522986.1;XP_011522987.1;XP_02 | 4272 | 608434 | . | . | 10.403165 | 11.599166 | 11.145742 | 11.752255 | 24.208591 | 26.811992 | 25.010724 | 26.870134 | 1.018752 | 1318.405996 | 0.131881709 | 0.203230385 | 0.838954964 | 0.998865404 | 10.351263 | 10.377773 | 10.318266 | 10.394502 | 10.357228 | 10.384855 | 1429 | 1480 | 1136 | 1270 |
| 28966 | NM_014035,N | SNX24 | sorting nexin 24 | protein_coding | NP_054754.1;XP_005272029.1;XP_006714655.1;XP_011541651.1;XP_01 | 21533 | . | . | . | 9.286205 | 9.495063 | 7.463608 | 8.522716 | 21.609378 | 21.948262 | 16.748122 | 19.486177 | -1.163972 | 552.975141 | 0.163613915 | -1.33886234 | 0.180615493 | 0.952541638 | 9.216638 | 8.997164 | 9.157488 | 9.159426 | 9.030687 | 9.087850 | 684 | 643 | 424 | 501 |
| 28968 | NM_014037,XI | SLC6A16 | solute carrier family 6 member 16 | protein_coding | NP_054756.2;XP_005258877.1;XP_006723231.1;XP_011525161.2;XP_01 | 13622 | 607972 | . | . | 0.041517 | 0.022912 | 0.007702 | 0.022769 | 0.096610 | 0.052962 | 0.017283 | 0.052057 | -1.829845 | 3.085216 | 1.747177154 | -0.49893145 | 0.61782768 | NA | 1.998355 | 1.121007 | 1.649647 | 1.638179 | 1.631048 | 1.640128 | 6 | 3 | 1 | 3 |
| 28969 | NM_00115976 | BZW2 | basic leucine zipper and W2 domains 2 | protein_coding | NP_001153239.1;NP_001349646.1;NP_001349647.1;NP_001349648.1;NI | 18808 | . | . | . | 63.451213 | 65.584099 | 63.822494 | 68.575465 | 147.653576 | 151.600585 | 143.215832 | 156.789649 | 1.010908 | 3679.492433 | 0.110544738 | 0.14158985 | 0.887403984 | 0.998865404 | 11.837499 | 11.853041 | 11.831193 | 11.850060 | 11.822822 | 11.875680 | 4166 | 3994 | 3114 | 3579 |
| 28970 | NM_00128606 | C11orf54 | chromosome 11 open reading frame 54 | protein_coding | NP_001272996.1;NP_001272997.1;NP_001272998.1;NP_001272999.1;NI | 30204 | 615810 | . | . | 3.012954 | 3.840719 | 3.663817 | 4.214094 | 7.011268 | 8.877994 | 8.221502 | 9.635029 | 1.181132 | 477.509473 | 0.173050044 | 1.387865715 | 0.16517794 | 0.940279399 | 8.774593 | 9.014238 | 8.806601 | 8.877820 | 8.935798 | 8.957903 | 475 | 500 | 442 | 493 |
| 28971 | NM_00131695 | AAMDC | adipogenesis associated Mth938 domain containing | protein_coding | NP_001303886.1;NP_001303887.1;NP_001303889.1;NP_001303890.1;NI | 30205 | . | . | . | 4.782803 | 5.233810 | 3.680403 | 6.119204 | 11.129778 | 12.098186 | 8.258717 | 13.990833 | -1.026172 | 93.440370 | 0.355277366 | -0.10490937 | 0.916447727 | 0.998865404 | 6.565899 | 6.525770 | 6.532473 | 6.560477 | 6.469053 | 6.604484 | 104 | 107 | 61 | 107 |
| 28972 | NM_014041 | SPCS1 | signal peptidase complex subunit 1 | protein_coding | NP_054760.4 | 23401 | 610358 | . | . | 22.527023 | 22.017214 | 23.703512 | 22.265596 | 52.421307 | 50.893772 | 53.189995 | 50.907639 | 1.017853 | 2669.325287 | 0.115514808 | 0.221002194 | 0.825090725 | 0.998865404 | 11.369355 | 11.395049 | 11.387391 | 11.361986 | 11.418286 | 11.359259 | 3094 | 2810 | 2427 | 2433 |
| 28973 | NM_014046,XI | MRPS18B | mitochondrial ribosomal protein S18B | protein_coding | NP_054765.1;XP_024302176.1 | 14516 | 611982 | . | . | 55.260885 | 50.466225 | 50.731731 | 60.600516 | 128.594340 | 116.654941 | 113.840527 | 138.555883 | 1.037608 | 2380.857610 | 0.129230078 | 0.41214214 | 0.680235243 | 0.998865404 | 11.190461 | 11.243581 | 11.239233 | 11.161539 | 11.162961 | 11.294489 | 2821 | 2400 | 1937 | 2458 |
| 28974 | NM_014047 | C19orf53 | chromosome 19 open reading frame 53 | protein_coding | NP_054766.1 | 24991 | . | . | . | 37.481953 | 41.640087 | 37.717632 | 46.188622 | 87.222046 | 96.252930 | 84.637276 | 105.604790 | 1.046147 | 1138.557834 | 0.147261576 | 0.441973484 | 0.658508394 | 0.998865404 | 10.120413 | 10.184851 | 10.097845 | 10.169783 | 10.093418 | 10.237488 | 1220 | 1259 | 915 | 1196 |
| 28976 | NM_014049,N | ACAD9 | acyl-CoA dehydrogenase family member 9 | protein_coding | NP_054768.2;XP_024309252.1;XP_024309253.1 | 21497 | 611103 | . | . | 15.945147 | 15.518290 | 16.787573 | 17.115386 | 37.105011 | 35.871224 | 37.670831 | 39.132297 | 1.060394 | 1276.376619 | 0.128781791 | 0.65692362 | 0.511230029 | 0.998865404 | 10.274874 | 10.359560 | 10.307762 | 10.281986 | 10.341089 | 10.337799 | 1451 | 1314 | 1145 | 1229 |
| 28977 | NM_014050,N | MRPL42 | mitochondrial ribosomal protein L42 | protein_coding | NP_054769.1;NP_751917.1 | 14493 | 611847 | . | . | 1.067383 | 1.008000 | 1.117357 | 0.916321 | 2.483845 | 2.330035 | 2.507319 | 2.095059 | -1.038388 | 500.375123 | 0.178809152 | -0.30393048 | 0.76118086 | 0.998865404 | 8.993286 | 8.939953 | 8.999018 | 8.954705 | 9.017154 | 8.885413 | 607 | 531 | 472 | 412 |
| 28978 | NM_014051,XI | TMEM14A | transmembrane protein 14A | protein_coding | NP_054770.1;XP_024302177.1 | 21076 | 616870 | . | . | 17.634345 | 18.176367 | 16.629484 | 15.957838 | 41.035846 | 42.015486 | 37.316081 | 36.485703 | -1.113007 | 526.631185 | 0.164634583 | -0.93821722 | 0.348132791 | 0.998865404 | 9.115741 | 8.961434 | 9.067142 | 9.081053 | 9.020570 | 8.990081 | 632 | 605 | 444 | 456 |
| 28981 | NM_00114377 | IFT81 | intraflagellar transport 81 | protein_coding | NP_001137251.1;NP_001334875.1;NP_001334876.1;NP_001334877.1;NI | 14313 | 605489 | . | . | 17.842876 | 16.948173 | 16.143816 | 15.424553 | 41.521113 | 39.176459 | 36.226256 | 35.266412 | -1.113514 | 1544.537008 | 0.12549751 | -1.23603367 | 0.216446049 | 0.982008277 | 10.668330 | 10.513432 | 10.653509 | 10.612206 | 10.568275 | 10.531105 | 1923 | 1710 | 1302 | 1337 |
| 28982 | NM_014053,XI | FLVCR1 | FLVCR heme transporter 1 | protein_coding | NP_054772.1;XP_011507748.1;XP_011507749.1;XP_011507750.1 | 24682 | 609144 | . | . | 2.017972 | 2.567952 | 2.364069 | 1.679511 | 4.695906 | 5.935939 | 5.304903 | 3.840002 | -1.042372 | 369.466726 | 0.211438191 | -0.28315669 | 0.77705673 | 0.998865404 | 8.558503 | 8.499495 | 8.493830 | 8.578463 | 8.601872 | 8.415800 | 403 | 436 | 366 | 284 |
| 28985 | NM_00113755 | MCTS1 | MCTS1 re-initiation and release factor | protein_coding | NP_001131026.1;NP_054779.1 | 23357 | 300587 | . | . | 2.556816 | 2.342646 | 2.536837 | 1.884334 | 5.949815 | 5.415130 | 5.692587 | 4.308306 | -1.124118 | 697.217309 | 0.176690053 | -0.95530775 | 0.339422089 | 0.998865404 | 9.526877 | 9.359182 | 9.514216 | 9.446863 | 9.498173 | 9.293564 | 890 | 758 | 658 | 522 |
| 28986 | NM_014061 | MAGEH1 | MAGE family member H1 | protein_coding | NP_054780.2 | 24092 | 300548 | . | . | 10.190746 | 10.342275 | 9.406115 | 9.960663 | 23.714287 | 23.906631 | 21.107048 | 22.773872 | -1.074472 | 447.936558 | 0.173535886 | -0.59715545 | 0.550403602 | 0.998865404 | 8.858145 | 8.754288 | 8.827557 | 8.830630 | 8.771329 | 8.798362 | 533 | 502 | 367 | 414 |
| 28987 | NM_014062,N | NOB1 | NIN1 (RPN12) binding protein 1 homolog | protein_coding | NP_054781.1 | 29540 | 613586 | . | . | 21.446923 | 21.296324 | 22.969165 | 25.747556 | 49.907873 | 49.227413 | 51.542143 | 58.868726 | 1.124376 | 1221.018178 | 0.133701634 | 1.264936915 | 0.205893943 | 0.970176829 | 10.166958 | 10.335841 | 10.213532 | 10.201005 | 10.255501 | 10.334212 | 1335 | 1230 | 1066 | 1274 |
| 28988 | NM_00101443 | DBNL | drebrin like | protein_coding | NP_001014436.1;NP_001116428.1;NP_001271242.1;NP_001271244.1;NI | 2696 | 610106 | . | . | 6.900588 | 7.140557 | 7.899443 | 7.306994 | 16.057956 | 16.505626 | 17.726118 | 16.706573 | 1.052778 | 2187.773649 | 0.120777375 | 0.614366538 | 0.538973142 | 0.998865404 | 11.057607 | 11.131931 | 11.056968 | 11.090360 | 11.152819 | 11.075786 | 2403 | 2348 | 2045 | 2003 |
| 28989 | NM_00128679 | NTMT1 | N-terminal Xaa-Pro-Lys N-methyltransferase 1 | protein_coding | NP_001273725.1;NP_001273726.1;NP_001273727.1;NP_001273728.1;NI | 23373 | 613560 | . | . | 13.121089 | 12.883716 | 12.570462 | 12.636872 | 30.533315 | 29.781287 | 28.207754 | 28.892709 | -1.067853 | 666.614777 | 0.153136737 | -0.61848289 | 0.536257075 | 0.998865404 | 9.427198 | 9.332673 | 9.421938 | 9.383052 | 9.360930 | 9.355330 | 815 | 722 | 563 | 602 |
| 28990 | NM_00128895 | ASTE1 | asteroid homolog 1 | protein_coding | NP_001275879.1;NP_054784.2;XP_016861755.1;XP_016861756.1;XP_01 | 25021 | . | . | . | 1.728575 | 2.029416 | 2.639551 | 2.237733 | 4.022464 | 4.691083 | 5.923078 | 5.116310 | 1.325243 | 186.034487 | 0.251691699 | 1.614103683 | 0.106504939 | 0.834803205 | 7.321972 | 7.728357 | 7.435375 | 7.498476 | 7.627452 | 7.549930 | 170 | 186 | 200 | 182 |
| 28991 | NM_00108100 | COMMD5 | COMM domain containing 5 | protein_coding | NP_001074472.1;NP_001074473.1;NP_001274166.1;NP_054785.2 | 17902 | 608216 | . | . | 8.166319 | 9.260326 | 7.490881 | 9.580970 | 19.003362 | 21.405656 | 16.809320 | 21.905753 | -1.033354 | 374.396325 | 0.197032001 | -0.24023798 | 0.81014577 | 0.998865404 | 8.572496 | 8.523940 | 8.531624 | 8.581123 | 8.470879 | 8.599871 | 420 | 428 | 284 | 384 |
| 28992 | NM_014067,XI | MACROD1 | mono-ADP ribosylhydrolase 1 | protein_coding | NP_054786.2;XP_005273996.1;XP_005273997.1;XP_006718584.1;XP_00 | 29598 | 610400 | . | . | 5.018327 | 4.508641 | 4.057017 | 5.721085 | 11.677853 | 10.421926 | 9.103831 | 13.080581 | 1.019857 | 216.799981 | 0.249808327 | 0.113556076 | 0.909589708 | 0.998865404 | 7.746527 | 7.773786 | 7.782189 | 7.722157 | 7.679715 | 7.837648 | 263 | 217 | 158 | 240 |
| 28996 | NM_00111323 | HIPK2 | homeodomain interacting protein kinase 2 | protein_coding | NP_001106710.1;NP_073577.3;XP_006715998.1;XP_011514379.1;XP_01 | 14402 | 606868 | . | . | 1.192019 | 1.476686 | 1.313194 | 1.342663 | 2.773876 | 3.413428 | 2.946768 | 3.069840 | -1.023114 | 636.417495 | 0.167817607 | -0.19644516 | 0.844261753 | 0.998865404 | 9.330506 | 9.296958 | 9.243453 | 9.391136 | 9.299599 | 9.309270 | 661 | 770 | 543 | 594 |
| 28998 | NM_014078 | MRPL13 | mitochondrial ribosomal protein L13 | protein_coding | NP_054797.2 | 14278 | 610200 | . | . | 30.548548 | 31.781441 | 28.395586 | 30.003992 | 71.087730 | 73.464226 | 63.718872 | 68.600563 | -1.082157 | 1185.306977 | 0.131643351 | -0.8652888 | 0.386880294 | 0.998865404 | 10.266973 | 10.152861 | 10.228289 | 10.251638 | 10.164197 | 10.196944 | 1397 | 1350 | 968 | 1091 |
| 28999 | NM_014079,XI | KLF15 | Kruppel like factor 15 | protein_coding | NP_054798.1;XP_005247457.1;XP_011511045.1 | 14536 | 606465 | . | . | 1.449963 | 1.020284 | 1.042906 | 1.114808 | 3.374124 | 2.358432 | 2.340250 | 2.548875 | -1.173507 | 92.481626 | 0.347956906 | -0.66337567 | 0.507089984 | 0.998865404 | 6.641413 | 6.411637 | 6.608235 | 6.493842 | 6.499185 | 6.513519 | 136 | 88 | 72 | 82 |

| | | | | | | | | | |
|---|---|---|---|---|---|---|---|---|---|
| 29015 | NM_00127820 | SLC43A3 | solute carrier family 43 member 3 | protein_coding | NP_001265130.1;NP_001265135.1;NP_054815.2;NP_060081.1;NP_9553( | 17466 | 618034 | . | . |
| 29028 | NM_014109,XI | ATAD2 | ATPase family AAA domain containing 2 | protein_coding | NP_054828.2;XP_011515296.1;XP_011515297.1;XP_011515298.1 | 30123 | 611941 | . | . |
| 29035 | NM_014117,XI | C16orf72 | chromosome 16 open reading frame 72 | protein_coding | NP_054836.2;XP_011520764.1 | 30103 | . | | . |
| 29057 | NM_00124248 | FAM156A | family with sequence similarity 156 member A | protein_coding | NP_001229418.1;NP_001229419.1;NP_001229420.1;NP_001229421.1;NI | 30114 | . | | . |
| 29058 | NM_00100992 | TMEM230 | transmembrane protein 230 | protein_coding | NP_001009923.1;NP_001009924.1;NP_001009925.1;NP_001317913.1;NI | 15876 | 617019 | . | . |
| 29062 | NM_00136273 | WDR91 | WD repeat domain 91 | protein_coding | NP_001349665.1;NP_001349666.1;NP_001349667.1;NP_054868.3;XP_01 | 24997 | 616303 | . | . |
| 29063 | NM_00131814 | ZCCHC4 | zinc finger CCHC-type containing 4 | protein_coding | NP_001305077.1;NP_079212.2;XP_011512137.1;XP_016863617.1;XP_01 | 22917 | 611792 | . | . |
| 29066 | NM_014153,XI | ZC3H7A | zinc finger CCCH-type containing 7A | protein_coding | NP_054872.2;XP_006720940.1;XP_011520765.1;XP_011520766.1;XP_01( | 30959 | . | | . |
| 29068 | NM_00130109 | ZBTB44 | zinc finger and BTB domain containing 44 | protein_coding | NP_001288027.1;NP_001288028.1;NP_001357148.1;NP_001357149.1;NI | 25001 | . | | . |
| 29070 | NM_00114230 | CCDC113 | coiled-coil domain containing 113 | protein_coding | NP_001135774.1;NP_054876.2;XP_011521340.1;XP_011521341.1;XP_01 | 25002 | 616070 | . | . |
| 29071 | NM_00101155 | C1GALT1C1 | C1GALT1 specific chaperone 1 | protein_coding | NP_001011551.1;NP_689905.1 | 24338 | 300611 | . | . |
| 29072 | NM_00134937 | SETD2 | SET domain containing 2, histone lysine methyltransferase | protein_coding | NP_001336299.1;NP_054878.5;XP_011531934.1;XP_011531936.1;XP_02 | 18420 | 612778 | . | . |
| 29074 | NM_00131881 | MRPL18 | mitochondrial ribosomal protein L18 | protein_coding | NP_001305746.1;NP_054880.2 | 14477 | 611831 | . | . |
| 29078 | NM_014165 | NDUFAF4 | NADH:ubiquinone oxidoreductase complex assembly factor 4 | protein_coding | NP_054884.1 | 21034 | 611776 | . | . |
| 29079 | NM_00127062 | MED4 | mediator complex subunit 4 | protein_coding | NP_001257558.1;NP_054885.1;XP_016876038.1 | 17903 | 605718 | . | . |
| 29080 | NM_014167,N | CCDC59 | coiled-coil domain containing 59 | protein_coding | NP_054886.2 | 25005 | . | | . |
| 29081 | NM_00129318 | METTL5 | methyltransferase like 5 | protein_coding | NP_001280115.1;NP_001280116.1;NP_054887.2;XP_011509325.1 | 25006 | 618628 | . | . |
| 29082 | NM_014169 | CHMP4A | charged multivesicular body protein 4A | protein_coding | NP_054888.3 | 20274 | 610051 | . | . |
| 29083 | NM_014170,N | GTPBP8 | GTP binding protein 8 (putative) | protein_coding | NP_054889.2;NP_612494.1 | 25007 | . | | . |
| 29085 | NM_00113586 | PHPT1 | phosphohistidine phosphatase 1 | protein_coding | NP_001129333.1;NP_001274271.1;NP_001274272.1;NP_054891.2 | 30033 | 610167 | . | . |
| 29086 | NM_00103354 | BABAM1 | BRISC and BRCA1 A complex member 1 | protein_coding | NP_001028721.1;NP_001275685.1;NP_001275686.1;NP_054892.2 | 25008 | 612766 | . | . |
| 29087 | NM_00103730 | THYN1 | thymocyte nuclear protein 1 | protein_coding | NP_001032381.1;NP_001032382.1;NP_054893.1;NP_954994.1;NP_9549( | 29560 | 613739 | . | . |
| 29088 | NM_014175,XI | MRPL15 | mitochondrial ribosomal protein L15 | protein_coding | NP_054894.1;XP_016868851.1 | 14054 | 611828 | . | . |
| 29089 | NM_00131032 | UBE2T | ubiquitin conjugating enzyme E2 T | protein_coding | NP_001297255.1;NP_054895.1 | 25009 | 610538 | . | . |
| 29090 | NM_014177,XI | TIMM21 | translocase of inner mitochondrial membrane 21 | protein_coding | NP_054896.2;XP_005266746.1 | 25010 | 615180 | . | . |
| 29091 | NM_00130447 | STXBP6 | syntaxin binding protein 6 | protein_coding | NP_001291405.1;NP_001291406.1;NP_001338869.1;NP_001338870.1;NI | 19666 | 607958 | . | . |
| 29092 | NR_023918,NF | LINC00339 | long intergenic non-protein coding RNA 339 | lncRNA | . | 25011 | . | | . |
| 29093 | NM_00101499 | MRPL22 | mitochondrial ribosomal protein L22 | protein_coding | NP_001014990.1;NP_054899.2 | 14480 | 611835 | . | . |
| 29094 | NM_014181 | LGALSL | galectin like | protein_coding | NP_054900.2 | 25012 | 617902 | . | . |
| 29095 | NM_014182 | ORMDL2 | ORMDL sphingolipid biosynthesis regulator 2 | protein_coding | NP_054901.1 | 16037 | 610074 | . | . |
| 29097 | NM_00127719 | CNIH4 | cornichon family AMPA receptor auxiliary protein 4 | protein_coding | NP_001264126.1;NP_001264127.1;NP_001264128.1;NP_001264129.1;NI | 25013 | 617483 | . | . |
| 29098 | NM_00117780 | RANGRF | RAN guanine nucleotide release factor | protein_coding | NP_001171272.1;NP_001171273.1;NP_001317056.1;NP_057576.2 | 17679 | 607954 | . | . |
| 29099 | NM_00110165 | COMMD9 | COMM domain containing 9 | protein_coding | NP_001095123.1;NP_001294861.1;NP_001294866.1;NP_054905.2;XP_01 | 25014 | 612299 | . | . |
| 29100 | NM_00131821 | TMEM208 | transmembrane protein 208 | protein_coding | NP_001305146.1;NP_054906.2 | 25015 | . | | . |
| 29101 | NM_014188 | SSU72 | SSU72 homolog, RNA polymerase II CTD phosphatase | protein_coding | NP_054907.1 | 25016 | 617680 | . | . |
| 29102 | NM_00110041 | DROSHA | drosha ribonuclease III | protein_coding | NP_001093882.1;NP_001369437.1;NP_037367.3;XP_005248349.1;XP_00 | 17904 | 608828 | . | . |
| 29103 | NM_013238 | DNAJC15 | DnaJ heat shock protein family (Hsp40) member C15 | protein_coding | NP_037370.2 | 20325 | 615339 | . | . |
| 29104 | NM_013240,N | N6AMT1 | N-6 adenine-specific DNA methyltransferase 1 | protein_coding | NP_037372.4;NP_877426.4 | 16021 | 614553 | . | . |
| 29105 | NM_013242,N | CFAP20 | cilia and flagella associated protein 20 | protein_coding | NP_037374.1 | 29523 | 617906 | . | . |
| 29106 | NM_00116525 | SCG3 | secretogranin III | protein_coding | NP_001158729.1;NP_037375.2 | 13707 | 611796 | . | . |
| 29107 | NM_013248 | NXT1 | nuclear transport factor 2 like export factor 1 | protein_coding | NP_037380.1 | 15913 | 605811 | . | . |
| 29108 | NM_013258,N | PYCARD | PYD and CARD domain containing | protein_coding | NP_037390.2;NP_660183.1 | 16608 | 606838 | . | . |
| 29109 | NM_00131820 | FHOD1 | formin homology 2 domain containing 1 | protein_coding | NP_001305131.1;NP_037373.2;XP_006721243.1;XP_011521345.1;XP_01 | 17905 | 606881 | . | . |
| 29110 | NM_013254,XI | TBK1 | TANK binding kinase 1 | protein_coding | NP_037386.1;XP_005268866.1;XP_005268867.1 | 11584 | 604834 | . | . |
| 29115 | NM_00130183 | SAP30BP | SAP30 binding protein | protein_coding | NP_001288768.1;NP_001288784.1;NP_037392.1;XP_011522995.1;XP_01 | 30785 | 610218 | . | . |
| 29116 | NM_013262,XI | MYLIP | myosin regulatory light chain interacting protein | protein_coding | NP_037394.2;XP_005249089.1;XP_005249090.1;XP_016866278.1 | 21155 | 610082 | . | . |
| 29117 | NM_00117398 | BRD7 | bromodomain containing 7 | protein_coding | NP_001167455.1;NP_037395.2;XP_011521348.1;XP_011521349.1;XP_01 | 14310 | 618489 | . | . |
| 29121 | NM_00100441 | CLEC2D | C-type lectin domain family 2 member D | protein_coding | NP_001004419.1;NP_001184246.1;NP_001184247.1;NP_001184248.1;NI | 14351 | 605659 | . | . |
| 29123 | NM_00125618 | ANKRD11 | ankyrin repeat domain 11 | protein_coding | NP_001243111.1;NP_001243112.1;NP_037407.4;XP_006721247.1;XP_01 | 21316 | 611192 | . | . |
| 29126 | NM_00126770 | CD274 | CD274 molecule | protein_coding | NP_001254635.1;NP_001300958.1;NP_054862.1 | 17635 | 605402 | . | . |
| 29127 | NM_00112610 | RACGAP1 | Rac GTPase activating protein 1 | protein_coding | NP_001119575.1;NP_001119576.1;NP_001306928.1;NP_001306929.1;NI | 9804 | 604980 | . | . |
| 29128 | NM_00104820 | UHRF1 | ubiquitin like with PHD and ring finger domains 1 | protein_coding | NP_001041666.1;NP_001276979.1;NP_001276980.1;NP_001276981.1;NI | 12556 | 607990 | . | . |
| 29761 | NM_00128304 | USP25 | ubiquitin specific peptidase 25 | protein_coding | NP_001269970.1;NP_001269971.1;NP_001339489.1;NP_001339490.1;NI | 12624 | 604736 | . | . |
| 29763 | NM_00118497 | PACSIN3 | protein kinase C and casein kinase substrate in neurons 3 | protein_coding | NP_001171903.1;NP_001171904.1;NP_057307.2 | 8572 | 606513 | . | . |
| 29766 | NM_014547,XI | TMOD3 | tropomodulin 3 | protein_coding | NP_055362.1;XP_016877576.1;XP_016877577.1 | 11873 | 605112 | . | . |
| 29767 | NM_00114288 | TMOD2 | tropomodulin 2 | protein_coding | NP_001136357.1;NP_055363.1;XP_016877578.1 | 11872 | 602928 | . | . |
| 29775 | NM_014550 | CARD10 | caspase recruitment domain family member 10 | protein_coding | NP_055365.2 | 16422 | 607209 | . | . |
| 29777 | NM_013375 | ABT1 | activator of basal transcription 1 | protein_coding | NP_037507.1 | 17369 | 618750 | . | . |
| 29780 | NM_00100382 | PARVB | parvin beta | protein_coding | NP_001003828.1;NP_001230314.1;NP_001230315.1;NP_037459.2;XP_00 | 14653 | 608121 | . | . |
| 29781 | NM_00118501 | NCAPH2 | non-SMC condensin II complex subunit H2 | protein_coding | NP_001171940.1;NP_055366.3;NP_689512.2;XP_005261969.1;XP_01152 | 25071 | 611230 | . | . |
| 29789 | NM_00101170 | OLA1 | Obg like ATPase 1 | protein_coding | NP_001011708.1;NP_001315617.1;NP_037473.3 | 28833 | 611175 | . | . |
| 29796 | NM_00100368 | UQCR10 | ubiquinol-cytochrome c reductase, complex III subunit X | protein_coding | NP_001003684.1;NP_037519.2 | 30863 | 610843 | . | . |
| 29798 | NR_149013,NF | C2orf27A | chromosome 2 open reading frame 27A | lncRNA | . | 25077 | . | | . |
| 29799 | NM_013313,N | YPEL1 | yippee like 1 | protein_coding | NP_037445.1 | 12845 | 608082 | . | . |
| 29800 | NM_00132362 | ZDHHC1 | zinc finger DHHC-type containing 1 | protein_coding | NP_001310556.1;NP_037436.1;XP_006721248.1;XP_011521362.1;XP_02 | 17916 | . | | . |
| 29801 | NM_00118502 | ZDHHC8 | zinc finger DHHC-type palmitoyltransferase 8 | protein_coding | NP_001171953.1;NP_037505.1;XP_006724302.1 | 18474 | 608784 | . | . |
| 29803 | NM_00109969 | REPIN1 | replication initiator 1 | protein_coding | NP_001093165.1;NP_001093166.1;NP_001349674.1;NP_001349675.1;NI | 17922 | . | | . |
| 29841 | NM_198182,XI | GRHL1 | grainyhead like transcription factor 1 | protein_coding | NP_937825.2;XP_005246216.1;XP_006711945.1;XP_006711947.1;XP_01 | 17923 | 609786 | . | . |
| 29842 | NM_014553,XI | TFCP2L1 | transcription factor CP2 like 1 | protein_coding | NP_055368.1;XP_016859391.1;XP_016859392.1;XP_016859393.1;XP_01( | 17925 | 609785 | . | . |
| 29843 | NM_00126759 | SENP1 | SUMO specific peptidase 1 | protein_coding | NP_001254523.1;NP_001254524.1;XP_006719424.1;XP_006719425.1;XP | 17927 | 612157 | . | . |
| 29844 | NM_00132179 | TFPT | TCF3 fusion partner | protein_coding | NP_001308721.1;NP_037474.1;XP_005278318.1 | 13630 | 609519 | . | . |
| 29851 | NM_012092 | ICOS | inducible T cell costimulator | protein_coding | NP_036224.1 | 5351 | 604558 | . | . |
| 29855 | NM_00107951 | UBN1 | ubinuclein 1 | protein_coding | NP_001072982.1;NP_001275585.1;XP_005255334.1;XP_005255335.1;XP | 12506 | 609771 | . | . |
| 29880 | NM_00114236 | ALG5 | ALG5 dolichyl-phosphate beta-glucosyltransferase | protein_coding | NP_001135836.1;NP_037470.1;XP_016876041.1 | 20266 | 604565 | . | . |
| 29882 | NM_013366,XI | ANAPC2 | anaphase promoting complex subunit 2 | protein_coding | NP_037498.1 | 19989 | 606946 | . | . |
| 29883 | NM_00132208 | CNOT7 | CCR4-NOT transcription complex subunit 7 | protein_coding | NP_001309016.1;NP_001309017.1;NP_001309018.1;NP_001309019.1;NI | 14101 | 604913 | . | . |
| 29886 | NM_013321,XI | SNX8 | sorting nexin 8 | protein_coding | NP_037453.1;XP_011513631.1;XP_011513632.1;XP_016867572.1;XP_01( | 14972 | 614905 | . | . |
| 29887 | NM_00119983 | SNX10 | sorting nexin 10 | protein_coding | NP_001186764.1;NP_001186766.1;NP_001186767.1;NP_001305127.1;NI | 14974 | 614780 | . | . |
| 29888 | NM_00103987 | STRN4 | striatin 4 | protein_coding | NP_001034966.1;NP_037535.2;XP_006723234.1;XP_006723235.1;XP_00 | 15721 | 614767 | . | . |
| 29889 | NM_00132362 | GNL2 | G protein nucleolar 2 | protein_coding | NP_001310552.1;NP_001310553.1;NP_037417.1;XP_024302359.1 | 29925 | 609365 | . | . |
| 29890 | NM_013286 | RBM15B | RNA binding motif protein 15B | protein_coding | NP_037418.3 | 24303 | 612602 | . | . |
| 29893 | NM_00125601 | PSMC3IP | PSMC3 interacting protein | protein_coding | NP_001242943.1;NP_001242944.1;NP_001242945.1;NP_037422.2;NP_0! | 17928 | 608665 | . | . |
| 29894 | NM_013291,XI | CPSF1 | cleavage and polyadenylation specific factor 1 | protein_coding | NP_037423.2;XP_006716611.1;XP_024302907.1;XP_024302908.1;XP_02 | 2324 | 606027 | . | . |
| 29896 | NM_00128275 | TRA2A | transformer 2 alpha homolog | protein_coding | NP_001269686.1;NP_001269687.1;NP_001269688.1;NP_001349688.1;NI | 16645 | 602718 | . | . |
| 29899 | NM_00132103 | GPSM2 | G protein signaling modulator 2 | protein_coding | NP_001307967.1;NP_001307968.1;NP_037428.3;XP_006710652.1;XP_01 | 29501 | 609245 | . | . |
| 29901 | NM_00136748 | SAC3D1 | SAC3 domain containing 1 | protein_coding | NP_001354414.1;NP_001354415.1;NP_001354416.1;NP_001354417.1;NI | 30179 | 618796 | . | . |
| 29902 | NM_013300,XI | FAM216A | family with sequence similarity 216 member A | protein_coding | NP_037432.2;XP_005253932.1;XP_011536548.1;XP_024304727.1 | 30180 | . | | . |
| 29903 | NM_00137046 | CCDC106 | coiled-coil domain containing 106 | protein_coding | NP_001357396.1;NP_001357397.1;NP_001357398.1;NP_001357399.1;NI | 30181 | 613478 | . | . |
| 29904 | NM_013302,XI | EEF2K | eukaryotic elongation factor 2 kinase | protein_coding | NP_037434.2;XP_016878682.1;XP_016878683.1 | 24615 | 606968 | . | . |
| 29907 | NM_013306,N | SNX15 | sorting nexin 15 | protein_coding | NP_037438.2;NP_680086.2 | 14978 | 605964 | . | . |
| 29911 | NM_00110017 | HOOK2 | hook microtubule tethering protein 2 | protein_coding | NP_001093646.1;NP_037444.2 | 19885 | 607824 | . | . |
| 29914 | NM_00133034 | UBIAD1 | UbiA prenyltransferase domain containing 1 | protein_coding | NP_001317278.1;NP_001317279.1;NP_037451.1 | 30791 | 611632 | . | . |
| 29915 | NM_013320,XI | HCFC2 | host cell factor C2 | protein_coding | NP_037452.1;XP_016874730.1;XP_016874731.1 | 24972 | 607926 | . | . |
| 29916 | NM_00133032 | SNX11 | sorting nexin 11 | protein_coding | NP_001317249.1;NP_037455.2;NP_689450.1;XP_005257317.1;XP_00525 | 14975 | 614906 | . | . |
| 29919 | NM_00127634 | RMC1 | regulator of MON1-CCZ1 | protein_coding | NP_001263271.1;NP_001305636.1;NP_001305637.1;NP_001305638.1;NI | 24326 | . | | . |
| 29920 | NM_00127168 | PYCR2 | pyrroline-5-carboxylate reductase 2 | protein_coding | NP_001258610.1;NP_037460.2 | 30262 | 616406 | . | . |
| 29922 | NM_013330,N | NME7 | NME/NM23 family member 7 | protein_coding | NP_037462.1;NP_932076.1 | 20461 | 613465 | . | . |
| 29923 | NM_00109878 | HILPDA | hypoxia inducible lipid droplet associated | protein_coding | NP_001092256.1;NP_037464.1 | 28859 | 617905 | . | . |
| 29924 | NM_00113007 | EPN1 | epsin 1 | protein_coding | NP_001123543.1;NP_001123544.1;NP_001308192.1;NP_037465.2 | 21604 | 607262 | . | . |
| 29925 | NM_013334,N | GMPPB | GDP-mannose pyrophosphorylase B | protein_coding | NP_037466.2;NP_068806.2 | 22932 | 615320 | . | . |
| 29926 | NM_00137429 | GMPPA | GDP-mannose pyrophosphorylase A | protein_coding | NP_001361223.1;NP_001361224.1;NP_037467.2;NP_995319.1 | 22923 | 615495 | . | . |
| 29927 | NM_013336 | SEC61A1 | SEC61 translocon subunit alpha 1 | protein_coding | NP_037468.1 | 18276 | 609213 | . | . |
| 29928 | NM_013337 | TIMM22 | translocase of inner mitochondrial membrane 22 | protein_coding | NP_037469.2 | 17317 | 607251 | . | . |
| 29929 | NM_013339 | ALG6 | ALG6 alpha-1,3-glucosyltransferase | protein_coding | NP_037471.2 | 23157 | 604566 | . | . |
| 29931 | NR_024065 | LINC00312 | long intergenic non-protein coding RNA 312 | lncRNA | . | 6662 | 610485 | . | . |
| 29934 | NM_00125618 | SNX12 | sorting nexin 12 | protein_coding | NP_001243114.1;NP_001243115.1;NP_001243116.1;NP_001243117.1;NI | 14976 | 300883 | . | . |
| 29935 | NM_013347 | RPA4 | replication protein A4 | protein_coding | NP_037479.1 | 30305 | 300767 | . | . |
| 29937 | NM_013349,N | NENF | neudesin neurotrophic factor | protein_coding | NP_037481.1 | 30384 | 611874 | . | . |
| 29940 | NM_00108097 | DSE | dermatan sulfate epimerase | protein_coding | NP_001074445.1;NP_001309866.1;NP_001309867.1;NP_001309868.1;NI | 21144 | 605942 | . | . |
| 29941 | NM_00131792 | PKN3 | protein kinase N3 | protein_coding | NP_001304855.1;NP_037487.2;XP_005252003.1;XP_006717143.1;XP_01 | 17999 | 610714 | . | . |
| 29942 | NM_00101550 | PURG | purine rich element binding protein G | protein_coding | NP_001015508.1;NP_001310240.1;NP_001310241.1;NP_037489.1 | 17930 | 618041 | . | . |
| 29944 | NM_00128253 | PNMA3 | PNMA family member 3 | protein_coding | NP_001269464.1;NP_037496.4 | 18742 | 300675 | . | . |
| 29945 | NM_00128675 | ANAPC4 | anaphase promoting complex subunit 4 | protein_coding | NP_001273685.1;NP_037499.2;XP_005248216.1;XP_011512140.1;XP_01 | 19990 | 606947 | . | . |
| 29946 | NM_013368,N | SERTAD3 | SERTA domain containing 3 | protein_coding | NP_037500.2;NP_976219.1;XP_005258889.1;XP_006723242.1 | 17931 | 612125 | . | . |
| 29948 | NM_182981 | OSGIN1 | oxidative stress induced growth inhibitor 1 | protein_coding | NP_892026.1 | 30093 | 607975 | . | . |
| 29950 | NM_013376 | SERTAD1 | SERTA domain containing 1 | protein_coding | NP_037508.2 | 17932 | 617850 | . | . |
| 29952 | NM_013379,XI | DPP7 | dipeptidyl peptidase 7 | protein_coding | NP_037511.2;XP_005266132.1;XP_006717146.1;XP_011516901.1;XP_01 | 14892 | 610537 | . | . |
| 29954 | NM_013382,XI | POMT2 | protein O-mannosyltransferase 2 | protein_coding | NP_037514.2;XP_011534977.1;XP_011534978.1;XP_011534979.1 | 19743 | 607439 | . | . |
| 29956 | NM_022075,N | CERS2 | ceramide synthase 2 | protein_coding | NP_071358.1;NP_859530.1;XP_011507753.1;XP_011507754.1 | 14076 | 606920 | . | . |
| 29957 | NM_013386,N | SLC25A24 | solute carrier family 25 member 24 | protein_coding | NP_037518.3;NP_998816.1 | 20662 | 608744 | . | . |
| 29959 | NM_00132135 | NRBP1 | nuclear receptor binding protein 1 | protein_coding | NP_001308286.1;NP_001308287.1;NP_001308288.1;NP_001308290.1;NI | 7993 | 606010 | . | . |
| 29960 | NM_013393 | MRM2 | mitochondrial rRNA methyltransferase 2 | protein_coding | NP_037525.1 | 16352 | 606906 | . | . |
| 29964 | NM_013397 | PRICKLE4 | prickle planar cell polarity protein 4 | protein_coding | NP_037529.3 | 16805 | 611389 | . | . |
| 29965 | NM_00119905 | CDIP1 | cell death inducing p53 target 1 | protein_coding | NP_001185983.1;NP_001185984.1;NP_001185985.1;NP_037531.2 | 13234 | 610503 | . | . |
| 29966 | NM_00108389 | STRN3 | striatin 3 | protein_coding | NP_001077362.1;NP_055389.3;XP_005267626.1;XP_005267627.1 | 15720 | 614766 | . | . |
| 29967 | NM_00113570 | LRP12 | LDL receptor related protein 12 | protein_coding | NP_001129175.1;NP_038465.1 | 31708 | 618299 | . | . |
| 29968 | NM_021154,N | PSAT1 | phosphoserine aminotransferase 1 | protein_coding | NP_066977.1;NP_478059.1 | 19129 | 610936 | . | . |
| 29969 | NM_00116634 | MDFIC | MyoD family inhibitor domain containing | protein_coding | NP_001159817.1;NP_001159818.3;NP_951038.1 | 28870 | 614511 | . | . |
| 29970 | NM_00119710 | SCHIP1 | schwannomin interacting protein 1 | protein_coding | NP_001184036.1;NP_001184037.1;NP_001184038.1;NP_055390.1 | 15678 | . | | . |
| 29978 | NM_013444 | UBQLN2 | ubiquilin 2 | protein_coding | NP_038472.2 | 12509 | 300264 | . | . |
| 29979 | NM_013438,N | UBQLN1 | ubiquilin 1 | protein_coding | NP_038466.2;NP_444295.1;XP_005252005.1 | 12508 | 605046 | . | . |
| 29980 | NM_017613 | DONSON | DNA replication fork stabilization factor DONSON | protein_coding | NP_060083.1 | 2993 | 611428 | . | . |
| 29982 | NM_00128240 | NRBF2 | nuclear receptor binding factor 2 | protein_coding | NP_001269334.1;NP_110386.2;XP_006717873.1 | 19692 | 616477 | . | . |
| 29984 | NM_00130088 | RHOD | ras homolog family member D | protein_coding | NP_001287815.1;NP_055393.1 | 670 | 605781 | . | . |
| 29985 | NM_144564,N | SLC39A3 | solute carrier family 39 member 3 | protein_coding | NP_653165.2;NP_998733.1 | 17128 | 612168 | . | . |
| 29990 | NM_00137193 | PILRB | paired immunoglobin like type 2 receptor beta | protein_coding | NP_001358860.1;NP_839956.1 | 18297 | 605342 | . | . |
| 29992 | NM_013439,N | PILRA | paired immunoglobin like type 2 receptor alpha | protein_coding | NP_038467.2;NP_840056.1;NP_840057.1;XP_024302507.1 | 20396 | 605341 | . | . |
| 29994 | NM_00128997 | BAZ2B | bromodomain adjacent to zinc finger domain 2B | protein_coding | NP_001276904.1;NP_001316786.1;NP_001316787.1;NP_038478.2;XP_00 | 963 | 605683 | . | . |
| 29995 | NM_00127823 | LMCD1 | LIM and cysteine rich domains 1 | protein_coding | NP_001265162.1;NP_001265163.1;NP_001265164.1;NP_055398.1 | 6633 | 604859 | . | . |
| 29997 | NM_015710 | NOP53 | NOP53 ribosome biogenesis factor | protein_coding | NP_056525.2 | 4333 | 605691 | . | . |
| 29998 | NM_015711,XI | BICRA | BRD4 interacting chromatin remodeling complex associated prote | protein_coding | NP_056526.3;XP_005258890.1;XP_006723243.1;XP_011525184.1;XP_01 | 4332 | 605690 | . | . |
| 30000 | NM_00113619 | TNPO2 | transportin 2 | protein_coding | NP_001129667.1;NP_001129668.1;NP_001369165.1;NP_001369166.1;NI | 19998 | 603002 | . | . |
| 30001 | NM_00138246 | ERO1A | endoplasmic reticulum oxidoreductase 1 alpha | protein_coding | NP_001369393.1;NP_001369394.1;NP_001369395.1;NP_001369396.1;NI | 13280 | 615435 | . | . |
| 30008 | NM_016938,N | EFEMP2 | EGF containing fibulin extracellular matrix protein 2 | protein_coding | NP_058634.4 | 3219 | 604633 | . | . |
| 30011 | NM_00102466 | SH3KBP1 | SH3 domain containing kinase binding protein 1 | protein_coding | NP_001019837.1;NP_001171889.1;NP_001340819.1;NP_001340820.1;NI | 13867 | 300374 | . | . |
| 30012 | NM_021025 | TLX3 | T cell leukemia homeobox 3 | protein_coding | NP_066305.2 | 13532 | 604640 | . | . |
| 30811 | NM_014586,XI | HUNK | hormonally up-regulated Neu-associated kinase | protein_coding | NP_055401.1;XP_011527839.1 | 13326 | 606532 | . | . |
| 30812 | NM_014587 | SOX8 | SRY-box transcription factor 8 | protein_coding | NP_055402.2 | 11203 | 605923 | . | . |
| 30815 | NM_00128699 | ST6GALNAC6 | ST6 N-acetylgalactosaminide alpha-2,6-sialyltransferase 6 | protein_coding | NP_001273928.1;NP_001273929.1;NP_001273930.1;NP_001273931.1;NI | 23364 | 610135 | . | . |
| 30817 | NM_00127105 | ADGRE2 | adhesion G protein-coupled receptor E2 | protein_coding | NP_001257981.1;NP_038475.2;XP_011526250.1;XP_011526251.1;XP_01 | 3337 | 606100 | . | . |
| 30818 | NM_00103491 | KCNIP3 | potassium voltage-gated channel interacting protein 3 | protein_coding | NP_001030086.1;NP_038462.1 | 15523 | 604662 | . | . |
| 30819 | NM_014591,N | KCNIP2 | potassium voltage-gated channel interacting protein 2 | protein_coding | NP_055406.2;NP_775283.1;NP_775284.1;NP_775285.1;NP_775286.1;NP | 15522 | 604661 | . | . |
| 30827 | NM_00110165 | CXXC1 | CXXC finger protein 1 | protein_coding | NP_001095124.1;NP_055408.2;XP_011524242.1;XP_011524243.1;XP_01 | 24343 | 609150 | . | . |
| 30832 | NM_014594,XI | ZNF354C | zinc finger protein 354C | protein_coding | NP_055409.1;XP_016864898.1 | 16736 | . | | . |
| 30833 | NM_00125237 | NT5C | 5', 3'-nucleotidase, cytosolic | protein_coding | NP_001239306.1;NP_055410.1;XP_011523002.2 | 17144 | 191720 | . | . |
| 30834 | NM_00127878 | POLR1H | RNA polymerase I subunit H | protein_coding | NP_001265714.1;NP_001265715.1;NP_055411.1;NP_740753.1 | 13182 | 607525 | . | . |
| 30836 | NM_014597 | DNTTIP2 | deoxynucleotidyltransferase terminal interacting protein 2 | protein_coding | NP_055412.2 | 24013 | 611199 | . | . |
| 30837 | NM_014598,XI | SOCS7 | suppressor of cytokine signaling 7 | protein_coding | NP_055413.2;XP_016880040.1;XP_016880041.1 | 29846 | 608788 | . | . |
| 30844 | NM_139265 | EHD4 | EH domain containing 4 | protein_coding | NP_644670.1 | 3245 | 605892 | . | . |
| 30845 | NM_014600,XI | EHD3 | EH domain containing 3 | protein_coding | NP_055415.1;XP_011531108.1 | 3244 | 605891 | . | . |
| 30846 | NM_014601 | EHD2 | EH domain containing 2 | protein_coding | NP_055416.2 | 3243 | 605890 | . | . |
| 30849 | NM_014602 | PIK3R4 | phosphoinositide-3-kinase regulatory subunit 4 | protein_coding | NP_055417.1 | 8982 | 602610 | . | . |

| | | | | | | | | | | | | | | | | | | | | | | | | | | | | | | | | | |
|---|---|---|---|---|---|---|---|---|---|---|---|---|---|---|---|---|---|---|---|---|---|---|---|---|---|---|---|---|---|---|---|---|---|
| 30850 | NM_014603,XI | CDR2L | cerebellar degeneration related protein 2 like | protein_coding | NP_055418.2;XP_006721915.1 | 29999 | . | . | . | 0.282348 | 0.318513 | 0.484967 | 0.273078 | 0.657034 | 0.736258 | 1.088252 | 0.624359 | 1.243695 | 37.816032 | 0.529043269 | 0.594719498 | 0.552030986 | 0.998865404 | 5.073577 | 5.390865 | 5.206631 | 5.219445 | 5.289618 | 5.198643 | 37 | 38 | 47 | 28 |
| 30851 | NM_00120469 | TAX1BP3 | Tax1 binding protein 3 | protein_coding | NP_001191627.1;NP_055419.1 | 30684 | 616484 | . | . | 15.569884 | 15.264493 | 12.350769 | 14.356475 | 36.231757 | 35.284558 | 27.714768 | 32.824371 | -1.170065 | 575.035698 | 0.164664548 | -1.37605993 | 0.16880308 | 0.945638095 | 9.276594 | 9.049494 | 9.226042 | 9.207120 | 9.068982 | 9.156008 | 724 | 660 | 429 | 531 |
| 30968 | NM_00128703 | STOML2 | stomatin like 2 | protein_coding | NP_001273960.1;NP_001273961.1;NP_001273962.1;NP_038470.1 | 14559 | 608292 | . | . | 157.285219 | 162.368517 | 142.177288 | 169.976291 | 366.009135 | 375.322105 | 319.041717 | 388.630620 | -1.040504 | 6697.388549 | 0.116637087 | -0.49111227 | 0.623347041 | 0.998865404 | 12.737802 | 12.680393 | 12.714902 | 12.735262 | 12.619179 | 12.759133 | 7771 | 7459 | 5234 | 6665 |
| 43847 | NM_00131118 | KLK14 | kallikrein related peptidase 14 | protein_coding | NP_001298111.2;NP_001356704.1;NP_071329.3;XP_016882309.1 | 6362 | 606135 | . | . | 0.461383 | 0.313266 | 0.385293 | 0.402395 | 1.073657 | 0.724129 | 0.864586 | 0.920027 | -1.042546 | 13.534431 | 0.83493929 | -0.07199468 | 0.942606139 | NA | 3.787711 | 3.728813 | 3.773090 | 3.748943 | 3.756028 | 3.761191 | 19 | 12 | 11 | 13 |
| 49854 | NM_00109840 | ZBTB21 | zinc finger and BTB domain containing 21 | protein_coding | NP_001091872.1;NP_001091873.1;NP_001307658.1;NP_001307660.1;NI | 13083 | 616485 | . | . | 3.569936 | 3.425489 | 4.792408 | 4.582452 | 8.307389 | 7.918173 | 10.754022 | 10.477234 | 1.309250 | 957.320676 | 0.139838057 | 2.779933198 | 0.005437008 | 0.225239728 | 9.695264 | 10.084305 | 9.804086 | 9.784671 | 10.018547 | 9.961563 | 968 | 882 | 980 | 979 |
| 49855 | NM_00114592 | SCAPER | S-phase cyclin A associated protein in the ER | protein_coding | NP_001139395.1;NP_001339938.1;NP_001339939.1;NP_001339940.1;NI | 13081 | 611611 | . | . | 3.026623 | 2.657216 | 2.778846 | 2.797823 | 7.043074 | 6.142275 | 6.235648 | 6.396890 | -1.025732 | 468.126249 | 0.17316062 | -0.21167584 | 0.832359939 | 0.998865404 | 8.888766 | 8.852514 | 8.909349 | 8.846509 | 8.868866 | 8.856474 | 573 | 486 | 406 | 429 |
| 49856 | NM_017818,XI | WRAP73 | WD repeat containing, antisense to TP73 | protein_coding | NP_060288.3;XP_005244811.1;XP_016856876.1 | 12759 | 606040 | . | . | 22.858528 | 23.327096 | 20.384000 | 24.928064 | 53.192733 | 53.921633 | 45.741104 | 56.995056 | -1.034872 | 1184.368545 | 0.144022179 | -0.34336161 | 0.73132644 | 0.998865404 | 10.234676 | 10.184696 | 10.217099 | 10.226249 | 10.119400 | 10.266870 | 1379 | 1308 | 912 | 1199 |
| 49861 | NM_00100134 | CLDN20 | claudin 20 | protein_coding | NP_001001346.1 | 2042 | . | . | . | 0.369777 | 0.281608 | 0.295007 | 0.155819 | 0.860487 | 0.650949 | 0.661988 | 0.356261 | -1.451513 | 10.946049 | 0.938919583 | -0.57252798 | 0.566964338 | NA | 3.694014 | 3.161867 | 3.473224 | 3.456695 | 3.458270 | 3.435946 | 17 | 12 | 10 | 6 |
| 50484 | NM_00117247 | RRM2B | ribonucleotide reductase regulatory TP53 inducible subunit M2B | protein_coding | NP_001165948.1;NP_001165949.1;NP_056528.2 | 17296 | 604712 | . | . | 4.070989 | 3.556867 | 3.818115 | 3.500740 | 9.473358 | 8.221859 | 8.567740 | 8.004027 | -1.056257 | 560.106966 | 0.16803682 | -0.46989926 | 0.63842699 | 0.998865404 | 9.168138 | 9.089919 | 9.190160 | 9.100728 | 9.140458 | 9.080373 | 707 | 579 | 496 | 487 |
| 50485 | NM_00112720 | SMARCAL1 | SWI/SNF related, matrix associated, actin dependent regulator of ( | protein_coding | NP_001120679.1;NP_054859.2 | 11102 | 606622 | . | . | 9.208166 | 10.857138 | 8.778940 | 11.195834 | 21.427782 | 25.096762 | 19.699687 | 25.597942 | -1.019243 | 984.414097 | 0.159044752 | -0.17289183 | 0.862736454 | 0.998865404 | 9.957217 | 9.928887 | 9.891969 | 10.001063 | 9.847551 | 10.013395 | 1060 | 1152 | 751 | 1018 |
| 50488 | NM_00102493 | MINK1 | misshapen like kinase 1 | protein_coding | NP_001020108.1;NP_001308165.1;NP_056531.1;NP_722549.2;NP_7337( | 17565 | 609426 | . | . | 19.229222 | 20.717205 | 22.273838 | 22.249441 | 44.747188 | 47.888748 | 49.981847 | 50.870702 | 1.101740 | 3299.519715 | 0.111813583 | 1.250151026 | 0.211244383 | 0.974817004 | 11.616489 | 11.756210 | 11.619058 | 11.673461 | 11.729788 | 11.722693 | 3495 | 3501 | 3021 | 3229 |
| 50489 | NM_015717,XI | CD207 | CD207 molecule | protein_coding | NP_056532.4;XP_011531176.1;XP_011531177.1;XP_011531178.1 | 17935 | 604862 | . | . | 0.223040 | 0.414528 | 0.304652 | 0.121530 | 0.519024 | 0.958200 | 0.683629 | 0.277863 | -1.501427 | 20.254881 | 0.702878824 | -0.83418957 | 0.404174165 | NA | 4.602488 | 4.019385 | 4.347206 | 4.368944 | 4.342682 | 4.304816 | 25 | 29 | 18 | 11 |
| 50509 | NM_015719,XI | COL5A3 | collagen type V alpha 3 chain | protein_coding | NP_056534.2;XP_011526344.1;XP_016882338.1 | 14864 | 120216 | . | . | 1.160728 | 1.254908 | 1.066443 | 1.088495 | 2.701061 | 2.900775 | 2.393066 | 2.488715 | -1.136750 | 222.280751 | 0.226780971 | -0.81538942 | 0.414849441 | 0.998865404 | 7.885882 | 7.700655 | 7.809096 | 7.842939 | 7.763410 | 7.767581 | 263 | 264 | 180 | 196 |
| 50512 | NM_015720 | PODXL2 | podocalyxin like 2 | protein_coding | NP_056535.1 | 17936 | 616627 | . | . | 36.887016 | 36.007900 | 29.719969 | 34.772163 | 85.837601 | 83.233871 | 66.690750 | 79.502411 | -1.146392 | 2328.332586 | 0.124508177 | -1.58303099 | 0.113414402 | 0.850211417 | 11.280377 | 11.083044 | 11.250333 | 11.223499 | 11.067420 | 11.183417 | 2911 | 2640 | 1748 | 2183 |
| 50515 | NM_00117398 | CHST11 | carbohydrate sulfotransferase 11 | protein_coding | NP_001167453.1;NP_060883.1;XP_016874858.1 | 17422 | 610128 | . | . | 22.465118 | 22.683895 | 19.173030 | 21.207043 | 52.277250 | 52.434838 | 43.023726 | 48.487386 | -1.133932 | 3815.539459 | 0.111969908 | -1.61949489 | 0.105340826 | 0.832940096 | 11.985551 | 11.804089 | 11.946716 | 11.947935 | 11.805395 | 11.879694 | 4666 | 4379 | 2968 | 3505 |
| 50516 | gene-PPP1R14 | PPP1R14BP1 | protein phosphatase 1 regulatory inhibitor subunit 14B pseudoger | pseudogene | . | 9058 | . | . | . | 0.287471 | 0.148264 | 0.207457 | 0.282873 | 0.668957 | 0.342718 | 0.465528 | 0.646756 | 1.086310 | 7.404942 | 1.127384735 | 0.105941068 | 0.91562911 | NA | 2.826455 | 2.947964 | 2.897949 | 2.875108 | 2.885755 | 2.892949 | 11 | 5 | 6 | 8 |
| 50615 | NM_021798,N | IL21R | interleukin 21 receptor | protein_coding | NP_068570.1;NP_851564.1;NP_851565.4;XP_011544159.1;XP_01154416 | 6006 | 605383 | . | . | 0.039580 | 0.035295 | 0.054997 | 0.036382 | 0.092103 | 0.081586 | 0.123411 | 0.083184 | 1.244255 | 7.026950 | 1.143009546 | 0.275834796 | 0.782674957 | NA | 2.644780 | 2.963455 | 2.806538 | 2.800097 | 2.816347 | 2.804103 | 8 | 6 | 8 | 6 |
| 50617 | NM_020632,N | ATP6V0A4 | ATPase H+ transporting V0 subunit a4 | protein_coding | NP_065683.2;NP_570855.2;NP_570856.2;XP_005250450.1;XP_00525045 | 866 | 605239 | . | . | 0.306234 | 0.316782 | 0.080638 | 0.133099 | 0.712620 | 0.732255 | 0.180950 | 0.304316 | -2.853079 | 23.304668 | 0.671655443 | -2.25192785 | 0.024326832 | NA | 5.110312 | 3.592943 | 4.588108 | 4.592912 | 4.486097 | 4.509326 | 39 | 38 | 8 | 14 |
| 50618 | NM_00134818 | ITSN2 | intersectin 2 | protein_coding | NP_001335110.1;NP_001335111.1;NP_001335112.1;NP_001335113.1;NI | 6184 | 604464 | . | . | 6.232240 | 6.293183 | 6.337817 | 6.333006 | 14.502679 | 14.546971 | 14.221881 | 14.479665 | -1.005463 | 1261.334160 | 0.128902458 | -0.06097257 | 0.951381055 | 0.998865404 | 10.304683 | 10.296776 | 10.293139 | 10.312341 | 10.299452 | 10.297835 | 1438 | 1382 | 1095 | 1178 |
| 50626 | NM_00112988 | CYHR1 | cysteine and histidine rich 1 | protein_coding | NP_001123360.1;NP_001317547.1;NP_116076.1;NP_612505.1 | 17806 | 616635 | . | . | 13.502576 | 12.932154 | 14.144034 | 14.421762 | 31.421048 | 29.893255 | 31.738803 | 32.973646 | 1.051125 | 656.124454 | 0.153173017 | 0.46963101 | 0.638618666 | 0.998865404 | 9.321491 | 9.393268 | 9.338806 | 9.342038 | 9.361563 | 9.386590 | 735 | 692 | 574 | 642 |
| 50628 | NM_015721,XI | GEMIN4 | gem nuclear organelle associated protein 4 | protein_coding | NP_056536.2;XP_005256724.1;XP_005256727.1;XP_011522212.1;XP_01: | 15717 | 606969 | . | . | 9.037648 | 9.062157 | 8.057099 | 9.466733 | 21.030977 | 20.947583 | 18.079894 | 21.644559 | -1.044056 | 1062.306147 | 0.140831387 | -0.44165441 | 0.658739303 | 0.998865404 | 10.083953 | 10.021336 | 10.069104 | 10.067599 | 9.983430 | 10.086658 | 1251 | 1170 | 840 | 1043 |
| 50631 | gene-YBX1P1 | YBX1P1 | Y-box binding protein 1 pseudogene 1 | pseudogene | . | 8015 | . | . | . | 7.041385 | 7.084807 | 6.207853 | 9.137536 | 16.385590 | 16.376848 | 13.930243 | 20.891890 | 1.071354 | 343.878744 | 0.21758559 | 0.456991538 | 0.647677128 | 0.998865404 | 8.375882 | 8.473964 | 8.402385 | 8.401060 | 8.329569 | 8.540682 | 383 | 358 | 252 | 395 |
| 50636 | NM_00100166 | ANO7 | anoctamin 7 | protein_coding | NP_001001666.2;NP_001357623.1;XP_011509565.1;XP_011509569.1;XP | 31677 | 605096 | . | . | 0.300104 | 0.372736 | 0.260746 | 0.398075 | 0.698355 | 0.861596 | 0.585106 | 0.910152 | -1.026169 | 47.913953 | 0.465821497 | -0.08000602 | 0.936232468 | 0.998865404 | 5.602488 | 5.561975 | 5.563293 | 5.602570 | 5.544045 | 5.611888 | 50 | 58 | 33 | 53 |
| 50640 | NM_00125600 | PNPLA8 | patatin like phospholipase domain containing 8 | protein_coding | NP_001242936.1;NP_001242937.1;NP_001242938.1;NP_001242939.1;NI | 28900 | 612123 | . | . | 6.504178 | 6.329495 | 6.944772 | 6.320951 | 15.135486 | 14.630914 | 15.583869 | 14.452104 | 1.034433 | 907.827926 | 0.142386798 | 0.343014694 | 0.731587406 | 0.998865404 | 9.801452 | 9.850678 | 9.831144 | 9.795411 | 9.866378 | 9.808728 | 1053 | 939 | 835 | 831 |
| 50649 | NM_00136749 | ARHGEF4 | Rho guanine nucleotide exchange factor 4 | protein_coding | NP_001354422.1;NP_001362829.1;NP_001362830.1;NP_001362831.1;NI | 684 | 605216 | . | . | 1.491006 | 2.149011 | 1.592893 | 1.737018 | 3.469634 | 4.967538 | 3.574405 | 3.971485 | 1.230960 | 351.915160 | 0.199201593 | 1.5049296 | 0.132342177 | 0.886721472 | 8.301006 | 8.601538 | 8.451168 | 8.331041 | 8.509692 | 8.512854 | 403 | 303 | 337 | 365 |
| 50650 | NM_00112861 | ARHGEF3 | Rho guanine nucleotide exchange factor 3 | protein_coding | NP_001122087.1;NP_001122088.1;NP_001276627.1;NP_001364336.1;NI | 683 | 612115 | . | . | 2.684850 | 2.546767 | 2.939706 | 3.093250 | 6.247757 | 5.886966 | 6.596613 | 7.072346 | 1.151072 | 313.038566 | 0.19883032 | 1.020861574 | 0.307320027 | 0.998865404 | 8.185110 | 8.388145 | 8.268888 | 8.228196 | 8.311202 | 8.339574 | 348 | 302 | 284 | 322 |
| 50717 | NM_015726,N | DCAF8 | DDB1 and CUL4 associated factor 8 | protein_coding | NP_056541.2 | 24891 | 615820 | . | . | 12.433036 | 13.551787 | 13.415832 | 13.618990 | 28.932186 | 31.325562 | 30.104736 | 31.138203 | 1.042071 | 1672.091553 | 0.124510621 | 0.47750096 | 0.633005437 | 0.998865404 | 10.677486 | 10.736781 | 10.657233 | 10.723315 | 10.720874 | 10.725029 | 1806 | 1842 | 1480 | 1604 |
| 50804 | NM_00130121 | MYEF2 | myelin expression factor 2 | protein_coding | NP_001288139.2;NP_057216.3;XP_005254479.2;XP_005254481.2;XP_00 | 17940 | . | . | . | 1.227757 | 1.047066 | 1.044178 | 1.031622 | 2.857039 | 2.420342 | 2.343106 | 2.358682 | -1.078844 | 261.299181 | 0.271513427 | -0.40324306 | 0.686769426 | 0.998865404 | 8.083217 | 7.973828 | 8.166604 | 7.892957 | 7.918809 | 8.075543 | 393 | 217 | 184 | 273 |
| 50807 | NM_00124799 | ASAP1 | ArfGAP with SH3 domain, ankyrin repeat and PH domain 1 | protein_coding | NP_001234925.1;NP_001349853.1;NP_001349854.1;NP_001349855.1;NI | 2720 | 605953 | . | . | 18.099017 | 20.410839 | 23.677404 | 20.801953 | 42.117155 | 47.180570 | 53.131411 | 47.561188 | 1.138621 | 4081.775938 | 0.121400845 | 1.542717283 | 0.122899387 | 0.867477288 | 11.898275 | 12.085610 | 11.891150 | 11.981823 | 12.101650 | 11.985799 | 4153 | 4346 | 4040 | 3794 |
| 50808 | NM_00119985 | AK3 | adenylate kinase 3 | protein_coding | NP_001186781.1;NP_001186782.1;NP_001186784.1;NP_001186785.1;NI | 17376 | 609290 | . | . | 4.519922 | 4.412460 | 4.622778 | 4.143150 | 10.518046 | 10.199599 | 10.373378 | 9.472820 | -1.033898 | 581.387731 | 0.163055252 | -0.29495196 | 0.768030576 | 0.998865404 | 9.206910 | 9.159413 | 9.206917 | 9.180369 | 9.210130 | 9.132229 | 694 | 625 | 527 | 504 |
| 50809 | NM_00137205 | HP1BP3 | heterochromatin protein 1 binding protein 3 | protein_coding | NP_001358981.1;NP_001363716.1;NP_001363717.1;NP_001363718.1;NI | 24973 | 616072 | . | . | 61.746284 | 65.767659 | 83.079034 | 59.314304 | 143.686139 | 152.024889 | 186.426930 | 135.615107 | 1.101306 | 12175.244704 | 0.138490073 | 1.005241157 | 0.314780742 | 0.998865404 | 13.500341 | 13.639625 | 13.501934 | 13.549224 | 13.733842 | 13.465753 | 12972 | 12852 | 12813 | 10116 |
| 50810 | NM_016073,XI | HDGFL3 | HDGF like 3 | protein_coding | NP_057157.1;XP_006720617.1 | 24937 | 616643 | . | . | 2.435338 | 2.254919 | 2.002578 | 2.202784 | 5.667131 | 5.212348 | 4.493726 | 5.036403 | -1.095402 | 809.502391 | 0.144593154 | -0.90917612 | 0.363257167 | 0.998865404 | 9.725133 | 9.593656 | 9.705915 | 9.678613 | 9.617846 | 9.638057 | 993 | 896 | 662 | 735 |
| 50813 | NM_00116409 | COPS7A | COP9 signalosome subunit 7A | protein_coding | NP_001157565.1;NP_001157566.1;NP_001157567.1;NP_057403.1;XP_0( | 16758 | 616009 | . | . | 20.935120 | 23.704914 | 20.663835 | 25.383858 | 48.716883 | 54.794972 | 46.369044 | 58.037180 | 1.027458 | 1368.023916 | 0.139842522 | 0.279452912 | 0.77989727 | 0.998865404 | 10.398480 | 10.437018 | 10.369134 | 10.442221 | 10.361060 | 10.488689 | 1480 | 1526 | 1105 | 1408 |
| 50814 | NM_00112976 | NSDHL | NAD(P) dependent steroid dehydrogenase-like | protein_coding | NP_001123237.1;NP_057006.1;XP_011529480.1;XP_016885053.1 | 13398 | 300275 | . | . | 12.530997 | 10.853606 | 18.067481 | 18.117382 | 29.160145 | 25.088594 | 40.542906 | 41.423243 | 1.531639 | 817.524264 | 0.151734163 | 4.053645461 | 5.04256E-05 | 0.007484138 | 9.334723 | 9.950333 | 9.551171 | 9.445566 | 9.810016 | 9.791985 | 803 | 641 | 871 | 916 |
| 50832 | NM_016944 | TAS2R4 | taste 2 receptor member 4 | protein_coding | NP_058640.1 | 14911 | 604869 | . | . | 0.043295 | 0.052686 | 0.064947 | 0.079939 | 0.100749 | 0.121787 | 0.145740 | 0.182771 | 1.516606 | 9.609503 | 0.982402426 | 0.611609121 | 0.540796402 | NA | 2.934350 | 3.532926 | 3.243064 | 3.249681 | 3.258672 | 3.269453 | 8 | 9 | 9 | 12 |
| 50840 | NM_023922 | TAS2R14 | taste 2 receptor member 14 | protein_coding | NP_076411.1 | 14920 | 604790 | . | . | 0.316349 | 0.255260 | 0.247731 | 0.245277 | 0.736158 | 0.590045 | 0.555902 | 0.560796 | -1.154122 | 8.352514 | 1.047924673 | -0.1973383 | 0.84356281 | NA | 3.161369 | 2.955730 | 3.070438 | 3.064820 | 3.063569 | 3.060959 | 11 | 9 | 7 | 7 |
| 50846 | NM_021044,XI | DHH | desert hedgehog signaling molecule | protein_coding | NP_066382.1;XP_016874869.1;XP_016874870.1 | 2865 | 605423 | . | . | 0.076232 | 0.040585 | 0.036290 | 0.041766 | 0.177394 | 0.093814 | 0.081434 | 0.095494 | -1.464598 | 7.481708 | 1.122513906 | -0.4904211 | 0.623835948 | NA | 3.151937 | 2.602870 | 2.928079 | 2.905948 | 2.903007 | 2.905732 | 13 | 7 | 5 | 6 |
| 50848 | NM_00134809 | F11R | F11 receptor | protein_coding | NP_001335020.1;NP_001369656.1;NP_001369659.1;NP_001369662.1;NI | 14685 | 605721 | . | . | 7.236984 | 7.228311 | 5.838896 | 6.660359 | 16.840759 | 16.708563 | 13.102312 | 15.228118 | -1.174284 | 975.526037 | 0.141855748 | -1.63392499 | 0.102274602 | 0.82130285 | 10.041458 | 9.809284 | 9.989013 | 9.982075 | 9.824587 | 9.910143 | 1219 | 1132 | 733 | 892 |
| 50853 | NM_00137026 | VILL | villin like | protein_coding | NP_001357193.1;NP_001357194.1;NP_056957.3 | 30906 | . | . | . | 1.403569 | 1.952297 | 1.656462 | 1.549699 | 3.266162 | 4.512821 | 3.717054 | 3.543206 | -1.043845 | 139.348541 | 0.289635609 | -0.21374415 | 0.830746579 | 0.998865404 | 7.153538 | 7.090890 | 7.067358 | 7.184572 | 7.125194 | 7.097856 | 139 | 177 | 123 | 123 |
| 50854 | NR_160945,NF | SNHG32 | small nucleolar RNA host gene 32 | lncRNA | . | 19078 | 605447 | . | . | 57.273806 | 55.487231 | 56.524649 | 64.991944 | 133.278513 | 128.261231 | 126.839674 | 148.596364 | 1.065840 | 1375.780633 | 0.132409416 | 0.694747559 | 0.487213518 | 0.998865404 | 10.379424 | 10.471186 | 10.414899 | 10.385965 | 10.401811 | 10.494632 | 1563 | 1410 | 1160 | 1411 |
| 50855 | NM_00103728 | PARD6A | par-6 family cell polarity regulator alpha | protein_coding | NP_001032358.1;NP_058644.1;XP_005256034.1;XP_011521397.1 | 15943 | 607484 | . | . | 1.303899 | 1.179034 | 0.837615 | 1.041840 | 3.034228 | 2.725391 | 1.879583 | 2.382044 | -1.311432 | 43.369085 | 0.475990774 | -0.82174499 | 0.411222046 | 0.998865404 | 5.621228 | 5.229445 | 5.474436 | 5.452254 | 5.407749 | 5.430202 | 60 | 50 | 30 | 38 |
| 50861 | NM_00127631 | STMN3 | stathmin 3 | protein_coding | NP_001263239.1;NP_056978.2 | 15926 | 608362 | . | . | 5.624543 | 5.553509 | 4.530894 | 6.937234 | 13.088541 | 12.837183 | 10.167196 | 15.861160 | 1.005476 | 401.149666 | 0.212707793 | 0.037039517 | 0.970453497 | 0.998865404 | 8.644708 | 8.651278 | 8.647504 | 8.636729 | 8.520353 | 8.756509 | 465 | 428 | 280 | 452 |
| 50862 | NM_016422 | RNF141 | ring finger protein 141 | protein_coding | NP_057506.2 | 21159 | 616641 | . | . | 3.470584 | 3.495770 | 3.341333 | 3.324728 | 8.076191 | 8.080628 | 7.497855 | 7.601594 | -1.059949 | 430.428384 | 0.175468395 | -0.47868633 | 0.632161794 | 0.998865404 | 8.790983 | 8.707056 | 8.767256 | 8.767282 | 8.736230 | 8.727633 | 510 | 478 | 366 | 389 |
| 50863 | NM_00104820 | NTM | neurotrimin | protein_coding | NP_001041674.1;NP_001137530.1;NP_001137531.1;NP_001338930.1;NI | 17941 | 607938 | . | . | 0.556276 | 0.678431 | 0.463557 | 0.353021 | 1.294474 | 1.568223 | 1.040207 | 0.807143 | -1.263965 | 62.319941 | 0.406051494 | -0.83230076 | 0.405239202 | 0.998865404 | 6.119956 | 5.783744 | 5.980270 | 5.992633 | 5.964045 | 5.906069 | 78 | 77 | 55 | 44 |
| 50865 | NM_015987 | HEBP1 | heme binding protein 1 | protein_coding | NP_057071.2 | 17176 | 605826 | . | . | 19.477703 | 20.549278 | 17.197252 | 23.065449 | 45.325417 | 47.500580 | 38.590134 | 52.736412 | -1.007756 | 724.683315 | 0.170652154 | -0.06531677 | 0.947921787 | 0.998865404 | 9.507230 | 9.495159 | 9.485788 | 9.517319 | 9.394297 | 9.588762 | 819 | 803 | 539 | 772 |
| 50937 | NM_00124359 | CDON | cell adhesion associated, oncogene regulated | protein_coding | NP_001230526.1;NP_001365893.1;NP_058648.4;XP_011541164.1;XP_01 | 17104 | 608707 | . | . | 0.713625 | 0.865650 | 0.920577 | 0.715414 | 1.660635 | 2.000987 | 2.065748 | 1.635711 | 1.018452 | 208.805156 | 0.248598356 | 0.10610489 | 0.915499131 | 0.998865404 | 7.692473 | 7.719428 | 7.653780 | 7.740414 | 7.763921 | 7.645555 | 216 | 244 | 207 | 172 |
| 50940 | NM_00107719 | PDE11A | phosphodiesterase 11A | protein_coding | NP_001070664.1;NP_001070665.1;NP_001070826.1;NP_058649.3 | 8773 | 604961 | . | . | 0.146600 | 0.184652 | 0.198184 | 0.188811 | 0.341143 | 0.426831 | 0.444720 | 0.431695 | 1.152093 | 52.268009 | 0.435354496 | 0.469173946 | 0.638945307 | 0.998865404 | 5.602488 | 5.806050 | 5.670095 | 5.709555 | 5.722443 | 5.711708 | 50 | 58 | 50 | 51 |
| 50943 | NM_00111437 | FOXP3 | forkhead box P3 | protein_coding | NP_001107849.1;NP_054728.2;XP_006724596.2;XP_016885056.1 | 6106 | 300292 | . | . | 0.174858 | 0.078106 | 0.213459 | 0.174600 | 0.406901 | 0.180546 | 0.478998 | 0.399204 | 1.562754 | 13.645914 | 0.856035986 | 0.752411192 | 0.451803818 | NA | 3.409440 | 4.058876 | 3.763668 | 3.726020 | 3.778484 | 3.764121 | 17 | 7 | 16 | 14 |
| 50999 | NM_00116783 | TMED5 | transmembrane p24 trafficking protein 5 | protein_coding | NP_001161302.1;NP_057124.3 | 24251 | 616876 | . | . | 10.350565 | 8.191661 | 8.603521 | 6.349195 | 24.086189 | 18.935391 | 19.306049 | 14.516678 | -1.261655 | 1517.813806 | 0.168505052 | -1.98995208 | 0.046596214 | 0.654401867 | 10.725397 | 10.390804 | 10.737195 | 10.546085 | 10.576698 | 10.342427 | 2182 | 1609 | 1349 | 1063 |
| 51000 | NM_00114254 | SLC35B3 | solute carrier family 35 member B3 | protein_coding | NP_001136012.1;NP_001136013.1;NP_001357405.1;NP_001357406.1;NI | 21601 | 610845 | . | . | 8.763272 | 7.724068 | 8.009524 | 5.884164 | 20.392495 | 17.854526 | 17.973141 | 13.453436 | -1.135068 | 567.554750 | 0.191035844 | -0.95677766 | 0.338679488 | 0.998865404 | 9.236492 | 9.055038 | 9.223218 | 9.144495 | 9.205707 | 8.986099 | 735 | 613 | 542 | 413 |
| 51001 | NM_00128664 | MTERF3 | mitochondrial transcription termination factor 3 | protein_coding | NP_001273572.1;NP_001349893.1;NP_057026.3;XP_011515356.1 | 24258 | 616930 | . | . | 12.552324 | 12.617860 | 12.624649 | 11.030048 | 29.209776 | 29.166750 | 28.329348 | 25.218895 | -1.076509 | 555.216182 | 0.166404073 | -0.63917092 | 0.522711745 | 0.998865404 | 9.168811 | 9.063064 | 9.143169 | 9.136546 | 9.136188 | 9.045801 | 666 | 618 | 498 | 466 |
| 51002 | NM_00133038 | TPRKB | TP53RK binding protein | protein_coding | NP_001317315.1;NP_001317316.1;NP_001317317.1;NP_001317318.1;NI | 24259 | 608680 | . | . | 23.295482 | 19.745804 | 19.309261 | 18.144467 | 54.209546 | 45.643309 | 43.329420 | 41.485165 | -1.131087 | 465.936801 | 0.178332699 | -0.99650611 | 0.319004301 | 0.998865404 | 8.949837 | 8.772711 | 8.952714 | 8.845298 | 8.831481 | 8.816470 | 617 | 489 | 385 | 405 |
| 51003 | NM_016060 | MED31 | mediator complex subunit 31 | protein_coding | NP_057144.1 | 24260 | . | . | . | 4.644070 | 4.148810 | 4.957336 | 4.575694 | 10.806942 | 9.590161 | 11.124118 | 10.461781 | 1.070686 | 233.090958 | 0.225536669 | 0.436895996 | 0.662186776 | 0.998865404 | 7.814164 | 7.913621 | 7.875452 | 7.816668 | 7.900981 | 7.857843 | 275 | 228 | 219 | 216 |
| 51004 | NM_182476,N | COQ6 | coenzyme Q6, monooxygenase | protein_coding | NP_872282.1;NP_872286.2;XP_006720219.1;XP_011535109.1;XP_01153 | 20233 | 614647 | . | . | 9.810240 | 9.559985 | 9.892674 | 9.049111 | 22.828833 | 22.098332 | 22.198873 | 20.689718 | -1.015997 | 591.819123 | 0.163394629 | -0.14013187 | 0.888555804 | 0.998865404 | 9.220124 | 9.197815 | 9.254855 | 9.170685 | 9.222789 | 9.183391 | 730 | 603 | 526 | 534 |
| 51005 | NM_00114581 | AMDHD2 | amidohydrolase domain containing 2 | protein_coding | NP_001139287.1;NP_001317378.1;NP_057028.2;XP_016878752.1;XP_01 | 24262 | . | . | . | 24.285925 | 25.568003 | 29.030372 | 26.456337 | 56.514348 | 59.101583 | 65.143310 | 60.489274 | 1.116380 | 2576.920841 | 0.119225406 | 1.332165432 | 0.182805833 | 0.954698793 | 11.249769 | 11.408720 | 11.268026 | 11.300688 | 11.417002 | 11.328543 | 2747 | 2681 | 2492 | 2411 |
| 51006 | NM_00128145 | SLC35C2 | solute carrier family 35 member C2 | protein_coding | NP_001268386.1;NP_001268387.1;NP_001268388.1;NP_001268389.1;NI | 17117 | . | . | . | 7.140363 | 6.614187 | 6.715605 | 7.220708 | 16.615919 | 15.288994 | 15.069620 | 16.509288 | -1.003113 | 770.263170 | 0.146110785 | -0.03068792 | 0.975518424 | 0.998865404 | 9.591509 | 9.586902 | 9.579633 | 9.600824 | 9.584176 | 9.591954 | 875 | 845 | 665 | 725 |
| 51008 | NM_00119879 | ASCC1 | activating signal cointegrator 1 complex subunit 1 | protein_coding | NP_001185727.1;NP_001185728.1;NP_001185729.1;NP_001356014.1;NI | 24268 | 614215 | . | . | 8.197684 | 8.497436 | 8.388263 | 9.939919 | 19.076347 | 19.642205 | 18.823017 | 22.726449 | 1.072336 | 709.041391 | 0.157832746 | 0.638375458 | 0.523229302 | 0.998865404 | 9.418745 | 9.518967 | 9.449159 | 9.438987 | 9.431671 | 9.549104 | 794 | 733 | 584 | 746 |
| 51009 | NM_00130477 | DERL2 | derlin 2 | protein_coding | NP_001291706.1;NP_001291708.1;NP_057125.2 | 17943 | 610304 | . | . | 10.870471 | 10.382061 | 11.338837 | 9.578067 | 25.296033 | 23.998600 | 25.444021 | 21.899113 | -1.032314 | 1398.022700 | 0.139194776 | -0.32962048 | 0.741686744 | 0.998865404 | 10.471679 | 10.426309 | 10.479635 | 10.440026 | 10.503767 | 10.362989 | 1677 | 1493 | 1307 | 1171 |
| 51010 | NM_00100226 | EXOSC3 | exosome component 3 | protein_coding | NP_001002269.1;NP_057126.2 | 17944 | 606489 | . | . | 7.094586 | 7.351647 | 7.298706 | 6.325859 | 16.509395 | 16.993662 | 16.378084 | 14.463326 | -1.076969 | 395.815012 | 0.185915365 | -0.57540419 | 0.565017972 | 0.998865404 | 8.680891 | 8.574513 | 8.641197 | 8.657947 | 8.647691 | 8.562203 | 466 | 449 | 356 | 331 |
| 51011 | NM_016044,XI | FAHD2A | fumarylacetoacetate hydrolase domain containing 2A | protein_coding | NP_057128.2;XP_011509584.1;XP_011509585.1;XP_011509586.1;XP_01: | 24252 | . | . | . | 15.385182 | 19.524467 | 15.987540 | 15.624665 | 35.801953 | 45.131678 | 35.875581 | 35.723941 | -1.015716 | 1802.706129 | 0.122335589 | -0.18389841 | 0.854093152 | 0.998865404 | 10.827262 | 10.804546 | 10.805904 | 10.837753 | 10.783440 | 10.835103 | 2050 | 2000 | 1504 | 1732 |
| 51012 | NM_00125640 | PRELID3B | PRELI domain containing 3B | protein_coding | NP_001243332.1;NP_057129.2 | 15892 | . | . | . | 31.629222 | 29.223947 | 29.494188 | 27.559218 | 73.602501 | 67.552460 | 66.184097 | 63.010876 | -1.080781 | 2298.058021 | 0.121306858 | -0.92388475 | 0.355546325 | 0.998865404 | 11.221023 | 11.109210 | 11.231096 | 11.160002 | 11.164891 | 11.101934 | 2869 | 2463 | 1997 | 1989 |
| 51013 | NM_00131836 | EXOSC1 | exosome component 1 | protein_coding | NP_001305291.1;NP_001305292.1;NP_001305293.1;NP_001305294.1;NI | 17286 | 606493 | . | . | 23.982992 | 20.965586 | 20.406986 | 29.343893 | 55.809409 | 48.462894 | 45.792685 | 67.091325 | -1.050770 | 1060.168502 | 0.141063492 | -0.50648988 | 0.612512794 | 0.998865404 | 10.085524 | 10.013738 | 10.081331 | 10.053964 | 9.979392 | 10.080181 | 1274 | 1151 | 837 | 1036 |
| 51014 | NM_181836 | TMED7 | transmembrane p24 trafficking protein 7 | protein_coding | NP_861974.1 | 24253 | . | . | . | 18.313456 | 11.341983 | 13.401562 | 8.235338 | 42.616165 | 26.217505 | 30.072714 | 18.829123 | -1.389055 | 1568.978841 | 0.488204198 | -0.971117 | 0.331490018 | 0.998865404 | 10.833214 | 10.359254 | 10.892744 | 10.499869 | 10.625462 | 10.259018 | 2603 | 1498 | 1420 | 932 |
| 51015 | NM_016048 | ISOC1 | isochorismatase domain containing 1 | protein_coding | NP_057132.2 | 24254 | . | . | . | 8.031786 | 7.326824 | 8.035509 | 8.028332 | 18.690294 | 16.936281 | 18.031450 | 18.355825 | 1.030617 | 475.416258 | 0.172161501 | 0.252717527 | 0.800486503 | 0.998865404 | 8.870944 | 8.914818 | 8.913809 | 8.851356 | 8.905367 | 8.898592 | 566 | 480 | 422 | 450 |
| 51016 | NM_00134687 | EMC9 | ER membrane protein complex subunit 9 | protein_coding | NP_001333803.1;NP_001333804.1;NP_001333805.1;NP_001333806.1;NI | 20273 | . | . | . | 7.053223 | 5.865227 | 6.389083 | 5.972553 | 16.413138 | 13.557736 | 14.336919 | 13.655537 | -1.078946 | 223.439364 | 0.228061154 | -0.48067158 | 0.630749933 | 0.998865404 | 7.857138 | 7.748288 | 7.845185 | 7.797258 | 7.802721 | 7.768027 | 280 | 238 | 194 | 194 |
| 51018 | NM_016052,XI | RRP15 | ribosomal RNA processing 15 homolog | protein_coding | NP_057136.2;XP_011507899.1 | 24255 | 611193 | . | . | 2.485187 | 2.429008 | 2.569842 | 2.876847 | 5.783131 | 5.614760 | 5.766650 | 6.577571 | 1.116962 | 526.240729 | 0.164769444 | 0.968501949 | 0.332793746 | 0.998865404 | 8.957622 | 9.117129 | 8.984545 | 9.019001 | 9.081036 | 9.064758 | 557 | 551 | 489 | 514 |
| 51019 | NM_00130110 | WASHC3 | WASH complex subunit 3 | protein_coding | NP_001288036.1;NP_057137.1;XP_011536725.1;XP_016874872.1;XP_02 | 24256 | . | . | . | 18.118064 | 16.737310 | 20.477065 | 19.816511 | 42.161480 | 38.689043 | 45.949935 | 45.308094 | 1.171079 | 518.647324 | 0.165835914 | 1.37387811 | 0.169479526 | 0.946181982 | 8.900014 | 9.128193 | 8.971924 | 8.956442 | 9.085566 | 9.044515 | 557 | 509 | 502 | 508 |
| 51020 | NM_016063,XI | HDDC2 | HD domain containing 2 | protein_coding | NP_057147.2;XP_024302218.1 | 21078 | . | . | . | 32.645700 | 31.444873 | 29.480720 | 36.610443 | 75.967888 | 72.686230 | 66.153879 | 83.705425 | 1.039429 | 1339.579421 | 0.131270936 | 0.425009174 | 0.670829987 | 0.998865404 | 10.359500 | 10.415093 | 10.384860 | 10.359504 | 10.362781 | 10.439170 | 1538 | 1394 | 1128 | 1344 |
| 51021 | NM_016065 | MRPS16 | mitochondrial ribosomal protein S16 | protein_coding | NP_057149.1 | 14048 | 609204 | . | . | 27.041906 | 27.285881 | 25.038427 | 29.607244 | 62.927624 | 63.072540 | 56.185505 | 67.693451 | -1.008202 | 2188.546606 | 0.126333168 | -0.09327723 | 0.92568332 | 0.998865404 | 11.101778 | 11.089712 | 11.098172 | 11.098562 | 11.026970 | 11.152662 | 2530 | 2372 | 1746 | 2204 |
| 51022 | NM_00124339 | GLRX2 | glutaredoxin 2 | protein_coding | NP_001230328.1;NP_057150.2;NP_932066.1 | 16065 | 606820 | . | . | 22.493489 | 20.215353 | 20.906633 | 16.122716 | 52.343276 | 46.728701 | 46.913878 | 36.862677 | -1.169719 | 425.173474 | 0.195513388 | -1.15675909 | 0.247370802 | 0.992733415 | 8.839918 | 8.615152 | 8.810521 | 8.737868 | 8.755872 | 8.598527 | 558 | 466 | 387 | 318 |
| 51023 | NM_00129776 | MRPS18C | mitochondrial ribosomal protein S18C | protein_coding | NP_001284696.1;NP_001284698.1;NP_001284699.1;NP_057151.1 | 16633 | 611983 | . | . | 13.604610 | 12.911321 | 13.316456 | 10.200894 | 31.658488 | 29.845094 | 29.881741 | 23.323132 | -1.203500 | 557.722647 | 0.184353442 | -1.44958635 | 0.147173905 | 0.90884489 | 9.250283 | 8.984282 | 9.202328 | 9.152291 | 9.147706 | 8.957672 | 727 | 633 | 507 | 403 |
| 51024 | NM_016068 | FIS1 | fission, mitochondrial 1 | protein_coding | NP_057152.2 | 21689 | 609003 | . | . | 57.776855 | 59.513718 | 55.018333 | 70.449683 | 134.449112 | 137.586634 | 123.459557 | 161.175430 | 1.055013 | 1644.838050 | 0.14205883 | 0.543868939 | 0.586531665 | 0.998865404 | 10.644818 | 10.721619 | 10.652381 | 10.668966 | 10.605410 | 10.791020 | 1824 | 1746 | 1295 | 1770 |
| 51025 | NM_016069 | PAM16 | presequence translocase associated motor 16 | protein_coding | NP_057153.8 | 29679 | 614336 | . | . | 14.521920 | 17.309322 | 15.080556 | 15.491077 | 33.793102 | 40.011276 | 33.840332 | 35.418510 | -1.052159 | 259.469419 | 0.217735029 | -0.33688868 | 0.73620082 | 0.998865404 | 8.055960 | 7.981932 | 7.987467 | 8.073725 | 8.001562 | 8.009735 | 281 | 311 | 218 | 239 |
| 51026 | NM_016072,XI | GOLT1B | golgi transport 1B | protein_coding | NP_057156.1;XP_024304767.1 | 20175 | 615078 | . | . | 13.889017 | 9.547066 | 11.420730 | 5.684161 | 32.320312 | 22.068470 | 25.627787 | 12.996159 | -1.388073 | 1029.967750 | 0.561298759 | -0.84283679 | 0.399319729 | 0.998865404 | 10.225537 | 9.752652 | 10.235592 | 9.939314 | 10.069534 | 9.583511 | 1640 | 1047 | 1005 | 534 |
| 51027 | NM_00132102 | BOLA1 | bolA family member 1 | protein_coding | NP_001307954.1;NP_001307955.1;NP_057158.1;XP_006711411.1 | 24263 | 613181 | . | . | 8.317229 | 11.007620 | 10.779837 | 12.106420 | 19.354534 | 25.444609 | 24.189642 | 27.679891 | 1.206529 | 307.770915 | 0.21820623 | 1.24131727 | 0.214488567 | 0.979353522 | 8.124786 | 8.394091 | 8.136839 | 8.272735 | 8.256428 | 8.358637 | 280 | 340 | 266 | 344 |
| 51028 | NM_00128216 | VPS36 | vacuolar protein sorting 36 homolog | protein_coding | NP_001269097.1;NP_001269098.1;NP_057159.2 | 20312 | 610903 | . | . | 7.944805 | 8.627531 | 8.024914 | 7.521275 | 18.487890 | 19.942926 | 18.007675 | 17.196501 | -1.084846 | 1113.673747 | 0.136457198 | -0.86100248 | 0.389236679 | 0.998865404 | 10.178625 | 10.061209 | 10.121960 | 10.177970 | 10.116262 | 10.062612 | 1285 | 1297 | 963 | 965 |
| 51029 | NM_00129774 | DESI2 | desumoylating isopeptidase 2 | protein_coding | NP_001284675.1;NP_057160.2;XP_011542505.1;XP_011542507.1;XP_01 | 24264 | 614638 | . | . | 8.534493 | 7.797814 | 8.271795 | 6.396407 | 19.860115 | 18.024994 | 18.561666 | 14.624623 | -1.149118 | 975.212020 | 0.163567671 | -1.22596077 | 0.220213438 | 0.982008277 | 10.025938 | 9.826305 | 10.018921 | 9.929690 | 9.968434 | 9.769977 | 1272 | 1058 | 899 | 733 |
| 51030 | NM_00131691 | TVP23B | trans-golgi network vesicle protein 23 homolog B | protein_coding | NP_001303848.1;NP_001303849.1;NP_001303850.1;NP_001303851.1;NI | 20399 | . | . | . | 24.468388 | 20.298554 | 23.419894 | 15.429870 | 56.938947 | 46.921014 | 52.553563 | 35.278564 | -1.157170 | 1245.786549 | 0.182319824 | -1.15511552 | 0.248043123 | 0.992733415 | 10.383908 | 10.174158 | 10.403189 | 10.249427 | 10.358599 | 10.056826 | 1691 | 1299 | 1208 | 864 |
| 51031 | NM_00136624 | GLOD4 | glyoxalase domain containing 4 | protein_coding | NP_001353176.1;NP_001353177.1;NP_001353178.1;NP_001353179.1;NI | 14111 | . | . | . | 24.622352 | 26.113258 | 23.387636 | 28.107299 | 57.297223 | 60.361963 | 52.481180 | 64.264003 | -1.002027 | 1484.952511 | 0.13675725 | -0.02136186 | 0.982956996 | 0.998865404 | 10.537901 | 10.534499 | 10.513242 | 10.558425 | 10.465487 | 10.598920 | 1662 | 1651 | 1178 | 1511 |
| 51035 | NM_00128607 | UBXN1 | UBX domain protein 1 | protein_coding | NP_001273006.1;NP_001273007.1;NP_056937.2;XP_005274090.1;XP_01 | 18402 | 616378 | . | . | 39.400488 | 36.139847 | 33.655787 | 40.467806 | 91.686553 | 83.538880 | 75.522615 | 92.524833 | -1.001069 | 1364.258440 | 0.135756979 | -0.01135837 | 0.990937529 | 0.999408937 | 10.414762 | 10.413040 | 10.445442 | 10.380864 | 10.361003 | 10.462085 | 1638 | 1411 | 1107 | 1363 |
| 51042 | NM_015871 | ZNF593 | zinc finger protein 593 | protein_coding | NP_056955.2 | 30943 | 616698 | . | . | 28.587372 | 30.591444 | 23.169502 | 29.181091 | 66.523987 | 70.713493 | 51.991695 | 66.719101 | -1.145114 | 554.640830 | 0.1744368 | -1.1206962 | 0.262417202 | 0.995798899 | 9.210314 | 9.013820 | 9.136968 | 9.176352 | 8.996756 | 9.135439 | 662 | 658 | 400 | 538 |
| 51043 | NM_00125240 | ZBTB7B | zinc finger and BTB domain containing 7B | protein_coding | NP_001239335.1;NP_001243384.1;NP_001364380.1;NP_001364381.1;NI | 18668 | 607646 | . | . | 5.158269 | 5.203094 | 5.326029 | 5.514395 | 12.003502 | 12.027183 | 11.951457 | 12.608009 | 1.050065 | 615.058275 | 0.15616045 | 0.451323697 | 0.651756268 | 0.998865404 | 9.228875 | 9.299421 | 9.241920 | 9.253398 | 9.290074 | 9.270432 | 685 | 653 | 556 | 582 |
| 51050 | NM_00132440 | PI15 | peptidase inhibitor 15 | protein_coding | NP_001311332.1;NP_056970.1 | 8946 | 607076 | . | . | 14.702672 | 11.990238 | 17.170397 | 11.773087 | 34.213722 | 27.715972 | 38.529872 | 26.917765 | 1.069405 | 2925.323666 | 0.16814305 | 0.575746448 | 0.564786575 | 0.998865404 | 11.464960 | 11.562164 | 11.562882 | 11.390008 | 11.681625 | 11.364129 | 3600 | 2727 | 3133 | 2292 |
| 51053 | NM_00125198 | GMNN | geminin DNA replication inhibitor | protein_coding | NP_001238918.1;NP_001238919.1;NP_001238920.1;NP_056979.1;XP_0( | 17493 | 602842 | . | . | 21.913147 | 24.114486 | 18.963382 | 22.936940 | 50.992798 | 55.741715 | 42.553278 | 52.442589 | -1.109566 | 870.374575 | 0.147189461 | -1.01905967 | 0.308174639 | 0.998865404 | 9.838809 | 9.688250 | 9.772156 | 9.829545 | 9.686491 | 9.765321 | 1013 | 1027 | 677 | 817 |
| 51054 | NR_037144 | PLEKHA8P1 | pleckstrin homology domain containing A8 pseudogene 1 | transcribed_ps | . | 30222 | . | . | . | 1.529669 | 1.752009 | 1.605609 | 1.432882 | 3.559602 | 4.049848 | 3.602941 | 3.276114 | -1.088556 | 92.671142 | 0.334280065 | -0.36620695 | 0.714210654 | 0.998865404 | 6.593851 | 6.471658 | 6.525963 | 6.565352 | 6.538473 | 6.503824 | 104 | 111 | 82 | 78 |
| 51056 | NM_015907 | LAP3 | leucine aminopeptidase 3 | protein_coding | NP_056991.2 | 18449 | 170250 | . | . | 20.990398 | 22.916658 | 20.418076 | 23.971767 | 48.845520 | 52.972881 | 45.817570 | 54.808601 | -1.003056 | 1444.362888 | 0.135790004 | -0.03241921 | 0.974137742 | 0.998865404 | 10.498657 | 10.493774 | 10.465568 | 10.526764 | 10.435083 | 10.549950 | 1600 | 1623 | 1160 | 1453 |
| 51057 | NM_00104269 | WDPCP | WD repeat containing planar cell polarity effector | protein_coding | NP_001036157.1;NP_001340973.1;NP_001340974.1;NP_056994.3;XP_0( | 28027 | 613580 | . | . | 1.934629 | 2.017932 | 1.722327 | 1.695954 | 4.501958 | 4.664541 | 3.864853 | 3.877596 | -1.015030 | 231.755108 | 0.223877412 | -0.09613563 | 0.923412857 | 0.998865404 | 7.866843 | 7.845996 | 7.869183 | 7.850128 | 7.875883 | 7.828710 | 274 | 247 | 210 | 205 |
| 51058 | NM_00124273 | ZNF691 | zinc finger protein 691 | protein_coding | NP_001229668.1;NP_056995.1;XP_006710734.1;XP_016856890.1;XP_01 | 28028 | . | . | . | 4.324876 | 3.911574 | 4.897411 | 4.351298 | 10.064162 | 9.041782 | 10.989649 | 9.948723 | 1.050381 | 269.840683 | 0.212492549 | 0.333721461 | 0.738589754 | 0.998865404 | 8.039657 | 8.111380 | 8.090512 | 8.032793 | 8.107632 | 8.066925 | 320 | 268 | 250 | 249 |
| 51059 | NM_00136296 | FAM135B | family with sequence similarity 135 member B | protein_coding | NP_001349894.1;NP_056996.2;XP_011515358.1;XP_011515360.1;XP_01 | 28029 | . | . | . | 0.212859 | 0.299792 | 0.250866 | 0.327205 | 0.495330 | 0.692984 | 0.562935 | 0.748113 | 1.103593 | 67.268318 | 0.400420948 | 0.355145835 | 0.722480341 | 0.998865404 | 6.000465 | 6.139879 | 6.015295 | 6.094131 | 6.049733 | 6.106115 | 61 | 81 | 54 | 74 |
| 51060 | NM_015913,N | TXNDC12 | thioredoxin domain containing 12 | protein_coding | NP_056997.1 | 24626 | 609448 | . | . | 64.909730 | 66.272138 | 67.976723 | 60.554452 | 151.047614 | 153.191022 | 152.537797 | 138.450568 | -1.011723 | 2941.903317 | 0.119857668 | -0.14028426 | 0.8884354 | 0.998865404 | 11.530825 | 11.514196 | 11.519617 | 11.533599 | 11.571587 | 11.459755 | 3378 | 3220 | 2718 | 2554 |
| 51061 | NM_00130344 | TXNDC11 | thioredoxin domain containing 11 | protein_coding | NP_001290376.1;NP_001310951.1;NP_001310953.1;NP_001310954.1;NI | 28030 | 617792 | . | . | 20.976080 | 19.412628 | 18.168817 | 20.484066 | 48.812203 | 44.873160 | 40.770295 | 46.834384 | -1.069838 | 1927.318428 | 0.126040218 | -0.77270325 | 0.439698021 | 0.998865404 | 10.960308 | 10.862803 | 10.969993 | 10.905231 | 10.841448 | 10.926017 | 2387 | 2063 | 1532 | 1840 |
| 51062 | NM_00112771 | ATL1 | atlastin GTPase 1 | protein_coding | NP_001121185.1;NP_056999.2;NP_853629.2 | 11231 | 606439 | . | . | 2.268110 | 2.134870 | 1.516494 | 1.756784 | 5.277986 | 4.934848 | 3.402969 | 4.016676 | -1.387291 | 157.550231 | 0.264887509 | -1.78291131 | 0.074600742 | 0.764716896 | 7.516765 | 7.044015 | 7.377522 | 7.343041 | 7.210578 | 7.250964 | 219 | 190 | 108 | 131 |
| 51063 | NM_015916,N | CALHM2 | calcium homeostasis modulator family member 2 | protein_coding | NP_057000.2;XP_006717943.1;XP_006717946.1;XP_006717947.1;XP_01: | 23493 | 612235 | . | . | 1.729808 | 1.287284 | 1.093958 | 1.377862 | 4.025334 | 2.975610 | 2.454809 | 3.150315 | -1.052537 | 96.296877 | 0.331701871 | -0.22270193 | 0.823767499 | 0.998865404 | 6.625875 | 6.552014 | 6.627180 | 6.565970 | 6.562059 | 6.601210 | 125 | 96 | 76 | 94 |
| 51065 | NM_015920 | RPS27L | ribosomal protein S27 like | protein_coding | NP_057004.1 | 18476 | 612055 | . | . | 2.994869 | 3.225842 | 3.139180 | 3.463942 | 6.969184 | 7.456679 | 7.044228 | 7.919891 | 1.047029 | 610.375266 | 0.159735254 | 0.415071461 | 0.67808959 | 0.998865404 | 9.220295 | 9.286061 | 9.215112 | 9.259402 | 9.238183 | 9.296935 | 664 | 665 | 519 | 611 |
| 51067 | NM_00104043 | YARS2 | tyrosyl-tRNA synthetase 2 | protein_coding | NP_001035526.1 | 24249 | 610957 | . | . | 7.300164 | 6.847554 | 7.094973 | 8.534021 | 16.987782 | 15.828429 | 15.920913 | 19.512023 | 1.063382 | 507.716688 | 0.176637486 | 0.501934949 | 0.61571328 | 0.998865404 | 8.943165 | 9.031251 | 8.985576 | 8.947122 | 8.940276 | 9.066920 | 584 | 515 | 410 | 539 |
| 51068 | NM_00132022 | NMD3 | NMD3 ribosome export adaptor | protein_coding | NP_001307156.1;NP_057022.2;XP_005247568.1;XP_005247569.1 | 24250 | 611021 | . | . | 17.198666 | 15.461389 | 15.942942 | 14.226867 | 40.022002 | 35.739695 | 35.775501 | 32.528045 | -1.111590 | 1510.784381 | 0.13511336 | -1.12960821 | 0.258641351 | 0.995760118 | 10.635161 | 10.482993 | 10.650285 | 10.546543 | 10.558527 | 10.475887 | 1952 | 1603 | 1312 | 1269 |
| 51069 | NM_00130084 | MRPL2 | mitochondrial ribosomal protein L2 | protein_coding | NP_001287777.1;NP_057034.2;XP_005249218.1 | 14056 | 611822 | . | . | 20.968099 | 20.217464 | 19.302780 | 23.616610 | 48.793627 | 46.733578 | 43.314875 | 53.996575 | 1.028565 | 792.290897 | 0.15691158 | 0.258954786 | 0.795670127 | 0.998865404 | 9.609683 | 9.649811 | 9.635323 | 9.601846 | 9.566954 | 9.705250 | 921 | 823 | 630 | 827 |
| 51070 | NM_00127096 | NOSIP | nitric oxide synthase interacting protein | protein_coding | NP_001257889.1;NP_001350578.1;NP_057037.1;XP_005259021.1;XP_01 | 17946 | 616759 | . | . | 34.150975 | 32.980879 | 26.455968 | 34.299986 | 79.470717 | 76.236776 | 59.366423 | 78.422838 | -1.131797 | 1072.673071 | 0.155109972 | -1.15154328 | 0.2495088 | 0.993339639 | 10.153812 | 9.974616 | 10.127282 | 10.091168 | 9.919568 | 10.106935 | 1343 | 1200 | 762 | 1065 |
| 51071 | NM_00130077 | DERA | deoxyribose-phosphate aldolase | protein_coding | NP_001287708.1;NP_057038.2;XP_024304768.1;XP_024304769.1 | 24269 | . | . | . | 17.624729 | 16.752950 | 17.946092 | 19.405243 | 41.013473 | 38.725192 | 40.270516 | 44.367783 | 1.068849 | 916.839172 | 0.1417667 | 0.677578966 | 0.498038686 | 0.998865404 | 9.791727 | 9.887726 | 9.837344 | 9.793210 | 9.839664 | 9.886962 | 1052 | 927 | 797 | 917 |
| 51072 | NM_00113760 | MEMO1 | mediator of cell motility 1 | protein_coding | NP_001131074.1;NP_001288762.1;NP_001288781.1;NP_001358841.1;NI | 14014 | 611786 | . | . | 3.271161 | 3.401337 | 3.622752 | 3.007570 | 7.612127 | 7.862344 | 8.129350 | 6.876452 | -1.044312 | 859.318755 | 0.149440103 | -0.4185809 | 0.675522449 | 0.998865404 | 9.777772 | 9.715659 | 9.750167 | 9.772709 | 9.784152 | 9.674178 | 995 | 962 | 788 | 727 |
| 51073 | NM_015956,N | MRPL4 | mitochondrial ribosomal protein L4 | protein_coding | NP_057040.2;NP_666499.1;NP_666500.1;XP_011526347.1 | 14276 | 611823 | . | . | 27.821713 | 29.516910 | 29.217250 | 32.320477 | 64.742269 | 68.229661 | 65.562659 | 73.896930 | 1.067083 | 1289.645068 | 0.132984795 | 0.704384229 | 0.481193525 | 0.998865404 | 10.285330 | 10.378677 | 10.302309 | 10.311809 | 10.311978 | 10.398739 | 1428 | 1355 | 1092 | 1319 |
| 51074 | NM_015957,XI | APIP | APAF1 interacting protein | protein_coding | NP_057041.2;XP_011518456.1;XP_016873364.1 | 17581 | 612491 | . | . | 6.978838 | 6.984983 | 5.760725 | 7.053182 | 16.240043 | 16.146104 | 12.926900 | 16.126262 | -1.099411 | 262.270672 | 0.218286858 | -0.62638094 | 0.531065107 | 0.998865404 | 8.102118 | 7.964423 | 8.061143 | 8.055659 | 7.961127 | 8.056725 | 318 | 295 | 196 | 257 |
| 51075 | NM_00114401 | TMX2 | thioredoxin related transmembrane protein 2 | protein_coding | NP_001137484.1;NP_001334819.1;NP_001334820.1;NP_001334821.1;NI | 30739 | 616715 | . | . | 87.792435 | 89.095389 | 84.277837 | 87.690470 | 204.296612 | 205.947990 | 189.117034 | 200.493851 | -1.041738 | 4478.637541 | 0.106516936 | -0.5538279 | 0.579696629 | 0.998865404 | 12.158059 | 12.099025 | 12.142865 | 12.148455 | 12.099567 | 12.123415 | 5245 | 4948 | 3760 | 4172 |
| 51076 | NM_015960 | CUTC | cutC copper transporter | protein_coding | NP_057044.2 | 24271 | 610101 | . | . | 9.098721 | 9.758477 | 8.647126 | 10.230900 | 21.173096 | 22.557158 | 19.403902 | 23.391739 | -1.012068 | 392.003570 | 0.187933604 | -0.09208558 | 0.926630041 | 0.998865404 | 8.623800 | 8.605589 | 8.598816 | 8.636217 | 8.563432 | 8.655416 | 440 | 439 | 312 | 394 |
| 51077 | NM_00131850 | FCF1 | FCF1 rRNA-processing protein | protein_coding | NP_001305437.1;NP_057046.1;XP_005267787.1;XP_011535117.1;XP_01 | 20220 | . | . | . | 6.681145 | 7.206942 | 6.147845 | 5.687474 | 15.547300 | 16.659169 | 13.795586 | 13.003732 | -1.179119 | 844.603016 | 0.142741398 | -1.66531047 | 0.095850831 | 0.813553488 | 9.836060 | 9.598424 | 9.769901 | 9.785680 | 9.671920 | 9.649415 | 1041 | 997 | 685 | 715 |
| 51078 | NM_00116435 | THAP4 | THAP domain containing 4 | protein_coding | NP_001157828.1;NP_057047.4;XP_005247073.3;XP_011509593.1;XP_01 | 23187 | 612533 | . | . | 23.162229 | 23.873975 | 20.332807 | 28.079110 | 53.899459 | 55.185768 | 45.626228 | 64.199545 | 1.018219 | 1551.282635 | 0.152924526 | 0.17033248 | 0.864748669 | 0.998865404 | 10.586438 | 10.611941 | 10.578262 | 10.600630 | 10.480369 | 10.713158 | 1745 | 1683 | 1157 | 1687 |
| 51079 | NM_015965 | NDUFA13 | NADH:ubiquinone oxidoreductase subunit A13 | protein_coding | NP_057049.5 | 17194 | 609435 | . | . | 173.787598 | 178.305664 | 147.384842 | 187.663345 | 404.410858 | 412.161499 | 330.727325 | 429.069977 | -1.065852 | 2788.506994 | 0.133204709 | -0.69072412 | 0.48973893 | 0.998865404 | 11.490700 | 11.398378 | 11.462119 | 11.476286 | 11.319745 | 11.506177 | 3285 | 3132 | 2077 | 2822 |
| 51081 | NM_015971 | MRPS7 | mitochondrial ribosomal protein S7 | protein_coding | NP_057055.2 | 14499 | 611974 | . | . | 51.780621 | 53.900261 | 51.470436 | 57.146694 | 120.495636 | 124.592865 | 115.498169 | 130.659134 | 1.013603 | 2009.472820 | 0.121854491 | 0.159966326 | 0.872907601 | 0.998865404 | 10.962944 | 10.982194 | 10.954243 | 10.979486 | 10.939387 | 11.014427 | 2262 | 2188 | 1676 | 1986 |
| 51082 | NM_00120655 | POLR1D | RNA polymerase I and III subunit D | protein_coding | NP_001193488.1;NP_001361336.1;NP_057056.1;NP_689918.1;XP_01687 | 20422 | 613715 | . | . | 56.429681 | 57.984984 | 47.771682 | 64.941702 | 131.314192 | 134.034894 | 107.198269 | 148.481487 | 1.002177 | 1917.147556 | 0.13905791 | 0.02255642 | 0.982004107 | 0.998865404 | 10.903384 | 10.906106 | 10.895264 | 10.909295 | 10.806765 | 10.992648 | 2186 | 2085 | 1473 | 2010 |
| 51084 | NM_00136364 | CRYL1 | crystallin lambda 1 | protein_coding | NP_001350576.1;NP_057058.2;XP_005266473.1;XP_016876113.1 | 18246 | 609877 | . | . | 10.561437 | 10.572014 | 10.632308 | 13.915843 | 24.576898 | 24.437683 | 23.858593 | 31.816918 | 1.153398 | 523.576964 | 0.179720953 | 1.145609671 | 0.251956694 | 0.994172589 | 8.926101 | 9.131136 | 8.981487 | 8.981957 | 8.988375 | 9.151528 | 560 | 525 | 427 | 591 |
| 51087 | NM_015982,XI | YBX2 | Y-box binding protein 2 | protein_coding | NP_057066.2;XP_016880202.1 | 17948 | 611447 | . | . | 11.767240 | 13.151300 | 13.006224 | 12.908353 | 27.382851 | 30.399820 | 29.185589 | 29.513416 | 1.024789 | 654.638188 | 0.156298346 | 0.2260184 | 0.82118711 | 0.998865404 | 9.336898 | 9.371997 | 9.309843 | 9.380309 | 9.367803 | 9.356561 | 706 | 734 | 581 | 616 |
| 51088 | NM_00100707 | KLHL5 | kelch like family member 5 | protein_coding | NP_001007076.1;NP_001165125.1;NP_057074.3;NP_950240.2;XP_01151 | 6356 | 608064 | . | . | 2.462113 | 2.633695 | 2.608234 | 2.683617 | 5.729440 | 6.087906 | 5.852802 | 6.135769 | -1.016959 | 580.596088 | 0.159394708 | -0.15221077 | 0.879020697 | 0.998865404 | 9.193501 | 9.169178 | 9.171784 | 9.201720 | 9.181475 | 9.170041 | 659 | 646 | 505 | 535 |
| 51090 | NM_015993 | PLLP | plasmolipin | protein_coding | NP_057077.1 | 18553 | 600340 | . | . | 3.303248 | 2.856720 | 2.317946 | 3.462827 | 7.686793 | 6.603436 | 5.201405 | 7.917344 | -1.083202 | 139.698803 | 0.298007714 | -0.38691243 | 0.698821039 | 0.998865404 | 7.183375 | 7.066615 | 7.167898 | 7.108915 | 7.037175 | 7.175884 | 180 | 145 | 94 | 150 |
| 51091 | NM_016955,XI | SEPSECS | Sep (O-phosphoserine) tRNA:Sec (selenocysteine) tRNA synthase | protein_coding | NP_058651.3;XP_011512148.1;XP_011512149.1;XP_011512150.1;XP_01( | 30605 | 613009 | . | . | 1.283102 | 1.149205 | 1.277892 | 1.068467 | 2.985826 | 2.656440 | 2.867554 | 2.442923 | -1.064874 | 209.734364 | 0.238258991 | -0.38060556 | 0.703495952 | 0.998865404 | 7.756420 | 7.667035 | 7.746689 | 7.704503 | 7.737386 | 7.654784 | 260 | 223 | 193 | 173 |
| 51092 | NM_00104045 | SIDT2 | SID1 transmembrane family member 2 | protein_coding | NP_001035545.1 | 24272 | 617551 | . | . | 3.472905 | 4.013243 | 3.748624 | 4.093752 | 8.081594 | 9.276792 | 8.411804 | 9.359878 | 1.034433 | 525.021368 | 0.170626159 | 0.286240452 | 0.774693963 | 0.998865404 | 9.011968 | 9.060095 | 8.979587 | 9.067218 | 9.020475 | 9.070736 | 554 | 595 | 446 | 520 |
| 51093 | NM_00114256 | RRNAD1 | ribosomal RNA adenine dimethylase domain containing 1 | protein_coding | NP_001136032.1;NP_057081.3;XP_005245291.1;XP_005245292.1;XP_00 | 24273 | . | . | . | 5.270550 | 4.684479 | 5.402141 | 5.631731 | 12.264786 | 10.828385 | 12.122250 | 12.876287 | 1.020760 | 322.414298 | 0.197861466 | 0.149819523 | 0.880907006 | 0.998865404 | 8.317746 | 8.347643 | 8.357250 | 8.295072 | 8.329680 | 8.346275 | 388 | 325 | 279 | 310 |
| 51094 | NM_00129055 | ADIPOR1 | adiponectin receptor 1 | protein_coding | NP_001277482.1;NP_001277486.1;NP_001277556.1;NP_057083.2;XP_01 | 24040 | 607945 | . | . | 63.029853 | 65.009250 | 63.324605 | 69.857961 | 146.673058 | 150.271795 | 142.098571 | 159.721929 | 1.030023 | 4279.306412 | 0.111356614 | 0.383238295 | 0.701543086 | 0.998865404 | 12.041730 | 12.084279 | 12.041178 | 12.059637 | 12.034928 | 12.113581 | 4801 | 4600 | 3598 | 4263 |
| 51095 | NM_00130294 | TRNT1 | tRNA nucleotidyl transferase 1 | protein_coding | NP_001289875.2;NP_001354250.1;NP_001354251.1;NP_001354252.1;NI | 17341 | 612907 | . | . | 7.661585 | 7.593116 | 8.951819 | 6.409187 | 17.828825 | 17.551827 | 20.087624 | 14.653846 | 1.004452 | 726.583098 | 0.186748357 | 0.034319181 | 0.972622629 | 0.998865404 | 9.501225 | 9.508733 | 9.531518 | 9.463812 | 9.625028 | 9.360941 | 875 | 744 | 754 | 553 |
| 51096 | NM_016001,XI | UTP18 | UTP18 small subunit processome component | protein_coding | NP_057085.2;XP_006721993.2;XP_011523172.1 | 24274 | 612816 | . | . | 26.929884 | 25.698304 | 23.373037 | 24.063495 | 62.666946 | 59.402778 | 52.448416 | 55.018322 | -1.126187 | 1463.422509 | 0.126348162 | -1.35693517 | 0.174801809 | 0.948677095 | 10.598302 | 10.426869 | 10.580414 | 10.537429 | 10.456260 | 10.478923 | 1834 | 1627 | 1179 | 1310 |
| 51097 | NM_016002 | SCCPDH | saccharopine dehydrogenase (putative) | protein_coding | NP_057086.2 | 24275 | . | . | . | 42.040474 | 39.097282 | 43.818989 | 41.888912 | 97.829910 | 90.375130 | 98.328537 | 95.774025 | 1.041398 | 2782.406172 | 0.116511075 | 0.502279465 | 0.615470952 | 0.998865404 | 11.412465 | 11.471172 | 11.457388 | 11.391546 | 11.480373 | 11.435282 | 3266 | 2822 | 2537 | 2588 |
| 51098 | NM_00130345 | IFT52 | intraflagellar transport 52 | protein_coding | NP_001290387.1;NP_001290388.1;NP_001310507.1;NP_001310508.1;NI | 15901 | 617094 | . | . | 7.877242 | 7.013569 | 8.439672 | 7.972527 | 18.330667 | 16.212181 | 18.938380 | 18.228232 | 1.090129 | 429.768819 | 0.181748403 | 0.685003814 | 0.493341514 | 0.998865404 | 8.683408 | 8.808706 | 8.758358 | 8.678088 | 8.797950 | 8.744487 | 505 | 414 | 407 | 402 |
| 51099 | NM_00135518 | ABHD5 | abhydrolase domain containing 5, lysophosphatidic acid acyltransf | protein_coding | NP_001342115.1;NP_001352578.1;NP_057090.2 | 21396 | 604780 | . | . | 2.669813 | 2.148062 | 2.365262 | 1.863383 | 6.212764 | 4.965341 | 5.307580 | 4.260404 | -1.187423 | 369.272202 | 0.215000386 | -1.15271417 | 0.249027723 | 0.992733415 | 8.646344 | 8.400256 | 8.637487 | 8.501617 | 8.554344 | 8.383730 | 513 | 384 | 339 | 268 |
| 51100 | NM_00120665 | SH3GLB1 | SH3 domain containing GRB2 like, endophilin B1 | protein_coding | NP_001193580.1;NP_001193581.1;NP_001193582.1;NP_057093.1;XP_0( | 10833 | 609287 | . | . | 8.028037 | 8.449322 | 7.771021 | 7.632905 | 18.681573 | 19.530987 | 17.437945 | 17.451734 | -1.085436 | 1569.483900 | 0.124174655 | -0.95248574 | 0.340850692 | 0.998865404 | 10.673980 | 10.555747 | 10.632642 | 10.661218 | 10.598603 | 10.567988 | 1847 | 1795 | 1335 | 1383 |
| 51101 | NM_00136296 | ZC2HC1A | zinc finger C2HC-type containing 1A | protein_coding | NP_001349898.1;NP_057094.2;XP_011515841.1 | 24277 | . | . | . | 1.874782 | 2.317099 | 2.344832 | 2.251665 | 4.362694 | 5.356078 | 5.261734 | 5.148167 | 1.071780 | 280.071048 | 0.217341937 | 0.460147224 | 0.64541055 | 0.998865404 | 8.079326 | 8.178276 | 8.045726 | 8.167903 | 8.130222 | 8.159194 | 275 | 326 | 245 | 279 |
| 51102 | NM_00102473 | MECR | mitochondrial trans-2-enoyl-CoA reductase | protein_coding | NP_001019903.3;NP_001336640.1;NP_001336641.1;NP_001336642.1;NI | 19691 | 608205 | . | . | 5.250775 | 5.220866 | 4.876370 | 5.724506 | 12.218766 | 12.068266 | 10.942438 | 13.088406 | 1.072809 | 373.843405 | 0.186004356 | 0.545113928 | 0.585675164 | 0.998865404 | 8.494952 | 8.595864 | 8.508685 | 8.540576 | 8.547106 | 8.583547 | 404 | 400 | 326 | 374 |
| 51103 | NM_016013,N | NDUFAF1 | NADH:ubiquinone oxidoreductase complex assembly factor 1 | protein_coding | NP_057097.2;XP_006720618.1;XP_011519960.1;XP_011519961.1;XP_02 | 18828 | 606934 | . | . | 8.518062 | 7.941023 | 7.068257 | 7.921515 | 19.821882 | 18.356031 | 15.860961 | 18.111601 | -1.116964 | 356.849748 | 0.190177191 | -0.839129 | 0.401396925 | 0.998865404 | 8.556805 | 8.397125 | 8.532283 | 8.487526 | 8.427815 | 8.465771 | 450 | 391 | 283 | 327 |
| 51104 | NM_00102578 | ABHD17B | abhydrolase domain containing 17B, depalmitoylase | protein_coding | NP_001020951.1;NP_057098.2;XP_006717197.2;XP_006717198.1;XP_01 | 24278 | 617943 | . | . | 4.210895 | 3.646870 | 4.044764 | 3.473194 | 9.798928 | 8.429905 | 9.076333 | 7.941046 | -1.064457 | 342.381496 | 0.205799914 | -0.43789187 | 0.661464669 | 0.998865404 | 8.463185 | 8.374370 | 8.496431 | 8.369372 | 8.439982 | 8.358001 | 451 | 339 | 310 | 288 |
| 51105 | NM_00127719 | PHF20L1 | PHD finger protein 20 like 1 | protein_coding | NP_001264125.1;NP_001349900.1;NP_057102.4;NP_940915.1;XP_0052! | 24280 | . | . | . | 7.093437 | 6.938532 | 7.147061 | 7.127375 | 16.506726 | 16.038729 | 16.037792 | 16.295892 | -1.003594 | 1307.283194 | 0.127778622 | -0.04050634 | 0.967689449 | 0.998865404 | 10.354917 | 10.349791 | 10.360781 | 10.346594 | 10.352488 | 10.349512 | 1522 | 1400 | 1137 | 1221 |
| 51106 | NM_00135050 | TFB1M | transcription factor B1, mitochondrial | protein_coding | NP_001337430.1;NP_001337431.1;NP_057104.2;XP_011534172.1;XP_01 | 17037 | 607033 | . | . | 6.198691 | 6.213819 | 7.696894 | 6.371654 | 14.424611 | 14.363522 | 17.271606 | 15.779697 | 1.038923 | 634.687673 | 0.164845333 | 0.334181391 | 0.738242686 | 0.998865404 | 9.281852 | 9.337419 | 9.273654 | 9.316430 | 9.380806 | 9.257607 | 694 | 693 | 614 | 551 |
| 51107 | NM_00107762 | APH1A | aph-1 homolog A, gamma-secretase subunit | protein_coding | NP_001071096.1;NP_001230700.1;NP_001230701.1;NP_057106.2;XP_01 | 29509 | 607629 | . | . | 50.210032 | 49.257767 | 47.809253 | 52.435911 | 116.840808 | 113.861533 | 107.282582 | 119.888475 | -1.010749 | 3027.740059 | 0.11457937 | -0.13462458 | 0.892908707 | 0.998865404 | 11.571782 | 11.556228 | 11.572358 | 11.564020 | 11.522399 | 11.595119 | 3523 | 3268 | 2502 | 2949 |
| 51108 | NM_00107718 | METTL9 | methyltransferase like 9 | protein_coding | NP_001070648.1;NP_001275588.1;NP_001275589.1;NP_057109.3;XP_01 | 24586 | 609388 | . | . | 55.579968 | 51.753913 | 50.329522 | 53.409810 | 129.336877 | 119.631483 | 112.937991 | 122.115181 | -1.049331 | 5270.909332 | 0.109003445 | -0.6373199 | 0.52391649 | 0.998865404 | 12.398151 | 12.328685 | 12.415458 | 12.350458 | 12.323125 | 12.362699 | 6455 | 5600 | 4367 | 4937 |
| 51109 | NM_00125265 | RDH11 | retinol dehydrogenase 11 | protein_coding | NP_001239579.1;NP_057110.3 | 17964 | 607849 | . | . | 42.005255 | 40.400776 | 46.481934 | 44.969241 | 97.747962 | 93.388207 | 104.304107 | 102.816821 | 1.094868 | 3434.873193 | 0.111184397 | 1.176033037 | 0.239581675 | 0.992733415 | 11.679122 | 11.809993 | 11.726574 | 11.687637 | 11.800295 | 11.763299 | 3866 | 3454 | 3192 | 3291 |
| 51110 | NM_016027 | LACTB2 | lactamase beta 2 | protein_coding | NP_057111.1 | 18512 | . | . | . | 13.380052 | 11.578749 | 11.795897 | 14.377578 | 31.135931 | 26.764797 | 26.469652 | 32.872627 | 1.034007 | 608.039968 | 0.172239287 | 0.280108221 | 0.779394477 | 0.998865404 | 9.223827 | 9.271818 | 9.284608 | 9.183001 | 9.192738 | 9.317930 | 741 | 596 | 487 | 634 |
| 51111 | NM_00130090 | KMT5B | lysine methyltransferase 5B | protein_coding | NP_001287836.1;NP_001287837.1;NP_001287838.1;NP_001350495.1;NI | 24283 | 610881 | . | . | 7.327372 | 6.601876 | 7.524230 | 7.439678 | 17.051099 | 15.260530 | 16.884161 | 17.009939 | 1.014173 | 846.696124 | 0.152794522 | 0.132878956 | 0.89428911 | 0.998865404 | 9.715207 | 9.736118 | 9.777696 | 9.657758 | 9.755584 | 9.703090 | 1049 | 831 | 769 | 770 |
| 51112 | NM_00132110 | TRAPPC12 | trafficking protein particle complex 12 | protein_coding | NP_001308031.1;NP_057114.5;XP_011508652.1;XP_011508654.1;XP_01 | 24284 | 614139 | . | . | 11.824802 | 9.270929 | 10.051034 | 12.211432 | 27.516800 | 21.430165 | 22.554230 | 27.919988 | 1.050835 | 1054.106611 | 0.1489137 | 0.48038253 | 0.630955414 | 0.998865404 | 10.005682 | 10.077046 | 10.069320 | 9.971982 | 9.995518 | 10.117528 | 1262 | 1036 | 862 | 1096 |
| 51114 | NM_00100822 | ZDHHC9 | zinc finger DHHC-type palmitoyltransferase 9 | protein_coding | NP_001008223.1;NP_057116.2;XP_011529650.1 | 18475 | 300646 | . | . | 4.194872 | 4.131028 | 5.020109 | 4.467401 | 9.761639 | 9.549055 | 11.264980 | 10.214181 | 1.130422 | 632.810541 | 0.158647737 | 1.114806447 | 0.264933373 | 0.996598404 | 9.214248 | 9.391567 | 9.263364 | 9.259548 | 9.383862 | 9.302422 | 686 | 639 | 619 | 592 |
| 51115 | NM_00128670 | RMDN1 | regulator of microtubule dynamics 1 | protein_coding | NP_001273636.1;NP_001273648.1;NP_001304736.1;NP_057117.2;XP_0( | 24285 | 611871 | . | . | 13.122112 | 12.536120 | 12.295426 | 11.230167 | 30.535691 | 28.977802 | 27.590584 | 25.676436 | -1.065135 | 1056.909484 | 0.138152245 | -0.65895805 | 0.509922705 | 0.998865404 | 10.090241 | 9.999608 | 10.091633 | 10.044174 | 10.052224 | 9.990258 | 1295 | 1139 | 928 | 919 |
| 51116 | NM_00137140 | MRPS2 | mitochondrial ribosomal protein S2 | protein_coding | NP_001358330.1;NP_057118.1 | 14495 | 611971 | . | . | 24.224401 | 24.585678 | 23.514803 | 26.315939 | 56.371173 | 56.830895 | 52.766540 | 60.168267 | 1.002499 | 1138.095190 | 0.13581572 | 0.026510163 | 0.978850428 | 0.998865404 | 10.150776 | 10.154033 | 10.144891 | 10.157517 | 10.113340 | 10.191141 | 1298 | 1237 | 939 | 1124 |
| 51117 | NM_00130594 | COQ4 | coenzyme Q4 | protein_coding | NP_001292871.2;NP_057119.3;XP_016870281.1;XP_016870282.1 | 19693 | 612898 | . | . | 15.754005 | 15.557426 | 15.125404 | 16.471451 | 36.660220 | 35.961685 | 33.940967 | 37.660014 | 1.003000 | 640.727318 | 0.155717794 | 0.027748542 | 0.977862708 | 0.998865404 | 9.321484 | 9.325646 | 9.335505 | 9.309243 | 9.301408 | 9.346957 | 751 | 677 | 539 | 622 |
| 51118 | NM_016037,XI | UTP11 | UTP11 small subunit processome component | protein_coding | NP_057121.2;XP_011539857.1 | 24329 | 609440 | . | . | 18.195737 | 18.435940 | 17.175940 | 21.217157 | 42.342227 | 42.615501 | 38.542309 | 48.510509 | 1.037771 | 1167.898721 | 0.143656203 | 0.37233909 | 0.709640398 | 0.998865404 | 10.162959 | 10.215954 | 10.172745 | 10.176318 | 10.127502 | 10.271815 | 1317 | 1240 | 935 | 1222 |
| 51119 | NM_016038 | SBDS | SBDS ribosome maturation factor | protein_coding | NP_057122.2 | 19440 | 607444 | . | . | 31.786549 | 28.148918 | 28.394520 | 26.397503 | 73.968605 | 65.067482 | 63.716484 | 60.354755 | -1.109394 | 1442.475107 | 0.134283885 | -1.1153406 | 0.264704492 | 0.996598404 | 10.567076 | 10.417722 | 10.581698 | 10.480297 | 10.483582 | 10.420444 | 1860 | 1531 | 1239 | 1229 |
| 51121 | NM_00131798 | RPL26L1 | ribosomal protein L26 like 1 | protein_coding | NP_001304909.1;NP_001304910.1;NP_001304911.1;NP_057177.1;XP_01 | 17050 | . | . | . | 57.818208 | 58.057316 | 49.990164 | 54.877011 | 134.545346 | 134.202090 | 112.176471 | 125.469766 | -1.115786 | 1284.624275 | 0.131494393 | -1.20202924 | 0.229352201 | 0.98402759 | 10.404116 | 10.245805 | 10.371540 | 10.362855 | 10.249002 | 10.317845 | 1569 | 1454 | 1007 | 1191 |
| 51122 | NM_016094,N | COMMD2 | COMM domain containing 2 | protein_coding | NP_057178.2 | 24993 | 616699 | . | . | 6.146444 | 6.267603 | 6.187204 | 5.747953 | 14.303028 | 14.487845 | 13.883905 | 13.142012 | -1.055051 | 702.708410 | 0.151935023 | -0.50885051 | 0.610857015 | 0.998865404 | 9.494772 | 9.417766 | 9.471620 | 9.477853 | 9.466379 | 9.408829 | 827 | 782 | 620 | 614 |
| 51123 | NM_00104251 | ZNF706 | zinc finger protein 706 | protein_coding | NP_001035975.1;NP_001254637.1;NP_001254638.1;NP_057180.1;XP_02 | 24992 | . | . | . | 25.001977 | 25.577873 | 26.943362 | 27.951141 | 58.180630 | 59.124400 | 60.460126 | 63.906959 | 1.044300 | 2377.578776 | 0.114933403 | 0.544117472 | 0.586364767 | 0.998865404 | 11.183631 | 11.246245 | 11.201207 | 11.193640 | 11.245516 | 11.219061 | 2691 | 2498 | 2147 | 2240 |
| 51124 | NM_016097 | IER3IP1 | immediate early response 3 interacting protein 1 | protein_coding | NP_057181.1 | 18550 | 609382 | . | . | 8.844623 | 7.596313 | 8.717962 | 5.771583 | 20.581800 | 17.559217 | 19.562855 | 13.196039 | -1.150116 | 887.667048 | 0.189529533 | -1.06463114 | 0.287042883 | 0.998865404 | 9.890738 | 9.690037 | 9.893545 | 9.774635 | 9.872014 | 9.577511 | 1181 | 942 | 868 | 613 |
| 51125 | NM_00100229 | GOLGA7 | golgin A7 | protein_coding | NP_001002296.1;NP_001167595.1;NP_001349908.1;NP_001349909.1;NI | 24876 | 609453 | . | . | 18.687823 | 16.160871 | 20.123789 | 19.321583 | 43.487334 | 37.356577 | 45.157201 | 44.176499 | 1.133236 | 1154.510654 | 0.137068449 | 1.316481576 | 0.188012458 | 0.960264078 | 10.079887 | 10.260590 | 10.168390 | 10.076282 | 10.215044 | 10.219076 | 1324 | 1095 | 1064 | 1153 |
| 51126 | NM_016100,N | NAA20 | N-alpha-acetyltransferase 20, NatB catalytic subunit | protein_coding | NP_057184.1;NP_852668.1;NP_852669.1 | 15908 | 610833 | . | . | 30.889961 | 31.118471 | 30.628222 | 31.597351 | 71.882207 | 71.931738 | 68.728845 | 72.243585 | -1.011936 | 1001.093546 | 0.136089527 | -0.12578884 | 0.899899076 | 0.998865404 | 9.975940 | 9.958730 | 9.968799 | 9.974666 | 9.955128 | 9.970838 | 1155 | 1091 | 856 | 943 |
| 51128 | NM_00103350 | SAR1B | secretion associated Ras related GTPase 1B | protein_coding | NP_001028675.1;NP_057187.1 | 10535 | 607690 | . | . | 9.196366 | 7.741014 | 8.106033 | 7.472701 | 21.400322 | 17.893699 | 18.189705 | 17.085442 | -1.103493 | 1655.477226 | 0.137174 | -1.03575113 | 0.300318262 | 0.998865404 | 10.762117 | 10.620471 | 10.797740 | 10.655405 | 10.686577 | 10.616721 | 2179 | 1706 | 1431 | 1409 |
| 51130 | NM_00120196 | ASB3 | ankyrin repeat and SOCS box containing 3 | protein_coding | NP_001188894.1;NP_057199.1;NP_665862.1 | 16013 | 605760 | . | . | 7.053329 | 6.926102 | 6.074376 | 6.187415 | 16.413385 | 16.009994 | 13.630722 | 14.146791 | -1.174135 | 448.173122 | 0.173214746 | -1.33705743 | 0.18120389 | 0.952877815 | 8.919039 | 8.687506 | 8.860677 | 8.849911 | 8.757842 | 8.755924 | 562 | 518 | 359 | 386 |
| 51132 | NM_016120,N | RLIM | ring finger protein, LIM domain interacting | protein_coding | NP_057204.2;NP_899196.1 | 13429 | 300379 | . | . | 3.337908 | 3.367901 | 2.936253 | 3.144979 | 7.767449 | 7.785054 | 6.588867 | 7.190620 | -1.116040 | 1055.706748 | 0.135443425 | -1.16941048 | 0.242238289 | 0.992733415 | 10.121111 | 9.962519 | 10.082502 | 10.084031 | 9.980602 | 10.023208 | 1281 | 1203 | 842 | 963 |
| 51133 | NM_00131929 | KCTD3 | potassium channel tetramerization domain containing 3 | protein_coding | NP_001306223.1;NP_001306224.1;NP_057205.2;XP_005273215.1 | 21305 | 613272 | . | . | 11.323563 | 12.377533 | 13.689649 | 14.321392 | 26.350399 | 28.611221 | 30.719179 | 32.744161 | 1.166531 | 1617.760718 | 0.12547943 | 1.77100871 | 0.076559253 | 0.767975096 | 10.544520 | 10.766513 | 10.564353 | 10.626716 | 10.701769 | 10.729902 | 1650 | 1677 | 1488 | 1662 |
| 51134 | NM_00104239 | CEP83 | centrosomal protein 83 | protein_coding | NP_001035858.1;NP_001333386.1;NP_001333387.1;NP_001333388.1;NI | 17966 | 615847 | . | . | 5.120985 | 4.651765 | 5.860465 | 3.712300 | 11.916745 | 10.752763 | 13.150715 | 8.487737 | 1.021415 | 582.985740 | 0.181912658 | 0.168044912 | 0.866547947 | 0.998865404 | 9.171416 | 9.203044 | 9.190809 | 9.162796 | 9.286024 | 9.086401 | 678 | 609 | 590 | 469 |
| 51135 | NM_00111418 | IRAK4 | interleukin 1 receptor associated kinase 4 | protein_coding | NP_001107654.1;NP_001138728.1;NP_001138729.1;NP_001138730.1;NI | 17967 | 606883 | . | . | 3.284647 | 3.883356 | 4.156742 | 3.520148 | 7.643507 | 8.976554 | 9.327608 | 8.048404 | 1.031110 | 481.540317 | 0.180390691 | 0.245017234 | 0.806443088 | 0.998865404 | 8.889158 | 8.933526 | 8.851564 | 8.945995 | 8.966289 | 8.869623 | 505 | 550 | 458 | 423 |
| 51136 | NM_016125 | RNFT1 | ring finger protein, transmembrane 1 | protein_coding | NP_057209.3 | 30206 | 615172 | . | . | 11.200776 | 9.377908 | 11.181849 | 7.702166 | 26.064665 | 21.677452 | 25.091747 | 17.610085 | -1.105338 | 653.265389 | 0.194920227 | -0.74126754 | 0.458531239 | 0.998865404 | 9.421340 | 9.278159 | 9.441557 | 9.312236 | 9.429446 | 9.177552 | 863 | 671 | 642 | 472 |
| 51138 | NM_00125800 | COPS4 | COP9 signalosome subunit 4 | protein_coding | NP_001244935.1;NP_001317656.1;NP_057213.2 | 16702 | 616008 | . | . | 13.377937 | 12.929175 | 12.663585 | 14.176164 | 31.131011 | 29.886367 | 28.416720 | 32.412116 | 1.005160 | 687.634974 | 0.154080674 | 0.048192277 | 0.961563005 | 0.998865404 | 9.421911 | 9.429078 | 9.437479 | 9.409332 | 9.392974 | 9.459847 | 806 | 725 | 570 | 678 |
| 51141 | NM_00132132 | INSIG2 | insulin induced gene 2 | protein_coding | NP_001308258.1;NP_001308259.1;NP_001308260.1;NP_001308261.1;NI | 20452 | 608660 | . | . | 7.794603 | 7.048728 | 10.231416 | 6.918361 | 18.138361 | 16.293452 | 22.958998 | 15.814230 | 1.080190 | 841.986813 | 0.181148962 | 0.614331073 | 0.538996573 | 0.998865404 | 9.660415 | 9.772708 | 9.730955 | 9.634905 | 9.861295 | 9.600666 | 995 | 814 | 898 | 672 |
| 51142 | NM_00132032 | CHCHD2 | coiled-coil-helix-coiled-coil-helix domain containing 2 | protein_coding | NP_001307256.1;NP_057223.1 | 21645 | 616244 | . | . | 359.315147 | 358.048389 | 293.186505 | 380.001887 | 836.141093 | 827.644858 | 657.902016 | 868.829104 | -1.080233 | 8659.408852 | 0.12793516 | -0.87030006 | 0.384136446 | 0.998865404 | 13.134703 | 13.023251 | 13.115724 | 13.104655 | 12.934602 | 13.143863 | 10423 | 9642 | 6345 | 8769 |
| 51143 | NM_00132913 | DYNC1LI1 | dynein cytoplasmic 1 light intermediate chain 1 | protein_coding | NP_001316064.1;NP_057225.2 | 18745 | 615890 | . | . | 10.837334 | 10.498343 | 9.446533 | 9.464863 | 25.218920 | 24.267389 | 21.197748 | 21.640285 | -1.144142 | 779.625976 | 0.145949762 | -1.33105005 | 0.183172545 | 0.954800306 | 9.700442 | 9.506308 | 9.666250 | 9.637660 | 9.560637 | 9.554833 | 977 | 880 | 635 | 679 |
| 51144 | NM_016142,XI | HSD17B12 | hydroxysteroid 17-beta dehydrogenase 12 | protein_coding | NP_057226.1;XP_011518458.1;XP_016873370.1;XP_024304339.1;XP_02 | 18646 | 609574 | . | . | 37.473090 | 33.092700 | 34.529683 | 29.514476 | 87.201418 | 76.495262 | 77.483612 | 67.481340 | -1.113476 | 2537.934741 | 0.131793741 | -1.1766174 | 0.239348251 | 0.992733415 | 11.384715 | 11.230018 | 11.402965 | 11.294102 | 11.327674 | 11.194193 | 3283 | 2694 | 2263 | 2064 |
| 51147 | NM_00112758 | ING4 | inhibitor of growth family member 4 | protein_coding | NP_001121054.1;NP_001121055.1;NP_001121056.1;NP_001121057.1;NI | 19423 | 608524 | . | . | 7.496979 | 6.865538 | 8.836354 | 8.083163 | 17.445778 | 15.869999 | 19.828521 | 18.481188 | 1.192649 | 386.179570 | 0.187869624 | 1.352901384 | 0.176087137 | 0.949193943 | 8.460001 | 8.715002 | 8.569973 | 8.501234 | 8.668239 | 8.610478 | 430 | 357 | 383 | 375 |
| 51148 | NM_00128676 | CERCAM | cerebral endothelial cell adhesion molecule | protein_coding | NP_001273689.1;NP_057258.3;XP_005252092.1;XP_011517064.1;XP_01 | 23723 | 616626 | . | . | 24.877317 | 24.784067 | 24.679719 | 26.307289 | 57.890540 | 57.289477 | 55.380574 | 60.148496 | 1.015876 | 1860.016742 | 0.120409589 | 0.188727913 | 0.850306073 | 0.998865404 | 10.849747 | 10.872365 | 10.858844 | 10.850669 | 10.845577 | 10.888304 | 2136 | 1981 | 1586 | 1805 |
| 51149 | NM_00101798 | MRNIP | MRN complex interacting protein | protein_coding | NP_001017987.1;NP_057259.2 | 30817 | 617154 | . | . | 5.356340 | 6.327590 | 5.606449 | 6.004184 | 12.464423 | 14.626509 | 12.580708 | 13.727854 | -1.074764 | 206.466639 | 0.238533035 | -0.43608089 | 0.662778042 | 0.998865404 | 7.741312 | 7.636307 | 7.665646 | 7.743937 | 7.657039 | 7.687618 | 226 | 250 | 167 | 193 |

| | | | | | | | | | | |
|---|---|---|---|---|---|---|---|---|---|---|
| 51150 | NM_016176,N | SDF4 | stromal cell derived factor 4 | protein_coding | NP_057260.3;NP_057631.2;XP_011539858.1;XP_024303241.1 | 24188 | 614282 | . | . | [illegible] |
| 51151 | NM_00101250 | SLC45A2 | solute carrier family 45 member 2 | protein_coding | NP_001012527.2;NP_001284346.2;NP_057264.4 | 16472 | 606202 | . | . | [illegible] |
| 51154 | NM_016183,XI | MRTO4 | MRT4 homolog, ribosome maturation factor | protein_coding | NP_057267.2;XP_006710738.1 | 18477 | . | . | . | [illegible] |
| 51155 | NM_00100203 | JPT1 | Jupiter microtubule associated homolog 1 | protein_coding | NP_001002032.1;NP_001002033.1;NP_001275538.1;NP_001275539.1;NI | 14569 | . | . | . | [illegible] |
| 51157 | NM_00116342 | ZNF580 | zinc finger protein 580 | protein_coding | NP_001156895.1;NP_057286.1;NP_996998.1 | 29473 | 617888 | . | . | [illegible] |
| 51160 | NM_016208,N | VPS28 | VPS28 subunit of ESCRT-I | protein_coding | NP_057292.1;NP_898880.1;XP_005272380.1;XP_005272381.1;XP_00527 | 18178 | 611952 | . | . | [illegible] |
| 51161 | NM_00117174 | C3orf18 | chromosome 3 open reading frame 18 | protein_coding | NP_001165211.1;NP_001165212.1;NP_001165214.2;NP_057294.2;XP_01 | 24837 | . | . | . | [illegible] |
| 51162 | NM_016215,N | EGFL7 | EGF like domain multiple 7 | protein_coding | NP_057299.1;NP_958854.1;XP_006717204.1;XP_011517066.1;XP_01151 | 20594 | 608582 | . | . | [illegible] |
| 51163 | NM_016216 | DBR1 | debranching RNA lariats 1 | protein_coding | NP_057300.2 | 15594 | 607024 | . | . | [illegible] |
| 51164 | NM_00113564 | DCTN4 | dynactin subunit 4 | protein_coding | NP_001129115.1;NP_001129116.1;NP_057305.1;XP_005268514.1;XP_01 | 15518 | 614758 | . | . | [illegible] |
| 51166 | NM_00128668 | AADAT | aminoadipate aminotransferase | protein_coding | NP_001273611.1;NP_001273612.1;NP_057312.1;NP_872603.1;XP_00671 | 17929 | 611754 | . | . | [illegible] |
| 51167 | NM_016230 | CYB5R4 | cytochrome b5 reductase 4 | protein_coding | NP_057314.2 | 20147 | 608343 | . | . | [illegible] |
| 51170 | NM_016245,XI | HSD17B11 | hydroxysteroid 17-beta dehydrogenase 11 | protein_coding | NP_057329.3;XP_011530323.1 | 22960 | 612831 | . | . | [illegible] |
| 51171 | NM_016246,XI | HSD17B14 | hydroxysteroid 17-beta dehydrogenase 14 | protein_coding | NP_057330.2;XP_005259026.1;XP_005259027.1 | 23238 | 612832 | . | . | [illegible] |
| 51172 | NM_016256,XI | NAGPA | N-acetylglucosamine-1-phosphodiester alpha-N-acetylglucosamini | protein_coding | NP_057340.2;XP_011520819.1 | 17378 | 607985 | . | . | [illegible] |
| 51174 | NM_00119360 | TUBD1 | tubulin delta 1 | protein_coding | NP_001180538.1;NP_001180539.1;NP_001180540.1;NP_001180541.1;NI | 16811 | 607344 | . | . | [illegible] |
| 51175 | NM_016262,XI | TUBE1 | tubulin epsilon 1 | protein_coding | NP_057346.1;XP_011534177.1;XP_016866407.1 | 20775 | 607345 | . | . | [illegible] |
| 51176 | NM_00113071 | LEF1 | lymphoid enhancer binding factor 1 | protein_coding | NP_001124185.1;NP_001124186.1;NP_001159591.1;NP_057353.1;XP_00 | 6551 | 153245 | . | . | [illegible] |
| 51177 | NM_00130472 | PLEKHO1 | pleckstrin homology domain containing O1 | protein_coding | NP_001291651.1;NP_001291652.1;NP_001291653.1;NP_057358.2;XP_01 | 24310 | 608335 | . | . | [illegible] |
| 51181 | NM_00119521 | DCXR | dicarbonyl and L-xylulose reductase | protein_coding | NP_001182147.1;NP_057370.1 | 18985 | 608347 | . | . | [illegible] |
| 51182 | NM_016299 | HSPA14 | heat shock protein family A (Hsp70) member 14 | protein_coding | NP_057383.2 | 29526 | 610369 | . | . | [illegible] |
| 51184 | NM_00116437 | GPN3 | GPN-loop GTPase 3 | protein_coding | NP_001157844.1;NP_001157845.1;NP_057385.3;XP_016874883.1 | 30186 | . | . | . | [illegible] |
| 51185 | NM_00117348 | CRBN | cereblon | protein_coding | NP_001166953.1;NP_057386.2;XP_005265259.1;XP_011532093.1;XP_01 | 30185 | 609262 | . | . | [illegible] |
| 51186 | NM_00100661 | TCEAL9 | transcription elongation factor A like 9 | protein_coding | NP_001006613.1;NP_001006614.1;NP_001006615.1;NP_057387.1 | 30084 | . | . | . | [illegible] |
| 51187 | NM_016304 | RSL24D1 | ribosomal L24 domain containing 1 | protein_coding | NP_057388.1 | 18479 | 613262 | . | . | [illegible] |
| 51188 | NM_00137030 | SS18L2 | SS18 like 2 | protein_coding | NP_001357229.1;NP_057389.1 | 15593 | 606473 | . | . | [illegible] |
| 51191 | NM_016323,XI | HERC5 | HECT and RLD domain containing E3 ubiquitin protein ligase 5 | protein_coding | NP_057407.2;XP_011530324.2 | 24368 | 608242 | . | . | [illegible] |
| 51192 | NM_00104013 | CKLF | chemokine like factor | protein_coding | NP_001035228.1;NP_057410.1;NP_058647.1;NP_857591.1;NP_857592.1 | 13253 | 616074 | . | . | [illegible] |
| 51193 | NM_00130342 | ZNF639 | zinc finger protein 639 | protein_coding | NP_001290354.1;NP_001290355.1;NP_001362729.1;NP_001362731.1;NI | 30950 | . | . | . | [illegible] |
| 51194 | NM_00113477 | IPO11 | importin 11 | protein_coding | NP_001128251.1;NP_057422.3 | 20628 | 610889 | . | . | [illegible] |
| 51195 | NM_00130353 | RAPGEFL1 | Rap guanine nucleotide exchange factor like 1 | protein_coding | NP_001290462.1;NP_001290463.1;NP_057423.2;XP_006721998.1;XP_00 | 17428 | . | . | . | [illegible] |
| 51196 | NM_00116597 | PLCE1 | phospholipase C epsilon 1 | protein_coding | NP_001159451.1;NP_001275918.1;NP_057425.3;XP_006717948.1;XP_00 | 17175 | 608414 | . | . | [illegible] |
| 51199 | NM_016350,N | NIN | ninein | protein_coding | NP_057434.4;NP_065972.4;NP_891989.2;NP_891991.1;XP_011535121.1 | 14906 | 608684 | . | . | [illegible] |
| 51200 | NM_00116344 | CPA4 | carboxypeptidase A4 | protein_coding | NP_001156918.1;NP_057436.2 | 15740 | 607635 | . | . | [illegible] |
| 51201 | NM_00136298 | ZDHHC2 | zinc finger DHHC-type palmitoyltransferase 2 | protein_coding | NP_001349917.1;NP_001349918.1;NP_057437.1;XP_011542846.1;XP_01 | 18469 | 618621 | . | . | [illegible] |
| 51202 | NM_016355,N | DDX47 | DEAD-box helicase 47 | protein_coding | NP_057439.2;NP_957518.1 | 18682 | 615428 | . | . | [illegible] |
| 51203 | NM_00124314 | NUSAP1 | nucleolar and spindle associated protein 1 | protein_coding | NP_001230071.1;NP_001230072.1;NP_001230073.1;NP_001288065.1;NI | 18538 | 612818 | . | . | [illegible] |
| 51204 | NM_016360 | TACO1 | translational activator of cytochrome c oxidase I | protein_coding | NP_057444.2 | 24316 | 612958 | . | . | [illegible] |
| 51205 | NM_00132362 | ACP6 | acid phosphatase 6, lysophosphatidic | protein_coding | NP_001310554.1;NP_057445.4;XP_011507903.1;XP_016856911.1;XP_02 | 29609 | 611471 | . | . | [illegible] |
| 51209 | NM_016370,N | RAB9B | RAB9B, member RAS oncogene family | protein_coding | NP_057454.1 | 14090 | 300285 | . | . | [illegible] |
| 51218 | NM_016417 | GLRX5 | glutaredoxin 5 | protein_coding | NP_057501.2 | 20134 | 609588 | . | . | [illegible] |
| 51222 | NM_00110167 | ZNF219 | zinc finger protein 219 | protein_coding | NP_001095142.1;NP_001095924.1;NP_057507.2;XP_005267796.1;XP_01 | 13011 | 605036 | . | . | [illegible] |
| 51226 | NM_016429,XI | COPZ2 | COPI coat complex subunit zeta 2 | protein_coding | NP_057513.1;XP_011523176.1 | 19356 | 615526 | . | . | [illegible] |
| 51227 | NM_00132048 | PIGP | phosphatidylinositol glycan anchor biosynthesis class P | protein_coding | NP_001307409.1;NP_057514.2;NP_710148.1;NP_710149.1;XP_0168838 | 3046 | 605938 | . | . | [illegible] |
| 51228 | NM_016433,XI | GLTP | glycolipid transfer protein | protein_coding | NP_057517.1;XP_011536736.1 | 24867 | 608949 | . | . | [illegible] |
| 51230 | NM_016436,XI | PHF20 | PHD finger protein 20 | protein_coding | NP_057520.2;XP_005260478.1;XP_011527145.1;XP_011527146.1;XP_01 | 16098 | 610335 | . | . | [illegible] |
| 51231 | NM_00102577 | VRK3 | VRK serine/threonine kinase 3 | protein_coding | NP_001020949.1;NP_001295349.1;NP_057524.3;XP_005259028.1;XP_00 | 18996 | . | . | . | [illegible] |
| 51232 | NM_016441,XI | CRIM1 | cysteine rich transmembrane BMP regulator 1 | protein_coding | NP_057525.1;XP_005264414.1;XP_011531200.1;XP_011531201.1;XP_01 | 2359 | 606189 | . | . | [illegible] |
| 51234 | NM_00128642 | EMC4 | ER membrane protein complex subunit 4 | protein_coding | NP_001273349.1;NP_001338302.1;NP_057538.1 | 28032 | 616245 | . | . | [illegible] |
| 51236 | NM_016458,XI | HGH1 | HGH1 homolog | protein_coding | NP_057542.2 | 24161 | . | . | . | [illegible] |
| 51239 | NM_016466 | ANKRD39 | ankyrin repeat domain 39 | protein_coding | NP_057550.3 | 28640 | . | . | . | [illegible] |
| 51241 | NM_00120409 | COX16 | cytochrome c oxidase assembly factor COX16 | protein_coding | NP_001191019.1;NP_057552.1 | 20213 | 618064 | . | . | [illegible] |
| 51244 | NM_016474,N | CCDC174 | coiled-coil domain containing 174 | protein_coding | NP_057558.3;XP_016862043.1;XP_016862044.1 | 28033 | 616735 | . | . | [illegible] |
| 51246 | NM_00127206 | SHISA5 | shisa family member 5 | protein_coding | NP_001258994.1;NP_001258995.1;NP_001258996.1;NP_001258997.1;NI | 30376 | 607290 | . | . | [illegible] |
| 51247 | NM_00103311 | PAIP2 | poly(A) binding protein interacting protein 2 | protein_coding | NP_001028284.1;NP_057564.3;XP_011541730.1;XP_016865011.1;XP_01 | 17970 | 605604 | . | . | [illegible] |
| 51248 | NM_00137017 | PDZD11 | PDZ domain containing 11 | protein_coding | NP_001357103.1;NP_001357104.1;NP_001357105.1;NP_001357106.1;NI | 28034 | 300632 | . | . | [illegible] |
| 51249 | NM_016486,XI | TMEM69 | transmembrane protein 69 | protein_coding | NP_057570.2;XP_024303276.1 | 28035 | . | . | . | [illegible] |
| 51250 | NM_00114246 | MTRES1 | mitochondrial transcription rescue factor 1 | protein_coding | NP_001135940.1;NP_001135942.1;NP_057571.1;XP_011534178.1;XP_01 | 17971 | 618583 | . | . | [illegible] |
| 51251 | NM_00100200 | NT5C3A | 5'-nucleotidase, cytosolic IIIA | protein_coding | NP_001002009.1;NP_001002010.2;NP_001159590.1;NP_001343925.1;NI | 17820 | 606224 | . | . | [illegible] |
| 51252 | NM_00112264 | FAM178B | family with sequence similarity 178 member B | protein_coding | NP_001116118.2;NP_001166138.1;NP_057574.2;XP_011509596.1;XP_01 | 28036 | . | . | . | [illegible] |
| 51253 | NM_00133060 | MRPL37 | mitochondrial ribosomal protein L37 | protein_coding | NP_001317531.1;NP_057575.2 | 14034 | 611843 | . | . | [illegible] |
| 51255 | NM_016494,XI | RNF181 | ring finger protein 181 | protein_coding | NP_057578.1;XP_005264416.1 | 28037 | 612490 | . | . | [illegible] |
| 51256 | NM_00114396 | TBC1D7 | TBC1 domain family member 7 | protein_coding | NP_001137436.1;NP_001137437.1;NP_001137438.1;NP_001245386.1;NI | 21066 | 612655 | . | . | [illegible] |
| 51257 | NM_00100541 | MARCHF2 | membrane associated ring-CH-type finger 2 | protein_coding | NP_001005415.1;NP_001005416.1;NP_001356705.1;NP_001356706.1;NI | 28038 | 613332 | . | . | [illegible] |
| 51258 | NM_016497 | MRPL51 | mitochondrial ribosomal protein L51 | protein_coding | NP_057581.2 | 14044 | 611855 | . | . | [illegible] |
| 51260 | NM_00130088 | PBDC1 | polysaccharide biosynthesis domain containing 1 | protein_coding | NP_001287817.1;NP_057584.2 | 28790 | . | . | . | [illegible] |
| 51263 | NM_145212,N | MRPL30 | mitochondrial ribosomal protein L30 | protein_coding | NP_660213.1 | 14036 | 611838 | . | . | [illegible] |
| 51264 | NM_016504 | MRPL27 | mitochondrial ribosomal protein L27 | protein_coding | NP_057588.1 | 14483 | 611837 | . | . | [illegible] |
| 51265 | NM_00111357 | CDKL3 | cyclin dependent kinase like 3 | protein_coding | NP_001107047.1;NP_001287782.1;NP_001336292.1;NP_001336293.1;NI | 15483 | 608459 | . | . | [illegible] |
| 51271 | NM_00117120 | UBAP1 | ubiquitin associated protein 1 | protein_coding | NP_001164672.1;NP_001164673.1;NP_001164674.1;NP_001164675.1;NI | 12461 | 609787 | . | . | [illegible] |
| 51272 | NM_00109878 | BET1L | Bet1 golgi vesicular membrane trafficking protein like | protein_coding | NP_001092257.1;NP_057610.2 | 19348 | 615417 | . | . | [illegible] |
| 51274 | NM_016531,XI | KLF3 | Kruppel like factor 3 | protein_coding | NP_057615.3;XP_016863768.1 | 16516 | 609392 | . | . | [illegible] |
| 51275 | NR_015404,NF | MAPKAPK5-AS | MAPKAPK5 antisense RNA 1 | lncRNA | . | 24091 | . | . | . | [illegible] |
| 51276 | NM_00129031 | ZNF571 | zinc finger protein 571 | protein_coding | NP_001277243.1;NP_001308201.1;NP_057620.3;XP_011525326.1;XP_01 | 25000 | . | . | . | [illegible] |
| 51277 | NM_00119855 | DNAJC27 | DnaJ heat shock protein family (Hsp40) member C27 | protein_coding | NP_001185488.1;NP_057628.1;XP_011531204.1 | 30290 | 613527 | . | . | [illegible] |
| 51278 | NM_016545 | IER5 | immediate early response 5 | protein_coding | NP_057629.2 | 5393 | 607177 | . | . | [illegible] |
| 51279 | NM_00129764 | C1RL | complement C1r subcomponent like | protein_coding | NP_001284569.1;NP_001284571.1;NP_001284572.1;NP_057630.2 | 21265 | 608974 | . | . | [illegible] |
| 51280 | NM_016548,N | GOLM1 | golgi membrane protein 1 | protein_coding | NP_057632.2;NP_808800.1 | 15451 | 606804 | . | . | [illegible] |
| 51281 | NM_00128277 | ANKMY1 | ankyrin repeat and MYND domain containing 1 | protein_coding | NP_001269700.1;NP_001269709.1;NP_001269710.1;NP_001295304.1;NI | 20987 | . | . | . | [illegible] |
| 51282 | NM_016558,N | SCAND1 | SCAN domain containing 1 | protein_coding | NP_057642.1;NP_361012.2 | 10566 | 610416 | . | . | [illegible] |
| 51283 | NM_00133050 | BFAR | bifunctional apoptosis regulator | protein_coding | NP_001317429.1;NP_057645.1;XP_005255407.1 | 17613 | . | . | . | [illegible] |
| 51287 | NM_016565,XI | COA4 | cytochrome c oxidase assembly factor 4 homolog | protein_coding | NP_057649.2;XP_016873372.1;XP_016873373.1 | 24604 | 608016 | . | . | [illegible] |
| 51290 | NM_016570,XI | ERGIC2 | ERGIC and golgi 2 | protein_coding | NP_057654.2;XP_024304777.1 | 30208 | 612236 | . | . | [illegible] |
| 51291 | NM_00128899 | GMIP | GEM interacting protein | protein_coding | NP_001275927.1;NP_001275928.1;NP_057657.2;XP_005259984.1;XP_01 | 24852 | 609694 | . | . | [illegible] |
| 51292 | NM_00100200 | GMPR2 | guanosine monophosphate reductase 2 | protein_coding | NP_001002000.1;NP_001002001.1;NP_001002002.1;NP_001269950.1;NI | 4377 | 610781 | . | . | [illegible] |
| 51293 | NM_00116589 | CD320 | CD320 molecule | protein_coding | NP_001159367.1;NP_057663.1 | 16692 | 606475 | . | . | [illegible] |
| 51295 | NM_00114246 | ECSIT | ECSIT signaling integrator | protein_coding | NP_001135936.1;NP_001135937.1;NP_001230133.1;NP_057665.2 | 29548 | 608388 | . | . | [illegible] |
| 51299 | NM_00127871 | NRN1 | neuritin 1 | protein_coding | NP_001265639.1;NP_001265640.1;NP_057672.1;XP_006715169.1;XP_00 | 17972 | 607409 | . | . | [illegible] |
| 51300 | NM_016589,XI | TIMMDC1 | translocase of inner mitochondrial membrane domain containing | protein_coding | NP_057673.2;XP_016862045.1 | 1321 | 615534 | . | . | [illegible] |
| 51302 | NM_00127873 | CYP39A1 | cytochrome P450 family 39 subfamily A member 1 | protein_coding | NP_001265667.1;NP_001265668.1;NP_057677.2;XP_005249228.1;XP_01 | 17449 | 605994 | . | . | [illegible] |
| 51303 | NM_00114378 | FKBP11 | FKBP prolyl isomerase 11 | protein_coding | NP_001137253.1;NP_001137254.1;NP_057678.1 | 18624 | 610571 | . | . | [illegible] |
| 51304 | NM_00113517 | ZDHHC3 | zinc finger DHHC-type palmitoyltransferase 3 | protein_coding | NP_001128651.1;NP_001317690.1;NP_001336305.1;NP_001336306.1;NI | 18470 | 617150 | . | . | [illegible] |
| 51305 | NM_00128253 | KCNK9 | potassium two pore domain channel subfamily K member 9 | protein_coding | NP_001269463.1;XP_011515403.1;XP_011515404.1;XP_011515405.1;XP_ | 6283 | 605874 | . | . | [illegible] |
| 51306 | NM_00110180 | FAM13B | family with sequence similarity 13 member B | protein_coding | NP_001095270.1;NP_001095271.1;NP_057687.2;XP_005272064.1;XP_00 | 1335 | 609371 | . | . | [illegible] |
| 51307 | NM_00113564 | FAM53C | family with sequence similarity 53 member C | protein_coding | NP_001129119.1;NP_001337123.1;NP_001337124.1;NP_057689.1;XP_02 | 1336 | 609372 | . | . | [illegible] |
| 51308 | NM_00127180 | REEP2 | receptor accessory protein 2 | protein_coding | NP_001258732.1;NP_057690.2 | 17975 | 609347 | . | . | [illegible] |
| 51309 | NM_016608 | ARMCX1 | armadillo repeat containing X-linked 1 | protein_coding | NP_057692.1 | 18073 | 300362 | . | . | [illegible] |
| 51310 | NM_00128905 | SLC22A17 | solute carrier family 22 member 17 | protein_coding | NP_001275979.1;NP_057693.3;NP_065105.2;XP_005267804.1;XP_00526 | 23095 | 611461 | . | . | [illegible] |
| 51312 | NM_00131781 | SLC25A37 | solute carrier family 25 member 37 | protein_coding | NP_001304741.1;NP_001304742.1;NP_001304743.1;NP_057696.2;XP_00 | 29786 | 610387 | . | . | [illegible] |
| 51313 | NM_00103170 | GASK1B | golgi associated kinase 1B | protein_coding | NP_001026870.2;NP_001121896.1;NP_057697.2;XP_024309846.1;XP_02 | 25312 | . | . | . | [illegible] |
| 51315 | NM_00130452 | KRCC1 | lysine rich coiled-coil 1 | protein_coding | NP_001291455.1;NP_057702.1;XP_016859781.1 | 28039 | . | . | . | [illegible] |
| 51317 | NM_00110180 | PHF21A | PHD finger protein 21A | protein_coding | NP_001095272.1;NP_001338954.1;NP_001338955.1;NP_001338956.1;NI | 24156 | 608325 | . | . | [illegible] |
| 51318 | NM_00136378 | MRPL35 | mitochondrial ribosomal protein L35 | protein_coding | NP_001350711.1;NP_057706.2;NP_663619.1 | 14489 | 611841 | . | . | [illegible] |
| 51319 | NM_00127183 | RSRC1 | arginine and serine rich coiled-coil 1 | protein_coding | NP_001258763.1;NP_001258767.1;NP_057709.2;XP_024309328.1;XP_02 | 24152 | 613352 | . | . | [illegible] |
| 51320 | NM_016626 | MEX3C | mex-3 RNA binding family member C | protein_coding | NP_057710.3 | 28040 | 611005 | . | . | [illegible] |
| 51321 | NM_00103356 | AMZ2 | archaelysin family metallopeptidase 2 | protein_coding | NP_001028741.1;NP_001028742.1;NP_001028743.1;NP_001028744.1;NI | 28041 | 615169 | . | . | [illegible] |
| 51322 | NM_016628,N | WAC | WW domain containing adaptor with coiled-coil | protein_coding | NP_057712.2;NP_567822.1;NP_567823.1;XP_016871804.1;XP_01687180 | 17327 | 615049 | . | . | [illegible] |
| 51324 | NM_00112788 | SPG21 | SPG21 abhydrolase domain containing, maspardin | protein_coding | NP_001121361.1;NP_001121362.1;NP_057714.1;XP_005254494.1;XP_01 | 20373 | 608181 | . | . | [illegible] |
| 51326 | NM_00111373 | ARL17A | ADP ribosylation factor like GTPase 17A | protein_coding | NP_001107210.1;NP_001275740.1;NP_001275741.1;NP_001275742.1;NI | 24096 | . | . | . | [illegible] |
| 51329 | NM_00100225 | ARL6IP4 | ADP ribosylation factor like GTPase 6 interacting protein 4 | protein_coding | NP_001002251.2;NP_001002252.2;NP_001265307.1;NP_001265308.1;NI | 18076 | 607668 | . | . | [illegible] |
| 51330 | NM_016639 | TNFRSF12A | TNF receptor superfamily member 12A | protein_coding | NP_057723.1 | 18152 | 605914 | . | . | [illegible] |
| 51332 | NM_016642,XI | SPTBN5 | spectrin beta, non-erythrocytic 5 | protein_coding | NP_057726.4;XP_016877788.1;XP_016877789.1;XP_016877790.1;XP_01 | 15680 | 605916 | . | . | [illegible] |
| 51333 | NM_00114230 | ZNF771 | zinc finger protein 771 | protein_coding | NP_001135777.1;NP_057727.2 | 29653 | . | . | . | [illegible] |
| 51334 | NM_00130078 | PRR16 | proline rich 16 | protein_coding | NP_001287712.1;NP_001295016.1;NP_001295017.1;NP_057728.1;XP_01 | 29654 | 615931 | . | . | [illegible] |
| 51335 | NM_00103308 | NGRN | neugrin, neurite outgrowth associated | protein_coding | NP_001028260.2 | 18077 | 616718 | . | . | [illegible] |
| 51337 | NM_00136300 | THEM6 | thioesterase superfamily member 6 | protein_coding | NP_001349929.1;NP_057731.1 | 29656 | . | . | . | [illegible] |
| 51339 | NM_00107952 | DACT1 | dishevelled binding antagonist of beta catenin 1 | protein_coding | NP_001072988.1;NP_057735.2 | 17748 | 607861 | . | . | [illegible] |
| 51340 | NM_00127862 | CRNKL1 | crooked neck pre-mRNA splicing factor 1 | protein_coding | NP_001265554.1;NP_001265555.1;NP_001265556.1;NP_001265557.1;NI | 15762 | 610952 | . | . | [illegible] |
| 51341 | NM_00131799 | ZBTB7A | zinc finger and BTB domain containing 7A | protein_coding | NP_001304919.1;NP_056982.1;XP_005259627.3;XP_005259628.1 | 18078 | 605878 | . | . | [illegible] |
| 51343 | NM_00113619 | FZR1 | fizzy and cell division cycle 20 related 1 | protein_coding | NP_001129669.1;NP_001129670.1;NP_057347.2;XP_005259630.1;XP_01 | 24824 | 603619 | . | . | [illegible] |
| 51347 | NM_00134648 | TAOK3 | TAO kinase 3 | protein_coding | NP_001333416.1;NP_001333417.1;NP_001333418.1;NP_001333419.1;NI | 18133 | 616711 | . | . | [illegible] |
| 51351 | NM_015852 | ZNF117 | zinc finger protein 117 | protein_coding | NP_056936.2 | 12897 | 194624 | . | . | [illegible] |
| 51360 | NM_015884 | MBTPS2 | membrane bound transcription factor peptidase, site 2 | protein_coding | NP_056968.1 | 15455 | 300294 | . | . | [illegible] |
| 51361 | NM_015888,XI | HOOK1 | hook microtubule tethering protein 1 | protein_coding | NP_056972.1;XP_006710739.1;XP_011539864.1;XP_011539865.1;XP_01 | 19884 | 607820 | . | . | [illegible] |
| 51362 | NM_015891 | CDC40 | cell division cycle 40 | protein_coding | NP_056975.1 | 17350 | 605585 | . | . | [illegible] |
| 51364 | NM_00130837 | ZMYND10 | zinc finger MYND-type containing 10 | protein_coding | NP_001295308.1;NP_056980.2;XP_005265273.1 | 19412 | 607070 | . | . | [illegible] |
| 51365 | NM_00120696 | PLA1A | phospholipase A1 member A | protein_coding | NP_001193889.1;NP_001193890.1;NP_001280154.1;NP_056984.1;XP_01 | 17661 | 607460 | . | . | [illegible] |
| 51366 | NM_00128287 | UBR5 | ubiquitin protein ligase E3 component n-recognin 5 | protein_coding | NP_001269802.1;NP_056986.2;XP_005251019.1;XP_011515406.2;XP_01 | 16806 | 608413 | . | . | [illegible] |
| 51367 | NM_015918,N | POP5 | POP5 homolog, ribonuclease P/MRP subunit | protein_coding | NP_057002.2;NP_937845.1;XP_011536743.1 | 17689 | 609992 | . | . | [illegible] |
| 51368 | NM_00112988 | TEX264 | testis expressed 264, ER-phagy receptor | protein_coding | NP_001123356.1;NP_001230654.1;NP_001230655.1;NP_001230656.1;NI | 30247 | . | . | . | [illegible] |
| 51371 | NM_015932 | POMP | proteasome maturation protein | protein_coding | NP_057016.1 | 20330 | 613386 | . | . | [illegible] |
| 51372 | NM_00132941 | TMA7 | translation machinery associated 7 homolog | protein_coding | NP_001316346.1;NP_057017.1 | 26932 | 615808 | . | . | [illegible] |
| 51373 | NM_015969 | MRPS17 | mitochondrial ribosomal protein S17 | protein_coding | NP_057053.1 | 14047 | 611980 | . | . | [illegible] |
| 51374 | NM_00117079 | ATRAID | all-trans retinoic acid induced differentiation factor | protein_coding | NP_001164266.1;NP_057169.2 | 24090 | . | . | . | [illegible] |
| 51375 | NM_00136490 | SNX7 | sorting nexin 7 | protein_coding | NP_001351832.1;NP_057060.2;NP_689424.2;XP_006710740.1;XP_01153 | 14971 | 614904 | . | . | [illegible] |
| 51377 | NM_00119926 | UCHL5 | ubiquitin C-terminal hydrolase L5 | protein_coding | NP_001186190.1;NP_001186191.1;NP_001186192.1;NP_001337769.1;NI | 19678 | 610667 | . | . | [illegible] |
| 51379 | NM_015986,N | CRLF3 | cytokine receptor like factor 3 | protein_coding | NP_057070.3 | 17177 | 614853 | . | . | [illegible] |
| 51380 | NM_00124470 | CSAD | cysteine sulfinic acid decarboxylase | protein_coding | NP_001231634.1;NP_001231635.1;NP_057073.4;XP_011536744.2;XP_01 | 18966 | 616569 | . | . | [illegible] |
| 51382 | NM_015994 | ATP6V1D | ATPase H+ transporting V1 subunit D | protein_coding | NP_057078.1 | 13527 | 609398 | . | . | [illegible] |
| 51385 | NM_016089 | ZNF589 | zinc finger protein 589 | protein_coding | NP_057173.2 | 16747 | 616702 | . | . | [illegible] |
| 51386 | NM_00124292 | EIF3L | eukaryotic translation initiation factor 3 subunit L | protein_coding | NP_001229852.1;NP_001350714.1;NP_057175.1;XP_006724323.1;XP_01 | 18138 | . | . | . | [illegible] |
| 51388 | NM_00119943 | NIP7 | nucleolar pre-rRNA processing protein NIP7 | protein_coding | NP_001186363.1;NP_057185.1 | 24328 | . | . | . | [illegible] |
| 51389 | NM_00100746 | RWDD1 | RWD domain containing 1 | protein_coding | NP_001007465.1;NP_057036.2;NP_057188.2 | 20993 | . | . | . | [illegible] |
| 51390 | NM_00128658 | AIG1 | androgen induced 1 | protein_coding | NP_001273516.1;NP_001273517.1;NP_001273518.1;NP_001350650.1;NI | 21607 | 608514 | . | . | [illegible] |
| 51393 | NM_016113,XI | TRPV2 | transient receptor potential cation channel subfamily V member 2 | protein_coding | NP_057197.2;XP_005256733.1;XP_005256734.1;XP_005256735.1;XP_00 | 18082 | 606676 | . | . | [illegible] |
| 51397 | NM_00130808 | COMMD10 | COMM domain containing 10 | protein_coding | NP_001295009.1;NP_057228.1 | 30201 | 616704 | . | . | [illegible] |
| 51398 | NM_016145,XI | WDR83OS | WD repeat domain 83 opposite strand | protein_coding | NP_057229.1;XP_016882353.1 | 30203 | 618474 | . | . | [illegible] |
| 51399 | NM_00131848 | TRAPPC4 | trafficking protein particle complex 4 | protein_coding | NP_001305415.1;NP_001305417.1;NP_001305418.1;NP_001305419.1;NI | 19943 | 610971 | . | . | [illegible] |
| 51400 | NM_00127159 | PPME1 | protein phosphatase methylesterase 1 | protein_coding | NP_001258522.1;NP_057231.1;XP_016873402.1 | 30178 | 611117 | . | . | [illegible] |
| 51406 | NM_00131772 | NOL7 | nucleolar protein 7 | protein_coding | NP_001304653.1;NP_057251.2 | 21040 | 611533 | . | . | [illegible] |
| 51409 | NM_00131785 | HEMK1 | HemK methyltransferase family member 1 | protein_coding | NP_001304780.1;NP_001364349.1;NP_001364350.1;NP_001364351.1;NI | 24923 | 618609 | . | . | [illegible] |
| 51421 | NM_00127868 | AMOTL2 | angiomotin like 2 | protein_coding | NP_001265612.1;NP_001265614.1;NP_001350872.1;NP_057285.3;XP_00 | 17812 | 614658 | . | . | [illegible] |
| 51422 | NM_00104063 | PRKAG2 | protein kinase AMP-activated non-catalytic subunit gamma 2 | protein_coding | NP_001035723.1;NP_001291456.1;NP_001291460.1;NP_001350627.1;NI | 9386 | 602743 | . | . | [illegible] |
| 51426 | NM_00134592 | POLK | DNA polymerase kappa | protein_coding | NP_001332850.1;NP_001332851.1;NP_057302.1;XP_005248591.1;XP_00 | 9183 | 605650 | . | . | [illegible] |
| 51427 | NM_00101374 | ZNF107 | zinc finger protein 107 | protein_coding | NP_001013768.1;NP_001269288.1;NP_001269289.1;NP_057304.1;XP_01 | 12887 | 603989 | . | . | [illegible] |
| 51428 | NM_00132173 | DDX41 | DEAD-box helicase 41 | protein_coding | NP_001308661.1;NP_001308759.1;NP_057306.2;XP_024301877.1 | 18674 | 608170 | . | . | [illegible] |
| 51429 | NM_016224,XI | SNX9 | sorting nexin 9 | protein_coding | NP_057308.1;XP_005267072.1;XP_011534188.1 | 14973 | 605952 | . | . | [illegible] |
| 51430 | NM_00128275 | SUCO | SUN domain containing ossification factor | protein_coding | NP_001269679.1;NP_001269680.1;NP_055098.1;NP_057311.3;XP_00671 | 1240 | . | . | . | [illegible] |
| 51433 | NM_00113755 | ANAPC5 | anaphase promoting complex subunit 5 | protein_coding | NP_001131031.1;NP_001317418.1;NP_057321.2;XP_016874912.1;XP_01 | 15713 | 606948 | . | . | [illegible] |
| 51434 | NM_00113766 | ANAPC7 | anaphase promoting complex subunit 7 | protein_coding | NP_001131136.1;NP_057322.2;XP_005253960.1;XP_011536756.1;XP_01 | 17380 | 606949 | . | . | [illegible] |
| 51439 | NM_016255 | FAM8A1 | family with sequence similarity 8 member A1 | protein_coding | NP_057339.1 | 16372 | 618409 | . | . | [illegible] |
| 51440 | NM_00128239 | HPCAL4 | hippocalcin like 4 | protein_coding | NP_001269325.1;NP_001269326.1;NP_057341.1;XP_016856936.1 | 18212 | . | . | . | [illegible] |

| | | | | | | | | | | | | | | | | | | | | | | | | | | | | | | | | | |
|---|---|---|---|---|---|---|---|---|---|---|---|---|---|---|---|---|---|---|---|---|---|---|---|---|---|---|---|---|---|---|---|---|---|
| 51441 | NM_00117282 | YTHDF2 | YTH N6-methyladenosine RNA binding protein 2 | protein_coding | NP_001166299.1;NP_001166599.1;NP_057342.2 | 31675 | 610640 | . | . | 28.407101 | 27.535823 | 29.581397 | 31.603067 | 66.104494 | 63.650283 | 66.379799 | 72.256655 | 1.078019 | 2502.538217 | 0.115827326 | 0.935730936 | 0.349411738 | 0.998865404 | 11.233991 | 11.342328 | 11.273726 | 11.241615 | 11.295220 | 11.341241 | 2829 | 2547 | 2194 | 2503 |
| 51444 | NM_00119132 | RNF138 | ring finger protein 138 | protein_coding | NP_001178253.2;NP_057355.2;NP_937761.1;XP_005258342.1;XP_00525 | 17765 | 616319 | . | . | 3.032212 | 2.230238 | 2.633137 | 2.581884 | 7.056084 | 5.155296 | 5.908685 | 5.903175 | -1.031676 | 287.397054 | 0.222710817 | -0.2020079 | 0.839910545 | 0.998865404 | 8.188658 | 8.144812 | 8.251280 | 8.087241 | 8.168536 | 8.143667 | 387 | 267 | 252 | 259 |
| 51447 | NM_00100590 | IP6K2 | inositol hexakisphosphate kinase 2 | protein_coding | NP_001005909.1;NP_001005910.1;NP_001005911.1;NP_001139650.1;NI | 17313 | 606992 | . | . | 11.650224 | 11.218460 | 12.446381 | 12.542430 | 27.110548 | 25.931974 | 27.929323 | 28.676778 | 1.141697 | 976.434008 | 0.140842624 | 1.357396851 | 0.174655148 | 0.948677095 | 9.832542 | 10.023876 | 9.860469 | 9.898843 | 10.010885 | 9.941174 | 1022 | 1010 | 948 | 930 |
| 51449 | NM_016297 | PCYOX1 | prenylcysteine oxidase 1 | protein_coding | NP_057381.3 | 20588 | 610995 | . | . | 23.868441 | 22.623884 | 23.818945 | 18.995848 | 55.542843 | 52.296120 | 53.449024 | 43.431751 | -1.100766 | 3716.352089 | 0.132245344 | -1.04735232 | 0.294937114 | 0.998865404 | 11.927132 | 11.788901 | 11.921841 | 11.870030 | 11.908991 | 11.715792 | 4623 | 4072 | 3439 | 2927 |
| 51450 | NM_016307,XI | PRRX2 | paired related homeobox 2 | protein_coding | NP_057391.1;XP_016870292.1 | 21338 | 604675 | . | . | 0.306394 | 0.121554 | 0.112521 | 0.191949 | 0.712990 | 0.280979 | 0.252494 | 0.438870 | -1.414211 | 7.930083 | 1.116860068 | -0.44768132 | 0.654383207 | NA | 3.216230 | 2.715150 | 3.017220 | 2.982976 | 2.979408 | 2.996523 | 15 | 6 | 4 | 8 |
| 51451 | NM_00103239 | LCMT1 | leucine carboxyl methyltransferase 1 | protein_coding | NP_001027563.1;NP_057393.2;XP_005255411.1;XP_011544164.1;XP_01 | 17557 | 610286 | . | . | 16.801810 | 20.176121 | 18.138438 | 22.004181 | 39.098504 | 46.638006 | 40.702130 | 50.309949 | 1.057875 | 828.992655 | 0.156995668 | 0.517013807 | 0.605146521 | 0.998865404 | 9.654461 | 9.734850 | 9.616122 | 9.727527 | 9.659793 | 9.763634 | 855 | 938 | 687 | 857 |
| 51454 | NM_00125266 | GULP1 | GULP PTB domain containing engulfment adaptor 1 | protein_coding | NP_001239597.1;NP_001239598.1;NP_001362854.1;NP_001362855.1;NI | 18649 | 608165 | . | . | 9.092872 | 8.917829 | 9.477647 | 8.794370 | 21.159485 | 20.613955 | 21.267560 | 20.107284 | -1.029511 | 977.965644 | 0.138079122 | -0.30388205 | 0.761217751 | 0.998865404 | 9.954366 | 9.912611 | 9.964908 | 9.922865 | 9.930594 | 9.915218 | 1175 | 1040 | 847 | 894 |
| 51455 | NM_00103787 | REV1 | REV1 DNA directed polymerase | protein_coding | NP_001032961.1;NP_001308383.1;NP_001308384.1;NP_001308387.1;NI | 14060 | 606134 | . | . | 7.098839 | 7.739190 | 8.300092 | 7.351500 | 16.519288 | 17.889483 | 18.625168 | 16.808329 | 1.033205 | 1172.437931 | 0.13951526 | 0.337790061 | 0.7355214 | 0.998865404 | 10.171435 | 10.218768 | 10.152180 | 10.210810 | 10.255719 | 10.154708 | 1276 | 1293 | 1106 | 1042 |
| 51460 | NM_016329,XI | SFMBT1 | Scm like with four mbt domains 1 | protein_coding | NP_057413.2;XP_005265278.1;XP_006713266.1;XP_006713267.1;XP_01 | 20255 | 607319 | . | . | 2.675047 | 3.665539 | 3.625525 | 3.183454 | 6.224945 | 8.473058 | 8.135573 | 7.278591 | 1.081086 | 667.328354 | 0.159866542 | 0.703592513 | 0.481686575 | 0.998865404 | 9.324833 | 9.437475 | 9.317848 | 9.388228 | 9.448881 | 9.362577 | 700 | 728 | 641 | 609 |
| 51463 | NM_00135018 | GPR89B | G protein-coupled receptor 89B | protein_coding | NP_001337109.1;NP_001337110.1;NP_001337111.1;NP_001337112.1;NI | 13840 | 612806 | . | . | 8.848518 | 8.875239 | 9.664485 | 7.346471 | 20.590866 | 20.515511 | 21.686822 | 16.796834 | -1.088068 | 595.556147 | 0.176604286 | -0.68950127 | 0.490507871 | 0.998865404 | 9.277197 | 9.156465 | 9.258691 | 9.227221 | 9.271164 | 9.096035 | 731 | 654 | 563 | 463 |
| 51465 | NM_016021,XI | UBE2J1 | ubiquitin conjugating enzyme E2 J1 | protein_coding | NP_057105.2;XP_011534189.1;XP_011534190.1 | 17598 | 616175 | . | . | 17.001244 | 16.144809 | 14.543647 | 16.656951 | 39.562593 | 37.319452 | 32.635524 | 38.084140 | -1.077879 | 2112.374911 | 0.123762066 | -0.87421645 | 0.382000391 | 0.998865404 | 11.097810 | 10.989457 | 11.095747 | 11.050737 | 10.966556 | 11.058514 | 2594 | 2299 | 1665 | 2017 |
| 51466 | NM_00133022 | EVL | Enah/Vasp-like | protein_coding | NP_001317150.1;NP_057421.1;XP_005267806.1;XP_011535130.1;XP_01 | 20234 | 616912 | . | . | 2.507208 | 2.244808 | 2.087577 | 2.684080 | 5.834378 | 5.188977 | 4.684461 | 6.136831 | -1.034585 | 191.872299 | 0.24737492 | -0.1982908 | 0.842817552 | 0.998865404 | 7.608660 | 7.558917 | 7.611574 | 7.570219 | 7.533168 | 7.617157 | 235 | 201 | 148 | 194 |
| 51474 | NM_00111354 | LIMA1 | LIM domain and actin binding 1 | protein_coding | NP_001107018.1;NP_001107019.1;NP_001230704.1;NP_057441.1;XP_01 | 24636 | 608364 | . | . | 63.629410 | 58.926378 | 50.481620 | 55.675816 | 148.068244 | 136.210969 | 113.279291 | 127.296133 | -1.169706 | 6490.997828 | 0.111036911 | -2.03667921 | **0.041682199** | 0.63293663 | 12.772885 | 12.546709 | 12.762767 | 12.688486 | 12.557258 | 12.629872 | 8409 | 7224 | 4975 | 5855 |
| 51477 | NM_00117093 | ISYNA1 | inositol-3-phosphate synthase 1 | protein_coding | NP_001164409.1;NP_001240318.1;NP_057452.1;XP_011526361.1;XP_02 | 29821 | 611670 | . | . | 13.295399 | 12.400578 | 12.618866 | 13.340296 | 30.938940 | 28.664494 | 28.316373 | 30.501004 | -1.005045 | 893.203519 | 0.145798838 | -0.04979149 | 0.960288545 | 0.998865404 | 9.806442 | 9.799239 | 9.844724 | 9.762029 | 9.777484 | 9.823224 | 1090 | 911 | 750 | 862 |
| 51478 | NM_00130451 | HSD17B7 | hydroxysteroid 17-beta dehydrogenase 7 | protein_coding | NP_001291441.1;NP_001291442.1;NP_057455.1 | 5215 | 606756 | . | . | 12.772547 | 12.712237 | 22.051937 | 16.901066 | 29.722244 | 29.384904 | 49.483911 | 38.642278 | 1.509444 | 752.475012 | 0.167768679 | 3.540692951 | **0.000399078** | **0.034002899** | 9.227683 | 9.822447 | 9.397880 | 9.376349 | 9.767186 | 9.571134 | 702 | 636 | 892 | 736 |
| 51479 | NM_00125799 | ANKFY1 | ankyrin repeat and FYVE domain containing 1 | protein_coding | NP_001244928.1;NP_001316992.1;NP_057460.3;XP_011522228.1;XP_01 | 20763 | 607927 | . | . | 11.251536 | 10.686690 | 10.818245 | 11.288944 | 26.182783 | 24.702763 | 24.275825 | 25.810826 | -1.009952 | 2596.189553 | 0.11450154 | -0.12477229 | 0.900703822 | 0.998865404 | 11.349303 | 11.335021 | 11.365069 | 11.326794 | 11.325630 | 11.350394 | 3075 | 2749 | 2212 | 2459 |
| 51480 | NM_016378 | VCX2 | variable charge X-linked 2 | protein_coding | NP_057462.2 | 18158 | 300532 | . | . | 0.366811 | 0.260357 | 0.446498 | 0.385035 | 0.853583 | 0.601826 | 1.001928 | 0.880338 | 1.304810 | 9.783309 | 0.971372325 | 0.395152209 | 0.69273056 | NA | 3.084644 | 3.470304 | 3.283724 | 3.273923 | 3.292145 | 3.288661 | 11 | 8 | 10 | 10 |
| 51481 | NM_016379 | VCX3A | variable charge X-linked 3A | protein_coding | NP_057463.2 | 18159 | 300533 | . | . | 2.455026 | 1.859526 | 1.435923 | 2.171279 | 5.712946 | 4.298378 | 3.222169 | 4.964373 | -1.209722 | 61.931322 | 0.420645246 | -0.65298609 | 0.513765247 | 0.998865404 | 6.084024 | 5.807986 | 6.005612 | 5.938980 | 5.891800 | 5.969807 | 89 | 63 | 39 | 63 |
| 51490 | NM_016390,XI | SPOUT1 | SPOUT domain containing methyltransferase 1 | protein_coding | NP_057474.2;XP_016870293.1 | 26933 | 617614 | . | . | 8.638996 | 8.332700 | 10.980415 | 9.665085 | 20.103299 | 19.261411 | 24.639731 | 22.098066 | 1.201491 | 1250.770891 | 0.136873021 | 1.934831084 | 0.053011027 | 0.681930298 | 10.149884 | 10.415144 | 10.230048 | 10.196463 | 10.404651 | 10.297298 | 1338 | 1198 | 1270 | 1191 |
| 51491 | NM_00125653 | NOP16 | NOP16 nucleolar protein | protein_coding | NP_001243468.2;NP_001243469.2;NP_001278234.1;NP_001278236.1;NI | 26934 | 612861 | . | . | 55.358873 | 57.320101 | 55.334386 | 63.781054 | 128.822377 | 132.497983 | 124.168755 | 145.827802 | 1.055850 | 1603.380533 | 0.129360211 | 0.606093235 | 0.544452843 | 0.998865404 | 10.607341 | 10.685406 | 10.613332 | 10.636113 | 10.615499 | 10.716011 | 1770 | 1708 | 1340 | 1643 |
| 51493 | NM_014306 | RTCB | RNA 2',3'-cyclic phosphate and 5'-OH ligase | protein_coding | NP_055121.1 | 26935 | 613901 | . | . | 68.957901 | 73.152252 | 66.854637 | 78.889450 | 160.467865 | 169.094696 | 150.019882 | 180.371353 | 1.011135 | 4525.142818 | 0.118643394 | 0.134646505 | 0.89289137 | 0.998865404 | 12.135832 | 12.151620 | 12.117014 | 12.159235 | 12.080806 | 12.210132 | 5051 | 4979 | 3650 | 4596 |
| 51495 | NM_016395,XI | HACD3 | 3-hydroxyacyl-CoA dehydratase 3 | protein_coding | NP_057479.2;XP_011519966.1 | 24175 | 615940 | . | . | 43.319144 | 41.583950 | 41.641628 | 34.053961 | 100.805428 | 96.123161 | 93.442599 | 77.860338 | -1.136938 | 4040.954430 | 0.127250557 | -1.45503146 | 0.145660552 | 0.906375892 | 12.069974 | 11.885066 | 12.050570 | 12.009016 | 12.006263 | 11.833536 | 5073 | 4525 | 3635 | 3173 |
| 51496 | NM_016396,XI | CTDSPL2 | CTD small phosphatase like 2 | protein_coding | NP_057480.2;XP_005254498.1;XP_006720628.1;XP_011519967.1;XP_01( | 26936 | 618739 | . | . | 3.132086 | 3.047267 | 2.764820 | 2.728405 | 7.288493 | 7.043896 | 6.204176 | 6.238172 | -1.121996 | 568.566974 | 0.160671214 | -1.03358417 | 0.301330603 | 0.998865404 | 9.231717 | 9.065894 | 9.203003 | 9.172342 | 9.120745 | 9.104251 | 708 | 634 | 472 | 496 |
| 51497 | NM_198976,XI | NELFCD | negative elongation factor complex member C/D | protein_coding | NP_945327.3 | 15934 | 605297 | . | . | 37.696104 | 40.286843 | 40.157196 | 46.160512 | 87.720389 | 93.124845 | 90.111588 | 105.540520 | 1.091354 | 2855.577355 | 0.120358904 | 1.047855468 | 0.294705202 | 0.998865404 | 11.415259 | 11.541130 | 11.418260 | 11.465571 | 11.460211 | 11.563954 | 3053 | 3033 | 2428 | 2970 |
| 51499 | NM_016399 | TRIAP1 | TP53 regulated inhibitor of apoptosis 1 | protein_coding | NP_057483.1 | 26937 | 614943 | . | . | 17.355179 | 16.018288 | 14.370918 | 13.610621 | 40.386215 | 37.026993 | 32.247925 | 31.119068 | -1.208445 | 551.111099 | 0.164989869 | -1.65556616 | 0.097809708 | 0.813553488 | 9.236081 | 8.963458 | 9.193608 | 9.133736 | 9.061232 | 9.021008 | 725 | 622 | 448 | 453 |
| 51501 | NM_00132240 | HIKESHI | heat shock protein nuclear import factor hikeshi | protein_coding | NP_001309333.1;NP_001309336.1;NP_001309338.1;NP_057485.2;XP_01 | 26938 | 614908 | . | . | 11.167218 | 9.710430 | 12.874701 | 12.366112 | 25.986576 | 22.446091 | 28.890458 | 28.273646 | 1.178371 | 420.855066 | 0.18495175 | 1.280303683 | 0.200438353 | 0.966220653 | 8.593479 | 8.831106 | 8.711923 | 8.611025 | 8.781776 | 8.741627 | 483 | 381 | 409 | 413 |
| 51503 | NM_00136337 | CWC15 | CWC15 spliceosome associated protein homolog | protein_coding | NP_001350300.1;NP_001350301.1;NP_057487.2 | 26939 | . | | . | 15.572357 | 15.795402 | 16.045627 | 17.203213 | 36.237514 | 36.511782 | 36.005919 | 39.333103 | 1.049281 | 771.353526 | 0.147028297 | 0.472029486 | 0.636905742 | 0.998865404 | 9.556264 | 9.625404 | 9.569871 | 9.577863 | 9.584907 | 9.629579 | 862 | 817 | 665 | 764 |
| 51504 | NM_00128608 | TRMT112 | tRNA methyltransferase subunit 11-2 | protein_coding | NP_001273011.1;NP_001273013.1;NP_001359000.1;NP_001359001.1;NI | 26940 | 618630 | . | . | 77.514001 | 75.597436 | 80.412271 | 85.595693 | 180.378274 | 174.746865 | 180.442816 | 195.704366 | 1.064908 | 2050.967227 | 0.119067498 | 0.761993967 | 0.446063605 | 0.998865404 | 10.956054 | 11.046700 | 10.986793 | 10.965694 | 11.001892 | 11.050208 | 2318 | 2115 | 1784 | 2043 |
| 51506 | NM_016406,XI | UFC1 | ubiquitin-fold modifier conjugating enzyme 1 | protein_coding | NP_057490.2;XP_005245311.1;XP_016856939.1;XP_016856940.1 | 26941 | 610554 | . | . | 53.392794 | 53.114638 | 56.533503 | 62.513975 | 124.247219 | 122.776866 | 126.859552 | 142.930778 | 1.103441 | 1581.841477 | 0.126192574 | 1.125337247 | 0.260446152 | 0.995760118 | 10.554733 | 10.696561 | 10.594018 | 10.580749 | 10.627810 | 10.698694 | 1747 | 1609 | 1378 | 1626 |
| 51507 | NM_00128303 | RTF2 | replication termination factor 2 | protein_coding | NP_001269964.1;NP_001269965.1;NP_001269966.1;NP_057491.2;XP_01 | 15890 | . | | . | 80.217070 | 79.018111 | 70.935261 | 84.926576 | 186.668415 | 182.653900 | 159.176667 | 194.174530 | -1.051553 | 4437.081682 | 0.121483462 | -0.59696657 | 0.550529698 | 0.998865404 | 12.151288 | 12.078587 | 12.134122 | 12.136014 | 12.014573 | 12.166052 | 5232 | 4914 | 3418 | 4425 |
| 51510 | NM_00119553 | CHMP5 | charged multivesicular body protein 5 | protein_coding | NP_001182465.1;NP_057494.3 | 26942 | 610900 | . | . | 17.099190 | 16.127814 | 15.921829 | 16.563145 | 39.790517 | 37.280162 | 35.728124 | 37.869662 | -1.037257 | 976.338890 | 0.138776811 | -0.38027347 | 0.703742429 | 0.998865404 | 9.957351 | 9.904644 | 9.969350 | 9.918813 | 9.907725 | 9.927620 | 1184 | 1036 | 822 | 911 |
| 51512 | NM_016426,XI | GTSE1 | G2 and S-phase expressed 1 | protein_coding | NP_057510.5;XP_005261684.1;XP_011528515.1;XP_016884305.1 | 13698 | 607477 | . | . | 19.540245 | 20.703615 | 16.406476 | 20.818136 | 45.470954 | 47.857338 | 36.815656 | 47.598192 | -1.087956 | 2725.197420 | 0.115072386 | -1.05690516 | 0.290554871 | 0.998865404 | 11.471742 | 11.349980 | 11.416997 | 11.472891 | 11.362957 | 11.390411 | 3159 | 3169 | 2231 | 2489 |
| 51514 | NM_00128622 | DTL | denticleless E3 ubiquitin protein ligase homolog | protein_coding | NP_001273158.1;NP_001273159.1;NP_057532.4;XP_011507916.1 | 30288 | 610617 | . | . | 11.204711 | 10.759604 | 9.665698 | 10.957291 | 26.073826 | 24.871305 | 21.689544 | 25.052544 | -1.078219 | 1462.096333 | 0.130875472 | -0.83018001 | 0.406437016 | 0.998865404 | 10.567226 | 10.458366 | 10.561326 | 10.522031 | 10.439013 | 10.526630 | 1791 | 1596 | 1153 | 1395 |
| 51517 | NM_016453,N | NCKIPSD | NCK interacting protein with SH3 domain | protein_coding | NP_057537.1;NP_909119.1;XP_016862083.1;XP_016862084.1;XP_01686 | 15486 | 606671 | . | . | 7.667407 | 9.113335 | 7.402363 | 9.219274 | 17.842368 | 21.065880 | 16.610686 | 21.078775 | -1.024280 | 781.908281 | 0.164852831 | -0.20994556 | 0.833710161 | 0.998865404 | 9.628520 | 9.592972 | 9.559363 | 9.672206 | 9.525544 | 9.669910 | 839 | 922 | 602 | 799 |
| 51520 | NM_00131796 | LARS1 | leucyl-tRNA synthetase 1 | protein_coding | NP_001304893.1;NP_001304894.1;NP_057544.2;NP_064502.9;XP_0115: | 6512 | 151350 | . | . | 33.755400 | 35.329778 | 33.949478 | 37.391123 | 78.550206 | 81.666360 | 76.181646 | 85.490361 | 1.018576 | 5209.696794 | 0.109351073 | 0.24282604 | 0.808140162 | 0.998865404 | 12.333702 | 12.360144 | 12.323355 | 12.354348 | 12.317590 | 12.389838 | 5834 | 5673 | 4375 | 5141 |
| 51522 | NM_00116525 | TMEM14C | transmembrane protein 14C | protein_coding | NP_001158730.1;NP_057546.1 | 20952 | 615318 | . | . | 53.037992 | 50.970071 | 55.949191 | 49.131909 | 123.421591 | 117.819598 | 125.548365 | 112.334273 | -1.002860 | 1665.610451 | 0.130485049 | -0.031579 | 0.974807792 | 0.998865404 | 10.703701 | 10.699968 | 10.721390 | 10.682533 | 10.753787 | 10.643342 | 1968 | 1755 | 1549 | 1449 |
| 51523 | NM_00131719 | CXXC5 | CXXC finger protein 5 | protein_coding | NP_001304128.1;NP_001304129.1;NP_001304130.1;NP_001304131.1;NI | 26943 | 612752 | . | . | 19.085956 | 18.044239 | 15.030590 | 16.561755 | 44.413800 | 41.710071 | 33.728208 | 37.866483 | -1.170168 | 1272.461520 | 0.132092533 | -1.71634153 | 0.086099552 | 0.793345872 | 10.422365 | 10.195542 | 10.393608 | 10.345540 | 10.223165 | 10.277643 | 1629 | 1435 | 982 | 1140 |
| 51524 | NM_00133028 | TMEM138 | transmembrane protein 138 | protein_coding | NP_001317210.1;NP_057548.1;XP_006718648.1;XP_006718649.1;XP_01 | 26944 | 614459 | . | . | 35.619386 | 40.283526 | 41.019822 | 36.262275 | 82.887774 | 93.117179 | 92.047296 | 82.909382 | -1.037189 | 1924.124601 | 0.129187151 | -0.4077705 | 0.683442172 | 0.998865404 | 10.935973 | 10.883524 | 10.901665 | 10.944691 | 10.953323 | 10.832224 | 2195 | 2171 | 1769 | 1635 |
| 51526 | NM_016470,XI | OSER1 | oxidative stress responsive serine rich 1 | protein_coding | NP_057554.4;XP_011527155.1;XP_011527156.1;XP_016883362.1;XP_02 | 16105 | . | | . | 20.565542 | 18.125351 | 23.101487 | 19.770638 | 47.856863 | 41.897564 | 51.839071 | 45.203209 | 1.109141 | 1699.584567 | 0.145503013 | 1.027080438 | 0.304382589 | 0.998865404 | 10.654031 | 10.804006 | 10.724722 | 10.646465 | 10.859718 | 10.666555 | 1949 | 1651 | 1744 | 1472 |
| 51527 | NM_00127190 | GSKIP | GSK3B interacting protein | protein_coding | NP_001258833.1;NP_001258834.1;NP_057556.2 | 20343 | 616605 | . | . | 5.670157 | 4.628591 | 5.525522 | 4.595477 | 13.194689 | 10.699198 | 12.399112 | 10.507011 | -1.032431 | 351.989927 | 0.205943697 | -0.22358529 | 0.823080011 | 0.998865404 | 8.481533 | 8.436902 | 8.525221 | 8.403025 | 8.499882 | 8.393334 | 455 | 345 | 330 | 294 |
| 51528 | NM_00109862 | JKAMP | JNK1/MAPK8 associated membrane protein | protein_coding | NP_001092095.1;NP_001271130.1;NP_001271131.1;NP_001271132.1;NI | 20184 | 611176 | . | . | 23.465560 | 19.941923 | 21.147645 | 16.816247 | 54.605321 | 46.096647 | 47.454704 | 38.448347 | -1.157245 | 1540.863853 | 0.15513847 | -1.3581012 | 0.174431576 | 0.948652105 | 10.690722 | 10.480688 | 10.712613 | 10.562705 | 10.618195 | 10.427533 | 2083 | 1615 | 1398 | 1172 |
| 51529 | NM_00100224 | ANAPC11 | anaphase promoting complex subunit 11 | protein_coding | NP_001002244.1;NP_001002245.1;NP_001002246.1;NP_001002247.1;NI | 14452 | 614534 | . | . | 30.659326 | 30.724266 | 23.539670 | 27.780988 | 71.345512 | 71.020512 | 52.822338 | 63.517924 | -1.236073 | 716.250603 | 0.155312211 | -1.96870416 | **0.048987073** | 0.663679701 | 9.629219 | 9.323228 | 9.584262 | 9.519401 | 9.368926 | 9.442220 | 953 | 816 | 526 | 634 |
| 51530 | NM_00128219 | ZC3HC1 | zinc finger C3HC-type containing 1 | protein_coding | NP_001269119.1;NP_001269120.1;NP_001350630.1;NP_057562.3;XP_0C | 29913 | . | | . | 15.975271 | 17.783986 | 16.943997 | 17.481464 | 37.175113 | 41.108476 | 38.021838 | 39.969288 | 1.010288 | 1033.510130 | 0.138367083 | 0.106717155 | 0.915013373 | 0.998865404 | 10.006043 | 10.020594 | 9.972326 | 10.044806 | 10.017271 | 10.016077 | 1126 | 1164 | 904 | 973 |
| 51531 | NM_00133072 | TRMO | tRNA methyltransferase O | protein_coding | NP_001317654.1;NP_001358586.1;NP_001358587.1;NP_001358588.1;NI | 30967 | . | | . | 3.690913 | 4.670114 | 4.040767 | 5.299504 | 8.588908 | 10.795177 | 9.067363 | 12.116688 | 1.009565 | 247.618906 | 0.232420387 | 0.059093319 | 0.95287778 | 0.998865404 | 7.945923 | 7.958007 | 7.888952 | 8.002025 | 7.902162 | 7.998828 | 252 | 296 | 195 | 256 |
| 51533 | NM_00127822 | PHF7 | PHD finger protein 7 | protein_coding | NP_001265150.1;NP_001308055.1;NP_001308056.1;NP_057567.3;XP_01 | 18458 | . | | . | 2.442512 | 2.409066 | 1.990505 | 2.770142 | 5.683824 | 5.568667 | 4.466635 | 6.333602 | 1.012762 | 170.801450 | 0.262465985 | 0.069706147 | 0.944427549 | 0.998865404 | 7.407589 | 7.424713 | 7.421898 | 7.402537 | 7.362506 | 7.469624 | 200 | 179 | 130 | 182 |
| 51534 | NM_00128637 | VTA1 | vesicle trafficking 1 | protein_coding | NP_001273300.1;NP_001273301.1;NP_057569.2 | 20954 | 610902 | . | . | 6.834039 | 4.781976 | 4.317454 | 4.045324 | 15.903089 | 11.053751 | 9.688242 | 9.249153 | -1.197499 | 987.689298 | 0.143590183 | -1.81087774 | 0.07015978 | 0.748967856 | 10.071838 | 9.812335 | 10.050428 | 9.966806 | 9.909398 | 9.845434 | 1308 | 1097 | 817 | 805 |
| 51535 | NM_00114378 | PPHLN1 | periphilin 1 | protein_coding | NP_001137259.1;NP_001137260.1;NP_001137261.1;NP_001351751.1;NI | 19369 | 608150 | . | . | 15.181679 | 20.752987 | 20.958295 | 25.248502 | 35.328387 | 47.971459 | 47.029814 | 57.727704 | 1.048450 | 2054.768215 | 0.118583628 | 0.575608468 | 0.564879856 | 0.998865404 | 10.970161 | 11.038548 | 10.987433 | 10.983314 | 11.045912 | 10.999773 | 2316 | 2159 | 1882 | 1915 |
| 51537 | NM_00100370 | MTFP1 | mitochondrial fission process 1 | protein_coding | NP_001003704.1;NP_057582.2 | 26945 | 610235 | . | . | 25.467490 | 28.520646 | 27.858145 | 30.467171 | 59.263893 | 65.926753 | 62.512879 | 69.659568 | 1.091482 | 960.273522 | 0.142598373 | 0.885626859 | 0.375818633 | 0.998865404 | 9.843004 | 9.968855 | 9.834323 | 9.911694 | 9.914778 | 9.959435 | 1001 | 1044 | 845 | 968 |
| 51538 | NM_00128256 | ZCCHC17 | zinc finger CCHC-type containing 17 | protein_coding | NP_001269495.1;NP_001269496.1;NP_001269497.1;NP_001269498.1;NI | 30246 | . | | . | 33.073632 | 33.830957 | 34.652496 | 39.408456 | 76.963699 | 78.201775 | 77.759193 | 90.102759 | 1.098768 | 1834.410314 | 0.124779359 | 1.089013132 | 0.276148095 | 0.998865404 | 10.771688 | 10.907327 | 10.799061 | 10.805394 | 10.830581 | 10.920341 | 2005 | 1894 | 1576 | 1902 |
| 51540 | NM_016510 | SCLY | selenocysteine lyase | protein_coding | NP_057594.5 | 18161 | 611056 | . | . | 3.648243 | 7.537778 | 8.711433 | 8.999060 | 8.489613 | 17.423912 | 19.548204 | 20.575281 | 1.563997 | 550.939448 | 0.500351091 | 1.28956964 | 0.197200122 | 0.96621494 | 8.747531 | 9.392481 | 8.689120 | 9.109814 | 9.204524 | 9.220677 | 325 | 623 | 578 | 637 |
| 51542 | NM_00100573 | VPS54 | VPS54 subunit of GARP complex | protein_coding | NP_001005739.1;NP_057600.2;XP_016859805.1 | 18652 | 614633 | . | . | 5.002554 | 4.554617 | 4.747445 | 4.525743 | 11.641148 | 10.528201 | 10.653126 | 10.347571 | 1.013627 | 676.286707 | 0.152633243 | 0.12792948 | 0.898204781 | 0.998865404 | 9.391579 | 9.411336 | 9.394439 | 9.398709 | 9.427288 | 9.383981 | 771 | 727 | 611 | 618 |
| 51545 | NM_016535,XI | ZNF581 | zinc finger protein 581 | protein_coding | NP_057619.1;XP_006723303.1;XP_016882356.1 | 25017 | . | | . | 8.847125 | 8.871121 | 9.048877 | 9.134708 | 20.587622 | 20.505995 | 20.305417 | 20.885425 | 1.002984 | 335.356129 | 0.191987234 | 0.022392023 | 0.982135244 | 0.998865404 | 8.387404 | 8.391693 | 8.390779 | 8.386635 | 8.388369 | 8.392377 | 387 | 360 | 291 | 316 |
| 51547 | NM_016538,XI | SIRT7 | sirtuin 7 | protein_coding | NP_057622.1;XP_011521882.1;XP_024306560.1;XP_024306561.1;XP_02 | 14935 | 606212 | . | . | 12.713300 | 11.204266 | 11.455407 | 11.238145 | 29.584366 | 25.899162 | 25.705602 | 25.694683 | 1.039317 | 714.555635 | 0.157292628 | 0.353707649 | 0.72355799 | 0.998865404 | 9.452454 | 9.508801 | 9.496728 | 9.434783 | 9.540303 | 9.443365 | 843 | 722 | 678 | 635 |
| 51548 | NM_00119328 | SIRT6 | sirtuin 6 | protein_coding | NP_001180214.1;NP_001307987.1;NP_001307988.1;NP_001307989.1;NI | 14934 | 606211 | . | . | 7.421220 | 7.519914 | 7.648489 | 7.680462 | 17.269483 | 17.382621 | 17.162985 | 17.560462 | 1.019323 | 372.201368 | 0.18480401 | 0.149406785 | 0.881232658 | 0.998865404 | 8.526102 | 8.553646 | 8.527980 | 8.540361 | 8.545699 | 8.544858 | 420 | 402 | 327 | 352 |
| 51550 | NM_00132004 | CINP | cyclin dependent kinase 2 interacting protein | protein_coding | NP_001306975.1;NP_116019.1 | 23789 | 613362 | . | . | 18.003246 | 19.250603 | 17.102831 | 23.478522 | 41.894294 | 44.498632 | 38.378259 | 53.680851 | 1.075471 | 576.771611 | 0.183372795 | 0.572434301 | 0.567027783 | 0.998865404 | 9.118981 | 9.222862 | 9.124294 | 9.163727 | 9.085052 | 9.288657 | 621 | 618 | 440 | 646 |
| 51552 | NM_016322 | RAB14 | RAB14, member RAS oncogene family | protein_coding | NP_057406.2 | 16524 | 612673 | . | . | 16.207811 | 14.480078 | 14.608705 | 12.493313 | 37.716244 | 33.471348 | 32.781509 | 28.564474 | -1.148372 | 1869.049634 | 0.135344372 | -1.47468007 | 0.140298577 | 0.900674731 | 10.964217 | 10.765096 | 10.966643 | 10.868908 | 10.872293 | 10.743233 | 2440 | 2025 | 1639 | 1496 |
| 51557 | NM_00114394 | LGSN | lengsin, lens protein with glutamine synthetase domain | protein_coding | NP_001137412.1;NP_057655.2;XP_011534191.1;XP_011534194.1;XP_01 | 21016 | 611470 | . | . | 0.065479 | 0.034013 | 0.097467 | 0.055453 | 0.152371 | 0.078622 | 0.218712 | 0.126786 | 1.462022 | 13.147983 | 0.868700518 | 0.630787266 | 0.52817963 | NA | 3.413393 | 3.967302 | 3.708055 | 3.678569 | 3.731422 | 3.698484 | 16 | 8 | 17 | 11 |
| 51559 | NM_00103170 | NT5DC3 | 5'-nucleotidase domain containing 3 | protein_coding | NP_001026871.1;XP_011536778.1;XP_016874943.1 | 30826 | 611076 | . | . | 10.577919 | 10.912547 | 11.586537 | 9.756682 | 24.615253 | 25.224840 | 25.999853 | 22.307497 | -1.037457 | 2066.533074 | 0.129658681 | -0.40915856 | 0.682423306 | 0.998865404 | 11.039118 | 10.986394 | 11.028093 | 11.024804 | 11.064933 | 10.924803 | 2428 | 2266 | 1920 | 1734 |
| 51560 | NM_00136395 | RAB6B | RAB6B, member RAS oncogene family | protein_coding | NP_001350882.1;NP_057661.3 | 14902 | 615852 | . | . | 3.186210 | 3.408249 | 3.202167 | 3.377036 | 7.414441 | 7.878318 | 7.185570 | 7.721192 | -1.015014 | 574.998347 | 0.16074683 | -0.13375126 | 0.893599273 | 0.998865404 | 9.178298 | 9.156447 | 9.152037 | 9.192026 | 9.148878 | 9.175656 | 647 | 644 | 486 | 546 |
| 51561 | NM_016584,XI | IL23A | interleukin 23 subunit alpha | protein_coding | NP_057668.1;XP_011536779.1 | 15488 | 605580 | . | . | 0.404622 | 0.557397 | 0.286631 | 0.397587 | 0.941571 | 1.288448 | 0.643192 | 0.909037 | -1.405287 | 16.560506 | 0.763701108 | -0.6427446 | 0.520389836 | NA | 4.275837 | 3.781347 | 4.052950 | 4.077791 | 4.032038 | 4.050853 | 19 | 24 | 10 | 15 |
| 51562 | NM_00114489 | MBIP | MAP3K12 binding inhibitory protein 1 | protein_coding | NP_001138363.1;NP_001295039.1;NP_057670.2;XP_005267811.1;XP_00 | 20427 | 609431 | . | . | 6.412594 | 6.099812 | 6.640176 | 6.975644 | 14.922373 | 14.099988 | 14.900363 | 15.948981 | 1.065502 | 331.701323 | 0.193850638 | 0.472181231 | 0.636797436 | 0.998865404 | 8.327281 | 8.418751 | 8.368765 | 8.341384 | 8.376091 | 8.404756 | 379 | 338 | 290 | 329 |
| 51564 | NM_00109841 | HDAC7 | histone deacetylase 7 | protein_coding | NP_001091886.1;NP_001295019.1;NP_001354975.1;NP_056216.2;XP_01 | 14067 | 606542 | . | . | 2.799926 | 2.906592 | 2.325135 | 1.874152 | 6.515541 | 6.718718 | 5.217535 | 4.285026 | -1.243740 | 340.343326 | 0.195123167 | -1.61275081 | 0.106798658 | 0.83501324 | 8.559371 | 8.245259 | 8.458187 | 8.478101 | 8.379851 | 8.310718 | 427 | 414 | 281 | 266 |
| 51566 | NM_016607,N | ARMCX3 | armadillo repeat containing X-linked 3 | protein_coding | NP_057691.1;NP_808816.1;NP_808817.1;XP_005262198.1 | 24065 | 300364 | . | . | 8.235038 | 6.503170 | 7.589498 | 5.534912 | 19.163273 | 15.032368 | 17.030614 | 12.654920 | -1.169862 | 805.757950 | 0.179987913 | -1.2575166 | 0.208566632 | 0.970482251 | 9.762424 | 9.537200 | 9.777696 | 9.621761 | 9.691096 | 9.482323 | 1107 | 837 | 744 | 591 |
| 51567 | NM_016614 | TDP2 | tyrosyl-DNA phosphodiesterase 2 | protein_coding | NP_057698.2 | 17768 | 605764 | . | . | 9.533115 | 8.634320 | 8.532765 | 8.663682 | 22.183952 | 19.958620 | 19.147280 | 19.808479 | -1.072270 | 533.711400 | 0.166423723 | -0.60488811 | 0.545253343 | 0.998865404 | 9.109230 | 9.008856 | 9.116577 | 9.046000 | 9.035679 | 9.038279 | 670 | 564 | 447 | 484 |
| 51569 | NM_00128670 | UFM1 | ubiquitin fold modifier 1 | protein_coding | NP_001273632.1;NP_001273633.1;NP_001273634.1;NP_001273635.1;NI | 20597 | 610553 | . | . | 11.148259 | 9.116413 | 9.162354 | 6.941690 | 25.942454 | 21.072996 | 20.560058 | 15.871350 | -1.274996 | 903.993675 | 0.173652559 | -2.01835908 | **0.043553877** | 0.642641319 | 9.984343 | 9.634884 | 9.971881 | 9.816377 | 9.816482 | 9.618378 | 1287 | 980 | 790 | 639 |
| 51571 | NM_00125676 | CYRIB | CYFIP related Rac1 interactor B | protein_coding | NP_001243692.1;NP_001317541.1;NP_001340171.1;NP_001340172.1;NI | 25216 | 617978 | . | . | 17.968969 | 16.733142 | 15.906077 | 15.525878 | 41.814542 | 38.679401 | 35.692775 | 35.498079 | -1.115811 | 1978.054957 | 0.119672037 | -1.3210445 | 0.186486527 | 0.95851159 | 11.026689 | 10.868723 | 11.015805 | 10.967027 | 10.911093 | 10.898530 | 2475 | 2183 | 1639 | 1739 |
| 51573 | NM_00132406 | GDE1 | glycerophosphodiester phosphodiesterase 1 | protein_coding | NP_001310995.1;NP_001310996.1;NP_057725.1 | 29644 | 605943 | . | . | 27.622576 | 26.595063 | 25.290689 | 29.008224 | 64.278866 | 61.475679 | 56.751570 | 66.323853 | -1.013137 | 2456.267140 | 0.122006765 | -0.15433358 | 0.877346718 | 0.998865404 | 11.271716 | 11.252725 | 11.285646 | 11.248064 | 11.204005 | 11.306323 | 2913 | 2606 | 1987 | 2433 |
| 51574 | NM_00126703 | LARP7 | La ribonucleoprotein 7, transcriptional regulator | protein_coding | NP_001253968.1;NP_001357903.1;NP_001357904.1;NP_001357905.1;NI | 24912 | 612026 | . | . | 12.975570 | 12.500595 | 12.216639 | 12.389187 | 30.194685 | 28.895689 | 27.413779 | 28.326403 | -1.040320 | 838.033380 | 0.143943747 | -0.39617438 | 0.691976388 | 0.998865404 | 9.738980 | 9.682189 | 9.747146 | 9.701763 | 9.704150 | 9.689132 | 1015 | 893 | 722 | 762 |
| 51575 | NM_00127638 | ESF1 | ESF1 nucleolar pre-rRNA processing protein homolog | protein_coding | NP_001263309.1;NP_057733.2;XP_016883363.1 | 15898 | 618765 | . | . | 7.069598 | 6.225559 | 7.227443 | 6.025212 | 16.451246 | 14.390658 | 16.218173 | 13.775931 | -1.008172 | 675.544874 | 0.166816154 | -0.07038663 | 0.943885935 | 0.998865404 | 9.405296 | 9.394499 | 9.449077 | 9.350162 | 9.455798 | 9.331393 | 837 | 679 | 639 | 573 |
| 51582 | NM_00130166 | AZIN1 | antizyme inhibitor 1 | protein_coding | NP_001288597.1;NP_001349939.1;NP_001349940.1;NP_001349941.1;NI | 16432 | 607909 | . | . | 28.581772 | 26.567936 | 26.802142 | 24.050293 | 66.510960 | 61.412972 | 60.143223 | 54.988140 | -1.115302 | 3546.067407 | 0.115876148 | -1.35864654 | 0.174258619 | 0.948096927 | 11.868481 | 11.711245 | 11.864651 | 11.804295 | 11.784260 | 11.704089 | 4461 | 3889 | 3059 | 2989 |
| 51585 | NM_00134641 | PCF11 | PCF11 cleavage and polyadenylation factor subunit | protein_coding | NP_001333342.1;NP_001333343.1;NP_001333344.1;NP_056969.2 | 30097 | 608876 | . | . | 6.243293 | 6.315595 | 7.041389 | 6.144098 | 14.528396 | 14.598782 | 15.800673 | 14.047752 | 1.035448 | 1539.750723 | 0.13269107 | 0.378736566 | 0.704883497 | 0.998865404 | 10.562961 | 10.613558 | 10.572100 | 10.575604 | 10.659429 | 10.538995 | 1738 | 1636 | 1468 | 1355 |
| 51586 | NM_00100389 | MED15 | mediator complex subunit 15 | protein_coding | NP_001003891.1;NP_001280163.1;NP_001280164.1;NP_001280165.1;NI | 14248 | 607372 | . | . | 42.378384 | 46.061594 | 47.317003 | 46.633905 | 98.616244 | 106.473430 | 106.177982 | 106.622887 | 1.024208 | 4909.707333 | 0.108024303 | 0.319450101 | 0.749385224 | 0.998865404 | 12.244085 | 12.278552 | 12.221115 | 12.280349 | 12.276673 | 12.265594 | 5388 | 5420 | 4349 | 4627 |
| 51588 | NM_015897,XI | PIAS4 | protein inhibitor of activated STAT 4 | protein_coding | NP_056981.2;XP_011526362.1;XP_016882357.1 | 17002 | 605989 | . | . | 4.753929 | 4.873649 | 5.586988 | 5.241264 | 11.062589 | 11.265658 | 12.537038 | 11.983528 | 1.111405 | 490.042155 | 0.169347391 | 0.899838317 | 0.3682063 | 0.998865404 | 8.858417 | 9.011071 | 8.896038 | 8.906553 | 8.992178 | 8.943060 | 530 | 505 | 466 | 465 |
| 51592 | NM_015906,N | TRIM33 | tripartite motif containing 33 | protein_coding | NP_056990.3;NP_148980.2;XP_005270993.1;XP_005270994.1;XP_01153 | 16290 | 605769 | . | . | 6.991435 | 6.866422 | 7.532182 | 6.030479 | 16.269358 | 15.872040 | 16.902000 | 13.787976 | -1.038810 | 1778.642349 | 0.139392882 | -0.39408187 | 0.693520594 | 0.998865404 | 10.823545 | 10.769063 | 10.820326 | 10.798849 | 10.865033 | 10.686919 | 2115 | 1929 | 1690 | 1450 |
| 51593 | NM_00112885 | SRRT | serrate, RNA effector molecule | protein_coding | NP_001122324.1;NP_001122325.1;NP_001122326.1;NP_056992.4;XP_0C | 24101 | 614469 | . | . | 43.504978 | 47.408175 | 50.458530 | 47.479516 | 101.237878 | 109.586120 | 113.227477 | 108.556276 | 1.067844 | 4442.821526 | 0.111216687 | 0.851494289 | 0.394494834 | 0.998865404 | 12.069129 | 12.163838 | 12.057276 | 12.119618 | 12.170073 | 12.116436 | 4762 | 4811 | 4118 | 4164 |
| 51594 | NM_015909,N | NBAS | NBAS subunit of NRZ tethering complex | protein_coding | NP_056993.2;XP_011508659.1;XP_011508660.1;XP_011508662.1;XP_01 | 15625 | 608025 | . | . | 5.249577 | 5.417952 | 4.880478 | 5.763409 | 12.215982 | 12.523840 | 10.951651 | 13.177350 | -1.015541 | 1202.310244 | 0.139270102 | -0.15982648 | 0.873017761 | 0.998865404 | 10.242905 | 10.220193 | 10.228958 | 10.244508 | 10.163898 | 10.282693 | 1381 | 1321 | 955 | 1207 |
| 51596 | NM_00101443 | CUTA | cutA divalent cation tolerance homolog | protein_coding | NP_001014433.2;NP_001014437.1;NP_001014838.1;NP_001014840.1;NI | 21101 | 616953 | . | . | 82.947903 | 78.819141 | 77.358801 | 80.907669 | 193.023187 | 182.193966 | 173.590907 | 184.985741 | -1.026616 | 2470.360048 | 0.11663176 | -0.32492428 | 0.745238377 | 0.998865404 | 11.289311 | 11.251452 | 11.309505 | 11.251428 | 11.247868 | 11.271358 | 2985 | 2604 | 2089 | 2319 |
| 51599 | NM_00126048 | LSR | lipolysis stimulated lipoprotein receptor | protein_coding | NP_001247418.1;NP_001247419.1;NP_057009.3;NP_991403.1;NP_9914( | 29572 | 616582 | . | . | 1.951252 | 2.083248 | 1.871593 | 2.129312 | 4.540644 | 4.815521 | 4.199800 | 4.868419 | -1.021445 | 142.924297 | 0.274256652 | -0.11161674 | 0.911127298 | 0.998865404 | 7.174760 | 7.143283 | 7.152957 | 7.173270 | 7.134053 | 7.175693 | 162 | 160 | 116 | 140 |
| 51601 | NM_00120483 | LIPT1 | lipoyltransferase 1 | protein_coding | NP_001191759.1;NP_057013.1;NP_660198.1;NP_660199.1;NP_660200.1 | 29569 | 610284 | . | . | 3.329273 | 2.940031 | 2.821824 | 2.458620 | 7.747353 | 6.796013 | 6.332090 | 5.621342 | -1.170722 | 128.204137 | 0.291085014 | -0.7812117 | 0.434677992 | 0.998865404 | 7.110889 | 6.884866 | 7.055114 | 7.003299 | 6.996940 | 6.950010 | 170 | 139 | 110 | 103 |
| 51602 | NM_015934 | NOP58 | NOP58 ribonucleoprotein | protein_coding | NP_057018.1 | 29926 | 616742 | . | . | 50.550659 | 53.176804 | 50.992317 | 48.586407 | 117.633461 | 122.920570 | 114.425285 | 111.087044 | -1.055895 | 3158.227221 | 0.11270026 | -0.6962376 | 0.486280046 | 0.998865404 | 11.663575 | 11.585158 | 11.629162 | 11.663834 | 11.625662 | 11.577933 | 3657 | 3574 | 2750 | 2796 |
| 51603 | NM_00100723 | EEF1AKNMT | eEF1A lysine and N-terminal methyltransferase | protein_coding | NP_001007240.1;NP_055770.1;NP_057019.3 | 24248 | 617987 | . | . | 10.734129 | 12.421337 | 11.399455 | 12.682572 | 24.978761 | 28.712479 | 25.580050 | 28.997193 | 1.018574 | 1233.780030 | 0.140157855 | 0.189434648 | 0.84975217 | 0.998865404 | 10.255793 | 10.281828 | 10.207034 | 10.312218 | 10.235253 | 10.312683 | 1310 | 1411 | 1027 | 1227 |
| 51604 | NM_00118472 | PIGT | phosphatidylinositol glycan anchor biosynthesis class T | protein_coding | NP_001171657.1;NP_001171658.1;NP_001171659.1;NP_057021.2 | 14938 | 610272 | . | . | 43.160746 | 43.710969 | 41.475718 | 45.617447 | 100.436827 | 101.039862 | 93.070306 | 104.298875 | -1.009875 | 2877.519686 | 0.114570194 | -0.12374154 | 0.901519917 | 0.998865404 | 11.497749 | 11.483435 | 11.493514 | 11.495409 | 11.452920 | 11.518813 | 3326 | 3124 | 2389 | 2793 |
| 51605 | NM_00128146 | TRMT6 | tRNA methyltransferase 6 | protein_coding | NP_001268396.1;NP_057023.2;XP_016883364.1 | 20900 | . | | . | 7.944538 | 7.769387 | 7.865122 | 8.416014 | 18.487266 | 17.959285 | 17.649104 | 19.242217 | -1.020020 | 712.110992 | 0.149550465 | -0.19121979 | 0.8483534 | 0.998865404 | 9.490125 | 9.461651 | 9.495247 | 9.470161 | 9.471250 | 9.467054 | 843 | 762 | 615 | 659 |
| 51606 | NM_015941,N | ATP6V1H | ATPase H+ transporting V1 subunit H | protein_coding | NP_057025.2;NP_998784.1;NP_998785.1;XP_006716518.1;XP_01151584 | 18303 | 608861 | . | . | 32.364259 | 32.037069 | 33.746379 | 34.845094 | 75.312964 | 74.055126 | 75.725893 | 79.669167 | 1.048705 | 2191.309662 | 0.116299156 | 0.589936404 | 0.555233287 | 0.998865404 | 11.062879 | 11.131461 | 11.085128 | 11.071093 | 11.105498 | 11.126779 | 2485 | 2288 | 1925 | 2131 |
| 51608 | NM_015949 | GET4 | guided entry of tail-anchored proteins factor 4 | protein_coding | NP_057033.2 | 21690 | 612056 | . | . | 23.731190 | 25.030807 | 24.651066 | 25.971203 | 55.223454 | 57.859837 | 55.316277 | 59.380070 | 1.024070 | 1618.126249 | 0.123776176 | 0.277232028 | 0.781601948 | 0.998865404 | 10.642942 | 10.677073 | 10.633038 | 10.667938 | 10.652685 | 10.685256 | 1800 | 1764 | 1394 | 1567 |
| 51611 | NM_00107739 | DPH5 | diphthamide biosynthesis 5 | protein_coding | NP_001070862.1;NP_001070863.1;NP_057042.2;XP_005270995.1;XP_00 | 24270 | 611075 | . | . | 9.371227 | 7.599216 | 7.382446 | 6.750905 | 21.807230 | 17.565925 | 16.565997 | 15.435144 | -1.094678 | 592.396059 | 0.1663681 | -0.78444114 | 0.432781305 | 0.998865404 | 9.274586 | 9.143263 | 9.216920 | 9.260767 | 9.124185 | 9.231117 | 690 | 690 | 451 | 574 |
| 51614 | NM_015966,N | ERGIC3 | ERGIC and golgi 3 | protein_coding | NP_057050.1;NP_938408.1 | 15927 | 616971 | . | . | 285.701286 | 292.520639 | 261.530769 | 273.585154 | 664.838711 | 676.174562 | 586.867457 | 625.519925 | -1.095824 | 11293.051099 | 0.100799668 | -1.30968799 | 0.190301409 | 0.961278413 | 13.527658 | 13.395617 | 13.494456 | 13.507416 | 13.407988 | 13.437446 | 13496 | 12839 | 9208 | 10279 |
| 51616 | NM_015975 | TAF9B | TATA-box binding protein associated factor 9b | protein_coding | NP_057059.2 | 17306 | 300754 | . | . | 2.945437 | 2.924576 | 2.390471 | 2.184514 | 6.854152 | 6.760289 | 5.364148 | 4.994632 | -1.298858 | 216.610079 | 0.230375981 | -1.63751466 | 0.101522978 | 0.819486355 | 7.935002 | 7.558404 | 7.820534 | 7.812190 | 7.715689 | 7.672819 | 284 | 262 | 172 | 168 |
| 51619 | NM_015983,XI | UBE2D4 | ubiquitin conjugating enzyme E2 D4 (putative) | protein_coding | NP_057067.1;XP_006715797.1;XP_011513722.1;XP_011513724.1;XP_01: | 21647 | . | | . | 1.559050 | 1.826156 | 1.446553 | 1.673446 | 3.627973 | 4.221242 | 3.246024 | 3.826138 | -1.105061 | 202.805435 | 0.242120948 | -0.59526254 | 0.551667992 | 0.998865404 | 7.734853 | 7.589387 | 7.649898 | 7.720210 | 7.607497 | 7.672013 | 227 | 247 | 155 | 194 |
| 51621 | NM_00130246 | KLF13 | Kruppel like factor 13 | protein_coding | NP_001289390.1;NP_057079.2 | 13672 | 605328 | . | . | 2.515270 | 2.582999 | 2.368244 | 2.521449 | 5.853137 | 5.970717 | 5.314271 | 5.764994 | -1.056819 | 532.788774 | 0.163975366 | -0.48622178 | 0.62680993 | 0.998865404 | 9.096868 | 9.016863 | 9.069450 | 9.080827 | 9.023313 | 9.055411 | 625 | 596 | 439 | 498 |
| 51622 | NM_015622 | CCZ1 | CCZ1 homolog, vacuolar protein trafficking and biogenesis associa | protein_coding | NP_056437.4 | 21691 | . | | . | 12.677227 | 12.910898 | 12.497275 | 12.561644 | 29.500431 | 29.844120 | 28.043524 | 28.720705 | -1.035731 | 939.041166 | 0.138175881 | -0.36655696 | 0.713949517 | 0.998865404 | 9.900151 | 9.849494 | 9.884226 | 9.891217 | 9.863861 | 9.860665 | 1095 | 1036 | 804 | 863 |
| 51626 | NM_00119346 | DYNC2LI1 | dynein cytoplasmic 2 light intermediate chain 1 | protein_coding | NP_001180393.1;NP_001335841.1;NP_001335842.1;NP_056337.1;NP_0! | 24595 | 617083 | . | . | 11.090508 | 10.415589 | 10.583458 | 9.939235 | 25.808067 | 24.076100 | 23.748974 | 22.724879 | -1.062294 | 461.608802 | 0.174621823 | -0.49926888 | 0.617589978 | 0.998865404 | 8.893168 | 8.806587 | 8.895540 | 8.841161 | 8.854029 | 8.808263 | 571 | 491 | 404 | 404 |
| 51629 | NM_00114378 | SLC25A39 | solute carrier family 25 member 39 | protein_coding | NP_001137252.1;NP_001308169.1;NP_001308170.1;NP_001353655.1;NI | 24279 | 610820 | . | . | 60.817622 | 61.947361 | 63.185775 | 68.547551 | 141.525105 | 143.194101 | 141.787039 | 156.725838 | 1.051106 | 3667.337436 | 0.109764033 | 0.655112119 | 0.51239557 | 0.998865404 | 11.804147 | 11.875993 | 11.824271 | 11.814595 | 11.838166 | 11.882565 | 4142 | 3836 | 3181 | 3618 |
| 51631 | NM_00124458 | LUC7L2 | LUC7 like 2, pre-mRNA splicing factor | protein_coding | NP_001231514.1;NP_001257572.1;NP_057103.2 | 21608 | 613056 | . | . | 33.562042 | 31.613848 | 31.231716 | 30.979353 | 78.100255 | 73.076826 | 70.083068 | 70.830608 | -1.062102 | 2644.223927 | 0.11503766 | -0.75559895 | 0.449889684 | 0.998865404 | 11.411379 | 11.324573 | 11.419391 | 11.364798 | 11.351119 | 11.336094 | 3241 | 2841 | 2251 | 2382 |
| 51633 | NM_00128674 | OTUD6B | OTU deubiquitinase 6B | protein_coding | NP_001273674.1;NP_057107.4;XP_011515431.2 | 24281 | 612021 | . | . | 3.598008 | 3.428467 | 3.764869 | 2.940359 | 8.372711 | 7.925057 | 8.448257 | 6.722784 | -1.061865 | 343.272357 | 0.205487733 | -0.42144026 | 0.673433628 | 0.998865404 | 8.465222 | 8.379938 | 8.453334 | 8.421870 | 8.474547 | 8.328227 | 418 | 371 | 328 | 271 |
| 51634 | NM_016024 | RBMX2 | RNA binding motif protein X-linked 2 | protein_coding | NP_057108.2 | 24282 | . | | . | 6.974409 | 7.670465 | 5.936490 | 7.432808 | 16.229736 | 17.730625 | 13.321312 | 16.994234 | -1.111335 | 398.163268 | 0.192205695 | -0.79234636 | 0.428158748 | 0.998865404 | 8.711914 | 8.558439 | 8.640119 | 8.692755 | 8.541246 | 8.663137 | 462 | 472 | 293 | 391 |
| 51635 | NM_00132228 | DHRS7 | dehydrogenase/reductase 7 | protein_coding | NP_001309209.1;NP_001309210.1;NP_001309211.1;NP_057113.1;XP_01 | 21524 | 612833 | . | . | 32.841251 | 29.799276 | 31.169096 | 28.850379 | 76.422939 | 68.882361 | 69.942551 | 65.962962 | -1.060070 | 1855.105136 | 0.12741214 | -0.6605261 | 0.508916276 | 0.998865404 | 10.898581 | 10.814773 | 10.918645 | 10.838265 | 10.870937 | 10.794337 | 2309 | 1956 | 1643 | 1606 |
| 51637 | NM_016039 | RTRAF | RNA transcription, translation and transport factor | protein_coding | NP_057123.1 | 23169 | 610858 | . | . | 8.921786 | 8.676450 | 8.466625 | 8.232480 | 20.761364 | 20.056002 | 18.998861 | 18.822588 | -1.068515 | 1872.790697 | 0.12020021 | -0.79539695 | 0.426382634 | 0.998865404 | 10.917914 | 10.822455 | 10.910850 | 10.882031 | 10.859217 | 10.829074 | 2268 | 2050 | 1605 | 1665 |
| 51639 | NM_016047 | SF3B6 | splicing factor 3b subunit 6 | protein_coding | NP_057131.1 | 30096 | 607835 | . | . | 125.141327 | 112.767677 | 107.787239 | 126.593742 | 291.209015 | 260.667542 | 241.871414 | 289.441528 | -1.029928 | 2394.998397 | 0.128933438 | -0.32996148 | 0.741429067 | 0.998865404 | 11.246980 | 11.204322 | 11.281222 | 11.189980 | 11.150848 | 11.271228 | 2956 | 2475 | 1898 | 2378 |
| 51642 | NM_00131849 | MRPL48 | mitochondrial ribosomal protein L48 | protein_coding | NP_001305427.1;NP_001305428.1;NP_001305429.1;NP_057139.1;XP_01 | 16653 | 611853 | . | . | 9.471154 | 10.690466 | 10.628991 | 10.007122 | 22.039765 | 24.711488 | 23.851148 | 22.880096 | -1.082584 | 437.406531 | 0.175975969 | -0.65053948 | 0.515343811 | 0.998865404 | 8.829144 | 8.714230 | 8.787571 | 8.805865 | 8.726619 | 8.769510 | 516 | 498 | 352 | 408 |
| 51643 | NM_00128260 | TMBIM4 | transmembrane BAX inhibitor motif containing 4 | protein_coding | NP_001269535.1;NP_001269538.1;NP_001269539.1;NP_057140.2 | 24257 | 616874 | . | . | 14.449588 | 12.123636 | 13.656416 | 9.188936 | 33.624785 | 28.024328 | 30.644602 | 21.009409 | -1.190593 | 1047.079123 | 0.189138916 | -1.33066485 | 0.183299313 | 0.954800306 | 10.152069 | 9.901363 | 10.149185 | 10.019185 | 10.103252 | 9.787397 | 1422 | 1123 | 1013 | 701 |
| 51645 | NM_016059 | PPIL1 | peptidylprolyl isomerase like 1 | protein_coding | NP_057143.1 | 9260 | 601301 | . | . | 17.767242 | 19.551481 | 20.954050 | 20.134041 | 41.345100 | 45.194126 | 47.020275 | 46.034092 | 1.085331 | 925.910684 | 0.142138709 | 0.831123516 | 0.405903856 | 0.998865404 | 9.794482 | 9.912561 | 9.791723 | 9.854464 | 9.900802 | 9.864524 | 978 | 1000 | 859 | 881 |
| 51646 | NM_00112739 | YPEL5 | yippee like 5 | protein_coding | NP_001120871.1;NP_001120872.1;NP_001120873.1;NP_057145.1;XP_01 | 18329 | 609726 | . | . | 11.443644 | 10.139446 | 9.824618 | 10.220283 | 26.629829 | 23.437781 | 22.046160 | 23.367463 | -1.100142 | 757.796047 | 0.150945916 | -0.91217831 | 0.361674862 | 0.998865404 | 9.632798 | 9.495257 | 9.636763 | 9.557412 | 9.519802 | 9.542682 | 965 | 809 | 617 | 688 |
| 51647 | NM_016062,N | CIAO2B | cytosolic iron-sulfur assembly component 2B | protein_coding | NP_057146.1 | 24261 | 614778 | . | . | 39.466737 | 37.442458 | 31.899312 | 40.090482 | 91.840717 | 86.549912 | 71.581132 | 91.662121 | -1.088418 | 797.313405 | 0.158711539 | -0.77015638 | 0.441207137 | 0.998865404 | 9.699082 | 9.576313 | 9.690020 | 9.643623 | 9.531839 | 9.677454 | 988 | 868 | 594 | 791 |
| 51649 | NM_016070 | MRPS23 | mitochondrial ribosomal protein S23 | protein_coding | NP_057154.2 | 14509 | 611985 | . | . | 8.341520 | 7.817363 | 6.951470 | 8.065682 | 19.411060 | 18.070185 | 15.598898 | 18.441225 | -1.091164 | 1362.658371 | 0.135701553 | -0.92753081 | 0.353651 | 0.998865404 | 10.473880 | 10.347782 | 10.472356 | 10.415087 | 10.324075 | 10.428002 | 1698 | 1478 | 1055 | 1306 |
| 51650 | NM_016071,N | MRPS33 | mitochondrial ribosomal protein S33 | protein_coding | NP_057155.1;NP_444263.1 | 16634 | 611993 | . | . | 6.474399 | 6.697042 | 6.067215 | 6.875036 | 15.066193 | 15.480513 | 13.614653 | 15.718954 | -1.031423 | 867.785375 | 0.145653651 | -0.30645419 | 0.759258851 | 0.998865404 | 9.783565 | 9.738472 | 9.762061 | 9.781763 | 9.708053 | 9.789715 | 1001 | 964 | 700 | 847 |
| 51651 | NM_00101550 | PTRH2 | peptidyl-tRNA hydrolase 2 | protein_coding | NP_001015509.1;NP_057161.1;XP_011523189.1 | 24265 | 608625 | . | . | 38.305270 | 35.129128 | 48.566885 | 40.295245 | 89.137935 | 81.202553 | 108.982677 | 92.130290 | 1.165905 | 1254.261461 | 0.144044533 | 1.537372362 | 0.12420215 | 0.87182553 | 10.177427 | 10.399296 | 10.220691 | 10.238042 | 10.422386 | 10.262856 | 1318 | 1264 | 1296 | 1135 |
| 51652 | NM_00100575 | CHMP3 | charged multivesicular body protein 3 | protein_coding | NP_001005753.1;NP_001180446.1;NP_057163.1 | 29865 | 610052 | . | . | 12.792207 | 12.547193 | 12.241296 | 12.230637 | 29.767991 | 29.003401 | 27.469116 | 27.963896 | -1.047743 | 1209.285902 | 0.130060492 | -0.51733111 | 0.604925044 | 0.998865404 | 10.273156 | 10.205940 | 10.264291 | 10.250286 | 10.225811 | 10.218624 | 1438 | 1323 | 1032 | 1102 |
| 51654 | NM_00127816 | CDK5RAP1 | CDK5 regulatory subunit associated protein 1 | protein_coding | NP_001265096.1;NP_001265097.1;NP_001265098.1;NP_001352657.1;NI | 15880 | 608200 | . | . | 14.683330 | 14.371385 | 15.300751 | 16.198973 | 34.168707 | 33.220098 | 34.334447 | 37.037025 | 1.067935 | 980.330199 | 0.137505439 | 0.689603631 | 0.490443483 | 0.998865404 | 9.888953 | 9.983738 | 9.924351 | 9.900145 | 9.946442 | 9.973986 | 1110 | 1006 | 864 | 967 |
| 51657 | NM_00131778 | STYXL1 | serine/threonine/tyrosine interacting like 1 | protein_coding | NP_001304714.1;NP_001304715.1;NP_001304716.1;NP_001304717.1;NI | 18165 | 616695 | . | . | 13.261986 | 11.763560 | 11.905270 | 14.426015 | 30.861188 | 27.191992 | 26.715083 | 32.983363 | 1.030082 | 495.440967 | 0.180943244 | 0.236309375 | 0.813192613 | 0.998865404 | 8.931044 | 8.973778 | 8.998479 | 8.881176 | 8.914147 | 9.004384 | 614 | 478 | 406 | 505 |
| 51659 | NM_016095 | GINS2 | GINS complex subunit 2 | protein_coding | NP_057179.1 | 24575 | 610609 | . | . | 6.635087 | 6.922948 | 6.243811 | 7.964548 | 15.440121 | 16.002705 | 14.010931 | 18.209993 | 1.032928 | 568.400160 | 0.174061412 | 0.268522301 | 0.788297311 | 0.998865404 | 9.127666 | 9.173497 | 9.127049 | 9.149733 | 9.080763 | 9.232631 | 633 | 614 | 444 | 604 |
| 51660 | NM_00127087 | MPC1 | mitochondrial pyruvate carrier 1 | protein_coding | NP_001257808.1;NP_001363494.1;NP_001363495.1;NP_001363496.1;NI | 21606 | 614738 | . | . | 15.722494 | 15.149301 | 14.225252 | 17.430958 | 36.586890 | 35.018288 | 31.921056 | 39.853815 | 1.028444 | 477.764895 | 0.176854259 | 0.228791241 | 0.819031175 | 0.998865404 | 8.880021 | 8.920016 | 8.910476 | 8.870054 | 8.855905 | 8.957413 | 560 | 492 | 387 | 491 |
| 51661 | NM_00113521 | FKBP7 | FKBP prolyl isomerase 7 | protein_coding | NP_001128684.1;NP_851939.1;XP_005246695.1;XP_011509650.1;XP_01 | 3723 | 607062 | . | . | 10.083394 | 8.869354 | 8.299205 | 8.898654 | 23.464474 | 20.501910 | 18.623177 | 20.345712 | -1.119205 | 823.469196 | 0.152497081 | -1.06542677 | 0.286682849 | 0.998865404 | 9.764451 | 9.602127 | 9.779526 | 9.664239 | 9.623262 | 9.663752 | 1081 | 864 | 656 | 750 |
| 51663 | NM_016107,N | ZFR | zinc finger RNA binding protein | protein_coding | NP_057191.2 | 17277 | 615635 | . | . | 13.741964 | 14.935925 | 13.487816 | 13.191212 | 31.978116 | 34.525056 | 30.266270 | 30.160140 | -1.089340 | 2033.878559 | 0.120286162 | -1.02633394 | 0.304734202 | 0.998865404 | 11.050441 | 10.926953 | 10.992816 | 11.052244 | 10.967814 | 10.941731 | 2352 | 2373 | 1721 | 1795 |
| 51665 | NM_00104044 | ASB1 | ankyrin repeat and SOCS box containing 1 | protein_coding | NP_001035535.1;NP_001317125.1 | 16011 | 605758 | . | . | 4.933779 | 4.773662 | 5.360795 | 5.072029 | 11.481105 | 11.034532 | 12.029468 | 11.596592 | 1.058368 | 1066.032459 | 0.136010971 | 0.601724926 | 0.547357255 | 0.998865404 | 10.016382 | 10.098520 | 10.054197 | 10.017895 | 10.100270 | 10.055768 | 1222 | 1090 | 982 | 997 |
| 51666 | NM_016116,N | ASB4 | ankyrin repeat and SOCS box containing 4 | protein_coding | NP_057200.1;NP_665879.1;XP_016867792.1 | 16009 | 605761 | . | . | 0.126007 | 0.123977 | 0.063870 | 0.172961 | 0.293223 | 0.286579 | 0.143321 | 0.395456 | -1.159468 | 14.160759 | 0.837290734 | -0.25494508 | 0.798765516 | NA | 3.928838 | 3.710567 | 3.833055 | 3.829409 | 3.800506 | 3.843324 | 18 | 16 | 7 | 17 |
| 51667 | NM_00124335 | NUB1 | negative regulator of ubiquitin like proteins 1 | protein_coding | NP_001230280.1;NP_001350458.1;NP_057202.3;XP_016867793.1;XP_01 | 17623 | 607981 | . | . | 6.483430 | 6.545454 | 5.210988 | 7.813736 | 15.087205 | 15.130112 | 11.693308 | 17.865180 | 1.000084 | 634.808466 | 0.189520765 | 0.000636348 | 0.999492268 | 0.999862634 | 9.310607 | 9.309748 | 9.315133 | 9.296932 | 9.170300 | 9.425761 | 741 | 676 | 447 | 708 |
| 51668 | NM_00131693 | HSPB11 | heat shock protein family B (small) member 11 | protein_coding | NP_001303864.1;NP_001369178.1;NP_001369179.1;NP_001369180.1;NI | 25019 | . | | . | 29.480618 | 31.680364 | 26.257891 | 27.126040 | 68.602611 | 73.230581 | 58.921948 | 62.020469 | -1.168754 | 627.080326 | 0.156526081 | -1.43727313 | 0.150640401 | 0.913627824 | 9.400702 | 9.175534 | 9.318852 | 9.365472 | 9.236997 | 9.238856 | 752 | 754 | 502 | 543 |
| 51669 | NM_00128423 | SARAF | store-operated calcium entry associated regulatory factor | protein_coding | NP_001271168.1;NP_057211.4;XP_016869056.1;XP_016869057.1;XP_01 | 28789 | 614768 | . | . | 69.837064 | 65.376277 | 65.651844 | 60.624820 | 162.513725 | 151.120181 | 147.320832 | 138.611440 | -1.086922 | 4112.797903 | 0.113880047 | -1.05591658 | 0.29100633 | 0.998865404 | 12.064695 | 11.944616 | 12.070261 | 12.006976 | 12.006373 | 11.932265 | 5119 | 4448 | 3582 | 3529 |
| 51676 | NM_00120242 | ASB2 | ankyrin repeat and SOCS box containing 2 | protein_coding | NP_001189358.1;NP_057234.2;XP_005267815.1;XP_011535136.1;XP_01 | 16012 | 605759 | . | . | 2.086988 | 2.044858 | 2.121487 | 1.784970 | 4.856507 | 4.726783 | 4.760552 | 4.081123 | -1.030087 | 166.546246 | 0.256259255 | -0.16688746 | 0.867458603 | 0.998865404 | 7.400689 | 7.358561 | 7.388763 | 7.382544 | 7.393857 | 7.352747 | 196 | 181 | 150 | 146 |
| 51678 | NM_00130303 | MPP6 | membrane palmitoylated protein 6 | protein_coding | NP_001289966.1;NP_057531.2;XP_005249832.1;XP_006715801.1;XP_00 | 18167 | 606959 | . | . | 2.207147 | 2.150880 | 1.845265 | 1.843212 | 5.136120 | 4.971852 | 4.140721 | 4.214285 | -1.194857 | 528.464639 | 0.16416936 | -1.56446667 | 0.117708014 | 0.857396579 | 9.168299 | 8.911626 | 9.110692 | 9.089360 | 8.990826 | 8.981001 | 673 | 611 | 422 | 448 |
| 51684 | NM_00117813 | SUFU | SUFU negative regulator of hedgehog signaling | protein_coding | NP_001171604.1;NP_057253.2;XP_011538160.1;XP_011538162.1;XP_01 | 16466 | 607035 | . | . | 3.728273 | 3.751312 | 2.774793 | 3.063820 | 8.675847 | 8.671325 | 6.226552 | 7.005061 | -1.263659 | 444.270144 | 0.178897343 | -1.88715694 | 0.05913922 | 0.718513409 | 8.954592 | 8.616197 | 8.850459 | 8.873398 | 8.677118 | 8.758649 | 561 | 545 | 317 | 394 |
| 51686 | NM_00113493 | OAZ3 | ornithine decarboxylase antizyme 3 | protein_coding | NP_001128411.1;NP_001288300.1;NP_057262.2 | 8097 | 605138 | . | . | 1.354932 | 1.349450 | 2.104314 | 2.257288 | 3.152982 | 3.119312 | 4.722019 | 5.161021 | 1.560272 | 47.623725 | 0.452263569 | 1.419078101 | 0.155876247 | 0.925004232 | 5.217479 | 5.859002 | 5.521539 | 5.520977 | 5.593107 | 5.605712 | 43 | 40 | 49 | 56 |
| 51690 | NM_016199,XI | LSM7 | LSM7 homolog, U6 small nuclear RNA and mRNA degradation ass | protein_coding | NP_057283.1;XP_016882358.1 | 20470 | 607287 | . | . | 44.927546 | 46.512769 | 43.098159 | 42.846524 | 104.548243 | 107.516348 | 96.711018 | 97.963474 | -1.070626 | 702.532150 | 0.149883862 | -0.65687167 | 0.511263438 | 0.998865404 | 9.504813 | 9.406349 | 9.473271 | 9.485809 | 9.436077 | 9.429265 | 829 | 791 | 593 | 633 |
| 51691 | NM_016200 | LSM8 | LSM8 homolog, U6 small nuclear RNA associated | protein_coding | NP_057284.1 | 20471 | 607288 | . | . | 3.793947 | 3.702238 | 3.565362 | 3.273721 | 8.828670 | 8.557889 | 8.000570 | 7.484975 | -1.111796 | 1383.627723 | 0.130122909 | -1.17498659 | 0.240000081 | 0.992733415 | 10.508508 | 10.355940 | 10.485930 | 10.460234 | 10.427487 | 10.355517 | 1705 | 1546 | 1194 | 1170 |
| 51692 | NM_00132183 | CPSF3 | cleavage and polyadenylation specific factor 3 | protein_coding | NP_001308762.1;NP_001308763.1;NP_001308764.1;NP_001308765.1;NI | 2326 | 606029 | . | . | 26.094031 | 27.778911 | 24.077637 | 25.505355 | 60.721885 | 64.212201 | 54.029518 | 58.314963 | -1.102121 | 1823.674509 | 0.121757049 | -1.15215467 | 0.24925752 | 0.992929739 | 10.901163 | 10.760684 | 10.848431 | 10.890505 | 10.774540 | 10.811186 | 2144 | 2118 | 1473 | 1665 |
| 51693 | NM_00131852 | TRAPPC2L | trafficking protein particle complex 2 like | protein_coding | NP_001305453.1;NP_001305454.1;NP_001305455.1;NP_001305456.1;NI | 30887 | 610970 | . | . | 13.239122 | 12.501755 | 12.997726 | 15.489122 | 30.807984 | 28.898368 | 29.166518 | 35.414035 | 1.095323 | 933.035404 | 0.148718492 | 0.883259266 | 0.377096207 | 0.998865404 | 9.798778 | 9.929823 | 9.855739 | 9.804830 | 9.835031 | 9.954532 | 1062 | 927 | 779 | 989 |
| 51696 | NM_016217 | HECA | hdc homolog, cell cycle regulator | protein_coding | NP_057301.1 | 21041 | 607977 | . | . | 1.245301 | 1.051770 | 1.475829 | 1.020615 | 2.897868 | 2.431213 | 3.311718 | 2.333514 | 1.070425 | 211.656112 | 0.258032882 | 0.380508502 | 0.703567987 | 0.998865404 | 7.674534 | 7.774878 | 7.743335 | 7.661891 | 7.818732 | 7.644007 | 256 | 201 | 226 | 167 |
| 51699 | NM_00128215 | VPS29 | VPS29 retromer complex component | protein_coding | NP_001269079.1;NP_001269080.1;NP_057310.1;NP_476528.1;XP_00671 | 14340 | 606932 | . | . | 38.875639 | 42.098050 | 37.052831 | 36.685001 | 90.465205 | 97.311526 | 83.145483 | 83.875903 | -1.058555 | 1474.336742 | 0.125639471 | -0.65342642 | 0.513481409 | 0.998865404 | 10.566242 | 10.484295 | 10.547868 | 10.546833 | 10.523774 | 10.482862 | 1747 | 1635 | 1279 | 1307 |
| 51700 | NM_00130282 | CYB5R2 | cytochrome b5 reductase 2 | protein_coding | NP_001289755.1;NP_001289756.1;NP_057313.2;XP_005253032.1;XP_00 | 24376 | 608342 | . | . | 0.752114 | 0.958885 | 0.921621 | 0.594636 | 1.750201 | 2.216505 | 2.068092 | 1.359564 | -1.041838 | 41.872791 | 0.48339306 | -0.12232613 | 0.902640738 | 0.998865404 | 5.417438 | 5.357829 | 5.372575 | 5.407910 | 5.389633 | 5.378005 | 44 | 51 | 37 | 37 |
| 51701 | NM_016231,XI | NLK | nemo like kinase | protein_coding | NP_057315.3;XP_005258045.1 | 29858 | 609476 | . | . | 5.642590 | 5.524942 | 6.895015 | 4.678802 | 13.130539 | 12.771153 | 15.472213 | 10.697524 | 1.020896 | 616.036231 | 0.191763332 | 0.155588253 | 0.876357585 | 0.998865404 | 9.251318 | 9.282259 | 9.264421 | 9.244866 | 9.389520 | 9.131500 | 712 | 648 | 646 | 471 |
| 51704 | NM_00130477 | GPRC5B | G protein-coupled receptor class C group 5 member B | protein_coding | NP_001291700.1;NP_057319.1;XP_006721114.1;XP_006721115.1 | 13308 | 605948 | . | . | 9.149069 | 8.615692 | 7.934196 | 8.930051 | 21.290261 | 19.915561 | 17.804108 | 20.417498 | -1.065336 | 1382.943710 | 0.132801327 | -0.68756145 | 0.491728994 | 0.998865404 | 10.478543 | 10.387060 | 10.483070 | 10.430135 | 10.364990 | 10.450083 | 1699 | 1487 | 1099 | 1326 |
| 51706 | NM_016243 | CYB5R1 | cytochrome b5 reductase 1 | protein_coding | NP_057327.2 | 13397 | 608341 | . | . | 34.339626 | 34.154095 | 38.232475 | 37.693478 | 79.909729 | 78.948723 | 85.792572 | 86.181664 | 1.093005 | 1826.586761 | 0.119774487 | 1.071185068 | 0.284086224 | 0.998865404 | 10.769317 | 10.897697 | 10.804077 | 10.793061 | 10.878192 | 10.859116 | 2024 | 1870 | 1679 | 1767 |
| 51710 | NM_00116427 | ZNF44 | zinc finger protein 44 | protein_coding | NP_001157748.1;NP_001340478.1;NP_001340479.1;NP_001340480.1;NI | 13110 | 194542 | . | . | 1.360737 | 0.641191 | 1.108199 | 0.996734 | 3.166491 | 1.482140 | 2.486765 | 2.278914 | -1.173481 | 133.732599 | 0.298458992 | -0.77328611 | 0.439353074 | 0.998865404 | 7.173012 | 6.944352 | 7.093284 | 7.084439 | 7.092211 | 6.965236 | 168 | 154 | 127 | 94 |
| 51714 | NM_016275 | SELENOT | selenoprotein T | protein_coding | NP_057359.2 | 18136 | 607912 | . | . | 18.924107 | 16.765327 | 13.227130 | 11.689106 | 44.037178 | 38.753803 | 29.681295 | 26.725752 | -1.452416 | 1625.497612 | 0.135582783 | -3.97140935 | **7.14487E-05** | **0.010039957** | 10.910713 | 10.372758 | 10.844028 | 10.738453 | 10.550845 | 10.451603 | 2360 | 1943 | 1230 | 1160 |
| 51715 | NM_00127866 | RAB23 | RAB23, member RAS oncogene family | protein_coding | NP_001265595.1;NP_001265596.1;NP_001265597.1;NP_057361.3;NP_8! | 14263 | 606144 | . | . | 2.104905 | 1.747328 | 1.900331 | 1.646885 | 4.898200 | 4.039028 | 4.264287 | 3.765407 | -1.101022 | 278.565246 | 0.218896543 | -0.63428678 | 0.525893682 | 0.998865404 | 8.188961 | 8.051509 | 8.196613 | 8.090426 | 8.134137 | 8.053953 | 369 | 284 | 249 | 229 |
| 51719 | NM_00113084 | CAB39 | calcium binding protein 39 | protein_coding | NP_001124321.1;NP_001124322.1;NP_057373.1;XP_011509652.1 | 20292 | 612174 | . | . | 14.244615 | 13.578317 | 13.354304 | 12.145591 | 33.147805 | 31.386888 | 29.966672 | 27.769448 | -1.107884 | 1599.856385 | 0.128170098 | -1.15321252 | 0.248823167 | 0.992733415 | 10.715573 | 10.568114 | 10.702774 | 10.660503 | 10.642044 | 10.560870 | 1988 | 1766 | 1390 | 1348 |
| 51720 | NM_00119929 | UIMC1 | ubiquitin interaction motif containing 1 | protein_coding | NP_001186226.1;NP_001186227.1;NP_001304890.1;NP_057374.3;XP_0C | 30298 | 609433 | . | . | 10.108681 | 11.034194 | 10.920814 | 12.013755 | 23.523318 | 25.506031 | 24.505990 | 27.468025 | 1.065124 | 852.665993 | 0.145122293 | 0.627203108 | 0.530526106 | 0.998865404 | 9.689822 | 9.780429 | 9.698039 | 9.726514 | 9.723177 | 9.790284 | 932 | 909 | 729 | 863 |
| 51726 | NM_00137845 | DNAJB11 | DnaJ heat shock protein family (Hsp40) member B11 | protein_coding | NP_001365380.1;NP_057390.1 | 14889 | 611341 | . | . | 91.997309 | 93.293654 | 77.853384 | 79.744907 | 214.081524 | 215.652469 | 174.700738 | 182.327256 | -1.192092 | 4382.868121 | 0.106414769 | -2.3821417 | **0.017212273** | 0.44170732 | 12.218857 | 11.965339 | 12.164313 | 12.168626 | 12.014467 | 12.025803 | 5475 | 5157 | 3453 | 3775 |
| 51727 | NM_00113614 | CMPK1 | cytidine/uridine monophosphate kinase 1 | protein_coding | NP_001129612.1;NP_001353064.1;NP_057392.1 | 18170 | 191710 | . | . | 16.729000 | 15.404987 | 15.199322 | 12.444580 | 38.929066 | 35.609317 | 34.106837 | 28.453054 | -1.179270 | 1368.999349 | 0.144421861 | -1.64721963 | 0.09951291 | 0.813730268 | 10.532670 | 10.295398 | 10.511921 | 10.440818 | 10.426284 | 10.269875 | 1783 | 1526 | 1207 | 1055 |
| 51728 | NM_016310 | POLR3K | RNA polymerase III subunit K | protein_coding | NP_057394.3 | 14121 | 606007 | . | . | 16.100336 | 16.578732 | 12.245569 | 16.899015 | 37.466145 | 38.322483 | 27.478701 | 38.637592 | -1.137021 | 661.035593 | 0.177015719 | -1.04657032 | 0.295297795 | 0.998865404 | 9.458718 | 9.272442 | 9.401243 | 9.415284 | 9.214216 | 9.417578 | 802 | 767 | 455 | 669 |
| 51729 | NM_016312 | WBP11 | WW domain binding protein 11 | protein_coding | NP_057396.1 | 16461 | 618083 | . | . | 21.167788 | 22.308018 | 21.304098 | 23.260391 | 49.258312 | 51.565983 | 47.805779 | 53.182121 | 1.010835 | 3140.649601 | 0.114088351 | 0.136271203 | 0.891606889 | 0.998865404 | 11.609136 | 11.624518 | 11.593990 | 11.630090 | 11.588644 | 11.652303 | 3518 | 3445 | 2639 | 3075 |
| 51734 | NM_00138226 | MSRB1 | methionine sulfoxide reductase B1 | protein_coding | NP_001369193.1;NP_001369194.1;NP_057416.1 | 14133 | 606216 | . | . | 10.005043 | 10.642897 | 9.311472 | 13.617554 | 23.282148 | 24.601535 | 20.894672 | 31.134917 | 1.094481 | 433.522898 | 0.204617092 | 0.636539787 | 0.524424657 | 0.998865404 | 8.694067 | 8.822983 | 8.710676 | 8.743494 | 8.663172 | 8.888625 | 464 | 459 | 322 | 502 |
| 51735 | NM_00116438 | RAPGEF6 | Rap guanine nucleotide exchange factor 6 | protein_coding | NP_001157858.1;NP_001157859.1;NP_001157860.1;NP_001157861.1;NI | 20655 | 610499 | . | . | 2.781128 | 2.734343 | 3.160949 | 2.596236 | 6.471799 | 6.320558 | 7.093077 | 5.935986 | 1.033593 | 726.299291 | 0.16282005 | 0.292762333 | 0.769703821 | 0.998865404 | 9.479981 | 9.528453 | 9.506499 | 9.474981 | 9.587545 | 9.434613 | 841 | 753 | 713 | 616 |
| 51741 | NM_00129199 | WWOX | WW domain containing oxidoreductase | protein_coding | NP_001278926.1;NP_057457.1;NP_570607.1 | 12799 | 605131 | . | . | 2.373829 | 2.832504 | 1.877798 | 2.311232 | 5.523998 | 6.547461 | 4.213724 | 5.284355 | -1.193503 | 141.961129 | 0.287915592 | -0.88637683 | 0.375414501 | 0.998865404 | 7.272304 | 7.014937 | 7.150838 | 7.209606 | 7.061701 | 7.163673 | 165 | 179 | 96 | 139 |
| 51742 | NM_00120679 | ARID4B | AT-rich interaction domain 4B | protein_coding | NP_001193723.1;NP_057458.4;NP_112739.2;XP_006711844.1;XP_01154 | 15550 | 609696 | . | . | 3.803484 | 3.386762 | 3.873423 | 3.474028 | 8.850864 | 7.828652 | 8.691848 | 7.942951 | 1.012887 | 677.535430 | 0.158621201 | 0.116457895 | 0.907289654 | 0.998865404 | 9.394542 | 9.413699 | 9.436972 | 9.359203 | 9.443629 | 9.370509 | 819 | 685 | 625 | 605 |
| 51747 | NM_00133033 | LUC7L3 | LUC7 like 3 pre-mRNA splicing factor | protein_coding | NP_001317259.1;NP_006098.2;NP_057508.2;XP_005257506.1;XP_00525 | 24309 | 609434 | . | . | 27.249651 | 21.873012 | 31.847625 | 24.744101 | 63.411056 | 50.560448 | 71.465147 | 56.574453 | 1.032801 | 3926.412531 | 0.14775672 | 0.315123557 | 0.752667843 | 0.998865404 | 11.915398 | 11.962215 | 11.933844 | 11.909584 | 12.079545 | 11.798672 | 4513 | 4107 | 4052 | 3113 |
| 51750 | NM_00128300 | RTEL1 | regulator of telomere elongation helicase 1 | protein_coding | NP_001269938.1;NP_001269939.1;NP_057518.1;NP_116575.3 | 15888 | 608833 | . | . | 16.474547 | 13.895420 | 15.914512 | 16.745492 | 38.336953 | 32.119886 | 35.711700 | 38.286579 | 1.054427 | 2226.352322 | 0.12922708 | 0.591662796 | 0.554076414 | 0.998865404 | 11.081653 | 11.158263 | 11.165980 | 11.024649 | 11.124605 | 11.155739 | 2716 | 2132 | 1949 | 2184 |
| 51752 | NM_00104045 | ERAP1 | endoplasmic reticulum aminopeptidase 1 | protein_coding | NP_001035548.1;NP_001185470.1;NP_001336173.1;NP_057526.3;XP_0C | 18173 | 606832 | . | . | 10.683470 | 10.535173 | 9.261049 | 8.738385 | 24.860876 | 24.352525 | 20.781525 | 19.979279 | -1.198781 | 1513.739108 | 0.124962449 | -2.09317194 | **0.036333808** | 0.593509535 | 10.688696 | 10.427279 | 10.646081 | 10.613143 | 10.500287 | 10.479271 | 1940 | 1744 | 1214 | 1273 |
| 51754 | NM_00104258 | TMEM8B | transmembrane protein 8B | protein_coding | NP_001036054.1;NP_001036055.2;NP_001350549.1;NP_001350550.1;NI | 21427 | 616888 | . | . | 3.917855 | 1.956377 | 3.040446 | 2.942592 | 9.117006 | 4.522255 | 6.822670 | 6.727887 | 1.048245 | 432.886307 | 0.182423215 | 0.372630605 | 0.709423391 | 0.998865404 | 8.723696 | 8.791203 | 8.762328 | 8.721923 | 8.720251 | 8.819196 | 503 | 441 | 354 | 449 |
| 51755 | NM_015083,N | CDK12 | cyclin dependent kinase 12 | protein_coding | NP_055898.1;NP_057591.2;XP_005257513.1;XP_005257515.1;XP_01152 | 24224 | 615514 | . | . | 9.142740 | 9.362771 | 9.789615 | 10.520612 | 21.275531 | 21.642461 | 21.967609 | 24.054127 | 1.075454 | 2485.797871 | 0.11590147 | 0.905473122 | 0.365214793 | 0.998865404 | 11.226140 | 11.330944 | 11.242873 | 11.255390 | 11.280234 | 11.334751 | 2737 | 2605 | 2165 | 2496 |
| 51759 | NM_016520,XI | C9orf78 | chromosome 9 open reading frame 78 | protein_coding | NP_057604.1;XP_011517082.1 | 24932 | . | | . | 20.424788 | 19.058609 | 17.559068 | 21.155942 | 47.529320 | 44.054825 | 39.402038 | 48.370542 | -1.034570 | 1093.081647 | 0.145782635 | -0.33633267 | 0.736620014 | 0.998865404 | 10.118659 | 10.069290 | 10.134522 | 10.076181 | 10.012655 | 10.143994 | 1330 | 1153 | 851 | 1097 |
| 51760 | NM_00130815 | SYT17 | synaptotagmin 17 | protein_coding | NP_001295086.1;NP_001317438.1;NP_057608.2;XP_005255419.1;XP_01 | 24119 | . | | . | 0.570597 | 0.727735 | 0.545949 | 1.489764 | 1.327801 | 1.682193 | 1.225096 | 3.406169 | 1.398501 | 61.133249 | 0.427587915 | 1.131652956 | 0.257780367 | 0.995760118 | 5.672908 | 6.154807 | 5.895040 | 5.874189 | 5.893133 | 6.017058 | 62 | 52 | 46 | 84 |
| 51762 | NM_016530,XI | RAB8B | RAB8B, member RAS oncogene family | protein_coding | NP_057614.1;XP_011519971.1;XP_016877801.1 | 30273 | 613532 | . | . | 2.747514 | 2.358201 | 2.477779 | 2.000243 | 6.393579 | 5.451086 | 5.560063 | 4.573318 | -1.150452 | 359.535076 | 0.204305932 | -0.98969506 | 0.322323191 | 0.998865404 | 8.586868 | 8.386132 | 8.575582 | 8.474997 | 8.507243 | 8.381666 | 480 | 380 | 324 | 278 |
| 51763 | NM_00113564 | INPP5K | inositol polyphosphate-5-phosphatase K | protein_coding | NP_001129114.1;NP_057616.2;NP_570122.1;XP_005256740.1;XP_00525 | 33882 | 607875 | . | . | 7.582190 | 7.440841 | 7.586446 | 8.017890 | 17.644065 | 17.199841 | 17.023766 | 18.331953 | 1.025485 | 662.507919 | 0.154535552 | 0.2349351 | 0.814259106 | 0.998865404 | 9.353638 | 9.389724 | 9.373420 | 9.352375 | 9.352523 | 9.406709 | 765 | 695 | 560 | 654 |
| 51765 | NM_00104245 | STK26 | serine/threonine kinase 26 | protein_coding | NP_001035917.1;NP_001035918.1;NP_057626.2;XP_011529651.1;XP_01 | 18174 | 300547 | . | . | 6.057014 | 6.213141 | 5.028222 | 5.335363 | 14.094921 | 14.361955 | 11.283184 | 12.198675 | -1.196318 | 571.364330 | 0.160559046 | -1.61062496 | 0.10726149 | 0.835490835 | 9.281913 | 9.023023 | 9.206840 | 9.221419 | 9.080886 | 9.112218 | 709 | 679 | 441 | 500 |
| 51768 | NM_016551,XI | TM7SF3 | transmembrane 7 superfamily member 3 | protein_coding | NP_057635.1;XP_005253448.1;XP_011518992.1;XP_011518993.1;XP_01( | 23049 | 605181 | . | . | 50.879519 | 51.204940 | 50.710731 | 53.336791 | 118.398735 | 118.362510 | 113.793409 | 121.948235 | 1.038366 | 7294.716527 | 0.103139245 | 0.526614151 | 0.598461563 | 0.998865404 | 12.805243 | 12.859519 | 12.811417 | 12.821419 | 12.831507 | 12.864795 | 8193 | 7769 | 6336 | 7104 |
| 51773 | NM_016578,XI | RSF1 | remodeling and spacing factor 1 | protein_coding | NP_057662.3;XP_005274108.1;XP_016873412.1 | 18118 | 608522 | . | . | 2.905864 | 3.140123 | 3.041097 | 3.410066 | 6.762065 | 7.258535 | 6.824133 | 7.796711 | 1.002062 | 791.431366 | 0.15171519 | 0.019583764 | 0.984375416 | 0.999197585 | 9.626525 | 9.630114 | 9.661756 | 9.590544 | 9.666210 | 9.589104 | 956 | 810 | 727 | 702 |
| 51776 | NM_016653,N | MAP3K20 | mitogen-activated protein kinase kinase kinase 20 | protein_coding | NP_057737.2;NP_598407.1;XP_005246697.1;XP_016859812.1;XP_01685 | 17797 | 609479 | . | . | 7.167043 | 7.162779 | 5.999710 | 7.300838 | 16.678005 | 16.557087 | 13.463175 | 16.692498 | -1.090390 | 1324.625075 | 0.137536834 | -0.90771306 | 0.36402984 | 0.998865404 | 10.432660 | 10.307357 | 10.396300 | 10.409336 | 10.272254 | 10.397466 | 1578 | 1504 | 1010 | 1287 |
| 51778 | NM_016599,XI | MYOZ2 | myozenin 2 | protein_coding | NP_057683.1;XP_006714297.1 | 1330 | 605602 | . | . | 0.201541 | 0.060849 | 0.080305 | 0.214052 | 0.468995 | 0.140655 | 0.180202 | 0.489403 | -1.062408 | 10.706846 | 0.990548455 | -0.08817088 | 0.929740864 | NA | 3.463417 | 3.376188 | 3.446543 | 3.396843 | 3.403989 | 3.434017 | 19 | 6 | 6 | 13 |
| 51780 | NM_016604,XI | KDM3B | lysine demethylase 3B | protein_coding | NP_057688.3;XP_005272075.1;XP_011541790.1;XP_011541791.1;XP_01( | 1337 | 609373 | . | . | 7.744243 | 8.118561 | 9.055147 | 9.374713 | 18.021171 | 18.766417 | 20.319489 | 21.434165 | 1.152528 | 1801.323183 | 0.121024492 | 1.692234383 | 0.090601293 | 0.805424371 | 10.708890 | 10.913541 | 10.740336 | 10.771247 | 10.852948 | 10.881868 | 1889 | 1842 | 1647 | 1841 |
| 51804 | NM_017420,XI | SIX4 | SIX homeobox 4 | protein_coding | NP_059116.3;XP_005267816.1 | 10890 | 606342 | . | . | 0.832501 | 0.948506 | 0.490832 | 0.711876 | 1.937265 | 2.192513 | 1.101412 | 1.627620 | -1.396375 | 138.653664 | 0.288047195 | -1.67224958 | 0.094475117 | 0.81228321 | 7.337028 | 6.853296 | 7.149663 | 7.194565 | 7.004600 | 7.091367 | 176 | 184 | 87 | 123 |

| | | | | | | | | | | | | | | | | | | | | | | | | | | | | | | | | | |
|---|---|---|---|---|---|---|---|---|---|---|---|---|---|---|---|---|---|---|---|---|---|---|---|---|---|---|---|---|---|---|---|---|---|
| 51805 | NM_017421,XI | COQ3 | coenzyme Q3, methyltransferase | protein_coding | NP_059117.3;XP_005267075.1;XP_006715563.1;XP_011534200.1;XP_01 | 18175 | 605196 | . | . | 3.212198 | 3.803831 | 2.730773 | 3.151076 | 7.474918 | 8.792726 | 6.127777 | 7.204563 | -1.220787 | 137.359641 | 0.285145117 | -1.00934928 | 0.312807153 | 0.998865404 | 7.239238 | 6.949909 | 7.099862 | 7.172872 | 7.039558 | 7.086560 | 158 | 178 | 100 | 124 |
| 51807 | NM_00119341 | TUBA8 | tubulin alpha 8 | protein_coding | NP_001180343.1;NP_061816.1 | 12410 | 605742 | . | . | 0.285467 | 0.262079 | 0.200186 | 0.208832 | 0.664293 | 0.605806 | 0.449211 | 0.477470 | -1.316611 | 15.358770 | 0.781663639 | -0.50767206 | 0.611683348 | NA | 4.125927 | 3.728813 | 3.958325 | 3.951346 | 3.934185 | 3.939170 | 21 | 18 | 11 | 13 |
| 51808 | NM_032177 | PHAX | phosphorylated adaptor for RNA export | protein_coding | NP_115553.2 | 10241 | 604924 | . | . | 8.222713 | 9.294602 | 7.360424 | 9.354887 | 19.134592 | 21.484890 | 16.516579 | 21.388836 | -1.061873 | 962.573098 | 0.157266498 | -0.55072628 | 0.58182133 | 0.998865404 | 9.953804 | 9.866376 | 9.887676 | 9.971677 | 9.801098 | 9.965495 | 1077 | 1131 | 719 | 975 |
| 51809 | NM_00137559 | GALNT7 | polypeptide N-acetylgalactosaminyltransferase 7 | protein_coding | NP_001362528.1;NP_001362529.1;NP_001362530.1;NP_059119.2;XP_01 | 4129 | 605005 | . | . | 9.032091 | 8.275541 | 7.017211 | 6.375184 | 21.018047 | 19.129285 | 15.746417 | 14.576100 | -1.313078 | 1023.295837 | 0.14255314 | -2.75653452 | 0.005841745 | 0.236462175 | 10.181910 | 9.789503 | 10.125333 | 10.052528 | 9.927086 | 9.851084 | 1401 | 1194 | 811 | 785 |
| 53335 | NM_00136386 | BCL11A | BAF chromatin remodeling complex subunit BCL11A | protein_coding | NP_001350793.1;NP_001352538.1;NP_060484.2;NP_075044.2;NP_6125( | 13221 | 606557 | . | . | 0.108197 | 0.111267 | 0.061931 | 0.069989 | 0.251779 | 0.257200 | 0.138971 | 0.160022 | -1.578339 | 15.708454 | 0.778360858 | -0.84588899 | 0.397614676 | NA | 4.264940 | 3.607807 | 4.001854 | 3.987541 | 3.966450 | 3.962232 | 24 | 19 | 11 | 11 |
| 53339 | NM_00101188 | BTBD1 | BTB domain containing 1 | protein_coding | NP_001011885.1;NP_079514.1 | 1120 | 608530 | . | . | 19.243610 | 17.124114 | 19.093044 | 17.102067 | 44.780673 | 39.583157 | 42.844236 | 39.101845 | -1.021284 | 1821.061232 | 0.132776866 | -0.22883516 | 0.81899704 | 0.998865404 | 10.845465 | 10.815507 | 10.889199 | 10.783487 | 10.869097 | 10.771094 | 2260 | 1853 | 1665 | 1584 |
| 53340 | NM_017425,XI | SPA17 | sperm autoantigenic protein 17 | protein_coding | NP_059121.1;XP_011541172.1;XP_024304351.1 | 11210 | 608621 | . | . | 1.231498 | 0.969676 | 0.704326 | 0.704359 | 2.865747 | 2.241448 | 1.580488 | 1.610432 | -1.379440 | 87.148555 | 0.345125271 | -1.34467843 | 0.178729103 | 0.951511445 | 6.658540 | 6.195232 | 6.473611 | 6.501870 | 6.419333 | 6.379013 | 111 | 114 | 69 | 63 |
| 53342 | NM_138284,XI | IL17D | interleukin 17D | protein_coding | NP_612141.1;XP_005266478.1;XP_005266479.1 | 5984 | 607587 | . | . | 0.877899 | 0.887475 | 2.009897 | 1.654624 | 2.042908 | 2.051438 | 4.510150 | 3.783099 | 1.990187 | 161.715117 | 0.273577086 | 3.62934024 | 0.000284147 | 0.026863524 | 6.756440 | 7.750298 | 7.149126 | 7.163478 | 7.495800 | 7.360746 | 122 | 119 | 214 | 173 |
| 53343 | NM_00124801 | NUDT9 | nudix hydrolase 9 | protein_coding | NP_001234940.1;NP_076952.1;NP_932155.1;XP_011530334.1 | 8056 | 606022 | . | . | 12.232012 | 11.215444 | 15.232638 | 12.715854 | 28.464391 | 25.925002 | 34.181599 | 29.073289 | 1.065596 | 1051.960052 | 0.140897379 | 0.650543985 | 0.515340899 | 0.998865404 | 9.992052 | 10.084205 | 10.048083 | 9.978758 | 10.097293 | 10.023453 | 1228 | 1047 | 991 | 967 |
| 53344 | NM_00103984 | CHIC1 | cysteine rich hydrophobic domain 1 | protein_coding | NP_001034929.2;NP_001287813.1;XP_016885071.1;XP_016885072.1;XP | 1934 | 300922 | . | . | 0.944742 | 0.755247 | 1.002823 | 0.693429 | 2.198453 | 1.745787 | 2.250305 | 1.585444 | -1.016496 | 183.579118 | 0.273271605 | -0.08637863 | 0.931165435 | 0.998865404 | 7.530814 | 7.509624 | 7.567184 | 7.467281 | 7.587852 | 7.428792 | 237 | 177 | 188 | 139 |
| 53349 | NM_00128173 | ZFYVE1 | zinc finger FYVE-type containing 1 | protein_coding | NP_001268663.1;NP_001268664.1;NP_067083.1;NP_848535.1;XP_01687 | 13180 | 605471 | . | . | 5.729325 | 5.777034 | 6.452047 | 6.716035 | 13.332374 | 13.353872 | 14.478206 | 15.355418 | 1.151914 | 831.045836 | 0.143835685 | 1.418517165 | 0.156039831 | 0.925053243 | 9.593175 | 9.797187 | 9.659944 | 9.630918 | 9.731063 | 9.760467 | 908 | 816 | 758 | 851 |
| 53354 | NM_138316,N | PANK1 | pantothenate kinase 1 | protein_coding | NP_612189.2;NP_683878.1;NP_683879.1;XP_016871822.1;XP_01687182 | 8598 | 606160 | . | . | 1.540135 | 1.981027 | 1.616727 | 1.640653 | 3.583960 | 4.579233 | 3.627890 | 3.751158 | -1.050028 | 242.116526 | 0.22274873 | -0.31617376 | 0.751870626 | 0.998865404 | 7.954694 | 7.883544 | 7.891981 | 7.968513 | 7.899056 | 7.914602 | 264 | 288 | 202 | 225 |
| 53358 | NM_016848,XI | SHC3 | SHC adaptor protein 3 | protein_coding | NP_058544.3;XP_011517087.1;XP_011517088.1;XP_016870298.1;XP_01( | 18181 | 605263 | . | . | 0.140653 | 0.119661 | 0.114025 | 0.131610 | 0.327306 | 0.276603 | 0.255869 | 0.300910 | -1.129888 | 46.350324 | 0.464151043 | -0.37957294 | 0.704262453 | 0.998865404 | 5.620373 | 5.443206 | 5.561002 | 5.532158 | 5.504776 | 5.545750 | 61 | 49 | 33 | 46 |
| 53371 | NM_00127860 | NUP54 | nucleoporin 54 | protein_coding | NP_001265532.1;NP_059122.2;XP_011530335.1;XP_016863784.1 | 17359 | 607607 | . | . | 11.098695 | 10.685919 | 10.274659 | 10.316154 | 25.827125 | 24.700981 | 23.056038 | 23.586657 | -1.076372 | 755.647224 | 0.147811199 | -0.71832522 | 0.472556785 | 0.998865404 | 9.613579 | 9.507614 | 9.606951 | 9.566090 | 9.540958 | 9.529975 | 928 | 821 | 638 | 677 |
| 53373 | NM_00114381 | TPCN1 | two pore segment channel 1 | protein_coding | NP_001137291.1;NP_001288143.1;NP_001338275.1;NP_001338276.1;NI | 18182 | 609666 | . | . | 13.099750 | 12.871175 | 12.931283 | 12.323024 | 30.483655 | 29.752301 | 29.017426 | 28.175130 | -1.031711 | 1998.887208 | 0.118849633 | -0.37895843 | 0.704718734 | 0.998865404 | 10.987256 | 10.942358 | 10.988242 | 10.966004 | 10.969764 | 10.934805 | 2370 | 2160 | 1749 | 1804 |
| 53383 | gene-MCUR1P: | MCUR1P1 | MCUR1 pseudogene 1 | pseudogene | . | 1217 | . | . | . | 0.074578 | 0.054299 | 0.144834 | 0.368356 | 0.173546 | 0.125516 | 0.325004 | 0.842203 | 3.425781 | 4.931346 | 1.431340562 | 1.241097224 | 0.214569835 | NA | 1.156660 | 2.932304 | 2.265910 | 2.262497 | 2.275602 | 2.302765 | 3 | 2 | 4 | 10 |
| 53407 | NM_00134628 | STX18 | syntaxin 18 | protein_coding | NP_001333210.1;NP_001333211.1;NP_001333229.1;NP_058626.1 | 15942 | 606046 | . | . | 8.881249 | 8.957324 | 7.808758 | 9.045211 | 20.667031 | 20.705254 | 17.522626 | 20.680802 | -1.073792 | 583.882557 | 0.163608042 | -0.62780953 | 0.530128719 | 0.998865404 | 9.240245 | 9.136976 | 9.213040 | 9.211764 | 9.120104 | 9.208649 | 697 | 652 | 456 | 564 |
| 53615 | NM_00128145 | MBD3 | methyl-CpG binding domain protein 3 | protein_coding | NP_001268382.1;NP_001268383.1 | 6918 | 603573 | . | . | 11.509391 | 11.008881 | 10.823437 | 13.266690 | 26.782826 | 25.447520 | 24.287479 | 30.332714 | 1.050774 | 2037.213494 | 0.133348016 | 0.53583432 | 0.59207306 | 0.998865404 | 10.956344 | 11.027541 | 10.993048 | 10.948630 | 10.929787 | 11.085220 | 2352 | 2084 | 1639 | 2147 |
| 53616 | NM_00132441 | ADAM22 | ADAM metallopeptidase domain 22 | protein_coding | NP_001311346.1;NP_001311347.1;NP_001311348.1;NP_001311349.1;NI | 201 | 603709 | . | . | 1.041141 | 0.995726 | 0.781053 | 1.036095 | 2.422780 | 2.301668 | 1.752662 | 2.368909 | -1.206426 | 231.497630 | 0.225142725 | -1.2025242 | 0.229160495 | 0.98402759 | 7.984166 | 7.712799 | 7.904808 | 7.892435 | 7.779733 | 7.835874 | 295 | 270 | 171 | 209 |
| 53635 | NM_00130510 | PTOV1 | PTOV1 extended AT-hook containing adaptor protein | protein_coding | NP_001292034.1;NP_001292037.1;NP_001351674.1;NP_001351676.1;NI | 9632 | 610195 | . | . | 22.592904 | 24.640001 | 21.910638 | 25.109758 | 52.574619 | 56.956465 | 49.166838 | 57.410473 | -1.015101 | 1273.898197 | 0.131657905 | -0.16424054 | 0.869541788 | 0.998865404 | 10.325963 | 10.304022 | 10.303363 | 10.337428 | 10.278224 | 10.339020 | 1445 | 1416 | 1055 | 1233 |
| 53637 | NM_00116621 | S1PR5 | sphingosine-1-phosphate receptor 5 | protein_coding | NP_001159687.1;NP_110387.1 | 14299 | 605146 | . | . | 2.699309 | 3.036836 | 3.279990 | 2.634771 | 6.281403 | 7.019783 | 7.360203 | 6.024090 | 1.033101 | 215.554198 | 0.242586243 | 0.193670668 | 0.846433761 | 0.998865404 | 7.727769 | 7.775647 | 7.715366 | 7.765848 | 7.816604 | 7.692016 | 231 | 241 | 216 | 178 |
| 53826 | NM_00116483 | FXYD6 | FXYD domain containing ion transport regulator 6 | protein_coding | NP_001158303.1;NP_001158304.1;NP_001158308.1;NP_001158309.1;NI | 4030 | 606683 | . | . | 1.716673 | 1.950558 | 0.847790 | 2.630406 | 3.994771 | 4.508803 | 1.902415 | 6.014111 | -1.001461 | 97.701861 | 0.399301219 | -0.0052754 | 0.995790862 | 0.999862634 | 6.613178 | 6.607445 | 6.582057 | 6.613329 | 6.464456 | 6.715254 | 107 | 111 | 49 | 130 |
| 53827 | NM_00116460 | FXYD5 | FXYD domain containing ion transport regulator 5 | protein_coding | NP_001158077.1;NP_001307841.1;NP_001307842.1;NP_054883.3;NP_6! | 4029 | 606669 | . | . | 174.848683 | 185.061893 | 182.374588 | 179.579623 | 406.880076 | 427.778849 | 409.243280 | 410.587461 | -1.018883 | 5140.726258 | 0.107456125 | -0.25115816 | 0.801691834 | 0.998865404 | 12.341188 | 12.314199 | 12.310942 | 12.359278 | 12.336105 | 12.303069 | 5803 | 5760 | 4516 | 4680 |
| 53831 | NM_020370,XI | GPR84 | G protein-coupled receptor 84 | protein_coding | NP_065103.1;XP_011536797.1 | 4535 | 606383 | . | . | 0.753861 | 0.626836 | 0.702764 | 0.577589 | 1.754265 | 1.448961 | 1.576982 | 1.320589 | -1.084231 | 31.803249 | 0.552350547 | -0.21122817 | 0.832709231 | 0.998865404 | 5.047242 | 4.932690 | 5.010493 | 4.982581 | 4.997345 | 4.975188 | 42 | 32 | 29 | 26 |
| 53833 | NM_144717,XI | IL20RB | interleukin 20 receptor subunit beta | protein_coding | NP_653318.2;XP_006713728.1;XP_011511212.1;XP_011511213.1;XP_01( | 6004 | 605621 | . | . | 0.273520 | 0.554960 | 0.369839 | 0.333384 | 0.636493 | 1.282814 | 0.829908 | 0.762243 | -1.111319 | 27.369672 | 0.598114161 | -0.25458804 | 0.7990413 | 0.998865404 | 4.849979 | 4.694866 | 4.761502 | 4.799111 | 4.757075 | 4.780223 | 28 | 36 | 20 | 27 |
| 53834 | NM_00100435 | FGFRL1 | fibroblast growth factor receptor like 1 | protein_coding | NP_001004356.1;NP_001004358.1;NP_001357225.1;NP_068742.2;XP_02 | 3693 | 605830 | . | . | 5.504812 | 5.654861 | 3.322858 | 3.994586 | 12.809924 | 13.071463 | 7.456395 | 9.133148 | -1.608898 | 461.592582 | 0.177307371 | -3.8694007 | 0.000109103 | 0.012952189 | 9.153418 | 8.466538 | 8.972456 | 8.977464 | 8.641646 | 8.721520 | 653 | 617 | 285 | 356 |
| 53838 | NM_00130091 | C11orf24 | chromosome 11 open reading frame 24 | protein_coding | NP_001287842.1;NP_071733.1;XP_005274110.1;XP_011543405.1 | 1174 | 610880 | . | . | 53.045172 | 55.272082 | 53.031240 | 54.773606 | 123.438302 | 127.763904 | 119.000557 | 125.233338 | -1.043352 | 3338.422973 | 0.111437248 | -0.54942355 | 0.582714814 | 0.998865404 | 11.735280 | 11.673971 | 11.696085 | 11.747412 | 11.682135 | 11.692109 | 3804 | 3793 | 2824 | 3082 |
| 53840 | NM_00100382 | TRIM34 | tripartite motif containing 34 | protein_coding | NP_001003827.1;NP_067629.2;NP_569074.2 | 10063 | 605684 | . | . | 2.137080 | 2.651034 | 2.202077 | 2.342373 | 4.973075 | 6.127984 | 4.941395 | 5.355558 | -1.089489 | 161.299038 | 0.265841584 | -0.46513363 | 0.641835761 | 0.998865404 | 7.394532 | 7.269968 | 7.303656 | 7.393820 | 7.311626 | 7.318426 | 173 | 201 | 133 | 146 |
| 53916 | NM_016154 | RAB4B | RAB4B, member RAS oncogene family | protein_coding | NP_057238.3 | 9782 | 612945 | . | . | 1.383071 | 1.049100 | 3.827275 | 8.002261 | 3.218462 | 2.425041 | 8.588293 | 18.296219 | 4.749280 | 128.274586 | 0.372202406 | 6.038942227 | 1.55128E-09 | 1.33827E-06 | 5.479531 | 7.727457 | 6.617369 | 6.568210 | 6.905030 | 7.256836 | 59 | 41 | 120 | 268 |
| 53917 | NM_00103167 | RAB24 | RAB24, member RAS oncogene family | protein_coding | NP_001026847.1;NP_570137.2 | 9765 | 612415 | . | . | 53.722145 | 53.731951 | 63.462627 | 54.543565 | 125.013631 | 124.203807 | 142.408295 | 124.707375 | 1.080652 | 2902.704343 | 0.123709346 | 0.904552617 | 0.365702447 | 0.998865404 | 11.446029 | 11.558156 | 11.469180 | 11.470021 | 11.599664 | 11.461744 | 3211 | 3012 | 2842 | 2590 |
| 53918 | NM_015946 | PELO | pelota mRNA surveillance and ribosome rescue factor | protein_coding | NP_057030.3 | 8829 | 605757 | . | . | 11.012439 | 12.122122 | 11.624143 | 11.939886 | 25.626398 | 28.020826 | 26.084238 | 27.299129 | 1.004589 | 1589.699942 | 0.125929628 | 0.05245456 | 0.958166501 | 0.998865404 | 10.631329 | 10.637741 | 10.598030 | 10.665877 | 10.629570 | 10.642399 | 1747 | 1786 | 1374 | 1506 |
| 53938 | NM_032472,N | PPIL3 | peptidylprolyl isomerase like 3 | protein_coding | NP_115861.1;NP_570981.1;XP_005246708.1;XP_005246709.1;XP_01150 | 9262 | 615811 | . | . | 13.977756 | 13.675754 | 13.735956 | 12.774514 | 32.526812 | 31.612121 | 30.823086 | 29.207408 | -1.058634 | 510.819329 | 0.166259928 | -0.49442711 | 0.621004574 | 0.998865404 | 9.037051 | 8.955125 | 9.019033 | 9.010159 | 8.994819 | 8.961640 | 609 | 563 | 443 | 453 |
| 53944 | NM_00132960 | CSNK1G1 | casein kinase 1 gamma 1 | protein_coding | NP_001316534.1;NP_001316535.1;NP_001316536.1;NP_071331.2 | 2454 | 606274 | . | . | 2.129706 | 2.274332 | 2.611903 | 2.577675 | 4.955913 | 5.257222 | 5.861035 | 5.893549 | 1.188231 | 586.936799 | 0.160583177 | 1.549449855 | 0.121273617 | 0.863030571 | 9.067360 | 9.316058 | 9.108145 | 9.162108 | 9.262156 | 9.236291 | 592 | 603 | 565 | 586 |
| 53947 | NM_00131803 | A4GALT | alpha 1,4-galactosyltransferase (P blood group) | protein_coding | NP_001304967.1;NP_059132.1;XP_005261701.1;XP_005261703.1;XP_00 | 18149 | 607922 | . | . | 1.582726 | 1.715868 | 0.483636 | 1.169906 | 3.683066 | 3.966304 | 1.085266 | 2.674852 | -1.744940 | 79.574285 | 0.40541949 | -1.98110264 | 0.047579764 | 0.659753773 | 6.662238 | 5.854651 | 6.379531 | 6.371281 | 6.157681 | 6.302771 | 118 | 108 | 32 | 74 |
| 53981 | NM_00132227 | CPSF2 | cleavage and polyadenylation specific factor 2 | protein_coding | NP_001309199.1;NP_001309200.1;NP_001309201.1;NP_059133.1 | 2325 | 606028 | . | . | 4.459729 | 4.522297 | 4.509960 | 4.686729 | 10.377973 | 10.453493 | 10.120218 | 10.715651 | 1.009453 | 1841.981369 | 0.119644201 | 0.113446472 | 0.909676598 | 0.998865404 | 10.840279 | 10.853775 | 10.841010 | 10.845581 | 10.839243 | 10.862014 | 2105 | 1984 | 1586 | 1760 |
| 54014 | NM_00100724 | BRWD1 | bromodomain and WD repeat domain containing 1 | protein_coding | NP_001007247.1;NP_061836.2;NP_387505.1;XP_011527913.1;XP_01152 | 12760 | 617824 | . | . | 1.957375 | 1.952515 | 2.047799 | 1.520225 | 4.554895 | 4.513325 | 4.595201 | 3.475814 | -1.079952 | 611.581344 | 0.187186348 | -0.59281944 | 0.553301994 | 0.998865404 | 9.310356 | 9.200349 | 9.260523 | 9.294059 | 9.335908 | 9.103404 | 713 | 702 | 602 | 454 |
| 54020 | NM_00132053 | SLC37A1 | solute carrier family 37 member 1 | protein_coding | NP_001307466.1;NP_061837.3;XP_011527916.1;XP_011527917.1;XP_01 | 11024 | 608094 | . | . | 0.154865 | 0.439943 | 0.088080 | 0.182322 | 0.360378 | 1.016949 | 0.197649 | 0.416858 | -1.503339 | 27.896922 | 0.620524134 | -0.9478608 | 0.343200323 | 0.998865404 | 5.066196 | 4.478414 | 4.778189 | 4.860336 | 4.791407 | 4.757029 | 27 | 47 | 23 | 17 |
| 54028 | gene-RPL23P2 | RPL23P2 | ribosomal protein L23 pseudogene 2 | pseudogene | . | 10324 | . | . | . | 0.579488 | 0.659294 | 1.282022 | 0.929374 | 1.348492 | 1.523987 | 2.876821 | 2.124902 | 1.744954 | 13.557489 | 0.8350029 | 0.961899962 | 0.336099858 | NA | 3.303566 | 4.107809 | 3.727565 | 3.730766 | 3.771042 | 3.751892 | 11 | 11 | 17 | 14 |
| 54029 | gene-RPL23AP4 | RPL23AP4 | ribosomal protein L23a pseudogene 4 | pseudogene | . | 10321 | . | . | . | 1.252804 | 1.240646 | 1.616184 | 1.567767 | 2.915327 | 2.867809 | 3.626669 | 3.584512 | 1.262115 | 23.320103 | 0.63513655 | 0.528773093 | 0.596962865 | NA | 4.365657 | 4.701810 | 4.527799 | 4.525994 | 4.551084 | 4.547880 | 24 | 22 | 23 | 24 |
| 54033 | NM_00132060 | RBM11 | RNA binding motif protein 11 | protein_coding | NP_001307531.1;NP_658983.3;XP_005261054.1;XP_016883873.1;XP_01 | 9897 | 617937 | . | . | 0.811445 | 1.138638 | 0.883446 | 0.722660 | 1.888265 | 2.632010 | 1.982426 | 1.652276 | -1.170940 | 76.607601 | 0.367983542 | -0.61868891 | 0.53612132 | 0.998865404 | 6.368205 | 6.141749 | 6.266773 | 6.290331 | 6.267358 | 6.208991 | 91 | 93 | 69 | 58 |
| 54053 | gene-EIF3FP1 | EIF3FP1 | eukaryotic translation initiation factor 3 subunit F pseudogene 1 | pseudogene | . | 3276 | . | . | . | 0.103142 | 0.130371 | 0.201163 | 0.089363 | 0.240016 | 0.301359 | 0.451404 | 0.204318 | 1.243167 | 4.551835 | 1.420730255 | 0.221026929 | 0.825071465 | NA | 2.019256 | 2.336262 | 2.171672 | 2.177214 | 2.187166 | 2.170688 | 4 | 5 | 6 | 3 |
| 54058 | NM_00128646 | C21orf58 | chromosome 21 open reading frame 58 | protein_coding | NP_001273391.1;NP_001273392.1;NP_001273405.1;NP_001273406.1;NI | 1300 | . | . | . | 4.004571 | 4.176311 | 4.882559 | 3.489279 | 9.318800 | 9.653730 | 10.956322 | 7.977821 | -1.382851 | 504.372203 | 0.180395857 | -2.59232786 | 0.009532888 | 0.310907798 | 9.193788 | 8.724983 | 9.050454 | 9.088816 | 8.793611 | 8.930507 | 655 | 650 | 325 | 443 |
| 54059 | NM_00100611 | YBEY | ybeY metalloendoribonuclease | protein_coding | NP_001006114.1;NP_001300951.1;NP_001300952.1;NP_001300953.1;NI | 1299 | 617461 | . | . | 5.512360 | 5.196810 | 4.772730 | 5.441525 | 12.827484 | 12.012662 | 10.709868 | 12.441400 | 1.057082 | 294.830139 | 0.202716721 | 0.395071737 | 0.692789946 | 0.998865404 | 8.163131 | 8.243238 | 8.204013 | 8.172068 | 8.206512 | 8.228945 | 340 | 300 | 258 | 290 |
| 54065 | NM_00137689 | SMIM11A | small integral membrane protein 11A | protein_coding | NP_001363828.1;NP_478062.1 | 1293 | . | . | . | 4.521414 | 4.463771 | 5.161025 | 5.541170 | 10.521518 | 10.318206 | 11.581192 | 12.669226 | 1.179882 | 132.829380 | 0.28118605 | 0.84870064 | 0.39604789 | 0.998865404 | 6.929183 | 7.167821 | 7.026235 | 7.013230 | 7.073410 | 7.086035 | 143 | 129 | 123 | 137 |
| 54067 | NR_024622,NF | C21orf62-AS1 | C21orf62 antisense RNA 1 | lncRNA | . | 1290 | . | . | . | 0.116368 | 0.304366 | 0.181722 | 0.266379 | 0.270793 | 0.703555 | 0.407780 | 0.609044 | 1.022352 | 8.706213 | 1.057091598 | 0.030169505 | 0.975931869 | NA | 3.109221 | 3.134756 | 3.101752 | 3.133322 | 3.110301 | 3.126756 | 6 | 13 | 6 | 10 |
| 54069 | NM_018944,XI | MIS18A | MIS18 kinetochore protein A | protein_coding | NP_061817.1;XP_016883889.1;XP_016883890.1;XP_016883891.1 | 1286 | 618137 | . | . | 8.240564 | 7.589199 | 9.200901 | 7.672568 | 19.176128 | 17.542773 | 20.646554 | 17.542415 | 1.053475 | 395.503292 | 0.190185354 | 0.395172333 | 0.69271571 | 0.998865404 | 8.588945 | 8.665141 | 8.633133 | 8.586274 | 8.694938 | 8.584013 | 461 | 399 | 386 | 345 |
| 54093 | NM_00100725 | SETD4 | SET domain containing 4 | protein_coding | NP_001007260.1;NP_001007262.1;NP_001273681.1;NP_059134.1;XP_01 | 1258 | . | . | . | 3.932119 | 4.515168 | 5.514303 | 4.416957 | 9.150205 | 10.437010 | 12.373933 | 10.098848 | 1.153817 | 484.702167 | 0.187680659 | 1.099814712 | 0.271412861 | 0.998865404 | 8.813736 | 9.020754 | 8.836917 | 8.901793 | 9.040819 | 8.872161 | 491 | 511 | 511 | 423 |
| 54101 | NM_020639 | RIPK4 | receptor interacting serine/threonine kinase 4 | protein_coding | NP_065690.2 | 496 | 605706 | . | . | 18.185667 | 16.961975 | 22.293344 | 20.200575 | 42.318794 | 39.208363 | 50.025620 | 46.186214 | 1.192003 | 2316.056885 | 0.122825113 | 2.062997594 | 0.039112857 | 0.616349735 | 11.045049 | 11.298724 | 11.131944 | 11.070347 | 11.286499 | 11.197424 | 2528 | 2191 | 2310 | 2233 |
| 54103 | NM_00135089 | GSAP | gamma-secretase activating protein | protein_coding | NP_001337825.1;NP_001337826.1;NP_001337827.1;NP_001337828.1;NI | 28042 | 613552 | . | . | 1.007116 | 1.051839 | 1.035033 | 0.917059 | 2.343602 | 2.431371 | 2.322584 | 2.096745 | -1.045015 | 106.581359 | 0.311728567 | -0.20377717 | 0.838527632 | 0.998865404 | 6.766824 | 6.704117 | 6.746937 | 6.738462 | 6.746710 | 6.710491 | 127 | 116 | 96 | 92 |
| 54107 | NM_00127825 | POLE3 | DNA polymerase epsilon 3, accessory subunit | protein_coding | NP_001265184.1;NP_059139.3 | 13546 | 607267 | . | . | 22.440946 | 24.854237 | 23.997227 | 24.207186 | 52.221006 | 57.451683 | 53.849083 | 55.346851 | -1.010999 | 1771.842520 | 0.124404836 | -0.12686206 | 0.899049574 | 0.998865404 | 10.799015 | 10.783010 | 10.757219 | 10.831538 | 10.773926 | 10.798670 | 1955 | 2013 | 1508 | 1678 |
| 54108 | NM_017444,N | CHRAC1 | chromatin accessibility complex subunit 1 | protein_coding | NP_059140.1 | 13544 | 607268 | . | . | 8.516246 | 8.566096 | 8.763435 | 10.111858 | 19.817654 | 19.800914 | 19.664892 | 23.119559 | 1.089138 | 691.585981 | 0.155254912 | 0.793446173 | 0.427517918 | 0.998865404 | 9.371096 | 9.493824 | 9.402415 | 9.403408 | 9.416244 | 9.504468 | 762 | 715 | 587 | 719 |
| 54145 | NM_017445 | H2BS1 | H2B.S histone 1 | protein_coding | NP_059141.1 | 4762 | . | . | . | 0.232410 | 0.117419 | 0.178257 | 0.114871 | 0.540827 | 0.271419 | 0.400003 | 0.262638 | 1.043482 | 2.727629 | 1.804920749 | 0.034021556 | 0.972859962 | NA | 1.413393 | 1.481107 | 1.449528 | 1.441809 | 1.449161 | 1.443240 | 4 | 2 | 3 | 2 |
| 54148 | NM_017446,N | MRPL39 | mitochondrial ribosomal protein L39 | protein_coding | NP_059142.3;NP_542984.3;XP_006724089.1;XP_011527953.1 | 14027 | 611845 | . | . | 18.149690 | 17.686412 | 14.961224 | 15.763555 | 42.235075 | 40.882935 | 33.572554 | 36.041494 | -1.169201 | 586.198888 | 0.159090673 | -1.41757469 | 0.156314973 | 0.925053243 | 9.303485 | 9.078224 | 9.253400 | 9.235607 | 9.155173 | 9.127946 | 737 | 673 | 480 | 496 |
| 54149 | NM_00110042 | C21orf91 | chromosome 21 open reading frame 91 | protein_coding | NP_001093890.1;NP_001093891.1;NP_059143.3 | 16459 | . | . | . | 3.403822 | 2.770157 | 3.143984 | 2.489150 | 7.920834 | 6.403339 | 7.055012 | 5.691145 | -1.111806 | 497.492582 | 0.191659936 | -0.7977929 | 0.424990684 | 0.998865404 | 9.032319 | 8.880765 | 9.055991 | 8.915137 | 8.992380 | 8.844409 | 668 | 504 | 459 | 389 |
| 54165 | NM_00130810 | DCUN1D1 | defective in cullin neddylation 1 domain containing 1 | protein_coding | NP_001295030.1;NP_065691.2;XP_011511214.1;XP_011511216.1;XP_01 | 18184 | 605905 | . | . | 2.685747 | 2.904371 | 3.471995 | 1.774351 | 6.249846 | 6.713587 | 7.791054 | 4.056842 | -1.105670 | 537.380250 | 0.201013732 | -0.72095184 | 0.470939149 | 0.998865404 | 9.139744 | 8.996290 | 9.171476 | 9.014850 | 9.132620 | 8.918941 | 726 | 537 | 519 | 398 |
| 54187 | NM_018946,XI | NANS | N-acetylneuraminate synthase | protein_coding | NP_061819.2;XP_011517089.1;XP_011517090.1;XP_016870300.1;XP_02( | 19237 | 605202 | . | . | 36.886264 | 34.576826 | 31.342530 | 37.036405 | 85.835848 | 79.925890 | 70.331727 | 84.679344 | -1.060957 | 1289.978661 | 0.137218262 | -0.62212198 | 0.533861663 | 0.998865404 | 10.375306 | 10.289687 | 10.380717 | 10.327898 | 10.254799 | 10.361684 | 1582 | 1384 | 1011 | 1257 |
| 54205 | NM_018947 | CYCS | cytochrome c, somatic | protein_coding | NP_061820.1 | 19986 | 123970 | . | . | 27.560442 | 23.484524 | 23.305445 | 21.274704 | 64.134285 | 54.285526 | 52.296741 | 48.642082 | -1.161353 | 4048.801978 | 0.124435167 | -1.73428818 | 0.082866964 | 0.78412713 | 12.087050 | 11.871462 | 12.109608 | 11.967570 | 11.957455 | 11.874691 | 5431 | 4300 | 3423 | 3335 |
| 54206 | NM_018948,XI | ERRFI1 | ERBB receptor feedback inhibitor 1 | protein_coding | NP_061821.1;XP_005263534.1;XP_006710760.1;XP_011539898.1 | 18185 | 608069 | . | . | 8.295379 | 8.867630 | 4.781417 | 6.569468 | 19.303687 | 20.497924 | 10.729361 | 15.020308 | -1.280353 | 791.531185 | 0.152461575 | -2.33856535 | 0.01935794 | 0.462516353 | 9.795708 | 9.439347 | 9.667464 | 9.746701 | 9.570785 | 9.496362 | 968 | 1011 | 637 | 615 |
| 54329 | NM_00114626 | GPR85 | G protein-coupled receptor 85 | protein_coding | NP_001139737.1;NP_001139738.1;NP_001139739.1;NP_061843.3;XP_0( | 4536 | 605188 | . | . | 0.966916 | 0.899347 | 1.126090 | 0.991572 | 2.250055 | 2.078882 | 2.526915 | 2.267110 | 1.135461 | 98.594229 | 0.323712454 | 0.566176425 | 0.571273854 | 0.998865404 | 6.528358 | 6.712625 | 6.605411 | 6.596151 | 6.661847 | 6.615941 | 108 | 98 | 98 | 91 |
| 54331 | NM_00124377 | GNG2 | G protein subunit gamma 2 | protein_coding | NP_001230702.1;NP_001230703.1;NP_444292.1;XP_006720236.1;XP_01 | 4404 | 606981 | . | . | 8.171383 | 7.358991 | 7.885517 | 8.203079 | 19.015143 | 17.010638 | 17.694876 | 18.755368 | 1.024407 | 894.036773 | 0.143156835 | 0.243011166 | 0.807996748 | 0.998865404 | 9.786624 | 9.821546 | 9.827844 | 9.761296 | 9.802521 | 9.822369 | 1064 | 909 | 776 | 860 |
| 54332 | NM_00104087 | GDAP1 | ganglioside induced differentiation associated protein 1 | protein_coding | NP_001035808.1;NP_001349858.1;NP_001349859.1;NP_001349860.1;NI | 15968 | 606598 | . | . | 5.390406 | 5.629766 | 5.538219 | 5.463110 | 12.543697 | 13.013456 | 12.427601 | 12.490749 | 1.020322 | 594.701638 | 0.157669859 | 0.18408123 | 0.853949731 | 0.998865404 | 9.201393 | 9.230505 | 9.207294 | 9.211010 | 9.230838 | 9.214091 | 676 | 637 | 529 | 556 |
| 54344 | NM_018973,N | DPM3 | dolichyl-phosphate mannosyltransferase subunit 3, regulatory | protein_coding | NP_061846.2;NP_714963.1;XP_016856987.1 | 3007 | 605951 | . | . | 19.609260 | 21.020455 | 20.760003 | 22.426881 | 45.631553 | 48.589725 | 46.584846 | 51.276397 | 1.085209 | 284.563862 | 0.207983868 | 0.567220642 | 0.570564285 | 0.998865404 | 8.092832 | 8.210007 | 8.103464 | 8.154650 | 8.148234 | 8.195794 | 299 | 309 | 246 | 290 |
| 54345 | NM_018419 | SOX18 | SRY-box transcription factor 18 | protein_coding | NP_060889.1 | 11194 | 601618 | . | . | 0.169299 | 0.245797 | 0.162050 | 0.231464 | 0.393966 | 0.568170 | 0.363634 | 0.529216 | -1.040090 | 12.366416 | 0.873536392 | -0.06491741 | 0.948239755 | NA | 3.658167 | 3.597915 | 3.619236 | 3.639124 | 3.619552 | 3.634615 | 12 | 16 | 9 | 13 |
| 54386 | NM_018975,N | TERF2IP | TERF2 interacting protein | protein_coding | NP_061848.2 | 19246 | 605061 | . | . | 21.428493 | 22.324367 | 20.991812 | 22.959624 | 49.864983 | 51.603775 | 47.105019 | 52.494451 | -1.008978 | 1459.278459 | 0.127834027 | -0.10086892 | 0.919654519 | 0.998865404 | 10.517609 | 10.504440 | 10.500942 | 10.527536 | 10.475797 | 10.537953 | 1659 | 1609 | 1212 | 1417 |
| 54407 | NM_00130793 | SLC38A2 | solute carrier family 38 member 2 | protein_coding | NP_001294865.1;NP_061849.2 | 13448 | 605180 | . | . | 21.802625 | 19.120710 | 20.222336 | 13.490650 | 50.735605 | 44.198376 | 45.378334 | 30.844767 | -1.229700 | 2786.552655 | 0.166902632 | -1.7873091 | 0.073887544 | 0.763883535 | 11.585528 | 11.287655 | 11.572154 | 11.458758 | 11.498937 | 11.172287 | 3779 | 3091 | 2616 | 1867 |
| 54414 | NM_00119992 | SIAE | sialic acid acetylesterase | protein_coding | NP_001186851.1;NP_733746.1;XP_016873419.1 | 18187 | 610079 | . | . | 2.669162 | 2.782524 | 2.188978 | 2.875230 | 6.211249 | 6.431928 | 4.912002 | 6.573872 | -1.085662 | 442.295807 | 0.190576925 | -0.62218867 | 0.533817813 | 0.998865404 | 8.847487 | 8.727765 | 8.803294 | 8.820728 | 8.677882 | 8.838834 | 523 | 504 | 319 | 451 |
| 54429 | NM_018980 | TAS2R5 | taste 2 receptor member 5 | protein_coding | NP_061853.1 | 14912 | 605062 | . | . | 0.162194 | 0.181205 | 0.298420 | 0.228960 | 0.377431 | 0.418864 | 0.669646 | 0.523491 | 1.484107 | 8.381533 | 1.050930299 | 0.54199103 | 0.587824685 | NA | 2.754145 | 3.324312 | 3.045586 | 3.051947 | 3.069860 | 3.057076 | 7 | 8 | 10 | 8 |
| 54431 | NM_00127158 | DNAJC10 | DnaJ heat shock protein family (Hsp40) member C10 | protein_coding | NP_001258510.1;NP_061854.1 | 24637 | 607987 | . | . | 9.787478 | 8.842728 | 9.429944 | 7.218228 | 22.775863 | 20.440359 | 21.160519 | 16.503621 | -1.137078 | 5547.313670 | 0.139347455 | -1.32999087 | 0.183521278 | 0.954800306 | 12.527164 | 12.342050 | 12.534628 | 12.435807 | 12.488262 | 12.255523 | 7191 | 6000 | 5145 | 4188 |
| 54432 | NM_018982,N | YIPF1 | Yip1 domain family member 1 | protein_coding | NP_061855.1 | 25231 | 617521 | . | . | 12.392251 | 13.042920 | 14.762568 | 14.112560 | 28.837279 | 30.149290 | 33.126775 | 32.266693 | 1.126583 | 747.097160 | 0.148674935 | 1.156576198 | 0.247445552 | 0.992733415 | 9.456583 | 9.628596 | 9.484887 | 9.517324 | 9.602969 | 9.564960 | 790 | 776 | 707 | 722 |
| 54433 | NM_018983,N | GAR1 | GAR1 ribonucleoprotein | protein_coding | NP_061856.1;NP_127460.1 | 14264 | 606468 | . | . | 11.263852 | 11.137978 | 9.881986 | 8.602223 | 26.211444 | 25.745936 | 22.174888 | 19.667961 | -1.227518 | 323.771579 | 0.199669397 | -1.48117064 | 0.138561109 | 0.900064053 | 8.478756 | 8.183866 | 8.398048 | 8.385781 | 8.317846 | 8.238664 | 415 | 381 | 272 | 252 |
| 54434 | NM_00116133 | SSH1 | slingshot protein phosphatase 1 | protein_coding | NP_001154802.1;NP_001154803.1;NP_061857.3;XP_005269041.1;XP_00 | 30579 | 606778 | . | . | 25.679960 | 28.623629 | 34.119387 | 34.315493 | 59.758323 | 66.164802 | 76.562919 | 78.458300 | 1.210098 | 8193.795729 | 0.106258642 | 2.589185538 | 0.009620324 | 0.312445503 | 12.856227 | 13.131312 | 12.875057 | 12.950857 | 13.081512 | 13.069487 | 8151 | 8364 | 7857 | 8353 |
| 54435 | NR_002139 | HCG4 | HLA complex group 4 | lncRNA | . | 21241 | . | . | . | 0.709303 | 0.838853 | 1.182259 | 0.553092 | 1.650578 | 1.939046 | 2.652954 | 1.264580 | 1.111140 | 52.719235 | 0.477710843 | 0.318269594 | 0.750280181 | 0.998865404 | 5.640659 | 5.795693 | 5.681812 | 5.711298 | 5.786847 | 5.641773 | 53 | 58 | 66 | 33 |
| 54436 | NM_00131848 | SH3TC1 | SH3 domain and tetratricopeptide repeats 1 | protein_coding | NP_001305409.2;NP_061859.4;XP_006713952.1;XP_006713954.1;XP_01 | 26009 | . | . | . | 5.228832 | 5.396326 | 6.206582 | 5.749418 | 12.167706 | 12.473847 | 13.927389 | 13.145361 | 1.080228 | 728.701328 | 0.150753181 | 0.738529943 | 0.460192478 | 0.998865404 | 9.452419 | 9.563800 | 9.457987 | 9.503627 | 9.556312 | 9.512196 | 780 | 781 | 679 | 687 |
| 54438 | NM_00124262 | GFOD1 | glucose-fructose oxidoreductase domain containing 1 | protein_coding | NP_001229557.1;NP_001229558.1;NP_001229559.1;NP_061861.1;XP_01 | 21096 | . | . | . | 0.771992 | 1.104393 | 0.861095 | 0.991500 | 1.796458 | 2.552854 | 1.932274 | 2.266948 | -1.019030 | 187.316939 | 0.255167184 | -0.10658568 | 0.915117681 | 0.998865404 | 7.563280 | 7.535259 | 7.488148 | 7.610141 | 7.549849 | 7.535503 | 188 | 232 | 164 | 171 |
| 54439 | NM_018989,XI | RBM27 | RNA binding motif protein 27 | protein_coding | NP_061862.1;XP_005268523.1;XP_005268524.1;XP_005268525.1;XP_00( | 29243 | . | . | . | 5.067839 | 5.046273 | 4.894281 | 5.263603 | 11.793068 | 11.664684 | 10.982627 | 12.034603 | -1.008597 | 1033.128121 | 0.136285062 | -0.09062242 | 0.927792614 | 0.998865404 | 10.019047 | 10.006533 | 10.021612 | 10.010193 | 9.986813 | 10.031821 | 1204 | 1111 | 867 | 994 |
| 54441 | NR_040583 | STAG3L1 | stromal antigen 3-like 1 (pseudogene) | transcribed_ps | . | 33852 | . | . | . | 4.679101 | 5.431763 | 5.968732 | 4.750385 | 10.888461 | 12.555763 | 13.393662 | 10.861190 | 1.048189 | 347.504911 | 0.205591277 | 0.330263817 | 0.74120063 | 0.998865404 | 8.406265 | 8.474702 | 8.383148 | 8.463575 | 8.516023 | 8.381374 | 363 | 392 | 346 | 294 |
| 54442 | NM_018992 | KCTD5 | potassium channel tetramerization domain containing 5 | protein_coding | NP_061865.1 | 21423 | 611285 | . | . | 23.864258 | 26.614914 | 24.553829 | 26.362103 | 55.533112 | 61.521564 | 55.098080 | 60.273808 | -1.005249 | 1922.128094 | 0.124389813 | -0.06071443 | 0.951586641 | 0.998865404 | 10.912398 | 10.904569 | 10.869548 | 10.949263 | 10.881535 | 10.929767 | 2108 | 2184 | 1616 | 1852 |
| 54443 | NM_00128430 | ANLN | anillin actin binding protein | protein_coding | NP_001271230.1;NP_001271231.1;NP_061155.2;XP_006715809.1;XP_00 | 14082 | 616027 | . | . | 46.311124 | 44.286502 | 32.133501 | 34.308142 | 107.767891 | 102.370236 | 72.106643 | 78.441493 | -1.384123 | 5780.739957 | 0.107159344 | -4.37640257 | 1.20654E-05 | 0.002345617 | 12.712564 | 12.243563 | 12.641626 | 12.594281 | 12.322844 | 12.366703 | 7940 | 7045 | 4103 | 4669 |
| 54454 | NM_00124233 | ATAD2B | ATPase family AAA domain containing 2B | protein_coding | NP_001229267.2;NP_001341036.1;NP_060022.2;XP_005264429.2;XP_00 | 29230 | 615347 | . | . | 1.478003 | 1.337492 | 1.516798 | 1.142636 | 3.439373 | 3.091674 | 3.403652 | 2.612501 | -1.069906 | 335.987546 | 0.221482855 | -0.44014214 | 0.659834163 | 0.998865404 | 8.439329 | 8.343611 | 8.444940 | 8.366303 | 8.463894 | 8.263565 | 427 | 349 | 334 | 249 |
| 54455 | NM_018994,XI | FBXO42 | F-box protein 42 | protein_coding | NP_061867.1;XP_006710761.1;XP_016856991.1;XP_016856992.1 | 29249 | 609109 | . | . | 5.251948 | 5.185564 | 6.097458 | 6.319239 | 12.221499 | 11.986664 | 13.682518 | 14.448189 | 1.165747 | 1107.612617 | 0.133008554 | 1.663459242 | 0.096220543 | 0.813553488 | 9.998409 | 10.219597 | 10.057094 | 10.048338 | 10.154242 | 10.178964 | 1185 | 1097 | 1020 | 1136 |
| 54458 | NM_00100535 | PRR13 | proline rich 13 | protein_coding | NP_001005354.1;NP_060927.1 | 24528 | 610459 | . | . | 61.197642 | 65.549471 | 59.822796 | 71.708916 | 142.409436 | 151.520553 | 134.240631 | 163.953908 | 1.023563 | 2220.017608 | 0.129156158 | 0.260147992 | 0.79474962 | 0.998865404 | 11.099638 | 11.132882 | 11.081635 | 11.129780 | 11.053672 | 11.190837 | 2451 | 2439 | 1786 | 2285 |
| 54460 | NM_018997,N | MRPS21 | mitochondrial ribosomal protein S21 | protein_coding | NP_061870.1;NP_114107.2 | 14046 | 611984 | . | . | 34.968657 | 34.054705 | 35.494265 | 35.499260 | 81.373497 | 78.718975 | 79.648105 | 81.164839 | 1.056285 | 1488.618641 | 0.125218717 | 0.630887413 | 0.528114143 | 0.998865404 | 10.499675 | 10.578758 | 10.536671 | 10.499015 | 10.558579 | 10.562068 | 1708 | 1524 | 1327 | 1437 |
| 54461 | NM_018998,XI | FBXW5 | F-box and WD repeat domain containing 5 | protein_coding | NP_061871.1;XP_005266146.2;XP_011517091.1;XP_016870301.1 | 13613 | 609072 | . | . | 22.055558 | 25.925987 | 23.475249 | 24.649966 | 51.324191 | 59.929080 | 52.677777 | 56.359219 | -1.004536 | 1829.321310 | 0.126039328 | -0.05180638 | 0.95868297 | 0.998865404 | 10.840479 | 10.833698 | 10.789958 | 10.884696 | 10.821800 | 10.847561 | 1985 | 2097 | 1561 | 1739 |
| 54462 | NM_00128424 | CCSER2 | coiled-coil serine rich protein 2 | protein_coding | NP_001271169.1;NP_001271170.1;NP_001271171.1;NP_001271172.1;NI | 29197 | . | . | . | 3.046121 | 3.372949 | 3.203373 | 2.445624 | 7.088450 | 7.796720 | 7.188277 | 5.591628 | -1.035307 | 650.893226 | 0.166412583 | -0.30080856 | 0.763560482 | 0.998865404 | 9.370638 | 9.321497 | 9.390576 | 9.320920 | 9.395581 | 9.266503 | 801 | 678 | 610 | 542 |
| 54463 | NM_00103485 | RETREG1 | reticulophagy regulator 1 | protein_coding | NP_001030022.1;NP_061873.2;XP_011512355.1;XP_011512356.1;XP_01 | 25964 | 613114 | . | . | 1.894019 | 1.573095 | 1.312236 | 1.305958 | 4.407460 | 3.636280 | 2.944620 | 2.985918 | -1.335004 | 156.371293 | 0.269058999 | -1.54926681 | 0.121317595 | 0.863030571 | 7.481886 | 7.065894 | 7.384542 | 7.298410 | 7.232996 | 7.223166 | 226 | 174 | 118 | 124 |
| 54464 | NM_00128285 | XRN1 | 5'-3' exoribonuclease 1 | protein_coding | NP_001269786.1;NP_001269788.1;NP_061874.3;XP_006713736.1;XP_01 | 30654 | 607994 | . | . | 2.046577 | 1.844302 | 2.035145 | 1.733537 | 4.762465 | 4.263186 | 4.566806 | 3.963523 | -1.032720 | 594.897218 | 0.166444964 | -0.27906449 | 0.780195337 | 0.998865404 | 9.239124 | 9.193508 | 9.257502 | 9.193530 | 9.253671 | 9.153628 | 729 | 621 | 548 | 507 |
| 54465 | NM_019002,XI | ETAA1 | ETAA1 activator of ATR kinase | protein_coding | NP_061875.2;XP_016859865.1;XP_016859866.1 | 24648 | 613196 | . | . | 4.221836 | 3.574795 | 3.971330 | 2.751652 | 9.824385 | 8.263298 | 8.911550 | 6.291326 | -1.175478 | 497.343247 | 0.196438792 | -1.18738087 | 0.235077423 | 0.990818043 | 9.069340 | 8.837557 | 9.064429 | 8.937835 | 8.989287 | 8.804233 | 678 | 524 | 458 | 364 |
| 54466 | NM_019003,XI | SPIN2A | spindlin family member 2A | protein_coding | NP_061876.3;XP_005262073.3;XP_005262074.3;XP_005262075.5;XP_00! | 20694 | 300621 | . | . | 1.061322 | 0.906694 | 1.408325 | 1.157499 | 2.469741 | 2.095864 | 3.160239 | 2.646485 | 1.277071 | 37.190223 | 0.517152288 | 0.682273377 | 0.495066104 | 0.998865404 | 5.028170 | 5.383686 | 5.202932 | 5.178840 | 5.250572 | 5.201601 | 41 | 32 | 42 | 33 |
| 54467 | NM_019004 | ANKIB1 | ankyrin repeat and IBR domain containing 1 | protein_coding | NP_061877.1 | 22215 | . | . | . | 9.103306 | 8.562199 | 9.437042 | 9.123675 | 21.183765 | 19.791906 | 21.176449 | 20.860199 | 1.036134 | 1789.113258 | 0.122856333 | 0.416832757 | 0.676800741 | 0.998865404 | 10.779087 | 10.830512 | 10.817604 | 10.762819 | 10.835583 | 10.801286 | 2094 | 1830 | 1618 | 1670 |
| 54468 | NM_00137007 | MIOS | meiosis regulator for oocyte development | protein_coding | NP_001357005.1;NP_001357006.1;NP_001357007.1;NP_001357009.1;NI | 21905 | 615359 | . | . | 4.278602 | 4.061965 | 4.376771 | 4.100328 | 9.956486 | 9.389412 | 9.821350 | 9.374914 | -1.011374 | 639.947854 | 0.156315139 | -0.10438371 | 0.916864854 | 0.998865404 | 9.329747 | 9.313831 | 9.349855 | 9.300263 | 9.334115 | 9.301311 | 768 | 669 | 567 | 582 |
| 54469 | NM_00124291 | ZFAND6 | zinc finger AN1-type containing 6 | protein_coding | NP_001229840.1;NP_001229841.1;NP_001229842.1;NP_001229843.1;NI | 30164 | 610183 | . | . | 23.931429 | 21.294724 | 19.217224 | 22.876774 | 55.689421 | 49.223713 | 43.122893 | 52.305023 | -1.084224 | 1135.430011 | 0.145924753 | -0.79947371 | 0.424015785 | 0.998865404 | 10.206297 | 10.089381 | 10.219328 | 10.133846 | 10.052720 | 10.177524 | 1437 | 1203 | 867 | 1108 |
| 54470 | NM_00100958 | ARMCX6 | armadillo repeat containing X-linked 6 | protein_coding | NP_001009584.1;NP_001171697.1;NP_061880.2;XP_011529278.1 | 26094 | 301048 | . | . | 9.064059 | 11.297567 | 9.905168 | 11.004354 | 21.092437 | 26.114834 | 22.226909 | 25.160147 | 1.042150 | 455.692590 | 0.179648953 | 0.331551102 | 0.740228248 | 0.998865404 | 8.802188 | 8.861046 | 8.764626 | 8.868261 | 8.828711 | 8.857355 | 471 | 522 | 395 | 446 |
| 54471 | NM_00130456 | MIEF1 | mitochondrial elongation factor 1 | protein_coding | NP_001291493.1;NP_061881.2;XP_011528538.1;XP_011528539.1;XP_01 | 25979 | 615497 | . | . | 7.699907 | 9.239186 | 7.788956 | 9.188991 | 17.918002 | 21.356792 | 17.478190 | 21.009533 | -1.015927 | 1502.683196 | 0.145703527 | -0.15646072 | 0.875669876 | 0.998865404 | 10.564973 | 10.541583 | 10.492113 | 10.619168 | 10.470033 | 10.615254 | 1602 | 1768 | 1174 | 1529 |
| 54472 | NM_00131851 | TOLLIP | toll interacting protein | protein_coding | NP_001305441.1;NP_001305443.1;NP_001305444.1;NP_001305445.1;NI | 16476 | 606277 | . | . | 13.406320 | 13.276809 | 13.250554 | 14.719562 | 31.197058 | 30.689943 | 29.733859 | 33.654529 | 1.008474 | 1796.989225 | 0.121821465 | 0.099937192 | 0.920394189 | 0.998865404 | 10.805339 | 10.817368 | 10.810760 | 10.805089 | 10.787549 | 10.840795 | 2068 | 1924 | 1516 | 1749 |
| 54475 | NM_00101444 | NLE1 | notchless homolog 1 | protein_coding | NP_001014445.1;NP_060566.2;XP_016880266.1 | 19889 | . | . | . | 5.221738 | 4.602916 | 5.193769 | 6.076764 | 12.151199 | 10.639846 | 11.654667 | 13.893799 | -1.003155 | 682.630570 | 0.160665575 | -0.02828777 | 0.977432634 | 0.998865404 | 9.417610 | 9.412308 | 9.366401 | 9.461614 | 9.372487 | 9.451551 | 733 | 789 | 558 | 676 |
| 54476 | NM_00137715 | RNF216 | ring finger protein 216 | protein_coding | NP_001364085.1;NP_996994.1;NP_996999.1;XP_005249842.1;XP_01151 | 21698 | 609948 | . | . | 9.938003 | 10.211871 | 11.264815 | 10.664261 | 23.126145 | 23.605195 | 25.277921 | 24.382563 | 1.055311 | 1906.296710 | 0.12011473 | 0.646619254 | 0.517878394 | 0.998865404 | 10.857157 | 10.934909 | 10.865714 | 10.883417 | 10.938747 | 10.895255 | 2112 | 2024 | 1749 | 1785 |
| 54477 | NM_00114382 | PLEKHA5 | pleckstrin homology domain containing A5 | protein_coding | NP_001137293.2;NP_001177789.2;NP_001243399.1;NP_001243716.1;NI | 30036 | 607770 | . | . | 6.658174 | 7.590937 | 7.286644 | 7.023957 | 15.493837 | 17.546794 | 16.351014 | 16.059439 | 1.003499 | 910.727477 | 0.142856949 | 0.035270908 | 0.971863721 | 0.998865404 | 9.828372 | 9.833375 | 9.796935 | 9.860474 | 9.850257 | 9.812811 | 1001 | 1024 | 814 | 833 |
| 54478 | NM_00119522 | PIMREG | PICALM interacting mitotic regulator | protein_coding | NP_001182157.1;NP_061886.2;XP_016880267.1 | 25483 | 617611 | . | . | 14.292650 | 17.797066 | 11.970310 | 14.746044 | 33.259584 | 41.138712 | 26.861030 | 33.715076 | -1.213852 | 868.610865 | 0.166204579 | -1.68221822 | 0.092526532 | 0.809520218 | 9.896024 | 9.615493 | 9.744667 | 9.899411 | 9.613667 | 9.754897 | 982 | 1136 | 612 | 811 |
| 54480 | NM_00128429 | CHPF2 | chondroitin polymerizing factor 2 | protein_coding | NP_001271224.1;NP_061888.1 | 29270 | 608037 | . | . | 13.153874 | 13.666909 | 15.201036 | 13.837292 | 30.609602 | 31.591669 | 34.110682 | 31.637322 | 1.051121 | 1666.645151 | 0.123808995 | 0.580968074 | 0.561261971 | 0.998865404 | 10.666262 | 10.738301 | 10.671171 | 10.693861 | 10.747871 | 10.694578 | 1844 | 1779 | 1536 | 1547 |
| 54482 | NM_019083,N | TRMT13 | tRNA methyltransferase 13 homolog | protein_coding | NP_061956.2;XP_005271002.1;XP_005271003.1;XP_016856993.1;XP_01( | 25502 | . | . | . | 2.337571 | 2.359825 | 2.831972 | 1.926413 | 5.439623 | 5.454843 | 6.354865 | 4.404515 | 1.021227 | 346.936336 | 0.234962133 | 0.128971178 | 0.897380455 | 0.998865404 | 8.422531 | 8.454348 | 8.416921 | 8.431638 | 8.566640 | 8.286822 | 390 | 375 | 380 | 248 |
| 54487 | NM_00119032 | DGCR8 | DGCR8 microprocessor complex subunit | protein_coding | NP_001177255.1;NP_073557.3;XP_006724331.1 | 2847 | 609030 | . | . | 9.782374 | 10.567594 | 12.006031 | 10.599631 | 22.763985 | 24.427467 | 26.941184 | 24.234799 | 1.093362 | 1495.316433 | 0.132274567 | 0.973514459 | 0.330297678 | 0.998865404 | 10.480320 | 10.609270 | 10.484203 | 10.534559 | 10.627753 | 10.527252 | 1591 | 1592 | 1446 | 1370 |
| 54491 | NM_019018,XI | OTULINL | OTU deubiquitinase with linear linkage specificity like | protein_coding | NP_061891.1;XP_011512358.1 | 25629 | . | . | . | 0.041250 | 0.053661 | 0.027828 | 0.025784 | 0.095990 | 0.124039 | 0.062445 | 0.058953 | -1.623735 | 8.725098 | 1.036979471 | -0.67437767 | 0.500071234 | NA | 3.432996 | 2.733338 | 3.135217 | 3.146418 | 3.123523 | 3.121135 | 11 | 13 | 6 | 6 |
| 54492 | NM_00114265 | NEURL1B | neuralized E3 ubiquitin protein ligase 1B | protein_coding | NP_001136123.1;NP_001295106.1;NP_001295107.1 | 35422 | 615893 | . | . | 2.041567 | 2.045743 | 1.114450 | 1.396756 | 4.750809 | 4.728827 | 2.500795 | 3.193516 | -1.541478 | 318.148000 | 0.207482516 | -3.00899719 | 0.002621115 | 0.135502831 | 8.592763 | 7.967020 | 8.390528 | 8.440445 | 8.126616 | 8.230469 | 426 | 434 | 194 | 260 |
| 54494 | NM_00127156 | C11orf71 | chromosome 11 open reading frame 71 | protein_coding | NP_001258491.1;NP_061894.2 | 25937 | . | . | . | 3.035736 | 2.414591 | 3.246837 | 2.619147 | 7.064282 | 5.581434 | 7.285810 | 5.988369 | 1.056674 | 175.104262 | 0.265655378 | 0.299375216 | 0.76465377 | 0.998865404 | 7.410754 | 7.492236 | 7.485046 | 7.385850 | 7.505998 | 7.411838 | 219 | 162 | 174 | 150 |
| 54495 | NM_00135051 | TMX3 | thioredoxin related transmembrane protein 3 | protein_coding | NP_001337441.1;NP_001337442.1;NP_001337443.1;NP_001337444.1;NI | 24718 | 616102 | . | . | 15.928605 | 13.098915 | 13.800692 | 9.246205 | 37.066518 | 30.278732 | 30.968352 | 21.140349 | -1.267767 | 1856.282337 | 0.17397313 | -1.96748351 | 0.049127495 | 0.663679701 | 11.018938 | 10.677282 | 11.014247 | 10.856791 | 10.893238 | 10.586091 | 2620 | 2023 | 1707 | 1230 |
| 54496 | NM_00118482 | PRMT7 | protein arginine methyltransferase 7 | protein_coding | NP_001171753.1;NP_001276947.1;NP_001338072.1;NP_001338073.1;NI | 25557 | 610087 | . | . | 15.234195 | 14.734990 | 13.561363 | 15.844172 | 35.450601 | 34.060588 | 30.431305 | 36.225820 | -1.004746 | 1674.554140 | 0.126484525 | -0.05400328 | 0.956932549 | 0.998865404 | 10.713129 | 10.705985 | 10.688962 | 10.732797 | 10.669309 | 10.743942 | 1879 | 1862 | 1383 | 1641 |
| 54497 | NM_019024,XI | HEATR5B | HEAT repeat containing 5B | protein_coding | NP_061897.1;XP_006712097.1;XP_006712098.1;XP_011531236.1;XP_01: | 29273 | . | . | . | 5.279532 | 6.048019 | 6.989952 | 6.895349 | 12.285691 | 13.980268 | 15.685246 | 15.765396 | 1.187568 | 1351.193112 | 0.134080742 | 1.849710916 | 0.064355226 | 0.731823074 | 10.270746 | 10.518653 | 10.281543 | 10.375026 | 10.483619 | 10.437047 | 1334 | 1416 | 1314 | 1334 |
| 54498 | NM_00127069 | SMOX | spermine oxidase | protein_coding | NP_001257620.1;NP_787033.1;NP_787034.1;NP_787035.1;NP_787036.1 | 15862 | 615854 | . | . | 4.697597 | 5.217333 | 4.831462 | 5.398979 | 10.931501 | 12.060099 | 10.841659 | 12.344125 | 1.025795 | 340.994843 | 0.19465997 | 0.188751969 | 0.850287218 | 0.998865404 | 8.395505 | 8.431483 | 8.379512 | 8.431360 | 8.393054 | 8.446210 | 370 | 380 | 286 | 339 |
| 54499 | NM_00125616 | TMCO1 | transmembrane and coiled-coil domains 1 | protein_coding | NP_001243093.1;NP_001243094.1;NP_061899.3 | 18188 | 614123 | . | . | 42.889407 | 41.309636 | 43.045316 | 33.596383 | 99.805407 | 95.489071 | 96.592442 | 76.814140 | -1.084610 | 2754.055123 | 0.131508064 | -0.89102265 | 0.372917019 | 0.998865404 | 11.484571 | 11.367748 | 11.498427 | 11.415925 | 11.467387 | 11.310702 | 3465 | 2937 | 2521 | 2237 |
| 54502 | NM_00109863 | RBM47 | RNA binding motif protein 47 | protein_coding | NP_001092104.1;NP_001358042.1;NP_001358043.1;NP_061900.2;XP_0( | 30358 | . | . | . | 2.991227 | 2.855865 | 2.778520 | 2.966131 | 6.960707 | 6.601462 | 6.234918 | 6.781702 | -1.035994 | 447.399615 | 0.175751039 | -0.29027295 | 0.77160743 | 0.998865404 | 8.830686 | 8.779704 | 8.842853 | 8.788741 | 8.778254 | 8.810391 | 547 | 470 | 372 | 423 |
| 54503 | NM_00100148 | ZDHHC13 | zinc finger DHHC-type palmitoyltransferase 13 | protein_coding | NP_001001483.1;NP_061901.2;XP_005253053.1;XP_011518495.1;XP_01 | 18413 | 612815 | . | . | 6.342905 | 6.296213 | 5.728870 | 4.873176 | 14.760201 | 14.553978 | 12.855419 | 11.141932 | -1.200894 | 433.074061 | 0.182711873 | -1.44549306 | 0.148319441 | 0.909660767 | 8.884103 | 8.620844 | 8.812853 | 8.808049 | 8.750249 | 8.645849 | 546 | 508 | 373 | 336 |
| 54504 | NM_00134805 | CPVL | carboxypeptidase vitellogenic like | protein_coding | NP_001334981.1;NP_001334983.1;NP_001358184.1;NP_001358185.1;NI | 14399 | 609780 | . | . | 14.654891 | 13.112074 | 12.032172 | 13.863295 | 34.102530 | 30.309146 | 26.999846 | 31.696779 | -1.066222 | 878.544564 | 0.151057056 | -0.61240386 | 0.54027059 | 0.998865404 | 9.824518 | 9.731941 | 9.849693 | 9.749399 | 9.713499 | 9.794543 | 1116 | 911 | 697 | 843 |
| 54505 | NM_00134596 | DHX29 | DExH-box helicase 29 | protein_coding | NP_001332893.1;NP_001332894.1;NP_061903.2 | 15815 | 612720 | . | . | 7.967861 | 7.300892 | 7.055333 | 6.472472 | 18.541540 | 16.876343 | 15.831962 | 14.798539 | -1.140620 | 1031.802456 | 0.138348168 | -1.37203247 | 0.17005333 | 0.946389416 | 10.102565 | 9.913122 | 10.081810 | 10.031834 | 9.985626 | 9.934792 | 1308 | 1147 | 868 | 872 |
| 54507 | NM_00128860 | ADAMTSL4 | ADAMTS like 4 | protein_coding | NP_001275536.1;NP_001275537.1;NP_001365525.1;NP_061905.2;NP_0: | 19706 | 610113 | . | . | 0.075911 | 0.128472 | 0.067604 | 0.150643 | 0.176648 | 0.296968 | 0.151701 | 0.344428 | 1.069876 | 15.875292 | 0.806387113 | 0.120839019 | 0.903818544 | NA | 3.942580 | 4.033413 | 3.963129 | 3.999556 | 3.960553 | 4.011257 | 13 | 21 | 9 | 21 |
| 54509 | NM_019034 | RHOF | ras homolog family member F, filopodia associated | protein_coding | NP_061907.2 | 15703 | 618867 | . | . | 0.505936 | 0.373537 | 0.532396 | 0.598115 | 1.177333 | 0.863449 | 1.194682 | 1.367518 | 1.286081 | 38.768685 | 0.505109793 | 0.718619839 | 0.472375187 | 0.998865404 | 5.083404 | 5.447344 | 5.272546 | 5.230768 | 5.281109 | 5.298111 | 45 | 31 | 36 | 43 |
| 54510 | NM_00130082 | PCDH18 | protocadherin 18 | protein_coding | NP_001287757.1;NP_061908.1;XP_006714302.1;XP_016863800.1 | 14268 | 608287 | . | . | 0.645941 | 0.560983 | 0.456910 | 0.500103 | 1.503133 | 1.296739 | 1.025292 | 1.143427 | -1.246227 | 99.892114 | 0.325769402 | -0.9748218 | 0.329648664 | 0.998865404 | 6.792272 | 6.474908 | 6.707442 | 6.642523 | 6.596983 | 6.620301 | 140 | 108 | 74 | 87 |
| 54512 | NM_019037,XI | EXOSC4 | exosome component 4 | protein_coding | NP_061910.1;XP_011515436.1 | 18189 | 606491 | . | . | 29.867781 | 30.162639 | 28.749350 | 35.612279 | 69.503555 | 69.722297 | 64.512705 | 81.423243 | 1.057073 | 816.018688 | 0.156027051 | 0.513216548 | 0.607799852 | 0.998865404 | 9.632184 | 9.711639 | 9.650514 | 9.651755 | 9.615351 | 9.760121 | 913 | 857 | 655 | 866 |
| 54516 | NM_00111418 | MTRF1L | mitochondrial translational release factor 1 like | protein_coding | NP_001107656.1;NP_001287976.1;NP_001288799.1;NP_001288800.1;NI | 21051 | 613542 | . | . | 5.506733 | 6.066195 | 5.497654 | 4.870797 | 12.814391 | 14.022282 | 12.336577 | 11.136496 | -1.126368 | 600.418718 | 0.15945421 | -1.07666242 | 0.281631105 | 0.998865404 | 9.312982 | 9.141580 | 9.257251 | 9.278321 | 9.216114 | 9.161085 | 721 | 697 | 512 | 507 |
| 54517 | NM_00131816 | PUS7 | pseudouridine synthase 7 | protein_coding | NP_001305092.1;NP_001305093.1;NP_061915.2;XP_005250519.1;XP_01 | 26033 | 616261 | . | . | 8.594353 | 7.806230 | 8.166259 | 7.755599 | 19.999411 | 18.044452 | 18.324850 | 17.732255 | -1.043301 | 886.796304 | 0.145115224 | -0.42142562 | 0.673444317 | 0.998865404 | 9.822495 | 9.761784 | 9.843618 | 9.769193 | 9.798285 | 9.754804 | 1096 | 927 | 778 | 789 |
| 54520 | NM_019044,XI | CCDC93 | coiled-coil domain containing 93 | protein_coding | NP_061917.3;XP_006712663.1;XP_011509661.1;XP_011509663.1;XP_01: | 25611 | . | . | . | 11.573503 | 11.797080 | 12.193396 | 10.967908 | 26.932016 | 27.269480 | 27.361625 | 25.076819 | -1.010598 | 2261.526162 | 0.120970898 | -0.12572165 | 0.899952264 | 0.998865404 | 11.150574 | 11.135548 | 11.138610 | 11.155018 | 11.181943 | 11.093054 | 2591 | 2478 | 2065 | 1992 |
| 54521 | NM_00118496 | WDR44 | WD repeat domain 44 | protein_coding | NP_001171894.1;NP_001171895.1;NP_061918.3;XP_011529655.1;XP_01 | 30512 | . | . | . | 2.730701 | 2.435631 | 2.311391 | 2.460290 | 6.354453 | 5.630071 | 5.186694 | 5.625161 | -1.099324 | 319.162747 | 0.199509339 | -0.68476277 | 0.493493633 | 0.998865404 | 8.384764 | 8.248308 | 8.377247 | 8.308935 | 8.278768 | 8.304382 | 408 | 339 | 258 | 292 |
| 54522 | NM_00100994 | ANKRD16 | ankyrin repeat domain 16 | protein_coding | NP_001009941.1;NP_001009943.1;NP_061919.1 | 23471 | 618017 | . | . | 2.398643 | 2.101403 | 2.430356 | 2.649470 | 5.581739 | 4.857487 | 5.453648 | 6.057700 | 1.043282 | 127.715879 | 0.286669713 | 0.213240937 | 0.831139039 | 0.998865404 | 6.966070 | 7.026877 | 6.993711 | 6.983716 | 6.989882 | 7.017420 | 146 | 133 | 109 | 127 |
| 54529 | NM_00135349 | ASNSD1 | asparagine synthetase domain containing 1 | protein_coding | NP_001340426.1;NP_061921.2 | 24910 | . | . | . | 17.368761 | 14.985261 | 16.892879 | 13.689218 | 40.417816 | 34.639101 | 37.907132 | 31.298769 | -1.073205 | 1183.384789 | 0.153838787 | -0.66254517 | 0.507621894 | 0.998865404 | 10.258409 | 10.157224 | 10.289889 | 10.171422 | 10.257839 | 10.094635 | 1520 | 1219 | 1102 | 953 |
| 54531 | NM_00134610 | MIER2 | MIER family member 2 | protein_coding | NP_001333034.1;NP_060020.1;XP_005259639.1;XP_006722832.1;XP_02 | 29210 | . | . | . | 5.699077 | 5.680528 | 6.237788 | 5.562793 | 13.261984 | 13.130794 | 13.997415 | 12.718663 | 1.027879 | 662.573131 | 0.163243937 | 0.243014927 | 0.807993834 | 0.998865404 | 9.351942 | 9.391656 | 9.308674 | 9.410776 | 9.420463 | 9.337148 | 696 | 758 | 620 | 591 |
| 54532 | NM_00137139 | USP53 | ubiquitin specific peptidase 53 | protein_coding | NP_001358324.1;NP_001358325.1;NP_001358326.1;NP_001358327.1;NI | 29255 | 617431 | . | . | 13.970483 | 14.295452 | 10.415494 | 10.349040 | 32.509891 | 33.044574 | 23.372068 | 23.661846 | -1.346783 | 2402.648438 | 0.11554971 | -3.71716513 | 0.000201471 | 0.020752992 | 11.429190 | 10.999796 | 11.342824 | 11.333437 | 11.117343 | 11.079304 | 3193 | 2959 | 1824 | 1873 |
| 54534 | NM_019051 | MRPL50 | mitochondrial ribosomal protein L50 | protein_coding | NP_061924.1 | 16654 | 611854 | . | . | 7.399160 | 6.691417 | 8.579819 | 8.016531 | 17.218147 | 15.467511 | 19.252865 | 18.328846 | 1.160536 | 797.377498 | 0.150098151 | 1.431005367 | 0.152428677 | 0.916353687 | 9.527441 | 9.742770 | 9.619574 | 9.544404 | 9.715180 | 9.659595 | 896 | 753 | 774 | 772 |
| 54535 | NM_00110556 | CCHCR1 | coiled-coil alpha-helical rod protein 1 | protein_coding | NP_001099033.1;NP_001099034.1;NP_061925.2;XP_011513004.1;XP_01 | 13930 | 605310 | . | . | 7.515328 | 8.026623 | 6.846809 | 7.835007 | 17.488479 | 18.553890 | 15.364039 | 17.913809 | -1.048442 | 626.270931 | 0.161398117 | -0.42285314 | 0.672402415 | 0.998865404 | 9.324701 | 9.255761 | 9.271849 | 9.338792 | 9.236660 | 9.310066 | 702 | 726 | 502 | 605 |
| 54536 | NM_00101384 | EXOC6 | exocyst complex component 6 | protein_coding | NP_001013870.1;NP_001306123.1;NP_001306124.1;NP_001306129.1;NI | 23196 | 609672 | . | . | 2.915419 | 2.349625 | 2.609813 | 2.243491 | 6.784298 | 5.431264 | 5.856343 | 5.129475 | -1.099645 | 282.469215 | 0.219958674 | -0.62301469 | 0.533274867 | 0.998865404 | 8.208153 | 8.072562 | 8.222568 | 8.102859 | 8.155022 | 8.072867 | 378 | 284 | 253 | 232 |
| 54537 | NM_00133011 | SHLD2 | shieldin complex subunit 2 | protein_coding | NP_001317041.1;NP_001364087.1;NP_001364088.1;NP_001364089.1;NI | 28773 | 618029 | . | . | 6.403201 | 5.725155 | 7.052618 | 6.233571 | 14.900516 | 13.233950 | 15.825867 | 14.252319 | 1.085663 | 716.546567 | 0.157857576 | 0.751160564 | 0.45255603 | 0.998865404 | 9.424030 | 9.543337 | 9.491647 | 9.415558 | 9.556303 | 9.463927 | 835 | 700 | 692 | 653 |
| 54539 | NM_00113599 | NDUFB11 | NADH:ubiquinone oxidoreductase subunit B11 | protein_coding | NP_001129470.1;NP_061929.2 | 20372 | 300403 | . | . | 55.123670 | 52.891977 | 57.025503 | 67.164355 | 128.275048 | 122.262175 | 127.963576 | 153.563322 | 1.118619 | 1158.162194 | 0.139941486 | 1.155618001 | 0.247837442 | 0.992733415 | 10.094643 | 10.256086 | 10.152185 | 10.113542 | 10.159437 | 10.271337 | 1292 | 1149 | 985 | 1231 |
| 54540 | NM_00119094 | FAM193B | family with sequence similarity 193 member B | protein_coding | NP_001177875.1;NP_001353427.1;NP_001353428.1;NP_001353429.1;XF | 25524 | 615813 | . | . | 10.168089 | 10.883289 | 12.314186 | 11.716099 | 23.661562 | 25.157209 | 27.632676 | 26.787468 | 1.074247 | 1236.120287 | 0.132613041 | 0.77914891 | 0.435892002 | 0.998865404 | 10.218923 | 10.322428 | 10.231538 | 10.254952 | 10.330689 | 10.263131 | 1351 | 1306 | 1163 | 1147 |
| 54541 | NM_019058 | DDIT4 | DNA damage inducible transcript 4 | protein_coding | NP_061931.1 | 24944 | 607729 | . | . | 36.449886 | 38.293808 | 42.804134 | 44.127842 | 84.820381 | 88.517853 | 96.051239 | 100.893074 | 1.147106 | 2195.116014 | 0.116765069 | 1.695696975 | 0.089943282 | 0.804059937 | 10.997743 | 11.195639 | 11.026194 | 11.058136 | 11.143801 | 11.160212 | 2307 | 2251 | 2019 | 2221 |
| 54542 | NM_00110058 | RC3H2 | ring finger and CCCH-type domains 2 | protein_coding | NP_001094058.1;NP_001341407.1;NP_001341408.1;NP_001341411.1;NI | 21461 | 615231 | . | . | 4.139510 | 4.261672 | 4.508837 | 3.878274 | 9.632806 | 9.851044 | 10.117700 | 8.867213 | -1.010461 | 1094.259606 | 0.1413392 | -0.10622387 | 0.915404733 | 0.998865404 | 10.103078 | 10.088363 | 10.077605 | 10.119410 | 10.143024 | 10.036380 | 1231 | 1220 | 1015 | 947 |
| 54543 | NM_019059,N | TOMM7 | translocase of outer mitochondrial membrane 7 | protein_coding | NP_061932.1 | 21648 | 607980 | . | . | 451.206143 | 433.501255 | 386.315187 | 474.928423 | 1049.975224 | 1002.057534 | 866.880071 | 1085.867401 | -1.042171 | 5899.099172 | 0.122394097 | -0.48688231 | 0.626341736 | 0.998865404 | 12.555825 | 12.496116 | 12.562641 | 12.521457 | 12.421968 | 12.586341 | 7102 | 6339 | 4532 | 5946 |
| 54545 | NM_00104044 | MTMR12 | myotubularin related protein 12 | protein_coding | NP_001035536.1;NP_001281272.1;NP_001281273.1 | 18191 | 606501 | . | . | 4.097031 | 3.797619 | 3.606948 | 4.431786 | 9.533958 | 8.778368 | 8.093886 | 10.132755 | 1.002998 | 634.336569 | 0.166771529 | 0.025897012 | 0.979339483 | 0.998865404 | 9.307173 | 9.311034 | 9.335806 | 9.277303 | 9.242273 | 9.371419 | 761 | 654 | 499 | 653 |
| 54552 | NM_00118481 | GNL3L | G protein nucleolar 3 like | protein_coding | NP_001171748.1;NP_061940.1 | 25553 | 300873 | . | . | 3.266061 | 3.429450 | 3.906877 | 3.520350 | 7.600259 | 7.927329 | 8.766920 | 8.048859 | 1.094630 | 948.643160 | 0.142755374 | 0.913756074 | 0.360845032 | 0.998865404 | 9.822886 | 9.953598 | 9.841730 | 9.867936 | 9.962074 | 9.877593 | 1024 | 995 | 912 | 876 |
| 54553 | NR_033797 | MIR4453HG | MIR4453 host gene | lncRNA | . | 25288 | . | . | . | 0.818662 | 0.959372 | 1.220122 | 0.738709 | 1.905061 | 2.217632 | 2.737918 | 1.688970 | 1.076280 | 87.773730 | 0.368273866 | 0.287972705 | 0.773367632 | 0.998865404 | 6.400715 | 6.508699 | 6.415798 | 6.457633 | 6.525830 | 6.384199 | 90 | 98 | 99 | 64 |
| 54554 | NM_019069 | WDR5B | WD repeat domain 5B | protein_coding | NP_061942.2 | 17826 | . | . | . | 1.714671 | 1.789423 | 1.595555 | 1.395602 | 3.990109 | 4.136333 | 3.580379 | 3.190879 | -1.190243 | 213.837225 | 0.233245993 | -1.07721723 | 0.281383231 | 0.998865404 | 7.860218 | 7.609654 | 7.770838 | 7.787348 | 7.729290 | 7.666280 | 263 | 255 | 182 | 170 |
| 54555 | NM_019070,N | DDX49 | DEAD-box helicase 49 | protein_coding | NP_061943.2;XP_011526385.1;XP_011526386.1 | 18684 | . | . | . | 22.264363 | 24.808556 | 23.176900 | 27.711081 | 51.810087 | 57.346087 | 52.008290 | 63.358094 | 1.068354 | 1377.285267 | 0.141105637 | 0.676020877 | 0.49902739 | 0.998865404 | 10.379416 | 10.474249 | 10.358875 | 10.440264 | 10.381979 | 10.515382 | 1452 | 1514 | 1130 | 1449 |
| 54556 | NM_019071,N | ING3 | inhibitor of growth family member 3 | protein_coding | NP_061944.2;NP_938008.1;XP_016867858.1 | 14587 | 607493 | . | . | 2.215075 | 2.504931 | 2.771907 | 2.125378 | 5.154571 | 5.790262 | 6.220081 | 4.859424 | 1.028192 | 188.376312 | 0.253217804 | 0.158399811 | 0.874141763 | 0.998865404 | 7.537192 | 7.577475 | 7.504440 | 7.590683 | 7.598977 | 7.520833 | 193 | 220 | 181 | 163 |
| 54557 | NM_019072,XI | SGTB | small glutamine rich tetratricopeptide repeat containing beta | protein_coding | NP_061945.1;XP_005248605.1;XP_016865086.1 | 23567 | . | . | . | 2.094041 | 2.090070 | 1.816481 | 1.494794 | 4.872919 | 4.831291 | 4.076131 | 3.417668 | -1.281928 | 319.296256 | 0.204112123 | -1.75548113 | 0.079177046 | 0.775642847 | 8.486349 | 8.129094 | 8.383013 | 8.379678 | 8.295916 | 8.192110 | 414 | 386 | 268 | 236 |
| 54558 | NM_00128623 | SPATA6 | spermatogenesis associated 6 | protein_coding | NP_001273167.1;NP_001273168.1;NP_061946.1;XP_006710762.1;XP_00 | 18309 | 613947 | . | . | 2.825569 | 2.778517 | 2.908866 | 2.943226 | 6.575214 | 6.422669 | 6.527413 | 6.729336 | 1.033194 | 386.936620 | 0.184717261 | 0.255041253 | 0.798691238 | 0.998865404 | 8.571921 | 8.619592 | 8.610591 | 8.560087 | 8.621841 | 8.587741 | 457 | 393 | 352 | 358 |
| 54566 | NM_018424,N | EPB41L4B | erythrocyte membrane protein band 4.1 like 4B | protein_coding | NP_060894.2;NP_061987.3;XP_011517092.1;XP_011517093.1;XP_01151 | 19818 | 610340 | . | . | 0.780320 | 0.684980 | 0.510193 | 0.505600 | 1.815837 | 1.583363 | 1.144858 | 1.155995 | -1.455085 | 78.130393 | 0.361761129 | -1.49574885 | 0.134719114 | 0.891755273 | 6.532860 | 5.992450 | 6.355069 | 6.313382 | 6.241652 | 6.235899 | 114 | 93 | 56 | 59 |
| 54581 | NR_003654,NF | SCAND2P | SCAN domain containing 2 pseudogene | transcribed_ps | . | 10567 | 610417 | . | . | 1.304429 | 1.344009 | 2.109160 | 1.207378 | 3.035460 | 3.106735 | 4.732894 | 2.760526 | 1.202909 | 191.300107 | 0.283061651 | 0.94158917 | 0.346403017 | 0.998865404 | 7.439234 | 7.707684 | 7.520792 | 7.528968 | 7.721782 | 7.476820 | 198 | 189 | 230 | 144 |
| 54583 | NM_00137726 | EGLN1 | egl-9 family hypoxia inducible factor 1 | protein_coding | NP_001364189.1;NP_001364190.1;NP_071334.1;XP_024303502.1 | 1232 | 606425 | . | . | 7.046001 | 6.724345 | 8.460405 | 7.502720 | 16.396332 | 15.543626 | 18.984905 | 17.154079 | 1.151281 | 966.976917 | 0.14610559 | 1.391051107 | 0.16420993 | 0.940065727 | 9.811855 | 10.015630 | 9.874993 | 9.848725 | 10.022192 | 9.904005 | 1054 | 952 | 972 | 893 |
| 54584 | NM_053004 | GNB1L | G protein subunit beta 1 like | protein_coding | NP_443730.1 | 4397 | 610778 | . | . | 1.326456 | 1.426582 | 1.250239 | 1.235527 | 3.086718 | 3.297607 | 2.805499 | 2.824886 | -1.122908 | 271.501025 | 0.209396737 | -0.79867481 | 0.424478996 | 0.998865404 | 8.166062 | 7.998715 | 8.096705 | 8.131355 | 8.060264 | 8.048484 | 320 | 320 | 225 | 237 |
| 54585 | NM_00127637 | LZTFL1 | leucine zipper transcription factor like 1 | protein_coding | NP_001263307.1;NP_001263308.1;NP_065080.1;XP_006713270.2;XP_01 | 6741 | 606568 | . | . | 2.911072 | 2.806118 | 2.856988 | 2.627710 | 6.774184 | 6.486469 | 6.410998 | 6.007945 | -1.057120 | 352.305806 | 0.190874339 | -0.4198536 | 0.674592408 | 0.998865404 | 8.499908 | 8.420365 | 8.487473 | 8.464661 | 8.470660 | 8.417902 | 425 | 383 | 312 | 306 |
| 54602 | NM_00116140 | NDFIP2 | Nedd4 family interacting protein 2 | protein_coding | NP_001154879.1;NP_061953.2;XP_006719903.1 | 18537 | 610041 | . | . | 8.091019 | 5.922707 | 6.519543 | 4.997637 | 18.828132 | 13.690604 | 14.629663 | 11.426501 | -1.238703 | 932.945898 | 0.184002311 | -1.67840272 | 0.093268506 | 0.809520218 | 10.011370 | 9.703540 | 10.045194 | 9.805720 | 9.868660 | 9.677096 | 1377 | 937 | 824 | 675 |
| 54606 | NM_00125718 | DDX56 | DEAD-box helicase 56 | protein_coding | NP_001244118.1;NP_061955.1 | 18193 | 608023 | . | . | 30.769945 | 33.152262 | 33.286083 | 34.794727 | 71.602928 | 76.632933 | 74.693008 | 79.554016 | 1.050768 | 2951.543486 | 0.113738888 | 0.628142731 | 0.529910443 | 0.998865404 | 11.491157 | 11.562469 | 11.479427 | 11.532836 | 11.532727 | 11.560825 | 3206 | 3208 | 2585 | 2884 |
| 54617 | NM_017553,N | INO80 | INO80 complex ATPase subunit | protein_coding | NP_060023.1;XP_011519987.1;XP_011519988.1 | 26956 | 610169 | . | . | 4.021764 | 4.246034 | 4.246047 | 4.579622 | 9.358810 | 9.814897 | 9.528006 | 10.470759 | 1.055837 | 847.572957 | 0.144153383 | 0.543773836 | 0.586597115 | 0.998865404 | 9.687647 | 9.765685 | 9.690067 | 9.723912 | 9.721856 | 9.769086 | 927 | 911 | 732 | 843 |
| 54619 | NM_00113437 | CCNJ | cyclin J | protein_coding | NP_001127847.1;NP_001127848.1;NP_061957.2;XP_005269978.1;XP_01 | 23434 | . | . | . | 2.058530 | 2.318892 | 1.966894 | 1.422635 | 4.790285 | 5.360223 | 4.413655 | 3.252685 | -1.283354 | 240.899363 | 0.240713532 | -1.49521881 | 0.134857345 | 0.891755273 | 8.080381 | 7.721992 | 7.946554 | 7.985677 | 7.921435 | 7.760597 | 300 | 303 | 216 | 163 |
| 54620 | NM_00109978 | FBXL19 | F-box and leucine rich repeat protein 19 | protein_coding | NP_001093254.2;NP_001269280.1;NP_001369708.1;NP_001369709.1;NI | 25300 | 609085 | . | . | 4.148808 | 3.820861 | 4.707778 | 4.995945 | 9.654445 | 8.832092 | 10.564118 | 11.422636 | 1.147710 | 585.808656 | 0.169470705 | 1.17282003 | 0.240867984 | 0.992733415 | 9.091652 | 9.290101 | 9.178077 | 9.110567 | 9.175447 | 9.292755 | 661 | 558 | 497 | 638 |
| 54621 | NM_019086,XI | VSIG10 | V-set and immunoglobulin domain containing 10 | protein_coding | NP_061959.2;XP_005253965.1;XP_006719530.1;XP_006719531.1;XP_01: | 26078 | . | . | . | 2.498979 | 2.410368 | 2.481622 | 2.751266 | 5.815227 | 5.571672 | 5.568689 | 6.290443 | 1.063731 | 384.768880 | 0.186473758 | 0.477993667 | 0.632654712 | 0.998865404 | 8.542646 | 8.631677 | 8.594908 | 8.541644 | 8.577247 | 8.631680 | 449 | 384 | 329 | 389 |
| 54622 | NM_019087,XI | ARL15 | ADP ribosylation factor like GTPase 15 | protein_coding | NP_061960.1;XP_011541800.1;XP_011541801.1;XP_011541802.1;XP_01( | 25945 | . | . | . | 3.849051 | 2.679279 | 3.271581 | 2.670529 | 8.956901 | 6.193274 | 7.341335 | 6.105846 | -1.118462 | 325.216728 | 0.228209412 | -0.70775426 | 0.479097877 | 0.998865404 | 8.422957 | 8.263134 | 8.468466 | 8.261998 | 8.362892 | 8.251100 | 466 | 304 | 295 | 258 |
| 54623 | NM_00125682 | PAF1 | PAF1 homolog, Paf1/RNA polymerase II complex component | protein_coding | NP_001243755.1;NP_061961.2 | 25459 | 610506 | . | . | 28.887098 | 30.056138 | 27.474403 | 31.434737 | 67.221462 | 69.476112 | 61.651758 | 71.871785 | -1.014012 | 2019.424653 | 0.124422945 | -0.16134569 | 0.871821139 | 0.998865404 | 10.989876 | 10.969510 | 10.972926 | 10.996620 | 10.923112 | 11.021868 | 2307 | 2227 | 1635 | 1997 |
| 54625 | NM_017554,XI | PARP14 | poly(ADP-ribose) polymerase family member 14 | protein_coding | NP_060024.2;XP_011511230.1;XP_011511231.1 | 29232 | 610028 | . | . | 4.418960 | 4.856700 | 6.032517 | 5.140340 | 10.283103 | 11.226480 | 13.536792 | 11.752775 | 1.170647 | 1216.468458 | 0.142737155 | 1.592479033 | 0.111277111 | 0.848126978 | 10.130166 | 10.357828 | 10.168062 | 10.199722 | 10.372248 | 10.228519 | 1263 | 1235 | 1248 | 1115 |
| 54626 | NM_019089 | HES2 | hes family bHLH transcription factor 2 | protein_coding | NP_061962.2 | 16005 | 609970 | . | . | 0.173162 | 0.090222 | 0.105938 | 0.043879 | 0.402956 | 0.208553 | 0.237721 | 0.100324 | -1.672001 | 14.211321 | 0.865096862 | -0.85721738 | 0.391324758 | NA | 4.149569 | 3.416142 | 3.870403 | 3.820166 | 3.832878 | 3.795859 | 27 | 13 | 13 | 6 |
| 54629 | NM_00104045 | MINDY2 | MINDY lysine 48 deubiquitinase 2 | protein_coding | NP_001035540.1;NP_001035543.1;XP_011519989.1;XP_011519990.1;XP | 26954 | 618408 | . | . | 1.745754 | 1.518022 | 1.555443 | 1.471767 | 4.062439 | 3.508978 | 3.490369 | 3.365020 | -1.095769 | 453.898166 | 0.179073067 | -0.73681532 | 0.461234666 | 0.998865404 | 8.890344 | 8.759123 | 8.899214 | 8.804997 | 8.816331 | 8.777139 | 587 | 474 | 389 | 393 |
| 54660 | NR_001281 | PCDHB18P | protocadherin beta 18 pseudogene | transcribed_ps | . | 14548 | . | . | . | 0.048516 | 0.072254 | 0.135180 | 0.110999 | 0.112900 | 0.167017 | 0.303341 | 0.253785 | 2.024529 | 9.539410 | 0.996598848 | 1.021058875 | 0.307226546 | NA | 2.658167 | 3.674293 | 3.215159 | 3.225738 | 3.253799 | 3.245180 | 6 | 8 | 12 | 11 |
| 54661 | NR_001280 | PCDHB17P | protocadherin beta 17 pseudogene | transcribed_ps | . | 14547 | . | . | . | 0.068101 | 0.062788 | 0.036614 | 0.040932 | 0.158475 | 0.145138 | 0.082160 | 0.093587 | -1.620123 | 4.355261 | 1.447443852 | -0.48091913 | 0.630573978 | NA | 2.429096 | 1.733338 | 2.134903 | 2.136464 | 2.126881 | 2.125664 | 6 | 6 | 3 | 3 |
| 54662 | NM_00128677 | TBC1D13 | TBC1 domain family member 13 | protein_coding | NP_001273701.1;NP_060671.3;XP_005252117.1 | 25571 | 616218 | . | . | 5.216062 | 6.065928 | 5.833577 | 6.420893 | 12.137992 | 14.021666 | 13.090377 | 14.680609 | 1.072057 | 706.557638 | 0.157843115 | 0.635957368 | 0.524804209 | 0.998865404 | 9.413941 | 9.513663 | 9.388507 | 9.487156 | 9.457274 | 9.515408 | 729 | 789 | 609 | 714 |
| 54663 | NM_00130797 | WDR74 | WD repeat domain 74 | protein_coding | NP_001294906.1;NP_001356376.1;NP_001356377.1;NP_001356378.1;NI | 25529 | 617947 | . | . | 14.574972 | 16.498334 | 16.815611 | 17.183419 | 33.916556 | 38.136639 | 37.733744 | 39.287849 | 1.077197 | 638.342490 | 0.157160366 | 0.682625124 | 0.494843754 | 0.998865404 | 9.263759 | 9.370636 | 9.259303 | 9.323387 | 9.330337 | 9.353166 | 674 | 695 | 566 | 631 |

| | | | | | | | | | | |
|---|---|---|---|---|---|---|---|---|---|---|
| 54664 | NM_00113423 | TMEM106B | transmembrane protein 106B | protein_coding | NP_001127704.1;NP_060844.2;XP_005249846.1 | 22407 | 613413 | . | . | [illegible] |
| 54665 | NM_018364,N | RSBN1 | round spermatid basic protein 1 | protein_coding | NP_060834.2;XP_016857007.1 | 25642 | 615858 | . | . | [illegible] |
| 54675 | NM_00112745 | CRLS1 | cardiolipin synthase 1 | protein_coding | NP_001120930.1;NP_001310490.1;NP_001310491.1;NP_001310492.1;NI | 16148 | 608188 | . | . | [illegible] |
| 54676 | NM_00128621 | GTPBP2 | GTP binding protein 2 | protein_coding | NP_001273145.1;NP_061969.3;XP_016866465.1;XP_024302243.1;XP_02 | 4670 | 607434 | . | . | [illegible] |
| 54677 | NM_00114393 | CROT | carnitine O-octanoyltransferase | protein_coding | NP_001137407.1;NP_001230674.1;NP_066974.2;XP_011514639.1;XP_01 | 2366 | 606090 | . | . | [illegible] |
| 54680 | NM_00117067 | ZNHIT6 | zinc finger HIT-type containing 6 | protein_coding | NP_001164141.1;NP_060423.3;XP_011539916.1;XP_024303504.1 | 26089 | . | . | . | [illegible] |
| 54682 | NM_00136361 | MANSC1 | MANSC domain containing 1 | protein_coding | NP_001350542.1;NP_060520.2 | 25505 | . | . | . | [illegible] |
| 54700 | NM_00130106 | RRN3 | RRN3 homolog, RNA polymerase I transcription factor | protein_coding | NP_001287993.1;NP_060897.3 | 30346 | 605121 | . | . | [illegible] |
| 54704 | NM_00116177 | PDP1 | pyruvate dehyrogenase phosphatase catalytic subunit 1 | protein_coding | NP_001155251.1;NP_001155252.1;NP_001155253.1;NP_060914.2;XP_01 | 9279 | 605993 | . | . | [illegible] |
| 54707 | NM_018066 | GPN2 | GPN-loop GTPase 2 | protein_coding | NP_060536.3 | 25513 | . | . | . | [illegible] |
| 54708 | NM_017824,XI | MARCHF5 | membrane associated ring-CH-type finger 5 | protein_coding | NP_060294.1;XP_011538188.1 | 26025 | 610637 | . | . | [illegible] |
| 54718 | NR_027795 | BTN2A3P | butyrophilin subfamily 2 member A3, pseudogene | transcribed_ps | . | 13229 | 613592 | . | . | [illegible] |
| 54726 | NM_00110265 | OTUD4 | OTU deubiquitinase 4 | protein_coding | NP_001096123.1;NP_001352986.1;NP_001352987.1;NP_059963.1;XP_01 | 24949 | 611744 | . | . | [illegible] |
| 54732 | NM_017510 | TMED9 | transmembrane p24 trafficking protein 9 | protein_coding | NP_059980.2 | 24878 | . | . | . | [illegible] |
| 54733 | NM_017515 | SLC35F2 | solute carrier family 35 member F2 | protein_coding | NP_059985.2 | 23615 | . | . | . | [illegible] |
| 54734 | NM_017516 | RAB39A | RAB39A, member RAS oncogene family | protein_coding | NP_059986.1 | 16521 | . | . | . | [illegible] |
| 54737 | NM_017520,XI | MPHOSPH8 | M-phase phosphoprotein 8 | protein_coding | NP_059990.2;XP_011533426.1 | 29810 | 611626 | . | . | [illegible] |
| 54741 | NM_00119868 | LEPROT | leptin receptor overlapping transcript | protein_coding | NP_001185610.1;NP_001185612.1;NP_059996.1 | 29477 | 613461 | . | . | [illegible] |
| 54742 | NM_00116035 | LY6K | lymphocyte antigen 6 family member K | protein_coding | NP_001153826.1;NP_001153827.1;NP_059997.3;XP_005251046.1;XP_00 | 24225 | 615093 | . | . | [illegible] |
| 54749 | NM_00124294 | EPDR1 | ependymin related 1 | protein_coding | NP_001229875.2;NP_001229877.1;NP_060019.2 | 17572 | . | . | . | [illegible] |
| 54751 | NM_00102421 | FBLIM1 | filamin binding LIM protein 1 | protein_coding | NP_001019386.1;NP_001019387.1;NP_001337080.1;NP_060026.2;XP_0C | 24686 | 607747 | . | . | [illegible] |
| 54756 | NM_00131886 | IL17RD | interleukin 17 receptor D | protein_coding | NP_001305793.1;NP_060033.3;XP_005265295.1;XP_011532151.2 | 17616 | 606807 | . | . | [illegible] |
| 54758 | NM_00118485 | KLHDC4 | kelch domain containing 4 | protein_coding | NP_001171783.1;NP_001171785.1;NP_001338866.1;NP_001338867.1;NI | 25272 | . | . | . | [illegible] |
| 54760 | NM_017573,XI | PCSK4 | proprotein convertase subtilisin/kexin type 4 | protein_coding | NP_060043.2;XP_005259643.1;XP_011526387.1;XP_011526388.1;XP_01 | 8746 | 600487 | . | . | [illegible] |
| 54762 | NM_00117210 | GRAMD1C | GRAM domain containing 1C | protein_coding | NP_001165576.1;NP_060047.3;XP_005247603.1;XP_005247604.1;XP_01 | 25252 | . | . | . | [illegible] |
| 54763 | NM_00131777 | ROPN1 | rhophilin associated tail protein 1 | protein_coding | NP_001304703.1;NP_001304704.1;NP_060048.2;XP_011511235.1;XP_01 | 17692 | 611757 | . | . | [illegible] |
| 54764 | NM_017580,XI | ZRANB1 | zinc finger RANBP2-type containing 1 | protein_coding | NP_060050.2;XP_006717970.1 | 18224 | 611749 | . | . | [illegible] |
| 54765 | NM_017583,XI | TRIM44 | tripartite motif containing 44 | protein_coding | NP_060053.2;XP_006718317.1 | 19016 | 612298 | . | . | [illegible] |
| 54776 | NM_00127161 | PPP1R12C | protein phosphatase 1 regulatory subunit 12C | protein_coding | NP_001258547.1;NP_060077.1;XP_005259070.1;XP_011525347.1 | 14947 | 613245 | . | . | [illegible] |
| 54778 | NM_00127052 | RNF111 | ring finger protein 111 | protein_coding | NP_001257457.1;NP_001257458.1;NP_001257459.1;NP_001317260.1;NI | 17384 | 605840 | . | . | [illegible] |
| 54780 | NM_00116786 | NSMCE4A | NSE4 homolog A, SMC5-SMC6 complex component | protein_coding | NP_001161337.1;NP_060085.2;XP_005269986.1;XP_005269987.1;XP_01 | 25935 | 612987 | . | . | [illegible] |
| 54784 | NM_017621,XI | ALKBH4 | alkB homolog 4, lysine demethylase | protein_coding | NP_060091.1;XP_005250521.1;XP_005250522.1 | 21900 | 613302 | . | . | [illegible] |
| 54785 | NM_017622 | BORCS6 | BLOC-1 related complex subunit 6 | protein_coding | NP_060092.2 | 25939 | 616599 | . | . | [illegible] |
| 54788 | NM_00100276 | DNAJB12 | DnaJ heat shock protein family (Hsp40) member B12 | protein_coding | NP_001002762.3;NP_001352009.1;NP_001352010.1;NP_060096.4 | 14891 | 608376 | . | . | [illegible] |
| 54790 | NM_00112720 | TET2 | tet methylcytosine dioxygenase 2 | protein_coding | NP_001120680.1;NP_060098.3;XP_005263139.1;XP_006714305.1;XP_01 | 25941 | 612839 | . | . | [illegible] |
| 54793 | NM_017634,XI | KCTD9 | potassium channel tetramerization domain containing 9 | protein_coding | NP_060104.2 | 22401 | 617265 | . | . | [illegible] |
| 54795 | NM_00119522 | TRPM4 | transient receptor potential cation channel subfamily M member 4 | protein_coding | NP_001182156.1;NP_001308210.1;NP_001308211.1;NP_001308212.1;NI | 17993 | 606936 | . | . | [illegible] |
| 54796 | NM_00131793 | BNC2 | basonuclin 2 | protein_coding | NP_001304868.1;NP_001304869.1;NP_060107.3 | 30988 | 608669 | . | . | [illegible] |
| 54797 | NM_00112735 | MED18 | mediator complex subunit 18 | protein_coding | NP_001120822.1;NP_060108.2;XP_005245971.1 | 25944 | 612384 | . | . | [illegible] |
| 54798 | NM_00114255 | DCHS2 | dachsous cadherin-related 2 | protein_coding | NP_001136024.1;NP_001345164.1;XP_011530348.1;XP_011530351.1 | 23111 | 612486 | . | . | [illegible] |
| 54799 | NM_017643,XI | MBTD1 | mbt domain containing 1 | protein_coding | NP_060113.2;XP_005257524.1;XP_005257525.1;XP_011523224.1;XP_01 | 19866 | 618705 | . | . | [illegible] |
| 54800 | NM_00134941 | KLHL24 | kelch like family member 24 | protein_coding | NP_001336342.1;NP_001336343.1;NP_001336344.1;NP_001336345.1;NI | 25947 | 611295 | . | . | [illegible] |
| 54801 | NM_00127089 | HAUS6 | HAUS augmin like complex subunit 6 | protein_coding | NP_001257819.1;NP_060115.3;XP_011516237.1 | 25948 | 613433 | . | . | [illegible] |
| 54802 | NM_00131269 | TRIT1 | tRNA isopentenyltransferase 1 | protein_coding | NP_001299620.1;NP_001299621.1;NP_060116.2;XP_005271011.1;XP_00 | 20286 | 617840 | . | . | [illegible] |
| 54805 | NM_017649,N | CNNM2 | cyclin and CBS domain divalent metal cation transport mediator 2 | protein_coding | NP_060119.3;NP_951058.1;NP_951059.1;XP_005269990.1;XP_00671797 | 103 | 607803 | . | . | [illegible] |
| 54806 | NM_00113483 | AHI1 | Abelson helper integration site 1 | protein_coding | NP_001128302.1;NP_001128303.1;NP_001128304.1;NP_001337432.1;NI | 21575 | 608894 | . | . | [illegible] |
| 54807 | NM_00107742 | ZNF586 | zinc finger protein 586 | protein_coding | NP_001070894.1;NP_001191743.1;NP_060122.2 | 25949 | . | . | . | [illegible] |
| 54808 | NM_00135321 | DYM | dymeclin | protein_coding | NP_001340139.1;NP_001340140.1;NP_001340141.1;NP_001340142.1;NI | 21317 | 607461 | . | . | [illegible] |
| 54809 | NM_00119330 | SAMD9 | sterile alpha motif domain containing 9 | protein_coding | NP_001180236.1;NP_060124.2 | 1348 | 610456 | . | . | [illegible] |
| 54811 | NM_00113003 | ZNF562 | zinc finger protein 562 | protein_coding | NP_001123503.1;NP_001123504.1;NP_001287814.1;NP_060126.2;XP_0C | 25950 | . | . | . | [illegible] |
| 54812 | NM_00100224 | AFTPH | aftiphilin | protein_coding | NP_001002243.1;NP_001362898.1;NP_001362899.1;NP_001362900.1;NI | 25951 | . | . | . | [illegible] |
| 54813 | NM_00130811 | KLHL28 | kelch like family member 28 | protein_coding | NP_001295041.1;NP_060128.2;XP_005267827.1;XP_006720237.1;XP_01 | 19741 | . | . | . | [illegible] |
| 54814 | NM_00116337 | QPCTL | glutaminyl-peptide cyclotransferase like | protein_coding | NP_001156849.1;NP_060129.2;XP_011525350.1;XP_016882389.1 | 25952 | . | . | . | [illegible] |
| 54815 | NM_00130094 | GATAD2A | GATA zinc finger domain containing 2A | protein_coding | NP_001287875.1;NP_001346560.1;NP_060130.3;XP_011526407.1;XP_01 | 29989 | 614997 | . | . | [illegible] |
| 54816 | NM_00100284 | ZNF280D | zinc finger protein 280D | protein_coding | NP_001002843.1;NP_001002844.1;NP_001275517.1;NP_001275518.1;NI | 25953 | . | . | . | [illegible] |
| 54819 | NM_00130081 | ZCCHC10 | zinc finger CCHC-type containing 10 | protein_coding | NP_001287745.1;NP_001287746.1;NP_001287747.1;NP_001287748.1;NI | 25954 | . | . | . | [illegible] |
| 54820 | NM_00114397 | NDE1 | nudE neurodevelopment protein 1 | protein_coding | NP_001137451.1;NP_060138.1;XP_005255453.1;XP_006720960.1;XP_00 | 17619 | 609449 | . | . | [illegible] |
| 54821 | NM_00100995 | ERCC6L | ERCC excision repair 6 like, spindle assembly checkpoint helicase | protein_coding | NP_001009954.1;NP_060139.2 | 20794 | 300687 | . | . | [illegible] |
| 54822 | NM_00130121 | TRPM7 | transient receptor potential cation channel subfamily M member 7 | protein_coding | NP_001288141.1;NP_060142.3;XP_005254543.1;XP_016877839.1;XP_01 | 17994 | 605692 | . | . | [illegible] |
| 54823 | NM_00110551 | SWT1 | SWT1 RNA endoribonuclease homolog | protein_coding | NP_001098988.1;NP_060143.4;XP_005245328.1;XP_005245330.1;XP_01 | 16785 | . | . | . | [illegible] |
| 54826 | NM_00131795 | GIN1 | gypsy retrotransposon integrase 1 | protein_coding | NP_001304883.1;NP_060146.2;XP_005272085.1;XP_016865088.1;XP_02 | 25959 | . | . | . | [illegible] |
| 54828 | NM_00109943 | BCAS3 | BCAS3 microtubule associated cell migration factor | protein_coding | NP_001092902.1;NP_001307399.1;NP_001317342.1;NP_001317343.1;NI | 14347 | 607470 | . | . | [illegible] |
| 54832 | NM_00101808 | VPS13C | vacuolar protein sorting 13 homolog C | protein_coding | NP_001018098.1;NP_060154.3;NP_060550.2;NP_065872.1;XP_01152001 | 23594 | 608879 | . | . | [illegible] |
| 54834 | NM_00113558 | GDAP2 | ganglioside induced differentiation associated protein 2 | protein_coding | NP_001129061.1;NP_060156.1 | 18010 | 618128 | . | . | [illegible] |
| 54838 | NM_00108391 | WBP1L | WW domain binding protein 1 like | protein_coding | NP_001077382.1;NP_060257.4;XP_011538215.1;XP_016871849.1 | 23510 | 611129 | . | . | [illegible] |
| 54839 | NM_00119901 | LRRC49 | leucine rich repeat containing 49 | protein_coding | NP_001185946.1;NP_001185947.1;NP_001271286.1;NP_001350661.1;NI | 25965 | . | . | . | [illegible] |
| 54840 | NM_00119524 | APTX | aprataxin | protein_coding | NP_001182177.2;NP_001182178.1;NP_001182179.2;NP_001182180.1;NI | 15984 | 606350 | . | . | [illegible] |
| 54841 | NM_00115959 | BIVM | basic, immunoglobulin-like variable motif containing | protein_coding | NP_001153068.1;NP_060163.2 | 16034 | . | . | . | [illegible] |
| 54842 | NM_00137598 | MFSD6 | major facilitator superfamily domain containing 6 | protein_coding | NP_001362915.1;NP_001362916.1;NP_001362917.1;NP_001362918.1;NI | 24711 | 613476 | . | . | [illegible] |
| 54843 | NM_00116295 | SYTL2 | synaptotagmin like 2 | protein_coding | NP_001156423.1;NP_001156424.1;NP_001156425.1;NP_001276537.1;NI | 15585 | 612880 | . | . | [illegible] |
| 54845 | NM_00103491 | ESRP1 | epithelial splicing regulatory protein 1 | protein_coding | NP_001030087.2;NP_001116297.1;NP_001116298.1;NP_001116299.1;NI | 25966 | 612959 | . | . | [illegible] |
| 54849 | NM_00124281 | DEF8 | differentially expressed in FDCP 8 homolog | protein_coding | NP_001229745.1;NP_001229746.1;NP_001229747.1;NP_001229748.1;NI | 25969 | . | . | . | [illegible] |
| 54850 | NM_00131693 | FBXL12 | F-box and leucine rich repeat protein 12 | protein_coding | NP_001303865.1;NP_001303866.1;NP_001303867.1;NP_001303868.1;NI | 13611 | 609079 | . | . | [illegible] |
| 54851 | NM_017704,XI | ANKRD49 | ankyrin repeat domain 49 | protein_coding | NP_060174.2;XP_016873430.1 | 25970 | . | . | . | [illegible] |
| 54853 | NM_017706,XI | WDR55 | WD repeat domain 55 | protein_coding | NP_060176.2;XP_005268526.1;XP_016865089.1 | 25971 | . | . | . | [illegible] |
| 54855 | NM_017709 | TENT5C | terminal nucleotidyltransferase 5C | protein_coding | NP_060179.2 | 24712 | 613952 | . | . | [illegible] |
| 54856 | NM_00128285 | GON4L | gon-4 like | protein_coding | NP_001269785.1;NP_001269787.1;NP_001269789.1;NP_001269790.1;NI | 25973 | 610393 | . | . | [illegible] |
| 54858 | NM_00130092 | PGPEP1 | pyroglutamyl-peptidase I | protein_coding | NP_001287856.1;NP_001295295.1;NP_001316400.1;NP_001316405.1;NI | 13568 | 610694 | . | . | [illegible] |
| 54859 | NM_00103170 | ELP6 | elongator acetyltransferase complex subunit 6 | protein_coding | NP_001026873.2;NP_001350886.1;XP_005265298.1;XP_011532153.1;XP | 25976 | 615020 | . | . | [illegible] |
| 54861 | NM_00110059 | SNRK | SNF related kinase | protein_coding | NP_001094064.1;NP_001317679.1;NP_060189.3;XP_005265302.1 | 30598 | 612760 | . | . | [illegible] |
| 54862 | NM_017721,XI | CC2D1A | coiled-coil and C2 domain containing 1A | protein_coding | NP_060191.3;XP_005260030.1;XP_005260031.1;XP_024307330.1;XP_02 | 30237 | 610055 | . | . | [illegible] |
| 54863 | NM_017723 | TOR4A | torsin family 4 member A | protein_coding | NP_060193.2 | 25981 | . | . | . | [illegible] |
| 54865 | NM_015590,N | GPATCH4 | G-patch domain containing 4 | protein_coding | NP_056405.2;NP_872620.1;XP_005245344.1 | 25982 | . | . | . | [illegible] |
| 54867 | NM_00108359 | TMEM214 | transmembrane protein 214 | protein_coding | NP_001077059.1;NP_060197.4;XP_005264438.1;XP_005264439.1;XP_00 | 25983 | 615301 | . | . | [illegible] |
| 54868 | NM_00132126 | TMEM104 | transmembrane protein 104 | protein_coding | NP_001308193.1;NP_060198.3;XP_006722026.1;XP_016880286.1;XP_01 | 25984 | . | . | . | [illegible] |
| 54869 | NM_017729,N | EPS8L1 | EPS8 like 1 | protein_coding | NP_060199.3;NP_573441.2;XP_005259077.1;XP_011525352.1;XP_01152 | 21295 | 614987 | . | . | [illegible] |
| 54870 | NM_00132058 | QRICH1 | glutamine rich 1 | protein_coding | NP_001307509.1;NP_001307510.1;NP_001307511.1;NP_001307512.1;NI | 24713 | 617387 | . | . | [illegible] |
| 54872 | NM_00112717 | PIGG | phosphatidylinositol glycan anchor biosynthesis class G | protein_coding | NP_001120650.1;NP_001275980.1;NP_001275981.1;NP_001275982.1;NI | 25985 | 616918 | . | . | [illegible] |
| 54874 | NM_00102494 | FNBP1L | formin binding protein 1 like | protein_coding | NP_001020119.1;NP_001157945.1;NP_060207.2;XP_011539927.1;XP_01 | 20851 | 608848 | . | . | [illegible] |
| 54875 | NM_00111439 | CNTLN | centlein | protein_coding | NP_001107867.1;NP_001273913.1;NP_001273914.1;NP_001351958.1;NI | 23432 | 611870 | . | . | [illegible] |
| 54876 | NM_00134588 | DCAF16 | DDB1 and CUL4 associated factor 16 | protein_coding | NP_001332809.1;NP_001332810.1;NP_001332811.1;NP_001332813.1;NI | 25987 | . | . | . | [illegible] |
| 54877 | NM_017742,N | ZCCHC2 | zinc finger CCHC-type containing 2 | protein_coding | NP_060212.4;XP_006722556.1;XP_011524346.1;XP_016881291.1 | 22916 | . | . | . | [illegible] |
| 54878 | NM_00132087 | DPP8 | dipeptidyl peptidase 8 | protein_coding | NP_001307804.1;NP_001307805.1;NP_060213.2;NP_569118.1;NP_93206 | 16490 | 606819 | . | . | [illegible] |
| 54879 | NM_00130826 | ST7L | suppression of tumorigenicity 7 like | protein_coding | NP_001295193.1;NP_060214.2;NP_620055.1;NP_620056.1;NP_620057.1 | 18441 | 617640 | . | . | [illegible] |
| 54880 | NM_00112338 | BCOR | BCL6 corepressor | protein_coding | NP_001116855.1;NP_001116856.1;NP_001116857.1;NP_060215.4;XP_0C | 20893 | 300485 | . | . | [illegible] |
| 54881 | NM_00116158 | TEX10 | testis expressed 10 | protein_coding | NP_001155056.1;NP_060216.2;XP_011517100.1;XP_016870337.1 | 25988 | 616717 | . | . | [illegible] |
| 54882 | NM_00119703 | ANKHD1 | ankyrin repeat and KH domain containing 1 | protein_coding | NP_001183959.1;NP_060217.1;NP_060448.1;NP_078944.2 | 24714 | 610500 | . | . | [illegible] |
| 54883 | NM_017748,N | CWC25 | CWC25 spliceosome associated protein homolog | protein_coding | NP_060218.1 | 25989 | . | . | . | [illegible] |
| 54884 | NM_017750 | RETSAT | retinol saturase | protein_coding | NP_060220.3 | 25991 | 617597 | . | . | [illegible] |
| 54885 | NM_017752,N | TBC1D8B | TBC1 domain family member 8B | protein_coding | NP_060222.2;NP_942582.1;XP_006724732.1;XP_006724733.1 | 24715 | 301027 | . | . | [illegible] |
| 54887 | NM_017754 | UHRF1BP1 | UHRF1 binding protein 1 | protein_coding | NP_060224.3 | 21216 | . | . | . | [illegible] |
| 54888 | NM_00119345 | NSUN2 | NOP2/Sun RNA methyltransferase 2 | protein_coding | NP_001180384.1;NP_060225.4 | 25994 | 610916 | . | . | [illegible] |
| 54890 | NM_017758 | ALKBH5 | alkB homolog 5, RNA demethylase | protein_coding | NP_060228.3 | 25996 | 613303 | . | . | [illegible] |
| 54891 | NM_017759,XI | INO80D | INO80 complex subunit D | protein_coding | NP_060229.3;XP_011509671.1;XP_011509672.1;XP_011509673.1;XP_01 | 25997 | . | . | . | [illegible] |
| 54892 | NM_00128193 | NCAPG2 | non-SMC condensin II complex subunit G2 | protein_coding | NP_001268861.1;NP_001268862.1;NP_060230.5;XP_005249604.1;XP_01 | 21904 | 608532 | . | . | [illegible] |
| 54893 | NM_017762,XI | MTMR10 | myotubularin related protein 10 | protein_coding | NP_060232.2;XP_005254565.1;XP_011520039.1;XP_011520040.1;XP_01 | 25999 | . | . | . | [illegible] |
| 54894 | NM_00130554 | RNF43 | ring finger protein 43 | protein_coding | NP_001292473.1;NP_001292474.1;NP_060233.3;XP_011523257.1;XP_01 | 18505 | 612482 | . | . | [illegible] |
| 54896 | NM_00104012 | SLC66A1 | solute carrier family 66 member 1 | protein_coding | NP_001035214.1;NP_001035215.1;NP_001274460.1;NP_060235.2;XP_0C | 26001 | 614760 | . | . | [illegible] |
| 54897 | NM_00107984 | CASZ1 | castor zinc finger 1 | protein_coding | NP_001073312.1;NP_060236.3;XP_005263536.1;XP_011539937.1;XP_01 | 26002 | 609895 | . | . | [illegible] |
| 54898 | NM_017770,XI | ELOVL2 | ELOVL fatty acid elongase 2 | protein_coding | NP_060240.3;XP_011513018.1;XP_011513019.1;XP_016866474.1 | 14416 | 611814 | . | . | [illegible] |
| 54899 | NM_00128909 | PXK | PX domain containing serine/threonine kinase like | protein_coding | NP_001276024.1;NP_001276025.1;NP_001276027.1;NP_001276028.1;NI | 23326 | 611450 | . | . | [illegible] |
| 54901 | NM_017774,XI | CDKAL1 | CDK5 regulatory subunit associated protein 1 like 1 | protein_coding | NP_060244.2;XP_011513020.1;XP_011513021.1;XP_016866475.1;XP_01 | 21050 | 611259 | . | . | [illegible] |
| 54902 | NM_00127142 | TTC19 | tetratricopeptide repeat domain 19 | protein_coding | NP_001258349.1;NP_060245.3;XP_016880290.2;XP_016880291.2;XP_02 | 26006 | 613814 | . | . | [illegible] |
| 54903 | NM_00116592 | MKS1 | MKS transition zone complex subunit 1 | protein_coding | NP_001159399.1;NP_001308197.1;NP_001308198.1;NP_001317326.1;NI | 7121 | 609883 | . | . | [illegible] |
| 54904 | NM_017778,N | NSD3 | nuclear receptor binding SET domain protein 3 | protein_coding | NP_060248.2;NP_075447.1 | 12767 | 607083 | . | . | [illegible] |
| 54906 | NM_00132178 | TASOR2 | transcription activation suppressor family member 2 | protein_coding | NP_001308712.1;NP_001308713.1;NP_001308714.1;NP_060252.4;XP_0C | 23484 | . | . | . | [illegible] |
| 54908 | NM_00132963 | SPDL1 | spindle apparatus coiled-coil protein 1 | protein_coding | NP_001316568.1;NP_001316569.1;NP_001316570.1;NP_001316571.1;NI | 26010 | 616401 | . | . | [illegible] |
| 54910 | NM_017789,XI | SEMA4C | semaphorin 4C | protein_coding | NP_060259.4;XP_006712669.2;XP_011509680.1;XP_011509681.1;XP_01 | 10731 | 604462 | . | . | [illegible] |
| 54913 | NM_017793 | RPP25 | ribonuclease P and MRP subunit p25 | protein_coding | NP_060263.2 | 30361 | . | . | . | [illegible] |
| 54914 | NM_00137556 | FOCAD | focadhesin | protein_coding | NP_001362496.1;NP_001362497.1;NP_001362499.1;NP_060264.4;XP_01 | 23377 | 614606 | . | . | [illegible] |
| 54915 | NM_017798,XI | YTHDF1 | YTH N6-methyladenosine RNA binding protein 1 | protein_coding | NP_060268.2;XP_024307682.1 | 15867 | 616529 | . | . | [illegible] |
| 54916 | NM_017799,XI | TMEM260 | transmembrane protein 260 | protein_coding | NP_060269.3;XP_006720239.1;XP_006720241.1;XP_011535153.1;XP_01 | 20185 | 617449 | . | . | [illegible] |
| 54918 | NM_017801 | CMTM6 | CKLF like MARVEL transmembrane domain containing 6 | protein_coding | NP_060271.1 | 19177 | 607889 | . | . | [illegible] |
| 54919 | NM_017802,N | DNAAF5 | dynein axonemal assembly factor 5 | protein_coding | NP_060272.3;XP_024302581.1;XP_024302582.1 | 26013 | 614864 | . | . | [illegible] |
| 54920 | NM_00127176 | DUS2 | dihydrouridine synthase 2 | protein_coding | NP_001258691.1;NP_001258692.1;NP_060273.1 | 26014 | 609707 | . | . | [illegible] |
| 54921 | NM_00103969 | CHTF8 | chromosome transmission fidelity factor 8 | protein_coding | NP_001034779.1;NP_001035236.1 | 24353 | 613202 | . | . | [illegible] |
| 54922 | NM_017805 | RASIP1 | Ras interacting protein 1 | protein_coding | NP_060275.2 | 24716 | 609623 | . | . | [illegible] |
| 54923 | NM_00130565 | LIME1 | Lck interacting transmembrane adaptor 1 | protein_coding | NP_001292583.1;NP_001292584.1;NP_060276.2 | 26016 | 609809 | . | . | [illegible] |
| 54925 | NM_00128452 | ZSCAN32 | zinc finger and SCAN domain containing 32 | protein_coding | NP_001271456.1;NP_001271457.1;NP_001271458.1;NP_001311269.1;NI | 20812 | . | . | . | [illegible] |
| 54926 | NM_017811,XI | UBE2R2 | ubiquitin conjugating enzyme E2 R2 | protein_coding | NP_060281.2;XP_005251553.1;XP_011516251.1;XP_016870349.1 | 19907 | 612506 | . | . | [illegible] |
| 54927 | NM_00131717 | CHCHD3 | coiled-coil-helix-coiled-coil-helix domain containing 3 | protein_coding | NP_001304106.1;NP_001304107.1;NP_060282.1 | 21906 | 613748 | . | . | [illegible] |
| 54928 | NM_017813,XI | BPNT2 | 3'(2'), 5'-bisphosphate nucleotidase 2 | protein_coding | NP_060283.3 | 26019 | 614010 | . | . | [illegible] |
| 54929 | NM_00125676 | TMEM161A | transmembrane protein 161A | protein_coding | NP_001243695.1;NP_060284.1;XP_011526419.1;XP_016882405.1 | 26020 | . | . | . | [illegible] |
| 54930 | NM_00116626 | HAUS4 | HAUS augmin like complex subunit 4 | protein_coding | NP_001159741.1;NP_001159742.1;NP_060285.2 | 20163 | 613431 | . | . | [illegible] |
| 54931 | NM_017819 | TRMT10C | tRNA methyltransferase 10C, mitochondrial RNase P subunit | protein_coding | NP_060289.2 | 26022 | 615423 | . | . | [illegible] |
| 54932 | NM_00128682 | EXD3 | exonuclease 3'-5' domain containing 3 | protein_coding | NP_001273752.1;NP_060290.3;XP_011517112.1;XP_011517116.1;XP_01 | 26023 | . | . | . | [illegible] |
| 54934 | NM_017822 | KANSL2 | KAT8 regulatory NSL complex subunit 2 | protein_coding | NP_060292.3 | 26024 | 615488 | . | . | [illegible] |
| 54935 | NM_00131965 | DUSP23 | dual specificity phosphatase 23 | protein_coding | NP_001306587.1;NP_001306588.1;NP_060293.2 | 21480 | 618361 | . | . | [illegible] |
| 54936 | NM_017825,XI | ADPRS | ADP-ribosylserine hydrolase | protein_coding | NP_060295.1;XP_011539938.1 | 21304 | 610624 | . | . | [illegible] |
| 54938 | NM_00114590 | SARS2 | seryl-tRNA synthetase 2, mitochondrial | protein_coding | NP_001139373.1;NP_060297.1 | 17697 | 612804 | . | . | [illegible] |
| 54939 | NM_00128437 | COMMD4 | COMM domain containing 4 | protein_coding | NP_001271306.1;NP_001271307.1;NP_001271308.1;NP_001308773.1;NI | 26027 | 616701 | . | . | [illegible] |
| 54940 | NM_00107983 | OCIAD1 | OCIA domain containing 1 | protein_coding | NP_001073308.1;NP_001073309.1;NP_001073310.1;NP_001073311.2;NI | 16074 | . | . | . | [illegible] |
| 54941 | NM_017831,XI | RNF125 | ring finger protein 125 | protein_coding | NP_060301.2;XP_011524347.1;XP_011524348.1;XP_011524349.1 | 21150 | 610432 | . | . | [illegible] |
| 54942 | NM_017832,XI | ABITRAM | actin binding transcription modulator | protein_coding | NP_060302.1;XP_011517118.1 | 1364 | . | . | . | [illegible] |
| 54943 | NM_00104019 | DNAJC28 | DnaJ heat shock protein family (Hsp40) member C28 | protein_coding | NP_001035282.1;NP_001307675.1;NP_060303.2 | 1297 | . | . | . | [illegible] |
| 54944 | NR_120386 | LINC01521 | long intergenic non-protein coding RNA 1521 | lncRNA | . | 26029 | . | . | . | [illegible] |
| 54946 | NM_00100848 | SLC41A3 | solute carrier family 41 member 3 | protein_coding | NP_001008485.2;NP_001008486.2;NP_001008487.1;NP_001157947.1;NI | 31046 | 610803 | . | . | [illegible] |
| 54947 | NM_017839,XI | LPCAT2 | lysophosphatidylcholine acyltransferase 2 | protein_coding | NP_060309.2;XP_005256063.1;XP_011521471.1;XP_016878865.1 | 26032 | 612040 | . | . | [illegible] |
| 54948 | NM_017840 | MRPL16 | mitochondrial ribosomal protein L16 | protein_coding | NP_060310.1 | 14476 | 611829 | . | . | [illegible] |
| 54949 | NM_017841 | SDHAF2 | succinate dehydrogenase complex assembly factor 2 | protein_coding | NP_060311.1 | 26034 | 613019 | . | . | [illegible] |
| 54951 | NM_00132966 | COMMD8 | COMM domain containing 8 | protein_coding | NP_001316597.1;NP_060315.1;XP_016863819.1 | 26036 | 616656 | . | . | [illegible] |
| 54952 | NM_017846,N | TRNAU1AP | tRNA selenocysteine 1 associated protein 1 | protein_coding | NP_060316.1 | 30813 | . | . | . | [illegible] |
| 54953 | NM_00116424 | ODR4 | odr-4 GPCR localization factor homolog | protein_coding | NP_001157717.1;NP_001157718.1;NP_060317.3;XP_011507962.1;XP_01 | 24299 | 609335 | . | . | [illegible] |
| 54954 | NM_00130078 | FAM120C | family with sequence similarity 120C | protein_coding | NP_001287717.1;NP_060318.4;NP_940858.2;XP_006724652.1 | 16949 | 300741 | . | . | [illegible] |
| 54955 | NM_00130303 | C1orf109 | chromosome 1 open reading frame 109 | protein_coding | NP_001289959.1;NP_001289960.1;NP_001337684.1;NP_001337685.1;NI | 26039 | 614799 | . | . | [illegible] |
| 54956 | NM_00131694 | PARP16 | poly(ADP-ribose) polymerase family member 16 | protein_coding | NP_001303872.1;NP_001303873.1;NP_060321.3;XP_006720652.1;XP_00 | 26040 | . | . | . | [illegible] |
| 54957 | NM_00114231 | TXNL4B | thioredoxin like 4B | protein_coding | NP_001135789.1;NP_001135790.1;NP_001311283.1;NP_001311284.1;NI | 26041 | 617722 | . | . | [illegible] |
| 54958 | NM_017854 | TMEM160 | transmembrane protein 160 | protein_coding | NP_060324.1 | 26042 | . | . | . | [illegible] |
| 54960 | NM_00104247 | GEMIN8 | gem nuclear organelle associated protein 8 | protein_coding | NP_001035944.1;NP_001035945.1;NP_060326.1;XP_005274612.1;XP_01 | 26044 | 300962 | . | . | [illegible] |
| 54961 | NM_017857,XI | SSH3 | slingshot protein phosphatase 3 | protein_coding | NP_060327.3;XP_016873431.1 | 30581 | 606780 | . | . | [illegible] |
| 54962 | NM_00128998 | TIPIN | TIMELESS interacting protein | protein_coding | NP_001276915.1;NP_060328.3;XP_005254578.2;XP_005254579.1;XP_00 | 30750 | 610716 | . | . | [illegible] |
| 54963 | NM_00119337 | UCKL1 | uridine-cytidine kinase 1 like 1 | protein_coding | NP_001180308.1;NP_001340404.1;NP_001340405.1;NP_001340406.1;NI | 15938 | 610866 | . | . | [illegible] |

| | | | | | | | | | | | | | | | | | | | | | | | | | | | | | | | | | |
|---|---|---|---|---|---|---|---|---|---|---|---|---|---|---|---|---|---|---|---|---|---|---|---|---|---|---|---|---|---|---|---|---|---|
| 54964 | NM_017860 | C1orf56 | chromosome 1 open reading frame 56 | protein_coding | NP_060330.2 | 26045 | . | . | . | 1.844219 | 1.920799 | 2.530224 | 1.853184 | 4.291574 | 4.440012 | 5.677750 | 4.237084 | 1.149699 | 132.651367 | 0.294917502 | 0.682415848 | 0.494976038 | 0.998865404 | 6.946570 | 7.149305 | 7.014667 | 7.025004 | 7.127919 | 7.009691 | 140 | 135 | 143 | 112 |
| 54965 | NM_00116630 | PIGX | phosphatidylinositol glycan anchor biosynthesis class X | protein_coding | NP_001159776.1;NP_060331.3 | 26046 | 610276 | . | . | 7.832862 | 6.871747 | 6.666574 | 6.965469 | 18.227392 | 15.884350 | 14.959597 | 15.925716 | -1.093733 | 672.707150 | 0.157700825 | -0.81966049 | 0.412409681 | 0.998865404 | 9.456911 | 9.327874 | 9.469267 | 9.375006 | 9.351760 | 9.372674 | 864 | 707 | 550 | 612 |
| 54968 | NM_00104061 | TMEM70 | transmembrane protein 70 | protein_coding | NP_001035703.1;NP_060336.3 | 26050 | 612418 | . | . | 8.864304 | 8.577422 | 8.705281 | 6.374686 | 20.627601 | 19.827096 | 19.534400 | 14.574962 | -1.165066 | 527.615071 | 0.187222867 | -1.17726659 | 0.239089118 | 0.992733415 | 9.148781 | 8.929585 | 9.102012 | 9.072491 | 9.081682 | 8.887818 | 668 | 599 | 490 | 386 |
| 54969 | NM_017867 | HPF1 | histone PARylation factor 1 | protein_coding | NP_060337.2 | 26051 | 616614 | . | . | 8.118122 | 8.146192 | 7.190289 | 5.013661 | 18.891205 | 18.830286 | 16.134800 | 11.463140 | -1.351536 | 270.395187 | 0.232741731 | -1.8673042 | 0.061859119 | 0.723723143 | 8.279154 | 7.846360 | 8.148939 | 8.144438 | 8.075572 | 7.901135 | 359 | 334 | 237 | 176 |
| 54970 | NM_00131853 | TTC12 | tetratricopeptide repeat domain 12 | protein_coding | NP_001305462.1;NP_001338966.1;NP_001364992.1;NP_001364993.1;NI | 23700 | 610732 | . | . | 3.084080 | 3.858126 | 4.652557 | 4.092205 | 7.176780 | 8.918231 | 10.440202 | 9.356340 | 1.136243 | 323.540810 | 0.201723528 | 0.913487778 | 0.360986059 | 0.998865404 | 8.242392 | 8.427296 | 8.286377 | 8.310581 | 8.418063 | 8.317951 | 341 | 334 | 326 | 294 |
| 54971 | NM_00117353 | BANP | BTG3 associated nuclear protein | protein_coding | NP_001167010.1;NP_001167011.1;NP_001167012.1;NP_001167013.1;NI | 13450 | 611564 | . | . | 4.892332 | 5.197192 | 6.322932 | 5.500255 | 11.384650 | 12.013545 | 14.188477 | 12.575674 | 1.141043 | 410.713247 | 0.18400252 | 1.034513957 | 0.300895955 | 0.998865404 | 8.583421 | 8.774248 | 8.625665 | 8.651691 | 8.762046 | 8.671674 | 432 | 423 | 409 | 380 |
| 54972 | NM_017870,N | TMEM132A | transmembrane protein 132A | protein_coding | NP_060340.2;NP_821174.1;XP_016873440.1;XP_016873441.1 | 31092 | 617363 | . | . | 3.923252 | 3.440921 | 3.832032 | 3.308655 | 9.129568 | 7.953843 | 8.598970 | 7.564846 | -1.146254 | 467.376443 | 0.18302411 | -1.07596132 | 0.281944554 | 0.998865404 | 8.963040 | 8.767202 | 8.957341 | 8.855771 | 8.870009 | 8.774437 | 620 | 496 | 409 | 376 |
| 54973 | NM_00125645 | INTS11 | integrator complex subunit 11 | protein_coding | NP_001243385.1;NP_001243389.1;NP_001243391.1;NP_001243392.1;NI | 26052 | 611354 | . | . | 31.837967 | 33.807933 | 31.298123 | 32.840243 | 74.088263 | 78.148553 | 70.232086 | 75.085310 | -1.040079 | 2147.507448 | 0.117775708 | -0.48137084 | 0.630252961 | 0.998865404 | 11.096586 | 11.039749 | 11.066386 | 11.101165 | 11.037340 | 11.067369 | 2467 | 2414 | 1796 | 2011 |
| 54974 | NM_00131782 | THG1L | tRNA-histidine guanylyltransferase 1 like | protein_coding | NP_001304753.1;NP_001304754.1;NP_001304755.1;NP_060342.2;XP_01 | 26053 | 618802 | . | . | 5.501860 | 5.304311 | 5.526482 | 5.173616 | 12.803051 | 12.261151 | 12.401263 | 11.828861 | -1.022820 | 484.718463 | 0.170288406 | -0.19116065 | 0.84839973 | 0.998865404 | 8.936959 | 8.904869 | 8.942013 | 8.913476 | 8.936154 | 8.890878 | 577 | 517 | 432 | 433 |
| 54976 | NM_00103914 | C20orf27 | chromosome 20 open reading frame 27 | protein_coding | NP_001034229.1;NP_001245358.1;NP_001245359.1;XP_011527568.1 | 15873 | . | . | . | 23.246544 | 25.997492 | 23.422275 | 26.653318 | 54.095664 | 60.094367 | 52.558906 | 60.939643 | -1.007556 | 990.825691 | 0.146515555 | -0.07412428 | 0.940911493 | 0.998865404 | 9.958219 | 9.946733 | 9.914172 | 9.993153 | 9.897337 | 9.997693 | 1084 | 1131 | 799 | 989 |
| 54977 | NM_00135479 | SLC25A38 | solute carrier family 25 member 38 | protein_coding | NP_001341727.1;NP_060345.2;XP_006713277.1;XP_011532171.1;XP_02 | 26054 | 610819 | . | . | 10.274476 | 10.543408 | 11.463110 | 10.468407 | 23.909127 | 24.371562 | 25.722888 | 23.934769 | 1.035915 | 702.160273 | 0.1527385 | 0.33328176 | 0.738921606 | 0.998865404 | 9.429816 | 9.481043 | 9.437399 | 9.448129 | 9.500335 | 9.432759 | 788 | 750 | 652 | 637 |
| 54978 | NM_017877 | SLC35F6 | solute carrier family 35 member F6 | protein_coding | NP_060347.2 | 26055 | . | . | . | 17.692350 | 18.290644 | 15.169074 | 17.823269 | 41.170826 | 42.279644 | 34.038963 | 40.750786 | -1.106054 | 2096.776246 | 0.12643459 | -1.15017127 | 0.250073336 | 0.993339639 | 11.104990 | 10.959245 | 11.062316 | 11.082200 | 10.930831 | 11.049306 | 2505 | 2406 | 1601 | 2007 |
| 54980 | NM_00134875 | C2orf42 | chromosome 2 open reading frame 42 | protein_coding | NP_001335687.1;NP_001335688.1;NP_001335689.1;NP_001335690.1;NI | 26056 | . | . | . | 3.097828 | 3.261246 | 3.344497 | 4.011898 | 7.208777 | 7.538520 | 7.504953 | 9.172727 | 1.144940 | 284.681947 | 0.211820583 | 0.921871674 | 0.356595516 | 0.998865404 | 8.052796 | 8.247083 | 8.105772 | 8.122081 | 8.132347 | 8.235300 | 301 | 291 | 239 | 312 |
| 54981 | NM_00112760 | NMRK1 | nicotinamide riboside kinase 1 | protein_coding | NP_001121075.1;NP_001317607.1;NP_001317608.1;NP_060351.1;XP_0C | 26057 | 608704 | . | . | 4.303678 | 5.228213 | 5.386449 | 4.670161 | 10.014838 | 12.085250 | 12.087036 | 10.677773 | 1.020453 | 471.358382 | 0.183841674 | 0.158882142 | 0.873761731 | 0.998865404 | 8.865700 | 8.895507 | 8.843485 | 8.899849 | 8.951321 | 8.813578 | 513 | 526 | 460 | 397 |
| 54982 | NM_017882 | CLN6 | CLN6 transmembrane ER protein | protein_coding | NP_060352.1 | 2077 | 606725 | . | . | 18.666235 | 19.624956 | 18.519033 | 17.174757 | 43.437096 | 45.363968 | 41.556175 | 39.268040 | -1.087233 | 1284.634822 | 0.131614877 | -0.91677508 | 0.359260515 | 0.998865404 | 10.386132 | 10.265638 | 10.341731 | 10.373619 | 10.325753 | 10.262040 | 1509 | 1474 | 1116 | 1105 |
| 54984 | NM_00128435 | PINX1 | PIN2 (TERF1) interacting telomerase inhibitor 1 | protein_coding | NP_001271285.1;NP_060354.4 | 30046 | 606505 | . | . | 9.370010 | 10.515502 | 10.183204 | 11.977344 | 21.804397 | 24.307056 | 22.850813 | 27.384774 | 1.100788 | 504.584734 | 0.174902968 | 0.792078886 | 0.428314681 | 0.998865404 | 8.908492 | 9.046132 | 8.911996 | 8.978855 | 8.957090 | 9.054278 | 524 | 546 | 425 | 533 |
| 54985 | NM_00100201 | HCFC1R1 | host cell factor C1 regulator 1 | protein_coding | NP_001002017.1;NP_001002018.1;NP_001275594.1;NP_001275595.1;NI | 21198 | 618818 | . | . | 31.436714 | 29.146851 | 26.223534 | 31.076307 | 73.154519 | 67.374259 | 58.844855 | 71.052285 | -1.024090 | 689.628782 | 0.157200393 | -0.21846466 | 0.827067092 | 0.998865404 | 9.446838 | 9.412308 | 9.469026 | 9.404636 | 9.380783 | 9.459705 | 841 | 718 | 558 | 676 |
| 54986 | NM_00132250 | ULK4 | unc-51 like kinase 4 | protein_coding | NP_001309429.1;NP_001309430.1;NP_060356.2;XP_011532182.1;XP_02 | 15784 | 617010 | . | . | 0.617743 | 0.589496 | 0.665214 | 0.749645 | 1.437514 | 1.362647 | 1.492720 | 1.713975 | 1.152909 | 88.693019 | 0.338702131 | 0.606075229 | 0.544464799 | 0.998865404 | 6.364637 | 6.569588 | 6.456595 | 6.438944 | 6.471412 | 6.504002 | 98 | 86 | 78 | 94 |
| 54987 | NM_00130475 | CZIB | CXXC motif containing zinc binding protein | protein_coding | NP_001291688.1;NP_001291689.1;NP_060357.1 | 26059 | . | . | . | 18.457744 | 19.483287 | 17.187851 | 19.996273 | 42.951932 | 45.036494 | 38.569042 | 45.719100 | -1.045365 | 607.086834 | 0.166550686 | -0.38430648 | 0.700751303 | 0.998865404 | 9.277821 | 9.212968 | 9.235961 | 9.282704 | 9.169659 | 9.286749 | 690 | 693 | 470 | 606 |
| 54989 | NM_014106,XI | ZNF770 | zinc finger protein 770 | protein_coding | NP_054825.2;XP_011520046.1 | 26061 | . | . | . | 4.524586 | 4.132361 | 5.020111 | 2.802740 | 10.528897 | 9.552137 | 11.264984 | 6.408133 | -1.141047 | 693.601549 | 0.218463191 | -0.87135329 | 0.383561284 | 0.998865404 | 9.529422 | 9.340312 | 9.505839 | 9.435831 | 9.553337 | 9.173069 | 893 | 758 | 722 | 437 |
| 54991 | NM_00133030 | C1orf159 | chromosome 1 open reading frame 159 | protein_coding | NP_001317235.1;NP_001350454.1;NP_060361.4 | 26062 | . | . | . | 7.283744 | 6.503475 | 9.604312 | 8.153026 | 16.949572 | 15.033075 | 21.551798 | 18.640927 | 1.236666 | 499.276746 | 0.178414967 | 1.717659997 | 0.08585865 | 0.792183786 | 8.801876 | 9.109182 | 8.897813 | 8.878824 | 9.089889 | 8.951683 | 522 | 474 | 531 | 463 |
| 54993 | NM_00100707 | ZSCAN2 | zinc finger and SCAN domain containing 2 | protein_coding | NP_001007073.1;NP_060364.4;NP_870992.2;XP_016877882.1;XP_02430 | 20994 | . | . | . | 2.164487 | 2.716239 | 2.746770 | 3.476327 | 5.036851 | 6.278710 | 6.163671 | 7.948210 | 1.193356 | 170.829387 | 0.258160597 | 0.987852848 | 0.323224721 | 0.998865404 | 7.283573 | 7.538045 | 7.391477 | 7.357782 | 7.408374 | 7.487037 | 187 | 161 | 147 | 190 |
| 54994 | NM_017896 | GID8 | GID complex subunit 8 homolog | protein_coding | NP_060366.1 | 15857 | 611625 | . | . | 11.200001 | 11.219130 | 10.867697 | 12.487904 | 26.062860 | 25.933519 | 24.386797 | 28.552107 | 1.026979 | 1557.707472 | 0.129714243 | 0.296084425 | 0.767165608 | 0.998865404 | 10.586024 | 10.624141 | 10.596934 | 10.591678 | 10.563572 | 10.663738 | 1776 | 1653 | 1284 | 1575 |
| 54995 | NM_00114539 | OXSM | 3-oxoacyl-ACP synthase, mitochondrial | protein_coding | NP_001138863.1;NP_060367.1;XP_006713279.1;XP_006713280.1;XP_01 | 26063 | 610324 | . | . | 5.922301 | 5.761164 | 6.111825 | 6.083449 | 13.781439 | 13.317188 | 13.714758 | 13.909082 | 1.140222 | 341.290538 | 0.194273841 | 0.974471482 | 0.329822492 | 0.998865404 | 8.316705 | 8.506754 | 8.393652 | 8.355231 | 8.478268 | 8.417975 | 380 | 332 | 333 | 323 |
| 54998 | NM_00112722 | AURKAIP1 | aurora kinase A interacting protein 1 | protein_coding | NP_001120701.1;NP_001120702.1;NP_060370.1;XP_011539972.1;XP_02 | 24114 | 609183 | . | . | 110.764111 | 106.384310 | 98.941407 | 117.314033 | 257.752641 | 245.912096 | 222.021652 | 268.224596 | 1.013523 | 3046.455489 | 0.119703819 | 0.16188739 | 0.871394533 | 0.998865404 | 11.563272 | 11.582496 | 11.581854 | 11.551588 | 11.520758 | 11.631773 | 3549 | 3205 | 2486 | 3070 |
| 55000 | NR_002323,NF | TUG1 | taurine up-regulated 1 | lncRNA | . | 26066 | 614971 | . | . | 13.530943 | 13.684392 | 16.208567 | 12.248019 | 31.487061 | 31.632087 | 36.371553 | 28.003642 | 1.034970 | 3272.273109 | 0.140667637 | 0.352525947 | 0.724443863 | 0.998865404 | 11.650934 | 11.700790 | 11.656195 | 11.661915 | 11.799343 | 11.561818 | 3690 | 3482 | 3310 | 2673 |
| 55002 | NM_00134974 | TMCO3 | transmembrane and coiled-coil domains 3 | protein_coding | NP_001336670.1;NP_001336671.1;NP_001336672.1;NP_001336673.1;NI | 20329 | 617134 | . | . | 14.851029 | 14.827148 | 13.244043 | 14.584577 | 34.558953 | 34.273621 | 29.719251 | 33.345909 | -1.073893 | 1352.535352 | 0.130165488 | -0.79015222 | 0.429438878 | 0.998865404 | 10.452079 | 10.348981 | 10.430859 | 10.425106 | 10.337555 | 10.408152 | 1619 | 1506 | 1081 | 1280 |
| 55003 | NM_017906,XI | PAK1IP1 | PAK1 interacting protein 1 | protein_coding | NP_060376.2;XP_005249261.1;XP_011513022.1;XP_011513023.1 | 20882 | 607811 | . | . | 15.016955 | 15.162065 | 14.704844 | 15.434656 | 34.945071 | 35.047792 | 32.997239 | 35.289503 | -1.020432 | 730.783006 | 0.148560401 | -0.19641908 | 0.844282164 | 0.998865404 | 9.527914 | 9.498535 | 9.512035 | 9.528552 | 9.494192 | 9.518187 | 840 | 806 | 618 | 690 |
| 55004 | NM_017907 | LAMTOR1 | late endosomal/lysosomal adaptor, MAPK and MTOR activator 1 | protein_coding | NP_060377.1 | 26068 | 613510 | . | . | 47.387344 | 51.384083 | 51.299812 | 56.295364 | 110.272293 | 118.776604 | 115.115295 | 128.712662 | 1.074048 | 1749.976160 | 0.125469443 | 0.821384488 | 0.411427296 | 0.998865404 | 10.720825 | 10.823584 | 10.715270 | 10.771821 | 10.766817 | 10.831926 | 1873 | 1887 | 1511 | 1769 |
| 55005 | NM_00127193 | RMND1 | required for meiotic nuclear division 1 homolog | protein_coding | NP_001258866.1;NP_060379.2;XP_005267097.1;XP_016866477.1 | 21176 | 614917 | . | . | 8.678061 | 8.621877 | 9.512125 | 8.377509 | 20.194204 | 19.929860 | 21.344935 | 19.154182 | 1.043183 | 526.321331 | 0.168327927 | 0.362345302 | 0.717094009 | 0.998865404 | 9.008706 | 9.070238 | 9.025969 | 9.024140 | 9.093509 | 9.009448 | 594 | 555 | 498 | 471 |
| 55006 | NM_017910,XI | TRMT61B | tRNA methyltransferase 61B | protein_coding | NP_060380.3;XP_005264450.1;XP_006712100.1;XP_016859890.1;XP_01 | 26070 | . | . | . | 9.947855 | 8.329581 | 9.926678 | 9.350639 | 23.149064 | 19.254202 | 22.275182 | 21.379126 | 1.041623 | 539.046261 | 0.172581211 | 0.340903905 | 0.733175925 | 0.998865404 | 9.044163 | 9.103752 | 9.116449 | 8.999131 | 9.106817 | 9.064764 | 663 | 518 | 494 | 499 |
| 55007 | NM_00110459 | FAM118A | family with sequence similarity 118 member A | protein_coding | NP_001098065.1;NP_001336842.1;NP_001336843.1;NP_001336844.1;NI | 1313 | . | . | . | 16.715278 | 19.377492 | 24.926826 | 17.076090 | 38.897141 | 44.791937 | 55.935072 | 39.042445 | 1.121481 | 2709.295039 | 0.143028218 | 1.156450317 | 0.247497011 | 0.992733415 | 11.318492 | 11.484159 | 11.332132 | 11.372036 | 11.556482 | 11.321363 | 2868 | 2824 | 2846 | 2303 |
| 55008 | NM_00116513 | HERC6 | HECT and RLD domain containing E3 ubiquitin protein ligase family | protein_coding | NP_001158608.1;NP_060382.3;XP_005263140.1;XP_011530355.1;XP_01 | 26072 | 609249 | . | . | 0.707070 | 0.541393 | 0.655809 | 0.715746 | 1.645382 | 1.251454 | 1.471615 | 1.636467 | 1.025213 | 81.877222 | 0.356160261 | 0.100862848 | 0.919659336 | 0.998865404 | 6.336973 | 6.373575 | 6.382205 | 6.319075 | 6.350034 | 6.365467 | 104 | 77 | 70 | 80 |
| 55009 | NM_017914 | FAM174C | family with sequence similarity 174 member C | protein_coding | NP_060384.3 | 26073 | . | . | . | 32.672649 | 34.753609 | 36.865398 | 39.094574 | 76.030602 | 80.334526 | 82.724884 | 89.385101 | 1.111070 | 971.789165 | 0.13875075 | 1.095126539 | 0.273461203 | 0.998865404 | 9.846675 | 9.998340 | 9.864794 | 9.903309 | 9.942955 | 9.978582 | 1032 | 1020 | 868 | 982 |
| 55010 | NM_00131998 | PARPBP | PARP1 binding protein | protein_coding | NP_001306917.1;NP_001306922.1;NP_001306923.1;NP_001306924.1;NI | 26074 | 613687 | . | . | 6.006948 | 5.628912 | 5.466718 | 4.405890 | 13.978413 | 13.011483 | 12.267155 | 10.073542 | -1.186593 | 472.609778 | 0.184644599 | -1.33675664 | 0.181302088 | 0.952975139 | 9.002111 | 8.756455 | 8.956014 | 8.910534 | 8.896263 | 8.752795 | 611 | 534 | 421 | 357 |
| 55011 | NM_017916,XI | PIH1D1 | PIH1 domain containing 1 | protein_coding | NP_060386.1;XP_024307336.1;XP_024307337.1;XP_024307338.1 | 26075 | 611480 | . | . | 23.994675 | 24.198271 | 27.080700 | 29.039472 | 55.836595 | 55.935386 | 60.768306 | 66.395296 | 1.160770 | 1138.841182 | 0.13711356 | 1.568642183 | 0.116731338 | 0.857396579 | 10.042029 | 10.256695 | 10.058792 | 10.127928 | 10.175510 | 10.234001 | 1157 | 1191 | 1023 | 1191 |
| 55012 | NM_00130515 | PPP2R3C | protein phosphatase 2 regulatory subunit B''gamma | protein_coding | NP_001292084.1;NP_001292085.1;NP_060387.2;XP_005267839.1;XP_01 | 17485 | 615902 | . | . | 6.595374 | 7.207663 | 7.378922 | 5.980650 | 15.347705 | 16.660838 | 16.558087 | 13.674044 | -1.055232 | 419.394465 | 0.187587096 | -0.41346428 | 0.679266491 | 0.998865404 | 8.750156 | 8.673144 | 8.698032 | 8.754531 | 8.749563 | 8.636010 | 473 | 486 | 389 | 346 |
| 55013 | NM_017918,XI | MCUB | mitochondrial calcium uniporter dominant negative subunit beta | protein_coding | NP_060388.2;XP_006714309.1 | 26076 | . | . | . | 3.427994 | 3.096196 | 2.401711 | 2.923497 | 7.977084 | 7.156996 | 5.389371 | 6.684230 | -1.246066 | 206.889566 | 0.240625109 | -1.31898431 | 0.187174358 | 0.959720529 | 7.843016 | 7.524919 | 7.765742 | 7.714788 | 7.599352 | 7.678954 | 278 | 235 | 145 | 189 |
| 55014 | NM_017919,XI | STX17 | syntaxin 17 | protein_coding | NP_060389.2;XP_011517122.1;XP_011517123.1;XP_011517125.1;XP_01 | 11432 | 604204 | . | . | 2.160846 | 2.183334 | 2.176742 | 1.823890 | 5.028376 | 5.046873 | 4.884547 | 4.170109 | -1.092755 | 444.131640 | 0.18558557 | -0.68954415 | 0.490480899 | 0.998865404 | 8.856976 | 8.729915 | 8.815296 | 8.824398 | 8.831175 | 8.695615 | 530 | 504 | 411 | 353 |
| 55015 | NM_017922 | PRPF39 | pre-mRNA processing factor 39 | protein_coding | NP_060392.3 | 20314 | 614907 | . | . | 10.038738 | 9.352677 | 11.968270 | 7.204311 | 23.360558 | 21.619133 | 26.856451 | 16.471804 | -1.025226 | 1062.124866 | 0.196867196 | -0.18257144 | 0.855134297 | 0.998865404 | 10.070174 | 10.035088 | 10.082612 | 10.023279 | 10.206027 | 9.829608 | 1287 | 1114 | 1143 | 735 |
| 55017 | NM_017924,XI | C14orf119 | chromosome 14 open reading frame 119 | protein_coding | NP_060394.1;XP_016876879.1 | 20270 | . | . | . | 15.030745 | 13.699246 | 15.633813 | 14.946081 | 34.977163 | 31.666423 | 35.081826 | 34.172439 | 1.027631 | 1250.286180 | 0.130414313 | 0.301515317 | 0.763021572 | 0.998865404 | 10.268146 | 10.307669 | 10.297463 | 10.256967 | 10.309596 | 10.286462 | 1458 | 1295 | 1119 | 1170 |
| 55020 | NM_017931,XI | TTC38 | tetratricopeptide repeat domain 38 | protein_coding | NP_060401.3;XP_011528561.1;XP_011528562.1 | 26082 | . | . | . | 8.459341 | 9.233518 | 8.815018 | 10.054460 | 19.685234 | 21.343695 | 19.780646 | 22.988328 | 1.047617 | 753.165043 | 0.151921937 | 0.441750118 | 0.658670038 | 0.998865404 | 9.523163 | 9.589714 | 9.510340 | 9.568320 | 9.536949 | 9.605896 | 812 | 827 | 637 | 758 |
| 55022 | NM_00110081 | PID1 | phosphotyrosine interaction domain containing 1 | protein_coding | NP_001094288.1;NP_001317085.1;NP_001317086.1;NP_001317087.1;NI | 26084 | 612930 | . | . | 1.839668 | 1.850718 | 1.075112 | 0.820511 | 4.280984 | 4.278015 | 2.412519 | 1.876002 | -1.939902 | 115.200803 | 0.312834512 | -3.05587779 | 0.002244027 | 0.120055447 | 7.247477 | 6.293683 | 6.940258 | 6.934513 | 6.758726 | 6.685334 | 176 | 163 | 78 | 63 |
| 55023 | NM_017934,XI | PHIP | pleckstrin homology domain interacting protein | protein_coding | NP_060404.4;XP_005248786.1;XP_011534220.1;XP_011534221.1;XP_01 | 15673 | 612870 | . | . | 2.825127 | 2.668842 | 2.893066 | 2.049482 | 6.574188 | 6.169146 | 6.491954 | 4.685898 | -1.111997 | 915.624876 | 0.16815563 | -0.9107775 | 0.362412618 | 0.998865404 | 9.912706 | 9.760508 | 9.924455 | 9.817814 | 9.892529 | 9.689398 | 1190 | 965 | 862 | 697 |
| 55027 | NM_00132972 | HEATR3 | HEAT repeat containing 3 | protein_coding | NP_001316658.1;NP_001316659.1;NP_001316660.1;NP_891552.1;XP_0C | 26087 | 614951 | . | . | 3.363261 | 3.530861 | 3.279910 | 3.356437 | 7.826447 | 8.161743 | 7.360025 | 7.674096 | -1.081580 | 461.422247 | 0.172768127 | -0.65487213 | 0.512550088 | 0.998865404 | 8.905610 | 8.792043 | 8.861019 | 8.886678 | 8.805622 | 8.844550 | 541 | 528 | 373 | 429 |
| 55028 | NM_00110062 | C17orf80 | chromosome 17 open reading frame 80 | protein_coding | NP_001094091.2;NP_001094092.2;NP_001275699.2;NP_001275700.2;NI | 29601 | . | . | . | 7.655552 | 8.236542 | 7.586409 | 8.679793 | 17.814780 | 19.039134 | 17.023682 | 19.845313 | 1.008221 | 898.769044 | 0.145159025 | 0.081372265 | 0.935145905 | 0.998865404 | 9.806143 | 9.817449 | 9.787274 | 9.829165 | 9.772561 | 9.854178 | 1001 | 994 | 740 | 894 |
| 55030 | NM_017943,N | FBXO34 | F-box protein 34 | protein_coding | NP_060413.2;NP_689417.1;XP_005267848.1;XP_006720248.1;XP_01687 | 20201 | 609104 | . | . | 5.175869 | 5.978392 | 4.836929 | 5.334184 | 12.044461 | 13.819323 | 10.853927 | 12.195978 | -1.119623 | 559.928661 | 0.167176918 | -0.9750955 | 0.329512892 | 0.998865404 | 9.208631 | 9.044925 | 9.120946 | 9.206058 | 9.063609 | 9.116732 | 638 | 679 | 440 | 516 |
| 55031 | NM_00128265 | USP47 | ubiquitin specific peptidase 47 | protein_coding | NP_001269588.1;NP_001317137.1;NP_001359020.1;NP_001359021.1;NI | 20076 | 614460 | . | . | 4.575239 | 4.028467 | 4.137114 | 3.971958 | 10.646767 | 9.311982 | 9.283566 | 9.081410 | -1.023460 | 1263.176114 | 0.131629702 | -0.25415342 | 0.799377037 | 0.998865404 | 10.319324 | 10.286166 | 10.336270 | 10.285726 | 10.315241 | 10.272008 | 1520 | 1333 | 1117 | 1137 |
| 55032 | NM_00134890 | SLC35A5 | solute carrier family 35 member A5 | protein_coding | NP_001335834.1;NP_001335835.1;NP_001335836.1;NP_001335837.1;NI | 20792 | . | . | . | 5.590988 | 4.657401 | 4.880036 | 4.293545 | 13.010459 | 10.765791 | 10.950660 | 9.816682 | -1.129854 | 660.311622 | 0.168068983 | -1.04800223 | 0.294637582 | 0.998865404 | 9.451961 | 9.276728 | 9.469433 | 9.336952 | 9.366691 | 9.276776 | 883 | 684 | 576 | 542 |
| 55033 | NM_017946,N | FKBP14 | FKBP prolyl isomerase 14 | protein_coding | NP_060416.1 | 18625 | 614505 | . | . | 13.893835 | 12.591345 | 11.913364 | 11.301300 | 32.331525 | 29.105457 | 26.733242 | 25.839079 | -1.156330 | 1930.964837 | 0.12508417 | -1.6752931 | 0.093876739 | 0.810751527 | 11.015949 | 10.806681 | 11.012087 | 10.924943 | 10.878416 | 10.829987 | 2513 | 2114 | 1606 | 1627 |
| 55034 | NM_017947 | MOCOS | molybdenum cofactor sulfurase | protein_coding | NP_060417.4 | 18234 | 613274 | . | . | 3.373160 | 3.661909 | 3.255057 | 3.436004 | 7.849482 | 8.464666 | 7.304254 | 7.856014 | -1.065976 | 661.511641 | 0.154728639 | -0.59571589 | 0.551365039 | 0.998865404 | 9.415154 | 9.322608 | 9.365027 | 9.416099 | 9.333299 | 9.361432 | 757 | 764 | 545 | 613 |
| 55035 | NM_00125639 | NOL8 | nucleolar protein 8 | protein_coding | NP_001243323.1;NP_001317651.1;NP_060418.4;XP_006717229.1;XP_00 | 23387 | 611534 | . | . | 7.583727 | 7.329324 | 9.297196 | 7.558398 | 17.647645 | 16.942057 | 20.862636 | 17.281376 | 1.131080 | 1130.971824 | 0.141293662 | 1.257670093 | 0.208511092 | 0.970482251 | 10.051480 | 10.229713 | 10.121978 | 10.065342 | 10.238901 | 10.130541 | 1269 | 1101 | 1119 | 1045 |
| 55036 | NM_00124334 | CCDC40 | coiled-coil domain containing 40 | protein_coding | NP_001230271.1;NP_001317437.1;NP_060420.2;XP_011523265.1;XP_01 | 26090 | 613799 | . | . | 1.245902 | 1.118414 | 1.496567 | 1.065875 | 2.899265 | 2.585262 | 3.358253 | 2.436997 | 1.119747 | 165.755200 | 0.277069001 | 0.588924279 | 0.55591207 | 0.998865404 | 7.287829 | 7.453253 | 7.393663 | 7.290624 | 7.447337 | 7.328926 | 203 | 147 | 174 | 141 |
| 55037 | NM_017952 | PTCD3 | pentatricopeptide repeat domain 3 | protein_coding | NP_060422.4 | 24717 | 614918 | . | . | 16.292555 | 16.553230 | 16.602043 | 15.518606 | 37.913445 | 38.263535 | 37.254505 | 35.481449 | -1.036665 | 3381.443242 | 0.112270009 | -0.462721 | 0.643564364 | 0.998865404 | 11.749111 | 11.697270 | 11.734817 | 11.740915 | 11.739669 | 11.675910 | 3949 | 3728 | 3000 | 2992 |
| 55038 | NM_017955,N | CDCA4 | cell division cycle associated 4 | protein_coding | NP_060425.2;NP_663747.1 | 14625 | 612270 | . | . | 9.942244 | 11.166952 | 9.202484 | 11.231747 | 23.136009 | 25.812910 | 20.650106 | 25.680056 | -1.046463 | 792.427756 | 0.158725842 | -0.41279194 | 0.679759059 | 0.998865404 | 9.662929 | 9.596579 | 9.605512 | 9.682577 | 9.545150 | 9.675851 | 883 | 922 | 610 | 794 |
| 55039 | NM_017956 | TRMT12 | tRNA methyltransferase 12 homolog | protein_coding | NP_060426.2 | 26091 | 611244 | . | . | 6.749320 | 6.088230 | 7.172541 | 6.404185 | 15.705944 | 14.073217 | 16.094975 | 14.642409 | 1.043094 | 454.545418 | 0.179167067 | 0.339733585 | 0.734057167 | 0.998865404 | 8.797082 | 8.858819 | 8.847874 | 8.778542 | 8.877879 | 8.800631 | 541 | 453 | 429 | 408 |
| 55041 | NM_00110062 | PLEKHB2 | pleckstrin homology domain containing B2 | protein_coding | NP_001094093.1;NP_001253991.1;NP_001253992.1;NP_001253993.1;NI | 19236 | 618452 | . | . | 44.181548 | 44.523370 | 40.441801 | 40.787719 | 102.812282 | 102.917765 | 90.750225 | 93.256275 | -1.099244 | 5731.421623 | 0.103915733 | -1.31367323 | 0.188956194 | 0.960264078 | 12.551322 | 12.414804 | 12.524100 | 12.522285 | 12.440274 | 12.447017 | 6919 | 6470 | 4729 | 5140 |
| 55048 | NM_017966,XI | VPS37C | VPS37C subunit of ESCRT-I | protein_coding | NP_060436.4;XP_005274134.1;XP_016873444.1 | 26097 | 610038 | . | . | 2.836041 | 2.980266 | 3.514444 | 3.082730 | 6.599583 | 6.889020 | 7.886309 | 7.048294 | 1.127836 | 261.612445 | 0.214944256 | 0.807450208 | 0.41940715 | 0.998865404 | 7.941653 | 8.115677 | 7.986548 | 8.008283 | 8.092009 | 8.024347 | 277 | 271 | 257 | 243 |
| 55049 | NM_00110041 | REX1BD | required for excision 1-B domain containing | protein_coding | NP_001093888.1;NP_001093889.1;XP_024307339.1 | 26098 | . | . | . | 17.793454 | 20.829072 | 17.877448 | 22.029382 | 41.406100 | 48.147334 | 40.116479 | 50.367562 | 1.018047 | 578.583787 | 0.166457363 | 0.155015506 | 0.876809093 | 0.998865404 | 9.163814 | 9.188842 | 9.131474 | 9.206435 | 9.141066 | 9.219873 | 623 | 654 | 477 | 580 |
| 55051 | NM_017970,XI | NRDE2 | NRDE-2, necessary for RNA interference, domain containing | protein_coding | NP_060440.2;XP_011535188.1;XP_016876883.1;XP_016876884.1;XP_01 | 20186 | 618631 | . | . | 1.290483 | 1.619653 | 1.424994 | 1.696620 | 3.003008 | 3.743900 | 3.197646 | 3.879121 | 1.073805 | 621.636009 | 0.161575012 | 0.635816218 | 0.524896216 | 0.998865404 | 9.227986 | 9.330062 | 9.214710 | 9.292919 | 9.271886 | 9.331394 | 650 | 685 | 535 | 630 |
| 55052 | NM_00131848 | MRPL20 | mitochondrial ribosomal protein L20 | protein_coding | NP_001305414.1;NP_060441.2 | 14478 | 611833 | . | . | 64.290242 | 63.150291 | 58.958279 | 70.049734 | 149.606039 | 145.974724 | 132.300675 | 160.160393 | 1.011330 | 1490.648564 | 0.132568697 | 0.122611841 | 0.902414477 | 0.998865404 | 10.533723 | 10.549682 | 10.547158 | 10.526285 | 10.487975 | 10.599734 | 1730 | 1578 | 1209 | 1507 |
| 55054 | NM_00119026 | ATG16L1 | autophagy related 16 like 1 | protein_coding | NP_001177195.1;NP_001177196.1;NP_001350671.1;NP_060444.3;NP_1 | 21498 | 610767 | . | . | 14.252391 | 15.680132 | 15.919681 | 15.454762 | 33.165899 | 36.245329 | 35.723304 | 35.335483 | 1.032429 | 1569.923767 | 0.126870588 | 0.362907147 | 0.716674248 | 0.998865404 | 10.593304 | 10.639287 | 10.568520 | 10.637428 | 10.643905 | 10.612474 | 1700 | 1741 | 1415 | 1465 |
| 55055 | NM_00128782 | ZWILCH | zwilch kinetochore protein | protein_coding | NP_001274750.1;NP_001274751.1;NP_001274752.1;NP_060445.3 | 25468 | 609984 | . | . | 10.844209 | 10.080569 | 10.156595 | 8.792811 | 25.234921 | 23.301690 | 22.791102 | 20.103716 | -1.124754 | 1101.010802 | 0.142704652 | -1.1885355 | 0.234622504 | 0.990818043 | 10.186651 | 10.017627 | 10.176501 | 10.114221 | 10.111129 | 10.003301 | 1397 | 1206 | 968 | 900 |
| 55056 | NR_026891 | GABPB1-IT1 | GABPB1 intronic transcript | lncRNA | . | 25469 | . | . | . | 0.701217 | 0.678043 | 0.740665 | 0.672123 | 1.631761 | 1.567327 | 1.662031 | 1.536730 | 1.013721 | 129.942694 | 0.285528438 | 0.068857101 | 0.945103366 | 0.998865404 | 7.011454 | 7.031937 | 7.026928 | 7.010171 | 7.042605 | 7.004037 | 152 | 136 | 120 | 116 |
| 55057 | NM_00103977 | CRYBG2 | crystallin beta-gamma domain containing 2 | protein_coding | NP_001034864.2;XP_005245975.1;XP_011539974.1;XP_011539975.1 | 17295 | . | . | . | 0.055735 | 0.039640 | 0.046251 | 0.094398 | 0.129698 | 0.091629 | 0.103785 | 0.215830 | 1.410380 | 10.737939 | 0.953853155 | 0.520084314 | 0.603004811 | NA | 3.156660 | 3.650563 | 3.416443 | 3.402712 | 3.406310 | 3.445186 | 12 | 8 | 7 | 16 |
| 55062 | NM_00132077 | WIPI1 | WD repeat domain, phosphoinositide interacting 1 | protein_coding | NP_001307701.1;NP_060453.3;XP_016880297.1 | 25471 | 609224 | . | . | 35.754185 | 35.228614 | 36.558362 | 43.207676 | 83.201463 | 81.432528 | 82.035915 | 98.789210 | 1.111984 | 2282.799461 | 0.126985134 | 1.205938394 | 0.227841245 | 0.982008277 | 11.078105 | 11.231022 | 11.125223 | 11.097426 | 11.127925 | 11.261012 | 2533 | 2292 | 1919 | 2438 |
| 55063 | NM_00125800 | ZCWPW1 | zinc finger CW-type and PWWP domain containing 1 | protein_coding | NP_001244937.1;NP_060454.3;XP_005250536.1;XP_005250537.1;XP_00 | 23486 | . | . | . | 5.509933 | 5.629345 | 5.216969 | 5.829280 | 12.821846 | 13.012477 | 11.706729 | 13.327951 | -1.043292 | 501.583925 | 0.170063921 | -0.35952581 | 0.719201771 | 0.998865404 | 9.000816 | 8.939222 | 8.989186 | 8.977975 | 8.919644 | 8.992296 | 595 | 548 | 402 | 487 |
| 55066 | NM_00132211 | PDPR | pyruvate dehydrogenase phosphatase regulatory subunit | protein_coding | NP_001309046.1;NP_001309047.1;NP_001309048.1;NP_060460.4;XP_0C | 30264 | 617835 | . | . | 11.375484 | 12.701086 | 14.658705 | 12.326825 | 26.471222 | 29.359132 | 32.893703 | 28.183822 | 1.117661 | 3485.944202 | 0.122229148 | 1.312961164 | 0.189196035 | 0.960264078 | 11.684814 | 11.845388 | 11.682870 | 11.753106 | 11.870513 | 11.746471 | 3628 | 3705 | 3438 | 3193 |
| 55068 | NM_00112761 | ENOX1 | ecto-NOX disulfide-thiol exchanger 1 | protein_coding | NP_001121087.1;NP_001229792.1;NP_001334892.1;NP_001334893.1;NI | 25474 | 610914 | . | . | 2.648410 | 2.446493 | 1.719745 | 2.391004 | 6.162959 | 5.655176 | 3.859058 | 5.466745 | -1.234660 | 235.529705 | 0.225872954 | -1.34639373 | 0.178175571 | 0.951511445 | 8.024188 | 7.719260 | 7.937839 | 7.919395 | 7.788240 | 7.862999 | 305 | 276 | 168 | 214 |
| 55069 | NM_017994,XI | TMEM248 | transmembrane protein 248 | protein_coding | NP_060464.1;XP_005250539.1;XP_024302587.1;XP_024302588.1;XP_02 | 25476 | . | . | . | 16.258959 | 16.666608 | 17.815271 | 17.965485 | 37.835271 | 38.525607 | 39.976952 | 41.075946 | 1.055046 | 2137.699977 | 0.121843916 | 0.634471754 | 0.525772995 | 0.998865404 | 11.022541 | 11.100104 | 11.054842 | 11.023876 | 11.123618 | 11.039450 | 2442 | 2201 | 2009 | 1950 |
| 55070 | NM_00114407 | DET1 | DET1 partner of COP1 E3 ubiquitin ligase | protein_coding | NP_001137546.1;NP_001308523.1;NP_001308525.1;NP_060466.2 | 25477 | 608727 | . | . | 2.836669 | 2.792213 | 2.372195 | 3.400755 | 6.601049 | 6.454326 | 5.323131 | 7.775422 | 1.029319 | 216.985999 | 0.246015183 | 0.169462779 | 0.865432647 | 0.998865404 | 7.741292 | 7.781347 | 7.743522 | 7.760381 | 7.686048 | 7.839086 | 242 | 235 | 160 | 240 |
| 55071 | NM_017998 | C9orf40 | chromosome 9 open reading frame 40 | protein_coding | NP_060468.2 | 23433 | . | . | . | 3.900439 | 3.728248 | 3.889252 | 3.806093 | 9.076482 | 8.618013 | 8.727369 | 8.702181 | -1.003897 | 281.242014 | 0.206142527 | -0.02721853 | 0.978285438 | 0.998865404 | 8.138291 | 8.133041 | 8.150492 | 8.122489 | 8.142798 | 8.126334 | 333 | 296 | 248 | 259 |
| 55072 | NM_00131033 | RNF31 | ring finger protein 31 | protein_coding | NP_001297261.1;NP_060469.4 | 16031 | 612487 | . | . | 7.915622 | 7.325438 | 7.727555 | 8.274211 | 18.419979 | 16.933077 | 17.340411 | 18.918001 | 1.035256 | 851.505816 | 0.145246257 | 0.344154459 | 0.730730129 | 0.998865404 | 9.708646 | 9.758666 | 9.751235 | 9.689830 | 9.726287 | 9.764772 | 1005 | 864 | 733 | 834 |
| 55073 | NR_002940 | LRRC37A4P | leucine rich repeat containing 37 member A4, pseudogene | transcribed_ps | . | 25479 | . | . | . | 0.905765 | 0.825569 | 1.168244 | 0.787638 | 2.107752 | 1.908339 | 2.621505 | 1.800840 | 1.113843 | 208.308957 | 0.259057194 | 0.600432361 | 0.548218126 | 0.998865404 | 7.621655 | 7.779208 | 7.692385 | 7.647676 | 7.809106 | 7.622418 | 238 | 202 | 229 | 165 |
| 55074 | NM_00119853 | OXR1 | oxidation resistance 1 | protein_coding | NP_001185461.1;NP_001185462.1;NP_001185463.1;NP_001185464.1;NI | 15822 | 605609 | . | . | 8.364381 | 6.502318 | 7.524741 | 6.345548 | 19.464255 | 15.030398 | 16.885304 | 14.508337 | -1.138053 | 826.083363 | 0.160862347 | -1.15979892 | 0.246130685 | 0.992733415 | 9.780062 | 9.594246 | 9.808343 | 9.650500 | 9.677214 | 9.603229 | 1122 | 846 | 708 | 686 |
| 55075 | NM_00100822 | UACA | uveal autoantigen with coiled-coil domains and ankyrin repeats | protein_coding | NP_001008225.1;NP_060473.2;XP_005254586.1;XP_011520054.2;XP_01 | 15947 | 612516 | . | . | 9.261310 | 9.022000 | 6.943337 | 6.930105 | 21.551446 | 20.854756 | 15.580649 | 15.844863 | -1.194349 | 1522.706249 | 0.129855009 | -1.97316211 | 0.048477095 | 0.660542298 | 10.694680 | 10.438836 | 10.657542 | 10.615416 | 10.541112 | 10.455945 | 1959 | 1741 | 1275 | 1228 |
| 55076 | NM_00136387 | TMEM45A | transmembrane protein 45A | protein_coding | NP_001350805.1;NP_060474.1;XP_005247626.1;XP_024309382.1;XP_02 | 25480 | 616928 | . | . | 4.398392 | 3.409925 | 3.723125 | 3.736659 | 10.235241 | 7.882195 | 8.354585 | 8.543884 | -1.058321 | 198.072785 | 0.247211191 | -0.33079691 | 0.740797897 | 0.998865404 | 7.669813 | 7.588824 | 7.696294 | 7.583592 | 7.612494 | 7.619695 | 263 | 193 | 166 | 182 |
| 55080 | NM_00135135 | TAPBPL | TAP binding protein like | protein_coding | NP_001338284.1;NP_060479.3;XP_005253757.1;XP_016875034.1;XP_01 | 30683 | 607081 | . | . | 2.448821 | 2.133518 | 1.740450 | 1.767277 | 5.698505 | 4.931720 | 3.905519 | 4.040668 | -1.080341 | 135.318796 | 0.280874163 | -0.39692878 | 0.691419977 | 0.998865404 | 7.134570 | 7.023738 | 7.116167 | 7.072630 | 7.072156 | 7.060495 | 171 | 143 | 115 | 120 |
| 55081 | NM_018010 | IFT57 | intraflagellar transport 57 | protein_coding | NP_060480.1 | 17367 | 606621 | . | . | 4.135880 | 4.105482 | 3.583207 | 4.227263 | 9.624364 | 9.490006 | 8.040614 | 9.665135 | -1.070029 | 382.740970 | 0.188196389 | -0.51887106 | 0.603850661 | 0.998865404 | 8.628545 | 8.530229 | 8.603947 | 8.595251 | 8.515487 | 8.602413 | 459 | 424 | 297 | 373 |
| 55082 | NM_018011,XI | ARGLU1 | arginine and glutamate rich 1 | protein_coding | NP_060481.3 | 25482 | 614046 | . | . | 23.278791 | 23.329391 | 27.264918 | 17.762862 | 54.170704 | 53.926935 | 61.181683 | 40.612673 | -1.048144 | 2553.906038 | 0.170063654 | -0.39889283 | 0.689972174 | 0.998865404 | 11.351820 | 11.284371 | 11.336673 | 11.325533 | 11.456579 | 11.098517 | 3030 | 2801 | 2646 | 1824 |
| 55083 | NM_018012 | KIF26B | kinesin family member 26B | protein_coding | NP_060482.2 | 25484 | 614026 | . | . | 0.168582 | 0.169886 | 0.157049 | 0.222072 | 0.392296 | 0.392700 | 0.352412 | 0.507741 | 1.106960 | 76.450110 | 0.372611393 | 0.39344827 | 0.693988424 | 0.998865404 | 6.182248 | 6.327016 | 6.241460 | 6.239323 | 6.221419 | 6.309156 | 84 | 78 | 58 | 88 |
| 55084 | NM_018013,XI | SOBP | sine oculis binding protein homolog | protein_coding | NP_060483.3;XP_005267098.1;XP_005267099.1;XP_011534222.1;XP_01 | 29256 | 613667 | . | . | 2.149829 | 2.029722 | 2.534200 | 2.876694 | 5.002739 | 4.691795 | 5.686669 | 6.577219 | 1.260186 | 430.159285 | 0.181196179 | 1.841303315 | 0.065577122 | 0.731993537 | 8.572473 | 8.905774 | 8.688707 | 8.652733 | 8.766074 | 8.854412 | 452 | 398 | 388 | 481 |
| 55086 | NM_00118478 | RADX | RPA1 related single stranded DNA binding protein, X-linked | protein_coding | NP_001171711.1;NP_060485.4 | 25486 | . | . | . | 2.398555 | 2.334061 | 2.794153 | 2.403039 | 5.581536 | 5.395288 | 6.269998 | 5.494261 | 1.083271 | 295.564320 | 0.206413476 | 0.559045772 | 0.576130482 | 0.998865404 | 8.148047 | 8.264269 | 8.190701 | 8.176672 | 8.267233 | 8.184192 | 331 | 302 | 288 | 266 |
| 55088 | NM_00132182 | CCDC186 | coiled-coil domain containing 186 | protein_coding | NP_001308758.1;NP_060487.2;NP_694981.1;XP_011538217.1;XP_01153 | 24349 | . | . | . | 1.743694 | 2.039295 | 1.982319 | 1.399416 | 4.057644 | 4.713919 | 4.448267 | 3.199598 | -1.126557 | 392.227505 | 0.203896012 | -0.84317817 | 0.399128804 | 0.998865404 | 8.698507 | 8.527524 | 8.612254 | 8.679415 | 8.660049 | 8.484782 | 452 | 473 | 370 | 293 |
| 55090 | NM_018019 | MED9 | mediator complex subunit 9 | protein_coding | NP_060489.1 | 25487 | 609878 | . | . | 5.244869 | 4.919261 | 4.060522 | 6.251281 | 12.205026 | 11.371091 | 9.111694 | 13.882478 | -1.000087 | 352.693343 | 0.223025395 | -0.00056444 | 0.99954964 | 0.999862634 | 8.462971 | 8.461569 | 8.478366 | 8.436685 | 8.333241 | 8.568392 | 421 | 367 | 243 | 399 |
| 55092 | NM_00113621 | TMEM51 | transmembrane protein 51 | protein_coding | NP_001129688.1;NP_001129689.1;NP_001129690.1;NP_001306594.1;NI | 25488 | . | . | . | 48.123834 | 48.383102 | 47.437391 | 47.765248 | 111.986138 | 111.839707 | 106.448134 | 109.209562 | -1.017380 | 2920.584179 | 0.111205751 | -0.22353882 | 0.823116176 | 0.998865404 | 11.524427 | 11.499548 | 11.516638 | 11.521468 | 11.501814 | 11.508136 | 3382 | 3188 | 2508 | 2725 |
| 55093 | NM_00128302 | NTAQ1 | N-terminal glutamine amidase 1 | protein_coding | NP_001269953.1;NP_001269956.1;NP_060494.1;XP_006716660.1;XP_01 | 25490 | . | . | . | 6.391240 | 6.101572 | 5.309559 | 6.074606 | 14.872678 | 14.104052 | 11.914500 | 13.888864 | -1.007069 | 334.645168 | 0.19687066 | -0.05162388 | 0.958828388 | 0.998865404 | 8.391754 | 8.381204 | 8.407359 | 8.368569 | 8.347425 | 8.419601 | 400 | 350 | 271 | 333 |
| 55094 | NM_018025,N | GPATCH1 | G-patch domain containing 1 | protein_coding | NP_060495.2;XP_006723318.1 | 24658 | . | . | . | 3.298569 | 3.857719 | 4.206820 | 3.907953 | 7.675907 | 8.917289 | 9.439883 | 8.935069 | 1.117603 | 380.161657 | 0.189885912 | 0.844758353 | 0.398245772 | 0.998865404 | 8.488120 | 8.648371 | 8.491403 | 8.576535 | 8.623325 | 8.575718 | 383 | 416 | 363 | 361 |
| 55095 | NM_00130361 | SAMD4B | sterile alpha motif domain containing 4B | protein_coding | NP_001290543.1;NP_060498.2;XP_011525359.1;XP_011525362.1;XP_01 | 25492 | . | . | . | 8.074291 | 9.134622 | 10.042158 | 11.023662 | 18.789207 | 21.115088 | 22.534314 | 25.204293 | 1.045704 | 1507.480110 | 0.139937662 | 0.460738046 | 0.644986555 | 0.998865404 | 10.525587 | 10.589551 | 10.472078 | 10.601780 | 10.524215 | 10.620327 | 1555 | 1724 | 1257 | 1534 |
| 55096 | NM_018029 | EBLN2 | endogenous Bornavirus like nucleoprotein 2 | protein_coding | NP_060499.3 | 25493 | 613250 | . | . | 2.421103 | 2.371048 | 2.923225 | 1.531572 | 5.634007 | 5.480785 | 6.559632 | 3.501757 | -1.091057 | 121.646643 | 0.339557566 | -0.37026547 | 0.711184702 | 0.998865404 | 6.986638 | 6.863856 | 6.937438 | 6.927520 | 7.000936 | 6.793029 | 148 | 135 | 133 | 75 |
| 55100 | NM_00134599 | WDR70 | WD repeat domain 70 | protein_coding | NP_001332927.1;NP_001332928.1;NP_060504.1;XP_011512363.1 | 25495 | 617233 | . | . | 4.600560 | 4.200814 | 4.253703 | 4.756085 | 10.705694 | 9.710369 | 9.545185 | 10.874221 | 1.018921 | 398.369654 | 0.186199089 | 0.14523353 | 0.884526494 | 0.998865404 | 8.624406 | 8.651396 | 8.667240 | 8.594920 | 8.612576 | 8.672200 | 482 | 400 | 332 | 396 |
| 55101 | NM_00116786 | DMAC2 | distal membrane arm assembly complex 2 | protein_coding | NP_001161339.1;NP_001161340.1;NP_001161341.1;NP_001161342.1;NI | 25496 | 617262 | . | . | 15.426286 | 18.210073 | 15.838061 | 18.266012 | 35.897600 | 42.093404 | 35.540150 | 41.763064 | 1.029324 | 948.443230 | 0.150509086 | 0.277037087 | 0.781751629 | 0.998865404 | 9.868753 | 9.909791 | 9.817313 | 9.934473 | 9.856286 | 9.938842 | 990 | 1090 | 789 | 953 |
| 55102 | NM_018036 | ATG2B | autophagy related 2B | protein_coding | NP_060506.6 | 20187 | 616226 | . | . | 1.923404 | 1.767450 | 2.506885 | 2.186549 | 4.475840 | 4.085540 | 5.625377 | 4.999284 | 1.253586 | 859.730072 | 0.151125369 | 2.157551198 | 0.030962742 | 0.558665013 | 9.575154 | 9.901868 | 9.690114 | 9.626155 | 9.872766 | 9.764963 | 919 | 785 | 893 | 831 |
| 55103 | NM_00128624 | RALGPS2 | Ral GEF with PH domain and SH3 binding motif 2 | protein_coding | NP_001273176.1;NP_689876.2;XP_006711473.1;XP_006711474.1;XP_01 | 30279 | 617819 | . | . | 3.121231 | 2.397597 | 2.478699 | 2.461111 | 7.263233 | 5.542153 | 5.562132 | 5.627038 | -1.127437 | 596.036492 | 0.17254497 | -1.00290527 | 0.31590657 | 0.998865404 | 9.302858 | 9.130512 | 9.330887 | 9.175547 | 9.190858 | 9.162583 | 810 | 604 | 498 | 514 |
| 55105 | NM_00129775 | GPATCH2 | G-patch domain containing 2 | protein_coding | NP_001284683.1;NP_060510.1;XP_011507991.1;XP_011507992.1;XP_01 | 25499 | 616836 | . | . | 3.160414 | 2.854634 | 3.974976 | 3.071459 | 7.354412 | 6.598617 | 8.919731 | 7.022525 | 1.161958 | 379.490614 | 0.199560024 | 1.085175824 | 0.27779379 | 0.998865404 | 8.454894 | 8.672732 | 8.547167 | 8.488177 | 8.677599 | 8.528945 | 425 | 359 | 399 | 335 |
| 55106 | NM_00128900 | SLFN12 | schlafen family member 12 | protein_coding | NP_001275938.1;NP_060512.3;XP_005258052.1;XP_011523269.1;XP_01 | 25500 | 614955 | . | . | 1.645568 | 1.689548 | 2.055653 | 1.792564 | 3.829303 | 3.905467 | 4.612824 | 4.098485 | 1.062200 | 194.267015 | 0.240366193 | 0.362179645 | 0.717217789 | 0.998865404 | 7.557487 | 7.644981 | 7.580972 | 7.593002 | 7.636499 | 7.590749 | 214 | 206 | 183 | 178 |
| 55108 | NM_00114388 | BSDC1 | BSD domain containing 1 | protein_coding | NP_001137360.1;NP_001137361.1;NP_001137362.1;NP_001287887.1;NI | 25501 | 617518 | . | . | 12.037676 | 13.452601 | 12.514781 | 13.746612 | 28.012165 | 31.096290 | 28.082805 | 31.429997 | 1.054200 | 1584.657812 | 0.127982788 | 0.594994545 | 0.551847118 | 0.998865404 | 10.591545 | 10.667370 | 10.575020 | 10.641143 | 10.619730 | 10.678465 | 1701 | 1736 | 1361 | 1582 |
| 55109 | NM_018046 | AGGF1 | angiogenic factor with G-patch and FHA domains 1 | protein_coding | NP_060516.2 | 24684 | 608464 | . | . | 7.061007 | 6.859643 | 6.828896 | 6.592309 | 16.431255 | 15.856374 | 15.323845 | 15.072532 | -1.051604 | 957.235910 | 0.13871465 | -0.52331395 | 0.600755797 | 0.998865404 | 9.938451 | 9.866104 | 9.934407 | 9.906841 | 9.900453 | 9.867672 | 1151 | 1039 | 830 | 855 |
| 55110 | NM_00130073 | MAGOHB | mago homolog B, exon junction complex subunit | protein_coding | NP_001287668.1;NP_001306914.1;NP_060518.1;XP_005253459.1 | 25504 | . | . | . | 3.407102 | 2.867112 | 2.914101 | 2.719598 | 7.928465 | 6.627456 | 6.539159 | 6.218039 | -1.191263 | 238.629196 | 0.226165191 | -1.11640589 | 0.264248434 | 0.996308385 | 8.018864 | 7.767451 | 7.978952 | 7.897877 | 7.876961 | 7.830229 | 322 | 258 | 199 | 194 |
| 55111 | NM_00130083 | PLEKHJ1 | pleckstrin homology domain containing J1 | protein_coding | NP_001287765.1;NP_060519.1;XP_016882414.1;XP_016882416.1;XP_01 | 18211 | 617834 | . | . | 19.984295 | 20.518585 | 18.413430 | 21.232516 | 46.504279 | 47.429630 | 41.319203 | 48.545625 | -1.042978 | 815.490288 | 0.149161762 | -0.4070026 | 0.684006075 | 0.998865404 | 9.701807 | 9.640592 | 9.679281 | 9.692597 | 9.608551 | 9.701424 | 950 | 907 | 648 | 798 |
| 55112 | NM_00135091 | DYNC2I1 | dynein 2 intermediate chain 1 | protein_coding | NP_001337843.1;NP_001337844.1;NP_001337845.1;NP_001337846.1;NI | 21862 | 615462 | . | . | 2.426354 | 2.381875 | 2.740175 | 2.651549 | 5.646227 | 5.505812 | 6.148873 | 6.062455 | 1.120984 | 307.491326 | 0.198840364 | 0.828633217 | 0.407311986 | 0.998865404 | 8.179509 | 8.344576 | 8.239306 | 8.223011 | 8.303336 | 8.282827 | 339 | 308 | 288 | 299 |
| 55113 | NM_018053,XI | XKR8 | XK related 8 | protein_coding | NP_060523.2;XP_011539981.1;XP_011539982.1 | 25508 | . | . | . | 1.235728 | 1.192862 | 1.008560 | 0.847416 | 2.875591 | 2.757355 | 2.263179 | 1.937516 | -1.402957 | 78.764817 | 0.363476425 | -1.34388671 | 0.178985024 | 0.951511445 | 6.522471 | 6.035606 | 6.338991 | 6.343305 | 6.281134 | 6.226485 | 105 | 100 | 64 | 54 |
| 55114 | NM_00100663 | ARHGAP17 | Rho GTPase activating protein 17 | protein_coding | NP_001006635.1;NP_060524.4;XP_011544175.1;XP_011544176.1;XP_01 | 18239 | 608293 | . | . | 10.288062 | 11.839348 | 11.745801 | 12.436377 | 23.940741 | 27.367182 | 26.357240 | 28.434298 | 1.061619 | 1254.151511 | 0.135017038 | 0.638927277 | 0.522870243 | 0.998865404 | 10.248906 | 10.334807 | 10.223565 | 10.312059 | 10.294709 | 10.332739 | 1319 | 1390 | 1095 | 1241 |
| 55116 | NM_00131967 | TMEM39B | transmembrane protein 39B | protein_coding | NP_001306606.1;NP_001306607.1;NP_001306608.1;NP_060526.2;XP_0C | 25510 | . | . | . | 6.714893 | 6.616620 | 7.111354 | 7.489331 | 15.625832 | 15.294613 | 15.957672 | 17.123464 | 1.049014 | 456.710532 | 0.173123689 | 0.398751348 | 0.690076429 | 0.998865404 | 8.800283 | 8.869167 | 8.831282 | 8.808127 | 8.829866 | 8.868389 | 523 | 472 | 394 | 452 |
| 55117 | NM_00114633 | SLC6A15 | solute carrier family 6 member 15 | protein_coding | NP_001139807.1;NP_060527.2;NP_877499.1;XP_011536827.1 | 13621 | 607971 | . | . | 5.033800 | 4.130975 | 5.115322 | 3.645325 | 11.713857 | 9.548935 | 11.478635 | 8.334602 | -1.059076 | 627.038002 | 0.193486475 | -0.42796636 | 0.668675612 | 0.998865404 | 9.332579 | 9.251089 | 9.375207 | 9.232268 | 9.375693 | 9.147314 | 820 | 623 | 621 | 473 |
| 55118 | NM_00120652 | CRTAC1 | cartilage acidic protein 1 | protein_coding | NP_001193457.1;NP_060528.3;XP_005269995.1;XP_011538219.1;XP_01 | 14882 | 606276 | . | . | 3.394480 | 3.977668 | 3.001889 | 3.675218 | 7.899096 | 9.194559 | 6.736153 | 8.402949 | -1.079431 | 284.000203 | 0.211964006 | -0.520236 | 0.6028991 | 0.998865404 | 8.204383 | 8.092963 | 8.145558 | 8.192142 | 8.084029 | 8.171655 | 325 | 332 | 217 | 278 |
| 55119 | NM_00134975 | PRPF38B | pre-mRNA processing factor 38B | protein_coding | NP_001336686.1;NP_001336687.1;NP_001336688.1;NP_001336690.1;NI | 25512 | . | . | . | 16.395146 | 16.404909 | 22.335353 | 15.191279 | 38.152179 | 37.920683 | 50.119889 | 34.733053 | 1.131772 | 2641.168454 | 0.159445368 | 1.120032212 | 0.262700035 | 0.995798899 | 11.274277 | 11.453650 | 11.312319 | 11.306693 | 11.556315 | 11.239849 | 2863 | 2664 | 2905 | 2127 |
| 55120 | NM_00111463 | FANCL | FA complementation group L | protein_coding | NP_001108108.1;NP_001361544.1;NP_060532.2;XP_005264452.1;XP_01 | 20748 | 608111 | . | . | 23.648988 | 21.342787 | 22.113982 | 18.182513 | 55.032164 | 49.334813 | 49.623133 | 41.572156 | -1.100841 | 1171.792583 | 0.153845591 | -0.90094134 | 0.36761951 | 0.998865404 | 10.261790 | 10.123920 | 10.264962 | 10.187952 | 10.243216 | 10.059126 | 1485 | 1258 | 1091 | 916 |
| 55122 | NM_018064 | AKIRIN2 | akirin 2 | protein_coding | NP_060534.1 | 21407 | 615165 | . | . | 12.578260 | 12.781018 | 11.155320 | 11.628698 | 29.270128 | 29.543896 | 25.032215 | 26.587637 | -1.128858 | 712.354672 | 0.14949938 | -1.16966281 | 0.242136691 | 0.992733415 | 9.561304 | 9.386296 | 9.514585 | 9.519252 | 9.423213 | 9.443572 | 867 | 818 | 573 | 637 |
| 55124 | NM_00113572 | PIWIL2 | piwi like RNA-mediated gene silencing 2 | protein_coding | NP_001129193.1;NP_001317409.1;NP_060538.2;XP_005273608.1 | 17644 | 610312 | . | . | 0.166619 | 0.175718 | 0.166971 | 0.137150 | 0.387730 | 0.406179 | 0.374678 | 0.313576 | -1.281572 | 23.936718 | 0.630136396 | -0.56799448 | 0.570038718 | NA | 4.749462 | 4.390584 | 4.597329 | 4.596961 | 4.566905 | 4.578330 | 31 | 29 | 17 | 21 |
| 55125 | NM_032142,XI | CEP192 | centrosomal protein 192 | protein_coding | NP_115518.3;XP_005258164.1;XP_005258166.1;XP_006722389.1;XP_00 | 25515 | 616426 | . | . | 4.515685 | 5.132177 | 4.825156 | 4.475042 | 10.508182 | 11.863260 | 10.827513 | 10.231649 | -1.039577 | 1163.492892 | 0.137133343 | -0.40833942 | 0.6830245 | 0.998865404 | 10.211978 | 10.155972 | 10.157034 | 10.237550 | 10.194804 | 10.142572 | 1293 | 1348 | 1027 | 1032 |
| 55127 | NM_018072 | HEATR1 | HEAT repeat containing 1 | protein_coding | NP_060542.4 | 25517 | . | . | . | 7.822259 | 8.091464 | 8.758227 | 7.467724 | 18.202721 | 18.703781 | 19.653208 | 17.074064 | 1.005958 | 2118.053031 | 0.128593177 | 0.066650215 | 0.946860169 | 0.998865404 | 11.044097 | 11.052936 | 11.035054 | 11.055081 | 11.115332 | 10.980128 | 2401 | 2307 | 2004 | 1823 |
| 55128 | NM_00130449 | TRIM68 | tripartite motif containing 68 | protein_coding | NP_001291425.1;NP_060543.5 | 21161 | 613184 | . | . | 2.648454 | 3.261603 | 2.981459 | 2.549657 | 6.163061 | 7.539343 | 6.690308 | 5.829488 | -1.145281 | 283.774066 | 0.213472119 | -0.91675205 | 0.359272586 | 0.998865404 | 8.243100 | 8.047471 | 8.135594 | 8.226276 | 8.140849 | 8.079643 | 320 | 354 | 244 | 233 |
| 55129 | NM_00120483 | ANO10 | anoctamin 10 | protein_coding | NP_001191760.1;NP_001191761.1;NP_001191762.1;NP_001191763.1;NI | 25519 | 613726 | . | . | 9.061390 | 8.664466 | 7.691909 | 8.309594 | 21.086225 | 20.028296 | 17.260420 | 18.998903 | -1.152762 | 756.542641 | 0.148285671 | -1.38310686 | 0.166632102 | 0.944905472 | 9.662181 | 9.457092 | 9.632129 | 9.588974 | 9.499428 | 9.523966 | 961 | 848 | 600 | 671 |
| 55131 | NM_00116613 | RBM28 | RNA binding motif protein 28 | protein_coding | NP_001159607.1;NP_060547.2;XP_011514672.1;XP_016867878.1;XP_01 | 21863 | 612074 | . | . | 5.590736 | 7.092972 | 6.150069 | 6.664571 | 13.009872 | 16.395722 | 13.800576 | 15.237750 | 1.036683 | 2514.069705 | 0.116719296 | 0.445298667 | 0.656103931 | 0.998865404 | 11.269508 | 11.321639 | 11.283388 | 11.278079 | 11.342972 | 11.275778 | 2852 | 2655 | 2318 | 2300 |
| 55132 | NM_00127860 | LARP1B | La ribonucleoprotein 1B | protein_coding | NP_001265533.1;NP_001337460.1;NP_060548.2;NP_115615.2;NP_8351 | 24704 | . | . | . | 3.384093 | 3.906205 | 3.705379 | 3.208663 | 7.874921 | 9.029366 | 8.314760 | 7.336230 | -1.091685 | 320.156772 | 0.201256258 | -0.6288321 | 0.529458979 | 0.998865404 | 8.384426 | 8.258078 | 8.313004 | 8.377056 | 8.328564 | 8.265161 | 363 | 381 | 282 | 270 |
| 55133 | NM_018079,XI | SRBD1 | S1 RNA binding domain 1 | protein_coding | NP_060549.4;XP_011531248.1;XP_016859907.1 | 25521 | . | . | . | 5.343777 | 5.454221 | 4.920259 | 6.121705 | 12.435189 | 12.607677 | 11.040920 | 13.996551 | 1.008732 | 623.156056 | 0.16653049 | 0.075323276 | 0.939957502 | 0.998865404 | 9.277543 | 9.289332 | 9.275109 | 9.283826 | 9.215398 | 9.350401 | 710 | 674 | 489 | 646 |
| 55135 | NM_00114399 | WRAP53 | WD repeat containing antisense to TP53 | protein_coding | NP_001137462.1;NP_001137463.1;NP_001137464.1;NP_060551.2;XP_01 | 25522 | 612661 | . | . | 8.232912 | 10.587859 | 9.413266 | 11.073137 | 19.158322 | 24.474306 | 21.123096 | 25.317412 | 1.095619 | 613.189533 | 0.171311727 | 0.769046846 | 0.441865501 | 0.998865404 | 9.193244 | 9.324165 | 9.150024 | 9.297756 | 9.261065 | 9.312956 | 599 | 702 | 536 | 624 |
| 55137 | NM_00132182 | FIGN | fidgetin, microtubule severing factor | protein_coding | NP_001308754.1;NP_060556.2;XP_011509691.1;XP_016859908.1 | 13285 | 605295 | . | . | 1.162962 | 1.046446 | 0.902950 | 0.877483 | 2.706260 | 2.418907 | 2.026194 | 2.006260 | -1.105665 | 420.356911 | 0.215828581 | -0.67143474 | 0.501943623 | 0.998865404 | 8.785515 | 8.641852 | 8.714282 | 8.763339 | 8.797535 | 8.543861 | 488 | 495 | 422 | 294 |
| 55139 | NM_00104241 | ANKZF1 | ankyrin repeat and zinc finger peptidyl tRNA hydrolase 1 | protein_coding | NP_001035869.1;NP_001269721.1;NP_060559.2;XP_005246720.1;XP_01 | 25527 | 617541 | . | . | 19.474890 | 19.530086 | 30.717548 | 25.324358 | 45.318872 | 45.144666 | 68.929290 | 57.901136 | 1.405634 | 1910.691193 | 0.134753631 | 3.645327297 | 0.000267052 | 0.025777026 | 10.633291 | 11.124804 | 10.741580 | 10.752570 | 11.096096 | 10.930353 | 1816 | 1726 | 2138 | 1882 |
| 55140 | NM_00128422 | ELP3 | elongator acetyltransferase complex subunit 3 | protein_coding | NP_001271149.1;NP_001271151.1;NP_001271153.1;NP_001271154.1;NI | 20696 | 612722 | . | . | 9.698822 | 9.885982 | 10.995145 | 10.479581 | 22.569556 | 22.851894 | 24.672784 | 23.960314 | 1.087664 | 973.457031 | 0.14000677 | 0.86590551 | 0.386541981 | 0.998865404 | 9.864974 | 9.986418 | 9.884003 | 9.905360 | 9.988790 | 9.922373 | 1058 | 1021 | 922 | 908 |
| 55142 | NM_00113044 | HAUS2 | HAUS augmin like complex subunit 2 | protein_coding | NP_001123919.1;NP_001310558.1;NP_001310559.1;NP_001310560.1;NI | 25530 | 613429 | . | . | 4.091955 | 4.186908 | 4.082616 | 3.833576 | 9.522148 | 9.678225 | 9.161276 | 8.765017 | -1.058877 | 487.576831 | 0.169257552 | -0.48763273 | 0.625810005 | 0.998865404 | 8.970084 | 8.887712 | 8.936681 | 8.957610 | 8.929289 | 8.892869 | 568 | 550 | 424 | 431 |
| 55143 | NM_00125687 | CDCA8 | cell division cycle associated 8 | protein_coding | NP_001243804.1;NP_060571.1 | 14629 | 609977 | . | . | 21.312541 | 23.606396 | 18.356985 | 19.045526 | 49.595160 | 54.567242 | 41.192539 | 43.545332 | -1.216484 | 1496.331221 | 0.128629837 | -2.19791328 | 0.027955287 | 0.539571956 | 10.681767 | 10.398808 | 10.577917 | 10.656196 | 10.455590 | 10.477110 | 1798 | 1860 | 1157 | 1284 |
| 55144 | NM_00113447 | LRRC8D | leucine rich repeat containing 8 VRAC subunit D | protein_coding | NP_001127951.1;NP_060573.2;XP_011539991.1;XP_016857090.1;XP_01 | 16992 | 612890 | . | . | 14.891844 | 14.011918 | 15.306057 | 13.225348 | 34.653933 | 32.389176 | 34.346349 | 30.238188 | -1.031906 | 1686.462690 | 0.132335579 | -0.34239624 | 0.732052717 | 0.998865404 | 10.742060 | 10.697160 | 10.748376 | 10.713331 | 10.767879 | 10.641354 | 2016 | 1807 | 1561 | 1430 |
| 55145 | NM_018105,N | THAP1 | THAP domain containing 1 | protein_coding | NP_060575.1;NP_945354.1 | 20856 | 609520 | . | . | 2.235861 | 2.478240 | 2.441484 | 2.861215 | 5.202940 | 5.728564 | 5.478620 | 6.541829 | 1.105863 | 168.704093 | 0.258328107 | 0.561969362 | 0.574136893 | 0.998865404 | 7.324570 | 7.468542 | 7.355797 | 7.393912 | 7.383189 | 7.448985 | 176 | 181 | 142 | 179 |
| 55146 | NM_00113438 | ZDHHC4 | zinc finger DHHC-type palmitoyltransferase 4 | protein_coding | NP_001127859.1;NP_001127860.1;NP_001127861.1;NP_001358221.1;NI | 18471 | . | . | . | 18.074313 | 18.167926 | 22.911339 | 23.211411 | 42.059670 | 41.995975 | 51.412382 | 53.070132 | 1.244527 | 999.186784 | 0.135985381 | 2.32081541 | 0.020296808 | 0.470997021 | 9.798179 | 10.113813 | 9.878796 | 9.876499 | 10.041194 | 10.034508 | 1027 | 959 | 968 | 1034 |
| 55147 | NM_00107735 | RBM23 | RNA binding motif protein 23 | protein_coding | NP_001070819.1;NP_001070820.1;NP_001294973.1;NP_001339691.1;NI | 20155 | . | . | . | 14.428841 | 14.584573 | 14.346982 | 17.768378 | 33.576507 | 33.712890 | 32.194210 | 40.625286 | 1.131102 | 1396.375239 | 0.130814248 | 1.358639645 | 0.174260806 | 0.948096927 | 10.356002 | 10.533485 | 10.402998 | 10.392179 | 10.447707 | 10.534423 | 1520 | 1404 | 1217 | 1467 |
| 55148 | NM_175748,N | UBR7 | ubiquitin protein ligase E3 component n-recognin 7 | protein_coding | NP_786924.2 | 20344 | 613816 | . | . | 10.437342 | 10.828377 | 9.699954 | 11.918483 | 24.288122 | 25.030277 | 21.766419 | 27.250195 | 1.004242 | 1169.320602 | 0.144538464 | 0.042251465 | 0.966298236 | 0.998865404 | 10.188675 | 10.194230 | 10.178173 | 10.199293 | 10.115166 | 10.263104 | 1325 | 1277 | 918 | 1207 |
| 55149 | NM_018109 | MTPAP | mitochondrial poly(A) polymerase | protein_coding | NP_060579.3 | 25532 | 613669 | . | . | 3.286854 | 3.354690 | 3.753920 | 3.725009 | 7.648643 | 7.754518 | 8.423687 | 8.516790 | 1.111222 | 611.575487 | 0.156227578 | 0.973877475 | 0.330117379 | 0.998865404 | 9.178291 | 9.330472 | 9.215253 | 9.223509 | 9.295779 | 9.283532 | 663 | 629 | 565 | 598 |
| 55150 | NM_018111 | C19orf73 | chromosome 19 open reading frame 73 | protein_coding | NP_060581.2 | 25534 | . | . | . | 0.848752 | 1.270724 | 0.984623 | 1.694330 | 1.975081 | 2.937335 | 2.209465 | 3.873884 | 1.262742 | 28.024043 | 0.609169305 | 0.552489887 | 0.580612761 | 0.998865404 | 4.633141 | 4.965003 | 4.765886 | 4.807196 | 4.780377 | 4.845410 | 23 | 32 | 20 | 37 |
| 55151 | NM_018112,XI | TMEM38B | transmembrane protein 38B | protein_coding | NP_060582.1;XP_005252132.1;XP_005252133.1;XP_005252134.1;XP_01 | 25535 | 611236 | . | . | 10.883723 | 10.243828 | 9.177424 | 7.955296 | 25.326873 | 23.679066 | 20.593873 | 18.188839 | -1.268278 | 996.057483 | 0.152330004 | -2.25084268 | 0.024395503 | 0.515584957 | 10.121016 | 9.778922 | 10.085641 | 9.989809 | 9.928735 | 9.797441 | 1363 | 1126 | 835 | 747 |
| 55152 | NM_00100999 | DALRD3 | DALR anticodon binding domain containing 3 | protein_coding | NP_001009996.1;NP_001263334.1;NP_060584.3;XP_005265326.1;XP_00 | 25536 | . | . | . | 8.864466 | 8.762784 | 9.417856 | 9.623490 | 20.627977 | 20.255565 | 21.133396 | 22.002965 | 1.077472 | 524.217687 | 0.164168915 | 0.655732414 | 0.511996309 | 0.998865404 | 8.979210 | 9.086828 | 9.014344 | 9.003752 | 9.050231 | 9.063605 | 586 | 540 | 468 | 515 |
| 55153 | NM_00128898 | SDAD1 | SDA1 domain containing 1 | protein_coding | NP_001275912.1;NP_001275913.1;NP_060585.2;XP_005263159.1;XP_00 | 25537 | . | . | . | 12.921514 | 11.561165 | 13.974223 | 12.772169 | 30.068895 | 26.724150 | 31.357750 | 29.202048 | 1.079680 | 1213.810434 | 0.141232222 | 0.783132048 | 0.433549573 | 0.998865404 | 10.188684 | 10.299830 | 10.264402 | 10.161714 | 10.314851 | 10.227888 | 1436 | 1173 | 1159 | 1114 |
| 55154 | NM_00125653 | MSTO1 | misato mitochondrial distribution and morphology regulator 1 | protein_coding | NP_001243461.1;NP_001243462.1;NP_001337701.1;NP_001337702.1;NI | 29678 | 617619 | . | . | 13.189753 | 14.971008 | 15.079067 | 15.499602 | 30.693096 | 34.606157 | 33.836991 | 35.438000 | 1.091874 | 1214.162962 | 0.13645357 | 0.929296622 | 0.352735385 | 0.998865404 | 10.181047 | 10.307669 | 10.161006 | 10.258232 | 10.288467 | 10.265298 | 1250 | 1334 | 1119 | 1170 |
| 55156 | NM_00128670 | ARMC1 | armadillo repeat containing 1 | protein_coding | NP_001273631.1;NP_060590.1;XP_005251321.1;XP_016869094.1 | 17684 | . | . | . | 10.683824 | 9.557286 | 9.237751 | 6.576964 | 24.861700 | 22.092093 | 20.729247 | 15.037448 | -1.302019 | 844.818728 | 0.179189866 | -2.12484689 | 0.033599391 | 0.575719787 | 9.899868 | 9.520223 | 9.847358 | 9.759252 | 9.731806 | 9.486659 | 1165 | 970 | 753 | 565 |
| 55157 | NM_00136521 | DARS2 | aspartyl-tRNA synthetase 2, mitochondrial | protein_coding | NP_001352141.1;NP_001352142.1;NP_060592.2 | 25538 | 610956 | . | . | 21.715110 | 22.935134 | 18.441346 | 22.656945 | 50.531951 | 53.015592 | 41.381847 | 51.802414 | -1.090254 | 2190.464618 | 0.130298906 | -0.95676084 | 0.338687983 | 0.998865404 | 11.158184 | 11.033151 | 11.112519 | 11.146665 | 10.985131 | 11.130564 | 2576 | 2518 | 1655 | 2145 |
| 55159 | NM_00137053 | RFWD3 | ring finger and WD repeat domain 3 | protein_coding | NP_001357463.1;NP_001357464.1;NP_001357465.1;NP_001357466.1;NI | 25539 | 614151 | . | . | 13.935816 | 13.837925 | 12.830605 | 13.787351 | 32.429219 | 31.986987 | 28.791501 | 31.523139 | -1.060971 | 2137.646572 | 0.118402042 | -0.72114971 | 0.470817407 | 0.998865404 | 11.103921 | 11.018428 | 11.093403 | 11.076449 | 11.011991 | 11.062682 | 2561 | 2350 | 1746 | 2007 |
| 55160 | NM_00101172 | ARHGEF10L | Rho guanine nucleotide exchange factor 10 like | protein_coding | NP_001011722.2;NP_001306766.1;NP_001306767.1;NP_001315053.1;NI | 25540 | 612494 | . | . | 1.531227 | 1.834752 | 0.966554 | 1.686293 | 3.563231 | 4.241114 | 2.168918 | 3.855507 | -1.321266 | 217.114045 | 0.275027735 | -1.461382 | 0.143910637 | 0.906161673 | 7.950485 | 7.545854 | 7.766030 | 7.865292 | 7.568674 | 7.791893 | 257 | 293 | 120 | 221 |
| 55161 | NM_018126,XI | TMEM33 | transmembrane protein 33 | protein_coding | NP_060596.2;XP_005248173.1;XP_005248174.1 | 25541 | 618515 | . | . | 12.402488 | 11.801909 | 13.862136 | 8.839723 | 28.861101 | 27.280642 | 31.106231 | 20.210976 | -1.176418 | 2115.579711 | 0.345265711 | -0.67889798 | 0.497202505 | 0.998865404 | 11.159216 | 10.924960 | 11.153497 | 11.046374 | 11.137133 | 10.773250 | 2806 | 2306 | 2079 | 1404 |
| 55163 | NM_018129,XI | PNPO | pyridoxamine 5'-phosphate oxidase | protein_coding | NP_060599.1;XP_005257557.1;XP_011523270.1;XP_016880302.1 | 30260 | 603287 | . | . | 5.530342 | 6.001535 | 5.905392 | 6.494507 | 12.869332 | 13.872818 | 13.251528 | 14.848918 | 1.076562 | 644.436251 | 0.157003729 | 0.677893468 | 0.497839242 | 0.998865404 | 9.277957 | 9.383887 | 9.280240 | 9.330658 | 9.329352 | 9.380768 | 688 | 695 | 559 | 650 |
| 55164 | NM_018130,XI | SHQ1 | SHQ1, H/ACA ribonucleoprotein assembly factor | protein_coding | NP_060600.2;XP_011532197.1;XP_011532198.1;XP_011532199.1;XP_01 | 25543 | 613663 | . | . | 18.872125 | 17.125802 | 17.101632 | 18.426432 | 43.916210 | 39.587057 | 38.375560 | 42.129839 | -1.028584 | 1567.351274 | 0.128880632 | -0.31548528 | 0.752393222 | 0.998865404 | 10.634299 | 10.593640 | 10.662650 | 10.585087 | 10.576144 | 10.628640 | 1921 | 1630 | 1298 | 1498 |
| 55165 | NM_00112718 | CEP55 | centrosomal protein 55 | protein_coding | NP_001120654.2;NP_060601.4;XP_011538220.1;XP_011538221.1;XP_01 | 1161 | 610000 | . | . | 20.577119 | 21.290844 | 18.547605 | 17.774208 | 47.883804 | 49.214739 | 41.620289 | 40.638616 | -1.181064 | 1600.002113 | 0.12492562 | -1.92183586 | 0.054626414 | 0.69238604 | 10.758851 | 10.518899 | 10.692442 | 10.717261 | 10.601540 | 10.548864 | 1964 | 1899 | 1320 | 1328 |
| 55166 | NM_018132,XI | CENPQ | centromere protein Q | protein_coding | NP_060602.2;XP_005249262.1;XP_011513025.2 | 21347 | 611506 | . | . | 4.291887 | 3.915643 | 3.885434 | 3.173334 | 9.987398 | 9.051183 | 8.718801 | 7.255453 | -1.182045 | 208.329102 | 0.239652076 | -1.00681111 | 0.314025553 | 0.998865404 | 7.817690 | 7.577789 | 7.761782 | 7.715753 | 7.703036 | 7.619960 | 272 | 232 | 182 | 162 |
| 55167 | NM_00114541 | MSL2 | MSL complex subunit 2 | protein_coding | NP_001138889.1;NP_060603.2;XP_005247628.1;XP_006713747.1;XP_01 | 25544 | 614802 | . | . | 2.846023 | 2.332863 | 2.407890 | 2.230281 | 6.622813 | 5.392517 | 5.403234 | 5.099273 | -1.083233 | 387.269222 | 0.187978696 | -0.6135969 | 0.539481732 | 0.998865404 | 8.653326 | 8.538787 | 8.658528 | 8.579395 | 8.596759 | 8.548458 | 494 | 406 | 337 | 334 |
| 55168 | NM_00119334 | MRPS18A | mitochondrial ribosomal protein S18A | protein_coding | NP_001180272.1;NP_060605.1;XP_006715197.1 | 14515 | 611981 | . | . | 29.444592 | 29.472649 | 25.208608 | 29.114686 | 68.518776 | 68.127349 | 56.567385 | 66.567268 | -1.096631 | 1036.397229 | 0.141610326 | -0.93974839 | 0.347346637 | 0.998865404 | 10.082565 | 9.949070 | 10.053959 | 10.046163 | 9.929758 | 10.031698 | 1255 | 1164 | 799 | 992 |
| 55170 | NM_018137 | PRMT6 | protein arginine methyltransferase 6 | protein_coding | NP_060607.2 | 18241 | 608274 | . | . | 6.273454 | 5.621276 | 6.951780 | 7.773007 | 14.598584 | 12.993831 | 15.599593 | 17.772058 | 1.219687 | 543.297412 | 0.168697529 | 1.698373053 | 0.089437375 | 0.802216049 | 8.935242 | 9.221753 | 9.051539 | 8.977450 | 9.109431 | 9.177065 | 597 | 497 | 493 | 588 |
| 55171 | NM_00113441 | TBCCD1 | TBCC domain containing 1 | protein_coding | NP_001127887.1;NP_001273678.1;NP_060608.1 | 25546 | . | . | . | 2.994557 | 3.642039 | 3.623329 | 2.902461 | 6.968458 | 8.418736 | 8.130646 | 6.636134 | -1.006082 | 280.669463 | 0.218613789 | -0.04001426 | 0.968081756 | 0.998865404 | 8.136994 | 8.128450 | 8.081346 | 8.180019 | 8.173716 | 8.081661 | 294 | 332 | 265 | 239 |
| 55172 | NM_00108390 | DNAAF2 | dynein axonemal assembly factor 2 | protein_coding | NP_001077377.1;NP_001365382.1;NP_060609.2 | 20188 | 612517 | . | . | 3.352387 | 2.983070 | 2.636469 | 2.556659 | 7.801141 | 6.895499 | 5.916159 | 5.845496 | -1.220726 | 264.972853 | 0.214500554 | -1.34143632 | 0.179778832 | 0.952541638 | 8.186106 | 7.899039 | 8.132636 | 8.062431 | 8.006877 | 7.985928 | 357 | 294 | 212 | 219 |
| 55173 | NM_018141,XI | MRPS10 | mitochondrial ribosomal protein S10 | protein_coding | NP_060611.2;XP_005249263.1;XP_011513026.1;XP_011513027.1;XP_01 | 14502 | 611976 | . | . | 15.505927 | 14.179271 | 12.211411 | 13.403571 | 36.082929 | 32.776020 | 27.402053 | 30.645671 | -1.173663 | 908.581634 | 0.144906385 | -1.59425909 | 0.110878017 | 0.848126978 | 9.938399 | 9.707301 | 9.916932 | 9.845850 | 9.737684 | 9.794990 | 1184 | 1008 | 697 | 816 |
| 55174 | NM_00135350 | INTS10 | integrator complex subunit 10 | protein_coding | NP_001340434.1;NP_001340435.1;NP_001340436.1;NP_001340437.1;NI | 25548 | 611353 | . | . | 19.440373 | 20.140165 | 24.027774 | 23.300792 | 45.238554 | 46.554890 | 53.917627 | 53.274492 | 1.127601 | 1803.905428 | 0.120491584 | 1.437913497 | 0.1504586 | 0.91327179 | 10.727657 | 10.900958 | 10.759093 | 10.775544 | 10.876192 | 10.847344 | 1932 | 1849 | 1693 | 1760 |
| 55175 | NM_018143,XI | KLHL11 | kelch like family member 11 | protein_coding | NP_060613.1 | 19008 | . | . | . | 0.886048 | 0.851070 | 0.979181 | 0.545125 | 2.061871 | 1.967287 | 2.197253 | 1.246363 | -1.155906 | 188.570211 | 0.289381468 | -0.72231454 | 0.470101115 | 0.998865404 | 7.658518 | 7.452015 | 7.592283 | 7.570690 | 7.630979 | 7.390670 | 238 | 213 | 196 | 117 |
| 55176 | NM_00114262 | SEC61A2 | SEC61 translocon subunit alpha 2 | protein_coding | NP_001136099.1;NP_001136100.1;NP_060614.2 | 17702 | 618271 | . | . | 8.239582 | 9.093904 | 8.315976 | 7.475405 | 19.173844 | 21.020964 | 18.660811 | 17.091625 | -1.109919 | 641.138522 | 0.160626656 | -0.9366674 | 0.34892967 | 0.998865404 | 9.397676 | 9.247397 | 9.322100 | 9.392415 | 9.323814 | 9.250348 | 737 | 765 | 558 | 538 |
| 55177 | NM_00130480 | RMDN3 | regulator of microtubule dynamics 3 | protein_coding | NP_001291731.1;NP_001310823.1;NP_001310824.1;NP_001310825.1;NI | 25550 | 611873 | . | . | 17.750646 | 17.464633 | 17.800875 | 16.302756 | 41.306484 | 40.370377 | 39.944649 | 37.274312 | -1.042272 | 1223.784482 | 0.133332916 | -0.4479902 | 0.654160268 | 0.998865404 | 10.286528 | 10.227128 | 10.281630 | 10.262733 | 10.276864 | 10.204058 | 1456 | 1331 | 1093 | 1069 |
| 55178 | NM_00131794 | MRM3 | mitochondrial rRNA methyltransferase 3 | protein_coding | NP_001304876.1;NP_060616.1 | 18485 | 612600 | . | . | 11.514798 | 13.357608 | 10.170861 | 12.392945 | 26.795406 | 30.876709 | 22.823116 | 28.334999 | -1.112472 | 630.299302 | 0.170464967 | -0.9020546 | 0.367027862 | 0.998865404 | 9.375212 | 9.220425 | 9.285710 | 9.378644 | 9.194149 | 9.324617 | 713 | 765 | 467 | 615 |
| 55179 | NM_00103303 | FAIM | Fas apoptotic inhibitory molecule | protein_coding | NP_001028202.1;NP_001028203.1;NP_001028204.1;NP_060617.1;XP_01 | 18703 | 617535 | . | . | 4.916425 | 5.406911 | 4.366918 | 5.129560 | 11.440722 | 12.498318 | 9.799237 | 11.728131 | -1.061594 | 171.347037 | 0.25671422 | -0.33590795 | 0.736940285 | 0.998865404 | 7.463830 | 7.376400 | 7.406582 | 7.458575 | 7.381110 | 7.433988 | 191 | 202 | 135 | 166 |
| 55180 | NM_00104061 | LINS1 | lines homolog 1 | protein_coding | NP_001035706.2;NP_001339436.1;NP_001339437.1;XP_005254998.1;XF | 30922 | 610350 | . | . | 7.091687 | 7.045649 | 10.860526 | 6.510362 | 16.502648 | 16.286333 | 24.370707 | 14.885169 | 1.185738 | 1158.895337 | 0.165996813 | 1.480660322 | 0.138697112 | 0.900064053 | 10.050073 | 10.296487 | 10.108742 | 10.103783 | 10.357265 | 10.094175 | 1225 | 1140 | 1281 | 977 |
| 55181 | NM_018149 | SMG8 | SMG8 nonsense mediated mRNA decay factor | protein_coding | NP_060619.4 | 25551 | 613175 | . | . | 9.517650 | 9.601418 | 9.394105 | 10.604425 | 22.147961 | 22.194107 | 21.080101 | 24.245758 | 1.031880 | 995.079408 | 0.140375019 | 0.3225275 | 0.747053108 | 0.998865404 | 9.936028 | 9.980958 | 9.946683 | 9.946686 | 9.928077 | 10.009408 | 1128 | 1057 | 830 | 1000 |
| 55182 | NM_00131995 | RNF220 | ring finger protein 220 | protein_coding | NP_001306885.1;NP_001306886.1;NP_001363415.1;NP_001363416.1;NI | 25552 | 616136 | . | . | 17.008162 | 17.230247 | 17.130047 | 17.135957 | 39.578691 | 39.828487 | 38.439330 | 39.179329 | 1.014863 | 1162.199706 | 0.132569868 | 0.160551434 | 0.872446709 | 0.998865404 | 10.172036 | 10.193171 | 10.182920 | 10.171019 | 10.164936 | 10.210532 | 1339 | 1235 | 987 | 1131 |
| 55183 | NM_00117766 | RIF1 | replication timing regulatory factor 1 | protein_coding | NP_001171134.1;NP_001171135.1;NP_001171136.1;NP_060621.3;XP_0C | 23207 | 608952 | . | . | 5.165588 | 4.313515 | 5.176558 | 2.876984 | 12.020536 | 9.970882 | 11.616044 | 6.577882 | -1.100832 | 1523.167048 | 0.166585351 | -0.83197512 | 0.405422987 | 0.998865404 | 10.640172 | 10.502249 | 10.646042 | 10.560909 | 10.661073 | 10.379772 | 1937 | 1629 | 1492 | 1111 |
| 55184 | NM_00109940 | DZANK1 | double zinc ribbon and ankyrin repeat domains 1 | protein_coding | NP_001092877.1;NP_001338612.1;NP_001338613.1;NP_001354540.1;NI | 15858 | . | . | . | 0.998016 | 1.137514 | 1.266450 | 1.050921 | 2.322427 | 2.629414 | 2.841877 | 2.402804 | 1.279994 | 130.311407 | 0.284570037 | 1.251490745 | 0.210755488 | 0.973906262 | 6.836465 | 7.193180 | 6.977690 | 6.970968 | 7.079451 | 7.045658 | 133 | 122 | 133 | 131 |
| 55186 | NM_00110464 | SLC25A36 | solute carrier family 25 member 36 | protein_coding | NP_001098117.1;NP_060625.2;XP_005247632.1 | 25554 | 616149 | . | . | 6.673155 | 5.393902 | 7.113750 | 4.302266 | 15.528706 | 12.468245 | 15.963050 | 9.836619 | -1.060971 | 1016.132749 | 0.397720194 | -0.21445758 | 0.830190241 | 0.998865404 | 10.030765 | 9.945729 | 10.073961 | 9.921904 | 10.128024 | 9.751336 | 1327 | 1014 | 1077 | 688 |
| 55187 | NM_015378,N | VPS13D | vacuolar protein sorting 13 homolog D | protein_coding | NP_056193.2;NP_060626.2 | 23595 | 608877 | . | . | 3.146421 | 3.363923 | 3.066251 | 3.291664 | 7.321853 | 7.775857 | 6.880578 | 7.525998 | -1.036932 | 1635.107293 | 0.124773149 | -0.41933504 | 0.67497129 | 0.998865404 | 10.701211 | 10.648649 | 10.665389 | 10.712163 | 10.638861 | 10.682044 | 1860 | 1850 | 1357 | 1547 |

| | | | | | | | | | | | | | | | | | | | | | | | | | | | | | | | | | |
|---|---|---|---|---|---|---|---|---|---|---|---|---|---|---|---|---|---|---|---|---|---|---|---|---|---|---|---|---|---|---|---|---|---|
| 55188 | NM_00133014 | RIC8B | RIC8 guanine nucleotide exchange factor B | protein_coding | NP_001317074.1;NP_001317075.1;NP_001317076.1;NP_001338290.1;NI | 25555 | 609147 | . | . | 4.423223 | 4.281860 | 5.298354 | 3.957317 | 10.293020 | 9.897713 | 11.889356 | 9.047934 | -1.007337 | 943.772130 | 0.160235701 | -0.06581811 | 0.947522638 | 0.998865404 | 9.887192 | 9.877381 | 9.890115 | 9.874605 | 9.971002 | 9.774079 | 1102 | 1011 | 929 | 762 |
| 55190 | NM_018159 | NUDT11 | nudix hydrolase 11 | protein_coding | NP_060629.2 | 18011 | 300528 | . | . | 1.926679 | 2.004849 | 1.526256 | 2.022249 | 4.483461 | 4.634298 | 3.424875 | 4.623632 | -1.121301 | 139.087479 | 0.284935323 | -0.57968687 | 0.562125803 | 0.998865404 | 7.200829 | 7.034052 | 7.135231 | 7.146170 | 7.050722 | 7.143164 | 167 | 161 | 99 | 139 |
| 55191 | NM_018161 | NADSYN1 | NAD synthetase 1 | protein_coding | NP_060631.2 | 29832 | 608285 | . | . | 8.990437 | 10.860827 | 9.145322 | 9.688640 | 20.921116 | 25.105289 | 20.521839 | 22.151922 | -1.067985 | 807.380400 | 0.154964118 | -0.6123399 | 0.540312895 | 0.998865404 | 9.704038 | 9.608594 | 9.612261 | 9.741524 | 9.615976 | 9.648395 | 874 | 981 | 663 | 749 |
| 55192 | NM_018163,XI | DNAJC17 | DnaJ heat shock protein family (Hsp40) member C17 | protein_coding | NP_060633.1;XP_016877890.1;XP_016877891.1;XP_024305749.1 | 25556 | 616844 | . | . | 2.289801 | 2.019282 | 2.614206 | 2.040467 | 5.328459 | 4.667661 | 5.866204 | 4.665286 | 1.188870 | 213.237550 | 0.231709306 | 1.077173588 | 0.281402723 | 0.998865404 | 7.606117 | 7.855735 | 7.672378 | 7.710104 | 7.789432 | 7.753759 | 215 | 219 | 209 | 209 |
| 55193 | NM_00135007 | PBRM1 | polybromo 1 | protein_coding | NP_001337003.1;NP_001337004.1;NP_001337005.1;NP_001337006.1;NI | 30064 | 606083 | . | . | 6.843971 | 6.375522 | 7.570773 | 7.062897 | 15.926203 | 14.737302 | 16.988594 | 16.148477 | 1.084769 | 1548.079091 | 0.131987586 | 0.889381781 | 0.373797933 | 0.998865404 | 10.536168 | 10.653955 | 10.582912 | 10.542292 | 10.679205 | 10.570013 | 1755 | 1560 | 1499 | 1405 |
| 55194 | NM_00130476 | EVA1B | eva-1 homolog B | protein_coding | NP_001291691.1;NP_060636.1 | 25558 | . | | . | 7.536514 | 6.478533 | 8.004125 | 8.537987 | 17.537779 | 14.975419 | 17.961023 | 19.521092 | 1.159278 | 232.735953 | 0.226471207 | 0.94151715 | 0.346439905 | 0.998865404 | 7.751764 | 7.965432 | 7.859872 | 7.784833 | 7.884481 | 7.904674 | 268 | 214 | 213 | 239 |
| 55196 | NM_018169,XI | RESF1 | retroelement silencing factor 1 | protein_coding | NP_060639.4;XP_005253462.1;XP_006719162.1;XP_011519024.1;XP_01( | 25559 | . | | . | 2.190221 | 2.158786 | 2.556736 | 1.702377 | 5.096735 | 4.990131 | 5.737240 | 3.892282 | -1.029302 | 422.034272 | 0.214374757 | -0.1943652 | 0.845889939 | 0.998865404 | 8.741183 | 8.700969 | 8.734247 | 8.713609 | 8.826492 | 8.570818 | 501 | 454 | 439 | 307 |
| 55197 | NM_00130341 | RPRD1A | regulation of nuclear pre-mRNA domain containing 1A | protein_coding | NP_001290340.1;NP_001290341.1;NP_001290342.1;NP_060640.2;XP_01 | 25560 | 610347 | . | . | 7.484269 | 8.152822 | 7.769000 | 6.901046 | 17.416201 | 18.845617 | 17.433410 | 15.778423 | -1.053108 | 939.478833 | 0.148267274 | -0.50350818 | 0.614607034 | 0.998865404 | 9.912393 | 9.838084 | 9.866611 | 9.919060 | 9.911892 | 9.796297 | 1070 | 1077 | 860 | 789 |
| 55198 | NM_00125190 | APPL2 | adaptor protein, phosphotyrosine interacting with PH domain and | protein_coding | NP_001238833.1;NP_001238834.1;NP_060641.2;XP_006719535.1;XP_01 | 18242 | 606231 | . | . | 8.118239 | 8.541087 | 9.841106 | 9.737190 | 18.891475 | 19.743106 | 22.083159 | 22.262929 | 1.136738 | 990.009574 | 0.137192085 | 1.347743355 | 0.177740942 | 0.951283903 | 9.855851 | 10.040822 | 9.892231 | 9.911265 | 10.011459 | 9.979286 | 1053 | 1013 | 936 | 966 |
| 55199 | XR_00464325( | FAM86C1P | family with sequence similarity 86 member C1, pseudogene | misc_RNA | . | 25561 | 616124 | . | . | 2.714515 | 2.707021 | 2.337174 | 3.006099 | 6.316788 | 6.257399 | 5.244552 | 6.873086 | -1.029211 | 218.641848 | 0.236878593 | -0.17535654 | 0.860799487 | 0.998865404 | 7.793619 | 7.750916 | 7.781268 | 7.775529 | 7.708387 | 7.818150 | 257 | 238 | 165 | 226 |
| 55201 | NM_00130836 | MAP1S | microtubule associated protein 1S | protein_coding | NP_001295292.1;NP_060644.4;XP_016882419.1;XP_016882420.1 | 15715 | 607573 | . | . | 6.513597 | 6.475780 | 5.493600 | 6.778999 | 15.157407 | 14.969055 | 12.327480 | 15.499375 | -1.071468 | 648.856303 | 0.16420815 | -0.60647807 | 0.544197337 | 0.998865404 | 9.391015 | 9.290754 | 9.370072 | 9.357710 | 9.249679 | 9.380369 | 780 | 718 | 491 | 645 |
| 55203 | NM_018176,XI | LGI2 | leucine rich repeat LGI family member 2 | protein_coding | NP_060646.2;XP_011512152.1;XP_016863845.1 | 18710 | 608301 | . | . | 0.356616 | 0.362775 | 0.382084 | 0.442071 | 0.829861 | 0.838570 | 0.857385 | 1.010742 | 1.161783 | 77.586642 | 0.360297501 | 0.600450222 | 0.548206226 | 0.998865404 | 6.166062 | 6.381384 | 6.246955 | 6.263254 | 6.276844 | 6.310062 | 80 | 80 | 68 | 83 |
| 55204 | NM_018178,XI | GOLPH3L | golgi phosphoprotein 3 like | protein_coding | NP_060648.2;XP_006711491.1 | 24882 | 612208 | . | . | 3.235853 | 3.482577 | 3.372322 | 3.055942 | 7.529963 | 8.050134 | 7.567392 | 6.987048 | -1.058777 | 320.727866 | 0.198425822 | -0.41526534 | 0.677947669 | 0.998865404 | 8.365671 | 8.283573 | 8.322731 | 8.357710 | 8.336513 | 8.280693 | 368 | 367 | 285 | 277 |
| 55205 | NM_00131872 | ZNF532 | zinc finger protein 532 | protein_coding | NP_001305655.1;NP_001305656.1;NP_001305657.1;NP_001340454.1;NI | 30940 | . | | . | 5.145024 | 5.541885 | 6.233269 | 6.536775 | 11.972682 | 12.810317 | 13.987279 | 14.945555 | 1.183478 | 1141.093083 | 0.133516566 | 1.82024474 | 0.068721743 | 0.741130004 | 10.029689 | 10.272507 | 10.063820 | 10.113469 | 10.207105 | 10.222274 | 1163 | 1166 | 1065 | 1171 |
| 55206 | NM_00116785 | SBNO1 | strawberry notch homolog 1 | protein_coding | NP_001161328.1;NP_060653.3 | 22973 | 614274 | . | . | 4.845536 | 5.112145 | 5.495858 | 5.173683 | 11.275762 | 11.816953 | 12.332548 | 11.829012 | 1.057098 | 1787.756831 | 0.122939438 | 0.651613821 | 0.51465033 | 0.998865404 | 10.763282 | 10.843473 | 10.763346 | 10.798919 | 10.852372 | 10.796859 | 1956 | 1918 | 1654 | 1662 |
| 55207 | NM_018184 | ARL8B | ADP ribosylation factor like GTPase 8B | protein_coding | NP_060654.1 | 25564 | 616596 | . | . | 23.188564 | 19.279705 | 22.029699 | 14.280307 | 53.960739 | 44.565903 | 49.434002 | 32.650223 | -1.185365 | 1802.361491 | 0.18081751 | -1.3567881 | 0.17484855 | 0.948677095 | 10.932839 | 10.688143 | 10.946872 | 10.791031 | 10.893824 | 10.553842 | 2468 | 1906 | 1748 | 1209 |
| 55208 | NM_00101428 | DCUN1D2 | defective in cullin neddylation 1 domain containing 2 | protein_coding | NP_001014305.1;XP_016876132.1;XP_016876133.1;XP_016876135.1;XP_ | 20328 | . | | . | 2.754449 | 2.941012 | 2.016652 | 2.562050 | 6.409718 | 6.798279 | 4.525311 | 5.857824 | -1.082617 | 368.087952 | 0.187003199 | -0.61241035 | 0.540266298 | 0.998865404 | 8.579932 | 8.465618 | 8.534778 | 8.557886 | 8.519866 | 8.480964 | 432 | 421 | 318 | 320 |
| 55209 | NM_00108051 | SETD5 | SET domain containing 5 | protein_coding | NP_001073986.1;NP_001278972.1;NP_001336380.1;XP_005265358.1;XP | 25566 | 615743 | . | . | 9.018475 | 9.590197 | 10.210278 | 9.673776 | 20.986360 | 22.168167 | 22.911568 | 22.117936 | 1.046581 | 2337.670325 | 0.119070342 | 0.551644434 | 0.581191987 | 0.998865404 | 11.157600 | 11.223362 | 11.150112 | 11.193108 | 11.240877 | 11.175027 | 2560 | 2531 | 2164 | 2150 |
| 55210 | NM_00117053 | ATAD3A | ATPase family AAA domain containing 3A | protein_coding | NP_001164006.1;NP_001164007.1;NP_060658.3;XP_024303866.1 | 25567 | 612316 | . | . | 33.767669 | 34.285129 | 31.363062 | 34.963071 | 78.578757 | 79.251608 | 70.377800 | 79.938904 | -1.043768 | 2597.912763 | 0.117138554 | -0.52759361 | 0.597781431 | 0.998865404 | 11.373791 | 11.311818 | 11.359365 | 11.360791 | 11.286487 | 11.362688 | 3053 | 2866 | 2106 | 2496 |
| 55212 | NM_018190,N | BBS7 | Bardet-Biedl syndrome 7 | protein_coding | NP_060660.2;NP_789794.1;XP_005263163.1;XP_011530381.1;XP_01153 | 18758 | 607590 | . | . | 4.780980 | 4.784840 | 4.706931 | 4.256046 | 11.125535 | 11.060371 | 10.562216 | 9.730940 | -1.055790 | 517.027683 | 0.168110219 | -0.46589837 | 0.641288247 | 0.998865404 | 9.052462 | 8.974685 | 9.058910 | 9.002224 | 9.017062 | 8.975281 | 638 | 548 | 452 | 456 |
| 55213 | NM_00135250 | RCBTB1 | RCC1 and BTB domain containing protein 1 | protein_coding | NP_001339429.1;NP_001339430.1;NP_001339431.1;NP_001339432.1;NI | 18243 | 607867 | . | . | 4.968771 | 4.405351 | 5.426713 | 4.843395 | 11.562538 | 10.183168 | 12.177388 | 11.073848 | 1.048700 | 679.333357 | 0.162416035 | 0.422382274 | 0.672746014 | 0.998865404 | 9.372847 | 9.442270 | 9.420213 | 9.358518 | 9.480770 | 9.360697 | 798 | 683 | 658 | 595 |
| 55214 | NM_00113441 | P3H2 | prolyl 3-hydroxylase 2 | protein_coding | NP_001127890.1;NP_060662.2;XP_011511257.1 | 19317 | 610341 | . | . | 4.161685 | 3.884229 | 2.935744 | 3.963873 | 9.684412 | 8.978569 | 6.587724 | 9.062925 | -1.190157 | 404.046859 | 0.197659214 | -1.27062984 | 0.203860366 | 0.969101675 | 8.778944 | 8.526812 | 8.729723 | 8.682255 | 8.520271 | 8.678576 | 525 | 456 | 276 | 394 |
| 55215 | NM_00111337 | FANCI | FA complementation group I | protein_coding | NP_001106849.1;NP_001363839.1;NP_001363840.1;NP_060663.2;XP_01 | 25568 | 611360 | . | . | 51.688682 | 52.535630 | 50.013515 | 55.514703 | 120.281686 | 121.438471 | 112.228886 | 126.927774 | -1.002410 | 7720.779299 | 0.106567458 | -0.03259105 | 0.974000707 | 0.998865404 | 12.916300 | 12.912759 | 12.912904 | 12.917725 | 12.873644 | 12.951386 | 8876 | 8365 | 6398 | 7561 |
| 55216 | NM_00108296 | NKAPD1 | NKAP domain containing 1 | protein_coding | NP_001076438.1;NP_001076439.1;NP_001287946.1;NP_001287948.1;NI | 25569 | . | | . | 4.967482 | 5.004880 | 4.815262 | 5.046087 | 11.559536 | 11.569003 | 10.805309 | 11.537276 | 1.011904 | 547.329283 | 0.161836013 | 0.105492079 | 0.915985355 | 0.998865404 | 9.087678 | 9.104801 | 9.099080 | 9.085566 | 9.098982 | 9.101091 | 633 | 581 | 478 | 517 |
| 55217 | NM_00118479 | TMLHE | trimethyllysine hydroxylase, epsilon | protein_coding | NP_001171726.1;NP_060666.1;XP_011529484.1;XP_016885109.1 | 18308 | 300777 | . | . | 1.649212 | 2.204623 | 1.337894 | 1.576486 | 3.837784 | 5.096082 | 3.002194 | 3.604447 | -1.098984 | 173.103391 | 0.258647449 | -0.52647088 | 0.598561083 | 0.998865404 | 7.502370 | 7.365359 | 7.405075 | 7.500553 | 7.413311 | 7.415246 | 186 | 217 | 143 | 155 |
| 55218 | NM_00119336 | EXD2 | exonuclease 3'-5' domain containing 2 | protein_coding | NP_001180289.1;NP_001180290.1;NP_001180291.1;NP_001180292.1;NI | 20217 | 616940 | . | . | 4.840665 | 4.682750 | 4.641913 | 4.952858 | 11.264427 | 10.824385 | 10.416319 | 11.324124 | 1.003267 | 749.028618 | 0.148124917 | 0.031764057 | 0.97466021 | 0.998865404 | 9.546570 | 9.551180 | 9.559705 | 9.535518 | 9.532380 | 9.567180 | 876 | 793 | 636 | 721 |
| 55219 | NM_00128256 | MACO1 | macoilin 1 | protein_coding | NP_001269493.1;NP_060672.2;XP_005245988.1;XP_011540006.1 | 25572 | 610301 | . | . | 6.856823 | 6.643872 | 6.767453 | 5.875286 | 15.956108 | 15.357609 | 15.185969 | 13.433143 | -1.100347 | 784.200437 | 0.148549727 | -0.92870556 | 0.353041697 | 0.998865404 | 9.682207 | 9.544674 | 9.653535 | 9.641324 | 9.615414 | 9.544161 | 954 | 879 | 683 | 664 |
| 55222 | NM_00127821 | LRRC20 | leucine rich repeat containing 20 | protein_coding | NP_001265140.1;NP_001265141.1;NP_001265142.1;NP_001265143.1;NI | 23421 | . | | . | 2.138698 | 1.929154 | 1.643304 | 2.395430 | 4.976839 | 4.459325 | 3.687527 | 5.476866 | -1.023579 | 193.648275 | 0.260624564 | -0.12900845 | 0.897350961 | 0.998865404 | 7.614636 | 7.579743 | 7.626424 | 7.572773 | 7.509685 | 7.663538 | 239 | 199 | 137 | 211 |
| 55223 | NM_00133048 | TRIM62 | tripartite motif containing 62 | protein_coding | NP_001317412.1;NP_060677.2;XP_016857118.1;XP_016857119.1;XP_01 | 25574 | 616755 | . | . | 5.524566 | 5.167672 | 4.030005 | 4.144972 | 12.855892 | 11.945305 | 9.043216 | 9.476985 | -1.305470 | 556.330843 | 0.161629669 | -2.37932355 | **0.017344444** | 0.44170732 | 9.299247 | 8.914818 | 9.210648 | 9.188510 | 9.030058 | 9.022806 | 737 | 669 | 422 | 450 |
| 55224 | NM_00129776 | ETNK2 | ethanolamine kinase 2 | protein_coding | NP_001284689.1;NP_001284690.1;NP_001284691.1;NP_060678.2;XP_01 | 25575 | 609859 | . | . | 9.252238 | 9.939825 | 10.120912 | 13.758297 | 21.530337 | 22.976352 | 22.711036 | 31.456711 | 1.213650 | 825.858789 | 0.168714925 | 1.655763521 | 0.097769718 | 0.813553488 | 9.543791 | 9.822292 | 9.588792 | 9.636489 | 9.637697 | 9.851604 | 830 | 833 | 672 | 973 |
| 55225 | NM_00136616 | RAVER2 | ribonucleoprotein, PTB binding 2 | protein_coding | NP_001353094.1;NP_060681.2;XP_011540008.1 | 25577 | 609953 | . | . | 6.063278 | 5.617331 | 5.755372 | 4.890026 | 14.109499 | 12.984711 | 12.914887 | 11.180467 | -1.082280 | 780.779258 | 0.158327221 | -0.72049209 | 0.471222069 | 0.998865404 | 9.664286 | 9.551034 | 9.671871 | 9.595781 | 9.639507 | 9.515170 | 984 | 829 | 711 | 640 |
| 55226 | NM_00114403 | NAT10 | N-acetyltransferase 10 | protein_coding | NP_001137502.2;NP_078938.3 | 29830 | 609221 | . | . | 8.991991 | 9.355456 | 9.651697 | 9.511686 | 20.924733 | 21.625556 | 21.658130 | 21.747339 | 1.029941 | 1160.475645 | 0.131736345 | 0.323083988 | 0.746631636 | 0.998865404 | 10.159059 | 10.201628 | 10.157404 | 10.180817 | 10.200704 | 10.181751 | 1296 | 1253 | 1037 | 1090 |
| 55227 | NM_018214,XI | LRRC1 | leucine rich repeat containing 1 | protein_coding | NP_060684.4;XP_011513028.1;XP_011513029.1;XP_016866486.1 | 14307 | 608195 | . | . | 2.317548 | 2.257452 | 2.201797 | 2.775013 | 5.393030 | 5.218199 | 4.940768 | 6.344739 | 1.066786 | 245.584667 | 0.227851488 | 0.409349938 | 0.682282873 | 0.998865404 | 7.893080 | 7.985591 | 7.943755 | 7.899571 | 7.894775 | 8.009753 | 286 | 245 | 195 | 265 |
| 55229 | NM_018216,XI | PANK4 | pantothenate kinase 4 (inactive) | protein_coding | NP_060686.3;XP_011540010.2 | 19366 | 606162 | . | . | 12.246872 | 13.195142 | 13.163910 | 13.512313 | 28.498975 | 30.501160 | 29.539436 | 30.894298 | 1.036085 | 1091.044711 | 0.134906921 | 0.379096676 | 0.704616075 | 0.998865404 | 10.065800 | 10.116740 | 10.052263 | 10.102968 | 10.098365 | 10.109955 | 1192 | 1196 | 958 | 1049 |
| 55230 | NM_00136547 | USP40 | ubiquitin specific peptidase 40 | protein_coding | NP_001352408.1;NP_001369224.1;NP_001369225.1;NP_001369226.1;NI | 20069 | 610570 | . | . | 7.645442 | 7.713737 | 8.913489 | 8.932004 | 17.791257 | 17.830652 | 20.001606 | 20.421966 | 1.154300 | 1424.110380 | 0.125418041 | 1.650630016 | 0.098814146 | 0.813553488 | 10.368602 | 10.575665 | 10.417890 | 10.417369 | 10.534108 | 10.521459 | 1523 | 1426 | 1338 | 1419 |
| 55231 | NM_018219 | CCDC87 | coiled-coil domain containing 87 | protein_coding | NP_060689.2 | 25579 | . | | . | 0.067760 | 0.138538 | 0.094843 | 0.134501 | 0.157681 | 0.320236 | 0.212825 | 0.307519 | 1.108102 | 10.480441 | 0.956215005 | 0.154871222 | 0.876922842 | NA | 3.316288 | 3.459419 | 3.368935 | 3.396610 | 3.379844 | 3.395454 | 8 | 14 | 8 | 12 |
| 55233 | NM_00131711 | MOB1A | MOB kinase activator 1A | protein_coding | NP_001304039.1;NP_001304040.1;NP_001304041.1;NP_060691.2;XP_02 | 16015 | 609281 | . | . | 14.841193 | 13.493912 | 13.673946 | 11.835795 | 34.536065 | 31.191781 | 30.683940 | 27.061137 | -1.124980 | 2056.401151 | 0.130758123 | -1.29933586 | 0.193828693 | 0.96621494 | 11.088164 | 10.918672 | 11.090205 | 11.008051 | 11.014584 | 10.894464 | 2634 | 2230 | 1812 | 1676 |
| 55234 | NM_018225,XI | SMU1 | SMU1 DNA replication regulator and spliceosomal factor | protein_coding | NP_060695.2;XP_005251560.1 | 18247 | 617811 | . | . | 4.978838 | 5.184133 | 4.851452 | 5.657167 | 11.585960 | 11.983357 | 10.886519 | 12.934441 | 1.017914 | 1069.850428 | 0.137443884 | 0.18636931 | 0.852155158 | 0.998865404 | 10.050466 | 10.075810 | 10.062294 | 10.050284 | 10.031265 | 10.106094 | 1231 | 1135 | 891 | 1063 |
| 55236 | NM_018227,XI | UBA6 | ubiquitin like modifier activating enzyme 6 | protein_coding | NP_060697.4;XP_016863848.1 | 25581 | 611361 | . | . | 4.523081 | 4.194210 | 5.464207 | 2.782079 | 10.525396 | 9.695104 | 12.261521 | 6.360894 | -1.119480 | 1124.796283 | 0.187526045 | -0.86830088 | 0.385229638 | 0.998865404 | 10.214166 | 10.052186 | 10.199004 | 10.138572 | 10.241405 | 9.900439 | 1422 | 1231 | 1136 | 766 |
| 55238 | NM_00130838 | SLC38A7 | solute carrier family 38 member 7 | protein_coding | NP_001295313.1;NP_001356537.1;NP_001356538.1;NP_001356539.1;NI | 25582 | 614236 | . | . | 8.584442 | 9.251610 | 10.797378 | 9.438795 | 19.976348 | 21.385512 | 24.229003 | 21.580685 | 1.104467 | 1199.483234 | 0.137138128 | 1.045294755 | 0.295886761 | 0.998865404 | 10.154636 | 10.298189 | 10.165769 | 10.210752 | 10.309468 | 10.215023 | 1273 | 1267 | 1163 | 1107 |
| 55239 | NM_00132435 | OGFOD1 | 2-oxoglutarate and iron dependent oxygenase domain containing | protein_coding | NP_001311286.1;NP_001311287.1;NP_001311288.1;NP_001311289.1;NI | 25585 | 615857 | . | . | 8.535214 | 8.707756 | 9.364189 | 9.538943 | 19.861793 | 20.128366 | 21.012969 | 21.809660 | 1.094477 | 1283.315814 | 0.128609302 | 1.01269269 | 0.311206971 | 0.998865404 | 10.259121 | 10.389266 | 10.277644 | 10.302358 | 10.355027 | 10.362008 | 1388 | 1344 | 1161 | 1263 |
| 55240 | NM_00100841 | STEAP3 | STEAP3 metalloreductase | protein_coding | NP_001008410.1;NP_060704.2;NP_619543.2;NP_878919.2;XP_00671267 | 24592 | 609671 | . | . | 3.216312 | 3.265026 | 3.459548 | 3.858273 | 7.484490 | 7.547258 | 7.763126 | 8.821483 | 1.101436 | 416.847666 | 0.179738369 | 0.775491238 | 0.438049441 | 0.998865404 | 8.632237 | 8.771172 | 8.671319 | 8.673810 | 8.699752 | 8.760548 | 456 | 429 | 361 | 430 |
| 55243 | NM_00128634 | KIRREL1 | kirre like nephrin family adhesion molecule 1 | protein_coding | NP_001273278.1;NP_060710.3;XP_005245362.1 | 15734 | 607428 | . | . | 43.956610 | 47.480545 | 43.072301 | 46.627813 | 102.288849 | 109.753410 | 96.652998 | 106.608950 | -1.031324 | 10543.426579 | 0.105668627 | -0.42110029 | 0.673681852 | 0.998865404 | 13.386164 | 13.341604 | 13.347012 | 13.405758 | 13.320676 | 13.379431 | 11905 | 11949 | 8715 | 10068 |
| 55244 | NM_018242 | SLC47A1 | solute carrier family 47 member 1 | protein_coding | NP_060712.2 | 25588 | 609832 | . | . | 7.489872 | 8.179064 | 8.806089 | 8.654042 | 17.429241 | 18.906271 | 19.760609 | 19.786438 | 1.098918 | 846.537821 | 0.144363422 | 0.94264455 | 0.345862745 | 0.998865404 | 9.655848 | 9.791812 | 9.661615 | 9.717800 | 9.766686 | 9.747780 | 892 | 905 | 781 | 820 |
| 55245 | NM_00118497 | UQCC1 | ubiquinol-cytochrome c reductase complex assembly factor 1 | protein_coding | NP_001171906.1;NP_060714.3;NP_955781.2;XP_011527179.1;XP_01152 | 15891 | 611797 | . | . | 22.381749 | 22.422570 | 21.961483 | 25.134761 | 52.083250 | 51.830776 | 49.280934 | 57.467645 | 1.055760 | 1635.278934 | 0.126600439 | 0.618337112 | 0.536353143 | 0.998865404 | 10.635756 | 10.713830 | 10.660934 | 10.645750 | 10.653765 | 10.735850 | 1850 | 1700 | 1384 | 1657 |
| 55246 | NM_00130452 | CCDC25 | coiled-coil domain containing 25 | protein_coding | NP_001291458.1;NP_001291459.1;NP_001291461.1;NP_060716.2;XP_01 | 25591 | . | | . | 4.191224 | 3.857681 | 3.393654 | 4.029199 | 9.753147 | 8.917203 | 7.615260 | 9.212284 | -1.096432 | 439.012166 | 0.182618841 | -0.72728645 | 0.467050497 | 0.998865404 | 8.843199 | 8.709960 | 8.834621 | 8.774803 | 8.700256 | 8.795217 | 554 | 472 | 335 | 424 |
| 55247 | NM_018248,XI | NEIL3 | nei like DNA glycosylase 3 | protein_coding | NP_060718.3;XP_016863849.1 | 24573 | 608934 | . | . | 4.901685 | 5.402773 | 3.978437 | 4.251705 | 11.406423 | 12.488753 | 8.927498 | 9.721019 | -1.272142 | 337.977631 | 0.194650798 | -1.78401288 | 0.074421572 | 0.764716896 | 8.564245 | 8.216407 | 8.435061 | 8.491740 | 8.315773 | 8.342079 | 415 | 428 | 252 | 286 |
| 55248 | NM_00119886 | PACC1 | proton activated chloride channel 1 | protein_coding | NP_001185791.1;NP_001364407.1;NP_001364408.1;NP_001364409.1;NI | 25593 | 618427 | . | . | 9.580089 | 9.734241 | 7.439304 | 7.622444 | 22.293261 | 22.501134 | 16.693582 | 17.427815 | -1.265769 | 649.401190 | 0.153754511 | -2.21141041 | **0.02700743** | 0.53637921 | 9.502965 | 9.162984 | 9.425074 | 9.408739 | 9.256307 | 9.259684 | 845 | 774 | 497 | 539 |
| 55249 | NM_00119889 | YY1AP1 | YY1 associated protein 1 | protein_coding | NP_001185828.1;NP_001185829.1;NP_001185830.1;NP_001185831.1;NI | 30935 | 607860 | . | . | 11.472299 | 11.700479 | 11.705875 | 12.237887 | 26.696511 | 27.046182 | 26.267641 | 27.980474 | 1.051800 | 1029.020485 | 0.135066842 | 0.539442151 | 0.589581803 | 0.998865404 | 9.970205 | 10.042989 | 9.984135 | 9.991832 | 10.019140 | 10.031087 | 1149 | 1088 | 910 | 997 |
| 55250 | NM_00124287 | ELP2 | elongator acetyltransferase complex subunit 2 | protein_coding | NP_001229804.1;NP_001229805.1;NP_001229806.1;NP_001229807.1;NI | 18248 | 616054 | . | . | 5.515454 | 5.506644 | 5.251774 | 5.253870 | 12.834689 | 12.728853 | 11.784832 | 12.012349 | -1.054043 | 1400.160462 | 0.126438571 | -0.60055595 | 0.548135786 | 0.998865404 | 10.488788 | 10.412968 | 10.484914 | 10.457385 | 10.436974 | 10.424850 | 1684 | 1523 | 1195 | 1268 |
| 55251 | NM_00110492 | PCMTD2 | protein-L-isoaspartate (D-aspartate) O-methyltransferase domain | protein_coding | NP_001098395.1;NP_060727.2 | 15882 | . | | . | 6.820392 | 6.141280 | 7.936462 | 7.076497 | 15.871331 | 14.195843 | 17.809191 | 16.179571 | 1.141460 | 840.635902 | 0.151328341 | 1.26136291 | 0.207178125 | 0.970482251 | 9.616394 | 9.807928 | 9.703416 | 9.624277 | 9.802681 | 9.712962 | 955 | 799 | 828 | 788 |
| 55252 | NM_00136934 | ASXL2 | ASXL transcriptional regulator 2 | protein_coding | NP_001356275.1;NP_001356276.1;NP_060733.4 | 23805 | 612991 | . | . | 3.480366 | 3.462098 | 3.974396 | 3.932367 | 8.098955 | 8.002795 | 8.918430 | 8.990891 | 1.122710 | 1485.728436 | 0.124483275 | 1.341429638 | 0.179781 | 0.952541638 | 10.451004 | 10.618071 | 10.495399 | 10.484934 | 10.587601 | 10.571376 | 1623 | 1500 | 1381 | 1458 |
| 55253 | NM_018264,N | TYW1 | tRNA-yW synthesizing protein 1 homolog | protein_coding | NP_060734.2;XP_011514674.1;XP_016867881.1 | 25598 | 611243 | . | . | 4.554341 | 4.276490 | 4.258996 | 4.201854 | 10.598138 | 9.885298 | 9.557065 | 9.607038 | -1.056410 | 450.279450 | 0.174365691 | -0.45404444 | 0.64979686 | 0.998865404 | 8.853534 | 8.774744 | 8.853900 | 8.808259 | 8.807382 | 8.787837 | 552 | 481 | 387 | 404 |
| 55254 | NM_018266,N | TMEM39A | transmembrane protein 39A | protein_coding | NP_060736.1;XP_005247635.1;XP_006713750.1;XP_011511260.1;XP_01( | 25600 | . | | . | 9.720362 | 9.602896 | 9.422052 | 7.684701 | 22.619682 | 22.197522 | 21.142813 | 17.570157 | -1.120328 | 1194.703564 | 0.141282014 | -1.1602415 | 0.245950495 | 0.992733415 | 10.301834 | 10.138415 | 10.266092 | 10.259170 | 10.240806 | 10.109767 | 1460 | 1356 | 1067 | 963 |
| 55255 | NM_018268,XI | WDR41 | WD repeat domain 41 | protein_coding | NP_060738.2;XP_005248608.1;XP_005248609.1;XP_011541807.1;XP_01 | 25601 | 617502 | . | . | 14.862988 | 14.548044 | 12.642925 | 13.142296 | 34.586782 | 33.628453 | 28.370356 | 30.048297 | -1.027273 | 1371.027682 | 0.129222779 | -0.3004061 | 0.763867408 | 0.998865404 | 10.440302 | 10.401522 | 10.459625 | 10.401778 | 10.396119 | 10.424771 | 1660 | 1443 | 1154 | 1292 |
| 55256 | NM_00130607 | ADI1 | acireductone dioxygenase 1 | protein_coding | NP_001293006.1;NP_060739.2 | 30576 | 613400 | . | . | 28.602895 | 29.357303 | 32.330390 | 34.649045 | 66.560113 | 67.860722 | 72.548459 | 79.220921 | 1.114496 | 2312.802249 | 0.115034892 | 1.359509175 | 0.173985297 | 0.948096927 | 11.095132 | 11.251486 | 11.122424 | 11.137625 | 11.215548 | 11.218719 | 2495 | 2383 | 2116 | 2290 |
| 55257 | NM_018270,N | MRGBP | MRG domain binding protein | protein_coding | NP_060740.1 | 15866 | 611157 | . | . | 18.365419 | 18.957894 | 18.381333 | 20.437963 | 42.737087 | 43.822025 | 41.247176 | 46.728973 | 1.026431 | 1642.523966 | 0.125762054 | 0.299273324 | 0.764731506 | 0.998865404 | 10.662890 | 10.700266 | 10.662818 | 10.679422 | 10.653601 | 10.727882 | 1847 | 1768 | 1378 | 1634 |
| 55258 | NM_00124467 | THNSL2 | threonine synthase like 2 | protein_coding | NP_001231605.1;NP_001231607.1;NP_060741.3;XP_005264457.1;XP_00 | 25602 | 611261 | . | . | 0.121735 | 0.149321 | 0.057322 | 0.114908 | 0.283284 | 0.345164 | 0.128628 | 0.262724 | -1.732242 | 14.175888 | 0.854710437 | -0.92737867 | 0.353729962 | NA | 4.170740 | 3.370388 | 3.837366 | 3.857943 | 3.795016 | 3.834092 | 18 | 22 | 5 | 14 |
| 55259 | NM_00108297 | CFAP94 | cilia and flagella associated protein 94 | protein_coding | NP_001076441.1;NP_001076442.2;NP_001191030.2;NP_001191031.2;NI | 29599 | 616906 | . | . | 0.131018 | 0.115011 | 0.096241 | 0.266505 | 0.304883 | 0.265853 | 0.215962 | 0.609331 | 1.594500 | 15.038491 | 0.844783511 | 0.796777554 | 0.425580233 | NA | 3.537347 | 4.206844 | 3.887335 | 3.874632 | 3.871527 | 3.949679 | 15 | 11 | 8 | 26 |
| 55260 | NM_00130353 | TMEM143 | transmembrane protein 143 | protein_coding | NP_001290467.1;NP_001290468.1;NP_001290469.1;NP_060743.2 | 25603 | . | | . | 3.731312 | 3.851256 | 3.896230 | 4.485701 | 8.682915 | 8.902351 | 8.743027 | 10.256022 | 1.097508 | 276.984820 | 0.210310278 | 0.638252479 | 0.52330934 | 0.998865404 | 8.045382 | 8.178858 | 8.080320 | 8.094707 | 8.099648 | 8.170918 | 300 | 289 | 235 | 290 |
| 55262 | NM_00130347 | MAP11 | microtubule associated protein 11 | protein_coding | NP_001290399.1;NP_060745.3 | 25604 | 618350 | . | . | 10.433587 | 10.753334 | 12.282952 | 10.398892 | 24.279387 | 24.856813 | 27.562587 | 23.775829 | 1.052600 | 871.425681 | 0.150929098 | 0.490015897 | 0.62412265 | 0.998865404 | 9.729528 | 9.803979 | 9.740117 | 9.754129 | 9.845580 | 9.718509 | 968 | 925 | 846 | 764 |
| 55266 | NM_018279 | TMEM19 | transmembrane protein 19 | protein_coding | NP_060749.2 | 25605 | . | | . | 5.566546 | 4.726930 | 5.254834 | 3.897806 | 12.953581 | 10.926510 | 11.791695 | 8.911871 | -1.141089 | 858.886108 | 0.174862022 | -1.08892416 | 0.276187333 | 0.998865404 | 9.837902 | 9.648534 | 9.848661 | 9.722989 | 9.795402 | 9.580527 | 1144 | 903 | 805 | 637 |
| 55267 | NR_165243 | PRR34 | PRR34 long non-coding RNA | lncRNA | . | 25606 | . | | . | 0.090074 | 0.075294 | 0.154407 | 0.195216 | 0.209606 | 0.174045 | 0.346484 | 0.446339 | **2.091978** | 15.893359 | 0.777840983 | 1.369004257 | 0.170997944 | NA | 3.361571 | 4.426810 | 3.951224 | 3.942333 | 3.987331 | 4.004035 | 13 | 10 | 17 | 22 |
| 55269 | NM_00104241 | PSPC1 | paraspeckle component 1 | protein_coding | NP_001035879.1;NP_001341837.1;NP_001341838.1;NP_001350589.1;XI | 20320 | 612408 | . | . | 20.878860 | 21.775352 | 23.439651 | 27.446709 | 48.585965 | 50.334700 | 52.597891 | 62.753641 | 1.072931 | 1804.076728 | 0.122694815 | 0.827721757 | 0.407828095 | 0.998865404 | 10.765492 | 10.866819 | 10.770831 | 10.805592 | 10.815560 | 10.871061 | 1960 | 1920 | 1567 | 1812 |
| 55270 | NM_00130474 | NUDT15 | nudix hydrolase 15 | protein_coding | NP_001291674.1;NP_060753.1 | 23063 | 615792 | . | . | 13.400406 | 10.128537 | 10.849531 | 10.250045 | 31.183296 | 23.412569 | 24.346034 | 23.435509 | -1.136515 | 673.232189 | 0.171813692 | -1.07451541 | 0.282591742 | 0.998865404 | 9.483977 | 9.300088 | 9.522226 | 9.339250 | 9.365655 | 9.329610 | 932 | 672 | 563 | 575 |
| 55272 | NM_018285 | IMP3 | IMP U3 small nucleolar ribonucleoprotein 3 | protein_coding | NP_060755.1 | 14497 | 612980 | . | . | 32.235016 | 31.731686 | 33.992420 | 36.557396 | 75.012207 | 73.349213 | 76.278008 | 83.584152 | 1.087981 | 1185.973908 | 0.132176007 | 0.920392419 | 0.357367729 | 0.998865404 | 10.149814 | 10.271341 | 10.187898 | 10.169913 | 10.218164 | 10.265711 | 1324 | 1211 | 1041 | 1195 |
| 55273 | NM_00109964 | TMEM100 | transmembrane protein 100 | protein_coding | NP_001093110.1;NP_060756.2;XP_016880304.1;XP_016880305.1;XP_02 | 25607 | 616334 | . | . | 0.048222 | 0.079512 | 0.088692 | 0.056906 | 0.112215 | 0.183796 | 0.199022 | 0.130109 | 1.235640 | 5.050188 | 1.346940393 | 0.226630872 | 0.820710785 | NA | 2.175403 | 2.481107 | 2.320361 | 2.330277 | 2.336017 | 2.324344 | 4 | 6 | 6 | 4 |
| 55274 | NM_018288,N | PHF10 | PHD finger protein 10 | protein_coding | NP_060758.2;NP_579866.2 | 18250 | 613069 | . | . | 5.290229 | 5.947156 | 5.242264 | 5.426265 | 12.310579 | 13.747117 | 11.763489 | 12.406510 | -1.065975 | 370.316192 | 0.188192315 | -0.48978222 | 0.624288015 | 0.998865404 | 8.578195 | 8.485545 | 8.517759 | 8.582589 | 8.506983 | 8.520212 | 416 | 435 | 308 | 340 |
| 55275 | NM_00112815 | VPS53 | VPS53 subunit of GARP complex | protein_coding | NP_001121631.1;NP_001353182.1;NP_001353183.1;NP_060759.2;XP_01 | 25608 | 615850 | . | . | 3.581748 | 3.511471 | 3.739135 | 4.459580 | 8.334874 | 8.116924 | 8.390512 | 10.196299 | 1.149927 | 1462.942817 | 0.130104264 | 1.549087268 | 0.121360743 | 0.863030571 | 10.410515 | 10.611786 | 10.458320 | 10.456718 | 10.518180 | 10.609999 | 1569 | 1467 | 1281 | 1553 |
| 55276 | NM_018290,XI | PGM2 | phosphoglucomutase 2 | protein_coding | NP_060760.2;XP_011512013.1 | 8906 | 172000 | . | . | 10.817434 | 9.488674 | 8.927390 | 9.344406 | 25.172611 | 21.933494 | 20.032806 | 21.364876 | -1.131545 | 967.715081 | 0.145001619 | -1.22960066 | 0.218846683 | 0.982008277 | 10.004745 | 9.826639 | 10.014071 | 9.905567 | 9.858963 | 9.883384 | 1269 | 1028 | 776 | 866 |
| 55277 | NM_00111341 | FGGY | FGGY carbohydrate kinase domain containing | protein_coding | NP_001106882.1;NP_001231643.1;NP_001265153.1;NP_001337719.1;NI | 25610 | 611370 | . | . | 2.207887 | 2.339641 | 1.632439 | 2.186543 | 5.137847 | 5.408186 | 3.663144 | 4.999270 | -1.141189 | 130.039520 | 0.287097425 | -0.6636689 | 0.50690225 | 0.998865404 | 7.115445 | 6.923808 | 7.035612 | 7.057574 | 6.975077 | 7.023086 | 155 | 154 | 98 | 122 |
| 55278 | NM_018292,XI | QRSL1 | glutaminyl-tRNA amidotransferase subunit QRSL1 | protein_coding | NP_060762.3;XP_011534226.1 | 21020 | 617209 | . | . | 4.116108 | 4.232870 | 4.126942 | 3.263829 | 9.578354 | 9.784470 | 9.260736 | 7.462356 | -1.144954 | 504.233551 | 0.180882286 | -1.07964958 | 0.280298255 | 0.998865404 | 9.071859 | 8.877496 | 9.011590 | 9.023414 | 9.004322 | 8.853573 | 614 | 586 | 459 | 387 |
| 55279 | NM_00135013 | ZNF654 | zinc finger protein 654 | protein_coding | NP_001337063.1;NP_001337064.1;NP_001337065.1;NP_001337066.1;NI | 25612 | . | | . | 3.482977 | 2.996364 | 3.440993 | 2.438175 | 8.105032 | 6.926231 | 7.721490 | 5.574597 | -1.206006 | 624.372543 | 0.19595546 | -1.3790728 | 0.167872308 | 0.945638095 | 9.414444 | 9.145574 | 9.393988 | 9.283417 | 9.332859 | 9.081377 | 848 | 678 | 588 | 428 |
| 55280 | NM_00130340 | CWF19L1 | CWF19 like cell cycle control factor 1 | protein_coding | NP_001290333.1;NP_001290334.1;NP_001290335.1;NP_001290336.1;NI | 25613 | 616120 | . | . | 8.314449 | 9.779397 | 7.854277 | 9.527093 | 19.348066 | 22.605517 | 17.624770 | 21.782567 | -1.055324 | 710.992562 | 0.165093398 | -0.47055886 | 0.637955791 | 0.998865404 | 9.512461 | 9.433849 | 9.435024 | 9.543198 | 9.391096 | 9.511769 | 776 | 849 | 548 | 706 |
| 55281 | NM_018295 | TMEM140 | transmembrane protein 140 | protein_coding | NP_060765.4 | 21870 | . | | . | 4.940283 | 6.977367 | 8.755100 | 7.756627 | 11.496242 | 16.128496 | 19.646191 | 17.734606 | 1.367942 | 441.802587 | 0.2024735 | 2.232426992 | **0.025586756** | 0.524911334 | 8.543945 | 8.995397 | 8.572867 | 8.767370 | 8.907680 | 8.823375 | 358 | 470 | 473 | 447 |
| 55282 | NM_00116157 | LRRC36 | leucine rich repeat containing 36 | protein_coding | NP_001155047.1;NP_060766.5;XP_005256082.1;XP_005256083.1;XP_00 | 25615 | . | | . | 0.119492 | 0.164575 | 0.162415 | 0.061477 | 0.278062 | 0.380424 | 0.364454 | 0.140561 | -1.229574 | 10.646765 | 0.969548094 | -0.30752283 | 0.75844545 | NA | 3.551764 | 3.257993 | 3.403655 | 3.423330 | 3.422094 | 3.385212 | 11 | 15 | 12 | 5 |
| 55283 | NM_00125369 | MCOLN3 | mucolipin 3 | protein_coding | NP_001240622.1;NP_060768.8;XP_005271060.1;XP_006710813.1;XP_01 | 13358 | 607400 | . | . | 11.961357 | 11.049966 | 16.028903 | 8.009816 | 27.834569 | 25.542490 | 35.968397 | 18.313496 | 1.150777 | 1666.249154 | 0.472222201 | 0.429052166 | 0.667885262 | 0.998865404 | 10.597478 | 10.800184 | 10.623560 | 10.636723 | 10.970478 | 10.443239 | 1769 | 1686 | 2053 | 1130 |
| 55284 | NM_00100148 | UBE2W | ubiquitin conjugating enzyme E2 W | protein_coding | NP_001001481.3;NP_001257944.2;NP_060769.5 | 25616 | 614277 | . | . | 1.625168 | 1.335930 | 1.450968 | 1.003491 | 3.781835 | 3.088064 | 3.255930 | 2.294364 | -1.221111 | 354.986650 | 0.22326442 | -1.29082209 | 0.196765377 | 0.96621494 | 8.607679 | 8.321383 | 8.580143 | 8.457061 | 8.502855 | 8.300274 | 495 | 378 | 330 | 244 |
| 55285 | NM_00117108 | RBM41 | RNA binding motif protein 41 | protein_coding | NP_001164551.1;NP_001311171.1;NP_001311172.1;NP_001311173.1;NI | 25617 | . | | . | 1.196675 | 1.005730 | 1.408677 | 0.968716 | 2.784712 | 2.324795 | 3.161032 | 2.214853 | 1.051953 | 238.581135 | 0.233493435 | 0.312944603 | 0.75432275 | 0.998865404 | 7.860561 | 7.935147 | 7.899476 | 7.865261 | 7.972477 | 7.837960 | 277 | 242 | 242 | 198 |
| 55286 | NM_00110462 | C4orf19 | chromosome 4 open reading frame 19 | protein_coding | NP_001098099.1;NP_060772.2;XP_011512014.1;XP_011512015.1 | 25618 | . | | . | 1.629503 | 2.205522 | 1.681403 | 1.745085 | 3.791921 | 5.098161 | 3.773018 | 3.989930 | -1.155224 | 260.011575 | 0.21751685 | -0.95703992 | 0.338547109 | 0.998865404 | 8.123262 | 7.914022 | 8.032610 | 8.082274 | 7.950217 | 8.017650 | 306 | 315 | 195 | 242 |
| 55288 | NM_00103356 | RHOT1 | ras homolog family member T1 | protein_coding | NP_001028738.1;NP_001028739.2;NP_001028740.1;NP_001275683.1;NI | 21168 | 613888 | . | . | 6.450583 | 7.648205 | 6.657323 | 6.142227 | 15.010774 | 17.679168 | 14.938834 | 14.043473 | -1.105467 | 687.286023 | 0.159192517 | -0.90868625 | 0.363515766 | 0.998865404 | 9.495314 | 9.350591 | 9.404557 | 9.507334 | 9.414636 | 9.361949 | 770 | 836 | 590 | 588 |
| 55290 | NM_018310 | BRF2 | BRF2 RNA polymerase III transcription initiation factor subunit | protein_coding | NP_060780.2 | 17298 | 607013 | . | . | 3.260086 | 2.750091 | 4.099334 | 2.735150 | 7.586353 | 6.356958 | 9.198787 | 6.253596 | 1.121257 | 256.707326 | 0.246960725 | 0.66859706 | 0.503752549 | 0.998865404 | 7.917978 | 8.085143 | 8.009293 | 7.921866 | 8.123737 | 7.914556 | 303 | 238 | 284 | 203 |
| 55291 | NM_00116416 | PPP6R3 | protein phosphatase 6 regulatory subunit 3 | protein_coding | NP_001157632.1;NP_001157633.1;NP_001157634.1;NP_001157635.1;NI | 1173 | 610879 | . | . | 21.889994 | 21.783824 | 21.514673 | 21.776625 | 50.938916 | 50.354285 | 48.278298 | 49.789673 | -1.023994 | 3450.590408 | 0.109641355 | -0.31198677 | 0.755050577 | 0.998865404 | 11.769601 | 11.735452 | 11.776117 | 11.748268 | 11.748833 | 11.736681 | 4088 | 3704 | 2987 | 3173 |
| 55293 | NM_00104069 | UEVLD | UEV and lactate/malate dehyrogenase domains | protein_coding | NP_001035787.1;NP_001248311.1;NP_001248312.1;NP_001248313.1;NI | 30866 | 610985 | . | . | 3.689512 | 3.773928 | 3.341546 | 2.829234 | 8.585645 | 8.723604 | 7.498332 | 6.468707 | -1.219859 | 439.999119 | 0.183004347 | -1.56670698 | 0.117183198 | 0.857396579 | 8.917232 | 8.631342 | 8.831195 | 8.843998 | 8.771071 | 8.658769 | 551 | 527 | 378 | 336 |
| 55294 | NM_00101341 | FBXW7 | F-box and WD repeat domain containing 7 | protein_coding | NP_001013433.1;NP_001243998.1;NP_001336727.1;NP_060785.2;NP_3( | 16712 | 606278 | . | . | 1.718816 | 1.933977 | 1.850330 | 1.643575 | 3.999756 | 4.470479 | 4.152088 | 3.757836 | -1.104801 | 270.217700 | 0.217104501 | -0.66229206 | 0.507784066 | 0.998865404 | 8.148078 | 8.004297 | 8.055772 | 8.146088 | 8.078614 | 8.021746 | 299 | 332 | 236 | 227 |
| 55295 | NM_00134598 | KLHL26 | kelch like family member 26 | protein_coding | NP_001332910.1;NP_001332911.1;NP_001332912.1;NP_001332913.1;NI | 25623 | . | | . | 0.697158 | 0.377612 | 0.598291 | 0.654486 | 1.622316 | 0.872866 | 1.342547 | 1.496407 | 1.096356 | 78.271591 | 0.393359398 | 0.337392839 | 0.735820783 | 0.998865404 | 6.221312 | 6.356363 | 6.343201 | 6.194062 | 6.297863 | 6.292515 | 111 | 57 | 72 | 76 |
| 55296 | NM_00129205 | TBC1D19 | TBC1 domain family member 19 | protein_coding | NP_001278983.1;NP_060787.2;XP_011512154.1;XP_016863852.1;XP_01 | 25624 | . | | . | 3.794481 | 2.964178 | 3.971804 | 2.448700 | 8.829917 | 6.851831 | 8.912612 | 5.598660 | 1.119018 | 328.585383 | 0.23136525 | 0.701198195 | 0.483179336 | 0.998865404 | 8.275737 | 8.439847 | 8.407458 | 8.225425 | 8.464214 | 8.295316 | 416 | 279 | 347 | 277 |
| 55297 | NM_00133036 | CCDC91 | coiled-coil domain containing 91 | protein_coding | NP_001317296.1;NP_001339007.1;NP_001339008.1;NP_001339009.1;NI | 24855 | 617366 | . | . | 6.169507 | 5.628803 | 5.056640 | 5.565661 | 14.356700 | 13.011225 | 11.346952 | 12.725221 | -1.114109 | 454.788121 | 0.177695494 | -0.87728703 | 0.380330754 | 0.998865404 | 8.904978 | 8.748905 | 8.893042 | 8.828471 | 8.762116 | 8.826165 | 580 | 491 | 354 | 425 |
| 55298 | NM_00130092 | RNF121 | ring finger protein 121 | protein_coding | NP_001287855.1;NP_060790.2;XP_006718692.1;XP_006718693.1;XP_00 | 21070 | . | | . | 10.322869 | 11.095037 | 12.323590 | 10.616722 | 24.021736 | 25.646671 | 27.653780 | 24.273876 | 1.051756 | 806.835243 | 0.154173724 | 0.472193579 | 0.636788623 | 0.998865404 | 9.619022 | 9.692308 | 9.624375 | 9.648512 | 9.733970 | 9.606678 | 890 | 863 | 784 | 706 |
| 55299 | NM_018321 | BRIX1 | biogenesis of ribosomes BRX1 | protein_coding | NP_060791.3 | 24170 | 618466 | . | . | 17.895243 | 18.597511 | 16.631712 | 16.312099 | 41.642967 | 42.988979 | 37.321079 | 37.295677 | -1.123111 | 861.076790 | 0.142061295 | -1.17907545 | 0.238368135 | 0.992733415 | 9.831301 | 9.663839 | 9.778940 | 9.801086 | 9.717738 | 9.696551 | 1033 | 998 | 716 | 749 |
| 55300 | NM_018323,N | PI4K2B | phosphatidylinositol 4-kinase type 2 beta | protein_coding | NP_060793.2;XP_005248231.1;XP_005248232.1 | 18215 | 612101 | . | . | 2.832701 | 2.427430 | 2.157013 | 2.571073 | 6.591813 | 5.611114 | 4.840276 | 5.878453 | -1.133316 | 277.291028 | 0.216457241 | -0.83411312 | 0.404217242 | 0.998865404 | 8.202783 | 8.022076 | 8.191418 | 8.099284 | 8.045350 | 8.115864 | 368 | 291 | 210 | 261 |
| 55308 | NM_00132052 | DDX19A | DEAD-box helicase 19A | protein_coding | NP_001307451.1;NP_001307454.1;NP_001307455.1;NP_001307456.1;NI | 25628 | . | | . | 13.983055 | 14.146533 | 15.559973 | 16.317968 | 32.539142 | 32.700346 | 34.916129 | 37.309095 | 1.071867 | 1439.655356 | 0.126614621 | 0.790794267 | 0.429064056 | 0.998865404 | 10.440681 | 10.540605 | 10.448320 | 10.479303 | 10.498991 | 10.535166 | 1568 | 1530 | 1265 | 1429 |
| 55311 | NM_00125379 | ZNF444 | zinc finger protein 444 | protein_coding | NP_001240721.1;NP_060807.2;XP_005259091.1;XP_005259092.1;XP_00 | 16052 | 607874 | . | . | 7.113846 | 8.271051 | 8.875548 | 8.892830 | 16.554212 | 19.118908 | 19.916474 | 20.332398 | 1.107313 | 548.731078 | 0.173075782 | 0.849700537 | 0.3954916 | 0.998865404 | 9.024868 | 9.171328 | 8.995169 | 9.127940 | 9.127925 | 9.131700 | 538 | 620 | 500 | 542 |
| 55312 | NM_018339 | RFK | riboflavin kinase | protein_coding | NP_060809.3 | 30324 | 613010 | . | . | 3.367689 | 3.717900 | 3.411400 | 3.253509 | 7.836752 | 8.594092 | 7.655082 | 7.438761 | -1.078352 | 278.635992 | 0.208601251 | -0.52170699 | 0.601874359 | 0.998865404 | 8.175633 | 8.066791 | 8.117115 | 8.165206 | 8.116934 | 8.087075 | 318 | 326 | 240 | 244 |
| 55313 | NM_00109945 | CPPED1 | calcineurin like phosphoesterase domain containing 1 | protein_coding | NP_001092925.1;NP_060810.2 | 25632 | 615603 | . | . | 3.366622 | 2.945494 | 3.087677 | 3.272893 | 7.834268 | 6.808641 | 6.928658 | 7.483082 | -1.005470 | 603.613494 | 0.163212203 | -0.0482216 | 0.961539639 | 0.998865404 | 9.241328 | 9.233624 | 9.283912 | 9.191454 | 9.216727 | 9.253234 | 745 | 608 | 509 | 580 |
| 55314 | NM_018342,XI | TMEM144 | transmembrane protein 144 | protein_coding | NP_060812.2;XP_005263167.1;XP_005263169.1;XP_006714317.1;XP_01( | 25633 | . | | . | 3.482693 | 3.013095 | 2.993208 | 2.688383 | 8.104370 | 6.964903 | 6.716674 | 6.146670 | -1.155426 | 307.118328 | 0.206806972 | -1.00782542 | 0.313538282 | 0.998865404 | 8.362594 | 8.155265 | 8.342097 | 8.255317 | 8.252743 | 8.189380 | 408 | 328 | 263 | 251 |
| 55315 | NM_00117409 | SLC29A3 | solute carrier family 29 member 3 | protein_coding | NP_001167569.1;NP_001350447.1;NP_060814.4;XP_016871866.1;XP_01 | 23096 | 612373 | . | . | 0.596224 | 1.196470 | 0.929109 | 0.557399 | 1.387438 | 2.765695 | 2.084894 | 1.274427 | -1.212198 | 61.016729 | 0.435540039 | -0.63742684 | 0.52384685 | 0.998865404 | 6.063054 | 5.785924 | 5.888055 | 5.998694 | 5.939740 | 5.865952 | 59 | 89 | 57 | 42 |
| 55316 | NM_018346,N | RSAD1 | radical S-adenosyl methionine domain containing 1 | protein_coding | NP_060816.1 | 25634 | . | | . | 19.199064 | 18.342923 | 20.733489 | 21.073399 | 44.677011 | 42.400488 | 46.525351 | 48.181824 | 1.102697 | 1508.878408 | 0.124389072 | 1.133837916 | 0.256862543 | 0.995760118 | 10.486985 | 10.628088 | 10.533440 | 10.506218 | 10.594785 | 10.596248 | 1682 | 1521 | 1375 | 1485 |
| 55317 | NM_00120444 | AP5S1 | adaptor related protein complex 5 subunit sigma 1 | protein_coding | NP_001191375.1;NP_001191376.1;NP_060817.1 | 15875 | 614824 | . | . | 3.348920 | 3.460681 | 4.893203 | 4.171737 | 7.793076 | 7.999521 | 10.980208 | 9.538178 | 1.197401 | 492.760113 | 0.167938893 | 1.547626991 | 0.121712129 | 0.864429515 | 8.809075 | 9.068740 | 8.874270 | 8.892292 | 8.985590 | 9.007948 | 509 | 491 | 459 | 512 |
| 55319 | NM_018352 | TMA16 | translation machinery associated 16 homolog | protein_coding | NP_060822.2 | 25638 | . | | . | 3.338371 | 4.557343 | 3.228350 | 2.588871 | 7.768526 | 10.534503 | 7.244324 | 5.919147 | -1.373244 | 187.270878 | 0.266268356 | -1.71852173 | 0.085701495 | 0.791792328 | 7.759569 | 7.302317 | 7.533603 | 7.678059 | 7.514714 | 7.424331 | 212 | 269 | 153 | 131 |
| 55320 | NM_018353,XI | MIS18BP1 | MIS18 binding protein 1 | protein_coding | NP_060823.3;XP_005267890.1;XP_011535225.1;XP_016876914.1 | 20190 | 618139 | . | . | 5.447559 | 4.477281 | 5.295053 | 4.666284 | 12.676692 | 10.349435 | 11.881947 | 10.668904 | -1.152382 | 874.881253 | 0.161478527 | -1.26715676 | 0.205099245 | 0.96993643 | 9.871208 | 9.667493 | 9.861916 | 9.775025 | 9.797613 | 9.631555 | 1142 | 951 | 791 | 672 |
| 55322 | NM_018356,N | C5orf22 | chromosome 5 open reading frame 22 | protein_coding | NP_060826.2;XP_006714542.1;XP_006714543.1;XP_011512364.1;XP_01( | 25639 | . | | . | 9.439998 | 9.384155 | 8.728734 | 8.975070 | 21.967261 | 21.691891 | 19.587025 | 20.520431 | -1.077873 | 1001.139605 | 0.136109381 | -0.79485898 | 0.426695536 | 0.998865404 | 10.020537 | 9.912287 | 9.996298 | 9.992538 | 9.931810 | 9.947080 | 1199 | 1118 | 829 | 913 |
| 55323 | NM_00128667 | LARP6 | La ribonucleoprotein 6, translational regulator | protein_coding | NP_001273608.1;NP_060827.2;NP_932062.1 | 24012 | 611300 | . | . | 17.040143 | 18.980767 | 12.454954 | 20.208363 | 39.653113 | 43.874892 | 27.948556 | 46.204023 | -1.103372 | 501.450988 | 0.198075428 | -0.71649417 | 0.473686268 | 0.998865404 | 9.039850 | 8.896521 | 8.961869 | 9.030728 | 8.816918 | 9.039432 | 574 | 598 | 340 | 527 |
| 55324 | NM_00135129 | ABCF3 | ATP binding cassette subfamily F member 3 | protein_coding | NP_001338227.1;NP_001338228.1;NP_001338229.1;NP_060828.2;XP_02 | 72 | . | | . | 18.845746 | 17.904418 | 21.134771 | 20.640300 | 43.854826 | 41.386863 | 47.425808 | 47.191593 | 1.119801 | 1578.093670 | 0.124832371 | 1.307692726 | 0.190977551 | 0.963121353 | 10.539932 | 10.703376 | 10.600266 | 10.554952 | 10.680294 | 10.651100 | 1765 | 1559 | 1477 | 1534 |
| 55325 | NM_018359,N | UFSP2 | UFM1 specific peptidase 2 | protein_coding | NP_060829.2 | 25640 | 611482 | . | . | 7.415489 | 6.633047 | 7.399313 | 6.923368 | 17.256148 | 15.332584 | 16.603846 | 15.829454 | 1.005400 | 523.281304 | 0.169493344 | 0.045843071 | 0.963435329 | 0.998865404 | 9.027220 | 9.035653 | 9.066133 | 8.990448 | 9.056919 | 9.007663 | 636 | 530 | 474 | 473 |
| 55326 | NM_018361,XI | AGPAT5 | 1-acylglycerol-3-phosphate O-acyltransferase 5 | protein_coding | NP_060831.2;XP_011533053.1 | 20886 | 614796 | . | . | 8.070962 | 7.469984 | 7.347085 | 6.377311 | 18.781458 | 17.267204 | 16.486645 | 14.580964 | -1.147753 | 1194.071186 | 0.141062666 | -1.4093869 | 0.158720796 | 0.931076558 | 10.317391 | 10.119151 | 10.302098 | 10.237432 | 10.221598 | 10.109070 | 1531 | 1319 | 1041 | 963 |
| 55327 | NM_018362 | LIN7C | lin-7 homolog C, crumbs cell polarity complex component | protein_coding | NP_060832.1 | 17789 | 612332 | . | . | 1.297577 | 1.376021 | 1.307964 | 1.125170 | 3.019516 | 3.180735 | 2.935034 | 2.572567 | -1.110257 | 193.212834 | 0.243860538 | -0.61877237 | 0.536066324 | 0.998865404 | 7.667191 | 7.516996 | 7.604357 | 7.628111 | 7.603471 | 7.535384 | 228 | 225 | 172 | 158 |
| 55328 | NM_00103170 | RNLS | renalase, FAD dependent amine oxidase | protein_coding | NP_001026879.2;NP_060833.1;XP_005270003.1;XP_005270004.1;XP_00 | 25641 | 609360 | . | . | 0.465475 | 0.493838 | 0.489629 | 0.794424 | 1.083181 | 1.141529 | 1.098713 | 1.816354 | 1.278222 | 48.971187 | 0.459188665 | 0.771225274 | 0.440573416 | 0.998865404 | 5.427143 | 5.779160 | 5.585681 | 5.582881 | 5.587936 | 5.668343 | 50 | 46 | 38 | 62 |
| 55329 | NM_018365 | MNS1 | meiosis specific nuclear structural 1 | protein_coding | NP_060835.1 | 29636 | 610766 | . | . | 5.377698 | 5.798958 | 4.441888 | 5.270149 | 12.514126 | 13.404552 | 9.967468 | 12.049568 | -1.167154 | 330.720758 | 0.19950247 | -1.11775743 | 0.263670608 | 0.996308385 | 8.477129 | 8.253124 | 8.391707 | 8.429243 | 8.281171 | 8.366104 | 397 | 397 | 244 | 309 |
| 55330 | NM_018366 | BLOC1S4 | biogenesis of lysosomal organelles complex 1 subunit 4 | protein_coding | NP_060836.1 | 24206 | 605695 | . | . | 5.564970 | 5.528575 | 4.950492 | 5.838945 | 12.949915 | 12.779551 | 11.108762 | 13.350054 | -1.040040 | 254.457769 | 0.218224574 | -0.25954359 | 0.795215852 | 0.998865404 | 8.019691 | 7.962303 | 8.004775 | 7.997250 | 7.942290 | 8.018682 | 301 | 278 | 200 | 252 |
| 55331 | NM_00130095 | ACER3 | alkaline ceramidase 3 | protein_coding | NP_001287882.1;NP_001287883.1;NP_001287884.1;NP_060837.3;XP_01 | 16066 | 617036 | . | . | 2.793200 | 2.548468 | 2.734525 | 1.871697 | 6.499892 | 5.890900 | 6.136195 | 4.279414 | -1.204868 | 555.777525 | 0.206443712 | -1.30241303 | 0.192775232 | 0.96621494 | 9.245878 | 8.978477 | 9.211374 | 9.125912 | 9.177273 | 8.900365 | 743 | 614 | 535 | 369 |
| 55332 | NM_018370,XI | DRAM1 | DNA damage regulated autophagy modulator 1 | protein_coding | NP_060840.2;XP_005269061.1;XP_005269062.1;XP_016875067.1 | 25645 | 610776 | . | . | 4.477560 | 4.296560 | 4.706734 | 4.726259 | 10.419465 | 9.931689 | 10.561774 | 10.806033 | 1.051040 | 470.323183 | 0.171317397 | 0.419208378 | 0.67506385 | 0.998865404 | 8.841019 | 8.913098 | 8.882208 | 8.839967 | 8.895009 | 8.889748 | 546 | 478 | 421 | 450 |
| 55333 | NM_018373 | SYNJ2BP | synaptojanin 2 binding protein | protein_coding | NP_060843.2 | 18955 | 609411 | . | . | 3.972324 | 4.281305 | 4.400569 | 4.203774 | 9.243762 | 9.896428 | 9.874750 | 9.611430 | 1.028156 | 926.650983 | 0.141287309 | 0.283528296 | 0.776771894 | 0.998865404 | 9.835702 | 9.875784 | 9.820789 | 9.869290 | 9.885573 | 9.844873 | 1017 | 1019 | 840 | 856 |
| 55334 | NM_00125214 | SLC39A9 | solute carrier family 39 member 9 | protein_coding | NP_001239077.1;NP_001239079.1;NP_001239080.1;NP_001239081.1;NI | 20182 | . | | . | 8.243485 | 7.581576 | 9.150654 | 8.073852 | 19.182928 | 17.525155 | 20.533801 | 18.459903 | 1.051301 | 1345.838838 | 0.135173742 | 0.533952795 | 0.593374194 | 0.998865404 | 10.357518 | 10.430148 | 10.410193 | 10.336392 | 10.454250 | 10.368483 | 1584 | 1347 | 1267 | 1221 |
| 55335 | NM_018376,N | NIPSNAP3B | nipsnap homolog 3B | protein_coding | NP_060846.2;XP_011517141.1 | 23641 | 608872 | . | . | 0.114197 | 0.054231 | 0.123541 | 0.139141 | 0.265742 | 0.125358 | 0.277222 | 0.318131 | 1.444724 | 20.091204 | 0.706785303 | 0.750996842 | 0.452654557 | NA | 4.037030 | 4.570852 | 4.326558 | 4.279759 | 4.331288 | 4.337866 | 25 | 12 | 20 | 23 |
| 55336 | NM_018378 | FBXL8 | F-box and leucine rich repeat protein 8 | protein_coding | NP_060848.2 | 17875 | 609077 | . | . | 0.861695 | 1.109505 | 0.953341 | 1.146826 | 2.005199 | 2.564671 | 2.139270 | 2.622081 | 1.063246 | 51.967724 | 0.439581291 | 0.201273409 | 0.840484794 | 0.998865404 | 5.655666 | 5.742126 | 5.667878 | 5.714160 | 5.688281 | 5.717459 | 51 | 61 | 43 | 54 |
| 55337 | NM_00130827 | SHFL | shiftless antiviral inhibitor of ribosomal frameshifting | protein_coding | NP_001295206.1;NP_060851.2;XP_011526423.1;XP_011526424.1 | 25649 | 616808 | . | . | 13.351565 | 13.820939 | 14.265902 | 14.730647 | 31.069642 | 31.947722 | 32.012272 | 33.679877 | 1.039361 | 859.762268 | 0.142506819 | 0.390835012 | 0.695919193 | 0.998865404 | 9.719803 | 9.775252 | 9.723273 | 9.743877 | 9.743123 | 9.778838 | 957 | 923 | 743 | 842 |
| 55339 | NM_00100662 | WDR33 | WD repeat domain 33 | protein_coding | NP_001006623.1;NP_001006624.1;NP_060853.3;XP_005263754.1;XP_01 | 25651 | 618082 | . | . | 12.167454 | 10.828798 | 11.868054 | 11.356974 | 28.314163 | 25.031255 | 26.631567 | 25.966374 | 1.030573 | 2692.329124 | 0.112636781 | 0.385719217 | 0.699704629 | 0.998865404 | 11.372732 | 11.416219 | 11.391344 | 11.373005 | 11.409253 | 11.404130 | 3088 | 2829 | 2384 | 2554 |
| 55341 | NM_018385,XI | LSG1 | large 60S subunit nuclear export GTPase 1 | protein_coding | NP_060855.2;XP_016862285.1 | 25652 | 610780 | . | . | 23.492861 | 24.547291 | 30.034017 | 26.541724 | 54.668853 | 56.742159 | 67.395464 | 60.684492 | 1.159032 | 2660.002205 | 0.120598349 | 1.765533499 | 0.077474145 | 0.767975096 | 11.266755 | 11.479810 | 11.295656 | 11.330828 | 11.483472 | 11.380468 | 2775 | 2717 | 2639 | 2510 |
| 55342 | NM_00117113 | STRBP | spermatid perinuclear RNA binding protein | protein_coding | NP_001164608.1;NP_001363035.1;NP_001363036.1;NP_001363038.1;NI | 16462 | 611138 | . | . | 3.567209 | 3.554632 | 2.452626 | 3.364193 | 8.301040 | 8.216691 | 5.503621 | 7.691827 | -1.198593 | 354.734501 | 0.195828267 | -1.3345444 | 0.182025507 | 0.953687323 | 8.595775 | 8.333514 | 8.520368 | 8.518745 | 8.363929 | 8.465882 | 446 | 417 | 254 | 331 |
| 55343 | NM_00114526 | SLC35C1 | solute carrier family 35 member C1 | protein_coding | NP_001138737.1;NP_001138738.1;NP_060859.4;XP_011518504.1;XP_01 | 20197 | 605881 | . | . | 8.692542 | 9.216179 | 8.676425 | 9.405056 | 20.227904 | 21.303609 | 19.469650 | 21.503542 | -1.010360 | 871.992766 | 0.142263237 | -0.10452028 | 0.916756474 | 0.998865404 | 9.775725 | 9.760580 | 9.757899 | 9.785547 | 9.746001 | 9.782383 | 990 | 964 | 735 | 834 |
| 55344 | NM_00137037 | PLCXD1 | phosphatidylinositol specific phospholipase C X domain containing | protein_coding | NP_001357299.1;NP_001357300.1;NP_001357301.1;NP_001357302.1;NI | 23148 | 300974 | . | . | 6.677280 | 7.003415 | 7.006418 | 6.386727 | 15.538300 | 16.188706 | 15.722201 | 14.602494 | -1.016658 | 1096.667086 | 0.138847574 | -0.17165815 | 0.863706291 | 0.998865404 | 10.110566 | 10.087159 | 10.113747 | 10.094990 | 10.138128 | 10.044280 | 1289 | 1178 | 1006 | 955 |
| 55345 | NM_00135039 | ZGRF1 | zinc finger GRF-type containing 1 | protein_coding | NP_001337326.1;NP_060862.3;XP_005263172.1;XP_011530393.1;XP_01 | 25654 | . | | . | 2.334784 | 2.431910 | 2.085083 | 1.503507 | 5.433140 | 5.621465 | 4.678865 | 3.437589 | -1.297299 | 417.448283 | 0.201722396 | -1.86152467 | 0.06267012 | 0.723723143 | 8.880378 | 8.506355 | 8.790309 | 8.758565 | 8.708662 | 8.519465 | 556 | 496 | 369 | 284 |
| 55346 | NM_00114554 | TCP11L1 | t-complex 11 like 1 | protein_coding | NP_001139013.1;NP_060863.3;XP_011518506.1;XP_011518507.1;XP_01 | 25655 | . | | . | 1.738025 | 2.365857 | 2.056796 | 2.462935 | 4.044454 | 5.468785 | 4.615391 | 5.631207 | -1.127502 | 323.060441 | 0.217449424 | -0.79618428 | 0.425924931 | 0.998865404 | 8.420200 | 8.245857 | 8.386713 | 8.344219 | 8.206891 | 8.383957 | 409 | 356 | 220 | 332 |
| 55347 | NM_00127206 | ABHD10 | abhydrolase domain containing 10, depalmitoylase | protein_coding | NP_001258998.1;NP_060864.1;XP_011511262.1 | 25656 | 618756 | . | . | 5.911826 | 5.955528 | 5.032299 | 6.297732 | 13.757061 | 13.766470 | 11.292335 | 14.399016 | -1.076187 | 467.340226 | 0.178058791 | -0.59490854 | 0.551904605 | 0.998865404 | 8.920680 | 8.814007 | 8.891536 | 8.889202 | 8.789804 | 8.896515 | 559 | 522 | 357 | 459 |
| 55351 | NM_00130608 | STK32B | serine/threonine kinase 32B | protein_coding | NP_001293011.1;NP_001332898.1;NP_060871.1;XP_005248039.1;XP_00 | 14217 | . | | . | 13.622675 | 13.894210 | 15.094638 | 15.512101 | 31.700523 | 32.117090 | 33.871929 | 35.466578 | 1.103283 | 1567.273447 | 0.123428192 | 1.148874816 | 0.250607603 | 0.993339639 | 10.541425 | 10.683177 | 10.579634 | 10.569043 | 10.638035 | 10.663068 | 1728 | 1597 | 1407 | 1566 |
| 55352 | NM_00133017 | COPRS | coordinator of PRMT5 and differentiation stimulator | protein_coding | NP_001317105.1;NP_060875.2 | 28848 | . | | . | 46.115839 | 47.074271 | 43.962036 | 50.592977 | 107.313456 | 108.814281 | 98.649532 | 115.674818 | 1.002974 | 1272.974547 | 0.134491766 | 0.031853827 | 0.97458862 | 0.998865404 | 10.312027 | 10.315946 | 10.307871 | 10.317040 | 10.265427 | 10.361522 | 1455 | 1380 | 1038 | 1271 |
| 55353 | NM_018407 | LAPTM4B | lysosomal protein transmembrane 4 beta | protein_coding | NP_060877.4 | 13646 | 613296 | . | . | 109.919044 | 99.524155 | 107.479424 | 84.293877 | 255.786118 | 230.054550 | 241.180710 | 192.727936 | -1.107133 | 7636.537909 | 0.132915132 | -1.10467781 | 0.269299222 | 0.998865404 | 12.970176 | 12.823504 | 12.983211 | 12.890833 | 12.952465 | 12.740180 | 9738 | 8192 | 7097 | 5940 |
| 55355 | NM_00128296 | HJURP | Holliday junction recognition protein | protein_coding | NP_001269891.1;NP_001269892.1;NP_060880.3;XP_011509739.1 | 25444 | 612667 | . | . | 41.406723 | 42.425908 | 40.667589 | 39.118553 | 96.355153 | 98.069388 | 91.256889 | 89.439926 | -1.058996 | 3931.611678 | 0.108100407 | -0.7649979 | 0.444272796 | 0.998865404 | 11.981656 | 11.898970 | 11.953226 | 11.975037 | 11.926822 | 11.906433 | 4596 | 4421 | 3362 | 3536 |
| 55356 | NM_018420,XI | SLC22A15 | solute carrier family 22 member 15 | protein_coding | NP_060890.2;XP_005271061.1;XP_024304006.1;XP_024304007.1 | 20301 | 608275 | . | . | 6.166491 | 5.842210 | 7.253403 | 4.680190 | 14.349679 | 13.504531 | 16.276423 | 10.700699 | 1.008668 | 672.895369 | 0.187474432 | 0.066416308 | 0.947046387 | 0.998865404 | 9.387460 | 9.400985 | 9.402008 | 9.370362 | 9.513330 | 9.255155 | 789 | 706 | 699 | 514 |
| 55357 | NM_00126757 | TBC1D2 | TBC1 domain family member 2 | protein_coding | NP_001254500.1;NP_001254501.1;NP_060891.3;XP_011517145.1;XP_01 | 18026 | 609871 | . | . | 1.760745 | 1.884544 | 1.807657 | 1.863846 | 4.097326 | 4.356205 | 4.056329 | 4.261461 | -1.046592 | 170.593253 | 0.255952183 | -0.256686 | 0.797421179 | 0.998865404 | 7.447399 | 7.380663 | 7.413264 | 7.434410 | 7.373981 | 7.434821 | 196 | 193 | 134 | 168 |
| 55359 | NM_018423,XI | STYK1 | serine/threonine/tyrosine kinase 1 | protein_coding | NP_060893.2;XP_005253474.1;XP_011519038.1;XP_011519039.1;XP_01 | 18889 | 611433 | . | . | 1.485320 | 1.713031 | 1.688082 | 1.367392 | 3.456401 | 3.959749 | 3.788006 | 3.126377 | -1.065837 | 139.431588 | 0.285423248 | -0.32227989 | 0.747240664 | 0.998865404 | 7.168403 | 7.076976 | 7.100929 | 7.164541 | 7.147432 | 7.070084 | 152 | 168 | 130 | 113 |
| 55361 | NM_018425 | PI4K2A | phosphatidylinositol 4-kinase type 2 alpha | protein_coding | NP_060895.1 | 30031 | 609763 | . | . | 10.400333 | 10.300102 | 11.535735 | 13.868845 | 24.202003 | 23.809147 | 25.885855 | 31.709465 | 1.209869 | 1503.309426 | 0.135597087 | 2.026970439 | **0.042665436** | 0.63734958 | 10.410144 | 10.684669 | 10.480835 | 10.467285 | 10.550325 | 10.687655 | 1581 | 1455 | 1307 | 1677 |
| 55362 | NM_00131879 | TMEM63B | transmembrane protein 63B | protein_coding | NP_001305721.1;NP_060896.1;XP_005249268.1;XP_005249270.1;XP_00 | 17735 | . | | . | 12.199840 | 12.960632 | 11.759791 | 11.743895 | 28.389527 | 29.959079 | 26.388626 | 26.851017 | -1.085058 | 1247.081046 | 0.129983998 | -0.90605097 | 0.364908875 | 0.998865404 | 10.342047 | 10.224227 | 10.296553 | 10.331977 | 10.258341 | 10.247064 | 1460 | 1433 | 1048 | 1113 |
| 55364 | NM_018439,XI | IMPACT | impact RWD domain protein | protein_coding | NP_060909.2;XP_016881321.1 | 20387 | 615319 | . | . | 5.816034 | 5.502519 | 6.326464 | 5.524973 | 13.534148 | 12.719321 | 14.196401 | 12.632194 | 1.039404 | 679.263076 | 0.158846376 | 0.351006108 | 0.725583766 | 0.998865404 | 9.379326 | 9.435775 | 9.414521 | 9.371598 | 9.469368 | 9.367789 | 791 | 696 | 647 | 601 |
| 55366 | NM_00134643 | LGR4 | leucine rich repeat containing G protein-coupled receptor 4 | protein_coding | NP_001333361.1;NP_060960.2 | 13299 | 606666 | . | . | 3.008478 | 2.256818 | 2.754560 | 1.976729 | 7.000852 | 5.216734 | 6.181154 | 4.519557 | -1.127892 | 409.776307 | 0.219360923 | -0.79152434 | 0.428638077 | 0.998865404 | 8.762075 | 8.590194 | 8.794363 | 8.612721 | 8.729562 | 8.532108 | 573 | 400 | 392 | 300 |
| 55367 | NM_145886,N | PIDD1 | p53-induced death domain protein 1 | protein_coding | NP_665893.2;NP_665894.2;XP_005253062.1;XP_005253063.1;XP_00525 | 16491 | 605247 | . | . | 15.125976 | 14.870946 | 19.650288 | 15.992014 | 35.198772 | 34.374859 | 44.094675 | 36.563839 | 1.120818 | 1629.250410 | 0.143869685 | 1.143761213 | 0.25272268 | 0.994320829 | 10.585124 | 10.750145 | 10.631782 | 10.610811 | 10.803182 | 10.608588 | 1793 | 1635 | 1682 | 1416 |
| 55370 | NR_003505 | PPP4R1L | protein phosphatase 4 regulatory subunit 1 like (pseudogene) | transcribed_ps | . | 15755 | . | | . | 2.134279 | 2.550526 | 2.233375 | 2.195010 | 4.966554 | 5.895655 | 5.011630 | 5.018629 | -1.074080 | 170.561515 | 0.257229457 | -0.40081718 | 0.688554729 | 0.998865404 | 7.465021 | 7.361416 | 7.390584 | 7.464556 | 7.404154 | 7.392714 | 186 | 207 | 145 | 152 |
| 55374 | NM_00130098 | TMCO6 | transmembrane and coiled-coil domains 6 | protein_coding | NP_001287909.1;NP_001287911.1;NP_060972.3;XP_005268534.1;XP_01 | 28814 | . | | . | 6.487130 | 7.111524 | 8.968960 | 8.358729 | 15.095818 | 16.438607 | 20.126086 | 19.111240 | 1.247798 | 473.726820 | 0.172913476 | 1.847077039 | 0.064735976 | 0.731823074 | 8.719365 | 9.038811 | 8.788386 | 8.834271 | 8.975572 | 8.923332 | 467 | 472 | 476 | 473 |
| 55379 | NM_018509 | LRRC59 | leucine rich repeat containing 59 | protein_coding | NP_060979.2 | 28817 | 614854 | . | . | 59.469955 | 62.359402 | 50.203381 | 58.663029 | 138.389023 | 144.146545 | 112.654930 | 134.126038 | -1.135037 | 5175.853344 | 0.115854886 | -1.57731251 | 0.114723636 | 0.855274326 | 12.426135 | 12.243234 | 12.371150 | 12.404134 | 12.219708 | 12.339863 | 6214 | 6054 | 3910 | 4875 |
| 55388 | NM_018518,N | MCM10 | minichromosome maintenance 10 replication initiation factor | protein_coding | NP_060988.3;NP_877428.1;XP_011517840.1 | 18043 | 609357 | . | . | 5.178921 | 5.615365 | 3.978738 | 5.199405 | 12.051564 | 12.980169 | 8.928172 | 11.887819 | -1.191139 | 708.600339 | 0.16875713 | -1.49529286 | 0.134838026 | 0.891755273 | 9.589945 | 9.336604 | 9.498826 | 9.549024 | 9.314737 | 9.485407 | 856 | 861 | 490 | 684 |
| 55421 | NM_00111411 | NCBP3 | nuclear cap binding subunit 3 | protein_coding | NP_001107590.1;XP_005256774.1;XP_006721619.1;XP_011522256.1;XP_ | 24612 | 616624 | . | . | 5.173450 | 4.656230 | 4.755304 | 4.201511 | 12.038830 | 10.763083 | 10.670765 | 9.606253 | 1.021043 | 1373.345035 | 0.126805165 | 0.236922469 | 0.812716938 | 0.998865404 | 10.408336 | 10.438464 | 10.412930 | 10.417719 | 10.444682 | 10.417721 | 1560 | 1471 | 1228 | 1278 |
| 55422 | NM_00107990 | ZNF331 | zinc finger protein 331 | protein_coding | NP_001073375.1;NP_001073376.1;NP_001240727.1;NP_001240728.1;NI | 15489 | 606043 | . | . | 8.380715 | 8.691823 | 9.899082 | 8.036174 | 19.502268 | 20.091537 | 22.213251 | 18.373760 | 1.037319 | 1372.542722 | 0.142507494 | 0.370928454 | 0.710690821 | 0.998865404 | 10.395742 | 10.449037 | 10.399156 | 10.414239 | 10.513431 | 10.349881 | 1536 | 1468 | 1345 | 1171 |
| 55425 | NM_00131695 | GPALPP1 | GPALPP motifs containing 1 | protein_coding | NP_001303880.1;NP_001303881.1;NP_061029.2;XP_005266501.1;XP_01 | 20298 | . | | . | 4.154206 | 3.523180 | 3.589848 | 4.510737 | 9.667006 | 8.143992 | 8.055512 | 10.313266 | -1.002157 | 383.954054 | 0.201613777 | -0.0154158 | 0.98770046 | 0.999197585 | 8.586540 | 8.583038 | 8.635827 | 8.527906 | 8.512808 | 8.646759 | 483 | 377 | 295 | 401 |
| 55432 | NM_00127632 | YOD1 | YOD1 deubiquitinase | protein_coding | NP_001263249.1;NP_061036.3 | 25035 | 612023 | . | . | 2.602319 | 2.218719 | 2.512354 | 1.738990 | 6.055703 | 5.128667 | 5.637650 | 3.975994 | -1.150074 | 443.662537 | 0.208971947 | -0.96532972 | 0.334379689 | 0.998865404 | 8.889901 | 8.689804 | 8.881581 | 8.774839 | 8.848791 | 8.627248 | 592 | 469 | 426 | 315 |
| 55435 | NM_00112842 | AP1AR | adaptor related protein complex 1 associated regulatory protein | protein_coding | NP_001121898.1;NP_061039.3;XP_016863867.1;XP_016863868.1;XP_01 | 28808 | 610851 | . | . | 2.396665 | 2.551332 | 2.573386 | 2.863455 | 5.577139 | 5.897519 | 5.774603 | 6.546947 | -1.034347 | 446.298523 | 0.178811165 | -0.27247019 | 0.785260507 | 0.998865404 | 8.825840 | 8.777485 | 8.789067 | 8.833187 | 8.830227 | 8.749619 | 504 | 507 | 407 | 384 |
| 55437 | NM_00120686 | STRADB | STE20 related adaptor beta | protein_coding | NP_001193793.1;NP_061041.2;XP_005246726.1;XP_016859927.1 | 13205 | 607333 | . | . | 17.757338 | 17.923813 | 21.285813 | 20.687773 | 41.322053 | 41.431693 | 47.764746 | 47.300133 | 1.161115 | 1342.976573 | 0.127482282 | 1.690518288 | 0.090928842 | 0.805877675 | 10.279391 | 10.494998 | 10.329687 | 10.331984 | 10.459399 | 10.429873 | 1429 | 1343 | 1279 | 1327 |
| 55449 | NR_023921,NF | DHRS4-AS1 | DHRS4 antisense RNA 1 | lncRNA | . | 23175 | 616925 | . | . | 3.467750 | 3.539353 | 3.490066 | 4.741301 | 8.181374 | 8.069597 | 7.831607 | 10.840422 | 1.148632 | 345.701771 | 0.204565705 | 0.977276631 | 0.328432219 | 0.998865404 | 8.330538 | 8.529384 | 8.388381 | 8.390640 | 8.388641 | 8.541311 | 370 | 348 | 279 | 392 |
| 55450 | NM_018584 | CAMK2N1 | calcium/calmodulin dependent protein kinase II inhibitor 1 | protein_coding | NP_061054.2 | 24190 | 614986 | . | . | 6.055874 | 5.367942 | 5.725611 | 5.595907 | 14.092267 | 12.408238 | 12.848104 | 12.794376 | -1.022120 | 412.535245 | 0.182503124 | -0.17295118 | 0.862689803 | 0.998865404 | 8.703788 | 8.672792 | 8.732787 | 8.654640 | 8.691627 | 8.671185 | 511 | 421 | 361 | 376 |
| 55454 | NM_00131965 | CSGALNACT2 | chondroitin sulfate N-acetylgalactosaminyltransferase 2 | protein_coding | NP_001306583.1;NP_001306585.1;NP_061060.3;XP_011538240.1;XP_01 | 24292 | 616616 | . | . | 7.722067 | 7.050301 | 7.365113 | 6.063758 | 17.969572 | 16.297358 | 16.527100 | 13.864066 | -1.104119 | 818.694965 | 0.162088651 | -0.88158882 | 0.377999201 | 0.998865404 | 9.746429 | 9.604443 | 9.755682 | 9.660610 | 9.711666 | 9.561420 | 1054 | 866 | 750 | 651 |
| 55466 | NM_00113018 | DNAJA4 | DnaJ heat shock protein family (Hsp40) member A4 | protein_coding | NP_001123654.1;NP_001123655.1;NP_061072.3;XP_005254592.1;XP_00 | 14885 | . | | . | 6.471678 | 7.369549 | 6.236276 | 6.480268 | 15.059860 | 17.035042 | 13.994023 | 14.816363 | -1.103897 | 631.353949 | 0.159767238 | -0.89258564 | 0.372079111 | 0.998865404 | 9.372062 | 9.229004 | 9.290119 | 9.376526 | 9.257988 | 9.277453 | 715 | 760 | 514 | 571 |
| 55471 | NM_00108394 | NDUFAF7 | NADH:ubiquinone oxidoreductase complex assembly factor 7 | protein_coding | NP_001077415.1;NP_001336953.1;NP_001336954.1;NP_001336956.1;NI | 28816 | 615898 | . | . | 8.191638 | 8.307306 | 11.206199 | 7.188071 | 19.062278 | 19.202707 | 25.146383 | 16.434673 | 1.122034 | 869.383749 | 0.16412589 | 1.012129872 | 0.311475961 | 0.998865404 | 9.678035 | 9.844845 | 9.715146 | 9.720174 | 9.897451 | 9.693045 | 940 | 887 | 912 | 741 |
| 55486 | NM_00103763 | PARL | presenilin associated rhomboid like | protein_coding | NP_001032728.1;NP_001311365.1;NP_001311366.1;NP_001311367.1;NI | 18253 | 607858 | . | . | 28.565325 | 28.155706 | 29.783972 | 33.936632 | 66.472683 | 65.083176 | 66.834371 | 77.592072 | 1.105229 | 1373.048996 | 0.132309659 | 1.090966517 | 0.275287616 | 0.998865404 | 10.349323 | 10.493415 | 10.392340 | 10.373396 | 10.411058 | 10.505567 | 1521 | 1390 | 1177 | 1434 |
| 55500 | NM_00103948 | ETNK1 | ethanolamine kinase 1 | protein_coding | NP_001034570.2;NP_061108.3;XP_005253477.1;XP_016875069.1;XP_01 | 24649 | 609858 | . | . | 7.156023 | 6.713092 | 8.124924 | 6.458509 | 16.652362 | 15.517618 | 18.232096 | 14.766612 | -1.055854 | 1329.872698 | 0.159266183 | -0.49231877 | 0.622494016 | 0.998865404 | 10.415402 | 10.337696 | 10.430942 | 10.355929 | 10.462651 | 10.229820 | 1649 | 1402 | 1298 | 1027 |
| 55501 | NM_00124379 | CHST12 | carbohydrate sulfotransferase 12 | protein_coding | NP_001230723.1;NP_001230724.1;NP_061111.1;XP_011513745.1;XP_01 | 17423 | 610129 | . | . | 3.616488 | 2.418301 | 2.769417 | 2.656992 | 8.415720 | 5.590010 | 6.214490 | 6.074896 | -1.085802 | 1119.191213 | 0.133238725 | -0.89133742 | 0.372748182 | 0.998865404 | 10.186334 | 10.067710 | 10.176227 | 10.139754 | 10.100682 | 10.093108 | 1374 | 1227 | 938 | 1001 |
| 55502 | NM_00114285 | HES6 | hes family bHLH transcription factor 6 | protein_coding | NP_001136325.1;NP_001269363.1;NP_061115.2 | 18254 | 610331 | . | . | 34.106488 | 35.655704 | 47.500654 | 50.501482 | 79.367201 | 82.419758 | 106.590088 | 115.465636 | 1.383252 | 1784.162097 | 0.121706055 | 3.845854165 | **0.000120133** | **0.013818322** | 10.548200 | 11.016091 | 10.644932 | 10.671681 | 10.892257 | 10.933950 | 1694 | 1644 | 1755 | 1991 |
| 55504 | NM_00120445 | TNFRSF19 | TNF receptor superfamily member 19 | protein_coding | NP_001191387.1;NP_001191388.1;NP_001341914.1;NP_061117.2;NP_6( | 11915 | 606122 | . | . | 16.534895 | 15.399570 | 17.374619 | 14.971103 | 38.477380 | 35.596796 | 38.988140 | 34.229643 | 1.005486 | 2122.763002 | 0.127577814 | 0.061866528 | 0.950669123 | 0.998865404 | 11.047596 | 11.055847 | 11.082710 | 11.013347 | 11.106704 | 10.996559 | 2543 | 2185 | 1979 | 1858 |
| 55505 | NM_018648 | NOP10 | NOP10 ribonucleoprotein | protein_coding | NP_061118.1 | 14378 | 606471 | . | . | 94.561127 | 99.250351 | 96.239395 | 107.934204 | 220.047623 | 229.421646 | 215.958420 | 246.778748 | 1.038641 | 1571.502123 | 0.127899242 | 0.42765853 | 0.668899746 | 0.998865404 | 10.590482 | 10.644863 | 10.587004 | 10.618010 | 10.590729 | 10.671585 | 1740 | 1697 | 1320 | 1579 |
| 55508 | NM_00135499 | SLC35E3 | solute carrier family 35 member E3 | protein_coding | NP_001341926.1;NP_001341927.1;NP_061126.2 | 20864 | . | | . | 1.701853 | 1.960254 | 2.815402 | 2.370109 | 3.960281 | 4.531214 | 6.317680 | 5.418972 | 1.275744 | 870.714721 | 0.157799534 | 2.226488212 | **0.025981498** | 0.527323808 | 9.579315 | 9.931372 | 9.688693 | 9.647847 | 9.923155 | 9.756185 | 907 | 801 | 945 | 812 |
| 55509 | NM_018664,XI | BATF3 | basic leucine zipper ATF-like transcription factor 3 | protein_coding | NP_061134.1;XP_016857172.1 | 28915 | 612470 | . | . | 2.970879 | 2.720308 | 1.399223 | 3.020058 | 6.913357 | 6.288113 | 3.139814 | 6.905006 | -1.271288 | 70.761042 | 0.425583953 | -0.81368431 | 0.415825826 | 0.998865404 | 6.309355 | 5.959220 | 6.174265 | 6.164071 | 6.029409 | 6.187698 | 93 | 84 | 34 | 80 |
| 55511 | NM_00138190 | SAGE1 | sarcoma antigen 1 | protein_coding | NP_001368831.1;NP_061136.2;XP_016885111.1;XP_016885112.1;XP_01 | 30369 | 300359 | . | . | 21.132437 | 20.803884 | 15.831450 | 13.712916 | 49.176047 | 48.089115 | 35.525313 | 31.352954 | -1.432260 | 1419.022576 | 0.131581959 | -3.9389386 | **8.18429E-05** | **0.01086228** | 10.706540 | 10.188582 | 10.582624 | 10.600085 | 10.360084 | 10.270381 | 1903 | 1823 | 1077 | 1027 |
| 55515 | NM_018674,N | ASIC4 | acid sensing ion channel subunit family member 4 | protein_coding | NP_061144.4;NP_878267.3;XP_016859928.1 | 21263 | 606715 | . | . | 0.081995 | 0.039942 | 0.127295 | 0.193772 | 0.190804 | 0.092329 | 0.285646 | 0.443036 | **2.469290** | 9.251073 | 1.034921466 | 1.260092034 | 0.207636164 | NA | 2.413393 | 3.719719 | 3.178517 | 3.163518 | 3.194032 | 3.218742 | 8 | 4 | 9 | 15 |

| | | | | | | | | | | | | | | | | | | | | | | | | | | | | | | | | | |
|---|---|---|---|---|---|---|---|---|---|---|---|---|---|---|---|---|---|---|---|---|---|---|---|---|---|---|---|---|---|---|---|---|---|
| 55520 | NM_018696 | ELAC1 | elaC ribonuclease Z 1 | protein_coding | NP_061166.1 | 14197 | 608079 | . | . | 1.134805 | 0.921394 | 1.104351 | 1.137445 | 2.640738 | 2.129844 | 2.478131 | 2.600633 | 1.076280 | 74.689258 | 0.369955756 | 0.286666907 | 0.774367379 | 0.998865404 | 6.168342 | 6.275333 | 6.238959 | 6.184063 | 6.228858 | 6.231665 | 92 | 69 | 67 | 73 |
| 55521 | NM_00101739 | TRIM36 | tripartite motif containing 36 | protein_coding | NP_001017397.1;NP_001017398.1;NP_001287681.1;NP_001287688.1;NI | 16280 | 609317 | . | . | 2.086876 | 1.507991 | 1.921766 | 2.544769 | 4.856244 | 3.485790 | 4.312387 | 5.818312 | 1.133449 | 213.180481 | 0.230226698 | 0.784961788 | 0.432475973 | 0.998865404 | 7.642510 | 7.823669 | 7.713701 | 7.694474 | 7.774208 | 7.750436 | 235 | 211 | 202 | 207 |
| 55526 | NM_018706 | DHTKD1 | dehydrogenase E1 and transketolase domain containing 1 | protein_coding | NP_061176.4 | 23537 | 614984 | . | . | 6.002871 | 6.122665 | 5.337533 | 6.467037 | 13.968925 | 14.152816 | 11.977269 | 14.786113 | -1.041439 | 964.482209 | 0.147959442 | -0.39590694 | 0.692173681 | 0.998865404 | 9.942879 | 9.883736 | 9.923388 | 9.931599 | 9.830229 | 9.961111 | 1127 | 1068 | 747 | 966 |
| 55527 | NM_018708 | FEM1A | fem-1 homolog A | protein_coding | NP_061178.1 | 16934 | 613538 | . | . | 1.917955 | 2.219803 | 2.098192 | 2.408867 | 4.463159 | 5.131174 | 4.708282 | 5.507588 | 1.075874 | 642.473583 | 0.163840152 | 0.64397396 | 0.51959232 | 0.998865404 | 9.274193 | 9.378894 | 9.253790 | 9.345953 | 9.305716 | 9.392404 | 664 | 714 | 542 | 664 |
| 55529 | NM_018710,XI | PIP4P2 | phosphatidylinositol-4,5-bisphosphate 4-phosphatase 2 | protein_coding | NP_061180.1;XP_011515454.1 | 25452 | 609864 | . | . | 8.210016 | 9.521045 | 8.678795 | 9.175881 | 19.105045 | 22.008322 | 19.474963 | 20.979560 | -1.006472 | 745.211922 | 0.152637215 | -0.06097131 | 0.951382058 | 0.998865404 | 9.546425 | 9.536572 | 9.505156 | 9.580053 | 9.511776 | 9.564856 | 815 | 850 | 621 | 723 |
| 55531 | NM_00113003 | ELMOD1 | ELMO domain containing 1 | protein_coding | NP_001123509.1;NP_001294947.1;NP_061182.3;XP_016873483.1;XP_01 | 25334 | 615456 | . | . | 1.150555 | 1.004783 | 0.973195 | 1.046286 | 2.677389 | 2.322600 | 2.183819 | 2.392210 | -1.035988 | 100.660752 | 0.321483651 | -0.1586625 | 0.873934785 | 0.998865404 | 6.678624 | 6.627640 | 6.680368 | 6.637291 | 6.634438 | 6.661861 | 126 | 103 | 82 | 97 |
| 55534 | NM_018717 | MAML3 | mastermind like transcriptional coactivator 3 | protein_coding | NP_061187.3 | 16272 | 608991 | . | . | 0.966379 | 1.167847 | 1.392967 | 1.661033 | 2.248803 | 2.699531 | 3.125777 | 3.797756 | 1.413409 | 276.568511 | 0.218915703 | 2.28023287 | 0.022593881 | 0.49659767 | 7.841241 | 8.339019 | 7.960969 | 8.049470 | 8.137585 | 8.231706 | 240 | 270 | 258 | 329 |
| 55536 | NM_00112737 | CDCA7L | cell division cycle associated 7 like | protein_coding | NP_001120842.1;NP_001120843.1;NP_061189.2 | 30777 | 609685 | . | . | 19.636356 | 20.261518 | 16.315985 | 21.015255 | 45.694612 | 46.835406 | 36.612599 | 48.048886 | -1.084019 | 1732.767473 | 0.141425361 | -0.82298224 | 0.410518094 | 0.998865404 | 10.816111 | 10.699248 | 10.779354 | 10.796926 | 10.627716 | 10.813360 | 2053 | 1967 | 1273 | 1745 |
| 55544 | NM_00129178 | RBM38 | RNA binding motif protein 38 | protein_coding | NP_001278709.1;NP_059965.2;NP_906270.1;XP_005260503.1;XP_01152 | 15818 | 612428 | . | . | 5.587805 | 6.458143 | 5.504719 | 6.459546 | 13.003054 | 14.928285 | 12.352432 | 14.768985 | -1.021987 | 448.746206 | 0.182303728 | -0.17211686 | 0.863345661 | 0.998865404 | 8.825850 | 8.793480 | 8.773506 | 8.855788 | 8.757212 | 8.844780 | 488 | 522 | 358 | 446 |
| 55552 | NM_00108049 | ZNF823 | zinc finger protein 823 | protein_coding | NP_001073962.1;NP_001284539.1;NP_059977.1;XP_016882430.1;XP_01 | 30936 | . | . | . | 1.245774 | 1.112570 | 0.949595 | 0.999639 | 2.898968 | 2.571757 | 2.130864 | 2.285555 | -1.168872 | 83.483906 | 0.348628083 | -0.64572112 | 0.518459979 | 0.998865404 | 6.491217 | 6.266926 | 6.420364 | 6.389807 | 6.370911 | 6.355443 | 109 | 92 | 69 | 70 |
| 55553 | NM_00114581 | SOX6 | SRY-box transcription factor 6 | protein_coding | NP_001139283.1;NP_001139291.2;NP_001354801.1;NP_001354802.1;NI | 16421 | 607257 | . | . | 3.387607 | 3.553616 | 3.763151 | 4.066728 | 7.883102 | 8.214342 | 8.444402 | 9.298091 | 1.112091 | 1024.535767 | 0.137221262 | 1.116988713 | 0.263999151 | 0.996308385 | 9.922237 | 10.075219 | 9.947146 | 9.972297 | 10.011991 | 10.062966 | 1098 | 1065 | 906 | 1046 |
| 55556 | NM_00131876 | ENOSF1 | enolase superfamily member 1 | protein_coding | NP_001305689.1;NP_001340994.1;NP_001340995.1;NP_001340996.1;NI | 30365 | 607427 | . | . | 9.076341 | 9.423739 | 12.208371 | 9.435399 | 21.121019 | 21.783392 | 27.395228 | 21.572923 | 1.147706 | 1192.513305 | 0.175544864 | 1.132208451 | 0.257546808 | 0.995760118 | 10.116584 | 10.316102 | 10.201862 | 10.115285 | 10.401403 | 10.103877 | 1353 | 1128 | 1323 | 964 |
| 55558 | NM_017514,XI | PLXNA3 | plexin A3 | protein_coding | NP_059984.3;XP_005274762.2;XP_005274763.2;XP_006724892.1;XP_01 | 9101 | 300022 | . | . | 3.080828 | 3.237310 | 2.840314 | 2.983805 | 7.169211 | 7.483187 | 6.373582 | 6.822117 | 1.060488 | 812.380607 | 0.155471293 | 0.544979212 | 0.585767815 | 0.998865404 | 9.622792 | 9.707974 | 9.623618 | 9.662258 | 9.749775 | 9.615410 | 883 | 874 | 796 | 710 |
| 55559 | NM_017518,N | HAUS7 | HAUS augmin like complex subunit 7 | protein_coding | NP_059988.3 | 32979 | 300540 | . | . | 5.259589 | 5.759214 | 4.199001 | 5.090809 | 12.239280 | 13.312681 | 9.422436 | 11.639528 | -1.191794 | 222.375757 | 0.235094335 | -1.0767381 | 0.281597285 | 0.998865404 | 7.918536 | 7.663959 | 7.817311 | 7.855197 | 7.703458 | 7.800729 | 268 | 271 | 157 | 211 |
| 55561 | NM_017525,XI | CDC42BPG | CDC42 binding protein kinase gamma | protein_coding | NP_059995.2;XP_011543457.1;XP_011543458.1;XP_011543459.1;XP_01 | 29829 | 613991 | . | . | 1.310543 | 1.300214 | 1.229679 | 1.161325 | 3.049688 | 3.005502 | 2.759365 | 2.655231 | -1.110204 | 236.063415 | 0.220690173 | -0.68342507 | 0.494338283 | 0.998865404 | 7.956353 | 7.805781 | 7.908400 | 7.908211 | 7.868197 | 7.846702 | 286 | 268 | 199 | 205 |
| 55565 | NM_00120155 | ZNF821 | zinc finger protein 821 | protein_coding | NP_001188481.1;NP_001188482.1;NP_001188483.1;NP_001188485.1;NI | 28043 | . | . | . | 1.623881 | 1.567547 | 1.258850 | 1.756270 | 3.778836 | 3.623457 | 2.824825 | 4.015501 | -1.071144 | 100.883372 | 0.329983847 | -0.30047607 | 0.763814045 | 0.998865404 | 6.706106 | 6.605219 | 6.671370 | 6.661817 | 6.597098 | 6.691689 | 122 | 111 | 71 | 106 |
| 55568 | NM_198321 | GALNT10 | polypeptide N-acetylgalactosaminyltransferase 10 | protein_coding | NP_938080.1 | 19873 | 608043 | . | . | 5.039785 | 5.318140 | 4.830897 | 5.112581 | 11.727787 | 12.293118 | 10.840391 | 11.689308 | -1.055951 | 942.448902 | 0.13947896 | -0.56311608 | 0.57335584 | 0.998865404 | 9.919135 | 9.840330 | 9.883264 | 9.915966 | 9.843759 | 9.876320 | 1090 | 1068 | 779 | 879 |
| 55571 | NM_017546 | CNOT11 | CCR4-NOT transcription complex subunit 11 | protein_coding | NP_060016.3 | 25217 | . | . | . | 23.212776 | 21.563547 | 21.950506 | 21.292978 | 54.017082 | 49.845104 | 49.256298 | 48.683865 | -1.049920 | 1724.857158 | 0.124359492 | -0.56513527 | 0.571981759 | 0.998865404 | 10.786862 | 10.716811 | 10.802823 | 10.737781 | 10.748285 | 10.716914 | 2118 | 1828 | 1493 | 1546 |
| 55572 | NM_017547,N | FOXRED1 | FAD dependent oxidoreductase domain containing 1 | protein_coding | NP_060017.1;XP_006718942.1;XP_006718944.1;XP_011541197.1;XP_01 | 26927 | 613622 | . | . | 9.016324 | 9.994776 | 10.193445 | 9.909589 | 20.981362 | 23.103376 | 22.873791 | 22.657099 | -1.032672 | 698.189737 | 0.152010782 | -0.30512665 | 0.760269696 | 0.998865404 | 9.470430 | 9.424150 | 9.440368 | 9.475765 | 9.457124 | 9.414810 | 796 | 785 | 616 | 624 |
| 55573 | NM_00113442 | CDV3 | CDV3 homolog | protein_coding | NP_001127894.1;NP_001127895.1;NP_001269691.1;NP_001269692.1;NI | 26928 | 618789 | . | . | 29.992450 | 30.732189 | 31.349177 | 28.279922 | 69.793667 | 71.038832 | 70.346645 | 64.658681 | -1.013849 | 2951.544014 | 0.11614662 | -0.17084748 | 0.864343692 | 0.998865404 | 11.537063 | 11.517378 | 11.530838 | 11.534114 | 11.562601 | 11.478287 | 3415 | 3213 | 2681 | 2606 |
| 55577 | NM_00133042 | NAGK | N-acetylglucosamine kinase | protein_coding | NP_001317354.1;NP_001317355.1;NP_001352395.1;NP_060037.4;XP_01 | 17174 | 606828 | . | . | 24.692499 | 26.130626 | 24.686963 | 26.574554 | 57.460461 | 60.402117 | 55.396832 | 60.759560 | 1.027769 | 1614.108483 | 0.124886336 | 0.31641667 | 0.751686266 | 0.998865404 | 10.636731 | 10.676045 | 10.643723 | 10.647396 | 10.635266 | 10.697434 | 1829 | 1722 | 1366 | 1595 |
| 55578 | NM_00101428 | SUPT20H | SPT20 homolog, SAGA complex component | protein_coding | NP_001014308.2;NP_001265409.1;NP_001265410.1;NP_001265411.1;NI | 20596 | 613417 | . | . | 8.651853 | 10.790880 | 10.205452 | 9.986712 | 20.133216 | 24.943613 | 22.900741 | 22.833421 | 1.026532 | 1026.441234 | 0.147550418 | 0.256034026 | 0.797924564 | 0.998865404 | 9.984529 | 10.022097 | 9.922768 | 10.057546 | 10.032339 | 9.990393 | 1060 | 1193 | 930 | 947 |
| 55582 | NM_00127192 | KIF27 | kinesin family member 27 | protein_coding | NP_001258856.1;NP_001258857.1;NP_001340998.1;NP_001340999.1;NI | 18632 | 611253 | . | . | 0.450367 | 0.637594 | 0.444835 | 0.258421 | 1.048025 | 1.473829 | 0.998198 | 0.590847 | 1.015408 | 91.803310 | 0.350448804 | 0.062946281 | 0.949809281 | 0.998865404 | 6.508315 | 6.532532 | 6.511280 | 6.515883 | 6.574651 | 6.457945 | 104 | 99 | 97 | 69 |
| 55585 | NM_017582 | UBE2Q1 | ubiquitin conjugating enzyme E2 Q1 | protein_coding | NP_060052.3 | 15698 | 617429 | . | . | 18.581047 | 18.291033 | 17.832273 | 20.172924 | 43.238861 | 42.280540 | 40.015106 | 46.122993 | 1.016315 | 1888.845191 | 0.124455273 | 0.187598109 | 0.851191709 | 0.998865404 | 10.871669 | 10.894816 | 10.886146 | 10.866741 | 10.843503 | 10.932969 | 2184 | 1997 | 1562 | 1886 |
| 55588 | NM_00131777 | MED29 | mediator complex subunit 29 | protein_coding | NP_001304699.2;NP_060062.2 | 23074 | 612914 | . | . | 9.652365 | 9.952582 | 8.987477 | 10.316493 | 22.461449 | 23.005840 | 20.167639 | 23.587435 | -1.031128 | 1087.780575 | 0.139243044 | -0.31760072 | 0.750787831 | 0.998865404 | 10.109326 | 10.064672 | 10.089889 | 10.106276 | 10.026490 | 10.121767 | 1258 | 1205 | 871 | 1069 |
| 55589 | NM_017593,N | BMP2K | BMP2 inducible kinase | protein_coding | NP_060063.2;NP_942595.1;XP_005263174.1;XP_011530404.1;XP_01686 | 18041 | 617648 | . | . | 3.245725 | 2.897764 | 3.112231 | 3.076161 | 7.552935 | 6.698314 | 6.983757 | 7.033275 | -1.202127 | 571.165135 | 0.160878998 | -1.65086222 | 0.098766712 | 0.813553488 | 9.284599 | 9.018695 | 9.198688 | 9.230873 | 9.081979 | 9.106737 | 701 | 689 | 442 | 496 |
| 55591 | NM_00135208 | VEZT | vezatin, adherens junctions transmembrane protein | protein_coding | NP_001339017.1;NP_001339018.1;NP_001339019.1;NP_001339020.1;NI | 18258 | . | . | . | 6.633407 | 6.375996 | 6.883101 | 5.873862 | 15.436209 | 14.738400 | 15.445478 | 13.429888 | -1.025012 | 946.489727 | 0.148423243 | -0.24012484 | 0.810233481 | 0.998865404 | 9.903855 | 9.868818 | 9.906712 | 9.881592 | 9.938323 | 9.810601 | 1122 | 1016 | 885 | 799 |
| 55592 | NR_024261,NF | GOLGA2P5 | GOLGA2 pseudogene 5 | transcribed_ps | . | 25315 | . | . | . | 0.944174 | 1.261426 | 1.923229 | 1.677711 | 2.197132 | 2.915842 | 4.315670 | 3.835886 | 1.606580 | 110.311012 | 0.318284249 | 2.149000037 | 0.031634397 | 0.563938326 | 6.403699 | 7.087004 | 6.647916 | 6.725545 | 6.866574 | 6.812782 | 84 | 104 | 127 | 118 |
| 55593 | NM_00113615 | OTUD5 | OTU deubiquitinase 5 | protein_coding | NP_001129629.1;NP_001129630.1;NP_001129631.1;NP_060072.1;XP_00 | 25402 | 300713 | . | . | 7.272938 | 8.991435 | 10.012604 | 10.131574 | 16.924423 | 20.784100 | 22.467995 | 23.164642 | 1.067190 | 1159.643430 | 0.132852381 | 0.70617666 | 0.480078283 | 0.998865404 | 10.131846 | 10.225563 | 10.129722 | 10.178172 | 10.208498 | 10.197229 | 1250 | 1250 | 1049 | 1114 |
| 55596 | NM_00135093 | ZCCHC8 | zinc finger CCHC-type containing 8 | protein_coding | NP_001337864.1;NP_001337865.1;NP_001337866.1;NP_001337867.1;NI | 25265 | 616381 | . | . | 4.764414 | 5.111416 | 6.513452 | 4.550084 | 11.086988 | 11.815266 | 14.615998 | 10.403225 | 1.065600 | 658.731908 | 0.180290783 | 0.508434518 | 0.611148651 | 0.998865404 | 9.316521 | 9.409090 | 9.334434 | 9.340383 | 9.486681 | 9.262477 | 731 | 691 | 688 | 533 |
| 55599 | NM_017619 | RNPC3 | RNA binding region (RNP1, RRM) containing 3 | protein_coding | NP_060089.1 | 18666 | 618016 | . | . | 5.599649 | 5.464359 | 5.802985 | 4.095681 | 13.030615 | 12.631111 | 13.021730 | 9.364287 | -1.132151 | 350.794920 | 0.214147165 | -0.83618544 | 0.403050587 | 0.998865404 | 8.540525 | 8.362984 | 8.495039 | 8.474914 | 8.507420 | 8.312468 | 436 | 395 | 337 | 254 |
| 55601 | NM_017631,XI | DDX60 | DExD/H-box helicase 60 | protein_coding | NP_060101.3;XP_011530405.1;XP_011530406.1;XP_016863872.1;XP_01 | 25942 | 613974 | . | . | 0.531260 | 0.629432 | 0.430236 | 0.486166 | 1.236264 | 1.454959 | 0.965437 | 1.111560 | -1.315238 | 100.450849 | 0.323800964 | -1.220885 | 0.222129565 | 0.982008277 | 6.835031 | 6.438500 | 6.663479 | 6.717145 | 6.595371 | 6.619240 | 121 | 133 | 72 | 85 |
| 55602 | NM_00131734 | CDKN2AIP | CDKN2A interacting protein | protein_coding | NP_001304272.1;NP_060102.1 | 24325 | 615914 | . | . | 2.860962 | 2.839958 | 3.031977 | 2.656950 | 6.657576 | 6.564691 | 6.803667 | 6.074801 | -1.015360 | 314.127675 | 0.200864879 | -0.10948001 | 0.912821776 | 0.998865404 | 8.305799 | 8.284537 | 8.302650 | 8.294652 | 8.327246 | 8.252248 | 367 | 339 | 290 | 272 |
| 55603 | NM_017633 | TENT5A | terminal nucleotidyltransferase 5A | protein_coding | NP_060103.2 | 18345 | 611357 | . | . | 4.428599 | 3.902669 | 4.308541 | 3.340567 | 10.305532 | 9.021194 | 9.668242 | 7.637810 | -1.104022 | 695.929181 | 0.173442948 | -0.82314671 | 0.410424572 | 0.998865404 | 9.511901 | 9.370215 | 9.519080 | 9.424909 | 9.489945 | 9.313188 | 897 | 735 | 651 | 539 |
| 55604 | NM_00117397 | CARMIL1 | capping protein regulator and myosin 1 linker 1 | protein_coding | NP_001167448.1;NP_060110.4;XP_005249275.1;XP_016866497.1;XP_01 | 21581 | 609593 | . | . | 4.410086 | 5.016287 | 5.419139 | 6.055783 | 10.262452 | 11.595375 | 12.160390 | 13.845828 | 1.198148 | 893.403689 | 0.148083857 | 1.761207755 | 0.078203246 | 0.769928249 | 9.667202 | 9.927417 | 9.689453 | 9.774191 | 9.824646 | 9.899162 | 880 | 930 | 804 | 959 |
| 55605 | NM_00117346 | KIF21A | kinesin family member 21A | protein_coding | NP_001166934.1;NP_001166935.1;NP_001166936.1;NP_001365368.1;NI | 19349 | 608283 | . | . | 3.135179 | 3.384596 | 3.996713 | 3.529422 | 7.295691 | 7.823647 | 8.968509 | 8.069606 | 1.149864 | 691.442832 | 0.158373947 | 1.272073231 | 0.203347101 | 0.968254058 | 9.329055 | 9.530828 | 9.357953 | 9.404697 | 9.527732 | 9.424365 | 715 | 718 | 691 | 642 |
| 55607 | NM_00116616 | PPP1R9A | protein phosphatase 1 regulatory subunit 9A | protein_coding | NP_001159632.1;NP_001159633.1;NP_001159634.1;NP_001159635.1;NI | 14946 | 602468 | . | . | 0.778333 | 0.872634 | 0.617299 | 0.723894 | 1.811212 | 2.017134 | 1.385202 | 1.655097 | -1.222966 | 248.616739 | 0.222921613 | -1.30262768 | 0.192701904 | 0.96621494 | 8.096201 | 7.804654 | 7.967415 | 8.041310 | 7.875005 | 7.933955 | 293 | 316 | 182 | 223 |
| 55608 | NM_00128672 | ANKRD10 | ankyrin repeat domain 10 | protein_coding | NP_001273650.1;NP_060134.2;XP_024305148.1;XP_024305149.1;XP_02 | 20265 | . | . | . | 15.514185 | 14.152683 | 17.645439 | 12.141798 | 36.102144 | 32.714559 | 39.595855 | 27.760775 | -1.055164 | 1773.815035 | 0.173949807 | -0.44534379 | 0.656071327 | 0.998865404 | 10.830571 | 10.753685 | 10.843138 | 10.770264 | 10.919700 | 10.583170 | 2193 | 1875 | 1822 | 1273 |
| 55609 | NM_017666,XI | ZNF280C | zinc finger protein 280C | protein_coding | NP_060136.1;XP_006724828.1 | 25955 | . | . | . | 1.035348 | 1.137413 | 1.347572 | 1.110250 | 2.409298 | 2.629181 | 3.023913 | 2.538454 | 1.120775 | 167.589226 | 0.26063848 | 0.631130701 | 0.527955069 | 0.998865404 | 7.303832 | 7.469014 | 7.345640 | 7.376784 | 7.448346 | 7.366944 | 175 | 177 | 169 | 150 |
| 55610 | NM_00125799 | VPS50 | VPS50 subunit of EARP/GARPII complex | protein_coding | NP_001244927.1;NP_060137.2;NP_078829.1;XP_011514697.1;XP_01151 | 25956 | 616465 | . | . | 4.754986 | 4.276412 | 4.736820 | 3.727604 | 11.065046 | 9.885119 | 10.629289 | 8.522726 | -1.155623 | 697.944700 | 0.17822397 | -1.17083182 | 0.241666383 | 0.992733415 | 9.547002 | 9.339479 | 9.535093 | 9.445233 | 9.488755 | 9.285241 | 916 | 756 | 649 | 515 |
| 55611 | NM_017670,N | OTUB1 | OTU deubiquitinase, ubiquitin aldehyde binding 1 | protein_coding | NP_060140.2 | 23077 | 608337 | . | . | 27.040380 | 29.065935 | 25.191229 | 31.991345 | 62.924073 | 67.187213 | 56.528388 | 73.144406 | 1.009981 | 2070.212198 | 0.137552761 | 0.104161223 | 0.917041406 | 0.998865404 | 11.008594 | 11.022498 | 10.986076 | 11.033404 | 10.924012 | 11.103714 | 2302 | 2289 | 1605 | 2170 |
| 55614 | NM_00119986 | KIF16B | kinesin family member 16B | protein_coding | NP_001186794.1;NP_001186795.1;NP_078980.3;XP_005260807.1;XP_00 | 15869 | 618171 | . | . | 3.894631 | 4.488909 | 4.091711 | 4.855503 | 9.062966 | 10.376312 | 9.181680 | 11.101533 | 1.065871 | 728.059274 | 0.160591274 | 0.573084384 | 0.56658756 | 0.998865404 | 9.461584 | 9.552799 | 9.433542 | 9.531894 | 9.475240 | 9.577826 | 754 | 815 | 605 | 756 |
| 55615 | NM_00101752 | PRR5 | proline rich 5 | protein_coding | NP_001017528.1;NP_001017529.1;NP_001017530.1;NP_001185650.1;NI | 31682 | 609406 | . | . | 7.378710 | 8.208661 | 8.972724 | 10.846824 | 17.170563 | 18.974684 | 20.134531 | 24.799974 | 1.253265 | 514.809614 | 0.175535275 | 1.855419136 | 0.063536397 | 0.729663894 | 8.836392 | 9.161160 | 8.897430 | 8.957185 | 9.011647 | 9.128542 | 501 | 517 | 454 | 584 |
| 55616 | NM_00114377 | ASAP3 | ArfGAP with SH3 domain, ankyrin repeat and PH domain 3 | protein_coding | NP_001137250.1;NP_060177.2;XP_011540057.1;XP_016857174.1;XP_01 | 14987 | 616594 | . | . | 6.899456 | 6.648416 | 8.339609 | 8.557322 | 16.055316 | 15.368110 | 18.713842 | 19.565299 | 1.230375 | 964.256350 | 0.137984983 | 2.167612302 | 0.030188197 | 0.554053715 | 9.755943 | 10.055120 | 9.847279 | 9.815577 | 9.977464 | 9.987623 | 1018 | 912 | 919 | 1004 |
| 55617 | NM_00132360 | TASP1 | taspase 1 | protein_coding | NP_001310531.1;NP_001310532.1;NP_001310533.1;NP_060184.2;XP_01 | 15859 | 608270 | . | . | 2.123856 | 2.162838 | 2.085011 | 2.161247 | 4.942300 | 4.999496 | 4.678706 | 4.941436 | -1.012769 | 158.603757 | 0.260303614 | -0.0703209 | 0.943938247 | 0.998865404 | 7.318551 | 7.299956 | 7.310213 | 7.313781 | 7.298176 | 7.315536 | 183 | 173 | 134 | 151 |
| 55619 | NM_00129026 | DOCK10 | dedicator of cytokinesis 10 | protein_coding | NP_001277192.1;NP_001350691.1;NP_055504.2;XP_006712680.1;XP_01 | 23479 | 611518 | . | . | 12.058870 | 12.418500 | 11.591163 | 12.753119 | 28.061480 | 28.705918 | 26.010235 | 29.158492 | -1.020784 | 2813.007103 | 0.115914736 | -0.256028 | 0.797929218 | 0.998865404 | 11.472751 | 11.442890 | 11.452279 | 11.479589 | 11.413119 | 11.484061 | 3218 | 3118 | 2315 | 2724 |
| 55621 | NM_00113603 | TRMT1 | tRNA methyltransferase 1 | protein_coding | NP_001129507.1;NP_001136026.1;NP_001338689.1;NP_001338690.1;NI | 25980 | 611669 | . | . | 29.585112 | 31.862934 | 35.835283 | 30.302581 | 68.845771 | 73.652600 | 80.413338 | 69.283247 | 1.064874 | 2129.614856 | 0.131938533 | 0.687309929 | 0.491887443 | 0.998865404 | 11.010218 | 11.101105 | 10.999012 | 11.058676 | 11.150151 | 11.003813 | 2286 | 2309 | 2084 | 1872 |
| 55622 | NM_00119350 | TTC27 | tetratricopeptide repeat domain 27 | protein_coding | NP_001180438.1;NP_060205.3;XP_005264473.1;XP_011531260.1 | 25986 | . | . | . | 4.206051 | 4.377007 | 4.027271 | 3.845011 | 9.787652 | 10.117649 | 9.037079 | 8.791159 | -1.102990 | 369.874524 | 0.186218347 | -0.75942769 | 0.447596756 | 0.998865404 | 8.599769 | 8.458561 | 8.551082 | 8.566226 | 8.518669 | 8.485145 | 441 | 424 | 315 | 320 |
| 55623 | NM_00130455 | THUMPD1 | THUMP domain containing 1 | protein_coding | NP_001291479.1;NP_060206.2;XP_011544193.1;XP_016878922.1 | 23807 | 616662 | . | . | 5.355345 | 5.251636 | 5.832041 | 5.206317 | 12.462109 | 12.139392 | 13.086931 | 11.903625 | 1.000662 | 686.662042 | 0.158277111 | 0.006033178 | 0.995186249 | 0.999862634 | 9.422641 | 9.424272 | 9.439305 | 9.404885 | 9.473422 | 9.369668 | 810 | 722 | 643 | 595 |
| 55624 | NM_00124376 | POMGNT1 | protein O-linked mannose N-acetylglucosaminyltransferase 1 (beta | protein_coding | NP_001230695.1;NP_001277058.1;NP_001277059.1;NP_060209.4;XP_00 | 19139 | 606822 | . | . | 55.994559 | 58.874339 | 59.301144 | 63.030874 | 130.301640 | 136.090670 | 133.070048 | 144.112602 | 1.051377 | 4999.716286 | 0.107338178 | 0.673379777 | 0.500705711 | 0.998865404 | 12.251586 | 12.323276 | 12.246936 | 12.285012 | 12.288354 | 12.327370 | 5485 | 5380 | 4353 | 4916 |
| 55625 | NM_00114554 | ZDHHC7 | zinc finger DHHC-type palmitoyltransferase 7 | protein_coding | NP_001139020.1;NP_060210.2;XP_011521524.1;XP_024306111.1 | 18459 | 614604 | . | . | 12.889543 | 13.894452 | 15.466486 | 16.014668 | 29.994496 | 32.117649 | 34.706345 | 36.615637 | 1.154974 | 1551.250401 | 0.125043239 | 1.662307662 | 0.096451102 | 0.813553488 | 10.491622 | 10.699340 | 10.514404 | 10.565113 | 10.649595 | 10.653954 | 1603 | 1605 | 1442 | 1563 |
| 55626 | NM_00126778 | AMBRA1 | autophagy and beclin 1 regulator 1 | protein_coding | NP_001254711.1;NP_001254712.1;NP_001287660.1;NP_001354397.1;NI | 25990 | 611359 | . | . | 3.807627 | 4.107678 | 4.676235 | 5.100658 | 8.860505 | 9.495081 | 10.493332 | 11.662047 | 1.203121 | 685.114260 | 0.154130272 | 1.730882898 | 0.083472652 | 0.785005898 | 9.280913 | 9.547217 | 9.329019 | 9.374357 | 9.450007 | 9.505259 | 692 | 694 | 625 | 729 |
| 55627 | NM_00117108 | SMPD4 | sphingomyelin phosphodiesterase 4 | protein_coding | NP_001164554.1;NP_060221.2;NP_060421.2;XP_011509745.1;XP_01150 | 32949 | 610457 | . | . | 42.069794 | 44.637670 | 43.915315 | 44.309724 | 97.898143 | 103.181978 | 98.544691 | 101.308919 | 1.018418 | 5802.040066 | 0.106286843 | 0.247729903 | 0.804343394 | 0.998865404 | 12.489146 | 12.515423 | 12.466271 | 12.521936 | 12.506110 | 12.513495 | 6401 | 6409 | 5069 | 5513 |
| 55628 | NM_00114618 | ZNF407 | zinc finger protein 407 | protein_coding | NP_001139661.1;NP_001139662.1;NP_060227.2;XP_005266783.1;XP_00 | 19904 | 615894 | . | . | 0.897202 | 1.044642 | 1.090534 | 1.304807 | 2.087825 | 2.414738 | 2.447128 | 2.983286 | 1.319975 | 485.350939 | 0.174704229 | 2.292507204 | 0.021876392 | 0.489944286 | 8.709225 | 9.108945 | 8.793211 | 8.855732 | 8.959698 | 9.037212 | 457 | 475 | 452 | 548 |
| 55629 | NM_017761,XI | PNRC2 | proline rich nuclear receptor coactivator 2 | protein_coding | NP_060231.1;XP_016857180.1 | 23158 | 611882 | . | . | 29.718590 | 26.281566 | 30.006336 | 22.532302 | 69.156380 | 60.751014 | 67.333344 | 51.517427 | -1.080750 | 2030.463650 | 0.151998701 | -0.73706104 | 0.461085228 | 0.998865404 | 11.042271 | 10.930762 | 11.066095 | 10.960100 | 11.062498 | 10.829867 | 2588 | 2126 | 1947 | 1561 |
| 55631 | NM_017768,XI | LRRC40 | leucine rich repeat containing 40 | protein_coding | NP_060238.3;XP_005271070.1;XP_011540065.1;XP_016857184.1;XP_01 | 26004 | . | . | . | 7.659313 | 6.397786 | 5.886504 | 3.971786 | 17.823537 | 14.788767 | 13.209144 | 9.081017 | -1.448333 | 530.638504 | 0.205459616 | -2.6009671 | 0.009296137 | 0.307003327 | 9.293509 | 8.760756 | 9.214797 | 9.083475 | 9.025224 | 8.788253 | 791 | 613 | 452 | 326 |
| 55632 | NM_00130809 | G2E3 | G2/M-phase specific E3 ubiquitin protein ligase | protein_coding | NP_001295026.1;NP_060239.2;XP_011535227.1;XP_011535228.1 | 20338 | 611299 | . | . | 3.236228 | 2.930992 | 2.649390 | 1.404110 | 7.530837 | 6.775118 | 5.945153 | 3.210333 | -1.503135 | 449.398811 | 0.240746142 | -2.44230202 | 0.014593932 | 0.403687453 | 9.075331 | 8.489259 | 8.940927 | 8.880509 | 8.817861 | 8.475883 | 649 | 556 | 407 | 235 |
| 55633 | NM_017772,N | TBC1D22B | TBC1 domain family member 22B | protein_coding | NP_060242.2;XP_011513040.1;XP_011513041.1 | 21602 | 616880 | . | . | 3.905245 | 4.044339 | 3.582643 | 4.315775 | 9.087667 | 9.348670 | 8.039348 | 9.867507 | -1.016671 | 414.253962 | 0.181469276 | -0.13144551 | 0.89542289 | 0.998865404 | 8.706509 | 8.682131 | 8.702911 | 8.695402 | 8.651675 | 8.725288 | 484 | 448 | 335 | 409 |
| 55634 | NM_00112989 | KRBOX4 | KRAB box domain containing 4 | protein_coding | NP_001123370.1;NP_001123371.1;NP_001123372.1;NP_060246.2 | 26007 | 300585 | . | . | 3.175652 | 3.063119 | 3.766679 | 3.718407 | 7.389873 | 7.080534 | 8.452318 | 8.501694 | 1.196131 | 262.739737 | 0.211324774 | 1.222646165 | 0.221463368 | 0.982008277 | 7.902371 | 8.161032 | 7.997048 | 7.977699 | 8.084915 | 8.072240 | 281 | 253 | 252 | 265 |
| 55635 | NM_00111412 | DEPDC1 | DEP domain containing 1 | protein_coding | NP_001107592.1;NP_060249.2 | 22949 | 612002 | . | . | 10.927401 | 9.784135 | 7.956038 | 5.382622 | 25.428510 | 22.616467 | 17.853119 | 12.306726 | -1.541785 | 1217.613473 | 0.176169622 | -3.54545665 | 0.000391933 | 0.033601598 | 10.528232 | 9.904565 | 10.428165 | 10.338253 | 10.190898 | 9.906849 | 1796 | 1504 | 1000 | 719 |
| 55636 | NM_00131669 | CHD7 | chromodomain helicase DNA binding protein 7 | protein_coding | NP_001303619.1;NP_060250.2;XP_011515855.1;XP_011515856.1;XP_01 | 20626 | 608892 | . | . | 3.465695 | 3.721675 | 3.542690 | 3.985080 | 8.064814 | 8.602822 | 7.949694 | 9.111415 | 1.023293 | 1257.259646 | 0.134573379 | 0.246844239 | 0.805028772 | 0.998865404 | 10.279571 | 10.312376 | 10.257156 | 10.315668 | 10.266689 | 10.340224 | 1376 | 1393 | 1052 | 1250 |
| 55638 | NM_00109974 | SYBU | syntabulin | protein_coding | NP_001093213.1;NP_001093214.1;NP_001093215.1;NP_001093216.1;NI | 26011 | 611568 | . | . | 0.248685 | 0.206110 | 0.119889 | 0.069856 | 0.578701 | 0.476431 | 0.269028 | 0.159718 | -1.730006 | 20.533760 | 0.693365128 | -1.14049143 | 0.254081623 | NA | 4.700498 | 3.913269 | 4.386889 | 4.386119 | 4.354467 | 4.326027 | 30 | 28 | 16 | 11 |
| 55640 | NM_00119528 | FLVCR2 | FLVCR heme transporter 2 | protein_coding | NP_001182212.1;NP_060261.2 | 20105 | 610865 | . | . | 5.050161 | 5.567796 | 2.922717 | 3.412561 | 11.751933 | 12.870208 | 6.558492 | 7.802414 | -1.668025 | 475.037049 | 0.176532244 | -4.18133646 | 2.89801E-05 | 0.004706022 | 9.214609 | 8.475550 | 9.005648 | 9.043346 | 8.667313 | 8.746356 | 665 | 659 | 287 | 358 |
| 55643 | NM_017797,XI | BTBD2 | BTB domain containing 2 | protein_coding | NP_060267.2;XP_011526429.1 | 15504 | 608531 | . | . | 5.016347 | 5.810308 | 4.528488 | 6.735380 | 11.673246 | 13.430788 | 10.161797 | 15.399645 | 1.025949 | 452.652337 | 0.208107145 | 0.177599221 | 0.859037736 | 0.998865404 | 8.804428 | 8.839874 | 8.765434 | 8.851586 | 8.700053 | 8.936308 | 479 | 516 | 322 | 512 |
| 55644 | NM_017807 | OSGEP | O-sialoglycoprotein endopeptidase | protein_coding | NP_060277.1 | 18028 | 610107 | . | . | 8.198997 | 8.458959 | 8.442229 | 8.111516 | 19.079401 | 19.553261 | 18.944118 | 18.546019 | -1.019515 | 511.745805 | 0.166182287 | -0.16778535 | 0.866752144 | 0.998865404 | 9.013082 | 8.985351 | 8.997795 | 9.012797 | 9.009051 | 8.976761 | 588 | 564 | 452 | 463 |
| 55646 | NM_00114572 | LYAR | Ly1 antibody reactive | protein_coding | NP_001139197.1;NP_060286.1;XP_011511807.1;XP_011511808.1 | 26021 | 617684 | . | . | 16.102861 | 17.739019 | 16.468375 | 13.909819 | 37.472016 | 41.004536 | 36.954561 | 31.803147 | -1.127082 | 778.791775 | 0.157389099 | -1.09659413 | 0.272818855 | 0.998865404 | 9.688616 | 9.516438 | 9.611934 | 9.675672 | 9.620204 | 9.497064 | 908 | 930 | 694 | 625 |
| 55647 | NM_017817 | RAB20 | RAB20, member RAS oncogene family | protein_coding | NP_060287.1 | 18260 | . | . | . | 4.051041 | 4.190175 | 3.950275 | 4.985698 | 9.426938 | 9.685777 | 8.864303 | 11.399206 | 1.068607 | 201.700850 | 0.243147209 | 0.393719285 | 0.693788302 | 0.998865404 | 7.608036 | 7.702563 | 7.633706 | 7.644467 | 7.617094 | 7.718557 | 222 | 213 | 161 | 217 |
| 55650 | NM_00120255 | PIGV | phosphatidylinositol glycan anchor biosynthesis class V | protein_coding | NP_001189483.1;NP_001361407.1;NP_001361409.1;NP_001361410.1;NI | 26031 | 610274 | . | . | 3.578056 | 3.932154 | 4.588599 | 4.000575 | 8.326284 | 9.089355 | 10.296684 | 9.146840 | 1.126026 | 593.658198 | 0.166642165 | 1.027590931 | 0.304142292 | 0.998865404 | 9.125131 | 9.296746 | 9.145929 | 9.194799 | 9.303203 | 9.192502 | 619 | 625 | 592 | 541 |
| 55651 | NM_00103483 | NHP2 | NHP2 ribonucleoprotein | protein_coding | NP_001030005.1;NP_060308.1 | 14377 | 606470 | . | . | 110.652182 | 115.956786 | 116.224271 | 136.562682 | 257.492145 | 268.039314 | 260.803879 | 312.234345 | 1.099864 | 2905.339007 | 0.122127423 | 1.12444519 | 0.26082421 | 0.995760118 | 11.434324 | 11.571402 | 11.448000 | 11.478861 | 11.477193 | 11.601403 | 3126 | 3043 | 2447 | 3068 |
| 55652 | NM_017842,XI | SLC48A1 | solute carrier family 48 member 1 | protein_coding | NP_060312.2;XP_005269073.1;XP_016875101.1;XP_016875103.1;XP_01 | 26035 | 612187 | . | . | 3.407451 | 3.564376 | 1.838424 | 1.789933 | 7.929278 | 8.239215 | 4.125370 | 4.092468 | -1.931260 | 247.493992 | 0.21961916 | -4.32358797 | 1.53512E-05 | 0.002788059 | 8.349236 | 7.399796 | 8.071106 | 8.090151 | 7.767012 | 7.753508 | 369 | 358 | 149 | 156 |
| 55654 | NM_00119330 | TMEM127 | transmembrane protein 127 | protein_coding | NP_001180233.1;NP_060319.1;XP_016859939.1;XP_016859941.1 | 26038 | 613403 | . | . | 12.956449 | 12.534118 | 13.147596 | 13.503478 | 30.150192 | 28.973179 | 29.502827 | 30.874097 | 1.030589 | 2541.875086 | 0.114233575 | 0.380523775 | 0.703556652 | 0.998865404 | 11.289766 | 11.333261 | 11.315715 | 11.282544 | 11.317237 | 11.330034 | 2942 | 2646 | 2226 | 2438 |
| 55656 | NM_017864,N | INTS8 | integrator complex subunit 8 | protein_coding | NP_060334.2;XP_016869105.1;XP_016869106.1;XP_016869107.1;XP_01 | 26048 | 611351 | . | . | 9.347061 | 9.728186 | 10.410665 | 8.138668 | 21.750995 | 22.487137 | 23.361231 | 18.608097 | -1.079014 | 1535.875770 | 0.144974888 | -0.75677792 | 0.449182924 | 0.998865404 | 10.638439 | 10.529185 | 10.601566 | 10.621587 | 10.643447 | 10.453396 | 1812 | 1742 | 1444 | 1214 |
| 55657 | NM_00113603 | ZNF692 | zinc finger protein 692 | protein_coding | NP_001129508.1;NP_001180257.1;NP_001337001.1;NP_001337002.1;NI | 26049 | 617758 | . | . | 8.098044 | 6.856158 | 10.401295 | 7.830995 | 18.844483 | 15.848317 | 23.340205 | 17.904638 | 1.209505 | 545.811654 | 0.183469486 | 1.495706067 | 0.134730269 | 0.891755273 | 8.947897 | 9.223481 | 9.064977 | 8.977846 | 9.228604 | 9.056121 | 608 | 496 | 587 | 488 |
| 55658 | NM_00136601 | RNF126 | ring finger protein 126 | protein_coding | NP_001352947.1;NP_919442.1 | 21151 | 615177 | . | . | 13.039146 | 15.063534 | 14.378797 | 15.021389 | 30.342628 | 34.820034 | 32.265606 | 34.344618 | 1.046740 | 755.771051 | 0.152049067 | 0.433436789 | 0.664697479 | 0.998865404 | 9.528667 | 9.594200 | 9.498332 | 9.589372 | 9.573340 | 9.579922 | 795 | 849 | 669 | 728 |
| 55659 | NM_00135340 | ZNF416 | zinc finger protein 416 | protein_coding | NP_001340334.1;NP_060349.1;XP_024307362.1 | 20645 | . | . | . | 0.153556 | 0.227204 | 0.321823 | 0.206123 | 0.357331 | 0.525192 | 0.722162 | 0.471276 | 1.351712 | 22.816633 | 0.663552335 | 0.655242079 | 0.512311906 | NA | 4.278006 | 4.713310 | 4.468225 | 4.501390 | 4.537160 | 4.489265 | 18 | 25 | 28 | 19 |
| 55660 | NM_00135443 | PRPF40A | pre-mRNA processing factor 40 homolog A | protein_coding | NP_001341360.1;NP_001341361.1;NP_001352525.1;NP_001352526.1;NI | 16463 | 612941 | . | . | 13.314545 | 13.757558 | 13.605668 | 10.890938 | 30.983492 | 31.801209 | 30.530725 | 24.900832 | -1.166350 | 2621.865357 | 0.120698206 | -1.83930485 | 0.06587036 | 0.73302766 | 11.462992 | 11.241251 | 11.428262 | 11.401589 | 11.340183 | 11.237979 | 3303 | 2997 | 2240 | 2124 |
| 55661 | NM_00134818 | DDX27 | DEAD-box helicase 27 | protein_coding | NP_001335116.1;NP_060365.7 | 15837 | 616621 | . | . | 60.822408 | 63.353829 | 62.509586 | 67.730767 | 141.536242 | 146.445201 | 140.269694 | 154.858360 | 1.035536 | 5367.599837 | 0.107733758 | 0.467616205 | 0.640059074 | 0.998865404 | 12.364700 | 12.414985 | 12.361858 | 12.388066 | 12.373760 | 12.433968 | 5978 | 5780 | 4579 | 5303 |
| 55662 | NM_017902,XI | HIF1AN | hypoxia inducible factor 1 subunit alpha inhibitor | protein_coding | NP_060372.2;XP_011538242.1;XP_011538243.1 | 17113 | 606615 | . | . | 5.058777 | 4.322406 | 5.653408 | 4.376887 | 11.771979 | 9.991434 | 12.686083 | 10.007230 | 1.076756 | 1700.179528 | 0.124514589 | 0.856854908 | 0.391525076 | 0.998865404 | 10.677070 | 10.783896 | 10.688262 | 10.714146 | 10.792006 | 10.725408 | 1854 | 1796 | 1598 | 1583 |
| 55663 | NM_00130445 | ZNF446 | zinc finger protein 446 | protein_coding | NP_001291382.1;NP_060378.1 | 21036 | . | . | . | 5.403899 | 4.617003 | 4.914913 | 5.747301 | 12.575093 | 10.672410 | 11.028922 | 13.140520 | 1.049483 | 330.338771 | 0.203432213 | 0.342518847 | 0.731960463 | 0.998865404 | 8.332585 | 8.402180 | 8.397781 | 8.305545 | 8.338600 | 8.418941 | 402 | 319 | 273 | 340 |
| 55664 | NM_017913 | CDC37L1 | cell division cycle 37 like 1 | protein_coding | NP_060383.2 | 17179 | 610346 | . | . | 0.877668 | 0.975440 | 0.964321 | 0.619519 | 2.042369 | 2.254773 | 2.163909 | 1.416457 | -1.185436 | 94.441390 | 0.350800822 | -0.69959282 | 0.48418163 | 0.998865404 | 6.677846 | 6.434609 | 6.566200 | 6.597624 | 6.591062 | 6.469679 | 112 | 116 | 92 | 63 |
| 55665 | NM_00107766 | URGCP | upregulator of cell proliferation | protein_coding | NP_001071131.1;NP_001071132.1;NP_001277004.1;NP_001277005.1;NI | 30890 | 610337 | . | . | 8.331458 | 8.357081 | 7.943304 | 9.578076 | 19.387646 | 19.317769 | 17.824543 | 21.899133 | 1.028015 | 991.344615 | 0.150477051 | 0.264903689 | 0.791083657 | 0.998865404 | 9.933468 | 9.972750 | 9.940466 | 9.943066 | 9.881940 | 10.035966 | 1124 | 1057 | 782 | 1041 |
| 55666 | NM_00136969 | NPLOC4 | NPL4 homolog, ubiquitin recognition factor | protein_coding | NP_001356627.1;NP_060391.2;XP_011523281.1;XP_011523282.1;XP_01 | 18261 | 606590 | . | . | 32.224641 | 34.596426 | 33.263658 | 37.903403 | 74.988062 | 79.971193 | 74.642689 | 86.661623 | 1.064014 | 4928.580015 | 0.114819298 | 0.779636221 | 0.43560503 | 0.998865404 | 12.221586 | 12.310943 | 12.198073 | 12.280004 | 12.249637 | 12.332048 | 5229 | 5407 | 4202 | 4997 |
| 55667 | NM_00133064 | DENND4C | DENN domain containing 4C | protein_coding | NP_001317569.1;NP_060395.5;XP_006716868.1;XP_006716870.1;XP_01 | 26079 | . | . | . | 1.447289 | 1.238902 | 1.513154 | 1.529014 | 3.367901 | 2.863779 | 3.395473 | 3.495909 | 1.118162 | 360.466438 | 0.192739674 | 0.835993154 | 0.403158754 | 0.998865404 | 8.410644 | 8.572274 | 8.502885 | 8.413072 | 8.519527 | 8.526797 | 423 | 338 | 329 | 359 |
| 55668 | NM_00132202 | GPATCH2L | G-patch domain containing 2 like | protein_coding | NP_001308955.1;NP_001308956.1;NP_001308957.1;NP_001308958.1;NI | 20210 | . | . | . | 4.298275 | 3.773172 | 4.607281 | 3.374283 | 10.002265 | 8.721857 | 10.338602 | 7.714896 | 1.051584 | 921.284097 | 0.158194029 | 0.458699851 | 0.646449721 | 0.998865404 | 9.810419 | 9.883656 | 9.828660 | 9.824413 | 9.950703 | 9.767326 | 1037 | 966 | 925 | 774 |
| 55669 | NM_033540,XI | MFN1 | mitofusin 1 | protein_coding | NP_284941.2;XP_005247653.2;XP_011511265.1 | 18262 | 608506 | . | . | 9.585559 | 9.904844 | 11.104523 | 9.012218 | 22.305990 | 22.895491 | 24.918229 | 20.605364 | -1.008952 | 1520.638307 | 0.140625533 | -0.09143456 | 0.927147305 | 0.998865404 | 10.576645 | 10.564251 | 10.576488 | 10.568092 | 10.645728 | 10.478564 | 1768 | 1639 | 1459 | 1266 |
| 55670 | NM_00112764 | PEX26 | peroxisomal biogenesis factor 26 | protein_coding | NP_001121121.1;NP_001186248.1;NP_060399.1 | 22965 | 608666 | . | . | 2.787217 | 2.686806 | 3.093546 | 3.272672 | 6.485968 | 6.210673 | 6.941827 | 7.482576 | 1.144451 | 1710.966668 | 0.122756144 | 1.585711178 | 0.112804842 | 0.850211417 | 10.639999 | 10.834633 | 10.702591 | 10.666941 | 10.771815 | 10.808903 | 1880 | 1682 | 1551 | 1752 |
| 55671 | NM_00128428 | PPP4R3A | protein phosphatase 4 regulatory subunit 3A | protein_coding | NP_001271209.1;NP_001271210.1;NP_001353361.1;XP_005267899.1 | 20219 | 610351 | . | . | 9.431591 | 9.455389 | 10.121586 | 8.464116 | 21.947697 | 21.856552 | 22.712541 | 19.352197 | -1.058501 | 1259.856229 | 0.146109611 | -0.56138132 | 0.574537618 | 0.998865404 | 10.339203 | 10.257734 | 10.322819 | 10.314209 | 10.357032 | 10.187203 | 1500 | 1390 | 1185 | 1018 |
| 55672 | NM_017940 | NBPF1 | NBPF member 1 | protein_coding | NP_060410.3 | 26088 | 610501 | . | . | 8.453694 | 9.321115 | 10.210115 | 9.277263 | 19.672092 | 21.546177 | 22.911203 | 21.211357 | 1.081608 | 1727.634938 | 0.128550025 | 0.880413012 | 0.37863561 | 0.998865404 | 10.696843 | 10.810100 | 10.686777 | 10.756354 | 10.824597 | 10.741008 | 1826 | 1871 | 1644 | 1594 |
| 55676 | NM_00119351 | SLC30A6 | solute carrier family 30 member 6 | protein_coding | NP_001180442.1;NP_001180443.1;NP_001180444.1;NP_001317405.1;NI | 19305 | 611148 | . | . | 7.048838 | 7.324772 | 8.060203 | 5.993322 | 16.402935 | 16.931538 | 18.086864 | 13.703017 | -1.028514 | 1069.881446 | 0.160513235 | -0.25270235 | 0.800498233 | 0.998865404 | 10.083027 | 10.043169 | 10.076606 | 10.064582 | 10.153884 | 9.934281 | 1259 | 1161 | 1054 | 842 |
| 55677 | NM_017969,XI | IWS1 | interacts with SUPT6H, CTD assembly factor 1 | protein_coding | NP_060439.2;XP_005263764.1;XP_006712689.1;XP_011509752.1;XP_01 | 25467 | . | . | . | 29.538362 | 28.658024 | 27.247976 | 29.544891 | 68.736987 | 66.244309 | 61.143668 | 67.550884 | -1.050723 | 2650.394678 | 0.117381067 | -0.6081244 | 0.543104973 | 0.998865404 | 11.407299 | 11.335799 | 11.406971 | 11.375855 | 11.313134 | 11.388160 | 3187 | 2875 | 2143 | 2536 |
| 55680 | NM_00104241 | RUFY2 | RUN and FYVE domain containing 2 | protein_coding | NP_001035882.1;NP_001265154.1;NP_001317032.1;NP_060457.4;XP_00 | 19761 | 610328 | . | . | 1.901615 | 1.932981 | 2.185338 | 1.201391 | 4.425136 | 4.468171 | 4.903839 | 2.746838 | -1.131619 | 251.020225 | 0.254383418 | -0.70125886 | 0.483141481 | 0.998865404 | 8.057190 | 7.880737 | 7.992383 | 7.997950 | 8.044309 | 7.808754 | 305 | 289 | 255 | 167 |
| 55681 | NM_00131778 | SCYL2 | SCY1 like pseudokinase 2 | protein_coding | NP_001304713.1;NP_001317182.1;NP_001317183.1;NP_001317185.1;NI | 19286 | 616365 | . | . | 8.083349 | 8.337340 | 7.523688 | 6.608308 | 18.810286 | 19.272134 | 16.882937 | 15.109109 | -1.179148 | 1392.176445 | 0.134375078 | -1.76926625 | 0.076849454 | 0.767975096 | 10.556962 | 10.319533 | 10.485990 | 10.518158 | 10.428441 | 10.320880 | 1699 | 1659 | 1191 | 1112 |
| 55683 | NM_00111501 | KANSL3 | KAT8 regulatory NSL complex subunit 3 | protein_coding | NP_001108488.1;NP_001336185.1;NP_001336186.1;NP_001336187.1;NI | 25473 | 617742 | . | . | 11.842182 | 12.292547 | 13.418648 | 14.379611 | 27.557243 | 28.414767 | 30.111058 | 32.877269 | 1.153514 | 2076.653511 | 0.119290375 | 1.7271814 | 0.08413509 | 0.787933172 | 10.913448 | 11.119304 | 10.947873 | 10.971926 | 11.046347 | 11.100197 | 2187 | 2113 | 1870 | 2155 |
| 55684 | NM_00117398 | RABL6 | RAB, member RAS oncogene family like 6 | protein_coding | NP_001167459.1;NP_001167460.1;NP_078994.3 | 24703 | 610615 | . | . | 28.304063 | 27.017465 | 26.672708 | 32.624970 | 65.864717 | 62.452086 | 59.852783 | 74.593119 | 1.057312 | 2693.184605 | 0.130209032 | 0.617482065 | 0.536916808 | 0.998865404 | 11.354449 | 11.434632 | 11.390351 | 11.350640 | 11.330028 | 11.495182 | 3089 | 2756 | 2164 | 2857 |
| 55686 | NM_00137218 | MREG | melanoregulin | protein_coding | NP_001359117.1;NP_001359118.1;NP_001359119.1;NP_060470.2;XP_01 | 25478 | 609207 | . | . | 19.938207 | 20.560286 | 21.473284 | 22.238883 | 46.397025 | 47.526020 | 48.185430 | 50.846566 | 1.107251 | 1922.450368 | 0.119301086 | 1.232029958 | 0.217937894 | 0.982008277 | 10.833376 | 10.980344 | 10.879313 | 10.854012 | 10.934624 | 10.959984 | 2135 | 1937 | 1729 | 1924 |
| 55687 | NM_00128278 | TRMU | tRNA 5-methylaminomethyl-2-thiouridylate methyltransferase | protein_coding | NP_001269711.1;NP_001269712.1;NP_001269713.1;NP_001269714.1;NI | 25481 | 610230 | . | . | 22.413769 | 23.533671 | 25.155926 | 20.863889 | 52.157764 | 54.399134 | 56.449169 | 47.702805 | 1.056305 | 1577.562213 | 0.129972939 | 0.60801949 | 0.543174548 | 0.998865404 | 10.583312 | 10.662563 | 10.585665 | 10.616092 | 10.692537 | 10.592309 | 1732 | 1688 | 1500 | 1422 |
| 55689 | NM_00135136 | YEATS2 | YEATS domain containing 2 | protein_coding | NP_001338298.1;NP_001338299.1;NP_001338300.1;NP_060493.3;XP_00 | 25489 | 613373 | . | . | 10.283531 | 11.175416 | 10.393796 | 11.762269 | 23.930205 | 25.832478 | 23.323375 | 26.893027 | 1.033432 | 2164.546279 | 0.12342147 | 0.384399743 | 0.700682191 | 0.998865404 | 11.056084 | 11.103229 | 11.033364 | 11.097717 | 11.050785 | 11.132056 | 2353 | 2390 | 1816 | 2167 |
| 55690 | NM_018026,XI | PACS1 | phosphofurin acidic cluster sorting protein 1 | protein_coding | NP_060496.2;XP_011543464.1;XP_011543465.1 | 30032 | 607492 | . | . | 5.692964 | 6.093381 | 5.094910 | 6.486051 | 13.247759 | 14.085122 | 11.432833 | 14.829585 | -1.037292 | 831.945559 | 0.160689162 | -0.32872487 | 0.742363652 | 0.998865404 | 9.726926 | 9.673266 | 9.685755 | 9.736951 | 9.598776 | 9.765364 | 941 | 947 | 626 | 856 |
| 55691 | NM_00131833 | FRMD4A | FERM domain containing 4A | protein_coding | NP_001305265.1;NP_001305266.1;NP_001305267.1;NP_060497.3;XP_00 | 25491 | 616305 | . | . | 14.574683 | 15.754820 | 18.514810 | 16.595055 | 33.915884 | 36.417973 | 41.546699 | 37.942621 | 1.199477 | 5459.873894 | 0.118882176 | 2.207272988 | 0.027294993 | 0.538018865 | 12.277474 | 12.539911 | 12.285770 | 12.376860 | 12.541249 | 12.425694 | 5402 | 5652 | 5529 | 5205 |
| 55692 | NM_00132022 | LUC7L | LUC7 like | protein_coding | NP_001307155.1;NP_001317349.1;NP_060502.1;NP_958815.1;XP_00525 | 6723 | 607782 | . | . | 12.975398 | 14.115466 | 14.883582 | 14.534850 | 30.194284 | 32.628533 | 33.398327 | 33.232202 | 1.043305 | 891.242520 | 0.141312869 | 0.432803938 | 0.665157212 | 0.998865404 | 9.768700 | 9.829997 | 9.775621 | 9.792042 | 9.834644 | 9.793711 | 993 | 952 | 814 | 829 |
| 55693 | NM_018039 | KDM4D | lysine demethylase 4D | protein_coding | NP_060509.2 | 25498 | 609766 | . | . | 0.724270 | 0.651023 | 0.922597 | 0.939310 | 1.685407 | 1.504869 | 2.070281 | 2.147621 | 1.331205 | 74.602089 | 0.366523579 | 1.12607319 | 0.260134541 | 0.995760118 | 5.999739 | 6.413048 | 6.190776 | 6.164586 | 6.248246 | 6.249539 | 78 | 65 | 74 | 80 |
| 55695 | NM_00116834 | NSUN5 | NOP2/Sun RNA methyltransferase 5 | protein_coding | NP_001161819.1;NP_001161820.1;NP_060514.1;NP_683759.1;XP_00527 | 16385 | 615732 | . | . | 10.175135 | 10.277576 | 7.024175 | 9.021774 | 23.677959 | 23.757109 | 15.762046 | 20.627216 | -1.098687 | 601.559260 | 0.164903298 | -0.82339647 | 0.41028257 | 0.998865404 | 9.299187 | 9.162713 | 9.259440 | 9.265741 | 9.143374 | 9.253557 | 722 | 683 | 456 | 583 |
| 55696 | NM_018047,XI | RBM22 | RNA binding motif protein 22 | protein_coding | NP_060517.1 | 25503 | 612430 | . | . | 20.891823 | 21.307896 | 21.878126 | 21.842469 | 48.616131 | 49.254160 | 49.093885 | 49.940209 | 1.023730 | 1515.415720 | 0.123833652 | 0.273232616 | 0.78467441 | 0.998865404 | 10.548475 | 10.582322 | 10.552444 | 10.560101 | 10.578912 | 10.569929 | 1716 | 1624 | 1341 | 1429 |
| 55697 | NM_00135115 | VAC14 | VAC14 component of PIKFYVE complex | protein_coding | NP_001338086.1;NP_060522.3;XP_005256095.1;XP_005256098.2;XP_01 | 25507 | 604632 | . | . | 7.677362 | 8.009277 | 6.104391 | 9.705532 | 17.865537 | 18.513799 | 13.698073 | 22.190546 | 1.021330 | 1046.374576 | 0.145999931 | 0.208558525 | 0.834792891 | 0.998865404 | 10.016172 | 10.046040 | 9.963309 | 10.078032 | 10.000670 | 10.073411 | 1100 | 1204 | 874 | 1040 |
| 55699 | NM_018060 | IARS2 | isoleucyl-tRNA synthetase 2, mitochondrial | protein_coding | NP_060530.3 | 29685 | 612801 | . | . | 43.208591 | 42.752205 | 37.860298 | 42.535130 | 100.548172 | 98.823631 | 84.957405 | 97.251518 | -1.084500 | 4574.546151 | 0.111833268 | -1.04647115 | 0.29534356 | 0.998865404 | 12.216793 | 12.099656 | 12.200126 | 12.183907 | 12.079481 | 12.167282 | 5534 | 5087 | 3614 | 4333 |
| 55700 | NM_00128636 | MAP7D1 | MAP7 domain containing 1 | protein_coding | NP_001273294.1;NP_001273295.1;NP_060537.3;XP_005271084.1;XP_00 | 25514 | . | . | . | 30.022973 | 29.613448 | 26.023012 | 33.230681 | 69.864685 | 68.452808 | 58.394882 | 75.978005 | -1.012055 | 3052.355478 | 0.129205167 | -0.13380508 | 0.893556712 | 0.998865404 | 11.584445 | 11.566916 | 11.587635 | 11.570954 | 11.479870 | 11.651823 | 3572 | 3280 | 2361 | 3143 |
| 55701 | NM_00127852 | ARHGEF40 | Rho guanine nucleotide exchange factor 40 | protein_coding | NP_001265458.1;NP_001265459.1;NP_060541.3;XP_005267901.1;XP_01 | 25516 | 610018 | . | . | 6.366776 | 7.076530 | 6.242586 | 7.749958 | 14.815750 | 16.357717 | 14.008183 | 17.719359 | 1.042081 | 1248.035541 | 0.145684472 | 0.40819771 | 0.683128529 | 0.998865404 | 10.255717 | 10.314569 | 10.233317 | 10.301825 | 10.219920 | 10.372712 | 1344 | 1379 | 996 | 1314 |
| 55702 | NM_018074 | YJU2 | YJU2 splicing factor homolog | protein_coding | NP_060544.2 | 25518 | . | . | . | 5.074492 | 5.178616 | 5.755830 | 7.131176 | 11.808553 | 11.970604 | 12.915915 | 16.304586 | 1.239798 | 258.016492 | 0.220502916 | 1.406352376 | 0.159619515 | 0.933525128 | 7.848482 | 8.157628 | 7.947329 | 7.952794 | 8.003257 | 8.111236 | 264 | 250 | 223 | 295 |
| 55703 | NM_00116070 | POLR3B | RNA polymerase III subunit B | protein_coding | NP_001154180.1;NP_060552.4;XP_016875110.1 | 30348 | 614366 | . | . | 2.761424 | 3.174976 | 2.957837 | 3.125772 | 6.425946 | 7.339098 | 6.637301 | 7.146709 | 1.007412 | 395.033203 | 0.18665816 | 0.057078759 | 0.954482457 | 0.998865404 | 8.620845 | 8.630798 | 8.578063 | 8.665701 | 8.612137 | 8.641780 | 420 | 456 | 336 | 381 |
| 55704 | NM_00113559 | CCDC88A | coiled-coil domain containing 88A | protein_coding | NP_001129069.1;NP_001241872.1;NP_001352409.1;NP_060554.3;XP_00 | 25523 | 609736 | . | . | 10.122094 | 9.493569 | 10.618865 | 8.929666 | 23.554527 | 21.944808 | 23.828426 | 20.416621 | -1.004091 | 2879.673667 | 0.131472825 | -0.04480287 | 0.964264436 | 0.998865404 | 11.494463 | 11.488911 | 11.535542 | 11.446016 | 11.564134 | 11.406944 | 3504 | 2943 | 2740 | 2435 |
| 55705 | NM_018085 | IPO9 | importin 9 | protein_coding | NP_060555.2 | 19425 | . | . | . | 11.124980 | 11.849649 | 12.038958 | 12.206974 | 25.888287 | 27.390995 | 27.015070 | 27.909792 | 1.040850 | 4197.815004 | 0.108316827 | 0.533275798 | 0.593842679 | 0.998865404 | 12.006280 | 12.063989 | 11.995579 | 12.040781 | 12.050261 | 12.053122 | 4608 | 4560 | 3717 | 4021 |
| 55706 | NM_00116855 | NDC1 | NDC1 transmembrane nucleoporin | protein_coding | NP_001162023.1;NP_060557.3;XP_011540068.1 | 25525 | 610115 | . | . | 14.130429 | 13.104650 | 12.503865 | 10.480455 | 32.882089 | 30.291987 | 28.058313 | 23.962316 | -1.211119 | 1768.666353 | 0.136960691 | -2.01766383 | 0.043626284 | 0.642661255 | 10.919783 | 10.643944 | 10.891098 | 10.822883 | 10.778793 | 10.630238 | 2325 | 2002 | 1526 | 1355 |
| 55707 | NM_00114527 | NECAP2 | NECAP endocytosis associated 2 | protein_coding | NP_001138749.1;NP_001138750.1;NP_060560.1;XP_011540069.1;XP_01 | 25528 | 611624 | . | . | 21.934948 | 21.514631 | 23.328199 | 26.770957 | 51.043527 | 49.732035 | 52.347801 | 61.208616 | 1.082426 | 1519.351101 | 0.130742436 | 0.873999008 | 0.382118794 | 0.998865404 | 10.511153 | 10.625078 | 10.528122 | 10.545999 | 10.545680 | 10.648361 | 1661 | 1593 | 1283 | 1578 |
| 55709 | NM_00131871 | KBTBD4 | kelch repeat and BTB domain containing 4 | protein_coding | NP_001305645.1;NP_001305646.1;NP_001305647.1;NP_001305648.1;NI | 23761 | 617645 | . | . | 3.056828 | 3.434838 | 3.561015 | 3.916718 | 7.113361 | 7.939781 | 7.990816 | 8.955112 | 1.161071 | 265.447455 | 0.214293337 | 1.005429234 | 0.314690209 | 0.998865404 | 7.940986 | 8.155604 | 7.977849 | 8.041516 | 8.067192 | 8.105488 | 265 | 282 | 239 | 277 |
| 55711 | NM_00127178 | FAR2 | fatty acyl-CoA reductase 2 | protein_coding | NP_001258712.1;NP_001258713.1;NP_060569.3;XP_011519049.1;XP_01 | 25531 | 616156 | . | . | 2.629579 | 2.657883 | 2.743508 | 2.937750 | 6.119138 | 6.143815 | 6.156353 | 6.716817 | 1.012827 | 318.378340 | 0.199616738 | 0.092114505 | 0.926607064 | 0.998865404 | 8.305120 | 8.324015 | 8.346046 | 8.273752 | 8.320198 | 8.315025 | 388 | 319 | 280 | 299 |
| 55713 | NM_00127049 | ZNF334 | zinc finger protein 334 | protein_coding | NP_001257426.1;NP_001340742.1;NP_001340743.1;NP_001340744.1;NI | 15806 | . | . | . | 0.962224 | 0.616730 | 0.807993 | 0.597930 | 2.239134 | 1.425600 | 1.813114 | 1.367095 | -1.240079 | 130.537389 | 0.291818871 | -1.06378239 | 0.287427289 | 0.998865404 | 7.174438 | 6.865716 | 7.094172 | 7.034655 | 7.018757 | 6.956357 | 179 | 144 | 111 | 99 |
| 55714 | NM_00108047 | TENM3 | teneurin transmembrane protein 3 | protein_coding | NP_001073946.1;XP_016863874.1;XP_016863875.1;XP_016863876.1;XP_ | 29944 | 610083 | . | . | 3.921405 | 4.412712 | 3.215408 | 3.478707 | 9.125270 | 10.200183 | 7.215284 | 7.953650 | -1.244003 | 1236.879056 | 0.132823253 | -2.37149801 | 0.017716142 | 0.446233408 | 10.421560 | 10.106221 | 10.331471 | 10.366116 | 10.152953 | 10.214326 | 1544 | 1513 | 918 | 1077 |
| 55715 | NM_00133055 | DOK4 | docking protein 4 | protein_coding | NP_001317485.1;NP_001356547.1;NP_001356548.1;NP_001356549.1;NI | 19868 | 608333 | . | . | 1.385101 | 1.728166 | 1.655916 | 1.789474 | 3.223187 | 3.994732 | 3.715827 | 4.091420 | 1.110994 | 143.261211 | 0.278879771 | 0.544505147 | 0.586093908 | 0.998865404 | 7.085329 | 7.235760 | 7.104376 | 7.173194 | 7.155091 | 7.202805 | 142 | 160 | 123 | 150 |
| 55716 | NM_00130075 | LMBR1L | limb development membrane protein 1 like | protein_coding | NP_001287679.1;NP_001287680.1;NP_001339090.1;NP_001339091.1;NI | 18268 | 610007 | . | . | 5.349388 | 6.035530 | 6.948142 | 6.443312 | 12.448250 | 13.951401 | 15.591433 | 14.731866 | 1.150372 | 552.546214 | 0.166276351 | 1.215451645 | 0.224193881 | 0.982008277 | 9.005225 | 9.207587 | 9.040126 | 9.080701 | 9.190734 | 9.111358 | 573 | 572 | 545 | 521 |
| 55717 | NM_018117,XI | WDR11 | WD repeat domain 11 | protein_coding | NP_060587.8;XP_005270020.1;XP_016871886.1;XP_016871887.1;XP_01 | 13831 | 606417 | . | . | 11.945298 | 14.154323 | 11.647352 | 13.671331 | 27.797199 | 32.718352 | 26.136317 | 31.257873 | -1.029476 | 1778.589389 | 0.133838515 | -0.31314382 | 0.754171399 | 0.998865404 | 10.817541 | 10.775184 | 10.754520 | 10.856651 | 10.729200 | 10.835813 | 1944 | 2073 | 1420 | 1755 |
| 55718 | NM_00125803 | POLR3E | RNA polymerase III subunit E | protein_coding | NP_001244962.1;NP_001244963.1;NP_001244964.1;NP_001244965.1;NI | 30347 | 617815 | . | . | 8.355358 | 8.364683 | 9.308068 | 8.713093 | 19.443262 | 19.335340 | 20.887036 | 19.921451 | 1.082594 | 981.025736 | 0.138275857 | 0.827997762 | 0.407671767 | 0.998865404 | 9.879629 | 9.994384 | 9.915997 | 9.899675 | 9.989616 | 9.941626 | 1097 | 1005 | 916 | 925 |
| 55719 | NM_00113612 | SLF2 | SMC5-SMC6 complex localization factor 2 | protein_coding | NP_001129595.1;NP_001230699.1;NP_060591.3;XP_005270022.1;XP_01 | 17814 | 610348 | . | . | 7.165476 | 6.137700 | 8.447824 | 6.593723 | 16.674357 | 14.187568 | 18.956674 | 15.075765 | 1.020299 | 1258.359819 | 0.145184785 | 0.199686704 | 0.841725613 | 0.998865404 | 10.282473 | 10.312033 | 10.304107 | 10.271492 | 10.381409 | 10.218794 | 1465 | 1315 | 1224 | 1064 |
| 55720 | NM_018128 | TSR1 | TSR1 ribosome maturation factor | protein_coding | NP_060598.3 | 25542 | 611214 | . | . | 22.287214 | 23.175310 | 23.084633 | 25.180870 | 51.863266 | 53.570766 | 51.801247 | 57.573071 | 1.046960 | 3098.136787 | 0.113704648 | 0.582263202 | 0.560389408 | 0.998865404 | 11.563779 | 11.629835 | 11.565409 | 11.590170 | 11.583357 | 11.646454 | 3433 | 3316 | 2650 | 3085 |
| 55721 | NM_00116004 | IQCC | IQ motif containing C | protein_coding | NP_001153514.1;NP_060604.2 | 25545 | . | . | . | 3.545164 | 4.567327 | 3.412521 | 4.217582 | 8.249743 | 10.557580 | 7.657598 | 9.643002 | -1.067085 | 249.210724 | 0.232596337 | -0.40273906 | 0.687140197 | 0.998865404 | 8.008095 | 7.912775 | 7.911728 | 8.034532 | 7.896323 | 7.986589 | 261 | 311 | 190 | 247 |
| 55722 | NM_018140,N | CEP72 | centrosomal protein 72 | protein_coding | NP_060610.2;XP_005248379.1;XP_011512365.1;XP_011512366.1;XP_01 | 25547 | 616475 | . | . | 1.910939 | 1.999680 | 1.871884 | 1.839709 | 4.446834 | 4.622349 | 4.200452 | 4.206280 | 1.007292 | 266.411719 | 0.21231266 | 0.049370573 | 0.960623979 | 0.998865404 | 8.051987 | 8.063019 | 8.082653 | 8.027093 | 8.066531 | 8.051505 | 322 | 271 | 236 | 247 |
| 55723 | NM_018154 | ASF1B | anti-silencing function 1B histone chaperone | protein_coding | NP_060624.1 | 20996 | 609190 | . | . | 13.375552 | 13.881699 | 12.115314 | 15.432268 | 31.125460 | 32.088165 | 27.186415 | 35.284050 | -1.002838 | 721.365925 | 0.163884541 | -0.02494587 | 0.98009814 | 0.998865404 | 9.497031 | 9.492140 | 9.484043 | 9.504194 | 9.409575 | 9.568331 | 820 | 791 | 554 | 753 |
| 55726 | NM_018164,XI | INTS13 | integrator complex subunit 13 | protein_coding | NP_060634.2;XP_005253479.1;XP_011519052.1;XP_011519053.1;XP_01 | 20174 | 615079 | . | . | 18.964907 | 18.885787 | 19.706754 | 17.801785 | 44.132123 | 43.655345 | 44.221383 | 40.701665 | -1.023241 | 1549.434058 | 0.128748851 | -0.25744397 | 0.796836066 | 0.998865404 | 10.613853 | 10.581011 | 10.610013 | 10.601750 | 10.632352 | 10.542031 | 1814 | 1682 | 1410 | 1352 |
| 55727 | NM_00100286 | BTBD7 | BTB domain containing 7 | protein_coding | NP_001002860.2;NP_001276062.1;NP_060637.1;XP_011535241.1;XP_01 | 18269 | 610386 | . | . | 2.398868 | 2.836231 | 3.122987 | 2.752949 | 5.582264 | 6.556074 | 7.007892 | 6.294290 | -1.011962 | 513.077524 | 0.171493766 | -0.10003621 | 0.920315581 | 0.998865404 | 9.011561 | 8.994455 | 8.966920 | 9.043504 | 9.028521 | 8.967270 | 559 | 590 | 465 | 455 |
| 55728 | NM_00131835 | N4BP2 | NEDD4 binding protein 2 | protein_coding | NP_001305288.1;NP_060647.2;XP_006714086.1;XP_011512018.1;XP_01 | 29851 | . | . | . | 0.607005 | 0.801965 | 0.715520 | 0.473753 | 1.412523 | 1.853777 | 1.605605 | 1.083177 | -1.139236 | 191.493804 | 0.277569021 | -0.67754891 | 0.498057746 | 0.998865404 | 7.671529 | 7.484736 | 7.545909 | 7.655768 | 7.633333 | 7.447602 | 207 | 246 | 190 | 131 |
| 55729 | NM_00128651 | ATF7IP | activating transcription factor 7 interacting protein | protein_coding | NP_001273443.1;NP_001273444.1;NP_060649.3;XP_851997.1;XP_00525 | 20092 | 613644 | . | . | 3.903211 | 4.459046 | 3.902874 | 3.705822 | 9.082935 | 10.307285 | 8.757934 | 8.472923 | -1.101786 | 1069.191728 | 0.14199435 | -0.98485939 | 0.324693145 | 0.998865404 | 10.130553 | 9.990667 | 10.045126 | 10.145259 | 10.050908 | 9.997719 | 1205 | 1290 | 917 | 919 |
| 55731 | NM_00107749 | FAM222B | family with sequence similarity 222 member B | protein_coding | NP_001070966.1;NP_001275560.1;NP_001275561.1;NP_001275562.1;NI | 25563 | . | . | . | 7.462813 | 8.368437 | 9.709768 | 9.921968 | 17.366273 | 19.344017 | 21.788440 | 22.685401 | 1.224788 | 1171.770952 | 0.134764918 | 2.170681563 | 0.029955252 | 0.553375672 | 10.040960 | 10.333214 | 10.074551 | 10.148491 | 10.257274 | 10.270911 | 1152 | 1194 | 1112 | 1220 |
| 55732 | NM_00132004 | C1orf112 | chromosome 1 open reading frame 112 | protein_coding | NP_001306976.1;NP_001306977.1;NP_001306979.1;NP_001306980.1;NI | 25565 | . | . | . | 6.591913 | 6.945996 | 6.859049 | 5.356394 | 15.339646 | 16.055980 | 15.391507 | 12.246756 | -1.078631 | 751.334857 | 0.163287333 | -0.66876959 | 0.503642465 | 0.998865404 | 9.606597 | 9.497984 | 9.552740 | 9.601181 | 9.600471 | 9.442725 | 867 | 870 | 701 | 600 |
| 55733 | NM_00112283 | HHAT | hedgehog acyltransferase | protein_coding | NP_001116306.1;NP_001164035.1;NP_001164051.1;NP_001164058.1;NI | 18270 | 605743 | . | . | 3.373263 | 3.430772 | 2.943108 | 3.216372 | 7.849723 | 7.930383 | 6.604246 | 7.353852 | -1.109641 | 364.314096 | 0.187795473 | -0.79923866 | 0.424152037 | 0.998865404 | 8.582324 | 8.431831 | 8.539483 | 8.537613 | 8.455727 | 8.501050 | 442 | 413 | 288 | 337 |
| 55734 | NM_00131914 | ZFP64 | ZFP64 zinc finger protein | protein_coding | NP_001306075.1;NP_060667.2;NP_071371.3;NP_955458.1;NP_955459.2 | 15940 | 618111 | . | . | 4.611292 | 4.814516 | 5.542888 | 5.617915 | 10.730666 | 11.128969 | 12.438079 | 12.844696 | 1.141180 | 475.691781 | 0.16994276 | 1.121120112 | 0.262236739 | 0.995760118 | 8.795598 | 8.985898 | 8.846625 | 8.853243 | 8.917909 | 8.946684 | 509 | 482 | 431 | 486 |
| 55735 | NM_018198,XI | DNAJC11 | DnaJ heat shock protein family (Hsp40) member C11 | protein_coding | NP_060668.2;XP_016857234.1 | 25570 | 614827 | . | . | 37.297883 | 38.302613 | 40.366057 | 42.517576 | 86.793704 | 88.538215 | 90.580265 | 97.211389 | 1.080407 | 3948.711840 | 0.1084348 | 1.02896193 | 0.303497565 | 0.998865404 | 11.890337 | 12.001841 | 11.906497 | 11.921695 | 11.960893 | 11.995242 | 4331 | 4134 | 3493 | 3924 |
| 55737 | NM_018206,XI | VPS35 | VPS35 retromer complex component | protein_coding | NP_060676.2;XP_005256102.1;XP_011521529.1 | 13487 | 601501 | . | . | 17.541121 | 19.624015 | 18.116574 | 14.750712 | 40.818907 | 45.361790 | 40.653067 | 33.725750 | -1.059425 | 2846.542012 | 0.113819211 | -0.73169377 | 0.464355497 | 0.998865404 | 11.515970 | 11.432822 | 11.520069 | 11.475328 | 11.466989 | 11.434666 | 3464 | 3074 | 2452 | 2540 |
| 55738 | NM_00128148 | ARFGAP1 | ADP ribosylation factor GTPase activating protein 1 | protein_coding | NP_001268411.1;NP_001268412.1;NP_001268413.1;NP_060679.1;NP_7( | 15852 | 608377 | . | . | 51.251363 | 51.581917 | 44.947038 | 47.005199 | 119.264031 | 119.233912 | 100.859854 | 107.471801 | -1.133872 | 4994.535586 | 0.105784315 | -1.71346401 | 0.086627209 | 0.795065303 | 12.373940 | 12.192637 | 12.335201 | 12.337677 | 12.216125 | 12.246328 | 6103 | 5736 | 3996 | 4469 |
| 55739 | NM_00124288 | NAXD | NAD(P)HX dehydratase | protein_coding | NP_001229810.1;NP_001229811.1;NP_001229812.1;NP_060680.2 | 25576 | 615910 | . | . | 10.608612 | 9.507233 | 10.482764 | 10.445219 | 24.686678 | 21.976396 | 23.523020 | 23.881751 | 1.025348 | 835.490149 | 0.146139102 | 0.247114226 | 0.804819824 | 0.998865404 | 9.688130 | 9.724597 | 9.731730 | 9.661151 | 9.725755 | 9.704005 | 997 | 846 | 747 | 781 |
| 55740 | NM_00100849 | ENAH | ENAH actin regulator | protein_coding | NP_001008493.1;NP_001364410.1;NP_001364411.1;NP_001364412.1;NI | 18271 | 609061 | . | . | 5.781753 | 5.657769 | 4.220802 | 4.602835 | 13.454374 | 13.078183 | 9.471354 | 10.523838 | -1.303030 | 2047.165834 | 0.118983882 | -3.20942731 | 0.001329997 | 0.083777674 | 11.177799 | 10.795815 | 11.103796 | 11.086779 | 10.862297 | 10.906737 | 2695 | 2474 | 1501 | 1715 |
| 55741 | NM_00114502 | EDEM2 | ER degradation enhancing alpha-mannosidase like protein 2 | protein_coding | NP_001138497.1;NP_060687.2 | 15877 | 610302 | . | . | 33.210934 | 31.907109 | 30.158978 | 32.652584 | 77.283210 | 73.754709 | 67.675874 | 74.656249 | -1.052472 | 1893.827650 | 0.120562107 | -0.61197762 | 0.540552567 | 0.998865404 | 10.923538 | 10.849696 | 10.922000 | 10.891678 | 10.845936 | 10.886646 | 2279 | 2056 | 1563 | 1775 |
| 55742 | NM_018222,XI | PARVA | parvin alpha | protein_coding | NP_060692.3;XP_005253072.1 | 14652 | 608120 | . | . | 12.171849 | 13.097819 | 11.801406 | 13.131233 | 28.324391 | 30.276196 | 26.482013 | 30.023003 | -1.027737 | 3385.051148 | 0.115930063 | -0.34047162 | 0.733501389 | 0.998865404 | 11.744660 | 11.704991 | 11.707903 | 11.762889 | 11.672560 | 11.752249 | 3820 | 3826 | 2762 | 3282 |
| 55743 | NM_00116134 | CHFR | checkpoint with forkhead and ring finger domains | protein_coding | NP_001154816.1;NP_001154817.1;NP_001154818.1;NP_001154819.1;NI | 20455 | 605209 | . | . | 3.541050 | 4.028684 | 3.588663 | 4.614614 | 8.240172 | 9.312483 | 8.052858 | 10.550766 | 1.053855 | 941.486152 | 0.141948985 | 0.533128804 | 0.593947882 | 0.998865404 | 9.840671 | 9.915939 | 9.845269 | 9.872514 | 9.859795 | 9.932772 | 1036 | 1008 | 798 | 951 |
| 55744 | NM_00132119 | COA1 | cytochrome c oxidase assembly factor 1 homolog | protein_coding | NP_001308126.1;NP_001308127.1;NP_001308128.1;NP_001308129.1;NI | 21868 | 614769 | . | . | 18.746278 | 22.528784 | 14.770744 | 26.772193 | 43.623357 | 52.076300 | 33.145129 | 61.211435 | 1.117231 | 2224.035771 | 0.122058226 | 1.310257336 | 0.190108795 | 0.961278413 | 11.036642 | 11.196842 | 11.104731 | 11.038533 | 11.195561 | 11.125295 | 2520 | 2172 | 2130 | 2105 |
| 55745 | NM_018229,N | AP5M1 | adaptor related protein complex 5 subunit mu 1 | protein_coding | NP_060699.3;XP_006720259.1;XP_011535242.1 | 20192 | 614368 | . | . | 1.817877 | 1.758405 | 2.083293 | 1.586160 | 4.230277 | 4.064632 | 4.674850 | 3.626565 | 1.013937 | 658.855231 | 0.1735843 | 0.115030265 | 0.908421133 | 0.998865404 | 9.353317 | 9.374243 | 9.371363 | 9.340937 | 9.455229 | 9.267852 | 770 | 690 | 657 | 536 |
| 55746 | NM_018230,XI | NUP133 | nucleoporin 133 | protein_coding | NP_060700.2 | 18016 | 607613 | . | . | 7.744530 | 8.038881 | 7.202933 | 7.938525 | 18.021839 | 18.582235 | 16.163174 | 18.150492 | -1.061541 | 1252.046357 | 0.131559035 | -0.65491762 | 0.512520793 | 0.998865404 | 10.332649 | 10.246201 | 10.300885 | 10.323397 | 10.237457 | 10.295627 | 1463 | 1412 | 1015 | 1183 |
| 55748 | NM_00116849 | CNDP2 | carnosine dipeptidase 2 | protein_coding | NP_001161971.1;NP_001357177.1;NP_001357178.1;NP_001357179.1;NI | 24437 | 169800 | . | . | 25.160765 | 25.391244 | 24.359989 | 28.856653 | 58.550129 | 58.692999 | 54.663111 | 65.977312 | 1.038165 | 4025.830824 | 0.119130299 | 0.453582782 | 0.650129149 | 0.998865404 | 11.947887 | 12.001751 | 11.958163 | 11.958961 | 11.921048 | 12.053864 | 4547 | 4265 | 3282 | 4151 |
| 55749 | NM_00128295 | CCAR1 | cell division cycle and apoptosis regulator 1 | protein_coding | NP_001269888.1;NP_001269889.1;NP_060707.2 | 24236 | 612569 | . | . | 17.021417 | 16.054669 | 17.803250 | 15.267654 | 39.609539 | 37.111088 | 39.949977 | 34.907677 | -1.015636 | 2386.439835 | 0.126871832 | -0.17642946 | 0.859956559 | 0.998865404 | 11.231626 | 11.209578 | 11.254704 | 11.196224 | 11.274307 | 11.148861 | 2869 | 2501 | 2220 | 2047 |
| 55750 | NM_00136494 | AGK | acylglycerol kinase | protein_coding | NP_001351877.1;NP_060708.1;XP_011514699.1;XP_024302603.1 | 21869 | 610345 | . | . | 5.934834 | 6.861533 | 5.430221 | 6.171601 | 13.810604 | 15.860740 | 12.185258 | 14.110632 | -1.075668 | 956.960699 | 0.148539044 | -0.70844873 | 0.478666639 | 0.998865404 | 9.954108 | 9.848595 | 9.962425 | 9.891457 | 9.816799 | 9.928933 | 1197 | 1019 | 739 | 932 |
| 55751 | NM_018241,XI | TMEM184C | transmembrane protein 184C | protein_coding | NP_060711.2 | 25587 | 613937 | . | . | 6.339785 | 6.221934 | 6.571589 | 5.182064 | 14.752941 | 14.382280 | 14.746454 | 11.848172 | -1.068913 | 849.678608 | 0.165473529 | -0.581029 | 0.561220909 | 0.998865404 | 9.777613 | 9.682362 | 9.776467 | 9.725766 | 9.797617 | 9.601136 | 1046 | 914 | 814 | 663 |
| 55752 | NM_00130614 | SEPTIN11 | septin 11 | protein_coding | NP_001293076.1;NP_060713.1;XP_006714322.1;XP_016863889.1;XP_01 | 25589 | 612887 | . | . | 21.144590 | 21.684103 | 22.033707 | 23.210458 | 49.204329 | 50.123770 | 49.443000 | 53.067954 | 1.028580 | 3655.631817 | 0.108612376 | 0.374297533 | 0.708182967 | 0.998865404 | 11.815445 | 11.856079 | 11.829825 | 11.818357 | 11.837669 | 11.857008 | 4179 | 3862 | 3186 | 3516 |
| 55753 | NM_00114399 | OGDHL | oxoglutarate dehydrogenase L | protein_coding | NP_001137468.1;NP_001137469.1;NP_001334748.1;NP_001334749.1;NI | 25590 | 617513 | . | . | 4.369422 | 4.305424 | 4.291862 | 4.530457 | 10.167824 | 9.952181 | 9.630814 | 10.358349 | -1.009593 | 500.453731 | 0.167599684 | -0.08218071 | 0.93450301 | 0.998865404 | 8.974069 | 8.960083 | 8.975598 | 8.964663 | 8.944921 | 8.982822 | 584 | 538 | 420 | 481 |
| 55754 | NM_00114395 | TMEM30A | transmembrane protein 30A | protein_coding | NP_001137430.1;NP_060717.1 | 16667 | 611028 | . | . | 31.803170 | 25.805081 | 25.834132 | 17.700944 | 74.007283 | 59.649602 | 57.971037 | 40.471107 | -1.341938 | 3485.500165 | 0.166938692 | -2.54175809 | 0.011029648 | 0.34058665 | 11.963606 | 11.539661 | 11.958613 | 11.773975 | 11.770657 | 11.461692 | 5097 | 3843 | 3085 | 2256 |
| 55755 | NM_00101164 | CDK5RAP2 | CDK5 regulatory subunit associated protein 2 | protein_coding | NP_001011649.1;NP_001258968.1;NP_060719.4;XP_006717245.1;XP_00 | 18672 | 608201 | . | . | 8.322341 | 8.114909 | 7.624971 | 8.446712 | 19.366428 | 18.757971 | 17.110216 | 19.312401 | -1.042259 | 1549.335058 | 0.12636535 | -0.47255386 | 0.636531504 | 0.998865404 | 10.627065 | 10.567180 | 10.622992 | 10.603272 | 10.548292 | 10.612756 | 1844 | 1685 | 1264 | 1482 |
| 55756 | NM_00114515 | INTS9 | integrator complex subunit 9 | protein_coding | NP_001138631.1;NP_001166033.1;NP_001349967.1;NP_060720.2;XP_00 | 25592 | 611352 | . | . | 7.539251 | 9.529600 | 10.537400 | 10.550224 | 17.544146 | 22.028101 | 23.645620 | 24.121837 | 1.233579 | 780.490540 | 0.162852816 | 1.859652898 | 0.06293465 | 0.724676532 | 9.449164 | 9.751500 | 9.439593 | 9.606341 | 9.681567 | 9.665626 | 707 | 846 | 758 | 799 |
| 55757 | NM_020121,XI | UGGT2 | UDP-glucose glycoprotein glucosyltransferase 2 | protein_coding | NP_064506.3;XP_011519396.1;XP_011519397.1;XP_011519398.1;XP_01 | 15664 | 605898 | . | . | 5.850541 | 4.714881 | 4.261240 | 3.704693 | 13.614452 | 10.898658 | 9.562100 | 8.470339 | -1.276762 | 710.195936 | 0.166145213 | -2.1215762 | 0.033873343 | 0.577226856 | 9.637197 | 9.285578 | 9.617871 | 9.473012 | 9.422123 | 9.336280 | 1009 | 773 | 578 | 547 |
| 55758 | NM_00113622 | RCOR3 | REST corepressor 3 | protein_coding | NP_001129695.1;NP_001129696.2;NP_001129697.1;NP_001336998.1;NI | 25594 | . | . | . | 3.828949 | 4.743534 | 5.576898 | 4.010654 | 8.910121 | 10.964891 | 12.514395 | 9.169887 | 1.157312 | 574.317428 | 0.182009786 | 1.15805727 | 0.246840666 | 0.992733415 | 9.056189 | 9.267490 | 9.070311 | 9.150461 | 9.294733 | 9.111716 | 575 | 610 | 609 | 499 |
| 55759 | NM_00137166 | WDR12 | WD repeat domain 12 | protein_coding | NP_001358593.1;NP_060726.3 | 14098 | 616620 | . | . | 10.228027 | 10.227634 | 9.718073 | 10.653599 | 23.801042 | 23.641638 | 21.807075 | 24.358189 | -1.018204 | 2565.434825 | 0.116352268 | -0.22368525 | 0.823002227 | 0.998865404 | 11.338010 | 11.311847 | 11.335697 | 11.328517 | 11.283854 | 11.349994 | 2994 | 2781 | 2120 | 2481 |
| 55760 | NM_018180 | DHX32 | DEAH-box helicase 32 (putative) | protein_coding | NP_060650.2 | 16717 | 607960 | . | . | 11.802980 | 12.922007 | 13.654986 | 14.226608 | 27.466021 | 29.869799 | 30.641396 | 32.527451 | 1.112941 | 1249.596012 | 0.13193116 | 1.170137469 | 0.241945652 | 0.992733415 | 10.208130 | 10.362248 | 10.213411 | 10.275187 | 10.313724 | 10.337359 | 1306 | 1329 | 1127 | 1253 |
| 55761 | NM_00130794 | TTC17 | tetratricopeptide repeat domain 17 | protein_coding | NP_001294872.1;NP_001363454.1;NP_001363455.1;NP_001363456.1;NI | 25596 | . | . | . | 12.366888 | 12.934621 | 12.954197 | 11.748283 | 28.778256 | 29.898959 | 29.068845 | 26.861054 | 1.025034 | 1667.050350 | 0.124325675 | 0.286919658 | 0.774173839 | 0.998865404 | 10.685167 | 10.720777 | 10.662985 | 10.722279 | 10.723715 | 10.700869 | 1824 | 1844 | 1489 | 1559 |
| 55762 | NM_00117265 | ZNF701 | zinc finger protein 701 | protein_coding | NP_001166126.1;NP_060730.2;XP_011525394.1 | 25597 | . | . | . | 0.139030 | 0.057004 | 0.055543 | 0.083834 | 0.323529 | 0.131768 | 0.124636 | 0.191676 | -1.193494 | 13.862867 | 0.846173711 | -0.30158302 | 0.762969952 | NA | 3.914964 | 3.660102 | 3.822887 | 3.779301 | 3.780878 | 3.800311 | 23 | 11 | 9 | 14 |
| 55763 | NM_00102492 | EXOC1 | exocyst complex component 1 | protein_coding | NP_001020095.1;NP_060731.2;NP_839955.1;XP_005265804.1;XP_00526 | 30380 | 607879 | . | . | 7.083780 | 7.442576 | 6.857920 | 5.958367 | 16.484244 | 17.203847 | 15.388974 | 13.623098 | -1.144628 | 748.615024 | 0.155225215 | -1.2554566 | 0.209313055 | 0.971746012 | 9.642001 | 9.447617 | 9.578360 | 9.606195 | 9.551611 | 9.442482 | 902 | 879 | 656 | 602 |
| 55764 | NM_00128054 | IFT122 | intraflagellar transport 122 | protein_coding | NP_001267470.1;NP_001267474.1;NP_001267475.1;NP_060732.2;NP_4( | 13556 | 606045 | . | . | 5.417594 | 5.708579 | 5.605319 | 5.780057 | 12.606966 | 13.195636 | 12.578175 | 13.215413 | 1.005309 | 687.433223 | 0.151887147 | 0.050297574 | 0.959885257 | 0.998865404 | 9.421379 | 9.428763 | 9.403302 | 9.442590 | 9.418348 | 9.434963 | 767 | 761 | 592 | 654 |

| | | | | | | | | | | | | | | | | | | | | | | | | | | | | | | | | | |
|---|---|---|---|---|---|---|---|---|---|---|---|---|---|---|---|---|---|---|---|---|---|---|---|---|---|---|---|---|---|---|---|---|---|
| 55766 | NM_177925,N | H2AJ | H2A.J histone | protein_coding | NP_808760.1 | 14456 | . | . | . | 8.281457 | 8.297606 | 7.941974 | 9.565072 | 19.271292 | 19.180286 | 17.821558 | 21.869402 | 1.041589 | 955.303772 | 0.147674787 | 0.398077181 | 0.690573294 | 0.998865404 | 9.870378 | 9.928664 | 9.885993 | 9.881717 | 9.847221 | 9.975512 | 1081 | 1007 | 773 | 994 |
| 55768 | NM_00114529 | NGLY1 | N-glycanase 1 | protein_coding | NP_001138765.1;NP_001138766.1;NP_001138767.1;NP_060767.2;XP_00 | 17646 | 610661 | . | . | 8.792782 | 9.591077 | 8.099480 | 7.948033 | 20.461170 | 22.170204 | 18.174996 | 18.172234 | -1.156470 | 672.806168 | 0.153262144 | -1.36842675 | 0.171178535 | 0.947127937 | 9.495127 | 9.285348 | 9.419342 | 9.462910 | 9.351590 | 9.333380 | 805 | 803 | 550 | 577 |
| 55769 | NM_00110554 | ZNF83 | zinc finger protein 83 | protein_coding | NP_001099019.1;NP_001099020.1;NP_001099021.1;NP_001099022.1;NI | 13158 | 194558 | . | . | 0.597542 | 0.881044 | 0.693505 | 0.585213 | 1.390504 | 2.036569 | 1.556204 | 1.338022 | -1.161746 | 68.387921 | 0.397180681 | -0.54457476 | 0.586046015 | 0.998865404 | 6.200031 | 5.983167 | 6.062256 | 6.155797 | 6.090750 | 6.059059 | 69 | 94 | 59 | 55 |
| 55770 | NM_018303,N | EXOC2 | exocyst complex component 2 | protein_coding | NP_060773.3;XP_016866507.1;XP_016866508.1;XP_016866509.1;XP_01 | 24968 | 615329 | . | . | 5.696097 | 5.529652 | 5.679157 | 5.974426 | 13.255055 | 12.782042 | 12.743865 | 13.659812 | 1.000409 | 763.465863 | 0.14882583 | 0.003961896 | 0.996838873 | 0.999862634 | 9.576323 | 9.576517 | 9.565621 | 9.586509 | 9.545547 | 9.606147 | 866 | 836 | 635 | 747 |
| 55771 | NM_018304,XI | PRR11 | proline rich 11 | protein_coding | NP_060774.2;XP_024306596.1 | 25619 | 615920 | . | . | 12.527036 | 12.559599 | 10.343775 | 12.898445 | 29.150928 | 29.032078 | 23.211133 | 29.490763 | -1.090080 | 2290.280406 | 0.127918142 | -0.97276315 | 0.330671032 | 0.998865404 | 11.222332 | 11.097590 | 11.192993 | 11.195416 | 11.052549 | 11.192311 | 2747 | 2582 | 1738 | 2235 |
| 55773 | NM_00119919 | TBC1D23 | TBC1 domain family member 23 | protein_coding | NP_001186127.1;NP_060779.2;XP_011511276.1;XP_016862330.1 | 25622 | 617687 | . | . | 3.968813 | 3.906057 | 4.037886 | 2.893641 | 9.235591 | 9.029028 | 9.060899 | 6.615966 | -1.152797 | 427.834801 | 0.201309788 | -1.01901784 | 0.308194497 | 0.998865404 | 8.839203 | 8.635427 | 8.786857 | 8.770859 | 8.788348 | 8.587500 | 534 | 488 | 405 | 309 |
| 55775 | NM_00100874 | TDP1 | tyrosyl-DNA phosphodiesterase 1 | protein_coding | NP_001008744.1;NP_001317134.1;NP_060789.2;XP_005267905.1;XP_00 | 18884 | 607198 | . | . | 3.223876 | 3.447207 | 3.166258 | 3.495661 | 7.502090 | 7.968376 | 7.104992 | 7.992411 | -1.032401 | 383.136252 | 0.183776948 | -0.25032079 | 0.802339279 | 0.998865404 | 8.604713 | 8.558342 | 8.584137 | 8.598067 | 8.555034 | 8.589458 | 443 | 425 | 318 | 364 |
| 55776 | NM_00130479 | SAYSD1 | SAYSVFN motif domain containing 1 | protein_coding | NP_001291722.1;NP_060792.1 | 21025 | . | . | . | 1.750421 | 1.724031 | 1.994176 | 2.251308 | 4.073302 | 3.985174 | 4.474874 | 5.147348 | 1.124259 | 190.195873 | 0.242251427 | 0.697516066 | 0.485479889 | 0.998865404 | 7.484237 | 7.653486 | 7.563474 | 7.522549 | 7.589303 | 7.600245 | 216 | 184 | 173 | 191 |
| 55777 | NM_00137812 | MBD5 | methyl-CpG binding domain protein 5 | protein_coding | NP_001365049.1;NP_060798.2;XP_011509772.1;XP_011509774.1;XP_02 | 20444 | 611472 | . | . | 0.597622 | 0.532444 | 0.599674 | 0.750687 | 1.390692 | 1.230767 | 1.345650 | 1.716357 | 1.180329 | 207.248567 | 0.242305821 | 0.98713951 | 0.323574251 | 0.998865404 | 7.570926 | 7.809647 | 7.680211 | 7.623423 | 7.678118 | 7.776143 | 233 | 192 | 175 | 232 |
| 55778 | NM_00126782 | ZNF839 | zinc finger protein 839 | protein_coding | NP_001254756.1;NP_001254757.1;NP_060805.3;XP_006720266.2;XP_01 | 20345 | . | . | . | 1.597364 | 2.601996 | 2.395047 | 2.040789 | 3.717132 | 6.014626 | 5.374416 | 4.666020 | 1.030379 | 245.763735 | 0.241089882 | 0.179085176 | 0.857870822 | 0.998865404 | 7.919789 | 7.962157 | 7.835409 | 8.014353 | 7.961975 | 7.924918 | 229 | 308 | 225 | 225 |
| 55779 | NM_00116449 | CFAP44 | cilia and flagella associated protein 44 | protein_coding | NP_001157968.1;NP_060808.2 | 25631 | 617559 | . | . | 0.547837 | 0.451832 | 0.698194 | 0.487274 | 1.274840 | 1.044431 | 1.566726 | 1.114093 | 1.168942 | 174.607357 | 0.2763784 | 0.814838438 | 0.415164795 | 0.998865404 | 7.329726 | 7.557253 | 7.446639 | 7.363915 | 7.549368 | 7.388520 | 204 | 156 | 194 | 144 |
| 55780 | NM_00127853 | ERMARD | ER membrane associated RNA degradation | protein_coding | NP_001265460.1;NP_001265461.1;NP_001265462.1;NP_060811.1;XP_01 | 21056 | 615532 | . | . | 3.873543 | 3.092159 | 3.568194 | 3.504817 | 9.013893 | 7.147660 | 8.006923 | 8.013348 | 1.048508 | 296.245194 | 0.206180108 | 0.331450236 | 0.740304424 | 0.998865404 | 8.175602 | 8.244862 | 8.223545 | 8.168835 | 8.252317 | 8.190742 | 350 | 296 | 279 | 268 |
| 55781 | NM_00115974 | RIOK2 | RIO kinase 2 | protein_coding | NP_001153221.1;NP_060813.2;XP_016865117.1 | 18999 | 617754 | . | . | 6.412323 | 5.623901 | 5.051954 | 4.960252 | 14.921741 | 12.999900 | 11.336435 | 11.341028 | -1.219044 | 664.602878 | 0.158114625 | -1.80723508 | 0.070725633 | 0.751131579 | 9.511958 | 9.226657 | 9.484575 | 9.391574 | 9.315038 | 9.294611 | 896 | 736 | 529 | 553 |
| 55783 | NM_00109964 | CMTR2 | cap methyltransferase 2 | protein_coding | NP_001093112.1;NP_001311303.1;NP_001311306.1;NP_001311307.1;NI | 25635 | 616190 | . | . | 2.348368 | 2.477182 | 2.876555 | 1.840109 | 5.464750 | 5.726115 | 6.454906 | 4.207191 | -1.039122 | 365.095719 | 0.225041848 | -0.24602109 | 0.80566591 | 0.998865404 | 8.538853 | 8.484904 | 8.503955 | 8.529310 | 8.613993 | 8.359453 | 419 | 410 | 381 | 261 |
| 55784 | NM_00115964 | MCTP2 | multiple C2 and transmembrane domain containing 2 | protein_coding | NP_001153115.1;NP_001153116.1;NP_060819.3;XP_005255012.1;XP_00 | 25636 | 616297 | . | . | 3.741237 | 3.820256 | 3.978439 | 3.823471 | 8.706015 | 8.830695 | 8.927503 | 8.741913 | -1.025914 | 765.784217 | 0.148592271 | -0.24839828 | 0.803826265 | 0.998865404 | 9.598956 | 9.562401 | 9.592599 | 9.586186 | 9.602259 | 9.540014 | 897 | 833 | 687 | 677 |
| 55785 | NM_018351 | FGD6 | FYVE, RhoGEF and PH domain containing 6 | protein_coding | NP_060821.3 | 21740 | 613520 | . | . | 0.432559 | 0.493420 | 0.431097 | 0.557353 | 1.006583 | 1.140563 | 0.967370 | 1.274320 | 1.056436 | 138.946312 | 0.285790022 | 0.277147588 | 0.781666782 | 0.998865404 | 7.079154 | 7.156575 | 7.082348 | 7.128864 | 7.079896 | 7.170567 | 146 | 155 | 109 | 150 |
| 55786 | NM_00113603 | ZNF415 | zinc finger protein 415 | protein_coding | NP_001129510.2;NP_001157781.1;NP_001317688.1;NP_001317695.1;NI | 20636 | . | . | . | 0.560329 | 0.448625 | 0.404902 | 0.370217 | 1.303909 | 1.037016 | 0.908588 | 0.846454 | -1.190737 | 38.240769 | 0.505906537 | -0.49782825 | 0.618605105 | 0.998865404 | 5.376728 | 5.126516 | 5.289074 | 5.253729 | 5.252941 | 5.238359 | 53 | 40 | 32 | 31 |
| 55787 | NM_00116868 | TXLNG | taxilin gamma | protein_coding | NP_001162154.1;NP_060830.2;XP_016885120.1;XP_024308166.1;XP_02 | 18578 | 300677 | . | . | 3.769708 | 3.942373 | 4.193873 | 3.307872 | 8.772263 | 9.112973 | 9.410930 | 7.563056 | -1.034803 | 526.811836 | 0.177055426 | -0.27876107 | 0.780428194 | 0.998865404 | 9.065250 | 9.016628 | 9.035552 | 9.064750 | 9.097337 | 8.954481 | 603 | 591 | 501 | 431 |
| 55788 | NM_00136372 | LMBRD1 | LMBR1 domain containing 1 | protein_coding | NP_001350651.1;NP_001354200.1;NP_001354201.1;NP_060838.3 | 23038 | 612625 | . | . | 4.650617 | 4.068025 | 4.883135 | 3.587965 | 10.822174 | 9.403422 | 10.957613 | 8.203455 | -1.044392 | 523.823459 | 0.192199274 | -0.32603562 | 0.744397404 | 0.998865404 | 9.063270 | 9.001953 | 9.088511 | 8.995177 | 9.110282 | 8.910361 | 660 | 536 | 516 | 405 |
| 55789 | NM_00114520 | DEPDC1B | DEP domain containing 1B | protein_coding | NP_001138680.1;NP_060839.2;XP_011541811.1;XP_011541812.1;XP_01 | 24902 | 616073 | . | . | 5.317401 | 5.534652 | 4.541731 | 5.697059 | 12.373815 | 12.793598 | 10.191515 | 13.025647 | -1.072634 | 414.291646 | 0.188553403 | -0.53649582 | 0.591615927 | 0.998865404 | 8.744707 | 8.642489 | 8.698228 | 8.730327 | 8.608707 | 8.731838 | 481 | 475 | 311 | 414 |
| 55790 | NM_00113051 | CSGALNACT1 | chondroitin sulfate N-acetylgalactosaminyltransferase 1 | protein_coding | NP_001123990.1;NP_001341404.1;NP_001341405.1;NP_001341406.1;NI | 24290 | 616615 | . | . | 4.590508 | 4.378768 | 4.359511 | 4.994001 | 10.682303 | 10.121719 | 9.782615 | 11.418196 | 1.123867 | 592.486740 | 0.160891085 | 1.047112184 | 0.29504784 | 0.998865404 | 9.123683 | 9.292650 | 9.197072 | 9.140992 | 9.266999 | 9.226230 | 670 | 576 | 562 | 570 |
| 55791 | NM_00100694 | LRIF1 | ligand dependent nuclear receptor interacting factor 1 | protein_coding | NP_001006946.1;NP_060842.3;XP_005271086.1;XP_011540070.1;XP_01 | 30299 | 615354 | . | . | 4.076239 | 3.983854 | 3.909839 | 3.618779 | 9.485577 | 9.208854 | 8.773567 | 8.273907 | -1.062816 | 345.320509 | 0.192506517 | -0.45656281 | 0.647985317 | 0.998865404 | 8.474767 | 8.387498 | 8.458951 | 8.438114 | 8.441989 | 8.385765 | 417 | 377 | 306 | 298 |
| 55793 | NM_00104021 | MINDY1 | MINDY lysine 48 deubiquitinase 1 | protein_coding | NP_001035307.2;NP_001156730.2;NP_001156731.2;NP_001156732.2;NI | 25648 | 618407 | . | . | 8.309236 | 9.833723 | 10.293793 | 11.562414 | 19.335933 | 22.731090 | 23.098974 | 26.436086 | 1.183392 | 815.652511 | 0.151669913 | 1.601688923 | 0.109224416 | 0.842810112 | 9.545587 | 9.787874 | 9.569301 | 9.644280 | 9.682368 | 9.768319 | 814 | 850 | 723 | 878 |
| 55794 | NM_018380 | DDX28 | DEAD-box helicase 28 | protein_coding | NP_060850.2 | 17330 | 607618 | . | . | 3.751062 | 3.640833 | 3.042952 | 4.151476 | 8.728875 | 8.415949 | 6.828296 | 9.491856 | -1.042438 | 263.682124 | 0.227141104 | -0.26398503 | 0.791791454 | 0.998865404 | 8.072936 | 8.011726 | 8.061564 | 8.041501 | 7.953096 | 8.101936 | 316 | 285 | 191 | 278 |
| 55795 | NM_00112720 | PCID2 | PCI domain containing 2 | protein_coding | NP_001120674.1;NP_001120675.1;NP_001245142.1;NP_001307584.1;NI | 25653 | 613713 | . | . | 15.541773 | 16.368664 | 14.663433 | 17.002891 | 36.166344 | 37.836906 | 32.904319 | 38.875088 | -1.030062 | 1155.227140 | 0.133933632 | -0.3190494 | 0.749689053 | 0.998865404 | 10.195320 | 10.152280 | 10.165086 | 10.204251 | 10.137880 | 10.186730 | 1315 | 1298 | 957 | 1102 |
| 55796 | NM_00117070 | MBNL3 | muscleblind like splicing regulator 3 | protein_coding | NP_001164172.1;NP_001164173.1;NP_001164174.1;NP_001164175.1;NI | 20564 | 300413 | . | . | 0.614003 | 0.729682 | 0.589859 | 0.578674 | 1.428810 | 1.686693 | 1.323627 | 1.323068 | -1.160724 | 214.318344 | 0.231171165 | -0.93015484 | 0.352290921 | 0.998865404 | 7.847040 | 7.632191 | 7.760439 | 7.793917 | 7.724638 | 7.691186 | 256 | 257 | 179 | 179 |
| 55798 | NM_018396 | METTL2B | methyltransferase like 2B | protein_coding | NP_060866.2 | 18272 | 607846 | . | . | 5.280332 | 4.852768 | 4.752627 | 5.209477 | 12.287550 | 11.217392 | 10.664758 | 11.910851 | -1.031952 | 914.479765 | 0.144270953 | -0.31452075 | 0.75312556 | 0.998865404 | 9.859344 | 9.813913 | 9.880904 | 9.813524 | 9.795698 | 9.853586 | 1119 | 956 | 751 | 878 |
| 55802 | NM_00129020 | DCP1A | decapping mRNA 1A | protein_coding | NP_001277133.1;NP_001277134.1;NP_001277135.1;NP_001277136.1;NI | 18714 | 607010 | . | . | 2.566391 | 2.760816 | 3.293727 | 2.726900 | 5.972097 | 6.381750 | 7.391028 | 6.234733 | 1.113286 | 525.538352 | 0.175431247 | 0.882538909 | 0.377485448 | 0.998865404 | 8.957876 | 9.113247 | 8.978650 | 9.020028 | 9.131960 | 9.001099 | 554 | 554 | 530 | 467 |
| 55805 | NM_00137744 | LRP2BP | LRP2 binding protein | protein_coding | NP_001364369.1;NP_001364370.1;NP_001364371.1;NP_001364372.1;NI | 25434 | . | . | . | 0.230188 | 0.156618 | 0.144331 | 0.129864 | 0.535657 | 0.362031 | 0.323873 | 0.296917 | -1.374652 | 33.554841 | 0.542457438 | -0.84627271 | 0.39740063 | 0.998865404 | 5.278976 | 4.821864 | 5.113270 | 5.069566 | 5.056238 | 5.043074 | 51 | 36 | 26 | 25 |
| 55806 | NM_005144,N | HR | HR lysine demethylase and nuclear receptor corepressor | protein_coding | NP_005135.2;NP_060881.2;XP_005273626.1;XP_006716430.1 | 5172 | 602302 | . | . | 9.096131 | 9.268242 | 7.800520 | 8.266496 | 21.167068 | 21.423958 | 17.504140 | 18.900364 | -1.158165 | 1437.094984 | 0.126439426 | -1.6754334 | 0.093849228 | 0.810751527 | 10.591043 | 10.379056 | 10.545854 | 10.539703 | 10.408918 | 10.450354 | 1783 | 1658 | 1126 | 1283 |
| 55809 | NM_00129757 | TRERF1 | transcriptional regulating factor 1 | protein_coding | NP_001284502.1;NP_277037.1;XP_011513043.1;XP_011513044.1;XP_01 | 18273 | 610322 | . | . | 0.375628 | 0.515982 | 0.489281 | 0.608832 | 0.874105 | 1.192715 | 1.097931 | 1.392022 | 1.263029 | 144.363692 | 0.279116667 | 1.20697893 | 0.227440258 | 0.982008277 | 6.996189 | 7.331500 | 7.091739 | 7.150662 | 7.174786 | 7.244369 | 135 | 149 | 128 | 164 |
| 55810 | NM_018416,XI | FOXJ2 | forkhead box J2 | protein_coding | NP_060886.1;XP_011519062.1;XP_011519063.1;XP_016875129.1 | 24818 | . | . | . | 1.870914 | 2.387948 | 2.123853 | 2.159991 | 4.353692 | 5.519851 | 4.765865 | 4.938563 | -1.012639 | 369.257032 | 0.196956654 | -0.09200263 | 0.926695946 | 0.998865404 | 8.537871 | 8.519031 | 8.462585 | 8.594001 | 8.521254 | 8.524830 | 380 | 446 | 318 | 345 |
| 55812 | NM_00104042 | SPATA7 | spermatogenesis associated 7 | protein_coding | NP_001035518.1;NP_060888.2;XP_005267908.1;XP_005267909.1;XP_00 | 20423 | 609868 | . | . | 1.664690 | 1.334110 | 1.854682 | 1.102195 | 3.873799 | 3.083854 | 4.161852 | 2.520040 | -1.045498 | 95.278759 | 0.362086507 | -0.17727768 | 0.859290284 | 0.998865404 | 6.604128 | 6.543399 | 6.616726 | 6.529230 | 6.628641 | 6.487847 | 128 | 90 | 101 | 66 |
| 55813 | NM_018428,XI | UTP6 | UTP6 small subunit processome component | protein_coding | NP_060898.2;XP_005258069.1;XP_011523299.1;XP_016880334.1 | 18279 | . | . | . | 24.207673 | 24.048491 | 18.789733 | 24.876047 | 56.332248 | 55.589163 | 42.163619 | 56.876125 | -1.074342 | 2092.244678 | 0.120028963 | -0.86190445 | 0.388740099 | 0.998865404 | 11.081718 | 10.978094 | 11.062016 | 11.055314 | 10.967095 | 11.034619 | 2506 | 2329 | 1679 | 1972 |
| 55814 | NM_018429,XI | BDP1 | B double prime 1, subunit of RNA polymerase III transcription initi | protein_coding | NP_060899.2;XP_011541813.1;XP_011541814.1;XP_016865119.1;XP_01 | 13652 | 607012 | . | . | 3.182305 | 3.192626 | 3.330146 | 2.904324 | 7.405354 | 7.379899 | 7.472751 | 6.640390 | -1.041985 | 1084.755444 | 0.140578979 | -0.42207628 | 0.672969342 | 0.998865404 | 10.112307 | 10.053400 | 10.097780 | 10.097249 | 10.119062 | 10.012439 | 1275 | 1194 | 991 | 924 |
| 55815 | NM_00128899 | TSNAXIP1 | translin associated factor X interacting protein 1 | protein_coding | NP_001275919.1;NP_001275920.1;NP_001275921.1;NP_001275922.1;NI | 18586 | 607720 | . | . | 1.154726 | 1.297523 | 2.306383 | 1.567736 | 2.687092 | 2.999279 | 5.175458 | 3.584443 | 1.545580 | 119.549358 | 0.308104834 | 2.038749314 | 0.041475053 | 0.632574762 | 6.553062 | 7.181880 | 6.787139 | 6.827075 | 7.006857 | 6.900066 | 101 | 108 | 144 | 117 |
| 55816 | NM_00129416 | DOK5 | docking protein 5 | protein_coding | NP_001281090.1;NP_060901.2;NP_808874.1;XP_011527206.1;XP_02430 | 16173 | 608334 | . | . | 4.445813 | 4.252459 | 2.943652 | 3.878174 | 10.345589 | 9.829750 | 6.605470 | 8.866983 | -1.303314 | 94.546500 | 0.336192397 | -1.13680261 | 0.255620811 | 0.995760118 | 6.742057 | 6.358418 | 6.611258 | 6.591066 | 6.489440 | 6.556955 | 127 | 112 | 62 | 87 |
| 55818 | NM_00114668 | KDM3A | lysine demethylase 3A | protein_coding | NP_001140160.1;NP_060903.2;XP_006712114.1;XP_016859983.1;XP_02 | 20815 | 611512 | . | . | 8.199738 | 8.115141 | 12.013371 | 9.899930 | 19.081129 | 18.758511 | 26.957657 | 22.635015 | 1.328950 | 1394.691546 | 0.140475698 | 2.920694661 | 0.003492519 | 0.166807075 | 10.225803 | 10.636524 | 10.331732 | 10.315729 | 10.617031 | 10.460702 | 1394 | 1278 | 1521 | 1345 |
| 55819 | NM_00128080 | RNF130 | ring finger protein 130 | protein_coding | NP_001267730.1;NP_060904.2;XP_011532895.1;XP_024301897.1 | 18280 | . | . | . | 60.128631 | 55.842657 | 64.435431 | 64.793727 | 139.921806 | 129.082811 | 144.591231 | 148.143159 | 1.107171 | 3768.255454 | 0.111840531 | 1.313277339 | 0.189089511 | 0.960264078 | 11.804311 | 11.951308 | 11.868764 | 11.801535 | 11.932840 | 11.906999 | 4287 | 3701 | 3497 | 3655 |
| 55823 | NM_00129018 | VPS11 | VPS11 core subunit of CORVET and HOPS complexes | protein_coding | NP_001277114.1;NP_001365147.1;NP_001365148.1;NP_001365149.1;NI | 14583 | 608549 | . | . | 5.860429 | 7.088227 | 7.246610 | 6.788792 | 13.637463 | 16.384757 | 16.261182 | 15.521769 | 1.087387 | 679.879071 | 0.161994121 | 0.746110378 | 0.455600741 | 0.998865404 | 9.347528 | 9.468217 | 9.317321 | 9.435268 | 9.462737 | 9.406820 | 685 | 764 | 641 | 637 |
| 55824 | NM_018440,XI | PAG1 | phosphoprotein membrane anchor with glycosphingolipid microdo | protein_coding | NP_060910.3;XP_016869129.1;XP_016869130.1;XP_016869131.1;XP_01 | 30043 | 605767 | . | . | 5.743085 | 5.339815 | 7.409848 | 5.863772 | 13.364393 | 12.343221 | 16.627483 | 13.406815 | 1.112640 | 1808.598734 | 0.137199857 | 1.122355215 | 0.261711438 | 0.995760118 | 10.741372 | 10.895810 | 10.803609 | 10.746041 | 10.936779 | 10.775922 | 2044 | 1779 | 1824 | 1606 |
| 55825 | NM_018441,XI | PECR | peroxisomal trans-2-enoyl-CoA reductase | protein_coding | NP_060911.2;XP_024308766.1;XP_024308767.1 | 18281 | 605843 | . | . | 6.255671 | 7.547201 | 5.205808 | 7.733503 | 14.557203 | 17.445694 | 11.681682 | 17.681739 | -1.095733 | 500.137408 | 0.190813299 | -0.69123122 | 0.489420248 | 0.998865404 | 9.031253 | 8.898034 | 8.956403 | 9.026393 | 8.835714 | 9.023283 | 570 | 595 | 352 | 515 |
| 55827 | NM_00101797 | DCAF6 | DDB1 and CUL4 associated factor 6 | protein_coding | NP_001017977.1;NP_001185885.1;NP_001185886.1;NP_001336702.1;NI | 30002 | 610494 | . | . | 9.585210 | 8.918760 | 8.431136 | 10.708835 | 22.305174 | 20.616113 | 18.919225 | 24.484477 | 1.015687 | 986.795563 | 0.152596714 | 0.147154865 | 0.883009785 | 0.998865404 | 9.935547 | 9.957584 | 9.967278 | 9.910871 | 9.870393 | 10.025207 | 1171 | 1016 | 773 | 1031 |
| 55829 | NM_018445,N | SELENOS | selenoprotein S | protein_coding | NP_060915.2;NP_982298.2 | 30396 | 607918 | . | . | 56.022412 | 56.086095 | 57.624946 | 55.091730 | 130.366453 | 129.645514 | 129.308715 | 125.960683 | -1.002612 | 2169.247384 | 0.117447444 | -0.03203999 | 0.974440163 | 0.998865404 | 11.084799 | 11.081156 | 11.085489 | 11.082290 | 11.103657 | 11.059571 | 2506 | 2339 | 1936 | 1975 |
| 55830 | NM_00101098 | GLT8D1 | glycosyltransferase 8 domain containing 1 | protein_coding | NP_001010983.1;NP_001265209.1;NP_001265210.1;NP_060916.1;NP_6 | 24870 | 618399 | . | . | 16.730818 | 15.915413 | 16.708475 | 17.344857 | 38.933304 | 36.789190 | 37.493332 | 39.656950 | 1.009553 | 1186.095734 | 0.13251037 | 0.103517301 | 0.917552419 | 0.998865404 | 10.205066 | 10.218911 | 10.234908 | 10.180862 | 10.212261 | 10.218576 | 1408 | 1228 | 1032 | 1122 |
| 55831 | NM_018447,N | EMC3 | ER membrane protein complex subunit 3 | protein_coding | NP_060917.1;XP_005265378.1 | 23999 | . | . | . | 27.500429 | 27.302417 | 30.069262 | 28.394237 | 63.994627 | 63.110755 | 67.474558 | 64.920049 | 1.056247 | 2292.785396 | 0.11701331 | 0.674685436 | 0.499875639 | 0.998865404 | 11.122794 | 11.201893 | 11.150209 | 11.130142 | 11.208175 | 11.159722 | 2600 | 2376 | 2110 | 2142 |
| 55832 | NM_00132967 | CAND1 | cullin associated and neddylation dissociated 1 | protein_coding | NP_001316603.1;NP_001316604.1;NP_001316605.1;NP_060918.2 | 30688 | 607727 | . | . | 8.728139 | 8.538679 | 8.321923 | 7.519104 | 20.310736 | 19.737542 | 18.674152 | 17.191539 | -1.105221 | 2883.885613 | 0.117606821 | -1.22726411 | 0.219723339 | 0.982008277 | 11.564060 | 11.419939 | 11.545450 | 11.520068 | 11.495156 | 11.405500 | 3540 | 3217 | 2515 | 2426 |
| 55833 | NM_00128259 | UBAP2 | ubiquitin associated protein 2 | protein_coding | NP_001269459.1;NP_001356988.1;NP_001356991.2;NP_001356993.1;NI | 14185 | . | . | . | 11.473142 | 13.347858 | 12.343956 | 14.437515 | 26.698474 | 30.854174 | 27.699480 | 33.009667 | 1.063182 | 1668.176372 | 0.136233504 | 0.648806976 | 0.516463148 | 0.998865404 | 10.659427 | 10.747346 | 10.619674 | 10.733837 | 10.673352 | 10.776262 | 1726 | 1873 | 1397 | 1717 |
| 55835 | NM_018451,N | CENPJ | centromere protein J | protein_coding | NP_060921.3;XP_011533451.1;XP_011533452.1;XP_016876162.1 | 17272 | 609279 | . | . | 8.107624 | 8.157810 | 8.386420 | 7.658675 | 18.866773 | 18.857147 | 18.818882 | 17.510651 | -1.055619 | 1224.842641 | 0.137074356 | -0.5696867 | 0.568890214 | 0.998865404 | 10.296696 | 10.219020 | 10.281295 | 10.274249 | 10.289430 | 10.181912 | 1455 | 1351 | 1111 | 1037 |
| 55837 | NM_00131891 | EAPP | E2F associated phosphoprotein | protein_coding | NP_001305845.1;NP_060923.2;XP_011535260.1;XP_016876949.1 | 19312 | 609486 | . | . | 8.902187 | 8.115099 | 8.819009 | 7.807399 | 20.715756 | 18.758417 | 19.789601 | 17.850689 | -1.039667 | 343.917788 | 0.19659579 | -0.28546725 | 0.775286192 | 0.998865404 | 8.453228 | 8.398085 | 8.463301 | 8.407912 | 8.452497 | 8.374378 | 423 | 360 | 314 | 294 |
| 55839 | NM_00110062 | CENPN | centromere protein N | protein_coding | NP_001094094.2;NP_001094095.2;NP_001257402.1;NP_001257403.1;NI | 30873 | 611509 | . | . | 10.553962 | 11.182269 | 10.187631 | 11.579311 | 24.559506 | 25.848317 | 22.860748 | 26.474716 | -1.065928 | 1264.176805 | 0.128945708 | -0.71432974 | 0.475023306 | 0.998865404 | 10.349291 | 10.257205 | 10.337872 | 10.317552 | 10.275782 | 10.283071 | 1522 | 1389 | 1059 | 1153 |
| 55840 | NM_00132004 | EAF2 | ELL associated factor 2 | protein_coding | NP_001306970.1;NP_060926.2;XP_005247675.1;XP_016862350.1;XP_01 | 23115 | 607659 | . | . | 2.420538 | 1.633082 | 1.670219 | 1.977998 | 5.632690 | 3.774941 | 3.747924 | 4.522457 | -1.129140 | 60.504862 | 0.419235019 | -0.41796144 | 0.675975311 | 0.998865404 | 6.003609 | 5.829074 | 5.974429 | 5.886687 | 5.891658 | 5.919160 | 88 | 56 | 46 | 57 |
| 55841 | NM_015691 | WWC3 | WWC family member 3 | protein_coding | NP_056506.2 | 29237 | . | . | . | 2.374039 | 2.652030 | 2.182147 | 2.532718 | 5.524487 | 6.130286 | 4.896674 | 5.790759 | -1.080318 | 527.784591 | 0.171437385 | -0.65012655 | 0.515610479 | 0.998865404 | 9.098864 | 8.986562 | 9.033422 | 9.100726 | 8.974033 | 9.059733 | 599 | 622 | 411 | 508 |
| 55843 | NM_018460,XI | ARHGAP15 | Rho GTPase activating protein 15 | protein_coding | NP_060930.3;XP_011509781.1;XP_011509783.1;XP_011509784.1;XP_01 | 21030 | 610578 | . | . | 0.141464 | 0.074569 | 0.111465 | 0.106336 | 0.329193 | 0.172369 | 0.250125 | 0.243126 | 1.019764 | 9.865904 | 0.983572027 | 0.028706916 | 0.977098341 | NA | 3.286426 | 3.318300 | 3.316221 | 3.286261 | 3.303814 | 3.300573 | 15 | 7 | 9 | 9 |
| 55844 | NM_00129131 | PPP2R2D | protein phosphatase 2 regulatory subunit Bdelta | protein_coding | NP_001278239.1;NP_060931.2;XP_006717977.1 | 23732 | 613992 | . | . | 5.337217 | 5.339701 | 5.166952 | 5.168475 | 12.419922 | 12.342958 | 11.594491 | 11.817103 | -1.061347 | 848.832942 | 0.142003596 | -0.60488645 | 0.545254449 | 0.998865404 | 9.771613 | 9.685784 | 9.754288 | 9.746312 | 9.711680 | 9.704104 | 1012 | 938 | 720 | 768 |
| 55845 | NM_018462 | BRK1 | BRICK1 subunit of SCAR/WAVE actin nucleating complex | protein_coding | NP_060932.2 | 23057 | 611183 | . | . | 41.233685 | 40.354759 | 41.064110 | 42.604565 | 95.952484 | 93.281830 | 92.146667 | 97.410278 | 1.010821 | 1480.432197 | 0.125017263 | 0.124207824 | 0.901150728 | 0.998865404 | 10.524025 | 10.539539 | 10.539127 | 10.515907 | 10.525540 | 10.546124 | 1721 | 1565 | 1277 | 1414 |
| 55846 | NM_018463,N | ITFG2 | integrin alpha FG-GAP repeat containing 2 | protein_coding | NP_060933.3;XP_005253764.1;XP_005253766.1;XP_016875130.1;XP_02 | 30879 | 617421 | . | . | 4.644301 | 6.016830 | 7.006897 | 5.565622 | 10.807481 | 13.908174 | 15.723273 | 12.725132 | 1.141110 | 477.514986 | 0.184701236 | 1.031055622 | 0.302514737 | 0.998865404 | 8.800774 | 8.991717 | 8.819346 | 8.885526 | 9.005638 | 8.860691 | 486 | 507 | 493 | 423 |
| 55847 | NM_018464 | CISD1 | CDGSH iron sulfur domain 1 | protein_coding | NP_060934.1 | 30880 | 611932 | . | . | 7.496463 | 7.289247 | 8.091448 | 7.493798 | 17.444578 | 16.849421 | 18.156975 | 17.133678 | 1.039605 | 562.121507 | 0.162781365 | 0.344239426 | 0.730666234 | 0.998865404 | 9.106213 | 9.162711 | 9.132924 | 9.109507 | 9.174677 | 9.118059 | 646 | 584 | 520 | 514 |
| 55848 | NM_018465,XI | PLGRKT | plasminogen receptor with a C-terminal lysine | protein_coding | NP_060935.2;XP_005251567.1;XP_005251569.1;XP_011516262.1 | 23633 | 618444 | . | . | 7.117230 | 5.396365 | 5.755979 | 7.140976 | 16.562086 | 12.473938 | 12.916248 | 16.326994 | 1.039582 | 200.178320 | 0.256631716 | 0.218224194 | 0.827254435 | 0.998865404 | 7.616735 | 7.673000 | 7.700909 | 7.560391 | 7.610545 | 7.689229 | 261 | 179 | 162 | 208 |
| 55850 | NM_018467,XI | USE1 | unconventional SNARE in the ER 1 | protein_coding | NP_060937.2;XP_016882465.1 | 30882 | 610675 | . | . | 14.063596 | 15.570922 | 17.375100 | 16.175409 | 32.726568 | 35.992884 | 38.989219 | 36.983150 | 1.125040 | 415.084416 | 0.181594664 | 0.936019841 | 0.349262972 | 0.998865404 | 8.609759 | 8.779758 | 8.629802 | 8.687371 | 8.754488 | 8.704303 | 428 | 442 | 398 | 395 |
| 55851 | NM_00128153 | PSENEN | presenilin enhancer, gamma-secretase subunit | protein_coding | NP_001268461.1;NP_758844.1 | 30100 | 607632 | . | . | 34.427409 | 35.371004 | 37.519877 | 40.052997 | 80.113994 | 81.761667 | 84.193513 | 91.576424 | 1.095303 | 1288.176680 | 0.129283094 | 1.015836362 | 0.309707317 | 0.998865404 | 10.264053 | 10.395197 | 10.288243 | 10.302075 | 10.343664 | 10.384336 | 1403 | 1339 | 1140 | 1296 |
| 55852 | NM_00128873 | TEX2 | testis expressed 2 | protein_coding | NP_001275661.1;NP_001275662.1;NP_060939.3;XP_011523300.1;XP_01 | 30884 | . | . | . | 11.270401 | 11.098224 | 10.932302 | 12.349527 | 26.226687 | 25.654040 | 24.531769 | 28.235727 | 1.026781 | 1867.956919 | 0.124293465 | 0.306756889 | 0.759028425 | 0.998865404 | 10.848160 | 10.886082 | 10.863513 | 10.849140 | 10.832647 | 10.919800 | 2142 | 1971 | 1555 | 1872 |
| 55853 | NR_024628,NF | IDI2-AS1 | IDI2 antisense RNA 1 | lncRNA | . | 30885 | 615391 | . | . | 0.741294 | 0.662336 | 0.545995 | 0.449044 | 1.725020 | 1.531020 | 1.225196 | 1.026685 | -1.287003 | 23.500846 | 0.637981235 | -0.57057376 | 0.568288611 | NA | 4.724393 | 4.362218 | 4.571528 | 4.567255 | 4.554931 | 4.534718 | 31 | 28 | 20 | 17 |
| 55854 | NM_018471,XI | ZC3H15 | zinc finger CCCH-type containing 15 | protein_coding | NP_060941.2;XP_016859991.1 | 29528 | . | . | . | 51.378993 | 47.850838 | 47.959090 | 39.846357 | 119.561026 | 110.609354 | 107.618804 | 91.103962 | -1.145623 | 2957.578275 | 0.129742172 | -1.51171301 | 0.130606886 | 0.882587711 | 11.624783 | 11.428980 | 11.614103 | 11.548475 | 11.547254 | 11.388257 | 3780 | 3273 | 2634 | 2330 |
| 55855 | NR_027141 | DENND10P1 | DENND10 pseudogene 1 | transcribed_ps | . | 30886 | . | . | . | 1.914543 | 1.477861 | 1.750620 | 1.210302 | 4.455220 | 3.416141 | 3.928340 | 2.767212 | -1.163146 | 124.633523 | 0.31640976 | -0.68908109 | 0.490772237 | 0.998865404 | 7.065122 | 6.849960 | 7.032412 | 6.931439 | 6.989721 | 6.867339 | 175 | 125 | 119 | 88 |
| 55856 | NM_00116009 | ACOT13 | acyl-CoA thioesterase 13 | protein_coding | NP_001153566.1;NP_060943.1 | 20999 | 615652 | . | . | 3.992487 | 3.589819 | 3.303645 | 4.155368 | 9.290681 | 8.298028 | 7.413285 | 9.500754 | -1.025800 | 476.249506 | 0.184150337 | -0.19955933 | 0.84182524 | 0.998865404 | 8.914097 | 8.876810 | 8.937509 | 8.865509 | 8.816145 | 8.951586 | 587 | 491 | 364 | 489 |
| 55857 | NM_00116302 | KIZ | kizuna centrosomal protein | protein_coding | NP_001156494.1;NP_001156495.1;NP_001263318.1;NP_001339363.1;NI | 15865 | 615757 | . | . | 3.935850 | 3.743725 | 4.591835 | 5.017474 | 9.158881 | 8.653787 | 10.303946 | 11.471860 | 1.178987 | 293.101987 | 0.203633113 | 1.166549863 | 0.243392201 | 0.992733415 | 8.071859 | 8.308931 | 8.142584 | 8.152339 | 8.211245 | 8.258339 | 307 | 293 | 264 | 310 |
| 55858 | NM_018475,N | TMEM165 | transmembrane protein 165 | protein_coding | NP_060945.2;XP_011532696.1;XP_016863901.1 | 30760 | 614726 | . | . | 27.458114 | 22.860844 | 24.925181 | 21.558904 | 63.896158 | 52.843860 | 55.931386 | 49.291873 | -1.095244 | 1642.099362 | 0.135051014 | -0.97187273 | 0.33111387 | 0.998865404 | 10.745303 | 10.614379 | 10.783481 | 10.641191 | 10.657286 | 10.630331 | 2154 | 1686 | 1387 | 1444 |
| 55859 | NM_018476 | BEX1 | brain expressed X-linked 1 | protein_coding | NP_060946.3 | 1036 | 300690 | . | . | 1.910859 | 1.005793 | 0.643932 | 1.028132 | 4.446649 | 2.324936 | 1.444966 | 2.350702 | -1.764923 | 28.838628 | 0.622068852 | -1.31754767 | 0.187655113 | 0.960264078 | 5.202709 | 4.381963 | 4.928083 | 4.832423 | 4.797040 | 4.835292 | 56 | 27 | 14 | 24 |
| 55860 | NM_018477,XI | ACTR10 | actin related protein 10 | protein_coding | NP_060947.1;XP_011535262.1;XP_011535263.1;XP_011535264.1;XP_01 | 17372 | . | . | . | 19.086833 | 17.232929 | 16.936245 | 19.070980 | 44.415849 | 39.834686 | 38.004444 | 43.603530 | -1.067338 | 1280.351344 | 0.130580649 | -0.71999282 | 0.471529418 | 0.998865404 | 10.368560 | 10.274557 | 10.371247 | 10.321113 | 10.282330 | 10.311621 | 1572 | 1380 | 1056 | 1184 |
| 55861 | NM_00104822 | DBNDD2 | dysbindin domain containing 2 | protein_coding | NP_001041686.1;NP_001041687.1;NP_001041688.1;NP_001041689.1;NI | 15881 | 611453 | . | . | 1.145535 | 0.571716 | 0.767443 | 0.713952 | 2.665709 | 1.321546 | 1.722119 | 1.632365 | -1.096714 | 30.137429 | 0.591641438 | -0.22511395 | 0.821890629 | 0.998865404 | 4.977288 | 4.846728 | 4.958338 | 4.874548 | 4.906009 | 4.906110 | 48 | 23 | 25 | 27 |
| 55862 | NM_00100203 | ECHDC1 | ethylmalonyl-CoA decarboxylase 1 | protein_coding | NP_001002030.1;NP_001099014.1;NP_001099015.1;NP_001132982.1;NI | 21489 | 612136 | . | . | 21.888174 | 19.635549 | 20.942239 | 20.507912 | 50.934678 | 45.388446 | 46.993774 | 46.888901 | -1.009116 | 1534.787167 | 0.127466858 | -0.10271103 | 0.918192318 | 0.998865404 | 10.590244 | 10.577374 | 10.623537 | 10.549156 | 10.588357 | 10.571712 | 1860 | 1584 | 1343 | 1417 |
| 55863 | NM_00119353 | TMEM126B | transmembrane protein 126B | protein_coding | NP_001180466.1;NP_001180467.1;NP_001243475.1;NP_001243476.1;NI | 30883 | 615533 | . | . | 19.501072 | 18.067722 | 24.043215 | 17.960940 | 45.379796 | 41.764349 | 53.952277 | 41.065552 | 1.049741 | 774.163892 | 0.179369771 | 0.390442448 | 0.696209405 | 0.998865404 | 9.560501 | 9.631613 | 9.633700 | 9.511938 | 9.713707 | 9.492128 | 945 | 744 | 798 | 627 |
| 55869 | NM_00116641 | HDAC8 | histone deacetylase 8 | protein_coding | NP_001159890.1;NP_001159891.1;NP_001159892.1;NP_001159894.1;NI | 13315 | 300269 | . | . | 5.279120 | 4.850048 | 5.418024 | 5.084285 | 12.284729 | 11.211102 | 12.157886 | 11.624612 | -1.029272 | 267.704833 | 0.211680858 | -0.19663825 | 0.844110633 | 0.998865404 | 8.084832 | 8.043877 | 8.082205 | 8.060851 | 8.081860 | 8.031227 | 319 | 287 | 241 | 235 |
| 55870 | NM_00136617 | ASH1L | ASH1 like histone lysine methyltransferase | protein_coding | NP_001353106.1;NP_060959.2;XP_005245394.1;XP_006711513.1;XP_00 | 19088 | 607999 | . | . | 2.376674 | 2.727138 | 2.987502 | 2.729774 | 5.530616 | 6.303903 | 6.703867 | 6.241303 | 1.081723 | 995.615870 | 0.148059043 | 0.765446203 | 0.444005889 | 0.998865404 | 9.901612 | 10.015050 | 9.880104 | 9.974463 | 10.035506 | 9.933657 | 1030 | 1099 | 962 | 903 |
| 55871 | NM_00114535 | CBWD1 | COBW domain containing 1 | protein_coding | NP_001138827.1;NP_001138828.1;NP_060961.3;XP_005251570.1;XP_00 | 17134 | 611078 | . | . | 7.680927 | 7.393976 | 9.029698 | 7.421782 | 17.873833 | 17.091509 | 20.262386 | 16.969023 | 1.099204 | 451.075594 | 0.181031109 | 0.753788464 | 0.450976252 | 0.998865404 | 8.746929 | 8.884255 | 8.798951 | 8.771248 | 8.898277 | 8.785892 | 506 | 453 | 448 | 403 |
| 55872 | NM_00127894 | PBK | PDZ binding kinase | protein_coding | NP_001265874.1;NP_001349969.1;NP_060962.2 | 18282 | 611210 | . | . | 38.602070 | 36.088891 | 30.760933 | 27.396813 | 89.828601 | 83.421086 | 69.026638 | 62.639562 | -1.299422 | 1894.407855 | 0.12760367 | -2.96127849 | 0.003063648 | 0.153945181 | 11.063963 | 10.686476 | 11.011308 | 10.950382 | 10.821426 | 10.725246 | 2557 | 2224 | 1523 | 1448 |
| 55876 | NM_00104247 | GSDMB | gasdermin B | protein_coding | NP_001035936.1;NP_001159430.1;NP_001159431.1;NP_001356331.1;NI | 23690 | 611221 | . | . | 1.150732 | 0.931184 | 1.314255 | 0.688531 | 2.677802 | 2.152474 | 2.949149 | 1.574243 | -1.034300 | 52.327058 | 0.470508313 | -0.10340748 | 0.917639574 | 0.998865404 | 5.731451 | 5.687180 | 5.730502 | 5.679562 | 5.755983 | 5.638116 | 69 | 50 | 59 | 33 |
| 55884 | NM_00127855 | WSB2 | WD repeat and SOCS box containing 2 | protein_coding | NP_001265486.1;NP_001265487.1;NP_061109.1;XP_016875131.1 | 19222 | . | . | . | 22.917736 | 22.476891 | 17.580251 | 23.404097 | 53.330511 | 51.956337 | 39.449569 | 53.510681 | -1.128377 | 1915.559848 | 0.145028634 | -1.20147695 | 0.229566242 | 0.98402759 | 10.988243 | 10.813574 | 10.958055 | 10.936725 | 10.739256 | 10.952946 | 2370 | 2163 | 1351 | 1918 |
| 55892 | NM_00118511 | MYNN | myoneurin | protein_coding | NP_001172047.1;NP_001172048.1;NP_061127.1 | 14955 | 606042 | . | . | 2.669284 | 3.135211 | 2.907921 | 2.515405 | 6.211533 | 7.247181 | 6.525290 | 5.751175 | -1.084916 | 436.193853 | 0.185983459 | -0.63222707 | 0.527238509 | 0.998865404 | 8.826313 | 8.708952 | 8.743140 | 8.837896 | 8.787011 | 8.695016 | 483 | 527 | 392 | 362 |
| 55893 | NM_018660 | ZNF395 | zinc finger protein 395 | protein_coding | NP_061130.1 | 18737 | 609494 | . | . | 2.192124 | 2.432479 | 2.560354 | 2.874399 | 5.101162 | 5.622785 | 5.745359 | 6.571972 | 1.159328 | 376.089315 | 0.188624087 | 1.130762065 | 0.25815525 | 0.995760118 | 8.444771 | 8.657274 | 8.480489 | 8.535205 | 8.562804 | 8.624319 | 382 | 394 | 333 | 398 |
| 55894 | NM_018661 | DEFB103B | defensin beta 103B | protein_coding | NP_061131.1 | 31702 | 606611 | . | . | 0.418617 | 0.258313 | 0.619616 | 0.347780 | 0.974139 | 0.597102 | 1.390399 | 0.795157 | 1.441305 | 7.082758 | 1.147930876 | 0.459414123 | 0.645936808 | NA | 2.533720 | 3.066070 | 2.813551 | 2.802767 | 2.828356 | 2.811121 | 8 | 5 | 9 | 6 |
| 55897 | NM_018670,XI | MESP1 | mesoderm posterior bHLH transcription factor 1 | protein_coding | NP_061140.1 | 29658 | 608689 | . | . | 4.569831 | 3.600586 | 2.296375 | 3.676526 | 10.634184 | 8.322914 | 5.152998 | 8.405940 | -1.394955 | 129.550537 | 0.315993193 | -1.51971017 | 0.128583836 | 0.877332024 | 7.238344 | 6.756322 | 7.113389 | 7.024580 | 6.879712 | 7.018510 | 193 | 145 | 73 | 124 |
| 55898 | NM_00103967 | UNC45A | unc-45 myosin chaperone A | protein_coding | NP_001034764.1;NP_001310548.1;NP_001310549.1;NP_001310550.1;NI | 30594 | 611219 | . | . | 35.655294 | 36.366464 | 34.107343 | 37.368286 | 82.971339 | 84.062709 | 76.535889 | 85.438144 | -1.013083 | 3760.408544 | 0.110832109 | -0.16919346 | 0.865644474 | 0.998865404 | 11.886055 | 11.867231 | 11.892634 | 11.871048 | 11.841575 | 11.899832 | 4415 | 4031 | 3134 | 3626 |
| 55900 | NM_00101232 | ZNF302 | zinc finger protein 302 | protein_coding | NP_001012320.1;NP_001276110.1;NP_001276111.1;NP_001276112.1;NI | 13848 | . | . | . | 8.147040 | 7.194981 | 9.286006 | 5.689850 | 18.958498 | 16.631517 | 20.837526 | 13.009164 | -1.018355 | 647.853057 | 0.207817183 | -0.12626862 | 0.899519294 | 0.998865404 | 9.351966 | 9.326971 | 9.372046 | 9.301944 | 9.476578 | 9.147907 | 791 | 669 | 694 | 456 |
| 55901 | NM_018676,N | THSD1 | thrombospondin type 1 domain containing 1 | protein_coding | NP_061146.1;NP_954872.1 | 17754 | 616821 | . | . | 4.675126 | 4.646135 | 3.484174 | 4.215318 | 10.879211 | 10.739750 | 7.818386 | 9.637825 | -1.228042 | 391.067503 | 0.186940582 | -1.58531632 | 0.112894483 | 0.850211417 | 8.752175 | 8.455107 | 8.672819 | 8.663813 | 8.504377 | 8.588563 | 500 | 462 | 282 | 354 |
| 55902 | NM_00107655 | ACSS2 | acyl-CoA synthetase short chain family member 2 | protein_coding | NP_001070020.2;NP_001229322.1;NP_061147.1;XP_005260512.1;XP_00 | 15814 | 605832 | . | . | 10.678658 | 10.564149 | 18.157358 | 19.165214 | 24.849675 | 24.419504 | 40.744584 | 43.818981 | 1.735780 | 1321.197929 | 0.1285366 | 6.18955443 | 6.03345E-10 | 6.40613E-07 | 9.915713 | 10.711243 | 10.122377 | 10.111741 | 10.517941 | 10.552690 | 1121 | 1034 | 1427 | 1605 |
| 55904 | NM_018682,N | KMT2E | lysine methyltransferase 2E (inactive) | protein_coding | NP_061152.3;NP_891847.1;XP_005250550.1;XP_011514702.1;XP_01686 | 18541 | 608444 | . | . | 6.520061 | 7.832916 | 9.548792 | 8.718442 | 15.172449 | 18.106140 | 21.427211 | 19.933684 | 1.223382 | 1890.143224 | 0.138200521 | 2.10472977 | 0.035314838 | 0.585116558 | 10.731491 | 11.022426 | 10.736745 | 10.854137 | 11.021460 | 10.886121 | 1809 | 1974 | 1961 | 1786 |
| 55905 | NM_018683 | RNF114 | ring finger protein 114 | protein_coding | NP_061153.1 | 13094 | 612451 | . | . | 40.203037 | 41.571190 | 43.486637 | 44.212509 | 93.554131 | 96.093666 | 97.582748 | 101.086647 | 1.057774 | 3229.259556 | 0.110109092 | 0.735914239 | 0.461782891 | 0.998865404 | 11.615923 | 11.696916 | 11.623137 | 11.643563 | 11.676935 | 11.682055 | 3569 | 3429 | 2878 | 3122 |
| 55906 | NM_00117803 | ZC4H2 | zinc finger C4H2-type containing | protein_coding | NP_001171503.1;NP_001171504.1;NP_001230733.1;NP_061154.1 | 24931 | 309605 | . | . | 0.912806 | 0.910960 | 0.976837 | 1.360659 | 2.124137 | 2.105723 | 2.191994 | 3.110985 | 1.190716 | 91.141865 | 0.344701135 | 0.73057135 | 0.465041017 | 0.998865404 | 6.379257 | 6.629950 | 6.494316 | 6.465102 | 6.483081 | 6.574654 | 101 | 85 | 73 | 107 |
| 55907 | NM_018686,N | CMAS | cytidine monophosphate N-acetylneuraminic acid synthetase | protein_coding | NP_061156.1 | 18290 | 603316 | . | . | 24.651326 | 21.418230 | 22.770751 | 20.092550 | 57.364649 | 49.509194 | 51.096907 | 45.939229 | -1.087688 | 1210.662256 | 0.143889506 | -0.84276616 | 0.399359237 | 0.998865404 | 10.300654 | 10.179985 | 10.325723 | 10.213413 | 10.257058 | 10.156652 | 1559 | 1261 | 1077 | 1014 |
| 55909 | NM_00136304 | BIN3 | bridging integrator 3 | protein_coding | NP_001349975.1;NP_061158.1;XP_011542888.1;XP_011542889.1 | 1054 | 606396 | . | . | 39.078505 | 37.733978 | 36.629138 | 45.025634 | 90.937290 | 87.223780 | 82.194722 | 102.945767 | 1.029674 | 2111.461335 | 0.130381908 | 0.323565425 | 0.746267068 | 0.998865404 | 11.022901 | 11.064846 | 11.051014 | 11.011053 | 10.980775 | 11.123291 | 2456 | 2189 | 1697 | 2187 |
| 55914 | NM_00100660 | ERBIN | erbb2 interacting protein | protein_coding | NP_001006600.1;NP_001240626.1;NP_001240627.1;NP_001240628.1;NI | 15842 | 606944 | . | . | 7.086464 | 6.689706 | 7.218472 | 6.200593 | 16.490496 | 15.463553 | 16.198044 | 14.176918 | 1.025653 | 1575.027955 | 0.137785883 | 0.265218072 | 0.790841474 | 0.998865404 | 10.602521 | 10.639565 | 10.643368 | 10.575066 | 10.697292 | 10.557206 | 1869 | 1604 | 1512 | 1361 |
| 55915 | NM_018697 | LANCL2 | LanC like 2 | protein_coding | NP_061167.1 | 6509 | 612919 | . | . | 2.685701 | 2.742370 | 3.105630 | 3.454127 | 6.249737 | 6.339111 | 6.968943 | 7.897451 | 1.190602 | 406.597535 | 0.181149165 | 1.389417497 | 0.164705833 | 0.940065727 | 8.536375 | 8.787616 | 8.607353 | 8.614493 | 8.685199 | 8.743876 | 425 | 403 | 366 | 434 |
| 55916 | NM_00124261 | NXT2 | nuclear transport factor 2 like export factor 2 | protein_coding | NP_001229546.1;NP_001229547.1;NP_061168.2;XP_011529291.1;XP_01 | 18151 | 300320 | . | . | 1.839810 | 1.226387 | 1.652446 | 1.175044 | 4.281316 | 2.834847 | 3.708043 | 2.686599 | -1.095497 | 120.688606 | 0.329130885 | -0.39979706 | 0.689305998 | 0.998865404 | 6.977990 | 6.849439 | 6.993326 | 6.850560 | 6.949522 | 6.833721 | 174 | 109 | 118 | 89 |
| 55917 | NM_018704,XI | CTTNBP2NL | CTTNBP2 N-terminal like | protein_coding | NP_061174.1;XP_011540083.1;XP_016857295.1 | 25330 | 615100 | . | . | 2.639480 | 2.764035 | 2.319128 | 2.822724 | 6.142178 | 6.389191 | 5.204056 | 6.453823 | -1.082446 | 473.099240 | 0.174534805 | -0.65485397 | 0.512561777 | 0.998865404 | 8.942328 | 8.827381 | 8.905153 | 8.915097 | 8.818103 | 8.901328 | 562 | 535 | 368 | 455 |
| 55920 | NM_00113620 | RCC2 | regulator of chromosome condensation 2 | protein_coding | NP_001129676.1;NP_061185.1 | 30297 | 609587 | . | . | 64.866507 | 69.722834 | 70.864955 | 76.540177 | 150.947029 | 161.167475 | 159.018906 | 175.000029 | 1.083164 | 8843.732208 | 0.105869856 | 1.088610864 | 0.276325524 | 0.998865404 | 13.051699 | 13.166881 | 13.047154 | 13.102643 | 13.115358 | 13.170182 | 9456 | 9463 | 7741 | 8898 |
| 55922 | NM_017544,N | NKRF | NFKB repressing factor | protein_coding | NP_060014.3 | 19374 | 300440 | . | . | 4.049265 | 4.186573 | 4.104199 | 3.385257 | 9.422806 | 9.677451 | 9.209704 | 7.739987 | -1.115511 | 465.818810 | 0.180862376 | -0.87195972 | 0.38323035 | 0.998865404 | 8.939950 | 8.783035 | 8.886896 | 8.903766 | 8.887241 | 8.764253 | 558 | 537 | 422 | 371 |
| 55924 | NM_019099,N | INKA2 | inka box actin regulator 2 | protein_coding | NP_061972.1;NP_945120.1;XP_011540085.1 | 28045 | . | . | . | 0.818076 | 0.995252 | 1.163150 | 1.137328 | 1.903696 | 2.300570 | 2.610075 | 2.600365 | 1.257822 | 192.350904 | 0.245542154 | 1.347742689 | 0.177741157 | 0.951283903 | 7.413029 | 7.743304 | 7.489135 | 7.567782 | 7.637026 | 7.624228 | 178 | 201 | 189 | 198 |
| 55929 | NM_00103402 | DMAP1 | DNA methyltransferase 1 associated protein 1 | protein_coding | NP_001029195.1;NP_001029196.1;NP_061973.1 | 18291 | 605077 | . | . | 11.733275 | 11.611221 | 11.012641 | 13.456148 | 27.303815 | 26.839859 | 24.712047 | 30.765882 | 1.039707 | 615.014421 | 0.166868299 | 0.336651983 | 0.736379264 | 0.998865404 | 9.236437 | 9.291982 | 9.259154 | 9.241025 | 9.207441 | 9.340251 | 704 | 642 | 491 | 646 |
| 55930 | NM_018728,XI | MYO5C | myosin VC | protein_coding | NP_061198.2;XP_011520083.1;XP_016877897.1;XP_016877898.1;XP_01 | 7604 | 610022 | . | . | 0.070439 | 0.032715 | 0.074791 | 0.026460 | 0.163914 | 0.075621 | 0.167828 | 0.060498 | 1.020254 | 11.673883 | 0.948651407 | 0.030494024 | 0.975673059 | NA | 3.526439 | 3.563745 | 3.555335 | 3.517973 | 3.560842 | 3.514159 | 18 | 8 | 15 | 6 |
| 55937 | NM_00125616 | APOM | apolipoprotein M | protein_coding | NP_001243098.1;NP_061974.2;XP_006715213.1 | 13916 | 606907 | . | . | 0.453951 | 0.687944 | 0.619858 | 1.207544 | 1.056362 | 1.590213 | 1.390943 | 2.760905 | 1.531480 | 17.663541 | 0.755761909 | 0.813651674 | 0.415844526 | NA | 3.805868 | 4.415590 | 4.101891 | 4.126339 | 4.119300 | 4.172357 | 13 | 18 | 13 | 26 |
| 55954 | NM_00100369 | ZMAT5 | zinc finger matrin-type 5 | protein_coding | NP_001003692.1;NP_001305058.1;NP_061976.1 | 28046 | . | . | . | 10.517997 | 8.101435 | 9.326219 | 11.573311 | 24.475810 | 18.726829 | 20.927766 | 26.461000 | 1.146855 | 295.697396 | 0.221813609 | 0.891212354 | 0.372815258 | 0.998865404 | 8.105723 | 8.303462 | 8.231022 | 8.096417 | 8.184519 | 8.291202 | 358 | 259 | 248 | 325 |
| 55957 | NM_00136978 | LIN37 | lin-37 DREAM MuvB core complex component | protein_coding | NP_001356709.1;NP_061977.1 | 33234 | . | . | . | 7.561857 | 8.403464 | 8.723360 | 6.472943 | 17.596752 | 19.424985 | 19.574965 | 14.799617 | -1.063016 | 228.399964 | 0.240686756 | -0.36629669 | 0.714143699 | 0.998865404 | 7.878272 | 7.791253 | 7.823078 | 7.870137 | 7.886484 | 7.743766 | 258 | 266 | 222 | 176 |
| 55958 | NM_018847 | KLHL9 | kelch like family member 9 | protein_coding | NP_061335.1 | 18732 | 611201 | . | . | 2.922833 | 3.316534 | 2.905331 | 2.783878 | 6.801552 | 7.666316 | 6.519478 | 6.365005 | -1.111054 | 533.822489 | 0.16778067 | -0.90552179 | 0.365189024 | 0.998865404 | 9.134235 | 8.982193 | 9.053175 | 9.130922 | 9.040882 | 9.008632 | 609 | 642 | 451 | 462 |
| 55964 | NM_00136384 | SEPTIN3 | septin 3 | protein_coding | NP_001350774.1;NP_061979.3;NP_663786.2 | 10750 | 608314 | . | . | 4.329642 | 4.017418 | 4.623442 | 5.474528 | 10.075254 | 9.286441 | 10.374868 | 12.516860 | 1.186935 | 643.149485 | 0.160857805 | 1.537018989 | 0.124288659 | 0.87182553 | 9.200321 | 9.447155 | 9.279423 | 9.253135 | 9.328062 | 9.432961 | 691 | 622 | 561 | 704 |
| 55967 | NM_00125833 | NDUFA12 | NADH:ubiquinone oxidoreductase subunit A12 | protein_coding | NP_001245267.1;NP_061326.1 | 23987 | 614530 | . | . | 49.577627 | 53.205343 | 52.737264 | 56.770224 | 115.369181 | 122.986556 | 118.340884 | 129.798361 | 1.051729 | 922.468481 | 0.141699704 | 0.513501542 | 0.607600532 | 0.998865404 | 9.812707 | 9.885096 | 9.808181 | 9.852209 | 9.841507 | 9.891457 | 1004 | 1000 | 794 | 917 |
| 55968 | NM_00120673 | NSFL1C | NSFL1 cofactor | protein_coding | NP_001193665.1;NP_057227.2;NP_061327.2;XP_006723655.1;XP_00672 | 15912 | 606610 | . | . | 18.691475 | 19.406485 | 18.835362 | 19.867438 | 43.495831 | 44.858964 | 42.266012 | 45.424532 | -1.026334 | 2035.469485 | 0.123214333 | -0.30435415 | 0.760858098 | 0.998865404 | 11.009905 | 10.972139 | 10.992477 | 11.009645 | 10.933574 | 11.025016 | 2349 | 2249 | 1646 | 1992 |
| 55969 | NM_00119953 | RAB5IF | RAB5 interacting factor | protein_coding | NP_001186463.1;NP_001361107.1;NP_061328.1;NP_955777.1 | 15870 | . | . | . | 113.415752 | 109.672795 | 116.674093 | 121.602776 | 263.923108 | 253.513583 | 261.813283 | 278.030293 | 1.054240 | 3864.886481 | 0.109148984 | 0.698154494 | 0.485080582 | 0.998865404 | 11.877606 | 11.953808 | 11.911378 | 11.875921 | 11.923813 | 11.951115 | 4426 | 3973 | 3393 | 3780 |
| 55970 | NM_018841,XI | GNG12 | G protein subunit gamma 12 | protein_coding | NP_061329.3;XP_016857298.1;XP_024304173.1 | 19663 | 615405 | . | . | 30.916640 | 28.876669 | 34.016039 | 28.365440 | 71.944289 | 66.749711 | 76.331002 | 64.854209 | 1.029040 | 4212.332136 | 0.125653722 | 0.328671328 | 0.742404128 | 0.998865404 | 12.019489 | 12.061019 | 12.060854 | 11.991902 | 12.127196 | 11.968413 | 4979 | 4294 | 4069 | 3625 |
| 55971 | NM_018842 | BAIAP2L1 | BAR/IMD domain containing adaptor protein 2 like 1 | protein_coding | NP_061330.2 | 21649 | 611877 | . | . | 20.254520 | 21.567823 | 17.903452 | 22.013357 | 47.133106 | 49.854984 | 40.174828 | 50.330925 | -1.062464 | 2320.633893 | 0.131470382 | -0.66489799 | 0.506115744 | 0.998865404 | 11.223528 | 11.135743 | 11.180927 | 11.224781 | 11.073061 | 11.229452 | 2679 | 2650 | 1765 | 2316 |
| 55972 | NM_018843 | SLC25A40 | solute carrier family 25 member 40 | protein_coding | NP_061331.2 | 29680 | 610821 | . | . | 5.093835 | 4.297507 | 4.450597 | 3.560977 | 11.853563 | 9.933880 | 9.987012 | 8.141750 | -1.188804 | 550.848761 | 0.183028205 | -1.36324044 | 0.172806746 | 0.947431715 | 9.224312 | 8.976045 | 9.214728 | 9.094365 | 9.114730 | 8.968334 | 750 | 588 | 489 | 417 |
| 55973 | NM_00100840 | BCAP29 | B cell receptor associated protein 29 | protein_coding | NP_001008405.1;NP_001350411.1;NP_001350412.1;NP_001358282.1;NI | 24131 | . | . | . | 23.191697 | 15.705700 | 16.212574 | 9.817526 | 53.968031 | 36.304432 | 36.380543 | 22.446610 | -1.526693 | 1399.793389 | 0.474191671 | -1.28726442 | 0.198002142 | 0.96621494 | 10.724089 | 10.113852 | 10.725608 | 10.404787 | 10.423802 | 10.046214 | 2330 | 1467 | 1212 | 771 |
| 55974 | NM_00112283 | SLC50A1 | solute carrier family 50 member 1 | protein_coding | NP_001116309.1;NP_001116311.1;NP_001274515.1;NP_001274516.1;NI | 30657 | 613683 | . | . | 26.546044 | 24.169875 | 31.915474 | 29.613739 | 61.779735 | 55.869752 | 71.617399 | 67.708291 | 1.200651 | 1105.587403 | 0.137804709 | 1.914426119 | 0.055565731 | 0.697883333 | 9.972437 | 10.236675 | 10.068184 | 10.005167 | 10.203718 | 10.141945 | 1207 | 1037 | 1092 | 1085 |
| 55975 | NM_00103171 | KLHL7 | kelch like family member 7 | protein_coding | NP_001026880.2;NP_001165899.1;NP_061334.4;XP_006715816.1;XP_00 | 15646 | 611119 | . | . | 15.243249 | 18.175085 | 14.204053 | 15.748920 | 35.471668 | 42.012520 | 31.873485 | 36.008034 | -1.019018 | 1418.550654 | 0.12602769 | -0.21566496 | 0.829248915 | 0.998865404 | 10.483665 | 10.456612 | 10.486342 | 10.468300 | 10.474143 | 10.451607 | 1668 | 1527 | 1240 | 1298 |
| 56005 | NM_019107,XI | MYDGF | myeloid derived growth factor | protein_coding | NP_061980.1;XP_016882476.1 | 16948 | 606746 | . | . | 114.604380 | 109.979399 | 110.874834 | 111.001087 | 266.689113 | 254.222297 | 248.799914 | 253.790802 | -1.026138 | 3412.561600 | 0.110427623 | -0.3370902 | 0.736048903 | 0.998865404 | 11.755097 | 11.717943 | 11.767240 | 11.726563 | 11.730835 | 11.720913 | 4078 | 3638 | 2947 | 3139 |
| 56006 | NM_019108,XI | SMG9 | SMG9 nonsense mediated mRNA decay factor | protein_coding | NP_061981.2;XP_005259114.1;XP_011525415.1;XP_011525416.1;XP_01 | 25763 | 613176 | . | . | 14.183221 | 12.916336 | 16.412575 | 12.951203 | 33.004938 | 29.856690 | 36.829344 | 29.611388 | 1.048111 | 1615.567125 | 0.123899614 | 0.547148246 | 0.584276889 | 0.998865404 | 10.623533 | 10.691320 | 10.618831 | 10.658725 | 10.689643 | 10.661298 | 1770 | 1746 | 1464 | 1522 |
| 56034 | NM_016205,N | PDGFC | platelet derived growth factor C | protein_coding | NP_057289.1;XP_011530426.1;XP_016863944.1;XP_016863945.1 | 8801 | 608452 | . | . | 0.762753 | 0.846185 | 0.471633 | 0.738094 | 1.774956 | 1.955993 | 1.058331 | 1.687565 | -1.351789 | 101.468188 | 0.339029525 | -1.28269088 | 0.199600397 | 0.96621494 | 6.867222 | 6.429464 | 6.691324 | 6.723842 | 6.556959 | 6.670763 | 128 | 132 | 59 | 98 |
| 56052 | NM_00133050 | ALG1 | ALG1 chitobiosyldiphosphodolichol beta-mannosyltransferase | protein_coding | NP_001317433.1;NP_061982.3;XP_016878946.1;XP_016878947.1 | 18294 | 605907 | . | . | 15.281924 | 12.829518 | 18.557578 | 12.073794 | 35.561662 | 29.656008 | 41.642668 | 27.605295 | 1.075149 | 1373.705669 | 0.139949389 | 0.746959188 | 0.455088197 | 0.998865404 | 10.370509 | 10.475302 | 10.362758 | 10.424120 | 10.515129 | 10.379280 | 1463 | 1486 | 1347 | 1217 |
| 56061 | NM_019116,XI | UBFD1 | ubiquitin family domain containing 1 | protein_coding | NP_061989.2;XP_011544196.1;XP_016878948.1;XP_016878949.1 | 30565 | . | . | . | 11.389360 | 13.308072 | 12.293124 | 12.916849 | 26.503509 | 30.762208 | 27.585409 | 29.532842 | 1.011856 | 1828.419301 | 0.128825731 | 0.131989736 | 0.894992411 | 0.998865404 | 10.827991 | 10.844723 | 10.771168 | 10.887562 | 10.829215 | 10.851074 | 1939 | 2106 | 1577 | 1748 |
| 56063 | NM_00136619 | TMEM234 | transmembrane protein 234 | protein_coding | NP_001353120.1;NP_061991.3;XP_011540097.1;XP_016857301.1;XP_01 | 28837 | . | . | . | 8.281756 | 7.650396 | 9.090249 | 8.417247 | 19.271986 | 17.684230 | 20.398258 | 19.245032 | 1.154184 | 599.048758 | 0.163784661 | 1.263083514 | 0.206559168 | 0.970482251 | 9.119041 | 9.326562 | 9.186657 | 9.162980 | 9.319332 | 9.218573 | 652 | 589 | 600 | 557 |
| 56097 | NM_018929,N | PCDHGC5 | protocadherin gamma subfamily C, 5 | protein_coding | NP_061752.1;NP_115783.1 | 8718 | 606306 | . | . | 0.057443 | 0.035536 | 0.042844 | 0.117454 | 0.133671 | 0.082143 | 0.096141 | 0.268543 | 1.962415 | 6.600210 | 1.20417508 | 0.807715203 | 0.41925455 | NA | 2.156660 | 3.127898 | 2.702652 | 2.695691 | 2.705480 | 2.731952 | 6 | 4 | 5 | 11 |
| 56098 | NM_018928,N | PCDHGC4 | protocadherin gamma subfamily C, 4 | protein_coding | NP_061751.1;NP_115782.1 | 8717 | 606305 | . | . | 0.047158 | 0.013363 | 0.032610 | 0.010224 | 0.109738 | 0.030888 | 0.073176 | 0.023375 | 1.080147 | 3.232233 | 1.683115441 | 0.066084235 | 0.947310762 | NA | 1.631267 | 1.751299 | 1.693964 | 1.682131 | 1.694716 | 1.683650 | 5 | 2 | 4 | 2 |
| 56101 | NM_018925,N | PCDHGB5 | protocadherin gamma subfamily B, 5 | protein_coding | NP_061748.1;NP_115270.1 | 8712 | 606302 | . | . | 0.346878 | 0.580731 | 0.225345 | 0.455676 | 0.807199 | 1.342385 | 0.505667 | 1.041849 | -1.585633 | 52.671026 | 0.558135985 | -1.19157154 | 0.23342929 | 0.988893316 | 6.016078 | 5.344290 | 5.675326 | 5.809764 | 5.565708 | 5.741135 | 53 | 90 | 15 | 60 |
| 56102 | NM_018924,N | PCDHGB3 | protocadherin gamma subfamily B, 3 | protein_coding | NP_061747.2;NP_115268.2 | 8710 | 606301 | . | . | 0.168344 | 0.123489 | 0.032147 | 0.064142 | 0.391744 | 0.285449 | 0.072138 | 0.146654 | -2.436503 | 10.385348 | 0.980273106 | -1.31066757 | 0.189970101 | NA | 3.881411 | 2.592943 | 3.415104 | 3.399174 | 3.362756 | 3.374881 | 19 | 14 | 4 | 7 |
| 56103 | NM_018923,N | PCDHGB2 | protocadherin gamma subfamily B, 2 | protein_coding | NP_061746.1;NP_115267.1 | 8709 | 606300 | . | . | 0.331559 | 0.252336 | 0.239509 | 0.405098 | 0.771551 | 0.583287 | 0.537450 | 0.926208 | -1.013110 | 28.759784 | 0.594416358 | -0.03161184 | 0.974781604 | 0.998865404 | 4.855648 | 4.836248 | 4.870060 | 4.821934 | 4.822469 | 4.866582 | 40 | 25 | 20 | 32 |
| 56104 | NM_018922,N | PCDHGB1 | protocadherin gamma subfamily B, 1 | protein_coding | NP_061745.1;NP_115266.1 | 8708 | 606299 | . | . | 0.282036 | 0.174407 | 0.233036 | 0.367443 | 0.656308 | 0.403149 | 0.522925 | 0.840114 | 1.896526 | 25.769626 | 0.675780398 | 1.366359985 | 0.171826002 | NA | 4.154300 | 5.076095 | 4.644654 | 4.610496 | 4.632051 | 4.753707 | 25 | 15 | 16 | 46 |
| 56105 | NM_018914,N | PCDHGA11 | protocadherin gamma subfamily A, 11 | protein_coding | NP_061737.1;NP_114480.1;NP_114481.1 | 8698 | 606298 | . | . | 0.148837 | 0.032460 | 0.251975 | 0.089733 | 0.346350 | 0.075032 | 0.565426 | 0.205165 | 2.236246 | 17.135416 | 0.893482257 | 1.299498249 | 0.193772994 | NA | 3.397517 | 4.568818 | 4.057900 | 3.995669 | 4.135702 | 4.028765 | 20 | 4 | 32 | 10 |
| 56106 | NM_018913,N | PCDHGA10 | protocadherin gamma subfamily A, 10 | protein_coding | NP_061736.1;NP_114479.1 | 8697 | 606297 | . | . | 0.292249 | 0.683330 | 0.174702 | 0.568788 | 0.680075 | 1.579548 | 0.392025 | 1.300465 | -1.261641 | 51.309344 | 1.168473668 | -0.28695703 | 0.774145221 | 0.998865404 | 5.839888 | 5.502761 | 5.577127 | 5.758563 | 5.509314 | 5.736025 | 38 | 88 | 13 | 71 |
| 56107 | NM_018921,N | PCDHGA9 | protocadherin gamma subfamily A, 9 | protein_coding | NP_061744.1;NP_114478.1 | 8707 | 606296 | . | . | 0.129936 | 0.190759 | 0.108810 | 0.137839 | 0.302366 | 0.440948 | 0.244166 | 0.315151 | 1.171878 | 13.678923 | 0.854998812 | 0.267628652 | 0.788985174 | NA | 3.658167 | 3.881002 | 3.754088 | 3.774066 | 3.749879 | 3.796912 | 12 | 16 | 8 | 19 |
| 56108 | NM_018920,N | PCDHGA7 | protocadherin gamma subfamily A, 7 | protein_coding | NP_061743.1;NP_114476.1 | 8705 | 606294 | . | . | 0.080231 | 0.108999 | 0.076290 | 0.085009 | 0.186699 | 0.251957 | 0.171192 | 0.194362 | -1.114503 | 11.117619 | 0.917119057 | -0.17053522 | 0.864589238 | NA | 3.551764 | 3.393446 | 3.467850 | 3.487543 | 3.471773 | 3.473087 | 11 | 15 | 9 | 10 |
| 56109 | NM_018919,N | PCDHGA6 | protocadherin gamma subfamily A, 6 | protein_coding | NP_061742.1;NP_114475.1 | 8704 | 606293 | . | . | 0.140650 | 0.263201 | 0.143015 | 0.153961 | 0.327298 | 0.608402 | 0.320921 | 0.352012 | -1.363533 | 18.341407 | 0.721793824 | -0.61977474 | 0.535406101 | NA | 4.403699 | 3.955730 | 4.198989 | 4.224133 | 4.190195 | 4.185050 | 21 | 26 | 14 | 14 |
| 56110 | NM_018918,N | PCDHGA5 | protocadherin gamma subfamily A, 5 | protein_coding | NP_061741.1;NP_114443.1 | 8703 | 606292 | . | . | 0.038672 | 0.074420 | 0.048330 | 0.055355 | 0.089991 | 0.172026 | 0.108450 | 0.126561 | -1.161769 | 7.155347 | 1.146950586 | -0.18860757 | 0.850400397 | NA | 2.945312 | 2.724273 | 2.829964 | 2.853019 | 2.832982 | 2.840495 | 6 | 11 | 5 | 7 |
| 56111 | NM_018917,N | PCDHGA4 | protocadherin gamma subfamily A, 4 | protein_coding | NP_061740.2;NP_114442.2 | 8702 | 606291 | . | . | 0.153009 | 0.239958 | 0.320356 | 0.212888 | 0.356060 | 0.554672 | 0.718870 | 0.486744 | 1.358603 | 25.979712 | 0.605721849 | 0.729912803 | 0.465443486 | NA | 4.461710 | 4.903263 | 4.665598 | 4.683596 | 4.711546 | 4.702939 | 23 | 26 | 27 | 27 |
| 56112 | NM_018916,N | PCDHGA3 | protocadherin gamma subfamily A, 3 | protein_coding | NP_061739.2;NP_114400.1 | 8701 | 606290 | . | . | 0.008735 | 0.026691 | 0.028411 | 0.035353 | 0.020328 | 0.061697 | 0.063753 | 0.080832 | 1.831615 | 3.844274 | 1.546577309 | 0.564547359 | 0.572381679 | NA | 1.444631 | 2.312263 | 1.918283 | 1.927806 | 1.931804 | 1.935203 | 2 | 4 | 4 | 5 |
| 56113 | NM_018915,N | PCDHGA2 | protocadherin gamma subfamily A, 2 | protein_coding | NP_061738.1;NP_114398.1 | 8700 | 606289 | . | . | 0.102991 | 0.083076 | 0.033446 | 0.046560 | 0.239665 | 0.192033 | 0.075052 | 0.106455 | -2.161932 | 10.455731 | 0.964248495 | -1.15356253 | 0.248679566 | NA | 3.838265 | 2.724273 | 3.419842 | 3.407756 | 3.376307 | 3.383444 | 18 | 14 | 5 | 7 |
| 56114 | NM_018912,N | PCDHGA1 | protocadherin gamma subfamily A, 1 | protein_coding | NP_061735.1;NP_114382.1 | 8696 | 606288 | . | . | 0.568957 | 0.430423 | 0.220109 | 0.359243 | 1.323987 | 0.994942 | 0.493917 | 0.821365 | -1.640719 | 34.116746 | 0.548630412 | -1.30202061 | 0.19290934 | 0.96621494 | 5.406572 | 4.690218 | 5.148376 | 5.106172 | 5.041869 | 5.081926 | 55 | 40 | 18 | 29 |
| 56121 | NM_018935 | PCDHB15 | protocadherin beta 15 | protein_coding | NP_061758.1 | 8686 | 606341 | . | . | 1.047215 | 0.807499 | 0.780583 | 0.961347 | 2.436912 | 1.866571 | 1.751607 | 2.198005 | -1.079670 | 111.786954 | 0.317463005 | -0.34835893 | 0.727570641 | 0.998865404 | 6.858913 | 6.748179 | 6.859563 | 6.770893 | 6.759051 | 6.822274 | 151 | 109 | 84 | 111 |
| 56122 | NM_018934 | PCDHB14 | protocadherin beta 14 | protein_coding | NP_061757.1 | 8685 | 606340 | . | . | 0.528073 | 0.401212 | 0.418809 | 0.474526 | 1.228847 | 0.927420 | 0.939795 | 1.084947 | -1.051294 | 56.615337 | 0.424040739 | -0.17018685 | 0.86486319 | 0.998865404 | 5.858551 | 5.786799 | 5.857116 | 5.798751 | 5.805930 | 5.830281 | 76 | 54 | 45 | 55 |
| 56123 | NM_018933 | PCDHB13 | protocadherin beta 13 | protein_coding | NP_061756.1 | 8684 | 606339 | . | . | 0.117125 | 0.197629 | 0.178737 | 0.234929 | 0.272554 | 0.456828 | 0.401081 | 0.537138 | 1.309073 | 29.174439 | 0.590557014 | 0.657930284 | 0.51058293 | 0.998865404 | 4.661494 | 5.046201 | 4.814782 | 4.867591 | 4.856262 | 4.890102 | 22 | 34 | 25 | 35 |
| 56124 | NM_018932 | PCDHB12 | protocadherin beta 12 | protein_coding | NP_061755.1 | 8683 | 606338 | . | . | 0.146014 | 0.053712 | 0.025418 | 0.121151 | 0.339781 | 0.124158 | 0.057038 | 0.276996 | -1.340983 | 10.776940 | 1.033442341 | -0.40959327 | 0.682104336 | NA | 3.627869 | 3.200321 | 3.465146 | 3.412183 | 3.400057 | 3.449551 | 21 | 7 | 3 | 14 |
| 56125 | NM_018931 | PCDHB11 | protocadherin beta 11 | protein_coding | NP_061754.1 | 8682 | 606337 | . | . | 0.067862 | 0.226444 | 0.151320 | 0.097002 | 0.157917 | 0.523434 | 0.339558 | 0.221783 | -1.210928 | 17.889566 | 0.79452347 | -0.3475205 | 0.728200322 | NA | 4.293102 | 4.015675 | 4.112379 | 4.200077 | 4.156838 | 4.128320 | 11 | 32 | 17 | 12 |
| 56126 | NM_018930 | PCDHB10 | protocadherin beta 10 | protein_coding | NP_061753.1 | 8681 | 606336 | . | . | 0.212549 | 0.227534 | 0.146868 | 0.248661 | 0.494611 | 0.525955 | 0.329568 | 0.568533 | -1.116774 | 22.093627 | 0.664225878 | -0.23988493 | 0.810419461 | NA | 4.544574 | 4.381963 | 4.469263 | 4.475668 | 4.442644 | 4.481525 | 26 | 26 | 14 | 24 |
| 56127 | NM_019119 | PCDHB9 | protocadherin beta 9 | protein_coding | NP_061992.3 | 8694 | 606335 | . | . | 0.294012 | 0.246803 | 0.221467 | 0.266050 | 0.684179 | 0.570497 | 0.496965 | 0.608292 | -1.115533 | 35.605580 | 0.522257004 | -0.30202235 | 0.762635026 | 0.998865404 | 5.230863 | 5.072877 | 5.176588 | 5.150394 | 5.138201 | 5.158241 | 47 | 37 | 27 | 34 |
| 56128 | NM_019120 | PCDHB8 | protocadherin beta 8 | protein_coding | NP_061993.3 | 8693 | 606334 | . | . | 0.513844 | 0.452500 | 0.226230 | 0.422258 | 1.195735 | 1.045974 | 0.507654 | 0.965442 | -1.502650 | 34.974628 | 0.550492221 | -1.06724268 | 0.285862262 | 0.998865404 | 5.393730 | 4.802613 | 5.170551 | 5.145682 | 5.069342 | 5.134969 | 52 | 42 | 17 | 34 |
| 56129 | NM_018940 | PCDHB7 | protocadherin beta 7 | protein_coding | NP_061763.1 | 8692 | 606333 | . | . | 0.098153 | 0.090194 | 0.159788 | 0.132254 | 0.228406 | 0.208489 | 0.358560 | 0.302383 | 1.525365 | 14.707174 | 0.798899894 | 0.762491726 | 0.445766579 | NA | 3.540965 | 4.151760 | 3.856295 | 3.851780 | 3.888049 | 3.873409 | 14 | 12 | 17 | 15 |
| 56130 | NM_00130314 | PCDHB6 | protocadherin beta 6 | protein_coding | NP_001290074.1;NP_061762.2 | 8691 | 606332 | . | . | 3.089481 | 3.230474 | 2.343804 | 3.098608 | 7.189348 | 7.467385 | 5.259429 | 7.084598 | -1.172220 | 299.153047 | 0.215099915 | -1.06575187 | 0.286535822 | 0.998865404 | 8.335466 | 8.104804 | 8.252673 | 8.276449 | 8.108062 | 8.245816 | 364 | 356 | 207 | 293 |
| 56131 | NM_018938 | PCDHB4 | protocadherin beta 4 | protein_coding | NP_061761.1 | 8689 | 606330 | . | . | 0.690121 | 0.709903 | 0.820452 | 0.483664 | 1.605940 | 1.640972 | 1.841070 | 1.105840 | -1.087574 | 80.994697 | 0.379087543 | -0.31948547 | 0.749358411 | 0.998865404 | 6.397726 | 6.279358 | 6.342642 | 6.348749 | 6.386650 | 6.256042 | 96 | 92 | 85 | 54 |
| 56132 | NM_018937 | PCDHB3 | protocadherin beta 3 | protein_coding | NP_061760.2 | 8688 | 606329 | . | . | 0.262991 | 0.279584 | 0.242159 | 0.136467 | 0.611992 | 0.646271 | 0.543397 | 0.312017 | -1.453040 | 24.636798 | 0.637023699 | -0.84623892 | 0.397419475 | NA | 4.865781 | 4.330299 | 4.640729 | 4.644610 | 4.626347 | 4.583410 | 33 | 32 | 22 | 14 |
| 56133 | NM_018936 | PCDHB2 | protocadherin beta 2 | protein_coding | NP_061759.1 | 8687 | 606328 | . | . | 1.155437 | 1.029809 | 0.899932 | 0.841769 | 2.688750 | 2.380450 | 2.019422 | 1.924604 | -1.268392 | 125.658481 | 0.29181704 | -1.17539525 | 0.239836621 | 0.992733415 | 7.134271 | 6.792232 | 7.037981 | 6.989073 | 6.942839 | 6.918406 | 172 | 142 | 100 | 100 |
| 56134 | NM_018899,N | PCDHAC2 | protocadherin alpha subfamily C, 2 | protein_coding | NP_061722.1;NP_114089.1 | 8677 | 606321 | . | . | 0.112578 | 0.137234 | 0.037791 | 0.114091 | 0.261973 | 0.317223 | 0.084802 | 0.260856 | -1.806615 | 16.714846 | 0.852873835 | -1.00048647 | 0.317075141 | NA | 4.431047 | 3.567825 | 4.080669 | 4.094166 | 4.009757 | 4.081206 | 23 | 25 | 3 | 19 |
| 56136 | NM_018904,N | PCDHA13 | protocadherin alpha 13 | protein_coding | NP_061727.1;NP_114071.1 | 8667 | 606319 | . | . | 0.077032 | 0.018337 | 0.037267 | 0.094445 | 0.179257 | 0.042387 | 0.083627 | 0.215938 | 1.374752 | 4.712294 | 1.432671916 | 0.320499751 | 0.748589519 | NA | 1.987791 | 2.448451 | 2.239018 | 2.219087 | 2.226962 | 2.244874 | 7 | 2 | 3 | 7 |
| 56138 | NM_018902,N | PCDHA11 | protocadherin alpha 11 | protein_coding | NP_061725.1;NP_114067.1 | 8665 | 606317 | . | . | 0.134396 | 0.022851 | 0.076318 | 0.076552 | 0.312745 | 0.052821 | 0.171256 | 0.175028 | -1.006392 | 10.523065 | 1.016367209 | -0.00904388 | 0.992784124 | NA | 3.397517 | 3.393446 | 3.422683 | 3.360415 | 3.390994 | 3.392364 | 20 | 4 | 9 | 10 |
| 56140 | NM_018911,N | PCDHA8 | protocadherin alpha 8 | protein_coding | NP_061734.1;NP_114062.1 | 8674 | 606314 | . | . | 0.067312 | 0.008290 | 0.052419 | 0.008982 | 0.156637 | 0.019163 | 0.117627 | 0.020536 | -1.093725 | 2.950990 | 1.832196642 | -0.07054345 | 0.943761118 | NA | 1.617627 | 1.502474 | 1.568892 | 1.548531 | 1.565424 | 1.549528 | 6 | 1 | 4 | 1 |
| 56141 | NM_018910,N | PCDHA7 | protocadherin alpha 7 | protein_coding | NP_061733.1;NP_114040.1 | 8673 | 606313 | . | . | 0.044836 | 0.015494 | 0.063194 | 0.089140 | 0.104335 | 0.035814 | 0.141805 | 0.203808 | 2.475486 | 4.635950 | 1.430217578 | 0.914344825 | 0.360535684 | NA | 1.413393 | 2.724273 | 2.189296 | 2.181675 | 2.199790 | 2.207625 | 4 | 2 | 5 | 7 |
| 56144 | NM_018907,N | PCDHA4 | protocadherin alpha 4 | protein_coding | NP_061730.1;NP_113688.1 | 8670 | 606310 | . | . | 0.077347 | 0.051239 | 0.061854 | 0.038630 | 0.179990 | 0.118440 | 0.138799 | 0.088324 | -1.307716 | 19.290238 | 0.718768829 | -0.5384891 | 0.590239429 | NA | 4.448287 | 4.066070 | 4.299854 | 4.263326 | 4.276386 | 4.243149 | 30 | 19 | 18 | 12 |
| 56145 | NM_018906,N | PCDHA3 | protocadherin alpha 3 | protein_coding | NP_061729.1;NP_113685.1 | 8669 | 606309 | . | . | 0.484439 | 0.486271 | 0.363669 | 0.613220 | 1.127309 | 1.124038 | 0.816063 | 1.402054 | -1.004950 | 37.972674 | 0.518821937 | -0.01373112 | 0.989044498 | 0.999197585 | 5.251870 | 5.241892 | 5.246805 | 5.245940 | 5.210496 | 5.279653 | 44 | 41 | 25 | 44 |
| 56146 | NM_018905,N | PCDHA2 | protocadherin alpha 2 | protein_coding | NP_061728.1;NP_113683.1;NP_113684.1 | 8668 | 606308 | . | . | 0.381997 | 0.379148 | 0.140123 | 0.346568 | 0.888922 | 0.876419 | 0.314433 | 0.792387 | -1.554157 | 27.941716 | 0.617631418 | -1.02995375 | 0.303031718 | 0.998865404 | 5.089593 | 4.448451 | 4.835115 | 4.829172 | 4.749710 | 4.814637 | 40 | 36 | 12 | 28 |
| 56147 | NM_018900,N | PCDHA1 | protocadherin alpha 1 | protein_coding | NP_061723.1;NP_113598.1;NP_113599.1 | 8663 | 606307 | . | . | 0.194986 | 0.092136 | 0.100748 | 0.110521 | 0.453742 | 0.212976 | 0.226075 | 0.252692 | -1.369981 | 11.127335 | 0.937678097 | -0.48434047 | 0.628144262 | NA | 3.684232 | 3.232645 | 3.508351 | 3.467638 | 3.472039 | 3.473612 | 20 | 9 | 8 | 9 |
| 56154 | NM_00135016 | TEX15 | testis expressed 15, meiosis and synapsis associated | protein_coding | NP_001337091.1 | 11738 | 605795 | . | . | 0.880723 | 0.780032 | 0.851772 | 0.588599 | 2.049478 | 1.803079 | 1.911352 | 1.345761 | -1.169040 | 277.834602 | 0.235004101 | -0.95881157 | 0.337653678 | 0.998865404 | 8.225437 | 8.002093 | 8.188018 | 8.116734 | 8.161433 | 7.973325 | 367 | 302 | 265 | 195 |
| 56155 | NM_00120145 | TEX14 | testis expressed 14, intercellular bridge forming factor | protein_coding | NP_001188386.1;NP_112562.3;NP_938207.2;XP_011523331.1;XP_01688 | 11737 | 605792 | . | . | 0.163325 | 0.127372 | 0.183019 | 0.189417 | 0.380063 | 0.294427 | 0.410690 | 0.433080 | 1.261240 | 26.264421 | 0.603429684 | 0.554899547 | 0.578963372 | NA | 4.537347 | 4.873224 | 4.709485 | 4.685417 | 4.720198 | 4.724874 | 30 | 22 | 25 | 28 |
| 56160 | NM_138704 | NSMCE3 | NSE3 homolog, SMC5-SMC6 complex component | protein_coding | NP_619649.1 | 7677 | 608243 | . | . | 4.244939 | 3.870672 | 3.972278 | 4.238739 | 9.878148 | 8.947232 | 8.913676 | 9.691373 | -1.001930 | 615.304708 | 0.159737979 | -0.01741012 | 0.986109438 | 0.999197585 | 9.266512 | 9.263802 | 9.298890 | 9.230974 | 9.246016 | 9.282120 | 745 | 631 | 520 | 592 |

| | | | | | | | |
|---|---|---|---|---|---|---|---|
| 56171 | NM_018897,XI | DNAH7 | dynein axonemal heavy chain 7 | protein_coding | NP_061720.2;XP_011509790.1;XP_011509791.1;XP_011509792.1;XP_01 | 18661 | 610061 |
| 56172 | NM_054027,XI | ANKH | ANKH inorganic pyrophosphate transport regulator | protein_coding | NP_473368.1;XP_011512369.1;XP_016865133.1 | 15492 | 605145 |
| 56180 | NM_00130618 | MOSPD1 | motile sperm domain containing 1 | protein_coding | NP_001293117.1;NP_062456.1;XP_011529668.1;XP_016885142.1;XP_01 | 25235 | 300674 |
| 56181 | NM_00109962 | MTFR1L | mitochondrial fission regulator 1 like | protein_coding | NP_001093095.1;NP_001093096.1;NP_001093097.1;NP_062457.3 | 28836 | . |
| 56203 | NM_00130441 | LMOD3 | leiomodin 3 | protein_coding | NP_001291347.1;NP_938012.2 | 6649 | 616112 |
| 56204 | NM_00128649 | FAM214A | family with sequence similarity 214 member A | protein_coding | NP_001273424.1;NP_062546.2;XP_005254604.1;XP_005254605.1;XP_01 | 25609 | . |
| 56241 | NM_019601 | SUSD2 | sushi domain containing 2 | protein_coding | NP_062547.1 | 30667 | 615825 |
| 56242 | NM_00133113 | ZNF253 | zinc finger protein 253 | protein_coding | NP_001318062.1;NP_001318063.1;NP_066385.2 | 13497 | 606954 |
| 56243 | NM_00109850 | KIAA1217 | KIAA1217 | protein_coding | NP_001091970.1;NP_001269696.1;NP_001269697.1;NP_001269698.1;NI | 25428 | 617367 |
| 56252 | NM_019589,XI | YLPM1 | YLP motif containing 1 | protein_coding | NP_062535.2;XP_005267917.1;XP_011535268.1;XP_011535269.1 | 17798 | . |
| 56254 | NM_019592,XI | RNF20 | ring finger protein 20 | protein_coding | NP_062538.5;XP_011517164.1 | 10062 | 607699 |
| 56255 | NM_021156 | TMX4 | thioredoxin related transmembrane protein 4 | protein_coding | NP_066979.2 | 25237 | 616766 |
| 56256 | NM_00135417 | SERTAD4 | SERTA domain containing 4 | protein_coding | NP_001341102.1;NP_001362357.1;NP_062551.1 | 25236 | . |
| 56257 | NM_00119499 | MEPCE | methylphosphate capping enzyme | protein_coding | NP_001181919.1;NP_001181920.1;NP_001181921.1;NP_001350415.1;NI | 20247 | 611478 |
| 56259 | NM_00128149 | CTNNBL1 | catenin beta like 1 | protein_coding | NP_001268424.1;NP_110517.2;XP_011527219.1;XP_024307715.1 | 15879 | 611537 |
| 56260 | NR_161216,NF | C8orf44 | chromosome 8 open reading frame 44 | lncRNA | . | 25646 | . |
| 56261 | NM_019593,XI | GPCPD1 | glycerophosphocholine phosphodiesterase 1 | protein_coding | NP_062539.1;XP_005260815.1;XP_005260818.1;XP_006723659.1;XP_02 | 26957 | 614124 |
| 56262 | NM_00112724 | LRRC8A | leucine rich repeat containing 8 VRAC subunit A | protein_coding | NP_001120716.1;NP_001120717.1;NP_062540.2;XP_005252152.1;XP_00 | 19027 | 608360 |
| 56267 | NM_00100866 | KYAT3 | kynurenine aminotransferase 3 | protein_coding | NP_001008661.1;NP_001008662.1;NP_001336376.1;NP_001336377.1;NI | 33238 | 610656 |
| 56270 | NM_019613,XI | WDR45B | WD repeat domain 45B | protein_coding | NP_062559.2;XP_005256434.1;XP_016880351.1;XP_016880352.1;XP_02 | 25072 | 609226 |
| 56271 | NM_00108042 | BEX4 | brain expressed X-linked 4 | protein_coding | NP_001073894.1;NP_001121160.1 | 25475 | 300692 |
| 56288 | NM_00118478 | PARD3 | par-3 family cell polarity regulator | protein_coding | NP_001171714.1;NP_001171715.1;NP_001171716.1;NP_001171717.1;NI | 16051 | 606745 |
| 56311 | NM_019644 | ANKRD7 | ankyrin repeat domain 7 | protein_coding | NP_062618.2 | 18588 | 610731 |
| 56339 | NM_019852,XI | METTL3 | methyltransferase like 3 | protein_coding | NP_062826.2;XP_006720269.1;XP_011535270.1 | 17563 | 612472 |
| 56342 | NM_00134613 | PPAN | peter pan homolog | protein_coding | NP_001333068.1;NP_001333070.1;NP_064615.3 | 9227 | 607793 |
| 56413 | NM_00116469 | LTB4R2 | leukotriene B4 receptor 2 | protein_coding | NP_001158164.1;NP_062813.2 | 19260 | 605773 |
| 56474 | NM_00114400 | CTPS2 | CTP synthase 2 | protein_coding | NP_001137474.1;NP_062831.3;NP_787055.1;XP_005274619.1;XP_00672 | 2520 | 300380 |
| 56477 | NM_00130187 | CCL28 | C-C motif chemokine ligand 28 | protein_coding | NP_001288802.1;NP_001288803.1;NP_001288804.1;NP_683513.1 | 17700 | 605240 |
| 56478 | NM_00116450 | EIF4ENIF1 | eukaryotic translation initiation factor 4E nuclear import factor 1 | protein_coding | NP_001157973.1;NP_001157974.1;NP_062817.2;XP_005261743.1;XP_00 | 16687 | 607445 |
| 56479 | NM_00116013 | KCNQ5 | potassium voltage-gated channel subfamily Q member 5 | protein_coding | NP_001153602.1;NP_001153604.1;NP_001153605.1;NP_001153606.1;NI | 6299 | 607357 |
| 56521 | NM_021800,N | DNAJC12 | DnaJ heat shock protein family (Hsp40) member C12 | protein_coding | NP_068572.1;NP_957714.1;XP_016871920.1;XP_016871921.1 | 28908 | 606060 |
| 56548 | NM_019886 | CHST7 | carbohydrate sulfotransferase 7 | protein_coding | NP_063939.2 | 13817 | 300375 |
| 56603 | NM_00127774 | CYP26B1 | cytochrome P450 family 26 subfamily B member 1 | protein_coding | NP_001264671.1;NP_063938.1;XP_005264490.1;XP_011531290.1 | 20581 | 605207 |
| 56604 | gene-TUBB7P | TUBB7P | tubulin beta 7 pseudogene | pseudogene | . | 12413 | . |
| 56605 | NM_019891,XI | ERO1B | endoplasmic reticulum oxidoreductase 1 beta | protein_coding | NP_063944.3;XP_016857323.1;XP_024304217.1 | 14355 | 615437 |
| 56616 | NM_00127830 | DIABLO | diablo IAP-binding mitochondrial protein | protein_coding | NP_001265231.1;NP_001265232.1;NP_001265233.1;NP_001265271.1;NI | 21528 | 605219 |
| 56623 | NM_00131850 | INPP5E | inositol polyphosphate-5-phosphatase E | protein_coding | NP_001305431.1;NP_063945.2;XP_016870415.1 | 21474 | 613037 |
| 56624 | NM_00114397 | ASAH2 | N-acylsphingosine amidohydrolase 2 | protein_coding | NP_001137446.1;NP_063946.2;XP_011538272.1;XP_011538273.1;XP_01 | 18860 | 611202 |
| 56647 | NM_016567,N | BCCIP | BRCA2 and CDKN1A interacting protein | protein_coding | NP_057651.1;NP_510868.1;NP_510869.1 | 978 | 611883 |
| 56648 | NM_020390 | EIF5A2 | eukaryotic translation initiation factor 5A2 | protein_coding | NP_065123.1 | 3301 | 605782 |
| 56650 | NM_00104018 | CLDND1 | claudin domain containing 1 | protein_coding | NP_001035271.1;NP_001035272.1;NP_001035273.1;NP_001035289.1;NI | 1322 | . |
| 56652 | NM_00116381 | TWNK | twinkle mtDNA helicase | protein_coding | NP_001157284.1;NP_001157285.1;NP_001157286.1;NP_001355204.1;NI | 1160 | 606075 |
| 56654 | NM_015392 | NPDC1 | neural proliferation, differentiation and control 1 | protein_coding | NP_056207.3 | 7899 | 605798 |
| 56655 | NM_019896 | POLE4 | DNA polymerase epsilon 4, accessory subunit | protein_coding | NP_063949.2 | 18755 | 607269 |
| 56658 | NM_00136952 | TRIM39 | tripartite motif containing 39 | protein_coding | NP_001356450.1;NP_001356451.1;NP_001356452.1;NP_067076.2;NP_7 | 10065 | 605700 |
| 56659 | NM_022054 | KCNK13 | potassium two pore domain channel subfamily K member 13 | protein_coding | NP_071337.2 | 6275 | 607367 |
| 56662 | NR_026705 | VTRNA1-3 | vault RNA 1-3 | vault_RNA | . | 12656 | 612697 |
| 56666 | NM_00116030 | PANX2 | pannexin 2 | protein_coding | NP_001153772.1;NP_443071.2 | 8600 | 608421 |
| 56672 | NM_00120664 | AKIP1 | A-kinase interacting protein 1 | protein_coding | NP_001193575.1;NP_001193576.1;NP_001193577.1;NP_065693.2;XP_01 | 1170 | 609191 |
| 56674 | NM_00128609 | TMEM9B | TMEM9 domain family member B | protein_coding | NP_001273023.1;NP_001273024.1;NP_065695.1 | 1168 | . |
| 56675 | NM_020645,XI | NRIP3 | nuclear receptor interacting protein 3 | protein_coding | NP_065696.1;XP_024304376.1;XP_024304377.1 | 1167 | 613125 |
| 56681 | NM_00114264 | SAR1A | secretion associated Ras related GTPase 1A | protein_coding | NP_001136120.1;NP_064535.1 | 10534 | 607691 |
| 56683 | NM_00135033 | CFAP298 | cilia and flagella associated protein 298 | protein_coding | NP_001337263.1;NP_001337264.1;NP_001337265.1;NP_001337266.1;NI | 1301 | 615494 |
| 56704 | NM_00131783 | JPH1 | junctophilin 1 | protein_coding | NP_001304759.1;NP_001349979.1;NP_001349980.1;NP_065698.1 | 14201 | 605266 |
| 56731 | NM_020062 | SLC2A4RG | SLC2A4 regulator | protein_coding | NP_064446.2 | 15930 | 609493 |
| 56776 | NM_00130542 | FMN2 | formin 2 | protein_coding | NP_001292353.1;NP_001335023.1;NP_064450.3;XP_011542539.1;XP_01 | 14074 | 606373 |
| 56829 | NM_00136349 | ZC3HAV1 | zinc finger CCCH-type containing, antiviral 1 | protein_coding | NP_001350420.1;NP_064504.2;NP_078901.3 | 23721 | 607312 |
| 56834 | NM_00117088 | GPR137 | G protein-coupled receptor 137 | protein_coding | NP_001164351.1;NP_001164352.1;NP_001170829.1;NP_001365005.1;NI | 24300 | . |
| 56848 | NM_00120415 | SPHK2 | sphingosine kinase 2 | protein_coding | NP_001191087.1;NP_001191088.1;NP_001191089.1;NP_001230805.1;NI | 18859 | 607092 |
| 56850 | NM_020137 | GRIPAP1 | GRIP1 associated protein 1 | protein_coding | NP_064522.4 | 18706 | 300408 |
| 56851 | NM_020154 | EMC7 | ER membrane protein complex subunit 7 | protein_coding | NP_064539.1 | 24301 | . |
| 56852 | NM_020165,XI | RAD18 | RAD18 E3 ubiquitin protein ligase | protein_coding | NP_064550.3;XP_016862362.1 | 18278 | 605256 |
| 56882 | NM_00103870 | CDC42SE1 | CDC42 small effector 1 | protein_coding | NP_001033796.1;NP_064624.1;XP_016857336.1 | 17719 | . |
| 56884 | NM_00112842 | FSTL5 | follistatin like 5 | protein_coding | NP_001121899.1;NP_001121900.1;NP_064501.2;XP_011530428.1 | 21386 | . |
| 56886 | NM_020120,N | UGGT1 | UDP-glucose glycoprotein glucosyltransferase 1 | protein_coding | NP_064505.1;XP_006712697.1;XP_006712698.1;XP_006712699.1;XP_01 | 15663 | 605897 |
| 56888 | NM_020122,XI | KCMF1 | potassium channel modulatory factor 1 | protein_coding | NP_064507.3;XP_006712115.1;XP_011531292.1;XP_016860000.1 | 20589 | 614719 |
| 56889 | NM_020123,XI | TM9SF3 | transmembrane 9 superfamily member 3 | protein_coding | NP_064508.3;XP_011538278.1;XP_011538279.1 | 21529 | 616872 |
| 56890 | NM_00120502 | MDM1 | Mdm1 nuclear protein | protein_coding | NP_001191957.1;NP_001191958.1;NP_001341898.1;NP_001341899.1;NI | 29917 | 613813 |
| 56893 | NM_00130434 | UBQLN4 | ubiquilin 4 | protein_coding | NP_001291271.1;NP_064516.2;XP_005245405.1;XP_024304237.1 | 1237 | 605440 |
| 56894 | NM_00103755 | AGPAT3 | 1-acylglycerol-3-phosphate O-acyltransferase 3 | protein_coding | NP_001032642.1;NP_001356807.1;NP_001356809.1;NP_001356810.1;NI | 326 | 614794 |
| 56895 | NM_020133,XI | AGPAT4 | 1-acylglycerol-3-phosphate O-acyltransferase 4 | protein_coding | NP_064518.1;XP_005267110.1;XP_006715577.1;XP_016866548.1 | 20885 | 614795 |
| 56896 | NM_00125372 | DPYSL5 | dihydropyrimidinase like 5 | protein_coding | NP_001240652.1;NP_001240653.1;NP_064519.2;XP_024308775.1 | 20637 | 608383 |
| 56897 | NM_020135,N | WRNIP1 | WRN helicase interacting protein 1 | protein_coding | NP_064520.2;NP_569079.1;XP_005249289.1;XP_016866549.1 | 20876 | 608196 |
| 56898 | NM_020139,XI | BDH2 | 3-hydroxybutyrate dehydrogenase 2 | protein_coding | NP_064524.3;XP_005263197.1;XP_006714337.1;XP_011530429.1;XP_01 | 32389 | . |
| 56899 | NM_00120406 | ANKS1B | ankyrin repeat and sterile alpha motif domain containing 1B | protein_coding | NP_001190994.1;NP_001190995.1;NP_001190996.1;NP_001190997.1;NI | 24600 | 607815 |
| 56900 | NM_00132224 | TMEM167B | transmembrane protein 167B | protein_coding | NP_001309177.1;NP_064526.1 | 30187 | . |
| 56902 | NM_00132991 | PNO1 | partner of NOB1 homolog | protein_coding | NP_001316845.1;NP_001316846.1;NP_064528.1 | 32790 | 618710 |
| 56904 | NM_00128704 | SH3GLB2 | SH3 domain containing GRB2 like, endophilin B2 | protein_coding | NP_001273974.1;NP_001273975.1;NP_001356842.1;NP_001356843.1;NI | 10834 | 609288 |
| 56905 | NM_015492 | C15orf39 | chromosome 15 open reading frame 39 | protein_coding | NP_056307.3 | 24497 | . |
| 56906 | NM_020147 | THAP10 | THAP domain containing 10 | protein_coding | NP_064532.1 | 23193 | 612538 |
| 56907 | NM_00112862 | SPIRE1 | spire type actin nucleation factor 1 | protein_coding | NP_001122098.1;NP_001122099.1;NP_064533.3;XP_005258179.1;XP_01 | 30622 | 609216 |
| 56910 | NM_020151,XI | STARD7 | StAR related lipid transfer domain containing 7 | protein_coding | NP_064536.2;XP_011509801.1 | 18063 | 616712 |
| 56912 | NM_00116861 | IFT46 | intraflagellar transport 46 | protein_coding | NP_001162089.1;NP_064538.3;XP_011541207.1;XP_011541208.1;XP_01 | 26146 | . |
| 56913 | NM_020156,XI | C1GALT1 | core 1 synthase, glycoprotein-N-acetylgalactosamine 3-beta-galac | protein_coding | NP_064541.1;XP_011513755.1;XP_011513757.1;XP_011513758.1;XP_01 | 24337 | 610555 |
| 56915 | NM_020158 | EXOSC5 | exosome component 5 | protein_coding | NP_064543.3 | 24662 | 606492 |
| 56916 | NM_00112842 | SMARCAD1 | SWI/SNF-related, matrix-associated actin-dependent regulator of | protein_coding | NP_001121901.1;NP_001121902.1;NP_001241878.1;NP_001362784.1;NI | 18398 | 612761 |
| 56917 | NM_00100981 | MEIS3 | Meis homeobox 3 | protein_coding | NP_001009813.1;NP_001287988.1;NP_001333077.1;NP_064545.1;XP_01 | 29537 | . |
| 56919 | NM_00119969 | DHX33 | DEAH-box helicase 33 | protein_coding | NP_001186628.1;NP_064547.2;XP_016880366.1 | 16718 | 614405 |
| 56920 | NM_020163,XI | SEMA3G | semaphorin 3G | protein_coding | NP_064548.1;XP_024309410.1 | 30400 | . |
| 56922 | NM_00129327 | MCCC1 | methylcrotonoyl-CoA carboxylase 1 | protein_coding | NP_001280202.1;NP_001350809.1;NP_064551.3;XP_011511294.1 | 6936 | 609010 |
| 56925 | NM_020169,XI | LXN | latexin | protein_coding | NP_064554.3;XP_016862365.1 | 13347 | 609305 |
| 56926 | NM_00132146 | NCLN | nicalin | protein_coding | NP_001308392.1;NP_064555.2 | 26923 | 609156 |
| 56927 | NM_00108045 | GPR108 | G protein-coupled receptor 108 | protein_coding | NP_001073921.1;NP_064556.1;XP_011526439.1;XP_016882501.1;XP_01 | 17829 | 618491 |
| 56928 | NM_00107723 | SPPL2B | signal peptide peptidase like 2B | protein_coding | NP_001070706.1;NP_694533.1;XP_011526440.1;XP_011526441.1;XP_01 | 30627 | 608239 |
| 56929 | NM_020177,XI | FEM1C | fem-1 homolog C | protein_coding | NP_064562.1;XP_005272092.1;XP_016865136.1 | 16933 | 608767 |
| 56931 | NM_00116161 | DUS3L | dihydrouridine synthase 3 like | protein_coding | NP_001155091.1;NP_064560.2;XP_016882509.1 | 26920 | . |
| 56935 | NM_020179,XI | SMCO4 | single-pass membrane protein with coiled-coil domains 4 | protein_coding | NP_064564.1;XP_011541209.1;XP_011541211.1;XP_011541212.1;XP_01 | 24810 | 609477 |
| 56937 | NM_00125597 | PMEPA1 | prostate transmembrane protein, androgen induced 1 | protein_coding | NP_001242905.1;NP_064567.2;NP_954638.1;NP_954639.1;NP_954640.1 | 14107 | 606564 |
| 56938 | NM_00124800 | ARNTL2 | aryl hydrocarbon receptor nuclear translocator like 2 | protein_coding | NP_001234931.1;NP_001234932.1;NP_001234933.1;NP_001234934.1;NI | 18984 | 614517 |
| 56940 | NM_00128655 | DUSP22 | dual specificity phosphatase 22 | protein_coding | NP_001273484.1;NP_064570.1;XP_011513059.1;XP_016866550.1;XP_01 | 16077 | 616778 |
| 56941 | NM_00100610 | HMCES | 5-hydroxymethylcytosine binding, ES cell specific | protein_coding | NP_001006109.1;NP_001350810.1;NP_001357272.1;NP_001357273.1;NI | 24446 | 618288 |
| 56942 | NM_00135196 | CMC2 | C-X9-C motif containing 2 | protein_coding | NP_001338896.1;NP_001338897.1;NP_001338899.1;NP_001338902.1;NI | 24447 | . |
| 56943 | NM_00119355 | ENY2 | ENY2 transcription and export complex 2 subunit | protein_coding | NP_001180486.1;NP_064574.1 | 24449 | . |
| 56944 | NM_00128635 | OLFML3 | olfactomedin like 3 | protein_coding | NP_001273281.1;NP_001273282.1;NP_064575.1;XP_016857337.1 | 24956 | 610088 |
| 56945 | NM_00136385 | MRPS22 | mitochondrial ribosomal protein S22 | protein_coding | NP_001350786.1;NP_001350822.1;NP_064576.1 | 14508 | 605810 |
| 56946 | NM_00130094 | EMSY | EMSY transcriptional repressor, BRCA2 interacting | protein_coding | NP_001287871.1;NP_001287872.1;NP_001287873.1;NP_064578.2;XP_00 | 18071 | 608574 |
| 56947 | NM_00127706 | MFF | mitochondrial fission factor | protein_coding | NP_001263990.1;NP_001263991.1;NP_001263992.1;NP_001263993.1;NI | 24858 | 614785 |
| 56948 | NM_00129029 | SDR39U1 | short chain dehydrogenase/reductase family 39U member 1 | protein_coding | NP_001277221.1;NP_001277222.1;NP_001295004.1;NP_064580.2;XP_01 | 20275 | 616162 |
| 56949 | NM_020196 | XAB2 | XPA binding protein 2 | protein_coding | NP_064581.2 | 14089 | 610850 |
| 56950 | NM_020197,XI | SMYD2 | SET and MYND domain containing 2 | protein_coding | NP_064582.2 | 20982 | 610663 |
| 56951 | NM_020199 | C5orf15 | chromosome 5 open reading frame 15 | protein_coding | NP_064584.1 | 20656 | . |
| 56953 | NM_020201,XI | NT5M | 5',3'-nucleotidase, mitochondrial | protein_coding | NP_064586.1;XP_005256788.1;XP_005256789.1;XP_005256790.1;XP_00 | 15769 | 605292 |
| 56954 | NM_020202 | NIT2 | nitrilase family member 2 | protein_coding | NP_064587.1 | 29878 | 616769 |
| 56957 | NM_020205,XI | OTUD7B | OTU deubiquitinase 7B | protein_coding | NP_064590.2;XP_011508084.1;XP_011508086.1;XP_011508087.1;XP_01 | 16683 | 611748 |
| 56963 | NM_00116628 | RGMA | repulsive guidance molecule BMP co-receptor a | protein_coding | NP_001159755.1;NP_001159758.1;NP_001159759.1;NP_001159760.1;NI | 30308 | 607362 |
| 56964 | NM_00128439 | WDR93 | WD repeat domain 93 | protein_coding | NP_001271324.1;NP_001271325.1;NP_064597.1;XP_006720677.1;XP_00 | 26924 | . |
| 56965 | NM_00132351 | PARP6 | poly(ADP-ribose) polymerase family member 6 | protein_coding | NP_001310444.1;NP_001310445.1;NP_001310448.1;NP_001310450.1;NI | 26921 | . |
| 56967 | NM_00125250 | C14orf132 | chromosome 14 open reading frame 132 | protein_coding | NP_001239436.1;NP_001269392.1;NP_001269393.1;NP_001276068.1;XF | 20346 | . |
| 56970 | NM_00138230 | ATXN7L3 | ataxin 7 like 3 | protein_coding | NP_001369237.1;NP_001369238.1;NP_001369239.1;NP_001369240.1;NI | 25416 | . |
| 56971 | NM_00112789 | CEACAM19 | CEA cell adhesion molecule 19 | protein_coding | NP_001121365.1;NP_064604.2 | 31951 | 606691 |
| 56975 | NM_020223,XI | FAM20C | FAM20C golgi associated secretory pathway kinase | protein_coding | NP_064608.2;XP_016867939.1;XP_016867940.1;XP_016867941.1;XP_01 | 22140 | 611061 |
| 56977 | NM_020225,N | STOX2 | storkhead box 2 | protein_coding | NP_064610.1;XP_011530431.1;XP_011530432.1;XP_011530433.1;XP_01 | 25450 | 617359 |
| 56978 | NM_00109940 | PRDM8 | PR/SET domain 8 | protein_coding | NP_001092873.1;NP_064611.3;XP_005263201.1;XP_005263202.1;XP_00 | 13993 | 616639 |
| 56980 | NM_00136789 | PRDM10 | PR/SET domain 10 | protein_coding | NP_001354819.1;NP_001354820.1;NP_001354821.1;NP_001354822.1;NI | 13995 | 618319 |
| 56981 | NM_00125669 | PRDM11 | PR/SET domain 11 | protein_coding | NP_001243624.1;NP_001243625.1;NP_001346562.1;XP_011518524.1;XP | 13996 | 616347 |
| 56983 | NM_152305,N | POGLUT1 | protein O-glucosyltransferase 1 | protein_coding | NP_689518.1;XP_006713768.1;XP_016862367.1;XP_016862368.1;XP_02 | 22954 | 615618 |
| 56984 | NM_020232,N | PSMG2 | proteasome assembly chaperone 2 | protein_coding | NP_064617.2;NP_671692.1 | 24929 | 609702 |
| 56985 | NM_020233 | ADPRM | ADP-ribose/CDP-alcohol diphosphatase, manganese dependent | protein_coding | NP_064618.3 | 30925 | . |
| 56986 | NM_00114495 | DTWD1 | DTW domain containing 1 | protein_coding | NP_001138427.1;NP_064619.2;XP_011520117.1;XP_016877908.1;XP_01 | 30926 | . |
| 56987 | NM_00114256 | BBX | BBX high mobility group box domain containing | protein_coding | NP_001136040.1;NP_001263215.1;NP_064620.2;XP_005247699.1;XP_00 | 14422 | . |
| 56990 | NM_00103870 | CDC42SE2 | CDC42 small effector 2 | protein_coding | NP_001033791.1;NP_001362562.1;NP_001362563.1;NP_001362564.1;NI | 18547 | . |
| 56992 | NM_020242,XI | KIF15 | kinesin family member 15 | protein_coding | NP_064627.1;XP_006713327.1;XP_011532266.1;XP_016862373.1;XP_01 | 17273 | 617569 |
| 56993 | NM_020243 | TOMM22 | translocase of outer mitochondrial membrane 22 | protein_coding | NP_064628.1 | 18002 | 607046 |
| 56994 | NM_020244,XI | CHPT1 | choline phosphotransferase 1 | protein_coding | NP_064629.2;XP_011536876.1;XP_011536877.1 | 17852 | 616747 |
| 56995 | NM_00100746 | TULP4 | TUB like protein 4 | protein_coding | NP_001007467.1;NP_064630.2;XP_011534248.1;XP_016866559.1 | 15530 | . |
| 56996 | NM_00126781 | SLC12A9 | solute carrier family 12 member 9 | protein_coding | NP_001254741.1;NP_001254743.1;NP_001350422.1;NP_001350423.1;NI | 17435 | 616861 |
| 56997 | NM_020247,XI | COQ8A | coenzyme Q8A | protein_coding | NP_064632.2;XP_005273258.1;XP_011542540.1;XP_011542541.1;XP_01 | 16812 | 606980 |
| 56998 | NM_00101232 | CTNNBIP1 | catenin beta interacting protein 1 | protein_coding | NP_001012329.1;NP_064633.1;XP_016857342.1;XP_016857343.1;XP_01 | 16913 | 607758 |
| 56999 | NM_00131878 | ADAMTS9 | ADAM metallopeptidase with thrombospondin type 1 motif 9 | protein_coding | NP_001305710.1;NP_891550.1 | 13202 | 605421 |
| 57000 | NR_103560 | GSN-AS1 | GSN antisense RNA 1 | lncRNA | . | 23372 | . |
| 57001 | NM_020186 | SDHAF3 | succinate dehydrogenase complex assembly factor 3 | protein_coding | NP_064571.1 | 21752 | 615773 |
| 57002 | NM_00128244 | YAE1 | YAE1 maturation factor of ABCE1 | protein_coding | NP_001269375.1;NP_064577.1 | 24857 | . |
| 57003 | NM_020198,XI | CCDC47 | coiled-coil domain containing 47 | protein_coding | NP_064583.2;XP_005257584.1 | 24856 | 618260 |
| 57016 | NM_020299,XI | AKR1B10 | aldo-keto reductase family 1 member B10 | protein_coding | NP_064695.3;XP_011514718.1;XP_011514719.1 | 382 | 604707 |
| 57017 | NM_020312 | COQ9 | coenzyme Q9 | protein_coding | NP_064708.1 | 25302 | 612837 |
| 57018 | NM_00130818 | CCNL1 | cyclin L1 | protein_coding | NP_001295114.1;NP_064703.1;XP_005247705.1;XP_006713773.1;XP_01 | 20569 | 613384 |
| 57019 | NM_00130834 | CIAPIN1 | cytokine induced apoptosis inhibitor 1 | protein_coding | NP_001295276.1;NP_001295287.1;NP_064709.2 | 28050 | 608943 |
| 57020 | NM_00130074 | VPS35L | VPS35 endosomal protein sorting factor like | protein_coding | NP_001287672.2;NP_001352222.1;NP_001352223.1;NP_001352224.1;NI | 24641 | . |
| 57026 | NM_020315 | PDXP | pyridoxal phosphatase | protein_coding | NP_064711.1 | 30259 | 609246 |
| 57030 | NM_020309 | SLC17A7 | solute carrier family 17 member 7 | protein_coding | NP_064705.1 | 16704 | 605208 |
| 57035 | NM_00132177 | RSRP1 | arginine and serine rich protein 1 | protein_coding | NP_001308701.1;NP_064713.3;XP_011540099.1 | 25234 | . |
| 57037 | NM_020319,XI | ANKMY2 | ankyrin repeat and MYND domain containing 2 | protein_coding | NP_064715.1;XP_011513760.1 | 25370 | . |
| 57038 | NM_00131878 | RARS2 | arginyl-tRNA synthetase 2, mitochondrial | protein_coding | NP_001305714.1;NP_001337434.1;NP_001337435.1;NP_001337436.1;NI | 21406 | 611524 |
| 57045 | NM_020648 | TWSG1 | twisted gastrulation BMP signaling modulator 1 | protein_coding | NP_065699.1 | 12429 | 605049 |
| 57048 | NM_00120157 | PLSCR3 | phospholipid scramblase 3 | protein_coding | NP_001188505.1;NP_001356336.1;NP_001356349.1;NP_001356350.1;NI | 16495 | 607611 |
| 57050 | NM_020368 | UTP3 | UTP3 small subunit processome component | protein_coding | NP_065101.1 | 24477 | 611614 |
| 57053 | NM_00130303 | CHRNA10 | cholinergic receptor nicotinic alpha 10 subunit | protein_coding | NP_001289963.1;NP_001289964.1;NP_065135.2;XP_011518536.1 | 13800 | 606372 |
| 57054 | NM_020364 | DAZ3 | deleted in azoospermia 3 | protein_coding | NP_065097.2 | 15965 | 400027 |
| 57055 | NM_00100578 | DAZ2 | deleted in azoospermia 2 | protein_coding | NP_001005785.1;NP_001005786.2;NP_065096.2 | 15964 | 400026 |
| 57060 | NM_00117410 | PCBP4 | poly(rC) binding protein 4 | protein_coding | NP_001167571.1;NP_001350814.1;NP_065151.2;NP_127501.1;NP_1275 | 8652 | 608503 |
| 57061 | NR_002768 | HYMAI | hydatidiform mole associated and imprinted | lncRNA | . | 5326 | 606546 |
| 57062 | NM_020414 | DDX24 | DEAD-box helicase 24 | protein_coding | NP_065147.1 | 13266 | 606181 |

| | | | | | | | | | | | | | | | | | | | | | | | | | | | | | | | | | |
|---|---|---|---|---|---|---|---|---|---|---|---|---|---|---|---|---|---|---|---|---|---|---|---|---|---|---|---|---|---|---|---|---|---|
| 57082 | NM_144508,N | KNL1 | kinetochore scaffold 1 | protein_coding | NP_653091.3;NP_733468.3;XP_016877921.1 | 24054 | 609173 | . | . | 2.530428 | 2.603548 | 2.551245 | 1.779593 | 5.888411 | 6.018218 | 5.724919 | 4.068828 | -1.202370 | 684.514556 | 0.184398901 | -1.44187694 | 0.149337093 | 0.910598055 | 9.545294 | 9.280443 | 9.466655 | 9.481852 | 9.462858 | 9.219629 | 851 | 815 | 640 | 476 |
| 57085 | NM_00104019 | AGTRAP | angiotensin II receptor associated protein | protein_coding | NP_001035284.1;NP_001035285.1;NP_001035286.1;NP_001035287.1;NI | 13539 | 608729 | . | . | 55.370263 | 53.409262 | 51.108512 | 56.990971 | 128.848889 | 123.457909 | 114.686019 | 130.303091 | -1.019416 | 1928.799550 | 0.123125521 | -0.2253225 | 0.821728396 | 0.998865404 | 10.927360 | 10.899480 | 10.938606 | 10.902709 | 10.867332 | 10.942372 | 2293 | 2054 | 1582 | 1876 |
| 57088 | NM_00112830 | PLSCR4 | phospholipid scramblase 4 | protein_coding | NP_001121776.1;NP_001121777.1;NP_001121778.1;NP_001170775.1;NI | 16497 | 607612 | . | . | 2.472409 | 2.068718 | 2.350392 | 2.187040 | 5.753397 | 4.781934 | 5.274212 | 5.000411 | -1.021473 | 237.549778 | 0.229567893 | -0.13351367 | 0.893787157 | 0.998865404 | 7.906704 | 7.877319 | 7.944429 | 7.844817 | 7.912321 | 7.858162 | 304 | 233 | 216 | 208 |
| 57089 | NM_00134996 | ENTPD7 | ectonucleoside triphosphate diphosphohydrolase 7 | protein_coding | NP_001336891.1;NP_001336892.1;NP_065087.1 | 19745 | 616753 | . | . | 2.487551 | 2.565698 | 2.585090 | 2.539919 | 5.788632 | 5.930725 | 5.800868 | 5.807223 | -1.002086 | 680.627205 | 0.151652158 | -0.01982658 | 0.984181715 | 0.999092463 | 9.412207 | 9.409233 | 9.401180 | 9.421396 | 9.419911 | 9.399837 | 773 | 746 | 600 | 628 |
| 57091 | NM_00116411 | CASS4 | Cas scaffold protein family member 4 | protein_coding | NP_001157586.1;NP_001157587.1;NP_001157588.1;NP_065089.2;XP_00 | 15878 | 618888 | . | . | 0.021151 | 0.072013 | 0.082655 | 0.056539 | 0.049219 | 0.166461 | 0.185476 | 0.129270 | 1.553418 | 8.712754 | 1.07117384 | 0.59322407 | 0.553031206 | NA | 2.772726 | 3.404839 | 3.081761 | 3.113365 | 3.123507 | 3.105754 | 4 | 11 | 11 | 8 |
| 57092 | NM_00132039 | PCNP | PEST proteolytic signal containing nuclear protein | protein_coding | NP_001307324.1;NP_001307326.1;NP_001307327.1;NP_001307328.1;NI | 30023 | 615210 | . | . | 14.709154 | 14.256584 | 14.796361 | 14.831816 | 34.228801 | 32.954732 | 33.202604 | 33.911183 | 1.005225 | 1058.740101 | 0.134804766 | 0.055777046 | 0.955519421 | 0.998865404 | 10.044303 | 10.051952 | 10.062514 | 10.029448 | 10.053107 | 10.046898 | 1243 | 1114 | 927 | 991 |
| 57095 | NM_020362,XI | PITHD1 | PITH domain containing 1 | protein_coding | NP_065095.2;XP_011540106.1 | 25022 | 618784 | . | . | 25.569110 | 27.188748 | 28.102686 | 27.445972 | 59.500369 | 62.848005 | 63.061614 | 62.751954 | 1.039982 | 1338.185331 | 0.128484874 | 0.440198498 | 0.659793347 | 0.998865404 | 10.357509 | 10.414061 | 10.351459 | 10.389507 | 10.413670 | 10.387272 | 1473 | 1451 | 1207 | 1257 |
| 57097 | NM_00128652 | PARP11 | poly(ADP-ribose) polymerase family member 11 | protein_coding | NP_001273450.1;NP_001273451.1;NP_065100.2;XP_011519272.1;XP_01 | 1186 | 616706 | . | . | 0.714017 | 0.596250 | 0.810058 | 0.743606 | 1.661547 | 1.378258 | 1.817748 | 1.700168 | -1.044964 | 91.904465 | 0.339068259 | -0.18714054 | 0.85155044 | 0.998865404 | 6.552567 | 6.490901 | 6.554468 | 6.499703 | 6.538745 | 6.490306 | 118 | 92 | 85 | 77 |
| 57099 | NM_020371,XI | AVEN | apoptosis and caspase activation inhibitor | protein_coding | NP_065104.1;XP_011520120.1;XP_011520121.1;XP_011520122.1;XP_02 | 13509 | 605265 | . | . | 2.312989 | 2.526907 | 2.784188 | 3.433957 | 5.382420 | 5.841058 | 6.247638 | 7.851336 | 1.250722 | 265.928075 | 0.219861507 | 1.468022023 | 0.142098241 | 0.903146012 | 7.883925 | 8.207729 | 7.992556 | 7.985723 | 8.162010 | 8.042089 | 274 | 253 | 286 | 246 |
| 57101 | NM_00127859 | ANO2 | anoctamin 2 | protein_coding | NP_001265525.1;NP_001265526.1;NP_001351720.1;XP_011519277.1;XP | 1183 | 610109 | . | . | 0.212109 | 0.161278 | 0.195578 | 0.203736 | 0.493586 | 0.372801 | 0.438872 | 0.465817 | 1.096223 | 40.681870 | 0.494479787 | 0.268040755 | 0.788667948 | 0.998865404 | 5.277893 | 5.411637 | 5.361809 | 5.311362 | 5.346853 | 5.355705 | 52 | 35 | 36 | 41 |
| 57102 | NM_00130481 | C12orf4 | chromosome 12 open reading frame 4 | protein_coding | NP_001291740.1;NP_001333082.1;NP_001333084.1;NP_001333085.1;NI | 1184 | 616082 | . | . | 5.695049 | 5.740083 | 5.829858 | 5.348858 | 13.252615 | 13.268458 | 13.082030 | 12.229529 | -1.031360 | 667.626309 | 0.154342527 | -0.28863305 | 0.772862198 | 0.998865404 | 9.404826 | 9.360629 | 9.393482 | 9.392631 | 9.401541 | 9.341910 | 781 | 731 | 597 | 589 |
| 57103 | NM_020375 | TIGAR | TP53 induced glycolysis regulatory phosphatase | protein_coding | NP_065108.1 | 1185 | 610775 | . | . | 3.947623 | 3.551722 | 3.788890 | 3.831876 | 9.186281 | 8.209964 | 8.502158 | 8.761128 | 1.001574 | 966.985015 | 0.141752894 | 0.016007635 | 0.987228301 | 0.999197585 | 9.916087 | 9.918611 | 9.957308 | 9.873721 | 9.917059 | 9.918117 | 1176 | 983 | 841 | 908 |
| 57104 | NM_020376,XI | PNPLA2 | patatin like phospholipase domain containing 2 | protein_coding | NP_065109.1;XP_006718328.1;XP_006718329.1;XP_016873517.1;XP_02 | 30802 | 609059 | . | . | 12.072299 | 12.609405 | 12.237929 | 12.218071 | 28.092736 | 29.147202 | 27.461558 | 27.935166 | -1.014411 | 785.705282 | 0.145652557 | -0.14172344 | 0.887298456 | 0.998865404 | 9.628131 | 9.607484 | 9.609345 | 9.636073 | 9.621606 | 9.603796 | 894 | 870 | 687 | 722 |
| 57106 | NM_020378 | NAT14 | N-acetyltransferase 14 (putative) | protein_coding | NP_065111.1 | 28918 | . | . | . | 20.160898 | 18.721657 | 19.147390 | 19.724335 | 46.915241 | 43.275948 | 42.966187 | 45.097351 | -1.014572 | 817.926579 | 0.145842563 | -0.14310327 | 0.886208619 | 0.998865404 | 9.686157 | 9.665424 | 9.709990 | 9.650708 | 9.663450 | 9.677653 | 988 | 852 | 699 | 769 |
| 57107 | NM_020381,XI | PDSS2 | decaprenyl diphosphate synthase subunit 2 | protein_coding | NP_065114.3;XP_011534258.1;XP_011534259.1;XP_011534260.1;XP_01 | 23041 | 610564 | . | . | 3.371936 | 4.424924 | 3.780523 | 4.081892 | 7.846635 | 10.228410 | 8.483382 | 9.332762 | -1.056450 | 345.772683 | 0.199981253 | -0.39616181 | 0.691985663 | 0.998865404 | 8.473160 | 8.393089 | 8.386344 | 8.505746 | 8.404594 | 8.428460 | 367 | 423 | 286 | 322 |
| 57108 | gene-KMT5AP1 | KMT5AP1 | KMT5A pseudogene 1 | pseudogene | . | 30088 | . | . | . | 0.134643 | 0.253714 | 0.142526 | 0.148025 | 0.313320 | 0.586471 | 0.319824 | 0.338441 | -1.349348 | 14.962447 | 0.809913041 | -0.53371494 | 0.593538768 | NA | 4.104339 | 3.669578 | 3.890600 | 3.934929 | 3.893554 | 3.893999 | 14 | 24 | 11 | 12 |
| 57109 | NM_00127934 | REXO4 | REX4 homolog, 3'-5' exonuclease | protein_coding | NP_001266278.1;NP_001266279.1;NP_001266280.1;NP_065118.2;XP_02 | 12820 | 602930 | . | . | 14.072231 | 15.664344 | 15.392965 | 17.345857 | 32.746660 | 36.208838 | 34.541363 | 39.659238 | 1.088239 | 1122.126446 | 0.140691927 | 0.86710583 | 0.385884027 | 0.998865404 | 10.070007 | 10.191476 | 10.063507 | 10.132666 | 10.114008 | 10.207004 | 1180 | 1214 | 954 | 1164 |
| 57110 | NM_00136611 | PLAAT1 | phospholipase A and acyltransferase 1 | protein_coding | NP_001353041.1;NP_065119.3;XP_011511336.1;XP_011511337.1 | 14922 | 606487 | . | . | 1.384205 | 1.303876 | 1.320060 | 1.364907 | 3.221101 | 3.013969 | 2.962175 | 3.120698 | -1.050331 | 43.670890 | 0.471247841 | -0.1503334 | 0.880501585 | 0.998865404 | 5.483297 | 5.413048 | 5.465171 | 5.441659 | 5.445156 | 5.445522 | 55 | 45 | 37 | 40 |
| 57116 | NM_00120422 | ZNF695 | zinc finger protein 695 | protein_coding | NP_001191150.2;NP_065127.5 | 30954 | 616348 | . | . | 1.649105 | 1.253543 | 1.586844 | 2.102634 | 3.837535 | 2.897620 | 3.560833 | 4.807421 | -1.158787 | 142.245795 | 0.285032358 | -0.74593422 | 0.45570715 | 0.998865404 | 7.254018 | 7.042738 | 7.232278 | 7.121417 | 7.133281 | 7.112264 | 200 | 142 | 118 | 120 |
| 57117 | NM_00114247 | INTS12 | integrator complex subunit 12 | protein_coding | NP_001135943.1;NP_065128.2;XP_005263205.1;XP_011530445.1;XP_01 | 25067 | 611355 | . | . | 7.594728 | 8.039774 | 7.639203 | 8.842274 | 17.673244 | 18.584301 | 17.142149 | 20.216806 | 1.015461 | 450.198432 | 0.175029127 | 0.126466974 | 0.89936229 | 0.998865404 | 8.803571 | 8.825182 | 8.796674 | 8.821990 | 8.791338 | 8.845438 | 504 | 492 | 377 | 444 |
| 57120 | NM_00101740 | GOPC | golgi associated PDZ and coiled-coil motif containing | protein_coding | NP_001017408.1;NP_065132.1;XP_024302264.1 | 17643 | 606845 | . | . | 6.762872 | 6.425808 | 6.644634 | 5.568926 | 15.737481 | 14.853542 | 14.910367 | 12.732687 | -1.095052 | 907.222646 | 0.151418706 | -0.86514783 | 0.386957654 | 0.998865404 | 9.888993 | 9.758692 | 9.878113 | 9.833360 | 9.854242 | 9.722916 | 1125 | 992 | 823 | 737 |
| 57121 | NM_00114296 | LPAR5 | lysophosphatidic acid receptor 5 | protein_coding | NP_001136433.1;NP_065133.1 | 13307 | 606926 | . | . | 0.298452 | 0.203504 | 0.162507 | 0.199646 | 0.694509 | 0.470409 | 0.364662 | 0.456468 | -1.414562 | 18.631944 | 0.723124965 | -0.69193423 | 0.488978634 | NA | 4.448287 | 3.947964 | 4.256271 | 4.219758 | 4.204208 | 4.217641 | 30 | 19 | 12 | 16 |
| 57122 | NM_00133019 | NUP107 | nucleoporin 107 | protein_coding | NP_001317121.1;NP_065134.1;XP_005269094.1 | 29914 | 607617 | . | . | 7.335991 | 7.760411 | 7.180735 | 7.203233 | 17.071151 | 17.938538 | 16.113359 | 16.469335 | -1.064300 | 1427.577863 | 0.126270764 | -0.71200168 | 0.476463739 | 0.998865404 | 10.523640 | 10.433665 | 10.484323 | 10.521016 | 10.457672 | 10.452180 | 1655 | 1626 | 1207 | 1292 |
| 57125 | NM_020405 | PLXDC1 | plexin domain containing 1 | protein_coding | NP_065138.2 | 20945 | 606826 | . | . | 0.161574 | 0.184161 | 0.345161 | 0.140396 | 0.375989 | 0.425696 | 0.774532 | 0.321000 | 1.396762 | 40.964307 | 0.542186431 | 0.889152262 | 0.373921255 | 0.998865404 | 5.093294 | 5.578664 | 5.298391 | 5.315152 | 5.430722 | 5.282331 | 37 | 39 | 59 | 26 |
| 57128 | NM_00116484 | LYRM4 | LYR motif containing 4 | protein_coding | NP_001158312.1;NP_001158313.1;NP_001305711.1;NP_001305712.1;NI | 21365 | 613311 | . | . | 10.177281 | 10.311989 | 9.749862 | 10.186642 | 23.682951 | 23.836622 | 21.878404 | 23.290546 | -1.038420 | 410.387763 | 0.181240076 | -0.3001003 | 0.764100649 | 0.998865404 | 8.708008 | 8.653159 | 8.696448 | 8.687643 | 8.636880 | 8.700944 | 485 | 448 | 331 | 398 |
| 57129 | NM_020409,N | MRPL47 | mitochondrial ribosomal protein L47 | protein_coding | NP_065142.2;NP_817125.1 | 16652 | 611852 | . | . | 28.576580 | 29.272658 | 24.225206 | 31.533839 | 66.498875 | 67.665062 | 54.360663 | 72.098384 | -1.053702 | 1191.934673 | 0.150786889 | -0.50048684 | 0.616732324 | 0.998865404 | 10.256628 | 10.180548 | 10.230520 | 10.243077 | 10.099909 | 10.286279 | 1397 | 1331 | 883 | 1224 |
| 57130 | NM_020410 | ATP13A1 | ATPase 13A1 | protein_coding | NP_065143.2 | 24215 | . | . | . | 20.889095 | 22.759899 | 28.441458 | 22.596226 | 48.609783 | 52.610523 | 63.821808 | 51.663582 | 1.153800 | 2837.795269 | 0.136287297 | 1.514397139 | 0.129925144 | 0.88082356 | 11.363566 | 11.570154 | 11.375442 | 11.437650 | 11.617993 | 11.420021 | 2917 | 2953 | 2960 | 2510 |
| 57132 | NM_020412 | CHMP1B | charged multivesicular body protein 1B | protein_coding | NP_065145.2 | 24287 | 606486 | . | . | 8.460228 | 7.374241 | 8.128768 | 8.014165 | 19.687298 | 17.045891 | 18.240721 | 18.323435 | 1.005360 | 755.779563 | 0.153518798 | 0.050238762 | 0.959932123 | 0.998865404 | 9.557737 | 9.565895 | 9.609012 | 9.506913 | 9.571389 | 9.554699 | 931 | 754 | 667 | 702 |
| 57134 | NM_00128901 | MAN1C1 | mannosidase alpha class 1C member 1 | protein_coding | NP_001275939.1;NP_065112.1;XP_016857349.1;XP_016857350.1 | 19080 | 616772 | . | . | 1.937971 | 2.204857 | 1.683203 | 1.831066 | 4.509738 | 5.096627 | 3.777057 | 4.186514 | -1.199800 | 264.623272 | 0.212822104 | -1.23480348 | 0.216903653 | 0.982008277 | 8.173465 | 7.910128 | 8.067243 | 8.117071 | 7.990082 | 8.008804 | 317 | 326 | 205 | 230 |
| 57135 | NM_00100537 | DAZ4 | deleted in azoospermia 4 | protein_coding | NP_001005375.1;NP_065153.1;XP_005262659.1;XP_011529811.1;XP_01 | 15966 | 400048 | . | . | 1.076074 | 0.658540 | 0.792256 | 0.625483 | 2.504069 | 1.522246 | 1.777799 | 1.430094 | -1.256581 | 89.614949 | 0.365875784 | -0.90058912 | 0.36780682 | 0.998865404 | 6.639727 | 6.313167 | 6.579201 | 6.441819 | 6.480607 | 6.414739 | 140 | 84 | 78 | 65 |
| 57136 | NM_020531,XI | APMAP | adipocyte plasma membrane associated protein | protein_coding | NP_065392.1;XP_005260820.1 | 13238 | 615884 | . | . | 127.027046 | 120.924484 | 100.294457 | 105.039734 | 295.597160 | 279.522349 | 225.057843 | 240.160859 | -1.224750 | 7778.530406 | 0.104516375 | -2.79848425 | **0.005134307** | 0.217389093 | 13.064127 | 12.771637 | 13.029543 | 12.979734 | 12.818371 | 12.848225 | 10147 | 8974 | 5971 | 6674 |
| 57139 | NM_00103522 | RGL3 | ral guanine nucleotide dissociation stimulator like 3 | protein_coding | NP_001030300.3;NP_001155088.2 | 30282 | 616743 | . | . | 3.337953 | 4.015111 | 4.355467 | 3.663651 | 7.767554 | 9.281110 | 9.773541 | 8.376503 | 1.074533 | 301.774338 | 0.213257212 | 0.48631245 | 0.626745653 | 0.998865404 | 8.184433 | 8.288349 | 8.165349 | 8.261316 | 8.301938 | 8.201875 | 305 | 342 | 297 | 266 |
| 57140 | NM_018226,XI | RNPEPL1 | arginyl aminopeptidase like 1 | protein_coding | NP_060696.4;XP_005247093.1 | 10079 | 605287 | . | . | 9.794928 | 10.516998 | 9.874754 | 8.089300 | 22.793199 | 24.310512 | 22.158662 | 18.495224 | -1.017174 | 1230.499875 | 0.136300064 | -0.18023878 | 0.856965115 | 0.998865404 | 10.277488 | 10.252463 | 10.241240 | 10.299880 | 10.214869 | 10.299371 | 1374 | 1391 | 1001 | 1208 |
| 57142 | NM_00132185 | RTN4 | reticulon 4 | protein_coding | NP_001308788.1;NP_001308789.1;NP_001308790.1;NP_001308791.1;NI | 14085 | 604475 | . | . | 164.884444 | 154.955831 | 137.671872 | 110.419178 | 383.692873 | 358.187334 | 308.931685 | 252.460322 | -1.320134 | 9768.711964 | 0.125466935 | -3.19354473 | **0.001405376** | 0.087623478 | 13.440375 | 13.039815 | 13.395413 | 13.321753 | 13.210168 | 13.027006 | 13340 | 11489 | 8128 | 7027 |
| 57143 | NM_00114254 | ADCK1 | aarF domain containing kinase 1 | protein_coding | NP_001136017.1;NP_001353414.1;NP_001353415.1;NP_001353416.1;NI | 19038 | . | . | . | 1.249526 | 1.197868 | 1.514818 | 1.487072 | 2.907698 | 2.768921 | 3.399211 | 3.400014 | 1.259565 | 133.425322 | 0.282738079 | 1.17750331 | 0.238994677 | 0.992733415 | 6.884299 | 7.216407 | 6.990615 | 7.031554 | 7.084971 | 7.105240 | 128 | 135 | 126 | 143 |
| 57146 | NM_00130176 | TMEM159 | transmembrane protein 159 | protein_coding | NP_001288698.1;NP_001288700.1;NP_001288702.1;NP_001288703.1;NI | 30136 | 611304 | . | . | 11.267601 | 10.133318 | 8.491792 | 9.562752 | 26.220172 | 23.423620 | 19.055336 | 21.864098 | -1.187683 | 556.198173 | 0.170146531 | -1.45845095 | 0.144716287 | 0.906189003 | 9.238289 | 8.989947 | 9.215310 | 9.126757 | 9.019912 | 9.099025 | 741 | 609 | 414 | 507 |
| 57147 | NM_020423,N | SCYL3 | SCY1 like pseudokinase 3 | protein_coding | NP_065156.5;NP_851607.2;XP_006711528.1;XP_011508103.1;XP_01150 | 19285 | 608192 | . | . | 1.638698 | 1.399791 | 1.684310 | 2.138725 | 3.813317 | 3.235679 | 3.779544 | 4.889941 | 1.092497 | 243.041918 | 0.234310032 | 0.54470493 | 0.585956474 | 0.998865404 | 7.860380 | 7.986966 | 7.918469 | 7.880586 | 7.866697 | 8.014554 | 278 | 241 | 188 | 273 |
| 57148 | NM_00128291 | RALGAPB | Ral GTPase activating protein non-catalytic subunit beta | protein_coding | NP_001269846.1;NP_001269847.1;NP_065069.1;XP_005260519.1;XP_00 | 29221 | 618833 | . | . | 5.881502 | 6.221197 | 6.445873 | 6.157316 | 13.686495 | 14.380575 | 14.464354 | 14.077968 | 1.041705 | 1654.641415 | 0.123203925 | 0.478449033 | 0.632330642 | 0.998865404 | 10.662477 | 10.721524 | 10.667764 | 10.683919 | 10.729781 | 10.685361 | 1847 | 1767 | 1510 | 1538 |
| 57149 | NM_00112830 | LYRM1 | LYR motif containing 1 | protein_coding | NP_001121773.1;NP_001121774.1;NP_001289764.1;NP_001289765.1;NI | 25074 | 614709 | . | . | 7.071024 | 6.874694 | 6.668705 | 7.161892 | 16.454562 | 15.891165 | 14.964384 | 16.374817 | -1.030613 | 346.243265 | 0.190834536 | -0.2279638 | 0.819674386 | 0.998865404 | 8.457409 | 8.413542 | 8.443231 | 8.445331 | 8.407281 | 8.446658 | 404 | 380 | 286 | 331 |
| 57150 | NM_00104249 | SMIM8 | small integral membrane protein 8 | protein_coding | NP_001035958.1;NP_001274374.1;NP_065158.3 | 21401 | . | . | . | 1.395382 | 1.573349 | 1.650559 | 1.393699 | 3.247111 | 3.636867 | 3.703807 | 3.186526 | 1.014830 | 114.041428 | 0.305616232 | 0.06949195 | 0.94459804 | 0.998865404 | 6.822529 | 6.844218 | 6.810217 | 6.848002 | 6.860297 | 6.807355 | 123 | 129 | 108 | 99 |
| 57153 | NM_00114505 | SLC44A2 | solute carrier family 44 member 2 | protein_coding | NP_001138528.1;NP_001350540.1;NP_065161.3;XP_005260056.1 | 17292 | 606106 | . | . | 3.928831 | 3.010678 | 2.329923 | 3.764592 | 9.142550 | 6.959318 | 5.228282 | 8.607293 | -1.151564 | 343.057166 | 0.241588601 | -0.84273007 | 0.399379426 | 0.998865404 | 8.520947 | 8.316420 | 8.531683 | 8.369656 | 8.242684 | 8.490404 | 481 | 342 | 214 | 367 |
| 57154 | NM_00119984 | SMURF1 | SMAD specific E3 ubiquitin protein ligase 1 | protein_coding | NP_001186776.1;NP_065162.1;NP_851994.1;XP_016867946.1 | 16807 | 605568 | . | . | 4.620594 | 5.358614 | 4.468542 | 5.491751 | 10.752314 | 12.386677 | 10.027278 | 12.556236 | -1.017089 | 881.051201 | 0.157989267 | -0.15473377 | 0.877031207 | 0.998865404 | 9.795665 | 9.770388 | 9.736541 | 9.836152 | 9.702874 | 9.842726 | 953 | 1025 | 685 | 899 |
| 57157 | NM_00112735 | PHTF2 | putative homeodomain transcription factor 2 | protein_coding | NP_001120829.1;NP_001120830.1;NP_001120831.1;NP_001120832.1;NI | 13411 | 616785 | . | . | 15.390545 | 12.445380 | 12.670394 | 9.516081 | 35.814428 | 28.768055 | 28.431993 | 21.757390 | -1.286198 | 1951.491317 | 0.166454763 | -2.18145286 | **0.029149938** | 0.54925494 | 11.100260 | 10.737750 | 11.101285 | 10.924735 | 10.940538 | 10.676938 | 2800 | 2114 | 1738 | 1328 |
| 57161 | NM_021255,XI | PELI2 | pellino E3 ubiquitin protein ligase family member 2 | protein_coding | NP_067078.1;XP_005267947.1;XP_006720274.1;XP_011535292.1;XP_01 | 8828 | 614798 | . | . | 0.763181 | 0.343641 | 0.761303 | 0.516251 | 1.775953 | 0.794343 | 1.708341 | 1.180347 | 1.005450 | 124.611294 | 0.302522734 | 0.025917828 | 0.97932288 | 0.998865404 | 6.956317 | 6.966247 | 6.996636 | 6.917187 | 6.997774 | 6.920019 | 159 | 119 | 121 | 104 |
| 57162 | NM_020651,XI | PELI1 | pellino E3 ubiquitin protein ligase 1 | protein_coding | NP_065702.2;XP_005264494.1;XP_011531296.1;XP_011531298.1;XP_01 | 8827 | 614797 | . | . | 4.474681 | 4.048740 | 4.854615 | 4.948536 | 10.412766 | 9.358845 | 10.893616 | 11.314239 | 1.124403 | 539.982542 | 0.166582728 | 1.015468589 | 0.309882512 | 0.998865404 | 8.989502 | 9.159057 | 9.074337 | 8.996647 | 9.113221 | 9.111115 | 621 | 515 | 498 | 535 |
| 57165 | NM_020435 | GJC2 | gap junction protein gamma 2 | protein_coding | NP_065168.2 | 17494 | 608803 | . | . | 0.175987 | 0.218045 | 0.127700 | 0.157228 | 0.409529 | 0.504020 | 0.286554 | 0.359482 | -1.356537 | 11.713143 | 0.894107252 | -0.49203145 | 0.622697113 | NA | 3.754145 | 3.312263 | 3.556446 | 3.568312 | 3.543706 | 3.549882 | 14 | 16 | 8 | 10 |
| 57168 | NM_020437 | ASPHD2 | aspartate beta-hydroxylase domain containing 2 | protein_coding | NP_065170.2 | 30437 | . | . | . | 3.126754 | 2.869291 | 2.870803 | 3.324924 | 7.276086 | 6.632496 | 6.441998 | 7.602043 | 1.019380 | 320.406377 | 0.201258292 | 0.13759403 | 0.890561274 | 0.998865404 | 8.310004 | 8.337384 | 8.344037 | 8.290836 | 8.290112 | 8.365115 | 383 | 326 | 262 | 324 |
| 57169 | NM_021035 | ZNFX1 | zinc finger NFX1-type containing 1 | protein_coding | NP_066363.1 | 29271 | . | . | . | 13.113062 | 13.957417 | 13.737682 | 15.086082 | 30.514635 | 32.263195 | 30.826963 | 34.492535 | 1.050052 | 3137.543434 | 0.11500271 | 0.612687074 | 0.540083271 | 0.998865404 | 11.579852 | 11.650132 | 11.572514 | 11.616506 | 11.599732 | 11.668514 | 3430 | 3392 | 2678 | 3139 |
| 57171 | NM_00113591 | DOLPP1 | dolichyldiphosphatase 1 | protein_coding | NP_001129389.1;NP_065171.2 | 29565 | 614516 | . | . | 8.638656 | 9.406329 | 9.435941 | 9.802923 | 20.102508 | 21.743148 | 21.173977 | 22.413221 | 1.049856 | 622.792304 | 0.156680117 | 0.447989333 | 0.654160895 | 0.998865404 | 9.247215 | 9.317152 | 9.242693 | 9.288136 | 9.293650 | 9.302667 | 676 | 678 | 551 | 602 |
| 57175 | NM_00101807 | CORO1B | coronin 1B | protein_coding | NP_001018080.1;NP_065174.1 | 2253 | 609849 | . | . | 16.847097 | 13.651693 | 25.179269 | 22.074941 | 39.203888 | 31.556499 | 56.501550 | 50.471729 | 1.055945 | 1916.269082 | 0.120276099 | 0.652951073 | 0.513787822 | 0.998865404 | 10.864367 | 10.942737 | 10.874688 | 10.889151 | 10.900240 | 10.949127 | 2127 | 2030 | 1659 | 1902 |
| 57176 | NM_00116773 | VARS2 | valyl-tRNA synthetase 2, mitochondrial | protein_coding | NP_001161205.1;NP_001161206.1;NP_065175.4 | 21642 | 612802 | . | . | 13.610658 | 15.114423 | 16.445566 | 16.984914 | 31.672558 | 34.937666 | 36.903373 | 38.833990 | 1.142463 | 1738.067523 | 0.124303493 | 1.545794075 | 0.122154308 | 0.86555232 | 10.664111 | 10.856047 | 10.672412 | 10.742519 | 10.802639 | 10.822265 | 1784 | 1830 | 1592 | 1759 |
| 57178 | NM_020338,XI | ZMIZ1 | zinc finger MIZ-type containing 1 | protein_coding | NP_065071.1;XP_005270044.1;XP_005270045.1;XP_006717986.1;XP_00 | 16493 | 607159 | . | . | 7.979708 | 8.809591 | 8.927204 | 9.231222 | 18.569108 | 20.363765 | 20.032389 | 21.106094 | 1.073217 | 2063.466359 | 0.12075784 | 0.844185845 | 0.398565563 | 0.998865404 | 10.959076 | 11.060838 | 10.946000 | 11.016096 | 11.028491 | 11.047191 | 2191 | 2243 | 1836 | 2026 |
| 57179 | NM_00107968 | KIAA1191 | KIAA1191 | protein_coding | NP_001073152.1;NP_001073153.1;NP_001274264.1;NP_001274265.1;NI | 29209 | . | . | . | 29.384580 | 30.442759 | 30.240624 | 33.569037 | 68.379123 | 70.369797 | 67.859084 | 76.751611 | 1.047845 | 2660.179037 | 0.117167735 | 0.575461681 | 0.564979099 | 0.998865404 | 11.343298 | 11.410534 | 11.347432 | 11.367902 | 11.355896 | 11.433803 | 2954 | 2839 | 2254 | 2674 |
| 57180 | NM_00104013 | ACTR3B | actin related protein 3B | protein_coding | NP_001035225.1;NP_001337869.1;NP_001337870.1;NP_001337871.1;NI | 17256 | . | . | . | 2.355157 | 2.382770 | 3.092884 | 3.500097 | 5.480547 | 5.507878 | 6.940342 | 8.002553 | 1.319984 | 193.931821 | 0.244143514 | 1.640513868 | 0.100898367 | 0.816490589 | 7.385783 | 7.785438 | 7.526857 | 7.529858 | 7.608868 | 7.692495 | 192 | 181 | 175 | 225 |
| 57181 | NM_00112725 | SLC39A10 | solute carrier family 39 member 10 | protein_coding | NP_001120729.1;NP_065075.1;XP_005246746.2;XP_011509806.1;XP_01 | 20861 | 608733 | . | . | 12.999427 | 11.298023 | 12.114306 | 9.967697 | 30.250202 | 26.115886 | 27.184154 | 22.789956 | -1.103277 | 1937.208312 | 0.142449106 | -0.99540445 | 0.319539597 | 0.998865404 | 10.988677 | 10.847392 | 11.011607 | 10.895901 | 10.958270 | 10.790056 | 2508 | 2035 | 1774 | 1542 |
| 57182 | NM_00116788 | ANKRD50 | ankyrin repeat domain 50 | protein_coding | NP_001161354.1;NP_065070.1;XP_016863960.1 | 29223 | . | . | . | 2.416204 | 2.368211 | 2.368026 | 2.281251 | 5.622604 | 5.474227 | 5.313780 | 5.215810 | -1.041179 | 654.014369 | 0.154253677 | -0.37741906 | 0.705862201 | 0.998865404 | 9.381837 | 9.323939 | 9.379771 | 9.353307 | 9.356888 | 9.321615 | 783 | 706 | 572 | 585 |
| 57184 | NM_00132192 | FAM219B | family with sequence similarity 219 member B | protein_coding | NP_001308849.1;NP_001308850.1;NP_001308851.1;NP_001308852.1;NI | 24695 | . | . | . | 10.393277 | 9.783434 | 11.545999 | 10.507642 | 24.185579 | 22.614845 | 25.908889 | 24.024478 | 1.080294 | 1021.829587 | 0.139972088 | 0.796040794 | 0.426008325 | 0.998865404 | 9.939941 | 10.051770 | 9.977832 | 9.956000 | 10.064729 | 9.981352 | 1148 | 1044 | 975 | 939 |
| 57185 | NM_00132285 | NIPAL3 | NIPA like domain containing 3 | protein_coding | NP_001309783.1;NP_001309784.1;NP_001309785.1;NP_001309786.1;NI | 25233 | . | . | . | 22.923096 | 25.769655 | 26.885281 | 27.123762 | 53.342985 | 59.567713 | 60.329794 | 62.015255 | 1.074558 | 3875.148913 | 0.111584977 | 0.929724135 | 0.352513935 | 0.998865404 | 11.867298 | 11.970914 | 11.857145 | 11.920352 | 11.930493 | 11.966666 | 4137 | 4186 | 3415 | 3845 |
| 57186 | NM_020343,XI | RALGAPA2 | Ral GTPase activating protein catalytic subunit alpha 2 | protein_coding | NP_065076.2;XP_005260825.1;XP_006723661.1;XP_006723662.1;XP_01 | 16207 | 618836 | . | . | 1.658470 | 1.633907 | 1.841773 | 1.549046 | 3.859328 | 3.776847 | 4.132884 | 3.541709 | 1.014999 | 514.370749 | 0.167373339 | 0.128323061 | 0.897893316 | 0.998865404 | 8.995760 | 9.017488 | 8.989481 | 9.013152 | 9.037856 | 8.983306 | 577 | 561 | 470 | 465 |
| 57187 | NM_00108155 | THOC2 | THO complex 2 | protein_coding | NP_001075019.1;XP_016885151.1;XP_016885152.1;XP_016885153.1;XP_ | 19073 | 300395 | . | . | 8.386081 | 8.301872 | 8.418559 | 7.695591 | 19.514753 | 19.190154 | 18.891003 | 17.595053 | -1.049950 | 1556.934443 | 0.127375195 | -0.55207936 | 0.580893985 | 0.998865404 | 10.639087 | 10.569048 | 10.629041 | 10.616474 | 10.623338 | 10.545281 | 1851 | 1707 | 1388 | 1352 |
| 57189 | NM_00108039 | DENND11 | DENN domain containing 11 | protein_coding | NP_001073861.1;XP_024302611.1 | 29472 | . | . | . | 1.884476 | 2.392780 | 1.922627 | 1.919229 | 4.385255 | 5.531019 | 4.314320 | 4.388090 | -1.116362 | 497.852173 | 0.183359353 | -0.8660858 | 0.386443113 | 0.998865404 | 9.037092 | 8.877652 | 8.906287 | 9.070024 | 8.921682 | 8.921552 | 529 | 638 | 409 | 441 |
| 57190 | NM_020451,N | SELENON | selenoprotein N | protein_coding | NP_065184.2;NP_996809.1 | 15999 | 606210 | . | . | 42.974688 | 44.395596 | 34.750165 | 39.608007 | 100.003867 | 102.622413 | 77.978361 | 90.558995 | -1.191251 | 5327.548832 | 0.111898128 | -2.25631852 | **0.02405069** | 0.512901603 | 12.500040 | 12.247428 | 12.437728 | 12.457981 | 12.251531 | 12.348291 | 6591 | 6325 | 3980 | 4826 |
| 57191 | NM_020633 | VN1R1 | vomeronasal 1 receptor 1 | protein_coding | NP_065684.1 | 13548 | 605234 | . | . | 0.020914 | 0.062466 | 0.033906 | 0.023446 | 0.048668 | 0.144392 | 0.076084 | 0.053606 | -1.225524 | 2.469287 | 1.896160988 | -0.15473285 | 0.877031932 | NA | 1.444631 | 1.148375 | 1.301291 | 1.310651 | 1.304063 | 1.303286 | 2 | 4 | 2 | 2 |
| 57192 | NM_020533 | MCOLN1 | mucolipin 1 | protein_coding | NP_065394.1 | 13356 | 605248 | . | . | 18.091297 | 19.974045 | 23.060762 | 20.209509 | 42.099194 | 46.170895 | 51.747681 | 46.206642 | 1.121958 | 1320.071354 | 0.137808656 | 1.204704996 | 0.228317207 | 0.982008277 | 10.280917 | 10.447101 | 10.284906 | 10.352934 | 10.460349 | 10.351139 | 1368 | 1403 | 1300 | 1216 |
| 57194 | NM_024490,XI | ATP10A | ATPase phospholipid transporting 10A (putative) | protein_coding | NP_077816.1;XP_005268318.1;XP_011520128.1;XP_011520130.1;XP_01 | 13542 | 605855 | . | . | 2.774206 | 2.792522 | 2.635965 | 3.118160 | 6.455692 | 6.455041 | 5.915027 | 7.129303 | 1.023390 | 470.276941 | 0.176283625 | 0.189219712 | 0.849920619 | 0.998865404 | 8.860933 | 8.893616 | 8.867813 | 8.871078 | 8.833600 | 8.931487 | 534 | 503 | 381 | 481 |
| 57198 | NM_00100585 | ATP8B2 | ATPase phospholipid transporting 8B2 | protein_coding | NP_001005855.1;NP_001354863.1;NP_001357525.1;NP_001357526.1;NI | 13534 | 605867 | . | . | 11.762310 | 12.545614 | 13.615383 | 14.218100 | 27.371382 | 28.999747 | 30.552524 | 32.507997 | 1.133843 | 2421.395203 | 0.116113067 | 1.560726868 | 0.118588209 | 0.858706505 | 11.148244 | 11.329323 | 11.166473 | 11.210068 | 11.275124 | 11.304023 | 2542 | 2516 | 2198 | 2455 |
| 57205 | NM_020453,XI | ATP10D | ATPase phospholipid transporting 10D (putative) | protein_coding | NP_065186.3;XP_005248176.1;XP_005248177.1;XP_011512024.1;XP_01 | 13549 | . | . | . | 4.059968 | 3.595309 | 3.369580 | 3.899546 | 9.447714 | 8.310718 | 7.561241 | 8.915845 | -1.126281 | 728.266004 | 0.15462503 | -1.10956972 | 0.267184485 | 0.998546485 | 9.591596 | 9.419945 | 9.585881 | 9.508139 | 9.433705 | 9.496172 | 937 | 787 | 568 | 672 |
| 57209 | NM_00126759 | ZNF248 | zinc finger protein 248 | protein_coding | NP_001254526.1;NP_001254534.1;NP_001254535.1;NP_001254536.1;NI | 13041 | . | . | . | 3.495980 | 3.431080 | 4.814471 | 3.440178 | 8.135289 | 7.931092 | 10.803534 | 7.865561 | 1.112502 | 466.584649 | 0.186742269 | 0.823635063 | 0.410146947 | 0.998865404 | 8.786543 | 8.941299 | 8.828978 | 8.827866 | 8.973638 | 8.809663 | 509 | 476 | 483 | 401 |
| 57210 | NM_00108043 | SLC45A4 | solute carrier family 45 member 4 | protein_coding | NP_001073900.1;NP_001273575.1;NP_001273577.1;XP_011515472.1;XP | 29196 | . | . | . | 1.177056 | 1.195018 | 1.040020 | 0.635595 | 2.739056 | 2.762336 | 2.333776 | 1.453211 | -1.463584 | 234.417456 | 0.228501804 | -2.40482021 | **0.016180424** | 0.427851323 | 8.121467 | 7.572494 | 7.923017 | 7.984900 | 7.803263 | 7.737712 | 302 | 318 | 178 | 165 |
| 57211 | NM_00103239 | ADGRG6 | adhesion G protein-coupled receptor G6 | protein_coding | NP_001027566.1;NP_001027567.1;NP_065188.4;NP_940971.2;XP_00526 | 13841 | 612243 | . | . | 16.310648 | 16.072774 | 16.276836 | 14.289739 | 37.955549 | 37.152938 | 36.524745 | 32.671787 | -1.072795 | 3372.651663 | 0.118350545 | -0.85655785 | 0.391689287 | 0.998865404 | 11.769363 | 11.668200 | 11.758874 | 11.735585 | 11.743751 | 11.632315 | 4076 | 3714 | 3023 | 2843 |
| 57212 | NR_033708,NF | TP73-AS1 | TP73 antisense RNA 1 | lncRNA | . | 29052 | . | . | . | 6.230443 | 6.396551 | 6.815457 | 5.829955 | 14.498496 | 14.785915 | 15.293689 | 13.329498 | -1.003534 | 1168.133438 | 0.143522698 | -0.03546186 | 0.971711456 | 0.998865404 | 10.192280 | 10.187696 | 10.192672 | 10.187333 | 10.256362 | 10.113716 | 1352 | 1258 | 1111 | 989 |
| 57213 | NM_00112748 | SPRYD7 | SPRY domain containing 7 | protein_coding | NP_001120954.1;NP_065189.1;XP_016876166.1 | 14297 | 607866 | . | . | 3.162093 | 2.431272 | 2.834749 | 2.486136 | 7.358321 | 5.619993 | 6.361096 | 5.684256 | -1.039301 | 257.792995 | 0.225825651 | -0.24627013 | 0.805473131 | 0.998865404 | 8.036937 | 7.982692 | 8.075399 | 7.957055 | 8.029749 | 7.966423 | 337 | 251 | 234 | 222 |
| 57214 | NM_00129329 | CEMIP | cell migration inducing hyaluronidase 1 | protein_coding | NP_001280227.1;NP_001280233.1;NP_061159.1;XP_024305769.1 | 29213 | 608366 | . | . | 4.360879 | 4.522950 | 5.204564 | 4.422884 | 10.147947 | 10.455001 | 11.678891 | 10.112398 | 1.075530 | 1030.963746 | 0.145080068 | 0.724067341 | 0.469024394 | 0.998865404 | 9.956108 | 10.061523 | 9.964243 | 9.997326 | 10.094803 | 9.970746 | 1118 | 1096 | 1007 | 918 |
| 57215 | NM_020457 | THAP11 | THAP domain containing 11 | protein_coding | NP_065190.2 | 23194 | 609119 | . | . | 10.901814 | 9.106770 | 10.076520 | 11.225660 | 25.368969 | 21.050707 | 22.611420 | 25.666138 | 1.050076 | 603.877356 | 0.169154676 | 0.416744672 | 0.676865176 | 0.998865404 | 9.202355 | 9.273004 | 9.280576 | 9.155880 | 9.219927 | 9.284381 | 742 | 576 | 512 | 608 |
| 57216 | NM_020335,XI | VANGL2 | VANGL planar cell polarity protein 2 | protein_coding | NP_065068.1;XP_005245414.1;XP_011508106.1;XP_011508107.1 | 15511 | 600533 | . | . | 1.953813 | 1.780756 | 1.414195 | 2.094681 | 4.546603 | 4.116297 | 3.173413 | 4.789240 | -1.078537 | 302.224058 | 0.228148119 | -0.47809155 | 0.632585046 | 0.998865404 | 8.293590 | 8.183251 | 8.283233 | 8.228757 | 8.111527 | 8.309780 | 379 | 322 | 205 | 324 |
| 57217 | NM_00128895 | TTC7A | tetratricopeptide repeat domain 7A | protein_coding | NP_001275880.1;NP_001275882.1;NP_001275884.1;NP_065191.2;XP_00 | 19750 | 609332 | . | . | 3.101519 | 2.978607 | 2.851270 | 2.724371 | 7.217364 | 6.885187 | 6.398169 | 6.228953 | -1.114466 | 449.584244 | 0.174197687 | -0.89755958 | 0.369420396 | 0.998865404 | 8.888332 | 8.732349 | 8.863772 | 8.825714 | 8.789997 | 8.766692 | 562 | 496 | 377 | 391 |
| 57221 | NM_020340,XI | ARFGEF3 | ARFGEF family member 3 | protein_coding | NP_065073.3 | 21213 | 617411 | . | . | 0.425971 | 0.397500 | 0.321555 | 0.397861 | 0.991252 | 0.918840 | 0.721562 | 0.909660 | -1.143444 | 164.926429 | 0.261984089 | -0.73815736 | 0.460418828 | 0.998865404 | 7.459536 | 7.265288 | 7.406304 | 7.379143 | 7.300544 | 7.374029 | 209 | 184 | 121 | 158 |
| 57222 | NM_00103171 | ERGIC1 | endoplasmic reticulum-golgi intermediate compartment 1 | protein_coding | NP_001026881.1;XP_011532899.1;XP_011532900.1;XP_016865143.1;XP_ | 29205 | 617946 | . | . | 64.958376 | 63.992448 | 61.864595 | 68.526043 | 151.160814 | 147.921415 | 138.822355 | 156.676643 | -1.002322 | 5852.069813 | 0.10856045 | -0.03082496 | 0.975409135 | 0.998865404 | 12.516439 | 12.513022 | 12.523973 | 12.506881 | 12.473900 | 12.551599 | 6815 | 6258 | 4848 | 5733 |
| 57223 | NM_00112296 | PPP4R3B | protein phosphatase 4 regulatory subunit 3B | protein_coding | NP_001116436.3;NP_001269779.1;NP_065196.1;XP_005264499.1;XP_00 | 29267 | 610352 | . | . | 11.103594 | 10.521525 | 11.125247 | 8.310145 | 25.838520 | 24.320977 | 24.964733 | 19.000154 | -1.141554 | 1684.187398 | 0.156815388 | -1.21798539 | 0.223229525 | 0.982008277 | 10.809891 | 10.619506 | 10.805137 | 10.721009 | 10.775631 | 10.530663 | 2176 | 1835 | 1585 | 1241 |
| 57224 | NM_00114406 | NHSL1 | NHS like 1 | protein_coding | NP_001137532.1;NP_065197.1;XP_005267119.1;XP_011534268.1;XP_01 | 21021 | . | . | . | 4.606360 | 5.290667 | 5.425821 | 5.783593 | 10.719188 | 12.229611 | 12.175387 | 13.223498 | 1.140105 | 1350.526909 | 0.131201785 | 1.441801427 | 0.1493584 | 0.910598055 | 10.301785 | 10.490652 | 10.308511 | 10.382745 | 10.430585 | 10.461840 | 1382 | 1429 | 1226 | 1376 |
| 57226 | NM_020466,N | LYRM2 | LYR motif containing 2 | protein_coding | NP_065199.1 | 25229 | . | . | . | 7.587138 | 7.633559 | 8.083601 | 7.203222 | 17.655578 | 17.645311 | 18.139366 | 16.469311 | -1.009317 | 1280.455317 | 0.134137502 | -0.09974008 | 0.920550682 | 0.998865404 | 10.328947 | 10.315906 | 10.325321 | 10.325407 | 10.364759 | 10.269937 | 1481 | 1388 | 1178 | 1120 |
| 57228 | NM_00103162 | SMAGP | small cell adhesion glycoprotein | protein_coding | NP_001026798.1;NP_001029045.1 | 26918 | . | . | . | 11.774791 | 13.950821 | 12.739451 | 12.646879 | 27.400422 | 32.247950 | 28.586959 | 28.915588 | -1.001596 | 718.459343 | 0.151061759 | -0.01523162 | 0.987847394 | 0.999197585 | 9.490085 | 9.487439 | 9.461863 | 9.515662 | 9.475549 | 9.500148 | 796 | 806 | 613 | 685 |
| 57231 | NM_00129761 | SNX14 | sorting nexin 14 | protein_coding | NP_001284543.1;NP_001291408.1;NP_001337461.1;NP_001337462.1;NI | 14977 | 616105 | . | . | 16.536340 | 15.347224 | 12.833721 | 9.630682 | 38.480750 | 35.475799 | 28.798499 | 22.019414 | -1.441749 | 1550.529847 | 0.153384413 | -3.44115638 | **0.000579234** | **0.045949208** | 10.838167 | 10.311039 | 10.753856 | 10.685165 | 10.539940 | 10.322338 | 2197 | 1892 | 1267 | 1016 |
| 57234 | NR_111950,NF | LINC00869 | long intergenic non-protein coding RNA 869 | lncRNA | . | 29050 | . | . | . | 2.229706 | 2.626379 | 1.452589 | 2.406339 | 5.188617 | 6.070993 | 3.259568 | 5.501809 | -1.267240 | 106.007389 | 0.329498662 | -1.03700049 | 0.299735628 | 0.998865404 | 6.889947 | 6.545595 | 6.741282 | 6.784509 | 6.633950 | 6.740074 | 128 | 136 | 66 | 104 |
| 57291 | NR_024031,NF | DANCR | differentiation antagonizing non-protein coding RNA | lncRNA | . | 28964 | 614625 | . | . | 54.986162 | 60.182829 | 55.787691 | 60.554271 | 127.955059 | 139.115310 | 125.185966 | 138.450153 | -1.004609 | 1657.492568 | 0.126582131 | -0.05241025 | 0.95820181 | 0.998865404 | 10.698251 | 10.691314 | 10.663324 | 10.728199 | 10.664673 | 10.719742 | 1834 | 1867 | 1387 | 1605 |
| 57325 | NM_020536,N | KAT14 | lysine acetyltransferase 14 | protein_coding | NP_065397.3 | 15904 | 617501 | . | . | 3.481775 | 4.141156 | 3.735999 | 4.535175 | 8.102236 | 9.572468 | 8.383473 | 10.369139 | 1.078011 | 451.883871 | 0.187570031 | 0.577769912 | 0.563419472 | 0.998865404 | 8.765222 | 8.872404 | 8.743709 | 8.841649 | 8.777841 | 8.898340 | 461 | 507 | 368 | 482 |
| 57326 | NM_00131773 | PBXIP1 | PBX homeobox interacting protein 1 | protein_coding | NP_001304663.1;NP_001304664.1;NP_065385.2 | 21199 | 618819 | . | . | 11.547400 | 11.298353 | 10.454353 | 12.350717 | 26.871275 | 26.116650 | 23.459266 | 28.238448 | -1.018282 | 1139.365534 | 0.140594966 | -0.18590596 | 0.852518511 | 0.998865404 | 10.167210 | 10.140698 | 10.171457 | 10.148744 | 10.086859 | 10.202891 | 1344 | 1222 | 905 | 1141 |
| 57332 | NM_020649 | CBX8 | chromobox 8 | protein_coding | NP_065700.1 | 15962 | 617354 | . | . | 3.846019 | 4.275796 | 3.833422 | 4.614681 | 8.949845 | 9.883694 | 8.602089 | 10.550920 | 1.026038 | 484.276530 | 0.178481096 | 0.207779906 | 0.835400823 | 0.998865404 | 8.901519 | 8.937629 | 8.879363 | 8.940446 | 8.869863 | 8.979804 | 524 | 541 | 389 | 500 |
| 57333 | NM_020650,XI | RCN3 | reticulocalbin 3 | protein_coding | NP_065701.2;XP_024307388.1 | 21145 | . | . | . | 55.950771 | 50.894843 | 46.231359 | 46.253570 | 130.199737 | 117.645714 | 103.741830 | 105.753290 | -1.171780 | 2298.133910 | 0.119778078 | -1.90938365 | 0.056212618 | 0.703410744 | 11.275993 | 11.047460 | 11.265585 | 11.185436 | 11.108244 | 11.089711 | 2995 | 2545 | 1862 | 1961 |
| 57335 | NM_00113084 | ZNF286A | zinc finger protein 286A | protein_coding | NP_001124314.1;NP_001275571.1;NP_001275572.1;NP_001275573.2;NI | 13501 | . | . | . | 2.741296 | 3.254173 | 3.898416 | 3.225806 | 6.379108 | 7.522162 | 8.747933 | 7.375422 | 1.183612 | 552.820749 | 0.179832609 | 1.352344914 | 0.176265003 | 0.949538805 | 8.983869 | 9.227217 | 8.989786 | 9.105366 | 9.222453 | 9.090442 | 530 | 596 | 573 | 506 |
| 57336 | NM_00134616 | ZNF287 | zinc finger protein 287 | protein_coding | NP_001330096.1;NP_001333097.1;NP_001333098.1;NP_001333099.1;NI | 13502 | . | . | . | 0.505478 | 0.476980 | 0.365599 | 0.694407 | 1.176267 | 1.102561 | 0.820394 | 1.587679 | 1.067745 | 123.049965 | 0.333896036 | 0.283245669 | 0.776988522 | 0.998865404 | 6.896242 | 6.988484 | 6.936069 | 6.911686 | 6.838666 | 7.044546 | 142 | 124 | 78 | 154 |
| 57337 | NM_00107720 | SENP7 | SUMO specific peptidase 7 | protein_coding | NP_001070671.1;NP_001269730.1;NP_001269731.1;NP_001269732.1;NI | 30402 | 612846 | . | . | 0.860033 | 0.691965 | 1.028234 | 0.676537 | 2.001331 | 1.599507 | 2.307328 | 1.546823 | 1.051871 | 137.731162 | 0.314229741 | 0.232179094 | 0.816398914 | 0.998865404 | 7.067319 | 7.143108 | 7.119780 | 7.054459 | 7.196408 | 7.007645 | 168 | 132 | 154 | 99 |
| 57343 | NM_00129031 | ZNF304 | zinc finger protein 304 | protein_coding | NP_001277247.1;NP_001277248.1;NP_001316385.1;NP_065708.2;XP_01 | 13505 | 613840 | . | . | 2.182340 | 2.539317 | 2.196874 | 2.395521 | 5.078394 | 5.869742 | 4.929723 | 5.477075 | -1.022191 | 328.360897 | 0.201262989 | -0.15733205 | 0.874983159 | 0.998865404 | 8.375402 | 8.342690 | 8.323163 | 8.403960 | 8.317993 | 8.384969 | 355 | 384 | 265 | 323 |
| 57380 | NM_00128626 | MRS2 | magnesium transporter MRS2 | protein_coding | NP_001273193.1;NP_001273194.1;NP_001273195.1;NP_065713.1;XP_00 | 13785 | . | . | . | 8.979629 | 8.414534 | 9.933626 | 9.129359 | 20.895966 | 19.450573 | 22.290767 | 20.873192 | 1.011104 | 893.824628 | 0.141279507 | 0.112763353 | 0.910218173 | 0.998865404 | 9.795730 | 9.811920 | 9.816119 | 9.782918 | 9.822586 | 9.792541 | 1047 | 937 | 798 | 825 |
| 57381 | NM_020663,XI | RHOJ | ras homolog family member J | protein_coding | NP_065714.1;XP_006720277.1;XP_011535295.1 | 688 | 607653 | . | . | 4.891162 | 4.506652 | 6.512869 | 4.841193 | 11.381932 | 10.417330 | 14.614686 | 11.068815 | 1.194884 | 535.767935 | 0.184025102 | 1.395846554 | 0.162760708 | 0.935298107 | 8.930717 | 9.188694 | 9.026817 | 8.973091 | 9.203599 | 9.019124 | 586 | 504 | 578 | 471 |
| 57396 | NM_020666 | CLK4 | CDC like kinase 4 | protein_coding | NP_065717.1 | 13659 | 607969 | . | . | 5.758798 | 5.209195 | 7.408366 | 5.167966 | 13.400959 | 12.041286 | 16.624159 | 11.815941 | 1.130323 | 459.762976 | 0.203691695 | 0.867661213 | 0.385579825 | 0.998865404 | 8.752935 | 8.931063 | 8.832146 | 8.765367 | 8.984513 | 8.753355 | 524 | 440 | 502 | 374 |
| 57403 | NM_020673 | RAB22A | RAB22A, member RAS oncogene family | protein_coding | NP_065724.1 | 9764 | 612966 | . | . | 5.308697 | 4.790108 | 5.840457 | 4.966356 | 12.353554 | 11.072550 | 13.105817 | 11.354984 | 1.055141 | 1409.259343 | 0.140856969 | 0.549748342 | 0.582491994 | 0.998865404 | 10.421200 | 10.499189 | 10.479951 | 10.394544 | 10.543824 | 10.410924 | 1667 | 1397 | 1367 | 1240 |
| 57404 | NM_00137169 | CYP20A1 | cytochrome P450 family 20 subfamily A member 1 | protein_coding | NP_001358624.1;NP_001358625.1;NP_001358626.1;NP_001358627.1;NI | 20576 | . | . | . | 2.000521 | 1.671488 | 1.748165 | 1.721182 | 4.655296 | 3.863720 | 3.922835 | 3.935275 | 1.037476 | 531.570809 | 0.169506293 | 0.313136768 | 0.754176756 | 0.998865404 | 9.026978 | 9.080757 | 9.081399 | 8.999222 | 9.090782 | 9.038499 | 639 | 527 | 490 | 487 |
| 57405 | NM_020675,XI | SPC25 | SPC25 component of NDC80 kinetochore complex | protein_coding | NP_065726.1;XP_011509818.1 | 24031 | 609395 | . | . | 16.804038 | 17.871683 | 12.690991 | 12.869711 | 39.103690 | 41.311201 | 28.478214 | 29.425063 | -1.381266 | 624.634868 | 0.156557756 | -2.97648271 | **0.002915755** | 0.147963858 | 9.501191 | 9.035058 | 9.364816 | 9.402608 | 9.168284 | 9.168154 | 812 | 803 | 456 | 492 |
| 57406 | NM_00132012 | ABHD6 | abhydrolase domain containing 6, acylglycerol lipase | protein_coding | NP_001307055.1;NP_065727.4;XP_005265391.1;XP_005265392.1 | 21398 | 616966 | . | . | 8.424058 | 8.196621 | 6.546060 | 8.126615 | 19.603127 | 18.946854 | 14.689166 | 18.580541 | -1.166614 | 558.447078 | 0.173100895 | -1.28438116 | 0.199008623 | 0.96621494 | 9.232479 | 9.009465 | 9.192213 | 9.152374 | 9.003661 | 9.135838 | 713 | 630 | 401 | 534 |
| 57407 | NM_00130514 | NMRAL1 | NmrA like redox sensor 1 | protein_coding | NP_001292070.1;NP_001292071.1;NP_001338923.1;NP_001338924.1;NI | 24987 | . | . | . | 26.470989 | 26.962915 | 25.033204 | 29.374513 | 61.599079 | 62.325988 | 56.173781 | 67.161333 | 1.017694 | 1071.864637 | 0.138511477 | 0.182679613 | 0.855049413 | 0.998865404 | 10.053381 | 10.078325 | 10.057446 | 10.060575 | 10.027600 | 10.114333 | 1221 | 1149 | 885 | 1073 |
| 57409 | NM_00124249 | MIF4GD | MIF4G domain containing | protein_coding | NP_001229427.1;NP_001229429.1;NP_001229430.1;NP_001350735.1;NI | 24030 | 612072 | . | . | 6.091149 | 4.534226 | 4.946370 | 5.846910 | 14.174355 | 10.481067 | 11.099511 | 13.368266 | -1.020450 | 241.028532 | 0.236347634 | -0.12356975 | 0.901655943 | 0.998865404 | 7.927784 | 7.898184 | 7.967460 | 7.861788 | 7.847596 | 7.961295 | 310 | 235 | 183 | 250 |
| 57410 | NM_00104821 | SCYL1 | SCY1 like pseudokinase 1 | protein_coding | NP_001041683.1;NP_065731.3;XP_005274175.1;XP_005274177.1;XP_00 | 14372 | 607982 | . | . | 20.160933 | 22.046719 | 22.104474 | 23.310979 | 46.915322 | 50.961977 | 49.601796 | 53.297782 | 1.073686 | 1771.922053 | 0.123457303 | 0.830834671 | 0.406067033 | 0.998865404 | 10.739026 | 10.841359 | 10.735160 | 10.788425 | 10.795630 | 10.839735 | 1901 | 1907 | 1551 | 1768 |
| 57414 | NM_00104045 | RHBDD2 | rhomboid domain containing 2 | protein_coding | NP_001035546.1;NP_001035547.1;NP_001333115.1;NP_001333116.1;NI | 23082 | 615203 | . | . | 16.037727 | 17.621824 | 18.633178 | 18.008845 | 37.320448 | 40.733634 | 41.812314 | 41.175081 | 1.087448 | 976.767430 | 0.140929356 | 0.858203503 | 0.390780102 | 0.998865404 | 9.870217 | 9.990998 | 9.860789 | 9.937142 | 9.970777 | 9.950793 | 1021 | 1063 | 897 | 941 |
| 57415 | NM_00129194 | C3orf14 | chromosome 3 open reading frame 14 | protein_coding | NP_001278870.1;NP_001278871.1;NP_001278872.1;NP_065736.1;XP_01 | 25024 | . | . | . | 19.997419 | 18.285913 | 14.541234 | 10.109010 | 46.534812 | 42.268707 | 32.630105 | 23.113051 | -1.212642 | 967.747792 | 0.159634581 | -1.74244119 | 0.081431273 | 0.782498736 | 10.050481 | 9.773190 | 10.044883 | 9.915361 | 9.913622 | 9.764714 | 1326 | 1046 | 841 | 734 |
| 57418 | NM_00137208 | WDR18 | WD repeat domain 18 | protein_coding | NP_001359014.1;NP_001359015.1;NP_077005.2 | 17956 | . | . | . | 14.099141 | 15.734033 | 14.558850 | 17.307239 | 32.809280 | 36.369924 | 32.669639 | 39.570947 | 1.002764 | 762.637713 | 0.154216222 | 0.025820816 | 0.979400258 | 0.998865404 | 9.573142 | 9.576564 | 9.572355 | 9.574159 | 9.516959 | 9.630003 | 876 | 823 | 610 | 774 |
| 57446 | NM_022477,N | NDRG3 | NDRG family member 3 | protein_coding | NP_071922.2;NP_114402.1;XP_006723900.1;XP_006723901.1;XP_01152 | 14462 | 605273 | . | . | 15.863704 | 15.527995 | 18.767279 | 19.236415 | 36.915486 | 35.893657 | 42.113230 | 43.981778 | 1.185431 | 1606.843080 | 0.122567583 | 2.002251864 | **0.04525765** | 0.649595749 | 10.522096 | 10.767507 | 10.586581 | 10.572525 | 10.705570 | 10.718065 | 1709 | 1572 | 1504 | 1647 |
| 57447 | NM_00128221 | NDRG2 | NDRG family member 2 | protein_coding | NP_001269140.1;NP_001269141.1;NP_001269142.1;NP_001269143.1;NI | 14460 | 605272 | . | . | 5.717281 | 5.248858 | 4.498563 | 4.891832 | 13.304345 | 12.132973 | 10.094644 | 11.184592 | -1.171533 | 353.130704 | 0.188734449 | -1.21015057 | 0.226221123 | 0.982008277 | 8.573798 | 8.345279 | 8.509486 | 8.505901 | 8.410531 | 8.424777 | 440 | 410 | 279 | 309 |
| 57448 | NM_00137812 | BIRC6 | baculoviral IAP repeat containing 6 | protein_coding | NP_001365054.1;NP_057336.3;XP_005264507.2;XP_005264508.2;XP_00 | 13516 | 605638 | . | . | 4.553877 | 4.466746 | 4.647442 | 4.801561 | 10.597059 | 10.325081 | 10.428727 | 10.978201 | 1.034653 | 2267.422351 | 0.115639082 | 0.425003405 | 0.670834193 | 0.998865404 | 11.122064 | 11.171193 | 11.140563 | 11.125253 | 11.150883 | 11.169575 | 2591 | 2382 | 1982 | 2187 |
| 57449 | NM_00104266 | PLEKHG5 | pleckstrin homology and RhoGEF domain containing G5 | protein_coding | NP_001036128.2;NP_001036129.1;NP_001036130.1;NP_001252521.2;NI | 29105 | 611101 | . | . | 2.978056 | 2.495779 | 2.160071 | 3.108510 | 6.930059 | 5.769107 | 4.847136 | 7.107239 | -1.093227 | 354.717054 | 0.195152948 | -0.6589331 | 0.509938727 | 0.998865404 | 8.533541 | 8.404630 | 8.526762 | 8.462497 | 8.404820 | 8.482290 | 450 | 378 | 274 | 340 |
| 57455 | NM_020695,XI | REXO1 | RNA exonuclease 1 homolog | protein_coding | NP_065746.3;XP_011526446.1;XP_011526448.1;XP_016882517.1;XP_01 | 24616 | 609614 | . | . | 4.598385 | 4.406952 | 5.236434 | 4.813027 | 10.700632 | 10.186867 | 11.750404 | 11.004415 | 1.107421 | 690.841939 | 0.153298392 | 0.960240934 | 0.336933969 | 0.998865404 | 9.356508 | 9.504140 | 9.406842 | 9.383317 | 9.495909 | 9.433441 | 768 | 695 | 660 | 650 |
| 57456 | NM_00132033 | KIAA1143 | KIAA1143 | protein_coding | NP_001307263.1;NP_065747.1 | 29198 | . | . | . | 2.922790 | 2.788635 | 3.148247 | 3.152867 | 6.801452 | 6.446055 | 7.064575 | 7.208656 | 1.088658 | 476.224605 | 0.170226589 | 0.719926794 | 0.47157007 | 0.998865404 | 8.832799 | 8.955586 | 8.885004 | 8.849813 | 8.923951 | 8.917483 | 540 | 478 | 434 | 463 |
| 57458 | NM_00130103 | TMCC3 | transmembrane and coiled-coil domain family 3 | protein_coding | NP_001287965.1;NP_065749.3 | 29199 | 617459 | . | . | 0.416701 | 0.346105 | 0.266454 | 0.332756 | 0.969680 | 0.800037 | 0.597915 | 0.760808 | -1.280527 | 62.871412 | 0.403971736 | -0.88307558 | 0.377195436 | 0.998865404 | 6.141971 | 5.784621 | 6.025541 | 5.978858 | 5.928448 | 5.967875 | 89 | 69 | 43 | 57 |
| 57459 | NM_020699,XI | GATAD2B | GATA zinc finger domain containing 2B | protein_coding | NP_065750.1;XP_005245421.1;XP_024304389.1 | 30778 | 614998 | . | . | 3.091913 | 3.354933 | 3.626428 | 3.473269 | 7.195010 | 7.755078 | 8.137600 | 7.941218 | 1.093607 | 778.641107 | 0.14897042 | 0.866580486 | 0.386171909 | 0.998865404 | 9.538803 | 9.667938 | 9.544031 | 9.598515 | 9.659627 | 9.608512 | 823 | 834 | 733 | 735 |
| 57460 | NM_020700,XI | PPM1H | protein phosphatase, Mg2+/Mn2+ dependent 1H | protein_coding | NP_065751.1;XP_011536880.1;XP_011536881.1;XP_016875165.1 | 18583 | 616016 | . | . | 2.462971 | 2.571805 | 2.689674 | 2.442413 | 5.731435 | 5.944843 | 6.035549 | 5.584288 | 1.028165 | 494.145863 | 0.169686058 | 0.236150188 | 0.813316131 | 0.998865404 | 8.928511 | 8.968794 | 8.924465 | 8.953994 | 8.983035 | 8.930132 | 548 | 538 | 454 | 450 |
| 57461 | NM_00119946 | ISY1 | ISY1 splicing factor homolog | protein_coding | NP_001186398.1;NP_065752.1 | 29201 | 612764 | . | . | 7.236445 | 7.638725 | 7.089917 | 8.309090 | 16.839504 | 17.657253 | 15.909568 | 18.997747 | 1.022951 | 852.514168 | 0.149164711 | 0.219471607 | 0.826282695 | 0.998865404 | 9.719409 | 9.751571 | 9.710641 | 9.742645 | 9.689567 | 9.793385 | 949 | 930 | 695 | 867 |
| 57462 | NM_020702,XI | MYORG | myogenesis regulating glycosidase (putative) | protein_coding | NP_065753.2;XP_011516268.1;XP_016870419.1 | 19918 | 618255 | . | . | 1.942213 | 1.681435 | 2.655343 | 3.049138 | 4.519610 | 3.886712 | 5.958512 | 6.971492 | 1.487489 | 477.250540 | 0.186218404 | 3.07638284 | **0.002095286** | 0.116149544 | 8.583704 | 9.156932 | 8.818223 | 8.683977 | 8.957911 | 9.043028 | 493 | 366 | 465 | 569 |
| 57463 | NM_020703,XI | AMIGO1 | adhesion molecule with Ig like domain 1 | protein_coding | NP_065754.2;XP_011540114.1 | 20824 | 615689 | . | . | 1.999344 | 1.931294 | 2.111522 | 2.290843 | 4.652556 | 4.464274 | 4.738195 | 5.237738 | 1.104559 | 333.267731 | 0.194035952 | 0.739399038 | 0.459664726 | 0.998865404 | 8.307113 | 8.450406 | 8.363378 | 8.338480 | 8.385395 | 8.427158 | 373 | 334 | 293 | 340 |
| 57464 | NM_00113433 | STRIP2 | striatin interacting protein 2 | protein_coding | NP_001127808.1;NP_065755.1;XP_006716132.1;XP_011514734.1;XP_01 | 22209 | 617919 | . | . | 9.358743 | 9.661947 | 12.351898 | 10.201886 | 21.778176 | 22.334025 | 27.717304 | 23.325402 | 1.194107 | 1690.348443 | 0.138169516 | 1.852305516 | 0.063981964 | 0.731304863 | 10.589287 | 10.845557 | 10.637755 | 10.656923 | 10.867175 | 10.698660 | 1752 | 1683 | 1769 | 1543 |
| 57465 | NM_00119910 | TBC1D24 | TBC1 domain family member 24 | protein_coding | NP_001186036.1;NP_065756.1;XP_016878982.1;XP_016878983.1;XP_01 | 29203 | 613577 | . | . | 1.771620 | 2.173557 | 2.240967 | 2.296396 | 4.122635 | 5.024274 | 5.028667 | 5.250435 | 1.106025 | 449.857993 | 0.173809595 | 0.836454825 | 0.402899081 | 0.998865404 | 8.738952 | 8.884053 | 8.760477 | 8.800114 | 8.837994 | 8.846809 | 476 | 476 | 408 | 446 |
| 57466 | NM_00114544 | SCAF4 | SR-related CTD associated factor 4 | protein_coding | NP_001138916.1;NP_001138917.1;NP_065757.1;XP_005261074.1;XP_00 | 19304 | 616023 | . | . | 6.765756 | 7.339770 | 7.752218 | 8.079071 | 15.744193 | 16.966205 | 17.395752 | 18.471835 | 1.094492 | 1048.730544 | 0.137141364 | 0.949833263 | 0.342196981 | 0.998865404 | 9.968003 | 10.097929 | 9.979553 | 10.019292 | 10.039981 | 10.091599 | 1122 | 1110 | 920 | 1063 |
| 57470 | NM_020710 | LRRC47 | leucine rich repeat containing 47 | protein_coding | NP_065761.1 | 29207 | . | . | . | 19.532032 | 19.346024 | 18.151417 | 20.189741 | 45.451843 | 44.719200 | 40.731255 | 46.161446 | -1.028397 | 2587.801445 | 0.117477326 | -0.34387623 | 0.730939369 | 0.998865404 | 11.357641 | 11.317096 | 11.355886 | 11.341157 | 11.286313 | 11.363849 | 3049 | 2806 | 2112 | 2507 |
| 57472 | NM_00137047 | CNOT6 | CCR4-NOT transcription complex subunit 6 | protein_coding | NP_001357401.1;NP_001357402.1;NP_001357403.1;XP_016865160.1;XP | 14099 | 608951 | . | . | 5.454825 | 5.330506 | 5.544262 | 5.306133 | 12.693603 | 12.321702 | 12.441164 | 12.131840 | -1.001704 | 1046.172904 | 0.136462906 | -0.01799612 | 0.985641945 | 0.999197585 | 10.031987 | 10.029823 | 10.048031 | 10.014289 | 10.051678 | 10.008130 | 1233 | 1104 | 936 | 951 |
| 57473 | NM_020713,XI | ZNF512B | zinc finger protein 512B | protein_coding | NP_065764.1;XP_011527232.1;XP_011527235.1;XP_024307718.1;XP_02 | 29212 | 617886 | . | . | 10.335202 | 10.600436 | 12.089625 | 10.219406 | 24.050436 | 24.503381 | 27.128771 | 23.365454 | 1.052317 | 2039.980899 | 0.126808716 | 0.580158287 | 0.56180788 | 0.998865404 | 10.956941 | 11.030794 | 10.973828 | 10.971706 | 11.072207 | 10.951181 | 2292 | 2142 | 1957 | 1813 |
| 57474 | NM_020714 | ZNF490 | zinc finger protein 490 | protein_coding | NP_065765.1 | 23705 | . | . | . | 0.490141 | 0.627720 | 0.585428 | 0.605949 | 1.140578 | 1.451002 | 1.313683 | 1.385430 | 1.049841 | 110.204473 | 0.312920419 | 0.224244854 | 0.822566784 | 0.998865404 | 6.749112 | 6.818140 | 6.733818 | 6.812011 | 6.785647 | 6.793188 | 109 | 130 | 97 | 107 |
| 57475 | NM_020715,XI | PLEKHH1 | pleckstrin homology, MyTH4 and FERM domain containing H1 | protein_coding | NP_065766.1;XP_011535311.1;XP_011535317.1;XP_016876988.1;XP_01 | 17733 | . | . | . | 6.727854 | 7.225320 | 11.261941 | 8.951394 | 15.655992 | 16.701653 | 25.271467 | 20.466298 | 1.437563 | 1775.492545 | 0.141717967 | 3.69483828 | **0.000220027** | **0.022007462** | 10.508408 | 11.032298 | 10.608498 | 10.652146 | 11.007406 | 10.815372 | 1629 | 1617 | 2035 | 1733 |
| 57477 | NM_020717,N | SHROOM4 | shroom family member 4 | protein_coding | NP_065768.2;XP_016885171.1;XP_016885172.1;XP_016885173.1;XP_01 | 29215 | 300579 | . | . | 1.493586 | 1.452183 | 1.496149 | 1.762387 | 3.475634 | 3.356784 | 3.357316 | 4.029487 | 1.132963 | 433.897346 | 0.179698339 | 1.002240066 | 0.316227661 | 0.998865404 | 8.668617 | 8.848217 | 8.727194 | 8.713495 | 8.752361 | 8.838421 | 474 | 434 | 373 | 462 |
| 57478 | NM_020718,XI | USP31 | ubiquitin specific peptidase 31 | protein_coding | NP_065769.3;XP_016878985.1;XP_016878986.1;XP_016878987.1;XP_01 | 20060 | . | . | . | 1.347436 | 1.273842 | 1.858377 | 1.148680 | 3.135540 | 2.944541 | 4.170145 | 2.626320 | 1.227374 | 468.319983 | 0.197724412 | 1.494881582 | 0.13494535 | 0.891755273 | 8.715378 | 9.012094 | 8.805871 | 8.788729 | 9.029778 | 8.808578 | 491 | 447 | 526 | 401 |
| 57479 | NM_020719,XI | PRR12 | proline rich 12 | protein_coding | NP_065770.1 | 29217 | 616633 | . | . | 4.047150 | 4.163811 | 3.697547 | 4.059084 | 9.417884 | 9.624835 | 8.297188 | 9.280614 | -1.072923 | 922.672335 | 0.141275238 | -0.71878603 | 0.472272768 | 0.998865404 | 9.899715 | 9.797836 | 9.867314 | 9.881854 | 9.793774 | 9.852606 | 1089 | 1041 | 742 | 869 |
| 57480 | NM_00102988 | PLEKHG1 | pleckstrin homology and RhoGEF domain containing G1 | protein_coding | NP_001025055.1;NP_001316727.1;NP_001316728.1;NP_001316729.1;NI | 20884 | . | . | . | 0.180050 | 0.151430 | 0.114205 | 0.164837 | 0.418983 | 0.350037 | 0.256271 | 0.376880 | -1.518687 | 31.866934 | 0.560484668 | -1.07554113 | 0.282132529 | 0.998865404 | 5.264940 | 4.660102 | 5.036706 | 5.010623 | 4.953122 | 4.989225 | 48 | 38 | 18 | 28 |
| 57482 | NM_020722,XI | CRACD | capping protein inhibiting regulator of actin dynamics | protein_coding | NP_065773.1;XP_005265809.1;XP_005265812.1;XP_005265813.1;XP_00 | 29219 | 618327 | . | . | 2.040970 | 2.061165 | 1.276533 | 1.979044 | 4.749421 | 4.764475 | 2.864501 | 4.524847 | -1.269589 | 398.932718 | 0.216749691 | -1.58875112 | 0.112116593 | 0.849366267 | 8.802630 | 8.456684 | 8.701966 | 8.702051 | 8.433617 | 8.673048 | 514 | 482 | 240 | 400 |
| 57484 | NM_020724,XI | RNF150 | ring finger protein 150 | protein_coding | NP_065775.1;XP_005263207.1;XP_011530449.1;XP_011530450.1;XP_01 | 23138 | . | . | . | 0.555997 | 0.388524 | 0.517147 | 0.451459 | 1.293828 | 0.898091 | 1.160463 | 1.032207 | 1.014888 | 159.017628 | 0.283569914 | 0.075186023 | 0.940066705 | 0.998865404 | 7.301378 | 7.324614 | 7.380043 | 7.225978 | 7.336193 | 7.287076 | 216 | 138 | 148 | 141 |
| 57486 | NM_020726,XI | NLN | neurolysin | protein_coding | NP_065777.1;XP_005248616.1;XP_006714724.1;XP_016865162.1 | 16058 | 611530 | . | . | 2.918424 | 3.009918 | 3.030632 | 2.628762 | 6.791293 | 6.957562 | 6.800650 | 6.010352 | -1.129420 | 717.636293 | 0.150824546 | -1.16414865 | 0.244363776 | 0.992733415 | 9.572331 | 9.396536 | 9.500151 | 9.554327 | 9.442445 | 9.444702 | 843 | 853 | 585 | 633 |
| 57488 | NM_00136777 | ESYT2 | extended synaptotagmin 2 | protein_coding | NP_001354702.1;NP_065779.2;XP_024302614.1;XP_024302615.1;XP_02 | 22211 | 616691 | . | . | 17.936926 | 17.059332 | 18.390901 | 18.483452 | 41.739961 | 39.433411 | 41.268653 | 42.260216 | 1.019337 | 3695.457411 | 0.108266566 | 0.255216768 | 0.79855568 | 0.998865404 | 11.837653 | 11.865289 | 11.850082 | 11.836958 | 11.855979 | 11.862765 | 4248 | 3918 | 3231 | 3512 |
| 57489 | NM_00100702 | ODF2L | outer dense fiber of sperm tails 2 like | protein_coding | NP_001007023.2;NP_001171694.1;NP_001171695.1;NP_001353708.1;NI | 29225 | . | . | . | 5.330986 | 4.372211 | 5.221534 | 4.166243 | 12.405422 | 10.106564 | 11.716971 | 9.525618 | -1.047963 | 896.366774 | 0.161474334 | -0.41856796 | 0.675531911 | 0.998865404 | 9.840913 | 9.774207 | 9.879950 | 9.761112 | 9.862280 | 9.709471 | 1142 | 909 | 843 | 733 |
| 57491 | NM_00137723 | AHRR | aryl-hydrocarbon receptor repressor | protein_coding | NP_001364165.1;NP_001364168.1 | 346 | 606517 | . | . | 4.516718 | 4.667144 | 6.262378 | 5.302779 | 10.510588 | 10.788314 | 14.052594 | 12.124174 | 1.241618 | 915.049071 | 0.149424466 | 2.089495434 | **0.036663149** | 0.596884093 | 9.672930 | 9.985576 | 9.742549 | 9.760626 | 9.971066 | 9.840898 | 928 | 892 | 959 | 867 |
| 57492 | NM_00136372 | ARID1B | AT-rich interaction domain 1B | protein_coding | NP_001350654.1;NP_001358585.1;NP_001361749.1;NP_001361757.1;NI | 18040 | 614556 | . | . | 4.266330 | 4.904539 | 5.060928 | 6.324588 | 9.927925 | 11.337062 | 11.356575 | 14.460418 | 1.065164 | 1501.851640 | 0.132328547 | 0.688252273 | 0.491293931 | 0.998865404 | 10.506468 | 10.597161 | 10.476896 | 10.574450 | 10.547411 | 10.602870 | 1569 | 1669 | 1299 | 1504 |
| 57493 | NM_020733,XI | HEG1 | heart development protein with EGF like domains 1 | protein_coding | NP_065784.1;XP_005247723.1 | 29227 | 614182 | . | . | 8.262047 | 8.534842 | 6.784506 | 6.027975 | 19.226122 | 19.728670 | 15.224235 | 13.782249 | -1.317975 | 2158.781026 | 0.123865794 | -3.21576102 | **0.001300991** | 0.083136952 | 11.261351 | 10.863279 | 11.164521 | 11.186334 | 11.004484 | 10.903050 | 2789 | 2684 | 1726 | 1632 |
| 57494 | NM_00129777 | RIMKLB | ribosomal modification protein rimK like family member B | protein_coding | NP_001284705.1;NP_001339196.1;NP_001339197.1;NP_001339198.1;NI | 29228 | 614054 | . | . | 4.328936 | 3.906147 | 4.111258 | 3.219384 | 10.073612 | 9.029235 | 9.225542 | 7.360737 | -1.141883 | 609.321947 | 0.180044262 | -1.06315483 | 0.287711742 | 0.998865404 | 9.343040 | 9.152816 | 9.353256 | 9.224197 | 9.277014 | 9.122297 | 818 | 635 | 554 | 470 |
| 57496 | NM_00130814 | MRTFB | myocardin related transcription factor B | protein_coding | NP_001295071.1;NP_001352340.1;NP_001352341.1;NP_001352342.1;NI | 29819 | 609463 | . | . | 2.648323 | 3.065237 | 3.473734 | 2.778002 | 6.162757 | 7.085432 | 7.794956 | 6.351572 | 1.100564 | 774.546780 | 0.154784055 | 0.893133707 | 0.371785574 | 0.998865404 | 9.526342 | 9.664756 | 9.523310 | 9.597273 | 9.675348 | 9.578754 | 804 | 838 | 754 | 709 |
| 57498 | NM_00134872 | KIDINS220 | kinase D interacting substrate 220 | protein_coding | NP_001335658.1;NP_001335660.1;NP_001335661.1;NP_001335663.1;NI | 29508 | 615759 | . | . | 10.246791 | 10.219210 | 9.989278 | 9.619415 | 23.844706 | 23.622162 | 22.415652 | 21.993649 | -1.058242 | 2285.136500 | 0.116315578 | -0.70213932 | 0.482592282 | 0.998865404 | 11.198265 | 11.116712 | 11.184612 | 11.176003 | 11.154542 | 11.114896 | 2720 | 2522 | 1979 | 2030 |
| 57504 | NM_00128275 | MTA3 | metastasis associated 1 family member 3 | protein_coding | NP_001269684.1;NP_001269685.1;NP_001317371.1;NP_001317372.1;NI | 23784 | 609050 | . | . | 10.134008 | 7.856961 | 10.259547 | 9.591706 | 23.582255 | 18.161713 | 23.022128 | 21.930299 | 1.111028 | 801.677587 | 0.146203381 | 1.03893029 | 0.298837158 | 0.998865404 | 9.568901 | 9.720856 | 9.631772 | 9.583343 | 9.668522 | 9.695491 | 905 | 791 | 720 | 806 |
| 57505 | NM_020745,XI | AARS2 | alanyl-tRNA synthetase 2, mitochondrial | protein_coding | NP_065796.2;XP_005249302.1;XP_011513066.1;XP_016866601.1 | 21022 | 612035 | . | . | 11.355710 | 12.620143 | 13.430120 | 13.412032 | 26.425205 | 29.172027 | 30.136800 | 30.665018 | 1.108756 | 1886.786510 | 0.122280059 | 1.218039742 | 0.223208871 | 0.982008277 | 10.805377 | 10.954217 | 10.804290 | 10.872527 | 10.928236 | 10.912847 | 1971 | 2015 | 1742 | 1842 |
| 57506 | NM_00120649 | MAVS | mitochondrial antiviral signaling protein | protein_coding | NP_001193420.1;NP_065797.2 | 29233 | 609676 | . | . | 6.884927 | 7.776194 | 7.355471 | 7.521207 | 16.021507 | 17.975021 | 16.505466 | 17.196346 | 1.001112 | 2696.233208 | 0.118486803 | 0.013526454 | 0.98920778 | 0.999197585 | 11.396005 | 11.397454 | 11.348912 | 11.440807 | 11.391257 | 11.402170 | 2928 | 3072 | 2330 | 2546 |
| 57507 | NM_020747,XI | ZNF608 | zinc finger protein 608 | protein_coding | NP_065798.2;XP_005272094.1;XP_005272095.1;XP_005272096.1;XP_00 | 29238 | . | . | . | 2.334677 | 2.546502 | 2.468554 | 2.523437 | 5.432889 | 5.886355 | 5.539365 | 5.769537 | 1.027337 | 452.599344 | 0.176752953 | 0.220138721 | 0.825763119 | 0.998865404 | 8.802744 | 8.841182 | 8.772201 | 8.852166 | 8.825114 | 8.833850 | 481 | 513 | 396 | 433 |
| 57508 | NM_00133041 | INTS2 | integrator complex subunit 2 | protein_coding | NP_001317346.2;NP_001338624.2;NP_065799.2;XP_011523360.1 | 29241 | 611346 | . | . | 3.502606 | 3.485775 | 3.736325 | 2.768386 | 8.150707 | 8.057525 | 8.384204 | 6.329586 | -1.108224 | 634.880822 | 0.175350662 | -0.84544597 | 0.397861888 | 0.998865404 | 9.382089 | 9.234839 | 9.350950 | 9.331700 | 9.361386 | 9.175037 | 779 | 710 | 598 | 485 |
| 57509 | NM_00100192 | MTUS1 | microtubule associated scaffold protein 1 | protein_coding | NP_001001924.1;NP_001001925.1;NP_001001931.1;NP_001159865.1;NI | 29789 | 609589 | . | . | 14.384259 | 14.400598 | 15.751027 | 16.590293 | 33.472765 | 33.287629 | 35.344849 | 37.931730 | 1.120672 | 2192.386854 | 0.116249346 | 1.413893364 | 0.157393215 | 0.926689685 | 11.013770 | 11.178125 | 11.040329 | 11.060904 | 11.148205 | 11.135404 | 2350 | 2260 | 2031 | 2155 |
| 57510 | NM_020750,N | XPO5 | exportin 5 | protein_coding | NP_065801.1 | 17675 | 607845 | . | . | 13.167658 | 14.042348 | 13.753313 | 15.467784 | 30.641684 | 32.459514 | 30.862036 | 35.365246 | 1.055702 | 2355.245180 | 0.121239873 | 0.645029121 | 0.518908314 | 0.998865404 | 11.162165 | 11.240106 | 11.156571 | 11.200856 | 11.177561 | 11.265552 | 2566 | 2541 | 1989 | 2391 |
| 57511 | NM_00114507 | COG6 | component of oligomeric golgi complex 6 | protein_coding | NP_001138551.1;NP_065802.1;XP_011533470.1 | 18621 | 606977 | . | . | 6.511170 | 6.518090 | 6.496688 | 6.844311 | 15.151762 | 15.066858 | 14.578379 | 15.648703 | 1.008759 | 738.338711 | 0.148055855 | 0.084981109 | 0.932276409 | 0.998865404 | 9.521899 | 9.534352 | 9.527929 | 9.522202 | 9.517247 | 9.544747 | 850 | 790 | 632 | 709 |
| 57512 | NM_020752,XI | GPR158 | G protein-coupled receptor 158 | protein_coding | NP_065803.2;XP_016871941.1 | 23689 | 614573 | . | . | 0.563959 | 0.729755 | 0.249759 | 0.382248 | 1.312355 | 1.686861 | 0.560452 | 0.873964 | **-2.285631** | 80.455112 | 0.379973404 | -3.13862167 | **0.001697444** | 0.099278924 | 6.807707 | 5.611258 | 6.376616 | 6.449464 | 6.170857 | 6.239276 | 115 | 134 | 34 | 55 |
| 57513 | NM_00114264 | CASKIN2 | CASK interacting protein 2 | protein_coding | NP_001136115.1;NP_065804.2 | 18200 | 612185 | . | . | 2.928403 | 3.304402 | 3.117081 | 3.315347 | 6.814515 | 7.638274 | 6.994639 | 7.580148 | 1.018530 | 490.381696 | 0.171867405 | 0.154119343 | 0.877515637 | 0.998865404 | 8.924744 | 8.950662 | 8.894964 | 8.966531 | 8.926249 | 8.959139 | 528 | 554 | 419 | 476 |
| 57514 | NM_020754,XI | ARHGAP31 | Rho GTPase activating protein 31 | protein_coding | NP_065805.2;XP_006713777.1;XP_016862444.1 | 29216 | 610911 | . | . | 5.282606 | 4.784384 | 6.964070 | 8.327429 | 12.292840 | 11.059319 | 15.627170 | 19.039675 | 1.435456 | 1654.714280 | 0.124498525 | 4.18888185 | **2.80332E-05** | **0.00460646** | 10.408311 | 10.929588 | 10.515315 | 10.549845 | 10.787271 | 10.841201 | 1529 | 1500 | 1640 | 1889 |
| 57515 | NM_020755 | SERINC1 | serine incorporator 1 | protein_coding | NP_065806.1 | 13464 | 614548 | . | . | 42.956264 | 37.016254 | 40.182060 | 31.003880 | 99.960991 | 85.564728 | 90.167374 | 70.886688 | -1.138902 | 3682.601090 | 0.143566101 | -1.30701974 | 0.191206006 | 0.963720425 | 11.937133 | 11.749810 | 11.958831 | 11.826863 | 11.891324 | 11.671892 | 4870 | 3899 | 3396 | 2796 |
| 57519 | NM_020759,XI | STARD9 | StAR related lipid transfer domain containing 9 | protein_coding | NP_065810.2;XP_011520133.1;XP_011520134.1;XP_011520135.1;XP_01 | 19162 | 614642 | . | . | 0.816855 | 0.876081 | 0.891420 | 0.925448 | 1.900856 | 2.025100 | 2.000321 | 2.115927 | 1.058224 | 427.659241 | 0.177161098 | 0.460851477 | 0.644905167 | 0.998865404 | 8.699104 | 8.780388 | 8.701116 | 8.743077 | 8.747926 | 8.765081 | 462 | 464 | 377 | 418 |
| 57520 | NM_00130484 | HECW2 | HECT, C2 and WW domain containing E3 ubiquitin protein ligase 2 | protein_coding | NP_001291769.1;NP_001335697.1;NP_065811.1;XP_006712709.1;XP_00 | 29853 | 617245 | . | . | 0.053256 | 0.052259 | 0.032425 | 0.031237 | 0.123930 | 0.120799 | 0.072761 | 0.071420 | -1.654398 | 16.887289 | 0.751230648 | -0.96682205 | 0.333633001 | NA | 4.395729 | 3.669578 | 4.104969 | 4.099000 | 4.066186 | 4.066458 | 25 | 22 | 11 | 12 |
| 57521 | NM_00116303 | RPTOR | regulatory associated protein of MTOR complex 1 | protein_coding | NP_001156506.1;NP_065812.1 | 30287 | 607130 | . | . | 6.946925 | 7.799695 | 8.107767 | 8.341186 | 16.165779 | 18.029342 | 18.193596 | 19.071130 | 1.102671 | 1658.425669 | 0.126097878 | 1.118200846 | 0.263481223 | 0.996308385 | 10.623488 | 10.764275 | 10.615339 | 10.694341 | 10.725319 | 10.738380 | 1724 | 1789 | 1500 | 1644 |
| 57522 | NM_00134620 | SRGAP1 | SLIT-ROBO Rho GTPase activating protein 1 | protein_coding | NP_001333130.1;NP_065813.1;XP_011536882.1;XP_011536883.1;XP_02 | 17382 | 606523 | . | . | 2.349842 | 1.301291 | 1.549156 | 1.488447 | 5.468181 | 3.007994 | 3.476260 | 3.403156 | 1.250168 | 881.887683 | 0.14390878 | 2.238373213 | **0.025196727** | 0.523780764 | 9.614455 | 9.936511 | 9.666506 | 9.726388 | 9.876540 | 9.837202 | 864 | 882 | 875 | 894 |
| 57524 | NM_020764,XI | CASKIN1 | CASK interacting protein 1 | protein_coding | NP_065815.1;XP_024306129.1 | 20879 | 612184 | . | . | 0.125048 | 0.065213 | 0.076904 | 0.088016 | 0.290992 | 0.150743 | 0.172569 | 0.201237 | -1.148713 | 16.611322 | 0.77270809 | -0.25885435 | 0.795747619 | NA | 4.149569 | 3.951852 | 4.085106 | 4.034960 | 4.048258 | 4.052581 | 27 | 13 | 13 | 15 |
| 57529 | NM_020769,XI | RTL9 | retrotransposon Gag like 9 | protein_coding | NP_065820.1;XP_011529300.1;XP_011529301.1;XP_016885183.1;XP_01 | 29245 | 300965 | . | . | 0.023219 | 0.119004 | 0.028984 | 0.058972 | 0.054032 | 0.275083 | 0.065040 | 0.134834 | -1.564503 | 11.984800 | 0.984989562 | -0.65554459 | 0.512117184 | NA | 3.873125 | 3.219802 | 3.542453 | 3.627290 | 3.552089 | 3.572936 | 6 | 26 | 6 | 11 |
| 57530 | NM_020770,XI | CGN | cingulin | protein_coding | NP_065821.1;XP_005245422.1 | 17429 | 609473 | . | . | 2.487769 | 2.240780 | 2.037920 | 1.980284 | 5.789142 | 5.179663 | 4.573035 | 4.527683 | -1.188747 | 347.807368 | 0.192724098 | -1.29429149 | 0.195564755 | 0.96621494 | 8.561230 | 8.312337 | 8.519993 | 8.456860 | 8.402152 | 8.380022 | 458 | 386 | 282 | 292 |
| 57531 | NM_00132108 | HACE1 | HECT domain and ankyrin repeat containing E3 ubiquitin protein li | protein_coding | NP_001308009.1;NP_001308012.1;NP_001308013.1;NP_001337483.1;NI | 21033 | 610876 | . | . | 2.512505 | 2.386455 | 2.322970 | 2.117306 | 5.846703 | 5.516395 | 5.212677 | 4.840971 | -1.129608 | 332.770128 | 0.194904968 | -0.90209376 | 0.367007064 | 0.998865404 | 8.463329 | 8.288119 | 8.427801 | 8.394674 | 8.365518 | 8.321144 | 418 | 370 | 283 | 281 |
| 57532 | NM_020772,XI | NUFIP2 | nuclear FMR1 interacting protein 2 | protein_coding | NP_065823.1;XP_016880385.1 | 17634 | 609356 | . | . | 9.561377 | 9.463005 | 10.443754 | 8.474998 | 22.249716 | 21.874159 | 23.435482 | 19.377078 | -1.009268 | 3183.563534 | 0.12933716 | -0.10290296 | 0.91803999 | 0.998865404 | 11.642931 | 11.629893 | 11.648838 | 11.628452 | 11.715111 | 11.539884 | 3728 | 3408 | 3050 | 2654 |

| | | | | | | | | | | | | | | | | | | | | | | | | | | | | | | | | | |
|---|---|---|---|---|---|---|---|---|---|---|---|---|---|---|---|---|---|---|---|---|---|---|---|---|---|---|---|---|---|---|---|---|---|
| 57533 | NM_00111336 | TBC1D14 | TBC1 domain family member 14 | protein_coding | NP_001106832.1;NP_001106834.1;NP_001273734.1;NP_001317567.1;NI | 29246 | 614855 | . | . | 24.808486 | 25.588844 | 28.848728 | 30.211801 | 57.730375 | 59.149762 | 64.735709 | 69.075692 | 1.164367 | 4169.773953 | 0.107751203 | 2.037528175 | 0.041597141 | 0.63293663 | 11.911843 | 12.131324 | 11.951073 | 11.966946 | 12.069216 | 12.101277 | 4394 | 4198 | 3827 | 4286 |
| 57534 | NM_020774,XI | MIB1 | mindbomb E3 ubiquitin protein ligase 1 | protein_coding | NP_065825.1;XP_011524400.1;XP_016881362.1;XP_016881363.1;XP_01 | 21086 | 608677 | . | . | 3.398464 | 2.704021 | 2.945042 | 2.510909 | 7.908363 | 6.250464 | 6.608587 | 5.740894 | -1.178252 | 866.427876 | 0.167046352 | -1.41665895 | 0.156582665 | 0.925053243 | 9.871993 | 9.636260 | 9.893897 | 9.722708 | 9.757995 | 9.627202 | 1204 | 894 | 757 | 676 |
| 57536 | NM_00132232 | KIAA1328 | KIAA1328 | protein_coding | NP_001309256.1;NP_001340847.1;NP_001340848.1;NP_001340849.1;NI | 29248 | 616480 | . | . | 1.598805 | 1.657599 | 1.105281 | 1.711182 | 3.720485 | 3.831615 | 2.480220 | 3.912411 | -1.000702 | 110.305628 | 0.312012239 | -0.00324564 | 0.997410355 | 0.999862634 | 6.786648 | 6.784076 | 6.773984 | 6.795451 | 6.747865 | 6.819860 | 123 | 123 | 85 | 115 |
| 57539 | NM_00100665 | WDR35 | WD repeat domain 35 | protein_coding | NP_001006658.1;NP_065830.2;XP_011531309.1 | 29250 | 613602 | . | . | 5.044262 | 5.022354 | 5.685098 | 5.011307 | 11.738205 | 11.609395 | 12.757196 | 11.457757 | 1.053420 | 1083.950338 | 0.139502248 | 0.538203007 | 0.590436904 | 0.998865404 | 10.043878 | 10.119303 | 10.055732 | 10.067380 | 10.148194 | 10.050030 | 1206 | 1148 | 1031 | 974 |
| 57541 | NM_020781,N | ZNF398 | zinc finger protein 398 | protein_coding | NP_065832.1;NP_733787.1;XP_011514741.1;XP_011514742.1;XP_01151 | 18373 | 618593 | . | . | 2.172159 | 2.530667 | 2.501000 | 2.542371 | 5.054702 | 5.849751 | 5.612172 | 5.812828 | 1.053538 | 414.278083 | 0.182958835 | 0.411256334 | 0.680884585 | 0.998865404 | 8.656583 | 8.731360 | 8.632978 | 8.720019 | 8.709281 | 8.708037 | 431 | 467 | 370 | 398 |
| 57542 | NM_020782,XI | KLHL42 | kelch like family member 42 | protein_coding | NP_065833.1;XP_016875187.1 | 29252 | . | . | . | 4.634166 | 4.401308 | 5.552757 | 4.273589 | 10.783894 | 10.173822 | 12.460227 | 9.771052 | 1.083526 | 959.369209 | 0.161987979 | 0.714462266 | 0.474941381 | 0.998865404 | 9.846524 | 9.963010 | 9.888186 | 9.857036 | 10.030453 | 9.821657 | 1083 | 972 | 991 | 803 |
| 57544 | NM_00116004 | TXNDC16 | thioredoxin domain containing 16 | protein_coding | NP_001153519.1;NP_065835.2;XP_016876994.1 | 19965 | 616179 | . | . | 1.064611 | 1.219370 | 1.119258 | 1.159750 | 2.477393 | 2.818629 | 2.511582 | 2.651631 | 1.003075 | 167.341907 | 0.257052483 | 0.017232756 | 0.98625093 | 0.999197585 | 7.384803 | 7.388504 | 7.357964 | 7.412110 | 7.379033 | 7.394140 | 180 | 192 | 143 | 160 |
| 57545 | NM_00108052 | CC2D2A | coiled-coil and C2 domain containing 2A | protein_coding | NP_001073991.2;NP_001158192.1;NP_001365544.1;NP_001365546.1;NI | 29253 | 612013 | . | . | 3.987851 | 3.522546 | 4.261862 | 4.355527 | 9.279893 | 8.142524 | 9.563497 | 9.958393 | 1.087826 | 319.461936 | 0.199691149 | 0.608181164 | 0.543067326 | 0.998865404 | 8.257321 | 8.379110 | 8.330504 | 8.257382 | 8.332171 | 8.349683 | 376 | 308 | 285 | 317 |
| 57546 | NM_00132992 | PDP2 | pyruvate dehyrogenase phosphatase catalytic subunit 2 | protein_coding | NP_001316857.1;NP_001316858.1;NP_001316859.1;NP_001316860.1;NI | 30263 | 615499 | . | . | 3.312869 | 3.421416 | 3.711647 | 3.966819 | 7.709185 | 7.908758 | 8.328829 | 9.069661 | 1.122470 | 794.402954 | 0.146086193 | 1.140948757 | 0.253891248 | 0.994807659 | 9.548144 | 9.714516 | 9.579424 | 9.602211 | 9.649757 | 9.693980 | 848 | 821 | 708 | 812 |
| 57547 | NM_020787,XI | ZNF624 | zinc finger protein 624 | protein_coding | NP_065838.2;XP_006721625.1;XP_011522272.1 | 29254 | . | . | . | 0.414570 | 0.564076 | 0.469061 | 0.323541 | 0.964721 | 1.303888 | 1.052560 | 0.739738 | -1.147793 | 76.573332 | 0.378332857 | -0.52562741 | 0.599147109 | 0.998865404 | 6.353987 | 6.156823 | 6.252236 | 6.293680 | 6.282890 | 6.191381 | 87 | 95 | 74 | 54 |
| 57551 | NM_020791,N | TAOK1 | TAO kinase 1 | protein_coding | NP_065842.1;NP_079418.1 | 29259 | 610266 | . | . | 5.441283 | 5.361899 | 5.337235 | 4.983918 | 12.662089 | 12.394271 | 11.976600 | 11.395136 | -1.061122 | 2081.053416 | 0.119831025 | -0.71425911 | 0.47506697 | 0.998865404 | 11.065161 | 10.979772 | 11.055455 | 11.036646 | 11.029425 | 10.967377 | 2496 | 2285 | 1825 | 1819 |
| 57552 | NM_00114627 | NCEH1 | neutral cholesterol ester hydrolase 1 | protein_coding | NP_001139748.2;NP_001139749.1;NP_001139750.1;NP_065843.4 | 29260 | 613234 | . | . | 10.990967 | 9.890142 | 9.731640 | 8.925940 | 25.576432 | 22.861506 | 21.837519 | 20.408102 | -1.129702 | 1395.304912 | 0.133754118 | -1.31541204 | 0.188371462 | 0.960264078 | 10.531454 | 10.355940 | 10.530879 | 10.448698 | 10.432684 | 10.360892 | 1794 | 1513 | 1194 | 1170 |
| 57553 | NM_00112273 | MICAL3 | microtubule associated monooxygenase, calponin and LIM domain | protein_coding | NP_001116203.1;NP_001129476.1;NP_056056.2 | 24694 | 608882 | . | . | 21.107142 | 22.399264 | 22.166067 | 21.266878 | 49.117190 | 51.776900 | 49.740015 | 48.624189 | -1.011810 | 6196.101289 | 0.105235779 | -0.16095051 | 0.87213238 | 0.998865404 | 12.605589 | 12.588652 | 12.581691 | 12.621849 | 12.606359 | 12.577581 | 7006 | 6885 | 5448 | 5676 |
| 57555 | NM_020795,XI | NLGN2 | neuroligin 2 | protein_coding | NP_065846.1;XP_005256801.1;XP_016880386.1 | 14290 | 606479 | . | . | 4.780018 | 4.757402 | 4.869050 | 5.019191 | 11.123300 | 10.996948 | 10.926010 | 11.475784 | 1.043417 | 733.136596 | 0.149009093 | 0.411486893 | 0.68071555 | 0.998865404 | 9.486865 | 9.548357 | 9.523250 | 9.481736 | 9.533132 | 9.531299 | 851 | 751 | 652 | 701 |
| 57556 | NM_00130078 | SEMA6A | semaphorin 6A | protein_coding | NP_001287709.1;NP_065847.1;XP_005272099.2;XP_006714726.1;XP_01 | 10738 | 605885 | . | . | 17.521539 | 18.206388 | 19.320499 | 17.828611 | 40.733343 | 42.084880 | 43.354639 | 40.763001 | 1.022929 | 3825.797309 | 0.113089333 | 0.289207642 | 0.772422484 | 0.998865404 | 11.885047 | 11.917856 | 11.880939 | 11.902324 | 11.951134 | 11.868188 | 4299 | 4134 | 3533 | 3446 |
| 57558 | NM_020798,XI | USP35 | ubiquitin specific peptidase 35 | protein_coding | NP_065849.1;XP_011543489.1;XP_011543490.1;XP_016873542.1 | 20061 | . | . | . | 4.999315 | 4.974803 | 4.814031 | 4.783900 | 11.633609 | 11.499478 | 10.802546 | 10.937823 | 1.011490 | 645.586503 | 0.15493249 | 0.106381682 | 0.915279526 | 0.998865404 | 9.326123 | 9.342762 | 9.320064 | 9.340437 | 9.358507 | 9.317176 | 728 | 703 | 582 | 590 |
| 57559 | NM_020799,XI | STAMBPL1 | STAM binding protein like 1 | protein_coding | NP_065850.1;XP_006717991.1;XP_006717993.1;XP_011538285.1;XP_01 | 24105 | 612352 | . | . | 6.272255 | 5.665281 | 4.978535 | 5.029041 | 14.595796 | 13.095553 | 11.171689 | 11.498303 | -1.160489 | 363.448835 | 0.189731624 | -1.13177325 | 0.257729777 | 0.995760118 | 8.609091 | 8.394126 | 8.568573 | 8.524805 | 8.435843 | 8.486405 | 466 | 406 | 279 | 330 |
| 57560 | NM_00119024 | IFT80 | intraflagellar transport 80 | protein_coding | NP_001177170.1;NP_001177171.1;NP_065851.1 | 29262 | 611177 | . | . | 8.471894 | 7.768269 | 9.299111 | 5.744917 | 19.714445 | 17.956701 | 20.866934 | 13.135073 | -1.082783 | 1013.756932 | 0.192946174 | -0.59469591 | 0.552046756 | 0.998865404 | 10.041286 | 9.927462 | 10.041525 | 9.970404 | 10.107389 | 9.759557 | 1271 | 1083 | 1048 | 696 |
| 57561 | NM_00132967 | ARRDC3 | arrestin domain containing 3 | protein_coding | NP_001316599.1;NP_001316600.1;NP_001316601.1;NP_065852.1 | 29263 | 612464 | . | . | 1.087793 | 1.128280 | 1.198689 | 0.762164 | 2.531337 | 2.608069 | 2.689824 | 1.742596 | -1.120086 | 140.586456 | 0.296588238 | -0.55164044 | 0.581194721 | 0.998865404 | 7.213830 | 7.052277 | 7.148021 | 7.156770 | 7.180370 | 7.034213 | 169 | 162 | 139 | 99 |
| 57562 | NM_00136354 | CEP126 | centrosomal protein 126 | protein_coding | NP_001350472.1;NP_065853.3 | 29264 | 614634 | . | . | 0.307180 | 0.433339 | 0.333345 | 0.263683 | 0.714820 | 1.001682 | 0.748016 | 0.602879 | -1.277490 | 80.352057 | 0.370517803 | -0.95356381 | 0.340304483 | 0.998865404 | 6.494068 | 6.140897 | 6.308335 | 6.402556 | 6.317678 | 6.266529 | 87 | 113 | 68 | 59 |
| 57563 | NM_00129200 | KLHL8 | kelch like family member 8 | protein_coding | NP_001278932.1;NP_001278935.1;NP_001278936.1;NP_065854.3 | 18644 | 611967 | . | . | 2.291864 | 2.446307 | 1.987179 | 1.684099 | 5.333263 | 5.654747 | 4.459171 | 3.850493 | -1.310078 | 352.172633 | 0.19536333 | -1.99450045 | 0.046097391 | 0.654058028 | 8.641806 | 8.252255 | 8.488779 | 8.570084 | 8.402584 | 8.350758 | 429 | 460 | 278 | 272 |
| 57565 | NM_020805,XI | KLHL14 | kelch like family member 14 | protein_coding | NP_065856.1;XP_011524411.2 | 29266 | 613772 | . | . | 0.127615 | 0.186805 | 0.125091 | 0.141402 | 0.296965 | 0.431808 | 0.280700 | 0.323300 | -1.161133 | 19.710143 | 0.698340349 | -0.30863656 | 0.757598009 | NA | 4.405684 | 4.187832 | 4.291006 | 4.324011 | 4.291212 | 4.299197 | 20 | 27 | 15 | 18 |
| 57567 | NM_020807,XI | ZNF319 | zinc finger protein 319 | protein_coding | NP_065858.1;XP_005256126.1;XP_024306130.1;XP_024306131.1 | 13644 | . | . | . | 1.213377 | 1.039592 | 0.909346 | 0.885500 | 2.823578 | 2.403064 | 2.040545 | 2.024590 | -1.279712 | 134.990340 | 0.285348011 | -1.24696669 | 0.212409712 | 0.975238634 | 7.243320 | 6.888321 | 7.157322 | 7.081063 | 7.033409 | 7.026486 | 191 | 148 | 104 | 110 |
| 57568 | NM_00137748 | SIPA1L2 | signal induced proliferation associated 1 like 2 | protein_coding | NP_001364417.1;NP_065859.3;XP_005273269.1;XP_005273270.1;XP_01 | 23800 | 611609 | . | . | 5.938764 | 6.616375 | 7.272733 | 8.039704 | 13.819749 | 15.294046 | 16.319802 | 18.381828 | 1.202093 | 1487.014868 | 0.130909913 | 2.028482712 | 0.042511006 | 0.637111201 | 10.399533 | 10.664706 | 10.423274 | 10.498727 | 10.570397 | 10.635939 | 1478 | 1530 | 1353 | 1585 |
| 57569 | NM_00125841 | ARHGAP20 | Rho GTPase activating protein 20 | protein_coding | NP_001245344.1;NP_001245345.1;NP_001245346.1;NP_001245347.1;NI | 18357 | 609568 | . | . | 0.313985 | 0.561105 | 0.477352 | 0.571701 | 0.730656 | 1.297020 | 1.071164 | 1.307127 | 1.182784 | 93.219983 | 0.36823263 | 0.657700652 | 0.510730503 | 0.998865404 | 6.417981 | 6.657245 | 6.425203 | 6.582327 | 6.528440 | 6.582571 | 71 | 118 | 80 | 103 |
| 57570 | NM_00135025 | TRMT5 | tRNA methyltransferase 5 | protein_coding | NP_001337182.1;NP_001337183.1;NP_065861.3 | 23141 | 611023 | . | . | 2.650641 | 2.476053 | 2.572184 | 2.232539 | 6.168150 | 5.723504 | 5.771906 | 5.104436 | -1.067174 | 417.765305 | 0.187600754 | -0.49997666 | 0.617091512 | 0.998865404 | 8.752165 | 8.659443 | 8.748412 | 8.699999 | 8.741630 | 8.626194 | 516 | 447 | 386 | 342 |
| 57572 | NM_00136783 | DOCK6 | dedicator of cytokinesis 6 | protein_coding | NP_001354759.1;NP_065863.2;XP_005260057.1;XP_006722867.1;XP_01 | 19189 | 614194 | . | . | 1.467202 | 1.456652 | 1.589506 | 1.765280 | 3.414241 | 3.367117 | 3.566806 | 4.036103 | 1.127110 | 322.254084 | 0.196239233 | 0.879685118 | 0.379029914 | 0.998865404 | 8.243349 | 8.415621 | 8.298251 | 8.295167 | 8.339712 | 8.385049 | 350 | 326 | 285 | 333 |
| 57575 | NM_020815,N | PCDH10 | protocadherin 10 | protein_coding | NP_065866.1;NP_116586.1;XP_011530452.1 | 13404 | 608286 | . | . | 0.160865 | 0.196324 | 0.145718 | 0.225085 | 0.374339 | 0.453811 | 0.326988 | 0.514629 | 1.150035 | 34.771113 | 0.562441564 | 0.358575541 | 0.719912647 | 0.998865404 | 5.017958 | 5.214957 | 5.087268 | 5.118130 | 5.070992 | 5.170934 | 33 | 39 | 21 | 47 |
| 57577 | NM_00135376 | CCDC191 | coiled-coil domain containing 191 | protein_coding | NP_001340695.1;NP_001340696.2;NP_065868.1;XP_005247738.1;XP_01 | 29272 | . | . | . | 0.414734 | 0.314315 | 0.409397 | 0.264063 | 0.965103 | 0.726555 | 0.918675 | 0.603750 | -1.055658 | 64.723595 | 0.422552065 | -0.18493028 | 0.853283722 | 0.998865404 | 6.052888 | 5.978596 | 6.060508 | 5.971144 | 6.051430 | 5.955437 | 91 | 58 | 67 | 46 |
| 57579 | NM_00110553 | FAM135A | family with sequence similarity 135 member A | protein_coding | NP_001099001.1;NP_001156001.1;NP_001317924.1;NP_001317925.1;NI | 21084 | . | . | . | 1.555101 | 1.820468 | 1.458561 | 1.185659 | 3.618786 | 4.208093 | 3.272973 | 2.710869 | -1.203076 | 264.546023 | 0.217486772 | -1.22640741 | 0.220045397 | 0.982008277 | 8.174484 | 7.907975 | 8.054290 | 8.127619 | 8.027373 | 7.966390 | 310 | 333 | 222 | 211 |
| 57580 | NM_020820,XI | PREX1 | phosphatidylinositol-3,4,5-trisphosphate dependent Rac exchange | protein_coding | NP_065871.3;XP_011527236.1 | 32594 | 606905 | . | . | 66.758265 | 69.543688 | 68.375629 | 80.949525 | 155.349231 | 160.753365 | 153.432933 | 185.081468 | 1.080789 | 14999.393345 | 0.112742472 | 0.994167998 | 0.320141085 | 0.998865404 | 13.815516 | 13.927543 | 13.823344 | 13.851631 | 13.833320 | 13.970535 | 16326 | 15815 | 12474 | 15767 |
| 57583 | NM_00137681 | TMEM181 | transmembrane protein 181 | protein_coding | NP_001363746.1;NP_001363779.1;NP_001363781.1;NP_001363783.1;NI | 20958 | 613209 | . | . | 14.285742 | 13.479281 | 15.183753 | 12.148818 | 33.243506 | 31.157958 | 34.071899 | 27.776829 | -1.030736 | 2203.962865 | 0.137696227 | -0.31718286 | 0.751104856 | 0.998865404 | 11.127351 | 11.084093 | 11.142017 | 11.089360 | 11.180139 | 10.995375 | 2660 | 2335 | 2106 | 1800 |
| 57584 | NM_00136744 | ARHGAP21 | Rho GTPase activating protein 21 | protein_coding | NP_001354376.1;NP_001354377.1;NP_001354378.1;NP_001354379.1;NI | 23725 | 609870 | . | . | 8.781458 | 9.106101 | 7.991330 | 10.208632 | 20.434816 | 21.049156 | 17.932314 | 23.340824 | 1.065190 | 2115.030790 | 0.128820023 | 0.707271945 | 0.479397496 | 0.998865404 | 11.000352 | 11.091146 | 11.013039 | 11.026429 | 10.998676 | 11.136860 | 2339 | 2229 | 1734 | 2221 |
| 57585 | NM_020825 | CRAMP1 | cramped chromatin regulator homolog 1 | protein_coding | NP_065876.3 | 14122 | . | . | . | 2.625128 | 2.556332 | 2.840527 | 2.528330 | 6.108781 | 5.909076 | 6.374059 | 5.780725 | 1.021993 | 655.606025 | 0.157903031 | 0.198764356 | 0.84244708 | 0.998865404 | 9.340617 | 9.372576 | 9.360591 | 9.336170 | 9.405294 | 9.319553 | 760 | 687 | 613 | 582 |
| 57587 | NM_00129203 | CFAP97 | cilia and flagella associated protein 97 | protein_coding | NP_001278962.1;NP_065878.1;XP_016863972.1;XP_016863973.1;XP_01 | 29276 | 616047 | . | . | 5.627517 | 4.929879 | 5.320578 | 4.284364 | 13.095464 | 11.395639 | 11.939225 | 9.795690 | -1.095508 | 859.899703 | 0.160145926 | -0.82174722 | 0.411220779 | 0.998865404 | 9.811924 | 9.681163 | 9.843932 | 9.707175 | 9.768442 | 9.654025 | 1132 | 879 | 772 | 707 |
| 57589 | NM_00113592 | RIC1 | RIC1 homolog, RAB6A GEF complex partner 1 | protein_coding | NP_001129392.2;NP_001193486.1;NP_065880.2;XP_005251580.1;XP_01 | 17686 | 610354 | . | . | 2.456873 | 2.292869 | 2.816177 | 2.361896 | 5.717244 | 5.300071 | 6.319419 | 5.400195 | 1.067951 | 517.271420 | 0.170568486 | 0.556057018 | 0.578171877 | 0.998865404 | 8.966239 | 9.061737 | 8.998638 | 8.985292 | 9.080536 | 8.985638 | 582 | 534 | 499 | 464 |
| 57590 | NM_020830 | WDFY1 | WD repeat and FYVE domain containing 1 | protein_coding | NP_065881.1 | 20451 | 618080 | . | . | 35.428810 | 33.530243 | 33.878761 | 31.441654 | 82.444298 | 77.506660 | 76.022949 | 71.887604 | -1.070518 | 4820.295928 | 0.11123784 | -0.88376868 | 0.376821097 | 0.998865404 | 12.283157 | 12.184985 | 12.288838 | 12.235212 | 12.240124 | 12.169662 | 5922 | 5207 | 4221 | 4180 |
| 57591 | NM_00128266 | MRTFA | myocardin related transcription factor A | protein_coding | NP_001269589.1;NP_001269590.2;NP_001269591.2;NP_001305068.1;NI | 14334 | 606078 | . | . | 6.103484 | 6.691790 | 6.660313 | 6.874665 | 14.203057 | 15.468375 | 14.945549 | 15.718104 | 1.030750 | 909.060316 | 0.140938374 | 0.310022948 | 0.756543506 | 0.998865404 | 9.806330 | 9.849806 | 9.792858 | 9.840497 | 9.832933 | 9.844505 | 997 | 998 | 796 | 872 |
| 57592 | NM_00130476 | ZNF687 | zinc finger protein 687 | protein_coding | NP_001291692.1;NP_001291693.1;NP_065883.1;XP_005245423.1;XP_01 | 29277 | 610568 | . | . | 2.656978 | 2.802991 | 3.042087 | 3.318175 | 6.182897 | 6.479238 | 6.826356 | 7.586615 | 1.122232 | 464.923237 | 0.17332061 | 0.959901275 | 0.337104905 | 0.998865404 | 8.775533 | 8.941400 | 8.813345 | 8.831992 | 8.863459 | 8.924149 | 497 | 480 | 407 | 483 |
| 57593 | NM_00111051 | EBF4 | EBF family member 4 | protein_coding | NP_001103984.1;XP_006723663.1;XP_016883472.1;XP_016883473.1;XP_ | 29278 | 609935 | . | . | 0.174484 | 0.175103 | 0.277500 | 0.138506 | 0.406032 | 0.404758 | 0.622702 | 0.316678 | 1.145183 | 17.820311 | 0.746258724 | 0.262077804 | 0.793261459 | NA | 4.052320 | 4.251698 | 4.141521 | 4.142398 | 4.177888 | 4.125938 | 19 | 18 | 22 | 12 |
| 57594 | NM_020834 | HOMEZ | homeobox and leucine zipper encoding | protein_coding | NP_065885.2 | 20164 | 608119 | . | . | 0.909605 | 0.797962 | 0.829387 | 1.035346 | 2.116689 | 1.844526 | 1.861120 | 2.367195 | 1.072422 | 139.033501 | 0.284062053 | 0.355110413 | 0.722506876 | 0.998865404 | 7.068260 | 7.168575 | 7.129026 | 7.078934 | 7.090227 | 7.168882 | 165 | 135 | 112 | 149 |
| 57595 | NM_00130351 | PDZD4 | PDZ domain containing 4 | protein_coding | NP_001290441.1;NP_001290442.1;NP_001290443.1;NP_001290444.1;NI | 21167 | 300634 | . | . | 5.261778 | 4.854141 | 4.490642 | 5.290815 | 12.244374 | 11.220567 | 10.076868 | 12.096819 | -1.051302 | 574.757821 | 0.165735858 | -0.43549304 | 0.663204588 | 0.998865404 | 9.202584 | 9.130127 | 9.210410 | 9.154185 | 9.107548 | 9.190349 | 707 | 609 | 456 | 559 |
| 57596 | NM_00115953 | BEGAIN | brain enriched guanylate kinase associated | protein_coding | NP_001153003.1;NP_065887.1;XP_005267976.1;XP_005267977.1;XP_00 | 24163 | 618597 | . | . | 0.188375 | 0.229788 | 0.388410 | 0.250772 | 0.438356 | 0.531166 | 0.871581 | 0.573361 | 1.582710 | 52.797933 | 0.45000352 | 1.471980858 | 0.141026047 | 0.902449014 | 5.352443 | 6.016603 | 5.650783 | 5.665388 | 5.794844 | 5.694407 | 45 | 46 | 69 | 47 |
| 57597 | NM_00129132 | BAHCC1 | BAH domain and coiled-coil containing 1 | protein_coding | NP_001278253.1;NP_001364377.1;XP_011523365.1;XP_016880387.1;XP | 29279 | 617646 | . | . | 4.918835 | 4.433726 | 5.584810 | 5.321585 | 11.446332 | 10.248758 | 12.532149 | 12.167171 | 1.149248 | 1703.509884 | 0.127682777 | 1.571785691 | 0.116000261 | 0.857396579 | 10.630296 | 10.831298 | 10.719322 | 10.631460 | 10.807462 | 10.762839 | 1929 | 1613 | 1629 | 1660 |
| 57599 | NM_00130340 | WDR48 | WD repeat domain 48 | protein_coding | NP_001290331.1;NP_001290332.1;NP_001333154.1;NP_001333155.1;NI | 30914 | 612167 | . | . | 4.659950 | 4.361776 | 4.091086 | 4.334322 | 10.843894 | 10.082441 | 9.180276 | 9.909914 | -1.081855 | 542.530692 | 0.16454079 | -0.6898444 | 0.490292044 | 0.998865404 | 9.139253 | 9.025632 | 9.128491 | 9.088292 | 9.036281 | 9.078604 | 669 | 590 | 438 | 505 |
| 57600 | NM_00132391 | FNIP2 | folliculin interacting protein 2 | protein_coding | NP_001310845.1;NP_001332972.1;NP_001353772.1;NP_065891.1;XP_0C | 29280 | 612768 | . | . | 5.935030 | 5.501036 | 7.653040 | 6.488723 | 13.811059 | 12.715894 | 17.173194 | 14.835697 | 1.224902 | 1343.058248 | 0.141028787 | 2.075226256 | 0.03796557 | 0.60652635 | 10.237301 | 10.530446 | 10.319728 | 10.292983 | 10.534430 | 10.382828 | 1415 | 1279 | 1411 | 1252 |
| 57602 | NM_00132129 | USP36 | ubiquitin specific peptidase 36 | protein_coding | NP_001308220.1;XP_005257599.1;XP_005257600.1;XP_011523367.1;XP_ | 20062 | 612543 | . | . | 9.559198 | 10.851718 | 11.217568 | 10.017824 | 22.244646 | 25.084235 | 25.171899 | 22.904566 | 1.019390 | 1873.133421 | 0.130270722 | 0.212682281 | 0.831574788 | 0.998865404 | 10.857260 | 10.885082 | 10.821802 | 10.901836 | 10.929044 | 10.823864 | 2028 | 2103 | 1754 | 1655 |
| 57604 | NM_00109967 | TRMT9B | tRNA methyltransferase 9B (putative) | protein_coding | NP_001093147.1;NP_065895.2;XP_005273641.1;XP_005273642.1;XP_00 | 26725 | 615666 | . | . | 0.777522 | 0.842966 | 0.876424 | 0.882321 | 1.809325 | 1.948555 | 1.966675 | 2.017322 | 1.062312 | 260.564993 | 0.212730464 | 0.40994436 | 0.681846763 | 0.998865404 | 7.981375 | 8.068315 | 7.989681 | 8.028068 | 8.040937 | 8.038497 | 280 | 283 | 234 | 251 |
| 57605 | NM_00130080 | PITPNM2 | phosphatidylinositol transfer protein membrane associated 2 | protein_coding | NP_001287730.1;NP_065896.1;XP_024304866.1;XP_024304867.1;XP_02 | 21044 | 608920 | . | . | 7.063292 | 7.800283 | 7.336069 | 6.946405 | 16.436571 | 18.030702 | 16.461928 | 15.882128 | -1.054527 | 1531.742701 | 0.128994349 | -0.59379774 | 0.552647399 | 0.998865404 | 10.618739 | 10.542161 | 10.564036 | 10.636202 | 10.586164 | 10.532447 | 1727 | 1775 | 1342 | 1349 |
| 57606 | NM_020846,XI | SLAIN2 | SLAIN motif family member 2 | protein_coding | NP_065897.1;XP_005248178.1 | 29282 | 610492 | . | . | 4.861641 | 5.154079 | 4.635724 | 5.417549 | 11.313239 | 11.913885 | 10.402431 | 12.386583 | 1.000016 | 854.851698 | 0.149191352 | 0.000153078 | 0.999877861 | 0.999950306 | 9.739773 | 9.739288 | 9.737862 | 9.740070 | 9.682669 | 9.792090 | 983 | 924 | 686 | 863 |
| 57608 | NM_00135000 | JCAD | junctional cadherin 5 associated | protein_coding | NP_001336930.1;NP_001336950.1;NP_001336951.1;NP_065899.1 | 29283 | 614398 | . | . | 0.056264 | 0.045108 | 0.034031 | 0.026961 | 0.130928 | 0.104268 | 0.076365 | 0.061644 | -1.658779 | 12.538279 | 0.871886991 | -0.83740431 | 0.402365343 | NA | 3.966708 | 3.239024 | 3.676411 | 3.660836 | 3.645509 | 3.637380 | 20 | 15 | 9 | 8 |
| 57609 | NM_173602 | DIP2B | disco interacting protein 2 homolog B | protein_coding | NP_775873.2 | 29284 | 611379 | . | . | 9.165947 | 8.253111 | 10.683633 | 10.833487 | 21.329535 | 19.077436 | 23.973763 | 24.769478 | 1.217783 | 2638.592197 | 0.118702646 | 2.394700479 | 0.016633947 | 0.431581156 | 11.216356 | 11.500756 | 11.323528 | 11.233416 | 11.437685 | 11.441014 | 2895 | 2422 | 2515 | 2722 |
| 57610 | NM_00132023 | RANBP10 | RAN binding protein 10 | protein_coding | NP_001307167.1;NP_001307168.1;NP_001307169.1;NP_065901.1;XP_0C | 29285 | 614031 | . | . | 5.212663 | 5.463250 | 6.809349 | 6.222213 | 12.130081 | 12.628550 | 15.279984 | 14.226349 | 1.206555 | 953.485357 | 0.140131157 | 1.933147502 | 0.05321803 | 0.683508073 | 9.755234 | 10.026196 | 9.804628 | 9.842011 | 9.984105 | 9.934895 | 969 | 957 | 937 | 945 |
| 57613 | NM_020853 | FAM234B | family with sequence similarity 234 member B | protein_coding | NP_065904.1 | 29288 | 617838 | . | . | 3.011336 | 2.760134 | 2.675340 | 2.758688 | 7.007501 | 6.380174 | 6.003386 | 6.307411 | -1.078359 | 411.557500 | 0.180642206 | -0.60250267 | 0.546839585 | 0.998865404 | 8.738242 | 8.629614 | 8.735874 | 8.677708 | 8.656335 | 8.667767 | 515 | 439 | 341 | 375 |
| 57614 | NM_00134622 | RELCH | RAB11 binding and LisH domain, coiled-coil and HEAT repeat cont | protein_coding | NP_001333158.1;NP_001333159.1;NP_001333160.1;NP_001333162.1;NI | 29289 | 618001 | . | . | 2.792863 | 2.917729 | 2.606491 | 2.743832 | 6.499106 | 6.744460 | 5.848889 | 6.273446 | -1.082089 | 732.142538 | 0.150058116 | -0.75850069 | 0.448151301 | 0.998865404 | 9.571944 | 9.457759 | 9.521293 | 9.563705 | 9.469564 | 9.506066 | 850 | 846 | 595 | 677 |
| 57615 | NM_020855 | ZNF492 | zinc finger protein 492 | protein_coding | NP_065906.1 | 23707 | . | . | . | 0.304364 | 0.178669 | 0.117764 | 0.211575 | 0.708266 | 0.413001 | 0.264259 | 0.483741 | -1.479113 | 27.186281 | 0.628082546 | -0.89913739 | 0.368579484 | 0.998865404 | 5.020416 | 4.453945 | 4.822029 | 4.753094 | 4.721278 | 4.768266 | 47 | 26 | 14 | 26 |
| 57616 | NM_020856,N | TSHZ3 | teashirt zinc finger homeobox 3 | protein_coding | NP_065907.2 | 30700 | 614119 | . | . | 1.681547 | 1.515391 | 1.628104 | 1.560935 | 3.913028 | 3.502896 | 3.653419 | 3.568892 | 1.051674 | 241.502588 | 0.218612633 | 0.332494426 | 0.739515948 | 0.998865404 | 7.878968 | 7.951900 | 7.900664 | 7.904595 | 7.940291 | 7.914416 | 270 | 255 | 221 | 226 |
| 57617 | NM_020857,XI | VPS18 | VPS18 core subunit of CORVET and HOPS complexes | protein_coding | NP_065908.1;XP_011520145.1;XP_011520146.1;XP_016877936.1 | 15972 | 608551 | . | . | 9.692810 | 10.793267 | 9.983954 | 11.567990 | 22.555566 | 24.949120 | 22.403703 | 26.448832 | 1.032206 | 1261.747929 | 0.139308638 | 0.328267821 | 0.742709172 | 0.998865404 | 10.278431 | 10.323631 | 10.251479 | 10.323633 | 10.256584 | 10.364592 | 1362 | 1404 | 1035 | 1287 |
| 57619 | NM_020859 | SHROOM3 | shroom family member 3 | protein_coding | NP_065910.3 | 30422 | 604570 | . | . | 0.623423 | 0.711559 | 0.568829 | 0.815054 | 1.450732 | 1.644801 | 1.276435 | 1.863524 | 1.023264 | 231.160364 | 0.245133461 | 0.135349007 | 0.89233594 | 0.998865404 | 7.837036 | 7.868295 | 7.813154 | 7.873342 | 7.768214 | 7.934295 | 247 | 262 | 168 | 257 |
| 57620 | NM_00116911 | STIM2 | stromal interaction molecule 2 | protein_coding | NP_001162588.1;NP_001162589.1;NP_065911.3 | 19205 | 610841 | . | . | 7.286006 | 6.862504 | 7.538637 | 6.019674 | 16.954836 | 15.862986 | 16.916483 | 13.763267 | -1.058787 | 1083.964350 | 0.152278823 | -0.54119048 | 0.588376299 | 0.998865404 | 10.122375 | 10.040672 | 10.126166 | 10.075214 | 10.141169 | 9.969260 | 1327 | 1162 | 1023 | 872 |
| 57621 | NM_020861,XI | ZBTB2 | zinc finger and BTB domain containing 2 | protein_coding | NP_065912.1;XP_005267133.1;XP_011534306.1;XP_016866626.1 | 20868 | 616595 | . | . | 5.228262 | 4.848228 | 4.005718 | 4.895141 | 12.166381 | 11.206895 | 8.988716 | 11.192157 | -1.126877 | 470.586199 | 0.178727399 | -0.96420416 | 0.334943578 | 0.998865404 | 8.962130 | 8.789329 | 8.941281 | 8.885810 | 8.788530 | 8.887873 | 599 | 515 | 353 | 449 |
| 57622 | NM_020862,XI | LRFN1 | leucine rich repeat and fibronectin type III domain containing 1 | protein_coding | NP_065913.1;XP_016882522.1 | 29290 | 612807 | . | . | 1.775332 | 1.653623 | 1.794806 | 2.171954 | 4.131270 | 3.822423 | 4.027493 | 4.965914 | 1.140722 | 206.513674 | 0.239237462 | 0.793970361 | 0.427212686 | 0.998865404 | 7.592304 | 7.781673 | 7.674043 | 7.638840 | 7.673346 | 7.758665 | 231 | 200 | 174 | 225 |
| 57623 | NM_00102993 | ZFAT | zinc finger and AT-hook domain containing | protein_coding | NP_001025110.2;NP_001161055.1;NP_001167628.1;NP_001167629.1;NI | 19899 | 610931 | . | . | 0.970621 | 1.428316 | 1.698254 | 2.061244 | 2.258677 | 3.301616 | 3.810835 | 4.712789 | 1.464941 | 217.656504 | 0.264992139 | 2.078714442 | 0.037643605 | 0.60652635 | 7.465760 | 8.014240 | 7.574230 | 7.728866 | 7.748217 | 7.912125 | 167 | 225 | 189 | 281 |
| 57630 | NM_020870 | SH3RF1 | SH3 domain containing ring finger 1 | protein_coding | NP_065921.2 | 17650 | 618642 | . | . | 1.058140 | 1.122759 | 0.831497 | 1.165277 | 2.462336 | 2.595307 | 1.865854 | 2.664207 | -1.102185 | 167.200500 | 0.269861666 | -0.52014381 | 0.602963343 | 0.998865404 | 7.454799 | 7.312567 | 7.393343 | 7.414540 | 7.298486 | 7.425132 | 197 | 194 | 116 | 173 |
| 57631 | NM_00124396 | LRCH2 | leucine rich repeats and calponin homology domain containing 2 | protein_coding | NP_001230892.1;NP_065922.3;XP_006724787.2;XP_016885185.1 | 29292 | . | . | . | 0.993407 | 0.581405 | 0.815326 | 0.761451 | 2.311699 | 1.343943 | 1.829570 | 1.740968 | -1.094323 | 115.815202 | 0.331481162 | -0.39229662 | 0.69483906 | 0.998865404 | 6.918445 | 6.790061 | 6.949852 | 6.774400 | 6.835035 | 6.836397 | 174 | 98 | 96 | 104 |
| 57634 | NM_015409 | EP400 | E1A binding protein p400 | protein_coding | NP_056224.3 | 11958 | 606265 | . | . | 3.802487 | 4.065164 | 4.376099 | 4.229229 | 8.848545 | 9.396809 | 9.819839 | 9.669631 | 1.079016 | 1576.682292 | 0.125044781 | 0.877410558 | 0.380263681 | 0.998865404 | 10.566775 | 10.676492 | 10.569385 | 10.614099 | 10.668356 | 10.633459 | 1696 | 1684 | 1455 | 1500 |
| 57636 | NM_00119941 | ARHGAP23 | Rho GTPase activating protein 23 | protein_coding | NP_001186346.1;XP_006722054.1;XP_006722055.1;XP_006722056.1;XP_ | 29293 | 610590 | . | . | 0.097416 | 0.079820 | 0.162118 | 0.108031 | 0.226692 | 0.184507 | 0.363787 | 0.246999 | 1.468632 | 21.544092 | 0.673793432 | 0.822913015 | 0.410557461 | NA | 4.123517 | 4.681336 | 4.406481 | 4.391946 | 4.455153 | 4.411821 | 22 | 17 | 27 | 19 |
| 57639 | NM_020879 | CCDC146 | coiled-coil domain containing 146 | protein_coding | NP_065930.2 | 29296 | . | . | . | 0.316755 | 0.363166 | 0.369533 | 0.345406 | 0.737103 | 0.839475 | 0.829222 | 0.789730 | 1.048758 | 36.807896 | 0.510815866 | 0.1344558 | 0.89304216 | 0.998865404 | 5.167233 | 5.235838 | 5.188500 | 5.205179 | 5.209094 | 5.198352 | 39 | 41 | 34 | 34 |
| 57643 | NM_020883,XI | ZSWIM5 | zinc finger SWIM-type containing 5 | protein_coding | NP_065934.1;XP_011540163.1;XP_016857402.1 | 29299 | . | . | . | 0.199535 | 0.226124 | 0.309023 | 0.281040 | 0.464327 | 0.522697 | 0.693438 | 0.642564 | 1.375059 | 46.936315 | 0.456310908 | 1.006975354 | 0.313946618 | 0.998865404 | 5.304630 | 5.764191 | 5.506539 | 5.524162 | 5.582010 | 5.559977 | 43 | 45 | 50 | 48 |
| 57644 | NM_020884,XI | MYH7B | myosin heavy chain 7B | protein_coding | NP_065935.3;XP_006723903.1;XP_011527243.1;XP_011527249.1;XP_01 | 15906 | 609928 | . | . | 0.204024 | 0.131203 | 0.166102 | 0.104909 | 0.474772 | 0.303282 | 0.372727 | 0.239862 | -1.199902 | 31.869617 | 0.575641796 | -0.45673554 | 0.64786114 | 0.998865404 | 5.117472 | 4.859205 | 5.034836 | 4.971158 | 5.006215 | 4.953676 | 49 | 29 | 31 | 21 |
| 57645 | NM_00131401 | POGK | pogo transposable element derived with KRAB domain | protein_coding | NP_001300943.1;NP_060012.3;XP_005245427.1;XP_011508116.1;XP_01 | 18800 | . | . | . | 8.445577 | 8.706694 | 9.610996 | 7.853804 | 19.653205 | 20.125912 | 21.566796 | 17.956791 | -1.005216 | 1285.800583 | 0.133378599 | -0.05627052 | 0.955126302 | 0.998865404 | 10.332101 | 10.324792 | 10.311070 | 10.348386 | 10.365703 | 10.284730 | 1448 | 1425 | 1175 | 1138 |
| 57646 | NM_00130102 | USP28 | ubiquitin specific peptidase 28 | protein_coding | NP_001287958.1;NP_001333181.1;NP_001333182.1;NP_001333183.1;NI | 12625 | 610748 | . | . | 5.105022 | 5.601962 | 6.487611 | 6.267139 | 11.879596 | 12.949188 | 14.558014 | 14.329068 | 1.159177 | 824.537680 | 0.145343525 | 1.466191457 | 0.142596135 | 0.903146012 | 9.576929 | 9.790088 | 9.611953 | 9.650760 | 9.758410 | 9.713470 | 855 | 847 | 794 | 804 |
| 57647 | NM_032656,XI | DHX37 | DEAH-box helicase 37 | protein_coding | NP_116045.2;XP_005253647.1;XP_011536900.1;XP_011536902.1 | 17210 | 617362 | . | . | 10.748844 | 10.771629 | 11.767508 | 11.596129 | 25.013000 | 24.899104 | 26.405946 | 26.513169 | 1.076043 | 1579.457722 | 0.122804446 | 0.861006106 | 0.38923468 | 0.998865404 | 10.571358 | 10.677131 | 10.596208 | 10.595449 | 10.658341 | 10.647209 | 1753 | 1641 | 1434 | 1524 |
| 57648 | NM_00119897 | KIAA1522 | KIAA1522 | protein_coding | NP_001185901.1;NP_001185902.1;NP_001356482.1;NP_065939.2;XP_01 | 29301 | . | . | . | 5.558771 | 6.310136 | 5.690490 | 7.131810 | 12.935488 | 14.586167 | 12.769293 | 16.306037 | 1.056170 | 1037.419260 | 0.151469651 | 0.520511524 | 0.602707098 | 0.998865404 | 9.979185 | 10.057323 | 9.962374 | 10.029769 | 9.956575 | 10.111083 | 1109 | 1139 | 830 | 1103 |
| 57649 | NM_00103356 | PHF12 | PHD finger protein 12 | protein_coding | NP_001028733.1;NP_001277060.1;NP_065940.1;XP_005258072.1;XP_00 | 20816 | 618645 | . | . | 7.198447 | 7.453558 | 7.289490 | 7.803861 | 16.751079 | 17.229233 | 16.357406 | 17.842602 | 1.036049 | 1098.444980 | 0.132832228 | 0.3846369 | 0.700506451 | 0.998865404 | 10.075460 | 10.126581 | 10.090520 | 10.085099 | 10.115560 | 10.112715 | 1247 | 1160 | 974 | 1046 |
| 57650 | NM_020890,XI | CIP2A | cellular inhibitor of PP2A | protein_coding | NP_065941.2;XP_006713779.1;XP_006713780.1;XP_011511358.1;XP_01 | 29302 | 610643 | . | . | 4.964400 | 4.997292 | 5.241925 | 3.208583 | 11.552364 | 11.551465 | 11.762727 | 7.336044 | -1.297520 | 732.834926 | 0.194891105 | -1.92803621 | 0.053850622 | 0.688240867 | 9.692476 | 9.317977 | 9.634908 | 9.551948 | 9.547628 | 9.259915 | 1007 | 842 | 675 | 469 |
| 57654 | NM_00131793 | UVSSA | UV stimulated scaffold protein A | protein_coding | NP_001304863.1;NP_001304864.1;NP_065945.2;XP_016863979.1;XP_01 | 29304 | 614632 | . | . | 5.319112 | 5.786368 | 4.948221 | 4.445855 | 12.377795 | 13.375448 | 11.103665 | 10.164920 | -1.118540 | 730.919664 | 0.149590414 | -1.08039547 | 0.279966112 | 0.998865404 | 9.592109 | 9.430506 | 9.531209 | 9.570516 | 9.486992 | 9.459882 | 864 | 856 | 612 | 634 |
| 57655 | NM_00113619 | GRAMD1A | GRAM domain containing 1A | protein_coding | NP_001129671.1;NP_001306963.1;NP_001306964.1;NP_001306965.1;NI | 29305 | . | . | . | 10.984604 | 11.400886 | 10.582290 | 12.782628 | 25.561621 | 26.353657 | 23.746355 | 29.225962 | 1.040029 | 954.853715 | 0.148260731 | 0.381918631 | 0.70252172 | 0.998865404 | 9.870828 | 9.926899 | 9.878845 | 9.888505 | 9.844077 | 9.975880 | 1071 | 1017 | 770 | 995 |
| 57657 | NM_020897,N | HCN3 | hyperpolarization activated cyclic nucleotide gated potassium cha | protein_coding | NP_065948.1;XP_011508118.1;XP_011508119.1;XP_011508120.1;XP_01 | 19183 | 609973 | . | . | 1.466807 | 1.701591 | 1.925815 | 1.661264 | 3.413320 | 3.933304 | 4.321474 | 3.798281 | 1.110755 | 193.088210 | 0.24890918 | 0.608818536 | 0.542644726 | 0.998865404 | 7.515205 | 7.667035 | 7.530482 | 7.599715 | 7.652751 | 7.570050 | 194 | 213 | 193 | 173 |
| 57658 | NM_00114368 | CALCOCO1 | calcium binding and coiled-coil domain 1 | protein_coding | NP_001137154.1;NP_065949.1;XP_011536903.1;XP_011536904.1;XP_01 | 29306 | . | . | . | 11.088559 | 10.853084 | 11.286738 | 13.806490 | 25.803531 | 25.087391 | 25.327116 | 31.566895 | 1.249671 | 1374.466710 | 0.130171044 | 2.470199611 | 0.013503768 | 0.383747314 | 10.254774 | 10.576625 | 10.346510 | 10.316862 | 10.537137 | 10.466454 | 1435 | 1292 | 1388 | 1367 |
| 57659 | NM_00112883 | ZBTB4 | zinc finger and BTB domain containing 4 | protein_coding | NP_001122305.1;NP_065950.2;XP_006721626.1;XP_006721627.1;XP_01 | 23847 | 612308 | . | . | 6.983382 | 7.950573 | 7.703621 | 8.303390 | 16.250617 | 18.378106 | 17.286702 | 18.984714 | 1.059555 | 1420.439055 | 0.133001277 | 0.627500203 | 0.530331403 | 0.998865404 | 10.429984 | 10.513063 | 10.400624 | 10.494722 | 10.467385 | 10.518137 | 1491 | 1580 | 1229 | 1415 |
| 57661 | NM_00128658 | PHRF1 | PHD and ring finger domains 1 | protein_coding | NP_001273510.1;NP_001273511.1;NP_001273512.1;NP_065952.2;XP_0C | 24351 | 611780 | . | . | 11.160050 | 11.000049 | 11.084844 | 12.276163 | 25.969895 | 25.427105 | 24.874071 | 28.067989 | 1.037452 | 1959.024218 | 0.121619672 | 0.436148621 | 0.662728903 | 0.998865404 | 10.909232 | 10.962122 | 10.928911 | 10.912802 | 10.914209 | 10.984568 | 2237 | 2054 | 1658 | 1953 |
| 57664 | NM_00116135 | PLEKHA4 | pleckstrin homology domain containing A4 | protein_coding | NP_001154826.1;NP_065955.2;XP_005259164.1;XP_006723364.1;XP_01 | 14339 | 607769 | . | . | 29.934758 | 30.724779 | 31.567149 | 33.344180 | 69.659417 | 71.021706 | 70.835770 | 76.237506 | 1.055744 | 2983.555335 | 0.111893567 | 0.699419033 | 0.484290197 | 0.998865404 | 11.503201 | 11.581373 | 11.513643 | 11.526559 | 11.546793 | 11.581756 | 3316 | 3157 | 2608 | 2934 |
| 57665 | NM_020905 | RDH14 | retinol dehydrogenase 14 | protein_coding | NP_065956.1 | 19979 | 616796 | . | . | 10.135124 | 8.125993 | 10.220820 | 8.425451 | 23.584852 | 18.783594 | 22.935225 | 19.263790 | 1.006880 | 449.313718 | 0.194435554 | 0.050873657 | 0.959426197 | 0.998865404 | 8.805957 | 8.817179 | 8.874558 | 8.733277 | 8.870979 | 8.746806 | 574 | 428 | 432 | 380 |
| 57666 | NM_00114264 | FBRSL1 | fibrosin like 1 | protein_coding | NP_001136113.1;NP_001354800.1;NP_001369668.1;NP_001369669.1;NI | 29308 | . | . | . | 3.962866 | 4.320118 | 4.286750 | 4.085770 | 9.221750 | 9.986147 | 9.619346 | 9.341630 | 1.017116 | 545.418793 | 0.163820031 | 0.149457779 | 0.881192423 | 0.998865404 | 9.078912 | 9.103425 | 9.064448 | 9.105692 | 9.113252 | 9.079034 | 603 | 602 | 491 | 502 |
| 57669 | NM_00118493 | EPB41L5 | erythrocyte membrane protein band 4.1 like 5 | protein_coding | NP_001171866.1;NP_001171867.1;NP_001171868.1;NP_001317236.1;NI | 19819 | 611730 | . | . | 2.649553 | 2.331063 | 2.327836 | 2.517215 | 6.165618 | 5.388358 | 5.223596 | 5.755316 | -1.034377 | 491.322723 | 0.17441549 | -0.27957638 | 0.779802533 | 0.998865404 | 8.964631 | 8.916014 | 8.995950 | 8.903111 | 8.908340 | 8.949859 | 617 | 500 | 406 | 468 |
| 57670 | NM_00116466 | KIAA1549 | KIAA1549 | protein_coding | NP_001158137.1;NP_065961.2;XP_011514744.1 | 22219 | 613344 | . | . | 1.361155 | 1.480612 | 1.487413 | 1.460730 | 3.167465 | 3.422501 | 3.337711 | 3.339786 | 1.023117 | 563.859988 | 0.161754705 | 0.203830356 | 0.83848607 | 0.998865404 | 9.122659 | 9.155540 | 9.108690 | 9.153434 | 9.155481 | 9.136885 | 620 | 622 | 503 | 527 |
| 57673 | NM_00108045 | BEND3 | BEN domain containing 3 | protein_coding | NP_001073919.1;NP_001354243.1;XP_005267137.1;XP_011534307.1 | 23040 | 616374 | . | . | 0.818711 | 0.980644 | 1.032080 | 0.952351 | 1.905174 | 2.266803 | 2.315958 | 2.177436 | 1.088841 | 196.731022 | 0.243782065 | 0.503703385 | 0.61446983 | 0.998865404 | 7.557510 | 7.680050 | 7.558613 | 7.636197 | 7.655399 | 7.616712 | 198 | 221 | 186 | 184 |
| 57674 | NM_00125607 | RNF213 | ring finger protein 213 | protein_coding | NP_001243000.2;NP_066005.2;XP_005257602.2;XP_005257603.2;XP_00 | 14539 | 613768 | . | . | 9.127979 | 9.490330 | 10.540655 | 11.139261 | 21.241181 | 21.937324 | 23.652929 | 25.468595 | 1.088892 | 5453.095712 | 0.105048884 | 1.169560369 | 0.242177934 | 0.992733415 | 12.350146 | 12.472961 | 12.369552 | 12.382339 | 12.432844 | 12.461804 | 5966 | 5677 | 4859 | 5421 |
| 57677 | NM_00129761 | ZFP14 | ZFP14 zinc finger protein | protein_coding | NP_001284548.1;NP_065968.1;XP_016882527.1;XP_024307393.1 | 29312 | . | . | . | 0.548308 | 0.429650 | 0.426648 | 0.427330 | 1.275937 | 0.993156 | 0.957384 | 0.977040 | -1.132821 | 109.743600 | 0.314741028 | -0.57164402 | 0.567563166 | 0.998865404 | 6.864661 | 6.685784 | 6.839807 | 6.753707 | 6.759161 | 6.755639 | 151 | 110 | 90 | 96 |
| 57678 | NM_00124494 | GPAM | glycerol-3-phosphate acyltransferase, mitochondrial | protein_coding | NP_001231878.1;NP_065969.3;XP_005270055.1;XP_024303857.1 | 24865 | 602395 | . | . | 2.822980 | 2.723934 | 2.372298 | 1.947821 | 6.569191 | 6.296495 | 5.323369 | 4.453462 | -1.212979 | 507.994629 | 0.167306072 | -1.664938 | 0.095925126 | 0.813553488 | 9.121066 | 8.842884 | 9.062943 | 9.030032 | 8.939016 | 8.909161 | 657 | 586 | 409 | 420 |
| 57679 | NM_00113574 | ALS2 | alsin Rho guanine nucleotide exchange factor ALS2 | protein_coding | NP_001129217.1;NP_065970.2;XP_006712717.1;XP_006712718.1;XP_01 | 443 | 606352 | . | . | 11.704598 | 12.380310 | 12.533179 | 12.462109 | 27.237076 | 28.617644 | 28.124090 | 28.493131 | 1.010077 | 2419.105148 | 0.116027655 | 0.124665115 | 0.900788676 | 0.998865404 | 11.233018 | 11.247462 | 11.213183 | 11.258412 | 11.255812 | 11.232300 | 2694 | 2670 | 2145 | 2246 |
| 57680 | NM_00117062 | CHD8 | chromodomain helicase DNA binding protein 8 | protein_coding | NP_001164100.1;NP_065971.2 | 20153 | 610528 | . | . | 7.046981 | 7.761696 | 7.602521 | 8.238511 | 16.398613 | 17.941508 | 17.059837 | 18.836374 | 1.038502 | 2004.114265 | 0.123515094 | 0.441272548 | 0.659015697 | 0.998865404 | 10.941384 | 10.995605 | 10.917694 | 10.987346 | 10.951827 | 11.013296 | 2165 | 2215 | 1707 | 1988 |
| 57683 | NM_00128554 | ZDBF2 | zinc finger DBF-type containing 2 | protein_coding | NP_001272478.1;NP_001356583.1;NP_065974.1;XP_005246769.1;XP_00 | 29313 | 617059 | . | . | 3.187827 | 3.354985 | 4.163961 | 3.158459 | 7.418204 | 7.755198 | 9.343806 | 7.221440 | 1.007706 | 983.841013 | 0.183228444 | 0.060443353 | 0.951802531 | 0.998865404 | 9.936324 | 9.948214 | 9.939128 | 9.927674 | 10.083208 | 9.771542 | 1137 | 1049 | 1041 | 730 |
| 57684 | NM_00130436 | ZBTB26 | zinc finger and BTB domain containing 26 | protein_coding | NP_001291292.1;NP_001291293.1;NP_065975.1 | 23383 | . | . | . | 1.592449 | 2.003560 | 1.620501 | 1.343477 | 3.705695 | 4.631318 | 3.636357 | 3.071700 | -1.145879 | 220.435525 | 0.235581954 | -0.83391164 | 0.404330779 | 0.998865404 | 7.878808 | 7.682977 | 7.784538 | 7.842911 | 7.792679 | 7.705390 | 255 | 269 | 196 | 174 |
| 57685 | NM_00129327 | CACHD1 | cache domain containing 1 | protein_coding | NP_001280203.1;NP_065976.3;XP_011540164.1;XP_011540165.1 | 29314 | . | . | . | 5.425019 | 4.987117 | 4.488278 | 4.788822 | 12.624242 | 11.527945 | 10.071564 | 10.949074 | -1.135948 | 910.372853 | 0.143204796 | -1.28415552 | 0.199087546 | 0.96621494 | 9.919330 | 9.735442 | 9.907032 | 9.840683 | 9.763125 | 9.801429 | 1165 | 998 | 721 | 821 |
| 57688 | NM_020928,XI | ZSWIM6 | zinc finger SWIM-type containing 6 | protein_coding | NP_065979.1;XP_016865166.1;XP_024301907.1;XP_024301908.1 | 29316 | 615951 | . | . | 1.782154 | 1.604407 | 1.780238 | 1.811167 | 4.147146 | 3.708659 | 3.994801 | 4.141019 | 1.062075 | 286.496348 | 0.204993635 | 0.423847671 | 0.671676909 | 0.998865404 | 8.118443 | 8.205004 | 8.148866 | 8.142402 | 8.157297 | 8.197060 | 322 | 298 | 247 | 287 |
| 57690 | NM_00114264 | TNRC6C | trinucleotide repeat containing adaptor 6C | protein_coding | NP_001136112.1;NP_061869.2;XP_006722059.3;XP_006722060.3;XP_00 | 29318 | 610741 | . | . | 0.507842 | 0.323336 | 0.568391 | 0.625890 | 1.181769 | 0.747406 | 1.275455 | 1.431023 | 1.061902 | 150.016671 | 0.268523992 | 0.322692232 | 0.746928335 | 0.998865404 | 7.185140 | 7.271525 | 7.193264 | 7.237712 | 7.246646 | 7.231441 | 158 | 166 | 137 | 142 |
| 57691 | NM_00128627 | KIAA1586 | KIAA1586 | protein_coding | NP_001273203.1;NP_001273204.1;NP_001273205.1;NP_065982.1;XP_01 | 21360 | . | . | . | 2.365942 | 2.407512 | 2.834818 | 1.482036 | 5.505643 | 5.565074 | 6.361252 | 3.388499 | -1.217220 | 148.972851 | 0.304707202 | -0.93069752 | 0.352010054 | 0.998865404 | 7.352390 | 7.071800 | 7.301175 | 7.195100 | 7.253231 | 7.086764 | 212 | 154 | 144 | 97 |
| 57692 | NM_020932 | MAGEE1 | MAGE family member E1 | protein_coding | NP_065983.1 | 24934 | 300759 | . | . | 0.471638 | 0.438169 | 0.484718 | 0.270698 | 1.097520 | 1.012847 | 1.087694 | 0.618919 | -1.219446 | 47.722044 | 0.476723535 | -0.60040279 | 0.548237831 | 0.998865404 | 5.710890 | 5.428479 | 5.599972 | 5.582204 | 5.600132 | 5.509537 | 63 | 54 | 48 | 29 |
| 57693 | NM_00119079 | ZNF317 | zinc finger protein 317 | protein_coding | NP_001177720.1;NP_065984.3;XP_024307394.1 | 13507 | 613864 | . | . | 7.123620 | 8.117811 | 8.573894 | 9.553997 | 16.576959 | 18.764682 | 19.239568 | 21.844078 | 1.173837 | 1045.653071 | 0.14286729 | 1.618510214 | 0.105552687 | 0.833478683 | 9.910232 | 10.140932 | 9.921043 | 10.010158 | 10.048102 | 10.119981 | 1040 | 1102 | 935 | 1109 |
| 57695 | NM_020935,XI | USP37 | ubiquitin specific peptidase 37 | protein_coding | NP_065986.3;XP_005246777.1;XP_005246778.1;XP_005246779.1;XP_00 | 20063 | . | . | . | 2.972706 | 3.102094 | 3.308506 | 2.804673 | 6.917606 | 7.170633 | 7.424191 | 6.412550 | -1.000238 | 742.073448 | 0.155622097 | -0.00220295 | 0.998242306 | 0.999862634 | 9.535367 | 9.535469 | 9.519221 | 9.549422 | 9.588268 | 9.477500 | 836 | 818 | 697 | 640 |
| 57696 | NM_020936,N | DDX55 | DEAD-box helicase 55 | protein_coding | NP_065987.1;XP_011536910.1;XP_016875199.1;XP_016875200.1;XP_01 | 20085 | . | . | . | 17.525624 | 18.889176 | 20.216700 | 20.582951 | 40.782845 | 43.663181 | 45.365693 | 47.060466 | 1.108342 | 1719.401597 | 0.123093248 | 1.205619467 | 0.227964249 | 0.982008277 | 10.671657 | 10.819917 | 10.676920 | 10.733416 | 10.783616 | 10.788659 | 1810 | 1824 | 1567 | 1700 |
| 57697 | NM_00130813 | FANCM | FA complementation group M | protein_coding | NP_001295062.1;NP_001295063.1;NP_065988.1;XP_011535336.1;XP_01 | 23168 | 609644 | . | . | 1.519662 | 1.572346 | 1.342499 | 0.998559 | 3.536318 | 3.634550 | 3.012528 | 2.283086 | -1.227194 | 276.471892 | 0.229365046 | -1.28774574 | 0.197834487 | 0.96621494 | 8.250323 | 7.956746 | 8.226006 | 8.085909 | 8.107245 | 7.993844 | 395 | 287 | 241 | 206 |
| 57698 | NM_00112721 | SHTN1 | shootin 1 | protein_coding | NP_001120683.1;NP_001245227.1;NP_001245228.1;NP_001245229.1;NI | 29319 | 611171 | . | . | 21.368850 | 22.733091 | 29.249509 | 21.818222 | 49.726192 | 52.548557 | 65.635048 | 49.884778 | 1.092308 | 5043.787528 | 0.141101843 | 0.902748742 | 0.366659261 | 0.998865404 | 12.235117 | 12.362649 | 12.233937 | 12.283525 | 12.453580 | 12.196644 | 5386 | 5357 | 5273 | 4190 |
| 57699 | NM_00131401 | CPNE5 | copine 5 | protein_coding | NP_001300946.1;NP_001300947.1;NP_001300948.1;NP_001300949.1;NI | 2318 | 604209 | . | . | 0.443047 | 0.142624 | 0.114703 | 0.548982 | 1.030989 | 0.329683 | 0.257391 | 1.255186 | -1.155855 | 23.504276 | 0.715149171 | -0.29219088 | 0.770140685 | NA | 4.656326 | 4.445696 | 4.600694 | 4.512743 | 4.505665 | 4.580056 | 41 | 16 | 11 | 29 |
| 57700 | NM_00113505 | FAM160B1 | family with sequence similarity 160 member B1 | protein_coding | NP_001128523.1;NP_065991.3 | 29320 | 617312 | . | . | 2.860338 | 2.581589 | 2.382166 | 2.149959 | 6.656125 | 5.967458 | 5.345512 | 4.915627 | -1.218402 | 449.485439 | 0.179213741 | -1.59022611 | 0.111783847 | 0.848562841 | 8.947225 | 8.663067 | 8.903935 | 8.832307 | 8.779820 | 8.714866 | 600 | 503 | 372 | 359 |
| 57701 | NM_00103780 | NCKAP5L | NCK associated protein 5 like | protein_coding | NP_001032895.2;NP_001354977.1;XP_005269107.1;XP_006719588.1;XP | 29321 | 615104 | . | . | 5.199176 | 5.688669 | 6.394834 | 6.444319 | 12.098696 | 13.149614 | 14.349822 | 14.734170 | 1.164033 | 896.573612 | 0.142044879 | 1.54269289 | 0.122905308 | 0.867477288 | 9.694676 | 9.913584 | 9.721168 | 9.777974 | 9.857644 | 9.860787 | 916 | 930 | 837 | 906 |
| 57703 | NM_00137602 | CWC22 | CWC22 spliceosome associated protein homolog | protein_coding | NP_001362958.1;NP_001362959.1;NP_001362961.1;NP_001362962.1;NI | 29322 | 615186 | . | . | 10.666301 | 10.109604 | 9.049847 | 9.045633 | 24.820922 | 23.368802 | 20.307595 | 20.681766 | -1.145237 | 964.008769 | 0.139424939 | -1.40323238 | 0.160547553 | 0.933525128 | 10.007276 | 9.811920 | 9.983687 | 9.935931 | 9.876389 | 9.846476 | 1223 | 1075 | 798 | 825 |
| 57704 | NM_00133066 | GBA2 | glucosylceramidase beta 2 | protein_coding | NP_001317589.1;NP_065995.1;XP_005251583.1;XP_006716872.1;XP_01 | 18986 | 609471 | . | . | 34.276829 | 35.341128 | 40.179283 | 36.606341 | 79.763593 | 81.692608 | 90.161147 | 83.696048 | 1.059214 | 4004.999430 | 0.109240463 | 0.759732895 | 0.447414261 | 0.998865404 | 11.925472 | 12.008506 | 11.930415 | 11.955525 | 12.013116 | 11.967965 | 4411 | 4261 | 3680 | 3758 |
| 57706 | NM_00135296 | DENND1A | DENN domain containing 1A | protein_coding | NP_001339893.1;NP_001339894.1;NP_001339895.1;NP_001339896.1;NI | 29324 | 613633 | . | . | 2.009248 | 2.592607 | 2.353423 | 2.752991 | 4.675603 | 5.992928 | 5.281013 | 6.294383 | 1.184195 | 325.503777 | 0.214911675 | 1.134914087 | 0.256411317 | 0.995760118 | 8.220190 | 8.462692 | 8.210069 | 8.368889 | 8.350435 | 8.420609 | 293 | 369 | 288 | 351 |
| 57707 | NM_020947,XI | MEAK7 | MTOR associated protein, eak-7 homolog | protein_coding | NP_065998.3;XP_005256132.1;XP_006721303.1;XP_011521550.1;XP_01 | 29325 | . | . | . | 3.395618 | 4.111782 | 3.958486 | 3.441996 | 7.901742 | 9.504567 | 8.882730 | 7.869714 | -1.036908 | 403.712252 | 0.189255552 | -0.2762823 | 0.782331252 | 0.998865404 | 8.683359 | 8.630524 | 8.604134 | 8.725487 | 8.648981 | 8.640868 | 426 | 488 | 347 | 369 |
| 57708 | NM_00107770 | MIER1 | MIER1 transcriptional regulator | protein_coding | NP_001071168.2;NP_001071169.1;NP_001071170.2;NP_001071171.2;NI | 29657 | 616848 | . | . | 3.707597 | 3.419666 | 4.150173 | 3.216560 | 8.627729 | 7.904717 | 9.312865 | 7.354284 | 1.020545 | 570.693818 | 0.181121592 | 0.161991168 | 0.871312809 | 0.998865404 | 9.141242 | 9.171743 | 9.186282 | 9.105734 | 9.243428 | 9.069859 | 690 | 572 | 568 | 469 |
| 57710 | NM_020950 | KIAA1614 | KIAA1614 | protein_coding | NP_066001.1 | 29327 | . | . | . | 0.106815 | 0.135080 | 0.129718 | 0.033837 | 0.248562 | 0.312243 | 0.291084 | 0.077364 | -1.508735 | 31.853817 | 0.63977231 | -0.92742241 | 0.353707261 | 0.998865404 | 5.257378 | 4.670041 | 4.991469 | 5.026321 | 5.015234 | 4.895356 | 39 | 46 | 35 | 10 |
| 57711 | NM_00114564 | ZNF529 | zinc finger protein 529 | protein_coding | NP_001139121.1;NP_001139122.1;NP_001308280.1;NP_001339201.1;NI | 29328 | . | . | . | 2.866750 | 2.739613 | 3.447704 | 2.028587 | 6.671047 | 6.332737 | 7.736549 | 4.638124 | -1.040898 | 460.485863 | 0.226449216 | -0.2553706 | 0.798436872 | 0.998865404 | 8.874863 | 8.818615 | 8.884327 | 8.815507 | 8.972611 | 8.653952 | 570 | 479 | 494 | 314 |
| 57713 | NM_00101803 | SFMBT2 | Scm like with four mbt domains 2 | protein_coding | NP_001018049.1;NP_001025051.1;XP_005252608.1;XP_006717553.1;XP | 20256 | 615392 | . | . | 1.174967 | 1.379531 | 1.707155 | 1.484902 | 2.734195 | 3.188848 | 3.830806 | 3.395051 | 1.231271 | 359.996807 | 0.196787494 | 1.525240923 | 0.127199017 | 0.874064017 | 8.333930 | 8.634159 | 8.382923 | 8.465069 | 8.586409 | 8.499369 | 344 | 374 | 371 | 345 |
| 57715 | NM_00120324 | SEMA4G | semaphorin 4G | protein_coding | NP_001190173.1;NP_060363.2;XP_024303858.1;XP_024303859.1 | 10735 | . | . | . | 2.158172 | 2.039235 | 2.009689 | 2.260883 | 5.022157 | 4.713784 | 4.509684 | 5.169241 | 1.002644 | 292.389689 | 0.204962144 | 0.018583833 | 0.9851731 | 0.999197585 | 8.189985 | 8.193510 | 8.206311 | 8.175213 | 8.165928 | 8.217901 | 346 | 306 | 242 | 288 |
| 57716 | NM_020956,N | PRX | periaxin | protein_coding | NP_066007.1;NP_870998.2;XP_011525473.1;XP_016882535.1;XP_01688 | 13797 | 605725 | . | . | 0.292808 | 0.398168 | 0.306509 | 0.296836 | 0.681377 | 0.920383 | 0.687798 | 0.678680 | -1.238163 | 54.548213 | 0.432720109 | -0.71224054 | 0.47631584 | 0.998865404 | 5.915807 | 5.606574 | 5.756098 | 5.822200 | 5.750425 | 5.744814 | 59 | 75 | 43 | 45 |
| 57717 | NM_020957 | PCDHB16 | protocadherin beta 16 | protein_coding | NP_066008.2 | 14546 | 606345 | . | . | 0.841033 | 0.768648 | 0.514717 | 0.613106 | 1.957118 | 1.776765 | 1.155010 | 1.401794 | -1.437225 | 97.516929 | 0.328536261 | -1.59278082 | 0.11120937 | 0.848126978 | 6.845925 | 6.321914 | 6.676044 | 6.641919 | 6.531692 | 6.571453 | 139 | 118 | 64 | 81 |
| 57718 | NM_00134814 | PPP4R4 | protein phosphatase 4 regulatory subunit 4 | protein_coding | NP_001335071.1;NP_001335072.1;NP_001335073.1;NP_001335074.1;NI | 23788 | 616790 | . | . | 1.038792 | 0.939553 | 0.677416 | 1.407240 | 2.417309 | 2.171818 | 1.520101 | 3.217483 | 1.204883 | 80.066647 | 0.353331914 | 0.761022523 | 0.446643615 | 0.998865404 | 6.182248 | 6.451470 | 6.295197 | 6.293204 | 6.349693 | 6.336351 | 84 | 78 | 78 | 80 |
| 57719 | NM_020959,XI | ANO8 | anoctamin 8 | protein_coding | NP_066010.1;XP_016882537.1 | 29329 | 610216 | . | . | 2.368414 | 2.618507 | 2.429404 | 2.520153 | 5.511396 | 6.052794 | 5.451512 | 5.762031 | -1.047391 | 368.069291 | 0.187344517 | -0.35656196 | 0.721419758 | 0.998865404 | 8.557052 | 8.489832 | 8.512768 | 8.560784 | 8.501902 | 8.518512 | 416 | 423 | 308 | 342 |
| 57720 | NM_00113655 | GPR107 | G protein-coupled receptor 107 | protein_coding | NP_001130029.1;NP_001130030.1;NP_001274275.1;NP_066011.2;XP_01 | 17830 | 618490 | . | . | 10.008824 | 10.954968 | 9.644519 | 10.560679 | 23.290944 | 25.322899 | 21.642019 | 24.145736 | -1.035340 | 2137.781697 | 0.122038994 | -0.41056008 | 0.681395141 | 0.998865404 | 11.086867 | 11.036491 | 11.046162 | 11.103669 | 11.011887 | 11.081883 | 2415 | 2431 | 1746 | 2056 |
| 57721 | NM_020961,XI | METTL14 | methyltransferase like 14 | protein_coding | NP_066012.1;XP_005263223.1;XP_011530464.1 | 29330 | 616504 | . | . | 2.776298 | 2.468955 | 2.476230 | 2.254474 | 6.460559 | 5.707099 | 5.556588 | 5.154587 | -1.155973 | 459.901606 | 0.177251496 | -1.17972596 | 0.238109225 | 0.992733415 | 8.945614 | 8.737231 | 8.926981 | 8.847116 | 8.822105 | 8.773911 | 603 | 499 | 386 | 384 |
| 57724 | NM_020964,XI | EPG5 | ectopic P-granules autophagy protein 5 homolog | protein_coding | NP_066015.2;XP_011524424.1;XP_016881378.1;XP_016881379.1;XP_01 | 29331 | 615068 | . | . | 4.486207 | 4.771726 | 5.151631 | 5.039745 | 10.439587 | 11.030060 | 11.560109 | 11.522776 | 1.101503 | 1876.918862 | 0.122564595 | 1.137953565 | 0.255139873 | 0.995760118 | 10.802760 | 10.942175 | 10.800900 | 10.866430 | 10.925368 | 10.895565 | 1971 | 2008 | 1743 | 1810 |
| 57727 | NM_00134814 | NCOA5 | nuclear receptor coactivator 5 | protein_coding | NP_001335077.1;NP_001335078.1;NP_001335079.1;NP_001335080.1;NI | 15909 | 616825 | . | . | 5.964404 | 6.618075 | 7.295664 | 7.876363 | 13.879413 | 15.297977 | 16.371258 | 18.008368 | 1.187996 | 699.405814 | 0.154287918 | 1.610821949 | 0.107218536 | 0.835490835 | 9.320657 | 9.568669 | 9.352383 | 9.418818 | 9.480130 | 9.527900 | 700 | 724 | 638 | 736 |
| 57728 | NM_00131792 | WDR19 | WD repeat domain 19 | protein_coding | NP_001304853.1;NP_079408.3;XP_011512026.1;XP_011512027.1;XP_01 | 18340 | 608151 | . | . | 6.131104 | 5.873806 | 8.057375 | 6.183579 | 14.267330 | 13.577564 | 18.080519 | 14.138020 | 1.154725 | 1104.970113 | 0.151068536 | 1.373876589 | 0.169479999 | 0.946181982 | 10.002001 | 10.210085 | 10.048697 | 10.053487 | 10.244041 | 10.064734 | 1177 | 1110 | 1152 | 979 |
| 57730 | NM_00135333 | ANKRD36B | ankyrin repeat domain 36B | protein_coding | NP_001340266.1;NP_079466.3;XP_005264052.1;XP_006712724.1;XP_01 | 29333 | . | . | . | 2.928346 | 3.188275 | 4.031177 | 1.906668 | 6.814382 | 7.369837 | 9.045842 | 4.359369 | 1.018551 | 443.567701 | 0.243444699 | 0.108931965 | 0.913256453 | 0.998865404 | 8.778954 | 8.806931 | 8.778632 | 8.770816 | 8.958801 | 8.577957 | 509 | 471 | 509 | 291 |
| 57731 | NM_020971,N | SPTBN4 | spectrin beta, non-erythrocytic 4 | protein_coding | NP_066022.2;NP_079489.2;XP_011525475.1;XP_016882538.1;XP_01688 | 14896 | 606214 | . | . | 0.067042 | 0.123122 | 0.038871 | 0.140903 | 0.156010 | 0.284601 | 0.087226 | 0.322158 | -1.060348 | 26.322114 | 0.69504225 | -0.12162973 | 0.90319227 | NA | 4.763465 | 4.671475 | 4.676591 | 4.744622 | 4.646934 | 4.759987 | 22 | 38 | 10 | 37 |
| 57732 | NM_00117265 | ZFYVE28 | zinc finger FYVE-type containing 28 | protein_coding | NP_001166127.1;NP_001166128.1;NP_001166129.1;NP_001166130.1;NI | 29334 | 614176 | . | . | 0.706166 | 0.464099 | 1.655723 | 0.709291 | 1.643279 | 1.072785 | 3.715394 | 1.621710 | 1.492521 | 72.938337 | 0.426338083 | 1.355148041 | 0.175370395 | 0.949120257 | 5.868587 | 6.450365 | 6.146150 | 6.078474 | 6.301526 | 6.111374 | 78 | 53 | 102 | 54 |
| 57758 | NM_00117069 | SCUBE2 | signal peptide, CUB domain and EGF like domain containing 2 | protein_coding | NP_001164161.1;NP_001317128.1;NP_001354906.1;NP_066025.2;XP_0C | 30425 | 611747 | . | . | 0.614616 | 0.806339 | 0.611430 | 1.059890 | 1.430237 | 1.863889 | 1.372028 | 2.423312 | 1.174809 | 110.874325 | 0.330529317 | 0.703193267 | 0.481935314 | 0.998865404 | 6.673363 | 6.903066 | 6.730011 | 6.786641 | 6.736807 | 6.880634 | 107 | 120 | 82 | 136 |
| 57761 | NM_00130118 | TRIB3 | tribbles pseudokinase 3 | protein_coding | NP_001288117.1;NP_001288119.1;NP_001288122.1;NP_001288125.1;NI | 16228 | 607898 | . | . | 10.189215 | 10.754601 | 9.939262 | 14.283021 | 23.710722 | 24.859744 | 22.303414 | 32.656430 | 1.135149 | 861.343870 | 0.177944205 | 1.027746294 | 0.304069185 | 0.998865404 | 9.656566 | 9.838587 | 9.683615 | 9.713119 | 9.646248 | 9.914430 | 910 | 889 | 650 | 1016 |
| 57763 | NM_023039,XI | ANKRA2 | ankyrin repeat family A member 2 | protein_coding | NP_075526.1;XP_005248617.1;XP_016865167.1 | 13208 | 605787 | . | . | 4.738820 | 4.920696 | 3.845731 | 4.806268 | 11.027428 | 11.374407 | 8.629710 | 10.988961 | -1.141519 | 300.642465 | 0.207373929 | -0.92082164 | 0.357143554 | 0.998865404 | 8.324688 | 8.132743 | 8.262995 | 8.269397 | 8.145340 | 8.242299 | 367 | 348 | 221 | 288 |
| 57786 | NM_00120445 | RBAK | RB associated KRAB zinc finger | protein_coding | NP_001191385.1;NP_066986.1 | 17680 | 608191 | . | . | 2.844388 | 2.978929 | 3.169534 | 2.554315 | 6.619008 | 6.885927 | 7.112342 | 5.840137 | -1.031156 | 595.268351 | 0.171181259 | -0.25857433 | 0.795963692 | 0.998865404 | 9.239007 | 9.195457 | 9.213323 | 9.237941 | 9.276477 | 9.129278 | 683 | 664 | 567 | 488 |
| 57787 | NM_00119986 | MARK4 | microtubule affinity regulating kinase 4 | protein_coding | NP_001186796.1;NP_113605.2;XP_006723370.1 | 13538 | 606495 | . | . | 3.016900 | 3.187525 | 2.566791 | 4.070529 | 7.020451 | 7.368107 | 5.759806 | 9.306780 | 1.057480 | 495.227658 | 0.20271951 | 0.397741385 | 0.690820828 | 0.998865404 | 8.911796 | 8.991013 | 8.886369 | 8.969870 | 8.836779 | 9.077501 | 518 | 554 | 359 | 567 |
| 57794 | NM_172231,XI | SUGP1 | SURP and G-patch domain containing 1 | protein_coding | NP_757386.2;XP_005260059.1;XP_011526457.1;XP_024307395.1;XP_02 | 18643 | 607992 | . | . | 9.937453 | 9.374071 | 10.142100 | 9.029816 | 23.124863 | 21.668586 | 22.758579 | 20.645601 | 1.043446 | 879.826291 | 0.14708675 | 0.417141247 | 0.676575097 | 0.998865404 | 9.749794 | 9.811692 | 9.782988 | 9.745986 | 9.841609 | 9.746855 | 1017 | 905 | 833 | 787 |
| 57798 | NM_021167,N | GATAD1 | GATA zinc finger domain containing 1 | protein_coding | NP_066990.3;XP_024302618.1 | 29941 | 614518 | . | . | 8.814133 | 7.928464 | 9.506765 | 8.193799 | 20.510852 | 18.327004 | 21.332906 | 18.734146 | -1.005171 | 1229.500738 | 0.141001196 | -0.05276757 | 0.9579171 | 0.998865404 | 10.267293 | 10.260412 | 10.298213 | 10.229967 | 10.318049 | 10.200465 | 1483 | 1270 | 1150 | 1060 |
| 57799 | NM_00117266 | RAB40C | RAB40C, member RAS oncogene family | protein_coding | NP_001166134.1;NP_001166135.1;NP_001166136.1;NP_001166137.1;NI | 18285 | . | . | . | 2.554590 | 2.766180 | 3.056087 | 3.057099 | 5.944633 | 6.394151 | 6.857770 | 6.989693 | 1.144831 | 241.650699 | 0.219335261 | 0.889662608 | 0.373647078 | 0.998865404 | 7.816051 | 8.010932 | 7.860698 | 7.899801 | 7.950519 | 7.943247 | 249 | 253 | 226 | 240 |
| 57801 | NM_00114246 | HES4 | hes family bHLH transcription factor 4 | protein_coding | NP_001135939.1;NP_066993.1;XP_005244828.1 | 24149 | 608060 | . | . | 2.921797 | 3.047039 | 2.926274 | 2.864427 | 6.799143 | 7.043368 | 6.566475 | 6.549169 | 1.016188 | 122.603958 | 0.292405066 | 0.079230221 | 0.936849507 | 0.998865404 | 6.926110 | 6.949518 | 6.924160 | 6.944369 | 6.954157 | 6.925404 | 136 | 135 | 112 | 111 |
| 57804 | NM_00125687 | POLD4 | DNA polymerase delta 4, accessory subunit | protein_coding | NP_001243799.1;NP_066996.3 | 14106 | 611525 | . | . | 7.609952 | 9.039523 | 8.938874 | 8.647843 | 17.708673 | 20.895261 | 20.058575 | 19.772266 | 1.045010 | 450.941361 | 0.181028132 | 0.350870535 | 0.725685478 | 0.998865404 | 8.784868 | 8.848033 | 8.745546 | 8.854608 | 8.842726 | 8.814252 | 463 | 518 | 410 | 422 |
| 57805 | NM_00136306 | CCAR2 | cell cycle and apoptosis regulator 2 | protein_coding | NP_001349997.1;NP_001349998.1;NP_066997.3;XP_011542906.1 | 23360 | 607359 | . | . | 21.973204 | 20.613918 | 21.736634 | 23.454022 | 51.132548 | 47.649994 | 48.776376 | 53.624836 | 1.111863 | 3147.965960 | 0.111709099 | 1.369440626 | 0.170861583 | 0.946389416 | 11.541730 | 11.694629 | 11.552415 | 11.596925 | 11.657384 | 11.666248 | 3339 | 3305 | 2867 | 3124 |
| 57819 | NM_021177 | LSM2 | LSM2 homolog, U6 small nuclear RNA and mRNA degradation ass | protein_coding | NP_067000.1 | 13940 | 607282 | . | . | 25.623711 | 24.030529 | 20.143785 | 28.997736 | 59.627426 | 55.547646 | 45.202072 | 66.299873 | -1.024752 | 660.289834 | 0.18430417 | -0.19139747 | 0.8482142 | 0.998865404 | 9.384896 | 9.348789 | 9.396612 | 9.346340 | 9.230487 | 9.466061 | 798 | 695 | 468 | 718 |
| 57820 | NM_021178,N | CCNB1IP1 | cyclin B1 interacting protein 1 | protein_coding | NP_067001.3;NP_878269.1;NP_878272.1 | 19437 | 608249 | . | . | 14.236253 | 14.282441 | 15.768946 | 15.566352 | 33.128346 | 33.014506 | 35.385058 | 35.590614 | 1.090226 | 684.668215 | 0.151088497 | 0.824864191 | 0.409448698 | 0.998865404 | 9.355559 | 9.480269 | 9.385342 | 9.391434 | 9.457214 | 9.437593 | 751 | 710 | 630 | 660 |
| 57823 | NM_00128258 | SLAMF7 | SLAM family member 7 | protein_coding | NP_001269517.1;NP_001269518.1;NP_001269519.1;NP_001269520.1;NI | 21394 | 606625 | . | . | 3.064644 | 3.777445 | 2.628188 | 2.622926 | 7.131555 | 8.731734 | 5.897579 | 5.997006 | -1.335202 | 257.951850 | 0.22100371 | -1.88710909 | 0.059145654 | 0.718513409 | 8.204683 | 7.787125 | 8.023457 | 8.128370 | 7.936611 | 7.926639 | 307 | 349 | 194 | 205 |
| 57826 | NM_00127118 | RAP2C | RAP2C, member of RAS oncogene family | protein_coding | NP_001258115.1;NP_001258116.1;NP_067006.3;XP_006724838.1;XP_01 | 21165 | 301016 | . | . | 2.870122 | 2.046312 | 2.754924 | 1.770578 | 6.678890 | 4.730143 | 6.181968 | 4.048217 | -1.089904 | 273.805331 | 0.258825918 | -0.47986472 | 0.631323587 | 0.998865404 | 8.156660 | 8.034780 | 8.196910 | 8.015343 | 8.170231 | 7.948614 | 384 | 256 | 278 | 192 |
| 57827 | NM_021184 | C6orf47 | chromosome 6 open reading frame 47 | protein_coding | NP_067007.3 | 19076 | . | . | . | 5.682075 | 5.843071 | 4.970684 | 6.295158 | 13.222422 | 13.506520 | 11.154071 | 14.393132 | -1.037050 | 440.744702 | 0.185833882 | -0.28243568 | 0.777609459 | 0.998865404 | 8.810283 | 8.756820 | 8.787941 | 8.799703 | 8.701646 | 8.836708 | 512 | 489 | 334 | 451 |
| 57828 | NM_00133049 | CATSPERG | cation channel sperm associated auxiliary subunit gamma | protein_coding | NP_001317425.1;NP_067008.3 | 25243 | 613452 | . | . | 0.119717 | 0.130682 | 0.092960 | 0.114257 | 0.278586 | 0.302078 | 0.208600 | 0.261235 | -1.201753 | 13.971148 | 0.817845099 | -0.32419405 | 0.745791128 | NA | 3.931597 | 3.664848 | 3.811208 | 3.815535 | 3.797213 | 3.807199 | 17 | 17 | 10 | 13 |
| 57835 | NM_021196,N | SLC4A5 | solute carrier family 4 member 5 | protein_coding | NP_067019.3;NP_597812.1 | 18168 | 606757 | . | . | 0.047957 | 0.027617 | 0.028139 | 0.041998 | 0.111597 | 0.063837 | 0.063142 | 0.096024 | -1.119017 | 7.566245 | 1.117329641 | -0.14519644 | 0.884555774 | NA | 2.998355 | 2.836248 | 2.936309 | 2.913836 | 2.915193 | 2.927363 | 12 | 6 | 5 | 8 |
| 57862 | NM_00124292 | ZNF410 | zinc finger protein 410 | protein_coding | NP_001229853.1;NP_001229855.1;NP_001229856.1;NP_001229857.1;NI | 20144 | . | . | . | 13.866080 | 15.730810 | 14.210327 | 16.532334 | 32.266942 | 36.362473 | 31.887559 | 37.799221 | 1.029103 | 1211.058871 | 0.141110183 | 0.293300754 | 0.769292277 | 0.998865404 | 10.221493 | 10.262325 | 10.184944 | 10.273522 | 10.200267 | 10.300502 | 1294 | 1364 | 997 | 1228 |
| 58155 | NM_00130098 | PTBP2 | polypyrimidine tract binding protein 2 | protein_coding | NP_001287914.1;NP_001287915.1;NP_001287916.1;NP_001287917.1;NI | 17662 | 608449 | . | . | 1.505636 | 1.135110 | 1.893221 | 1.602594 | 3.503676 | 2.623855 | 4.248332 | 3.664140 | -1.004609 | 232.190004 | 0.248813738 | -0.02666375 | 0.97872793 | 0.998865404 | 7.861447 | 7.856874 | 7.891715 | 7.817560 | 7.938252 | 7.760360 | 288 | 232 | 239 | 177 |
| 58190 | NM_00120687 | CTDSP1 | CTD small phosphatase 1 | protein_coding | NP_001193807.1;NP_067021.1;NP_872580.1;XP_011509871.1;XP_01150 | 21614 | 605323 | . | . | 30.190183 | 29.321222 | 33.756624 | 31.685102 | 70.253799 | 67.777320 | 75.748886 | 72.444216 | 1.087822 | 2358.057217 | 0.117285798 | 1.03543747 | 0.300464654 | 0.998865404 | 11.141293 | 11.262910 | 11.181892 | 11.154225 | 11.263271 | 11.207886 | 2646 | 2395 | 2210 | 2225 |
| 58191 | NM_00110081 | CXCL16 | C-X-C motif chemokine ligand 16 | protein_coding | NP_001094282.1;NP_071342.2 | 16642 | 605398 | . | . | 6.038474 | 6.098109 | 7.380072 | 5.890216 | 14.051780 | 14.096051 | 16.560667 | 13.467278 | 1.084201 | 456.697893 | 0.185716213 | 0.628013211 | 0.529995286 | 0.998865404 | 8.775149 | 8.892652 | 8.802411 | 8.810987 | 8.929151 | 8.779063 | 501 | 476 | 463 | 392 |
| 58472 | NM_00127121 | SQOR | sulfide quinone oxidoreductase | protein_coding | NP_001258142.1;NP_067022.1 | 20390 | 617658 | . | . | 1.721904 | 2.072962 | 1.543315 | 1.613334 | 4.006940 | 4.791744 | 3.463153 | 3.688697 | -1.216121 | 92.546868 | 0.335070087 | -0.84246994 | 0.399524958 | 0.998865404 | 6.666799 | 6.383545 | 6.532947 | 6.586496 | 6.497963 | 6.509168 | 107 | 119 | 71 | 80 |
| 58473 | NM_00113003 | PLEKHB1 | pleckstrin homology domain containing B1 | protein_coding | NP_001123505.1;NP_001123506.1;NP_001123507.1;NP_001123508.1;NI | 19079 | 607651 | . | . | 3.102120 | 4.656492 | 2.896007 | 3.677804 | 7.218760 | 10.763690 | 6.498557 | 8.408862 | 1.074346 | 290.556956 | 0.212919269 | 0.485903687 | 0.627035454 | 0.998865404 | 8.130571 | 8.232967 | 8.158618 | 8.164311 | 8.137006 | 8.258237 | 321 | 304 | 232 | 314 |
| 58475 | NM_021201,N | MS4A7 | membrane spanning 4-domains A7 | protein_coding | NP_067024.1;NP_996821.1;NP_996822.1;NP_996823.1 | 13378 | 606502 | . | . | 0.153916 | 0.205510 | 0.220380 | 0.112601 | 0.358169 | 0.475047 | 0.494526 | 0.257448 | -1.093345 | 16.397841 | 0.777936288 | -0.16549986 | 0.868550559 | NA | 4.096985 | 3.971139 | 4.024097 | 4.044811 | 4.049164 | 4.006771 | 17 | 21 | 18 | 10 |
| 58476 | NM_00132942 | TP53INP2 | tumor protein p53 inducible nuclear protein 2 | protein_coding | NP_001316358.1;NP_001316359.1;NP_001316360.1;NP_067025.1;XP_0C | 16104 | 617549 | . | . | 5.018768 | 5.413572 | 5.608300 | 6.055035 | 11.678881 | 12.513713 | 12.584862 | 13.844116 | 1.101084 | 706.290600 | 0.152287487 | 0.912252527 | 0.361635799 | 0.998865404 | 9.393206 | 9.531707 | 9.403120 | 9.454734 | 9.474979 | 9.515346 | 745 | 753 | 625 | 714 |
| 58477 | NM_00137931 | SRPRB | SRP receptor subunit beta | protein_coding | NP_001366242.1;NP_067026.3 | 24085 | 616883 | . | . | 32.787433 | 32.932462 | 31.163208 | 31.781993 | 76.297707 | 76.124866 | 69.929338 | 72.665749 | -1.059742 | 2598.776298 | 0.113007876 | -0.74077105 | 0.458832274 | 0.998865404 | 11.384876 | 11.301142 | 11.367465 | 11.365995 | 11.314843 | 11.324544 | 3082 | 2883 | 2181 | 2380 |
| 58478 | NM_00129201 | ENOPH1 | enolase-phosphatase 1 | protein_coding | NP_001278946.1;NP_067027.1 | 24599 | . | . | . | 19.762790 | 17.500600 | 13.933731 | 16.675868 | 45.988876 | 40.453420 | 31.266891 | 38.127392 | -1.226667 | 1095.132772 | 0.14766071 | -1.99608776 | 0.045924371 | 0.653499062 | 10.236872 | 9.941852 | 10.212876 | 10.118408 | 9.952611 | 10.073795 | 1475 | 1222 | 783 | 1000 |
| 58480 | NM_021205,N | RHOU | ras homolog family member U | protein_coding | NP_067028.1 | 17794 | 606366 | . | . | 0.447086 | 0.356212 | 0.267834 | 0.330507 | 1.040389 | 0.823401 | 0.601012 | 0.755666 | -1.333406 | 43.774154 | 0.477261442 | -0.86978677 | 0.384416942 | 0.998865404 | 5.644780 | 5.229445 | 5.498833 | 5.456567 | 5.419251 | 5.441666 | 64 | 48 | 30 | 38 |
| 58485 | NM_00116662 | TRAPPC1 | trafficking protein particle complex 1 | protein_coding | NP_001160093.1;NP_067033.1 | 19894 | 610969 | . | . | 59.294494 | 59.696794 | 59.734239 | 69.493619 | 137.980734 | 137.991814 | 134.041908 | 158.888913 | 1.079904 | 1547.563620 | 0.132288568 | 0.838345753 | 0.401836546 | 0.998865404 | 10.539435 | 10.650013 | 10.566723 | 10.562043 | 10.562714 | 10.681671 | 1719 | 1601 | 1291 | 1621 |
| 58486 | NM_00114366 | ZBED5 | zinc finger BED-type containing 5 | protein_coding | NP_001137139.1;NP_067034.2 | 30803 | 615251 | . | . | 10.289472 | 7.874184 | 10.585316 | 5.966566 | 23.944024 | 18.201531 | 23.753142 | 13.641842 | -1.110814 | 729.488445 | 0.450999263 | -0.3361813 | 0.736734154 | 0.998865404 | 9.584405 | 9.433113 | 9.627094 | 9.431752 | 9.635733 | 9.249004 | 1005 | 715 | 770 | 466 |
| 58487 | NM_00103961 | CREBZF | CREB/ATF bZIP transcription factor | protein_coding | NP_001034707.1;XP_006718705.1;XP_011543497.1;XP_016873575.1;XP_ | 24905 | 606444 | . | . | 8.882420 | 7.436150 | 10.101254 | 7.598868 | 20.669753 | 17.188993 | 22.666919 | 17.373911 | 1.065267 | 1622.170640 | 0.180593636 | 0.505084354 | 0.61349959 | 0.998865404 | 10.617044 | 10.708914 | 10.709233 | 10.546821 | 10.832623 | 10.502457 | 1997 | 1518 | 1759 | 1242 |
| 58488 | NM_00110240 | PCTP | phosphatidylcholine transfer protein | protein_coding | NP_001095872.1;NP_001317306.1;NP_001317307.1;NP_067036.2;XP_01 | 8752 | 606055 | . | . | 5.826054 | 4.578628 | 5.322814 | 5.543591 | 13.557469 | 10.583707 | 11.944241 | 12.674761 | 1.019046 | 357.934817 | 0.194933558 | 0.139636221 | 0.888947421 | 0.998865404 | 8.469568 | 8.497404 | 8.527727 | 8.423776 | 8.489303 | 8.486657 | 445 | 348 | 315 | 338 |

| | | | | | | | | | | |
|---|---|---|---|---|---|---|---|---|---|---|
| 58489 | NM_021214 | ABHD17C | abhydrolase domain containing 17C, depalmitoylase | protein_coding | NP_067037.1 | 26925 | 617944 | . | . | [illegible] |
| 58490 | NM_021215,XI | RPRD1B | regulation of nuclear pre-mRNA domain containing 1B | protein_coding | NP_067038.1;XP_005260534.1;XP_005260537.1;XP_011527256.1;XP_01 | 16209 | 614694 | . | . | [illegible] |
| 58491 | NM_00137021 | ZNF71 | zinc finger protein 71 | protein_coding | NP_001357143.1;NP_001357144.1;NP_067039.1 | 13141 | 194545 | . | . | [illegible] |
| 58492 | NM_021217,XI | ZNF77 | zinc finger protein 77 | protein_coding | NP_067040.1;XP_016882570.1;XP_016882571.1 | 13150 | 194551 | . | . | [illegible] |
| 58493 | NM_00132958 | INIP | INTS3 and NABP interacting protein | protein_coding | NP_001316514.1;NP_001316515.1;NP_001316516.1;NP_001316517.1;NI | 24994 | 613273 | . | . | [illegible] |
| 58494 | NM_00127040 | JAM2 | junctional adhesion molecule 2 | protein_coding | NP_001257336.1;NP_001257337.1;NP_067042.1;XP_024307891.1;XP_02 | 14686 | 606870 | . | . | [illegible] |
| 58495 | NM_00130346 | OVOL2 | ovo like zinc finger 2 | protein_coding | NP_001290390.1;NP_001290391.1;NP_067043.2 | 15804 | 616441 | . | . | [illegible] |
| 58496 | NM_021221 | LY6G5B | lymphocyte antigen 6 family member G5B | protein_coding | NP_067044.2 | 13931 | 610433 | . | . | [illegible] |
| 58497 | NM_00130322 | PRUNE1 | prune exopolyphosphatase 1 | protein_coding | NP_001290158.1;NP_001290171.1;NP_001290172.1;NP_067045.1;XP_00 | 13420 | 617413 | . | . | [illegible] |
| 58499 | NM_00134799 | ZNF462 | zinc finger protein 462 | protein_coding | NP_001334926.1;NP_067047.4;XP_006717272.1;XP_006717274.1;XP_00 | 21684 | 617371 | . | . | [illegible] |
| 58500 | NM_00110968 | ZNF250 | zinc finger protein 250 | protein_coding | NP_001103159.1;NP_001350027.1;NP_001350028.1;NP_001350029.1;NI | 13044 | . | . | . | [illegible] |
| 58504 | NM_00125602 | ARHGAP22 | Rho GTPase activating protein 22 | protein_coding | NP_001242953.1;NP_001242954.1;NP_001242955.1;NP_001334664.1;NI | 30320 | 610585 | . | . | [illegible] |
| 58505 | NM_00126781 | OSTC | oligosaccharyltransferase complex non-catalytic subunit | protein_coding | NP_001254746.1;NP_001254747.1;NP_067050.1 | 24448 | . | . | . | [illegible] |
| 58506 | NM_021228,XI | SCAF1 | SR-related CTD associated factor 1 | protein_coding | NP_067051.2;XP_005259179.1;XP_011525496.1;XP_016882572.1 | 30403 | 617264 | . | . | [illegible] |
| 58508 | NM_170606,XI | KMT2C | lysine methyltransferase 2C | protein_coding | NP_733751.2;XP_005250082.1;XP_005250083.1;XP_005250084.1;XP_00 | 13726 | 606833 | . | . | [illegible] |
| 58509 | NM_00108054 | CACTIN | cactin, spliceosome C complex subunit | protein_coding | NP_001074012.1;NP_067054.1;XP_011526462.1;XP_011526463.1 | 29938 | 618536 | . | . | [illegible] |
| 58513 | NM_00125837 | EPS15L1 | epidermal growth factor receptor pathway substrate 15 like 1 | protein_coding | NP_001245303.1;NP_001245304.1;NP_001245305.1;NP_067058.1;XP_01 | 24634 | 616826 | . | . | [illegible] |
| 58515 | NM_021237 | SELENOK | selenoprotein K | protein_coding | NP_067060.2 | 30394 | 607916 | . | . | [illegible] |
| 58516 | NM_00113581 | SINHCAF | SIN3-HDAC complex associated factor | protein_coding | NP_001129283.1;NP_001129284.1;NP_067061.1;XP_011519105.1;XP_01 | 30702 | 615027 | . | . | [illegible] |
| 58517 | NM_021239,XI | RBM25 | RNA binding motif protein 25 | protein_coding | NP_067062.1;XP_011535346.1;XP_024305448.1 | 23244 | 612427 | . | . | [illegible] |
| 58525 | NM_00133039 | WIZ | WIZ zinc finger | protein_coding | NP_001317324.2;NP_001358518.1;NP_001358532.2;NP_067064.2;XP_00 | 30917 | . | . | . | [illegible] |
| 58526 | NM_00109879 | MID1IP1 | MID1 interacting protein 1 | protein_coding | NP_001092260.1;NP_001092261.1;NP_067065.1 | 20715 | 300961 | . | . | [illegible] |
| 58527 | NM_021243,XI | ABRACL | ABRA C-terminal like | protein_coding | NP_067066.1;XP_016866640.1 | 21230 | . | . | . | [illegible] |
| 58528 | NM_021244,XI | RRAGD | Ras related GTP binding D | protein_coding | NP_067067.1;XP_005248812.1;XP_011534324.1 | 19903 | 608268 | . | . | [illegible] |
| 58533 | NM_00136651 | SNX6 | sorting nexin 6 | protein_coding | NP_001353448.1;NP_067072.3;NP_689419.3 | 14970 | 606098 | . | . | [illegible] |
| 58538 | NM_033066,XI | MPP4 | membrane palmitoylated protein 4 | protein_coding | NP_149055.2;XP_016860109.1 | 13680 | 606575 | . | . | [illegible] |
| 58985 | NM_021258,XI | IL22RA1 | interleukin 22 receptor subunit alpha 1 | protein_coding | NP_067081.2;XP_011540184.1 | 13700 | 605457 | . | . | [illegible] |
| 58986 | NM_021259,XI | PGAP6 | post-glycosylphosphatidylinositol attachment to proteins 6 | protein_coding | NP_067082.2;XP_024306134.1 | 17205 | . | . | . | [illegible] |
| 59084 | NM_00129007 | ENPP5 | ectonucleotide pyrophosphatase/phosphodiesterase family memb | protein_coding | NP_001277001.1;NP_001277002.1;NP_067547.1;XP_005249317.1;XP_01 | 13717 | 617001 | . | . | [illegible] |
| 59269 | NM_00112771 | HIVEP3 | HIVEP zinc finger 3 | protein_coding | NP_001121186.1;NP_078779.2;XP_011540186.1;XP_011540187.1;XP_01 | 13561 | 606649 | . | . | [illegible] |
| 59271 | NM_00128655 | EVA1C | eva-1 homolog C | protein_coding | NP_001273485.1;NP_001307673.1;NP_001307674.1;NP_478067.2;XP_00 | 13239 | . | . | . | [illegible] |
| 59274 | NM_022566 | TLNRD1 | talin rod domain containing 1 | protein_coding | NP_072088.1 | 13519 | 615466 | . | . | [illegible] |
| 59277 | NM_00132970 | NTN4 | netrin 4 | protein_coding | NP_001316629.1;NP_001316630.1;NP_001316631.1;NP_067052.2 | 13658 | 610401 | . | . | [illegible] |
| 59286 | NM_00104824 | UBL5 | ubiquitin like 5 | protein_coding | NP_001041706.1;NP_077268.1 | 13736 | 606849 | . | . | [illegible] |
| 59307 | NM_00113505 | SIGIRR | single Ig and TIR domain containing | protein_coding | NP_001128525.1;NP_001128526.1;NP_068577.2;XP_005253101.1;XP_00 | 30575 | 605478 | . | . | [illegible] |
| 59336 | NM_021620 | PRDM13 | PR/SET domain 13 | protein_coding | NP_067633.2 | 13998 | 616741 | . | . | [illegible] |
| 59338 | NM_00100197 | PLEKHA1 | pleckstrin homology domain containing A1 | protein_coding | NP_001001974.1;NP_001182537.1;NP_001317107.1;NP_001364159.1;NI | 14335 | 607772 | . | . | [illegible] |
| 59339 | NM_021623,XI | PLEKHA2 | pleckstrin homology domain containing A2 | protein_coding | NP_067636.1;XP_011542907.1;XP_011542908.1;XP_011542909.1;XP_01 | 14336 | 607773 | . | . | [illegible] |
| 59341 | NM_00117742 | TRPV4 | transient receptor potential cation channel subfamily V member 4 | protein_coding | NP_001170899.1;NP_001170902.1;NP_001170904.1;NP_067638.3;NP_67 | 18083 | 605427 | . | . | [illegible] |
| 59342 | NM_021626,XI | SCPEP1 | serine carboxypeptidase 1 | protein_coding | NP_067639.1;XP_005257614.1 | 29507 | . | . | . | [illegible] |
| 59343 | NM_021627,XI | SENP2 | SUMO specific peptidase 2 | protein_coding | NP_067640.2;XP_005247747.2;XP_005247748.1;XP_016862462.1 | 23116 | 608261 | . | . | [illegible] |
| 59345 | NM_021629,XI | GNB4 | G protein subunit beta 4 | protein_coding | NP_067642.1;XP_005247749.1;XP_006713784.1 | 20731 | 610863 | . | . | [illegible] |
| 59348 | NM_021632,XI | ZNF350 | zinc finger protein 350 | protein_coding | NP_067645.3;XP_011525498.1;XP_016882583.1;XP_016882584.1;XP_01 | 16656 | 605422 | . | . | [illegible] |
| 59349 | NM_00130305 | KLHL12 | kelch like family member 12 | protein_coding | NP_001289980.1;NP_001290038.1;NP_067646.1;XP_011508137.1;XP_01 | 19360 | 614522 | . | . | [illegible] |
| 60312 | NM_00113464 | AFAP1 | actin filament associated protein 1 | protein_coding | NP_001128119.1;NP_001358019.1;NP_001358020.1;NP_940997.1;XP_00 | 24017 | 608252 | . | . | [illegible] |
| 60313 | NM_021639,XI | GPBP1L1 | GC-rich promoter binding protein 1 like 1 | protein_coding | NP_067652.1;XP_016857508.1;XP_024304574.1;XP_024304576.1;XP_02 | 28843 | . | . | . | [illegible] |
| 60314 | NM_021640 | MYG1 | MYG1 exonuclease | protein_coding | NP_067653.4 | 17590 | 611366 | . | . | [illegible] |
| 60343 | NM_00117113 | FAM3A | FAM3 metabolism regulating signaling molecule A | protein_coding | NP_001164603.1;NP_001164604.1;NP_001164605.1;NP_001269240.1;NI | 13749 | 300492 | . | . | [illegible] |
| 60370 | NM_021732,XI | AVPI1 | arginine vasopressin induced 1 | protein_coding | NP_068378.2;XP_016871983.1 | 30898 | 618537 | . | . | [illegible] |
| 60386 | NM_00112612 | SLC25A19 | solute carrier family 25 member 19 | protein_coding | NP_001119593.1;NP_001119594.1;NP_068380.3;XP_005257616.1;XP_00 | 14409 | 606521 | . | . | [illegible] |
| 60412 | NM_00103712 | EXOC4 | exocyst complex component 4 | protein_coding | NP_001032203.1;NP_068579.3;XP_005250580.1;XP_016867983.1 | 30389 | 608185 | . | . | [illegible] |
| 60436 | NM_00119951 | TGIF2 | TGFB induced factor homeobox 2 | protein_coding | NP_001186442.1;NP_001186443.1;NP_001186444.1;NP_068581.1 | 15764 | 607294 | . | . | [illegible] |
| 60437 | NM_00134820 | CDH26 | cadherin 26 | protein_coding | NP_001335133.1;NP_068582.2;NP_817089.1;XP_011527271.1;XP_01152 | 15902 | 617685 | . | . | [illegible] |
| 60468 | NM_00117079 | BACH2 | BTB domain and CNC homolog 2 | protein_coding | NP_001164265.1;NP_068585.1;XP_005248815.1;XP_005248816.1;XP_01 | 14078 | 605394 | . | . | [illegible] |
| 60481 | NM_00124282 | ELOVL5 | ELOVL fatty acid elongase 5 | protein_coding | NP_001229757.1;NP_001229759.1;NP_001229760.1;NP_001288785.1;NI | 21308 | 611805 | . | . | [illegible] |
| 60485 | NM_021818,XI | SAV1 | salvador family WW domain containing protein 1 | protein_coding | NP_068590.1;XP_011535359.1;XP_016877057.1 | 17795 | 607203 | . | . | [illegible] |
| 60487 | NM_00103171 | TRMT11 | tRNA methyltransferase 11 homolog | protein_coding | NP_001026882.2;NP_001337509.1;NP_001337510.1;NP_001337511.1;NI | 21080 | . | . | . | [illegible] |
| 60488 | NM_00119086 | MRPS35 | mitochondrial ribosomal protein S35 | protein_coding | NP_001177793.1;NP_068593.2;XP_016875269.1 | 16635 | 611995 | . | . | [illegible] |
| 60489 | NM_00134943 | APOBEC3G | apolipoprotein B mRNA editing enzyme catalytic subunit 3G | protein_coding | NP_001336365.1;NP_001336366.1;NP_001336367.1;NP_068594.1;XP_01 | 17357 | 607113 | . | . | [illegible] |
| 60490 | NM_00130110 | PPCDC | phosphopantothenoylcysteine decarboxylase | protein_coding | NP_001288030.1;NP_001288031.1;NP_001288032.1;NP_001288033.1;NI | 28107 | 609854 | . | . | [illegible] |
| 60491 | NM_00113603 | NIF3L1 | NGG1 interacting factor 3 like 1 | protein_coding | NP_001129511.1;NP_001135827.1;NP_001135828.1;NP_001356370.1;NI | 13390 | 605778 | . | . | [illegible] |
| 60492 | NM_00128611 | CCDC90B | coiled-coil domain containing 90B | protein_coding | NP_001273045.1;NP_001273046.1;NP_001273047.1;NP_001273048.1;NI | 28108 | . | . | . | [illegible] |
| 60493 | NM_021826 | FASTKD5 | FAST kinase domains 5 | protein_coding | NP_068598.1 | 25790 | 614272 | . | . | [illegible] |
| 60496 | NM_015423,XI | AASDHPPT | aminoadipate-semialdehyde dehydrogenase-phosphopantetheiny | protein_coding | NP_056238.2;XP_024304409.1 | 14235 | 607756 | . | . | [illegible] |
| 60509 | NM_00103550 | AGBL5 | ATP/GTP binding protein like 5 | protein_coding | NP_001030584.1;NP_068603.4;XP_005264534.1;XP_011531313.1;XP_01 | 26147 | 615900 | . | . | [illegible] |
| 60526 | NM_00128271 | LDAH | lipid droplet associated hydrolase | protein_coding | NP_001269648.1;NP_001269649.1;NP_001269650.1;NP_001269651.1;NI | 26145 | 613570 | . | . | [illegible] |
| 60528 | NM_00116596 | ELAC2 | elaC ribonuclease Z 2 | protein_coding | NP_001159434.1;NP_060597.4;NP_776065.1;XP_024306618.1;XP_02430 | 14198 | 605367 | . | . | [illegible] |
| 60558 | NM_00134586 | GUF1 | GUF1 homolog, GTPase | protein_coding | NP_001332796.1;NP_001332797.1;NP_001332798.1;NP_068746.2;XP_02 | 25799 | 617064 | . | . | [illegible] |
| 60559 | NM_021928 | SPCS3 | signal peptidase complex subunit 3 | protein_coding | NP_068747.1 | 26212 | 618854 | . | . | [illegible] |
| 60560 | NM_00132188 | NAA35 | N-alpha-acetyltransferase 35, NatC auxiliary subunit | protein_coding | NP_001308810.1;NP_001308811.1;NP_078911.3;XP_005252184.1;XP_01 | 24340 | . | . | . | [illegible] |
| 60561 | NM_00134659 | RINT1 | RAD50 interactor 1 | protein_coding | NP_001333528.1;NP_001333529.1;NP_001333530.1;NP_001333532.1;NI | 21876 | 610089 | . | . | [illegible] |
| 60592 | NM_00115344 | SCOC | short coiled-coil protein | protein_coding | NP_001146918.1;NP_001146956.1;NP_001147024.1;NP_001147057.1;NI | 20335 | . | . | . | [illegible] |
| 60625 | NM_00119080 | DHX35 | DEAH-box helicase 35 | protein_coding | NP_001177738.1;NP_068750.2;XP_006723911.1;XP_011527274.1;XP_01 | 15861 | . | . | . | [illegible] |
| 60626 | NM_00128613 | RIC8A | RIC8 guanine nucleotide exchange factor A | protein_coding | NP_001273063.1;NP_068751.4;XP_005253109.1;XP_016873596.1 | 29550 | 609146 | . | . | [illegible] |
| 60672 | NM_021933,XI | MIIP | migration and invasion inhibitory protein | protein_coding | NP_068752.2;XP_005263544.1;XP_011540197.1;XP_011540198.1 | 25715 | 608772 | . | . | [illegible] |
| 60673 | NM_00109867 | ATG101 | autophagy related 101 | protein_coding | NP_001092143.1;NP_068753.2;XP_024304888.1 | 25679 | 615089 | . | . | [illegible] |
| 60674 | NR_002578,NF | GAS5 | growth arrest specific 5 | lncRNA | . | 16355 | 608280 | . | . | [illegible] |
| 60677 | NM_00117268 | CELF6 | CUGBP Elav-like family member 6 | protein_coding | NP_001166155.1;NP_001166156.1;NP_443072.3 | 14059 | 612681 | . | . | [illegible] |
| 60678 | NM_021937,XI | EEFSEC | eukaryotic elongation factor, selenocysteine-tRNA specific | protein_coding | NP_068756.2;XP_005247753.1;XP_011511368.1;XP_024309460.1;XP_02 | 24614 | 607695 | . | . | [illegible] |
| 60681 | NM_021939,XI | FKBP10 | FKBP prolyl isomerase 10 | protein_coding | NP_068758.3;XP_011523401.1;XP_011523402.1 | 18169 | 607063 | . | . | [illegible] |
| 60682 | NM_00104430 | SMAP1 | small ArfGAP 1 | protein_coding | NP_001037770.1;NP_001268368.1;NP_001268369.1;NP_068759.2;XP_00 | 19651 | 611372 | . | . | [illegible] |
| 60684 | NM_021942,N | TRAPPC11 | trafficking protein particle complex 11 | protein_coding | NP_068761.4;NP_951008.1;XP_016864026.1;XP_016864027.1;XP_02430 | 25751 | 614138 | . | . | [illegible] |
| 60685 | NM_021943,XI | ZFAND3 | zinc finger AN1-type containing 3 | protein_coding | NP_068762.1;XP_005249326.1;XP_011513092.1;XP_016866660.1 | 18019 | 607455 | . | . | [illegible] |
| 60686 | NM_00113070 | C14orf93 | chromosome 14 open reading frame 93 | protein_coding | NP_001124178.1;NP_001124180.1;NP_001269897.1;NP_001269898.1;NI | 20162 | . | . | . | [illegible] |
| 63027 | NM_00128645 | SLC22A23 | solute carrier family 22 member 23 | protein_coding | NP_001273384.1;NP_001273385.1;NP_001369246.1;NP_001369247.1;NI | 21106 | 611697 | . | . | [illegible] |
| 63035 | NM_00118477 | BCORL1 | BCL6 corepressor like 1 | protein_coding | NP_001171701.1;NP_001366379.1;NP_001366380.1;NP_068765.3;XP_00 | 25657 | 300688 | . | . | [illegible] |
| 63826 | NM_00130480 | SRR | serine racemase | protein_coding | NP_001291732.1;NP_068766.1;XP_006721628.1;XP_006721629.1;XP_01 | 14398 | 606477 | . | . | [illegible] |
| 63827 | NM_021948,N | BCAN | brevican | protein_coding | NP_068767.3;NP_940819.1;XP_011508168.1;XP_016857536.1 | 23059 | 600347 | . | . | [illegible] |
| 63874 | NM_022060,XI | ABHD4 | abhydrolase domain containing 4, N-acyl phospholipase B | protein_coding | NP_071343.2;XP_005268043.1 | 20154 | . | . | . | [illegible] |
| 63875 | NM_022061 | MRPL17 | mitochondrial ribosomal protein L17 | protein_coding | NP_071344.1 | 14053 | 611830 | . | . | [illegible] |
| 63876 | NM_00138232 | PKNOX2 | PBX/knotted 1 homeobox 2 | protein_coding | NP_001369252.1;NP_001369253.1;NP_001369254.1;NP_001369255.1;NI | 16714 | 613066 | . | . | [illegible] |
| 63877 | NM_00113467 | FAM204A | family with sequence similarity 204 member A | protein_coding | NP_001128144.1;NP_071346.1;XP_005270081.1 | 25794 | . | . | . | [illegible] |
| 63891 | NM_022064,N | RNF123 | ring finger protein 123 | protein_coding | NP_071347.2;XP_006713348.1;XP_011532297.1;XP_016862507.1 | 21148 | 614472 | . | . | [illegible] |
| 63892 | NM_00108395 | THADA | THADA armadillo repeat containing | protein_coding | NP_001077422.1;NP_001258572.1;NP_001258573.1;NP_001332852.1;NI | 19217 | 611800 | . | . | [illegible] |
| 63893 | NM_022066,XI | UBE2O | ubiquitin conjugating enzyme E2 O | protein_coding | NP_071349.3;XP_005257624.1;XP_016880424.1;XP_024306633.1;XP_02 | 29554 | 617649 | . | . | [illegible] |
| 63894 | NM_00119331 | VIPAS39 | VPS33B interacting protein, apical-basolateral polarity regulator, s | protein_coding | NP_001180243.1;NP_001180244.1;NP_001180245.1;NP_001180246.1;NI | 20347 | 613401 | . | . | [illegible] |
| 63895 | NM_00137818 | PIEZO2 | piezo type mechanosensitive ion channel component 2 | protein_coding | NP_001365112.1;NP_071351.2;XP_011524025.1;XP_011524026.1;XP_01 | 26270 | 613629 | . | . | [illegible] |
| 63897 | NM_022070 | HEATR6 | HEAT repeat containing 6 | protein_coding | NP_071353.4 | 24076 | . | . | . | [illegible] |
| 63899 | NM_022072,XI | NSUN3 | NOP2/Sun RNA methyltransferase 3 | protein_coding | NP_071355.1;XP_024309467.1 | 26208 | 617491 | . | . | [illegible] |
| 63901 | NM_00114251 | FAM111A | family with sequence similarity 111 member A | protein_coding | NP_001135991.1;NP_001135992.1;NP_001135993.1;NP_001299838.1;NI | 24725 | 615292 | . | . | [illegible] |
| 63905 | NM_00100389 | MANBAL | mannosidase beta like | protein_coding | NP_001003897.1;NP_001356671.1;NP_001356672.1;NP_001363458.1;NI | 15799 | . | . | . | [illegible] |
| 63906 | NM_022078 | GPATCH3 | G-patch domain containing 3 | protein_coding | NP_071361.2 | 25720 | 617486 | . | . | [illegible] |
| 63908 | NM_00128301 | NAPB | NSF attachment protein beta | protein_coding | NP_001269947.1;NP_001269949.1;NP_001269955.1;NP_071363.1;XP_01 | 15751 | 611270 | . | . | [illegible] |
| 63910 | NM_00130264 | SLC17A9 | solute carrier family 17 member 9 | protein_coding | NP_001289572.2;NP_071365.4;XP_011527280.1 | 16192 | 612107 | . | . | [illegible] |
| 63915 | NM_00119932 | BLOC1S5 | biogenesis of lysosomal organelles complex 1 subunit 5 | protein_coding | NP_001186251.1;NP_001186252.1;NP_958437.1 | 18561 | 607289 | . | . | [illegible] |
| 63916 | NM_00131825 | ELMO2 | engulfment and cell motility 2 | protein_coding | NP_001305182.1;NP_573403.1;NP_877496.1;XP_005260553.1;XP_00526 | 17233 | 606421 | . | . | [illegible] |
| 63917 | NM_00130451 | GALNT11 | polypeptide N-acetylgalactosaminyltransferase 11 | protein_coding | NP_001291443.1;NP_001358387.1;NP_001358388.1;NP_001358389.1;NI | 19875 | 615130 | . | . | [illegible] |
| 63920 | NM_00130325 | ZBED8 | zinc finger BED-type containing 8 | protein_coding | NP_001290180.1;NP_071373.2 | 30804 | 615253 | . | . | [illegible] |
| 63922 | NM_022092,XI | CHTF18 | chromosome transmission fidelity factor 18 | protein_coding | NP_071375.1;XP_005255527.1;XP_005255528.1;XP_011520874.1;XP_01 | 18435 | 613201 | . | . | [illegible] |
| 63925 | NM_022095,XI | ZNF335 | zinc finger protein 335 | protein_coding | NP_071378.1;XP_005260561.1;XP_011527281.1;XP_016883501.1 | 15807 | 610827 | . | . | [illegible] |
| 63926 | NM_00130347 | ANKEF1 | ankyrin repeat and EF-hand domain containing 1 | protein_coding | NP_001290401.1;NP_071379.3;NP_942093.1;XP_005260849.1 | 15803 | . | . | . | [illegible] |
| 63929 | NM_00120482 | XPNPEP3 | X-prolyl aminopeptidase 3 | protein_coding | NP_001191756.1;NP_071381.1 | 28052 | 613553 | . | . | [illegible] |
| 63931 | NM_022100,N | MRPS14 | mitochondrial ribosomal protein S14 | protein_coding | NP_071383.1 | 14049 | 611978 | . | . | [illegible] |
| 63932 | NM_00117056 | CXorf56 | chromosome X open reading frame 56 | protein_coding | NP_001164040.1;NP_001164041.1;NP_071384.1 | 26239 | 301012 | . | . | [illegible] |
| 63933 | NM_00103171 | MCUR1 | mitochondrial calcium uniporter regulator 1 | protein_coding | NP_001026883.1;XP_011513104.1 | 21097 | 616952 | . | . | [illegible] |
| 63935 | NM_022104,XI | PCIF1 | PDX1 C-terminal inhibiting factor 1 | protein_coding | NP_071387.1;XP_011527282.1;XP_011527283.1;XP_016883502.1;XP_01 | 16200 | 618626 | . | . | [illegible] |
| 63939 | NM_00119082 | FAM217B | family with sequence similarity 217 member B | protein_coding | NP_001177755.1;NP_001177756.1;NP_071389.1;XP_011527287.1;XP_01 | 16170 | . | . | . | [illegible] |
| 63940 | NM_00127650 | GPSM3 | G protein signaling modulator 3 | protein_coding | NP_001263430.1;NP_071390.1 | 13945 | 618558 | . | . | [illegible] |
| 63941 | NM_031231,N | NECAB3 | N-terminal EF-hand calcium binding protein 3 | protein_coding | NP_112508.3;NP_112509.3;XP_005260567.1;XP_011527293.1;XP_01152 | 15851 | 612478 | . | . | [illegible] |
| 63943 | NM_022110 | FKBPL | FKBP prolyl isomerase like | protein_coding | NP_071393.2 | 13949 | 617076 | . | . | [illegible] |
| 63951 | NM_022160 | DMRTA1 | DMRT like family A1 | protein_coding | NP_071443.2 | 13826 | 614803 | . | . | [illegible] |
| 63967 | NM_00119048 | CLSPN | claspin | protein_coding | NP_001177410.1;NP_001317419.1;NP_071394.2;XP_011540240.1;XP_01 | 19715 | 605434 | . | . | [illegible] |
| 63970 | NM_00119519 | TP53AIP1 | tumor protein p53 regulated apoptosis inducing protein 1 | protein_coding | NP_001182123.1;NP_001182124.1;NP_001238893.1;NP_071395.2;XP_01 | 29984 | 605426 | . | . | [illegible] |
| 63971 | NM_00110556 | KIF13A | kinesin family member 13A | protein_coding | NP_001099036.1;NP_001099037.1;NP_001099038.1;NP_001230352.1;NI | 14566 | 605433 | . | . | [illegible] |
| 63973 | NM_024019 | NEUROG2 | neurogenin 2 | protein_coding | NP_076924.1 | 13805 | 606624 | . | . | [illegible] |
| 63977 | NM_00104042 | PRDM15 | PR/SET domain 15 | protein_coding | NP_001035514.2;NP_001269863.2;NP_071398.5;XP_006724102.1;XP_00 | 13999 | 617692 | . | . | [illegible] |
| 63979 | NM_00104276 | FIGNL1 | fidgetin like 1 | protein_coding | NP_001036227.1;NP_001274421.1;NP_001274422.1;NP_001274423.1;NI | 13286 | 615383 | . | . | [illegible] |
| 64061 | NM_022117,XI | TSPYL2 | TSPY like 2 | protein_coding | NP_071400.1;XP_006724655.1;XP_016885215.1;XP_016885216.1 | 24358 | 300564 | . | . | [illegible] |
| 64062 | NM_00128663 | RBM26 | RNA binding motif protein 26 | protein_coding | NP_001273560.1;NP_001273561.1;NP_001353664.1;NP_071401.3;XP_00 | 20327 | . | . | . | [illegible] |
| 64064 | NM_022120 | OXCT2 | 3-oxoacid CoA-transferase 2 | protein_coding | NP_071403.1 | 18606 | 610289 | . | . | [illegible] |
| 64065 | NM_022121,XI | PERP | p53 apoptosis effector related to PMP22 | protein_coding | NP_071404.2;XP_024302288.1 | 17637 | 609301 | . | . | [illegible] |
| 64077 | NM_00116788 | LHPP | phospholysine phosphohistidine inorganic pyrophosphate phosph | protein_coding | NP_001161352.1;NP_001305260.1;NP_001305261.1;NP_071409.3;XP_00 | 30042 | 617231 | . | . | [illegible] |
| 64080 | NM_00128758 | RBKS | ribokinase | protein_coding | NP_001274509.1;NP_071411.1 | 30325 | 611132 | . | . | [illegible] |
| 64081 | NM_00103308 | PBLD | phenazine biosynthesis like protein domain containing | protein_coding | NP_001028255.1;NP_071412.2;XP_005270085.1;XP_011538362.1;XP_01 | 23301 | 612189 | . | . | [illegible] |
| 64083 | NM_022130,XI | GOLPH3 | golgi phosphoprotein 3 | protein_coding | NP_071413.1;XP_016865175.1 | 15452 | 612207 | . | . | [illegible] |
| 64084 | NM_022131,XI | CLSTN2 | calsyntenin 2 | protein_coding | NP_071414.2;XP_016862511.1 | 17448 | 611323 | . | . | [illegible] |
| 64087 | NM_00136314 | MCCC2 | methylcrotonoyl-CoA carboxylase 2 | protein_coding | NP_001350076.1;NP_071415.1;XP_011541831.1;XP_016865177.1 | 6937 | 609014 | . | . | [illegible] |
| 64089 | NM_00134818 | SNX16 | sorting nexin 16 | protein_coding | NP_001335118.1;NP_071416.2;NP_690049.1;NP_690050.1;XP_0052513 | 14980 | 614903 | . | . | [illegible] |
| 64097 | NM_00134788 | EPB41L4A | erythrocyte membrane protein band 4.1 like 4A | protein_coding | NP_001334816.1;NP_001334817.1;NP_071423.4;XP_011541833.1;XP_01 | 13278 | 612141 | . | . | [illegible] |
| 64101 | NM_022143,XI | LRRC4 | leucine rich repeat containing 4 | protein_coding | NP_071426.1;XP_011514763.1 | 15586 | 610486 | . | . | [illegible] |
| 64105 | NM_00126703 | CENPK | centromere protein K | protein_coding | NP_001253967.1;NP_001336296.1;NP_001336297.1;NP_001336298.1;NI | 29479 | 611502 | . | . | [illegible] |
| 64108 | NM_022147 | RTP4 | receptor transporter protein 4 | protein_coding | NP_071430.2 | 23992 | 609350 | . | . | [illegible] |
| 64110 | NM_022149 | MAGEF1 | MAGE family member F1 | protein_coding | NP_071432.2 | 29639 | 609267 | . | . | [illegible] |
| 64112 | NM_022151 | MOAP1 | modulator of apoptosis 1 | protein_coding | NP_071434.2 | 16658 | 609485 | . | . | [illegible] |
| 64114 | NM_00132142 | TMBIM1 | transmembrane BAX inhibitor motif containing 1 | protein_coding | NP_001308356.1;NP_001308357.1;NP_001308358.1;NP_001308359.1;NI | 23410 | 610364 | . | . | [illegible] |
| 64116 | NM_00113514 | SLC39A8 | solute carrier family 39 member 8 | protein_coding | NP_001128618.1;NP_001128619.1;NP_001128620.1;NP_071437.3;XP_00 | 20862 | 608732 | . | . | [illegible] |
| 64118 | NM_022156,XI | DUS1L | dihydrouridine synthase 1 like | protein_coding | NP_071439.3;XP_005256450.1;XP_005256451.1;XP_006722351.1;XP_00 | 30086 | . | . | . | [illegible] |
| 64121 | NM_00127185 | RRAGC | Ras related GTP binding C | protein_coding | NP_001258780.1;NP_071440.1 | 19902 | 608267 | . | . | [illegible] |
| 64122 | NM_022158,XI | FN3K | fructosamine 3 kinase | protein_coding | NP_071441.1;XP_024306640.1 | 24822 | 608425 | . | . | [illegible] |
| 64129 | NM_00120441 | TINAGL1 | tubulointerstitial nephritis antigen like 1 | protein_coding | NP_001191343.1;NP_001191344.1;NP_071447.1;XP_005271163.1;XP_00 | 19168 | 616064 | . | . | [illegible] |
| 64130 | NM_00130841 | LIN7B | lin-7 homolog B, crumbs cell polarity complex component | protein_coding | NP_001295348.1;NP_071448.1;XP_006723386.1;XP_016882620.1 | 17788 | 612331 | . | . | [illegible] |
| 64131 | NM_022166,XI | XYLT1 | xylosyltransferase 1 | protein_coding | NP_071449.1;XP_016879028.1;XP_016879029.1 | 15516 | 608124 | . | . | [illegible] |
| 64132 | NM_022167,N | XYLT2 | xylosyltransferase 2 | protein_coding | NP_071450.2;XP_005257629.1 | 15517 | 608125 | . | . | [illegible] |
| 64135 | NM_022168 | IFIH1 | interferon induced with helicase C domain 1 | protein_coding | NP_071451.2 | 18873 | 606951 | . | . | [illegible] |
| 64137 | NM_00114250 | ABCG4 | ATP binding cassette subfamily G member 4 | protein_coding | NP_001135977.1;NP_001335120.1;NP_001335121.1;NP_071452.2;XP_01 | 13884 | 607784 | . | . | [illegible] |
| 64145 | NM_00130237 | RBSN | rabenosyn, RAB effector | protein_coding | NP_001289307.1;NP_071735.2;XP_005265441.1;XP_005265442.1;XP_01 | 20759 | 609511 | . | . | [illegible] |
| 64146 | NM_022341 | PDF | peptide deformylase, mitochondrial | protein_coding | NP_071736.1 | 30012 | 618720 | . | . | [illegible] |
| 64147 | NM_00113487 | KIF9 | kinesin family member 9 | protein_coding | NP_001128350.1;NP_001364403.1;NP_001364404.1;NP_001364405.1;NI | 16666 | 607910 | . | . | [illegible] |
| 64149 | NM_022344,XI | C17orf75 | chromosome 17 open reading frame 75 | protein_coding | NP_071739.2;XP_005258079.1;XP_016880429.1;XP_016880430.1 | 30173 | . | . | . | [illegible] |
| 64151 | NM_022346,N | NCAPG | non-SMC condensin I complex subunit G | protein_coding | NP_071741.2;XP_011512178.1;XP_016864032.1;XP_016864033.1 | 24304 | 606280 | . | . | [illegible] |
| 64167 | NM_00113014 | ERAP2 | endoplasmic reticulum aminopeptidase 2 | protein_coding | NP_001123612.1;NP_001316158.1;NP_001316162.1;NP_071745.1;XP_01 | 29499 | 609497 | . | . | [illegible] |
| 64172 | NM_00135434 | OSGEPL1 | O-sialoglycoprotein endopeptidase like 1 | protein_coding | NP_001341276.2;NP_001363006.1;NP_001363007.1;NP_001363008.1;NI | 23075 | . | . | . | [illegible] |

| | | | | | | | | | | | | | | | | | | | | | | | | | | | | | | | | | |
|---|---|---|---|---|---|---|---|---|---|---|---|---|---|---|---|---|---|---|---|---|---|---|---|---|---|---|---|---|---|---|---|---|---|
| 64175 | NM_00114628 | P3H1 | prolyl 3-hydroxylase 1 | protein_coding | NP_001139761.1;NP_001230175.1;NP_071751.3;XP_005271167.1;XP_01 | 19316 | 610339 | . | . | 46.647566 | 47.978075 | 41.855647 | 44.227577 | 108.550805 | 110.903471 | 93.922860 | 101.121101 | -1.121429 | 3651.216215 | 0.112115756 | -1.47470944 | 0.140290678 | 0.900674731 | 11.914529 | 11.749051 | 11.871389 | 11.887609 | 11.749577 | 11.818837 | 4402 | 4206 | 2866 | 3364 |
| 64207 | NM_024496 | IRF2BPL | interferon regulatory factor 2 binding protein like | protein_coding | NP_078772.1 | 14282 | 611720 | . | . | 6.849594 | 6.718671 | 5.749326 | 6.461344 | 15.939285 | 15.530510 | 12.901322 | 14.773095 | -1.127177 | 837.134176 | 0.14639494 | -1.17978289 | 0.238086577 | 0.992733415 | 9.793240 | 9.620205 | 9.760488 | 9.739990 | 9.627664 | 9.701228 | 1036 | 944 | 648 | 777 |
| 64208 | NM_022361,N | POPDC3 | popeye domain containing 3 | protein_coding | NP_071756.2;XP_011534369.1;XP_016866683.1;XP_016866684.1 | 17649 | 605824 | . | . | 16.024245 | 17.308585 | 21.620763 | 15.460304 | 37.289079 | 40.009569 | 48.516365 | 35.348147 | 1.122642 | 1018.010400 | 0.173255275 | 0.963303004 | 0.335395486 | 0.998865404 | 9.905352 | 10.072862 | 9.920647 | 9.963218 | 10.154666 | 9.882107 | 1072 | 1065 | 1107 | 826 |
| 64210 | NM_00128940 | MMS19 | MMS19 homolog, cytosolic iron-sulfur assembly component | protein_coding | NP_001276332.1;NP_001276333.1;NP_001276334.1;NP_001317057.1;NI | 13824 | 614777 | . | . | 22.850979 | 23.723119 | 24.806511 | 25.763419 | 53.175166 | 54.837049 | 55.665090 | 58.904992 | 1.048622 | 2747.887843 | 0.112243829 | 0.610226341 | 0.541711883 | 0.998865404 | 11.389460 | 11.457942 | 11.409358 | 11.399444 | 11.436578 | 11.449428 | 3108 | 2877 | 2427 | 2658 |
| 64215 | NM_022365,XI | DNAJC1 | DnaJ heat shock protein family (Hsp40) member C1 | protein_coding | NP_071760.2;XP_011517916.1;XP_016872025.1 | 20090 | 611207 | . | . | 13.974408 | 13.313698 | 11.336664 | 11.982846 | 32.519023 | 30.775208 | 25.439147 | 27.397354 | -1.194309 | 822.171397 | 0.14464472 | -1.77107456 | 0.076548304 | 0.767975096 | 9.805705 | 9.549529 | 9.762070 | 9.723057 | 9.606130 | 9.628148 | 1058 | 940 | 641 | 714 |
| 64216 | NM_022366,XI | TFB2M | transcription factor B2, mitochondrial | protein_coding | NP_071761.1;XP_011542550.1;XP_016857544.1 | 18559 | 607055 | . | . | 12.378942 | 11.971766 | 15.490996 | 11.788468 | 28.806307 | 27.673277 | 34.761346 | 26.952929 | 1.107700 | 712.202899 | 0.171457919 | 0.86066216 | 0.38942414 | 0.998865404 | 9.400005 | 9.548467 | 9.451028 | 9.421194 | 9.600864 | 9.403607 | 795 | 713 | 743 | 603 |
| 64219 | NM_00103239 | PJA1 | praja ring finger ubiquitin ligase 1 | protein_coding | NP_001027568.1;NP_001369704.1;NP_001369705.1;NP_071763.2;NP_6( | 16648 | 300420 | . | . | 6.397849 | 5.857514 | 5.203829 | 6.437530 | 14.888055 | 13.539906 | 11.677237 | 14.718644 | -1.059374 | 443.549126 | 0.185179537 | -0.44936077 | 0.653171427 | 0.998865404 | 8.834163 | 8.750525 | 8.846292 | 8.771339 | 8.717226 | 8.827924 | 557 | 463 | 340 | 441 |
| 64221 | NM_00137035 | ROBO3 | roundabout guidance receptor 3 | protein_coding | NP_001357285.1;NP_001357286.1;NP_001357287.1;NP_001357288.1;NI | 13433 | 608630 | . | . | 8.310016 | 8.064900 | 9.994795 | 9.610717 | 19.337750 | 18.642377 | 22.428028 | 21.973761 | 1.154983 | 481.131872 | 0.171281559 | 1.213628707 | 0.22488954 | 0.982008277 | 8.802337 | 9.010722 | 8.883902 | 8.838920 | 8.972283 | 8.931389 | 533 | 464 | 463 | 468 |
| 64222 | NM_022371 | TOR3A | torsin family 3 member A | protein_coding | NP_071766.2 | 11997 | 607555 | . | . | 21.696863 | 20.926437 | 19.091187 | 21.271284 | 50.489487 | 48.372398 | 42.840069 | 48.634266 | -1.070906 | 1313.280250 | 0.131971084 | -0.74889182 | 0.45392241 | 0.998865404 | 10.407620 | 10.308599 | 10.401381 | 10.366952 | 10.294809 | 10.368119 | 1599 | 1433 | 1049 | 1247 |
| 64223 | NM_00119917 | MLST8 | MTOR associated protein, LST8 homolog | protein_coding | NP_001186102.1;NP_001186103.1;NP_001186104.1;NP_001338986.1;NI | 24825 | 612190 | . | . | 18.961198 | 21.666951 | 20.464226 | 22.153585 | 44.123487 | 50.084127 | 45.921130 | 50.651538 | 1.005969 | 1241.346213 | 0.138387927 | 0.062039586 | 0.950531308 | 0.998865404 | 10.273646 | 10.281722 | 10.239368 | 10.308930 | 10.228830 | 10.326971 | 1360 | 1397 | 1012 | 1243 |
| 64224 | NM_022373,XI | HERPUD2 | HERPUD family member 2 | protein_coding | NP_071768.3;XP_006715828.1;XP_016867995.1 | 21915 | . | . | . | 5.360759 | 5.586533 | 5.789320 | 6.894637 | 12.474705 | 12.913522 | 12.991064 | 15.763769 | 1.104598 | 619.270484 | 0.162736638 | 0.881924055 | 0.377817878 | 0.998865404 | 9.201178 | 9.344133 | 9.240053 | 9.237798 | 9.249760 | 9.358046 | 679 | 634 | 520 | 658 |
| 64225 | NM_00113567 | ATL2 | atlastin GTPase 2 | protein_coding | NP_001129145.1;NP_001295005.1;NP_001317387.1;NP_001317388.1;NI | 24047 | 609368 | . | . | 18.680050 | 17.278805 | 18.214820 | 12.000705 | 43.469244 | 39.940730 | 40.873525 | 27.438186 | -1.175868 | 2140.060820 | 0.17307585 | -1.35042686 | 0.176879101 | 0.949538805 | 11.175335 | 10.942122 | 11.150035 | 11.086172 | 11.148440 | 10.800219 | 2767 | 2398 | 2087 | 1439 |
| 64236 | NM_00136812 | PDLIM2 | PDZ and LIM domain 2 | protein_coding | NP_001355049.1;NP_067643.3;NP_789847.1;NP_932159.1 | 13992 | 609722 | . | . | 8.280740 | 7.525806 | 6.942325 | 7.389231 | 19.269620 | 17.396236 | 15.578373 | 16.894600 | -1.085328 | 522.328187 | 0.168639406 | -0.70049207 | 0.483620054 | 0.998865404 | 9.086588 | 8.968627 | 9.092395 | 9.014827 | 8.989459 | 9.014177 | 663 | 552 | 427 | 479 |
| 64240 | NM_022436,XI | ABCG5 | ATP binding cassette subfamily G member 5 | protein_coding | NP_071881.1;XP_005264537.1;XP_006712136.1;XP_006712137.1;XP_01 | 13886 | 605459 | . | . | 0.086925 | 0.240913 | 0.165382 | 0.049767 | 0.202279 | 0.556882 | 0.371113 | 0.113787 | -1.460763 | 14.576974 | 0.899516642 | -0.60779496 | 0.54332347 | NA | 4.111656 | 3.568818 | 3.831349 | 3.899494 | 3.870125 | 3.813855 | 11 | 27 | 16 | 5 |
| 64282 | NM_00104028 | TENT4B | terminal nucleotidyltransferase 4B | protein_coding | NP_001035374.2;NP_001035375.2;NP_001352252.1;NP_001352253.1;XF | 30758 | 605540 | . | . | 1.488755 | 0.920187 | 0.916991 | 0.851939 | 3.464393 | 2.127052 | 2.057701 | 1.947856 | -1.174578 | 249.931434 | 0.224440011 | -1.03431613 | 0.300988399 | 0.998865404 | 8.076319 | 7.845212 | 8.054768 | 7.949016 | 7.939833 | 7.906224 | 342 | 262 | 207 | 208 |
| 64283 | NM_00108047 | ARHGEF28 | Rho guanine nucleotide exchange factor 28 | protein_coding | NP_001073948.2;NP_001171164.1;NP_001231293.1;XP_005248625.1;XF | 30322 | 612790 | . | . | 0.660113 | 0.688014 | 0.506527 | 0.607900 | 1.536110 | 1.590375 | 1.136632 | 1.389893 | -1.225828 | 122.967745 | 0.292926463 | -1.00283481 | 0.315940571 | 0.998865404 | 7.081918 | 6.787344 | 6.970122 | 6.984208 | 6.890337 | 6.923135 | 153 | 149 | 91 | 109 |
| 64284 | NM_022449,N | RAB17 | RAB17, member RAS oncogene family | protein_coding | NP_071894.1;XP_006712752.1;XP_016860182.1;XP_016860183.1 | 16523 | 602206 | . | . | 30.810859 | 33.519382 | 51.938187 | 41.870849 | 71.698135 | 77.481550 | 116.547780 | 95.732720 | 1.451825 | 2424.675666 | 0.137719682 | 3.905522503 | 9.4022E-05 | 0.012046912 | 10.949598 | 11.487672 | 11.047514 | 11.094178 | 11.467221 | 11.266572 | 2209 | 2198 | 2796 | 2370 |
| 64285 | NM_022450,XI | RHBDF1 | rhomboid 5 homolog 1 | protein_coding | NP_071895.3;XP_005255551.1;XP_005255555.1;XP_006720984.1;XP_01( | 20561 | 614403 | . | . | 26.217937 | 25.906966 | 29.610265 | 25.455873 | 61.010212 | 59.885113 | 66.444574 | 58.201830 | 1.034486 | 2398.647778 | 0.124586548 | 0.392606845 | 0.694609883 | 0.998865404 | 11.203195 | 11.252397 | 11.227149 | 11.199246 | 11.299761 | 11.178113 | 2762 | 2500 | 2281 | 2115 |
| 64288 | NM_00113521 | ZSCAN31 | zinc finger and SCAN domain containing 31 | protein_coding | NP_001128687.1;NP_001128688.1;NP_001230170.1;NP_001230171.1;NI | 14097 | 610794 | . | . | 0.236220 | 0.129480 | 0.105396 | 0.259486 | 0.549689 | 0.299299 | 0.236506 | 0.593283 | -1.045643 | 26.641315 | 0.602386568 | -0.10689104 | 0.914875421 | NA | 4.766397 | 4.704117 | 4.755024 | 4.721011 | 4.739420 | 4.727075 | 36 | 25 | 24 | 23 |
| 64318 | NM_022451,XI | NOC3L | NOC3 like DNA replication regulator | protein_coding | NP_071896.8;XP_016872033.1 | 24034 | 610769 | . | . | 8.882169 | 7.714953 | 8.643078 | 7.958199 | 20.669174 | 17.833458 | 19.394815 | 18.195479 | 1.005817 | 919.883528 | 0.150135314 | 0.055731737 | 0.955555516 | 0.998865404 | 9.840821 | 9.849780 | 9.900806 | 9.779711 | 9.874361 | 9.817644 | 1144 | 907 | 834 | 831 |
| 64319 | NM_00110507 | FBRS | fibrosin | protein_coding | NP_001098549.2;XP_011544218.1;XP_011544219.1;XP_011544221.1 | 20442 | 608601 | . | . | 9.473791 | 10.593102 | 11.162800 | 11.776967 | 22.045905 | 24.486428 | 25.049000 | 26.926635 | 1.126971 | 1738.706561 | 0.125817668 | 1.370634459 | 0.170488935 | 0.946389416 | 10.675141 | 10.847323 | 10.673633 | 10.753503 | 10.790484 | 10.825334 | 1786 | 1855 | 1567 | 1765 |
| 64320 | NM_022453,XI | RNF25 | ring finger protein 25 | protein_coding | NP_071898.2;XP_016860184.1 | 14662 | 616014 | . | . | 25.327220 | 25.905304 | 25.716061 | 31.143248 | 58.937481 | 59.881270 | 57.706092 | 71.205343 | 1.078681 | 1252.177644 | 0.14105505 | 0.774653006 | 0.438544727 | 0.998865404 | 10.234819 | 10.343579 | 10.244057 | 10.275061 | 10.245728 | 10.383355 | 1359 | 1327 | 1028 | 1328 |
| 64324 | NM_00136568 | NSD1 | nuclear receptor binding SET domain protein 1 | protein_coding | NP_001352613.1;NP_071900.2;NP_758859.1;XP_024301918.1;XP_02430 | 14234 | 606681 | . | . | 6.668877 | 6.910203 | 6.772353 | 7.625604 | 15.518749 | 15.973244 | 15.196968 | 17.435042 | 1.074746 | 2675.232903 | 0.12067217 | 0.861801514 | 0.388796752 | 0.998865404 | 11.332644 | 11.436389 | 11.311922 | 11.396103 | 11.378475 | 11.446921 | 2815 | 2928 | 2309 | 2707 |
| 64326 | NM_00100174 | COP1 | COP1 E3 ubiquitin ligase | protein_coding | NP_001001740.1;NP_001273573.1;NP_071902.2;XP_005245504.1;XP_00 | 17440 | 608067 | . | . | 6.946309 | 7.153230 | 5.958108 | 6.647105 | 16.164351 | 16.535012 | 13.369825 | 15.197818 | -1.067410 | 748.645116 | 0.150047912 | -0.6272331 | 0.530506452 | 0.998865404 | 9.594585 | 9.500146 | 9.576588 | 9.564218 | 9.491026 | 9.557733 | 898 | 827 | 599 | 712 |
| 64327 | NM_00135095 | LMBR1 | limb development membrane protein 1 | protein_coding | NP_001337882.1;NP_001337883.1;NP_001337884.1;NP_001337885.1;NI | 13243 | 605522 | . | . | 4.351082 | 4.928817 | 3.856809 | 3.619090 | 10.125148 | 11.393182 | 8.654568 | 8.274619 | -1.179733 | 968.134951 | 0.14264826 | -1.67166507 | 0.094590387 | 0.812444344 | 10.033143 | 9.795184 | 9.993921 | 9.957176 | 9.896319 | 9.812903 | 1236 | 1103 | 818 | 784 |
| 64328 | NM_00137206 | XPO4 | exportin 4 | protein_coding | NP_001358990.1;NP_071904.4;XP_005266559.1;XP_005266561.1;XP_01 | 17796 | 611449 | . | . | 4.640602 | 4.745635 | 5.188573 | 4.361343 | 10.798870 | 10.969747 | 11.643010 | 9.971694 | -1.042241 | 1256.364475 | 0.141304053 | -0.422416 | 0.672721401 | 0.998865404 | 10.324324 | 10.265148 | 10.318938 | 10.299609 | 10.345919 | 10.205603 | 1495 | 1366 | 1170 | 1046 |
| 64332 | NM_00100547 | NFKBIZ | NFKB inhibitor zeta | protein_coding | NP_001005474.1;NP_113607.1 | 29805 | 608004 | . | . | 5.315955 | 5.711269 | 5.927160 | 4.180979 | 12.370446 | 13.201854 | 13.300376 | 9.559312 | -1.101041 | 645.899136 | 0.185713972 | -0.7477522 | 0.454609639 | 0.998865404 | 9.402492 | 9.264541 | 9.340946 | 9.383873 | 9.409088 | 9.172796 | 755 | 753 | 630 | 474 |
| 64342 | NM_022460,XI | HS1BP3 | HCLS1 binding protein 3 | protein_coding | NP_071905.3;XP_016860185.1;XP_016860186.1;XP_016860187.1;XP_01( | 24979 | 609359 | . | . | 12.791513 | 12.336351 | 11.767943 | 16.467878 | 29.766377 | 28.516030 | 26.406926 | 37.651849 | 1.089484 | 964.575490 | 0.161483007 | 0.765687563 | 0.443862229 | 0.998865404 | 9.850919 | 9.973959 | 9.891523 | 9.866233 | 9.831849 | 10.040227 | 1082 | 979 | 751 | 1076 |
| 64343 | NM_00113443 | AZI2 | 5-azacytidine induced 2 | protein_coding | NP_001127904.1;NP_001127905.1;NP_001258579.1;NP_071906.1;XP_0C | 24002 | 609916 | . | . | 7.132965 | 8.120672 | 7.216938 | 7.334180 | 16.598702 | 18.771292 | 16.194603 | 16.768730 | 1.005818 | 562.140825 | 0.16042953 | 0.052169583 | 0.958393569 | 0.998865404 | 9.130646 | 9.138917 | 9.128325 | 9.137381 | 9.132336 | 9.140905 | 641 | 609 | 487 | 532 |
| 64359 | NM_00120531 | NXN | nucleoredoxin | protein_coding | NP_001192248.1;NP_071908.2;XP_005256813.1;XP_005256815.1;XP_01 | 18008 | 612895 | . | . | 3.011919 | 3.609685 | 2.317391 | 3.601376 | 7.008859 | 8.343949 | 5.200159 | 8.234120 | -1.149467 | 295.326750 | 0.237140765 | -0.84745151 | 0.396743504 | 0.998865404 | 8.304112 | 8.101088 | 8.187550 | 8.285402 | 8.053294 | 8.262155 | 332 | 371 | 190 | 310 |
| 64374 | NM_00103763 | SIL1 | SIL1 nucleotide exchange factor | protein_coding | NP_001032722.1;NP_071909.1;XP_011541872.1;XP_024301932.1 | 24624 | 608005 | . | . | 30.926526 | 29.266862 | 24.229211 | 27.269041 | 71.967302 | 67.651656 | 54.369648 | 62.347420 | -1.186691 | 1655.758946 | 0.128154484 | -1.92692512 | 0.053988962 | 0.689370625 | 10.811549 | 10.564436 | 10.781827 | 10.729592 | 10.580647 | 10.662467 | 2137 | 1876 | 1246 | 1496 |
| 64375 | NM_00135108 | IKZF4 | IKAROS family zinc finger 4 | protein_coding | NP_001338018.1;NP_001338019.1;NP_001338020.1;NP_001338021.1;NI | 13179 | 606239 | . | . | 0.697115 | 0.599756 | 0.730847 | 0.704618 | 1.622214 | 1.386362 | 1.639999 | 1.611024 | -1.008895 | 118.844058 | 0.299976516 | -0.04259084 | 0.966027701 | 0.998865404 | 6.899027 | 6.886799 | 6.925303 | 6.861863 | 6.884915 | 6.897234 | 150 | 117 | 101 | 113 |
| 64376 | NM_00127184 | IKZF5 | IKAROS family zinc finger 5 | protein_coding | NP_001258769.1;NP_001359052.1;NP_001359053.1;NP_001359054.1;NI | 14283 | 606238 | . | . | 1.978767 | 1.730922 | 2.270077 | 1.798779 | 4.604674 | 4.001104 | 5.093990 | 4.112694 | 1.069369 | 279.551758 | 0.223063391 | 0.433775624 | 0.664451384 | 0.998865404 | 8.076942 | 8.175324 | 8.149812 | 8.059213 | 8.202857 | 8.074656 | 338 | 266 | 282 | 238 |
| 64393 | NM_00137582 | ZMAT3 | zinc finger matrin-type 3 | protein_coding | NP_001362753.1;NP_001362754.1;NP_001362755.1;NP_001362756.1;NI | 29983 | 606452 | . | . | 3.178310 | 2.656437 | 2.957412 | 2.626129 | 7.396058 | 6.140472 | 6.636347 | 6.004329 | -1.054756 | 810.421362 | 0.160794212 | -0.47830718 | 0.632431587 | 0.998865404 | 9.700067 | 9.623986 | 9.744712 | 9.609455 | 9.691317 | 9.589782 | 1048 | 813 | 736 | 686 |
| 64395 | NM_178439,XI | GMCL1 | germ cell-less 1, spermatogenesis associated | protein_coding | NP_848526.1;XP_011531335.1;XP_011531336.1;XP_016860193.1;XP_01( | 23843 | 618627 | . | . | 5.142414 | 3.767816 | 5.038178 | 4.536284 | 11.966609 | 8.709475 | 11.305527 | 10.371674 | 1.064243 | 597.826821 | 0.184839127 | 0.485977671 | 0.626982997 | 0.998865404 | 9.177446 | 9.268291 | 9.296272 | 9.090839 | 9.276549 | 9.202567 | 770 | 528 | 566 | 546 |
| 64397 | NM_00128430 | ZNF106 | zinc finger protein 106 | protein_coding | NP_001271235.1;NP_001271236.1;NP_001353773.1;NP_001353774.1;NI | 12886 | . | . | . | 20.930018 | 21.281914 | 26.013237 | 27.904798 | 48.705018 | 49.194097 | 58.372950 | 63.801001 | 1.257947 | 7582.260571 | 0.103164726 | 3.209148766 | 0.001331286 | 0.083777674 | 12.713419 | 13.044457 | 12.782201 | 12.784295 | 12.957740 | 12.997196 | 7724 | 7256 | 7179 | 8101 |
| 64398 | NM_00125655 | MPP5 | membrane palmitoylated protein 5 | protein_coding | NP_001243479.1;NP_071919.2;XP_005268060.1;XP_011535388.1;XP_01 | 18669 | 606958 | . | . | 3.523165 | 3.184573 | 2.836146 | 2.747842 | 8.198550 | 7.361282 | 6.364231 | 6.282614 | -1.226352 | 520.859819 | 0.168640186 | -1.74557129 | 0.080885475 | 0.780742805 | 9.164215 | 8.870349 | 9.122237 | 9.048341 | 8.969202 | 8.941683 | 697 | 585 | 416 | 429 |
| 64400 | NM_00101239 | AKTIP | AKT interacting protein | protein_coding | NP_001012398.1;NP_001295254.1;NP_071921.1;XP_005256152.1;XP_00 | 16710 | 608483 | . | . | 3.861359 | 3.329180 | 2.888849 | 3.085306 | 8.985541 | 7.695547 | 6.482493 | 7.054185 | -1.080801 | 245.234099 | 0.217099061 | -0.51635754 | 0.605604718 | 0.998865404 | 7.993056 | 7.880792 | 7.949788 | 7.964853 | 7.916752 | 7.921042 | 289 | 279 | 204 | 222 |
| 64403 | NM_022478,N | CDH24 | cadherin 24 | protein_coding | NP_071923.2;NP_659422.2;XP_011535391.1 | 14265 | 618599 | . | . | 4.859216 | 4.759203 | 4.566577 | 4.441024 | 11.307594 | 11.001111 | 10.247263 | 10.153874 | -1.084555 | 507.365813 | 0.16662222 | -0.70281015 | 0.482174071 | 0.998865404 | 9.044108 | 8.927293 | 9.020971 | 9.003459 | 8.974854 | 8.946364 | 616 | 562 | 433 | 446 |
| 64410 | NM_022480 | KLHL25 | kelch like family member 25 | protein_coding | NP_071925.2 | 25732 | . | . | . | 1.406903 | 1.243436 | 0.925795 | 1.255619 | 3.273921 | 2.874258 | 2.077457 | 2.870824 | -1.227219 | 137.963743 | 0.29067453 | -1.0162326 | 0.309518634 | 0.998865404 | 7.248832 | 6.952243 | 7.171301 | 7.116672 | 7.017990 | 7.116633 | 187 | 153 | 92 | 133 |
| 64411 | NM_022481,XI | ARAP3 | ArfGAP with RhoGAP domain, ankyrin repeat and PH domain 3 | protein_coding | NP_071926.4;XP_005268554.1;XP_005268555.1;XP_005268556.1;XP_00! | 24097 | 606647 | . | . | 4.876170 | 5.088207 | 6.684888 | 5.775673 | 11.347048 | 11.761618 | 15.000694 | 13.205393 | 1.230990 | 895.133003 | 0.148974479 | 2.012556034 | 0.044161356 | 0.645513836 | 9.648108 | 9.948228 | 9.703183 | 9.744226 | 9.929367 | 9.814469 | 897 | 891 | 926 | 854 |
| 64412 | NM_00131701 | GZF1 | GDNF inducible zinc finger protein 1 | protein_coding | NP_001303941.1;NP_001303948.1;NP_071927.1;XP_005260857.1;XP_00 | 15808 | 613842 | . | . | 5.310611 | 4.644459 | 4.259046 | 4.224232 | 12.358010 | 10.735877 | 9.557177 | 9.658206 | -1.279428 | 768.679668 | 0.156927326 | -2.26537124 | 0.023489908 | 0.50908701 | 9.752827 | 9.397872 | 9.727873 | 9.597439 | 9.507978 | 9.477126 | 1082 | 848 | 600 | 618 |
| 64417 | NM_00137739 | TMEM267 | transmembrane protein 267 | protein_coding | NP_001364323.1;NP_001364324.1;NP_001364325.1;NP_001364326.1;NI | 26139 | . | . | . | 3.837729 | 3.913799 | 3.851178 | 2.941518 | 8.930555 | 9.046922 | 8.641931 | 6.725430 | -1.151608 | 366.115894 | 0.202761485 | -1.00437945 | 0.315195753 | 0.998865404 | 8.613800 | 8.411422 | 8.557070 | 8.549136 | 8.547577 | 8.390114 | 454 | 420 | 338 | 274 |
| 64418 | NM_00128749 | TMEM168 | transmembrane protein 168 | protein_coding | NP_001274426.1;NP_071929.3;XP_016868012.1;XP_016868013.1;XP_02 | 25826 | . | . | . | 3.104763 | 2.577760 | 2.807756 | 2.000579 | 7.224911 | 5.958605 | 6.300524 | 4.574084 | -1.258604 | 563.532516 | 0.185956214 | -1.78442078 | 0.074355317 | 0.764716896 | 9.294172 | 8.963649 | 9.273141 | 9.132357 | 9.129664 | 8.974787 | 797 | 608 | 488 | 410 |
| 64419 | NM_00107752 | MTMR14 | myotubularin related protein 14 | protein_coding | NP_001070993.1;NP_001070994.1;NP_071930.2;XP_005265457.1;XP_01 | 26190 | 611089 | . | . | 9.877258 | 11.225089 | 11.842860 | 12.357462 | 22.984788 | 25.947292 | 26.575028 | 28.253876 | 1.115863 | 816.451661 | 0.146714178 | 1.078015945 | 0.28102664 | 0.998865404 | 9.592169 | 9.749966 | 9.598725 | 9.664649 | 9.697368 | 9.721940 | 847 | 872 | 736 | 821 |
| 64421 | NM_00103385 | DCLRE1C | DNA cross-link repair 1C | protein_coding | NP_001029027.1;NP_001029029.1;NP_001029030.1;NP_001276005.1;NI | 17642 | 605988 | . | . | 4.925728 | 4.340468 | 5.477764 | 5.290823 | 11.462373 | 10.033188 | 12.291947 | 12.096835 | 1.112312 | 873.037724 | 0.153584026 | 0.999855457 | 0.317380463 | 0.998865404 | 9.690701 | 9.844977 | 9.782477 | 9.671631 | 9.849542 | 9.758513 | 1026 | 822 | 850 | 808 |
| 64422 | NM_00127871 | ATG3 | autophagy related 3 | protein_coding | NP_001265641.1;NP_071933.2;XP_011511376.1 | 20962 | 609606 | . | . | 18.507258 | 19.021350 | 19.526067 | 21.764134 | 43.067149 | 43.968706 | 43.815929 | 49.761104 | 1.054567 | 1030.933472 | 0.135763347 | 0.564586467 | 0.572355072 | 0.998865404 | 9.970997 | 10.047461 | 9.983450 | 9.995760 | 10.011457 | 10.045600 | 1146 | 1092 | 899 | 1015 |
| 64423 | NM_00103171 | INF2 | inverted formin 2 | protein_coding | NP_001026884.3;NP_071934.3;NP_116103.1 | 23791 | 610982 | . | . | 5.802657 | 5.759987 | 4.935354 | 6.025458 | 13.503018 | 13.314467 | 11.074791 | 13.776494 | -1.096772 | 602.963695 | 0.164234017 | -0.81142357 | 0.417122471 | 0.998865404 | 9.301382 | 9.167362 | 9.245160 | 9.285327 | 9.153465 | 9.251971 | 705 | 701 | 462 | 580 |
| 64425 | NM_00128276 | POLR1E | RNA polymerase I subunit E | protein_coding | NP_001269695.1;NP_071935.1 | 17631 | . | . | . | 14.536502 | 16.159968 | 13.528698 | 14.546399 | 33.827038 | 37.354492 | 30.358007 | 33.258611 | -1.108467 | 849.727488 | 0.146407964 | -1.01473975 | 0.310229899 | 0.998865404 | 9.803440 | 9.654427 | 9.730354 | 9.800906 | 9.670861 | 9.714094 | 979 | 1011 | 679 | 779 |
| 64426 | NM_022491,XI | SUDS3 | SDS3 homolog, SIN3A corepressor complex component | protein_coding | NP_071936.2;XP_011536973.1;XP_016875306.1;XP_016875307.1;XP_01( | 29545 | 608250 | . | . | 7.696061 | 7.741006 | 9.089088 | 7.588753 | 17.909051 | 17.893680 | 20.395651 | 17.350782 | 1.015897 | 1238.709479 | 0.137531676 | 0.165446899 | 0.868592242 | 0.998865404 | 10.262996 | 10.286156 | 10.269371 | 10.266077 | 10.335293 | 10.221049 | 1418 | 1323 | 1168 | 1082 |
| 64427 | NM_00137612 | TTC31 | tetratricopeptide repeat domain 31 | protein_coding | NP_001363058.1;NP_001363059.1;NP_001363061.1;NP_001363062.1;NI | 25759 | . | . | . | 14.707779 | 15.061730 | 16.613068 | 14.862261 | 34.225604 | 34.815862 | 37.279239 | 33.980789 | 1.039397 | 1394.352383 | 0.131257824 | 0.424713918 | 0.671045237 | 0.998865404 | 10.417101 | 10.473114 | 10.423665 | 10.435764 | 10.503189 | 10.413757 | 1562 | 1487 | 1308 | 1255 |
| 64428 | NM_00130479 | CIAO3 | cytosolic iron-sulfur assembly component 3 | protein_coding | NP_001291728.1;NP_071938.1 | 14179 | 611118 | . | . | 7.317877 | 7.709674 | 7.605847 | 8.174653 | 17.029001 | 17.821258 | 17.067299 | 18.690372 | 1.031227 | 504.699472 | 0.167344212 | 0.265095898 | 0.790935587 | 0.998865404 | 8.957115 | 9.001111 | 8.958755 | 8.979478 | 8.969097 | 9.007787 | 563 | 545 | 432 | 495 |
| 64429 | NM_00130313 | ZDHHC6 | zinc finger DHHC-type palmitoyltransferase 6 | protein_coding | NP_001290063.1;NP_001338011.1;NP_001338012.1;NP_001338013.1;NI | 19160 | 618715 | . | . | 17.477558 | 17.963851 | 20.675024 | 15.057556 | 40.670991 | 41.524244 | 46.394151 | 34.427317 | 1.022493 | 1318.405849 | 0.15564892 | 0.206173039 | 0.836655747 | 0.998865404 | 10.348179 | 10.380795 | 10.340702 | 10.365069 | 10.477011 | 10.250629 | 1477 | 1429 | 1331 | 1065 |
| 64430 | NM_00133017 | PCNX4 | pecanex 4 | protein_coding | NP_001317106.1;NP_071940.4 | 20349 | . | . | . | 6.065588 | 5.459625 | 6.234516 | 5.881517 | 14.114873 | 12.620167 | 13.990074 | 13.447389 | -1.058421 | 1701.068826 | 0.162341158 | -0.50457336 | 0.613858527 | 0.998865404 | 10.772322 | 10.690983 | 10.773121 | 10.725886 | 10.837312 | 10.553987 | 2074 | 1831 | 1698 | 1269 |
| 64431 | NM_022496,N | ACTR6 | actin related protein 6 | protein_coding | NP_071941.1 | 24025 | . | . | . | 5.983701 | 4.774496 | 6.227215 | 3.029752 | 13.924319 | 11.036463 | 13.973693 | 6.927169 | -1.147073 | 347.691203 | 0.584211584 | -0.338845 | 0.734726501 | 0.998865404 | 8.537007 | 8.339568 | 8.534751 | 8.380835 | 8.571607 | 8.147707 | 483 | 349 | 389 | 188 |
| 64432 | NM_022497,N | MRPS25 | mitochondrial ribosomal protein S25 | protein_coding | NP_071942.1 | 14511 | 611987 | . | . | 10.629651 | 11.857170 | 13.159180 | 11.101275 | 24.735631 | 27.408377 | 29.528817 | 25.381748 | 1.068659 | 2301.835776 | 0.147139226 | 0.651096971 | 0.514983892 | 0.998865404 | 11.119785 | 11.215758 | 11.073531 | 11.198898 | 11.291290 | 11.081192 | 2362 | 2589 | 2336 | 1941 |
| 64434 | NM_00135336 | NOM1 | nucleolar protein with MIF4G domain 1 | protein_coding | NP_001340295.1;NP_612409.1 | 13244 | 611269 | . | . | 3.583430 | 3.515384 | 3.817023 | 3.073782 | 8.338790 | 8.125966 | 8.565291 | 7.027835 | -1.058654 | 636.993996 | 0.168192336 | -0.48890752 | 0.624907174 | 0.998865404 | 9.355231 | 9.273895 | 9.348449 | 9.316021 | 9.365172 | 9.217924 | 772 | 690 | 598 | 516 |
| 64493 | NR_024121 | LINC00235 | long intergenic non-protein coding RNA 235 | lncRNA | . | 14138 | . | . | . | 0.354178 | 0.108392 | 0.432111 | 0.506456 | 0.824186 | 0.250552 | 0.969645 | 1.157952 | 1.808964 | 6.844399 | 1.184816085 | 0.72176924 | 0.470436364 | NA | 2.282109 | 3.141582 | 2.763539 | 2.744001 | 2.768298 | 2.775291 | 8 | 3 | 7 | 9 |
| 64506 | NM_00107953 | CPEB1 | cytoplasmic polyadenylation element binding protein 1 | protein_coding | NP_001073001.1;NP_001073002.1;NP_001073003.1;NP_001275748.1;NI | 21744 | 607342 | . | . | 9.973615 | 10.738325 | 10.130591 | 10.607818 | 23.209014 | 24.822115 | 22.732755 | 24.253517 | -1.008919 | 1108.850217 | 0.135944803 | -0.09423514 | 0.924922374 | 0.998865404 | 10.121388 | 10.108280 | 10.084346 | 10.150407 | 10.098131 | 10.124120 | 1226 | 1255 | 943 | 1053 |
| 64518 | NM_031898,XI | TEKT3 | tektin 3 | protein_coding | NP_114104.1;XP_011522290.1;XP_011522291.1;XP_011522292.1;XP_01( | 14293 | 612683 | . | . | 0.314980 | 0.507438 | 0.603451 | 0.503763 | 0.732972 | 1.172965 | 1.354126 | 1.151795 | -1.164916 | 37.751844 | 0.521622417 | -0.42219444 | 0.672883103 | 0.998865404 | 5.345328 | 5.123071 | 5.209683 | 5.283828 | 5.223788 | 5.226287 | 36 | 54 | 30 | 33 |
| 64582 | NM_022571,XI | GPR135 | G protein-coupled receptor 135 | protein_coding | NP_072093.2;XP_016877085.1;XP_016877086.1;XP_016877087.1;XP_01( | 19991 | 607970 | . | . | 0.516080 | 0.716658 | 0.803151 | 0.465714 | 1.200941 | 1.656590 | 1.802248 | 1.064802 | 1.031885 | 155.306194 | 0.322969839 | 0.140205421 | 0.888497689 | 0.998865404 | 7.255482 | 7.302085 | 7.190399 | 7.322645 | 7.382566 | 7.145865 | 147 | 192 | 178 | 104 |
| 64598 | NM_00104009 | MOSPD3 | motile sperm domain containing 3 | protein_coding | NP_001035186.1;NP_001035187.1;NP_001035188.1;NP_001350344.1;NI | 25078 | 609125 | . | . | 7.881175 | 8.269984 | 8.242338 | 9.609932 | 18.339819 | 19.116436 | 18.495567 | 21.971965 | 1.085025 | 303.398595 | 0.203768789 | 0.577753983 | 0.563430228 | 0.998865404 | 8.185414 | 8.302358 | 8.212963 | 8.229734 | 8.225789 | 8.303367 | 330 | 319 | 255 | 317 |
| 64599 | NM_00137575 | GIGYF1 | GRB10 interacting GYF protein 1 | protein_coding | NP_001362688.1;NP_001362689.1;NP_001362690.1;NP_001362691.1;NI | 9126 | 612064 | . | . | 13.742686 | 14.761323 | 18.317856 | 15.633174 | 31.979796 | 34.121460 | 41.104737 | 35.743394 | 1.159847 | 3170.050927 | 0.127313207 | 1.680375573 | 0.092884266 | 0.809520218 | 11.519280 | 11.733365 | 11.536848 | 11.591966 | 11.762065 | 11.604981 | 3265 | 3275 | 3240 | 2891 |
| 64601 | NM_022575,N | VPS16 | VPS16 core subunit of CORVET and HOPS complexes | protein_coding | NP_072097.2;NP_536338.1 | 14584 | 608550 | . | . | 20.996202 | 22.737105 | 22.323703 | 23.086270 | 48.859027 | 52.557838 | 50.093742 | 52.784018 | 1.024402 | 1868.296904 | 0.121988461 | 0.285122436 | 0.775550342 | 0.998865404 | 10.850100 | 10.884708 | 10.825773 | 10.888611 | 10.868375 | 10.884857 | 2041 | 2071 | 1627 | 1791 |
| 64645 | NM_033055,XI | MFSD14A | major facilitator superfamily domain containing 14A | protein_coding | NP_149044.2;XP_005271175.1;XP_011540259.1;XP_016857573.1 | 23363 | . | . | . | 10.873209 | 10.206387 | 12.664753 | 10.030870 | 25.302401 | 23.592521 | 28.419339 | 22.934392 | 1.062666 | 949.313312 | 0.157465566 | 0.556870402 | 0.577615978 | 0.998865404 | 9.845838 | 9.934287 | 9.891570 | 9.839161 | 9.995048 | 9.817473 | 1098 | 957 | 954 | 806 |
| 64651 | NM_00132055 | CSRNP1 | cysteine and serine rich nuclear protein 1 | protein_coding | NP_001307488.1;NP_001307489.1;NP_149016.2;XP_016862538.1 | 14300 | 606458 | . | . | 1.497085 | 1.320307 | 1.440087 | 1.640737 | 3.483780 | 3.051949 | 3.231514 | 3.751349 | 1.081049 | 146.143758 | 0.274403382 | 0.40973134 | 0.682003037 | 0.998865404 | 7.133972 | 7.246330 | 7.200709 | 7.147875 | 7.180635 | 7.227823 | 173 | 141 | 124 | 151 |
| 64682 | NM_022662,XI | ANAPC1 | anaphase promoting complex subunit 1 | protein_coding | NP_073153.1;XP_011509935.1;XP_011509936.1;XP_016860199.1;XP_01( | 19988 | 608473 | . | . | 13.343121 | 14.325153 | 13.478971 | 14.256815 | 31.049994 | 33.113234 | 30.246418 | 32.596513 | -1.007055 | 3353.950982 | 0.112221426 | -0.09037583 | 0.92798856 | 0.998865404 | 11.716771 | 11.706502 | 11.687929 | 11.740177 | 11.691415 | 11.725404 | 3753 | 3747 | 2846 | 3198 |
| 64689 | NM_00127878 | GORASP1 | golgi reassembly stacking protein 1 | protein_coding | NP_001265718.1;NP_001265719.1;NP_114105.1;XP_006713364.1;XP_01 | 16769 | 606867 | . | . | 9.288905 | 9.753021 | 9.192889 | 8.511111 | 21.615665 | 22.544539 | 20.628574 | 19.459643 | -1.033029 | 984.566192 | 0.136917995 | -0.34239656 | 0.732052474 | 0.998865404 | 9.966663 | 9.919642 | 9.946069 | 9.964204 | 9.922591 | 9.940106 | 1138 | 1093 | 832 | 919 |
| 64708 | NM_00128294 | COPS7B | COP9 signalosome subunit 7B | protein_coding | NP_001269878.1;NP_001269879.1;NP_001269880.1;NP_001269881.1;NI | 16760 | 616010 | . | . | 20.455166 | 20.348706 | 17.285230 | 19.495432 | 47.600012 | 47.036944 | 38.787557 | 44.573992 | -1.088019 | 1657.265476 | 0.122935639 | -0.98997794 | 0.322184902 | 0.998865404 | 10.754239 | 10.632366 | 10.716355 | 10.736872 | 10.642050 | 10.679131 | 1963 | 1888 | 1347 | 1524 |
| 64710 | NM_022731,XI | NUCKS1 | nuclear casein kinase and cyclin dependent kinase substrate 1 | protein_coding | NP_073568.2;XP_005245510.1 | 29923 | 611912 | . | . | 26.601899 | 23.322197 | 20.399888 | 21.781064 | 61.903712 | 53.910301 | 45.776757 | 49.799812 | -1.204628 | 4514.120921 | 0.117583031 | -2.28423849 | 0.022357518 | 0.494552608 | 12.268267 | 11.999712 | 12.271469 | 12.149376 | 12.034254 | 12.079250 | 6088 | 4941 | 3463 | 3945 |
| 64718 | NM_00103712 | UNKL | unk like zinc finger | protein_coding | NP_001032202.1;NP_001180317.2;NP_001180318.1;NP_001263343.1;NI | 14184 | 617463 | . | . | 4.094464 | 5.129951 | 5.760340 | 5.612977 | 9.527987 | 11.858112 | 12.926033 | 12.833405 | 1.218205 | 522.088954 | 0.171428858 | 1.661078012 | 0.09669778 | 0.813553488 | 8.878719 | 9.163549 | 8.922228 | 8.993549 | 9.122570 | 9.045495 | 511 | 537 | 528 | 506 |
| 64743 | NM_00116642 | WDR13 | WD repeat domain 13 | protein_coding | NP_001159898.1;NP_001334146.1;NP_001334148.1;NP_060353.2;XP_01 | 14352 | 300512 | . | . | 5.987385 | 5.838143 | 5.961632 | 6.352850 | 13.932889 | 13.495127 | 13.377729 | 14.525037 | 1.026945 | 1021.313748 | 0.137422624 | 0.279132469 | 0.780143166 | 0.998865404 | 9.976859 | 10.015306 | 9.968492 | 10.003207 | 10.028192 | 9.982427 | 1133 | 1113 | 928 | 940 |
| 64744 | NM_00119897 | SMAP2 | small ArfGAP2 | protein_coding | NP_001185907.1;NP_001185908.1;NP_001185909.1;NP_073570.1;XP_01 | 25082 | 616916 | . | . | 13.588902 | 15.276736 | 15.467444 | 16.072505 | 31.621933 | 35.312862 | 34.708496 | 36.747875 | 1.074041 | 1362.064604 | 0.130982351 | 0.786738581 | 0.431434914 | 0.998865404 | 10.359268 | 10.462061 | 10.342793 | 10.421846 | 10.429631 | 10.445511 | 1434 | 1491 | 1214 | 1336 |
| 64745 | NM_00102999 | METTL17 | methyltransferase like 17 | protein_coding | NP_001025162.1;NP_073571.1;XP_006720299.1;XP_016877090.1 | 19280 | 616091 | . | . | 34.436473 | 32.952449 | 36.391198 | 35.272600 | 80.135093 | 76.171061 | 81.660795 | 80.646607 | 1.047523 | 1746.589843 | 0.123010252 | 0.544519891 | 0.586083764 | 0.998865404 | 10.736338 | 10.803530 | 10.775753 | 10.726644 | 10.805478 | 10.770235 | 2026 | 1783 | 1589 | 1638 |
| 64746 | NM_022735 | ACBD3 | acyl-CoA binding domain containing 3 | protein_coding | NP_073572.2 | 15453 | 606809 | . | . | 10.059605 | 9.158257 | 8.731421 | 9.431151 | 23.409113 | 21.169724 | 19.593054 | 21.563208 | -1.072821 | 1044.230992 | 0.139863562 | -0.72505992 | 0.468415272 | 0.998865404 | 10.078036 | 9.976632 | 10.090735 | 10.014519 | 9.977066 | 10.025353 | 1308 | 1107 | 847 | 976 |
| 64747 | NM_00116790 | MFSD1 | major facilitator superfamily domain containing 1 | protein_coding | NP_001161375.2;NP_001276335.1;NP_001276336.1;NP_073573.3;XP_0C | 25874 | . | . | . | 50.255387 | 44.558601 | 46.757866 | 37.314215 | 116.946360 | 102.999205 | 104.923290 | 85.314520 | -1.142776 | 3126.925012 | 0.138516433 | -1.39003664 | 0.16451775 | 0.940065727 | 11.703425 | 11.511237 | 11.712286 | 11.605979 | 11.643843 | 11.449293 | 4085 | 3369 | 2843 | 2408 |
| 64748 | NM_00117063 | PLPPR2 | phospholipid phosphatase related 2 | protein_coding | NP_001164106.1;NP_073574.2;XP_005260074.1;XP_016882636.1;XP_01 | 29566 | . | . | . | 5.503754 | 5.246547 | 5.536054 | 6.288148 | 12.807461 | 12.127627 | 12.422743 | 14.377102 | 1.094281 | 475.080913 | 0.173645356 | 0.748554843 | 0.454125558 | 0.998865404 | 8.825728 | 8.955417 | 8.879154 | 8.844853 | 8.881601 | 8.953950 | 537 | 476 | 407 | 492 |
| 64750 | NM_022739,XI | SMURF2 | SMAD specific E3 ubiquitin protein ligase 2 | protein_coding | NP_073576.1;XP_005257642.1 | 16809 | 605532 | . | . | 3.457872 | 3.480626 | 2.566768 | 2.508245 | 8.046610 | 8.045622 | 5.759752 | 5.734804 | -1.346095 | 576.086116 | 0.160018575 | -2.67956659 | 0.007371754 | 0.269900048 | 9.368567 | 8.940004 | 9.266955 | 9.250495 | 9.074281 | 9.053791 | 770 | 705 | 434 | 453 |
| 64753 | NM_00120137 | CCDC136 | coiled-coil domain containing 136 | protein_coding | NP_001188301.1;NP_001350352.1;NP_001350353.1;NP_001354690.1;NI | 22225 | 611902 | . | . | 4.784263 | 5.495318 | 6.239295 | 5.344191 | 11.133176 | 12.702670 | 14.000799 | 12.218858 | 1.118957 | 680.745652 | 0.161624698 | 1.003278434 | 0.315726539 | 0.998865404 | 9.327571 | 9.489814 | 9.322818 | 9.413746 | 9.491005 | 9.398805 | 690 | 740 | 667 | 629 |
| 64754 | NM_00116774 | SMYD3 | SET and MYND domain containing 3 | protein_coding | NP_001161212.1;NP_001362891.1;NP_001362892.1;NP_001362894.1;NI | 15513 | 608783 | . | . | 6.073856 | 6.352891 | 5.564522 | 6.833045 | 14.134111 | 14.684991 | 12.486627 | 15.622944 | -1.014768 | 302.247340 | 0.207803699 | -0.10177983 | 0.918931433 | 0.998865404 | 8.250642 | 8.228444 | 8.233894 | 8.251666 | 8.180917 | 8.285954 | 345 | 334 | 235 | 309 |
| 64755 | NM_022744,XI | RUSF1 | RUS family member 1 | protein_coding | NP_073581.2;XP_016879059.1 | 25848 | . | . | . | 24.837505 | 23.739384 | 21.363612 | 26.259765 | 57.797894 | 54.874648 | 47.939326 | 60.039832 | -1.042488 | 2058.284407 | 0.132893771 | -0.45171756 | 0.651472467 | 0.998865404 | 11.037064 | 10.976759 | 11.043339 | 11.001269 | 10.909797 | 11.063040 | 2483 | 2208 | 1583 | 2072 |
| 64756 | NM_00104254 | ATPAF1 | ATP synthase mitochondrial F1 complex assembly factor 1 | protein_coding | NP_001036011.2;NP_001230657.1;NP_001243347.1;NP_073582.3 | 18803 | 608917 | . | . | 8.562384 | 8.548601 | 6.722698 | 8.274377 | 19.925018 | 19.760474 | 15.085541 | 18.918376 | -1.159016 | 456.662774 | 0.180956584 | -1.17652965 | 0.239383292 | 0.992733415 | 8.937903 | 8.724159 | 8.879834 | 8.876733 | 8.727733 | 8.843014 | 566 | 528 | 332 | 435 |
| 64759 | NM_022748,XI | TNS3 | tensin 3 | protein_coding | NP_073585.8;XP_011513778.1;XP_011513779.1;XP_011513780.1;XP_01 | 21616 | 606825 | . | . | 23.271797 | 23.585454 | 19.570737 | 25.092847 | 54.154429 | 54.518829 | 43.916169 | 57.371809 | -1.016724 | 5408.863888 | 0.116988003 | -0.20454034 | 0.837931274 | 0.998865404 | 12.413090 | 12.389030 | 12.411028 | 12.403564 | 12.323835 | 12.458423 | 6308 | 5859 | 4290 | 5432 |
| 64760 | NM_00135425 | FAM160B2 | family with sequence similarity 160 member B2 | protein_coding | NP_001341179.1;NP_001341180.1;NP_073586.5;XP_005273659.1;XP_00 | 16492 | . | . | . | 7.798099 | 7.670159 | 9.220045 | 7.425638 | 18.146494 | 17.729917 | 20.689512 | 16.977840 | 1.082790 | 916.889055 | 0.144789777 | 0.792555805 | 0.428036667 | 0.998865404 | 9.781957 | 9.896959 | 9.790947 | 9.828876 | 9.910507 | 9.822904 | 987 | 975 | 879 | 840 |
| 64761 | NM_022750,N | PARP12 | poly(ADP-ribose) polymerase family member 12 | protein_coding | NP_073587.1;XP_005250095.1;XP_005250096.1;XP_005250097.1;XP_01 | 21919 | 612481 | . | . | 3.485245 | 3.839476 | 4.234331 | 4.628242 | 8.110307 | 8.875123 | 9.501717 | 10.581928 | 1.199954 | 384.210880 | 0.185651049 | 1.416525486 | 0.156621708 | 0.925053243 | 8.448603 | 8.710992 | 8.498059 | 8.555770 | 8.618172 | 8.649681 | 382 | 396 | 355 | 403 |
| 64762 | NM_00124240 | GAREM1 | GRB2 associated regulator of MAPK1 subtype 1 | protein_coding | NP_001229338.1;NP_073588.1;XP_016881408.1;XP_024307002.1;XP_02 | 26136 | 617998 | . | . | 0.813815 | 0.743666 | 0.543873 | 0.695815 | 1.893780 | 1.719015 | 1.220436 | 1.590900 | -1.070498 | 148.989757 | 0.277924077 | -0.35362719 | 0.723618292 | 0.998865404 | 7.268246 | 7.168157 | 7.200223 | 7.261258 | 7.162051 | 7.245745 | 164 | 179 | 111 | 150 |
| 64763 | NM_00133051 | ZNF574 | zinc finger protein 574 | protein_coding | NP_001317448.1;NP_073589.4;XP_005259216.1;XP_011525530.1;XP_01 | 26166 | . | . | . | 3.097618 | 4.279122 | 3.619169 | 3.939096 | 7.208288 | 9.891383 | 8.121312 | 9.006275 | 1.014776 | 366.857835 | 0.20232738 | 0.10458813 | 0.916702634 | 0.998865404 | 8.508929 | 8.529155 | 8.430713 | 8.587722 | 8.511136 | 8.529943 | 363 | 446 | 316 | 352 |
| 64764 | NM_00125377 | CREB3L2 | cAMP responsive element binding protein 3 like 2 | protein_coding | NP_001240704.1;NP_001305175.1;NP_919047.2;XP_005250598.1;XP_01 | 23720 | 608834 | . | . | 11.364681 | 11.786157 | 8.721488 | 8.933916 | 26.446081 | 27.244232 | 19.570768 | 20.426337 | -1.306481 | 2330.354716 | 0.116713197 | -3.30455877 | 0.00095126 | 0.065898069 | 11.366367 | 10.980588 | 11.258675 | 11.308711 | 11.066109 | 11.072201 | 2948 | 2935 | 1750 | 1902 |
| 64766 | NM_00125612 | S100PBP | S100P binding protein | protein_coding | NP_001243050.1;NP_073590.2;XP_011540263.1;XP_011540264.1;XP_01 | 25768 | 611889 | . | . | 6.093268 | 5.666806 | 5.907638 | 5.469483 | 14.179287 | 13.099075 | 13.256563 | 12.505324 | -1.043021 | 765.751345 | 0.150728294 | -0.40316054 | 0.686830124 | 0.998865404 | 9.610538 | 9.550297 | 9.611093 | 9.578269 | 9.604018 | 9.525472 | 921 | 824 | 689 | 663 |
| 64768 | NM_022755,XI | IPPK | inositol-pentakisphosphate 2-kinase | protein_coding | NP_073592.1;XP_016870530.1 | 14645 | . | . | . | 2.825837 | 3.134480 | 3.368743 | 2.541677 | 6.575839 | 7.245491 | 7.559360 | 5.811241 | -1.028464 | 352.278264 | 0.227153446 | -0.17825711 | 0.85852106 | 0.998865404 | 8.479792 | 8.441092 | 8.499507 | 8.423500 | 8.560197 | 8.318094 | 438 | 360 | 367 | 256 |
| 64769 | NM_00127087 | MEAF6 | MYST/Esa1 associated factor 6 | protein_coding | NP_001257804.1;NP_001257805.1;NP_073593.2 | 25674 | 611001 | . | . | 4.715672 | 4.229246 | 4.733397 | 5.185863 | 10.973565 | 9.776089 | 10.621602 | 11.856858 | 1.088002 | 695.647714 | 0.155925887 | 0.780375107 | 0.435170116 | 0.998865404 | 9.380071 | 9.501789 | 9.449619 | 9.372074 | 9.442240 | 9.495973 | 811 | 678 | 606 | 706 |
| 64770 | NM_00130831 | CCDC14 | coiled-coil domain containing 14 | protein_coding | NP_001295246.1;NP_001353264.1;NP_001353265.1;NP_001353266.1;NI | 25766 | 617147 | . | . | 30.633473 | 32.247744 | 37.663830 | 27.970953 | 71.285350 | 74.542100 | 84.516535 | 63.952260 | -1.011389 | 5021.880392 | 0.154808347 | -0.10553704 | 0.915949682 | 0.998865404 | 12.302063 | 12.285915 | 12.282151 | 12.305839 | 12.434076 | 12.109727 | 5733 | 5526 | 5177 | 3782 |
| 64771 | NM_022758,N | ILRUN | inflammation and lipid regulator with UBA-like and NBR1-like dom | protein_coding | NP_073595.2;NP_077270.1;XP_005249355.1 | 21215 | 612217 | . | . | 12.994097 | 13.718812 | 12.988642 | 17.188760 | 30.237800 | 31.711647 | 29.146136 | 39.300056 | 1.092269 | 1880.167816 | 0.136837461 | 0.930507533 | 0.352108366 | 0.998865404 | 10.811800 | 10.938702 | 10.818120 | 10.859756 | 10.821208 | 10.989552 | 2015 | 1991 | 1527 | 2034 |
| 64772 | NM_00104257 | ENGASE | endo-beta-N-acetylglucosaminidase | protein_coding | NP_001036038.1;XP_006722080.1;XP_006722081.1;XP_006722082.1;XP_ | 24622 | 611898 | . | . | 8.933347 | 9.278169 | 7.538156 | 7.980243 | 20.788265 | 21.446901 | 16.915403 | 18.245875 | -1.146144 | 1260.856915 | 0.129061823 | -1.52475889 | 0.127319246 | 0.874064017 | 10.395277 | 10.198389 | 10.341815 | 10.357418 | 10.239496 | 10.253402 | 1535 | 1468 | 1012 | 1112 |
| 64773 | NM_00127116 | PCED1A | PC-esterase domain containing 1A | protein_coding | NP_001258097.1;NP_073597.2;XP_005260861.1;XP_005260862.1;XP_00 | 16212 | . | . | . | 35.235411 | 37.400818 | 46.538581 | 38.682824 | 81.994253 | 86.453667 | 104.431227 | 88.443681 | 1.158574 | 2374.483951 | 0.128726661 | 1.649619836 | 0.099020715 | 0.813553488 | 11.103210 | 11.315764 | 11.129943 | 11.168989 | 11.338557 | 11.192933 | 2471 | 2432 | 2414 | 2177 |
| 64776 | NM_00133037 | C11orf1 | chromosome 11 open reading frame 1 | protein_coding | NP_001317300.1;NP_073598.1;XP_011541259.1;XP_011541260.1 | 1163 | . | . | . | 10.337171 | 9.977882 | 9.911844 | 10.232592 | 24.055025 | 23.064320 | 22.241890 | 23.395604 | 1.082516 | 507.989149 | 0.168619894 | 0.678381609 | 0.497529767 | 0.998865404 | 8.930468 | 9.044584 | 8.974538 | 8.949492 | 8.980957 | 9.043129 | 573 | 516 | 437 | 519 |
| 64777 | NM_00128879 | RMND5B | required for meiotic nuclear division 5 homolog B | protein_coding | NP_001275723.1;NP_001275724.1;NP_073599.2;XP_005266026.1;XP_00 | 26181 | . | . | . | 9.289126 | 9.061557 | 9.939930 | 10.320687 | 21.616178 | 20.946195 | 22.304915 | 23.597023 | 1.070112 | 988.778493 | 0.140678803 | 0.694931668 | 0.487098126 | 0.998865404 | 9.899975 | 9.997388 | 9.922846 | 9.924419 | 9.930972 | 10.013569 | 1099 | 1032 | 839 | 1012 |
| 64778 | NM_00113509 | FNDC3B | fibronectin type III domain containing 3B | protein_coding | NP_001128567.1;NP_073600.3;XP_016862551.1;XP_016862552.1;XP_01 | 24670 | 611909 | . | . | 10.960954 | 10.514877 | 9.835740 | 11.115913 | 25.506590 | 24.305608 | 22.071115 | 25.415212 | -1.031144 | 2587.108615 | 0.121556646 | -0.36399029 | 0.715865259 | 0.998865404 | 11.359146 | 11.314762 | 11.372682 | 11.324645 | 11.272521 | 11.373100 | 3114 | 2751 | 2077 | 2537 |
| 64779 | NM_00115937 | MTHFSD | methenyltetrahydrofolate synthetase domain containing | protein_coding | NP_001152849.1;NP_001152850.1;NP_001152851.1;NP_001152852.1;NI | 25778 | 616820 | . | . | 2.722018 | 3.265798 | 3.013143 | 2.950849 | 6.334244 | 7.549038 | 6.761407 | 6.746765 | 1.008260 | 275.698465 | 0.211926747 | 0.05599853 | 0.955342978 | 0.998865404 | 8.101232 | 8.112641 | 8.062835 | 8.143236 | 8.112961 | 8.103106 | 292 | 319 | 243 | 257 |
| 64780 | NM_00115929 | MICAL1 | microtubule associated monooxygenase, calponin and LIM domai | protein_coding | NP_001152763.1;NP_001273542.1;NP_073602.3 | 20619 | 607129 | . | . | 66.795697 | 71.292272 | 91.555855 | 77.977614 | 155.436325 | 164.795285 | 205.448687 | 178.286550 | 1.208853 | 8210.605139 | 0.117559702 | 2.32766113 | 0.019930105 | 0.467054728 | 12.859941 | 13.133643 | 12.892741 | 12.939279 | 13.148973 | 13.000406 | 8322 | 8245 | 8487 | 7701 |
| 64781 | NM_022766,XI | CERK | ceramide kinase | protein_coding | NP_073603.2;XP_016884398.1 | 19256 | 610307 | . | . | 18.280132 | 19.009355 | 14.843194 | 20.054947 | 42.538620 | 43.940974 | 33.307698 | 45.853249 | -1.081059 | 2492.674157 | 0.144090715 | -0.78037609 | 0.435169535 | 0.998865404 | 11.338790 | 11.225962 | 11.301766 | 11.321707 | 11.125591 | 11.358506 | 2946 | 2829 | 1781 | 2571 |
| 64782 | NM_022767,XI | AEN | apoptosis enhancing nuclease | protein_coding | NP_073604.3;XP_005255023.1;XP_005255024.1;XP_006720708.2;XP_01 | 25722 | 610177 | . | . | 31.116284 | 31.200825 | 22.969038 | 28.825263 | 72.408868 | 72.122101 | 51.541856 | 65.905538 | -1.203198 | 2911.003423 | 0.126365968 | -2.11191814 | 0.034693471 | 0.581842449 | 11.634716 | 11.367532 | 11.556631 | 11.597285 | 11.350902 | 11.496102 | 3572 | 3515 | 2089 | 2702 |
| 64783 | NM_00120154 | RBM15 | RNA binding motif protein 15 | protein_coding | NP_001188474.1;NP_073605.4 | 14959 | 606077 | . | . | 6.387861 | 6.054997 | 6.283029 | 6.089429 | 14.864815 | 13.996397 | 14.098937 | 13.922757 | -1.014753 | 658.243719 | 0.153900073 | -0.13728523 | 0.890805343 | 0.998865404 | 9.372854 | 9.352027 | 9.383810 | 9.350532 | 9.371250 | 9.343481 | 782 | 698 | 580 | 600 |
| 64784 | NM_00104257 | CRTC3 | CREB regulated transcription coactivator 3 | protein_coding | NP_001036039.1;NP_073606.3;XP_005255025.1;XP_011520208.1;XP_02 | 26148 | 608986 | . | . | 7.640837 | 8.845885 | 11.371540 | 8.967945 | 17.780545 | 20.447658 | 25.517409 | 20.504142 | 1.213281 | 1535.535504 | 0.148561674 | 1.877424763 | 0.0604599 | 0.719735192 | 10.438229 | 10.717345 | 10.449004 | 10.551532 | 10.740403 | 10.554178 | 1490 | 1598 | 1636 | 1393 |
| 64785 | NM_00112612 | GINS3 | GINS complex subunit 3 | protein_coding | NP_001119601.1;NP_001119602.1;NP_073607.2 | 25851 | 610610 | . | . | 5.969106 | 6.427646 | 6.490628 | 6.751437 | 13.890353 | 14.857791 | 14.564780 | 15.436357 | 1.067772 | 437.570814 | 0.176037389 | 0.537403936 | 0.590988625 | 0.998865404 | 8.725496 | 8.819711 | 8.729053 | 8.775049 | 8.782893 | 8.801513 | 469 | 474 | 387 | 430 |
| 64786 | NM_00114621 | TBC1D15 | TBC1 domain family member 15 | protein_coding | NP_001139685.2;NP_001139686.1;NP_073608.4;XP_006719627.1;XP_00 | 25694 | 612662 | . | . | 4.860029 | 5.629484 | 4.947929 | 4.822461 | 11.309490 | 13.012808 | 11.103004 | 11.025982 | -1.037592 | 857.006197 | 0.149182861 | -0.35687649 | 0.721184272 | 0.998865404 | 9.769269 | 9.716574 | 9.761242 | 9.749498 | 9.782257 | 9.673204 | 1013 | 934 | 788 | 728 |
| 64787 | NM_022772,XI | EPS8L2 | EPS8 like 2 | protein_coding | NP_073609.2;XP_016873620.1;XP_016873621.1 | 21296 | 614988 | . | . | 0.046658 | 0.041047 | 0.108075 | 0.099149 | 0.108575 | 0.094882 | 0.242517 | 0.226693 | 2.249791 | 8.002698 | 1.085041828 | 1.07810672 | 0.280986133 | NA | 2.299300 | 3.470304 | 2.968181 | 2.965674 | 2.996520 | 2.993186 | 6 | 5 | 10 | 10 |
| 64788 | NM_00135201 | LMF1 | lipase maturation factor 1 | protein_coding | NP_001338946.1;NP_001338947.1;NP_001338948.1;NP_001338949.1;NI | 14154 | 611761 | . | . | 0.599380 | 0.512748 | 0.409928 | 0.813241 | 1.394781 | 1.185241 | 0.919867 | 1.859379 | 1.121487 | 49.804359 | 0.464295363 | 0.356265392 | 0.721641824 | 0.998865404 | 5.554358 | 5.717436 | 5.632644 | 5.613533 | 5.594559 | 5.691807 | 57 | 48 | 34 | 62 |
| 64789 | NM_00134694 | EXO5 | exonuclease 5 | protein_coding | NP_001333875.1;NP_001333876.1;NP_001333877.1;NP_001333878.1;NI | 26115 | 618601 | . | . | 4.210556 | 5.701236 | 4.853072 | 4.536686 | 9.798135 | 13.178658 | 10.890152 | 10.372592 | 1.011510 | 309.682279 | 0.213343735 | 0.077386279 | 0.938316256 | 0.998865404 | 8.266637 | 8.282609 | 8.193928 | 8.338514 | 8.293137 | 8.256427 | 308 | 376 | 280 | 282 |
| 64792 | NM_00113082 | IFT22 | intraflagellar transport 22 | protein_coding | NP_001124292.1;NP_001124293.1;NP_001124294.1;NP_001274454.1;NI | 21895 | . | . | . | 4.538277 | 4.344767 | 3.772761 | 4.266496 | 10.560756 | 10.043124 | 8.465966 | 9.754835 | -1.173413 | 611.208139 | 0.1675938 | -1.37661085 | 0.168632596 | 0.945638095 | 9.366607 | 9.135161 | 9.313700 | 9.297432 | 9.131201 | 9.262407 | 769 | 704 | 439 | 581 |
| 64793 | NM_00128151 | CEP85 | centrosomal protein 85 | protein_coding | NP_001268446.1;NP_001268447.1;NP_001306873.1;NP_073615.2;XP_01 | 25309 | . | . | . | 12.399696 | 13.996479 | 13.447007 | 12.827793 | 28.854602 | 32.353487 | 30.174693 | 29.329221 | -1.017118 | 1605.572794 | 0.128896978 | -0.18997588 | 0.849328033 | 0.998865404 | 10.661100 | 10.636540 | 10.608796 | 10.699067 | 10.660933 | 10.622081 | 1757 | 1848 | 1419 | 1455 |
| 64794 | NM_00132234 | DDX31 | DEAD-box helicase 31 | protein_coding | NP_001309269.1;NP_001309270.1;NP_001309271.1;NP_001309272.1;NI | 16715 | 616533 | . | . | 5.949177 | 5.442192 | 5.536351 | 5.961271 | 13.843980 | 12.579869 | 12.423408 | 13.629737 | -1.009421 | 764.228157 | 0.149507969 | -0.09048052 | 0.927905376 | 0.998865404 | 9.584598 | 9.571089 | 9.610643 | 9.550301 | 9.554833 | 9.593540 | 922 | 793 | 643 | 733 |
| 64795 | NM_022780 | RMND5A | required for meiotic nuclear division 5 homolog A | protein_coding | NP_073617.1 | 25850 | . | . | . | 3.822754 | 3.777218 | 3.953586 | 4.246686 | 8.895705 | 8.731208 | 8.871733 | 9.709543 | 1.063340 | 761.016209 | 0.147596798 | 0.600305027 | 0.548302969 | 0.998865404 | 9.526889 | 9.615323 | 9.556515 | 9.542585 | 9.570503 | 9.613871 | 858 | 788 | 661 | 758 |
| 64798 | NM_00128301 | DEPTOR | DEP domain containing MTOR interacting protein | protein_coding | NP_001269941.1;NP_073620.2 | 22953 | 612974 | . | . | 0.849782 | 0.735724 | 1.316124 | 1.322418 | 1.977477 | 1.700659 | 2.953344 | 3.023551 | 1.663036 | 84.611957 | 0.347625139 | 2.110951285 | 0.034776499 | 0.581842449 | 5.989254 | 6.723816 | 6.329163 | 6.299968 | 6.451648 | 6.452050 | 78 | 64 | 92 | 99 |
| 64799 | NM_00103171 | IQCH | IQ motif containing H | protein_coding | NP_001026885.2;NP_001271276.2;NP_001271277.2;NP_001271278.1;NI | 25721 | 612523 | . | . | 0.274990 | 0.446195 | 0.516643 | 0.329144 | 0.639914 | 1.031401 | 1.159333 | 0.752546 | -1.161817 | 40.711159 | 0.500679956 | -0.43217732 | 0.665612539 | 0.998865404 | 5.452235 | 5.234243 | 5.321455 | 5.391835 | 5.335107 | 5.332553 | 40 | 57 | 33 | 35 |
| 64800 | NM_022785,N | EFCAB6 | EF-hand calcium binding domain 6 | protein_coding | NP_073622.2;NP_942153.1;XP_005261761.1;XP_005261762.1;XP_01152 | 24204 | . | . | . | 0.194470 | 0.180960 | 0.190932 | 0.169717 | 0.452539 | 0.418297 | 0.428445 | 0.388038 | -1.050801 | 35.844229 | 0.518781566 | -0.13780332 | 0.89039586 | 0.998865404 | 5.198253 | 5.128235 | 5.173243 | 5.161170 | 5.171664 | 5.148675 | 44 | 38 | 33 | 30 |
| 64801 | NM_00134699 | ARV1 | ARV1 homolog, fatty acid homeostasis modulator | protein_coding | NP_001333921.1;NP_073623.1;XP_024304970.1 | 29561 | 611647 | . | . | 20.760966 | 19.581357 | 20.919674 | 19.974258 | 48.311616 | 45.263186 | 46.943142 | 45.668768 | 1.002213 | 920.502870 | 0.141273537 | 0.022579238 | 0.981985905 | 0.998865404 | 9.844522 | 9.848033 | 9.869831 | 9.819972 | 9.863441 | 9.830086 | 1095 | 958 | 820 | 844 |
| 64802 | NM_00129777 | NMNAT1 | nicotinamide nucleotide adenylyltransferase 1 | protein_coding | NP_001284707.1;NP_001284708.1;NP_073624.2;XP_011540273.1;XP_01 | 17877 | 608700 | . | . | 7.133320 | 6.995333 | 7.481025 | 9.807803 | 16.599528 | 16.170023 | 16.787201 | 22.424374 | 1.095257 | 754.262493 | 0.147759086 | 0.888402817 | 0.374324111 | 0.998865404 | 9.491937 | 9.622936 | 9.511351 | 9.539979 | 9.573905 | 9.604092 | 812 | 793 | 671 | 755 |
| 64834 | NM_00125639 | ELOVL1 | ELOVL fatty acid elongase 1 | protein_coding | NP_001243328.1;NP_001243330.1;NP_001243331.1;NP_073732.1 | 14418 | 611813 | . | . | 65.638005 | 69.404853 | 68.961754 | 63.796920 | 152.742333 | 160.432420 | 154.748177 | 145.864083 | -1.041615 | 3133.351163 | 0.114066212 | -0.51568853 | 0.606071968 | 0.998865404 | 11.642546 | 11.583838 | 11.621772 | 11.637618 | 11.631098 | 11.560407 | 3646 | 3483 | 2785 | 2753 |
| 64837 | NM_00113477 | KLC2 | kinesin light chain 2 | protein_coding | NP_001128246.1;NP_001128247.1;NP_001128248.1;NP_001305663.1;NI | 20716 | 611729 | . | . | 6.913944 | 7.764641 | 8.799770 | 9.007656 | 16.089032 | 17.948312 | 19.746428 | 20.594933 | 1.202112 | 748.528292 | 0.150491852 | 1.76468747 | 0.077616306 | 0.767975096 | 9.409212 | 9.674441 | 9.442348 | 9.513623 | 9.601921 | 9.611366 | 742 | 772 | 706 | 771 |
| 64838 | NM_022823,XI | FNDC4 | fibronectin type III domain containing 4 | protein_coding | NP_073734.1;XP_005264556.1 | 20239 | 611905 | . | . | 2.626429 | 2.928915 | 3.800623 | 4.294485 | 6.111808 | 6.770320 | 8.528489 | 9.818831 | 1.433724 | 175.356818 | 0.253184295 | 2.05292001 | 0.040080336 | 0.625824522 | 7.171585 | 7.690332 | 7.342895 | 7.385214 | 7.490656 | 7.540644 | 157 | 164 | 170 | 204 |
| 64839 | NM_00116331 | FBXL17 | F-box and leucine rich repeat protein 17 | protein_coding | NP_001156787.2;XP_005272105.1;XP_005272107.1;XP_011541876.1;XP_ | 13615 | 609083 | . | . | 1.635425 | 1.528504 | 1.888105 | 1.566658 | 3.805699 | 3.533206 | 4.236854 | 3.581975 | 1.082872 | 242.368596 | 0.223833212 | 0.513163505 | 0.607836952 | 0.998865404 | 7.862008 | 7.977787 | 7.900825 | 7.896917 | 7.982017 | 7.892408 | 269 | 250 | 239 | 215 |
| 64840 | NM_00128216 | PORCN | porcupine O-acyltransferase | protein_coding | NP_001269096.1;NP_073736.2;NP_982299.1;NP_982300.1;NP_982301.1 | 17652 | 300651 | . | . | 5.948550 | 7.863551 | 7.186366 | 7.125806 | 13.842519 | 18.176951 | 16.125997 | 16.292309 | 1.030059 | 409.352023 | 0.194392383 | 0.219796755 | 0.826029447 | 0.998865404 | 8.655979 | 8.698110 | 8.585701 | 8.739869 | 8.694531 | 8.672001 | 405 | 491 | 368 | 382 |
| 64841 | NM_198066,XI | GNPNAT1 | glucosamine-phosphate N-acetyltransferase 1 | protein_coding | NP_932332.1;XP_005268069.1;XP_006720301.1 | 19980 | 616510 | . | . | 5.875884 | 5.500069 | 5.945498 | 4.444870 | 13.673427 | 12.713653 | 13.341524 | 10.162667 | -1.107295 | 658.105019 | 0.176011364 | -0.83539512 | 0.403495277 | 0.998865404 | 9.433310 | 9.287348 | 9.418724 | 9.366474 | 9.415755 | 9.223263 | 826 | 718 | 622 | 501 |
| 64844 | NM_00128280 | MARCHF7 | membrane associated ring-CH-type finger 7 | protein_coding | NP_001269734.1;NP_001269735.1;NP_001269736.1;NP_001363163.1;NI | 17393 | 613334 | . | . | 15.646802 | 15.184570 | 15.409566 | 12.611627 | 36.410748 | 35.099822 | 34.578618 | 28.834983 | -1.029034 | 1767.949642 | 0.123120948 | -0.33536895 | 0.737346786 | 0.998865404 | 10.808289 | 10.767141 | 10.830574 | 10.766066 | 10.778225 | 10.774552 | 2149 | 1856 | 1519 | 1630 |
| 64847 | NM_00125837 | SPATA20 | spermatogenesis associated 20 | protein_coding | NP_001245301.1;NP_001245302.1;NP_073738.2 | 26125 | 613939 | . | . | 34.609755 | 35.416394 | 39.500446 | 39.581275 | 80.538325 | 81.866583 | 88.637852 | 90.497878 | 1.115008 | 3141.247053 | 0.110308407 | 1.423767303 | 0.154513844 | 0.922573785 | 11.536460 | 11.693511 | 11.564374 | 11.576948 | 11.664122 | 11.655629 | 3394 | 3230 | 2895 | 3089 |
| 64848 | NM_00134597 | YTHDC2 | YTH domain containing 2 | protein_coding | NP_001332904.1;NP_001332905.1;NP_073739.3;XP_011541883.1;XP_01 | 24721 | 616530 | . | . | 6.404654 | 6.043313 | 6.446946 | 5.094960 | 14.903894 | 13.969390 | 14.466760 | 11.649020 | -1.083680 | 1156.252361 | 0.151390385 | -0.76582795 | 0.44377868 | 0.998865404 | 10.231713 | 10.116467 | 10.228339 | 10.176197 | 10.227464 | 10.050112 | 1432 | 1253 | 1081 | 916 |
| 64849 | NM_00101155 | SLC13A3 | solute carrier family 13 member 3 | protein_coding | NP_001011554.1;NP_001180268.1;NP_001180269.1;NP_001180271.1;NI | 14430 | 606411 | . | . | 0.538279 | 0.572871 | 0.595252 | 0.429235 | 1.252598 | 1.324216 | 1.335727 | 0.981394 | -1.103536 | 68.478317 | 0.390911845 | -0.36359657 | 0.716159289 | 0.998865404 | 6.166062 | 6.025675 | 6.099330 | 6.115340 | 6.120237 | 6.046978 | 80 | 80 | 66 | 51 |
| 64852 | NM_00136790 | TUT1 | terminal uridylyl transferase 1, U6 snRNA-specific | protein_coding | NP_001354835.1;NP_073741.3 | 26184 | 610641 | . | . | 5.863098 | 5.903640 | 5.727519 | 6.343793 | 13.643671 | 13.646528 | 12.852386 | 14.504328 | 1.010567 | 502.878153 | 0.168577279 | 0.089961692 | 0.928317657 | 0.998865404 | 8.966659 | 8.981433 | 8.971535 | 8.969538 | 8.947200 | 9.006196 | 577 | 539 | 419 | 496 |
| 64853 | NM_022831,XI | AIDA | axin interactor, dorsalization associated | protein_coding | NP_073742.2 | 25761 | 612375 | . | . | 15.174031 | 14.472950 | 14.785942 | 13.020382 | 35.310590 | 33.454872 | 33.179222 | 29.769553 | -1.081400 | 1338.034038 | 0.135103359 | -0.83566038 | 0.403345992 | 0.998865404 | 10.441025 | 10.328583 | 10.438829 | 10.389257 | 10.403325 | 10.303866 | 1652 | 1452 | 1192 | 1126 |
| 64854 | NM_00113422 | USP46 | ubiquitin specific peptidase 46 | protein_coding | NP_001127695.1;NP_001273696.1;NP_001273697.1;NP_073743.2;XP_01 | 20075 | 612849 | . | . | 1.987793 | 2.020576 | 1.945958 | 1.606862 | 4.625675 | 4.670649 | 4.366673 | 3.673900 | -1.145404 | 468.782276 | 0.179998673 | -1.0881011 | 0.276550479 | 0.998865404 | 8.966995 | 8.771967 | 8.908811 | 8.915834 | 8.886306 | 8.766191 | 573 | 543 | 418 | 369 |
| 64855 | NM_00103553 | NIBAN2 | niban apoptosis regulator 2 | protein_coding | NP_001030611.1;NP_073744.2;XP_005252192.2;XP_011517227.1 | 25282 | 614045 | . | . | 47.162188 | 48.022888 | 40.063386 | 51.794126 | 109.748349 | 111.007053 | 89.901073 | 118.421117 | -1.051261 | 5742.628266 | 0.129426981 | -0.55723143 | 0.577369316 | 0.998865404 | 12.523187 | 12.450899 | 12.502530 | 12.510306 | 12.356483 | 12.561899 | 6747 | 6381 | 4273 | 5891 |
| 64856 | NM_022834,N | VWA1 | von Willebrand factor A domain containing 1 | protein_coding | NP_073745.2;NP_954572.2 | 30910 | 611901 | . | . | 2.580557 | 3.180596 | 2.506229 | 3.183769 | 6.005063 | 7.352089 | 5.623905 | 7.279310 | -1.020959 | 399.983024 | 0.199991141 | -0.1496319 | 0.881055041 | 0.998865404 | 8.659383 | 8.628037 | 8.586752 | 8.704825 | 8.566220 | 8.699380 | 420 | 479 | 305 | 413 |
| 64857 | NM_00135169 | PLEKHG2 | pleckstrin homology and RhoGEF domain containing G2 | protein_coding | NP_001338622.1;NP_001338623.1;NP_073746.2;XP_005259220.1;XP_00 | 29515 | 611893 | . | . | 2.089979 | 2.724885 | 2.746176 | 2.663423 | 4.863465 | 6.298693 | 6.162337 | 6.089602 | 1.090470 | 633.619835 | 0.17141268 | 0.728943905 | 0.466035976 | 0.998865404 | 9.243940 | 9.368327 | 9.189091 | 9.354264 | 9.338096 | 9.328156 | 612 | 735 | 579 | 615 |
| 64858 | NM_00131994 | DCLRE1B | DNA cross-link repair 1B | protein_coding | NP_001306875.1;NP_001306876.1;NP_001350619.1;NP_001350620.1;NI | 17641 | 609683 | . | . | 4.370234 | 4.180548 | 3.988929 | 5.020378 | 10.169714 | 9.663526 | 8.951041 | 11.478498 | 1.045458 | 507.935715 | 0.178931173 | 0.358430754 | 0.72002098 | 0.998865404 | 8.956429 | 9.019878 | 8.989096 | 8.955814 | 8.926986 | 9.069605 | 587 | 522 | 401 | 541 |
| 64859 | NM_00103171 | NABP1 | nucleic acid binding protein 1 | protein_coding | NP_001026886.1;NP_001241665.1 | 26232 | 612103 | . | . | 8.872986 | 8.141736 | 11.281362 | 7.260426 | 20.647801 | 18.819987 | 25.315049 | 16.600101 | 1.067575 | 1609.695282 | 0.17723323 | 0.532277518 | 0.594533801 | 0.998865404 | 10.604318 | 10.699264 | 10.657768 | 10.580292 | 10.828636 | 10.484512 | 1882 | 1596 | 1760 | 1220 |
| 64860 | NM_00116847 | ARMCX5 | armadillo repeat containing X-linked 5 | protein_coding | NP_001161950.1;NP_001161951.1;NP_001161952.1;NP_001161954.1;NI | 25772 | 301047 | . | . | 5.334590 | 5.258178 | 5.307593 | 5.660181 | 12.413810 | 12.154513 | 11.910085 | 12.941333 | 1.020104 | 465.387472 | 0.171182571 | 0.167748533 | 0.866781114 | 0.998865404 | 8.847934 | 8.876502 | 8.859957 | 8.851945 | 8.855019 | 8.881401 | 534 | 494 | 400 | 450 |
| 64863 | NM_00130840 | METTL4 | methyltransferase like 4 | protein_coding | NP_001295330.1;NP_073751.3;XP_005258189.1;XP_005258190.1;XP_01 | 24726 | . | . | . | 2.504740 | 2.163182 | 2.933489 | 1.814396 | 5.828634 | 5.000294 | 6.582664 | 4.148403 | 1.048659 | 253.470311 | 0.271310007 | 0.252647778 | 0.800540406 | 0.998865404 | 7.949940 | 8.020543 | 7.974526 | 7.948805 | 8.125758 | 7.823140 | 292 | 260 | 293 | 171 |
| 64864 | NM_00136807 | RFX7 | regulatory factor X7 | protein_coding | NP_001355002.1;NP_001355003.1;NP_001357483.1;NP_001357490.1;NI | 25777 | 612660 | . | . | 1.209847 | 1.201191 | 1.639976 | 1.333428 | 2.815364 | 2.776606 | 3.680059 | 3.048724 | 1.208233 | 454.945209 | 0.185084683 | 1.474455685 | 0.140358941 | 0.900674731 | 8.686167 | 8.959960 | 8.768542 | 8.759478 | 8.951649 | 8.805067 | 478 | 441 | 483 | 413 |
| 64895 | NM_022894,XI | PAPOLG | poly(A) polymerase gamma | protein_coding | NP_075045.2;XP_005264557.1;XP_005264558.1;XP_011531343.1;XP_01( | 14982 | 616865 | . | . | 2.395396 | 2.179416 | 1.786838 | 1.622202 | 5.574186 | 5.037819 | 4.009611 | 3.708969 | -1.134911 | 394.960378 | 0.181965022 | -1.00337735 | 0.315678829 | 0.998865404 | 8.713787 | 8.531593 | 8.671449 | 8.652231 | 8.603532 | 8.570691 | 491 | 446 | 331 | 337 |
| 64897 | NM_00128619 | C12orf43 | chromosome 12 open reading frame 43 | protein_coding | NP_001273120.1;NP_001273121.1;NP_001273124.1;NP_001273125.1;NI | 25719 | . | . | . | 4.889578 | 4.551813 | 4.857753 | 5.251117 | 11.378250 | 10.521717 | 10.900657 | 12.006053 | 1.011459 | 490.592060 | 0.176136734 | 0.093328031 | 0.925642964 | 0.998865404 | 8.930286 | 8.946429 | 8.966443 | 8.900405 | 8.893223 | 8.986794 | 591 | 499 | 397 | 497 |
| 64900 | NM_00130186 | LPIN3 | lipin 3 | protein_coding | NP_001288789.1;NP_075047.1;XP_005260573.1;XP_006723926.1;XP_01 | 14451 | 605520 | . | . | 12.474802 | 15.059096 | 17.402765 | 14.503828 | 29.029375 | 34.809777 | 39.051301 | 33.161281 | 1.099349 | 1796.227841 | 0.139530706 | 0.979348568 | 0.327407781 | 0.998865404 | 10.740774 | 10.877497 | 10.706504 | 10.830460 | 10.908662 | 10.776663 | 1814 | 1993 | 1772 | 1617 |
| 64901 | NM_022897,XI | RANBP17 | RAN binding protein 17 | protein_coding | NP_075048.1;XP_011532929.1;XP_011532938.1;XP_011532939.1;XP_01( | 14428 | 606141 | . | . | 1.090355 | 0.478414 | 0.467769 | 0.488160 | 2.537299 | 1.105875 | 1.049659 | 1.116119 | -1.416034 | 78.190504 | 0.387994867 | -1.29345908 | 0.19585233 | 0.96621494 | 6.517382 | 6.017349 | 6.399542 | 6.250126 | 6.241377 | 6.235423 | 132 | 74 | 57 | 60 |
| 64921 | NM_00136342 | CASD1 | CAS1 domain containing 1 | protein_coding | NP_001350355.1;NP_001350356.1;NP_001350357.1;NP_075051.4;XP_0C | 16014 | 611686 | . | . | 2.785949 | 1.629455 | 2.449625 | 1.172154 | 6.483017 | 3.766559 | 5.496885 | 2.679993 | -1.311214 | 241.368129 | 0.598313414 | -0.65334183 | 0.513535933 | 0.998865404 | 8.097010 | 7.706878 | 8.091220 | 7.805321 | 7.959916 | 7.666055 | 399 | 217 | 240 | 133 |
| 64924 | NM_00125196 | SLC30A5 | solute carrier family 30 member 5 | protein_coding | NP_001238898.1;NP_075053.2;NP_076960.1;XP_005248626.1;XP_00671 | 19089 | 607819 | . | . | 23.527210 | 22.247523 | 22.486017 | 17.627872 | 54.748780 | 51.426147 | 50.457972 | 40.304036 | -1.207818 | 2196.291550 | 0.157777249 | -1.72650681 | 0.084256275 | 0.787933172 | 11.230401 | 10.958516 | 11.217561 | 11.115258 | 11.141836 | 10.873695 | 2939 | 2431 | 2025 | 1548 |
| 64925 | NM_022903 | CCDC71 | coiled-coil domain containing 71 | protein_coding | NP_075054.3 | 25760 | . | . | . | 5.015772 | 5.092930 | 4.313864 | 5.215535 | 11.671907 | 11.772537 | 9.680185 | 11.924702 | -1.075394 | 274.223561 | 0.213706421 | -0.4906973 | 0.623640553 | 0.998865404 | 8.151142 | 8.045336 | 8.115498 | 8.119741 | 8.033495 | 8.124126 | 326 | 308 | 209 | 270 |
| 64927 | NM_00128861 | TTC23 | tetratricopeptide repeat domain 23 | protein_coding | NP_001275544.1;NP_001275545.1;NP_001340798.1;XP_011520229.1;XF | 25730 | . | . | . | 5.801894 | 5.516671 | 5.905114 | 6.076087 | 13.501243 | 12.752028 | 13.250906 | 13.892260 | 1.094195 | 676.013855 | 0.152034738 | 0.854212065 | 0.392987499 | 0.998865404 | 9.334483 | 9.464411 | 9.382857 | 9.354609 | 9.424171 | 9.436187 | 759 | 682 | 609 | 668 |
| 64928 | NM_00131876 | MRPL14 | mitochondrial ribosomal protein L14 | protein_coding | NP_001305696.1;NP_001305697.1;NP_001305698.1;NP_001305699.1;NI | 14279 | 611827 | . | . | 31.358781 | 30.157203 | 24.745595 | 35.538174 | 72.973170 | 69.709723 | 55.528396 | 81.253807 | -1.034581 | 910.258025 | 0.17251753 | -0.28429957 | 0.776180818 | 0.998865404 | 9.854802 | 9.805032 | 9.856465 | 9.821150 | 9.683155 | 9.930467 | 1092 | 975 | 644 | 983 |
| 64940 | NR_040585,NF | STAG3L4 | stromal antigen 3-like 4 (pseudogene) | transcribed_ps | . | 33887 | . | . | . | 6.055851 | 6.022022 | 7.242018 | 6.463517 | 14.092215 | 13.920173 | 16.250878 | 14.778064 | 1.122871 | 432.574506 | 0.17960289 | 0.930895967 | 0.351907381 | 0.998865404 | 8.670465 | 8.838268 | 8.722111 | 8.715050 | 8.830142 | 8.746564 | 472 | 437 | 425 | 400 |
| 64943 | NM_00113423 | NT5DC2 | 5'-nucleotidase domain containing 2 | protein_coding | NP_001127703.1;NP_075059.1;XP_005265463.1;XP_006713366.1;XP_00 | 25717 | . | . | . | 28.923937 | 31.651330 | 28.042941 | 30.063532 | 67.307189 | 73.163465 | 62.927546 | 68.736692 | -1.052559 | 1741.030862 | 0.12506605 | -0.59089367 | 0.554591669 | 0.998865404 | 10.802338 | 10.728160 | 10.752715 | 10.816823 | 10.719904 | 10.769479 | 1973 | 2005 | 1428 | 1641 |
| 64946 | NM_022909 | CENPH | centromere protein H | protein_coding | NP_075060.1 | 17268 | 605607 | . | . | 15.307781 | 12.259569 | 11.625823 | 10.900349 | 35.621838 | 28.338541 | 26.088009 | 24.922350 | -1.241923 | 529.083675 | 0.179267526 | -1.74362496 | 0.081224507 | 0.782373954 | 9.194813 | 8.883085 | 9.185195 | 9.030704 | 8.994741 | 8.948321 | 752 | 560 | 426 | 426 |
| 64949 | NM_030811 | MRPS26 | mitochondrial ribosomal protein S26 | protein_coding | NP_110438.1 | 14045 | 611988 | . | . | 26.236286 | 25.205391 | 21.590118 | 27.475376 | 61.052917 | 58.263393 | 48.447598 | 62.819183 | -1.062858 | 795.827342 | 0.160472604 | -0.54806005 | 0.583650674 | 0.998865404 | 9.679912 | 9.591353 | 9.673097 | 9.638953 | 9.530403 | 9.689540 | 967 | 864 | 594 | 806 |
| 64951 | NM_032014 | MRPS24 | mitochondrial ribosomal protein S24 | protein_coding | NP_114403.1 | 14510 | 611986 | . | . | 71.267197 | 73.024773 | 66.095787 | 80.014488 | 165.841690 | 168.800049 | 148.317032 | 182.943619 | -1.001427 | 1508.591037 | 0.136072098 | -0.01512345 | 0.987933696 | 0.999197585 | 10.560226 | 10.557745 | 10.552580 | 10.564733 | 10.485826 | 10.624381 | 1725 | 1642 | 1193 | 1540 |
| 64960 | NM_031280 | MRPS15 | mitochondrial ribosomal protein S15 | protein_coding | NP_112570.2 | 14504 | 611979 | . | . | 60.988457 | 65.297882 | 60.757751 | 75.928642 | 141.922638 | 150.938995 | 136.338638 | 173.601807 | 1.067109 | 1950.978882 | 0.136587001 | 0.686065531 | 0.492671785 | 0.998865404 | 10.882588 | 10.975870 | 10.878250 | 10.925452 | 10.865450 | 11.034657 | 2109 | 2098 | 1566 | 2088 |
| 64963 | NM_00132197 | MRPS11 | mitochondrial ribosomal protein S11 | protein_coding | NP_001308899.1;NP_001308901.1;NP_001308902.1;NP_001308903.1;NI | 14050 | 611977 | . | . | 22.499260 | 22.009386 | 19.839340 | 23.381360 | 52.356702 | 50.875680 | 44.518905 | 53.458701 | -1.028347 | 2113.885415 | 0.126337961 | -0.31919653 | 0.749577489 | 0.998865404 | 11.065848 | 11.025229 | 11.053765 | 11.058548 | 10.971281 | 11.092614 | 2461 | 2320 | 1674 | 2103 |
| 64965 | NM_182640,XI | MRPS9 | mitochondrial ribosomal protein S9 | protein_coding | NP_872578.1;XP_011509946.1 | 14501 | 611975 | . | . | 34.528628 | 34.259504 | 32.718205 | 36.002070 | 80.349545 | 79.192381 | 73.418702 | 82.314456 | -1.016472 | 1548.585921 | 0.125439788 | -0.18789993 | 0.850955095 | 0.998865404 | 10.608536 | 10.584838 | 10.612971 | 10.592863 | 10.563597 | 10.616234 | 1821 | 1663 | 1290 | 1489 |
| 64968 | NM_032476 | MRPS6 | mitochondrial ribosomal protein S6 | protein_coding | NP_115865.1 | 14051 | 611973 | . | . | 75.268623 | 66.941170 | 81.638306 | 80.492401 | 175.153168 | 154.737518 | 183.194000 | 184.036301 | 1.124004 | 2206.771744 | 0.12300846 | 1.371016234 | 0.170369896 | 0.946389416 | 11.020820 | 11.189685 | 11.106904 | 11.007203 | 11.161811 | 11.142080 | 2543 | 2101 | 2056 | 2163 |
| 64969 | NM_00132199 | MRPS5 | mitochondrial ribosomal protein S5 | protein_coding | NP_001308924.1;NP_001308925.1;NP_001308926.1;NP_114108.1 | 14498 | 611972 | . | . | 27.421525 | 27.720065 | 26.158141 | 29.264442 | 63.811018 | 64.076174 | 58.698114 | 66.909673 | -1.007985 | 2835.963992 | 0.116607435 | -0.0983963 | 0.921617615 | 0.998865404 | 11.475433 | 11.463790 | 11.471665 | 11.473533 | 11.423445 | 11.506970 | 3275 | 3076 | 2330 | 2784 |
| 64975 | NM_032477 | MRPL41 | mitochondrial ribosomal protein L41 | protein_coding | NP_115866.1 | 14492 | 611846 | . | . | 54.518177 | 57.719147 | 44.096943 | 58.976875 | 126.866035 | 133.420395 | 98.952255 | 134.843613 | -1.104660 | 976.399087 | 0.160694128 | -0.89364058 | 0.371514229 | 0.998865404 | 10.001723 | 9.857320 | 9.946437 | 9.982309 | 9.786523 | 9.986976 | 1152 | 1133 | 694 | 991 |
| 64976 | NM_00131815 | MRPL40 | mitochondrial ribosomal protein L40 | protein_coding | NP_001305080.1;NP_001305081.1;NP_003767.2 | 14491 | 605089 | . | . | 31.275888 | 29.375118 | 27.508480 | 32.451879 | 72.780275 | 67.901899 | 61.728230 | 74.197359 | -1.021245 | 1268.935486 | 0.142808866 | -0.21237646 | 0.831813351 | 0.998865404 | 10.324677 | 10.293965 | 10.336374 | 10.295761 | 10.223858 | 10.371293 | 1516 | 1347 | 985 | 1292 |
| 64978 | NM_032478 | MRPL38 | mitochondrial ribosomal protein L38 | protein_coding | NP_115867.2 | 14033 | 611844 | . | . | 39.990101 | 43.336136 | 39.183647 | 45.889236 | 93.058601 | 100.173416 | 87.926971 | 104.920288 | 1.006869 | 1781.027017 | 0.1308001 | 0.075505192 | 0.939812767 | 0.998865404 | 10.793750 | 10.803222 | 10.766474 | 10.822848 | 10.740602 | 10.856702 | 1971 | 1984 | 1439 | 1799 |
| 64979 | NM_032479,XI | MRPL36 | mitochondrial ribosomal protein L36 | protein_coding | NP_115868.1;XP_011512381.1;XP_011512382.1;XP_016865239.1;XP_01( | 14490 | 611842 | . | . | 38.811200 | 40.835820 | 42.890723 | 40.552453 | 90.315257 | 94.393826 | 96.245541 | 92.718364 | 1.035983 | 802.939832 | 0.145910917 | 0.349526654 | 0.726693964 | 0.998865404 | 9.623373 | 9.674470 | 9.620285 | 9.651718 | 9.682115 | 9.639626 | 888 | 870 | 732 | 743 |

| | | | | | | | | | | | | | | | | | | | | | | | | | | | | | | | | | |
|---|---|---|---|---|---|---|---|---|---|---|---|---|---|---|---|---|---|---|---|---|---|---|---|---|---|---|---|---|---|---|---|---|---|
| 64981 | NM_00138234 | MRPL34 | mitochondrial ribosomal protein L34 | protein_coding | NP_001369271.1;NP_001369272.1;NP_076426.1 | 14488 | 611840 | . | . | 42.838243 | 43.905493 | 43.845602 | 55.934373 | 99.686359 | 101.489519 | 98.388263 | 127.887295 | 1.140152 | 1127.499301 | 0.150020709 | 1.261333713 | 0.20718864 | 0.970482251 | 10.041502 | 10.230157 | 10.078709 | 10.092812 | 10.091466 | 10.267997 | 1202 | 1148 | 923 | 1257 |
| 64983 | NM_031903,N | MRPL32 | mitochondrial ribosomal protein L32 | protein_coding | NP_114109.1 | 14035 | 611839 | . | . | 46.930560 | 48.808244 | 52.665449 | 48.142522 | 109.209341 | 112.822444 | 118.179745 | 110.072152 | 1.036640 | 1320.022460 | 0.131338142 | 0.395274961 | 0.692639976 | 0.998865404 | 10.340047 | 10.392176 | 10.342460 | 10.361375 | 10.416177 | 10.341310 | 1474 | 1416 | 1226 | 1198 |
| 65003 | NM_016050,N | MRPL11 | mitochondrial ribosomal protein L11 | protein_coding | NP_057134.1;NP_733934.1;NP_733935.1;XP_016873624.1;XP_02430442 | 14042 | 611826 | . | . | 17.269601 | 18.365373 | 16.782022 | 17.626116 | 40.187068 | 42.452383 | 37.658369 | 40.300022 | -1.054546 | 864.088357 | 0.142777076 | -0.53665121 | 0.591508562 | 0.998865404 | 9.792970 | 9.716075 | 9.754098 | 9.793037 | 9.721784 | 9.749443 | 994 | 983 | 717 | 804 |
| 65005 | NM_00130073 | MRPL9 | mitochondrial ribosomal protein L9 | protein_coding | NP_001287662.1;NP_113608.1 | 14277 | 611824 | . | . | 30.744857 | 28.959850 | 27.020620 | 34.615826 | 71.544548 | 66.941987 | 60.633485 | 79.144975 | 1.015609 | 1137.320599 | 0.151005387 | 0.147976738 | 0.882361127 | 0.998865404 | 10.140438 | 10.162325 | 10.165750 | 10.121629 | 10.063587 | 10.239584 | 1338 | 1182 | 879 | 1201 |
| 65008 | NM_020236,XI | MRPL1 | mitochondrial ribosomal protein L1 | protein_coding | NP_064621.3 | 14275 | 611821 | . | . | 8.213205 | 6.969725 | 6.667468 | 6.390442 | 19.112468 | 16.110832 | 14.961606 | 14.610985 | -1.158372 | 271.297807 | 0.213983714 | -0.99118882 | 0.321593389 | 0.998865404 | 8.185528 | 7.974207 | 8.160943 | 8.077606 | 8.056481 | 8.032178 | 361 | 290 | 224 | 230 |
| 65009 | NM_00113048 | NDRG4 | NDRG family member 4 | protein_coding | NP_001123959.1;NP_001229762.1;NP_001229763.1;NP_001229764.1;NI | 14466 | 614463 | . | . | 0.050733 | 0.120361 | 0.103772 | 0.116279 | 0.118058 | 0.278220 | 0.232860 | 0.265857 | 1.220014 | 11.070004 | 0.93942254 | 0.305397594 | 0.760063354 | NA | 3.320504 | 3.602870 | 3.437352 | 3.473337 | 3.462413 | 3.468114 | 7 | 15 | 10 | 12 |
| 65010 | NM_00104045 | SLC26A6 | solute carrier family 26 member 6 | protein_coding | NP_001035544.1;NP_001268661.1;NP_001268662.1;NP_075062.2;NP_5 | 14472 | 610068 | . | . | 22.812082 | 24.482290 | 28.082731 | 22.223386 | 53.084653 | 56.591907 | 63.016832 | 50.811130 | 1.042967 | 2007.559994 | 0.141962828 | 0.427528224 | 0.668994632 | 0.998865404 | 10.940402 | 11.001408 | 10.927850 | 10.975446 | 11.081258 | 10.879893 | 2195 | 2184 | 2007 | 1680 |
| 65018 | NM_032409 | PINK1 | PTEN induced kinase 1 | protein_coding | NP_115785.1 | 14581 | 608309 | . | . | 17.428072 | 15.956366 | 14.987281 | 20.321987 | 40.555840 | 36.883854 | 33.631023 | 46.463806 | 1.042350 | 1420.922654 | 0.155679918 | 0.384381171 | 0.700695954 | 0.998865404 | 10.442581 | 10.502031 | 10.490030 | 10.415753 | 10.366257 | 10.592090 | 1680 | 1429 | 1077 | 1558 |
| 65055 | NM_00116473 | REEP1 | receptor accessory protein 1 | protein_coding | NP_001158202.1;NP_001158203.1;NP_001158204.1;NP_001358208.1;NI | 25786 | 609139 | . | . | 0.079978 | 0.107530 | 0.089560 | 0.138992 | 0.186112 | 0.248561 | 0.200969 | 0.317789 | 1.222710 | 14.491847 | 0.811481531 | 0.357472563 | 0.720738062 | NA | 3.706954 | 3.993209 | 3.838021 | 3.854076 | 3.844797 | 3.872580 | 13 | 16 | 11 | 18 |
| 65056 | NM_00112723 | GPBP1 | GC-rich promoter binding protein 1 | protein_coding | NP_001120707.2;NP_001120708.1;NP_001190175.1;NP_001317966.1;NI | 29520 | 608412 | . | . | 13.065818 | 12.571053 | 12.455689 | 10.536579 | 30.404694 | 29.058553 | 27.950207 | 24.090639 | -1.129063 | 1697.913843 | 0.134186659 | -1.30509483 | 0.191860559 | 0.965106155 | 10.814246 | 10.639565 | 10.790790 | 10.757184 | 10.746265 | 10.607533 | 2117 | 1902 | 1512 | 1361 |
| 65057 | NM_00108248 | ACD | ACD shelterin complex subunit and telomerase recruitment factor | protein_coding | NP_001075955.2;NP_075065.3;XP_005256172.1 | 25070 | 609377 | . | . | 11.970162 | 12.027989 | 12.142991 | 12.790742 | 27.855058 | 27.803233 | 27.248517 | 29.244509 | 1.037862 | 586.038630 | 0.159574174 | 0.335982552 | 0.736884024 | 0.998865404 | 9.167941 | 9.221270 | 9.181826 | 9.182562 | 9.183442 | 9.229341 | 662 | 621 | 501 | 579 |
| 65059 | NM_00132972 | RAPH1 | Ras association (RalGDS/AF-6) and pleckstrin homology domains 1 | protein_coding | NP_001316657.1;NP_976241.1;NP_998754.1;XP_005246834.1;XP_00524 | 14436 | 609035 | . | . | 3.868967 | 3.991954 | 3.739278 | 3.181156 | 9.003243 | 9.227584 | 8.390834 | 7.273337 | -1.151417 | 1083.687769 | 0.143728634 | -1.4152361 | 0.156999283 | 0.925698889 | 10.179666 | 9.976666 | 10.112742 | 10.148425 | 10.085652 | 9.963362 | 1304 | 1281 | 950 | 865 |
| 65061 | NM_00126143 | CDK15 | cyclin dependent kinase 15 | protein_coding | NP_001248364.1;NP_001248365.1;NP_001353315.1;NP_631897.1;XP_00 | 14434 | 616147 | . | . | 0.124668 | 0.075703 | 0.104236 | 0.076258 | 0.290105 | 0.174993 | 0.233901 | 0.174355 | -1.259134 | 11.220619 | 0.911715278 | -0.36462256 | 0.715393171 | NA | 3.644780 | 3.312263 | 3.504853 | 3.492254 | 3.484077 | 3.490299 | 16 | 12 | 8 | 10 |
| 65062 | NM_00104438 | TMEM237 | transmembrane protein 237 | protein_coding | NP_001037850.1;NP_689601.2 | 14432 | 614423 | . | . | 9.839714 | 8.698694 | 8.327259 | 8.426646 | 22.897419 | 20.107420 | 18.686130 | 19.266525 | -1.122719 | 1467.603916 | 0.131748731 | -1.26753732 | 0.204963229 | 0.96993643 | 10.600228 | 10.433448 | 10.612521 | 10.508360 | 10.472325 | 10.472865 | 1906 | 1564 | 1202 | 1297 |
| 65065 | NM_00111413 | NBEAL1 | neurobeachin like 1 | protein_coding | NP_001107604.1;NP_001364955.1;XP_005246844.1;XP_005246845.1;XP | 20681 | 609816 | . | . | 1.496564 | 1.370311 | 1.268827 | 1.065375 | 3.482566 | 3.167537 | 2.847212 | 2.435854 | -1.030956 | 591.403231 | 0.182583041 | -0.24088816 | 0.809641803 | 0.998865404 | 9.229252 | 9.186427 | 9.292998 | 9.130284 | 9.256708 | 9.129250 | 775 | 569 | 556 | 493 |
| 65078 | NM_023004 | RTN4R | reticulon 4 receptor | protein_coding | NP_075380.1 | 18601 | 605566 | . | . | 7.267065 | 7.669859 | 8.001625 | 8.205587 | 16.910759 | 17.729225 | 17.955416 | 18.761101 | 1.070594 | 471.884027 | 0.170465862 | 0.577307298 | 0.563731885 | 0.998865404 | 8.832357 | 8.930549 | 8.845845 | 8.874043 | 8.897583 | 8.907473 | 513 | 503 | 421 | 461 |
| 65080 | NM_022915,XI | MRPL44 | mitochondrial ribosomal protein L44 | protein_coding | NP_075066.1;XP_011509970.1 | 16650 | 611849 | . | . | 41.952746 | 39.475871 | 39.773015 | 49.724266 | 97.625767 | 91.250241 | 89.249492 | 113.688634 | 1.083172 | 2248.534079 | 0.134832855 | 0.854856468 | 0.392630613 | 0.998865404 | 11.076108 | 11.191138 | 11.127447 | 11.072306 | 11.075003 | 11.246472 | 2572 | 2249 | 1818 | 2424 |
| 65082 | NM_00135101 | VPS33A | VPS33A core subunit of CORVET and HOPS complexes | protein_coding | NP_001337947.1;NP_001337948.1;NP_001337949.1;NP_001337950.1;NI | 18179 | 610034 | . | . | 7.682492 | 7.613166 | 7.598536 | 8.049469 | 17.877475 | 17.598175 | 17.050895 | 18.404154 | 1.006480 | 1104.339175 | 0.133825034 | 0.069629413 | 0.944488626 | 0.998865404 | 10.104400 | 10.113521 | 10.102009 | 10.110976 | 10.090760 | 10.131322 | 1260 | 1195 | 937 | 1067 |
| 65083 | NM_022917,N | NOL6 | nucleolar protein 6 | protein_coding | NP_075068.2;NP_631981.2;XP_016870533.1 | 19910 | 611532 | . | . | 10.580765 | 11.669918 | 11.944659 | 10.730397 | 24.621874 | 26.975539 | 26.803468 | 24.533776 | 1.003194 | 1777.346576 | 0.130324966 | 0.03530201 | 0.97183892 | 0.998865404 | 10.793167 | 10.797847 | 10.749528 | 10.835410 | 10.840486 | 10.749288 | 1932 | 2019 | 1638 | 1572 |
| 65084 | NM_00116872 | TMEM135 | transmembrane protein 135 | protein_coding | NP_001162195.1;NP_075069.3;XP_016873630.1;XP_016873631.1 | 26167 | 616360 | . | . | 2.136134 | 1.678593 | 1.769256 | 1.267354 | 4.970872 | 3.880144 | 3.970159 | 2.897655 | -1.302979 | 474.537418 | 0.203261715 | -1.8784352 | 0.060321654 | 0.719735192 | 9.068008 | 8.687766 | 9.041274 | 8.875995 | 8.881144 | 8.705289 | 695 | 507 | 412 | 329 |
| 65094 | NM_00116146 | JMJD4 | jumonji domain containing 4 | protein_coding | NP_001154937.1;NP_075383.2;XP_011542564.1 | 25724 | . | . | . | 6.890280 | 6.684750 | 6.568868 | 7.774884 | 16.033963 | 15.452100 | 14.740351 | 17.776348 | 1.042165 | 561.308065 | 0.166686295 | 0.357458603 | 0.720748512 | 0.998865404 | 9.102799 | 9.161894 | 9.128389 | 9.108021 | 9.095227 | 9.192711 | 643 | 584 | 461 | 577 |
| 65095 | NM_023008,XI | KRI1 | KRI1 homolog | protein_coding | NP_075384.4;XP_011526492.1 | 25769 | . | . | . | 18.205421 | 18.721470 | 20.008223 | 19.257848 | 42.364765 | 43.275513 | 44.897875 | 44.030785 | 1.049019 | 1772.631469 | 0.121304628 | 0.569155871 | 0.569250365 | 0.998865404 | 10.756702 | 10.825824 | 10.764706 | 10.779900 | 10.829071 | 10.790521 | 1973 | 1885 | 1617 | 1660 |
| 65108 | NM_023009,N | MARCKSL1 | MARCKS like 1 | protein_coding | NP_075385.1 | 7142 | 602940 | . | . | 83.647629 | 89.095868 | 91.195145 | 111.114520 | 194.651456 | 205.949092 | 204.639265 | 254.050148 | 1.154106 | 4507.895367 | 0.123211471 | 1.67821481 | 0.093305172 | 0.809520218 | 12.031255 | 12.237833 | 12.051256 | 12.096687 | 12.111656 | 12.269721 | 4687 | 4641 | 3808 | 4951 |
| 65109 | NM_023010,N | UPF3B | UPF3B regulator of nonsense mediated mRNA decay | protein_coding | NP_075386.1;NP_542199.1;XP_016885226.1;XP_016885227.1;XP_01688 | 20439 | 300298 | . | . | 5.595717 | 4.764888 | 5.314112 | 4.413672 | 13.021465 | 11.014254 | 11.924713 | 10.091336 | -1.080915 | 370.253734 | 0.20023209 | -0.56061191 | 0.57506213 | 0.998865404 | 8.586759 | 8.475851 | 8.602608 | 8.499794 | 8.561673 | 8.450767 | 481 | 379 | 340 | 301 |
| 65110 | NM_00135364 | UPF3A | UPF3A regulator of nonsense mediated mRNA decay | protein_coding | NP_001340573.1;NP_001340574.1;NP_001340575.1;NP_001340576.1;NI | 20332 | 605530 | . | . | 19.090284 | 20.506989 | 18.659836 | 16.971269 | 44.423870 | 47.402824 | 41.872139 | 38.802788 | -1.125158 | 1345.651607 | 0.135734282 | -1.25338428 | 0.21006589 | 0.972673425 | 10.476545 | 10.306633 | 10.405620 | 10.466802 | 10.395062 | 10.296033 | 1576 | 1598 | 1174 | 1109 |
| 65117 | NM_023012,N | RSRC2 | arginine and serine rich coiled-coil 2 | protein_coding | NP_075388.2;XP_005253658.1;XP_005253659.1;XP_005253661.1;XP_01 | 30559 | . | . | . | 14.412924 | 13.662217 | 16.038566 | 15.047139 | 33.539468 | 31.580826 | 35.990079 | 34.403494 | 1.122719 | 1357.115976 | 0.127504962 | 1.30972702 | 0.190288199 | 0.961278413 | 10.320347 | 10.487472 | 10.360248 | 10.359609 | 10.466441 | 10.429954 | 1473 | 1379 | 1278 | 1314 |
| 65123 | NM_00132447 | INTS3 | integrator complex subunit 3 | protein_coding | NP_001311404.1;NP_075391.3 | 26153 | 611347 | . | . | 47.095619 | 50.579338 | 55.691461 | 55.655809 | 109.593445 | 116.916401 | 124.970020 | 127.250384 | 1.121195 | 8528.806062 | 0.103533848 | 1.594036998 | 0.110927748 | 0.848126978 | 12.973266 | 13.138275 | 12.981009 | 13.033171 | 13.106980 | 13.102231 | 8973 | 8946 | 7862 | 8429 |
| 65124 | NM_023016 | SOWAHC | sosondowah ankyrin repeat domain family member C | protein_coding | NP_075392.2 | 26149 | . | . | . | 3.736991 | 3.609709 | 3.635176 | 3.286959 | 8.696133 | 8.344004 | 8.157230 | 7.515241 | -1.076738 | 514.661016 | 0.168842172 | -0.63175403 | 0.52754762 | 0.998865404 | 9.059532 | 8.953477 | 9.044679 | 9.015703 | 9.017359 | 8.947889 | 628 | 563 | 455 | 439 |
| 65125 | NM_00118498 | WNK1 | WNK lysine deficient protein kinase 1 | protein_coding | NP_001171914.1;NP_055638.2;NP_061852.3;NP_998820.3;XP_0067190 | 14540 | 605232 | . | . | 11.111144 | 11.628845 | 12.475046 | 12.932808 | 25.856091 | 26.880593 | 27.993643 | 29.569330 | 1.099761 | 3869.788602 | 0.110783351 | 1.23836411 | 0.215581082 | 0.982008277 | 11.847878 | 11.984946 | 11.852326 | 11.901309 | 11.934289 | 11.976873 | 4118 | 4096 | 3435 | 3897 |
| 65220 | NM_00119899 | NADK | NAD kinase | protein_coding | NP_001185922.1;NP_001185923.1;NP_001185924.1;NP_001340570.1;NI | 29831 | 611616 | . | . | 22.775762 | 22.511099 | 22.413935 | 27.667829 | 53.000132 | 52.035413 | 50.296220 | 63.259205 | 1.060940 | 2350.061152 | 0.123167053 | 0.692902591 | 0.488370684 | 0.998865404 | 11.155238 | 11.240469 | 11.201790 | 11.144521 | 11.171552 | 11.268440 | 2720 | 2373 | 1978 | 2404 |
| 65243 | NM_00136956 | ZFP69B | ZFP69 zinc finger protein B | protein_coding | NP_001356494.1;NP_075558.2;XP_005271193.1;XP_005271195.1;XP_00 | 28053 | . | . | . | 0.378932 | 0.231503 | 0.826880 | 0.399327 | 0.881792 | 0.535129 | 1.855495 | 0.913013 | 1.052524 | 47.632963 | 0.523717768 | 0.141017612 | 0.887856029 | 0.998865404 | 5.533720 | 5.612968 | 5.588944 | 5.517795 | 5.637106 | 5.482631 | 64 | 40 | 61 | 26 |
| 65244 | NM_00129328 | SPATS2 | spermatogenesis associated serine rich 2 | protein_coding | NP_001280214.1;NP_001280215.1;NP_075559.2;XP_011536993.1;XP_01 | 18650 | 611667 | . | . | 11.366813 | 12.267678 | 12.830693 | 14.147458 | 26.451039 | 28.357286 | 28.791701 | 32.346482 | 1.156370 | 1315.896523 | 0.132330512 | 1.583933885 | 0.113208765 | 0.850211417 | 10.253394 | 10.462683 | 10.262952 | 10.341534 | 10.398098 | 10.428651 | 1332 | 1386 | 1203 | 1349 |
| 65249 | NM_00136783 | ZSWIM4 | zinc finger SWIM-type containing 4 | protein_coding | NP_001354763.1;NP_075560.2;XP_016882642.1;XP_016882643.1;XP_01 | 25704 | . | . | . | 0.574333 | 0.469956 | 0.299853 | 0.643580 | 1.336497 | 1.086325 | 0.672862 | 1.471471 | -1.177220 | 70.133027 | 0.438110889 | -0.53727024 | 0.591080958 | 0.998865404 | 6.246341 | 6.007859 | 6.176240 | 6.113151 | 6.016486 | 6.188632 | 98 | 72 | 34 | 84 |
| 65250 | NM_023073,XI | CPLANE1 | ciliogenesis and planar polarity effector 1 | protein_coding | NP_075561.3;XP_005248402.1;XP_005248403.1;XP_005248404.1;XP_00 | 25801 | 614571 | . | . | 2.031614 | 2.132606 | 2.404125 | 1.774251 | 4.727654 | 4.929614 | 5.394788 | 4.056618 | 1.058933 | 759.810270 | 0.160878365 | 0.513501012 | 0.607600903 | 0.998865404 | 9.527356 | 9.610438 | 9.520652 | 9.572722 | 9.658115 | 9.510883 | 818 | 826 | 751 | 656 |
| 65251 | NM_023074 | ZNF649 | zinc finger protein 649 | protein_coding | NP_075562.2 | 25741 | 611903 | . | . | 1.510953 | 1.286290 | 1.844690 | 1.366657 | 3.516049 | 2.973316 | 4.139430 | 3.124699 | 1.132225 | 149.797704 | 0.28438811 | 0.629988742 | 0.52870195 | 0.998865404 | 7.133373 | 7.314678 | 7.227322 | 7.164767 | 7.305006 | 7.181047 | 175 | 139 | 160 | 126 |
| 65258 | NM_00124290 | MPPE1 | metallophosphoesterase 1 | protein_coding | NP_001229833.1;NP_001306083.1;NP_001317492.1;NP_075563.3;XP_00 | 15988 | 611900 | . | . | 6.953402 | 7.046249 | 6.907308 | 6.831380 | 16.180851 | 16.287719 | 15.499795 | 15.619144 | -1.037811 | 634.943773 | 0.155264922 | -0.34485461 | 0.730203673 | 0.998865404 | 9.337144 | 9.283324 | 9.313066 | 9.332706 | 9.282103 | 9.313486 | 734 | 708 | 529 | 598 |
| 65260 | NM_023077 | COA7 | cytochrome c oxidase assembly factor 7 (putative) | protein_coding | NP_075565.2 | 25716 | 615623 | . | . | 7.525035 | 8.707881 | 8.290633 | 8.012651 | 17.511066 | 20.128658 | 18.603941 | 18.319975 | -1.008999 | 1010.890569 | 0.142920845 | -0.09043089 | 0.927944807 | 0.998865404 | 9.987940 | 9.974853 | 9.932008 | 10.033444 | 9.993031 | 9.961444 | 1089 | 1171 | 894 | 923 |
| 65263 | NM_00132986 | PYCR3 | pyrroline-5-carboxylate reductase 3 | protein_coding | NP_001316795.2;NP_075566.3 | 25846 | 616408 | . | . | 4.147462 | 4.452697 | 3.931855 | 4.275950 | 9.651313 | 10.292609 | 8.822968 | 9.776452 | -1.063048 | 438.442944 | 0.176659799 | -0.49930154 | 0.617566968 | 0.998865404 | 8.819935 | 8.731191 | 8.774909 | 8.814058 | 8.735188 | 8.778852 | 504 | 503 | 356 | 413 |
| 65264 | NM_023079 | UBE2Z | ubiquitin conjugating enzyme E2 Z | protein_coding | NP_075567.2 | 25847 | 611362 | . | . | 27.755234 | 29.694332 | 30.081821 | 33.592918 | 64.587563 | 68.639778 | 67.502739 | 76.806213 | 1.093067 | 2908.215579 | 0.117610051 | 1.091585637 | 0.275015272 | 0.998865404 | 11.440413 | 11.568579 | 11.443863 | 11.491569 | 11.498520 | 11.580943 | 3106 | 3087 | 2509 | 2990 |
| 65265 | NM_023080,XI | C8orf33 | chromosome 8 open reading frame 33 | protein_coding | NP_075568.1;XP_024303002.1 | 26104 | . | . | . | 13.103166 | 11.163687 | 12.369320 | 13.402265 | 30.491603 | 25.805362 | 27.756397 | 30.642686 | 1.082859 | 1174.800962 | 0.134711834 | 0.852529304 | 0.393920382 | 0.998865404 | 10.139581 | 10.254531 | 10.201973 | 10.130771 | 10.212909 | 10.240399 | 1361 | 1159 | 1043 | 1166 |
| 65266 | NM_00132129 | WNK4 | WNK lysine deficient protein kinase 4 | protein_coding | NP_001308228.1;NP_115763.2;XP_016880451.1;XP_016880455.1 | 14544 | 601844 | . | . | 3.018905 | 3.183648 | 2.287279 | 2.866639 | 7.025112 | 7.359146 | 5.132588 | 6.554231 | -1.140796 | 375.735537 | 0.189784873 | -1.00134741 | 0.31665888 | 0.998865404 | 8.645747 | 8.455107 | 8.603406 | 8.577566 | 8.470657 | 8.555346 | 471 | 423 | 282 | 354 |
| 65267 | NM_00100283 | WNK3 | WNK lysine deficient protein kinase 3 | protein_coding | NP_001002838.1;NP_065973.2;XP_006724656.1;XP_011529104.1;XP_01 | 14543 | 300358 | . | . | 0.422192 | 0.474428 | 0.459046 | 0.528581 | 0.982459 | 1.096661 | 1.030085 | 1.208537 | 1.078519 | 166.579083 | 0.259018907 | 0.421017545 | 0.67374227 | 0.998865404 | 7.325095 | 7.433014 | 7.338753 | 7.386003 | 7.367697 | 7.419008 | 174 | 183 | 141 | 172 |
| 65268 | NM_00128239 | WNK2 | WNK lysine deficient protein kinase 2 | protein_coding | NP_001269323.1;NP_006639.3;XP_005252194.1;XP_005252197.1;XP_00 | 14542 | 606249 | . | . | 0.024828 | 0.026161 | 0.014445 | 0.014070 | 0.057776 | 0.060471 | 0.032415 | 0.032168 | -1.495083 | 9.669752 | 0.988498354 | -0.58697702 | 0.557219142 | NA | 3.533720 | 2.955730 | 3.299158 | 3.278082 | 3.272959 | 3.270237 | 16 | 10 | 7 | 7 |
| 65977 | NM_019091,XI | PLEKHA3 | pleckstrin homology domain containing A3 | protein_coding | NP_061964.3;XP_011509991.1 | 14338 | 607774 | . | . | 1.346309 | 1.038772 | 1.254396 | 1.073357 | 3.132914 | 2.401168 | 2.814828 | 2.454102 | -1.037877 | 509.448628 | 0.190041617 | -0.28222828 | 0.777768476 | 0.998865404 | 9.018765 | 8.966344 | 9.084318 | 8.911187 | 9.030092 | 8.922884 | 677 | 485 | 472 | 429 |
| 65979 | NM_00104818 | PHACTR4 | phosphatase and actin regulator 4 | protein_coding | NP_001041648.1;NP_001337087.1;NP_001337088.1;NP_001337089.1;NI | 25793 | 608726 | . | . | 6.784022 | 7.410311 | 7.642149 | 7.539847 | 15.786701 | 17.129267 | 17.148764 | 17.238966 | 1.055420 | 1408.016083 | 0.128515186 | 0.605508531 | 0.544841158 | 0.998865404 | 10.420072 | 10.497778 | 10.406132 | 10.468887 | 10.486428 | 10.472216 | 1513 | 1539 | 1269 | 1343 |
| 65980 | NM_00100987 | BRD9 | bromodomain containing 9 | protein_coding | NP_001009877.2;NP_001304880.1;NP_001362790.1;NP_001362791.1;NI | 25818 | 618465 | . | . | 8.893155 | 9.895518 | 9.855738 | 9.707279 | 20.694738 | 22.873934 | 22.115990 | 22.194534 | 1.041379 | 823.310989 | 0.145424481 | 0.402237783 | 0.687509038 | 0.998865404 | 9.655795 | 9.714201 | 9.642571 | 9.697289 | 9.710161 | 9.687734 | 893 | 904 | 742 | 775 |
| 65981 | NM_00100225 | CAPRIN2 | caprin family member 2 | protein_coding | NP_001002259.1;NP_001193785.1;NP_001306771.1;NP_001306772.1;NI | 21259 | 610375 | . | . | 21.064847 | 21.686846 | 22.218308 | 20.404447 | 49.018771 | 50.130102 | 49.857244 | 46.652344 | -1.006775 | 2863.091458 | 0.122097627 | -0.07978069 | 0.936411683 | 0.998865404 | 11.488150 | 11.478550 | 11.459595 | 11.510400 | 11.535543 | 11.420996 | 3205 | 3196 | 2656 | 2487 |
| 65985 | NM_00131983 | AACS | acetoacetyl-CoA synthetase | protein_coding | NP_001306768.1;NP_001306769.1;NP_076417.2;XP_011536994.1 | 21298 | 614364 | . | . | 7.544967 | 7.309313 | 6.988834 | 8.404709 | 17.557450 | 16.895802 | 15.682742 | 19.216365 | 1.016772 | 749.067741 | 0.155841011 | 0.153981854 | 0.877624043 | 0.998865404 | 9.537156 | 9.560653 | 9.552481 | 9.532115 | 9.493202 | 9.611004 | 868 | 790 | 601 | 768 |
| 65986 | NM_00110553 | ZBTB10 | zinc finger and BTB domain containing 10 | protein_coding | NP_001099009.1;NP_001264074.1;NP_076418.3 | 30953 | 618576 | . | . | 1.275173 | 1.240598 | 1.082452 | 1.043024 | 2.967381 | 2.867699 | 2.428990 | 2.384750 | -1.204797 | 359.611305 | 0.188223174 | -1.42803937 | 0.153280528 | 0.917721153 | 8.618268 | 8.349855 | 8.551199 | 8.531036 | 8.448805 | 8.420501 | 460 | 417 | 291 | 298 |
| 65987 | NM_00128240 | KCTD14 | potassium channel tetramerization domain containing 14 | protein_coding | NP_001269335.1;NP_076419.2 | 23295 | . | . | . | 6.012803 | 6.225429 | 5.819800 | 6.107304 | 13.992041 | 14.390358 | 13.059462 | 13.963624 | -1.040125 | 315.363879 | 0.196976948 | -0.2881434 | 0.773236974 | 0.998865404 | 8.329113 | 8.272070 | 8.303760 | 8.319716 | 8.280594 | 8.299716 | 365 | 352 | 264 | 295 |
| 65988 | NM_00130501 | ZNF747 | zinc finger protein 747 | protein_coding | NP_001291947.1;NP_001291948.1;NP_001291949.1;NP_076420.1 | 28350 | . | . | . | 3.286229 | 3.097267 | 4.181936 | 3.912007 | 7.647191 | 7.159472 | 9.384145 | 8.944338 | 1.252913 | 395.557251 | 0.183555277 | 1.772142394 | 0.076370923 | 0.767975096 | 8.455613 | 8.781507 | 8.574265 | 8.533284 | 8.711159 | 8.662519 | 419 | 365 | 398 | 396 |
| 65989 | NM_00128665 | DLK2 | delta like non-canonical Notch ligand 2 | protein_coding | NP_001273584.1;NP_001273585.1;NP_076421.2;NP_996262.1;XP_00524 | 21113 | . | . | . | 2.299496 | 2.336252 | 2.727032 | 3.394853 | 5.351024 | 5.400351 | 6.119382 | 7.761929 | 1.176750 | 167.397958 | 0.258719877 | 0.907577288 | 0.364101598 | 0.998865404 | 7.265516 | 7.499300 | 7.342051 | 7.356639 | 7.379496 | 7.452149 | 174 | 169 | 144 | 184 |
| 65990 | NM_00127128 | ANTKMT | adenine nucleotide translocase lysine methyltransferase | protein_coding | NP_001258214.1;NP_076422.1 | 14152 | 618566 | . | . | 9.138713 | 8.157702 | 8.418649 | 10.980661 | 21.266161 | 18.856893 | 18.891202 | 25.105978 | 1.103274 | 289.077179 | 0.21234051 | 0.667753401 | 0.504291017 | 0.998865404 | 8.103020 | 8.244152 | 8.165229 | 8.127695 | 8.142801 | 8.251582 | 328 | 286 | 237 | 313 |
| 65991 | NM_023934 | FUNDC2 | FUN14 domain containing 2 | protein_coding | NP_076423.2 | 24925 | 301042 | . | . | 3.315978 | 3.471948 | 3.256381 | 3.799361 | 7.716417 | 8.025564 | 7.307223 | 8.686789 | 1.025797 | 679.148171 | 0.157676144 | 0.233045544 | 0.815726041 | 0.998865404 | 9.389405 | 9.425534 | 9.385489 | 9.410735 | 9.365259 | 9.463215 | 758 | 737 | 555 | 691 |
| 65992 | NM_023935 | DDRGK1 | DDRGK domain containing 1 | protein_coding | NP_076424.1 | 16110 | 616177 | . | . | 46.913330 | 48.189243 | 47.006020 | 49.016285 | 109.169235 | 111.391594 | 105.480148 | 112.069908 | -1.004256 | 1939.874506 | 0.118793739 | -0.0515796 | 0.958863673 | 0.998865404 | 10.924858 | 10.918630 | 10.916093 | 10.930696 | 10.907623 | 10.932262 | 2218 | 2117 | 1657 | 1843 |
| 65993 | NM_00130090 | MRPS34 | mitochondrial ribosomal protein S34 | protein_coding | NP_001287829.1;NP_076425.1 | 16618 | 611994 | . | . | 56.847807 | 55.750805 | 56.117005 | 59.174876 | 132.287183 | 128.870479 | 125.924915 | 135.296320 | 1.010205 | 1776.382814 | 0.121252698 | 0.120801314 | 0.903848409 | 0.998865404 | 10.787414 | 10.802003 | 10.801372 | 10.779817 | 10.781000 | 10.815929 | 2063 | 1881 | 1518 | 1711 |
| 65996 | NR_026052 | CENPBD1P1 | CENPB DNA-binding domains containing 1 pseudogene 1 | transcribed_ps | . | 28421 | . | . | . | 4.427346 | 4.431936 | 4.496805 | 5.442520 | 10.302616 | 10.244619 | 10.090699 | 12.443676 | 1.106265 | 489.441386 | 0.175938643 | 0.828112217 | 0.407606951 | 0.998865404 | 8.860661 | 9.005680 | 8.902640 | 8.898783 | 8.905472 | 9.019838 | 537 | 500 | 407 | 525 |
| 65997 | NM_023940 | RASL11B | RAS like family 11 member B | protein_coding | NP_076429.1 | 23804 | 612404 | . | . | 0.257953 | 0.404214 | 0.365999 | 0.426487 | 0.600267 | 0.934358 | 0.821290 | 0.975112 | 1.198539 | 22.797168 | 0.653266077 | 0.399954452 | 0.689190065 | NA | 4.375822 | 4.634191 | 4.477674 | 4.514765 | 4.506114 | 4.521137 | 19 | 27 | 20 | 25 |
| 65998 | NM_00114493 | C11orf95 | chromosome 11 open reading frame 95 | protein_coding | NP_001138408.1;XP_024304430.1 | 28449 | 615699 | . | . | 1.333761 | 1.311832 | 1.358266 | 1.815620 | 3.103717 | 3.032359 | 3.047909 | 4.151200 | 1.063276 | 244.996563 | 0.218376977 | 0.405334488 | 0.685231659 | 0.998865404 | 7.891859 | 7.980029 | 7.926779 | 7.914235 | 7.928940 | 7.972287 | 277 | 253 | 210 | 247 |
| 65999 | NM_00114292 | LRRC61 | leucine rich repeat containing 61 | protein_coding | NP_001136400.1;NP_001350362.1;NP_001350363.1;NP_076431.1 | 21704 | . | . | . | 19.438550 | 17.083461 | 11.913895 | 15.129008 | 45.234303 | 39.489189 | 26.734434 | 34.590682 | -1.376453 | 905.656626 | 0.161457992 | -2.85495274 | 0.004304323 | 0.193526306 | 10.035160 | 9.573744 | 9.978386 | 9.866801 | 9.613471 | 9.768761 | 1297 | 1049 | 587 | 796 |
| 66000 | NM_00113646 | TMEM108 | transmembrane protein 108 | protein_coding | NP_001129941.1;NP_001269794.1;NP_076432.1 | 28451 | 617361 | . | . | 2.726149 | 2.977800 | 2.023210 | 2.314945 | 6.343861 | 6.883317 | 4.540026 | 5.292849 | -1.324438 | 296.505020 | 0.206430346 | -1.96376074 | 0.049557845 | 0.664973499 | 8.400827 | 7.994480 | 8.260178 | 8.301616 | 8.098445 | 8.163138 | 375 | 378 | 207 | 255 |
| 66005 | NM_00114267 | CHID1 | chitinase domain containing 1 | protein_coding | NP_001136146.1;NP_001136147.1;NP_001136148.1;NP_001136149.1;NI | 28474 | 615692 | . | . | 27.365875 | 28.799367 | 26.291403 | 24.991558 | 63.681517 | 66.571028 | 58.997143 | 57.140225 | -1.100185 | 1881.557131 | 0.122161944 | -1.12757042 | 0.259501385 | 0.995760118 | 10.945041 | 10.807085 | 10.912062 | 10.916314 | 10.804097 | 10.872083 | 2264 | 2133 | 1490 | 1753 |
| 66008 | NM_015049,XI | TRAK2 | trafficking kinesin protein 2 | protein_coding | NP_055864.2;XP_011509992.1;XP_016860261.1 | 13206 | 607334 | . | . | 15.588736 | 17.156029 | 19.133694 | 19.253855 | 36.275633 | 39.656930 | 42.935454 | 44.021647 | 1.156713 | 3539.879592 | 0.112470333 | 1.867436097 | 0.061840713 | 0.723723143 | 11.680697 | 11.890641 | 11.690015 | 11.760769 | 11.847608 | 11.844536 | 3616 | 3696 | 3308 | 3553 |
| 66035 | NM_00102493 | SLC2A11 | solute carrier family 2 member 11 | protein_coding | NP_001020109.1;NP_001020110.1;NP_001269793.1;NP_110434.3 | 14239 | 610367 | . | . | 18.137355 | 19.394001 | 24.176785 | 20.598771 | 42.206372 | 44.830103 | 54.252007 | 47.096638 | 1.142074 | 1823.827096 | 0.132783221 | 1.443371871 | 0.148915744 | 0.909660767 | 10.733611 | 10.925519 | 10.757773 | 10.794885 | 10.949664 | 10.808146 | 1914 | 1881 | 1841 | 1662 |
| 66036 | NM_015458,XI | MTMR9 | myotubularin related protein 9 | protein_coding | NP_056273.2;XP_011542132.1;XP_011542133.1;XP_016869242.1 | 14596 | 606260 | . | . | 1.784318 | 1.537646 | 1.519242 | 1.450453 | 4.152181 | 3.554338 | 3.409134 | 3.316289 | -1.130667 | 348.082802 | 0.196385367 | -0.90217815 | 0.366962237 | 0.998865404 | 8.528790 | 8.352395 | 8.523475 | 8.427294 | 8.422037 | 8.391831 | 460 | 366 | 292 | 298 |
| 78987 | NM_00103171 | CRELD1 | cysteine rich with EGF like domains 1 | protein_coding | NP_001026887.2;NP_001070883.2;NP_001361245.1;NP_001361246.1;NI | 14630 | 607170 | . | . | 8.802179 | 9.699824 | 9.109698 | 9.641428 | 20.483029 | 22.421576 | 20.441898 | 22.043982 | -1.009185 | 680.737210 | 0.154689384 | -0.08527223 | 0.932044968 | 0.998865404 | 9.417726 | 9.404150 | 9.377704 | 9.448559 | 9.397970 | 9.416753 | 747 | 776 | 581 | 644 |
| 78988 | NM_024026,XI | MRPL57 | mitochondrial ribosomal protein L57 | protein_coding | NP_076931.1;XP_016876229.1 | 14514 | 611997 | . | . | 14.463712 | 13.375145 | 13.125489 | 14.886146 | 33.657653 | 30.917245 | 29.453217 | 34.035403 | -1.048390 | 895.625253 | 0.14334969 | -0.47559065 | 0.634366041 | 0.998865404 | 9.840570 | 9.772121 | 9.833235 | 9.813548 | 9.753862 | 9.823659 | 1070 | 976 | 723 | 860 |
| 78990 | NM_023112 | OTUB2 | OTU deubiquitinase, ubiquitin aldehyde binding 2 | protein_coding | NP_075601.1 | 20351 | 608338 | . | . | 0.454114 | 0.701787 | 0.841746 | 0.938568 | 1.056743 | 1.622211 | 1.888854 | 2.145924 | 1.520828 | 89.800101 | 0.355734257 | 1.700306035 | 0.089073376 | 0.802216049 | 6.156194 | 6.758652 | 6.358312 | 6.461899 | 6.515079 | 6.544218 | 65 | 93 | 90 | 106 |
| 78991 | NM_00130105 | PCYOX1L | prenylcysteine oxidase 1 like | protein_coding | NP_001287983.1;NP_001287986.1;NP_076933.3;XP_011535984.1;XP_02 | 28477 | . | . | . | 10.480208 | 10.754990 | 8.863059 | 9.075415 | 24.387874 | 24.860640 | 19.888449 | 20.749860 | -1.196611 | 777.884193 | 0.145623221 | -1.77825045 | 0.07536273 | 0.766002767 | 9.727114 | 9.468099 | 9.657922 | 9.667979 | 9.535431 | 9.539255 | 969 | 921 | 614 | 666 |
| 78992 | NM_00132143 | YIPF2 | Yip1 domain family member 2 | protein_coding | NP_001308368.1;NP_001308369.1;NP_076934.1;XP_011526572.1;XP_01 | 28476 | 617522 | . | . | 20.008618 | 19.578856 | 19.380135 | 19.135514 | 46.560878 | 45.257406 | 43.488462 | 43.751082 | -1.048077 | 1271.372952 | 0.129156562 | -0.52451496 | 0.599920419 | 0.998865404 | 10.345573 | 10.277978 | 10.335693 | 10.323667 | 10.308908 | 10.279213 | 1510 | 1393 | 1101 | 1141 |
| 78994 | NM_00132046 | PRR14 | proline rich 14 | protein_coding | NP_001307393.1;NP_076936.1 | 28458 | 617423 | . | . | 8.540188 | 8.222788 | 9.601704 | 9.077449 | 19.873368 | 19.007343 | 21.545946 | 20.754507 | 1.121143 | 563.076601 | 0.163252618 | 1.010519232 | 0.31224659 | 0.998865404 | 9.052074 | 9.217558 | 9.113687 | 9.080564 | 9.201436 | 9.141954 | 627 | 558 | 541 | 533 |
| 78995 | NM_00117125 | HROB | homologous recombination factor with OB-fold | protein_coding | NP_001164722.1;NP_001308239.1;NP_001308240.1;NP_076937.2;XP_01 | 28460 | 618611 | . | . | 4.450274 | 3.881045 | 4.140607 | 4.576391 | 10.355970 | 8.971207 | 9.291404 | 10.463376 | -1.004041 | 339.864591 | 0.195823696 | -0.02970804 | 0.976299902 | 0.998865404 | 8.411755 | 8.405871 | 8.440769 | 8.377398 | 8.380685 | 8.433239 | 414 | 347 | 281 | 333 |
| 78996 | NM_00124374 | CYREN | cell cycle regulator of NHEJ | protein_coding | NP_001230678.1;NP_001230680.1;NP_001230681.1;NP_001230682.1;NI | 22432 | 616980 | . | . | 8.710562 | 11.019936 | 10.015912 | 11.873635 | 20.269831 | 25.473077 | 22.475414 | 27.147655 | 1.079639 | 1368.308342 | 0.129309784 | 0.854914947 | 0.392598235 | 0.998865404 | 10.361914 | 10.472329 | 10.400999 | 10.374314 | 10.415670 | 10.476000 | 1542 | 1395 | 1187 | 1384 |
| 78998 | NR_026785 | RHPN1-AS1 | RHPN1 antisense RNA 1 (head to head) | lncRNA | . | 28457 | . | . | . | 1.152736 | 1.063974 | 1.014512 | 1.643095 | 2.682465 | 2.459424 | 2.276535 | 3.756741 | 1.180639 | 77.067137 | 0.381940746 | 0.62723855 | 0.530502877 | 0.998865404 | 6.144352 | 6.381963 | 6.250841 | 6.229698 | 6.220142 | 6.341911 | 85 | 73 | 56 | 96 |
| 78999 | NM_00136352 | LRFN4 | leucine rich repeat and fibronectin type III domain containing 4 | protein_coding | NP_001350453.1;NP_076941.2 | 28456 | 612810 | . | . | 4.256877 | 4.654748 | 3.967642 | 4.569702 | 9.905928 | 10.759659 | 8.903273 | 10.448081 | -1.086922 | 382.682546 | 0.188359261 | -0.63840157 | 0.523212306 | 0.998865404 | 8.639288 | 8.518179 | 8.574327 | 8.632143 | 8.520738 | 8.587700 | 437 | 451 | 300 | 364 |
| 79000 | NM_00128749 | AUNIP | aurora kinase A and ninein interacting protein | protein_coding | NP_001274419.1;NP_076942.1 | 28363 | . | . | . | 5.145334 | 6.338264 | 4.646528 | 5.889065 | 11.973403 | 14.651181 | 10.426674 | 13.464647 | -1.075871 | 361.782265 | 0.19973191 | -0.52823041 | 0.597339428 | 0.998865404 | 8.551393 | 8.444587 | 8.478105 | 8.557192 | 8.417426 | 8.531769 | 403 | 432 | 272 | 360 |
| 79001 | NM_00131131 | VKORC1 | vitamin K epoxide reductase complex subunit 1 | protein_coding | NP_001298240.1;NP_076869.1;NP_996560.1 | 23663 | 608547 | . | . | 134.596923 | 139.424694 | 121.049399 | 127.188887 | 313.212570 | 322.286421 | 271.631350 | 290.802297 | -1.112199 | 3541.364867 | 0.110155101 | -1.39271605 | 0.163705673 | 0.939554848 | 11.864805 | 11.711293 | 11.823080 | 11.841528 | 11.724410 | 11.764469 | 4247 | 4069 | 2844 | 3221 |
| 79002 | NM_00132973 | TRIR | telomerase RNA component interacting RNase | protein_coding | NP_001316667.1;NP_001316668.1;NP_076943.1 | 28424 | . | . | . | 104.174035 | 102.140070 | 92.483441 | 110.486174 | 242.417260 | 236.101336 | 207.530162 | 252.613515 | -1.028662 | 2819.403286 | 0.123831141 | -0.32922718 | 0.741983978 | 0.998865404 | 11.481518 | 11.440539 | 11.483322 | 11.460240 | 11.380106 | 11.513082 | 3339 | 3042 | 2219 | 2819 |
| 79003 | NM_00125821 | MIS12 | MIS12 kinetochore complex component | protein_coding | NP_001245146.1;NP_001245147.1;NP_001245148.1;NP_001245149.1;NI | 24967 | 609178 | . | . | 12.069296 | 11.399753 | 10.885186 | 8.922960 | 28.085746 | 26.351036 | 24.426042 | 20.401288 | -1.142379 | 927.142861 | 0.154014044 | -1.2469088 | 0.21243094 | 0.975238634 | 9.949140 | 9.757817 | 9.912195 | 9.890295 | 9.880406 | 9.723922 | 1155 | 1051 | 836 | 722 |
| 79004 | NM_024040 | CUEDC2 | CUE domain containing 2 | protein_coding | NP_076945.2 | 28352 | 614142 | . | . | 23.108561 | 21.828121 | 21.569443 | 26.391397 | 53.774574 | 50.456684 | 48.401207 | 60.340797 | 1.051803 | 796.096455 | 0.156739497 | 0.464877026 | 0.642019524 | 0.998865404 | 9.600131 | 9.672558 | 9.640220 | 9.594138 | 9.583230 | 9.718201 | 923 | 810 | 642 | 838 |
| 79005 | NM_00120485 | SCNM1 | sodium channel modifier 1 | protein_coding | NP_001191785.1;NP_076946.1 | 23136 | 608095 | . | . | 6.623476 | 8.342095 | 7.806001 | 7.341711 | 15.413101 | 19.283125 | 17.516440 | 16.785950 | 1.003149 | 453.750625 | 0.185792311 | 0.024410661 | 0.980525045 | 0.998865404 | 8.823653 | 8.827856 | 8.751597 | 8.889320 | 8.847402 | 8.801393 | 464 | 543 | 410 | 410 |
| 79006 | NM_024042 | METRN | meteorin, glial cell differentiation regulator | protein_coding | NP_076947.1 | 14151 | 610998 | . | . | 4.722682 | 5.021099 | 3.480573 | 4.309271 | 10.989875 | 11.606495 | 7.810306 | 9.852635 | -1.267583 | 454.577852 | 0.182782703 | -1.87151076 | 0.06127432 | 0.722879863 | 8.989777 | 8.646635 | 8.878696 | 8.912562 | 8.688789 | 8.806852 | 570 | 563 | 313 | 414 |
| 79007 | NM_00104261 | DBNDD1 | dysbindin domain containing 1 | protein_coding | NP_001036075.1;NP_001275637.1;NP_001275638.2;NP_001358510.1;NI | 28455 | . | . | . | 7.819863 | 7.272656 | 7.022628 | 9.172295 | 18.197139 | 16.811070 | 15.758574 | 20.971363 | 1.062005 | 498.965163 | 0.183766799 | 0.4722855 | 0.636723019 | 0.998865404 | 8.919117 | 9.005190 | 8.963125 | 8.918806 | 8.894876 | 9.057155 | 577 | 504 | 390 | 543 |
| 79008 | NM_024044,N | SLX1B | SLX1 homolog B, structure-specific endonuclease subunit | protein_coding | NP_076949.1;NP_835145.1 | 28748 | 615823 | . | . | 11.440882 | 12.634350 | 9.856953 | 13.398513 | 26.623400 | 29.204866 | 22.118717 | 30.634107 | -1.044754 | 425.586888 | 0.197129638 | -0.32041374 | 0.748654715 | 0.998865404 | 8.765212 | 8.700686 | 8.714238 | 8.773290 | 8.626804 | 8.800744 | 477 | 492 | 309 | 447 |
| 79009 | NM_024045,XI | DDX50 | DExD-box helicase 50 | protein_coding | NP_076950.1;XP_005270205.1;XP_011538446.1;XP_016872115.1;XP_01 | 17906 | 610373 | . | . | 10.332404 | 10.239135 | 9.355171 | 9.225593 | 24.043929 | 23.668215 | 20.992730 | 21.093224 | -1.081008 | 848.751520 | 0.142769454 | -0.78711995 | 0.43121165 | 0.998865404 | 9.784313 | 9.671894 | 9.768973 | 9.743764 | 9.688766 | 9.712971 | 1033 | 935 | 697 | 778 |
| 79016 | NM_024050,XI | DDA1 | DET1 and DDB1 associated 1 | protein_coding | NP_076955.1;XP_024307469.1 | 28360 | . | . | . | 8.062732 | 8.050893 | 7.857852 | 9.635373 | 18.762308 | 18.610001 | 17.632792 | 22.030134 | 1.070243 | 1058.190284 | 0.146677289 | 0.667717897 | 0.504313683 | 0.998865404 | 9.997838 | 10.095284 | 10.023967 | 10.016974 | 9.995767 | 10.140228 | 1183 | 1098 | 859 | 1125 |
| 79017 | NM_00119981 | GGCT | gamma-glutamylcyclotransferase | protein_coding | NP_001186744.1;NP_001186745.1;NP_001186746.1;NP_076956.1 | 21705 | 137170 | . | . | 38.749691 | 34.921736 | 37.111567 | 38.386554 | 90.172122 | 80.723162 | 83.277277 | 87.766294 | 1.004882 | 1302.732150 | 0.133202732 | 0.052746891 | 0.957933577 | 0.998865404 | 10.343726 | 10.350914 | 10.388260 | 10.299922 | 10.340308 | 10.357213 | 1583 | 1321 | 1123 | 1238 |
| 79018 | NM_024052,XI | GID4 | GID complex subunit 4 homolog | protein_coding | NP_076957.3;XP_005256856.1;XP_005256857.1;XP_011522310.1 | 28453 | 617699 | . | . | 2.897171 | 3.181217 | 2.726239 | 3.119719 | 6.741835 | 7.353526 | 6.117600 | 7.132865 | -1.041846 | 358.202736 | 0.19521185 | -0.30296138 | 0.761919291 | 0.998865404 | 8.514388 | 8.454251 | 8.481716 | 8.509540 | 8.419221 | 8.522350 | 411 | 404 | 277 | 359 |
| 79019 | NM_00100287 | CENPM | centromere protein M | protein_coding | NP_001002876.1;NP_001103685.1;NP_001291299.1;NP_001291300.1;NI | 18352 | 610152 | . | . | 15.918066 | 18.089512 | 18.245179 | 20.407849 | 37.041995 | 41.814716 | 40.941651 | 46.660120 | 1.081368 | 545.076612 | 0.169307057 | 0.666584775 | 0.505037397 | 0.998865404 | 9.033173 | 9.145280 | 9.017958 | 9.106267 | 9.083421 | 9.142989 | 562 | 604 | 470 | 555 |
| 79020 | NM_00109985 | C7orf25 | chromosome 7 open reading frame 25 | protein_coding | NP_001093328.1;NP_001350365.1;NP_076959.2;XP_016868085.1;XP_01 | 21703 | . | . | . | 7.338877 | 8.037313 | 6.018469 | 7.637002 | 17.077867 | 18.578608 | 13.505269 | 17.461098 | -1.127385 | 429.402482 | 0.187373394 | -0.92318802 | 0.355909231 | 0.998865404 | 8.830623 | 8.656500 | 8.751216 | 8.809048 | 8.645924 | 8.765765 | 500 | 514 | 315 | 417 |
| 79022 | NM_00114384 | TMEM106C | transmembrane protein 106C | protein_coding | NP_001137313.1;NP_001137314.1;NP_001137315.1;NP_076961.1;XP_02 | 28775 | . | . | . | 80.504665 | 79.791493 | 66.690088 | 74.159651 | 187.337675 | 184.441607 | 149.650621 | 169.557233 | -1.154941 | 3541.600757 | 0.113979552 | -1.82330506 | 0.068257201 | 0.741130004 | 11.890420 | 11.682464 | 11.852516 | 11.840405 | 11.684975 | 11.768523 | 4403 | 4067 | 2711 | 3240 |
| 79023 | NM_024057,XI | NUP37 | nucleoporin 37 | protein_coding | NP_076962.2 | 29929 | 609264 | . | . | 11.834617 | 11.403241 | 9.851073 | 10.155317 | 27.539639 | 26.359103 | 22.105522 | 23.218924 | -1.170231 | 793.505599 | 0.145399197 | -1.55980013 | 0.118807121 | 0.858706505 | 9.741036 | 9.514255 | 9.699235 | 9.670283 | 9.567107 | 9.581628 | 1005 | 905 | 629 | 693 |
| 79026 | NM_00134644 | AHNAK | AHNAK nucleoprotein | protein_coding | NP_001333374.1;NP_001333375.1;NP_001611.1;NP_076965.2;XP_01687 | 347 | 103390 | . | . | 45.419220 | 47.073317 | 46.826759 | 43.616810 | 105.692391 | 108.812073 | 105.077891 | 99.724654 | -1.034985 | 26405.272236 | 0.101901712 | -0.48684185 | 0.62637041 | 0.998865404 | 14.713123 | 14.663527 | 14.690849 | 14.714941 | 14.707123 | 14.638405 | 30497 | 29386 | 23469 | 23357 |
| 79027 | NM_00100995 | ZNF655 | zinc finger protein 655 | protein_coding | NP_001009958.1;NP_001009960.1;NP_001077425.1;NP_001078835.1;NI | 30899 | 617891 | . | . | 18.218201 | 19.467684 | 21.524460 | 16.931572 | 42.394495 | 45.000430 | 48.300265 | 38.712030 | -1.027672 | 1463.413432 | 0.179259353 | -0.21967985 | 0.826120498 | 0.998865404 | 10.534372 | 10.495611 | 10.525979 | 10.512053 | 10.657218 | 10.309627 | 1723 | 1586 | 1537 | 1050 |
| 79029 | NM_00132364 | SPATA5L1 | spermatogenesis associated 5 like 1 | protein_coding | NP_001310569.1;NP_076968.2 | 28762 | . | . | . | 2.760363 | 2.713873 | 2.499962 | 2.288207 | 6.423478 | 6.273241 | 5.609845 | 5.231716 | -1.159099 | 208.349766 | 0.234978809 | -0.90648243 | 0.364680558 | 0.998865404 | 7.805057 | 7.592876 | 7.740498 | 7.730934 | 7.696587 | 7.638584 | 260 | 239 | 179 | 169 |
| 79031 | NM_024065,XI | PDCL3 | phosducin like 3 | protein_coding | NP_076970.1;XP_016860379.1 | 28860 | 611678 | . | . | 29.903475 | 29.601304 | 25.071346 | 33.445798 | 69.586615 | 68.424736 | 56.259374 | 76.469842 | -1.007971 | 1039.785308 | 0.154778763 | -0.07400665 | 0.941005091 | 0.998865404 | 10.028075 | 10.016040 | 10.033514 | 10.012891 | 9.920458 | 10.105697 | 1219 | 1111 | 787 | 1093 |
| 79033 | NM_00130169 | ERI3 | ERI1 exoribonuclease family member 3 | protein_coding | NP_001288627.1;NP_001288628.1;NP_001288629.1;NP_001288630.1;NI | 17276 | 609917 | . | . | 20.574161 | 22.603853 | 21.309440 | 24.426482 | 47.876920 | 52.249819 | 47.817767 | 55.848245 | 1.048942 | 1112.388307 | 0.142659393 | 0.483215367 | 0.628942825 | 0.998865404 | 10.084848 | 10.153231 | 10.077255 | 10.123106 | 10.074291 | 10.193943 | 1212 | 1208 | 911 | 1153 |
| 79034 | NM_00130303 | C7orf26 | chromosome 7 open reading frame 26 | protein_coding | NP_001289968.1;NP_076972.2;XP_005249902.1;XP_011513820.1;XP_01 | 21702 | . | . | . | 7.449066 | 7.492429 | 7.714381 | 7.377224 | 17.334284 | 17.319088 | 17.310846 | 16.867147 | 1.004634 | 503.503117 | 0.167678691 | 0.039781839 | 0.968267056 | 0.998865404 | 8.972320 | 8.979385 | 8.985799 | 8.962276 | 8.997172 | 8.956682 | 589 | 532 | 453 | 458 |
| 79035 | NM_024068,XI | NABP2 | nucleic acid binding protein 2 | protein_coding | NP_076973.1;XP_005269204.1;XP_005269206.1;XP_024304952.1 | 28412 | 612104 | . | . | 18.783083 | 18.681305 | 17.031962 | 21.147734 | 43.709002 | 43.182672 | 38.219228 | 48.351781 | 1.004900 | 841.985308 | 0.155911047 | 0.04522695 | 0.963926413 | 0.998865404 | 9.714438 | 9.720857 | 9.715924 | 9.713527 | 9.640081 | 9.790490 | 969 | 905 | 656 | 875 |
| 79036 | NM_00117194 | KXD1 | KxDL motif containing 1 | protein_coding | NP_001165419.1;NP_001165420.1;NP_076974.2;XP_005260130.1;XP_00 | 28420 | 615178 | . | . | 34.048041 | 36.210548 | 34.112821 | 40.085084 | 79.231197 | 83.702306 | 76.548189 | 91.649777 | 1.069209 | 1586.652468 | 0.129989352 | 0.74270646 | 0.457659418 | 0.998865404 | 10.582876 | 10.679062 | 10.589436 | 10.619386 | 10.604286 | 10.706085 | 1732 | 1687 | 1333 | 1637 |
| 79037 | NM_024070,XI | PVRIG | PVR related immunoglobulin domain containing | protein_coding | NP_076975.2;XP_011514877.1 | 32190 | 617012 | . | . | 2.021276 | 2.658754 | 3.456566 | 2.314163 | 4.703592 | 6.145828 | 7.756434 | 5.291060 | 1.223458 | 129.187479 | 0.315760856 | 0.921469508 | 0.356805354 | 0.998865404 | 6.860075 | 7.151843 | 6.918281 | 7.009265 | 7.110885 | 6.957112 | 116 | 142 | 149 | 106 |
| 79038 | NM_00119895 | ZFYVE21 | zinc finger FYVE-type containing 21 | protein_coding | NP_001185882.1;NP_076976.1 | 20760 | 613504 | . | . | 5.690085 | 5.845481 | 6.643105 | 6.154723 | 13.241062 | 13.512092 | 14.906938 | 14.072041 | 1.093129 | 264.817442 | 0.212250545 | 0.605245088 | 0.545016161 | 0.998865404 | 7.983021 | 8.111815 | 8.014703 | 8.033376 | 8.093819 | 8.044858 | 286 | 278 | 252 | 247 |
| 79039 | NM_00111132 | DDX54 | DEAD-box helicase 54 | protein_coding | NP_001104792.1;NP_076977.3 | 20084 | 611665 | . | . | 14.302632 | 15.368896 | 14.201119 | 15.074653 | 33.282812 | 35.525893 | 31.866898 | 34.466402 | -1.027716 | 2010.752567 | 0.120397243 | -0.32759149 | 0.743220557 | 0.998865404 | 10.993205 | 10.953563 | 10.958600 | 11.009107 | 10.942703 | 10.981648 | 2273 | 2269 | 1682 | 1905 |
| 79041 | NM_024074 | TMEM38A | transmembrane protein 38A | protein_coding | NP_076979.1 | 28462 | 611235 | . | . | 2.065239 | 1.759302 | 1.819093 | 1.994035 | 4.805897 | 4.066705 | 4.081992 | 4.559124 | -1.018827 | 157.270128 | 0.266395961 | -0.10101136 | 0.91954144 | 0.998865404 | 7.310388 | 7.283691 | 7.333913 | 7.265970 | 7.277142 | 7.308780 | 198 | 157 | 130 | 152 |
| 79042 | NM_00107744 | TSEN34 | tRNA splicing endonuclease subunit 34 | protein_coding | NP_001070914.1;NP_001269261.1;NP_001269262.1;NP_076980.2;XP_00 | 15506 | 608754 | . | . | 25.420176 | 22.717227 | 20.903015 | 26.335917 | 59.153792 | 52.511888 | 46.905758 | 60.213947 | -1.035069 | 1511.427722 | 0.145442945 | -0.34189576 | 0.73242934 | 0.998865404 | 10.586436 | 10.536525 | 10.630360 | 10.512626 | 10.467640 | 10.619635 | 1904 | 1534 | 1164 | 1530 |
| 79047 | NM_00112999 | KCTD15 | potassium channel tetramerization domain containing 15 | protein_coding | NP_001123466.1;NP_001123467.1;NP_076981.2;XP_011525598.2;XP_01 | 23297 | 615240 | . | . | 12.910237 | 12.483692 | 11.104499 | 12.756516 | 30.042648 | 28.856616 | 24.918175 | 29.166258 | -1.084448 | 1518.865889 | 0.129318605 | -0.90444204 | 0.365761056 | 0.998865404 | 10.626193 | 10.508985 | 10.609864 | 10.588237 | 10.491266 | 10.579409 | 1845 | 1682 | 1194 | 1445 |
| 79048 | NM_00128268 | SECISBP2 | SECIS binding protein 2 | protein_coding | NP_001269617.1;NP_001269618.1;NP_001269619.1;NP_001341625.1;NI | 30972 | 607693 | . | . | 5.768408 | 6.556502 | 6.081407 | 6.798998 | 13.423324 | 15.155647 | 13.646498 | 15.545100 | 1.071600 | 729.815423 | 0.153722696 | 0.649005882 | 0.516334574 | 0.998865404 | 9.460928 | 9.560142 | 9.445734 | 9.524830 | 9.508572 | 9.558456 | 765 | 804 | 633 | 733 |
| 79050 | NM_024078,XI | NOC4L | nucleolar complex associated 4 homolog | protein_coding | NP_076983.1;XP_016875449.1 | 28461 | 612819 | . | . | 10.608284 | 12.036952 | 11.981681 | 13.335962 | 24.685913 | 27.823954 | 26.886545 | 30.491094 | 1.102175 | 616.469079 | 0.162524002 | 0.863585829 | 0.387815451 | 0.998865404 | 9.196364 | 9.336027 | 9.192626 | 9.272145 | 9.264961 | 9.329993 | 635 | 671 | 535 | 635 |
| 79053 | NM_00100702 | ALG8 | ALG8 alpha-1,3-glucosyltransferase | protein_coding | NP_001007028.1;NP_076984.2;XP_005274304.1;XP_011543554.1;XP_01 | 23161 | 608103 | . | . | 34.086708 | 30.481991 | 33.629584 | 25.067970 | 79.321173 | 70.460487 | 75.463810 | 57.314932 | -1.073986 | 1703.121793 | 0.147424661 | -0.69849334 | 0.484868722 | 0.998865404 | 10.784292 | 10.681821 | 10.766732 | 10.750876 | 10.805019 | 10.589814 | 2051 | 1884 | 1626 | 1327 |
| 79054 | NM_024080,XI | TRPM8 | transient receptor potential cation channel subfamily M member 8 | protein_coding | NP_076985.4;XP_011510112.1;XP_016860380.1;XP_024308900.1;XP_02 | 17961 | 606678 | . | . | 2.334027 | 2.610008 | 1.470662 | 2.286981 | 5.431374 | 6.033149 | 3.300122 | 5.228909 | -1.348408 | 429.370482 | 0.193800051 | -2.22527 | 0.026063118 | 0.527491527 | 8.946134 | 8.513747 | 8.842098 | 8.811981 | 8.563870 | 8.722167 | 581 | 520 | 273 | 391 |
| 79056 | NM_024081,XI | PRRG4 | proline rich and Gla domain 4 | protein_coding | NP_076986.1;XP_006718376.3;XP_006718377.1 | 30799 | 611690 | . | . | 0.584370 | 0.618887 | 0.400771 | 0.266194 | 1.359853 | 1.430586 | 0.899318 | 0.608620 | -1.801683 | 64.327064 | 0.399425147 | -2.12641938 | 0.033468355 | 0.575719787 | 6.370187 | 5.521454 | 6.089498 | 6.043780 | 5.939161 | 5.937550 | 103 | 82 | 40 | 43 |
| 79058 | NM_00125188 | ASPSCR1 | ASPSCR1 tether for SLC2A4, UBX domain containing | protein_coding | NP_001238817.1;NP_001317457.1;NP_076988.1;XP_011521903.1;XP_01 | 13825 | 606236 | . | . | 8.664412 | 8.257516 | 9.613567 | 9.020521 | 20.162443 | 19.087620 | 21.572561 | 20.624347 | 1.105724 | 538.832320 | 0.163017024 | 0.889427302 | 0.373773478 | 0.998865404 | 8.999354 | 9.144387 | 9.030319 | 9.048447 | 9.114725 | 9.093911 | 581 | 560 | 500 | 522 |
| 79064 | NM_00108050 | TMEM223 | transmembrane protein 223 | protein_coding | NP_001073970.1 | 28464 | . | . | . | 11.658187 | 11.110785 | 12.754323 | 13.850275 | 27.129080 | 25.683075 | 28.620333 | 31.667007 | 1.151297 | 489.479949 | 0.170048209 | 1.195310915 | 0.231965577 | 0.986726781 | 8.829957 | 9.033110 | 8.905019 | 8.869912 | 8.952574 | 8.999472 | 539 | 477 | 439 | 509 |
| 79065 | NM_00107719 | ATG9A | autophagy related 9A | protein_coding | NP_001070666.1;NP_076990.4 | 22408 | 612204 | . | . | 23.791705 | 24.600047 | 27.025906 | 27.907263 | 55.364272 | 56.864114 | 60.645348 | 63.806640 | 1.119492 | 3018.102348 | 0.111317457 | 1.462881954 | 0.143499683 | 0.905681811 | 11.475745 | 11.638518 | 11.501007 | 11.521506 | 11.593240 | 11.613773 | 3238 | 3112 | 2738 | 3026 |
| 79066 | NM_024086,XI | METTL16 | methyltransferase like 16 | protein_coding | NP_076991.3;XP_005256858.1;XP_016880535.1;XP_024306696.1 | 28484 | . | . | . | 4.353979 | 3.846209 | 4.706441 | 4.481866 | 10.131889 | 8.890684 | 10.561119 | 10.247252 | 1.103942 | 774.836943 | 0.151793402 | 0.939861548 | 0.347288583 | 0.998865404 | 9.524378 | 9.667568 | 9.604043 | 9.515669 | 9.650835 | 9.609306 | 902 | 744 | 728 | 740 |
| 79068 | NM_00108043 | FTO | FTO alpha-ketoglutarate dependent dioxygenase | protein_coding | NP_001073901.1;NP_001350820.1;NP_001350823.1;NP_001350825.1;NI | 24678 | 610966 | . | . | 2.551655 | 2.259698 | 2.284027 | 3.307089 | 5.937802 | 5.223394 | 5.125288 | 7.561264 | 1.022162 | 712.830493 | 0.161311896 | 0.196039009 | 0.844579633 | 0.998865404 | 9.461857 | 9.492807 | 9.469466 | 9.467803 | 9.410166 | 9.552112 | 813 | 760 | 562 | 745 |
| 79070 | NM_00131873 | POGLUT2 | protein O-glucosyltransferase 2 | protein_coding | NP_001305661.1;NP_076994.2;XP_005254132.1 | 19350 | 611613 | . | . | 9.382045 | 8.530045 | 8.117768 | 8.480724 | 21.832404 | 19.717582 | 18.216034 | 19.390167 | -1.094847 | 549.770900 | 0.165244962 | -0.79112257 | 0.428872467 | 0.998865404 | 9.166471 | 9.035952 | 9.167225 | 9.094082 | 9.064975 | 9.079977 | 698 | 586 | 451 | 498 |
| 79071 | NM_00113072 | ELOVL6 | ELOVL fatty acid elongase 6 | protein_coding | NP_001124193.1;NP_076995.1;XP_011530535.1;XP_011530536.1 | 15829 | 611546 | . | . | 2.529818 | 2.363115 | 3.133235 | 3.199747 | 5.886990 | 5.462445 | 7.030889 | 7.315841 | 1.280376 | 571.925030 | 0.160857968 | 2.216658745 | 0.02664642 | 0.532994094 | 8.970318 | 9.327050 | 9.091115 | 9.048960 | 9.226625 | 9.238032 | 597 | 523 | 554 | 607 |
| 79072 | NM_024091,N | FASTKD3 | FAST kinase domains 3 | protein_coding | NP_076996.2;XP_006714561.1 | 28758 | 617530 | . | . | 4.167734 | 4.066214 | 4.567961 | 3.052801 | 9.698487 | 9.399234 | 10.250374 | 6.979868 | -1.104812 | 297.321151 | 0.230509397 | -0.62384237 | 0.532731108 | 0.998865404 | 8.285191 | 8.143066 | 8.241723 | 8.232940 | 8.282828 | 8.074670 | 362 | 334 | 295 | 212 |
| 79073 | NM_024092 | TMEM109 | transmembrane protein 109 | protein_coding | NP_076997.1 | 28771 | . | . | . | 35.728561 | 34.621758 | 31.732594 | 30.792080 | 83.141838 | 80.029739 | 71.207024 | 70.402428 | -1.141094 | 2204.749489 | 0.117296682 | -1.62338925 | 0.104506221 | 0.830227047 | 11.198402 | 11.008127 | 11.172989 | 11.140432 | 11.067035 | 11.035318 | 2760 | 2485 | 1827 | 1892 |
| 79074 | NM_00128653 | C2orf49 | chromosome 2 open reading frame 49 | protein_coding | NP_001273466.1;NP_076998.1;XP_016860381.1;XP_024308903.1;XP_02 | 28772 | . | . | . | 6.446389 | 6.598730 | 6.580987 | 6.286621 | 15.001012 | 15.253263 | 14.767547 | 14.373613 | -1.057214 | 915.566214 | 0.140248481 | -0.57232418 | 0.567102373 | 0.998865404 | 9.878039 | 9.797888 | 9.848403 | 9.867980 | 9.836341 | 9.799534 | 1069 | 1029 | 794 | 813 |
| 79075 | NM_024094,XI | DSCC1 | DNA replication and sister chromatid cohesion 1 | protein_coding | NP_076999.2;XP_005251122.1 | 24453 | 613203 | . | . | 11.632330 | 12.232625 | 11.051756 | 9.996675 | 27.068911 | 28.276263 | 24.799820 | 22.856211 | -1.146854 | 768.951591 | 0.150260194 | -1.31559633 | 0.188309566 | 0.960264078 | 9.682064 | 9.484689 | 9.616408 | 9.648033 | 9.573160 | 9.498655 | 925 | 906 | 657 | 635 |
| 79077 | NM_024096,N | DCTPP1 | dCTP pyrophosphatase 1 | protein_coding | NP_077001.1 | 28777 | 615840 | . | . | 39.783278 | 35.556405 | 29.548899 | 40.938096 | 92.577313 | 82.190225 | 66.306875 | 93.600091 | -1.084830 | 1342.133515 | 0.16124454 | -0.72851186 | 0.466300309 | 0.998865404 | 10.448060 | 10.330157 | 10.462912 | 10.369949 | 10.229889 | 10.466833 | 1705 | 1417 | 944 | 1396 |
| 79078 | NM_024097,N | C1orf50 | chromosome 1 open reading frame 50 | protein_coding | NP_077002.2 | 28795 | . | . | . | 1.439176 | 1.541421 | 1.472718 | 1.644993 | 3.349020 | 3.563064 | 3.304736 | 3.761082 | 1.030772 | 226.409165 | 0.226317553 | 0.193200245 | 0.846802147 | 0.998865404 | 7.801142 | 7.844115 | 7.800180 | 7.829510 | 7.805691 | 7.852482 | 249 | 248 | 190 | 226 |
| 79080 | NM_024098 | CCDC86 | coiled-coil domain containing 86 | protein_coding | NP_077003.1 | 28359 | 611293 | . | . | 22.159525 | 22.587524 | 24.558310 | 24.774942 | 51.566124 | 52.212074 | 55.108135 | 56.644962 | 1.087659 | 1356.421883 | 0.126649414 | 0.957181616 | 0.338475594 | 0.998865404 | 10.343714 | 10.464921 | 10.368079 | 10.376582 | 10.437071 | 10.435957 | 1488 | 1410 | 1230 | 1324 |
| 79081 | NM_00136794 | LBHD1 | LBH domain containing 1 | protein_coding | NP_001354869.1;NP_001354870.1;NP_077004.2 | 28351 | . | . | . | 10.708493 | 11.823798 | 15.123773 | 12.108376 | 24.919106 | 27.331237 | 33.937310 | 27.684363 | 1.137736 | 636.520952 | 0.1675762 | 1.110935163 | 0.266596259 | 0.998274218 | 9.217776 | 9.404326 | 9.233121 | 9.298305 | 9.418013 | 9.283966 | 652 | 674 | 647 | 573 |
| 79083 | NM_00104246 | MLPH | melanophilin | protein_coding | NP_001035932.1;NP_001268402.1;NP_001268403.1;NP_077006.1;XP_00 | 29643 | 606526 | . | . | 18.300662 | 18.486963 | 15.770938 | 18.881698 | 42.586394 | 42.733444 | 35.389524 | 43.170757 | -1.078358 | 1520.225362 | 0.133066853 | -0.81790216 | 0.413413056 | 0.998865404 | 10.623629 | 10.514445 | 10.603778 | 10.592279 | 10.478320 | 10.597306 | 1830 | 1690 | 1173 | 1478 |
| 79084 | NM_00131706 | WDR77 | WD repeat domain 77 | protein_coding | NP_001303991.1;NP_001303992.1;NP_001303993.1;NP_077007.1 | 29652 | 611734 | . | . | 28.090671 | 28.127666 | 26.309931 | 26.593355 | 65.368151 | 65.018359 | 59.038723 | 60.802547 | -1.080304 | 2095.730561 | 0.11677014 | -0.95432747 | 0.339917906 | 0.998865404 | 11.087872 | 10.976453 | 11.068366 | 11.058260 | 11.001731 | 11.001743 | 2522 | 2334 | 1752 | 1889 |
| 79085 | NM_024103,XI | SLC25A23 | solute carrier family 25 member 23 | protein_coding | NP_077008.2;XP_011526576.1;XP_011526577.1;XP_011526578.1;XP_01 | 19375 | 608746 | . | . | 4.723212 | 5.500155 | 5.339807 | 7.680054 | 10.991110 | 12.713854 | 11.982375 | 17.559530 | 1.255238 | 610.583966 | 0.184238942 | 1.780084335 | 0.07506216 | 0.765332165 | 9.081391 | 9.408228 | 9.128848 | 9.205249 | 9.205967 | 9.421154 | 587 | 619 | 499 | 736 |
| 79086 | NM_00130092 | SMIM7 | small integral membrane protein 7 | protein_coding | NP_001287854.1;NP_077009.2 | 28419 | . | . | . | 35.049149 | 33.775064 | 30.142486 | 35.118689 | 81.560808 | 78.072569 | 67.638866 | 80.294712 | -1.057507 | 1368.290984 | 0.130676285 | -0.61730136 | 0.537035969 | 0.998865404 | 10.458060 | 10.377123 | 10.442014 | 10.436014 | 10.355848 | 10.434959 | 1626 | 1512 | 1096 | 1312 |
| 79087 | NM_024105,XI | ALG12 | ALG12 alpha-1,6-mannosyltransferase | protein_coding | NP_077010.1;XP_011528673.1;XP_016884425.1;XP_016884426.1 | 19358 | 607144 | . | . | 6.793071 | 7.744853 | 7.516249 | 7.332082 | 15.807754 | 17.902575 | 16.866246 | 16.763933 | 1.005303 | 1131.657453 | 0.137920522 | 0.055322678 | 0.955881395 | 0.998865404 | 10.140429 | 10.148005 | 10.098903 | 10.182667 | 10.168186 | 10.122438 | 1227 | 1286 | 1017 | 1031 |
| 79088 | NM_00130088 | ZNF426 | zinc finger protein 426 | protein_coding | NP_001287812.1;NP_001304984.1;NP_001304985.1;NP_077011.1;XP_01 | 20725 | . | . | . | 1.041378 | 1.435420 | 1.583975 | 2.145096 | 2.423330 | 3.318038 | 3.554394 | 4.904506 | 1.165683 | 254.766005 | 0.224928001 | 0.983317719 | 0.325451092 | 0.998865404 | 7.878827 | 8.098850 | 7.889394 | 8.005700 | 8.009910 | 8.042356 | 239 | 284 | 231 | 265 |
| 79089 | NM_00107667 | TMUB2 | transmembrane and ubiquitin like domain containing 2 | protein_coding | NP_001070142.1;NP_001317164.1;NP_001340102.1;NP_001340103.1;NI | 28459 | . | . | . | 19.974723 | 19.651080 | 23.144443 | 20.725548 | 46.482005 | 45.424354 | 51.935464 | 47.386500 | 1.065901 | 1487.552650 | 0.131653602 | 0.699359084 | 0.484327652 | 0.998865404 | 10.491785 | 10.584186 | 10.514001 | 10.511321 | 10.614202 | 10.507143 | 1661 | 1551 | 1427 | 1340 |
| 79090 | NM_00127089 | TRAPPC6A | trafficking protein particle complex 6A | protein_coding | NP_001257820.1;NP_001257821.1;NP_001257822.1;NP_077013.1;XP_01 | 23069 | 610396 | . | . | 6.024421 | 6.580138 | 5.677122 | 7.061024 | 14.019077 | 15.210286 | 12.739298 | 16.144191 | 1.093316 | 169.032528 | 0.25841163 | 0.498083819 | 0.618424965 | 0.998865404 | 7.335987 | 7.463510 | 7.366991 | 7.393806 | 7.379797 | 7.453719 | 180 | 180 | 140 | 180 |
| 79091 | NM_024109,N | METTL22 | methyltransferase like 22 | protein_coding | NP_077014.4;XP_005255627.1;XP_005255628.1;XP_011520959.1;XP_01 | 28368 | 615261 | . | . | 3.826060 | 3.978862 | 4.179818 | 2.649341 | 8.903396 | 9.197314 | 9.379386 | 6.057405 | -1.052232 | 392.142110 | 0.182869008 | -0.40166628 | 0.687929645 | 0.998865404 | 8.651489 | 8.578042 | 8.610303 | 8.649303 | 8.613288 | 8.586469 | 448 | 448 | 340 | 350 |
| 79092 | NM_00125797 | CARD14 | caspase recruitment domain family member 14 | protein_coding | NP_001244899.1;NP_001353314.1;NP_077015.2;NP_438170.1;XP_01152 | 16446 | 607211 | . | . | 3.922798 | 4.159316 | 4.491311 | 4.306238 | 9.128511 | 9.614443 | 10.078373 | 9.845700 | 1.116887 | 536.727421 | 0.163925106 | 0.972904951 | 0.330600544 | 0.998865404 | 8.986158 | 9.145534 | 9.010388 | 9.049661 | 9.108731 | 9.094287 | 566 | 564 | 498 | 525 |
| 79094 | NM_00114277 | CHAC1 | ChaC glutathione specific gamma-glutamylcyclotransferase 1 | protein_coding | NP_001136248.2;NP_077016.3;XP_024305813.1;XP_024305814.1;XP_02 | 28680 | 614587 | . | . | 6.136635 | 4.706861 | 3.516566 | 6.986771 | 14.280202 | 10.880120 | 7.891071 | 15.974421 | -1.060079 | 264.315251 | 0.279297063 | -0.30137071 | 0.76313183 | 0.998865404 | 8.088341 | 8.002617 | 8.100363 | 7.997275 | 7.847533 | 8.167819 | 344 | 265 | 152 | 317 |
| 79095 | NM_024112,XI | C9orf16 | chromosome 9 open reading frame 16 | protein_coding | NP_077017.1;XP_011517306.1 | 17823 | . | . | . | 22.170301 | 19.700910 | 16.057494 | 22.208537 | 51.591201 | 45.539536 | 36.032552 | 50.777178 | -1.108353 | 438.570886 | 0.199759001 | -0.74298018 | 0.45749368 | 0.998865404 | 8.849365 | 8.700109 | 8.845277 | 8.762460 | 8.644444 | 8.830520 | 566 | 465 | 305 | 451 |
| 79096 | NM_00100367 | C11orf49 | chromosome 11 open reading frame 49 | protein_coding | NP_001003676.1;NP_001003677.1;NP_001003678.1;NP_001265151.1;NI | 28720 | . | . | . | 5.040895 | 5.556768 | 3.697231 | 5.736514 | 11.730370 | 12.844716 | 8.296479 | 13.115855 | -1.048367 | 281.387457 | 0.223364529 | -0.30507835 | 0.760306478 | 0.998865404 | 8.170855 | 8.101131 | 8.125830 | 8.167273 | 8.042136 | 8.195751 | 319 | 323 | 203 | 296 |
| 79101 | NM_024116,N | TAF1D | TATA-box binding protein associated factor, RNA polymerase I sub | protein_coding | NP_077021.1 | 28759 | 612823 | . | . | 22.549130 | 21.247585 | 25.475336 | 17.054582 | 52.472750 | 49.114743 | 57.165919 | 38.993275 | -1.041877 | 1135.541990 | 0.17843022 | -0.33169922 | 0.740116392 | 0.998865404 | 10.178062 | 10.119679 | 10.184432 | 10.132327 | 10.269992 | 9.963534 | 1380 | 1207 | 1166 | 829 |
| 79102 | NM_032015 | RNF26 | ring finger protein 26 | protein_coding | NP_114404.1 | 14646 | 606130 | . | . | 8.552259 | 9.412458 | 7.413167 | 8.541697 | 19.901459 | 21.757318 | 16.634932 | 19.529573 | -1.141603 | 735.517214 | 0.1552997 | -1.23027223 | 0.218595179 | 0.982008277 | 9.615307 | 9.423556 | 9.535056 | 9.596664 | 9.429230 | 9.517485 | 864 | 883 | 558 | 686 |
| 79109 | NM_00100661 | MAPKAP1 | MAPK associated protein 1 | protein_coding | NP_001006618.1;NP_001006619.1;NP_001006620.1;NP_001006621.1;NI | 18752 | 610558 | . | . | 15.356949 | 15.491795 | 13.688115 | 18.111353 | 35.736252 | 35.809977 | 30.715735 | 41.409457 | 1.037905 | 1513.339423 | 0.15147244 | 0.354352423 | 0.723074785 | 0.998865404 | 10.536718 | 10.589833 | 10.528831 | 10.563678 | 10.458009 | 10.680140 | 1672 | 1639 | 1149 | 1651 |
| 79132 | NM_024119,XI | DHX58 | DExH-box helicase 58 | protein_coding | NP_077024.2;XP_016880548.1 | 29517 | 608588 | . | . | 2.141723 | 2.067808 | 1.967664 | 1.929376 | 4.983877 | 4.779830 | 4.415381 | 4.411289 | -1.124646 | 158.803806 | 0.260247846 | -0.65119098 | 0.514923212 | 0.998865404 | 7.393255 | 7.223986 | 7.335877 | 7.333833 | 7.293804 | 7.281823 | 194 | 181 | 132 | 138 |
| 79133 | NM_00103937 | NDUFAF5 | NADH:ubiquinone oxidoreductase complex assembly factor 5 | protein_coding | NP_001034464.1;NP_001339332.1;NP_001339335.1;NP_001339336.1;NI | 15899 | 612360 | . | . | 2.571225 | 2.319515 | 2.535809 | 2.521004 | 5.983345 | 5.361660 | 5.690281 | 5.763978 | 1.071287 | 341.146545 | 0.191876498 | 0.517753214 | 0.604630465 | 0.998865404 | 8.363501 | 8.463270 | 8.402029 | 8.385155 | 8.450833 | 8.413756 | 385 | 350 | 317 | 320 |
| 79134 | NM_024121 | TMEM185B | transmembrane protein 185B | protein_coding | NP_077026.2 | 18896 | . | . | . | 14.634709 | 15.048407 | 14.991498 | 14.475836 | 34.055569 | 34.785065 | 33.640488 | 33.097279 | -1.021207 | 2728.586442 | 0.113367874 | -0.26705056 | 0.789430231 | 0.998865404 | 11.428968 | 11.398750 | 11.414378 | 11.430226 | 11.423456 | 11.386808 | 3144 | 3004 | 2401 | 2474 |
| 79135 | NM_024122,N | APOO | apolipoprotein O | protein_coding | NP_077027.1;XP_011543886.1;XP_011543887.1;XP_011543888.1;XP_01 | 28727 | 300753 | . | . | 18.113222 | 19.540790 | 20.600444 | 20.785428 | 42.150212 | 45.169409 | 46.226797 | 47.523413 | 1.117194 | 688.563434 | 0.1532614 | 1.043184671 | 0.296862772 | 0.998865404 | 9.345519 | 9.504969 | 9.371793 | 9.402768 | 9.434311 | 9.491064 | 732 | 718 | 606 | 709 |
| 79137 | NM_00132110 | RETREG2 | reticulophagy regulator family member 2 | protein_coding | NP_001308038.1;NP_001308039.1;NP_077269.3;XP_005246905.1 | 28450 | . | . | . | 31.624211 | 30.391877 | 29.633583 | 32.348902 | 73.590840 | 70.252183 | 66.496896 | 73.961919 | 1.016772 | 4240.472636 | 0.109420774 | 0.219300532 | 0.826415947 | 0.998865404 | 12.038007 | 12.061913 | 12.042488 | 12.043316 | 12.025800 | 12.086697 | 4840 | 4540 | 3582 | 4155 |
| 79139 | NM_00113467 | DERL1 | derlin 1 | protein_coding | NP_001128143.1;NP_001317530.1;NP_001350892.1;NP_077271.1 | 28454 | 608813 | . | . | 25.801580 | 26.050498 | 25.091950 | 22.657875 | 60.041334 | 60.216892 | 56.305610 | 51.804537 | -1.100946 | 2497.094884 | 0.119215003 | -1.16381558 | 0.244498756 | 0.992733415 | 11.353643 | 11.215102 | 11.322946 | 11.323625 | 11.289044 | 11.200312 | 3012 | 2825 | 2182 | 2105 |
| 79140 | NM_00130101 | CCDC28B | coiled-coil domain containing 28B | protein_coding | NP_001287940.1;NP_077272.2;XP_011540415.1;XP_016857795.1;XP_01 | 28163 | 610162 | . | . | 9.991530 | 9.696177 | 8.390555 | 10.088610 | 23.250701 | 22.413144 | 18.828164 | 23.066405 | -1.073588 | 315.404212 | 0.201368128 | -0.50872384 | 0.610945807 | 0.998865404 | 8.351800 | 8.248466 | 8.303295 | 8.336609 | 8.243355 | 8.317500 | 365 | 363 | 246 | 305 |
| 79142 | NM_00128451 | PHF23 | PHD finger protein 23 | protein_coding | NP_001271446.1;NP_001271447.1;NP_077273.2;XP_024306706.1;XP_02 | 28428 | 612910 | . | . | 27.825287 | 29.905349 | 28.476235 | 34.101818 | 64.750587 | 69.127550 | 63.899844 | 77.969757 | 1.066576 | 1830.463692 | 0.130162716 | 0.714385713 | 0.474988704 | 0.998865404 | 10.790943 | 10.883558 | 10.793233 | 10.828766 | 10.797738 | 10.922232 | 1994 | 1955 | 1514 | 1910 |
| 79143 | NM_00114605 | MBOAT7 | membrane bound O-acyltransferase domain containing 7 | protein_coding | NP_001139528.1;NP_001139554.1;NP_001139555.1;NP_077274.3;XP_01 | 15505 | 606048 | . | . | 38.095604 | 38.925099 | 35.712127 | 38.489387 | 88.650040 | 89.977107 | 80.136977 | 88.001402 | -1.048630 | 2674.206251 | 0.113835708 | -0.60179454 | 0.54731091 | 0.998865404 | 11.418796 | 11.350178 | 11.396708 | 11.410965 | 11.345147 | 11.385005 | 3125 | 2980 | 2214 | 2508 |
| 79144 | NM_00135342 | PPDPF | pancreatic progenitor cell differentiation and proliferation factor | protein_coding | NP_001340352.1;NP_077275.1 | 16142 | . | . | . | 30.562256 | 32.545918 | 26.083520 | 37.119426 | 71.119626 | 75.231346 | 58.530661 | 84.869161 | -1.023680 | 794.447585 | 0.179318478 | -0.18829725 | 0.85064364 | 0.998865404 | 9.651070 | 9.616337 | 9.621394 | 9.654889 | 9.486682 | 9.740795 | 904 | 888 | 559 | 869 |
| 79145 | NM_00101166 | CHCHD7 | coiled-coil-helix-coiled-coil-helix domain containing 7 | protein_coding | NP_001011667.1;NP_001011668.1;NP_001011669.1;NP_001011670.1;NI | 28314 | 611238 | . | . | 5.708900 | 4.999548 | 5.095964 | 5.273647 | 13.284846 | 11.556682 | 11.435194 | 12.057566 | -1.062769 | 313.950052 | 0.200876224 | -0.43722784 | 0.66194612 | 0.998865404 | 8.337377 | 8.250085 | 8.346941 | 8.272999 | 8.282049 | 8.272851 | 397 | 326 | 267 | 283 |
| 79147 | NM_00103988 | FKRP | fukutin related protein | protein_coding | NP_001034974.1;NP_077277.1;XP_005259304.1;XP_005259305.1;XP_00 | 17997 | 606596 | . | . | 6.178636 | 6.383288 | 6.023933 | 5.334061 | 14.377943 | 14.755254 | 13.517534 | 12.195697 | -1.131778 | 642.065428 | 0.15908409 | -1.12262097 | 0.261598506 | 0.995760118 | 9.412879 | 9.234781 | 9.360264 | 9.372301 | 9.326091 | 9.238064 | 778 | 742 | 559 | 527 |
| 79149 | NM_00132206 | ZSCAN5A | zinc finger and SCAN domain containing 5A | protein_coding | NP_001308990.1;NP_001308991.1;NP_001308993.1;NP_001308994.1;NI | 23710 | . | . | . | 1.416281 | 1.621579 | 2.061771 | 1.764701 | 3.295747 | 3.748352 | 4.626551 | 4.034780 | 1.022489 | 147.760227 | 0.293131176 | 0.109455853 | 0.912840935 | 0.998865404 | 7.191197 | 7.222858 | 7.121703 | 7.267121 | 7.240650 | 7.169297 | 137 | 187 | 142 | 127 |
| 79153 | NM_024307 | GDPD3 | glycerophosphodiester phosphodiesterase domain containing 3 | protein_coding | NP_077283.2 | 28638 | 616318 | . | . | 0.389395 | 0.622393 | 0.520403 | 0.471105 | 0.906138 | 1.438690 | 1.167770 | 1.077125 | -1.022815 | 17.402450 | 0.742701522 | -0.04382081 | 0.965047235 | NA | 4.137922 | 4.104320 | 4.102966 | 4.135656 | 4.122792 | 4.112725 | 16 | 23 | 16 | 15 |
| 79154 | NM_024308,XI | DHRS11 | dehydrogenase/reductase 11 | protein_coding | NP_077284.2;XP_005257715.1;XP_011523535.1 | 28639 | 616159 | . | . | 9.111363 | 9.769228 | 7.852044 | 9.807158 | 21.202515 | 22.582009 | 17.619760 | 22.422904 | -1.088194 | 405.875554 | 0.190136447 | -0.64130547 | 0.521324239 | 0.998865404 | 8.725099 | 8.602066 | 8.670745 | 8.706504 | 8.574680 | 8.697968 | 473 | 470 | 302 | 403 |
| 79155 | NM_00116152 | TNIP2 | TNFAIP3 interacting protein 2 | protein_coding | NP_001154999.1;NP_001278945.1;NP_077285.3 | 19118 | 610669 | . | . | 6.768523 | 7.493509 | 5.623513 | 8.126122 | 15.750631 | 17.321583 | 12.619000 | 18.579411 | -1.061329 | 420.631056 | 0.205837534 | -0.41718285 | 0.676544672 | 0.998865404 | 8.759441 | 8.672059 | 8.698165 | 8.762377 | 8.584549 | 8.794016 | 473 | 492 | 292 | 450 |
| 79156 | NM_024310,XI | PLEKHF1 | pleckstrin homology and FYVE domain containing 1 | protein_coding | NP_077286.3;XP_005259313.1;XP_011525611.1 | 20764 | 615200 | . | . | 19.883202 | 18.886481 | 23.376355 | 26.140013 | 46.269030 | 43.656947 | 52.455862 | 59.766035 | 1.161403 | 1369.814300 | 0.132032555 | 1.634964562 | 0.102056477 | 0.821228746 | 10.307943 | 10.523539 | 10.362515 | 10.355083 | 10.425269 | 10.518961 | 1467 | 1361 | 1203 | 1463 |
| 79157 | NM_00124253 | MFSD11 | major facilitator superfamily domain containing 11 | protein_coding | NP_001229461.1;NP_001229462.1;NP_001229463.1;NP_001229464.1;NI | 25458 | . | . | . | 11.814943 | 10.528258 | 11.810076 | 11.186542 | 27.493858 | 24.336543 | 26.501473 | 25.576695 | 1.037771 | 1017.109403 | 0.140651031 | 0.38028656 | 0.703732711 | 0.998865404 | 9.963049 | 10.016966 | 10.012608 | 9.937662 | 10.027353 | 9.978411 | 1208 | 1022 | 931 | 939 |
| 79158 | NM_024312,XI | GNPTAB | N-acetylglucosamine-1-phosphate transferase subunits alpha and | protein_coding | NP_077288.2;XP_006719656.1;XP_011537033.1;XP_016875450.1;XP_01 | 29670 | 607840 | . | . | 24.022756 | 23.899483 | 23.040703 | 22.687334 | 55.901943 | 55.244727 | 51.702671 | 51.871893 | -1.066873 | 4017.275668 | 0.107551973 | -0.86831193 | 0.385223594 | 0.998865404 | 12.017934 | 11.924558 | 11.991175 | 12.005330 | 11.952991 | 11.936086 | 4735 | 4513 | 3415 | 3607 |
| 79159 | NM_024313 | NOL12 | nucleolar protein 12 | protein_coding | NP_077289.1 | 28585 | . | . | . | 9.090141 | 9.366312 | 10.237385 | 9.945251 | 21.153131 | 21.650650 | 22.972395 | 22.738634 | 1.078706 | 852.103726 | 0.142378773 | 0.767687192 | 0.442673047 | 0.998865404 | 9.679159 | 9.788539 | 9.699172 | 9.714277 | 9.774614 | 9.746818 | 934 | 894 | 784 | 813 |

| | | | | | | | | | | | | | | | | | | | | | | | | | | | | | | | | | |
|---|---|---|---|---|---|---|---|---|---|---|---|---|---|---|---|---|---|---|---|---|---|---|---|---|---|---|---|---|---|---|---|---|---|
| 79160 | NR_125359 | LINC01711 | long intergenic non-protein coding RNA 1711 | lncRNA | . | 28663 | . | . | . | 0.307082 | 0.258822 | 0.243113 | 0.225331 | 0.714593 | 0.598279 | 0.545539 | 0.515193 | -1.154122 | 8.352514 | 1.047924673 | -0.1973383 | 0.84356281 | NA | 3.161369 | 2.955730 | 3.070438 | 3.064820 | 3.063569 | 3.060959 | 11 | 9 | 7 | 7 |
| 79161 | NM_00132947 | TMEM243 | transmembrane protein 243 | protein_coding | NP_001316401.1;NP_001316402.1;NP_001316403.1;NP_001316404.1;NI | 21707 | 616993 | . | . | 16.111650 | 15.999559 | 12.759784 | 10.829474 | 37.492471 | 36.983696 | 28.632590 | 24.760303 | -1.337124 | 561.652826 | 0.168413683 | -2.48871479 | 0.012820577 | 0.374881745 | 9.327616 | 8.909226 | 9.216431 | 9.221423 | 9.081428 | 8.975561 | 737 | 696 | 454 | 412 |
| 79165 | NM_024316 | LENG1 | leukocyte receptor cluster member 1 | protein_coding | NP_077292.2 | 15502 | . | . | . | 4.351251 | 5.085390 | 4.750657 | 5.296151 | 10.125541 | 11.755106 | 10.660337 | 12.109020 | 1.054269 | 209.946933 | 0.236616261 | 0.322223285 | 0.747283546 | 0.998865404 | 7.675812 | 7.750971 | 7.665250 | 7.733781 | 7.701068 | 7.747182 | 218 | 237 | 178 | 212 |
| 79169 | NM_024319,N | C1orf35 | chromosome 1 open reading frame 35 | protein_coding | NP_077295.1 | 19032 | . | . | . | 11.749890 | 10.709018 | 11.091508 | 11.999970 | 27.342476 | 24.754374 | 24.889025 | 27.436505 | 1.014266 | 459.344170 | 0.175311405 | 0.116571429 | 0.90719968 | 0.998865404 | 8.833159 | 8.853632 | 8.869924 | 8.806354 | 8.826877 | 8.867536 | 552 | 467 | 389 | 448 |
| 79171 | NM_00131911 | RBM42 | RNA binding motif protein 42 | protein_coding | NP_001306042.1;NP_077297.2 | 28117 | 613232 | . | . | 31.251730 | 31.771486 | 30.564713 | 34.846219 | 72.724060 | 73.441208 | 68.586326 | 79.671749 | 1.023761 | 1672.680733 | 0.127740226 | 0.265222347 | 0.790838181 | 0.998865404 | 10.691056 | 10.724641 | 10.694688 | 10.701809 | 10.666735 | 10.763970 | 1895 | 1792 | 1380 | 1685 |
| 79172 | NM_00119980 | CENPO | centromere protein O | protein_coding | NP_001186732.1;NP_001309030.1;NP_077298.1 | 28152 | 611504 | . | . | 11.819125 | 13.121611 | 11.997820 | 13.247634 | 27.503588 | 30.331189 | 26.922757 | 30.289141 | 1.000084 | 1609.462626 | 0.128765354 | 0.000940394 | 0.999249674 | 0.999862634 | 10.652479 | 10.652247 | 10.616139 | 10.686467 | 10.617185 | 10.685380 | 1770 | 1815 | 1338 | 1575 |
| 79173 | NM_00134584 | C19orf57 | chromosome 19 open reading frame 57 | protein_coding | NP_001332772.1;NP_001332773.1;NP_001332775.1;NP_001332776.1;NI | 28153 | . | . | . | 6.878807 | 8.437778 | 7.270691 | 9.166806 | 16.007270 | 19.504299 | 16.315221 | 20.958812 | 1.047066 | 695.782300 | 0.173313886 | 0.382847412 | 0.701832905 | 0.998865404 | 9.409472 | 9.474774 | 9.354055 | 9.489030 | 9.378166 | 9.526530 | 706 | 806 | 552 | 739 |
| 79174 | NM_00113510 | CRELD2 | cysteine rich with EGF like domains 2 | protein_coding | NP_001128573.1;NP_001271246.1;NP_001271247.1;NP_077300.3;XP_0C | 28150 | 607171 | . | . | 64.457235 | 66.146081 | 53.649626 | 54.211722 | 149.994634 | 152.899622 | 120.388211 | 123.948658 | -1.227747 | 2679.441639 | 0.112644893 | -2.62784792 | 0.008592692 | 0.293930247 | 11.528154 | 11.232120 | 11.459002 | 11.471517 | 11.298517 | 11.298979 | 3375 | 3211 | 2091 | 2256 |
| 79175 | NM_00128249 | ZNF343 | zinc finger protein 343 | protein_coding | NP_001269424.1;NP_001269425.1;NP_001269426.1;NP_001269427.1;NI | 16017 | . | . | . | 2.779506 | 3.130784 | 3.295621 | 3.494535 | 6.468027 | 7.236946 | 7.395274 | 7.989840 | 1.123083 | 343.646469 | 0.192544653 | 0.869745213 | 0.384439658 | 0.998865404 | 8.338834 | 8.505895 | 8.360912 | 8.417234 | 8.452786 | 8.457074 | 354 | 367 | 315 | 342 |
| 79176 | NM_024326,XI | FBXL15 | F-box and leucine rich repeat protein 15 | protein_coding | NP_077302.3;XP_005270206.1;XP_005270207.1;XP_005270208.1;XP_01 | 28155 | 610287 | . | . | 2.298402 | 2.666237 | 2.327652 | 2.832931 | 5.348477 | 6.163127 | 5.223183 | 6.477161 | 1.067518 | 182.716182 | 0.250806498 | 0.375828242 | 0.70704459 | 0.998865404 | 7.466186 | 7.559235 | 7.483540 | 7.511594 | 7.485245 | 7.564783 | 197 | 197 | 149 | 193 |
| 79177 | NM_00114534 | ZNF576 | zinc finger protein 576 | protein_coding | NP_001138819.1;NP_077303.1 | 28357 | . | . | . | 4.603835 | 5.071578 | 4.862281 | 5.333316 | 10.713315 | 11.723180 | 10.910818 | 12.193992 | 1.016096 | 383.201899 | 0.186180552 | 0.123736061 | 0.901524258 | 0.998865404 | 8.570739 | 8.593096 | 8.552511 | 8.600544 | 8.560027 | 8.611406 | 420 | 427 | 321 | 378 |
| 79178 | NM_00112633 | THTPA | thiamine triphosphatase | protein_coding | NP_001119811.1;NP_001242991.1;NP_001243250.1;NP_001243251.1;NI | 18987 | 611612 | . | . | 4.226875 | 4.799290 | 4.876860 | 4.975689 | 9.836110 | 11.093773 | 10.943531 | 11.376318 | 1.071069 | 307.311962 | 0.20048805 | 0.494050148 | 0.621270764 | 0.998865404 | 8.213278 | 8.312148 | 8.218337 | 8.268152 | 8.289708 | 8.271674 | 326 | 335 | 281 | 293 |
| 79180 | NM_024329,XI | EFHD2 | EF-hand domain family member D2 | protein_coding | NP_077305.2;XP_005246057.1 | 28670 | 616450 | . | . | 30.576636 | 31.018181 | 28.902896 | 33.901770 | 71.153090 | 71.699919 | 64.857260 | 77.512363 | 1.005771 | 2328.926421 | 0.124197024 | 0.066840753 | 0.94670848 | 0.998865404 | 11.181422 | 11.189465 | 11.181254 | 11.183819 | 11.126069 | 11.244653 | 2670 | 2510 | 1881 | 2351 |
| 79183 | NM_00103919 | TTPAL | alpha tocopherol transfer protein like | protein_coding | NP_001034288.1;NP_001248768.1;NP_077307.2;XP_011527345.1;XP_01 | 16114 | . | . | . | 2.929164 | 2.785447 | 3.106345 | 2.774253 | 6.816284 | 6.438687 | 6.970548 | 6.342998 | 1.004421 | 560.283705 | 0.164449937 | 0.038703155 | 0.969127058 | 0.998865404 | 9.126529 | 9.133490 | 9.148585 | 9.106953 | 9.163169 | 9.097777 | 664 | 584 | 513 | 500 |
| 79184 | NM_00101805 | BRCC3 | BRCA1/BRCA2-containing complex subunit 3 | protein_coding | NP_001018065.1;NP_001229569.1;NP_077308.1;XP_005274808.1;XP_01 | 24185 | 300617 | . | . | 4.199959 | 3.858155 | 4.444778 | 4.217439 | 9.773478 | 8.918299 | 9.973955 | 9.642672 | -1.062406 | 518.186535 | 0.167600669 | -0.52108886 | 0.602304869 | 0.998865404 | 9.060076 | 8.973274 | 9.059661 | 9.012153 | 9.020288 | 8.974382 | 637 | 555 | 453 | 454 |
| 79187 | NM_00133042 | FSD1 | fibronectin type III and SPRY domain containing 1 | protein_coding | NP_001317358.1;NP_077309.1 | 13745 | 609828 | . | . | 6.870514 | 7.140293 | 8.687351 | 8.416321 | 15.987970 | 16.505107 | 19.494163 | 19.242917 | 1.201706 | 442.861646 | 0.174169145 | 1.521990058 | 0.128011572 | 0.876241045 | 8.652105 | 8.917281 | 8.722812 | 8.736512 | 8.854639 | 8.829338 | 458 | 439 | 429 | 444 |
| 79188 | NM_024334,XI | TMEM43 | transmembrane protein 43 | protein_coding | NP_077310.1;XP_016862665.1 | 28472 | 612048 | . | . | 26.202132 | 27.020994 | 24.714504 | 21.417129 | 60.973442 | 62.460239 | 55.458628 | 48.967722 | -1.168780 | 2503.907304 | 0.123979137 | -1.8148534 | 0.069546442 | 0.744724233 | 11.397973 | 11.173212 | 11.339595 | 11.358263 | 11.282097 | 11.160298 | 3072 | 2945 | 2162 | 1999 |
| 79190 | NM_024335,XI | IRX6 | iroquois homeobox 6 | protein_coding | NP_077311.2;XP_005256194.1 | 14675 | 606196 | . | . | 3.715747 | 4.819794 | 6.503573 | 6.209128 | 8.646696 | 11.141169 | 14.593829 | 14.196432 | 1.426957 | 473.236408 | 0.182686003 | 2.807777776 | 0.004988464 | 0.213175742 | 8.607531 | 9.120027 | 8.692956 | 8.819140 | 9.005033 | 8.953339 | 401 | 466 | 503 | 501 |
| 79191 | NM_024336,XI | IRX3 | iroquois homeobox 3 | protein_coding | NP_077312.2;XP_005256196.1 | 14360 | 612985 | . | . | 2.450916 | 2.737581 | 2.021872 | 2.894181 | 5.703383 | 6.328042 | 4.537021 | 6.617202 | -1.061127 | 212.738739 | 0.248456282 | -0.34451872 | 0.730456221 | 0.998865404 | 7.776023 | 7.688529 | 7.716904 | 7.771411 | 7.642924 | 7.785694 | 238 | 250 | 151 | 224 |
| 79192 | NM_024337,XI | IRX1 | iroquois homeobox 1 | protein_coding | NP_077313.3;XP_016865334.1 | 14358 | 606197 | . | . | 0.049075 | 0.087986 | 0.076758 | 0.102943 | 0.114200 | 0.203383 | 0.172242 | 0.235366 | 1.331319 | 5.794381 | 1.262673799 | 0.326969752 | 0.743690763 | NA | 2.316288 | 2.724273 | 2.516343 | 2.530618 | 2.527190 | 2.534872 | 4 | 7 | 5 | 7 |
| 79228 | NM_00114235 | THOC6 | THO complex 6 | protein_coding | NP_001135822.1;NP_001334632.1;NP_001334633.1;NP_077315.2 | 28369 | 615403 | . | . | 9.818156 | 12.375768 | 12.127684 | 11.943974 | 22.847251 | 28.607143 | 27.214172 | 27.308479 | 1.071793 | 508.428892 | 0.178895199 | 0.559137072 | 0.576068176 | 0.998865404 | 8.939301 | 9.038789 | 8.891549 | 9.033792 | 9.017013 | 9.001351 | 502 | 589 | 463 | 487 |
| 79230 | NM_00104438 | ZNF557 | zinc finger protein 557 | protein_coding | NP_001037852.1;NP_001037853.1;NP_077317.2 | 28632 | . | . | . | 0.902482 | 0.951661 | 1.169365 | 1.138054 | 2.100114 | 2.199804 | 2.624023 | 2.602026 | 1.230849 | 187.672198 | 0.242585341 | 1.235253119 | 0.216736316 | 0.982008277 | 7.394506 | 7.694110 | 7.490599 | 7.511387 | 7.598820 | 7.584527 | 189 | 186 | 183 | 191 |
| 79269 | NM_00128681 | DCAF10 | DDB1 and CUL4 associated factor 10 | protein_coding | NP_001273739.1;NP_077321.3;XP_005251634.1;XP_005251635.1;XP_01 | 23686 | . | . | . | 1.318393 | 1.377476 | 1.649648 | 1.732262 | 3.067955 | 3.184097 | 3.701761 | 3.960611 | -1.049931 | 306.967252 | 0.198901116 | -0.35341113 | 0.723780244 | 0.998865404 | 8.296669 | 8.226356 | 8.265873 | 8.284391 | 8.254050 | 8.243486 | 356 | 345 | 263 | 278 |
| 79339 | NM_033179 | OR51B4 | olfactory receptor family 51 subfamily B member 4 | protein_coding | NP_149419.2 | 14708 | . | . | . | 0.102971 | 0.115551 | 0.047490 | 0.112174 | 0.239617 | 0.267101 | 0.106566 | 0.256472 | -1.090094 | 3.436841 | 1.614238885 | -0.07709675 | 0.93854658 | NA | 1.844134 | 1.715150 | 1.784541 | 1.785633 | 1.779321 | 1.788114 | 4 | 4 | 2 | 4 |
| 79363 | NM_030907,XI | CPLANE2 | ciliogenesis and planar polarity effector 2 | protein_coding | NP_112169.2;XP_011540428.1 | 28127 | . | . | . | 1.797286 | 2.455040 | 2.375750 | 2.048030 | 4.182359 | 5.674937 | 5.331114 | 4.682577 | 1.029676 | 105.567583 | 0.323933743 | 0.13024397 | 0.896373408 | 0.998865404 | 6.700945 | 6.742798 | 6.668135 | 6.757527 | 6.745839 | 6.699704 | 103 | 128 | 100 | 93 |
| 79364 | NM_00104065 | ZXDC | ZXD family zinc finger C | protein_coding | NP_001035743.1;NP_079388.3;XP_005247814.1;XP_006713804.1;XP_01 | 28160 | 615746 | . | . | 5.128878 | 5.830388 | 7.274711 | 6.010040 | 11.935112 | 13.477204 | 16.324242 | 13.741239 | 1.194709 | 710.681879 | 0.160319906 | 1.600923212 | 0.109393928 | 0.843131948 | 9.338899 | 9.595800 | 9.370088 | 9.440158 | 9.584444 | 9.469056 | 707 | 735 | 728 | 666 |
| 79365 | NM_030762 | BHLHE41 | basic helix-loop-helix family member e41 | protein_coding | NP_110389.1 | 16617 | 606200 | . | . | 7.628429 | 7.284781 | 10.941516 | 10.858171 | 17.751667 | 16.839096 | 24.552444 | 24.825918 | 1.440971 | 1069.420802 | 0.135001961 | 3.903952347 | 9.46344E-05 | 0.012046912 | 9.775043 | 10.302280 | 9.927721 | 9.890239 | 10.186016 | 10.172351 | 1036 | 920 | 1108 | 1173 |
| 79366 | NM_030763 | HMGN5 | high mobility group nucleosome binding domain 5 | protein_coding | NP_110390.1 | 8013 | 300385 | . | . | 1.755131 | 1.623149 | 1.626460 | 2.084959 | 4.084263 | 3.751981 | 3.649729 | 4.767012 | 1.088002 | 116.340304 | 0.30601926 | 0.397627841 | 0.690904535 | 0.998865404 | 6.800556 | 6.921331 | 6.860278 | 6.831229 | 6.833367 | 6.913351 | 134 | 115 | 93 | 127 |
| 79369 | NM_00133049 | B3GNT4 | UDP-GlcNAc:betaGal beta-1,3-N-acetylglucosaminyltransferase 4 | protein_coding | NP_001317421.1;NP_110392.1 | 15683 | 605864 | . | . | 0.351853 | 0.184975 | 0.403732 | 0.435663 | 0.818776 | 0.427577 | 0.905964 | 0.996093 | 1.464468 | 30.680765 | 0.579972834 | 0.948970277 | 0.34263573 | 0.998865404 | 4.636349 | 5.189480 | 4.934936 | 4.873461 | 4.948083 | 4.955033 | 37 | 19 | 31 | 35 |
| 79400 | NM_00118477 | NOX5 | NADPH oxidase 5 | protein_coding | NP_001171708.1;NP_001171709.1;NP_078781.3 | 14874 | 606572 | . | . | 0.026050 | 0.029788 | 0.009492 | 0.017159 | 0.060620 | 0.068855 | 0.021299 | 0.039233 | -1.722740 | 6.718762 | 1.18741495 | -0.66085132 | 0.508707666 | NA | 3.089593 | 2.300113 | 2.767405 | 2.765702 | 2.744013 | 2.756872 | 10 | 9 | 3 | 6 |
| 79411 | NM_00128642 | GLB1L | galactosidase beta 1 like | protein_coding | NP_001273352.1;NP_001273356.1;NP_078782.3;XP_016860384.1;XP_02 | 28129 | . | . | . | 4.087412 | 4.380711 | 4.508797 | 5.360751 | 9.511573 | 10.126208 | 10.117612 | 12.256721 | 1.056961 | 392.201135 | 0.186242395 | 0.429127109 | 0.667830725 | 0.998865404 | 8.575343 | 8.654472 | 8.580003 | 8.615577 | 8.590150 | 8.669329 | 426 | 424 | 327 | 403 |
| 79412 | NM_00125372 | KREMEN2 | kringle containing transmembrane protein 2 | protein_coding | NP_001240654.1;NP_001240655.1;NP_078783.1;NP_757384.1 | 18797 | 609899 | . | . | 4.414379 | 3.640500 | 3.636802 | 4.062859 | 10.272439 | 8.415180 | 8.160878 | 9.289245 | -1.057515 | 239.168151 | 0.22774013 | -0.35425754 | 0.723145889 | 0.998865404 | 7.941617 | 7.861020 | 7.959933 | 7.867718 | 7.860886 | 7.913414 | 309 | 241 | 191 | 230 |
| 79414 | NM_024509 | LRFN3 | leucine rich repeat and fibronectin type III domain containing 3 | protein_coding | NP_078785.1 | 28370 | 612809 | . | . | 2.360125 | 2.270132 | 2.414789 | 2.845010 | 5.492107 | 5.247511 | 5.418716 | 6.504776 | 1.122068 | 361.647674 | 0.19246541 | 0.86332602 | 0.387958242 | 0.998865404 | 8.413237 | 8.578893 | 8.476131 | 8.449067 | 8.482892 | 8.572744 | 402 | 359 | 307 | 386 |
| 79415 | NM_00103304 | CYBC1 | cytochrome b-245 chaperone 1 | protein_coding | NP_001028218.1;NP_001093877.1;NP_001093878.1;NP_001180582.1;NI | 28672 | 618334 | . | . | 16.957957 | 19.322176 | 20.767633 | 20.250720 | 39.461860 | 44.664072 | 46.601963 | 46.300864 | 1.029927 | 1357.719660 | 0.128099906 | 0.332102652 | 0.739811747 | 0.998865404 | 10.385472 | 10.428152 | 10.390219 | 10.400430 | 10.440540 | 10.394777 | 1530 | 1453 | 1234 | 1253 |
| 79441 | NM_00130314 | HAUS3 | HAUS augmin like complex subunit 3 | protein_coding | NP_001290072.1;NP_078787.2 | 28719 | 613430 | . | . | 2.438099 | 2.106318 | 2.476610 | 1.517690 | 5.673557 | 4.868848 | 5.557440 | 3.470017 | -1.170894 | 389.506932 | 0.23096374 | -0.98547909 | 0.324388796 | 0.998865404 | 8.713967 | 8.488222 | 8.689149 | 8.593603 | 8.678308 | 8.402192 | 521 | 418 | 388 | 255 |
| 79443 | NM_024513,XI | FYCO1 | FYVE and coiled-coil domain autophagy adaptor 1 | protein_coding | NP_078789.2;XP_006713396.1;XP_006713397.1;XP_011532413.1 | 14673 | 607182 | . | . | 3.990393 | 4.281955 | 4.738458 | 5.591656 | 9.285808 | 9.897928 | 10.632961 | 12.784658 | 1.236560 | 1205.738167 | 0.140141877 | 2.185872112 | 0.028824963 | 0.547128317 | 10.074696 | 10.380528 | 10.125203 | 10.171254 | 10.246233 | 10.366580 | 1203 | 1200 | 1069 | 1347 |
| 79444 | NM_022161,N | BIRC7 | baculoviral IAP repeat containing 7 | protein_coding | NP_071444.1;NP_647478.1 | 13702 | 605737 | . | . | 114.722408 | 112.442230 | 118.444527 | 122.280697 | 266.963760 | 259.915245 | 265.786080 | 279.580284 | 1.043209 | 4787.436997 | 0.106030425 | 0.575574105 | 0.564903088 | 0.998865404 | 12.194203 | 12.255227 | 12.217465 | 12.196505 | 12.232853 | 12.251841 | 5464 | 4993 | 4203 | 4635 |
| 79446 | NM_00116147 | WDR25 | WD repeat domain 25 | protein_coding | NP_001154948.1;NP_001337876.1;NP_001337877.1;NP_001337878.1;NI | 21064 | 618059 | . | . | 2.074220 | 2.157800 | 2.616770 | 2.144611 | 4.826794 | 4.987855 | 5.871958 | 4.903399 | 1.053439 | 131.337761 | 0.286825887 | 0.261854229 | 0.793433829 | 0.998865404 | 6.999082 | 7.074216 | 7.052483 | 7.000156 | 7.026168 | 7.064342 | 158 | 128 | 111 | 133 |
| 79447 | NM_024516 | PAGR1 | PAXIP1 associated glutamate rich protein 1 | protein_coding | NP_078792.1 | 28707 | 612033 | . | . | 9.360117 | 9.573256 | 9.003547 | 9.299251 | 21.781378 | 22.129009 | 20.203697 | 21.261631 | -1.048820 | 1124.284241 | 0.132378201 | -0.51947925 | 0.60342658 | 0.998865404 | 10.168815 | 10.099946 | 10.147011 | 10.157760 | 10.108854 | 10.124759 | 1316 | 1251 | 944 | 1040 |
| 79465 | NM_024518 | ULBP3 | UL16 binding protein 3 | protein_coding | NP_078794.1 | 14895 | 605699 | . | . | 6.877308 | 7.374770 | 6.626508 | 5.853148 | 16.003777 | 17.047112 | 14.869690 | 13.382529 | -1.157717 | 648.708116 | 0.159667134 | -1.32327036 | 0.185745486 | 0.958175316 | 9.443026 | 9.232124 | 9.366731 | 9.409095 | 9.332396 | 9.244416 | 777 | 774 | 558 | 526 |
| 79469 | NR_002771 | DLEU2L | deleted in lymphocytic leukemia 2 like | ncRNA_pseudc | . | 13225 | . | . | . | 0.557886 | 0.753449 | 1.043969 | 0.322434 | 1.298223 | 1.741632 | 2.342636 | 0.737207 | 1.022286 | 35.123129 | 0.604761703 | 0.052580179 | 0.95806641 | 0.998865404 | 5.116387 | 5.152091 | 5.094044 | 5.136235 | 5.187569 | 5.043824 | 34 | 43 | 47 | 16 |
| 79541 | NM_030908 | OR2A4 | olfactory receptor family 2 subfamily A member 4 | protein_coding | NP_112170.1 | 14729 | . | . | . | 0.039279 | 0.016877 | 0.034161 | 0.028400 | 0.091403 | 0.039012 | 0.076655 | 0.064934 | -1.007340 | 2.223025 | 1.975467281 | -0.0053412 | 0.995738358 | NA | 1.156660 | 1.148375 | 1.155078 | 1.151489 | 1.153584 | 1.152831 | 3 | 2 | 2 | 2 |
| 79567 | NM_00119352 | RIPOR1 | RHO family interacting cell polarization regulator 1 | protein_coding | NP_001180451.1;NP_001180452.1;NP_001180453.1;NP_078795.2;XP_01 | 25836 | . | . | . | 9.727434 | 10.821479 | 10.680050 | 11.902993 | 22.636139 | 25.014336 | 23.965723 | 27.214780 | 1.087947 | 1380.780329 | 0.133868723 | 0.908411057 | 0.363661086 | 0.998865404 | 10.369407 | 10.490585 | 10.360118 | 10.433226 | 10.420716 | 10.501041 | 1450 | 1496 | 1186 | 1419 |
| 79568 | NM_00136939 | MAIP1 | matrix AAA peptidase interacting protein 1 | protein_coding | NP_001356328.1;NP_078796.2 | 26198 | 617267 | . | . | 13.014774 | 14.999044 | 12.795020 | 13.582575 | 30.285911 | 34.670965 | 28.711653 | 31.054944 | -1.079202 | 636.975491 | 0.160229136 | -0.68630081 | 0.492523438 | 0.998865404 | 9.369287 | 9.258786 | 9.293241 | 9.384271 | 9.272575 | 9.303898 | 711 | 761 | 520 | 588 |
| 79570 | NM_024522,XI | NKAIN1 | sodium/potassium transporting ATPase interacting 1 | protein_coding | NP_078798.2;XP_016857809.1 | 25743 | 612871 | . | . | 0.119995 | 0.084660 | 0.039667 | 0.060001 | 0.279232 | 0.195696 | 0.089011 | 0.137186 | -1.861825 | 7.874698 | 1.102496961 | -0.81335153 | 0.416016539 | NA | 3.357474 | 2.459419 | 3.007358 | 2.989838 | 2.973521 | 2.981213 | 14 | 9 | 4 | 6 |
| 79571 | NM_024523 | GCC1 | GRIP and coiled-coil domain containing 1 | protein_coding | NP_078799.2 | 19095 | 607418 | . | . | 4.318034 | 4.950739 | 4.321742 | 4.813055 | 10.048242 | 11.443855 | 9.697864 | 11.004478 | -1.027492 | 592.028416 | 0.165063552 | -0.23704084 | 0.812625109 | 0.998865404 | 9.229310 | 9.189460 | 9.174174 | 9.259634 | 9.167414 | 9.231003 | 647 | 689 | 483 | 574 |
| 79572 | NM_00136754 | ATP13A3 | ATPase 13A3 | protein_coding | NP_001354478.1;NP_001361765.1;NP_078800.3;XP_005269414.1;XP_01 | 24113 | 610232 | . | . | 11.153533 | 9.089155 | 9.981090 | 7.762465 | 25.954731 | 21.009986 | 22.397278 | 17.747954 | -1.158198 | 2218.301024 | 0.15368743 | -1.37865701 | 0.168000531 | 0.945638095 | 11.217061 | 11.005683 | 11.250345 | 11.077789 | 11.149464 | 10.939891 | 3030 | 2298 | 2025 | 1672 |
| 79573 | NM_00112283 | TTC13 | tetratricopeptide repeat domain 13 | protein_coding | NP_001116307.2;NP_001363436.1;NP_001363437.1;NP_001363438.1;NI | 26204 | . | . | . | 6.563774 | 6.162070 | 5.660203 | 5.391582 | 15.274170 | 14.243903 | 12.701333 | 12.327217 | -1.077640 | 792.334733 | 0.16932023 | -0.63710684 | 0.524055255 | 0.998865404 | 9.682519 | 9.575426 | 9.642073 | 9.663506 | 9.698433 | 9.491846 | 932 | 900 | 762 | 609 |
| 79575 | NM_024527 | ABHD8 | abhydrolase domain containing 8 | protein_coding | NP_078803.4 | 23759 | . | . | . | 4.250745 | 3.984992 | 3.052675 | 4.544200 | 9.891658 | 9.211488 | 6.850114 | 10.389775 | -1.099255 | 252.156626 | 0.240457196 | -0.56777511 | 0.570187685 | 0.998865404 | 8.045525 | 7.907529 | 8.017006 | 7.980053 | 7.854823 | 8.039121 | 315 | 275 | 169 | 268 |
| 79576 | NM_024528,XI | NKAP | NFKB activating protein | protein_coding | NP_078804.2;XP_016885331.1 | 29873 | 300766 | . | . | 1.744654 | 1.785726 | 1.623427 | 1.813625 | 4.059881 | 4.127786 | 3.642923 | 4.146638 | -1.042811 | 322.763511 | 0.196957038 | -0.30705782 | 0.758799365 | 0.998865404 | 8.364523 | 8.303499 | 8.342225 | 8.349625 | 8.295378 | 8.349345 | 377 | 358 | 261 | 311 |
| 79577 | NM_024529,XI | CDC73 | cell division cycle 73 | protein_coding | NP_078805.3;XP_006711600.1 | 16783 | 607393 | . | . | 6.162655 | 6.218368 | 5.371434 | 5.430158 | 14.340751 | 14.374036 | 12.053343 | 12.415414 | -1.164209 | 1054.936244 | 0.134304542 | -1.63322636 | 0.102421399 | 0.821931724 | 10.148455 | 9.929095 | 10.100060 | 10.092903 | 9.981279 | 9.987427 | 1313 | 1219 | 844 | 918 |
| 79581 | NM_00125381 | SLC52A2 | solute carrier family 52 member 2 | protein_coding | NP_001240744.1;NP_001240745.1;NP_001350047.1;NP_001350049.1;NI | 30224 | 607882 | . | . | 34.420893 | 33.344299 | 30.024910 | 28.401716 | 80.098834 | 77.076842 | 67.375026 | 64.937147 | -1.155421 | 2037.595162 | 0.119056147 | -1.75058839 | 0.080016837 | 0.779933816 | 11.093024 | 10.884771 | 11.055579 | 11.038955 | 10.955389 | 10.908935 | 2541 | 2333 | 1693 | 1720 |
| 79582 | NM_00102543 | SPAG16 | sperm associated antigen 16 | protein_coding | NP_001020607.1;NP_078808.3;XP_006712809.1;XP_011510116.1;XP_01 | 23225 | 612173 | . | . | 8.251374 | 5.684004 | 6.519534 | 5.851484 | 19.201290 | 13.138832 | 14.629648 | 13.378726 | -1.186315 | 291.594787 | 0.236682959 | -1.04142331 | 0.297679128 | 0.998865404 | 8.305078 | 8.060184 | 8.330996 | 8.109495 | 8.171789 | 8.097929 | 436 | 274 | 249 | 232 |
| 79583 | NM_00107741 | TMEM231 | transmembrane protein 231 | protein_coding | NP_001070884.2;NP_001070886.1 | 37234 | 614949 | . | . | 1.634496 | 1.679149 | 1.795380 | 1.627623 | 3.803541 | 3.881428 | 4.028781 | 3.721364 | 1.022151 | 200.158017 | 0.238864526 | 0.132324942 | 0.89472728 | 0.998865404 | 7.628640 | 7.661168 | 7.657135 | 7.620771 | 7.674157 | 7.623372 | 237 | 205 | 186 | 179 |
| 79585 | NM_00120147 | CORO7 | coronin 7 | protein_coding | NP_001188401.1;NP_001188402.1;NP_001338658.1;NP_078811.3 | 26161 | 611668 | . | . | 3.530668 | 3.636232 | 3.279518 | 3.175808 | 8.216010 | 8.405311 | 7.359142 | 7.261107 | -1.125564 | 368.443439 | 0.186117771 | -0.91688466 | 0.359203083 | 0.998865404 | 8.608036 | 8.437528 | 8.551684 | 8.565598 | 8.502511 | 8.478171 | 444 | 426 | 308 | 318 |
| 79586 | NM_00119573 | CHPF | chondroitin polymerizing factor | protein_coding | NP_001182660.1;NP_078812.3;XP_011510140.1 | 24291 | 610405 | . | . | 58.411266 | 58.276132 | 52.895237 | 52.754051 | 135.925417 | 134.707888 | 118.695379 | 120.615863 | -1.103712 | 5139.614757 | 0.10496874 | -1.35624376 | 0.175021623 | 0.949120257 | 12.396863 | 12.254516 | 12.364826 | 12.369961 | 12.291959 | 12.277504 | 6191 | 5837 | 4286 | 4541 |
| 79587 | NM_00135225 | CARS2 | cysteinyl-tRNA synthetase 2, mitochondrial | protein_coding | NP_001339181.1;NP_001339182.1;NP_078813.1;XP_006720016.1;XP_01 | 25695 | 612800 | . | . | 25.865335 | 29.309912 | 28.324019 | 30.110294 | 60.189700 | 67.751171 | 63.558278 | 68.843613 | 1.050099 | 1736.381636 | 0.124655116 | 0.565763999 | 0.571554222 | 0.998865404 | 10.726288 | 10.796592 | 10.704930 | 10.777759 | 10.768625 | 10.791784 | 1860 | 1913 | 1524 | 1692 |
| 79589 | NM_024539,N | RNF128 | ring finger protein 128 | protein_coding | NP_078815.3;NP_919445.1 | 21153 | 300439 | . | . | 6.268598 | 5.790821 | 5.849549 | 5.001824 | 14.587285 | 13.385742 | 13.126218 | 11.436077 | -1.127336 | 501.127727 | 0.175577178 | -0.98485408 | 0.324695751 | 0.998865404 | 9.052462 | 8.880485 | 9.032931 | 8.975919 | 8.979150 | 8.876321 | 638 | 548 | 444 | 405 |
| 79590 | NM_024540,N | MRPL24 | mitochondrial ribosomal protein L24 | protein_coding | NP_078816.2;NP_663781.1;XP_011508283.1;XP_011508284.1 | 14037 | 611836 | . | . | 57.301235 | 55.425207 | 52.756174 | 57.539395 | 133.342329 | 128.117842 | 118.383329 | 131.556989 | -1.032999 | 1523.354685 | 0.126568798 | -0.37006962 | 0.711330615 | 0.998865404 | 10.596337 | 10.549352 | 10.597699 | 10.572912 | 10.529626 | 10.589895 | 1811 | 1644 | 1252 | 1460 |
| 79591 | NM_024541,XI | ARMH3 | armadillo like helical domain containing 3 | protein_coding | NP_078817.2;XP_006718032.1;XP_011538453.1;XP_011538454.1;XP_01 | 25788 | . | . | . | 6.105645 | 5.918812 | 6.253973 | 7.133311 | 14.208089 | 13.681599 | 14.033732 | 16.309465 | 1.074702 | 816.881550 | 0.149252878 | 0.696380032 | 0.486190864 | 0.998865404 | 9.621298 | 9.724812 | 9.650277 | 9.643984 | 9.648593 | 9.745209 | 911 | 846 | 686 | 847 |
| 79594 | NM_024544,XI | MUL1 | mitochondrial E3 ubiquitin protein ligase 1 | protein_coding | NP_078820.2;XP_011540439.1 | 25762 | 612037 | . | . | 30.507726 | 31.597244 | 29.628007 | 31.731854 | 70.992731 | 73.038443 | 66.484394 | 72.551118 | -1.026200 | 2342.070012 | 0.116520445 | -0.32021342 | 0.748806551 | 0.998865404 | 11.212176 | 11.174718 | 11.194964 | 11.212634 | 11.159139 | 11.206271 | 2702 | 2588 | 1951 | 2231 |
| 79595 | NM_00114592 | SAP130 | Sin3A associated protein 130 | protein_coding | NP_001139400.1;NP_001317228.1;NP_001317229.1;NP_001317230.1;NI | 29813 | 609697 | . | . | 10.977299 | 11.992016 | 12.168456 | 10.785170 | 25.544627 | 27.720082 | 27.305664 | 24.659013 | -1.017652 | 1460.310125 | 0.13324783 | -0.18945307 | 0.849737735 | 0.998865404 | 10.524554 | 10.499455 | 10.484040 | 10.550870 | 10.549935 | 10.457114 | 1623 | 1658 | 1335 | 1275 |
| 79596 | NM_024546,XI | OBI1 | ORC ubiquitin ligase 1 | protein_coding | NP_078822.3;XP_011533527.1;XP_024305178.1 | 20308 | 615906 | . | . | 4.007451 | 3.724607 | 3.458450 | 2.638221 | 9.325504 | 8.609595 | 7.760662 | 6.031980 | -1.293131 | 377.318297 | 0.200848677 | -1.84650666 | 0.064818674 | 0.731823074 | 8.732560 | 8.363131 | 8.650963 | 8.602094 | 8.548801 | 8.401027 | 509 | 441 | 325 | 267 |
| 79598 | NM_00130340 | CEP97 | centrosomal protein 97 | protein_coding | NP_001290330.1;NP_078824.2;XP_006713806.1;XP_011511429.1;XP_01 | 26244 | 615864 | . | . | 1.406900 | 1.321391 | 1.284534 | 1.191591 | 3.273914 | 3.054455 | 2.882456 | 2.724433 | -1.117428 | 311.483983 | 0.200010691 | -0.80086651 | 0.423208931 | 0.998865404 | 8.360575 | 8.201046 | 8.332477 | 8.292029 | 8.272761 | 8.231128 | 392 | 342 | 266 | 265 |
| 79600 | NM_00108253 | TCTN1 | tectonic family member 1 | protein_coding | NP_001076006.1;NP_001076007.1;NP_001167446.1;NP_001167447.1;NI | 26113 | 609863 | . | . | 15.311182 | 16.908556 | 16.214450 | 14.677339 | 35.629755 | 39.084886 | 36.384755 | 33.557996 | -1.049949 | 967.871891 | 0.13932529 | -0.50471326 | 0.613760246 | 0.998865404 | 9.953282 | 9.883212 | 9.933081 | 9.938914 | 9.928953 | 9.871368 | 1137 | 1074 | 854 | 850 |
| 79602 | NM_00137536 | ADIPOR2 | adiponectin receptor 2 | protein_coding | NP_001362292.1;NP_001362293.1;NP_001362294.1;NP_078827.2;XP_0C | 24041 | 607946 | . | . | 29.771010 | 30.516325 | 30.294574 | 33.404691 | 69.278368 | 70.539850 | 67.980144 | 76.375857 | 1.045569 | 3866.302066 | 0.112450371 | 0.57170544 | 0.567521549 | 0.998865404 | 11.884305 | 11.948459 | 11.891019 | 11.904328 | 11.893654 | 11.973912 | 4318 | 4112 | 3271 | 3884 |
| 79603 | NM_024552,XI | CERS4 | ceramide synthase 4 | protein_coding | NP_078828.2;XP_011526592.1;XP_011526593.1;XP_011526594.1;XP_01 | 23747 | 615334 | . | . | 4.512192 | 4.083867 | 3.987985 | 4.392916 | 10.500058 | 9.440042 | 8.948922 | 10.043879 | -1.029956 | 241.727545 | 0.221035735 | -0.1926519 | 0.847231598 | 0.998865404 | 7.938399 | 7.895762 | 7.948288 | 7.900132 | 7.890993 | 7.928610 | 296 | 252 | 199 | 232 |
| 79605 | NM_00125831 | PGBD5 | piggyBac transposable element derived 5 | protein_coding | NP_001245240.1 | 19405 | 616791 | . | . | 0.078723 | 0.036525 | 0.057239 | 0.046752 | 0.183193 | 0.084428 | 0.128444 | 0.106893 | -1.150081 | 19.078535 | 0.734242283 | -0.27475286 | 0.78350612 | NA | 4.349244 | 4.151760 | 4.290183 | 4.225856 | 4.254576 | 4.239687 | 32 | 14 | 17 | 15 |
| 79607 | NM_00133044 | FAM118B | family with sequence similarity 118 member B | protein_coding | NP_001317375.1;NP_078832.1;XP_011541279.1;XP_011541280.1;XP_01 | 26110 | 616587 | . | . | 4.642214 | 4.678065 | 4.639315 | 5.486147 | 10.802625 | 10.813557 | 10.410490 | 12.543424 | 1.069598 | 327.644991 | 0.201540949 | 0.481635517 | 0.630064891 | 0.998865404 | 8.306981 | 8.403388 | 8.349589 | 8.321438 | 8.319458 | 8.423655 | 374 | 333 | 267 | 347 |
| 79609 | NM_00104066 | VCPKMT | valosin containing protein lysine methyltransferase | protein_coding | NP_001035752.1;NP_078834.2;XP_005268117.1;XP_016877129.1;XP_01 | 20352 | 615260 | . | . | 3.768355 | 2.960691 | 3.826831 | 2.965598 | 8.769117 | 6.843767 | 8.587298 | 6.780489 | 1.014723 | 209.626880 | 0.270801443 | 0.077862577 | 0.937937368 | 0.998865404 | 7.699849 | 7.723414 | 7.767849 | 7.628826 | 7.796437 | 7.610054 | 275 | 191 | 221 | 158 |
| 79612 | NM_00111079 | NAA16 | N-alpha-acetyltransferase 16, NatA auxiliary subunit | protein_coding | NP_001104268.1;NP_060997.2;NP_078837.3;XP_005266580.1;XP_00671 | 26164 | . | . | . | 3.036279 | 2.963024 | 3.147682 | 2.707663 | 7.065542 | 6.849159 | 7.063305 | 6.190751 | -1.032189 | 389.444657 | 0.202527686 | -0.22568231 | 0.821448522 | 0.998865404 | 8.627269 | 8.582939 | 8.631846 | 8.590584 | 8.677347 | 8.500913 | 471 | 412 | 384 | 305 |
| 79613 | NM_024562,XI | TANGO6 | transport and golgi organization 6 homolog | protein_coding | NP_078838.1;XP_011521629.1 | 25749 | . | . | . | 3.120771 | 2.709684 | 3.186869 | 3.881860 | 7.262161 | 6.263555 | 7.151242 | 8.875411 | 1.187236 | 456.571479 | 0.188998993 | 1.310094648 | 0.190163819 | 0.961278413 | 8.705682 | 8.953115 | 8.828195 | 8.719974 | 8.816180 | 8.943150 | 524 | 410 | 388 | 511 |
| 79616 | NM_00130817 | CCNJL | cyclin J like | protein_coding | NP_001295102.1;NP_078841.3;XP_006714980.1;XP_011532948.1;XP_01 | 25876 | . | . | . | 5.423625 | 4.937889 | 6.982858 | 6.013071 | 12.621000 | 11.414148 | 15.669329 | 13.748174 | 1.185512 | 794.807842 | 0.148905769 | 1.64876109 | 0.099196589 | 0.813553488 | 9.506490 | 9.752002 | 9.538284 | 9.600274 | 9.716169 | 9.662333 | 800 | 820 | 778 | 778 |
| 79618 | NM_00113572 | HMBOX1 | homeobox containing 1 | protein_coding | NP_001129198.1;NP_001311311.1;NP_001311312.1;NP_001311313.1;NI | 26137 | 618610 | . | . | 2.443599 | 2.730144 | 5.206999 | 3.130018 | 5.686356 | 6.310850 | 11.684348 | 7.156414 | 1.202419 | 1157.843538 | 0.150689551 | 1.764817559 | 0.077594433 | 0.767975096 | 10.037939 | 10.304239 | 10.072411 | 10.128966 | 10.323565 | 10.147369 | 1164 | 1178 | 1226 | 1049 |
| 79621 | NM_00114227 | RNASEH2B | ribonuclease H2 subunit B | protein_coding | NP_001135751.1;NP_078846.2;XP_006719930.1;XP_011533531.1;XP_01 | 25671 | 610326 | . | . | 9.703279 | 8.681810 | 8.134716 | 7.634595 | 22.579931 | 20.068393 | 18.254066 | 17.455592 | -1.168085 | 436.881788 | 0.182132082 | -1.23067678 | 0.218443776 | 0.982008277 | 8.878413 | 8.655156 | 8.857212 | 8.772569 | 8.753331 | 8.688107 | 578 | 474 | 370 | 357 |
| 79622 | NM_024571 | SNRNP25 | small nuclear ribonucleoprotein U11/U12 subunit 25 | protein_coding | NP_078847.2 | 14161 | . | . | . | 33.189346 | 33.535149 | 30.112558 | 35.185310 | 77.232979 | 77.517998 | 67.571709 | 80.447021 | -1.035793 | 1118.059421 | 0.139651481 | -0.36330138 | 0.71637976 | 0.998865404 | 10.152133 | 10.100976 | 10.138620 | 10.140029 | 10.058228 | 10.164984 | 1309 | 1229 | 886 | 1104 |
| 79623 | NM_00125382 | GALNT14 | polypeptide N-acetylgalactosaminyltransferase 14 | protein_coding | NP_001240755.1;NP_001240756.1;NP_001316024.1;NP_001316025.1;NI | 22946 | 608225 | . | . | 0.856922 | 0.728491 | 0.557625 | 1.056593 | 1.994094 | 1.683939 | 1.251293 | 2.415774 | -1.143003 | 69.303617 | 0.40917047 | -0.47126827 | 0.637449173 | 0.998865404 | 6.209604 | 6.013451 | 6.124972 | 6.132051 | 6.031737 | 6.158106 | 84 | 81 | 40 | 78 |
| 79624 | NM_00128656 | ARMT1 | acidic residue methyltransferase 1 | protein_coding | NP_001273491.1;NP_078849.1 | 17872 | 616332 | . | . | 13.680701 | 11.753764 | 12.289593 | 10.269346 | 31.835555 | 27.169351 | 27.577493 | 23.479641 | -1.143574 | 897.917552 | 0.158806848 | -1.21877227 | 0.22293064 | 0.982008277 | 9.903573 | 9.710875 | 9.912811 | 9.793360 | 9.823164 | 9.689348 | 1192 | 950 | 798 | 711 |
| 79625 | NM_024574,XI | NDNF | neuron derived neurotrophic factor | protein_coding | NP_078850.3;XP_024309980.1;XP_024309981.1 | 26256 | 616506 | . | . | 0.285467 | 0.554526 | 0.151667 | 0.182758 | 0.664294 | 1.281813 | 0.340337 | 0.417854 | -1.739486 | 64.248616 | 0.398907804 | -2.00211919 | 0.045271914 | 0.649595749 | 6.350898 | 5.550733 | 6.059933 | 6.064966 | 5.926833 | 5.956943 | 93 | 89 | 37 | 48 |
| 79627 | NM_00132426 | OGFRL1 | opioid growth factor receptor like 1 | protein_coding | NP_001311195.1;NP_078852.3;XP_011534419.1;XP_011534420.1;XP_01 | 21378 | . | . | . | 0.734625 | 0.674845 | 0.517299 | 0.726570 | 1.709504 | 1.559936 | 1.160803 | 1.661215 | -1.155813 | 177.325981 | 0.26647034 | -0.78398154 | 0.433050944 | 0.998865404 | 7.571589 | 7.361273 | 7.521250 | 7.476074 | 7.368019 | 7.502185 | 230 | 195 | 119 | 180 |
| 79628 | NM_024577 | SH3TC2 | SH3 domain and tetratricopeptide repeats 2 | protein_coding | NP_078853.2 | 29427 | 608206 | . | . | 3.455471 | 3.726340 | 4.590867 | 4.133312 | 8.041023 | 8.613601 | 10.301771 | 9.450327 | 1.198393 | 3277.300169 | 0.118034668 | 2.21207256 | 0.026961652 | 0.536241613 | 11.541802 | 11.802978 | 11.570935 | 11.625190 | 11.794149 | 11.697971 | 3318 | 3325 | 3286 | 3157 |
| 79629 | NM_024578,XI | OCEL1 | occludin/ELL domain containing 1 | protein_coding | NP_078854.1;XP_005260136.1;XP_006722962.1;XP_016882795.1 | 26221 | . | . | . | 6.058664 | 5.483251 | 6.483636 | 8.690166 | 14.098759 | 12.674780 | 14.549093 | 19.869035 | 1.254264 | 216.377333 | 0.240128119 | 1.361109335 | 0.17347914 | 0.94763772 | 7.585140 | 7.911278 | 7.718051 | 7.674540 | 7.741141 | 7.860482 | 232 | 197 | 184 | 253 |
| 79630 | NM_00130103 | C1orf54 | chromosome 1 open reading frame 54 | protein_coding | NP_001287968.1;NP_001287969.1;NP_001287970.1;NP_001287971.1;NI | 26258 | . | . | . | 0.200077 | 0.363950 | 0.568973 | 0.631147 | 0.465587 | 0.841287 | 1.276758 | 1.443042 | 1.966421 | 10.037763 | 0.973910933 | 1.001706005 | 0.316485606 | NA | 2.760365 | 3.733338 | 3.287109 | 3.301999 | 3.325885 | 3.321887 | 6 | 9 | 12 | 12 |
| 79631 | NM_00104061 | EFL1 | elongation factor like GTPase 1 | protein_coding | NP_001035700.1;NP_001309773.1;NP_001309774.1;NP_078856.4;XP_01 | 25789 | 617538 | . | . | 3.555865 | 3.699480 | 3.283650 | 4.433195 | 8.274642 | 8.551512 | 7.368416 | 10.135974 | 1.076966 | 420.454267 | 0.196214927 | 0.545178205 | 0.58563096 | 0.998865404 | 8.661823 | 8.767840 | 8.702436 | 8.677752 | 8.642482 | 8.819471 | 476 | 428 | 324 | 468 |
| 79632 | NM_00110041 | FAM184A | family with sequence similarity 184 member A | protein_coding | NP_001093881.1;NP_001275505.1;NP_078857.5 | 20991 | . | . | . | 1.180519 | 1.316354 | 1.044443 | 0.769327 | 2.747115 | 3.042812 | 2.343699 | 1.758976 | -1.394980 | 139.228910 | 0.290583188 | -1.6526934 | 0.098393279 | 0.813553488 | 7.340897 | 6.862206 | 7.155019 | 7.197655 | 7.103628 | 7.000453 | 177 | 184 | 117 | 92 |
| 79634 | NM_00119352 | SCRN3 | secernin 3 | protein_coding | NP_001180457.1;NP_078859.2;XP_005246910.1;XP_005246911.1;XP_00 | 30382 | 614967 | . | . | 6.351689 | 6.033672 | 6.122869 | 5.812278 | 14.780640 | 13.947102 | 13.739539 | 13.289083 | -1.063911 | 728.852280 | 0.154782058 | -0.5774376 | 0.563643884 | 0.998865404 | 9.553206 | 9.464393 | 9.533070 | 9.526640 | 9.536650 | 9.435039 | 869 | 807 | 660 | 613 |
| 79635 | NM_00114268 | CCDC121 | coiled-coil domain containing 121 | protein_coding | NP_001136155.1;NP_078860.2;XP_005264617.1 | 25833 | . | . | . | 1.268972 | 1.050627 | 0.969276 | 0.823840 | 2.952953 | 2.428571 | 2.175026 | 1.883612 | -1.212141 | 86.438783 | 0.35002127 | -0.79297416 | 0.427792877 | 0.998865404 | 6.564965 | 6.289080 | 6.494938 | 6.422986 | 6.421466 | 6.391426 | 122 | 90 | 72 | 69 |
| 79637 | NM_00130427 | ARMC7 | armadillo repeat containing 7 | protein_coding | NP_001291200.1;NP_001291201.1;NP_001291202.1;NP_078861.1 | 26168 | . | . | . | 3.656535 | 4.335259 | 3.385398 | 3.911892 | 8.508906 | 10.021143 | 7.596735 | 8.944077 | -1.155656 | 215.101635 | 0.236889585 | -0.88104968 | 0.378290933 | 0.998865404 | 7.849960 | 7.640170 | 7.732396 | 7.827117 | 7.696852 | 7.729483 | 240 | 273 | 167 | 194 |
| 79639 | NM_00130074 | TMEM53 | transmembrane protein 53 | protein_coding | NP_001287675.1;NP_001287676.1;NP_001287677.1;NP_078863.2;XP_01 | 26186 | . | . | . | 1.822417 | 1.714063 | 2.128367 | 2.362283 | 4.240840 | 3.962134 | 4.775994 | 5.401081 | 1.238576 | 141.329587 | 0.275015934 | 1.122414427 | 0.261686273 | 0.995760118 | 6.980358 | 7.289005 | 7.110649 | 7.083585 | 7.162691 | 7.192145 | 151 | 131 | 131 | 152 |
| 79640 | NR_160905 | C22orf46 | chromosome 22 open reading frame 46 | transcribed_ps | . | 26294 | . | . | . | 5.357155 | 5.964275 | 7.416899 | 6.307847 | 12.466320 | 13.786689 | 16.643307 | 14.422144 | 1.195633 | 978.139420 | 0.149641436 | 1.72261769 | 0.084957684 | 0.788085288 | 9.799133 | 10.057140 | 9.826338 | 9.897442 | 10.055102 | 9.927035 | 975 | 1009 | 1006 | 913 |
| 79641 | NM_024589,N | ROGDI | rogdi atypical leucine zipper | protein_coding | NP_078865.1;XP_006721010.1;XP_006721011.1 | 29478 | 614574 | . | . | 6.005826 | 5.564344 | 6.026812 | 6.123375 | 13.975801 | 12.862230 | 13.523992 | 14.000367 | 1.047081 | 275.946820 | 0.208094058 | 0.318953803 | 0.749761545 | 0.998865404 | 8.074481 | 8.141240 | 8.114772 | 8.075976 | 8.124734 | 8.114297 | 322 | 280 | 248 | 262 |
| 79642 | NM_00135421 | ARSJ | arylsulfatase family member J | protein_coding | NP_001341139.1;NP_001341140.1;NP_078866.3;XP_011530540.1;XP_01 | 26286 | 610010 | . | . | 2.087351 | 2.069748 | 2.111069 | 2.289599 | 4.857350 | 4.784314 | 4.737176 | 5.234896 | 1.027536 | 324.710980 | 0.197702563 | 0.198218729 | 0.842873935 | 0.998865404 | 8.323507 | 8.362258 | 8.345648 | 8.324239 | 8.316364 | 8.382480 | 376 | 339 | 269 | 327 |
| 79643 | NM_024591,XI | CHMP6 | charged multivesicular body protein 6 | protein_coding | NP_078867.2;XP_005257725.1 | 25675 | 610901 | . | . | 7.653073 | 6.757368 | 6.091403 | 7.900967 | 17.809018 | 15.619958 | 13.668932 | 18.064623 | -1.046550 | 360.786645 | 0.203986894 | -0.321791 | 0.747611035 | 0.998865404 | 8.527777 | 8.461466 | 8.544234 | 8.466913 | 8.406821 | 8.548427 | 453 | 372 | 268 | 372 |
| 79644 | NM_024592,XI | SRD5A3 | steroid 5 alpha-reductase 3 | protein_coding | NP_078868.1;XP_005265823.1;XP_005265824.1;XP_016864090.1 | 25812 | 611715 | . | . | 6.163732 | 7.321216 | 6.217726 | 5.958882 | 14.343258 | 16.923318 | 13.952398 | 13.624274 | 1.006788 | 731.520811 | 0.151087147 | 0.06460203 | 0.948490869 | 0.998865404 | 9.509874 | 9.519620 | 9.482954 | 9.540191 | 9.535636 | 9.497316 | 805 | 819 | 656 | 669 |
| 79646 | NM_024594 | PANK3 | pantothenate kinase 3 | protein_coding | NP_078870.1 | 19365 | 606161 | . | . | 6.933860 | 6.857621 | 7.608144 | 6.478593 | 16.135378 | 15.851698 | 17.072454 | 14.812533 | 1.007467 | 2231.513661 | 0.127815399 | 0.083969989 | 0.933080292 | 0.998865404 | 11.118273 | 11.129320 | 11.127681 | 11.111827 | 11.191913 | 11.055262 | 2585 | 2375 | 2114 | 1921 |
| 79647 | NM_00113627 | AKIRIN1 | akirin 1 | protein_coding | NP_001129747.1;NP_078871.1 | 25744 | 615164 | . | . | 27.444444 | 26.262999 | 25.480959 | 26.704848 | 63.864350 | 60.708099 | 57.178532 | 61.057462 | -1.046003 | 2202.883720 | 0.118109992 | -0.54937504 | 0.582748099 | 0.998865404 | 11.137252 | 11.072374 | 11.146930 | 11.098160 | 11.072080 | 11.101365 | 2672 | 2357 | 1838 | 2056 |
| 79648 | NM_00117257 | MCPH1 | microcephalin 1 | protein_coding | NP_001166045.1;NP_001166046.1;NP_001308971.2;NP_001308972.1;NI | 6954 | 607117 | . | . | 2.047627 | 2.068295 | 2.102843 | 2.042090 | 4.764913 | 4.780953 | 4.718718 | 4.668999 | -1.064392 | 372.246177 | 0.185752857 | -0.48467699 | 0.62790549 | 0.998865404 | 8.584385 | 8.494440 | 8.575526 | 8.540532 | 8.515542 | 8.528453 | 455 | 402 | 310 | 342 |
| 79649 | NM_00117351 | MAP7D3 | MAP7 domain containing 3 | protein_coding | NP_001166558.1;NP_001166988.1;NP_078873.2;XP_005262529.1;XP_01 | 25742 | 300930 | . | . | 4.049558 | 4.899959 | 3.684481 | 4.351849 | 9.423489 | 11.326474 | 8.267868 | 9.949988 | -1.114917 | 571.053309 | 0.171007607 | -0.9177141 | 0.358768563 | 0.998865404 | 9.234248 | 9.076400 | 9.137894 | 9.241713 | 9.076046 | 9.161361 | 640 | 700 | 438 | 540 |
| 79650 | NM_00119530 | USB1 | U6 snRNA biogenesis phosphodiesterase 1 | protein_coding | NP_001182231.1;NP_001191840.1;NP_001317497.1;NP_001317498.1;NI | 25792 | 613276 | . | . | 15.732659 | 17.004139 | 13.811232 | 17.598795 | 36.610545 | 39.305827 | 30.992005 | 40.237554 | -1.054250 | 1178.538458 | 0.147365662 | -0.51719518 | 0.605019918 | 0.998865404 | 10.240708 | 10.163835 | 10.187703 | 10.253291 | 10.101924 | 10.254743 | 1333 | 1362 | 895 | 1186 |
| 79651 | NM_00100549 | RHBDF2 | rhomboid 5 homolog 2 | protein_coding | NP_001005498.2;NP_001363157.1;NP_001363158.1;NP_001363159.1;NI | 20788 | 614404 | . | . | 6.708845 | 6.208887 | 6.361227 | 6.374008 | 15.611755 | 14.352119 | 14.274411 | 14.573411 | -1.044257 | 777.589991 | 0.149429439 | -0.41810266 | 0.675872058 | 0.998865404 | 9.633640 | 9.571421 | 9.655972 | 9.577789 | 9.585676 | 9.588965 | 965 | 810 | 660 | 715 |
| 79654 | NM_024602,XI | HECTD3 | HECT domain E3 ubiquitin protein ligase 3 | protein_coding | NP_078878.3;XP_024305571.1 | 26117 | 618638 | . | . | 12.588914 | 13.044060 | 12.453896 | 14.655802 | 29.294917 | 30.151931 | 27.946183 | 33.508751 | 1.045089 | 1475.712762 | 0.133724644 | 0.47580038 | 0.634216599 | 0.998865404 | 10.495238 | 10.558462 | 10.497968 | 10.520270 | 10.482091 | 10.600660 | 1638 | 1580 | 1211 | 1522 |
| 79656 | NM_00130208 | BEND5 | BEN domain containing 5 | protein_coding | NP_001289011.1;NP_001336722.1;NP_001336723.1;NP_001336724.1;NI | 25668 | . | . | . | 0.180753 | 0.508709 | 0.768800 | 0.442745 | 0.420621 | 1.175903 | 1.725167 | 1.012283 | 2.042971 | 31.736613 | 0.596833033 | 1.726895912 | 0.084186359 | 0.787933172 | 4.385915 | 5.411637 | 4.876002 | 4.953077 | 4.998821 | 5.008517 | 14 | 32 | 36 | 41 |
| 79657 | NM_00114607 | RPAP3 | RNA polymerase II associated protein 3 | protein_coding | NP_001139547.1;NP_001139548.1;NP_078880.2 | 26151 | 611477 | . | . | 5.958593 | 5.691680 | 5.723504 | 5.305026 | 13.865889 | 13.156574 | 12.843377 | 12.129312 | -1.083535 | 745.547047 | 0.148301645 | -0.78047278 | 0.435112643 | 0.998865404 | 9.598739 | 9.483262 | 9.575912 | 9.562866 | 9.532901 | 9.494155 | 901 | 829 | 640 | 652 |
| 79658 | NM_024605,XI | ARHGAP10 | Rho GTPase activating protein 10 | protein_coding | NP_078881.3;XP_005263272.1;XP_016864091.1 | 26099 | 609746 | . | . | 2.174069 | 2.420910 | 1.965424 | 2.427623 | 5.059147 | 5.596042 | 4.410355 | 5.550471 | -1.049572 | 228.348949 | 0.232535089 | -0.30017208 | 0.764045897 | 0.998865404 | 7.870262 | 7.799052 | 7.819128 | 7.871792 | 7.773902 | 7.868787 | 255 | 266 | 174 | 230 |
| 79659 | NM_00108046 | DYNC2H1 | dynein cytoplasmic 2 heavy chain 1 | protein_coding | NP_001073932.1;NP_001368.2;XP_006718966.1;XP_016873780.1;XP_01 | 2962 | 603297 | . | . | 1.864727 | 1.873230 | 2.230067 | 1.442255 | 4.339294 | 4.330054 | 5.004207 | 3.297544 | -1.029462 | 791.274260 | 0.19156841 | -0.21866894 | 0.826907949 | 0.998865404 | 9.648359 | 9.607419 | 9.631842 | 9.631901 | 9.754251 | 9.445634 | 924 | 866 | 827 | 571 |
| 79660 | NM_00120132 | PPP1R3B | protein phosphatase 1 regulatory subunit 3B | protein_coding | NP_001188258.1;NP_078883.2;XP_006716316.1;XP_016869324.1 | 14942 | 610541 | . | . | 2.906897 | 2.970636 | 2.710883 | 2.945408 | 6.764467 | 6.866758 | 6.083142 | 6.734326 | -1.054154 | 498.893808 | 0.168597022 | -0.45128737 | 0.651782449 | 0.998865404 | 9.000332 | 8.923832 | 8.974076 | 8.984270 | 8.919923 | 8.970855 | 585 | 557 | 405 | 474 |
| 79661 | NM_00125655 | NEIL1 | nei like DNA glycosylase 1 | protein_coding | NP_001243481.1;NP_001339448.1;NP_001339449.1;NP_078884.2;XP_0C | 18448 | 608844 | . | . | 0.306744 | 0.267706 | 0.297187 | 0.142781 | 0.713805 | 0.618813 | 0.666877 | 0.326452 | -1.509087 | 25.993195 | 0.647763272 | -0.9165009 | 0.359404237 | NA | 4.964015 | 4.376754 | 4.741501 | 4.697867 | 4.708920 | 4.643262 | 42 | 28 | 25 | 12 |
| 79663 | NM_00132072 | HSPBAP1 | HSPB1 associated protein 1 | protein_coding | NP_001307657.1;NP_078886.2;XP_005247819.1;XP_011511432.1;XP_01 | 16389 | 608263 | . | . | 5.981608 | 5.105151 | 5.941260 | 6.059591 | 13.919449 | 11.800787 | 13.332012 | 13.854532 | 1.064833 | 367.711575 | 0.196002053 | 0.462377908 | 0.643810339 | 0.998865404 | 8.476148 | 8.567275 | 8.560779 | 8.440128 | 8.526434 | 8.550690 | 453 | 344 | 323 | 363 |
| 79664 | NM_00101808 | ICE2 | interactor of little elongation complex ELL subunit 2 | protein_coding | NP_001018099.1;NP_001263314.1;NP_078887.2;XP_005254718.1;XP_00 | 29885 | 610835 | . | . | 2.762673 | 3.060389 | 3.012612 | 2.426186 | 6.428854 | 7.074227 | 6.760216 | 5.547186 | -1.067760 | 506.623170 | 0.178937954 | -0.52860536 | 0.597079242 | 0.998865404 | 9.030834 | 8.937184 | 9.008713 | 8.998489 | 9.031081 | 8.888082 | 607 | 560 | 475 | 407 |
| 79665 | NM_00116630 | DHX40 | DEAH-box helicase 40 | protein_coding | NP_001159773.1;NP_078888.4;XP_006722146.1;XP_011523555.1;XP_01 | 18018 | 607570 | . | . | 15.142762 | 14.741192 | 13.225802 | 16.498710 | 35.237830 | 34.074927 | 29.678315 | 37.722338 | -1.038369 | 1604.288584 | 0.13226246 | -0.41069094 | 0.681299173 | 0.998865404 | 10.674731 | 10.620190 | 10.687438 | 10.635086 | 10.574944 | 10.686446 | 1945 | 1705 | 1270 | 1582 |
| 79666 | NM_024613 | PLEKHF2 | pleckstrin homology and FYVE domain containing 2 | protein_coding | NP_078889.1 | 20757 | 615208 | . | . | 2.913050 | 2.772715 | 3.028818 | 2.230958 | 6.778788 | 6.409254 | 6.796580 | 5.100821 | -1.097479 | 247.961726 | 0.233850851 | -0.57384041 | 0.566075799 | 0.998865404 | 8.018718 | 7.886186 | 7.988350 | 7.959295 | 8.000552 | 7.849428 | 307 | 272 | 238 | 187 |
| 79668 | NM_00117805 | PARP8 | poly(ADP-ribose) polymerase family member 8 | protein_coding | NP_001171526.1;NP_001171527.1;NP_001317957.1;NP_078891.2;XP_0C | 26124 | . | . | . | 4.330368 | 4.821687 | 3.260790 | 3.822894 | 10.076948 | 11.145546 | 7.317119 | 8.740591 | -1.319740 | 926.597729 | 0.152579269 | -2.62325063 | 0.008709518 | 0.293930247 | 10.042428 | 9.641390 | 9.903475 | 9.986993 | 9.682756 | 9.804426 | 1147 | 1201 | 633 | 815 |
| 79669 | NM_00117174 | C3orf52 | chromosome 3 open reading frame 52 | protein_coding | NP_001165218.1;NP_078892.3 | 26255 | 611956 | . | . | 2.510273 | 2.625273 | 2.358778 | 1.848590 | 5.841509 | 6.068435 | 5.293028 | 4.226581 | -1.115989 | 294.767815 | 0.217455838 | -0.72806927 | 0.466571183 | 0.998865404 | 8.279831 | 8.122767 | 8.223752 | 8.235564 | 8.242278 | 8.094942 | 354 | 339 | 277 | 224 |
| 79670 | NM_00118505 | TUT7 | terminal uridylyl transferase 7 | protein_coding | NP_001171988.1;NP_001172003.1;NP_001317647.1;NP_078893.2;XP_0C | 25817 | 613467 | . | . | 2.435636 | 2.630211 | 2.716890 | 2.582342 | 5.667824 | 6.079849 | 6.096621 | 5.904219 | 1.067538 | 495.170741 | 0.169893547 | 0.554981114 | 0.578907578 | 0.998865404 | 8.903874 | 8.998151 | 8.903728 | 8.955297 | 8.989955 | 8.951964 | 529 | 538 | 458 | 465 |
| 79671 | NM_00128214 | NLRX1 | NLR family member X1 | protein_coding | NP_001269072.1;NP_001269073.1;NP_001269287.1;NP_078894.2;XP_0C | 29890 | 611947 | . | . | 1.362742 | 1.545233 | 1.255162 | 1.397643 | 3.171155 | 3.571875 | 2.816546 | 3.195545 | -1.115190 | 168.289585 | 0.259189833 | -0.6068485 | 0.543951456 | 0.998865404 | 7.471858 | 7.313397 | 7.383619 | 7.448800 | 7.350686 | 7.392074 | 189 | 206 | 131 | 157 |
| 79672 | NM_024619,N | FN3KRP | fructosamine 3 kinase related protein | protein_coding | NP_078895.2;XP_024306716.1 | 25700 | 611683 | . | . | 14.841663 | 14.350416 | 14.526585 | 17.079659 | 34.537162 | 33.171630 | 32.597236 | 39.050612 | 1.063441 | 868.551773 | 0.148142019 | 0.599019396 | 0.54915995 | 0.998865404 | 9.717612 | 9.805971 | 9.750524 | 9.727798 | 9.729654 | 9.834019 | 985 | 894 | 722 | 900 |
| 79673 | NM_024620,XI | ZNF329 | zinc finger protein 329 | protein_coding | NP_078896.3;XP_006723444.1;XP_006723445.1;XP_006723446.1;XP_00 | 14209 | . | . | . | 0.189903 | 0.286599 | 0.340182 | 0.238249 | 0.441912 | 0.662486 | 0.763358 | 0.544728 | 1.049294 | 31.438019 | 0.563311376 | 0.123234501 | 0.901921406 | 0.998865404 | 4.939842 | 5.008225 | 4.940175 | 4.992934 | 4.981967 | 4.963911 | 28 | 40 | 30 | 28 |
| 79674 | NM_00116791 | VEPH1 | ventricular zone expressed PH domain containing 1 | protein_coding | NP_001161383.1;NP_001161384.1;NP_001161387.1;NP_001161388.1;NI | 25735 | 609594 | . | . | 2.564983 | 2.830118 | 3.610577 | 3.273506 | 5.968821 | 6.541944 | 8.102031 | 7.484484 | 1.220976 | 434.249283 | 0.177443851 | 1.62324167 | 0.104537753 | 0.830227047 | 8.611254 | 8.899165 | 8.668570 | 8.722056 | 8.831747 | 8.802392 | 430 | 441 | 425 | 437 |
| 79675 | NM_00128147 | FASTKD1 | FAST kinase domains 1 | protein_coding | NP_001268405.1;NP_001308975.1;NP_001308977.1;NP_001308978.1;NI | 26150 | 617529 | . | . | 11.332608 | 11.228035 | 12.335776 | 8.833268 | 26.371448 | 25.954104 | 27.681124 | 20.196218 | -1.096785 | 1448.654287 | 0.162764453 | -0.81885615 | 0.412868492 | 0.998865404 | 10.565302 | 10.432645 | 10.540155 | 10.521930 | 10.586067 | 10.316304 | 1765 | 1616 | 1414 | 1067 |
| 79676 | NM_00130483 | OGFOD2 | 2-oxoglutarate and iron dependent oxygenase domain containing | protein_coding | NP_001291762.1;NP_001291763.1;NP_001291764.1;NP_001291765.1;NI | 25823 | . | . | . | 9.961055 | 10.769490 | 8.708217 | 9.331930 | 23.179782 | 24.894160 | 19.540986 | 21.336350 | -1.150857 | 619.033861 | 0.158165312 | -1.28162121 | 0.199975555 | 0.96621494 | 9.371865 | 9.168740 | 9.292566 | 9.344508 | 9.208915 | 9.240855 | 734 | 742 | 488 | 553 |
| 79677 | NM_00114228 | SMC6 | structural maintenance of chromosomes 6 | protein_coding | NP_001135758.1;NP_078900.1;XP_011531409.1;XP_011531410.1;XP_01 | 20466 | 609387 | . | . | 7.512866 | 7.031065 | 8.094591 | 5.220131 | 17.482745 | 16.252618 | 18.164031 | 11.935212 | -1.094719 | 1115.265539 | 0.178413432 | -0.73178949 | 0.464297064 | 0.998865404 | 10.186583 | 10.056844 | 10.173517 | 10.127018 | 10.223515 | 9.920529 | 1383 | 1219 | 1116 | 794 |
| 79680 | NM_024627 | RTL10 | retrotransposon Gag like 10 | protein_coding | NP_078903.3 | 26112 | . | . | . | 2.908504 | 3.451867 | 3.358446 | 2.931134 | 6.768209 | 7.979146 | 7.536255 | 6.701690 | -1.024460 | 656.035385 | 0.166516184 | -0.20937179 | 0.834158017 | 0.998865404 | 9.374902 | 9.340148 | 9.306548 | 9.418504 | 9.395818 | 9.297279 | 702 | 775 | 605 | 563 |
| 79682 | NM_024629,N | CENPU | centromere protein U | protein_coding | NP_078905.2;XP_005263275.2 | 21348 | 611511 | . | . | 17.334965 | 17.104203 | 14.152345 | 15.828271 | 40.339174 | 39.537134 | 31.757449 | 36.189464 | -1.162014 | 1256.962763 | 0.132796746 | -1.63127256 | 0.102832821 | 0.823247517 | 10.400101 | 10.183207 | 10.357213 | 10.342150 | 10.198170 | 10.272559 | 1571 | 1444 | 960 | 1145 |
| 79683 | NM_024630,N | ZDHHC14 | zinc finger DHHC-type palmitoyltransferase 14 | protein_coding | NP_078906.2;NP_714968.1;XP_005267203.1;XP_011534422.1;XP_01153 | 20341 | . | . | . | 3.408521 | 3.150182 | 3.466329 | 3.764146 | 7.931767 | 7.281788 | 7.778341 | 8.606273 | 1.223538 | 437.054124 | 0.176141594 | 1.652414647 | 0.098450052 | 0.813553488 | 8.618589 | 8.910052 | 8.709834 | 8.698696 | 8.851739 | 8.802039 | 457 | 420 | 435 | 433 |
| 79684 | NM_00130802 | MSANTD2 | Myb/SANT DNA binding domain containing 2 | protein_coding | NP_001294956.1;NP_001299848.1;NP_001299849.1;NP_001299850.1;NI | 26266 | . | . | . | 2.759645 | 2.535629 | 2.948541 | 2.307002 | 6.421806 | 5.861221 | 6.616440 | 5.274687 | 1.030229 | 391.691936 | 0.21847918 | 0.196655438 | 0.844097185 | 0.998865404 | 8.591357 | 8.635457 | 8.569557 | 8.626036 | 8.732117 | 8.484682 | 423 | 436 | 418 | 295 |
| 79685 | NM_00113106 | SAP30L | SAP30 like | protein_coding | NP_001124534.1;NP_001124535.1;NP_078908.1 | 25663 | 610398 | . | . | 2.310775 | 2.144206 | 2.859641 | 2.242994 | 5.377266 | 4.956426 | 6.416953 | 5.128340 | 1.076376 | 378.358112 | 0.198962517 | 0.533679183 | 0.593563514 | 0.998865404 | 8.508869 | 8.616347 | 8.570610 | 8.505577 | 8.649553 | 8.509149 | 443 | 371 | 382 | 324 |
| 79690 | NM_024637 | GAL3ST4 | galactose-3-O-sulfotransferase 4 | protein_coding | NP_078913.3 | 24145 | 608235 | . | . | 0.881323 | 0.778058 | 0.994104 | 0.747352 | 2.050874 | 1.798517 | 2.230740 | 1.708730 | 1.030348 | 64.099180 | 0.401375476 | 0.107459061 | 0.914424802 | 0.998865404 | 5.979359 | 6.024752 | 6.009488 | 5.982484 | 6.035027 | 5.969410 | 77 | 64 | 65 | 52 |
| 79691 | NM_00125683 | QTRT2 | queuine tRNA-ribosyltransferase accessory subunit 2 | protein_coding | NP_001243764.1;NP_001243765.1;NP_001243766.1;NP_078914.1;XP_0C | 25771 | . | . | . | 5.423469 | 5.729866 | 6.619832 | 6.570296 | 12.620636 | 13.244842 | 14.854712 | 15.022202 | 1.163449 | 753.074337 | 0.147257197 | 1.483170033 | 0.138029241 | 0.899796454 | 9.443322 | 9.661718 | 9.488650 | 9.512072 | 9.614410 | 9.597194 | 788 | 764 | 713 | 750 |
| 79692 | NM_00124279 | ZNF322 | zinc finger protein 322 | protein_coding | NP_001229726.1;NP_001229727.1;NP_001229728.1;NP_078915.2 | 23640 | 610847 | . | . | 2.522469 | 3.049436 | 2.930428 | 2.909006 | 5.869889 | 7.048908 | 6.575796 | 6.651096 | 1.034346 | 428.484678 | 0.183983611 | 0.264801333 | 0.79116251 | 0.998865404 | 8.718787 | 8.767010 | 8.674403 | 8.785245 | 8.756748 | 8.746685 | 441 | 496 | 382 | 405 |
| 79693 | NM_024640 | YRDC | yrdC N6-threonylcarbamoyltransferase domain containing | protein_coding | NP_078916.3 | 28905 | 612276 | . | . | 10.529799 | 11.586545 | 10.123520 | 12.345788 | 24.503277 | 26.782816 | 22.716887 | 28.227177 | 1.002631 | 641.762610 | 0.165848802 | 0.022853198 | 0.981767373 | 0.998865404 | 9.324439 | 9.327351 | 9.293822 | 9.353057 | 9.260343 | 9.386642 | 706 | 722 | 506 | 659 |
| 79694 | NM_024641,XI | MANEA | mannosidase endo-alpha | protein_coding | NP_078917.2;XP_005267204.1 | 21072 | 612327 | . | . | 2.234209 | 1.891451 | 1.800848 | 1.441116 | 5.199098 | 4.372175 | 4.041050 | 3.294940 | -1.291605 | 263.461896 | 0.226793748 | -1.62775405 | 0.103577047 | 0.826207034 | 8.213517 | 7.846047 | 8.147668 | 8.050513 | 8.023072 | 7.914508 | 372 | 292 | 223 | 191 |
| 79695 | NM_024642,XI | GALNT12 | polypeptide N-acetylgalactosaminyltransferase 12 | protein_coding | NP_078918.3;XP_006717350.1;XP_011517320.1;XP_011517322.1;XP_01 | 19877 | 610290 | . | . | 4.744124 | 4.662173 | 3.729921 | 4.545651 | 11.039771 | 10.776821 | 8.369834 | 10.393092 | -1.152196 | 407.921057 | 0.185591544 | -1.10126598 | 0.270780913 | 0.998865404 | 8.771036 | 8.565976 | 8.723775 | 8.701496 | 8.579043 | 8.674390 | 512 | 463 | 302 | 385 |
| 79696 | NM_00104243 | ZC2HC1C | zinc finger C2HC-type containing 1C | protein_coding | NP_001035895.1;NP_001317120.1;NP_078919.2;XP_005268119.1;XP_00 | 20354 | . | . | . | 1.685124 | 1.916450 | 1.945218 | 2.114027 | 3.921353 | 4.429959 | 4.365015 | 4.833470 | 1.082884 | 152.342564 | 0.265243166 | 0.4331092 | 0.66493544 | 0.998865404 | 7.192575 | 7.307488 | 7.223524 | 7.244160 | 7.274331 | 7.255836 | 164 | 162 | 141 | 145 |
| 79697 | NM_024644 | RIOX1 | ribosomal oxygenase 1 | protein_coding | NP_078920.2 | 20968 | 611919 | . | . | 5.317885 | 5.784895 | 5.339259 | 6.088154 | 12.374939 | 13.372044 | 11.981141 | 13.919841 | 1.015671 | 432.382980 | 0.180116354 | 0.124550191 | 0.900879663 | 0.998865404 | 8.745295 | 8.766956 | 8.726851 | 8.774274 | 8.723047 | 8.795941 | 475 | 481 | 356 | 433 |
| 79699 | NM_024646,XI | ZYG11B | zyg-11 family member B, cell cycle regulator | protein_coding | NP_078922.1;XP_006710961.1;XP_016857825.1 | 25820 | 618673 | . | . | 2.660947 | 2.289321 | 2.292780 | 2.234839 | 6.192133 | 5.291869 | 5.144932 | 5.109694 | -1.114090 | 598.828676 | 0.164765099 | -0.94598642 | 0.344155515 | 0.998865404 | 9.301547 | 9.146277 | 9.310759 | 9.206864 | 9.204664 | 9.172312 | 782 | 629 | 505 | 518 |
| 79701 | NM_024648,N | OGFOD3 | 2-oxoglutarate and iron dependent oxygenase domain containing | protein_coding | NP_078924.1;NP_787098.3 | 26174 | . | . | . | 7.193339 | 6.763386 | 7.202831 | 7.284040 | 16.739196 | 15.633869 | 16.162944 | 16.654095 | 1.024209 | 917.106443 | 0.140393003 | 0.245811087 | 0.805828474 | 0.998865404 | 9.823563 | 9.858121 | 9.852377 | 9.811336 | 9.839411 | 9.859209 | 1073 | 950 | 796 | 882 |
| 79703 | NM_00130208 | C11orf80 | chromosome 11 open reading frame 80 | protein_coding | NP_001289013.1;NP_078926.3 | 26197 | 616109 | . | . | 4.451770 | 3.584811 | 4.789628 | 3.808540 | 10.359452 | 8.286451 | 10.747788 | 8.707774 | 1.024450 | 275.570738 | 0.226877773 | 0.153607247 | 0.877919423 | 0.998865404 | 8.087878 | 8.124448 | 8.151107 | 8.038678 | 8.169726 | 8.044698 | 347 | 262 | 272 | 230 |
| 79705 | NM_024652,XI | LRRK1 | leucine rich repeat kinase 1 | protein_coding | NP_078928.3;XP_005255036.1;XP_011520314.1;XP_011520315.1;XP_01 | 18608 | 610986 | . | . | 2.514348 | 2.621676 | 2.403715 | 3.001751 | 5.850991 | 6.060117 | 5.393865 | 6.863145 | 1.099059 | 1063.890475 | 0.141734956 | 0.961436782 | 0.336332598 | 0.998865404 | 9.985595 | 10.121475 | 9.962913 | 10.070401 | 10.077953 | 10.097607 | 1082 | 1174 | 956 | 1058 |
| 79706 | NM_024653 | PRKRIP1 | PRKR interacting protein 1 | protein_coding | NP_078929.1 | 21894 | 617458 | . | . | 8.189288 | 8.070831 | 8.840524 | 9.770225 | 19.056808 | 18.656088 | 19.837879 | 22.338459 | 1.128717 | 586.650677 | 0.160536643 | 1.088124742 | 0.276540042 | 0.998865404 | 9.106518 | 9.280930 | 9.161289 | 9.146559 | 9.203640 | 9.263152 | 642 | 588 | 517 | 609 |
| 79707 | NM_024654,XI | NOL9 | nucleolar protein 9 | protein_coding | NP_078930.4;XP_005263550.1;XP_011540449.1;XP_016857826.1;XP_01 | 26265 | . | . | . | 6.251735 | 6.725105 | 7.138420 | 6.431278 | 14.548047 | 15.545381 | 16.018408 | 14.704352 | 1.032467 | 1282.589708 | 0.134099046 | 0.343742825 | 0.731039702 | 0.998865404 | 10.301585 | 10.347734 | 10.277290 | 10.345447 | 10.368376 | 10.302947 | 1388 | 1423 | 1182 | 1169 |
| 79709 | NM_024656,XI | COLGALT1 | collagen beta(1-O)galactosyltransferase 1 | protein_coding | NP_078932.2;XP_005260137.1;XP_011526599.1;XP_011526600.1;XP_01 | 26182 | 617531 | . | . | 101.529623 | 106.949767 | 82.906789 | 94.997586 | 236.263597 | 247.219172 | 186.040429 | 217.200687 | -1.188998 | 10925.455054 | 0.109512657 | -2.28052624 | 0.022576496 | 0.49659767 | 13.534917 | 13.285092 | 13.464743 | 13.503673 | 13.283258 | 13.387293 | 13360 | 13095 | 8143 | 9937 |
| 79710 | NM_00108535 | MORC4 | MORC family CW-type zinc finger 4 | protein_coding | NP_001078823.1;NP_078933.3;XP_005262247.1;XP_006724754.1;XP_01 | 23485 | 300970 | . | . | 3.085142 | 3.029427 | 2.574059 | 3.452468 | 7.179252 | 7.002656 | 5.776114 | 7.893660 | -1.038036 | 366.638358 | 0.199520043 | -0.26992794 | 0.787215694 | 0.998865404 | 8.545338 | 8.490570 | 8.542379 | 8.512782 | 8.433227 | 8.573923 | 441 | 393 | 274 | 379 |
| 79711 | NM_024658,N | IPO4 | importin 4 | protein_coding | NP_078934.3 | 19426 | . | . | . | 19.217581 | 20.230523 | 18.276821 | 19.811106 | 44.720099 | 46.763764 | 41.012658 | 45.295740 | -1.050962 | 2128.009313 | 0.1192168 | -0.60151297 | 0.547498378 | 0.998865404 | 11.090795 | 11.018886 | 11.054862 | 11.094210 | 11.010379 | 11.058970 | 2450 | 2411 | 1749 | 2005 |
| 79712 | NM_00100663 | GTDC1 | glycosyltransferase like domain containing 1 | protein_coding | NP_001006637.1;NP_001158101.1;NP_001271162.1;NP_001271163.1;NI | 20887 | 610165 | . | . | 1.431271 | 1.778545 | 1.703585 | 2.089533 | 3.330626 | 4.111186 | 3.822793 | 4.777471 | -1.091514 | 338.792924 | 0.193328259 | -0.65344933 | 0.513466647 | 0.998865404 | 8.466305 | 8.339427 | 8.408127 | 8.448457 | 8.362498 | 8.395778 | 393 | 395 | 273 | 313 |
| 79713 | NM_00134600 | IGFLR1 | IGF like family receptor 1 | protein_coding | NP_001332932.1;NP_001332933.1;NP_001332934.1;NP_001332935.1;NI | 23620 | 614143 | . | . | 3.994550 | 3.004862 | 3.548134 | 3.874707 | 9.295483 | 6.945875 | 7.961910 | 8.859053 | 1.054309 | 185.710796 | 0.25666731 | 0.297265334 | 0.766263945 | 0.998865404 | 7.497907 | 7.574894 | 7.581451 | 7.460745 | 7.530545 | 7.561178 | 237 | 168 | 160 | 185 |
| 79714 | NM_00125696 | CCDC51 | coiled-coil domain containing 51 | protein_coding | NP_001243893.1;NP_001243894.1;NP_001243895.1;NP_001243896.1;NI | 25714 | 618585 | . | . | 13.295889 | 13.400308 | 10.531435 | 13.676759 | 30.940084 | 30.975410 | 23.632231 | 31.270286 | -1.130871 | 627.752518 | 0.16883372 | -1.05094004 | 0.293286129 | 0.998865404 | 9.380463 | 9.202232 | 9.337921 | 9.327838 | 9.173496 | 9.321647 | 773 | 714 | 454 | 615 |
| 79716 | NM_00120487 | NPEPL1 | aminopeptidase like 1 | protein_coding | NP_001191801.1;NP_001191802.1;NP_078939.3 | 16244 | . | . | . | 10.239643 | 10.083372 | 9.153823 | 11.199261 | 23.828070 | 23.308164 | 20.540915 | 25.605780 | -1.013306 | 693.934940 | 0.161283673 | -0.11823516 | 0.905881328 | 0.998865404 | 9.448464 | 9.428782 | 9.450428 | 9.434092 | 9.364693 | 9.497189 | 814 | 745 | 541 | 709 |
| 79717 | NM_00107744 | PPCS | phosphopantothenoylcysteine synthetase | protein_coding | NP_001070915.1;NP_001274435.1;NP_001274436.1;NP_001274437.1;NI | 25686 | 609853 | . | . | 13.106707 | 13.428213 | 14.444933 | 12.741827 | 30.499850 | 31.039912 | 32.414007 | 29.132673 | 1.009666 | 927.714839 | 0.143543662 | 0.096684004 | 0.922977343 | 0.998865404 | 9.850432 | 9.864608 | 9.841028 | 9.865641 | 9.901708 | 9.817474 | 1045 | 1013 | 858 | 823 |
| 79718 | NM_00132119 | TBL1XR1 | TBL1X receptor 1 | protein_coding | NP_001308122.1;NP_001308123.1;NP_001308124.1;NP_001361256.1;NI | 29529 | 608628 | . | . | 6.643928 | 5.652060 | 6.613598 | 4.915404 | 15.460695 | 13.064991 | 14.840721 | 11.238491 | -1.120061 | 1424.385023 | 0.15539693 | -1.05264239 | 0.292504922 | 0.998865404 | 10.555272 | 10.392380 | 10.562132 | 10.465968 | 10.529233 | 10.316045 | 1839 | 1524 | 1333 | 1083 |
| 79719 | NM_00127188 | AAGAB | alpha and gamma adaptin binding protein | protein_coding | NP_001258814.1;NP_001258815.1;NP_078942.3;XP_011520322.1;XP_02 | 25662 | 614888 | . | . | 11.431663 | 10.680582 | 10.594695 | 12.019878 | 26.601950 | 24.688644 | 23.774190 | 27.482028 | 1.006989 | 1085.489336 | 0.139039524 | 0.072263264 | 0.9423924 | 0.998865404 | 10.079160 | 10.089082 | 10.110194 | 10.051198 | 10.048749 | 10.122420 | 1295 | 1121 | 900 | 1072 |
| 79720 | NM_024667,XI | VPS37B | VPS37B subunit of ESCRT-I | protein_coding | NP_078943.1;XP_005253679.1;XP_006719665.1;XP_011537045.1;XP_01 | 25754 | 610037 | . | . | 17.109274 | 19.395964 | 20.478191 | 22.097677 | 39.813983 | 44.834645 | 45.952472 | 50.523714 | 1.129890 | 1670.021211 | 0.127271506 | 1.384299841 | 0.166266663 | 0.944092297 | 10.615042 | 10.790903 | 10.618654 | 10.690584 | 10.721059 | 10.779316 | 1722 | 1771 | 1484 | 1722 |
| 79723 | NM_00119342 | SUV39H2 | suppressor of variegation 3-9 homolog 2 | protein_coding | NP_001180353.1;NP_001180354.1;NP_001180355.1;NP_001180356.1;NI | 17287 | 606503 | . | . | 4.991148 | 5.158663 | 4.816602 | 4.127984 | 11.614606 | 11.924479 | 10.808316 | 9.438144 | -1.145029 | 451.737252 | 0.179113939 | -1.0908396 | 0.27534347 | 0.998865404 | 8.913427 | 8.718687 | 8.849408 | 8.868095 | 8.822661 | 8.726163 | 547 | 528 | 396 | 363 |
| 79724 | NM_024671,XI | ZNF768 | zinc finger protein 768 | protein_coding | NP_078947.3;XP_016879154.1;XP_016879155.1 | 26273 | 618032 | . | . | 17.945798 | 17.981574 | 15.626543 | 20.028048 | 41.760609 | 41.565216 | 35.065508 | 45.791752 | -1.020192 | 1312.405663 | 0.146825184 | -0.19642545 | 0.844277182 | 0.998865404 | 10.372610 | 10.343237 | 10.362738 | 10.365331 | 10.254305 | 10.435081 | 1524 | 1433 | 995 | 1363 |
| 79725 | NM_00131777 | THAP9 | THAP domain containing 9 | protein_coding | NP_001304705.1;NP_078948.3;XP_005263277.1;XP_011530548.1;XP_01 | 23192 | 612537 | . | . | 0.864638 | 0.942533 | 0.796562 | 0.343387 | 2.012045 | 2.178708 | 1.787462 | 0.785114 | -1.525558 | 91.885213 | 0.380993149 | -1.59933963 | 0.109745158 | 0.843391194 | 6.793070 | 6.187331 | 6.555504 | 6.584822 | 6.528775 | 6.364143 | 122 | 125 | 85 | 45 |
| 79726 | NM_00132417 | WDR59 | WD repeat domain 59 | protein_coding | NP_001311100.1;NP_001311101.1;NP_085058.3;XP_005256203.1;XP_01 | 25706 | 617418 | . | . | 4.987903 | 5.020129 | 5.704205 | 5.276730 | 11.607054 | 11.604256 | 12.800068 | 12.064616 | 1.079939 | 777.021966 | 0.146638526 | 0.756620407 | 0.449277312 | 0.998865404 | 9.545356 | 9.656141 | 9.552058 | 9.594879 | 9.627893 | 9.627236 | 834 | 831 | 702 | 756 |
| 79728 | NM_024675,XI | PALB2 | partner and localizer of BRCA2 | protein_coding | NP_078951.2;XP_011544248.1;XP_011544249.1;XP_011544250.1;XP_01 | 26144 | 610355 | . | . | 7.313785 | 7.824564 | 8.003258 | 7.443589 | 17.019479 | 18.086832 | 17.959081 | 17.018881 | 1.008551 | 976.425133 | 0.138845059 | 0.088475233 | 0.929498969 | 0.998865404 | 9.925163 | 9.937541 | 9.911124 | 9.944523 | 9.956371 | 9.911544 | 1094 | 1073 | 879 | 891 |
| 79729 | NM_00116253 | SH3D21 | SH3 domain containing 21 | protein_coding | NP_001156002.1;NP_078952.4;XP_011540455.1;XP_016857829.1;XP_01 | 26236 | . | . | . | 6.266862 | 5.739884 | 7.230441 | 7.113402 | 14.583248 | 13.267997 | 16.224902 | 16.263949 | 1.188633 | 538.190420 | 0.168795367 | 1.476952718 | 0.139688312 | 0.900674731 | 8.941549 | 9.191577 | 9.033753 | 8.987562 | 9.165389 | 9.078261 | 587 | 511 | 542 | 512 |
| 79731 | NM_00124325 | NARS2 | asparaginyl-tRNA synthetase 2, mitochondrial | protein_coding | NP_001230180.1;NP_078954.4;XP_011543555.1;XP_016873791.1;XP_01 | 26274 | 612803 | . | . | 16.071669 | 17.334183 | 19.820102 | 17.828976 | 37.399434 | 40.068739 | 44.475732 | 40.763834 | 1.139229 | 1462.439609 | 0.125061939 | 1.51080096 | 0.13083917 | 0.882587711 | 10.416548 | 10.605587 | 10.463982 | 10.458104 | 10.573942 | 10.549794 | 1580 | 1469 | 1376 | 1438 |
| 79733 | NM_00125637 | E2F8 | E2F transcription factor 8 | protein_coding | NP_001243300.1;NP_001243301.1;NP_078956.2;XP_011518669.1 | 24727 | 612047 | . | . | 3.239156 | 4.111261 | 3.651129 | 4.189278 | 7.537649 | 9.503364 | 8.193027 | 9.578287 | 1.098992 | 431.543000 | 0.191829754 | 0.648064539 | 0.516943209 | 0.998865404 | 8.690460 | 8.813633 | 8.650376 | 8.792058 | 8.737030 | 8.813376 | 420 | 498 | 367 | 448 |
| 79734 | NM_00128268 | KCTD17 | potassium channel tetramerization domain containing 17 | protein_coding | NP_001269613.1;NP_001269614.1;NP_001269615.1;NP_078957.2;XP_0C | 25705 | 616386 | . | . | 8.419730 | 7.739131 | 7.772561 | 8.423785 | 19.593057 | 17.889352 | 17.441404 | 19.259985 | 1.009492 | 441.842446 | 0.175712048 | 0.077565086 | 0.938174029 | 0.998865404 | 8.780561 | 8.794184 | 8.805531 | 8.762669 | 8.774519 | 8.805788 | 524 | 458 | 376 | 427 |
| 79735 | NM_00116822 | TBC1D17 | TBC1 domain family member 17 | protein_coding | NP_001161694.1;NP_078958.2;XP_011525619.1 | 25699 | 616659 | . | . | 10.361850 | 11.408004 | 12.758287 | 11.914956 | 24.112453 | 26.370113 | 28.629226 | 27.242133 | 1.135054 | 771.255325 | 0.150094608 | 1.217634328 | 0.223362963 | 0.982008277 | 9.496810 | 9.679537 | 9.509933 | 9.576359 | 9.655865 | 9.608384 | 792 | 817 | 737 | 743 |
| 79736 | NM_024683,XI | TEFM | transcription elongation factor, mitochondrial | protein_coding | NP_078959.3;XP_006722147.1 | 26223 | 616422 | . | . | 6.325461 | 6.807561 | 6.674761 | 7.587563 | 14.719606 | 15.735983 | 14.977969 | 17.348060 | 1.130319 | 314.970006 | 0.202256771 | 0.873790579 | 0.382232312 | 0.998865404 | 8.208355 | 8.384444 | 8.217664 | 8.305034 | 8.326042 | 8.332412 | 313 | 345 | 289 | 315 |

| | | | | | | | | | | | | | | | | | | | | | | | |
|---|---|---|---|---|---|---|---|---|---|---|---|---|---|---|---|---|---|---|---|---|---|---|---|
| 79738 | NM_024685 | BBS10 | Bardet-Biedl syndrome 10 | protein_coding | NP_078961.3 | 26291 | 610148 | . | . | 2.144894 | 2.044276 | 2.718162 | 1.362087 | 4.991257 | 4.725437 | 6.099476 | 3.114250 | -1.038234 | 230.562346 | 0.290197962 | -0.18653171 | 0.852027818 | [illegible] |
| 79739 | NM_00135021 | TTLL7 | tubulin tyrosine ligase like 7 | protein_coding | NP_001337143.1;NP_001337144.1;NP_078962.4;XP_011540461.1;XP_01 | 26242 | 618813 | . | . | 2.649818 | 2.712219 | 3.278506 | 2.380776 | 6.166235 | 6.269413 | 7.356873 | 5.443362 | 1.063249 | 700.036832 | 0.179588992 | 0.49267524 | 0.622242075 | [illegible] |
| 79741 | NM_00102638 | CCDC7 | coiled-coil domain containing 7 | protein_coding | NP_001021554.1;NP_001308044.1;NP_659460.3;XP_005252650.1;XP_00 | 26533 | . | . | . | 0.658679 | 0.829928 | 0.586843 | 0.378523 | 1.532773 | 1.918418 | 1.316859 | 0.865449 | -1.549367 | 86.637868 | 0.363786644 | -1.73640038 | 0.082493065 | [illegible] |
| 79744 | NM_00109849 | ZNF419 | zinc finger protein 419 | protein_coding | NP_001091961.1;NP_001091962.1;NP_001091963.1;NP_001091964.1;NI | 20648 | 617410 | . | . | 1.483755 | 1.427984 | 1.320563 | 1.457659 | 3.452760 | 3.300850 | 2.963305 | 3.332763 | -1.013495 | 158.385091 | 0.263565905 | -0.07337683 | 0.941506254 | [illegible] |
| 79745 | NM_00128752 | CLIP4 | CAP-Gly domain containing linker protein family member 4 | protein_coding | NP_001274456.1;NP_001274457.1;NP_078968.3;XP_005264619.1;XP_00 | 26108 | . | . | . | 21.764578 | 20.121563 | 23.074433 | 21.608083 | 50.647063 | 46.511891 | 51.778359 | 49.404309 | 1.049340 | 3019.331025 | 0.119079816 | 0.583493653 | 0.559561028 | [illegible] |
| 79746 | NM_024693,XI | ECHDC3 | enoyl-CoA hydratase domain containing 3 | protein_coding | NP_078969.3;XP_011517991.1 | 23489 | . | . | . | 2.803353 | 2.626470 | 1.764773 | 3.113498 | 6.523516 | 6.071203 | 3.960098 | 7.118642 | -1.124377 | 131.027673 | 0.320618109 | -0.5274983 | 0.597847593 | [illegible] |
| 79751 | NM_00119106 | SLC25A22 | solute carrier family 25 member 22 | protein_coding | NP_001177989.1;NP_001177990.1;NP_078974.1;XP_011518671.1;XP_01 | 19954 | 609302 | . | . | 9.891380 | 10.342518 | 9.690054 | 12.056161 | 23.017643 | 23.907194 | 21.744200 | 27.564976 | 1.057428 | 879.196893 | 0.153608226 | 0.524450078 | 0.599965533 | [illegible] |
| 79752 | NM_00117079 | ZFAND1 | zinc finger AN1-type containing 1 | protein_coding | NP_001164267.1;NP_001164268.1;NP_078975.2 | 25858 | . | . | . | 8.030193 | 7.245233 | 8.378330 | 7.613138 | 18.686591 | 16.747679 | 18.800731 | 17.406536 | 1.013309 | 523.614822 | 0.173791013 | 0.109752566 | 0.91260561 | [illegible] |
| 79753 | NM_024700 | SNIP1 | Smad nuclear interacting protein 1 | protein_coding | NP_078976.2 | 30587 | 608241 | . | . | 2.661885 | 2.759532 | 2.810954 | 3.170338 | 6.194316 | 6.378782 | 6.307699 | 7.248600 | 1.089186 | 404.723889 | 0.182194224 | 0.676475319 | 0.498738911 | [illegible] |
| 79754 | NM_024701,N | ASB13 | ankyrin repeat and SOCS box containing 13 | protein_coding | NP_078977.2 | 19765 | 615055 | . | . | 4.846443 | 4.861732 | 4.608037 | 6.243338 | 11.277871 | 11.238111 | 10.340301 | 14.274652 | 1.108016 | 437.832968 | 0.193732204 | 0.763827963 | 0.444969771 | [illegible] |
| 79758 | NM_00103171 | DHRS12 | dehydrogenase/reductase 12 | protein_coding | NP_001026889.1;NP_001257353.1;NP_001364462.1;NP_001364858.1;NI | 25832 | 616163 | . | . | 2.089333 | 2.055108 | 1.961569 | 1.872845 | 4.861964 | 4.750473 | 4.401702 | 4.282035 | -1.086663 | 98.491919 | 0.327634585 | -0.36596853 | 0.714388559 | [illegible] |
| 79759 | NM_00117266 | ZNF668 | zinc finger protein 668 | protein_coding | NP_001166139.1;NP_001166140.1;NP_001166141.1;NP_078982.3 | 25821 | 617103 | . | . | 3.817829 | 4.140253 | 3.809847 | 4.035292 | 8.884246 | 9.570377 | 8.549186 | 9.226218 | -1.024829 | 332.517831 | 0.19454111 | -0.18187744 | 0.855678913 | [illegible] |
| 79760 | NM_00100726 | GEMIN7 | gem nuclear organelle associated protein 7 | protein_coding | NP_001007270.1;NP_001007271.1;NP_001305983.1;NP_001305984.1;NI | 20045 | 607419 | . | . | 5.984892 | 6.404256 | 6.369841 | 7.147109 | 13.927088 | 14.803723 | 14.293739 | 16.341014 | 1.107218 | 299.810621 | 0.202880554 | 0.724263121 | 0.468904213 | [illegible] |
| 79762 | NM_024709 | C1orf115 | chromosome 1 open reading frame 115 | protein_coding | NP_078985.3 | 25873 | . | . | . | 0.094785 | 0.118478 | 0.182016 | 0.226229 | 0.220568 | 0.273868 | 0.408438 | 0.517247 | 1.897416 | 14.330412 | 0.815320936 | 1.133340238 | 0.2570714 | [illegible] |
| 79763 | NM_00113620 | ISOC2 | isochorismatase domain containing 2 | protein_coding | NP_001129673.1;NP_001129674.1;NP_078986.1 | 26278 | 612928 | . | . | 36.124075 | 35.667042 | 34.181182 | 39.527467 | 84.062210 | 82.445967 | 76.701580 | 90.374858 | 1.012825 | 1094.301890 | 0.142502109 | 0.129015201 | 0.897345621 | [illegible] |
| 79767 | NM_024712,XI | ELMO3 | engulfment and cell motility 3 | protein_coding | NP_078988.3;XP_024306215.1 | 17289 | 606422 | . | . | 0.846459 | 1.278421 | 0.859341 | 0.825419 | 1.969745 | 2.955126 | 1.928336 | 1.887223 | -1.276350 | 74.414252 | 0.384539182 | -0.91544483 | 0.359958155 | [illegible] |
| 79768 | NM_024713,XI | KATNBL1 | katanin regulatory subunit B1 like 1 | protein_coding | NP_078989.1;XP_005254725.1;XP_016878060.1;XP_016878061.1;XP_01 | 26199 | 616235 | . | . | 3.482358 | 2.286876 | 2.822425 | 1.978944 | 8.103591 | 5.286215 | 6.333441 | 4.524621 | -1.213334 | 215.549061 | 0.271355796 | -1.02808253 | 0.303911008 | [illegible] |
| 79770 | NM_00135073 | TXNDC15 | thioredoxin domain containing 15 | protein_coding | NP_001337664.1;NP_078991.3 | 20652 | 617778 | . | . | 17.566523 | 16.508480 | 16.158672 | 14.101821 | 40.878022 | 38.160090 | 36.259590 | 32.242138 | -1.140416 | 1552.196519 | 0.133726473 | -1.41752443 | 0.156329657 | [illegible] |
| 79772 | NM_00100279 | MCTP1 | multiple C2 and transmembrane domain containing 1 | protein_coding | NP_001002796.1;NP_001284706.1;NP_078993.4;XP_005272139.1;XP_00 | 26183 | 616296 | . | . | 4.188442 | 4.458462 | 3.969317 | 4.281633 | 9.746682 | 10.305933 | 8.907030 | 9.789447 | -1.051661 | 579.759302 | 0.162136179 | -0.4482055 | 0.654004891 | [illegible] |
| 79777 | NM_00113570 | ACBD4 | acyl-CoA binding domain containing 4 | protein_coding | NP_001129177.1;NP_001129178.1;NP_001129179.1;NP_001308281.1;NI | 23337 | . | . | . | 3.340973 | 2.697146 | 2.219982 | 3.808410 | 7.774578 | 6.234578 | 4.981576 | 8.707476 | -1.015582 | 216.511908 | 0.257718429 | -0.08655493 | 0.931025292 | [illegible] |
| 79778 | NM_182924,XI | MICALL2 | MICAL like 2 | protein_coding | NP_891554.1;XP_011513822.1;XP_011513824.1;XP_011513826.1;XP_02 | 29672 | . | . | . | 6.116656 | 5.427798 | 5.866318 | 5.967885 | 14.233708 | 12.546602 | 13.163847 | 13.644859 | -1.020456 | 612.476061 | 0.158851957 | -0.18390582 | 0.854087337 | [illegible] |
| 79780 | NM_00131873 | CCDC82 | coiled-coil domain containing 82 | protein_coding | NP_001305665.1;NP_001305666.1;NP_001350523.1;NP_079001.2;XP_00 | 26282 | . | . | . | 11.607702 | 9.795267 | 12.440706 | 7.606191 | 27.011603 | 22.642199 | 27.916587 | 17.390656 | -1.019904 | 1241.696519 | 0.426470308 | -0.06667028 | 0.946844197 | [illegible] |
| 79783 | NM_00119331 | SUGCT | succinyl-CoA:glutarate-CoA transferase | protein_coding | NP_001180240.2;NP_001180241.2;NP_001180242.2;NP_079004.2;XP_00 | 16001 | 609187 | . | . | 2.096699 | 1.962355 | 1.976286 | 2.537901 | 4.879104 | 4.536071 | 4.434729 | 5.802611 | 1.073178 | 112.358049 | 0.308812295 | 0.329940669 | 0.741444794 | [illegible] |
| 79784 | NM_00107718 | MYH14 | myosin heavy chain 14 | protein_coding | NP_001070654.1;NP_001139281.1;NP_079005.3;XP_006723449.1;XP_01 | 23212 | 608568 | . | . | 6.243675 | 6.542382 | 5.581665 | 6.737713 | 14.529289 | 15.123008 | 12.525095 | 15.404981 | -1.052407 | 1332.628175 | 0.138793069 | -0.53095464 | 0.595450214 | [illegible] |
| 79786 | NM_00130345 | KLHL36 | kelch like family member 36 | protein_coding | NP_001290380.1;NP_079007.2;XP_005256206.1 | 17844 | . | . | . | 3.919504 | 3.695346 | 3.669802 | 3.656672 | 9.120844 | 8.541961 | 8.234927 | 8.360543 | -1.043108 | 965.935022 | 0.13874136 | -0.43886674 | 0.660758097 | [illegible] |
| 79789 | NM_024734,XI | CLMN | calmin | protein_coding | NP_079010.2;XP_011535460.1;XP_011535461.1;XP_016877135.1;XP_01 | 19972 | 611121 | . | . | 4.011587 | 4.393268 | 4.004016 | 4.223156 | 9.335130 | 10.155237 | 8.984896 | 9.655741 | -1.032544 | 1645.238213 | 0.125603323 | -0.36784562 | 0.712988349 | [illegible] |
| 79791 | NM_00128268 | FBXO31 | F-box protein 31 | protein_coding | NP_001269612.1;NP_079011.3 | 16510 | 609102 | . | . | 3.930050 | 3.770718 | 4.168557 | 4.611557 | 9.145387 | 8.716183 | 9.354121 | 10.543776 | 1.123618 | 764.305327 | 0.14957676 | 1.124186778 | 0.260933797 | [illegible] |
| 79792 | NM_00116623 | GSDMD | gasdermin D | protein_coding | NP_001159709.1;NP_079012.3;XP_011515603.1 | 25697 | 617042 | . | . | 13.582291 | 14.659700 | 14.188477 | 15.524936 | 31.606549 | 33.886555 | 31.838532 | 35.495923 | 1.067763 | 917.250247 | 0.14107421 | 0.670507966 | 0.50253403 | [illegible] |
| 79794 | NM_00135362 | SPRING1 | SREBF pathway regulator in golgi 1 | protein_coding | NP_001340552.1;NP_001340553.1;NP_001340554.1;NP_079014.1;XP_02 | 26128 | . | . | . | 8.425441 | 8.472947 | 9.258997 | 8.849324 | 19.606351 | 19.585597 | 20.776922 | 20.232926 | 1.032776 | 2207.086573 | 0.115850299 | 0.401620576 | 0.687963285 | [illegible] |
| 79796 | NM_00107769 | ALG9 | ALG9 alpha-1,2-mannosyltransferase | protein_coding | NP_001071158.1;NP_001071159.1;NP_001071160.1;NP_001339338.1;NI | 15672 | 606941 | . | . | 2.785827 | 3.850813 | 3.303322 | 3.497735 | 6.482733 | 8.901333 | 7.412554 | 7.997156 | -1.034173 | 575.261299 | 0.16452684 | -0.29464762 | 0.768263079 | [illegible] |
| 79797 | NM_00118475 | ZNF408 | zinc finger protein 408 | protein_coding | NP_001171680.1;NP_079017.1 | 20041 | 616454 | . | . | 2.493078 | 2.848449 | 2.440737 | 3.046908 | 5.801495 | 6.584318 | 5.476943 | 6.966394 | 1.013824 | 188.968371 | 0.250916083 | 0.078939345 | 0.937080868 | [illegible] |
| 79798 | NM_00110524 | ARMC5 | armadillo repeat containing 5 | protein_coding | NP_001098717.1;NP_001275696.1;NP_001288749.1;NP_079018.1;XP_00 | 25781 | 615549 | . | . | 2.852292 | 2.662363 | 2.400937 | 3.173377 | 6.637400 | 6.154174 | 5.387632 | 7.255550 | 1.096409 | 318.214924 | 0.196415637 | 0.67604626 | 0.499011275 | [illegible] |
| 79800 | NM_00110458 | CARF | calcium responsive transcription factor | protein_coding | NP_001098056.1;NP_001269839.1;NP_001269840.1;NP_001269841.1;NI | 14435 | 607586 | . | . | 0.857755 | 0.668097 | 0.801388 | 0.648117 | 1.996027 | 1.544335 | 1.798293 | 1.481842 | 1.000801 | 160.409300 | 0.277543411 | 0.004162141 | 0.996679101 | [illegible] |
| 79801 | NM_00132431 | SHCBP1 | SHC binding and spindle associated 1 | protein_coding | NP_001311247.1;NP_001311248.1;NP_079021.4;XP_011521637.1 | 29547 | 611027 | . | . | 11.571752 | 11.278831 | 9.506409 | 9.900511 | 26.927945 | 26.071521 | 21.332106 | 22.636342 | -1.196551 | 1693.506556 | 0.122246438 | -2.11770547 | **0.03420002** | [illegible] |
| 79802 | NM_024746,XI | HHIPL2 | HHIP like 2 | protein_coding | NP_079022.2;XP_011508288.1;XP_016857839.1;XP_024305582.1 | 25842 | . | . | . | 2.244858 | 2.345603 | 1.006753 | 1.398177 | 5.223876 | 5.421968 | 2.259123 | 3.196766 | -1.566261 | 118.506290 | 0.298702264 | -2.16712282 | **0.030225491** | [illegible] |
| 79803 | NM_024747 | HPS6 | HPS6 biogenesis of lysosomal organelles complex 2 subunit 3 | protein_coding | NP_079023.2 | 18817 | 607522 | . | . | 4.327193 | 3.193839 | 3.599045 | 4.592849 | 10.069556 | 7.382701 | 8.076153 | 10.501003 | 1.073754 | 329.332988 | 0.222855097 | 0.460672178 | 0.645033819 | [illegible] |
| 79805 | NM_00113647 | VASH2 | vasohibin 2 | protein_coding | NP_001129946.1;NP_001129947.1;NP_001287985.1;NP_079025.2 | 25723 | 610471 | . | . | 0.157943 | 0.169242 | 0.180350 | 0.131335 | 0.367541 | 0.391211 | 0.404700 | 0.300281 | -1.054796 | 22.295222 | 0.655398255 | -0.11743087 | 0.906518625 | [illegible] |
| 79807 | NM_00103172 | GSTCD | glutathione S-transferase C-terminal domain containing | protein_coding | NP_001026890.2;NP_001357110.1;NP_079027.2;XP_011530550.1;XP_01 | 25806 | 615912 | . | . | 3.109329 | 3.261813 | 2.879497 | 3.165729 | 7.235535 | 7.539827 | 6.461508 | 7.238063 | -1.077803 | 414.964160 | 0.180255776 | -0.59967062 | 0.54872577 | [illegible] |
| 79809 | NM_024753,XI | TTC21B | tetratricopeptide repeat domain 21B | protein_coding | NP_079029.3;XP_006712824.1;XP_011510173.1;XP_011510174.1;XP_01 | 25660 | 612014 | . | . | 3.241666 | 3.440699 | 3.538798 | 2.523412 | 7.543490 | 7.953332 | 7.940963 | 5.769482 | 1.023459 | 596.446130 | 0.16162253 | 0.206981856 | 0.836024029 | [illegible] |
| 79810 | NM_00128440 | PTCD2 | pentatricopeptide repeat domain 2 | protein_coding | NP_001271332.1;NP_001271333.1;NP_001271334.1;NP_079030.3;XP_00 | 25734 | 615484 | . | . | 2.263665 | 1.993754 | 2.438692 | 2.473452 | 5.267643 | 4.608647 | 5.472356 | 5.655251 | 1.124136 | 590.503111 | 0.163223893 | 1.034261158 | 0.30101409 | [illegible] |
| 79811 | NM_00101384 | SLTM | SAFB like transcription modulator | protein_coding | NP_001013865.1;NP_079031.2;XP_006720749.3;XP_006720753.1;XP_01 | 20709 | . | . | . | 17.575271 | 17.035024 | 17.850475 | 15.952766 | 40.898381 | 39.377224 | 40.055947 | 36.474107 | -1.040830 | 2362.814698 | 0.121309709 | -0.47592068 | 0.63413089 | [illegible] |
| 79813 | NM_00114552 | EHMT1 | euchromatic histone lysine methyltransferase 1 | protein_coding | NP_001138999.1;NP_001341188.1;NP_001341192.1;NP_001341540.1;NI | 24650 | 607001 | . | . | 5.939656 | 5.916682 | 5.288853 | 6.100389 | 13.821822 | 13.676676 | 11.868034 | 13.947817 | -1.046644 | 909.594684 | 0.145122885 | -0.45320459 | 0.650401424 | [illegible] |
| 79815 | NM_00132163 | NIPAL2 | NIPA like domain containing 2 | protein_coding | NP_001308564.1;NP_001308565.1;NP_079035.1;XP_011515604.2 | 25854 | . | . | . | 0.330827 | 0.306585 | 0.375659 | 0.157694 | 0.769846 | 0.708685 | 0.842969 | 0.360549 | -1.085180 | 48.228168 | 0.463920332 | -0.25421111 | 0.799332471 | [illegible] |
| 79816 | NM_00114398 | TLE6 | TLE family member 6, subcortical maternal complex member | protein_coding | NP_001137458.1;NP_079036.1;XP_005259702.1;XP_011526602.1;XP_01 | 30788 | 612399 | . | . | 2.308698 | 1.997156 | 1.391693 | 2.991199 | 5.372434 | 4.616515 | 3.122920 | 6.839021 | -1.028119 | 100.639570 | 0.351378497 | -0.11385911 | 0.909349479 | [illegible] |
| 79817 | NM_024761 | MOB3B | MOB kinase activator 3B | protein_coding | NP_079037.3 | 23825 | 617652 | . | . | 0.653822 | 0.629537 | 0.888620 | 0.794036 | 1.521471 | 1.455202 | 1.994038 | 1.815469 | 1.292415 | 149.990061 | 0.268972056 | 1.375864541 | 0.168863575 | [illegible] |
| 79818 | NM_024762,XI | ZNF552 | zinc finger protein 552 | protein_coding | NP_079038.2;XP_005259324.1 | 26135 | . | . | . | 3.427202 | 3.051963 | 2.970968 | 2.941674 | 7.975239 | 7.054748 | 6.666764 | 6.725787 | -1.042138 | 241.776715 | 0.219066335 | -0.27182015 | 0.78576031 | [illegible] |
| 79819 | NM_024763,N | DNAI4 | dynein axonemal intermediate chain 4 | protein_coding | NP_079039.4;NP_996897.2;XP_005271269.1;XP_011540463.1;XP_01154 | 26252 | . | . | . | 0.307019 | 0.303567 | 0.492972 | 0.272172 | 0.714448 | 0.701708 | 1.106215 | 0.622288 | -1.052582 | 37.562649 | 0.506531015 | -0.14595938 | 0.883953452 | [illegible] |
| 79822 | NM_00101000 | ARHGAP28 | Rho GTPase activating protein 28 | protein_coding | NP_001010000.1;NP_001353159.1;NP_001353160.1;XP_005258203.1 | 25509 | 610592 | . | . | 1.072317 | 0.909719 | 1.099970 | 1.921780 | 2.495324 | 2.102858 | 2.468301 | 4.393921 | -1.111873 | 158.807755 | 0.272983549 | -0.56044347 | 0.57517699 | [illegible] |
| 79823 | NM_024766,XI | CAMKMT | calmodulin-lysine N-methyltransferase | protein_coding | NP_079042.1;XP_011531413.1;XP_011531415.1;XP_016860459.1;XP_01 | 26276 | 609559 | . | . | 3.335531 | 3.026839 | 3.610754 | 3.351987 | 7.761914 | 6.996670 | 8.102426 | 7.663920 | 1.053762 | 165.136268 | 0.256484729 | 0.294554111 | 0.768334522 | [illegible] |
| 79828 | NM_00132115 | METTL8 | methyltransferase like 8 | protein_coding | NP_001308083.1;NP_001308084.1;NP_001308085.1;NP_001308086.1;NI | 25856 | 609525 | . | . | 2.783470 | 3.530303 | 3.876777 | 3.650726 | 6.477254 | 8.160453 | 8.699374 | 8.346948 | 1.174859 | 937.654850 | 0.150350057 | 1.546311809 | 0.122029281 | [illegible] |
| 79829 | NM_00130080 | NAA40 | N-alpha-acetyltransferase 40, NatD catalytic subunit | protein_coding | NP_001287729.1;NP_079047.2;XP_006718752.1;XP_011543556.1;XP_01 | 25845 | . | . | . | 10.611314 | 10.453756 | 11.377437 | 11.276917 | 24.692964 | 24.164324 | 25.530638 | 25.783333 | 1.063210 | 1244.978343 | 0.129455993 | 0.683060554 | 0.49456858 | [illegible] |
| 79830 | NM_00128908 | ZMYM1 | zinc finger MYM-type containing 1 | protein_coding | NP_001276017.1;NP_001276018.1;NP_001276019.1;NP_001276020.1;NI | 26253 | . | . | . | 3.508762 | 2.395342 | 2.516127 | 1.879117 | 8.165035 | 5.536940 | 5.646119 | 4.296375 | -1.299833 | 388.664839 | 0.215301457 | -1.75719355 | 0.078884824 | [illegible] |
| 79831 | NM_00114534 | KDM8 | lysine demethylase 8 | protein_coding | NP_001138820.1;NP_079049.2;XP_016879165.1;XP_016879166.1;XP_01 | 25840 | 611917 | . | . | 0.432482 | 0.480119 | 0.794981 | 0.488154 | 1.006404 | 1.109815 | 1.783914 | 1.116107 | 1.382839 | 51.892279 | 0.450914432 | 1.037077918 | 0.299699545 | [illegible] |
| 79832 | NM_00107678 | QSER1 | glutamine and serine rich 1 | protein_coding | NP_001070254.2;XP_016873816.1;XP_016873818.1;XP_016873819.1 | 26154 | . | . | . | 2.342406 | 2.580523 | 2.120304 | 2.725249 | 5.450876 | 5.964995 | 4.757902 | 6.230957 | -1.116972 | 674.221534 | 0.152150404 | -1.04891769 | 0.294216003 | [illegible] |
| 79833 | NM_024775 | GEMIN6 | gem nuclear organelle associated protein 6 | protein_coding | NP_079051.9 | 20044 | 607006 | . | . | 5.447568 | 5.511208 | 5.693999 | 5.847097 | 12.676714 | 12.739405 | 12.777168 | 13.368692 | 1.038698 | 649.866025 | 0.153294107 | 0.3573264 | 0.720847469 | [illegible] |
| 79834 | NM_024776,XI | PEAK1 | pseudopodium enriched atypical kinase 1 | protein_coding | NP_079052.2;XP_005254727.1;XP_005254728.1;XP_005254730.1;XP_00 | 29431 | 614248 | . | . | 3.177409 | 4.015591 | 3.826543 | 3.283076 | 7.393959 | 9.282219 | 8.586655 | 7.506366 | 1.006091 | 1429.516439 | 0.145606253 | 0.060169042 | 0.952021002 | [illegible] |
| 79836 | NM_00103185 | LONRF3 | LON peptidase N-terminal domain and ring finger 3 | protein_coding | NP_001027026.1;NP_001276038.1;NP_079054.3;XP_005262533.1;XP_01 | 21152 | . | . | . | 2.319554 | 2.155609 | 2.687681 | 2.400112 | 5.397696 | 4.982787 | 6.031080 | 5.487571 | 1.128026 | 480.040301 | 0.177028499 | 0.981767998 | 0.326214157 | [illegible] |
| 79837 | NM_00114625 | PIP4K2C | phosphatidylinositol-5-phosphate 4-kinase type 2 gamma | protein_coding | NP_001139730.1;NP_001139731.1;NP_001139732.1;NP_079055.3;XP_00 | 23786 | 617104 | . | . | 14.306626 | 13.842766 | 15.037636 | 16.807095 | 33.292106 | 31.998172 | 33.744020 | 38.427423 | 1.119892 | 1469.139399 | 0.129103006 | 1.265339404 | 0.205749687 | [illegible] |
| 79839 | NM_00109372 | CCDC102B | coiled-coil domain containing 102B | protein_coding | NP_001087198.2;NP_079057.3;XP_011524476.1;XP_011524477.1;XP_01 | 26295 | . | . | . | 0.576607 | 0.507591 | 0.406788 | 0.548809 | 1.341786 | 1.173317 | 0.912821 | 1.254789 | 1.400112 | 40.243207 | 0.491833526 | 0.987208291 | 0.323540538 | [illegible] |
| 79840 | NM_00137749 | NHEJ1 | non-homologous end joining factor 1 | protein_coding | NP_001364427.1;NP_001364428.1;NP_079058.1 | 25737 | 611290 | . | . | 3.179437 | 3.224749 | 3.755369 | 3.874482 | 7.398682 | 7.454152 | 8.426941 | 8.858544 | 1.175507 | 877.367905 | 0.140765717 | 1.657243623 | 0.09747023 | [illegible] |
| 79841 | NM_024783,XI | AGBL2 | ATP/GTP binding protein like 2 | protein_coding | NP_079059.2;XP_005253195.1;XP_005253196.1;XP_005253197.1;XP_00 | 26296 | 617345 | . | . | 0.119204 | 0.117028 | 0.148067 | 0.166755 | 0.277392 | 0.270515 | 0.332258 | 0.381265 | 1.328719 | 14.097206 | 0.812334325 | 0.504762194 | 0.613725873 | [illegible] |
| 79842 | NM_00136310 | ZBTB3 | zinc finger and BTB domain containing 3 | protein_coding | NP_001350037.1;NP_001350038.1;NP_001357738.1 | 22918 | . | . | . | 0.486650 | 0.562320 | 0.588319 | 0.638943 | 1.132454 | 1.299828 | 1.320169 | 1.460867 | 1.163542 | 52.947649 | 0.4299815 | 0.50821437 | 0.611303014 | [illegible] |
| 79844 | NM_024786,XI | ZDHHC11 | zinc finger DHHC-type containing 11 | protein_coding | NP_079062.1;XP_016865358.1;XP_016865359.1;XP_016865360.1;XP_01 | 19158 | . | . | . | 1.775506 | 1.865900 | 1.227583 | 1.383791 | 4.131675 | 4.313108 | 2.754659 | 3.163873 | -1.318644 | 143.465812 | 0.274001999 | -1.4563966 | 0.145283016 | [illegible] |
| 79845 | NM_024787,XI | RNF122 | ring finger protein 122 | protein_coding | NP_079063.2;XP_016869335.1 | 21147 | . | . | . | 1.019554 | 1.105981 | 1.132591 | 1.433426 | 2.372545 | 2.556525 | 2.541500 | 3.277359 | 1.194142 | 68.890406 | 0.38392882 | 0.666724038 | 0.504948422 | [illegible] |
| 79846 | NM_00103970 | CFAP69 | cilia and flagella associated protein 69 | protein_coding | NP_001034795.2;NP_001153610.1;NP_001350367.1;XP_005250657.1;XP | 26107 | 617949 | . | . | 0.560658 | 0.446992 | 0.738042 | 0.505509 | 1.304676 | 1.033242 | 1.656144 | 1.155783 | 1.149854 | 89.500047 | 0.352086941 | 0.57216297 | 0.567211572 | [illegible] |
| 79847 | NM_024789,XI | MFSD13A | major facilitator superfamily domain containing 13A | protein_coding | NP_079065.2;XP_006718036.1;XP_006718037.1;XP_011538462.1;XP_01 | 26196 | . | . | . | 2.409166 | 2.342187 | 2.400979 | 2.709301 | 5.606225 | 5.414071 | 5.387727 | 6.194492 | 1.186849 | 223.105202 | 0.225845521 | 1.094272144 | 0.273835639 | [illegible] |
| 79848 | NM_00129133 | CSPP1 | centrosome and spindle pole associated protein 1 | protein_coding | NP_001278268.1;NP_001350060.1;NP_001350061.1;NP_001350062.1;NI | 26193 | 611654 | . | . | 3.573032 | 3.133248 | 4.227723 | 3.897694 | 8.314588 | 7.242644 | 9.486890 | 8.911614 | 1.149593 | 639.076541 | 0.163434471 | 1.23060106 | 0.218472109 | [illegible] |
| 79850 | NM_00131800 | TLCD3A | TLC domain containing 3A | protein_coding | NP_001304935.1;NP_001304936.1;NP_001304937.1;NP_079068.1 | 29646 | 611627 | . | . | 10.525243 | 10.253806 | 11.451997 | 12.401639 | 24.492673 | 23.702134 | 25.697951 | 28.354875 | 1.130657 | 772.310002 | 0.147448338 | 1.201510364 | 0.229553289 | [illegible] |
| 79854 | NR_024321 | LINC00115 | long intergenic non-protein coding RNA 115 | lncRNA | . | 26211 | . | . | . | 0.951942 | 1.013876 | 1.152217 | 1.045191 | 2.215209 | 2.343619 | 2.585542 | 2.389705 | 1.101227 | 43.386050 | 0.472389742 | 0.294485979 | 0.768386576 | [illegible] |
| 79856 | NM_024798,N | SNX22 | sorting nexin 22 | protein_coding | NP_079074.2;XP_005254734.1;XP_016878070.1 | 16315 | . | . | . | 0.171092 | 0.190259 | 0.230883 | 0.110403 | 0.398138 | 0.439793 | 0.518095 | 0.252423 | -1.078481 | 20.366364 | 0.70560259 | -0.15447979 | 0.877231444 | [illegible] |
| 79858 | NM_00114600 | NEK11 | NIMA related kinase 11 | protein_coding | NP_001139475.1;NP_001308149.1;NP_001308150.1;NP_001308151.1;NI | 18593 | 609779 | . | . | 1.230291 | 1.506495 | 1.832126 | 1.486127 | 2.862941 | 3.482335 | 4.111238 | 3.397858 | 1.127129 | 134.339376 | 0.292906967 | 0.589444508 | 0.555563127 | [illegible] |
| 79862 | NM_00114257 | ZNF669 | zinc finger protein 669 | protein_coding | NP_001136044.1;NP_079080.2 | 25736 | . | . | . | 0.107656 | 0.210391 | 0.269533 | 0.188555 | 0.250520 | 0.486328 | 0.604824 | 0.431109 | 1.456019 | 12.226229 | 0.893121095 | 0.606893716 | 0.543921448 | [illegible] |
| 79863 | NM_00117196 | RBFA | ribosome binding factor A | protein_coding | NP_001165438.1;NP_079081.2 | 26120 | . | . | . | 5.995985 | 6.689207 | 6.550005 | 7.269229 | 13.952903 | 15.462403 | 14.698023 | 16.620229 | 1.074081 | 1153.813131 | 0.138343546 | 0.745263026 | 0.456112728 | [illegible] |
| 79864 | NM_00136308 | JHY | junctional cadherin complex regulator | protein_coding | NP_001350016.1;NP_001350017.1;NP_001350018.1;NP_079082.2;NP_9 | 26288 | 617594 | . | . | 0.707300 | 1.122355 | 0.899461 | 1.247125 | 1.645916 | 2.594374 | 2.018363 | 2.851404 | 1.043323 | 114.479731 | 0.316926366 | 0.19306064 | 0.846911478 | [illegible] |
| 79866 | NM_00128674 | BORA | BORA aurora kinase A activator | protein_coding | NP_001273675.2;NP_001273676.1;NP_001353593.1;NP_079084.4 | 24724 | 610510 | . | . | 3.114701 | 3.678754 | 3.163526 | 2.655538 | 7.248039 | 8.503606 | 7.098859 | 6.071573 | -1.193401 | 268.409472 | 0.21790681 | -1.17058566 | 0.241765363 | [illegible] |
| 79867 | NM_00114385 | TCTN2 | tectonic family member 2 | protein_coding | NP_001137322.1;NP_079085.2;XP_005253680.1;XP_006719668.1;XP_01 | 25774 | 613846 | . | . | 12.943205 | 13.048144 | 11.896932 | 12.037672 | 30.119372 | 30.161368 | 26.696372 | 27.522705 | -1.095491 | 1110.472213 | 0.132930169 | -0.98982636 | 0.322258999 | [illegible] |
| 79868 | NM_00103921 | ALG13 | ALG13 UDP-N-acetylglucosaminyltransferase subunit | protein_coding | NP_001034299.3;NP_001093392.1;NP_001161857.1;NP_001244159.1;NI | 30881 | 300776 | . | . | 3.778780 | 3.748946 | 3.586669 | 4.035376 | 8.793374 | 8.665862 | 8.048381 | 9.226410 | 1.086915 | 472.636865 | 0.178074074 | 0.675222124 | 0.499534653 | [illegible] |
| 79869 | NM_00113604 | CPSF7 | cleavage and polyadenylation specific factor 7 | protein_coding | NP_001129512.1;NP_001136037.1;NP_079087.3;XP_005274355.1;XP_00 | 30098 | . | . | . | 20.319180 | 21.282376 | 26.267816 | 22.519651 | 47.283562 | 49.195165 | 58.944219 | 51.488502 | 1.166288 | 2498.465082 | 0.125777251 | 1.764424697 | 0.077660503 | [illegible] |
| 79871 | NM_024813,XI | RPAP2 | RNA polymerase II associated protein 2 | protein_coding | NP_079089.2;XP_005271280.1;XP_011540469.1;XP_016857851.1;XP_01 | 25791 | 611476 | . | . | 2.328479 | 2.849001 | 3.011577 | 3.345893 | 5.418465 | 6.585592 | 6.757894 | 7.649988 | 1.112065 | 606.804088 | 0.172836739 | 0.886620645 | 0.375283175 | [illegible] |
| 79872 | NM_00128429 | CBLL1 | Cbl proto-oncogene like 1 | protein_coding | NP_001271220.1;NP_079090.2;XP_011514882.1;XP_016868132.1;XP_01 | 21225 | 606872 | . | . | 5.716356 | 4.142408 | 4.779308 | 4.324556 | 13.302196 | 9.575360 | 10.724627 | 9.887579 | -1.122883 | 750.399403 | 0.157643249 | -1.06067259 | 0.288838722 | [illegible] |
| 79873 | NM_024815,XI | NUDT18 | nudix hydrolase 18 | protein_coding | NP_079091.3;XP_011542952.1 | 26194 | 615791 | . | . | 1.618230 | 2.360150 | 1.589605 | 1.859728 | 3.765687 | 5.455592 | 3.567027 | 4.252047 | -1.205548 | 98.953452 | 0.32212295 | -0.83722572 | 0.402465701 | [illegible] |
| 79874 | NM_024816 | RABEP2 | rabaptin, RAB GTPase binding effector protein 2 | protein_coding | NP_079092.2 | 24817 | 611869 | . | . | 5.875323 | 5.890541 | 7.372721 | 7.623567 | 13.672118 | 13.616248 | 16.544172 | 17.430382 | 1.255987 | 480.876103 | 0.16897548 | 1.945972051 | 0.051658097 | [illegible] |
| 79875 | NM_00128642 | THSD4 | thrombospondin type 1 domain containing 4 | protein_coding | NP_001273358.1;NP_079093.2;XP_006720755.1;XP_011520345.1;XP_01 | 25835 | 614476 | . | . | 0.182652 | 0.160387 | 0.126056 | 0.124867 | 0.425038 | 0.370741 | 0.282866 | 0.285494 | -1.392982 | 43.059535 | 0.476511923 | -1.0034926 | 0.315623245 | [illegible] |
| 79876 | NM_00132021 | UBA5 | ubiquitin like modifier activating enzyme 5 | protein_coding | NP_001307139.1;NP_001308167.1;NP_001308168.1;NP_079094.1;NP_9 | 23230 | 610552 | . | . | 5.257831 | 5.357235 | 4.870381 | 3.691397 | 12.235193 | 12.383490 | 10.928994 | 8.439942 | -1.251349 | 683.746375 | 0.173622443 | -1.86314628 | 0.062441688 | [illegible] |
| 79877 | NM_00112863 | DCAKD | dephospho-CoA kinase domain containing | protein_coding | NP_001122103.1;NP_001275583.1;NP_001275584.1;NP_001308255.1;NI | 26238 | . | . | . | 5.160317 | 5.818361 | 5.060158 | 5.564630 | 12.008270 | 13.449401 | 11.354849 | 12.722860 | -1.036038 | 353.865583 | 0.192996165 | -0.26464899 | 0.791279879 | [illegible] |
| 79879 | NM_00130479 | CCDC134 | coiled-coil domain containing 134 | protein_coding | NP_001291726.1;NP_001369275.1;NP_079097.1 | 26185 | 618788 | . | . | 6.119103 | 6.260692 | 6.804433 | 5.948167 | 14.239402 | 14.471869 | 15.268947 | 13.599777 | 1.014683 | 1396.883778 | 0.134010214 | 0.156919252 | 0.875308487 | [illegible] |
| 79882 | NM_00116010 | ZC3H14 | zinc finger CCCH-type containing 14 | protein_coding | NP_001153575.1;NP_001153576.1;NP_001313224.1;NP_001313225.1;NI | 20509 | 613279 | . | . | 4.639591 | 3.968615 | 4.808066 | 4.125273 | 10.796519 | 9.173626 | 10.789163 | 9.431948 | -1.035078 | 1874.433136 | 0.119110782 | -0.4175921 | 0.676245371 | [illegible] |
| 79883 | NM_00114625 | PODNL1 | podocan like 1 | protein_coding | NP_001139726.1;NP_001139727.1;NP_001357024.1;NP_079101.4;XP_00 | 26275 | . | . | . | 0.066123 | 0.138602 | 0.077607 | 0.189916 | 0.153871 | 0.320387 | 0.174147 | 0.434221 | 1.301728 | 13.021341 | 0.90932892 | 0.418360864 | 0.675683296 | [illegible] |
| 79885 | NM_00113604 | HDAC11 | histone deacetylase 11 | protein_coding | NP_001129513.1;NP_001317565.1;NP_079103.2;XP_005265548.1;XP_00 | 19086 | 607226 | . | . | 2.021910 | 2.742441 | 2.083802 | 2.722861 | 4.705067 | 6.339276 | 4.675991 | 6.225494 | 1.021593 | 204.532298 | 0.2515793 | 0.122507158 | 0.902497377 | [illegible] |
| 79886 | NM_00116757 | CAAP1 | caspase activity and apoptosis inhibitor 1 | protein_coding | NP_001161047.1;NP_079104.3 | 25834 | . | . | . | 2.197950 | 2.110934 | 1.935026 | 2.435547 | 5.114720 | 4.879519 | 4.342143 | 5.568591 | 1.014702 | 192.641202 | 0.247634779 | 0.085027516 | 0.932239515 | [illegible] |
| 79888 | NM_024830,XI | LPCAT1 | lysophosphatidylcholine acyltransferase 1 | protein_coding | NP_079106.3;XP_005248430.1;XP_011512434.1;XP_011512436.1;XP_01 | 25718 | 610472 | . | . | 16.242498 | 15.184701 | 14.897497 | 16.582571 | 37.796959 | 35.100118 | 33.429550 | 37.914080 | -1.012644 | 1933.369705 | 0.124722324 | -0.14533758 | 0.884444349 | [illegible] |
| 79890 | NM_00131998 | RIN3 | Ras and Rab interactor 3 | protein_coding | NP_001306916.1;NP_079108.3;XP_011535466.1;XP_011535467.1;XP_01 | 18751 | 610223 | . | . | 1.788975 | 2.316329 | 1.583538 | 1.936494 | 4.163018 | 5.354297 | 3.553413 | 4.427565 | -1.173589 | 227.872801 | 0.240633032 | -0.95966561 | 0.337223536 | [illegible] |
| 79891 | NM_00132137 | ZNF671 | zinc finger protein 671 | protein_coding | NP_001308304.1;NP_001308305.1;NP_079109.2;XP_016882803.1 | 26279 | . | . | . | 1.143039 | 1.237651 | 1.205709 | 1.116764 | 2.659898 | 2.860884 | 2.705578 | 2.553349 | 1.005843 | 101.000039 | 0.322133549 | 0.026090112 | 0.979185464 | [illegible] |
| 79892 | NM_00125637 | MCMBP | minichromosome maintenance complex binding protein | protein_coding | NP_001243307.1;NP_001243308.1;NP_079110.1;XP_005270214.1;XP_01 | 25782 | 610909 | . | . | 18.791854 | 18.315616 | 15.776309 | 16.366525 | 43.729416 | 42.337366 | 35.401579 | 37.420115 | -1.165346 | 2275.602515 | 0.115677419 | -1.90840056 | 0.056339464 | [illegible] |
| 79893 | NM_024835,XI | GGNBP2 | gametogenetin binding protein 2 | protein_coding | NP_079111.1;XP_005257746.1;XP_011523567.1;XP_011523568.1;XP_01 | 19357 | 612275 | . | . | 18.402640 | 18.445003 | 18.030656 | 17.461951 | 42.823701 | 42.636447 | 40.460272 | 39.924676 | -1.052883 | 1589.938428 | 0.123402638 | -0.60246194 | 0.546866693 | [illegible] |
| 79894 | NM_024836,XI | ZNF672 | zinc finger protein 672 | protein_coding | NP_079112.1;XP_005270393.1 | 26179 | . | . | . | 1.688623 | 1.771005 | 1.587281 | 2.176447 | 3.929495 | 4.093758 | 3.561813 | 4.976188 | 1.083758 | 170.947751 | 0.266068159 | 0.436137911 | 0.662736673 | [illegible] |
| 79896 | NM_024838,XI | THNSL1 | threonine synthase like 1 | protein_coding | NP_079114.3;XP_005252654.1;XP_016872154.1 | 26160 | 611260 | . | . | 2.934317 | 2.714876 | 3.033148 | 2.595472 | 6.828277 | 6.275558 | 6.806296 | 5.934240 | -1.017003 | 331.316088 | 0.200541501 | -0.12129471 | 0.903457611 | [illegible] |
| 79897 | NM_00119912 | RPP21 | ribonuclease P/MRP subunit p21 | protein_coding | NP_001186049.1;NP_001186050.1;NP_079115.1 | 21300 | 612524 | . | . | 18.983234 | 16.713117 | 16.651500 | 18.202357 | 44.174769 | 38.633114 | 37.365481 | 41.617526 | -1.037853 | 299.534219 | 0.205565384 | -0.26075514 | 0.794281339 | [illegible] |
| 79898 | NM_00103172 | ZNF613 | zinc finger protein 613 | protein_coding | NP_001026891.2;NP_079116.2;XP_005259326.1;XP_011525635.1;XP_01 | 25827 | . | . | . | 0.698111 | 0.611067 | 0.681987 | 0.894441 | 1.624533 | 1.412508 | 1.530358 | 2.045035 | 1.171179 | 70.409336 | 0.385489395 | 0.591355572 | 0.554282202 | [illegible] |
| 79901 | NM_00112738 | CYBRD1 | cytochrome b reductase 1 | protein_coding | NP_001120855.1;NP_001243838.1;NP_079119.3 | 20797 | 605745 | . | . | 15.163322 | 14.345442 | 13.594397 | 12.777507 | 35.285676 | 33.160130 | 30.505432 | 29.214252 | -1.135289 | 1838.123994 | 0.123777853 | -1.47893736 | 0.139157057 | [illegible] |
| 79902 | NM_00130327 | NUP85 | nucleoporin 85 | protein_coding | NP_001290205.1;NP_001317401.1;NP_079120.1;XP_005257747.1;XP_01 | 8734 | 170285 | . | . | 30.093532 | 30.982086 | 27.625782 | 29.599699 | 70.028889 | 71.616481 | 61.991448 | 67.676197 | -1.068517 | 2079.111170 | 0.117246686 | -0.81545812 | 0.414810129 | [illegible] |
| 79903 | NM_00108360 | NAA60 | N-alpha-acetyltransferase 60, NatF catalytic subunit | protein_coding | NP_001077069.1;NP_001077070.1;NP_001304022.1;NP_001304023.1;NI | 25875 | 614246 | . | . | 6.415368 | 8.136129 | 7.846107 | 8.458551 | 14.928823 | 18.807027 | 17.606435 | 19.339471 | 1.104530 | 621.500973 | 0.170435255 | 0.841566667 | 0.400030554 | [illegible] |
| 79905 | NM_00116036 | TMC7 | transmembrane channel like 7 | protein_coding | NP_001153836.1;NP_001287661.1;NP_001311192.1;NP_001311194.1;NI | 23000 | 617198 | . | . | 0.197471 | 0.201689 | 0.149002 | 0.211998 | 0.459524 | 0.466212 | 0.334355 | 0.484709 | -1.131587 | 27.009300 | 0.596658272 | -0.29890958 | 0.76500904 | [illegible] |
| 79906 | NM_00130106 | MORN1 | MORN repeat containing 1 | protein_coding | NP_001287989.1;NP_079124.1 | 25852 | . | . | . | 3.941819 | 4.215983 | 4.533787 | 4.532342 | 9.172775 | 9.745433 | 10.173685 | 10.362661 | 1.088640 | 385.990859 | 0.197021062 | 0.621900262 | 0.534007453 | [illegible] |
| 79912 | NM_00135091 | PYROXD1 | pyridine nucleotide-disulphide oxidoreductase domain 1 | protein_coding | NP_001337841.1;NP_001337842.1;NP_079130.2;XP_006719216.1;XP_01 | 26162 | 617220 | . | . | 2.499823 | 3.093841 | 2.277448 | 1.465391 | 5.817189 | 7.151550 | 5.110527 | 3.350443 | -1.477716 | 283.986417 | 0.242186191 | -2.32618313 | **0.020008783** | [illegible] |
| 79913 | NM_024855 | ACTR5 | actin related protein 5 | protein_coding | NP_079131.3 | 14671 | . | . | . | 4.903540 | 5.012317 | 4.288688 | 5.012547 | 11.410739 | 11.586194 | 9.623693 | 11.460594 | -1.080802 | 381.923060 | 0.187846677 | -0.59677203 | 0.550659593 | [illegible] |
| 79915 | NM_024857,XI | ATAD5 | ATPase family AAA domain containing 5 | protein_coding | NP_079133.3;XP_011523571.1;XP_011523572.1;XP_011523575.1;XP_01 | 25752 | 609534 | . | . | 2.498718 | 2.633015 | 2.812464 | 1.404344 | 5.814620 | 6.086330 | 6.311087 | 3.210863 | -1.228104 | 513.983809 | 0.231482882 | -1.28058058 | 0.200341024 | [illegible] |
| 79917 | NM_00109968 | MAGIX | MAGI family member, X-linked | protein_coding | NP_001093151.2;NP_001093152.2;NP_079135.3;XP_005278119.1;XP_00 | 30006 | . | . | . | 1.198400 | 0.800505 | 0.842213 | 1.159407 | 2.788727 | 1.850404 | 1.889903 | 2.650845 | -1.103515 | 58.568909 | 0.430096928 | -0.33040604 | 0.741093181 | [illegible] |
| 79918 | NM_00116030 | SETD6 | SET domain containing 6, protein lysine methyltransferase | protein_coding | NP_001153777.1;NP_079136.2;XP_011521638.1 | 26116 | 616424 | . | . | 2.742805 | 2.846558 | 3.813608 | 2.457136 | 6.382622 | 6.579947 | 8.557627 | 5.617949 | 1.032380 | 429.303039 | 0.188542391 | 0.243836579 | 0.807357389 | [illegible] |
| 79921 | NM_00100693 | TCEAL4 | transcription elongation factor A like 4 | protein_coding | NP_001006936.1;NP_001006938.1;NP_001287830.1;NP_001292769.1;NI | 26121 | . | . | . | 31.417774 | 31.345203 | 27.781290 | 35.168493 | 73.110452 | 72.455837 | 62.340409 | 80.408585 | -1.011254 | 1265.915308 | 0.145831199 | -0.11070971 | 0.911846545 | [illegible] |
| 79922 | NM_024864,XI | MRM1 | mitochondrial rRNA methyltransferase 1 | protein_coding | NP_079140.2;XP_005257751.1;XP_011523577.1;XP_011523578.1;XP_01 | 26202 | 618099 | . | . | 5.721120 | 6.810246 | 5.812945 | 5.813679 | 13.313283 | 15.742190 | 13.044079 | 13.292287 | -1.066163 | 342.727631 | 0.196658021 | -0.46999038 | 0.638361894 | [illegible] |
| 79924 | NM_00125384 | ADM2 | adrenomedullin 2 | protein_coding | NP_001240774.1;NP_001356811.1 | 28898 | 608682 | . | . | 0.922973 | 0.883600 | 0.551448 | 0.874711 | 2.147795 | 2.042482 | 1.237434 | 1.999922 | -1.281300 | 108.386014 | 0.331747983 | -1.07795036 | 0.281055909 | [illegible] |
| 79929 | NM_024871,XI | MAP6D1 | MAP6 domain containing 1 | protein_coding | NP_079147.1 | 25753 | 610593 | . | . | 1.893572 | 2.248576 | 2.215204 | 2.116813 | 4.406420 | 5.197685 | 4.970854 | 4.839841 | 1.035951 | 137.687864 | 0.280042069 | 0.181959932 | 0.85561417 | [illegible] |
| 79930 | NM_00114487 | DOK3 | docking protein 3 | protein_coding | NP_001138347.1;NP_001138348.1;NP_001295164.1;NP_001295165.1;NI | 24583 | 611435 | . | . | 1.088853 | 1.617190 | 2.026126 | 0.744254 | 2.533804 | 3.738206 | 4.546568 | 1.701648 | 1.103215 | 138.802659 | 0.373212607 | 0.379712436 | 0.704158888 | [illegible] |
| 79932 | NM_024874,XI | KIAA0319L | KIAA0319 like | protein_coding | NP_079150.3;XP_006710970.1;XP_006710972.1;XP_006710973.1;XP_01 | 30071 | 613535 | . | . | 17.249621 | 16.273074 | 16.457286 | 19.002607 | 40.140581 | 37.615945 | 36.929669 | 43.447201 | 1.053752 | 2551.044288 | 0.121596215 | 0.621195659 | 0.534470895 | [illegible] |
| 79934 | NM_00114255 | COQ8B | coenzyme Q8B | protein_coding | NP_001136027.1;NP_079152.3 | 19041 | 615567 | . | . | 13.858541 | 13.296738 | 14.353123 | 16.099091 | 32.249396 | 30.736008 | 32.207993 | 36.808660 | 1.105858 | 1091.906916 | 0.137196476 | 1.058088721 | 0.290014996 | [illegible] |
| 79937 | NM_033655 | CNTNAP3 | contactin associated protein family member 3 | protein_coding | NP_387504.2 | 13834 | 610517 | . | . | 1.060532 | 1.015854 | 1.073193 | 0.904008 | 2.467901 | 2.348191 | 2.408214 | 2.066908 | -1.064165 | 415.904305 | 0.185649677 | -0.48328323 | 0.628894645 | [illegible] |
| 79939 | NM_024881 | SLC35E1 | solute carrier family 35 member E1 | protein_coding | NP_079157.3 | 20803 | . | . | . | 11.622670 | 11.771057 | 12.542004 | 11.317102 | 27.046432 | 27.209328 | 28.143892 | 25.875210 | 1.005906 | 1886.994144 | 0.124230058 | 0.068381301 | 0.945482107 | [illegible] |
| 79940 | NR_026807,NF | LINC00472 | long intergenic non-protein coding RNA 472 | lncRNA | . | 21380 | . | . | . | 0.267070 | 0.246097 | 0.273316 | 0.219567 | 0.621483 | 0.568865 | 0.613313 | 0.502014 | -1.057097 | 75.404889 | 0.36921447 | -0.21696827 | 0.828233075 | [illegible] |
| 79943 | NM_030895 | ZNF696 | zinc finger protein 696 | protein_coding | NP_112157.2 | 25872 | . | . | . | 1.146685 | 1.115872 | 1.518904 | 1.225677 | 2.668382 | 2.579387 | 3.408377 | 2.802365 | 1.195636 | 182.094993 | 0.253553719 | 1.016661003 | 0.309314723 | [illegible] |
| 79944 | NM_024884,XI | L2HGDH | L-2-hydroxyglutarate dehydrogenase | protein_coding | NP_079160.1;XP_005268132.1;XP_011535468.1;XP_011535469.1;XP_01 | 20499 | 609584 | . | . | 2.665554 | 2.624935 | 2.489588 | 2.538209 | 6.202856 | 6.067654 | 5.586563 | 5.803313 | -1.163937 | 374.817700 | 0.190854275 | -1.14753747 | 0.251159555 | [illegible] |
| 79947 | NM_00124356 | DHDDS | dehydrodolichyl diphosphate synthase subunit | protein_coding | NP_001230493.1;NP_001230494.1;NP_001306888.1;NP_079163.2;NP_9 | 20603 | 608172 | . | . | 10.911366 | 10.626824 | 11.994248 | 11.412064 | 25.391194 | 24.564378 | 26.914745 | 26.092329 | 1.072661 | 1158.643631 | 0.132890004 | 0.761490318 | 0.446364259 | [illegible] |
| 79949 | NM_00119343 | PLEKHS1 | pleckstrin homology domain containing S1 | protein_coding | NP_001180363.1;NP_001180364.1;NP_079165.3;NP_872407.1;XP_00527 | 26285 | . | . | . | 1.878990 | 2.255598 | 1.259711 | 1.029747 | 4.372486 | 5.213915 | 2.826753 | 2.354395 | -1.382491 | 159.499175 | 0.27007343 | -1.73015817 | 0.083602019 | [illegible] |
| 79954 | NM_00126139 | NOL10 | nucleolar protein 10 | protein_coding | NP_001248321.1;NP_001248323.1;NP_079170.2;XP_011508700.1;XP_01 | 25862 | 616197 | . | . | 10.075368 | 12.083675 | 11.544132 | 10.500232 | 23.445797 | 27.931958 | 25.904698 | 24.007536 | -1.025593 | 1141.736407 | 0.137889481 | -0.26440665 | 0.791466589 | [illegible] |
| 79955 | NM_00119526 | PDZD7 | PDZ domain containing 7 | protein_coding | NP_001182192.1;NP_001337973.1;NP_079171.1;XP_005270222.1;XP_01 | 26257 | 612971 | . | . | 1.352409 | 0.996916 | 0.959568 | 1.302183 | 3.147109 | 2.304416 | 2.153242 | 2.977284 | -1.088234 | 122.946745 | 0.311288771 | -0.39188156 | 0.695145726 | [illegible] |
| 79956 | NM_024896,XI | ERMP1 | endoplasmic reticulum metallopeptidase 1 | protein_coding | NP_079172.2;XP_005251644.1;XP_016870628.1 | 23703 | 611156 | . | . | 8.196775 | 8.180419 | 7.108111 | 7.085551 | 19.074232 | 18.909406 | 15.950393 | 16.200270 | -1.164907 | 1259.163797 | 0.128836293 | -1.70925792 | 0.087403184 | [illegible] |
| 79957 | NM_00127210 | PAQR6 | progestin and adipoQ receptor family member 6 | protein_coding | NP_001259033.1;NP_001259034.1;NP_001259035.1;NP_001259036.1;NI | 30132 | 614579 | . | . | 0.969799 | 1.397848 | 0.917334 | 1.202855 | 2.256761 | 3.231186 | 2.058470 | 2.750184 | -1.106462 | 76.757470 | 0.376439811 | -0.38772293 | 0.698221087 | [illegible] |
| 79959 | NM_00127198 | CEP76 | centrosomal protein 76 | protein_coding | NP_001258918.1;NP_079175.2;XP_005258206.1;XP_016881470.1;XP_01 | 25727 | . | . | . | 3.740122 | 3.029111 | 2.939500 | 2.759349 | 8.703421 | 7.001926 | 6.596151 | 6.308925 | -1.169165 | 271.038235 | 0.214292302 | -1.05220247 | 0.292706666 | [illegible] |
| 79960 | NM_00128743 | JADE1 | jade family PHD finger 1 | protein_coding | NP_001274366.1;NP_001274368.1;NP_001274369.1;NP_001274370.1;NI | 30027 | 610514 | . | . | 3.022653 | 2.766080 | 2.595943 | 2.415104 | 7.033841 | 6.393917 | 5.825220 | 5.521848 | -1.188878 | 447.242102 | 0.17566603 | -1.42087938 | 0.155351827 | [illegible] |
| 79961 | NM_00127183 | DENND2D | DENN domain containing 2D | protein_coding | NP_001258762.1;NP_079177.2;XP_006710984.2;XP_006710985.1;XP_00 | 26192 | 615111 | . | . | 0.883620 | 0.647898 | 1.127702 | 0.902707 | 2.056220 | 1.497646 | 2.530531 | 2.063931 | 1.201685 | 69.061197 | 0.379497666 | 0.698444797 | 0.484899069 | [illegible] |
| 79962 | NM_00130494 | DNAJC22 | DnaJ heat shock protein family (Hsp40) member C22 | protein_coding | NP_001291873.1;NP_079178.2;XP_005269212.1;XP_005269213.1;XP_00 | 25802 | . | . | . | 4.258515 | 4.270946 | 4.907134 | 4.379124 | 9.909738 | 9.872484 | 11.011465 | 10.012345 | 1.089260 | 421.250535 | 0.181216202 | 0.680669784 | 0.49608046 | [illegible] |
| 79963 | NR_002451 | ABCA11P | ATP binding cassette subfamily A member 11, pseudogene | transcribed_ps | . | 31 | . | . | . | 0.699183 | 0.526047 | 0.724767 | 0.855167 | 1.627027 | 1.215981 | 1.626356 | 1.955238 | 1.266307 | 41.973840 | 0.487178019 | 0.699183128 | 0.484437593 | [illegible] |
| 79966 | NM_00103758 | SCD5 | stearoyl-CoA desaturase 5 | protein_coding | NP_001032671.2;NP_079182.2 | 21088 | 608370 | . | . | 4.445177 | 4.250237 | 3.423124 | 3.606687 | 10.344107 | 9.824614 | 7.681392 | 8.246262 | -1.241707 | 367.521344 | 0.187158933 | -1.66876872 | 0.095163225 | [illegible] |
| 79968 | NM_00116794 | WDR76 | WD repeat domain 76 | protein_coding | NP_001161413.1;NP_079184.2;XP_005254737.1 | 25773 | . | . | . | 6.086058 | 6.417810 | 6.144689 | 6.523611 | 14.162507 | 14.835052 | 13.788503 | 14.915461 | 1.009906 | 768.600359 | 0.147270839 | 0.096567627 | 0.923069766 | [illegible] |
| 79969 | NM_00103172 | ATAT1 | alpha tubulin acetyltransferase 1 | protein_coding | NP_001026892.1;NP_001177653.1;NP_001241881.1;NP_001305691.1;NI | 21186 | 615556 | . | . | 5.053618 | 5.576948 | 6.688382 | 5.731555 | 11.759981 | 12.891365 | 15.008536 | 13.104518 | 1.126857 | 410.510621 | 0.193280433 | 0.891475717 | 0.372674013 | [illegible] |
| 79970 | NR_027788,NF | ZNF767P | zinc finger family member 767, pseudogene | transcribed_ps | . | 21884 | . | . | . | 3.244241 | 3.778452 | 3.949129 | 2.982373 | 7.549483 | 8.734060 | 8.861730 | 6.818842 | -1.057475 | 356.381320 | 0.20745468 | -0.38863431 | 0.697546682 | [illegible] |
| 79971 | NM_00100229 | WLS | Wnt ligand secretion mediator | protein_coding | NP_001002292.3;NP_001180263.1;NP_079187.3;XP_011540493.1;XP_01 | 30238 | 611514 | . | . | 22.870064 | 21.706415 | 20.796591 | 20.707186 | 53.219576 | 50.175349 | 46.666945 | 47.344521 | -1.089451 | 1773.099232 | 0.12103568 | -1.02119207 | 0.307163451 | [illegible] |
| 79974 | NM_00110553 | CPED1 | cadherin like and PC-esterase domain containing 1 | protein_coding | NP_001099003.1;NP_079189.4;XP_016868138.1;XP_024302709.1 | 26159 | . | . | . | 1.252987 | 1.525516 | 1.356685 | 0.957897 | 2.915753 | 3.526299 | 3.044362 | 2.190118 | -1.120377 | 137.243483 | 0.279057311 | -0.58763495 | 0.556777348 | [illegible] |
| 79979 | NM_00116797 | TRMT2B | tRNA methyltransferase 2 homolog B | protein_coding | NP_001161442.1;NP_001161443.1;NP_001161444.1;NP_079193.2;XP_00 | 25748 | . | . | . | 2.912843 | 2.498510 | 2.646272 | 2.827697 | 6.778306 | 5.775422 | 5.938156 | 6.465193 | 1.111761 | 378.205136 | 0.185422203 | 0.824317742 | 0.40975904 | [illegible] |
| 79980 | NM_00114531 | DSN1 | DSN1 component of MIS12 kinetochore complex | protein_coding | NP_001138787.1;NP_001138788.1;NP_001138789.1;NP_001138790.1;NI | 16165 | 609175 | . | . | 20.323508 | 20.418324 | 18.863957 | 20.431021 | 47.293635 | 47.197871 | 42.330167 | 46.713104 | -1.054395 | 1388.942508 | 0.128646896 | -0.59399598 | 0.552514802 | [illegible] |
| 79982 | NM_00103172 | DNAJB14 | DnaJ heat shock protein family (Hsp40) member B14 | protein_coding | NP_001026893.1;NP_001265239.1;NP_001265240.1;XP_011530564.1;XP | 25881 | 617487 | . | . | 6.023619 | 4.330615 | 5.310354 | 3.896801 | 14.017206 | 10.010410 | 11.916281 | 8.909572 | -1.153289 | 824.572755 | 0.198728713 | -1.03535032 | 0.300505337 | [illegible] |
| 79989 | NM_00114492 | TTC26 | tetratricopeptide repeat domain 26 | protein_coding | NP_001138392.1;NP_001138395.1;NP_001274441.1;NP_001274442.1;NI | 21882 | 617453 | . | . | 2.911440 | 3.391944 | 2.808913 | 2.842287 | 6.775040 | 7.840632 | 6.303118 | 6.498552 | -1.149139 | 383.361683 | 0.191074479 | -1.04960944 | 0.293897714 | [illegible] |
| 79990 | NM_024927,N | PLEKHH3 | pleckstrin homology, MyTH4 and FERM domain containing H3 | protein_coding | NP_079203.4;XP_016880602.1;XP_016880603.1;XP_016880604.1;XP_01 | 26105 | . | . | . | 3.493755 | 3.253345 | 3.321271 | 3.832586 | 8.130110 | 7.520250 | 7.452836 | 8.762752 | 1.048519 | 330.634463 | 0.19762351 | 0.345872498 | 0.729438538 | [illegible] |
| 79991 | NM_024928 | STN1 | STN1 subunit of CST complex | protein_coding | NP_079204.2 | 26200 | 613128 | . | . | 1.952041 | 1.666628 | 1.685260 | 1.907811 | 4.542479 | 3.852484 | 3.781675 | 4.361982 | -1.021555 | 358.338276 | 0.19656575 | -0.15651876 | 0.875624137 | [illegible] |
| 79992 | NR_024277 | AGPAT4-IT1 | AGPAT4 intronic transcript 1 | lncRNA | . | 20988 | . | . | . | 0.136831 | 0.163774 | 0.027038 | 0.140268 | 0.318410 | 0.378572 | 0.060672 | 0.320706 | -1.731145 | 7.427975 | 1.16491468 | -0.67964358 | 0.496730166 | [illegible] |
| 79993 | NM_00110455 | ELOVL7 | ELOVL fatty acid elongase 7 | protein_coding | NP_001098028.1;NP_001284546.1;NP_001284547.1;NP_079206.2;XP_00 | 26292 | 614451 | . | . | 2.733184 | 2.511445 | 2.206860 | 2.470384 | 6.360232 | 5.805316 | 4.952133 | 5.648240 | -1.141545 | 301.008014 | 0.204851095 | -0.93232655 | 0.351167793 | [illegible] |
| 80000 | NM_00114296 | GREB1L | GREB1 like retinoic acid receptor coactivator | protein_coding | NP_001136438.1;XP_006722610.1;XP_011524481.1;XP_016881477.1;XP | 31042 | 617782 | . | . | 1.588077 | 1.914957 | 1.851696 | 1.825805 | 3.695522 | 4.426511 | 4.155153 | 4.174485 | 1.054621 | 532.109408 | 0.180477335 | 0.425119578 | 0.670749507 | [illegible] |
| 80003 | NM_00132860 | PCNX2 | pecanex 2 | protein_coding | NP_001315536.1;NP_055616.3;XP_006711879.1;XP_011542580.2;XP_01 | 8736 | 617656 | . | . | 1.854816 | 2.107745 | 2.379027 | 2.667025 | 4.316229 | 4.872149 | 5.338465 | 6.097834 | 1.205892 | 415.582392 | 0.178808293 | 1.5105589 | 0.130900872 | [illegible] |
| 80004 | NM_00136526 | ESRP2 | epithelial splicing regulatory protein 2 | protein_coding | NP_001352193.1;NP_001352194.1;NP_079215.2;XP_005256210.1;XP_00 | 26152 | 612960 | . | . | 0.048028 | 0.062175 | 0.059221 | 0.024670 | 0.111762 | 0.143720 | 0.132888 | 0.056404 | -1.305321 | 10.344696 | 0.973914161 | -0.39470056 | 0.693063888 | [illegible] |
| 80005 | NM_00132281 | DOCK5 | dedicator of cytokinesis 5 | protein_coding | NP_001309739.1;NP_079216.4 | 23476 | 616904 | . | . | 2.644177 | 2.927675 | 3.155114 | 2.524577 | 6.153109 | 6.767451 | 7.079983 | 5.772146 | -1.028902 | 805.639087 | 0.163660904 | -0.25116514 | 0.801686437 | [illegible] |
| 80006 | NM_00109375 | TRAPPC13 | trafficking protein particle complex 13 | protein_coding | NP_001087224.1;NP_001087225.1;NP_001230666.1;NP_001352271.1;NI | 25828 | . | . | . | 4.555254 | 3.972369 | 3.113810 | 3.250591 | 10.600262 | 9.182307 | 6.987299 | 7.432091 | -1.351613 | 352.518228 | 0.194733675 | -2.23218821 | **0.025602526** | [illegible] |
| 80007 | NM_024942,XI | C10orf88 | chromosome 10 open reading frame 88 | protein_coding | NP_079218.2 | 25822 | . | . | . | 4.111600 | 4.159101 | 4.102764 | 3.658373 | 9.567860 | 9.613949 | 9.206483 | 8.364436 | -1.070819 | 382.402720 | 0.18643524 | -0.52948849 | 0.59646663 | [illegible] |
| 80010 | NM_00135829 | RMI1 | RecQ mediated genome instability 1 | protein_coding | NP_001345220.1;NP_001345221.1;NP_001345222.1;NP_001345223.1;NI | 25764 | 610404 | . | . | 3.369530 | 3.037834 | 3.569434 | 1.984527 | 7.841035 | 7.022087 | 8.009705 | 4.537383 | -1.175068 | 316.711415 | 0.253803524 | -0.91702443 | 0.359129841 | [illegible] |
| 80011 | NM_00135407 | PSME3IP1 | proteasome activator subunit 3 interacting protein 1 | protein_coding | NP_001341007.1;NP_001341008.1;NP_001341009.1;NP_001341010.1;NI | 29856 | 617766 | . | . | 15.383783 | 15.756751 | 15.735646 | 17.316239 | 35.798696 | 36.422435 | 35.310332 | 39.591525 | 1.097022 | 1609.105917 | 0.123323484 | 1.083265608 | 0.278690546 | [illegible] |
| 80012 | NM_00130811 | PHC3 | polyhomeotic homolog 3 | protein_coding | NP_001295045.1;NP_079223.3;XP_005247846.1;XP_005247848.1;XP_00 | 15682 | . | . | . | 2.141290 | 2.101356 | 2.197149 | 1.983457 | 4.982869 | 4.857379 | 4.930341 | 4.534936 | -1.065693 | 777.300596 | 0.165698607 | -0.55396716 | 0.579601318 | [illegible] |
| 80013 | NM_00131833 | MINDY3 | MINDY lysine 48 deubiquitinase 3 | protein_coding | NP_001305259.1;NP_079224.1;XP_005252657.1;XP_005252659.1;XP_00 | 23578 | 611649 | . | . | 12.821056 | 12.068755 | 11.560843 | 9.574513 | 29.835124 | 27.897469 | 25.942197 | 21.890984 | -1.199165 | 959.729937 | 0.15109605 | -1.73419473 | 0.082883539 | [illegible] |
| 80014 | NM_024949,XI | WWC2 | WW and C2 domain containing 2 | protein_coding | NP_079225.5;XP_011530571.1;XP_024309993.1 | 24148 | . | . | . | 2.743197 | 2.733440 | 2.589132 | 2.508809 | 6.383534 | 6.318467 | 5.809937 | 5.736095 | -1.103762 | 700.498330 | 0.151178332 | -0.94213036 | 0.3461259 | [illegible] |
| 80017 | NM_00110236 | DGLUCY | D-glutamate cyclase | protein_coding | NP_001095836.1;NP_001095837.1;NP_001095838.1;NP_001095839.1;NI | 20498 | . | . | . | 3.221505 | 3.124553 | 3.124647 | 3.407115 | 7.496576 | 7.222544 | 7.011617 | 7.789964 | 1.005217 | 301.103708 | 0.200492089 | 0.037440268 | 0.970133966 | [illegible] |
| 80018 | NM_024953,XI | NAA25 | N-alpha-acetyltransferase 25, NatB auxiliary subunit | protein_coding | NP_079229.2;XP_005253995.1;XP_006719669.1;XP_016875466.1 | 25783 | 612755 | . | . | 6.144558 | 6.365798 | 6.948833 | 5.197391 | 14.298640 | 14.714827 | 15.592979 | 11.883214 | -1.068882 | 1112.408572 | 0.16770727 | -0.57303813 | 0.56661888 | [illegible] |
| 80019 | NM_024954 | UBTD1 | ubiquitin domain containing 1 | protein_coding | NP_079230.1 | 25683 | 616388 | . | . | 11.229111 | 11.012754 | 9.989038 | 12.612515 | 26.130600 | 25.456474 | 22.415112 | 28.837019 | 1.002115 | 575.407812 | 0.172382355 | 0.017680801 | 0.985893497 | [illegible] |
| 80020 | NM_00110237 | FOXRED2 | FAD dependent oxidoreductase domain containing 2 | protein_coding | NP_001095841.1;NP_001349970.1;NP_001349971.1;NP_079231.4 | 26264 | 613777 | . | . | 30.380047 | 32.630268 | 30.423340 | 32.346735 | 70.695616 | 75.426322 | 68.269090 | 73.956961 | -1.015515 | 4779.917354 | 0.108652153 | -0.20443413 | 0.838014261 | [illegible] |
| 80021 | NM_00134700 | TMEM62 | transmembrane protein 62 | protein_coding | NP_001333933.1;NP_001333934.1;NP_001333935.1;NP_001333936.1;NI | 26269 | . | . | . | 5.418507 | 5.376310 | 5.509719 | 4.734572 | 12.609089 | 12.427586 | 12.363647 | 10.825037 | -1.032236 | 440.478354 | 0.18133869 | -0.25241769 | 0.800718227 | [illegible] |

| | | | | | | | | | | |
|---|---|---|---|---|---|---|---|---|---|---|
| 80022 | NM_00130924 | MYO15B | myosin XVB | protein_coding | NP_001296171.1;XP_016880609.1;XP_016880610.1;XP_016880611.1;XP | 14083 | . | . | . | [illegible] |
| 80023 | NM_00132367 | NRSN2 | neurensin 2 | protein_coding | NP_001310608.1;NP_001310609.1;NP_001310610.1;NP_001310611.1;NI | 16229 | 610666 | . | . | [illegible] |
| 80024 | NM_00133046 | SLC8B1 | solute carrier family 8 member B1 | protein_coding | NP_001317395.1;NP_001345274.1;NP_079235.2;XP_011537054.1;XP_01 | 26175 | 609841 | . | . | [illegible] |
| 80025 | NM_00132419 | PANK2 | pantothenate kinase 2 | protein_coding | NP_001311120.1;NP_001311121.1;NP_001311122.1;NP_079236.3;NP_7( | 15894 | 606157 | . | . | [illegible] |
| 80028 | NM_00132121 | FBXL18 | F-box and leucine rich repeat protein 18 | protein_coding | NP_001308142.1;NP_001350370.1;NP_001350371.1;NP_001354709.1;NI | 21874 | 609084 | . | . | [illegible] |
| 80031 | NM_00119899 | SEMA6D | semaphorin 6D | protein_coding | NP_001185928.1;NP_001345280.1;NP_001345281.1;NP_065909.1;NP_0 | 16770 | 609295 | . | . | [illegible] |
| 80036 | NM_00100747 | TRPM3 | transient receptor potential cation channel subfamily M member 3 | protein_coding | NP_001007471.1;NP_001007472.2;NP_001353070.1;NP_001353071.1;NI | 17992 | 608961 | . | . | [illegible] |
| 80039 | NM_024974 | FAM106A | family with sequence similarity 106 member A | protein_coding | NP_079250.1 | 25682 | . | . | . | [illegible] |
| 80045 | NM_024980,XI | GPR157 | G protein-coupled receptor 157 | protein_coding | NP_079256.4;XP_005263553.1;XP_005263554.1 | 23687 | . | . | . | [illegible] |
| 80055 | NM_00132109 | PGAP1 | post-GPI attachment to proteins inositol deacylase 1 | protein_coding | NP_001308028.1;NP_001308029.1;NP_079265.2;XP_016860481.1;XP_01 | 25712 | 611655 | . | . | [illegible] |
| 80060 | gene-GNL3LP1 | GNL3LP1 | G protein nucleolar 3 like pseudogene 1 | pseudogene | . | 25733 | . | . | . | [illegible] |
| 80063 | NM_00125616 | ATF7IP2 | activating transcription factor 7 interacting protein 2 | protein_coding | NP_001243089.1;NP_001339049.1;NP_079273.2;XP_006721016.1;XP_00 | 20397 | 613645 | . | . | [illegible] |
| 80067 | NM_00116482 | DCAF17 | DDB1 and CUL4 associated factor 17 | protein_coding | NP_001158293.1;NP_079276.2;XP_006712830.1;XP_006712831.1;XP_01 | 25784 | 612515 | . | . | [illegible] |
| 80071 | NM_025004,XI | CCDC15 | coiled-coil domain containing 15 | protein_coding | NP_079280.2;XP_016873835.1;XP_016873836.1;XP_016873837.1;XP_01 | 25798 | . | . | . | [illegible] |
| 80072 | NR_027262 | HEXA-AS1 | HEXA antisense RNA 1 | lncRNA | . | 25810 | . | . | . | [illegible] |
| 80094 | NR_027715 | BIN3-IT1 | BIN3 intronic transcript 1 | lncRNA | . | 25868 | . | . | . | [illegible] |
| 80097 | NM_00133028 | MZT2B | mitotic spindle organizing protein 2B | protein_coding | NP_001317211.1;NP_001317213.1;NP_079305.2 | 25886 | 613450 | . | . | [illegible] |
| 80110 | NM_025040 | ZNF614 | zinc finger protein 614 | protein_coding | NP_079316.2 | 24722 | . | . | . | [illegible] |
| 80114 | NM_00108051 | BICC1 | BicC family RNA binding protein 1 | protein_coding | NP_001073981.1;XP_005270226.1;XP_011538487.1;XP_011538492.1;XP | 19351 | 614295 | . | . | [illegible] |
| 80115 | NM_025045,XI | BAIAP2L2 | BAR/IMD domain containing adaptor protein 2 like 2 | protein_coding | NP_079321.3;XP_005261808.1;XP_011528681.1;XP_011528682.1;XP_01 | 26203 | 617536 | . | . | [illegible] |
| 80119 | NM_00128649 | PIF1 | PIF1 5'-to-3' DNA helicase | protein_coding | NP_001273425.1;NP_001273426.1;NP_001273428.1;NP_079325.2;XP_01 | 26220 | 610953 | . | . | [illegible] |
| 80124 | NM_025054 | VCPIP1 | valosin containing protein interacting protein 1 | protein_coding | NP_079330.2 | 30897 | 611745 | . | . | [illegible] |
| 80127 | NM_025057,XI | BBOF1 | basal body orientation factor 1 | protein_coding | NP_079333.2;XP_005268149.1;XP_005268155.1;XP_011535472.1;XP_01 | 19855 | . | . | . | [illegible] |
| 80128 | NM_00125659 | TRIM46 | tripartite motif containing 46 | protein_coding | NP_001243528.1;NP_001243530.1;NP_001269307.1;NP_001269308.1;NI | 19019 | 600986 | . | . | [illegible] |
| 80131 | NM_00126828 | LRRC8E | leucine rich repeat containing 8 VRAC subunit E | protein_coding | NP_001255213.1;NP_001255214.1;NP_079337.2;XP_011526621.1 | 26272 | 612891 | . | . | [illegible] |
| 80135 | NM_025065 | RPF1 | ribosome production factor 1 homolog | protein_coding | NP_079341.2 | 30350 | . | . | . | [illegible] |
| 80139 | NM_025069 | ZNF703 | zinc finger protein 703 | protein_coding | NP_079345.1 | 25883 | 617045 | . | . | [illegible] |
| 80142 | NM_00125633 | PTGES2 | prostaglandin E synthase 2 | protein_coding | NP_001243264.1;NP_079348.1;NP_945176.1;XP_005252296.1;XP_01151 | 17822 | 608152 | . | . | [illegible] |
| 80143 | NM_00110239 | SIKE1 | suppressor of IKBKE 1 | protein_coding | NP_001095866.1;NP_079349.2 | 26119 | 611656 | . | . | [illegible] |
| 80144 | NM_00116613 | FRAS1 | Fraser extracellular matrix complex subunit 1 | protein_coding | NP_001159605.1;NP_079350.5 | 19185 | 607830 | . | . | [illegible] |
| 80145 | NM_00128538 | THOC7 | THO complex 7 | protein_coding | NP_001272316.1;NP_001272333.1;NP_079351.2;XP_006713402.1 | 29874 | 611965 | . | . | [illegible] |
| 80146 | NM_00125387 | UXS1 | UDP-glucuronate decarboxylase 1 | protein_coding | NP_001240804.1;NP_001240805.1;NP_001364433.1;NP_001364434.1;NI | 17729 | 609749 | . | . | [illegible] |
| 80148 | NM_00114634 | SLC66A2 | solute carrier family 66 member 2 | protein_coding | NP_001139815.1;NP_001139817.1;NP_079354.2;XP_005266827.1;XP_01 | 26188 | . | . | . | [illegible] |
| 80149 | NM_00132355 | ZC3H12A | zinc finger CCCH-type containing 12A | protein_coding | NP_001310479.1;NP_001310480.1;NP_079355.2;XP_011540500.1 | 26259 | 610562 | . | . | [illegible] |
| 80150 | NM_00108392 | ASRGL1 | asparaginase and isoaspartyl peptidase 1 | protein_coding | NP_001077395.1;NP_079356.3;XP_005274362.1;XP_005274363.1;XP_01 | 16448 | 609212 | . | . | [illegible] |
| 80152 | NM_025082,XI | CENPT | centromere protein T | protein_coding | NP_079358.3;XP_011521647.1;XP_016879204.1;XP_016879206.1;XP_02 | 25787 | 611510 | . | . | [illegible] |
| 80153 | NM_00114244 | EDC3 | enhancer of mRNA decapping 3 | protein_coding | NP_001135915.1;NP_001135916.1;NP_001338307.1;NP_001338308.1;NI | 26114 | 609842 | . | . | [illegible] |
| 80154 | NR_026811,NF | GOLGA2P10 | GOLGA2 pseudogene 10 | transcribed_ps | . | 26229 | . | . | . | [illegible] |
| 80155 | NM_057175,XI | NAA15 | N-alpha-acetyltransferase 15, NatA auxiliary subunit | protein_coding | NP_476516.1;XP_005263293.1 | 30782 | 608000 | . | . | [illegible] |
| 80161 | NR_026710,NF | ASMTL-AS1 | ASMTL antisense RNA 1 | lncRNA | . | 25811 | . | . | . | [illegible] |
| 80162 | NM_025092,XI | PGGHG | protein-glucosylgalactosylhydroxylysine glucosidase | protein_coding | NP_079368.3;XP_011518685.1;XP_011518686.1;XP_011518687.1;XP_01 | 26210 | 617032 | . | . | [illegible] |
| 80167 | NM_00103971 | ABHD18 | abhydrolase domain containing 18 | protein_coding | NP_001034806.1;NP_001306234.1;NP_001306235.1;NP_001306236.1;NI | 26111 | . | . | . | [illegible] |
| 80169 | NM_025099,N | CTC1 | CST telomere replication complex component 1 | protein_coding | NP_079375.3;XP_006721640.1;XP_006721641.1;XP_011522312.1;XP_01 | 26169 | 613129 | . | . | [illegible] |
| 80173 | NM_00109922 | IFT74 | intraflagellar transport 74 | protein_coding | NP_001092692.1;NP_001092693.1;NP_001092694.1;NP_001336857.1;NI | 21424 | 608040 | . | . | [illegible] |
| 80174 | NM_025104,N | DBF4B | DBF4 zinc finger B | protein_coding | NP_079380.2;NP_663696.1;XP_006722161.1;XP_006722165.1;XP_01152 | 17883 | 611661 | . | . | [illegible] |
| 80176 | NM_025106 | SPSB1 | splA/ryanodine receptor domain and SOCS box containing 1 | protein_coding | NP_079382.2 | 30628 | 611657 | . | . | [illegible] |
| 80178 | NM_025108,XI | TEDC2 | tubulin epsilon and delta complex 2 | protein_coding | NP_079384.2;XP_011520969.1 | 25849 | . | . | . | [illegible] |
| 80179 | NM_00103358 | MYO19 | myosin XIX | protein_coding | NP_001028752.1;NP_001157207.1;NP_079385.2;XP_011523588.1;XP_01 | 26234 | 617379 | . | . | [illegible] |
| 80183 | NM_00128676 | RUBCNL | rubicon like autophagy enhancer | protein_coding | NP_001273690.1;NP_001273691.1;NP_001273692.1;NP_001273693.1;NI | 20420 | . | . | . | [illegible] |
| 80184 | NM_025114,XI | CEP290 | centrosomal protein 290 | protein_coding | NP_079390.3;XP_011537058.1;XP_011537059.1;XP_011537060.1;XP_01 | 29021 | 610142 | . | . | [illegible] |
| 80185 | NM_00110240 | TTI2 | TELO2 interacting protein 2 | protein_coding | NP_001095871.1;NP_001252510.1;NP_001317434.1;NP_079391.2 | 26262 | 614426 | . | . | [illegible] |
| 80194 | NM_00107865 | TMEM134 | transmembrane protein 134 | protein_coding | NP_001072118.1;NP_001072119.1;NP_079400.1;XP_006718756.1;XP_01 | 26142 | . | . | . | [illegible] |
| 80195 | NM_00127036 | TMEM254 | transmembrane protein 254 | protein_coding | NP_001257296.1;NP_001257297.1;NP_001257298.1;NP_001257299.1;NI | 25804 | . | . | . | [illegible] |
| 80196 | NM_00125685 | RNF34 | ring finger protein 34 | protein_coding | NP_001243787.1;NP_079402.2;NP_919247.1;XP_024304958.1;XP_02430 | 17297 | 608299 | . | . | [illegible] |
| 80198 | NM_00135028 | MUS81 | MUS81 structure-specific endonuclease subunit | protein_coding | NP_001337212.1;NP_079404.3;XP_011543571.1;XP_011543572.1 | 29814 | 606591 | . | . | [illegible] |
| 80199 | NM_00117193 | FUZ | fuzzy planar cell polarity protein | protein_coding | NP_001165408.1;NP_001339191.1;NP_001350592.1;NP_079405.2;XP_00 | 26219 | 610622 | . | . | [illegible] |
| 80201 | NM_025130,XI | HKDC1 | hexokinase domain containing 1 | protein_coding | NP_079406.4;XP_011538497.1 | 23302 | 617221 | . | . | [illegible] |
| 80204 | NM_00119027 | FBXO11 | F-box protein 11 | protein_coding | NP_001177203.1;NP_001361254.1;NP_079409.3;XP_005264629.1;XP_00 | 13590 | 607871 | . | . | [illegible] |
| 80205 | NM_00130831 | CHD9 | chromodomain helicase DNA binding protein 9 | protein_coding | NP_001295248.1;NP_001339056.1;NP_001339085.1;NP_001339086.1;NI | 25701 | 616936 | . | . | [illegible] |
| 80206 | NM_00128173 | FHOD3 | formin homology 2 domain containing 3 | protein_coding | NP_001268668.1;NP_001268669.1;NP_079411.2;XP_005258406.1;XP_00 | 26178 | 609691 | . | . | [illegible] |
| 80207 | NM_00101798 | OPA3 | outer mitochondrial membrane lipid metabolism regulator OPA3 | protein_coding | NP_001017989.2;NP_079412.1;XP_006723466.1 | 8142 | 606580 | . | . | [illegible] |
| 80208 | NM_00116022 | SPG11 | SPG11 vesicle trafficking associated, spatacsin | protein_coding | NP_001153699.1;NP_079413.3;XP_006720763.1;XP_006720764.1;XP_01 | 11226 | 610844 | . | . | [illegible] |
| 80209 | NM_025138,N | PROSER1 | proline and serine rich 1 | protein_coding | NP_079414.3;NP_733837.2;XP_011533541.1 | 20291 | . | . | . | [illegible] |
| 80210 | NM_00127146 | ARMC9 | armadillo repeat containing 9 | protein_coding | NP_001258395.2;NP_001278585.2;NP_001339683.2;NP_001339684.2;NI | 20730 | 617612 | . | . | [illegible] |
| 80212 | NM_00130495 | CCDC92 | coiled-coil domain containing 92 | protein_coding | NP_001291886.1;NP_001291887.1;NP_001291888.1;NP_001291889.1;NI | 29563 | . | . | . | [illegible] |
| 80213 | NM_00130796 | TM2D3 | TM2 domain containing 3 | protein_coding | NP_001294889.1;NP_001294955.1;NP_079417.2;NP_510883.2 | 24128 | 610014 | . | . | [illegible] |
| 80216 | NM_00110240 | ALPK1 | alpha kinase 1 | protein_coding | NP_001095876.1;NP_001240813.1;NP_079420.3 | 20917 | 607347 | . | . | [illegible] |
| 80217 | NM_025145,XI | CFAP43 | cilia and flagella associated protein 43 | protein_coding | NP_079421.5;XP_005270228.1;XP_005270229.1;XP_011538498.1;XP_01 | 26684 | 617558 | . | . | [illegible] |
| 80218 | NM_00130844 | NAA50 | N-alpha-acetyltransferase 50, NatE catalytic subunit | protein_coding | NP_001295374.1;NP_079422.1 | 29533 | 610834 | . | . | [illegible] |
| 80219 | NM_00132081 | COQ10B | coenzyme Q10B | protein_coding | NP_001307747.1;NP_001307748.1;NP_001307749.1;NP_079423.1 | 25819 | . | . | . | [illegible] |
| 80221 | NM_00128896 | ACSF2 | acyl-CoA synthetase family member 2 | protein_coding | NP_001275897.1;NP_001275898.1;NP_001275899.1;NP_001275900.1;NI | 26101 | 610465 | . | . | [illegible] |
| 80222 | NM_00127189 | TARS2 | threonyl-tRNA synthetase 2, mitochondrial | protein_coding | NP_001258824.1;NP_001258825.1;NP_079426.2;XP_006711618.1;XP_01 | 30740 | 612805 | . | . | [illegible] |
| 80223 | NM_00100281 | RAB11FIP1 | RAB11 family interacting protein 1 | protein_coding | NP_001002814.2;NP_079427.4;XP_016869358.1 | 30265 | 608737 | . | . | [illegible] |
| 80224 | NM_00120157 | NUBPL | nucleotide binding protein like | protein_coding | NP_001188502.1;NP_001188503.1;NP_079428.2;XP_011535483.1;XP_01 | 20278 | 613621 | . | . | [illegible] |
| 80227 | NM_00126780 | PAAF1 | proteasomal ATPase associated factor 1 | protein_coding | NP_001254732.1;NP_001254733.1;NP_001254734.1;NP_001254735.1;NI | 25687 | . | . | . | [illegible] |
| 80228 | NM_00112634 | ORAI2 | ORAI calcium release-activated calcium modulator 2 | protein_coding | NP_001119812.1;NP_001258747.1;NP_001258748.1;NP_116220.1 | 21667 | 610929 | . | . | [illegible] |
| 80230 | NM_00104045 | RUFY1 | RUN and FYVE domain containing 1 | protein_coding | NP_001035541.1;NP_001035542.1;NP_079434.3;XP_005266050.1;XP_00 | 19760 | 610327 | . | . | [illegible] |
| 80232 | NM_00111511 | WDR26 | WD repeat domain 26 | protein_coding | NP_001108585.2;NP_001366332.1;NP_079436.4 | 21208 | 617424 | . | . | [illegible] |
| 80233 | NM_025161,N | FAAP100 | FA core complex associated protein 100 | protein_coding | NP_079437.5;XP_006722174.1;XP_006722175.1;XP_016880651.1;XP_01 | 26171 | 611301 | . | . | [illegible] |
| 80235 | NM_025163,XI | PIGZ | phosphatidylinositol glycan anchor biosynthesis class Z | protein_coding | NP_079439.2;XP_011511492.1;XP_011511494.1;XP_016862729.1;XP_01 | 30596 | 611671 | . | . | [illegible] |
| 80237 | NM_025165 | ELL3 | elongation factor for RNA polymerase II 3 | protein_coding | NP_079441.1 | 23113 | 609885 | . | . | [illegible] |
| 80254 | NM_00104238 | CEP63 | centrosomal protein 63 | protein_coding | NP_001035842.1;NP_001035843.1;NP_001035859.1;NP_001340037.1;NI | 25815 | 614724 | . | . | [illegible] |
| 80255 | NM_00133031 | SLC35F5 | solute carrier family 35 member F5 | protein_coding | NP_001317243.1;NP_001317244.1;NP_001317245.1;NP_001317246.1;NI | 23617 | . | . | . | [illegible] |
| 80256 | NM_00131799 | FAM214B | family with sequence similarity 214 member B | protein_coding | NP_001304920.1;NP_079458.2;XP_005251645.1;XP_005251647.1;XP_00 | 25666 | . | . | . | [illegible] |
| 80262 | NM_00132054 | C16orf70 | chromosome 16 open reading frame 70 | protein_coding | NP_001307469.1;NP_001307470.1;NP_001307471.1;NP_001307472.1;NI | 29564 | . | . | . | [illegible] |
| 80263 | NM_00114563 | TRIM45 | tripartite motif containing 45 | protein_coding | NP_001139107.1;NP_079464.2;XP_011540501.1 | 19018 | 609318 | . | . | [illegible] |
| 80264 | NM_00117267 | ZNF430 | zinc finger protein 430 | protein_coding | NP_001166142.1;NP_079465.3 | 20808 | . | . | . | [illegible] |
| 80267 | NM_00131996 | EDEM3 | ER degradation enhancing alpha-mannosidase like protein 3 | protein_coding | NP_001306889.1;NP_079467.3;XP_005245556.1;XP_011508314.1;XP_01 | 16787 | 610214 | . | . | [illegible] |
| 80270 | NM_00114277 | HSD3B7 | hydroxy-delta-5-steroid dehydrogenase, 3 beta- and steroid delta- | protein_coding | NP_001136249.1;NP_001136250.1;NP_079469.2;XP_005255658.2;XP_01 | 18324 | 607764 | . | . | [illegible] |
| 80271 | NM_025194,XI | ITPKC | inositol-trisphosphate 3-kinase C | protein_coding | NP_079470.1;XP_016882813.1 | 14897 | 606476 | . | . | [illegible] |
| 80273 | NM_025196 | GRPEL1 | GrpE like 1, mitochondrial | protein_coding | NP_079472.1 | 19696 | 606173 | . | . | [illegible] |
| 80279 | NM_00127819 | CDK5RAP3 | CDK5 regulatory subunit associated protein 3 | protein_coding | NP_001265126.1;NP_001265127.1;NP_001265145.1;NP_001265146.1;NI | 18673 | 608202 | . | . | [illegible] |
| 80298 | NM_00103305 | MTERF2 | mitochondrial transcription termination factor 2 | protein_coding | NP_001028222.1;NP_079474.2;XP_011537069.1;XP_016875474.1;XP_01 | 30779 | 616929 | . | . | [illegible] |
| 80301 | NM_00119505 | PLEKHO2 | pleckstrin homology domain containing O2 | protein_coding | NP_001181988.1;NP_079477.2 | 30026 | . | . | . | [illegible] |
| 80303 | NM_00124325 | EFHD1 | EF-hand domain family member D1 | protein_coding | NP_001230181.1;NP_001295324.1;NP_079478.1 | 29556 | 611617 | . | . | [illegible] |
| 80304 | NM_00114231 | WDCP | WD repeat and coiled coil containing | protein_coding | NP_001135791.1;NP_079479.1;XP_016860518.1 | 26157 | 616234 | . | . | [illegible] |
| 80305 | NM_00132048 | TRABD | TraB domain containing | protein_coding | NP_001307413.1;NP_001307414.1;NP_001307416.1;NP_001307417.1;NI | 28805 | . | . | . | [illegible] |
| 80306 | NM_025205 | MED28 | mediator complex subunit 28 | protein_coding | NP_079481.2 | 24628 | 610311 | . | . | [illegible] |
| 80307 | NR_119376 | FER1L4 | fer-1 like family member 4 (pseudogene) | transcribed_ps | . | 15801 | . | . | . | [illegible] |
| 80308 | NM_00118489 | FLAD1 | flavin adenine dinucleotide synthetase 1 | protein_coding | NP_001171820.1;NP_001171821.1;NP_079483.3;NP_958800.1 | 24671 | 610595 | . | . | [illegible] |
| 80310 | NM_025208,N | PDGFD | platelet derived growth factor D | protein_coding | NP_079484.1;NP_149126.1 | 30620 | 609673 | . | . | [illegible] |
| 80311 | NM_030624 | KLHL15 | kelch like family member 15 | protein_coding | NP_085127.2 | 29347 | 300980 | . | . | [illegible] |
| 80312 | NM_030625,XI | TET1 | tet methylcytosine dioxygenase 1 | protein_coding | NP_085128.2;XP_011538506.1;XP_011538507.1;XP_011538508.1;XP_01 | 29484 | 607790 | . | . | [illegible] |
| 80313 | NM_00114375 | LRRC27 | leucine rich repeat containing 27 | protein_coding | NP_001137229.1;NP_001137230.1;NP_001137231.1;NP_001296403.1;NI | 29346 | . | . | . | [illegible] |
| 80314 | NM_00127200 | EPC1 | enhancer of polycomb homolog 1 | protein_coding | NP_001258933.1;NP_001258948.1;NP_001269320.1;NP_001369682.1;NI | 19876 | 610999 | . | . | [illegible] |
| 80315 | NM_00130818 | CPEB4 | cytoplasmic polyadenylation element binding protein 4 | protein_coding | NP_001295118.1;NP_001295120.1;NP_001295121.1;NP_001295122.1;NI | 21747 | 610607 | . | . | [illegible] |
| 80317 | NM_00124289 | ZKSCAN3 | zinc finger with KRAB and SCAN domains 3 | protein_coding | NP_001229823.1;NP_001229824.1;NP_077819.2;XP_005249480.1;XP_00 | 13853 | 612791 | . | . | [illegible] |
| 80318 | NM_00113595 | GKAP1 | G kinase anchoring protein 1 | protein_coding | NP_001129425.1;NP_079487.2;XP_005252298.1;XP_006717356.1;XP_01 | 17496 | 611356 | . | . | [illegible] |
| 80321 | NM_00128896 | CEP70 | centrosomal protein 70 | protein_coding | NP_001275893.1;NP_001275894.1;NP_001275895.1;NP_001275896.1;NI | 29972 | 614310 | . | . | [illegible] |
| 80323 | NM_00114382 | CCDC68 | coiled-coil domain containing 68 | protein_coding | NP_001137301.1;NP_079490.1;XP_006722615.1;XP_011524500.1;XP_01 | 24350 | 616909 | . | . | [illegible] |
| 80324 | NM_00100201 | PUS1 | pseudouridine synthase 1 | protein_coding | NP_001002019.1;NP_001002020.1;NP_079491.2;XP_011537070.1;XP_01 | 15508 | 608109 | . | . | [illegible] |
| 80325 | NM_032548,N | ABTB1 | ankyrin repeat and BTB domain containing 1 | protein_coding | NP_115937.1;NP_742024.1;XP_006713832.1;XP_011511515.1;XP_01686 | 18275 | 608308 | . | . | [illegible] |
| 80328 | NM_025217,XI | ULBP2 | UL16 binding protein 2 | protein_coding | NP_079493.1;XP_016866810.1 | 14894 | 605698 | . | . | [illegible] |
| 80331 | NM_025219,XI | DNAJC5 | DnaJ heat shock protein family (Hsp40) member C5 | protein_coding | NP_079495.1;XP_011527350.1;XP_011527351.1 | 16235 | 611203 | . | . | [illegible] |
| 80335 | NM_025222,XI | WDR82 | WD repeat domain 82 | protein_coding | NP_079498.2;XP_011532438.1 | 28826 | 611059 | . | . | [illegible] |
| 80336 | NM_00137217 | PABPC1L | poly(A) binding protein cytoplasmic 1 like | protein_coding | NP_001359108.1 | 15797 | . | . | . | [illegible] |
| 80339 | NM_025225 | PNPLA3 | patatin like phospholipase domain containing 3 | protein_coding | NP_079501.2 | 18590 | 609567 | . | . | [illegible] |
| 80344 | NM_00116348 | DCAF11 | DDB1 and CUL4 associated factor 11 | protein_coding | NP_001156956.1;NP_079506.3;NP_852002.1 | 20258 | 613317 | . | . | [illegible] |
| 80345 | NM_00132055 | ZSCAN16 | zinc finger and SCAN domain containing 16 | protein_coding | NP_001307484.1;NP_001307485.1;NP_001307486.1;NP_001307487.1;NI | 20813 | 618544 | . | . | [illegible] |
| 80346 | NM_00131696 | REEP4 | receptor accessory protein 4 | protein_coding | NP_001303893.1;NP_001303894.1;NP_079508.2 | 26176 | 609349 | . | . | [illegible] |
| 80347 | NM_00104252 | COASY | Coenzyme A synthase | protein_coding | NP_001035994.1;NP_001035997.2;NP_079509.5;XP_006722179.1;XP_01 | 29932 | 609855 | . | . | [illegible] |
| 80349 | NM_00130324 | WDR61 | WD repeat domain 61 | protein_coding | NP_001290176.1;NP_001290177.1;NP_079510.1;XP_011520396.1;XP_01 | 30300 | 609540 | . | . | [illegible] |
| 80350 | NR_028092,NF | LPAL2 | lipoprotein(a) like 2, pseudogene | transcribed_ps | . | 21210 | 611682 | . | . | [illegible] |
| 80351 | NM_025235,XI | TNKS2 | tankyrase 2 | protein_coding | NP_079511.1;XP_005270242.2;XP_011538515.1;XP_011538517.1;XP_01 | 15677 | 607128 | . | . | [illegible] |
| 80381 | NM_00102473 | CD276 | CD276 molecule | protein_coding | NP_001019907.1;NP_001316557.1;NP_001316558.1;NP_079516.1;XP_00 | 19137 | 605715 | . | . | [illegible] |
| 80700 | NM_00117109 | UBXN6 | UBX domain protein 6 | protein_coding | NP_001164562.1;NP_079517.1;XP_016882814.1 | 14928 | 611946 | . | . | [illegible] |
| 80704 | NM_00137141 | SLC19A3 | solute carrier family 19 member 3 | protein_coding | NP_001358340.1;NP_001358341.1;NP_001358342.1;NP_001358343.1;NI | 16266 | 606152 | . | . | [illegible] |
| 80705 | NM_00134901 | TSGA10 | testis specific 10 | protein_coding | NP_001335941.1;NP_001335942.1;NP_001335943.1;NP_079520.1;NP_8 | 14927 | 607166 | . | . | [illegible] |
| 80709 | NM_00131795 | AKNA | AT-hook transcription factor | protein_coding | NP_001304879.1;NP_001304881.1;NP_110394.3;XP_005252301.1;XP_00 | 24108 | 605729 | . | . | [illegible] |
| 80714 | NM_025245,N | PBX4 | PBX homeobox 4 | protein_coding | NP_079521.1;XP_006722974.1;XP_011526622.1;XP_011526624.1;XP_01 | 13403 | 608127 | . | . | [illegible] |
| 80724 | NM_00113653 | ACAD10 | acyl-CoA dehydrogenase family member 10 | protein_coding | NP_001130010.1;NP_079523.3 | 21597 | 611181 | . | . | [illegible] |
| 80725 | NM_025248,XI | SRCIN1 | SRC kinase signaling inhibitor 1 | protein_coding | NP_079524.2;XP_016880658.1;XP_016880659.1;XP_016880660.1;XP_01 | 29506 | 610786 | . | . | [illegible] |
| 80727 | NM_025250,XI | TTYH3 | tweety family member 3 | protein_coding | NP_079526.1;XP_011513837.1;XP_016868144.1;XP_016868145.1 | 22222 | 608919 | . | . | [illegible] |
| 80728 | NM_00130820 | ARHGAP39 | Rho GTPase activating protein 39 | protein_coding | NP_001295136.1;NP_001295137.1;NP_079527.1;XP_011515610.1;XP_01 | 29351 | 615880 | . | . | [illegible] |
| 80742 | NM_00107749 | PRR3 | proline rich 3 | protein_coding | NP_001070965.1;NP_079539.2 | 21149 | . | . | . | [illegible] |
| 80745 | NM_00132146 | THUMPD2 | THUMP domain containing 2 | protein_coding | NP_001308397.1;NP_001308398.1;NP_001308399.1;NP_001308403.1;NI | 14890 | 611751 | . | . | [illegible] |
| 80746 | NM_00114539 | TSEN2 | tRNA splicing endonuclease subunit 2 | protein_coding | NP_001138864.1;NP_001138865.1;NP_001138866.1;NP_001308206.1;NI | 28422 | 608753 | . | . | [illegible] |
| 80755 | NM_00126143 | AARSD1 | alanyl-tRNA synthetase domain containing 1 | protein_coding | NP_001248363.1 | 28417 | 613212 | . | . | [illegible] |
| 80757 | NM_00133123 | TMEM121 | transmembrane protein 121 | protein_coding | NP_001318167.1;NP_079544.1;XP_006720324.1 | 20511 | . | . | . | [illegible] |
| 80758 | NM_00117410 | PRR7 | proline rich 7, synaptic | protein_coding | NP_001167572.1;NP_001167573.1;NP_001362522.1;NP_001362523.1;NI | 28130 | 618306 | . | . | [illegible] |
| 80760 | NM_00100185 | ITIH5 | inter-alpha-trypsin inhibitor heavy chain 5 | protein_coding | NP_001001851.1;NP_085046.5;NP_116206.4;XP_011518015.1;XP_01151 | 21449 | 609783 | . | . | [illegible] |
| 80762 | NM_030571 | NDFIP1 | Nedd4 family interacting protein 1 | protein_coding | NP_085048.1 | 17592 | 612050 | . | . | [illegible] |
| 80763 | NM_030572,N | SPX | spexin hormone | protein_coding | NP_085049.1 | 28139 | . | . | . | [illegible] |
| 80764 | NM_00100869 | THAP7 | THAP domain containing 7 | protein_coding | NP_001008695.1;NP_085050.2 | 23190 | 609518 | . | . | [illegible] |
| 80765 | NM_181900,N | STARD5 | StAR related lipid transfer domain containing 5 | protein_coding | NP_871629.1 | 18065 | 607050 | . | . | [illegible] |
| 80772 | NM_00102988 | CPTP | ceramide-1-phosphate transfer protein | protein_coding | NP_001025056.1;XP_005244858.1;XP_005244859.1;XP_011540502.1 | 28116 | 615467 | . | . | [illegible] |
| 80774 | NM_030576,XI | LIMD2 | LIM domain containing 2 | protein_coding | NP_085053.1;XP_005257760.1;XP_005257762.1;XP_006722187.1 | 28142 | . | . | . | [illegible] |
| 80775 | NM_00110519 | TMEM177 | transmembrane protein 177 | protein_coding | NP_001098668.1;NP_001098669.1;NP_085054.2 | 28143 | . | . | . | [illegible] |
| 80776 | NM_030578,XI | B9D2 | B9 domain containing 2 | protein_coding | NP_085055.2;XP_011525651.1;XP_011525652.1 | 28636 | 611951 | . | . | [illegible] |
| 80777 | NM_030579 | CYB5B | cytochrome b5 type B | protein_coding | NP_085056.2 | 24374 | 611964 | . | . | [illegible] |
| 80778 | NM_00128676 | ZNF34 | zinc finger protein 34 | protein_coding | NP_001273698.1;NP_001273699.1;NP_001364956.1;NP_001364957.1;NI | 13098 | 194526 | . | . | [illegible] |
| 80781 | NM_00137950 | COL18A1 | collagen type XVIII alpha 1 chain | protein_coding | NP_001366429.1;NP_085059.2;NP_569711.2 | 2195 | 120328 | . | . | [illegible] |
| 80789 | NM_030628 | INTS5 | integrator complex subunit 5 | protein_coding | NP_085131.1 | 29352 | 611349 | . | . | [illegible] |
| 80790 | NM_030629,N | CMIP | c-Maf inducing protein | protein_coding | NP_085132.1;NP_938204.2;XP_005256236.4;XP_005256238.1;XP_00525 | 24319 | 610112 | . | . | [illegible] |
| 80817 | NM_00104015 | CEP44 | centrosomal protein 44 | protein_coding | NP_001035247.1;NP_001138786.1;XP_011530587.1;XP_011530588.1;XP | 29356 | . | . | . | [illegible] |
| 80818 | NM_00107719 | ZNF436 | zinc finger protein 436 | protein_coding | NP_001070663.1;NP_001357581.1;NP_085137.1 | 20814 | 611703 | . | . | [illegible] |
| 80820 | NM_030636,XI | EEPD1 | endonuclease/exonuclease/phosphatase family domain containing | protein_coding | NP_085139.2;XP_016868146.1;XP_016868147.1 | 22223 | 617192 | . | . | [illegible] |
| 80821 | NM_00116014 | DDHD1 | DDHD domain containing 1 | protein_coding | NP_001153619.1;NP_001153620.1;NP_085140.2;XP_005268159.1;XP_00 | 19714 | 614603 | . | . | [illegible] |
| 80824 | NM_030640,XI | DUSP16 | dual specificity phosphatase 16 | protein_coding | NP_085143.1;XP_005253545.1;XP_006719218.1;XP_011519158.1;XP_01 | 17909 | 607175 | . | . | [illegible] |
| 80829 | NM_00119705 | ZFP91 | ZFP91 zinc finger protein, atypical E3 ubiquitin ligase | protein_coding | NP_001183980.1;NP_444251.1 | 14983 | . | . | . | [illegible] |
| 80830 | NM_030641,XI | APOL6 | apolipoprotein L6 | protein_coding | NP_085144.1;XP_011528694.1 | 14870 | 607256 | . | . | [illegible] |
| 80851 | NM_00132246 | SH3BP5L | SH3 binding domain protein 5 like | protein_coding | NP_001309391.1;NP_001309392.1;NP_001309393.1;NP_085148.1;XP_01 | 29360 | . | . | . | [illegible] |

| | | | | | | | | | | |
|---|---|---|---|---|---|---|---|---|---|---|
| 80852 | NM_00108042 | GRIP2 | glutamate receptor interacting protein 2 | protein_coding | NP_001073892.3;XP_011532443.1;XP_011532444.1;XP_011532445.1;XP_ | 23841 | . | . | . | [illegible] |
| 80853 | NM_030647,XI | KDM7A | lysine demethylase 7A | protein_coding | NP_085150.1;XP_011514889.1 | 22224 | . | . | . | [illegible] |
| 80854 | NM_00130619 | SETD7 | SET domain containing 7, histone lysine methyltransferase | protein_coding | NP_001293128.1;NP_001293129.1;NP_085151.1;XP_016864150.1 | 30412 | 606594 | . | . | [illegible] |
| 80856 | NM_00130500 | LNPK | lunapark, ER junction formation factor | protein_coding | NP_001291937.1;NP_001291938.1;NP_001291939.1;NP_001291940.1;NI | 21610 | 610236 | . | . | [illegible] |
| 80862 | NR_026751,NF | ZNRD1ASP | zinc ribbon domain containing 1 antisense, pseudogene | transcribed_ps | . | 13924 | 615714 | . | . | [illegible] |
| 80863 | NM_00136378 | PRRT1 | proline rich transmembrane protein 1 | protein_coding | NP_001350709.1;NP_085154.3;XP_024302328.1 | 13943 | 618297 | . | . | [illegible] |
| 80864 | NM_030652,N | EGFL8 | EGF like domain multiple 8 | protein_coding | NP_085155.1 | 13944 | 609897 | . | . | [illegible] |
| 80895 | NM_030768,XI | ILKAP | ILK associated serine/threonine phosphatase | protein_coding | NP_110395.1;XP_005246163.1;XP_006712847.1;XP_011510248.1;XP_01 | 15566 | . | . | . | [illegible] |
| 80896 | NM_00120005 | NPL | N-acetylneuraminate pyruvate lyase | protein_coding | NP_001186979.1;NP_001186980.1;NP_001186981.1;NP_001186985.1;NI | 16781 | 611412 | . | . | [illegible] |
| 80975 | NM_00128874 | TMPRSS5 | transmembrane serine protease 5 | protein_coding | NP_001275678.1;NP_001275679.1;NP_001275680.1;NP_001275681.1;NI | 14908 | 606751 | . | . | [illegible] |
| 81029 | NM_030775,N | WNT5B | Wnt family member 5B | protein_coding | NP_110402.2;NP_116031.1;XP_024304971.1;XP_024304972.1;XP_02430 | 16265 | 606361 | . | . | [illegible] |
| 81031 | NM_030777,XI | SLC2A10 | solute carrier family 2 member 10 | protein_coding | NP_110404.1;XP_011527362.1;XP_011527363.1;XP_011527364.1;XP_01 | 13444 | 606145 | . | . | [illegible] |
| 81034 | NM_030780,N | SLC25A32 | solute carrier family 25 member 32 | protein_coding | NP_110407.2 | 29683 | 610815 | . | . | [illegible] |
| 81035 | NM_130386,XI | COLEC12 | collectin subfamily member 12 | protein_coding | NP_569057.2;XP_011524043.1 | 16016 | 607621 | . | . | [illegible] |
| 81037 | NM_030782,XI | CLPTM1L | CLPTM1 like | protein_coding | NP_110409.2;XP_011512446.1;XP_024301989.1;XP_024301990.1 | 24308 | 612585 | . | . | [illegible] |
| 81399 | NM_00100527 | OR4F16 | olfactory receptor family 4 subfamily F member 16 | protein_coding | NP_001005277.1;XP_016857897.1;XP_016857898.1;XP_016857899.1;XP_ | 15079 | . | . | . | [illegible] |
| 81488 | NM_00101810 | POLR2M | RNA polymerase II subunit M | protein_coding | NP_001018112.1;NP_056347.1 | 14862 | 606485 | . | . | [illegible] |
| 81490 | NM_00132954 | PTDSS2 | phosphatidylserine synthase 2 | protein_coding | NP_001316473.1;NP_001316474.1;NP_001316477.1;NP_110410.1;XP_01 | 15463 | 612793 | . | . | [illegible] |
| 81491 | NM_00114395 | GPR63 | G protein-coupled receptor 63 | protein_coding | NP_001137429.1;NP_110411.1;XP_006715633.1;XP_011534455.1;XP_01 | 13302 | 606915 | . | . | [illegible] |
| 81493 | NM_00116170 | SYNC | syncoilin, intermediate filament protein | protein_coding | NP_001155180.2;NP_110413.3;XP_024305778.1;XP_024305779.1;XP_02 | 28897 | 611750 | . | . | [illegible] |
| 81502 | NM_030789,N | HM13 | histocompatibility minor 13 | protein_coding | NP_110416.1;NP_848695.1;NP_848696.1;NP_848697.1 | 16435 | 607106 | . | . | [illegible] |
| 81532 | NM_00117222 | MOB2 | MOB kinase activator 2 | protein_coding | NP_001165694.1;NP_443731.2 | 24904 | 611969 | . | . | [illegible] |
| 81533 | NM_00130500 | ITFG1 | integrin alpha FG-GAP repeat containing 1 | protein_coding | NP_001291931.1;NP_110417.2 | 30697 | 611803 | . | . | [illegible] |
| 81537 | NM_030791,XI | SGPP1 | sphingosine-1-phosphate phosphatase 1 | protein_coding | NP_110418.1;XP_016877167.1 | 17720 | 612826 | . | . | [illegible] |
| 81542 | NM_030755 | TMX1 | thioredoxin related transmembrane protein 1 | protein_coding | NP_110382.3 | 15487 | 610527 | . | . | [illegible] |
| 81543 | NM_030891 | LRRC3 | leucine rich repeat containing 3 | protein_coding | NP_112153.1 | 14965 | 617620 | . | . | [illegible] |
| 81544 | NM_00135116 | GDPD5 | glycerophosphodiester phosphodiesterase domain containing 5 | protein_coding | NP_001338096.1;NP_001338097.1;NP_110419.5;XP_006718760.1;XP_01 | 28804 | 609632 | . | . | [illegible] |
| 81545 | NM_00127172 | FBXO38 | F-box protein 38 | protein_coding | NP_001258652.1;NP_110420.3;NP_995308.1;XP_006714860.1;XP_01153 | 28844 | 608533 | . | . | [illegible] |
| 81550 | NM_00114607 | TDRD3 | tudor domain containing 3 | protein_coding | NP_001139542.1;NP_001139543.1;NP_110421.1;XP_005266613.1;XP_00 | 20612 | 614392 | . | . | [illegible] |
| 81552 | NM_00128428 | VOPP1 | VOPP1 WW domain binding protein | protein_coding | NP_001271211.1;NP_001271212.1;NP_001271213.1;NP_001308171.1;NI | 34518 | 611915 | . | . | [illegible] |
| 81554 | NM_00128144 | RCC1L | RCC1 like | protein_coding | NP_001268370.1;NP_001350376.1;NP_110425.2;NP_683682.1 | 14948 | . | . | . | [illegible] |
| 81555 | NM_00102494 | YIPF5 | Yip1 domain family member 5 | protein_coding | NP_001020118.1;NP_001258661.1;NP_110426.4 | 24877 | 611483 | . | . | [illegible] |
| 81556 | NM_00113604 | INTS14 | integrator complex subunit 14 | protein_coding | NP_001129515.1;NP_001193987.2;NP_001193988.2;NP_001353289.1;NI | 25372 | . | . | . | [illegible] |
| 81557 | NM_00124236 | MAGED4B | MAGE family member D4B | protein_coding | NP_001229291.1;NP_110428.2;NP_803879.1;NP_803881.1 | 22880 | 300765 | . | . | [illegible] |
| 81558 | NM_030802,XI | FAM117A | family with sequence similarity 117 member A | protein_coding | NP_110429.1;XP_016880671.1 | 24179 | . | . | . | [illegible] |
| 81559 | NM_145214,XI | TRIM11 | tripartite motif containing 11 | protein_coding | NP_660215.1;XP_011542587.1;XP_016857901.1 | 16281 | 607868 | . | . | [illegible] |
| 81562 | NM_00114229 | LMAN2L | lectin, mannose binding 2 like | protein_coding | NP_001135764.1;NP_001309275.1;NP_001309276.1;NP_001309279.1;NI | 19263 | 609552 | . | . | [illegible] |
| 81563 | NM_030806 | C1orf21 | chromosome 1 open reading frame 21 | protein_coding | NP_110433.1 | 15494 | . | . | . | [illegible] |
| 81565 | NM_00102557 | NDEL1 | nudE neurodevelopment protein 1 like 1 | protein_coding | NP_001020750.1;NP_001317058.1;NP_110435.1;XP_016880672.1;XP_01 | 17620 | 607538 | . | . | [illegible] |
| 81566 | NM_030809,N | CSRNP2 | cysteine and serine rich nuclear protein 2 | protein_coding | NP_110436.1;XP_006719684.1;XP_016875480.1;XP_024304978.1;XP_02 | 16006 | . | . | . | [illegible] |
| 81567 | NM_00114554 | TXNDC5 | thioredoxin domain containing 5 | protein_coding | NP_001139021.1;NP_110437.2 | 21073 | 616412 | . | . | [illegible] |
| 81569 | NM_030812,XI | ACTL8 | actin like 8 | protein_coding | NP_110439.2;XP_011540514.1 | 24018 | . | . | . | [illegible] |
| 81570 | NM_00125839 | CLPB | caseinolytic mitochondrial matrix peptidase chaperone subunit B | protein_coding | NP_001245321.1;NP_001245322.1;NP_001245323.1;NP_110440.1;XP_00 | 30664 | 616254 | . | . | [illegible] |
| 81571 | NR_026677 | MIR600HG | MIR600 host gene | lncRNA | . | 23642 | . | . | . | [illegible] |
| 81572 | NM_030815 | PDRG1 | p53 and DNA damage regulated 1 | protein_coding | NP_110442.1 | 16119 | 610789 | . | . | [illegible] |
| 81573 | NM_030816,XI | ANKRD13C | ankyrin repeat domain 13C | protein_coding | NP_110443.3;XP_005271291.1;XP_005271292.1;XP_006710992.1;XP_01 | 25374 | 615125 | . | . | [illegible] |
| 81575 | NM_00113041 | APOLD1 | apolipoprotein L domain containing 1 | protein_coding | NP_001123887.1;NP_110444.3 | 25268 | 612456 | . | . | [illegible] |
| 81576 | NM_00132056 | CCDC130 | coiled-coil domain containing 130 | protein_coding | NP_001307490.1;NP_001307493.1;NP_001307494.1;NP_001307495.1;NI | 28118 | . | . | . | [illegible] |
| 81577 | NM_00124365 | GFOD2 | glucose-fructose oxidoreductase domain containing 2 | protein_coding | NP_001230579.1;NP_110446.3;XP_006721351.1 | 28159 | . | . | . | [illegible] |
| 81579 | NM_030821 | PLA2G12A | phospholipase A2 group XIIA | protein_coding | NP_110448.2 | 18554 | 611652 | . | . | [illegible] |
| 81602 | NM_00119347 | CDADC1 | cytidine and dCMP deaminase domain containing 1 | protein_coding | NP_001180407.1;NP_112173.1;XP_006719934.1;XP_011533552.1;XP_01 | 20299 | . | . | . | [illegible] |
| 81603 | NM_00134595 | TRIM8 | tripartite motif containing 8 | protein_coding | NP_001332879.1;NP_112174.2 | 15579 | 606125 | . | . | [illegible] |
| 81605 | NM_00113594 | URM1 | ubiquitin related modifier 1 | protein_coding | NP_001129419.1;NP_001252511.1;NP_112176.1 | 28378 | 612693 | . | . | [illegible] |
| 81606 | NM_030915,XI | LBH | LBH regulator of WNT signaling pathway | protein_coding | NP_112177.2;XP_016860548.1 | 29532 | 611763 | . | . | [illegible] |
| 81608 | NM_00113493 | FIP1L1 | factor interacting with PAPOLA and CPSF1 | protein_coding | NP_001128409.1;NP_001128410.1;NP_001363673.1;NP_001363674.1;NI | 19124 | 607686 | . | . | [illegible] |
| 81609 | NM_00133072 | SNX27 | sorting nexin 27 | protein_coding | NP_001317652.1;NP_112180.4;XP_005245567.1;XP_005245568.1;XP_01 | 20073 | 611541 | . | . | [illegible] |
| 81610 | NM_030919,XI | FAM83D | family with sequence similarity 83 member D | protein_coding | NP_112181.3;XP_016883577.1 | 16122 | 618380 | . | . | [illegible] |
| 81611 | NM_00113647 | ANP32E | acidic nuclear phosphoprotein 32 family member E | protein_coding | NP_001129950.1;NP_001129951.1;NP_001267488.1;NP_001267489.1;NI | 16673 | 609611 | . | . | [illegible] |
| 81614 | NM_00100886 | NIPA2 | NIPA magnesium transporter 2 | protein_coding | NP_001008860.1;NP_001008892.1;NP_001008894.1;NP_001171817.1;NI | 17044 | 608146 | . | . | [illegible] |
| 81615 | NM_030923,XI | TMEM163 | transmembrane protein 163 | protein_coding | NP_112185.1;XP_011510252.1;XP_016860549.1;XP_016860550.1;XP_01 | 25380 | . | . | . | [illegible] |
| 81617 | NM_00107967 | CAB39L | calcium binding protein 39 like | protein_coding | NP_001073138.1;NP_001274266.1;NP_001274267.1;NP_001274268.1;NI | 20290 | 612175 | . | . | [illegible] |
| 81618 | NM_00101251 | ITM2C | integral membrane protein 2C | protein_coding | NP_001012532.1;NP_001012534.1;NP_001274169.1;NP_001274170.1;NI | 6175 | 609554 | . | . | [illegible] |
| 81619 | NM_00112830 | TSPAN14 | tetraspanin 14 | protein_coding | NP_001121781.1;NP_001338195.1;NP_001338196.1;NP_001338197.1;NI | 23303 | . | . | . | [illegible] |
| 81620 | NM_030928 | CDT1 | chromatin licensing and DNA replication factor 1 | protein_coding | NP_112190.2 | 24576 | 605525 | . | . | [illegible] |
| 81621 | NM_00131930 | KAZALD1 | Kazal type serine peptidase inhibitor domain 1 | protein_coding | NP_001306232.1;NP_112191.2;XP_005270251.1;XP_016872204.1;XP_02 | 25460 | 609208 | . | . | [illegible] |
| 81622 | NM_030930,XI | UNC93B1 | unc-93 homolog B1, TLR signaling regulator | protein_coding | NP_112192.2;XP_011543592.1;XP_011543593.1 | 13481 | 608204 | . | . | [illegible] |
| 81624 | NM_00104251 | DIAPH3 | diaphanous related formin 3 | protein_coding | NP_001035982.1;NP_001245295.1;NP_001245296.1;NP_001245297.1;NI | 15480 | 614567 | . | . | [illegible] |
| 81627 | NM_00120242 | TRMT1L | tRNA methyltransferase 1 like | protein_coding | NP_001189352.1;NP_112196.3;XP_016857908.1 | 16782 | 611673 | . | . | [illegible] |
| 81628 | NM_00130304 | TSC22D4 | TSC22 domain family member 4 | protein_coding | NP_001289972.1;NP_112197.1 | 21696 | 611914 | . | . | [illegible] |
| 81631 | NM_022818 | MAP1LC3B | microtubule associated protein 1 light chain 3 beta | protein_coding | NP_073729.1 | 13352 | 609604 | . | . | [illegible] |
| 81669 | NM_00103957 | CCNL2 | cyclin L2 | protein_coding | NP_001034666.1;NP_001307082.1;NP_001307084.1;NP_001337426.1;NI | 20570 | 613482 | . | . | [illegible] |
| 81671 | NM_00132939 | VMP1 | vacuole membrane protein 1 | protein_coding | NP_001316323.1;NP_001316324.1;NP_001316325.1;NP_001316326.1;NI | 29559 | 611753 | . | . | [illegible] |
| 81688 | NM_030939,XI | C6orf62 | chromosome 6 open reading frame 62 | protein_coding | NP_112201.1;XP_005249490.1;XP_011513230.1 | 20998 | . | . | . | [illegible] |
| 81689 | NM_030940 | ISCA1 | iron-sulfur cluster assembly 1 | protein_coding | NP_112202.2 | 28660 | 611006 | . | . | [illegible] |
| 81691 | NM_00114492 | REXO5 | RNA exonuclease 5 | protein_coding | NP_001138396.1;NP_001185982.1;NP_112203.2;XP_005255661.1;XP_01 | 24661 | . | . | . | [illegible] |
| 81704 | NM_00119045 | DOCK8 | dedicator of cytokinesis 8 | protein_coding | NP_001177387.1;NP_001180465.1;NP_982272.2;XP_011516347.1;XP_01 | 19191 | 611432 | . | . | [illegible] |
| 81706 | NM_030949,XI | PPP1R14C | protein phosphatase 1 regulatory inhibitor subunit 14C | protein_coding | NP_112211.1;XP_005267209.1 | 14952 | 613242 | . | . | [illegible] |
| 81786 | NM_033342,N | TRIM7 | tripartite motif containing 7 | protein_coding | NP_203128.1;NP_976038.1;NP_976039.1;NP_976040.1;NP_976041.1;NP_ | 16278 | 609315 | . | . | [illegible] |
| 81788 | NM_030952,XI | NUAK2 | NUAK family kinase 2 | protein_coding | NP_112214.3;XP_005245572.1 | 29558 | 608131 | . | . | [illegible] |
| 81789 | NM_00124325 | TIGD6 | tigger transposable element derived 6 | protein_coding | NP_001230182.1;NP_112215.1 | 18332 | . | . | . | [illegible] |
| 81790 | NM_00116022 | RNF170 | ring finger protein 170 | protein_coding | NP_001153695.1;NP_001153696.1;NP_001153697.1;NP_112216.3;XP_00 | 25358 | 614649 | . | . | [illegible] |
| 81796 | NM_00114600 | SLCO5A1 | solute carrier organic anion transporter family member 5A1 | protein_coding | NP_001139480.1;NP_001139481.1;NP_112220.2;XP_005251370.1;XP_01 | 19046 | 613543 | . | . | [illegible] |
| 81831 | NM_00120147 | NETO2 | neuropilin and tolloid like 2 | protein_coding | NP_001188406.1;NP_060562.3;XP_006721352.1;XP_006721354.1;XP_00 | 14644 | 607974 | . | . | [illegible] |
| 81839 | NM_00117241 | VANGL1 | VANGL planar cell polarity protein 1 | protein_coding | NP_001165882.1;NP_001165883.1;NP_620409.1 | 15512 | 610132 | . | . | [illegible] |
| 81844 | NM_030961,XI | TRIM56 | tripartite motif containing 56 | protein_coding | NP_112223.1;XP_011514891.1 | 19028 | 616996 | . | . | [illegible] |
| 81846 | NM_030962,XI | SBF2 | SET binding factor 2 | protein_coding | NP_112224.1;XP_005253211.1;XP_005253212.1;XP_011518696.1;XP_01 | 2135 | 607697 | . | . | [illegible] |
| 81847 | NM_00124284 | RNF146 | ring finger protein 146 | protein_coding | NP_001229773.1;NP_001229774.1;NP_001229775.1;NP_001229776.1;NI | 21336 | 612137 | . | . | [illegible] |
| 81848 | NM_00112749 | SPRY4 | sprouty RTK signaling antagonist 4 | protein_coding | NP_001120968.1;NP_001280218.1;NP_001280219.1;NP_112226.2;XP_01 | 15533 | 607984 | . | . | [illegible] |
| 81853 | NM_00112771 | TMEM14B | transmembrane protein 14B | protein_coding | NP_001121183.1;NP_001273413.1;NP_001273417.1;NP_001273418.1;NI | 21384 | . | . | . | [illegible] |
| 81855 | NM_030971,N | SFXN3 | sideroflexin 3 | protein_coding | NP_112233.2 | 16087 | 615571 | . | . | [illegible] |
| 81856 | NM_00116149 | ZNF611 | zinc finger protein 611 | protein_coding | NP_001154971.1;NP_001154972.1;NP_001154973.1;NP_112234.3 | 28766 | . | . | . | [illegible] |
| 81857 | NM_00137835 | MED25 | mediator complex subunit 25 | protein_coding | NP_001365284.1;NP_112235.2 | 28845 | 610197 | . | . | [illegible] |
| 81858 | NM_030974,N | SHARPIN | SHANK associated RH domain interactor | protein_coding | NP_112236.3;XP_016869376.1;XP_016869377.1 | 25321 | 611885 | . | . | [illegible] |
| 81873 | NM_030978,XI | ARPC5L | actin related protein 2/3 complex subunit 5 like | protein_coding | NP_112240.1;XP_011517369.1 | 23366 | . | . | . | [illegible] |
| 81875 | NM_00130309 | ISG20L2 | interferon stimulated exonuclease gene 20 like 2 | protein_coding | NP_001290024.1;NP_001357079.1;NP_001357080.1;NP_001357081.1;XF | 25745 | 611930 | . | . | [illegible] |
| 81876 | NM_030981,XI | RAB1B | RAB1B, member RAS oncogene family | protein_coding | NP_112243.1;XP_016873867.1 | 18370 | 612565 | . | . | [illegible] |
| 81887 | NM_00117064 | LAS1L | LAS1 like ribosome biogenesis factor | protein_coding | NP_001164120.1;NP_001164121.1;NP_001362257.1;NP_001362258.1;NI | 25726 | 300964 | . | . | [illegible] |
| 81888 | NM_00119088 | HYI | hydroxypyruvate isomerase (putative) | protein_coding | NP_001177809.1;NP_001230455.1;NP_001317455.1;NP_112484.3;XP_00 | 26948 | . | . | . | [illegible] |
| 81889 | NM_00101810 | FAHD1 | fumarylacetoacetate hydrolase domain containing 1 | protein_coding | NP_001018114.1;NP_001135870.1;NP_112485.1 | 14169 | 616320 | . | . | [illegible] |
| 81890 | NM_031209 | QTRT1 | queuine tRNA-ribosyltransferase catalytic subunit 1 | protein_coding | NP_112486.1 | 23797 | 609615 | . | . | [illegible] |
| 81892 | NM_00126786 | SLIRP | SRA stem-loop interacting RNA binding protein | protein_coding | NP_001254792.1;NP_001254793.1;NP_112487.1 | 20495 | 610211 | . | . | [illegible] |
| 81893 | gene-SLC7A5P1 | SLC7A5P1 | solute carrier family 7 member 5 pseudogene 1 | pseudogene | . | 29458 | . | . | . | [illegible] |
| 81894 | NM_031212,XI | SLC25A28 | solute carrier family 25 member 28 | protein_coding | NP_112489.3;XP_005270265.1;XP_005270268.1;XP_006718068.1;XP_00 | 23472 | 609767 | . | . | [illegible] |
| 81926 | NM_00113011 | ABHD17A | abhydrolase domain containing 17A, depalmitoylase | protein_coding | NP_001123583.1;NP_112490.3;XP_011526640.1 | 28756 | 617942 | . | . | [illegible] |
| 81928 | NM_031215 | CABLES2 | Cdk5 and Abl enzyme substrate 2 | protein_coding | NP_112492.2 | 16143 | 618772 | . | . | [illegible] |
| 81929 | NM_00101343 | SEH1L | SEH1 like nucleoporin | protein_coding | NP_001013455.1;NP_112493.2;XP_005258209.1;XP_011524044.1;XP_01 | 30379 | 609263 | . | . | [illegible] |
| 81930 | NM_031217,XI | KIF18A | kinesin family member 18A | protein_coding | NP_112494.3;XP_016873868.1;XP_016873869.1 | 29441 | 611271 | . | . | [illegible] |
| 81931 | NM_031218 | ZNF93 | zinc finger protein 93 | protein_coding | NP_112495.2 | 13169 | 603975 | . | . | [illegible] |
| 81932 | NM_00130450 | HDHD3 | haloacid dehalogenase like hydrolase domain containing 3 | protein_coding | NP_001291438.1;NP_001291439.1;NP_001291440.1;NP_001358852.1;NI | 28171 | . | . | . | [illegible] |
| 83394 | NM_00116596 | PITPNM3 | PITPNM family member 3 | protein_coding | NP_001159438.1;NP_112497.2;XP_011522317.1;XP_011522318.1;XP_01 | 21043 | 608921 | . | . | [illegible] |
| 83401 | NM_152310,XI | ELOVL3 | ELOVL fatty acid elongase 3 | protein_coding | NP_689523.1;XP_011538547.1 | 18047 | 611815 | . | . | [illegible] |
| 83439 | NM_031283,XI | TCF7L1 | transcription factor 7 like 1 | protein_coding | NP_112573.1;XP_006712172.1 | 11640 | 604652 | . | . | [illegible] |
| 83440 | NM_00136522 | ADPGK | ADP dependent glucokinase | protein_coding | NP_001352152.1;NP_001352153.1;NP_001352154.1;NP_001352155.1;NI | 25250 | 611861 | . | . | [illegible] |
| 83442 | NM_031286 | SH3BGRL3 | SH3 domain binding glutamate rich protein like 3 | protein_coding | NP_112576.1 | 15568 | 615679 | . | . | [illegible] |
| 83443 | NM_031287 | SF3B5 | splicing factor 3b subunit 5 | protein_coding | NP_112577.1 | 21083 | 617847 | . | . | [illegible] |
| 83444 | NM_031288 | INO80B | INO80 complex subunit B | protein_coding | NP_112578.2 | 13324 | 616456 | . | . | [illegible] |
| 83448 | NM_00109861 | PUS7L | pseudouridine synthase 7 like | protein_coding | NP_001092084.1;NP_001092085.1;NP_001258755.1;NP_112582.3;XP_00 | 25276 | . | . | . | [illegible] |
| 83450 | NM_00113009 | DRC3 | dynein regulatory complex subunit 3 | protein_coding | NP_001123562.1;NP_001123563.1;NP_001123564.1;NP_112584.3;XP_01 | 25384 | 618758 | . | . | [illegible] |
| 83451 | NM_00114536 | ABHD11 | abhydrolase domain containing 11 | protein_coding | NP_001138836.1;NP_001287987.1;NP_001308311.1;NP_001308312.1;NI | 16407 | . | . | . | [illegible] |
| 83452 | NM_031296,XI | RAB33B | RAB33B, member RAS oncogene family | protein_coding | NP_112586.1;XP_011530601.1 | 16075 | 605950 | . | . | [illegible] |
| 83460 | NM_00101476 | EMC6 | ER membrane protein complex subunit 6 | protein_coding | NP_001014764.1;NP_112588.1 | 28430 | . | . | . | [illegible] |
| 83461 | NM_00129760 | CDCA3 | cell division cycle associated 3 | protein_coding | NP_001284531.1;NP_001284532.1;NP_001284533.1;NP_001317948.1;NI | 14624 | 607749 | . | . | [illegible] |
| 83463 | NM_00114293 | MXD3 | MAX dimerization protein 3 | protein_coding | NP_001136407.1;NP_112590.1 | 14008 | 609450 | . | . | [illegible] |
| 83464 | NM_00114564 | APH1B | aph-1 homolog B, gamma-secretase subunit | protein_coding | NP_001139118.1;NP_112591.2;XP_011520407.1;XP_024305853.1;XP_02 | 24080 | 607630 | . | . | [illegible] |
| 83473 | NM_00135389 | KATNAL2 | katanin catalytic subunit A1 like 2 | protein_coding | NP_001340828.1;NP_001340829.1;NP_001340830.1;NP_001340831.1;NI | 25387 | 614697 | . | . | [illegible] |
| 83475 | NM_00114516 | DOHH | deoxyhypusine hydroxylase | protein_coding | NP_001138637.1;NP_112594.1;XP_011526643.1 | 28662 | 611262 | . | . | [illegible] |
| 83478 | NM_00102561 | ARHGAP24 | Rho GTPase activating protein 24 | protein_coding | NP_001020787.2;NP_001036134.1;NP_001274734.1;NP_001333022.1;NI | 25361 | 610586 | . | . | [illegible] |
| 83479 | NM_00103172 | DDX59 | DEAD-box helicase 59 | protein_coding | NP_001026895.2;NP_001307110.1;NP_001307111.1;NP_001336728.1;NI | 25360 | 615464 | . | . | [illegible] |
| 83480 | NM_00127198 | PUS3 | pseudouridine synthase 3 | protein_coding | NP_001258914.1;NP_112597.4;XP_005271745.1 | 25461 | 616283 | . | . | [illegible] |
| 83481 | NM_031308,XI | EPPK1 | epiplakin 1 | protein_coding | NP_112598.3;XP_016869379.1;XP_016869381.1 | 15577 | 607553 | . | . | [illegible] |
| 83530 | gene-CYP51A1 | CYP51A1P2 | cytochrome P450 family 51 subfamily A member 1 pseudogene 2 | pseudogene | . | 20251 | . | . | . | [illegible] |
| 83538 | NM_00135031 | TTC25 | tetratricopeptide repeat domain 25 | protein_coding | NP_001337248.1;NP_113609.1 | 25280 | 617095 | . | . | [illegible] |
| 83540 | NM_031423,N | NUF2 | NUF2 component of NDC80 kinetochore complex | protein_coding | NP_113611.2;NP_663735.2;XP_011508338.1;XP_024305880.1;XP_02430 | 14621 | 611772 | . | . | [illegible] |
| 83541 | NM_00104235 | FAM110A | family with sequence similarity 110 member A | protein_coding | NP_001035812.1;NP_001276074.1;NP_001276075.1;NP_001276076.1;NI | 16188 | 611393 | . | . | [illegible] |
| 83543 | NM_00118509 | AIF1L | allograft inflammatory factor 1 like | protein_coding | NP_001172024.1;NP_001172025.1;NP_113614.1;XP_016870666.1 | 28904 | . | . | . | [illegible] |
| 83544 | NM_00120136 | DNAL1 | dynein axonemal light chain 1 | protein_coding | NP_001188295.1;NP_113615.2;XP_016877168.1;XP_024305483.1 | 23247 | 610062 | . | . | [illegible] |
| 83547 | NM_031430,XI | RILP | Rab interacting lysosomal protein | protein_coding | NP_113618.2;XP_005256868.1;XP_016880684.1 | 30266 | 607848 | . | . | [illegible] |
| 83548 | NM_031431,XI | COG3 | component of oligomeric golgi complex 3 | protein_coding | NP_113619.3;XP_011533568.1;XP_011533569.1 | 18619 | 606975 | . | . | [illegible] |
| 83549 | NM_00113595 | UCK1 | uridine-cytidine kinase 1 | protein_coding | NP_001129426.1;NP_001248379.1;NP_001248380.1;NP_001305448.1;NI | 14859 | 609328 | . | . | [illegible] |
| 83552 | NM_031433 | MFRP | membrane frizzled-related protein | protein_coding | NP_113621.1 | 18121 | 606227 | . | . | [illegible] |
| 83590 | NM_00113604 | TMUB1 | transmembrane and ubiquitin like domain containing 1 | protein_coding | NP_001129516.1;NP_113622.1 | 21709 | 614792 | . | . | [illegible] |
| 83591 | NM_031435 | THAP2 | THAP domain containing 2 | protein_coding | NP_113623.1 | 20854 | 612531 | . | . | [illegible] |
| 83592 | NM_00104017 | AKR1E2 | aldo-keto reductase family 1 member E2 | protein_coding | NP_001035267.1;NP_001257950.1;NP_001257954.1;XP_011518017.1;XF | 23437 | 617451 | . | . | [illegible] |
| 83593 | NM_182663,N | RASSF5 | Ras association domain family member 5 | protein_coding | NP_872604.1;NP_872605.1;NP_872606.1 | 17609 | 607020 | . | . | [illegible] |
| 83594 | NM_00130074 | NUDT12 | nudix hydrolase 12 | protein_coding | NP_001287670.1;NP_113626.1;XP_005272152.1;XP_005272154.1 | 18826 | 609232 | . | . | [illegible] |
| 83596 | NM_00104066 | BCL2L12 | BCL2 like 12 | protein_coding | NP_001035758.1;NP_001269445.1;NP_001269446.1;NP_001269448.1;NI | 13787 | 610837 | . | . | [illegible] |
| 83604 | NM_031442 | TMEM47 | transmembrane protein 47 | protein_coding | NP_113630.1 | 18515 | 300698 | . | . | [illegible] |
| 83605 | NM_00102983 | CCM2 | CCM2 scaffold protein | protein_coding | NP_001025006.1;NP_001161406.1;NP_001161407.1;NP_001350387.1;NI | 21708 | 607929 | . | . | [illegible] |
| 83606 | NM_00128425 | GUCD1 | guanylyl cyclase domain containing 1 | protein_coding | NP_001271180.1;NP_001271181.1;NP_001271182.1;NP_001271183.1;NI | 14237 | . | . | . | [illegible] |
| 83607 | NM_00119914 | AMMECR1L | AMMECR1 like | protein_coding | NP_001186069.1;NP_113633.2;XP_005263861.1;XP_005263863.1;XP_00 | 28658 | . | . | . | [illegible] |
| 83608 | NM_00120147 | C18orf21 | chromosome 18 open reading frame 21 | protein_coding | NP_001188403.1;NP_001188404.1;NP_001188405.1;NP_113634.3;XP_00 | 28802 | . | . | . | [illegible] |
| 83636 | NM_00103172 | C19orf12 | chromosome 19 open reading frame 12 | protein_coding | NP_001026896.2;NP_001242975.1;NP_001242976.1;NP_001269858.1;NI | 25443 | 614297 | . | . | [illegible] |
| 83637 | NM_00130095 | ZMIZ2 | zinc finger MIZ-type containing 2 | protein_coding | NP_001287888.1;NP_113637.3;NP_777589.2;XP_005249923.1;XP_00524 | 22229 | 611196 | . | . | [illegible] |
| 83638 | NM_00113563 | C11orf68 | chromosome 11 open reading frame 68 | protein_coding | NP_001129107.1;NP_113638.2 | 28801 | . | . | . | [illegible] |
| 83640 | NM_031452 | RAMAC | RNA guanine-7 methyltransferase activating subunit | protein_coding | NP_113640.1 | 31022 | 614547 | . | . | [illegible] |
| 83641 | NM_00128269 | FAM107B | family with sequence similarity 107 member B | protein_coding | NP_001269624.1;NP_001269625.1;NP_001269626.1;NP_001269627.1;NI | 23726 | . | . | . | [illegible] |
| 83642 | NM_031454 | SELENOO | selenoprotein O | protein_coding | NP_113642.1 | 30395 | 607917 | . | . | [illegible] |
| 83643 | NM_00128265 | CCDC3 | coiled-coil domain containing 3 | protein_coding | NP_001269587.1;NP_113643.1 | 23813 | . | . | . | [illegible] |
| 83658 | NM_00131915 | DYNLRB1 | dynein light chain roadblock-type 1 | protein_coding | NP_001306086.1;NP_001369294.1;NP_001369295.1;NP_001369296.1;NI | 15468 | 607167 | . | . | [illegible] |
| 83660 | NM_015059,XI | TLN2 | talin 2 | protein_coding | NP_055874.2;XP_005254765.1;XP_005254767.1;XP_005254768.1;XP_00 | 15447 | 607349 | . | . | [illegible] |
| 83666 | NM_00114610 | PARP9 | poly(ADP-ribose) polymerase family member 9 | protein_coding | NP_001139574.1;NP_001139575.1;NP_001139576.1;NP_001139577.1;NI | 24118 | 612065 | . | . | [illegible] |
| 83667 | NM_031459,XI | SESN2 | sestrin 2 | protein_coding | NP_113647.1 | 20746 | 607767 | . | . | [illegible] |

| | | | | | | | | |
|---|---|---|---|---|---|---|---|---|
| 83690 | NM_00128677 | CRISPLD1 | cysteine rich secretory protein LCCL domain containing 1 | protein_coding | NP_001273706.1;NP_001273707.1;NP_113649.1 | 18206 | . | [illegible] |
| 83692 | NM_00118480 | CD99L2 | CD99 molecule like 2 | protein_coding | NP_001171737.1;NP_001229543.1;NP_113650.2;NP_604394.1;NP_6043 | 18237 | 300846 | [illegible] |
| 83693 | NM_00114605 | HSDL1 | hydroxysteroid dehydrogenase like 1 | protein_coding | NP_001139523.1;NP_113651.4;XP_005256246.1 | 16475 | . | [illegible] |
| 83694 | NM_00137025 | RPS6KL1 | ribosomal protein S6 kinase like 1 | protein_coding | NP_001357181.1;NP_001357182.1;NP_001357184.1;NP_001357185.1;NI | 20222 | . | [illegible] |
| 83695 | NM_00125249 | RHNO1 | RAD9-HUS1-RAD1 interacting nuclear orphan 1 | protein_coding | NP_001239428.1;NP_001239429.1;NP_001244026.1;NP_001244027.1 | 28206 | 614085 | [illegible] |
| 83696 | NM_00116037 | TRAPPC9 | trafficking protein particle complex 9 | protein_coding | NP_001153844.1;NP_001308575.1;NP_001361611.1;NP_001361612.1;NI | 30832 | 611966 | [illegible] |
| 83699 | NM_031469,XI | SH3BGRL2 | SH3 domain binding glutamate rich protein like 2 | protein_coding | NP_113657.1;XP_011534467.1 | 15567 | 615678 | [illegible] |
| 83700 | NM_00120532 | JAM3 | junctional adhesion molecule 3 | protein_coding | NP_001192258.1;NP_116190.3 | 15532 | 606871 | [illegible] |
| 83706 | NM_00138236 | FERMT3 | fermitin family member 3 | protein_coding | NP_001369290.1;NP_001369291.1;NP_001369292.1;NP_001369293.1;NI | 23151 | 607901 | [illegible] |
| 83707 | NM_00103367 | TRPT1 | tRNA phosphotransferase 1 | protein_coding | NP_001028850.2;NP_001153861.1;NP_001153862.1;NP_001153864.1;NI | 20316 | 610470 | [illegible] |
| 83716 | NM_031476,XI | CRISPLD2 | cysteine rich secretory protein LCCL domain containing 2 | protein_coding | NP_113664.1;XP_005256247.1;XP_024306231.1 | 25248 | 612434 | [illegible] |
| 83719 | NM_00114552 | YPEL3 | yippee like 3 | protein_coding | NP_001138996.1;NP_113665.3 | 18327 | 609724 | [illegible] |
| 83729 | NM_031479 | INHBE | inhibin subunit beta E | protein_coding | NP_113667.1 | 24029 | 612031 | [illegible] |
| 83732 | NM_00134819 | RIOK1 | RIO kinase 1 | protein_coding | NP_001335123.1;NP_113668.2;XP_011513235.2 | 18656 | 617753 | [illegible] |
| 83734 | NM_00113102 | ATG10 | autophagy related 10 | protein_coding | NP_001124500.1;NP_113670.1;XP_005248667.1;XP_005248668.1;XP_00 | 20315 | 610800 | [illegible] |
| 83737 | NM_00125713 | ITCH | itchy E3 ubiquitin protein ligase | protein_coding | NP_001244066.1;NP_001244067.1;NP_001311126.1;NP_001311127.1;NI | 13890 | 606409 | [illegible] |
| 83742 | NM_031484 | MARVELD1 | MARVEL domain containing 1 | protein_coding | NP_113672.1 | 28674 | 616970 | [illegible] |
| 83743 | NM_031485 | GRWD1 | glutamate rich WD repeat containing 1 | protein_coding | NP_113673.3 | 21270 | 610597 | [illegible] |
| 83744 | NM_00100710 | ZNF484 | zinc finger protein 484 | protein_coding | NP_001007102.1;NP_001248387.1;NP_001248388.1;NP_001248389.1;NI | 23385 | . | [illegible] |
| 83746 | NM_031488,XI | L3MBTL2 | L3MBTL histone methyl-lysine binding protein 2 | protein_coding | NP_113676.2;XP_011528722.1;XP_011528723.1;XP_016884465.1;XP_01 | 18594 | 611865 | [illegible] |
| 83752 | NM_00130094 | LONP2 | lon peptidase 2, peroxisomal | protein_coding | NP_001287877.1;NP_001335007.1;NP_113678.2;XP_016879244.1;XP_01 | 20598 | 617774 | [illegible] |
| 83759 | NM_00128613 | RBM4B | RNA binding motif protein 4B | protein_coding | NP_001273064.1;NP_113680.1;XP_011543599.1;XP_016873891.1 | 28842 | . | [illegible] |
| 83786 | NM_00130083 | FRMD8 | FERM domain containing 8 | protein_coding | NP_001287761.1;NP_001287762.1;NP_114110.1 | 25462 | 618337 | [illegible] |
| 83787 | NM_00116100 | ARMC10 | armadillo repeat containing 10 | protein_coding | NP_001154481.1;NP_001154482.1;NP_001154483.1;NP_001154484.1;NI | 21706 | 611864 | [illegible] |
| 83852 | NM_00116030 | SETDB2 | SET domain bifurcated histone lysine methyltransferase 2 | protein_coding | NP_001153780.1;NP_001307628.1;NP_114121.2 | 20263 | 607865 | [illegible] |
| 83853 | NM_00120146 | ROPN1L | rhophilin associated tail protein 1 like | protein_coding | NP_001188395.1;NP_114122.2;XP_006714567.1;XP_016865435.1;XP_01 | 24060 | 611756 | [illegible] |
| 83854 | NM_00132141 | ANGPTL6 | angiopoietin like 6 | protein_coding | NP_001308340.1;NP_114123.2;XP_011526649.1;XP_011526650.1;XP_01 | 23140 | 609336 | [illegible] |
| 83855 | NM_031918 | KLF16 | Kruppel like factor 16 | protein_coding | NP_114124.1 | 16857 | 606139 | [illegible] |
| 83856 | NM_00114531 | FSD1L | fibronectin type III and SPRY domain containing 1 like | protein_coding | NP_001138785.1;NP_001274120.1;NP_001274121.1;NP_001317668.1;NI | 13753 | 609829 | [illegible] |
| 83857 | NM_00119345 | TMTC1 | transmembrane O-mannosyltransferase targeting cadherins 1 | protein_coding | NP_001180380.1;NP_001354804.1;NP_787057.2;XP_016875492.1;XP_01 | 24099 | 615855 | [illegible] |
| 83858 | NM_00131723 | ATAD3B | ATPase family AAA domain containing 3B | protein_coding | NP_001304167.1;NP_114127.3;XP_005244863.1;XP_011540543.1;XP_01 | 24007 | 612317 | [illegible] |
| 83860 | NM_031923,XI | TAF3 | TATA-box binding protein associated factor 3 | protein_coding | NP_114129.1;XP_011518043.2 | 17303 | 606576 | [illegible] |
| 83861 | NM_00134641 | RSPH3 | radial spoke head 3 | protein_coding | NP_001333347.1;NP_114130.4;XP_016866836.1;XP_024302334.1 | 21054 | 615876 | [illegible] |
| 83862 | NM_00131780 | TMEM120A | transmembrane protein 120A | protein_coding | NP_001304732.1;NP_001350391.1;NP_114131.1;XP_024302721.1 | 21697 | 616550 | [illegible] |
| 83871 | NM_00114262 | RAB34 | RAB34, member RAS oncogene family | protein_coding | NP_001136096.2;NP_001136097.2;NP_001138414.1;NP_001138415.1;NI | 16519 | 610917 | [illegible] |
| 83872 | NM_031935,XI | HMCN1 | hemicentin 1 | protein_coding | NP_114141.2;XP_011508340.1;XP_011508343.1;XP_016857926.1;XP_02 | 19194 | 608548 | [illegible] |
| 83874 | NM_00120424 | TBC1D10A | TBC1 domain family member 10A | protein_coding | NP_001191169.1;NP_114143.1 | 23609 | 610020 | [illegible] |
| 83875 | NM_00103729 | BCO2 | beta-carotene oxygenase 2 | protein_coding | NP_001032367.3;NP_001243326.2;NP_001243327.2;NP_001243329.2;NI | 18503 | 611740 | [illegible] |
| 83876 | NM_00112717 | MRO | maestro | protein_coding | NP_001120646.1;NP_001120647.1;NP_001120648.1;NP_001356437.1;NI | 24121 | 608080 | [illegible] |
| 83877 | NM_00102438 | TM2D2 | TM2 domain containing 2 | protein_coding | NP_001019551.1;NP_001019552.1;NP_114146.3;NP_510882.1;XP_0052 | 24127 | 610081 | [illegible] |
| 83879 | NM_031942,N | CDCA7 | cell division cycle associated 7 | protein_coding | NP_114148.3;NP_665809.1 | 14628 | 609937 | [illegible] |
| 83880 | gene-EIF3FP2 | EIF3FP2 | eukaryotic translation initiation factor 3 subunit F pseudogene 2 | pseudogene | . | 28863 | . | [illegible] |
| 83882 | NM_00129021 | TSPAN10 | tetraspanin 10 | protein_coding | NP_001277141.1;NP_114151.3 | 29942 | . | [illegible] |
| 83888 | NM_031950 | FGFBP2 | fibroblast growth factor binding protein 2 | protein_coding | NP_114156.1 | 29451 | 607713 | [illegible] |
| 83890 | NM_00134930 | SPATA9 | spermatogenesis associated 9 | protein_coding | NP_001336232.1;NP_114158.2;XP_011541966.1;XP_011541968.1;XP_01 | 22988 | 608039 | [illegible] |
| 83891 | NM_00131778 | SNX25 | sorting nexin 25 | protein_coding | NP_001304710.1;NP_001364961.1;NP_001364963.1;NP_001364964.1;NI | 21883 | . | [illegible] |
| 83892 | NM_00131739 | KCTD10 | potassium channel tetramerization domain containing 10 | protein_coding | NP_001304324.1;NP_001304328.1;NP_114160.1 | 23236 | 613421 | [illegible] |
| 83903 | NM_031965 | HASPIN | histone H3 associated protein kinase | protein_coding | NP_114171.2 | 19682 | 609240 | [illegible] |
| 83930 | NM_00136333 | STARD3NL | STARD3 N-terminal like | protein_coding | NP_001350268.1;NP_001350269.1;NP_001350272.1;NP_001350273.1;NI | 19169 | 611759 | [illegible] |
| 83931 | NM_00128254 | STK40 | serine/threonine kinase 40 | protein_coding | NP_001269475.1;NP_001269476.1;NP_114406.1 | 21373 | 609437 | [illegible] |
| 83932 | NM_00101098 | SPRTN | SprT-like N-terminal domain | protein_coding | NP_001010984.1;NP_001248391.1;NP_114407.3;XP_006711881.1;XP_01 | 25356 | 616086 | [illegible] |
| 83933 | NM_00115928 | HDAC10 | histone deacetylase 10 | protein_coding | NP_001152758.1;NP_114408.3 | 18128 | 608544 | [illegible] |
| 83937 | NM_032023,XI | RASSF4 | Ras association domain family member 4 | protein_coding | NP_114412.2;XP_005271890.1;XP_005271892.1;XP_006718076.1;XP_00 | 20793 | 610559 | [illegible] |
| 83938 | NM_00130558 | LRMDA | leucine rich melanocyte differentiation associated | protein_coding | NP_001292510.1;NP_114413.1 | 23405 | 614537 | [illegible] |
| 83939 | NM_00131904 | EIF2A | eukaryotic translation initiation factor 2A | protein_coding | NP_001305972.1;NP_001305973.1;NP_001305974.1;NP_001305975.1;NI | 3254 | 609234 | [illegible] |
| 83940 | NM_00114616 | TATDN1 | TatD DNase domain containing 1 | protein_coding | NP_001139632.1;NP_001304818.1;NP_001304819.1;NP_001304820.1;NI | 24220 | . | [illegible] |
| 83941 | NM_032027,N | TM2D1 | TM2 domain containing 1 | protein_coding | NP_114416.1 | 24142 | 610080 | [illegible] |
| 83943 | NM_00124460 | IMMP2L | inner mitochondrial membrane peptidase subunit 2 | protein_coding | NP_001231535.1;NP_001337888.1;NP_001337889.1;NP_001337890.1;NI | 14598 | 605977 | [illegible] |
| 83959 | NM_00117408 | SLC4A11 | solute carrier family 4 member 11 | protein_coding | NP_001167560.1;NP_001167561.1;NP_001350674.1;NP_114423.1;XP_00 | 16438 | 610206 | [illegible] |
| 83982 | NM_032036 | IFI27L2 | interferon alpha inducible protein 27 like 2 | protein_coding | NP_114425.1 | 19753 | 611319 | [illegible] |
| 83983 | NM_032037 | TSSK6 | testis specific serine kinase 6 | protein_coding | NP_114426.1 | 30410 | 610712 | [illegible] |
| 83985 | NM_00114244 | SPNS1 | sphingolipid transporter 1 (putative) | protein_coding | NP_001135920.1;NP_001135921.1;NP_001135922.1;NP_001135923.1;NI | 30621 | 612583 | [illegible] |
| 83986 | NM_00128449 | FAM234A | family with sequence similarity 234 member A | protein_coding | NP_001271426.1;NP_114428.1;XP_005255679.1;XP_005255680.1;XP_00 | 14163 | . | [illegible] |
| 83988 | NM_00104062 | NCALD | neurocalcin delta | protein_coding | NP_001035714.1;NP_001035715.1;NP_001035716.1;NP_001035717.1;NI | 7655 | 606722 | [illegible] |
| 83989 | NM_00116341 | FAM172A | family with sequence similarity 172 member A | protein_coding | NP_001156889.1;NP_001156890.1;NP_114431.2;XP_005272160.1;XP_00 | 25365 | . | [illegible] |
| 83990 | NM_032043,XI | BRIP1 | BRCA1 interacting protein C-terminal helicase 1 | protein_coding | NP_114432.2;XP_011523634.1;XP_011523635.1;XP_011523636.1;XP_01 | 20473 | 605882 | [illegible] |
| 83992 | NM_00136334 | CTTNBP2 | cortactin binding protein 2 | protein_coding | NP_001350278.1;NP_001350279.1;NP_001350280.1;NP_219499.1;XP_01 | 15679 | 609772 | [illegible] |
| 83999 | NM_00103957 | KREMEN1 | kringle containing transmembrane protein 1 | protein_coding | NP_001034659.2;NP_114434.3;XP_011528731.1;XP_011528732.1;XP_01 | 17550 | 609898 | [illegible] |
| 84002 | NM_032047,XI | B3GNT5 | UDP-GlcNAc:betaGal beta-1,3-N-acetylglucosaminyltransferase 5 | protein_coding | NP_114436.1;XP_005247880.1;XP_005247881.1;XP_005247882.1;XP_01 | 15684 | 615333 | [illegible] |
| 84033 | NM_00109862 | OBSCN | obscurin, cytoskeletal calmodulin and titin-interacting RhoGEF | protein_coding | NP_001092093.2;NP_001258152.2;NP_443075.3 | 15719 | 608616 | [illegible] |
| 84034 | NM_032048,XI | EMILIN2 | elastin microfibril interfacer 2 | protein_coding | NP_114437.2;XP_016881527.1 | 19881 | 608928 | [illegible] |
| 84056 | NM_00101438 | KATNAL1 | katanin catalytic subunit A1 like 1 | protein_coding | NP_001014402.1;NP_115492.1;XP_005266631.1;XP_016876280.1;XP_01 | 28361 | 614764 | [illegible] |
| 84057 | NM_00125386 | MND1 | meiotic nuclear divisions 1 | protein_coding | NP_001240790.1;NP_115493.1;XP_005263332.1;XP_016864177.1;XP_02 | 24839 | 611422 | [illegible] |
| 84058 | NM_00132082 | WDR54 | WD repeat domain 54 | protein_coding | NP_001307752.1;NP_001307753.1;NP_001307754.1;NP_115494.1;XP_01 | 25770 | . | [illegible] |
| 84060 | NM_00136336 | RBM48 | RNA binding motif protein 48 | protein_coding | NP_001350295.1;NP_001350296.1;NP_115496.2;XP_005250693.1;XP_00 | 21785 | . | [illegible] |
| 84061 | NM_00136791 | MAGT1 | magnesium transporter 1 | protein_coding | NP_001354845.1;NP_115497.4 | 28880 | 300715 | [illegible] |
| 84062 | NM_00127166 | DTNBP1 | dystrobrevin binding protein 1 | protein_coding | NP_001258596.1;NP_001258597.1;NP_001258598.1;NP_115498.2;NP_8 | 17328 | 607145 | [illegible] |
| 84063 | NM_00132953 | KIRREL2 | kirre like nephrin family adhesion molecule 2 | protein_coding | NP_001316459.1;NP_001350596.1;NP_115499.5;NP_954648.3;NP_9546 | 18816 | 607762 | [illegible] |
| 84064 | NM_00131876 | HDHD2 | haloacid dehalogenase like hydrolase domain containing 2 | protein_coding | NP_001305694.1;NP_115500.1;XP_005258428.1;XP_006722620.1;XP_01 | 25364 | . | [illegible] |
| 84065 | NM_032125,N | TMEM222 | transmembrane protein 222 | protein_coding | NP_115501.2 | 25363 | . | [illegible] |
| 84067 | NM_00109879 | FAM160A2 | family with sequence similarity 160 member A2 | protein_coding | NP_001092264.1;NP_115503.2;XP_006718406.1;XP_011518699.1;XP_01 | 25378 | . | [illegible] |
| 84068 | NM_00102999 | SLC10A7 | solute carrier family 10 member 7 | protein_coding | NP_001025169.1;NP_001287771.1;NP_001304745.1;NP_001304746.1;NI | 23088 | 611459 | [illegible] |
| 84069 | NM_00116018 | PLEKHN1 | pleckstrin homology domain containing N1 | protein_coding | NP_001153656.1;NP_001354481.1;NP_115505.2;XP_011540550.2;XP_01 | 25284 | . | [illegible] |
| 84070 | NM_032130,N | FAM186B | family with sequence similarity 186 member B | protein_coding | NP_115506.1;XP_006719688.1;XP_006719689.1;XP_006719690.1;XP_01 | 25296 | . | [illegible] |
| 84071 | NM_00128660 | ARMC2 | armadillo repeat containing 2 | protein_coding | NP_001273538.1;NP_115507.4;XP_005267211.1;XP_005267212.1;XP_00 | 23045 | 618424 | [illegible] |
| 84073 | NM_00136629 | MYCBPAP | MYCBP associated protein | protein_coding | NP_001353223.1;NP_115509.5;XP_005257783.1;XP_011523644.1;XP_01 | 19677 | 609835 | [illegible] |
| 84074 | NM_032134,N | QRICH2 | glutamine rich 2 | protein_coding | NP_115510.1;XP_005257785.1;XP_006722199.1;XP_011523646.1;XP_01 | 25326 | 618304 | [illegible] |
| 84078 | NM_032138 | KBTBD7 | kelch repeat and BTB domain containing 7 | protein_coding | NP_115514.2 | 25266 | 617739 | [illegible] |
| 84079 | NM_032139,XI | ANKRD27 | ankyrin repeat domain 27 | protein_coding | NP_115515.2;XP_006723475.1;XP_016882838.1 | 25310 | . | [illegible] |
| 84080 | NM_032140,N | ENKD1 | enkurin domain containing 1 | protein_coding | NP_115516.1;XP_024306237.1;XP_024306238.1 | 25246 | . | [illegible] |
| 84081 | NM_00126146 | NSRP1 | nuclear speckle splicing regulatory protein 1 | protein_coding | NP_001248396.1;NP_115517.1;XP_011523647.1 | 25305 | 616173 | [illegible] |
| 84083 | NM_00128656 | ZRANB3 | zinc finger RANBP2-type containing 3 | protein_coding | NP_001273497.1;NP_001273498.1;NP_115519.2;XP_005263866.1;XP_00 | 25249 | 615655 | [illegible] |
| 84084 | NM_032144 | RAB6C | RAB6C, member RAS oncogene family | protein_coding | NP_115520.2 | 16525 | 612909 | [illegible] |
| 84085 | NM_00134809 | FBXO30 | F-box protein 30 | protein_coding | NP_001335021.1;NP_115521.3;XP_011534479.1;XP_016866842.1 | 15600 | 609101 | [illegible] |
| 84100 | NM_00127829 | ARL6 | ADP ribosylation factor like GTPase 6 | protein_coding | NP_001265222.1;NP_001310442.1;NP_001310443.1;NP_115522.1;NP_8 | 13210 | 608845 | [illegible] |
| 84102 | NM_00135216 | SLC41A2 | solute carrier family 41 member 2 | protein_coding | NP_001339098.1;NP_001339099.1;NP_001339100.1;NP_001339101.1;NI | 31045 | 610802 | [illegible] |
| 84105 | NM_032151 | PCBD2 | pterin-4 alpha-carbinolamine dehydratase 2 | protein_coding | NP_115527.3 | 24474 | 609836 | [illegible] |
| 84107 | NM_00116837 | ZIC4 | Zic family member 4 | protein_coding | NP_001161850.1;NP_001161851.1;NP_001230185.1;NP_115529.2;XP_01 | 20393 | 608948 | [illegible] |
| 84108 | NM_00101166 | PCGF6 | polycomb group ring finger 6 | protein_coding | NP_001011663.1;NP_115530.2 | 21156 | 607816 | [illegible] |
| 84124 | NM_00134596 | ZNF394 | zinc finger protein 394 | protein_coding | NP_001332896.1;NP_001332897.1;NP_115540.2 | 18832 | . | [illegible] |
| 84125 | NM_00107991 | LRRIQ1 | leucine rich repeats and IQ motif containing 1 | protein_coding | NP_001073379.1;XP_011537119.1;XP_011537120.1;XP_011537121.1;XP | 25708 | . | [illegible] |
| 84126 | NM_00127102 | ATRIP | ATR interacting protein | protein_coding | NP_001257951.1;NP_001257952.1;NP_115542.2;NP_569055.1 | 33499 | 606605 | [illegible] |
| 84128 | NM_00130309 | WDR75 | WD repeat domain 75 | protein_coding | NP_001290025.1;NP_115544.1;XP_024308936.1 | 25725 | . | [illegible] |
| 84129 | NM_032169,N | ACAD11 | acyl-CoA dehydrogenase family member 11 | protein_coding | NP_115545.3 | 30211 | 614288 | [illegible] |
| 84131 | NM_00109880 | CEP78 | centrosomal protein 78 | protein_coding | NP_001092272.1;NP_001317620.1;NP_001317622.1;NP_001317623.1;NI | 25740 | 617110 | [illegible] |
| 84132 | NM_00136576 | USP42 | ubiquitin specific peptidase 42 | protein_coding | NP_001352693.1;NP_115548.1;XP_005249940.1;XP_006715853.1;XP_00 | 20068 | . | [illegible] |
| 84133 | NM_00120699 | ZNRF3 | zinc and ring finger 3 | protein_coding | NP_001193927.1;NP_115549.2;XP_011528737.1;XP_011528738.1;XP_01 | 18126 | 612062 | [illegible] |
| 84134 | NM_00128637 | TOMM40L | translocase of outer mitochondrial membrane 40 like | protein_coding | NP_001273302.1;NP_001273303.1;NP_115550.2;XP_006711635.1;XP_01 | 25756 | . | [illegible] |
| 84135 | NM_00128443 | UTP15 | UTP15 small subunit processome component | protein_coding | NP_001271359.1;NP_001271360.1;NP_115551.2;XP_011541982.1 | 25758 | 616194 | [illegible] |
| 84138 | NM_032178,XI | SLC7A6OS | solute carrier family 7 member 6 opposite strand | protein_coding | NP_115554.2;XP_011521674.1 | 25807 | . | [illegible] |
| 84140 | NM_00120154 | FAM161A | FAM161 centrosomal protein A | protein_coding | NP_001188472.1;NP_115556.2;XP_016860561.1;XP_016860562.1;XP_01 | 25808 | 613596 | [illegible] |
| 84141 | NM_00113503 | EVA1A | eva-1 homolog A, regulator of programmed cell death | protein_coding | NP_001128504.1;NP_001356453.1;NP_001356454.1;NP_115557.1 | 25816 | . | [illegible] |
| 84142 | NM_00134596 | ABRAXAS1 | abraxas 1, BRCA1 A complex subunit | protein_coding | NP_001332891.1;NP_620775.2;XP_005263337.1 | 25829 | 611143 | [illegible] |
| 84144 | NM_032184,XI | SYDE2 | synapse defective Rho GTPase homolog 2 | protein_coding | NP_115560.1;XP_006711009.1;XP_016857972.1;XP_016857973.1;XP_01 | 25841 | . | [illegible] |
| 84146 | NM_016620,N | ZNF644 | zinc finger protein 644 | protein_coding | NP_057704.2;NP_115562.3;NP_958357.1;XP_005271314.1;XP_00527131 | 29222 | 614159 | [illegible] |
| 84148 | NM_032188,N | KAT8 | lysine acetyltransferase 8 | protein_coding | NP_115564.2;NP_892003.2;XP_011544272.1;XP_011544273.1 | 17933 | 609912 | [illegible] |
| 84153 | NM_032193 | RNASEH2C | ribonuclease H2 subunit C | protein_coding | NP_115569.2 | 24116 | 610330 | [illegible] |
| 84154 | NM_00128911 | RPF2 | ribosome production factor 2 homolog | protein_coding | NP_001276040.1;NP_115570.1;XP_016866843.1 | 20870 | 618471 | [illegible] |
| 84159 | NM_00124463 | ARID5B | AT-rich interaction domain 5B | protein_coding | NP_001231567.1;NP_115575.1;XP_011538564.1;XP_024303998.1;XP_02 | 17362 | 608538 | [illegible] |
| 84162 | NM_015312,XI | KIAA1109 | KIAA1109 | protein_coding | NP_056127.2;XP_005263339.1;XP_005263344.1;XP_006714407.1;XP_01 | 26953 | 611565 | [illegible] |
| 84163 | NM_00128144 | GTF2IRD2 | GTF2I repeat domain containing 2 | protein_coding | NP_001268376.1;NP_001355229.1;NP_775808.4 | 30775 | 608899 | [illegible] |
| 84164 | NM_00124290 | ASCC2 | activating signal cointegrator 1 complex subunit 2 | protein_coding | NP_001229835.1;NP_001356849.1;NP_001356850.1;NP_001356851.1;NI | 24103 | 614216 | [illegible] |
| 84166 | NM_00133055 | NLRC5 | NLR family CARD domain containing 5 | protein_coding | NP_001317481.1;NP_115582.4;XP_005256250.1;XP_005256251.1;XP_00 | 29933 | 613537 | [illegible] |
| 84167 | NM_00128883 | C19orf44 | chromosome 19 open reading frame 44 | protein_coding | NP_001275763.1;NP_115583.1;XP_006722983.1;XP_011526656.1;XP_01 | 26141 | . | [illegible] |
| 84168 | NM_018153,N | ANTXR1 | ANTXR cell adhesion molecule 1 | protein_coding | NP_060623.2;NP_115584.1;NP_444262.1;XP_016860564.1;XP_0168605 | 21014 | 606410 | [illegible] |
| 84172 | NM_00113760 | POLR1B | RNA polymerase I subunit B | protein_coding | NP_001131076.1;NP_001269701.1;NP_001269703.1;NP_001269705.1;NI | 20454 | 602000 | [illegible] |
| 84173 | NM_00113502 | ELMOD3 | ELMO domain containing 3 | protein_coding | NP_001128493.1;NP_001128494.1;NP_001128495.1;NP_001316720.1;NI | 26158 | 615427 | [illegible] |
| 84179 | NM_00129434 | SLC49A3 | solute carrier family 49 member 3 | protein_coding | NP_001281270.1;NP_001281271.1;NP_001364988.1;NP_001364989.1;NI | 26177 | . | [illegible] |
| 84181 | NM_032221,XI | CHD6 | chromodomain helicase DNA binding protein 6 | protein_coding | NP_115597.3;XP_005260630.1;XP_005260633.1;XP_011527382.1;XP_01 | 19057 | 616114 | [illegible] |
| 84182 | NM_032222 | MINDY4 | MINDY lysine 48 deubiquitinase 4 | protein_coding | NP_115598.2 | 21916 | . | [illegible] |
| 84186 | NM_00128911 | ZCCHC7 | zinc finger CCHC-type containing 7 | protein_coding | NP_001276048.1;NP_001276049.1;NP_001276050.1;NP_115602.2;XP_00 | 26209 | . | [illegible] |
| 84187 | NM_00135384 | TMEM164 | transmembrane protein 164 | protein_coding | NP_001340778.1;NP_001340779.1;NP_001340780.1;NP_060168.2;NP_1 | 26217 | . | [illegible] |
| 84188 | NM_032228,XI | FAR1 | fatty acyl-CoA reductase 1 | protein_coding | NP_115604.1;XP_011518702.1 | 26222 | 616107 | [illegible] |
| 84189 | NM_032229 | SLITRK6 | SLIT and NTRK like family member 6 | protein_coding | NP_115605.2 | 23503 | 609681 | [illegible] |
| 84190 | NM_00131967 | METTL25 | methyltransferase like 25 | protein_coding | NP_001306604.1;NP_001334863.1;NP_115606.2;XP_005269243.2;XP_01 | 26228 | . | [illegible] |
| 84191 | NM_00101481 | CIAO2A | cytosolic iron-sulfur assembly component 2A | protein_coding | NP_001014812.1;NP_001276037.1;NP_115607.1 | 26235 | 618382 | [illegible] |
| 84193 | NM_032233,N | SETD3 | SET domain containing 3, actin histidine methyltransferase | protein_coding | NP_115609.2;NP_954574.1;XP_005268184.1;XP_011535533.2;XP_01153 | 20493 | 615671 | [illegible] |
| 84196 | NM_00103273 | USP48 | ubiquitin specific peptidase 48 | protein_coding | NP_001027902.1;NP_001317323.1;NP_001337093.1;NP_001337095.1;NI | 18533 | 617445 | [illegible] |
| 84197 | NM_00127797 | POMK | protein O-mannose kinase | protein_coding | NP_001264900.1;NP_115613.1 | 26267 | 615247 | [illegible] |
| 84206 | NM_032246 | MEX3B | mex-3 RNA binding family member B | protein_coding | NP_115622.2 | 25297 | 611008 | [illegible] |
| 84210 | NM_032250 | ANKRD20A1 | ankyrin repeat domain 20 family member A1 | protein_coding | NP_115626.2 | 23665 | . | [illegible] |
| 84214 | XR_00295650 | LOC84214 | uncharacterized LOC84214 | lncRNA | . | . | . | [illegible] |
| 84216 | NM_00128621 | TMEM117 | transmembrane protein 117 | protein_coding | NP_001273140.1;NP_001273141.1;NP_001273142.1;NP_115632.1;XP_01 | 25308 | . | [illegible] |
| 84217 | NM_00114619 | ZMYND12 | zinc finger MYND-type containing 12 | protein_coding | NP_001139664.1;NP_115633.3 | 21192 | . | [illegible] |
| 84218 | NM_032258 | TBC1D3F | TBC1 domain family member 3F | protein_coding | NP_115634.3 | 18257 | 610809 | [illegible] |
| 84219 | NM_032259,XI | WDR24 | WD repeat domain 24 | protein_coding | NP_115635.1;XP_011521001.1 | 20852 | . | [illegible] |
| 84220 | NM_005054,N | RGPD5 | RANBP2 like and GRIP domain containing 5 | protein_coding | NP_005045.2;NP_115636.1;XP_006712857.2;XP_016860569.1;XP_01686 | 32418 | 612708 | [illegible] |
| 84221 | NM_00114285 | SPATC1L | spermatogenesis and centriole associated 1 like | protein_coding | NP_001136326.1;NP_115637.3;XP_005261245.1;XP_011528058.1;XP_01 | 1298 | 612412 | [illegible] |
| 84222 | NR_026815 | TMEM191A | transmembrane protein 191A (pseudogene) | transcribed_ps | . | 25317 | . | [illegible] |
| 84223 | NM_00113443 | IQCG | IQ motif containing G | protein_coding | NP_001127907.1;NP_001309956.1;NP_001309957.1;NP_001309958.1;NI | 25251 | 612477 | [illegible] |
| 84224 | NM_00125641 | NBPF3 | NBPF member 3 | protein_coding | NP_001243345.1;NP_001243346.1;NP_001317310.1;NP_001364420.1;NI | 25076 | 612992 | [illegible] |
| 84225 | NM_00113604 | ZMYND15 | zinc finger MYND-type containing 15 | protein_coding | NP_001129518.1;NP_001254751.1;NP_115641.1;XP_016880707.1 | 20997 | 614312 | [illegible] |
| 84230 | NM_032270,XI | LRRC8C | leucine rich repeat containing 8 VRAC subunit C | protein_coding | NP_115646.3;XP_006711023.1;XP_011540584.1;XP_011540585.1;XP_01 | 25075 | 612889 | [illegible] |
| 84231 | NM_032271,XI | TRAF7 | TNF receptor associated factor 7 | protein_coding | NP_115647.2;XP_005255684.1;XP_011521002.1 | 20456 | 606692 | [illegible] |
| 84232 | NM_032272 | MAF1 | MAF1 homolog, negative regulator of RNA polymerase III | protein_coding | NP_115648.2 | 24966 | 610210 | [illegible] |
| 84233 | NM_00124473 | TMEM126A | transmembrane protein 126A | protein_coding | NP_001231664.1;NP_115649.1 | 25382 | 612988 | [illegible] |
| 84236 | NM_00116760 | RHBDD1 | rhomboid domain containing 1 | protein_coding | NP_001161080.1;NP_001335998.1;NP_001336000.1;NP_001336001.1;NI | 23081 | 617515 | [illegible] |
| 84240 | NM_00113103 | ZCCHC9 | zinc finger CCHC-type containing 9 | protein_coding | NP_001124507.1;NP_001124508.1;NP_115656.1;XP_005248673.1 | 25424 | . | [illegible] |
| 84243 | NM_032283 | ZDHHC18 | zinc finger DHHC-type palmitoyltransferase 18 | protein_coding | NP_115659.1 | 20712 | . | [illegible] |
| 84245 | NM_00103172 | MRI1 | methylthioribose-1-phosphate isomerase 1 | protein_coding | NP_001026897.1;NP_001316501.1;NP_115661.1;XP_011526658.1;XP_01 | 28469 | 615105 | [illegible] |
| 84246 | NM_032286 | MED10 | mediator complex subunit 10 | protein_coding | NP_115662.2 | 28760 | 612382 | [illegible] |
| 84247 | NM_032287 | RTL6 | retrotransposon Gag like 6 | protein_coding | NP_115663.2 | 13343 | . | [illegible] |

| | | | | | | | | | | | | | | | | | | | | | | | | | | | | | | | | | |
|---|---|---|---|---|---|---|---|---|---|---|---|---|---|---|---|---|---|---|---|---|---|---|---|---|---|---|---|---|---|---|---|---|---|
| 84248 | NM_00101153 | FYTTD1 | forty-two-three domain containing 1 | protein_coding | NP_001011537.2;NP_115664.2 | 25407 | 616933 | . | . | 6.165752 | 5.483456 | 5.697821 | 5.512083 | 14.347959 | 12.675255 | 12.785742 | 12.602723 | -1.052655 | 1198.555237 | 0.136226851 | -0.54345028 | 0.586819815 | 0.998865404 | 10.263453 | 10.189767 | 10.290129 | 10.199116 | 10.221984 | 10.192082 | 1500 | 1247 | 1036 | 1073 |
| 84249 | NM_032289,XI | PSD2 | pleckstrin and Sec7 domain containing 2 | protein_coding | NP_115665.1;XP_011535998.1;XP_016865465.1;XP_016865466.1 | 19092 | . | . | . | 0.298738 | 0.434774 | 0.149673 | 0.217836 | 0.695175 | 1.004998 | 0.335862 | 0.498056 | -1.639011 | 50.853553 | 0.449155988 | -1.58703408 | 0.112504927 | 0.850211417 | 5.982023 | 5.266773 | 5.714503 | 5.712590 | 5.599199 | 5.645820 | 73 | 68 | 28 | 42 |
| 84250 | NM_032290,XI | SLF1 | SMC5-SMC6 complex localization factor 1 | protein_coding | NP_115666.2;XP_016865468.1;XP_016865470.1;XP_016865472.1;XP_01 | 25408 | 618467 | . | . | 2.942686 | 3.126824 | 2.200034 | 1.473907 | 6.847750 | 7.227797 | 4.936811 | 3.369914 | -1.326590 | 476.192539 | 0.175041792 | -2.32928973 | 0.019843722 | 0.466614813 | 9.084693 | 8.677468 | 8.957340 | 8.993439 | 8.838316 | 8.759048 | 608 | 602 | 380 | 358 |
| 84253 | NM_00128677 | GARNL3 | GTPase activating Rap/RanGAP domain like 3 | protein_coding | NP_001273708.1;NP_115669.3;XP_005252324.1;XP_005252325.1;XP_01 | 25425 | . | . | . | 0.206873 | 0.263500 | 0.094609 | 0.281080 | 0.481402 | 0.609092 | 0.212300 | 0.642657 | -1.236201 | 26.816452 | 0.652313976 | -0.46896605 | 0.639093905 | NA | 4.892781 | 4.580439 | 4.739803 | 4.770040 | 4.685268 | 4.776386 | 30 | 36 | 11 | 33 |
| 84254 | NM_032294,N | CAMKK1 | calcium/calmodulin dependent protein kinase kinase 1 | protein_coding | NP_115670.1;NP_757343.2;NP_757344.2;XP_005256874.1;XP_00672165 | 1469 | 611411 | . | . | 1.970512 | 1.840318 | 2.196003 | 1.628002 | 4.585460 | 4.253978 | 4.927769 | 3.722233 | 1.037215 | 218.082040 | 0.235267123 | 0.224065013 | 0.822706716 | 0.998865404 | 7.741468 | 7.795481 | 7.778126 | 7.737693 | 7.819961 | 7.728758 | 257 | 221 | 212 | 188 |
| 84255 | NM_00128749 | SLC37A3 | solute carrier family 37 member 3 | protein_coding | NP_001274427.1;NP_001350302.1;NP_001350303.1;NP_001350304.1;NI | 20651 | . | . | . | 13.013892 | 12.043991 | 12.803248 | 12.430197 | 30.283859 | 27.840223 | 28.730116 | 28.420168 | -1.005121 | 1319.395372 | 0.13025961 | -0.05657299 | 0.954885349 | 0.998865404 | 10.369221 | 10.362092 | 10.397034 | 10.336799 | 10.375088 | 10.351882 | 1583 | 1371 | 1162 | 1215 |
| 84256 | NM_00130806 | FLYWCH1 | FLYWCH-type zinc finger 1 | protein_coding | NP_001294997.1;NP_065963.1;NP_115672.2;XP_006721025.1;XP_01152 | 25404 | . | . | . | 6.873401 | 7.832950 | 7.320528 | 8.074567 | 15.994687 | 18.106211 | 16.427050 | 18.461539 | 1.022130 | 1111.502428 | 0.142775042 | 0.221180389 | 0.824951978 | 0.998865404 | 10.102708 | 10.133716 | 10.065239 | 10.151373 | 10.077073 | 10.171399 | 1193 | 1255 | 915 | 1120 |
| 84258 | NM_00116032 | SYT3 | synaptotagmin 3 | protein_coding | NP_001153800.1;NP_001153801.1;NP_115674.1 | 11511 | 600327 | . | . | 0.263100 | 0.250159 | 0.209410 | 0.345528 | 0.612244 | 0.578254 | 0.469910 | 0.790011 | 1.066941 | 24.990330 | 0.627975186 | 0.148859452 | 0.881664538 | NA | 4.597285 | 4.687889 | 4.638885 | 4.637990 | 4.619796 | 4.669918 | 28 | 26 | 17 | 30 |
| 84259 | NM_00131873 | DCUN1D5 | defective in cullin neddylation 1 domain containing 5 | protein_coding | NP_001305668.1;NP_001305669.1;NP_001305670.1;NP_115675.1;XP_01 | 28409 | 616522 | . | . | 6.234983 | 5.717659 | 5.674664 | 7.464046 | 14.509061 | 13.216622 | 12.733780 | 17.065653 | -1.080551 | 1046.817660 | 0.151423768 | -0.7381113 | 0.460446816 | 0.998865404 | 10.086243 | 9.975210 | 10.097545 | 10.016953 | 10.072258 | 9.924545 | 1319 | 1110 | 964 | 848 |
| 84260 | NM_00114385 | TCHP | trichoplein keratin filament binding | protein_coding | NP_001137324.1;NP_115676.1;XP_011537138.1;XP_011537139.1;XP_01 | 28135 | 612654 | . | . | 7.822499 | 7.687095 | 6.895869 | 7.412375 | 18.203279 | 17.769064 | 15.474130 | 16.947514 | -1.076865 | 740.911544 | 0.148316517 | -0.72033778 | 0.47131705 | 0.998865404 | 9.585584 | 9.478754 | 9.573779 | 9.542913 | 9.497783 | 9.516222 | 904 | 811 | 612 | 678 |
| 84261 | NM_032301,XI | FBXW9 | F-box and WD repeat domain containing 9 | protein_coding | NP_115677.2;XP_005260153.1 | 28136 | 609074 | . | . | 4.464065 | 4.782612 | 4.589004 | 5.464270 | 10.388062 | 11.055222 | 10.297592 | 12.493402 | 1.069671 | 264.167768 | 0.218133557 | 0.445447421 | 0.655996449 | 0.998865404 | 7.996501 | 8.092523 | 8.008623 | 8.043571 | 8.010914 | 8.107862 | 284 | 285 | 215 | 280 |
| 84262 | NM_00113434 | PSMG3 | proteasome assembly chaperone 3 | protein_coding | NP_001127812.1;NP_115678.1;XP_024302744.1 | 22420 | 617528 | . | . | 33.117164 | 32.689177 | 34.766365 | 35.722986 | 77.065002 | 75.562500 | 78.014707 | 81.676369 | 1.032355 | 1463.435388 | 0.124994049 | 0.367536245 | 0.713219061 | 0.998865404 | 10.492015 | 10.537907 | 10.507590 | 10.497700 | 10.516480 | 10.537774 | 1669 | 1544 | 1275 | 1413 |
| 84263 | NM_00119582 | HSDL2 | hydroxysteroid dehydrogenase like 2 | protein_coding | NP_001182751.1;NP_115679.2;XP_011517393.1;XP_016870692.1;XP_01 | 18572 | . | . | . | 7.349951 | 6.618165 | 6.854396 | 6.718638 | 17.103635 | 15.298184 | 15.381067 | 15.361365 | -1.056798 | 675.124731 | 0.155338103 | -0.51307511 | 0.607898782 | 0.998865404 | 9.438130 | 9.358800 | 9.454126 | 9.379133 | 9.386904 | 9.372572 | 842 | 708 | 577 | 609 |
| 84264 | NM_00129013 | HAGHL | hydroxyacylglutathione hydrolase like | protein_coding | NP_001277066.1;NP_001277068.1;NP_001310564.1;NP_001310565.1;NI | 14177 | . | . | . | 12.449878 | 11.828973 | 10.911484 | 12.916757 | 28.971376 | 27.343198 | 24.485055 | 29.532630 | -1.038761 | 524.239320 | 0.17191913 | -0.31912848 | 0.749629086 | 0.998865404 | 9.061485 | 9.006148 | 9.066417 | 9.024504 | 8.968781 | 9.070992 | 635 | 558 | 411 | 521 |
| 84265 | NM_00133068 | POLR3GL | RNA polymerase III subunit GL | protein_coding | NP_001317614.1;NP_115681.1;XP_005277488.1 | 28466 | 617457 | . | . | 9.747574 | 9.797262 | 9.291336 | 12.298435 | 22.683005 | 22.646812 | 20.849487 | 28.118914 | 1.095590 | 376.624051 | 0.19998449 | 0.658593661 | 0.510156735 | 0.998865404 | 8.490190 | 8.620817 | 8.525523 | 8.528608 | 8.497117 | 8.656919 | 413 | 389 | 296 | 419 |
| 84266 | NM_032306,XI | ALKBH7 | alkB homolog 7 | protein_coding | NP_115682.1;XP_005259715.1;XP_016882844.1 | 21306 | 613305 | . | . | 10.690521 | 9.619542 | 9.626283 | 10.398572 | 24.877280 | 22.236004 | 21.601102 | 23.775097 | -1.034156 | 297.654466 | 0.205200677 | -0.23613224 | 0.813330061 | 0.998865404 | 8.241683 | 8.192894 | 8.247216 | 8.205021 | 8.175944 | 8.239688 | 362 | 314 | 239 | 291 |
| 84267 | NM_00131799 | C9orf64 | chromosome 9 open reading frame 64 | protein_coding | NP_001304926.1;NP_115683.3;XP_005252328.1 | 28144 | 611342 | . | . | 6.500209 | 6.378186 | 5.584718 | 6.774925 | 15.126253 | 14.743464 | 12.531946 | 15.490060 | -1.085284 | 470.057588 | 0.176858721 | -0.66761327 | 0.504380486 | 0.998865404 | 8.934830 | 8.816116 | 8.908761 | 8.893979 | 8.799593 | 8.898576 | 570 | 522 | 360 | 457 |
| 84268 | NM_00103300 | RPAIN | RPA interacting protein | protein_coding | NP_001028174.2;NP_001153715.1;NP_001153716.1;NP_001153718.1;NI | 28641 | 617299 | . | . | 15.987404 | 14.219359 | 16.797984 | 17.215274 | 37.203346 | 32.868681 | 37.694189 | 39.360679 | 1.134079 | 789.599728 | 0.146038939 | 1.24296193 | 0.213881863 | 0.978201242 | 9.531236 | 9.712998 | 9.598188 | 9.557654 | 9.674579 | 9.658765 | 877 | 775 | 738 | 778 |
| 84269 | NM_00130435 | CHCHD5 | coiled-coil-helix-coiled-coil-helix domain containing 5 | protein_coding | NP_001291282.1;NP_001291283.1;NP_001358251.1;NP_001358252.1;NI | 17840 | 616978 | . | . | 22.706903 | 23.848550 | 19.187697 | 23.294672 | 52.839891 | 55.126989 | 43.056635 | 53.260501 | -1.009399 | 395.085350 | 0.184330232 | -0.07321869 | 0.941632098 | 0.998865404 | 8.633085 | 8.618921 | 8.616503 | 8.640478 | 8.588358 | 8.656330 | 448 | 437 | 322 | 390 |
| 84270 | NM_00131801 | CARD19 | caspase recruitment domain family member 19 | protein_coding | NP_001304939.1;NP_001304940.1;NP_115686.3 | 28148 | 617726 | . | . | 19.288189 | 16.486889 | 16.908112 | 18.651869 | 44.884413 | 38.110186 | 37.941312 | 42.645280 | 1.008062 | 710.269356 | 0.150983813 | 0.076725695 | 0.938841764 | 0.998865404 | 9.466596 | 9.477827 | 9.454173 | 9.484227 | 9.453463 | 9.495596 | 797 | 780 | 601 | 689 |
| 84271 | NM_00127865 | POLDIP3 | DNA polymerase delta interacting protein 3 | protein_coding | NP_001265586.1;NP_001349981.1;NP_115687.2;NP_835237.1;XP_0168 | 23782 | 611520 | . | . | 24.119843 | 26.320384 | 25.127505 | 28.437007 | 56.127863 | 60.840747 | 56.385393 | 65.017836 | 1.038977 | 2692.224913 | 0.121448328 | 0.45421589 | 0.649673447 | 0.998865404 | 11.366875 | 11.421770 | 11.348743 | 11.406919 | 11.358249 | 11.457713 | 2932 | 2952 | 2240 | 2729 |
| 84272 | NM_032312,XI | YIPF4 | Yip1 domain family member 4 | protein_coding | NP_115688.1;XP_005264656.1;XP_011531427.1;XP_024308941.1 | 28145 | 617534 | . | . | 4.611151 | 4.122233 | 4.824796 | 3.818240 | 10.730338 | 9.528723 | 10.826701 | 8.729951 | -1.028917 | 1356.668268 | 0.150855214 | -0.27262669 | 0.785140193 | 0.998865404 | 10.425945 | 10.385476 | 10.455660 | 10.371112 | 10.482485 | 10.294178 | 1671 | 1403 | 1307 | 1099 |
| 84273 | NM_032313 | NOA1 | nitric oxide associated 1 | protein_coding | NP_115689.1 | 28473 | 614919 | . | . | 6.374622 | 6.696937 | 5.582583 | 6.005766 | 14.834006 | 15.480270 | 12.527154 | 13.731471 | -1.142761 | 426.429531 | 0.177540147 | -1.08439198 | 0.278191036 | 0.998865404 | 8.829421 | 8.636458 | 8.762433 | 8.788095 | 8.675957 | 8.712885 | 513 | 501 | 335 | 385 |
| 84274 | NM_032314,XI | COQ5 | coenzyme Q5, methyltransferase | protein_coding | NP_115690.3;XP_006719702.1;XP_024304992.1 | 28722 | 616359 | . | . | 11.589950 | 11.636416 | 9.453599 | 12.644631 | 26.970288 | 26.898101 | 21.213601 | 28.910449 | -1.061149 | 537.094835 | 0.179094256 | -0.47811289 | 0.632569858 | 0.998865404 | 9.111665 | 9.025103 | 9.087575 | 9.085438 | 8.963409 | 9.126088 | 638 | 596 | 395 | 551 |
| 84275 | NM_032315,XI | SLC25A33 | solute carrier family 25 member 33 | protein_coding | NP_115691.1;XP_005263560.1;XP_011540598.1 | 29681 | 610816 | . | . | 7.025096 | 8.178512 | 7.471614 | 7.845461 | 16.347686 | 18.904997 | 16.766084 | 17.937712 | 1.024765 | 903.115132 | 0.143175209 | 0.246499412 | 0.80529566 | 0.998865404 | 9.801194 | 9.836127 | 9.774963 | 9.842906 | 9.812552 | 9.841278 | 979 | 1008 | 779 | 874 |
| 84276 | NM_032316 | NICN1 | nicolin 1 | protein_coding | NP_115692.1 | 18317 | 611516 | . | . | 3.294698 | 2.822213 | 3.217849 | 3.280790 | 7.666897 | 6.523673 | 7.220760 | 7.501136 | 1.047725 | 317.318046 | 0.201911977 | 0.333113399 | 0.739048686 | 0.998865404 | 8.275494 | 8.343281 | 8.339642 | 8.249876 | 8.318602 | 8.324217 | 386 | 307 | 281 | 306 |
| 84277 | NM_032317 | DNAJC30 | DnaJ heat shock protein family (Hsp40) member C30 | protein_coding | NP_115693.2 | 16410 | 618202 | . | . | 4.143147 | 4.448745 | 3.919302 | 5.461052 | 9.641273 | 10.283474 | 8.794800 | 12.486047 | 1.075771 | 355.211141 | 0.209822198 | 0.502189352 | 0.615534332 | 0.998865404 | 8.419612 | 8.523582 | 8.429911 | 8.465455 | 8.393314 | 8.577281 | 382 | 381 | 269 | 400 |
| 84278 | NM_00135522 | MFSD14C | major facilitator superfamily domain containing 14C | protein_coding | NP_001342157.1;NP_001342158.1;NP_001342159.1;XP_024303466.1;XP | 23672 | . | . | . | 7.056340 | 7.615301 | 8.341530 | 7.335693 | 16.420392 | 17.603107 | 18.718148 | 16.772190 | 1.063921 | 287.970448 | 0.210408022 | 0.424844797 | 0.670949819 | 0.998865404 | 8.123834 | 8.214302 | 8.171987 | 8.130077 | 8.227854 | 8.138512 | 334 | 289 | 280 | 255 |
| 84279 | NM_032319,N | PRADC1 | protease associated domain containing 1 | protein_coding | NP_115695.1 | 16047 | . | . | . | 16.036271 | 16.191228 | 14.733418 | 17.286274 | 37.317064 | 37.426752 | 33.061362 | 39.523011 | -1.021938 | 546.293689 | 0.16767807 | -0.18670969 | 0.85188826 | 0.998865404 | 9.109398 | 9.077491 | 9.100192 | 9.100174 | 9.037356 | 9.132381 | 636 | 596 | 435 | 544 |
| 84280 | NM_00129774 | BTBD10 | BTB domain containing 10 | protein_coding | NP_001284670.1;NP_001284671.1;NP_115696.2;XP_016873894.1;XP_01 | 21445 | 615933 | . | . | 8.038242 | 7.099522 | 7.108607 | 8.121352 | 18.705322 | 16.410864 | 15.951510 | 18.568506 | -1.004304 | 575.504265 | 0.166984578 | -0.03710778 | 0.970399071 | 0.998865404 | 9.171838 | 9.165520 | 9.208417 | 9.128905 | 9.127542 | 9.204197 | 704 | 585 | 470 | 570 |
| 84282 | NM_00118499 | RNF135 | ring finger protein 135 | protein_coding | NP_001171921.1;NP_115698.3;NP_922921.1;XP_016880712.1;XP_02430 | 21158 | 611358 | . | . | 3.172884 | 4.948716 | 4.206260 | 5.611379 | 7.383434 | 11.439178 | 9.438725 | 12.829751 | 1.169220 | 298.954973 | 0.23414576 | 0.963275174 | 0.335409449 | 0.998865404 | 8.107634 | 8.331277 | 8.083874 | 8.254880 | 8.178866 | 8.329354 | 266 | 346 | 242 | 343 |
| 84283 | NM_032323,N | TMEM79 | transmembrane protein 79 | protein_coding | NP_115699.1 | 28196 | 615531 | . | . | 1.834239 | 2.929555 | 3.192043 | 2.375584 | 4.268348 | 6.771798 | 7.162852 | 5.431491 | 1.149597 | 180.055502 | 0.289324574 | 0.695166032 | 0.486951257 | 0.998865404 | 7.388348 | 7.589258 | 7.352302 | 7.539664 | 7.575186 | 7.440759 | 150 | 221 | 193 | 153 |
| 84284 | NM_00132945 | NTPCR | nucleoside-triphosphatase, cancer-related | protein_coding | NP_001316381.1;NP_001316382.1;NP_115700.1 | 28204 | . | . | . | 3.470462 | 3.605161 | 3.132360 | 3.983392 | 8.075909 | 8.333496 | 7.028926 | 9.107554 | -1.013103 | 586.042569 | 0.177059931 | -0.10607006 | 0.915526768 | 0.998865404 | 9.204732 | 9.184924 | 9.179016 | 9.215207 | 9.097848 | 9.271729 | 661 | 654 | 439 | 618 |
| 84285 | NM_00124248 | EIF1AD | eukaryotic translation initiation factor 1A domain containing | protein_coding | NP_001229410.1;NP_001229411.1;NP_001229412.1;NP_001229413.1;NI | 28147 | 618473 | . | . | 7.111034 | 8.053794 | 7.750419 | 7.992143 | 16.547668 | 18.616708 | 17.391718 | 18.273087 | 1.036173 | 668.885715 | 0.155640851 | 0.329381252 | 0.741867537 | 0.998865404 | 9.359904 | 9.410877 | 9.334642 | 9.409578 | 9.398097 | 9.395691 | 715 | 748 | 593 | 637 |
| 84286 | NM_00129742 | TMEM175 | transmembrane protein 175 | protein_coding | NP_001284352.1;NP_001284353.1;NP_001284354.1;NP_001284355.1;NI | 28709 | 616660 | . | . | 11.081364 | 11.229176 | 11.155899 | 11.306454 | 25.786791 | 25.956744 | 25.033518 | 25.850864 | 1.008462 | 671.383668 | 0.152951823 | 0.079484765 | 0.936647049 | 0.998865404 | 9.384784 | 9.397177 | 9.386284 | 9.389438 | 9.414999 | 9.371866 | 768 | 723 | 605 | 612 |
| 84287 | NM_00128780 | ZDHHC16 | zinc finger DHHC-type palmitoyltransferase 16 | protein_coding | NP_001274732.1;NP_001274733.1;NP_115703.2;NP_932160.1;NP_9321 | 20714 | 616750 | . | . | 28.018927 | 28.168883 | 27.502124 | 30.378915 | 65.201196 | 65.113630 | 61.713970 | 69.457780 | 1.019632 | 1633.066373 | 0.12460463 | 0.225102787 | 0.821899313 | 0.998865404 | 10.659379 | 10.687222 | 10.662489 | 10.668627 | 10.649300 | 10.711069 | 1855 | 1752 | 1377 | 1607 |
| 84288 | NM_00114394 | EFCAB2 | EF-hand calcium binding domain 2 | protein_coding | NP_001137415.1;NP_001277256.1;NP_115704.1;XP_011542602.1;XP_01 | 28166 | . | . | . | 1.100109 | 1.606551 | 1.707267 | 1.194761 | 2.560000 | 3.713615 | 3.831058 | 2.731676 | 1.563901 | 102.867622 | 0.321116436 | 2.009082763 | 0.044528357 | 0.646973597 | 6.325592 | 6.971909 | 6.601662 | 6.592909 | 6.768952 | 6.700083 | 94 | 85 | 120 | 106 |
| 84289 | NM_00133016 | ING5 | inhibitor of growth family member 5 | protein_coding | NP_001317090.1;NP_001317091.1;NP_115705.2;XP_016860586.1;XP_01 | 19421 | 608525 | . | . | 15.716434 | 16.369568 | 16.727675 | 16.147015 | 36.572788 | 37.838989 | 37.536418 | 36.918227 | 1.045360 | 1685.319579 | 0.126948538 | 0.504139821 | 0.614163126 | 0.998865404 | 10.686290 | 10.750607 | 10.717535 | 10.683312 | 10.777340 | 10.691654 | 1939 | 1739 | 1580 | 1527 |
| 84290 | NM_032330 | CAPNS2 | calpain small subunit 2 | protein_coding | NP_115706.1 | 16371 | 616767 | . | . | 0.538250 | 0.289953 | 0.440953 | 0.215318 | 1.252530 | 0.670240 | 0.989487 | 0.492300 | -1.258203 | 11.975662 | 0.912598723 | -0.36309972 | 0.716530393 | NA | 3.735321 | 3.410501 | 3.606486 | 3.569983 | 3.589926 | 3.562499 | 20 | 10 | 12 | 7 |
| 84292 | NM_00109973 | WDR83 | WD repeat domain 83 | protein_coding | NP_001093207.1;NP_115708.1 | 32672 | 616850 | . | . | 4.979106 | 5.156003 | 5.638138 | 5.498310 | 11.586585 | 11.918334 | 12.651821 | 12.571232 | 1.020752 | 224.034945 | 0.230606259 | 0.128499023 | 0.897754072 | 0.998865404 | 7.793260 | 7.821759 | 7.778164 | 7.824853 | 7.775421 | 7.846213 | 243 | 251 | 182 | 228 |
| 84293 | NM_00124377 | PRXL2A | peroxiredoxin like 2A | protein_coding | NP_001230707.1;NP_001230708.1;NP_001230709.1;NP_001230710.1;NI | 28651 | 617165 | . | . | 17.015183 | 16.510851 | 20.103548 | 22.093281 | 39.595031 | 38.165570 | 45.111780 | 50.513662 | 1.232029 | 3312.991824 | 0.112735353 | 2.670293493 | 0.007578497 | 0.272411452 | 11.535578 | 11.836587 | 11.621451 | 11.581130 | 11.746175 | 11.799181 | 3504 | 3123 | 3080 | 3537 |
| 84294 | NM_032334 | UTP23 | UTP23 small subunit processome component | protein_coding | NP_115710.2 | 28224 | . | . | . | 3.191478 | 2.635010 | 3.129722 | 2.753923 | 7.426701 | 6.090943 | 7.023006 | 6.296519 | -1.006679 | 335.527660 | 0.204634838 | -0.04692923 | 0.96256963 | 0.998865404 | 8.394467 | 8.386097 | 8.445073 | 8.332429 | 8.424243 | 8.347827 | 426 | 327 | 311 | 292 |
| 84295 | NM_00101587 | PHF6 | PHD finger protein 6 | protein_coding | NP_001015877.1;NP_115711.2;NP_115834.1 | 18145 | 300414 | . | . | 5.876263 | 5.692549 | 5.515707 | 5.072787 | 13.674305 | 13.158582 | 12.377087 | 11.598324 | -1.117633 | 751.982273 | 0.151931534 | -1.05604666 | 0.290946897 | 0.998865404 | 9.632279 | 9.472403 | 9.614161 | 9.568529 | 9.551441 | 9.475724 | 943 | 829 | 652 | 629 |
| 84296 | NM_032336,XI | GINS4 | GINS complex subunit 4 | protein_coding | NP_115712.1;XP_005273716.1 | 28226 | 610611 | . | . | 12.727048 | 13.194720 | 11.613110 | 12.802313 | 29.616363 | 30.500185 | 26.059479 | 29.270969 | -1.074475 | 1472.881412 | 0.127861034 | -0.81050384 | 0.417650659 | 0.998865404 | 10.575447 | 10.471533 | 10.540245 | 10.562461 | 10.462283 | 10.528480 | 1732 | 1670 | 1181 | 1389 |
| 84298 | NM_032338 | LLPH | LLP homolog, long-term synaptic facilitation factor | protein_coding | NP_115714.1 | 28229 | 616998 | . | . | 3.399275 | 3.683286 | 3.386356 | 3.793105 | 7.910253 | 8.514080 | 7.598883 | 8.672484 | -1.000060 | 858.470609 | 0.146660591 | -0.0005853 | 0.999532999 | 0.999862634 | 9.745927 | 9.745323 | 9.718984 | 9.770879 | 9.707893 | 9.780948 | 953 | 960 | 708 | 846 |
| 84299 | NM_00133020 | MIEN1 | migration and invasion enhancer 1 | protein_coding | NP_001317135.1;NP_115715.3 | 28230 | 611802 | . | . | 16.602040 | 19.751681 | 16.840585 | 19.594870 | 38.633629 | 45.656896 | 37.789781 | 44.801342 | -1.007055 | 810.580117 | 0.158180652 | -0.06411874 | 0.948875679 | 0.998865404 | 9.668282 | 9.657319 | 9.615848 | 9.709931 | 9.595747 | 9.718202 | 875 | 936 | 641 | 823 |
| 84300 | NM_032340 | UQCC2 | ubiquinol-cytochrome c reductase complex assembly factor 2 | protein_coding | NP_115716.1 | 21237 | 614461 | . | . | 10.361919 | 12.376674 | 12.581131 | 13.518539 | 24.112614 | 28.609236 | 28.231691 | 30.908531 | 1.132387 | 488.282927 | 0.176305817 | 1.017363797 | 0.308980398 | 0.998865404 | 8.839507 | 9.018115 | 8.835215 | 8.940104 | 8.947202 | 8.986773 | 483 | 536 | 437 | 501 |
| 84301 | NM_032341 | DDI2 | DNA damage inducible 1 homolog 2 | protein_coding | NP_115717.3 | 24578 | . | . | . | 6.363389 | 6.907380 | 7.058570 | 7.299120 | 14.807868 | 15.966718 | 15.839228 | 16.688572 | 1.067019 | 2294.788862 | 0.117803466 | 0.794424935 | 0.426948093 | 0.998865404 | 11.116671 | 11.210108 | 11.107465 | 11.166193 | 11.179778 | 11.198714 | 2463 | 2484 | 2036 | 2247 |
| 84303 | NM_00132061 | CHCHD6 | coiled-coil-helix-coiled-coil-helix domain containing 6 | protein_coding | NP_001307539.1;NP_115719.1;XP_011511536.1;XP_011511537.1;XP_01 | 28184 | 615634 | . | . | 9.410832 | 8.524275 | 8.336486 | 9.842446 | 21.899391 | 19.704247 | 18.706831 | 22.503585 | 1.002017 | 297.841317 | 0.207022091 | 0.014041034 | 0.988797244 | 0.999197585 | 8.217040 | 8.219759 | 8.248957 | 8.184611 | 8.184593 | 8.251018 | 363 | 302 | 243 | 297 |
| 84304 | NM_00112861 | NUDT22 | nudix hydrolase 22 | protein_coding | NP_001122084.1;NP_001122085.1;NP_001258760.1;NP_115720.2;XP_02 | 28189 | . | . | . | 12.861640 | 15.027213 | 11.933243 | 15.502173 | 29.929564 | 34.736076 | 26.777854 | 35.443874 | -1.004074 | 460.758303 | 0.187094884 | -0.03134746 | 0.97499244 | 0.998865404 | 8.851398 | 8.844326 | 8.804322 | 8.888662 | 8.775309 | 8.909568 | 496 | 532 | 356 | 478 |
| 84305 | NM_00114385 | PYM1 | PYM homolog 1, exon junction complex associated factor | protein_coding | NP_001137325.1;NP_115721.1 | 30258 | . | . | . | 4.260743 | 3.777763 | 4.974609 | 6.400619 | 9.914924 | 8.732469 | 11.162879 | 14.634255 | 1.421576 | 172.199501 | 0.264025027 | 1.922133282 | 0.054588989 | 0.69238604 | 7.152294 | 7.659269 | 7.365392 | 7.313924 | 7.428544 | 7.542119 | 175 | 143 | 154 | 213 |
| 84306 | NM_00135343 | PDCD2L | programmed cell death 2 like | protein_coding | NP_001340362.1;NP_115722.1 | 28194 | 615661 | . | . | 6.100037 | 5.457826 | 5.751872 | 6.765800 | 14.195037 | 12.616009 | 12.907035 | 15.469198 | 1.065761 | 217.660449 | 0.234315102 | 0.392137198 | 0.694956844 | 0.998865404 | 7.719433 | 7.810986 | 7.776585 | 7.721518 | 7.741667 | 7.815558 | 257 | 214 | 180 | 227 |
| 84307 | NM_00113517 | ZNF397 | zinc finger protein 397 | protein_coding | NP_001128650.1;NP_115723.1;XP_006722621.1;XP_011524531.1;XP_01 | 18818 | 609601 | . | . | 1.899057 | 1.746011 | 2.130301 | 1.629129 | 4.419183 | 4.035980 | 4.780335 | 3.724810 | -1.063109 | 425.207604 | 0.191699165 | -0.46056561 | 0.645110288 | 0.998865404 | 8.775265 | 8.687446 | 8.702344 | 8.791812 | 8.772894 | 8.645296 | 468 | 507 | 397 | 345 |
| 84309 | NM_00119345 | NUDT16L1 | nudix hydrolase 16 like 1 | protein_coding | NP_001180381.1;NP_001357514.1;NP_001357515.1;NP_001357516.1;NI | 28154 | 617338 | . | . | 7.886254 | 6.216859 | 6.880822 | 6.608541 | 18.351639 | 14.370551 | 15.440364 | 15.109644 | -1.080389 | 382.847436 | 0.189723297 | -0.58796757 | 0.556554062 | 0.998865404 | 8.635592 | 8.523482 | 8.620891 | 8.583963 | 8.509945 | 8.602435 | 472 | 416 | 294 | 373 |
| 84310 | NM_00113439 | C7orf50 | chromosome 7 open reading frame 50 | protein_coding | NP_001127867.1;NP_001127868.1;NP_001305181.1;NP_001337897.1;NI | 22421 | . | . | . | 32.872058 | 35.681535 | 27.414282 | 35.190567 | 76.494628 | 82.479467 | 61.516857 | 80.459054 | -1.098846 | 1386.029616 | 0.143653165 | -0.94665143 | 0.343816426 | 0.998865404 | 10.503409 | 10.366846 | 10.449638 | 10.490971 | 10.312429 | 10.477216 | 1627 | 1608 | 1023 | 1373 |
| 84311 | NM_00127827 | MRPL45 | mitochondrial ribosomal protein L45 | protein_coding | NP_001265208.2;NP_115727.5 | 16651 | 611850 | . | . | 33.357645 | 32.815671 | 35.261347 | 40.855434 | 77.624611 | 75.854887 | 79.125440 | 93.411091 | 1.133982 | 1720.768680 | 0.128572297 | 1.410866466 | 0.158283992 | 0.929304099 | 10.655417 | 10.836575 | 10.706043 | 10.687107 | 10.739619 | 10.847788 | 1880 | 1719 | 1482 | 1831 |
| 84312 | NM_032352,XI | BRMS1L | BRMS1 like transcriptional repressor | protein_coding | NP_115728.2;XP_005268185.1;XP_006720337.1;XP_006720338.1;XP_01 | 20512 | 618514 | . | . | 2.508708 | 2.520261 | 2.310626 | 2.325277 | 5.837868 | 5.825696 | 5.184978 | 5.316470 | -1.097210 | 199.629070 | 0.236023593 | -0.56706181 | 0.570672189 | 0.998865404 | 7.706509 | 7.572748 | 7.665249 | 7.659327 | 7.621815 | 7.619249 | 242 | 224 | 166 | 178 |
| 84313 | NM_032353 | VPS25 | vacuolar protein sorting 25 homolog | protein_coding | NP_115729.1 | 28122 | 610907 | . | . | 43.197632 | 44.733768 | 40.583729 | 56.348785 | 100.522675 | 103.404106 | 91.068710 | 128.834808 | 1.086532 | 1565.371335 | 0.155344026 | 0.770743549 | 0.440858954 | 0.998865404 | 10.551459 | 10.670658 | 10.565536 | 10.586206 | 10.508837 | 10.756853 | 1705 | 1641 | 1194 | 1769 |
| 84314 | NM_00135127 | TMEM107 | transmembrane protein 107 | protein_coding | NP_001338207.1;NP_001338208.1;NP_001338209.1;NP_115730.2;NP_8 | 28128 | 616183 | . | . | 5.659988 | 5.367963 | 4.855841 | 4.748543 | 13.171023 | 12.408287 | 10.896365 | 10.856979 | -1.186058 | 355.836016 | 0.18922221 | -1.30098453 | 0.193263749 | 0.96621494 | 8.592740 | 8.346941 | 8.539758 | 8.503460 | 8.434094 | 8.415060 | 458 | 404 | 288 | 300 |
| 84315 | NM_00114250 | MON1A | MON1 homolog A, secretory trafficking associated | protein_coding | NP_001135973.1;NP_115731.2;XP_006713408.1;XP_011532462.1 | 28207 | 611464 | . | . | 2.870558 | 3.474891 | 3.951706 | 4.114772 | 6.679909 | 8.032367 | 8.867514 | 9.407939 | 1.263584 | 245.927533 | 0.226835492 | 1.487956752 | 0.1367623 | 0.8955076 | 7.763985 | 8.100603 | 7.818471 | 7.929569 | 7.991763 | 7.990956 | 221 | 262 | 239 | 257 |
| 84316 | NM_00132092 | NAA38 | N-alpha-acetyltransferase 38, NatC auxiliary subunit | protein_coding | NP_001307853.1;NP_001307854.1;NP_001317040.1;NP_115732.2 | 28212 | 617990 | . | . | 44.766035 | 43.655996 | 38.421808 | 46.054350 | 104.172400 | 100.912792 | 86.217430 | 105.297800 | -1.060189 | 1206.687838 | 0.14016946 | -0.60156845 | 0.547461433 | 0.998865404 | 10.278541 | 10.193891 | 10.278799 | 10.236400 | 10.152386 | 10.271578 | 1470 | 1303 | 937 | 1186 |
| 84317 | NM_00132111 | CCDC115 | coiled-coil domain containing 115 | protein_coding | NP_001308047.1;NP_001308048.2;NP_115733.2 | 28178 | 613734 | . | . | 24.332336 | 24.154410 | 24.682771 | 27.389018 | 56.622345 | 55.834000 | 55.387421 | 62.621730 | 1.087660 | 1496.636029 | 0.129729319 | 0.934462784 | 0.350065225 | 0.998865404 | 10.485733 | 10.606745 | 10.525465 | 10.501473 | 10.532586 | 10.622275 | 1677 | 1523 | 1278 | 1546 |
| 84318 | NM_00113014 | CCDC77 | coiled-coil domain containing 77 | protein_coding | NP_001123618.1;NP_001123619.1;NP_001123620.1;NP_115734.1 | 28203 | . | . | . | 12.248060 | 12.176016 | 11.390164 | 12.236998 | 28.501738 | 28.145401 | 25.559196 | 27.978443 | -1.045174 | 854.232159 | 0.142676245 | -0.44677114 | 0.655040312 | 0.998865404 | 9.770083 | 9.706178 | 9.758878 | 9.749384 | 9.703623 | 9.741175 | 1012 | 936 | 707 | 804 |
| 84319 | NM_00116792 | CMSS1 | cms1 ribosomal small subunit homolog | protein_coding | NP_001161396.1;NP_115735.2 | 28666 | . | . | . | 2.989372 | 3.392490 | 2.851891 | 3.145058 | 6.956391 | 7.841894 | 6.399560 | 7.190801 | -1.072438 | 413.452585 | 0.182696824 | -0.55225176 | 0.580775874 | 0.998865404 | 8.741498 | 8.639868 | 8.678004 | 8.744758 | 8.644543 | 8.694602 | 466 | 487 | 332 | 390 |
| 84320 | NM_032360,XI | ACBD6 | acyl-CoA binding domain containing 6 | protein_coding | NP_115736.1 | 23339 | 616352 | . | . | 10.491552 | 10.490638 | 9.017890 | 11.164238 | 24.414273 | 24.249577 | 20.235883 | 25.525704 | 1.110650 | 826.205973 | 0.143328345 | 1.056348869 | 0.290808857 | 0.998865404 | 9.612677 | 9.764068 | 9.660225 | 9.642442 | 9.717034 | 9.734464 | 913 | 834 | 747 | 825 |
| 84321 | NM_00137690 | THOC3 | THO complex 3 | protein_coding | NP_001363831.1;NP_115737.1;XP_016865474.1 | 19072 | 606929 | . | . | 28.399686 | 28.580176 | 25.529275 | 28.966128 | 66.087231 | 66.064357 | 57.286954 | 66.227614 | -1.059721 | 1392.266408 | 0.131050767 | -0.6385687 | 0.523103547 | 0.998865404 | 10.484601 | 10.400616 | 10.465875 | 10.463794 | 10.375858 | 10.462859 | 1652 | 1544 | 1108 | 1340 |
| 84324 | NM_033082,N | SARNP | SAP domain containing ribonucleoprotein | protein_coding | NP_149073.1 | 24432 | 610049 | . | . | 20.254405 | 18.849487 | 19.797912 | 24.864781 | 47.132832 | 43.571432 | 44.425943 | 56.850368 | 1.067446 | 621.078250 | 0.166489798 | 0.565580404 | 0.571679052 | 0.998865404 | 9.231172 | 9.324579 | 9.247171 | 9.262695 | 9.229137 | 9.363440 | 684 | 656 | 502 | 661 |
| 84326 | NM_00104016 | METTL26 | methyltransferase like 26 | protein_coding | NP_001035250.1;NP_001035251.1;NP_001035252.1;NP_001035255.1;NI | 14141 | . | . | . | 21.136827 | 21.606441 | 23.245676 | 23.201093 | 49.186264 | 49.944252 | 52.162626 | 53.046543 | 1.098546 | 488.645599 | 0.16862302 | 0.804129733 | 0.421322055 | 0.998865404 | 8.863273 | 8.998833 | 8.886979 | 8.915597 | 8.969202 | 8.951846 | 524 | 514 | 451 | 473 |
| 84327 | NM_00132956 | ZBED3 | zinc finger BED-type containing 3 | protein_coding | NP_001316493.1;NP_115743.1 | 20711 | 615250 | . | . | 1.679806 | 2.276472 | 2.224924 | 2.395487 | 3.908979 | 5.262166 | 4.992666 | 5.476996 | 1.156840 | 385.909914 | 0.204806391 | 1.026280453 | 0.304759408 | 0.998865404 | 8.483946 | 8.692746 | 8.459666 | 8.620090 | 8.579072 | 8.675602 | 355 | 440 | 331 | 419 |
| 84328 | NM_00131697 | LZIC | leucine zipper and CTNNBIP1 domain containing | protein_coding | NP_001303902.1;NP_001303903.1;NP_001303904.1;NP_001303905.1;NI | 17497 | 610458 | . | . | 16.285951 | 15.181328 | 17.727716 | 16.688522 | 37.898077 | 35.092320 | 39.780482 | 38.156322 | 1.003967 | 2328.663487 | 0.119957418 | 0.047621476 | 0.962017916 | 0.998865404 | 11.182309 | 11.188258 | 11.214150 | 11.151458 | 11.215368 | 11.156911 | 2780 | 2410 | 2103 | 2108 |
| 84330 | NM_00114617 | ZNF414 | zinc finger protein 414 | protein_coding | NP_001139647.1;NP_115746.2 | 20630 | . | . | . | 2.814993 | 2.506956 | 3.042394 | 3.858274 | 6.550606 | 5.794943 | 6.827043 | 8.821486 | 1.161246 | 126.997915 | 0.288040755 | 0.748759737 | 0.454002031 | 0.998865404 | 6.877199 | 7.092125 | 6.950219 | 6.967487 | 6.994011 | 7.029028 | 132 | 130 | 113 | 134 |
| 84331 | NM_00133122 | MCRIP2 | MAPK regulated corepressor interacting protein 2 | protein_coding | NP_001318158.1;NP_001318159.1;NP_612427.2 | 14142 | . | . | . | 11.909486 | 13.673307 | 12.244321 | 13.168855 | 27.713862 | 31.606463 | 27.475904 | 30.109024 | -1.014498 | 380.721640 | 0.188056878 | -0.11042105 | 0.912075461 | 0.998865404 | 8.583323 | 8.561782 | 8.542733 | 8.608830 | 8.541834 | 8.593075 | 417 | 437 | 314 | 370 |
| 84333 | NM_00125654 | PCGF5 | polycomb group ring finger 5 | protein_coding | NP_001243478.1;NP_001244030.1;NP_115749.2;XP_006718093.1;XP_01 | 28264 | 617407 | . | . | 9.712055 | 9.798524 | 9.576876 | 9.237654 | 22.600349 | 22.649724 | 21.490233 | 21.120800 | -1.022231 | 676.185397 | 0.154165306 | -0.20576367 | 0.83697552 | 0.998865404 | 9.416962 | 9.385415 | 9.439475 | 9.376706 | 9.387162 | 9.400061 | 823 | 704 | 576 | 633 |
| 84334 | NM_00130265 | COA8 | cytochrome c oxidase assembly factor 8 | protein_coding | NP_001289581.2;NP_001289582.2;NP_001357524.1 | 20492 | 616003 | . | . | 12.025951 | 14.722999 | 13.121770 | 14.857998 | 27.984882 | 34.032873 | 29.444871 | 33.971049 | 1.019238 | 529.595916 | 0.175735095 | 0.156437909 | 0.875687858 | 0.998865404 | 9.035410 | 9.061964 | 8.981335 | 9.097524 | 9.014095 | 9.090794 | 550 | 617 | 437 | 531 |
| 84335 | NM_00109863 | AKT1S1 | AKT1 substrate 1 | protein_coding | NP_001092102.1;NP_001092103.1;NP_001265088.1;NP_001265089.1;NI | 28426 | 610221 | . | . | 30.085931 | 30.776991 | 36.180489 | 36.444662 | 70.011198 | 71.142391 | 81.187966 | 83.326392 | 1.218733 | 2115.005342 | 0.117364579 | 2.431584312 | 0.015032949 | 0.411705946 | 10.896719 | 11.182084 | 10.947119 | 10.975964 | 11.128490 | 11.110478 | 2155 | 2095 | 2043 | 2154 |
| 84336 | NM_00130481 | TMEM101 | transmembrane protein 101 | protein_coding | NP_001291742.1;NP_001291743.1;NP_115752.1;XP_011523655.1;XP_02 | 28653 | . | . | . | 22.974815 | 23.349950 | 21.209906 | 23.636510 | 53.463338 | 53.974456 | 47.594418 | 54.042074 | -1.061468 | 1104.168876 | 0.135918884 | -0.6331814 | 0.526615192 | 0.998865404 | 10.151290 | 10.064907 | 10.125016 | 10.135875 | 10.051217 | 10.119645 | 1300 | 1236 | 888 | 1051 |
| 84337 | NM_00136367 | ELOF1 | elongation factor 1 homolog | protein_coding | NP_001350602.1;NP_001350603.1;NP_001350604.1;NP_115753.1;XP_01 | 28691 | . | . | . | 23.810812 | 24.936677 | 21.464274 | 25.271355 | 55.408741 | 57.642241 | 48.165217 | 57.779948 | -1.048858 | 775.677668 | 0.153161884 | -0.44932725 | 0.6531956 | 0.998865404 | 9.633609 | 9.564183 | 9.605502 | 9.625128 | 9.526425 | 9.634252 | 902 | 869 | 607 | 765 |
| 84340 | NM_00128130 | GFM2 | GTP dependent ribosome recycling factor mitochondrial 2 | protein_coding | NP_001268231.1;NP_115756.2;NP_733781.1;NP_733792.1;XP_0115419 | 29682 | 606544 | . | . | 16.275728 | 15.677847 | 14.870063 | 15.473233 | 37.874289 | 36.240048 | 33.367990 | 35.377713 | -1.071432 | 1451.349346 | 0.126392934 | -0.78754714 | 0.430961646 | 0.998865404 | 10.552058 | 10.452586 | 10.551529 | 10.506259 | 10.470730 | 10.481367 | 1779 | 1573 | 1210 | 1323 |
| 84342 | NM_00137487 | COG8 | component of oligomeric golgi complex 8 | protein_coding | NP_001361800.1;NP_001366190.1;NP_001366191.1;NP_001366192.1;NI | 18623 | 606979 | . | . | 4.905047 | 4.972248 | 4.713779 | 5.638936 | 11.414248 | 11.493572 | 10.577583 | 12.892757 | 1.027146 | 702.641368 | 0.156162741 | 0.247445838 | 0.804563204 | 0.998865404 | 9.437488 | 9.475550 | 9.438575 | 9.454839 | 9.413765 | 9.513810 | 789 | 757 | 574 | 716 |
| 84343 | NM_00130825 | HPS3 | HPS3 biogenesis of lysosomal organelles complex 2 subunit 1 | protein_coding | NP_001295187.1;NP_115759.2;XP_005247891.1;XP_016862812.1 | 15597 | 606118 | . | . | 10.919348 | 10.748212 | 9.603871 | 9.632534 | 25.409768 | 24.844976 | 21.550811 | 22.023646 | -1.143247 | 1390.349327 | 0.126156543 | -1.53093476 | 0.125785517 | 0.874064017 | 10.534542 | 10.341466 | 10.494534 | 10.486165 | 10.393131 | 10.382692 | 1717 | 1592 | 1136 | 1208 |
| 84364 | NM_00124283 | ARFGAP2 | ADP ribosylation factor GTPase activating protein 2 | protein_coding | NP_001229761.1;NP_115765.2;XP_005253223.1;XP_005253224.1;XP_00 | 13504 | 606908 | . | . | 8.515620 | 8.189469 | 8.563456 | 9.586232 | 19.816199 | 18.930327 | 19.216147 | 21.917778 | 1.071370 | 757.149807 | 0.150444414 | 0.661088129 | 0.508555798 | 0.998865404 | 9.513973 | 9.613191 | 9.554862 | 9.523403 | 9.550505 | 9.622831 | 861 | 771 | 646 | 772 |
| 84365 | NM_032390 | NIFK | nucleolar protein interacting with the FHA domain of MKI67 | protein_coding | NP_115766.3 | 17838 | 611970 | . | . | 31.264608 | 31.746975 | 31.091162 | 31.649700 | 72.754028 | 73.384560 | 69.767670 | 72.363281 | -1.018544 | 1651.858709 | 0.121726457 | -0.21777037 | 0.827608034 | 0.998865404 | 10.703083 | 10.676544 | 10.694351 | 10.699727 | 10.679842 | 10.685522 | 1913 | 1805 | 1418 | 1540 |
| 84376 | NM_032410 | HOOK3 | hook microtubule tethering protein 3 | protein_coding | NP_115786.1 | 23576 | 607825 | . | . | 1.398533 | 1.417302 | 1.593266 | 1.261362 | 3.254444 | 3.276158 | 3.575243 | 2.883954 | 1.000807 | 634.418423 | 0.170210273 | 0.006833326 | 0.994547837 | 0.999862634 | 9.308308 | 9.310273 | 9.305006 | 9.308759 | 9.385512 | 9.223253 | 728 | 686 | 619 | 523 |
| 84418 | NM_032412 | CYSTM1 | cysteine rich transmembrane module containing 1 | protein_coding | NP_115788.1 | 30239 | . | . | . | 58.701557 | 62.412205 | 59.695053 | 70.757278 | 136.600937 | 144.268600 | 133.953964 | 161.778091 | 1.062300 | 1547.434763 | 0.134008026 | 0.650645224 | 0.51527553 | 0.998865404 | 10.551630 | 10.638391 | 10.550372 | 10.590899 | 10.553742 | 10.677687 | 1683 | 1662 | 1276 | 1613 |
| 84436 | NM_032423,XI | ZNF528 | zinc finger protein 528 | protein_coding | NP_115799.2;XP_005259380.1;XP_006723481.1;XP_011525694.1;XP_01 | 29384 | 615580 | . | . | 1.264583 | 1.465248 | 1.627202 | 1.264394 | 2.942737 | 3.386986 | 3.651395 | 2.890886 | 1.054983 | 178.706590 | 0.261062291 | 0.295788443 | 0.76739165 | 0.998865404 | 7.441897 | 7.519946 | 7.437459 | 7.493931 | 7.544750 | 7.429994 | 187 | 200 | 181 | 149 |
| 84437 | NM_00131874 | MSANTD4 | Myb/SANT DNA binding domain containing 4 with coiled-coils | protein_coding | NP_001305676.1;NP_001305677.1;NP_001305678.1;NP_001305679.1;NI | 29383 | . | . | . | 7.899373 | 7.298316 | 7.421080 | 6.446466 | 18.382166 | 16.870380 | 16.652687 | 14.739078 | -1.116719 | 838.498240 | 0.147510967 | -1.07969176 | 0.280279464 | 0.998865404 | 9.788850 | 9.630114 | 9.770726 | 9.727214 | 9.708505 | 9.631758 | 1049 | 926 | 727 | 702 |
| 84440 | NM_00130354 | RAB11FIP4 | RAB11 family interacting protein 4 | protein_coding | NP_001290471.2;NP_001333676.1;NP_001333677.1;NP_001333678.1;NI | 30267 | 611999 | . | . | 8.434636 | 8.243800 | 7.678846 | 9.124939 | 19.627744 | 19.055916 | 17.231109 | 20.863088 | -1.012863 | 2186.707111 | 0.127269263 | -0.14488307 | 0.884803195 | 0.998865404 | 11.103863 | 11.085165 | 11.108304 | 11.089450 | 11.021339 | 11.151724 | 2564 | 2347 | 1735 | 2203 |
| 84441 | NM_032427,XI | MAML2 | mastermind like transcriptional coactivator 2 | protein_coding | NP_115803.1;XP_011541325.1;XP_011541326.1;XP_011541327.1 | 16259 | 607537 | . | . | 1.919446 | 1.754627 | 1.808752 | 1.890648 | 4.466631 | 4.055901 | 4.058789 | 4.322740 | -1.020369 | 401.371795 | 0.183730284 | -0.15833245 | 0.874194838 | 0.998865404 | 8.663047 | 8.634402 | 8.691465 | 8.615986 | 8.644983 | 8.640062 | 496 | 410 | 347 | 371 |
| 84444 | NM_032482,XI | DOT1L | DOT1 like histone lysine methyltransferase | protein_coding | NP_115871.1;XP_005259716.1;XP_005259717.1;XP_011526661.1;XP_01 | 24948 | 607375 | . | . | 5.345263 | 5.519456 | 5.771978 | 5.715730 | 12.438645 | 12.758471 | 12.952154 | 13.068337 | 1.031405 | 1383.573484 | 0.128277035 | 0.347768927 | 0.728013727 | 0.998865404 | 10.411754 | 10.456270 | 10.394462 | 10.448743 | 10.452031 | 10.439137 | 1513 | 1522 | 1232 | 1306 |
| 84445 | NM_00131809 | LZTS2 | leucine zipper tumor suppressor 2 | protein_coding | NP_001305028.1;NP_001305029.1;NP_001305030.1;NP_115805.1;XP_00 | 29381 | 610454 | . | . | 29.523318 | 29.448413 | 31.111409 | 32.614960 | 68.701973 | 68.071327 | 69.813097 | 74.570231 | 1.072074 | 2771.485827 | 0.112259009 | 0.894394358 | 0.37111093 | 0.998865404 | 11.385392 | 11.485751 | 11.407493 | 11.407194 | 11.453107 | 11.474685 | 3081 | 2886 | 2461 | 2724 |
| 84446 | NM_032430,XI | BRSK1 | BR serine/threonine kinase 1 | protein_coding | NP_115806.1;XP_005259384.1;XP_011525697.2;XP_024307507.1 | 18994 | 609235 | . | . | 2.628405 | 2.421793 | 2.647938 | 2.643273 | 6.116407 | 5.598082 | 5.941896 | 6.043528 | -1.023964 | 254.823156 | 0.217714429 | -0.15692847 | 0.875301221 | 0.998865404 | 8.010475 | 7.976024 | 8.021483 | 7.976348 | 7.957628 | 8.015982 | 310 | 266 | 206 | 250 |
| 84447 | NM_032431,N | SYVN1 | synoviolin 1 | protein_coding | NP_115807.1;NP_757385.1;XP_011543605.1 | 20738 | 608046 | . | . | 14.495316 | 15.690070 | 15.333826 | 13.774231 | 33.731196 | 36.268300 | 34.408658 | 31.493139 | -1.051198 | 1403.153896 | 0.133059261 | -0.54137125 | 0.588251721 | 0.998865404 | 10.489939 | 10.418082 | 10.444766 | 10.499882 | 10.478361 | 10.388573 | 1597 | 1607 | 1260 | 1207 |
| 84448 | NM_00113008 | ABLIM2 | actin binding LIM protein family member 2 | protein_coding | NP_001123555.1;NP_001123556.1;NP_001123557.1;NP_001123558.1;NI | 19195 | 612544 | . | . | 1.681715 | 1.693412 | 1.333124 | 2.050108 | 3.913420 | 3.914398 | 2.991491 | 4.687328 | -1.008734 | 204.759481 | 0.259857797 | -0.04828058 | 0.96149263 | 0.998865404 | 7.684952 | 7.670585 | 7.676378 | 7.676159 | 7.576087 | 7.760010 | 237 | 222 | 141 | 230 |
| 84449 | NM_00130091 | ZNF333 | zinc finger protein 333 | protein_coding | NP_001287841.1;NP_001339168.1;NP_001339169.1;NP_001339170.1;NI | 15624 | 611811 | . | . | 0.436477 | 0.421371 | 0.332063 | 0.389787 | 1.015700 | 0.974017 | 0.745140 | 0.891199 | 1.019336 | 80.963793 | 0.352790326 | 0.078315905 | 0.937576766 | 0.998865404 | 6.325592 | 6.352691 | 6.341261 | 6.331733 | 6.326501 | 6.356408 | 94 | 85 | 67 | 81 |
| 84450 | NM_00127128 | ZNF512 | zinc finger protein 512 | protein_coding | NP_001258215.1;NP_001258216.1;NP_001258217.1;NP_001258218.1;NI | 29380 | . | . | . | 8.122945 | 8.507250 | 9.891994 | 9.353855 | 18.902425 | 19.664888 | 22.197346 | 21.386478 | 1.143736 | 990.558907 | 0.137980056 | 1.404219748 | 0.160253423 | 0.933525128 | 9.851926 | 10.045766 | 9.886281 | 9.913607 | 10.019571 | 9.976429 | 1044 | 1016 | 946 | 962 |
| 84451 | NM_032435,XI | MAP3K21 | mitogen-activated protein kinase kinase kinase 21 | protein_coding | NP_115811.2;XP_011542607.1 | 29798 | 614793 | . | . | 0.604384 | 0.650350 | 0.611997 | 0.640732 | 1.406425 | 1.503313 | 1.373304 | 1.464958 | -1.016156 | 114.848330 | 0.300463221 | -0.07695508 | 0.938659277 | 0.998865404 | 6.855361 | 6.831714 | 6.835190 | 6.857281 | 6.834208 | 6.847637 | 129 | 129 | 97 | 109 |
| 84455 | NM_032437,XI | EFCAB7 | EF-hand calcium binding domain 7 | protein_coding | NP_115813.2;XP_005271331.1;XP_006711039.1;XP_006711040.1;XP_01 | 29379 | 617632 | . | . | 7.405686 | 6.373061 | 7.140851 | 5.708038 | 17.233334 | 14.731619 | 16.023863 | 13.050748 | -1.072239 | 513.207663 | 0.199449049 | -0.50452558 | 0.613892092 | 0.998865404 | 9.052129 | 8.952965 | 9.085463 | 8.951282 | 9.074608 | 8.867243 | 674 | 514 | 502 | 388 |
| 84456 | NM_00100710 | L3MBTL3 | L3MBTL histone methyl-lysine binding protein 3 | protein_coding | NP_001007103.1;NP_001333479.1;NP_001333480.1;NP_115814.1;XP_00 | 23035 | 618844 | . | . | 1.437367 | 1.097383 | 1.162774 | 1.661255 | 3.344810 | 2.536650 | 2.609232 | 3.798262 | 1.081950 | 186.947026 | 0.262840334 | 0.432329637 | 0.66550185 | 0.998865404 | 7.488878 | 7.601881 | 7.568732 | 7.480087 | 7.492516 | 7.622905 | 228 | 174 | 144 | 209 |
| 84458 | NM_00117076 | LCOR | ligand dependent nuclear receptor corepressor | protein_coding | NP_001164236.1;NP_001164237.1;NP_001333445.1;NP_115816.2 | 29503 | 607698 | . | . | 1.586816 | 1.726598 | 1.591184 | 1.261099 | 3.692586 | 3.991107 | 3.570571 | 2.883353 | -1.087496 | 600.127224 | 0.170579391 | -0.70940288 | 0.478074499 | 0.998865404 | 9.287931 | 9.167819 | 9.265295 | 9.244070 | 9.269415 | 9.123920 | 731 | 664 | 556 | 479 |
| 84460 | NM_00101165 | ZMAT1 | zinc finger matrin-type 1 | protein_coding | NP_001011657.2;NP_001269329.1;NP_001269330.1;XP_005262269.2;XP | 29377 | 301007 | . | . | 0.048430 | 0.079724 | 0.067833 | 0.046074 | 0.112698 | 0.184286 | 0.152215 | 0.105344 | -1.191560 | 10.388001 | 0.958855041 | -0.26370124 | 0.79201014 | NA | 3.497331 | 3.245375 | 3.367585 | 3.391263 | 3.380210 | 3.362603 | 10 | 15 | 10 | 7 |
| 84461 | NM_00100540 | NEURL4 | neuralized E3 ubiquitin protein ligase 4 | protein_coding | NP_001005408.1;NP_115818.2 | 34410 | 615865 | . | . | 12.268061 | 12.319175 | 12.711718 | 11.489993 | 28.548283 | 28.476325 | 28.524727 | 26.270504 | -1.030214 | 1979.048059 | 0.12317439 | -0.34864052 | 0.727359201 | 0.998865404 | 10.971784 | 10.929081 | 10.963754 | 10.959989 | 10.981429 | 10.893335 | 2318 | 2162 | 1790 | 1726 |
| 84464 | NM_032444,XI | SLX4 | SLX4 structure-specific endonuclease subunit | protein_coding | NP_115820.2;XP_011521017.1;XP_016879264.1;XP_024306239.1 | 23845 | 613278 | . | . | 2.033939 | 1.775775 | 2.259935 | 1.614804 | 4.733058 | 4.104783 | 5.071230 | 3.692060 | 1.045696 | 504.204905 | 0.185104329 | 0.34825207 | 0.727650886 | 0.998865404 | 8.944882 | 9.010114 | 8.938283 | 8.981936 | 9.071533 | 8.898971 | 548 | 550 | 509 | 418 |
| 84466 | NM_00125654 | MEGF10 | multiple EGF like domains 10 | protein_coding | NP_001243474.1;NP_001295048.1;NP_001295050.1;NP_115822.1;XP_01 | 29634 | 612453 | . | . | 0.046131 | 0.210151 | 0.143738 | 0.056700 | 0.107348 | 0.485775 | 0.322544 | 0.129638 | -1.290005 | 23.081074 | 0.735167847 | -0.4997188 | 0.617273091 | NA | 4.700668 | 4.333284 | 4.463703 | 4.581806 | 4.525948 | 4.473460 | 14 | 43 | 23 | 13 |
| 84498 | NM_00128637 | FAM120B | family with sequence similarity 120B | protein_coding | NP_001273308.1;NP_001273309.1;NP_001273310.1;NP_115824.1;XP_01 | 21109 | 612266 | . | . | 4.165777 | 4.391646 | 3.571531 | 4.450005 | 9.693934 | 10.151485 | 8.014415 | 10.174404 | -1.069434 | 632.471971 | 0.163485227 | -0.59239365 | 0.553587016 | 0.998865404 | 9.352816 | 9.255250 | 9.320204 | 9.332657 | 9.220667 | 9.338319 | 746 | 712 | 484 | 624 |
| 84501 | NM_032451 | SPIRE2 | spire type actin nucleation factor 2 | protein_coding | NP_115827.1 | 30623 | 609217 | . | . | 2.507277 | 2.748956 | 2.637103 | 3.213298 | 5.834537 | 6.354335 | 5.917583 | 7.346824 | 1.098522 | 282.973381 | 0.212784059 | 0.637095714 | 0.5240625 | 0.998865404 | 8.075745 | 8.210170 | 8.095830 | 8.138384 | 8.116734 | 8.214077 | 298 | 303 | 234 | 303 |
| 84503 | NM_032453,XI | ZNF527 | zinc finger protein 527 | protein_coding | NP_115829.1;XP_005259385.1;XP_005259386.1;XP_016882869.1;XP_01 | 29385 | . | . | . | 0.417070 | 0.421413 | 0.633152 | 0.468173 | 0.970538 | 0.974115 | 1.420775 | 1.070425 | 1.271721 | 80.001864 | 0.36395222 | 0.952822182 | 0.340680173 | 0.998865404 | 6.137262 | 6.485677 | 6.280960 | 6.281633 | 6.386616 | 6.300088 | 81 | 76 | 90 | 71 |
| 84513 | NM_00110255 | PLPP5 | phospholipid phosphatase 5 | protein_coding | NP_001096029.1;NP_001096030.1;NP_001341381.1;NP_001341383.1;NI | 25026 | 610626 | . | . | 12.010542 | 13.899023 | 14.359255 | 11.551731 | 27.949024 | 32.128216 | 32.221753 | 26.411662 | -1.013308 | 896.714745 | 0.161861981 | -0.11783615 | 0.906197485 | 0.998865404 | 9.817715 | 9.799237 | 9.788700 | 9.832195 | 9.892474 | 9.701217 | 1008 | 1003 | 878 | 724 |
| 84514 | NM_00114262 | GHDC | GH3 domain containing | protein_coding | NP_001136095.1;NP_115873.1 | 24438 | 608587 | . | . | 11.537816 | 11.294864 | 12.713868 | 13.106857 | 26.848973 | 26.108585 | 28.529550 | 29.967269 | 1.113337 | 921.833081 | 0.139427355 | 1.110904794 | 0.266609332 | 0.998274218 | 9.768835 | 9.923733 | 9.819986 | 9.795269 | 9.876020 | 9.894504 | 1022 | 925 | 834 | 922 |
| 84515 | NM_00128152 | MCM8 | minichromosome maintenance 8 homologous recombination repa | protein_coding | NP_001268449.1;NP_001268450.1;NP_001268451.1;NP_115874.3;NP_8 | 16147 | 608187 | . | . | 7.777465 | 9.230167 | 9.214055 | 10.609660 | 18.098478 | 21.335941 | 20.676074 | 24.257731 | -1.002859 | 1823.922852 | 0.130265494 | -0.03162042 | 0.974774758 | 0.998865404 | 10.834814 | 10.830842 | 10.797030 | 10.867805 | 10.883113 | 10.775741 | 2009 | 2059 | 1692 | 1591 |
| 84516 | NM_00119901 | DCTN5 | dynactin subunit 5 | protein_coding | NP_001185940.1;NP_001186672.1;NP_115875.1 | 24594 | 612962 | . | . | 8.785667 | 9.622222 | 9.477438 | 9.154791 | 20.444608 | 22.242192 | 21.267095 | 20.931343 | -1.013258 | 2931.870367 | 0.11503987 | -0.16517606 | 0.868805407 | 0.998865404 | 11.527073 | 11.508076 | 11.492808 | 11.551744 | 11.533350 | 11.490088 | 3276 | 3299 | 2600 | 2658 |
| 84517 | NM_032487 | ACTRT3 | actin related protein T3 | protein_coding | NP_115876.3 | 24022 | 608534 | . | . | 0.094080 | 0.046762 | 0.079110 | 0.103908 | 0.218929 | 0.108092 | 0.177520 | 0.237573 | 1.375174 | 4.747697 | 1.389328296 | 0.330817656 | 0.740782224 | NA | 1.998355 | 2.459419 | 2.244817 | 2.233361 | 2.241959 | 2.250038 | 6 | 3 | 4 | 6 |
| 84518 | NM_032488,XI | CNFN | cornifelin | protein_coding | NP_115877.2;XP_005259389.1;XP_011525698.1;XP_011525699.1 | 30183 | 611764 | . | . | 2.554530 | 2.866213 | 0.939720 | 2.794933 | 5.944495 | 6.625379 | 2.108704 | 6.390282 | -1.443202 | 45.928128 | 0.529323847 | -0.99990522 | 0.317356376 | 0.998865404 | 5.764158 | 5.229131 | 5.536931 | 5.559756 | 5.415345 | 5.551354 | 59 | 62 | 17 | 52 |
| 84520 | NM_032490 | GON7 | GON7 subunit of KEOPS complex | protein_coding | NP_115879.2 | 20356 | 617436 | . | . | 13.506061 | 11.563228 | 11.712596 | 10.427131 | 31.429159 | 26.728918 | 26.282722 | 23.840397 | -1.148568 | 414.590480 | 0.18817999 | -1.06194097 | 0.288262483 | 0.998865404 | 8.791523 | 8.592721 | 8.784310 | 8.682155 | 8.687682 | 8.614117 | 552 | 439 | 357 | 339 |
| 84522 | NM_00136389 | JAGN1 | jagunal homolog 1 | protein_coding | NP_001350819.1;NP_115881.3 | 26926 | 616012 | . | . | 19.248427 | 19.652397 | 20.089382 | 19.472876 | 44.791879 | 45.427397 | 45.079990 | 44.522417 | 1.002577 | 1002.683403 | 0.136468423 | 0.027212195 | 0.978290489 | 0.998865404 | 9.967747 | 9.971551 | 9.962799 | 9.974362 | 9.984571 | 9.956257 | 1144 | 1089 | 890 | 923 |
| 84524 | NM_032494,XI | ZC3H8 | zinc finger CCCH-type containing 8 | protein_coding | NP_115883.2;XP_016860596.1;XP_024308942.1;XP_024308943.1 | 30941 | . | . | . | 3.295521 | 2.990646 | 3.988307 | 3.261532 | 7.668810 | 6.913012 | 8.949644 | 7.457105 | 1.131781 | 562.846241 | 0.16726763 | 1.067717847 | 0.285647805 | 0.998865404 | 9.044188 | 9.223442 | 9.100101 | 9.085224 | 9.224960 | 9.119867 | 615 | 563 | 561 | 516 |
| 84527 | NM_00120240 | ZNF559 | zinc finger protein 559 | protein_coding | NP_001189335.1;NP_001189336.1;NP_001189337.1;NP_001189338.1;NI | 28197 | . | . | . | 2.085766 | 2.155041 | 1.755736 | 1.501377 | 4.853660 | 4.981472 | 3.939823 | 3.432720 | -1.200484 | 183.847783 | 0.250318776 | -1.05311986 | 0.29228606 | 0.998865404 | 7.647741 | 7.385054 | 7.550116 | 7.569446 | 7.518398 | 7.442142 | 226 | 221 | 159 | 142 |
| 84529 | NM_00113001 | CDIN1 | CDAN1 interacting nuclease 1 | protein_coding | NP_001123482.1;NP_001277161.1;NP_001277162.1;NP_001308685.1;NI | 26929 | 615626 | . | . | 2.065469 | 2.409935 | 2.859280 | 2.524845 | 4.806430 | 5.570670 | 6.416143 | 5.772757 | 1.118607 | 217.601725 | 0.236527106 | 0.683655031 | 0.494193026 | 0.998865404 | 7.682336 | 7.844218 | 7.698318 | 7.770800 | 7.825548 | 7.748533 | 218 | 239 | 216 | 198 |
| 84532 | NM_00125267 | ACSS1 | acyl-CoA synthetase short chain family member 1 | protein_coding | NP_001239604.1;NP_001239605.1;NP_001239606.1;NP_115890.2;XP_00 | 16091 | 614355 | . | . | 18.100898 | 17.820090 | 19.296868 | 20.023477 | 42.121533 | 41.191937 | 43.301614 | 45.781296 | 1.056385 | 2086.033130 | 0.117062019 | 0.676011395 | 0.499033411 | 0.998865404 | 10.986433 | 11.065574 | 11.012881 | 10.995349 | 11.042393 | 11.053362 | 2362 | 2165 | 1851 | 2023 |
| 84536 | NR_027128,NF | LINC01547 | long intergenic non-protein coding RNA 1547 | lncRNA | . | 15707 | . | . | . | 0.526356 | 0.544936 | 0.566520 | 0.701414 | 1.224852 | 1.259644 | 1.271254 | 1.603699 | 1.231195 | 33.483478 | 0.535952529 | 0.559861591 | 0.575573849 | 0.998865404 | 4.908124 | 5.207163 | 5.048593 | 5.045724 | 5.060821 | 5.088630 | 35 | 32 | 29 | 38 |
| 84541 | NM_032505,XI | KBTBD8 | kelch repeat and BTB domain containing 8 | protein_coding | NP_115894.2;XP_005264828.1 | 30691 | 616607 | . | . | 0.902878 | 0.905310 | 1.021125 | 0.529598 | 2.101034 | 2.092666 | 2.291374 | 1.210863 | -1.218513 | 117.736397 | 0.333542477 | -0.85482872 | 0.392645974 | 0.998865404 | 7.013734 | 6.731299 | 6.896327 | 6.914692 | 6.921672 | 6.748651 | 145 | 143 | 118 | 72 |
| 84542 | NM_00112999 | KIAA1841 | KIAA1841 | protein_coding | NP_001123465.1;NP_001317361.1;NP_001317362.1;NP_001317363.1;NI | 29387 | . | . | . | 2.390789 | 2.442763 | 2.630947 | 2.227513 | 5.563463 | 5.646556 | 5.903767 | 5.092943 | -1.010815 | 342.631805 | 0.196886627 | -0.07882178 | 0.937174377 | 0.998865404 | 8.427876 | 8.413117 | 8.418751 | 8.426086 | 8.463443 | 8.367242 | 394 | 374 | 322 | 292 |
| 84545 | NM_00130839 | MRPL43 | mitochondrial ribosomal protein L43 | protein_coding | NP_001295325.1;NP_115488.2;NP_789762.1;NP_789763.1;NP_789764.1 | 14517 | 611848 | . | . | 26.977111 | 28.657835 | 27.506357 | 30.998884 | 62.776849 | 66.243870 | 61.723465 | 70.875269 | 1.041595 | 889.329052 | 0.145101164 | 0.405192445 | 0.685336058 | 0.998865404 | 9.767124 | 9.825434 | 9.762895 | 9.799229 | 9.769440 | 9.849715 | 978 | 964 | 745 | 898 |
| 84546 | NR_001285 | SNORD35B | small nucleolar RNA, C/D box 35B | snoRNA | . | 17365 | . | . | . | 10.141864 | 13.359252 | 14.555841 | 7.584228 | 23.600534 | 30.880508 | 32.662884 | 17.340435 | -1.069257 | 31.827705 | 0.57656913 | -0.16755668 | 0.866932054 | 0.998865404 | 5.038444 | 4.944447 | 4.973298 | 5.007661 | 5.018603 | 4.944163 | 33 | 40 | 35 | 20 |
| 84547 | NM_00118474 | PGBD1 | piggyBac transposable element derived 1 | protein_coding | NP_001171672.1;NP_115896.1;XP_006715291.1;XP_011513244.1;XP_01 | 19398 | . | . | . | 2.035875 | 1.659589 | 2.318770 | 1.601570 | 4.737566 | 3.836212 | 5.203251 | 3.661802 | 1.059807 | 216.301489 | 0.256919644 | 0.326178129 | 0.744289588 | 0.998865404 | 7.713255 | 7.799263 | 7.794171 | 7.676918 | 7.839788 | 7.683087 | 272 | 198 | 226 | 174 |
| 84548 | NM_00117409 | TMEM185A | transmembrane protein 185A | protein_coding | NP_001167563.1;NP_001269231.1;NP_115897.1 | 17125 | 300031 | . | . | 4.599038 | 4.429829 | 4.447521 | 5.027713 | 10.702150 | 10.239749 | 9.980110 | 11.495267 | 1.060038 | 363.608936 | 0.187796836 | 0.447912229 | 0.654216542 | 0.998865404 | 8.463818 | 8.547458 | 8.476408 | 8.500661 | 8.501676 | 8.542080 | 398 | 389 | 314 | 363 |
| 84549 | NM_032509 | MAK16 | MAK16 homolog | protein_coding | NP_115898.2 | 13703 | . | . | . | 6.597566 | 7.000147 | 6.772288 | 6.151836 | 15.352807 | 16.181154 | 15.196818 | 14.065441 | -1.066138 | 736.576725 | 0.151958785 | -0.60802443 | 0.543171273 | 0.998865404 | 9.570020 | 9.477893 | 9.528068 | 9.563816 | 9.537410 | 9.464760 | 853 | 841 | 653 | 633 |
| 84552 | NM_032510 | PARD6G | par-6 family cell polarity regulator gamma | protein_coding | NP_115899.1 | 16076 | 608976 | . | . | 4.661775 | 4.090998 | 3.712584 | 4.010947 | 10.848141 | 9.456526 | 8.330931 | 9.170554 | -1.148726 | 492.839284 | 0.173936518 | -1.15004342 | 0.250125989 | 0.993339639 | 9.041477 | 8.841549 | 9.031280 | 8.940201 | 8.879524 | 8.918694 | 649 | 529 | 386 | 444 |
| 84553 | NM_00134653 | FAXC | failed axon connections homolog, metaxin like GST domain contai | protein_coding | NP_001333459.1;NP_001333460.1;NP_001333461.1;NP_001333462.1;NI | 20742 | . | . | . | 1.662357 | 1.673932 | 1.194246 | 1.712758 | 3.868372 | 3.869367 | 2.679855 | 3.916019 | -1.209161 | 537.093483 | 0.191436955 | -1.43131611 | 0.152339639 | 0.916288189 | 9.200121 | 8.924824 | 9.108955 | 9.139079 | 8.887783 | 9.103422 | 662 | 649 | 349 | 535 |
| 84557 | NM_032514,N | MAP1LC3A | microtubule associated protein 1 light chain 3 alpha | protein_coding | NP_115903.1;NP_852610.1;XP_011527385.1;XP_011527386.1;XP_01152 | 6838 | 601242 | . | . | 1.131078 | 1.020530 | 0.664944 | 0.779368 | 2.632064 | 2.359001 | 1.492115 | 1.781932 | -1.499371 | 27.603304 | 0.591242966 | -0.98835401 | 0.322979301 | 0.998865404 | 5.049783 | 4.464871 | 4.820273 | 4.806800 | 4.763228 | 4.774018 | 40 | 34 | 18 | 22 |
| 84561 | NM_00119548 | SLC12A8 | solute carrier family 12 member 8 | protein_coding | NP_001182412.2;NP_078904.4 | 15595 | 611316 | . | . | 0.254057 | 0.465712 | 0.217399 | 0.153208 | 0.591201 | 1.076514 | 0.487837 | 0.350292 | -1.933040 | 30.211421 | 0.600545459 | -1.58334662 | 0.113342482 | 0.850211417 | 5.315232 | 4.365137 | 4.904313 | 4.992868 | 4.886789 | 4.856964 | 33 | 55 | 21 | 16 |
| 84570 | NM_00125607 | COL25A1 | collagen type XXV alpha 1 chain | protein_coding | NP_001243003.1;NP_115907.2;NP_942014.1;XP_011530635.1;XP_01153 | 18603 | 610004 | . | . | 0.382507 | 0.417841 | 0.093095 | 0.146929 | 0.890111 | 0.965857 | 0.208902 | 0.335937 | -2.108999 | 49.897535 | 0.450406597 | -2.39019212 | 0.016839561 | 0.436090928 | 6.080986 | 5.004486 | 5.701598 | 5.700517 | 5.571126 | 5.569149 | 78 | 73 | 28 | 30 |
| 84572 | NM_032520,XI | GNPTG | N-acetylglucosamine-1-phosphate transferase subunit gamma | protein_coding | NP_115909.1;XP_016879271.1;XP_016879272.1 | 23026 | 607838 | . | . | 28.721165 | 28.090209 | 27.957108 | 29.621994 | 66.835327 | 64.931776 | 62.734940 | 67.727171 | 1.014838 | 1711.864080 | 0.121610163 | 0.174733816 | 0.861288795 | 0.998865404 | 10.730701 | 10.751926 | 10.746379 | 10.724493 | 10.735176 | 10.758740 | 1984 | 1808 | 1477 | 1641 |
| 84612 | NM_032521 | PARD6B | par-6 family cell polarity regulator beta | protein_coding | NP_115910.1 | 16245 | 608975 | . | . | 0.366570 | 0.405768 | 0.298850 | 0.362926 | 0.853024 | 0.937951 | 0.670610 | 0.829788 | -1.186504 | 51.277239 | 0.439505086 | -0.56135159 | 0.574557878 | 0.998865404 | 5.799086 | 5.550733 | 5.687879 | 5.707659 | 5.649947 | 5.681210 | 61 | 63 | 37 | 48 |
| 84614 | NM_00112277 | ZBTB37 | zinc finger and BTB domain containing 37 | protein_coding | NP_001116242.1;NP_001333044.1;NP_001356775.1;NP_115911.1;XP_00 | 28365 | . | . | . | 0.480244 | 0.562179 | 0.552475 | 0.437001 | 1.117547 | 1.299502 | 1.239741 | 0.999152 | -1.042203 | 317.475614 | 0.215770123 | -0.27638932 | 0.782249059 | 0.998865404 | 8.339613 | 8.280791 | 8.280234 | 8.355364 | 8.369942 | 8.216011 | 348 | 373 | 309 | 250 |
| 84617 | NM_00130352 | TUBB6 | tubulin beta 6 class V | protein_coding | NP_001290453.1;NP_001290454.1;NP_001290455.1;NP_001290456.1;NI | 20776 | 615103 | . | . | 69.986459 | 66.457927 | 57.568256 | 68.633936 | 162.861377 | 153.620480 | 129.181493 | 156.923332 | -1.097305 | 3853.090555 | 0.121497755 | -1.10261181 | 0.270195781 | 0.998865404 | 11.977298 | 11.843187 | 11.971989 | 11.923944 | 11.801494 | 11.936590 | 4776 | 4226 | 2932 | 3728 |
| 84619 | NM_00108311 | ZGPAT | zinc finger CCCH-type and G-patch domain containing | protein_coding | NP_001076582.1;NP_001182582.1;NP_001182583.1;NP_115916.3;NP_8 | 15948 | . | . | . | 14.284021 | 15.144377 | 17.144895 | 17.651919 | 33.239504 | 35.006905 | 38.472647 | 40.359017 | 1.168068 | 959.550114 | 0.13831413 | 1.62039791 | 0.105146829 | 0.832886861 | 9.789905 | 10.013842 | 9.828730 | 9.863010 | 9.951452 | 9.966547 | 995 | 978 | 890 | 979 |
| 84622 | NM_032530,XI | ZNF594 | zinc finger protein 594 | protein_coding | NP_115919.1;XP_005256884.1 | 29392 | . | . | . | 1.376800 | 1.668729 | 2.068822 | 1.730531 | 3.203869 | 3.857342 | 4.642375 | 3.956655 | 1.273953 | 278.965385 | 0.216129457 | 1.616219223 | 0.106046924 | 0.833478683 | 7.938606 | 8.288157 | 8.016192 | 8.085324 | 8.224998 | 8.128186 | 263 | 283 | 296 | 267 |
| 84624 | NM_032532,XI | FNDC1 | fibronectin type III domain containing 1 | protein_coding | NP_115921.2;XP_011534492.1;XP_011534493.1 | 21184 | 609991 | . | . | 0.099994 | 0.118212 | 0.089141 | 0.136461 | 0.232690 | 0.273251 | 0.200031 | 0.312001 | 1.008202 | 22.999413 | 0.650588943 | 0.018113185 | 0.98554856 | NA | 4.519313 | 4.527725 | 4.513400 | 4.531319 | 4.504039 | 4.542029 | 24 | 27 | 16 | 26 |
| 84626 | NM_00129018 | KRBA1 | KRAB-A domain containing 1 | protein_coding | NP_001277116.1;NP_115923.2;XP_011514938.1;XP_011514939.1;XP_01 | 22228 | . | . | . | 0.893909 | 0.879551 | 1.249587 | 1.234058 | 2.080163 | 2.033120 | 2.804033 | 2.821526 | 1.426980 | 132.783601 | 0.281530799 | 1.822058142 | 0.068446164 | 0.741130004 | 6.773719 | 7.286773 | 6.973219 | 6.979030 | 7.110732 | 7.098561 | 125 | 119 | 138 | 144 |
| 84629 | NM_00108049 | TNRC18 | trinucleotide repeat containing 18 | protein_coding | NP_001073964.2;XP_016868217.1;XP_016868219.1;XP_016868220.1;XP_ | 11962 | . | . | . | 7.538104 | 8.127577 | 7.070462 | 8.713448 | 17.541480 | 18.787261 | 15.865911 | 19.922261 | -1.006228 | 2541.666932 | 0.13040801 | -0.0686827 | 0.94524219 | 0.998865404 | 11.316202 | 11.306902 | 11.288230 | 11.336964 | 11.225738 | 11.383692 | 2848 | 2834 | 1988 | 2607 |
| 84631 | NM_00114400 | SLITRK2 | SLIT and NTRK like family member 2 | protein_coding | NP_001137475.1;NP_001137476.1;NP_001137477.1;NP_001137478.1;NI | 13449 | 300561 | . | . | 1.633614 | 1.548258 | 2.079090 | 2.203863 | 3.801488 | 3.578869 | 4.665420 | 5.038868 | 1.406373 | 456.760864 | 0.17534479 | 2.805783476 | 0.005019441 | 0.213837476 | 8.568315 | 9.060499 | 8.699736 | 8.729752 | 8.957776 | 8.891673 | 426 | 420 | 488 | 475 |
| 84632 | NM_00100193 | AFAP1L2 | actin filament associated protein 1 like 2 | protein_coding | NP_001001936.1;NP_001274753.1;NP_001337993.1;NP_001337994.1;NI | 25901 | 612420 | . | . | 2.249469 | 2.189128 | 2.046998 | 2.666940 | 5.234604 | 5.060268 | 4.593406 | 6.097641 | 1.057758 | 278.151514 | 0.218768234 | 0.370295781 | 0.711162117 | 0.998865404 | 8.079293 | 8.159059 | 8.095366 | 8.110298 | 8.063312 | 8.196422 | 307 | 296 | 217 | 302 |
| 84640 | NM_00129032 | USP38 | ubiquitin specific peptidase 38 | protein_coding | NP_001277254.1;NP_001277255.1;NP_115946.2;XP_011530662.1 | 20067 | 618322 | . | . | 1.857927 | 2.092292 | 1.918189 | 1.798472 | 4.323473 | 4.836426 | 4.304359 | 4.111994 | -1.105923 | 409.268836 | 0.182555013 | -0.79574596 | 0.426179708 | 0.998865404 | 8.747830 | 8.602312 | 8.668000 | 8.742542 | 8.652738 | 8.638660 | 465 | 492 | 342 | 360 |
| 84641 | NM_032558,XI | MFSD14B | major facilitator superfamily domain containing 14B | protein_coding | NP_115947.2;XP_011517416.1;XP_016870706.1;XP_016870707.1;XP_01 | 23376 | . | . | . | 21.768699 | 20.503262 | 18.902506 | 18.863325 | 50.656652 | 47.394207 | 42.416674 | 43.128754 | -1.132671 | 2099.299336 | 0.118407915 | -1.51788063 | 0.129044493 | 0.878737612 | 11.122696 | 10.943103 | 11.104920 | 11.061088 | 10.994498 | 10.973392 | 2637 | 2341 | 1735 | 1821 |
| 84645 | NM_00120706 | C22orf23 | chromosome 22 open reading frame 23 | protein_coding | NP_001193991.1;NP_115950.3;XP_005261838.1;XP_005261839.1;XP_00 | 18589 | . | . | . | 1.103656 | 1.145690 | 0.647965 | 1.519694 | 2.568255 | 2.648315 | 1.454013 | 3.474599 | -1.143461 | 58.746832 | 0.436933844 | -0.44264512 | 0.658022442 | 0.998865404 | 5.971407 | 5.774776 | 5.891644 | 5.886778 | 5.804380 | 5.914936 | 73 | 67 | 34 | 66 |
| 84649 | NM_00125389 | DGAT2 | diacylglycerol O-acyltransferase 2 | protein_coding | NP_001240820.1;NP_115953.2;XP_011543606.1 | 16940 | 606983 | . | . | 3.340675 | 3.300400 | 4.099990 | 3.308796 | 7.773887 | 7.629021 | 9.200259 | 7.565168 | 1.105376 | 262.930979 | 0.22036125 | 0.655909176 | 0.511882564 | 0.998865404 | 7.963900 | 8.109508 | 8.011861 | 8.006762 | 8.116624 | 8.001173 | 289 | 268 | 267 | 230 |
| 84650 | NM_00127863 | EBPL | EBP like | protein_coding | NP_001265565.1;NP_115954.1 | 18061 | 617335 | . | . | 12.763756 | 11.229305 | 11.133763 | 10.501081 | 29.701786 | 25.957042 | 24.983837 | 24.009474 | -1.123835 | 348.801051 | 0.194683648 | -0.86514987 | 0.386956534 | 0.998865404 | 8.527663 | 8.359989 | 8.517540 | 8.437045 | 8.429767 | 8.393925 | 454 | 371 | 295 | 298 |
| 84656 | NM_00130809 | GLYR1 | glyoxylate reductase 1 homolog | protein_coding | NP_001295025.1;NP_001311025.2;NP_001311026.2;NP_001311027.2;NI | 24434 | 610660 | . | . | 12.899648 | 15.056562 | 15.605977 | 15.614924 | 30.018012 | 34.803913 | 35.019363 | 35.701670 | 1.107981 | 1757.722185 | 0.129722173 | 1.140386485 | 0.254125321 | 0.994807659 | 10.703758 | 10.851447 | 10.676086 | 10.794217 | 10.815975 | 10.818683 | 1775 | 1936 | 1604 | 1735 |
| 84657 | NR_026829 | LINC00852 | long intergenic non-protein coding RNA 852 | lncRNA | . | 29904 | . | . | . | 0.207164 | 0.167450 | 0.271881 | 0.231259 | 0.482080 | 0.387068 | 0.610093 | 0.528747 | 1.308909 | 9.299531 | 0.995064628 | 0.39029116 | 0.696321259 | NA | 3.008843 | 3.399154 | 3.208049 | 3.202245 | 3.220355 | 3.212226 | 10 | 8 | 10 | 9 |
| 84660 | NM_201435,N | CCDC62 | coiled-coil domain containing 62 | protein_coding | NP_958843.2;XP_006719706.1;XP_006719707.1;XP_011537158.1;XP_01 | 30723 | 613481 | . | . | 0.229351 | 0.196986 | 0.157568 | 0.115733 | 0.533709 | 0.455343 | 0.353578 | 0.264610 | -1.636150 | 21.991343 | 0.667783524 | -1.06367523 | 0.287475849 | NA | 4.769484 | 4.062477 | 4.494299 | 4.475107 | 4.452391 | 4.428282 | 34 | 27 | 17 | 13 |
| 84661 | NM_00132120 | DPY30 | dpy-30 histone methyltransferase complex regulatory subunit | protein_coding | NP_001308138.1;NP_001308139.1;NP_115963.1;XP_006712180.1;XP_00 | 24590 | 612032 | . | . | 42.827850 | 35.272883 | 37.668875 | 32.835854 | 99.662161 | 81.534847 | 84.527860 | 75.075289 | -1.099826 | 829.105469 | 0.159925141 | -0.85837069 | 0.390687807 | 0.998865404 | 9.762042 | 9.625555 | 9.795987 | 9.652044 | 9.702964 | 9.613403 | 1099 | 844 | 731 | 693 |
| 84662 | NM_00131891 | GLIS2 | GLIS family zinc finger 2 | protein_coding | NP_001305847.1;NP_115964.2 | 29450 | 608539 | . | . | 0.150909 | 0.193844 | 0.289020 | 0.309708 | 0.351171 | 0.448079 | 0.648554 | 0.708111 | 1.498719 | 30.911481 | 0.559492619 | 1.043320954 | 0.296799669 | 0.998865404 | 4.629747 | 5.212035 | 4.900323 | 4.926561 | 4.962097 | 4.964617 | 25 | 30 | 32 | 35 |
| 84665 | NM_00125626 | MYPN | myopalladin | protein_coding | NP_001243196.1;NP_001243197.1;NP_115967.2;XP_016872322.1;XP_01 | 23246 | 608517 | . | . | 0.186966 | 0.105510 | 0.206764 | 0.093802 | 0.435078 | 0.243891 | 0.463974 | 0.214468 | 1.022293 | 28.811347 | 0.631365326 | 0.050381257 | 0.959818573 | 0.998865404 | 4.829417 | 4.867463 | 4.870180 | 4.803012 | 4.882451 | 4.794798 | 42 | 22 | 35 | 17 |
| 84667 | NM_00116596 | HES7 | hes family bHLH transcription factor 7 | protein_coding | NP_001159439.1;NP_115969.2;XP_016880721.1;XP_024306775.1 | 15977 | 608059 | . | . | 0.941327 | 1.051722 | 1.240146 | 1.198349 | 2.190506 | 2.431102 | 2.782851 | 2.739883 | -1.018045 | 74.934422 | 0.368141153 | -0.0700844 | 0.944126484 | 0.998865404 | 6.239736 | 6.215274 | 6.249804 | 6.208504 | 6.237632 | 6.212008 | 94 | 75 | 68 | 66 |
| 84668 | NM_00136346 | FAM126A | family with sequence similarity 126 member A | protein_coding | NP_001350395.1;NP_001350396.1;NP_115970.2;XP_011513891.1;XP_01 | 24587 | 610531 | . | . | 10.519945 | 9.922856 | 9.844623 | 9.378726 | 24.480345 | 22.937126 | 22.091051 | 21.443343 | -1.081845 | 4104.646751 | 0.111499117 | -1.01788538 | 0.308732432 | 0.998865404 | 12.058608 | 11.945251 | 12.063813 | 12.004463 | 11.994174 | 11.943838 | 5086 | 4440 | 3534 | 3584 |
| 84669 | NM_032582,XI | USP32 | ubiquitin specific peptidase 32 | protein_coding | NP_115971.2;XP_011523673.1;XP_011523674.1;XP_011523675.1;XP_01 | 19143 | 607740 | . | . | 12.754109 | 12.967233 | 15.413658 | 16.242442 | 29.679340 | 29.974341 | 34.587802 | 37.136410 | 1.211677 | 3086.430105 | 0.111761249 | 2.478545962 | 0.013191911 | 0.380410307 | 11.446629 | 11.723550 | 11.504911 | 11.510443 | 11.642959 | 11.685845 | 3203 | 3022 | 2865 | 3252 |
| 84674 | NM_032587,XI | CARD6 | caspase recruitment domain family member 6 | protein_coding | NP_115976.2;XP_016865478.1;XP_016865479.1 | 16394 | 609986 | . | . | 1.144444 | 1.033055 | 1.253342 | 1.594141 | 2.663169 | 2.387954 | 2.812464 | 3.644814 | 1.237386 | 154.683119 | 0.267876882 | 1.147153296 | 0.25131827 | 0.993682637 | 7.111497 | 7.418541 | 7.245432 | 7.204513 | 7.278563 | 7.338982 | 168 | 141 | 138 | 172 |
| 84676 | NM_032588,XI | TRIM63 | tripartite motif containing 63 | protein_coding | NP_115977.2;XP_016858048.1 | 16007 | 606131 | . | . | 16.381939 | 16.221126 | 20.206448 | 18.652373 | 38.121441 | 37.495858 | 45.342683 | 42.646433 | 1.177496 | 982.905415 | 0.139344355 | 1.691654855 | 0.090711801 | 0.805424371 | 9.818073 | 10.054102 | 9.882713 | 9.871048 | 10.028408 | 9.963317 | 1048 | 966 | 965 | 953 |
| 84677 | NR_026838,NF | DSCR8 | Down syndrome critical region 8 | lncRNA | . | 16707 | 613396 | . | . | 5.144669 | 3.714060 | 3.696835 | 4.403309 | 11.971857 | 8.585217 | 8.295591 | 10.067642 | -1.084536 | 77.725182 | 0.364188933 | -0.32147568 | 0.747849942 | 0.998865404 | 6.337494 | 6.220766 | 6.319454 | 6.262088 | 6.259637 | 6.280956 | 103 | 78 | 62 | 73 |
| 84678 | NM_00100536 | KDM2B | lysine demethylase 2B | protein_coding | NP_001005366.1;NP_115979.3;XP_005254012.1;XP_005254013.1;XP_00 | 13610 | 609078 | . | . | 4.413592 | 4.661466 | 5.683701 | 4.882957 | 10.270609 | 10.775188 | 12.754058 | 11.164303 | 1.144348 | 868.141453 | 0.151562691 | 1.283466748 | 0.199328602 | 0.96621494 | 9.660951 | 9.856032 | 9.713042 | 9.705876 | 9.870012 | 9.738068 | 937 | 869 | 879 | 790 |
| 84679 | NM_00125729 | SLC9A7 | solute carrier family 9 member A7 | protein_coding | NP_001244220.1;NP_115980.1;XP_005272738.1;XP_005272739.1;XP_00 | 17123 | 300368 | . | . | 0.693061 | 0.643870 | 0.543642 | 0.637537 | 1.612780 | 1.488335 | 1.219917 | 1.457652 | -1.116314 | 184.882184 | 0.244210433 | -0.65002763 | 0.515674376 | 0.998865404 | 7.607820 | 7.448720 | 7.551365 | 7.557701 | 7.496242 | 7.517285 | 223 | 212 | 148 | 168 |
| 84680 | NM_00112721 | ACCS | 1-aminocyclopropane-1-carboxylate synthase homolog (inactive) | protein_coding | NP_001120691.1;NP_115981.1;XP_005253227.2;XP_005253229.1;XP_00 | 23989 | 608405 | . | . | 5.844664 | 5.368952 | 6.544154 | 7.284713 | 13.600773 | 12.410576 | 14.684891 | 16.655634 | 1.178023 | 557.092083 | 0.161862736 | 1.460298275 | 0.144208116 | 0.906161673 | 8.998870 | 9.235021 | 9.044442 | 9.083826 | 9.169643 | 9.172470 | 571 | 569 | 523 | 566 |

| | | | | | | | | | | | | | | | | | | | | | | | | | | | | | | | | | |
|---|---|---|---|---|---|---|---|---|---|---|---|---|---|---|---|---|---|---|---|---|---|---|---|---|---|---|---|---|---|---|---|---|---|
| 84681 | NM_032593,XI | HINT2 | histidine triad nucleotide binding protein 2 | protein_coding | NP_115982.1;XP_024303470.1;XP_024303471.1;XP_024303472.1 | 18344 | 609997 | . | . | 46.223838 | 44.915521 | 38.524027 | 49.089111 | 107.564764 | 103.824234 | 86.446790 | 112.236419 | -1.051612 | 910.621481 | 0.156931977 | -0.4626353 | 0.643625801 | 0.998865404 | 9.866843 | 9.793644 | 9.861288 | 9.832909 | 9.722272 | 9.892130 | 1096 | 988 | 680 | 931 |
| 84684 | NM_032594 | INSM2 | INSM transcriptional repressor 2 | protein_coding | NP_115983.3 | 17539 | 614027 | . | . | 0.034931 | 0.070416 | 0.018493 | 0.043204 | 0.081285 | 0.162770 | 0.041498 | 0.098782 | -1.509307 | 4.131900 | 1.501259357 | -0.39559188 | 0.692406128 | NA | 2.316288 | 1.715150 | 2.049618 | 2.063685 | 2.044653 | 2.053357 | 4 | 7 | 2 | 4 |
| 84687 | NM_032595 | PPP1R9B | protein phosphatase 1 regulatory subunit 9B | protein_coding | NP_115984.3 | 9298 | 603325 | . | . | 11.113606 | 11.776610 | 10.436264 | 12.296549 | 25.861820 | 27.222164 | 23.418673 | 28.114595 | -1.020964 | 1496.746769 | 0.133994576 | -0.22338634 | 0.823234834 | 0.998865404 | 10.562720 | 10.532350 | 10.535744 | 10.573782 | 10.478036 | 10.596095 | 1699 | 1672 | 1189 | 1495 |
| 84688 | NM_00125219 | C9orf24 | chromosome 9 open reading frame 24 | protein_coding | NP_001239124.1;NP_115985.2;NP_671697.1;NP_671698.1;XP_00525167 | 19919 | . | . | . | 0.145143 | 0.076198 | 0.228416 | 0.227334 | 0.337754 | 0.176136 | 0.512559 | 0.519772 | 1.371510 | 19.055340 | 0.707792904 | 0.643924861 | 0.51962416 | NA | 4.006229 | 4.462148 | 4.242206 | 4.223148 | 4.247829 | 4.268551 | 21 | 15 | 17 | 23 |
| 84690 | NM_00117069 | SPATA22 | spermatogenesis associated 22 | protein_coding | NP_001164166.1;NP_001164167.1;NP_001164168.1;NP_001164169.1;NI | 30705 | 617673 | . | . | 2.064218 | 2.182918 | 3.343909 | 1.976965 | 4.803520 | 5.045912 | 7.503633 | 4.520097 | 1.244084 | 130.510867 | 0.322139276 | 0.978099688 | 0.328025022 | 0.998865404 | 6.860761 | 7.177898 | 6.963745 | 6.983640 | 7.150113 | 6.944077 | 130 | 129 | 160 | 99 |
| 84693 | NM_032601,XI | MCEE | methylmalonyl-CoA epimerase | protein_coding | NP_115990.3;XP_005264670.1 | 16732 | 608419 | . | . | 4.846911 | 5.978793 | 4.736357 | 5.241165 | 11.278962 | 13.820251 | 10.628248 | 11.983300 | -1.106970 | 140.426841 | 0.283097277 | -0.5179006 | 0.604527625 | 0.998865404 | 7.205680 | 7.057886 | 7.104159 | 7.196458 | 7.104513 | 7.121903 | 150 | 178 | 113 | 128 |
| 84695 | NM_00128916 | LOXL3 | lysyl oxidase like 3 | protein_coding | NP_001276093.1;NP_001276094.1;NP_115992.1;XP_011531436.1;XP_01 | 13869 | 607163 | . | . | 218.993264 | 226.342918 | 212.181708 | 233.156777 | 509.606334 | 523.201782 | 476.129610 | 533.085264 | -1.054566 | 24099.076742 | 0.101098965 | -0.75816452 | 0.448352496 | 0.998865404 | 14.594517 | 14.517845 | 14.568407 | 14.589483 | 14.502723 | 14.562863 | 28140 | 27020 | 19684 | 22764 |
| 84698 | NM_00128654 | CAPS2 | calcyphosine 2 | protein_coding | NP_001273476.1;NP_001273477.1;NP_001341952.1;NP_001341953.1;NI | 16471 | 607724 | . | . | 0.212319 | 0.373257 | 0.184232 | 0.367680 | 0.494074 | 0.862800 | 0.413411 | 0.840660 | -1.092046 | 45.850549 | 0.514956236 | -0.24668745 | 0.805150119 | 0.998865404 | 5.583502 | 5.451201 | 5.466974 | 5.567819 | 5.451152 | 5.553594 | 40 | 66 | 26 | 54 |
| 84705 | NM_00112885 | GTPBP3 | GTP binding protein 3, mitochondrial | protein_coding | NP_001122327.1;NP_001182351.1;NP_116009.2;NP_598399.2 | 14880 | 608536 | . | . | 8.016479 | 8.064427 | 7.462230 | 7.652334 | 18.654675 | 18.641284 | 16.745027 | 17.496150 | -1.079347 | 625.261141 | 0.155157404 | -0.70998387 | 0.477714142 | 0.998865404 | 9.342361 | 9.232165 | 9.315014 | 9.312117 | 9.258770 | 9.266070 | 749 | 699 | 520 | 567 |
| 84706 | NM_00114246 | GPT2 | glutamic--pyruvic transaminase 2 | protein_coding | NP_001135938.1;NP_597700.1;XP_016879279.1 | 18062 | 138210 | . | . | 7.484407 | 7.474947 | 6.372619 | 7.623326 | 17.416522 | 17.278673 | 14.299971 | 17.429827 | -1.082412 | 901.889405 | 0.148927869 | -0.76714745 | 0.442993855 | 0.998865404 | 9.873053 | 9.758278 | 9.847685 | 9.840556 | 9.724871 | 9.845438 | 1085 | 1007 | 689 | 881 |
| 84717 | NM_00100152 | HDGFL2 | HDGF like 2 | protein_coding | NP_001001520.1;NP_001335098.1;NP_116020.1;XP_011526678.1;XP_01 | 14680 | 617884 | . | . | 22.130188 | 22.175801 | 20.487901 | 23.681905 | 51.497856 | 51.260356 | 45.974257 | 54.145860 | -1.014073 | 1577.531109 | 0.13063908 | -0.15432791 | 0.87735119 | 0.998865404 | 10.633656 | 10.613176 | 10.630354 | 10.626418 | 10.561421 | 10.670539 | 1834 | 1710 | 1266 | 1572 |
| 84719 | NR_046213 | LINC00260 | long intergenic non-protein coding RNA 260 | lncRNA | . | 28770 | . | . | . | 0.307341 | 0.517812 | 0.524186 | 0.206638 | 0.715196 | 1.196946 | 1.176258 | 0.472453 | -1.132979 | 16.181152 | 0.809269957 | -0.22257188 | 0.823868726 | NA | 4.101892 | 3.925185 | 3.994413 | 4.030987 | 4.031962 | 3.975545 | 15 | 23 | 19 | 8 |
| 84720 | NM_00120148 | PIGO | phosphatidylinositol glycan anchor biosynthesis class O | protein_coding | NP_001188413.1;NP_116023.2;NP_690577.2;XP_005251676.1;XP_01687 | 23215 | 614730 | . | . | 24.905779 | 22.846202 | 20.876337 | 24.388379 | 57.956772 | 52.810021 | 46.845896 | 55.761127 | -1.078206 | 2909.466931 | 0.111668768 | -0.97281242 | 0.330646539 | 0.998865404 | 11.559818 | 11.451217 | 11.548478 | 11.524407 | 11.474494 | 11.475677 | 3527 | 3210 | 2433 | 2627 |
| 84722 | NM_00100529 | PSRC1 | proline and serine rich coiled-coil 1 | protein_coding | NP_001005290.1;NP_001027462.1;NP_001337166.1;NP_001337167.1;NI | 24472 | 613126 | . | . | 24.228353 | 23.747851 | 24.231928 | 24.369591 | 56.380370 | 54.894218 | 54.375749 | 55.718176 | 1.008653 | 1396.205049 | 0.126019097 | 0.098630505 | 0.921431649 | 0.998865404 | 10.441034 | 10.453529 | 10.445483 | 10.442370 | 10.458440 | 10.442610 | 1604 | 1497 | 1232 | 1301 |
| 84725 | NM_00119702 | PLEKHA8 | pleckstrin homology domain containing A8 | protein_coding | NP_001183955.1;NP_001183956.1;NP_001337902.1;NP_001337903.1;NI | 30037 | 608639 | . | . | 3.860724 | 3.364550 | 3.066444 | 3.266384 | 8.984064 | 7.777307 | 6.881010 | 7.468200 | -1.125530 | 640.265762 | 0.160696645 | -1.06165653 | 0.288391639 | 0.998865404 | 9.404993 | 9.235060 | 9.399891 | 9.319374 | 9.311840 | 9.249108 | 826 | 689 | 549 | 538 |
| 84726 | NM_013318 | PRRC2B | proline rich coiled-coil 2B | protein_coding | NP_037450.2 | 28121 | . | . | . | 10.219721 | 10.853910 | 11.464067 | 12.114975 | 23.781712 | 25.089300 | 25.725037 | 27.699448 | 1.103350 | 3845.237244 | 0.110101462 | 1.288722692 | 0.19749451 | 0.96621494 | 11.836225 | 11.978007 | 11.845184 | 11.887456 | 11.928940 | 11.966536 | 4102 | 4047 | 3429 | 3867 |
| 84727 | NM_00114631 | SPSB2 | splA/ryanodine receptor domain and SOCS box containing 2 | protein_coding | NP_001139788.1;NP_001306599.1;NP_116030.1 | 29522 | 611658 | . | . | 3.478534 | 2.977140 | 2.506807 | 2.988433 | 8.094691 | 6.881793 | 5.625200 | 6.832697 | -1.132490 | 130.079226 | 0.29005333 | -0.618846 | 0.536017815 | 0.998865404 | 7.110280 | 6.930624 | 7.075822 | 7.014253 | 6.978353 | 7.023124 | 172 | 137 | 99 | 122 |
| 84733 | NM_005189,N | CBX2 | chromobox 2 | protein_coding | NP_005180.1;NP_116036.1;XP_011523684.1;XP_011523685.1 | 1552 | 602770 | . | . | 5.608639 | 6.267633 | 4.821307 | 6.053955 | 13.051534 | 14.487911 | 10.818874 | 13.841646 | -1.058044 | 798.197391 | 0.158427816 | -0.51379699 | 0.607393933 | 0.998865404 | 9.681096 | 9.598938 | 9.645979 | 9.672060 | 9.541788 | 9.691045 | 928 | 902 | 602 | 805 |
| 84734 | NM_032648 | FAM167B | family with sequence similarity 167 member B | protein_coding | NP_116037.2 | 28133 | . | . | . | 44.948288 | 43.741833 | 58.251591 | 68.833214 | 104.596512 | 101.111198 | 130.714890 | 157.378967 | 1.412060 | 1589.641988 | 0.132844684 | 3.747240162 | 0.000178791 | 0.018983466 | 10.364356 | 10.861925 | 10.495927 | 10.469528 | 10.685704 | 10.812138 | 1545 | 1397 | 1492 | 1881 |
| 84747 | NM_00108053 | UNC119B | unc-119 lipid binding chaperone B | protein_coding | NP_001074002.1 | 16488 | . | . | . | 11.545753 | 11.221097 | 10.969574 | 12.715413 | 26.867443 | 25.938068 | 24.615408 | 29.072283 | 1.026050 | 1584.843637 | 0.130667733 | 0.283935116 | 0.776460105 | 0.998865404 | 10.611581 | 10.648433 | 10.633758 | 10.604961 | 10.584575 | 10.691401 | 1835 | 1657 | 1300 | 1608 |
| 84749 | NM_00130117 | USP30 | ubiquitin specific peptidase 30 | protein_coding | NP_001288104.1;NP_116052.2;XP_005254019.1;XP_005254022.1;XP_00 | 20065 | 612492 | . | . | 3.039493 | 2.410759 | 2.266402 | 2.855975 | 7.073025 | 5.572575 | 5.085739 | 6.529846 | -1.080918 | 310.804592 | 0.218281849 | -0.51427514 | 0.607059638 | 0.998865404 | 8.335035 | 8.222499 | 8.361452 | 8.231365 | 8.196173 | 8.312892 | 415 | 308 | 231 | 311 |
| 84750 | NM_032664,XI | FUT10 | fucosyltransferase 10 | protein_coding | NP_116053.3;XP_011542978.1;XP_011542979.1;XP_011542980.1;XP_01 | 19234 | 616931 | . | . | 1.647457 | 1.705613 | 1.766139 | 1.368579 | 3.833698 | 3.942602 | 3.963166 | 3.129095 | -1.092597 | 179.147041 | 0.258075812 | -0.49505596 | 0.62056062 | 0.998865404 | 7.546801 | 7.420434 | 7.499248 | 7.504516 | 7.518010 | 7.407256 | 214 | 203 | 169 | 139 |
| 84752 | NM_033309 | B3GNT9 | UDP-GlcNAc:betaGal beta-1,3-N-acetylglucosaminyltransferase 9 | protein_coding | NP_171608.2 | 28714 | . | . | . | 0.308217 | 0.387677 | 0.561362 | 0.078858 | 0.717234 | 0.896132 | 1.259681 | 0.180298 | -1.085239 | 29.012525 | 0.722435747 | -0.16335323 | 0.870240327 | 0.998865404 | 4.913716 | 4.801303 | 4.826923 | 4.853817 | 4.909481 | 4.744673 | 31 | 36 | 42 | 7 |
| 84759 | NM_032673,XI | PCGF1 | polycomb group ring finger 1 | protein_coding | NP_116062.2;XP_024308949.1 | 17615 | 610231 | . | . | 16.788334 | 18.099405 | 19.534260 | 20.757313 | 39.067147 | 41.837586 | 43.834313 | 47.459131 | 1.139069 | 533.219155 | 0.164947369 | 1.13887994 | 0.254753233 | 0.995760118 | 8.961819 | 9.149266 | 8.992427 | 9.034881 | 9.081827 | 9.113242 | 555 | 556 | 482 | 545 |
| 84765 | NM_00113559 | ZNF577 | zinc finger protein 577 | protein_coding | NP_001129062.1;NP_001357376.1;NP_001357377.1;NP_001357378.1;NI | 28673 | . | . | . | 0.391952 | 0.362863 | 0.789799 | 0.534955 | 0.912087 | 0.838776 | 1.772286 | 1.223113 | 1.848281 | 118.272111 | 0.308870738 | 2.869109054 | 0.004116298 | 0.190661973 | 6.375262 | 7.262390 | 6.742427 | 6.758371 | 7.008605 | 6.899429 | 93 | 92 | 152 | 124 |
| 84766 | NM_00114495 | CRACR2A | calcium release activated channel regulator 2A | protein_coding | NP_001138430.1;NP_116069.1;XP_006719084.1;XP_011519336.1;XP_01 | 28657 | 614178 | . | . | 2.988458 | 2.833179 | 2.666661 | 3.528795 | 6.954265 | 6.549020 | 5.983913 | 8.068170 | 1.075764 | 389.701143 | 0.19267062 | 0.54684697 | 0.584483871 | 0.998865404 | 8.553173 | 8.657393 | 8.541586 | 8.621651 | 8.573025 | 8.674801 | 403 | 433 | 321 | 411 |
| 84767 | NM_032681 | TRIM51 | tripartite motif-containing 51 | protein_coding | NP_116070.2 | 19023 | . | . | . | 6.320346 | 6.414896 | 5.662038 | 6.302497 | 14.707705 | 14.828321 | 12.705451 | 14.409909 | -1.078454 | 314.057984 | 0.19884888 | -0.54797653 | 0.58370802 | 0.998865404 | 8.348591 | 8.239106 | 8.313494 | 8.317409 | 8.248169 | 8.299458 | 374 | 353 | 250 | 297 |
| 84769 | NM_032683 | MPV17L2 | MPV17 mitochondrial inner membrane protein like 2 | protein_coding | NP_116072.2 | 28177 | 616133 | . | . | 11.950247 | 9.802999 | 11.621984 | 13.069688 | 27.808716 | 22.660072 | 26.079397 | 29.882292 | 1.118039 | 571.191105 | 0.174030826 | 0.924951825 | 0.354990967 | 0.998865404 | 9.075002 | 9.236159 | 9.182310 | 9.048814 | 9.154243 | 9.226508 | 685 | 522 | 496 | 596 |
| 84771 | NR_024004,NF | DDX11L2 | DEAD/H-box helicase 11 like 2 (pseudogene) | transcribed_ps | . | 37103 | . | . | . | 3.051548 | 2.814792 | 4.494418 | 3.865501 | 7.101078 | 6.506520 | 10.085343 | 8.838009 | 1.293139 | 204.256661 | 0.239408603 | 1.549140173 | 0.121348028 | 0.863030571 | 7.476539 | 7.848084 | 7.597615 | 7.615177 | 7.764772 | 7.680529 | 201 | 196 | 218 | 197 |
| 84775 | NM_00117267 | ZNF607 | zinc finger protein 607 | protein_coding | NP_001166148.1;NP_001362824.1;NP_116078.4;XP_006723498.1 | 28192 | . | . | . | 0.384610 | 0.505704 | 0.469342 | 0.411901 | 0.895003 | 1.168957 | 1.053188 | 0.941763 | -1.075537 | 67.319856 | 0.390867523 | -0.26877978 | 0.78809915 | 0.998865404 | 6.124786 | 6.019204 | 6.048191 | 6.110790 | 6.073312 | 6.052845 | 70 | 85 | 59 | 58 |
| 84777 | NR_119377 | DLGAP1-AS2 | DLGAP1 antisense RNA 2 | lncRNA | . | 28146 | . | . | . | 2.360571 | 2.857683 | 2.901309 | 3.036234 | 5.493146 | 6.605662 | 6.510454 | 6.941989 | 1.121781 | 80.575952 | 0.355372596 | 0.466528237 | 0.640837441 | 0.998865404 | 6.247506 | 6.412343 | 6.291658 | 6.335958 | 6.340311 | 6.348299 | 80 | 89 | 73 | 81 |
| 84787 | NM_032701,XI | KMT5C | lysine methyltransferase 5C | protein_coding | NP_116090.2;XP_005259395.1;XP_006723505.1;XP_011525717.1;XP_01 | 28405 | 613198 | . | . | 3.437895 | 3.000286 | 2.878731 | 3.099761 | 8.000123 | 6.935297 | 6.459786 | 7.087237 | -1.097876 | 237.001071 | 0.222651145 | -0.6050485 | 0.545146773 | 0.998865404 | 7.954659 | 7.819685 | 7.929649 | 7.892928 | 7.843220 | 7.888168 | 296 | 258 | 187 | 222 |
| 84789 | NR_026877 | MGC2889 | uncharacterized protein MGC2889 | lncRNA | . | . | . | . | . | 0.053616 | 0.083012 | 0.157556 | 0.048484 | 0.124766 | 0.191887 | 0.353551 | 0.110853 | 1.452912 | 6.648959 | 1.206571117 | 0.44667684 | 0.655108406 | NA | 2.436885 | 2.978782 | 2.705941 | 2.716154 | 2.737965 | 2.706153 | 5 | 7 | 10 | 4 |
| 84790 | NM_00130311 | TUBA1C | tubulin alpha 1c | protein_coding | NP_001290043.1;NP_001290044.1;NP_001290045.1;NP_001290046.1;NI | 20768 | . | . | . | 183.742535 | 187.103579 | 145.071344 | 171.861279 | 427.576393 | 432.498334 | 325.535892 | 392.940380 | -1.187476 | 15119.070062 | 0.112297678 | -2.20751019 | 0.027278436 | 0.538018865 | 14.002740 | 13.754783 | 13.948606 | 13.956680 | 13.735757 | 13.870347 | 18814 | 17795 | 11064 | 13990 |
| 84791 | NR_026761 | LINC00467 | long intergenic non-protein coding RNA 467 | lncRNA | . | 28227 | . | . | . | 6.242304 | 5.761002 | 5.462819 | 7.080369 | 14.526097 | 13.316813 | 12.258407 | 16.188423 | 1.027227 | 145.165196 | 0.279944447 | 0.138439585 | 0.889893014 | 0.998865404 | 7.162544 | 7.200313 | 7.191353 | 7.158577 | 7.137121 | 7.231307 | 172 | 148 | 112 | 155 |
| 84792 | NM_00103716 | FAM220A | family with sequence similarity 220 member A | protein_coding | NP_001032240.1 | 22422 | 616628 | . | . | 4.912879 | 5.134788 | 4.153887 | 4.085559 | 11.432470 | 11.869290 | 9.321202 | 9.341146 | -1.235916 | 316.915034 | 0.197009388 | -1.55109739 | 0.120878347 | 0.862265544 | 8.452653 | 8.147104 | 8.352386 | 8.373331 | 8.254749 | 8.239719 | 396 | 385 | 250 | 262 |
| 84793 | NR_026878 | FOXD2-AS1 | FOXD2 adjacent opposite strand RNA 1 | lncRNA | . | 44256 | . | . | . | 2.142872 | 2.058521 | 1.626011 | 2.212852 | 4.986551 | 4.758364 | 3.648723 | 5.059422 | -1.109181 | 157.943882 | 0.273421723 | -0.54675319 | 0.584548307 | 0.998865404 | 7.376755 | 7.225837 | 7.333232 | 7.312802 | 7.223977 | 7.336037 | 196 | 175 | 111 | 161 |
| 84795 | NM_032709,XI | PYROXD2 | pyridine nucleotide-disulphide oxidoreductase domain 2 | protein_coding | NP_116098.2;XP_011538595.1;XP_011538603.1;XP_016872324.1;XP_01 | 23517 | 617889 | . | . | 5.365439 | 5.142111 | 3.628949 | 3.551850 | 12.485598 | 11.886216 | 8.143255 | 8.120879 | -1.387050 | 364.754317 | 0.187401152 | -2.51876521 | 0.011776716 | 0.354930366 | 8.727451 | 8.255721 | 8.607462 | 8.584332 | 8.417124 | 8.399641 | 497 | 449 | 270 | 282 |
| 84798 | NM_00129014 | C19orf48 | chromosome 19 open reading frame 48 | protein_coding | NP_001277078.1;NP_001277079.1;NP_001277080.1;NP_001277081.1;NI | 29667 | 618864 | . | . | 46.788410 | 49.669292 | 44.428871 | 46.591693 | 108.878550 | 114.812797 | 99.697096 | 106.526370 | -1.056488 | 2434.661257 | 0.115928353 | -0.68384373 | 0.49407385 | 0.998865404 | 11.288663 | 11.209255 | 11.248510 | 11.293430 | 11.214784 | 11.238759 | 2801 | 2774 | 2028 | 2253 |
| 84803 | NM_00125642 | GPAT3 | glycerol-3-phosphate acyltransferase 3 | protein_coding | NP_001243350.1;NP_001243351.1;NP_116106.2;XP_011530685.1;XP_01 | 28157 | 610958 | . | . | 49.738148 | 43.512343 | 44.087599 | 35.789271 | 115.742713 | 100.580735 | 98.931290 | 81.827914 | -1.150912 | 4060.971151 | 0.12069977 | -1.68001299 | 0.092954789 | 0.809520218 | 12.085332 | 11.882782 | 12.081798 | 12.001188 | 11.982166 | 11.865859 | 5243 | 4465 | 3517 | 3289 |
| 84804 | NM_00132208 | MFSD9 | major facilitator superfamily domain containing 9 | protein_coding | NP_001309009.1;NP_001309010.1;NP_116107.3;XP_011510307.1;XP_01 | 28158 | . | . | . | 3.648887 | 3.232577 | 3.416653 | 2.570485 | 8.491109 | 7.472245 | 7.666872 | 5.877106 | -1.121341 | 513.508259 | 0.185787743 | -0.88932362 | 0.37382918 | 0.998865404 | 9.083894 | 8.919938 | 9.079530 | 8.992837 | 9.041966 | 8.880007 | 666 | 547 | 476 | 395 |
| 84807 | NM_00132183 | NFKBID | NFKB inhibitor delta | protein_coding | NP_001308760.1;NP_001352634.1;NP_001352635.1;NP_116110.2;NP_6 | 15671 | 618887 | . | . | 3.744936 | 3.432105 | 5.141891 | 4.392240 | 8.714621 | 7.933463 | 11.538251 | 10.042336 | 1.354030 | 305.329908 | 0.205357279 | 2.129264862 | 0.033232353 | 0.575540992 | 8.018556 | 8.456748 | 8.175644 | 8.144218 | 8.368410 | 8.277236 | 308 | 271 | 330 | 303 |
| 84809 | NR_026752 | CROCCP2 | CROCC pseudogene 2 | transcribed_ps | . | 28170 | . | . | . | 13.795753 | 13.229536 | 20.281862 | 15.143636 | 32.103283 | 30.580666 | 45.511909 | 34.624123 | 1.292131 | 858.654939 | 0.168054112 | 2.200198257 | 0.027792832 | 0.53804272 | 9.548800 | 9.919342 | 9.658779 | 9.624554 | 9.929962 | 9.708400 | 884 | 788 | 968 | 772 |
| 84811 | NM_00115973 | BUD13 | BUD13 homolog | protein_coding | NP_001153208.1;NP_116114.1;XP_011541337.1 | 28199 | . | . | . | 3.238887 | 2.997149 | 3.363613 | 3.394016 | 7.537020 | 6.928043 | 7.547850 | 7.760016 | -1.012661 | 229.587861 | 0.223165327 | -0.08133875 | 0.93517256 | 0.998865404 | 7.851926 | 7.833823 | 7.836569 | 7.855016 | 7.848371 | 7.831135 | 261 | 254 | 202 | 210 |
| 84812 | NM_032726,XI | PLCD4 | phospholipase C delta 4 | protein_coding | NP_116115.1;XP_005246970.1;XP_011510314.1;XP_016860603.1;XP_01 | 9062 | 605939 | . | . | 0.149369 | 0.267606 | 0.147021 | 0.197761 | 0.347588 | 0.618583 | 0.329911 | 0.452158 | -1.048886 | 22.061432 | 0.67249147 | -0.10239284 | 0.918444866 | NA | 4.499192 | 4.426810 | 4.436230 | 4.488743 | 4.450417 | 4.466435 | 19 | 31 | 17 | 22 |
| 84815 | NR_026880 | MGC12916 | uncharacterized protein MGC12916 | lncRNA | . | . | . | . | . | 0.021089 | 0.019769 | 0.055311 | 0.019159 | 0.049075 | 0.045697 | 0.124116 | 0.043804 | 1.870564 | 2.580901 | 1.868819978 | 0.483445811 | 0.628779227 | NA | 0.844134 | 1.751299 | 1.345797 | 1.346419 | 1.358911 | 1.347831 | 2 | 2 | 4 | 2 |
| 84816 | NM_00131874 | RTN4IP1 | reticulon 4 interacting protein 1 | protein_coding | NP_001305675.1;NP_116119.2;XP_011534494.1;XP_016866865.1 | 18647 | 610502 | . | . | 4.088998 | 4.037845 | 3.365207 | 4.281745 | 9.515265 | 9.333659 | 7.551426 | 9.789699 | -1.144012 | 330.642660 | 0.194220109 | -0.99938975 | 0.317605924 | 0.998865404 | 8.463103 | 8.268605 | 8.402469 | 8.408967 | 8.312666 | 8.348789 | 404 | 383 | 259 | 299 |
| 84817 | NM_032731 | TXNDC17 | thioredoxin domain containing 17 | protein_coding | NP_116120.1 | 28218 | 616967 | . | . | 22.424517 | 21.310869 | 18.055735 | 22.512491 | 52.182770 | 49.261028 | 40.516548 | 51.472141 | -1.093410 | 1259.085076 | 0.145057648 | -0.88816205 | 0.374453592 | 0.998865404 | 10.361339 | 10.232087 | 10.352631 | 10.306598 | 10.180759 | 10.337532 | 1560 | 1377 | 936 | 1246 |
| 84818 | NM_00120326 | IL17RC | interleukin 17 receptor C | protein_coding | NP_001190192.2;NP_001190193.1;NP_001190194.2;NP_001354207.1;NI | 18358 | 610925 | . | . | 9.550840 | 9.031997 | 10.462123 | 9.247174 | 22.225194 | 20.877867 | 23.476699 | 21.142565 | 1.059355 | 727.531495 | 0.156115215 | 0.532849158 | 0.594138002 | 0.998865404 | 9.464335 | 9.548179 | 9.503361 | 9.466784 | 9.577306 | 9.470581 | 835 | 742 | 701 | 648 |
| 84820 | NR_003655 | POLR2J4 | RNA polymerase II subunit J4, pseudogene | transcribed_ps | . | 28195 | . | . | . | 1.386324 | 1.320140 | 1.181440 | 1.374091 | 3.226033 | 3.051563 | 2.651118 | 3.141696 | -1.073599 | 267.266602 | 0.214229896 | -0.47824856 | 0.632473307 | 0.998865404 | 8.112713 | 8.009721 | 8.095462 | 8.064636 | 8.007298 | 8.078772 | 328 | 290 | 208 | 259 |
| 84823 | NM_032737 | LMNB2 | lamin B2 | protein_coding | NP_116126.3 | 6638 | 150341 | . | . | 17.849113 | 18.962265 | 15.576599 | 17.675152 | 41.535622 | 43.832127 | 34.953434 | 40.412136 | -1.122499 | 2529.390866 | 0.120699618 | -1.38123371 | 0.167207114 | 0.945199193 | 11.385650 | 11.218670 | 11.327679 | 11.370204 | 11.208246 | 11.300836 | 3001 | 2962 | 1952 | 2364 |
| 84824 | NM_00118486 | FCRLA | Fc receptor like A | protein_coding | NP_001171795.2;NP_001171796.2;NP_001171799.2;NP_001171800.2;NI | 18504 | 606891 | . | . | 2.608654 | 2.142793 | 2.890035 | 3.100784 | 6.070444 | 4.953161 | 6.485155 | 7.089576 | 1.249589 | 173.013019 | 0.256584596 | 1.252818334 | 0.210271828 | 0.972954982 | 7.264940 | 7.586637 | 7.414160 | 7.347521 | 7.452902 | 7.496426 | 192 | 152 | 159 | 189 |
| 84826 | NM_032740 | SFT2D3 | SFT2 domain containing 3 | protein_coding | NP_116129.3 | 28767 | . | . | . | 1.093092 | 0.812444 | 0.870015 | 0.804578 | 2.543671 | 1.878001 | 1.952289 | 1.839572 | -1.153590 | 105.000080 | 0.325538638 | -0.63319942 | 0.526603422 | 0.998865404 | 6.812836 | 6.608423 | 6.784769 | 6.683162 | 6.704790 | 6.675753 | 149 | 103 | 89 | 87 |
| 84833 | NM_00120642 | ATP5MD | ATP synthase membrane subunit DAPIT | protein_coding | NP_001193355.1;NP_001193356.1;NP_116136.1;XP_024304005.1 | 30889 | 615204 | . | . | 113.454060 | 112.760192 | 89.986059 | 87.010396 | 264.012282 | 260.650237 | 201.926117 | 198.938927 | -1.272625 | 1293.850111 | 0.128365674 | -2.70950867 | 0.006738294 | 0.260528509 | 10.500863 | 10.153146 | 10.428635 | 10.414856 | 10.245551 | 10.232232 | 1683 | 1550 | 999 | 1058 |
| 84836 | NM_00114631 | ABHD14B | abhydrolase domain containing 14B | protein_coding | NP_001139786.1;NP_001241682.1;NP_116139.1 | 28235 | . | . | . | 13.963834 | 14.524014 | 13.792824 | 14.452556 | 32.494418 | 33.572907 | 30.950697 | 33.044053 | 1.002215 | 1018.897336 | 0.138441678 | 0.02306128 | 0.981601392 | 0.998865404 | 9.991376 | 9.994209 | 9.977143 | 10.006225 | 9.961558 | 10.024092 | 1149 | 1120 | 849 | 997 |
| 84837 | NR_027263 | ARHGAP5-AS1 | ARHGAP5 antisense RNA 1 (head to head) | lncRNA | . | 20279 | . | . | . | 4.345234 | 5.135445 | 1.411112 | 4.570241 | 10.111538 | 11.870811 | 3.166494 | 10.449315 | -1.609820 | 154.444531 | 0.781168757 | -0.87932306 | 0.379226136 | 0.998865404 | 7.574263 | 6.886401 | 7.298796 | 7.370809 | 6.960485 | 7.308727 | 202 | 222 | 49 | 169 |
| 84838 | NM_00132973 | ZNF496 | zinc finger protein 496 | protein_coding | NP_001316662.1;NP_116141.1 | 23713 | 613911 | . | . | 3.317866 | 3.797386 | 3.040569 | 3.591941 | 7.720811 | 8.777828 | 6.822948 | 8.212547 | -1.068326 | 531.391188 | 0.175365344 | -0.54373171 | 0.586626109 | 0.998865404 | 9.101002 | 9.004651 | 9.025358 | 9.119395 | 8.979189 | 9.079844 | 587 | 635 | 411 | 520 |
| 84842 | NM_032756 | HPDL | 4-hydroxyphenylpyruvate dioxygenase like | protein_coding | NP_116145.1 | 28242 | . | . | . | 0.152750 | 0.029787 | 0.034259 | 0.172821 | 0.355455 | 0.068854 | 0.076877 | 0.395134 | 1.130033 | 5.609466 | 1.380702669 | 0.127735908 | 0.898357972 | NA | 2.397517 | 2.572884 | 2.500867 | 2.468911 | 2.471906 | 2.504165 | 10 | 2 | 2 | 9 |
| 84844 | NM_032758 | PHF5A | PHD finger protein 5A | protein_coding | NP_116147.1 | 18000 | 617846 | . | . | 42.200134 | 40.738163 | 40.601009 | 41.269966 | 98.201454 | 94.168091 | 91.107483 | 94.358879 | -1.027759 | 1351.960196 | 0.127563092 | -0.30966619 | 0.756814815 | 0.998865404 | 10.420418 | 10.380986 | 10.428103 | 10.394012 | 10.387987 | 10.392639 | 1613 | 1446 | 1156 | 1254 |
| 84848 | NR_024607 | MIR503HG | MIR503 host gene | lncRNA | . | 28258 | . | . | . | 0.976411 | 1.340573 | 1.488965 | 0.713352 | 2.272149 | 3.098795 | 3.341193 | 1.630995 | -1.051971 | 28.152926 | 0.615794958 | -0.11869977 | 0.905513215 | 0.998865404 | 4.849979 | 4.779589 | 4.791871 | 4.829574 | 4.841664 | 4.767068 | 28 | 36 | 32 | 17 |
| 84851 | NM_00134604 | TRIM52 | tripartite motif containing 52 | protein_coding | NP_001332977.1;NP_001332978.1;NP_001332979.1;NP_001332980.1;NI | 19024 | . | . | . | 4.537037 | 4.749879 | 7.246107 | 3.785636 | 10.557870 | 10.979557 | 16.260056 | 8.655409 | 1.094831 | 518.384202 | 0.230505284 | 0.567048378 | 0.570681314 | 0.998865404 | 8.950365 | 9.082372 | 8.986390 | 8.965519 | 9.205725 | 8.828110 | 579 | 525 | 607 | 362 |
| 84852 | NR_024124,NF | ATP1A1-AS1 | ATP1A1 antisense RNA 1 | lncRNA | . | 28262 | 618305 | . | . | 0.315967 | 0.424473 | 0.552921 | 0.316291 | 0.735265 | 0.981189 | 1.240738 | 0.723163 | 1.026891 | 16.369919 | 0.754183935 | 0.05076162 | 0.959515474 | NA | 4.014059 | 4.051647 | 4.029178 | 4.033848 | 4.031352 | 4.035433 | 18 | 18 | 14 | 16 |
| 84856 | NR_026827 | LINC00839 | long intergenic non-protein coding RNA 839 | lncRNA | . | 28269 | . | . | . | 8.719822 | 8.865949 | 11.013806 | 9.721675 | 20.291386 | 20.494036 | 24.714661 | 22.227455 | 1.162002 | 688.395635 | 0.156097649 | 1.387673403 | 0.165236519 | 0.940279399 | 9.314489 | 9.531543 | 9.368569 | 9.374543 | 9.520357 | 9.426373 | 730 | 690 | 687 | 647 |
| 84858 | NM_032772,N | ZNF503 | zinc finger protein 503 | protein_coding | NP_116161.2 | 23589 | 613902 | . | . | 13.008135 | 12.204968 | 12.812662 | 12.069788 | 30.270465 | 28.212329 | 28.751241 | 27.596136 | -1.020535 | 1237.457265 | 0.132629415 | -0.22111084 | 0.825006131 | 0.998865404 | 10.287591 | 10.258589 | 10.305933 | 10.254410 | 10.289449 | 10.240473 | 1488 | 1303 | 1100 | 1111 |
| 84859 | NM_00136388 | LRCH3 | leucine rich repeats and calponin homology domain containing 3 | protein_coding | NP_001350816.1;NP_001352644.1;NP_001352645.1;NP_001352646.1;NI | 28637 | . | . | . | 4.075775 | 4.543851 | 4.606136 | 3.796313 | 9.484497 | 10.503317 | 10.336037 | 8.679817 | -1.005846 | 888.092166 | 0.150659346 | -0.05581837 | 0.955486497 | 0.998865404 | 9.798666 | 9.790454 | 9.754837 | 9.834549 | 9.841058 | 9.739078 | 969 | 1014 | 825 | 771 |
| 84861 | NM_032775,N | KLHL22 | kelch like family member 22 | protein_coding | NP_116164.2;XP_016884507.1;XP_016884508.1;XP_016884509.1;XP_01 | 25888 | 618020 | . | . | 5.004834 | 4.330110 | 5.230014 | 5.903780 | 11.646454 | 10.009244 | 11.735999 | 13.498290 | 1.064508 | 501.682852 | 0.175722989 | 0.513231592 | 0.60778933 | 0.998865404 | 8.925245 | 9.014634 | 8.942536 | 8.955699 | 8.926144 | 9.047578 | 554 | 530 | 407 | 531 |
| 84864 | NM_00104253 | RIOX2 | ribosomal oxygenase 2 | protein_coding | NP_001035998.1;NP_001248758.1;NP_116167.3;NP_694822.2;XP_0052 | 19441 | 612049 | . | . | 4.484363 | 4.537169 | 4.831458 | 4.359479 | 10.435298 | 10.487869 | 10.841653 | 9.967431 | -1.046049 | 673.795257 | 0.151927208 | -0.42750972 | 0.669008109 | 0.998865404 | 9.428303 | 9.363298 | 9.400629 | 9.422060 | 9.383429 | 9.378015 | 781 | 755 | 575 | 615 |
| 84865 | NM_00136557 | CCDC142 | coiled-coil domain containing 142 | protein_coding | NP_001352504.1;NP_116168.3 | 25889 | . | . | . | 3.629052 | 4.039811 | 4.370894 | 3.941333 | 8.444954 | 9.338203 | 9.808160 | 9.011390 | 1.068535 | 513.679341 | 0.17112527 | 0.558851489 | 0.576263079 | 0.998865404 | 8.956032 | 9.051827 | 8.950310 | 9.012324 | 9.060218 | 8.986685 | 544 | 562 | 488 | 469 |
| 84868 | NM_032782 | HAVCR2 | hepatitis A virus cellular receptor 2 | protein_coding | NP_116171.3 | 18437 | 606652 | . | . | 0.084185 | 0.158894 | 0.080865 | 0.218280 | 0.195903 | 0.367290 | 0.181459 | 0.499071 | 1.271587 | 9.737478 | 1.022005525 | 0.339167037 | 0.7344839 | NA | 3.104339 | 3.442936 | 3.258823 | 3.282181 | 3.258587 | 3.303033 | 7 | 12 | 5 | 15 |
| 84869 | NM_032783,XI | CBR4 | carbonyl reductase 4 | protein_coding | NP_116172.2;XP_005263372.1;XP_005263373.1;XP_006714454.1;XP_00 | 25891 | . | . | . | 6.389535 | 5.635666 | 6.634154 | 6.429599 | 14.868711 | 13.027092 | 14.886849 | 14.700511 | 1.083097 | 676.906264 | 0.155968599 | 0.738366722 | 0.460291631 | 0.998865404 | 9.343823 | 9.459484 | 9.411027 | 9.335467 | 9.446025 | 9.410709 | 790 | 662 | 628 | 643 |
| 84872 | NM_00130312 | ZC3H10 | zinc finger CCCH-type containing 10 | protein_coding | NP_001290053.1;NP_001290054.1;NP_116175.1 | 25893 | . | . | . | 0.818000 | 0.888283 | 0.941800 | 0.711082 | 1.903520 | 2.053306 | 2.113373 | 1.625805 | -1.041369 | 187.924216 | 0.256635875 | -0.22787889 | 0.819740393 | 0.998865404 | 7.582336 | 7.525111 | 7.544326 | 7.575541 | 7.604046 | 7.476714 | 213 | 214 | 184 | 147 |
| 84874 | NM_00131800 | ZNF514 | zinc finger protein 514 | protein_coding | NP_001304934.1;NP_116177.1;XP_006712869.1 | 25894 | . | . | . | 3.061067 | 3.471802 | 4.002391 | 3.220231 | 7.123228 | 8.025228 | 8.981249 | 7.362675 | 1.084339 | 661.396637 | 0.166882566 | 0.699990992 | 0.48393293 | 0.998865404 | 9.309575 | 9.426789 | 9.303402 | 9.373025 | 9.460750 | 9.322318 | 693 | 720 | 659 | 580 |
| 84875 | NM_00131789 | PARP10 | poly(ADP-ribose) polymerase family member 10 | protein_coding | NP_001304824.1;NP_116178.2;XP_011515638.1 | 25895 | 609564 | . | . | 8.509972 | 7.930136 | 9.805050 | 9.144092 | 19.803055 | 18.330866 | 22.002249 | 20.906878 | 1.144327 | 968.444638 | 0.141963012 | 1.370069776 | 0.170665121 | 0.946389416 | 9.818757 | 10.013713 | 9.894990 | 9.838469 | 9.997728 | 9.932063 | 1081 | 936 | 939 | 926 |
| 84876 | NM_032790 | ORAI1 | ORAI calcium release-activated calcium modulator 1 | protein_coding | NP_116179.2 | 25896 | 610277 | . | . | 16.210930 | 17.325380 | 15.659823 | 19.338739 | 37.723503 | 40.048393 | 35.140190 | 44.215729 | 1.029207 | 800.207093 | 0.157326084 | 0.263995332 | 0.791783516 | 0.998865404 | 9.623693 | 9.664478 | 9.611954 | 9.653050 | 9.579669 | 9.721128 | 882 | 876 | 635 | 837 |
| 84878 | NM_00131697 | ZBTB45 | zinc finger and BTB domain containing 45 | protein_coding | NP_001303907.1;NP_001303908.1;NP_001303909.1;NP_001303910.1;NI | 23715 | . | . | . | 2.028211 | 2.450465 | 2.059465 | 2.288520 | 4.719731 | 5.664361 | 4.621381 | 5.232427 | 1.015983 | 155.742163 | 0.265856598 | 0.08604846 | 0.931427895 | 0.998865404 | 7.272032 | 7.293916 | 7.251632 | 7.305697 | 7.269410 | 7.300865 | 166 | 178 | 131 | 153 |
| 84879 | NM_00113649 | MFSD2A | major facilitator superfamily domain containing 2A | protein_coding | NP_001129965.1;NP_001274737.1;NP_001274738.1;NP_001336750.1;NI | 25897 | 614397 | . | . | 0.148299 | 0.145659 | 0.068237 | 0.124247 | 0.345098 | 0.336697 | 0.153123 | 0.284075 | -1.440453 | 8.724676 | 1.035395902 | -0.50852299 | 0.61108662 | NA | 3.365657 | 2.836248 | 3.139850 | 3.138706 | 3.119166 | 3.131200 | 12 | 11 | 5 | 8 |
| 84881 | NM_00114482 | RPUSD4 | RNA pseudouridine synthase D4 | protein_coding | NP_001138299.1;NP_001350445.1;NP_116184.2;XP_011541341.1 | 25898 | 617488 | . | . | 5.752261 | 6.940346 | 6.234067 | 5.522478 | 13.385747 | 16.042920 | 13.989064 | 12.626490 | -1.093025 | 471.378950 | 0.182069128 | -0.70482007 | 0.480922216 | 0.998865404 | 8.943457 | 8.815180 | 8.848466 | 8.961055 | 8.889471 | 8.810709 | 517 | 578 | 417 | 395 |
| 84883 | NM_00119869 | AIFM2 | apoptosis inducing factor mitochondria associated 2 | protein_coding | NP_001185625.1;NP_116186.1 | 21411 | 605159 | . | . | 13.149763 | 12.894262 | 15.946331 | 16.206642 | 30.600039 | 29.805663 | 35.783105 | 37.054561 | 1.214501 | 1432.093055 | 0.125430171 | 2.235217489 | 0.025403074 | 0.524911334 | 10.336910 | 10.617306 | 10.412493 | 10.394748 | 10.550107 | 10.555994 | 1507 | 1379 | 1361 | 1478 |
| 84885 | NM_00131801 | ZDHHC12 | zinc finger DHHC-type palmitoyltransferase 12 | protein_coding | NP_001304944.2;NP_001304945.1;NP_001304949.2;NP_001304952.2;NI | 19159 | . | . | . | 23.287082 | 24.942019 | 20.903779 | 22.201416 | 54.189995 | 57.654593 | 46.907478 | 50.760905 | -1.131692 | 829.980945 | 0.1460438 | -1.22210581 | 0.221667618 | 0.982008277 | 9.783618 | 9.604708 | 9.716858 | 9.760931 | 9.624459 | 9.677460 | 984 | 980 | 651 | 758 |
| 84886 | NM_00113649 | C1orf198 | chromosome 1 open reading frame 198 | protein_coding | NP_001129966.1;NP_001129967.1;NP_116189.1;XP_016858088.1 | 25900 | . | . | . | 4.095616 | 4.489289 | 2.743240 | 4.122525 | 9.530667 | 10.377193 | 6.155752 | 9.425662 | -1.266904 | 449.203118 | 0.206624124 | -1.65182535 | 0.098570158 | 0.813553488 | 8.972505 | 8.629617 | 8.848880 | 8.901095 | 8.609726 | 8.837099 | 555 | 564 | 277 | 444 |
| 84888 | NM_032802,XI | SPPL2A | signal peptide peptidase like 2A | protein_coding | NP_116191.2;XP_005254779.1;XP_016878169.1;XP_016878170.1 | 30227 | 608238 | . | . | 5.505146 | 5.795819 | 5.623874 | 3.795449 | 12.810702 | 13.397293 | 12.619810 | 8.677845 | -1.166341 | 1153.750699 | 0.175074428 | -1.26797155 | 0.204808112 | 0.96993643 | 10.278477 | 10.057285 | 10.235768 | 10.207926 | 10.242942 | 9.949846 | 1456 | 1316 | 1111 | 800 |
| 84890 | NM_032804 | ADO | 2-aminoethanethiol dioxygenase | protein_coding | NP_116193.2 | 23506 | 611392 | . | . | 7.992302 | 6.472334 | 6.583338 | 5.413985 | 18.598415 | 14.961090 | 14.772820 | 12.378432 | -1.223112 | 776.502680 | 0.170725126 | -1.70189783 | 0.088774523 | 0.802216049 | 9.738369 | 9.448822 | 9.740800 | 9.581520 | 9.590355 | 9.451488 | 1091 | 821 | 670 | 588 |
| 84892 | NM_032806,XI | POMGNT2 | protein O-linked mannose N-acetylglucosaminyltransferase 2 (beta | protein_coding | NP_116195.2;XP_005265572.1;XP_011532465.1;XP_016862842.1 | 25902 | 614828 | . | . | 8.952164 | 9.585496 | 8.357192 | 7.858460 | 20.832057 | 22.157306 | 18.753296 | 17.967438 | -1.167621 | 693.597752 | 0.153095821 | -1.46034129 | 0.1441963 | 0.906161673 | 9.545356 | 9.321912 | 9.466585 | 9.509678 | 9.405417 | 9.358844 | 834 | 831 | 576 | 579 |
| 84893 | NM_00125845 | FBH1 | F-box DNA helicase 1 | protein_coding | NP_001245381.1;NP_001245382.1;NP_116196.3;NP_835363.1;XP_01151 | 13620 | 607222 | . | . | 14.513513 | 15.807086 | 14.985708 | 15.594904 | 33.773538 | 36.538789 | 33.627492 | 35.655896 | -1.043295 | 1777.939192 | 0.123412389 | -0.49546865 | 0.620269347 | 0.998865404 | 10.826347 | 10.764981 | 10.781201 | 10.842427 | 10.762685 | 10.794811 | 2010 | 2035 | 1482 | 1665 |
| 84894 | NM_00130118 | LINGO1 | leucine rich repeat and Ig domain containing 1 | protein_coding | NP_001288115.1;NP_001288116.1;NP_001288118.1;NP_001288120.1;NI | 21205 | 609791 | . | . | 0.770334 | 1.296686 | 1.057796 | 1.053113 | 1.792597 | 2.997348 | 2.373665 | 2.407817 | -1.008790 | 114.594721 | 0.326335696 | -0.03868923 | 0.96913816 | 0.998865404 | 6.847463 | 6.833296 | 6.757499 | 6.909229 | 6.836061 | 6.830549 | 103 | 152 | 100 | 106 |
| 84895 | NM_00132999 | MIGA2 | mitoguardin 2 | protein_coding | NP_001316919.1;NP_116198.3;XP_011517420.1 | 23621 | 616774 | . | . | 6.358505 | 6.752330 | 6.377045 | 6.995320 | 14.796504 | 15.608314 | 14.309904 | 15.993969 | 1.011668 | 806.056543 | 0.145578312 | 0.114962257 | 0.908475039 | 0.998865404 | 9.646493 | 9.662935 | 9.629608 | 9.670904 | 9.644564 | 9.672389 | 896 | 890 | 691 | 775 |
| 84896 | NM_00132196 | ATAD1 | ATPase family AAA domain containing 1 | protein_coding | NP_001308896.1;NP_001308897.1;NP_001308898.1;NP_116199.2;XP_0C | 25903 | 614452 | . | . | 13.549807 | 14.039820 | 12.567030 | 10.483065 | 31.530957 | 32.453676 | 28.200052 | 23.968280 | -1.165123 | 1690.076947 | 0.131648273 | -1.67477823 | 0.093977752 | 0.810751527 | 10.828723 | 10.608638 | 10.786390 | 10.771240 | 10.719829 | 10.595230 | 2114 | 1944 | 1468 | 1345 |
| 84897 | NM_032811,N | TBRG1 | transforming growth factor beta regulator 1 | protein_coding | NP_116200.2;XP_005271766.1;XP_016873931.1 | 29551 | 610614 | . | . | 2.938889 | 2.901574 | 3.219271 | 3.106766 | 6.838913 | 6.707120 | 7.223952 | 7.103250 | 1.053517 | 547.812207 | 0.164339469 | 0.457674443 | 0.647186351 | 0.998865404 | 9.059225 | 9.134859 | 9.081365 | 9.077781 | 9.145097 | 9.080900 | 616 | 574 | 513 | 501 |
| 84898 | NM_00128273 | PLXDC2 | plexin domain containing 2 | protein_coding | NP_001269665.1;NP_116201.7;XP_011518052.1 | 21013 | 606827 | . | . | 0.061301 | 0.138633 | 0.039467 | 0.052396 | 0.142650 | 0.320457 | 0.088564 | 0.120416 | -1.456225 | 22.352945 | 0.65701226 | -0.82530192 | 0.409200202 | NA | 4.729157 | 4.184531 | 4.494613 | 4.512932 | 4.460664 | 4.477569 | 28 | 31 | 14 | 19 |
| 84899 | NM_00107966 | TMTC4 | transmembrane O-mannosyltransferase targeting cadherins 4 | protein_coding | NP_001073137.1;NP_001273382.1;NP_001337500.1;NP_001337501.1;NI | 25904 | 618203 | . | . | 3.738903 | 3.821803 | 4.099203 | 3.733305 | 8.700583 | 8.834269 | 9.198496 | 8.535760 | 1.031813 | 455.602054 | 0.173675687 | 0.260150119 | 0.794747979 | 0.998865404 | 8.808699 | 8.854203 | 8.817266 | 8.825336 | 8.865747 | 8.815048 | 513 | 487 | 419 | 416 |
| 84900 | NM_00110990 | RNFT2 | ring finger protein, transmembrane 2 | protein_coding | NP_001103373.1;NP_001369195.1;NP_116203.2 | 25905 | . | . | . | 1.794905 | 1.479923 | 1.509497 | 1.559916 | 4.176817 | 3.420909 | 3.387267 | 3.566563 | -1.039795 | 271.510756 | 0.214438238 | -0.26254001 | 0.792905147 | 0.998865404 | 8.112393 | 8.056802 | 8.138623 | 8.048010 | 8.079625 | 8.068851 | 346 | 273 | 234 | 247 |
| 84901 | NM_032815,XI | NFATC2IP | nuclear factor of activated T cells 2 interacting protein | protein_coding | NP_116204.3 | 25906 | 614525 | . | . | 13.737849 | 13.637599 | 16.066380 | 15.459476 | 31.968540 | 31.523918 | 36.052490 | 35.346253 | 1.127862 | 1772.131293 | 0.121928426 | 1.423708259 | 0.154530942 | 0.922573785 | 10.701776 | 10.875535 | 10.747641 | 10.735438 | 10.860832 | 10.810987 | 1933 | 1783 | 1685 | 1706 |
| 84902 | NM_032816,XI | CEP89 | centrosomal protein 89 | protein_coding | NP_116205.3;XP_005259401.1;XP_016882887.1;XP_024307513.1 | 25907 | 615470 | . | . | 2.294267 | 1.956744 | 2.318911 | 2.828661 | 5.338854 | 4.523102 | 5.203568 | 6.467398 | 1.149099 | 376.823754 | 0.185141723 | 1.082970114 | 0.278821689 | 0.998865404 | 8.453827 | 8.654680 | 8.534006 | 8.493953 | 8.601042 | 8.589301 | 418 | 365 | 354 | 374 |
| 84904 | NM_032818,XI | ARHGEF39 | Rho guanine nucleotide exchange factor 39 | protein_coding | NP_116207.2;XP_011516359.1;XP_016870714.1;XP_016870715.1;XP_01 | 25909 | . | . | . | 28.155929 | 31.238198 | 29.800989 | 30.137770 | 65.520006 | 72.208495 | 66.872553 | 68.906428 | 1.012698 | 2892.307069 | 0.112486313 | 0.161836474 | 0.87143463 | 0.998865404 | 11.488888 | 11.507064 | 11.473951 | 11.511193 | 11.509463 | 11.496546 | 3234 | 3172 | 2551 | 2707 |
| 84905 | NM_00128293 | ZNF341 | zinc finger protein 341 | protein_coding | NP_001269862.1;NP_001269864.1;NP_116208.3 | 15992 | 618269 | . | . | 0.959667 | 1.054173 | 1.070902 | 0.839081 | 2.233184 | 2.436768 | 2.403072 | 1.918459 | -1.049615 | 107.033785 | 0.317157058 | -0.22027161 | 0.825659625 | 0.998865404 | 6.776023 | 6.706996 | 6.729094 | 6.766588 | 6.766472 | 6.697630 | 119 | 125 | 101 | 87 |
| 84908 | NM_00132975 | FAM136A | family with sequence similarity 136 member A | protein_coding | NP_001316681.1;NP_001316682.1;NP_001316684.1;NP_001316686.1;NI | 25911 | 616275 | . | . | 23.327812 | 23.303601 | 23.875789 | 26.338019 | 54.284778 | 53.867313 | 53.576577 | 60.218744 | 1.084151 | 1445.913189 | 0.127255555 | 0.915991791 | 0.3596712 | 0.998865404 | 10.438385 | 10.554798 | 10.472219 | 10.458701 | 10.498717 | 10.555746 | 1612 | 1484 | 1260 | 1462 |
| 84909 | NM_00119332 | AOPEP | aminopeptidase O (putative) | protein_coding | NP_001180258.1;NP_001180260.1;NP_116212.3;XP_005252341.1;XP_00 | 1361 | . | . | . | 8.735983 | 8.104371 | 8.184755 | 9.003910 | 20.328992 | 18.733611 | 18.366353 | 20.586367 | 1.096949 | 959.397001 | 0.137336852 | 0.972035246 | 0.331033014 | 0.998865404 | 9.837701 | 9.971181 | 9.870417 | 9.871417 | 9.936634 | 9.939853 | 1053 | 988 | 871 | 943 |
| 84910 | NM_00132991 | TMEM87B | transmembrane protein 87B | protein_coding | NP_001316843.1;NP_116213.1;XP_005263884.1;XP_016860610.1 | 25913 | 617203 | . | . | 16.520626 | 15.832797 | 17.565860 | 12.128598 | 38.444176 | 36.598217 | 39.417278 | 27.730599 | -1.107762 | 2408.445617 | 0.168670105 | -0.87536308 | 0.381376383 | 0.998865404 | 11.305604 | 11.158417 | 11.307802 | 11.226686 | 11.339917 | 11.003651 | 3059 | 2596 | 2403 | 1695 |
| 84911 | NM_00125683 | ZNF382 | zinc finger protein 382 | protein_coding | NP_001243767.1;NP_116214.2 | 17409 | 609516 | . | . | 0.572408 | 0.719969 | 0.687897 | 0.760674 | 1.332018 | 1.664241 | 1.543620 | 1.739191 | 1.108385 | 178.725212 | 0.256127864 | 0.579628343 | 0.562165275 | 0.998865404 | 7.406154 | 7.553294 | 7.409059 | 7.501180 | 7.480245 | 7.519434 | 174 | 203 | 156 | 184 |
| 84912 | NM_032826,XI | SLC35B4 | solute carrier family 35 member B4 | protein_coding | NP_116215.1;XP_011514947.1 | 20584 | 610923 | . | . | 12.741570 | 12.624599 | 15.738158 | 12.735408 | 29.650154 | 29.182327 | 35.315970 | 29.117997 | 1.110600 | 2834.688512 | 0.131873812 | 1.147609344 | 0.251129871 | 0.993339639 | 11.391170 | 11.542797 | 11.431470 | 11.414135 | 11.595872 | 11.412993 | 3126 | 2867 | 2882 | 2487 |
| 84913 | NM_032827,XI | ATOH8 | atonal bHLH transcription factor 8 | protein_coding | NP_116216.2;XP_006712185.1;XP_011531441.1 | 24126 | . | . | . | 2.988459 | 3.477726 | 1.818609 | 3.285817 | 6.954265 | 8.038921 | 4.080907 | 7.512631 | -1.309815 | 678.744247 | 0.194421009 | -2.00267745 | 0.045211921 | 0.649595749 | 9.588859 | 9.198228 | 9.464687 | 9.507587 | 9.154565 | 9.431010 | 860 | 856 | 402 | 668 |
| 84914 | NM_00120481 | ZNF587 | zinc finger protein 587 | protein_coding | NP_001191746.1;NP_116217.1 | 30955 | . | . | . | 5.542952 | 5.884098 | 6.651489 | 5.103365 | 12.898678 | 13.601357 | 14.925749 | 11.668236 | 1.015352 | 1244.052706 | 0.153849594 | 0.142870999 | 0.886392063 | 0.998865404 | 10.269551 | 10.292026 | 10.252795 | 10.291537 | 10.381977 | 10.175619 | 1385 | 1366 | 1239 | 1015 |
| 84915 | NM_032829,XI | FAM222A | family with sequence similarity 222 member A | protein_coding | NP_116218.2;XP_006719717.1;XP_016875544.1;XP_024304997.1 | 25915 | . | . | . | 3.071101 | 3.453288 | 2.852427 | 3.544750 | 7.146577 | 7.982430 | 6.400763 | 8.104649 | -1.033653 | 370.071059 | 0.197445001 | -0.24184672 | 0.808898938 | 0.998865404 | 8.555946 | 8.506955 | 8.507720 | 8.571211 | 8.458798 | 8.578876 | 410 | 428 | 283 | 377 |
| 84916 | NM_00131839 | UTP4 | UTP4 small subunit processome component | protein_coding | NP_001305320.1;NP_116219.2;XP_005256262.2 | 1983 | 607456 | . | . | 27.894191 | 30.032491 | 29.421223 | 30.919229 | 64.910927 | 69.421450 | 66.020373 | 70.693139 | 1.023014 | 2004.840526 | 0.120937571 | 0.271427976 | 0.786061888 | 0.998865404 | 10.952871 | 10.985489 | 10.930084 | 10.988918 | 10.959682 | 10.995850 | 2198 | 2218 | 1723 | 1944 |
| 84918 | NM_032832,XI | LRP11 | LDL receptor related protein 11 | protein_coding | NP_116221.3;XP_011534496.1;XP_016866869.1 | 16936 | . | . | . | 12.526011 | 11.184485 | 11.747608 | 9.961667 | 29.148541 | 25.853439 | 26.361290 | 22.776168 | -1.105737 | 1280.543106 | 0.144383707 | -1.00432757 | 0.315220748 | 0.998865404 | 10.392926 | 10.248543 | 10.403722 | 10.310178 | 10.346312 | 10.213244 | 1642 | 1363 | 1152 | 1039 |
| 84919 | NM_032833,XI | PPP1R15B | protein phosphatase 1 regulatory subunit 15B | protein_coding | NP_116222.4;XP_005245608.2 | 14951 | 613257 | . | . | 10.826979 | 11.150814 | 12.197234 | 11.346135 | 25.194826 | 25.775603 | 27.370239 | 25.941589 | 1.042290 | 1826.152848 | 0.122813597 | 0.486565637 | 0.626566181 | 0.998865404 | 10.804314 | 10.864247 | 10.813365 | 10.821885 | 10.884367 | 10.815152 | 2048 | 1940 | 1692 | 1671 |
| 84920 | NM_032834,XI | ALG10 | ALG10 alpha-1,2-glucosyltransferase | protein_coding | NP_116223.3;XP_024304998.1;XP_024304999.1 | 23162 | 618355 | . | . | 2.674491 | 2.382040 | 2.585055 | 1.820690 | 6.223650 | 5.506192 | 5.800790 | 4.162791 | -1.167788 | 344.248442 | 0.226364928 | -0.98857316 | 0.322872023 | 0.998865404 | 8.533983 | 8.312108 | 8.511650 | 8.414036 | 8.479997 | 8.261256 | 462 | 367 | 332 | 238 |
| 84922 | NM_032836,XI | FIZ1 | FLT3 interacting zinc finger 1 | protein_coding | NP_116225.2;XP_005259409.1;XP_011525728.1 | 25917 | 609133 | . | . | 2.450720 | 2.313799 | 2.568396 | 3.001108 | 5.702926 | 5.348451 | 5.763405 | 6.861678 | 1.183059 | 248.401290 | 0.217237093 | 1.116395075 | 0.264253061 | 0.996308385 | 7.829864 | 8.072965 | 7.924920 | 7.895764 | 8.010895 | 7.976985 | 270 | 238 | 242 | 244 |
| 84923 | NM_00109883 | FAM104A | family with sequence similarity 104 member A | protein_coding | NP_001092302.1;NP_001276339.1;NP_001276340.1;NP_001276341.1;NI | 25918 | . | . | . | 13.172021 | 12.395147 | 12.479999 | 12.011849 | 30.651834 | 28.651937 | 28.004753 | 27.463666 | -1.060359 | 1113.850177 | 0.135610332 | -0.62350272 | 0.532954214 | 0.998865404 | 10.162837 | 10.078613 | 10.169584 | 10.114767 | 10.119716 | 10.077925 | 1368 | 1192 | 967 | 985 |
| 84924 | NM_00114534 | ZNF566 | zinc finger protein 566 | protein_coding | NP_001138815.1;NP_001138816.1;NP_001138817.1;NP_001287899.1;NI | 25919 | . | . | . | 0.985366 | 0.949047 | 1.170867 | 0.773890 | 2.292987 | 2.193763 | 2.627393 | 1.769408 | -1.004860 | 163.207511 | 0.285418573 | -0.02450768 | 0.98044766 | 0.998865404 | 7.352931 | 7.348192 | 7.358380 | 7.332972 | 7.428924 | 7.250899 | 194 | 171 | 173 | 119 |
| 84925 | NM_032839,XI | SLC49A4 | solute carrier family 49 member 4 | protein_coding | NP_116228.1 | 16628 | 602773 | . | . | 5.912718 | 5.169292 | 4.999894 | 4.036149 | 13.759136 | 11.949049 | 11.219619 | 9.228175 | -1.092785 | 688.187709 | 0.155775684 | -0.82175305 | 0.411217457 | 0.998865404 | 9.488961 | 9.361543 | 9.472228 | 9.439061 | 9.435459 | 9.353344 | 847 | 757 | 607 | 579 |
| 84926 | NM_032840 | SPRYD3 | SPRY domain containing 3 | protein_coding | NP_116229.1 | 25920 | . | . | . | 2.761376 | 2.794430 | 2.062086 | 2.679861 | 6.425836 | 6.459449 | 4.627261 | 6.127183 | -1.187377 | 233.235160 | 0.232413432 | -1.06611076 | 0.286373575 | 0.998865404 | 7.984831 | 7.735712 | 7.904053 | 7.906210 | 7.763115 | 7.877559 | 291 | 274 | 162 | 225 |
| 84928 | NM_00130116 | TMEM209 | transmembrane protein 209 | protein_coding | NP_001288092.1;NP_001350407.1;NP_116231.2;XP_005250712.1 | 21898 | . | . | . | 5.758808 | 5.813145 | 4.388743 | 5.874137 | 13.400984 | 13.437343 | 9.848214 | 13.430516 | -1.138584 | 593.818508 | 0.181428117 | -1.03203593 | 0.302055286 | 0.998865404 | 9.304942 | 9.116677 | 9.255665 | 9.250493 | 9.062512 | 9.261693 | 731 | 680 | 408 | 601 |
| 84930 | NM_00117230 | MASTL | microtubule associated serine/threonine kinase like | protein_coding | NP_001165774.1;NP_001165775.1;NP_001307685.1;NP_001307686.1;NI | 19042 | 608221 | . | . | 6.418278 | 5.597509 | 5.948836 | 5.507703 | 14.935596 | 12.938893 | 13.349016 | 12.592706 | -1.064644 | 755.685918 | 0.155239883 | -0.58214064 | 0.560471956 | 0.998865404 | 9.605813 | 9.516077 | 9.631497 | 9.530636 | 9.567845 | 9.508850 | 961 | 781 | 664 | 657 |
| 84932 | NM_00116338 | RAB2B | RAB2B, member RAS oncogene family | protein_coding | NP_001156852.1;NP_116235.2;XP_016877199.1;XP_016877201.1 | 20246 | 607466 | . | . | 5.262403 | 5.022186 | 5.639501 | 4.481581 | 12.245825 | 11.609010 | 12.654876 | 10.246605 | -1.031624 | 461.067687 | 0.184518771 | -0.24343228 | 0.80767054 | 0.998865404 | 8.870594 | 8.826743 | 8.873976 | 8.838882 | 8.905126 | 8.765000 | 554 | 491 | 440 | 377 |
| 84933 | NM_032847 | C8orf76 | chromosome 8 open reading frame 76 | protein_coding | NP_116236.1 | 25924 | . | . | . | 10.065070 | 9.786599 | 8.779999 | 9.808312 | 23.421831 | 22.622162 | 19.702061 | 22.425543 | -1.084526 | 392.659932 | 0.18394711 | -0.63640544 | 0.524512199 | 0.998865404 | 8.674659 | 8.557225 | 8.651681 | 8.629772 | 8.563470 | 8.621391 | 479 | 433 | 311 | 371 |
| 84934 | NM_00128621 | RITA1 | RBPJ interacting and tubulin associated 1 | protein_coding | NP_001273144.1;NP_116237.1;XP_005254029.1 | 25925 | . | . | . | 10.613924 | 12.124040 | 10.635865 | 14.541372 | 24.699036 | 28.025262 | 23.866570 | 33.247118 | 1.098177 | 704.852554 | 0.175848753 | 0.768334667 | 0.442288384 | 0.998865404 | 9.392594 | 9.526649 | 9.378600 | 9.468594 | 9.384682 | 9.583447 | 723 | 773 | 550 | 790 |
| 84936 | NM_00107726 | ZFYVE19 | zinc finger FYVE-type containing 19 | protein_coding | NP_001070736.1;NP_001245349.1;NP_001245350.1;NP_116239.3;XP_01 | 20758 | . | . | . | 11.435965 | 13.696645 | 14.581580 | 12.927548 | 26.611958 | 31.660407 | 32.720641 | 29.557305 | 1.072408 | 1017.432179 | 0.144792569 | 0.696536318 | 0.486093021 | 0.998865404 | 9.939459 | 10.040216 | 9.908286 | 10.015339 | 10.042674 | 9.985645 | 1048 | 1137 | 951 | 949 |
| 84937 | NM_032268,XI | ZNRF1 | zinc and ring finger 1 | protein_coding | NP_115644.1;XP_011521694.1;XP_016879282.1;XP_016879283.1 | 18452 | 612060 | . | . | 4.451735 | 4.787555 | 3.183746 | 4.364223 | 10.359372 | 11.066649 | 7.144235 | 9.978278 | -1.214491 | 590.081957 | 0.182286943 | -1.5379715 | 0.124055584 | 0.87182553 | 9.338704 | 9.057121 | 9.229751 | 9.295166 | 9.035331 | 9.224621 | 710 | 732 | 394 | 574 |
| 84938 | NM_032852,N | ATG4C | autophagy related 4C cysteine peptidase | protein_coding | NP_116241.2;NP_835739.1;XP_005271345.1;XP_011540615.1;XP_01685 | 16040 | 611339 | . | . | 5.028538 | 4.281807 | 5.578316 | 3.727225 | 11.701614 | 9.897587 | 12.517580 | 8.521857 | -1.014071 | 403.378547 | 0.216036717 | -0.09330859 | 0.925658406 | 0.998865404 | 8.665202 | 8.646720 | 8.704210 | 8.601992 | 8.754469 | 8.525051 | 507 | 401 | 416 | 303 |
| 84939 | NM_00136978 | PWWP3A | PWWP domain containing 3A, DNA repair factor | protein_coding | NP_001356718.1;NP_001356719.1;NP_001356721.1;NP_001356722.1;NI | 29641 | . | . | . | 9.647288 | 8.500003 | 9.390914 | 9.309352 | 22.449632 | 19.648135 | 21.072931 | 21.284727 | 1.029624 | 1072.197847 | 0.136912467 | 0.307627176 | 0.758366039 | 0.998865404 | 10.045087 | 10.087315 | 10.087709 | 10.021437 | 10.065713 | 10.087680 | 1271 | 1089 | 932 | 1035 |
| 84942 | NM_032856,N | WDR73 | WD repeat domain 73 | protein_coding | NP_116245.2 | 25928 | 616144 | . | . | 9.731278 | 11.771078 | 13.055751 | 10.220146 | 22.645085 | 27.209370 | 29.296727 | 23.367147 | 1.084747 | 1177.807291 | 0.133754651 | 0.877412304 | 0.380262732 | 0.998865404 | 10.141973 | 10.259413 | 10.149059 | 10.190412 | 10.257608 | 10.203644 | 1265 | 1253 | 1104 | 1108 |
| 84945 | NM_032859,XI | ABHD13 | abhydrolase domain containing 13 | protein_coding | NP_116248.2;XP_011519430.1 | 20293 | . | . | . | 2.153646 | 1.901250 | 1.489569 | 1.659839 | 5.011624 | 4.394824 | 3.342549 | 3.795025 | -1.329564 | 305.005810 | 0.204154603 | -2.01295004 | 0.044119886 | 0.645513836 | 8.443659 | 8.032536 | 8.355197 | 8.290832 | 8.146910 | 8.192827 | 423 | 355 | 217 | 257 |
| 84946 | NM_00132995 | LTV1 | LTV1 ribosome biogenesis factor | protein_coding | NP_001316882.1;NP_116249.2 | 21173 | . | . | . | 25.779031 | 25.373071 | 28.445324 | 22.736385 | 59.988863 | 58.650991 | 63.830480 | 51.984045 | -1.012092 | 1485.008047 | 0.144188487 | -0.1202601 | 0.904277109 | 0.998865404 | 10.544643 | 10.527818 | 10.548426 | 10.529409 | 10.617183 | 10.434358 | 1741 | 1592 | 1436 | 1220 |
| 84947 | NM_032861,N | SERAC1 | serine active site containing 1 | protein_coding | NP_116250.3;XP_006715649.1;XP_011534498.1;XP_011534500.1;XP_02 | 21061 | 614725 | . | . | 1.878989 | 2.176547 | 2.185180 | 2.248999 | 4.372484 | 5.031188 | 4.903480 | 5.142071 | 1.076098 | 257.898031 | 0.216419483 | 0.488910843 | 0.624904819 | 0.998865404 | 7.957048 | 8.062345 | 7.954179 | 8.025105 | 8.028430 | 8.026170 | 266 | 287 | 233 | 250 |
| 84948 | NM_032862 | TIGD5 | tigger transposable element derived 5 | protein_coding | NP_116251.4 | 18336 | . | . | . | 1.709810 | 1.487184 | 1.458222 | 1.859420 | 3.978798 | 3.437693 | 3.272207 | 4.251342 | 1.022907 | 274.088477 | 0.221883692 | 0.147263677 | 0.882923902 | 0.998865404 | 8.082351 | 8.114466 | 8.128321 | 8.051776 | 8.040771 | 8.160587 | 335 | 271 | 213 | 290 |
| 84950 | NM_032864,XI | PRPF38A | pre-mRNA processing factor 38A | protein_coding | NP_116253.2;XP_011540617.1 | 25930 | 617031 | . | . | 7.082909 | 7.392535 | 7.184676 | 6.384014 | 16.482220 | 17.088176 | 16.122204 | 14.596289 | -1.049751 | 1102.841396 | 0.139163239 | -0.50334559 | 0.614721318 | 0.998865404 | 10.141452 | 10.071725 | 10.111665 | 10.136577 | 10.135199 | 10.038930 | 1279 | 1239 | 997 | 943 |
| 84952 | NM_00125233 | CGNL1 | cingulin like 1 | protein_coding | NP_001239264.1;NP_116255.2;XP_005254783.1;XP_005254784.1;XP_01 | 25931 | 607856 | . | . | 0.024209 | 0.055140 | 0.025723 | 0.036376 | 0.056335 | 0.127459 | 0.057723 | 0.083168 | -1.206262 | 9.120029 | 1.034534385 | -0.26151191 | 0.793697765 | NA | 3.320504 | 3.044381 | 3.173786 | 3.209537 | 3.178369 | 3.190083 | 7 | 15 | 6 | 9 |
| 84954 | NM_00115984 | MPND | MPN domain containing | protein_coding | NP_001153318.1;NP_001287791.1;NP_116257.2;XP_006722989.1 | 25934 | . | . | . | 3.440826 | 3.238496 | 2.533663 | 3.433158 | 8.006943 | 7.485932 | 5.685467 | 7.849508 | -1.134940 | 158.727787 | 0.272615604 | -0.67070202 | 0.502410373 | 0.998865404 | 7.399444 | 7.215520 | 7.349185 | 7.320310 | 7.229238 | 7.336041 | 201 | 176 | 111 | 159 |
| 84955 | NM_00112821 | NUDCD1 | NudC domain containing 1 | protein_coding | NP_001121683.1;NP_116258.2;XP_016869399.1;XP_016869400.1 | 24306 | 606109 | . | . | 7.589139 | 7.039858 | 6.457848 | 5.974304 | 17.660235 | 16.272946 | 14.491223 | 13.659535 | -1.195130 | 827.842402 | 0.14774188 | -1.74065473 | 0.081744118 | 0.782498736 | 9.815840 | 9.559185 | 9.781807 | 9.722283 | 9.656367 | 9.595659 | 1080 | 933 | 685 | 676 |
| 84957 | NM_032871,N | RELT | RELT TNF receptor | protein_coding | NP_116260.2;NP_689408.1;XP_011543607.1 | 13764 | 611211 | . | . | 6.054550 | 7.003268 | 7.273139 | 6.286497 | 14.089187 | 16.188368 | 16.320712 | 14.373328 | 1.027668 | 708.417260 | 0.161713465 | 0.243480587 | 0.807633122 | 0.998865404 | 9.448549 | 9.488091 | 9.408530 | 9.503837 | 9.527105 | 9.422422 | 749 | 806 | 672 | 622 |
| 84959 | NM_00136336 | UBASH3B | ubiquitin associated and SH3 domain containing B | protein_coding | NP_001350294.1;NP_116262.2;XP_005271769.1;XP_011541343.1 | 29884 | 609201 | . | . | 5.082369 | 5.069340 | 3.984225 | 3.822327 | 11.826880 | 11.718006 | 8.940486 | 8.739296 | -1.318464 | 962.461866 | 0.138550673 | -2.87878626 | 0.003992088 | 0.185531278 | 10.096177 | 9.697545 | 10.006787 | 9.997410 | 9.819489 | 9.782659 | 1268 | 1174 | 741 | 758 |
| 84961 | NM_00118490 | FBXL20 | F-box and leucine rich repeat protein 20 | protein_coding | NP_001171835.1;NP_001357137.1;NP_001357138.1;NP_116264.2 | 24679 | 609086 | . | . | 1.003821 | 0.877092 | 1.111937 | 1.157051 | 2.335931 | 2.027435 | 2.495154 | 2.645460 | 1.191414 | 336.723503 | 0.196668223 | 1.284778855 | 0.198869573 | 0.96621494 | 8.263313 | 8.516440 | 8.380860 | 8.300557 | 8.433312 | 8.446121 | 380 | 307 | 315 | 347 |
| 84962 | NM_00128909 | AJUBA | ajuba LIM protein | protein_coding | NP_001276026.1;NP_116265.1;NP_932352.1 | 20250 | 609066 | . | . | 1.862549 | 1.968565 | 1.657666 | 1.875099 | 4.334228 | 4.550425 | 3.719756 | 4.287190 | -1.070760 | 227.511213 | 0.22686045 | -0.43478467 | 0.663718732 | 0.998865404 | 7.878611 | 7.779267 | 7.848094 | 7.844305 | 7.781329 | 7.844471 | 272 | 253 | 178 | 220 |
| 84964 | NM_00129770 | ALKBH6 | alkB homolog 6 | protein_coding | NP_001284630.1;NP_116267.3;NP_942567.1;XP_005259414.1;XP_02430 | 28243 | 613304 | . | . | 16.154832 | 15.591520 | 19.171692 | 18.098904 | 37.592958 | 36.040497 | 43.020721 | 41.380996 | 1.146597 | 627.453584 | 0.154912477 | 1.274000346 | 0.202663293 | 0.968254058 | 9.191280 | 9.388701 | 9.245399 | 9.243907 | 9.341003 | 9.331642 | 674 | 630 | 587 | 624 |
| 84967 | NM_032881 | LSM10 | LSM10, U7 small nuclear RNA associated | protein_coding | NP_116270.1 | 17562 | 617909 | . | . | 28.657425 | 31.009773 | 30.411209 | 34.372898 | 66.687012 | 71.680481 | 68.241875 | 78.589546 | 1.070971 | 833.612010 | 0.148147071 | 0.667712793 | 0.504316942 | 0.998865404 | 9.653203 | 9.751585 | 9.652725 | 9.701885 | 9.685696 | 9.765138 | 895 | 899 | 708 | 853 |
| 84968 | NM_032882 | PNMA6A | PNMA family member 6A | protein_coding | NP_116271.3 | 28248 | 300917 | . | . | 0.931172 | 0.997368 | 1.021243 | 1.475834 | 2.166877 | 2.305461 | 2.291640 | 3.374320 | 1.275035 | 77.533163 | 0.372741239 | 0.94043073 | 0.346996666 | 0.998865404 | 6.092061 | 6.440441 | 6.229529 | 6.245403 | 6.249226 | 6.348033 | 76 | 76 | 62 | 96 |
| 84969 | NM_00109879 | TOX2 | TOX high mobility group box family member 2 | protein_coding | NP_001092266.1;NP_001092267.1;NP_001092268.1;NP_116272.1;XP_0C | 16095 | 611163 | . | . | 1.671812 | 2.550490 | 1.452023 | 2.103441 | 3.890376 | 5.895573 | 3.258297 | 4.809266 | -1.162102 | 146.400638 | 0.308244219 | -0.70313205 | 0.481973459 | 0.998865404 | 7.299384 | 7.079827 | 7.127638 | 7.296309 | 7.096026 | 7.215999 | 144 | 205 | 98 | 148 |
| 84971 | NM_00128150 | ATG4D | autophagy related 4D cysteine peptidase | protein_coding | NP_001268433.1;NP_116274.3;XP_006722990.1 | 20789 | 611340 | . | . | 10.900381 | 12.683425 | 11.922676 | 11.450983 | 25.365634 | 29.318307 | 26.754141 | 26.181315 | -1.019166 | 713.515729 | 0.156049251 | -0.17551615 | 0.860674084 | 0.998865404 | 9.492533 | 9.464938 | 9.433442 | 9.533220 | 9.486295 | 9.456309 | 770 | 833 | 628 | 648 |
| 84973 | NR_003672,NF | SNHG7 | small nucleolar RNA host gene 7 | lncRNA | . | 28254 | . | . | . | 25.484660 | 29.732160 | 27.578708 | 29.075356 | 59.303851 | 68.727219 | 61.885818 | 66.477345 | -1.021605 | 1001.312845 | 0.141978971 | -0.21719775 | 0.828054236 | 0.998865404 | 9.983200 | 9.951986 | 9.927366 | 10.020478 | 9.947072 | 9.971206 | 1092 | 1161 | 847 | 944 |
| 84975 | NM_00117079 | MFSD5 | major facilitator superfamily domain containing 5 | protein_coding | NP_001164261.1;NP_116278.3;XP_005269254.1;XP_005269255.1 | 28156 | . | . | . | 16.043646 | 18.128500 | 15.534423 | 15.545464 | 37.334224 | 41.904842 | 34.858797 | 35.542855 | -1.094697 | 973.766749 | 0.142057795 | -0.91885903 | 0.358169312 | 0.998865404 | 9.991347 | 9.860554 | 9.913867 | 10.002606 | 9.892806 | 9.893420 | 1102 | 1164 | 808 | 872 |
| 84976 | NM_00135063 | DISP1 | dispatched RND transporter family member 1 | protein_coding | NP_001337559.1;NP_001356523.1;NP_001364157.1;NP_001364158.1;NI | 19711 | 607502 | . | . | 9.677084 | 10.053332 | 8.344906 | 7.888686 | 22.518970 | 23.238733 | 18.725729 | 18.036545 | -1.052068 | 330.002911 | 0.194236211 | -0.37700677 | 0.706168573 | 0.998865404 | 8.402555 | 8.329182 | 8.361758 | 8.398600 | 8.356674 | 8.347492 | 377 | 377 | 282 | 299 |
| 84978 | NM_00128649 | FRMD5 | FERM domain containing 5 | protein_coding | NP_001273419.1;NP_001273420.1;NP_001309878.1;NP_001309879.1;NI | 28214 | 616309 | . | . | 6.095164 | 7.194382 | 6.739656 | 7.315091 | 14.183699 | 16.630137 | 15.123594 | 16.725088 | 1.049017 | 1003.558500 | 0.138648037 | 0.497936239 | 0.618528984 | 0.998865404 | 9.936042 | 10.004954 | 9.921033 | 9.983321 | 9.996236 | 9.979057 | 1080 | 1102 | 904 | 952 |
| 84981 | NR_028502,NF | MIR22HG | MIR22 host gene | lncRNA | . | 28219 | . | . | . | 6.869655 | 7.254562 | 9.690657 | 6.111493 | 15.985970 | 16.769246 | 21.745557 | 13.973202 | 1.082395 | 598.842782 | 0.202557064 | 0.563924962 | 0.572805201 | 0.998865404 | 9.167298 | 9.282471 | 9.168362 | 9.214434 | 9.382476 | 9.084769 | 639 | 642 | 661 | 455 |
| 84983 | NR_026661,NF | FAM222A-AS1 | FAM222A antisense RNA 1 | lncRNA | . | 28223 | . | . | . | 0.247776 | 0.567857 | 0.283920 | 0.572755 | 0.576585 | 1.312627 | 0.637108 | 1.309535 | 1.060519 | 11.189653 | 0.945532652 | 0.089653034 | 0.928562939 | NA | 3.444631 | 3.522505 | 3.459757 | 3.495720 | 3.465919 | 3.495598 | 8 | 16 | 7 | 14 |
| 84984 | NM_00137946 | CEP19 | centrosomal protein 19 | protein_coding | NP_001366397.1;NP_001366398.1;NP_001366399.1;NP_116287.3 | 28209 | 615586 | . | . | 3.944474 | 4.048530 | 3.705260 | 4.366649 | 9.178953 | 9.358361 | 8.314498 | 9.983825 | 1.000132 | 269.855270 | 0.2135706 | 0.000891946 | 0.99928833 | 0.999862634 | 8.076352 | 8.075732 | 8.074258 | 8.076945 | 8.034375 | 8.115162 | 310 | 292 | 216 | 273 |
| 84986 | NM_00120430 | ARHGAP19 | Rho GTPase activating protein 19 | protein_coding | NP_001191229.1;NP_001243352.1;NP_116289.4 | 23724 | 611587 | . | . | 3.330709 | 3.175249 | 3.042825 | 3.303604 | 7.750697 | 7.339729 | 6.828011 | 7.553300 | -1.037757 | 543.964684 | 0.164323569 | -0.32539033 | 0.744885671 | 0.998865404 | 9.113920 | 9.060320 | 9.117672 | 9.080563 | 9.052705 | 9.097311 | 656 | 581 | 448 | 518 |
| 84987 | NM_00125713 | COX14 | cytochrome c oxidase assembly factor COX14 | protein_coding | NP_001244062.1;NP_001244063.1;NP_116290.1 | 28216 | 614478 | . | . | 19.680890 | 23.389752 | 18.943517 | 23.303616 | 45.798242 | 54.066464 | 42.508704 | 53.280949 | -1.022526 | 333.124110 | 0.202995843 | -0.15831785 | 0.874206343 | 0.998865404 | 8.396504 | 8.363135 | 8.346246 | 8.422281 | 8.322612 | 8.419824 | 362 | 388 | 261 | 336 |
| 84988 | NM_00132944 | PPP1R16A | protein phosphatase 1 regulatory subunit 16A | protein_coding | NP_001316371.1;NP_001316372.1;NP_001316373.1;NP_001316374.1;NI | 14941 | 609172 | . | . | 10.981837 | 12.693771 | 13.384341 | 13.334321 | 25.555186 | 29.342221 | 30.034073 | 30.487341 | 1.107970 | 1130.687639 | 0.138275915 | 1.069737813 | 0.284737341 | 0.998865404 | 10.067251 | 10.214941 | 10.053186 | 10.149518 | 10.185766 | 10.172045 | 1155 | 1233 | 1043 | 1104 |
| 84989 | NR_027182 | JMJD1C-AS1 | JMJD1C antisense RNA 1 | lncRNA | . | 28222 | . | . | . | 0.107694 | 0.153942 | 0.246317 | 0.242034 | 0.250607 | 0.355843 | 0.552729 | 0.553382 | 1.798531 | 8.178576 | 1.065322664 | 0.794894419 | 0.426674921 | NA | 2.548173 | 3.393446 | 3.004191 | 3.010363 | 3.027591 | 3.029195 | 6 | 7 | 9 | 10 |
| 84991 | NM_00114554 | RBM17 | RNA binding motif protein 17 | protein_coding | NP_001139019.1;NP_116294.1 | 16944 | 606935 | . | . | 13.006929 | 13.981069 | 13.027428 | 12.116446 | 30.267658 | 32.317868 | 29.233172 | 27.702816 | -1.088599 | 1342.763006 | 0.131055685 | -0.93451223 | 0.350039729 | 0.998865404 | 10.450867 | 10.328518 | 10.399082 | 10.444800 | 10.388554 | 10.325505 | 1564 | 1555 | 1165 | 1155 |
| 84992 | NM_00104261 | PIGY | phosphatidylinositol glycan anchor biosynthesis class Y | protein_coding | NP_001036081.1 | 28213 | 610662 | . | . | 7.514238 | 5.011446 | 10.158925 | 8.119757 | 17.485941 | 11.584183 | 22.796333 | 18.564859 | 1.437534 | 315.078443 | 0.237171677 | 2.207665988 | 0.027267565 | 0.538018865 | 8.013014 | 8.538527 | 8.276545 | 8.070270 | 8.443079 | 8.306875 | 358 | 222 | 361 | 308 |
| 84993 | NM_00128673 | UBL7 | ubiquitin like 7 | protein_coding | NP_001273668.1;NP_001273669.1;NP_001273670.1;NP_001273671.1;NI | 28221 | 609748 | . | . | 32.221138 | 32.056705 | 28.528254 | 39.397324 | 74.979916 | 74.100513 | 64.016573 | 90.077293 | 1.009899 | 1550.895079 | 0.145339562 | 0.097775652 | 0.922110441 | 0.998865404 | 10.591955 | 10.605782 | 10.615594 | 10.570360 | 10.501308 | 10.691403 | 1829 | 1617 | 1189 | 1640 |
| 84996 | NR_026845 | URB1-AS1 | URB1 antisense RNA 1 (head to head) | lncRNA | . | 23128 | . | . | . | 4.086052 | 3.979551 | 4.548120 | 5.046017 | 9.508411 | 9.198911 | 10.205849 | 11.537119 | 1.170067 | 114.244472 | 0.301581023 | 0.751343229 | 0.452446119 | 0.998865404 | 6.718456 | 6.944648 | 6.810993 | 6.802081 | 6.841888 | 6.874993 | 123 | 112 | 102 | 121 |
| 85001 | NR_026914 | MGC16275 | uncharacterized protein MGC16275 | lncRNA | . | . | . | . | . | 0.704302 | 0.521200 | 0.696657 | 0.765903 | 1.638940 | 1.204777 | 1.563279 | 1.751146 | 1.191937 | 52.822296 | 0.437967303 | 0.578372233 | 0.563012837 | 0.998865404 | 5.590318 | 5.844638 | 5.729435 | 5.673238 | 5.727463 | 5.743726 | 64 | 44 | 48 | 56 |
| 85002 | NM_00108353 | FAM86B1 | family with sequence similarity 86 member B1 | protein_coding | NP_001077006.1;XP_006716319.1;XP_006716320.1;XP_006716323.1;XP_ | 28268 | 616122 | . | . | 0.725337 | 0.677867 | 0.848334 | 0.514777 | 1.687890 | 1.566922 | 1.903637 | 1.176975 | 1.025362 | 52.262764 | 0.444136313 | 0.081356347 | 0.935158564 | 0.998865404 | 5.688401 | 5.726767 | 5.700228 | 5.702733 | 5.742366 | 5.668792 | 59 | 56 | 55 | 40 |
| 85007 | NM_00127834 | PHYKPL | 5-phosphohydroxy-L-lysine phospho-lyase | protein_coding | NP_001265275.1;NP_699204.1;XP_006714988.1;XP_011532971.1;XP_01 | 28249 | 614683 | . | . | 7.426667 | 8.604554 | 8.845999 | 9.302415 | 17.282159 | 19.889810 | 19.850167 | 21.268870 | 1.070616 | 1088.882108 | 0.1424252 | 0.691177744 | 0.489453848 | 0.998865404 | 10.038499 | 10.137081 | 10.023633 | 10.098080 | 10.158863 | 10.063244 | 1150 | 1192 | 1042 | 988 |
| 85009 | NR_026664 | MGC16025 | uncharacterized LOC85009 | lncRNA | . | . | . | . | . | 1.977369 | 1.779527 | 1.555467 | 2.057403 | 4.601420 | 4.113457 | 3.490423 | 4.704007 | -1.051669 | 122.614844 | 0.302070923 | -0.24060608 | 0.809860439 | 0.998865404 | 6.974386 | 6.900652 | 6.966309 | 6.925635 | 6.882990 | 6.972419 | 153 | 128 | 90 | 127 |
| 85012 | NM_00100693 | TCEAL3 | transcription elongation factor A like 3 | protein_coding | NP_001006934.1;NP_116315.1 | 28247 | . | . | . | 12.660677 | 12.314141 | 11.156153 | 12.496599 | 29.461916 | 28.464690 | 25.034081 | 28.571988 | -1.071285 | 406.709233 | 0.181964336 | -0.54594768 | 0.585101903 | 0.998865404 | 8.716868 | 8.617116 | 8.697627 | 8.678633 | 8.615112 | 8.677940 | 492 | 447 | 323 | 388 |
| 85013 | NM_00129755 | TMEM128 | transmembrane protein 128 | protein_coding | NP_001284480.1;NP_001284481.1;NP_116316.1;XP_005248091.1;XP_02 | 28201 | . | . | . | 11.739698 | 8.843746 | 9.610925 | 8.376284 | 27.318759 | 20.442716 | 21.566637 | 19.151376 | -1.114988 | 450.359618 | 0.181654916 | -0.81932842 | 0.41259906 | 0.998865404 | 8.890769 | 8.734890 | 8.920638 | 8.763794 | 8.817269 | 8.737977 | 614 | 449 | 395 | 373 |
| 85014 | NM_032928 | TMEM141 | transmembrane protein 141 | protein_coding | NP_116317.1 | 28211 | . | . | . | 25.803190 | 26.262043 | 21.589838 | 30.763601 | 60.045086 | 60.705891 | 48.446972 | 70.337318 | -1.008904 | 682.330309 | 0.180682585 | -0.07078399 | 0.943569671 | 0.998865404 | 9.421188 | 9.407432 | 9.411165 | 9.417151 | 9.285400 | 9.517343 | 786 | 743 | 490 | 745 |
| 85015 | NM_00108048 | USP45 | ubiquitin specific peptidase 45 | protein_coding | NP_001073950.1;NP_001332950.1;NP_001332951.1;NP_001332952.1;NI | 20080 | 618439 | . | . | 2.661938 | 2.828777 | 3.544704 | 1.934926 | 6.194443 | 6.538847 | 7.954213 | 4.423976 | -1.092176 | 423.567786 | 0.22010502 | -0.57792863 | 0.563312304 | 0.998865404 | 8.787846 | 8.662323 | 8.786760 | 8.703375 | 8.816647 | 8.551477 | 543 | 445 | 431 | 295 |

| | | | | | | | | | | | | | | | | | | | | | | | | | | | | | | | | | |
|---|---|---|---|---|---|---|---|---|---|---|---|---|---|---|---|---|---|---|---|---|---|---|---|---|---|---|---|---|---|---|---|---|---|
| 85019 | NM_00131883 | TMEM241 | transmembrane protein 241 | protein_coding | NP_001305763.1;NP_116322.3;XP_011524535.1;XP_016881532.1;XP_01 | 31723 | 615430 | . | . | 1.316189 | 1.919887 | 1.870943 | 1.911312 | 3.062829 | 4.437901 | 4.198343 | 4.369989 | -1.146358 | 213.714592 | 0.229807804 | -0.85748815 | 0.391175161 | 0.998865404 | 7.834827 | 7.637515 | 7.763875 | 7.778090 | 7.702205 | 7.712840 | 259 | 250 | 171 | 189 |
| 85021 | NM_00112861 | REPS1 | RALBP1 associated Eps domain containing 1 | protein_coding | NP_001122089.1;NP_001273540.1;NP_001273541.1;NP_114128.3;XP_00 | 15578 | 614825 | . | . | 6.946531 | 8.115597 | 7.332086 | 7.400994 | 16.164863 | 18.759566 | 16.452990 | 16.921491 | -1.026908 | 884.330715 | 0.140795064 | -0.27207757 | 0.785562373 | 0.998865404 | 9.807443 | 9.769188 | 9.790306 | 9.805438 | 9.789314 | 9.768283 | 1021 | 977 | 770 | 806 |
| 85025 | NM_032936 | TMEM60 | transmembrane protein 60 | protein_coding | NP_116325.1 | 21754 | . | . | . | 36.091606 | 32.459942 | 37.443531 | 26.738115 | 83.986656 | 75.032608 | 84.022202 | 61.133522 | -1.083449 | 907.706412 | 0.174860434 | -0.66127398 | 0.508436628 | 0.998865404 | 9.882272 | 9.767614 | 9.891425 | 9.807174 | 9.907645 | 9.662699 | 1149 | 960 | 888 | 677 |
| 85026 | NR_122035,NF | ARRDC1-AS1 | ARRDC1 antisense RNA 1 | lncRNA | . | 23395 | . | . | . | 5.873941 | 5.448259 | 5.881774 | 6.273315 | 13.668901 | 12.593893 | 13.198530 | 14.343189 | 1.071769 | 269.802720 | 0.211959702 | 0.47175636 | 0.637100703 | 0.998865404 | 8.024817 | 8.124968 | 8.085381 | 8.027052 | 8.077757 | 8.106749 | 317 | 265 | 236 | 269 |
| 85027 | NM_032947 | SMIM3 | small integral membrane protein 3 | protein_coding | NP_116565.3 | 30248 | 608324 | . | . | 2.384696 | 2.529278 | 2.018409 | 2.909360 | 5.549285 | 5.846539 | 4.529251 | 6.651906 | -1.008752 | 179.027743 | 0.265661459 | -0.04731998 | 0.962258206 | 0.998865404 | 7.491242 | 7.476801 | 7.472616 | 7.494351 | 7.404050 | 7.550147 | 202 | 199 | 128 | 196 |
| 85028 | NR_024127,NF | SNHG12 | small nucleolar RNA host gene 12 | lncRNA | . | 30062 | . | . | . | 34.254473 | 36.770496 | 45.073268 | 35.754916 | 79.711565 | 84.996647 | 101.143109 | 81.749367 | 1.133730 | 1164.899258 | 0.157123404 | 1.152449733 | 0.249136315 | 0.992733415 | 10.092412 | 10.273865 | 10.093838 | 10.171848 | 10.324917 | 10.120549 | 1191 | 1240 | 1221 | 1005 |
| 85236 | NM_00131265 | H2BC12 | H2B clustered histone 12 | protein_coding | NP_001299582.1;NP_542160.1 | 13954 | 615045 | . | . | 13.690473 | 12.668498 | 15.209299 | 11.547282 | 31.858293 | 29.283800 | 34.129224 | 26.401486 | -1.000378 | 347.064050 | 0.208005204 | -0.0026237 | 0.997906591 | 0.999862634 | 8.438615 | 8.439501 | 8.460875 | 8.410736 | 8.513287 | 8.352535 | 417 | 358 | 345 | 279 |
| 85294 | NM_033184 | KRTAP2-4 | keratin associated protein 2-4 | protein_coding | NP_149440.1 | 18891 | . | . | . | 0.072162 | 0.015957 | 0.024204 | 0.073129 | 0.167925 | 0.036884 | 0.054314 | 0.167202 | -1.076094 | 1.703845 | 2.268349467 | -0.04664372 | 0.962797187 | NA | 0.820514 | 0.715150 | 0.778739 | 0.770724 | 0.771939 | 0.776369 | 3 | 1 | 1 | 2 |
| 85301 | NM_032888,XI | COL27A1 | collagen type XXVII alpha 1 chain | protein_coding | NP_116277.2;XP_006717371.1;XP_006717373.1;XP_011517440.1;XP_01 | 22986 | 608461 | . | . | 8.350287 | 10.267751 | 9.340007 | 7.111594 | 19.431464 | 23.734363 | 20.958699 | 16.259818 | 1.005235 | 1801.713704 | 0.143936764 | 0.05233242 | 0.958263821 | 0.998865404 | 10.811194 | 10.819103 | 10.793883 | 10.827439 | 10.911464 | 10.708605 | 2025 | 1980 | 1773 | 1476 |
| 85302 | NM_00131919 | FBF1 | Fas binding factor 1 | protein_coding | NP_001306122.1 | 24674 | 616807 | . | . | 5.131147 | 5.125266 | 5.741563 | 5.197556 | 11.940390 | 11.847282 | 12.883901 | 11.883595 | 1.051943 | 772.745289 | 0.149466545 | 0.488779417 | 0.624997872 | 0.998865404 | 9.556649 | 9.630114 | 9.578131 | 9.571989 | 9.648580 | 9.571279 | 871 | 809 | 727 | 702 |
| 85313 | NM_139126 | PPIL4 | peptidylprolyl isomerase like 4 | protein_coding | NP_624311.1 | 15702 | 607609 | . | . | 9.152815 | 9.435603 | 10.839732 | 9.858031 | 21.298977 | 21.810820 | 24.324045 | 22.539219 | 1.097875 | 757.487305 | 0.150134025 | 0.897287832 | 0.369565348 | 0.998865404 | 9.495989 | 9.631009 | 9.522580 | 9.538408 | 9.632349 | 9.557861 | 822 | 788 | 726 | 704 |
| 85315 | NM_133367 | PAQR8 | progestin and adipoQ receptor family member 8 | protein_coding | NP_588608.1 | 15708 | 607780 | . | . | 2.985916 | 2.308996 | 2.978172 | 2.981227 | 6.948349 | 5.337348 | 6.682933 | 6.816222 | 1.109292 | 413.486572 | 0.194227131 | 0.770431157 | 0.441044177 | 0.998865404 | 8.614506 | 8.764966 | 8.732201 | 8.575268 | 8.721789 | 8.716526 | 511 | 367 | 380 | 406 |
| 85319 | NM_182482 | BAGE2 | BAGE family member 2 | protein_coding | NP_872288.2 | 15723 | 617776 | . | . | 0.560523 | 0.724677 | 0.611486 | 0.551862 | 1.304360 | 1.675124 | 1.372157 | 1.261767 | -1.107901 | 36.823025 | 0.515733644 | -0.28663821 | 0.774389351 | 0.998865404 | 5.274751 | 5.126516 | 5.190020 | 5.228485 | 5.201928 | 5.187420 | 39 | 47 | 32 | 31 |
| 85352 | NM_00109929 | SHISAL1 | shisa like 1 | protein_coding | NP_001092764.1;XP_005261847.1 | 29335 | . | . | . | 0.166395 | 0.178449 | 0.168051 | 0.169493 | 0.387208 | 0.412492 | 0.377101 | 0.387526 | -1.031768 | 37.115926 | 0.507910641 | -0.0888326 | 0.929214949 | 0.998865404 | 5.236451 | 5.191126 | 5.211592 | 5.221348 | 5.212984 | 5.210904 | 42 | 42 | 32 | 34 |
| 85358 | NM_00137204 | SHANK3 | SH3 and multiple ankyrin repeat domains 3 | protein_coding | NP_001358973.1 | 14294 | 606230 | . | . | 0.771557 | 0.816733 | 0.960278 | 1.064992 | 1.795446 | 1.887914 | 2.154837 | 2.434978 | 1.257790 | 215.709129 | 0.230450339 | 1.435843849 | 0.151046779 | 0.913627824 | 7.578404 | 7.908628 | 7.682249 | 7.703146 | 7.775822 | 7.822883 | 215 | 211 | 199 | 236 |
| 85359 | NM_033257 | DGCR6L | DiGeorge syndrome critical region gene 6 like | protein_coding | NP_150282.2 | 18551 | 609459 | . | . | 23.286337 | 22.194672 | 22.894354 | 28.453011 | 54.188267 | 51.303974 | 51.374268 | 65.054428 | 1.113114 | 883.865754 | 0.154708365 | 0.999307619 | 0.317645696 | 0.998865404 | 9.708550 | 9.862701 | 9.766195 | 9.725649 | 9.745031 | 9.895459 | 991 | 877 | 726 | 963 |
| 85360 | NM_00130091 | SYDE1 | synapse defective Rho GTPase homolog 1 | protein_coding | NP_001287839.1;NP_149014.3;XP_024307518.1 | 25824 | 617377 | . | . | 6.503446 | 7.238724 | 6.628346 | 7.911094 | 15.133786 | 16.732636 | 14.873818 | 18.087777 | 1.056723 | 704.413615 | 0.16113554 | 0.493978528 | 0.621321344 | 0.998865404 | 9.420349 | 9.499134 | 9.406926 | 9.471324 | 9.414883 | 9.536324 | 752 | 774 | 574 | 738 |
| 85363 | NM_033034,N | TRIM5 | tripartite motif containing 5 | protein_coding | NP_149023.2;NP_149083.2;NP_149084.2;XP_005253240.1;XP_00525324 | 16276 | 608487 | . | . | 4.660530 | 4.387801 | 5.077153 | 4.660769 | 10.845244 | 10.142600 | 11.392983 | 10.656295 | 1.045565 | 484.767095 | 0.17197923 | 0.373782892 | 0.708565849 | 0.998865404 | 8.888332 | 8.953234 | 8.924716 | 8.887137 | 8.965753 | 8.901691 | 562 | 496 | 453 | 441 |
| 85364 | NM_033089 | ZCCHC3 | zinc finger CCHC-type containing 3 | protein_coding | NP_149080.2 | 16230 | 618326 | . | . | 7.953079 | 8.235328 | 9.161004 | 8.701130 | 18.507141 | 19.036329 | 20.557028 | 19.894098 | 1.088449 | 729.731357 | 0.149456244 | 0.818120231 | 0.413288538 | 0.998865404 | 9.448711 | 9.571136 | 9.471232 | 9.489506 | 9.560505 | 9.517312 | 794 | 764 | 682 | 691 |
| 85365 | NM_033087,N | ALG2 | ALG2 alpha-1,3/1,6-mannosyltransferase | protein_coding | NP_149078.1 | 23159 | 607905 | . | . | 10.241775 | 9.509425 | 9.298783 | 8.722430 | 23.833032 | 21.982462 | 20.866200 | 19.942801 | -1.109018 | 842.068458 | 0.147515355 | -1.01197737 | 0.311548872 | 0.998865404 | 9.790261 | 9.641493 | 9.786282 | 9.718663 | 9.708798 | 9.649570 | 1067 | 911 | 724 | 717 |
| 85366 | NM_033118 | MYLK2 | myosin light chain kinase 2 | protein_coding | NP_149109.1 | 16243 | 606566 | . | . | 0.080446 | 0.152598 | 0.108063 | 0.184382 | 0.187201 | 0.352737 | 0.242490 | 0.421569 | 1.217804 | 12.016732 | 0.898243391 | 0.31648611 | 0.751633566 | NA | 3.440763 | 3.719719 | 3.561587 | 3.589446 | 3.573861 | 3.598149 | 9 | 15 | 9 | 15 |
| 85369 | NM_00127076 | STRIP1 | striatin interacting protein 1 | protein_coding | NP_001257697.1;NP_149079.2;XP_006711058.1;XP_016858118.1 | 25916 | 617918 | . | . | 10.937973 | 11.795400 | 12.556405 | 13.408971 | 25.453117 | 27.265596 | 28.176211 | 30.658019 | 1.117835 | 1212.176136 | 0.133032629 | 1.20802518 | 0.227037577 | 0.982008277 | 10.160963 | 10.321349 | 10.175110 | 10.222964 | 10.257646 | 10.307855 | 1276 | 1275 | 1076 | 1239 |
| 85371 | NR_046312 | KIAA1656 | KIAA1656 protein | lncRNA | . | . | . | . | . | 1.096272 | 0.965845 | 1.372597 | 1.145621 | 2.551070 | 2.232595 | 3.080067 | 2.619326 | 1.204314 | 271.803369 | 0.218788221 | 1.22589754 | 0.220237235 | 0.982008277 | 7.945358 | 8.214909 | 8.064792 | 7.997642 | 8.177561 | 8.075932 | 303 | 248 | 283 | 252 |
| 85376 | NM_015672 | RIMBP3 | RIMS binding protein 3 | protein_coding | NP_056487.1 | 29344 | 612699 | . | . | 0.026675 | 0.030996 | 0.010577 | 0.071243 | 0.062073 | 0.071648 | 0.023733 | 0.162888 | 1.390100 | 6.966122 | 1.230146203 | 0.386286283 | 0.699284658 | NA | 2.548173 | 3.014946 | 2.783506 | 2.789562 | 2.771226 | 2.822034 | 6 | 7 | 2 | 13 |
| 85377 | NM_033386,XI | MICALL1 | MICAL like 1 | protein_coding | NP_203744.1;XP_005261848.1;XP_005261849.1;XP_011528773.1;XP_01 | 29804 | . | . | . | 8.488858 | 8.761080 | 9.141177 | 10.739817 | 19.753922 | 20.251627 | 20.512534 | 24.555314 | 1.141562 | 1309.633198 | 0.136382312 | 1.400546317 | 0.161349784 | 0.933525128 | 10.256490 | 10.447112 | 10.297201 | 10.305344 | 10.341015 | 10.459397 | 1401 | 1327 | 1121 | 1409 |
| 85378 | NM_020461,XI | TUBGCP6 | tubulin gamma complex associated protein 6 | protein_coding | NP_065194.3 | 18127 | 610053 | . | . | 15.482655 | 18.137744 | 18.535924 | 16.790733 | 36.028775 | 41.926208 | 41.594077 | 38.390013 | 1.076630 | 3156.021057 | 0.119535909 | 0.891134762 | 0.372856878 | 0.998865404 | 11.569597 | 11.676216 | 11.563479 | 11.619853 | 11.702782 | 11.599965 | 3379 | 3393 | 3023 | 2877 |
| 85379 | NM_00114520 | KIAA1671 | KIAA1671 | protein_coding | NP_001138678.1;XP_005261850.1;XP_005261851.1;XP_006724409.1 | 29345 | . | . | . | 0.808766 | 0.695179 | 0.753717 | 0.884766 | 1.882032 | 1.606937 | 1.691318 | 2.022912 | 1.076264 | 264.562174 | 0.219940266 | 0.482094762 | 0.629738631 | 0.998865404 | 7.993517 | 8.099465 | 8.068099 | 7.983294 | 8.025575 | 8.101236 | 318 | 252 | 221 | 276 |
| 85388 | NR_001452 | SNORD14B | small nucleolar RNA, C/D box 14B | snoRNA | . | 31803 | . | . | . | 3.173196 | 2.750338 | 7.895103 | 1.591276 | 7.384158 | 6.357528 | 17.716381 | 3.638263 | 1.576129 | 11.556596 | 0.975066712 | 0.673170394 | 0.500838894 | NA | 3.161369 | 3.824402 | 3.496306 | 3.490866 | 3.553436 | 3.477928 | 11 | 9 | 20 | 5 |
| 85389 | NR_001453 | SNORD14C | small nucleolar RNA, C/D box 14C | snoRNA | . | 30352 | . | . | . | 2.555625 | 3.965715 | 5.721635 | 4.542738 | 5.947045 | 9.166928 | 12.839185 | 10.386431 | 1.526860 | 11.959140 | 0.88820073 | 0.687420746 | 0.491817628 | NA | 3.243129 | 3.852980 | 3.548928 | 3.564332 | 3.585634 | 3.571772 | 9 | 12 | 14 | 12 |
| 85390 | NR_001454 | SNORD14D | small nucleolar RNA, C/D box 14D | snoRNA | . | 30353 | . | . | . | 7.259334 | 6.756403 | 6.241976 | 3.265769 | 16.892769 | 15.617729 | 14.006811 | 7.466794 | -1.438713 | 16.339670 | 0.774428772 | -0.67763401 | 0.498003774 | NA | 4.267126 | 3.746830 | 4.048534 | 4.042069 | 4.036261 | 4.003528 | 23 | 20 | 15 | 9 |
| 85391 | NR_003125 | SNORD14E | small nucleolar RNA, C/D box 14E | snoRNA | . | 30354 | . | . | . | 2.483671 | 3.741726 | 3.712197 | 2.629329 | 5.779604 | 8.649166 | 8.330063 | 6.011648 | 1.039188 | 8.739542 | 1.032096721 | 0.053731412 | 0.957149156 | NA | 3.099441 | 3.155137 | 3.115324 | 3.130342 | 3.130634 | 3.118057 | 8 | 11 | 9 | 7 |
| 85395 | NM_00131698 | FAM207A | family with sequence similarity 207 member A | protein_coding | NP_001303912.1;NP_001303913.1;NP_001303914.1;NP_001303915.1;NI | 15811 | . | . | . | 12.533369 | 11.494234 | 11.727405 | 13.955900 | 29.165668 | 26.569436 | 26.315960 | 31.908507 | 1.062686 | 351.802443 | 0.196123048 | 0.447248558 | 0.654695605 | 0.998865404 | 8.414334 | 8.501591 | 8.462832 | 8.416260 | 8.425722 | 8.520797 | 409 | 353 | 289 | 368 |
| 85403 | NM_033083,XI | EAF1 | ELL associated factor 1 | protein_coding | NP_149074.3;XP_005265575.1;XP_011532467.1;XP_011532468.1 | 20907 | 608315 | . | . | 5.923405 | 6.005246 | 5.812019 | 5.977241 | 13.784007 | 13.881397 | 13.042003 | 13.666253 | -1.024777 | 817.689334 | 0.143491378 | -0.24608013 | 0.805620201 | 0.998865404 | 9.693000 | 9.657601 | 9.681029 | 9.687165 | 9.660426 | 9.673030 | 949 | 897 | 696 | 764 |
| 85406 | NM_032364 | DNAJC14 | DnaJ heat shock protein family (Hsp40) member C14 | protein_coding | NP_115740.5 | 24581 | 606092 | . | . | 11.081956 | 11.030939 | 10.177268 | 12.655602 | 25.788166 | 25.498510 | 22.837494 | 28.935528 | 1.017877 | 1165.668581 | 0.145472571 | 0.175720864 | 0.860513243 | 0.998865404 | 10.174352 | 10.199423 | 10.183792 | 10.174311 | 10.111950 | 10.266642 | 1339 | 1239 | 917 | 1216 |
| 85407 | NM_033119,XI | NKD1 | NKD inhibitor of WNT signaling pathway 1 | protein_coding | NP_149110.1;XP_011521701.1;XP_011521702.1;XP_016879293.1 | 17045 | 607851 | . | . | 0.028633 | 0.023882 | 0.036844 | 0.038080 | 0.066631 | 0.055205 | 0.082678 | 0.087065 | 1.373666 | 19.111570 | 0.70141178 | 0.653012784 | 0.513748037 | NA | 4.008843 | 4.467590 | 4.241980 | 4.230889 | 4.260886 | 4.263403 | 20 | 16 | 19 | 21 |
| 85414 | NM_033102 | SLC45A3 | solute carrier family 45 member 3 | protein_coding | NP_149093.1 | 8642 | 605097 | . | . | 4.471849 | 3.851582 | 4.329100 | 4.517621 | 10.406177 | 8.903105 | 9.714375 | 10.329002 | 1.046938 | 454.069833 | 0.178804154 | 0.370101083 | 0.711307175 | 0.998865404 | 8.793125 | 8.859649 | 8.858804 | 8.761738 | 8.829068 | 8.850432 | 551 | 441 | 397 | 443 |
| 85415 | NM_033103 | RHPN2 | rhophilin Rho GTPase binding protein 2 | protein_coding | NP_149094.3 | 19974 | 617932 | . | . | 3.235646 | 3.129120 | 3.369714 | 3.518558 | 7.529481 | 7.233101 | 7.561543 | 8.044764 | 1.066816 | 361.600134 | 0.187294103 | 0.498208726 | 0.618339633 | 0.998865404 | 8.450851 | 8.544144 | 8.490152 | 8.467316 | 8.505868 | 8.525932 | 411 | 370 | 319 | 356 |
| 85416 | NM_033132,N | ZIC5 | Zic family member 5 | protein_coding | NP_149123.2 | 20322 | 617896 | . | . | 0.645449 | 0.788302 | 0.594604 | 0.665404 | 1.501986 | 1.822195 | 1.334276 | 1.521366 | -1.236625 | 89.358814 | 0.340531415 | -0.89979223 | 0.368230829 | 0.998865404 | 6.626345 | 6.320558 | 6.495212 | 6.526184 | 6.471305 | 6.430452 | 108 | 112 | 75 | 69 |
| 85417 | NM_033031,N | CCNB3 | cyclin B3 | protein_coding | NP_149020.2;NP_391990.1;XP_016885402.1;XP_016885403.1;XP_01688 | 18709 | 300456 | . | . | 0.345791 | 0.354013 | 0.353263 | 0.511558 | 0.804670 | 0.818317 | 0.792712 | 1.169617 | 1.227581 | 58.390790 | 0.420986371 | 0.702680188 | 0.482255077 | 0.998865404 | 5.713297 | 6.007108 | 5.838915 | 5.841867 | 5.843026 | 5.921759 | 60 | 57 | 46 | 71 |
| 85437 | NM_033114,XI | ZCRB1 | zinc finger CCHC-type and RNA binding motif containing 1 | protein_coding | NP_149105.3;XP_016875613.1 | 29620 | 610750 | . | . | 17.655871 | 18.761505 | 15.727159 | 17.363510 | 41.085938 | 43.368059 | 35.291291 | 39.699601 | -1.127076 | 964.887680 | 0.141223386 | -1.22207177 | 0.221680489 | 0.982008277 | 9.998134 | 9.825116 | 9.934683 | 9.977046 | 9.836083 | 9.900568 | 1144 | 1135 | 753 | 889 |
| 85439 | NM_00125643 | STON2 | stonin 2 | protein_coding | NP_001243359.1;NP_001353778.1;NP_001353779.1;XP_011535542.1;XP | 30652 | 608467 | . | . | 0.350071 | 0.373314 | 0.387024 | 0.382838 | 0.814631 | 0.862932 | 0.868471 | 0.875314 | -1.050431 | 118.009987 | 0.306243723 | -0.23178151 | 0.816707722 | 0.998865404 | 6.918518 | 6.846104 | 6.897449 | 6.883778 | 6.827063 | 6.918746 | 142 | 128 | 86 | 123 |
| 85440 | NM_00127199 | DOCK7 | dedicator of cytokinesis 7 | protein_coding | NP_001258928.1;NP_001258929.1;NP_001258930.1;NP_001258931.1;NI | 19190 | 615730 | . | . | 9.198922 | 8.785887 | 7.894297 | 7.927309 | 21.406267 | 20.308971 | 17.714571 | 18.124850 | -1.144302 | 1855.281591 | 0.119660199 | -1.62516677 | 0.104127028 | 0.828395026 | 10.951357 | 10.756944 | 10.921224 | 10.895157 | 10.802684 | 10.801321 | 2317 | 2102 | 1506 | 1621 |
| 85441 | NM_00103733 | HELZ2 | helicase with zinc finger 2 | protein_coding | NP_001032412.2;NP_208384.3;XP_024307774.1;XP_024307775.1 | 30021 | 611265 | . | . | 1.187082 | 1.334066 | 1.288399 | 1.158526 | 2.762390 | 3.083753 | 2.891131 | 2.648832 | -1.018756 | 383.902502 | 0.186963447 | -0.14338869 | 0.885983213 | 0.998865404 | 8.597839 | 8.571230 | 8.563560 | 8.614475 | 8.608210 | 8.547868 | 427 | 436 | 348 | 338 |
| 85442 | NM_00134786 | KNDC1 | kinase non-catalytic C-lobe domain containing 1 | protein_coding | NP_001334793.1;NP_001334794.1;NP_001334795.1;NP_689856.6;XP_01 | 29374 | 616237 | . | . | 0.171597 | 0.076737 | 0.100553 | 0.124851 | 0.399313 | 0.177381 | 0.225638 | 0.285457 | -1.060495 | 10.777651 | 0.946234618 | -0.08955241 | 0.9286429 | NA | 3.471023 | 3.387716 | 3.449526 | 3.416098 | 3.424660 | 3.435571 | 17 | 8 | 8 | 11 |
| 85444 | NM_00134963 | LRRCC1 | leucine rich repeat and coiled-coil centrosomal protein 1 | protein_coding | NP_001336565.1;NP_001336566.1;NP_001336567.1;NP_001336568.1;NI | 29373 | 617791 | . | . | 2.602553 | 2.749577 | 1.920968 | 1.654811 | 6.056261 | 6.355769 | 4.310598 | 3.783529 | -1.413322 | 247.718965 | 0.221444945 | -2.25379073 | 0.024209336 | 0.514295174 | 8.180234 | 7.682100 | 8.030001 | 8.029478 | 7.894643 | 7.821963 | 334 | 313 | 193 | 177 |
| 85446 | NM_033400,XI | ZFHX2 | zinc finger homeobox 2 | protein_coding | NP_207646.2;XP_005268192.1;XP_011535547.1;XP_011535548.1;XP_01 | 20152 | 617828 | . | . | 0.033146 | 0.043636 | 0.043206 | 0.048523 | 0.077131 | 0.100866 | 0.096954 | 0.110942 | 1.270090 | 15.304772 | 0.781816737 | 0.441191489 | 0.659074375 | NA | 3.754145 | 4.097315 | 3.916360 | 3.928502 | 3.933929 | 3.942636 | 14 | 16 | 14 | 17 |
| 85450 | NM_00127201 | ITPRIP | inositol 1,4,5-trisphosphate receptor interacting protein | protein_coding | NP_001258941.1;NP_001258942.1;NP_203755.1;XP_005270314.1 | 29370 | . | . | . | 4.166650 | 4.259092 | 3.028389 | 3.755339 | 9.695965 | 9.845083 | 6.795618 | 8.586135 | -1.258648 | 781.568420 | 0.157277579 | -2.11012532 | 0.034847563 | 0.582325556 | 9.766974 | 9.434354 | 9.679885 | 9.688756 | 9.453043 | 9.588171 | 997 | 946 | 541 | 714 |
| 85451 | NM_00108041 | UNK | unk zinc finger | protein_coding | NP_001073888.2;XP_006722208.1;XP_006722211.1;XP_011523702.1;XP_ | 29369 | 616375 | . | . | 10.844774 | 8.455302 | 7.323968 | 7.437409 | 25.236234 | 19.544812 | 16.434771 | 17.004749 | 1.152212 | 1034.346458 | 0.143143038 | 1.427989904 | 0.153294765 | 0.917721153 | 9.908527 | 10.113226 | 9.939972 | 9.976385 | 10.116342 | 10.006320 | 1079 | 1063 | 1034 | 962 |
| 85455 | NM_033510,XI | DISP2 | dispatched RND transporter family member 2 | protein_coding | NP_277045.1;XP_011520427.1;XP_016878176.1 | 19712 | 607503 | . | . | 0.317034 | 0.343159 | 0.339676 | 0.396578 | 0.737752 | 0.793226 | 0.762224 | 0.906729 | 1.099900 | 137.540257 | 0.280244843 | 0.49018661 | 0.624001855 | 0.998865404 | 7.033958 | 7.170244 | 7.071944 | 7.095844 | 7.090969 | 7.146610 | 146 | 146 | 116 | 145 |
| 85456 | NM_033396,XI | TNKS1BP1 | tankyrase 1 binding protein 1 | protein_coding | NP_203754.2;XP_006718788.1;XP_011543627.1 | 19081 | 607104 | . | . | 6.564563 | 7.072332 | 5.992956 | 6.825377 | 15.276007 | 16.348013 | 13.448020 | 15.605414 | -1.078020 | 1197.495833 | 0.136676231 | -0.79300236 | 0.42777645 | 0.998865404 | 10.279199 | 10.170359 | 10.229452 | 10.276173 | 10.151828 | 10.239189 | 1387 | 1382 | 943 | 1144 |
| 85457 | NM_033426 | CIPC | CLOCK interacting pacemaker | protein_coding | NP_219494.2 | 20365 | 616995 | . | . | 4.586248 | 5.030948 | 5.057243 | 5.062799 | 10.672389 | 11.629263 | 11.348308 | 11.575489 | 1.037992 | 665.106833 | 0.154392355 | 0.348435967 | 0.727512795 | 0.998865404 | 9.350397 | 9.403990 | 9.335160 | 9.392409 | 9.392757 | 9.386150 | 720 | 734 | 592 | 632 |
| 85458 | NM_00103795 | DIXDC1 | DIX domain containing 1 | protein_coding | NP_001033043.1;NP_001265471.1;NP_219493.1;XP_016873955.1;XP_01 | 23695 | 610493 | . | . | 1.999702 | 2.103673 | 2.091398 | 1.978206 | 4.653387 | 4.862733 | 4.693037 | 4.522932 | 1.007214 | 361.175839 | 0.189070366 | 0.054848328 | 0.956259298 | 0.998865404 | 8.491242 | 8.501854 | 8.479357 | 8.508322 | 8.522691 | 8.472921 | 404 | 398 | 329 | 325 |
| 85459 | NM_033395,XI | CEP295 | centrosomal protein 295 | protein_coding | NP_203753.1;XP_005274423.1;XP_005274424.1;XP_005274425.1;XP_01 | 29366 | 617728 | . | . | 3.263662 | 3.496048 | 3.923833 | 3.125902 | 7.594675 | 8.081270 | 8.804972 | 7.147001 | 1.028957 | 871.150898 | 0.158606781 | 0.259649313 | 0.795134294 | 0.998865404 | 9.745780 | 9.787477 | 9.730273 | 9.777643 | 9.855026 | 9.687800 | 956 | 957 | 858 | 732 |
| 85460 | NM_00137581 | ZNF518B | zinc finger protein 518B | protein_coding | NP_001362745.1;NP_001362746.1;NP_444270.2;XP_016864275.1;XP_02 | 29365 | 617734 | . | . | 3.648491 | 3.210283 | 3.519131 | 3.587010 | 8.490189 | 7.420713 | 7.896828 | 8.201271 | -1.010391 | 831.371609 | 0.148147033 | -0.1006667 | 0.919815044 | 0.998865404 | 9.706544 | 9.692120 | 9.741029 | 9.662374 | 9.718209 | 9.671773 | 1015 | 852 | 743 | 750 |
| 85461 | NM_00114590 | TANC1 | tetratricopeptide repeat, ankyrin repeat and coiled-coil containing | protein_coding | NP_001139381.1;NP_001336991.1;NP_001336992.1;NP_001336993.1;NI | 29364 | 611397 | . | . | 19.965750 | 21.046713 | 18.740958 | 20.318885 | 46.461120 | 48.650419 | 42.054168 | 46.456716 | -1.021257 | 4080.395719 | 0.10848988 | -0.27970688 | 0.779702396 | 0.998865404 | 12.009625 | 11.979201 | 11.985048 | 12.020864 | 11.973084 | 11.997990 | 4646 | 4545 | 3458 | 3842 |
| 85462 | NM_00137111 | FHDC1 | FH2 domain containing 1 | protein_coding | NP_001358045.1;NP_203751.2;XP_005263376.1;XP_011530691.1 | 29363 | . | . | . | 2.178989 | 2.302609 | 2.069163 | 2.334058 | 5.070596 | 5.322584 | 4.643141 | 5.336547 | -1.029661 | 455.765964 | 0.175096982 | -0.24083169 | 0.809685565 | 0.998865404 | 8.853372 | 8.810610 | 8.828524 | 8.853378 | 8.790502 | 8.854228 | 522 | 509 | 370 | 443 |
| 85463 | NM_033390,XI | ZC3H12C | zinc finger CCCH-type containing 12C | protein_coding | NP_203748.1;XP_005271772.1;XP_011541357.1;XP_016873966.1 | 29362 | 615001 | . | . | 2.537718 | 2.512123 | 2.699595 | 2.204508 | 5.905376 | 5.806884 | 6.057813 | 5.040348 | -1.015590 | 699.385597 | 0.164847267 | -0.13538376 | 0.892308463 | 0.998865404 | 9.460670 | 9.439138 | 9.458210 | 9.448527 | 9.518393 | 9.361462 | 817 | 755 | 673 | 576 |
| 85464 | NM_00128212 | SSH2 | slingshot protein phosphatase 2 | protein_coding | NP_001269058.1;NP_001269059.1;NP_001269060.1;NP_203747.2;XP_00 | 30580 | 606779 | . | . | 3.292907 | 3.240926 | 3.294643 | 3.426989 | 7.662732 | 7.491546 | 7.393085 | 7.835407 | -1.021060 | 840.729654 | 0.143429522 | -0.20963651 | 0.833951384 | 0.998865404 | 9.730558 | 9.700280 | 9.723466 | 9.722341 | 9.686928 | 9.728674 | 979 | 916 | 702 | 803 |
| 85465 | NM_033505,N | SELENOI | selenoprotein I | protein_coding | NP_277040.1 | 29361 | 607915 | . | . | 11.535550 | 10.108605 | 11.594296 | 8.330700 | 26.843698 | 23.366493 | 26.017264 | 19.047155 | -1.101124 | 2613.069696 | 0.15622209 | -0.88961341 | 0.373673505 | 0.998865404 | 11.419138 | 11.280598 | 11.440939 | 11.326313 | 11.434195 | 11.162109 | 3373 | 2749 | 2528 | 1939 |
| 85474 | NM_00100981 | LBX2 | ladybird homeobox 2 | protein_coding | NP_001009812.1;NP_001269359.1 | 15525 | 607164 | . | . | 2.454012 | 2.615143 | 1.899725 | 4.082993 | 5.710586 | 6.045020 | 4.262927 | 9.335278 | 1.120846 | 117.068174 | 0.346033092 | 0.475641627 | 0.634329714 | 0.998865404 | 6.788171 | 6.949719 | 6.819552 | 6.862770 | 6.766513 | 6.982154 | 119 | 127 | 74 | 152 |
| 85476 | NM_00130816 | GFM1 | G elongation factor mitochondrial 1 | protein_coding | NP_001295093.1;NP_001295095.1;NP_001361284.1;NP_001361285.1;NI | 13780 | 606639 | . | . | 11.004706 | 9.327592 | 10.188088 | 10.241416 | 25.608400 | 21.561143 | 22.861772 | 23.415777 | 1.025358 | 1703.951377 | 0.127429623 | 0.283506946 | 0.776788258 | 0.998865404 | 10.716566 | 10.752547 | 10.747642 | 10.700461 | 10.700033 | 10.785922 | 1996 | 1761 | 1418 | 1706 |
| 85477 | NM_00111270 | SCIN | scinderin | protein_coding | NP_001106177.1;NP_149119.1;XP_024302748.1 | 21695 | 613416 | . | . | 0.619859 | 0.518549 | 0.668398 | 0.630706 | 1.442438 | 1.198650 | 1.499867 | 1.442036 | 1.114564 | 183.448965 | 0.25130808 | 0.622660353 | 0.533507742 | 0.998865404 | 7.438239 | 7.595925 | 7.530772 | 7.452999 | 7.559637 | 7.519261 | 218 | 170 | 176 | 173 |
| 85480 | NM_033035,N | TSLP | thymic stromal lymphopoietin | protein_coding | NP_149024.1;NP_612561.2;XP_011542000.1 | 30743 | 607003 | . | . | 0.439265 | 0.257053 | 0.148622 | 0.254039 | 1.022187 | 0.594189 | 0.333502 | 0.580830 | -1.706633 | 20.790547 | 0.708230522 | -1.08884419 | 0.276222599 | NA | 4.713213 | 3.940155 | 4.433324 | 4.375806 | 4.343043 | 4.374917 | 38 | 21 | 10 | 18 |
| 85495 | NR_002312 | RPPH1 | ribonuclease P RNA component H1 | RNase_P_RNA | . | 19273 | 608513 | . | . | 0.553382 | 1.491181 | 0.940697 | 1.738857 | 1.287744 | 3.446932 | 2.110898 | 3.975690 | 1.305127 | 13.077570 | 0.898923649 | 0.427389605 | 0.669095578 | NA | 3.508460 | 3.885075 | 3.665498 | 3.713906 | 3.686232 | 3.724007 | 7 | 18 | 9 | 18 |
| 85865 | NM_00104271 | GTPBP10 | GTP binding protein 10 | protein_coding | NP_001036182.1;NP_149098.2 | 25106 | 610920 | . | . | 2.809540 | 2.605762 | 2.489109 | 2.901127 | 6.537914 | 6.023335 | 5.585488 | 6.633082 | -1.018852 | 630.685443 | 0.162615168 | -0.16569087 | 0.868400232 | 0.998865404 | 9.314343 | 9.287082 | 9.334643 | 9.277447 | 9.244688 | 9.340743 | 764 | 658 | 504 | 628 |
| 86781 | gene-RPS2P7 | RPS2P7 | ribosomal protein S2 pseudogene 7 | pseudogene | . | 15838 | . | . | . | 0.113069 | 0.123895 | 0.072776 | 0.190863 | 0.263117 | 0.286390 | 0.163308 | 0.436385 | 1.216734 | 3.970175 | 1.523074266 | 0.185817133 | 0.852588168 | NA | 1.844134 | 2.121007 | 1.984946 | 1.986080 | 1.979899 | 1.998241 | 4 | 4 | 2 | 6 |
| 87178 | NM_033109,XI | PNPT1 | polyribonucleotide nucleotidyltransferase 1 | protein_coding | NP_149100.2;XP_005264686.1;XP_016860661.1 | 23166 | 610316 | . | . | 11.858906 | 11.198257 | 11.198368 | 8.969288 | 27.596160 | 25.885273 | 25.128814 | 20.507213 | -1.182348 | 1512.934262 | 0.139009886 | -1.73839569 | 0.082141116 | 0.783811235 | 10.678660 | 10.437544 | 10.649484 | 10.595988 | 10.563970 | 10.418252 | 1951 | 1709 | 1322 | 1175 |
| 87688 | gene-RPL7AP5 | RPL7AP50 | ribosomal protein L7a pseudogene 50 | pseudogene | . | 36117 | . | . | . | 0.188194 | 0.179684 | 0.083865 | 0.153286 | 0.437936 | 0.415349 | 0.188192 | 0.350470 | -1.507010 | 4.839039 | 1.373456333 | -0.43080322 | 0.666611457 | NA | 2.540965 | 1.947964 | 2.288759 | 2.286269 | 2.276818 | 2.280358 | 7 | 6 | 3 | 4 |
| 87769 | NM_00119508 | GGACT | gamma-glutamylamine cyclotransferase | protein_coding | NP_001182016.1;NP_149101.1;XP_005254140.1;XP_011519431.1 | 25100 | 613378 | . | . | 0.289876 | 0.323839 | 0.346981 | 0.841201 | 0.674553 | 0.748567 | 0.778616 | 1.923304 | 1.952292 | 39.688717 | 0.534691356 | 1.805095367 | 0.07105976 | 0.751131579 | 4.751025 | 5.712861 | 5.226983 | 5.235094 | 5.263620 | 5.395686 | 30 | 30 | 30 | 66 |
| 88455 | NM_033121,XI | ANKRD13A | ankyrin repeat domain 13A | protein_coding | NP_149112.1;XP_005254037.1;XP_005254038.1;XP_005254039.1;XP_00 | 21268 | 615123 | . | . | 5.727158 | 6.523670 | 6.509692 | 7.436688 | 13.327337 | 15.079754 | 14.607565 | 17.003104 | 1.128817 | 831.627616 | 0.151169221 | 1.15639609 | 0.24751918 | 0.992733415 | 9.610082 | 9.784252 | 9.614194 | 9.691944 | 9.695672 | 9.781694 | 850 | 890 | 721 | 876 |
| 88745 | NM_00132970 | RRP36 | ribosomal RNA processing 36 | protein_coding | NP_001316633.1;NP_149103.1 | 21374 | 613475 | . | . | 27.131507 | 26.338677 | 27.358696 | 29.813045 | 63.136132 | 60.883037 | 61.392124 | 68.163981 | 1.056777 | 1026.066475 | 0.137659953 | 0.578753278 | 0.562755661 | 0.998865404 | 9.962598 | 10.042123 | 9.995835 | 9.967746 | 9.993991 | 10.050149 | 1171 | 1056 | 882 | 1026 |
| 89122 | NM_033017,N | TRIM4 | tripartite motif containing 4 | protein_coding | NP_148977.2;NP_149082.1 | 16275 | . | . | . | 5.799493 | 5.315495 | 6.244349 | 5.831786 | 13.495659 | 12.287004 | 14.012137 | 13.333684 | 1.070269 | 600.210415 | 0.16121642 | 0.607717697 | 0.543374723 | 0.998865404 | 9.179211 | 9.277755 | 9.235024 | 9.174859 | 9.276445 | 9.224032 | 698 | 597 | 561 | 559 |
| 89761 | gene-NUFIP1P1 | NUFIP1P1 | NUFIP1 pseudogene 1 | pseudogene | . | 8058 | . | . | . | 0.018046 | 0.030594 | 0.022893 | 0.006179 | 0.041994 | 0.070720 | 0.051372 | 0.014127 | -1.404040 | 2.707224 | 1.839081235 | -0.26621147 | 0.790076352 | NA | 1.658167 | 1.175234 | 1.437226 | 1.442423 | 1.441014 | 1.430231 | 3 | 4 | 3 | 1 |
| 89781 | NM_00134989 | HPS4 | HPS4 biogenesis of lysosomal organelles complex 3 subunit 2 | protein_coding | NP_001336825.1;NP_001336827.1;NP_001336828.1;NP_001336829.1;NI | 15844 | 606682 | . | . | 44.077396 | 48.513222 | 66.116134 | 57.786299 | 102.569915 | 112.140483 | 148.362687 | 132.121514 | 1.317984 | 7784.744045 | 0.115026569 | 3.462961147 | 0.000534266 | 0.043125539 | 12.713523 | 13.111927 | 12.785695 | 12.808447 | 13.098084 | 12.959806 | 7628 | 7347 | 8311 | 7639 |
| 89782 | NM_00113604 | LMLN | leishmanolysin like peptidase | protein_coding | NP_001129521.2;NP_149018.2 | 15991 | 609380 | . | . | 4.223305 | 3.978310 | 4.673115 | 4.350758 | 9.827806 | 9.196041 | 10.486335 | 9.947491 | 1.086688 | 954.645867 | 0.142238749 | 0.843217235 | 0.399106961 | 0.998865404 | 9.837370 | 9.957763 | 9.892957 | 9.840186 | 9.960124 | 9.893750 | 1092 | 951 | 904 | 890 |
| 89795 | NM_00102438 | NAV3 | neuron navigator 3 | protein_coding | NP_001019554.1;NP_055718.4;XP_005269272.1;XP_011537242.1;XP_01 | 15998 | 611629 | . | . | 0.702808 | 0.751896 | 0.457666 | 0.519047 | 1.635465 | 1.738040 | 1.026989 | 1.186740 | -1.548368 | 190.025883 | 0.244688821 | -2.57775807 | 0.009944359 | 0.320705593 | 7.851561 | 7.219960 | 7.649751 | 7.660103 | 7.442773 | 7.485896 | 263 | 252 | 122 | 148 |
| 89796 | NM_00116773 | NAV1 | neuron navigator 1 | protein_coding | NP_001161210.1;NP_065176.3 | 15989 | 611628 | . | . | 8.692354 | 9.464557 | 8.901486 | 9.638792 | 20.227465 | 21.877748 | 19.974678 | 22.037952 | 1.021955 | 3694.381145 | 0.111746476 | 0.280386729 | 0.779180818 | 0.998865404 | 11.835436 | 11.866629 | 11.821834 | 11.861518 | 11.830741 | 11.887987 | 4107 | 4038 | 3135 | 3622 |
| 89797 | NM_00111101 | NAV2 | neuron navigator 2 | protein_coding | NP_001104488.1;NP_001104489.1;NP_001231892.1;NP_660093.2;NP_8 | 15997 | 607026 | . | . | 14.720487 | 14.552234 | 11.472169 | 12.788062 | 34.255174 | 33.638144 | 25.743218 | 29.238388 | -1.223577 | 4805.791363 | 0.111122472 | -2.61967604 | 0.008801333 | 0.294943244 | 12.368828 | 12.077619 | 12.318365 | 12.299126 | 12.097314 | 12.181218 | 6159 | 5643 | 3566 | 4260 |
| 89801 | NM_00118474 | PPP1R3F | protein phosphatase 1 regulatory subunit 3F | protein_coding | NP_001171674.1;NP_149992.3;XP_005272744.1;XP_016885421.1;XP_01 | 14944 | . | . | . | 1.730357 | 1.588399 | 1.970512 | 1.387165 | 4.026612 | 3.671656 | 4.421772 | 3.171586 | 1.031622 | 240.105381 | 0.239741086 | 0.187344027 | 0.851390905 | 0.998865404 | 7.884103 | 7.930571 | 7.897196 | 7.893705 | 7.992483 | 7.822097 | 273 | 254 | 249 | 189 |
| 89832 | NM_139320,N | CHRFAM7A | CHRNA7 (exons 5-10) and FAM7A (exons A-E) fusion | protein_coding | NP_647536.1;NP_683709.1;XP_005254807.1;XP_011520455.1;XP_01152 | 15781 | 609756 | . | . | 1.297943 | 1.427662 | 1.636409 | 1.171058 | 3.020366 | 3.300104 | 3.672055 | 2.677485 | 1.054244 | 151.800001 | 0.280696581 | 0.271498882 | 0.786007359 | 0.998865404 | 7.206790 | 7.284227 | 7.212353 | 7.249798 | 7.312247 | 7.187926 | 162 | 167 | 157 | 123 |
| 89837 | NR_026859 | ULK4P3 | ULK4 pseudogene 3 | transcribed_ps | . | 15777 | . | . | . | 0.393066 | 0.491269 | 0.610757 | 0.338183 | 0.914681 | 1.135589 | 1.370522 | 0.773216 | 1.039004 | 16.986982 | 0.756898184 | 0.072930427 | 0.941861481 | NA | 4.057381 | 4.114763 | 4.071296 | 4.088028 | 4.101246 | 4.063335 | 17 | 20 | 19 | 12 |
| 89838 | NR_026858 | ULK4P1 | ULK4 pseudogene 1 | transcribed_ps | . | 15775 | . | . | . | 0.327794 | 0.287385 | 0.497512 | 0.293133 | 0.762789 | 0.664303 | 1.116403 | 0.670213 | 1.245033 | 10.056648 | 0.9653496 | 0.327533579 | 0.74326435 | NA | 3.161369 | 3.481107 | 3.320581 | 3.315061 | 3.338879 | 3.316187 | 11 | 9 | 12 | 8 |
| 89839 | NM_00103984 | ARHGAP11B | Rho GTPase activating protein 11B | protein_coding | NP_001034930.1 | 15782 | 616310 | . | . | 4.266335 | 4.387471 | 3.701885 | 4.578084 | 9.927939 | 10.141835 | 8.306922 | 10.467245 | -1.175147 | 301.153354 | 0.206007964 | -1.13025247 | 0.258369854 | 0.995760118 | 8.345634 | 8.113768 | 8.285179 | 8.265609 | 8.230886 | 8.145065 | 381 | 345 | 261 | 238 |
| 89845 | NM_00119893 | ABCC10 | ATP binding cassette subfamily C member 10 | protein_coding | NP_001185863.1;NP_001337447.1;NP_258261.2;XP_011513276.2;XP_01 | 52 | 612509 | . | . | 5.581691 | 5.904768 | 7.444389 | 6.472888 | 12.988827 | 13.649140 | 16.704995 | 14.799491 | 1.175108 | 928.503885 | 0.147603133 | 1.577153451 | 0.114760221 | 0.855274326 | 9.737534 | 9.970592 | 9.770474 | 9.819775 | 9.966217 | 9.855488 | 949 | 953 | 938 | 870 |
| 89846 | NM_00108353 | FGD3 | FYVE, RhoGEF and PH domain containing 3 | protein_coding | NP_001077005.1;NP_001273922.1;NP_001356880.1;NP_001356881.1;NI | 16027 | 617554 | . | . | 0.084695 | 0.281644 | 0.095814 | 0.132278 | 0.197088 | 0.651032 | 0.215004 | 0.302438 | -1.617372 | 16.586324 | 0.830633697 | -0.8350877 | 0.403668334 | NA | 4.359738 | 3.660102 | 4.011607 | 4.107336 | 4.017243 | 4.036545 | 11 | 34 | 9 | 14 |
| 89848 | NM_033449,XI | FCHSD1 | FCH and double SH3 domains 1 | protein_coding | NP_258260.1;XP_005268581.1;XP_006714866.1;XP_011536000.1;XP_01 | 25463 | 617555 | . | . | 14.330949 | 13.643528 | 15.709014 | 15.443304 | 33.348701 | 31.537624 | 35.250572 | 35.309277 | 1.098598 | 1441.798796 | 0.126900315 | 1.069053124 | 0.285045732 | 0.998865404 | 10.424190 | 10.559927 | 10.480121 | 10.431158 | 10.520140 | 10.536435 | 1633 | 1435 | 1299 | 1430 |
| 89849 | NM_00131876 | ATG16L2 | autophagy related 16 like 2 | protein_coding | NP_001305695.1;NP_203746.1;XP_005274433.1;XP_006718795.1;XP_00 | 25464 | 618716 | . | . | 7.904384 | 8.261091 | 7.527249 | 7.246618 | 18.393826 | 19.095880 | 16.890927 | 16.568531 | -1.115875 | 707.292176 | 0.151562497 | -1.04362926 | 0.296656952 | 0.998865404 | 9.543177 | 9.384804 | 9.474373 | 9.530174 | 9.428048 | 9.426308 | 825 | 837 | 582 | 626 |
| 89853 | NM_00101170 | MVB12B | multivesicular body subunit 12B | protein_coding | NP_001011703.1;NP_258257.1;XP_005252354.1;XP_016870764.1;XP_01 | 23368 | . | . | . | 1.423406 | 1.533255 | 1.076032 | 1.308330 | 3.312324 | 3.544186 | 2.414584 | 2.991339 | -1.256436 | 181.376880 | 0.248127326 | -1.32729334 | 0.184411667 | 0.956844295 | 7.658310 | 7.328585 | 7.562850 | 7.537120 | 7.433458 | 7.471015 | 239 | 212 | 133 | 158 |
| 89857 | NM_130446,XI | KLHL6 | kelch like family member 6 | protein_coding | NP_569713.2;XP_011511575.1;XP_011511576.1 | 18653 | 614214 | . | . | 0.425805 | 0.278787 | 0.265013 | 0.241550 | 0.990865 | 0.644426 | 0.594681 | 0.552275 | -1.293877 | 55.269795 | 0.442246749 | -0.84048092 | 0.400638794 | 0.998865404 | 5.961318 | 5.591942 | 5.862188 | 5.760031 | 5.769872 | 5.752260 | 88 | 52 | 44 | 43 |
| 89874 | NM_00117117 | SLC25A21 | solute carrier family 25 member 21 | protein_coding | NP_001164641.1;NP_085134.1;XP_011535589.1;XP_011535590.1;XP_01 | 14411 | 607571 | . | . | 0.095716 | 0.121901 | 0.131315 | 0.143776 | 0.222736 | 0.281778 | 0.294668 | 0.328727 | 1.303258 | 17.598179 | 0.730033783 | 0.523432248 | 0.600673488 | NA | 3.934350 | 4.315285 | 4.114651 | 4.127232 | 4.139299 | 4.142475 | 16 | 18 | 17 | 19 |
| 89884 | NM_033343,XI | LHX4 | LIM homeobox 4 | protein_coding | NP_203129.1;XP_011508407.1;XP_011508408.1;XP_011508410.1;XP_01 | 21734 | 602146 | . | . | 0.140354 | 0.079003 | 0.092283 | 0.060340 | 0.326609 | 0.182833 | 0.207081 | 0.137961 | -1.361631 | 17.753526 | 0.755013178 | -0.58983847 | 0.555298946 | NA | 4.353365 | 3.913269 | 4.186859 | 4.138045 | 4.153532 | 4.125229 | 30 | 16 | 16 | 11 |
| 89886 | NM_00114617 | SLAMF9 | SLAM family member 9 | protein_coding | NP_001139644.1;NP_001139645.1;NP_254273.2;XP_016858245.1 | 18430 | 608589 | . | . | 0.225903 | 0.159129 | 0.080204 | 0.454485 | 0.525685 | 0.367833 | 0.179976 | 1.039127 | 1.108961 | 9.369203 | 1.014237293 | 0.147114652 | 0.883041525 | NA | 3.151937 | 3.300113 | 3.236019 | 3.213954 | 3.216493 | 3.242132 | 13 | 7 | 6 | 12 |
| 89887 | NM_033113,XI | ZNF628 | zinc finger protein 628 | protein_coding | NP_149104.3;XP_005259428.1;XP_024307520.1 | 28054 | 610671 | . | . | 1.109899 | 1.235543 | 0.957438 | 1.398979 | 2.582780 | 2.856011 | 2.148463 | 3.198603 | 1.082993 | 141.659655 | 0.279858735 | 0.411008632 | 0.681066205 | 0.998865404 | 7.088390 | 7.201946 | 7.107978 | 7.149610 | 7.122545 | 7.194111 | 148 | 155 | 116 | 151 |
| 89890 | NM_152903 | KBTBD6 | kelch repeat and BTB domain containing 6 | protein_coding | NP_690867.3 | 25340 | 617738 | . | . | 2.051713 | 2.248548 | 2.125571 | 1.900747 | 4.774419 | 5.197619 | 4.769721 | 4.345833 | -1.081762 | 339.988114 | 0.195724085 | -0.57930251 | 0.562385071 | 0.998865404 | 8.464770 | 8.351695 | 8.406372 | 8.454023 | 8.418520 | 8.353191 | 390 | 397 | 301 | 288 |
| 89891 | NM_052844,XI | DYNC2I2 | dynein 2 intermediate chain 2 | protein_coding | NP_443076.2;XP_011517481.1 | 28296 | 613363 | . | . | 29.443854 | 31.443173 | 26.949640 | 33.802270 | 68.517056 | 72.682306 | 60.474210 | 77.284876 | -1.016470 | 1713.323395 | 0.139763118 | -0.16862672 | 0.866090263 | 0.998865404 | 10.754558 | 10.730506 | 10.725396 | 10.769197 | 10.645005 | 10.816961 | 1934 | 1916 | 1315 | 1768 |
| 89894 | NM_00119345 | TMEM116 | transmembrane protein 116 | protein_coding | NP_001180382.1;NP_001180460.1;NP_001281243.1;NP_612350.1;XP_00 | 25084 | . | . | . | 3.860986 | 4.270447 | 3.706218 | 4.143769 | 8.984675 | 9.871326 | 8.316642 | 9.474236 | -1.046171 | 200.078802 | 0.238311935 | -0.27325067 | 0.784660534 | 0.998865404 | 7.677024 | 7.611071 | 7.639516 | 7.669517 | 7.610715 | 7.657148 | 228 | 228 | 161 | 193 |
| 89910 | NM_00127044 | UBE3B | ubiquitin protein ligase E3B | protein_coding | NP_001257378.1;NP_001257379.1;NP_001257380.1;NP_569733.2;NP_9 | 13478 | 608047 | . | . | 6.823610 | 7.183245 | 7.509261 | 8.378548 | 15.878821 | 16.604392 | 16.850562 | 19.156561 | 1.132821 | 1312.781054 | 0.130643636 | 1.377184512 | 0.168455209 | 0.945638095 | 10.265744 | 10.445483 | 10.316468 | 10.300247 | 10.369196 | 10.436192 | 1433 | 1314 | 1160 | 1364 |
| 89927 | NM_00114246 | BMERB1 | bMERB domain containing 1 | protein_coding | NP_001135941.1;NP_149978.1 | 19213 | . | . | . | 3.075870 | 3.760069 | 2.813082 | 3.425410 | 7.157676 | 8.691569 | 6.312475 | 7.831792 | -1.109628 | 215.349803 | 0.241147505 | -0.62234092 | 0.533717718 | 0.998865404 | 7.824386 | 7.672706 | 7.725460 | 7.818410 | 7.682277 | 7.764828 | 236 | 268 | 161 | 209 |
| 89932 | NM_00136590 | PAPLN | papilin, proteoglycan like sulfated glycoprotein | protein_coding | NP_001352835.1;NP_001352836.1;NP_775733.3;XP_011535593.1;XP_01 | 19262 | 617785 | . | . | 1.000763 | 1.199128 | 0.912591 | 1.011501 | 2.328818 | 2.771837 | 2.047826 | 2.312675 | -1.165163 | 189.148315 | 0.247156515 | -0.89227647 | 0.372244761 | 0.998865404 | 7.669878 | 7.448376 | 7.556404 | 7.633312 | 7.515333 | 7.541504 | 215 | 238 | 147 | 169 |
| 89941 | NM_00135227 | RHOT2 | ras homolog family member T2 | protein_coding | NP_001339204.1;NP_001339205.1;NP_001339206.1;NP_001339207.1;NI | 21169 | 613889 | . | . | 29.508739 | 31.917239 | 30.146383 | 30.070210 | 68.668046 | 73.778129 | 67.647607 | 68.751967 | -1.030133 | 2535.354695 | 0.115569601 | -0.37060359 | 0.710932804 | 0.998865404 | 11.329266 | 11.286359 | 11.291985 | 11.346784 | 11.297962 | 11.293440 | 2863 | 2870 | 2178 | 2335 |
| 89944 | NM_00137046 | GLB1L2 | galactosidase beta 1 like 2 | protein_coding | NP_001357389.1;NP_001357390.1;NP_001357391.1;NP_001357392.1;NI | 25129 | . | . | . | 18.665823 | 17.592485 | 14.722613 | 17.707523 | 43.436139 | 40.665816 | 33.037116 | 40.486150 | -1.136536 | 1704.074757 | 0.135199035 | -1.36571138 | 0.172029572 | 0.947431715 | 10.824275 | 10.639349 | 10.810257 | 10.752747 | 10.610512 | 10.748044 | 2163 | 1886 | 1263 | 1629 |
| 89953 | NM_00128903 | KLC4 | kinesin light chain 4 | protein_coding | NP_001275963.1;NP_001275964.1;NP_612352.1;NP_958929.1;NP_9589 | 21624 | . | . | . | 4.822331 | 4.401160 | 3.893534 | 4.528256 | 11.221766 | 10.173477 | 8.736978 | 10.353320 | -1.089815 | 337.730265 | 0.19551132 | -0.63466271 | 0.525648418 | 0.998865404 | 8.460479 | 8.336306 | 8.452056 | 8.393937 | 8.351706 | 8.398165 | 426 | 361 | 269 | 316 |
| 89958 | NM_178448,XI | SAPCD2 | suppressor APC domain containing 2 | protein_coding | NP_848543.2;XP_011517482.1 | 28055 | 612057 | . | . | 6.429273 | 6.078388 | 4.117047 | 5.768823 | 14.961184 | 14.050465 | 9.238535 | 13.189729 | -1.284549 | 668.106923 | 0.179559881 | -2.01193171 | 0.044227136 | 0.645513836 | 9.553698 | 9.191499 | 9.481705 | 9.437642 | 9.179437 | 9.389071 | 892 | 786 | 426 | 637 |
| 89970 | NM_00130516 | RSPRY1 | ring finger and SPRY domain containing 1 | protein_coding | NP_001292092.1;NP_001292093.1;NP_001292111.1;NP_588609.1;XP_00 | 29420 | 616585 | . | . | 5.638937 | 4.794277 | 5.089349 | 5.592650 | 13.122038 | 11.082186 | 11.420354 | 12.786930 | -1.026497 | 573.120785 | 0.167398403 | -0.22538888 | 0.821676764 | 0.998865404 | 9.181436 | 9.143708 | 9.215695 | 9.123414 | 9.121514 | 9.184407 | 715 | 583 | 468 | 556 |
| 89978 | NM_00114197 | DPH6 | diphthamine biosynthesis 6 | protein_coding | NP_001135444.1;NP_542381.1;XP_011520462.1;XP_016878196.1;XP_01 | 30543 | 618391 | . | . | 1.867300 | 1.455882 | 1.046682 | 1.157258 | 4.345282 | 3.365336 | 2.348725 | 2.645933 | -1.432845 | 90.083125 | 0.337811957 | -1.53601054 | 0.124535795 | 0.87301553 | 6.729512 | 6.210413 | 6.555165 | 6.529187 | 6.432699 | 6.449757 | 127 | 110 | 62 | 72 |
| 90007 | NM_177401,XI | MIDN | midnolin | protein_coding | NP_796375.3;XP_005259728.1;XP_005259729.1;XP_024307521.1;XP_02 | 16298 | 606700 | . | . | 6.631571 | 7.253409 | 6.768308 | 6.843324 | 15.431938 | 16.766581 | 15.187887 | 15.646447 | -1.023203 | 827.111566 | 0.144939541 | -0.22832246 | 0.819395561 | 0.998865404 | 9.708521 | 9.675162 | 9.673274 | 9.725885 | 9.675713 | 9.691365 | 928 | 936 | 703 | 775 |
| 90024 | NR_033874 | FLJ20021 | uncharacterized LOC90024 | lncRNA | . | . | . | . | . | 1.703323 | 2.102601 | 1.231082 | 1.740810 | 3.963702 | 4.860257 | 2.762513 | 3.980155 | -1.302999 | 42.490636 | 0.487978734 | -0.78248485 | 0.433929686 | 0.998865404 | 5.588663 | 5.203906 | 5.413632 | 5.449229 | 5.366886 | 5.410647 | 50 | 57 | 27 | 40 |
| 90025 | NM_00130443 | UBE3D | ubiquitin protein ligase E3D | protein_coding | NP_001291366.1;NP_001337531.1;NP_001337532.1;NP_001337533.1;NI | 21381 | 612495 | . | . | 1.482973 | 1.142298 | 1.663066 | 1.466315 | 3.450940 | 2.640473 | 3.731875 | 3.352558 | 1.195317 | 94.640698 | 0.336053573 | 0.765930271 | 0.443717793 | 0.998865404 | 6.429045 | 6.688117 | 6.568138 | 6.496008 | 6.599759 | 6.570449 | 112 | 81 | 94 | 92 |
| 90060 | NM_00116332 | CCDC120 | coiled-coil domain containing 120 | protein_coding | NP_001156793.2;NP_001156794.1;NP_001156795.1;NP_001258764.1;NI | 28910 | 300947 | . | . | 0.746755 | 0.702646 | 0.701161 | 0.562989 | 1.737730 | 1.624198 | 1.573386 | 1.287208 | -1.115903 | 85.805103 | 0.349127925 | -0.45316368 | 0.650430884 | 0.998865404 | 6.499094 | 6.342650 | 6.448193 | 6.426523 | 6.437887 | 6.375811 | 108 | 94 | 79 | 67 |
| 90075 | NM_00109943 | ZNF30 | zinc finger protein 30 | protein_coding | NP_001092907.1;NP_001092908.1;NP_919306.2;XP_011525745.1;XP_01 | 13090 | . | . | . | 1.716169 | 1.609280 | 1.501032 | 1.408209 | 3.993599 | 3.719925 | 3.368273 | 3.219702 | -1.128366 | 133.930499 | 0.282766216 | -0.616179 | 0.537776386 | 0.998865404 | 7.149366 | 6.976118 | 7.100236 | 7.074027 | 7.063751 | 7.022448 | 169 | 148 | 116 | 111 |
| 90102 | NM_00113443 | PHLDB2 | pleckstrin homology like domain family B member 2 | protein_coding | NP_001127909.1;NP_001127910.1;NP_001127911.1;NP_665696.1 | 29573 | 610298 | . | . | 0.685331 | 0.538397 | 0.391693 | 0.540055 | 1.594794 | 1.244528 | 0.878947 | 1.234771 | -1.364914 | 100.935433 | 0.335413465 | -1.33807845 | 0.180870866 | 0.952877815 | 6.864661 | 6.415024 | 6.744250 | 6.657483 | 6.569953 | 6.642662 | 151 | 110 | 64 | 91 |
| 90120 | NM_00125652 | TMEM250 | transmembrane protein 250 | protein_coding | NP_001243455.1;NP_690046.3;XP_024303480.1 | 31009 | . | . | . | 8.172789 | 7.696665 | 7.467298 | 8.993788 | 19.018415 | 17.791186 | 16.756401 | 20.563225 | 1.019044 | 710.511460 | 0.159235355 | 0.170920591 | 0.864286206 | 0.998865404 | 9.459237 | 9.486066 | 9.490885 | 9.438966 | 9.418529 | 9.534484 | 841 | 731 | 571 | 729 |
| 90121 | NM_00134678 | TSR2 | TSR2 ribosome maturation factor | protein_coding | NP_001333718.1;NP_001333719.1;NP_001333720.1;NP_001333721.1;NI | 25455 | 300945 | . | . | 7.761062 | 8.156759 | 8.210657 | 8.905388 | 18.060311 | 18.854717 | 18.424475 | 20.361109 | 1.062449 | 1051.473703 | 0.136692564 | 0.639341898 | 0.522600538 | 0.998865404 | 9.994000 | 10.081080 | 9.999090 | 10.030736 | 10.033367 | 10.085276 | 1149 | 1124 | 909 | 1051 |
| 90133 | gene-KRT8P12 | KRT8P12 | keratin 8 pseudogene 12 | pseudogene | . | 28057 | . | . | . | 3.304977 | 3.070225 | 3.990635 | 3.608668 | 7.690817 | 7.096963 | 8.954869 | 8.250790 | 1.172585 | 184.878657 | 0.246597653 | 0.931444172 | 0.351623851 | 0.998865404 | 7.410533 | 7.641132 | 7.508711 | 7.474300 | 7.584534 | 7.536407 | 204 | 176 | 183 | 177 |
| 90135 | NM_033271,XI | BTBD6 | BTB domain containing 6 | protein_coding | NP_150374.2;XP_005268245.1 | 19897 | . | . | . | 13.301628 | 14.935786 | 14.591312 | 15.829456 | 30.953436 | 34.524738 | 32.742481 | 36.192169 | 1.072352 | 980.559597 | 0.14131469 | 0.713151896 | 0.475751769 | 0.998865404 | 9.886415 | 9.986763 | 9.876960 | 9.943479 | 9.935184 | 9.987406 | 1040 | 1068 | 851 | 985 |
| 90141 | NM_00128426 | EFCAB11 | EF-hand calcium binding domain 11 | protein_coding | NP_001271195.1;NP_001271196.1;NP_001271197.1;NP_001271198.1;NI | 20357 | . | . | . | 1.295530 | 1.646260 | 1.453349 | 1.929644 | 3.014752 | 3.805406 | 3.261273 | 4.411902 | 1.163752 | 123.826156 | 0.305498949 | 0.716152233 | 0.473897354 | 0.998865404 | 6.839773 | 7.056444 | 6.899587 | 6.940278 | 6.912797 | 7.029787 | 124 | 131 | 97 | 145 |
| 90161 | NM_00107718 | HS6ST2 | heparan sulfate 6-O-sulfotransferase 2 | protein_coding | NP_001070656.1;NP_671704.3;XP_005262548.1;XP_011529708.1;XP_01 | 19133 | 300545 | . | . | 0.248868 | 0.362062 | 0.328089 | 0.267737 | 0.579127 | 0.836922 | 0.736222 | 0.612149 | -1.001010 | 42.799826 | 0.486799413 | -0.00299103 | 0.997613509 | 0.999862634 | 5.420387 | 5.418678 | 5.387445 | 5.443866 | 5.432073 | 5.399442 | 41 | 54 | 41 | 36 |
| 90196 | NM_00109979 | SYS1 | SYS1 golgi trafficking protein | protein_coding | NP_001093261.1;NP_001184058.1;NP_291020.1 | 16162 | 612979 | . | . | 9.851795 | 9.808553 | 12.075107 | 9.623995 | 22.925535 | 22.672912 | 27.096189 | 22.004121 | 1.088082 | 885.153753 | 0.157372305 | 0.773882947 | 0.439000015 | 0.998865404 | 9.727258 | 9.849713 | 9.760542 | 9.751874 | 9.899081 | 9.727901 | 982 | 909 | 897 | 763 |
| 90203 | NM_00104263 | SNX21 | sorting nexin family member 21 | protein_coding | NP_001036097.1;NP_001036098.1;NP_219489.1;NP_690857.1;XP_0052 | 16154 | . | . | . | 3.451709 | 3.635548 | 4.300290 | 3.703599 | 8.032269 | 8.403732 | 9.649726 | 8.467841 | 1.158492 | 360.214734 | 0.191611133 | 1.107699894 | 0.267991445 | 0.998865404 | 8.382411 | 8.595178 | 8.438539 | 8.454115 | 8.571270 | 8.488785 | 379 | 365 | 360 | 337 |
| 90204 | NM_080603,XI | ZSWIM1 | zinc finger SWIM-type containing 1 | protein_coding | NP_542170.3;XP_005260667.1;XP_005260668.1;XP_011527402.1 | 16155 | . | . | . | 1.613292 | 1.615609 | 2.151763 | 1.374676 | 3.754195 | 3.734552 | 4.828494 | 3.143034 | -1.071065 | 135.714053 | 0.284399269 | -0.34826168 | 0.727643669 | 0.998865404 | 7.132505 | 7.034690 | 7.097144 | 7.094987 | 7.107493 | 7.032928 | 162 | 151 | 126 | 110 |
| 90226 | NM_033199 | UCN2 | urocortin 2 | protein_coding | NP_149976.1 | 18414 | 605902 | . | . | 229.389587 | 237.976532 | 206.540390 | 243.734329 | 533.798950 | 550.093445 | 463.470703 | 557.269592 | -1.052857 | 10704.946688 | 0.113084844 | -0.65711666 | 0.511105905 | 0.998865404 | 13.422706 | 13.348315 | 13.394702 | 13.418894 | 13.294848 | 13.426679 | 12466 | 12016 | 8366 | 10535 |
| 90231 | NM_138346 | KIAA2013 | KIAA2013 | protein_coding | NP_612355.1 | 28513 | . | . | . | 51.838882 | 54.012619 | 44.961998 | 48.085953 | 120.631218 | 124.852592 | 100.893410 | 109.942825 | -1.153418 | 4368.861867 | 0.108610221 | -1.89591059 | 0.057971868 | 0.713843573 | 12.192375 | 11.986360 | 12.135762 | 12.163093 | 12.007071 | 12.053651 | 5302 | 5132 | 3428 | 3912 |
| 90233 | NM_00127093 | ZNF551 | zinc finger protein 551 | protein_coding | NP_001257867.1;NP_612356.2 | 25108 | . | . | . | 0.746430 | 1.068774 | 1.096775 | 0.763191 | 1.736974 | 2.470517 | 2.461131 | 1.744946 | 1.027739 | 130.255651 | 0.314900001 | 0.125352617 | 0.900244398 | 0.998865404 | 7.005022 | 7.045104 | 6.952181 | 7.069719 | 7.086254 | 6.954446 | 124 | 161 | 136 | 101 |
| 90268 | NM_138348,XI | OTULIN | OTU deubiquitinase with linear linkage specificity | protein_coding | NP_612357.4;XP_011512453.1;XP_011512454.1;XP_011512456.1;XP_01 | 25118 | 615712 | . | . | 2.105064 | 2.213146 | 2.114272 | 2.025292 | 4.898572 | 5.115784 | 4.744368 | 4.630589 | -1.028160 | 543.441491 | 0.164506245 | -0.24354355 | 0.807584355 | 0.998865404 | 9.105771 | 9.065916 | 9.079639 | 9.109390 | 9.102410 | 9.050301 | 620 | 608 | 485 | 482 |
| 90271 | NR_026762 | OLMALINC | oligodendrocyte maturation-associated long intergenic non-coding | lncRNA | . | 28060 | 616240 | . | . | 2.767154 | 2.801095 | 2.863134 | 3.729069 | 6.439282 | 6.474856 | 6.424790 | 8.526073 | 1.166239 | 94.159904 | 0.335231316 | 0.661823345 | 0.508084443 | 0.998865404 | 6.442597 | 6.663070 | 6.529890 | 6.531012 | 6.536316 | 6.613260 | 100 | 94 | 77 | 107 |
| 90288 | NM_207307,XI | EFCAB12 | EF-hand calcium binding domain 12 | protein_coding | NP_997190.1;XP_011511595.1;XP_011511596.1;XP_011511597.1;XP_01 | 28061 | . | . | . | 0.201059 | 0.176115 | 0.174169 | 0.123444 | 0.467874 | 0.407097 | 0.390831 | 0.282240 | -1.077786 | 17.223785 | 0.73925161 | -0.14618959 | 0.88377172 | NA | 4.159017 | 4.051647 | 4.120122 | 4.101540 | 4.103329 | 4.107328 | 23 | 17 | 14 | 16 |
| 90293 | NM_00116829 | KLHL13 | kelch like family member 13 | protein_coding | NP_001161771.1;NP_001161772.1;NP_001161773.1;NP_001161774.1;NI | 22931 | 300655 | . | . | 1.187278 | 1.152974 | 0.830915 | 1.944194 | 2.762843 | 2.665150 | 1.864549 | 4.445167 | 1.144588 | 146.531596 | 0.328615393 | 0.592874717 | 0.553264999 | 0.998865404 | 7.095724 | 7.288010 | 7.151995 | 7.159369 | 7.064684 | 7.333418 | 156 | 149 | 90 | 196 |
| 90313 | NM_00134607 | TP53I13 | tumor protein p53 inducible protein 13 | protein_coding | NP_001333006.1;NP_001333007.1;NP_001333008.1;NP_001333010.1;NI | 25102 | . | . | . | 28.290321 | 28.653854 | 23.556070 | 27.811188 | 65.832733 | 66.234668 | 52.859139 | 63.586977 | -1.114809 | 1268.517268 | 0.137727555 | -1.13845249 | 0.254931587 | 0.995760118 | 10.385409 | 10.228163 | 10.344112 | 10.352247 | 10.204787 | 10.323209 | 1532 | 1451 | 960 | 1214 |
| 90317 | NM_178523,N | ZNF616 | zinc finger protein 616 | protein_coding | NP_848618.2 | 28062 | . | . | . | 0.862239 | 1.218254 | 1.537161 | 1.420869 | 2.006465 | 2.816048 | 3.449346 | 3.248650 | 1.404332 | 172.724486 | 0.266388265 | 1.838986788 | 0.065917129 | 0.73302766 | 7.167264 | 7.656170 | 7.284929 | 7.403993 | 7.504765 | 7.471406 | 140 | 179 | 182 | 182 |
| 90321 | NM_00101085 | ZNF766 | zinc finger protein 766 | protein_coding | NP_001010851.1;XP_011525762.1;XP_016882929.1;XP_016882930.1 | 28063 | . | . | . | 1.185303 | 1.338928 | 1.232681 | 1.200999 | 2.758246 | 3.094993 | 2.766099 | 2.745940 | 1.053702 | 227.102602 | 0.229614775 | 0.32866456 | 0.742409244 | 0.998865404 | 7.789313 | 7.864119 | 7.769147 | 7.854708 | 7.838683 | 7.836937 | 232 | 260 | 203 | 218 |
| 90324 | NM_00134610 | CCDC97 | coiled-coil domain containing 97 | protein_coding | NP_001333029.1;NP_443080.1;XP_016882931.1 | 28289 | . | . | . | 7.127766 | 7.741533 | 7.670789 | 8.159113 | 16.586604 | 17.894899 | 17.213029 | 18.654842 | 1.046714 | 796.625171 | 0.147340663 | 0.447040365 | 0.654845915 | 0.998865404 | 9.604616 | 9.670154 | 9.588701 | 9.652278 | 9.643545 | 9.662472 | 856 | 878 | 699 | 774 |
| 90326 | NM_00119575 | THAP3 | THAP domain containing 3 | protein_coding | NP_001182681.1;NP_001182682.1;NP_612359.2;XP_005263589.1;XP_01 | 20855 | 612532 | . | . | 11.223477 | 12.957482 | 10.311479 | 11.392328 | 26.117493 | 29.951800 | 23.138661 | 26.047200 | -1.084761 | 557.734971 | 0.172722468 | -0.6795707 | 0.496776327 | 0.998865404 | 9.181401 | 9.063044 | 9.115533 | 9.180347 | 9.032950 | 9.153869 | 636 | 657 | 421 | 549 |
| 90333 | NM_00100880 | ZNF468 | zinc finger protein 468 | protein_coding | NP_001008801.1;NP_001264049.1;XP_016882932.1;XP_016882933.1 | 33105 | 616841 | . | . | 1.527719 | 1.503542 | 1.667028 | 1.228355 | 3.555063 | 3.475504 | 3.740762 | 2.808487 | -1.053618 | 210.220830 | 0.247051586 | -0.30500309 | 0.760363801 | 0.998865404 | 7.752175 | 7.678406 | 7.728323 | 7.721542 | 7.769026 | 7.627885 | 250 | 231 | 206 | 162 |
| 90338 | NM_00110260 | ZNF160 | zinc finger protein 160 | protein_coding | NP_001096073.1;NP_001309054.1;NP_001309055.1;NP_001309057.1;NI | 12948 | 600398 | . | . | 2.819399 | 2.929197 | 4.232471 | 3.314012 | 6.560854 | 6.770969 | 9.497545 | 7.577093 | 1.288044 | 437.303095 | 0.195228275 | 1.870535223 | 0.06140953 | 0.723723143 | 8.577853 | 8.943962 | 8.676503 | 8.690171 | 8.928347 | 8.738851 | 435 | 417 | 492 | 393 |
| 90353 | NM_145232 | CTU1 | cytosolic thiouridylase subunit 1 | protein_coding | NP_660275.2 | 29590 | 612694 | . | . | 2.049036 | 2.406428 | 2.988556 | 2.772901 | 4.768190 | 5.562567 | 6.706234 | 6.339908 | 1.274301 | 169.651197 | 0.257237087 | 1.359470043 | 0.173997689 | 0.948096927 | 7.221148 | 7.570598 | 7.318194 | 7.377095 | 7.465996 | 7.431020 | 159 | 173 | 172 | 171 |
| 90355 | NM_00131696 | MACIR | macrophage immunometabolism regulator | protein_coding | NP_001303897.1;NP_001303898.1;NP_001364212.1;NP_001364213.1;NI | 25052 | 616608 | . | . | 7.020840 | 6.982013 | 8.230440 | 7.628333 | 16.337782 | 16.139238 | 18.468866 | 17.441276 | 1.111194 | 701.325165 | 0.152290359 | 0.998824165 | 0.317879878 | 0.998865404 | 9.375692 | 9.528161 | 9.419073 | 9.411974 | 9.518856 | 9.456337 | 769 | 713 | 671 | 661 |
| 90362 | NM_00137798 | FAM110B | family with sequence similarity 110 member B | protein_coding | NP_001364918.1;NP_001364926.1;NP_001364927.1;NP_671722.1;XP_01 | 28587 | 611394 | . | . | 0.495416 | 0.480326 | 0.512130 | 0.632624 | 1.152853 | 1.110295 | 1.149205 | 1.446418 | 1.167962 | 56.319514 | 0.419160989 | 0.534384866 | 0.593075287 | 0.998865404 | 5.699689 | 5.922818 | 5.800225 | 5.792921 | 5.808749 | 5.846975 | 61 | 55 | 48 | 62 |
| 90378 | NM_138352 | SAMD1 | sterile alpha motif domain containing 1 | protein_coding | NP_612361.1 | 17958 | . | . | . | 12.808674 | 12.877872 | 12.530153 | 12.986441 | 29.806311 | 29.767776 | 28.117300 | 29.691956 | -1.021484 | 876.060436 | 0.140919582 | -0.21761866 | 0.827726245 | 0.998865404 | 9.790180 | 9.759429 | 9.783593 | 9.781479 | 9.758109 | 9.776359 | 1021 | 954 | 744 | 823 |
| 90379 | NM_138353 | DCAF15 | DDB1 and CUL4 associated factor 15 | protein_coding | NP_612362.2 | 25095 | . | . | . | 12.551233 | 14.037372 | 15.777679 | 14.423716 | 29.207237 | 32.448013 | 35.404655 | 32.978111 | 1.120511 | 1015.617628 | 0.142597338 | 1.151190307 | 0.249653952 | 0.993339639 | 9.903703 | 10.067911 | 9.905388 | 9.980443 | 10.063454 | 9.989398 | 1046 | 1087 | 980 | 956 |
| 90381 | NM_00130802 | TICRR | TOPBP1 interacting checkpoint and replication regulator | protein_coding | NP_001294954.1;NP_689472.3 | 28704 | 613298 | . | . | 4.659656 | 5.187524 | 6.223433 | 6.109320 | 10.843212 | 11.991195 | 13.965205 | 13.968233 | 1.232624 | 1167.907498 | 0.134962368 | 2.235680313 | 0.02537272 | 0.524911334 | 10.031052 | 10.332636 | 10.067620 | 10.140389 | 10.272538 | 10.249987 | 1145 | 1185 | 1138 | 1191 |
| 90390 | NM_00128298 | MED30 | mediator complex subunit 30 | protein_coding | NP_001269915.1;NP_001350111.1;NP_542382.1 | 23032 | 610237 | . | . | 5.180893 | 5.027816 | 3.105684 | 3.989904 | 12.056150 | 11.622023 | 6.969063 | 9.122440 | -1.437876 | 138.495567 | 0.283407348 | -1.84871572 | 0.064498872 | 0.731823074 | 7.352416 | 6.827487 | 7.193503 | 7.158135 | 7.010258 | 7.072826 | 196 | 169 | 89 | 117 |
| 90407 | NM_080652,XI | TMEM41A | transmembrane protein 41A | protein_coding | NP_542383.1;XP_016862926.1 | 30544 | . | . | . | 6.369791 | 6.051260 | 5.468038 | 5.269085 | 14.822763 | 13.987759 | 12.270119 | 12.047138 | -1.163824 | 511.752994 | 0.167499687 | -1.306707 | 0.191312241 | 0.963729349 | 9.104388 | 8.885960 | 9.066414 | 9.024225 | 8.966393 | 8.931491 | 654 | 575 | 423 | 431 |
| 90410 | NM_00126777 | IFT20 | intraflagellar transport 20 | protein_coding | NP_001254703.1;NP_001254704.1;NP_001254705.1;NP_001254706.1;NI | 30989 | 614394 | . | . | 10.172813 | 8.721878 | 8.263113 | 10.030009 | 23.672557 | 20.161012 | 18.542188 | 22.932424 | -1.051239 | 287.985104 | 0.213534253 | -0.3376102 | 0.735656955 | 0.998865404 | 8.205621 | 8.133171 | 8.219812 | 8.143154 | 8.110148 | 8.199437 | 364 | 296 | 223 | 286 |
| 90411 | NM_00117150 | MCFD2 | multiple coagulation factor deficiency 2, ER cargo receptor comple | protein_coding | NP_001164977.1;NP_001164978.1;NP_001164979.1;NP_001164980.1;NI | 18451 | 607788 | . | . | 43.499381 | 40.626687 | 37.896455 | 34.802383 | 101.224846 | 93.910411 | 85.038545 | 79.571516 | -1.171193 | 5045.339993 | 0.112727918 | -2.02237764 | 0.043137353 | 0.639862814 | 12.410158 | 12.182329 | 12.395680 | 12.328974 | 12.268921 | 12.190813 | 6513 | 5643 | 4233 | 4151 |
| 90416 | NM_00108079 | CCDC32 | coiled-coil domain containing 32 | protein_coding | NP_001074260.2;NP_001074261.1;NP_001276061.1;NP_001369363.1;NI | 28295 | . | . | . | 3.721337 | 4.259713 | 3.883493 | 3.783414 | 8.659703 | 9.846520 | 8.714446 | 8.650328 | -1.084852 | 191.106380 | 0.242018263 | -0.48549553 | 0.627324882 | 0.998865404 | 7.635826 | 7.518244 | 7.575857 | 7.615738 | 7.569692 | 7.550392 | 219 | 224 | 163 | 168 |
| 90417 | NM_00114276 | KNSTRN | kinetochore localized astrin (SPAG5) binding protein | protein_coding | NP_001136233.1;NP_001136234.1;NP_150628.3 | 30767 | 614718 | . | . | 15.741477 | 16.161513 | 13.653838 | 15.808004 | 36.631061 | 37.358064 | 30.638817 | 36.143124 | -1.096836 | 845.209586 | 0.147270293 | -0.90546615 | 0.365218485 | 0.998865404 | 9.788532 | 9.654696 | 9.747647 | 9.762321 | 9.644410 | 9.731818 | 1008 | 964 | 657 | 803 |
| 90423 | NM_00131806 | ATP6V1E2 | ATPase H+ transporting V1 subunit E2 | protein_coding | NP_001304992.1;NP_001358210.1;NP_001358211.1;NP_001358212.1;XI | 18125 | 617385 | . | . | 3.168647 | 3.286295 | 2.142667 | 3.661616 | 7.373569 | 7.596416 | 4.808084 | 8.371850 | -1.111909 | 251.413343 | 0.256114112 | -0.59753999 | 0.550146916 | 0.998865404 | 8.049324 | 7.894351 | 7.998120 | 7.991202 | 7.813142 | 8.055321 | 307 | 284 | 155 | 279 |
| 90427 | NM_00100394 | BMF | Bcl2 modifying factor | protein_coding | NP_001003940.1;NP_001003942.1;NP_001003943.1;NP_277038.1 | 24132 | 606266 | . | . | 0.810249 | 1.089179 | 0.997763 | 1.091084 | 1.885483 | 2.517686 | 2.238952 | 2.494634 | 1.112595 | 145.423310 | 0.279570013 | 0.55059113 | 0.581913991 | 0.998865404 | 7.105803 | 7.258394 | 7.112126 | 7.205431 | 7.188312 | 7.212762 | 139 | 167 | 129 | 148 |
| 90441 | NM_033414 | ZNF622 | zinc finger protein 622 | protein_coding | NP_219482.1 | 30958 | 608694 | . | . | 16.247303 | 15.314066 | 16.904776 | 18.041412 | 37.808140 | 35.399151 | 37.933826 | 41.249542 | 1.091314 | 884.576754 | 0.142889201 | 0.882268899 | 0.377631411 | 0.998865404 | 9.724437 | 9.850497 | 9.779759 | 9.731768 | 9.798866 | 9.838281 | 1007 | 882 | 781 | 889 |
| 90459 | NM_00135463 | ERI1 | exoribonuclease 1 | protein_coding | NP_001341564.1;NP_001341565.1;NP_001341567.1;NP_699163.2;XP_00 | 23994 | 608739 | . | . | 3.478646 | 2.988085 | 3.097380 | 2.759820 | 8.094953 | 6.907095 | 6.950434 | 6.309999 | -1.174567 | 417.418684 | 0.196233784 | -1.18292125 | 0.236840357 | 0.992733415 | 8.816105 | 8.585383 | 8.798827 | 8.697159 | 8.719629 | 8.581060 | 561 | 447 | 374 | 317 |
| 90480 | NM_052850 | GADD45GIP1 | GADD45G interacting protein 1 | protein_coding | NP_443082.2 | 29996 | 605162 | . | . | 19.184725 | 20.532598 | 17.521929 | 20.387411 | 44.643642 | 47.462025 | 39.318699 | 46.613396 | -1.062228 | 1070.102962 | 0.141747785 | -0.61442791 | 0.538932599 | 0.998865404 | 10.106683 | 10.019055 | 10.062578 | 10.106959 | 9.987610 | 10.090142 | 1232 | 1225 | 839 | 1041 |
| 90488 | NM_00131966 | TMEM263 | transmembrane protein 263 | protein_coding | NP_001306590.1;NP_001306591.1;NP_001306592.1;NP_001306593.1;NI | 28281 | . | . | . | 10.602174 | 9.382449 | 9.653479 | 7.432184 | 24.671696 | 21.687952 | 21.662127 | 16.992807 | -1.184136 | 961.151425 | 0.163590282 | -1.49052241 | 0.136086928 | 0.894235858 | 10.024974 | 9.782052 | 10.013134 | 9.912649 | 9.931803 | 9.741923 | 1280 | 1049 | 868 | 715 |
| 90506 | NM_033413,XI | LRRC46 | leucine rich repeat containing 46 | protein_coding | NP_219481.1;XP_005257832.1 | 25047 | . | . | . | 1.101701 | 1.545932 | 1.362320 | 1.257099 | 2.563704 | 3.573491 | 3.057007 | 2.874206 | -1.060125 | 76.628556 | 0.3751775 | -0.22451912 | 0.822353392 | 0.998865404 | 6.301490 | 6.216890 | 6.219855 | 6.306409 | 6.269144 | 6.228931 | 76 | 99 | 70 | 64 |
| 90507 | NM_00114502 | SCRN2 | secernin 2 | protein_coding | NP_001138495.1;NP_612364.2;XP_005257833.1;XP_016880775.1;XP_02 | 30381 | 614966 | . | . | 13.010886 | 13.121373 | 12.988492 | 12.379924 | 30.276865 | 30.330644 | 29.145802 | 28.305223 | -1.045025 | 701.929490 | 0.151271306 | -0.42002277 | 0.674468821 | 0.998865404 | 9.486467 | 9.423205 | 9.470062 | 9.469980 | 9.463441 | 9.415687 | 826 | 774 | 618 | 621 |
| 90522 | NM_00103967 | YIF1B | Yip1 interacting factor homolog B, membrane trafficking protein | protein_coding | NP_001034760.1;NP_001034761.1;NP_001034762.1;NP_001138933.1;NI | 30511 | . | . | . | 32.465254 | 35.714692 | 25.050711 | 29.940374 | 75.547978 | 82.556106 | 56.213073 | 68.455107 | -1.049266 | 2085.773020 | 0.118525521 | -0.58535788 | 0.558307102 | 0.998865404 | 11.060668 | 10.991229 | 11.059004 | 11.031235 | 10.986511 | 11.026799 | 2502 | 2265 | 1725 | 1957 |
| 90523 | NM_00128174 | MLIP | muscular LMNA interacting protein | protein_coding | NP_001268675.1;NP_001268676.1;NP_612636.2;XP_005249533.1;XP_00 | 21355 | 614106 | . | . | 1.590503 | 1.955028 | 1.248878 | 2.325413 | 3.701166 | 4.519135 | 2.802447 | 5.316780 | -1.168189 | 159.148090 | 0.295019225 | -0.76020119 | 0.447134334 | 0.998865404 | 7.423305 | 7.196221 | 7.273475 | 7.401633 | 7.198952 | 7.349604 | 168 | 213 | 102 | 165 |
| 90525 | NM_00130116 | SHF | Src homology 2 domain containing F | protein_coding | NP_001288097.2;NP_001288098.2;NP_001288099.2;NP_001288100.1;NI | 25116 | 617313 | . | . | 0.980922 | 1.138744 | 0.698399 | 1.869020 | 2.282649 | 2.632262 | 1.567187 | 4.273292 | 1.193488 | 75.698757 | 0.430791056 | 0.592362099 | 0.553608141 | 0.998865404 | 6.110920 | 6.362519 | 6.196116 | 6.212088 | 6.133362 | 6.353179 | 77 | 77 | 42 | 109 |
| 90529 | NM_00119901 | STPG1 | sperm tail PG-rich repeat containing 1 | protein_coding | NP_001185941.1;NP_001185942.1;NP_001185943.1;NP_835223.1;XP_00 | 28070 | 615826 | . | . | 4.314523 | 5.191589 | 3.689807 | 5.556855 | 10.040075 | 12.000587 | 8.279821 | 12.705088 | -1.055253 | 407.384800 | 0.214003595 | -0.36255788 | 0.716935182 | 0.998865404 | 8.709336 | 8.630072 | 8.627092 | 8.733429 | 8.533315 | 8.753446 | 440 | 491 | 280 | 441 |
| 90550 | NM_00127067 | MCU | mitochondrial calcium uniporter | protein_coding | NP_001257608.1;NP_001257609.1;NP_612366.1;XP_016872371.1 | 23526 | 614197 | . | . | 6.126981 | 4.732822 | 4.859843 | 5.863746 | 14.257738 | 10.940129 | 10.905345 | 13.406758 | -1.026990 | 496.948766 | 0.191062091 | -0.20109728 | 0.840622511 | 0.998865404 | 8.976022 | 8.937629 | 9.043423 | 8.874891 | 8.889592 | 8.999116 | 656 | 472 | 389 | 500 |
| 90557 | NM_00125830 | CCDC74A | coiled-coil domain containing 74A | protein_coding | NP_001245233.1;NP_001245234.1;NP_001245235.1;NP_001335970.1;NI | 25197 | . | . | . | 0.089050 | 0.049270 | 0.028782 | 0.057560 | 0.207224 | 0.113891 | 0.064585 | 0.131605 | -1.512468 | 4.803636 | 1.399572964 | -0.42649054 | 0.669750437 | NA | 2.533720 | 1.932304 | 2.284120 | 2.273272 | 2.263046 | 2.276435 | 8 | 5 | 2 | 5 |
| 90576 | NM_00108082 | ZNF799 | zinc finger protein 799 | protein_coding | NP_001074290.1;NP_001309426.1;NP_001309427.1 | 28071 | . | . | . | 0.404924 | 0.340037 | 0.496539 | 0.357482 | 0.942275 | 0.786011 | 1.114219 | 0.817341 | 1.133933 | 32.961416 | 0.544671375 | 0.3329271 | 0.739189313 | 0.998865404 | 4.947897 | 5.131667 | 5.038689 | 5.021176 | 5.066245 | 5.026263 | 38 | 31 | 35 | 28 |
| 90580 | NM_138358 | TIMM29 | translocase of inner mitochondrial membrane 29 | protein_coding | NP_612367.1 | 25152 | 617380 | . | . | 5.571478 | 7.261045 | 4.660360 | 7.642117 | 12.965059 | 16.784233 | 10.457712 | 17.472792 | -1.055856 | 264.068224 | 0.258869066 | -0.30290674 | 0.761960932 | 0.998865404 | 8.084570 | 8.003834 | 7.982353 | 8.115509 | 7.896891 | 8.135591 | 273 | 330 | 170 | 298 |

| | | | | | | | | | | | | | | | | | | | | | | | | | | | | | | | | | |
|---|---|---|---|---|---|---|---|---|---|---|---|---|---|---|---|---|---|---|---|---|---|---|---|---|---|---|---|---|---|---|---|---|---|
| 90589 | NM_145233,N | ZNF625 | zinc finger protein 625 | protein_coding | NP_660276.2 | 30571 | . | . | . | 0.226971 | 0.480071 | 0.186221 | 0.222439 | 0.528170 | 1.109706 | 0.417875 | 0.508580 | -1.760752 | 15.264648 | 0.816115177 | -1.00009385 | 0.317265094 | NA | 4.284495 | 3.464871 | 3.922513 | 3.978758 | 3.911851 | 3.917426 | 15 | 28 | 9 | 11 |
| 90592 | NM_00127184 | ZNF700 | zinc finger protein 700 | protein_coding | NP_001258777.1;NP_653167.1 | 25292 | . | . | . | 4.220613 | 4.844615 | 6.995748 | 4.472582 | 9.821541 | 11.198545 | 15.698257 | 10.226029 | 1.249868 | 457.015058 | 0.21829636 | 1.474032262 | 0.140472905 | 0.900674731 | 8.665763 | 8.988428 | 8.713447 | 8.789555 | 9.026583 | 8.733337 | 438 | 466 | 542 | 368 |
| 90594 | NM_00134871 | ZNF439 | zinc finger protein 439 | protein_coding | NP_001335647.1;NP_001335648.1;NP_001335649.1;NP_001335650.1;NI | 20873 | . | . | . | 0.051750 | 0.105055 | 0.087593 | 0.075760 | 0.120425 | 0.242839 | 0.196556 | 0.173217 | 1.013201 | 7.751160 | 1.104644003 | 0.017128543 | 0.986334068 | NA | 2.945312 | 2.963455 | 2.937299 | 2.960450 | 2.955531 | 2.943237 | 6 | 11 | 8 | 6 |
| 90624 | NM_00129373 | LYRM7 | LYR motif containing 7 | protein_coding | NP_001280664.1;NP_859056.2 | 28072 | 615831 | . | . | 2.741432 | 2.632596 | 2.773932 | 2.350799 | 6.379424 | 6.085364 | 6.224622 | 5.374822 | -1.064333 | 509.306746 | 0.174117132 | -0.51660308 | 0.605433264 | 0.998865404 | 9.036250 | 8.947157 | 9.026346 | 8.995249 | 9.025009 | 8.914769 | 619 | 553 | 467 | 422 |
| 90627 | NM_00124346 | STARD13 | StAR related lipid transfer domain containing 13 | protein_coding | NP_001230395.1;NP_001230403.1;NP_001230405.1;NP_443083.1;NP_8 | 19164 | 609866 | . | . | 2.404681 | 2.295609 | 3.411286 | 3.149073 | 5.595792 | 5.306401 | 7.654829 | 7.199984 | 1.323076 | 536.594694 | 0.168047052 | 2.403470295 | 0.016240286 | 0.428496267 | 8.851269 | 9.255838 | 8.999401 | 8.932114 | 9.174823 | 9.118561 | 561 | 471 | 554 | 549 |
| 90632 | NR_026860,NF | LINC00473 | long intergenic non-protein coding RNA 473 | lncRNA | . | 21160 | . | . | . | 5.085664 | 5.572052 | 5.818796 | 5.023025 | 11.834548 | 12.880049 | 13.057209 | 11.484550 | 1.016146 | 286.466363 | 0.212525521 | 0.108731635 | 0.913415349 | 0.998865404 | 8.150270 | 8.174076 | 8.137853 | 8.174093 | 8.215175 | 8.111510 | 316 | 317 | 276 | 244 |
| 90634 | NM_00107969 | N4BP2L1 | NEDD4 binding protein 2 like 1 | protein_coding | NP_001073159.1;NP_001273388.1;NP_001273389.1;NP_001273390.1;NI | 25037 | . | . | . | 0.308195 | 0.696068 | 0.703935 | 0.552403 | 0.717183 | 1.608993 | 1.579610 | 1.263004 | 1.205958 | 28.024368 | 0.627500428 | 0.430565361 | 0.666784432 | 0.998865404 | 4.668126 | 4.936619 | 4.741748 | 4.825348 | 4.825481 | 4.785856 | 18 | 38 | 31 | 24 |
| 90637 | NM_00136538 | ZFAND2A | zinc finger AN1-type containing 2A | protein_coding | NP_001352310.1;NP_001352312.1;NP_872297.2 | 28073 | 610699 | . | . | 8.657451 | 8.911089 | 9.325004 | 10.812557 | 20.146243 | 20.598381 | 20.925038 | 24.721625 | 1.088441 | 362.158347 | 0.190980842 | 0.640186646 | 0.522051264 | 0.998865404 | 8.438289 | 8.560094 | 8.434119 | 8.512593 | 8.524057 | 8.521462 | 372 | 400 | 329 | 353 |
| 90639 | NM_00103161 | COX19 | cytochrome c oxidase assembly factor COX19 | protein_coding | NP_001026788.1 | 28074 | 610429 | . | . | 2.214186 | 1.995409 | 2.236458 | 1.982766 | 5.152503 | 4.612476 | 5.018548 | 4.533360 | -1.012617 | 318.088000 | 0.202180542 | -0.08946452 | 0.928712747 | 0.998865404 | 8.321799 | 8.304715 | 8.346347 | 8.284235 | 8.343966 | 8.272847 | 389 | 326 | 293 | 277 |
| 90649 | NM_052852 | ZNF486 | zinc finger protein 486 | protein_coding | NP_443084.2 | 20807 | . | . | . | 1.500034 | 1.188602 | 1.343931 | 1.342925 | 3.490640 | 2.747507 | 3.015742 | 3.070438 | -1.016858 | 163.492930 | 0.265532329 | -0.090828 | 0.927629262 | 0.998865404 | 7.364610 | 7.341466 | 7.402955 | 7.305542 | 7.351428 | 7.345905 | 212 | 157 | 142 | 151 |
| 90668 | NM_138360 | CARMIL3 | capping protein regulator and myosin 1 linker 3 | protein_coding | NP_612369.3 | 20272 | 614716 | . | . | 0.635193 | 1.052377 | 1.022509 | 0.788727 | 1.478121 | 2.432617 | 2.294479 | 1.803330 | 1.061769 | 125.290260 | 0.327740477 | 0.263834544 | 0.791907415 | 0.998865404 | 6.925491 | 7.011489 | 6.859524 | 7.030710 | 7.015297 | 6.922217 | 106 | 163 | 127 | 105 |
| 90673 | NM_00127631 | PPP1R3E | protein phosphatase 1 regulatory subunit 3E | protein_coding | NP_001263247.1 | 14943 | . | . | . | 2.366696 | 2.367709 | 2.950030 | 2.166756 | 5.507400 | 5.473066 | 6.619781 | 4.954030 | 1.065280 | 366.787850 | 0.206686659 | 0.441405939 | 0.658919144 | 0.998865404 | 8.471833 | 8.564290 | 8.497754 | 8.494528 | 8.620372 | 8.437101 | 410 | 382 | 381 | 299 |
| 90678 | NM_00100537 | LRSAM1 | leucine rich repeat and sterile alpha motif containing 1 | protein_coding | NP_001005373.1;NP_001005374.1;NP_001177652.1;NP_612370.3;XP_00 | 25135 | 610933 | . | . | 4.012155 | 4.107363 | 4.134057 | 4.273292 | 9.336450 | 9.494353 | 9.276703 | 9.770374 | 1.022351 | 428.498958 | 0.175708612 | 0.181499056 | 0.855975874 | 0.998865404 | 8.727164 | 8.758957 | 8.731260 | 8.741216 | 8.746851 | 8.752417 | 484 | 461 | 375 | 408 |
| 90693 | NM_138771,XI | CCDC126 | coiled-coil domain containing 126 | protein_coding | NP_620126.2;XP_016868264.1;XP_016868265.1;XP_024302761.1 | 22398 | . | . | . | 2.238385 | 1.612583 | 1.103151 | 1.576655 | 5.208815 | 3.727557 | 2.475441 | 3.604834 | -1.470159 | 128.448630 | 0.315171314 | -1.76403154 | 0.077726669 | 0.767975096 | 7.256742 | 6.699964 | 7.131155 | 6.993047 | 6.879162 | 6.975156 | 206 | 137 | 76 | 113 |
| 90701 | NM_00130794 | SEC11C | SEC11 homolog C, signal peptidase complex subunit | protein_coding | NP_001294870.1;NP_150596.1;XP_011524562.1;XP_016881562.1 | 23400 | . | . | . | 68.289980 | 65.386275 | 82.039535 | 64.460078 | 158.913591 | 151.143309 | 184.094340 | 147.380314 | 1.081816 | 1722.335513 | 0.143980882 | 0.78798295 | 0.430706679 | 0.998865404 | 10.692057 | 10.805995 | 10.734705 | 10.697086 | 10.872439 | 10.673832 | 1951 | 1742 | 1752 | 1468 |
| 90736 | NM_00116669 | FAM104B | family with sequence similarity 104 member B | protein_coding | NP_001160171.1;NP_001160172.1;NP_001160173.1;NP_001160174.1;NI | 25085 | . | . | . | 2.733129 | 2.827899 | 2.835413 | 3.175800 | 6.360103 | 6.536812 | 6.362584 | 7.261087 | 1.180087 | 158.808158 | 0.267274142 | 0.893812841 | 0.371422037 | 0.998865404 | 7.186873 | 7.425549 | 7.235081 | 7.308319 | 7.361954 | 7.315075 | 152 | 172 | 158 | 152 |
| 90737 | NM_00101343 | PAGE5 | PAGE family member 5 | protein_coding | NP_001013453.1;NP_569734.2 | 29992 | 301009 | . | . | 0.984131 | 1.004510 | 0.650921 | 1.376913 | 2.290112 | 2.321969 | 1.460649 | 3.148149 | 1.002652 | 17.485853 | 0.754160818 | 0.005067104 | 0.995957053 | NA | 4.128334 | 4.127898 | 4.127081 | 4.128134 | 4.102551 | 4.152517 | 20 | 19 | 10 | 22 |
| 90768 | NR_027107,NF | TENM3-AS1 | TENM3 antisense RNA 1 | lncRNA | . | 28076 | . | . | . | 2.061578 | 2.234311 | 2.258315 | 2.244554 | 4.797377 | 5.164710 | 5.067593 | 5.131907 | 1.046332 | 358.441659 | 0.187731923 | 0.34805007 | 0.727802581 | 0.998865404 | 8.452533 | 8.517916 | 8.462803 | 8.481021 | 8.507231 | 8.488483 | 397 | 384 | 324 | 338 |
| 90780 | NM_138300 | PYGO2 | pygopus family PHD finger 2 | protein_coding | NP_612157.1 | 30257 | 606903 | . | . | 6.485283 | 7.030508 | 6.428954 | 7.825779 | 15.091520 | 16.251333 | 14.426387 | 17.892715 | 1.040337 | 688.024526 | 0.162134538 | 0.351877186 | 0.724930369 | 0.998865404 | 9.397919 | 9.454165 | 9.386550 | 9.435609 | 9.372195 | 9.500322 | 749 | 754 | 553 | 719 |
| 90799 | NM_00131699 | CEP95 | centrosomal protein 95 | protein_coding | NP_001303919.1;NP_612372.1;XP_005257836.1;XP_005257837.1;XP_00 | 25141 | . | . | . | 10.158108 | 9.371717 | 10.813459 | 7.991606 | 23.638336 | 21.663140 | 24.265089 | 18.271856 | -1.022195 | 1052.828128 | 0.151041374 | -0.20968326 | 0.83391489 | 0.998865404 | 10.055500 | 10.024441 | 10.049758 | 10.042893 | 10.110752 | 9.942952 | 1231 | 1143 | 1009 | 865 |
| 90806 | NM_00130075 | ANGEL2 | angel homolog 2 | protein_coding | NP_001287682.1;NP_001287684.1;NP_001287686.1;NP_001287687.1;NI | 30534 | . | . | . | 3.449033 | 3.422166 | 4.404052 | 3.678616 | 8.026043 | 7.910487 | 9.882565 | 8.410719 | 1.151515 | 551.303728 | 0.170264793 | 1.195390108 | 0.231934649 | 0.986726781 | 9.000980 | 9.205206 | 9.059667 | 9.053283 | 9.206872 | 9.086350 | 593 | 550 | 560 | 503 |
| 90809 | NM_00110081 | PIP4P1 | phosphatidylinositol-4,5-bisphosphate 4-phosphatase 1 | protein_coding | NP_001094284.1;NP_653169.2;XP_024305507.1;XP_024305508.1 | 19299 | 609865 | . | . | 12.960039 | 12.578001 | 14.438834 | 14.644165 | 30.158541 | 29.074612 | 32.400329 | 33.482144 | 1.126537 | 776.575542 | 0.145831671 | 1.178716043 | 0.238551265 | 0.992733415 | 9.512431 | 9.684411 | 9.569182 | 9.544630 | 9.638928 | 9.642080 | 856 | 774 | 714 | 773 |
| 90826 | NM_00130445 | PRMT9 | protein arginine methyltransferase 9 | protein_coding | NP_001291387.1;NP_001337070.1;NP_001337071.1;NP_001337072.1;NI | 25099 | 616125 | . | . | 2.414414 | 2.380613 | 2.662754 | 1.910455 | 5.618440 | 5.502895 | 5.975143 | 4.368027 | -1.039707 | 280.409889 | 0.214994575 | -0.26129437 | 0.793865507 | 0.998865404 | 8.158769 | 8.103489 | 8.128435 | 8.151339 | 8.173649 | 8.062681 | 322 | 315 | 265 | 230 |
| 90835 | NM_00101497 | CCDC189 | coiled-coil domain containing 189 | protein_coding | NP_001014979.2;NP_001182549.1;XP_011544278.1;XP_011544280.1;XP | 28078 | 618318 | . | . | 5.079566 | 5.776260 | 5.295076 | 6.237361 | 11.820360 | 13.352085 | 11.881998 | 14.260987 | 1.066794 | 319.763921 | 0.204053789 | 0.457139412 | 0.647570844 | 0.998865404 | 8.274057 | 8.366198 | 8.264387 | 8.336600 | 8.293331 | 8.377955 | 333 | 356 | 265 | 333 |
| 90843 | NM_00100668 | TCEAL8 | transcription elongation factor A like 8 | protein_coding | NP_001006685.1;NP_699164.1 | 28683 | . | . | . | 22.746817 | 20.427223 | 20.404334 | 20.314691 | 52.932779 | 47.218442 | 45.786734 | 46.447126 | -1.072908 | 765.242018 | 0.150442476 | -0.67484891 | 0.499771763 | 0.998865404 | 9.629479 | 9.528292 | 9.643354 | 9.562281 | 9.559779 | 9.549213 | 964 | 806 | 647 | 687 |
| 90850 | NM_178167 | ZNF598 | zinc finger protein 598, E3 ubiquitin ligase | protein_coding | NP_835461.2 | 28079 | 617508 | . | . | 13.887463 | 14.617249 | 14.399678 | 15.267395 | 32.316696 | 33.788425 | 32.312458 | 34.907082 | 1.026491 | 1525.441507 | 0.125335895 | 0.300962397 | 0.763443169 | 0.998865404 | 10.556130 | 10.593648 | 10.548127 | 10.580915 | 10.565764 | 10.603534 | 1697 | 1659 | 1311 | 1484 |
| 90853 | NM_00128198 | SPOCD1 | SPOC domain containing 1 | protein_coding | NP_001268916.1;NP_001268917.1;NP_653170.3;XP_011540715.1;XP_01 | 26338 | . | . | . | 0.623103 | 0.719524 | 0.378945 | 0.509684 | 1.449986 | 1.663210 | 0.850341 | 1.165332 | -1.457553 | 67.745147 | 0.391030906 | -1.39003974 | 0.16451681 | 0.940065727 | 6.329258 | 5.783530 | 6.108289 | 6.140286 | 6.014294 | 6.061048 | 87 | 92 | 42 | 58 |
| 90859 | gene-NT5C3AP | NT5C3AP1 | NT5C3A pseudogene 1 | pseudogene | . | 18530 | . | . | . | 0.060887 | 0.070364 | 0.025077 | 0.025654 | 0.141687 | 0.162650 | 0.056271 | 0.058654 | -1.620127 | 2.903507 | 1.757623934 | -0.39605015 | 0.692068032 | NA | 1.844134 | 1.148375 | 1.548992 | 1.550042 | 1.543603 | 1.542786 | 4 | 4 | 2 | 2 |
| 90861 | NM_144570 | JPT2 | Jupiter microtubule associated homolog 2 | protein_coding | NP_653171.1 | 14137 | . | . | . | 21.491137 | 20.739300 | 20.805300 | 21.693150 | 50.010754 | 47.939823 | 46.686489 | 49.598808 | -1.007773 | 2438.440211 | 0.115398448 | -0.09680546 | 0.922880885 | 0.998865404 | 11.257318 | 11.246146 | 11.272528 | 11.236753 | 11.235817 | 11.261122 | 2881 | 2583 | 2079 | 2313 |
| 90864 | NM_00132408 | SPSB3 | splA/ryanodine receptor domain and SOCS box containing 3 | protein_coding | NP_001311010.1;NP_543137.2;XP_005255730.1 | 30629 | 611659 | . | . | 9.625312 | 8.483404 | 8.111127 | 9.988857 | 22.398499 | 19.609769 | 18.201135 | 22.838335 | -1.020540 | 438.587938 | 0.184234141 | -0.15921321 | 0.873500898 | 0.998865404 | 8.791533 | 8.761759 | 8.812842 | 8.750363 | 8.714768 | 8.821630 | 536 | 454 | 344 | 443 |
| 90871 | NM_00131805 | DMAC1 | distal membrane arm assembly complex 1 | protein_coding | NP_001304987.1;NP_001304988.1;NP_219500.1 | 30536 | 617261 | . | . | 8.563116 | 8.539397 | 7.251916 | 7.842279 | 19.926722 | 19.739199 | 16.273089 | 17.930440 | -1.147722 | 617.299045 | 0.157138377 | -1.26495558 | 0.205887253 | 0.970176829 | 9.365912 | 9.166880 | 9.318108 | 9.310279 | 9.200284 | 9.243589 | 764 | 708 | 483 | 557 |
| 90874 | NM_00108047 | ZNF697 | zinc finger protein 697 | protein_coding | NP_001073939.1;XP_005271372.1;XP_011540718.1 | 32034 | . | . | . | 18.475662 | 17.409716 | 19.489494 | 17.694897 | 42.993625 | 40.243338 | 43.733862 | 40.457281 | 1.018889 | 3251.806360 | 0.127569871 | 0.211623824 | 0.832400521 | 0.998865404 | 11.653316 | 11.680606 | 11.703788 | 11.609889 | 11.740610 | 11.601420 | 3921 | 3277 | 3096 | 2817 |
| 90933 | NM_033549,N | TRIM41 | tripartite motif containing 41 | protein_coding | NP_291027.3;NP_963921.1;XP_006714992.1;XP_006714993.1;XP_00671 | 19013 | 610530 | . | . | 21.104688 | 21.253460 | 22.145466 | 22.040212 | 49.111476 | 49.128322 | 49.693783 | 50.392324 | 1.063085 | 2087.001367 | 0.123758544 | 0.713137088 | 0.475760932 | 0.998865404 | 10.982318 | 11.070758 | 10.988026 | 11.015001 | 11.098394 | 11.000195 | 2290 | 2220 | 1985 | 1893 |
| 90956 | NM_052853,XI | ADCK2 | aarF domain containing kinase 2 | protein_coding | NP_443085.2;XP_006716233.1;XP_011514977.1 | 19039 | . | . | . | 8.044543 | 8.039503 | 6.436257 | 7.766642 | 18.719981 | 18.583673 | 14.442775 | 17.757504 | -1.147091 | 606.483658 | 0.165039261 | -1.19958953 | 0.230298795 | 0.985678841 | 9.340235 | 9.141582 | 9.291165 | 9.284320 | 9.138272 | 9.251287 | 750 | 696 | 448 | 576 |
| 90957 | NM_00132996 | DHX57 | DExH-box helicase 57 | protein_coding | NP_001316892.1;NP_945314.1;XP_011531456.1;XP_011531457.1;XP_01 | 20086 | . | . | . | 5.145774 | 5.418044 | 5.387113 | 5.694377 | 11.974431 | 12.524052 | 12.088525 | 13.019514 | 1.035991 | 820.471776 | 0.144159593 | 0.353849858 | 0.723451406 | 0.998865404 | 9.654700 | 9.705473 | 9.656587 | 9.678210 | 9.676652 | 9.707981 | 914 | 883 | 710 | 800 |
| 90987 | NM_138367,XI | ZNF251 | zinc finger protein 251 | protein_coding | NP_612376.1;XP_005272413.1;XP_006716746.1;XP_011515663.1;XP_02 | 13045 | . | . | . | 4.939204 | 4.190771 | 4.571505 | 5.487145 | 11.493733 | 9.687153 | 10.258324 | 12.545707 | 1.100092 | 380.481761 | 0.19091645 | 0.720860945 | 0.470995074 | 0.998865404 | 8.501277 | 8.638813 | 8.579153 | 8.502465 | 8.559609 | 8.633471 | 445 | 365 | 325 | 397 |
| 90990 | NM_00136976 | KIFC2 | kinesin family member C2 | protein_coding | NP_001356698.1;NP_665697.1;XP_011515664.1 | 29530 | 615216 | . | . | 17.777673 | 18.052456 | 23.161265 | 20.335494 | 41.369375 | 41.729061 | 51.973210 | 46.494692 | 1.200942 | 2028.143793 | 0.126344144 | 2.090847921 | 0.036541699 | 0.596199845 | 10.847688 | 11.112102 | 10.904711 | 10.909943 | 11.110428 | 10.992408 | 2115 | 1995 | 2065 | 1924 |
| 91010 | NM_00136783 | FMNL3 | formin like 3 | protein_coding | NP_001354764.1;NP_783863.4;NP_944489.2;XP_011537270.1;XP_01153 | 23698 | 616288 | . | . | 3.519731 | 3.930006 | 4.271528 | 4.355057 | 8.190556 | 9.084384 | 9.585186 | 9.957316 | 1.139643 | 1591.602428 | 0.127191195 | 1.482664182 | 0.138163656 | 0.899796454 | 10.539004 | 10.727380 | 10.541178 | 10.622186 | 10.681746 | 10.686418 | 1623 | 1690 | 1470 | 1594 |
| 91012 | NM_00128173 | CERS5 | ceramide synthase 5 | protein_coding | NP_001268660.1;NP_001317998.1;NP_001317999.1;NP_001318000.1;NI | 23749 | 615335 | . | . | 11.004139 | 10.143633 | 12.756940 | 10.987387 | 25.607087 | 23.447461 | 28.626206 | 25.121355 | 1.043424 | 863.404935 | 0.145906853 | 0.420308352 | 0.674260209 | 0.998865404 | 9.722657 | 9.784468 | 9.765335 | 9.709456 | 9.799634 | 9.735973 | 1011 | 876 | 801 | 790 |
| 91039 | NM_00136598 | DPP9 | dipeptidyl peptidase 9 | protein_coding | NP_001352916.1;NP_631898.3;XP_005259730.1;XP_011526707.1;XP_01 | 18648 | 608258 | . | . | 15.179227 | 16.042215 | 16.805124 | 17.869970 | 35.322687 | 37.082302 | 37.710211 | 40.857563 | 1.107701 | 2147.926902 | 0.12050449 | 1.224583948 | 0.220732019 | 0.982008277 | 10.993182 | 11.140517 | 11.006485 | 11.045103 | 11.072897 | 11.141284 | 2290 | 2253 | 1880 | 2206 |
| 91050 | NM_00113072 | CCDC149 | coiled-coil domain containing 149 | protein_coding | NP_001124198.1;NP_001317572.1;NP_001317573.1;NP_775734.2;XP_01 | 25405 | . | . | . | 0.520386 | 0.624017 | 0.361654 | 0.328768 | 1.210962 | 1.442443 | 0.811541 | 0.751689 | -1.023321 | 72.930033 | 0.374803238 | -0.0887374 | 0.929290607 | 0.998865404 | 6.205110 | 6.171577 | 6.162756 | 6.216065 | 6.196676 | 6.171375 | 76 | 88 | 66 | 64 |
| 91056 | NM_138368 | AP5B1 | adaptor related protein complex 5 subunit beta 1 | protein_coding | NP_612377.4 | 25104 | 614367 | . | . | 2.207104 | 2.384395 | 2.141498 | 1.914519 | 5.136021 | 5.511636 | 4.805460 | 4.377319 | -1.146032 | 470.709001 | 0.175185037 | -1.12251542 | 0.261643355 | 0.995760118 | 8.973501 | 8.777211 | 8.896847 | 8.940823 | 8.870735 | 8.796409 | 559 | 561 | 405 | 386 |
| 91057 | NM_030771,N | CCDC34 | coiled-coil domain containing 34 | protein_coding | NP_110398.1;NP_542385.1 | 25079 | 612324 | . | . | 10.599846 | 9.163912 | 9.749492 | 9.866351 | 24.666277 | 21.182794 | 21.877577 | 22.558239 | -1.035021 | 466.170572 | 0.176067034 | -0.28205433 | 0.777901855 | 0.998865404 | 8.889112 | 8.839897 | 8.919072 | 8.828707 | 8.854438 | 8.852715 | 585 | 475 | 399 | 429 |
| 91107 | NM_033452,XI | TRIM47 | tripartite motif containing 47 | protein_coding | NP_258411.2;XP_005257844.1;XP_005257845.1 | 19020 | 611041 | . | . | 16.764232 | 15.093897 | 18.892892 | 19.036848 | 39.011057 | 34.890221 | 42.395102 | 43.525491 | 1.181047 | 1213.699967 | 0.133036645 | 1.804509201 | 0.071151518 | 0.751131579 | 10.120083 | 10.360319 | 10.211312 | 10.143884 | 10.298710 | 10.308399 | 1340 | 1146 | 1136 | 1240 |
| 91120 | NM_00107734 | ZNF682 | zinc finger protein 682 | protein_coding | NP_001070817.1;NP_149973.1;XP_016882944.1;XP_016882945.1 | 28857 | . | . | . | 1.021359 | 0.970142 | 1.604672 | 0.745703 | 2.376746 | 2.242525 | 3.600839 | 1.704961 | 1.168787 | 112.340358 | 0.366097653 | 0.614621752 | 0.538804546 | 0.998865404 | 6.693244 | 6.921224 | 6.775242 | 6.760271 | 6.937077 | 6.685385 | 122 | 109 | 144 | 72 |
| 91133 | NM_00133055 | L3MBTL4 | L3MBTL histone methyl-lysine binding protein 4 | protein_coding | NP_001317488.1;NP_001352694.1;NP_001352695.1;NP_001352696.1;NI | 26677 | 617135 | . | . | 1.926322 | 1.729556 | 2.249196 | 2.445388 | 4.482630 | 3.997944 | 5.047135 | 5.591089 | 1.216692 | 221.149357 | 0.233516433 | 1.211750836 | 0.225607773 | 0.982008277 | 7.640615 | 7.923313 | 7.764265 | 7.708376 | 7.788633 | 7.868691 | 244 | 202 | 194 | 246 |
| 91137 | NM_00130324 | SLC25A46 | solute carrier family 25 member 46 | protein_coding | NP_001290178.1;NP_001290179.1;NP_620128.1 | 25198 | 610826 | . | . | 7.398521 | 6.394777 | 6.656183 | 5.390409 | 17.216663 | 14.781813 | 14.936283 | 12.324530 | -1.159202 | 956.522489 | 0.159386132 | -1.33720433 | 0.181155948 | 0.952877815 | 10.003889 | 9.791620 | 10.007064 | 9.891052 | 9.918227 | 9.762706 | 1274 | 1022 | 855 | 740 |
| 91147 | NM_00114230 | TMEM67 | transmembrane protein 67 | protein_coding | NP_001135773.1;NP_714915.3;XP_006716749.1;XP_011515665.1;XP_02 | 28396 | 609884 | . | . | 3.921202 | 3.225250 | 3.353825 | 2.776878 | 9.124796 | 7.455311 | 7.525887 | 6.349005 | -1.138416 | 463.539970 | 0.189361814 | -0.98767785 | 0.323310446 | 0.998865404 | 8.946449 | 8.760675 | 8.955548 | 8.826704 | 8.868482 | 8.754120 | 625 | 479 | 413 | 368 |
| 91151 | NM_033208 | TIGD7 | tigger transposable element derived 7 | protein_coding | NP_149985.2 | 18331 | 612969 | . | . | 1.448213 | 1.765896 | 1.765783 | 1.342470 | 3.370050 | 4.081947 | 3.962366 | 3.069398 | -1.045679 | 168.959502 | 0.270666639 | -0.23808056 | 0.811818609 | 0.998865404 | 7.432022 | 7.368343 | 7.365003 | 7.445894 | 7.443343 | 7.328947 | 180 | 204 | 164 | 133 |
| 91156 | NM_00116458 | IGFN1 | immunoglobulin like and fibronectin type III domain containing 1 | protein_coding | NP_001158058.1;NP_001354770.1;XP_005245637.1;XP_006711681.1;XP | 24607 | 617309 | . | . | 0.322259 | 0.332408 | 0.480427 | 0.190836 | 0.749909 | 0.768377 | 1.078064 | 0.436324 | 1.014831 | 124.388776 | 0.375419208 | 0.056575683 | 0.954883206 | 0.998865404 | 6.946570 | 6.970754 | 6.931659 | 6.941824 | 7.081421 | 6.779415 | 140 | 135 | 156 | 67 |
| 91179 | NM_153334,N | SCARF2 | scavenger receptor class F member 2 | protein_coding | NP_699165.3;NP_878315.2;XP_016884554.1 | 19869 | 613619 | . | . | 6.298129 | 6.115776 | 5.123250 | 5.592832 | 14.656002 | 14.136890 | 11.496423 | 12.787347 | -1.187608 | 607.436809 | 0.162235326 | -1.52900562 | 0.12626305 | 0.874064017 | 9.365530 | 9.116957 | 9.310265 | 9.291123 | 9.140612 | 9.230022 | 770 | 702 | 449 | 557 |
| 91252 | NM_00112822 | SLC39A13 | solute carrier family 39 member 13 | protein_coding | NP_001121697.2;NP_001317174.1;NP_689477.2;XP_006718444.1;XP_00 | 20859 | 608735 | . | . | 9.554042 | 10.064105 | 10.744274 | 9.971053 | 22.232644 | 23.263626 | 24.109839 | 22.797621 | 1.039752 | 730.661034 | 0.150698322 | 0.373195451 | 0.709002981 | 0.998865404 | 9.484549 | 9.541016 | 9.487062 | 9.510418 | 9.555576 | 9.495276 | 811 | 786 | 676 | 668 |
| 91272 | NM_00115965 | BOD1 | biorientation of chromosomes in cell division 1 | protein_coding | NP_001153123.1;NP_612378.1;XP_016865506.1 | 25114 | 616745 | . | . | 32.469008 | 33.039075 | 32.998375 | 35.613222 | 75.556715 | 76.371296 | 74.047393 | 81.425398 | 1.039210 | 2020.880813 | 0.119751437 | 0.463355845 | 0.643109323 | 0.998865404 | 10.952838 | 11.008168 | 10.963894 | 10.966280 | 10.966260 | 11.024153 | 2279 | 2142 | 1726 | 2001 |
| 91283 | NM_00119880 | MSANTD3 | Myb/SANT DNA binding domain containing 3 | protein_coding | NP_001185734.1;NP_001185735.1;NP_001185736.1;NP_542386.1 | 23370 | . | . | . | 23.068954 | 25.964655 | 28.461486 | 28.463589 | 53.682404 | 60.018460 | 63.866748 | 65.078609 | 1.156699 | 1485.294632 | 0.129265873 | 1.624659789 | 0.10423507 | 0.828776884 | 10.427800 | 10.637643 | 10.434034 | 10.517749 | 10.595575 | 10.582537 | 1499 | 1568 | 1397 | 1481 |
| 91289 | NM_00136381 | LMF2 | lipase maturation factor 2 | protein_coding | NP_001350745.1;NP_149977.2;XP_006724489.1;XP_006724490.1;XP_01 | 25096 | . | . | . | 56.992111 | 57.958205 | 62.885546 | 55.555922 | 132.622985 | 133.972990 | 141.113341 | 127.022008 | 1.017621 | 5124.721143 | 0.115317688 | 0.218528427 | 0.827017412 | 0.998865404 | 12.310564 | 12.335841 | 12.288164 | 12.341531 | 12.383179 | 12.273609 | 5663 | 5656 | 4789 | 4530 |
| 91298 | NM_00100989 | C12orf29 | chromosome 12 open reading frame 29 | protein_coding | NP_001009894.2 | 25322 | . | . | . | 2.327715 | 2.087960 | 2.642642 | 2.085334 | 5.416689 | 4.826414 | 5.930012 | 4.767869 | 1.057195 | 202.187863 | 0.246770186 | 0.325169397 | 0.745052869 | 0.998865404 | 7.618043 | 7.699902 | 7.668988 | 7.617841 | 7.723420 | 7.611708 | 239 | 200 | 203 | 171 |
| 91300 | NM_138774,XI | R3HDM4 | R3H domain containing 4 | protein_coding | NP_620129.2;XP_011526718.1;XP_024307539.1;XP_024307540.1 | 28270 | . | . | . | 7.015031 | 7.966180 | 7.565012 | 7.505877 | 16.324266 | 18.414181 | 16.975669 | 17.161296 | -1.015347 | 415.551447 | 0.182399056 | -0.12046376 | 0.904115783 | 0.998865404 | 8.710028 | 8.687651 | 8.660066 | 8.743845 | 8.695998 | 8.691184 | 449 | 483 | 360 | 385 |
| 91304 | NM_00103302 | TMEM259 | transmembrane protein 259 | protein_coding | NP_001028198.1;NP_219488.1;XP_005259732.1;XP_005259733.1;XP_00 | 17039 | 611011 | . | . | 51.260231 | 51.096274 | 50.570333 | 50.049394 | 119.284652 | 118.111322 | 113.478357 | 114.431997 | -1.030077 | 4255.294405 | 0.10685341 | -0.40010085 | 0.689082241 | 0.998865404 | 12.076243 | 12.033528 | 12.072364 | 12.062093 | 12.050857 | 12.034284 | 5003 | 4631 | 3682 | 3891 |
| 91316 | NR_024448 | GUSBP11 | GUSB pseudogene 11 | transcribed_ps | . | 42325 | . | . | . | 4.798696 | 4.858290 | 7.040123 | 4.849235 | 11.166761 | 11.230156 | 15.797832 | 11.087198 | 1.215315 | 573.816721 | 0.193498094 | 1.453917012 | 0.145969316 | 0.906757185 | 9.016391 | 9.298712 | 9.088248 | 9.091282 | 9.338246 | 9.082025 | 595 | 560 | 651 | 479 |
| 91319 | NM_00100286 | DERL3 | derlin 3 | protein_coding | NP_001002862.1;NP_001129223.1;NP_001350001.1;NP_940842.2;XP_01 | 14236 | 610305 | . | . | 0.571292 | 0.646755 | 0.850909 | 0.575329 | 1.329417 | 1.495004 | 1.909417 | 1.315419 | 1.036594 | 49.928836 | 0.466409013 | 0.111169341 | 0.911482063 | 0.998865404 | 5.614466 | 5.668628 | 5.616785 | 5.645963 | 5.685715 | 5.588046 | 52 | 57 | 56 | 35 |
| 91351 | NM_00101296 | DDX60L | DExD/H-box 60 like | protein_coding | NP_001012985.2;NP_001278439.1;NP_001332856.1;NP_001364998.1;NI | 26429 | 616725 | . | . | 3.205588 | 2.721020 | 3.098623 | 2.432864 | 7.459533 | 6.289759 | 6.953219 | 5.562456 | -1.065078 | 524.921259 | 0.197606602 | -0.4603028 | 0.645298892 | 0.998865404 | 9.080038 | 8.990487 | 9.097343 | 9.002983 | 9.117477 | 8.891898 | 668 | 542 | 521 | 392 |
| 91355 | NM_00113577 | LRP5L | LDL receptor related protein 5 like | protein_coding | NP_001129244.1;NP_872298.1;XP_006724435.1;XP_016884572.1;XP_01 | 25323 | . | . | . | 0.722898 | 0.976883 | 1.102813 | 0.818493 | 1.682214 | 2.258111 | 2.474679 | 1.871388 | 1.040570 | 120.879450 | 0.309322058 | 0.185483702 | 0.852849662 | 0.998865404 | 6.888565 | 6.945719 | 6.852925 | 6.956593 | 6.950549 | 6.887979 | 116 | 147 | 117 | 105 |
| 91368 | NM_080656,XI | CDKN2AIPNL | CDKN2A interacting protein N-terminal like | protein_coding | NP_542387.1;XP_011542011.1 | 30545 | . | . | . | 25.638564 | 26.816142 | 21.012771 | 26.937756 | 59.661989 | 61.986711 | 47.152048 | 61.589519 | -1.107204 | 967.918664 | 0.154564953 | -0.95054671 | 0.341834534 | 0.998865404 | 9.990678 | 9.843030 | 9.939691 | 9.966907 | 9.791612 | 9.959441 | 1150 | 1118 | 705 | 962 |
| 91369 | NM_052855,XI | ANKRD40 | ankyrin repeat domain 40 | protein_coding | NP_443087.1;XP_016880785.1 | 28233 | . | . | . | 7.427212 | 7.423142 | 7.629668 | 8.126282 | 17.283429 | 17.158925 | 17.120754 | 18.579779 | 1.048896 | 991.939056 | 0.137176317 | 0.502066568 | 0.615620696 | 0.998865404 | 9.919343 | 9.988054 | 9.937445 | 9.934885 | 9.951641 | 9.990090 | 1117 | 1043 | 860 | 977 |
| 91370 | NR_148946 | LOC91370 | uncharacterized LOC91370 | lncRNA | . | . | . | . | . | 0.297809 | 0.551861 | 0.877898 | 0.308143 | 0.693015 | 1.275652 | 1.969979 | 0.704532 | 1.347665 | 12.758333 | 0.912427518 | 0.471776492 | 0.637086332 | NA | 3.440763 | 3.873625 | 3.631881 | 3.659879 | 3.692682 | 3.631459 | 9 | 15 | 19 | 7 |
| 91373 | NM_207309,XI | UAP1L1 | UDP-N-acetylglucosamine pyrophosphorylase 1 like 1 | protein_coding | NP_997192.2;XP_006717380.1;XP_011517484.1 | 28082 | . | . | . | 23.015208 | 23.231314 | 27.107367 | 25.103678 | 53.557336 | 53.700221 | 60.828148 | 57.396572 | 1.119443 | 2509.797135 | 0.116540085 | 1.396782012 | 0.162479133 | 0.935295382 | 11.209600 | 11.372513 | 11.243437 | 11.247598 | 11.369010 | 11.303567 | 2720 | 2562 | 2398 | 2386 |
| 91380 | NR_001293 | SNORD107 | small nucleolar RNA, C/D box 107 | snoRNA | . | 32771 | . | . | . | 0.000189 | 0.000007 | 0.000128 | 0.000034 | 0.000440 | 0.000017 | 0.000287 | 0.000077 | 1.234655 | 1.002957 | 2.868339486 | 0.106022226 | 0.915564717 | NA | -0.155866 | 0.148375 | -0.001261 | -0.000972 | 0.000053 | -0.000315 | 1 | 1 | 1 | 1 |
| 91404 | NM_178123,XI | SESTD1 | SEC14 and spectrin domain containing 1 | protein_coding | NP_835224.3;XP_011510441.1;XP_011510443.1;XP_016860739.1;XP_01 | 18379 | . | . | . | 1.554775 | 1.569954 | 1.338031 | 1.429655 | 3.618025 | 3.629019 | 3.002503 | 3.268737 | -1.133999 | 486.674267 | 0.169032882 | -1.07328026 | 0.283145391 | 0.998865404 | 9.014774 | 8.833138 | 8.969026 | 8.961526 | 8.870002 | 8.902224 | 599 | 555 | 386 | 439 |
| 91405 | gene-PLEKHA3 | PLEKHA3P1 | pleckstrin homology domain containing A3 pseudogene 1 | pseudogene | . | 19648 | . | . | . | 0.169154 | 0.204263 | 0.149029 | 0.297856 | 0.393627 | 0.472164 | 0.334416 | 0.681011 | 1.167108 | 11.180907 | 0.926923417 | 0.240514078 | 0.809931751 | NA | 3.369732 | 3.587955 | 3.472407 | 3.479634 | 3.466447 | 3.500689 | 11 | 12 | 7 | 15 |
| 91408 | NM_00113649 | BTF3L4 | basic transcription factor 3 like 4 | protein_coding | NP_001129969.1;NP_001230696.1;NP_689478.1 | 30547 | . | . | . | 5.376393 | 5.042749 | 4.788811 | 3.868484 | 12.511087 | 11.656541 | 10.745956 | 8.844826 | -1.218759 | 674.802016 | 0.166623283 | -1.71292247 | 0.086726804 | 0.795065303 | 9.533547 | 9.249096 | 9.485182 | 9.434200 | 9.395976 | 9.249852 | 884 | 771 | 588 | 507 |
| 91419 | NM_00132040 | ATP23 | ATP23 metallopeptidase and ATP synthase assembly factor homol | protein_coding | NP_001307337.1;NP_001307338.1;NP_001307339.1;NP_150592.1;XP_01 | 29452 | . | . | . | 1.988938 | 2.073738 | 0.863934 | 1.662955 | 4.628340 | 4.793539 | 1.938643 | 3.802148 | -1.340998 | 117.920021 | 0.310274357 | -1.36429805 | 0.17247378 | 0.947431715 | 7.078499 | 6.653668 | 6.948680 | 6.907801 | 6.782888 | 6.874503 | 164 | 138 | 74 | 109 |
| 91433 | NM_00101791 | RCCD1 | RCC1 domain containing 1 | protein_coding | NP_001017919.1;NP_291022.2 | 30457 | 617997 | . | . | 16.599565 | 15.409376 | 15.022192 | 15.588108 | 38.627870 | 35.619464 | 33.709366 | 35.640358 | -1.090150 | 1653.269217 | 0.123047059 | -1.01202381 | 0.311526668 | 0.998865404 | 10.752051 | 10.627473 | 10.737802 | 10.709776 | 10.639841 | 10.672914 | 2021 | 1828 | 1346 | 1515 |
| 91442 | NM_00130097 | FAAP24 | FA core complex associated protein 24 | protein_coding | NP_001287907.1;NP_689479.1;XP_005259450.1 | 28467 | 610884 | . | . | 4.164376 | 4.231509 | 4.018374 | 5.125065 | 9.690674 | 9.781322 | 9.017115 | 11.717853 | 1.104485 | 297.698469 | 0.208527471 | 0.687551091 | 0.491735517 | 0.998865404 | 8.144829 | 8.287082 | 8.153845 | 8.221885 | 8.201433 | 8.279988 | 305 | 325 | 252 | 314 |
| 91445 | NM_00113582 | RNF185 | ring finger protein 185 | protein_coding | NP_001129297.1;NP_689480.2 | 26783 | . | . | . | 14.383269 | 15.528573 | 17.686111 | 17.540594 | 33.470460 | 35.894992 | 39.687120 | 40.104480 | 1.155416 | 1606.686921 | 0.124123706 | 1.679072052 | 0.093138002 | 0.809520218 | 10.541942 | 10.750258 | 10.565534 | 10.614550 | 10.707003 | 10.698510 | 1662 | 1660 | 1506 | 1606 |
| 91450 | NR_026998 | LOC91450 | uncharacterized LOC91450 | lncRNA | . | . | . | . | . | 0.106369 | 0.059757 | 0.163732 | 0.046651 | 0.247525 | 0.138131 | 0.367410 | 0.106662 | 1.254522 | 3.534302 | 1.608430777 | 0.203389261 | 0.838830793 | NA | 1.644780 | 1.978782 | 1.811584 | 1.808305 | 1.821681 | 1.805476 | 4 | 3 | 5 | 2 |
| 91452 | NM_00104247 | ACBD5 | acyl-CoA binding domain containing 5 | protein_coding | NP_001035938.1;NP_001258441.1;NP_001288180.1;NP_001288181.1;NI | 23338 | 616618 | . | . | 6.512082 | 6.193069 | 5.982158 | 6.554726 | 15.153880 | 14.315552 | 13.423791 | 14.986596 | -1.036868 | 787.621316 | 0.145784596 | -0.35828217 | 0.720132162 | 0.998865404 | 9.647205 | 9.595040 | 9.651325 | 9.616386 | 9.602878 | 9.614239 | 946 | 844 | 667 | 731 |
| 91461 | NM_138370 | PKDCC | protein kinase domain containing, cytoplasmic | protein_coding | NP_612379.2 | 25123 | 614150 | . | . | 3.458440 | 3.936339 | 1.481798 | 1.823892 | 8.047932 | 9.099025 | 3.325112 | 4.170112 | -2.258975 | 208.593090 | 0.245062464 | -4.79742403 | 1.60719E-06 | 0.000418567 | 8.176360 | 6.998858 | 7.809090 | 7.874877 | 7.445573 | 7.511962 | 313 | 331 | 100 | 132 |
| 91526 | NM_00119514 | ANKRD44 | ankyrin repeat domain 44 | protein_coding | NP_001182073.1;NP_001354424.1;NP_001354425.1;NP_001354426.1;NI | 25259 | . | . | . | 6.889540 | 6.563538 | 6.359035 | 6.641863 | 16.032243 | 15.171914 | 14.269488 | 15.185831 | 1.037258 | 1420.061274 | 0.129223062 | 0.408400332 | 0.68297979 | 0.998865404 | 10.445071 | 10.497920 | 10.490881 | 10.422308 | 10.470804 | 10.499561 | 1677 | 1437 | 1234 | 1381 |
| 91544 | NM_00107726 | UBXN11 | UBX domain protein 11 | protein_coding | NP_001070730.1;NP_663320.2;NP_892120.2 | 30600 | 609151 | . | . | 15.686807 | 17.934648 | 19.067302 | 18.638276 | 36.503845 | 41.456741 | 42.786476 | 42.614202 | 1.109241 | 1019.757858 | 0.140116337 | 1.06749127 | 0.285750054 | 0.998865404 | 9.917376 | 10.066780 | 9.912005 | 9.993810 | 10.041183 | 10.018444 | 1051 | 1102 | 947 | 990 |
| 91574 | NM_00114390 | C12orf65 | chromosome 12 open reading frame 65 | protein_coding | NP_001137377.1;NP_001181924.1;NP_689482.1;XP_005253687.1;XP_01 | 26784 | 613541 | . | . | 5.167952 | 4.997610 | 4.444975 | 4.769612 | 12.026034 | 11.552201 | 9.974394 | 10.905152 | -1.120140 | 240.281762 | 0.220326646 | -0.74289359 | 0.457546109 | 0.998865404 | 7.988112 | 7.824413 | 7.954704 | 7.917379 | 7.868549 | 7.892577 | 303 | 264 | 192 | 218 |
| 91582 | NM_194326,N | RPS19BP1 | ribosomal protein S19 binding protein 1 | protein_coding | NP_919307.1 | 28749 | 610225 | . | . | 73.050154 | 70.460799 | 74.655122 | 82.070462 | 169.990699 | 162.873308 | 167.523929 | 187.644374 | 1.079383 | 1925.436526 | 0.122519256 | 0.899507758 | 0.368382265 | 0.998865404 | 10.854870 | 10.964970 | 10.897234 | 10.861509 | 10.904902 | 10.973947 | 2181 | 1953 | 1663 | 1955 |
| 91603 | NM_052857 | ZNF830 | zinc finger protein 830 | protein_coding | NP_443089.3 | 28291 | . | . | . | 4.178111 | 4.645766 | 4.893097 | 4.905036 | 9.722635 | 10.738895 | 10.979969 | 11.214781 | 1.095502 | 325.058388 | 0.196323176 | 0.670282734 | 0.502677572 | 0.998865404 | 8.277422 | 8.408703 | 8.290498 | 8.343675 | 8.370576 | 8.365167 | 340 | 351 | 297 | 317 |
| 91607 | NM_00110458 | SLFN11 | schlafen family member 11 | protein_coding | NP_001098057.1;NP_001098058.1;NP_001098059.1;NP_001098060.1;NI | 26633 | 614953 | . | . | 0.105109 | 0.055461 | 0.202923 | 0.059144 | 0.244593 | 0.128201 | 0.455354 | 0.135227 | 1.611486 | 19.191441 | 0.790977401 | 0.870305457 | 0.384133498 | NA | 3.872825 | 4.568818 | 4.233737 | 4.193708 | 4.304791 | 4.197176 | 22 | 11 | 32 | 10 |
| 91612 | NM_00120406 | CHURC1 | churchill domain containing 1 | protein_coding | NP_001190992.1;NP_001190993.1;NP_660148.3 | 20099 | 608577 | . | . | 12.191891 | 12.146067 | 10.782623 | 11.067282 | 28.371032 | 28.076176 | 24.195891 | 25.304025 | -1.128084 | 1297.953005 | 0.128066975 | -1.35768706 | 0.174563006 | 0.948677095 | 10.426355 | 10.252453 | 10.390795 | 10.381482 | 10.288573 | 10.300691 | 1594 | 1476 | 1052 | 1153 |
| 91614 | NM_00107724 | DEPDC7 | DEP domain containing 7 | protein_coding | NP_001070710.1;NP_631899.2 | 29899 | 612294 | . | . | 1.896094 | 2.098479 | 1.648147 | 2.128429 | 4.412288 | 4.850728 | 3.698393 | 4.866400 | -1.073714 | 105.740194 | 0.319268345 | -0.32139048 | 0.747914496 | 0.998865404 | 6.775639 | 6.671232 | 6.719177 | 6.751239 | 6.676068 | 6.748342 | 120 | 124 | 78 | 107 |
| 91646 | NM_00111082 | TDRD12 | tudor domain containing 12 | protein_coding | NP_001104292.1;NP_001353031.1;XP_011525773.1;XP_011525775.1;XP | 25044 | . | . | . | 0.016676 | 0.031233 | 0.029463 | 0.015725 | 0.038807 | 0.072196 | 0.066115 | 0.035953 | 1.121093 | 5.281874 | 1.323243912 | 0.124622788 | 0.900822187 | NA | 2.316288 | 2.481107 | 2.385328 | 2.399568 | 2.401049 | 2.389365 | 4 | 7 | 6 | 4 |
| 91647 | NM_145691,XI | ATPAF2 | ATP synthase mitochondrial F1 complex assembly factor 2 | protein_coding | NP_663729.1;XP_005256905.1;XP_011522367.1;XP_016880791.1;XP_01 | 18802 | 608918 | . | . | 6.339794 | 5.337813 | 6.831414 | 6.854508 | 14.752959 | 12.338595 | 15.329496 | 15.672020 | 1.201850 | 338.519818 | 0.200873035 | 1.320521958 | 0.18666081 | 0.95851159 | 8.263983 | 8.529968 | 8.400878 | 8.287250 | 8.448043 | 8.449995 | 391 | 297 | 321 | 347 |
| 91653 | NM_00130186 | BOC | BOC cell adhesion associated, oncogene regulated | protein_coding | NP_001288790.1;NP_001288796.1;NP_001365002.1;NP_001365003.1;NI | 17173 | 608708 | . | . | 0.502156 | 0.673011 | 0.932881 | 1.166147 | 1.168539 | 1.555697 | 2.093360 | 2.666256 | 1.996249 | 90.858820 | 0.353394217 | 2.822037502 | 0.00477196 | 0.205194266 | 5.924070 | 6.918850 | 6.350376 | 6.394958 | 6.507633 | 6.622591 | 63 | 72 | 88 | 132 |
| 91661 | NM_00104018 | ZNF765 | zinc finger protein 765 | protein_coding | NP_001035275.1;NP_001337424.1 | 25092 | . | . | . | 1.455386 | 1.841953 | 1.449925 | 1.023995 | 3.386745 | 4.257757 | 3.253589 | 2.341244 | -1.346621 | 205.099675 | 0.260092372 | -1.65073517 | 0.098792662 | 0.813553488 | 7.878470 | 7.450230 | 7.681978 | 7.791940 | 7.671471 | 7.529920 | 241 | 282 | 178 | 136 |
| 91663 | NM_00102081 | MYADM | myeloid associated differentiation marker | protein_coding | NP_001018654.1;NP_001018655.1;NP_001018656.1;NP_001018657.1;NI | 7544 | 609959 | . | . | 6.388719 | 7.429169 | 4.924197 | 5.862834 | 14.866809 | 17.172856 | 11.049755 | 13.404673 | -1.297648 | 616.477070 | 0.172798011 | -2.17536839 | 0.029602521 | 0.553061869 | 9.444120 | 9.067129 | 9.293491 | 9.396569 | 9.106327 | 9.234412 | 743 | 807 | 419 | 554 |
| 91664 | NM_00132152 | ZNF845 | zinc finger protein 845 | protein_coding | NP_001308451.1;NP_001308452.1;NP_001308453.1;NP_612383.1 | 25112 | . | . | . | 1.112676 | 1.079083 | 1.102967 | 0.872505 | 2.589244 | 2.494347 | 2.475026 | 1.994879 | -1.117623 | 205.912315 | 0.243428369 | -0.65906113 | 0.509856517 | 0.998865404 | 7.763171 | 7.604227 | 7.722429 | 7.696099 | 7.711151 | 7.603230 | 257 | 228 | 190 | 160 |
| 91687 | NM_00112718 | CENPL | centromere protein L | protein_coding | NP_001120653.1;NP_001164653.1;NP_201576.1 | 17879 | 611503 | . | . | 4.434167 | 3.959841 | 3.930697 | 3.844446 | 10.318489 | 9.153350 | 8.820369 | 8.789870 | -1.088508 | 316.570767 | 0.197978815 | -0.61800537 | 0.536571797 | 0.998865404 | 8.366154 | 8.244031 | 8.353014 | 8.305220 | 8.279786 | 8.286200 | 396 | 341 | 262 | 286 |
| 91689 | NM_033318,N | SMDT1 | single-pass membrane protein with aspartate rich tail 1 | protein_coding | NP_201575.3;XP_011528811.1 | 25055 | 615588 | . | . | 10.450558 | 10.467806 | 9.962280 | 13.288930 | 24.318878 | 24.196802 | 22.355067 | 30.383557 | 1.110229 | 534.328065 | 0.182199057 | 0.827982997 | 0.407680129 | 0.998865404 | 8.984664 | 9.134605 | 9.024618 | 9.023283 | 8.996397 | 9.178150 | 584 | 546 | 421 | 600 |
| 91694 | NM_00132997 | LONRF1 | LON peptidase N-terminal domain and ring finger 1 | protein_coding | NP_001316905.1;NP_689484.3;XP_011542996.1 | 26302 | . | . | . | 9.655401 | 9.916663 | 12.509593 | 11.268044 | 22.468515 | 22.922809 | 28.071168 | 25.763045 | 1.197446 | 1217.158511 | 0.135094243 | 1.924286403 | 0.054318694 | 0.691661377 | 10.113329 | 10.373555 | 10.169291 | 10.182737 | 10.356432 | 10.265406 | 1264 | 1206 | 1222 | 1170 |
| 91695 | NR_002184 | RRP7BP | ribosomal RNA processing 7 homolog B, pseudogene | transcribed_ps | . | 30454 | . | . | . | 8.760641 | 10.126302 | 11.264487 | 9.222528 | 20.386372 | 23.407402 | 25.277184 | 21.086214 | 1.070638 | 715.526207 | 0.166530037 | 0.591309785 | 0.554312875 | 0.998865404 | 9.432618 | 9.531413 | 9.408671 | 9.501566 | 9.573673 | 9.427889 | 742 | 796 | 711 | 621 |
| 91746 | NM_00103173 | YTHDC1 | YTH domain containing 1 | protein_coding | NP_001026902.1;NP_001317627.1;NP_588611.2;XP_005265764.1;XP_00 | 30626 | 617283 | . | . | 10.510077 | 9.536126 | 9.995225 | 7.334790 | 24.457382 | 22.043180 | 22.428993 | 16.770130 | -1.026520 | 1504.865188 | 0.136572129 | -0.27650153 | 0.782162881 | 0.998865404 | 10.573961 | 10.536635 | 10.570842 | 10.557741 | 10.612316 | 10.471264 | 1770 | 1631 | 1410 | 1265 |
| 91748 | NM_00104331 | MIDEAS | mitotic deacetylase associated SANT domain protein | protein_coding | NP_001036783.1;NP_001354639.1;NP_919254.2;XP_005268262.1;XP_00 | 19853 | . | . | . | 4.263946 | 4.306839 | 4.061451 | 4.242581 | 9.922376 | 9.955450 | 9.113774 | 9.700156 | -1.036998 | 1021.186097 | 0.135732933 | -0.38615297 | 0.699383381 | 0.998865404 | 10.022067 | 9.969515 | 10.006675 | 10.011968 | 9.969298 | 9.995693 | 1193 | 1126 | 856 | 957 |
| 91749 | NM_153369,XI | MFSD4B | major facilitator superfamily domain containing 4B | protein_coding | NP_699200.2;XP_016866972.1 | 21053 | 617331 | . | . | 1.549440 | 1.624975 | 1.827915 | 1.764013 | 3.605610 | 3.756202 | 4.101787 | 4.033205 | 1.044974 | 719.893383 | 0.16006821 | 0.396495762 | 0.691739334 | 0.998865404 | 9.459184 | 9.523381 | 9.487639 | 9.461298 | 9.570135 | 9.435728 | 826 | 745 | 702 | 623 |
| 91750 | NM_00102467 | LIN52 | lin-52 DREAM MuvB core complex component | protein_coding | NP_001019845.2;NP_001358934.1;NP_001358935.1;XP_011535623.1;XP | 19856 | . | . | . | 2.846031 | 2.874720 | 2.635864 | 3.410941 | 6.622831 | 6.645043 | 5.914802 | 7.798712 | 1.061649 | 273.092062 | 0.215506797 | 0.400485634 | 0.688798863 | 0.998865404 | 8.049959 | 8.135268 | 8.072393 | 8.079828 | 8.053503 | 8.157815 | 303 | 288 | 220 | 290 |
| 91752 | NM_194250 | ZNF804A | zinc finger protein 804A | protein_coding | NP_919226.1 | 21711 | 612282 | . | . | 0.042806 | 0.044774 | 0.032275 | 0.030623 | 0.099611 | 0.103497 | 0.072424 | 0.070015 | -1.516150 | 5.575329 | 1.282129046 | -0.46829369 | 0.639574578 | NA | 2.747897 | 2.148375 | 2.492751 | 2.490531 | 2.482050 | 2.480459 | 8 | 7 | 4 | 4 |
| 91754 | NM_00132923 | NEK9 | NIMA related kinase 9 | protein_coding | NP_001316166.1;NP_001316167.1;NP_149107.4;XP_005268266.1;XP_02 | 18591 | 609798 | . | . | 6.652378 | 6.774577 | 7.071357 | 7.826578 | 15.480359 | 15.659739 | 15.867921 | 17.894539 | 1.098113 | 1788.353673 | 0.122984885 | 1.097914163 | 0.272241973 | 0.998865404 | 10.735417 | 10.870266 | 10.767964 | 10.764122 | 10.805521 | 10.872640 | 1968 | 1835 | 1559 | 1829 |
| 91768 | NM_00110061 | CABLES1 | Cdk5 and Abl enzyme substrate 1 | protein_coding | NP_001094089.1;NP_001243367.1;NP_612384.1 | 25097 | 609194 | . | . | 17.276238 | 17.599179 | 20.987741 | 23.187881 | 40.202517 | 40.681289 | 47.095884 | 53.016335 | 1.255021 | 3071.321010 | 0.114464624 | 2.862996413 | 0.004196553 | 0.192048182 | 11.411574 | 11.739157 | 11.482793 | 11.484820 | 11.631351 | 11.706703 | 3133 | 2943 | 2839 | 3349 |
| 91775 | NM_00113445 | NXPE3 | neurexophilin and PC-esterase domain family member 3 | protein_coding | NP_001127928.1;NP_001335919.1;NP_001335920.1;NP_001335921.1;NI | 28238 | . | . | . | 3.628035 | 3.272490 | 3.951233 | 3.499759 | 8.442586 | 7.564507 | 8.866453 | 8.001782 | 1.068027 | 972.571322 | 0.146740787 | 0.647043528 | 0.517603773 | 0.998865404 | 9.877086 | 9.972652 | 9.936122 | 9.862152 | 9.997355 | 9.896169 | 1138 | 963 | 934 | 877 |
| 91782 | NM_00131789 | CHMP7 | charged multivesicular body protein 7 | protein_coding | NP_001304828.1;NP_001350112.1;NP_689485.1;XP_016869450.1;XP_01 | 28439 | 611130 | . | . | 11.692189 | 11.493718 | 11.348653 | 13.596179 | 27.208207 | 26.568243 | 25.466049 | 31.086048 | -1.019511 | 1113.081324 | 0.145525328 | -0.19156174 | 0.848085516 | 0.998865404 | 10.134478 | 10.106069 | 10.124616 | 10.128441 | 10.036445 | 10.182348 | 1291 | 1216 | 864 | 1135 |
| 91801 | NM_00130101 | ALKBH8 | alkB homolog 8, tRNA methyltransferase | protein_coding | NP_001287939.2;NP_001365062.1;NP_620130.2;XP_005271785.1;XP_00 | 25189 | 613306 | . | . | 3.681314 | 3.484018 | 3.731220 | 3.331830 | 8.566569 | 8.053464 | 8.372749 | 7.617835 | -1.023409 | 442.610133 | 0.177401495 | -0.18817502 | 0.850739455 | 0.998865404 | 8.806175 | 8.773425 | 8.806695 | 8.786069 | 8.817511 | 8.746025 | 524 | 475 | 403 | 386 |
| 91833 | NM_00124241 | WDR20 | WD repeat domain 20 | protein_coding | NP_001229343.1;NP_001229344.1;NP_001229345.1;NP_001229346.1;NI | 19667 | 617741 | . | . | 5.410579 | 5.377400 | 6.064152 | 6.220484 | 12.590638 | 12.430112 | 13.607788 | 14.222399 | 1.062717 | 456.146938 | 0.177370802 | 0.494767652 | 0.620764141 | 0.998865404 | 8.788417 | 8.876935 | 8.846108 | 8.778837 | 8.880832 | 8.819726 | 537 | 451 | 429 | 419 |
| 91860 | NM_00103173 | CALML4 | calmodulin like 4 | protein_coding | NP_001026903.2;NP_001273623.1;NP_001273624.1;NP_219501.2 | 18445 | . | . | . | 0.994764 | 0.935524 | 1.167207 | 0.985320 | 2.314858 | 2.162507 | 2.619178 | 2.252817 | 1.095129 | 146.017243 | 0.274624858 | 0.477383044 | 0.633089386 | 0.998865404 | 7.122341 | 7.254618 | 7.177978 | 7.161302 | 7.239548 | 7.168331 | 164 | 147 | 145 | 130 |
| 91869 | NM_052859,XI | RFT1 | RFT1 homolog | protein_coding | NP_443091.1;XP_005265594.1;XP_006713447.1;XP_011532516.1;XP_01 | 30220 | 611908 | . | . | 5.904347 | 5.705994 | 4.875245 | 5.419224 | 13.739661 | 13.189664 | 10.939913 | 12.390413 | -1.117802 | 776.346481 | 0.146286651 | -1.09828954 | 0.272078077 | 0.998865404 | 9.678748 | 9.517884 | 9.637205 | 9.639731 | 9.543740 | 9.576815 | 942 | 886 | 622 | 704 |
| 91875 | NM_138376 | TTC5 | tetratricopeptide repeat domain 5 | protein_coding | NP_612385.2 | 19274 | . | . | . | 2.844053 | 2.822780 | 2.950890 | 3.188827 | 6.618227 | 6.524982 | 6.621711 | 7.290874 | 1.067979 | 435.596799 | 0.175799691 | 0.539720385 | 0.589389879 | 0.998865404 | 8.718755 | 8.813392 | 8.750673 | 8.741115 | 8.764990 | 8.806462 | 489 | 451 | 378 | 436 |
| 91893 | NM_138378,N | FDXACB1 | ferredoxin-fold anticodon binding domain containing 1 | protein_coding | NP_612387.1 | 25110 | . | . | . | 1.079031 | 1.273480 | 1.529422 | 1.431698 | 2.510951 | 2.943705 | 3.431978 | 3.273408 | 1.244093 | 114.964144 | 0.30334873 | 1.038720791 | 0.298934609 | 0.998865404 | 6.679078 | 6.993866 | 6.777562 | 6.825902 | 6.887531 | 6.861607 | 109 | 119 | 115 | 115 |
| 91942 | NM_174889 | NDUFAF2 | NADH:ubiquinone oxidoreductase complex assembly factor 2 | protein_coding | NP_777549.1 | 28086 | 609653 | . | . | 26.499846 | 26.437325 | 22.595728 | 28.200756 | 61.666233 | 61.111061 | 50.704155 | 64.477676 | -1.056053 | 511.267608 | 0.176479711 | -0.44584088 | 0.655712193 | 0.998865404 | 9.037131 | 8.957644 | 9.017608 | 9.010946 | 8.911529 | 9.042933 | 608 | 564 | 387 | 515 |
| 91947 | NM_183376 | ARRDC4 | arrestin domain containing 4 | protein_coding | NP_899232.2 | 28087 | . | . | . | 11.293909 | 9.921844 | 9.063746 | 8.652942 | 26.281391 | 22.934790 | 20.338785 | 19.783924 | -1.214590 | 1233.360904 | 0.137833528 | -2.03484025 | 0.041866949 | 0.633602228 | 10.401627 | 10.121560 | 10.388720 | 10.281441 | 10.210440 | 10.169006 | 1665 | 1359 | 996 | 1015 |
| 91949 | NM_153603,XI | COG7 | component of oligomeric golgi complex 7 | protein_coding | NP_705831.1;XP_016879359.1 | 18622 | 606978 | . | . | 5.597338 | 6.626816 | 5.993855 | 6.116792 | 13.025235 | 15.318182 | 13.450034 | 13.985317 | -1.024204 | 565.254787 | 0.166367346 | -0.20739239 | 0.835703427 | 0.998865404 | 9.160100 | 9.125204 | 9.099686 | 9.197642 | 9.136006 | 9.132217 | 610 | 663 | 487 | 522 |
| 91966 | NM_00117190 | EOLA1 | endothelium and lymphocyte associated ASCH domain 1 | protein_coding | NP_001165378.1;NP_001165379.1;NP_001165380.1;NP_001311203.1;NI | 28089 | 300954 | . | . | 2.651443 | 2.287176 | 2.525671 | 2.783643 | 6.170017 | 5.286911 | 5.667530 | 6.364467 | 1.045316 | 219.610266 | 0.23528366 | 0.271755167 | 0.78581028 | 0.998865404 | 7.746723 | 7.810182 | 7.794529 | 7.738553 | 7.742938 | 7.830701 | 262 | 218 | 177 | 230 |
| 91975 | NM_00117283 | ZNF300 | zinc finger protein 300 | protein_coding | NP_001166302.1;NP_001166303.1;NP_443092.1;XP_011536004.1 | 13091 | 612429 | . | . | 3.614094 | 4.738147 | 5.817170 | 3.969170 | 8.410145 | 10.952439 | 13.053559 | 9.075037 | 1.119033 | 480.492606 | 0.202887982 | 0.799715941 | 0.423875395 | 0.998865404 | 8.824562 | 8.987577 | 8.821766 | 8.907564 | 9.046373 | 8.812913 | 486 | 523 | 522 | 389 |
| 91978 | NM_033513 | TPGS1 | tubulin polyglutamylase complex subunit 1 | protein_coding | NP_277048.2 | 25058 | . | . | . | 2.903504 | 2.829188 | 2.365408 | 3.524449 | 6.756574 | 6.539795 | 5.307908 | 8.058233 | 1.015035 | 101.509014 | 0.333793393 | 0.06450096 | 0.948571343 | 0.998865404 | 6.655623 | 6.675239 | 6.666121 | 6.655878 | 6.606311 | 6.721563 | 118 | 107 | 72 | 114 |
| 92002 | NM_00113099 | CCNQ | cyclin Q | protein_coding | NP_001124469.1;NP_689487.2;XP_005277977.1;XP_005277978.1;XP_01 | 28434 | 300708 | . | . | 8.539193 | 8.960135 | 8.210769 | 8.843888 | 19.871057 | 20.711754 | 18.424728 | 20.220502 | -1.026936 | 366.638033 | 0.188606874 | -0.20330957 | 0.838893076 | 0.998865404 | 8.537496 | 8.498668 | 8.530146 | 8.521502 | 8.478285 | 8.541566 | 431 | 398 | 297 | 358 |
| 92014 | NM_033412,N | SLC25A51 | solute carrier family 25 member 51 | protein_coding | NP_219480.1 | 23323 | . | . | . | 3.139325 | 3.293477 | 3.770523 | 3.315250 | 7.305338 | 7.613021 | 8.460944 | 7.579924 | 1.095415 | 212.868960 | 0.229798242 | 0.57214621 | 0.567222925 | 0.998865404 | 7.666570 | 7.798078 | 7.705074 | 7.716829 | 7.759162 | 7.747426 | 231 | 222 | 196 | 206 |
| 92017 | NM_00137649 | SNX29 | sorting nexin 29 | protein_coding | NP_001363419.1;NP_115543.3;XP_005255739.1;XP_005255740.1;XP_01 | 30542 | . | . | . | 1.202844 | 1.106050 | 1.057783 | 1.771682 | 2.799066 | 2.556682 | 2.373636 | 4.050743 | 1.097775 | 308.310797 | 0.226957408 | 0.592984827 | 0.553191307 | 0.998865404 | 8.199956 | 8.333441 | 8.264675 | 8.209533 | 8.179632 | 8.386172 | 356 | 301 | 228 | 359 |
| 92070 | NR_033339 | CTBP1-DT | CTBP1 divergent transcript | lncRNA | . | 28307 | . | . | . | 2.049124 | 2.425548 | 1.621086 | 2.585622 | 4.768397 | 5.606762 | 3.637669 | 5.911716 | -1.076192 | 198.421475 | 0.270938196 | -0.39099379 | 0.695801824 | 0.998865404 | 7.685587 | 7.577228 | 7.606014 | 7.680823 | 7.509179 | 7.703484 | 218 | 240 | 129 | 219 |
| 92092 | NM_080660,XI | ZC3HAV1L | zinc finger CCCH-type containing, antiviral 1 like | protein_coding | NP_542391.2;XP_006716239.1;XP_011514990.1;XP_011514991.1 | 22423 | . | . | . | 5.978093 | 6.596297 | 8.636093 | 7.187136 | 13.911269 | 15.247634 | 19.379145 | 16.432532 | 1.070152 | 517.916567 | 0.176618276 | 0.553824479 | 0.579698974 | 0.998865404 | 8.966500 | 9.064972 | 8.977630 | 9.006538 | 9.101588 | 8.965317 | 563 | 552 | 515 | 449 |
| 92104 | NM_152275 | TTC30A | tetratricopeptide repeat domain 30A | protein_coding | NP_689488.3 | 25853 | . | . | . | 1.843345 | 1.641060 | 1.696275 | 1.880669 | 4.289541 | 3.793383 | 3.806392 | 4.299927 | 1.011665 | 316.362094 | 0.20153349 | 0.083018685 | 0.933836682 | 0.998865404 | 8.297042 | 8.313775 | 8.335896 | 8.266287 | 8.282770 | 8.332700 | 385 | 318 | 264 | 312 |
| 92105 | NM_033547,XI | INTS4 | integrator complex subunit 4 | protein_coding | NP_291025.3;XP_011543654.1;XP_011543655.1;XP_016874049.1 | 25048 | 611348 | . | . | 6.154790 | 7.030288 | 6.813948 | 6.865379 | 14.322449 | 16.250821 | 15.290301 | 15.696871 | 1.030062 | 688.502123 | 0.156096946 | 0.273743403 | 0.784281821 | 0.998865404 | 9.406017 | 9.448307 | 9.370421 | 9.460345 | 9.428049 | 9.444950 | 730 | 780 | 600 | 663 |
| 92106 | NM_00133067 | OXNAD1 | oxidoreductase NAD binding domain containing 1 | protein_coding | NP_001317599.1;NP_001317600.1;NP_001339906.1;NP_001339907.1;NI | 25128 | . | . | . | 3.877751 | 5.015841 | 3.676327 | 4.906421 | 9.023682 | 11.594339 | 8.249577 | 11.217947 | -1.023664 | 563.616636 | 0.179606723 | -0.18786388 | 0.850983356 | 0.998865404 | 9.155803 | 9.121130 | 9.055760 | 9.225018 | 9.098304 | 9.157650 | 573 | 694 | 462 | 546 |
| 92126 | NM_032160 | DSEL | dermatan sulfate epimerase like | protein_coding | NP_115536.2 | 18144 | 611125 | . | . | 0.050604 | 0.069968 | 0.106899 | 0.067473 | 0.117757 | 0.161734 | 0.239879 | 0.154269 | 1.423742 | 21.834081 | 0.671068453 | 0.759516717 | 0.447543518 | NA | 4.170740 | 4.681336 | 4.408795 | 4.429977 | 4.473005 | 4.429651 | 18 | 22 | 27 | 19 |
| 92140 | NM_00136313 | MTDH | metadherin | protein_coding | NP_001350065.1;NP_001350066.1;NP_001350067.1;NP_001350068.1;NI | 29608 | 610323 | . | . | 32.898822 | 29.466788 | 30.962596 | 25.847439 | 76.556906 | 68.113798 | 69.479167 | 59.097097 | -1.119224 | 7037.631500 | 0.1255899 | -1.29388593 | 0.195704825 | 0.96621494 | 12.859761 | 12.697423 | 12.874837 | 12.772848 | 12.808120 | 12.645127 | 9069 | 7543 | 6337 | 5622 |
| 92154 | NM_138383,XI | MTSS2 | MTSS I-BAR domain containing 2 | protein_coding | NP_612392.1;XP_005256293.3;XP_011521752.1;XP_011521755.3;XP_01 | 25094 | 616951 | . | . | 2.049676 | 2.209933 | 2.675869 | 2.484077 | 4.769682 | 5.108357 | 6.004574 | 5.679548 | 1.191595 | 354.871768 | 0.190798501 | 1.325449045 | 0.185022259 | 0.958025663 | 8.338950 | 8.592253 | 8.410295 | 8.422032 | 8.547800 | 8.483788 | 369 | 353 | 354 | 342 |
| 92162 | NM_00131994 | TMEM88 | transmembrane protein 88 | protein_coding | NP_001306870.1;NP_981956.1;XP_005256913.1 | 32371 | 617813 | . | . | 0.668718 | 0.981224 | 0.992810 | 1.133072 | 1.556134 | 2.268142 | 2.227838 | 2.590635 | 1.347446 | 22.214146 | 0.658076653 | 0.653764727 | 0.513263395 | NA | 4.243129 | 4.671937 | 4.435999 | 4.465154 | 4.482055 | 4.474573 | 18 | 24 | 23 | 23 |
| 92170 | NM_138384 | MTG1 | mitochondrial ribosome associated GTPase 1 | protein_coding | NP_612393.2 | 32159 | . | . | . | 16.935459 | 17.333069 | 20.598814 | 17.985991 | 39.409512 | 40.066166 | 46.223137 | 41.122829 | 1.110626 | 1942.809184 | 0.127173455 | 1.190286364 | 0.233933858 | 0.988970612 | 10.846124 | 10.997751 | 10.873978 | 10.885647 | 11.019944 | 10.902962 | 2104 | 2001 | 1908 | 1777 |
| 92181 | NM_152277,XI | UBTD2 | ubiquitin domain containing 2 | protein_coding | NP_689490.2;XP_016865511.1;XP_016865512.1 | 24463 | 610174 | . | . | 10.757312 | 10.700438 | 11.454078 | 10.770886 | 25.032708 | 24.734543 | 25.702620 | 24.626353 | 1.023029 | 1018.397345 | 0.137115381 | 0.239558031 | 0.810672904 | 0.998865404 | 9.975439 | 10.008541 | 9.988258 | 9.978404 | 10.025431 | 9.974228 | 1167 | 1079 | 927 | 932 |
| 92211 | NM_00117197 | CDHR1 | cadherin related family member 1 | protein_coding | NP_001165442.1;NP_149091.1;XP_011538639.1;XP_011538640.1;XP_01 | 14550 | 609502 | . | . | 0.016594 | 0.043114 | 0.142646 | 0.077927 | 0.038614 | 0.099660 | 0.320093 | 0.178171 | 3.551348 | 16.504750 | 0.827092455 | 2.210595279 | 0.027063876 | NA | 2.861598 | 4.686013 | 3.941251 | 3.969615 | 4.067527 | 4.005374 | 5 | 11 | 29 | 17 |
| 92249 | NR_015353,NF | LINC01278 | long intergenic non-protein coding RNA 1278 | lncRNA | . | 28090 | . | . | . | 7.006257 | 7.083553 | 7.376005 | 8.141767 | 16.303849 | 16.373950 | 16.551541 | 18.615182 | 1.106298 | 591.802254 | 0.158174539 | 0.92138118 | 0.356847808 | 0.998865404 | 9.134369 | 9.279905 | 9.165578 | 9.182099 | 9.229900 | 9.251234 | 639 | 614 | 532 | 592 |
| 92255 | NM_00100752 | LMBRD2 | LMBR1 domain containing 2 | protein_coding | NP_001007528.1;XP_011512464.1;XP_016865513.1 | 25287 | . | . | . | 1.079838 | 0.977903 | 0.989431 | 0.735986 | 2.512825 | 2.260467 | 2.220256 | 1.682745 | -1.210734 | 240.374211 | 0.237074258 | -1.16369416 | 0.244547978 | 0.992733415 | 8.039657 | 7.765627 | 7.980321 | 7.921577 | 7.925699 | 7.785059 | 320 | 268 | 218 | 173 |
| 92259 | NM_033281 | MRPS36 | mitochondrial ribosomal protein S36 | protein_coding | NP_150597.1 | 16631 | 611996 | . | . | 11.683620 | 13.439312 | 9.751094 | 11.828238 | 27.188267 | 31.065571 | 21.881172 | 27.043858 | -1.180140 | 478.726291 | 0.181614847 | -1.31574313 | 0.188260274 | 0.960264078 | 9.018103 | 8.778035 | 8.908161 | 8.993036 | 8.791651 | 8.900671 | 558 | 596 | 347 | 449 |
| 92270 | NM_00101797 | ATP6AP1L | ATPase H+ transporting accessory protein 1 like | protein_coding | NP_001017971.1;NP_001336300.1;NP_001336301.1;NP_001336302.1;NI | 28091 | . | . | . | 1.165901 | 1.722278 | 2.107529 | 1.349722 | 2.713100 | 3.981120 | 4.729230 | 3.085982 | 1.127030 | 197.481756 | 0.283150681 | 0.609307698 | 0.542320505 | 0.998865404 | 7.536310 | 7.709639 | 7.516233 | 7.652012 | 7.740800 | 7.528488 | 181 | 231 | 224 | 151 |
| 92283 | NM_00129762 | ZNF461 | zinc finger protein 461 | protein_coding | NP_001284552.1;NP_001309750.1;NP_001309752.1;NP_001309754.1;NI | 21629 | 608640 | . | . | 0.743007 | 0.695848 | 0.896451 | 0.707340 | 1.729004 | 1.608485 | 2.011608 | 1.617251 | 1.098823 | 94.979989 | 0.333946336 | 0.407129922 | 0.683912568 | 0.998865404 | 6.499094 | 6.636728 | 6.562608 | 6.541296 | 6.615322 | 6.543026 | 108 | 94 | 97 | 82 |
| 92285 | NM_152279 | ZNF585B | zinc finger protein 585B | protein_coding | NP_689492.3 | 30948 | . | . | . | 0.619250 | 0.678867 | 0.790994 | 0.696262 | 1.441021 | 1.569232 | 1.774969 | 1.591920 | 1.130427 | 134.864903 | 0.282498218 | 0.626082537 | 0.531260806 | 0.998865404 | 6.984018 | 7.161283 | 7.035025 | 7.063559 | 7.119466 | 7.068435 | 140 | 142 | 133 | 125 |
| 92305 | NM_00112726 | TMEM129 | transmembrane protein 129, E3 ubiquitin ligase | protein_coding | NP_001120738.1;NP_612394.1;XP_005248095.1;XP_005248096.1;XP_01 | 25137 | 615975 | . | . | 10.016621 | 9.240733 | 8.890985 | 9.469589 | 23.309089 | 21.360367 | 19.951110 | 21.651088 | -1.052426 | 812.940604 | 0.146717207 | -0.50245241 | 0.61534932 | 0.998865404 | 9.703395 | 9.629675 | 9.712710 | 9.655945 | 9.629355 | 9.667486 | 998 | 864 | 670 | 763 |
| 92312 | NM_00109372 | MEX3A | mex-3 RNA binding family member A | protein_coding | NP_001087194.1 | 33482 | 611007 | . | . | 0.695268 | 0.617108 | 0.644198 | 0.796865 | 1.617916 | 1.426472 | 1.445562 | 1.821938 | 1.079050 | 141.825262 | 0.281018674 | 0.390587417 | 0.696102228 | 0.998865404 | 7.092337 | 7.201538 | 7.154768 | 7.107916 | 7.120886 | 7.198156 | 167 | 138 | 115 | 152 |
| 92335 | NM_00100378 | STRADA | STE20 related adaptor alpha | protein_coding | NP_001003786.1;NP_001003787.1;NP_001003788.1;NP_001159441.1;NI | 30172 | 608626 | . | . | 12.215719 | 13.017491 | 13.022903 | 13.639756 | 28.426484 | 30.090509 | 29.223019 | 31.185689 | 1.052880 | 953.898236 | 0.138160431 | 0.538081034 | 0.590521106 | 0.998865404 | 9.860128 | 9.934302 | 9.867381 | 9.889345 | 9.905630 | 9.925948 | 1054 | 1018 | 839 | 930 |
| 92342 | NM_00132019 | METTL18 | methyltransferase like 18 | protein_coding | NP_001307128.1;NP_001307130.1;NP_219486.1;XP_006711690.1 | 28793 | 615255 | . | . | 2.534057 | 2.635762 | 1.871911 | 1.826873 | 5.896856 | 6.092681 | 4.200516 | 4.176928 | -1.284044 | 107.780742 | 0.313945815 | -1.14890805 | 0.250593898 | 0.993339639 | 6.920602 | 6.560952 | 6.781490 | 6.798747 | 6.738102 | 6.682448 | 136 | 134 | 90 | 80 |
| 92344 | NM_00114603 | GORAB | golgin, RAB6 interacting | protein_coding | NP_001139511.2;NP_001307181.1;NP_689494.3;XP_006711691.1;XP_01 | 25676 | 607983 | . | . | 3.954386 | 4.213228 | 4.886126 | 3.034768 | 9.202019 | 9.739070 | 10.964327 | 6.938639 | -1.039113 | 338.711740 | 0.22468417 | -0.24635442 | 0.80540789 | 0.998865404 | 8.430534 | 8.376794 | 8.418854 | 8.399403 | 8.496368 | 8.266075 | 404 | 366 | 349 | 247 |
| 92345 | NM_00112893 | NAF1 | nuclear assembly factor 1 ribonucleoprotein | protein_coding | NP_001122403.1;NP_612395.2;XP_011530712.1;XP_011530713.1 | 25126 | 617868 | . | . | 2.771686 | 2.595670 | 2.333137 | 3.579223 | 6.449824 | 6.000009 | 5.235494 | 8.183468 | 1.004381 | 278.788533 | 0.206680477 | 0.030517176 | 0.975654595 | 0.998865404 | 8.119789 | 8.126259 | 8.136040 | 8.107399 | 8.122047 | 8.126199 | 329 | 292 | 242 | 263 |
| 92370 | NM_00103717 | PXYLP1 | 2-phosphoxylose phosphatase 1 | protein_coding | NP_001032249.1;NP_001269657.1;NP_689495.1;XP_005247960.1;XP_01 | 26303 | . | . | . | 7.063636 | 6.734311 | 5.776539 | 6.618206 | 16.437371 | 15.566658 | 12.962386 | 15.131741 | -1.144349 | 674.114197 | 0.157782783 | -1.23288281 | 0.21761949 | 0.982008277 | 9.491054 | 9.296060 | 9.452911 | 9.428436 | 9.301111 | 9.394399 | 843 | 763 | 508 | 631 |
| 92399 | NM_00117351 | MRRF | mitochondrial ribosome recycling factor | protein_coding | NP_001166983.1;NP_001333268.1;NP_001333270.1;NP_001333272.1;NI | 7234 | 604602 | . | . | 5.483193 | 5.199856 | 5.653589 | 5.998807 | 12.759613 | 12.019701 | 12.686489 | 13.715559 | 1.067591 | 969.525536 | 0.138836411 | 0.679640175 | 0.496732325 | 0.998865404 | 9.873053 | 9.967666 | 9.899730 | 9.892809 | 9.969969 | 9.917414 | 1085 | 1007 | 902 | 905 |
| 92400 | NM_033117,N | RBM18 | RNA binding motif protein 18 | protein_coding | NP_149108.1;XP_006717382.1 | 28413 | . | . | . | 5.569884 | 5.160501 | 5.257239 | 4.806123 | 12.961349 | 11.928728 | 11.797092 | 10.988629 | -1.025406 | 693.666131 | 0.156009257 | -0.23200547 | 0.816533764 | 0.998865404 | 9.455786 | 9.420189 | 9.481715 | 9.409071 | 9.458474 | 9.398656 | 851 | 718 | 622 | 614 |
| 92427 | gene-RPS2P4 | RPS2P4 | ribosomal protein S2 pseudogene 4 | pseudogene | . | 19814 | . | . | . | 0.029273 | 0.042318 | 0.007591 | 0.109218 | 0.068121 | 0.097821 | 0.017034 | 0.249714 | 1.216986 | 1.985087 | 2.102786642 | 0.134731979 | 0.892823788 | NA | 0.844134 | 1.121007 | 0.986974 | 0.987547 | 0.984432 | 0.993706 | 2 | 2 | 1 | 3 |
| 92482 | NM_00119530 | BBIP1 | BBSome interacting protein 1 | protein_coding | NP_001182233.1;NP_001182234.1;NP_001182235.1;NP_001182236.1;NI | 28093 | 613605 | . | . | 4.729148 | 3.557778 | 4.085738 | 4.380363 | 11.004923 | 8.223962 | 9.168281 | 10.015179 | 1.002361 | 269.877253 | 0.226644729 | 0.015009509 | 0.988024594 | 0.999197585 | 8.074136 | 8.078180 | 8.147151 | 7.993386 | 8.056901 | 8.091697 | 356 | 248 | 227 | 262 |
| 92521 | NM_00103355 | SPECC1 | sperm antigen with calponin homology and coiled-coil domains 1 | protein_coding | NP_001028725.1;NP_001028726.1;NP_001028727.1;NP_001230367.1;NI | 30615 | 608793 | . | . | 16.027655 | 15.673139 | 14.610330 | 17.248145 | 37.297008 | 36.229164 | 32.785158 | 39.435836 | -1.035501 | 2986.544844 | 0.126033122 | -0.39933272 | 0.689648063 | 0.998865404 | 11.569323 | 11.518758 | 11.563680 | 11.551087 | 11.449798 | 11.602148 | 3526 | 3254 | 2313 | 3008 |
| 92558 | NM_00136788 | BICDL1 | BICD family like cargo adaptor 1 | protein_coding | NP_001354815.1;NP_997194.2;XP_006719757.1;XP_006719759.1;XP_00 | 28095 | 617002 | . | . | 11.839156 | 11.969335 | 10.903489 | 12.087947 | 27.550202 | 27.667653 | 24.467113 | 27.637655 | -1.032539 | 1233.955406 | 0.131449316 | -0.35143547 | 0.725261673 | 0.998865404 | 10.292109 | 10.245666 | 10.275344 | 10.286653 | 10.228919 | 10.283870 | 1433 | 1363 | 1017 | 1180 |
| 92565 | NM_00135093 | FANK1 | fibronectin type III and ankyrin repeat domains 1 | protein_coding | NP_001337868.1;NP_001350478.1;NP_660278.3;XP_005270337.1;XP_00 | 23527 | 611640 | . | . | 0.582167 | 0.876011 | 0.487576 | 0.518468 | 1.354727 | 2.024937 | 1.094108 | 1.185415 | -1.353678 | 34.517670 | 0.542451748 | -0.8053881 | 0.420595756 | 0.998865404 | 5.312060 | 4.873224 | 5.094888 | 5.161888 | 5.085936 | 5.089956 | 36 | 52 | 25 | 28 |
| 92579 | NM_00131994 | G6PC3 | glucose-6-phosphatase catalytic subunit 3 | protein_coding | NP_001306874.1;NP_612396.1;XP_011523775.1;XP_011523776.1;XP_01 | 24861 | 611045 | . | . | 48.736553 | 51.024498 | 52.300076 | 52.483999 | 113.411962 | 117.945413 | 117.359852 | 119.998425 | 1.022341 | 2359.826553 | 0.115110868 | 0.276916535 | 0.781844196 | 0.998865404 | 11.188442 | 11.220312 | 11.183275 | 11.207424 | 11.220649 | 11.205707 | 2647 | 2556 | 2094 | 2216 |
| 92595 | NM_00117267 | ZNF764 | zinc finger protein 764 | protein_coding | NP_001166150.1;NP_219363.2 | 28200 | . | . | . | 2.832407 | 2.877079 | 2.907388 | 2.784190 | 6.591126 | 6.650496 | 6.524094 | 6.365721 | -1.016395 | 253.543201 | 0.214573505 | -0.10933867 | 0.912933878 | 0.998865404 | 7.997636 | 7.974446 | 7.987829 | 7.992303 | 7.996077 | 7.967600 | 293 | 277 | 225 | 229 |
| 92597 | NM_00124476 | MOB1B | MOB kinase activator 1B | protein_coding | NP_001231695.1;NP_001231696.1;NP_775739.1;XP_005265766.1;XP_01 | 29801 | 609282 | . | . | 5.071653 | 4.360130 | 5.506229 | 3.614026 | 11.801943 | 10.078637 | 12.355818 | 8.263038 | -1.068271 | 989.007893 | 0.188255403 | -0.50610786 | 0.612780931 | 0.998865404 | 9.996220 | 9.901916 | 10.024705 | 9.905776 | 10.062113 | 9.751886 | 1270 | 1014 | 1009 | 706 |
| 92609 | NM_00100156 | TIMM50 | translocase of inner mitochondrial membrane 50 | protein_coding | NP_001001563.2;NP_001316488.1;XP_011525793.1 | 23656 | 607381 | . | . | 36.848247 | 37.918349 | 40.978136 | 46.283164 | 85.747387 | 87.649966 | 91.953753 | 105.820961 | 1.149444 | 6337.792015 | 0.110239534 | 1.822718845 | 0.068345985 | 0.741130004 | 12.525854 | 12.726691 | 12.559337 | 12.575993 | 12.635409 | 12.732606 | 6723 | 6427 | 5559 | 6716 |
| 92610 | NM_052864 | TIFA | TRAF interacting protein with forkhead associated domain | protein_coding | NP_443096.1 | 19075 | 609028 | . | . | 1.534300 | 1.790131 | 1.625621 | 1.849491 | 3.500567 | 4.137967 | 3.647846 | 4.228641 | 1.039743 | 219.964134 | 0.234146544 | 0.240132904 | 0.810227226 | 0.998865404 | 7.753365 | 7.808360 | 7.730551 | 7.808583 | 7.759348 | 7.816994 | 228 | 252 | 183 | 223 |
| 92659 | NR_015454 | MAFG-DT | MAFG divergent transcript | lncRNA | . | 43649 | . | . | . | 2.600837 | 2.533593 | 1.872856 | 2.329747 | 6.052254 | 5.856514 | 4.202634 | 5.326690 | -1.236535 | 138.128675 | 0.282877981 | -1.08281139 | 0.278892148 | 0.998865404 | 7.255424 | 6.947964 | 7.155456 | 7.141127 | 7.033386 | 7.104721 | 179 | 162 | 96 | 128 |
| 92667 | NM_00131033 | MGME1 | mitochondrial genome maintenance exonuclease 1 | protein_coding | NP_001297267.1;NP_001297268.1;NP_001350667.1;NP_443097.1;XP_00 | 16205 | 615076 | . | . | 19.878376 | 18.895784 | 19.991281 | 21.244606 | 46.257795 | 43.678450 | 44.859856 | 48.573269 | 1.036699 | 1407.536413 | 0.128286484 | 0.40531849 | 0.685243417 | 0.998865404 | 10.432738 | 10.484707 | 10.467287 | 10.421634 | 10.451150 | 10.493133 | 1639 | 1447 | 1212 | 1380 |
| 92675 | NM_00131804 | DTD1 | D-aminoacyl-tRNA deacylase 1 | protein_coding | NP_001304972.1;NP_543010.3;XP_005260929.1 | 16219 | 610996 | . | . | 2.643436 | 2.543161 | 2.225631 | 2.884274 | 6.151383 | 5.878631 | 4.994252 | 6.594551 | -1.036115 | 311.910678 | 0.209957167 | -0.24378464 | 0.807397614 | 0.998865404 | 8.310799 | 8.258709 | 8.310649 | 8.276065 | 8.206995 | 8.336945 | 377 | 332 | 234 | 322 |
| 92689 | NM_00133076 | FAM114A1 | family with sequence similarity 114 member A1 | protein_coding | NP_001317693.1;NP_001337560.1;NP_001337561.1;NP_001337562.1;NI | 25087 | . | . | . | 8.402631 | 8.346561 | 6.566659 | 8.203903 | 19.553267 | 19.293451 | 14.735393 | 18.757250 | -1.161590 | 965.538079 | 0.15389954 | -1.40416855 | 0.160268666 | 0.933525128 | 10.019488 | 9.802758 | 9.974691 | 9.957241 | 9.771519 | 9.935357 | 1209 | 1107 | 687 | 934 |
| 92691 | NM_00114231 | TMEM169 | transmembrane protein 169 | protein_coding | NP_001135782.1;NP_001135783.1;NP_001135784.1;NP_612399.1;XP_00 | 25130 | . | . | . | 1.545902 | 1.643667 | 1.535110 | 1.868140 | 3.597378 | 3.799409 | 3.444744 | 4.271281 | 1.041208 | 181.112974 | 0.251670007 | 0.231485574 | 0.816937592 | 0.998865404 | 7.471833 | 7.529091 | 7.492173 | 7.489758 | 7.465683 | 7.549083 | 205 | 191 | 145 | 190 |
| 92703 | NM_00132295 | TMEM183A | transmembrane protein 183A | protein_coding | NP_001309884.1;NP_001309885.1;NP_001309886.1;NP_001309887.1;NI | 20173 | . | . | . | 12.965019 | 12.824018 | 13.184603 | 15.374706 | 30.170132 | 29.643293 | 29.585857 | 35.152440 | 1.093516 | 1313.131927 | 0.137077832 | 0.940879481 | 0.346766624 | 0.998865404 | 10.293034 | 10.421691 | 10.335397 | 10.310037 | 10.328547 | 10.449410 | 1469 | 1331 | 1099 | 1387 |

| | | | | | | | | | | | | | | | | | | | | | | | | | | | | | | | | | |
|---|---|---|---|---|---|---|---|---|---|---|---|---|---|---|---|---|---|---|---|---|---|---|---|---|---|---|---|---|---|---|---|---|---|
| 92714 | NM_00131796 | ARRDC1 | arrestin domain containing 1 | protein_coding | NP_001304897.1;NP_689498.1;XP_005266176.1;XP_006717383.1;XP_00 | 28633 | . | . | . | 3.877062 | 3.637415 | 5.637217 | 4.447752 | 9.022083 | 8.408048 | 12.649751 | 10.169255 | 1.212571 | 292.768554 | 0.232637081 | 1.195290965 | 0.231973368 | 0.986726781 | 8.046940 | 8.326748 | 8.160512 | 8.101789 | 8.335417 | 8.122298 | 322 | 269 | 333 | 243 |
| 92715 | NM_00134637 | DPH7 | diphthamide biosynthesis 7 | protein_coding | NP_001333299.1;NP_001333300.1;NP_001333301.1;NP_001333302.1;NI | 25199 | 613210 | . | . | 5.252925 | 5.517059 | 5.025510 | 5.115341 | 12.223775 | 12.752929 | 11.277098 | 11.695621 | 1.044044 | 462.927283 | 0.171416485 | 0.362756428 | 0.716786843 | 0.998865404 | 8.823160 | 8.885452 | 8.835233 | 8.846115 | 8.879048 | 8.855721 | 517 | 493 | 419 | 435 |
| 92737 | NM_139072,XI | DNER | delta/notch like EGF repeat containing | protein_coding | NP_620711.3;XP_005247007.1 | 24456 | 607299 | . | . | 1.054685 | 0.993109 | 0.885687 | 0.649874 | 2.454294 | 2.295615 | 1.987455 | 1.485858 | -1.362274 | 91.315466 | 0.344981319 | -1.29287124 | 0.196055596 | 0.96621494 | 6.717657 | 6.273933 | 6.564276 | 6.543510 | 6.505165 | 6.425034 | 125 | 110 | 78 | 61 |
| 92755 | gene-TUBBP1 | TUBBP1 | tubulin beta pseudogene 1 | pseudogene | . | 12414 | . | . | . | 1.011648 | 1.123333 | 0.645827 | 1.059414 | 2.354147 | 2.596633 | 1.449216 | 2.422223 | -1.269969 | 50.513581 | 0.456186626 | -0.75581666 | 0.449759124 | 0.998865404 | 5.822157 | 5.474099 | 5.671414 | 5.691297 | 5.599954 | 5.672895 | 62 | 64 | 30 | 51 |
| 92797 | NM_00137028 | HELB | DNA helicase B | protein_coding | NP_001357214.1;NP_387467.2;XP_016875719.1 | 17196 | 614539 | . | . | 0.678672 | 0.540460 | 0.730370 | 0.550961 | 1.579297 | 1.249298 | 1.638930 | 1.259707 | 1.020022 | 70.422457 | 0.387791396 | 0.073750213 | 0.941209144 | 0.998865404 | 6.122310 | 6.153450 | 6.161879 | 6.102714 | 6.167956 | 6.106039 | 90 | 66 | 70 | 58 |
| 92799 | NM_138392,XI | SHKBP1 | SH3KBP1 binding protein 1 | protein_coding | NP_612401.2;XP_006723537.1;XP_006723538.1;XP_011525794.1;XP_01 | 19214 | 617322 | . | . | 20.047250 | 19.855800 | 19.206262 | 23.580040 | 46.650775 | 45.897568 | 43.098296 | 53.912960 | 1.057437 | 1502.235284 | 0.137437473 | 0.586244797 | 0.557711022 | 0.998865404 | 10.512233 | 10.592443 | 10.539440 | 10.519988 | 10.496871 | 10.643580 | 1703 | 1556 | 1216 | 1584 |
| 92806 | NM_145039 | CENPBD1 | CENPB DNA-binding domain containing 1 | protein_coding | NP_659476.2 | 28272 | . | . | . | 1.561787 | 1.987468 | 1.859288 | 2.257014 | 3.634342 | 4.594120 | 4.172188 | 5.160395 | 1.146048 | 182.880266 | 0.258575166 | 0.760580705 | 0.446907548 | 0.998865404 | 7.414015 | 7.608918 | 7.430859 | 7.527337 | 7.496951 | 7.579489 | 174 | 205 | 154 | 200 |
| 92815 | NM_033445 | H2AW | H2A.W histone | protein_coding | NP_254280.1 | 20507 | 615015 | . | . | 3.570043 | 4.241616 | 4.354725 | 5.061687 | 8.307636 | 9.804685 | 9.771878 | 11.572946 | 1.194400 | 68.996613 | 0.382383302 | 0.670233161 | 0.502709168 | 0.998865404 | 5.975424 | 6.230246 | 6.065729 | 6.099809 | 6.108283 | 6.140995 | 67 | 73 | 61 | 75 |
| 92822 | NM_00111352 | ZNF276 | zinc finger protein 276 | protein_coding | NP_001106997.1;NP_689500.2;XP_005256381.1;XP_005256385.1;XP_01 | 23330 | 608460 | . | . | 7.986830 | 8.277413 | 7.944786 | 6.955884 | 18.585684 | 19.133612 | 17.827868 | 15.903802 | -1.076133 | 980.701007 | 0.146667963 | -0.72174405 | 0.470451852 | 0.998865404 | 9.989354 | 9.884064 | 9.970227 | 9.955531 | 9.970124 | 9.844995 | 1182 | 1086 | 893 | 809 |
| 92840 | NM_00132955 | REEP6 | receptor accessory protein 6 | protein_coding | NP_001316485.1;NP_612402.1 | 30078 | 609346 | . | . | 3.836747 | 4.220475 | 4.374022 | 5.048483 | 8.928269 | 9.755817 | 9.815179 | 11.542757 | 1.153457 | 185.551274 | 0.247433644 | 0.832402748 | 0.405181651 | 0.998865404 | 7.429584 | 7.634494 | 7.482593 | 7.519193 | 7.531982 | 7.593262 | 190 | 194 | 161 | 199 |
| 92856 | NM_00132030 | IMP4 | IMP U3 small nucleolar ribonucleoprotein 4 | protein_coding | NP_001307233.1;NP_001307234.1;NP_001307235.1;NP_001307236.1;NI | 30856 | 612981 | . | . | 41.333529 | 42.881410 | 39.276250 | 41.391925 | 96.184819 | 99.122302 | 88.134770 | 94.637718 | -1.022158 | 3887.205993 | 0.109560854 | -0.28858589 | 0.772898294 | 0.998865404 | 11.940284 | 11.908577 | 11.910361 | 11.956023 | 11.903551 | 11.926816 | 4401 | 4357 | 3296 | 3655 |
| 92906 | NM_00114265 | HNRNPLL | heterogeneous nuclear ribonucleoprotein L like | protein_coding | NP_001136122.1;NP_612403.2;XP_005264696.2;XP_005264697.2;XP_01 | 25127 | 611208 | . | . | 9.641964 | 11.477490 | 8.245040 | 8.145266 | 22.437247 | 26.530728 | 18.501630 | 18.623181 | -1.096881 | 959.782989 | 0.137463099 | -0.9704963 | 0.331799166 | 0.998865404 | 9.971701 | 9.838348 | 9.942495 | 9.936290 | 9.875734 | 9.868552 | 1161 | 1079 | 800 | 854 |
| 92912 | NM_00114533 | UBE2Q2 | ubiquitin conjugating enzyme E2 Q2 | protein_coding | NP_001138807.1;NP_001271311.1;NP_775740.1;XP_005254844.1;XP_00 | 19248 | 612501 | . | . | 4.036169 | 4.332750 | 4.211645 | 4.212273 | 9.392331 | 10.015343 | 9.450807 | 9.630862 | 1.005598 | 390.878173 | 0.181829837 | 0.044295841 | 0.964668587 | 0.998865404 | 8.606537 | 8.614602 | 8.599690 | 8.617793 | 8.619159 | 8.604938 | 442 | 427 | 345 | 363 |
| 92922 | NM_033212,XI | CCDC102A | coiled-coil domain containing 102A | protein_coding | NP_149989.2;XP_011521771.1 | 28097 | . | . | . | 1.960680 | 1.936596 | 1.312926 | 2.313954 | 4.562584 | 4.476527 | 2.946168 | 5.290581 | 1.029599 | 150.926670 | 0.276289644 | 0.152311517 | 0.878941242 | 0.998865404 | 7.217196 | 7.257924 | 7.234240 | 7.226655 | 7.187681 | 7.293113 | 173 | 159 | 115 | 163 |
| 92935 | NM_138395 | MARS2 | methionyl-tRNA synthetase 2, mitochondrial | protein_coding | NP_612404.1 | 25133 | 609728 | . | . | 6.182556 | 6.162097 | 5.995049 | 6.487512 | 14.387063 | 14.243964 | 13.452717 | 14.832926 | -1.002526 | 585.508060 | 0.159641678 | -0.02279762 | 0.98181171 | 0.998865404 | 9.195489 | 9.191599 | 9.198166 | 9.190510 | 9.169882 | 9.214866 | 679 | 629 | 491 | 567 |
| 92960 | NM_00127053 | PEX11G | peroxisomal biogenesis factor 11 gamma | protein_coding | NP_001257468.1;NP_001287810.1;NP_542393.1;XP_005272563.1;XP_01 | 20208 | 607583 | . | . | 0.335343 | 0.491064 | 0.420875 | 0.588755 | 0.780356 | 1.135118 | 0.944431 | 1.346120 | 1.187941 | 16.231095 | 0.764493508 | 0.325004436 | 0.745177715 | NA | 3.892781 | 4.138173 | 4.002381 | 4.018803 | 4.011454 | 4.033839 | 15 | 18 | 13 | 19 |
| 92979 | NM_138396 | MARCHF9 | membrane associated ring-CH-type finger 9 | protein_coding | NP_612405.2 | 25139 | 613336 | . | . | 3.622262 | 3.291730 | 3.531514 | 3.890828 | 8.429153 | 7.608982 | 7.924615 | 8.895916 | 1.058578 | 333.064489 | 0.196513836 | 0.417924233 | 0.676002514 | 0.998865404 | 8.338027 | 8.420121 | 8.391231 | 8.333368 | 8.369952 | 8.418432 | 392 | 331 | 285 | 335 |
| 92999 | NM_145166 | ZBTB47 | zinc finger and BTB domain containing 47 | protein_coding | NP_660149.2 | 26955 | . | . | . | 1.023367 | 1.100064 | 1.073960 | 1.045853 | 2.381416 | 2.542846 | 2.409934 | 2.391220 | -1.014245 | 181.820184 | 0.246243288 | -0.08287247 | 0.933952946 | 0.998865404 | 7.516559 | 7.496105 | 7.494884 | 7.523347 | 7.511217 | 7.494964 | 204 | 204 | 160 | 166 |
| 93058 | NM_00109933 | COQ10A | coenzyme Q10A | protein_coding | NP_001092807.1;NP_653177.3 | 26515 | . | . | . | 4.993302 | 4.663856 | 4.520390 | 4.892939 | 11.619620 | 10.780714 | 10.143624 | 11.187123 | -1.015000 | 220.214608 | 0.226545295 | -0.09481266 | 0.924463629 | 0.998865404 | 7.793429 | 7.772024 | 7.791196 | 7.781608 | 7.779231 | 7.779559 | 258 | 237 | 190 | 205 |
| 93081 | NM_00128677 | TEX30 | testis expressed 30 | protein_coding | NP_001273704.1;NP_001273705.1;NP_620134.3;XP_005254154.1;XP_01 | 25188 | . | . | . | 8.817599 | 8.813011 | 8.295209 | 8.392870 | 20.518915 | 20.371666 | 18.614205 | 19.189301 | -1.070355 | 395.330584 | 0.180919259 | -0.54217221 | 0.587699878 | 0.998865404 | 8.675070 | 8.577098 | 8.652856 | 8.641322 | 8.610143 | 8.603409 | 475 | 437 | 334 | 356 |
| 93082 | NM_00128548 | NEURL3 | neuralized E3 ubiquitin protein ligase 3 | protein_coding | NP_001272414.1;NP_001272415.1;XP_011510480.1;XP_011510481.1 | 25162 | 617206 | . | . | 0.053288 | 0.043090 | 0.127989 | 0.079147 | 0.124004 | 0.099605 | 0.287202 | 0.180960 | 2.134481 | 4.909290 | 1.376808346 | 0.79450788 | 0.426899824 | NA | 1.644780 | 2.742346 | 2.269661 | 2.266474 | 2.290372 | 2.278458 | 4 | 3 | 7 | 5 |
| 93100 | NM_00128682 | NAPRT | nicotinate phosphoribosyltransferase | protein_coding | NP_001273758.1;NP_001350074.1;NP_001350075.1;NP_660202.3;XP_01 | 30450 | 611552 | . | . | 23.672450 | 27.269662 | 25.921750 | 25.806788 | 55.086761 | 63.035047 | 58.167653 | 59.004152 | -1.008411 | 1388.907675 | 0.134163753 | -0.09006637 | 0.928234474 | 0.998865404 | 10.445857 | 10.433587 | 10.386089 | 10.495503 | 10.445478 | 10.425983 | 1492 | 1612 | 1218 | 1280 |
| 93109 | NM_00101165 | TMEM44 | transmembrane protein 44 | protein_coding | NP_001011655.1;NP_001159777.1;NP_001159778.1;NP_612408.3;XP_00 | 25120 | . | . | . | 4.945053 | 4.381319 | 5.294329 | 4.402995 | 11.507336 | 10.127614 | 11.880319 | 10.066925 | 1.005030 | 385.523836 | 0.194940741 | 0.037135341 | 0.970377094 | 0.998865404 | 8.586442 | 8.594898 | 8.620565 | 8.552728 | 8.648808 | 8.528309 | 468 | 391 | 371 | 325 |
| 93129 | NM_152288 | ORAI3 | ORAI calcium release-activated calcium modulator 3 | protein_coding | NP_689501.1 | 28185 | 610930 | . | . | 4.500635 | 4.452332 | 4.681814 | 6.086263 | 10.473164 | 10.291766 | 10.505855 | 13.915518 | 1.186967 | 338.525334 | 0.205024093 | 1.206102071 | 0.227778137 | 0.982008277 | 8.274708 | 8.521031 | 8.354244 | 8.344143 | 8.369333 | 8.516176 | 360 | 331 | 279 | 388 |
| 93134 | NM_00133036 | ZNF561 | zinc finger protein 561 | protein_coding | NP_001317294.1;NP_689502.2;XP_005260207.1;XP_006723001.1;XP_01 | 28684 | . | . | . | 3.232957 | 3.148437 | 3.336394 | 3.476133 | 7.523223 | 7.277753 | 7.486770 | 7.947765 | 1.091709 | 496.402781 | 0.169130019 | 0.748465327 | 0.454179531 | 0.998865404 | 8.890451 | 9.017488 | 8.934440 | 8.917250 | 9.008405 | 8.953788 | 554 | 505 | 470 | 465 |
| 93144 | gene-RPS10P5 | RPS10P5 | ribosomal protein S10 pseudogene 5 | pseudogene | . | 15795 | . | . | . | 0.158893 | 0.229311 | 0.115382 | 0.148539 | 0.369751 | 0.530064 | 0.258914 | 0.339617 | -1.470152 | 3.401861 | 1.628545485 | -0.34138746 | 0.732811914 | NA | 2.019256 | 1.459419 | 1.773648 | 1.779066 | 1.768429 | 1.772398 | 4 | 5 | 2 | 3 |
| 93145 | NM_00130434 | OLFM2 | olfactomedin 2 | protein_coding | NP_001291276.1;NP_001291277.1;NP_477512.1 | 17189 | 617492 | . | . | 1.956082 | 2.861349 | 1.325724 | 2.058648 | 4.551881 | 6.614138 | 2.974887 | 4.706855 | -1.434846 | 131.934456 | 0.321107221 | -1.62218777 | 0.10476315 | 0.83153868 | 7.282037 | 6.757988 | 7.016656 | 7.172410 | 6.906255 | 7.025756 | 145 | 200 | 76 | 121 |
| 93164 | NR_002774 | HTR7P1 | 5-hydroxytryptamine receptor 7 pseudogene 1 | transcribed_ps | . | 30411 | . | . | . | 0.627573 | 0.622250 | 0.602048 | 0.654152 | 1.460389 | 1.438359 | 1.350978 | 1.495641 | -1.009659 | 86.090747 | 0.342002856 | -0.04055114 | 0.967653736 | 0.998865404 | 6.434891 | 6.420647 | 6.430439 | 6.428281 | 6.417650 | 6.435673 | 100 | 93 | 72 | 83 |
| 93166 | NM_00113623 | PRDM6 | PR/SET domain 6 | protein_coding | NP_001129711.1;XP_011542028.1 | 9350 | 616982 | . | . | 0.642897 | 0.510999 | 0.806644 | 0.456176 | 1.496046 | 1.181198 | 1.810086 | 1.042994 | 1.224851 | 121.570641 | 0.302173195 | 0.968341249 | 0.33287397 | 0.998865404 | 6.771836 | 7.064636 | 6.851775 | 6.913612 | 6.987943 | 6.916472 | 114 | 129 | 128 | 113 |
| 93183 | NM_145167 | PIGM | phosphatidylinositol glycan anchor biosynthesis class M | protein_coding | NP_660150.1 | 18858 | 610273 | . | . | 2.170882 | 2.157805 | 2.557934 | 1.969583 | 5.051733 | 4.987864 | 5.739929 | 4.503218 | 1.030869 | 485.795742 | 0.185871032 | 0.235977908 | 0.813449813 | 0.998865404 | 8.901589 | 8.946474 | 8.916626 | 8.906780 | 9.012609 | 8.842363 | 555 | 512 | 487 | 400 |
| 93185 | NM_00120666 | IGSF8 | immunoglobulin superfamily member 8 | protein_coding | NP_001193594.1;NP_001307176.1;NP_443100.1;XP_006711694.1;XP_01 | 17813 | 606644 | . | . | 66.351451 | 66.996179 | 40.158704 | 49.090176 | 154.402550 | 154.864664 | 90.114965 | 112.238850 | -1.523072 | 4002.857305 | 0.125049789 | -4.85393794 | 1.21034E-06 | 0.000340944 | 12.238714 | 11.631485 | 12.119312 | 12.110582 | 11.683279 | 11.843019 | 5593 | 5189 | 2481 | 3274 |
| 93210 | NM_00129172 | PGAP3 | post-GPI attachment to proteins phospholipase 3 | protein_coding | NP_001278655.1;NP_001278657.1;NP_001278659.1;NP_001278661.1;NI | 23719 | 611801 | . | . | 2.421981 | 3.325228 | 2.479762 | 2.507573 | 5.636049 | 7.686414 | 5.564514 | 5.733268 | -1.112202 | 236.321053 | 0.232356261 | -0.66027394 | 0.50907805 | 0.998865404 | 7.959584 | 7.805513 | 7.843166 | 7.969221 | 7.864710 | 7.847820 | 251 | 302 | 198 | 206 |
| 93233 | NM_00136417 | CCDC114 | coiled-coil domain containing 114 | protein_coding | NP_001351100.1;NP_653178.3 | 26560 | 615038 | . | . | 0.339729 | 0.222843 | 0.280844 | 0.189077 | 0.790562 | 0.515112 | 0.630205 | 0.432301 | -1.173910 | 24.774225 | 0.634281438 | -0.36469984 | 0.715335481 | NA | 4.740050 | 4.512525 | 4.661590 | 4.615971 | 4.637322 | 4.607690 | 37 | 23 | 23 | 18 |
| 93323 | NM_00101169 | HAUS8 | HAUS augmin like complex subunit 8 | protein_coding | NP_001011699.1;NP_219485.1;XP_005260211.1 | 30532 | 613434 | . | . | 11.171531 | 12.808345 | 8.819087 | 12.451633 | 25.996612 | 29.607063 | 19.789777 | 28.469177 | -1.143198 | 500.041319 | 0.194904242 | -0.9906147 | 0.321873756 | 0.998865404 | 9.059872 | 8.865385 | 8.961933 | 9.043967 | 8.813006 | 9.014470 | 576 | 612 | 339 | 509 |
| 93343 | NM_00130454 | MVB12A | multivesicular body subunit 12A | protein_coding | NP_001291476.1;NP_612410.1 | 25153 | . | . | . | 20.111893 | 18.231009 | 18.857487 | 22.329375 | 46.801205 | 42.141792 | 42.315651 | 51.053462 | 1.059154 | 774.525793 | 0.156545721 | 0.52963375 | 0.59636589 | 0.998865404 | 9.555247 | 9.637908 | 9.611623 | 9.537956 | 9.558417 | 9.669933 | 912 | 769 | 638 | 806 |
| 93349 | NM_00130816 | SP140L | SP140 nuclear body protein like | protein_coding | NP_001295091.1;NP_001295092.1;NP_001339821.1;NP_001339822.1;NI | 25105 | 617747 | . | . | 3.956915 | 4.490079 | 4.595061 | 4.679695 | 9.207905 | 10.379020 | 10.311180 | 10.699567 | 1.073867 | 366.654506 | 0.190583378 | 0.539477392 | 0.589557492 | 0.998865404 | 8.466198 | 8.568542 | 8.453817 | 8.536408 | 8.538449 | 8.535583 | 378 | 409 | 331 | 355 |
| 93380 | NM_00133000 | MMGT1 | membrane magnesium transporter 1 | protein_coding | NP_001316929.1;NP_775741.1 | 28100 | . | . | . | 5.608097 | 4.608273 | 4.757034 | 3.620876 | 13.050273 | 10.652230 | 10.674645 | 8.278702 | -1.232810 | 756.135315 | 0.178516899 | -1.6914376 | 0.090753257 | 0.805424371 | 9.705078 | 9.404272 | 9.699050 | 9.549552 | 9.571467 | 9.382273 | 1060 | 808 | 672 | 546 |
| 93408 | NM_138403,XI | MYL10 | myosin light chain 10 | protein_coding | NP_612412.2;XP_016868282.1 | 29825 | 617177 | . | . | 3.555335 | 4.413870 | 2.962659 | 2.966952 | 8.273412 | 10.202859 | 6.648119 | 6.783583 | -1.253642 | 97.737242 | 0.328197096 | -0.9936875 | 0.32037503 | 0.998865404 | 6.765319 | 6.437807 | 6.616837 | 6.670455 | 6.560796 | 6.591728 | 115 | 127 | 71 | 86 |
| 93432 | NM_00129362 | MGAM2 | maltase-glucoamylase 2 (putative) | protein_coding | NP_001280555.1;XP_011514994.1;XP_011514995.1;XP_011514996.1;XP_ | 28101 | . | . | . | 3.719995 | 3.735454 | 5.059136 | 4.654278 | 8.656581 | 8.634670 | 11.352553 | 10.641455 | 1.199557 | 1047.006557 | 0.152080192 | 1.726073479 | 0.084334194 | 0.788061307 | 9.894543 | 10.157592 | 9.957291 | 9.958787 | 10.176721 | 10.000753 | 1095 | 1028 | 1109 | 946 |
| 93436 | NM_00119919 | ARMC6 | armadillo repeat containing 6 | protein_coding | NP_001186125.1;NP_219483.1;XP_005260214.1;XP_016882973.1;XP_01 | 25049 | . | . | . | 22.224953 | 23.300900 | 23.323875 | 23.299837 | 51.718379 | 53.861073 | 52.338099 | 53.272308 | 1.023006 | 1759.179725 | 0.120898994 | 0.271421131 | 0.786067152 | 0.998865404 | 10.764218 | 10.796970 | 10.756699 | 10.786277 | 10.789876 | 10.789001 | 1965 | 1912 | 1548 | 1667 |
| 93474 | NM_00120422 | ZNF670 | zinc finger protein 670 | protein_coding | NP_001191149.1;NP_149990.1 | 28167 | . | . | . | 0.440194 | 0.438209 | 0.438844 | 0.334188 | 1.024350 | 1.012939 | 0.984754 | 0.764081 | -1.161553 | 54.667761 | 0.428992123 | -0.5036343 | 0.614518383 | 0.998865404 | 5.875767 | 5.661523 | 5.788613 | 5.785988 | 5.782296 | 5.732557 | 68 | 63 | 50 | 41 |
| 93487 | NM_144578,XI | MAPK1IP1L | mitogen-activated protein kinase 1 interacting protein 1 like | protein_coding | NP_653179.1;XP_011535664.2;XP_016877278.1 | 19840 | 617226 | . | . | 7.329494 | 6.882021 | 8.916997 | 6.430446 | 17.056034 | 15.908101 | 20.009484 | 14.702452 | 1.043675 | 1431.111318 | 0.152881169 | 0.403400674 | 0.686653491 | 0.998865404 | 10.451442 | 10.513726 | 10.491276 | 10.434460 | 10.599367 | 10.381780 | 1672 | 1455 | 1450 | 1178 |
| 93550 | NM_00112832 | ZFAND4 | zinc finger AN1-type containing 4 | protein_coding | NP_001121796.1;NP_001269834.1;NP_001269835.1;NP_777550.2;XP_00 | 23504 | . | . | . | 0.149914 | 0.242511 | 0.139833 | 0.211210 | 0.348856 | 0.560571 | 0.313782 | 0.482908 | -1.013635 | 27.720876 | 0.597746226 | -0.0326877 | 0.973923633 | 0.998865404 | 4.804363 | 4.781347 | 4.773219 | 4.810818 | 4.772604 | 4.808469 | 27 | 35 | 20 | 30 |
| 93556 | NR_021485 | EGFEM1P | EGF like and EMI domain containing 1, pseudogene | transcribed_ps | . | 25149 | . | . | . | 0.491984 | 0.466886 | 1.009549 | 0.761500 | 1.144866 | 1.079229 | 2.265398 | 1.741080 | 1.823155 | 82.254478 | 0.361574395 | 2.396290476 | 0.016561959 | 0.431581156 | 5.863615 | 6.731753 | 6.260272 | 6.248864 | 6.457855 | 6.365351 | 69 | 61 | 106 | 85 |
| 93587 | NM_00113466 | TRMT10A | tRNA methyltransferase 10A | protein_coding | NP_001128137.1;NP_001128138.1;NP_001362809.1;NP_001362810.1;NI | 28403 | 616013 | . | . | 1.312506 | 1.151860 | 1.210623 | 1.106347 | 3.054256 | 2.662576 | 2.716605 | 2.529531 | -1.065366 | 134.454532 | 0.283497876 | -0.32222354 | 0.74728335 | 0.998865404 | 7.115414 | 7.025123 | 7.108938 | 7.055054 | 7.073976 | 7.044368 | 171 | 139 | 118 | 117 |
| 93594 | NM_00114508 | TBC1D31 | TBC1 domain family member 31 | protein_coding | NP_001138560.1;NP_001317535.1;NP_001350077.1;NP_001350078.1;NI | 30888 | . | . | . | 2.664636 | 2.848762 | 2.676922 | 2.064696 | 6.200717 | 6.585041 | 6.006933 | 4.720682 | -1.142142 | 270.710548 | 0.21857397 | -0.87723855 | 0.380357079 | 0.998865404 | 8.172882 | 7.982025 | 8.094077 | 8.130508 | 8.096942 | 7.988261 | 321 | 322 | 244 | 211 |
| 93611 | NM_00101476 | FBXO44 | F-box protein 44 | protein_coding | NP_001014765.1;NP_001291719.1;NP_001291720.1;NP_001317284.1;NI | 24847 | 609111 | . | . | 45.441852 | 46.671211 | 39.772300 | 43.295668 | 105.745055 | 107.882594 | 89.247892 | 98.990403 | -1.106631 | 3669.697904 | 0.108484776 | -1.34741544 | 0.17784647 | 0.951478615 | 11.912699 | 11.766489 | 11.873500 | 11.890227 | 11.793748 | 11.802617 | 4395 | 4202 | 3013 | 3284 |
| 93621 | NM_00127205 | MRFAP1 | Morf4 family associated protein 1 | protein_coding | NP_001258982.1;NP_001258983.1;NP_150638.1 | 24549 | 616905 | . | . | 54.424000 | 55.311244 | 56.339357 | 54.853949 | 126.646880 | 127.854421 | 126.423891 | 125.417036 | 1.011582 | 3716.962490 | 0.10888044 | 0.152577287 | 0.878731638 | 0.998865404 | 11.851544 | 11.868225 | 11.861577 | 11.848476 | 11.879768 | 11.849200 | 4289 | 3956 | 3311 | 3440 |
| 93622 | NR_015433,NF | LOC93622 | Morf4 family associated protein 1 like 1 pseudogene | transcribed_ps | . | . | . | | . | 4.446314 | 3.480720 | 3.499731 | 3.111101 | 10.346756 | 8.045840 | 7.853295 | 7.113162 | -1.189633 | 187.485338 | 0.254597726 | -0.9839722 | 0.325129182 | 0.998865404 | 7.669813 | 7.420716 | 7.641081 | 7.527713 | 7.532224 | 7.487824 | 263 | 193 | 157 | 152 |
| 93624 | NM_152293,XI | TADA2B | transcriptional adaptor 2B | protein_coding | NP_689506.2;XP_011511897.1 | 30781 | 608790 | . | . | 2.355036 | 2.591301 | 2.422504 | 2.764351 | 5.480266 | 5.989907 | 5.436029 | 6.320360 | 1.032670 | 346.227969 | 0.194815428 | 0.238071249 | 0.811825833 | 0.998865404 | 8.412645 | 8.458153 | 8.400103 | 8.450485 | 8.408293 | 8.477835 | 375 | 384 | 287 | 350 |
| 93627 | NM_00116343 | TBCK | TBC1 domain containing kinase | protein_coding | NP_001156907.2;NP_001156908.2;NP_001156909.2;NP_001277697.2;NI | 28261 | 616899 | . | . | 0.960752 | 0.941036 | 0.690919 | 0.956702 | 2.235708 | 2.175244 | 1.550403 | 2.187383 | -1.152037 | 198.783177 | 0.243423053 | -0.83881629 | 0.401572407 | 0.998865404 | 7.734042 | 7.528766 | 7.669761 | 7.662386 | 7.557698 | 7.645216 | 247 | 228 | 144 | 191 |
| 93643 | NM_00114601 | TJAP1 | tight junction associated protein 1 | protein_coding | NP_001139488.1;NP_001139489.1;NP_001139490.1;NP_001139491.1;NI | 17949 | 612658 | . | . | 12.369811 | 13.532121 | 15.670879 | 14.314007 | 28.785059 | 31.280101 | 35.164997 | 32.727278 | 1.146545 | 1195.865993 | 0.135253825 | 1.458685866 | 0.144651588 | 0.906189003 | 10.121799 | 10.319138 | 10.138652 | 10.198740 | 10.302894 | 10.239841 | 1231 | 1251 | 1157 | 1148 |
| 93653 | NR_002330 | ST7-AS1 | ST7 antisense RNA 1 | lncRNA | . | 16000 | . | . | . | 0.752821 | 1.002744 | 0.843670 | 0.964857 | 1.751846 | 2.317888 | 1.893170 | 2.206031 | 1.026816 | 52.900720 | 0.436510844 | 0.087459683 | 0.930306131 | 0.998865404 | 5.706954 | 5.743249 | 5.693725 | 5.746781 | 5.715067 | 5.736265 | 52 | 64 | 44 | 53 |
| 93655 | NR_002332 | ST7-OT3 | ST7 overlapping transcript 3 | lncRNA | . | 16045 | . | . | . | 0.001560 | 0.089815 | 0.048274 | 0.077928 | 0.003630 | 0.207611 | 0.108327 | 0.178172 | 1.209283 | 2.518843 | 1.905639286 | 0.143863333 | 0.88560839 | NA | 1.193905 | 1.459419 | 1.317363 | 1.330972 | 1.324370 | 1.328457 | 1 | 4 | 2 | 3 |
| 93663 | NM_033515,XI | ARHGAP18 | Rho GTPase activating protein 18 | protein_coding | NP_277050.2;XP_005267269.2;XP_005267270.1;XP_016866983.1 | 21035 | 613351 | . | . | 5.761071 | 5.458178 | 7.097900 | 6.049163 | 13.406247 | 12.616824 | 15.927484 | 13.830691 | 1.155461 | 837.650156 | 0.152908934 | 1.363352235 | 0.172771527 | 0.947431715 | 9.601853 | 9.810983 | 9.676153 | 9.632153 | 9.817446 | 9.693553 | 923 | 812 | 848 | 770 |
| 93664 | NM_00100957 | CADPS2 | calcium dependent secretion activator 2 | protein_coding | NP_001009571.2;NP_001161412.1;NP_001350318.1;NP_001350319.1;NI | 16018 | 609978 | . | . | 2.076614 | 2.121439 | 2.246955 | 2.147936 | 4.832363 | 4.903800 | 5.042101 | 4.911001 | 1.025754 | 388.814114 | 0.182296481 | 0.201237598 | 0.840512794 | 0.998865404 | 8.584396 | 8.621242 | 8.591033 | 8.599085 | 8.622920 | 8.597033 | 439 | 417 | 350 | 361 |
| 93953 | NM_052957,XI | GCNA | germ cell nuclear acidic peptidase | protein_coding | NP_443189.1;XP_006724779.1;XP_011529365.1;XP_011529366.1;XP_01 | 15805 | 300369 | . | . | 0.244888 | 0.246242 | 0.461363 | 0.311638 | 0.569865 | 0.569198 | 1.035285 | 0.712524 | 1.570465 | 39.219195 | 0.508672278 | 1.280178941 | 0.20048221 | 0.966220653 | 4.930218 | 5.583440 | 5.241691 | 5.242939 | 5.344625 | 5.274639 | 35 | 33 | 50 | 36 |
| 93973 | NM_022899,XI | ACTR8 | actin related protein 8 | protein_coding | NP_075050.3;XP_005265644.1;XP_011532551.1 | 14672 | . | . | . | 3.674481 | 4.801063 | 4.623788 | 4.027775 | 8.550669 | 11.097873 | 10.375646 | 9.209031 | 1.133548 | 884.491202 | 0.142997171 | 1.264678788 | 0.205986497 | 0.970176829 | 9.695376 | 9.876360 | 9.725444 | 9.756217 | 9.858640 | 9.802429 | 934 | 914 | 849 | 847 |
| 93974 | NM_016311,N | ATP5IF1 | ATP synthase inhibitory factor subunit 1 | protein_coding | NP_057395.1;NP_835497.1;NP_835498.1 | 871 | 614981 | . | . | 71.985384 | 64.488530 | 64.664010 | 86.972586 | 167.512953 | 149.068125 | 145.104159 | 198.852475 | 1.156978 | 1614.624436 | 0.131917617 | 1.594637909 | 0.11079323 | 0.848126978 | 10.548011 | 10.758299 | 10.634724 | 10.556571 | 10.661887 | 10.755494 | 1812 | 1533 | 1417 | 1720 |
| 93995 | gene-NDUFB3F | NDUFB3P4 | NADH:ubiquinone oxidoreductase subunit B3 pseudogene 4 | pseudogene | . | 13676 | . | . | . | 0.681130 | 0.425496 | 0.527997 | 0.646026 | 1.585018 | 0.983554 | 1.184809 | 1.477061 | 1.043018 | 8.639585 | 1.037842612 | 0.058548908 | 0.953311406 | NA | 3.079678 | 3.141582 | 3.119425 | 3.101284 | 3.108631 | 3.115409 | 12 | 7 | 7 | 9 |
| 94005 | NM_033198 | PIGS | phosphatidylinositol glycan anchor biosynthesis class S | protein_coding | NP_149975.1 | 14937 | 610271 | . | . | 32.503113 | 30.163450 | 26.018805 | 28.562368 | 75.636078 | 69.724167 | 58.385441 | 65.304459 | -1.164391 | 2311.059214 | 0.121156475 | -1.8123352 | 0.069934419 | 0.747718654 | 11.279982 | 11.060343 | 11.265341 | 11.197554 | 11.076790 | 11.141899 | 2982 | 2572 | 1780 | 2085 |
| 94015 | NM_00133045 | TTYH2 | tweety family member 2 | protein_coding | NP_001317382.1;NP_116035.5;NP_443101.1 | 13877 | 608855 | . | . | 26.009676 | 27.795638 | 35.455777 | 32.210786 | 60.525586 | 64.250857 | 79.561736 | 73.646132 | 1.238382 | 3218.361670 | 0.116484598 | 2.648046546 | 0.008095838 | 0.285797567 | 11.489617 | 11.798141 | 11.533664 | 11.579974 | 11.774486 | 11.688968 | 3216 | 3192 | 3255 | 3168 |
| 94032 | NM_033259 | CAMK2N2 | calcium/calmodulin dependent protein kinase II inhibitor 2 | protein_coding | NP_150284.1 | 24197 | 608721 | . | . | 1.653563 | 1.721876 | 1.600923 | 1.110014 | 3.847908 | 3.980193 | 3.592425 | 2.537914 | -1.268010 | 69.444484 | 0.390782063 | -0.87661585 | 0.380695325 | 0.998865404 | 6.277949 | 5.937600 | 6.140253 | 6.147898 | 6.125964 | 6.048716 | 88 | 85 | 63 | 47 |
| 94039 | NM_00130094 | ZNF101 | zinc finger protein 101 | protein_coding | NP_001287878.1;NP_149981.2;XP_024307553.1;XP_024307554.1;XP_02 | 12881 | 603983 | . | . | 1.664510 | 1.826574 | 2.143287 | 1.744361 | 3.873384 | 4.222208 | 4.809476 | 3.988275 | 1.116989 | 221.810386 | 0.229216578 | 0.696348242 | 0.486210768 | 0.998865404 | 7.710911 | 7.871016 | 7.753728 | 7.774222 | 7.847632 | 7.784337 | 236 | 231 | 217 | 205 |
| 94056 | NM_032796,N | SYAP1 | synapse associated protein 1 | protein_coding | NP_116185.2 | 16273 | . | . | . | 5.756810 | 5.300726 | 5.378467 | 6.140482 | 13.396334 | 12.252865 | 12.069126 | 14.039482 | 1.027177 | 1082.601202 | 0.141330647 | 0.273719345 | 0.78430031 | 0.998865404 | 10.060894 | 10.099421 | 10.102244 | 10.035855 | 10.043344 | 10.133497 | 1285 | 1101 | 896 | 1091 |
| 94059 | NM_00130178 | LENG9 | leukocyte receptor cluster member 9 | protein_coding | NP_001288711.1 | 16306 | . | . | . | 1.191988 | 1.093880 | 1.516632 | 0.674671 | 2.773804 | 2.528552 | 3.403279 | 1.542555 | -1.063691 | 71.747116 | 0.429837926 | -0.20723946 | 0.835822855 | 0.998865404 | 6.206760 | 6.121684 | 6.172979 | 6.149225 | 6.232707 | 6.051041 | 89 | 76 | 84 | 40 |
| 94081 | NM_00132297 | SFXN1 | sideroflexin 1 | protein_coding | NP_001309906.1;NP_001309907.1;NP_001309909.1;NP_001309910.1;NI | 16085 | 615569 | . | . | 37.476671 | 36.437982 | 35.382034 | 37.464103 | 87.209757 | 84.228020 | 79.396266 | 85.657219 | -1.029473 | 4497.969089 | 0.109836472 | -0.38152826 | 0.702811306 | 0.998865404 | 12.155883 | 12.113928 | 12.165086 | 12.128332 | 12.091923 | 12.152464 | 5370 | 4816 | 3714 | 4307 |
| 94097 | NM_00133040 | SFXN5 | sideroflexin 5 | protein_coding | NP_001317329.1;NP_001317330.1;NP_001317331.1;NP_001317332.1;NI | 16073 | 615572 | . | . | 4.657548 | 5.333119 | 5.461031 | 4.673256 | 10.838306 | 12.327741 | 12.254395 | 10.684845 | -1.002371 | 655.301447 | 0.160427311 | -0.02129369 | 0.98301138 | 0.998865404 | 9.357612 | 9.354416 | 9.322416 | 9.388139 | 9.396828 | 9.309833 | 719 | 742 | 606 | 574 |
| 94101 | NM_00112815 | ORMDL1 | ORMDL sphingolipid biosynthesis regulator 1 | protein_coding | NP_001121622.1;NP_001358313.1;NP_001358314.1;NP_001358315.1;NI | 16036 | 610073 | . | . | 34.252047 | 32.326349 | 37.603224 | 29.270898 | 79.705919 | 74.723804 | 84.380543 | 66.924426 | -1.011849 | 2259.496904 | 0.143281511 | -0.11860473 | 0.905588519 | 0.998865404 | 11.150047 | 11.133477 | 11.169017 | 11.118323 | 11.237688 | 11.020146 | 2699 | 2375 | 2218 | 1821 |
| 94103 | NM_00132080 | ORMDL3 | ORMDL sphingolipid biosynthesis regulator 3 | protein_coding | NP_001307730.1;NP_001307731.1;NP_001307732.1;NP_644809.1 | 16038 | 610075 | . | . | 15.513851 | 15.496430 | 17.966540 | 17.065524 | 36.101371 | 35.820694 | 40.316397 | 39.018293 | 1.113006 | 1086.928697 | 0.134560286 | 1.147901128 | 0.251009383 | 0.993339639 | 10.006635 | 10.161305 | 10.048266 | 10.040386 | 10.146209 | 10.100875 | 1191 | 1104 | 1026 | 1041 |
| 94104 | NM_013329,N | PAXBP1 | PAX3 and PAX7 binding protein 1 | protein_coding | NP_037461.2;NP_057715.2;XP_006724129.1;XP_011528106.1 | 13579 | 617621 | . | . | 10.386152 | 10.048283 | 11.164479 | 10.504412 | 24.169002 | 23.227061 | 25.052769 | 24.017090 | -1.031906 | 1130.814206 | 0.152323658 | -0.29747036 | 0.766107431 | 0.998865404 | 10.165274 | 10.120674 | 10.185598 | 10.118648 | 10.213026 | 10.038375 | 1381 | 1184 | 1082 | 921 |
| 94107 | NM_053045 | TMEM203 | transmembrane protein 203 | protein_coding | NP_444273.1 | 28217 | 616499 | . | . | 13.923829 | 11.378901 | 10.085619 | 13.154377 | 32.401321 | 26.302835 | 22.631836 | 30.075918 | -1.104366 | 592.908908 | 0.18472371 | -0.77531027 | 0.438156342 | 0.998865404 | 9.281675 | 9.138087 | 9.309876 | 9.166957 | 9.090086 | 9.253572 | 792 | 601 | 428 | 595 |
| 94120 | NM_00100999 | SYTL3 | synaptotagmin like 3 | protein_coding | NP_001009991.2;NP_001229313.1;NP_001229323.1;NP_001229324.1;NI | 15587 | . | . | . | 0.334457 | 0.333906 | 0.376993 | 0.652162 | 0.778294 | 0.771841 | 0.845960 | 1.491090 | 1.637154 | 55.922856 | 0.464981881 | 1.529500611 | 0.126140388 | 0.874064017 | 5.407564 | 6.116854 | 5.749258 | 5.714958 | 5.747819 | 5.908144 | 54 | 41 | 40 | 87 |
| 94121 | NM_00112989 | SYTL4 | synaptotagmin like 4 | protein_coding | NP_001123368.1;NP_001167539.1;NP_001357089.1;NP_001357090.1;NI | 15588 | 300723 | . | . | 1.314764 | 1.464426 | 1.129436 | 1.259491 | 3.059511 | 3.385088 | 2.534420 | 2.879677 | -1.200772 | 157.546919 | 0.263354435 | -1.00230884 | 0.316194455 | 0.998865404 | 7.425946 | 7.161201 | 7.315926 | 7.354435 | 7.249386 | 7.275468 | 189 | 194 | 120 | 139 |
| 94134 | NM_00127069 | ARHGAP12 | Rho GTPase activating protein 12 | protein_coding | NP_001257624.1;NP_001257625.1;NP_001257626.1;NP_001257627.1;NI | 16348 | 610577 | . | . | 5.238625 | 5.378259 | 5.679212 | 4.223340 | 12.190495 | 12.432087 | 12.743987 | 9.656167 | -1.090114 | 802.443438 | 0.169988387 | -0.7322804 | 0.463997437 | 0.998865404 | 9.708762 | 9.585101 | 9.668564 | 9.681276 | 9.715055 | 9.502291 | 955 | 911 | 770 | 610 |
| 94161 | NR_000024 | SNORD46 | small nucleolar RNA, C/D box 46 | snoRNA | . | 10186 | . | . | . | 1.779621 | 1.279940 | 1.629148 | 1.155480 | 4.141251 | 2.958638 | 3.655761 | 2.641867 | -1.072000 | 5.182339 | 1.322372303 | -0.07585255 | 0.939536404 | NA | 2.421266 | 2.324312 | 2.378555 | 2.371759 | 2.376811 | 2.370200 | 7 | 5 | 5 | 4 |
| 94162 | NR_001456 | SNORD38A | small nucleolar RNA, C/D box 38A | snoRNA | . | 30355 | . | . | . | 6.915530 | 4.484961 | 5.033750 | 8.085596 | 16.092724 | 10.367188 | 11.295589 | 18.486755 | 1.147903 | 13.864808 | 0.834799194 | 0.238381516 | 0.811585202 | NA | 3.690761 | 3.889137 | 3.799305 | 3.774857 | 3.781332 | 3.809525 | 18 | 11 | 10 | 17 |
| 94163 | NR_001457 | SNORD38B | small nucleolar RNA, C/D box 38B | snoRNA | . | 30356 | . | . | . | 3.496680 | 5.555833 | 8.390406 | 7.176153 | 8.136917 | 12.842556 | 18.827827 | 16.407421 | 1.733975 | 13.565814 | 0.839667871 | 0.94571099 | 0.344296016 | NA | 3.312060 | 4.104320 | 3.720195 | 3.739904 | 3.767001 | 3.757201 | 9 | 13 | 16 | 15 |
| 94235 | NM_033258,XI | GNG8 | G protein subunit gamma 8 | protein_coding | NP_150283.1;XP_011525826.1;XP_016882992.1;XP_016882993.1;XP_01 | 19664 | . | . | . | 0.146014 | 0.158661 | 0.078058 | 0.119206 | 0.339779 | 0.366751 | 0.175161 | 0.272550 | -1.275067 | 11.272671 | 0.923479912 | -0.37962202 | 0.704226011 | NA | 3.661494 | 3.306201 | 3.488764 | 3.516648 | 3.482859 | 3.498668 | 11 | 17 | 7 | 11 |
| 94239 | NM_012412,N | H2AZ2 | H2A.Z variant histone 2 | protein_coding | NP_036544.1;NP_619541.1;NP_958844.1;NP_958924.1;NP_958925.1 | 20664 | . | . | . | 27.992217 | 26.468934 | 23.589291 | 26.332008 | 65.139041 | 61.184127 | 52.933684 | 60.205009 | -1.077899 | 2603.158205 | 0.117672185 | -0.91968935 | 0.357735122 | 0.998865404 | 11.399220 | 11.290839 | 11.382063 | 11.368896 | 11.273834 | 11.353883 | 3135 | 2891 | 2070 | 2466 |
| 94241 | NM_00113573 | TP53INP1 | tumor protein p53 inducible nuclear protein 1 | protein_coding | NP_001129205.1;NP_150601.1;XP_011515688.1 | 18022 | 606185 | . | . | 1.544247 | 1.230862 | 1.402971 | 1.364754 | 3.593527 | 2.845193 | 3.148227 | 3.120347 | -1.025362 | 242.682657 | 0.228463635 | -0.15815633 | 0.874333614 | 0.998865404 | 7.940419 | 7.905221 | 7.985014 | 7.867141 | 7.921434 | 7.909585 | 316 | 234 | 211 | 222 |
| 95681 | NM_00125715 | CEP41 | centrosomal protein 41 | protein_coding | NP_001244087.1;NP_001244088.1;NP_001244089.1;NP_061188.1;XP_01 | 12370 | 610523 | . | . | 4.883151 | 4.595505 | 4.883959 | 4.845207 | 11.363293 | 10.622717 | 10.959462 | 11.077990 | -1.011508 | 843.736469 | 0.147160408 | -0.11217244 | 0.91068668 | 0.998865404 | 9.728670 | 9.712583 | 9.764541 | 9.682270 | 9.731022 | 9.701170 | 1033 | 863 | 745 | 770 |
| 96459 | NM_00100873 | FNIP1 | folliculin interacting protein 1 | protein_coding | NP_001008738.3;NP_001333042.2;NP_001333043.1;NP_588613.3 | 29418 | 610594 | . | . | 4.840177 | 5.043303 | 5.558625 | 4.345681 | 11.263288 | 11.657822 | 12.473393 | 9.935884 | -1.015740 | 1003.402438 | 0.155534233 | -0.1448627 | 0.884819279 | 0.998865404 | 9.981616 | 9.959670 | 9.962616 | 9.985678 | 10.050092 | 9.867620 | 1146 | 1108 | 974 | 817 |
| 96610 | NR_027293 | BMS1P20 | BMS1 pseudogene 20 | transcribed_ps | . | 49153 | . | . | . | 7.712014 | 7.904648 | 10.415963 | 10.322939 | 17.946173 | 18.271948 | 23.373119 | 23.602173 | 1.309720 | 569.159276 | 0.159941483 | 2.433754728 | 0.014943117 | 0.410059331 | 8.944958 | 9.334239 | 9.049386 | 9.059702 | 9.237879 | 9.224494 | 563 | 536 | 567 | 599 |
| 96626 | NM_033514,N | LIMS3 | LIM zinc finger domain containing 3 | protein_coding | NP_277049.1 | 30047 | . | . | . | 0.234035 | 0.307042 | 0.126256 | 0.157986 | 0.544608 | 0.709742 | 0.283316 | 0.361216 | -1.800416 | 14.672590 | 0.810698608 | -1.04641868 | 0.295367772 | NA | 4.238681 | 3.387716 | 3.893548 | 3.906447 | 3.858467 | 3.868961 | 20 | 22 | 8 | 11 |
| 96764 | NM_00131790 | TGS1 | trimethylguanosine synthase 1 | protein_coding | NP_001304831.1;NP_001350113.1;NP_079107.6;XP_006716548.1;XP_01 | 17843 | 606461 | . | . | 6.950542 | 6.603565 | 8.066250 | 7.346778 | 16.174198 | 15.264439 | 18.100431 | 16.797534 | 1.120979 | 950.883560 | 0.138792286 | 1.18709469 | 0.235190273 | 0.990818043 | 9.808272 | 9.973263 | 9.862240 | 9.836488 | 9.948163 | 9.916537 | 1051 | 950 | 893 | 922 |
| 103910 | NM_00114494 | MYL12B | myosin light chain 12B | protein_coding | NP_001138416.1;NP_001138417.1;NP_291024.1 | 29827 | 609211 | . | . | 98.244151 | 96.058666 | 79.476599 | 87.935399 | 228.618152 | 222.043910 | 178.343176 | 201.053839 | -1.176791 | 3276.501969 | 0.114041938 | -2.05939605 | 0.039456314 | 0.61817878 | 11.790648 | 11.555677 | 11.753330 | 11.728439 | 11.570312 | 11.642767 | 4140 | 3766 | 2500 | 2949 |
| 112398 | NM_053046,N | EGLN2 | egl-9 family hypoxia inducible factor 2 | protein_coding | NP_444274.1;NP_542770.2 | 14660 | 606424 | . | . | 26.518177 | 26.320362 | 27.382951 | 28.895539 | 61.708885 | 60.840696 | 61.446553 | 66.066220 | 1.050472 | 1785.183603 | 0.12087804 | 0.587685861 | 0.556743289 | 0.998865404 | 10.765938 | 10.836903 | 10.788224 | 10.775793 | 10.803433 | 10.837742 | 2021 | 1864 | 1556 | 1752 |
| 112399 | NM_00130810 | EGLN3 | egl-9 family hypoxia inducible factor 3 | protein_coding | NP_001295032.1;NP_071356.1 | 14661 | 606426 | . | . | 0.829787 | 0.578468 | 0.553472 | 0.785219 | 1.930947 | 1.337154 | 1.241975 | 1.795311 | -1.061871 | 58.403007 | 0.432171171 | -0.20040262 | 0.841165713 | 0.998865404 | 5.910849 | 5.823779 | 5.912059 | 5.831989 | 5.826511 | 5.893032 | 82 | 53 | 41 | 62 |
| 112464 | NM_145040 | CAVIN3 | caveolae associated protein 3 | protein_coding | NP_659477.2 | 9400 | 618303 | . | . | 11.543720 | 13.372617 | 12.728477 | 13.850771 | 26.862713 | 30.911402 | 28.562332 | 31.668144 | 1.051765 | 412.449796 | 0.184051184 | 0.395605773 | 0.692395877 | 0.998865404 | 8.651604 | 8.723646 | 8.629219 | 8.712326 | 8.679207 | 8.723800 | 431 | 464 | 354 | 411 |
| 112476 | NM_00125644 | PRRT2 | proline rich transmembrane protein 2 | protein_coding | NP_001243371.1;NP_001243372.1;NP_660282.2;XP_011544017.1;XP_01 | 30500 | 614386 | . | . | 1.766374 | 1.826450 | 2.003000 | 1.692435 | 4.110423 | 4.221918 | 4.494674 | 3.869552 | 1.005738 | 155.105081 | 0.270525658 | 0.030511598 | 0.975659044 | 0.998865404 | 7.272276 | 7.281912 | 7.280797 | 7.267287 | 7.320675 | 7.230098 | 181 | 164 | 151 | 129 |
| 112479 | NM_00114272 | ERI2 | ERI1 exoribonuclease family member 2 | protein_coding | NP_001136197.1;NP_542394.2 | 30541 | . | . | . | 4.568108 | 3.473544 | 3.134345 | 3.044165 | 10.630175 | 8.029251 | 7.033379 | 6.960121 | -1.324040 | 379.775110 | 0.201036184 | -2.01429896 | 0.043978155 | 0.645513836 | 8.756875 | 8.352947 | 8.720281 | 8.557436 | 8.499245 | 8.457876 | 563 | 406 | 295 | 295 |
| 112483 | NM_00132084 | SAT2 | spermidine/spermine N1-acetyltransferase family member 2 | protein_coding | NP_001307774.1;NP_001307775.1;NP_001307776.1;NP_597998.1;XP_01 | 23160 | 611463 | . | . | 39.527989 | 40.451994 | 42.191590 | 44.982601 | 91.983255 | 93.506594 | 94.676708 | 102.847369 | 1.081919 | 1192.995360 | 0.130744239 | 0.868811918 | 0.384950013 | 0.998865404 | 10.162522 | 10.275993 | 10.185904 | 10.193574 | 10.235993 | 10.261596 | 1313 | 1243 | 1060 | 1182 |
| 112487 | NM_080664 | DTD2 | D-aminoacyl-tRNA deacylase 2 | protein_coding | NP_542395.1 | 20277 | . | . | . | 3.641057 | 2.794640 | 3.583292 | 3.222605 | 8.472890 | 6.459936 | 8.040804 | 7.368103 | 1.041431 | 280.331841 | 0.222718343 | 0.262964081 | 0.792578265 | 0.998865404 | 8.100708 | 8.160653 | 8.186093 | 8.043558 | 8.168025 | 8.108550 | 359 | 256 | 263 | 253 |
| 112495 | NM_138408 | GTF3C6 | general transcription factor IIIC subunit 6 | protein_coding | NP_612417.1 | 20872 | 611784 | . | . | 41.102707 | 42.050632 | 43.911953 | 48.495800 | 95.647697 | 97.201920 | 98.537155 | 110.879883 | 1.096216 | 1077.125772 | 0.136396932 | 0.971660992 | 0.331219229 | 0.998865404 | 10.005339 | 10.137574 | 10.032065 | 10.042694 | 10.071547 | 10.138131 | 1175 | 1117 | 936 | 1103 |
| 112574 | NM_00110257 | SNX18 | sorting nexin 18 | protein_coding | NP_001096045.1;NP_001138899.1;NP_443102.2;XP_016864486.1 | 19245 | . | . | . | 10.568592 | 10.406931 | 9.736903 | 9.805905 | 24.593548 | 24.056084 | 21.849326 | 22.420038 | -1.096264 | 1633.327784 | 0.122266693 | -1.08447166 | 0.278155726 | 0.998865404 | 10.738351 | 10.605803 | 10.718412 | 10.698351 | 10.637359 | 10.636272 | 1992 | 1820 | 1356 | 1460 |
| 112597 | NR_024204,NF | CYTOR | cytoskeleton regulator RNA | lncRNA | . | 28717 | . | . | . | 78.556166 | 71.076959 | 68.818765 | 81.211970 | 182.803408 | 164.297567 | 154.427307 | 185.681513 | -1.026079 | 1331.757463 | 0.137591255 | -0.26994761 | 0.787200556 | 0.998865404 | 10.397678 | 10.360311 | 10.419147 | 10.356206 | 10.311208 | 10.422682 | 1617 | 1396 | 1059 | 1323 |
| 112609 | NM_00134654 | MRAP2 | melanocortin 2 receptor accessory protein 2 | protein_coding | NP_001333470.1;NP_001333471.1;NP_001333472.1;NP_001333473.1;NI | 21232 | 615410 | . | . | 0.570397 | 0.734283 | 0.519189 | 0.591276 | 1.327338 | 1.697328 | 1.165046 | 1.351882 | -1.127324 | 51.720074 | 0.437211767 | -0.3954656 | 0.692499302 | 0.998865404 | 5.777174 | 5.602870 | 5.688124 | 5.721129 | 5.672698 | 5.691356 | 58 | 64 | 40 | 48 |
| 112611 | NM_00132233 | RWDD2A | RWD domain containing 2A | protein_coding | NP_001309264.1;NP_001309265.1;NP_001309266.1;NP_001309268.1;NI | 21385 | . | . | . | 1.123167 | 0.878544 | 1.019976 | 0.935694 | 2.613655 | 2.030792 | 2.288798 | 2.139353 | -1.032849 | 120.438813 | 0.303075009 | -0.15385319 | 0.877725495 | 0.998865404 | 6.934622 | 6.889336 | 6.960906 | 6.870194 | 6.914945 | 6.896064 | 159 | 115 | 106 | 108 |
| 112616 | NM_138410,N | CMTM7 | CKLF like MARVEL transmembrane domain containing 7 | protein_coding | NP_612419.1;NP_852137.1;XP_011531621.1;XP_016861135.1 | 19178 | 607890 | . | . | 16.399064 | 17.101575 | 17.274192 | 18.446507 | 38.161300 | 39.531057 | 38.762786 | 42.175747 | 1.038500 | 945.629914 | 0.140167618 | 0.38882394 | 0.69740639 | 0.998865404 | 9.857792 | 9.911963 | 9.852837 | 9.888789 | 9.874811 | 9.921260 | 1042 | 1026 | 811 | 932 |
| 112724 | NM_00114597 | RDH13 | retinol dehydrogenase 13 | protein_coding | NP_001139443.1;NP_612421.1;XP_005258530.1;XP_011524710.1;XP_02 | 19978 | . | . | . | 6.209306 | 7.144482 | 5.529678 | 7.791148 | 14.449309 | 16.514789 | 12.408437 | 17.813534 | 1.028599 | 444.765233 | 0.19356417 | 0.210163943 | 0.833539721 | 0.998865404 | 8.777068 | 8.816464 | 8.755747 | 8.814679 | 8.712986 | 8.885277 | 481 | 496 | 337 | 482 |
| 112752 | NM_00110256 | IFT43 | intraflagellar transport 43 | protein_coding | NP_001096034.1;NP_001242924.1;NP_443105.2 | 29669 | 614068 | . | . | 10.903539 | 13.798150 | 11.117431 | 13.518523 | 25.372980 | 31.895040 | 24.947192 | 30.908498 | -1.003352 | 324.132508 | 0.208201845 | -0.02318486 | 0.981502819 | 0.998865404 | 8.343489 | 8.337384 | 8.277812 | 8.397585 | 8.299321 | 8.374201 | 334 | 388 | 262 | 324 |
| 112755 | NM_052874,XI | STX1B | syntaxin 1B | protein_coding | NP_443106.1;XP_016878382.1 | 18539 | 601485 | . | . | 0.142772 | 0.231410 | 0.114462 | 0.154098 | 0.332236 | 0.534915 | 0.256849 | 0.352328 | -1.408846 | 23.912558 | 0.647988571 | -0.76315281 | 0.445372271 | NA | 4.807371 | 4.309235 | 4.570773 | 4.623273 | 4.553180 | 4.574069 | 25 | 37 | 15 | 21 |
| 112770 | NM_00125660 | GLMP | glycosylated lysosomal membrane protein | protein_coding | NP_001243533.1;NP_001243534.1;NP_001243537.1;NP_001243538.1;NI | 29436 | . | . | . | 120.560767 | 113.950591 | 125.046696 | 124.830437 | 280.549855 | 263.401900 | 280.601172 | 285.409883 | 1.062764 | 5950.446821 | 0.104344743 | 0.841649898 | 0.39998395 | 0.998865404 | 12.494185 | 12.582042 | 12.527136 | 12.497716 | 12.568382 | 12.559176 | 6760 | 6116 | 5361 | 5717 |
| 112812 | NM_00103173 | FDX2 | ferredoxin 2 | protein_coding | NP_001026904.2 | 30546 | 614585 | . | . | 11.435858 | 12.244893 | 11.000557 | 13.808262 | 26.611710 | 28.304619 | 24.684931 | 31.570950 | 1.035488 | 382.587683 | 0.19401501 | 0.259312453 | 0.795394172 | 0.998865404 | 8.554829 | 8.604045 | 8.550034 | 8.585417 | 8.523274 | 8.649122 | 420 | 418 | 302 | 404 |
| 112840 | NM_00100872 | WDR89 | WD repeat domain 89 | protein_coding | NP_001008726.1;NP_001245201.1;NP_001369352.1;NP_001369353.1;NI | 20489 | . | . | . | 2.219660 | 2.264130 | 2.063145 | 1.584496 | 5.165241 | 5.233635 | 4.629640 | 3.622763 | -1.177187 | 209.377341 | 0.24357864 | -0.96619023 | 0.333949002 | 0.998865404 | 7.822138 | 7.588322 | 7.751098 | 7.738065 | 7.726951 | 7.608851 | 264 | 241 | 190 | 156 |
| 112846 | gene-RBM8B | RBM8B | RNA binding motif protein 8B pseudogene | pseudogene | . | 13426 | . | . | . | 1.204570 | 1.267924 | 1.043196 | 1.189450 | 2.803084 | 2.930863 | 2.340902 | 2.719537 | -1.114742 | 103.373099 | 0.316309452 | -0.49543253 | 0.62029484 | 0.998865404 | 6.768368 | 6.610764 | 6.702764 | 6.715823 | 6.658279 | 6.692484 | 123 | 120 | 80 | 97 |
| 112849 | NM_00133115 | L3HYPDH | trans-L-3-hydroxyproline dehydratase | protein_coding | NP_001318087.1;NP_001318088.1;NP_001318093.1;NP_653182.1;XP_02 | 20488 | 614811 | . | . | 8.848182 | 7.744524 | 8.351009 | 7.385581 | 20.590085 | 17.901812 | 18.739420 | 16.886252 | 1.075033 | 613.446822 | 0.162024301 | 0.644231932 | 0.519425048 | 0.998865404 | 9.207322 | 9.312356 | 9.272518 | 9.195991 | 9.312193 | 9.253693 | 720 | 601 | 577 | 570 |
| 112858 | NM_033550 | TP53RK | TP53 regulating kinase | protein_coding | NP_291028.3 | 16197 | 608679 | . | . | 3.926414 | 4.152299 | 3.990075 | 4.731977 | 9.136928 | 9.598226 | 8.953613 | 10.819104 | 1.066140 | 416.580943 | 0.183982833 | 0.502207046 | 0.615521887 | 0.998865404 | 8.655946 | 8.747507 | 8.666769 | 8.696459 | 8.670795 | 8.767140 | 453 | 446 | 344 | 435 |
| 112869 | NM_138414,XI | SGF29 | SAGA complex associated factor 29 | protein_coding | NP_612423.1;XP_016878383.1 | 25156 | 613374 | . | . | 4.222060 | 4.880551 | 4.956094 | 5.033527 | 9.824907 | 11.281610 | 11.121331 | 11.508562 | 1.115160 | 180.115649 | 0.250272244 | 0.628318075 | 0.529795593 | 0.998865404 | 7.412288 | 7.569017 | 7.433282 | 7.498615 | 7.517956 | 7.508319 | 181 | 198 | 167 | 176 |
| 112885 | NM_00113586 | PHF21B | PHD finger protein 21B | protein_coding | NP_001129334.1;NP_001229379.1;NP_001271225.1;NP_612424.1;XP_00 | 25161 | 616727 | . | . | 1.548210 | 2.389046 | 2.275591 | 1.638867 | 3.602751 | 5.522391 | 5.106359 | 3.747074 | -1.058118 | 203.932251 | 0.26982339 | -0.30204938 | 0.76261442 | 0.998865404 | 7.711956 | 7.630795 | 7.591506 | 7.757019 | 7.720699 | 7.578026 | 199 | 266 | 200 | 156 |
| 112936 | NM_052875,XI | VPS26B | VPS26, retromer complex component B | protein_coding | NP_443107.1;XP_011540867.1 | 28119 | 610027 | . | . | 7.545832 | 7.918794 | 7.326224 | 7.656086 | 17.559463 | 18.304648 | 16.439835 | 17.504732 | -1.050377 | 846.342053 | 0.143411636 | -0.49443418 | 0.62099958 | 0.998865404 | 9.760251 | 9.689065 | 9.726714 | 9.757855 | 9.690749 | 9.723618 | 977 | 956 | 701 | 792 |
| 112937 | NM_00108040 | GLB1L3 | galactosidase beta 1 like 3 | protein_coding | NP_001073876.2;XP_011540868.1;XP_011540869.1;XP_011540870.1;XP_ | 25147 | . | . | . | 0.039392 | 0.063819 | 0.093315 | 0.083214 | 0.091666 | 0.147521 | 0.209396 | 0.190260 | 1.623411 | 14.800457 | 0.805764849 | 0.867533531 | 0.385649748 | NA | 3.497331 | 4.194412 | 3.845786 | 3.869830 | 3.893627 | 3.883523 | 10 | 15 | 17 | 16 |
| 112939 | NM_052876,XI | NACC1 | nucleus accumbens associated 1 | protein_coding | NP_443108.1;XP_005259778.1 | 20967 | 610672 | . | . | 12.703776 | 13.492782 | 12.238707 | 12.948299 | 29.562210 | 31.189171 | 27.463302 | 29.604749 | -1.056521 | 1819.236258 | 0.122017608 | -0.65008491 | 0.515637376 | 0.998865404 | 10.868334 | 10.788805 | 10.828491 | 10.871707 | 10.786783 | 10.826708 | 2094 | 2072 | 1499 | 1701 |
| 112942 | NM_00128276 | CFAP36 | cilia and flagella associated protein 36 | protein_coding | NP_001269690.1;NP_542398.3;XP_011530801.1;XP_016858743.1;XP_01 | 30540 | . | . | . | 37.965735 | 37.125867 | 39.921674 | 45.437267 | 88.347819 | 85.818104 | 89.583081 | 103.886914 | 1.116788 | 1504.776042 | 0.129086144 | 1.234490744 | 0.217020094 | 0.982008277 | 10.473564 | 10.632715 | 10.521409 | 10.499291 | 10.551981 | 10.637740 | 1661 | 1512 | 1305 | 1570 |
| 112950 | NM_00100165 | MED8 | mediator complex subunit 8 | protein_coding | NP_001001653.1;NP_443109.2;NP_963836.2 | 19971 | 607956 | . | . | 18.338240 | 18.818446 | 18.232121 | 19.460159 | 42.673838 | 43.499675 | 40.912355 | 44.493345 | 1.004630 | 1123.783785 | 0.133068048 | 0.050078697 | 0.960059675 | 0.998865404 | 10.130909 | 10.137381 | 10.122993 | 10.141677 | 10.119380 | 10.151879 | 1275 | 1225 | 958 | 1079 |
| 112970 | NM_138417 | KTI12 | KTI12 chromatin associated homolog | protein_coding | NP_612426.1 | 25160 | . | . | . | 6.462159 | 6.006504 | 4.102075 | 6.203657 | 15.037708 | 13.884301 | 9.204937 | 14.183924 | -1.226326 | 303.448551 | 0.229484776 | -1.28262181 | 0.199624605 | 0.96621494 | 8.385644 | 8.089835 | 8.317797 | 8.270369 | 8.078826 | 8.279730 | 401 | 346 | 190 | 306 |
| 113000 | NM_00132408 | RPUSD1 | RNA pseudouridine synthase domain containing 1 | protein_coding | NP_001311015.1;NP_001311339.1;NP_001311340.1;NP_001311341.1;NI | 14173 | . | . | . | 13.499879 | 13.647613 | 12.945976 | 12.788317 | 31.414774 | 31.547073 | 29.050396 | 29.238970 | -1.047532 | 806.184631 | 0.144504585 | -0.46361483 | 0.64292373 | 0.998865404 | 9.687993 | 9.621166 | 9.672838 | 9.669491 | 9.652805 | 9.623849 | 952 | 888 | 699 | 723 |
| 113026 | NM_133373,XI | PLCD3 | phospholipase C delta 3 | protein_coding | NP_588614.1;XP_011522555.1;XP_024306322.1 | 9061 | 608795 | . | . | 1.542737 | 1.361471 | 1.129857 | 1.817251 | 3.590010 | 3.147101 | 2.535367 | 4.154927 | -1.060343 | 244.368635 | 0.223576809 | -0.37808383 | 0.705368318 | 0.998865404 | 7.975072 | 7.889490 | 7.940502 | 7.953481 | 7.872264 | 7.961652 | 286 | 275 | 187 | 243 |
| 113115 | NM_00109928 | MTFR2 | mitochondrial fission regulator 2 | protein_coding | NP_001092756.1;NP_001305667.1;NP_612428.2;XP_011533712.1;XP_01 | 21115 | . | . | . | 5.943533 | 5.799942 | 4.819602 | 5.580735 | 13.830844 | 13.406826 | 10.815048 | 12.759688 | -1.108854 | 305.860394 | 0.200925787 | -0.74191552 | 0.458138524 | 0.998865404 | 8.329492 | 8.180101 | 8.295839 | 8.272765 | 8.205168 | 8.251281 | 378 | 340 | 241 | 284 |
| 113130 | NM_080668,XI | CDCA5 | cell division cycle associated 5 | protein_coding | NP_542399.1;XP_005273790.1;XP_005273791.1;XP_011543045.1;XP_01 | 14626 | 609374 | . | . | 18.888589 | 19.089754 | 16.929811 | 21.063871 | 43.954525 | 44.126823 | 37.990004 | 48.160035 | -1.017432 | 1460.410041 | 0.140066827 | -0.17800514 | 0.858718941 | 0.998865404 | 10.524796 | 10.499408 | 10.516555 | 10.518769 | 10.421966 | 10.580338 | 1694 | 1592 | 1129 | 1497 |
| 113146 | NM_00135092 | AHNAK2 | AHNAK nucleoprotein 2 | protein_coding | NP_001337858.1;NP_612429.2;XP_024305231.1 | 20125 | 608570 | . | . | 0.698062 | 0.715811 | 0.719085 | 0.598297 | 1.624419 | 1.654629 | 1.613606 | 1.367934 | -1.090776 | 391.210739 | 0.18996204 | -0.6598968 | 0.509320061 | 0.998865404 | 8.672723 | 8.548194 | 8.630805 | 8.640737 | 8.638799 | 8.527579 | 466 | 444 | 357 | 317 |
| 113157 | NR_002775 | RPLP0P2 | ribosomal protein lateral stalk subunit P0 pseudogene 2 | transcribed_ps | . | 17960 | . | . | . | 0.125587 | 0.079502 | 0.142526 | 0.122168 | 0.292246 | 0.183772 | 0.319825 | 0.279323 | 1.244186 | 13.499319 | 0.838803626 | 0.375776302 | 0.707083207 | NA | 3.586822 | 3.905270 | 3.755469 | 3.730639 | 3.760663 | 3.751320 | 17 | 10 | 14 | 13 |
| 113174 | NM_138421 | SAAL1 | serum amyloid A like 1 | protein_coding | NP_612430.2 | 25158 | . | . | . | 13.487001 | 12.705584 | 12.456331 | 11.326755 | 31.384808 | 29.369526 | 27.951647 | 25.897278 | -1.116543 | 613.036567 | 0.16051938 | -0.99077512 | 0.321795402 | 0.998865404 | 9.336871 | 9.178441 | 9.318188 | 9.271444 | 9.255000 | 9.187349 | 770 | 674 | 530 | 515 |
| 113177 | NM_00103173 | IZUMO4 | IZUMO family member 4 | protein_coding | NP_001026905.2;NP_001034935.1;NP_001350517.1;XP_024307110.1;XP | 26950 | . | . | . | 0.330836 | 0.393927 | 0.452857 | 0.360464 | 0.769869 | 0.910581 | 1.016198 | 0.824157 | 1.111545 | 18.986223 | 0.704690589 | 0.216500822 | 0.828597385 | NA | 4.168403 | 4.321310 | 4.232339 | 4.245430 | 4.255070 | 4.239771 | 19 | 21 | 19 | 17 |
| 113178 | NM_00132953 | SCAMP4 | secretory carrier membrane protein 4 | protein_coding | NP_001316468.1;NP_001316469.1;NP_524558.1 | 30385 | 613764 | . | . | 15.831176 | 17.576701 | 18.801798 | 17.172590 | 36.839796 | 40.629331 | 42.190689 | 39.263086 | 1.062959 | 1348.731481 | 0.134071629 | 0.657004601 | 0.511177958 | 0.998865404 | 10.352636 | 10.440792 | 10.335931 | 10.408291 | 10.456900 | 10.380385 | 1434 | 1478 | 1269 | 1238 |
| 113179 | NM_00132953 | ADAT3 | adenosine deaminase tRNA specific 3 | protein_coding | NP_001316462.1;NP_612431.2 | 25151 | 615302 | . | . | 4.278241 | 3.800102 | 4.816988 | 4.682261 | 9.955643 | 8.784106 | 10.809183 | 10.705434 | 1.113524 | 213.316099 | 0.232121073 | 0.668323827 | 0.503926907 | 0.998865404 | 7.656852 | 7.812643 | 7.735869 | 7.682382 | 7.767867 | 7.751208 | 246 | 205 | 199 | 207 |
| 113189 | NM_130468 | CHST14 | carbohydrate sulfotransferase 14 | protein_coding | NP_569735.1 | 24464 | 608429 | . | . | 12.867656 | 13.360864 | 10.613968 | 11.590799 | 29.943562 | 30.884233 | 23.817436 | 26.500984 | -1.197172 | 821.453644 | 0.145536617 | -1.78394936 | 0.074431894 | 0.764716896 | 9.806185 | 9.546188 | 9.731751 | 9.752705 | 9.588131 | 9.640747 | 1016 | 980 | 625 | 728 |
| 113201 | NM_138423,N | GOLM2 | golgi membrane protein 2 | protein_coding | NP_612432.2;NP_816929.1;XP_016877369.1 | 24892 | . | . | . | 10.015708 | 9.454938 | 8.774893 | 9.234049 | 23.306963 | 21.855514 | 19.690607 | 21.112555 | -1.092340 | 1149.486514 | 0.133751049 | -0.95268425 | 0.340750076 | 0.998865404 | 10.229071 | 10.101665 | 10.223828 | 10.173256 | 10.118604 | 10.146915 | 1428 | 1252 | 937 | 1050 |
| 113220 | NM_138424,XI | KIF12 | kinesin family member 12 | protein_coding | NP_612433.1;XP_005251740.1;XP_006717010.1;XP_024303172.1;XP_02 | 21495 | 611278 | . | . | 0.483893 | 0.879059 | 0.711272 | 0.836216 | 1.126041 | 2.031983 | 1.596073 | 1.911906 | -1.016400 | 50.161433 | 0.462308597 | -0.05076367 | 0.959513844 | 0.998865404 | 5.661494 | 5.635401 | 5.597545 | 5.691204 | 5.633200 | 5.651704 | 44 | 68 | 41 | 49 |
| 113230 | NM_00129129 | MISP3 | MISP family member 3 | protein_coding | NP_001278220.1 | 26963 | . | . | . | 0.540723 | 0.665361 | 0.453580 | 0.451268 | 1.258285 | 1.538013 | 1.017820 | 1.031770 | -1.352648 | 25.403470 | 0.613795832 | -0.70998578 | 0.477712957 | NA | 4.868663 | 4.432387 | 4.676055 | 4.694905 | 4.655387 | 4.652973 | 31 | 34 | 19 | 20 |
| 113235 | NM_00124236 | SLC46A1 | solute carrier family 46 member 1 | protein_coding | NP_001229295.1;NP_542400.2;XP_005277843.1;XP_016879599.1 | 30521 | 611672 | . | . | 3.565593 | 3.784601 | 3.530248 | 3.727431 | 8.297282 | 8.748275 | 7.921773 | 8.522328 | -1.026641 | 739.086777 | 0.149028199 | -0.25452615 | 0.799089105 | 0.998865404 | 9.548593 | 9.510353 | 9.520344 | 9.556635 | 9.504846 | 9.535547 | 840 | 829 | 620 | 699 |
| 113246 | NM_00130183 | C12orf57 | chromosome 12 open reading frame 57 | protein_coding | NP_001288763.1;NP_001288765.1;NP_001288766.1;NP_001288767.1;NI | 29521 | 615140 | . | . | 28.916777 | 30.091620 | 31.356507 | 35.604915 | 67.290529 | 69.558126 | 70.363094 | 81.406412 | 1.103372 | 570.091513 | 0.163398072 | 0.868550201 | 0.385093203 | 0.998865404 | 9.082608 | 9.224027 | 9.115319 | 9.126475 | 9.146228 | 9.222941 | 619 | 590 | 490 | 593 |
| 113251 | NM_00117080 | LARP4 | La ribonucleoprotein 4 | protein_coding | NP_001164274.1;NP_001164275.1;NP_001164279.1;NP_001317344.1;NI | 24320 | 618657 | . | . | 9.787266 | 9.643697 | 9.079416 | 7.981366 | 22.775377 | 22.291839 | 20.373949 | 18.248445 | -1.160529 | 1728.058265 | 0.126466808 | -1.69833218 | 0.089445084 | 0.802216049 | 10.858183 | 10.643727 | 10.825141 | 10.794714 | 10.735563 | 10.649521 | 2178 | 1965 | 1469 | 1416 |
| 113263 | NM_138426 | GLCCI1 | glucocorticoid induced 1 | protein_coding | NP_612435.1 | 18713 | 614283 | . | . | 2.065663 | 1.796075 | 2.750208 | 1.873640 | 4.806882 | 4.151709 | 6.171388 | 4.283855 | 1.179096 | 314.035555 | 0.229688846 | 1.034796233 | 0.300764083 | 0.998865404 | 8.170095 | 8.409549 | 8.278123 | 8.200223 | 8.433760 | 8.215015 | 356 | 288 | 353 | 257 |
| 113277 | NM_00129158 | TMEM106A | transmembrane protein 106A | protein_coding | NP_001278515.1;NP_001278516.1;NP_001278517.1;NP_659478.1 | 28288 | . | . | . | 2.845175 | 3.314490 | 3.138748 | 3.484711 | 6.620839 | 7.661592 | 7.043258 | 7.967378 | 1.068394 | 319.492585 | 0.199071156 | 0.479446408 | 0.631621085 | 0.998865404 | 8.271488 | 8.366234 | 8.269060 | 8.330161 | 8.318893 | 8.353487 | 336 | 352 | 278 | 319 |
| 113402 | NM_145169,N | SFT2D1 | SFT2 domain containing 1 | protein_coding | NP_660152.1 | 21102 | . | . | . | 23.660785 | 22.791267 | 22.113762 | 19.680962 | 55.059618 | 52.683037 | 49.622638 | 44.998185 | -1.128535 | 712.429973 | 0.154850639 | -1.12657829 | 0.259920824 | 0.995760118 | 9.560908 | 9.387067 | 9.532684 | 9.499600 | 9.475854 | 9.388528 | 890 | 796 | 620 | 587 |
| 113419 | NM_144582 | TEX261 | testis expressed 261 | protein_coding | NP_653183.2 | 30712 | 618562 | . | . | 35.963493 | 39.445690 | 40.627159 | 39.933010 | 83.688522 | 91.180473 | 91.166176 | 91.302078 | 1.053853 | 4086.893790 | 0.111039527 | 0.681504443 | 0.495552356 | 0.998865404 | 11.958477 | 12.034109 | 11.939099 | 12.008361 | 12.029125 | 12.006388 | 4391 | 4474 | 3697 | 3878 |
| 113444 | NM_00116482 | SMIM12 | small integral membrane protein 12 | protein_coding | NP_001158296.1;NP_001158297.1;NP_001307190.1;NP_612437.3;XP_00 | 25154 | . | . | . | 5.016953 | 5.122651 | 5.494313 | 6.085346 | 11.674656 | 11.841233 | 12.329082 | 13.913416 | 1.115897 | 967.119230 | 0.14132247 | 1.119455896 | 0.262945696 | 0.995917097 | 9.836470 | 9.994315 | 9.872089 | 9.878720 | 9.913658 | 9.995204 | 1048 | 991 | 838 | 1009 |
| 113451 | NM_00129356 | AZIN2 | antizyme inhibitor 2 | protein_coding | NP_001280491.1;NP_001288752.1;NP_001288753.1;NP_001288754.1;NI | 29957 | 608353 | . | . | 0.831493 | 1.068025 | 1.161073 | 1.108499 | 1.934919 | 2.468788 | 2.605415 | 2.534453 | -1.024690 | 113.128806 | 0.304994068 | -0.11536911 | 0.908152564 | 0.998865404 | 6.839000 | 6.804438 | 6.851585 | 6.799359 | 6.817171 | 6.818418 | 142 | 114 | 97 | 105 |
| 113452 | NM_00132972 | TMEM54 | transmembrane protein 54 | protein_coding | NP_001316651.1;NP_001316652.1;NP_001316653.1;NP_001316654.1;NI | 24143 | . | . | . | 17.012974 | 19.101953 | 16.625479 | 18.550474 | 39.589891 | 44.155015 | 37.307093 | 42.413458 | -1.036609 | 573.823603 | 0.165385987 | -0.313644 | 0.753791434 | 0.998865404 | 9.190511 | 9.137937 | 9.139718 | 9.210477 | 9.118549 | 9.184311 | 637 | 664 | 465 | 555 |
| 113510 | NM_00129775 | HELQ | helicase, POLQ like | protein_coding | NP_001284684.2;NP_001284685.1;NP_001284686.1;NP_001284687.2;NI | 18536 | 606769 | . | . | 3.013217 | 3.483325 | 3.426340 | 3.007625 | 7.011878 | 8.051862 | 7.688606 | 6.876579 | -1.072374 | 316.541816 | 0.203003451 | -0.49658575 | 0.619481204 | 0.998865404 | 8.355772 | 8.254973 | 8.282807 | 8.363969 | 8.312453 | 8.258650 | 350 | 379 | 279 | 272 |
| 113540 | NM_052999,N | CMTM1 | CKLF like MARVEL transmembrane domain containing 1 | protein_coding | NP_443725.3;NP_851785.2;NP_851786.1;NP_851787.1;NP_851788.1;NP_ | 19172 | 607884 | . | . | 0.633250 | 0.671765 | 0.373032 | 0.651455 | 1.473599 | 1.552814 | 0.837072 | 1.489475 | -1.276831 | 19.982349 | 0.702574919 | -0.5018218 | 0.615792877 | NA | 4.487990 | 4.131331 | 4.331447 | 4.337550 | 4.293588 | 4.333941 | 25 | 25 | 11 | 21 |
| 113612 | NM_183075,XI | CYP2U1 | cytochrome P450 family 2 subfamily U member 1 | protein_coding | NP_898898.1;XP_005262774.1;XP_005262777.1 | 20582 | 610670 | . | . | 0.770131 | 0.656797 | 0.894482 | 0.703697 | 1.792128 | 1.518213 | 2.007191 | 1.608919 | 1.168956 | 110.311706 | 0.313161274 | 0.719184273 | 0.472027388 | 0.998865404 | 6.667583 | 6.894396 | 6.765010 | 6.744169 | 6.845180 | 6.763882 | 121 | 106 | 116 | 98 |
| 113622 | NM_138430,N | ADPRHL1 | ADP-ribosylhydrolase like 1 | protein_coding | NP_612439.2;NP_954631.1 | 21303 | 610620 | . | . | 0.277321 | 0.278965 | 0.173826 | 0.215805 | 0.645336 | 0.644840 | 0.390060 | 0.493411 | -1.354427 | 15.569629 | 0.779297694 | -0.56163773 | 0.574362867 | NA | 4.163717 | 3.724273 | 3.977515 | 3.974605 | 3.948597 | 3.962973 | 21 | 19 | 10 | 14 |
| 113655 | NM_138431,N | MFSD3 | major facilitator superfamily domain containing 3 | protein_coding | NP_612440.1;XP_011515108.1;XP_016868494.1 | 25157 | . | . | . | 12.165022 | 10.372784 | 12.247310 | 12.698883 | 28.308505 | 23.977152 | 27.482608 | 29.034489 | 1.184285 | 696.278842 | 0.156954844 | 1.55469144 | 0.120019574 | 0.859972379 | 9.316580 | 9.560191 | 9.391376 | 9.370056 | 9.443475 | 9.547343 | 746 | 677 | 608 | 760 |
| 113675 | NM_00130499 | SDSL | serine dehydratase like | protein_coding | NP_001291922.1;NP_612441.1;XP_005253888.1;XP_011536148.1;XP_01 | 30404 | . | . | . | 2.191755 | 2.433516 | 2.565515 | 2.654811 | 5.100305 | 5.625181 | 5.756943 | 6.069910 | 1.091999 | 103.312876 | 0.315657396 | 0.402245999 | 0.687502992 | 0.998865404 | 6.626345 | 6.752644 | 6.659481 | 6.690780 | 6.696566 | 6.708646 | 108 | 112 | 92 | 103 |
| 113691 | NR_003608 | TUBA3FP | tubulin alpha 3f pseudogene | transcribed_ps | . | 24067 | . | . | . | 1.649264 | 2.071764 | 1.332926 | 1.533018 | 3.837904 | 4.788976 | 2.991047 | 3.505063 | -1.319042 | 101.203788 | 0.326750393 | -1.22261618 | 0.221474699 | 0.982008277 | 6.847790 | 6.446655 | 6.663018 | 6.738021 | 6.599453 | 6.633484 | 118 | 138 | 71 | 87 |
| 113763 | NM_138434 | ZBED6CL | ZBED6 C-terminal like | protein_coding | NP_612443.1 | 21720 | 615252 | . | . | 4.162789 | 4.626182 | 4.279760 | 3.534737 | 9.686982 | 10.693626 | 9.603657 | 8.081758 | -1.137662 | 372.156766 | 0.195640806 | -0.95109173 | 0.341557815 | 0.998865404 | 8.629512 | 8.444066 | 8.545122 | 8.603067 | 8.554087 | 8.442536 | 434 | 448 | 333 | 294 |
| 113791 | NM_00113591 | PIK3IP1 | phosphoinositide-3-kinase interacting protein 1 | protein_coding | NP_001129383.1;NP_443112.2 | 24942 | . | . | . | 0.127781 | 0.102815 | 0.079744 | 0.065793 | 0.297352 | 0.237661 | 0.178942 | 0.150427 | -1.536839 | 7.748797 | 1.097646628 | -0.56481378 | 0.572200436 | NA | 3.229742 | 2.612728 | 2.972974 | 2.963332 | 2.956958 | 2.949909 | 12 | 9 | 6 | 5 |
| 113802 | NM_00110259 | HENMT1 | HEN methyltransferase 1 | protein_coding | NP_001096062.1;NP_653185.2;XP_005270468.1;XP_016855668.1 | 26400 | 612178 | . | . | 29.304754 | 28.914099 | 32.243675 | 36.254202 | 68.193369 | 66.836236 | 72.353874 | 82.890928 | 1.163456 | 1668.334327 | 0.123906736 | 1.762745749 | 0.077943381 | 0.768496883 | 10.590934 | 10.809204 | 10.646160 | 10.636745 | 10.729586 | 10.789014 | 1787 | 1654 | 1502 | 1745 |
| 113829 | NM_080670 | SLC35A4 | solute carrier family 35 member A4 | protein_coding | NP_542401.1 | 20753 | . | . | . | 23.790680 | 26.401690 | 24.459406 | 28.574133 | 55.361889 | 61.028690 | 54.886196 | 65.331360 | 1.042156 | 2357.229258 | 0.127347294 | 0.467789198 | 0.639935346 | 0.998865404 | 11.172958 | 11.232186 | 11.146094 | 11.222667 | 11.157871 | 11.275234 | 2532 | 2610 | 1940 | 2419 |
| 113835 | NM_00131699 | ZNF257 | zinc finger protein 257 | protein_coding | NP_001303925.1;NP_001303926.1;NP_001303927.1;NP_258429.2 | 13498 | 606957 | . | . | 1.700307 | 1.665008 | 2.162364 | 1.596805 | 3.956683 | 3.848742 | 4.852280 | 3.650906 | 1.089242 | 218.708297 | 0.242540439 | 0.508469965 | 0.611123797 | 0.998865404 | 7.709103 | 7.833926 | 7.755919 | 7.740387 | 7.857105 | 7.715045 | 245 | 222 | 228 | 182 |
| 113878 | NM_00110259 | DTX2 | deltex E3 ubiquitin ligase 2 | protein_coding | NP_001096064.1;NP_001096065.1;NP_001096066.1;NP_065943.2;XP_00 | 15973 | 613141 | . | . | 4.186678 | 3.611310 | 4.808500 | 4.586084 | 9.742571 | 8.347702 | 10.790132 | 10.485541 | 1.192954 | 350.678876 | 0.190154864 | 1.338587653 | 0.180704948 | 0.952541638 | 8.320892 | 8.575870 | 8.416628 | 8.381583 | 8.512005 | 8.487068 | 380 | 334 | 339 | 350 |
| 114034 | NM_025077,XI | TOE1 | target of EGR1, exonuclease | protein_coding | NP_079353.3;XP_005270469.1;XP_005270470.1;XP_011538871.1;XP_02 | 15954 | 613931 | . | . | 11.273174 | 13.336790 | 13.037832 | 12.182295 | 26.233141 | 30.828596 | 29.256516 | 27.853369 | -1.003461 | 808.827975 | 0.15432516 | -0.03229992 | 0.974232877 | 0.998865404 | 9.662162 | 9.657212 | 9.607375 | 9.709452 | 9.696377 | 9.616576 | 866 | 937 | 742 | 714 |
| 114036 | NR_027266,NF | LINC00310 | long intergenic non-protein coding RNA 310 | lncRNA | . | 16414 | . | . | . | 0.452940 | 0.631178 | 0.819141 | 0.599705 | 1.054011 | 1.458996 | 1.838128 | 1.371156 | 1.231656 | 16.182672 | 0.792375978 | 0.379365099 | 0.704416764 | NA | 3.858702 | 4.158506 | 3.974771 | 4.019339 | 4.028339 | 3.995012 | 11 | 21 | 19 | 13 |
| 114043 | NR_026547,NF | TSPEAR-AS2 | TSPEAR antisense RNA 2 | lncRNA | . | 16428 | . | . | . | 0.323861 | 0.683718 | 0.411969 | 0.595826 | 0.753637 | 1.580444 | 0.924445 | 1.362287 | 1.015475 | 16.401144 | 0.778659852 | 0.028452673 | 0.977301114 | NA | 4.027016 | 4.044381 | 4.008836 | 4.053310 | 4.021036 | 4.043742 | 13 | 23 | 12 | 18 |
| 114044 | NR_002776,NF | MCM3AP-AS1 | MCM3AP antisense RNA 1 | lncRNA | . | 16417 | . | . | . | 0.267009 | 0.507395 | 0.854914 | 0.612829 | 0.621340 | 1.172867 | 1.918402 | 1.401161 | 1.828135 | 43.489915 | 0.5063119 | 1.719044255 | 0.085609314 | 0.791792328 | 4.943947 | 5.812480 | 5.327822 | 5.403861 | 5.493138 | 5.435070 | 25 | 43 | 56 | 45 |
| 114049 | NM_00120256 | BUD23 | BUD23 rRNA methyltransferase and ribosome maturation factor | protein_coding | NP_001189489.1;NP_059998.2;XP_006715910.1;XP_011514080.1;XP_01 | 16405 | 615733 | . | . | 47.854359 | 50.363711 | 39.542175 | 43.318020 | 111.359063 | 116.417981 | 88.731497 | 99.041505 | -1.171053 | 2003.393008 | 0.11891844 | -1.91565387 | 0.055409168 | 0.697001222 | 11.077693 | 10.849774 | 11.007394 | 11.047436 | 10.908639 | 10.908856 | 2427 | 2390 | 1591 | 1745 |
| 114088 | NM_015163,N | TRIM9 | tripartite motif containing 9 | protein_coding | NP_055978.4;NP_443210.1;XP_005267358.1;XP_005267359.1;XP_00672 | 16288 | 606555 | . | . | 2.780913 | 3.241090 | 2.805180 | 3.149306 | 6.471298 | 7.491927 | 6.294744 | 7.200513 | -1.011714 | 463.439100 | 0.182241151 | -0.09219683 | 0.926541656 | 0.998865404 | 8.865088 | 8.847330 | 8.799746 | 8.914119 | 8.817622 | 8.883590 | 488 | 549 | 379 | 455 |
| 114112 | NM_00117351 | TXNRD3 | thioredoxin reductase 3 | protein_coding | NP_001166984.1;NP_443115.1 | 20667 | 606235 | . | . | 2.066501 | 2.081579 | 1.523576 | 2.077239 | 4.808834 | 4.811663 | 3.418860 | 4.749360 | -1.158462 | 175.296618 | 0.262792699 | -0.8075204 | 0.419366727 | 0.998865404 | 7.556593 | 7.342802 | 7.485868 | 7.480586 | 7.359806 | 7.478785 | 218 | 202 | 120 | 175 |
| 114134 | NM_052885,XI | SLC2A13 | solute carrier family 2 member 13 | protein_coding | NP_443117.3;XP_011536149.1;XP_011536151.1;XP_011536152.1;XP_01 | 15956 | 611036 | . | . | 1.252718 | 1.251520 | 1.259665 | 0.960145 | 2.915126 | 2.892942 | 2.826652 | 2.195255 | -1.188360 | 255.813112 | 0.235464202 | -1.05736773 | 0.290343793 | 0.998865404 | 8.117232 | 7.870089 | 8.054470 | 8.020043 | 8.030832 | 7.863579 | 330 | 290 | 239 | 181 |

| | | | | | | | | | | | | | | | | | | | | | | | | | | | | | | | | | | |
|---|---|---|---|---|---|---|---|---|---|---|---|---|---|---|---|---|---|---|---|---|---|---|---|---|---|---|---|---|---|---|---|---|---|---|
| 114294 | NM_00128858 | LACTB | lactamase beta | protein_coding | NP_001275514.1;NP_116246.2;NP_741982.1 | 16468 | 608440 | . | . | . | 3.837781 | 3.290639 | 4.057509 | 3.009090 | 8.930674 | 7.606460 | 9.104933 | 6.879927 | -1.039538 | 242.602527 | 0.2374091 | -0.23563866 | 0.813713071 | 0.998865404 | 7.949241 | 7.895153 | 7.976270 | 7.879013 | 7.973110 | 7.842329 | 312 | 241 | 235 | 193 |
| 114327 | NM_00117242 | EFHC1 | EF-hand domain containing 1 | protein_coding | NP_001165891.1;NP_060570.2 | 16406 | 608815 | . | . | . | 3.784169 | 3.883820 | 4.657324 | 3.876700 | 8.805918 | 8.977625 | 10.450898 | 8.863614 | 1.083773 | 980.850863 | 0.153905924 | 0.754115187 | 0.450780059 | 0.998865404 | 9.878392 | 9.995031 | 9.900807 | 9.909995 | 10.046888 | 9.874639 | 1077 | 1022 | 991 | 845 |
| 114548 | NM_00107982 | NLRP3 | NLR family pyrin domain containing 3 | protein_coding | NP_001073289.2;NP_001120933.2;NP_001120934.2;NP_001230062.1;NI | 16400 | 606416 | . | . | . | 0.432440 | 0.617957 | 0.218608 | 0.379801 | 1.006306 | 1.428435 | 0.490549 | 0.868370 | -1.851682 | 53.348789 | 0.45218826 | -1.96563291 | 0.049341029 | 0.66444315 | 6.115780 | 5.223024 | 5.756490 | 5.822230 | 5.645417 | 5.700747 | 69 | 85 | 26 | 42 |
| 114599 | NR_000025 | SNORD15B | small nucleolar RNA, C/D box 15B | snoRNA | . | 16649 | . | . | . | . | 2.506851 | 3.281624 | 2.375184 | 2.190471 | 5.833547 | 7.585621 | 5.329844 | 5.008252 | -1.258718 | 12.519816 | 0.864965412 | -0.38377818 | 0.701142859 | NA | 3.802857 | 3.470304 | 3.646807 | 3.662833 | 3.644132 | 3.640418 | 14 | 17 | 10 | 10 |
| 114609 | NM_00103966 | TIRAP | TIR domain containing adaptor protein | protein_coding | NP_001034750.1;NP_001305705.1;NP_001305706.1;NP_683708.1 | 17192 | 606252 | . | . | . | 2.438139 | 2.321824 | 2.521175 | 2.297674 | 5.673650 | 5.367001 | 5.657445 | 5.253359 | -1.004481 | 150.265753 | 0.272680417 | -0.02365693 | 0.981126264 | 0.998865404 | 7.233939 | 7.228801 | 7.239115 | 7.222368 | 7.268153 | 7.188485 | 177 | 159 | 144 | 126 |
| 114625 | NM_00101792 | ERMAP | erythroblast membrane associated protein (Scianna blood group) | protein_coding | NP_001017922.1;NP_061008.2;XP_006710376.1;XP_011538872.1;XP_01 | 15743 | 609017 | . | . | . | 0.989146 | 0.953107 | 1.250882 | 0.961004 | 2.301783 | 2.203148 | 2.806944 | 2.197224 | 1.136014 | 113.438997 | 0.312188016 | 0.589324402 | 0.555643679 | 0.998865404 | 6.729909 | 6.915662 | 6.809248 | 6.789561 | 6.888862 | 6.792073 | 126 | 111 | 120 | 97 |
| 114659 | NM_00132135 | LRRC37B | leucine rich repeat containing 37B | protein_coding | NP_001308279.1;NP_443120.2 | 29070 | 616558 | . | . | . | 1.526323 | 1.437638 | 1.812811 | 1.535231 | 3.551818 | 3.323165 | 4.067897 | 3.510122 | 1.113086 | 174.119312 | 0.252390461 | 0.612406331 | 0.540268954 | 0.998865404 | 7.364126 | 7.519554 | 7.436683 | 7.400678 | 7.483641 | 7.443972 | 198 | 170 | 167 | 164 |
| 114757 | NM_134268,XI | CYGB | cytoglobin | protein_coding | NP_599030.1;XP_005257062.1;XP_016879605.1 | 16505 | 608759 | . | . | . | 0.300223 | 0.301184 | 0.138696 | 0.332506 | 0.698633 | 0.696201 | 0.311229 | 0.760236 | -1.333661 | 17.225569 | 0.765583589 | -0.5425827 | 0.587417154 | NA | 4.301434 | 3.881002 | 4.122535 | 4.120151 | 4.076776 | 4.123183 | 23 | 21 | 8 | 19 |
| 114769 | NM_00101753 | CARD16 | caspase recruitment domain family member 16 | protein_coding | NP_001017534.1;NP_443121.1;XP_011540885.1 | 33701 | 615680 | . | . | . | 4.102877 | 6.152632 | 4.477069 | 5.796845 | 9.547564 | 14.222085 | 10.046414 | 13.253798 | -1.009336 | 131.321901 | 0.302846096 | -0.04426873 | 0.964690196 | 0.998865404 | 7.044796 | 7.029089 | 6.975151 | 7.091600 | 6.991792 | 7.070019 | 128 | 165 | 101 | 136 |
| 114780 | NM_00107678 | PKD1L2 | polycystin 1 like 2 (gene/pseudogene) | protein_coding | NP_001070248.1;NP_001265352.1;NP_001265354.1 | 21715 | 607894 | . | . | . | 0.190443 | 0.210297 | 0.074089 | 0.024264 | 0.443168 | 0.486111 | 0.166253 | 0.055476 | -1.895579 | 17.897763 | 0.778826278 | -1.18465269 | 0.236154796 | NA | 4.548173 | 3.632244 | 4.174182 | 4.195651 | 4.163338 | 4.111830 | 24 | 28 | 16 | 6 |
| 114781 | NM_00109927 | BTBD9 | BTB domain containing 9 | protein_coding | NP_001092742.1;NP_001165889.1;NP_443125.1;NP_689946.2;XP_01151 | 21228 | 611237 | . | . | . | 0.684423 | 0.526357 | 0.736848 | 0.695334 | 1.592678 | 1.216698 | 1.653466 | 1.589799 | 1.174925 | 180.827376 | 0.252827036 | 0.919872782 | 0.357639244 | 0.998865404 | 7.376981 | 7.610517 | 7.500830 | 7.415974 | 7.533746 | 7.523405 | 211 | 161 | 172 | 181 |
| 114785 | NM_052897,XI | MBD6 | methyl-CpG binding domain protein 6 | protein_coding | NP_443129.3;XP_005268674.1;XP_005268675.1;XP_005268677.1;XP_00 | 20445 | . | . | . | . | 9.402037 | 10.425410 | 12.298872 | 10.875889 | 21.878928 | 24.098803 | 27.598309 | 24.866425 | 1.160182 | 1402.002738 | 0.13495808 | 1.588278868 | 0.112223294 | 0.849708242 | 10.342054 | 10.556529 | 10.357799 | 10.425252 | 10.554480 | 10.455334 | 1428 | 1463 | 1393 | 1322 |
| 114787 | NM_052899 | GPRIN1 | G protein regulated inducer of neurite outgrowth 1 | protein_coding | NP_443131.2 | 24835 | 611239 | . | . | . | 7.207786 | 7.432099 | 6.540643 | 8.506350 | 16.772814 | 17.179630 | 14.677014 | 19.448759 | 1.013412 | 979.043638 | 0.156384973 | 0.122910856 | 0.902177689 | 0.998865404 | 9.925933 | 9.944467 | 9.920581 | 9.936429 | 9.842195 | 10.025760 | 1108 | 1061 | 749 | 1040 |
| 114789 | NM_00100664 | SLC25A25 | solute carrier family 25 member 25 | protein_coding | NP_001006642.1;NP_001006643.1;NP_001252543.1;NP_001317917.1;NI | 20663 | 608745 | . | . | . | 4.468606 | 4.378883 | 4.912312 | 3.915872 | 10.398631 | 10.121984 | 11.023086 | 8.953177 | -1.027889 | 473.556116 | 0.180258927 | -0.22015451 | 0.825750825 | 0.998865404 | 8.906644 | 8.867879 | 8.904037 | 8.884796 | 8.938927 | 8.812039 | 561 | 510 | 448 | 393 |
| 114790 | NM_052902,XI | STK11IP | serine/threonine kinase 11 interacting protein | protein_coding | NP_443134.3;XP_011508828.1;XP_011508829.1;XP_011508830.1;XP_01 | 19184 | 607172 | . | . | . | 15.235485 | 15.608211 | 16.481547 | 16.853014 | 35.453598 | 36.079082 | 36.984110 | 38.532411 | 1.070094 | 1791.841293 | 0.119881572 | 0.815283644 | 0.414909969 | 0.998865404 | 10.757545 | 10.855255 | 10.776869 | 10.782750 | 10.830670 | 10.835539 | 1986 | 1875 | 1606 | 1742 |
| 114791 | NM_00110261 | TUBGCP5 | tubulin gamma complex associated protein 5 | protein_coding | NP_001096080.1;NP_001341301.1;NP_001341302.1;NP_001341303.1;NI | 18600 | 608147 | . | . | . | 7.540040 | 7.840744 | 8.275966 | 7.271600 | 17.545985 | 18.124230 | 18.571026 | 16.625649 | 1.000131 | 908.131186 | 0.146453427 | 0.001289129 | 0.998971424 | 0.999862634 | 9.826479 | 9.827035 | 9.812258 | 9.839296 | 9.875857 | 9.773729 | 1026 | 998 | 846 | 791 |
| 114792 | NM_00128625 | KLHL32 | kelch like family member 32 | protein_coding | NP_001273179.1;NP_001273180.1;NP_001273181.1;NP_001273183.1;NI | 21221 | . | . | . | . | 0.186565 | 0.134298 | 0.299207 | 0.177875 | 0.434145 | 0.310435 | 0.671413 | 0.406689 | 1.382397 | 28.696521 | 0.587332769 | 0.795412408 | 0.426373644 | 0.998865404 | 4.588571 | 5.058876 | 4.833784 | 4.794848 | 4.865319 | 4.838594 | 33 | 21 | 32 | 28 |
| 114793 | NM_052905,XI | FMNL2 | formin like 2 | protein_coding | NP_443137.2;XP_005246320.1;XP_005246322.1;XP_011508832.1;XP_01 | 18267 | 616285 | . | . | . | 33.067709 | 34.405637 | 38.832376 | 34.067123 | 76.949924 | 79.530175 | 87.138723 | 77.890432 | 1.066724 | 6207.140868 | 0.113680162 | 0.819727814 | 0.41237129 | 0.998865404 | 12.552323 | 12.645596 | 12.557913 | 12.584242 | 12.683033 | 12.564912 | 6808 | 6583 | 5960 | 5589 |
| 114795 | NM_00128621 | TMEM132B | transmembrane protein 132B | protein_coding | NP_001273148.1;NP_001353783.1;NP_443139.2;XP_011536156.1;XP_01 | 29397 | . | . | . | . | 0.043524 | 0.031680 | 0.042487 | 0.018553 | 0.101283 | 0.073231 | 0.095339 | 0.042419 | -1.202843 | 13.541526 | 0.845786464 | -0.31502968 | 0.752739119 | NA | 3.884262 | 3.622519 | 3.768535 | 3.760721 | 3.769843 | 3.737340 | 18 | 15 | 14 | 8 |
| 114796 | NR_021487,NF | PSMG3-AS1 | PSMG3 antisense RNA 1 (head to head) | lncRNA | . | 22230 | . | . | . | . | 0.714128 | 0.793562 | 0.978063 | 0.876840 | 1.661804 | 1.834356 | 2.194743 | 2.004789 | 1.110803 | 133.946712 | 0.28036956 | 0.540724942 | 0.588697184 | 0.998865404 | 6.987964 | 7.139110 | 7.038947 | 7.049758 | 7.071676 | 7.093544 | 144 | 139 | 119 | 136 |
| 114799 | NM_052911,XI | ESCO1 | establishment of sister chromatid cohesion N-acetyltransferase 1 | protein_coding | NP_443143.2;XP_011524100.1;XP_011524101.1 | 24645 | 609674 | . | . | . | 3.712133 | 3.495475 | 3.718862 | 2.191218 | 8.638286 | 8.079945 | 8.345019 | 5.009960 | -1.222668 | 473.332014 | 0.232676309 | -1.24650553 | 0.212578859 | 0.975238634 | 9.023694 | 8.735338 | 8.971466 | 8.899772 | 8.966273 | 8.626725 | 632 | 531 | 475 | 287 |
| 114803 | NM_00108548 | MYSM1 | Myb like, SWIRM and MPN domains 1 | protein_coding | NP_001078956.1;XP_006710377.1;XP_011538875.1;XP_011538876.1 | 29401 | 612176 | . | . | . | 5.600864 | 6.115646 | 7.227168 | 4.746087 | 13.033443 | 14.136590 | 16.217557 | 10.851365 | 1.018701 | 1432.975208 | 0.180616703 | 0.147997764 | 0.882344533 | 0.998865404 | 10.471111 | 10.498356 | 10.439363 | 10.501180 | 10.644272 | 10.296667 | 1569 | 1593 | 1539 | 1053 |
| 114804 | NM_00133050 | RNF157 | ring finger protein 157 | protein_coding | NP_001317430.1;NP_443148.1;XP_005257064.1;XP_011522575.1;XP_01 | 29402 | . | . | . | . | 6.985653 | 7.389381 | 11.262533 | 12.304949 | 16.255904 | 17.080884 | 25.272796 | 28.133805 | 1.595295 | 1480.576222 | 0.129725572 | 5.194222118 | 2.05578E-07 | 8.10739E-05 | 10.156247 | 10.829741 | 10.300690 | 10.341997 | 10.640274 | 10.720340 | 1277 | 1266 | 1505 | 1790 |
| 114814 | NR_002328,NF | GNRHR2 | gonadotropin releasing hormone receptor 2 (pseudogene) | transcribed_ps | . | 16341 | 612875 | . | . | . | 1.446551 | 0.554619 | 1.029580 | 0.653709 | 3.366182 | 1.282026 | 2.310347 | 1.494629 | -1.378838 | 30.240983 | 0.597098726 | -0.77617503 | 0.437645646 | 0.998865404 | 5.130611 | 4.669578 | 4.984059 | 4.887434 | 4.897476 | 4.898315 | 54 | 25 | 22 | 24 |
| 114815 | NM_00101303 | SORCS1 | sortilin related VPS10 domain containing receptor 1 | protein_coding | NP_001013049.1;NP_001193498.1;NP_001193499.1;NP_001193500.1;NI | 16697 | 606283 | . | . | . | 3.695510 | 3.352152 | 4.055686 | 4.522764 | 8.599604 | 7.748648 | 9.100842 | 10.340762 | 1.163273 | 807.769581 | 0.152234833 | 1.43324772 | 0.151787057 | 0.914688406 | 9.544719 | 9.762658 | 9.626461 | 9.573398 | 9.653833 | 9.757706 | 893 | 775 | 701 | 873 |
| 114817 | gene-CSPG4P5 | CSPG4P5 | chondroitin sulfate proteoglycan 4 pseudogene 5 | pseudogene | . | 29403 | . | . | . | . | 0.026838 | 0.025562 | 0.010424 | 0.013627 | 0.062453 | 0.059087 | 0.023391 | 0.031157 | -1.620127 | 2.903507 | 1.757623934 | -0.39605015 | 0.692068032 | NA | 1.844134 | 1.148375 | 1.548992 | 1.550042 | 1.543603 | 1.542786 | 4 | 4 | 2 | 2 |
| 114818 | NM_052920,XI | KLHL29 | kelch like family member 29 | protein_coding | NP_443152.1;XP_006711992.1;XP_011530803.1;XP_016858753.1 | 29404 | . | . | . | . | 4.795471 | 5.571619 | 7.149829 | 7.009292 | 11.159259 | 12.879046 | 16.044011 | 16.025913 | 1.302049 | 968.489256 | 0.140476086 | 2.710665296 | 0.006714837 | 0.260350829 | 9.716789 | 10.097369 | 9.778517 | 9.847049 | 10.012027 | 9.999114 | 922 | 952 | 961 | 1018 |
| 114819 | NR_023386 | CROCCP3 | CROCC pseudogene 3 | transcribed_ps | . | 29405 | . | . | . | . | 1.175132 | 1.028734 | 1.363601 | 1.017802 | 2.734581 | 2.377966 | 3.059882 | 2.327083 | 1.063225 | 192.100701 | 0.257026974 | 0.34411355 | 0.730760893 | 0.998865404 | 7.539835 | 7.630188 | 7.597667 | 7.537071 | 7.659367 | 7.526459 | 229 | 187 | 198 | 158 |
| 114822 | NM_052924,XI | RHPN1 | rhophilin Rho GTPase binding protein 1 | protein_coding | NP_443156.2;XP_005250829.1;XP_005250830.1;XP_006716565.1;XP_01 | 19973 | 601031 | . | . | . | 9.061429 | 8.429827 | 9.783017 | 8.085562 | 21.086317 | 19.485922 | 21.952807 | 18.486677 | 1.007273 | 1113.072970 | 0.146045251 | 0.071590326 | 0.942927939 | 0.998865404 | 10.114785 | 10.125859 | 10.141720 | 10.089978 | 10.189237 | 10.049495 | 1321 | 1155 | 1063 | 949 |
| 114823 | NM_00137563 | LENG8 | leukocyte receptor cluster member 8 | protein_coding | NP_001362567.1;NP_001362568.1;NP_001362569.1;NP_001362570.1;NI | 15500 | 616575 | . | . | . | 34.551948 | 40.650367 | 44.935656 | 33.684863 | 80.403799 | 93.965140 | 100.834312 | 77.016444 | 1.042711 | 6747.873320 | 0.147480783 | 0.409131733 | 0.682442989 | 0.998865404 | 12.689689 | 12.750113 | 12.630563 | 12.761887 | 12.860381 | 12.589313 | 7016 | 7683 | 6942 | 5433 |
| 114825 | NM_00113086 | PWWP2A | PWWP domain containing 2A | protein_coding | NP_001124336.1;NP_001253964.1;NP_001336661.1;NP_001336662.1;NI | 29406 | 617823 | . | . | . | 3.362666 | 2.717585 | 3.967899 | 3.019928 | 7.825061 | 6.281823 | 8.903847 | 6.904705 | 1.095671 | 392.128046 | 0.205439209 | 0.641620768 | 0.521119445 | 0.998865404 | 8.547002 | 8.680283 | 8.619199 | 8.545119 | 8.725487 | 8.540095 | 458 | 378 | 412 | 325 |
| 114826 | NM_052928,XI | SMYD4 | SET and MYND domain containing 4 | protein_coding | NP_443160.2;XP_011521934.1;XP_011521935.1;XP_016879610.1;XP_02 | 21067 | . | . | . | . | 4.442340 | 5.217512 | 4.984244 | 5.869199 | 10.337505 | 12.060515 | 11.184500 | 13.419224 | 1.113351 | 706.986534 | 0.161562604 | 0.958817882 | 0.337650499 | 0.998865404 | 9.386432 | 9.540533 | 9.370434 | 9.477802 | 9.453448 | 9.543128 | 710 | 779 | 606 | 743 |
| 114827 | NM_052929,XI | FHAD1 | forkhead associated phosphopeptide binding domain 1 | protein_coding | NP_443161.1;XP_011538877.1;XP_011538878.1;XP_011538879.1;XP_01 | 29408 | . | . | . | . | 0.223763 | 0.188388 | 0.164462 | 0.335840 | 0.520709 | 0.435468 | 0.369048 | 0.767860 | 1.193910 | 36.716229 | 0.542174255 | 0.471609284 | 0.637205699 | 0.998865404 | 5.066064 | 5.319511 | 5.194485 | 5.163284 | 5.156639 | 5.250443 | 43 | 32 | 24 | 49 |
| 114876 | NM_00124250 | OSBPL1A | oxysterol binding protein like 1A | protein_coding | NP_001229437.1;NP_060500.3;NP_542164.2;XP_006722443.1;XP_00672 | 16398 | 606730 | . | . | . | 9.546117 | 9.827098 | 10.583289 | 11.004163 | 22.214207 | 22.715777 | 23.748594 | 25.159710 | 1.131166 | 1215.674819 | 0.131686554 | 1.350257028 | 0.176933551 | 0.949538805 | 10.155900 | 10.333708 | 10.218974 | 10.177987 | 10.274166 | 10.308520 | 1350 | 1197 | 1097 | 1237 |
| 114879 | NM_00114406 | OSBPL5 | oxysterol binding protein like 5 | protein_coding | NP_001137535.1;NP_065947.1;NP_663613.1;XP_011518175.1;XP_01687 | 16392 | 606733 | . | . | . | 2.286393 | 2.092934 | 1.965145 | 1.378139 | 5.320531 | 4.837910 | 4.409727 | 3.150952 | -1.312806 | 242.468952 | 0.234057864 | -1.67759351 | 0.09342648 | 0.809520218 | 8.103938 | 7.712967 | 7.990624 | 7.972737 | 7.914503 | 7.777436 | 322 | 292 | 210 | 167 |
| 114880 | NM_00120148 | OSBPL6 | oxysterol binding protein like 6 | protein_coding | NP_001188409.1;NP_001188410.1;NP_001188411.1;NP_115912.1;NP_6 | 16388 | 606734 | . | . | . | 4.071096 | 3.852870 | 4.390774 | 4.754691 | 9.473606 | 8.906075 | 9.852771 | 10.871033 | 1.057798 | 1053.184412 | 0.134768195 | 0.601509812 | 0.547500478 | 0.998865404 | 9.999514 | 10.080436 | 10.016225 | 10.022176 | 10.045943 | 10.075150 | 1174 | 1109 | 923 | 1035 |
| 114881 | NM_145798,XI | OSBPL7 | oxysterol binding protein like 7 | protein_coding | NP_665741.1;XP_016879611.1;XP_016879612.1;XP_016879613.1;XP_01 | 16387 | 606735 | . | . | . | 1.527183 | 1.607599 | 1.696820 | 1.558241 | 3.553817 | 3.716036 | 3.807615 | 3.562734 | 1.028979 | 181.634908 | 0.249675939 | 0.165069566 | 0.868889227 | 0.998865404 | 7.483792 | 7.525699 | 7.488567 | 7.505899 | 7.543077 | 7.474995 | 202 | 197 | 173 | 159 |
| 114882 | NM_00100371 | OSBPL8 | oxysterol binding protein like 8 | protein_coding | NP_001003712.1;NP_001306581.1;NP_001306582.1;NP_001306584.1;NI | 16396 | 606736 | . | . | . | 3.382571 | 3.130595 | 3.539124 | 2.840155 | 7.871382 | 7.236510 | 7.941691 | 6.493680 | -1.037310 | 723.857883 | 0.165476264 | -0.31935974 | 0.74945374 | 0.998865404 | 9.525291 | 9.473367 | 9.540602 | 9.478687 | 9.560567 | 9.403358 | 886 | 760 | 689 | 590 |
| 114883 | NM_00133058 | OSBPL9 | oxysterol binding protein like 9 | protein_coding | NP_001317509.1;NP_001337137.1;NP_001337138.1;NP_001337139.1;NI | 16386 | 606737 | . | . | . | 29.593718 | 32.347031 | 36.399844 | 32.771691 | 68.865798 | 74.771612 | 81.680196 | 74.928576 | 1.122100 | 3403.387665 | 0.115887019 | 1.434162119 | 0.151526004 | 0.914154052 | 11.647189 | 11.813531 | 11.678510 | 11.687061 | 11.825481 | 11.727639 | 3674 | 3479 | 3314 | 3176 |
| 114884 | NM_00117406 | OSBPL10 | oxysterol binding protein like 10 | protein_coding | NP_001167531.1;NP_060254.2;XP_005264900.1;XP_005264901.1;XP_00 | 16395 | 606738 | . | . | . | 0.725882 | 0.904826 | 1.042140 | 0.715932 | 1.689157 | 2.091544 | 2.338533 | 1.636894 | 1.172112 | 105.227671 | 0.326550995 | 0.701605924 | 0.482924956 | 0.998865404 | 6.597285 | 6.828224 | 6.686350 | 6.683833 | 6.790363 | 6.676773 | 112 | 104 | 116 | 88 |
| 114885 | NM_022776,XI | OSBPL11 | oxysterol binding protein like 11 | protein_coding | NP_073613.2 | 16397 | 606739 | . | . | . | 4.615548 | 4.194945 | 4.836427 | 4.961513 | 10.740568 | 9.696803 | 10.852802 | 11.343910 | 1.086002 | 663.399073 | 0.155768033 | 0.764126696 | 0.444791745 | 0.998865404 | 9.312896 | 9.432109 | 9.378256 | 9.309045 | 9.390281 | 9.410245 | 770 | 651 | 592 | 657 |
| 114897 | NM_030968,N | C1QTNF1 | C1q and TNF related 1 | protein_coding | NP_112230.1;NP_699203.1;NP_940995.1;NP_940996.1;XP_006721726.1 | 14324 | 610365 | . | . | . | 2.009996 | 1.488401 | 1.013584 | 1.712569 | 4.677342 | 3.440505 | 2.274454 | 3.915585 | -1.303389 | 137.115385 | 0.320209836 | -1.193805 | 0.232554257 | 0.988283993 | 7.278435 | 6.894604 | 7.201104 | 7.076648 | 6.958126 | 7.121252 | 206 | 142 | 78 | 139 |
| 114899 | NM_030945,N | C1QTNF3 | C1q and TNF related 3 | protein_coding | NP_112207.1;NP_852100.3 | 14326 | 612045 | . | . | . | 0.232154 | 0.304937 | 0.280763 | 0.402470 | 0.540232 | 0.704876 | 0.630023 | 0.920199 | 1.254629 | 41.047909 | 0.496990794 | 0.658484662 | 0.51022675 | 0.998865404 | 5.188027 | 5.512267 | 5.316850 | 5.351799 | 5.341212 | 5.398391 | 37 | 44 | 33 | 50 |
| 114904 | NM_00136587 | C1QTNF6 | C1q and TNF related 6 | protein_coding | NP_001352807.1;NP_114116.3;NP_872292.1;XP_011528159.1;XP_01688 | 14343 | 614910 | . | . | . | 0.702717 | 0.695885 | 0.500525 | 0.404748 | 1.635251 | 1.608569 | 1.123162 | 0.925407 | -1.344709 | 104.110822 | 0.323488517 | -1.32089303 | 0.186537034 | 0.95851159 | 6.899027 | 6.473688 | 6.776990 | 6.712631 | 6.680700 | 6.625171 | 150 | 117 | 85 | 75 |
| 114907 | NM_00124246 | FBXO32 | F-box protein 32 | protein_coding | NP_001229392.1;NP_478136.1;NP_680482.1 | 16731 | 606604 | . | . | . | 7.002460 | 7.015465 | 8.108276 | 8.530948 | 16.295013 | 16.216561 | 18.194738 | 19.504998 | 1.173055 | 1607.293183 | 0.122961684 | 1.872703392 | 0.061109355 | 0.722512417 | 10.530736 | 10.760926 | 10.586517 | 10.582535 | 10.693293 | 10.723722 | 1708 | 1592 | 1480 | 1658 |
| 114908 | NM_052932 | TMEM123 | transmembrane protein 123 | protein_coding | NP_443164.2 | 30138 | 606356 | . | . | . | 45.078072 | 38.333111 | 41.995087 | 34.455544 | 104.898529 | 88.608704 | 94.235756 | 78.778503 | -1.106335 | 4116.792651 | 0.135562171 | -1.0754344 | 0.282180289 | 0.998865404 | 12.078228 | 11.932714 | 12.114786 | 11.971519 | 12.043637 | 11.872710 | 5403 | 4269 | 3752 | 3285 |
| 114915 | NR_015370 | EPB41L4A-AS1 | EPB41L4A antisense RNA 1 | lncRNA | . | 30749 | . | . | . | . | 6.145174 | 6.273575 | 7.199638 | 6.776632 | 14.300072 | 14.501650 | 16.155779 | 15.493964 | 1.109557 | 245.971142 | 0.21744584 | 0.689753834 | 0.490349005 | 0.998865404 | 7.865240 | 8.015538 | 7.911590 | 7.917202 | 7.983489 | 7.948097 | 267 | 253 | 233 | 234 |
| 114926 | NM_00113567 | SMIM19 | small integral membrane protein 19 | protein_coding | NP_001129146.1;NP_001129147.1;NP_001129148.1;NP_001350115.1;NI | 25166 | . | . | . | . | 1.969910 | 1.674604 | 1.883310 | 1.894266 | 4.584063 | 3.870923 | 4.226095 | 4.331012 | 1.055575 | 206.867865 | 0.23684084 | 0.329455349 | 0.741811538 | 0.998865404 | 7.652700 | 7.731360 | 7.714338 | 7.642108 | 7.704184 | 7.703283 | 250 | 200 | 185 | 199 |
| 114928 | NM_00100405 | GPRASP2 | G protein-coupled receptor associated sorting protein 2 | protein_coding | NP_001004051.1;NP_001171803.1;NP_001171804.1;NP_001171805.1;NI | 25169 | 300969 | . | . | . | 1.005246 | 1.234944 | 0.962972 | 1.637247 | 2.339250 | 2.854628 | 2.160883 | 3.743371 | 1.146592 | 134.050847 | 0.318020326 | 0.620564681 | 0.534886084 | 0.998865404 | 6.966106 | 7.160616 | 6.996944 | 7.066381 | 6.982672 | 7.173136 | 130 | 148 | 93 | 168 |
| 114932 | NM_203462,XI | MRFAP1L1 | Morf4 family associated protein 1 like 1 | protein_coding | NP_982287.1;XP_005247989.1 | 28796 | . | . | . | . | 17.114520 | 17.171691 | 17.045735 | 15.957127 | 39.826191 | 39.693134 | 38.250135 | 36.484070 | -1.053912 | 827.424880 | 0.144898903 | -0.5228102 | 0.601106343 | 0.998865404 | 9.729721 | 9.654262 | 9.712414 | 9.708950 | 9.700469 | 9.645853 | 980 | 914 | 728 | 726 |
| 114971 | NM_00114398 | PTPMT1 | protein tyrosine phosphatase mitochondrial 1 | protein_coding | NP_001137456.1;NP_783859.1 | 26965 | 609538 | . | . | . | 7.770425 | 9.449762 | 8.919148 | 8.122605 | 18.082099 | 21.843548 | 20.014308 | 18.571370 | -1.020062 | 664.725727 | 0.164936063 | -0.17374924 | 0.862062534 | 0.998865404 | 9.390923 | 9.362165 | 9.316770 | 9.442907 | 9.401514 | 9.333930 | 702 | 791 | 601 | 586 |
| 114984 | NM_00114249 | FLYWCH2 | FLYWCH family member 2 | protein_coding | NP_001135971.1;NP_001135972.1;NP_612448.1;XP_005255135.1 | 25178 | . | . | . | . | 7.892647 | 7.996968 | 7.155763 | 8.475201 | 18.366513 | 18.485348 | 16.057325 | 19.377541 | -1.043978 | 252.212265 | 0.217950793 | -0.2848897 | 0.775728646 | 0.998865404 | 8.009513 | 7.946794 | 7.994935 | 7.983571 | 7.933977 | 8.000237 | 300 | 275 | 200 | 247 |
| 114987 | NM_00100661 | WDR31 | WD repeat domain 31 | protein_coding | NP_001006616.1;NP_001012361.1;NP_660284.1;XP_011516496.1 | 21421 | . | . | . | . | 0.708854 | 0.868321 | 0.618558 | 0.793408 | 1.649532 | 2.007162 | 1.388028 | 1.814034 | -1.141066 | 113.401120 | 0.310712866 | -0.61272787 | 0.540056294 | 0.998865404 | 6.918207 | 6.725977 | 6.815056 | 6.876551 | 6.772106 | 6.832559 | 127 | 142 | 83 | 109 |
| 114990 | NM_138440 | VASN | vasorin | protein_coding | NP_612449.2 | 18517 | 608843 | . | . | . | 0.311506 | 0.221778 | 0.216252 | 0.135624 | 0.724886 | 0.512650 | 0.485263 | 0.310088 | -1.493754 | 20.132024 | 0.701258966 | -0.82557611 | 0.409044589 | NA | 4.590318 | 4.015675 | 4.366823 | 4.335077 | 4.331607 | 4.303043 | 32 | 22 | 17 | 12 |
| 114991 | NM_00131804 | ZNF618 | zinc finger protein 618 | protein_coding | NP_001304969.1;NP_001304970.1;NP_001304971.1;NP_588615.2;XP_00 | 29416 | 617077 | . | . | . | 1.365742 | 1.711652 | 1.502056 | 1.858267 | 3.178139 | 3.956495 | 3.370570 | 4.248709 | 1.075731 | 475.241583 | 0.185434846 | 0.567951596 | 0.570067835 | 0.998865404 | 8.839507 | 8.943648 | 8.813032 | 8.917993 | 8.849874 | 8.970970 | 483 | 536 | 387 | 506 |
| 115004 | NM_138441 | CGAS | cyclic GMP-AMP synthase | protein_coding | NP_612450.2 | 21367 | 613973 | . | . | . | 1.736151 | 1.780894 | 1.406167 | 1.775207 | 4.040094 | 4.116616 | 3.155399 | 4.058801 | -1.122880 | 167.379326 | 0.261501338 | -0.63939804 | 0.522564025 | 0.998865404 | 7.468775 | 7.300262 | 7.406367 | 7.414562 | 7.318785 | 7.404628 | 202 | 193 | 122 | 164 |
| 115024 | NM_052935,N | NT5C3B | 5'-nucleotidase, cytosolic IIIB | protein_coding | NP_443167.4;XP_006721732.1;XP_006721733.1;XP_011522578.1;XP_01 | 28300 | . | . | . | . | 23.319238 | 24.934444 | 24.419931 | 27.386371 | 54.264820 | 57.637088 | 54.797615 | 62.615681 | 1.076352 | 1227.096622 | 0.13218639 | 0.803031162 | 0.421956724 | 0.998865404 | 10.207134 | 10.312991 | 10.217109 | 10.247100 | 10.258922 | 10.315516 | 1334 | 1301 | 1065 | 1237 |
| 115098 | NM_00113620 | CCDC124 | coiled-coil domain containing 124 | protein_coding | NP_001129675.1;NP_612451.1 | 25171 | . | . | . | . | 50.538483 | 50.892148 | 42.014803 | 54.185865 | 117.605130 | 117.639489 | 94.279998 | 123.889551 | -1.067333 | 1524.123691 | 0.143638539 | -0.65449481 | 0.512793072 | 0.998865404 | 10.620238 | 10.525744 | 10.597683 | 10.596309 | 10.450729 | 10.633882 | 1814 | 1697 | 1129 | 1547 |
| 115106 | NM_138443,N | HAUS1 | HAUS augmin like complex subunit 1 | protein_coding | NP_612452.1 | 25174 | 608775 | . | . | . | 17.101527 | 16.483839 | 13.236797 | 14.427400 | 39.795957 | 38.103131 | 29.702989 | 32.986536 | -1.229991 | 513.281108 | 0.167556165 | -1.78237445 | 0.074688189 | 0.764716896 | 9.145318 | 8.846442 | 9.081098 | 9.050245 | 8.910629 | 8.956543 | 667 | 597 | 385 | 448 |
| 115110 | NR_037844 | TNFRSF14-AS1 | TNFRSF14 antisense RNA 1 | lncRNA | . | 26966 | . | . | . | . | 5.760369 | 6.833846 | 7.697083 | 6.285117 | 13.404615 | 15.796742 | 17.272030 | 14.370173 | 1.095823 | 956.278439 | 0.159236315 | 0.829046213 | 0.407078258 | 0.998865404 | 9.833658 | 9.965887 | 9.804118 | 9.920745 | 10.005910 | 9.849697 | 966 | 1064 | 962 | 838 |
| 115123 | NM_178450,XI | MARCHF3 | membrane associated ring-CH-type finger 3 | protein_coding | NP_848545.1;XP_011541430.1;XP_011541431.1;XP_011541432.1;XP_01 | 28728 | 613333 | . | . | . | 2.149614 | 2.238881 | 1.618559 | 1.415068 | 5.002240 | 5.175275 | 3.631998 | 3.235382 | -1.425863 | 227.367467 | 0.227646183 | -2.24838043 | 0.024551942 | 0.516601303 | 8.061887 | 7.550858 | 7.904375 | 7.906211 | 7.765271 | 7.707273 | 307 | 289 | 174 | 164 |
| 115196 | NM_00110265 | ZNF554 | zinc finger protein 554 | protein_coding | NP_001096121.1;XP_006722698.1;XP_011525962.1;XP_011525964.1;XP_ | 26629 | . | . | . | . | 0.771606 | 0.466157 | 0.440613 | 0.512053 | 1.795559 | 1.077541 | 0.988725 | 1.170747 | -1.072336 | 57.871228 | 0.4152913 | -0.24261628 | 0.808302668 | 0.998865404 | 5.903915 | 5.803900 | 5.884690 | 5.840175 | 5.845944 | 5.850524 | 76 | 58 | 48 | 53 |
| 115201 | NM_00132128 | ATG4A | autophagy related 4A cysteine peptidase | protein_coding | NP_001308216.1;NP_001308217.1;NP_001308218.1;NP_001308219.1;NI | 16489 | 300663 | . | . | . | 3.984157 | 3.668826 | 3.865156 | 4.328489 | 9.271298 | 8.480655 | 8.673295 | 9.896573 | 1.054666 | 279.632901 | 0.210616754 | 0.364577383 | 0.7154269 | 0.998865404 | 8.088667 | 8.165102 | 8.133010 | 8.091360 | 8.104685 | 8.174448 | 326 | 282 | 233 | 287 |
| 115207 | NM_138444 | KCTD12 | potassium channel tetramerization domain containing 12 | protein_coding | NP_612453.1 | 14678 | 610521 | . | . | . | 0.501578 | 0.472010 | 0.481312 | 0.562267 | 1.167192 | 1.091071 | 1.080049 | 1.285556 | 1.059977 | 98.693702 | 0.323739256 | 0.259571362 | 0.795194429 | 0.998865404 | 6.582532 | 6.666032 | 6.624962 | 6.605131 | 6.610782 | 6.654129 | 114 | 100 | 82 | 102 |
| 115209 | NM_145243 | OMA1 | OMA1 zinc metallopeptidase | protein_coding | NP_660286.1 | 29661 | 617081 | . | . | . | 7.019075 | 8.032249 | 8.353386 | 6.003242 | 16.333677 | 18.566904 | 18.744755 | 13.725698 | -1.061056 | 426.848384 | 0.203961051 | -0.41919867 | 0.675070941 | 0.998865404 | 8.779261 | 8.694659 | 8.711307 | 8.789367 | 8.810821 | 8.609792 | 474 | 504 | 421 | 323 |
| 115286 | NM_00116479 | SLC25A26 | solute carrier family 25 member 26 | protein_coding | NP_001158268.1;NP_001337920.1;NP_001337921.1;NP_001337922.1;NI | 20661 | 611037 | . | . | . | 11.444798 | 10.038658 | 10.947836 | 14.022139 | 26.632514 | 23.204810 | 24.566633 | 32.059961 | 1.119091 | 639.033795 | 0.159750246 | 1.016128262 | 0.309568312 | 0.998865404 | 9.236213 | 9.398711 | 9.321057 | 9.235829 | 9.336349 | 9.373628 | 739 | 609 | 571 | 650 |
| 115290 | NM_024907,N | FBXO17 | F-box protein 17 | protein_coding | NP_079183.4;NP_680474.1 | 18754 | 609094 | . | . | . | 7.739287 | 7.091080 | 7.500976 | 8.216055 | 18.009641 | 16.391349 | 16.831974 | 18.785032 | 1.046509 | 529.466592 | 0.167849954 | 0.39073831 | 0.695990679 | 0.998865404 | 9.015245 | 9.080802 | 9.063920 | 9.001018 | 9.036001 | 9.087727 | 625 | 531 | 452 | 528 |
| 115294 | NM_00128678 | PCMTD1 | protein-L-isoaspartate (D-aspartate) O-methyltransferase domain | protein_coding | NP_001273711.1;NP_001273712.1;NP_001350122.1;NP_443169.2 | 30483 | . | . | . | . | 3.327820 | 3.319794 | 3.213351 | 2.628094 | 7.743973 | 7.673851 | 7.210665 | 6.008827 | -1.120149 | 367.453985 | 0.206308382 | -0.79342815 | 0.427528415 | 0.998865404 | 8.600300 | 8.437976 | 8.557248 | 8.543425 | 8.569328 | 8.393028 | 452 | 414 | 349 | 274 |
| 115330 | NM_00130347 | GPR146 | G protein-coupled receptor 146 | protein_coding | NP_001290402.1;NP_001290403.1;NP_612454.1;XP_005249670.2;XP_00 | 21718 | . | . | . | . | 0.233724 | 0.169634 | 0.587102 | 0.350892 | 0.543886 | 0.392117 | 1.317440 | 0.802273 | 2.214108 | 28.761100 | 0.602635072 | 1.902851912 | 0.057059871 | 0.708295137 | 4.159017 | 5.309532 | 4.778564 | 4.760671 | 4.894910 | 4.820751 | 23 | 17 | 43 | 28 |
| 115353 | NM_00125640 | LRRC42 | leucine rich repeat containing 42 | protein_coding | NP_001243338.1;NP_443172.1;XP_006710390.1;XP_006710391.1 | 28792 | . | . | . | . | 17.064615 | 16.152235 | 13.301846 | 18.420882 | 39.710060 | 37.336616 | 29.848957 | 42.117158 | -1.062413 | 885.793298 | 0.168712653 | -0.51771246 | 0.604658904 | 0.998865404 | 9.834173 | 9.746137 | 9.830786 | 9.787845 | 9.648657 | 9.871033 | 1082 | 956 | 630 | 931 |
| 115399 | NM_198075,XI | LRRC56 | leucine rich repeat containing 56 | protein_coding | NP_932341.1;XP_005252832.1;XP_005252833.1;XP_006718195.1;XP_00 | 25430 | 618227 | . | . | . | 1.513572 | 1.263042 | 1.410081 | 1.128867 | 3.522142 | 2.919578 | 3.164180 | 2.581022 | -1.018591 | 140.716360 | 0.293947318 | -0.09040898 | 0.927962225 | 0.998865404 | 7.148742 | 7.124448 | 7.190509 | 7.079687 | 7.173225 | 7.084366 | 187 | 131 | 136 | 115 |
| 115416 | NM_138446,XI | MALSU1 | mitochondrial assembly of ribosomal large subunit 1 | protein_coding | NP_612455.1;XP_016867220.1 | 21721 | 614624 | . | . | . | 16.515663 | 15.746496 | 13.380499 | 13.656195 | 38.432629 | 36.398735 | 30.025454 | 31.223264 | -1.210369 | 1342.738707 | 0.128416822 | -2.14494185 | 0.031957496 | 0.56480067 | 10.522090 | 10.246715 | 10.480998 | 10.436812 | 10.310071 | 10.317582 | 1741 | 1542 | 1051 | 1145 |
| 115426 | NM_152896,N | UHRF2 | ubiquitin like with PHD and ring finger domains 2 | protein_coding | NP_690856.1;XP_011516006.1;XP_011516007.1;XP_016869742.1;XP_01 | 12557 | 615211 | . | . | . | 4.236756 | 4.427248 | 4.400998 | 3.643504 | 9.859109 | 10.233783 | 9.875712 | 8.330441 | -1.087216 | 470.636236 | 0.180640337 | -0.66783928 | 0.504236191 | 0.998865404 | 8.937140 | 8.817313 | 8.896547 | 8.908701 | 8.912036 | 8.785389 | 559 | 534 | 433 | 379 |
| 115509 | NM_138447,N | ZNF689 | zinc finger protein 689 | protein_coding | NP_612456.1 | 25173 | 618033 | . | . | . | 7.826121 | 8.471130 | 5.736402 | 7.224190 | 18.211704 | 19.581398 | 12.872321 | 16.517254 | -1.270863 | 732.511137 | 0.162243577 | -2.13141536 | 0.033054939 | 0.575268292 | 9.679677 | 9.332955 | 9.564372 | 9.619773 | 9.354775 | 9.493133 | 908 | 919 | 502 | 668 |
| 115548 | NM_00114603 | FCHO2 | FCH and mu domain containing endocytic adaptor 2 | protein_coding | NP_001139504.1;NP_620137.2;XP_016864505.1;XP_016864506.1;XP_01 | 25180 | 613438 | . | . | . | 1.825668 | 2.284758 | 2.492426 | 2.253751 | 4.248405 | 5.281322 | 5.592931 | 5.152934 | 1.169043 | 326.563510 | 0.200846101 | 1.121891208 | 0.261908699 | 0.995760118 | 8.234248 | 8.459415 | 8.260393 | 8.344130 | 8.415047 | 8.363698 | 320 | 350 | 320 | 315 |
| 115557 | NM_00111127 | ARHGEF25 | Rho guanine nucleotide exchange factor 25 | protein_coding | NP_001104740.2;NP_001334862.2;NP_891992.3 | 30275 | 610215 | . | . | . | 3.830196 | 3.837758 | 3.644903 | 3.950191 | 8.913024 | 8.871151 | 8.179056 | 9.031641 | -1.020345 | 304.884114 | 0.199902075 | -0.14535808 | 0.884428158 | 0.998865404 | 8.266731 | 8.237354 | 8.258252 | 8.257118 | 8.229559 | 8.263663 | 355 | 332 | 254 | 292 |
| 115560 | NM_00125828 | ZNF501 | zinc finger protein 501 | protein_coding | NP_001245209.1;NP_659481.2 | 23717 | . | . | . | . | 0.106714 | 0.166897 | 0.098508 | 0.127003 | 0.248328 | 0.385790 | 0.221050 | 0.290376 | -1.190085 | 13.278585 | 0.849433612 | -0.29556768 | 0.767560259 | NA | 3.852893 | 3.597915 | 3.723024 | 3.751203 | 3.719652 | 3.734607 | 13 | 19 | 9 | 13 |
| 115650 | NM_052945 | TNFRSF13C | TNF receptor superfamily member 13C | protein_coding | NP_443177.1 | 17755 | 606269 | . | . | . | 0.280442 | 0.177226 | 0.157096 | 0.191283 | 0.652601 | 0.409666 | 0.352519 | 0.437347 | -1.325082 | 24.998949 | 0.631336678 | -0.64320866 | 0.520088717 | NA | 4.832372 | 4.426810 | 4.687099 | 4.634718 | 4.624422 | 4.640143 | 40 | 24 | 17 | 22 |
| 115701 | NM_052947 | ALPK2 | alpha kinase 2 | protein_coding | NP_443179.3 | 20565 | . | . | . | . | 0.025212 | 0.024095 | 0.028607 | 0.036081 | 0.058670 | 0.055696 | 0.064194 | 0.082496 | 1.229559 | 6.999873 | 1.140440974 | 0.261425975 | 0.793764028 | NA | 2.651489 | 2.947964 | 2.799278 | 2.801239 | 2.803171 | 2.810422 | 7 | 7 | 6 | 8 |
| 115703 | NM_00117263 | ARHGAP33 | Rho GTPase activating protein 33 | protein_coding | NP_001166101.1;NP_001353107.1;NP_443180.2;XP_006723062.1;XP_00 | 23085 | 614902 | . | . | . | 3.012151 | 3.521004 | 3.174367 | 2.655762 | 7.009394 | 8.138959 | 7.123189 | 6.072084 | -1.115646 | 398.206994 | 0.190892515 | -0.82706089 | 0.408252548 | 0.998865404 | 8.713908 | 8.556553 | 8.634812 | 8.699226 | 8.655125 | 8.547540 | 458 | 477 | 358 | 320 |
| 115704 | NM_00115994 | EVI5L | ecotropic viral integration site 5 like | protein_coding | NP_001153416.1;NP_660288.1;XP_016881728.1 | 30464 | . | . | . | . | 1.139577 | 0.838381 | 0.940799 | 1.492972 | 2.651842 | 1.937955 | 2.111125 | 3.413502 | 1.221993 | 134.775497 | 0.316019705 | 0.915248517 | 0.36006118 | 0.998865404 | 6.922958 | 7.211471 | 7.078788 | 6.972574 | 7.013794 | 7.180726 | 161 | 111 | 101 | 169 |
| 115708 | NM_152307 | TRMT61A | tRNA methyltransferase 61A | protein_coding | NP_689520.2 | 23790 | . | . | . | . | 4.767000 | 4.835799 | 4.522501 | 5.349695 | 11.093006 | 11.178167 | 10.148360 | 12.231441 | 1.013825 | 488.371664 | 0.174162461 | 0.113736974 | 0.909446302 | 0.998865404 | 8.922232 | 8.941376 | 8.925048 | 8.928743 | 8.885114 | 8.983335 | 557 | 525 | 394 | 497 |
| 115752 | NM_00114368 | DIS3L | DIS3 like exosome 3'-5' exoribonuclease | protein_coding | NP_001137160.1;NP_001310865.1;NP_001310866.1;NP_001310867.1;NI | 28698 | 614183 | . | . | . | 4.664493 | 4.362465 | 5.076208 | 4.630168 | 10.854464 | 10.084030 | 11.390864 | 10.586329 | 1.048995 | 571.365551 | 0.16482654 | 0.418666427 | 0.675459935 | 0.998865404 | 9.123012 | 9.192687 | 9.162444 | 9.119738 | 9.212235 | 9.131837 | 663 | 582 | 540 | 515 |
| 115811 | NM_00133045 | IQCD | IQ motif containing D | protein_coding | NP_001317381.1;NP_612460.1;XP_005253890.1;XP_005253891.1;XP_01 | 25168 | . | . | . | . | 1.042257 | 0.763232 | 0.662303 | 0.439739 | 2.425375 | 1.764246 | 1.486189 | 1.005412 | -1.603662 | 51.964821 | 0.439107318 | -1.55171626 | 0.120730131 | 0.861666629 | 5.999739 | 5.319806 | 5.756612 | 5.729315 | 5.665135 | 5.644899 | 78 | 65 | 37 | 35 |
| 115817 | NM_00113605 | DHRS1 | dehydrogenase/reductase 1 | protein_coding | NP_001129522.1;NP_612461.1 | 16445 | 610410 | . | . | . | 10.274178 | 9.512557 | 9.226344 | 10.681200 | 23.908433 | 21.988702 | 20.703650 | 24.421297 | -1.008575 | 441.296163 | 0.1801964 | -0.06836145 | 0.945497908 | 0.998865404 | 8.791912 | 8.779267 | 8.813598 | 8.761200 | 8.740312 | 8.822923 | 532 | 458 | 356 | 440 |
| 115825 | NM_052950,XI | WDFY2 | WD repeat and FYVE domain containing 2 | protein_coding | NP_443182.1;XP_011533216.1;XP_011533217.1;XP_016875875.1;XP_02 | 20482 | 610418 | . | . | . | 4.849471 | 6.638703 | 5.832529 | 6.286073 | 11.284919 | 15.345658 | 13.088025 | 14.372356 | 1.000740 | 1381.099097 | 0.149473392 | 0.007135314 | 0.994306892 | 0.999862634 | 10.430852 | 10.432350 | 10.405705 | 10.451648 | 10.523367 | 10.326460 | 1542 | 1534 | 1356 | 1129 |
| 115827 | NM_00131791 | RAB3C | RAB3C, member RAS oncogene family | protein_coding | NP_001304844.1;NP_612462.1 | 30269 | 612829 | . | . | . | 0.030931 | 0.040267 | 0.015152 | 0.038845 | 0.071978 | 0.093079 | 0.034001 | 0.088815 | -1.316037 | 9.483445 | 1.017584921 | -0.38935344 | 0.697014713 | NA | 3.432996 | 3.029739 | 3.250215 | 3.261470 | 3.228749 | 3.259706 | 11 | 13 | 4 | 11 |
| 115861 | NM_138454 | NXNL1 | nucleoredoxin like 1 | protein_coding | NP_612463.1 | 25179 | 608791 | . | . | . | 0.398693 | 0.276477 | 0.485853 | 0.312865 | 0.927775 | 0.639088 | 1.090239 | 0.715329 | 1.193864 | 11.262140 | 0.918190323 | 0.278415578 | 0.780693363 | NA | 3.357474 | 3.617632 | 3.491675 | 3.474255 | 3.503192 | 3.480276 | 14 | 9 | 13 | 9 |
| 115908 | NM_00125609 | CTHRC1 | collagen triple helix repeat containing 1 | protein_coding | NP_001243028.1;NP_612464.1;XP_011515126.1 | 18831 | 610635 | . | . | . | 140.595862 | 129.621032 | 122.491782 | 122.197521 | 327.172373 | 299.624844 | 274.868004 | 279.390116 | -1.120012 | 4923.653196 | 0.109277306 | -1.49632431 | 0.134569163 | 0.891755273 | 12.344920 | 12.181481 | 12.346680 | 12.273811 | 12.221367 | 12.210794 | 6248 | 5372 | 4065 | 4327 |
| 115939 | NM_00100141 | TSR3 | TSR3 ribosome maturation factor | protein_coding | NP_001001410.1 | 14175 | 617058 | . | . | . | 18.769556 | 19.490644 | 17.325184 | 18.301529 | 43.677525 | 45.053497 | 38.877213 | 41.844269 | -1.088827 | 696.498511 | 0.151020689 | -0.81296952 | 0.416235536 | 0.998865404 | 9.504182 | 9.381149 | 9.462418 | 9.482700 | 9.399277 | 9.428909 | 824 | 795 | 567 | 639 |
| 115948 | NM_00130245 | CCDC151 | coiled-coil domain containing 151 | protein_coding | NP_001289382.1;NP_001289383.1;NP_659482.3;XP_016881730.1 | 28303 | 615956 | . | . | . | 0.251462 | 0.370413 | 0.240443 | 0.377085 | 0.585161 | 0.856226 | 0.539548 | 0.862160 | -1.013451 | 21.022651 | 0.686831886 | -0.02806562 | 0.977609811 | NA | 4.405684 | 4.381963 | 4.376161 | 4.409245 | 4.371985 | 4.411033 | 20 | 27 | 14 | 24 |
| 115950 | NM_138783 | ZNF653 | zinc finger protein 653 | protein_coding | NP_620138.2 | 25196 | 611371 | . | . | . | 1.250616 | 1.173985 | 1.363108 | 1.425618 | 2.910236 | 2.713718 | 3.058774 | 3.259506 | 1.134143 | 87.934672 | 0.338704406 | 0.536170026 | 0.591841047 | 0.998865404 | 6.364637 | 6.546365 | 6.447100 | 6.429416 | 6.468871 | 6.478148 | 98 | 86 | 80 | 89 |
| 115992 | NM_00117181 | RNF166 | ring finger protein 166 | protein_coding | NP_001165286.1;NP_001165287.1;NP_849163.1;XP_011521147.1;XP_01 | 28856 | 617178 | . | . | . | 4.318040 | 4.000668 | 4.777718 | 4.970200 | 10.048257 | 9.247721 | 10.721063 | 11.363771 | 1.078016 | 371.697689 | 0.187904093 | 0.576777579 | 0.564089717 | 0.998865404 | 8.482839 | 8.591102 | 8.537033 | 8.492288 | 8.533082 | 8.583165 | 428 | 371 | 321 | 377 |
| 116028 | NM_152308,N | RMI2 | RecQ mediated genome instability 2 | protein_coding | NP_689521.1 | 28349 | 612426 | . | . | . | 12.677752 | 11.193594 | 11.796771 | 12.282066 | 29.501646 | 25.874494 | 26.471610 | 28.081486 | -1.010018 | 528.117334 | 0.167614907 | -0.0857956 | 0.931628906 | 0.998865404 | 9.051749 | 9.037646 | 9.085010 | 9.008717 | 9.035650 | 9.046780 | 647 | 539 | 453 | 497 |
| 116039 | NM_00114246 | OSR2 | odd-skipped related transciption factor 2 | protein_coding | NP_001135934.1;NP_001273770.1;NP_443727.2;XP_005250835.1;XP_01 | 15830 | 611297 | . | . | . | 0.564960 | 0.488305 | 0.472074 | 0.887226 | 1.314686 | 1.128739 | 1.059323 | 2.028538 | 1.241744 | 36.634055 | 0.531385349 | 0.587836873 | 0.556641792 | 0.998865404 | 5.032032 | 5.341619 | 5.173973 | 5.171871 | 5.162837 | 5.243727 | 38 | 35 | 26 | 48 |
| 116064 | NM_00109967 | LRRC58 | leucine rich repeat containing 58 | protein_coding | NP_001093148.1 | 26968 | . | . | . | . | 7.692887 | 5.732738 | 6.623256 | 5.087140 | 17.901663 | 13.251481 | 14.862393 | 11.631140 | -1.162839 | 1552.093427 | 0.170606482 | -1.27575246 | 0.202043038 | 0.967656221 | 10.704412 | 10.487437 | 10.762292 | 10.524047 | 10.631850 | 10.425611 | 2210 | 1530 | 1418 | 1163 |
| 116068 | NM_00128681 | LYSMD3 | LysM domain containing 3 | protein_coding | NP_001273741.1;NP_938014.1;XP_005248478.1 | 26969 | . | . | . | . | 4.341305 | 3.213887 | 2.942113 | 2.615424 | 10.102393 | 7.429044 | 6.602017 | 5.979857 | -1.358275 | 444.071558 | 0.199264177 | -2.21703526 | 0.026620682 | 0.532994094 | 8.998160 | 8.557638 | 8.968037 | 8.775759 | 8.731046 | 8.645841 | 675 | 471 | 352 | 327 |
| 116071 | NM_00130080 | BATF2 | basic leucine zipper ATF-like transcription factor 2 | protein_coding | NP_001287736.1;NP_001287737.1;NP_612465.3 | 25163 | 614983 | . | . | . | 0.258178 | 0.245129 | 0.188055 | 0.344798 | 0.600790 | 0.566627 | 0.421991 | 0.788341 | 1.031684 | 17.274994 | 0.750091642 | 0.059993845 | 0.952160537 | NA | 4.089593 | 4.131331 | 4.110615 | 4.107598 | 4.090869 | 4.131618 | 20 | 18 | 11 | 21 |
| 116092 | NM_052951,XI | DNTTIP1 | deoxynucleotidyltransferase terminal interacting protein 1 | protein_coding | NP_443183.1;XP_024307591.1 | 16160 | 611388 | . | . | . | 15.401679 | 15.655441 | 13.915935 | 16.669727 | 35.840336 | 36.188251 | 31.226954 | 38.113353 | -1.029801 | 607.012165 | 0.164402663 | -0.2576982 | 0.796639832 | 0.998865404 | 9.266936 | 9.223907 | 9.252501 | 9.256999 | 9.177900 | 9.289183 | 707 | 667 | 476 | 608 |
| 116113 | NM_00101242 | FOXP4 | forkhead box P4 | protein_coding | NP_001012426.1;NP_001012427.1;NP_612466.1;XP_006715054.1;XP_01 | 20842 | 608924 | . | . | . | 1.850594 | 2.101268 | 1.903812 | 1.807461 | 4.306408 | 4.857173 | 4.272099 | 4.132543 | -1.075889 | 284.716703 | 0.208800737 | -0.50540869 | 0.613271816 | 0.998865404 | 8.205523 | 8.099289 | 8.134929 | 8.207866 | 8.121757 | 8.145043 | 317 | 340 | 233 | 263 |
| 116115 | NM_00131403 | ZNF526 | zinc finger protein 526 | protein_coding | NP_001300962.1;NP_597701.1 | 29415 | 614387 | . | . | . | 2.948098 | 2.927899 | 2.595456 | 3.046157 | 6.860345 | 6.767971 | 5.824129 | 6.964677 | -1.059090 | 362.551909 | 0.191598163 | -0.43228708 | 0.665532773 | 0.998865404 | 8.543221 | 8.459656 | 8.516119 | 8.520301 | 8.441761 | 8.526640 | 428 | 404 | 283 | 355 |
| 116138 | NM_057161,N | KLHDC3 | kelch domain containing 3 | protein_coding | NP_476502.1;XP_024302088.1 | 20704 | 611248 | . | . | . | 18.581022 | 19.014152 | 19.732525 | 22.742200 | 43.238800 | 43.952061 | 44.279213 | 51.997338 | 1.111074 | 1170.177996 | 0.136936504 | 1.109673469 | 0.267139759 | 0.998546485 | 10.114721 | 10.266323 | 10.149823 | 10.151904 | 10.179686 | 10.277461 | 1275 | 1198 | 1002 | 1229 |
| 116143 | NM_00125647 | WDR92 | WD repeat domain 92 | protein_coding | NP_001243405.1;NP_612467.1 | 25176 | 610729 | . | . | . | 5.028213 | 5.122339 | 5.165616 | 5.949151 | 11.700859 | 11.840517 | 11.591492 | 13.602026 | 1.068624 | 407.353507 | 0.181818736 | 0.526649852 | 0.598436766 | 0.998865404 | 8.621666 | 8.717033 | 8.657582 | 8.640811 | 8.655515 | 8.721048 | 460 | 419 | 346 | 416 |
| 116149 | gene-RPL12P4 | RPL12P4 | ribosomal protein L12 pseudogene 4 | pseudogene | . | 16587 | . | . | . | . | 0.931382 | 0.508314 | 0.447690 | 1.620023 | 2.167364 | 1.174990 | 1.004604 | 3.703991 | 1.408478 | 17.624932 | 0.81512597 | 0.60620885 | 0.544376077 | NA | 3.872825 | 4.364566 | 4.132447 | 4.092588 | 4.090120 | 4.185121 | 22 | 11 | 8 | 30 |
| 116150 | NM_138459 | NUS1 | NUS1 dehydrodolichyl diphosphate synthase subunit | protein_coding | NP_612468.1 | 21042 | 610463 | . | . | . | 8.566962 | 7.801977 | 7.744164 | 5.447994 | 19.935675 | 18.034620 | 17.377680 | 12.456192 | -1.258131 | 1106.097521 | 0.171390259 | -1.93290975 | 0.053247317 | 0.683508073 | 10.267020 | 9.936631 | 10.224594 | 10.146077 | 10.136721 | 9.877838 | 1491 | 1262 | 1005 | 754 |
| 116151 | NM_080821 | FAM210B | family with sequence similarity 210 member B | protein_coding | NP_543011.2 | 16102 | 618722 | . | . | . | 22.741621 | 21.571495 | 14.445845 | 18.839979 | 52.920685 | 49.863480 | 32.416058 | 43.075375 | -1.350675 | 1806.971662 | 0.143763033 | -3.01663839 | 0.002555945 | 0.133671005 | 11.020146 | 10.586036 | 10.948846 | 10.898452 | 10.586649 | 10.778913 | 2465 | 2172 | 1167 | 1624 |
| 116224 | NM_138333 | FAM122A | family with sequence similarity 122A | protein_coding | NP_612206.5 | 23490 | 617249 | . | . | . | 1.418923 | 1.340272 | 1.481843 | 1.197506 | 3.301893 | 3.098098 | 3.325213 | 2.737955 | -1.043602 | 233.818679 | 0.230450892 | -0.26717865 | 0.78933161 | 0.998865404 | 7.899046 | 7.838819 | 7.894441 | 7.861334 | 7.908369 | 7.803847 | 284 | 249 | 221 | 191 |
| 116225 | NM_138462,XI | ZMYND19 | zinc finger MYND-type containing 19 | protein_coding | NP_612471.1;XP_005266109.1;XP_016869744.1 | 21146 | 611424 | . | . | . | 16.861042 | 15.194732 | 16.782515 | 16.714249 | 39.236341 | 35.123305 | 37.659476 | 38.215147 | 1.037596 | 731.692322 | 0.152818052 | 0.348419046 | 0.727525501 | 0.998865404 | 9.488077 | 9.541613 | 9.543461 | 9.457801 | 9.523866 | 9.530501 | 878 | 727 | 645 | 702 |
| 116228 | NM_00131287 | COX20 | cytochrome c oxidase assembly factor COX20 | protein_coding | NP_001299800.1;NP_001299801.1;NP_001299802.1;NP_001299803.1;NI | 26970 | 614698 | . | . | . | 20.610780 | 19.687705 | 23.058373 | 20.885957 | 47.962136 | 45.509011 | 51.742316 | 47.753262 | 1.076223 | 1523.543524 | 0.129117258 | 0.820784961 | 0.411768766 | 0.998865404 | 10.519076 | 10.625395 | 10.563749 | 10.522547 | 10.640013 | 10.559104 | 1735 | 1541 | 1445 | 1404 |
| 116236 | NM_198147 | ABHD15 | abhydrolase domain containing 15 | protein_coding | NP_937790.2 | 26971 | . | . | . | . | 3.991575 | 4.104265 | 3.969832 | 4.124293 | 9.288559 | 9.487192 | 8.908188 | 9.429707 | -1.014719 | 440.737505 | 0.174244925 | -0.12098355 | 0.903704067 | 0.998865404 | 8.794377 | 8.773096 | 8.781952 | 8.794628 | 8.770679 | 8.787914 | 506 | 484 | 375 | 416 |
| 116238 | NM_00116040 | TLCD1 | TLC domain containing 1 | protein_coding | NP_001153879.1;NP_612472.1;XP_006721734.1;XP_011522580.1 | 25177 | . | . | . | . | 12.726669 | 14.331910 | 14.745323 | 15.736210 | 29.615482 | 33.128850 | 33.088079 | 35.978976 | 1.083073 | 518.084575 | 0.167342436 | 0.687996354 | 0.491455077 | 0.998865404 | 8.958553 | 9.073256 | 8.959759 | 9.019485 | 9.030699 | 9.051269 | 546 | 562 | 461 | 513 |
| 116254 | NM_138785 | GINM1 | glycoprotein integral membrane 1 | protein_coding | NP_620140.1 | 21074 | . | . | . | . | 13.194445 | 11.837652 | 9.712940 | 10.153872 | 30.704016 | 27.363264 | 21.795557 | 23.215624 | -1.277831 | 680.827645 | 0.156121591 | -2.26552578 | 0.023480434 | 0.50908701 | 9.577120 | 9.223564 | 9.529167 | 9.445434 | 9.310221 | 9.331889 | 931 | 776 | 511 | 570 |
| 116349 | NR_126522 | EXOC3-AS1 | EXOC3 antisense RNA 1 | lncRNA | . | 25175 | . | . | . | . | 2.107908 | 2.026611 | 2.334358 | 2.625842 | 4.905187 | 4.684602 | 5.238233 | 6.003675 | 1.182000 | 94.311521 | 0.329179606 | 0.732822796 | 0.463666516 | 0.998865404 | 6.433918 | 6.674766 | 6.537933 | 6.524385 | 6.563250 | 6.596818 | 102 | 91 | 84 | 101 |
| 116372 | NM_00107742 | LYPD1 | LY6/PLAUR domain containing 1 | protein_coding | NP_001070895.1;NP_001308163.1;NP_001308164.1;NP_653187.3;XP_01 | 28431 | 610450 | . | . | . | 2.624337 | 3.004747 | 0.963909 | 1.222343 | 6.106943 | 6.945608 | 2.162984 | 2.794742 | -2.530347 | 204.580428 | 0.248125237 | -5.39781844 | 6.74561E-08 | 4.00991E-05 | 8.196579 | 6.855373 | 7.787906 | 7.858953 | 7.388089 | 7.446989 | 316 | 337 | 91 | 119 |
| 116412 | NM_138466,N | ZNF837 | zinc finger protein 837 | protein_coding | NP_612475.1;XP_016881731.1 | 25164 | . | . | . | . | 0.271831 | 0.139640 | 0.162902 | 0.188417 | 0.632561 | 0.322785 | 0.365547 | 0.430793 | -1.179055 | 15.602114 | 0.793234555 | -0.29957279 | 0.764503044 | NA | 4.077189 | 3.840449 | 3.992937 | 3.950239 | 3.953651 | 3.967711 | 25 | 13 | 11 | 15 |
| 116447 | NM_00125844 | TOP1MT | DNA topoisomerase I mitochondrial | protein_coding | NP_001245375.1;NP_001245376.1;NP_443195.1;XP_005250838.1 | 29787 | 606387 | . | . | . | 12.348772 | 11.661677 | 11.433638 | 13.546441 | 28.736102 | 26.956489 | 25.656752 | 30.972329 | 1.034364 | 747.431128 | 0.154578554 | 0.315331667 | 0.752509844 | 0.998865404 | 9.521378 | 9.569809 | 9.556576 | 9.509218 | 9.506009 | 9.604841 | 875 | 766 | 614 | 763 |
| 116461 | NM_00112739 | TSEN15 | tRNA splicing endonuclease subunit 15 | protein_coding | NP_001120866.1;NP_001287693.1;NP_001287695.1;NP_001350572.1;NI | 16791 | 608756 | . | . | . | 10.959364 | 9.414852 | 7.924834 | 7.040285 | 25.502890 | 21.762854 | 17.783099 | 16.096773 | -1.384842 | 525.014388 | 0.175024925 | -2.68373732 | 0.007280428 | 0.268886682 | 9.251642 | 8.782872 | 9.178520 | 9.075543 | 8.959444 | 8.881696 | 755 | 608 | 408 | 386 |
| 116496 | NM_052966,XI | NIBAN1 | niban apoptosis regulator 1 | protein_coding | NP_443198.1;XP_011507442.1;XP_011507443.1 | 16784 | . | . | . | . | 3.513978 | 3.229318 | 1.856776 | 2.207247 | 8.177171 | 7.464713 | 4.166551 | 5.046608 | -1.683106 | 581.144269 | 0.168935348 | -4.44623571 | 8.73881E-06 | 0.001773849 | 9.510190 | 8.758486 | 9.357317 | 9.289513 | 8.935537 | 9.029679 | 879 | 750 | 346 | 439 |
| 116540 | NM_053050 | MRPL53 | mitochondrial ribosomal protein L53 | protein_coding | NP_444278.1 | 16684 | 611857 | . | . | . | 63.410755 | 61.120930 | 56.438950 | 66.810043 | 147.559433 | 141.283768 | 126.647362 | 152.753220 | -1.024413 | 957.339125 | 0.146472611 | -0.23757272 | 0.81221251 | 0.998865404 | 9.920372 | 9.885186 | 9.927449 | 9.894288 | 9.834347 | 9.949120 | 1141 | 1022 | 757 | 957 |
| 116541 | NM_172251 | MRPL54 | mitochondrial ribosomal protein L54 | protein_coding | NP_758455.1 | 16685 | 611858 | . | . | . | 12.934590 | 13.313265 | 10.383865 | 13.694092 | 30.099321 | 30.774208 | 23.301092 | 31.309916 | -1.103985 | 238.772032 | 0.231729277 | -0.61589329 | 0.537964949 | 0.998865404 | 7.969748 | 7.825634 | 7.917767 | 7.926971 | 7.809340 | 7.934507 | 286 | 273 | 171 | 241 |
| 116828 | NR_026928 | N4BP2L2-IT2 | N4BPL2 intronic transcript 2 | lncRNA | . | 25038 | . | . | . | . | 0.328298 | 0.247621 | 0.476340 | 0.132135 | 0.763964 | 0.572386 | 1.068892 | 0.302110 | 1.034984 | 45.487416 | 0.567590262 | 0.087402837 | 0.930351314 | 0.998865404 | 5.479531 | 5.534732 | 5.508407 | 5.456619 | 5.580856 | 5.386588 | 59 | 41 | 63 | 19 |
| 116832 | NM_052969 | RPL39L | ribosomal protein L39 like | protein_coding | NP_443201.1 | 17094 | 607547 | . | . | . | 27.648594 | 29.569107 | 19.582148 | 27.974630 | 64.339417 | 68.350311 | 43.941772 | 63.960667 | -1.220268 | 595.400010 | 0.186781373 | -1.53761416 | 0.124142984 | 0.87182553 | 9.354738 | 9.066301 | 9.256743 | 9.296584 | 9.029746 | 9.248206 | 732 | 727 | 386 | 589 |
| 116840 | NM_00103714 | CNTROB | centrobin, centriole duplication and spindle assembly protein | protein_coding | NP_001032221.1;NP_001317053.1;NP_001340131.1;NP_001340132.1;NI | 29616 | 611425 | . | . | . | 12.363146 | 12.529127 | 13.174027 | 11.124596 | 28.769544 | 28.961637 | 29.562136 | 25.435068 | -1.024314 | 1483.198783 | 0.135483797 | -0.25581033 | 0.798097293 | 0.998865404 | 10.551506 | 10.517283 | 10.552922 | 10.532352 | 10.587652 | 10.456740 | 1751 | 1598 | 1383 | 1257 |
| 116841 | NM_00132393 | SNAP47 | synaptosome associated protein 47 | protein_coding | NP_001310859.1;NP_001310860.1;NP_001310861.1;NP_001310862.1;NI | 30669 | . | . | . | . | 8.298502 | 7.857474 | 7.199809 | 10.686843 | 19.310952 | 18.162905 | 16.156164 | 24.434193 | 1.039584 | 620.436995 | 0.173134894 | 0.323479897 | 0.74633183 | 0.998865404 | 9.249185 | 9.304566 | 9.286828 | 9.236641 | 9.201726 | 9.367395 | 727 | 632 | 482 | 666 |
| 116842 | NM_052971 | LEAP2 | liver enriched antimicrobial peptide 2 | protein_coding | NP_443203.1 | 29571 | 611373 | . | . | . | 0.766806 | 0.375640 | 0.388726 | 0.633678 | 1.784389 | 0.868309 | 0.872289 | 1.448829 | -1.086855 | 14.900804 | 0.821298991 | -0.14630416 | 0.883681283 | NA | 3.955908 | 3.836248 | 3.924416 | 3.877088 | 3.883801 | 3.907320 | 24 | 11 | 10 | 16 |
| 116843 | NM_052831,XI | SLC18B1 | solute carrier family 18 member B1 | protein_coding | NP_439896.1;XP_016865725.1;XP_024302089.1;XP_024302090.1 | 21573 | 613361 | . | . | . | 19.214491 | 17.558321 | 26.361200 | 21.521829 | 44.712910 | 40.586844 | 59.153770 | 49.207102 | 1.264485 | 1689.642791 | 0.146358485 | 2.313155445 | 0.020714094 | 0.476527738 | 10.543041 | 10.882090 | 10.648082 | 10.591409 | 10.902165 | 10.695461 | 1782 | 1550 | 1852 | 1542 |
| 116931 | NM_053002,XI | MED12L | mediator complex subunit 12L | protein_coding | NP_443728.3;XP_006713550.1;XP_011510692.1;XP_011510696.1;XP_01 | 16050 | 611318 | . | . | . | 0.344297 | 0.285440 | 0.244310 | 0.319834 | 0.801192 | 0.659808 | 0.548224 | 0.731264 | -1.129992 | 101.486961 | 0.329802187 | -0.53460127 | 0.592925606 | 0.998865404 | 6.750984 | 6.573886 | 6.714633 | 6.651486 | 6.607712 | 6.682934 | 136 | 105 | 72 | 101 |
| 116935 | gene-RPL3P2 | RPL3P2 | ribosomal protein L3 pseudogene 2 | pseudogene | . | 17834 | . | . | . | . | 0.233905 | 0.153452 | 0.135878 | 0.289524 | 0.544307 | 0.354711 | 0.304906 | 0.661961 | 1.091767 | 8.380821 | 1.066487015 | 0.118768521 | 0.905458748 | NA | 3.003609 | 3.127898 | 3.072376 | 3.058230 | 3.055505 | 3.081633 | 11 | 7 | 5 | 11 |
| 116936 | NR_000026 | SNORD139 | small nucleolar RNA, C/D box 139 | snoRNA | . | 50421 | 611069 | . | . | . | 19.020802 | 19.788519 | 23.788689 | 18.585842 | 44.262188 | 45.742046 | 53.381130 | 42.494320 | 1.089740 | 35.172360 | 0.525431913 | 0.235965466 | 0.813459468 | 0.998865404 | 5.072328 | 5.197690 | 5.124318 | 5.129649 | 5.156707 | 5.120768 | 38 | 37 | 36 | 30 |
| 116937 | NR_000027 | SNORD83A | small nucleolar RNA, C/D box 83A | snoRNA | . | 17131 | 611070 | . | . | . | 11.862437 | 8.086039 | 10.638628 | 11.764368 | 27.604380 | 18.691240 | 23.872774 | 26.897831 | 1.127918 | 31.775196 | 0.557839909 | 0.311312267 | 0.755563243 | 0.998865404 | 4.899695 | 5.074662 | 5.002876 | 4.955297 | 4.988776 | 5.000578 | 41 | 26 | 28 | 33 |
| 116938 | NR_000028 | SNORD83B | small nucleolar RNA, C/D box 83B | snoRNA | . | 17132 | 611071 | . | . | . | 9.690513 | 8.193601 | 9.792487 | 13.891735 | 22.550224 | 18.939875 | 21.974058 | 31.761801 | 1.314256 | 30.509165 | 0.56799446 | 0.694101961 | 0.487618269 | 0.998865404 | 4.721207 | 5.114423 | 4.917388 | 4.898406 | 4.918205 | 4.964199 | 33 | 26 | 25 | 38 |
| 116966 | NM_00135072 | WDR17 | WD repeat domain 17 | protein_coding | NP_001337656.1;NP_001365032.1;NP_001365033.1;NP_001365034.1;NI | 16661 | 609005 | . | . | . | 0.534831 | 0.565736 | 0.524941 | 0.664444 | 1.244573 | 1.307723 | 1.177953 | 1.519172 | 1.068196 | 134.489443 | 0.286839107 | 0.331812269 | 0.740031019 | 0.998865404 | 7.023720 | 7.117456 | 7.047949 | 7.066710 | 7.038671 | 7.122510 | 146 | 144 | 107 | 145 |
| 116983 | NM_030649,XI | ACAP3 | ArfGAP with coiled-coil, ankyrin repeat and PH domains 3 | protein_coding | NP_085152.2;XP_005244772.1;XP_005244774.1;XP_011538908.1;XP_01 | 16754 | . | . | . | . | 27.906028 | 26.936004 | 32.145529 | 28.299398 | 64.938476 | 62.263783 | 72.133636 | 64.703212 | 1.109135 | 3566.699518 | 0.116403945 | 1.283762961 | 0.199224907 | 0.96621494 | 11.723647 | 11.873225 | 11.750058 | 11.760720 | 11.892435 | 11.786788 | 3869 | 3673 | 3469 | 3294 |
| 116984 | NM_015230,N | ARAP2 | ArfGAP with RhoGAP domain, ankyrin repeat and PH domain 2 | protein_coding | NP_056045.2;XP_016863188.1;XP_016863189.1;XP_016863190.1;XP_01 | 16924 | 606645 | . | . | . | 0.755308 | 0.741769 | 1.011201 | 0.775517 | 1.757633 | 1.714633 | 2.269106 | 1.773126 | 1.171645 | 195.771456 | 0.249321286 | 0.91663289 | 0.359335043 | 0.998865404 | 7.493553 | 7.723359 | 7.572368 | 7.570611 | 7.701189 | 7.579893 | 208 | 194 | 208 | 172 |
| 116985 | NM_00104011 | ARAP1 | ArfGAP with RhoGAP domain, ankyrin repeat and PH domain 1 | protein_coding | NP_001035207.1;NP_001128662.1;NP_001356418.1;NP_056057.2 | 16925 | 606646 | . | . | . | 13.411044 | 14.033874 | 15.149553 | 13.813128 | 31.208052 | 32.439928 | 33.995158 | 31.582077 | 1.034282 | 2200.096164 | 0.120396119 | 0.403913901 | 0.686276034 | 0.998865404 | 11.078774 | 11.127516 | 11.071247 | 11.106961 | 11.152128 | 11.079150 | 2435 | 2386 | 2034 | 2002 |
| 116986 | NM_00112277 | AGAP2 | ArfGAP with GTPase domain, ankyrin repeat and PH domain 2 | protein_coding | NP_001116244.1;NP_055585.1;XP_005268682.1;XP_005268683.1;XP_01 | 16921 | 605476 | . | . | . | 0.066366 | 0.079298 | 0.117947 | 0.069215 | 0.154435 | 0.183301 | 0.264670 | 0.158251 | 1.364778 | 16.453094 | 0.768229953 | 0.584026578 | 0.559202431 | NA | 3.796818 | 4.248540 | 4.017670 | 4.017862 | 4.060795 | 4.018090 | 16 | 15 | 21 | 13 |
| 116987 | NM_00103713 | AGAP1 | ArfGAP with GTPase domain, ankyrin repeat and PH domain 1 | protein_coding | NP_001032208.1;NP_001231817.1;NP_055729.2;XP_005246116.1;XP_00 | 16922 | 608651 | . | . | . | 3.719388 | 3.973011 | 3.268443 | 4.409750 | 8.655168 | 9.183796 | 7.334293 | 10.082366 | -1.042380 | 1198.679961 | 0.160708755 | -0.37260861 | 0.709439763 | 0.998865404 | 10.257216 | 10.196609 | 10.215170 | 10.263943 | 10.085599 | 10.318103 | 1364 | 1363 | 862 | 1271 |
| 116988 | NM_00104253 | AGAP3 | ArfGAP with GTPase domain, ankyrin repeat and PH domain 3 | protein_coding | NP_001036000.1;NP_001268229.1;NP_001295233.1;NP_001295234.1;NI | 16923 | 616813 | . | . | . | 32.724682 | 31.295689 | 36.102123 | 39.412271 | 76.151681 | 72.341384 | 81.012117 | 90.111477 | 1.134192 | 3345.905646 | 0.11311669 | 1.605999429 | 0.108274035 | 0.838332591 | 11.614529 | 11.796123 | 11.667705 | 11.639637 | 11.723558 | 11.790039 | 3673 | 3325 | 2969 | 3467 |
| 117143 | NM_053053 | TADA1 | transcriptional adaptor 1 | protein_coding | NP_444281.1 | 30631 | 612763 | . | . | . | 2.937783 | 2.223777 | 2.602340 | 2.662137 | 6.836342 | 5.140361 | 5.839575 | 6.086660 | 1.005303 | 170.988239 | 0.26471368 | 0.02882742 | 0.977002232 | 0.998865404 | 7.413445 | 7.422049 | 7.472704 | 7.353408 | 7.413929 | 7.420204 | 224 | 158 | 148 | 162 |
| 117145 | NM_053055,XI | THEM4 | thioesterase superfamily member 4 | protein_coding | NP_444283.2;XP_011507444.1 | 17947 | 606388 | . | . | . | 24.749881 | 22.871226 | 31.240338 | 24.915507 | 57.593990 | 52.867859 | 70.102414 | 56.966345 | 1.158620 | 2790.723924 | 0.114096207 | 1.861653887 | 0.062651892 | 0.723723143 | 11.336250 | 11.548777 | 11.386704 | 11.379332 | 11.530349 | 11.474787 | 2991 | 2777 | 2693 | 2714 |
| 117151 | gene-SLC31A1I | SLC31A1P1 | solute carrier family 31 member 1 pseudogene 1 | pseudogene | . | 17124 | . | . | . | . | 0.246430 | 0.240851 | 0.179690 | 0.217930 | 0.573452 | 0.556738 | 0.403219 | 0.498271 | -1.210401 | 33.538029 | 0.536205606 | -0.51376682 | 0.607415026 | 0.998865404 | 5.199399 | 4.922818 | 5.085707 | 5.080719 | 5.047220 | 5.068417 | 43 | 39 | 24 | 31 |
| 117155 | NM_00128230 | CATSPER2 | cation channel sperm associated 2 | protein_coding | NP_001269238.1;NP_001269239.1;NP_742093.1 | 18810 | 607249 | . | . | . | 0.406380 | 0.427851 | 0.765020 | 0.222818 | 0.945663 | 0.988998 | 1.716683 | 0.509446 | 1.112521 | 30.444550 | 0.620591433 | 0.247880252 | 0.80422706 | 0.998865404 | 4.847059 | 5.004852 | 4.894650 | 4.917510 | 4.976316 | 4.858389 | 30 | 34 | 41 | 16 |
| 117177 | NM_00102464 | RAB3IP | RAB3A interacting protein | protein_coding | NP_001019818.1;NP_001265331.1;NP_071901.2;NP_783322.1;NP_7833 | 16508 | 608686 | . | . | . | 0.833763 | 0.864272 | 0.807683 | 0.826536 | 1.940201 | 1.997806 | 1.812417 | 1.889777 | -1.057959 | 246.476513 | 0.216735166 | -0.37503899 | 0.70763147 | 0.998865404 | 7.985494 | 7.903967 | 7.951041 | 7.967805 | 7.926270 | 7.936690 | 287 | 278 | 206 | 227 |
| 117178 | NM_00116629 | SSX2IP | SSX family member 2 interacting protein | protein_coding | NP_001159765.1;NP_001159766.1;NP_001159767.1;NP_001159889.1;NI | 16509 | 608690 | . | . | . | 4.920450 | 5.048142 | 4.793182 | 4.379064 | 11.450090 | 11.669007 | 10.755763 | 10.012210 | -1.088553 | 803.182381 | 0.148870609 | -0.82227236 | 0.410921902 | 0.998865404 | 9.709254 | 9.587339 | 9.688108 | 9.668501 | 9.659335 | 9.576222 | 977 | 891 | 709 | 678 |
| 117196 | NM_054032 | MRGPRX4 | MAS related GPR family member X4 | protein_coding | NP_473373.2 | 17617 | 607230 | . | . | . | 0.343999 | 0.217266 | 0.490444 | 0.376649 | 0.800499 | 0.502220 | 1.100542 | 0.861162 | 1.496536 | 16.690188 | 0.764254231 | 0.761039121 | 0.446633701 | NA | 3.738475 | 4.324312 | 4.049025 | 4.020813 | 4.075808 | 4.051445 | 19 | 11 | 20 | 16 |
| 117246 | NM_017647,XI | FTSJ3 | FtsJ RNA 2'-O-methyltransferase 3 | protein_coding | NP_060117.3;XP_024306330.1 | 17136 | 618411 | . | . | . | 22.626345 | 23.296605 | 22.292935 | 22.551521 | 52.652439 | 53.851147 | 50.024699 | 51.561376 | -1.034669 | 2464.150235 | 0.114139533 | -0.43077896 | 0.666629102 | 0.998865404 | 11.291277 | 11.242052 | 11.270948 | 11.289977 | 11.247919 | 11.257986 | 2852 | 2736 | 2093 | 2285 |
| 117247 | NM_018593,XI | SLC16A10 | solute carrier family 16 member 10 | protein_coding | NP_061063.2;XP_006715392.1;XP_011533724.1;XP_016865726.1 | 17027 | 607550 | . | . | . | 0.782863 | 1.102201 | 0.326908 | 1.027345 | 1.821753 | 2.547785 | 0.733572 | 2.348901 | -1.229156 | 111.349826 | 0.315223717 | -0.9443066 | 0.345012993 | 0.998865404 | 6.940077 | 6.642518 | 6.874851 | 6.786210 | 6.744262 | 6.784021 | 159 | 116 | 81 | 100 |
| 117248 | NM_00131905 | GALNT15 | polypeptide N-acetylgalactosaminyltransferase 15 | protein_coding | NP_001305980.1;NP_001305981.1;NP_473451.3;XP_005264909.1;XP_01 | 21531 | 615131 | . | . | . | 0.014087 | 0.009468 | 0.024101 | 0.006132 | 0.032782 | 0.021885 | 0.054082 | 0.014020 | 1.258236 | 6.065647 | 1.263497175 | 0.26228978 | 0.79309804 | NA | 2.421266 | 2.760196 | 2.588903 | 2.582140 | 2.607479 | 2.575878 | 7 | 5 | 9 | 3 |
| 117583 | NM_00130276 | PARD3B | par-3 family cell polarity regulator beta | protein_coding | NP_001289698.1;NP_476518.4;NP_689739.4;NP_995585.2;XP_0115088 | 14446 | . | . | . | . | 0.505556 | 0.632981 | 0.439913 | 0.609302 | 1.176449 | 1.463165 | 0.987150 | 1.393097 | -1.070770 | 149.665670 | 0.272406089 | -0.36213665 | 0.717249918 | 0.998865404 | 7.274722 | 7.174745 | 7.224453 | 7.252821 | 7.177006 | 7.246084 | 172 | 173 | 114 | 148 |
| 117584 | NM_00101736 | RFFL | ring finger and FYVE like domain containing E3 ubiquitin protein lig | protein_coding | NP_001017368.1 | 24821 | 609735 | . | . | . | 1.364470 | 1.851458 | 1.855132 | 1.804382 | 3.175177 | 4.279728 | 4.162863 | 4.125505 | 1.124981 | 386.611228 | 0.20161951 | 0.842679889 | 0.399407498 | 0.998865404 | 8.507671 | 8.676852 | 8.466715 | 8.636178 | 8.635858 | 8.612220 | 358 | 450 | 363 | 376 |
| 117608 | NM_058230 | ZNF354B | zinc finger protein 354B | protein_coding | NP_478137.1 | 17197 | . | . | . | . | 4.826308 | 4.428199 | 5.790845 | 3.071184 | 11.231018 | 10.235981 | 12.994490 | 7.021896 | -1.059465 | 395.284517 | 0.251115821 | -0.33186503 | 0.739991181 | 0.998865404 | 8.666962 | 8.585380 | 8.659522 | 8.602437 | 8.763903 | 8.397421 | 490 | 418 | 438 | 248 |
| 117854 | NM_00100381 | TRIM6 | tripartite motif containing 6 | protein_coding | NP_001003818.1;NP_001185573.1;NP_001185574.1;NP_477514.1 | 16277 | 607564 | . | . | . | 0.197782 | 0.306594 | 0.091278 | 0.182076 | 0.460248 | 0.708705 | 0.204826 | 0.416296 | -1.855819 | 21.155910 | 0.704301933 | -1.26658233 | 0.205304673 | NA | 4.783296 | 3.885075 | 4.406995 | 4.455613 | 4.359707 | 4.396444 | 25 | 36 | 9 | 18 |
| 118424 | NM_058167,N | UBE2J2 | ubiquitin conjugating enzyme E2 J2 | protein_coding | NP_477515.2;NP_919296.1;NP_919439.1;NP_919440.1;XP_005244775.1 | 19268 | . | . | . | . | 19.178718 | 18.561308 | 19.694822 | 21.971370 | 44.629663 | 42.905294 | 44.194611 | 50.234928 | 1.102022 | 1492.512377 | 0.128711788 | 1.088893589 | 0.276200814 | 0.998865404 | 10.471827 | 10.611832 | 10.521241 | 10.486922 | 10.540791 | 10.616281 | 1672 | 1498 | 1295 | 1538 |
| 118426 | NM_00130074 | BORCS5 | BLOC-1 related complex subunit 5 | protein_coding | NP_001287671.1;NP_001317285.1;NP_477517.1;XP_011518853.1;XP_01 | 17950 | 616598 | . | . | . | 1.377544 | 1.334756 | 1.373345 | 1.604844 | 3.205604 | 3.085346 | 3.081747 | 3.669284 | 1.053327 | 245.714396 | 0.219932112 | 0.34080537 | 0.733250107 | 0.998865404 | 7.903268 | 7.977456 | 7.898219 | 7.954589 | 7.936045 | 7.968580 | 260 | 273 | 212 | 244 |
| 118429 | NM_00114579 | ANTXR2 | ANTXR cell adhesion molecule 2 | protein_coding | NP_001139266.1;NP_001273709.1;NP_001273710.1;NP_477520.2;XP_01 | 21732 | 608041 | . | . | . | 2.065993 | 2.377271 | 1.800464 | 2.085785 | 4.807651 | 5.495167 | 4.040188 | 4.768896 | -1.319833 | 443.719496 | 0.17982262 | -2.22638832 | 0.025988182 | 0.527323808 | 8.980143 | 8.579081 | 8.827196 | 8.912942 | 8.681531 | 8.720655 | 543 | 581 | 321 | 371 |
| 118433 | NR_000029,NF | RPL23AP7 | ribosomal protein L23a pseudogene 7 | transcribed_ps | . | 17336 | . | . | . | . | 12.061186 | 12.431338 | 15.155288 | 13.389403 | 28.066874 | 28.735597 | 34.008024 | 30.613280 | 1.176745 | 456.331672 | 0.17220364 | 1.363513928 | 0.172720598 | 0.947431715 | 8.711703 | 8.946622 | 8.776160 | 8.783320 | 8.891683 | 8.868608 | 480 | 455 | 437 | 454 |
| 118460 | NM_058219 | EXOSC6 | exosome component 6 | protein_coding | NP_478126.1 | 19055 | 606490 | . | . | . | 4.497417 | 4.230018 | 4.498470 | 4.076275 | 10.465675 | 9.777876 | 10.094437 | 9.319919 | -1.032802 | 696.456246 | 0.155050914 | -0.30031044 | 0.763940374 | 0.998865404 | 9.466696 | 9.420716 | 9.478134 | 9.429972 | 9.468184 | 9.395229 | 843 | 737 | 628 | 608 |
| 118472 | NM_145806,N | ZNF511 | zinc finger protein 511 | protein_coding | NP_665805.2 | 28445 | . | . | . | . | 14.174953 | 15.186407 | 13.112643 | 17.496822 | 32.985700 | 35.104065 | 29.424389 | 40.004404 | 1.029170 | 506.551926 | 0.185232034 | 0.223943212 | 0.822821491 | 0.998865404 | 8.964246 | 9.004605 | 8.953842 | 8.991897 | 8.899505 | 9.075281 | 558 | 555 | 385 | 548 |
| 118487 | NM_203298 | CHCHD1 | coiled-coil-helix-coiled-coil-helix domain containing 1 | protein_coding | NP_976043.1 | 23518 | 608842 | . | . | . | 20.748205 | 21.088964 | 21.553200 | 22.577560 | 48.281929 | 48.748089 | 48.364750 | 51.620914 | 1.040285 | 569.451082 | 0.160022538 | 0.356068524 | 0.721789249 | 0.998865404 | 9.124746 | 9.181551 | 9.137221 | 9.142999 | 9.154259 | 9.177521 | 640 | 605 | 496 | 554 |
| 118490 | NM_00102459 | MSS51 | MSS51 mitochondrial translational activator | protein_coding | NP_001019764.1 | 21000 | 614773 | . | . | . | 0.285434 | 0.577814 | 0.710002 | 0.577805 | 0.664217 | 1.335644 | 1.593225 | 1.321082 | 1.460185 | 41.211347 | 0.519623326 | 1.05105146 | 0.293234956 | 0.998865404 | 5.067450 | 5.611499 | 5.271603 | 5.361718 | 5.395829 | 5.358432 | 26 | 48 | 47 | 41 |
| 118491 | NM_00135093 | CFAP70 | cilia and flagella associated protein 70 | protein_coding | NP_001337862.1;NP_001337863.1;NP_001354730.1;XP_005269546.1;XF | 30726 | 618661 | . | . | . | 0.184487 | 0.210063 | 0.229451 | 0.228995 | 0.429308 | 0.485572 | 0.514881 | 0.523571 | 1.194711 | 32.430964 | 0.543751392 | 0.472019899 | 0.636912585 | 0.998865404 | 4.885686 | 5.141582 | 5.007652 | 5.000899 | 5.015175 | 5.039331 | 35 | 31 | 28 | 36 |
| 118611 | NM_00100429 | C10orf90 | chromosome 10 open reading frame 90 | protein_coding | NP_001004298.2;NP_001337850.1;NP_001337851.1;NP_001337852.1;XF | 26563 | 617735 | . | . | . | 12.046164 | 13.000316 | 15.366745 | 17.401015 | 28.031923 | 30.050815 | 34.482530 | 39.785353 | 1.323863 | 1527.960638 | 0.128594218 | 3.147527346 | 0.001646577 | 0.097126935 | 10.361046 | 10.765485 | 10.420628 | 10.492539 | 10.651362 | 10.696657 | 1442 | 1487 | 1471 | 1678 |

| | | | | | | | | | | | | | | | | | | | | | | | | | | | | | | | | | |
|---|---|---|---|---|---|---|---|---|---|---|---|---|---|---|---|---|---|---|---|---|---|---|---|---|---|---|---|---|---|---|---|---|---|
| 118663 | NM_00131818 | BTBD16 | BTB domain containing 16 | protein_coding | NP_001305118.1;NP_653188.2;XP_011537541.1;XP_011537542.1;XP_01 | 26340 | . | . | . | 0.018752 | 0.085929 | 0.072116 | 0.048437 | 0.043637 | 0.198628 | 0.161826 | 0.110745 | 1.218078 | 5.577692 | 1.33690256 | 0.212885119 | 0.83141657 | NA | 2.333078 | 2.612728 | 2.449103 | 2.480332 | 2.473303 | 2.466398 | 2 | 9 | 6 | 5 |
| 118672 | NM_00136353 | PSTK | phosphoseryl-tRNA kinase | protein_coding | NP_001350460.1;NP_699167.2;XP_016871130.1 | 28578 | 611310 | . | . | 1.710939 | 2.232766 | 2.114193 | 2.317748 | 3.981425 | 5.161139 | 4.744187 | 5.299256 | 1.056061 | 81.789901 | 0.354426795 | 0.222027975 | 0.824292106 | 0.998865404 | 6.314648 | 6.392016 | 6.319163 | 6.369587 | 6.348031 | 6.370244 | 83 | 94 | 70 | 82 |
| 118738 | NM_00134693 | ZNF488 | zinc finger protein 488 | protein_coding | NP_001333861.1;NP_001333862.1;NP_001333863.1;NP_001333864.1;NI | 23535 | . | . | . | 0.578912 | 0.389409 | 0.425768 | 0.407054 | 1.347151 | 0.900137 | 0.955409 | 0.930681 | -1.316706 | 59.768739 | 0.414358177 | -0.95794834 | 0.338088809 | 0.998865404 | 6.085264 | 5.690446 | 5.957083 | 5.902785 | 5.889242 | 5.855102 | 87 | 65 | 49 | 44 |
| 118812 | NM_00109883 | MORN4 | MORN repeat containing 4 | protein_coding | NP_001092301.1;NP_849154.1;XP_011537552.1;XP_011537553.1 | 24001 | 617736 | . | . | 3.293891 | 4.328353 | 3.882854 | 3.873465 | 7.665020 | 10.005183 | 8.713012 | 8.856217 | 1.002529 | 282.624073 | 0.220042907 | 0.016563332 | 0.986784978 | 0.999197585 | 8.141305 | 8.144175 | 8.064816 | 8.209002 | 8.146988 | 8.134381 | 281 | 346 | 249 | 262 |
| 118813 | NM_00100226 | ZFYVE27 | zinc finger FYVE-type containing 27 | protein_coding | NP_001002261.1;NP_001002262.1;NP_001167590.1;NP_001167591.1;NI | 26559 | 610243 | . | . | 23.931651 | 23.211078 | 23.275608 | 21.360204 | 55.689933 | 53.653439 | 52.229791 | 48.837565 | -1.041771 | 2200.067956 | 0.121218601 | -0.48703586 | 0.626232917 | 0.998865404 | 11.132426 | 11.073640 | 11.132449 | 11.105578 | 11.127550 | 11.043578 | 2628 | 2382 | 1973 | 1914 |
| 118881 | NM_144589,XI | COMTD1 | catechol-O-methyltransferase domain containing 1 | protein_coding | NP_653190.2;XP_024303559.1 | 26309 | . | . | . | 4.699423 | 5.584490 | 6.426383 | 5.265510 | 10.935750 | 12.908801 | 14.420619 | 12.038963 | 1.142925 | 202.306376 | 0.242422268 | 0.795022363 | 0.426600494 | 0.998865404 | 7.560832 | 7.753534 | 7.587148 | 7.662584 | 7.716152 | 7.655316 | 199 | 221 | 200 | 189 |
| 118924 | NM_00134771 | FRA10AC1 | FRA10A associated CGG repeat 1 | protein_coding | NP_001334641.1;NP_001334642.1;NP_001334643.1;NP_001334644.1;NI | 1162 | 608866 | . | . | 3.161061 | 2.569622 | 2.107469 | 1.644386 | 7.355918 | 5.939796 | 4.729098 | 3.759694 | -1.524497 | 236.926610 | 0.237304345 | -2.56351416 | 0.010361847 | 0.328038024 | 8.159899 | 7.553420 | 8.029049 | 7.920556 | 7.820419 | 7.722198 | 362 | 278 | 182 | 156 |
| 118980 | NM_00135098 | SFXN2 | sideroflexin 2 | protein_coding | NP_001337918.1;NP_849189.1;XP_011537563.1;XP_011537565.1;XP_01 | 16086 | 615570 | . | . | 1.124627 | 1.004462 | 1.251362 | 1.840572 | 2.617054 | 2.321858 | 2.808021 | 4.208249 | 1.199404 | 231.241118 | 0.228796266 | 1.146513326 | 0.251582817 | 0.994159066 | 7.716258 | 7.978362 | 7.832252 | 7.775448 | 7.853953 | 7.928629 | 257 | 213 | 203 | 254 |
| 118987 | NM_173791,XI | PDZD8 | PDZ domain containing 8 | protein_coding | NP_776152.1;XP_005269575.1;XP_011537567.1;XP_011537568.1 | 26974 | 614235 | . | . | 3.127641 | 3.340707 | 2.857129 | 2.228453 | 7.278150 | 7.722192 | 6.411317 | 5.095093 | -1.273717 | 831.656553 | 0.15699981 | -2.22321529 | 0.026201286 | 0.529511499 | 9.863485 | 9.515239 | 9.790337 | 9.764840 | 9.677193 | 9.530010 | 1091 | 988 | 705 | 612 |
| 119016 | NM_00127634 | AGAP4 | ArfGAP with GTPase domain, ankyrin repeat and PH domain 4 | protein_coding | NP_001263272.2;NP_001278308.1;NP_597703.2 | 23459 | . | . | . | 1.064939 | 1.389085 | 1.522954 | 1.002277 | 2.478157 | 3.210933 | 3.417464 | 2.291587 | 1.020367 | 101.040656 | 0.348423008 | 0.083483767 | 0.933466885 | 0.998865404 | 6.643606 | 6.673820 | 6.603592 | 6.690697 | 6.717882 | 6.586323 | 99 | 123 | 108 | 75 |
| 119032 | NM_00113620 | BORCS7 | BLOC-1 related complex subunit 7 | protein_coding | NP_001129672.1;NP_653192.2 | 23516 | 616600 | . | . | 3.034972 | 2.222815 | 3.011999 | 2.344172 | 7.062503 | 5.138134 | 6.758838 | 5.359671 | -1.006427 | 187.278154 | 0.269697805 | -0.03426935 | 0.972662364 | 0.998865404 | 7.552543 | 7.545526 | 7.616764 | 7.469117 | 7.596000 | 7.486857 | 252 | 169 | 183 | 153 |
| 119369 | NR_002779 | NUDT9P1 | nudix hydrolase 9 pseudogene 1 | transcribed_ps | . | 28523 | . | . | . | 0.124811 | 0.215530 | 0.179225 | 0.393558 | 0.290441 | 0.498206 | 0.402177 | 0.899824 | 1.774596 | 8.130672 | 1.093387016 | 0.756813702 | 0.449161482 | NA | 2.555346 | 3.376188 | 2.991920 | 3.006511 | 3.004432 | 3.034979 | 5 | 8 | 6 | 13 |
| 119385 | NM_133447 | AGAP11 | ArfGAP with GTPase domain, ankyrin repeat and PH domain 11 | protein_coding | NP_597704.1 | 29421 | . | . | . | 0.164771 | 0.144715 | 0.161475 | 0.098610 | 0.383428 | 0.334515 | 0.362345 | 0.225459 | -1.190631 | 16.480014 | 0.763248966 | -0.32980913 | 0.741544187 | NA | 4.161369 | 3.913269 | 4.054928 | 4.044179 | 4.051241 | 4.022970 | 22 | 18 | 16 | 11 |
| 119391 | NM_00119101 | GSTO2 | glutathione S-transferase omega 2 | protein_coding | NP_001177942.1;NP_001177943.1;NP_001177944.1;NP_899062.1;XP_0C | 23064 | 612314 | . | . | 0.643759 | 0.420294 | 1.034992 | 0.696116 | 1.498053 | 0.971527 | 2.322493 | 1.591585 | 1.554042 | 59.720876 | 0.43066069 | 1.476860654 | 0.139712994 | 0.900674731 | 5.545380 | 6.184693 | 5.879228 | 5.791578 | 5.950688 | 5.907159 | 67 | 38 | 69 | 62 |
| 119392 | NM_00100275 | SFR1 | SWI5 dependent homologous recombination repair protein 1 | protein_coding | NP_001002759.1;NP_660290.3;XP_005269578.1;XP_016871161.1 | 29574 | 616527 | . | . | 4.359388 | 4.610412 | 4.833843 | 5.261605 | 10.144475 | 10.657173 | 10.847001 | 12.030035 | 1.115178 | 238.540870 | 0.219943977 | 0.715063276 | 0.474569945 | 0.998865404 | 7.817523 | 7.974399 | 7.864186 | 7.874596 | 7.907212 | 7.937882 | 257 | 246 | 212 | 243 |
| 119504 | NM_00124254 | ANAPC16 | anaphase promoting complex subunit 16 | protein_coding | NP_001229475.1;NP_001229476.1;NP_001229477.1;NP_001353720.1;NI | 26976 | 613427 | . | . | 9.002671 | 8.764077 | 9.620795 | 9.986006 | 20.949583 | 20.258556 | 21.588782 | 22.831817 | 1.089983 | 993.324482 | 0.137910782 | 0.901347916 | 0.367403365 | 0.998865404 | 9.892647 | 10.016921 | 9.941260 | 9.904959 | 9.968118 | 10.004168 | 1122 | 1000 | 879 | 995 |
| 119548 | NM_00101170 | PNLIPRP3 | pancreatic lipase related protein 3 | protein_coding | NP_001011709.2;XP_011537578.1;XP_011537579.1;XP_011537580.1;XP_ | 23492 | . | . | . | 0.575390 | 0.975764 | 0.885054 | 0.561834 | 1.338956 | 2.255522 | 1.986035 | 1.284567 | -1.088033 | 52.930329 | 0.464333716 | -0.262143 | 0.7932112 | 0.998865404 | 5.785585 | 5.663895 | 5.672006 | 5.778263 | 5.742608 | 5.676917 | 47 | 75 | 52 | 39 |
| 119559 | NM_213649,N | SFXN4 | sideroflexin 4 | protein_coding | NP_998814.1;XP_005269582.1;XP_005269583.1;XP_005269584.1;XP_01 | 16088 | 615564 | . | . | 18.468293 | 17.705268 | 19.740321 | 19.332279 | 42.976478 | 40.926518 | 44.296705 | 44.200960 | 1.054153 | 1085.849756 | 0.134544411 | 0.565493539 | 0.571738118 | 0.998865404 | 10.046015 | 10.122197 | 10.053743 | 10.074852 | 10.123115 | 10.083501 | 1200 | 1157 | 995 | 1017 |
| 119710 | NM_00127672 | IFTAP | intraflagellar transport associated protein | protein_coding | NP_001263651.1;NP_001263652.1;NP_001263653.1;NP_001263654.1;NI | 25142 | . | . | . | 9.506696 | 8.152552 | 8.271492 | 9.144113 | 22.122474 | 18.844992 | 18.560989 | 20.906928 | -1.066259 | 252.532568 | 0.221422364 | -0.4180168 | 0.675934834 | 0.998865404 | 8.025607 | 7.933577 | 8.042683 | 7.946529 | 7.957750 | 7.969499 | 328 | 255 | 210 | 232 |
| 120103 | NM_00128613 | SLC36A4 | solute carrier family 36 member 4 | protein_coding | NP_001273068.1;NP_689526.2;XP_016872663.1;XP_016872664.1;XP_01 | 19660 | 613760 | . | . | 2.029730 | 1.479497 | 2.827694 | 1.345278 | 4.723265 | 3.419924 | 6.345265 | 3.075819 | 1.033295 | 324.448121 | 0.278844438 | 0.16945952 | 0.86543521 | 0.998865404 | 8.316928 | 8.366337 | 8.401271 | 8.222052 | 8.501183 | 8.133056 | 426 | 289 | 381 | 208 |
| 120227 | NM_00137721 | CYP2R1 | cytochrome P450 family 2 subfamily R member 1 | protein_coding | NP_001364143.1;NP_001364144.1;NP_001364145.1;NP_001364146.1;NI | 20580 | 608713 | . | . | 5.253887 | 5.112969 | 6.235133 | 4.813347 | 12.226007 | 11.818856 | 13.991458 | 11.005145 | 1.068890 | 457.612749 | 0.192934473 | 0.498163114 | 0.618369079 | 0.998865404 | 8.788522 | 8.885805 | 8.826171 | 8.800064 | 8.943851 | 8.756842 | 520 | 467 | 473 | 377 |
| 120379 | NM_00108261 | PIH1D2 | PIH1 domain containing 2 | protein_coding | NP_001076088.1;NP_620144.1;XP_016872690.1;XP_016872691.1;XP_01 | 25210 | . | . | . | 1.828619 | 1.363411 | 2.637693 | 1.910349 | 4.255271 | 3.151584 | 5.918906 | 4.367785 | 1.374316 | 76.427459 | 0.379443605 | 1.208910357 | 0.226697287 | 0.982008277 | 6.006883 | 6.468402 | 6.229538 | 6.174670 | 6.328809 | 6.235091 | 83 | 61 | 90 | 69 |
| 120526 | NM_181706,XI | DNAJC24 | DnaJ heat shock protein family (Hsp40) member C24 | protein_coding | NP_859057.4 | 26979 | 611072 | . | . | 3.257162 | 1.975893 | 2.183887 | 1.522552 | 7.579549 | 4.567365 | 4.900580 | 3.481135 | -1.361104 | 165.387284 | 0.281550513 | -1.57974091 | 0.114166213 | 0.853186917 | 7.573973 | 7.131667 | 7.487787 | 7.356894 | 7.351807 | 7.242681 | 251 | 176 | 140 | 112 |
| 120534 | NM_152316,XI | ARL14EP | ADP ribosylation factor like GTPase 14 effector protein | protein_coding | NP_689529.1;XP_005252849.1 | 26798 | 612295 | . | . | 2.903384 | 3.533470 | 3.071699 | 3.087991 | 6.756293 | 8.167774 | 6.892803 | 7.060323 | -1.031441 | 263.589945 | 0.213639289 | -0.20905056 | 0.834408767 | 0.998865404 | 8.064560 | 8.019390 | 8.018404 | 8.080041 | 8.030263 | 8.037400 | 290 | 306 | 224 | 245 |
| 120824 | gene-TRIM51G | TRIM51GP | tripartite motif-containing 51G, pseudogene | pseudogene | . | 43972 | . | . | . | 0.257014 | 0.226854 | 0.161400 | 0.287321 | 0.598082 | 0.524384 | 0.362177 | 0.656925 | -1.073129 | 12.518972 | 0.869925311 | -0.1170486 | 0.906821541 | NA | 3.697260 | 3.592943 | 3.654535 | 3.646187 | 3.634161 | 3.658682 | 16 | 13 | 8 | 14 |
| 120863 | NM_00131931 | DEPDC4 | DEP domain containing 4 | protein_coding | NP_001306239.1;NP_001306240.1;NP_001351747.1;NP_689530.1;XP_01 | 22952 | . | . | . | 0.782640 | 0.833938 | 0.706482 | 1.060380 | 1.821235 | 1.927684 | 1.585329 | 2.424435 | 1.174312 | 49.015076 | 0.453474954 | 0.511198448 | 0.609212105 | 0.998865404 | 5.494437 | 5.726544 | 5.620176 | 5.572848 | 5.609097 | 5.642660 | 59 | 42 | 42 | 54 |
| 120892 | NM_198578,XI | LRRK2 | leucine rich repeat kinase 2 | protein_coding | NP_940980.4;XP_005268686.1;XP_011536179.1;XP_011536183.1;XP_01 | 18618 | 609007 | . | . | 2.289753 | 2.067290 | 2.307305 | 2.000732 | 5.328351 | 4.778633 | 5.177524 | 4.574436 | -1.021852 | 621.927081 | 0.163832397 | -0.1903535 | 0.849032129 | 0.998865404 | 9.295719 | 9.265325 | 9.318346 | 9.254200 | 9.319536 | 9.223206 | 758 | 646 | 574 | 535 |
| 121053 | NM_152318 | C12orf45 | chromosome 12 open reading frame 45 | protein_coding | NP_689531.2 | 28628 | . | . | . | 0.442057 | 0.420424 | 0.332530 | 0.377249 | 1.028684 | 0.971829 | 0.746188 | 0.862535 | -1.231075 | 288.932960 | 0.207033978 | -1.44864625 | 0.147436397 | 0.90884489 | 8.316969 | 8.016608 | 8.243440 | 8.210177 | 8.087886 | 8.145465 | 378 | 334 | 212 | 257 |
| 121227 | NM_00113605 | LRIG3 | leucine rich repeats and immunoglobulin like domains 3 | protein_coding | NP_001129523.1;NP_700356.2;XP_016874279.1 | 30991 | 608870 | . | . | 2.064265 | 2.283116 | 2.003955 | 2.274159 | 4.803629 | 5.277524 | 4.496817 | 5.199596 | -1.023297 | 262.256575 | 0.213771543 | -0.15542024 | 0.876490031 | 0.998865404 | 8.051703 | 8.017768 | 8.017308 | 8.063197 | 8.009979 | 8.047084 | 292 | 299 | 217 | 252 |
| 121260 | NM_145648,XI | SLC15A4 | solute carrier family 15 member 4 | protein_coding | NP_663623.1;XP_011536197.1;XP_016874280.1;XP_016874281.1;XP_01 | 23090 | 615806 | . | . | 18.831009 | 18.004541 | 19.308763 | 18.993736 | 43.820531 | 41.618305 | 43.328302 | 43.426925 | 1.006866 | 1613.486080 | 0.124406264 | 0.079353259 | 0.936751645 | 0.998865404 | 10.650923 | 10.660990 | 10.677076 | 10.628531 | 10.671673 | 10.645133 | 1909 | 1681 | 1432 | 1492 |
| 121268 | NM_00130312 | RHEBL1 | RHEB like 1 | protein_coding | NP_001290055.1;NP_653194.1 | 21166 | . | . | . | 4.451364 | 3.623935 | 4.225696 | 6.868684 | 10.358507 | 8.376890 | 9.482341 | 15.704431 | 1.330294 | 176.876042 | 0.287858489 | 1.430374399 | 0.152609592 | 0.916437617 | 7.246618 | 7.657426 | 7.434783 | 7.343992 | 7.400797 | 7.611812 | 195 | 145 | 135 | 233 |
| 121274 | NM_00117268 | ZNF641 | zinc finger protein 641 | protein_coding | NP_001166152.1;NP_001166153.1;NP_001352732.1;NP_001352733.1;NI | 31834 | 613906 | . | . | 0.812076 | 0.967860 | 0.865891 | 0.791074 | 1.889735 | 2.237251 | 1.943036 | 1.808698 | -1.096523 | 194.101941 | 0.242878393 | -0.54733531 | 0.58414839 | 0.998865404 | 7.665556 | 7.532729 | 7.592813 | 7.647155 | 7.600475 | 7.557979 | 220 | 232 | 169 | 165 |
| 121441 | NM_00113517 | NEDD1 | NEDD1 gamma-tubulin ring complex targeting factor | protein_coding | NP_001128647.1;NP_001128648.1;NP_001128649.1;NP_690869.1;XP_0C | 7723 | 600372 | . | . | 6.043499 | 5.414463 | 4.918915 | 4.967564 | 14.063474 | 12.515771 | 11.037903 | 11.357746 | -1.177030 | 641.943713 | 0.156277746 | -1.50470032 | 0.132401141 | 0.886721472 | 9.438918 | 9.204147 | 9.404236 | 9.352124 | 9.282965 | 9.254545 | 829 | 721 | 524 | 541 |
| 121457 | NM_153687,N | IKBIP | IKBKB interacting protein | protein_coding | NP_710154.1;NP_963906.1;NP_963907.1;XP_024304604.1 | 26430 | 609861 | . | . | 32.698896 | 27.244248 | 24.271163 | 20.457159 | 76.091678 | 62.976301 | 54.463782 | 46.772864 | -1.335970 | 2036.252976 | 0.139815259 | -2.98885923 | 0.002800211 | 0.142624776 | 11.185352 | 10.767967 | 11.148945 | 11.035519 | 10.939065 | 10.782087 | 2866 | 2340 | 1671 | 1468 |
| 121512 | NM_00130448 | FGD4 | FYVE, RhoGEF and PH domain containing 4 | protein_coding | NP_001291409.1;NP_001291410.1;NP_001291412.1;NP_001291413.1;NI | 19125 | 611104 | . | . | 3.091382 | 2.601727 | 3.034269 | 2.687760 | 7.193774 | 6.014008 | 6.808814 | 6.145244 | -1.009693 | 729.807108 | 0.1611525 | -0.08635697 | 0.931182651 | 0.998865404 | 9.517895 | 9.504818 | 9.561464 | 9.462518 | 9.558954 | 9.451662 | 904 | 735 | 681 | 628 |
| 121536 | NM_00111417 | AEBP2 | AE binding protein 2 | protein_coding | NP_001107648.1;NP_001253972.1;NP_001350665.1;NP_694939.2;XP_01 | 24051 | 617934 | . | . | 2.462144 | 2.238943 | 2.011542 | 2.281898 | 5.729512 | 5.175417 | 4.513843 | 5.217288 | -1.053953 | 391.921008 | 0.187778122 | -0.40372489 | 0.686415035 | 0.998865404 | 8.651874 | 8.575965 | 8.666191 | 8.591191 | 8.566915 | 8.628930 | 492 | 407 | 314 | 377 |
| 121601 | NM_00128661 | ANO4 | anoctamin 4 | protein_coding | NP_001273544.1;NP_001273545.1;NP_849148.2;XP_011536213.1;XP_01 | 23837 | 610111 | . | . | 7.318445 | 7.206404 | 6.877593 | 7.786080 | 17.030326 | 16.657925 | 15.433123 | 17.801948 | -1.008693 | 952.101100 | 0.141015653 | -0.0885477 | 0.929441378 | 0.998865404 | 9.901325 | 9.888589 | 9.907750 | 9.888045 | 9.856569 | 9.925435 | 1116 | 1018 | 785 | 931 |
| 121642 | NM_00100165 | ALKBH2 | alkB homolog 2, alpha-ketoglutarate dependent dioxygenase | protein_coding | NP_001001655.1;NP_001138846.1;NP_001138847.1;NP_001192108.1;NI | 32487 | 610602 | . | . | 21.245069 | 19.384036 | 18.706198 | 24.274758 | 49.438147 | 44.807066 | 41.976160 | 55.501354 | 1.060853 | 738.574163 | 0.165377187 | 0.515334188 | 0.606319513 | 0.998865404 | 9.485639 | 9.570316 | 9.530946 | 9.479852 | 9.459575 | 9.626908 | 856 | 745 | 582 | 798 |
| 121643 | NM_213596,XI | FOXN4 | forkhead box N4 | protein_coding | NP_998761.2;XP_011536224.1;XP_011536225.1;XP_016874307.1;XP_01 | 21399 | 609429 | . | . | 0.236876 | 0.191910 | 0.267502 | 0.215592 | 0.551220 | 0.443609 | 0.600266 | 0.492925 | 1.022840 | 25.758297 | 0.610068635 | 0.053404186 | 0.95740987 | NA | 4.669607 | 4.704117 | 4.698354 | 4.671201 | 4.693380 | 4.681090 | 33 | 24 | 24 | 23 |
| 121665 | NM_139015,XI | SPPL3 | signal peptide peptidase like 3 | protein_coding | NP_620584.2;XP_011536227.1 | 30424 | 608240 | . | . | 9.383608 | 8.407763 | 9.593728 | 8.369522 | 21.836040 | 19.434921 | 21.528048 | 19.135918 | -1.048966 | 1127.671572 | 0.138704608 | -0.49723299 | 0.619024756 | 0.998865404 | 10.172997 | 10.104451 | 10.199842 | 10.110550 | 10.144053 | 10.096675 | 1408 | 1172 | 988 | 999 |
| 122060 | NM_00104015 | SLAIN1 | SLAIN motif family member 1 | protein_coding | NP_001035243.2;NP_001229797.1;NP_001229798.1;NP_001229799.1;NI | 26387 | 610491 | . | . | 2.589934 | 2.710751 | 3.907681 | 3.850638 | 6.026882 | 6.266023 | 8.768722 | 8.804024 | 1.408043 | 303.441211 | 0.199770301 | 2.471293823 | 0.013462516 | 0.383747314 | 7.977393 | 8.471117 | 8.135212 | 8.143416 | 8.329481 | 8.317457 | 288 | 274 | 312 | 329 |
| 122416 | NM_00134865 | ANKRD9 | ankyrin repeat domain 9 | protein_coding | NP_001335580.1;NP_001335581.1;NP_689539.1;XP_005267367.1 | 20096 | 618605 | . | . | 2.105364 | 1.812558 | 1.865158 | 1.943350 | 4.899269 | 4.189809 | 4.185358 | 4.443238 | -1.028778 | 350.508733 | 0.195953811 | -0.20888758 | 0.834535998 | 0.998865404 | 8.473453 | 8.432874 | 8.508597 | 8.410483 | 8.432484 | 8.456663 | 444 | 351 | 294 | 331 |
| 122481 | NM_00135088 | AK7 | adenylate kinase 7 | protein_coding | NP_001337817.1;NP_001337819.1;NP_001337820.1;NP_001337821.1;NI | 20091 | 615364 | . | . | 0.185666 | 0.269834 | 0.135951 | 0.165973 | 0.432050 | 0.623733 | 0.305071 | 0.379477 | -1.504778 | 22.130005 | 0.666691731 | -0.8842921 | 0.376538552 | NA | 4.733907 | 4.141582 | 4.468162 | 4.509121 | 4.445349 | 4.457844 | 25 | 34 | 14 | 18 |
| 122509 | NM_145249,N | IFI27L1 | interferon alpha inducible protein 27 like 1 | protein_coding | NP_660292.1;NP_996832.1;XP_005267371.1;XP_011534707.1;XP_01153 | 19754 | 611320 | . | . | 12.566707 | 12.265306 | 12.612166 | 12.093241 | 29.243243 | 28.351801 | 28.301337 | 27.649759 | -1.004790 | 407.574128 | 0.184676934 | -0.03732968 | 0.970222142 | 0.998865404 | 8.673938 | 8.667893 | 8.691362 | 8.651080 | 8.710208 | 8.625469 | 486 | 426 | 379 | 354 |
| 122525 | NM_00101792 | C14orf28 | chromosome 14 open reading frame 28 | protein_coding | NP_001017923.1;XP_011534710.1;XP_024305236.1 | 19834 | . | . | . | 0.453079 | 0.520333 | 0.253896 | 0.527796 | 1.054335 | 1.202774 | 0.569736 | 1.206744 | -1.106380 | 35.791013 | 0.540087198 | -0.27004238 | 0.787127646 | 0.998865404 | 5.234218 | 5.084975 | 5.171155 | 5.166495 | 5.116188 | 5.191684 | 44 | 40 | 21 | 41 |
| 122553 | NM_00107953 | TRAPPC6B | trafficking protein particle complex 6B | protein_coding | NP_001073005.1;NP_803235.1;XP_011534711.1;XP_016876453.1 | 23066 | 610397 | . | . | 5.578220 | 4.242579 | 5.953611 | 4.379353 | 12.980750 | 9.806912 | 13.359728 | 10.012867 | 1.040990 | 493.409421 | 0.212197767 | 0.27312499 | 0.784757137 | 0.998865404 | 8.916577 | 8.976092 | 9.015468 | 8.829386 | 9.058045 | 8.834189 | 638 | 445 | 515 | 389 |
| 122616 | NM_00136417 | CLBA1 | clathrin binding box of aftiphilin containing 1 | protein_coding | NP_001351099.1;NP_777551.2;XP_005267375.1 | 20126 | . | . | . | 3.220239 | 3.381674 | 4.803974 | 3.734899 | 7.493628 | 7.816892 | 10.779978 | 8.539402 | 1.281857 | 279.544579 | 0.219020318 | 1.635624555 | 0.101918188 | 0.820756563 | 7.936204 | 8.295368 | 8.042565 | 8.060732 | 8.248838 | 8.109421 | 277 | 269 | 308 | 257 |
| 122622 | NM_00132042 | ADSS1 | adenylosuccinate synthase 1 | protein_coding | NP_001307353.1;NP_689541.1;NP_954634.1;XP_006720089.1;XP_01153 | 20093 | 612498 | . | . | 0.594492 | 0.643605 | 0.192058 | 0.401565 | 1.383407 | 1.487722 | 0.430972 | 0.918131 | -1.882280 | 26.083373 | 0.629974411 | -1.44844236 | 0.147493376 | NA | 5.092061 | 4.174582 | 4.737053 | 4.745583 | 4.653748 | 4.697653 | 38 | 38 | 11 | 22 |
| 122704 | NM_00131887 | MRPL52 | mitochondrial ribosomal protein L52 | protein_coding | NP_001305799.1;NP_848026.1;NP_851313.1;NP_851821.1;NP_851822.1 | 16655 | 611856 | . | . | 22.664825 | 20.726803 | 19.518277 | 25.329232 | 52.741981 | 47.910933 | 43.798446 | 57.932277 | 1.024389 | 771.589267 | 0.162077277 | 0.214491064 | 0.830164135 | 0.998865404 | 9.574504 | 9.608672 | 9.598274 | 9.564633 | 9.513624 | 9.676590 | 899 | 803 | 600 | 817 |
| 122769 | NM_152329,N | LRR1 | leucine rich repeat protein 1 | protein_coding | NP_689542.2;NP_982292.1 | 19742 | 609193 | . | . | 11.902717 | 8.282176 | 8.514823 | 7.577107 | 27.698112 | 19.144621 | 19.107016 | 17.324154 | -1.323860 | 381.905429 | 0.2156311 | -1.87705104 | 0.060511098 | 0.719735192 | 8.764754 | 8.361269 | 8.756773 | 8.525822 | 8.514414 | 8.452793 | 593 | 383 | 302 | 291 |
| 122786 | NM_00104248 | FRMD6 | FERM domain containing 6 | protein_coding | NP_001035946.1;NP_001253975.1;NP_001253976.1;NP_689543.1;XP_0C | 19839 | 614555 | . | . | 41.456334 | 40.860542 | 31.979959 | 36.691986 | 96.470587 | 94.450977 | 71.762100 | 83.891867 | -1.220799 | 5663.542920 | 0.112334261 | -2.56222266 | 0.010400462 | 0.328507034 | 12.604282 | 12.316354 | 12.558116 | 12.532063 | 12.325250 | 12.427121 | 7279 | 6617 | 4161 | 5077 |
| 122809 | NM_080867,N | SOCS4 | suppressor of cytokine signaling 4 | protein_coding | NP_543143.1;NP_955453.1;XP_011534727.1;XP_011534728.1 | 19392 | 616337 | . | . | 4.506833 | 3.712341 | 4.770032 | 2.743474 | 10.487586 | 8.581245 | 10.703816 | 6.272627 | -1.105796 | 844.539671 | 0.417014257 | -0.34791473 | 0.727904223 | 0.998865404 | 9.792599 | 9.647813 | 9.820777 | 9.666520 | 9.847116 | 9.466318 | 1129 | 856 | 885 | 550 |
| 122830 | NM_00101171 | NAA30 | N-alpha-acetyltransferase 30, NatC catalytic subunit | protein_coding | NP_001011713.2 | 19844 | 617989 | . | . | 1.929316 | 1.525679 | 1.691869 | 1.235132 | 4.489598 | 3.526676 | 3.796506 | 2.823983 | -1.197497 | 378.406547 | 0.216825878 | -1.1992191 | 0.230442764 | 0.985768201 | 8.687145 | 8.428900 | 8.680531 | 8.532288 | 8.590181 | 8.410149 | 532 | 391 | 348 | 271 |
| 122945 | NM_00111347 | NOXRED1 | NADP dependent oxidoreductase domain containing 1 | protein_coding | NP_001106946.1;XP_005267387.1;XP_011534729.1;XP_011534730.1;XP_ | 20487 | . | . | . | 0.075839 | 0.200741 | 0.131995 | 0.356703 | 0.176480 | 0.464021 | 0.296193 | 0.815558 | 1.622707 | 13.695440 | 0.901085727 | 0.775068277 | 0.438299316 | NA | 3.389932 | 4.079653 | 3.718534 | 3.767030 | 3.743563 | 3.799158 | 6 | 17 | 9 | 22 |
| 122953 | NM_00113504 | JDP2 | Jun dimerization protein 2 | protein_coding | NP_001128519.1;NP_001128520.1;NP_001128521.1;NP_569736.1;XP_0C | 17546 | 608657 | . | . | 0.773499 | 0.544300 | 0.339024 | 0.536982 | 1.799962 | 1.258175 | 0.760759 | 1.227745 | -1.415490 | 69.158581 | 0.388920802 | -1.28895604 | 0.197413371 | 0.96621494 | 6.341648 | 5.838350 | 6.157272 | 6.147423 | 6.039789 | 6.101232 | 95 | 86 | 42 | 62 |
| 122961 | NM_00127200 | ISCA2 | iron-sulfur cluster assembly 2 | protein_coding | NP_001258936.1;NP_919255.2 | 19857 | 615317 | . | . | 4.476339 | 4.030500 | 4.210022 | 4.681979 | 10.416625 | 9.316681 | 9.447166 | 10.704790 | -1.077835 | 314.611204 | 0.214058099 | -0.50517144 | 0.613424625 | 0.998865404 | 8.351014 | 8.241771 | 8.332464 | 8.299365 | 8.194754 | 8.349223 | 387 | 342 | 225 | 325 |
| 123016 | NM_00128878 | TTC8 | tetratricopeptide repeat domain 8 | protein_coding | NP_001275710.1;NP_001275711.1;NP_001275712.1;NP_001353464.1;NI | 20087 | 608132 | . | . | 9.395422 | 7.437336 | 9.297591 | 6.552299 | 21.863530 | 17.191734 | 20.863527 | 14.981053 | 1.119182 | 805.301840 | 0.144360131 | 1.125273742 | 0.260473053 | 0.995760118 | 9.569959 | 9.732251 | 9.607441 | 9.615834 | 9.678722 | 9.703256 | 870 | 825 | 727 | 811 |
| 123036 | NM_00112859 | TC2N | tandem C2 domains, nuclear | protein_coding | NP_001122067.2;NP_001122068.2;NP_001276063.2;NP_689545.2 | 19859 | . | . | . | 0.833561 | 0.744761 | 0.598278 | 0.600943 | 1.939732 | 1.721548 | 1.342519 | 1.373985 | -1.321588 | 112.425407 | 0.305672039 | -1.31602596 | 0.188165329 | 0.960264078 | 6.999739 | 6.598037 | 6.878473 | 6.835400 | 6.769131 | 6.763228 | 156 | 130 | 85 | 90 |
| 123096 | NM_00103935 | SLC25A29 | solute carrier family 25 member 29 | protein_coding | NP_001034444.1;NP_001278742.1;NP_001278743.1;NP_001339749.1;NI | 20116 | 615064 | . | . | 11.623946 | 11.896871 | 11.772377 | 11.377347 | 27.049404 | 27.500151 | 26.416870 | 26.012952 | -1.062631 | 894.930760 | 0.141061859 | -0.6212946 | 0.534405809 | 0.998865404 | 9.848851 | 9.761078 | 9.806440 | 9.847367 | 9.783843 | 9.782915 | 1032 | 1023 | 755 | 813 |
| 123169 | NM_00128643 | LEO1 | LEO1 homolog, Paf1/RNA polymerase II complex component | protein_coding | NP_001273359.1;NP_001310832.1;NP_001310833.1;NP_620147.1;XP_01 | 30401 | 610507 | . | . | 8.745706 | 9.953016 | 8.311950 | 10.610828 | 20.351619 | 23.006844 | 18.651577 | 24.260398 | -1.003736 | 637.787517 | 0.172481766 | -0.03119182 | 0.97511656 | 0.998865404 | 9.320137 | 9.313720 | 9.274891 | 9.356115 | 9.232697 | 9.388547 | 692 | 731 | 489 | 666 |
| 123207 | NM_00116011 | C15orf40 | chromosome 15 open reading frame 40 | protein_coding | NP_001153585.1;NP_001153586.1;NP_001153587.1;NP_001153588.1;NI | 28443 | . | . | . | 3.125224 | 3.344736 | 2.913813 | 3.059860 | 7.272525 | 7.731510 | 6.538511 | 6.996006 | -1.006332 | 603.371898 | 0.16479548 | -0.05526252 | 0.955929325 | 0.998865404 | 9.241467 | 9.232326 | 9.281605 | 9.193123 | 9.199494 | 9.267616 | 743 | 610 | 496 | 593 |
| 123228 | NM_00116634 | SENP8 | SUMO peptidase family member, NEDD8 specific | protein_coding | NP_001159812.1;NP_001165580.1;NP_001165581.1;NP_001165582.1;NI | 22992 | 608659 | . | . | 0.372487 | 0.335591 | 0.521406 | 0.365990 | 0.866793 | 0.775735 | 1.170019 | 0.836792 | 1.230574 | 52.458362 | 0.441901966 | 0.677371644 | 0.498170185 | 0.998865404 | 5.554358 | 5.856096 | 5.694819 | 5.675832 | 5.759368 | 5.689736 | 57 | 48 | 59 | 45 |
| 123263 | NM_139242,XI | MTFMT | mitochondrial methionyl-tRNA formyltransferase | protein_coding | NP_640335.2;XP_005254215.2 | 29666 | 611766 | . | . | 5.740169 | 6.004058 | 7.078345 | 7.702654 | 13.357609 | 13.878649 | 15.883601 | 17.611202 | 1.160667 | 538.452535 | 0.165622446 | 1.297853278 | 0.194337758 | 0.96621494 | 8.961482 | 9.175909 | 9.004153 | 9.037532 | 9.086476 | 9.146949 | 559 | 552 | 480 | 567 |
| 123283 | NM_152334,XI | TARS3 | threonyl-tRNA synthetase 3 | protein_coding | NP_689547.2;XP_006720449.1;XP_016877401.1;XP_024305604.1 | 24728 | . | . | . | 3.662758 | 3.229707 | 3.131498 | 3.957903 | 8.523391 | 7.465613 | 7.026988 | 9.049275 | 1.013676 | 377.865538 | 0.19666345 | 0.099645678 | 0.920625629 | 0.998865404 | 8.552130 | 8.571265 | 8.593333 | 8.517909 | 8.503292 | 8.621600 | 460 | 379 | 297 | 393 |
| 123355 | NM_00128440 | LRRC28 | leucine rich repeat containing 28 | protein_coding | NP_001271329.1;NP_001308604.1;NP_001308605.1;NP_001308606.1;NI | 28355 | . | . | . | 3.349689 | 3.765359 | 3.808419 | 4.229578 | 7.794866 | 8.703790 | 8.545981 | 9.670428 | 1.044666 | 585.076519 | 0.160688181 | 0.392321613 | 0.694820596 | 0.998865404 | 9.160820 | 9.223463 | 9.149924 | 9.204167 | 9.193546 | 9.218563 | 632 | 643 | 510 | 571 |
| 123591 | NM_152335,XI | TMEM266 | transmembrane protein 266 | protein_coding | NP_689548.2;XP_005254217.1;XP_016877404.1;XP_016877405.1;XP_01 | 26763 | 618691 | . | . | 0.189798 | 0.044035 | 0.080054 | 0.059837 | 0.441667 | 0.101790 | 0.179639 | 0.136810 | -1.517625 | 8.397633 | 1.130821995 | -0.53219313 | 0.594592239 | NA | 3.336810 | 2.742346 | 3.107769 | 3.049374 | 3.069495 | 3.057434 | 19 | 4 | 7 | 5 |
| 123606 | NM_00114227 | NIPA1 | NIPA magnesium transporter 1 | protein_coding | NP_001135747.1;NP_653200.2 | 17043 | 608145 | . | . | 7.730029 | 7.226210 | 6.944967 | 6.598620 | 17.988094 | 16.703709 | 15.584305 | 15.086960 | -1.120148 | 1455.756687 | 0.128692851 | -1.27193763 | 0.203395279 | 0.968254058 | 10.586937 | 10.423546 | 10.578542 | 10.519350 | 10.485197 | 10.438629 | 1838 | 1597 | 1231 | 1248 |
| 123688 | NM_00101361 | HYKK | hydroxylysine kinase | protein_coding | NP_001013641.2;NP_001077081.1 | 34403 | 614681 | . | . | 2.271003 | 0.465587 | 0.683937 | 2.448911 | 5.284717 | 1.076226 | 1.534734 | 5.599141 | 1.161663 | 84.251695 | 0.366494126 | 0.589889403 | 0.555264799 | 0.998865404 | 6.283786 | 6.501291 | 6.425220 | 6.312484 | 6.397480 | 6.424811 | 109 | 66 | 75 | 89 |
| 123720 | NM_00108043 | WHAMM | WASP homolog associated with actin, golgi membranes and micro | protein_coding | NP_001073904.1;XP_005272478.1;XP_005272479.1;XP_005272480.1;XP_ | 30493 | 612393 | . | . | 2.280628 | 2.864979 | 3.203499 | 2.960207 | 5.307114 | 6.622526 | 7.188558 | 6.768159 | 1.174658 | 392.200790 | 0.187695064 | 1.237333393 | 0.215963335 | 0.982008277 | 8.494615 | 8.726939 | 8.528812 | 8.596388 | 8.692539 | 8.622241 | 391 | 412 | 390 | 374 |
| 123775 | NM_00110087 | C16orf46 | chromosome 16 open reading frame 46 | protein_coding | NP_001094343.1;NP_689550.2;XP_005255849.1;XP_011521150.2;XP_01 | 26525 | . | . | . | 0.374475 | 0.788181 | 1.095225 | 0.661454 | 0.871420 | 1.821916 | 2.457653 | 1.512335 | 1.357966 | 41.496477 | 0.499023384 | 0.884622602 | 0.376360212 | 0.998865404 | 5.137922 | 5.578418 | 5.307873 | 5.368359 | 5.405567 | 5.368317 | 32 | 46 | 46 | 40 |
| 123803 | NM_00127076 | NTAN1 | N-terminal asparagine amidase | protein_coding | NP_001257695.1;NP_001257696.1;NP_775745.1;XP_011520657.1 | 29909 | 615367 | . | . | 22.722099 | 22.321655 | 24.829280 | 26.749733 | 52.875257 | 51.597503 | 55.716184 | 61.160086 | 1.125959 | 903.150382 | 0.141498117 | 1.209584933 | 0.226438202 | 0.982008277 | 9.730800 | 9.901782 | 9.782111 | 9.765377 | 9.832617 | 9.885165 | 990 | 906 | 802 | 929 |
| 123811 | NM_00130449 | CEP20 | centrosomal protein 20 | protein_coding | NP_001291426.1;NP_001291427.1;NP_001291428.1;NP_001291429.1;NI | 26435 | 617149 | . | . | 16.496589 | 14.084796 | 14.997123 | 15.550810 | 38.388237 | 32.557636 | 33.653109 | 35.555078 | -1.016242 | 1073.948059 | 0.143356096 | -0.16214482 | 0.871191815 | 0.998865404 | 10.080150 | 10.057175 | 10.133343 | 10.010652 | 10.054459 | 10.069458 | 1349 | 1072 | 917 | 1009 |
| 123872 | NM_00131875 | DNAAF1 | dynein axonemal assembly factor 1 | protein_coding | NP_001305685.1;NP_848547.4;XP_006721192.1;XP_011521155.1;XP_01 | 30539 | 613190 | . | . | 0.042517 | 0.077446 | 0.039421 | 0.088904 | 0.098939 | 0.179020 | 0.088460 | 0.203269 | 1.033353 | 9.596712 | 1.053975012 | 0.044909302 | 0.964179602 | NA | 3.243129 | 3.281693 | 3.251053 | 3.266217 | 3.232571 | 3.286998 | 9 | 12 | 3 | 15 |
| 123879 | NM_173475 | DCUN1D3 | defective in cullin neddylation 1 domain containing 3 | protein_coding | NP_775746.1 | 28734 | 616167 | . | . | 1.984680 | 1.594780 | 1.862893 | 1.682757 | 4.618432 | 3.686407 | 4.180278 | 3.847424 | -1.023562 | 341.008994 | 0.204877513 | -0.16399 | 0.869739016 | 0.998865404 | 8.429777 | 8.397373 | 8.480935 | 8.351536 | 8.436648 | 8.373071 | 442 | 330 | 310 | 298 |
| 123920 | NM_00136391 | CMTM3 | CKLF like MARVEL transmembrane domain containing 3 | protein_coding | NP_001350847.1;NP_001350852.1;NP_653202.1;NP_853531.1;XP_01152 | 19174 | 607886 | . | . | 9.193065 | 10.039554 | 8.349925 | 8.598844 | 21.392641 | 23.206880 | 18.736989 | 19.660236 | -1.029909 | 572.779312 | 0.162661624 | -0.26138316 | 0.793797041 | 0.998865404 | 9.183196 | 9.140154 | 9.144760 | 9.197103 | 9.128255 | 9.174833 | 643 | 652 | 473 | 548 |
| 124044 | NM_152339,XI | SPATA2L | spermatogenesis associated 2 like | protein_coding | NP_689552.2;XP_005256336.1 | 28393 | . | . | . | 2.100216 | 2.000176 | 1.698108 | 2.255246 | 4.887287 | 4.623496 | 3.810506 | 5.156351 | -1.061114 | 150.524522 | 0.274044194 | -0.31228382 | 0.754824833 | 0.998865404 | 7.276594 | 7.189811 | 7.250727 | 7.239507 | 7.176704 | 7.265267 | 181 | 165 | 112 | 153 |
| 124045 | NM_00127190 | SPATA33 | spermatogenesis associated 33 | protein_coding | NP_001258836.1;NP_001258837.1;NP_001258838.1;NP_001258839.1;NI | 26463 | 615409 | . | . | 3.244857 | 4.167746 | 4.141660 | 3.662508 | 7.550918 | 9.633932 | 9.293764 | 8.373888 | 1.027785 | 188.420040 | 0.25517864 | 0.154945772 | 0.876864108 | 0.998865404 | 7.537871 | 7.577475 | 7.497806 | 7.596837 | 7.598848 | 7.520704 | 190 | 223 | 181 | 163 |
| 124093 | NM_00103173 | CCDC78 | coiled-coil domain containing 78 | protein_coding | NP_001026907.2;NP_001364959.1;NP_001364960.1;NP_001364962.1;XI | 14153 | 614666 | . | . | 7.982873 | 8.282750 | 9.966232 | 8.950716 | 18.576473 | 19.145949 | 22.363937 | 20.464745 | 1.144930 | 502.033516 | 0.168496668 | 1.158835462 | 0.24652326 | 0.992733415 | 8.870513 | 9.066140 | 8.925453 | 8.925534 | 9.039882 | 8.982217 | 539 | 505 | 487 | 480 |
| 124152 | NM_00130512 | IQCK | IQ motif containing K | protein_coding | NP_001292050.1;NP_001292051.1;NP_694940.1;XP_011544038.1 | 28556 | . | . | . | 0.418454 | 0.605204 | 0.353622 | 0.477902 | 0.973760 | 1.398954 | 0.793518 | 1.092667 | -1.226943 | 43.146435 | 0.473476626 | -0.62319455 | 0.533156686 | 0.998865404 | 5.571170 | 5.276108 | 5.447554 | 5.442291 | 5.417974 | 5.415029 | 54 | 52 | 34 | 36 |
| 124222 | NM_00128451 | PAQR4 | progestin and adipoQ receptor family member 4 | protein_coding | NP_001271440.1;NP_001271441.1;NP_001271442.1;NP_001311047.1;NI | 26386 | 614578 | . | . | 19.932199 | 21.211313 | 16.606724 | 20.857662 | 46.383050 | 49.030901 | 37.265005 | 47.688571 | -1.115561 | 1640.785555 | 0.140113888 | -1.12600862 | 0.260161871 | 0.995760118 | 10.757142 | 10.598860 | 10.698426 | 10.741236 | 10.547588 | 10.714734 | 1939 | 1918 | 1202 | 1612 |
| 124245 | NM_00129434 | ZC3H18 | zinc finger CCCH-type containing 18 | protein_coding | NP_001281269.1;NP_653205.3;XP_011521165.1;XP_011521167.1;XP_01 | 25091 | . | . | . | 12.917879 | 15.301632 | 14.581445 | 16.176385 | 30.060436 | 35.370409 | 32.720341 | 36.985381 | 1.070730 | 1698.827955 | 0.134372457 | 0.733739156 | 0.46310773 | 0.998865404 | 10.680403 | 10.778576 | 10.638116 | 10.761718 | 10.717057 | 10.791910 | 1740 | 1911 | 1455 | 1725 |
| 124359 | NM_152342,XI | CDYL2 | chromodomain Y like 2 | protein_coding | NP_689555.2;XP_011521168.1;XP_011521169.1;XP_024305919.1 | 23030 | 618816 | . | . | 0.023642 | 0.011859 | 0.020316 | 0.067243 | 0.055015 | 0.027412 | 0.045588 | 0.153744 | 2.243011 | 8.634432 | 1.103401353 | 1.056221779 | 0.290866903 | NA | 2.413393 | 3.577925 | 3.082422 | 3.067406 | 3.077700 | 3.131539 | 8 | 4 | 5 | 17 |
| 124401 | NM_00124292 | ANKS3 | ankyrin repeat and sterile alpha motif domain containing 3 | protein_coding | NP_001229858.1;NP_001295018.1;NP_001311058.1;NP_001311059.1;NI | 29422 | 617310 | . | . | 3.216378 | 2.777892 | 3.694380 | 3.553023 | 7.484645 | 6.421224 | 8.290084 | 8.123566 | 1.238707 | 262.766551 | 0.21219442 | 1.4554324 | 0.145549592 | 0.906189003 | 7.874693 | 8.184033 | 7.990671 | 7.963157 | 8.100433 | 8.071199 | 278 | 246 | 260 | 265 |
| 124402 | NM_00133046 | UBALD1 | UBA like domain containing 1 | protein_coding | NP_001317396.1;NP_660296.1 | 29576 | . | . | . | 6.631458 | 6.893565 | 7.162379 | 7.976783 | 15.431676 | 15.934785 | 16.072172 | 18.237967 | 1.105001 | 301.131193 | 0.201171862 | 0.716042567 | 0.473965065 | 0.998865404 | 8.160665 | 8.304260 | 8.198225 | 8.212918 | 8.239355 | 8.278716 | 325 | 313 | 265 | 307 |
| 124411 | NM_00113091 | ZNF720 | zinc finger protein 720 | protein_coding | NP_001124385.1 | 26987 | . | . | . | 1.614365 | 1.207431 | 1.053012 | 1.390754 | 3.756695 | 2.791031 | 2.362928 | 3.179793 | -1.170632 | 114.148224 | 0.321747442 | -0.70641643 | 0.479929205 | 0.998865404 | 6.944145 | 6.716406 | 6.910656 | 6.804171 | 6.764003 | 6.847484 | 163 | 113 | 79 | 112 |
| 124446 | NM_00108361 | TMEM219 | transmembrane protein 219 | protein_coding | NP_001077082.1;NP_001356617.1;NP_001356618.1;NP_001356619.1;NI | 25201 | . | . | . | 32.851375 | 36.498951 | 31.292790 | 33.784156 | 76.446500 | 84.368959 | 70.220115 | 77.243459 | 1.025625 | 1524.502717 | 0.125471114 | 0.290926063 | 0.771107868 | 0.998865404 | 10.555801 | 10.592215 | 10.538574 | 10.588889 | 10.588523 | 10.578643 | 1677 | 1677 | 1351 | 1438 |
| 124454 | NM_00108361 | EARS2 | glutamyl-tRNA synthetase 2, mitochondrial | protein_coding | NP_001077083.1;NP_001295140.1;XP_011544040.1 | 29419 | 612799 | . | . | 8.841512 | 10.326350 | 10.253290 | 10.524580 | 20.574564 | 23.869822 | 23.008084 | 24.063201 | 1.091596 | 1467.795579 | 0.130004064 | 0.972575278 | 0.330764434 | 0.998865404 | 10.454958 | 10.581154 | 10.438816 | 10.527262 | 10.546679 | 10.556283 | 1523 | 1602 | 1324 | 1445 |
| 124491 | NM_00130499 | TMEM170A | transmembrane protein 170A | protein_coding | NP_001291925.1;NP_001291926.1;NP_001291927.1;NP_660297.1;XP_01 | 29577 | . | . | . | 2.185793 | 2.219796 | 2.731000 | 1.891816 | 5.086428 | 5.131160 | 6.128285 | 4.325413 | 1.034471 | 355.523804 | 0.216723974 | 0.225598154 | 0.821513976 | 0.998865404 | 8.448470 | 8.497401 | 8.454141 | 8.463464 | 8.580960 | 8.364198 | 399 | 380 | 374 | 275 |
| 124512 | NM_00108051 | METTL23 | methyltransferase like 23 | protein_coding | NP_001073979.3;NP_001193912.1;NP_001193913.1;NP_001193914.1;NI | 26988 | 615262 | . | . | 23.859869 | 21.494946 | 24.276817 | 23.591049 | 55.522894 | 49.686527 | 54.476469 | 53.938132 | 1.021656 | 828.887443 | 0.14722497 | 0.209949784 | 0.833706806 | 0.998865404 | 9.679291 | 9.710604 | 9.724469 | 9.647782 | 9.716042 | 9.688086 | 995 | 837 | 743 | 770 |
| 124540 | NM_00132225 | MSI2 | musashi RNA binding protein 2 | protein_coding | NP_001309179.1;NP_001309180.1;NP_620412.1;NP_733839.1;XP_00525 | 18585 | 607897 | . | . | 19.700607 | 18.463350 | 19.169798 | 18.323987 | 45.844122 | 42.678865 | 43.016473 | 41.895619 | -1.070471 | 3375.053328 | 0.11488021 | -0.85520373 | 0.392438373 | 0.998865404 | 11.768898 | 11.670824 | 11.783344 | 11.710802 | 11.716466 | 11.666464 | 4194 | 3601 | 2922 | 2963 |
| 124565 | NM_00103798 | SLC38A10 | solute carrier family 38 member 10 | protein_coding | NP_001033073.1;NP_612637.1;XP_005257076.1;XP_011522590.1;XP_01 | 28237 | 616525 | . | . | 32.325189 | 32.486492 | 31.059021 | 35.079520 | 75.222040 | 75.093985 | 69.695542 | 80.205155 | 1.002382 | 3888.141279 | 0.113020351 | 0.030364636 | 0.975776248 | 0.998865404 | 11.923209 | 11.926519 | 11.927473 | 11.919685 | 11.881564 | 11.967653 | 4493 | 4171 | 3209 | 3839 |
| 124583 | NM_00115977 | CANT1 | calcium activated nucleotidase 1 | protein_coding | NP_001153244.1;NP_001153245.1;NP_620148.1;XP_005257078.1;XP_00 | 19721 | 613165 | . | . | 16.376622 | 16.365766 | 14.516481 | 16.036867 | 38.109077 | 37.830200 | 32.574562 | 36.666391 | -1.089423 | 1676.589911 | 0.123882239 | -0.99743663 | 0.318552616 | 0.998865404 | 10.771851 | 10.648125 | 10.750418 | 10.737232 | 10.647452 | 10.705388 | 2030 | 1871 | 1343 | 1561 |
| 124637 | NM_00133011 | CYB5D1 | cytochrome b5 domain containing 1 | protein_coding | NP_001317039.1;NP_653208.2 | 26516 | . | . | . | 8.647881 | 9.072650 | 9.820135 | 9.944269 | 20.123975 | 20.971835 | 22.036099 | 22.736391 | 1.109603 | 1012.585922 | 0.13605051 | 1.102850609 | 0.270092046 | 0.998865404 | 9.906916 | 10.056848 | 9.929673 | 9.957940 | 10.018324 | 10.022041 | 1084 | 1056 | 924 | 1001 |
| 124641 | NM_080822 | OVCA2 | OVCA2 serine hydrolase domain containing | protein_coding | NP_543012.1 | 24203 | 607896 | . | . | 17.162367 | 17.449247 | 21.130913 | 23.922054 | 39.937531 | 40.334721 | 47.417156 | 54.694935 | 1.284177 | 639.506426 | 0.157829352 | 2.286290489 | 0.02223727 | 0.492741101 | 9.129378 | 9.489800 | 9.224743 | 9.230855 | 9.355811 | 9.434784 | 642 | 607 | 590 | 712 |
| 124739 | NM_00126757 | USP43 | ubiquitin specific peptidase 43 | protein_coding | NP_001254505.1;NP_694942.3;XP_011521941.1;XP_011521942.1;XP_01 | 20072 | . | . | . | 0.114324 | 0.134349 | 0.191099 | 0.193012 | 0.266038 | 0.310554 | 0.428820 | 0.441300 | 1.532060 | 21.034888 | 0.67021104 | 0.918326964 | 0.358447715 | NA | 4.054853 | 4.669578 | 4.360471 | 4.369508 | 4.401636 | 4.403221 | 18 | 19 | 22 | 24 |
| 124751 | NM_00130494 | KRBA2 | KRAB-A domain containing 2 | protein_coding | NP_001291876.1;NP_998762.1 | 26989 | . | . | . | 0.514093 | 0.570107 | 0.603338 | 0.579378 | 1.196316 | 1.317827 | 1.353872 | 1.324680 | 1.056874 | 193.610180 | 0.250702064 | 0.31831718 | 0.750244354 | 0.998865404 | 7.556170 | 7.636728 | 7.554829 | 7.607050 | 7.658382 | 7.550578 | 204 | 215 | 194 | 164 |
| 124790 | NM_00130343 | HEXIM2 | HEXIM P-TEFb complex subunit 2 | protein_coding | NP_001290365.1;NP_001290366.1;NP_001290367.1;NP_001290368.1;NI | 28591 | 615695 | . | . | 5.434077 | 5.772381 | 5.367964 | 6.466499 | 12.645320 | 13.343121 | 12.045556 | 14.784883 | -1.002272 | 261.205256 | 0.21894698 | -0.01495707 | 0.988066432 | 0.999197585 | 8.031245 | 8.026832 | 8.014666 | 8.042808 | 7.978174 | 8.074644 | 293 | 290 | 205 | 268 |
| 124801 | NM_00136948 | LSM12 | LSM12 homolog | protein_coding | NP_001356413.1;NP_001356414.1;NP_001356415.1;NP_001358374.1;NI | 26407 | 611793 | . | . | 9.825952 | 10.827755 | 9.931737 | 11.110703 | 22.865397 | 25.028841 | 22.286529 | 25.403302 | 1.003623 | 825.964398 | 0.148670929 | 0.03509859 | 0.972001126 | 0.998865404 | 9.687601 | 9.692266 | 9.659615 | 9.716001 | 9.652213 | 9.727616 | 912 | 925 | 681 | 817 |
| 124808 | NM_00109922 | CCDC43 | coiled-coil domain containing 43 | protein_coding | NP_001092695.1;NP_653210.2 | 26472 | . | . | . | 16.216144 | 15.656783 | 18.902440 | 17.479868 | 37.735633 | 36.191355 | 42.416529 | 39.965641 | 1.127082 | 1120.313569 | 0.136054606 | 1.268554609 | 0.204599962 | 0.96993643 | 10.040620 | 10.213574 | 10.098782 | 10.065861 | 10.205774 | 10.136511 | 1240 | 1111 | 1080 | 1062 |
| 124923 | NM_00117410 | RSKR | ribosomal protein S6 kinase related | protein_coding | NP_001167574.1 | 26314 | . | . | . | 1.373988 | 1.409049 | 2.013336 | 1.042345 | 3.197327 | 3.257079 | 4.517868 | 2.383198 | 1.081507 | 172.361087 | 0.308563905 | 0.366352706 | 0.714101905 | 0.998865404 | 7.370468 | 7.485808 | 7.395587 | 7.402449 | 7.560084 | 7.285139 | 189 | 180 | 206 | 114 |
| 124930 | NM_152345,XI | ANKRD13B | ankyrin repeat domain 13B | protein_coding | NP_689558.4;XP_005257977.1;XP_011522621.1;XP_011522622.1;XP_01 | 26363 | 615124 | . | . | 3.777950 | 3.767025 | 4.822992 | 4.306778 | 8.791450 | 8.707646 | 10.822654 | 9.846937 | 1.163858 | 438.436800 | 0.180284121 | 1.214279585 | 0.224640979 | 0.982008277 | 8.662238 | 8.881862 | 8.732346 | 8.718491 | 8.865382 | 8.768857 | 472 | 432 | 442 | 408 |
| 124935 | NM_00128449 | SLC43A2 | solute carrier family 43 member 2 | protein_coding | NP_001271427.1;NP_001271428.1;NP_001308293.1;NP_001308294.1;NI | 23087 | 610791 | . | . | 0.392070 | 0.319204 | 0.349589 | 0.405135 | 0.912382 | 0.737854 | 0.784468 | 0.926293 | 1.064231 | 96.852425 | 0.333531501 | 0.270897871 | 0.786469242 | 0.998865404 | 6.552149 | 6.641917 | 6.616420 | 6.556234 | 6.581430 | 6.628736 | 119 | 91 | 80 | 101 |
| 124936 | NM_00125475 | CYB5D2 | cytochrome b5 domain containing 2 | protein_coding | NP_001241684.1;NP_001241685.1;NP_653212.1;XP_006721510.2;XP_01 | 28471 | . | . | . | 8.735106 | 8.499988 | 7.857040 | 9.941525 | 20.326949 | 19.648102 | 17.630969 | 22.730115 | 1.030018 | 505.820843 | 0.177909936 | 0.239841276 | 0.810453307 | 0.998865404 | 8.961295 | 9.003363 | 8.992602 | 8.950321 | 8.922342 | 9.054178 | 593 | 520 | 400 | 531 |
| 124944 | NM_00114279 | C17orf49 | chromosome 17 open reading frame 49 | protein_coding | NP_001136270.1;NP_001136271.1;NP_777553.1 | 28737 | 617215 | . | . | 8.829457 | 6.877278 | 5.815524 | 7.639357 | 20.546509 | 15.897134 | 13.049865 | 17.466481 | -1.178176 | 193.151196 | 0.26002944 | -0.90972456 | 0.36296779 | 0.998865404 | 7.707272 | 7.470171 | 7.682433 | 7.567749 | 7.498593 | 7.607030 | 270 | 198 | 134 | 188 |
| 124961 | NM_153018 | ZFP3 | ZFP3 zinc finger protein | protein_coding | NP_694563.1 | 12861 | 194480 | . | . | 0.050843 | 0.047772 | 0.051914 | 0.027729 | 0.118313 | 0.110426 | 0.116493 | 0.063400 | -1.269016 | 7.622896 | 1.114870804 | -0.30829561 | 0.757857407 | NA | 3.089593 | 2.751299 | 2.936041 | 2.934392 | 2.937917 | 2.916116 | 10 | 9 | 8 | 4 |
| 124974 | gene-TXNP4 | TXNP4 | thioredoxin pseudogene 4 | pseudogene | . | 49484 | . | . | . | 0.307535 | 0.021626 | 0.027707 | 0.088417 | 0.715648 | 0.049990 | 0.062175 | 0.202155 | -1.869654 | 2.355176 | 2.022867145 | -0.44628297 | 0.655392853 | NA | 1.617627 | 0.715150 | 1.263891 | 1.243670 | 1.245056 | 1.249391 | 6 | 1 | 1 | 2 |
| 124989 | NM_00119519 | EFCAB13 | EF-hand calcium binding domain 13 | protein_coding | NP_001182121.1;NP_689560.3 | 26864 | . | . | . | 0.621526 | 0.445567 | 0.352402 | 0.404781 | 1.446317 | 1.029948 | 0.790781 | 0.925485 | -1.408814 | 57.186705 | 0.427968089 | -1.15541583 | 0.247920181 | 0.992733415 | 6.063551 | 5.569588 | 5.911473 | 5.831177 | 5.793088 | 5.811671 | 90 | 60 | 39 | 47 |
| 124995 | NM_145255,N | MRPL10 | mitochondrial ribosomal protein L10 | protein_coding | NP_660298.2;NP_683685.1;XP_024306343.1 | 14055 | 611825 | . | . | 19.821220 | 20.103604 | 20.129417 | 22.872478 | 46.124794 | 46.470381 | 45.169828 | 52.295205 | 1.058353 | 1133.602497 | 0.136863785 | 0.597823548 | 0.549957677 | 0.998865404 | 10.105385 | 10.186863 | 10.123148 | 10.126103 | 10.121801 | 10.210026 | 1266 | 1191 | 954 | 1157 |
| 124997 | NM_00116367 | WDR81 | WD repeat domain 81 | protein_coding | NP_001157145.1;NP_001157281.1;NP_001157283.1;NP_689561.2;XP_01 | 26600 | 614218 | . | . | 5.907977 | 5.780273 | 7.063266 | 6.367167 | 13.748106 | 13.361359 | 15.849763 | 14.557768 | 1.157460 | 1313.007866 | 0.129117172 | 1.633882464 | 0.102283532 | 0.82130285 | 10.249230 | 10.460371 | 10.301427 | 10.297773 | 10.435067 | 10.386228 | 1405 | 1310 | 1264 | 1279 |
| 125050 | NR_001445 | RN7SK | RNA component of 7SK nuclear ribonucleoprotein | snRNA | . | 10037 | 606515 | . | . | 8.024332 | 5.015541 | 5.927339 | 6.597740 | 18.672951 | 11.593649 | 13.300776 | 15.084949 | -1.052308 | 66.857159 | 0.406759779 | -0.18083735 | 0.856495247 | 0.998865404 | 6.098699 | 6.026416 | 6.120288 | 6.009991 | 6.044356 | 6.065244 | 97 | 57 | 54 | 64 |
| 125058 | NM_00127184 | TBC1D16 | TBC1 domain family member 16 | protein_coding | NP_001258773.1;NP_001258774.1;NP_001258775.1;NP_061893.2;XP_0C | 28356 | 616637 | . | . | 36.171323 | 39.205365 | 41.006703 | 44.944099 | 84.172151 | 90.624959 | 92.017852 | 102.759338 | 1.084393 | 11354.231571 | 0.106645694 | 1.096035646 | 0.273063175 | 0.998865404 | 13.411348 | 13.528173 | 13.413084 | 13.456424 | 13.461879 | 13.544799 | 12225 | 12056 | 9778 | 11609 |
| 125061 | NM_00101098 | AFMID | arylformamidase | protein_coding | NP_001010982.2;NP_001138998.1;XP_011522628.1;XP_011522629.1;XP | 20910 | . | . | . | 8.700511 | 8.453502 | 9.920022 | 11.919617 | 20.246449 | 19.540651 | 22.260241 | 27.252788 | 1.065466 | 444.590994 | 0.179099313 | 0.51080596 | 0.609486936 | 0.998865404 | 8.750243 | 8.840999 | 8.765810 | 8.785231 | 8.767224 | 8.859358 | 488 | 472 | 369 | 462 |
| 125144 | NR_027158,NF | SNHG29 | small nucleolar RNA host gene 29 | lncRNA | . | 28619 | . | . | . | 250.770853 | 251.993988 | 245.605275 | 259.873729 | 583.554085 | 582.495403 | 551.131107 | 594.170391 | 1.026496 | 8271.711605 | 0.106392113 | 0.354607196 | 0.722883885 | 0.998865404 | 12.995009 | 13.032685 | 13.010459 | 12.994486 | 12.984839 | 13.063039 | 9489 | 8726 | 6952 | 8217 |
| 125150 | NM_00104269 | ZSWIM7 | zinc finger SWIM-type containing 7 | protein_coding | NP_001036162.1;NP_001036163.1 | 26993 | 614535 | . | . | 5.627319 | 5.568179 | 6.380550 | 5.822800 | 13.095003 | 12.871096 | 14.317769 | 13.313138 | 1.077990 | 355.896638 | 0.190535832 | 0.568624415 | 0.569611053 | 0.998865404 | 8.419845 | 8.528730 | 8.454487 | 8.449864 | 8.526597 | 8.462780 | 396 | 368 | 339 | 327 |
| 125170 | NM_00114490 | MIEF2 | mitochondrial elongation factor 2 | protein_coding | NP_001138372.1;NP_631901.2;NP_683684.2;XP_016879679.1 | 17920 | 615498 | . | . | 2.529028 | 2.418877 | 2.476027 | 2.636973 | 5.885153 | 5.591343 | 5.556133 | 6.029125 | 1.027294 | 243.578306 | 0.217373635 | 0.178723219 | 0.858155036 | 0.998865404 | 7.908649 | 7.947572 | 7.921789 | 7.920737 | 7.937572 | 7.931548 | 277 | 259 | 216 | 230 |
| 125208 | gene-RPS2P46 | RPS2P46 | ribosomal protein S2 pseudogene 46 | pseudogene | . | 35839 | . | . | . | 17.429705 | 16.209589 | 12.504566 | 19.230906 | 40.559639 | 37.469189 | 28.059885 | 43.969181 | -1.075734 | 480.608895 | 0.207254599 | -0.50817531 | 0.611330405 | 0.998865404 | 8.960958 | 8.854518 | 8.952380 | 8.900403 | 8.744831 | 9.000679 | 597 | 516 | 319 | 524 |
| 125228 | NM_00109880 | FAM210A | family with sequence similarity 210 member A | protein_coding | NP_001092271.1;NP_689565.2;XP_024306850.1;XP_024306851.1 | 28346 | 617975 | . | . | 15.953990 | 14.862399 | 16.004900 | 14.062846 | 37.125591 | 34.355100 | 35.914535 | 32.153025 | -1.040937 | 2013.674676 | 0.12816844 | -0.45161466 | 0.651546604 | 0.998865404 | 11.004084 | 10.946573 | 11.021900 | 10.958154 | 11.012479 | 10.902041 | 2459 | 2128 | 1836 | 1721 |
| 125371 | gene-RPL22P1 | RPL22P1 | ribosomal protein L22 pseudogene 1 | pseudogene | . | 17998 | . | . | . | 47.637230 | 39.508495 | 41.259235 | 46.679230 | 110.853790 | 91.325653 | 92.584534 | 106.726501 | -1.005432 | 822.773539 | 0.158866391 | -0.04919989 | 0.960760002 | 0.998865404 | 9.688216 | 9.680477 | 9.752343 | 9.612523 | 9.640171 | 9.720264 | 1043 | 803 | 673 | 813 |
| 125476 | NM_00109881 | INO80C | INO80 complex subunit C | protein_coding | NP_001092287.1;NP_001294993.1;NP_919257.2;XP_005258253.1;XP_00 | 26994 | . | . | . | 4.829550 | 6.056279 | 7.108015 | 7.652654 | 11.238561 | 13.999360 | 15.950179 | 17.496886 | 1.376992 | 195.638904 | 0.247532582 | 1.864481361 | 0.062254137 | 0.723723143 | 7.363669 | 7.823883 | 7.479471 | 7.574286 | 7.652785 | 7.687727 | 168 | 198 | 190 | 220 |
| 125488 | NM_00113599 | TTC39C | tetratricopeptide repeat domain 39C | protein_coding | NP_001129465.1;NP_001230354.1;NP_001278959.1;NP_694943.2;XP_0C | 26595 | . | . | . | 2.100031 | 3.428000 | 2.717248 | 3.513731 | 4.886857 | 7.923974 | 6.097427 | 8.033729 | 1.078737 | 228.524844 | 0.231061345 | 0.473220792 | 0.636055665 | 0.998865404 | 7.781154 | 7.889236 | 7.796220 | 7.835120 | 7.803629 | 7.898443 | 243 | 247 | 186 | 244 |
| 125704 | NM_00104436 | DIPK1C | divergent protein kinase domain 1C | protein_coding | NP_001037834.2;XP_011524118.1;XP_016881040.1 | 31729 | 614544 | . | . | 0.116728 | 0.078069 | 0.282355 | 0.182089 | 0.271630 | 0.180460 | 0.633597 | 0.416325 | 2.101640 | 11.109030 | 0.932154181 | 1.149504514 | 0.250348007 | NA | 2.838265 | 3.913269 | 3.437115 | 3.431181 | 3.483292 | 3.455164 | 9 | 7 | 16 | 11 |
| 125875 | NM_152353,XI | CLDND2 | claudin domain containing 2 | protein_coding | NP_689566.1;XP_011524727.1;XP_011524730.1;XP_016881733.1;XP_01 | 28511 | . | . | . | 0.874750 | 0.480369 | 0.751334 | 0.641478 | 2.035579 | 1.110395 | 1.685973 | 1.466663 | -1.016252 | 39.681148 | 0.523257503 | -0.04444756 | 0.964547651 | 0.998865404 | 5.320284 | 5.300412 | 5.354493 | 5.258397 | 5.316874 | 5.293115 | 60 | 30 | 37 | 34 |
| 125893 | NM_00103166 | ZNF816 | zinc finger protein 816 | protein_coding | NP_001026835.1;NP_001189385.1;NP_001189386.1 | 26995 | . | . | . | 0.937153 | 1.060278 | 1.015772 | 0.964054 | 2.180793 | 2.450880 | 2.279362 | 2.204196 | -1.013458 | 80.299573 | 0.354274618 | -0.05544051 | 0.956584203 | 0.998865404 | 6.337028 | 6.317547 | 6.313781 | 6.343239 | 6.331176 | 6.318990 | 88 | 92 | 71 | 73 |
| 125919 | NM_213598 | ZNF543 | zinc finger protein 543 | protein_coding | NP_998763.2 | 25281 | 616847 | . | . | 0.493347 | 0.756125 | 0.719171 | 0.847813 | 1.148038 | 1.747817 | 1.613799 | 1.938424 | 1.232245 | 81.522808 | 0.37117501 | 0.811842369 | 0.416882087 | 0.998865404 | 6.192060 | 6.490770 | 6.259656 | 6.356250 | 6.345239 | 6.390253 | 67 | 95 | 72 | 91 |
| 125950 | NM_00136617 | RAVER1 | ribonucleoprotein, PTB binding 1 | protein_coding | NP_001353103.1;NP_597709.3 | 30296 | 609950 | . | . | 5.189447 | 6.546482 | 7.672338 | 7.136312 | 12.076056 | 15.132485 | 17.216504 | 16.316329 | 1.245094 | 719.150762 | 0.164940516 | 1.917383412 | 0.05518924 | 0.697001222 | 9.323548 | 9.476634 | 9.328495 | 9.447024 | 9.586718 | 9.526661 | 656 | 768 | 723 | 716 |
| 125972 | NM_145046 | CALR3 | calreticulin 3 | protein_coding | NP_659483.2 | 20407 | 611414 | . | . | 0.209506 | 0.134667 | 0.298920 | 0.257561 | 0.487530 | 0.311290 | 0.670768 | 0.588882 | 1.632808 | 9.390319 | 1.000323333 | 0.707126565 | 0.479487829 | NA | 2.832372 | 3.543272 | 3.216475 | 3.202059 | 3.233738 | 3.225368 | 10 | 6 | 11 | 10 |
| 125988 | NM_00130824 | MICOS13 | mitochondrial contact site and cristae organizing system subunit 1 | protein_coding | NP_001295169.1;NP_001352690.1;NP_991330.1;XP_011525977.1 | 33702 | 616658 | . | . | 41.328622 | 42.422802 | 37.609087 | 52.628884 | 96.173407 | 98.062204 | 84.393692 | 120.329686 | 1.064764 | 746.569320 | 0.166876712 | 0.542515962 | 0.587463114 | 0.998865404 | 9.498525 | 9.588342 | 9.527126 | 9.512060 | 9.466338 | 9.651654 | 842 | 772 | 581 | 817 |
| 126003 | NM_00104246 | TRAPPC5 | trafficking protein particle complex 5 | protein_coding | NP_001035926.1;NP_001035927.1;NP_777554.1 | 23067 | . | . | . | 2.531316 | 2.464374 | 1.801176 | 2.624970 | 5.890475 | 5.696512 | 4.041788 | 6.001683 | -1.138564 | 409.268286 | 0.205942113 | -0.90906934 | 0.363313526 | 0.998865404 | 8.768068 | 8.579586 | 8.722765 | 8.700368 | 8.524965 | 8.732178 | 511 | 462 | 273 | 423 |
| 126006 | NM_00127183 | PCP2 | Purkinje cell protein 2 | protein_coding | NP_001258759.1;NP_777555.1;XP_006722702.1;XP_016881738.1;XP_01 | 30209 | . | . | . | 2.262118 | 1.993336 | 2.676086 | 2.308983 | 5.264046 | 4.607684 | 6.005060 | 5.279215 | 1.143240 | 115.134096 | 0.310638683 | 0.62171358 | 0.53413022 | 0.998865404 | 6.746331 | 6.941421 | 6.841176 | 6.797440 | 6.908410 | 6.818026 | 132 | 108 | 121 | 100 |
| 126014 | NM_00128234 | OSCAR | osteoclast associated Ig-like receptor | protein_coding | NP_001269278.1;NP_001269279.1;NP_570127.3;NP_573398.2;NP_5733 | 29960 | 606862 | . | . | 0.071441 | 0.087999 | 0.173241 | 0.054390 | 0.166246 | 0.203413 | 0.388747 | 0.124357 | 1.343963 | 6.866069 | 1.1835297 | 0.360357131 | 0.71858008 | NA | 2.548173 | 2.978782 | 2.758016 | 2.764094 | 2.785940 | 2.754143 | 6 | 7 | 10 | 4 |
| 126017 | NM_00100430 | ZNF813 | zinc finger protein 813 | protein_coding | NP_001004301.2 | 33257 | . | . | . | 0.103186 | 0.139139 | 0.113172 | 0.141908 | 0.240119 | 0.321627 | 0.253955 | 0.324456 | 1.041006 | 24.325266 | 0.62942412 | 0.092113081 | 0.926608195 | NA | 4.576468 | 4.631770 | 4.587778 | 4.613560 | 4.592855 | 4.616668 | 24 | 29 | 19 | 26 |
| 126068 | NM_152355 | ZNF441 | zinc finger protein 441 | protein_coding | NP_689568.2 | 20875 | . | . | . | 1.150200 | 0.922282 | 1.404558 | 0.976506 | 2.676562 | 2.131897 | 3.151788 | 2.232664 | 1.130312 | 154.773362 | 0.28910967 | 0.611261764 | 0.5410263 | 0.998865404 | 7.181638 | 7.360828 | 7.284678 | 7.197451 | 7.361852 | 7.213977 | 186 | 139 | 169 | 126 |
| 126070 | NM_152357,XI | ZNF440 | zinc finger protein 440 | protein_coding | NP_689570.2;XP_005259788.1;XP_016881743.1 | 20874 | . | . | . | 1.489061 | 1.889047 | 2.021712 | 2.101485 | 3.465107 | 4.366619 | 4.536664 | 4.804794 | 1.223220 | 264.962733 | 0.219422259 | 1.324770006 | 0.185247446 | 0.958025663 | 7.897514 | 8.187254 | 7.936648 | 8.043114 | 8.086688 | 8.101671 | 245 | 285 | 250 | 277 |
| 126074 | NM_175871 | SWSAP1 | SWIM-type zinc finger 7 associated protein 1 | protein_coding | NP_787067.3 | 26638 | 614536 | . | . | 1.839098 | 2.123596 | 2.207843 | 2.130762 | 4.279656 | 4.908787 | 4.954336 | 4.871734 | 1.078871 | 59.764293 | 0.406287211 | 0.269570348 | 0.787490815 | 0.998865404 | 5.845597 | 5.954762 | 5.877524 | 5.904429 | 5.911313 | 5.903002 | 62 | 66 | 55 | 57 |
| 126075 | NM_00108050 | CCDC159 | coiled-coil domain containing 159 | protein_coding | NP_001073972.2;XP_006722706.1;XP_011525980.2;XP_016881744.1;XP_ | 26996 | . | . | . | 2.436033 | 2.628932 | 3.074812 | 2.988124 | 5.668747 | 6.076895 | 6.899791 | 6.831988 | 1.184385 | 107.877370 | 0.310391227 | 0.786548277 | 0.431546348 | 0.998865404 | 6.625920 | 6.870245 | 6.708557 | 6.734359 | 6.792496 | 6.758448 | 109 | 111 | 107 | 104 |
| 126119 | NM_00127063 | JOSD2 | Josephin domain containing 2 | protein_coding | NP_001257568.1;NP_001257569.1;NP_001257570.1;NP_001257615.1;NI | 28853 | 615324 | . | . | 10.782580 | 11.222891 | 9.684851 | 11.976811 | 25.091506 | 25.942216 | 21.732523 | 27.383553 | -1.037035 | 296.063019 | 0.209623387 | -0.25028048 | 0.802370452 | 0.998865404 | 8.236312 | 8.182711 | 8.202588 | 8.234700 | 8.146754 | 8.249046 | 337 | 335 | 228 | 299 |
| 126129 | NM_00113605 | CPT1C | carnitine palmitoyltransferase 1C | protein_coding | NP_001129524.1;NP_001186681.1;NP_001186682.1;NP_001365411.1;NI | 18540 | 608846 | . | . | 0.017727 | 0.083368 | 0.049100 | 0.068628 | 0.041252 | 0.192708 | 0.110178 | 0.156911 | 1.213142 | 6.040643 | 1.264676042 | 0.220410909 | 0.82555115 | NA | 2.452336 | 2.724273 | 2.570124 | 2.597147 | 2.585216 | 2.592871 | 3 | 9 | 5 | 7 |
| 126133 | NM_00114539 | ALDH16A1 | aldehyde dehydrogenase 16 family member A1 | protein_coding | NP_001138868.1;NP_699160.2;XP_011524743.1;XP_011524744.1 | 28114 | 613358 | . | . | 6.169998 | 6.608198 | 6.954077 | 7.087007 | 14.357837 | 15.275146 | 15.604745 | 16.203597 | 1.073460 | 643.861619 | 0.154904979 | 0.66019972 | 0.509125672 | 0.998865404 | 9.278694 | 9.380716 | 9.284595 | 9.326307 | 9.347853 | 9.358922 | 693 | 691 | 575 | 630 |
| 126208 | NM_00100283 | ZNF787 | zinc finger protein 787 | protein_coding | NP_001002836.2;NP_001338611.1;XP_011524747.1 | 26998 | . | . | . | 11.105787 | 12.009232 | 10.375149 | 11.955739 | 25.843625 | 27.759882 | 23.281536 | 27.335378 | -1.048633 | 695.943173 | 0.154612906 | -0.4431908 | 0.65762773 | 0.998865404 | 9.476952 | 9.407872 | 9.438591 | 9.478449 | 9.389126 | 9.461614 | 797 | 791 | 559 | 671 |
| 126231 | NM_00117268 | ZNF573 | zinc finger protein 573 | protein_coding | NP_001166160.1;NP_001166161.1;NP_001166162.1;NP_001166163.1;NI | 26420 | . | . | . | 0.594111 | 0.562680 | 0.687763 | 0.613690 | 1.382520 | 1.300658 | 1.543320 | 1.403131 | 1.149994 | 47.178272 | 0.454927088 | 0.443206959 | 0.657616046 | 0.998865404 | 5.455014 | 5.657956 | 5.552403 | 5.535775 | 5.581464 | 5.555929 | 53 | 45 | 47 | 44 |
| 126272 | NM_152361 | EID2B | EP300 interacting inhibitor of differentiation 2B | protein_coding | NP_689574.1 | 26796 | 617355 | . | . | 2.050053 | 2.288972 | 2.084859 | 2.207807 | 4.770558 | 5.291061 | 4.678362 | 5.047888 | -1.020970 | 125.748838 | 0.289482575 | -0.10342558 | 0.917625211 | 0.998865404 | 6.989620 | 6.959021 | 6.959732 | 6.995784 | 6.963655 | 6.977494 | 139 | 144 | 106 | 119 |
| 126282 | NM_00116794 | TNFAIP8L1 | TNF alpha induced protein 8 like 1 | protein_coding | NP_001161414.1;NP_689575.2;XP_005259544.1;XP_011525982.1 | 28279 | 615869 | . | . | 4.216839 | 4.945208 | 3.502744 | 4.420818 | 9.812758 | 11.431070 | 7.860057 | 10.107674 | -1.170521 | 508.582707 | 0.183603472 | -1.23718145 | 0.216019726 | 0.982008277 | 9.100014 | 8.871635 | 8.985362 | 9.085293 | 8.865773 | 9.001918 | 584 | 637 | 362 | 488 |
| 126295 | NM_00131908 | ZNF57 | zinc finger protein 57 | protein_coding | NP_001306012.1;NP_775751.1;XP_011525984.1 | 13125 | . | . | . | 1.562132 | 1.590461 | 1.271795 | 2.046749 | 3.635145 | 3.676422 | 2.853870 | 4.679648 | 1.035452 | 99.841801 | 0.343498117 | 0.146320436 | 0.883668434 | 0.998865404 | 6.617423 | 6.665324 | 6.632374 | 6.632733 | 6.571153 | 6.710431 | 113 | 106 | 68 | 117 |
| 126298 | NM_00100756 | IRGQ | immunity related GTPase Q | protein_coding | NP_001007562.1;XP_005258572.1 | 24868 | . | . | . | 3.654571 | 3.695869 | 3.497031 | 3.524410 | 8.504337 | 8.543168 | 7.847237 | 8.058145 | -1.041614 | 1074.488492 | 0.134193219 | -0.43833266 | 0.661145152 | 0.998865404 | 10.098587 | 10.039680 | 10.069171 | 10.099365 | 10.054756 | 10.053561 | 1237 | 1207 | 916 | 986 |
| 126299 | NM_182498 | ZNF428 | zinc finger protein 428 | protein_coding | NP_872304.2 | 20804 | . | . | . | 14.109343 | 12.360574 | 11.856668 | 14.657518 | 32.833019 | 28.572021 | 26.606020 | 33.512672 | -1.012783 | 479.983715 | 0.183584379 | -0.09981794 | 0.920488867 | 0.998865404 | 8.916169 | 8.897453 | 8.952549 | 8.864346 | 8.837134 | 8.961824 | 595 | 485 | 373 | 492 |
| 126308 | NM_130807,XI | MOB3A | MOB kinase activator 3A | protein_coding | NP_570719.1;XP_011525985.1;XP_024307128.1 | 29802 | . | . | . | 3.366939 | 3.941573 | 4.417081 | 4.341831 | 7.835004 | 9.111122 | 9.911860 | 9.927078 | 1.183969 | 422.993499 | 0.181857892 | 1.339682517 | 0.180348584 | 0.952541638 | 8.597525 | 8.840814 | 8.625051 | 8.709946 | 8.781432 | 8.759119 | 414 | 448 | 406 | 422 |
| 126321 | NM_00128752 | MFSD12 | major facilitator superfamily domain containing 12 | protein_coding | NP_001274458.1;NP_778148.2;XP_005259547.1;XP_006722710.1;XP_01 | 28299 | 617745 | . | . | 448.620082 | 443.830047 | 392.065920 | 418.954163 | 1043.957328 | 1025.933033 | 879.784582 | 957.888939 | -1.119914 | 28206.758375 | 0.099862926 | -1.63612547 | 0.10181333 | 0.820390774 | 14.863141 | 14.699742 | 14.839300 | 14.821745 | 14.707300 | 14.757845 | 34663 | 31834 | 22456 | 25686 |
| 126328 | NM_00119337 | NDUFA11 | NADH:ubiquinone oxidoreductase subunit A11 | protein_coding | NP_001180304.1;NP_783313.1 | 20371 | 612638 | . | . | 115.619182 | 111.943993 | 83.317070 | 114.075496 | 269.050592 | 258.763565 | 186.961084 | 260.820071 | -1.173323 | 1930.759391 | 0.115160239 | -1.52553856 | 0.127124824 | 0.874064017 | 11.025850 | 10.794816 | 10.989705 | 10.954494 | 10.717395 | 10.959241 | 2453 | 2201 | 1305 | 1924 |
| 126374 | NM_00108043 | WTIP | WT1 interacting protein | protein_coding | NP_001073905.1;XP_006723077.1;XP_011524754.1;XP_011524755.1 | 20964 | 614790 | . | . | 0.050152 | 0.039308 | 0.037416 | 0.010539 | 0.116706 | 0.090863 | 0.083960 | 0.024095 | -1.514891 | 13.436584 | 0.880079431 | -0.68086306 | 0.495958146 | NA | 4.014059 | 3.421760 | 3.757619 | 3.762089 | 3.758895 | 3.712300 | 18 | 18 | 14 | 5 |
| 126375 | NM_175872 | ZNF792 | zinc finger protein 792 | protein_coding | NP_787068.3 | 24751 | . | . | . | 0.507753 | 0.645932 | 0.333509 | 0.594594 | 1.181563 | 1.493102 | 0.748384 | 1.359470 | -1.255831 | 66.207297 | 0.419306462 | -0.7837763 | 0.43317138 | 0.998865404 | 6.205680 | 5.873023 | 6.041819 | 6.101278 | 5.962806 | 6.074146 | 75 | 89 | 37 | 70 |
| 126382 | NM_00130094 | NR2C2AP | nuclear receptor 2C2 associated protein | protein_coding | NP_001287874.1;NP_795361.1 | 30763 | 608719 | . | . | 16.240359 | 16.446785 | 13.418523 | 17.039254 | 37.791982 | 38.017481 | 30.110777 | 38.958229 | -1.088694 | 621.519060 | 0.17087997 | -0.71745694 | 0.473092194 | 0.998865404 | 9.340040 | 9.216631 | 9.312415 | 9.299966 | 9.167127 | 9.325111 | 753 | 693 | 455 | 625 |
| 126393 | NM_144617 | HSPB6 | heat shock protein family B (small) member 6 | protein_coding | NP_653218.1 | 26511 | 610695 | . | . | 0.052156 | 0.109253 | 0.228489 | 0.044185 | 0.121369 | 0.252544 | 0.512724 | 0.101025 | 1.666171 | 5.449718 | 1.380840378 | 0.533397113 | 0.593758716 | NA | 2.029593 | 2.769039 | 2.406587 | 2.420775 | 2.446680 | 2.405118 | 3 | 6 | 10 | 2 |
| 126526 | NM_00125644 | C19orf47 | chromosome 19 open reading frame 47 | protein_coding | NP_001243369.1;NP_001243370.1;XP_005258577.1;XP_011524762.1;XP | 26723 | . | . | . | 4.315794 | 3.856660 | 3.677661 | 5.284693 | 10.043033 | 8.914838 | 8.252566 | 12.082820 | 1.073303 | 610.728111 | 0.187899344 | 0.543147435 | 0.587028295 | 0.998865404 | 9.202949 | 9.304053 | 9.233328 | 9.218954 | 9.145025 | 9.387173 | 686 | 629 | 451 | 699 |
| 126549 | NM_00127844 | ANKLE1 | ankyrin repeat and LEM domain containing 1 | protein_coding | NP_001265372.2;NP_001265373.2;NP_001265374.2;NP_689576.6 | 26812 | . | . | . | 1.552628 | 1.658302 | 1.926168 | 1.533536 | 3.613030 | 3.833238 | 4.322266 | 3.506249 | 1.066469 | 153.980528 | 0.27233542 | 0.340909961 | 0.733171365 | 0.998865404 | 7.218891 | 7.312790 | 7.240154 | 7.261081 | 7.323008 | 7.226972 | 167 | 165 | 155 | 131 |
| 126626 | NM_00132390 | GABPB2 | GA binding protein transcription factor subunit beta 2 | protein_coding | NP_001310835.1;NP_001310836.1;NP_001310837.1;NP_001310838.1;NI | 28441 | . | . | . | 0.578046 | 0.557325 | 0.400518 | 0.201359 | 1.345132 | 1.288283 | 0.898749 | 0.460382 | -1.106499 | 85.358862 | 0.396500459 | -0.36822688 | 0.712704066 | 0.998865404 | 6.484657 | 6.342795 | 6.474312 | 6.361948 | 6.467884 | 6.309291 | 123 | 78 | 92 | 53 |
| 126661 | NM_00110260 | CCDC163 | coiled-coil domain containing 163 | protein_coding | NP_001096071.1;NP_001345335.1;NP_001345336.1;XP_024308812.1;XP | 27003 | . | . | . | 0.640821 | 0.859380 | 0.767456 | 0.558321 | 1.491219 | 1.986494 | 1.722148 | 1.276534 | -1.150789 | 49.693237 | 0.459681087 | -0.44079113 | 0.659364212 | 0.998865404 | 5.732325 | 5.530583 | 5.614520 | 5.674172 | 5.647434 | 5.589861 | 52 | 66 | 47 | 36 |
| 126731 | NM_145257,XI | CCSAP | centriole, cilia and spindle associated protein | protein_coding | NP_660300.3;XP_024308823.1 | 29578 | 616762 | . | . | 4.899353 | 3.930890 | 5.135451 | 4.736460 | 11.400997 | 9.086429 | 11.523803 | 10.829353 | 1.096445 | 755.406130 | 0.164167151 | 0.80913254 | 0.418438913 | 0.998865404 | 9.492764 | 9.626362 | 9.602179 | 9.442903 | 9.619023 | 9.560287 | 925 | 688 | 716 | 710 |
| 126789 | NM_00134611 | PUSL1 | pseudouridine synthase like 1 | protein_coding | NP_001333045.1;NP_699170.1;XP_005244777.1;XP_024308825.1 | 26914 | . | . | . | 20.225284 | 19.335886 | 19.463220 | 22.756854 | 47.065069 | 44.695763 | 43.674899 | 52.030841 | 1.062895 | 784.972972 | 0.150469326 | 0.584829868 | 0.558662119 | 0.998865404 | 9.571982 | 9.659684 | 9.610248 | 9.577438 | 9.592286 | 9.679400 | 897 | 802 | 660 | 805 |
| 126792 | NM_080605 | B3GALT6 | beta-1,3-galactosyltransferase 6 | protein_coding | NP_542172.2 | 17978 | 615291 | . | . | 11.794799 | 12.256722 | 13.292960 | 12.486471 | 27.446981 | 28.331963 | 29.829018 | 28.548830 | 1.057337 | 1088.955148 | 0.135388805 | 0.594110733 | 0.552438702 | 0.998865404 | 10.047871 | 10.128461 | 10.056245 | 10.078017 | 10.135467 | 10.081021 | 1201 | 1159 | 1009 | 1011 |
| 126820 | NM_00128856 | DNAI3 | dynein axonemal intermediate chain 3 | protein_coding | NP_001275492.1;NP_660155.2 | 30711 | 617968 | . | . | 1.364928 | 1.286536 | 1.128184 | 1.047502 | 3.176244 | 2.973886 | 2.531613 | 2.394990 | -1.239412 | 112.897262 | 0.304115139 | -1.0182208 | 0.308573038 | 0.998865404 | 6.965061 | 6.656170 | 6.866603 | 6.840747 | 6.794910 | 6.771969 | 149 | 130 | 91 | 91 |
| 126826 | gene-TGIF2P1 | TGIF2P1 | TGFB induced factor homeobox 2 pseudogene 1 | pseudogene | . | 33317 | . | . | . | 0.058862 | 0.095502 | 0.078580 | 0.061118 | 0.136973 | 0.220758 | 0.176332 | 0.139739 | -1.017609 | 2.237600 | 1.972119472 | -0.01276997 | 0.989811317 | NA | 1.175403 | 1.148375 | 1.158018 | 1.163002 | 1.160711 | 1.159957 | 2 | 3 | 2 | 2 |
| 126868 | NM_152367,XI | MAB21L3 | mab-21 like 3 | protein_coding | NP_689580.2;XP_011538925.1;XP_011538927.1;XP_011538928.1 | 26787 | . | . | . | 0.071255 | 0.098889 | 0.053867 | 0.105312 | 0.165812 | 0.228587 | 0.120877 | 0.240784 | -1.093106 | 8.606679 | 1.046956266 | -0.12267363 | 0.902365544 | NA | 3.170740 | 3.037079 | 3.103483 | 3.114392 | 3.094763 | 3.116202 | 9 | 11 | 5 | 10 |

| | | | | | | | | | | | | | | | | | | | | | | | | | | | | | | | | | |
|---|---|---|---|---|---|---|---|---|---|---|---|---|---|---|---|---|---|---|---|---|---|---|---|---|---|---|---|---|---|---|---|---|---|
| 126917 | NM_00113626 | IFFO2 | intermediate filament family orphan 2 | protein_coding | NP_001129737.1;XP_011538932.1 | 27006 | . | . | . | 2.835414 | 2.952858 | 1.868079 | 2.662069 | 6.598125 | 6.825665 | 4.191913 | 6.086505 | -1.290684 | 496.541020 | 0.193261086 | -1.90486183 | 0.056798034 | 0.708205297 | 9.128710 | 8.759237 | 9.018019 | 9.037505 | 8.760587 | 8.962266 | 635 | 613 | 313 | 475 |
| 126969 | NM_00111410 | SLC44A3 | solute carrier family 44 member 3 | protein_coding | NP_001107578.1;NP_001245269.1;NP_001245270.1;NP_001245271.1;NI | 28689 | . | . | . | 2.336222 | 2.173920 | 1.289792 | 1.458251 | 5.436481 | 5.025113 | 2.894259 | 3.334119 | -1.595708 | 132.682601 | 0.284462376 | -2.37007317 | **0.017784565** | 0.447140899 | 7.350125 | 6.675354 | 7.130193 | 7.124445 | 6.947613 | 6.972564 | 189 | 175 | 85 | 100 |
| 127002 | NM_00135017 | ATXN7L2 | ataxin 7 like 2 | protein_coding | NP_001337103.1;NP_001337104.1;NP_001337106.1;NP_699171.3;XP_01 | 28713 | . | . | . | 2.344073 | 2.671313 | 2.383702 | 2.840814 | 5.454755 | 6.174855 | 5.348962 | 6.495183 | 1.123640 | 209.838159 | 0.233688692 | 0.719672329 | 0.471726767 | 0.998865404 | 7.626771 | 7.794616 | 7.655815 | 7.710175 | 7.744104 | 7.730133 | 214 | 226 | 196 | 205 |
| 127018 | NM_00130076 | LYPLAL1 | lysophospholipase like 1 | protein_coding | NP_001287698.1;NP_001287699.1;NP_001287700.1;NP_001287701.1;NI | 20440 | 616548 | . | . | 6.166529 | 5.235517 | 5.780162 | 3.494288 | 14.349764 | 12.102127 | 12.970516 | 7.989275 | -1.147501 | 340.973514 | 0.241845117 | -0.82075308 | 0.411786932 | 0.998865404 | 8.508395 | 8.311955 | 8.502268 | 8.379609 | 8.490022 | 8.223012 | 463 | 352 | 344 | 225 |
| 127253 | NM_00116291 | TYW3 | tRNA-yW synthesizing protein 3 homolog | protein_coding | NP_001156388.1;NP_612476.1;XP_006710410.1 | 24757 | 611245 | . | . | 10.784854 | 10.275142 | 10.357786 | 10.302000 | 25.096798 | 23.751451 | 23.242574 | 23.554295 | -1.038119 | 1011.560523 | 0.136847535 | -0.39439483 | 0.693289562 | 0.998865404 | 10.008987 | 9.955246 | 10.002417 | 9.989259 | 9.989123 | 9.947411 | 1197 | 1102 | 888 | 904 |
| 127255 | NM_00110565 | LRRIQ3 | leucine rich repeats and IQ motif containing 3 | protein_coding | NP_001099129.1;NP_001309244.1;XP_016855776.1;XP_024308953.1;XP | 28318 | 617957 | . | . | 2.566561 | 2.270661 | 2.018978 | 2.248947 | 5.972495 | 5.248734 | 4.530530 | 5.141951 | -1.166077 | 85.573645 | 0.346365367 | -0.63996817 | 0.522193292 | 0.998865404 | 6.525622 | 6.304070 | 6.462874 | 6.418668 | 6.389045 | 6.408729 | 114 | 92 | 66 | 77 |
| 127262 | NM_182752 | TPRG1L | tumor protein p63 regulated 1 like | protein_coding | NP_877429.2 | 27007 | 611460 | . | . | 21.866695 | 21.761055 | 22.211977 | 26.828350 | 50.884701 | 50.301651 | 49.843029 | 61.339836 | 1.107903 | 1732.125878 | 0.133141713 | 1.110332941 | 0.266855583 | 0.998484037 | 10.682690 | 10.830197 | 10.720623 | 10.710538 | 10.722694 | 10.864240 | 1905 | 1762 | 1442 | 1859 |
| 127281 | NM_00119573 | PRXL2B | peroxiredoxin like 2B | protein_coding | NP_001182665.4;NP_001182666.4;NP_001182667.4;NP_001182669.4;NI | 28390 | . | . | . | 15.235552 | 15.135696 | 18.412388 | 18.612108 | 35.453757 | 34.986839 | 41.316863 | 42.554373 | 1.202320 | 1437.221117 | 0.125191832 | 2.123309615 | **0.033727916** | 0.576885276 | 10.350029 | 10.615874 | 10.419187 | 10.407184 | 10.553323 | 10.556494 | 1515 | 1397 | 1362 | 1474 |
| 127343 | NM_147192,N | DMBX1 | diencephalon/mesencephalon homeobox 1 | protein_coding | NP_671725.1;NP_757379.1;XP_011538970.1;XP_016855778.1 | 19026 | 607410 | . | . | 0.544358 | 0.529543 | 0.676463 | 0.458642 | 1.266745 | 1.224062 | 1.517962 | 1.048629 | 1.075970 | 71.698592 | 0.373396079 | 0.282910763 | 0.777245244 | 0.998865404 | 6.109703 | 6.216082 | 6.151280 | 6.154521 | 6.186939 | 6.154137 | 79 | 75 | 69 | 65 |
| 127396 | NM_152373 | ZNF684 | zinc finger protein 684 | protein_coding | NP_689586.3 | 28418 | . | . | . | 1.710313 | 1.954086 | 1.665877 | 1.807829 | 3.979969 | 4.516956 | 3.738180 | 4.133386 | -1.068033 | 112.598782 | 0.305153453 | -0.31117604 | 0.755666796 | 0.998865404 | 6.862208 | 6.766293 | 6.805478 | 6.846014 | 6.792118 | 6.816354 | 126 | 133 | 91 | 106 |
| 127428 | NM_153035,N | TCEANC2 | transcription elongation factor A N-terminal and central domain co | protein_coding | NP_694580.1 | 26494 | . | . | . | 1.346320 | 1.214011 | 1.336790 | 1.342065 | 3.132942 | 2.806241 | 2.999718 | 3.068472 | 1.025208 | 424.224375 | 0.179465392 | 0.200129667 | 0.841379172 | 0.998865404 | 8.710388 | 8.746750 | 8.753558 | 8.686482 | 8.741235 | 8.730150 | 509 | 427 | 377 | 399 |
| 127544 | NM_00112736 | RNF19B | ring finger protein 19B | protein_coding | NP_001120833.1;NP_001287755.1;NP_699172.2;XP_006710419.1;XP_00 | 26886 | 610872 | . | . | 9.443704 | 10.702143 | 8.500621 | 11.139861 | 21.975891 | 24.738486 | 19.075147 | 25.469967 | -1.054724 | 876.951511 | 0.164584941 | -0.46702706 | 0.640480518 | 0.998865404 | 9.814707 | 9.736952 | 9.761702 | 9.823350 | 9.652310 | 9.847518 | 993 | 1013 | 640 | 910 |
| 127602 | NM_00114515 | DNAH14 | dynein axonemal heavy chain 14 | protein_coding | NP_001138626.1;NP_001336840.1;NP_001336841.1;NP_001354408.1;NI | 2945 | 603341 | . | . | 5.870128 | 7.079632 | 4.194642 | 4.304769 | 13.660030 | 16.364885 | 9.412660 | 9.842343 | -1.369151 | 364.946737 | 0.210624196 | -2.15208575 | **0.031390601** | 0.561754658 | 8.719722 | 8.268173 | 8.642196 | 8.535755 | 8.486390 | 8.327626 | 527 | 416 | 308 | 246 |
| 127665 | NM_00100999 | ZNF648 | zinc finger protein 648 | protein_coding | NP_001009992.1;XP_024309026.1;XP_024309028.1;XP_024309031.1 | 18190 | . | . | . | 0.030587 | 0.027276 | 0.036899 | 0.063567 | 0.071177 | 0.063049 | 0.082798 | 0.145338 | 1.419968 | 10.289565 | 0.955825975 | 0.529236468 | 0.596641422 | NA | 3.089593 | 3.592943 | 3.349012 | 3.347548 | 3.351748 | 3.376609 | 10 | 9 | 8 | 14 |
| 127687 | NM_00114272 | C1orf122 | chromosome 1 open reading frame 122 | protein_coding | NP_001136198.1;NP_940848.2 | 24789 | . | . | . | 19.651694 | 21.687611 | 22.157390 | 24.749013 | 45.730298 | 50.131888 | 49.720539 | 56.585679 | 1.118062 | 821.407885 | 0.149345516 | 1.078042349 | 0.281014858 | 0.998865404 | 9.599509 | 9.759943 | 9.609471 | 9.669600 | 9.679079 | 9.757093 | 855 | 873 | 713 | 857 |
| 127700 | NM_00133049 | OSCP1 | organic solute carrier partner 1 | protein_coding | NP_001317422.1;NP_659484.4;NP_996668.1;XP_005270519.1;XP_00527 | 29971 | 608854 | . | . | 3.683808 | 3.701987 | 3.233941 | 3.496651 | 8.572371 | 8.557309 | 7.256869 | 7.994676 | -1.135879 | 139.227859 | 0.278075297 | -0.66100662 | 0.508608068 | 0.998865404 | 7.210740 | 7.025955 | 7.129978 | 7.160089 | 7.078042 | 7.116950 | 164 | 166 | 107 | 129 |
| 127703 | NM_00134869 | C1orf216 | chromosome 1 open reading frame 216 | protein_coding | NP_001335620.1;NP_689587.1 | 26800 | . | . | . | 16.193667 | 16.382642 | 16.312782 | 19.303648 | 37.683331 | 37.869212 | 36.605409 | 44.135496 | 1.088399 | 1428.532996 | 0.130075609 | 0.939509383 | 0.347469277 | 0.998865404 | 10.418079 | 10.539984 | 10.439719 | 10.452841 | 10.468108 | 10.552795 | 1562 | 1489 | 1224 | 1472 |
| 127829 | NM_00125612 | ARL8A | ADP ribosylation factor like GTPase 8A | protein_coding | NP_001243058.1;NP_620150.1 | 25192 | 616597 | . | . | 10.812808 | 11.452909 | 11.208751 | 13.950442 | 25.161848 | 26.473912 | 25.152113 | 31.896027 | 1.115400 | 660.882500 | 0.165284269 | 0.953276035 | 0.340450231 | 0.998865404 | 9.287743 | 9.444500 | 9.311369 | 9.344482 | 9.328032 | 9.471020 | 702 | 691 | 543 | 721 |
| 127833 | NM_00113650 | SYT2 | synaptotagmin 2 | protein_coding | NP_001129976.1;NP_796376.2;XP_011507494.1;XP_016855798.1;XP_01 | 11510 | 600104 | . | . | 0.014627 | 0.015580 | 0.027123 | 0.016929 | 0.034038 | 0.036013 | 0.060862 | 0.038705 | 1.346629 | 6.326063 | 1.206906701 | 0.355745883 | 0.72203088 | NA | 2.429096 | 2.861274 | 2.646414 | 2.648129 | 2.664223 | 2.647230 | 6 | 6 | 8 | 5 |
| 127933 | NM_00118476 | UHMK1 | U2AF homology motif kinase 1 | protein_coding | NP_001171692.1;NP_653225.2;NP_787062.1 | 19683 | 608849 | . | . | 7.300484 | 5.950571 | 7.097295 | 5.181415 | 16.988527 | 13.755013 | 15.926122 | 11.846689 | -1.093985 | 1697.046018 | 0.165597797 | -0.78257559 | 0.43387638 | 0.998865404 | 10.791845 | 10.662895 | 10.839335 | 10.670342 | 10.805690 | 10.555100 | 2258 | 1710 | 1637 | 1275 |
| 127943 | NM_00100290 | FCRLB | Fc receptor like B | protein_coding | NP_001002901.1;NP_001275758.1;NP_001275759.1;NP_001275760.1;NI | 26431 | 609251 | . | . | 0.505387 | 0.577211 | 0.739626 | 0.483081 | 1.176056 | 1.334250 | 1.659701 | 1.104508 | 1.126415 | 36.322756 | 0.528446916 | 0.324986908 | 0.74519098 | 0.998865404 | 5.093294 | 5.267079 | 5.159240 | 5.175658 | 5.216715 | 5.151024 | 37 | 39 | 41 | 28 |
| 128061 | NM_00130083 | C1orf131 | chromosome 1 open reading frame 131 | protein_coding | NP_001287759.1;NP_689592.2 | 25332 | . | . | . | 4.528891 | 4.224932 | 3.817599 | 6.094607 | 10.538915 | 9.766121 | 8.566583 | 13.934594 | 1.118285 | 208.538637 | 0.26405027 | 0.610821285 | 0.541317901 | 0.998865404 | 7.622082 | 7.781839 | 7.689541 | 7.653114 | 7.613422 | 7.821964 | 236 | 204 | 149 | 252 |
| 128077 | NM_153713,XI | LIX1L | limb and CNS expressed 1 like | protein_coding | NP_714924.1;XP_016855805.1 | 28715 | . | . | . | 5.954875 | 5.948135 | 5.793512 | 5.745606 | 13.857239 | 13.749380 | 13.000473 | 13.136646 | -1.047123 | 728.686374 | 0.148327764 | -0.44786623 | 0.654249742 | 0.998865404 | 9.541930 | 9.475616 | 9.528795 | 9.521152 | 9.500764 | 9.485451 | 863 | 800 | 626 | 660 |
| 128153 | NM_00137565 | SPATA17 | spermatogenesis associated 17 | protein_coding | NP_001362584.1;NP_620151.1;XP_006711228.1;XP_011507496.1 | 25184 | 611032 | . | . | 0.105117 | 0.278991 | 0.099712 | 0.321827 | 0.244611 | 0.644902 | 0.223750 | 0.735817 | 1.023417 | 18.086266 | 0.722235267 | 0.046236841 | 0.963121479 | NA | 4.159017 | 4.194412 | 4.184342 | 4.165801 | 4.182041 | 4.171717 | 23 | 17 | 17 | 16 |
| 128178 | NM_080738,N | EDARADD | EDAR associated death domain | protein_coding | NP_542776.1;NP_665860.2 | 14341 | 606603 | . | . | 0.412672 | 0.319573 | 0.295035 | 0.519357 | 0.960306 | 0.738708 | 0.662049 | 1.187446 | 1.092088 | 37.802203 | 0.529636141 | 0.239956538 | 0.81036395 | 0.998865404 | 5.176444 | 5.301637 | 5.248790 | 5.211465 | 5.202004 | 5.283010 | 47 | 34 | 25 | 47 |
| 128229 | NM_00130481 | TSACC | TSSK6 activating cochaperone | protein_coding | NP_001291746.1;NP_001291747.1;NP_001291748.1;NP_001291749.1;NI | 30636 | . | . | . | 0.512054 | 0.223870 | 0.255757 | 0.179633 | 1.191569 | 0.517485 | 0.573909 | 0.410709 | -1.598138 | 7.185890 | 1.173392453 | -0.57644179 | 0.564316607 | NA | 3.142442 | 2.470304 | 2.878372 | 2.839789 | 2.845336 | 2.843344 | 15 | 5 | 5 | 5 |
| 128239 | NM_178229,XI | IQGAP3 | IQ motif containing GTPase activating protein 3 | protein_coding | NP_839943.3;XP_011507500.1;XP_011507502.1;XP_011507503.1;XP_02 | 20669 | . | . | . | 9.843505 | 11.445500 | 10.278492 | 9.407846 | 22.906242 | 26.456787 | 23.064632 | 21.509922 | -1.108124 | 1870.869459 | 0.12883291 | -1.14970569 | 0.25026511 | 0.993339639 | 10.941690 | 10.793492 | 10.849094 | 10.963696 | 10.848155 | 10.805082 | 2103 | 2274 | 1586 | 1618 |
| 128240 | NM_144772,XI | NAXE | NAD(P)HX epimerase | protein_coding | NP_658985.2;XP_016855808.1 | 18453 | 608862 | . | . | 62.192356 | 58.550967 | 55.622277 | 71.716956 | 144.724168 | 135.343192 | 124.814775 | 163.972293 | 1.036582 | 2168.675002 | 0.139319857 | 0.372053182 | 0.709853253 | 0.998865404 | 11.056590 | 11.108146 | 11.092354 | 11.039308 | 10.993731 | 11.186923 | 2534 | 2222 | 1689 | 2318 |
| 128272 | NM_153213,XI | ARHGEF19 | Rho guanine nucleotide exchange factor 19 | protein_coding | NP_694945.2;XP_011539008.1 | 26604 | 612496 | . | . | 3.619414 | 3.546672 | 3.305184 | 3.982796 | 8.422523 | 8.198291 | 7.416736 | 9.106188 | 1.000342 | 386.719617 | 0.188752431 | 0.002614108 | 0.997914246 | 0.999862634 | 8.595232 | 8.595057 | 8.602003 | 8.586903 | 8.544474 | 8.642318 | 451 | 412 | 308 | 393 |
| 128308 | NM_00132128 | MRPL55 | mitochondrial ribosomal protein L55 | protein_coding | NP_001308213.1;NP_852106.1;NP_852119.1;NP_852120.1;NP_852121.1 | 16686 | 611859 | . | . | 28.638945 | 28.049908 | 27.774490 | 32.996921 | 66.643998 | 64.838618 | 62.325149 | 75.443538 | 1.099912 | 853.585592 | 0.148305155 | 0.926388068 | 0.354244343 | 0.998865404 | 9.667283 | 9.804251 | 9.705700 | 9.697144 | 9.716491 | 9.819511 | 942 | 872 | 722 | 898 |
| 128338 | NM_00134988 | DRAM2 | DNA damage regulated autophagy modulator 2 | protein_coding | NP_001336810.1;NP_001336811.1;NP_001336813.1;NP_001336814.1;NI | 28769 | 613360 | . | . | 19.928571 | 16.865280 | 18.113745 | 12.711129 | 46.374606 | 38.984850 | 40.646718 | 29.062484 | -1.222800 | 1116.886616 | 0.175945212 | -1.64931082 | 0.099083973 | 0.813553488 | 10.262666 | 9.973392 | 10.247771 | 10.129689 | 10.166355 | 9.897372 | 1523 | 1224 | 1036 | 768 |
| 128346 | NM_00130083 | C1orf162 | chromosome 1 open reading frame 162 | protein_coding | NP_001287763.1;NP_001287764.1;NP_777556.1;XP_016855812.1;XP_01 | 28344 | . | . | . | 0.387379 | 0.504823 | 0.351386 | 0.363358 | 0.901447 | 1.166920 | 0.788500 | 0.830775 | -1.352043 | 86.503023 | 0.350700428 | -1.24077786 | 0.214687821 | 0.979615203 | 6.636482 | 6.199984 | 6.433604 | 6.512449 | 6.383939 | 6.397795 | 100 | 121 | 62 | 71 |
| 128387 | NM_00104255 | TATDN3 | TatD DNase domain containing 3 | protein_coding | NP_001036017.1;NP_001036018.1;NP_001139641.1;NP_001139642.1;NI | 27010 | . | . | . | 2.968667 | 3.189911 | 2.389583 | 3.195565 | 6.908211 | 7.373624 | 5.362156 | 7.306280 | -1.116440 | 235.548968 | 0.233286684 | -0.68116041 | 0.495769998 | 0.998865404 | 7.957876 | 7.797432 | 7.888657 | 7.921372 | 7.788369 | 7.910833 | 277 | 277 | 168 | 236 |
| 128439 | NR_003239 | SNHG11 | small nucleolar RNA host gene 11 | lncRNA | . | 25046 | . | . | . | 4.953683 | 4.906709 | 7.514871 | 6.127161 | 11.527424 | 11.342078 | 16.863153 | 14.009025 | 1.363904 | 199.259942 | 0.243938753 | 1.835467293 | 0.066436481 | 0.733835894 | 7.396728 | 7.845525 | 7.560073 | 7.552233 | 7.744839 | 7.644173 | 196 | 180 | 221 | 193 |
| 128486 | NM_00108047 | FITM2 | fat storage inducing transmembrane protein 2 | protein_coding | NP_001073941.1 | 16135 | 612029 | . | . | 7.852424 | 7.736693 | 7.471260 | 7.509096 | 18.272913 | 17.883713 | 16.765289 | 17.168655 | -1.055508 | 1102.287265 | 0.132980141 | -0.58608832 | 0.557816167 | 0.998865404 | 10.144694 | 10.066825 | 10.135029 | 10.117125 | 10.087853 | 10.084117 | 1319 | 1207 | 935 | 1003 |
| 128497 | NM_080608,XI | SPATA25 | spermatogenesis associated 25 | protein_coding | NP_542175.1;XP_024307594.1 | 16158 | . | . | . | 0.151967 | 0.143446 | 0.120311 | 0.056834 | 0.353632 | 0.331581 | 0.269974 | 0.129943 | -1.556221 | 12.145712 | 0.896925013 | -0.71137149 | 0.476854064 | NA | 3.884262 | 3.251698 | 3.621660 | 3.613768 | 3.608076 | 3.580801 | 18 | 15 | 11 | 6 |
| 128553 | NM_00119342 | TSHZ2 | teashirt zinc finger homeobox 2 | protein_coding | NP_001180350.1;NP_775756.3;XP_016883129.1;XP_016883130.1 | 13010 | 614118 | . | . | 0.063303 | 0.125077 | 0.093132 | 0.090469 | 0.147309 | 0.289120 | 0.208986 | 0.206846 | -1.063034 | 36.021211 | 0.547760555 | -0.16099732 | 0.872095508 | 0.998865404 | 5.215335 | 5.124794 | 5.119629 | 5.215607 | 5.163134 | 5.157219 | 29 | 53 | 31 | 32 |
| 128637 | NM_144628,N | TBC1D20 | TBC1 domain family member 20 | protein_coding | NP_653229.1;XP_005260718.1;XP_006723603.1;XP_016883134.1 | 16133 | 611663 | . | . | 14.315272 | 16.360678 | 17.965237 | 17.173682 | 33.312228 | 37.818436 | 40.313471 | 39.265583 | 1.078005 | 1999.367143 | 0.124217812 | 0.872373486 | 0.38300466 | 0.998865404 | 10.910143 | 11.018479 | 10.891866 | 10.974554 | 11.019590 | 10.967206 | 2100 | 2185 | 1863 | 1881 |
| 128710 | NM_00100960 | SLX4IP | SLX4 interacting protein | protein_coding | NP_001009608.1;XP_016883136.1;XP_016883140.1;XP_024307595.1;XP | 16225 | 615958 | . | . | 0.871534 | 0.999665 | 0.976302 | 0.795955 | 2.028094 | 2.310771 | 2.190794 | 1.819859 | 1.043953 | 208.315604 | 0.257316186 | 0.241169619 | 0.809423657 | 0.998865404 | 7.670248 | 7.734295 | 7.707465 | 7.666381 | 7.792431 | 7.613801 | 245 | 210 | 221 | 161 |
| 128853 | NM_00101264 | DUSP15 | dual specificity phosphatase 15 | protein_coding | NP_001012662.1;NP_001307407.1;NP_001307408.1;NP_542178.2;NP_8 | 16236 | 616776 | . | . | 4.069813 | 2.851534 | 4.483631 | 4.516140 | 9.470619 | 6.591450 | 10.061137 | 10.325618 | 1.294584 | 178.023133 | 0.269351016 | 1.382912658 | 0.166691648 | 0.944905472 | 7.276808 | 7.650856 | 7.482250 | 7.332589 | 7.525703 | 7.514750 | 212 | 136 | 177 | 186 |
| 128866 | NM_176812 | CHMP4B | charged multivesicular body protein 4B | protein_coding | NP_789782.1 | 16171 | 610897 | . | . | 40.203129 | 43.230640 | 39.227703 | 48.964203 | 93.554337 | 99.929565 | 88.025826 | 111.950821 | 1.042354 | 2140.847198 | 0.135138379 | 0.442843534 | 0.657878912 | 0.998865404 | 11.033938 | 11.093382 | 11.019781 | 11.070943 | 10.990692 | 11.159809 | 2337 | 2335 | 1700 | 2264 |
| 128869 | NM_080476,XI | PIGU | phosphatidylinositol glycan anchor biosynthesis class U | protein_coding | NP_536724.1;XP_011526844.1;XP_016883153.1;XP_016883154.1 | 15791 | 608528 | . | . | 17.798748 | 18.585508 | 13.485369 | 14.412126 | 41.418423 | 42.961233 | 30.260778 | 32.951611 | -1.319856 | 821.502919 | 0.144908294 | -2.76298899 | **0.00572747** | 0.233204322 | 9.868599 | 9.467913 | 9.758328 | 9.784865 | 9.557080 | 9.593311 | 1057 | 1027 | 599 | 682 |
| 128977 | NM_00116624 | C22orf39 | chromosome 22 open reading frame 39 | protein_coding | NP_001159714.2;NP_776154.4 | 27012 | . | . | . | 5.370889 | 5.817781 | 5.061361 | 6.386802 | 12.498279 | 13.448060 | 11.357547 | 14.602664 | 1.043734 | 538.937638 | 0.167513606 | 0.36865513 | 0.712384798 | 0.998865404 | 9.043089 | 9.104213 | 9.048111 | 9.070377 | 9.043206 | 9.128697 | 597 | 579 | 447 | 550 |
| 128989 | NM_00128310 | TANGO2 | transport and golgi organization 2 homolog | protein_coding | NP_001270035.1;NP_001270045.1;NP_001270058.1;NP_001270077.1;NI | 25439 | 616830 | . | . | 18.446843 | 20.054815 | 23.117053 | 21.710477 | 42.926556 | 46.357608 | 51.873996 | 49.638423 | 1.105576 | 1761.072069 | 0.124743902 | 1.16076868 | 0.245735983 | 0.992733415 | 10.707940 | 10.852897 | 10.728143 | 10.753320 | 10.857794 | 10.780203 | 1895 | 1834 | 1687 | 1649 |
| 129025 | NM_080740 | ZNF280A | zinc finger protein 280A | protein_coding | NP_542778.2 | 18597 | . | . | . | 1.197440 | 1.083262 | 0.903017 | 1.669704 | 2.786492 | 2.504008 | 2.026344 | 3.817581 | 1.111852 | 81.682103 | 0.386588224 | 0.395679979 | 0.692341127 | 0.998865404 | 6.274694 | 6.425275 | 6.344715 | 6.316373 | 6.276682 | 6.435133 | 94 | 79 | 53 | 104 |
| 129049 | NM_00103994 | SGSM1 | small G protein signaling modulator 1 | protein_coding | NP_001035037.1;NP_001091967.1;NP_001091968.1;NP_597711.1 | 29410 | 611417 | . | . | 0.129370 | 0.051983 | 0.116266 | 0.131215 | 0.301049 | 0.120161 | 0.260898 | 0.300007 | 1.299801 | 23.434762 | 0.674687453 | 0.560691132 | 0.575008113 | NA | 4.347179 | 4.728813 | 4.564751 | 4.492619 | 4.549880 | 4.559945 | 33 | 13 | 22 | 26 |
| 129080 | NM_00126789 | EMID1 | EMI domain containing 1 | protein_coding | NP_001254824.1;NP_597712.2;XP_005261385.1;XP_005261386.1;XP_01 | 18036 | 608926 | . | . | 0.584482 | 0.708067 | 0.311843 | 0.620520 | 1.360111 | 1.636728 | 0.699768 | 1.418745 | -1.274171 | 43.253583 | 0.478523245 | -0.73049411 | 0.465088214 | 0.998865404 | 5.599889 | 5.248230 | 5.447240 | 5.465812 | 5.399365 | 5.434250 | 53 | 55 | 29 | 40 |
| 129138 | NM_00134985 | ANKRD54 | ankyrin repeat domain 54 | protein_coding | NP_001336782.1;NP_001350768.1;NP_620152.1;XP_006724199.1;XP_00 | 25185 | 613383 | . | . | 24.869318 | 25.628355 | 25.754275 | 29.074070 | 57.871930 | 59.241095 | 57.791849 | 66.474402 | 1.056957 | 1787.581664 | 0.123385522 | 0.647697461 | 0.517180647 | 0.998865404 | 10.763398 | 10.843088 | 10.773981 | 10.788539 | 10.789369 | 10.859035 | 1983 | 1893 | 1527 | 1798 |
| 129285 | NM_00113562 | PPP1R21 | protein phosphatase 1 regulatory subunit 21 | protein_coding | NP_001129101.1;NP_001180404.1;NP_694539.1;XP_016858790.1;XP_01 | 30595 | 618159 | . | . | 6.024893 | 5.500594 | 6.225698 | 6.600104 | 14.020173 | 12.714869 | 13.970288 | 15.090354 | 1.095390 | 609.216487 | 0.159613758 | 0.823517988 | 0.410213492 | 0.998865404 | 9.183527 | 9.315096 | 9.250235 | 9.186332 | 9.264879 | 9.293937 | 702 | 597 | 542 | 610 |
| 129303 | NM_00103173 | TMEM150A | transmembrane protein 150A | protein_coding | NP_001026908.1;NP_001356846.1;NP_699173.2;XP_006711993.1;XP_00 | 24677 | 616757 | . | . | 8.863969 | 9.075016 | 12.578830 | 8.316439 | 20.626825 | 20.977305 | 28.226529 | 19.014552 | 1.171323 | 566.184863 | 0.204962283 | 1.113080709 | 0.265673768 | 0.997577536 | 9.025955 | 9.255206 | 9.086395 | 9.078597 | 9.325912 | 9.024711 | 604 | 559 | 650 | 445 |
| 129401 | NM_00128758 | NUP35 | nucleoporin 35 | protein_coding | NP_001274513.1;NP_001274514.1;NP_612142.2;XP_006712317.1;XP_01 | 29797 | 608140 | . | . | 18.645088 | 17.147032 | 15.846539 | 16.031672 | 43.387885 | 39.636137 | 35.559175 | 36.654511 | -1.124872 | 820.775390 | 0.146038317 | -1.16244468 | 0.245054885 | 0.992733415 | 9.763117 | 9.593593 | 9.752718 | 9.688234 | 9.640015 | 9.635407 | 1045 | 896 | 674 | 722 |
| 129450 | NM_00103969 | TYW5 | tRNA-yW synthesizing protein 5 | protein_coding | NP_001034782.1;XP_016858799.1 | 26754 | . | . | . | 2.676560 | 2.418483 | 3.020018 | 2.283477 | 6.228465 | 5.590433 | 6.776834 | 5.220899 | 1.026170 | 415.054433 | 0.198415022 | 0.187835179 | 0.851005858 | 0.998865404 | 8.677711 | 8.716344 | 8.718003 | 8.652828 | 8.783885 | 8.612237 | 497 | 418 | 418 | 338 |
| 129530 | NM_00137138 | LYG1 | lysozyme g1 | protein_coding | NP_001358309.1;NP_777558.1;XP_005263927.1;XP_005263928.1;XP_01 | 27014 | . | . | . | 1.126737 | 1.384931 | 1.390946 | 0.560243 | 2.621963 | 3.201330 | 3.121245 | 1.280929 | -1.222813 | 42.970881 | 0.517462449 | -0.56082095 | 0.574919604 | 0.998865404 | 5.561500 | 5.274861 | 5.417053 | 5.452799 | 5.454759 | 5.346686 | 49 | 56 | 46 | 23 |
| 129531 | NM_00132041 | MITD1 | microtubule interacting and trafficking domain containing 1 | protein_coding | NP_001307346.1;NP_001307347.1;NP_001307348.1;NP_620153.1;XP_01 | 25207 | . | . | . | 16.178480 | 14.156350 | 18.464673 | 14.920856 | 37.647986 | 32.723036 | 41.434190 | 34.114764 | 1.095643 | 511.391226 | 0.184330368 | 0.714897997 | 0.474672074 | 0.998865404 | 8.930286 | 9.063220 | 8.997287 | 8.931554 | 9.103116 | 8.937066 | 591 | 499 | 525 | 437 |
| 129563 | NM_00125728 | DIS3L2 | DIS3 like 3'-5' exoribonuclease 2 | protein_coding | NP_001244210.1;NP_001244211.1;NP_689596.4 | 28648 | 614184 | . | . | 6.728957 | 7.334122 | 6.714011 | 8.220332 | 15.658560 | 16.953151 | 15.066046 | 18.794809 | 1.035492 | 770.682609 | 0.15475697 | 0.325130268 | 0.745082482 | 0.998865404 | 9.564956 | 9.614603 | 9.557951 | 9.595204 | 9.542093 | 9.656716 | 849 | 839 | 626 | 795 |
| 129642 | NM_00132126 | MBOAT2 | membrane bound O-acyltransferase domain containing 2 | protein_coding | NP_001308194.1;NP_001308195.1;NP_001308196.1;NP_620154.2;XP_01 | 25193 | 611949 | . | . | 7.101716 | 6.047723 | 6.094718 | 4.495737 | 16.525986 | 13.979585 | 13.676370 | 10.278968 | -1.263960 | 1393.211516 | 0.161781266 | -2.08893731 | **0.036713366** | 0.596884093 | 10.602967 | 10.265773 | 10.584026 | 10.459020 | 10.451436 | 10.224060 | 1932 | 1546 | 1234 | 978 |
| 129685 | NM_138572,XI | TAF8 | TATA-box binding protein associated factor 8 | protein_coding | NP_612639.2;XP_005248906.1;XP_011512598.1;XP_016865730.1;XP_01 | 17300 | 609514 | . | . | 2.278400 | 1.992234 | 2.364779 | 2.346142 | 5.301928 | 4.605140 | 5.306495 | 5.364173 | 1.027755 | 375.556319 | 0.189046245 | 0.208926883 | 0.834505318 | 0.998865404 | 8.532746 | 8.572748 | 8.585059 | 8.501440 | 8.561650 | 8.558579 | 457 | 371 | 332 | 356 |
| 129787 | NM_00135268 | TMEM18 | transmembrane protein 18 | protein_coding | NP_001339609.1;NP_001339610.1;NP_690047.2 | 25257 | 613220 | . | . | 5.921613 | 6.229682 | 6.476054 | 5.732533 | 13.779835 | 14.400187 | 14.532079 | 13.106754 | -1.009567 | 1183.390479 | 0.138116939 | -0.09945424 | 0.920777621 | 0.998865404 | 10.215415 | 10.201974 | 10.194835 | 10.227913 | 10.254779 | 10.151535 | 1339 | 1311 | 1095 | 1028 |
| 129831 | NM_00136557 | RBM45 | RNA binding motif protein 45 | protein_coding | NP_001352507.1;NP_001352508.1;NP_694453.2;XP_016858809.1;XP_01 | 24468 | 608888 | . | . | 5.956820 | 5.765487 | 6.609209 | 4.975670 | 13.861765 | 13.327182 | 14.830874 | 11.376278 | -1.015021 | 335.484005 | 0.206680658 | -0.1040687 | 0.917114826 | 0.998865404 | 8.400191 | 8.379938 | 8.402011 | 8.381599 | 8.454313 | 8.307753 | 396 | 358 | 328 | 271 |
| 129880 | NM_152384 | BBS5 | Bardet-Biedl syndrome 5 | protein_coding | NP_689597.1 | 970 | 603650 | . | . | 12.291134 | 10.979717 | 13.914604 | 10.979037 | 28.601974 | 25.380112 | 31.223969 | 25.102264 | 1.054557 | 1185.744848 | 0.154538377 | 0.495910025 | 0.619957893 | 0.998865404 | 10.172381 | 10.249738 | 10.232772 | 10.142285 | 10.317481 | 10.131404 | 1409 | 1170 | 1189 | 1001 |
| 130026 | NM_00128862 | ICA1L | islet cell autoantigen 1 like | protein_coding | NP_001275551.1;NP_001275552.1;NP_001275553.1;NP_612477.3 | 14442 | . | . | . | 0.183463 | 0.286477 | 0.424273 | 0.277451 | 0.426925 | 0.662205 | 0.952056 | 0.634360 | 1.065140 | 62.971329 | 0.451081385 | 0.20183355 | 0.840046852 | 0.998865404 | 5.929600 | 6.022162 | 5.906080 | 6.002010 | 6.039441 | 5.893464 | 55 | 80 | 75 | 41 |
| 130074 | NM_00100999 | FAM168B | family with sequence similarity 168 member B | protein_coding | NP_001009993.2;NP_001308672.1;NP_001308673.1;NP_001308674.1;NI | 27016 | . | . | . | 27.072231 | 26.792414 | 24.304620 | 26.775686 | 62.998192 | 61.931868 | 54.538863 | 61.219425 | -1.072335 | 4583.567495 | 0.10754865 | -0.93683776 | 0.348842016 | 0.998865404 | 12.211784 | 12.110965 | 12.193393 | 12.187917 | 12.108982 | 12.155251 | 5480 | 5102 | 3739 | 4263 |
| 130162 | NM_00113559 | CLHC1 | clathrin heavy chain linker domain containing 1 | protein_coding | NP_001129070.1;NP_001340708.1;NP_001340709.1;NP_001340710.1;NI | 26453 | . | . | . | 0.482554 | 0.555152 | 0.704164 | 0.434162 | 1.122923 | 1.283257 | 1.580123 | 0.992661 | 1.100765 | 95.838618 | 0.352094888 | 0.393377355 | 0.694040792 | 0.998865404 | 6.510623 | 6.651032 | 6.543035 | 6.576602 | 6.657204 | 6.514686 | 99 | 104 | 108 | 72 |
| 130271 | NM_172069,XI | PLEKHH2 | pleckstrin homology, MyTH4 and FERM domain containing H2 | protein_coding | NP_742066.2;XP_016858840.1;XP_016858841.1;XP_016858842.1;XP_02 | 30506 | 612723 | . | . | 2.445181 | 2.414388 | 2.229591 | 2.392476 | 5.690034 | 5.580967 | 5.003139 | 5.470111 | -1.108572 | 505.044103 | 0.168294281 | -0.88358465 | 0.376920468 | 0.998865404 | 9.052867 | 8.903817 | 9.018134 | 9.006404 | 8.919015 | 8.973733 | 617 | 568 | 398 | 469 |
| 130340 | NM_00103956 | AP1S3 | adaptor related protein complex 1 subunit sigma 3 | protein_coding | NP_001034658.1;XP_011508902.1 | 18971 | 615781 | . | . | 4.219182 | 3.810168 | 3.690050 | 3.829449 | 9.818210 | 8.807376 | 8.280366 | 8.755580 | -1.084143 | 483.532028 | 0.172081034 | -0.67732888 | 0.498197311 | 0.998865404 | 8.974469 | 8.858121 | 8.976955 | 8.906315 | 8.883214 | 8.899846 | 611 | 513 | 398 | 441 |
| 130355 | NM_00101792 | C2orf76 | chromosome 2 open reading frame 76 | protein_coding | NP_001017927.2;NP_001309258.1;NP_001309259.1;NP_001309260.1;NI | 27017 | . | . | . | 5.785082 | 5.861686 | 7.614556 | 6.324879 | 13.462122 | 13.549548 | 17.086841 | 14.461085 | 1.033245 | 185.874209 | 0.269501846 | 0.175072208 | 0.861022896 | 0.998865404 | 7.513340 | 7.562605 | 7.541026 | 7.509303 | 7.624743 | 7.446683 | 218 | 190 | 198 | 141 |
| 130399 | NM_00111103 | ACVR1C | activin A receptor type 1C | protein_coding | NP_001104501.1;NP_001104502.1;NP_001104503.1;NP_660302.2 | 18123 | 608981 | . | . | 0.826503 | 0.672548 | 0.695541 | 0.571621 | 1.923307 | 1.554627 | 1.560774 | 1.306942 | -1.204516 | 190.235059 | 0.253776324 | -1.05783623 | 0.29013011 | 0.998865404 | 7.698859 | 7.432107 | 7.657044 | 7.559614 | 7.565160 | 7.489050 | 264 | 201 | 164 | 147 |
| 130502 | NM_00100823 | TTC32 | tetratricopeptide repeat domain 32 | protein_coding | NP_001008238.1 | 32954 | . | . | . | 1.967365 | 1.918628 | 2.189714 | 2.076358 | 4.578138 | 4.434994 | 4.913656 | 4.747347 | 1.082754 | 60.196019 | 0.403663335 | 0.284160426 | 0.776287441 | 0.998865404 | 5.852816 | 5.968075 | 5.906008 | 5.896892 | 5.924524 | 5.912281 | 68 | 61 | 56 | 57 |
| 130507 | NM_172070,XI | UBR3 | ubiquitin protein ligase E3 component n-recognin 3 | protein_coding | NP_742067.3;XP_005246356.1;XP_005246357.1;XP_006712331.1;XP_00 | 30467 | 613831 | . | . | 4.377717 | 4.476289 | 4.505095 | 4.201635 | 10.187125 | 10.347143 | 10.109300 | 9.606538 | -1.033955 | 1090.365200 | 0.135047225 | -0.35671036 | 0.721308649 | 0.998865404 | 10.114383 | 10.066409 | 10.097765 | 10.107608 | 10.104173 | 10.051106 | 1268 | 1204 | 966 | 969 |
| 130535 | NM_00132154 | KCTD18 | potassium channel tetramerization domain containing 18 | protein_coding | NP_001308476.1;NP_001308477.1;NP_001308479.1;NP_689600.2 | 26446 | . | . | . | 4.283846 | 4.764422 | 4.518409 | 5.340901 | 9.968686 | 11.013176 | 10.139176 | 12.211335 | 1.087846 | 424.732310 | 0.183016628 | 0.663730643 | 0.506862723 | 0.998865404 | 8.668834 | 8.789465 | 8.686611 | 8.719113 | 8.703708 | 8.801542 | 456 | 451 | 354 | 448 |
| 130540 | NM_00112739 | FLACC1 | flagellum associated containing coiled-coil domains 1 | protein_coding | NP_001120863.1;NP_001276922.1;NP_631902.2;XP_011508908.1;XP_01 | 14439 | . | . | . | 0.564680 | 0.486337 | 0.948118 | 0.850845 | 1.314033 | 1.124190 | 2.127550 | 1.945355 | 1.655251 | 49.304959 | 0.447339369 | 1.625275185 | 0.104103936 | 0.828395026 | 5.214084 | 5.942303 | 5.572648 | 5.550577 | 5.658431 | 5.646903 | 46 | 37 | 55 | 56 |
| 130557 | NM_00120145 | ZNF513 | zinc finger protein 513 | protein_coding | NP_001188388.1;NP_653232.3;XP_005264199.1;XP_005264200.1 | 26498 | 613598 | . | . | 6.696084 | 6.539705 | 8.074044 | 6.637850 | 15.582061 | 15.116823 | 18.117921 | 15.176655 | 1.093302 | 455.537063 | 0.181385067 | 0.709496747 | 0.478016269 | 0.998865404 | 8.765192 | 8.894750 | 8.810622 | 8.791220 | 8.914889 | 8.794028 | 509 | 462 | 454 | 403 |
| 130574 | NM_00119568 | LYPD6 | LY6/PLAUR domain containing 6 | protein_coding | NP_001182614.1;NP_919298.1;XP_024308466.1;XP_024308467.1;XP_02 | 28751 | 613359 | . | . | 2.601461 | 2.956601 | 2.467266 | 2.436313 | 6.053706 | 6.834316 | 5.536473 | 5.570341 | -1.150071 | 329.143200 | 0.196780774 | -1.02511537 | 0.305308752 | 0.998865404 | 8.460039 | 8.258040 | 8.365936 | 8.432814 | 8.327953 | 8.316492 | 382 | 402 | 269 | 284 |
| 130589 | NM_138801,XI | GALM | galactose mutarotase | protein_coding | NP_620156.1;XP_011530842.1 | 24063 | 137030 | . | . | 1.508550 | 1.411663 | 1.671704 | 1.805636 | 3.510458 | 3.263122 | 3.751256 | 4.128371 | 1.200937 | 111.417598 | 0.304567016 | 0.867331147 | 0.385760596 | 0.998865404 | 6.661823 | 6.925786 | 6.772026 | 6.759459 | 6.813193 | 6.837585 | 119 | 107 | 102 | 118 |
| 130612 | NM_00100520 | TMEM198 | transmembrane protein 198 | protein_coding | NP_001005209.1;NP_001290027.1;XP_016858856.1 | 33704 | . | . | . | 6.399172 | 7.065085 | 7.446741 | 7.388177 | 14.891137 | 16.331262 | 16.710271 | 16.892190 | 1.080341 | 490.427867 | 0.170677706 | 0.653196968 | 0.513629305 | 0.998865404 | 8.881200 | 8.992450 | 8.879163 | 8.943809 | 8.967887 | 8.953420 | 515 | 535 | 448 | 472 |
| 130617 | NM_00127099 | ZFAND2B | zinc finger AN1-type containing 2B | protein_coding | NP_001257927.1;NP_001257928.1;NP_620157.1;XP_006712348.1;XP_00 | 25206 | 613474 | . | . | 11.911377 | 13.161387 | 14.268226 | 13.952218 | 27.718263 | 30.423134 | 32.017486 | 31.900088 | 1.125355 | 520.468665 | 0.166570113 | 1.022870363 | 0.306369144 | 0.998865404 | 8.936128 | 9.106198 | 8.954847 | 9.011077 | 9.057155 | 9.060728 | 539 | 552 | 478 | 518 |
| 130749 | NM_173077,XI | CPO | carboxypeptidase O | protein_coding | NP_775100.1;XP_016858861.1 | 21011 | 609563 | . | . | 0.010216 | 0.026704 | 0.024221 | 0.019227 | 0.023772 | 0.061727 | 0.054352 | 0.043961 | 1.184288 | 15.384323 | 0.820466217 | 0.297416664 | 0.766148423 | NA | 3.817846 | 4.058876 | 3.897905 | 3.954506 | 3.945106 | 3.930619 | 9 | 22 | 16 | 14 |
| 130752 | NM_00103984 | MDH1B | malate dehydrogenase 1B | protein_coding | NP_001034934.1;NP_001269869.1;NP_001317152.1;NP_001317153.1;NI | 17836 | . | . | . | 1.928506 | 1.478610 | 1.406159 | 1.551690 | 4.487712 | 3.417874 | 3.155380 | 3.547755 | -1.108592 | 124.265442 | 0.291923004 | -0.50948029 | 0.610415608 | 0.998865404 | 7.029459 | 6.881302 | 6.998712 | 6.951942 | 6.941386 | 6.938017 | 160 | 132 | 103 | 110 |
| 130814 | NM_00128271 | SLC66A3 | solute carrier family 66 member 3 | protein_coding | NP_001269639.1;NP_001269640.1;NP_001269641.1;NP_689604.1;XP_01 | 28503 | . | . | . | 4.795071 | 4.830377 | 4.225292 | 5.021872 | 11.158328 | 11.165633 | 9.481432 | 11.481913 | -1.003505 | 327.985455 | 0.193832356 | -0.02604263 | 0.97922334 | 0.998865404 | 8.360076 | 8.354895 | 8.360968 | 8.356063 | 8.347456 | 8.365572 | 380 | 353 | 280 | 312 |
| 130827 | NM_00132134 | TMEM182 | transmembrane protein 182 | protein_coding | NP_001308272.2;NP_001308273.2;NP_001308274.2;NP_001308275.2;NI | 26391 | . | . | . | 2.088017 | 1.851655 | 1.796064 | 2.307704 | 4.858901 | 4.280184 | 4.030313 | 5.276294 | 1.035530 | 179.637090 | 0.249192694 | 0.202128419 | 0.839816331 | 0.998865404 | 7.463805 | 7.513647 | 7.488710 | 7.473024 | 7.468848 | 7.522045 | 207 | 187 | 149 | 182 |
| 130872 | NR_152210,NF | AHSA2P | activator of HSP90 ATPase homolog 2, pseudogene | transcribed_ps | . | 20437 | . | . | . | 21.931970 | 22.630347 | 30.149068 | 22.404174 | 51.036596 | 52.311061 | 67.653635 | 51.224478 | 1.154842 | 2388.209213 | 0.148763269 | 1.396149934 | 0.16266935 | 0.935295382 | 11.113949 | 11.321985 | 11.154349 | 11.160065 | 11.394442 | 11.136284 | 2543 | 2400 | 2576 | 2023 |
| 130888 | NM_174899,XI | FBXO36 | F-box protein 36 | protein_coding | NP_777559.3;XP_005246374.1 | 27020 | 609105 | . | . | 0.554907 | 0.446953 | 0.391699 | 0.497036 | 1.291292 | 1.033152 | 0.878962 | 1.136415 | -1.160713 | 43.998614 | 0.484143677 | -0.44410686 | 0.65696533 | 0.998865404 | 5.563187 | 5.347534 | 5.500808 | 5.443788 | 5.424461 | 5.470799 | 63 | 43 | 30 | 44 |
| 130916 | NM_00133017 | MTERF4 | mitochondrial transcription termination factor 4 | protein_coding | NP_001317108.1;NP_001317109.1;NP_872307.2;XP_006712355.1;XP_01 | 28785 | 615393 | . | . | 7.545326 | 6.849340 | 7.458662 | 6.492857 | 17.558287 | 15.832556 | 16.737023 | 14.845143 | 1.078809 | 935.896430 | 0.140704226 | 0.777801691 | 0.436685931 | 0.998865404 | 9.814375 | 9.923955 | 9.826842 | 9.855763 | 9.923014 | 9.869114 | 1016 | 991 | 876 | 877 |
| 130940 | NM_00130168 | CCDC148 | coiled-coil domain containing 148 | protein_coding | NP_001288613.1;NP_001288614.1;NP_620158.3;XP_005246376.1;XP_01 | 25191 | . | . | . | 0.482752 | 0.371667 | 0.417232 | 0.239257 | 1.123386 | 0.859127 | 0.936255 | 0.547033 | -1.027621 | 32.851067 | 0.561291668 | -0.0700319 | 0.944168268 | 0.998865404 | 5.056118 | 5.019385 | 5.022709 | 5.046154 | 5.065313 | 4.995269 | 35 | 39 | 36 | 22 |
| 130951 | NM_00128129 | M1AP | meiosis 1 associated protein | protein_coding | NP_001268224.1;NP_001268225.1;NP_001308668.1;NP_620159.2;XP_00 | 25183 | . | . | . | 0.489679 | 0.920975 | 0.254025 | 0.556049 | 1.139502 | 2.128877 | 0.570025 | 1.271339 | -1.752973 | 40.961283 | 0.501142568 | -1.61591493 | 0.106112707 | 0.833478683 | 5.706148 | 4.893187 | 5.375127 | 5.421155 | 5.295205 | 5.329010 | 53 | 63 | 22 | 32 |
| 131076 | NM_00101792 | CCDC58 | coiled-coil domain containing 58 | protein_coding | NP_001017928.1;NP_001295255.1;XP_011510712.1;XP_016861186.1 | 31136 | . | . | . | 22.173991 | 19.542838 | 20.306407 | 23.549474 | 51.599789 | 45.174145 | 45.566993 | 53.843073 | 1.030283 | 481.708570 | 0.180085264 | 0.239000087 | 0.811105515 | 0.998865404 | 8.890433 | 8.933283 | 8.945786 | 8.854766 | 8.869996 | 8.967585 | 586 | 475 | 392 | 494 |
| 131096 | NM_144633,XI | KCNH8 | potassium voltage-gated channel subfamily H member 8 | protein_coding | NP_653234.2;XP_016861187.1;XP_016861188.1;XP_016861189.1;XP_01 | 18864 | 608260 | . | . | 0.270841 | 0.400642 | 0.276027 | 0.293548 | 0.630257 | 0.926103 | 0.619397 | 0.671162 | -1.206956 | 49.450411 | 0.455977315 | -0.59514664 | 0.551745453 | 0.998865404 | 5.758036 | 5.484875 | 5.606646 | 5.679727 | 5.605659 | 5.613349 | 51 | 69 | 38 | 43 |
| 131118 | NM_00119023 | DNAJC19 | DnaJ heat shock protein family (Hsp40) member C19 | protein_coding | NP_001177162.1;NP_660304.1 | 30528 | 608977 | . | . | 10.006485 | 9.648782 | 9.202403 | 10.747921 | 23.285505 | 22.303594 | 20.649925 | 24.573843 | 1.005313 | 438.763294 | 0.179782643 | 0.042522119 | 0.966082479 | 0.998865404 | 8.773709 | 8.780881 | 8.789783 | 8.760418 | 8.733839 | 8.821005 | 516 | 461 | 355 | 442 |
| 131408 | NM_00117109 | FAM131A | family with sequence similarity 131 member A | protein_coding | NP_001164564.1;NP_001353062.1;NP_001353063.1;NP_653236.3;XP_00 | 28308 | . | . | . | 3.818146 | 3.818746 | 3.254078 | 3.518427 | 8.884985 | 8.827200 | 7.302057 | 8.044464 | -1.146090 | 276.995379 | 0.207651624 | -0.94735608 | 0.343457365 | 0.998865404 | 8.208893 | 8.011818 | 8.149155 | 8.148005 | 8.061259 | 8.093842 | 341 | 319 | 217 | 250 |
| 131474 | NM_00109850 | CHCHD4 | coiled-coil-helix-coiled-coil-helix domain containing 4 | protein_coding | NP_001091972.1;NP_653237.1 | 26467 | 611077 | . | . | 6.049161 | 6.324333 | 7.231460 | 6.565529 | 14.076646 | 14.618978 | 16.227188 | 15.011302 | 1.101428 | 298.347594 | 0.203642782 | 0.684408274 | 0.493717392 | 0.998865404 | 8.149233 | 8.289080 | 8.186186 | 8.199012 | 8.275808 | 8.212162 | 323 | 310 | 288 | 276 |
| 131544 | NM_153605,XI | CRYBG3 | crystallin beta-gamma domain containing 3 | protein_coding | NP_705833.3;XP_005247174.1 | 34427 | . | . | . | 1.379319 | 1.398426 | 2.006656 | 1.596215 | 3.209731 | 3.232523 | 4.502878 | 3.649557 | 1.275551 | 536.348448 | 0.177522722 | 1.97788885 | **0.047941247** | 0.659753773 | 8.880387 | 9.232263 | 8.975002 | 8.981423 | 9.211786 | 9.053044 | 540 | 511 | 585 | 497 |
| 131566 | NM_080927,XI | DCBLD2 | discoidin, CUB and LCCL domain containing 2 | protein_coding | NP_563615.3;XP_011510721.1;XP_024309115.1;XP_024309116.1 | 24627 | 608698 | . | . | 17.503928 | 15.531355 | 14.447162 | 12.141891 | 40.732362 | 35.901423 | 32.419015 | 27.760991 | -1.243359 | 2764.580782 | 0.133656618 | -2.35112358 | **0.018716817** | 0.458064219 | 11.581259 | 11.267393 | 11.555112 | 11.469632 | 11.416101 | 11.249467 | 3709 | 3137 | 2368 | 2069 |
| 131583 | NM_153690 | FAM43A | family with sequence similarity 43 member A | protein_coding | NP_710157.2 | 26888 | . | . | . | 8.985006 | 8.188409 | 11.215697 | 9.902950 | 20.908478 | 18.927874 | 25.167698 | 22.641918 | 1.212227 | 940.870171 | 0.147509357 | 1.882318207 | 0.059792824 | 0.719735192 | 9.732016 | 10.010290 | 9.836895 | 9.765050 | 9.989970 | 9.890481 | 1029 | 871 | 957 | 902 |
| 131601 | NM_00113605 | TPRA1 | transmembrane protein adipocyte associated 1 | protein_coding | NP_001129525.1;NP_001136118.1;NP_001339930.1;NP_001339931.1;NI | 30413 | 608336 | . | . | 10.508422 | 9.458718 | 8.570626 | 9.453816 | 24.453528 | 21.864251 | 19.232233 | 21.615026 | -1.074313 | 798.878425 | 0.146419313 | -0.70628504 | 0.480010892 | 0.998865404 | 9.692497 | 9.589323 | 9.659092 | 9.673717 | 9.642731 | 9.588545 | 943 | 902 | 696 | 694 |
| 131616 | NM_144638 | TMEM42 | transmembrane protein 42 | protein_coding | NP_653239.1 | 28444 | . | . | . | 3.295909 | 4.248146 | 3.425983 | 4.791325 | 7.669716 | 9.819780 | 7.687806 | 10.954795 | 1.081285 | 120.266694 | 0.314658694 | 0.358316302 | 0.72010662 | 0.998865404 | 6.853946 | 6.964135 | 6.849706 | 6.932468 | 6.861148 | 6.972986 | 117 | 140 | 91 | 136 |
| 131870 | NM_00117190 | NUDT16 | nudix hydrolase 16 | protein_coding | NP_001165376.1;NP_001165377.1;NP_689608.2 | 26442 | 617381 | . | . | 4.411381 | 3.959244 | 5.105679 | 4.913141 | 10.265467 | 9.151969 | 11.456995 | 11.233313 | 1.071314 | 685.901766 | 0.15229842 | 0.65254668 | 0.51404857 | 0.998865404 | 9.371293 | 9.470710 | 9.379524 | 9.414574 | 9.460833 | 9.427502 | 743 | 733 | 632 | 649 |
| 131961 | gene-IMPDH1F | IMPDH1P8 | inosine monophosphate dehydrogenase 1 pseudogene 8 | pseudogene | . | 33963 | . | . | . | 0.259693 | 0.307014 | 0.345180 | 0.190024 | 0.604316 | 0.709676 | 0.774575 | 0.434467 | -1.044590 | 21.131383 | 0.684669846 | -0.0919234 | 0.926758901 | NA | 4.431047 | 4.370958 | 4.392514 | 4.406287 | 4.418981 | 4.371494 | 23 | 25 | 23 | 14 |
| 131965 | NM_00130179 | METTL6 | methyltransferase like 6 | protein_coding | NP_001288719.1;NP_001288720.1;NP_001288721.1;NP_001317591.1;NI | 28343 | . | . | . | 5.441097 | 4.576577 | 6.414754 | 5.445641 | 12.661661 | 10.578964 | 14.394524 | 12.450809 | 1.142134 | 469.516623 | 0.173648832 | 1.104137613 | 0.269533447 | 0.998865404 | 8.775821 | 8.967858 | 8.817120 | 8.843297 | 8.946812 | 8.878338 | 494 | 483 | 459 | 444 |
| 132001 | NM_00128440 | TAMM41 | TAM41 mitochondrial translocator assembly and maintenance ho | protein_coding | NP_001271330.1;NP_001308223.1;NP_001308224.1;NP_001352960.1;NI | 25187 | 614948 | . | . | 4.243758 | 4.134718 | 4.875983 | 4.231865 | 9.875396 | 9.557591 | 10.941568 | 9.675657 | 1.094716 | 234.979586 | 0.22455107 | 0.581413024 | 0.560962121 | 0.998865404 | 7.809149 | 7.940639 | 7.873648 | 7.830103 | 7.924087 | 7.867444 | 270 | 231 | 226 | 217 |
| 132014 | NM_00119338 | IL17RE | interleukin 17 receptor E | protein_coding | NP_001180309.1;NP_705613.1;NP_705614.1;NP_705616.2;XP_0067130 | 18439 | 614995 | . | . | 0.166683 | 0.225050 | 0.383001 | 0.448076 | 0.387879 | 0.520212 | 0.859444 | 1.024474 | **2.108174** | 27.279454 | 0.602936487 | 1.784589623 | 0.074327906 | 0.764716896 | 4.135531 | 5.208789 | 4.692292 | 4.717827 | 4.775665 | 4.795730 | 17 | 22 | 30 | 37 |
| 132158 | NM_00114495 | GLYCTK | glycerate kinase | protein_coding | NP_001138423.1;NP_660305.2;XP_016861219.1;XP_024309119.1;XP_02 | 24247 | 610516 | . | . | 1.790260 | 1.934358 | 2.245499 | 1.860086 | 4.166009 | 4.471355 | 5.038835 | 4.252867 | 1.097079 | 170.389377 | 0.27476528 | 0.486478475 | 0.626627963 | 0.998865404 | 7.343462 | 7.478751 | 7.377845 | 7.395667 | 7.503759 | 7.340662 | 183 | 179 | 185 | 135 |
| 132160 | NM_00112287 | PPM1M | protein phosphatase, Mg2+/Mn2+ dependent 1M | protein_coding | NP_001116342.1;NP_653242.3;XP_005264936.1;XP_005264937.1;XP_00 | 26506 | 608979 | . | . | 4.136199 | 3.501847 | 3.021704 | 3.304893 | 9.625107 | 8.094674 | 6.780616 | 7.556245 | -1.213045 | 232.472857 | 0.228247278 | -1.22075136 | 0.222180176 | 0.982008277 | 7.993351 | 7.715089 | 7.949279 | 7.859449 | 7.803178 | 7.821105 | 319 | 251 | 179 | 201 |
| 132241 | NR_003111 | RPL32P3 | ribosomal protein L32 pseudogene 3 | transcribed_ps | . | 27024 | . | . | . | 1.448148 | 1.408167 | 1.875133 | 1.524913 | 3.369901 | 3.255041 | 4.207745 | 3.486532 | 1.173791 | 101.826751 | 0.322226766 | 0.717430827 | 0.473108304 | 0.998865404 | 6.549025 | 6.781561 | 6.645929 | 6.634196 | 6.723362 | 6.654515 | 110 | 99 | 106 | 92 |
| 132320 | NM_00130089 | SCLT1 | sodium channel and clathrin linker 1 | protein_coding | NP_001287826.1;NP_001287827.1;NP_653244.2;XP_016863205.1;XP_01 | 26406 | 611399 | . | . | 4.301910 | 4.010726 | 3.893273 | 3.955018 | 10.010720 | 9.270976 | 8.736395 | 9.042678 | -1.079152 | 297.602292 | 0.203454605 | -0.5401591 | 0.589087317 | 0.998865404 | 8.271187 | 8.161201 | 8.259415 | 8.215157 | 8.178044 | 8.215139 | 370 | 320 | 240 | 278 |
| 132321 | NM_00109978 | C4orf33 | chromosome 4 open reading frame 33 | protein_coding | NP_001093253.1;NP_775758.2;XP_005262791.1;XP_016863211.1;XP_01 | 27025 | . | . | . | 2.078807 | 1.698297 | 1.204678 | 1.288059 | 4.837471 | 3.925687 | 2.703264 | 2.944993 | -1.195547 | 198.393008 | 0.2410898 | -1.06877417 | 0.285171443 | 0.998865404 | 7.755252 | 7.497704 | 7.703757 | 7.637404 | 7.577366 | 7.604511 | 266 | 217 | 152 | 175 |
| 132660 | NM_00111500 | LIN54 | lin-54 DREAM MuvB core complex component | protein_coding | NP_001108479.1;NP_001108480.1;NP_001275925.1;NP_001275926.1;NI | 25397 | 613367 | . | . | 3.025291 | 2.887731 | 3.193183 | 3.401784 | 7.039978 | 6.675119 | 7.165409 | 7.777769 | 1.072019 | 544.427961 | 0.163250147 | 0.61458422 | 0.538829338 | 0.998865404 | 9.037603 | 9.137851 | 9.078666 | 9.051347 | 9.092821 | 9.127174 | 618 | 555 | 477 | 542 |
| 132720 | NM_152400 | FAM241A | family with sequence similarity 241 member A | protein_coding | NP_689613.2 | 26813 | . | . | . | 0.395756 | 0.342882 | 0.282837 | 0.335188 | 0.920941 | 0.792588 | 0.634677 | 0.766368 | -1.210396 | 93.919618 | 0.334315448 | -0.82401108 | 0.409933262 | 0.998865404 | 6.684726 | 6.408811 | 6.604338 | 6.558344 | 6.504766 | 6.546770 | 127 | 103 | 68 | 86 |
| 132789 | NM_00127088 | GNPDA2 | glucosamine-6-phosphate deaminase 2 | protein_coding | NP_001257809.1;NP_001257810.1;NP_612208.1 | 21526 | 613222 | . | . | 4.608510 | 3.307118 | 3.235271 | 2.804347 | 10.724190 | 7.644549 | 7.259852 | 6.411806 | -1.320756 | 242.671576 | 0.242428002 | -1.65560137 | 0.097802572 | 0.813553488 | 8.108894 | 7.709239 | 8.065292 | 7.893187 | 7.883128 | 7.808041 | 369 | 250 | 197 | 180 |
| 132864 | NM_00117738 | CPEB2 | cytoplasmic polyadenylation element binding protein 2 | protein_coding | NP_001170852.1;NP_001170853.1;NP_001170854.1;NP_001170855.1;NI | 21745 | 610605 | . | . | 3.547592 | 3.182626 | 4.445595 | 3.269431 | 8.255393 | 7.356783 | 9.975788 | 7.475168 | 1.124293 | 783.578710 | 0.1778327 | 0.950432116 | 0.341892732 | 0.998865404 | 9.526420 | 9.696442 | 9.605400 | 9.525641 | 9.760578 | 9.523546 | 898 | 750 | 841 | 649 |
| 132884 | NM_00116613 | EVC2 | EvC ciliary complex subunit 2 | protein_coding | NP_001159608.1;NP_667338.3;XP_016863225.1;XP_016863226.1;XP_01 | 19747 | 607261 | . | . | 1.876070 | 1.617337 | 1.686320 | 1.745731 | 4.365690 | 3.738547 | 3.784052 | 3.991406 | 1.024730 | 242.173968 | 0.225816678 | 0.156069846 | 0.875977967 | 0.998865404 | 7.902177 | 7.937408 | 7.952542 | 7.871480 | 7.894145 | 7.954639 | 298 | 237 | 200 | 244 |
| 132949 | NM_00128666 | AASDH | aminoadipate-semialdehyde dehydrogenase | protein_coding | NP_001273597.1;NP_001273598.1;NP_001273599.1;NP_001273600.1;NI | 23993 | 614365 | . | . | 2.442676 | 2.295373 | 2.967794 | 2.073255 | 5.684209 | 5.305856 | 6.659644 | 4.740253 | 1.081359 | 281.378749 | 0.22187914 | 0.508591291 | 0.611038735 | 0.998865404 | 8.078032 | 8.192439 | 8.144313 | 8.078052 | 8.220442 | 8.080578 | 331 | 273 | 288 | 238 |
| 133015 | NM_00113072 | PACRGL | parkin coregulated like | protein_coding | NP_001124199.1;NP_001245274.1;NP_001245275.1;NP_001304778.1;NI | 28442 | . | . | . | 2.870532 | 2.842374 | 3.147662 | 3.514833 | 6.679848 | 6.570278 | 7.063258 | 8.036252 | -1.000758 | 355.818301 | 0.188879975 | -0.00578596 | 0.995383498 | 0.999862634 | 8.475358 | 8.474636 | 8.478019 | 8.471886 | 8.493905 | 8.454897 | 412 | 382 | 320 | 322 |
| 133022 | NM_152402 | TRAM1L1 | translocation associated membrane protein 1 like 1 | protein_coding | NP_689615.2 | 28371 | 617505 | . | . | 0.714099 | 1.269703 | 1.119732 | 0.921718 | 1.661739 | 2.934975 | 2.512647 | 2.107399 | 1.018427 | 63.888164 | 0.426080762 | 0.061826918 | 0.950700667 | 0.998865404 | 5.984751 | 6.010091 | 5.923104 | 6.047456 | 6.014458 | 5.967597 | 53 | 87 | 62 | 54 |
| 133283 | gene-EEF1A1P | EEF1A1P13 | eukaryotic translation elongation factor 1 alpha 1 pseudogene 13 | pseudogene | . | 3196 | . | . | . | 15.082689 | 13.939396 | 12.074406 | 11.852961 | 35.098038 | 32.221539 | 27.094616 | 27.100386 | -1.229396 | 721.397571 | 0.151549581 | -1.96601881 | **0.049296438** | 0.66444315 | 9.635798 | 9.338182 | 9.592227 | 9.529462 | 9.428624 | 9.409063 | 956 | 821 | 571 | 598 |
| 133308 | NM_00130075 | SLC9B2 | solute carrier family 9 member B2 | protein_coding | NP_001287683.1;NP_001287685.1;NP_001357128.1;NP_001357129.1;NI | 25143 | 611789 | . | . | 0.655079 | 0.835236 | 0.803046 | 0.647328 | 1.524397 | 1.930687 | 1.802014 | 1.480038 | 1.019332 | 137.453296 | 0.28302589 | 0.097604304 | 0.922246507 | 0.998865404 | 7.088699 | 7.116760 | 7.073765 | 7.120304 | 7.135228 | 7.072614 | 147 | 156 | 131 | 119 |
| 133383 | NM_00117199 | SETD9 | SET domain containing 9 | protein_coding | NP_001165461.1;NP_001309947.1;NP_001309948.1;NP_001309949.1;NI | 28508 | . | . | . | 6.143001 | 5.565538 | 5.702381 | 5.843905 | 14.295016 | 12.864990 | 12.795976 | 13.361391 | -1.000437 | 266.456290 | 0.211230829 | -0.00298206 | 0.997620662 | 0.999862634 | 8.057805 | 8.057705 | 8.074649 | 8.040273 | 8.071137 | 8.043578 | 317 | 278 | 238 | 243 |
| 133418 | NM_198449,XI | EMB | embigin | protein_coding | NP_940851.1;XP_011541448.1 | 30465 | 615669 | . | . | 8.629312 | 7.120453 | 5.554077 | 4.538226 | 20.080763 | 16.459246 | 12.463187 | 10.376115 | -1.581886 | 848.118117 | 0.166795525 | -3.9668084 | **7.28415E-05** | **0.010039957** | 10.020949 | 9.360317 | 9.937499 | 9.784473 | 9.605958 | 9.466453 | 1316 | 1009 | 632 | 551 |
| 133491 | NM_00114495 | C5orf47 | chromosome 5 open reading frame 47 | protein_coding | NP_001138426.1;XP_011532733.1;XP_016864517.1 | 27026 | . | . | . | 0.141751 | 0.102598 | 0.229292 | 0.281511 | 0.329860 | 0.237161 | 0.514526 | 0.643641 | **2.280201** | 14.673291 | 0.810874808 | 1.466515938 | 0.142507782 | NA | 3.161369 | 4.350483 | 3.831087 | 3.825880 | 3.870130 | 3.887296 | 11 | 9 | 16 | 21 |
| 133522 | NM_00117269 | PPARGC1B | PPARG coactivator 1 beta | protein_coding | NP_001166169.1;NP_001166170.1;NP_573570.3;XP_005268429.1;XP_01 | 30022 | 608886 | . | . | 1.327812 | 1.219363 | 1.402502 | 1.247296 | 3.089877 | 2.818614 | 3.147176 | 2.851793 | 1.167238 | 406.403389 | 0.185211811 | 1.204557898 | 0.228374018 | 0.982008277 | 8.550472 | 8.774386 | 8.631030 | 8.600816 | 8.754176 | 8.660637 | 443 | 394 | 410 | 379 |
| 133584 | NM_00120530 | EGFLAM | EGF like, fibronectin type III and laminin G domains | protein_coding | NP_001192230.1;NP_689616.2;NP_877950.1;NP_877953.1 | 26810 | 617683 | . | . | 0.060293 | 0.136476 | 0.105131 | 0.221263 | 0.140305 | 0.315469 | 0.235911 | 0.505892 | 1.650026 | 19.776893 | 0.74574836 | 0.968810945 | 0.332639525 | NA | 3.904062 | 4.619599 | 4.241663 | 4.290381 | 4.272452 | 4.341889 | 11 | 22 | 14 | 31 |
| 133619 | NM_00128680 | PRRC1 | proline rich coiled-coil 1 | protein_coding | NP_001273737.1;NP_570721.1;XP_006714592.1 | 28164 | . | . | . | 11.375594 | 10.892772 | 9.217063 | 9.564636 | 26.471474 | 25.179130 | 20.682820 | 21.868404 | -1.209506 | 1493.007422 | 0.125541848 | -2.18586722 | **0.028825321** | 0.547128317 | 10.674682 | 10.400303 | 10.633115 | 10.592003 | 10.459767 | 10.472575 | 1931 | 1718 | 1165 | 1278 |
| 133686 | NM_00108541 | NADK2 | NAD kinase 2, mitochondrial | protein_coding | NP_001078880.1;NP_001274269.1;NP_001274270.1;NP_694558.1;XP_00 | 26404 | 615787 | . | . | 2.354298 | 2.295524 | 1.795677 | 1.647510 | 5.478548 | 5.306208 | 4.029446 | 3.766836 | -1.353915 | 245.903628 | 0.218891114 | -1.99705241 | **0.04581949** | 0.653353744 | 8.143773 | 7.707228 | 8.015050 | 8.006879 | 7.882271 | 7.840615 | 328 | 303 | 190 | 187 |
| 133746 | NM_152405,XI | JMY | junction mediating and regulatory protein, p53 cofactor | protein_coding | NP_689618.4;XP_005248487.1;XP_011541457.1 | 28916 | 604279 | . | . | 1.052416 | 0.785633 | 0.756725 | 0.829940 | 2.449016 | 1.816025 | 1.698070 | 1.897559 | -1.176740 | 243.517170 | 0.235595495 | -0.99660229 | 0.318957593 | 0.998865404 | 8.040262 | 7.805997 | 8.038932 | 7.884463 | 7.866201 | 7.902269 | 348 | 242 | 187 | 218 |
| 133957 | NM_145265 | CCDC127 | coiled-coil domain containing 127 | protein_coding | NP_660308.1 | 30520 | . | . | . | 1.938771 | 1.800260 | 2.170580 | 1.794077 | 4.511600 | 4.161383 | 4.870719 | 4.101944 | 1.045220 | 557.394890 | 0.172998222 | 0.368828742 | 0.71225538 | 0.998865404 | 9.089817 | 9.154569 | 9.132170 | 9.079095 | 9.198136 | 9.067757 | 653 | 564 | 545 | 481 |
| 134145 | NM_00125838 | ATPSCKMT | ATP synthase c subunit lysine N-methyltransferase | protein_coding | NP_001245317.1;NP_001245318.1;NP_954584.2;XP_011512266.1 | 27029 | 618568 | . | . | 5.265275 | 4.125289 | 4.184372 | 3.070074 | 12.252512 | 9.535793 | 9.389609 | 7.019358 | -1.256696 | 332.494113 | 0.215769221 | -1.52772479 | 0.126580883 | 0.874064017 | 8.531837 | 8.203942 | 8.486450 | 8.378285 | 8.378948 | 8.228926 | 465 | 363 | 293 | 237 |
| 134147 | NM_138809 | CMBL | carboxymethylenebutenolidase homolog | protein_coding | NP_620164.1 | 25090 | 613379 | . | . | 4.223804 | 4.326403 | 5.031582 | 5.528181 | 9.828964 | 10.000673 | 11.290726 | 12.639529 | 1.217598 | 579.184399 | 0.160868902 | 1.765649975 | 0.07745459 | 0.767975096 | 9.029081 | 9.312754 | 9.102523 | 9.112498 | 9.208271 | 9.264898 | 597 | 568 | 530 | 621 |
| 134218 | NM_00101233 | DNAJC21 | DnaJ heat shock protein family (Hsp40) member C21 | protein_coding | NP_001012339.2;NP_001335349.1;NP_919259.3;XP_005248307.2;XP_01 | 27030 | 617048 | . | . | 4.813114 | 4.157289 | 4.650824 | 3.923634 | 11.200316 | 9.609760 | 10.436316 | 8.970922 | -1.056589 | 750.511282 | 0.159527743 | -0.49781007 | 0.618617915 | 0.998865404 | 9.590505 | 9.511884 | 9.622934 | 9.512631 | 9.575775 | 9.484537 | 957 | 767 | 677 | 639 |
| 134265 | NM_00114633 | AFAP1L1 | actin filament associated protein 1 like 1 | protein_coding | NP_001139809.1;NP_001309991.1;NP_001309992.1;NP_689619.1;XP_01 | 26714 | 614410 | . | . | 7.789847 | 8.498131 | 7.560759 | 9.118680 | 18.127294 | 19.643812 | 16.966128 | 20.848778 | 1.058327 | 1456.468995 | 0.136271972 | 0.600166915 | 0.548395 | 0.998865404 | 10.467006 | 10.548366 | 10.476572 | 10.493406 | 10.458989 | 10.593755 | 1612 | 1544 | 1189 | 1526 |
| 134266 | NM_152407 | GRPEL2 | GrpE like 2, mitochondrial | protein_coding | NP_689620.2 | 21060 | 618545 | . | . | 6.600922 | 6.512994 | 6.451318 | 5.038219 | 15.360617 | 15.055078 | 14.476570 | 11.519289 | -1.156731 | 771.476876 | 0.164429752 | -1.27746579 | 0.201437852 | 0.967111885 | 9.692289 | 9.483092 | 9.646055 | 9.630240 | 9.620132 | 9.445239 | 963 | 883 | 702 | 585 |
| 134353 | NM_173491 | LSM11 | LSM11, U7 small nuclear RNA associated | protein_coding | NP_775762.1 | 30860 | 617910 | . | . | 1.280743 | 1.328816 | 1.670554 | 1.349840 | 2.980341 | 3.071618 | 3.748675 | 3.086249 | 1.139294 | 288.435529 | 0.213391412 | 0.88166462 | 0.377958199 | 0.998865404 | 8.074497 | 8.263525 | 8.125459 | 8.140397 | 8.260454 | 8.140191 | 306 | 295 | 297 | 256 |
| 134359 | NM_00109927 | POC5 | POC5 centriolar protein | protein_coding | NP_001092741.1;NP_689621.2;XP_005248493.1;XP_011541460.1;XP_01 | 26658 | 617880 | . | . | 5.945965 | 5.425762 | 6.026772 | 4.683029 | 13.836505 | 12.541891 | 13.523901 | 10.707189 | -1.056951 | 369.126981 | 0.199048297 | -0.40145163 | 0.688087641 | 0.998865404 | 8.566752 | 8.488124 | 8.567822 | 8.515265 | 8.574675 | 8.440975 | 456 | 391 | 349 | 297 |
| 134429 | NM_00130805 | STARD4 | StAR related lipid transfer domain containing 4 | protein_coding | NP_001294985.1;NP_001294986.1;NP_001294987.1;NP_001294988.1;NI | 18058 | 607049 | . | . | 7.204502 | 4.476039 | 7.141389 | 5.449924 | 16.765175 | 10.346563 | 16.025068 | 12.460600 | 1.144432 | 1015.104397 | 0.188187151 | 1.034245338 | 0.301021484 | 0.998865404 | 9.886329 | 10.081875 | 9.989378 | 9.862609 | 10.173793 | 9.856608 | 1185 | 932 | 1141 | 802 |
| 134430 | NM_139281 | WDR36 | WD repeat domain 36 | protein_coding | NP_644810.1 | 30696 | 609669 | . | . | 5.870602 | 5.709049 | 6.499417 | 5.170691 | 13.661132 | 13.196722 | 14.584503 | 11.822171 | -1.005848 | 1194.438088 | 0.149393918 | -0.05630499 | 0.955098843 | 0.998865404 | 10.226010 | 10.218212 | 10.235873 | 10.208572 | 10.303488 | 10.124960 | 1404 | 1269 | 1159 | 984 |
| 134466 | NR_026867 | ZNF300P1 | zinc finger protein 300 pseudogene 1 | transcribed_ps | . | 27032 | . | . | . | 0.170300 | 0.061058 | 0.245435 | 0.151862 | 0.396295 | 0.141137 | 0.550748 | 0.347214 | 1.761900 | 12.235814 | 0.923952705 | 0.884387462 | 0.376487089 | NA | 3.142442 | 3.967302 | 3.598862 | 3.560287 | 3.625935 | 3.593013 | 15 | 5 | 17 | 11 |
| 134492 | NM_00132999 | NUDCD2 | NudC domain containing 2 | protein_coding | NP_001316920.1;NP_660309.1 | 30535 | . | . | . | 2.672288 | 2.175537 | 2.121705 | 2.504298 | 6.218522 | 5.028852 | 4.761043 | 5.725780 | -1.065742 | 607.064666 | 0.174463023 | -0.52652093 | 0.598526312 | 0.998865404 | 9.290906 | 9.199044 | 9.334574 | 9.190105 | 9.173958 | 9.268816 | 798 | 604 | 474 | 591 |
| 134510 | NM_145049 | UBLCP1 | ubiquitin like domain containing CTD phosphatase 1 | protein_coding | NP_659486.2 | 28110 | 609867 | . | . | 14.465589 | 13.447558 | 12.905278 | 9.974044 | 33.662018 | 31.084631 | 28.959074 | 22.804466 | -1.236478 | 863.529717 | 0.163544858 | -1.8724926 | 0.061138485 | 0.722512417 | 9.898693 | 9.593389 | 9.852414 | 9.791759 | 9.758746 | 9.575127 | 1144 | 988 | 761 | 628 |
| 134526 | NM_130767,XI | ACOT12 | acyl-CoA thioesterase 12 | protein_coding | NP_570123.1;XP_006714594.1;XP_006714595.1;XP_016864534.1;XP_01 | 24436 | 614315 | . | . | 0.651125 | 0.435143 | 0.193040 | 0.278824 | 1.515194 | 1.005852 | 0.433176 | 0.637498 | -1.800621 | 28.755224 | 0.599803089 | -1.41462153 | 0.157179492 | 0.926365724 | 5.207270 | 4.362218 | 4.912522 | 4.845915 | 4.823169 | 4.802584 | 52 | 31 | 20 | 17 |
| 134549 | NM_00117270 | SHROOM1 | shroom family member 1 | protein_coding | NP_001166171.1;NP_597713.2;XP_005271942.1;XP_005271943.1;XP_00 | 24084 | 611179 | . | . | 3.592684 | 3.533519 | 4.407521 | 4.387237 | 8.360324 | 8.167888 | 9.890351 | 10.030895 | 1.256285 | 483.026279 | 0.169842552 | 1.938054651 | 0.052616555 | 0.678753563 | 8.741861 | 9.071290 | 8.857698 | 8.816252 | 8.981901 | 8.978932 | 510 | 446 | 469 | 503 |
| 134553 | NM_00113558 | C5orf24 | chromosome 5 open reading frame 24 | protein_coding | NP_001129058.1;NP_001287823.1;NP_689622.2;XP_005271946.1;XP_01 | 26746 | . | . | . | 8.350742 | 7.167862 | 7.633080 | 7.831508 | 19.432522 | 16.568837 | 17.128412 | 17.905811 | -1.022345 | 1162.637102 | 0.141855529 | -0.22475265 | 0.822171703 | 0.998865404 | 10.198879 | 10.167319 | 10.253546 | 10.123377 | 10.174106 | 10.174015 | 1470 | 1159 | 1000 | 1078 |
| 134637 | NM_00128625 | ADAT2 | adenosine deaminase tRNA specific 2 | protein_coding | NP_001273188.1;NP_872309.2;XP_011533741.1;XP_011533745.1;XP_01 | 21172 | 615388 | . | . | 2.346947 | 2.389065 | 3.184177 | 1.821390 | 5.461440 | 5.522432 | 7.145198 | 4.164394 | -1.018284 | 643.748146 | 0.196309265 | -0.13315742 | 0.894068881 | 0.998865404 | 9.342808 | 9.317789 | 9.339798 | 9.320955 | 9.451755 | 9.164826 | 758 | 691 | 673 | 471 |
| 134728 | NM_00101084 | IRAK1BP1 | interleukin 1 receptor associated kinase 1 binding protein 1 | protein_coding | NP_001010844.1;XP_006715399.1;XP_011533749.1;XP_011533750.1;XP | 17368 | 615375 | . | . | 0.308297 | 0.463072 | 0.250558 | 0.362307 | 0.717422 | 1.070415 | 0.562245 | 0.828371 | -1.079657 | 54.941035 | 0.434363343 | -0.25456206 | 0.799061365 | 0.998865404 | 5.835399 | 5.721998 | 5.768494 | 5.805045 | 5.738885 | 5.803524 | 60 | 67 | 38 | 58 |
| 134957 | NM_00112771 | STXBP5 | syntaxin binding protein 5 | protein_coding | NP_001121187.1;NP_640337.3;XP_006715402.1;XP_011533755.1;XP_01 | 19665 | 604586 | . | . | 3.168214 | 3.094083 | 3.869306 | 3.224996 | 7.372563 | 7.152114 | 8.682611 | 7.373570 | 1.118433 | 952.584323 | 0.149695591 | 1.078711795 | 0.280716226 | 0.998865404 | 9.812360 | 9.974493 | 9.878794 | 9.823783 | 9.996125 | 9.866280 | 1075 | 933 | 952 | 860 |
| 134997 | gene-PPIAP29 | PPIAP29 | peptidylprolyl isomerase A pseudogene 29 | pseudogene | . | 43021 | . | . | . | 15.392525 | 14.492810 | 12.128347 | 17.356354 | 35.819038 | 33.500778 | 27.215660 | 39.683243 | -1.027970 | 375.337151 | 0.209956346 | -0.18955766 | 0.84965577 | 0.998865404 | 8.572363 | 8.531434 | 8.576733 | 8.537076 | 8.436448 | 8.635634 | 453 | 397 | 266 | 407 |
| 135112 | NM_00112284 | NCOA7 | nuclear receptor coactivator 7 | protein_coding | NP_001116314.1;NP_001186548.1;NP_001186549.1;NP_001186550.1;NI | 21081 | 609752 | . | . | 3.492300 | 3.621242 | 3.787071 | 3.989888 | 8.126727 | 8.370663 | 8.498077 | 9.122403 | 1.098172 | 689.453389 | 0.15322498 | 0.881732587 | 0.377921434 | 0.998865404 | 9.360403 | 9.495074 | 9.379431 | 9.411550 | 9.431052 | 9.487337 | 739 | 726 | 602 | 704 |
| 135114 | NM_138571 | HINT3 | histidine triad nucleotide binding protein 3 | protein_coding | NP_612638.3 | 18468 | 609998 | . | . | 2.951511 | 2.372494 | 2.776506 | 2.285071 | 6.868289 | 5.484126 | 6.230399 | 5.224543 | -1.067067 | 269.677061 | 0.226675209 | -0.41314902 | 0.679497436 | 0.998865404 | 8.120362 | 8.028349 | 8.147664 | 8.026778 | 8.105304 | 8.003777 | 357 | 266 | 250 | 220 |
| 135138 | NM_00108037 | PACRG | parkin coregulated | protein_coding | NP_001073847.1;NP_001073848.1;NP_689623.2;XP_005266882.1;XP_00 | 19152 | 608427 | . | . | 0.150737 | 0.193137 | 0.155013 | 0.095171 | 0.350771 | 0.446448 | 0.347843 | 0.217597 | 1.081909 | 11.656781 | 0.892408172 | 0.127272696 | 0.89872457 | NA | 3.486115 | 3.597915 | 3.540625 | 3.539888 | 3.536724 | 3.551867 | 13 | 12 | 9 | 13 |
| 135154 | NM_145267 | SDHAF4 | succinate dehydrogenase complex assembly factor 4 | protein_coding | NP_660310.2 | 20957 | . | . | . | 1.843610 | 1.775452 | 1.328232 | 2.082640 | 4.290156 | 4.104038 | 2.980514 | 4.761708 | -1.069773 | 65.380793 | 0.405999428 | -0.23966871 | 0.810587096 | 0.998865404 | 6.079743 | 5.980127 | 6.044256 | 6.031227 | 5.974594 | 6.068147 | 80 | 71 | 43 | 72 |
| 135228 | NM_00115958 | CD109 | CD109 molecule | protein_coding | NP_001153059.1;NP_001153060.1;NP_598000.2 | 21685 | 608859 | . | . | 52.862671 | 47.036173 | 43.991796 | 44.412377 | 123.013605 | 108.726225 | 98.716320 | 101.543619 | -1.146251 | 13256.957281 | 0.108132386 | -1.82112655 | 0.068587622 | 0.741130004 | 13.789551 | 13.592661 | 13.802587 | 13.693888 | 13.633082 | 13.632533 | 17327 | 14322 | 10746 | 11578 |
| 135293 | NM_00101085 | PM20D2 | peptidase M20 domain containing 2 | protein_coding | NP_001010853.1;XP_005248718.1;XP_005248719.1;XP_011533783.1;XP | 21408 | 615913 | . | . | 3.162800 | 2.764113 | 3.142050 | 2.309521 | 7.359965 | 6.389369 | 7.050669 | 5.280445 | -1.089473 | 415.317592 | 0.200014776 | -0.61810363 | 0.53650703 | 0.998865404 | 8.757857 | 8.635700 | 8.766905 | 8.671340 | 8.747401 | 8.583911 | 537 | 431 | 394 | 321 |
| 135295 | NM_00137689 | SRSF12 | serine and arginine rich splicing factor 12 | protein_coding | NP_001363825.1;NP_001363826.1;NP_001363827.1;NP_542781.3;XP_01 | 21220 | . | . | . | 1.612132 | 1.514618 | 1.842301 | 1.466631 | 3.751498 | 3.501107 | 4.134070 | 3.353280 | 1.032629 | 185.273892 | 0.251671689 | 0.184054945 | 0.853970351 | 0.998865404 | 7.509446 | 7.557191 | 7.543318 | 7.505104 | 7.582894 | 7.491933 | 219 | 188 | 181 | 158 |
| 135398 | NM_00114565 | C6orf141 | chromosome 6 open reading frame 141 | protein_coding | NP_001139124.2;XP_005248907.1;XP_005248909.1;XP_005248910.1;XP | 21351 | . | . | . | 0.008448 | 0.009093 | 0.042568 | 0.046858 | 0.019659 | 0.021018 | 0.095520 | 0.107135 | **2.469696** | 1.557117 | 2.386421563 | 0.546564555 | 0.584677926 | NA | -0.155866 | 1.148375 | 0.616459 | 0.616793 | 0.623168 | 0.622477 | 1 | 1 | 2 | 2 |
| 135458 | NM_148959 | HUS1B | HUS1 checkpoint clamp component B | protein_coding | NP_683762.2 | 16485 | 609713 | . | . | 0.282241 | 0.062130 | 0.331922 | 0.383276 | 0.656787 | 0.143616 | 0.744823 | 0.876315 | 1.961976 | 9.158211 | 1.055183342 | 0.921458199 | 0.356811255 | NA | 2.624463 | 3.602870 | 3.181619 | 3.150072 | 3.189840 | 3.195740 | 11 | 3 | 10 | 12 |
| 136051 | NM_152411 | ZNF786 | zinc finger protein 786 | protein_coding | NP_689624.2 | 21806 | . | . | . | 1.481941 | 1.661844 | 1.623354 | 1.714001 | 3.448537 | 3.841428 | 3.642760 | 3.918860 | 1.046835 | 162.200245 | 0.25952283 | 0.254442051 | 0.799154068 | 0.998865404 | 7.308587 | 7.373937 | 7.309791 | 7.352385 | 7.340637 | 7.358953 | 173 | 180 | 141 | 159 |
| 136259 | NM_138693 | KLF14 | Kruppel like factor 14 | protein_coding | NP_619638.2 | 23025 | 609393 | . | . | 0.394899 | 0.431055 | 0.444198 | 0.596761 | 0.918945 | 0.996403 | 0.996769 | 1.364423 | 1.239929 | 51.749216 | 0.440397313 | 0.704493841 | 0.481125284 | 0.998865404 | 5.531089 | 5.839400 | 5.659771 | 5.675002 | 5.680118 | 5.735487 | 51 | 52 | 43 | 61 |
| 136319 | NM_145808 | MTPN | myotrophin | protein_coding | NP_665807.1 | 15667 | 606484 | . | . | 32.694397 | 32.011253 | 28.018492 | 31.627035 | 76.081207 | 73.995445 | 62.872684 | 72.311462 | -1.100209 | 3663.959400 | 0.114686848 | -1.20131812 | 0.229627825 | 0.98402759 | 11.906492 | 11.768590 | 11.889396 | 11.864608 | 11.751852 | 11.842365 | 4486 | 4081 | 2866 | 3452 |
| 136332 | NM_00136570 | LRGUK | leucine rich repeats and guanylate kinase domain containing | protein_coding | NP_001352629.1;NP_001352630.1;NP_653249.1;XP_016867238.1;XP_02 | 21964 | 616478 | . | . | 1.729020 | 1.578750 | 3.041004 | 1.799161 | 4.023498 | 3.649351 | 6.823925 | 4.113567 | 1.350869 | 296.555422 | 0.238372631 | 1.820207469 | 0.068727416 | 0.741130004 | 7.978227 | 8.413397 | 8.100278 | 8.126322 | 8.392482 | 8.140819 | 283 | 279 | 362 | 249 |
| 136647 | NM_138701 | MPLKIP | M-phase specific PLK1 interacting protein | protein_coding | NP_619646.1 | 16002 | 609188 | . | . | 2.927646 | 2.310537 | 2.382865 | 2.519468 | 6.812753 | 5.340909 | 5.347080 | 5.760463 | -1.084097 | 603.145118 | 0.174059924 | -0.66927116 | 0.503322518 | 0.998865404 | 9.293233 | 9.177155 | 9.340946 | 9.174145 | 9.192568 | 9.221901 | 811 | 594 | 492 | 555 |
| 136853 | NM_080744,XI | SSC4D | scavenger receptor cysteine rich family member with 4 domains | protein_coding | NP_542782.1;XP_016867239.1;XP_024302432.1 | 14461 | 607639 | . | . | 0.317089 | 0.203640 | 0.183092 | 0.271922 | 0.737879 | 0.470723 | 0.410854 | 0.621717 | -1.150022 | 22.461221 | 0.667290908 | -0.30220846 | 0.762493159 | NA | 4.586822 | 4.384843 | 4.520939 | 4.474048 | 4.469671 | 4.499398 | 34 | 20 | 15 | 23 |
| 136895 | NM_00137135 | C7orf31 | chromosome 7 open reading frame 31 | protein_coding | NP_001358280.1;NP_001358281.1;NP_620166.3;XP_011513427.1 | 21722 | 616071 | . | . | 1.557344 | 1.329836 | 1.382829 | 1.625771 | 3.624003 | 3.073976 | 3.103027 | 3.717131 | 1.003810 | 161.355643 | 0.264606259 | 0.020731207 | 0.983460074 | 0.998865404 | 7.331555 | 7.336641 | 7.357943 | 7.306927 | 7.300935 | 7.366546 | 197 | 163 | 129 | 164 |
| 137075 | NM_194284 | CLDN23 | claudin 23 | protein_coding | NP_919260.2 | 17591 | 609203 | . | . | 0.768250 | 0.654289 | 0.677781 | 0.590650 | 1.787749 | 1.512417 | 1.520920 | 1.350450 | -1.141713 | 45.731092 | 0.464454695 | -0.41166463 | 0.680585251 | 0.998865404 | 5.606719 | 5.417273 | 5.543424 | 5.510973 | 5.516736 | 5.492250 | 61 | 48 | 40 | 37 |
| 137196 | NR_130917,NF | CCDC26 | CCDC26 long non-coding RNA | lncRNA | . | 28416 | 613040 | . | . | 0.058780 | 0.103960 | 0.117657 | 0.059920 | 0.136784 | 0.240309 | 0.264018 | 0.137000 | 1.012637 | 5.015207 | 1.366080212 | 0.013262587 | 0.989418297 | NA | 2.316288 | 2.336262 | 2.312346 | 2.326561 | 2.327972 | 2.311481 | 4 | 7 | 6 | 3 |
| 137209 | NM_152412 | ZNF572 | zinc finger protein 572 | protein_coding | NP_689625.2 | 26758 | . | . | . | 0.285587 | 0.270188 | 0.510849 | 0.332151 | 0.664572 | 0.624552 | 1.146330 | 0.759424 | 1.515399 | 36.090384 | 0.530262475 | 1.130944665 | 0.258078382 | 0.995760118 | 4.841203 | 5.443477 | 5.131866 | 5.125061 | 5.221363 | 5.151305 | 34 | 30 | 46 | 32 |

| | | | | | | | | | | | | | | | | | | | | | | | | | | | | | | | | | |
|---|---|---|---|---|---|---|---|---|---|---|---|---|---|---|---|---|---|---|---|---|---|---|---|---|---|---|---|---|---|---|---|---|---|
| 137392 | NM_00128303 | CIBAR1 | CBY1 interacting BAR domain containing 1 | protein_coding | NP_001269963.1;NP_660312.2 | 30452 | 617273 | . | . | 4.545783 | 3.980476 | 4.958521 | 3.802639 | 10.578223 | 9.201047 | 11.126773 | 8.694281 | -1.020910 | 517.027991 | 0.18565025 | -0.1608136 | 0.872240213 | 0.998865404 | 9.028331 | 8.999724 | 9.073186 | 8.959472 | 9.076104 | 8.928310 | 654 | 514 | 497 | 424 |
| 137492 | NM_00114515 | VPS37A | VPS37A subunit of ESCRT-I | protein_coding | NP_001138624.1;NP_001350096.1;NP_001350097.1;NP_001350098.1;NI | 24928 | 609927 | . | . | 5.564278 | 6.056470 | 5.676010 | 5.284422 | 12.948297 | 13.999803 | 12.736806 | 12.082202 | -1.101730 | 774.309958 | 0.15153349 | -0.92237141 | 0.356334878 | 0.998865404 | 9.664883 | 9.525276 | 9.596826 | 9.660514 | 9.596404 | 9.525159 | 893 | 915 | 674 | 655 |
| 137529 | gene-CKS1BP7 | CKS1BP7 | CDC28 protein kinase regulatory subunit 1B pseudogene 7 | pseudogene | . | 1999 | . | | . | 0.501321 | 0.259535 | 0.218627 | 0.324937 | 1.166595 | 0.599928 | 0.490593 | 0.742930 | -1.305229 | 8.232152 | 1.077187389 | -0.35676483 | 0.721267865 | NA | 3.220748 | 2.836248 | 3.065048 | 3.039004 | 3.036255 | 3.048339 | 14 | 7 | 5 | 8 |
| 137682 | NM_00133058 | NDUFAF6 | NADH:ubiquinone oxidoreductase complex assembly factor 6 | protein_coding | NP_001317511.1;NP_001341443.1;NP_001341444.1;NP_001341445.1;NI | 28625 | 612392 | . | . | 5.571482 | 5.581567 | 4.522451 | 5.280638 | 12.965065 | 12.902043 | 10.148253 | 12.073553 | -1.099349 | 354.883933 | 0.190408319 | -0.71766796 | 0.472962041 | 0.998865404 | 8.538175 | 8.400971 | 8.493719 | 8.501983 | 8.415351 | 8.471342 | 425 | 404 | 279 | 333 |
| 137695 | NM_00128665 | TMEM68 | transmembrane protein 68 | protein_coding | NP_001273586.1;NP_001273589.1;NP_001273590.1;NP_001350105.1;NI | 26510 | . | | . | 9.814958 | 10.118138 | 10.427757 | 8.192185 | 22.839809 | 23.388527 | 23.399581 | 18.730457 | -1.086790 | 774.760069 | 0.162889774 | -0.73714043 | 0.461036955 | 0.998865404 | 9.656045 | 9.536699 | 9.617240 | 9.630993 | 9.647042 | 9.477904 | 920 | 879 | 725 | 611 |
| 137872 | NM_144650 | ADHFE1 | alcohol dehydrogenase iron containing 1 | protein_coding | NP_653251.2 | 16354 | 611083 | . | . | 0.492161 | 0.577235 | 0.834221 | 0.483044 | 1.145279 | 1.334304 | 1.871969 | 1.104424 | 1.208894 | 37.255620 | 0.529811053 | 0.516576892 | 0.60545155 | 0.998865404 | 5.074826 | 5.350772 | 5.185083 | 5.205193 | 5.263253 | 5.180603 | 36 | 39 | 45 | 28 |
| 137886 | NM_00107761 | UBXN2B | UBX domain protein 2B | protein_coding | NP_001071087.1;NP_001317464.1;NP_001350110.1 | 27035 | 610686 | . | . | 3.534704 | 3.206026 | 3.691879 | 2.811031 | 8.225401 | 7.410873 | 8.284471 | 6.427088 | -1.031357 | 496.866145 | 0.187008552 | -0.23818821 | 0.811735117 | 0.998865404 | 8.978201 | 8.934901 | 8.999853 | 8.926213 | 9.024037 | 8.857927 | 614 | 513 | 482 | 398 |
| 137964 | NM_00136319 | GPAT4 | glycerol-3-phosphate acyltransferase 4 | protein_coding | NP_001350126.1;NP_001350127.1;NP_848934.1 | 20880 | 608143 | . | . | 36.765494 | 36.298594 | 43.210587 | 41.730122 | 85.554810 | 83.905826 | 96.963305 | 95.410967 | 1.149696 | 7781.938923 | 0.103259721 | 1.94898758 | 0.051296908 | 0.672475152 | 12.821758 | 13.023054 | 12.871898 | 12.855604 | 13.001603 | 12.961969 | 8418 | 7736 | 7406 | 7623 |
| 137994 | NM_00119965 | LETM2 | leucine zipper and EF-hand containing transmembrane protein 2 | protein_coding | NP_001186588.1;NP_001186589.1;NP_001273716.1;NP_001273748.1;NI | 14648 | . | | . | 0.949577 | 0.987018 | 0.930495 | 1.149574 | 2.209707 | 2.281538 | 2.088002 | 2.628367 | -1.314099 | 99.743710 | 0.326609674 | -1.2065586 | 0.227602179 | 0.982008277 | 6.824386 | 6.428904 | 6.647120 | 6.711501 | 6.583306 | 6.610824 | 118 | 134 | 71 | 85 |
| 138050 | NM_00136322 | HGSNAT | heparan-alpha-glucosaminide N-acetyltransferase | protein_coding | NP_001350156.1;NP_001350157.1;NP_001350158.1;NP_689632.2;XP_0C | 26527 | 610453 | . | . | 4.077033 | 4.020469 | 4.044529 | 3.401858 | 9.487426 | 9.293494 | 9.075807 | 7.777944 | -1.105084 | 629.482669 | 0.165663518 | -0.87017051 | 0.384207229 | 0.998865404 | 9.367937 | 9.224547 | 9.332304 | 9.326465 | 9.325836 | 9.195019 | 764 | 710 | 572 | 505 |
| 138151 | NM_144653,XI | NACC2 | NACC family member 2 | protein_coding | NP_653254.1;XP_011516523.1 | 23846 | 615786 | . | . | 9.406976 | 9.181165 | 6.638629 | 6.946957 | 21.890419 | 21.222673 | 14.896891 | 15.883392 | -1.388321 | 1733.586793 | 0.121829166 | -3.8852846 | 0.00010221 | 0.01259648 | 10.976908 | 10.503520 | 10.887327 | 10.860320 | 10.604381 | 10.629435 | 2359 | 2139 | 1241 | 1384 |
| 138162 | NM_00104826 | C9orf116 | chromosome 9 open reading frame 116 | protein_coding | NP_001041730.1;NP_653255.1 | 28435 | 614502 | . | . | 2.116326 | 1.870904 | 1.929728 | 2.077508 | 4.924778 | 4.324677 | 4.330252 | 4.749974 | -1.025387 | 41.981199 | 0.4800657 | -0.07534052 | 0.939943782 | 0.998865404 | 5.409544 | 5.373575 | 5.404654 | 5.383876 | 5.385771 | 5.394789 | 52 | 43 | 35 | 40 |
| 138199 | NM_00132049 | CARNMT1 | carnosine N-methyltransferase 1 | protein_coding | NP_001307426.1;NP_689633.1;XP_005251758.2;XP_011516525.1;XP_01 | 23435 | 616552 | . | . | 3.023848 | 2.362644 | 2.941544 | 1.888657 | 7.036616 | 5.461358 | 6.600739 | 4.318189 | -1.133581 | 341.476430 | 0.238270497 | -0.75917043 | 0.447750614 | 0.998865404 | 8.502297 | 8.323448 | 8.510863 | 8.368914 | 8.486865 | 8.241804 | 468 | 344 | 341 | 233 |
| 138241 | NM_00136505 | C9orf85 | chromosome 9 open reading frame 85 | protein_coding | NP_001351982.1;NP_001351984.1;NP_001351986.1;NP_872311.2 | 28784 | . | | . | 4.653893 | 4.355226 | 4.003243 | 3.020260 | 10.829801 | 10.067303 | 8.983160 | 6.905464 | -1.186438 | 195.720053 | 0.251560923 | -0.98042454 | 0.3268766 | 0.998865404 | 7.729909 | 7.485006 | 7.662263 | 7.632726 | 7.629754 | 7.509726 | 252 | 222 | 178 | 144 |
| 138311 | NM_152421,XI | DIPK1B | divergent protein kinase domain 1B | protein_coding | NP_689634.2;XP_006717017.1;XP_011516540.1;XP_016869762.1 | 28290 | 614543 | . | . | 10.000150 | 11.222205 | 10.380830 | 11.904162 | 23.270761 | 25.940631 | 23.294282 | 27.217450 | 1.048781 | 803.166530 | 0.150550141 | 0.456411508 | 0.648094092 | 0.998865404 | 9.615093 | 9.683214 | 9.604075 | 9.658739 | 9.621843 | 9.706805 | 868 | 879 | 672 | 817 |
| 138428 | NM_00100291 | PTRH1 | peptidyl-tRNA hydrolase 1 homolog | protein_coding | NP_001002913.1;NP_001332906.1;NP_001332907.1;NP_001332908.1;NI | 27039 | . | | . | 6.327540 | 6.302381 | 5.126467 | 6.904208 | 14.724445 | 14.568234 | 11.503641 | 15.785653 | -1.055127 | 190.693565 | 0.253629605 | -0.30523478 | 0.760187349 | 0.998865404 | 7.614014 | 7.535132 | 7.586681 | 7.584901 | 7.498358 | 7.621936 | 226 | 211 | 138 | 199 |
| 138429 | NM_00113521 | PIP5KL1 | phosphatidylinositol-4-phosphate 5-kinase like 1 | protein_coding | NP_001128691.1;NP_775763.1;XP_016869768.1 | 28711 | 612865 | . | . | 0.813434 | 0.907723 | 0.795001 | 0.572660 | 1.892894 | 2.098241 | 1.783960 | 1.309318 | -1.220666 | 44.935660 | 0.465991921 | -0.61732542 | 0.537020103 | 0.998865404 | 5.626345 | 5.338943 | 5.498446 | 5.517213 | 5.482101 | 5.466243 | 54 | 56 | 37 | 36 |
| 138639 | NM_00125382 | PTPDC1 | protein tyrosine phosphatase domain containing 1 | protein_coding | NP_001240758.1;NP_001240759.1;NP_689635.3;NP_818931.1;XP_0052 | 30184 | . | | . | 3.136240 | 3.191236 | 3.276186 | 3.317703 | 7.298159 | 7.376686 | 7.351666 | 7.585533 | 1.046827 | 451.361904 | 0.173064905 | 0.381496678 | 0.702834736 | 0.998865404 | 8.784838 | 8.850692 | 8.807298 | 8.799796 | 8.817041 | 8.846161 | 511 | 473 | 392 | 443 |
| 138649 | NR_026868 | ANKRD19P | ankyrin repeat domain 19, pseudogene | transcribed_ps | . | 22567 | . | | . | 3.553436 | 3.721610 | 4.282186 | 2.852641 | 8.268991 | 8.602668 | 9.609101 | 6.522224 | -1.033815 | 259.602891 | 0.242005467 | -0.1982494 | 0.842849939 | 0.998865404 | 8.043169 | 7.996784 | 8.013422 | 8.032920 | 8.104494 | 7.897859 | 298 | 290 | 268 | 190 |
| 138715 | NM_00101736 | ARID3C | AT-rich interaction domain 3C | protein_coding | NP_001017363.1;NP_001358874.1;XP_016869772.1 | 21209 | . | | . | 0.161066 | 0.099823 | 0.147037 | 0.151051 | 0.374808 | 0.230745 | 0.329946 | 0.345360 | 1.150487 | 19.628386 | 0.702893975 | 0.287731844 | 0.773552012 | NA | 4.189060 | 4.393446 | 4.304723 | 4.266728 | 4.296212 | 4.298940 | 26 | 15 | 18 | 20 |
| 138716 | NM_148178,N | RPP25L | ribonuclease P/MRP subunit p25 like | protein_coding | NP_680544.1;NP_680545.1 | 19909 | . | | . | 14.321759 | 12.736379 | 15.663589 | 16.564452 | 33.327321 | 29.440713 | 35.148640 | 37.872647 | 1.157965 | 435.828526 | 0.180377847 | 1.173044696 | 0.240777882 | 0.992733415 | 8.657778 | 8.869681 | 8.761667 | 8.674725 | 8.794947 | 8.822090 | 499 | 404 | 398 | 448 |
| 139221 | NM_00117102 | PWWP3B | PWWP domain containing 3B | protein_coding | NP_001164491.1;NP_689636.3;XP_005262134.1;XP_011529158.1;XP_01 | 26583 | . | | . | 0.902479 | 0.793795 | 0.605291 | 0.607408 | 2.100105 | 1.834895 | 1.358257 | 1.388765 | -1.372414 | 97.080924 | 0.326783729 | -1.39760857 | 0.162230645 | 0.935295382 | 6.811378 | 6.354895 | 6.666971 | 6.626577 | 6.549150 | 6.556985 | 137 | 114 | 70 | 78 |
| 139231 | NM_207318,XI | FAM199X | family with sequence similarity 199, X-linked | protein_coding | NP_997201.1;XP_005262136.1 | 25195 | . | | . | 3.166290 | 2.706941 | 3.000410 | 2.185266 | 7.368089 | 6.257215 | 6.732833 | 4.996352 | -1.144542 | 654.561166 | 0.185243085 | -1.05143474 | 0.293058966 | 0.998865404 | 9.447903 | 9.254380 | 9.450687 | 9.334627 | 9.403284 | 9.191911 | 871 | 691 | 616 | 481 |
| 139285 | NM_152424 | AMER1 | APC membrane recruitment protein 1 | protein_coding | NP_689637.3 | 26837 | 300647 | . | . | 0.311239 | 0.453162 | 0.378294 | 0.421142 | 0.724266 | 1.047503 | 0.848880 | 0.962892 | 1.034664 | 102.704144 | 0.333334626 | 0.147486785 | 0.88274781 | 0.998865404 | 6.658583 | 6.705733 | 6.613717 | 6.728705 | 6.667033 | 6.699534 | 95 | 129 | 86 | 103 |
| 139322 | NM_198450,XI | APOOL | apolipoprotein O like | protein_coding | NP_940852.3;XP_011529161.1;XP_016884760.1;XP_016884761.1;XP_01 | 24009 | 300955 | . | . | 1.078266 | 1.172717 | 1.150543 | 1.269639 | 2.509168 | 2.710789 | 2.581785 | 2.902879 | -1.040227 | 214.707237 | 0.23625318 | -0.24083581 | 0.809682375 | 0.998865404 | 7.773911 | 7.718002 | 7.748364 | 7.759754 | 7.781825 | 7.687507 | 249 | 239 | 202 | 177 |
| 139324 | NM_00117747 | HDX | highly divergent homeobox | protein_coding | NP_001170949.1;NP_001170950.1;NP_653258.2;XP_006724682.1;XP_01 | 26411 | 300994 | . | . | 0.297793 | 0.347586 | 0.410900 | 0.321984 | 0.692977 | 0.803460 | 0.922047 | 0.736179 | 1.084655 | 68.382534 | 0.388144476 | 0.302044331 | 0.762618269 | 0.998865404 | 6.035242 | 6.153450 | 6.067978 | 6.095869 | 6.131540 | 6.069624 | 71 | 75 | 70 | 58 |
| 139341 | NM_173794 | FUNDC1 | FUN14 domain containing 1 | protein_coding | NP_776155.1 | 28746 | 300871 | . | . | 13.834669 | 12.522563 | 13.597821 | 11.694516 | 32.193844 | 28.946465 | 30.513113 | 26.738123 | -1.056300 | 424.432692 | 0.185234725 | -0.42659288 | 0.669675885 | 0.998865404 | 8.767865 | 8.689864 | 8.777406 | 8.710797 | 8.758386 | 8.662767 | 529 | 445 | 388 | 356 |
| 139596 | NM_00130794 | UPRT | uracil phosphoribosyltransferase homolog | protein_coding | NP_001294873.1;NP_001350750.1;NP_659489.1;XP_011529169.1 | 28334 | 300656 | . | . | 1.863901 | 2.085702 | 1.751988 | 2.197571 | 4.337373 | 4.821193 | 3.931411 | 5.024484 | 1.003462 | 154.683917 | 0.271644021 | 0.018354192 | 0.985356296 | 0.999197585 | 7.271488 | 7.274869 | 7.248699 | 7.293428 | 7.229260 | 7.314150 | 168 | 176 | 120 | 161 |
| 139716 | NM_00108157 | GAB3 | GRB2 associated binding protein 3 | protein_coding | NP_001075042.1;NP_001269212.1;NP_542179.1;XP_005274705.1;XP_00 | 17515 | 300482 | . | . | 0.293354 | 0.304281 | 0.237075 | 0.220638 | 0.682649 | 0.703359 | 0.531988 | 0.504463 | -1.178567 | 37.433125 | 0.511530145 | -0.46338238 | 0.643090309 | 0.998865404 | 5.340146 | 5.102572 | 5.219066 | 5.257616 | 5.219690 | 5.209513 | 41 | 49 | 31 | 31 |
| 139728 | NM_00103958 | PNCK | pregnancy up-regulated nonubiquitous CaM kinase | protein_coding | NP_001034671.3;NP_001129212.1;NP_001353904.1;NP_001353905.1;NI | 13415 | 300680 | . | . | 0.602827 | 0.766478 | 1.429417 | 1.101463 | 1.402805 | 1.771748 | 3.207571 | 2.518364 | 1.759312 | 57.008794 | 0.419538136 | 1.942640329 | 0.052059631 | 0.675217033 | 5.368714 | 6.183866 | 5.745436 | 5.763814 | 5.886329 | 5.850283 | 45 | 47 | 68 | 63 |
| 139818 | NM_144658,XI | DOCK11 | dedicator of cytokinesis 11 | protein_coding | NP_653259.3;XP_005262425.1;XP_005262426.1;XP_011529578.1;XP_01 | 23483 | 300681 | . | . | 0.833768 | 0.658754 | 0.875825 | 0.763548 | 1.940212 | 1.522739 | 1.965325 | 1.745762 | 1.080898 | 169.221843 | 0.26547772 | 0.422748933 | 0.67247845 | 0.998865404 | 7.344702 | 7.458595 | 7.432180 | 7.331749 | 7.445318 | 7.385007 | 210 | 154 | 164 | 153 |
| 139886 | NM_00101296 | SPIN4 | spindlin family member 4 | protein_coding | NP_001012986.2 | 27040 | . | | . | 1.725149 | 1.349778 | 1.221300 | 0.929870 | 4.014493 | 3.120072 | 2.740561 | 2.126037 | -1.445198 | 167.821854 | 0.27599936 | -1.92488714 | 0.054243478 | 0.691341398 | 7.631144 | 7.102220 | 7.513440 | 7.396518 | 7.354812 | 7.254113 | 257 | 187 | 136 | 111 |
| 140459 | NM_00120240 | ASB6 | ankyrin repeat and SOCS box containing 6 | protein_coding | NP_001189332.1;NP_060343.1;NP_821066.1 | 17181 | 615051 | . | . | 7.748431 | 7.959702 | 7.640927 | 7.657993 | 18.030918 | 18.399209 | 17.146017 | 17.509093 | -1.040097 | 1112.732462 | 0.132590374 | -0.4277712 | 0.668817709 | 0.998865404 | 10.147981 | 10.091243 | 10.127222 | 10.141554 | 10.107815 | 10.102468 | 1294 | 1236 | 952 | 1019 |
| 140460 | NM_024708,N | ASB7 | ankyrin repeat and SOCS box containing 7 | protein_coding | NP_078984.2;NP_937886.1 | 17182 | 615052 | . | . | 4.608783 | 5.004618 | 5.139334 | 4.917491 | 10.724829 | 11.568399 | 11.532516 | 11.243262 | 1.013482 | 716.667224 | 0.150824372 | 0.128098056 | 0.898071375 | 0.998865404 | 9.475364 | 9.494889 | 9.470249 | 9.489955 | 9.514956 | 9.463282 | 808 | 779 | 651 | 651 |
| 140461 | NM_00131929 | ASB8 | ankyrin repeat and SOCS box containing 8 | protein_coding | NP_001306225.1;NP_001306226.1;NP_001306227.1;NP_001306228.1;NI | 17183 | 615053 | . | . | 3.753376 | 3.684774 | 4.304449 | 4.581802 | 8.734260 | 8.517521 | 9.659060 | 10.475746 | 1.174732 | 326.468051 | 0.194728277 | 1.193105194 | 0.232828178 | 0.98883918 | 8.230052 | 8.462214 | 8.308950 | 8.295948 | 8.375223 | 8.407735 | 350 | 320 | 298 | 340 |
| 140462 | NM_00103173 | ASB9 | ankyrin repeat and SOCS box containing 9 | protein_coding | NP_001026909.1;NP_001162002.1;NP_001162003.1;NP_076992.1;XP_0C | 17184 | 300890 | . | . | 8.530019 | 7.660953 | 8.044478 | 7.345181 | 19.849707 | 17.708639 | 18.051576 | 16.793883 | -1.045723 | 432.173907 | 0.178479273 | -0.36139312 | 0.717805594 | 0.998865404 | 8.787095 | 8.723133 | 8.800221 | 8.736126 | 8.755406 | 8.727629 | 535 | 452 | 376 | 387 |
| 140465 | NM_00119962 | MYL6B | myosin light chain 6B | protein_coding | NP_001186558.1;NP_002466.1 | 29823 | 609930 | . | . | 35.449384 | 41.744998 | 35.531770 | 50.129815 | 82.492178 | 96.495430 | 79.732263 | 114.615863 | 1.097388 | 1060.741232 | 0.168996969 | 0.793350544 | 0.427573616 | 0.998865404 | 9.982710 | 10.115930 | 9.950926 | 10.068414 | 9.956801 | 10.189958 | 1072 | 1179 | 815 | 1202 |
| 140467 | NM_018083,XI | ZNF358 | zinc finger protein 358 | protein_coding | NP_060553.4;XP_005272517.1;XP_011525997.1 | 16838 | . | | . | 4.456735 | 4.453388 | 5.163720 | 6.297919 | 10.371003 | 10.294207 | 11.587238 | 14.399444 | 1.231579 | 307.477418 | 0.211381063 | 1.421647175 | 0.155128702 | 0.92370092 | 8.106823 | 8.406333 | 8.202916 | 8.198127 | 8.239965 | 8.381870 | 319 | 296 | 258 | 358 |
| 140469 | NM_00108361 | MYO3B | myosin IIIB | protein_coding | NP_001077084.2;NP_620482.3;XP_006712362.1;XP_011508956.1;XP_01 | 15576 | 610040 | . | . | 0.139556 | 0.159308 | 0.092023 | 0.167400 | 0.324753 | 0.368245 | 0.206497 | 0.382741 | -1.131042 | 27.459748 | 0.607845726 | -0.29226676 | 0.770082675 | 0.998865404 | 4.867222 | 4.685555 | 4.781301 | 4.792805 | 4.743550 | 4.802208 | 32 | 33 | 16 | 31 |
| 140545 | NM_00118499 | RNF32 | ring finger protein 32 | protein_coding | NP_001171925.1;NP_001171926.1;NP_001295202.1;NP_001295203.1;NI | 17118 | 610241 | . | . | 0.924319 | 0.636272 | 0.729708 | 0.716907 | 2.150925 | 1.470773 | 1.637445 | 1.639124 | -1.094493 | 51.590575 | 0.450576241 | -0.28910258 | 0.772502877 | 0.998865404 | 5.751724 | 5.623500 | 5.737711 | 5.653743 | 5.684238 | 5.674682 | 75 | 46 | 44 | 45 |
| 140564 | NM_00136378 | APOBEC3D | apolipoprotein B mRNA editing enzyme catalytic subunit 3D | protein_coding | NP_001350710.1;NP_689639.2;XP_016884085.1 | 17354 | 609900 | . | . | 0.935065 | 0.968270 | 0.666508 | 0.977662 | 2.175934 | 2.238201 | 1.495625 | 2.235308 | -1.199560 | 73.153044 | 0.385088952 | -0.68167355 | 0.495445393 | 0.998865404 | 6.319341 | 6.054185 | 6.209369 | 6.220505 | 6.123727 | 6.214720 | 90 | 88 | 46 | 75 |
| 140576 | NM_00131700 | S100A16 | S100 calcium binding protein A16 | protein_coding | NP_001303936.1;NP_001303937.1;NP_525127.1 | 20441 | 617437 | . | . | 2.539766 | 2.312930 | 2.128716 | 2.509732 | 5.910141 | 5.346442 | 4.776778 | 5.738205 | -1.015198 | 97.254293 | 0.328652284 | -0.06621448 | 0.947207069 | 0.998865404 | 6.614850 | 6.592443 | 6.622438 | 6.589040 | 6.573453 | 6.628496 | 119 | 100 | 76 | 99 |
| 140609 | NM_133494,XI | NEK7 | NIMA related kinase 7 | protein_coding | NP_598001.1;XP_011507511.1;XP_016855833.1;XP_016855834.1;XP_01 | 13386 | 606848 | . | . | 6.611925 | 5.350840 | 6.573755 | 5.273276 | 15.386222 | 12.368707 | 14.751315 | 12.056719 | -1.024344 | 757.394696 | 0.172751195 | -0.20087046 | 0.840799868 | 0.998865404 | 9.581637 | 9.547969 | 9.642406 | 9.492377 | 9.628443 | 9.472727 | 976 | 739 | 724 | 623 |
| 140612 | NM_00130844 | ZFP28 | ZFP28 zinc finger protein | protein_coding | NP_001295369.1;NP_065879.1;XP_011524764.1;XP_011524765.2 | 17801 | 616798 | . | . | 0.019500 | 0.072223 | 0.046281 | 0.050748 | 0.045378 | 0.166948 | 0.103853 | 0.116029 | 1.350197 | 10.122854 | 0.979371846 | 0.442293424 | 0.658276889 | NA | 3.109221 | 3.538109 | 3.306372 | 3.338119 | 3.335274 | 3.336426 | 6 | 13 | 10 | 11 |
| 140623 | gene-FAUP1 | FAUP1 | FAU pseudogene 1 | pseudogene | . | 17984 | . | | . | 0.052307 | 0.056299 | 0.094499 | 0.247427 | 0.121722 | 0.130137 | 0.212054 | 0.565713 | 1.833800 | 2.539248 | 1.876543826 | 0.466195476 | 0.641075587 | NA | 0.844134 | 1.715150 | 1.330022 | 1.330639 | 1.332827 | 1.341772 | 2 | 2 | 2 | 4 |
| 140680 | NM_080571,N | C20orf96 | chromosome 20 open reading frame 96 | protein_coding | NP_542138.1;NP_695001.2;XP_006723606.1 | 16227 | . | | . | 5.073126 | 4.597031 | 5.740224 | 5.821120 | 11.805371 | 10.626244 | 12.880895 | 13.309299 | 1.184705 | 260.176456 | 0.213437907 | 1.145660611 | 0.251935608 | 0.994172589 | 7.895713 | 8.140601 | 7.998660 | 7.953109 | 8.061155 | 8.063253 | 287 | 245 | 245 | 265 |
| 140685 | NM_00136974 | ZBTB46 | zinc finger and BTB domain containing 46 | protein_coding | NP_001356670.1;NP_079500.2;XP_005260252.1;XP_005260254.1;XP_00 | 16094 | 614639 | . | . | 0.617581 | 0.442052 | 0.332274 | 0.542598 | 1.437136 | 1.021823 | 0.745613 | 1.240585 | -1.174397 | 68.901917 | 0.385654059 | -0.60137001 | 0.547593567 | 0.998865404 | 6.218609 | 5.984879 | 6.121231 | 6.131426 | 6.060620 | 6.115748 | 84 | 82 | 48 | 67 |
| 140686 | NM_080614,XI | WFDC3 | WAP four-disulfide core domain 3 | protein_coding | NP_542181.1;XP_011526855.1;XP_016883157.1 | 15957 | . | | . | 0.128899 | 0.035774 | 0.089189 | 0.189917 | 0.299953 | 0.082694 | 0.200138 | 0.434223 | 1.588295 | 4.011406 | 1.523437328 | 0.438140037 | 0.661284773 | NA | 1.631267 | 2.300113 | 1.998914 | 1.987132 | 1.994836 | 2.008092 | 5 | 2 | 3 | 6 |
| 140688 | NM_00125679 | NOL4L | nucleolar protein 4 like | protein_coding | NP_001243727.1;NP_001338609.1;NP_542183.2;XP_005260340.1;XP_00 | 16106 | . | | . | 0.905713 | 1.007009 | 0.652316 | 0.862852 | 2.107631 | 2.327746 | 1.463779 | 1.972808 | -1.179774 | 170.797160 | 0.258255893 | -0.92354143 | 0.355725121 | 0.998865404 | 7.531089 | 7.291233 | 7.430917 | 7.468002 | 7.345422 | 7.414870 | 204 | 208 | 124 | 160 |
| 140699 | NM_152503,N | MROH8 | maestro heat like repeat family member 8 | protein_coding | NP_689716.4;NP_998796.1;NP_998797.2 | 16125 | . | | . | 0.204839 | 0.154414 | 0.252105 | 0.509616 | 0.476669 | 0.356935 | 0.565717 | 1.165176 | 1.509169 | 25.213427 | 0.618734128 | 0.959628267 | 0.337242337 | NA | 4.328680 | 4.922818 | 4.635420 | 4.614121 | 4.656224 | 4.678284 | 26 | 19 | 24 | 31 |
| 140700 | NM_080621,XI | SAMD10 | sterile alpha motif domain containing 10 | protein_coding | NP_542188.1;XP_005260256.1;XP_006723768.1;XP_011526867.1;XP_01 | 16129 | . | | . | 0.122917 | 0.077780 | 0.175141 | 0.180884 | 0.286032 | 0.179792 | 0.393012 | 0.413571 | 1.831582 | 14.524622 | 0.811148743 | 1.076362482 | 0.281765175 | NA | 3.357474 | 4.232645 | 3.835449 | 3.818237 | 3.862401 | 3.866130 | 14 | 9 | 16 | 18 |
| 140707 | NM_080626,XI | BRI3BP | BRI3 binding protein | protein_coding | NP_542193.3;XP_011536242.1 | 14251 | 615627 | . | . | 7.058292 | 6.943432 | 6.162206 | 5.725414 | 16.424936 | 16.050053 | 13.827810 | 13.090480 | -1.192684 | 1343.641408 | 0.131056345 | -1.93971097 | 0.052414826 | 0.677417458 | 10.513289 | 10.259423 | 10.469401 | 10.440421 | 10.355850 | 10.284403 | 1714 | 1548 | 1117 | 1094 |
| 140710 | NM_080627,N | SOGA1 | suppressor of glucose, autophagy associated 1 | protein_coding | NP_542194.2;NP_954650.2 | 16111 | . | | . | 7.377311 | 7.873693 | 7.461883 | 8.058858 | 17.167307 | 18.200392 | 16.744247 | 18.425620 | 1.002370 | 3391.391243 | 0.112441688 | 0.03036692 | 0.975774426 | 0.998865404 | 11.726024 | 11.729297 | 11.702635 | 11.749855 | 11.704903 | 11.751386 | 3789 | 3760 | 2869 | 3273 |
| 140733 | NM_00103308 | MACROD2 | mono-ADP ribosylhydrolase 2 | protein_coding | NP_001028259.1;NP_001338590.1;NP_001338592.1;NP_001338593.1;NI | 16126 | 611567 | . | . | 0.032257 | 0.122737 | 0.088790 | 0.101282 | 0.075062 | 0.283712 | 0.199243 | 0.231568 | 1.212803 | 13.617998 | 0.8938911 | 0.311386638 | 0.755506711 | NA | 3.624820 | 3.897226 | 3.716001 | 3.780737 | 3.755680 | 3.765153 | 6 | 21 | 12 | 15 |
| 140735 | NM_080677 | DYNLL2 | dynein light chain LC8-type 2 | protein_coding | NP_542408.1 | 24596 | 608942 | . | . | 8.499810 | 8.541859 | 8.376468 | 8.449662 | 19.779409 | 19.744888 | 18.796549 | 19.319149 | -1.026912 | 1795.801839 | 0.119821457 | -0.31975115 | 0.749156985 | 0.998865404 | 10.829437 | 10.791134 | 10.822223 | 10.819406 | 10.800501 | 10.799334 | 2099 | 1960 | 1542 | 1660 |
| 140739 | NM_00127830 | UBE2F | ubiquitin conjugating enzyme E2 F (putative) | protein_coding | NP_001265234.1;NP_001265235.1;NP_001265236.1;NP_001265237.1;NI | 12480 | 617700 | . | . | 16.424715 | 17.314178 | 15.644629 | 17.676046 | 38.220989 | 40.022496 | 35.106096 | 40.414178 | -1.019046 | 1125.198048 | 0.137145866 | -0.19847047 | 0.842676986 | 0.998865404 | 10.149714 | 10.122080 | 10.125878 | 10.159213 | 10.087122 | 10.168253 | 1279 | 1253 | 916 | 1102 |
| 140775 | NM_144775 | SMCR8 | SMCR8-C9orf72 complex subunit | protein_coding | NP_658988.2 | 17921 | 617074 | . | . | 8.111012 | 9.258043 | 9.104870 | 9.436469 | 18.874660 | 21.400383 | 20.431065 | 21.575365 | 1.053245 | 2320.035717 | 0.121988851 | 0.613502047 | 0.53954443 | 0.998865404 | 11.142132 | 11.216766 | 11.107503 | 11.206400 | 11.189323 | 11.210135 | 2442 | 2589 | 2043 | 2260 |
| 140809 | NM_080725 | SRXN1 | sulfiredoxin 1 | protein_coding | NP_542763.1 | 16132 | 617583 | . | . | 41.046841 | 42.373009 | 46.203728 | 50.796753 | 95.517685 | 97.947098 | 103.679825 | 116.140739 | 1.146669 | 3550.499209 | 0.11247254 | 1.755528301 | 0.079168984 | 0.775642847 | 11.691778 | 11.889093 | 11.724772 | 11.743740 | 11.811268 | 11.881884 | 3765 | 3611 | 3159 | 3706 |
| 140823 | NM_080748,XI | ROMO1 | reactive oxygen species modulator 1 | protein_coding | NP_542786.1;XP_016883167.1 | 16185 | . | | . | 145.657250 | 134.122633 | 120.778094 | 144.535431 | 338.950424 | 310.030479 | 271.022548 | 330.463066 | -1.065865 | 1641.489488 | 0.136666101 | -0.67335313 | 0.500722661 | 0.998865404 | 10.726185 | 10.633919 | 10.739342 | 10.667769 | 10.583945 | 10.720176 | 2039 | 1745 | 1260 | 1621 |
| 140825 | NM_00127853 | NEURL2 | neuralized E3 ubiquitin protein ligase 2 | protein_coding | NP_001265464.1;NP_542787.1 | 16156 | 608597 | . | . | 0.081331 | 0.074601 | 0.010996 | 0.078490 | 0.189260 | 0.172444 | 0.024674 | 0.179459 | -1.477586 | 3.366458 | 1.667253268 | -0.33782609 | 0.735494249 | NA | 2.008843 | 1.437399 | 1.766095 | 1.763060 | 1.751648 | 1.765522 | 5 | 4 | 1 | 4 |
| 140831 | NM_080752,N | ZSWIM3 | zinc finger SWIM-type containing 3 | protein_coding | NP_542790.2 | 16157 | . | | . | 0.689619 | 0.470087 | 0.382936 | 0.750933 | 1.604772 | 1.086627 | 0.859297 | 1.716920 | -1.042871 | 51.468496 | 0.475039265 | -0.12748486 | 0.898558655 | 0.998865404 | 5.716416 | 5.654148 | 5.722504 | 5.648095 | 5.623755 | 5.729635 | 72 | 46 | 31 | 61 |
| 140838 | NM_152667 | NANP | N-acetylneuraminic acid phosphatase | protein_coding | NP_689880.1 | 16140 | 610763 | . | . | 2.705697 | 2.362567 | 2.701852 | 2.343258 | 6.296269 | 5.461179 | 6.062877 | 5.357580 | -1.018843 | 300.123487 | 0.209236529 | -0.12871303 | 0.897584724 | 0.998865404 | 8.242224 | 8.216486 | 8.271957 | 8.192675 | 8.262061 | 8.182839 | 374 | 303 | 278 | 258 |
| 140849 | NR_040415 | LINC00266-1 | long intergenic non-protein coding RNA 266-1 | lncRNA | . | 16202 | . | | . | 0.043890 | 0.089355 | 0.074935 | 0.030458 | 0.102133 | 0.206549 | 0.168153 | 0.069638 | -1.345114 | 1.970934 | 2.116122047 | -0.20212829 | 0.839816435 | NA | 1.175403 | 0.751299 | 0.979035 | 0.983991 | 0.981632 | 0.976028 | 2 | 3 | 2 | 1 |
| 140862 | NM_080826,XI | ISM1 | isthmin 1 | protein_coding | NP_543016.1;XP_016883169.1 | 16213 | 615793 | . | . | 3.553887 | 3.600278 | 2.826021 | 3.710726 | 8.270041 | 8.322204 | 6.341510 | 8.484135 | -1.110366 | 329.645431 | 0.206457398 | -0.73155409 | 0.464440781 | 0.998865404 | 8.438870 | 8.286660 | 8.391464 | 8.392986 | 8.263281 | 8.399837 | 399 | 375 | 237 | 330 |
| 140883 | NM_080764,N | ZNF280B | zinc finger protein 280B | protein_coding | NP_542942.2;XP_011528199.1 | 23022 | . | | . | 1.009037 | 1.132366 | 1.093014 | 0.768306 | 2.348070 | 2.617514 | 2.452692 | 1.756639 | -1.150710 | 165.895672 | 0.277065839 | -0.73096066 | 0.464803183 | 0.998865404 | 7.471147 | 7.270121 | 7.372106 | 7.424200 | 7.410264 | 7.266502 | 192 | 203 | 159 | 118 |
| 140885 | NM_00104002 | SIRPA | signal regulatory protein alpha | protein_coding | NP_001035111.1;NP_001035112.1;NP_001317657.1;NP_542970.1;XP_0C | 9662 | 602461 | . | . | 79.589056 | 85.781503 | 103.825699 | 95.612935 | 185.207018 | 198.287792 | 232.981852 | 218.607613 | 1.172708 | 12574.021619 | 0.104191415 | 2.205982388 | 0.027385233 | 0.538018865 | 13.498667 | 13.728506 | 13.513825 | 13.577060 | 13.703122 | 13.660887 | 12829 | 12957 | 12091 | 12415 |
| 140886 | NM_080832 | PABPC5 | poly(A) binding protein cytoplasmic 5 | protein_coding | NP_543022.1 | 13629 | 300407 | . | . | 0.210359 | 0.208177 | 0.230379 | 0.328372 | 0.489515 | 0.481210 | 0.516963 | 0.750784 | 1.304846 | 24.773087 | 0.625074814 | 0.61413338 | 0.539127192 | NA | 4.427143 | 4.809059 | 4.611802 | 4.610290 | 4.617858 | 4.658314 | 25 | 23 | 20 | 31 |
| 140890 | NM_00107719 | SREK1 | splicing regulatory glutamic acid and lysine rich protein 1 | protein_coding | NP_001070667.1;NP_001257421.1;NP_001310456.1;NP_001310458.1;NI | 17882 | 609268 | . | . | 10.019599 | 10.493639 | 15.744339 | 9.209957 | 23.316021 | 24.256519 | 35.329841 | 21.057468 | 1.127250 | 2112.999173 | 0.174473096 | 0.990455483 | 0.321951538 | 0.998865404 | 10.955870 | 11.129086 | 10.985328 | 10.991056 | 11.252126 | 10.883229 | 2279 | 2151 | 2387 | 1626 |
| 140893 | NM_080833 | RBBP8NL | RBBP8 N-terminal like | protein_coding | NP_543023.2 | 16144 | . | | . | 0.115549 | 0.136593 | 0.265317 | 0.238569 | 0.268888 | 0.315741 | 0.595363 | 0.545460 | 2.024678 | 16.987947 | 0.750556858 | 1.355916313 | 0.175125796 | NA | 3.489863 | 4.507249 | 4.037436 | 4.045243 | 4.096703 | 4.085600 | 12 | 13 | 21 | 20 |
| 140894 | NM_00120707 | CNBD2 | cyclic nucleotide binding domain containing 2 | protein_coding | NP_001194005.1;NP_001291296.1;NP_001352638.1;NP_543024.2;XP_01 | 16145 | . | | . | 0.207392 | 0.189839 | 0.165142 | 0.240067 | 0.482608 | 0.438822 | 0.370574 | 0.548884 | -1.170629 | 12.406936 | 0.909118187 | -0.25000547 | 0.802583116 | NA | 3.744764 | 3.512009 | 3.646363 | 3.634112 | 3.607194 | 3.655636 | 17 | 13 | 5 | 16 |
| 140901 | NM_080836,XI | STK35 | serine/threonine kinase 35 | protein_coding | NP_543026.2;XP_011527476.1 | 16254 | 609370 | . | . | 3.160879 | 3.533635 | 3.427491 | 3.656781 | 7.355496 | 8.168155 | 7.691191 | 8.360796 | -1.006809 | 848.992516 | 0.146607213 | -0.0667746 | 0.946761142 | 0.998865404 | 9.734700 | 9.724498 | 9.691325 | 9.770402 | 9.711485 | 9.741641 | 927 | 970 | 720 | 810 |
| 140921 | gene-PPP2R5C | PPP2R5CP | protein phosphatase 2 regulatory subunit B'gamma pseudogene | pseudogene | . | 17835 | . | | . | 0.113444 | 0.099114 | 0.030494 | 0.118733 | 0.263988 | 0.229106 | 0.068427 | 0.271468 | -1.284927 | 4.369415 | 1.452511177 | -0.24900756 | 0.80335493 | NA | 2.299300 | 1.932304 | 2.138343 | 2.135598 | 2.125231 | 2.138692 | 6 | 5 | 2 | 5 |
| 142678 | NM_00117068 | MIB2 | mindbomb E3 ubiquitin protein ligase 2 | protein_coding | NP_001164157.2;NP_001164158.2;NP_001164159.1;NP_001164160.1;NI | 30577 | 611141 | . | . | 12.923764 | 11.033069 | 13.787913 | 13.244195 | 30.074133 | 25.503429 | 30.939677 | 30.281276 | 1.131903 | 1357.913245 | 0.136814636 | 1.306515262 | 0.191377392 | 0.963729349 | 10.314812 | 10.493980 | 10.364261 | 10.348556 | 10.512872 | 10.385755 | 1482 | 1360 | 1357 | 1241 |
| 142679 | NM_00114231 | DUSP19 | dual specificity phosphatase 19 | protein_coding | NP_001135786.1;NP_001308448.1;NP_543152.1 | 18894 | 611437 | . | . | 0.801761 | 0.550900 | 0.743171 | 0.461333 | 1.865731 | 1.273430 | 1.667653 | 1.054782 | -1.127240 | 104.398239 | 0.355741352 | -0.48572988 | 0.627158695 | 0.998865404 | 6.788092 | 6.618855 | 6.777463 | 6.652963 | 6.748316 | 6.605552 | 151 | 97 | 106 | 70 |
| 142891 | NM_00117415 | SAMD8 | sterile alpha motif domain containing 8 | protein_coding | NP_001167627.1;NP_653261.1;XP_005269598.1;XP_011537614.1;XP_01 | 26320 | 611575 | . | . | 4.062101 | 3.583181 | 3.832605 | 2.637940 | 9.452678 | 8.282683 | 8.600254 | 6.031337 | -1.195446 | 744.213932 | 0.186820576 | -1.37859186 | 0.168020629 | 0.945638095 | 9.662066 | 9.405708 | 9.640296 | 9.544249 | 9.588932 | 9.333939 | 995 | 816 | 701 | 516 |
| 142910 | NM_00101093 | LIPJ | lipase family member J | protein_coding | NP_001010939.2;XP_005269599.1;XP_006717698.1;XP_011537616.1;XP_ | 21773 | 613921 | . | . | 0.009842 | 0.074625 | 0.067895 | 0.064752 | 0.022904 | 0.172500 | 0.152356 | 0.148048 | 1.828067 | 5.184703 | 1.41408068 | 0.615466311 | 0.538246811 | NA | 1.878854 | 2.742346 | 2.331847 | 2.358790 | 2.365285 | 2.353344 | 1 | 7 | 7 | 5 |
| 142913 | NR_028492 | CFL1P1 | cofilin 1 pseudogene 1 | transcribed_ps | . | 28560 | . | | . | 0.138783 | 0.128172 | 0.088957 | 0.110712 | 0.322953 | 0.296276 | 0.199617 | 0.253130 | -1.309600 | 8.296706 | 1.053272774 | -0.36944474 | 0.711796253 | NA | 3.234218 | 2.844638 | 3.063360 | 3.061974 | 3.051505 | 3.053883 | 11 | 10 | 6 | 7 |
| 142940 | NM_139169 | TRUB1 | TruB pseudouridine synthase family member 1 | protein_coding | NP_631908.1 | 16060 | 610726 | . | . | 5.032474 | 4.893515 | 5.185448 | 4.921767 | 11.710772 | 11.311581 | 11.635996 | 11.253036 | 1.004404 | 531.739025 | 0.164689942 | 0.038494125 | 0.969293716 | 0.998865404 | 9.051211 | 9.057930 | 9.064636 | 9.041001 | 9.075672 | 9.035542 | 622 | 562 | 478 | 484 |
| 143098 | NM_00131817 | MPP7 | membrane palmitoylated protein 7 | protein_coding | NP_001305099.1;NP_775767.2;XP_005252426.1;XP_011517639.1;XP_01 | 26542 | 610973 | . | . | 0.277552 | 0.262025 | 0.118253 | 0.101173 | 0.645876 | 0.605681 | 0.265358 | 0.231322 | -1.705186 | 27.020836 | 0.600520002 | -1.28210438 | 0.199806033 | 0.96621494 | 5.089593 | 4.321310 | 4.791066 | 4.785090 | 4.738505 | 4.722729 | 40 | 36 | 19 | 17 |
| 143187 | NM_00131820 | VTI1A | vesicle transport through interaction with t-SNAREs 1A | protein_coding | NP_001305132.1;NP_001305134.1;NP_001352639.1;NP_001352640.1;NI | 17792 | 614316 | . | . | 5.690842 | 5.538411 | 5.487929 | 4.991372 | 13.242821 | 12.802284 | 12.314754 | 11.412176 | -1.072887 | 580.206904 | 0.161261867 | -0.62939595 | 0.529089867 | 0.998865404 | 9.230129 | 9.128944 | 9.194687 | 9.211134 | 9.185722 | 9.126854 | 683 | 656 | 509 | 501 |
| 143244 | NM_00109969 | EIF5AL1 | eukaryotic translation initiation factor 5A like 1 | protein_coding | NP_001093162.1 | 17419 | . | | . | 1.013244 | 0.932704 | 0.841208 | 1.003289 | 2.357860 | 2.155987 | 1.887646 | 2.293901 | -1.066590 | 114.029835 | 0.305407928 | -0.30453077 | 0.760723553 | 0.998865404 | 6.879307 | 6.785711 | 6.860580 | 6.827785 | 6.796372 | 6.848869 | 142 | 121 | 88 | 112 |
| 143279 | NM_00128427 | HECTD2 | HECT domain E3 ubiquitin protein ligase 2 | protein_coding | NP_001271203.2;NP_001335294.1;NP_775768.4;NP_877497.4;XP_01687 | 26736 | . | | . | 1.789087 | 1.542037 | 2.199773 | 1.341263 | 4.163279 | 3.564488 | 4.936227 | 3.066639 | 1.103921 | 220.345611 | 0.270301963 | 0.527697084 | 0.597709598 | 0.998865404 | 7.709505 | 7.854121 | 7.753282 | 7.745745 | 7.915153 | 7.661807 | 243 | 224 | 254 | 160 |
| 143282 | NM_152429 | FGFBP3 | fibroblast growth factor binding protein 3 | protein_coding | NP_689642.3 | 23428 | . | | . | 1.951141 | 1.680615 | 2.439080 | 1.723373 | 4.540384 | 3.884816 | 5.473224 | 3.940288 | 1.129413 | 155.344593 | 0.284315621 | 0.617527679 | 0.536886731 | 0.998865404 | 7.187679 | 7.365502 | 7.278023 | 7.217412 | 7.366443 | 7.221480 | 181 | 145 | 169 | 127 |
| 143341 | gene-EIF3LP2 | EIF3LP2 | eukaryotic translation initiation factor 3 subunit L pseudogene 2 | pseudogene | . | 24657 | . | | . | 0.450559 | 0.400386 | 0.571368 | 0.431384 | 1.048470 | 0.925511 | 1.282133 | 0.986308 | 1.199131 | 14.783940 | 0.796216733 | 0.329043052 | 0.742123148 | NA | 3.747897 | 4.011955 | 3.876845 | 3.872787 | 3.895926 | 3.876910 | 16 | 14 | 16 | 13 |
| 143371 | gene-YWHAZP: | YWHAZP3 | tyrosine 3-monooxygenase/tryptophan 5-monooxygenase activati | pseudogene | . | 31101 | . | | . | 1.371063 | 0.969485 | 0.946333 | 1.037267 | 3.190519 | 2.241008 | 2.123544 | 2.371589 | -1.195741 | 98.253239 | 0.33872691 | -0.76139332 | 0.446422174 | 0.998865404 | 6.741230 | 6.484204 | 6.699814 | 6.584121 | 6.579850 | 6.599718 | 145 | 95 | 75 | 87 |
| 143384 | NM_153810 | CACUL1 | CDK2 associated cullin domain 1 | protein_coding | NP_722517.3 | 23727 | 618764 | . | . | 2.935040 | 2.802308 | 2.939864 | 2.524546 | 6.829957 | 6.477661 | 6.596969 | 5.772074 | -1.064574 | 963.572989 | 0.14694588 | -0.61434966 | 0.538984293 | 0.998865404 | 9.956386 | 9.866721 | 9.953511 | 9.913447 | 9.945219 | 9.827957 | 1175 | 1043 | 878 | 804 |
| 143458 | NM_00130426 | LDLRAD3 | low density lipoprotein receptor class A domain containing 3 | protein_coding | NP_001291192.1;NP_001291193.1;NP_777562.1 | 27046 | 617986 | . | . | 22.351863 | 22.328457 | 23.671994 | 22.119015 | 52.013700 | 51.613230 | 53.119274 | 50.572500 | 1.011423 | 2687.427116 | 0.115429264 | 0.14196382 | 0.887108583 | 0.998865404 | 11.383719 | 11.400254 | 11.391123 | 11.383099 | 11.427911 | 11.363897 | 3092 | 2869 | 2443 | 2434 |
| 143570 | NM_00127038 | XRRA1 | X-ray radiation resistance associated 1 | protein_coding | NP_001257309.1;NP_001257310.1;NP_001365086.1;NP_001365087.1;NI | 18868 | 609788 | . | . | 1.434935 | 1.704771 | 2.071123 | 1.590091 | 3.339153 | 3.940660 | 4.647540 | 3.635555 | 1.126845 | 239.629111 | 0.245236617 | 0.702541968 | 0.482341239 | 0.998865404 | 7.815510 | 7.988619 | 7.823153 | 7.910401 | 8.009194 | 7.836221 | 236 | 265 | 259 | 197 |
| 143666 | NR_026967 | LOC143666 | uncharacterized LOC143666 | lncRNA | . | . | . | | . | 0.276871 | 0.624268 | 0.671984 | 0.256394 | 0.644291 | 1.443024 | 1.507913 | 0.586214 | 1.028620 | 29.944937 | 0.651266682 | 0.06250893 | 0.950157551 | 0.998865404 | 4.882987 | 4.925185 | 4.834932 | 4.929510 | 4.938271 | 4.832359 | 21 | 44 | 38 | 16 |
| 143684 | NM_00133035 | FAM76B | family with sequence similarity 76 member B | protein_coding | NP_001317286.1;NP_653265.3;XP_005273833.1;XP_011540912.1;XP_01 | 28492 | . | | . | 2.225257 | 1.606401 | 1.850710 | 1.530577 | 5.178264 | 3.713267 | 4.152941 | 3.499484 | -1.141944 | 224.672938 | 0.251756553 | -0.76062162 | 0.446883101 | 0.998865404 | 7.903370 | 7.713770 | 7.920822 | 7.748805 | 7.817934 | 7.730793 | 323 | 214 | 200 | 178 |
| 143686 | NM_00127159 | SESN3 | sestrin 3 | protein_coding | NP_001258523.1;NP_653266.2;XP_011540916.1;XP_024304123.1 | 23060 | 607768 | . | . | 4.285842 | 3.642218 | 5.308998 | 4.132906 | 9.973332 | 8.419149 | 11.913237 | 9.449398 | 1.174109 | 1294.574670 | 0.158241245 | 1.463371805 | 0.143365669 | 0.905249924 | 10.217453 | 10.449733 | 10.331956 | 10.204328 | 10.488078 | 10.286805 | 1488 | 1175 | 1374 | 1141 |
| 143689 | NM_152431 | PIWIL4 | piwi like RNA-mediated gene silencing 4 | protein_coding | NP_689644.2 | 18444 | 610315 | . | . | 0.582728 | 0.382941 | 0.268310 | 0.369269 | 1.356032 | 0.885186 | 0.602079 | 0.844290 | -1.532906 | 39.724560 | 0.516708469 | -1.192683 | 0.232993545 | 0.988893316 | 5.587697 | 4.970764 | 5.382744 | 5.304994 | 5.260900 | 5.298549 | 67 | 41 | 23 | 34 |
| 143872 | NM_00136794 | ARHGAP42 | Rho GTPase activating protein 42 | protein_coding | NP_001354874.1;NP_689645.2;XP_011540917.1;XP_011540918.1;XP_01 | 26545 | 615936 | . | . | 1.638160 | 1.501625 | 1.773853 | 1.513476 | 3.812067 | 3.471075 | 3.980474 | 3.460384 | 1.032548 | 409.099624 | 0.187429662 | 0.246540809 | 0.805263618 | 0.998865404 | 8.652502 | 8.699729 | 8.693876 | 8.635358 | 8.733335 | 8.632890 | 486 | 413 | 392 | 357 |
| 143879 | NM_00133035 | KBTBD3 | kelch repeat and BTB domain containing 3 | protein_coding | NP_001317288.1;NP_689646.2;NP_940841.1;XP_006718830.1;XP_01154 | 22934 | . | | . | 0.785708 | 0.558712 | 0.433884 | 0.624686 | 1.828373 | 1.291487 | 0.973624 | 1.428270 | -1.348254 | 96.644756 | 0.344302981 | -1.25207223 | 0.210543547 | 0.973252704 | 6.793749 | 6.363536 | 6.696869 | 6.569486 | 6.537400 | 6.555684 | 152 | 97 | 69 | 80 |
| 143884 | NM_152434,XI | CWF19L2 | CWF19 like cell cycle control factor 2 | protein_coding | NP_689647.2;XP_011540922.1;XP_016872728.1 | 26508 | . | | . | 2.717151 | 2.387440 | 2.530135 | 2.828904 | 6.322920 | 5.518675 | 5.677551 | 6.467949 | 1.021271 | 260.886043 | 0.215861832 | 0.14066969 | 0.888130891 | 0.998865404 | 8.011925 | 8.042465 | 8.054764 | 7.986736 | 8.015889 | 8.047927 | 317 | 260 | 222 | 255 |
| 143888 | NM_00136350 | POGLUT3 | protein O-glucosyltransferase 3 | protein_coding | NP_001350431.1;NP_001350432.1;NP_714916.3;XP_011540923.1 | 28496 | 618503 | . | . | 20.485264 | 19.433501 | 16.993176 | 17.033273 | 47.670049 | 44.921411 | 38.132192 | 38.944555 | -1.193405 | 2446.930010 | 0.115970122 | -2.19956938 | 0.027837461 | 0.538151934 | 11.378621 | 11.123642 | 11.348865 | 11.297855 | 11.186897 | 11.175633 | 3161 | 2784 | 1954 | 2077 |
| 144097 | NM_138471,XI | SPINDOC | spindlin interactor and repressor of chromatin binding | protein_coding | NP_612480.1;XP_005273839.1;XP_005273840.1;XP_006718500.1;XP_01 | 25115 | . | | . | 10.653101 | 12.721724 | 11.403651 | 11.183330 | 24.790204 | 29.406838 | 25.589463 | 25.569354 | -1.057018 | 707.843183 | 0.159857146 | -0.50044831 | 0.616759445 | 0.998865404 | 9.506907 | 9.426546 | 9.418925 | 9.547130 | 9.454269 | 9.439203 | 761 | 857 | 605 | 638 |
| 144100 | NM_00132963 | PLEKHA7 | pleckstrin homology domain containing A7 | protein_coding | NP_001316559.1;NP_001316560.1;NP_778228.3;XP_016872730.1;XP_01 | 27049 | 612686 | . | . | 1.463682 | 1.491252 | 0.458133 | 0.490054 | 3.406049 | 3.447099 | 1.028038 | 1.120450 | -2.860613 | 198.302274 | 0.248428591 | -6.10366246 | 1.03665E-09 | 9.53926E-07 | 8.199413 | 6.681692 | 7.770596 | 7.804040 | 7.320328 | 7.358851 | 328 | 327 | 83 | 103 |
| 144106 | gene-ST13P5 | ST13P5 | ST13, Hsp70 interacting protein pseudogene 5 | pseudogene | . | 18556 | . | | . | 0.253067 | 0.165857 | 0.162654 | 0.283957 | 0.588898 | 0.383385 | 0.364992 | 0.649234 | 1.083480 | 11.606803 | 0.905378403 | 0.127762222 | 0.898337147 | NA | 3.478589 | 3.592943 | 3.542932 | 3.525865 | 3.526163 | 3.550810 | 15 | 10 | 8 | 14 |
| 144108 | NM_194285,XI | SPTY2D1 | SPT2 chromatin protein domain containing 1 | protein_coding | NP_919261.2;XP_011518221.1 | 26818 | . | | . | 3.891852 | 3.840329 | 3.071414 | 3.661227 | 9.056500 | 8.877091 | 6.892163 | 8.370962 | -1.164333 | 631.777327 | 0.162244852 | -1.35291371 | 0.1760832 | 0.949193943 | 9.409130 | 9.189028 | 9.358999 | 9.344429 | 9.195080 | 9.301142 | 791 | 726 | 466 | 592 |
| 144110 | NM_153347 | TMEM86A | transmembrane protein 86A | protein_coding | NP_699178.1 | 26890 | . | | . | 1.323492 | 1.004164 | 1.243627 | 1.283621 | 3.079819 | 2.321170 | 2.790663 | 2.934847 | 1.067636 | 137.082194 | 0.288417656 | 0.327371028 | 0.743387277 | 0.998865404 | 7.050385 | 7.145832 | 7.134491 | 7.031372 | 7.104962 | 7.112044 | 174 | 123 | 122 | 134 |
| 144132 | NM_144666,N | DNHD1 | dynein heavy chain domain 1 | protein_coding | NP_653267.2;NP_775860.3 | 26532 | 617277 | . | . | 0.616813 | 0.686984 | 0.619572 | 0.574355 | 1.435349 | 1.587995 | 1.390300 | 1.313194 | -1.171002 | 262.490837 | 0.214939709 | -1.05956942 | 0.289340532 | 0.998865404 | 8.145259 | 7.918051 | 8.061462 | 8.087473 | 8.025443 | 7.962240 | 318 | 313 | 224 | 212 |
| 144165 | NM_00114488 | PRICKLE1 | prickle planar cell polarity protein 1 | protein_coding | NP_001138353.1;NP_001138354.1;NP_001138355.1;NP_694571.2;XP_01 | 17019 | 608500 | . | . | 0.651159 | 0.700258 | 0.565694 | 0.708937 | 1.515273 | 1.618678 | 1.269403 | 1.620902 | -1.112587 | 118.411336 | 0.298651135 | -0.51537666 | 0.606289836 | 0.998865404 | 6.963075 | 6.808091 | 6.894569 | 6.917558 | 6.848694 | 6.890840 | 139 | 139 | 91 | 112 |
| 144193 | NM_152435 | AMDHD1 | amidohydrolase domain containing 1 | protein_coding | NP_689648.2 | 28577 | . | | . | 0.558981 | 0.464521 | 0.642057 | 0.386151 | 1.300772 | 1.073761 | 1.440757 | 0.882888 | -1.014898 | 35.975018 | 0.536915568 | -0.03973519 | 0.968304246 | 0.998865404 | 5.177607 | 5.160188 | 5.180809 | 5.150546 | 5.198011 | 5.128068 | 46 | 35 | 39 | 25 |
| 144195 | NM_00128623 | SLC2A14 | solute carrier family 2 member 14 | protein_coding | NP_001273162.1;NP_001273163.1;NP_001273164.1;NP_001273165.1;NI | 18301 | 611039 | . | . | 0.591640 | 0.591440 | 0.827286 | 0.718119 | 1.376771 | 1.367139 | 1.856405 | 1.641894 | 1.286401 | 90.228949 | 0.337057571 | 1.077976219 | 0.281044369 | 0.998865404 | 6.301967 | 6.666148 | 6.452304 | 6.451854 | 6.546099 | 6.500524 | 91 | 85 | 95 | 88 |
| 144203 | NR_153414 | OVOS2 | alpha-2-macroglobulin like 1 pseudogene | transcribed_ps | . | . | . | | . | 0.684178 | 0.696499 | 0.799001 | 0.652512 | 1.592110 | 1.609987 | 1.792934 | 1.491891 | 1.033266 | 126.773653 | 0.292767753 | 0.161260015 | 0.871888615 | 0.998865404 | 6.961726 | 7.010091 | 6.976920 | 6.979607 | 7.026472 | 6.951869 | 143 | 135 | 124 | 108 |
| 144233 | NM_181708 | BCDIN3D | BCDIN3 domain containing RNA methyltransferase | protein_coding | NP_859059.1 | 27050 | . | | . | 0.924743 | 0.740870 | 0.968375 | 1.265428 | 2.151915 | 1.712555 | 2.173005 | 2.893251 | 1.317687 | 98.335262 | 0.335493103 | 1.186337446 | 0.235489064 | 0.990979804 | 6.407068 | 6.804870 | 6.600236 | 6.536758 | 6.609997 | 6.691972 | 109 | 81 | 85 | 118 |
| 144245 | NM_00101362 | ALG10B | ALG10 alpha-1,2-glucosyltransferase B | protein_coding | NP_001013642.2;NP_001295269.2;XP_005268722.1 | 31088 | 603313 | . | . | 1.597306 | 1.210899 | 1.381883 | 1.036638 | 3.716998 | 2.799047 | 3.100907 | 2.370150 | -1.287220 | 354.948147 | 0.218294277 | -1.66865677 | 0.095185421 | 0.813553488 | 8.641330 | 8.278896 | 8.602235 | 8.461156 | 8.470177 | 8.303808 | 513 | 381 | 312 | 246 |
| 144321 | NM_00127039 | GLIPR1L2 | GLIPR1 like 2 | protein_coding | NP_001257325.1;NP_689649.1;XP_011536251.1;XP_011536252.1;XP_01 | 28592 | 610394 | . | . | 0.154814 | 0.182435 | 0.344437 | 0.071617 | 0.360257 | 0.421707 | 0.772907 | 0.163744 | 1.059743 | 11.542153 | 0.9731451 | 0.086025198 | 0.931446387 | NA | 3.482357 | 3.573873 | 3.518883 | 3.509938 | 3.549661 | 3.483808 | 14 | 11 | 17 | 4 |
| 144348 | NM_00120429 | ZNF664 | zinc finger protein 664 | protein_coding | NP_001191227.1;NP_689650.1 | 25406 | 617890 | . | . | 21.254969 | 22.273444 | 21.707557 | 21.039764 | 49.461188 | 51.486067 | 48.711126 | 48.104920 | -1.029759 | 2886.715576 | 0.113049441 | -0.37422743 | 0.708235118 | 0.998865404 | 11.516199 | 11.473918 | 11.490374 | 11.523281 | 11.500158 | 11.465649 | 3305 | 3224 | 2526 | 2610 |
| 144363 | NM_00100166 | ETFRF1 | electron transfer flavoprotein regulatory factor 1 | protein_coding | NP_001001660.2;XP_005253376.1;XP_005253377.1;XP_016874339.1 | 27052 | . | | . | 5.037495 | 3.599230 | 2.766294 | 2.078098 | 11.722458 | 8.319780 | 6.207483 | 4.751324 | -1.591528 | 156.071535 | 0.302515743 | -2.21612307 | 0.026683075 | 0.532994094 | 7.581412 | 6.914266 | 7.418359 | 7.299772 | 7.255529 | 7.094322 | 249 | 180 | 130 | 86 |
| 144402 | NM_153634,XI | CPNE8 | copine 8 | protein_coding | NP_705898.1;XP_011536253.1;XP_011536254.1;XP_016874341.1 | 23498 | . | | . | 11.095562 | 11.816333 | 13.987668 | 11.183900 | 25.819827 | 27.313985 | 31.387923 | 25.570655 | 1.090496 | 1299.821415 | 0.147769972 | 0.845801998 | 0.397663211 | 0.998865404 | 10.280037 | 10.405434 | 10.287742 | 10.327581 | 10.460790 | 10.278913 | 1394 | 1377 | 1319 | 1121 |
| 144404 | NM_00108082 | TMEM120B | transmembrane protein 120B | protein_coding | NP_001074294.2;XP_024304619.1;XP_024304620.1 | 32008 | 616551 | . | . | 5.534360 | 5.513416 | 6.506925 | 5.840092 | 12.878682 | 12.744509 | 14.601352 | 13.352678 | 1.041786 | 1067.344856 | 0.134057113 | 0.44055515 | 0.659535078 | 0.998865404 | 10.030003 | 10.089015 | 10.033845 | 10.054627 | 10.076785 | 10.072339 | 1187 | 1144 | 950 | 1018 |
| 144406 | NM_00117800 | CFAP251 | cilia and flagella associated protein 251 | protein_coding | NP_001171474.1;NP_653269.3 | 28506 | 618146 | . | . | 0.280830 | 0.230119 | 0.523276 | 0.272364 | 0.653505 | 0.531931 | 1.174217 | 0.622728 | 1.485629 | 38.388391 | 0.536824137 | 1.063800006 | 0.287419309 | 0.998865404 | 4.946534 | 5.521710 | 5.225594 | 5.200828 | 5.326215 | 5.214711 | 39 | 30 | 53 | 29 |
| 144455 | NM_203394,XI | E2F7 | E2F transcription factor 7 | protein_coding | NP_976328.2;XP_011536268.1;XP_011536271.1 | 23820 | 612046 | . | . | 3.527203 | 4.068950 | 4.970785 | 4.582507 | 8.207945 | 9.405561 | 11.154299 | 10.477357 | 1.246993 | 757.780392 | 0.155859423 | 2.043207785 | 0.041031873 | 0.628593728 | 9.397683 | 9.716060 | 9.430756 | 9.528204 | 9.669862 | 9.598193 | 721 | 780 | 768 | 749 |
| 144486 | NR_027035 | CEP83-DT | CEP83 divergent transcript | lncRNA | . | 27055 | . | | . | 0.181467 | 0.267432 | 0.159794 | 0.222762 | 0.422281 | 0.618180 | 0.358573 | 0.509319 | -1.155958 | 17.234606 | 0.749080038 | -0.27912711 | 0.780147277 | NA | 4.209663 | 3.996978 | 4.096646 | 4.129336 | 4.093365 | 4.111470 | 17 | 24 | 12 | 17 |
| 144535 | NM_00130608 | CFAP54 | cilia and flagella associated protein 54 | protein_coding | NP_001293013.1;NP_001354814.1 | 26456 | . | | . | 0.051488 | 0.042129 | 0.026662 | 0.045289 | 0.119814 | 0.097384 | 0.059829 | 0.103548 | -1.257475 | 13.633676 | 0.837575182 | -0.39462631 | 0.693118693 | NA | 3.926074 | 3.592943 | 3.786563 | 3.774901 | 3.754915 | 3.779281 | 19 | 15 | 8 | 14 |
| 144577 | NM_00130094 | C12orf66 | chromosome 12 open reading frame 66 | protein_coding | NP_001287869.1;NP_001287870.2;NP_689653.4 | 26517 | 617420 | . | . | 2.065416 | 2.316200 | 1.996589 | 2.251889 | 4.806307 | 5.354001 | 4.480289 | 5.148678 | 1.039167 | 227.711695 | 0.224918827 | 0.246434073 | 0.805346233 | 0.998865404 | 7.803383 | 7.858225 | 7.813325 | 7.829177 | 7.818581 | 7.860249 | 253 | 245 | 193 | 227 |
| 144581 | gene-RPL14P1 | RPL14P1 | ribosomal protein L14 pseudogene 1 | pseudogene | . | 31384 | . | | . | 2.994490 | 2.814083 | 3.027110 | 4.173790 | 6.968302 | 6.504879 | 6.792747 | 9.542877 | 1.221714 | 73.587702 | 0.376788686 | 0.766761413 | 0.443223382 | 0.998865404 | 6.050418 | 6.338054 | 6.179150 | 6.162998 | 6.181340 | 6.258452 | 79 | 69 | 60 | 87 |
| 144608 | NM_175874,XI | C12orf60 | chromosome 12 open reading frame 60 | protein_coding | NP_787070.2;XP_005253379.1;XP_011518870.1;XP_011518871.1;XP_01 | 28726 | . | | . | 0.817374 | 1.302435 | 1.137110 | 1.196971 | 1.902064 | 3.010636 | 2.551641 | 2.736733 | 1.055842 | 60.717664 | 0.421699513 | 0.185899066 | 0.852523914 | 0.998865404 | 5.885760 | 5.961339 | 5.867795 | 5.957066 | 5.904318 | 5.945527 | 54 | 77 | 49 | 64 |
| 144699 | NM_152441,XI | FBXL14 | F-box and leucine rich repeat protein 14 | protein_coding | NP_689654.1;XP_016874364.1;XP_016874365.1 | 28624 | 609081 | . | . | 0.628947 | 2.028978 | 0.759385 | 0.620762 | 1.463586 | 4.690074 | 1.704038 | 1.419299 | -1.053876 | 140.325294 | 0.287669435 | -0.26316565 | 0.792422908 | 0.998865404 | 7.169864 | 7.094412 | 7.094858 | 7.182595 | 7.156558 | 7.082201 | 147 | 173 | 131 | 115 |
| 144712 | gene-HMGA1P | HMGA1P3 | high mobility group AT-hook 1 pseudogene 3 | pseudogene | . | 13323 | . | | . | 1.091455 | 0.972088 | 0.959923 | 1.163772 | 2.539861 | 2.247026 | 2.154039 | 2.660826 | 1.015054 | 58.596536 | 0.412405708 | 0.052270288 | 0.958313328 | 0.998865404 | 5.862170 | 5.883240 | 5.883694 | 5.858229 | 5.856669 | 5.891361 | 71 | 59 | 47 | 60 |
| 144715 | NM_00128653 | RAD9B | RAD9 checkpoint clamp component B | protein_coding | NP_001273460.1;NP_001273461.1;NP_001273462.1;NP_001273463.1;NI | 21700 | 608368 | . | . | 0.663262 | 0.624117 | 0.554472 | 0.364655 | 1.543440 | 1.442672 | 1.244218 | 0.833740 | -1.392785 | 79.229871 | 0.371031974 | -1.28822508 | 0.197667624 | 0.96621494 | 6.526078 | 6.050918 | 6.366915 | 6.328007 | 6.301699 | 6.220881 | 113 | 93 | 68 | 51 |
| 144717 | NM_00117799 | PHETA1 | PH domain containing endocytic trafficking adaptor 1 | protein_coding | NP_001171467.1;NP_001171468.1;NP_653272.2;XP_006719320.1;XP_01 | 26509 | 614239 | . | . | 2.336894 | 2.325221 | 2.125068 | 2.443134 | 5.438048 | 5.374853 | 4.768589 | 5.585935 | -1.028395 | 222.864439 | 0.228783919 | -0.1765647 | 0.859850327 | 0.998865404 | 7.820484 | 7.779267 | 7.800491 | 7.812688 | 7.761086 | 7.824414 | 257 | 247 | 178 | 220 |
| 144742 | NR_024246 | LINC00934 | long intergenic non-protein coding RNA 934 | lncRNA | . | 27059 | . | | . | 0.048129 | 0.093218 | 0.050988 | 0.030854 | 0.111999 | 0.215476 | 0.114415 | 0.070545 | -1.395494 | 4.671906 | 1.413361316 | -0.34016497 | 0.733732297 | NA | 2.444631 | 1.963455 | 2.219269 | 2.237685 | 2.224653 | 2.218328 | 4 | 8 | 4 | 3 |
| 144766 | NR_145420 | LINC00355 | long intergenic non-protein coding RNA 355 | lncRNA | . | 27061 | . | | . | 0.626520 | 0.459405 | 0.445544 | 0.571904 | 1.457938 | 1.061936 | 0.999788 | 1.307591 | -1.094472 | 31.165360 | 0.561905818 | -0.23177398 | 0.81671357 | 0.998865404 | 5.025590 | 4.895208 | 4.988307 | 4.949213 | 4.944594 | 4.970545 | 43 | 30 | 23 | 31 |
| 144811 | NM_00112830 | LACC1 | laccase domain containing 1 | protein_coding | NP_001121775.1;NP_001337567.1;NP_001337568.1;NP_001337569.1;NI | 26789 | 613409 | . | . | 2.317717 | 1.722296 | 1.743524 | 1.543267 | 5.393426 | 3.981164 | 3.912415 | 3.528498 | -1.163729 | 197.424833 | 0.248217577 | -0.88130166 | 0.378154568 | 0.998865404 | 7.729690 | 7.512460 | 7.703537 | 7.608051 | 7.621357 | 7.555819 | 269 | 206 | 171 | 158 |
| 144983 | NM_00101172 | HNRNPA1L2 | heterogeneous nuclear ribonucleoprotein A1 like 2 | protein_coding | NP_001011724.1;NP_001011725.1 | 27067 | . | | . | 4.887490 | 4.723427 | 4.517016 | 4.771016 | 11.373390 | 10.918413 | 10.136053 | 10.908361 | -1.057290 | 330.706843 | 0.193117002 | -0.41617635 | 0.677280961 | 0.998865404 | 8.409075 | 8.328622 | 8.380061 | 8.389923 | 8.352954 | 8.355011 | 388 | 370 | 279 | 302 |
| 145165 | NR_002183 | ST13P4 | ST13, Hsp70 interacting protein pseudogene 4 | transcribed_ps | . | 18487 | . | | . | 0.450717 | 0.533037 | 0.536281 | 0.703260 | 1.048837 | 1.232138 | 1.203399 | 1.607920 | 1.249741 | 28.280733 | 0.583577279 | 0.551133403 | 0.581542229 | 0.998865404 | 4.653161 | 4.972679 | 4.796303 | 4.811021 | 4.814164 | 4.844290 | 27 | 29 | 24 | 33 |
| 145173 | NM_194318,XI | B3GLCT | beta 3-glucosyltransferase | protein_coding | NP_919299.3;XP_006719831.1;XP_011533238.1;XP_011533240.1;XP_01 | 20207 | 610308 | . | . | 3.912144 | 3.686544 | 3.534618 | 3.304796 | 9.103719 | 8.521610 | 7.931581 | 7.556023 | -1.126125 | 476.554029 | 0.172323164 | -0.99445409 | 0.320001848 | 0.998865404 | 8.979368 | 8.808572 | 8.952984 | 8.911160 | 8.883018 | 8.832816 | 600 | 527 | 406 | 403 |
| 145200 | NR_026774 | LINC00239 | long intergenic non-protein coding RNA 239 | lncRNA | . | 20119 | . | | . | 0.379212 | 0.673078 | 0.207619 | 0.723699 | 0.882443 | 1.555850 | 0.465890 | 1.654652 | -1.105053 | 10.312597 | 1.00436681 | -0.14348946 | 0.885903631 | NA | 3.440763 | 3.287859 | 3.353418 | 3.381084 | 3.340239 | 3.384948 | 9 | 15 | 4 | 14 |
| 145241 | NR_003951 | ADAM21P1 | ADAM metallopeptidase domain 21 pseudogene 1 | transcribed_ps | . | 19822 | . | | . | 0.031036 | 0.014337 | 0.021446 | 0.059116 | 0.072222 | 0.033139 | 0.048123 | 0.135162 | 1.855457 | 2.524672 | 1.904053786 | 0.468355742 | 0.639530212 | NA | 0.820514 | 1.715150 | 1.327318 | 1.319311 | 1.325917 | 1.334856 | 3 | 1 | 2 | 4 |
| 145258 | NM_173849 | GSC | goosecoid homeobox | protein_coding | NP_776248.1 | 4612 | 138890 | . | . | 0.233428 | 0.096115 | 0.200759 | 0.159832 | 0.543196 | 0.222174 | 0.450498 | 0.365437 | 1.161735 | 6.710305 | 1.18695546 | 0.182214621 | 0.855414299 | NA | 2.631267 | 2.852980 | 2.751965 | 2.728724 | 2.748553 | 2.741337 | 10 | 4 | 7 | 6 |
| 145259 | gene-RPSAP4 | RPSAP4 | ribosomal protein SA pseudogene 4 | pseudogene | . | 20018 | . | | . | 0.064746 | 0.022452 | 0.010742 | 0.037989 | 0.150666 | 0.051898 | 0.024106 | 0.086856 | -1.076094 | 1.703845 | 2.268349467 | -0.04664372 | 0.962797187 | NA | 0.820514 | 0.715150 | 0.778739 | 0.770724 | 0.771939 | 0.776369 | 3 | 1 | 1 | 2 |
| 145270 | NM_178013,XI | PRIMA1 | proline rich membrane anchor 1 | protein_coding | NP_821092.1;XP_011534758.1;XP_024305250.1 | 18319 | 613851 | . | . | 0.212413 | 0.307405 | 0.267054 | 0.233862 | 0.494293 | 0.710579 | 0.599262 | 0.534697 | 1.052518 | 31.379716 | 0.553781846 | 0.133345396 | 0.893920227 | 0.998865404 | 4.934350 | 5.008225 | 4.954447 | 4.977529 | 4.981486 | 4.963434 | 32 | 36 | 30 | 28 |
| 145282 | NM_00119529 | MIPOL1 | mirror-image polydactyly 1 | protein_coding | NP_001182225.1;NP_001182226.1;NP_620059.1;XP_011534765.1;XP_01 | 21460 | 606850 | . | . | 1.073754 | 0.999555 | 1.233218 | 0.933342 | 2.498673 | 2.310517 | 2.767305 | 2.133978 | 1.000145 | 157.259893 | 0.261753104 | 0.000798773 | 0.999362672 | 0.999862634 | 7.296655 | 7.297359 | 7.307853 | 7.285640 | 7.306096 | 7.287615 | 186 | 165 | 140 | 144 |
| 145375 | gene-NUP50P1 | NUP50P1 | nucleoporin 50 pseudogene 1 | pseudogene | . | 20025 | . | | . | 0.041911 | 0.035394 | 0.042661 | 0.033048 | 0.097528 | 0.081815 | 0.095731 | 0.075561 | -1.099830 | 6.416983 | 1.191012301 | -0.11526346 | 0.908236298 | NA | 2.747897 | 2.612728 | 2.685324 | 2.683157 | 2.684940 | 2.677987 | 8 | 7 | 6 | 5 |
| 145376 | NM_172365,XI | PPP1R36 | protein phosphatase 1 regulatory subunit 36 | protein_coding | NP_758953.1;XP_005267411.1;XP_005267412.1;XP_011534770.1;XP_02 | 20097 | . | | . | 0.316672 | 0.256874 | 0.267201 | 0.112269 | 0.736910 | 0.593777 | 0.599592 | 0.256690 | -1.492706 | 10.462272 | 0.966567945 | -0.59792006 | 0.549893273 | NA | 3.644780 | 3.073227 | 3.405835 | 3.393181 | 3.394634 | 3.367601 | 16 | 12 | 10 | 5 |
| 145389 | NM_00117270 | SLC38A6 | solute carrier family 38 member 6 | protein_coding | NP_001166173.1;NP_722518.2;XP_006720113.1;XP_011534771.1;XP_01 | 19863 | 616518 | . | . | 13.790476 | 9.851883 | 15.567438 | 7.216633 | 32.091001 | 22.773072 | 34.932882 | 16.499969 | -1.036363 | 629.126072 | 0.626504421 | -0.08224989 | 0.934448005 | 0.998865404 | 9.322612 | 9.271343 | 9.394258 | 9.163247 | 9.496776 | 8.952219 | 861 | 575 | 751 | 349 |
| 145407 | NM_00100187 | ARMH4 | armadillo like helical domain containing 4 | protein_coding | NP_001001872.2;NP_001307102.1;XP_011534773.1;XP_016876518.1;XP | 19846 | . | | . | 0.974488 | 0.934858 | 0.940106 | 1.361568 | 2.267673 | 2.160966 | 2.109567 | 3.113063 | 1.066052 | 178.871613 | 0.257995872 | 0.357671915 | 0.720588853 | 0.998865404 | 7.436323 | 7.527789 | 7.489184 | 7.444209 | 7.440744 | 7.545055 | 210 | 177 | 141 | 194 |
| 145438 | NR_037676 | FRMD6-AS1 | FRMD6 antisense RNA 1 | lncRNA | . | 20129 | . | | . | 0.644063 | 0.812740 | 0.484796 | 0.549736 | 1.498761 | 1.878685 | 1.087869 | 1.256906 | -1.417950 | 44.096212 | 0.475337049 | -1.05989404 | 0.289192806 | 0.998865404 | 5.693286 | 5.187832 | 5.470774 | 5.513474 | 5.427413 | 5.441743 | 53 | 62 | 30 | 36 |
| 145482 | NM_00114615 | PTGR2 | prostaglandin reductase 2 | protein_coding | NP_001139626.1;NP_001139627.1;NP_001358254.1;NP_001358255.1;NI | 20149 | 608642 | . | . | 2.885904 | 3.479331 | 2.898943 | 2.638642 | 6.715616 | 8.042630 | 6.505147 | 6.032942 | -1.191034 | 223.707571 | 0.234385828 | -1.07606559 | 0.281897922 | 0.998865404 | 7.925955 | 7.673999 | 7.801985 | 7.884506 | 7.794543 | 7.726622 | 257 | 284 | 191 | 177 |
| 145483 | NM_152445,XI | FAM161B | FAM161 centrosomal protein B | protein_coding | NP_689658.3;XP_011534777.1 | 19854 | . | | . | 0.889614 | 0.910891 | 0.742084 | 1.083159 | 2.070168 | 2.105564 | 1.665215 | 2.476515 | 1.011230 | 112.690764 | 0.317870763 | 0.050684152 | 0.959577205 | 0.998865404 | 6.809168 | 6.823248 | 6.806818 | 6.817829 | 6.756829 | 6.871840 | 127 | 123 | 81 | 125 |
| 145501 | NM_182509,N | ISM2 | isthmin 2 | protein_coding | NP_872315.2;NP_954993.1;XP_011534791.1 | 23176 | 612684 | . | . | 0.316081 | 0.322278 | 0.123744 | 0.283375 | 0.735533 | 0.744960 | 0.277679 | 0.647904 | -1.591442 | 24.786118 | 0.656089486 | -1.02171144 | 0.306917497 | NA | 4.930218 | 4.254232 | 4.658232 | 4.658675 | 4.580105 | 4.642198 | 35 | 33 | 10 | 25 |
| 145508 | NM_152446,N | CEP128 | centrosomal protein 128 | protein_coding | NP_689659.2;XP_011534792.1;XP_011534793.1;XP_011534794.1;XP_01 | 20359 | . | | . | 2.447386 | 2.468074 | 2.331962 | 2.202326 | 5.695168 | 5.705063 | 5.232856 | 5.035357 | -1.100892 | 350.617452 | 0.189276341 | -0.73265102 | 0.463771304 | 0.998865404 | 8.521327 | 8.382860 | 8.490331 | 8.471068 | 8.431466 | 8.420908 | 430 | 390 | 293 | 310 |
| 145553 | NM_00119982 | MDP1 | magnesium dependent phosphatase 1 | protein_coding | NP_001186750.1;NP_001186751.1;NP_612485.2 | 28781 | . | | . | 4.361480 | 2.645411 | 2.427738 | 5.297290 | 10.149342 | 6.114986 | 5.447775 | 12.111625 | 1.070321 | 85.615060 | 0.414908251 | 0.236300214 | 0.813199721 | 0.998865404 | 6.370642 | 6.467324 | 6.460870 | 6.330565 | 6.315145 | 6.508635 | 118 | 68 | 50 | 112 |
| 145567 | NM_00101085 | TTC7B | tetratricopeptide repeat domain 7B | protein_coding | NP_001010854.1;NP_001307350.1;XP_005267424.1;XP_011534799.1;XP | 19858 | . | | . | 0.592347 | 1.158147 | 0.548403 | 1.023769 | 1.378417 | 2.677105 | 1.230598 | 2.340726 | 1.012288 | 330.744702 | 0.203935836 | 0.086396478 | 0.931151247 | 0.998865404 | 8.361255 | 8.377846 | 8.303954 | 8.421636 | 8.349381 | 8.392461 | 339 | 392 | 278 | 324 |
| 145624 | NR_022009 | PWAR1 | Prader Willi/Angelman region RNA 1 | lncRNA | . | 30089 | 600161 | . | . | 0.034286 | 0.013397 | 0.132537 | 0.132307 | 0.079785 | 0.030967 | 0.297409 | 0.302504 | 3.930318 | 6.585924 | 1.246307381 | 1.58439739 | 0.113103316 | NA | 1.413393 | 3.393446 | 2.673832 | 2.666336 | 2.705178 | 2.706950 | 4 | 2 | 9 | 10 |
| 145694 | NR_132969,NF | LOC145694 | uncharacterized LOC145694 | lncRNA | . | . | . | | . | 0.574362 | 0.585563 | 0.709444 | 0.374706 | 1.336565 | 1.353556 | 1.591973 | 0.856721 | -1.052082 | 33.276963 | 0.561447985 | -0.13046134 | 0.896201438 | 0.998865404 | 5.090828 | 5.021237 | 5.054036 | 5.055694 | 5.086669 | 5.007991 | 39 | 37 | 37 | 21 |
| 145748 | NM_00128441 | LYSMD4 | LysM domain containing 4 | protein_coding | NP_001271346.1;NP_001271347.1;NP_001271348.1;NP_001271349.1;NI | 26571 | . | | . | 3.254026 | 3.310655 | 3.356514 | 3.681759 | 7.572249 | 7.652730 | 7.531921 | 8.417904 | 1.071390 | 296.145854 | 0.201615841 | 0.493434195 | 0.621705826 | 0.998865404 | 8.159636 | 8.258982 | 8.195108 | 8.186413 | 8.216605 | 8.238493 | 332 | 306 | 261 | 293 |
| 145767 | gene-RPS3AP6 | RPS3AP6 | RPS3A pseudogene 6 | pseudogene | . | 18630 | . | | . | 10.488760 | 10.871557 | 8.894998 | 8.817133 | 24.407776 | 25.130091 | 19.960115 | 20.159328 | -1.221066 | 261.887481 | 0.211835576 | -1.36020902 | 0.173763788 | 0.947973539 | 8.169687 | 7.881557 | 8.074167 | 8.088686 | 7.984923 | 7.975472 | 327 | 315 | 207 | 219 |
| 145773 | NM_152450,XI | FAM81A | family with sequence similarity 81 member A | protein_coding | NP_689663.2;XP_005254223.1;XP_006720461.1;XP_011519549.1;XP_01 | 28379 | . | | . | 1.861947 | 1.603846 | 1.914513 | 1.828864 | 4.332827 | 3.707361 | 4.296113 | 4.181478 | 1.052664 | 202.897691 | 0.238987377 | 0.309827886 | 0.756691844 | 0.998865404 | 7.626726 | 7.701522 | 7.687684 | 7.614215 | 7.679936 | 7.670059 | 246 | 196 | 183 | 193 |
| 145783 | NR_015419 | LOC145783 | uncharacterized LOC145783 | lncRNA | . | . | . | | . | 1.583284 | 1.473186 | 1.323780 | 1.748016 | 3.684368 | 3.405337 | 2.970525 | 3.996630 | -1.008685 | 144.379481 | 0.282145636 | -0.04421728 | 0.964731206 | 0.998865404 | 7.180509 | 7.166903 | 7.190377 | 7.158004 | 7.119045 | 7.220382 | 174 | 150 | 108 | 153 |
| 145788 | NM_00119878 | C15orf65 | chromosome 15 open reading frame 65 | protein_coding | NP_001185713.1 | 44654 | . | | . | 1.006511 | 0.958770 | 0.452246 | 0.760211 | 2.342193 | 2.216241 | 1.014828 | 1.738132 | -1.604440 | 22.162201 | 0.67039017 | -1.01742221 | 0.308952623 | NA | 4.772565 | 4.086744 | 4.498896 | 4.494762 | 4.435878 | 4.471521 | 32 | 29 | 11 | 20 |
| 145853 | NM_00114393 | C15orf61 | chromosome 15 open reading frame 61 | protein_coding | NP_001137408.1 | 34453 | . | | . | 0.729867 | 0.956856 | 0.542247 | 1.101666 | 1.698431 | 2.211815 | 1.216787 | 2.518828 | -1.039707 | 109.383651 | 0.364609451 | -0.15407452 | 0.87755098 | 0.998865404 | 6.802857 | 6.743029 | 6.726184 | 6.812405 | 6.647125 | 6.857433 | 112 | 136 | 62 | 134 |
| 145864 | NM_00130795 | HAPLN3 | hyaluronan and proteoglycan link protein 3 | protein_coding | NP_001294881.1;NP_839946.1;XP_011519563.1;XP_016877423.1;XP_01 | 21446 | . | | . | 48.128396 | 46.588595 | 26.421689 | 33.208108 | 111.996749 | 107.691633 | 59.289504 | 75.926387 | -1.607134 | 2446.443769 | 0.129181935 | -5.29865244 | 1.1666E-07 | 5.70364E-05 | 11.558627 | 10.873795 | 11.430312 | 11.401062 | 10.959183 | 11.103230 | 3533 | 3199 | 1477 | 1926 |
| 145957 | NM_138573,XI | NRG4 | neuregulin 4 | protein_coding | NP_612640.1;XP_016877426.1;XP_016877427.1;XP_016877428.1;XP_01 | 29862 | 610894 | . | . | 0.025384 | 0.053144 | 0.054753 | 0.014606 | 0.059068 | 0.122843 | 0.122865 | 0.033395 | -1.484232 | 12.435279 | 0.874530454 | -0.65145387 | 0.514753547 | NA | 3.892781 | 3.324312 | 3.642508 | 3.658643 | 3.635870 | 3.622842 | 15 | 18 | 10 | 8 |
| 145978 | NR_026869 | LINC00052 | long intergenic non-protein coding RNA 52 | lncRNA | . | 26455 | . | | . | 0.156943 | 0.400996 | 0.361059 | 0.078005 | 0.365213 | 0.926920 | 0.810206 | 0.178348 | -1.260353 | 14.688588 | 0.89875115 | -0.37143467 | 0.710313808 | NA | 4.032166 | 3.702263 | 3.839953 | 3.900314 | 3.888276 | 3.822459 | 11 | 25 | 18 | 5 |
| 145989 | gene-FSCN1P1 | FSCN1P1 | fascin actin-bundling protein 1 pseudogene 1 | pseudogene | . | 49669 | . | | . | 0.013156 | 0.006921 | 0.026103 | 0.022734 | 0.030613 | 0.015999 | 0.058575 | 0.051978 | 1.659343 | 1.774228 | 2.21400588 | 0.329995761 | 0.741403165 | NA | 0.413393 | 1.148375 | 0.817239 | 0.813367 | 0.819776 | 0.819065 | 2 | 1 | 2 | 2 |
| 146050 | NM_00137208 | ZSCAN29 | zinc finger and SCAN domain containing 29 | protein_coding | NP_001359009.1;NP_689668.3;XP_016877438.1 | 26673 | . | | . | 2.858342 | 2.728303 | 3.087054 | 2.751663 | 6.651478 | 6.306592 | 6.927260 | 6.291351 | -1.009136 | 491.428051 | 0.178729765 | -0.07341343 | 0.941477133 | 0.998865404 | 8.946893 | 8.934755 | 8.964021 | 8.919743 | 8.997088 | 8.872257 | 588 | 514 | 468 | 413 |
| 146053 | gene-RPS3AP4 | RPS3AP47 | RPS3A pseudogene 47 | pseudogene | . | 36744 | . | | . | 0.807592 | 0.767608 | 0.719625 | 0.865145 | 1.879299 | 1.774359 | 1.614817 | 1.978052 | -1.019772 | 21.727716 | 0.659198515 | -0.04284894 | 0.965821953 | NA | 4.455972 | 4.426810 | 4.445973 | 4.440515 | 4.433415 | 4.449450 | 26 | 23 | 17 | 22 |
| 146057 | NM_173500,XI | TTBK2 | tau tubulin kinase 2 | protein_coding | NP_775771.3;XP_005254228.1;XP_005254230.1;XP_006720465.1;XP_00 | 19141 | 611695 | . | . | 1.090006 | 0.966583 | 0.764008 | 0.847912 | 2.536487 | 2.234299 | 1.714411 | 1.938649 | -1.151342 | 299.186207 | 0.201878641 | -1.00712119 | 0.313876541 | 0.998865404 | 8.322981 | 8.119661 | 8.278842 | 8.244501 | 8.176185 | 8.196427 | 380 | 335 | 237 | 266 |

| | | | | | | | | | | | | | | | | | | | | | | | | | | | | | | | | | | |
|---|---|---|---|---|---|---|---|---|---|---|---|---|---|---|---|---|---|---|---|---|---|---|---|---|---|---|---|---|---|---|---|---|---|---|
| 146059 | NM_138477,X | CDAN1 | codanin 1 | protein_coding | NP_612486.2;XP_005254233.1;XP_011519572.1;XP_011519573.1;XP_01 | 1713 | 607465 | . | . | . | 3.721540 | 3.902049 | 4.637544 | 4.237716 | 8.660178 | 9.019762 | 10.406515 | 9.689035 | 1.146668 | 595.599065 | 0.159962699 | 1.234336608 | 0.2170775 | 0.982008277 | 9.115968 | 9.313660 | 9.159336 | 9.179714 | 9.289795 | 9.229956 | 629 | 608 | 578 | 570 |
| 146110 | gene-RPL7P47 | RPL7P47 | ribosomal protein L7 pseudogene 47 | pseudogene | . | 35913 | . | . | . | . | 0.013533 | 0.075494 | 0.018066 | 0.078736 | 0.031491 | 0.174508 | 0.040541 | 0.180020 | 1.212988 | 1.501310 | 2.393488391 | 0.116384722 | 0.907347643 | NA | 0.444631 | 0.715150 | 0.579258 | 0.584004 | 0.580748 | 0.585221 | 1 | 2 | 1 | 2 |
| 146198 | NM_00130520 | ZFP90 | ZFP90 zinc finger protein | protein_coding | NP_001292132.1;NP_001292133.1;NP_001292135.1;NP_001292136.1;NI | 23329 | 609451 | . | . | . | 2.501572 | 2.199684 | 2.689034 | 2.510667 | 5.821260 | 5.084667 | 6.034115 | 5.740341 | 1.081424 | 355.018865 | 0.1913039 | 0.590329503 | 0.554969763 | 0.998865404 | 8.413840 | 8.527428 | 8.476540 | 8.418135 | 8.510457 | 8.474296 | 413 | 349 | 331 | 335 |
| 146206 | NM_00101383 | CARMIL2 | capping protein regulator and myosin 1 linker 2 | protein_coding | NP_001013860.1;NP_001303955.1;XP_011521176.1;XP_011521177.2;XP | 27089 | 610859 | . | . | . | 0.075691 | 0.044922 | 0.035440 | 0.049146 | 0.176137 | 0.103840 | 0.079526 | 0.112365 | -1.345430 | 7.748375 | 1.103070971 | -0.38806825 | 0.697965529 | NA | 3.151937 | 2.724273 | 2.976462 | 2.954339 | 2.951465 | 2.958893 | 13 | 7 | 5 | 7 |
| 146223 | NM_178818,N | CMTM4 | CKLF like MARVEL transmembrane domain containing 4 | protein_coding | NP_848933.1;NP_852662.1;XP_011521184.1;XP_016878443.1 | 19175 | 607887 | . | . | . | 2.805122 | 2.933598 | 2.571812 | 2.658649 | 6.527634 | 6.781143 | 5.771072 | 6.078684 | -1.052420 | 659.548309 | 0.154395518 | -0.47741024 | 0.633070024 | 0.998865404 | 9.401864 | 9.327856 | 9.384349 | 9.380645 | 9.321429 | 9.373495 | 781 | 728 | 537 | 626 |
| 146330 | NM_153350,X | FBXL16 | F-box and leucine rich repeat protein 16 | protein_coding | NP_699181.2;XP_024305928.1 | 14150 | 609082 | . | . | . | 0.256764 | 0.379035 | 0.275668 | 0.408413 | 0.597500 | 0.876156 | 0.618591 | 0.933788 | 1.058330 | 41.722695 | 0.502730743 | 0.162690373 | 0.870762227 | 0.998865404 | 5.343257 | 5.421211 | 5.343390 | 5.403354 | 5.351274 | 5.413629 | 38 | 52 | 30 | 48 |
| 146434 | NM_152457 | ZNF597 | zinc finger protein 597 | protein_coding | NP_689670.1 | 26573 | 614685 | . | . | . | 0.465222 | 0.397479 | 0.403785 | 0.324868 | 1.082590 | 0.918792 | 0.906082 | 0.742773 | -1.201602 | 68.452342 | 0.387888478 | -0.68308034 | 0.494556077 | 0.998865404 | 6.222498 | 5.959598 | 6.139082 | 6.097157 | 6.098529 | 6.051850 | 93 | 74 | 60 | 52 |
| 146481 | NR_026980 | FRG2DP | FSHD region gene 2 family member D, pseudogene | transcribed_ps | . | 27093 | . | . | . | . | 2.208491 | 2.758569 | 3.732871 | 2.327335 | 5.139248 | 6.376555 | 8.376453 | 5.321176 | 1.204611 | 156.783618 | 0.29898747 | 0.898255693 | 0.369049248 | 0.998865404 | 7.151469 | 7.421204 | 7.199791 | 7.283120 | 7.409665 | 7.214189 | 146 | 170 | 184 | 123 |
| 146512 | NR_026674 | FLJ30679 | uncharacterized protein FLJ30679 | lncRNA | . | . | . | . | . | . | 0.057067 | 0.028337 | 0.068798 | 0.096877 | 0.132798 | 0.065503 | 0.154382 | 0.221498 | 1.771459 | 4.298900 | 1.471491047 | 0.560613802 | 0.575060841 | NA | 1.631267 | 2.459419 | 2.094189 | 2.082410 | 2.095305 | 2.103449 | 5 | 2 | 4 | 6 |
| 146540 | NM_152458,X | ZNF785 | zinc finger protein 785 | protein_coding | NP_689671.2;XP_011544054.1;XP_011544055.1;XP_016878456.1;XP_02 | 26496 | . | . | . | . | 3.809096 | 3.820137 | 4.823466 | 4.104462 | 8.863920 | 8.830418 | 10.823719 | 9.384366 | 1.155784 | 381.413583 | 0.188923154 | 1.105588517 | 0.268904662 | 0.998865404 | 8.466649 | 8.676175 | 8.526624 | 8.530665 | 8.659751 | 8.564919 | 406 | 383 | 384 | 353 |
| 146542 | NM_00102468 | ZNF688 | zinc finger protein 688 | protein_coding | NP_001019854.1;NP_660314.1;XP_005255196.1;XP_005255197.1;XP_02 | 30489 | . | . | . | . | 3.988798 | 5.050096 | 4.739795 | 5.981993 | 9.282094 | 11.673523 | 10.635960 | 13.677115 | 1.240572 | 253.851529 | 0.232971662 | 1.334950869 | 0.181892427 | 0.953687323 | 7.824942 | 8.134199 | 7.871413 | 7.975573 | 7.967932 | 8.094246 | 233 | 271 | 215 | 295 |
| 146547 | NM_00125829 | PRSS36 | serine protease 36 | protein_coding | NP_001245219.1;NP_001245220.1;NP_775773.2;XP_011544060.1;XP_01 | 26906 | 610560 | . | . | . | 0.157563 | 0.098207 | 0.079874 | 0.052825 | 0.366657 | 0.227010 | 0.179236 | 0.120777 | -2.165375 | 8.813945 | 1.054495344 | -1.05701424 | 0.29050509 | NA | 3.590318 | 2.481107 | 3.172185 | 3.155292 | 3.140820 | 3.128944 | 16 | 11 | 6 | 4 |
| 146562 | NM_139170,X | C16orf71 | chromosome 16 open reading frame 71 | protein_coding | NP_631909.2;XP_005255201.1;XP_016878464.1;XP_016878465.1 | 25081 | . | . | . | . | 0.130077 | 0.163871 | 0.086345 | 0.045538 | 0.302696 | 0.378795 | 0.193755 | 0.104117 | -2.261583 | 9.089358 | 1.039529996 | -1.13256287 | 0.25739787 | NA | 3.654832 | 2.481107 | 3.202215 | 3.213719 | 3.182723 | 3.170818 | 13 | 15 | 6 | 4 |
| 146664 | NM_00119917 | MGAT5B | alpha-1,6-mannosylglycoprotein 6-beta-N-acetylglucosaminyltrans | protein_coding | NP_001186101.1;NP_653278.2;NP_945193.1;XP_006721770.1;XP_01152 | 24140 | 612441 | . | . | . | 1.790688 | 1.892863 | 1.628945 | 1.804565 | 4.167003 | 4.375438 | 3.655303 | 4.125922 | -1.048143 | 246.717332 | 0.218158163 | -0.31094727 | 0.75584071 | 0.998865404 | 7.980542 | 7.912076 | 7.948273 | 7.968576 | 7.913061 | 7.956627 | 285 | 278 | 200 | 236 |
| 146691 | NM_00103355 | TOM1L2 | target of myb1 like 2 membrane trafficking protein | protein_coding | NP_001028723.1;NP_001076437.1;NP_001275715.1;NP_001275716.1;NI | 11984 | 615519 | . | . | . | 5.672594 | 5.555065 | 5.145319 | 6.241073 | 13.200361 | 12.840782 | 11.545946 | 14.269473 | -1.000183 | 1011.092695 | 0.147134143 | -0.00179212 | 0.998570093 | 0.999862634 | 9.982060 | 9.981339 | 9.992487 | 9.969125 | 9.911018 | 10.045780 | 1183 | 1074 | 798 | 1035 |
| 146705 | NM_00136376 | TEPSIN | TEPSIN adaptor related protein complex 4 accessory protein | protein_coding | NP_001350693.1;NP_653280.1;XP_005257123.1;XP_006721772.1;XP_00 | 26458 | . | . | . | . | 8.909755 | 9.006631 | 10.657508 | 8.412937 | 20.733368 | 20.819236 | 23.915142 | 19.235179 | 1.175774 | 894.615072 | 0.169824039 | 1.37560264 | 0.168944689 | 0.945638095 | 9.683230 | 9.917515 | 9.729561 | 9.748641 | 9.968218 | 9.729806 | 934 | 899 | 977 | 760 |
| 146712 | NM_00100990 | B3GNTL1 | UDP-GlcNAc:betaGal beta-1,3-N-acetylglucosaminyltransferase lik | protein_coding | NP_001009905.1;NP_001307671.1;NP_001307672.1;XP_006722335.1;XP | 21727 | 615337 | . | . | . | 0.960394 | 1.122808 | 0.748519 | 1.537528 | 2.234878 | 2.595420 | 1.679655 | 3.515375 | -1.102017 | 82.117467 | 0.362508934 | -0.38660291 | 0.699050205 | 0.998865404 | 6.429096 | 6.286622 | 6.363043 | 6.381222 | 6.301204 | 6.389311 | 96 | 96 | 56 | 86 |
| 146754 | NM_00130327 | DNAH2 | dynein axonemal heavy chain 2 | protein_coding | NP_001290199.1;NP_065928.2;XP_011521965.1;XP_011521969.1;XP_01 | 2948 | 603333 | . | . | . | 0.009485 | 0.022559 | 0.002621 | 0.003211 | 0.022071 | 0.052147 | 0.005881 | 0.007343 | -3.679980 | 5.191218 | 1.427763039 | -1.31653347 | 0.187995052 | NA | 3.029593 | 1.148375 | 2.386814 | 2.413713 | 2.374063 | 2.373032 | 6 | 12 | 2 | 2 |
| 146760 | NM_178568 | RTN4RL1 | reticulon 4 receptor like 1 | protein_coding | NP_848663.1 | 21329 | 610461 | . | . | . | 0.280861 | 0.347078 | 0.324338 | 0.194549 | 0.653574 | 0.802287 | 0.727805 | 0.444813 | -1.217872 | 32.795392 | 0.558783576 | -0.50889598 | 0.610825136 | 0.998865404 | 5.169572 | 4.887505 | 5.030524 | 5.061465 | 5.048562 | 4.991578 | 37 | 43 | 32 | 21 |
| 146857 | NM_144682,X | SLFN13 | schlafen family member 13 | protein_coding | NP_653283.3;XP_005257979.1;XP_011522685.1;XP_016879721.1;XP_01 | 26481 | 614957 | . | . | . | 0.011732 | 0.007031 | 0.010966 | 0.019599 | 0.027301 | 0.016252 | 0.024609 | 0.044811 | 1.647268 | 3.527629 | 1.605996642 | 0.448366454 | 0.653888748 | NA | 1.413393 | 2.134756 | 1.810493 | 1.802817 | 1.810399 | 1.818922 | 4 | 2 | 3 | 5 |
| 146880 | NR_027487 | ARHGAP27P1 | Rho GTPase activating protein 27 pseudogene 1 | transcribed_ps | . | 28630 | . | . | . | . | 1.170475 | 1.477853 | 1.195503 | 1.444926 | 2.723745 | 3.416124 | 2.682673 | 3.303650 | -1.021321 | 118.804321 | 0.306022975 | -0.09945674 | 0.920775636 | 0.998865404 | 6.908510 | 6.876196 | 6.855073 | 6.932260 | 6.857102 | 6.916416 | 123 | 144 | 93 | 120 |
| 146909 | NM_00126457 | KIF18B | kinesin family member 18B | protein_coding | NP_001251503.1;NP_001252506.1;XP_011522687.1;XP_011522688.1;XP | 27102 | 614570 | . | . | . | 13.461708 | 15.703326 | 13.763319 | 11.587453 | 31.325947 | 36.298939 | 30.884491 | 26.493332 | -1.181213 | 1706.456142 | 0.138299489 | -1.73730931 | 0.082332589 | 0.78412713 | 10.851851 | 10.611745 | 10.739734 | 10.851195 | 10.727416 | 10.601920 | 1980 | 2133 | 1472 | 1347 |
| 146923 | NM_00132138 | RUNDC1 | RUN domain containing 1 | protein_coding | NP_001308310.2;NP_775102.3;XP_005257135.1;XP_005257137.1 | 25418 | . | . | . | . | 4.016133 | 4.076941 | 4.693159 | 4.381547 | 9.345706 | 9.424032 | 10.531312 | 10.017886 | 1.118918 | 668.998499 | 0.153534704 | 1.05581166 | 0.29105427 | 0.998865404 | 9.302574 | 9.464598 | 9.322788 | 9.367520 | 9.432448 | 9.411063 | 703 | 704 | 624 | 652 |
| 146956 | NM_00116613 | EME1 | essential meiotic structure-specific endonuclease 1 | protein_coding | NP_001159603.1;NP_689676.2;XP_005257138.1;XP_011522697.1;XP_01 | 24965 | 610885 | . | . | . | 10.651876 | 12.261106 | 11.583086 | 11.689473 | 24.787352 | 28.342095 | 25.992108 | 26.726592 | -1.010774 | 934.473620 | 0.142762083 | -0.10829269 | 0.913763518 | 0.998865404 | 9.875876 | 9.860101 | 9.830313 | 9.911115 | 9.857258 | 9.870008 | 1022 | 1070 | 801 | 879 |
| 147007 | NM_152464 | TMEM199 | transmembrane protein 199 | protein_coding | NP_689677.1 | 18085 | 616815 | . | . | . | 12.177614 | 12.479924 | 13.033716 | 13.324325 | 28.337811 | 28.847904 | 29.247278 | 30.464487 | 1.054233 | 1224.707155 | 0.129629297 | 0.587783398 | 0.556677689 | 0.998865404 | 10.219656 | 10.295782 | 10.231709 | 10.243838 | 10.273357 | 10.281829 | 1362 | 1297 | 1087 | 1185 |
| 147011 | NM_00130494 | PROCA1 | protein interacting with cyclin A1 | protein_coding | NP_001291878.1;NP_001291880.1;NP_001291881.1;NP_001291882.1;NI | 28600 | 617376 | . | . | . | 0.725886 | 0.853209 | 0.870335 | 0.642452 | 1.689166 | 1.972233 | 1.953007 | 1.468889 | -1.019264 | 43.175744 | 0.476754658 | -0.05774025 | 0.953955535 | 0.998865404 | 5.445495 | 5.418678 | 5.421538 | 5.443009 | 5.445381 | 5.412740 | 47 | 50 | 41 | 36 |
| 147015 | NM_144683 | DHRS13 | dehydrogenase/reductase 13 | protein_coding | NP_653284.2 | 28326 | 616157 | . | . | . | 11.152884 | 11.450113 | 12.726103 | 12.602526 | 25.953218 | 26.467449 | 28.557009 | 28.814177 | 1.104980 | 720.265334 | 0.148796798 | 0.967897678 | 0.333095474 | 0.998865404 | 9.418567 | 9.562608 | 9.450077 | 9.462345 | 9.531893 | 9.518410 | 781 | 745 | 664 | 702 |
| 147040 | NM_00100291 | KCTD11 | potassium channel tetramerization domain containing 11 | protein_coding | NP_001002914.1;NP_001350571.1 | 21302 | 609848 | . | . | . | 1.227635 | 0.578669 | 1.225567 | 0.536967 | 2.856757 | 1.337621 | 2.750139 | 1.227712 | -1.035549 | 81.598601 | 0.45428835 | -0.11093236 | 0.911669986 | 0.998865404 | 6.373128 | 6.327456 | 6.426465 | 6.231026 | 6.420148 | 6.216862 | 129 | 58 | 97 | 46 |
| 147081 | NR_026680,N | LINC02210 | long intergenic non-protein coding RNA 2210 | lncRNA | . | 26327 | . | . | . | . | 3.528212 | 2.963670 | 3.152710 | 4.025385 | 8.210293 | 6.850654 | 7.074590 | 9.203563 | 1.029100 | 286.215400 | 0.205376942 | 0.201499531 | 0.840307996 | 0.998865404 | 8.140066 | 8.181551 | 8.172635 | 8.132999 | 8.158786 | 8.177439 | 337 | 293 | 248 | 277 |
| 147138 | NM_152468,X | TMC8 | transmembrane channel like 8 | protein_coding | NP_689681.2;XP_016879731.1;XP_016879732.1;XP_016879733.1;XP_02 | 20474 | 605829 | . | . | . | 0.264673 | 0.378942 | 0.286377 | 0.102338 | 0.615906 | 0.875940 | 0.642620 | 0.233984 | -1.646898 | 31.890581 | 0.592786922 | -1.21418227 | 0.22467813 | 0.982008277 | 5.308880 | 4.593431 | 4.998539 | 5.044123 | 4.995990 | 4.920957 | 39 | 49 | 29 | 14 |
| 147150 | gene-UPF3AP2 | UPF3AP2 | UPF3A pseudogene 2 | pseudogene | . | 30567 | . | . | . | . | 0.325534 | 0.319168 | 0.338148 | 0.169745 | 0.757530 | 0.737770 | 0.758795 | 0.388101 | -1.267843 | 19.656861 | 0.713902223 | -0.47958466 | 0.631522756 | NA | 4.455972 | 4.118228 | 4.309030 | 4.303437 | 4.309883 | 4.262675 | 26 | 23 | 20 | 11 |
| 147166 | NM_00103733 | TRIM16L | tripartite motif containing 16 like | protein_coding | NP_001032407.1;NP_001340148.1;NP_001340149.1;NP_001340150.1;NI | 32670 | . | . | . | . | 18.600987 | 20.903033 | 29.681925 | 28.586565 | 43.285259 | 48.318297 | 66.605374 | 65.359784 | 1.449773 | 1545.722179 | 0.128861591 | 4.158162138 | 3.20818E-05 | 0.005089948 | 10.301418 | 10.837244 | 10.399056 | 10.460649 | 10.750291 | 10.684556 | 1393 | 1418 | 1656 | 1645 |
| 147172 | gene-LRRC37B | LRRC37BP1 | leucine rich repeat containing 37B pseudogene 1 | pseudogene | . | 25390 | . | . | . | . | 0.406550 | 0.790879 | 0.491681 | 0.231645 | 0.946058 | 1.828152 | 1.103319 | 0.529628 | -1.571470 | 12.233166 | 0.911705848 | -0.71526861 | 0.474443083 | NA | 3.901250 | 3.251698 | 3.603172 | 3.643318 | 3.613214 | 3.585907 | 12 | 21 | 11 | 6 |
| 147179 | NM_133264,X | WIPF2 | WAS/WASL interacting protein family member 2 | protein_coding | NP_573571.1;XP_005257140.1;XP_005257141.1;XP_011522714.1;XP_01 | 30923 | 609692 | . | . | . | 3.400387 | 3.215033 | 4.200469 | 4.402697 | 7.912842 | 7.431693 | 9.425730 | 10.066241 | 1.285237 | 886.394666 | 0.141980184 | 2.549892991 | 0.010775599 | 0.334237281 | 9.599433 | 9.961519 | 9.713234 | 9.671925 | 9.859016 | 9.886726 | 920 | 812 | 851 | 952 |
| 147184 | NR_160886,N | TMEM99 | transmembrane protein 99 (putative) | lncRNA | . | 28305 | . | . | . | . | 9.106609 | 8.241158 | 9.200462 | 8.599485 | 21.191454 | 19.049805 | 20.645568 | 19.661700 | 1.011240 | 560.880876 | 0.165368552 | 0.097510902 | 0.922320677 | 0.998865404 | 9.123155 | 9.139897 | 9.162253 | 9.090916 | 9.158762 | 9.109940 | 677 | 569 | 509 | 509 |
| 147226 | gene-UPF3AP1 | UPF3AP1 | UPF3A pseudogene 1 | pseudogene | . | 30568 | . | . | . | . | 0.583880 | 0.500278 | 0.326834 | 0.455967 | 1.358713 | 1.156416 | 0.733406 | 1.042515 | -1.387266 | 31.523501 | 0.561559645 | -0.84095106 | 0.40037535 | 0.998865404 | 5.195961 | 4.721998 | 5.015616 | 4.988922 | 4.943380 | 4.979144 | 46 | 36 | 19 | 29 |
| 147339 | NM_00100823 | C18orf25 | chromosome 18 open reading frame 25 | protein_coding | NP_001008240.1;NP_659492.1;XP_011524124.1;XP_016881044.1 | 28172 | . | . | . | . | 4.274722 | 4.242721 | 4.157790 | 3.501177 | 9.947453 | 9.807241 | 9.329962 | 8.005026 | -1.144649 | 667.322784 | 0.161700108 | -1.2053489 | 0.228068641 | 0.982008277 | 9.476060 | 9.281893 | 9.432161 | 9.418444 | 9.392670 | 9.271128 | 828 | 761 | 591 | 530 |
| 147407 | NM_00103417 | SLC25A52 | solute carrier family 25 member 52 | protein_coding | NP_001029344.4 | 23324 | 616153 | . | . | . | 0.081122 | 0.084707 | 0.089782 | 0.044183 | 0.188773 | 0.195804 | 0.201467 | 0.101018 | -1.070014 | 3.478495 | 1.609933276 | -0.06064235 | 0.951644048 | NA | 1.844134 | 1.751299 | 1.796180 | 1.797278 | 1.801166 | 1.790082 | 4 | 4 | 4 | 2 |
| 147463 | NM_00130823 | ANKRD29 | ankyrin repeat domain 29 | protein_coding | NP_001295167.1;NP_775776.2;XP_011524132.1;XP_011524133.1;XP_01 | 27110 | . | . | . | . | 0.246013 | 0.309058 | 0.225973 | 0.211167 | 0.572483 | 0.714402 | 0.507077 | 0.482809 | -1.257590 | 45.653619 | 0.467328206 | -0.70755872 | 0.479219338 | 0.998865404 | 5.668867 | 5.337457 | 5.507380 | 5.556889 | 5.497042 | 5.489439 | 51 | 62 | 36 | 37 |
| 147495 | NM_153000,X | APCDD1 | APC down-regulated 1 | protein_coding | NP_694545.1;XP_011523905.1 | 15718 | 607479 | . | . | . | 1.358202 | 1.258293 | 1.247516 | 1.273198 | 3.160590 | 2.908600 | 2.799390 | 2.911016 | -1.036104 | 160.653561 | 0.259464453 | -0.1972086 | 0.843664305 | 0.998865404 | 7.352931 | 7.302242 | 7.347738 | 7.322335 | 7.328733 | 7.312879 | 194 | 171 | 140 | 145 |
| 147525 | NR_026849 | LINC00526 | long intergenic non-protein coding RNA 526 | lncRNA | . | 28278 | . | . | . | . | 0.835929 | 1.051766 | 1.093227 | 1.121387 | 1.945241 | 2.431204 | 2.453169 | 2.563919 | 1.159633 | 42.252593 | 0.481035192 | 0.444183846 | 0.656909671 | 0.998865404 | 5.290842 | 5.503280 | 5.367090 | 5.402253 | 5.406743 | 5.410721 | 40 | 47 | 39 | 43 |
| 147645 | NM_00116392 | VSIG10L | V-set and immunoglobulin domain containing 10 like | protein_coding | NP_001157394.1 | 27111 | 617740 | . | . | . | 1.759325 | 1.828094 | 2.118576 | 1.989088 | 4.094021 | 4.225721 | 4.754024 | 4.547814 | 1.130886 | 220.808237 | 0.227215237 | 0.780994097 | 0.434805967 | 0.998865404 | 7.695071 | 7.872761 | 7.747657 | 7.761922 | 7.829614 | 7.795969 | 235 | 227 | 211 | 212 |
| 147650 | NM_00131697 | SPACA6 | sperm acrosome associated 6 | protein_coding | NP_001303901.1;NP_001303923.1;XP_016881788.1;XP_016881789.1;XP | 27113 | . | . | . | . | 1.972578 | 2.260133 | 3.257369 | 2.070192 | 4.590272 | 5.224400 | 7.309443 | 4.733247 | 1.337193 | 149.263325 | 0.302839087 | 1.384258776 | 0.166279232 | 0.944092297 | 6.996189 | 7.416707 | 7.117579 | 7.176910 | 7.355053 | 7.156020 | 135 | 149 | 184 | 122 |
| 147657 | NM_00129762 | ZNF480 | zinc finger protein 480 | protein_coding | NP_001284553.1;NP_001284554.1;NP_653285.2;XP_011524767.1 | 23305 | 613910 | . | . | . | 1.953488 | 1.547783 | 1.655191 | 1.693462 | 4.545848 | 3.577770 | 3.714198 | 3.871903 | -1.053484 | 255.586426 | 0.224065043 | -0.33547788 | 0.737264631 | 0.998865404 | 8.034389 | 7.959986 | 8.067974 | 7.947615 | 7.982262 | 7.984525 | 337 | 250 | 216 | 234 |
| 147687 | NM_00129773 | ZNF417 | zinc finger protein 417 | protein_coding | NP_001284663.1;NP_689688.2;XP_011524775.1 | 20646 | . | . | . | . | 1.511057 | 1.791490 | 1.553620 | 1.521172 | 3.516290 | 4.141110 | 3.486279 | 3.477977 | -1.107932 | 255.636681 | 0.218280546 | -0.67742934 | 0.498133587 | 0.998865404 | 8.070171 | 7.921924 | 7.982373 | 8.061553 | 7.978336 | 7.963783 | 286 | 312 | 214 | 224 |
| 147694 | NM_00117277 | ZNF548 | zinc finger protein 548 | protein_coding | NP_001166244.1;NP_690873.2;XP_016881805.1 | 26561 | . | . | . | . | 2.398996 | 2.187832 | 2.361243 | 1.870878 | 5.582561 | 5.057272 | 5.298561 | 4.277539 | -1.152455 | 356.341296 | 0.198952916 | -1.02894096 | 0.30350742 | 0.998865404 | 8.575331 | 8.371723 | 8.518632 | 8.510912 | 8.494907 | 8.368999 | 442 | 409 | 321 | 275 |
| 147699 | NM_00108040 | PPM1N | protein phosphatase, Mg2+/Mn2+ dependent 1N (putative) | protein_coding | NP_001073870.1 | 26845 | . | . | . | . | 1.825322 | 1.576892 | 1.328022 | 1.814057 | 4.247598 | 3.645057 | 2.980042 | 4.147627 | -1.138374 | 119.658318 | 0.304058948 | -0.61492922 | 0.538601462 | 0.998865404 | 6.993517 | 6.805945 | 6.954449 | 6.893841 | 6.846204 | 6.913629 | 159 | 126 | 87 | 116 |
| 147700 | NM_177417,X | KLC3 | kinesin light chain 3 | protein_coding | NP_803136.2;XP_024307137.1;XP_024307138.1;XP_024307139.1 | 20717 | 601334 | . | . | . | 0.562629 | 0.292650 | 0.388627 | 0.548855 | 1.309260 | 0.676472 | 0.872067 | 1.254892 | 1.014711 | 29.528981 | 0.595534441 | 0.035378553 | 0.971777886 | 0.998865404 | 4.872825 | 4.895208 | 4.917187 | 4.843199 | 4.869459 | 4.895582 | 44 | 22 | 23 | 31 |
| 147727 | NR_024333 | ILF3-DT | ILF3 divergent transcript | lncRNA | . | 27115 | . | . | . | . | 3.125676 | 3.513978 | 3.694805 | 3.502920 | 7.273577 | 8.122719 | 8.291036 | 8.009009 | 1.067597 | 217.277538 | 0.230458873 | 0.409476548 | 0.682189974 | 0.998865404 | 7.715499 | 7.809752 | 7.720892 | 7.771288 | 7.791112 | 7.762821 | 229 | 239 | 201 | 204 |
| 147804 | NR_003148 | TPM3P9 | tropomyosin 3 pseudogene 9 | transcribed_ps | . | 44142 | . | . | . | . | 4.881268 | 5.019325 | 5.391665 | 4.937892 | 11.358911 | 11.602395 | 12.098740 | 11.289904 | 1.028103 | 462.559465 | 0.173606201 | 0.230319202 | 0.81784374 | 0.998865404 | 8.833178 | 8.873530 | 8.837991 | 8.850308 | 8.891504 | 8.830442 | 520 | 497 | 428 | 418 |
| 147807 | NM_153219,X | ZNF524 | zinc finger protein 524 | protein_coding | NP_694951.1;XP_011524789.1 | 28322 | . | . | . | . | 2.837245 | 3.601857 | 2.970127 | 4.267846 | 6.602388 | 8.325855 | 6.664880 | 9.757923 | 1.087270 | 129.404112 | 0.304197707 | 0.396814652 | 0.691504145 | 0.998865404 | 6.955372 | 7.073682 | 6.963591 | 7.027811 | 6.961913 | 7.086531 | 130 | 146 | 97 | 148 |
| 147808 | NM_203374 | ZNF784 | zinc finger protein 784 | protein_coding | NP_976308.1 | 33111 | . | . | . | . | 0.888571 | 0.651380 | 1.298325 | 0.891445 | 2.067740 | 1.505694 | 2.913403 | 2.038183 | 1.395183 | 58.297598 | 0.432059251 | 1.112010057 | 0.266133826 | 0.998274218 | 5.603263 | 6.087090 | 5.846140 | 5.787140 | 5.927136 | 5.842834 | 65 | 44 | 70 | 52 |
| 147837 | NM_00136360 | ZNF563 | zinc finger protein 563 | protein_coding | NP_001350537.1;NP_660319.1;XP_005259807.1;XP_006722713.1;XP_00 | 30498 | . | . | . | . | 0.228553 | 0.259514 | 0.273810 | 0.438543 | 0.531851 | 0.599879 | 0.614421 | 1.002675 | 1.223527 | 22.922225 | 0.65789131 | 0.442391509 | 0.658205922 | NA | 4.367696 | 4.655341 | 4.500854 | 4.506806 | 4.495236 | 4.550363 | 23 | 23 | 16 | 30 |
| 147841 | NM_00131703 | SPC24 | SPC24 component of NDC80 kinetochore complex | protein_coding | NP_001303960.1;NP_001303961.1;NP_001303962.1;NP_872319.1;XP_00 | 26913 | 609394 | . | . | . | 9.262169 | 10.787766 | 7.413212 | 8.296591 | 21.553452 | 24.936405 | 16.635035 | 18.969170 | -1.257912 | 533.543607 | 0.164620743 | -2.01087194 | 0.044338985 | 0.645513836 | 9.215656 | 8.884283 | 9.126074 | 9.130953 | 8.960155 | 9.001054 | 682 | 644 | 397 | 458 |
| 147872 | NM_144688,X | KASH5 | KASH domain containing 5 | protein_coding | NP_653289.3;XP_005258606.1;XP_011524790.1;XP_011524791.1;XP_01 | 26520 | 618125 | . | . | . | 0.358957 | 0.186381 | 0.379775 | 0.463576 | 0.835308 | 0.430828 | 0.852205 | 1.059911 | 1.234066 | 28.460368 | 0.580771573 | 0.522442178 | 0.601362496 | 0.998865404 | 4.671258 | 4.974591 | 4.824136 | 4.804751 | 4.828075 | 4.849462 | 32 | 25 | 25 | 32 |
| 147906 | NM_00130104 | DACT3 | dishevelled binding antagonist of beta catenin 3 | protein_coding | NP_001287975.1;NP_659493.2;XP_011524801.1;XP_011524802.1;XP_01 | 30745 | 611112 | . | . | . | 2.040167 | 2.529537 | 1.630674 | 2.261745 | 4.747551 | 5.847137 | 3.659185 | 5.171212 | -1.229822 | 191.109602 | 0.257202676 | -1.16036766 | 0.245899146 | 0.992733415 | 7.720672 | 7.420227 | 7.572674 | 7.665418 | 7.475826 | 7.579281 | 219 | 250 | 130 | 181 |
| 147912 | NM_175875 | SIX5 | SIX homeobox 5 | protein_coding | NP_787071.3 | 10891 | 600963 | . | . | . | 4.160442 | 4.404843 | 4.412393 | 4.356626 | 9.681520 | 10.181993 | 9.901283 | 9.960908 | 1.009778 | 451.765495 | 0.173189285 | 0.08105496 | 0.935398244 | 0.998865404 | 8.812427 | 8.826400 | 8.800901 | 8.831301 | 8.828753 | 8.815593 | 505 | 497 | 399 | 421 |
| 147923 | NM_00132951 | ZNF420 | zinc finger protein 420 | protein_coding | NP_001316444.1;NP_001316445.1;NP_001316446.1;NP_001316447.1;NI | 20649 | 617216 | . | . | . | 1.025077 | 1.168585 | 1.116762 | 1.176064 | 2.385396 | 2.701237 | 2.505982 | 2.688933 | 1.035668 | 116.699512 | 0.3022612 | 0.167278771 | 0.867150707 | 0.998865404 | 6.840469 | 6.892374 | 6.874454 | 6.843575 | 6.898657 | 6.842459 | 138 | 118 | 112 | 102 |
| 147929 | NM_00104247 | ZNF565 | zinc finger protein 565 | protein_coding | NP_001035939.1;NP_001353117.1;NP_001353118.1;NP_001353119.1;NI | 26726 | 614275 | . | . | . | 0.451911 | 0.829500 | 0.640897 | 0.488971 | 1.051614 | 1.917427 | 1.438154 | 1.117974 | 1.028498 | 43.879417 | 0.482551381 | 0.084008343 | 0.933049798 | 0.998865404 | 5.434941 | 5.475715 | 5.421856 | 5.474808 | 5.475679 | 5.430522 | 42 | 54 | 44 | 36 |
| 147947 | NR_003127,N | ZNF542P | zinc finger protein 542, pseudogene | transcribed_ps | . | 25393 | . | . | . | . | 5.221861 | 6.332343 | 7.359695 | 5.233079 | 12.151484 | 14.637498 | 16.514939 | 11.964812 | 1.071067 | 684.965280 | 0.18889509 | 0.524357602 | 0.60002984 | 0.998865404 | 9.369230 | 9.468826 | 9.329707 | 9.447164 | 9.552134 | 9.307775 | 696 | 775 | 725 | 547 |
| 147948 | NM_00132037 | ZNF582 | zinc finger protein 582 | protein_coding | NP_001307300.1;NP_653291.1 | 26421 | 615600 | . | . | . | 0.164329 | 0.171302 | 0.154800 | 0.146404 | 0.382401 | 0.395973 | 0.347367 | 0.334736 | -1.151868 | 15.499536 | 0.775656103 | -0.26297222 | 0.792571988 | NA | 4.052320 | 3.848815 | 3.960653 | 3.961383 | 3.953889 | 3.949205 | 19 | 18 | 13 | 13 |
| 147949 | NM_00115986 | ZNF583 | zinc finger protein 583 | protein_coding | NP_001153332.1;NP_001153333.1;NP_689691.2;XP_005258615.1;XP_01 | 26427 | . | . | . | . | 0.565970 | 0.569225 | 0.796906 | 0.443973 | 1.317035 | 1.315791 | 1.788233 | 1.015092 | 1.041919 | 62.134497 | 0.409727185 | 0.14459318 | 0.885032082 | 0.998865404 | 5.926074 | 5.987909 | 5.967957 | 5.930462 | 5.992446 | 5.923033 | 76 | 60 | 64 | 50 |
| 147965 | NM_00135167 | FAM98C | family with sequence similarity 98 member C | protein_coding | NP_001338604.1;NP_777565.3;XP_016881843.1 | 27119 | . | . | . | . | 5.203527 | 6.247364 | 4.450929 | 6.437824 | 12.108820 | 14.441059 | 9.987755 | 14.719320 | -1.061885 | 218.971910 | 0.251563308 | -0.34435722 | 0.730577656 | 0.998865404 | 7.818289 | 7.729551 | 7.741848 | 7.827307 | 7.675796 | 7.832299 | 237 | 265 | 153 | 233 |
| 147968 | NM_144691,X | CAPN12 | calpain 12 | protein_coding | NP_653292.2;XP_016881844.1;XP_016881845.1;XP_016881846.1;XP_01 | 13249 | 608839 | . | . | . | 0.019196 | 0.022952 | 0.011167 | 0.057666 | 0.044669 | 0.053054 | 0.025061 | 0.131845 | 2.045617 | 4.096365 | 1.495516147 | 0.690421152 | 0.489929379 | NA | 1.429096 | 2.459419 | 2.012929 | 2.013872 | 2.022329 | 2.030512 | 3 | 3 | 4 | 6 |
| 147991 | NM_00117277 | DPY19L3 | dpy-19 like C-mannosyltransferase 3 | protein_coding | NP_001166245.1;NP_997208.2;XP_006723108.1;XP_011524828.1;XP_01 | 27120 | 613894 | . | . | . | 6.408241 | 5.325703 | 8.155368 | 5.057166 | 14.912241 | 12.310603 | 18.300410 | 11.562609 | 1.110776 | 1236.630828 | 0.400188353 | 0.37874103 | 0.704880182 | 0.998865404 | 10.194322 | 10.346087 | 10.285666 | 10.151140 | 10.483001 | 10.077170 | 1470 | 1151 | 1428 | 901 |
| 148014 | NM_152479,X | TTC9B | tetratricopeptide repeat domain 9B | protein_coding | NP_689692.2;XP_016881857.1 | 26395 | . | . | . | . | 0.362219 | 0.218694 | 0.255455 | 0.189017 | 0.842898 | 0.505521 | 0.573234 | 0.432164 | -1.285088 | 9.508449 | 0.995061748 | -0.36366315 | 0.716109564 | NA | 3.417335 | 3.058876 | 3.269146 | 3.247728 | 3.251730 | 3.244066 | 15 | 9 | 8 | 7 |
| 148022 | NM_182919 | TICAM1 | toll like receptor adaptor molecule 1 | protein_coding | NP_891549.1 | 18348 | 607601 | . | . | . | 2.469293 | 3.437960 | 3.495945 | 3.326973 | 5.746145 | 7.946999 | 7.844799 | 7.606728 | 1.138792 | 266.433123 | 0.229955114 | 0.815393326 | 0.414847203 | 0.998865404 | 7.961273 | 8.147952 | 7.934018 | 8.095806 | 8.100075 | 8.069191 | 241 | 312 | 254 | 258 |
| 148046 | NR_027271 | CIRBP-AS1 | CIRBP antisense RNA 1 | lncRNA | . | 28588 | . | . | . | . | 0.574644 | 0.297006 | 0.756134 | 0.566590 | 1.337221 | 0.686543 | 1.696742 | 1.295441 | 1.481900 | 23.722967 | 0.659569454 | 0.860330991 | 0.389606614 | NA | 4.253961 | 4.826107 | 4.562332 | 4.509057 | 4.589676 | 4.559317 | 29 | 14 | 28 | 23 |
| 148103 | NM_00100724 | ZNF599 | zinc finger protein 599 | protein_coding | NP_001007249.1;XP_016881858.1 | 26408 | . | . | . | . | 1.157146 | 0.714285 | 0.789124 | 0.848373 | 2.692728 | 1.651101 | 1.770771 | 1.939704 | -1.147738 | 86.616309 | 0.368339566 | -0.53970212 | 0.589402477 | 0.998865404 | 6.531903 | 6.334482 | 6.520887 | 6.378860 | 6.406269 | 6.421996 | 132 | 76 | 68 | 78 |
| 148137 | NM_00103988 | PROSER3 | proline and serine rich 3 | protein_coding | NP_001034976.2;NP_001354785.1;XP_011524830.1;XP_011524831.1;XP | 25204 | . | . | . | . | 3.367685 | 4.134028 | 4.734471 | 3.907123 | 7.836742 | 9.555992 | 10.624012 | 8.933170 | 1.118953 | 581.316117 | 0.180009818 | 0.900784196 | 0.367703072 | 0.998865404 | 9.099715 | 9.262077 | 9.077509 | 9.200943 | 9.285364 | 9.139939 | 572 | 648 | 592 | 513 |
| 148156 | NM_00130435 | ZNF558 | zinc finger protein 558 | protein_coding | NP_001291279.1;NP_653294.1;XP_011526005.1;XP_016881859.1;XP_01 | 26422 | . | . | . | . | 3.794364 | 2.564130 | 3.193906 | 3.451249 | 8.829641 | 5.927104 | 7.167034 | 7.890868 | -1.101530 | 446.413389 | 0.174122902 | -0.8012122 | 0.423008811 | 0.998865404 | 8.870154 | 8.730963 | 8.836225 | 8.826308 | 8.788970 | 8.754825 | 543 | 501 | 380 | 387 |
| 148189 | NR_027301 | LINC00662 | long intergenic non-protein coding RNA 662 | lncRNA | . | 27122 | . | . | . | . | 1.350100 | 1.717565 | 1.220517 | 1.667255 | 3.141738 | 3.970228 | 2.738804 | 3.811980 | -1.075326 | 95.448393 | 0.342286862 | -0.30609843 | 0.759529703 | 0.998865404 | 6.629322 | 6.521980 | 6.549751 | 6.620080 | 6.521219 | 6.605186 | 101 | 119 | 68 | 99 |
| 148198 | NM_00109862 | ZNF98 | zinc finger protein 98 | protein_coding | NP_001092096.1 | 13174 | 603980 | . | . | . | 0.510316 | 0.430584 | 0.319589 | 0.480223 | 1.187525 | 0.995313 | 0.717150 | 1.097974 | -1.173424 | 32.267562 | 0.555543967 | -0.41531269 | 0.677913014 | 0.998865404 | 5.123517 | 4.891163 | 5.038086 | 5.010830 | 4.982118 | 5.025388 | 44 | 34 | 21 | 33 |
| 148203 | NM_00135523 | ZNF738 | zinc finger protein 738 | protein_coding | NP_001342166.1;NP_001342168.1;NP_001342169.1;NP_001342170.1 | 32469 | . | . | . | . | 5.017811 | 5.766079 | 5.305738 | 5.582104 | 11.676651 | 13.328549 | 11.905923 | 12.762818 | 1.011131 | 470.873471 | 0.170604865 | 0.093609513 | 0.925419353 | 0.998865404 | 8.871142 | 8.887205 | 8.867116 | 8.883851 | 8.894779 | 8.869944 | 532 | 512 | 420 | 435 |
| 148206 | NM_182515,N | ZNF714 | zinc finger protein 714 | protein_coding | NP_872321.2 | 27124 | . | . | . | . | 1.639076 | 1.804293 | 1.870165 | 1.280600 | 3.814197 | 4.170704 | 4.196595 | 2.927940 | -1.131643 | 355.849744 | 0.198878426 | -0.89712668 | 0.369651325 | 0.998865404 | 8.561142 | 8.383651 | 8.497151 | 8.518555 | 8.497462 | 8.373631 | 427 | 415 | 323 | 278 |
| 148213 | NM_138286 | ZNF681 | zinc finger protein 681 | protein_coding | NP_612143.2 | 26457 | . | . | . | . | 0.601985 | 0.532261 | 0.535327 | 0.452946 | 1.400845 | 1.230346 | 1.201259 | 1.035607 | -1.160555 | 107.915541 | 0.314945788 | -0.68207055 | 0.495194341 | 0.998865404 | 6.856412 | 6.643238 | 6.798680 | 6.754586 | 6.756880 | 6.701204 | 142 | 117 | 95 | 85 |
| 148223 | NM_152482,X | C19orf25 | chromosome 19 open reading frame 25 | protein_coding | NP_689695.2;XP_005259563.1;XP_006722716.1 | 26711 | . | . | . | . | 5.509046 | 5.775462 | 6.501353 | 6.366910 | 12.819776 | 13.350238 | 14.588849 | 14.557184 | 1.123244 | 423.853754 | 0.176803438 | 0.948345424 | 0.342953634 | 0.998865404 | 8.641164 | 8.808814 | 8.678554 | 8.701345 | 8.767994 | 8.752405 | 451 | 439 | 395 | 415 |
| 148229 | NM_00117800 | ATP8B3 | ATPase phospholipid transporting 8B3 | protein_coding | NP_001171473.1;NP_620168.1;XP_006722717.1;XP_006722718.1;XP_00 | 13535 | 605866 | . | . | . | 0.251207 | 0.287902 | 0.213314 | 0.259269 | 0.584569 | 0.665499 | 0.478671 | 0.592787 | -1.129743 | 42.335452 | 0.480515518 | -0.36626178 | 0.71416975 | 0.998865404 | 5.489863 | 5.312263 | 5.402194 | 5.427234 | 5.384091 | 5.406007 | 48 | 52 | 32 | 40 |
| 148252 | NM_145173,X | DIRAS1 | DIRAS family GTPase 1 | protein_coding | NP_660156.1;XP_011526015.1 | 19127 | 607862 | . | . | . | 1.586227 | 1.810456 | 1.339046 | 1.796689 | 3.691215 | 4.184950 | 3.004780 | 4.107916 | -1.087005 | 176.606661 | 0.257876173 | -0.46672759 | 0.64069479 | 0.998865404 | 7.524071 | 7.402146 | 7.462618 | 7.499701 | 7.397482 | 7.492008 | 203 | 207 | 130 | 177 |
| 148254 | NM_00117277 | ZNF555 | zinc finger protein 555 | protein_coding | NP_001166246.1;NP_690004.4;XP_011526018.1;XP_016881864.1 | 28382 | . | . | . | . | 0.334624 | 0.217070 | 0.221701 | 0.265535 | 0.778685 | 0.501768 | 0.497491 | 0.607115 | -1.124122 | 70.258729 | 0.398797544 | -0.42326814 | 0.672099633 | 0.998865404 | 6.216289 | 6.048019 | 6.200232 | 6.090485 | 6.097191 | 6.140039 | 104 | 63 | 52 | 68 |
| 148266 | NM_00133048 | ZNF569 | zinc finger protein 569 | protein_coding | NP_001317411.1;NP_689697.2;XP_006723109.1;XP_006723110.1;XP_00 | 24737 | 613904 | . | . | . | 0.473951 | 0.436856 | 0.251917 | 0.325349 | 1.102904 | 1.009811 | 0.565296 | 0.743872 | -1.546238 | 50.516371 | 0.446965641 | -1.406736 | 0.159505686 | 0.933525128 | 5.939770 | 5.309235 | 5.703840 | 5.695097 | 5.600726 | 5.638477 | 72 | 65 | 30 | 42 |
| 148268 | NM_00130099 | ZNF570 | zinc finger protein 570 | protein_coding | NP_001287922.1;NP_001308920.1;NP_001308921.1;NP_001308922.1;NI | 26416 | . | . | . | . | 0.568673 | 0.327833 | 0.465446 | 0.610924 | 1.323324 | 0.757801 | 1.044447 | 1.396803 | -1.029038 | 74.069994 | 0.400680246 | -0.10306505 | 0.917911346 | 0.998865404 | 6.230244 | 6.191126 | 6.279500 | 6.134040 | 6.204032 | 6.200487 | 111 | 58 | 64 | 68 |
| 148281 | NM_00125377 | SYT6 | synaptotagmin 6 | protein_coding | NP_001240701.1;NP_001257734.1;NP_001353152.1;NP_001353153.1;NI | 18638 | 607718 | . | . | . | 0.207360 | 0.226303 | 0.454767 | 0.377550 | 0.482535 | 0.523109 | 1.020485 | 0.863222 | 1.833398 | 57.885429 | 0.416653815 | 2.098912214 | 0.03582464 | 0.590080555 | 5.352443 | 6.227198 | 5.762247 | 5.777075 | 5.910878 | 5.874599 | 45 | 46 | 70 | 65 |
| 148304 | NM_152485 | C1orf74 | chromosome 1 open reading frame 74 | protein_coding | NP_689698.1 | 26319 | . | . | . | . | 1.074592 | 0.616049 | 0.673345 | 0.790092 | 2.500621 | 1.424024 | 1.510967 | 1.806451 | -1.173151 | 115.694261 | 0.340228362 | -0.67715927 | 0.498304909 | 0.998865404 | 6.964317 | 6.734921 | 6.967727 | 6.771394 | 6.798516 | 6.842681 | 183 | 98 | 86 | 107 |
| 148327 | NM_00125597 | CREB3L4 | cAMP responsive element binding protein 3 like 4 | protein_coding | NP_001242907.1;NP_001242908.1;NP_001242909.1;NP_001242910.1;NI | 18854 | 607138 | . | . | . | 3.698253 | 4.270785 | 3.185979 | 4.825061 | 8.605983 | 9.872111 | 7.149242 | 11.031928 | -1.003111 | 221.439245 | 0.250961104 | -0.01785695 | 0.985752973 | 0.999197585 | 7.794018 | 7.787509 | 7.756348 | 7.818660 | 7.697234 | 7.868370 | 239 | 255 | 157 | 245 |
| 148362 | NM_00128857 | BROX | BRO1 domain and CAAX motif containing | protein_coding | NP_001275508.1;NP_001275509.1;NP_001275510.1;NP_001362590.1;NI | 26512 | . | . | . | . | 7.626435 | 7.153250 | 7.377022 | 6.411876 | 17.747029 | 16.535055 | 16.553824 | 14.659997 | -1.086260 | 735.872428 | 0.17046556 | -0.70025704 | 0.483766798 | 0.998865404 | 9.581270 | 9.462927 | 9.590354 | 9.505346 | 9.577057 | 9.398141 | 935 | 777 | 694 | 575 |
| 148398 | NM_152486 | SAMD11 | sterile alpha motif domain containing 11 | protein_coding | NP_689699.3 | 28706 | 616765 | . | . | . | 0.064014 | 0.102437 | 0.093629 | 0.052916 | 0.148963 | 0.236789 | 0.210102 | 0.120986 | -1.101817 | 6.466961 | 1.204349184 | -0.11615004 | 0.90753363 | NA | 2.760365 | 2.622519 | 2.683953 | 2.698531 | 2.696778 | 2.680008 | 6 | 9 | 7 | 4 |
| 148413 | NR_015434 | MRPL20-AS1 | MRPL20 antisense RNA 1 | lncRNA | . | 27125 | . | . | . | . | 8.550274 | 9.999168 | 9.459409 | 8.890458 | 19.896839 | 23.113525 | 21.226639 | 20.326977 | -1.024273 | 503.116365 | 0.173559378 | -0.19935767 | 0.841982973 | 0.998865404 | 8.992008 | 8.957279 | 8.932789 | 9.027734 | 8.988759 | 8.942972 | 543 | 590 | 448 | 449 |
| 148418 | NM_00101097 | SAMD13 | sterile alpha motif domain containing 13 | protein_coding | NP_001010971.1;NP_001128135.1;NP_001128136.1;XP_016855866.1 | 24582 | . | . | . | . | 0.789423 | 0.871061 | 0.919015 | 1.199258 | 1.837021 | 2.013497 | 2.062244 | 2.741960 | 1.285916 | 43.585083 | 0.476458094 | 0.761445498 | 0.446391021 | 0.998865404 | 5.254076 | 5.614942 | 5.411594 | 5.425312 | 5.436014 | 5.483477 | 42 | 43 | 37 | 52 |
| 148423 | NM_198077,N | C1orf52 | chromosome 1 open reading frame 52 | protein_coding | NP_932343.1;XP_024309131.1 | 24871 | . | . | . | . | 4.508191 | 4.059143 | 4.176016 | 3.876065 | 10.490746 | 9.382889 | 9.370861 | 8.862161 | -1.065724 | 608.401760 | 0.177355166 | -0.51779771 | 0.604599417 | 0.998865404 | 9.293516 | 9.202820 | 9.315254 | 9.216732 | 9.301998 | 9.141703 | 775 | 628 | 575 | 485 |
| 148479 | NM_153812,X | PHF13 | PHD finger protein 13 | protein_coding | NP_722519.2;XP_011539064.1 | 22983 | . | . | . | . | 11.092625 | 12.040112 | 11.810459 | 12.925171 | 25.812994 | 27.831257 | 26.502328 | 29.551869 | 1.055566 | 1354.315786 | 0.131355458 | 0.593931517 | 0.552557918 | 0.998865404 | 10.363994 | 10.441658 | 10.353113 | 10.409850 | 10.391895 | 10.453280 | 1461 | 1475 | 1161 | 1356 |
| 148523 | NM_00130083 | CIART | circadian associated repressor of transcription | protein_coding | NP_001287767.1;NP_001287768.1;NP_001287769.1;NP_001287770.1;NI | 25200 | 615782 | . | . | . | 1.805014 | 1.991444 | 1.704880 | 1.540472 | 4.200344 | 4.603313 | 3.825702 | 3.522107 | -1.018507 | 93.721546 | 0.329030668 | -0.08040718 | 0.935913415 | 0.998865404 | 6.563678 | 6.536815 | 6.553544 | 6.553096 | 6.538270 | 6.557547 | 109 | 102 | 78 | 90 |
| 148534 | NM_00119967 | TLCD4 | TLC domain containing 4 | protein_coding | NP_001186608.1;NP_689700.1 | 26477 | . | . | . | . | 1.130599 | 0.795370 | 0.761405 | 0.753334 | 2.630949 | 1.838533 | 1.708570 | 1.722409 | -1.293650 | 183.409901 | 0.267066664 | -1.39084016 | 0.164273902 | 0.940065727 | 7.692185 | 7.321987 | 7.649614 | 7.479870 | 7.462878 | 7.451710 | 281 | 183 | 141 | 148 |
| 148641 | NM_00130084 | SLC35F3 | solute carrier family 35 member F3 | protein_coding | NP_001287774.1;NP_775779.1 | 23616 | . | . | . | . | 0.054795 | 0.098885 | 0.095071 | 0.161909 | 0.127510 | 0.228576 | 0.213336 | 0.370186 | 1.625102 | 10.655766 | 0.950881492 | 0.736716694 | 0.461294653 | NA | 3.024434 | 3.719719 | 3.380286 | 3.399551 | 3.400314 | 3.424809 | 7 | 11 | 9 | 15 |
| 148645 | NR_103534 | LINC00337 | long intergenic non-protein coding RNA 337 | lncRNA | . | 28620 | . | . | . | . | 1.896222 | 2.465266 | 1.861253 | 2.869064 | 4.412586 | 5.698573 | 4.176598 | 6.559773 | 1.070653 | 116.403146 | 0.327189228 | 0.301021584 | 0.763398036 | 0.998865404 | 6.814409 | 6.909981 | 6.801613 | 6.887571 | 6.796219 | 6.935906 | 113 | 137 | 83 | 136 |
| 148709 | NR_002929 | ACTG1P25 | actin gamma 1 pseudogene 25 | transcribed_ps | . | 54872 | . | . | . | . | 0.997650 | 0.953539 | 1.005785 | 0.988623 | 2.321573 | 2.204147 | 2.256953 | 2.260369 | 1.011835 | 72.097121 | 0.371120816 | 0.0457355 | 0.963521068 | 0.998865404 | 6.163131 | 6.180556 | 6.177622 | 6.162741 | 6.176586 | 6.169562 | 85 | 75 | 64 | 67 |
| 148756 | NR_125716 | OVAAL | ovarian adenocarcinoma amplified long non-coding RNA | lncRNA | . | 49422 | . | . | . | . | 0.622927 | 0.589877 | 0.268437 | 0.412415 | 1.449576 | 1.363528 | 0.602365 | 0.942939 | -1.796133 | 22.294774 | 0.666698851 | -1.26728047 | 0.205055022 | NA | 4.841203 | 3.993209 | 4.514440 | 4.506896 | 4.444243 | 4.471014 | 34 | 30 | 11 | 18 |
| 148789 | NM_00127715 | B3GALNT2 | beta-1,3-N-acetylgalactosaminyltransferase 2 | protein_coding | NP_001264084.1;NP_689703.1;XP_006711812.1;XP_016855883.1;XP_01 | 28596 | 610194 | . | . | . | 5.325909 | 5.027785 | 6.146236 | 5.237965 | 12.393607 | 11.621948 | 13.791975 | 11.975986 | 1.086071 | 799.027120 | 0.151018123 | 0.788766311 | 0.430248603 | 0.998865404 | 9.581007 | 9.700712 | 9.631432 | 9.590119 | 9.715752 | 9.620795 | 908 | 802 | 772 | 728 |
| 148867 | NM_00114488 | SLC30A7 | solute carrier family 30 member 7 | protein_coding | NP_001138356.1;NP_598003.2;XP_011539081.1;XP_016855889.1;XP_01 | 19306 | 611149 | . | . | . | 2.600533 | 1.904104 | 2.365653 | 1.246121 | 6.051547 | 4.401422 | 5.308455 | 2.849110 | -1.267909 | 502.622897 | 0.538710603 | -0.63568782 | 0.524979914 | 0.998865404 | 9.134240 | 8.792200 | 9.130693 | 8.912102 | 9.060483 | 8.648659 | 752 | 508 | 515 | 276 |
| 148898 | NR_033690,N | ZNF436-AS1 | ZNF436 antisense RNA 1 | lncRNA | . | 25122 | . | . | . | . | 1.611518 | 2.273062 | 2.653188 | 2.272953 | 3.750069 | 5.254285 | 5.953676 | 5.196835 | 1.264105 | 195.421189 | 0.261431338 | 1.293326076 | 0.195898306 | 0.96621494 | 7.431804 | 7.769380 | 7.471786 | 7.616675 | 7.690631 | 7.610567 | 165 | 218 | 206 | 187 |
| 148932 | NM_145279,N | MOB3C | MOB kinase activator 3C | protein_coding | NP_660322.3;NP_958805.1 | 29800 | . | . | . | . | 2.178027 | 2.663484 | 2.719337 | 2.632866 | 5.068358 | 6.156763 | 6.102114 | 6.019735 | 1.094912 | 224.662625 | 0.231828634 | 0.564272055 | 0.572568997 | 0.998865404 | 7.745021 | 7.875273 | 7.740043 | 7.831998 | 7.839092 | 7.821260 | 223 | 254 | 208 | 216 |
| 149013 | NM_00127814 | NBPF12 | NBPF member 12 | protein_coding | NP_001265070.1;XP_024309211.1 | 24297 | 608607 | . | . | . | 3.128889 | 3.471780 | 4.182889 | 3.153563 | 7.281053 | 8.025176 | 9.386282 | 7.210247 | 1.063193 | 759.328760 | 0.165684179 | 0.533570097 | 0.593639002 | 0.998865404 | 9.523449 | 9.612344 | 9.510223 | 9.576715 | 9.668503 | 9.498322 | 807 | 832 | 763 | 645 |
| 149041 | NM_00130085 | RC3H1 | ring finger and CCCH-type domains 1 | protein_coding | NP_001287779.1;NP_001287780.1;NP_001287781.1;NP_742068.1;XP_00 | 29434 | 609424 | . | . | . | 1.199232 | 1.045647 | 1.278328 | 1.008333 | 2.790660 | 2.417058 | 2.868532 | 2.305431 | -1.009043 | 396.103683 | 0.194981194 | -0.0666111 | 0.946891308 | 0.998865404 | 8.635581 | 8.623864 | 8.668770 | 8.590239 | 8.686319 | 8.560037 | 488 | 401 | 380 | 330 |
| 149076 | NM_00137021 | ZNF362 | zinc finger protein 362 | protein_coding | NP_001357141.1;NP_689706.2;XP_005270564.1;XP_005270565.1 | 18079 | . | . | . | . | 3.545655 | 3.582405 | 2.670619 | 3.419483 | 8.250887 | 8.280891 | 5.992792 | 7.818241 | -1.099380 | 352.283435 | 0.193465604 | -0.70653565 | 0.479855088 | 0.998865404 | 8.527686 | 8.390226 | 8.482923 | 8.490949 | 8.391040 | 8.473104 | 422 | 401 | 270 | 338 |
| 149111 | NM_00132230 | CNIH3 | cornichon family AMPA receptor auxiliary protein 3 | protein_coding | NP_001309231.1;NP_001309232.1;NP_001309233.1;NP_001309234.1;NI | 26802 | . | . | . | . | 3.511458 | 3.335151 | 3.474320 | 3.701096 | 8.171308 | 7.709351 | 7.796271 | 8.462118 | 1.029250 | 240.890354 | 0.219277916 | 0.189684118 | 0.849556667 | 0.998865404 | 7.891328 | 7.932839 | 7.916672 | 7.892852 | 7.906248 | 7.931565 | 280 | 250 | 207 | 235 |
| 149134 | NR_015422 | LINC01341 | long intergenic non-protein coding RNA 1341 | lncRNA | . | 49457 | . | . | . | . | 0.218248 | 0.343981 | 0.398140 | 0.262169 | 0.507871 | 0.795126 | 0.893415 | 0.599419 | 1.180711 | 11.335019 | 0.920140033 | 0.260456076 | 0.794511993 | NA | 3.377846 | 3.617632 | 3.478204 | 3.501909 | 3.509841 | 3.486900 | 9 | 14 | 13 | 9 |
| 149175 | NM_00103174 | MANEAL | mannosidase endo-alpha like | protein_coding | NP_001026910.1;NP_001106954.1;NP_689709.1;XP_005270567.1;XP_01 | 26452 | . | . | . | . | 14.312482 | 15.451535 | 15.195861 | 14.947908 | 33.305733 | 35.716916 | 34.099071 | 34.176615 | -1.003337 | 1181.405297 | 0.133743634 | -0.03593272 | 0.971336009 | 0.998865404 | 10.208731 | 10.203842 | 10.174832 | 10.238668 | 10.216610 | 10.192689 | 1305 | 1331 | 1042 | 1088 |
| 149281 | NM_00113610 | METTL11B | methyltransferase like 11B | protein_coding | NP_001129579.1;XP_011507534.1;XP_011507535.1;XP_011507536.1 | 31932 | . | . | . | . | 0.156617 | 0.109701 | 0.144218 | 0.051390 | 0.364455 | 0.253577 | 0.323621 | 0.117497 | -1.226319 | 8.112925 | 1.092336459 | -0.26945414 | 0.787580226 | NA | 3.156660 | 2.869520 | 3.030631 | 3.016729 | 3.029575 | 3.002793 | 12 | 8 | 9 | 4 |
| 149297 | NM_00101796 | FAM78B | family with sequence similarity 78 member B | protein_coding | NP_001017961.1;NP_001307231.1;XP_016855906.1 | 13495 | . | . | . | . | 1.240665 | 1.071049 | 1.255506 | 1.355502 | 2.887078 | 2.475777 | 2.817318 | 3.099192 | 1.028191 | 242.517143 | 0.219768225 | 0.182505159 | 0.855186306 | 0.998865404 | 7.901844 | 7.941766 | 7.930046 | 7.899128 | 7.907959 | 7.948399 | 284 | 250 | 205 | 240 |
| 149345 | NM_198149,N | SHISA4 | shisa family member 4 | protein_coding | NP_937792.2 | 27139 | 617326 | . | . | . | 20.885720 | 22.490723 | 26.777246 | 29.475330 | 48.601933 | 51.988314 | 60.087369 | 67.391847 | 1.283043 | 1100.678740 | 0.136824348 | 2.627965448 | 0.008589724 | 0.293930247 | 9.913431 | 10.272624 | 9.981532 | 10.023634 | 10.153587 | 10.219966 | 1078 | 1071 | 1029 | 1210 |
| 149371 | NM_175876 | EXOC8 | exocyst complex component 8 | protein_coding | NP_787072.2 | 24659 | 615283 | . | . | . | 3.211647 | 2.831411 | 3.211583 | 2.380063 | 7.473635 | 6.544933 | 7.206699 | 5.441731 | -1.096359 | 462.971616 | 0.196335669 | -0.67598873 | 0.499047799 | 0.998865404 | 8.918934 | 8.787639 | 8.921472 | 8.835226 | 8.910906 | 8.725363 | 595 | 487 | 443 | 351 |
| 149420 | NM_00124353 | PDIK1L | PDLIM1 interacting kinase 1 like | protein_coding | NP_001230461.1;NP_001230462.1;NP_690048.1;XP_005245800.1;XP_00 | 18981 | 610785 | . | . | . | 0.930938 | 0.727571 | 0.668392 | 0.624366 | 2.166330 | 1.681813 | 1.499852 | 1.427540 | -1.248079 | 96.557333 | 0.334638272 | -0.95538883 | 0.339381099 | 0.998865404 | 6.743587 | 6.425549 | 6.665541 | 6.582651 | 6.574616 | 6.543527 | 139 | 101 | 79 | 76 |
| 149465 | NM_00116796 | CFAP57 | cilia and flagella associated protein 57 | protein_coding | NP_001161437.1;NP_001182760.2;NP_001365118.1;NP_689711.2;XP_00 | 26485 | 614259 | . | . | . | 0.048973 | 0.104823 | 0.052617 | 0.063080 | 0.113962 | 0.242302 | 0.118071 | 0.144224 | -1.254355 | 9.357966 | 1.01197084 | -0.32307864 | 0.746635683 | NA | 3.381886 | 3.051647 | 3.214883 | 3.246572 | 3.220457 | 3.222470 | 8 | 15 | 7 | 8 |
| 149473 | NM_00134912 | CCDC24 | coiled-coil domain containing 24 | protein_coding | NP_001336056.1;NP_001336057.1;NP_001336058.1;NP_689712.1 | 28688 | . | . | . | . | 3.563654 | 3.824680 | 3.808182 | 4.216476 | 8.292769 | 8.840922 | 8.545448 | 9.640472 | 1.133615 | 168.420644 | 0.255514746 | 0.708101164 | 0.478882437 | 0.998865404 | 7.303033 | 7.483196 | 7.358590 | 7.375907 | 7.395359 | 7.442321 | 178 | 174 | 147 | 177 |
| 149478 | NM_00113653 | BTBD19 | BTB domain containing 19 | protein_coding | NP_001130009.1;XP_006710449.1;XP_011539105.1;XP_011539107.1;XP | 27145 | . | . | . | . | 8.670681 | 8.466282 | 9.128362 | 8.887272 | 20.177032 | 19.570187 | 20.483780 | 20.319689 | 1.020353 | 543.053700 | 0.166655317 | 0.17441795 | 0.861537009 | 0.998865404 | 9.070069 | 9.099682 | 9.079747 | 9.075365 | 9.131704 | 9.048032 | 621 | 578 | 508 | 481 |
| 149483 | NM_00111493 | CCDC17 | coiled-coil domain containing 17 | protein_coding | NP_001108410.2;NP_001177111.1;XP_011539114.1;XP_011539116.1;XP | 26574 | . | . | . | . | 0.487533 | 0.579076 | 0.717491 | 0.581227 | 1.134512 | 1.338560 | 1.610027 | 1.328906 | 1.315632 | 54.197447 | 0.431438787 | 0.917293866 | 0.358988671 | 0.998865404 | 5.549072 | 5.944256 | 5.700586 | 5.747012 | 5.791427 | 5.764241 | 47 | 57 | 57 | 54 |
| 149603 | NM_00101085 | RNF187 | ring finger protein 187 | protein_coding | NP_001010858.2 | 27146 | 613754 | . | . | . | 20.418127 | 21.996233 | 20.406223 | 25.270384 | 47.513821 | 50.845276 | 45.790974 | 57.777733 | 1.061990 | 2138.360466 | 0.134161313 | 0.646763186 | 0.517785222 | 0.998865404 | 11.018456 | 11.104830 | 11.009788 | 11.062217 | 10.999479 | 11.162650 | 2310 | 2312 | 1721 | 2274 |
| 149840 | NM_00130347 | SHLD1 | shieldin complex subunit 1 | protein_coding | NP_001290406.1;NP_001290407.1;NP_001290408.1;NP_689717.2;XP_00 | 26318 | 618028 | . | . | . | 0.740711 | 1.416931 | 0.933647 | 1.215467 | 1.723665 | 3.275297 | 2.095077 | 2.779021 | 1.019888 | 33.292317 | 0.560810133 | 0.050659631 | 0.959596746 | 0.998865404 | 5.044829 | 5.069302 | 5.014879 | 5.085957 | 5.034796 | 5.072286 | 28 | 45 | 25 | 36 |
| 149844 | gene-LOC1498 | LOC149844 | synaptotagmin binding cytoplasmic RNA interacting protein pseud | pseudogene | . | . | . | . | . | . | 0.104365 | 0.122356 | 0.087712 | 0.104439 | 0.242862 | 0.282832 | 0.196824 | 0.238787 | -1.109250 | 6.410732 | 1.191181247 | -0.12557716 | 0.900066641 | NA | 2.754145 | 2.602870 | 2.680554 | 2.686746 | 2.679228 | 2.682068 | 7 | 8 | 5 | 6 |
| 149951 | NM_00109933 | COMMD7 | COMM domain containing 7 | protein_coding | NP_001092809.1;NP_444269.2;XP_005260356.1;XP_005260357.1;XP_01 | 16223 | 616703 | . | . | . | 12.564647 | 12.097005 | 10.012457 | 14.274816 | 29.238455 | 27.962768 | 22.467661 | 32.637672 | -1.001972 | 532.250628 | 0.186033644 | -0.01528155 | 0.987807565 | 0.999197585 | 9.057805 | 9.054116 | 9.075473 | 9.032369 | 8.952116 | 9.144344 | 634 | 556 | 393 | 573 |
| 149986 | NM_144703,X | LSM14B | LSM family member 14B | protein_coding | NP_653304.2;XP_005260359.1;XP_011526907.1;XP_011526908.1;XP_01 | 15887 | . | . | . | . | 22.495723 | 22.880180 | 21.727125 | 22.257966 | 52.348471 | 52.888564 | 48.755039 | 50.890195 | -1.046064 | 1820.530929 | 0.12022649 | -0.54040782 | 0.588915817 | 0.998865404 | 10.862300 | 10.797254 | 10.851915 | 10.843419 | 10.797212 | 10.826900 | 2155 | 1998 | 1518 | 1700 |
| 150051 | NM_00132102 | LOC150051 | uncharacterized LOC150051 | protein_coding | NP_001307952.1 | . | . | . | . | . | 0.937586 | 0.884792 | 0.772764 | 0.478095 | 2.181802 | 2.045237 | 1.734060 | 1.093108 | -1.447212 | 20.463245 | 0.692529029 | -0.77004162 | 0.441275204 | NA | 4.595547 | 4.066070 | 4.377542 | 4.368731 | 4.358143 | 4.324853 | 29 | 25 | 18 | 12 |
| 150082 | NM_152505,X | LCA5L | lebercilin LCA5 like | protein_coding | NP_689718.1;XP_005260983.1;XP_006724030.1;XP_006724035.1;XP_00 | 1255 | . | . | . | . | 0.203317 | 0.119208 | 0.303017 | 0.097810 | 0.473129 | 0.275554 | 0.679962 | 0.223631 | 1.125223 | 17.522419 | 0.79441701 | 0.214258419 | 0.830345542 | NA | 4.039590 | 4.217207 | 4.133251 | 4.094295 | 4.159498 | 4.084475 | 24 | 13 | 24 | 9 |
| 150094 | NM_173354,X | SIK1 | salt inducible kinase 1 | protein_coding | NP_775490.2;XP_011527776.1 | 11142 | 605705 | . | . | . | 0.476867 | 0.432327 | 0.303663 | 0.297494 | 1.109690 | 0.999344 | 0.681410 | 0.680184 | -1.546690 | 56.383792 | 0.420613809 | -1.49587052 | 0.1346874 | 0.891755273 | 6.097535 | 5.468947 | 5.873414 | 5.847633 | 5.776207 | 5.771356 | 83 | 70 | 39 | 41 |
| 150135 | NR_027272 | LINC00479 | long intergenic non-protein coding RNA 479 | lncRNA | . | 19727 | . | . | . | . | 0.832084 | 0.730707 | 1.016960 | 1.310794 | 1.936293 | 1.689062 | 2.282028 | 2.996974 | 1.449874 | 61.909830 | 0.406153204 | 1.319520915 | 0.186995023 | 0.959513866 | 5.659745 | 6.195071 | 5.912462 | 5.888173 | 5.950632 | 6.010383 | 62 | 51 | 56 | 77 |
| 150142 | NR_027273,N | ZNF295-AS1 | ZNF295 antisense RNA 1 | lncRNA | . | 23130 | . | . | . | . | 1.835331 | 2.412287 | 5.183632 | 4.435609 | 4.270890 | 5.576110 | 11.631920 | 10.141492 | 2.243299 | 108.747948 | 0.322235581 | 3.617296641 | 0.000297696 | 0.027953052 | 6.068012 | 7.232724 | 6.558236 | 6.622337 | 6.872901 | 6.812544 | 67 | 82 | 141 | 130 |
| 150165 | NM_00131825 | XKR3 | XK related 3 | protein_coding | NP_001305180.1;NP_787074.2;XP_016884089.1 | 28778 | 611674 | . | . | . | 0.358192 | 0.540519 | 0.484412 | 0.280012 | 0.833528 | 1.249434 | 1.087006 | 0.640213 | -1.170418 | 23.019854 | 0.663267889 | -0.34228156 | 0.732139008 | NA | 4.633141 | 4.407673 | 4.508235 | 4.549298 | 4.535193 | 4.492178 | 23 | 32 | 23 | 15 |
| 150197 | NR_026919 | LINC00896 | long intergenic non-protein coding RNA 896 | lncRNA | . | 26519 | . | . | . | . | 0.108257 | 0.062907 | 0.117690 | 0.158816 | 0.251918 | 0.145413 | 0.264094 | 0.363115 | 1.679391 | 8.345709 | 1.062615916 | 0.703865361 | 0.481516625 | NA | 2.638039 | 3.387716 | 3.048351 | 3.033593 | 3.054513 | 3.065735 | 9 | 5 | 8 | 11 |
| 150209 | NM_00101806 | AIFM3 | apoptosis inducing factor mitochondria associated 3 | protein_coding | NP_001018070.1;NP_001139760.1;NP_653305.1 | 26398 | 617298 | . | . | . | 0.465071 | 0.437764 | 0.301125 | 0.242305 | 1.082240 | 1.011911 | 0.675717 | 0.554001 | -1.657946 | 27.308330 | 0.598000392 | -1.2197273 | 0.22256827 | 0.982008277 | 5.089593 | 4.362218 | 4.804884 | 4.798923 | 4.757008 | 4.736594 | 40 | 36 | 20 | 17 |
| 150221 | NM_00112863 | RIMBP3C | RIMS binding protein 3C | protein_coding | NP_001122105.1 | 33892 | 612701 | . | . | . | 0.014736 | 0.018540 | 0.023927 | 0.032895 | 0.034292 | 0.042856 | 0.053691 | 0.075211 | 1.534310 | 4.545162 | 1.410259528 | 0.437926456 | 0.661439597 | NA | 1.844134 | 2.459419 | 2.171168 | 2.172348 | 2.176527 | 2.184641 | 4 | 4 | 4 | 6 |
| 150223 | NM_00101796 | YDJC | YdjC chitooligosaccharide deacetylase homolog | protein_coding | NP_001017964.1;NP_001358279.1 | 27158 | . | . | . | . | 12.812587 | 16.443698 | 15.249363 | 17.378641 | 29.815417 | 38.010347 | 34.219130 | 39.734199 | 1.091881 | 663.328970 | 0.170453004 | 0.743987996 | 0.456883737 | 0.998865404 | 9.309264 | 9.435152 | 9.263237 | 9.412161 | 9.354514 | 9.443435 | 650 | 760 | 563 | 691 |
| 150274 | NM_00131831 | HSCB | HscB mitochondrial iron-sulfur cluster cochaperone | protein_coding | NP_001305243.1;NP_001305244.1;NP_001305245.1;NP_001350785.1;NI | 28913 | 608142 | . | . | . | 7.574948 | 9.834033 | 6.125258 | 7.979297 | 17.627216 | 22.731810 | 13.744900 | 18.243711 | -1.188930 | 251.226207 | 0.229430527 | -1.08818779 | 0.276512215 | 0.998865404 | 8.093002 | 7.841760 | 7.957008 | 8.065664 | 7.881759 | 7.968754 | 282 | 325 | 181 | 235 |
| 150275 | NM_00128426 | CCDC117 | coiled-coil domain containing 117 | protein_coding | NP_001271192.1;NP_001271193.1;NP_001271194.1;NP_775781.1 | 26599 | . | . | . | . | 4.526976 | 3.914433 | 3.812237 | 3.879879 | 10.534459 | 9.048388 | 8.554550 | 8.870882 | -1.116525 | 497.695739 | 0.173459681 | -0.9167303 | 0.359283986 | 0.998865404 | 9.036233 | 8.877652 | 9.041424 | 8.941209 | 8.922955 | 8.922818 | 651 | 523 | 409 | 441 |
| 150290 | NM_00130479 | DUSP18 | dual specificity phosphatase 18 | protein_coding | NP_001291723.1;NP_001291724.1;NP_001291725.1;NP_689724.3;XP_00 | 18484 | 611446 | . | . | . | 2.198663 | 2.261615 | 2.163106 | 1.857584 | 5.116379 | 5.227825 | 4.853948 | 4.247146 | -1.179293 | 246.906312 | 0.219610119 | -1.08338383 | 0.278638091 | 0.998865404 | 8.061903 | 7.823935 | 7.958395 | 8.014903 | 7.919596 | 7.890823 | 291 | 304 | 203 | 206 |
| 150350 | NM_152512,X | ENTHD1 | ENTH domain containing 1 | protein_coding | NP_689725.2;XP_006724212.1;XP_006724213.1;XP_006724214.1;XP_00 | 26352 | . | . | . | . | 0.339194 | 0.394105 | 0.354397 | 0.303295 | 0.789318 | 0.910992 | 0.795255 | 0.693448 | -1.357345 | 22.204144 | 0.653699455 | -0.67429673 | 0.500122683 | NA | 4.676202 | 4.235838 | 4.488986 | 4.492016 | 4.465424 | 4.459289 | 29 | 28 | 17 | 17 |
| 150368 | NM_00100203 | PHETA2 | PH domain containing endocytic trafficking adaptor 2 | protein_coding | NP_001002034.2;XP_005261430.1 | 27161 | 614240 | . | . | . | 0.297398 | 0.240238 | 0.154924 | 0.175878 | 0.692058 | 0.555322 | 0.347646 | 0.402124 | -1.585342 | 17.139381 | 0.746061503 | -0.89107111 | 0.372891022 | NA | 4.393730 | 3.728813 | 4.128957 | 4.115256 | 4.086494 | 4.091310 | 26 | 21 | 11 | 13 |
| 150381 | NR_027034 | PRR34-AS1 | PRR34 antisense RNA 1 | lncRNA | . | 50499 | . | . | . | . | 2.006844 | 1.995475 | 1.929179 | 2.417328 | 4.670008 | 4.612629 | 4.329021 | 5.526934 | 1.075572 | 106.929870 | 0.315100512 | 0.333558777 | 0.738712529 | 0.998865404 | 6.687585 | 6.791583 | 6.730661 | 6.723814 | 6.715378 | 6.784235 | 120 | 110 | 86 | 115 |
| 150383 | NM_207327,X | CDPF1 | cysteine rich DPF motif domain containing 1 | protein_coding | NP_997210.3;XP_011528262.1;XP_011528265.1;XP_011528266.1;XP_01 | 33710 | . | . | . | . | 4.617405 | 5.070187 | 4.493436 | 6.228270 | 10.744889 | 11.719966 | 10.083138 | 14.240199 | 1.072344 | 265.116051 | 0.216273374 | 0.465927618 | 0.641267312 | 0.998865404 | 7.999774 | 8.099465 | 8.005032 | 8.056030 | 8.027911 | 8.103552 | 280 | 290 | 221 | 276 |
| 150384 | NR_024009 | GTSE1-DT | GTSE1 divergent transcript | lncRNA | . | 27163 | . | . | . | . | 0.008667 | 0.001842 | 0.008408 | 0.008823 | 0.020168 | 0.004258 | 0.018868 | 0.020172 | 1.234655 | 1.002957 | 2.868339486 | 0.106022226 | 0.915564717 | NA | -0.155866 | 0.148375 | -0.001261 | -0.000972 | 0.000053 | -0.000315 | 1 | 1 | 1 | 1 |
| 150417 | gene-LOC1504 | LOC150417 | small nuclear ribonucleoprotein polypeptide A' pseudogene | pseudogene | . | . | . | . | . | . | 0.056199 | 0.165449 | 0.051591 | 0.127791 | 0.130778 | 0.382443 | 0.115768 | 0.292179 | -1.236989 | 3.683103 | 1.580948665 | -0.19408121 | 0.846112291 | NA | 2.029593 | 1.715150 | 1.878176 | 1.892094 | 1.877195 | 1.885962 | 3 | 6 | 2 | 4 |
| 150465 | NM_00137171 | TTL | tubulin tyrosine ligase | protein_coding | NP_001358641.1;NP_714923.1;XP_005263656.1;XP_011508967.1 | 21586 | 608291 | . | . | . | 13.790115 | 13.172400 | 13.170660 | 13.593173 | 32.090165 | 30.448593 | 29.554577 | 31.079175 | 1.125734 | 4284.807587 | 0.108987452 | 1.567753958 | 0.116938564 | 0.857396579 | 11.977103 | 12.147876 | 12.006651 | 12.020251 | 12.083626 | 12.139614 | 4604 | 4386 | 3815 | 4396 |
| 150468 | NM_00130436 | CKAP2L | cytoskeleton associated protein 2 like | protein_coding | NP_001291290.1;NP_689728.3;XP_011508968.1 | 26877 | 616174 | . | . | . | 12.357130 | 12.158604 | 9.736903 | 10.458884 | 28.755547 | 28.105155 | 21.849331 | 23.912995 | -1.224605 | 1626.330434 | 0.12404088 | -2.35661692 | 0.018442261 | 0.457491143 | 10.806222 | 10.513795 | 10.753528 | 10.728454 | 10.560800 | 10.605138 | 2094 | 1902 | 1234 | 1411 |
| 150472 | NM_00133033 | CBWD2 | COBW domain containing 2 | protein_coding | NP_001317265.1;NP_001317266.1;NP_001317268.1;NP_001317269.1;NI | 17907 | 611079 | . | . | . | 8.474399 | 8.630572 | 9.726130 | 8.240603 | 19.720278 | 19.949952 | 21.825151 | 18.841157 | 1.088171 | 552.430496 | 0.170699128 | 0.714154562 | 0.47513161 | 0.998865404 | 9.047036 | 9.169657 | 9.081456 | 9.077539 | 9.194656 | 9.070890 | 611 | 569 | 548 | 489 |
| 150478 | gene-SNRPA1P | SNRPA1P1 | SNRPA1 pseudogene 1 | pseudogene | . | 54744 | . | . | . | . | 0.084536 | 0.171045 | 0.171390 | 0.129227 | 0.196719 | 0.395379 | 0.384595 | 0.295462 | 1.109089 | 4.762694 | 1.38020612 | 0.108226518 | 0.91381601 | NA | 2.175403 | 2.324312 | 2.240500 | 2.250382 | 2.250998 | 2.244409 | 4 | 6 | 5 | 4 |
| 150554 | gene-YWHAZP | YWHAZP2 | tyrosine 3-monooxygenase/tryptophan 5-monooxygenase activati | pseudogene | . | 31078 | . | . | . | . | 0.751769 | 0.808622 | 0.581393 | 0.952193 | 1.749397 | 1.869165 | 1.304629 | 2.177077 | -1.023907 | 70.012987 | 0.397638165 | -0.08571783 | 0.931690726 | 0.998865404 | 6.147915 | 6.110950 | 6.122231 | 6.138154 | 6.068944 | 6.177359 | 79 | 79 | 46 | 80 |
| 150568 | NR_015399 | LINC01102 | long intergenic non-protein coding RNA 1102 | lncRNA | . | 27165 | . | . | . | . | 0.375575 | 0.165128 | 0.457913 | 0.370503 | 0.873978 | 0.381701 | 1.027544 | 0.847110 | 1.499240 | 16.640209 | 0.779350692 | 0.749638281 | 0.453472588 | NA | 3.732160 | 4.321310 | 4.052659 | 4.008273 | 4.066990 | 4.051845 | 21 | 9 | 19 | 17 |
| 150572 | NM_00133036 | SMYD1 | SET and MYND domain containing 1 | protein_coding | NP_001317293.1;NP_938015.1 | 20986 | 606846 | . | . | . | 0.102014 | 0.079653 | 0.127935 | 0.113859 | 0.237392 | 0.184121 | 0.287083 | 0.260325 | 1.326825 | 15.071012 | 0.79020018 | 0.516296472 | 0.60564736 | NA | 3.694014 | 4.104320 | 3.905875 | 3.889566 | 3.921117 | 3.911218 | 17 | 12 | 16 | 15 |
| 150590 | NM_00131799 | C2orf15 | chromosome 2 open reading frame 15 | protein_coding | NP_001304921.2;NP_653307.2 | 28436 | . | . | . | . | 0.708046 | 0.401907 | 0.601027 | 0.514582 | 1.647653 | 0.929025 | 1.348688 | 1.176530 | 1.001112 | 29.381023 | 0.591069148 | 0.002713136 | 0.997835234 | 0.999862634 | 4.874259 | 4.879362 | 4.906733 | 4.840150 | 4.884955 | 4.863091 | 43 | 23 | 28 | 25 |
| 150678 | NM_00116342 | COPS9 | COP9 signalosome subunit 9 | protein_coding | NP_001156896.1;NP_612209.1;XP_016858900.1 | 21314 | . | . | . | . | 63.513064 | 59.307461 | 56.459840 | 69.950092 | 147.797501 | 137.091854 | 126.694237 | 159.932597 | 1.014392 | 818.522125 | 0.156574174 | 0.131668141 | 0.895246786 | 0.998865404 | 9.666761 | 9.686924 | 9.694658 | 9.645981 | 9.608487 | 9.747978 | 968 | 847 | 646 | 849 |
| 150684 | NM_00132178 | COMMD1 | copper metabolism domain containing 1 | protein_coding | NP_001308710.1;NP_001308711.1;NP_001358694.1;NP_689729.1;XP_01 | 23024 | 607238 | . | . | . | 12.046281 | 13.353594 | 10.480104 | 19.154984 | 28.032193 | 30.867436 | 23.517049 | 43.795599 | 1.038970 | 472.269643 | 0.175788328 | 0.313751887 | 0.753709489 | 0.998865404 | 8.855915 | 8.910502 | 8.823005 | 8.915694 | 8.885957 | 8.902274 | 494 | 537 | 413 | 457 |
| 150696 | NM_00116597 | PROM2 | prominin 2 | protein_coding | NP_001159449.1;NP_001159450.1;NP_001307999.1;NP_653308.2;XP_00 | 20685 | 617160 | . | . | . | 0.109528 | 0.151939 | 0.205268 | 0.163049 | 0.254877 | 0.351214 | 0.460615 | 0.372792 | 1.371326 | 24.030028 | 0.634827257 | 0.717631307 | 0.472984649 | NA | 4.341184 | 4.796563 | 4.548049 | 4.573511 | 4.605022 | 4.579218 | 20 | 25 | 27 | 23 |
| 150709 | NM_00137806 | ANKAR | ankyrin and armadillo repeat containing | protein_coding | NP_001364997.1;NP_653309.3;XP_011508975.1;XP_011508977.1;XP_01 | 26350 | 609803 | . | . | . | 0.361899 | 0.349475 | 0.471811 | 0.146318 | 0.842154 | 0.807826 | 1.058731 | 0.334536 | -1.153255 | 44.782194 | 0.531407679 | -0.38710587 | 0.698677832 | 0.998865404 | 5.581654 | 5.381087 | 5.495998 | 5.476769 | 5.532123 | 5.386778 | 58 | 49 | 53 | 21 |
| 150726 | NM_00108041 | FBXO41 | F-box protein 41 | protein_coding | NP_001073879.2;NP_001358318.1;XP_005264215.1;XP_011530865.1;XP | 29409 | 609108 | . | . | . | 4.864455 | 4.828973 | 3.755293 | 4.377961 | 11.319788 | 11.162386 | 8.426771 | 10.009689 | 1.026660 | 1062.010415 | 0.13749809 | 0.276067054 | 0.782496565 | 0.998865404 | 10.033308 | 10.071602 | 10.047958 | 10.036531 | 10.097676 | 10.024654 | 1216 | 1122 | 982 | 959 |
| 150737 | NM_152517 | TTC30B | tetratricopeptide repeat domain 30B | protein_coding | NP_689730.2 | 26425 | . | . | . | . | 2.256402 | 1.836067 | 2.153095 | 2.087291 | 5.250739 | 4.244152 | 4.831482 | 4.772343 | 1.021335 | 247.089574 | 0.225081516 | 0.13531124 | 0.892365798 | 0.998865404 | 7.933092 | 7.964517 | 7.993416 | 7.888207 | 7.963575 | 7.941837 | 311 | 236 | 222 | 229 |
| 150759 | NR_103734 | LINC00342 | long intergenic non-protein coding RNA 342 | lncRNA | . | 42470 | . | . | . | . | 0.614850 | 0.947791 | 0.935831 | 0.659042 | 1.430781 | 2.190861 | 2.099979 | 1.506822 | 1.016309 | 77.365848 | 0.390270296 | 0.059801845 | 0.952313456 | 0.998865404 | 6.261770 | 6.285383 | 6.207914 | 6.316439 | 6.312276 | 6.221237 | 70 | 100 | 80 | 60 |
| 150776 | NR_026922 | SMPD4BP | sphingomyelin phosphodiesterase 4B, pseudogene | transcribed_ps | . | 24761 | . | . | . | . | 11.229659 | 11.885018 | 13.192008 | 12.056573 | 26.131882 | 27.472752 | 29.602488 | 27.565920 | 1.077387 | 1427.076392 | 0.130191655 | 0.825987524 | 0.408811169 | 0.998865404 | 10.423995 | 10.531690 | 10.430263 | 10.466984 | 10.544287 | 10.466522 | 1544 | 1518 | 1352 | 1318 |
| 150786 | NM_00107763 | RAB6D | RAB6D, member RAS oncogene family | protein_coding | NP_001071105.1 | 30272 | . | . | . | . | 0.168866 | 0.100822 | 0.097248 | 0.076757 | 0.392957 | 0.233054 | 0.218221 | 0.175495 | -1.521168 | 13.280399 | 0.855321952 | -0.70754558 | 0.479227496 | NA | 4.000985 | 3.399154 | 3.765846 | 3.730592 | 3.728586 | 3.720173 | 23 | 13 | 10 | 9 |
| 150864 | NM_173511 | FAM117B | family with sequence similarity 117 member B | protein_coding | NP_775782.2 | 14440 | . | . | . | . | 0.521498 | 0.586679 | 0.808153 | 0.679746 | 1.213547 | 1.356134 | 1.813471 | 1.554158 | 1.320850 | 121.449593 | 0.298377987 | 1.345496017 | 0.178465107 | 0.951511445 | 6.709325 | 7.111205 | 6.848901 | 6.883729 | 6.994396 | 6.928724 | 114 | 119 | 131 | 118 |
| 150946 | NM_00116824 | GAREM2 | GRB2 associated regulator of MAPK1 subtype 2 | protein_coding | NP_001161713.1;NP_001177962.1;XP_006712014.1;XP_011530866.1;XP | 27172 | 617999 | . | . | . | 0.530686 | 0.407214 | 0.526660 | 0.402805 | 1.234926 | 0.941294 | 1.181809 | 0.920966 | -1.058231 | 63.522251 | 0.405617587 | -0.20130955 | 0.840456534 | 0.998865404 | 6.028170 | 5.949127 | 6.014587 | 5.972282 | 6.013381 | 5.947749 | 82 | 64 | 62 | 49 |
| 150962 | NM_00132212 | PUS10 | pseudouridine synthase 10 | protein_coding | NP_001309052.1;NP_001309053.1;NP_001309056.1;NP_653310.2;XP_01 | 26505 | 612787 | . | . | . | 0.741247 | 1.056377 | 0.468815 | 0.535100 | 1.724911 | 2.441864 | 1.052006 | 1.223442 | -1.578783 | 100.497208 | 0.323785309 | -2.03472069 | 0.041878985 | 0.633602228 | 6.943533 | 6.283691 | 6.692258 | 6.735592 | 6.566931 | 6.586109 | 133 | 141 | 65 | 76 |
| 150967 | NR_037701 | LINC01963 | long intergenic non-protein coding RNA 1963 | lncRNA | . | 25283 | . | . | . | . | 1.314017 | 1.371161 | 1.684183 | 1.077617 | 3.057771 | 3.169499 | 3.779258 | 2.463843 | 1.014718 | 140.629654 | 0.301398832 | 0.069937729 | 0.944243223 | 0.998865404 | 7.124151 | 7.147270 | 7.120621 | 7.133090 | 7.212523 | 7.045306 | 158 | 153 | 151 | 103 |
| 151011 | NM_00132149 | SEPTIN10 | septin 10 | protein_coding | NP_001308425.1;NP_001308427.1;NP_001308428.1;NP_001308429.1;NI | 14349 | 611737 | . | . | . | 18.255608 | 16.172015 | 15.527213 | 14.985429 | 42.481552 | 37.382333 | 34.842619 | 34.262406 | -1.111113 | 1826.627869 | 0.125342476 | -1.21272536 | 0.225234841 | 0.982008277 | 10.908849 | 10.757096 | 10.917246 | 10.830320 | 10.807678 | 10.775545 | 2334 | 1962 | 1536 | 1589 |

| | | | | | | | | | | |
|---|---|---|---|---|---|---|---|---|---|---|
| 151050 | NM_00130797 | KANSL1L | KAT8 regulatory NSL complex subunit 1 like | protein_coding | NP_001294905.1;NP_689732.2;XP_005246385.1;XP_005246386.1;XP_00 | 26310 | 613833 | . | . | [illegible] |
| 151056 | NM_00117058 | PLB1 | phospholipase B1 | protein_coding | NP_001164056.1;NP_694566.4;XP_011530881.1;XP_011530883.1;XP_01 | 30041 | 610179 | . | . | [illegible] |
| 151176 | NM_00129183 | ERFE | erythroferrone | protein_coding | NP_001278761.1 | 26727 | 615099 | . | . | [illegible] |
| 151188 | NM_00135006 | ARL6IP6 | ADP ribosylation factor like GTPase 6 interacting protein 6 | protein_coding | NP_001336997.1;NP_001358901.1;NP_689735.1 | 24048 | 616495 | . | . | [illegible] |
| 151194 | NM_00112739 | METTL21A | methyltransferase like 21A | protein_coding | NP_001120867.1;NP_001294950.1;NP_001317059.1;NP_001317060.1;NI | 30476 | 615257 | . | . | [illegible] |
| 151195 | NM_00114230 | CCNYL1 | cyclin Y like 1 | protein_coding | NP_001135772.1;NP_001317147.1;NP_689736.1 | 26868 | . | . | . | [illegible] |
| 151230 | NM_144711 | KLHL23 | kelch like family member 23 | protein_coding | NP_653312.2 | 27506 | . | . | . | [illegible] |
| 151246 | NM_00116003 | SGO2 | shugoshin 2 | protein_coding | NP_001153505.1;NP_001153518.1;NP_689737.4;XP_005246401.2;XP_00 | 30812 | 612425 | . | . | [illegible] |
| 151278 | NR_161172 | CCDC140 | CCDC140 long non-coding RNA | lncRNA | . | 26514 | . | . | . | [illegible] |
| 151295 | NM_00114488 | SLC23A3 | solute carrier family 23 member 3 | protein_coding | NP_001138361.1;NP_001138362.1;NP_653313.3 | 20601 | . | . | . | [illegible] |
| 151313 | NM_00132084 | FAHD2B | fumarylacetoacetate hydrolase domain containing 2B | protein_coding | NP_001307777.1;NP_001307778.1;NP_955368.1;XP_011509047.1;XP_01 | 25318 | . | . | . | [illegible] |
| 151354 | NM_00136936 | LRATD1 | LRAT domain containing 1 | protein_coding | NP_001356293.1;NP_660158.2 | 20743 | 611234 | . | . | [illegible] |
| 151393 | NM_00117079 | RMDN2 | regulator of microtubule dynamics 2 | protein_coding | NP_001164262.1;NP_001164263.1;NP_001164264.1;NP_001309140.1;NI | 26567 | 611872 | . | . | [illegible] |
| 151457 | gene-DRG1P1 | DRG1P1 | DRG1 pseudogene 1 | pseudogene | . | 54701 | . | . | . | [illegible] |
| 151477 | NR_024079 | LINC00471 | long intergenic non-protein coding RNA 471 | lncRNA | . | 28668 | . | . | . | [illegible] |
| 151507 | NR_024322 | MSL3P1 | MSL complex subunit 3 pseudogene 1 | transcribed_ps | . | 17837 | . | . | . | [illegible] |
| 151516 | NM_152792 | ASPRV1 | aspartic peptidase retroviral like 1 | protein_coding | NP_690005.2 | 26321 | 611765 | . | . | [illegible] |
| 151525 | NM_00112821 | WDSUB1 | WD repeat, sterile alpha motif and U-box domain containing 1 | protein_coding | NP_001121684.1;NP_001121685.1;NP_001294923.1;NP_001317203.1;NI | 26697 | . | . | . | [illegible] |
| 151534 | NR_024606,NF | LBX2-AS1 | LBX2 antisense RNA 1 | lncRNA | . | 25136 | . | . | . | [illegible] |
| 151556 | NM_00103304 | GPR155 | G protein-coupled receptor 155 | protein_coding | NP_001028217.1;NP_001253979.1;NP_001253980.1;NP_689742.4;XP_01 | 22951 | . | . | . | [illegible] |
| 151579 | gene-BZW1P2 | BZW1P2 | basic leucine zipper and W2 domains 1 pseudogene 2 | pseudogene | . | 33954 | . | . | . | [illegible] |
| 151613 | NM_00104260 | TTC14 | tetratricopeptide repeat domain 14 | protein_coding | NP_001036066.1;NP_001275511.1;NP_597719.1;XP_016861228.1;XP_01 | 24697 | . | . | . | [illegible] |
| 151636 | NM_138287 | DTX3L | deltex E3 ubiquitin ligase 3L | protein_coding | NP_612144.1 | 30323 | 613143 | . | . | [illegible] |
| 151645 | gene-PSMC1P1 | PSMC1P1 | proteasome 26S subunit, ATPase 1 pseudogene 1 | pseudogene | . | 30147 | . | . | . | [illegible] |
| 151648 | NM_00101240 | SGO1 | shugoshin 1 | protein_coding | NP_001012409.1;NP_001012410.1;NP_001012411.1;NP_001012412.1;NI | 25088 | 609168 | . | . | [illegible] |
| 151742 | NM_00131791 | PPM1L | protein phosphatase, Mg2+/Mn2+ dependent 1L | protein_coding | NP_001304840.1;NP_001304841.1;NP_640338.2;XP_011510742.1 | 16381 | 611931 | . | . | [illegible] |
| 151877 | NR_145422 | MAGI1-IT1 | MAGI1 intronic transcript 1 | lncRNA | . | 42436 | . | . | . | [illegible] |
| 151887 | NM_199511,N | CCDC80 | coiled-coil domain containing 80 | protein_coding | NP_955805.1;NP_955806.1 | 30649 | 608298 | . | . | [illegible] |
| 151903 | NM_00127707 | CCDC12 | coiled-coil domain containing 12 | protein_coding | NP_001264003.1;NP_653317.2;XP_011531692.1;XP_011531695.1;XP_01 | 28332 | . | . | . | [illegible] |
| 151963 | NM_178496 | MB21D2 | Mab-21 domain containing 2 | protein_coding | NP_848591.2 | 30438 | . | . | . | [illegible] |
| 151987 | NM_00131802 | PPP4R2 | protein phosphatase 4 regulatory subunit 2 | protein_coding | NP_001304954.1;NP_001304955.1;NP_001304956.1;NP_001304957.1;NI | 18296 | 613822 | . | . | [illegible] |
| 152002 | NM_00130806 | XXYLT1 | xyloside xylosyltransferase 1 | protein_coding | NP_001294998.1;NP_689744.3;XP_005269343.1;XP_011510750.1;XP_01 | 26639 | 614552 | . | . | [illegible] |
| 152006 | NM_022781,N | RNF38 | ring finger protein 38 | protein_coding | NP_073618.3;NP_919309.1;NP_919310.1;NP_919311.1;NP_919313.1;XP_ | 18052 | 612488 | . | . | [illegible] |
| 152007 | NM_00128701 | GLIPR2 | GLI pathogenesis related 2 | protein_coding | NP_001273939.1;NP_001273940.1;NP_001273941.1;NP_001273942.1;NI | 18007 | 607141 | . | . | [illegible] |
| 152015 | NM_00101233 | ROPN1B | rhophilin associated tail protein 1B | protein_coding | NP_001012337.1;NP_001295242.1;XP_005247195.1;XP_006713576.1 | 31927 | . | . | . | [illegible] |
| 152048 | NR_149029 | LOC152048 | uncharacterized FLJ31715 | lncRNA | . | . | . | . | . | [illegible] |
| 152098 | NM_00104043 | ZCWPW2 | zinc finger CW-type and PWWP domain containing 2 | protein_coding | NP_001035522.1;NP_001311098.1;NP_001311099.1;XP_016861244.1;XP | 23574 | . | . | . | [illegible] |
| 152100 | NM_00133118 | CMC1 | C-X9-C motif containing 1 | protein_coding | NP_001318114.1;NP_001318115.1;NP_001318116.1;NP_001318118.1;NI | 28783 | 615166 | . | . | [illegible] |
| 152110 | NM_00103174 | NEK10 | NIMA related kinase 10 | protein_coding | NP_001026911.1;NP_001291313.1;NP_689747.3;NP_955379.2;XP_00671 | 18592 | 618726 | . | . | [illegible] |
| 152137 | NM_174908,N | CCDC50 | coiled-coil domain containing 50 | protein_coding | NP_777568.1;NP_848018.1;XP_011510762.1 | 18111 | 611051 | . | . | [illegible] |
| 152185 | NM_00133107 | SPICE1 | spindle and centriole associated protein 1 | protein_coding | NP_001318007.1;NP_001318008.1;NP_653319.1 | 25083 | 613447 | . | . | [illegible] |
| 152189 | NM_00132030 | CMTM8 | CKLF like MARVEL transmembrane domain containing 8 | protein_coding | NP_001307237.1;NP_849199.2;XP_011531718.1;XP_016861268.1;XP_01 | 19179 | 607891 | . | . | [illegible] |
| 152217 | NM_00135524 | NCBP2AS2 | NCBP2 antisense 2 (head to head) | protein_coding | NP_001342172.1 | 25121 | . | . | . | [illegible] |
| 152225 | NR_026934 | LINC02085 | long intergenic non-protein coding RNA 2085 | lncRNA | . | 27190 | . | . | . | [illegible] |
| 152302 | NR_002786 | CIDECP1 | cell death inducing DFFA like effector c pseudogene 1 | transcribed_ps | . | 24230 | . | . | . | [illegible] |
| 152330 | NM_00120695 | CNTN4 | contactin 4 | protein_coding | NP_001193884.1;NP_001193885.1;NP_001337024.1;NP_783200.1;NP_78 | 2174 | 607280 | . | . | [illegible] |
| 152404 | NM_00101588 | IGSF11 | immunoglobulin superfamily member 11 | protein_coding | NP_001015887.1;NP_001340247.1;NP_001340248.1;NP_001340249.1;NI | 16669 | 608351 | . | . | [illegible] |
| 152485 | NM_00130621 | ZNF827 | zinc finger protein 827 | protein_coding | NP_001293144.1;NP_849157.2;XP_011529936.1;XP_011529937.1;XP_01 | 27193 | 617962 | . | . | [illegible] |
| 152503 | NM_00100955 | SH3D19 | SH3 domain containing 19 | protein_coding | NP_001009555.3;NP_001122395.1;NP_001122396.1;NP_001230278.1;NI | 30418 | 608674 | . | . | [illegible] |
| 152518 | NM_00127862 | NFXL1 | nuclear transcription factor, X-box binding like 1 | protein_coding | NP_001265552.1;NP_001265553.1;NP_694540.3 | 18726 | . | . | . | [illegible] |
| 152519 | NM_207330,XI | NIPAL1 | NIPA like domain containing 1 | protein_coding | NP_997213.1;XP_016863273.1 | 27194 | . | . | . | [illegible] |
| 152559 | NM_00104020 | PAQR3 | progestin and adipoQ receptor family member 3 | protein_coding | NP_001035292.1;NP_001337034.1;NP_001337035.1;XP_005262826.1;XP | 30130 | 614577 | . | . | [illegible] |
| 152579 | NM_152540,XI | SCFD2 | sec1 family domain containing 2 | protein_coding | NP_689753.2;XP_011532677.1;XP_011532678.1;XP_011532680.1;XP_01 | 30676 | . | . | . | [illegible] |
| 152667 | gene-FAM192E | FAM192BP | family with sequence similarity 192 member B, pseudogene | pseudogene | . | 37194 | . | . | . | [illegible] |
| 152687 | NM_00128605 | ZNF595 | zinc finger protein 595 | protein_coding | NP_001272981.1;NP_001272982.1;NP_001272983.1;NP_872330.1;XP_01 | 27196 | . | . | . | [illegible] |
| 152709 | gene-ARL4AP2 | ARL4AP2 | ADP ribosylation factor like GTPase 4A pseudogene 2 | pseudogene | . | 32379 | . | . | . | [illegible] |
| 152756 | NR_160935 | FAM218A | family with sequence similarity 218 member A | lncRNA | . | 26466 | . | . | . | [illegible] |
| 152815 | NM_00131779 | THAP6 | THAP domain containing 6 | protein_coding | NP_001304720.1;NP_653322.1;XP_005262829.1;XP_005262831.1;XP_00 | 23189 | 612535 | . | . | [illegible] |
| 152877 | NM_00101362 | FAM53A | family with sequence similarity 53 member A | protein_coding | NP_001013644.1;NP_001167541.1;NP_001284364.1;XP_005248005.1;XP | 31860 | 617229 | . | . | [illegible] |
| 152905 | gene-RTN3P1 | RTN3P1 | reticulon 3 pseudogene 1 | pseudogene | . | 19212 | . | . | . | [illegible] |
| 152926 | NM_152542,XI | PPM1K | protein phosphatase, Mg2+/Mn2+ dependent 1K | protein_coding | NP_689755.3;XP_005262832.1;XP_006714174.1;XP_016863292.1;XP_01 | 25415 | 611065 | . | . | [illegible] |
| 152992 | NM_00135023 | TRMT44 | tRNA methyltransferase 44 homolog | protein_coding | NP_001337162.1;NP_689757.2;XP_011511708.1;XP_011511709.1;XP_01 | 26653 | 614309 | . | . | [illegible] |
| 153090 | NM_032552,N | DAB2IP | DAB2 interacting protein | protein_coding | NP_115941.2;NP_619723.1;XP_005251776.1;XP_005251778.1;XP_01151 | 17294 | 609205 | . | . | [illegible] |
| 153129 | NM_00125828 | SLC38A9 | solute carrier family 38 member 9 | protein_coding | NP_001245215.1;NP_001245216.1;NP_001269358.1;NP_001336311.1;NI | 26907 | 616203 | . | . | [illegible] |
| 153222 | NM_00116839 | CREBRF | CREB3 regulatory factor | protein_coding | NP_001161865.1;NP_001161866.1;NP_705835.2;XP_005265878.1;XP_00 | 24050 | 617109 | . | . | [illegible] |
| 153241 | NM_00116622 | CEP120 | centrosomal protein 120 | protein_coding | NP_001159698.1;NP_001362334.1;NP_001362335.1;NP_001362336.1;NI | 26690 | 613446 | . | . | [illegible] |
| 153328 | NM_00134933 | SLC25A48 | solute carrier family 25 member 48 | protein_coding | NP_001336264.1;NP_001336265.1;NP_001336274.1;NP_660325.4;XP_00 | 30451 | 616150 | . | . | [illegible] |
| 153339 | NM_174909 | TMEM167A | transmembrane protein 167A | protein_coding | NP_777569.1 | 28330 | . | . | . | [illegible] |
| 153364 | NM_203406 | MBLAC2 | metallo-beta-lactamase domain containing 2 | protein_coding | NP_981951.2 | 33711 | . | . | . | [illegible] |
| 153396 | NM_00128900 | TMEM161B | transmembrane protein 161B | protein_coding | NP_001275936.1;NP_001275937.1;NP_001336329.1;NP_001336331.1;NI | 28483 | . | . | . | [illegible] |
| 153443 | NM_152546,XI | SRFBP1 | serum response factor binding protein 1 | protein_coding | NP_689759.2;XP_016864600.2 | 26333 | 610479 | . | . | [illegible] |
| 153527 | NM_144723 | ZMAT2 | zinc finger matrin-type 2 | protein_coding | NP_653324.1 | 26433 | . | . | . | [illegible] |
| 153571 | NM_00129433 | C5orf38 | chromosome 5 open reading frame 38 | protein_coding | NP_001281266.1;NP_001293078.1;NP_001293079.1;NP_001352614.1;NI | 24226 | 610522 | . | . | [illegible] |
| 153572 | NM_00113422 | IRX2 | iroquois homeobox 2 | protein_coding | NP_001127694.1;NP_150366.1;XP_011512281.1;XP_024310147.1 | 14359 | 606198 | . | . | [illegible] |
| 153579 | NM_00130824 | BTNL9 | butyrophilin like 9 | protein_coding | NP_001295174.1;NP_689760.2;XP_006714886.1;XP_006714887.1;XP_00 | 24176 | . | . | . | [illegible] |
| 153642 | NM_198150,XI | ARSK | arylsulfatase family member K | protein_coding | NP_937793.1;XP_005271961.1 | 25239 | 610011 | . | . | [illegible] |
| 153657 | NM_00131794 | TTC23L | tetratricopeptide repeat domain 23 like | protein_coding | NP_001304878.1;NP_653326.3;XP_011512286.1;XP_011512287.1;XP_01 | 26355 | 616344 | . | . | [illegible] |
| 153684 | NR_015447 | LOC153684 | uncharacterized LOC153684 | lncRNA | . | . | . | . | . | [illegible] |
| 153733 | NM_00104044 | CCDC112 | coiled-coil domain containing 112 | protein_coding | NP_001035530.1;NP_001362825.1;NP_001362826.1;NP_001362827.1;NI | 28599 | . | . | . | [illegible] |
| 153743 | NM_206858 | PPP1R2B | PPP1R2 family member B | protein_coding | NP_996740.2 | 16318 | . | . | . | [illegible] |
| 153768 | NM_138492,N | PRELID2 | PRELI domain containing 2 | protein_coding | NP_612501.3;NP_892005.1;NP_995318.1;XP_016864616.1;XP_01686461 | 28306 | . | . | . | [illegible] |
| 153770 | NM_00102986 | PLAC8L1 | PLAC8 like 1 | protein_coding | NP_001025040.1;XP_005268438.1;XP_011535873.1 | 31746 | . | . | . | [illegible] |
| 153830 | NM_00119938 | RNF145 | ring finger protein 145 | protein_coding | NP_001186309.1;NP_001186310.1;NP_001186311.1;NP_001186312.1;NI | 20853 | . | . | . | [illegible] |
| 154007 | NM_152551,XI | SNRNP48 | small nuclear ribonucleoprotein U11/U12 subunit 48 | protein_coding | NP_689764.3;XP_011512614.1 | 21368 | . | . | . | [illegible] |
| 154043 | NM_00136811 | CNKSR3 | CNKSR family member 3 | protein_coding | NP_001355045.1;NP_001355046.1;NP_001355047.1;NP_001355048.1;NI | 23034 | 617476 | . | . | [illegible] |
| 154091 | NM_145176,XI | SLC2A12 | solute carrier family 2 member 12 | protein_coding | NP_660159.1;XP_006715412.1;XP_016865800.1 | 18067 | 610372 | . | . | [illegible] |
| 154141 | NM_00108048 | MBOAT1 | membrane bound O-acyltransferase domain containing 1 | protein_coding | NP_001073949.1;XP_006715062.1;XP_006715063.1;XP_011512615.1 | 21579 | 611732 | . | . | [illegible] |
| 154197 | NM_00127186 | PNLDC1 | PARN like, ribonuclease domain containing 1 | protein_coding | NP_001258791.1;NP_775787.1;XP_011533793.1;XP_011533795.1;XP_01 | 21185 | . | . | . | [illegible] |
| 154214 | NM_00128639 | RNF217 | ring finger protein 217 | protein_coding | NP_001273327.1;NP_689766.1;XP_011533796.1;XP_011533797.1;XP_01 | 21487 | 618592 | . | . | [illegible] |
| 154313 | NM_00103174 | CFAP206 | cilia and flagella associated protein 206 | protein_coding | NP_001026913.1 | 21405 | . | . | . | [illegible] |
| 154442 | NR_037157 | BVES-AS1 | BVES antisense RNA 1 | lncRNA | . | 21223 | . | . | . | [illegible] |
| 154467 | NM_138493 | CCDC167 | coiled-coil domain containing 167 | protein_coding | NP_612502.1 | 21239 | . | . | . | [illegible] |
| 154743 | NM_152556 | BMT2 | base methyltransferase of 25S rRNA 2 homolog | protein_coding | NP_689769.2 | 26475 | 617855 | . | . | [illegible] |
| 154761 | NR_015421 | LOC154761 | family with sequence similarity 115, member C pseudogene | transcribed_ps | . | . | . | . | . | [illegible] |
| 154791 | NM_197964,N | FMC1 | formation of mitochondrial complex V assembly factor 1 homolog | protein_coding | NP_932068.2 | 26946 | . | . | . | [illegible] |
| 154796 | NM_00111349 | AMOT | angiomotin | protein_coding | NP_001106962.1;NP_573572.1;XP_005262144.1;XP_005262147.1;XP_01 | 17810 | 300410 | . | . | [illegible] |
| 154807 | NM_00128434 | VKORC1L1 | vitamin K epoxide reductase complex subunit 1 like 1 | protein_coding | NP_001271271.1;NP_775788.2;XP_011514133.1 | 21492 | 608838 | . | . | [illegible] |
| 154810 | NM_00130100 | AMOTL1 | angiomotin like 1 | protein_coding | NP_001287936.1;NP_570899.1;XP_005273855.1;XP_005273858.1;XP_00 | 17811 | 614657 | . | . | [illegible] |
| 154860 | NR_036484 | FEZF1-AS1 | FEZF1 antisense RNA 1 | lncRNA | . | 41001 | . | . | . | [illegible] |
| 154865 | NM_00128285 | IQUB | IQ motif and ubiquitin domain containing | protein_coding | NP_001269784.1;NP_001308222.1;NP_849149.3;XP_005250218.1;XP_00 | 21995 | . | . | . | [illegible] |
| 154881 | NM_00116796 | KCTD7 | potassium channel tetramerization domain containing 7 | protein_coding | NP_001161433.1;NP_694578.1 | 21957 | 611725 | . | . | [illegible] |
| 155019 | gene-ST13P7 | ST13P7 | ST13, Hsp70 interacting protein pseudogene 7 | pseudogene | . | 18558 | . | . | . | [illegible] |
| 155054 | NM_00100166 | ZNF425 | zinc finger protein 425 | protein_coding | NP_001001661.1 | 20690 | . | . | . | [illegible] |
| 155060 | NR_036573 | LOC155060 | AI894139 pseudogene | transcribed_ps | . | . | . | . | . | [illegible] |
| 155061 | NM_00116347 | ZNF746 | zinc finger protein 746 | protein_coding | NP_001156946.1;NP_001350446.1;NP_689770.3;XP_005250012.1;XP_00 | 21948 | 613914 | . | . | [illegible] |
| 155066 | NM_00110059 | ATP6V0E2 | ATPase H+ transporting V0 subunit e2 | protein_coding | NP_001094062.1;NP_001276919.1;NP_001354717.1;NP_001354718.1;NI | 21723 | 611019 | . | . | [illegible] |
| 155368 | NM_152559,XI | METTL27 | methyltransferase like 27 | protein_coding | NP_689772.2;XP_016867266.1;XP_016867267.1 | 19068 | 612546 | . | . | [illegible] |
| 155370 | NR_001588,NF | SBDSP1 | SBDS pseudogene 1 | transcribed_ps | . | 21646 | . | . | . | [illegible] |
| 155382 | NM_00107762 | VPS37D | VPS37D subunit of ESCRT-I | protein_coding | NP_001071089.1;XP_016867268.1 | 18287 | 610039 | . | . | [illegible] |
| 155400 | NR_033322,NF | NSUN5P1 | NSUN5 pseudogene 1 | transcribed_ps | . | 19146 | . | . | . | [illegible] |
| 155435 | NM_053043,XI | RBM33 | RNA binding motif protein 33 | protein_coding | NP_444271.2;XP_005249588.1;XP_005249589.1;XP_005249590.1;XP_00 | 27223 | . | . | . | [illegible] |
| 157273 | NR_040039 | LOC157273 | uncharacterized LOC157273 | lncRNA | . | . | . | . | . | [illegible] |
| 157285 | NM_00108082 | PRAG1 | PEAK1 related, kinase-activating pseudokinase 1 | protein_coding | NP_001074295.2;NP_001356688.1 | 25438 | 617344 | . | . | [illegible] |
| 157313 | NM_00131790 | CDCA2 | cell division cycle associated 2 | protein_coding | NP_001304835.1;NP_001304836.1;NP_689775.2;XP_006716357.1;XP_01 | 14623 | 618785 | . | . | [illegible] |
| 157378 | NM_194291 | TMEM65 | transmembrane protein 65 | protein_coding | NP_919267.2 | 25203 | 616609 | . | . | [illegible] |
| 157489 | NR_144550 | SDAD1P1 | SDA1 domain containing 1 pseudogene 1 | transcribed_ps | . | 31403 | . | . | . | [illegible] |
| 157506 | NM_172037 | RDH10 | retinol dehydrogenase 10 | protein_coding | NP_742034.1 | 19975 | 607599 | . | . | [illegible] |
| 157530 | gene-ST13P6 | ST13P6 | ST13, Hsp70 interacting protein pseudogene 6 | pseudogene | . | 18557 | . | . | . | [illegible] |
| 157567 | NM_00127037 | ANKRD46 | ankyrin repeat domain 46 | protein_coding | NP_001257306.1;NP_001257307.1;NP_001257308.1;NP_940683.1 | 27229 | . | . | . | [illegible] |
| 157570 | NM_00101742 | ESCO2 | establishment of sister chromatid cohesion N-acetyltransferase 2 | protein_coding | NP_001017420.1;XP_011542723.1;XP_011542724.1 | 27230 | 609353 | . | . | [illegible] |
| 157574 | NM_00125821 | FBXO16 | F-box protein 16 | protein_coding | NP_001245140.1;NP_758954.1 | 13618 | 608519 | . | . | [illegible] |
| 157638 | NM_174911,N | LRATD2 | LRAT domain containing 2 | protein_coding | NP_777571.1;XP_016868596.1;XP_016868597.1 | 24166 | 609483 | . | . | [illegible] |
| 157657 | NM_00136326 | C8orf37 | chromosome 8 open reading frame 37 | protein_coding | NP_001350189.1;NP_808880.1 | 27232 | 614477 | . | . | [illegible] |
| 157680 | NM_015243,N | VPS13B | vacuolar protein sorting 13 homolog B | protein_coding | NP_056058.2;NP_060360.3;NP_689777.3;NP_858047.2 | 2183 | 607817 | . | . | [illegible] |
| 157695 | NM_00125611 | TDRP | testis development related protein | protein_coding | NP_001243042.1;NP_001356645.1;NP_778250.2;XP_006716295.1 | 26951 | . | . | . | [illegible] |
| 157697 | NM_00130310 | ERICH1 | glutamate rich 1 | protein_coding | NP_001290029.1;NP_997215.1;XP_006716297.2;XP_011533033.1;XP_01 | 27234 | . | . | . | [illegible] |
| 157713 | gene-LYPLA1P3 | LYPLA1P3 | LYPLA1 pseudogene 3 | pseudogene | . | 44007 | . | . | . | [illegible] |
| 157753 | NM_153015,N | TMEM74 | transmembrane protein 74 | protein_coding | NP_694560.1 | 26409 | 613935 | . | . | [illegible] |
| 157769 | NM_00131791 | FAM91A1 | family with sequence similarity 91 member A1 | protein_coding | NP_001304846.2;NP_001304847.1;NP_659400.3 | 26306 | . | . | . | [illegible] |
| 157773 | NM_00100709 | C8orf48 | chromosome 8 open reading frame 48 | protein_coding | NP_001007091.2 | 26345 | . | . | . | [illegible] |
| 157777 | NM_00113616 | MCMDC2 | minichromosome maintenance domain containing 2 | protein_coding | NP_001129632.1;NP_001129633.1;NP_775789.3;XP_005251231.1;XP_00 | 26368 | 617545 | . | . | [illegible] |
| 157922 | NM_015447,XI | CAMSAP1 | calmodulin regulated spectrin associated protein 1 | protein_coding | NP_056262.3;XP_005263453.1;XP_005263454.1;XP_011516574.1;XP_01 | 19946 | 613774 | . | . | [illegible] |
| 157956 | gene-CDC20P1 | CDC20P1 | cell division cycle 20 pseudogene 1 | pseudogene | . | 29487 | . | . | . | [illegible] |
| 158038 | NM_00125828 | LINGO2 | leucine rich repeat and Ig domain containing 2 | protein_coding | NP_001245211.1;NP_001341503.1;NP_001341504.1;NP_689783.1;XP_01 | 21207 | 609793 | . | . | [illegible] |
| 158055 | NM_152571 | C9orf163 | chromosome 9 open reading frame 163 | protein_coding | NP_689784.1 | 26718 | . | . | . | [illegible] |
| 158056 | NM_206920 | MAMDC4 | MAM domain containing 4 | protein_coding | NP_996803.2 | 24083 | 617208 | . | . | [illegible] |
| 158067 | NM_00131795 | AK8 | adenylate kinase 8 | protein_coding | NP_001304887.1;NP_001304888.1;NP_001358700.1;NP_001358701.1;NI | 26526 | 615365 | . | . | [illegible] |
| 158078 | gene-EEF1A1P5 | EEF1A1P5 | eukaryotic translation elongation factor 1 alpha 1 pseudogene 5 | pseudogene | . | 3200 | . | . | . | [illegible] |
| 158104 | gene-RPS10P3 | RPS10P3 | ribosomal protein S10 pseudogene 3 | pseudogene | . | 23684 | . | . | . | [illegible] |
| 158135 | NM_00113944 | TTLL11 | tubulin tyrosine ligase like 11 | protein_coding | NP_001132914.1;NP_919228.2;XP_005251785.1;XP_005251786.1;XP_00 | 18113 | . | . | . | [illegible] |
| 158158 | NM_152573,XI | RASEF | RAS and EF-hand domain containing | protein_coding | NP_689786.2;XP_005251787.1;XP_005251788.1 | 26464 | 611344 | . | . | [illegible] |
| 158160 | NR_003086 | HSD17B7P2 | hydroxysteroid 17-beta dehydrogenase 7 pseudogene 2 | transcribed_ps | . | 28120 | . | . | . | [illegible] |
| 158219 | NM_00116833 | TTC39B | tetratricopeptide repeat domain 39B | protein_coding | NP_001161811.1;NP_001161812.1;NP_001161813.1;NP_001161814.1;NI | 23704 | 613574 | . | . | [illegible] |
| 158222 | gene-LDHAP4 | LDHAP4 | lactate dehydrogenase A pseudogene 4 | pseudogene | . | 6539 | . | . | . | [illegible] |
| 158234 | NM_00128695 | TRMT10B | tRNA methyltransferase 10B | protein_coding | NP_001273879.1;NP_001273880.1;NP_001273881.1;NP_001273882.1;NI | 26454 | . | . | . | [illegible] |
| 158293 | NM_00132222 | FAM120AOS | family with sequence similarity 120A opposite strand | protein_coding | NP_001309153.1;NP_942138.2 | 23389 | . | . | . | [illegible] |
| 158326 | NM_00117770 | FREM1 | FRAS1 related extracellular matrix 1 | protein_coding | NP_001171175.1;NP_001356987.1;NP_001356989.1;NP_001356990.1;NI | 23399 | 608944 | . | . | [illegible] |
| 158345 | gene-RPL4P5 | RPL4P5 | ribosomal protein L4 pseudogene 5 | pseudogene | . | 35699 | . | . | . | [illegible] |
| 158358 | NM_00101796 | KIAA2026 | KIAA2026 | protein_coding | NP_001017969.2;XP_005251442.1;XP_011516061.1;XP_011516062.1;XP_ | 23378 | . | . | . | [illegible] |
| 158373 | gene-YBX1P10 | YBX1P10 | Y-box binding protein 1 pseudogene 10 | pseudogene | . | 42432 | . | . | . | [illegible] |
| 158383 | gene-YWHAZP6 | YWHAZP6 | tyrosine 3-monooxygenase/tryptophan 5-monooxygenase activati | pseudogene | . | 30849 | . | . | . | [illegible] |
| 158399 | NM_00100716 | ZNF483 | zinc finger protein 483 | protein_coding | NP_001007170.1;NP_597721.2;XP_011516602.1;XP_016869826.1;XP_01 | 23384 | . | . | . | [illegible] |
| 158405 | NM_00128703 | KIAA1958 | KIAA1958 | protein_coding | NP_001273965.1;NP_001273967.1;NP_597722.1;XP_011516613.1 | 23427 | 617390 | . | . | [illegible] |
| 158427 | NM_139246 | TSTD2 | thiosulfate sulfurtransferase like domain containing 2 | protein_coding | NP_640339.4 | 30087 | . | . | . | [illegible] |
| 158431 | NM_00100166 | ZNF782 | zinc finger protein 782 | protein_coding | NP_001001662.1;NP_001333920.1;NP_001333922.1;NP_001333924.1;XP | 33110 | . | . | . | [illegible] |
| 158435 | NR_033838 | LOC158435 | uncharacterized LOC158435 | lncRNA | . | . | . | . | . | [illegible] |
| 158471 | NM_00130804 | PRUNE2 | prune homolog 2 with BCH domain | protein_coding | NP_001294976.1;NP_001294977.1;NP_001294978.1;NP_001294979.1;NI | 25209 | 610691 | . | . | [illegible] |
| 158511 | NM_00110257 | CSAG1 | chondrosarcoma associated gene 1 | protein_coding | NP_001096046.2;NP_705611.2 | 24294 | 300944 | . | . | [illegible] |
| 158572 | NR_026742 | USP27X-AS1 | USP27X antisense RNA 1 (head to head) | lncRNA | . | 27249 | . | . | . | [illegible] |
| 158586 | NM_007157 | ZXDB | zinc finger X-linked duplicated B | protein_coding | NP_009088.1 | 13199 | 300236 | . | . | [illegible] |
| 158747 | NM_00117747 | MOSPD2 | motile sperm domain containing 2 | protein_coding | NP_001170946.1;NP_001317170.1;NP_689794.1 | 28381 | . | . | . | [illegible] |
| 158787 | NM_00103174 | RIBC1 | RIB43A domain with coiled-coils 1 | protein_coding | NP_001026915.1;NP_001253982.1;NP_659405.1;XP_005262045.1;XP_00 | 26537 | . | . | . | [illegible] |

| | | | | | | | | | | | | | | | | | | | | | | | | | | | | | | | | | |
|---|---|---|---|---|---|---|---|---|---|---|---|---|---|---|---|---|---|---|---|---|---|---|---|---|---|---|---|---|---|---|---|---|---|
| 158801 | NR_027131 | NKAPP1 | NFKB activating protein pseudogene 1 | transcribed_ps | . | 26706 | . | . | . | 0.378433 | 0.492503 | 0.334310 | 0.418453 | 0.880628 | 1.138443 | 0.750181 | 0.956742 | -1.176281 | 31.205753 | 0.560787953 | -0.41768406 | 0.676178127 | 0.998865404 | 5.077319 | 4.840449 | 4.957643 | 4.991977 | 4.941457 | 4.967763 | 34 | 41 | 22 | 30 |
| 158880 | NM_201286,XI | USP51 | ubiquitin specific peptidase 51 | protein_coding | NP_958443.1;XP_016884788.1;XP_016884789.1;XP_016884790.1 | 23086 | . | . | . | 1.732333 | 1.945215 | 1.744196 | 2.150382 | 4.031210 | 4.496453 | 3.913925 | 4.916592 | 1.055431 | 278.220119 | 0.216299974 | 0.359831948 | 0.71897281 | 0.998865404 | 8.081313 | 8.157838 | 8.073667 | 8.133527 | 8.079403 | 8.182107 | 294 | 309 | 224 | 294 |
| 158960 | NR_103768 | CH17-340M24. | uncharacterized protein BC009467 | lncRNA | . | . | . | . | . | 3.485574 | 1.934975 | 2.387549 | 2.648164 | 8.111073 | 4.472780 | 5.357590 | 6.054713 | -1.098759 | 49.246767 | 0.472905334 | -0.28732101 | 0.773866537 | 0.998865404 | 5.687500 | 5.553294 | 5.681221 | 5.572078 | 5.603185 | 5.618484 | 76 | 40 | 39 | 46 |
| 159013 | NM_00133045 | CXorf38 | chromosome X open reading frame 38 | protein_coding | NP_001317384.1;NP_659407.1;XP_005272646.2;XP_006724590.2;XP_00 | 28589 | . | . | . | 2.726662 | 3.340006 | 3.259410 | 2.846768 | 6.345052 | 7.720575 | 7.314017 | 6.508796 | 1.006811 | 408.020695 | 0.187251777 | 0.052300484 | 0.958289268 | 0.998865404 | 8.667512 | 8.677468 | 8.628064 | 8.708369 | 8.712237 | 8.632371 | 437 | 468 | 380 | 358 |
| 159090 | NM_00116659 | FAM122B | family with sequence similarity 122B | protein_coding | NP_001160071.1;NP_001160072.1;NP_001164227.1;NP_001164228.1;NI | 30490 | . | . | . | 5.581668 | 5.100269 | 5.985261 | 4.101237 | 12.988774 | 11.789503 | 13.430751 | 9.376988 | -1.097619 | 610.569801 | 0.190923256 | -0.70383255 | 0.481537059 | 0.998865404 | 9.319064 | 9.185889 | 9.298621 | 9.260442 | 9.334009 | 9.086544 | 756 | 670 | 602 | 443 |
| 159091 | NM_00117077 | FAM122C | family with sequence similarity 122C | protein_coding | NP_001164250.1;NP_001164251.1;NP_001164252.1;NP_001164253.1;NI | 25202 | . | . | . | 0.809320 | 0.693640 | 0.713907 | 1.116178 | 1.883319 | 1.603378 | 1.601986 | 2.552006 | 1.218609 | 56.593662 | 0.426047567 | 0.669491546 | 0.503181972 | 0.998865404 | 5.673732 | 5.957476 | 5.804229 | 5.789964 | 5.800614 | 5.872873 | 61 | 53 | 45 | 68 |
| 159195 | NM_00132043 | USP54 | ubiquitin specific peptidase 54 | protein_coding | NP_001307366.1;NP_001307370.1;NP_001337924.1;NP_001365137.1;NI | 23513 | . | . | . | 4.666485 | 5.023094 | 4.752009 | 5.119202 | 10.859103 | 11.611105 | 10.663371 | 11.704448 | 1.087256 | 1192.472854 | 0.136048578 | 0.887120033 | 0.375014279 | 0.998865404 | 10.158306 | 10.278666 | 10.140906 | 10.230505 | 10.238010 | 10.260801 | 1237 | 1307 | 1064 | 1182 |
| 159371 | NM_00113465 | SLC35G1 | solute carrier family 35 member G1 | protein_coding | NP_001128130.1;NP_001332922.1;NP_694958.1;XP_011537675.1 | 26607 | 617167 | . | . | 6.164900 | 6.447045 | 4.755425 | 4.052114 | 14.345974 | 14.902632 | 10.671035 | 9.264679 | -1.467814 | 552.508727 | 0.16840184 | -3.28778801 | **0.001009778** | 0.068323392 | 9.360019 | 8.807016 | 9.207902 | 9.228943 | 9.016268 | 8.921911 | 745 | 720 | 420 | 387 |
| 160335 | NM_00132032 | TMTC2 | transmembrane O-mannosyltransferase targeting cadherins 2 | protein_coding | NP_001307250.1;NP_001307251.1;NP_689801.1;XP_016874373.1;XP_02 | 25440 | 615856 | . | . | 10.072437 | 9.264673 | 9.983823 | 9.227912 | 23.438970 | 21.415710 | 22.403411 | 21.098525 | -1.063456 | 1612.458786 | 0.128387329 | -0.69134954 | 0.489345907 | 0.998865404 | 10.698576 | 10.610162 | 10.711398 | 10.643613 | 10.663113 | 10.596409 | 1996 | 1716 | 1418 | 1402 |
| 160418 | NM_00136657 | TMTC3 | transmembrane O-mannosyltransferase targeting cadherins 3 | protein_coding | NP_001353503.1;NP_001353508.1;NP_001353509.1;NP_001353512.1;NI | 26899 | 617218 | . | . | 2.189867 | 1.595280 | 1.995585 | 1.137208 | 5.095912 | 3.687561 | 4.478034 | 2.600091 | -1.226598 | 388.835229 | 0.25438246 | -1.15834252 | 0.246724287 | 0.992733415 | 8.741918 | 8.449300 | 8.743737 | 8.541360 | 8.672856 | 8.356743 | 573 | 387 | 389 | 236 |
| 160428 | NM_00103417 | ALDH1L2 | aldehyde dehydrogenase 1 family member L2 | protein_coding | NP_001029345.2;XP_011536288.1;XP_011536290.1;XP_011536291.1;XP_ | 26777 | 613584 | . | . | 3.124431 | 3.375141 | 1.781056 | 2.925614 | 7.270679 | 7.801790 | 3.996635 | 6.689072 | -1.393200 | 615.179737 | 0.197948037 | -2.41681021 | **0.015657178** | 0.418829521 | 9.484901 | 9.005121 | 9.342213 | 9.376334 | 8.995605 | 9.263361 | 805 | 792 | 351 | 585 |
| 160518 | NM_00130833 | DENND5B | DENN domain containing 5B | protein_coding | NP_001295268.1;NP_001353819.1;NP_001353820.1;NP_001353821.1;NI | 28338 | 617279 | . | . | 1.932714 | 1.722268 | 1.763208 | 1.866073 | 4.497506 | 3.981099 | 3.956589 | 4.266557 | -1.050082 | 456.704698 | 0.182959423 | -0.38533875 | 0.699986459 | 0.998865404 | 8.870433 | 8.798917 | 8.831616 | 8.866395 | 8.753633 | 8.879983 | 524 | 519 | 347 | 461 |
| 160622 | NM_00127185 | TAMALIN | trafficking regulator and scaffold protein tamalin | protein_coding | NP_001258785.1;NP_859062.1;XP_005268748.1;XP_011536298.1 | 18707 | 612027 | . | . | 7.762357 | 7.441836 | 8.367324 | 8.771848 | 18.063322 | 17.202136 | 18.776033 | 20.055788 | 1.086947 | 522.107537 | 0.165070127 | 0.728670389 | 0.46620331 | 0.998865404 | 8.966734 | 9.087160 | 9.017498 | 8.982570 | 9.050908 | 9.056402 | 592 | 525 | 471 | 512 |
| 160760 | NM_139283,N | PPTC7 | protein phosphatase targeting COQ7 | protein_coding | NP_644812.1;XP_024304637.1;XP_024304638.1 | 30695 | 609668 | . | . | 6.737235 | 6.645858 | 7.466398 | 6.509933 | 15.677823 | 15.362198 | 16.754382 | 14.884187 | 1.027907 | 1059.359354 | 0.140781366 | 0.282071586 | 0.77788862 | 0.998865404 | 10.028755 | 10.068918 | 10.047122 | 10.028526 | 10.107758 | 10.006450 | 1218 | 1113 | 998 | 938 |
| 160851 | NM_00120450 | DGKH | diacylglycerol kinase eta | protein_coding | NP_001191433.1;NP_001191434.1;NP_001191435.1;NP_001284358.1;NI | 2854 | 604071 | . | . | 0.494173 | 0.436750 | 0.591793 | 0.353994 | 1.149961 | 1.009566 | 1.327966 | 0.809365 | -1.095680 | 244.530907 | 0.304166018 | -0.43340232 | 0.664722518 | 0.998865404 | 7.997125 | 7.867720 | 7.970015 | 7.907426 | 8.062986 | 7.690067 | 312 | 259 | 281 | 135 |
| 160857 | NM_00135061 | CCDC122 | coiled-coil domain containing 122 | protein_coding | NP_001337546.1;NP_001337547.1;NP_659411.2;XP_016875886.1;XP_01 | 26478 | 613408 | . | . | 1.544838 | 1.303221 | 1.203155 | 0.891092 | 3.594901 | 3.012456 | 2.699846 | 2.037374 | -1.328608 | 83.649554 | 0.364023612 | -1.12606855 | 0.260136505 | 0.995760118 | 6.575540 | 6.168402 | 6.432969 | 6.409494 | 6.391370 | 6.296286 | 114 | 99 | 75 | 54 |
| 160897 | NM_180989 | GPR180 | G protein-coupled receptor 180 | protein_coding | NP_851320.1 | 28899 | 607787 | . | . | 4.807949 | 4.080453 | 4.416659 | 2.676831 | 11.188295 | 9.432151 | 9.910856 | 6.120257 | -1.269387 | 1107.872554 | 0.381183867 | -0.90279749 | 0.366633382 | 0.998865404 | 10.275291 | 9.931419 | 10.251759 | 10.118303 | 10.174329 | 9.812131 | 1550 | 1222 | 1061 | 687 |
| 161145 | NM_00134854 | TMEM229B | transmembrane protein 229B | protein_coding | NP_001335470.1;NP_001335471.1;NP_001335472.1;NP_001335473.1;NI | 20130 | . | . | . | 8.434035 | 8.762281 | 9.399007 | 10.958551 | 19.626344 | 20.254405 | 21.091098 | 25.055429 | 1.165276 | 1179.433356 | 0.138822696 | 1.589593715 | 0.111926415 | 0.848857309 | 10.089567 | 10.309793 | 10.129497 | 10.155097 | 10.199577 | 10.311500 | 1233 | 1196 | 1023 | 1277 |
| 161176 | NM_00136369 | SYNE3 | spectrin repeat containing nuclear envelope family member 3 | protein_coding | NP_001350621.1;NP_689805.3;XP_005267434.1;XP_006720126.1;XP_01 | 19861 | 610861 | . | . | 1.926145 | 2.275991 | 1.689995 | 1.562289 | 4.482220 | 5.261056 | 3.792298 | 3.571989 | -1.222049 | 650.771575 | 0.155223935 | -1.86377153 | 0.062353795 | 0.723723143 | 9.483563 | 9.193941 | 9.385323 | 9.433209 | 9.263951 | 9.284599 | 796 | 799 | 501 | 558 |
| 161357 | NM_00111349 | MDGA2 | MAM domain containing glycosylphosphatidylinositol anchor 2 | protein_coding | NP_001106970.3;NP_878250.2;XP_011534821.1;XP_011534824.1;XP_01 | 19835 | 611128 | . | . | 1.026263 | 1.173727 | 1.584866 | 1.035423 | 2.388158 | 2.713124 | 3.556394 | 2.367373 | 1.095686 | 243.823291 | 0.230875618 | 0.571019074 | 0.567986713 | 0.998865404 | 7.861447 | 7.994854 | 7.939275 | 7.865621 | 8.003569 | 7.890120 | 288 | 232 | 247 | 212 |
| 161394 | NM_00101086 | SAMD15 | sterile alpha motif domain containing 15 | protein_coding | NP_001010860.1 | 18631 | . | . | . | 0.443995 | 0.796133 | 0.451432 | 0.833762 | 1.033195 | 1.840297 | 1.013001 | 1.906297 | 1.029469 | 42.146487 | 0.527627758 | 0.07941341 | 0.936703803 | 0.998865404 | 5.378857 | 5.415590 | 5.341716 | 5.429818 | 5.342730 | 5.438116 | 35 | 57 | 26 | 52 |
| 161424 | NM_00128636 | NOP9 | NOP9 nucleolar protein | protein_coding | NP_001273296.1;NP_777573.1;XP_005267442.1;XP_011534828.1;XP_01 | 19826 | 618308 | . | . | 5.357031 | 5.783123 | 5.603266 | 5.407954 | 12.466033 | 13.367952 | 12.573566 | 12.364645 | -1.029887 | 1016.323743 | 0.139069872 | -0.30550463 | 0.759981843 | 0.998865404 | 10.010260 | 9.967714 | 9.965616 | 10.032398 | 9.993899 | 9.961665 | 1134 | 1163 | 890 | 918 |
| 161497 | NM_153700 | STRC | stereocilin | protein_coding | NP_714544.1 | 16035 | 606440 | . | . | 0.028555 | 0.034897 | 0.078815 | 0.034760 | 0.066449 | 0.080666 | 0.176858 | 0.079475 | 1.723658 | 7.974390 | 1.098550482 | 0.715008815 | 0.474603596 | NA | 2.548173 | 3.336262 | 2.963653 | 2.969818 | 3.001815 | 2.969661 | 6 | 7 | 12 | 6 |
| 161582 | NM_00103355 | DNAAF4 | dynein axonemal assembly factor 4 | protein_coding | NP_001028731.1;NP_001028732.1;NP_570722.2 | 21493 | 608706 | . | . | 1.593702 | 2.022398 | 1.711255 | 1.833233 | 3.708609 | 4.674862 | 3.840006 | 4.191470 | -1.040883 | 108.870560 | 0.313301525 | -0.18451275 | 0.853611226 | 0.998865404 | 6.795722 | 6.736612 | 6.737939 | 6.805037 | 6.746556 | 6.771547 | 115 | 132 | 89 | 104 |
| 161725 | NM_00132990 | OTUD7A | OTU deubiquitinase 7A | protein_coding | NP_001316836.1;NP_001369566.1;NP_570971.1 | 20718 | 612024 | . | . | 0.379271 | 0.236537 | 0.094111 | 0.150337 | 0.882579 | 0.546765 | 0.211181 | 0.343728 | -1.546955 | 54.616467 | 0.463138915 | -1.35905592 | 0.17412887 | 0.948096927 | 6.053654 | 5.419808 | 5.748350 | 5.862330 | 5.686433 | 5.755863 | 58 | 89 | 29 | 49 |
| 161742 | NM_152594,XI | SPRED1 | sprouty related EVH1 domain containing 1 | protein_coding | NP_689807.1;XP_005254259.1;XP_011519591.1 | 20249 | 609291 | . | . | 7.103028 | 6.335662 | 6.539591 | 6.504223 | 16.529036 | 14.645167 | 14.674652 | 14.871132 | -1.043867 | 1581.028813 | 0.128801531 | -0.48087439 | 0.630605778 | 0.998865404 | 10.657214 | 10.595421 | 10.688279 | 10.595067 | 10.601142 | 10.617605 | 1970 | 1639 | 1331 | 1466 |
| 161779 | NM_152595 | PGBD4 | piggyBac transposable element derived 4 | protein_coding | NP_689808.2 | 19401 | . | . | . | 1.199839 | 1.145180 | 1.159165 | 1.023348 | 2.792074 | 2.647134 | 2.601132 | 2.339763 | -1.091248 | 233.118996 | 0.225264629 | -0.55925055 | 0.575990737 | 0.998865404 | 7.926092 | 7.801045 | 7.899591 | 7.871024 | 7.874333 | 7.811185 | 288 | 255 | 207 | 195 |
| 161823 | NM_00101296 | ADAL | adenosine deaminase like | protein_coding | NP_001012987.1;NP_001152752.1;NP_001311293.1;NP_001311294.1;NI | 31853 | . | . | . | 5.204588 | 4.582468 | 5.192568 | 4.239573 | 12.111288 | 10.592579 | 11.651975 | 9.693280 | 1.026194 | 552.238826 | 0.165907098 | 0.2248424 | 0.82210188 | 0.998865404 | 9.090048 | 9.127999 | 9.122927 | 9.076267 | 9.151497 | 9.080925 | 650 | 567 | 513 | 496 |
| 161829 | NM_00128644 | EXD1 | exonuclease 3'-5' domain containing 1 | protein_coding | NP_001273370.1;NP_689809.2;XP_006720478.1;XP_011519600.1;XP_01 | 28507 | . | . | . | 0.117390 | 0.065428 | 0.107670 | 0.059647 | 0.273171 | 0.151239 | 0.241609 | 0.136375 | -1.172565 | 10.321374 | 0.972889031 | -0.2360674 | 0.813380373 | NA | 3.474811 | 3.251698 | 3.383280 | 3.357954 | 3.376890 | 3.349685 | 16 | 9 | 11 | 6 |
| 161835 | NM_00132433 | FSIP1 | fibrous sheath interacting protein 1 | protein_coding | NP_001311267.1;NP_689810.3;XP_011519607.1;XP_011519608.1;XP_01 | 21674 | 615795 | . | . | 0.206155 | 0.183232 | 0.111472 | 0.217356 | 0.479732 | 0.423550 | 0.250140 | 0.496958 | -1.144675 | 30.621598 | 0.571483246 | -0.34110959 | 0.733021082 | 0.998865404 | 5.032032 | 4.834143 | 4.947736 | 4.945310 | 4.903238 | 4.955895 | 38 | 35 | 19 | 33 |
| 161882 | NM_153813,XI | ZFPM1 | zinc finger protein, FOG family member 1 | protein_coding | NP_722520.2;XP_011521214.1;XP_011521216.1;XP_011521219.1;XP_01( | 19762 | 601950 | . | . | 0.995830 | 0.919205 | 0.846726 | 0.967578 | 2.317336 | 2.124786 | 1.900029 | 2.212252 | -1.050806 | 151.166167 | 0.268749914 | -0.26603173 | 0.790214772 | 0.998865404 | 7.275508 | 7.203578 | 7.264756 | 7.234905 | 7.206220 | 7.254353 | 185 | 161 | 120 | 147 |
| 162073 | NM_00103484 | ITPRIPL2 | ITPRIP like 2 | protein_coding | NP_001030013.1 | 27257 | . | . | . | 5.986825 | 6.089069 | 6.481808 | 5.520809 | 13.931588 | 14.075158 | 14.544990 | 12.622674 | -1.023416 | 1442.538260 | 0.135203732 | -0.2469781 | 0.804925172 | 0.998865404 | 10.510807 | 10.477792 | 10.499049 | 10.505549 | 10.545667 | 10.419695 | 1673 | 1581 | 1342 | 1227 |
| 162239 | NM_00131846 | ZFP1 | ZFP1 zinc finger protein | protein_coding | NP_001305398.1;NP_001305400.1;NP_001305401.1;NP_001305402.1;NI | 23328 | 617230 | . | . | 1.864339 | 1.405056 | 1.724330 | 1.707438 | 4.338393 | 3.247851 | 3.869344 | 3.903855 | 1.089552 | 179.157708 | 0.262085828 | 0.472117745 | 0.636842747 | 0.998865404 | 7.421021 | 7.546428 | 7.520405 | 7.401756 | 7.526896 | 7.471551 | 225 | 159 | 173 | 164 |
| 162394 | NM_00133018 | SLFN5 | schlafen family member 5 | protein_coding | NP_001317112.1;NP_659412.3 | 28286 | 614952 | . | . | 4.223873 | 4.285821 | 2.956212 | 3.665146 | 9.829125 | 9.906867 | 6.633653 | 8.379920 | -1.299730 | 1243.307853 | 0.143457022 | -2.63641341 | **0.008378758** | 0.289854633 | 10.456963 | 10.078181 | 10.367894 | 10.371115 | 10.094694 | 10.242811 | 1615 | 1520 | 844 | 1117 |
| 162417 | NM_153006,XI | NAGS | N-acetylglutamate synthase | protein_coding | NP_694551.1;XP_011522740.1;XP_011522741.1 | 17996 | 608300 | . | . | 0.525221 | 0.541102 | 0.515846 | 0.556267 | 1.222209 | 1.250780 | 1.157543 | 1.271839 | -1.013619 | 35.628770 | 0.51773476 | -0.03769414 | 0.969931549 | 0.998865404 | 5.164890 | 5.144983 | 5.155173 | 5.157324 | 5.150956 | 5.157739 | 41 | 39 | 30 | 34 |
| 162427 | NM_178126,N | RETREG3 | reticulophagy regulator family member 3 | protein_coding | NP_835227.1;XP_011522742.1 | 27258 | 616498 | . | . | 10.565285 | 10.663277 | 11.458972 | 11.086907 | 24.585854 | 24.648643 | 25.713602 | 25.348895 | 1.045233 | 1278.732716 | 0.129035218 | 0.494626201 | 0.620864004 | 0.998865404 | 10.288170 | 10.352119 | 10.302790 | 10.303682 | 10.353649 | 10.319732 | 1439 | 1350 | 1162 | 1198 |
| 162461 | NM_00116821 | TMEM92 | transmembrane protein 92 | protein_coding | NP_001161687.1;NP_694961.2;XP_006721793.1 | 26579 | . | . | . | 0.041132 | 0.051914 | 0.066435 | 0.113099 | 0.095716 | 0.120003 | 0.149078 | 0.258588 | 1.833563 | 6.348119 | 1.216150005 | 0.719195651 | 0.472020379 | NA | 2.166062 | 3.037079 | 2.644685 | 2.646202 | 2.651570 | 2.673388 | 5 | 5 | 5 | 10 |
| 162466 | NM_00114380 | PHOSPHO1 | phosphoethanolamine/phosphocholine phosphatase 1 | protein_coding | NP_001137276.1;NP_848595.1;XP_016879760.1 | 16815 | . | . | . | 0.309858 | 0.464821 | 0.567765 | 0.651325 | 0.721051 | 1.074455 | 1.274048 | 1.489176 | 1.497287 | 33.192386 | 0.546679393 | 1.065251639 | 0.286762071 | 0.998865404 | 4.733907 | 5.313775 | 4.992712 | 5.034456 | 5.059534 | 5.073941 | 25 | 34 | 33 | 39 |
| 162494 | NM_00133018 | RHBDL3 | rhomboid like 3 | protein_coding | NP_001317110.1;NP_001350763.1;NP_001350764.1;NP_001350765.1;NI | 16502 | . | . | . | 0.299438 | 0.217110 | 0.373945 | 0.316393 | 0.696804 | 0.501860 | 0.839123 | 0.723396 | 1.323704 | 47.239776 | 0.461196552 | 0.877241351 | 0.38035556 | 0.998865404 | 5.344184 | 5.751080 | 5.549669 | 5.507448 | 5.594910 | 5.561046 | 53 | 38 | 51 | 46 |
| 162514 | NM_00125820 | TRPV3 | transient receptor potential cation channel subfamily V member 3 | protein_coding | NP_001245134.1;NP_659505.1 | 18084 | 607066 | . | . | 0.165275 | 0.223494 | 0.285580 | 0.223970 | 0.384601 | 0.516617 | 0.640833 | 0.512082 | 1.295171 | 41.866566 | 0.490369158 | 0.760942009 | 0.446691705 | 0.998865404 | 5.189181 | 5.562225 | 5.336264 | 5.378484 | 5.419271 | 5.378078 | 36 | 45 | 46 | 39 |
| 162632 | NR_003190 | USP32P1 | ubiquitin specific peptidase 32 pseudogene 1 | transcribed_ps | . | 43570 | . | . | . | 5.203266 | 5.489029 | 6.864041 | 4.819418 | 12.108213 | 12.688136 | 15.402709 | 11.019027 | 1.077764 | 680.233280 | 0.184250815 | 0.586381086 | 0.557619452 | 0.998865404 | 9.354416 | 9.463301 | 9.363439 | 9.393691 | 9.546624 | 9.298411 | 737 | 722 | 724 | 543 |
| 162655 | NM_145287,N | ZNF519 | zinc finger protein 519 | protein_coding | NP_660330.2 | 30574 | . | . | . | 0.824099 | 0.711778 | 0.889660 | 0.753038 | 1.917713 | 1.645307 | 1.996370 | 1.721731 | 1.055425 | 169.673979 | 0.261356079 | 0.297769543 | 0.765879062 | 0.998865404 | 7.366395 | 7.445756 | 7.425138 | 7.359712 | 7.451502 | 7.378057 | 205 | 164 | 165 | 149 |
| 162681 | NM_00128898 | C18orf54 | chromosome 18 open reading frame 54 | protein_coding | NP_001275909.1;NP_001275910.1;NP_001275911.1;NP_001357238.1;NI | 13796 | 613258 | . | . | 1.815454 | 1.634536 | 1.896054 | 0.986617 | 4.224637 | 3.778304 | 4.254689 | 2.255781 | -1.217460 | 302.676821 | 0.263796938 | -1.07610853 | 0.28187872 | 0.998865404 | 8.375616 | 8.093925 | 8.308791 | 8.253059 | 8.321726 | 8.003561 | 401 | 341 | 311 | 177 |
| 162962 | NM_00110265 | ZNF836 | zinc finger protein 836 | protein_coding | NP_001096127.1;XP_011524860.1;XP_011524861.1;XP_016881878.1 | 34333 | . | . | . | 0.755758 | 0.560828 | 0.387567 | 0.511794 | 1.758680 | 1.296378 | 0.869688 | 1.170159 | -1.466053 | 71.283463 | 0.39223317 | -1.40716495 | 0.159378481 | 0.933525128 | 6.405581 | 5.852776 | 6.239653 | 6.156487 | 6.078384 | 6.135618 | 112 | 78 | 43 | 62 |
| 162966 | NM_00132186 | ZNF600 | zinc finger protein 600 | protein_coding | NP_001308795.1;NP_001308796.1;NP_940859.3;XP_016881879.1;XP_01 | 30951 | . | . | . | 1.709890 | 1.641826 | 2.225115 | 1.594299 | 3.978983 | 3.795154 | 4.993096 | 3.645175 | 1.069978 | 242.979148 | 0.230009273 | 0.424251187 | 0.671382634 | 0.998865404 | 7.874334 | 7.973345 | 7.922222 | 7.886674 | 7.998579 | 7.873824 | 280 | 244 | 246 | 206 |
| 162967 | NM_00135177 | ZNF320 | zinc finger protein 320 | protein_coding | NP_001338702.1;NP_001338703.1;NP_001338704.1;NP_001338705.1;NI | 13842 | 606427 | . | . | 0.817811 | 0.889878 | 1.078911 | 1.070080 | 1.903079 | 2.056992 | 2.421046 | 2.446610 | 1.179797 | 177.539824 | 0.255257609 | 0.934502639 | 0.350044675 | 0.998865404 | 7.347314 | 7.586760 | 7.423288 | 7.438981 | 7.547255 | 7.453711 | 184 | 179 | 185 | 161 |
| 162968 | NM_00120700 | ZNF497 | zinc finger protein 497 | protein_coding | NP_001193938.1;NP_940860.2 | 23714 | . | . | . | 0.386729 | 0.247068 | 0.351308 | 0.266394 | 0.899934 | 0.571109 | 0.788326 | 0.609078 | -1.042743 | 34.044605 | 0.549646005 | -0.10985984 | 0.912520531 | 0.998865404 | 5.117472 | 5.060678 | 5.121125 | 5.057575 | 5.101784 | 5.066225 | 49 | 29 | 33 | 27 |
| 162972 | NM_00127709 | ZNF550 | zinc finger protein 550 | protein_coding | NP_001264019.1;NP_001264020.1;NP_001264021.1;NP_001264022.1;XI | 28643 | . | . | . | 0.183492 | 0.323718 | 0.297904 | 0.259057 | 0.426993 | 0.748288 | 0.668489 | 0.592305 | 1.111962 | 42.876750 | 0.497104195 | 0.307999286 | 0.758082876 | 0.998865404 | 5.343257 | 5.496901 | 5.374814 | 5.434884 | 5.454909 | 5.389229 | 38 | 52 | 47 | 34 |
| 162979 | NM_145288 | ZNF296 | zinc finger protein 296 | protein_coding | NP_660331.1 | 15981 | 613226 | . | . | 0.168675 | 0.069722 | 0.202214 | 0.181063 | 0.392513 | 0.161165 | 0.453763 | 0.413980 | 1.604367 | 8.085293 | 1.089716797 | 0.625854332 | 0.531410491 | NA | 2.631267 | 3.318300 | 3.007707 | 2.984515 | 3.014688 | 3.011599 | 10 | 4 | 9 | 9 |
| 162989 | NM_00127061 | DEDD2 | death effector domain containing 2 | protein_coding | NP_001257543.1;NP_001257544.1;NP_579874.1;XP_011524871.1;XP_01 | 24450 | 617078 | . | . | 9.060828 | 10.872260 | 9.253621 | 10.681347 | 21.084919 | 25.131717 | 20.764856 | 24.421632 | -1.030355 | 544.048776 | 0.167328445 | -0.25782269 | 0.796543755 | 0.998865404 | 9.109338 | 9.065513 | 9.064379 | 9.128283 | 9.045408 | 9.108255 | 605 | 625 | 443 | 527 |
| 162993 | NM_00107762 | ZNF846 | zinc finger protein 846 | protein_coding | NP_001071092.1;NP_001340727.1;NP_001340728.1;NP_001340729.1 | 27260 | . | . | . | 1.117994 | 1.101279 | 1.123653 | 1.208620 | 2.601618 | 2.545656 | 2.521447 | 2.763365 | 1.033998 | 76.110601 | 0.361915547 | 0.133271261 | 0.893978855 | 0.998865404 | 6.225873 | 6.273780 | 6.247781 | 6.243004 | 6.245262 | 6.262383 | 87 | 80 | 65 | 75 |
| 163033 | NM_152600,XI | ZNF579 | zinc finger protein 579 | protein_coding | NP_689813.2;XP_016881898.1;XP_016881899.1;XP_016881900.1 | 26646 | . | . | . | 2.420975 | 2.478418 | 2.773078 | 2.530616 | 5.633708 | 5.728975 | 6.222706 | 5.785953 | 1.090964 | 221.896417 | 0.226770958 | 0.553879015 | 0.579661647 | 0.998865404 | 7.729533 | 7.855216 | 7.758922 | 7.783586 | 7.825779 | 7.799169 | 238 | 235 | 207 | 211 |
| 163049 | NM_153358,XI | ZNF791 | zinc finger protein 791 | protein_coding | NP_699189.2;XP_016881901.1 | 26895 | . | . | . | 2.172884 | 2.060672 | 2.394351 | 1.548139 | 5.056389 | 4.763336 | 5.372855 | 3.539637 | -1.036468 | 396.798747 | 0.21910387 | -0.23584649 | 0.813551789 | 0.998865404 | 8.657060 | 8.607034 | 8.668922 | 8.603501 | 8.732602 | 8.483448 | 490 | 412 | 411 | 288 |
| 163050 | NM_144976 | ZNF564 | zinc finger protein 564 | protein_coding | NP_659413.1 | 31106 | . | . | . | 1.073087 | 1.072288 | 1.383378 | 1.300849 | 2.497117 | 2.478642 | 3.104261 | 2.974236 | 1.230015 | 108.976523 | 0.307346653 | 0.971788843 | 0.331155608 | 0.998865404 | 6.610598 | 6.909677 | 6.730285 | 6.728062 | 6.806592 | 6.784729 | 113 | 105 | 108 | 109 |
| 163051 | NM_152601 | ZNF709 | zinc finger protein 709 | protein_coding | NP_689814.1 | 20629 | . | . | . | 0.176845 | 0.294057 | 0.191879 | 0.119683 | 0.411526 | 0.679725 | 0.430570 | 0.273641 | -1.525727 | 30.308328 | 0.595253704 | -1.02392821 | 0.305869183 | 0.998865404 | 5.193784 | 4.585944 | 4.906845 | 4.977703 | 4.915790 | 4.867703 | 32 | 49 | 26 | 17 |
| 163059 | NM_00108041 | ZNF433 | zinc finger protein 433 | protein_coding | NP_001073880.1;NP_001295275.1;NP_001295277.1;NP_001295280.1;NI | 20811 | . | . | . | 0.564241 | 0.481647 | 0.637254 | 0.888870 | 1.313011 | 1.113349 | 1.429979 | 2.032299 | 1.435596 | 53.784987 | 0.438166502 | 1.190529509 | 0.233838339 | 0.988893316 | 5.464470 | 5.986776 | 5.729861 | 5.675349 | 5.752478 | 5.794002 | 59 | 40 | 50 | 65 |
| 163071 | NM_00133109 | ZNF114 | zinc finger protein 114 | protein_coding | NP_001318026.1;NP_001318027.1;NP_001356740.1;NP_001356741.1;NI | 12894 | 603996 | . | . | 0.389978 | 0.233565 | 0.047841 | 0.240509 | 0.907494 | 0.539896 | 0.107354 | 0.549896 | **-2.117965** | 18.616789 | 0.807355951 | -1.34101747 | 0.179914782 | NA | 4.662982 | 3.572884 | 4.278284 | 4.220608 | 4.162115 | 4.223393 | 37 | 20 | 4 | 18 |
| 163081 | NM_00130097 | ZNF567 | zinc finger protein 567 | protein_coding | NP_001287908.1;NP_001309840.1;NP_001309841.1;NP_001309842.1;NI | 28696 | . | . | . | 2.092597 | 2.610678 | 2.984071 | 1.826096 | 4.869559 | 6.034702 | 6.696166 | 4.175150 | 1.004592 | 214.294911 | 0.268803688 | 0.024587791 | 0.980383758 | 0.998865404 | 7.739563 | 7.747334 | 7.680723 | 7.785035 | 7.835666 | 7.626310 | 219 | 256 | 229 | 156 |
| 163087 | NM_00134594 | ZNF383 | zinc finger protein 383 | protein_coding | NP_001332876.1;NP_001332877.1;NP_001332878.1;NP_689817.1;XP_0( | 18609 | . | . | . | 0.798297 | 0.728689 | 0.917143 | 1.109784 | 1.857668 | 1.684400 | 2.058041 | 2.537388 | 1.266544 | 115.695539 | 0.300297637 | 1.135198447 | 0.25629218 | 0.995760118 | 6.673732 | 7.014563 | 6.820090 | 6.796859 | 6.872988 | 6.901419 | 122 | 106 | 108 | 126 |
| 163115 | NM_152605 | ZNF781 | zinc finger protein 781 | protein_coding | NP_689818.2 | 26745 | . | . | . | 0.318043 | 0.260452 | 0.327584 | 0.180095 | 0.740099 | 0.602046 | 0.735088 | 0.411766 | -1.157519 | 27.103457 | 0.613382344 | -0.34405175 | 0.730807365 | 0.998865404 | 4.860000 | 4.653422 | 4.779478 | 4.753619 | 4.778394 | 4.721618 | 37 | 28 | 28 | 17 |
| 163126 | NM_153232 | EID2 | EP300 interacting inhibitor of differentiation 2 | protein_coding | NP_694964.3 | 28292 | 609773 | . | . | 7.846153 | 7.458997 | 6.942409 | 6.746346 | 18.258320 | 17.241804 | 15.578565 | 15.424717 | -1.132299 | 329.404611 | 0.1946463 | -0.92092986 | 0.357087048 | 0.998865404 | 8.450370 | 8.271525 | 8.414837 | 8.379694 | 8.339096 | 8.317811 | 415 | 366 | 274 | 284 |
| 163131 | NM_00100585 | ZNF780B | zinc finger protein 780B | protein_coding | NP_001005851.1;XP_005258647.1;XP_005258648.1;XP_005258649.1;XP_ | 33109 | . | . | . | 0.600215 | 0.644028 | 0.885128 | 0.346223 | 1.396727 | 1.488698 | 1.986199 | 0.791595 | 1.081732 | 161.684266 | 0.351341322 | 0.322599372 | 0.74699867 | 0.998865404 | 7.277949 | 7.393797 | 7.292032 | 7.304383 | 7.493203 | 7.135398 | 176 | 170 | 209 | 90 |
| 163154 | NM_00113431 | PRR22 | proline rich 22 | protein_coding | NP_001127788.1 | 28354 | . | . | . | 1.156130 | 1.596727 | 1.584755 | 1.516775 | 2.690361 | 3.690906 | 3.556145 | 3.467925 | 1.118311 | 62.965983 | 0.404321823 | 0.398993531 | 0.689897971 | 0.998865404 | 5.894196 | 6.054362 | 5.925855 | 5.992205 | 5.988866 | 5.979625 | 58 | 74 | 59 | 61 |
| 163183 | NM_00103987 | SYNE4 | spectrin repeat containing nuclear envelope family member 4 | protein_coding | NP_001034965.1;NP_001284664.1 | 26703 | 615535 | . | . | 0.176008 | 0.405329 | 0.277364 | 0.290812 | 0.409577 | 0.936935 | 0.622398 | 0.664906 | 1.003492 | 12.718463 | 0.881301093 | 0.005707104 | 0.995446414 | NA | 3.668126 | 3.669578 | 3.641058 | 3.685335 | 3.663343 | 3.663991 | 9 | 19 | 11 | 12 |
| 163223 | NM_00100141 | ZNF676 | zinc finger protein 676 | protein_coding | NP_001001411.2 | 20429 | . | . | . | 0.262833 | 0.201422 | 0.206585 | 0.161205 | 0.611623 | 0.465596 | 0.463571 | 0.368576 | -1.297424 | 19.782340 | 0.697878584 | -0.53827396 | 0.590387929 | NA | 4.480474 | 4.107809 | 4.331981 | 4.307238 | 4.307679 | 4.288215 | 29 | 21 | 17 | 14 |
| 163227 | NM_00135166 | ZNF100 | zinc finger protein 100 | protein_coding | NP_001338598.1;NP_001338599.1;NP_001338600.1;NP_001338601.1;NI | 12880 | 603982 | . | . | 1.599678 | 1.530250 | 1.544983 | 1.491878 | 3.722516 | 3.537241 | 3.466896 | 3.411003 | -1.040440 | 274.556977 | 0.208292124 | -0.27458204 | 0.783637366 | 0.998865404 | 8.129070 | 8.072295 | 8.125368 | 8.096939 | 8.101470 | 8.079914 | 331 | 294 | 239 | 247 |
| 163255 | NM_00117222 | ZNF540 | zinc finger protein 540 | protein_coding | NP_001165696.1;NP_001165697.1;NP_689819.1 | 25331 | 613903 | . | . | 0.444079 | 0.492602 | 0.489776 | 0.356477 | 1.033391 | 1.138671 | 1.099044 | 0.815041 | -1.115115 | 44.731342 | 0.474083644 | -0.33157242 | 0.740212149 | 0.998865404 | 5.558826 | 5.403420 | 5.483668 | 5.498721 | 5.499127 | 5.445834 | 52 | 53 | 43 | 33 |
| 163259 | NM_00125640 | DENND2C | DENN domain containing 2C | protein_coding | NP_001243333.1;NP_940861.3 | 24748 | . | . | . | 1.062584 | 0.919624 | 0.769163 | 0.839321 | 2.472676 | 2.125753 | 1.725980 | 1.919007 | -1.221993 | 171.288743 | 0.255085393 | -1.1338776 | 0.256845895 | 0.995760118 | 7.557687 | 7.268410 | 7.485743 | 7.435681 | 7.363109 | 7.391771 | 229 | 192 | 129 | 150 |
| 163404 | NM_00101086 | PLPPR5 | phospholipid phosphatase related 5 | protein_coding | NP_001010861.1;NP_001032394.1;XP_011539138.1;XP_011539139.1;XP | 31703 | 617287 | . | . | 0.043417 | 0.088319 | 0.029510 | 0.111116 | 0.101033 | 0.204154 | 0.066220 | 0.254054 | 1.206112 | 9.933762 | 1.0274418 | 0.263143132 | 0.792440261 | NA | 3.175403 | 3.437399 | 3.291608 | 3.310877 | 3.282324 | 3.336283 | 8 | 12 | 4 | 16 |
| 163486 | NM_00119521 | DENND1B | DENN domain containing 1B | protein_coding | NP_001182144.1;NP_001182145.1;NP_001287787.1;NP_659414.2;XP_0C | 28404 | 613292 | . | . | 0.385163 | 0.363392 | 0.463880 | 0.520383 | 0.896293 | 0.839998 | 1.040934 | 1.189795 | 1.078493 | 129.490878 | 0.302113836 | 0.360847404 | 0.718213524 | 0.998865404 | 6.960003 | 7.071266 | 7.048448 | 6.946174 | 7.065182 | 6.984657 | 164 | 115 | 130 | 112 |
| 163590 | NM_00119926 | TOR1AIP2 | torsin 1A interacting protein 2 | protein_coding | NP_001186189.1;NP_001336860.1;NP_001336862.1;NP_001336863.1;NI | 24055 | 614513 | . | . | 8.732359 | 8.755933 | 9.394589 | 8.348837 | 20.320561 | 20.239727 | 21.081186 | 19.088623 | -1.001036 | 2264.206678 | 0.122469664 | -0.01219592 | 0.990269305 | 0.999289967 | 11.145416 | 11.144163 | 11.146670 | 11.142559 | 11.193868 | 11.091328 | 2615 | 2438 | 2094 | 1986 |
| 163688 | NM_00133031 | CALML6 | calmodulin like 6 | protein_coding | NP_001317242.1;NP_619650.2;XP_005244786.2 | 24193 | 610171 | . | . | 0.781100 | 0.686225 | 1.278929 | 0.573756 | 1.817653 | 1.586240 | 2.869879 | 1.311826 | 1.240634 | 57.729319 | 0.466379351 | 0.66700579 | 0.504768435 | 0.998865404 | 5.685135 | 6.000166 | 5.822987 | 5.798615 | 5.936687 | 5.766946 | 63 | 52 | 77 | 37 |
| 163786 | NM_00130482 | SASS6 | SAS-6 centriolar assembly protein | protein_coding | NP_001291758.1;NP_919268.1;XP_016855975.1 | 25403 | 609321 | . | . | 2.866930 | 2.538117 | 2.454641 | 1.545446 | 6.671465 | 5.866971 | 5.508145 | 3.533481 | -1.222476 | 303.912598 | 0.230618004 | -1.25664968 | 0.208880515 | 0.971093888 | 8.384261 | 8.096433 | 8.334120 | 8.252905 | 8.280172 | 8.082435 | 412 | 335 | 284 | 207 |
| 163859 | NM_152608 | SDE2 | SDE2 telomere maintenance homolog | protein_coding | NP_689821.3 | 26643 | . | . | . | 6.243593 | 6.044226 | 5.877507 | 6.221318 | 14.529098 | 13.971498 | 13.188956 | 14.224304 | -1.030564 | 757.833114 | 0.147708854 | -0.29405142 | 0.768718614 | 0.998865404 | 9.587323 | 9.543822 | 9.590606 | 9.561585 | 9.539045 | 9.571052 | 904 | 813 | 634 | 716 |
| 163882 | NM_00113945 | CNST | consortin, connexin sorting protein | protein_coding | NP_001132931.1;NP_689822.2;XP_005273138.2;XP_005273140.1;XP_01 | 26486 | 613439 | . | . | 6.197658 | 6.885599 | 8.373029 | 7.368620 | 14.422206 | 15.916370 | 18.788836 | 16.847472 | 1.183434 | 1138.639830 | 0.140731437 | 1.726547314 | 0.084248995 | 0.787933172 | 10.026412 | 10.269548 | 10.052795 | 10.116314 | 10.260390 | 10.158913 | 1149 | 1174 | 1145 | 1080 |
| 163933 | NM_207334 | FAM43B | family with sequence similarity 43 member B | protein_coding | NP_997217.1 | 31791 | . | . | . | 0.154750 | 0.244917 | 0.027972 | 0.127697 | 0.360109 | 0.566137 | 0.062769 | 0.291965 | **-2.417710** | 11.146612 | 1.003566476 | -1.26911499 | 0.204400054 | NA | 3.982758 | 2.696731 | 3.489561 | 3.521512 | 3.446838 | 3.482905 | 14 | 21 | 2 | 10 |
| 164045 | NM_00101797 | HFM1 | helicase for meiosis 1 | protein_coding | NP_001017975.5;XP_011539151.1;XP_011539152.1;XP_011539153.1;XP_ | 20193 | 615684 | . | . | 0.455941 | 0.274956 | 0.323590 | 0.162362 | 1.060993 | 0.635574 | 0.726127 | 0.371221 | -1.491384 | 48.730413 | 0.501414191 | -1.15005077 | 0.250122962 | 0.993339639 | 5.864262 | 5.293067 | 5.687483 | 5.578763 | 5.609387 | 5.513927 | 84 | 47 | 45 | 25 |
| 164091 | NM_178422,XI | PAQR7 | progestin and adipoQ receptor family member 7 | protein_coding | NP_848509.1;XP_005245803.1;XP_011539163.1;XP_011539164.1 | 23146 | 607779 | . | . | 4.115920 | 4.408126 | 3.972797 | 4.047029 | 9.577914 | 10.189584 | 8.914841 | 9.253049 | -1.075509 | 346.821469 | 0.190628815 | -0.55091343 | 0.581693023 | 0.998865404 | 8.489734 | 8.384444 | 8.440237 | 8.476397 | 8.414303 | 8.420170 | 401 | 400 | 289 | 315 |
| 164395 | NM_00100840 | TTLL9 | tubulin tyrosine ligase like 9 | protein_coding | NP_001008409.1;NP_001354549.1 | 16118 | . | . | . | 0.038587 | 0.089537 | 0.049980 | 0.080543 | 0.089795 | 0.206969 | 0.112153 | 0.184153 | -1.177521 | 8.333761 | 1.07968134 | -0.21835404 | 0.827153276 | NA | 3.175403 | 2.932304 | 3.054287 | 3.073417 | 3.044502 | 3.070989 | 8 | 12 | 4 | 10 |
| 164592 | NM_00133106 | CCDC116 | coiled-coil domain containing 116 | protein_coding | NP_001317995.1;NP_689825.2;XP_006724222.1;XP_011528286.1;XP_01 | 26688 | . | . | . | 0.082259 | 0.109595 | 0.111609 | 0.088254 | 0.191419 | 0.253333 | 0.250447 | 0.201782 | -1.080752 | 8.669159 | 1.03176668 | -0.1085864 | 0.913530547 | NA | 3.170740 | 3.058876 | 3.111468 | 3.122397 | 3.117732 | 3.110122 | 9 | 11 | 8 | 7 |
| 164684 | NM_152613,XI | WBP2NL | WBP2 N-terminal like | protein_coding | NP_689826.2 | 28389 | 610981 | . | . | 0.039721 | 0.067606 | 0.029249 | 0.016228 | 0.092433 | 0.156274 | 0.065633 | 0.037103 | **-2.025975** | 8.438026 | 1.077993677 | -0.94491871 | 0.34470038 | NA | 3.497331 | 2.481107 | 3.083431 | 3.106879 | 3.075659 | 3.063793 | 10 | 15 | 6 | 4 |
| 164832 | NM_00137178 | LONRF2 | LON peptidase N-terminal domain and ring finger 2 | protein_coding | NP_001358712.1;NP_940863.3 | 24788 | . | . | . | 1.143126 | 1.161171 | 1.055668 | 0.899408 | 2.660100 | 2.684100 | 2.368887 | 2.056389 | -1.198870 | 502.178416 | 0.173769812 | -1.50587059 | 0.132100399 | 0.885996989 | 9.096638 | 8.835691 | 9.023392 | 9.028194 | 8.961404 | 8.857973 | 628 | 593 | 431 | 392 |
| 165055 | NM_00130310 | CCDC138 | coiled-coil domain containing 138 | protein_coding | NP_001290034.1;NP_001290035.1;NP_001338473.1;NP_001338474.1;NI | 26531 | . | . | . | 4.891182 | 3.751921 | 4.296665 | 3.430165 | 11.381980 | 8.672732 | 9.641593 | 7.842663 | -1.088756 | 355.385450 | 0.209430587 | -0.58578437 | 0.558020431 | 0.998865404 | 8.532519 | 8.411422 | 8.538073 | 8.447994 | 8.521486 | 8.363666 | 459 | 369 | 338 | 274 |
| 165186 | NM_00132153 | TOGARAM2 | TOG array regulator of axonemal microtubules 2 | protein_coding | NP_001308467.1;NP_001308468.1;NP_954974.2;XP_011530930.1;XP_01 | 33715 | . | . | . | 0.551046 | 0.456049 | 0.622942 | 0.902437 | 1.282307 | 1.054175 | 1.397864 | 2.063316 | 1.264132 | 64.249333 | 0.397280521 | 0.851153714 | 0.394683971 | 0.998865404 | 5.826533 | 6.164892 | 5.994281 | 5.952277 | 6.008686 | 6.042758 | 72 | 55 | 58 | 72 |
| 165215 | NM_177454 | FAM171B | family with sequence similarity 171 member B | protein_coding | NP_803237.3 | 29412 | . | . | . | 2.798791 | 2.384806 | 2.012834 | 2.311466 | 6.512903 | 5.512586 | 4.516740 | 5.284894 | -1.215368 | 424.997021 | 0.187330872 | -1.50211583 | 0.133067196 | 0.888450802 | 8.865211 | 8.583695 | 8.838173 | 8.731621 | 8.630685 | 8.704792 | 582 | 461 | 312 | 383 |
| 165324 | NM_181713,XI | UBXN2A | UBX domain protein 2A | protein_coding | NP_859064.2;XP_005264223.1;XP_005264224.1;XP_005264225.1;XP_00! | 27265 | . | . | . | 8.113643 | 7.420351 | 7.661987 | 7.990078 | 18.880780 | 17.152475 | 17.193274 | 18.268365 | 1.051115 | 1234.836452 | 0.130404351 | 0.551522457 | 0.581275576 | 0.998865404 | 10.233733 | 10.305581 | 10.261289 | 10.240152 | 10.270647 | 10.305828 | 1406 | 1281 | 1075 | 1214 |
| 165530 | NM_00125802 | CLEC4F | C-type lectin domain family 4 member F | protein_coding | NP_001244956.1;NP_001308237.1;NP_775806.2;XP_011530937.1;XP_01 | 25357 | . | . | . | 0.049073 | 0.082918 | 0.077963 | 0.059087 | 0.114195 | 0.191669 | 0.174947 | 0.135096 | -1.061742 | 7.449091 | 1.113902058 | -0.07759548 | 0.938149836 | NA | 2.939842 | 2.852980 | 2.889028 | 2.903806 | 2.898069 | 2.890790 | 7 | 10 | 7 | 6 |
| 165545 | NM_133637,XI | DQX1 | DEAQ-box RNA dependent ATPase 1 | protein_coding | NP_598376.2;XP_011530947.1;XP_016859009.1 | 20410 | . | . | . | 0.104884 | 0.075031 | 0.207183 | 0.213608 | 0.244070 | 0.173437 | 0.464913 | 0.488390 | **2.258645** | 12.371981 | 0.880699704 | 1.334686177 | 0.18197908 | NA | 2.923304 | 4.100822 | 3.591897 | 3.581996 | 3.629544 | 3.629204 | 10 | 7 | 15 | 16 |
| 165679 | NM_00104010 | SPTSSB | serine palmitoyltransferase small subunit B | protein_coding | NP_001035189.1;NP_001307608.1;XP_016861282.1;XP_016861283.1;XP | 24045 | 610412 | . | . | 0.057423 | 0.046832 | 0.053991 | 0.098840 | 0.133626 | 0.108256 | 0.121153 | 0.225985 | 1.223603 | 6.943643 | 1.167521803 | 0.249362087 | 0.803080709 | NA | 2.644780 | 2.932304 | 2.793275 | 2.786846 | 2.782988 | 2.809694 | 8 | 6 | 4 | 10 |
| 165904 | NM_00119862 | XIRP1 | xin actin binding repeat containing 1 | protein_coding | NP_001185550.1;NP_001338306.1;NP_919269.2;XP_005264966.1 | 14301 | 609777 | . | . | 0.040426 | 0.046603 | 0.054296 | 0.054544 | 0.094074 | 0.107724 | 0.121838 | 0.124708 | 1.231487 | 10.527924 | 0.935327616 | 0.32117231 | 0.748079815 | NA | 3.238681 | 3.538109 | 3.382905 | 3.389927 | 3.395595 | 3.396705 | 10 | 11 | 10 | 11 |
| 165918 | NM_152617 | RNF168 | ring finger protein 168 | protein_coding | NP_689830.2 | 26661 | 612688 | . | . | 2.277904 | 1.871703 | 2.403211 | 2.215505 | 5.300775 | 4.326525 | 5.392736 | 5.065490 | 1.096610 | 365.520578 | 0.197146148 | 0.674882028 | 0.499750719 | 0.998865404 | 8.445203 | 8.579299 | 8.540334 | 8.424707 | 8.564057 | 8.510290 | 442 | 338 | 348 | 342 |
| 166336 | NM_00137052 | PRICKLE2 | prickle planar cell polarity protein 2 | protein_coding | NP_001357457.1;NP_942559.1;XP_011531734.1;XP_011531735.1;XP_01 | 20340 | 608501 | . | . | 2.241849 | 2.397902 | 2.275298 | 2.429844 | 5.216873 | 5.542856 | 5.105702 | 5.555548 | 1.006741 | 718.579120 | 0.153022902 | 0.063342507 | 0.949493768 | 0.998865404 | 9.484385 | 9.493607 | 9.449613 | 9.521639 | 9.472114 | 9.509098 | 782 | 813 | 610 | 694 |
| 166378 | NM_00131779 | SPATA5 | spermatogenesis associated 5 | protein_coding | NP_001304728.1;NP_001332785.1;NP_660208.2;XP_011529980.1;XP_01 | 18119 | 613940 | . | . | 1.267779 | 1.147138 | 1.210620 | 1.240470 | 2.950173 | 2.651664 | 2.716596 | 2.836188 | 1.048812 | 314.978415 | 0.206042278 | 0.33369716 | 0.738608093 | 0.998865404 | 8.263983 | 8.333401 | 8.339972 | 8.225458 | 8.307537 | 8.313466 | 391 | 297 | 279 | 304 |
| 166379 | NM_00117800 | BBS12 | Bardet-Biedl syndrome 12 | protein_coding | NP_001171478.1;NP_689831.2;XP_011529982.1;XP_016863320.1 | 26648 | 610683 | . | . | 1.448945 | 1.284618 | 1.501832 | 1.238831 | 3.371753 | 2.969453 | 3.370068 | 2.832440 | -1.014375 | 138.676147 | 0.283825692 | -0.07254847 | 0.942165433 | 0.998865404 | 7.125024 | 7.106066 | 7.140691 | 7.092136 | 7.147949 | 7.073617 | 171 | 141 | 132 | 116 |
| 166614 | NM_00104026 | DCLK2 | doublecortin like kinase 2 | protein_coding | NP_001035350.2;NP_001035351.4;XP_005262842.1;XP_005262843.1;XP | 19002 | 613166 | . | . | 0.460468 | 0.449084 | 0.397415 | 0.389643 | 1.071529 | 1.038077 | 0.891789 | 0.890873 | -1.157819 | 55.531182 | 0.419710599 | -0.50370264 | 0.614470352 | 0.998865404 | 5.896948 | 5.685784 | 5.815911 | 5.806990 | 5.783268 | 5.781050 | 70 | 63 | 45 | 48 |
| 166647 | NM_145290,XI | ADGRA3 | adhesion G protein-coupled receptor A3 | protein_coding | NP_660333.2;XP_005248194.1;XP_011512113.1 | 13839 | 612303 | . | . | 14.006391 | 14.169332 | 13.253530 | 13.661249 | 32.593449 | 32.753048 | 29.740537 | 31.234822 | -1.061550 | 1955.493938 | 0.118416915 | -0.72770749 | 0.466792662 | 0.998865404 | 10.975794 | 10.889552 | 10.955930 | 10.957284 | 10.897919 | 10.920397 | 2317 | 2175 | 1626 | 1804 |
| 166785 | NM_00137564 | MMAA | metabolism of cobalamin associated A | protein_coding | NP_001362573.1;NP_758454.1;XP_011529986.1;XP_011529988.1 | 18871 | 607481 | . | . | 0.394042 | 0.397762 | 0.429879 | 0.460630 | 0.916953 | 0.919446 | 0.964636 | 1.053175 | 1.131570 | 80.690611 | 0.352404633 | 0.506025097 | 0.612839031 | 0.998865404 | 6.242516 | 6.420647 | 6.320917 | 6.310406 | 6.339056 | 6.357412 | 89 | 80 | 72 | 83 |
| 166793 | NM_00133062 | ZBTB49 | zinc finger and BTB domain containing 49 | protein_coding | NP_001317554.1;NP_660334.3;XP_005248008.1;XP_011511718.1;XP_02 | 19883 | 616238 | . | . | 1.238545 | 1.349884 | 1.452626 | 1.249747 | 2.882146 | 3.120319 | 3.259651 | 2.857398 | -1.042341 | 107.723919 | 0.311425564 | -0.19210902 | 0.847656812 | 0.998865404 | 6.781345 | 6.720401 | 6.743089 | 6.772533 | 6.725293 | 6.763617 | 121 | 124 | 86 | 105 |
| 166815 | NM_00138238 | TIGD2 | tigger transposable element derived 2 | protein_coding | NP_001369309.1;NP_663761.1 | 18333 | 612973 | . | . | 1.685301 | 1.081534 | 1.480542 | 1.637951 | 3.921765 | 2.500013 | 3.322295 | 3.744980 | 1.092440 | 145.359720 | 0.299412307 | 0.426015971 | 0.670096209 | 0.998865404 | 7.117775 | 7.246330 | 7.240437 | 7.074212 | 7.170677 | 7.218043 | 195 | 117 | 124 | 151 |
| 166863 | NM_00127717 | RBM46 | RNA binding motif protein 46 | protein_coding | NP_001264100.1;NP_001264102.1;NP_659416.1;XP_006714182.1;XP_00 | 28401 | . | . | . | 0.049267 | 0.016107 | 0.034789 | 0.031483 | 0.114646 | 0.037233 | 0.078066 | 0.071981 | -1.002052 | 4.431473 | 1.437315308 | -0.0020573 | 0.998358513 | NA | 2.147197 | 2.148375 | 2.156352 | 2.140801 | 2.149297 | 2.147812 | 7 | 3 | 4 | 4 |
| 166929 | NM_00113625 | SGMS2 | sphingomyelin synthase 2 | protein_coding | NP_001129729.1;NP_001129730.1;NP_001362834.1;NP_001362835.1;NI | 28395 | 611574 | . | . | 1.674229 | 1.877032 | 1.719740 | 1.313862 | 3.895999 | 4.338841 | 3.859047 | 3.003988 | -1.179423 | 320.502354 | 0.212698047 | -1.11934064 | 0.262994843 | 0.995917097 | 8.437926 | 8.200717 | 8.329568 | 8.399616 | 8.342335 | 8.201843 | 375 | 397 | 290 | 239 |
| 166968 | NM_00129759 | MIER3 | MIER family member 3 | protein_coding | NP_001284527.1;NP_001284528.1;NP_689835.3;XP_011541518.1;XP_01 | 26678 | . | . | . | 3.577088 | 3.235978 | 3.516882 | 2.560915 | 8.324031 | 7.480109 | 7.891781 | 5.855229 | -1.137148 | 525.044329 | 0.191039695 | -0.97058383 | 0.331755562 | 0.998865404 | 9.125400 | 8.941323 | 9.109009 | 9.034864 | 9.085567 | 8.885127 | 679 | 569 | 496 | 387 |
| 166979 | NM_00114573 | CDC20B | cell division cycle 20B | protein_coding | NP_001139206.2;NP_001163873.1;NP_689836.2;XP_011541520.1 | 24222 | . | . | . | 0.271833 | 0.209050 | 0.611294 | 0.383045 | 0.632568 | 0.483227 | 1.371724 | 0.875788 | 1.955796 | 33.690927 | 0.555531883 | 1.74203487 | 0.081502343 | 0.782498736 | 4.508267 | 5.479761 | 5.018053 | 4.990439 | 5.125626 | 5.055871 | 30 | 21 | 47 | 33 |
| 167153 | NM_00111439 | TENT2 | terminal nucleotidyltransferase 2 | protein_coding | NP_001107865.1;NP_001107866.1;NP_001284673.1;NP_001284674.1;NI | 26776 | 614121 | . | . | 9.214589 | 8.283016 | 9.314510 | 7.245552 | 21.442728 | 19.146564 | 20.901486 | 16.566095 | -1.067933 | 1074.464453 | 0.151236712 | -0.62697473 | 0.530675798 | 0.998865404 | 10.115683 | 10.021587 | 10.130430 | 10.051078 | 10.116500 | 9.963858 | 1345 | 1134 | 998 | 873 |
| 167227 | NM_00124237 | DCP2 | decapping mRNA 2 | protein_coding | NP_001229306.1;NP_689837.2;XP_016864652.1 | 24452 | 609844 | . | . | 2.644572 | 2.479479 | 2.445615 | 1.451883 | 6.154028 | 5.731427 | 5.487891 | 3.319559 | -1.348263 | 686.182934 | 0.204423144 | -2.10887069 | **0.034955745** | 0.583157098 | 9.621394 | 9.191615 | 9.534771 | 9.480293 | 9.449848 | 9.133148 | 941 | 818 | 632 | 415 |
| 167359 | NM_153361,N | NIM1K | NIM1 serine/threonine protein kinase | protein_coding | NP_699192.1 | 28646 | . | . | . | 0.254749 | 0.118966 | 0.104695 | 0.176721 | 0.592813 | 0.274995 | 0.234934 | 0.404052 | -1.278847 | 15.012551 | 0.8218869 | -0.4317422 | 0.665928791 | NA | 4.074695 | 3.719719 | 3.944180 | 3.893571 | 3.891257 | 3.915092 | 26 | 12 | 9 | 15 |
| 167555 | NM_205548,XI | FAM151B | family with sequence similarity 151 member B | protein_coding | NP_991111.2;XP_006714627.2;XP_011541536.1;XP_011541537.1;XP_01: | 33716 | . | . | . | 0.784787 | 0.910621 | 1.295326 | 1.025044 | 1.826230 | 2.104941 | 2.906676 | 2.343640 | 1.369866 | 49.033861 | 0.452729894 | 1.002883105 | 0.315917265 | 0.998865404 | 5.370748 | 5.825047 | 5.560186 | 5.592214 | 5.655975 | 5.610007 | 43 | 49 | 55 | 47 |
| 167681 | NM_00117042 | PRSS35 | serine protease 35 | protein_coding | NP_001163894.1;NP_699193.2 | 21387 | . | . | . | 4.486950 | 4.929324 | 5.622516 | 3.670854 | 10.441319 | 11.394353 | 12.616763 | 8.392971 | -1.028451 | 361.454529 | 0.225555914 | -0.17943402 | 0.857596921 | 0.998865404 | 8.517133 | 8.477941 | 8.472732 | 8.524980 | 8.600440 | 8.350544 | 403 | 413 | 378 | 261 |
| 167691 | NM_00112276 | LCA5 | lebercilin LCA5 | protein_coding | NP_001116241.1;NP_859065.2;XP_005248722.1;XP_011533806.1 | 31923 | 611408 | . | . | 0.537670 | 0.486041 | 0.474822 | 0.488145 | 1.251180 | 1.123505 | 1.065486 | 1.116087 | -1.078574 | 72.467342 | 0.371187392 | -0.29399097 | 0.768764804 | 0.998865404 | 6.232541 | 6.123933 | 6.202375 | 6.176117 | 6.172294 | 6.169430 | 91 | 77 | 61 | 65 |
| 168374 | NM_00128753 | ZNF92 | zinc finger protein 92 | protein_coding | NP_001274461.1;NP_001274462.1;NP_001274463.1;NP_009070.2;NP_6! | 13168 | 603974 | . | . | 2.675275 | 1.962049 | 2.424914 | 1.441431 | 6.225477 | 4.535364 | 5.441437 | 3.295661 | -1.217711 | 208.545626 | 0.284428377 | -0.99909703 | 0.317747688 | 0.998865404 | 7.838062 | 7.556678 | 7.813350 | 7.657757 | 7.751123 | 7.532655 | 305 | 208 | 205 | 132 |
| 168433 | NM_139175 | RNF133 | ring finger protein 133 | protein_coding | NP_631914.1 | 21154 | . | . | . | 0.082491 | 0.042906 | 0.052955 | 0.067736 | 0.191960 | 0.099179 | 0.118830 | 0.154869 | -1.127667 | 2.923912 | 1.754679452 | -0.09878791 | 0.921306672 | NA | 1.631267 | 1.459419 | 1.559607 | 1.547801 | 1.550083 | 1.554103 | 5 | 2 | 2 | 3 |
| 168451 | NM_00113047 | THAP5 | THAP domain containing 5 | protein_coding | NP_001123947.1;NP_001274527.1;NP_001274528.1;NP_001274530.1;NI | 23188 | 612534 | . | . | 5.061137 | 4.754200 | 4.647090 | 3.360949 | 11.777471 | 10.989545 | 10.427935 | 7.684408 | -1.240012 | 470.655147 | 0.196246188 | -1.58145487 | 0.113774071 | 0.851639642 | 9.024745 | 8.715800 | 8.958877 | 8.916643 | 8.899221 | 8.699420 | 619 | 544 | 427 | 328 |
| 168455 | NM_175884 | CCDC71L | coiled-coil domain containing 71 like | protein_coding | NP_787080.2 | 26685 | . | . | . | 1.643727 | 1.496522 | 1.633155 | 1.438533 | 3.825020 | 3.459277 | 3.664752 | 3.289036 | -1.035362 | 329.617493 | 0.199606344 | -0.25116789 | 0.801684306 | 0.998865404 | 8.389009 | 8.339870 | 8.402326 | 8.343640 | 8.390918 | 8.316188 | 406 | 343 | 301 | 283 |
| 168507 | NM_138295,XI | PKD1L1 | polycystin 1 like 1, transient receptor potential channel interacting | protein_coding | NP_612152.1;XP_016867287.1 | 18053 | 609721 | . | . | 0.421566 | 0.359618 | 0.430456 | 0.371687 | 0.981001 | 0.831274 | 0.965931 | 0.849818 | 1.012480 | 196.259618 | 0.246123402 | 0.072700879 | 0.942044149 | 0.998865404 | 7.606935 | 7.626240 | 7.648765 | 7.574235 | 7.651251 | 7.583275 | 243 | 193 | 185 | 171 |
| 168544 | NM_00132985 | ZNF467 | zinc finger protein 467 | protein_coding | NP_001316785.1;NP_997219.1;XP_005250016.1;XP_005250017.1;XP_00 | 23154 | 614040 | . | . | 0.110984 | 0.072128 | 0.073118 | 0.135915 | 0.258264 | 0.166726 | 0.164074 | 0.310753 | -1.193444 | 9.011748 | 1.026415006 | -0.24856527 | 0.803697073 | NA | 3.295021 | 3.037079 | 3.186948 | 3.173411 | 3.162357 | 3.183718 | 13 | 9 | 5 | 10 |
| 168667 | NM_00136530 | BMPER | BMP binding endothelial regulator | protein_coding | NP_001352237.1;NP_597725.1;XP_005249690.1;XP_016867289.1;XP_01 | 24154 | 608699 | . | . | 0.048064 | 0.044732 | 0.067831 | 0.071616 | 0.111846 | 0.103400 | 0.152212 | 0.163742 | 1.494663 | 10.618712 | 0.932728582 | 0.621638983 | 0.534179282 | NA | 3.089593 | 3.669578 | 3.390839 | 3.389395 | 3.408476 | 3.409318 | 10 | 9 | 11 | 12 |
| 168850 | NM_176814,XI | ZNF800 | zinc finger protein 800 | protein_coding | NP_789784.2;XP_005250238.1;XP_006715930.1;XP_006715931.1;XP_00 | 27267 | . | . | . | 2.306702 | 2.049414 | 2.087440 | 1.863521 | 5.367790 | 4.737314 | 4.684155 | 4.260721 | -1.035818 | 624.887340 | 0.189626 | -0.2677438 | 0.788896529 | 0.998865404 | 9.312050 | 9.262428 | 9.312054 | 9.276583 | 9.388161 | 9.138735 | 751 | 668 | 634 | 468 |
| 169044 | NM_152888,XI | COL22A1 | collagen type XXII alpha 1 chain | protein_coding | NP_690848.1;XP_011515185.1;XP_011515186.1;XP_011515187.1;XP_01: | 22989 | 610026 | . | . | 1.671065 | 1.552039 | 1.656952 | 2.032412 | 3.888634 | 3.587607 | 3.718152 | 4.646869 | 1.105270 | 346.119799 | 0.200803766 | 0.719102295 | 0.472077893 | 0.998865404 | 8.361329 | 8.505334 | 8.437720 | 8.368949 | 8.404543 | 8.513644 | 402 | 333 | 286 | 373 |
| 169200 | NM_00100849 | TMEM64 | transmembrane protein 64 | protein_coding | NP_001008495.2;NP_001139745.1 | 25441 | . | . | . | 11.172979 | 8.890012 | 10.425069 | 8.762705 | 25.999981 | 20.549663 | 23.393552 | 20.034884 | -1.050548 | 1497.531809 | 0.151371702 | -0.46998237 | 0.638367616 | 0.998865404 | 10.583214 | 10.512666 | 10.652373 | 10.468550 | 10.586695 | 10.462734 | 1974 | 1461 | 1373 | 1259 |
| 169270 | NM_00104241 | ZNF596 | zinc finger protein 596 | protein_coding | NP_001035880.1;NP_001035881.1;NP_001274183.1;NP_001274184.1;NI | 27268 | . | . | . | 0.332063 | 0.252853 | 0.507665 | 0.281510 | 0.772724 | 0.584481 | 1.139187 | 0.643641 | 1.208710 | 37.220218 | 0.524974641 | 0.520916637 | 0.602424845 | 0.998865404 | 5.073577 | 5.349298 | 5.188600 | 5.201374 | 5.258973 | 5.184759 | 37 | 38 | 44 | 29 |
| 169436 | NM_153710,N | STKLD1 | serine/threonine kinase like domain containing 1 | protein_coding | NP_714921.4 | 28669 | 618530 | . | . | 0.815700 | 0.922091 | 0.824365 | 0.790285 | 1.898165 | 2.131453 | 1.849850 | 1.806894 | -1.074194 | 88.470511 | 0.339588864 | -0.30405956 | 0.761082519 | 0.998865404 | 6.517937 | 6.414458 | 6.458867 | 6.495082 | 6.463702 | 6.450347 | 99 | 105 | 76 | 78 |
| 169611 | NM_00128271 | OLFML2A | olfactomedin like 2A | protein_coding | NP_001269644.1;NP_872293.2;XP_005251817.1;XP_006717052.1 | 27270 | 615899 | . | . | 3.510538 | 3.814924 | 2.937637 | 2.830155 | 8.169167 | 8.818366 | 6.591974 | 6.470812 | -1.284511 | 665.396890 | 0.154651807 | -2.33569379 | **0.019507207** | 0.462516353 | 9.547425 | 9.186160 | 9.429089 | 9.482391 | 9.297651 | 9.275797 | 829 | 838 | 515 | 537 |
| 169714 | NM_181701 | QSOX2 | quiescin sulfhydryl oxidase 2 | protein_coding | NP_859052.3 | 30249 | 612860 | . | . | 16.215981 | 17.000881 | 15.377844 | 15.999539 | 37.735252 | 39.298298 | 34.507435 | 36.581047 | -1.073462 | 2264.980542 | 0.116400284 | -0.87861043 | 0.379612539 | 0.998865404 | 11.195568 | 11.093182 | 11.157185 | 11.188714 | 11.104520 | 11.127655 | 2648 | 2580 | 1872 | 2078 |
| 169834 | NM_00110133 | ZNF883 | zinc finger protein 883 | protein_coding | NP_001094808.1 | 27271 | . | . | . | 1.052029 | 1.053647 | 1.265715 | 1.315019 | 2.448114 | 2.435552 | 2.840228 | 3.006634 | 1.212817 | 87.291533 | 0.339392151 | 0.820176559 | 0.412115463 | 0.998865404 | 6.301967 | 6.580180 | 6.416724 | 6.416174 | 6.465666 | 6.474514 | 91 | 85 | 82 | 91 |
| 169841 | NM_00130127 | ZNF169 | zinc finger protein 169 | protein_coding | NP_001288204.1;NP_003439.2;NP_919301.2;XP_016869853.1;XP_02430 | 12957 | 603404 | . | . | 0.280081 | 0.310828 | 0.521612 | 0.260929 | 0.651759 | 0.718494 | 1.170484 | 0.596583 | 1.383928 | 39.269853 | 0.523671124 | 0.895158335 | 0.370702454 | 0.998865404 | 5.039590 | 5.512525 | 5.293217 | 5.221487 | 5.333624 | 5.280152 | 48 | 26 | 46 | 36 |
| 169981 | NM_00101086 | SPIN3 | spindlin family member 3 | protein_coding | NP_001010862.2;XP_006724640.1;XP_006724641.1;XP_016884804.1 | 27272 | . | . | . | 3.310490 | 3.435653 | 4.475258 | 3.248357 | 7.703644 | 7.941668 | 10.042352 | 7.426984 | 1.112486 | 481.245413 | 0.199449985 | 0.771057129 | 0.44067307 | 0.998865404 | 8.831167 | 8.985942 | 8.853123 | 8.887461 | 9.046997 | 8.814922 | 510 | 505 | 521 | 389 |
| 170082 | NM_00129756 | TCEANC | transcription elongation factor A N-terminal and central domain co | protein_coding | NP_001284492.1;NP_001284493.1;NP_689847.2;XP_016884805.1;XP_01 | 28277 | . | . | . | 0.398067 | 0.464603 | 0.366144 | 0.317848 | 0.926317 | 1.073950 | 0.821617 | 0.726723 | -1.278189 | 40.440055 | 0.491351033 | -0.72066876 | 0.471113336 | 0.998865404 | 5.503736 | 5.150069 | 5.347172 | 5.368699 | 5.329511 | 5.310468 | 49 | 52 | 33 | 31 |
| 170384 | NM_00128419 | FUT11 | fucosyltransferase 11 | protein_coding | NP_001271123.1;NP_775811.2;XP_006717719.1 | 19233 | 616932 | . | . | 7.424012 | 7.099986 | 8.740218 | 7.404670 | 17.275979 | 16.411935 | 19.612797 | 16.929898 | 1.099159 | 485.976688 | 0.175659909 | 0.776503181 | 0.437451943 | 0.998865404 | 8.854534 | 8.991694 | 8.901516 | 8.882989 | 9.004112 | 8.896946 | 541 | 492 | 480 | 437 |
| 170394 | NM_00109863 | PWWP2B | PWWP domain containing 2B | protein_coding | NP_001092107.1;NP_612508.3;XP_006717722.1;XP_011537689.1 | 25150 | . | . | . | 0.968693 | 1.035235 | 1.274062 | 0.950222 | 2.254190 | 2.392992 | 2.858957 | 2.172569 | 1.009677 | 81.616680 | 0.350905235 | 0.039592279 | 0.968418183 | 0.998865404 | 6.343721 | 6.357829 | 6.341561 | 6.356154 | 6.361226 | 6.341573 | 91 | 90 | 74 | 74 |
| 170425 | NR_038220 | LINC00858 | long intergenic non-protein coding RNA 858 | lncRNA | . | 27276 | . | . | . | 0.155830 | 0.058970 | 0.054416 | 0.104924 | 0.362623 | 0.136312 | 0.122108 | 0.239896 | -1.368704 | 8.413861 | 1.09520999 | -0.41344612 | 0.679279793 | NA | 3.282109 | 2.827809 | 3.103393 | 3.065267 | 3.061823 | 3.083564 | 16 | 6 | 4 | 9 |
| 170463 | NM_00100999 | SSBP4 | single stranded DNA binding protein 4 | protein_coding | NP_001009998.1;NP_116016.1;XP_005259847.1;XP_006722728.1;XP_00 | 15676 | 607391 | . | . | 14.224986 | 15.026707 | 15.965532 | 16.856714 | 33.102127 | 34.734908 | 35.826191 | 38.540874 | 1.138644 | 935.195874 | 0.139521024 | 1.34256881 | 0.179411635 | 0.952086703 | 9.772484 | 9.959696 | 9.798426 | 9.839848 | 9.918492 | 9.908340 | 978 | 971 | 872 | 927 |
| 170485 | gene-ARF4P2 | ARF4P2 | ADP ribosylation factor 4 pseudogene 2 | pseudogene | . | 16599 | . | . | . | 0.214654 | 0.084736 | 0.106911 | 0.137666 | 0.499509 | 0.195871 | 0.239906 | 0.314757 | -1.108322 | 4.627757 | 1.421079843 | -0.1044113 | 0.916842956 | NA | 2.282109 | 2.134756 | 2.225376 | 2.205801 | 2.209265 | 2.217619 | 8 | 3 | 3 | 5 |
| 170487 | NM_00102467 | ACTL10 | actin like 10 | protein_coding | NP_001019846.1 | 16127 | . | . | . | 0.229655 | 0.915044 | 0.213974 | 0.556225 | 0.534417 | 2.115165 | 0.480152 | 1.271742 | -1.459537 | 24.200443 | 0.752244529 | -0.72517687 | 0.468343531 | NA | 4.847212 | 4.293999 | 4.526020 | 4.666148 | 4.526851 | 4.594009 | 14 | 49 | 10 | 26 |
| 170506 | NM_00111439 | DHX36 | DEAH-box helicase 36 | protein_coding | NP_001107869.1;NP_065916.2 | 14410 | 612767 | . | . | 7.960033 | 8.262343 | 8.817252 | 7.146727 | 18.523324 | 19.098775 | 19.785658 | 16.340143 | -1.116651 | 1558.994368 | 0.129624691 | -1.22799563 | 0.219448607 | 0.982008277 | 10.683629 | 10.524802 | 10.656396 | 10.637900 | 10.608817 | 10.512304 | 1916 | 1754 | 1362 | 1294 |
| 170556 | gene-UBE2V1P | UBE2V1P1 | ubiquitin conjugating enzyme E2 V1 pseudogene 1 | pseudogene | . | 16581 | . | . | . | 0.057135 | 0.114633 | 0.061719 | 0.111618 | 0.132956 | 0.264979 | 0.138495 | 0.255201 | 1.113346 | 5.738573 | 1.27824251 | 0.121183096 | 0.903546013 | NA | 2.444631 | 2.592943 | 2.505961 | 2.524494 | 2.511714 | 2.524445 | 4 | 8 | 4 | 7 |
| 170622 | NM_00128739 | COMMD6 | COMM domain containing 6 | protein_coding | NP_001274321.1;NP_001274322.1;NP_001274323.1;NP_987091.1;NP_9! | 24015 | 612377 | . | . | 9.411807 | 8.688354 | 8.971113 | 6.507384 | 21.901660 | 20.083520 | 20.130918 | 14.878362 | -1.186012 | 448.275536 | 0.198150498 | -1.2420817 | 0.214206421 | 0.979036831 | 8.925400 | 8.680722 | 8.882234 | 8.826742 | 8.842117 | 8.649240 | 584 | 502 | 415 | 322 |
| 170685 | NM_00130496 | NUDT10 | nudix hydrolase 10 | protein_coding | NP_001291892.1;NP_694853.1 | 17621 | 300527 | . | . | 0.837224 | 1.259720 | 1.250164 | 0.903400 | 1.948256 | 2.911899 | 2.805332 | 2.065518 | 1.023948 | 64.382369 | 0.416061132 | 0.082060285 | 0.934598774 | 0.998865404 | 5.991308 | 6.025675 | 5.954018 | 6.042532 | 6.040473 | 5.967284 | 59 | 82 | 66 | 51 |
| 170688 | NR_104005 | NUDT4P2 | nudix hydrolase 4 pseudogene 2 | transcribed_ps | . | 18013 | . | . | . | 3.656442 | 2.723126 | 1.715041 | 1.984368 | 8.508693 | 6.294627 | 3.848501 | 4.537022 | -1.752632 | 302.352128 | 0.224135459 | -3.61175808 | **0.000304128** | **0.028364066** | 8.588801 | 7.779267 | 8.449343 | 8.265081 | 8.031920 | 8.092551 | 510 | 353 | 178 | 220 |
| 170689 | NM_139055,XI | ADAMTS15 | ADAM metallopeptidase with thrombospondin type 1 motif 15 | protein_coding | NP_620686.1;XP_005271476.1 | 16305 | 607509 | . | . | 0.088602 | 0.059427 | 0.052673 | 0.030088 | 0.206180 | 0.137367 | 0.118196 | 0.068794 | -1.762607 | 11.541573 | 0.920299995 | -0.88852649 | 0.374257613 | NA | 3.878555 | 3.066070 | 3.560941 | 3.537041 | 3.529722 | 3.512285 | 20 | 13 | 9 | 6 |
| 170690 | NM_139056,N | ADAMTS16 | ADAM metallopeptidase with thrombospondin type 1 motif 16 | protein_coding | NP_620687.2 | 17108 | 607510 | . | . | 0.014800 | 0.116523 | 0.032875 | 0.038111 | 0.034441 | 0.269349 | 0.073770 | 0.087137 | -1.733180 | 8.322524 | 1.176995117 | -0.67410797 | 0.500242673 | NA | 3.401919 | 2.602870 | 3.020588 | 3.093681 | 3.036459 | 3.039136 | 3 | 20 | 5 | 6 |
| 170691 | NM_139057,XI | ADAMTS17 | ADAM metallopeptidase with thrombospondin type 1 motif 17 | protein_coding | NP_620688.2;XP_005254929.1;XP_011519614.1;XP_016877462.1;XP_01( | 17109 | 607511 | . | . | 4.200004 | 4.039426 | 4.126778 | 4.192862 | 9.773579 | 9.337317 | 9.260375 | 9.586477 | 1.002500 | 942.807713 | 0.138948179 | 0.02592471 | 0.979317391 | 0.998865404 | 9.878959 | 9.882678 | 9.898326 | 9.860988 | 9.880874 | 9.882476 | 1112 | 990 | 820 | 886 |
| 170850 | NM_133329,N | KCNG3 | potassium voltage-gated channel modifier subfamily G member 3 | protein_coding | NP_579875.1;NP_758847.1 | 18306 | 606767 | . | . | 1.295753 | 1.301288 | 0.755563 | 0.750058 | 3.015272 | 3.007985 | 1.695463 | 1.714918 | -1.727793 | 118.563627 | 0.299054106 | -2.63808447 | **0.008337581** | 0.289154856 | 7.230612 | 6.441960 | 6.970303 | 6.969192 | 6.793512 | 6.782217 | 173 | 162 | 77 | 80 |
| 170954 | NM_00113487 | PPP1R18 | protein phosphatase 1 regulatory subunit 18 | protein_coding | NP_001128342.1;NP_597728.1 | 29413 | 610990 | . | . | 24.490601 | 26.060280 | 27.133941 | 29.319518 | 56.990637 | 60.239501 | 60.887771 | 67.035598 | 1.110137 | 2704.300462 | 0.116106418 | 1.2982727 | 0.194193644 | 0.96621494 | 11.323799 | 11.474354 | 11.332264 | 11.380620 | 11.414220 | 11.468065 | 2863 | 2849 | 2393 | 2755 |
| 170958 | NM_00134815 | ZNF525 | zinc finger protein 525 | protein_coding | NP_001335085.1;NP_001335086.1 | 29423 | . | . | . | 0.582282 | 0.775151 | 1.143719 | 0.747869 | 1.354995 | 1.791796 | 2.566472 | 1.709914 | 1.158646 | 131.267699 | 0.333067605 | 0.637826827 | 0.523586414 | 0.998865404 | 6.925454 | 7.139360 | 6.948421 | 7.038927 | 7.150314 | 6.933957 | 122 | 148 | 158 | 94 |
| 170959 | NM_00131912 | ZNF431 | zinc finger protein 431 | protein_coding | NP_001306053.1;NP_001306055.1;NP_001306056.1;NP_597730.2;XP_01 | 20809 | . | . | . | 1.463803 | 1.356285 | 1.738038 | 0.932460 | 3.406329 | 3.135113 | 3.900108 | 2.131961 | 1.033174 | 523.645848 | 0.217678937 | 0.21629987 | 0.828754011 | 0.998865404 | 9.008052 | 9.056437 | 9.022929 | 9.003131 | 9.183905 | 8.858307 | 602 | 547 | 580 | 373 |
| 170960 | NM_133474 | ZNF721 | zinc finger protein 721 | protein_coding | NP_597731.2 | 29425 | . | . | . | 2.417682 | 2.474066 | 3.198050 | 2.258466 | 5.626046 | 5.718915 | 7.176332 | 5.163713 | 1.101622 | 377.141186 | 0.210345136 | 0.663811228 | 0.506811138 | 0.998865404 | 8.486818 | 8.627667 | 8.520197 | 8.528972 | 8.680891 | 8.470702 | 410 | 390 | 405 | 305 |
| 170961 | NM_133475,XI | ANKRD24 | ankyrin repeat domain 24 | protein_coding | NP_597732.1;XP_011526058.1;XP_011526059.1;XP_011526060.1;XP_01: | 29424 | . | . | . | 0.241421 | 0.199559 | 0.190660 | 0.115817 | 0.561797 | 0.461289 | 0.427836 | 0.264803 | -1.372330 | 25.132656 | 0.631981217 | -0.72253365 | 0.469966442 | NA | 4.860000 | 4.407673 | 4.681647 | 4.655694 | 4.657508 | 4.614479 | 37 | 28 | 23 | 15 |
| 171017 | NM_00103992 | ZNF384 | zinc finger protein 384 | protein_coding | NP_001035009.1;NP_001129206.1;NP_597733.2;XP_006719032.1;XP_01 | 11955 | 609951 | . | . | 12.838972 | 13.944532 | 15.671887 | 15.634744 | 29.876815 | 32.233411 | 35.167257 | 35.746984 | 1.155653 | 1495.701091 | 0.12744441 | 1.637645014 | 0.101495767 | 0.819486355 | 10.438553 | 10.647128 | 10.452848 | 10.520619 | 10.605403 | 10.592597 | 1527 | 1564 | 1406 | 1491 |
| 171022 | NR_026690 | ABHD11-AS1 | ABHD11 antisense RNA 1 (tail to tail) | lncRNA | . | 18289 | 612545 | . | . | 0.968818 | 0.684397 | 0.184254 | 0.700794 | 2.254480 | 1.582015 | 0.413460 | 1.602282 | -1.759943 | 9.768575 | 1.024876057 | -0.79573385 | 0.426186746 | NA | 3.641413 | 2.819320 | 3.320007 | 3.299294 | 3.270070 | 3.301289 | 17 | 11 | 3 | 10 |
| 171023 | NM_00116460 | ASXL1 | ASXL transcriptional regulator 1 | protein_coding | NP_001158075.1;NP_001350663.1;NP_056153.2;XP_006723790.1;XP_00 | 18318 | 612990 | . | . | 10.972694 | 11.324420 | 12.321657 | 11.891070 | 25.533907 | 26.176900 | 27.649437 | 27.187522 | 1.074552 | 2501.856521 | 0.11456241 | 0.905496677 | 0.365202319 | 0.998865404 | 11.235960 | 11.339741 | 11.250277 | 11.267252 | 11.333435 | 11.300154 | 2748 | 2630 | 2300 | 2380 |
| 171024 | NM_00112893 | SYNPO2 | synaptopodin 2 | protein_coding | NP_001122405.1;NP_001122406.1;NP_001273683.1;NP_001273684.1;NI | 17732 | . | . | . | 0.033599 | 0.032049 | 0.025840 | 0.009682 | 0.078186 | 0.074082 | 0.057983 | 0.022137 | -1.234551 | 10.675917 | 0.982673254 | -0.30934667 | 0.757057831 | NA | 3.558919 | 3.257993 | 3.396832 | 3.432822 | 3.423351 | 3.386453 | 9 | 17 | 12 | 5 |
| 171177 | NM_133639 | RHOV | ras homolog family member V | protein_coding | NP_598378.3 | 18313 | . | . | . | 0.106360 | 0.072638 | 0.074569 | 0.065717 | 0.247505 | 0.167906 | 0.167332 | 0.150254 | -1.209732 | 4.894846 | 1.359067692 | -0.20211421 | 0.839827441 | NA | 2.421266 | 2.148375 | 2.300838 | 2.294032 | 2.293956 | 2.292427 | 7 | 5 | 4 | 4 |
| 171220 | NR_033796 | DSTNP2 | DSTN pseudogene 2 | transcribed_ps | . | 34546 | . | . | . | 1.863340 | 1.712562 | 1.622148 | 1.806699 | 4.336068 | 3.958663 | 3.640054 | 4.130803 | -1.052568 | 60.665063 | 0.403660588 | -0.18310631 | 0.854714604 | 0.998865404 | 5.959362 | 5.885275 | 5.939613 | 5.918640 | 5.908602 | 5.927697 | 75 | 64 | 49 | 58 |
| 171391 | NR_126342,NF | LOC171391 | uncharacterized LOC171391 | lncRNA | . | . | . | . | . | 2.393513 | 2.839598 | 2.566849 | 2.031585 | 5.569803 | 6.563860 | 5.759936 | 4.644977 | -1.120056 | 61.292137 | 0.409908664 | -0.39904323 | 0.689861348 | 0.998865404 | 6.016660 | 5.854019 | 5.927972 | 5.968321 | 5.950574 | 5.896520 | 68 | 76 | 57 | 47 |
| 171392 | NM_138330 | ZNF675 | zinc finger protein 675 | protein_coding | NP_612203.2 | 30768 | . | . | . | 3.127190 | 2.600083 | 3.279183 | 2.332656 | 7.277100 | 6.010210 | 7.358392 | 5.333341 | -1.037346 | 204.668558 | 0.258662175 | -0.20450525 | 0.837958688 | 0.998865404 | 7.702275 | 7.651571 | 7.725379 | 7.634838 | 7.738731 | 7.584348 | 263 | 203 | 205 | 156 |
| 171423 | NR_002305 | PDIA3P1 | protein disulfide isomerase family A member 3 pseudogene 1 | transcribed_ps | . | 4607 | . | . | . | 2.382196 | 2.545285 | 2.097144 | 2.237455 | 5.543469 | 5.883539 | 4.705931 | 5.115676 | -1.150541 | 152.215224 | 0.265925098 | -0.76078562 | 0.446785124 | 0.998865404 | 7.347831 | 7.144983 | 7.265204 | 7.289595 | 7.213534 | 7.231584 | 182 | 181 | 120 | 136 |
| 171425 | NM_206808,N | CLYBL | citramalyl-CoA lyase | protein_coding | NP_996531.1 | 18355 | 609686 | . | . | 1.473811 | 1.184567 | 1.284130 | 2.105483 | 3.429618 | 2.738180 | 2.881549 | 4.813937 | 1.247635 | 67.680020 | 0.413809483 | 0.77136069 | 0.440493169 | 0.998865404 | 5.912945 | 6.230889 | 6.075156 | 6.014923 | 6.033066 | 6.157358 | 79 | 56 | 49 | 88 |
| 171523 | NR_144551 | CYP2T1P | cytochrome P450 family 2 subfamily T member 1, pseudogene | transcribed_ps | . | 18852 | . | . | . | 0.052483 | 0.022298 | 0.041336 | 0.161353 | 0.122129 | 0.051542 | 0.092758 | 0.368915 | **2.622026** | 5.611407 | 1.364666219 | 1.019063836 | 0.308172661 | NA | 1.631267 | 3.022362 | 2.461824 | 2.450136 | 2.458272 | 2.499726 | 5 | 2 | 3 | 12 |
| 171546 | NM_138288 | SPTSSA | serine palmitoyltransferase small subunit A | protein_coding | NP_612145.2 | 20361 | 613540 | . | . | 4.694370 | 3.917381 | 3.757451 | 3.801765 | 10.923992 | 9.055202 | 8.431611 | 8.692286 | -1.154759 | 336.165833 | 0.200238127 | -1.03672644 | 0.299863365 | 0.998865404 | 8.492828 | 8.285810 | 8.485046 | 8.374513 | 8.350223 | 8.351424 | 454 | 352 | 271 | 293 |
| 171558 | NM_00124316 | PTCRA | pre T cell antigen receptor alpha | protein_coding | NP_001230097.1;NP_001230098.1;NP_001230099.1;NP_612153.2;XP_02 | 21290 | 606817 | . | . | 0.412890 | 0.450778 | 0.171371 | 0.467285 | 0.960814 | 1.041993 | 0.384554 | 1.068392 | -1.324072 | 16.489279 | 0.789631174 | -0.51287445 | 0.608039148 | NA | 4.234218 | 3.823571 | 4.058653 | 4.056013 | 4.011541 | 4.062949 | 22 | 20 | 7 | 19 |
| 171568 | NM_00101805 | POLR3H | RNA polymerase III subunit H | protein_coding | NP_001018060.1;NP_001018062.1;NP_001269813.1;NP_001269814.1;NI | 30349 | . | . | . | 14.856058 | 14.281605 | 15.332407 | 16.616181 | 34.570659 | 33.012569 | 34.405478 | 37.990924 | 1.021235 | 1788.828036 | 0.121247864 | 0.250026133 | 0.802567139 | 0.998865404 | 10.789575 | 10.819864 | 10.776749 | 10.815525 | 10.822364 | 10.803230 | 1988 | 1957 | 1591 | 1674 |
| 171586 | NM_00130825 | ABHD3 | abhydrolase domain containing 3, phospholipase | protein_coding | NP_001295185.1;NP_001295186.1;NP_612213.2;XP_011524149.1;XP_01 | 18718 | 612197 | . | . | 10.647097 | 10.381369 | 9.806160 | 8.905793 | 24.776235 | 23.997002 | 22.004740 | 20.362039 | -1.120840 | 676.204124 | 0.155487412 | -1.05848309 | 0.289835254 | 0.998865404 | 9.481041 | 9.316925 | 9.438732 | 9.438400 | 9.399121 | 9.321296 | 823 | 771 | 587 | 563 |

| | | | | | | | | | | | | | | | | | | | | | | | | | | | | | | | | | |
|---|---|---|---|---|---|---|---|---|---|---|---|---|---|---|---|---|---|---|---|---|---|---|---|---|---|---|---|---|---|---|---|---|---|
| 192111 | NM_00117054 | PGAM5 | PGAM family member 5, mitochondrial serine/threonine protein p | protein_coding | NP_001164014.1;NP_001164015.1;NP_612642.2 | 28763 | 614939 | . | . | 14.817652 | 13.932915 | 12.618557 | 13.220461 | 34.481284 | 32.206557 | 28.315675 | 30.227014 | -1.124605 | 1174.631331 | 0.133782598 | -1.26636913 | 0.205380957 | 0.970176829 | 10.280219 | 10.110794 | 10.267751 | 10.212092 | 10.134271 | 10.170058 | 1484 | 1293 | 938 | 1062 |
| 192146 | gene-DDX50P1 | DDX50P1 | DEAD-box helicase 50 pseudogene 1 | pseudogene | . | 18974 | . | . | . | 0.044509 | 0.047647 | 0.088045 | 0.024435 | 0.103574 | 0.110138 | 0.197571 | 0.055867 | 1.252703 | 4.053482 | 1.517094015 | 0.21425438 | 0.830348692 | NA | 1.844134 | 2.175234 | 2.004894 | 2.006037 | 2.020228 | 1.998913 | 4 | 4 | 6 | 2 |
| 192217 | NR_132962 | OXCT2P1 | 3-oxoacid CoA-transferase 2 pseudogene 1 | transcribed_ps | . | 21627 | . | . | . | 0.517948 | 0.532230 | 0.763324 | 0.444811 | 1.205287 | 1.230274 | 1.712878 | 1.017007 | 1.150629 | 32.352160 | 0.561206219 | 0.360692793 | 0.718329113 | 0.998865404 | 4.909524 | 5.114763 | 4.996519 | 5.001017 | 5.051236 | 4.981236 | 34 | 33 | 38 | 24 |
| 192286 | NM_138820 | HIGD2A | HIG1 hypoxia inducible domain family member 2A | protein_coding | NP_620175.1 | 28311 | . | . | . | 92.693184 | 104.603905 | 92.702499 | 123.182304 | 215.700836 | 241.796600 | 208.021713 | 281.641693 | 1.078995 | 2103.401225 | 0.14697673 | 0.746293089 | 0.455490385 | 0.998865404 | 10.982867 | 11.092083 | 10.959182 | 11.048118 | 10.949586 | 11.169148 | 2200 | 2306 | 1640 | 2325 |
| 192669 | NM_024852,N | AGO3 | argonaute RISC catalytic component 3 | protein_coding | NP_079128.2;NP_803171.1;XP_005270632.1;XP_005270633.1;XP_01153 | 18421 | 607355 | . | . | 1.368850 | 1.687623 | 2.225891 | 1.399458 | 3.185370 | 3.901018 | 4.994832 | 3.199695 | 1.093890 | 592.022998 | 0.175318378 | 0.738468344 | 0.460229897 | 0.998865404 | 9.143052 | 9.273040 | 9.142041 | 9.208745 | 9.311982 | 9.152758 | 618 | 641 | 602 | 511 |
| 192670 | NM_017629,N | AGO4 | argonaute RISC component 4 | protein_coding | NP_060099.2;XP_005270635.1;XP_005270636.1;XP_011539185.1;XP_01 | 18424 | 607356 | . | . | 2.182737 | 2.391480 | 2.741141 | 2.499577 | 5.079318 | 5.528011 | 6.151042 | 5.714985 | 1.070104 | 448.230304 | 0.175884289 | 0.555766326 | 0.578370608 | 0.998865404 | 8.758276 | 8.856253 | 8.768932 | 8.802422 | 8.856135 | 8.798019 | 485 | 480 | 422 | 414 |
| 192683 | NM_00117811 | SCAMP5 | secretory carrier membrane protein 5 | protein_coding | NP_001171582.1;NP_001171583.1;NP_620417.1;XP_006720483.1;XP_00 | 30386 | 613766 | . | . | 3.986352 | 4.564547 | 3.579742 | 4.579242 | 9.276405 | 10.551158 | 8.032838 | 10.469893 | -1.048326 | 421.852480 | 0.189334539 | -0.35961371 | 0.719136031 | 0.998865404 | 8.754806 | 8.685552 | 8.702922 | 8.763247 | 8.639927 | 8.765865 | 473 | 489 | 320 | 427 |
| 195827 | NM_153698,XI | PRXL2C | peroxiredoxin like 2C | protein_coding | NP_714542.1;XP_005251840.1;XP_005251841.1 | 16881 | . | . | . | 2.980350 | 3.400122 | 3.195166 | 3.973404 | 6.935397 | 7.859534 | 7.169859 | 9.084714 | 1.119033 | 251.946520 | 0.228281758 | 0.710758383 | 0.477233974 | 0.998865404 | 7.894336 | 8.055133 | 7.924872 | 7.964544 | 7.932052 | 8.065851 | 263 | 267 | 201 | 282 |
| 195828 | NM_153695,XI | ZNF367 | zinc finger protein 367 | protein_coding | NP_710162.1;XP_011516647.1 | 18320 | 610160 | . | . | 3.545517 | 3.063312 | 2.580694 | 2.394451 | 8.250564 | 7.080982 | 5.791004 | 5.474629 | -1.354020 | 327.358850 | 0.202002652 | -2.16457115 | 0.030420542 | 0.555416333 | 8.556458 | 8.120135 | 8.478545 | 8.380431 | 8.284605 | 8.241587 | 469 | 373 | 252 | 250 |
| 196047 | NR_002791,NF | EMX2OS | EMX2 opposite strand/antisense RNA | antisense_RNA | . | 18511 | 607637 | . | . | 0.337667 | 0.420809 | 0.389221 | 0.238939 | 0.785765 | 0.972719 | 0.873400 | 0.546306 | -1.229235 | 78.369363 | 0.385490209 | -0.77241994 | 0.439865745 | 0.998865404 | 6.432509 | 6.136800 | 6.283376 | 6.342752 | 6.315610 | 6.201755 | 89 | 103 | 76 | 50 |
| 196051 | NM_00103005 | PLPP4 | phospholipid phosphatase 4 | protein_coding | NP_001025230.1;NP_001305095.1;NP_001305096.1;NP_001305097.1;NI | 23531 | . | . | . | 6.112990 | 6.736030 | 1.951760 | 2.508936 | 14.225181 | 15.570632 | 4.379693 | 5.736381 | -2.267221 | 409.925463 | 0.192282722 | -6.14160753 | 8.16905E-10 | 8.0541E-07 | 9.152547 | 7.970092 | 8.833106 | 8.879086 | 8.324876 | 8.427707 | 633 | 635 | 194 | 261 |
| 196074 | NM_00111352 | METTL15 | methyltransferase like 15 | protein_coding | NP_001107000.1;NP_001284704.1;NP_689849.2;XP_011518237.1;XP_01 | 26606 | 618711 | . | . | 1.853460 | 1.899833 | 1.732613 | 1.531105 | 4.313076 | 4.391550 | 3.887934 | 3.500693 | -1.119019 | 215.165725 | 0.233518439 | -0.69473716 | 0.487220035 | 0.998865404 | 7.827664 | 7.666444 | 7.782836 | 7.766224 | 7.757944 | 7.685147 | 266 | 241 | 191 | 175 |
| 196264 | NM_00128615 | MPZL3 | myelin protein zero like 3 | protein_coding | NP_001273081.1;NP_938016.1;XP_006718838.1;XP_016872793.1 | 27279 | 611707 | . | . | 0.441607 | 0.522537 | 0.488719 | 0.452856 | 1.027636 | 1.207868 | 1.096672 | 1.035402 | 1.007031 | 63.522089 | 0.394680746 | 0.025610739 | 0.97956782 | 0.998865404 | 5.984018 | 5.994337 | 5.978602 | 5.996518 | 5.999187 | 5.979024 | 70 | 71 | 58 | 57 |
| 196294 | NM_00130427 | IMMP1L | inner mitochondrial membrane peptidase subunit 1 | protein_coding | NP_001291203.1;NP_659418.1;XP_005252869.1;XP_011518244.1;XP_01 | 26317 | 612323 | . | . | 2.129860 | 2.552598 | 2.501375 | 2.010635 | 4.956268 | 5.900446 | 5.613015 | 4.597078 | -1.000239 | 61.580729 | 0.403393645 | -0.00085529 | 0.999317573 | 0.999862634 | 5.944558 | 5.944256 | 5.924249 | 5.961266 | 5.956494 | 5.929071 | 65 | 72 | 57 | 54 |
| 196383 | NM_145058,N | RILPL2 | Rab interacting lysosomal protein like 2 | protein_coding | NP_659495.1;XP_011536314.1 | 28787 | 614093 | . | . | 5.065820 | 5.079085 | 6.038243 | 6.227130 | 11.788372 | 11.740534 | 13.549644 | 14.237595 | 1.181243 | 309.453895 | 0.198076873 | 1.213193203 | 0.225055962 | 0.982008277 | 8.148492 | 8.388683 | 8.226338 | 8.222550 | 8.306126 | 8.323505 | 328 | 305 | 287 | 319 |
| 196385 | NM_00137210 | DNAH10 | dynein axonemal heavy chain 10 | protein_coding | NP_001359035.1;NP_997320.2;XP_011536316.1;XP_011536317.1;XP_01 | 2941 | 605884 | . | . | 0.336853 | 0.349696 | 0.200957 | 0.357281 | 0.783872 | 0.808337 | 0.450943 | 0.816880 | -1.242232 | 138.325025 | 0.313860012 | -0.99705158 | 0.318739476 | 0.998865404 | 7.261164 | 6.945433 | 7.140938 | 7.152612 | 6.969651 | 7.154433 | 174 | 168 | 78 | 147 |
| 196394 | NM_00111340 | AMN1 | antagonist of mitotic exit network 1 homolog | protein_coding | NP_001106873.1;NP_001265340.1;NP_001265341.1;XP_006719117.1;XF | 27281 | . | . | . | 0.886630 | 0.849312 | 0.824431 | 0.997825 | 2.063225 | 1.963224 | 1.849999 | 2.281407 | 1.111799 | 67.253773 | 0.382925156 | 0.399285293 | 0.689683002 | 0.998865404 | 5.993221 | 6.145832 | 6.052884 | 6.061390 | 6.079387 | 6.083982 | 72 | 70 | 61 | 67 |
| 196410 | NM_152637 | METTL7B | methyltransferase like 7B | protein_coding | NP_689850.2 | 28276 | . | . | . | 0.321700 | 0.306034 | 0.195260 | 0.289182 | 0.748610 | 0.707411 | 0.438158 | 0.661179 | -1.355891 | 11.663164 | 0.898821498 | -0.48868621 | 0.625063868 | NA | 3.747897 | 3.306201 | 3.559740 | 3.555491 | 3.534189 | 3.549933 | 16 | 14 | 7 | 11 |
| 196441 | NM_144982 | ZFC3H1 | zinc finger C3H1-type containing | protein_coding | NP_659419.3 | 28328 | . | . | . | 6.087925 | 5.990495 | 7.278994 | 5.020598 | 14.166852 | 13.847297 | 16.333851 | 11.479000 | 1.004568 | 1337.011622 | 0.168446707 | 0.039035663 | 0.968861956 | 0.998865404 | 10.381181 | 10.388402 | 10.387889 | 10.368628 | 10.516205 | 10.227398 | 1555 | 1421 | 1385 | 1020 |
| 196463 | NM_00115972 | PLBD2 | phospholipase B domain containing 2 | protein_coding | NP_001153199.1;NP_775813.2;XP_011536325.1;XP_016874466.1 | 27283 | . | . | . | 35.336431 | 38.474759 | 39.465754 | 39.180496 | 82.229328 | 88.936126 | 88.560007 | 89.581550 | 1.051673 | 5650.016093 | 0.107898751 | 0.673653007 | 0.500531945 | 0.998865404 | 12.427257 | 12.499907 | 12.407458 | 12.475910 | 12.493265 | 12.475811 | 6083 | 6186 | 5089 | 5374 |
| 196483 | NM_00128902 | EEF2KMT | eukaryotic elongation factor 2 lysine methyltransferase | protein_coding | NP_001275958.1;NP_958802.1;NP_963892.1;XP_005255214.1;XP_00525 | 32221 | 615263 | . | . | 3.674134 | 4.588020 | 4.030183 | 5.273715 | 8.549860 | 10.605413 | 9.043612 | 12.057722 | 1.113193 | 326.882143 | 0.2169435 | 0.713106192 | 0.475780048 | 0.998865404 | 8.274072 | 8.427125 | 8.255046 | 8.376662 | 8.302203 | 8.446786 | 317 | 371 | 261 | 364 |
| 196513 | NM_00131929 | DCP1B | decapping mRNA 1B | protein_coding | NP_001306221.1;NP_689853.3;XP_011519229.1 | 24451 | 609843 | . | . | 1.650802 | 2.399533 | 1.950687 | 2.883778 | 3.841486 | 5.546628 | 4.377285 | 6.593415 | 1.197667 | 145.515974 | 0.29951595 | 0.868824727 | 0.384943006 | 0.998865404 | 7.050484 | 7.308098 | 7.074404 | 7.203472 | 7.147885 | 7.268676 | 126 | 168 | 117 | 171 |
| 196527 | NM_00102535 | ANO6 | anoctamin 6 | protein_coding | NP_001020527.2;NP_001136150.1;NP_001136151.1;NP_001191732.1;XF | 25240 | 608663 | . | . | 10.347956 | 9.869139 | 9.360874 | 9.242701 | 24.080118 | 22.812958 | 21.005529 | 21.132339 | -1.102442 | 1818.752432 | 0.12128704 | -1.16008003 | 0.246016224 | 0.992733415 | 10.897296 | 10.756749 | 10.887388 | 10.843866 | 10.800146 | 10.777969 | 2256 | 2002 | 1526 | 1599 |
| 196528 | NM_00134783 | ARID2 | AT-rich interaction domain 2 | protein_coding | NP_001334768.1;NP_689854.2;XP_006719335.1;XP_011536327.1 | 18037 | 609539 | . | . | 2.013236 | 2.119424 | 2.268541 | 1.920844 | 4.684882 | 4.899143 | 5.090540 | 4.391780 | 1.006064 | 552.829273 | 0.169161169 | 0.051560864 | 0.958878604 | 0.998865404 | 9.105992 | 9.115373 | 9.105956 | 9.109069 | 9.166015 | 9.054094 | 633 | 596 | 524 | 475 |
| 196743 | NM_152911,N | PAOX | polyamine oxidase | protein_coding | NP_690875.1;NP_997010.1;NP_997011.1 | 20837 | 615853 | . | . | 0.662270 | 0.450842 | 0.412808 | 0.585967 | 1.541130 | 1.042142 | 0.926329 | 1.339744 | -1.075383 | 29.692779 | 0.578413318 | -0.18127374 | 0.856152717 | 0.998865404 | 4.943804 | 4.838350 | 4.917216 | 4.877332 | 4.870981 | 4.906203 | 41 | 28 | 21 | 31 |
| 196792 | NM_00120436 | FAM24B | family with sequence similarity 24 member B | protein_coding | NP_001191293.1;NP_689857.2 | 23475 | . | . | . | 2.770055 | 3.795692 | 3.981708 | 3.723353 | 6.446034 | 8.773912 | 8.934840 | 8.513006 | 1.164733 | 81.925417 | 0.362000418 | 0.607731312 | 0.543365692 | 0.998865404 | 6.242574 | 6.461599 | 6.286657 | 6.370757 | 6.382152 | 6.358204 | 73 | 95 | 80 | 79 |
| 196872 | NR_024396 | LINC00638 | long intergenic non-protein coding RNA 638 | lncRNA | . | 28325 | . | . | . | 0.207137 | 0.301573 | 0.357605 | 0.369989 | 0.482016 | 0.697099 | 0.802454 | 0.845936 | 1.419610 | 24.536283 | 0.628931534 | 0.803735833 | 0.421549555 | NA | 4.343257 | 4.846728 | 4.572041 | 4.605328 | 4.623946 | 4.624490 | 19 | 26 | 25 | 27 |
| 196913 | NR_131171 | LINC01599 | long intergenic non-protein coding RNA 1599 | lncRNA | . | 27285 | . | . | . | 0.252153 | 0.506475 | 0.286400 | 0.332232 | 0.586771 | 1.170739 | 0.642673 | 0.759610 | -1.183219 | 10.859276 | 0.944037536 | -0.25710514 | 0.797097607 | NA | 3.558919 | 3.312263 | 3.425738 | 3.461691 | 3.432698 | 3.438969 | 9 | 17 | 8 | 10 |
| 196951 | NM_00133029 | FAM227B | family with sequence similarity 227 member B | protein_coding | NP_001317222.1;NP_689860.2;XP_005254270.1;XP_005254271.1;XP_00 | 26543 | . | . | . | 0.253250 | 0.270880 | 0.276764 | 0.244305 | 0.589326 | 0.626150 | 0.621053 | 0.558572 | -1.010710 | 26.734598 | 0.595003877 | -0.0258305 | 0.979392535 | NA | 4.747897 | 4.733338 | 4.745103 | 4.737351 | 4.744062 | 4.736044 | 32 | 28 | 24 | 24 |
| 196968 | NR_003260 | DNM1P46 | dynamin 1 pseudogene 46 | transcribed_ps | . | 35199 | . | . | . | 0.223773 | 0.194534 | 0.348310 | 0.330513 | 0.520730 | 0.449674 | 0.781599 | 0.755679 | 1.579371 | 33.475310 | 0.536916487 | 1.228030934 | 0.219435356 | 0.982008277 | 4.697260 | 5.357829 | 5.029508 | 5.014345 | 5.087556 | 5.076439 | 32 | 26 | 37 | 37 |
| 197021 | NM_00127856 | LCTL | lactase like | protein_coding | NP_001265491.1;NP_997221.2;XP_024305637.1;XP_024305638.1 | 15583 | 617060 | . | . | 0.816750 | 0.926228 | 0.789977 | 0.635238 | 1.900611 | 2.141018 | 1.772687 | 1.452397 | -1.226583 | 62.940855 | 0.399633994 | -0.73728712 | 0.460947766 | 0.998865404 | 6.115110 | 5.821864 | 6.017882 | 5.980102 | 5.965812 | 5.942807 | 86 | 69 | 52 | 50 |
| 197131 | NM_174916 | UBR1 | ubiquitin protein ligase E3 component n-recognin 1 | protein_coding | NP_777576.1 | 16808 | 605981 | . | . | 3.868223 | 3.547092 | 3.831881 | 3.382057 | 9.001513 | 8.199262 | 8.598631 | 7.732672 | -1.042623 | 878.032728 | 0.149404389 | -0.40305045 | 0.686911109 | 0.998865404 | 9.807611 | 9.748036 | 9.825995 | 9.756932 | 9.808890 | 9.713454 | 1081 | 921 | 798 | 752 |
| 197135 | NM_00114511 | PATL2 | PAT1 homolog 2 | protein_coding | NP_001138584.1;NP_001317212.1;XP_011519638.2;XP_011519639.2;XP | 33630 | 614661 | . | . | 0.070265 | 0.092727 | 0.051727 | 0.049716 | 0.163509 | 0.214342 | 0.116074 | 0.113670 | -1.348352 | 5.877398 | 1.249150276 | -0.34519275 | 0.729949469 | NA | 2.754145 | 2.324312 | 2.559657 | 2.565804 | 2.558155 | 2.551490 | 7 | 8 | 5 | 4 |
| 197187 | NR_024399,NF | SNAI3-AS1 | SNAI3 antisense RNA 1 | lncRNA | . | 28327 | . | . | . | 0.824876 | 1.058818 | 1.381381 | 1.216933 | 1.919518 | 2.447505 | 3.099779 | 2.782373 | 1.390378 | 75.842725 | 0.371023503 | 1.281527691 | 0.20000838 | 0.96621494 | 5.987999 | 6.462962 | 6.162692 | 6.219520 | 6.293130 | 6.255385 | 64 | 77 | 82 | 77 |
| 197257 | NM_153486,N | LDHD | lactate dehydrogenase D | protein_coding | NP_705690.2;NP_919417.1;XP_024305942.1;XP_024305943.1;XP_02430 | 19708 | 607490 | . | . | 4.281925 | 3.577018 | 3.570715 | 4.934178 | 9.964217 | 8.268436 | 8.012584 | 11.281410 | 1.062777 | 274.408680 | 0.231325491 | 0.379722654 | 0.704151302 | 0.998865404 | 8.056068 | 8.142987 | 8.116155 | 8.043425 | 8.021545 | 8.195084 | 328 | 267 | 205 | 309 |
| 197258 | NM_145059,XI | FCSK | fucose kinase | protein_coding | NP_659496.2;XP_006721225.1;XP_011521230.1;XP_011521231.1;XP_01 | 29500 | 608675 | . | . | 4.041898 | 4.063935 | 4.240698 | 3.388165 | 9.405663 | 9.393968 | 9.516005 | 7.746636 | 1.031902 | 552.833151 | 0.167768515 | 0.270048024 | 0.787123308 | 0.998865404 | 9.087524 | 9.133511 | 9.108083 | 9.090299 | 9.168656 | 9.068528 | 635 | 579 | 526 | 486 |
| 197259 | NM_00114249 | MLKL | mixed lineage kinase domain like pseudokinase | protein_coding | NP_001135969.1;NP_689862.1;XP_005255891.1;XP_011521238.1 | 26617 | 615153 | . | . | 0.527834 | 0.663046 | 0.919326 | 0.790731 | 1.228291 | 1.532663 | 2.062940 | 1.807912 | 1.417019 | 58.678345 | 0.413501255 | 1.216100179 | 0.223946763 | 0.982008277 | 5.601622 | 6.104320 | 5.810487 | 5.843326 | 5.915206 | 5.883365 | 51 | 57 | 64 | 60 |
| 197320 | NM_00120140 | ZNF778 | zinc finger protein 778 | protein_coding | NP_001188336.1;NP_001365810.1;NP_872337.2;XP_011521242.1;XP_01 | 26479 | . | . | . | 1.683171 | 1.583065 | 2.115999 | 1.560527 | 3.916802 | 3.659329 | 4.748239 | 3.567959 | 1.009893 | 528.578502 | 0.164122362 | 0.08653982 | 0.931037303 | 0.998865404 | 9.038794 | 9.053118 | 9.056877 | 9.028331 | 9.047772 | 9.050354 | 619 | 555 | 461 | 499 |
| 197322 | NM_00112721 | ACSF3 | acyl-CoA synthetase family member 3 | protein_coding | NP_001120686.1;NP_001230208.1;NP_001271245.1;NP_777577.2;XP_00 | 27288 | 614245 | . | . | 3.215096 | 3.147972 | 1.664849 | 3.656002 | 7.481661 | 7.276680 | 3.735870 | 8.359011 | 1.010006 | 318.910673 | 0.21258986 | 0.067568007 | 0.946129524 | 0.998865404 | 8.310269 | 8.323717 | 8.335489 | 8.288519 | 8.239250 | 8.390412 | 381 | 328 | 240 | 342 |
| 197331 | gene-TUBB8P7 | TUBB8P7 | tubulin beta 8 class VIII pseudogene 7 | pseudogene | . | 42345 | . | . | . | 1.006258 | 0.875275 | 0.852025 | 0.592978 | 2.341603 | 2.023238 | 1.911919 | 1.355774 | -1.328368 | 40.814321 | 0.497501019 | -0.82342446 | 0.410266659 | 0.998865404 | 5.540061 | 5.133380 | 5.384978 | 5.361849 | 5.353366 | 5.304669 | 57 | 47 | 36 | 27 |
| 197335 | NM_145294,XI | WDR90 | WD repeat domain 90 | protein_coding | NP_660337.3;XP_016878512.1;XP_016878513.1 | 26960 | 618290 | . | . | 6.955847 | 7.552480 | 9.298571 | 6.801342 | 16.186542 | 17.457894 | 20.865724 | 15.550459 | 1.122469 | 1307.440610 | 0.162473429 | 1.025866504 | 0.304954513 | 0.998865404 | 10.266619 | 10.433611 | 10.244369 | 10.357597 | 10.501189 | 10.266673 | 1312 | 1429 | 1385 | 1100 |
| 197342 | NM_00125737 | EME2 | essential meiotic structure-specific endonuclease subunit 2 | protein_coding | NP_001244299.1;XP_016878514.1;XP_016878515.1 | 27289 | 610886 | . | . | 3.862662 | 4.522782 | 5.788489 | 3.973976 | 8.988574 | 10.454615 | 12.989201 | 9.086024 | 1.148150 | 178.208974 | 0.271689149 | 0.733600381 | 0.463192329 | 0.998865404 | 7.373768 | 7.574133 | 7.406352 | 7.471918 | 7.575597 | 7.415396 | 176 | 193 | 196 | 146 |
| 197370 | NM_145080,XI | NSMCE1 | NSE1 homolog, SMC5-SMC6 complex component | protein_coding | NP_659547.2;XP_006721086.1;XP_011544064.1;XP_016878529.1;XP_01 | 29897 | 617263 | . | . | 17.530565 | 16.029470 | 12.944653 | 16.525301 | 40.794347 | 37.052840 | 29.047428 | 37.783138 | -1.090764 | 595.639920 | 0.168805041 | -0.74250448 | 0.457781741 | 0.998865404 | 9.280041 | 9.153791 | 9.219895 | 9.270620 | 9.124094 | 9.247795 | 689 | 696 | 448 | 585 |
| 197407 | NM_00121490 | ZNF48 | zinc finger protein 48 | protein_coding | NP_001201835.1;NP_001201836.1;NP_001201838.1;NP_001311423.1;NI | 13114 | . | . | . | 0.128319 | 0.092952 | 0.116663 | 0.323401 | 0.298604 | 0.214862 | 0.261788 | 0.739417 | 1.900932 | 17.451416 | 0.786966326 | 1.177568881 | 0.238968522 | NA | 3.590318 | 4.514640 | 4.089035 | 4.072501 | 4.084777 | 4.169576 | 16 | 11 | 11 | 31 |
| 199221 | NM_00117053 | DZIP1L | DAZ interacting zinc finger protein 1 like | protein_coding | NP_001164009.1;NP_775814.2;XP_005247255.1;XP_005247257.1;XP_00 | 26551 | 617570 | . | . | 4.801705 | 5.650269 | 5.507268 | 5.249163 | 11.173759 | 13.060849 | 12.358148 | 12.001587 | 1.033743 | 578.486839 | 0.166693043 | 0.287221205 | 0.77394295 | 0.998865404 | 9.152045 | 9.199840 | 9.115340 | 9.210191 | 9.213094 | 9.157525 | 608 | 658 | 533 | 528 |
| 199223 | NM_00110551 | TTC21A | tetratricopeptide repeat domain 21A | protein_coding | NP_001098983.2;NP_001353828.1;NP_001353829.1;NP_665698.2;XP_00 | 30761 | 611430 | . | . | 1.277750 | 0.946295 | 1.212071 | 0.686014 | 2.973375 | 2.187403 | 2.719851 | 1.568491 | -1.302870 | 129.462559 | 0.313600327 | -1.21713299 | 0.223553621 | 0.982008277 | 7.193377 | 6.814617 | 7.112655 | 6.997874 | 7.020670 | 6.901039 | 193 | 135 | 116 | 86 |
| 199692 | NM_00129008 | ZNF627 | zinc finger protein 627 | protein_coding | NP_001277012.1;NP_001277013.1;NP_001277014.1;NP_660338.1;XP_01 | 30570 | 612248 | . | . | 1.062564 | 1.092110 | 1.107485 | 1.159014 | 2.472632 | 2.524460 | 2.485165 | 2.649949 | 1.065644 | 99.116555 | 0.320734628 | 0.285984799 | 0.774889764 | 0.998865404 | 6.584725 | 6.675942 | 6.616137 | 6.624573 | 6.629534 | 6.649606 | 109 | 105 | 86 | 99 |
| 199704 | NM_00128880 | ZNF585A | zinc finger protein 585A | protein_coding | NP_001275729.1;NP_689868.1;NP_954577.1 | 26305 | . | . | . | 2.954691 | 1.703895 | 1.939433 | 3.147218 | 6.875686 | 3.938630 | 4.352032 | 7.195738 | -1.207008 | 121.855114 | 0.296130936 | -0.91660725 | 0.359348482 | 0.998865404 | 7.058926 | 6.786255 | 6.945339 | 6.976629 | 6.873389 | 6.918970 | 147 | 150 | 89 | 111 |
| 199731 | NM_145296,XI | CADM4 | cell adhesion molecule 4 | protein_coding | NP_660339.1;XP_005258677.1;XP_016881941.1 | 30825 | 609744 | . | . | 47.644320 | 44.043610 | 30.538765 | 34.212069 | 110.870285 | 101.808778 | 68.528108 | 78.221825 | -1.437564 | 2648.261237 | 0.121432168 | -4.31209096 | 1.61718E-05 | 0.00289895 | 11.609061 | 11.085325 | 11.537259 | 11.459805 | 11.164436 | 11.244293 | 3762 | 3216 | 1792 | 2142 |
| 199745 | NM_00133110 | THAP8 | THAP domain containing 8 | protein_coding | NP_001318031.1;NP_001318032.1;NP_001318033.1;NP_689871.1 | 23191 | 612536 | . | . | 3.203599 | 3.420408 | 3.772166 | 3.183460 | 7.454906 | 7.906426 | 8.464633 | 7.278604 | 1.066744 | 152.030940 | 0.267064682 | 0.349031138 | 0.727065933 | 0.998865404 | 7.200543 | 7.294374 | 7.230591 | 7.237021 | 7.283292 | 7.234431 | 168 | 160 | 145 | 138 |
| 199746 | NM_00104042 | U2AF1L4 | U2 small nuclear RNA auxiliary factor 1 like 4 | protein_coding | NP_001035515.1;NP_001356753.1;NP_659424.2 | 23020 | 601080 | . | . | 12.566042 | 11.879150 | 15.489786 | 12.497580 | 29.241698 | 27.459191 | 34.758631 | 28.574229 | 1.131780 | 319.027451 | 0.205256805 | 0.870098615 | 0.384246514 | 0.998865404 | 8.224889 | 8.404592 | 8.295542 | 8.262840 | 8.403501 | 8.288357 | 355 | 313 | 325 | 285 |
| 199777 | NM_00107667 | ZNF626 | zinc finger protein 626 | protein_coding | NP_001070143.1;NP_660340.1 | 30461 | . | . | . | 2.200145 | 1.560709 | 1.400997 | 2.447270 | 5.119828 | 3.607647 | 3.143797 | 5.595394 | 1.045007 | 118.838904 | 0.301651018 | 0.210549922 | 0.833238493 | 0.998865404 | 6.860723 | 6.924303 | 6.914682 | 6.852767 | 6.877659 | 6.920211 | 146 | 114 | 99 | 121 |
| 199786 | NM_00109852 | NIBAN3 | niban apoptosis regulator 3 | protein_coding | NP_001091994.1;NP_001308755.2;NP_001308756.2;NP_001308757.1;NI | 24130 | 609967 | . | . | 0.130839 | 0.115231 | 0.233024 | 0.152655 | 0.304467 | 0.266362 | 0.522899 | 0.349029 | 1.515259 | 18.045192 | 0.731483197 | 0.819655177 | 0.412412709 | NA | 3.841203 | 4.443477 | 4.145995 | 4.142364 | 4.194907 | 4.156513 | 17 | 15 | 23 | 16 |
| 199800 | NM_00110134 | ADM5 | adrenomedullin 5 (putative) | protein_coding | NP_001094810.1 | 27293 | . | . | . | 0.843217 | 0.822296 | 0.511133 | 0.607486 | 1.962201 | 1.900774 | 1.146967 | 1.388944 | -1.479960 | 46.539813 | 0.460685468 | -1.22764401 | 0.219580632 | 0.982008277 | 5.796061 | 5.229445 | 5.579510 | 5.572546 | 5.497499 | 5.519671 | 65 | 59 | 30 | 38 |
| 199857 | NM_00130524 | ALG14 | ALG14 UDP-N-acetylglucosaminyltransferase subunit | protein_coding | NP_001292171.1;NP_659425.1;XP_005270639.1;XP_011539199.1 | 28287 | 612866 | . | . | 5.185771 | 2.207780 | 3.010781 | 1.106519 | 12.067503 | 5.103383 | 6.756107 | 2.529925 | 1.012268 | 250.146253 | 0.232985794 | 0.075505991 | 0.939812131 | 0.998865404 | 7.957654 | 7.975547 | 8.028879 | 7.888652 | 7.933065 | 8.000427 | 326 | 231 | 204 | 252 |
| 199870 | NM_00114391 | FAM76A | family with sequence similarity 76 member A | protein_coding | NP_001137384.1;NP_001137385.1;NP_001137386.1;NP_001137387.1;NI | 28530 | . | . | . | 1.748081 | 1.710822 | 1.971797 | 1.763242 | 4.067854 | 3.954642 | 4.424655 | 4.031445 | 1.042192 | 196.765193 | 0.241482997 | 0.246896936 | 0.804987987 | 0.998865404 | 7.589709 | 7.650317 | 7.619964 | 7.599053 | 7.661552 | 7.593584 | 227 | 203 | 188 | 174 |
| 199953 | NM_00101086 | TMEM201 | transmembrane protein 201 | protein_coding | NP_001010866.1;NP_001124396.2;XP_006710480.1;XP_011539212.1;XP | 33719 | . | . | . | 25.646371 | 26.366236 | 26.794554 | 25.527833 | 59.680155 | 60.946734 | 60.126205 | 58.366357 | -1.005298 | 3117.480641 | 0.112648128 | -0.06767876 | 0.946041359 | 0.998865404 | 11.609935 | 11.602385 | 11.597802 | 11.618398 | 11.629886 | 11.577140 | 3554 | 3415 | 2791 | 2821 |
| 199990 | NM_00114631 | FAAP20 | FA core complex associated protein 20 | protein_coding | NP_001139782.1;NP_001243874.2;NP_001243875.2;NP_001243876.2;NI | 26428 | 615183 | . | . | 32.622374 | 33.519944 | 33.296887 | 40.481873 | 75.913609 | 77.482847 | 74.717251 | 92.556993 | 1.111496 | 1042.011608 | 0.143037435 | 1.066170336 | 0.286346649 | 0.998865404 | 9.947115 | 10.099191 | 9.985212 | 9.982331 | 10.001352 | 10.118540 | 1139 | 1063 | 879 | 1109 |
| 200010 | NM_00101154 | SLC5A9 | solute carrier family 5 member 9 | protein_coding | NP_001011547.2;NP_001128653.1;XP_011539226.1;XP_011539227.1;XP | 22146 | . | . | . | 0.069128 | 0.027570 | 0.068715 | 0.035439 | 0.160863 | 0.063728 | 0.154194 | 0.081028 | -1.020811 | 7.901643 | 1.116606962 | -0.02661261 | 0.978768719 | NA | 2.993083 | 2.971139 | 2.994616 | 2.963802 | 2.989489 | 2.967490 | 13 | 5 | 9 | 5 |
| 200014 | NM_00133058 | CC2D1B | coiled-coil and C2 domain containing 1B | protein_coding | NP_001317514.1;NP_115825.1;XP_005270647.1;XP_005270650.1;XP_00 | 29386 | . | . | . | 12.260380 | 11.654941 | 16.866731 | 13.923114 | 28.530406 | 26.940913 | 37.848454 | 31.833547 | 1.193586 | 2074.341184 | 0.118841512 | 2.148260244 | 0.031693087 | 0.563938326 | 10.885237 | 11.140372 | 10.926991 | 10.959294 | 11.063207 | 11.104455 | 2133 | 2083 | 1913 | 2170 |
| 200030 | NM_00110166 | NBPF11 | NBPF member 11 | protein_coding | NP_001095133.3;NP_899228.4;XP_024309626.1;XP_024309627.1 | 31993 | 614001 | . | . | 4.743923 | 5.246630 | 6.478611 | 5.647830 | 11.039305 | 12.127818 | 14.537815 | 12.913094 | 1.167813 | 860.785738 | 0.157815575 | 1.418171242 | 0.156140776 | 0.925053243 | 9.632979 | 9.857328 | 9.675797 | 9.700942 | 9.879620 | 9.712605 | 898 | 872 | 899 | 770 |
| 200035 | NM_00101275 | NUDT17 | nudix hydrolase 17 | protein_coding | NP_001012776.1;XP_011507562.1;XP_011507564.1;XP_016856049.1 | 26618 | . | . | . | 0.857218 | 0.820690 | 0.881768 | 1.083660 | 1.994783 | 1.897061 | 1.978660 | 2.477760 | 1.031123 | 54.089659 | 0.441902115 | 0.100058112 | 0.92029819 | 0.998865404 | 5.736193 | 5.778065 | 5.757466 | 5.747015 | 5.713662 | 5.801036 | 63 | 56 | 37 | 63 |
| 200081 | NM_00137685 | TXLNA | taxilin alpha | protein_coding | NP_001363786.1;NP_001363787.1;NP_001363788.1;NP_787048.1;XP_01 | 30685 | 608676 | . | . | 26.228876 | 28.568598 | 29.004920 | 31.164743 | 61.035671 | 66.037599 | 65.086204 | 71.254481 | 1.083693 | 4450.045802 | 0.111130458 | 1.043429046 | 0.296749627 | 0.998865404 | 12.060529 | 12.176355 | 12.052969 | 12.115790 | 12.125382 | 12.177474 | 4732 | 4784 | 3900 | 4474 |
| 200185 | NM_173852 | KRTCAP2 | keratinocyte associated protein 2 | protein_coding | NP_776251.2 | 28942 | . | . | . | 60.122288 | 55.825562 | 65.426277 | 58.492336 | 139.907043 | 129.043289 | 146.814667 | 133.735779 | 1.053708 | 977.619802 | 0.144127204 | 0.523668631 | 0.600509038 | 0.998865404 | 9.894618 | 9.970640 | 9.942082 | 9.882447 | 9.994916 | 9.905176 | 1141 | 985 | 926 | 883 |
| 200186 | NM_181715,XI | CRTC2 | CREB regulated transcription coactivator 2 | protein_coding | NP_859066.1;XP_005245003.1;XP_005245006.1;XP_016856064.1;XP_01 | 27301 | 608972 | . | . | 10.103575 | 10.910101 | 12.723910 | 10.951022 | 23.511437 | 25.219185 | 28.552084 | 25.038214 | 1.111278 | 908.754969 | 0.148144753 | 1.027503511 | 0.304183433 | 0.998865404 | 9.749472 | 9.901994 | 9.764111 | 9.809107 | 9.916849 | 9.806340 | 960 | 958 | 895 | 829 |
| 200197 | NR_027136 | TMEM51-AS1 | TMEM51 antisense RNA 1 | lncRNA | . | 26301 | . | . | . | 1.529344 | 1.603065 | 1.658249 | 1.697813 | 3.558846 | 3.705555 | 3.721062 | 3.881847 | 1.054681 | 353.001786 | 0.188701218 | 0.407026543 | 0.683988493 | 0.998865404 | 8.424723 | 8.501324 | 8.436492 | 8.458682 | 8.474054 | 8.481823 | 388 | 378 | 313 | 342 |
| 200205 | NM_00101086 | IBA57 | iron-sulfur cluster assembly factor IBA57 | protein_coding | NP_001010867.1;NP_001297256.1 | 27302 | 615316 | . | . | 0.656213 | 0.581584 | 0.682129 | 0.662263 | 1.527034 | 1.344358 | 1.530677 | 1.514186 | 1.070319 | 157.965924 | 0.26328463 | 0.372377474 | 0.709611823 | 0.998865404 | 7.253220 | 7.352027 | 7.313689 | 7.262468 | 7.323823 | 7.307379 | 187 | 154 | 145 | 150 |
| 200232 | NM_00101297 | FAM209A | family with sequence similarity 209 member A | protein_coding | NP_001012989.2 | 16100 | . | . | . | 0.191010 | 0.073523 | 0.111568 | 0.179690 | 0.444489 | 0.169951 | 0.250355 | 0.410839 | 1.233450 | 3.976426 | 1.50608734 | 0.200983681 | 0.840711334 | NA | 1.832372 | 2.134756 | 1.991200 | 1.983822 | 1.987121 | 1.995574 | 5 | 3 | 3 | 5 |
| 200312 | NM_00101798 | RNF215 | ring finger protein 215 | protein_coding | NP_001017981.1 | 33434 | . | . | . | 11.590402 | 10.600803 | 11.140136 | 9.617458 | 26.971342 | 24.504230 | 24.998140 | 21.989176 | -1.084030 | 673.545693 | 0.161384108 | -0.721289 | 0.470731724 | 0.998865404 | 9.452274 | 9.336675 | 9.454864 | 9.386966 | 9.417462 | 9.313304 | 846 | 719 | 606 | 559 |
| 200315 | NM_00127040 | APOBEC3A | apolipoprotein B mRNA editing enzyme catalytic subunit 3A | protein_coding | NP_001257335.1;NP_663745.1 | 17343 | 607109 | . | . | 1.974092 | 1.846831 | 1.452156 | 1.716218 | 4.593793 | 4.269033 | 3.258596 | 3.923929 | -1.216659 | 74.628150 | 0.368648092 | -0.76746629 | 0.442804327 | 0.998865404 | 6.356501 | 6.072877 | 6.257901 | 6.236565 | 6.181101 | 6.215675 | 98 | 85 | 54 | 68 |
| 200316 | NM_00100666 | APOBEC3F | apolipoprotein B mRNA editing enzyme catalytic subunit 3F | protein_coding | NP_001006667.1;NP_660341.2;XP_016884131.1;XP_016884132.1;XP_02 | 17356 | 608993 | . | . | 2.324706 | 1.996447 | 2.040955 | 2.679780 | 5.409687 | 4.614876 | 4.579844 | 6.126999 | 1.075781 | 269.635780 | 0.221982554 | 0.474743418 | 0.634969867 | 0.998865404 | 8.021439 | 8.126389 | 8.091720 | 8.013706 | 8.033942 | 8.145190 | 322 | 259 | 217 | 290 |
| 200407 | NM_153836,XI | CREG2 | cellular repressor of E1A stimulated genes 2 | protein_coding | NP_722578.1;XP_011509079.1;XP_016859054.1 | 14272 | 618540 | . | . | 0.093771 | 0.067525 | 0.085281 | 0.085117 | 0.218208 | 0.156088 | 0.191369 | 0.194609 | 1.021367 | 7.672031 | 1.092509516 | 0.027918962 | 0.977726785 | NA | 2.923304 | 2.955730 | 2.944137 | 2.933951 | 2.941056 | 2.938506 | 10 | 7 | 7 | 7 |
| 200424 | NM_00128749 | TET3 | tet methylcytosine dioxygenase 3 | protein_coding | NP_001274420.1;NP_001352951.1;XP_011530984.1;XP_011530985.1;XP | 28313 | 613555 | . | . | 1.589391 | 1.648779 | 1.707374 | 1.561943 | 3.698581 | 3.811225 | 3.831298 | 3.571195 | 1.025787 | 611.115478 | 0.157836099 | 0.232719924 | 0.815978897 | 0.998865404 | 9.236723 | 9.273643 | 9.235978 | 9.256567 | 9.286269 | 9.239881 | 684 | 661 | 557 | 560 |
| 200539 | NM_144994 | ANKRD23 | ankyrin repeat domain 23 | protein_coding | NP_659431.5 | 24470 | 610736 | . | . | 4.455617 | 3.993266 | 5.054217 | 3.565282 | 10.368405 | 9.230613 | 11.341517 | 8.151593 | 1.006584 | 344.085072 | 0.218351282 | 0.043360343 | 0.96541429 | 0.998865404 | 8.421046 | 8.432175 | 8.453175 | 8.385739 | 8.519488 | 8.318372 | 418 | 348 | 354 | 266 |
| 200558 | NM_173545 | APLF | aprataxin and PNKP like factor | protein_coding | NP_775816.1 | 28724 | 611035 | . | . | 0.707207 | 0.754754 | 1.071447 | 0.796911 | 1.645700 | 1.744648 | 2.404299 | 1.822042 | 1.251778 | 99.577650 | 0.331301202 | 0.977898827 | 0.328124365 | 0.998865404 | 6.465948 | 6.791253 | 6.586622 | 6.602665 | 6.710720 | 6.612737 | 99 | 98 | 111 | 88 |
| 200576 | NM_00117800 | PIKFYVE | phosphoinositide kinase, FYVE-type zinc finger containing | protein_coding | NP_001171471.1;NP_055855.2;NP_689884.1;XP_011509080.1;XP_01150 | 23785 | 609414 | . | . | 7.557799 | 7.477732 | 8.037759 | 7.479834 | 17.587306 | 17.285107 | 18.036500 | 17.101754 | 1.019966 | 2311.134322 | 0.117833775 | 0.242045594 | 0.808744835 | 0.998865404 | 11.159963 | 11.188665 | 11.176790 | 11.155304 | 11.211828 | 11.151407 | 2670 | 2436 | 2106 | 2106 |
| 200728 | NM_198276,XI | TMEM17 | transmembrane protein 17 | protein_coding | NP_938017.2;XP_011530995.1;XP_011530996.1;XP_024308517.1;XP_02 | 26623 | 614950 | . | . | 3.361906 | 3.181263 | 3.218609 | 3.166637 | 7.823293 | 7.353632 | 7.222466 | 7.240138 | -1.037139 | 168.052674 | 0.254471028 | -0.20674101 | 0.836212129 | 0.998865404 | 7.418616 | 7.366452 | 7.413313 | 7.387461 | 7.392485 | 7.378331 | 203 | 179 | 146 | 152 |
| 200734 | NM_00112821 | SPRED2 | sprouty related EVH1 domain containing 2 | protein_coding | NP_001121682.1;NP_861449.2;XP_005264257.2;XP_005264258.1;XP_00 | 17722 | 609292 | . | . | 17.031734 | 17.616104 | 19.161065 | 18.979203 | 39.633544 | 40.720411 | 42.996870 | 43.393692 | 1.098555 | 2511.612825 | 0.114324417 | 1.186163264 | 0.235557831 | 0.990979804 | 11.225009 | 11.360603 | 11.240219 | 11.269504 | 11.339200 | 11.322600 | 2706 | 2630 | 2310 | 2440 |
| 200765 | NM_145702 | TIGD1 | tigger transposable element derived 1 | protein_coding | NP_663748.1 | 14523 | 612972 | . | . | 1.597187 | 1.745243 | 2.156895 | 1.492274 | 3.716720 | 4.034207 | 4.840011 | 3.411908 | 1.078276 | 363.996964 | 0.21604899 | 0.503249964 | 0.614788541 | 0.998865404 | 8.451825 | 8.561651 | 8.456449 | 8.502593 | 8.626542 | 8.408538 | 387 | 393 | 390 | 288 |
| 200844 | NM_00135153 | CFAP20DC | CFAP20 domain containing | protein_coding | NP_001338459.1;NP_001338460.1;NP_001338461.1;NP_001338462.1;NI | 24763 | . | . | . | 7.114840 | 6.769144 | 7.427120 | 8.503592 | 16.556530 | 15.647179 | 16.666242 | 19.442454 | 1.203208 | 676.802356 | 0.157729693 | 1.692049585 | 0.09063652 | 0.805424371 | 9.263122 | 9.529757 | 9.355818 | 9.311898 | 9.413599 | 9.504750 | 731 | 641 | 601 | 738 |
| 200845 | NM_00112821 | KCTD6 | potassium channel tetramerization domain containing 6 | protein_coding | NP_001121686.1;NP_699162.3;XP_005264994.1 | 22235 | 618791 | . | . | 1.697679 | 1.952977 | 1.631614 | 1.646405 | 3.950568 | 4.514392 | 3.661295 | 3.764309 | 1.010241 | 104.463440 | 0.323839536 | 0.045390176 | 0.963796311 | 0.998865404 | 6.699327 | 6.714342 | 6.667928 | 6.735336 | 6.737647 | 6.672069 | 107 | 124 | 101 | 88 |
| 200879 | NM_139248,XI | LIPH | lipase H | protein_coding | NP_640341.1;XP_006713592.1;XP_011510832.1;XP_011510833.1;XP_01 | 18483 | 607365 | . | . | 0.169652 | 0.138839 | 0.129780 | 0.105318 | 0.394788 | 0.320932 | 0.291222 | 0.240798 | -1.265767 | 17.608872 | 0.734789114 | -0.46273411 | 0.643554965 | NA | 4.297162 | 3.959598 | 4.159001 | 4.141011 | 4.139538 | 4.125132 | 25 | 19 | 15 | 13 |
| 200894 | NM_00117415 | ARL13B | ADP ribosylation factor like GTPase 13B | protein_coding | NP_001167621.1;NP_001167622.1;NP_001308257.1;NP_659433.2;NP_87 | 25419 | 608922 | . | . | 3.055938 | 2.128439 | 2.651646 | 2.542698 | 7.111294 | 4.919983 | 5.950216 | 5.813580 | -1.006025 | 319.245967 | 0.220939821 | -0.03922542 | 0.968710665 | 0.998865404 | 8.322283 | 8.314756 | 8.409577 | 8.214228 | 8.323380 | 8.303287 | 433 | 285 | 282 | 293 |
| 200895 | NM_00119564 | DHFR2 | dihydrofolate reductase 2 | protein_coding | NP_001182572.1;NP_789785.1;XP_011510839.1 | 27309 | 616588 | . | . | 1.898217 | 1.420949 | 1.329366 | 1.852693 | 4.417229 | 3.284588 | 2.983058 | 4.235962 | -1.065292 | 193.734076 | 0.268563606 | -0.33976458 | 0.734033822 | 0.998865404 | 7.643098 | 7.551311 | 7.673586 | 7.538366 | 7.508753 | 7.646672 | 264 | 184 | 137 | 204 |
| 200916 | NM_00109964 | RPL22L1 | ribosomal protein L22 like 1 | protein_coding | NP_001093115.1;NP_001307380.1 | 27610 | . | . | . | 8.860878 | 7.830526 | 7.778929 | 8.089291 | 20.619628 | 18.100612 | 17.455694 | 18.495201 | -1.066421 | 493.740679 | 0.172158225 | -0.53890576 | 0.589951885 | 0.998865404 | 8.993113 | 8.900625 | 9.008531 | 8.924422 | 8.919988 | 8.933539 | 625 | 514 | 411 | 453 |
| 200933 | NM_00110557 | FBXO45 | F-box protein 45 | protein_coding | NP_001099043.1 | 29148 | 609112 | . | . | 2.291899 | 2.341925 | 2.365928 | 2.201334 | 5.333344 | 5.413465 | 5.309074 | 5.033088 | -1.027337 | 425.245552 | 0.177850137 | -0.21877651 | 0.826824142 | 0.998865404 | 8.751307 | 8.712740 | 8.736090 | 8.743988 | 8.748307 | 8.698686 | 493 | 468 | 380 | 377 |
| 200942 | NM_173546,XI | KLHDC8B | kelch domain containing 8B | protein_coding | NP_775817.1;XP_005264995.1;XP_005264997.1;XP_006713078.1;XP_00 | 28557 | 613169 | . | . | 4.870652 | 5.294577 | 4.017580 | 4.850021 | 11.334205 | 12.238652 | 9.015332 | 11.088995 | -1.163796 | 294.903475 | 0.209082647 | -1.04665941 | 0.295256692 | 0.998865404 | 8.309899 | 8.089922 | 8.222235 | 8.262992 | 8.116144 | 8.205524 | 352 | 355 | 216 | 278 |
| 200958 | NM_00128250 | MUC20 | mucin 20, cell surface associated | protein_coding | NP_001269435.1 | 23282 | 610360 | . | . | 0.007664 | 0.092436 | 0.064825 | 0.079465 | 0.017834 | 0.213670 | 0.145466 | 0.181688 | 1.475830 | 5.108069 | 1.424544847 | 0.394179565 | 0.693448471 | NA | 2.050049 | 2.602870 | 2.317072 | 2.348240 | 2.340304 | 2.343296 | 1 | 8 | 5 | 6 |
| 201134 | NM_00103732 | CEP112 | centrosomal protein 112 | protein_coding | NP_001032402.1;NP_001186094.1;NP_001289820.1;NP_001340056.1;NI | 28514 | . | . | . | 0.960689 | 1.002445 | 0.796359 | 1.115145 | 2.235562 | 2.317196 | 1.787004 | 2.549646 | -1.029705 | 112.213370 | 0.318431914 | -0.13262342 | 0.894491212 | 0.998865404 | 6.832081 | 6.787781 | 6.807764 | 6.819008 | 6.745455 | 6.858873 | 129 | 125 | 79 | 122 |
| 201158 | NM_00113503 | TVP23C | trans-golgi network vesicle protein 23 homolog C | protein_coding | NP_001128508.1;NP_660344.2 | 30453 | . | . | . | 3.366126 | 3.163599 | 3.370800 | 3.270252 | 7.833113 | 7.312801 | 7.563976 | 7.477042 | -1.055920 | 285.509795 | 0.205781063 | -0.38147677 | 0.702849506 | 0.998865404 | 8.195976 | 8.117757 | 8.194421 | 8.148543 | 8.142025 | 8.144222 | 352 | 303 | 241 | 261 |
| 201161 | NM_181716 | CENPV | centromere protein V | protein_coding | NP_859067.2 | 29920 | 608139 | . | . | 1.108822 | 1.700014 | 2.063814 | 1.896199 | 2.580274 | 3.929660 | 4.631139 | 4.335433 | 1.396098 | 59.979616 | 0.420745857 | 1.144159066 | 0.252557676 | 0.994320829 | 5.646548 | 6.126516 | 5.818702 | 5.895961 | 5.941011 | 5.916392 | 46 | 65 | 64 | 62 |
| 201163 | NM_00135322 | FLCN | folliculin | protein_coding | NP_001340158.1;NP_001340159.1;NP_001340160.1;NP_653207.1;NP_65 | 27310 | 607273 | . | . | 8.135962 | 9.242651 | 9.973406 | 8.967784 | 18.932718 | 21.364808 | 22.380034 | 20.503774 | 1.111390 | 995.670330 | 0.145523175 | 1.047015442 | 0.295092456 | 0.998865404 | 9.881195 | 10.033819 | 9.890969 | 9.944355 | 10.049500 | 9.938374 | 1046 | 1055 | 980 | 909 |
| 201164 | NM_178836,XI | PLD6 | phospholipase D family member 6 | protein_coding | NP_849158.2;XP_016879799.2 | 30447 | 614960 | . | . | 0.800266 | 0.846060 | 0.511068 | 0.658515 | 1.862251 | 1.955706 | 1.146821 | 1.505617 | -1.266055 | 59.777842 | 0.421909122 | -0.80666686 | 0.419858444 | 0.998865404 | 6.062988 | 5.719719 | 5.920817 | 5.932697 | 5.836142 | 5.916146 | 75 | 74 | 36 | 60 |
| 201176 | NM_00128229 | ARHGAP27 | Rho GTPase activating protein 27 | protein_coding | NP_001269219.1;NP_777579.2;NP_954976.1;XP_005257185.1;XP_00672 | 31813 | 610591 | . | . | 4.582134 | 4.953124 | 5.036624 | 4.771334 | 10.662813 | 11.449366 | 11.302041 | 10.909093 | 1.038061 | 561.979289 | 0.161446306 | 0.333805361 | 0.738526437 | 0.998865404 | 9.107212 | 9.161032 | 9.102878 | 9.140070 | 9.154090 | 9.137897 | 617 | 612 | 504 | 530 |
| 201229 | NM_00107668 | LYRM9 | LYR motif containing 9 | protein_coding | NP_001070148.1;XP_011522793.1;XP_016879814.1;XP_024306404.1;XP | 27314 | . | . | . | 1.875446 | 1.040149 | 0.795935 | 1.924128 | 4.364241 | 2.404351 | 1.786054 | 4.399292 | -1.095671 | 79.260857 | 0.396655789 | -0.3323137 | 0.7396524 | 0.998865404 | 6.373688 | 6.240303 | 6.362685 | 6.267239 | 6.223433 | 6.356654 | 112 | 74 | 48 | 90 |
| 201232 | NM_201566 | SLC16A13 | solute carrier family 16 member 13 | protein_coding | NP_963860.1 | 31037 | . | . | . | 3.351876 | 3.314370 | 3.684581 | 3.626582 | 7.799953 | 7.661314 | 8.268093 | 8.291749 | 1.080029 | 196.713787 | 0.2374006 | 0.46785904 | 0.639885396 | 0.998865404 | 7.563233 | 7.674529 | 7.605863 | 7.596947 | 7.641739 | 7.630012 | 220 | 202 | 180 | 189 |
| 201254 | NM_00127100 | CENPX | centromere protein X | protein_coding | NP_001257935.1;NP_001257936.1;NP_001317465.1;NP_659435.2;XP_01 | 11422 | 615128 | . | . | 29.937639 | 33.301165 | 34.041386 | 37.929976 | 69.666113 | 76.977136 | 76.387880 | 86.722380 | 1.109480 | 820.355859 | 0.146369948 | 1.024005582 | 0.305832638 | 0.998865404 | 9.603433 | 9.752909 | 9.614578 | 9.667455 | 9.692271 | 9.736846 | 862 | 871 | 727 | 834 |
| 201255 | NM_144999,XI | LRRC45 | leucine rich repeat containing 45 | protein_coding | NP_659436.1 | 28302 | . | . | . | 12.823079 | 12.880116 | 12.023171 | 13.204943 | 29.839834 | 29.772964 | 26.979648 | 30.191535 | -1.032452 | 1079.767281 | 0.13609875 | -0.33854184 | 0.734954899 | 0.998865404 | 10.099496 | 10.053142 | 10.086815 | 10.089522 | 10.032241 | 10.095612 | 1261 | 1186 | 884 | 1039 |
| 201266 | NM_00115977 | SLC39A11 | solute carrier family 39 member 11 | protein_coding | NP_001153242.1;NP_001339620.1;NP_001339621.1;NP_001339622.1;NI | 14463 | 616508 | . | . | 5.534007 | 4.766143 | 6.358349 | 5.514039 | 12.877863 | 11.017158 | 14.267955 | 12.607196 | 1.081658 | 402.528001 | 0.188570081 | 0.600542795 | 0.548144549 | 0.998865404 | 8.594676 | 8.708952 | 8.662000 | 8.590911 | 8.719099 | 8.626616 | 472 | 392 | 392 | 362 |
| 201283 | NR_026903 | AMZ2P1 | archaelysin family metallopeptidase 2 pseudogene 1 | transcribed_ps | . | 26491 | . | . | . | 1.622836 | 1.497203 | 1.685332 | 2.042455 | 3.776407 | 3.460852 | 3.781837 | 4.669831 | 1.176802 | 183.656679 | 0.250892963 | 0.936143071 | 0.349199529 | 0.998865404 | 7.398946 | 7.633284 | 7.499645 | 7.462603 | 7.510226 | 7.591604 | 203 | 174 | 157 | 203 |
| 201292 | NM_00125612 | TRIM65 | tripartite motif containing 65 | protein_coding | NP_001243053.1;NP_775818.2;XP_006721823.1;XP_006721824.1;XP_01 | 27316 | . | . | . | 7.598029 | 8.143790 | 9.969737 | 9.481235 | 17.680925 | 18.824735 | 22.371800 | 21.677714 | 1.216733 | 917.866467 | 0.14307556 | 1.978060241 | 0.047921911 | 0.659753773 | 9.693666 | 9.976751 | 9.740250 | 9.789902 | 9.937229 | 9.875533 | 920 | 925 | 913 | 905 |
| 201294 | NM_199242 | UNC13D | unc-13 homolog D | protein_coding | NP_954712.1 | 23147 | 608897 | . | . | 0.087967 | 0.115559 | 0.141942 | 0.164254 | 0.204702 | 0.267119 | 0.318513 | 0.375547 | 1.474343 | 16.679759 | 0.751824489 | 0.744950708 | 0.456301519 | NA | 3.754145 | 4.312263 | 4.030638 | 4.042874 | 4.058186 | 4.070655 | 14 | 16 | 16 | 20 |
| 201299 | NM_00103483 | RDM1 | RAD52 motif containing 1 | protein_coding | NP_001030008.1;NP_001156592.1;NP_001156593.1;NP_001156594.1;NI | 19950 | 612896 | . | . | 4.955964 | 3.674831 | 2.933455 | 4.282647 | 11.532728 | 8.494537 | 6.582589 | 9.791766 | -1.169044 | 143.239360 | 0.295712427 | -0.76198763 | 0.446067387 | 0.998865404 | 7.270886 | 7.044836 | 7.242255 | 7.132551 | 7.073780 | 7.183104 | 202 | 144 | 97 | 143 |
| 201456 | NM_00114295 | FBXO15 | F-box protein 15 | protein_coding | NP_001136430.1;NP_689889.1;XP_011524158.1;XP_011524160.1;XP_01 | 13617 | 609093 | . | . | 0.192233 | 0.389385 | 0.113950 | 0.139788 | 0.447333 | 0.900081 | 0.255699 | 0.319609 | -1.920005 | 24.197272 | 0.69417212 | -1.35572967 | 0.175185196 | NA | 4.993359 | 4.048019 | 4.556149 | 4.675958 | 4.548054 | 4.560792 | 20 | 50 | 13 | 17 |
| 201475 | NM_00102530 | RAB12 | RAB12, member RAS oncogene family | protein_coding | NP_001020471.2;XP_006722363.1 | 31332 | 616448 | . | . | 9.550668 | 9.247719 | 11.206800 | 8.654201 | 22.224796 | 21.376517 | 25.147734 | 19.786805 | 1.052608 | 697.389924 | 0.172405749 | 0.429039894 | 0.667894192 | 0.998865404 | 9.407899 | 9.482773 | 9.435863 | 9.413491 | 9.554039 | 9.356779 | 795 | 721 | 711 | 575 |
| 201514 | NM_00131800 | ZNF584 | zinc finger protein 584 | protein_coding | NP_001304931.1;NP_001350609.1;NP_775819.1;XP_005258685.1;XP_00 | 27318 | . | . | . | 4.204608 | 5.147356 | 3.988098 | 4.778595 | 9.784297 | 11.898345 | 8.949176 | 10.925690 | -1.006859 | 429.411138 | 0.180730514 | -0.05456613 | 0.956484126 | 0.998865404 | 8.751328 | 8.741086 | 8.703111 | 8.790156 | 8.748047 | 8.738618 | 461 | 498 | 375 | 398 |
| 201562 | NM_00132978 | HACD2 | 3-hydroxyacyl-CoA dehydratase 2 | protein_coding | NP_001316712.1;NP_001316713.1;NP_001316715.1;NP_001316716.1;NI | 9640 | 615939 | . | . | 9.045430 | 8.289797 | 6.608323 | 5.806874 | 21.049088 | 19.162238 | 14.828888 | 13.276726 | -1.406598 | 931.579350 | 0.148545852 | -3.31352023 | 0.000921294 | 0.064386964 | 10.088469 | 9.596934 | 10.014787 | 9.927834 | 9.769390 | 9.680522 | 1325 | 1108 | 717 | 679 |
| 201595 | NM_178862,XI | STT3B | STT3 oligosaccharyltransferase complex catalytic subunit B | protein_coding | NP_849193.1;XP_011531767.1;XP_016861346.1;XP_016861347.1;XP_01 | 30611 | 608605 | . | . | 38.947294 | 34.227438 | 33.220520 | 24.984129 | 90.631964 | 79.118262 | 74.545878 | 57.123234 | -1.267292 | 4230.769840 | 0.144327001 | -2.36787649 | 0.017890507 | 0.447807832 | 12.207354 | 11.865899 | 12.191129 | 12.073806 | 12.048760 | 11.811796 | 5820 | 4752 | 3717 | 2991 |
| 201625 | NM_00136602 | DNAH12 | dynein axonemal heavy chain 12 | protein_coding | NP_001352957.1;NP_940966.2;XP_011531769.1;XP_011531770.1;XP_01 | 2943 | 603340 | . | . | 0.014023 | 0.003344 | 0.022469 | 0.002702 | 0.032633 | 0.007729 | 0.050420 | 0.006178 | 1.416785 | 4.816428 | 1.456223032 | 0.345153616 | 0.729978887 | NA | 1.987791 | 2.502474 | 2.258002 | 2.237981 | 2.271199 | 2.239775 | 7 | 2 | 8 | 2 |
| 201626 | NM_00132217 | PDE12 | phosphodiesterase 12 | protein_coding | NP_001309105.1;NP_001309106.1;NP_808881.3 | 25386 | 616519 | . | . | 2.320106 | 2.392378 | 2.747644 | 2.550232 | 5.398982 | 5.530090 | 6.165634 | 5.830803 | -1.028839 | 635.059758 | 0.154640008 | -0.26524177 | 0.790823222 | 0.998865404 | 9.331125 | 9.290080 | 9.310475 | 9.329996 | 9.305592 | 9.296711 | 731 | 705 | 548 | 583 |
| 201627 | NM_152678,XI | DENND6A | DENN domain containing 6A | protein_coding | NP_689891.1;XP_006713082.1;XP_006713083.1;XP_016861353.1 | 26635 | . | . | . | 2.139709 | 1.991931 | 1.946322 | 1.821207 | 4.979190 | 4.604438 | 4.367490 | 4.163975 | -1.143810 | 293.410351 | 0.204664396 | -0.94714585 | 0.343564461 | 0.998865404 | 8.290129 | 8.096962 | 8.250645 | 8.211816 | 8.181210 | 8.138548 | 373 | 326 | 248 | 246 |
| 201725 | NM_00100839 | C4orf46 | chromosome 4 open reading frame 46 | protein_coding | NP_001008394.1 | 27320 | 616210 | . | . | 3.974065 | 3.564353 | 3.397489 | 2.872290 | 9.247810 | 8.239161 | 7.623867 | 6.567149 | -1.216775 | 362.921207 | 0.196472877 | -1.4407202 | 0.149663746 | 0.911252178 | 8.637578 | 8.355701 | 8.590159 | 8.520268 | 8.490476 | 8.393875 | 485 | 405 | 310 | 280 |
| 201780 | NM_152679 | SLC10A4 | solute carrier family 10 member 4 | protein_coding | NP_689892.1 | 22980 | 618563 | . | . | 0.626067 | 0.429134 | 0.254084 | 0.258490 | 1.456885 | 0.991963 | 0.570157 | 0.591006 | -2.079339 | 26.399758 | 0.621854264 | -1.69834767 | 0.089442163 | NA | 5.155481 | 4.100822 | 4.790118 | 4.733605 | 4.686964 | 4.685443 | 49 | 31 | 15 | 16 |
| 201798 | NM_145720,XI | TIGD4 | tigger transposable element derived 4 | protein_coding | NP_663772.1;XP_005262864.1 | 18335 | . | . | . | 0.382883 | 0.289118 | 0.376556 | 0.315660 | 0.890985 | 0.668310 | 0.844980 | 0.721720 | 1.082904 | 32.061459 | 0.559331427 | 0.205432811 | 0.837233988 | 0.998865404 | 4.942436 | 5.060678 | 5.017136 | 4.969935 | 5.021589 | 4.986093 | 42 | 27 | 33 | 27 |
| 201853 | NR_126435 | LINC00504 | long intergenic non-protein coding RNA 504 | lncRNA | . | 43555 | . | . | . | 7.976449 | 7.948690 | 5.520326 | 7.638489 | 18.561523 | 18.373755 | 12.387453 | 17.464497 | -1.227135 | 1058.941821 | 0.162585777 | -1.8162351 | 0.069334317 | 0.744184737 | 10.188847 | 9.892808 | 10.121241 | 10.112014 | 9.847675 | 10.070603 | 1352 | 1252 | 698 | 1030 |
| 201895 | NM_00131789 | SMIM14 | small integral membrane protein 14 | protein_coding | NP_001304825.1;NP_001304826.1;NP_777581.1 | 27321 | . | . | . | 0.786623 | 0.717324 | 0.589515 | 0.837317 | 1.830504 | 1.658129 | 1.322855 | 1.914426 | -1.084065 | 144.461919 | 0.288571602 | -0.40354497 | 0.686547362 | 0.998865404 | 7.232262 | 7.114423 | 7.210863 | 7.165490 | 7.093656 | 7.217788 | 183 | 153 | 100 | 152 |
| 201931 | NM_00110038 | TMEM192 | transmembrane protein 192 | protein_coding | NP_001093859.1;XP_011530019.1;XP_011530020.1 | 26775 | . | . | . | 3.245187 | 3.123318 | 3.945077 | 3.452694 | 7.551685 | 7.219688 | 8.852637 | 7.894174 | 1.093300 | 1002.666215 | 0.14620031 | 0.880227548 | 0.378736053 | 0.998865404 | 9.903548 | 10.032809 | 9.951497 | 9.917116 | 10.060704 | 9.935535 | 1129 | 1009 | 988 | 899 |
| 201965 | NM_00130792 | RWDD4 | RWD domain containing 4 | protein_coding | NP_001294851.1;NP_689895.2 | 23750 | . | . | . | 9.308798 | 8.421670 | 8.554982 | 6.928572 | 21.661955 | 19.467066 | 19.197133 | 15.841356 | -1.157857 | 676.641651 | 0.168171883 | -1.25738952 | 0.208612624 | 0.970482251 | 9.503642 | 9.293186 | 9.485518 | 9.409487 | 9.419344 | 9.272395 | 881 | 741 | 606 | 523 |
| 201973 | NM_00130076 | PRIMPOL | primase and DNA directed polymerase | protein_coding | NP_001287696.1;NP_001287697.1;NP_001332820.1;NP_001332821.1;NI | 26575 | 615421 | . | . | 5.083345 | 5.140570 | 4.660597 | 4.595358 | 11.829156 | 11.882659 | 10.458247 | 10.506738 | -1.149810 | 333.248179 | 0.192789726 | -1.04463511 | 0.296191648 | 0.998865404 | 8.477589 | 8.276302 | 8.416435 | 8.420340 | 8.347474 | 8.333706 | 409 | 386 | 273 | 287 |
| 202018 | NM_153365,XI | TAPT1 | transmembrane anterior posterior transformation 1 | protein_coding | NP_699196.2;XP_005248196.1;XP_005248197.1;XP_011512114.1;XP_01 | 26887 | 612758 | . | . | 4.350692 | 4.974126 | 5.485182 | 3.753025 | 10.124238 | 11.497916 | 12.308589 | 8.580849 | -1.057409 | 597.010393 | 0.175279983 | -0.4594554 | 0.645907176 | 0.998865404 | 9.260966 | 9.181155 | 9.214966 | 9.259678 | 9.280526 | 9.114087 | 683 | 684 | 569 | 475 |
| 202020 | NR_027696,NF | TAPT1-AS1 | TAPT1 antisense RNA 1 (head to head) | lncRNA | . | 26832 | . | . | . | 0.478888 | 0.270344 | 0.469987 | 0.358930 | 1.114392 | 0.624911 | 1.054638 | 0.820650 | 1.085507 | 32.017732 | 0.571362566 | 0.207169875 | 0.835877193 | 0.998865404 | 4.938327 | 5.060678 | 5.024507 | 4.955063 | 5.018431 | 4.982944 | 45 | 24 | 33 | 27 |
| 202051 | NM_194296,XI | SPATA24 | spermatogenesis associated 24 | protein_coding | NP_919272.1;XP_005271973.1;XP_011541554.1;XP_011541555.1;XP_01 | 27322 | . | . | . | 1.608470 | 1.491433 | 1.776922 | 2.201977 | 3.742977 | 3.447514 | 3.987360 | 5.034557 | 1.220251 | 42.718877 | 0.476279615 | 0.602961546 | 0.546534271 | 0.998865404 | 5.266033 | 5.553294 | 5.405530 | 5.387160 | 5.420866 | 5.436652 | 47 | 39 | 39 | 46 |
| 202052 | NM_152686 | DNAJC18 | DnaJ heat shock protein family (Hsp40) member C18 | protein_coding | NP_689899.1 | 28429 | . | . | . | 2.320207 | 2.409300 | 2.867753 | 2.937829 | 5.399216 | 5.569205 | 6.435154 | 6.716997 | 1.210197 | 418.821461 | 0.177308974 | 1.55232967 | 0.120583362 | 0.861496971 | 8.566109 | 8.841182 | 8.638978 | 8.656440 | 8.757862 | 8.766977 | 430 | 415 | 396 | 433 |
| 202181 | NR_026921 | LOC202181 | SUMO interacting motifs containing 1 pseudogene | transcribed_ps | . | . | . | . | . | 1.646422 | 1.791175 | 1.972937 | 1.937187 | 3.831292 | 4.140382 | 4.427213 | 4.429149 | 1.122470 | 163.860810 | 0.257381731 | 0.647585335 | 0.517253185 | 0.998865404 | 7.270671 | 7.437181 | 7.314650 | 7.345489 | 7.383461 | 7.371073 | 171 | 173 | 153 | 160 |
| 202243 | NM_00129769 | CCDC125 | coiled-coil domain containing 125 | protein_coding | NP_001284625.1;NP_001284626.1;NP_789786.2;XP_005248515.1;XP_00 | 28924 | 613781 | . | . | 2.624207 | 2.763231 | 2.610487 | 2.732065 | 6.106637 | 6.387334 | 5.857857 | 6.246540 | 1.014972 | 277.858166 | 0.210018578 | 0.102084903 | 0.918689284 | 0.998865404 | 8.107754 | 8.128580 | 8.078838 | 8.147191 | 8.114058 | 8.128400 | 297 | 317 | 240 | 266 |
| 202333 | NM_153610,XI | CMYA5 | cardiomyopathy associated 5 | protein_coding | NP_705838.3;XP_016864701.1 | 14305 | 612193 | . | . | 0.067211 | 0.089990 | 0.116042 | 0.079067 | 0.156403 | 0.208016 | 0.260395 | 0.180777 | 1.237244 | 35.770538 | 0.532395484 | 0.576883951 | 0.564017853 | 0.998865404 | 4.998493 | 5.306499 | 5.118970 | 5.153479 | 5.194459 | 5.137188 | 32 | 39 | 41 | 30 |
| 202374 | NM_00111272 | STK32A | serine/threonine kinase 32A | protein_coding | NP_001106195.1;NP_001274669.1;NP_659438.1;XP_011535878.1;XP_01 | 28317 | . | . | . | 8.053118 | 6.633939 | 10.097391 | 9.451319 | 18.739933 | 15.334648 | 22.658256 | 21.609315 | 1.178709 | 1097.768492 | 0.138534262 | 1.712262248 | 0.08684835 | 0.795087717 | 9.976707 | 10.214242 | 10.078822 | 9.988722 | 10.163503 | 10.150522 | 1232 | 1020 | 1042 | 1104 |
| 202500 | NM_182539,XI | TCTE1 | t-complex-associated-testis-expressed 1 | protein_coding | NP_872345.2;XP_005248929.1;XP_005248930.1;XP_005248931.1;XP_00 | 11693 | 186975 | . | . | 0.260758 | 0.261001 | 0.259541 | 0.037695 | 0.606796 | 0.603314 | 0.582402 | 0.086184 | -1.781104 | 16.493492 | 0.838636943 | -0.99300644 | 0.320706818 | NA | 4.397726 | 3.573873 | 4.060904 | 4.062659 | 4.054307 | 3.988352 | 24 | 23 | 17 | 4 |
| 202781 | NR_028090 | PAXIP1-AS1 | PAXIP1 antisense RNA 1 (head to head) | lncRNA | . | 27328 | . | . | . | 1.961473 | 1.886588 | 2.149864 | 1.610548 | 4.564428 | 4.360932 | 4.824232 | 3.682327 | -1.035324 | 135.269723 | 0.291139306 | -0.17202314 | 0.863419336 | 0.998865404 | 7.103632 | 7.055355 | 7.092308 | 7.074934 | 7.123227 | 7.015881 | 162 | 145 | 133 | 106 |
| 203054 | NM_174922,XI | ADCK5 | aarF domain containing kinase 5 | protein_coding | NP_777582.4;XP_006716590.1;XP_011515209.1;XP_011515211.1;XP_01 | 21738 | . | . | . | 4.841825 | 5.455900 | 6.673098 | 5.771885 | 11.267125 | 12.611559 | 14.974236 | 13.196731 | 1.203879 | 359.352442 | 0.197831315 | 1.353123271 | 0.176016253 | 0.949193943 | 8.348992 | 8.617083 | 8.398445 | 8.459336 | 8.590498 | 8.477987 | 355 | 371 | 374 | 333 |
| 203062 | NM_00129193 | TSNARE1 | t-SNARE domain containing 1 | protein_coding | NP_001278860.1;NP_001350669.1;NP_001353830.1;NP_001353831.1;NI | 26437 | . | . | . | 1.237635 | 0.957529 | 1.153068 | 1.247154 | 2.880027 | 2.213370 | 2.587451 | 2.851469 | 1.177675 | 133.413662 | 0.284633169 | 0.828932504 | 0.407142602 | 0.998865404 | 6.936684 | 7.173162 | 7.055081 | 6.995949 | 7.077771 | 7.093178 | 153 | 121 | 123 | 138 |
| 203068 | NM_00129321 | TUBB | tubulin beta class I | protein_coding | NP_001280141.1;NP_001280142.1;NP_001280143.1;NP_001280144.1;NI | 20778 | 191130 | . | . | 511.356543 | 535.635459 | 436.557942 | 519.022147 | 1189.947752 | 1238.145343 | 979.623315 | 1186.682402 | -1.111814 | 39046.807089 | 0.11184503 | -1.36720481 | 0.171561119 | 0.947194954 | 15.327361 | 15.174421 | 15.279279 | 15.312859 | 15.132664 | 15.271834 | 46431 | 45219 | 29548 | 37480 |
| 203069 | NM_00113610 | R3HCC1 | R3H domain and coiled-coil containing 1 | protein_coding | NP_001129580.2;NP_001288579.1 | 27329 | . | . | . | 5.881114 | 5.511346 | 5.308165 | 6.103498 | 13.685593 | 12.739722 | 11.911367 | 13.954922 | -1.014085 | 279.673431 | 0.210129523 | -0.096028 | 0.923498336 | 0.998865404 | 8.137843 | 8.117282 | 8.149953 | 8.111774 | 8.090190 | 8.156115 | 336 | 293 | 226 | 277 |
| 203190 | NM_139278 | LGI3 | leucine rich repeat LGI family member 3 | protein_coding | NP_644807.1 | 18711 | 608302 | . | . | 5.579378 | 5.934954 | 10.503659 | 9.101480 | 12.983440 | 13.718912 | 23.569906 | 20.809454 | 1.680039 | 784.770515 | 0.153823802 | 4.86592172 | 1.13925E-06 | 0.000327604 | 9.193717 | 9.942461 | 9.380709 | 9.416387 | 9.817476 | 9.703081 | 656 | 649 | 921 | 852 |
| 203197 | NM_00133076 | TMEM268 | transmembrane protein 268 | protein_coding | NP_001317689.1;NP_694590.2;XP_011516652.1;XP_011516653.1;XP_01 | 24513 | . | . | . | 5.046346 | 5.001139 | 5.391615 | 5.585673 | 11.743054 | 11.560359 | 12.098630 | 12.770974 | 1.103694 | 736.289583 | 0.147885232 | 0.962502509 | 0.335797244 | 0.998865404 | 9.451151 | 9.593593 | 9.497545 | 9.478936 | 9.558917 | 9.554580 | 817 | 745 | 674 | 722 |
| 203228 | NM_00125605 | C9orf72 | C9orf72-SMCR8 complex subunit | protein_coding | NP_001242983.1;NP_060795.1;NP_659442.2 | 28337 | 614260 | . | . | 1.149685 | 0.856975 | 0.782203 | 0.895933 | 2.675364 | 1.980938 | 1.755241 | 2.048445 | -1.208580 | 90.157386 | 0.346022813 | -0.78986972 | 0.429603855 | 0.998865404 | 6.624597 | 6.351219 | 6.559491 | 6.480943 | 6.445234 | 6.488654 | 128 | 93 | 65 | 83 |
| 203238 | NM_00134800 | CCDC171 | coiled-coil domain containing 171 | protein_coding | NP_001334931.1;NP_001342476.1;NP_775821.2;XP_005251454.1;XP_00 | 29828 | . | . | . | 0.462499 | 0.442778 | 0.451485 | 0.534961 | 1.076254 | 1.023500 | 1.013121 | 1.223126 | 1.093816 | 107.298175 | 0.311763352 | 0.414962381 | 0.678169442 | 0.998865404 | 6.679445 | 6.808628 | 6.745259 | 6.713526 | 6.739699 | 6.776184 | 124 | 105 | 92 | 111 |
| 203245 | NM_197956,XI | NAIF1 | nuclear apoptosis inducing factor 1 | protein_coding | NP_931045.1 | 25446 | 610673 | . | . | 2.440055 | 2.572145 | 2.511238 | 2.849408 | 5.678109 | 5.945627 | 5.635146 | 6.514833 | 1.064724 | 246.420269 | 0.21900267 | 0.41314566 | 0.679499898 | 0.998865404 | 7.899416 | 7.989143 | 7.912015 | 7.944234 | 7.934235 | 7.983703 | 266 | 266 | 210 | 250 |
| 203259 | NM_00118494 | FAM219A | family with sequence similarity 219 member A | protein_coding | NP_001171869.1;NP_001171870.1;NP_001171871.1;NP_001171872.1;NI | 19920 | . | . | . | 18.173683 | 19.429086 | 22.323061 | 20.788369 | 42.290907 | 44.911204 | 50.092301 | 47.530138 | 1.130684 | 2275.695319 | 0.119787546 | 1.479252251 | 0.139072909 | 0.900674731 | 11.060742 | 11.238000 | 11.076430 | 11.123326 | 11.231203 | 11.165110 | 2387 | 2373 | 2186 | 2172 |
| 203260 | NM_00119520 | CCDC107 | coiled-coil domain containing 107 | protein_coding | NP_001182129.1;NP_001182130.1;NP_001182146.1;NP_777583.2;XP_00 | 28465 | . | . | . | 28.042476 | 28.110645 | 28.918466 | 32.876530 | 65.255987 | 64.979017 | 64.892196 | 75.168282 | 1.099829 | 1007.145336 | 0.139666363 | 0.982908977 | 0.325652239 | 0.998865404 | 9.905903 | 10.042955 | 9.951381 | 9.927345 | 9.971000 | 10.045981 | 1123 | 1018 | 871 | 1039 |
| 203274 | NR_146625,NF | LINC00537 | long intergenic non-protein coding RNA 537 | lncRNA | . | 43654 | . | . | . | 0.311301 | 0.339035 | 0.371525 | 0.314728 | 0.724410 | 0.783694 | 0.833692 | 0.719588 | 1.034520 | 44.753266 | 0.466718216 | 0.104904753 | 0.916451388 | 0.998865404 | 5.458844 | 5.508570 | 5.473709 | 5.484843 | 5.499373 | 5.470479 | 49 | 49 | 43 | 39 |
| 203286 | NM_173551,XI | ANKS6 | ankyrin repeat and sterile alpha motif domain containing 6 | protein_coding | NP_775822.3;XP_005251850.1;XP_005251851.1;XP_006717061.1;XP_00 | 26724 | 615370 | . | . | 3.337888 | 3.150540 | 3.635226 | 3.441878 | 7.767400 | 7.282614 | 8.157343 | 7.869444 | 1.088856 | 595.675810 | 0.158263033 | 0.776008537 | 0.437743945 | 0.998865404 | 9.155682 | 9.278473 | 9.202019 | 9.175606 | 9.232739 | 9.257666 | 670 | 603 | 530 | 593 |
| 203414 | NR_126059 | LINC01560 | long intergenic non-protein coding RNA 1560 | lncRNA | . | 27333 | . | . | . | 0.476634 | 0.462913 | 0.793337 | 0.674852 | 1.109146 | 1.070045 | 1.780226 | 1.542969 | 1.518022 | 47.411635 | 0.455378984 | 1.322399535 | 0.186035144 | 0.958175316 | 5.234218 | 5.837506 | 5.519233 | 5.515175 | 5.607327 | 5.573149 | 44 | 40 | 54 | 49 |
| 203427 | NM_145305,XI | SLC25A43 | solute carrier family 25 member 43 | protein_coding | NP_660348.2;XP_016884829.1;XP_016884830.1 | 30557 | 300641 | . | . | 5.145879 | 3.999737 | 5.517197 | 5.178741 | 11.974673 | 9.245571 | 12.380431 | 11.840575 | 1.160606 | 383.597956 | 0.198568536 | 1.082134213 | 0.279192895 | 0.998865404 | 8.471440 | 8.687389 | 8.605105 | 8.458462 | 8.642458 | 8.600926 | 461 | 334 | 371 | 373 |
| 203522 | NM_00135160 | INTS6L | integrator complex subunit 6 like | protein_coding | NP_001338530.1;NP_001338532.1;NP_001338533.1;NP_001338534.1;NI | 27334 | . | . | . | 0.154958 | 0.166739 | 0.067549 | 0.076976 | 0.360594 | 0.385425 | 0.151577 | 0.175996 | -1.913530 | 24.208249 | 0.635185328 | -1.47395759 | 0.140493009 | NA | 4.990578 | 4.055266 | 4.638125 | 4.627780 | 4.573196 | 4.567380 | 38 | 33 | 15 | 15 |
| 203523 | NM_152695,XI | ZNF449 | zinc finger protein 449 | protein_coding | NP_689908.3;XP_011529614.1;XP_011529615.1;XP_011529616.1;XP_01 | 21039 | 300627 | . | . | 1.132450 | 1.199704 | 1.293728 | 0.886763 | 2.635257 | 2.773170 | 2.903087 | 2.027478 | -1.053870 | 138.375639 | 0.294771121 | -0.25679825 | 0.797334521 | 0.998865404 | 7.149100 | 7.074836 | 7.096017 | 7.140331 | 7.160886 | 7.032956 | 154 | 162 | 138 | 104 |
| 203547 | NM_00101798 | VMA21 | vacuolar ATPase assembly factor VMA21 | protein_coding | NP_001017980.1;NP_001350739.1 | 22082 | 300913 | . | . | 5.457629 | 4.815476 | 5.569780 | 3.665016 | 12.700130 | 11.131187 | 12.498425 | 8.379624 | -1.128325 | 720.903907 | 0.195165262 | -0.89248813 | 0.372131352 | 0.998865404 | 9.577555 | 9.404590 | 9.574526 | 9.478871 | 9.578091 | 9.290608 | 939 | 769 | 714 | 501 |
| 203859 | NM_00114264 | ANO5 | anoctamin 5 | protein_coding | NP_001136121.1;NP_998764.1;XP_005252877.2;XP_005252878.2;XP_00 | 27337 | 608662 | . | . | 0.293658 | 0.247596 | 0.370442 | 0.346598 | 0.683354 | 0.572329 | 0.831261 | 0.792456 | 1.305570 | 66.520974 | 0.388427475 | 0.990351093 | 0.322002541 | 0.998865404 | 5.849903 | 6.235838 | 6.035178 | 6.000165 | 6.087908 | 6.069759 | 72 | 57 | 68 | 68 |
| 204851 | NM_00136980 | HIPK1 | homeodomain interacting protein kinase 1 | protein_coding | NP_001356735.1;NP_001356736.1;NP_689909.2;NP_852003.1;NP_93800 | 19006 | 608003 | . | . | 4.800234 | 4.907331 | 4.848112 | 4.800634 | 11.170343 | 11.343513 | 10.879024 | 10.976082 | -1.023242 | 1219.631168 | 0.129816896 | -0.25533973 | 0.79846072 | 0.998865404 | 10.268699 | 10.235570 | 10.255225 | 10.266442 | 10.249674 | 10.237360 | 1410 | 1341 | 1057 | 1121 |
| 204962 | NM_00113005 | SLC44A5 | solute carrier family 44 member 5 | protein_coding | NP_001123530.1;NP_001307212.1;NP_001307214.1;NP_001307216.1;NI | 28524 | . | . | . | 0.047997 | 0.050737 | 0.042330 | 0.060255 | 0.111692 | 0.117282 | 0.094988 | 0.137765 | 1.059158 | 13.375202 | 0.836205898 | 0.099160269 | 0.92101102 | NA | 3.700498 | 3.781347 | 3.739479 | 3.739269 | 3.733112 | 3.752308 | 15 | 14 | 10 | 15 |
| 205251 | NM_00135525 | MTLN | mitoregulin | protein_coding | NP_001342180.1;XP_024308519.1 | 27339 | . | . | . | 8.586743 | 8.289351 | 6.310302 | 9.370080 | 19.981703 | 19.161209 | 14.160135 | 21.423574 | -1.092060 | 520.344306 | 0.197695231 | -0.64266486 | 0.520441587 | 0.998865404 | 9.085985 | 8.957815 | 9.061145 | 9.033246 | 8.864015 | 9.103315 | 639 | 574 | 350 | 555 |
| 205327 | NM_153689 | C2orf69 | chromosome 2 open reading frame 69 | protein_coding | NP_710156.3 | 26799 | . | . | . | 4.923256 | 4.042000 | 4.356715 | 3.165166 | 11.456619 | 9.343263 | 9.776341 | 7.236775 | -1.208740 | 475.024159 | 0.202318587 | -1.35184938 | 0.176423503 | 0.949538805 | 9.021430 | 8.749488 | 9.008704 | 8.873859 | 8.917983 | 8.723168 | 660 | 503 | 435 | 338 |
| 205428 | NM_00113447 | DIPK2A | divergent protein kinase domain 2A | protein_coding | NP_001127942.1;NP_001350873.1;NP_775823.1;XP_011510845.1 | 28490 | 612200 | . | . | 1.138494 | 1.169871 | 1.591085 | 0.693662 | 2.649321 | 2.704208 | 3.570349 | 1.585976 | -1.025205 | 155.175758 | 0.344136416 | -0.10435457 | 0.91688797 | 0.998865404 | 7.294274 | 7.261054 | 7.261507 | 7.269269 | 7.401201 | 7.084635 | 179 | 171 | 187 | 86 |
| 205564 | NM_00130804 | SENP5 | SUMO specific peptidase 5 | protein_coding | NP_001294974.1;NP_689912.2;XP_005269367.1;XP_011510846.1 | 28407 | 612845 | . | . | 4.464760 | 4.492813 | 4.342328 | 4.230732 | 10.389680 | 10.385340 | 9.744058 | 9.673063 | -1.074482 | 845.653380 | 0.142936472 | -0.72508938 | 0.468397198 | 0.998865404 | 9.774717 | 9.671274 | 9.750697 | 9.747261 | 9.715554 | 9.679912 | 1011 | 943 | 727 | 745 |
| 205717 | NM_00100989 | USF3 | upstream transcription factor family member 3 | protein_coding | NP_001009899.3;XP_005247265.2;XP_016861360.1;XP_016861361.1;XP | 30494 | 617568 | . | . | 0.427687 | 0.377180 | 0.331778 | 0.473686 | 0.995248 | 0.871868 | 0.744501 | 1.083027 | -1.029506 | 186.413998 | 0.248154471 | -0.16905953 | 0.865749819 | 0.998865404 | 7.563678 | 7.520735 | 7.548833 | 7.548156 | 7.495851 | 7.574137 | 218 | 204 | 145 | 188 |
| 206358 | NM_00130815 | SLC36A1 | solute carrier family 36 member 1 | protein_coding | NP_001295079.1;NP_001295080.1;NP_001336669.1;NP_510968.2;XP_00 | 18761 | 606561 | . | . | 8.932566 | 9.828518 | 12.377668 | 9.046659 | 20.786443 | 22.719061 | 27.775127 | 20.684110 | 1.106582 | 1654.510443 | 0.149374623 | 0.978151456 | 0.327999421 | 0.998865404 | 10.617086 | 10.763575 | 10.625886 | 10.670061 | 10.832190 | 10.610881 | 1757 | 1743 | 1723 | 1402 |
| 206412 | NM_00101086 | C6orf163 | chromosome 6 open reading frame 163 | protein_coding | NP_001010868.2 | 21403 | . | . | . | 0.358222 | 0.243267 | 0.304881 | 0.141189 | 0.833597 | 0.562323 | 0.684145 | 0.322811 | -1.330428 | 11.738147 | 0.917123408 | -0.44911177 | 0.653351033 | NA | 3.741623 | 3.336262 | 3.571640 | 3.551255 | 3.562753 | 3.530620 | 18 | 12 | 12 | 6 |
| 207063 | NM_145177,N | DHRSX | dehydrogenase/reductase X-linked | protein_coding | NP_660160.2 | 18399 | 301034 | . | . | 3.245035 | 3.674066 | 3.490968 | 3.663180 | 7.551331 | 8.492769 | 7.833630 | 8.375425 | 1.020791 | 283.573417 | 0.20771365 | 0.142925524 | 0.886349001 | 0.998865404 | 8.132970 | 8.162040 | 8.111189 | 8.170952 | 8.141517 | 8.162849 | 305 | 320 | 244 | 274 |
| 219285 | NM_00130349 | SAMD9L | sterile alpha motif domain containing 9 like | protein_coding | NP_001290425.1;NP_001290426.1;NP_001290427.1;NP_001290429.1;NI | 1349 | 611170 | . | . | 0.942170 | 1.046454 | 0.792700 | 0.713008 | 2.192471 | 2.418928 | 1.778796 | 1.630208 | -1.299153 | 192.524403 | 0.247120494 | -1.52788311 | 0.126541562 | 0.874064017 | 7.765319 | 7.387857 | 7.602354 | 7.680673 | 7.550050 | 7.503121 | 230 | 254 | 154 | 148 |
| 219293 | NM_00103921 | ATAD3C | ATPase family AAA domain containing 3C | protein_coding | NP_001034300.2 | 32151 | 617227 | . | . | 0.929974 | 1.124541 | 0.789887 | 0.953069 | 2.164088 | 2.599428 | 1.772484 | 2.179079 | -1.191385 | 114.894660 | 0.30852741 | -0.81885489 | 0.412869207 | 0.998865404 | 6.965769 | 6.711366 | 6.839728 | 6.903690 | 6.786655 | 6.840883 | 131 | 147 | 83 | 107 |
| 219333 | NM_182488,XI | USP12 | ubiquitin specific peptidase 12 | protein_coding | NP_872294.2;XP_005266339.1;XP_011533258.1;XP_016875905.1 | 20485 | 603091 | . | . | 8.084537 | 7.802453 | 6.969697 | 6.798774 | 18.813047 | 18.035718 | 15.639800 | 15.544589 | -1.152010 | 1012.232024 | 0.136203384 | -1.4988893 | 0.133902353 | 0.891584264 | 10.081716 | 9.877728 | 10.040921 | 10.024973 | 9.941469 | 9.917002 | 1261 | 1157 | 832 | 867 |
| 219347 | NR_027428,NF | TMEM254-AS1 | TMEM254 antisense RNA 1 | lncRNA | . | 27340 | . | . | . | 0.423626 | 0.622109 | 0.741316 | 0.591603 | 0.985794 | 1.438034 | 1.663491 | 1.352631 | 1.284243 | 74.629884 | 0.381054363 | 0.947155365 | 0.343559616 | 0.998865404 | 6.030170 | 6.390725 | 6.141309 | 6.217171 | 6.271236 | 6.212602 | 63 | 82 | 81 | 70 |

| | | | | | | | | | |
|---|---|---|---|---|---|---|---|---|---|
| 219402 | NM_00116626 | MTIF3 | mitochondrial translational initiation factor 3 | protein_coding | NP_001159733.1;NP_001159734.1;NP_001159735.1;NP_690876.3;XP_00 | 29788 | . | . | [illegible] |
| 219539 | NM_00136348 | YPEL4 | yippee like 4 | protein_coding | NP_001350416.1;NP_659445.1 | 18328 | 609725 | . | [illegible] |
| 219541 | NM_00131707 | MED19 | mediator complex subunit 19 | protein_coding | NP_001304007.2;NP_703151.3 | 29600 | 612385 | . | [illegible] |
| 219595 | NM_153696,XI | FOLH1B | folate hydrolase 1B | protein_coding | NP_710163.1;XP_016872808.1 | 13636 | 609020 | . | [illegible] |
| 219654 | NM_153367,XI | ZCCHC24 | zinc finger CCHC-type containing 24 | protein_coding | NP_699198.2;XP_011537754.1 | 26911 | . | . | [illegible] |
| 219670 | NM_00127038 | ENKUR | enkurin, TRPC channel interacting protein | protein_coding | NP_001257312.1;NP_659447.1 | 28388 | 611025 | . | [illegible] |
| 219699 | NM_00124488 | UNC5B | unc-5 netrin receptor B | protein_coding | NP_001231818.1;NP_734465.2;XP_011537755.1;XP_016871323.1;XP_01 | 12568 | 607870 | . | [illegible] |
| 219736 | NM_00113015 | STOX1 | storkhead box 1 | protein_coding | NP_001123631.1;NP_001123632.1;NP_001123633.1;NP_689922.3;XP_01 | 23508 | 609397 | . | [illegible] |
| 219738 | NM_145306,XI | FAM241B | family with sequence similarity 241 member B | protein_coding | NP_660349.1;XP_005269663.1;XP_005269665.1;XP_011537757.1;XP_01 | 23519 | . | . | [illegible] |
| 219743 | NM_00104027 | TYSND1 | trypsin domain containing 1 | protein_coding | NP_001035363.1;NP_775826.2 | 28531 | 611017 | . | [illegible] |
| 219749 | NM_00132964 | ZNF25 | zinc finger protein 25 | protein_coding | NP_001316576.1;NP_001316577.1;NP_001316578.1;NP_001316579.1;NI | 13043 | 194528 | . | [illegible] |
| 219771 | NM_00128285 | CCNY | cyclin Y | protein_coding | NP_001269781.1;NP_001269782.1;NP_001269783.1;NP_659449.3;NP_8! | 23354 | 612786 | . | [illegible] |
| 219790 | NM_00128294 | RTKN2 | rhotekin 2 | protein_coding | NP_001269870.1;NP_660350.2;XP_011537758.1;XP_011537759.1;XP_01 | 19364 | 618450 | . | [illegible] |
| 219844 | NM_00113479 | HYLS1 | HYLS1 centriolar and ciliogenesis associated | protein_coding | NP_001128265.1;NP_001364198.1;NP_001364199.1;NP_659451.1;XP_01 | 26558 | 610693 | . | [illegible] |
| 219854 | NM_00108054 | TMEM218 | transmembrane protein 218 | protein_coding | NP_001074015.1;NP_001245167.1;NP_001245168.1;NP_001245169.1;NI | 27344 | . | . | [illegible] |
| 219855 | NM_00114529 | SLC37A2 | solute carrier family 37 member 2 | protein_coding | NP_001138762.1;NP_938018.1 | 20644 | . | . | [illegible] |
| 219899 | NM_00113004 | TBCEL | tubulin folding cofactor E like | protein_coding | NP_001123519.1;NP_001350573.1;NP_689928.3 | 28115 | 610451 | . | [illegible] |
| 219902 | NM_00119867 | TLCD5 | TLC domain containing 5 | protein_coding | NP_001185599.1;NP_001185600.1;NP_001185601.1;NP_001185602.1;NI | 28280 | . | . | [illegible] |
| 219927 | NM_181514,N | MRPL21 | mitochondrial ribosomal protein L21 | protein_coding | NP_852615.1;NP_852616.1;XP_005273880.1 | 14479 | 611834 | . | [illegible] |
| 219931 | NM_139075 | TPCN2 | two pore segment channel 2 | protein_coding | NP_620714.2 | 20820 | 612163 | . | [illegible] |
| 219968 | NM_00100521 | OR5B21 | olfactory receptor family 5 subfamily B member 21 | protein_coding | NP_001005218.1 | 19616 | . | . | [illegible] |
| 219988 | NM_152716 | PATL1 | PAT1 homolog 1, processing body mRNA decay factor | protein_coding | NP_689929.2 | 26721 | 614660 | . | [illegible] |
| 220001 | NM_152718,XI | VWCE | von Willebrand factor C and EGF domains | protein_coding | NP_689931.2;XP_006718523.1;XP_006718525.1;XP_011543119.1;XP_01 | 26487 | 611115 | . | [illegible] |
| 220002 | NM_00116145 | CYB561A3 | cytochrome b561 family member A3 | protein_coding | NP_001154924.1;NP_001154926.1;NP_001287692.1;NP_705839.3;XP_01 | 23014 | 618757 | . | [illegible] |
| 220042 | NM_00136348 | DDIAS | DNA damage induced apoptosis suppressor | protein_coding | NP_001350410.1;NP_659455.3;XP_011543138.1;XP_011543139.1;XP_02 | 26351 | 618045 | . | [illegible] |
| 220064 | NM_153451,XI | LTO1 | LTO1 maturation factor of ABCE1 | protein_coding | NP_703152.1;XP_006718533.1 | 17589 | 607224 | . | [illegible] |
| 220074 | NM_00114530 | LRTOMT | leucine rich transmembrane and O-methyltransferase domain con | protein_coding | NP_001138779.1;NP_001138780.1;NP_001138781.1;NP_001138782.1;NI | 25033 | 612414 | . | [illegible] |
| 220108 | NM_00124231 | FAM124A | family with sequence similarity 124 member A | protein_coding | NP_001229241.1;NP_001317451.1;NP_659456.3;XP_011533280.1;XP_01 | 26413 | . | . | [illegible] |
| 220112 | gene-CTAGE3P | CTAGE3P | CTAGE family member 3, pseudogene | pseudogene | . | 24348 | 608857 | . | [illegible] |
| 220134 | NM_00103953 | SKA1 | spindle and kinetochore associated complex subunit 1 | protein_coding | NP_001034624.1;NP_659497.1 | 28109 | 616673 | . | [illegible] |
| 220136 | NM_145020,XI | CFAP53 | cilia and flagella associated protein 53 | protein_coding | NP_659457.2;XP_024306868.1 | 26530 | 614759 | . | [illegible] |
| 220202 | NM_145178 | ATOH7 | atonal bHLH transcription factor 7 | protein_coding | NP_660161.1 | 13907 | 609875 | . | [illegible] |
| 220213 | NM_00114537 | OTUD1 | OTU deubiquitinase 1 | protein_coding | NP_001138845.1 | 27346 | 612022 | . | [illegible] |
| 220323 | NM_178507 | OAF | out at first homolog | protein_coding | NP_848602.1 | 28752 | . | . | [illegible] |
| 220388 | NM_152723 | CCDC89 | coiled-coil domain containing 89 | protein_coding | NP_689936.1 | 26762 | . | . | [illegible] |
| 220416 | NM_00128246 | LRRC63 | leucine rich repeat containing 63 | protein_coding | NP_001269389.1;XP_011533291.1;XP_011533292.1;XP_011533294.1;XP | 34296 | . | . | [illegible] |
| 220433 | gene-RPS4XP1( | RPS4XP16 | ribosomal protein S4X pseudogene 16 | pseudogene | . | 36365 | . | . | [illegible] |
| 220441 | NM_173557,XI | RNF152 | ring finger protein 152 | protein_coding | NP_775828.1;XP_005266707.1;XP_005266709.1;XP_011524180.1;XP_01 | 26811 | 616512 | . | [illegible] |
| 220594 | NR_003554 | USP32P2 | ubiquitin specific peptidase 32 pseudogene 2 | transcribed_ps | . | 30751 | . | . | [illegible] |
| 220717 | gene-RPLP0P6 | RPLP0P6 | ribosomal protein lateral stalk subunit P0 pseudogene 6 | pseudogene | . | 36404 | . | . | [illegible] |
| 220729 | NR_003266 | LOC220729 | succinate dehydrogenase complex flavoprotein subunit A pseudog | transcribed_ps | . | . | . | . | [illegible] |
| 220869 | NM_00102491 | CBWD5 | COBW domain containing 5 | protein_coding | NP_001020087.2;NP_001273764.1;NP_001273765.1;NP_001317597.1;NI | 24584 | . | . | [illegible] |
| 220885 | gene-RPSAP15 | RPSAP15 | ribosomal protein SA pseudogene 15 | pseudogene | . | 31464 | . | . | [illegible] |
| 220906 | NR_033805 | WAC-AS1 | WAC antisense RNA 1 (head to head) | lncRNA | . | 27347 | . | . | [illegible] |
| 220929 | NM_00114376 | ZNF438 | zinc finger protein 438 | protein_coding | NP_001137238.1;NP_001137239.1;NP_001137240.1;NP_001137241.1;NI | 21029 | . | . | [illegible] |
| 220930 | NR_024284,NF | ZEB1-AS1 | ZEB1 antisense RNA 1 | lncRNA | . | 42354 | 616915 | . | [illegible] |
| 220972 | NM_00100226 | MARCHF8 | membrane associated ring-CH-type finger 8 | protein_coding | NP_001002266.1;NP_001269795.1;NP_659458.2;XP_005271861.1;XP_00 | 23356 | 613335 | . | [illegible] |
| 220988 | NM_00133024 | HNRNPA3 | heterogeneous nuclear ribonucleoprotein A3 | protein_coding | NP_001317176.1;NP_001317177.1;NP_001317178.1;NP_001317179.1;NI | 24941 | 605372 | . | [illegible] |
| 220992 | NM_00131814 | ZNF485 | zinc finger protein 485 | protein_coding | NP_001305069.1;NP_001305070.1;NP_001305071.1;NP_001305072.1;NI | 23440 | . | . | [illegible] |
| 221035 | NM_00100133 | REEP3 | receptor accessory protein 3 | protein_coding | NP_001001330.1;XP_011537803.1;XP_016871385.1 | 23711 | 609348 | . | [illegible] |
| 221037 | NM_00128294 | JMJD1C | jumonji domain containing 1C | protein_coding | NP_001269877.1;NP_001305082.1;NP_001305083.1;NP_001309181.1;NI | 12313 | 604503 | . | [illegible] |
| 221060 | NR_160789 | RPP38-DT | RPP38 divergent transcript | lncRNA | . | 28582 | . | . | [illegible] |
| 221061 | NM_00101092 | FAM171A1 | family with sequence similarity 171 member A1 | protein_coding | NP_001010924.1;XP_011517680.1;XP_016871393.1;XP_016871394.1 | 23522 | . | . | [illegible] |
| 221074 | NM_00114519 | SLC39A12 | solute carrier family 39 member 12 | protein_coding | NP_001138667.1;NP_001269662.1;NP_001269663.1;NP_689938.2 | 20860 | 608734 | . | [illegible] |
| 221078 | NM_00135111 | NSUN6 | NOP2/Sun RNA methyltransferase 6 | protein_coding | NP_001338044.1;NP_001338045.1;NP_001338046.1;NP_001338047.1;NI | 23529 | 617199 | . | [illegible] |
| 221079 | NM_178815,XI | ARL5B | ADP ribosylation factor like GTPase 5B | protein_coding | NP_848930.1;XP_005252457.1;XP_005252458.1 | 23052 | 608909 | . | [illegible] |
| 221092 | NM_00107955 | HNRNPUL2 | heterogeneous nuclear ribonucleoprotein U like 2 | protein_coding | NP_001073027.1 | 25451 | . | . | [illegible] |
| 221120 | NM_139178 | ALKBH3 | alkB homolog 3, alpha-ketoglutarate dependent dioxygenase | protein_coding | NP_631917.1 | 30141 | 610603 | . | [illegible] |
| 221143 | NM_00131893 | EEF1AKMT1 | EEF1A lysine methyltransferase 1 | protein_coding | NP_001305868.1;NP_777588.1;XP_016875921.1 | 27351 | 617793 | . | [illegible] |
| 221150 | NM_00116601 | SKA3 | spindle and kinetochore associated complex subunit 3 | protein_coding | NP_001159489.1;NP_659498.4;XP_005266345.1;XP_011533296.1 | 20262 | . | . | [illegible] |
| 221154 | NM_152726,XI | MICU2 | mitochondrial calcium uptake 2 | protein_coding | NP_689939.1;XP_016875922.1 | 31830 | 610632 | . | [illegible] |
| 221178 | NM_00116627 | SPATA13 | spermatogenesis associated 13 | protein_coding | NP_001159743.1;NP_001273721.1;NP_001273722.1;NP_001273723.1;NI | 23222 | 613324 | . | [illegible] |
| 221184 | NM_152727,XI | CPNE2 | copine 2 | protein_coding | NP_689940.3 | 2315 | 604206 | . | [illegible] |
| 221264 | NM_00114512 | AK9 | adenylate kinase 9 | protein_coding | NP_001138600.2;NP_001316531.1;NP_001316532.1;NP_659462.1;XP_00 | 33814 | 615358 | . | [illegible] |
| 221294 | NM_152729,XI | NT5DC1 | 5'-nucleotidase domain containing 1 | protein_coding | NP_689942.2;XP_006715441.1 | 21556 | . | . | [illegible] |
| 221302 | NM_00136118 | ZUP1 | zinc finger containing ubiquitin peptidase 1 | protein_coding | NP_001348118.1;NP_001348119.1;NP_001348120.1;NP_659499.2 | 21224 | . | . | [illegible] |
| 221322 | NM_00136775 | TBC1D32 | TBC1 domain family member 32 | protein_coding | NP_001354688.1;NP_001354689.1;NP_689943.4;XP_011533871.1;XP_01 | 21485 | 615867 | . | [illegible] |
| 221336 | NM_00131853 | BEND6 | BEN domain containing 6 | protein_coding | NP_001305468.1;NP_689944.2;XP_005248946.1;XP_011512648.1;XP_01 | 20871 | . | . | [illegible] |
| 221409 | NM_00137208 | SPATS1 | spermatogenesis associated serine rich 1 | protein_coding | NP_001359010.1;NP_659463.1 | 22957 | . | . | [illegible] |
| 221421 | NM_00119334 | RSPH9 | radial spoke head component 9 | protein_coding | NP_001180270.1;NP_689945.2;XP_005248958.1 | 21057 | 612648 | . | [illegible] |
| 221424 | NM_00101297 | LRRC73 | leucine rich repeat containing 73 | protein_coding | NP_001012992.1;NP_001258811.1 | 21375 | . | . | [illegible] |
| 221442 | NR_026938 | ADCY10P1 | ADCY10 pseudogene 1 | transcribed_ps | . | 44143 | . | . | [illegible] |
| 221443 | NM_00132968 | OARD1 | O-acyl-ADP-ribose deacylase 1 | protein_coding | NP_001316613.1;NP_001316614.1;NP_001316615.1;NP_001316617.1;NI | 21257 | 614393 | . | [illegible] |
| 221458 | NM_00128902 | KIF6 | kinesin family member 6 | protein_coding | NP_001275949.1;NP_001275950.1;NP_001275953.1;NP_001338432.1;NI | 21202 | 613919 | . | [illegible] |
| 221468 | NM_00116290 | TMEM217 | transmembrane protein 217 | protein_coding | NP_001156372.1;NP_001273330.1;NP_001358484.1;NP_660359.2;XP_01 | 21238 | . | . | [illegible] |
| 221477 | NM_00128663 | C6orf89 | chromosome 6 open reading frame 89 | protein_coding | NP_001273564.1;NP_001273565.1;NP_001273566.1;NP_689947.2;XP_01 | 21114 | 616642 | . | [illegible] |
| 221481 | NM_00128657 | ARMC12 | armadillo repeat containing 12 | protein_coding | NP_001273503.1;NP_001273505.1;NP_659465.2;XP_011512683.1;XP_01 | 21099 | . | . | [illegible] |
| 221491 | NM_00100870 | SMIM29 | small integral membrane protein 29 | protein_coding | NP_001008703.2;NP_001008704.2;NP_001274325.2;NP_848603.3 | 1340 | 611419 | . | [illegible] |
| 221496 | NM_00114394 | LEMD2 | LEM domain nuclear envelope protein 2 | protein_coding | NP_001137416.1;NP_001335638.1;NP_001335639.1;NP_851853.1;XP_01 | 21244 | 616312 | . | [illegible] |
| 221504 | NM_152735 | ZBTB9 | zinc finger and BTB domain containing 9 | protein_coding | NP_689948.1 | 28323 | . | . | [illegible] |
| 221527 | NM_181842,XI | ZBTB12 | zinc finger and BTB domain containing 12 | protein_coding | NP_862825.1;XP_011512685.2 | 19066 | . | . | [illegible] |
| 221545 | NM_00110993 | C6orf136 | chromosome 6 open reading frame 136 | protein_coding | NP_001103408.1;NP_001154848.1;NP_659466.2;XP_011512686.1 | 21301 | . | . | [illegible] |
| 221547 | gene-RANP1 | RANP1 | RAN pseudogene 1 | pseudogene | . | 21631 | . | . | [illegible] |
| 221656 | NM_00136461 | KDM1B | lysine demethylase 1B | protein_coding | NP_001351543.1;NP_694587.3;XP_005248982.1;XP_005248985.1;XP_01 | 21577 | 613081 | . | [illegible] |
| 221662 | NM_00114394 | RBM24 | RNA binding motif protein 24 | protein_coding | NP_001137413.1;NP_001137414.1;NP_694565.1;XP_005248989.1;XP_01 | 21539 | 617603 | . | [illegible] |
| 221687 | NM_00116503 | RNF182 | ring finger protein 182 | protein_coding | NP_001158504.1;NP_001158505.1;NP_001158506.1;NP_689950.1;XP_01 | 28522 | . | . | [illegible] |
| 221692 | NM_00124264 | PHACTR1 | phosphatase and actin regulator 1 | protein_coding | NP_001229577.1;NP_001309237.1;NP_001309238.1;NP_001309239.1;NI | 20990 | 608723 | . | [illegible] |
| 221710 | NM_00113557 | SMIM13 | small integral membrane protein 13 | protein_coding | NP_001129047.1 | 27356 | . | . | [illegible] |
| 221718 | NR_027793 | LINC00518 | long intergenic non-protein coding RNA 518 | lncRNA | . | 28626 | . | . | [illegible] |
| 221749 | NM_183373,XI | PXDC1 | PX domain containing 1 | protein_coding | NP_899229.2;XP_011512695.1 | 21361 | . | . | [illegible] |
| 221785 | NM_00135097 | ZSCAN25 | zinc finger and SCAN domain containing 25 | protein_coding | NP_001337908.1;NP_001337909.1;NP_001337910.1;NP_001337911.1;NI | 21961 | . | . | [illegible] |
| 221786 | NM_145111,XI | FAM200A | family with sequence similarity 200 member A | protein_coding | NP_659802.1;XP_011514215.1;XP_024302449.1 | 25401 | . | . | [illegible] |
| 221830 | NM_00100292 | POLR1F | RNA polymerase I subunit F | protein_coding | NP_001002926.1 | 18027 | 608312 | . | [illegible] |
| 221833 | NM_182700,N | SP8 | Sp8 transcription factor | protein_coding | NP_874359.2;NP_945194.1 | 19196 | 608306 | . | [illegible] |
| 221838 | gene-EEF1A1P( | EEF1A1P6 | eukaryotic translation elongation factor 1 alpha 1 pseudogene 6 | pseudogene | . | 3201 | . | . | [illegible] |
| 221883 | NR_002795 | HOXA11-AS | HOXA11 antisense RNA | lncRNA | . | 24957 | 607530 | . | [illegible] |
| 221895 | NM_175061,XI | JAZF1 | JAZF zinc finger 1 | protein_coding | NP_778231.2;XP_006715719.1 | 28917 | 606246 | . | [illegible] |
| 221908 | NM_00134693 | PPP1R35 | protein phosphatase 1 regulatory subunit 35 | protein_coding | NP_001333867.1;NP_659467.1;XP_011514216.1 | 28320 | . | . | [illegible] |
| 221914 | NM_152742 | GPC2 | glypican 2 | protein_coding | NP_689955.1 | 4450 | 618446 | . | [illegible] |
| 221927 | NM_00135062 | BRAT1 | BRCA1 associated ATM activator 1 | protein_coding | NP_001337555.1;NP_001337556.1;NP_689956.2;XP_011513479.1;XP_01 | 21701 | 614506 | . | [illegible] |
| 221937 | NM_00103716 | FOXK1 | forkhead box K1 | protein_coding | NP_001032242.1 | 23480 | 616302 | . | [illegible] |
| 221955 | NM_00114293 | DAGLB | diacylglycerol lipase beta | protein_coding | NP_001136408.1;NP_631918.3 | 28923 | 614016 | . | [illegible] |
| 221960 | NM_198097 | CCZ1B | CCZ1 homolog B, vacuolar protein trafficking and biogenesis assoc | protein_coding | NP_932765.1 | 21717 | . | . | [illegible] |
| 222068 | NM_00130305 | TMED4 | transmembrane p24 trafficking protein 4 | protein_coding | NP_001289987.1;NP_001289988.1;NP_001289989.1;NP_001289990.1;NI | 22301 | 612038 | . | [illegible] |
| 222161 | NR_002186 | DKFZP586I142( | uncharacterized protein DKFZp586I1420 | transcribed_ps | . | . | . | . | [illegible] |
| 222166 | NM_152793,XI | MTURN | maturin, neural progenitor differentiation regulator homolog | protein_coding | NP_690006.2;XP_005249709.1 | 25457 | . | . | [illegible] |
| 222183 | NM_00111019 | SRRM3 | serine/arginine repetitive matrix 3 | protein_coding | NP_001103669.1;NP_001278760.1 | 26729 | . | . | [illegible] |
| 222194 | NM_198467 | RSBN1L | round spermatid basic protein 1 like | protein_coding | NP_940869.2 | 24765 | . | . | [illegible] |
| 222223 | NM_00114274 | ELAPOR2 | endosome-lysosome associated apoptosis and autophagy regulat( | protein_coding | NP_001136221.1;NP_001278919.1;NP_001278920.1;NP_689961.3;XP_00 | 21945 | 614048 | . | [illegible] |
| 222229 | NM_00131772 | LRWD1 | leucine rich repeats and WD repeat domain containing 1 | protein_coding | NP_001304650.1;NP_690852.1 | 21769 | 615167 | . | [illegible] |
| 222234 | NM_00114526 | FAM185A | family with sequence similarity 185 member A | protein_coding | NP_001138740.2;NP_001138741.1;NP_001337916.1 | 22412 | . | . | [illegible] |
| 222235 | NM_00111103 | FBXL13 | F-box and leucine rich repeat protein 13 | protein_coding | NP_001104508.1;NP_001274079.1;NP_659469.3;XP_005250262.1;XP_00 | 21658 | 609080 | . | [illegible] |
| 222236 | NM_00112283 | NAPEPLD | N-acyl phosphatidylethanolamine phospholipase D | protein_coding | NP_001116310.1;NP_945341.3;XP_005250271.3;XP_005250272.1;XP_00 | 21683 | 612334 | . | [illegible] |
| 222255 | NM_00131822 | ATXN7L1 | ataxin 7 like 1 | protein_coding | NP_001305158.1;NP_065776.1;NP_612504.1;NP_689962.1;XP_0052502 | 22210 | . | . | [illegible] |
| 222256 | NM_00130116 | CDHR3 | cadherin related family member 3 | protein_coding | NP_001288090.1;NP_689963.2;XP_005250281.1;XP_005250282.1;XP_00 | 26308 | 615610 | . | [illegible] |
| 222389 | NM_00110091 | BEND7 | BEN domain containing 7 | protein_coding | NP_001094382.1;NP_001356792.1;NP_001357004.2;NP_001365078.1;NI | 23514 | . | . | [illegible] |
| 222484 | NM_153371,XI | LNX2 | ligand of numb-protein X 2 | protein_coding | NP_699202.1;XP_011533297.1;XP_016875923.1 | 20421 | 609733 | . | [illegible] |
| 222553 | NM_00102985 | SLC35F1 | solute carrier family 35 member F1 | protein_coding | NP_001025029.2;XP_005266922.1 | 21483 | . | . | [illegible] |
| 222643 | NM_173561 | UNC5CL | unc-5 family C-terminal like | protein_coding | NP_775832.2 | 21203 | 617464 | . | [illegible] |
| 222658 | NM_00128657 | KCTD20 | potassium channel tetramerization domain containing 20 | protein_coding | NP_001273508.1;NP_001273509.1;NP_775833.2;XP_005248997.1;XP_00 | 21052 | 615932 | . | [illegible] |
| 222663 | NM_00130313 | SCUBE3 | signal peptide, CUB domain and EGF like domain containing 3 | protein_coding | NP_001290065.1;NP_689966.2;XP_005249000.1;XP_005249004.1;XP_01 | 13655 | 614708 | . | [illegible] |
| 222696 | NM_00101245 | ZSCAN23 | zinc finger and SCAN domain containing 23 | protein_coding | NP_001012458.1;XP_005249007.1;XP_011512708.1;XP_011512710.1;XP | 21193 | . | . | [illegible] |
| 222699 | NR_002936 | TOB2P1 | transducer of ERBB2, 2 pseudogene 1 | transcribed_ps | . | 13986 | . | . | [illegible] |
| 222950 | NM_173564 | NYAP1 | neuronal tyrosine phosphorylated phosphoinositide-3-kinase adap | protein_coding | NP_775835.2 | 22009 | 615477 | . | [illegible] |
| 222962 | NM_00104066 | SLC29A4 | solute carrier family 29 member 4 | protein_coding | NP_001035751.1;NP_001287776.1;NP_694979.2;XP_005249715.1;XP_00 | 23097 | 609149 | . | [illegible] |
| 223082 | NM_147128,XI | ZNRF2 | zinc and ring finger 2 | protein_coding | NP_667339.1 | 22316 | 612061 | . | [illegible] |
| 223117 | NM_152754,XI | SEMA3D | semaphorin 3D | protein_coding | NP_689967.2;XP_011514262.1;XP_011514263.1;XP_016867362.1 | 10726 | 609907 | . | [illegible] |
| 245812 | NM_152755 | CNPY4 | canopy FGF signaling regulator 4 | protein_coding | NP_689968.1 | 28631 | 610047 | . | [illegible] |
| 245972 | NM_152565 | ATP6V0D2 | ATPase H+ transporting V0 subunit d2 | protein_coding | NP_689778.1 | 18266 | 618072 | . | [illegible] |
| 245973 | NM_00103936 | ATP6V1C2 | ATPase H+ transporting V1 subunit C2 | protein_coding | NP_001034451.1;NP_653184.2;XP_011508641.1;XP_011508642.1;XP_01 | 18264 | 618070 | . | [illegible] |
| 246126 | NR_045128,NF | TXLNGY | taxilin gamma pseudogene, Y-linked | transcribed_ps | . | 18473 | 400031 | . | [illegible] |
| 246175 | NM_00128679 | CNOT6L | CCR4-NOT transcription complex subunit 6 like | protein_coding | NP_001273719.1;NP_001351935.1;NP_001351936.1;NP_653172.2;XP_01 | 18042 | 618069 | . | [illegible] |
| 246181 | NR_040288,NF | AKR7L | aldo-keto reductase family 7 like (gene/pseudogene) | transcribed_ps | . | 24056 | 608478 | . | [illegible] |
| 246184 | NM_139286,XI | CDC26 | cell division cycle 26 | protein_coding | NP_644815.1;XP_016870062.1;XP_016870063.1;XP_016870064.1 | 17839 | 614533 | . | [illegible] |
| 246243 | NM_00128683 | RNASEH1 | ribonuclease H1 | protein_coding | NP_001273763.1;NP_001273766.1;NP_001365200.1;NP_001365201.1;NI | 18466 | 604123 | . | [illegible] |
| 246269 | NM_00132300 | AFG1L | AFG1 like ATPase | protein_coding | NP_001309934.1;NP_660358.2;XP_005266942.1;XP_011533959.1;XP_01 | 16411 | 617469 | . | [illegible] |
| 246312 | NR_038870,NF | C21orf91-OT1 | C21orf91 overlapping transcript 1 | lncRNA | . | 16729 | . | . | [illegible] |
| 246330 | NM_00109851 | PELI3 | pellino E3 ubiquitin protein ligase family member 3 | protein_coding | NP_001091980.1;NP_001230064.1;NP_001230065.1;NP_659502.2;XP_01 | 30010 | 609827 | . | [illegible] |
| 246714 | gene-CKS1BP2 | CKS1BP2 | CDC28 protein kinase regulatory subunit 1B pseudogene 2 | pseudogene | . | 24232 | . | . | [illegible] |
| 246715 | gene-CKS1BP3 | CKS1BP3 | CDC28 protein kinase regulatory subunit 1B pseudogene 3 | pseudogene | . | 24233 | . | . | [illegible] |
| 246716 | gene-YWHAQP | YWHAQP2 | YWHAQ pseudogene 2 | pseudogene | . | 23211 | . | . | [illegible] |
| 246719 | gene-UBE2SP1 | UBE2SP1 | ubiquitin conjugating enzyme E2 S pseudogene 1 | pseudogene | . | 30864 | . | . | [illegible] |
| 246720 | gene-DYNLL1P: | DYNLL1P1 | dynein light chain LC8-type 1 pseudogene 1 | pseudogene | . | 20723 | . | . | [illegible] |
| 246721 | NM_032959 | POLR2J2 | RNA polymerase II subunit J2 | protein_coding | NP_116581.3 | 23208 | 609881 | . | [illegible] |
| 246737 | gene-YWHAZP4 | YWHAZP4 | tyrosine 3-monooxygenase/tryptophan 5-monooxygenase activati | pseudogene | . | 30851 | . | . | [illegible] |
| 246777 | NM_145658 | SPESP1 | sperm equatorial segment protein 1 | protein_coding | NP_663633.1 | 15570 | 609399 | . | [illegible] |
| 246784 | gene-SMU1P1 | SMU1P1 | SMU1 pseudogene 1 | pseudogene | . | 54527 | . | . | [illegible] |
| 252839 | NM_00128856 | TMEM9 | transmembrane protein 9 | protein_coding | NP_001275493.1;NP_001275494.1;NP_001275495.1;NP_001275496.1;NI | 18823 | 616877 | . | [illegible] |
| 252884 | NM_00132228 | ZNF396 | zinc finger protein 396 | protein_coding | NP_001309215.1;NP_001309219.1;NP_001309220.1;NP_665699.1;XP_00 | 18824 | 609600 | . | [illegible] |
| 252969 | NM_00113574 | NEIL2 | nei like DNA glycosylase 2 | protein_coding | NP_001129218.1;NP_001129219.1;NP_001129220.1;NP_001336368.1;NI | 18956 | 608933 | . | [illegible] |
| 252983 | NM_178509,XI | STXBP4 | syntaxin binding protein 4 | protein_coding | NP_848604.3;XP_005257244.1;XP_006721860.1;XP_006721861.1;XP_01 | 19694 | 610415 | . | [illegible] |
| 253018 | NR_026791 | HCG27 | HLA complex group 27 | lncRNA | . | 27366 | . | . | [illegible] |
| 253039 | NR_024408,NF | CUTALP | cutA divalent cation tolerance like, pseudogene | transcribed_ps | . | 27367 | . | . | [illegible] |
| 253143 | NM_173566 | PRR14L | proline rich 14 like | protein_coding | NP_775837.2 | 28738 | . | . | [illegible] |
| 253260 | NM_00128543 | RICTOR | RPTOR independent companion of MTOR complex 2 | protein_coding | NP_001272368.1;NP_001272369.1;NP_689969.2;XP_006714526.1;XP_01 | 28611 | 609022 | . | [illegible] |
| 253272 | gene-PCGF7P | PCGF7P | polycomb group ring finger 7 pseudogene | pseudogene | . | 30341 | . | . | [illegible] |
| 253430 | NM_152230,XI | IPMK | inositol polyphosphate multikinase | protein_coding | NP_689416.1;XP_011537867.1;XP_016871502.1 | 20739 | 609851 | . | [illegible] |
| 253461 | NM_00108041 | ZBTB38 | zinc finger and BTB domain containing 38 | protein_coding | NP_001073881.2;NP_001337028.1;NP_001337029.1;NP_001363041.1;NI | 26636 | 612218 | . | [illegible] |
| 253512 | NM_00101087 | SLC25A30 | solute carrier family 25 member 30 | protein_coding | NP_001010875.1;NP_001273735.1;NP_001273736.1;XP_005266378.1;XP | 27371 | 610793 | . | [illegible] |
| 253558 | NM_00100225 | LCLAT1 | lysocardiolipin acyltransferase 1 | protein_coding | NP_001002257.1;NP_001291374.1;NP_001291375.1;NP_872357.2;XP_00 | 26756 | 614241 | . | [illegible] |
| 253635 | NM_00127850 | GPATCH11 | G-patch domain containing 11 | protein_coding | NP_001265434.1;NP_001309178.3;NP_001358785.2;NP_001358787.2;NI | 26768 | . | . | [illegible] |
| 253639 | NM_00125616 | ZNF620 | zinc finger protein 620 | protein_coding | NP_001243096.1;NP_001243097.1;NP_787084.1;XP_005265068.1;XP_00 | 28742 | . | . | [illegible] |

| | | | | | | | | |
|---|---|---|---|---|---|---|---|---|
| 253650 | NM_00133110 | ANKRD18A | ankyrin repeat domain 18A | protein_coding | NP_001318029.1;NP_671728.2;XP_011516141.1;XP_016870065.1;XP_01 | 23643 | . | [illegible] |
| 253714 | NM_00135059 | MMS22L | MMS22 like, DNA repair protein | protein_coding | NP_001337528.1;NP_001337529.1;NP_940870.2;XP_006715495.1;XP_01 | 21475 | 615614 | [illegible] |
| 253724 | NR_027249 | TTC41P | tetratricopeptide repeat domain 41, pseudogene | transcribed_ps | . | 49210 | . | [illegible] |
| 253725 | NM_00116910 | WASHC2C | WASH complex subunit 2C | protein_coding | NP_001162577.1;NP_001162578.1;NP_001317003.1;NP_001354322.1;NI | 23414 | 613631 | [illegible] |
| 253769 | NM_00120255 | WDR27 | WD repeat domain 27 | protein_coding | NP_001189479.1;NP_001337552.1;NP_001337553.1;NP_001337554.1;NI | 21248 | . | [illegible] |
| 253782 | NM_00125612 | CERS6 | ceramide synthase 6 | protein_coding | NP_001243055.1;NP_982288.1;XP_005246497.1;XP_016859238.1;XP_02 | 23826 | 615336 | [illegible] |
| 253827 | NM_00103167 | MSRB3 | methionine sulfoxide reductase B3 | protein_coding | NP_001026849.1;NP_001180389.1;NP_001180390.1;NP_932346.1;XP_02 | 27375 | 613719 | [illegible] |
| 253832 | NM_00128663 | ZDHHC20 | zinc finger DHHC-type palmitoyltransferase 20 | protein_coding | NP_001273567.1;NP_001316988.1;NP_694983.2;XP_006719859.1;XP_00 | 20749 | 617972 | [illegible] |
| 253943 | NM_00127781 | YTHDF3 | YTH N6-methyladenosine RNA binding protein 3 | protein_coding | NP_001264742.1;NP_001264743.1;NP_001264744.1;NP_001264745.1;NI | 26465 | 618669 | [illegible] |
| 253959 | NM_00128304 | RALGAPA1 | Ral GTPase activating protein catalytic subunit alpha 1 | protein_coding | NP_001269972.1;NP_001269973.1;NP_001317004.1;NP_001333172.1;NI | 17770 | 608884 | [illegible] |
| 253980 | NM_178863,N | KCTD13 | potassium channel tetramerization domain containing 13 | protein_coding | NP_849194.1;XP_011544085.1;XP_011544086.1;XP_016878593.1;XP_01 | 22234 | 608947 | [illegible] |
| 253982 | NM_181718,XI | ASPHD1 | aspartate beta-hydroxylase domain containing 1 | protein_coding | NP_859069.2;XP_016878596.1;XP_024305990.1;XP_024305991.1 | 27380 | . | [illegible] |
| 254013 | NM_00113586 | ETFBKMT | electron transfer flavoprotein subunit beta lysine methyltransferas | protein_coding | NP_001129335.1;NP_001129336.1;NP_776163.1;XP_024304691.1;XP_02 | 28739 | 615256 | [illegible] |
| 254042 | NM_00132227 | METAP1D | methionyl aminopeptidase type 1D, mitochondrial | protein_coding | NP_001309207.1;NP_001309208.1;NP_954697.1;XP_016859239.1 | 32583 | 610267 | [illegible] |
| 254048 | NM_173569,XI | UBN2 | ubinuclein 2 | protein_coding | NP_775840.3;XP_005250306.2;XP_006715979.2;XP_011514304.1;XP_01 | 21931 | 613841 | [illegible] |
| 254065 | NM_153252,XI | BRWD3 | bromodomain and WD repeat domain containing 3 | protein_coding | NP_694984.5;XP_005262170.1;XP_016884873.1;XP_016884874.1 | 17342 | 300553 | [illegible] |
| 254100 | NR_038923 | ZNRD2-AS1 | ZNRD2 antisense RNA 1 (head to head) | lncRNA | . | 27384 | . | [illegible] |
| 254102 | NM_00109940 | EHBP1L1 | EH domain binding protein 1 like 1 | protein_coding | NP_001092879.1;NP_001338016.1;XP_005273924.1;XP_005273925.1;XP | 30682 | . | [illegible] |
| 254128 | NR_037856,NF | NIFK-AS1 | NIFK antisense RNA 1 | lncRNA | . | 27385 | . | [illegible] |
| 254170 | NM_203301 | FBXO33 | F-box protein 33 | protein_coding | NP_976046.1 | 19833 | 609103 | [illegible] |
| 254225 | NM_00109863 | RNF169 | ring finger protein 169 | protein_coding | NP_001092108.1;XP_011543191.1 | 26961 | 618650 | [illegible] |
| 254251 | NM_00116613 | LCORL | ligand dependent nuclear receptor corepressor like | protein_coding | NP_001159611.1;NP_001352587.1;NP_001352588.1;NP_001352589.1;NI | 30776 | 611799 | [illegible] |
| 254263 | NM_182553,N | CNIH2 | cornichon family AMPA receptor auxiliary protein 2 | protein_coding | NP_872359.1 | 28744 | 611288 | [illegible] |
| 254359 | NM_00134857 | ZDHHC24 | zinc finger DHHC-type containing 24 | protein_coding | NP_001335500.1;NP_997223.1;XP_005273931.1;XP_011543195.1;XP_01 | 27387 | . | [illegible] |
| 254394 | NM_00137835 | MCM9 | minichromosome maintenance 9 homologous recombination repa | protein_coding | NP_001365285.1;NP_001365286.1;NP_001365287.1;NP_001365288.1;NI | 21484 | 610098 | [illegible] |
| 254427 | NM_153256,XI | PROSER2 | proline and serine rich 2 | protein_coding | NP_694988.3;XP_011517740.1;XP_016871517.1 | 23728 | . | [illegible] |
| 254428 | NM_173854,XI | SLC41A1 | solute carrier family 41 member 1 | protein_coding | NP_776253.3;XP_005245126.1 | 19429 | 610801 | [illegible] |
| 254528 | NM_00116356 | MEIOB | meiosis specific with OB-fold | protein_coding | NP_001157032.1;NP_689977.2 | 28569 | 617670 | [illegible] |
| 254531 | NM_153613,XI | LPCAT4 | lysophosphatidylcholine acyltransferase 4 | protein_coding | NP_705841.2;XP_006720517.1;XP_011519724.1 | 30059 | 612039 | [illegible] |
| 254552 | NM_00124375 | NUDT8 | nudix hydrolase 8 | protein_coding | NP_001230679.1;NP_862826.1;XP_016872963.1 | 8055 | . | [illegible] |
| 254559 | NR_015411,NF | MIR9-3HG | MIR9-3 host gene | lncRNA | . | 27388 | . | [illegible] |
| 254700 | gene-USP12PX | USP12PX | USP12 pseudogene X-linked | pseudogene | . | 21410 | . | [illegible] |
| 254778 | NM_152765 | VXN | vexin | protein_coding | NP_689978.2 | 28498 | . | [illegible] |
| 254863 | NM_152766 | TMEM256 | transmembrane protein 256 | protein_coding | NP_689979.1 | 28618 | 617779 | [illegible] |
| 254887 | NM_00132046 | ZDHHC23 | zinc finger DHHC-type palmitoyltransferase 23 | protein_coding | NP_001307395.1;NP_001307396.1;NP_001307397.1;NP_001350881.1;NI | 28654 | 617334 | [illegible] |
| 254896 | NR_046173 | LOC254896 | uncharacterized LOC254896 | lncRNA | . | . | . | [illegible] |
| 254948 | gene-RPL9P8 | RPL9P8 | ribosomal protein L9 pseudogene 8 | pseudogene | . | 30334 | . | [illegible] |
| 255043 | NM_00137201 | TMEM86B | transmembrane protein 86B | protein_coding | NP_001358942.1;NP_776165.3 | 28448 | 617806 | [illegible] |
| 255057 | NM_152769,XI | CBARP | CACN subunit beta associated regulatory protein | protein_coding | NP_689982.3;XP_016882044.1;XP_016882045.1;XP_016882046.1;XP_01 | 28617 | . | [illegible] |
| 255082 | NR_026939,NF | CASC2 | cancer susceptibility 2 | lncRNA | . | 22933 | 608598 | [illegible] |
| 255101 | NM_00127829 | CFAP65 | cilia and flagella associated protein 65 | protein_coding | NP_001265224.1;NP_001265225.1;NP_689602.2;NP_919278.2;XP_0115C | 25325 | 614270 | [illegible] |
| 255104 | NM_00134911 | TMCO4 | transmembrane and coiled-coil domains 4 | protein_coding | NP_001336041.1;NP_001336042.1;NP_001336043.1;NP_001336044.1;NI | 27393 | . | [illegible] |
| 255180 | NR_136885 | GOLGA2P11 | GOLGA2 pseudogene 11 | transcribed_ps | . | 26788 | . | [illegible] |
| 255187 | NR_122044 | SH3TC2-DT | SH3TC2 divergent transcript | lncRNA | . | 52905 | . | [illegible] |
| 255231 | NM_00133064 | MCOLN2 | mucolipin 2 | protein_coding | NP_001317576.1;NP_694991.2;XP_005270776.1;XP_011539489.1;XP_01 | 13357 | 607399 | [illegible] |
| 255252 | NM_153260,XI | LRRC57 | leucine rich repeat containing 57 | protein_coding | NP_694992.2;XP_011519725.1;XP_011519726.1 | 26719 | . | [illegible] |
| 255308 | NM_00135773 | EIF2S3B | eukaryotic translation initiation factor 2 subunit gamma B | protein_coding | NP_001344660.1;NP_001344663.1 | 43863 | . | [illegible] |
| 255374 | NM_203397,XI | MBLAC1 | metallo-beta-lactamase domain containing 1 | protein_coding | NP_981942.1;XP_005250307.1 | 22180 | . | [illegible] |
| 255394 | NM_00128626 | TCP11L2 | t-complex 11 like 2 | protein_coding | NP_001273191.1;NP_689985.1;XP_005268824.1;XP_005268825.1;XP_01 | 28627 | . | [illegible] |
| 255403 | NM_00103912 | ZNF718 | zinc finger protein 718 | protein_coding | NP_001034216.2;NP_001276859.1;NP_001276860.1;XP_016863468.1;XP | 26889 | . | [illegible] |
| 255488 | NM_182757,XI | RNF144B | ring finger protein 144B | protein_coding | NP_877434.2;XP_005249043.1 | 21578 | 618869 | [illegible] |
| 255512 | NR_029409 | TOLLIP-AS1 | TOLLIP antisense RNA 1 (head to head) | lncRNA | . | 27403 | . | [illegible] |
| 255520 | NM_153702,XI | ELMOD2 | ELMO domain containing 2 | protein_coding | NP_714913.1;XP_005262942.1;XP_005262943.1;XP_011530120.1;XP_01 | 28111 | 610196 | [illegible] |
| 255631 | NM_00134995 | COL24A1 | collagen type XXIV alpha 1 chain | protein_coding | NP_001336884.1;NP_690850.2;XP_016856413.1;XP_016856414.1;XP_01 | 20821 | 610025 | [illegible] |
| 255758 | NM_00135162 | TCTEX1D2 | Tctex1 domain containing 2 | protein_coding | NP_001338557.1;NP_689986.2 | 28482 | 617353 | [illegible] |
| 255783 | NM_178511,N | INAFM1 | InaF motif containing 1 | protein_coding | NP_848606.3 | 27406 | . | [illegible] |
| 255809 | NM_00113648 | C19orf38 | chromosome 19 open reading frame 38 | protein_coding | NP_001129954.1;XP_005259903.1;XP_005259904.1;XP_024307216.1;XP | 34073 | . | [illegible] |
| 255812 | NR_003264 | SDHAP1 | succinate dehydrogenase complex flavoprotein subunit A pseudog | transcribed_ps | . | 32455 | . | [illegible] |
| 255919 | NM_00128178 | CNEP1R1 | CTD nuclear envelope phosphatase 1 regulatory subunit 1 | protein_coding | NP_001268718.1;NP_694993.2 | 26759 | 616869 | [illegible] |
| 255928 | NM_00114626 | SYT14 | synaptotagmin 14 | protein_coding | NP_001139733.1;NP_001139734.1;NP_001139736.1;NP_001242935.1;NI | 23143 | 610949 | [illegible] |
| 255967 | NM_175854,XI | PAN3 | poly(A) specific ribonuclease subunit PAN3 | protein_coding | NP_787050.6;XP_005266389.1;XP_005266390.1;XP_005266391.1;XP_01 | 29991 | 617448 | [illegible] |
| 256006 | NM_00116444 | ANKRD31 | ankyrin repeat domain 31 | protein_coding | NP_001157915.1;NP_001358982.1;XP_011541600.1;XP_011541602.1;XP | 26853 | 618423 | [illegible] |
| 256051 | NM_00119929 | ZNF549 | zinc finger protein 549 | protein_coding | NP_001186224.2;NP_694995.2 | 26632 | . | [illegible] |
| 256076 | NM_00127829 | COL6A5 | collagen type VI alpha 5 chain | protein_coding | NP_001265227.1;NP_694996.5;XP_011510923.1;XP_011510924.1;XP_01 | 26674 | 611916 | [illegible] |
| 256126 | NM_00110557 | SYCE2 | synaptonemal complex central element protein 2 | protein_coding | NP_001099048.1;XP_005259905.1;XP_011526184.1;XP_011526185.1 | 27411 | 611487 | [illegible] |
| 256227 | NM_00116446 | STEAP1B | STEAP family member 1B | protein_coding | NP_001157932.1;NP_001369376.1;NP_997225.1 | 41907 | . | [illegible] |
| 256281 | NM_00131838 | NUDT14 | nudix hydrolase 14 | protein_coding | NP_001305309.1;NP_803877.2 | 20141 | 609219 | [illegible] |
| 256302 | NM_152914 | NATD1 | N-acetyltransferase domain containing 1 | protein_coding | NP_690878.2 | 30770 | . | [illegible] |
| 256329 | NM_173573,XI | LMNTD2 | lamin tail domain containing 2 | protein_coding | NP_775844.2;XP_011518267.2;XP_011518269.2;XP_016872965.1;XP_01 | 28561 | . | [illegible] |
| 256355 | NR_026676 | RPS2P32 | ribosomal protein S2 pseudogene 32 | transcribed_ps | . | 30518 | . | [illegible] |
| 256356 | NM_00103954 | GK5 | glycerol kinase 5 | protein_coding | NP_001034636.1;XP_011510926.1;XP_016861577.1;XP_016861578.1;XP | 28635 | . | [illegible] |
| 256364 | NM_00130079 | EML3 | EMAP like 3 | protein_coding | NP_001287722.1;NP_001287723.1;NP_694997.2;XP_005273935.1;XP_00 | 26666 | 618118 | [illegible] |
| 256380 | NM_00128640 | SCML4 | Scm polycomb group protein like 4 | protein_coding | NP_001273337.1;NP_001273338.1;NP_932347.2;XP_011534005.1;XP_01 | 21397 | . | [illegible] |
| 256435 | NM_00116001 | ST6GALNAC3 | ST6 N-acetylgalactosaminide alpha-2,6-sialyltransferase 3 | protein_coding | NP_001153483.1;NP_001336034.1;NP_001336035.1;NP_001336036.1;NI | 19343 | 610133 | [illegible] |
| 256471 | NM_00136352 | MFSD8 | major facilitator superfamily domain containing 8 | protein_coding | NP_001350449.1;NP_001350450.1;NP_001358519.1;NP_001358520.1;NI | 28486 | 611124 | [illegible] |
| 256586 | NM_00114391 | LYSMD2 | LysM domain containing 2 | protein_coding | NP_001137389.1;NP_001350898.1;NP_699205.1 | 28571 | . | [illegible] |
| 256643 | NM_00136777 | BCLAF3 | BCLAF1 and THRAP3 family member 3 | protein_coding | NP_001354703.1;NP_938020.2;XP_005274530.1;XP_005274531.1;XP_00 | 27413 | . | [illegible] |
| 256691 | NM_00134799 | MAMDC2 | MAM domain containing 2 | protein_coding | NP_001334919.1;NP_694999.3;XP_011516773.1;XP_011516777.1 | 23673 | 612879 | [illegible] |
| 256764 | NM_00127717 | WDR72 | WD repeat domain 72 | protein_coding | NP_001264105.1;NP_877435.3;XP_011519735.1;XP_011519737.1;XP_01 | 26790 | 613214 | [illegible] |
| 256815 | NM_00135130 | C10orf67 | chromosome 10 open reading frame 67 | protein_coding | NP_001338235.1;NP_001352791.1;NP_001358838.1;NP_714925.2;XP_01 | 28716 | . | [illegible] |
| 256949 | NM_198471,XI | KANK3 | KN motif and ankyrin repeat domains 3 | protein_coding | NP_940873.2;XP_006722781.1;XP_011526186.1;XP_016882052.1 | 24796 | 614611 | [illegible] |
| 256987 | NM_00117407 | SERINC5 | serine incorporator 5 | protein_coding | NP_001167542.1;NP_001167543.1;NP_840060.1;XP_011541606.1;XP_01 | 18825 | 614551 | [illegible] |
| 257000 | NR_027064 | TINCR | TINCR ubiquitin domain containing | lncRNA | . | 14607 | 615241 | [illegible] |
| 257019 | NM_00124495 | FRMD3 | FERM domain containing 3 | protein_coding | NP_001231888.1;NP_001231889.1;NP_001231890.1;NP_001231891.1;NI | 24125 | 607619 | [illegible] |
| 257044 | NM_00113095 | CATSPERE | catsper channel auxiliary subunit epsilon | protein_coding | NP_001124429.1;NP_001229269.1;NP_776168.1;XP_011542440.1;XP_01 | 28491 | 617510 | [illegible] |
| 257160 | NM_00107723 | RNF214 | ring finger protein 214 | protein_coding | NP_001070707.1;NP_001265178.1;NP_997226.2 | 25335 | . | [illegible] |
| 257169 | NM_00127862 | C9orf43 | chromosome 9 open reading frame 43 | protein_coding | NP_001265558.1;NP_001265559.1;NP_689999.1;XP_005251942.1;XP_01 | 23570 | . | [illegible] |
| 257203 | NR_026719 | DSCR9 | Down syndrome critical region 9 | lncRNA | . | 16301 | . | [illegible] |
| 257218 | NM_00104268 | SHPRH | SNF2 histone linker PHD RING helicase | protein_coding | NP_001036148.2;NP_001357256.1;NP_001357257.1;NP_775105.1;XP_00 | 19336 | 608048 | [illegible] |
| 257236 | NM_153376 | CCDC96 | coiled-coil domain containing 96 | protein_coding | NP_699207.1 | 26900 | . | [illegible] |
| 257358 | NR_026945,NF | LINC01366 | long intergenic non-protein coding RNA 1366 | lncRNA | . | 27416 | . | [illegible] |
| 257364 | NM_00131814 | SNX33 | sorting nexin 33 | protein_coding | NP_001305075.1;NP_695003.1 | 28468 | . | [illegible] |
| 257396 | NR_034107,NF | LOC257396 | uncharacterized LOC257396 | lncRNA | . | . | . | [illegible] |
| 257397 | NM_152787,XI | TAB3 | TGF-beta activated kinase 1 (MAP3K7) binding protein 3 | protein_coding | NP_690000.3;XP_005274540.2;XP_005274541.2;XP_011543790.1;XP_01 | 30681 | 300480 | [illegible] |
| 257407 | NM_00114499 | C2orf72 | chromosome 2 open reading frame 72 | protein_coding | NP_001138466.1 | 27418 | . | [illegible] |
| 257415 | NM_00104005 | FAM133B | family with sequence similarity 133 member B | protein_coding | NP_001035146.1;NP_001275513.1;NP_690002.2;XP_011514306.1;XP_01 | 28629 | . | [illegible] |
| 257468 | gene-MEIS3P2 | MEIS3P2 | Meis homeobox 3 pseudogene 2 | pseudogene | . | 17638 | . | [illegible] |
| 258010 | NM_00132034 | SVIP | small VCP interacting protein | protein_coding | NP_001307269.1;NP_001307270.1;NP_001307271.1;NP_683691.1;XP_02 | 25238 | . | [illegible] |
| 259173 | NM_00119070 | ALS2CL | ALS2 C-terminal like | protein_coding | NP_001177636.1;NP_667340.2;XP_005265082.1;XP_006713154.1;XP_00 | 20605 | 612402 | [illegible] |
| 259217 | NM_00133016 | HSPA12A | heat shock protein family A (Hsp70) member 12A | protein_coding | NP_001317093.1;NP_079291.2;XP_005269730.1;XP_011537881.1;XP_01 | 19022 | 610701 | [illegible] |
| 259230 | NM_147156,XI | SGMS1 | sphingomyelin synthase 1 | protein_coding | NP_671512.1;XP_005269732.1;XP_011537885.1;XP_011537886.1 | 29799 | 611573 | [illegible] |
| 259266 | NM_00120684 | ASPM | assembly factor for spindle microtubules | protein_coding | NP_001193775.1;NP_060606.3 | 19048 | 605481 | [illegible] |
| 259282 | NM_148894,XI | BOD1L1 | biorientation of chromosomes in cell division 1 like 1 | protein_coding | NP_683692.2;XP_005248207.1;XP_005248208.1;XP_006714021.1;XP_01 | 31792 | 616746 | [illegible] |
| 259290 | NM_176885 | TAS2R31 | taste 2 receptor member 31 | protein_coding | NP_795366.2 | 19113 | 612669 | [illegible] |
| 259294 | NM_176888 | TAS2R19 | taste 2 receptor member 19 | protein_coding | NP_795369.1 | 19108 | 613961 | [illegible] |
| 259295 | NM_176889 | TAS2R20 | taste 2 receptor member 20 | protein_coding | NP_795370.2 | 19109 | 613962 | [illegible] |
| 260294 | NR_033323 | NSUN5P2 | NSUN5 pseudogene 2 | transcribed_ps | . | 16609 | . | [illegible] |
| 260334 | NM_00135868 | TUBB8B | tubulin beta 8B | protein_coding | NP_001345618.1;XP_024306911.1;XP_024306912.1;XP_024306913.1;XP | 24983 | . | [illegible] |
| 260425 | NM_00114278 | MAGI3 | membrane associated guanylate kinase, WW and PDZ domain cor | protein_coding | NP_001136254.1;NP_690864.2;XP_005270794.1;XP_011539510.1;XP_01 | 29647 | 615943 | [illegible] |
| 260429 | NM_152891,XI | PRSS33 | serine protease 33 | protein_coding | NP_690851.2;XP_011520753.1;XP_011520754.1 | 30405 | 613797 | [illegible] |
| 261726 | NM_00103180 | TIPRL | TOR signaling pathway regulator | protein_coding | NP_001026970.1;NP_690866.1 | 30231 | 611807 | [illegible] |
| 261729 | NM_00104066 | STEAP2 | STEAP2 metalloreductase | protein_coding | NP_001035755.1;NP_001035756.1;NP_001231873.1;NP_001231874.1;NI | 17885 | 605094 | [illegible] |
| 261734 | NM_00129159 | NPHP4 | nephrocystin 4 | protein_coding | NP_001278522.1;NP_001278523.1;NP_055917.1;XP_006710626.1;XP_01 | 19104 | 607215 | [illegible] |
| 261735 | gene-TVP23CP | TVP23CP2 | TVP23C pseudogene 2 | pseudogene | . | 44647 | . | [illegible] |
| 266655 | NM_00135525 | BRD3OS | BRD3 opposite strand | protein_coding | NP_001342185.1 | 24742 | . | [illegible] |
| 266657 | gene-TAS2R15 | TAS2R15P | taste 2 receptor member 15 pseudogene | pseudogene | . | 19106 | . | [illegible] |
| 266699 | gene-FABP5P1 | FABP5P11 | fatty acid binding protein 5 pseudogene 11 | pseudogene | . | 19328 | . | [illegible] |
| 266723 | gene-PSMC6P1 | PSMC6P1 | proteasome 26S subunit, ATPase, 6 pseudogene 1 | pseudogene | . | 20634 | . | [illegible] |
| 266724 | gene-HSPA9P1 | HSPA9P1 | heat shock protein family A (Hsp70) member 9 pseudogene 1 | pseudogene | . | 24915 | . | [illegible] |
| 266727 | NM_153487,XI | MDGA1 | MAM domain containing glycosylphosphatidylinositol anchor 1 | protein_coding | NP_705691.1;XP_006715119.1;XP_016866223.1 | 19267 | 609626 | [illegible] |
| 266740 | NM_00132140 | MAGEA2B | MAGE family member A2B | protein_coding | NP_001308329.1;NP_001308330.1;NP_001308331.1;NP_001308332.1;NI | 19340 | 300549 | [illegible] |
| 266785 | gene-NUP50P2 | NUP50P2 | nucleoporin 50 pseudogene 2 | pseudogene | . | 29928 | . | [illegible] |
| 266786 | gene-NUP50P3 | NUP50P3 | nucleoporin 50 pseudogene 3 | pseudogene | . | 29927 | . | [illegible] |
| 266812 | NM_153757 | NAP1L5 | nucleosome assembly protein 1 like 5 | protein_coding | NP_715638.1 | 19968 | 612203 | [illegible] |
| 266920 | gene-AK2P1 | AK2P1 | adenylate kinase 2 pseudogene 1 | pseudogene | . | 24037 | . | [illegible] |
| 266971 | NR_002319 | PIPSL | PIP5K1A and PSMD4 like (pseudogene) | transcribed_ps | . | 23733 | . | [illegible] |
| 267002 | NM_00101743 | PGBD2 | piggyBac transposable element derived 2 | protein_coding | NP_001017434.1;NP_733843.1;XP_005270390.1;XP_011542461.1;XP_01 | 19399 | . | [illegible] |
| 267004 | NM_170753 | PGBD3 | piggyBac transposable element derived 3 | protein_coding | NP_736609.2 | 19400 | . | [illegible] |
| 280636 | NM_00132133 | SELENOH | selenoprotein H | protein_coding | NP_001308264.1;NP_734467.1 | 18251 | 607914 | [illegible] |
| 280661 | gene-EIF1AXP1 | EIF1AXP1 | EIF1AX pseudogene 1 | pseudogene | . | 19900 | . | [illegible] |
| 282679 | NM_00136347 | AQP11 | aquaporin 11 | protein_coding | NP_001350406.1;NP_766627.1 | 19940 | 609914 | [illegible] |
| 282763 | NM_00100556 | OR51B5 | olfactory receptor family 51 subfamily B member 5 | protein_coding | NP_001005567.2 | 19599 | . | [illegible] |
| 282809 | NM_00119977 | POC1B | POC1 centriolar protein B | protein_coding | NP_001186706.1;NP_758440.1 | 30836 | 614784 | [illegible] |
| 282890 | NM_00101087 | ZNF311 | zinc finger protein 311 | protein_coding | NP_001010877.2;NP_001337566.1;NP_001369289.1;XP_005249076.1;XP | 13847 | . | [illegible] |
| 282969 | NM_00109848 | FUOM | fucose mutarotase | protein_coding | NP_001091953.1;NP_001288756.1;NP_001288757.1;NP_940874.2;XP_01 | 24733 | 617725 | [illegible] |
| 282973 | NM_00110552 | JAKMIP3 | Janus kinase and microtubule interacting protein 3 | protein_coding | NP_001098991.1;NP_001310015.1;NP_001310016.1;NP_001310017.1;NI | 23523 | 611198 | [illegible] |
| 282974 | NM_00131887 | STK32C | serine/threonine kinase 32C | protein_coding | NP_001305807.1;NP_001305808.1;NP_001305809.1;NP_001305810.1;NI | 21332 | . | [illegible] |
| 282991 | NM_00100134 | BLOC1S2 | biogenesis of lysosomal organelles complex 1 subunit 2 | protein_coding | NP_001001342.1;NP_001269365.1;NP_001269366.1;NP_001269367.1;NI | 20984 | 609768 | [illegible] |
| 282996 | NM_00113436 | RBM20 | RNA binding motif protein 20 | protein_coding | NP_001127835.2;XP_016871592.1;XP_016871593.1 | 27424 | 613171 | [illegible] |
| 282997 | NR_026932 | PDCD4-AS1 | PDCD4 antisense RNA 1 | lncRNA | . | 27425 | . | [illegible] |
| 283008 | NM_00135526 | NUTM2E | NUT family member 2E | protein_coding | NP_001342192.1;XP_024303718.1;XP_024303719.1;XP_024303720.1 | 23448 | . | [illegible] |
| 283011 | NR_026835 | FLJ37201 | tigger transposable element derived 2 pseudogene | transcribed_ps | . | . | . | [illegible] |
| 283014 | gene-EIF2S2P3 | EIF2S2P3 | eukaryotic translation initiation factor 2 subunit 2 beta pseudogen | pseudogene | . | 31664 | . | [illegible] |
| 283050 | NR_015429,NF | ZMIZ1-AS1 | ZMIZ1 antisense RNA 1 | lncRNA | . | 27433 | . | [illegible] |
| 283104 | NR_036485 | SBF2-AS1 | SBF2 antisense RNA 1 | lncRNA | . | 27438 | . | [illegible] |
| 283106 | NM_00125668 | CSNK2A3 | casein kinase 2 alpha 3 | protein_coding | NP_001243615.1 | 2458 | . | [illegible] |
| 283114 | gene-RPS25P1 | RPS25P1 | ribosomal protein S25 pseudogene 1 | pseudogene | . | 34027 | . | [illegible] |
| 283120 | NR_002196,NF | H19 | H19 imprinted maternally expressed transcript | lncRNA | . | 4713 | 103280 | [illegible] |
| 283130 | NM_00107724 | SLC25A45 | solute carrier family 25 member 45 | protein_coding | NP_001070709.3;NP_001265179.3;NP_001265180.2;NP_001287749.2;NI | 27442 | 610825 | [illegible] |
| 283131 | NR_028272,NF | NEAT1 | nuclear paraspeckle assembly transcript 1 | lncRNA | . | 30815 | 612769 | [illegible] |
| 283140 | NR_126004 | LINC02762 | long intergenic non-protein coding RNA 2762 | lncRNA | . | 27443 | . | [illegible] |
| 283149 | NM_00137821 | BCL9L | BCL9 like | protein_coding | NP_001365142.1;NP_001365143.1;NP_872363.1;XP_005271570.1;XP_00 | 23688 | 609004 | [illegible] |
| 283152 | NM_00114501 | CCDC153 | coiled-coil domain containing 153 | protein_coding | NP_001138490.1 | 27446 | . | [illegible] |
| 283209 | NM_173582,XI | PGM2L1 | phosphoglucomutase 2 like 1 | protein_coding | NP_775853.2;XP_011543255.1 | 20898 | 611610 | [illegible] |
| 283219 | NM_00102985 | KCTD21 | potassium channel tetramerization domain containing 21 | protein_coding | NP_001025030.1;XP_005273985.1;XP_006718579.1;XP_006718580.1;XP | 27452 | 618790 | [illegible] |
| 283232 | NM_00104246 | TMEM80 | transmembrane protein 80 | protein_coding | NP_001035928.2;NP_001263182.1;NP_001263203.1;NP_777600.3;XP_00 | 27453 | . | [illegible] |
| 283237 | NM_00131881 | TTC9C | tetratricopeptide repeat domain 9C | protein_coding | NP_001305741.1;NP_001305742.1;NP_001305743.1;NP_001305744.1;NI | 28432 | . | [illegible] |
| 283248 | NM_00136364 | RCOR2 | REST corepressor 2 | protein_coding | NP_001350577.1;NP_775858.2 | 27455 | 616019 | [illegible] |
| 283254 | NM_173811,XI | HARBI1 | harbinger transposase derived 1 | protein_coding | NP_776172.1;XP_011518327.1;XP_011518328.1;XP_011518329.1;XP_01 | 26522 | 615086 | [illegible] |
| 283267 | NR_015451 | LINC00294 | long intergenic non-protein coding RNA 294 | lncRNA | . | 27456 | 612296 | [illegible] |
| 283314 | NR_026947 | C1RL-AS1 | C1RL antisense RNA 1 | lncRNA | . | 27461 | . | [illegible] |
| 283335 | NR_033854 | LOC283335 | uncharacterized LOC283335 | lncRNA | . | . | . | [illegible] |

| | | | | | | | | | | | | | | | | | | | | | | | | | | | | | | | | | |
|---|---|---|---|---|---|---|---|---|---|---|---|---|---|---|---|---|---|---|---|---|---|---|---|---|---|---|---|---|---|---|---|---|---|
| 283337 | NM_00100430 | ZNF740 | zinc finger protein 740 | protein_coding | NP_001004304.1;XP_006719407.1;XP_011536496.1;XP_016874670.1 | 27465 | . | . | . | 3.149671 | 2.916599 | 3.265514 | 4.106548 | 7.329413 | 6.741847 | 7.327719 | 9.389135 | 1.094549 | 742.069039 | 0.153392564 | 0.849693912 | 0.395495284 | 0.998865404 | 9.469006 | 9.598891 | 9.505469 | 9.498890 | 9.514377 | 9.613007 | 820 | 761 | 627 | 778 |
| 283345 | NR_002803 | RPL13P5 | ribosomal protein L13 pseudogene 5 | transcribed_ps | . | 30363 | . | . | . | 1.275973 | 1.288105 | 1.402374 | 1.570120 | 2.969243 | 2.977513 | 3.146886 | 3.589894 | 1.148039 | 27.921205 | 0.582134449 | 0.342141066 | 0.732244734 | 0.998865404 | 4.700498 | 4.899241 | 4.792935 | 4.792606 | 4.804096 | 4.812884 | 30 | 28 | 25 | 29 |
| 283349 | NM_178169,N | RASSF3 | Ras association domain family member 3 | protein_coding | NP_835463.1;XP_006719408.1;XP_011536497.1;XP_016874671.1 | 14271 | 607019 | . | . | 21.887824 | 20.815040 | 22.858522 | 22.086873 | 50.933864 | 48.114894 | 51.293860 | 50.499010 | 1.036432 | 2388.037350 | 0.116986273 | 0.441289859 | 0.659003167 | 0.998865404 | 11.195481 | 11.247274 | 11.230167 | 11.182793 | 11.254223 | 11.216724 | 2780 | 2456 | 2163 | 2226 |
| 283352 | NR_104172,NF | TMEM132D-AS | TMEM132D antisense RNA 1 | lncRNA | . | 27466 | . | . | . | 17.566978 | 20.648926 | 18.869392 | 26.197564 | 40.879082 | 47.730917 | 42.342371 | 59.897617 | 1.160759 | 320.762071 | 0.218331159 | 0.985054905 | 0.324597102 | 0.998865404 | 8.214695 | 8.428138 | 8.235719 | 8.318155 | 8.270077 | 8.442314 | 316 | 345 | 254 | 372 |
| 283358 | NM_173593 | B4GALNT3 | beta-1,4-N-acetyl-galactosaminyltransferase 3 | protein_coding | NP_775864.3 | 24137 | 612220 | . | . | 1.645237 | 1.493170 | 0.964826 | 1.362856 | 3.828535 | 3.451528 | 2.165042 | 3.116008 | -1.362565 | 234.655088 | 0.241445478 | -1.84855593 | 0.064521961 | 0.731823074 | 8.080970 | 7.633224 | 7.969061 | 7.913579 | 7.717008 | 7.861797 | 328 | 277 | 144 | 217 |
| 283373 | NM_173595,XI | ANKRD52 | ankyrin repeat domain 52 | protein_coding | NP_775866.2;XP_011536499.1;XP_016874672.1;XP_016874673.1 | 26614 | . | . | . | 9.899008 | 10.884519 | 10.080308 | 10.788782 | 23.035397 | 25.160050 | 22.619917 | 24.667270 | -1.012169 | 2799.926686 | 0.116906815 | -0.1492602 | 0.881348322 | 0.998865404 | 11.459961 | 11.442332 | 11.417109 | 11.493063 | 11.427632 | 11.463791 | 3093 | 3181 | 2366 | 2667 |
| 283377 | NM_207344 | SPRYD4 | SPRY domain containing 4 | protein_coding | NP_997227.1 | 27468 | . | . | . | 1.807011 | 1.584742 | 2.049159 | 1.865600 | 4.204988 | 3.663202 | 4.598252 | 4.265473 | 1.137403 | 613.031660 | 0.164507953 | 1.129084238 | 0.2588623 | 0.995760118 | 9.163555 | 9.350057 | 9.257606 | 9.166893 | 9.333887 | 9.263578 | 706 | 576 | 597 | 580 |
| 283383 | NM_00133049 | ADGRD1 | adhesion G protein-coupled receptor D1 | protein_coding | NP_001317426.1;NP_942122.2;XP_005253623.1;XP_011536506.1;XP_01 | 19893 | 613639 | . | . | 0.032985 | 0.030388 | 0.039729 | 0.018639 | 0.076756 | 0.070243 | 0.089151 | 0.042617 | -1.121359 | 5.954032 | 1.257226038 | -0.13143838 | 0.895428529 | NA | 2.651489 | 2.491830 | 2.571877 | 2.573760 | 2.580426 | 2.558880 | 7 | 7 | 7 | 3 |
| 283385 | NM_00136368 | MORN3 | MORN repeat containing 3 | protein_coding | NP_001350614.1;NP_776254.3;XP_011536515.1;XP_011536516.1 | 29807 | . | . | . | 0.199058 | 0.202069 | 0.378952 | 0.094290 | 0.463218 | 0.467091 | 0.850358 | 0.215582 | -1.156357 | 23.015706 | 0.687238709 | -0.30496982 | 0.760389141 | NA | 4.622936 | 4.418954 | 4.527961 | 4.523204 | 4.550573 | 4.470961 | 29 | 26 | 27 | 11 |
| 283416 | NR_121682 | LINC01465 | long intergenic non-protein coding RNA 1465 | lncRNA | . | 26364 | . | . | . | 0.096109 | 0.131702 | 0.104376 | 0.061130 | 0.223650 | 0.304435 | 0.234216 | 0.139766 | -1.413385 | 5.393621 | 1.313018945 | -0.38015811 | 0.703828049 | NA | 2.658167 | 2.161867 | 2.432816 | 2.443027 | 2.435244 | 2.423809 | 6 | 8 | 5 | 3 |
| 283417 | NM_173812,XI | DPY19L2 | dpy-19 like 2 | protein_coding | NP_776173.3;XP_006719415.1;XP_011536517.1;XP_011536520.1;XP_01( | 19414 | 613893 | . | . | 0.056933 | 0.147112 | 0.356040 | 0.051549 | 0.132486 | 0.340056 | 0.798943 | 0.117860 | 1.153861 | 27.803954 | 0.652305798 | 0.316521513 | 0.751606699 | 0.998865404 | 4.689302 | 4.897621 | 4.738566 | 4.802791 | 4.836528 | 4.735571 | 21 | 36 | 37 | 16 |
| 283431 | NM_00130313 | GAS2L3 | growth arrest specific 2 like 3 | protein_coding | NP_001290059.1;NP_001290060.1;NP_001350601.1;NP_777602.1;XP_0C | 27475 | 617224 | . | . | 8.319452 | 8.291791 | 9.597331 | 8.384067 | 19.359708 | 19.166845 | 21.536133 | 19.169175 | 1.037400 | 1323.800667 | 0.139667912 | 0.379276314 | 0.704482687 | 0.998865404 | 10.343631 | 10.396820 | 10.319126 | 10.389633 | 10.442221 | 10.320228 | 1427 | 1467 | 1266 | 1163 |
| 283440 | NR_146317,NF | ITFG2-AS1 | ITFG2 antisense RNA 1 | lncRNA | . | 53128 | . | . | . | 0.246470 | 0.059841 | 0.352086 | 0.277750 | 0.573546 | 0.138326 | 0.790071 | 0.635042 | 1.723175 | 4.824331 | 1.399958301 | 0.560780247 | 0.574947354 | NA | 1.820514 | 2.612728 | 2.259945 | 2.244053 | 2.267235 | 2.260393 | 6 | 2 | 6 | 5 |
| 283450 | NM_00110966 | HECTD4 | HECT domain E3 ubiquitin protein ligase 4 | protein_coding | NP_001103132.4 | 26611 | . | . | . | 1.748283 | 1.773857 | 1.941357 | 2.116929 | 4.068326 | 4.100350 | 4.356348 | 4.840107 | 1.136336 | 931.438337 | 0.140414787 | 1.313178867 | 0.189122683 | 0.960264078 | 9.768315 | 9.952447 | 9.812076 | 9.816739 | 9.878675 | 9.934409 | 1001 | 944 | 829 | 964 |
| 283455 | NM_173598,XI | KSR2 | kinase suppressor of ras 2 | protein_coding | NP_775869.4;XP_011536526.1;XP_011536527.1;XP_011536528.1;XP_01 | 18610 | 610737 | . | . | 0.030703 | 0.032407 | 0.019114 | 0.023643 | 0.071446 | 0.074909 | 0.042891 | 0.054057 | -1.450036 | 15.549224 | 0.78044181 | -0.68690426 | 0.492143061 | NA | 4.202828 | 3.664848 | 3.971910 | 3.980886 | 3.946832 | 3.956652 | 20 | 21 | 10 | 13 |
| 283459 | NM_176818,N | GATC | glutamyl-tRNA amidotransferase subunit C | protein_coding | NP_789788.1 | 25068 | 617210 | . | . | 8.002808 | 7.579817 | 7.867893 | 8.216042 | 18.622863 | 17.521085 | 17.655324 | 18.785003 | 1.016718 | 1048.683250 | 0.136141152 | 0.175697491 | 0.860531606 | 0.998865404 | 10.022337 | 10.046290 | 10.050320 | 10.005278 | 10.027798 | 10.052636 | 1234 | 1088 | 904 | 1008 |
| 283464 | NM_00109965 | GXYLT1 | glucoside xylosyltransferase 1 | protein_coding | NP_001093120.1;NP_775872.1;XP_016874700.1 | 27482 | 613321 | . | . | 7.873058 | 6.126935 | 6.243983 | 4.069411 | 18.320929 | 14.162682 | 14.011318 | 9.304225 | -1.376957 | 1411.626846 | 0.362837738 | -1.27187175 | 0.203418692 | 0.968254058 | 10.675416 | 10.214158 | 10.663373 | 10.456736 | 10.468026 | 10.145151 | 2125 | 1538 | 1258 | 871 |
| 283487 | NR_027701 | PRECSIT | p53 regulated carcinoma associated Stat3 activating long intergen | lncRNA | . | 27492 | . | . | . | 0.039804 | 0.027508 | 0.073214 | 0.021005 | 0.092626 | 0.063585 | 0.164290 | 0.048026 | 1.341347 | 8.365306 | 1.104338578 | 0.383652395 | 0.7012361 | NA | 2.832372 | 3.264261 | 3.050556 | 3.036053 | 3.077260 | 3.030682 | 10 | 6 | 13 | 4 |
| 283489 | NM_00116414 | CHAMP1 | chromosome alignment maintaining phosphoprotein 1 | protein_coding | NP_001157616.1;NP_001157617.1;NP_115812.1 | 20311 | 616327 | . | . | 12.811396 | 13.903145 | 14.643852 | 14.525507 | 29.812647 | 32.137742 | 32.860378 | 33.210848 | 1.077475 | 1651.801534 | 0.124039392 | 0.867905375 | 0.385446136 | 0.998865404 | 10.635035 | 10.742609 | 10.631573 | 10.686861 | 10.725719 | 10.709869 | 1766 | 1777 | 1505 | 1590 |
| 283507 | NR_003365 | SUGT1P3 | SUGT1 pseudogene 3 | transcribed_ps | . | 20513 | . | . | . | 0.049854 | 0.038397 | 0.024160 | 0.029037 | 0.116011 | 0.088756 | 0.054215 | 0.066390 | -1.620137 | 1.451754 | 2.440017378 | -0.28529139 | 0.775420906 | NA | 0.844134 | 0.148375 | 0.548030 | 0.548560 | 0.545314 | 0.544903 | 2 | 2 | 1 | 1 |
| 283521 | NM_00135100 | TMEM272 | transmembrane protein 272 | protein_coding | NP_001337932.1;NP_001337934.1;NP_001337935.1 | 26737 | . | . | . | 1.789460 | 2.171805 | 2.437729 | 1.901795 | 4.164146 | 5.020224 | 5.470192 | 4.348229 | 1.083127 | 268.906947 | 0.22784774 | 0.505609572 | 0.61313076 | 0.998865404 | 8.011976 | 8.127633 | 7.995680 | 8.093035 | 8.150367 | 8.018768 | 269 | 305 | 274 | 229 |
| 283537 | NM_00113591 | SLC46A3 | solute carrier family 46 member 3 | protein_coding | NP_001129391.1;NP_001334889.1;NP_861450.1;XP_005266418.1 | 27501 | 616764 | . | . | 4.960047 | 4.626923 | 4.967004 | 3.857218 | 11.542232 | 10.695337 | 11.145813 | 8.819070 | -1.115264 | 458.172391 | 0.193023391 | -0.81536828 | 0.414861536 | 0.998865404 | 8.915656 | 8.759620 | 8.902410 | 8.835981 | 8.885322 | 8.712225 | 585 | 494 | 431 | 348 |
| 283551 | NR_126496,NF | LINC01588 | long intergenic non-protein coding RNA 1588 | lncRNA | . | 27503 | . | . | . | 0.816258 | 0.731105 | 0.575318 | 0.654646 | 1.899467 | 1.689982 | 1.290999 | 1.496772 | -1.032506 | 59.333640 | 0.421745463 | -0.10942687 | 0.912863922 | 0.998865404 | 5.915110 | 5.866029 | 5.861968 | 5.921569 | 5.854063 | 5.915957 | 60 | 74 | 43 | 63 |
| 283554 | NM_00109965 | GPR137C | G protein-coupled receptor 137C | protein_coding | NP_001093122.1;NP_001340290.1;XP_024305309.1 | 25445 | . | . | . | 0.319123 | 0.422860 | 0.286744 | 0.175127 | 0.742611 | 0.977460 | 0.643446 | 0.400407 | -1.373812 | 52.309123 | 0.453119006 | -1.01117974 | 0.311930409 | 0.998865404 | 5.918518 | 5.463777 | 5.741290 | 5.732380 | 5.716836 | 5.634566 | 71 | 64 | 48 | 31 |
| 283556 | gene-UBE2L1 | UBE2L1 | ubiquitin conjugating enzyme E2 L1 (pseudogene) | pseudogene | . | 12486 | 600012 | . | . | 9.203346 | 9.161076 | 9.472075 | 12.379803 | 21.416565 | 21.176237 | 21.255062 | 28.304947 | 1.173200 | 233.192979 | 0.232950936 | 0.989261526 | 0.322535203 | 0.998865404 | 7.746156 | 7.975500 | 7.824864 | 7.818369 | 7.833227 | 7.960679 | 249 | 230 | 191 | 266 |
| 283578 | NM_00134613 | TMED8 | transmembrane p24 trafficking protein family member 8 | protein_coding | NP_001333060.1;NP_001333062.1;NP_001333063.1;NP_998766.1;XP_01 | 18633 | . | . | . | 4.323588 | 4.387792 | 4.868684 | 4.319460 | 10.061167 | 10.142580 | 10.925186 | 9.875931 | 1.040000 | 1083.104511 | 0.139304559 | 0.406188784 | 0.684603895 | 0.998865404 | 10.052202 | 10.109150 | 10.063237 | 10.067596 | 10.141881 | 10.045083 | 1219 | 1149 | 1023 | 968 |
| 283596 | NR_001459,NF | SNHG10 | small nucleolar RNA host gene 10 | lncRNA | . | 27510 | . | . | . | 4.396457 | 4.650476 | 4.914678 | 3.936243 | 10.230737 | 10.749784 | 11.028395 | 8.999751 | -1.159278 | 214.810380 | 0.239881036 | -0.8888861 | 0.374064295 | 0.998865404 | 7.849029 | 7.637030 | 7.770577 | 7.786942 | 7.764608 | 7.652321 | 261 | 253 | 195 | 163 |
| 283603 | gene-RPL26P4 | RPL26P4 | ribosomal protein L26 pseudogene 4 | pseudogene | . | 19807 | . | . | . | 0.314402 | 0.411380 | 0.076696 | 0.226178 | 0.731627 | 0.950923 | 0.172103 | 0.517130 | -1.774075 | 4.566121 | 1.430744364 | -0.57806756 | 0.563218508 | NA | 2.548173 | 1.715150 | 2.204250 | 2.210104 | 2.191236 | 2.199876 | 6 | 7 | 2 | 4 |
| 283624 | NR_038970,NF | LINC00641 | long intergenic non-protein coding RNA 641 | lncRNA | . | 27511 | . | . | . | 1.143629 | 1.343105 | 1.294655 | 0.976857 | 2.661272 | 3.104648 | 2.905167 | 2.233466 | -1.062904 | 192.324049 | 0.259304227 | -0.33941411 | 0.734297792 | 0.998865404 | 7.630149 | 7.543336 | 7.568533 | 7.625015 | 7.638868 | 7.496562 | 214 | 227 | 189 | 146 |
| 283635 | NM_00107951 | FAM177A1 | family with sequence similarity 177 member A1 | protein_coding | NP_001072987.1;NP_001275951.1;NP_775878.2 | 19829 | . | . | . | 8.200577 | 8.113163 | 7.684164 | 7.108878 | 19.083081 | 18.753940 | 17.243042 | 16.253604 | -1.122004 | 737.989431 | 0.150207397 | -1.10565847 | 0.268874372 | 0.998865404 | 9.607922 | 9.442236 | 9.571847 | 9.559906 | 9.516063 | 9.455280 | 906 | 835 | 632 | 623 |
| 283638 | NM_00111272 | CEP170B | centrosomal protein 170B | protein_coding | NP_001106197.1;NP_055820.2;XP_005267607.1;XP_011534967.1;XP_01 | 20362 | . | . | . | 9.670016 | 10.494725 | 11.897092 | 11.523676 | 22.502527 | 24.259027 | 26.696734 | 26.347513 | 1.174292 | 2227.290709 | 0.116275141 | 1.993469613 | 0.046210048 | 0.654058028 | 11.000516 | 11.232330 | 11.043363 | 11.062191 | 11.194104 | 11.168655 | 2331 | 2237 | 2125 | 2220 |
| 283643 | NM_00113487 | TEDC1 | tubulin epsilon and delta complex 1 | protein_coding | NP_001128347.1;NP_001128348.1;NP_001128349.1;NP_001185912.1;NI | 20127 | . | . | . | 8.303261 | 7.529768 | 8.102060 | 8.448990 | 19.322030 | 17.405395 | 18.180787 | 19.317614 | 1.024193 | 437.280828 | 0.176441725 | 0.195458613 | 0.845033934 | 0.998865404 | 8.754970 | 8.789654 | 8.790892 | 8.737910 | 8.773286 | 8.785417 | 519 | 446 | 381 | 419 |
| 283652 | NM_205850,XI | SLC24A5 | solute carrier family 24 member 5 | protein_coding | NP_995322.1;XP_016877568.1;XP_016877569.1;XP_024305669.1 | 20611 | 609802 | . | . | 87.130036 | 83.456622 | 98.885991 | 82.516539 | 202.755178 | 192.913708 | 221.897308 | 188.664264 | 1.049566 | 5166.877161 | 0.122733085 | 0.568656319 | 0.569589397 | 0.998865404 | 12.299666 | 12.369639 | 12.334740 | 12.291092 | 12.431252 | 12.270014 | 5957 | 5298 | 5042 | 4487 |
| 283659 | NM_173814,XI | PRTG | protogenin | protein_coding | NP_776175.2;XP_011519761.1;XP_011519762.1;XP_016877570.1;XP_01( | 26373 | 613261 | . | . | 1.416293 | 1.721089 | 1.111547 | 1.940195 | 3.295774 | 3.978376 | 2.494278 | 4.436027 | -1.267056 | 315.013386 | 0.205975507 | -1.65786782 | 0.097344149 | 0.813553488 | 8.460372 | 8.117889 | 8.367394 | 8.351514 | 8.175059 | 8.282463 | 411 | 375 | 216 | 288 |
| 283663 | NR_024433 | LINC00926 | long intergenic non-protein coding RNA 926 | lncRNA | . | 27514 | . | . | . | 0.083637 | 0.134502 | 0.145833 | 0.059191 | 0.194626 | 0.310908 | 0.327246 | 0.135332 | -1.075401 | 8.725388 | 1.05292567 | -0.09960326 | 0.920659302 | NA | 3.175403 | 3.073227 | 3.111890 | 3.131110 | 3.132117 | 3.105098 | 8 | 12 | 10 | 5 |
| 283673 | NR_026949 | EWSAT1 | Ewing sarcoma associated transcript 1 | lncRNA | . | 26596 | 616492 | . | . | 0.258566 | 0.131359 | 0.393717 | 0.339612 | 0.601694 | 0.303643 | 0.883490 | 0.776482 | 1.800334 | 22.419882 | 0.673020057 | 1.260385349 | 0.207530384 | NA | 3.998355 | 4.850899 | 4.463105 | 4.420141 | 4.504526 | 4.487464 | 24 | 12 | 27 | 25 |
| 283683 | NR_040057 | LOC283683 | uncharacterized LOC283683 | lncRNA | . | . | . | . | . | 4.498827 | 4.079285 | 4.653713 | 4.882478 | 10.468954 | 9.429450 | 10.442798 | 11.163205 | 1.096189 | 204.067486 | 0.235881582 | 0.561707846 | 0.574315088 | 0.998865404 | 7.605015 | 7.737739 | 7.673643 | 7.626341 | 7.683003 | 7.701150 | 236 | 199 | 182 | 204 |
| 283687 | NR_028330 | ST20-AS1 | ST20 antisense RNA 1 | lncRNA | . | 27521 | . | . | . | 1.433160 | 1.789877 | 1.881428 | 1.712969 | 3.335021 | 4.137381 | 4.221869 | 3.916501 | 1.106918 | 111.352664 | 0.310896154 | 0.471374369 | 0.637373416 | 0.998865404 | 6.724036 | 6.870245 | 6.744351 | 6.811784 | 6.828723 | 6.794563 | 109 | 126 | 107 | 104 |
| 283692 | NR_046096 | CPEB1-AS1 | CPEB1 antisense RNA 1 | lncRNA | . | 27523 | . | . | . | 0.191930 | 0.262168 | 0.273553 | 0.056319 | 0.446629 | 0.606013 | 0.613846 | 0.128767 | -1.357990 | 13.325259 | 0.912980924 | -0.48355126 | 0.62870437 | NA | 3.937099 | 3.502474 | 3.728753 | 3.749042 | 3.751287 | 3.690304 | 15 | 19 | 16 | 4 |
| 283693 | NR_036446 | ACTG1P17 | actin gamma 1 pseudogene 17 | transcribed_ps | . | 51497 | . | . | . | 0.136302 | 0.153562 | 0.237033 | 0.133969 | 0.317180 | 0.354967 | 0.531896 | 0.306305 | 1.242687 | 9.068267 | 1.015311313 | 0.308735998 | 0.757522358 | NA | 3.014059 | 3.330299 | 3.167140 | 3.169643 | 3.188344 | 3.165874 | 9 | 9 | 11 | 7 |
| 283726 | NM_00100822 | SAXO2 | stabilizer of axonemal microtubules 2 | protein_coding | NP_001008227.1;NP_001335628.1;NP_001335629.1;NP_001335630.1;NI | 33727 | . | . | . | 0.710475 | 0.459005 | 1.143969 | 0.878325 | 1.653303 | 1.061011 | 2.567033 | 2.008186 | 1.487777 | 57.303255 | 0.434493428 | 1.319142489 | 0.187121481 | 0.959720529 | 5.523795 | 6.100117 | 5.802641 | 5.762827 | 5.910951 | 5.814934 | 59 | 44 | 72 | 51 |
| 283742 | NM_173611 | FAM98B | family with sequence similarity 98 member B | protein_coding | NP_775882.2 | 26773 | 616142 | . | . | 5.174802 | 4.543488 | 4.670189 | 3.746728 | 12.041978 | 10.502476 | 10.479771 | 8.566448 | -1.169882 | 620.817827 | 0.174494795 | -1.29724997 | 0.194545192 | 0.96621494 | 9.386260 | 9.161009 | 9.372842 | 9.276731 | 9.293028 | 9.143940 | 824 | 672 | 555 | 475 |
| 283747 | gene-ELOCP2 | ELOCP2 | elongin C pseudogene 2 | pseudogene | . | 17762 | . | . | . | 1.835574 | 1.790982 | 0.869758 | 2.295316 | 4.271455 | 4.139936 | 1.951711 | 5.247969 | -1.161756 | 35.370657 | 0.572831307 | -0.37761018 | 0.705720202 | 0.998865404 | 5.250766 | 5.029739 | 5.156898 | 5.148707 | 5.074983 | 5.185910 | 45 | 40 | 16 | 44 |
| 283755 | NR_036432 | HERC2P3 | hect domain and RLD 2 pseudogene 3 | transcribed_ps | . | 4871 | . | . | . | 8.126633 | 9.171988 | 11.617077 | 10.168394 | 18.911009 | 21.201462 | 26.068384 | 23.248825 | 1.242969 | 1302.126629 | 0.139838726 | 2.243946495 | 0.024835844 | 0.518622021 | 10.181188 | 10.495080 | 10.213429 | 10.296738 | 10.475600 | 10.363998 | 1262 | 1323 | 1345 | 1256 |
| 283807 | NM_00136780 | FBXL22 | F-box and leucine rich repeat protein 22 | protein_coding | NP_001354736.1;NP_001354737.1;NP_001354738.1;NP_976307.2;XP_0C | 27537 | 609088 | . | . | 0.041704 | 0.058420 | 0.034774 | 0.080182 | 0.097046 | 0.135041 | 0.078032 | 0.183326 | 1.639830 | 8.787833 | 1.059950178 | 0.673188623 | 0.500827298 | NA | 2.735321 | 3.448451 | 3.125119 | 3.106364 | 3.117377 | 3.152389 | 10 | 5 | 6 | 14 |
| 283820 | NM_00100406 | NOMO2 | NODAL modulator 2 | protein_coding | NP_001004060.1;NP_775885.1;XP_016878645.1 | 22652 | 609158 | . | . | 74.008477 | 75.006516 | 66.021166 | 72.572838 | 172.220758 | 173.380913 | 148.149584 | 165.929166 | -1.089806 | 9485.246650 | 0.104637961 | -1.1857183 | 0.235733566 | 0.991416943 | 13.272198 | 13.148072 | 13.243538 | 13.250114 | 13.138363 | 13.207626 | 11348 | 10716 | 7570 | 8859 |
| 283869 | NM_00109945 | NPW | neuropeptide W | protein_coding | NP_001092926.2 | 30509 | 607997 | . | . | 0.061782 | 0.343824 | 0.327177 | 0.215207 | 0.143769 | 0.794765 | 0.734175 | 0.492044 | 1.332059 | 5.865186 | 1.310607899 | 0.315623328 | 0.752288429 | NA | 2.333078 | 2.742346 | 2.517622 | 2.548901 | 2.546968 | 2.535001 | 2 | 9 | 7 | 5 |
| 283870 | NM_182563 | BRICD5 | BRICHOS domain containing 5 | protein_coding | NP_872369.2 | 28309 | . | . | . | 1.003726 | 0.883314 | 1.490384 | 1.266461 | 2.335712 | 2.041820 | 3.344378 | 2.895612 | 1.437341 | 37.640539 | 0.508738945 | 1.028822381 | 0.303563148 | 0.998865404 | 4.947897 | 5.473012 | 5.205730 | 5.188494 | 5.264150 | 5.235687 | 38 | 31 | 42 | 38 |
| 283871 | NM_00104237 | PGP | phosphoglycolate phosphatase | protein_coding | NP_001035830.1 | 8909 | 172280 | . | . | 10.054364 | 10.885708 | 11.884220 | 12.807542 | 23.396915 | 25.162802 | 26.667845 | 29.282923 | 1.163107 | 1124.131410 | 0.135495396 | 1.608790809 | 0.107662091 | 0.835804188 | 10.021675 | 10.239315 | 10.048654 | 10.100107 | 10.162157 | 10.211444 | 1155 | 1161 | 1017 | 1170 |
| 283876 | NR_033904 | LINC00921 | long intergenic non-protein coding RNA 921 | lncRNA | . | 26830 | . | . | . | 0.220199 | 0.149223 | 0.433036 | 0.280498 | 0.512413 | 0.344936 | 0.971720 | 0.641325 | 1.853302 | 21.024780 | 0.691301299 | 1.287568313 | 0.197896275 | NA | 3.878555 | 4.773010 | 4.357983 | 4.334561 | 4.423298 | 4.375376 | 20 | 13 | 29 | 20 |
| 283887 | NR_135196,NF | LINC02175 | long intergenic non-protein coding RNA 2175 | lncRNA | . | 27550 | . | . | . | 0.151504 | 0.231572 | 0.224103 | 0.244298 | 0.352556 | 0.535290 | 0.502880 | 0.558558 | 1.180703 | 13.278453 | 0.843526847 | 0.284100435 | 0.776333413 | NA | 3.607672 | 3.844638 | 3.708910 | 3.732904 | 3.728183 | 3.733108 | 11 | 16 | 12 | 14 |
| 283899 | NM_00130456 | INO80E | INO80 complex subunit E | protein_coding | NP_001291491.1;NP_001291492.1;NP_775889.1;XP_011544111.1;XP_01 | 26905 | . | . | . | 31.070122 | 34.753716 | 33.921171 | 37.555438 | 72.301451 | 80.334772 | 76.118127 | 85.866049 | 1.049801 | 1509.964508 | 0.131141011 | 0.53465954 | 0.592885304 | 0.998865404 | 10.525010 | 10.594745 | 10.498979 | 10.580569 | 10.544732 | 10.610268 | 1607 | 1674 | 1288 | 1511 |
| 283902 | NR_027756 | HCCAT5 | hepatocellular carcinoma associated transcript 5 | lncRNA | . | 48612 | 615613 | . | . | 6.094276 | 6.932276 | 5.815176 | 7.592703 | 14.181630 | 16.024267 | 13.049086 | 17.359812 | 1.015351 | 222.579955 | 0.239354703 | 0.09182135 | 0.926839981 | 0.998865404 | 7.787981 | 7.808307 | 7.762979 | 7.821305 | 7.737768 | 7.857969 | 239 | 253 | 170 | 237 |
| 283922 | NR_026950 | LOC283922 | pyruvate dehydrogenase phosphatase regulatory subunit pseudog | transcribed_ps | . | . | . | . | . | 3.166164 | 3.906951 | 4.737091 | 4.276803 | 7.367794 | 9.031093 | 10.629893 | 9.778402 | 1.254669 | 290.516772 | 0.213495254 | 1.533088188 | 0.125254128 | 0.873614312 | 8.009711 | 8.336750 | 8.061245 | 8.163646 | 8.264553 | 8.202592 | 267 | 306 | 297 | 286 |
| 283927 | NM_00110566 | NUDT7 | nudix hydrolase 7 | protein_coding | NP_001099133.1;NP_001230586.1;NP_001230589.1;NP_001230590.1 | 8054 | 609231 | . | . | 0.793797 | 1.202160 | 0.881320 | 1.413019 | 1.847197 | 2.778847 | 1.977655 | 3.230699 | 1.115473 | 39.050452 | 0.522918449 | 0.301491552 | 0.763039691 | 0.998865404 | 5.208527 | 5.361932 | 5.246305 | 5.299168 | 5.251026 | 5.326930 | 35 | 47 | 27 | 48 |
| 283932 | NR_024348 | FBXL19-AS1 | FBXL19 antisense RNA 1 | lncRNA | . | 27557 | . | . | . | 0.657522 | 0.857768 | 1.044298 | 1.002680 | 1.530080 | 1.982770 | 2.343373 | 2.292508 | 1.330818 | 110.135371 | 0.314054103 | 1.312872217 | 0.18922601 | 0.960264078 | 6.562835 | 6.974207 | 6.687620 | 6.764179 | 6.827222 | 6.810808 | 95 | 115 | 112 | 115 |
| 283951 | NM_00127205 | C16orf91 | chromosome 16 open reading frame 91 | protein_coding | NP_001258980.1 | 27558 | . | . | . | 9.446921 | 7.777985 | 6.840904 | 7.806584 | 21.983372 | 17.979160 | 15.350792 | 17.848827 | -1.193655 | 228.300112 | 0.233863544 | -1.09203102 | 0.274819467 | 0.998865404 | 7.956825 | 7.701465 | 7.925141 | 7.822185 | 7.762633 | 7.816968 | 315 | 241 | 170 | 207 |
| 283955 | gene-PKD1P2 | PKD1P2 | polycystin 1, transient receptor potential channel interacting pseu | pseudogene | . | 30066 | . | . | . | 2.339661 | 2.702257 | 2.846057 | 2.267925 | 5.444488 | 6.246388 | 6.386469 | 5.185340 | 1.001312 | 784.226904 | 0.166192469 | 0.011379983 | 0.990920284 | 0.999408937 | 9.613983 | 9.616271 | 9.562828 | 9.658966 | 9.692411 | 9.526990 | 841 | 903 | 763 | 649 |
| 283970 | NR_003610 | PDXDC2P-NPIP | nuclear pore complex-interacting protein | transcribed_ps | . | . | . | . | . | 7.450424 | 7.551733 | 8.580222 | 7.562080 | 17.337442 | 17.456169 | 19.253769 | 17.289799 | 1.061311 | 1014.920808 | 0.141873401 | 0.60509872 | 0.545113405 | 0.998865404 | 9.943387 | 10.029627 | 9.962106 | 9.965708 | 10.056880 | 9.956121 | 1131 | 1065 | 971 | 913 |
| 283971 | NM_173619,XI | CLEC18C | C-type lectin domain family 18 member C | protein_coding | NP_775890.2;XP_011521338.1;XP_024306007.1;XP_024306008.1;XP_02( | 28538 | 616573 | . | . | 0.035656 | 0.067566 | 0.049262 | 0.029012 | 0.082972 | 0.156183 | 0.110542 | 0.066333 | -1.158570 | 4.208534 | 1.467917282 | -0.14465707 | 0.884981639 | NA | 2.175403 | 1.963455 | 2.069335 | 2.079156 | 2.074568 | 2.068277 | 4 | 6 | 4 | 3 |
| 283981 | NR_027231,NF | LINC00685 | long intergenic non-protein coding RNA 685 | lncRNA | . | 27560 | . | . | . | 0.748439 | 0.588454 | 0.741742 | 0.594382 | 1.741649 | 1.360236 | 1.664446 | 1.358985 | -1.009019 | 46.270544 | 0.465079453 | -0.02785315 | 0.977779276 | 0.998865404 | 5.537347 | 5.526677 | 5.553063 | 5.509531 | 5.546363 | 5.513283 | 60 | 44 | 44 | 39 |
| 283989 | NM_207346 | TSEN54 | tRNA splicing endonuclease subunit 54 | protein_coding | NP_997229.2 | 27561 | 608755 | . | . | 7.602048 | 8.618430 | 7.579002 | 7.924871 | 17.690279 | 19.921886 | 17.007061 | 18.119278 | -1.061305 | 478.861782 | 0.173065908 | -0.49559454 | 0.619898263 | 0.998865404 | 8.945991 | 8.859649 | 8.883930 | 8.957867 | 8.873635 | 8.894658 | 535 | 563 | 397 | 443 |
| 283991 | NM_182565 | UBALD2 | UBA like domain containing 2 | protein_coding | NP_872371.1 | 28438 | . | . | . | 19.651966 | 20.654894 | 21.682177 | 25.161289 | 45.730934 | 47.744713 | 48.654175 | 57.528294 | 1.146178 | 980.943606 | 0.143713513 | 1.369610977 | 0.170808371 | 0.946389416 | 9.836520 | 10.032858 | 9.869750 | 9.899689 | 9.931700 | 10.034472 | 1031 | 1007 | 848 | 1050 |
| 284001 | NM_00131632 | CCDC57 | coiled-coil domain containing 57 | protein_coding | NP_001303250.1;NP_001354757.1;NP_932348.2;XP_011521847.2;XP_01 | 27564 | . | . | . | 3.592577 | 4.773156 | 3.671671 | 3.674144 | 8.360075 | 11.033361 | 8.239118 | 8.400491 | 1.022165 | 785.665730 | 0.160073067 | 0.197585849 | 0.843369107 | 0.998865404 | 9.601893 | 9.633478 | 9.539895 | 9.672090 | 9.665639 | 9.579612 | 811 | 917 | 733 | 699 |
| 284004 | NM_00133054 | HEXD | hexosaminidase D | protein_coding | NP_001317471.1;NP_001356416.1;NP_001356417.1;NP_775891.2;XP_0C | 26307 | 616864 | . | . | 4.916745 | 4.994296 | 7.131783 | 7.513060 | 11.441470 | 11.544540 | 16.003513 | 17.177723 | 1.380448 | 448.300622 | 0.177272161 | 2.623855749 | 0.00869406 | 0.293930247 | 8.556693 | 9.022513 | 8.718298 | 8.672286 | 8.925385 | 8.860712 | 451 | 390 | 475 | 463 |
| 284009 | NR_028335 | LOC284009 | uncharacterized LOC284009 | lncRNA | . | . | . | . | . | 0.326963 | 0.307532 | 0.273367 | 0.228762 | 0.760856 | 0.710874 | 0.613428 | 0.523037 | -1.164080 | 5.414026 | 1.29429889 | -0.16935084 | 0.865520687 | NA | 2.540965 | 2.324312 | 2.441780 | 2.439324 | 2.440117 | 2.433489 | 7 | 6 | 5 | 4 |
| 284018 | NM_00138235 | C17orf58 | chromosome 17 open reading frame 58 | protein_coding | NP_001369288.1;NP_858041.2;NP_858042.2 | 27568 | . | . | . | 5.944473 | 6.416402 | 6.599421 | 5.703791 | 13.833032 | 14.831802 | 14.808908 | 13.041039 | -1.012295 | 277.960002 | 0.211080458 | -0.08352142 | 0.933436946 | 0.998865404 | 8.127297 | 8.110118 | 8.102805 | 8.138842 | 8.149854 | 8.078226 | 311 | 312 | 257 | 241 |
| 284021 | NM_00108542 | MILR1 | mast cell immunoglobulin like receptor 1 | protein_coding | NP_001078892.1;NP_001278245.1;NP_001278246.1;NP_001356422.1;XF | 27570 | . | . | . | 0.267899 | 0.255497 | 0.181094 | 0.164699 | 0.623412 | 0.590591 | 0.406370 | 0.376564 | -1.612045 | 11.620281 | 0.903804319 | -0.76221375 | 0.445932443 | NA | 3.841203 | 3.155137 | 3.558558 | 3.554526 | 3.538952 | 3.526203 | 17 | 15 | 9 | 7 |
| 284023 | NM_00135869 | RNF227 | ring finger protein 227 | protein_coding | NP_001345628.1 | 27571 | . | . | . | 2.278488 | 2.574999 | 4.009853 | 2.531413 | 5.302137 | 5.952226 | 8.997994 | 5.787777 | 1.332201 | 236.640848 | 0.249570539 | 1.658093436 | 0.097298609 | 0.813553488 | 7.664106 | 8.079243 | 7.787529 | 7.813725 | 8.040613 | 7.829705 | 227 | 225 | 283 | 202 |
| 284047 | NR_036647 | CCDC144B | coiled-coil domain containing 144B (pseudogene) | transcribed_ps | . | 26704 | . | . | . | 0.398650 | 0.550185 | 0.391755 | 0.368713 | 0.927675 | 1.271777 | 0.879088 | 0.843019 | -1.258487 | 116.426000 | 0.313112076 | -1.05933453 | 0.289447454 | 0.998865404 | 7.019904 | 6.687534 | 6.838772 | 6.953560 | 6.832565 | 6.809211 | 126 | 162 | 93 | 93 |
| 284058 | NM_00119346 | KANSL1 | KAT8 regulatory NSL complex subunit 1 | protein_coding | NP_001180394.1;NP_001180395.1;NP_001366127.1;NP_056258.1;XP_0C | 24565 | 612452 | . | . | 5.787530 | 5.243503 | 6.359247 | 6.004582 | 13.467817 | 12.120593 | 14.269966 | 13.728762 | 1.161052 | 1104.960065 | 0.140737873 | 1.530732688 | 0.125835471 | 0.874064017 | 9.997792 | 10.213695 | 10.063936 | 10.035882 | 10.214918 | 10.104576 | 1200 | 1082 | 1108 | 1032 |
| 284069 | NM_198475,XI | FAM171A2 | family with sequence similarity 171 member A2 | protein_coding | NP_940877.2;XP_016879979.1 | 30480 | . | . | . | 1.051392 | 1.074456 | 0.868084 | 0.870276 | 2.446631 | 2.483654 | 1.947955 | 1.989782 | -1.225886 | 94.841667 | 0.327478048 | -0.89723685 | 0.369592548 | 0.998865404 | 6.706912 | 6.413048 | 6.597253 | 6.598504 | 6.538071 | 6.538356 | 120 | 113 | 74 | 80 |
| 284071 | NM_00114508 | MEIOC | meiosis specific with coiled-coil domain | protein_coding | NP_001138552.2;XP_005257293.1 | 26670 | 616934 | . | . | 1.239487 | 0.883182 | 0.978849 | 1.029122 | 2.884338 | 2.041514 | 2.196509 | 2.352966 | -1.076590 | 147.575406 | 0.284757567 | -0.37389234 | 0.708484416 | 0.998865404 | 7.257017 | 7.151677 | 7.281226 | 7.149641 | 7.187286 | 7.190997 | 205 | 138 | 123 | 134 |
| 284076 | NM_00113091 | TTLL6 | tubulin tyrosine ligase like 6 | protein_coding | NP_001124390.1;NP_001353243.1;NP_775894.2;XP_016879980.1;XP_01 | 26664 | 610849 | . | . | 0.021911 | 0.089772 | 0.013262 | 0.074076 | 0.050987 | 0.207511 | 0.029761 | 0.169366 | -1.156534 | 7.641198 | 1.18628939 | -0.17686018 | 0.859618217 | NA | 3.039857 | 2.819320 | 2.909303 | 2.953152 | 2.910608 | 2.942261 | 4 | 14 | 3 | 10 |
| 284086 | NM_178170 | NEK8 | NIMA related kinase 8 | protein_coding | NP_835464.1 | 13387 | 609799 | . | . | 1.482721 | 1.396680 | 2.311842 | 1.532184 | 3.450353 | 3.228489 | 5.187706 | 3.503157 | 1.312518 | 187.589205 | 0.270635729 | 1.449688695 | 0.14714535 | 0.90884489 | 7.340870 | 7.735142 | 7.490763 | 7.463449 | 7.688886 | 7.494230 | 193 | 169 | 224 | 158 |
| 284098 | NM_00134675 | PIGW | phosphatidylinositol glycan anchor biosynthesis class W | protein_coding | NP_001333683.1;NP_001333684.1;NP_848612.2 | 23213 | 610275 | . | . | 7.161700 | 6.406213 | 8.240939 | 5.627675 | 16.665569 | 14.808249 | 18.492432 | 12.867010 | 1.015701 | 607.707876 | 0.196342238 | 0.114475995 | 0.908860475 | 0.998865404 | 9.235313 | 9.259057 | 9.275922 | 9.194861 | 9.370238 | 9.106282 | 736 | 611 | 638 | 461 |
| 284106 | NM_00113649 | CISD3 | CDGSH iron sulfur domain 3 | protein_coding | NP_001129970.1 | 27578 | 611933 | . | . | 8.422079 | 8.524703 | 8.381017 | 9.401041 | 19.598524 | 19.705235 | 18.806761 | 21.494364 | 1.035216 | 690.662722 | 0.153744859 | 0.324772998 | 0.745352882 | 0.998865404 | 9.406859 | 9.456391 | 9.418028 | 9.420940 | 9.406674 | 9.478232 | 780 | 734 | 579 | 693 |
| 284114 | NM_00132044 | TMEM102 | transmembrane protein 102 | protein_coding | NP_001307373.1;NP_848613.1 | 26722 | 613936 | . | . | 3.216266 | 4.531474 | 4.027320 | 4.556308 | 7.484382 | 10.474705 | 9.037191 | 10.417456 | 1.099648 | 264.838439 | 0.232445784 | 0.589563694 | 0.555483198 | 0.998865404 | 7.979595 | 8.115159 | 7.932698 | 8.099292 | 8.041449 | 8.093734 | 243 | 317 | 229 | 273 |
| 284119 | NM_012232,XI | CAVIN1 | caveolae associated protein 1 | protein_coding | NP_036364.2;XP_005257299.1 | 9688 | 603198 | . | . | 32.229127 | 33.138237 | 29.357997 | 36.100529 | 74.998505 | 76.600528 | 65.878493 | 82.539561 | -1.019457 | 3578.340550 | 0.126654522 | -0.21949827 | 0.826261924 | 0.998865404 | 11.819028 | 11.790986 | 11.801694 | 11.821784 | 11.708481 | 11.876271 | 4110 | 3946 | 2768 | 3660 |
| 284129 | NM_00116634 | SLC26A11 | solute carrier family 26 member 11 | protein_coding | NP_001159819.1;NP_001159820.1;NP_001159821.1;NP_775897.3;XP_0C | 14471 | 610117 | . | . | 6.072936 | 6.871833 | 7.559793 | 7.147517 | 14.131972 | 15.884553 | 16.963956 | 16.341946 | 1.114360 | 613.869353 | 0.160235856 | 0.974909354 | 0.329605226 | 0.998865404 | 9.181619 | 9.337736 | 9.187332 | 9.257818 | 9.314756 | 9.275710 | 633 | 660 | 579 | 589 |
| 284131 | NM_00116463 | ENDOV | endonuclease V | protein_coding | NP_001158109.1;NP_001158110.1;NP_001339689.1;NP_001339690.1;NI | 26640 | . | . | . | 2.665231 | 2.767245 | 2.880939 | 3.419850 | 6.202100 | 6.396609 | 6.464745 | 7.819078 | 1.084378 | 137.128773 | 0.287219842 | 0.406892465 | 0.68408697 | 0.998865404 | 7.040296 | 7.156154 | 7.097335 | 7.066326 | 7.062373 | 7.159121 | 158 | 136 | 108 | 151 |
| 284161 | NM_00116599 | GDPD1 | glycerophosphodiester phosphodiesterase domain containing 1 | protein_coding | NP_001159465.1;NP_001159466.1;NP_872375.2;XP_016880010.1 | 20883 | 616317 | . | . | 3.928406 | 3.060342 | 4.029623 | 3.000987 | 9.141563 | 7.074118 | 9.042359 | 6.861401 | -1.066175 | 338.165059 | 0.20852162 | -0.44332934 | 0.657527538 | 0.998865404 | 8.446397 | 8.355334 | 8.481395 | 8.348433 | 8.425055 | 8.335501 | 448 | 333 | 308 | 282 |
| 284167 | gene-POLRMTF | POLRMTP1 | RNA polymerase mitochondrial pseudogene 1 | pseudogene | . | 28865 | . | . | . | 0.028710 | 0.014107 | 0.031427 | 0.019017 | 0.066810 | 0.032609 | 0.070521 | 0.043481 | 1.458369 | 3.281789 | 1.655938297 | 0.328729604 | 0.742360076 | NA | 1.413393 | 1.963455 | 1.706670 | 1.698968 | 1.711598 | 1.705370 | 4 | 2 | 4 | 3 |
| 284184 | NM_00108652 | NDUFAF8 | NADH:ubiquinone oxidoreductase complex assembly factor 8 | protein_coding | NP_001079990.1;NP_001340331.1;NP_001340332.1 | 33551 | 618461 | . | . | 33.069272 | 28.110576 | 26.765470 | 39.731095 | 76.953555 | 64.978859 | 60.060939 | 90.840427 | 1.090884 | 620.271138 | 0.190453266 | 0.658939004 | 0.509934938 | 0.998865404 | 9.212958 | 9.337851 | 9.292082 | 9.189698 | 9.175739 | 9.411471 | 736 | 591 | 465 | 712 |
| 284185 | NR_038080 | LINC00482 | long intergenic non-protein coding RNA 482 | lncRNA | . | 26816 | . | . | . | 0.094009 | 0.116118 | 0.056962 | 0.053378 | 0.218764 | 0.268412 | 0.127822 | 0.122042 | -1.865387 | 7.939252 | 1.090650899 | -0.82471366 | 0.409534177 | NA | 3.369732 | 2.470304 | 3.004872 | 3.011845 | 2.987942 | 2.985879 | 11 | 12 | 5 | 5 |
| 284207 | NM_00100443 | METRNL | meteorin like, glial cell differentiation regulator | protein_coding | NP_001004431.1;NP_001350782.1;XP_016880012.1 | 27584 | 616241 | . | . | 4.998110 | 3.766215 | 3.534040 | 4.441418 | 11.630807 | 8.705776 | 7.930284 | 10.154773 | -1.109779 | 222.694911 | 0.247490429 | -0.60718622 | 0.543727336 | 0.998865404 | 7.872163 | 7.721769 | 7.888538 | 7.748227 | 7.721763 | 7.818622 | 308 | 217 | 164 | 219 |
| 284217 | NM_005559 | LAMA1 | laminin subunit alpha 1 | protein_coding | NP_005550.2 | 6481 | 150320 | . | . | 37.767952 | 38.872173 | 36.731506 | 41.102287 | 87.887581 | 89.854767 | 82.424446 | 93.975494 | 1.001383 | 11598.065882 | 0.106006789 | 0.018805935 | 0.98499592 | 0.999197585 | 13.500627 | 13.502565 | 13.492007 | 13.509308 | 13.457298 | 13.544556 | 13211 | 12633 | 9577 | 11436 |
| 284252 | NM_00113620 | KCTD1 | potassium channel tetramerization domain containing 1 | protein_coding | NP_001129677.1;NP_001136202.1;NP_001245150.1;NP_001245151.1;NI | 18249 | 613420 | . | . | 3.238082 | 3.505642 | 3.748296 | 4.319899 | 7.535151 | 8.103448 | 8.411067 | 9.876936 | 1.143554 | 352.631520 | 0.190319019 | 1.016840276 | 0.309229419 | 0.998865404 | 8.362237 | 8.555341 | 8.420958 | 8.420930 | 8.469913 | 8.523948 | 379 | 355 | 312 | 369 |
| 284273 | NM_00130609 | ZADH2 | zinc binding alcohol dehydrogenase domain containing 2 | protein_coding | NP_001293022.1;NP_787103.1;XP_024306934.1 | 28697 | . | . | . | 6.472916 | 5.967400 | 6.736572 | 6.003412 | 15.062740 | 13.793914 | 15.116672 | 13.726086 | 1.025512 | 1437.061285 | 0.132482684 | 0.274334208 | 0.783827799 | 0.998865404 | 10.470411 | 10.507167 | 10.503624 | 10.452481 | 10.543286 | 10.450361 | 1688 | 1480 | 1342 | 1282 |
| 284306 | NM_173631 | ZNF547 | zinc finger protein 547 | protein_coding | NP_775902.2 | 26432 | . | . | . | 0.716750 | 0.885179 | 0.867579 | 0.944869 | 1.667906 | 2.046130 | 1.946821 | 2.160329 | 1.118469 | 73.815669 | 0.371838933 | 0.434394587 | 0.664001928 | 0.998865404 | 6.123580 | 6.283691 | 6.163293 | 6.213527 | 6.206016 | 6.226756 | 72 | 83 | 65 | 76 |
| 284307 | NM_00101087 | ZIK1 | zinc finger protein interacting with K protein 1 | protein_coding | NP_001010879.2;NP_001308074.1;NP_001308075.1;NP_001308076.1;XF | 33104 | . | . | . | 0.068029 | 0.139234 | 0.060532 | 0.224915 | 0.158305 | 0.321846 | 0.135832 | 0.514241 | 1.205721 | 18.437018 | 0.785674411 | 0.343521249 | 0.731206358 | NA | 4.067450 | 4.329714 | 4.161104 | 4.209645 | 4.159292 | 4.240141 | 13 | 24 | 9 | 28 |
| 284309 | NM_00134800 | ZNF776 | zinc finger protein 776 | protein_coding | NP_001334936.1;NP_775903.3 | 26765 | . | . | . | 1.754198 | 1.721660 | 1.983128 | 1.701539 | 4.082091 | 3.979691 | 4.450081 | 3.890366 | 1.044467 | 253.253919 | 0.215784629 | 0.29087976 | 0.771143281 | 0.998865404 | 7.952658 | 8.015538 | 7.956008 | 7.988727 | 8.011912 | 7.976466 | 276 | 276 | 233 | 234 |
| 284323 | NM_00101088 | ZNF780A | zinc finger protein 780A | protein_coding | NP_001010880.2;NP_001136049.1;NP_001136050.1;NP_001136051.1;XF | 27603 | . | . | . | 2.166907 | 1.795706 | 2.383042 | 1.664950 | 5.042481 | 4.150854 | 5.347477 | 3.806708 | 1.017623 | 370.141454 | 0.218691144 | 0.115244834 | 0.908251063 | 0.998865404 | 8.518474 | 8.545267 | 8.543979 | 8.497794 | 8.640603 | 8.409030 | 439 | 380 | 390 | 280 |
| 284325 | NM_00135380 | C19orf54 | chromosome 19 open reading frame 54 | protein_coding | NP_001340734.1;NP_001340735.1;NP_001340736.1;NP_001340737.1;NI | 24758 | . | . | . | 7.454688 | 7.852576 | 7.790584 | 8.390127 | 17.347367 | 18.151578 | 17.481843 | 19.183029 | 1.054031 | 699.649625 | 0.155253582 | 0.488987268 | 0.624850711 | 0.998865404 | 9.412337 | 9.487658 | 9.405217 | 9.456982 | 9.428839 | 9.504547 | 755 | 763 | 590 | 710 |
| 284338 | NM_199285,XI | PRR19 | proline rich 19 | protein_coding | NP_954979.2;XP_005258836.2;XP_005258837.1;XP_006723217.1;XP_01: | 33728 | . | . | . | 0.780639 | 0.732099 | 1.714833 | 1.028049 | 1.816579 | 1.692280 | 3.848035 | 2.350511 | 1.776779 | 51.035653 | 0.462161546 | 1.794316917 | 0.072762599 | 0.759139955 | 5.198253 | 6.030279 | 5.598982 | 5.587618 | 5.754157 | 5.642012 | 44 | 38 | 71 | 46 |
| 284339 | NM_00136691 | TMEM145 | transmembrane protein 145 | protein_coding | NP_001353839.1;NP_775904.2;XP_011525093.1;XP_011525094.1 | 26912 | . | . | . | 1.261701 | 1.228691 | 1.457488 | 1.206186 | 2.936031 | 2.840174 | 3.270561 | 2.757803 | 1.224175 | 81.896586 | 0.361270526 | 0.807732257 | 0.41924473 | 0.998865404 | 6.201686 | 6.494904 | 6.312217 | 6.328912 | 6.418464 | 6.328542 | 82 | 82 | 91 | 71 |
| 284346 | NM_174945,XI | ZNF575 | zinc finger protein 575 | protein_coding | NP_777605.1;XP_005258840.1;XP_011525095.1 | 27606 | . | . | . | 0.838452 | 0.538027 | 0.571214 | 0.604928 | 1.951111 | 1.243675 | 1.281787 | 1.383095 | -1.159415 | 34.017237 | 0.544040277 | -0.39224584 | 0.694876575 | 0.998865404 | 5.190212 | 4.978409 | 5.129743 | 5.067120 | 5.077238 | 5.080765 | 51 | 31 | 27 | 30 |
| 284349 | NM_00129775 | ZNF283 | zinc finger protein 283 | protein_coding | NP_001284681.1;NP_862828.1;XP_005258841.1;XP_005258842.1;XP_00 | 13077 | . | . | . | 1.122282 | 1.119349 | 1.121529 | 0.975614 | 2.611598 | 2.587423 | 2.516680 | 2.230625 | -1.019681 | 324.004688 | 0.222803289 | -0.12619843 | 0.899574851 | 0.998865404 | 8.353060 | 8.326560 | 8.356086 | 8.324741 | 8.428326 | 8.220898 | 387 | 343 | 332 | 244 |
| 284352 | NM_019121,XI | PPP1R37 | protein phosphatase 1 regulatory subunit 37 | protein_coding | NP_061994.1;XP_006723222.1 | 27607 | . | . | . | 25.349637 | 26.173702 | 27.444263 | 30.748072 | 58.989646 | 60.501682 | 61.584131 | 70.301813 | 1.116261 | 2485.741630 | 0.119360027 | 1.329374263 | 0.183724525 | 0.954800306 | 11.198057 | 11.356515 | 11.221720 | 11.243819 | 11.276551 | 11.364820 | 2668 | 2570 | 2157 | 2591 |
| 284358 | NM_00113091 | MAMSTR | MEF2 activating motif and SAP domain containing transcriptional r | protein_coding | NP_001124387.1;NP_001284682.1;NP_872380.1;XP_011525109.1;XP_01 | 26689 | 610349 | . | . | 0.413223 | 0.381895 | 0.345159 | 0.442162 | 0.961588 | 0.882767 | 0.774526 | 1.010953 | -1.159112 | 30.824133 | 0.570483285 | -0.37340358 | 0.708848096 | 0.998865404 | 5.049783 | 4.834143 | 4.963719 | 4.950424 | 4.912181 | 4.964805 | 40 | 34 | 19 | 33 |
| 284361 | NM_175063,N | EMC10 | ER membrane protein complex subunit 10 | protein_coding | NP_778233.4;NP_996261.1;XP_006723226.1;XP_011525120.1 | 27609 | 614545 | . | . | 4.295660 | 4.005115 | 3.634184 | 4.065878 | 9.996178 | 9.258003 | 8.155001 | 9.296147 | -1.096976 | 1160.955468 | 0.138046299 | -0.96730263 | 0.333392771 | 0.998865404 | 10.246401 | 10.112696 | 10.244622 | 10.183296 | 10.103120 | 10.185340 | 1455 | 1258 | 909 | 1096 |
| 284370 | NM_00119932 | ZNF615 | zinc finger protein 615 | protein_coding | NP_001186253.1;NP_001308246.1;NP_001308247.1;NP_001308248.1;NI | 24740 | . | . | . | 0.807490 | 0.847175 | 0.983459 | 0.822972 | 1.879061 | 1.958283 | 2.206852 | 1.881628 | 1.106643 | 111.513575 | 0.308569901 | 0.473765361 | 0.635667238 | 0.998865404 | 6.725585 | 6.872811 | 6.778799 | 6.783507 | 6.846566 | 6.780966 | 121 | 115 | 112 | 99 |
| 284371 | NM_00113649 | ZNF841 | zinc finger protein 841 | protein_coding | NP_001129971.1;NP_001308278.1;NP_001339227.1;NP_001339228.1;NI | 27611 | . | . | . | 2.507791 | 2.561555 | 3.007228 | 2.175202 | 5.835734 | 5.921149 | 6.748130 | 4.973341 | 1.003720 | 315.492361 | 0.216689622 | 0.024721352 | 0.980277224 | 0.998865404 | 8.298126 | 8.304789 | 8.294662 | 8.299070 | 8.382282 | 8.208372 | 361 | 341 | 319 | 249 |
| 284385 | NR_122038 | ZNF561-AS1 | ZNF561 antisense RNA 1 (head to head) | lncRNA | . | 27613 | . | . | . | 0.458605 | 0.696190 | 0.490975 | 0.749899 | 1.067193 | 1.609275 | 1.101733 | 1.714555 | 1.057877 | 34.778629 | 0.546616915 | 0.14849992 | 0.881948249 | 0.998865404 | 5.081051 | 5.158176 | 5.083307 | 5.139734 | 5.092548 | 5.146152 | 31 | 44 | 25 | 40 |
| 284390 | NM_00101275 | ZNF763 | zinc finger protein 763 | protein_coding | NP_001012771.1;NP_001354101.1;NP_001354102.1;NP_001354103.1 | 27614 | . | . | . | 0.019077 | 0.044419 | 0.048402 | 0.043576 | 0.044392 | 0.102677 | 0.108613 | 0.099631 | 1.396326 | 3.809293 | 1.552872665 | 0.310158077 | 0.756440749 | NA | 1.671431 | 2.148375 | 1.908976 | 1.922846 | 1.922466 | 1.921035 | 2 | 5 | 4 | 4 |
| 284393 | gene-RPL10P1( | RPL10P16 | ribosomal protein L10 pseudogene 16 | pseudogene | . | 36882 | . | . | . | 4.117768 | 4.674871 | 3.342678 | 4.777880 | 9.582215 | 10.806174 | 7.500872 | 10.924054 | -1.095511 | 99.169943 | 0.335970784 | -0.39171215 | 0.69527091 | 0.998865404 | 6.697302 | 6.563247 | 6.624330 | 6.664184 | 6.565815 | 6.665272 | 112 | 119 | 68 | 104 |
| 284402 | NM_00102559 | SCGB2B2 | secretoglobin family 2B member 2 | protein_coding | NP_001020762.1 | 27616 | 615063 | . | . | 0.046625 | 0.108336 | 0.100382 | 0.080521 | 0.108497 | 0.250423 | 0.225254 | 0.184103 | 1.158197 | 8.803674 | 1.04131996 | 0.203472614 | 0.838765649 | NA | 3.029593 | 3.239024 | 3.113781 | 3.141231 | 3.137321 | 3.129458 | 6 | 12 | 9 | 8 |
| 284403 | NM_00108396 | WDR62 | WD repeat domain 62 | protein_coding | NP_001077430.1;NP_775907.4;XP_005258866.1;XP_011525139.1;XP_01 | 24502 | 613583 | . | . | 13.446705 | 14.576899 | 13.683694 | 13.268942 | 31.291038 | 33.695156 | 30.705811 | 30.337860 | -1.025165 | 1911.002392 | 0.122529605 | -0.29263563 | 0.769800677 | 0.998865404 | 10.917932 | 10.882073 | 10.877726 | 10.940477 | 10.910753 | 10.868640 | 2140 | 2170 | 1685 | 1722 |
| 284406 | NM_00132191 | ZFP82 | ZFP82 zinc finger protein | protein_coding | NP_001308846.1;NP_001308847.1;NP_001308848.1;NP_597723.1 | 28682 | . | . | . | 0.338988 | 0.268342 | 0.329203 | 0.406442 | 0.788838 | 0.620284 | 0.738721 | 0.929281 | 1.253180 | 61.445070 | 0.406640436 | 0.800691409 | 0.423310317 | 0.998865404 | 5.769484 | 6.094680 | 5.927482 | 5.894467 | 5.934442 | 5.984719 | 68 | 54 | 53 | 71 |
| 284439 | NM_00132154 | SLC25A42 | solute carrier family 25 member 42 | protein_coding | NP_001308473.1;NP_848621.2;XP_005259918.1;XP_011526239.1;XP_01 | 28380 | 610823 | . | . | 1.174889 | 1.214502 | 1.122995 | 1.290518 | 2.734016 | 2.807376 | 2.519969 | 2.950617 | 1.001262 | 124.780018 | 0.290507634 | 0.006262046 | 0.995003643 | 0.999862634 | 6.962738 | 6.963748 | 6.953844 | 6.971845 | 6.945565 | 6.980711 | 140 | 138 | 103 | 123 |
| 284440 | NR_026956 | LINC00663 | long intergenic non-protein coding RNA 663 | lncRNA | . | 28609 | 617117 | . | . | 0.110806 | 0.061055 | 0.072403 | 0.163615 | 0.257851 | 0.141131 | 0.162471 | 0.374086 | 1.393804 | 7.967005 | 1.106547812 | 0.43290302 | 0.665085226 | NA | 2.735321 | 3.213338 | 2.991783 | 2.972989 | 2.978762 | 3.009626 | 10 | 5 | 5 | 12 |
| 284443 | NM_00107667 | ZNF493 | zinc finger protein 493 | protein_coding | NP_001070146.1;NP_663299.2;NP_787106.4 | 23708 | . | . | . | 0.916976 | 0.545809 | 1.024619 | 0.560477 | 2.133840 | 1.261662 | 2.299214 | 1.281465 | -1.205014 | 87.254167 | 0.430464447 | -0.62502319 | 0.53195584 | 0.998865404 | 6.573287 | 6.308923 | 6.524054 | 6.382874 | 6.506325 | 6.288843 | 135 | 79 | 98 | 43 |
| 284454 | NR_036515 | LOC284454 | uncharacterized LOC284454 | lncRNA | . | . | . | . | . | 1.330907 | 1.575333 | 1.825868 | 1.208776 | 3.097074 | 3.641452 | 4.097195 | 2.763722 | 1.030617 | 389.420458 | 0.221190022 | 0.196698272 | 0.844063662 | 0.998865404 | 8.582786 | 8.627241 | 8.540614 | 8.635681 | 8.721612 | 8.477901 | 406 | 447 | 415 | 294 |
| 284459 | NM_00132976 | ZNF875 | zinc finger protein 875 | protein_coding | NP_001316690.1;NP_001316691.1;NP_001316692.1;NP_001316693.1;NI | 4928 | 165250 | . | . | 3.595743 | 3.522764 | 3.565129 | 3.539381 | 8.367440 | 8.143021 | 8.000046 | 8.092375 | -1.088074 | 337.929497 | 0.200212281 | -0.60823666 | 0.543030524 | 0.998865404 | 8.459655 | 8.338979 | 8.439168 | 8.405951 | 8.427334 | 8.320877 | 417 | 369 | 309 | 274 |
| 284467 | NM_00100444 | TAFA3 | TAFA chemokine like family member 3 | protein_coding | NP_001004440.1;NP_877436.1;XP_005270823.1;XP_006710645.1 | 21590 | 617497 | . | . | 36.010813 | 38.491319 | 36.534161 | 40.339714 | 83.798647 | 88.974406 | 81.981601 | 92.231958 | 1.019368 | 1626.458623 | 0.126809328 | 0.218241693 | 0.827240801 | 0.998865404 | 10.653772 | 10.681136 | 10.636233 | 10.682576 | 10.639114 | 10.708719 | 1800 | 1790 | 1364 | 1608 |
| 284561 | XR_00173773( | LINC02806 | long intergenic non-protein coding RNA 2806 | lncRNA | . | 27628 | . | . | . | 0.626391 | 0.642943 | 0.304134 | 0.665967 | 1.457637 | 1.486192 | 0.682469 | 1.522654 | -1.324033 | 171.451805 | 0.321675359 | -1.25884329 | 0.20808694 | 0.970482251 | 7.611243 | 7.203338 | 7.463240 | 7.471498 | 7.201697 | 7.479814 | 223 | 213 | 81 | 189 |
| 284565 | NM_00117075 | NBPF15 | NBPF member 15 | protein_coding | NP_001164226.1;NP_775909.2 | 28791 | 614005 | . | . | 9.510128 | 9.496821 | 11.967702 | 10.452179 | 22.130459 | 21.952326 | 26.855174 | 23.897665 | 1.153886 | 1457.977979 | 0.142428587 | 1.449850541 | 0.147100202 | 0.90884489 | 10.402648 | 10.609454 | 10.423165 | 10.475630 | 10.639540 | 10.474130 | 1505 | 1511 | 1502 | 1310 |
| 284581 | NR_046097,NF | LOC284581 | uncharacterized LOC284581 | lncRNA | . | . | . | . | . | 1.779568 | 1.569624 | 2.089936 | 1.244547 | 4.141130 | 3.628256 | 4.689755 | 2.845507 | -1.012290 | 133.082843 | 0.31543414 | -0.05586768 | 0.955447221 | 0.998865404 | 7.063584 | 7.048739 | 7.075161 | 7.026984 | 7.132802 | 6.953010 | 164 | 135 | 144 | 93 |
| 284600 | XR_00295852( | LOC284600 | uncharacterized LOC284600 | lncRNA | . | . | . | . | . | 0.045913 | 0.088554 | 0.043661 | 0.065640 | 0.106841 | 0.204695 | 0.097975 | 0.150078 | -1.199790 | 7.870810 | 1.09717914 | -0.23950663 | 0.810712754 | NA | 3.104339 | 2.836248 | 2.969231 | 2.992421 | 2.968394 | 2.980548 | 7 | 12 | 5 | 8 |
| 284611 | NM_00101088 | FAM102B | family with sequence similarity 102 member B | protein_coding | NP_001010883.2 | 27637 | . | . | . | 15.068141 | 13.599968 | 13.768321 | 13.130177 | 35.064182 | 31.436935 | 30.895714 | 30.020590 | -1.081065 | 2417.921069 | 0.121280029 | -0.92722391 | 0.353810294 | 0.998865404 | 11.294569 | 11.182352 | 11.313997 | 11.223435 | 11.229667 | 11.183688 | 3053 | 2560 | 2079 | 2116 |
| 284612 | NM_00104070 | SYPL2 | synaptophysin like 2 | protein_coding | NP_001035799.1;XP_011539585.1;XP_011539586.1;XP_011539587.1 | 27638 | . | . | . | 0.361893 | 0.608932 | 0.764212 | 0.533849 | 0.842140 | 1.407574 | 1.714869 | 1.220583 | 1.320896 | 66.081930 | 0.422885143 | 0.949469134 | 0.342382063 | 0.998865404 | 5.831789 | 6.232802 | 5.943805 | 6.050349 | 6.105066 | 6.017179 | 49 | 77 | 77 | 58 |
| 284613 | NM_00113440 | CYB561D1 | cytochrome b561 family member D1 | protein_coding | NP_001127872.1;NP_001127874.1;NP_001127875.1;NP_001127876.1;NI | 26804 | . | . | . | 2.999708 | 3.262434 | 3.250770 | 3.273202 | 6.980443 | 7.541263 | 7.294632 | 7.483787 | 1.021042 | 497.932455 | 0.168007177 | 0.178817365 | 0.85808111 | 0.998865404 | 8.944796 | 8.974662 | 8.932007 | 8.973327 | 8.968194 | 8.963840 | 549 | 549 | 439 | 470 |
| 284615 | NM_00103988 | ANKRD34A | ankyrin repeat domain 34A | protein_coding | NP_001034977.1 | 27639 | . | . | . | 1.329620 | 1.588990 | 1.687018 | 1.655113 | 3.094081 | 3.673021 | 3.785619 | 3.784219 | 1.131555 | 176.294143 | 0.253531558 | 0.703293639 | 0.481872774 | 0.998865404 | 7.370240 | 7.547971 | 7.395324 | 7.467555 | 7.490300 | 7.478536 | 174 | 194 | 165 | 173 |
| 284618 | NR_145424,NF | RUSC1-AS1 | RUSC1 antisense RNA 1 | lncRNA | . | 26680 | . | . | . | 0.744916 | 0.822070 | 1.057404 | 0.932066 | 1.733450 | 1.900251 | 2.372783 | 2.131058 | 1.247831 | 48.948612 | 0.446537929 | 0.715332406 | 0.474403669 | 0.998865404 | 5.444530 | 5.764191 | 5.579619 | 5.594363 | 5.638167 | 5.616039 | 48 | 49 | 50 | 48 |
| 284649 | NR_040063 | SMG7-AS1 | SMG7 antisense RNA 1 | lncRNA | . | 24518 | . | . | . | 0.138799 | 0.182641 | 0.209632 | 0.074608 | 0.322991 | 0.422183 | 0.470408 | 0.170583 | -1.164414 | 12.154326 | 0.908447368 | -0.24173626 | 0.808984529 | NA | 3.706954 | 3.491830 | 3.595225 | 3.611125 | 3.615795 | 3.573902 | 13 | 16 | 14 | 6 |
| 284672 | gene-PTGES3P: | PTGES3P1 | prostaglandin E synthase 3 pseudogene 1 | pseudogene | . | 43824 | . | . | . | 0.229911 | 0.095262 | 0.135545 | 0.031920 | 0.535012 | 0.220203 | 0.304159 | 0.072982 | -1.776656 | 6.239162 | 1.281458894 | -0.64704736 | 0.517601292 | NA | 2.993083 | 2.175234 | 2.669596 | 2.638808 | 2.649113 | 2.627830 | 13 | 5 | 6 | 2 |
| 284677 | NR_147122,NF | SNHG28 | small nucleolar RNA host gene 28 | lncRNA | . | 27647 | . | . | . | 0.199360 | 0.208909 | 0.121957 | 0.172555 | 0.463918 | 0.482902 | 0.273668 | 0.394527 | -1.384599 | 13.129494 | 0.84826286 | -0.55344699 | 0.579957371 | NA | 3.931597 | 3.459419 | 3.727430 | 3.731697 | 3.703392 | 3.718659 | 17 | 17 | 8 | 12 |
| 284685 | gene-LOC2846 | LOC284685 | EWS RNA binding protein 1 pseudogene | pseudogene | . | . | . | . | . | 0.053511 | 0.073204 | 0.077581 | 0.093370 | 0.124522 | 0.169213 | 0.174090 | 0.213480 | 1.228632 | 5.513138 | 1.280223022 | 0.232032061 | 0.816513112 | NA | 2.307819 | 2.602870 | 2.452722 | 2.458507 | 2.459322 | 2.462261 | 5 | 6 | 5 | 6 |
| 284695 | NM_00132018 | ZNF326 | zinc finger protein 326 | protein_coding | NP_001307114.1;NP_861446.2;NP_892020.1;NP_892021.1;XP_00527083 | 14104 | 614601 | . | . | 4.000093 | 3.799341 | 4.167454 | 3.165677 | 9.308380 | 8.782349 | 9.351645 | 7.237944 | -1.081467 | 1050.411604 | 0.145581173 | -0.77612688 | 0.437674074 | 0.998865404 | 10.091857 | 9.979432 | 10.071264 | 10.056451 | 10.070597 | 9.937543 | 1269 | 1166 | 958 | 860 |
| 284697 | NM_00137613 | BTBD8 | BTB domain containing 8 | protein_coding | NP_001363060.1;NP_056052.3;XP_016856289.1 | 21019 | 617945 | . | . | 0.278138 | 0.315449 | 0.277858 | 0.203861 | 0.647240 | 0.729175 | 0.623506 | 0.466103 | -1.260034 | 51.547697 | 0.443783525 | -0.75140733 | 0.452407553 | 0.998865404 | 5.844134 | 5.512525 | 5.702482 | 5.715921 | 5.692147 | 5.638550 | 64 | 64 | 46 | 36 |
| 284716 | NM_173642,XI | RIMKLA | ribosomal modification protein rimK like family member A | protein_coding | NP_775913.2;XP_006710648.1 | 28725 | . | . | . | 0.378691 | 0.365385 | 0.476418 | 0.517731 | 0.881229 | 0.844604 | 1.069067 | 1.183730 | 1.272502 | 146.065072 | 0.270736845 | 1.284154113 | 0.199088037 | 0.96621494 | 7.006263 | 7.353793 | 7.147110 | 7.128271 | 7.214525 | 7.244875 | 152 | 135 | 137 | 159 |
| 284723 | NM_207348,XI | SLC25A34 | solute carrier family 25 member 34 | protein_coding | NP_997231.1;XP_011539594.1;XP_011539595.1;XP_016856572.1 | 27653 | 610817 | . | . | 0.173570 | 0.325732 | 0.168983 | 0.215160 | 0.403904 | 0.752944 | 0.379192 | 0.491938 | -1.315299 | 30.648259 | 0.590050953 | -0.6700952 | 0.502797105 | 0.998865404 | 5.123643 | 4.724273 | 4.907591 | 4.993193 | 4.904043 | 4.931211 | 28 | 49 | 20 | 28 |
| 284739 | NM_00134809 | C20orf204 | chromosome 20 open reading frame 204 | protein_coding | NP_001335019.1;XP_024307644.1 | 27655 | . | . | . | 0.661824 | 0.410643 | 0.670767 | 0.340314 | 1.540092 | 0.949220 | 1.505181 | 0.778087 | -1.069518 | 43.971759 | 0.519543851 | -0.18662655 | 0.851953447 | 0.998865404 | 5.503639 | 5.411913 | 5.497885 | 5.411836 | 5.494710 | 5.391398 | 65 | 37 | 49 | 27 |
| 284751 | NR_034124 | LINC01270 | long intergenic non-protein coding RNA 1270 | lncRNA | . | 27658 | . | . | . | 0.168575 | 0.354881 | 0.307351 | 0.253214 | 0.392281 | 0.820323 | 0.689686 | 0.578945 | 1.050783 | 19.522474 | 0.724452145 | 0.098646853 | 0.921418668 | NA | 4.251986 | 4.321310 | 4.246707 | 4.307131 | 4.289143 | 4.273806 | 14 | 28 | 19 | 17 |
| 284764 | gene-FTLP3 | FTLP3 | ferritin light chain pseudogene 3 | pseudogene | . | 4000 | . | . | . | 0.103511 | 0.136255 | 0.074835 | 0.168222 | 0.240875 | 0.314959 | 0.167927 | 0.384621 | -1.057136 | 3.935194 | 1.523609365 | -0.05261253 | 0.958040637 | NA | 2.019256 | 1.932304 | 1.976409 | 1.981877 | 1.971361 | 1.984902 | 4 | 5 | 2 | 5 |
| 284802 | NR_003579,NF | FRG1BP | FSHD region gene 1 family member B, pseudogene | transcribed_ps | . | 15792 | . | . | . | 7.548857 | 9.680941 | 8.635316 | 11.460142 | 17.566499 | 22.377928 | 19.377398 | 26.202254 | 1.237325 | 339.798642 | 0.196361426 | 1.564587199 | 0.117679732 | 0.857396579 | 8.247062 | 8.553743 | 8.349926 | 8.334330 | 8.416368 | 8.505287 | 355 | 323 | 302 | 379 |
| 284821 | gene-RPL13AP7 | RPL13AP7 | ribosomal protein L13a pseudogene 7 | pseudogene | . | 23755 | . | . | . | 1.387793 | 1.165108 | 0.836796 | 1.496195 | 3.229450 | 2.693200 | 1.877746 | 3.420871 | -1.108033 | 43.342683 | 0.496564462 | -0.29805067 | 0.765664492 | 0.998865404 | 5.511036 | 5.360470 | 5.460910 | 5.430746 | 5.386438 | 5.469976 | 57 | 45 | 26 | 49 |
| 284889 | NR_038911 | MIF-AS1 | MIF antisense RNA 1 | lncRNA | . | 27669 | . | . | . | 0.836704 | 1.103375 | 0.987228 | 1.309747 | 1.947045 | 2.550500 | 2.215311 | 2.994581 | 1.167585 | 72.640137 | 0.383856806 | 0.58231893 | 0.560351877 | 0.998865404 | 6.068012 | 6.288929 | 6.126192 | 6.188991 | 6.161690 | 6.229149 | 67 | 82 | 59 | 83 |
| 284900 | NR_026962,NF | TTC28-AS1 | TTC28 antisense RNA 1 | lncRNA | . | 29336 | . | . | . | 2.647014 | 3.283578 | 2.909326 | 2.256599 | 6.159710 | 7.590138 | 6.528442 | 5.159446 | -1.086915 | 113.137462 | 0.315390668 | -0.38123882 | 0.703026048 | 0.998865404 | 6.880415 | 6.760979 | 6.800317 | 6.864545 | 6.847090 | 6.762230 | 123 | 139 | 107 | 88 |
| 284930 | NR_122046 | LOC284930 | uncharacterized LOC284930 | lncRNA | . | . | . | . | . | 0.735786 | 0.870264 | 1.066576 | 0.879503 | 1.712203 | 2.011655 | 2.393365 | 2.010880 | 1.199340 | 50.919967 | 0.442060399 | 0.593222809 | 0.55303205 | 0.998865404 | 5.532907 | 5.795481 | 5.632528 | 5.661522 | 5.700015 | 5.661920 | 49 | 54 | 53 | 47 |
| 284942 | NR_026981,NF | RPL23AP82 | ribosomal protein L23a pseudogene 82 | transcribed_ps | . | 33730 | . | . | . | 6.277795 | 7.164935 | 6.431185 | 6.584522 | 14.608686 | 16.562069 | 14.431393 | 15.054728 | -1.039131 | 304.101902 | 0.202964806 | -0.27284031 | 0.784975975 | 0.998865404 | 8.276080 | 8.220201 | 8.222304 | 8.293066 | 8.234860 | 8.240167 | 334 | 356 | 258 | 281 |
| 284948 | NM_201594,XI | SH2D6 | SH2 domain containing 6 | protein_coding | NP_963888.2;XP_011531085.1;XP_011531098.1;XP_016859323.1;XP_01( | 30439 | . | . | . | 0.169043 | 0.368192 | 0.187378 | 0.151008 | 0.393369 | 0.851092 | 0.420471 | 0.345261 | -1.484632 | 22.006600 | 0.694591664 | -0.82077774 | 0.411772883 | NA | 4.718182 | 4.144983 | 4.431549 | 4.514574 | 4.436279 | 4.439611 | 19 | 39 | 15 | 17 |
| 284988 | gene-RHOQP3 | RHOQP3 | ras homolog family member Q pseudogene 3 | pseudogene | . | 37837 | . | . | . | 0.304460 | 0.254536 | 0.164561 | 0.171094 | 0.708490 | 0.588372 | 0.369271 | 0.391185 | -1.644280 | 21.213399 | 0.673995042 | -1.06448277 | 0.287110054 | NA | 4.721207 | 4.004486 | 4.443593 | 4.424091 | 4.390953 | 4.390099 | 33 | 26 | 14 | 15 |
| 284992 | NM_00108053 | CCDC150 | coiled-coil domain containing 150 | protein_coding | NP_001074008.1;NP_001340268.1;NP_001340269.1;XP_006712501.1;XF | 26834 | . | . | . | 3.418906 | 3.832907 | 4.750295 | 4.332437 | 7.955936 | 8.859941 | 10.659525 | 9.905603 | 1.161113 | 482.880421 | 0.182560722 | 1.18047249 | 0.237812346 | 0.992733415 | 8.803487 | 9.019481 | 8.832048 | 8.893233 | 9.023900 | 8.885455 | 489 | 506 | 500 | 434 |
| 284996 | NM_173647,XI | RNF149 | ring finger protein 149 | protein_coding | NP_775918.2;XP_005263977.1;XP_005263978.1;XP_011509292.1;XP_02( | 23137 | . | . | . | 21.250562 | 18.849751 | 21.878622 | 14.711042 | 49.450933 | 43.572046 | 49.094995 | 33.635048 | -1.123367 | 1665.922898 | 0.1661205 | -1.0102881 | 0.31235728 | 0.998865404 | 10.783278 | 10.616094 | 10.796048 | 10.683241 | 10.780521 | 10.501140 | 2173 | 1767 | 1611 | 1206 |
| 285000 | NR_038891 | LOC285000 | uncharacterized LOC285000 | lncRNA | . | . | . | . | . | 8.888757 | 9.075753 | 2.719232 | 6.212429 | 20.684502 | 20.979012 | 6.101877 | 14.203979 | -2.036092 | 297.691497 | 0.286627921 | -3.57886572 | 0.000345089 | 0.030730707 | 8.642206 | 7.613535 | 8.351010 | 8.358285 | 7.801694 | 8.132174 | 457 | 434 | 105 | 254 |
| 285003 | gene-MRPL36F | MRPL36P1 | mitochondrial ribosomal protein L36 pseudogene 1 | pseudogene | . | 29708 | . | . | . | 0.056668 | 0.301723 | 0.062686 | 0.105851 | 0.131868 | 0.697446 | 0.140665 | 0.242015 | -2.487616 | 2.876852 | 1.856726397 | -0.70810844 | 0.478877916 | NA | 2.039857 | 0.715150 | 1.527342 | 1.549566 | 1.525134 | 1.529429 | 2 | 7 | 1 | 2 |
| 285025 | NM_00131674 | CCDC141 | coiled-coil domain containing 141 | protein_coding | NP_001303674.1;NP_775919.3;XP_011509293.1;XP_011509294.1;XP_01 | 26821 | 616031 | . | . | 0.092865 | 0.126821 | 0.107437 | 0.110037 | 0.216100 | 0.293153 | 0.241084 | 0.251585 | -1.018539 | 32.986522 | 0.541084841 | -0.0489777 | 0.960937066 | 0.998865404 | 5.057381 | 5.030099 | 5.028489 | 5.059253 | 5.044217 | 5.038939 | 34 | 40 | 29 | 30 |
| 285033 | NR_046322 | STARD7-AS1 | STARD7 antisense RNA 1 | lncRNA | . | 40827 | . | . | . | 1.156200 | 1.191574 | 1.437867 | 1.082925 | 2.690525 | 2.754377 | 3.226532 | 2.475979 | 1.058069 | 94.652108 | 0.3368166 | 0.241773799 | 0.808955442 | 0.998865404 | 6.522471 | 6.605461 | 6.550306 | 6.555146 | 6.611267 | 6.525576 | 105 | 100 | 97 | 78 |
| 285053 | gene-RPL18AP( | RPL18AP6 | ribosomal protein L18a pseudogene 6 | pseudogene | . | 36266 | . | . | . | 1.642119 | 0.942847 | 0.911802 | 1.312135 | 3.821280 | 2.179434 | 2.046057 | 3.000040 | -1.151656 | 23.666712 | 0.655979815 | -0.31054328 | 0.75614785 | NA | 4.662982 | 4.459419 | 4.601256 | 4.543638 | 4.544252 | 4.573475 | 37 | 20 | 16 | 24 |
| 285074 | NR_026846 | ANAPC1P2 | ANAPC1 pseudogene 2 | transcribed_ps | . | 54708 | . | . | . | 4.571288 | 4.500733 | 4.312224 | 4.706377 | 10.637574 | 10.403646 | 9.676507 | 10.760572 | -1.019379 | 310.247740 | 0.19906086 | -0.13910351 | 0.889368358 | 0.998865404 | 8.291203 | 8.263215 | 8.289150 | 8.275987 | 8.251535 | 8.292271 | 365 | 334 | 257 | 299 |
| 285097 | NR_149023 | LOC285097 | uncharacterized FLJ38379 | lncRNA | . | . | . | . | . | 0.010111 | 0.006114 | 0.030788 | 0.014426 | 0.023528 | 0.014132 | 0.069088 | 0.032983 | 2.914566 | 2.615882 | 1.896469644 | 0.813765289 | 0.415779423 | NA | 0.413393 | 1.963455 | 1.359184 | 1.355355 | 1.372337 | 1.366135 | 2 | 1 | 4 | 3 |
| 285116 | NR_077058 | AHCTF1P1 | AT-hook containing transcription factor 1 pseudogene 1 | transcribed_ps | . | 24620 | . | . | . | 0.341225 | 0.333684 | 0.319997 | 0.279803 | 0.794045 | 0.771326 | 0.718064 | 0.639737 | -1.139032 | 88.387788 | 0.340119969 | -0.55218414 | 0.580822202 | 0.998865404 | 6.556147 | 6.369362 | 6.488916 | 6.479649 | 6.466435 | 6.428915 | 110 | 100 | 77 | 72 |
| 285141 | NM_00128994 | ERICH2 | glutamate rich 2 | protein_coding | NP_001276876.1;NP_001276959.1;NP_001276960.1;XP_011509303.1;XF | 44395 | . | . | . | 2.915835 | 2.249452 | 2.634516 | 2.312786 | 6.785266 | 5.199707 | 5.911777 | 5.287913 | -1.088579 | 114.450354 | 0.307686284 | -0.39795943 | 0.690660091 | 0.998865404 | 6.897970 | 6.776636 | 6.891215 | 6.811782 | 6.829733 | 6.818521 | 153 | 114 | 97 | 101 |
| 285148 | NM_00103961 | IAH1 | isoamyl acetate hydrolyzing esterase 1 (putative) | protein_coding | NP_001034702.1;NP_001307787.1;NP_001307788.1;NP_001307789.1;NI | 27696 | . | . | . | 16.165067 | 16.109690 | 13.338847 | 17.502469 | 37.616772 | 37.238269 | 29.931983 | 40.017315 | -1.022117 | 1125.868489 | 0.154879206 | -0.20377466 | 0.838529594 | 0.998865404 | 10.152831 | 10.120635 | 10.140178 | 10.145759 | 10.021776 | 10.221500 | 1306 | 1233 | 838 | 1184 |
| 285172 | NM_00132161 | FAM126B | family with sequence similarity 126 member B | protein_coding | NP_001308547.1;NP_001308548.1;NP_001308550.1;NP_001308551.1;NI | 28593 | . | . | . | 2.165861 | 2.011129 | 2.140373 | 1.700492 | 5.040046 | 4.648818 | 4.802938 | 3.887971 | -1.104798 | 583.077664 | 0.176430263 | -0.81495512 | 0.415098003 | 0.998865404 | 9.257122 | 9.114440 | 9.249897 | 9.183763 | 9.227835 | 9.069814 | 739 | 628 | 541 | 456 |
| 285175 | NM_00137198 | UNC80 | unc-80 homolog, NALCN channel complex subunit | protein_coding | NP_001358915.1;NP_115893.1;NP_872393.3;XP_005246533.1;XP_01150 | 26582 | 612636 | . | . | 0.534699 | 0.497467 | 0.465374 | 0.461531 | 1.244267 | 1.149915 | 1.044283 | 1.055235 | -1.133524 | 204.821258 | 0.234633218 | -0.77062647 | 0.440928364 | 0.998865404 | 7.765665 | 7.585135 | 7.724501 | 7.688429 | 7.649781 | 7.649367 | 260 | 226 | 167 | 180 |
| 285190 | NM_182588,XI | RGPD4 | RANBP2 like and GRIP domain containing 4 | protein_coding | NP_872394.2;XP_011509319.1;XP_011509320.1;XP_011509321.1;XP_01: | 32417 | 612707 | . | . | 0.673987 | 0.437756 | 0.830879 | 0.476778 | 1.568395 | 1.011892 | 1.864469 | 1.090096 | 1.171074 | 121.820138 | 0.317263438 | 0.718115831 | 0.47268587 | 0.998865404 | 6.808754 | 7.039266 | 6.930181 | 6.853387 | 7.012420 | 6.877229 | 144 | 107 | 136 | 100 |
| 285193 | NM_00103357 | DUSP28 | dual specificity phosphatase 28 | protein_coding | NP_001028747.1;NP_001357394.1 | 33237 | . | . | . | 1.549665 | 1.905199 | 2.188100 | 1.788022 | 3.606133 | 4.403953 | 4.910034 | 4.088100 | 1.120600 | 250.497943 | 0.237438855 | 0.691844531 | 0.489034968 | 0.998865404 | 7.883962 | 8.048650 | 7.877705 | 7.986589 | 8.058346 | 7.917994 | 242 | 283 | 262 | 214 |
| 285195 | NM_173653,XI | SLC9A9 | solute carrier family 9 member A9 | protein_coding | NP_775924.1;XP_011511005.1;XP_011511006.1;XP_016861691.1;XP_01( | 20653 | 608396 | . | . | 0.293117 | 0.225348 | 0.297290 | 0.116539 | 0.682096 | 0.520904 | 0.667110 | 0.266453 | -1.279364 | 24.318369 | 0.674776379 | -0.52673208 | 0.598379656 | NA | 4.767942 | 4.418954 | 4.625184 | 4.598454 | 4.626072 | 4.546482 | 35 | 26 | 27 | 11 |
| 285203 | NM_00127868 | EOGT | EGF domain specific O-linked N-acetylglucosamine transferase | protein_coding | NP_001265618.1;NP_775925.1;XP_005264800.1;XP_011531901.1;XP_01 | 28526 | 614789 | . | . | 14.020162 | 13.255562 | 14.486373 | 11.135641 | 32.625489 | 30.640825 | 32.506999 | 25.460320 | -1.062685 | 1830.432427 | 0.147916228 | -0.59299885 | 0.553181922 | 0.998865404 | 10.880906 | 10.793714 | 10.886588 | 10.830326 | 10.914673 | 10.695706 | 2248 | 1963 | 1759 | 1432 |
| 285237 | NM_173824,XI | C3orf38 | chromosome 3 open reading frame 38 | protein_coding | NP_776185.2;XP_005264802.1 | 28384 | . | . | . | 5.159620 | 4.021753 | 4.758106 | 4.452185 | 12.006648 | 9.296460 | 10.677051 | 10.179391 | -1.013209 | 345.678791 | 0.204922889 | -0.09238407 | 0.926392892 | 0.998865404 | 8.442196 | 8.424325 | 8.504702 | 8.360153 | 8.446738 | 8.408335 | 451 | 328 | 309 | 311 |

| | | | | | | | | | | | | | | | | | | | | | | | | | | | | | | | | | |
|---|---|---|---|---|---|---|---|---|---|---|---|---|---|---|---|---|---|---|---|---|---|---|---|---|---|---|---|---|---|---|---|---|---|
| 285266 | NR_040100 | ENTPD3-AS1 | ENTPD3 antisense RNA 1 | lncRNA | . | 26710 | . | . | . | 0.414684 | 0.282375 | 0.321107 | 0.427456 | 0.964986 | 0.652723 | 0.720556 | 0.977327 | 1.065506 | 23.472370 | 0.64609336 | 0.141679387 | 0.887333258 | NA | 4.506418 | 4.597915 | 4.566512 | 4.530594 | 4.540555 | 4.569101 | 31 | 20 | 18 | 26 |
| 285267 | NM_00114508 | ZNF619 | zinc finger protein 619 | protein_coding | NP_001138554.1;NP_001138555.1;NP_001138565.1;NP_001138566.1;NI | 26910 | . | . | . | 0.611647 | 0.672612 | 0.556958 | 0.614233 | 1.423329 | 1.554775 | 1.249797 | 1.404370 | -1.036726 | 92.850741 | 0.330920224 | -0.15724217 | 0.875053996 | 0.998865404 | 6.562788 | 6.510419 | 6.548548 | 6.536696 | 6.520527 | 6.543671 | 111 | 100 | 76 | 89 |
| 285268 | NM_00109841 | ZNF621 | zinc finger protein 621 | protein_coding | NP_001091884.1;NP_001274174.1;NP_940886.1;XP_005265136.1;XP_00 | 24787 | . | . | . | 2.791030 | 3.077272 | 3.602873 | 3.141126 | 6.494843 | 7.113251 | 8.084743 | 7.181814 | 1.092844 | 810.908633 | 0.150680094 | 0.850060336 | 0.395291542 | 0.998865404 | 9.597839 | 9.726102 | 9.599239 | 9.659513 | 9.734829 | 9.648667 | 854 | 872 | 781 | 746 |
| 285282 | NM_00136396 | RABL3 | RAB, member of RAS oncogene family like 3 | protein_coding | NP_001350893.1;NP_001350894.1;NP_776186.2 | 18072 | 618542 | . | . | 1.320775 | 1.323957 | 1.564822 | 1.067019 | 3.073499 | 3.060384 | 3.511415 | 2.439614 | -1.045920 | 219.122290 | 0.260945657 | -0.2482242 | 0.803960942 | 0.998865404 | 7.806748 | 7.743749 | 7.767012 | 7.792530 | 7.858980 | 7.647476 | 251 | 248 | 229 | 155 |
| 285313 | NM_00117814 | IGSF10 | immunoglobulin superfamily member 10 | protein_coding | NP_001171616.1;NP_001171617.1;NP_849144.2;XP_011511010.1;XP_01 | 26384 | 617351 | . | . | 0.090019 | 0.092981 | 0.072504 | 0.089608 | 0.209478 | 0.214929 | 0.162696 | 0.204879 | -1.134171 | 28.264770 | 0.581256828 | -0.31249203 | 0.754666616 | 0.998865404 | 4.909524 | 4.726544 | 4.828177 | 4.832446 | 4.806591 | 4.825039 | 34 | 33 | 21 | 27 |
| 285315 | NM_00130822 | C3orf33 | chromosome 3 open reading frame 33 | protein_coding | NP_001295158.1;NP_775928.1;XP_005247436.1;XP_011511012.1;XP_01 | 26434 | . | . | . | 1.653476 | 1.528929 | 0.839071 | 1.178947 | 3.847707 | 3.534190 | 1.882852 | 2.695525 | -1.328294 | 83.682949 | 0.34760293 | -1.17828159 | 0.238684366 | 0.992733415 | 6.577303 | 6.167403 | 6.425503 | 6.424861 | 6.343653 | 6.354378 | 110 | 103 | 61 | 69 |
| 285329 | gene-UBQLN4F | UBQLN4P1 | ubiquilin 4 pseudogene 1 | pseudogene | . | 31133 | . | . | . | 0.440347 | 0.414405 | 0.492301 | 0.432800 | 1.024705 | 0.957916 | 1.104710 | 0.989546 | 1.075934 | 49.759305 | 0.443159693 | 0.238265378 | 0.811675272 | 0.998865404 | 5.582532 | 5.689282 | 5.634112 | 5.622127 | 5.654420 | 5.628638 | 57 | 50 | 48 | 45 |
| 285331 | NM_00101250 | CCDC66 | coiled-coil domain containing 66 | protein_coding | NP_001012524.4;NP_001135419.1;NP_001340076.1;NP_001340077.1;NI | 27709 | . | . | . | 3.059014 | 3.386722 | 4.179051 | 3.063636 | 7.118455 | 7.828559 | 9.377670 | 7.004644 | 1.114292 | 350.826915 | 0.215900555 | 0.723141698 | 0.469592834 | 0.998865404 | 8.373881 | 8.531070 | 8.394706 | 8.438952 | 8.576815 | 8.370352 | 367 | 372 | 379 | 285 |
| 285343 | NM_00102983 | TCAIM | T cell activation inhibitor, mitochondrial | protein_coding | NP_001025010.1;NP_001025011.1;NP_001269842.1;NP_001269843.1;NI | 25241 | . | . | . | 3.179406 | 3.319589 | 3.532757 | 2.839825 | 7.398607 | 7.673380 | 7.927403 | 6.492924 | -1.073441 | 264.584093 | 0.214898653 | -0.47577191 | 0.634236882 | 0.998865404 | 8.097398 | 7.995986 | 8.067680 | 8.062120 | 8.066191 | 7.989974 | 317 | 294 | 239 | 221 |
| 285346 | NM_00128734 | ZNF852 | zinc finger protein 852 | protein_coding | NP_001274278.1 | 27713 | . | . | . | 0.668818 | 0.906715 | 0.773989 | 0.650744 | 1.556368 | 2.095912 | 1.736808 | 1.487849 | -1.118831 | 51.187874 | 0.447100947 | -0.36231784 | 0.717114528 | 0.998865404 | 5.756481 | 5.594432 | 5.656114 | 5.715768 | 5.681914 | 5.648430 | 53 | 67 | 46 | 41 |
| 285352 | NR_033373 | KIF9-AS1 | KIF9 antisense RNA 1 | lncRNA | . | 26822 | . | . | . | 0.102973 | 0.143806 | 0.103863 | 0.084209 | 0.239622 | 0.332414 | 0.233066 | 0.192533 | -1.243982 | 9.349642 | 0.998672426 | -0.31538425 | 0.752469925 | NA | 3.373794 | 3.058876 | 3.223337 | 3.238631 | 3.225778 | 3.218117 | 10 | 13 | 8 | 7 |
| 285359 | NR_002941 | PDCL3P4 | PDCL3 pseudogene 4 | transcribed_ps | . | 44505 | . | . | . | 0.315597 | 0.239667 | 0.207865 | 0.110308 | 0.734407 | 0.554002 | 0.466442 | 0.252207 | -1.602909 | 7.250444 | 1.142362217 | -0.59586413 | 0.551265998 | NA | 3.156660 | 2.481107 | 2.879310 | 2.865391 | 2.863129 | 2.851354 | 12 | 8 | 6 | 4 |
| 285362 | NM_00116467 | SUMF1 | sulfatase modifying factor 1 | protein_coding | NP_001158146.1;NP_001158147.1;NP_877437.2;XP_011531926.1;XP_01 | 20376 | 607939 | . | . | 15.505362 | 16.166059 | 14.766719 | 15.389493 | 36.081616 | 37.368572 | 33.136091 | 35.186248 | -1.065476 | 1037.300866 | 0.135448291 | -0.67552038 | 0.499345208 | 0.998865404 | 10.063727 | 9.972055 | 10.029852 | 10.053260 | 9.983713 | 10.005881 | 1213 | 1173 | 860 | 956 |
| 285367 | NM_00114254 | RPUSD3 | RNA pseudouridine synthase D3 | protein_coding | NP_001136019.1;NP_001338665.1;NP_001338666.1;NP_001338667.1;NI | 28437 | 617759 | . | . | 24.050001 | 28.000836 | 26.759203 | 31.750480 | 55.965343 | 64.725187 | 60.046870 | 72.593703 | 1.106555 | 1112.369983 | 0.148119472 | 0.986194212 | 0.324037819 | 0.998865404 | 10.044886 | 10.190294 | 10.026829 | 10.128131 | 10.088970 | 10.215088 | 1133 | 1218 | 931 | 1187 |
| 285368 | NM_00131887 | PRRT3 | proline rich transmembrane protein 3 | protein_coding | NP_001305800.1;NP_997234.3;XP_011531930.1;XP_011531931.1 | 26591 | . | . | . | 0.496863 | 0.699598 | 0.526512 | 0.745362 | 1.156219 | 1.617153 | 1.181476 | 1.704181 | 1.033286 | 68.244330 | 0.405494124 | 0.116500236 | 0.907256099 | 0.998865404 | 6.070517 | 6.114423 | 6.039009 | 6.126990 | 6.051398 | 6.130375 | 63 | 86 | 50 | 76 |
| 285381 | NM_00104743 | DPH3 | diphthamide biosynthesis 3 | protein_coding | NP_001040899.1;NP_996662.1 | 27717 | 608959 | . | . | 4.731622 | 4.504848 | 5.153918 | 4.478823 | 11.010677 | 10.413157 | 11.565244 | 10.240297 | 1.029045 | 586.351903 | 0.165240361 | 0.249976199 | 0.802605755 | 0.998865404 | 9.174453 | 9.216486 | 9.203302 | 9.166248 | 9.252096 | 9.153518 | 684 | 606 | 556 | 516 |
| 285382 | NM_00102526 | C3orf70 | chromosome 3 open reading frame 70 | protein_coding | NP_001020437.1 | 33731 | . | . | . | 0.511321 | 0.531134 | 0.712318 | 0.545462 | 1.189866 | 1.227740 | 1.598422 | 1.247133 | 1.186914 | 128.570675 | 0.295325531 | 0.837093683 | 0.402539911 | 0.998865404 | 6.876841 | 7.125309 | 6.963195 | 6.975293 | 7.079565 | 6.978606 | 133 | 129 | 138 | 113 |
| 285401 | NR_027104 | LINC00698 | long intergenic non-protein coding RNA 698 | lncRNA | . | 27720 | . | . | . | 60.623631 | 63.710415 | 59.678242 | 58.242889 | 141.073685 | 147.269470 | 133.916245 | 133.165436 | -1.068945 | 3465.008028 | 0.110405802 | -0.87121615 | 0.383636144 | 0.998865404 | 11.805930 | 11.709753 | 11.768207 | 11.802370 | 11.744530 | 11.716290 | 4038 | 3943 | 2963 | 3086 |
| 285429 | NM_00102995 | DCAF4L1 | DDB1 and CUL4 associated factor 4 like 1 | protein_coding | NP_001025126.2 | 27723 | . | . | . | 0.150083 | 0.084946 | 0.158058 | 0.038831 | 0.349249 | 0.196357 | 0.354678 | 0.088783 | -1.170770 | 16.525845 | 0.835969882 | -0.27208859 | 0.785553901 | NA | 4.151937 | 3.933075 | 4.064758 | 4.022400 | 4.069312 | 3.994238 | 26 | 14 | 21 | 6 |
| 285440 | NM_207352,XI | CYP4V2 | cytochrome P450 family 4 subfamily V member 2 | protein_coding | NP_997235.3;XP_005262992.1;XP_016863526.1 | 23198 | 608614 | . | . | 3.906912 | 3.385074 | 3.556167 | 3.870506 | 9.091546 | 7.824750 | 7.979935 | 8.849454 | 1.018735 | 529.447161 | 0.172040399 | 0.155657132 | 0.876303289 | 0.998865404 | 9.034799 | 9.061761 | 9.092537 | 8.987595 | 9.030073 | 9.076184 | 653 | 520 | 448 | 519 |
| 285442 | gene-LOC2854 | LOC285442 | mitochondrial translational release factor 1 like pseudogene | pseudogene | . | . | . | . | . | 2.104688 | 2.115526 | 2.253736 | 2.256704 | 4.897696 | 4.890134 | 5.057318 | 5.159686 | 1.055500 | 184.293770 | 0.243808302 | 0.319623186 | 0.749253996 | 0.998865404 | 7.486349 | 7.564324 | 7.514410 | 7.512318 | 7.537801 | 7.535762 | 207 | 193 | 165 | 177 |
| 285500 | XR_00174192E | LOC285500 | uncharacterized LOC285500, transcript variant X6 | lncRNA | . | . | . | . | . | 0.149568 | 0.091490 | 0.149054 | 0.072339 | 0.348052 | 0.211483 | 0.334474 | 0.165395 | -1.015359 | 6.191548 | 1.229892724 | -0.01787898 | 0.985735398 | NA | 2.638039 | 2.622519 | 2.635786 | 2.620908 | 2.636257 | 2.619518 | 9 | 5 | 7 | 4 |
| 285512 | NR_002806 | FAM13A-AS1 | FAM13A antisense RNA 1 | lncRNA | . | 19370 | 613300 | . | . | 0.418461 | 0.328567 | 0.685612 | 0.301986 | 0.973775 | 0.759498 | 1.538494 | 0.690455 | 1.298080 | 43.695401 | 0.524994 | 0.716920383 | 0.473423225 | 0.998865404 | 5.246341 | 5.627390 | 5.426946 | 5.390321 | 5.521178 | 5.377730 | 49 | 36 | 60 | 28 |
| 285513 | NM_198281,XI | GPRIN3 | GPRIN family member 3 | protein_coding | NP_938022.2;XP_005262993.1;XP_005262994.1;XP_016863532.1;XP_01 | 27733 | 611241 | . | . | 0.049865 | 0.041971 | 0.055971 | 0.065407 | 0.116037 | 0.097017 | 0.125597 | 0.149545 | 1.283593 | 25.521879 | 0.611481059 | 0.589042156 | 0.555832995 | NA | 4.482357 | 4.842545 | 4.663671 | 4.646976 | 4.672505 | 4.690607 | 28 | 22 | 23 | 29 |
| 285521 | NM_00129773 | COX18 | cytochrome c oxidase assembly factor COX18 | protein_coding | NP_001284661.1;NP_001284662.1;NP_001287658.1;NP_776188.1;XP_00 | 26801 | 610428 | . | . | 1.927895 | 2.045298 | 2.026134 | 1.521962 | 4.486293 | 4.727795 | 4.546584 | 3.479786 | -1.017183 | 323.548278 | 0.20459746 | -0.12013429 | 0.904376766 | 0.998865404 | 8.349467 | 8.326113 | 8.375375 | 8.305450 | 8.379306 | 8.281915 | 399 | 330 | 304 | 274 |
| 285527 | NM_015030,XI | FRYL | FRY like transcription coactivator | protein_coding | NP_055845.1;XP_005248147.1;XP_005248150.1;XP_011511982.2;XP_01 | 29127 | . | . | . | 2.528740 | 2.579572 | 2.690615 | 2.273851 | 5.884481 | 5.962797 | 6.037663 | 5.198889 | -1.039777 | 915.704799 | 0.147041698 | -0.38270989 | 0.701934878 | 0.998865404 | 9.866357 | 9.810581 | 9.851993 | 9.851686 | 9.878297 | 9.766244 | 1075 | 1007 | 842 | 776 |
| 285540 | NR_037934 | SEPSECS-AS1 | SEPSECS antisense RNA 1 (head to head) | lncRNA | . | 27737 | . | . | . | 0.580405 | 0.757583 | 0.460589 | 0.420180 | 1.350626 | 1.751187 | 1.033549 | 0.960692 | -1.537186 | 19.687797 | 0.699446717 | -0.88683176 | 0.375169487 | NA | 4.576468 | 3.955730 | 4.307457 | 4.332921 | 4.287675 | 4.282426 | 24 | 29 | 14 | 14 |
| 285550 | NM_00114519 | FAM200B | family with sequence similarity 200 member B | protein_coding | NP_001138663.1;XP_016863537.1;XP_024309764.1;XP_024309765.1;XP | 27740 | . | . | . | 5.312273 | 4.298329 | 5.145169 | 3.241417 | 12.361875 | 9.935785 | 11.545610 | 7.411118 | -1.162157 | 575.176480 | 0.209842234 | -1.03318008 | 0.301519633 | 0.998865404 | 9.271506 | 9.056189 | 9.274081 | 9.137316 | 9.240802 | 8.958146 | 783 | 600 | 565 | 389 |
| 285556 | NM_00135443 | C4orf54 | chromosome 4 open reading frame 54 | protein_coding | NP_001341364.1;XP_006714493.1;XP_373030.5 | 27741 | 617881 | . | . | 0.028974 | 0.035471 | 0.056114 | 0.029144 | 0.067423 | 0.081992 | 0.125919 | 0.066635 | 1.294062 | 12.400034 | 0.88384443 | 0.420783224 | 0.673913383 | NA | 3.432996 | 3.807334 | 3.608245 | 3.619766 | 3.646563 | 3.609095 | 11 | 13 | 16 | 9 |
| 285590 | NM_00101799 | SH3PXD2B | SH3 and PX domains 2B | protein_coding | NP_001017995.1;NP_001295104.1;XP_016864840.1 | 29242 | 613293 | . | . | 7.482526 | 8.107469 | 6.009822 | 8.603494 | 17.412147 | 18.740780 | 13.485865 | 19.670867 | -1.073055 | 1817.732868 | 0.159004044 | -0.63976025 | 0.522328477 | 0.998865404 | 10.878158 | 10.775879 | 10.820788 | 10.881291 | 10.651120 | 10.922456 | 2085 | 2108 | 1261 | 1928 |
| 285600 | NM_00114567 | KIAA0825 | KIAA0825 | protein_coding | NP_001139150.1;NP_775936.1;XP_006714651.1;XP_011541626.1;XP_01 | 28532 | 617266 | . | . | 0.029791 | 0.035091 | 0.017522 | 0.035208 | 0.069325 | 0.081114 | 0.039318 | 0.080499 | -1.228540 | 13.424890 | 0.851964671 | -0.34854071 | 0.727434144 | NA | 3.889947 | 3.587955 | 3.752315 | 3.760512 | 3.727326 | 3.761204 | 16 | 17 | 7 | 15 |
| 285605 | NM_00130808 | DTWD2 | DTW domain containing 2 | protein_coding | NP_001295010.1;NP_775937.1;XP_011541640.3;XP_011541642.1 | 19334 | . | . | . | 1.372895 | 1.096003 | 1.274736 | 1.207043 | 3.194783 | 2.533459 | 2.860470 | 2.759758 | -1.010506 | 222.738206 | 0.235350282 | -0.06406772 | 0.948916308 | 0.998865404 | 7.806145 | 7.792232 | 7.851800 | 7.745999 | 7.812840 | 7.777709 | 286 | 215 | 200 | 200 |
| 285613 | NM_00113002 | RELL2 | RELT like 2 | protein_coding | NP_001123501.1;NP_776189.3;XP_005268471.1;XP_011535926.1;XP_01 | 26902 | 611213 | . | . | 15.741957 | 15.931647 | 20.643423 | 17.551197 | 36.632181 | 36.826716 | 46.323238 | 40.128730 | 1.188356 | 1338.721090 | 0.138544188 | 1.797020203 | 0.072332413 | 0.758556671 | 10.256593 | 10.505927 | 10.309478 | 10.320927 | 10.513282 | 10.376142 | 1398 | 1330 | 1376 | 1243 |
| 285636 | NM_175921,XI | C5orf51 | chromosome 5 open reading frame 51 | protein_coding | NP_787117.3;XP_005248346.1;XP_011512334.1 | 27750 | . | . | . | 6.428701 | 5.798324 | 6.620439 | 4.584949 | 14.959853 | 13.403089 | 14.856073 | 10.482942 | -1.108193 | 961.884606 | 0.177816302 | -0.83349643 | 0.404564813 | 0.998865404 | 9.981469 | 9.834215 | 9.981885 | 9.898848 | 9.992048 | 9.723332 | 1229 | 1030 | 942 | 696 |
| 285638 | NR_149040 | LOC285638 | uncharacterized LOC285638 | lncRNA | . | . | . | . | . | 0.039112 | 0.128226 | 0.093945 | 0.047978 | 0.091014 | 0.296400 | 0.210810 | 0.109696 | -1.184718 | 13.455597 | 0.90462772 | -0.27032503 | 0.786910211 | NA | 3.867373 | 3.622519 | 3.709338 | 3.777796 | 3.750398 | 3.717850 | 8 | 24 | 14 | 8 |
| 285647 | gene-SUDS3P1 | SUDS3P1 | SDS3 homolog, SIN3A corepressor complex component pseudogei | pseudogene | . | 44000 | . | . | . | 0.016426 | 0.055594 | 0.094803 | 0.030911 | 0.038223 | 0.128507 | 0.212736 | 0.070675 | 1.767034 | 4.405107 | 1.48099413 | 0.554580323 | 0.579181751 | NA | 1.671431 | 2.491830 | 2.105000 | 2.118966 | 2.133964 | 2.112379 | 2 | 5 | 7 | 3 |
| 285672 | NM_173829,XI | SREK1IP1 | SREK1 interacting protein 1 | protein_coding | NP_776190.1;XP_011541646.1 | 26716 | . | . | . | 1.346967 | 1.198275 | 1.304628 | 1.065498 | 3.134447 | 2.769867 | 2.927547 | 2.436134 | -1.090640 | 264.705586 | 0.222160322 | -0.56344639 | 0.573130955 | 0.998865404 | 8.108773 | 7.985065 | 8.101430 | 8.034352 | 8.076147 | 7.969558 | 338 | 279 | 244 | 212 |
| 285704 | NM_00101276 | RGMB | repulsive guidance molecule BMP co-receptor b | protein_coding | NP_001012779.2;NP_001353437.1;NP_001353438.1;NP_001353439.1;NI | 26896 | 612687 | . | . | 3.282680 | 3.145288 | 3.423778 | 3.435376 | 7.638929 | 7.270474 | 7.682861 | 7.854577 | 1.051745 | 451.726062 | 0.173227666 | 0.420169049 | 0.674361963 | 0.998865404 | 8.782367 | 8.855320 | 8.819918 | 8.786046 | 8.833046 | 8.835425 | 521 | 462 | 402 | 435 |
| 285735 | NR_026969 | LINC00326 | long intergenic non-protein coding RNA 326 | lncRNA | . | 41926 | . | . | . | 6.150387 | 5.839805 | 5.751121 | 6.072547 | 14.312203 | 13.498971 | 12.905348 | 13.884158 | -1.027652 | 225.346257 | 0.225430493 | -0.17456054 | 0.86142496 | 0.998865404 | 7.835544 | 7.796186 | 7.837360 | 7.807804 | 7.799307 | 7.819957 | 271 | 239 | 189 | 213 |
| 285753 | NM_00108353 | CEP57L1 | centrosomal protein 57 like 1 | protein_coding | NP_001077004.1;NP_001258781.1;NP_001258782.1;NP_001337581.1;NI | 21561 | . | . | . | 0.925688 | 0.787172 | 0.764614 | 0.627832 | 2.154115 | 1.819586 | 1.715774 | 1.435462 | -1.138287 | 249.949122 | 0.223122621 | -0.8374947 | 0.402314555 | 0.998865404 | 8.055469 | 7.869525 | 7.985143 | 8.006485 | 7.978773 | 7.881693 | 300 | 293 | 224 | 197 |
| 285755 | NM_00111129 | PPIL6 | peptidylprolyl isomerase like 6 | protein_coding | NP_001104768.2;NP_001273289.1;NP_001273290.1;NP_775943.1;XP_01 | 21557 | . | . | . | 0.422099 | 0.816279 | 0.261136 | 0.439253 | 0.982243 | 1.886865 | 0.585980 | 1.004301 | -1.633515 | 38.479784 | 0.517602629 | -1.36780631 | 0.171372711 | 0.947127937 | 5.578230 | 4.867059 | 5.271057 | 5.330722 | 5.214343 | 5.244213 | 46 | 60 | 22 | 31 |
| 285761 | NM_00136645 | DCBLD1 | discoidin, CUB and LCCL domain containing 1 | protein_coding | NP_001353387.1;NP_001353388.1;NP_001353389.1;NP_775945.1;XP_01 | 21479 | . | . | . | 3.417573 | 3.196991 | 3.168781 | 2.907979 | 7.952829 | 7.389986 | 7.110655 | 6.648753 | -1.088853 | 347.773537 | 0.193858096 | -0.63350221 | 0.526405737 | 0.998865404 | 8.501741 | 8.379687 | 8.497816 | 8.431769 | 8.438708 | 8.395343 | 441 | 369 | 301 | 300 |
| 285785 | gene-RPL39P3 | RPL39P3 | ribosomal protein L39 pseudogene 3 | pseudogene | . | 21383 | . | . | . | 24.284147 | 21.427917 | 19.472603 | 23.842791 | 56.510208 | 49.531586 | 43.695957 | 54.513702 | -1.069386 | 268.755537 | 0.219635117 | -0.44065241 | 0.659464653 | 0.998865404 | 8.117974 | 8.020688 | 8.120856 | 8.050934 | 8.002525 | 8.099040 | 341 | 280 | 204 | 267 |
| 285829 | gene-SUMO2P | SUMO2P1 | SUMO2 pseudogene 1 | pseudogene | . | 13985 | . | . | . | 0.570995 | 0.546850 | 0.470238 | 0.691018 | 1.328730 | 1.264068 | 1.055199 | 1.579930 | 1.018830 | 18.515888 | 0.71875036 | 0.037444564 | 0.97013054 | NA | 4.198253 | 4.223024 | 4.213472 | 4.206918 | 4.196089 | 4.227055 | 22 | 19 | 13 | 21 |
| 285830 | NR_026972,NF | HLA-F-AS1 | HLA-F antisense RNA 1 | lncRNA | . | 26645 | . | . | . | 1.019677 | 1.449623 | 1.574038 | 1.215668 | 2.372831 | 3.350868 | 3.532095 | 2.779480 | 1.121150 | 78.402809 | 0.376386525 | 0.438325821 | 0.661150111 | 0.998865404 | 6.207958 | 6.372992 | 6.227701 | 6.312040 | 6.336098 | 6.263287 | 71 | 93 | 82 | 67 |
| 285834 | NR_003948,NF | HCG22 | HLA complex group 22 | transcribed_ps | . | 27780 | 613918 | . | . | 0.535485 | 0.708622 | 0.848280 | 0.586313 | 1.246095 | 1.638013 | 1.903517 | 1.340535 | 1.134048 | 46.143963 | 0.476032882 | 0.381238232 | 0.703026483 | 0.998865404 | 5.433969 | 5.616407 | 5.486222 | 5.532268 | 5.565518 | 5.495822 | 43 | 53 | 51 | 37 |
| 285849 | gene-COX6A1P | COX6A1P2 | cytochrome c oxidase subunit 6A1 pseudogene 2 | pseudogene | . | 35239 | . | . | . | 7.688824 | 7.435545 | 8.110274 | 8.863235 | 17.892208 | 17.187595 | 18.199221 | 20.264732 | 1.107974 | 134.568470 | 0.280365542 | 0.527611538 | 0.597768983 | 0.998865404 | 6.996484 | 7.144133 | 7.058743 | 7.044392 | 7.073738 | 7.104188 | 150 | 135 | 118 | 138 |
| 285855 | NM_00136648 | RPL7L1 | ribosomal protein L7 like 1 | protein_coding | NP_001353410.1;NP_940888.3 | 21370 | 617417 | . | . | 17.863054 | 19.281994 | 18.931698 | 21.403066 | 41.568063 | 44.575194 | 42.482181 | 48.935567 | 1.075020 | 2541.792916 | 0.117668869 | 0.886919987 | 0.37512198 | 0.998865404 | 11.258613 | 11.362769 | 11.258433 | 11.303646 | 11.307200 | 11.371244 | 2742 | 2718 | 2200 | 2566 |
| 285872 | gene-THAP5P1 | THAP5P1 | THAP domain containing 5 pseudogene 1 | pseudogene | . | 22340 | . | . | . | 0.017558 | 0.073439 | 0.102042 | 0.011038 | 0.040858 | 0.169758 | 0.228979 | 0.025236 | 1.489070 | 2.847990 | 1.860722199 | 0.308703778 | 0.757546869 | NA | 1.193905 | 1.769039 | 1.478348 | 1.492059 | 1.500901 | 1.479767 | 1 | 4 | 5 | 1 |
| 285905 | NR_146905,NF | INTS4P1 | integrator complex subunit 4 pseudogene 1 | transcribed_ps | . | 21925 | . | . | . | 0.281120 | 0.361519 | 0.293704 | 0.307097 | 0.654177 | 0.835666 | 0.659063 | 0.702141 | -1.071814 | 37.819310 | 0.507478392 | -0.19716046 | 0.843701974 | 0.998865404 | 5.290842 | 5.189480 | 5.228450 | 5.263408 | 5.232698 | 5.239047 | 40 | 47 | 31 | 35 |
| 285908 | NR_026873 | LINC00174 | long intergenic non-protein coding RNA 174 | lncRNA | . | 27788 | . | . | . | 0.760338 | 0.806768 | 1.406798 | 0.871197 | 1.769338 | 1.864881 | 3.156814 | 1.991889 | 1.431466 | 133.191714 | 0.312531786 | 1.655809502 | 0.097760403 | 0.813553488 | 6.774487 | 7.293758 | 6.963450 | 6.980026 | 7.192754 | 7.002236 | 123 | 121 | 169 | 112 |
| 285943 | NR_122069 | HOXA-AS2 | HOXA cluster antisense RNA 2 | lncRNA | . | 43745 | 616068 | . | . | 0.421064 | 0.533075 | 0.434213 | 0.516154 | 0.979832 | 1.232226 | 0.974362 | 1.180124 | -1.009904 | 16.097001 | 0.762642509 | -0.01864239 | 0.985126385 | NA | 4.016660 | 4.000737 | 4.002621 | 4.015233 | 4.003809 | 4.012715 | 17 | 19 | 13 | 16 |
| 285958 | NR_003697,NF | SNHG15 | small nucleolar RNA host gene 15 | lncRNA | . | 27797 | . | . | . | 14.376870 | 18.957346 | 20.596833 | 18.599278 | 33.455569 | 43.820753 | 46.218696 | 42.525038 | 1.151187 | 493.855791 | 0.186714609 | 1.087874336 | 0.276650587 | 0.998865404 | 8.842982 | 9.045788 | 8.814772 | 8.974357 | 9.022894 | 8.949058 | 461 | 559 | 486 | 467 |
| 285962 | NR_015392 | WEE2-AS1 | WEE2 antisense RNA 1 | lncRNA | . | 48669 | . | . | . | 0.113540 | 0.094153 | 0.090818 | 0.145631 | 0.264213 | 0.217638 | 0.203793 | 0.332967 | 1.092667 | 8.416224 | 1.048899662 | 0.121893463 | 0.902983393 | NA | 3.008843 | 3.134756 | 3.074026 | 3.068175 | 3.066198 | 3.082670 | 10 | 8 | 6 | 10 |
| 285966 | NM_00113002 | TCAF2 | TRPM8 channel associated factor 2 | protein_coding | NP_001123498.2;NP_001350467.1;NP_001352356.1;NP_001352357.1;NI | 26878 | 616252 | . | . | 0.347570 | 0.390701 | 0.673298 | 0.530816 | 0.808807 | 0.903124 | 1.510862 | 1.213648 | 1.650870 | 70.924714 | 0.38326118 | 1.887033723 | 0.05915579 | 0.718513409 | 5.741705 | 6.465005 | 6.049258 | 6.086341 | 6.216285 | 6.157503 | 56 | 63 | 85 | 74 |
| 285971 | NM_173680 | ZNF775 | zinc finger protein 775 | protein_coding | NP_775951.2 | 28501 | . | . | . | 2.030384 | 1.411877 | 1.123576 | 1.923555 | 4.724786 | 3.263617 | 2.521273 | 4.397981 | -1.145716 | 114.716635 | 0.347336607 | -0.56501186 | 0.572065699 | 0.998865404 | 6.937334 | 6.739769 | 6.921416 | 6.791089 | 6.729518 | 6.890167 | 167 | 108 | 69 | 126 |
| 285973 | NM_00131705 | ATG9B | autophagy related 9B | protein_coding | NP_001303985.1;XP_011514367.2;XP_011514368.2;XP_024302487.1;XP | 21899 | 612205 | . | . | 0.128996 | 0.183371 | 0.112030 | 0.196268 | 0.300179 | 0.423869 | 0.251392 | 0.448745 | -1.029907 | 21.738114 | 0.677956483 | -0.06270955 | 0.949997793 | NA | 4.465522 | 4.418403 | 4.425339 | 4.458512 | 4.417539 | 4.460833 | 21 | 28 | 14 | 25 |
| 285987 | NR_015448 | DLX6-AS1 | DLX6 antisense RNA 1 | lncRNA | . | 37151 | . | . | . | 0.514729 | 0.489791 | 0.719120 | 0.625588 | 1.197794 | 1.132175 | 1.613685 | 1.430334 | 1.308121 | 36.708097 | 0.512680802 | 0.755824163 | 0.449754628 | 0.998865404 | 4.990578 | 5.379361 | 5.176328 | 5.166667 | 5.221889 | 5.197958 | 38 | 33 | 39 | 36 |
| 285989 | NM_00101325 | ZNF789 | zinc finger protein 789 | protein_coding | NP_001013276.1;NP_001337928.1;NP_001337929.1;NP_001337930.1;NI | 27801 | . | . | . | 5.442558 | 5.514778 | 6.579392 | 5.470825 | 12.665056 | 12.747659 | 14.763965 | 12.508393 | 1.013592 | 419.105238 | 0.19358815 | 0.100612669 | 0.91985794 | 0.998865404 | 8.701014 | 8.721252 | 8.678432 | 8.729582 | 8.786467 | 8.632888 | 459 | 468 | 414 | 345 |
| 286006 | NM_00113446 | LSMEM1 | leucine rich single-pass membrane protein 1 | protein_coding | NP_001127940.1;NP_872403.1;XP_011514376.1;XP_011514377.1;XP_01 | 22036 | . | . | . | 0.319640 | 0.301248 | 0.563419 | 0.387755 | 0.743815 | 0.696348 | 1.264296 | 0.886555 | 1.261246 | 75.037413 | 0.437613855 | 0.76517183 | 0.444169232 | 0.998865404 | 6.050418 | 6.388855 | 6.184740 | 6.168481 | 6.343773 | 6.112737 | 79 | 69 | 104 | 45 |
| 286016 | NR_002187 | TPI1P2 | triosephosphate isomerase 1 pseudogene 2 | transcribed_ps | . | 38069 | . | . | . | 0.784152 | 0.928749 | 0.403765 | 0.845012 | 1.824754 | 2.146844 | 0.906038 | 1.932019 | -1.370236 | 47.366018 | 0.490150408 | -0.92711169 | 0.353868546 | 0.998865404 | 5.777174 | 5.318005 | 5.575471 | 5.608391 | 5.485198 | 5.585738 | 58 | 64 | 23 | 50 |
| 286042 | NR_024361,NF | FAM86B3P | family with sequence similarity 86 member B3, pseudogene | transcribed_ps | . | 44371 | . | . | . | 1.323151 | 1.644249 | 2.159148 | 2.487499 | 3.079029 | 3.800755 | 4.845065 | 5.687372 | 1.527968 | 84.610829 | 0.348375258 | 1.75561951 | 0.079153399 | 0.775642847 | 6.065502 | 6.675942 | 6.311000 | 6.349025 | 6.434364 | 6.455393 | 71 | 78 | 86 | 99 |
| 286046 | NM_173683,N | XKR6 | XK related 6 | protein_coding | NP_775954.2;XP_011542123.1;XP_024302897.1 | 27806 | . | . | . | 1.411849 | 0.860464 | 1.666023 | 1.647070 | 3.285429 | 1.989001 | 3.738509 | 3.765831 | 1.183045 | 94.555739 | 0.337821441 | 0.717850368 | 0.472849553 | 0.998865404 | 6.435812 | 6.680050 | 6.572641 | 6.491905 | 6.595411 | 6.569464 | 114 | 80 | 93 | 92 |
| 286053 | NM_00134948 | NSMCE2 | NSE2 (MMS21) homolog, SMC5-SMC6 complex SUMO ligase | protein_coding | NP_001336414.1;NP_001336415.1;NP_001336416.1;NP_775956.1;XP_01 | 26513 | 617246 | . | . | 12.070113 | 11.950606 | 12.226887 | 12.591234 | 28.087651 | 27.624360 | 27.436778 | 28.788363 | 1.018822 | 475.522538 | 0.169967977 | 0.158279434 | 0.874236613 | 0.998865404 | 8.879941 | 8.906675 | 8.890301 | 8.884505 | 8.884988 | 8.912878 | 545 | 506 | 408 | 460 |
| 286059 | NR_038873 | LOC286059 | TNF receptor superfamily member 10d pseudogene | transcribed_ps | . | . | . | . | . | 0.118039 | 0.022511 | 0.056261 | 0.059750 | 0.274681 | 0.052035 | 0.126249 | 0.136612 | -1.112364 | 5.839500 | 1.323286516 | -0.1160969 | 0.907575744 | NA | 2.617627 | 2.470304 | 2.567512 | 2.527554 | 2.545802 | 2.543931 | 12 | 2 | 5 | 5 |
| 286075 | NM_00110059 | ZNF707 | zinc finger protein 707 | protein_coding | NP_001094068.1;NP_001094069.1;NP_001275734.1;NP_001275735.1;NI | 27815 | . | . | . | 3.113928 | 3.444971 | 4.123358 | 4.261214 | 7.246236 | 7.963205 | 9.252694 | 9.742760 | 1.283432 | 272.438662 | 0.208491647 | 1.726723039 | 0.084217416 | 0.787933172 | 7.898535 | 8.258628 | 8.010388 | 8.024282 | 8.157528 | 8.134946 | 271 | 261 | 272 | 281 |
| 286076 | NR_015445 | LINC02878 | long intergenic non-protein coding RNA 2878 | lncRNA | . | 54524 | . | . | . | 0.306979 | 0.225741 | 0.358145 | 0.208556 | 0.714353 | 0.521811 | 0.803666 | 0.476840 | 1.060208 | 9.200419 | 1.008755252 | 0.083614656 | 0.933362814 | NA | 3.156660 | 3.245375 | 3.204430 | 3.190586 | 3.208645 | 3.191112 | 12 | 8 | 10 | 7 |
| 286077 | NM_198488 | FAM83H | family with sequence similarity 83 member H | protein_coding | NP_940890.4 | 24797 | 611927 | . | . | 3.626026 | 3.600866 | 3.992986 | 3.882956 | 8.437912 | 8.323564 | 8.960146 | 8.877918 | 1.074999 | 662.702567 | 0.152843251 | 0.682627214 | 0.494842433 | 0.998865404 | 9.319005 | 9.423538 | 9.351637 | 9.341562 | 9.409152 | 9.382161 | 741 | 684 | 609 | 631 |
| 286097 | NM_00134981 | MICU3 | mitochondrial calcium uptake family member 3 | protein_coding | NP_001336739.1;NP_859074.1;XP_005273534.1;XP_006716391.1;XP_00 | 27820 | 610633 | . | . | 0.497642 | 0.383614 | 0.407773 | 0.238098 | 1.158029 | 0.886741 | 0.915030 | 0.544383 | -1.425462 | 77.141102 | 0.39712603 | -1.28782596 | 0.197806553 | 0.96621494 | 6.500904 | 5.993579 | 6.350763 | 6.261711 | 6.273189 | 6.158048 | 120 | 83 | 70 | 44 |
| 286101 | NR_023392 | ZNF252P | zinc finger protein 252, pseudogene | transcribed_ps | . | 13046 | . | . | . | 1.725725 | 1.647378 | 1.493670 | 2.109028 | 4.015833 | 3.807987 | 3.351753 | 4.822041 | 1.054618 | 295.302096 | 0.22292622 | 0.344154308 | 0.730730242 | 0.998865404 | 8.167787 | 8.243320 | 8.202627 | 8.174120 | 8.124297 | 8.301609 | 340 | 302 | 220 | 331 |
| 286103 | NR_026974 | ZNF252P-AS1 | ZNF252P antisense RNA 1 | lncRNA | . | 27821 | . | . | . | 0.156527 | 0.133937 | 0.161364 | 0.228731 | 0.364244 | 0.309601 | 0.362096 | 0.522967 | 1.340077 | 17.779466 | 0.731382266 | 0.577422158 | 0.563654309 | NA | 3.926074 | 4.347534 | 4.140792 | 4.129369 | 4.144271 | 4.170218 | 19 | 15 | 15 | 22 |
| 286128 | NM_00127115 | ZFP41 | ZFP41 zinc finger protein | protein_coding | NP_001258085.3;NP_776193.3 | 26786 | . | . | . | 1.376838 | 1.593610 | 1.532091 | 1.615836 | 3.203958 | 3.683700 | 3.437967 | 3.694418 | 1.069436 | 185.618201 | 0.25062895 | 0.38643027 | 0.699178036 | 0.998865404 | 7.487546 | 7.583256 | 7.474881 | 7.561688 | 7.533042 | 7.563435 | 186 | 213 | 161 | 186 |
| 286133 | NM_173833 | SCARA5 | scavenger receptor class A member 5 | protein_coding | NP_776194.2 | 28701 | 611306 | . | . | 0.074711 | 0.086107 | 0.063905 | 0.142321 | 0.173855 | 0.199041 | 0.143400 | 0.325399 | 1.271024 | 11.714241 | 0.914953247 | 0.378151449 | 0.70531809 | NA | 3.369732 | 3.710567 | 3.534761 | 3.542026 | 3.528966 | 3.572233 | 11 | 12 | 7 | 17 |
| 286140 | NR_003129 | RNF5P1 | ring finger protein 5 pseudogene 1 | transcribed_ps | . | 31053 | . | . | . | 0.238217 | 0.268578 | 0.161830 | 0.095985 | 0.554341 | 0.620829 | 0.363141 | 0.219458 | -1.795029 | 6.992102 | 1.159355054 | -0.72799709 | 0.466615369 | NA | 3.166062 | 2.324312 | 2.821606 | 2.824141 | 2.808355 | 2.801608 | 10 | 10 | 5 | 4 |
| 286144 | NM_00117179 | TRIQK | triple QxxK/R motif containing | protein_coding | NP_001165266.1;NP_001165267.1;NP_001165268.1;NP_001165269.1;NI | 27828 | . | . | . | 3.493554 | 2.918779 | 2.988654 | 2.530076 | 8.129643 | 6.746888 | 6.706457 | 5.784715 | -1.176169 | 347.856045 | 0.205224748 | -1.14067949 | 0.254003326 | 0.994807659 | 8.554010 | 8.321310 | 8.540771 | 8.424857 | 8.441951 | 8.340993 | 475 | 366 | 304 | 272 |
| 286148 | NM_181787,XI | DPY19L4 | dpy-19 like 4 | protein_coding | NP_861452.2;XP_005250951.2;XP_005250952.1;XP_011515285.1;XP_01 | 27829 | 613895 | . | . | 4.545037 | 4.117126 | 4.365549 | 2.461304 | 10.576487 | 9.516921 | 9.796167 | 5.627481 | -1.298460 | 682.806744 | 0.200743605 | -1.87702781 | 0.060514283 | 0.719735192 | 9.590892 | 9.215412 | 9.527386 | 9.452800 | 9.451176 | 9.151926 | 934 | 789 | 635 | 430 |
| 286151 | NM_00102986 | FBXO43 | F-box protein 43 | protein_coding | NP_001025031.2;XP_011515289.1;XP_011515291.1;XP_011515292.1;XP | 28521 | 609110 | . | . | 2.705585 | 3.299979 | 4.326994 | 2.466236 | 6.296007 | 7.628046 | 9.709652 | 5.638754 | 1.184973 | 314.106849 | 0.245242216 | 0.998418347 | 0.31807654 | 0.998865404 | 8.166632 | 8.413079 | 8.236885 | 8.233513 | 8.464440 | 8.171075 | 332 | 309 | 373 | 237 |
| 286205 | NM_00114487 | SCAI | suppressor of cancer cell invasion | protein_coding | NP_001138349.1;NP_775961.2 | 26709 | . | . | . | 0.557377 | 0.504466 | 0.397371 | 0.324384 | 1.297038 | 1.166097 | 0.891690 | 0.741666 | 1.143105 | 119.054196 | 0.315497664 | 0.611598474 | 0.540803448 | 0.998865404 | 6.794547 | 6.989801 | 6.881022 | 6.849217 | 6.976574 | 6.840505 | 134 | 114 | 132 | 96 |
| 286223 | NM_00100193 | C9orf47 | chromosome 9 open reading frame 47 | protein_coding | NP_001001938.1;NP_001135885.1 | 23669 | . | . | . | 0.099804 | 0.049362 | 0.121093 | 0.091838 | 0.232247 | 0.114103 | 0.271730 | 0.209977 | 1.389037 | 14.347224 | 0.827840528 | 0.572677164 | 0.566863302 | NA | 3.583318 | 4.062477 | 3.839020 | 3.806075 | 3.854884 | 3.831132 | 18 | 9 | 17 | 13 |
| 286257 | NM_00132967 | PAXX | PAXX non-homologous end joining factor | protein_coding | NP_001316607.1;NP_899064.1 | 27849 | 616315 | . | . | 15.600182 | 17.437117 | 20.438039 | 16.094771 | 36.302265 | 40.306678 | 45.862365 | 36.798779 | 1.081947 | 428.725710 | 0.189604844 | 0.599298845 | 0.548973618 | 0.998865404 | 8.685678 | 8.799885 | 8.686662 | 8.745636 | 8.830785 | 8.693579 | 451 | 466 | 431 | 371 |
| 286262 | NM_00112822 | TPRN | taperin | protein_coding | NP_001121700.2 | 26894 | 613354 | . | . | 3.026977 | 2.750420 | 2.509446 | 2.997302 | 7.043900 | 6.357719 | 5.631124 | 6.852974 | -1.063619 | 233.992454 | 0.227795028 | -0.39062193 | 0.696076711 | 0.998865404 | 7.914358 | 7.824891 | 7.910154 | 7.859033 | 7.815728 | 7.892780 | 292 | 247 | 181 | 230 |
| 286297 | NR_046175 | LOC286297 | methylenetetrahydrofolate dehydrogenase (NADP+ dependent) 1 | transcribed_ps | . | . | . | . | . | 0.036178 | 0.086258 | 0.083307 | 0.077102 | 0.084188 | 0.199390 | 0.186938 | 0.176285 | 1.224379 | 6.046894 | 1.240318181 | 0.235463994 | 0.813848622 | NA | 2.444631 | 2.733338 | 2.576456 | 2.595045 | 2.592431 | 2.590278 | 4 | 8 | 6 | 6 |
| 286319 | NM_00100412 | TUSC1 | tumor suppressor candidate 1 | protein_coding | NP_001004125.1 | 31010 | 610529 | . | . | 2.628755 | 1.980063 | 1.592385 | 2.540226 | 6.117222 | 4.577003 | 3.573265 | 5.807926 | -1.127367 | 168.761468 | 0.29371733 | -0.58885737 | 0.555956954 | 0.998865404 | 7.483272 | 7.309161 | 7.479736 | 7.352569 | 7.276186 | 7.452844 | 236 | 165 | 107 | 182 |
| 286333 | NR_024366 | FAM225A | family with sequence similarity 225 member A | lncRNA | . | 27855 | . | . | . | 0.265193 | 0.272290 | 0.255231 | 0.259457 | 0.617114 | 0.629409 | 0.572731 | 0.593216 | -1.048763 | 48.189818 | 0.448221131 | -0.15324806 | 0.878202663 | 0.998865404 | 5.624641 | 5.555851 | 5.592886 | 5.598187 | 5.587430 | 5.586566 | 56 | 54 | 41 | 44 |
| 286336 | NM_033387,XI | FAM78A | family with sequence similarity 78 member A | protein_coding | NP_203745.2;XP_011516869.1;XP_011516870.1 | 25465 | . | . | . | 4.028373 | 4.271209 | 4.257669 | 4.321692 | 9.374189 | 9.873089 | 9.554086 | 9.881034 | 1.011812 | 491.300989 | 0.169670121 | 0.099844558 | 0.920467732 | 0.998865404 | 8.932113 | 8.948765 | 8.908424 | 8.963636 | 8.942722 | 8.944770 | 538 | 550 | 429 | 464 |
| 286343 | NM_203403,XI | LURAP1L | leucine rich adaptor protein 1 like | protein_coding | NP_981948.1;XP_005251500.1 | 31452 | 616130 | . | . | 0.755191 | 0.740043 | 0.937437 | 1.124026 | 1.757361 | 1.710641 | 2.103582 | 2.569952 | 1.278025 | 56.123098 | 0.418056228 | 0.84657487 | 0.397232126 | 0.998865404 | 5.622936 | 5.976501 | 5.783890 | 5.775960 | 5.818220 | 5.840468 | 58 | 52 | 52 | 62 |
| 286380 | NM_199135 | FOXD4L3 | forkhead box D4 like 3 | protein_coding | NP_954586.4 | 18523 | 611086 | . | . | 0.008614 | 0.037878 | 0.026911 | 0.031251 | 0.020044 | 0.087558 | 0.060387 | 0.071452 | 1.219513 | 2.020490 | 2.092167648 | 0.136846221 | 0.891152349 | NA | 0.867373 | 1.148375 | 1.001080 | 1.010279 | 1.007918 | 1.007184 | 1 | 3 | 2 | 2 |
| 286410 | NM_00101098 | ATP11C | ATPase phospholipid transporting 11C | protein_coding | NP_001010986.2;NP_001340739.2;NP_001340740.2;NP_001340741.2;NI | 13554 | 300516 | . | . | 1.574738 | 1.461570 | 1.483683 | 1.250103 | 3.664484 | 3.378484 | 3.329341 | 2.858214 | -1.121049 | 314.804543 | 0.203565214 | -0.80980905 | 0.41804993 | 0.998865404 | 8.377880 | 8.214100 | 8.351421 | 8.304019 | 8.307318 | 8.222280 | 399 | 344 | 279 | 256 |
| 286437 | NR_039980 | LOC286437 | uncharacterized LOC286437 | lncRNA | . | . | . | . | . | 0.680100 | 0.591551 | 0.724033 | 0.847538 | 1.582620 | 1.367397 | 1.624709 | 1.937795 | 1.210583 | 51.906372 | 0.436055274 | 0.632263004 | 0.527215034 | 0.998865404 | 5.553462 | 5.829074 | 5.688340 | 5.662511 | 5.697656 | 5.725838 | 58 | 47 | 46 | 57 |
| 286444 | gene-RPS2P55 | RPS2P55 | ribosomal protein S2 pseudogene 55 | pseudogene | . | 36140 | . | . | . | 2.097135 | 2.146902 | 1.607149 | 2.269920 | 4.880120 | 4.962659 | 3.606395 | 5.189901 | -1.114761 | 59.938881 | 0.414598317 | -0.37804002 | 0.705400866 | 0.998865404 | 5.982688 | 5.823779 | 5.915107 | 5.920035 | 5.860794 | 5.927153 | 72 | 69 | 41 | 62 |
| 286451 | NM_00119521 | YIPF6 | Yip1 domain family member 6 | protein_coding | NP_001182143.1;NP_776195.2;XP_011529225.1;XP_011529226.1;XP_01 | 28304 | 300996 | . | . | 5.580526 | 4.737508 | 5.032348 | 4.314464 | 12.986113 | 10.950961 | 11.292443 | 9.864510 | -1.140518 | 893.895778 | 0.154297283 | -1.22937254 | 0.218932159 | 0.982008277 | 9.895350 | 9.706393 | 9.907682 | 9.784632 | 9.798587 | 9.706571 | 1188 | 942 | 774 | 732 |
| 286467 | NR_026975,NF | FIRRE | firre intergenic repeating RNA element | lncRNA | . | 49627 | 300999 | . | . | 0.573660 | 0.391245 | 0.369497 | 0.360509 | 1.334929 | 0.904381 | 0.829141 | 0.824261 | -1.338690 | 38.182598 | 0.51295343 | -0.82038995 | 0.411993845 | 0.998865404 | 5.449250 | 5.030099 | 5.307071 | 5.247950 | 5.237703 | 5.232070 | 59 | 39 | 29 | 30 |
| 286480 | NR_110506 | UBE2E4P | ubiquitin conjugating enzyme E2 E4 pseudogene | transcribed_ps | . | 32199 | . | . | . | 0.681267 | 0.818412 | 0.633793 | 0.908188 | 1.585337 | 1.891797 | 1.422214 | 2.076463 | 1.024458 | 27.726705 | 0.593848748 | 0.0587039 | 0.953187953 | 0.998865404 | 4.777174 | 4.809059 | 4.781372 | 4.800140 | 4.773326 | 4.813400 | 29 | 32 | 20 | 31 |
| 286495 | NR_030737 | TTC3P1 | tetratricopeptide repeat domain 3 pseudogene 1 | transcribed_ps | . | 23318 | . | . | . | 0.061844 | 0.043104 | 0.056469 | 0.047786 | 0.143914 | 0.099638 | 0.126715 | 0.109257 | -1.007286 | 4.446049 | 1.421176525 | -0.00736951 | 0.994120033 | NA | 2.156660 | 2.148375 | 2.156736 | 2.149644 | 2.153787 | 2.152299 | 6 | 4 | 4 | 4 |
| 286499 | NM_00117110 | FAM133A | family with sequence similarity 133 member A | protein_coding | NP_001164580.1;NP_001164581.1;NP_001164582.1;NP_775969.1;XP_00 | 26748 | . | . | . | 0.499930 | 0.451671 | 0.397306 | 0.575537 | 1.163357 | 1.044059 | 0.891545 | 1.315897 | -1.069670 | 59.276729 | 0.414684922 | -0.23431187 | 0.814742875 | 0.998865404 | 5.937713 | 5.839400 | 5.910902 | 5.883148 | 5.855286 | 5.909839 | 75 | 62 | 43 | 61 |
| 286527 | NM_00135021 | TMSB15B | thymosin beta 15B | protein_coding | NP_001337140.1;NP_001337141.1;NP_001337142.1;NP_919305.2 | 28612 | 301011 | . | . | 0.439127 | 0.676180 | 0.909765 | 0.592231 | 1.021865 | 1.563022 | 2.041154 | 1.354066 | 1.616324 | 40.659158 | 0.511060616 | 1.355449297 | 0.175274452 | 0.949120257 | 4.957404 | 5.651032 | 5.269560 | 5.304434 | 5.403718 | 5.317423 | 31 | 38 | 54 | 36 |
| 286826 | NM_00127040 | LIN9 | lin-9 DREAM MuvB core complex component | protein_coding | NP_001257338.1;NP_001257339.1;NP_001353166.1;NP_001353167.1;NI | 30830 | 609375 | . | . | 3.852549 | 4.941272 | 4.338107 | 3.172729 | 8.965041 | 11.421973 | 9.734585 | 7.254068 | -1.175377 | 390.393014 | 0.214113779 | -1.08878192 | 0.276250065 | 0.998865404 | 8.720383 | 8.487822 | 8.569582 | 8.723327 | 8.637286 | 8.463677 | 425 | 512 | 360 | 285 |
| 317648 | NR_015453 | NOP14-AS1 | NOP14 antisense RNA 1 | lncRNA | . | 20205 | . | . | . | 1.271540 | 1.035960 | 1.489524 | 1.341194 | 2.958926 | 2.394668 | 3.342449 | 3.066480 | 1.204501 | 155.221117 | 0.270993957 | 0.99056058 | 0.321900194 | 0.998865404 | 7.136932 | 7.406826 | 7.274168 | 7.195049 | 7.332308 | 7.285554 | 179 | 136 | 156 | 150 |
| 317649 | NM_00113464 | EIF4E3 | eukaryotic translation initiation factor 4E family member 3 | protein_coding | NP_001128121.1;NP_001128122.1;NP_001128123.1;NP_001269815.1;NI | 31837 | 609896 | . | . | 1.231804 | 0.937554 | 1.034946 | 0.887982 | 2.866459 | 2.167199 | 2.322390 | 2.030264 | -1.185416 | 301.153836 | 0.209695246 | -1.17023989 | 0.241904444 | 0.992733415 | 8.351502 | 8.106852 | 8.334264 | 8.217107 | 8.191088 | 8.181020 | 415 | 316 | 242 | 256 |
| 317662 | NM_173348,XI | FAM149B1 | family with sequence similarity 149 member B1 | protein_coding | NP_775483.1;XP_005269801.1;XP_005269802.1;XP_005269804.1;XP_01 | 29162 | 618413 | . | . | 2.294971 | 2.159540 | 1.548027 | 1.949452 | 5.340489 | 4.991870 | 3.473728 | 4.457193 | 1.076543 | 218.212215 | 0.245928215 | 0.432672252 | 0.665252891 | 0.998865404 | 7.716321 | 7.820962 | 7.681814 | 7.810276 | 7.739195 | 7.824366 | 209 | 258 | 179 | 231 |
| 317671 | NM_00113106 | RFESD | Rieske Fe-S domain containing | protein_coding | NP_001124537.1;NP_001124538.1;NP_001362323.1;NP_775498.1;XP_01 | 29587 | . | . | . | 1.313042 | 1.055778 | 1.397265 | 1.248640 | 3.055504 | 2.440480 | 3.135423 | 2.854864 | 1.119806 | 102.284131 | 0.324920255 | 0.502428989 | 0.615365791 | 0.998865404 | 6.591627 | 6.756540 | 6.692523 | 6.616585 | 6.706801 | 6.673582 | 125 | 91 | 99 | 96 |
| 317726 | gene-HMGN2P | HMGN2P2 | high mobility group nucleosomal binding domain 2 pseudogene 2 | pseudogene | . | 20016 | . | . | . | 0.237041 | 0.162131 | 0.170741 | 0.121969 | 0.551604 | 0.374774 | 0.383138 | 0.278868 | -1.314443 | 7.068314 | 1.145786524 | -0.34426245 | 0.730648922 | NA | 3.003609 | 2.612728 | 2.837418 | 2.823196 | 2.825148 | 2.818151 | 11 | 7 | 6 | 5 |
| 317749 | NM_00119363 | DHRS4L2 | dehydrogenase/reductase 4 like 2 | protein_coding | NP_001180564.1;NP_001180565.1;NP_001180566.1;NP_932349.2 | 19731 | 615196 | . | . | 6.636721 | 5.875674 | 6.552277 | 6.870432 | 15.443926 | 13.581887 | 14.703117 | 15.708426 | 1.038242 | 259.370025 | 0.217000804 | 0.249505336 | 0.802969914 | 0.998865404 | 7.991387 | 8.045835 | 8.045787 | 7.969623 | 8.014586 | 8.040202 | 315 | 254 | 224 | 254 |
| 317762 | NM_00114499 | CCDC85C | coiled-coil domain containing 85C | protein_coding | NP_001138467.1;XP_011535007.1;XP_011535008.1;XP_011535009.1 | 35459 | . | . | . | 1.658283 | 1.680267 | 1.908942 | 1.702758 | 3.858892 | 3.884013 | 4.283609 | 3.893152 | 1.023229 | 847.759435 | 0.143017741 | 0.23164009 | 0.816817567 | 0.998865404 | 9.710755 | 9.744075 | 9.717573 | 9.720426 | 9.755521 | 9.714920 | 963 | 906 | 767 | 781 |
| 317780 | gene-CKS1BP1 | CKS1BP1 | CDC28 protein kinase regulatory subunit 1B pseudogene 1 | pseudogene | . | 20004 | . | . | . | 2.499839 | 1.631550 | 1.108363 | 1.323424 | 5.817229 | 3.771400 | 2.487135 | 3.025851 | -1.703708 | 40.277201 | 0.511296775 | -1.50338946 | 0.132738644 | 0.887687926 | 5.665470 | 4.897226 | 5.412186 | 5.328969 | 5.282166 | 5.298657 | 71 | 43 | 24 | 30 |
| 317781 | NM_175066,XI | DDX51 | DEAD-box helicase 51 | protein_coding | NP_778236.2;XP_011536558.1 | 20082 | . | . | . | 6.785444 | 7.744539 | 8.705142 | 7.125937 | 15.790008 | 17.901851 | 19.534088 | 16.292611 | 1.085083 | 1011.689194 | 0.152055281 | 0.774751194 | 0.438486694 | 0.998865404 | 9.922385 | 10.040307 | 9.891776 | 10.004529 | 10.068553 | 9.946987 | 1031 | 1128 | 991 | 906 |
| 319093 | gene-RHOQP1 | RHOQP1 | ras homolog family member Q pseudogene 1 | pseudogene | . | 20003 | . | . | . | 0.780576 | 0.942696 | 0.751981 | 0.735157 | 1.816431 | 2.179084 | 1.687425 | 1.680850 | -1.163730 | 31.758262 | 0.550106484 | -0.3976619 | 0.690879423 | 0.998865404 | 5.094525 | 4.875273 | 4.988060 | 5.011520 | 4.982889 | 4.978370 | 36 | 40 | 26 | 27 |
| 319128 | gene-PPIAP6 | PPIAP6 | peptidylprolyl isomerase A pseudogene 6 | pseudogene | . | 20028 | . | . | . | 0.305169 | 0.223680 | 0.167235 | 0.214264 | 0.710142 | 0.517046 | 0.375271 | 0.489889 | -1.346981 | 5.827420 | 1.254842717 | -0.34245694 | 0.732007044 | NA | 2.741623 | 2.312263 | 2.558676 | 2.548136 | 2.544005 | 2.547171 | 9 | 6 | 4 | 5 |
| 319129 | gene-RPSAP5 | RPSAP5 | ribosomal protein SA pseudogene 5 | pseudogene | . | 20139 | . | . | . | 0.053482 | 0.027759 | 0.017954 | 0.053415 | 0.124455 | 0.064167 | 0.040288 | 0.122126 | 1.233060 | 1.486734 | 2.400765875 | 0.125894475 | 0.899815456 | NA | 0.413393 | 0.715150 | 0.572744 | 0.568868 | 0.570025 | 0.574493 | 2 | 1 | 1 | 2 |
| 319131 | gene-HMGN1P | HMGN1P1 | high mobility group nucleosome binding domain 1 pseudogene 1 | pseudogene | . | 20015 | . | . | . | 0.772451 | 0.973535 | 1.237709 | 0.602278 | 1.797525 | 2.250371 | 2.777382 | 1.377037 | 1.026201 | 28.748318 | 0.607484803 | 0.06142337 | 0.951022041 | 0.998865404 | 4.825128 | 4.865403 | 4.822352 | 4.848819 | 4.877328 | 4.798397 | 29 | 34 | 34 | 18 |
| 319138 | gene-BNIP3P1 | BNIP3P1 | BCL2 interacting protein 3 pseudogene 1 | pseudogene | . | 19922 | . | . | . | 0.285313 | 0.420645 | 0.494057 | 0.548041 | 0.663935 | 0.972339 | 1.108649 | 1.253031 | 1.410943 | 21.723696 | 0.666224531 | 0.745484623 | 0.455978804 | NA | 4.173073 | 4.667215 | 4.401384 | 4.430436 | 4.441983 | 4.452482 | 17 | 23 | 21 | 25 |
| 326300 | gene-HSPE1P2 | HSPE1P2 | heat shock protein family E (Hsp10) member 1 pseudogene 2 | pseudogene | . | 20052 | . | . | . | 0.581449 | 0.552377 | 0.242282 | 0.396758 | 1.353055 | 1.276843 | 0.543673 | 0.907141 | -1.545142 | 7.006256 | 1.151784559 | -0.54501426 | 0.585743712 | NA | 3.089593 | 2.459419 | 2.824631 | 2.822946 | 2.806355 | 2.814151 | 10 | 9 | 4 | 6 |
| 326302 | gene-NEK2P1 | NEK2P1 | NEK2 pseudogene 1 | pseudogene | . | 19272 | . | . | . | 0.040259 | 0.042851 | 0.142569 | 0.066014 | 0.093685 | 0.099052 | 0.319920 | 0.150933 | 2.179725 | 5.715963 | 1.288961093 | 0.872133322 | 0.383135649 | NA | 1.844134 | 2.971139 | 2.481796 | 2.483067 | 2.512810 | 2.490792 | 4 | 4 | 9 | 5 |
| 326307 | gene-RPL3P4 | RPL3P4 | ribosomal protein L3 pseudogene 4 | pseudogene | . | 19805 | . | . | . | 3.669467 | 4.043907 | 2.885902 | 4.956582 | 8.539001 | 9.347672 | 6.475881 | 11.332635 | 1.006993 | 58.511126 | 0.4328513 | 0.023228198 | 0.981468246 | 0.998865404 | 5.867222 | 5.874048 | 5.856811 | 5.877182 | 5.815048 | 5.919328 | 64 | 66 | 38 | 69 |
| 326310 | gene-RPL7AP6 | RPL7AP6 | ribosomal protein L7a pseudogene 6 | pseudogene | . | 19785 | . | . | . | 5.143043 | 6.123898 | 4.558880 | 6.738016 | 11.968072 | 14.155665 | 10.229996 | 15.405674 | -1.008516 | 155.193609 | 0.286289588 | -0.04273231 | 0.965914925 | 0.998865404 | 7.285219 | 7.270594 | 7.242719 | 7.308508 | 7.196685 | 7.343448 | 165 | 182 | 109 | 172 |
| 326591 | gene-MORF4L1 | MORF4L1P1 | mortality factor 4 like 1 pseudogene 1 | pseudogene | . | 20400 | . | . | . | 0.733363 | 0.483226 | 0.557381 | 0.760059 | 1.706566 | 1.116998 | 1.250748 | 1.737784 | 1.058569 | 22.469413 | 0.660783073 | 0.124268866 | 0.901102398 | NA | 4.448287 | 4.530328 | 4.504261 | 4.467927 | 4.477007 | 4.506027 | 30 | 19 | 17 | 25 |
| 326594 | gene-MORF4L1 | MORF4L1P2 | mortality factor 4 like 1 pseudogene 2 | pseudogene | . | 20401 | . | . | . | 0.350438 | 0.374861 | 0.484839 | 0.163867 | 0.815484 | 0.866507 | 1.087965 | 0.374663 | -1.118676 | 17.882923 | 0.774934845 | -0.20878221 | 0.83461826 | NA | 4.236451 | 4.080349 | 4.155123 | 4.160419 | 4.183733 | 4.113321 | 21 | 21 | 22 | 8 |
| 326615 | gene-MED15P: | MED15P1 | mediator complex subunit 15 pseudogene 1 | pseudogene | . | 19271 | . | . | . | 0.456085 | 0.397205 | 0.424407 | 0.355254 | 1.061328 | 0.918158 | 0.952356 | 0.812245 | -1.100376 | 19.215546 | 0.700471104 | -0.19700505 | 0.843823585 | NA | 4.330772 | 4.194412 | 4.276326 | 4.262465 | 4.266791 | 4.256388 | 25 | 20 | 17 | 16 |
| 326625 | NM_052845,N | MMAB | metabolism of cobalamin associated B | protein_coding | NP_443077.1;XP_011536569.1;XP_011536570.1;XP_011536571.1;XP_02 | 19331 | 607568 | . | . | 4.436745 | 5.087935 | 6.484007 | 7.634972 | 10.324488 | 11.760991 | 14.549924 | 17.456456 | 1.283595 | 545.649373 | 0.166540583 | 2.162775266 | 0.030558469 | 0.555729309 | 8.900806 | 9.260493 | 8.999256 | 9.005019 | 9.119373 | 9.206465 | 548 | 518 | 499 | 612 |
| 333926 | NM_005167 | PPM1J | protein phosphatase, Mg2+/Mn2+ dependent 1J | protein_coding | NP_005158.5 | 20785 | 609957 | . | . | 2.508767 | 2.514070 | 2.153952 | 1.858254 | 5.838003 | 5.811386 | 4.833406 | 4.248678 | -1.271390 | 120.978124 | 0.296731435 | -1.16740653 | 0.243046236 | 0.992733415 | 7.080986 | 6.735595 | 6.958714 | 6.957850 | 6.900743 | 6.851152 | 156 | 146 | 100 | 92 |
| 337867 | NM_00114407 | UBAC2 | UBA domain containing 2 | protein_coding | NP_001137544.1;NP_808882.1;XP_006720011.1;XP_011519384.1;XP_01 | 20486 | . | . | . | 19.450482 | 20.567504 | 17.271285 | 18.828260 | 45.262072 | 47.542705 | 38.756268 | 43.048579 | -1.080634 | 1381.683862 | 0.127372467 | -0.87835227 | 0.37975258 | 0.998865404 | 10.487144 | 10.375105 | 10.442409 | 10.479637 | 10.392732 | 10.410772 | 1613 | 1586 | 1141 | 1260 |
| 337873 | NR_036461 | H2BC20P | H2B clustered histone 20, pseudogene | transcribed_ps | . | 20516 | . | . | . | 0.715388 | 0.649091 | 1.249873 | 1.243154 | 1.664738 | 1.500403 | 2.804678 | 2.842324 | 1.748761 | 17.835563 | 0.728648892 | 1.106613879 | 0.268460904 | NA | 3.697260 | 4.504604 | 4.125342 | 4.117316 | 4.164211 | 4.162163 | 16 | 13 | 20 | 21 |
| 337874 | gene-H2BC19P | H2BC19P | H2B clustered histone 19, pseudogene | pseudogene | . | 20517 | . | . | . | 0.849598 | 0.963696 | 1.390188 | 1.702320 | 1.977050 | 2.227625 | 3.119542 | 3.892152 | 1.664402 | 18.538524 | 0.716504079 | 1.025819412 | 0.304976714 | NA | 3.799841 | 4.532926 | 4.175508 | 4.183903 | 4.209104 | 4.229579 | 15 | 16 | 18 | 24 |
| 337875 | NR_027337,NF | H2BP1 | H2B histone pseudogene 1 | transcribed_ps | . | 20560 | . | . | . | 1.220094 | 1.231445 | 2.442343 | 1.474809 | 2.839208 | 2.846539 | 5.480548 | 3.371977 | 1.155568 | 47.636430 | 0.509490138 | 0.40943204 | 0.68222263 | 0.998865404 | 5.464570 | 5.675700 | 5.517085 | 5.570477 | 5.640553 | 5.493769 | 43 | 55 | 62 | 29 |
| 337876 | NM_175856,XI | CHSY3 | chondroitin sulfate synthase 3 | protein_coding | NP_787052.3;XP_005272039.1;XP_005272040.1;XP_011541665.1;XP_01 | 24293 | 609963 | . | . | 0.138880 | 0.282169 | 0.146112 | 0.287154 | 0.323180 | 0.652247 | 0.327870 | 0.656543 | -1.189149 | 30.259026 | 0.568546403 | -0.43959365 | 0.660231443 | 0.998865404 | 5.039724 | 4.787887 | 4.909368 | 4.950947 | 4.904748 | 4.913889 | 32 | 41 | 23 | 27 |
| 337878 | NM_181606 | KRTAP7-1 | keratin associated protein 7-1 | protein_coding | NP_853637.2 | 18934 | . | . | . | 0.233562 | 0.351612 | 0.282892 | 0.352661 | 0.543509 | 0.812766 | 0.634800 | 0.806317 | 1.072468 | 7.463245 | 1.108817788 | 0.091028672 | 0.927469809 | NA | 2.849979 | 2.947964 | 2.891451 | 2.901989 | 2.895497 | 2.902695 | 7 | 9 | 6 | 8 |
| 337882 | NM_181607 | KRTAP19-1 | keratin associated protein 19-1 | protein_coding | NP_853638.1 | 18936 | . | . | . | 0.141825 | 0.214771 | 0.471836 | 0.181420 | 0.330033 | 0.496452 | 1.058787 | 0.414795 | 1.799479 | 5.680287 | 1.296005817 | 0.653993247 | 0.513116157 | NA | 2.019256 | 2.869520 | 2.475517 | 2.481160 | 2.506490 | 2.479661 | 4 | 5 | 9 | 4 |
| 338005 | NR_027457 | LINC00221 | long intergenic non-protein coding RNA 221 | lncRNA | . | 20169 | . | . | . | 2.399985 | 2.156563 | 2.772094 | 2.602555 | 5.584863 | 4.984992 | 6.220497 | 5.950433 | 1.162697 | 136.022907 | 0.280579413 | 0.77509128 | 0.438285724 | 0.998865404 | 6.974386 | 7.192770 | 7.076087 | 7.036002 | 7.125702 | 7.097535 | 153 | 128 | 132 | 132 |
| 338069 | NR_002329 | ST7-OT4 | ST7 overlapping transcript 4 | lncRNA | . | 18835 | . | . | . | 0.177693 | 0.130981 | 0.018351 | 0.137833 | 0.413500 | 0.302769 | 0.041180 | 0.315140 | -1.954371 | 6.395866 | 1.282195051 | -0.75394474 | 0.450882402 | NA | 3.084644 | 2.107126 | 2.704147 | 2.694204 | 2.666569 | 2.694279 | 11 | 8 | 1 | 7 |
| 338376 | NM_176891 | IFNE | interferon epsilon | protein_coding | NP_795372.1 | 18163 | 615223 | . | . | 0.020354 | 0.080459 | 0.048411 | 0.061649 | 0.047365 | 0.185985 | 0.108632 | 0.140954 | 1.014447 | 2.735953 | 1.804031249 | 0.011470344 | 0.99084819 | NA | 1.444631 | 1.459419 | 1.445375 | 1.454766 | 1.448241 | 1.452296 | 2 | 4 | 2 | 3 |
| 338382 | NM_00116452 | RAB7B | RAB7B, member RAS oncogene family | protein_coding | NP_001157994.1;NP_001291768.1;NP_796377.3;XP_006711351.1 | 30513 | . | . | . | 10.139837 | 11.676253 | 8.032379 | 9.835549 | 23.595818 | 26.990180 | 18.024425 | 22.487814 | -1.254075 | 842.502464 | 0.160098011 | -2.04014976 | 0.041335412 | 0.631142361 | 9.873027 | 9.545498 | 9.751662 | 9.830602 | 9.552489 | 9.703230 | 1022 | 1066 | 578 | 778 |
| 338391 | NR_160913,NF | H2BP2 | H2B histone pseudogene 2 | transcribed_ps | . | 20654 | . | . | . | 0.162342 | 0.145574 | 0.066888 | 0.092349 | 0.377777 | 0.336502 | 0.150094 | 0.211144 | -1.494583 | 9.619774 | 1.008092634 | -0.57508929 | 0.565230908 | NA | 3.526439 | 2.947964 | 3.299423 | 3.262124 | 3.260475 | 3.267416 | 18 | 8 | 6 | 8 |
| 338427 | NR_001292 | SNORD108 | small nucleolar RNA, C/D box 108 | snoRNA | . | 32772 | . | . | . | 0.006732 | 0.021232 | 0.007200 | 0.007995 | 0.015665 | 0.049080 | 0.016156 | 0.018280 | 1.234655 | 1.002957 | 2.868339486 | 0.106022226 | 0.915564717 | NA | -0.155866 | 0.148375 | -0.001261 | -0.000972 | 0.000053 | -0.000315 | 1 | 1 | 1 | 1 |
| 338428 | NR_001295 | SNORD109A | small nucleolar RNA, C/D box 109A | snoRNA | . | 32773 | . | . | . | 1.662027 | 5.160472 | 3.519745 | 5.167623 | 3.867605 | 11.928661 | 7.898206 | 11.815156 | 1.215452 | 8.544910 | 1.067617019 | 0.263664497 | 0.792038455 | NA | 2.950762 | 3.226238 | 3.067552 | 3.099103 | 3.085188 | 3.096688 | 5 | 12 | 7 | 10 |
| 338596 | NM_00100447 | ST8SIA6 | ST8 alpha-N-acetyl-neuraminide alpha-2,8-sialyltransferase 6 | protein_coding | NP_001004470.1;NP_001332890.1;XP_016871663.1;XP_016871664.1;XP | 23317 | 610139 | . | . | 0.528810 | 0.445888 | 0.738330 | 0.469457 | 1.230561 | 1.030691 | 1.656792 | 1.073358 | 1.202659 | 114.338852 | 0.327297583 | 0.813409847 | 0.415983115 | 0.998865404 | 6.696449 | 6.965380 | 6.823717 | 6.767734 | 6.929250 | 6.780112 | 130 | 102 | 132 | 92 |
| 338657 | NM_198489,N | CCDC84 | coiled-coil domain containing 84 | protein_coding | NP_940891.1;XP_011541101.1;XP_011541102.1 | 30460 | . | . | . | 2.602517 | 3.346414 | 4.115368 | 3.244182 | 6.056168 | 7.735384 | 9.234769 | 7.417438 | 1.225992 | 135.718247 | 0.292421732 | 1.005223265 | 0.314789355 | 0.998865404 | 6.929909 | 7.224064 | 6.999352 | 7.076359 | 7.156438 | 7.068133 | 125 | 146 | 145 | 124 |
| 338692 | NM_00134790 | ANKRD13D | ankyrin repeat domain 13D | protein_coding | NP_001334830.1;NP_997237.2;XP_011543287.1;XP_011543288.1;XP_01 | 27880 | 615126 | . | . | 8.370865 | 8.530767 | 10.258814 | 8.350435 | 19.479348 | 19.719249 | 23.020482 | 19.092277 | 1.081659 | 615.876566 | 0.169562482 | 0.667873798 | 0.504214154 | 0.998865404 | 9.208403 | 9.322342 | 9.229207 | 9.245247 | 9.362495 | 9.211672 | 673 | 646 | 619 | 533 |
| 338699 | NM_00130097 | ANKRD42 | ankyrin repeat domain 42 | protein_coding | NP_001287901.1;NP_001287902.1;NP_001287904.1;NP_001287905.1;NI | 26752 | . | . | . | 3.799433 | 3.742065 | 4.715494 | 4.325593 | 8.841439 | 8.649950 | 10.581431 | 9.889953 | 1.134710 | 471.039211 | 0.170440266 | 1.069718637 | 0.284745975 | 0.998865404 | 8.785591 | 8.968051 | 8.850718 | 8.824728 | 8.915766 | 8.917392 | 519 | 466 | 435 | 470 |
| 338707 | NM_178537,XI | B4GALNT4 | beta-1,4-N-acetyl-galactosaminyltransferase 4 | protein_coding | NP_848632.2;XP_016873143.1 | 26315 | 618560 | . | . | 1.012991 | 0.669374 | 1.645892 | 1.723856 | 2.357273 | 1.547287 | 3.693332 | 3.941392 | 1.977271 | 147.315582 | 0.29226086 | 3.365181398 | 0.000764934 | 0.056765469 | 6.627533 | 7.612914 | 7.092988 | 6.969762 | 7.282076 | 7.300767 | 137 | 85 | 167 | 187 |
| 338758 | NR_028138 | ATP2B1-AS1 | ATP2B1 antisense RNA 1 | lncRNA | . | 27883 | . | . | . | 0.152562 | 0.130856 | 0.207884 | 0.213508 | 0.355018 | 0.302480 | 0.466486 | 0.488161 | 1.434933 | 16.888545 | 0.744767028 | 0.699525378 | 0.484223759 | NA | 3.793789 | 4.315285 | 4.060709 | 4.052867 | 4.081237 | 4.084477 | 17 | 14 | 17 | 19 |
| 338761 | NM_00100822 | C1QL4 | complement C1q like 4 | protein_coding | NP_001008224.1;XP_011536572.1 | 31416 | 615229 | . | . | 2.295655 | 1.954942 | 1.584070 | 1.678501 | 5.342084 | 4.518935 | 3.554606 | 3.837693 | -1.317995 | 121.105935 | 0.298154675 | -1.33603427 | 0.181538073 | 0.953435029 | 7.105434 | 6.707459 | 6.997527 | 6.932754 | 6.865305 | 6.877263 | 172 | 136 | 89 | 100 |
| 338799 | NR_002809,NF | LINC01089 | long intergenic non-protein coding RNA 1089 | lncRNA | . | 27886 | . | . | . | 7.415075 | 7.701446 | 9.468629 | 8.258169 | 17.255182 | 17.802236 | 21.247332 | 18.881321 | 1.159826 | 367.481725 | 0.192867654 | 1.109092769 | 0.267390164 | 0.998546485 | 8.410175 | 8.624900 | 8.478705 | 8.469480 | 8.611301 | 8.505774 | 395 | 364 | 374 | 337 |
| 338811 | NM_178539,XI | TAFA2 | TAFA chemokine like family member 2 | protein_coding | NP_848634.1;XP_011536574.1;XP_024304730.1;XP_024304731.1;XP_02 | 21589 | 617496 | . | . | 0.020389 | 0.062417 | 0.036259 | 0.060120 | 0.047445 | 0.144279 | 0.081363 | 0.137455 | 1.103039 | 5.254796 | 1.342398413 | 0.105395886 | 0.916061681 | NA | 2.324708 | 2.459419 | 2.375509 | 2.398185 | 2.385265 | 2.393292 | 3 | 8 | 4 | 6 |
| 339005 | NR_003521 | WHAMMP3 | WAS protein homolog associated with actin, golgi membranes anc | transcribed_ps | . | 27892 | . | . | . | 0.687727 | 0.791823 | 0.737496 | 0.542791 | 1.600368 | 1.830333 | 1.654920 | 1.241028 | -1.164961 | 77.681363 | 0.369954575 | -0.59542897 | 0.551556768 | 0.998865404 | 6.384803 | 6.165894 | 6.279306 | 6.314877 | 6.294000 | 6.220922 | 90 | 96 | 72 | 57 |
| 339039 | gene-PKD1P3 | PKD1P3 | polycystin 1, transient receptor potential channel interacting pseu | pseudogene | . | 30067 | . | . | . | 3.835698 | 3.926261 | 4.507664 | 3.348020 | 8.925828 | 9.075727 | 10.115067 | 7.654850 | -1.002124 | 785.931549 | 0.170660212 | -0.01793613 | 0.985689808 | 0.999197585 | 9.619391 | 9.617128 | 9.610528 | 9.620988 | 9.715928 | 9.501788 | 899 | 855 | 787 | 624 |
| 339044 | NR_036447 | PKD1P1 | polycystin 1, transient receptor potential channel interacting pseu | transcribed_ps | . | 30065 | . | . | . | 10.855294 | 12.423504 | 14.047252 | 11.042461 | 25.260715 | 28.717487 | 31.521627 | 25.247274 | 1.063733 | 2371.272443 | 0.142977918 | 0.623424264 | 0.533005757 | 0.998865404 | 11.166090 | 11.255419 | 11.131395 | 11.232769 | 11.329537 | 11.126889 | 2479 | 2636 | 2391 | 2006 |
| 339105 | NM_00103950 | PRSS53 | serine protease 53 | protein_coding | NP_001034592.1;XP_011544118.1;XP_011544119.1;XP_011544120.1;XP | 34407 | 610561 | . | . | 0.904972 | 0.925872 | 1.009371 | 0.805493 | 2.105907 | 2.140194 | 2.265000 | 1.841665 | -1.013883 | 93.992294 | 0.33473657 | -0.05942192 | 0.952616055 | 0.998865404 | 6.563678 | 6.545204 | 6.555107 | 6.554663 | 6.584777 | 6.515796 | 109 | 102 | 91 | 77 |
| 339122 | NM_00120488 | RAB43 | RAB43, member RAS oncogene family | protein_coding | NP_001191812.1;NP_001191813.1;NP_001191814.1;NP_001191815.1;NI | 19983 | . | . | . | 7.066915 | 6.131410 | 7.567779 | 9.433402 | 16.445001 | 14.173026 | 16.981879 | 21.568351 | 1.268820 | 1053.717341 | 0.155039294 | 2.21548554 | 0.026726756 | 0.532994094 | 9.859425 | 10.202742 | 9.995108 | 9.888845 | 10.036646 | 10.195636 | 1149 | 928 | 918 | 1220 |
| 339123 | NM_00100592 | JMJD8 | jumonji domain containing 8 | protein_coding | NP_001005920.3;NP_001310847.2;NP_001310848.2;NP_001310849.2;NI | 14148 | . | . | . | 24.739842 | 25.289557 | 26.094344 | 25.145640 | 57.570629 | 58.457945 | 58.554952 | 57.492513 | 1.018117 | 1534.212404 | 0.124562786 | 0.207949443 | 0.835268443 | 0.998865404 | 10.570236 | 10.596212 | 10.566893 | 10.585263 | 10.607642 | 10.572161 | 1730 | 1660 | 1377 | 1418 |
| 339175 | NM_181725,XI | METTL2A | methyltransferase like 2A | protein_coding | NP_859076.3;XP_016880061.1 | 25755 | . | . | . | 3.904443 | 3.734262 | 4.052500 | 3.606456 | 9.085798 | 8.631916 | 9.093692 | 8.245732 | -1.102281 | 631.747185 | 0.155753163 | -0.90201398 | 0.367049443 | 0.998865404 | 9.371598 | 9.231405 | 9.341567 | 9.328434 | 9.288859 | 9.250295 | 770 | 708 | 538 | 547 |
| 339201 | NR_049729,NF | ASB16-AS1 | ASB16 antisense RNA 1 | lncRNA | . | 25442 | . | . | . | 3.665678 | 2.879885 | 3.563261 | 2.438569 | 8.530182 | 6.656984 | 7.995854 | 5.575498 | -1.022892 | 198.239785 | 0.262062744 | -0.12460073 | 0.900839649 | 0.998865404 | 7.646250 | 7.615795 | 7.667393 | 7.594412 | 7.699755 | 7.536638 | 249 | 199 | 202 | 150 |
| 339210 | NM_00108543 | C17orf67 | chromosome 17 open reading frame 67 | protein_coding | NP_001078899.2 | 27900 | . | . | . | 0.410803 | 0.381725 | 0.199721 | 0.366558 | 0.955956 | 0.882375 | 0.448168 | 0.838091 | -1.390577 | 22.302545 | 0.666794553 | -0.71338801 | 0.475605691 | NA | 4.698880 | 4.219802 | 4.503027 | 4.494800 | 4.448455 | 4.488084 | 31 | 27 | 12 | 22 |
| 339229 | NM_00103984 | OXLD1 | oxidoreductase like domain containing 1 | protein_coding | NP_001034931.1;NP_001291923.1;NP_001291924.1;NP_001291928.1 | 27901 | . | . | . | 17.669380 | 19.241820 | 19.394163 | 21.931092 | 41.117378 | 44.478327 | 43.519940 | 50.142837 | 1.110744 | 835.442386 | 0.146060423 | 1.037419944 | 0.299540187 | 0.998865404 | 9.628875 | 9.779964 | 9.642302 | 9.691239 | 9.713394 | 9.768948 | 880 | 884 | 735 | 856 |
| 339230 | NM_199287,XI | CCDC137 | coiled-coil domain containing 137 | protein_coding | NP_954981.1;XP_011523040.1;XP_016880062.1 | 33451 | 614271 | . | . | 19.730303 | 21.473531 | 21.412286 | 24.962140 | 45.913227 | 49.637026 | 48.048547 | 57.072972 | 1.105900 | 1392.320652 | 0.136674957 | 1.062523614 | 0.287998044 | 0.998865404 | 10.369086 | 10.513836 | 10.378620 | 10.425458 | 10.416265 | 10.538750 | 1476 | 1471 | 1171 | 1479 |
| 339231 | NM_00104002 | ARL16 | ADP ribosylation factor like GTPase 16 | protein_coding | NP_001035114.1;NP_001316537.1;NP_001316538.1 | 27902 | . | . | . | 16.478591 | 16.016019 | 20.657883 | 19.261241 | 38.346358 | 37.021742 | 46.355689 | 44.038537 | 1.218043 | 850.614308 | 0.148470621 | 1.916643497 | 0.055283238 | 0.697001222 | 9.582812 | 9.867854 | 9.661745 | 9.649400 | 9.846705 | 9.743463 | 891 | 820 | 870 | 814 |
| 339263 | NR_160801 | LINC02693 | long intergenic non-protein coding RNA 2693 | lncRNA | . | 27904 | . | . | . | 3.217390 | 3.229096 | 3.083550 | 3.543883 | 7.487000 | 7.464199 | 6.919398 | 8.102666 | 1.013848 | 806.069461 | 0.149226109 | 0.132961588 | 0.894223759 | 0.998865404 | 9.645023 | 9.664433 | 9.651042 | 9.647881 | 9.612472 | 9.703609 | 924 | 862 | 660 | 810 |
| 339287 | NM_00101224 | MSL1 | MSL complex subunit 1 | protein_coding | NP_001012241.1;NP_001352848.1;NP_001352849.1;NP_001352850.1 | 27905 | 614801 | . | . | 21.603729 | 20.883279 | 22.396298 | 23.037197 | 50.272765 | 48.272635 | 50.256646 | 52.671811 | 1.045128 | 2627.074929 | 0.112973062 | 0.563674403 | 0.572975741 | 0.998865404 | 11.327026 | 11.390753 | 11.347989 | 11.333926 | 11.382026 | 11.371542 | 2984 | 2748 | 2350 | 2501 |
| 339290 | NR_015389 | LINC00667 | long intergenic non-protein coding RNA 667 | lncRNA | . | 27906 | . | . | . | 5.293446 | 5.141591 | 5.507035 | 5.798495 | 12.318068 | 11.885017 | 12.357627 | 13.257570 | 1.067929 | 674.036777 | 0.152493946 | 0.621767667 | 0.53409465 | 0.998865404 | 9.348520 | 9.443291 | 9.386282 | 9.360463 | 9.404068 | 9.432161 | 765 | 690 | 593 | 666 |
| 339318 | NM_00102999 | ZNF181 | zinc finger protein 181 | protein_coding | NP_001025168.2;NP_001139137.1;XP_005258906.1;XP_005258907.1;XP | 12971 | 606741 | . | . | 1.827303 | 2.098829 | 1.749527 | 1.527984 | 4.252212 | 4.851538 | 3.925889 | 3.493555 | -1.166247 | 233.572692 | 0.233159131 | -0.95159795 | 0.341300924 | 0.998865404 | 7.973734 | 7.753310 | 7.910206 | 7.893964 | 7.885926 | 7.766166 | 294 | 267 | 212 | 176 |
| 339324 | NM_00101275 | ZNF260 | zinc finger protein 260 | protein_coding | NP_001012774.1;NP_001159508.1;NP_001159509.1;NP_001159510.1;NI | 13499 | 613749 | . | . | 2.690019 | 2.224611 | 2.837274 | 2.350567 | 6.259785 | 5.142288 | 6.366762 | 5.374290 | 1.022560 | 415.003966 | 0.195352697 | 0.164755447 | 0.869136473 | 0.998865404 | 8.680141 | 8.713627 | 8.747899 | 8.623364 | 8.755316 | 8.643920 | 521 | 397 | 399 | 357 |
| 339327 | NM_00129776 | ZNF546 | zinc finger protein 546 | protein_coding | NP_001284692.1;NP_848639.2 | 28671 | . | . | . | 0.333556 | 0.393849 | 0.456907 | 0.242439 | 0.776198 | 0.910399 | 1.025286 | 0.554308 | -1.046599 | 89.280325 | 0.377911976 | -0.17387505 | 0.86196366 | 0.998865404 | 6.511546 | 6.448302 | 6.455143 | 6.500158 | 6.544988 | 6.377966 | 97 | 106 | 99 | 57 |
| 339344 | NM_00101264 | MYPOP | Myb related transcription factor, partner of profilin | protein_coding | NP_001012661.1;XP_016882231.1;XP_016882232.1;XP_024307252.1 | 20178 | 617861 | . | . | 3.512935 | 3.640968 | 4.631004 | 4.072030 | 8.174746 | 8.416261 | 10.391841 | 9.310209 | 1.252581 | 271.288820 | 0.208561579 | 1.55782949 | 0.119273673 | 0.858706505 | 7.912127 | 8.236995 | 8.010968 | 8.026892 | 8.140310 | 8.129963 | 273 | 264 | 265 | 280 |

| | | | | | | | | | | | | | | | | | | | | | | | | | | | | | | | | | |
|---|---|---|---|---|---|---|---|---|---|---|---|---|---|---|---|---|---|---|---|---|---|---|---|---|---|---|---|---|---|---|---|---|---|
| 339366 | NM_00136719 | ADAMTSL5 | ADAMTS like 5 | protein_coding | NP_001354126.1;NP_998769.2;XP_011526263.1;XP_011526265.1;XP_01 | 27912 | . | . | . | 2.663175 | 2.773766 | 3.105428 | 2.839235 | 6.197314 | 6.411683 | 6.968490 | 6.491574 | 1.073627 | 231.209004 | 0.22363966 | 0.458292565 | 0.646742265 | 0.998865404 | 7.800765 | 7.903514 | 7.823468 | 7.845105 | 7.888837 | 7.848030 | 251 | 246 | 217 | 215 |
| 339400 | NR_103778,NF | FLG-AS1 | FLG antisense RNA 1 | lncRNA | . | 27913 | . | . | . | 0.154320 | 0.221391 | 0.169556 | 0.203032 | 0.359109 | 0.511755 | 0.380479 | 0.464208 | -1.019829 | 24.521972 | 0.628692058 | -0.0450578 | 0.964061239 | NA | 4.631445 | 4.600394 | 4.599046 | 4.632546 | 4.604178 | 4.623656 | 24 | 31 | 19 | 25 |
| 339416 | NM_198493,N | ANKRD45 | ankyrin repeat domain 45 | protein_coding | NP_940895.1;XP_011507768.1;XP_011507772.1;XP_011507773.1;XP_01( | 24786 | 618712 | . | . | 0.165314 | 0.096742 | 0.108291 | 0.045691 | 0.384692 | 0.223624 | 0.243001 | 0.104468 | -1.540771 | 7.763373 | 1.094837485 | -0.56962974 | 0.568928854 | NA | 3.234218 | 2.612728 | 2.971266 | 2.969843 | 2.959246 | 2.952194 | 11 | 10 | 6 | 5 |
| 339448 | NM_207356,XI | C1orf174 | chromosome 1 open reading frame 174 | protein_coding | NP_997239.2;XP_005244800.1;XP_011539625.1 | 27915 | . | . | . | 15.882367 | 15.675299 | 17.267532 | 15.700284 | 36.958919 | 36.234158 | 38.747838 | 35.896838 | 1.026084 | 876.022518 | 0.145549272 | 0.255231907 | 0.798543989 | 0.998865404 | 9.755954 | 9.793451 | 9.757833 | 9.771728 | 9.825444 | 9.739464 | 986 | 942 | 818 | 782 |
| 339451 | NM_198317,XI | KLHL17 | kelch like family member 17 | protein_coding | NP_938073.1;XP_006710663.1;XP_006710664.1 | 24023 | . | . | . | 14.651371 | 16.120841 | 18.482802 | 15.163992 | 34.094341 | 37.264045 | 41.474869 | 34.670666 | 1.079256 | 1351.906846 | 0.143916872 | 0.764587566 | 0.444517175 | 0.998865404 | 10.344487 | 10.454959 | 10.360175 | 10.377608 | 10.508707 | 10.338986 | 1480 | 1419 | 1355 | 1171 |
| 339483 | NR_026850 | MTMR9LP | myotubularin related protein 9 like, pseudogene | transcribed_ps | . | 27920 | . | . | . | 8.113587 | 8.959136 | 11.260099 | 9.636781 | 18.880650 | 20.709444 | 25.267338 | 22.033352 | 1.207602 | 687.443263 | 0.161003054 | 1.690313567 | 0.090967981 | 0.805877675 | 9.282460 | 9.554891 | 9.323678 | 9.383587 | 9.539360 | 9.423818 | 685 | 702 | 708 | 647 |
| 339487 | NM_00130813 | ZBTB8OS | zinc finger and BTB domain containing 8 opposite strand | protein_coding | NP_001295064.1;NP_001295065.1;NP_001295066.1;NP_001295067.1;NI | 24094 | 615891 | . | . | 16.185684 | 15.977378 | 15.637968 | 17.252557 | 37.664759 | 36.932429 | 35.091139 | 39.445917 | 1.041003 | 960.556753 | 0.141172292 | 0.410660608 | 0.681321415 | 0.998865404 | 9.878611 | 9.936267 | 9.895207 | 9.889836 | 9.882443 | 9.959493 | 1088 | 1012 | 807 | 967 |
| 339488 | NM_178548,XI | TFAP2E | transcription factor AP-2 epsilon | protein_coding | NP_848643.2;XP_016856628.1 | 30774 | 614428 | . | . | 0.163936 | 0.450141 | 0.127200 | 0.126758 | 0.381486 | 1.040521 | 0.285433 | 0.289816 | -1.350339 | 33.187247 | 0.538169413 | -0.80517724 | 0.420717406 | 0.998865404 | 5.252974 | 4.819737 | 5.072014 | 5.078051 | 5.037948 | 5.033557 | 43 | 42 | 25 | 26 |
| 339500 | NM_00136790 | ZNF678 | zinc finger protein 678 | protein_coding | NP_001354838.1;NP_001354839.1;NP_001354840.1;NP_848644.2 | 28652 | . | . | . | 0.588506 | 0.547653 | 0.834416 | 0.477353 | 1.369476 | 1.265924 | 1.872403 | 1.091410 | 1.137729 | 141.816738 | 0.319454307 | 0.582734103 | 0.560072312 | 0.998865404 | 7.050418 | 7.239180 | 7.123334 | 7.097557 | 7.267279 | 7.038340 | 158 | 138 | 170 | 100 |
| 339535 | NR_015407 | LINC01139 | long intergenic non-protein coding RNA 1139 | lncRNA | . | 27924 | . | . | . | 0.184866 | 0.091523 | 0.229374 | 0.398727 | 0.430190 | 0.211560 | 0.514708 | 0.911641 | 2.160940 | 11.221585 | 0.947620331 | 1.173105618 | 0.240753454 | NA | 2.826455 | 3.940155 | 3.462569 | 3.439919 | 3.471439 | 3.504596 | 11 | 5 | 10 | 18 |
| 339559 | NM_00132017 | ZFP69 | ZFP69 zinc finger protein | protein_coding | NP_001307107.1;NP_001307108.1;NP_940896.2;XP_005270866.1;XP_00 | 24708 | 617939 | . | . | 1.755018 | 1.687032 | 2.005597 | 1.701656 | 4.083999 | 3.899650 | 4.500503 | 3.890636 | 1.053758 | 152.451007 | 0.267585953 | 0.282313503 | 0.777703133 | 0.998865404 | 7.213547 | 7.289848 | 7.238111 | 7.242354 | 7.289007 | 7.231823 | 170 | 161 | 146 | 136 |
| 339665 | NM_00100147 | SLC35E4 | solute carrier family 35 member E4 | protein_coding | NP_001001479.1;NP_001305299.1;NP_001305300.1;XP_016884283.1;XF | 17058 | . | . | . | 2.936392 | 2.912900 | 2.275587 | 2.367608 | 6.833106 | 6.733299 | 5.106350 | 5.413257 | -1.274234 | 213.183601 | 0.230407139 | -1.51744531 | 0.129154291 | 0.878949649 | 7.900278 | 7.550475 | 7.793189 | 7.785882 | 7.669888 | 7.685019 | 277 | 256 | 160 | 179 |
| 339674 | NR_024355 | LINC00634 | long intergenic non-protein coding RNA 634 | lncRNA | . | 27930 | . | . | . | 0.546949 | 0.528801 | 0.468308 | 0.651652 | 1.272773 | 1.222348 | 1.050869 | 1.489924 | 1.057146 | 26.252051 | 0.604895345 | 0.13254358 | 0.894554355 | NA | 4.674556 | 4.753092 | 4.712812 | 4.708491 | 4.700651 | 4.732438 | 30 | 27 | 20 | 29 |
| 339745 | NM_00100166 | SPOPL | speckle type BTB/POZ protein like | protein_coding | NP_001001664.1;XP_005263712.1;XP_005263713.1;XP_005263714.1 | 27934 | . | . | . | 2.241063 | 2.276354 | 2.042307 | 1.357935 | 5.215044 | 5.261895 | 4.582876 | 3.104758 | -1.343813 | 368.958765 | 0.217150083 | -1.96330546 | 0.049610692 | 0.664973499 | 8.724105 | 8.299380 | 8.596108 | 8.605695 | 8.533057 | 8.316643 | 483 | 460 | 329 | 236 |
| 339761 | NM_00100166 | CYP27C1 | cytochrome P450 family 27 subfamily C member 1 | protein_coding | NP_001001665.3;NP_001354430.1;NP_001354431.1;XP_024308606.1 | 33480 | . | . | . | 0.625355 | 0.501266 | 0.441626 | 0.468124 | 1.455227 | 1.158698 | 0.990997 | 1.070310 | -1.255772 | 81.046947 | 0.358878315 | -0.91555887 | 0.359898313 | 0.998865404 | 6.495370 | 6.167403 | 6.404389 | 6.336058 | 6.305315 | 6.316216 | 116 | 86 | 61 | 69 |
| 339799 | gene-EIF3FP3 | EIF3FP3 | eukaryotic translation initiation factor 3 subunit F pseudogene 3 | pseudogene | . | 37625 | . | . | . | 4.046139 | 3.873693 | 4.180312 | 3.904322 | 9.415531 | 8.954214 | 9.380499 | 8.926768 | 1.007821 | 155.632682 | 0.263356241 | 0.042678254 | 0.965958014 | 0.998865404 | 7.276051 | 7.287927 | 7.290195 | 7.269801 | 7.297789 | 7.268359 | 183 | 163 | 141 | 141 |
| 339803 | NR_036496 | LOC339803 | uncharacterized LOC339803 | lncRNA | . | . | . | . | . | 3.027430 | 3.018100 | 2.966855 | 3.353456 | 7.044954 | 6.976473 | 6.657536 | 7.667278 | 1.032312 | 246.835188 | 0.218486095 | 0.209986972 | 0.833677841 | 0.998865404 | 7.924553 | 7.969899 | 7.941205 | 7.936843 | 7.927099 | 7.981669 | 281 | 261 | 206 | 248 |
| 339804 | NM_00114395 | C2orf74 | chromosome 2 open reading frame 74 | protein_coding | NP_001137431.1;NP_001137432.1;NP_001303246.1;NP_001353998.1;NI | 34439 | . | . | . | 11.395822 | 9.709845 | 11.298048 | 11.794979 | 26.518547 | 22.444743 | 25.352490 | 26.966675 | 1.077056 | 337.593321 | 0.198408567 | 0.5397588 | 0.589363384 | 0.998865404 | 8.344327 | 8.451951 | 8.424044 | 8.326486 | 8.413426 | 8.422910 | 407 | 320 | 302 | 331 |
| 339834 | NM_00113519 | IHO1 | interactor of HORMAD1 1 | protein_coding | NP_001128669.1;NP_835467.2;XP_006713187.1;XP_011531972.1;XP_01 | 27945 | . | . | . | 1.655120 | 2.157185 | 1.766988 | 2.072544 | 3.851532 | 4.986430 | 3.965068 | 4.738625 | 1.025135 | 159.786868 | 0.269252593 | 0.133012434 | 0.894183547 | 0.998865404 | 7.302795 | 7.337013 | 7.270110 | 7.354197 | 7.292359 | 7.352605 | 163 | 188 | 130 | 163 |
| 339879 | gene-ALG1L6P | ALG1L6P | ALG1 like 6, pseudogene | pseudogene | . | 44375 | . | . | . | 0.050494 | 0.032616 | 0.050766 | 0.052668 | 0.117501 | 0.075394 | 0.113917 | 0.120418 | 1.240863 | 5.000210 | 1.34320253 | 0.231792155 | 0.816699451 | NA | 2.156660 | 2.470304 | 2.318848 | 2.311782 | 2.321143 | 2.319344 | 6 | 4 | 5 | 5 |
| 339883 | NM_00130283 | C3orf35 | chromosome 3 open reading frame 35 | protein_coding | NP_001289760.1;NP_001289761.1;NP_848029.2;NP_848032.1;NP_8480: | 24082 | 611429 | . | . | 0.153816 | 0.048691 | 0.360768 | 0.115274 | 0.357936 | 0.112551 | 0.809553 | 0.263562 | 1.328783 | 19.710331 | 0.760004006 | 0.539609371 | 0.589466452 | NA | 4.077189 | 4.494499 | 4.291404 | 4.248777 | 4.337112 | 4.248566 | 25 | 13 | 29 | 11 |
| 339929 | NR_036497 | LPP-AS2 | LPP antisense RNA 2 | lncRNA | . | 27952 | . | . | . | 1.057584 | 1.032279 | 0.605673 | 0.781591 | 2.461041 | 2.386160 | 1.359114 | 1.787014 | -1.511064 | 78.899271 | 0.363778596 | -1.63716169 | 0.10159669 | 0.819602054 | 6.569843 | 5.972679 | 6.357676 | 6.348194 | 6.221345 | 6.272177 | 111 | 101 | 48 | 66 |
| 339942 | NR_026991 | H1-10-AS1 | H1-10 antisense RNA 1 | lncRNA | . | 27953 | . | . | . | 0.622403 | 0.765042 | 0.411902 | 0.632778 | 1.448357 | 1.768429 | 0.924296 | 1.446771 | -1.324609 | 42.182316 | 0.494793812 | -0.81966675 | 0.412406111 | 0.998865404 | 5.588663 | 5.179565 | 5.403989 | 5.439565 | 5.348289 | 5.404947 | 50 | 57 | 25 | 41 |
| 339966 | gene-LOC3399 | LOC339966 | ubiquitin family domain containing 1 pseudogene | pseudogene | . | . | . | . | . | 0.012449 | 0.036783 | 0.029307 | 0.040604 | 0.028969 | 0.085025 | 0.065765 | 0.092836 | 1.212988 | 1.501310 | 2.393488391 | 0.116384722 | 0.907347643 | NA | 0.444631 | 0.715150 | 0.579258 | 0.584004 | 0.580748 | 0.585221 | 1 | 2 | 1 | 2 |
| 339970 | gene-GCOM2 | GCOM2 | GRINL1B complex locus 2 (pseudogene) | pseudogene | . | 15712 | 608311 | . | . | 0.213994 | 0.180191 | 0.169414 | 0.154021 | 0.497972 | 0.416520 | 0.380160 | 0.352150 | -1.211063 | 23.268738 | 0.639554116 | -0.43197835 | 0.66575715 | NA | 4.671258 | 4.396303 | 4.562113 | 4.542426 | 4.537138 | 4.530471 | 32 | 25 | 19 | 19 |
| 339977 | NM_00102461 | LRRC66 | leucine rich repeat containing 66 | protein_coding | NP_001019782.1 | 34299 | . | . | . | 0.078900 | 0.131186 | 0.078247 | 0.095900 | 0.183603 | 0.303241 | 0.175585 | 0.219264 | -1.180781 | 10.844701 | 0.936062724 | -0.25611655 | 0.797860843 | NA | 3.555346 | 3.312263 | 3.429290 | 3.457097 | 3.432211 | 3.438482 | 10 | 16 | 8 | 10 |
| 339983 | NM_178557 | NAT8L | N-acetyltransferase 8 like | protein_coding | NP_848652.2 | 26742 | 610647 | . | . | 5.489038 | 4.836460 | 4.568419 | 4.648741 | 12.773218 | 11.179692 | 10.251401 | 10.628795 | -1.135835 | 922.874005 | 0.145192176 | -1.26558991 | 0.205659942 | 0.970176829 | 9.938819 | 9.755331 | 9.942385 | 9.842902 | 9.799525 | 9.804139 | 1206 | 988 | 749 | 813 |
| 340037 | NR_038915,NF | PRR7-AS1 | PRR7 antisense RNA 1 | lncRNA | . | 27961 | . | . | . | 2.477239 | 2.401221 | 2.787817 | 2.320653 | 5.764636 | 5.550530 | 6.255782 | 5.305898 | 1.013699 | 122.276915 | 0.293844726 | 0.066799597 | 0.946741244 | 0.998865404 | 6.923687 | 6.944256 | 6.938776 | 6.922752 | 6.957320 | 6.913268 | 143 | 128 | 114 | 108 |
| 340061 | NM_00130173 | STING1 | stimulator of interferon response cGAMP interactor 1 | protein_coding | NP_001288667.1;NP_001354187.1;NP_938023.1;XP_011535941.1 | 27962 | 612374 | . | . | 0.519059 | 0.778166 | 0.526665 | 0.422128 | 1.207873 | 1.798766 | 1.181820 | 0.965146 | -1.323305 | 36.809426 | 0.525324328 | -0.76932608 | 0.441699757 | 0.998865404 | 5.389828 | 4.986014 | 5.192220 | 5.248391 | 5.196204 | 5.165005 | 40 | 53 | 31 | 26 |
| 340075 | NM_00101230 | ARSI | arylsulfatase family member I | protein_coding | NP_001012301.1 | 32521 | 610009 | . | . | 0.258377 | 0.205903 | 0.147627 | 0.269620 | 0.601254 | 0.475955 | 0.331270 | 0.616453 | -1.113982 | 22.014498 | 0.674714701 | -0.23080353 | 0.817467442 | NA | 4.537347 | 4.379078 | 4.479189 | 4.454900 | 4.432906 | 4.480870 | 30 | 22 | 13 | 25 |
| 340156 | NM_00101241 | MYLK4 | myosin light chain kinase family member 4 | protein_coding | NP_001012418.2;NP_001334801.1;XP_005249135.2;XP_005249136.1;XP | 27972 | . | . | . | 0.023766 | 0.045752 | 0.052787 | 0.051697 | 0.055305 | 0.105756 | 0.118452 | 0.118199 | 1.586465 | 9.946975 | 0.980641876 | 0.678959493 | 0.497163528 | NA | 2.945312 | 3.607807 | 3.278889 | 3.302247 | 3.312690 | 3.308944 | 6 | 11 | 11 | 11 |
| 340252 | NM_00113002 | ZNF680 | zinc finger protein 680 | protein_coding | NP_001123494.1;NP_848653.2;XP_006716022.1;XP_016867615.1;XP_01 | 26897 | . | . | . | 2.676686 | 1.726357 | 1.939236 | 1.433358 | 6.228755 | 3.990554 | 4.351592 | 3.277205 | -1.286599 | 187.685026 | 0.28841944 | -1.26053426 | 0.207476697 | 0.970482251 | 7.721365 | 7.360463 | 7.693116 | 7.488915 | 7.554274 | 7.412768 | 295 | 179 | 168 | 127 |
| 340273 | NM_00116394 | ABCB5 | ATP binding cassette subfamily B member 5 | protein_coding | NP_001157413.1;NP_001157414.1;NP_001157465.1;NP_848654.3;XP_01 | 46 | 611785 | . | . | 1.683184 | 1.295866 | 1.623929 | 1.737438 | 3.916840 | 2.995453 | 3.644049 | 3.972445 | 1.096907 | 111.437586 | 0.312845896 | 0.426538903 | 0.669715207 | 0.998865404 | 6.731451 | 6.865615 | 6.822633 | 6.740326 | 6.801279 | 6.823747 | 138 | 100 | 98 | 113 |
| 340277 | NM_00112736 | FAM221A | family with sequence similarity 221 member A | protein_coding | NP_001120836.1;NP_001120837.1;NP_001287861.1;NP_954587.2;XP_01 | 27977 | . | . | . | 1.283197 | 1.442210 | 1.049425 | 1.010658 | 2.986054 | 3.333734 | 2.354881 | 2.310751 | -1.301269 | 59.886997 | 0.405906656 | -0.9359762 | 0.34928544 | 0.998865404 | 6.081607 | 5.701810 | 5.930368 | 5.936047 | 5.880969 | 5.874444 | 77 | 74 | 46 | 48 |
| 340307 | NM_178561 | CTAGE6 | CTAGE family member 6 | protein_coding | NP_848656.2 | 28644 | . | . | . | 0.086374 | 0.096831 | 0.155835 | 0.074266 | 0.200996 | 0.223829 | 0.349688 | 0.169801 | 1.176060 | 8.801600 | 1.036046648 | 0.225821441 | 0.821340301 | NA | 3.014059 | 3.251698 | 3.125457 | 3.127941 | 3.146573 | 3.119373 | 9 | 9 | 11 | 6 |
| 340348 | NM_178562,XI | TSPAN33 | tetraspanin 33 | protein_coding | NP_848657.1;XP_006716023.1 | 28743 | 610120 | . | . | 3.314293 | 2.976638 | 2.238305 | 2.667951 | 7.712495 | 6.880632 | 5.022692 | 6.099951 | -1.399437 | 250.146400 | 0.218891728 | -2.2150045 | 0.026759756 | 0.532994094 | 8.188688 | 7.704061 | 8.073456 | 8.011180 | 7.870711 | 7.881346 | 355 | 297 | 179 | 198 |
| 340351 | NM_00134585 | AGBL3 | ATP/GTP binding protein like 3 | protein_coding | NP_001332779.1;NP_001332780.1;NP_001332781.1;NP_001332782.1;NI | 27981 | 617346 | . | . | 2.696319 | 3.004201 | 2.355339 | 2.602273 | 6.274443 | 6.944344 | 5.285311 | 5.949791 | -1.222516 | 127.238759 | 0.287455759 | -1.00834119 | 0.313290694 | 0.998865404 | 7.129236 | 6.838978 | 7.016756 | 7.036332 | 6.950972 | 6.960579 | 157 | 155 | 98 | 109 |
| 340357 | NR_015383,NF | LOC340357 | uncharacterized LOC340357 | lncRNA | . | . | . | . | . | 3.100171 | 2.645930 | 5.428519 | 4.614123 | 7.214225 | 6.116186 | 12.181439 | 10.549644 | 1.791022 | 150.062848 | 0.273280219 | 3.076634746 | 0.002093517 | 0.116149544 | 6.747897 | 7.589821 | 7.096994 | 7.076022 | 7.348018 | 7.271691 | 128 | 112 | 182 | 165 |
| 340371 | NM_178564,XI | NRBP2 | nuclear receptor binding protein 2 | protein_coding | NP_848659.2;XP_016868867.1;XP_016868868.1;XP_016868869.1;XP_01( | 19339 | 615563 | . | . | 18.515012 | 19.238319 | 21.125268 | 18.935485 | 43.085193 | 44.470232 | 47.404490 | 43.293731 | 1.039291 | 2264.406488 | 0.123172631 | 0.451399085 | 0.651701943 | 0.998865404 | 11.116760 | 11.172536 | 11.112408 | 11.144410 | 11.209917 | 11.106564 | 2506 | 2444 | 2136 | 2025 |
| 340385 | NM_00131793 | ZNF517 | zinc finger protein 517 | protein_coding | NP_001304865.1;NP_998770.2;XP_011515316.1;XP_011515317.1;XP_01 | 27984 | . | . | . | 0.297202 | 0.206257 | 0.375832 | 0.300531 | 0.691603 | 0.476775 | 0.843354 | 0.687128 | -1.098229 | 42.203849 | 0.488186347 | -0.27690009 | 0.781856822 | 0.998865404 | 5.465422 | 5.330007 | 5.431579 | 5.383490 | 5.375447 | 5.409700 | 58 | 41 | 31 | 42 |
| 340390 | NM_00131630 | WDR97 | WD repeat domain 97 | protein_coding | NP_001303238.1;XP_024302913.1 | 26959 | . | . | . | 0.022494 | 0.055472 | 0.050830 | 0.051104 | 0.052345 | 0.128226 | 0.114060 | 0.116844 | 1.350197 | 10.122854 | 0.979371846 | 0.442293424 | 0.658276889 | NA | 3.109221 | 3.538109 | 3.306372 | 3.338119 | 3.335274 | 3.336426 | 6 | 13 | 10 | 11 |
| 340393 | NM_00125240 | TMEM249 | transmembrane protein 249 | protein_coding | NP_001239331.1;NP_001239333.1;NP_001267490.1 | 44155 | . | . | . | 0.488433 | 0.502851 | 0.752535 | 0.746980 | 1.136605 | 1.162362 | 1.688667 | 1.707880 | 1.459220 | 30.369822 | 0.56142373 | 0.971098215 | 0.331499373 | 0.998865404 | 4.626345 | 5.171577 | 4.885487 | 4.896447 | 4.944014 | 4.929699 | 27 | 28 | 33 | 32 |
| 340481 | NM_00135411 | ZDHHC21 | zinc finger DHHC-type palmitoyltransferase 21 | protein_coding | NP_001341047.1;NP_001341048.1;NP_001341049.1;NP_001341050.1;NI | 20750 | 614605 | . | . | 0.834161 | 0.716215 | 0.833699 | 0.493965 | 1.941126 | 1.655565 | 1.870796 | 1.129393 | -1.148196 | 215.927410 | 0.271633768 | -0.73396306 | 0.46297125 | 0.998865404 | 7.849557 | 7.652526 | 7.794096 | 7.762673 | 7.824912 | 7.589062 | 274 | 241 | 221 | 139 |
| 340485 | NM_00101088 | ACER2 | alkaline ceramidase 2 | protein_coding | NP_001010887.2;XP_005251505.1;XP_011516160.1;XP_011516161.1;XP_ | 23675 | 613492 | . | . | 0.168514 | 0.186916 | 0.126311 | 0.050753 | 0.392139 | 0.432065 | 0.283437 | 0.116040 | -1.537063 | 20.614150 | 0.725364899 | -0.85498587 | 0.392558973 | NA | 4.639727 | 4.026777 | 4.405823 | 4.359200 | 4.373807 | 4.317434 | 35 | 21 | 20 | 9 |
| 340526 | NM_00102445 | RTL5 | retrotransposon Gag like 5 | protein_coding | NP_001019626.1 | 29430 | . | . | . | 0.455575 | 0.426632 | 0.405806 | 0.485035 | 1.060141 | 0.986179 | 0.910617 | 1.108974 | -1.004891 | 60.294552 | 0.405941094 | -0.01734139 | 0.986164264 | 0.999197585 | 5.917822 | 5.910079 | 5.922174 | 5.907185 | 5.896418 | 5.930617 | 72 | 63 | 48 | 61 |
| 340544 | NR_038978 | MORF4L2-AS1 | MORF4L2 antisense RNA 1 | lncRNA | . | 27991 | . | . | . | 0.212757 | 0.129451 | 0.243613 | 0.235198 | 0.495094 | 0.299231 | 0.546660 | 0.537753 | 1.345999 | 18.801176 | 0.718562198 | 0.596577263 | 0.550789655 | NA | 4.000985 | 4.432887 | 4.231147 | 4.196123 | 4.238437 | 4.236550 | 23 | 13 | 19 | 20 |
| 340554 | NM_00101088 | ZC3H12B | zinc finger CCCH-type containing 12B | protein_coding | NP_001010888.3;XP_011529241.1;XP_016884967.1;XP_016884968.1;XP_ | 17407 | 300889 | . | . | 0.045783 | 0.107385 | 0.099993 | 0.045967 | 0.106538 | 0.248224 | 0.224383 | 0.105099 | -1.257921 | 18.165528 | 0.72227145 | -0.4583337 | 0.64671272 | NA | 4.339107 | 4.008225 | 4.185372 | 4.202636 | 4.181283 | 4.171383 | 21 | 24 | 15 | 14 |
| 340591 | NR_026551,NF | CA5BP1 | carbonic anhydrase 5B pseudogene 1 | transcribed_ps | . | 29544 | . | . | . | 7.578420 | 7.543447 | 7.318892 | 8.318407 | 17.635294 | 17.437016 | 16.423382 | 19.019051 | 1.033697 | 328.250562 | 0.195592025 | 0.244453912 | 0.806879292 | 0.998865404 | 8.334815 | 8.382105 | 8.344770 | 8.353160 | 8.339949 | 8.393929 | 369 | 351 | 276 | 328 |
| 340719 | NM_199461 | NANOS1 | nanos C2HC-type zinc finger 1 | protein_coding | NP_955631.1 | 23044 | 608226 | . | . | 1.607158 | 1.589166 | 1.403875 | 1.790200 | 3.739922 | 3.673429 | 3.150255 | 4.093081 | -1.015419 | 200.581385 | 0.244244505 | -0.09038398 | 0.927982084 | 0.998865404 | 7.659142 | 7.635946 | 7.654753 | 7.645831 | 7.590843 | 7.693190 | 235 | 216 | 153 | 208 |
| 340780 | gene-IMPDH1F | IMPDH1P5 | inosine monophosphate dehydrogenase 1 pseudogene 5 | pseudogene | . | 33960 | . | . | . | 0.065508 | 0.050956 | 0.021383 | 0.019154 | 0.152440 | 0.117788 | 0.047983 | 0.043792 | -2.014346 | 3.337728 | 1.664788034 | -0.60687087 | 0.543936609 | NA | 2.156660 | 1.148375 | 1.760619 | 1.753493 | 1.747193 | 1.746336 | 6 | 4 | 2 | 2 |
| 340784 | NM_00110557 | HMX3 | H6 family homeobox 3 | protein_coding | NP_001099044.1 | 5019 | 613380 | . | . | 0.097783 | 0.141565 | 0.016206 | 0.122215 | 0.227546 | 0.327233 | 0.036366 | 0.279430 | -1.593563 | 9.140144 | 1.082312593 | -0.62112955 | 0.53451439 | NA | 3.493602 | 2.810781 | 3.200074 | 3.215423 | 3.168444 | 3.209470 | 11 | 14 | 2 | 11 |
| 340900 | NM_00119993 | EBLN1 | endogenous Bornavirus like nucleoprotein 1 | protein_coding | NP_001186867.1 | 39430 | 613249 | . | . | 0.024960 | 0.152151 | 0.062283 | 0.065696 | 0.058083 | 0.351704 | 0.139760 | 0.150205 | -1.487750 | 3.431012 | 1.66373865 | -0.34448434 | 0.730482069 | NA | 2.039857 | 1.459419 | 1.771966 | 1.794306 | 1.775046 | 1.779018 | 2 | 7 | 2 | 3 |
| 341019 | NM_00136797 | DCDC1 | doublecortin domain containing 1 | protein_coding | NP_001354908.1;NP_065920.2;NP_861523.2;XP_024304238.1;XP_02430 | 20625 | 608062 | . | . | 0.056293 | 0.033152 | 0.083144 | 0.042250 | 0.130995 | 0.076633 | 0.186572 | 0.096601 | 1.405659 | 12.910467 | 0.866937412 | 0.566646563 | 0.570954333 | NA | 3.421266 | 3.917252 | 3.675379 | 3.662298 | 3.706604 | 3.669016 | 14 | 10 | 17 | 10 |
| 341350 | NM_00135317 | OVCH1 | ovochymase 1 | protein_coding | NP_001340108.1;XP_011518940.1;XP_011518942.1;XP_011518943.1;XP_ | 23080 | . | . | . | 0.029718 | 0.036986 | 0.022361 | 0.030035 | 0.069156 | 0.085495 | 0.050177 | 0.068671 | -1.290188 | 7.574992 | 1.103464968 | -0.33311562 | 0.739047006 | NA | 3.094525 | 2.724273 | 2.926937 | 2.933555 | 2.917883 | 2.925340 | 9 | 10 | 5 | 7 |
| 341640 | NM_207361,XI | FREM2 | FRAS1 related extracellular matrix 2 | protein_coding | NP_997244.4;XP_016876043.1 | 25396 | 608945 | . | . | 0.534062 | 0.557460 | 0.366162 | 0.518987 | 1.242785 | 1.288593 | 0.821657 | 1.186602 | -1.281097 | 233.969935 | 0.229618685 | -1.55640761 | 0.119611197 | 0.858998979 | 8.038234 | 7.679935 | 7.944163 | 7.908293 | 7.760006 | 7.851045 | 313 | 274 | 160 | 212 |
| 341676 | NM_00136555 | NEK5 | NIMA related kinase 5 | protein_coding | NP_001352481.1;NP_954983.1;XP_006719871.2;XP_011533368.1;XP_01 | 7748 | 616731 | . | . | 0.374872 | 0.259218 | 0.314016 | 0.136084 | 0.872343 | 0.599194 | 0.704643 | 0.311141 | -1.809397 | 38.139979 | 0.515081294 | -1.66092076 | 0.096729442 | 0.813553488 | 5.617806 | 4.764191 | 5.320698 | 5.270713 | 5.217079 | 5.203776 | 64 | 46 | 25 | 24 |
| 342132 | NM_00100430 | ZNF774 | zinc finger protein 774 | protein_coding | NP_001004309.2;XP_016877616.1;XP_016877617.1;XP_016877618.1 | 33108 | . | . | . | 0.891013 | 1.078593 | 0.914578 | 0.980852 | 2.073424 | 2.493214 | 2.052287 | 2.242600 | -1.044480 | 97.219607 | 0.327872782 | -0.19149359 | 0.848138899 | 0.998865404 | 6.634790 | 6.570852 | 6.580561 | 6.636923 | 6.586780 | 6.605335 | 104 | 117 | 80 | 92 |
| 342184 | NM_00110318 | FMN1 | formin 1 | protein_coding | NP_001096654.1;NP_001264242.1;NP_001264243.1;XP_011519806.1;XF | 3768 | 136535 | . | . | 9.393719 | 10.040419 | 10.329068 | 10.342866 | 21.859565 | 23.208880 | 23.178129 | 23.647735 | 1.051523 | 4115.884764 | 0.108714639 | 0.666700494 | 0.504963463 | 0.998865404 | 11.970295 | 12.042769 | 11.965263 | 12.005538 | 12.040413 | 12.014070 | 4508 | 4435 | 3727 | 3893 |
| 342357 | NM_00101298 | ZKSCAN2 | zinc finger with KRAB and SCAN domains 2 | protein_coding | NP_001012999.3;XP_016878689.1 | 25677 | . | . | . | 1.512255 | 1.468448 | 1.413137 | 1.160202 | 3.519080 | 3.394382 | 3.171039 | 2.652665 | -1.175410 | 324.294539 | 0.203091114 | -1.14807415 | 0.250937954 | 0.993339639 | 8.452520 | 8.220482 | 8.396832 | 8.369011 | 8.344823 | 8.240340 | 413 | 369 | 285 | 252 |
| 342371 | NM_00113767 | ATXN1L | ataxin 1 like | protein_coding | NP_001131147.1 | 33279 | 614301 | . | . | 3.457920 | 3.932567 | 3.557965 | 4.131833 | 8.046721 | 9.090304 | 7.983969 | 9.446945 | 1.026612 | 927.427186 | 0.14961335 | 0.253257178 | 0.800069485 | 0.998865404 | 9.838359 | 9.875581 | 9.802121 | 9.888989 | 9.813657 | 9.914438 | 991 | 1047 | 760 | 942 |
| 342574 | NM_181537 | KRT27 | keratin 27 | protein_coding | NP_853515.2 | 30841 | 616676 | . | . | 1.030867 | 0.841032 | 1.000935 | 1.257697 | 2.398871 | 1.944083 | 2.246068 | 2.875575 | 1.199295 | 53.070210 | 0.435377786 | 0.602204001 | 0.547038353 | 0.998865404 | 5.592935 | 5.854854 | 5.725667 | 5.690071 | 5.721300 | 5.764623 | 61 | 47 | 45 | 60 |
| 342865 | NM_00114633 | VSTM2B | V-set and transmembrane domain containing 2B | protein_coding | NP_001139811.1;XP_005258913.1;XP_011525204.1;XP_011525205.1;XP_ | 33595 | . | . | . | 0.799067 | 1.369727 | 0.803065 | 1.256002 | 1.859459 | 3.166186 | 1.802055 | 2.871699 | -1.074743 | 59.537019 | 0.41871725 | -0.24835888 | 0.803856744 | 0.998865404 | 5.947968 | 5.841498 | 5.866546 | 5.935800 | 5.866856 | 5.905888 | 60 | 77 | 45 | 59 |
| 342892 | NM_00119355 | ZNF850 | zinc finger protein 850 | protein_coding | NP_001180481.1;NP_001254708.1 | 27994 | . | . | . | 0.356488 | 0.446914 | 0.339321 | 0.340228 | 0.829561 | 1.033061 | 0.761426 | 0.777891 | -1.199400 | 89.497055 | 0.342292332 | -0.76633996 | 0.443474049 | 0.998865404 | 6.609351 | 6.346202 | 6.474246 | 6.541888 | 6.458403 | 6.456160 | 100 | 117 | 71 | 76 |
| 342908 | NM_00103371 | ZNF404 | zinc finger protein 404 | protein_coding | NP_001028891.2 | 19417 | . | . | . | 0.570956 | 0.385903 | 0.399852 | 0.118587 | 1.328637 | 0.892032 | 0.897257 | 0.271136 | -1.883349 | 21.876003 | 0.725716625 | -1.25848029 | 0.208218111 | NA | 4.833848 | 3.929136 | 4.499456 | 4.454500 | 4.453527 | 4.387919 | 39 | 25 | 20 | 7 |
| 342909 | NM_00103781 | ZNF284 | zinc finger protein 284 | protein_coding | NP_001032902.1;XP_011525209.1;XP_011525210.1;XP_024307254.1;XP_ | 13078 | . | . | . | 1.083198 | 1.237115 | 1.326518 | 1.287801 | 2.520645 | 2.859647 | 2.976670 | 2.944404 | 1.205338 | 157.051544 | 0.26228448 | 1.027275146 | 0.304290922 | 0.998865404 | 7.153772 | 7.423800 | 7.266913 | 7.239166 | 7.337507 | 7.317471 | 170 | 148 | 153 | 157 |
| 342918 | NM_00119507 | C19orf81 | chromosome 19 open reading frame 81 | protein_coding | NP_001182005.1 | 40041 | . | . | . | 1.072247 | 0.850912 | 0.593787 | 0.865907 | 2.495163 | 1.966921 | 1.332441 | 1.979793 | -1.278718 | 20.681163 | 0.685358713 | -0.51753702 | 0.604781331 | NA | 4.537347 | 4.181222 | 4.397189 | 4.372826 | 4.350532 | 4.376792 | 30 | 22 | 13 | 20 |
| 342945 | NM_00132111 | ZSCAN22 | zinc finger and SCAN domain containing 22 | protein_coding | NP_001308045.1;NP_001308046.1;NP_862829.1;XP_006723255.1;XP_01 | 4929 | 165260 | . | . | 1.043655 | 1.215409 | 1.494131 | 1.148118 | 2.428628 | 2.809473 | 3.352786 | 2.625035 | 1.077958 | 199.240762 | 0.246554078 | 0.439257767 | 0.660474772 | 0.998865404 | 7.582775 | 7.691900 | 7.597852 | 7.637538 | 7.703875 | 7.596863 | 211 | 216 | 201 | 171 |
| 343099 | NM_00130607 | CCDC18 | coiled-coil domain containing 18 | protein_coding | NP_001293005.1;NP_001365133.1;NP_996769.3;XP_011539663.2;XP_01 | 30370 | . | . | . | 2.811996 | 2.360753 | 3.125239 | 1.761021 | 6.543626 | 5.456986 | 7.012945 | 4.026362 | -1.239921 | 446.682737 | 0.227706591 | -1.36249165 | 0.173042777 | 0.947431715 | 8.949183 | 8.640556 | 8.865947 | 8.847830 | 8.869022 | 8.556883 | 577 | 526 | 439 | 275 |
| 343171 | NM_00100195 | OR2W3 | olfactory receptor family 2 subfamily W member 3 | protein_coding | NP_001001957.2 | 15021 | 616729 | . | . | 1.071218 | 1.337603 | 0.545208 | 1.376927 | 2.492768 | 3.091929 | 1.223432 | 3.148181 | -1.265862 | 32.153843 | 0.596616561 | -0.57008194 | 0.568622126 | 0.998865404 | 5.169572 | 4.823571 | 5.004880 | 5.035849 | 4.940860 | 5.037497 | 37 | 43 | 14 | 38 |
| 343472 | NM_020063 | BARHL2 | BarH like homeobox 2 | protein_coding | NP_064447.1 | 954 | 605212 | . | . | 0.427423 | 0.507176 | 0.298107 | 0.695518 | 0.994630 | 1.172360 | 0.668944 | 1.590219 | 1.072077 | 31.143949 | 0.595470594 | 0.168620035 | 0.866095521 | 0.998865404 | 4.912320 | 5.007859 | 4.941274 | 4.960547 | 4.909223 | 5.007891 | 32 | 35 | 17 | 42 |
| 344148 | NM_207363,N | NCKAP5 | NCK associated protein 5 | protein_coding | NP_997246.2;NP_997364.3;XP_005263716.1;XP_005263717.1;XP_01150 | 29847 | 608789 | . | . | 0.915150 | 0.927477 | 0.760434 | 0.940538 | 2.129596 | 2.143905 | 1.706392 | 2.150427 | -1.144550 | 216.432437 | 0.235807987 | -0.82601244 | 0.408797038 | 0.998865404 | 7.852271 | 7.656649 | 7.804914 | 7.771794 | 7.680205 | 7.768394 | 275 | 241 | 158 | 208 |
| 344167 | NM_00113564 | FOXI3 | forkhead box I3 | protein_coding | NP_001129121.1 | 35123 | 612351 | . | . | 0.155151 | 0.227891 | 0.270804 | 0.404500 | 0.361042 | 0.526781 | 0.607676 | 0.924841 | 1.767414 | 23.674901 | 0.652402985 | 1.259405508 | 0.207883901 | NA | 4.099441 | 4.916862 | 4.505623 | 4.534701 | 4.550649 | 4.599294 | 16 | 22 | 21 | 34 |
| 344405 | NR_027258 | PRORSD1P | prolyl-tRNA synthetase associated domain containing 1, pseudoge | transcribed_ps | . | 34379 | . | . | . | 0.781160 | 0.701525 | 0.789660 | 0.782186 | 1.817792 | 1.621606 | 1.771974 | 1.788376 | 1.039647 | 51.568224 | 0.435317501 | 0.128858644 | 0.897469501 | 0.998865404 | 5.659745 | 5.716518 | 5.696348 | 5.671582 | 5.692505 | 5.690307 | 62 | 51 | 46 | 49 |
| 344595 | NR_028301,NF | DUBR | DPPA2 upstream binding RNA | lncRNA | . | 48569 | 616619 | . | . | 1.163199 | 1.175567 | 1.209723 | 1.144545 | 2.706813 | 2.717376 | 2.714584 | 2.616866 | -1.008964 | 119.642280 | 0.294788079 | -0.04367341 | 0.965164736 | 0.998865404 | 6.908824 | 6.896316 | 6.903031 | 6.904847 | 6.911872 | 6.889828 | 138 | 130 | 107 | 108 |
| 344697 | gene-HNRNPA: | HNRNPA1P21 | heterogeneous nuclear ribonucleoprotein A1 pseudogene 21 | pseudogene | . | 39539 | . | . | . | 0.132828 | 0.045971 | 0.031171 | 0.029210 | 0.309096 | 0.106263 | 0.069948 | 0.066785 | -3.196093 | 2.320196 | 2.070611742 | -0.80957213 | 0.418186129 | NA | 1.820514 | 0.148375 | 1.248353 | 1.232544 | 1.229567 | 1.229073 | 6 | 2 | 1 | 1 |
| 344838 | NM_00137530 | PAQR9 | progestin and adipoQ receptor family member 9 | protein_coding | NP_001362229.1;NP_001362230.1;NP_940906.1 | 30131 | 614580 | . | . | 0.093409 | 0.073132 | 0.060225 | 0.026840 | 0.217366 | 0.169048 | 0.135143 | 0.061366 | -1.879027 | 18.504946 | 0.745540598 | -1.22057124 | 0.2222484 | NA | 4.592063 | 3.688346 | 4.248854 | 4.224672 | 4.207431 | 4.170010 | 31 | 23 | 15 | 8 |
| 344887 | NR_151491 | NMRAL2P | NmrA like redox sensor 2, pseudogene | transcribed_ps | . | 52332 | . | . | . | 7.333138 | 7.016976 | 12.899807 | 8.686258 | 17.064514 | 16.220053 | 28.946795 | 19.860098 | 1.483165 | 502.987751 | 0.203726775 | 2.79137946 | 0.00524839 | 0.220864422 | 8.661511 | 9.231342 | 8.839437 | 8.807715 | 9.200575 | 8.929401 | 479 | 425 | 627 | 451 |
| 345462 | NM_00113611 | ZNF879 | zinc finger protein 879 | protein_coding | NP_001129588.1;NP_001340301.1;NP_001340302.1;NP_001340303.1;XF | 37273 | . | . | . | 1.079912 | 1.464646 | 1.595113 | 0.983405 | 2.512999 | 3.385595 | 3.579387 | 2.248438 | 1.102927 | 164.956501 | 0.291713752 | 0.484507897 | 0.628025461 | 0.998865404 | 7.292693 | 7.435650 | 7.309816 | 7.361778 | 7.472491 | 7.270110 | 169 | 180 | 187 | 123 |
| 345630 | NM_00135527 | FBLL1 | fibrillarin like 1 | protein_coding | NP_001342203.1 | 35458 | . | . | . | 0.405497 | 0.501595 | 0.515412 | 0.225494 | 0.943609 | 1.159459 | 1.156569 | 0.515564 | -1.233696 | 17.533371 | 0.763630072 | -0.39677233 | 0.69153536 | NA | 4.273664 | 3.974966 | 4.128199 | 4.145317 | 4.146618 | 4.094877 | 20 | 23 | 19 | 9 |
| 345643 | NM_00119078 | MCIDAS | multiciliate differentiation and DNA synthesis associated cell cycle | protein_coding | NP_001177716.1;XP_016864928.1 | 40050 | 614086 | . | . | 2.924078 | 3.627726 | 3.485299 | 3.225498 | 6.804451 | 8.385651 | 7.820911 | 7.374718 | 1.049440 | 273.348661 | 0.223405075 | 0.311626892 | 0.755324094 | 0.998865404 | 8.059258 | 8.129094 | 8.024565 | 8.129741 | 8.154382 | 8.049913 | 276 | 317 | 268 | 236 |
| 345651 | NM_00101799 | ACTBL2 | actin beta like 2 | protein_coding | NP_001017992.1 | 17780 | 614835 | . | . | 0.162645 | 0.054356 | 0.057274 | 0.075531 | 0.378482 | 0.125647 | 0.128521 | 0.172692 | -1.539454 | 8.385131 | 1.090265562 | -0.57088765 | 0.568075802 | NA | 3.345111 | 2.724273 | 3.103095 | 3.061084 | 3.062649 | 3.070002 | 17 | 6 | 5 | 7 |
| 345757 | NM_198507,XI | FAM174A | family with sequence similarity 174 member A | protein_coding | NP_940909.1;XP_006714663.1 | 24943 | . | . | . | 4.644032 | 5.182721 | 6.000048 | 4.117006 | 10.806852 | 11.980093 | 13.463935 | 9.413045 | 1.023213 | 202.674045 | 0.258586384 | 0.128028298 | 0.898126579 | 0.998865404 | 7.645664 | 7.680165 | 7.631325 | 7.673372 | 7.744669 | 7.574237 | 220 | 226 | 212 | 156 |
| 345778 | NM_00101089 | MTX3 | metaxin 3 | protein_coding | NP_001010891.4;NP_001161213.1;NP_001350747.1;XP_016864929.1 | 24812 | . | . | . | 4.215145 | 4.300205 | 4.615711 | 3.251602 | 9.808816 | 9.940117 | 10.357521 | 7.434401 | -1.098555 | 1006.340090 | 0.171252811 | -0.79185005 | 0.428448115 | 0.998865404 | 10.040748 | 9.905907 | 9.999524 | 10.009165 | 10.056732 | 9.797980 | 1205 | 1144 | 983 | 739 |
| 345895 | NM_00101089 | RSPH4A | radial spoke head component 4A | protein_coding | NP_001010892.1;NP_001155136.1;XP_016866315.1 | 21558 | 612647 | . | . | 0.050779 | 0.112493 | 0.101875 | 0.176895 | 0.118164 | 0.260033 | 0.228605 | 0.404449 | 1.566468 | 18.547271 | 0.729686338 | 0.887388115 | 0.37486998 | NA | 3.855800 | 4.499300 | 4.163735 | 4.200375 | 4.209249 | 4.225345 | 12 | 20 | 18 | 23 |
| 346085 | gene-GAPDHP: | GAPDHP72 | glyceraldehyde-3-phosphate dehydrogenase pseudogene 72 | pseudogene | . | 22955 | . | . | . | 0.422266 | 0.201714 | 0.406000 | 0.326739 | 0.982631 | 0.466272 | 0.911051 | 0.747049 | 1.206374 | 14.206457 | 0.833865924 | 0.324604575 | 0.745480364 | NA | 3.684232 | 3.959598 | 3.837066 | 3.796400 | 3.835589 | 3.821400 | 20 | 9 | 15 | 13 |
| 346157 | NM_00107678 | ZNF391 | zinc finger protein 391 | protein_coding | NP_001070249.1;NP_001309217.1;NP_001309218.1;NP_001309222.1;XF | 18779 | . | . | . | 0.422836 | 0.444543 | 0.316307 | 0.500507 | 0.983956 | 1.027579 | 0.709785 | 1.144350 | -1.084961 | 53.838122 | 0.441990777 | -0.26616656 | 0.790110939 | 0.998865404 | 5.809543 | 5.689054 | 5.753193 | 5.763221 | 5.700826 | 5.782059 | 63 | 62 | 35 | 59 |
| 346653 | NM_00101245 | FAM71F2 | family with sequence similarity 71 member F2 | protein_coding | NP_001012457.3;NP_001122398.1;NP_001277183.1;NP_001277184.1;XF | 27998 | . | . | . | 0.272895 | 0.606418 | 0.847164 | 0.569649 | 0.635039 | 1.401764 | 1.901011 | 1.302435 | 2.211122 | 69.136908 | 0.43966449 | 2.6037543 | 0.009220883 | 0.30595157 | 5.430174 | 6.572240 | 5.907460 | 6.036022 | 6.198533 | 6.111375 | 31 | 64 | 95 | 76 |
| 347240 | NM_194313,XI | KIF24 | kinesin family member 24 | protein_coding | NP_919289.2;XP_006716827.1;XP_011516163.1;XP_011516165.1;XP_01( | 19916 | 613747 | . | . | 3.428577 | 4.230245 | 4.169033 | 4.067114 | 7.978442 | 9.778399 | 9.355190 | 9.298971 | 1.106279 | 760.118614 | 0.154292313 | 0.9444098 | 0.344960276 | 0.998865404 | 9.495547 | 9.640952 | 9.476491 | 9.585172 | 9.612515 | 9.592880 | 766 | 840 | 704 | 738 |
| 347273 | NM_00101811 | CAVIN4 | caveolae associated protein 4 | protein_coding | NP_001018126.1 | 33742 | 617714 | . | . | 0.467013 | 0.620322 | 0.912507 | 0.601995 | 1.086759 | 1.433901 | 2.047638 | 1.376389 | 1.374500 | 63.103471 | 0.419104117 | 1.094971782 | 0.273528999 | 0.998865404 | 5.731534 | 6.191287 | 5.902051 | 5.955629 | 6.049403 | 5.946540 | 53 | 65 | 77 | 54 |
| 347344 | NM_00137815 | ZNF81 | zinc finger protein 81 | protein_coding | NP_001365081.1;NP_001365082.1;NP_001365083.1;NP_001365084.1;NI | 13156 | 314998 | . | . | 0.774821 | 0.963256 | 1.216602 | 1.612616 | 1.803042 | 2.226613 | 2.730018 | 3.687056 | 1.183511 | 275.053197 | 0.237250082 | 1.024545525 | 0.30557768 | 0.998865404 | 7.976241 | 8.220561 | 8.030452 | 8.068926 | 8.238848 | 8.025094 | 279 | 282 | 311 | 224 |
| 347365 | NM_198510 | ITIH6 | inter-alpha-trypsin inhibitor heavy chain family member 6 | protein_coding | NP_940912.1 | 28907 | . | . | . | 4.371182 | 5.019901 | 4.559235 | 5.992342 | 10.171922 | 11.603727 | 10.230791 | 13.700779 | 1.108246 | 770.712322 | 0.169615175 | 0.874204069 | 0.38200713 | 0.998865404 | 9.514515 | 9.661824 | 9.504130 | 9.593565 | 9.524214 | 9.710649 | 788 | 840 | 612 | 859 |
| 347404 | NM_00117033 | LANCL3 | LanC like 3 | protein_coding | NP_001163802.1;NP_940913.1;XP_011542206.1 | 24767 | . | . | . | 0.046702 | 0.039950 | 0.059883 | 0.047272 | 0.108677 | 0.092347 | 0.134375 | 0.108082 | 1.203277 | 15.772321 | 0.772956117 | 0.345386241 | 0.72980402 | NA | 3.838265 | 4.107809 | 3.973720 | 3.961901 | 3.990005 | 3.970751 | 18 | 14 | 17 | 14 |
| 347475 | NM_00110135 | CCDC160 | coiled-coil domain containing 160 | protein_coding | NP_001094827.1;NP_001340382.1;XP_005262467.1 | 37286 | . | . | . | 0.054322 | 0.115868 | 0.101238 | 0.083865 | 0.126409 | 0.267833 | 0.227175 | 0.191748 | 1.006277 | 4.994380 | 1.354912675 | 0.006662934 | 0.994683787 | NA | 2.316288 | 2.324312 | 2.309109 | 2.323311 | 2.319678 | 2.313073 | 4 | 7 | 5 | 4 |
| 347686 | NR_001294 | SNORD64 | small nucleolar RNA, C/D box 64 | snoRNA | . | 32725 | . | . | . | 0.001574 | 0.001815 | 0.004823 | 0.010573 | 0.003663 | 0.004196 | 0.010822 | 0.024174 | 1.234655 | 1.002957 | 2.868339486 | 0.106022226 | 0.915564717 | NA | -0.155866 | 0.148375 | -0.001261 | -0.000972 | 0.000053 | -0.000315 | 1 | 1 | 1 | 1 |
| 347688 | NM_177987,XI | TUBB8 | tubulin beta 8 class VIII | protein_coding | NP_817124.1;XP_011517761.1;XP_011517762.1;XP_016871681.1;XP_01( | 20773 | 616768 | . | . | 0.089915 | 0.098897 | 0.054550 | 0.090690 | 0.209237 | 0.228605 | 0.122408 | 0.207352 | -1.327971 | 5.120281 | 1.336933271 | -0.30609102 | 0.759535345 | NA | 2.548173 | 2.134756 | 2.362177 | 2.368086 | 2.354481 | 2.362709 | 6 | 7 | 3 | 5 |
| 347716 | NR_146076 | USP32P3 | ubiquitin specific peptidase 32 pseudogene 3 | transcribed_ps | . | 43576 | . | . | . | 0.870401 | 0.900459 | 1.173715 | 0.775325 | 2.025460 | 2.081452 | 2.633781 | 1.772687 | 1.084972 | 129.612668 | 0.307358113 | 0.382803915 | 0.701865158 | 0.998865404 | 6.957030 | 7.076618 | 6.991481 | 7.001865 | 7.099177 | 6.946995 | 141 | 136 | 142 | 100 |
| 347733 | NM_178012 | TUBB2B | tubulin beta 2B class IIb | protein_coding | NP_821080.1 | 30829 | 612850 | . | . | 27.075468 | 27.950586 | 21.129953 | 23.633162 | 63.005722 | 64.609032 | 47.415005 | 54.034416 | -1.246033 | 1495.043919 | 0.128478944 | -2.46999108 | 0.013511642 | 0.383747314 | 10.696078 | 10.378415 | 10.615347 | 10.633764 | 10.416295 | 10.491709 | 1888 | 1811 | 1099 | 1311 |
| 347734 | NM_00128650 | SLC35B2 | solute carrier family 35 member B2 | protein_coding | NP_001273438.1;NP_001273439.1;NP_001273440.1;NP_001273441.1;NI | 16872 | 610788 | . | . | 77.959866 | 77.239235 | 69.484286 | 73.534855 | 181.415807 | 178.541950 | 155.920725 | 168.128715 | -1.100725 | 4686.655780 | 0.106941322 | -1.2946788 | 0.195431055 | 0.96621494 | 12.261931 | 12.123432 | 12.239851 | 12.226255 | 12.133642 | 12.171907 | 5701 | 5257 | 3790 | 4280 |
| 347735 | NM_00119903 | SERINC2 | serine incorporator 2 | protein_coding | NP_001185966.1;NP_001185967.1;NP_001185968.1;NP_061035.2;NP_8‹ | 23231 | 614549 | . | . | 6.111066 | 5.895320 | 6.248010 | 6.564402 | 14.220702 | 13.627297 | 14.020353 | 15.008728 | 1.057188 | 369.588520 | 0.18598872 | 0.43138061 | 0.666191646 | 0.998865404 | 8.489137 | 8.569302 | 8.524374 | 8.501455 | 8.532162 | 8.557975 | 422 | 380 | 323 | 364 |
| 347744 | NM_00114502 | C6orf52 | chromosome 6 open reading frame 52 | protein_coding | NP_001138492.1;NP_001341286.1;XP_006715151.1;XP_011512874.1;XP | 20881 | . | . | . | 0.393855 | 0.422678 | 0.410743 | 0.286298 | 0.916519 | 0.977040 | 0.921693 | 0.654586 | -1.368259 | 10.499326 | 0.94139711 | -0.48050013 | 0.630871807 | NA | 3.600756 | 3.148375 | 3.400130 | 3.407636 | 3.390954 | 3.387906 | 13 | 14 | 8 | 8 |
| 347746 | NR_022011 | PWARSN | Prader Willi/Angelman region RNA, SNRPN neighbor | lncRNA | . | 30001 | . | . | . | 0.649913 | 1.079732 | 1.042816 | 1.069588 | 1.512375 | 2.495850 | 2.340049 | 2.445487 | 1.203649 | 54.222553 | 0.444921312 | 0.601038035 | 0.547814654 | 0.998865404 | 5.622171 | 5.887307 | 5.687625 | 5.778458 | 5.768541 | 5.772661 | 43 | 66 | 51 | 56 |
| 347862 | NM_00131881 | GATD1 | glutamine amidotransferase like class 1 domain containing 1 | protein_coding | NP_001305747.1;NP_001305749.1;NP_001305750.1;NP_001305751.1;NI | 26616 | . | . | . | 15.091077 | 15.992305 | 18.168472 | 17.295220 | 35.117558 | 36.966932 | 40.769528 | 39.543466 | 1.131591 | 2029.499247 | 0.119548103 | 1.49189478 | 0.135726731 | 0.893814919 | 10.894892 | 11.073406 | 10.939951 | 10.930368 | 11.061157 | 11.005277 | 2206 | 2042 | 1939 | 1950 |
| 347902 | NM_00114366 | AMIGO2 | adhesion molecule with Ig like domain 2 | protein_coding | NP_001137140.1;NP_001357228.1;NP_862830.1 | 24073 | 615690 | . | . | 0.462747 | 0.375648 | 0.410233 | 0.180353 | 1.076830 | 0.868328 | 0.920552 | 0.412355 | -1.422996 | 43.427031 | 0.518242958 | -0.98203202 | 0.326084074 | 0.998865404 | 5.670433 | 5.166894 | 5.485062 | 5.442848 | 5.457152 | 5.352867 | 65 | 49 | 43 | 21 |
| 347918 | NR_003290 | EP400P1 | EP400 pseudogene 1 | transcribed_ps | . | 26602 | . | . | . | 1.208415 | 1.134370 | 1.840322 | 1.679373 | 2.812032 | 2.622148 | 4.129629 | 3.839688 | 1.481001 | 70.570014 | 0.376885415 | 1.503302272 | 0.132761115 | 0.887687926 | 5.829494 | 6.397016 | 6.089131 | 6.073731 | 6.187996 | 6.161433 | 68 | 59 | 77 | 75 |
| 348013 | NM_00134866 | TMEM255B | transmembrane protein 255B | protein_coding | NP_001335592.1;NP_872420.1;XP_005272470.1;XP_005272472.1;XP_00 | 28297 | . | . | . | 1.872577 | 1.846506 | 1.507963 | 1.452711 | 4.357561 | 4.268281 | 3.383825 | 3.321453 | -1.120590 | 264.310372 | 0.211624049 | -0.77617889 | 0.437643371 | 0.998865404 | 8.126094 | 7.961386 | 8.070533 | 8.079265 | 7.999302 | 8.033959 | 319 | 304 | 209 | 242 |
| 348093 | NM_194272,N | RBPMS2 | RNA binding protein, mRNA processing factor 2 | protein_coding | NP_919248.1 | 19098 | . | . | . | 2.566254 | 2.597756 | 3.682362 | 3.464939 | 5.971778 | 6.004830 | 8.263113 | 7.922172 | 1.364224 | 193.478130 | 0.239997586 | 1.867019323 | 0.06189882 | 0.723723143 | 7.354474 | 7.802877 | 7.514949 | 7.517457 | 7.670018 | 7.637327 | 188 | 177 | 201 | 202 |
| 348094 | NM_182703 | ANKDD1A | ankyrin repeat and death domain containing 1A | protein_coding | NP_874362.3 | 28002 | . | . | . | 0.668444 | 0.528898 | 0.475348 | 0.627309 | 1.555497 | 1.222752 | 1.066667 | 1.434268 | -1.103392 | 56.174575 | 0.429223348 | -0.33070176 | 0.740869776 | 0.998865404 | 5.881411 | 5.738754 | 5.847698 | 5.796237 | 5.775865 | 5.828287 | 76 | 56 | 40 | 57 |
| 348110 | NM_00128238 | ARPIN | actin related protein 2/3 complex inhibitor | protein_coding | NP_001269309.1;NP_872422.1 | 28782 | 615543 | . | . | 4.242242 | 4.190272 | 4.132770 | 4.997560 | 9.871872 | 9.686000 | 9.273817 | 11.426327 | 1.068147 | 1024.830472 | 0.148944432 | 0.638559843 | 0.523109312 | 0.998865404 | 9.953112 | 10.047678 | 9.974718 | 9.975311 | 9.944575 | 10.096568 | 1141 | 1070 | 826 | 1094 |
| 348156 | gene-PKD1P5 | PKD1P5 | polycystin 1, transient receptor potential channel interacting pseu | pseudogene | . | 30069 | . | . | . | 1.309475 | 1.462816 | 1.482065 | 1.291384 | 3.047201 | 3.381365 | 3.325710 | 2.952595 | -1.013271 | 450.195428 | 0.180760639 | -0.10522016 | 0.916201116 | 0.998865404 | 8.823726 | 8.805028 | 8.784818 | 8.848623 | 8.853666 | 8.762648 | 495 | 514 | 418 | 388 |
| 348174 | NM_00113621 | CLEC18A | C-type lectin domain family 18 member A | protein_coding | NP_001129686.1;NP_001258126.1;NP_001357452.1;NP_872425.2;XP_01 | 30388 | 616571 | . | . | 0.007587 | 0.094784 | 0.097343 | 0.057903 | 0.017655 | 0.219098 | 0.218434 | 0.132387 | -1.042183 | 6.293578 | 1.301712603 | -0.04579238 | 0.963475734 | NA | 2.684574 | 2.622519 | 2.619438 | 2.663377 | 2.648850 | 2.632065 | 2 | 12 | 7 | 4 |
| 348180 | NM_00101275 | CTU2 | cytosolic thiouridylase subunit 2 | protein_coding | NP_001012777.1;NP_001012780.1;NP_001305436.1;NP_001305442.1;XF | 28005 | 617057 | . | . | 9.008125 | 9.759386 | 8.867934 | 9.640175 | 20.962276 | 22.559257 | 19.899387 | 22.041114 | -1.008938 | 506.252036 | 0.167549706 | -0.07662079 | 0.938925219 | 0.998865404 | 8.990273 | 8.977121 | 8.962443 | 9.009792 | 8.974585 | 8.986708 | 564 | 569 | 434 | 477 |
| 348235 | NM_00110059 | SKA2 | spindle and kinetochore associated complex subunit 2 | protein_coding | NP_001094065.1;NP_001317328.1;NP_872426.1 | 28006 | 616674 | . | . | 17.833087 | 14.073046 | 16.488573 | 14.441929 | 41.498325 | 32.530474 | 36.999881 | 33.019755 | -1.046416 | 1377.043790 | 0.152735327 | -0.42856111 | 0.668224649 | 0.998865404 | 10.459418 | 10.394571 | 10.532158 | 10.343072 | 10.460113 | 10.351748 | 1819 | 1334 | 1254 | 1172 |
| 348254 | NR_023380 | CCDC144CP | coiled-coil domain containing 144C, pseudogene | transcribed_ps | . | 29073 | . | . | . | 0.186212 | 0.195884 | 0.178128 | 0.175367 | 0.433322 | 0.452795 | 0.399713 | 0.400955 | -1.073911 | 24.687452 | 0.618118973 | -0.16643249 | 0.86781661 | NA | 4.676202 | 4.573379 | 4.628770 | 4.631952 | 4.624495 | 4.621657 | 29 | 28 | 21 | 22 |
| 348262 | NM_00109376 | MCRIP1 | MAPK regulated corepressor interacting protein 1 | protein_coding | NP_001087236.1;NP_001275727.1;NP_001275728.1;NP_997251.2;XP_01 | 28007 | 616514 | . | . | 13.238848 | 13.531180 | 9.217429 | 13.834030 | 30.807342 | 31.277925 | 20.683642 | 31.629866 | -1.180273 | 480.439817 | 0.204193117 | -1.17105165 | 0.241578019 | 0.992733415 | 9.023477 | 8.782931 | 8.940634 | 8.967028 | 8.721216 | 8.963947 | 587 | 573 | 306 | 496 |
| 348327 | NM_00132198 | ZNF530 | zinc finger protein 530 | protein_coding | NP_001308910.1;NP_065931.3;XP_006723257.1;XP_011525221.1;XP_01 | 29297 | . | . | . | 0.581343 | 0.508451 | 0.662366 | 0.472087 | 1.352810 | 1.175307 | 1.486331 | 1.079371 | -1.228797 | 71.894270 | 0.376711883 | -0.78905497 | 0.430079877 | 0.998865404 | 6.309355 | 6.010840 | 6.198963 | 6.188875 | 6.120351 | 6.167260 | 93 | 84 | 50 | 67 |
| 348654 | NM_00113000 | GEN1 | GEN1 Holliday junction 5' flap endonuclease | protein_coding | NP_001123481.3;NP_872431.5;XP_005262670.1;XP_006712068.1;XP_01 | 26881 | 612449 | . | . | 3.364186 | 2.905965 | 3.632429 | 2.099503 | 7.828601 | 6.717269 | 8.151066 | 4.800266 | -1.118805 | 867.822403 | 0.528977589 | -0.3061726 | 0.759473235 | 0.998865404 | 9.839858 | 9.678124 | 9.857441 | 9.704098 | 9.922723 | 9.421108 | 1165 | 886 | 961 | 500 |
| 348751 | NM_00135085 | FTCDNL1 | formiminotransferase cyclodeaminase N-terminal like | protein_coding | NP_001337782.1;NP_001337783.1;NP_001337784.1;NP_001350815.1;XF | 48661 | 614308 | . | . | 1.293074 | 1.103731 | 1.209871 | 1.714621 | 3.009039 | 2.551326 | 2.714915 | 3.920278 | 1.170265 | 101.053570 | 0.329363098 | 0.688708494 | 0.49100673 | 0.998865404 | 6.541868 | 6.767288 | 6.637384 | 6.622675 | 6.627488 | 6.726760 | 110 | 98 | 80 | 118 |
| 348761 | NR_033879 | SPATA3-AS1 | SPATA3 antisense RNA 1 (head to head) | lncRNA | . | 28013 | . | . | . | 0.156727 | 0.197819 | 0.323712 | 0.253176 | 0.364711 | 0.457267 | 0.726400 | 0.578858 | 1.600200 | 25.748316 | 0.615279297 | 1.102348271 | 0.270310291 | NA | 4.307819 | 4.986014 | 4.635728 | 4.657334 | 4.711028 | 4.680263 | 20 | 24 | 31 | 26 |
| 348793 | NM_00134590 | WDR53 | WD repeat domain 53 | protein_coding | NP_001332835.1;NP_001332836.1;NP_001332837.1;NP_001332838.1;NI | 28786 | 615110 | . | . | 6.088725 | 6.010795 | 7.104586 | 7.414210 | 14.168709 | 13.894222 | 15.942486 | 16.951712 | 1.181868 | 359.546362 | 0.187101264 | 1.288440006 | 0.19759284 | 0.96621494 | 8.364523 | 8.605495 | 8.434352 | 8.442013 | 8.530726 | 8.536997 | 377 | 358 | 337 | 367 |
| 348801 | NM_00108545 | LNP1 | leukemia NUP98 fusion partner 1 | protein_coding | NP_001078920.1 | 28014 | . | . | . | 0.206748 | 0.359158 | 0.295462 | 0.212260 | 0.481111 | 0.830211 | 0.663009 | 0.485308 | -1.113571 | 15.880735 | 0.786138846 | -0.19741315 | 0.843504241 | NA | 4.064940 | 3.909275 | 3.969183 | 4.009667 | 3.991678 | 3.972784 | 14 | 23 | 15 | 12 |
| 348926 | NR_130740,NF | FAM86EP | family with sequence similarity 86 member E, pseudogene | transcribed_ps | . | 28017 | . | . | . | 0.918918 | 0.854364 | 0.875964 | 0.887958 | 2.138358 | 1.974900 | 1.965638 | 2.030211 | -1.028040 | 72.532185 | 0.371196556 | -0.10748129 | 0.914407167 | 0.998865404 | 6.199971 | 6.160863 | 6.184783 | 6.182668 | 6.191315 | 6.162420 | 85 | 79 | 66 | 63 |
| 348995 | NM_198887,N | NUP43 | nucleoporin 43 | protein_coding | NP_942590.1;XP_005267017.1;XP_005267018.1;XP_005267019.1;XP_01: | 21182 | 608141 | . | . | 10.837079 | 10.484501 | 11.211228 | 11.006222 | 25.218330 | 24.235395 | 25.157669 | 25.164418 | -1.068830 | 1431.219618 | 0.12835205 | -0.74819725 | 0.454341189 | 0.998865404 | 10.530159 | 10.434308 | 10.540866 | 10.473898 | 10.460482 | 10.453139 | 1776 | 1527 | 1209 | 1291 |
| 349075 | NM_00136679 | ZNF713 | zinc finger protein 713 | protein_coding | NP_001353725.1;NP_872439.2 | 22043 | 616181 | . | . | 0.351716 | 0.425940 | 0.488948 | 0.330780 | 0.818459 | 0.984580 | 1.097187 | 0.756290 | 1.051493 | 54.916544 | 0.438899572 | 0.165047058 | 0.868906943 | 0.998865404 | 5.741705 | 5.815684 | 5.750179 | 5.786585 | 5.818328 | 5.737033 | 56 | 63 | 59 | 42 |
| 349114 | NR_026999 | LINC00265 | long intergenic non-protein coding RNA 265 | lncRNA | . | 28019 | . | . | . | 1.048306 | 1.337656 | 1.620726 | 1.302824 | 2.439452 | 3.092051 | 3.636861 | 2.978751 | 1.209267 | 192.398036 | 0.256955876 | 1.066847131 | 0.286040872 | 0.998865404 | 7.444314 | 7.718572 | 7.485882 | 7.586357 | 7.672458 | 7.567941 | 177 | 210 | 204 | 175 |
| 349149 | NM_181538 | GJC3 | gap junction protein gamma 3 | protein_coding | NP_853516.1 | 17495 | 611925 | . | . | 0.127075 | 0.072402 | 0.045895 | 0.443367 | 0.295709 | 0.167361 | 0.102986 | 1.013705 | 2.294420 | 6.572711 | 1.302857996 | 0.919616565 | 0.357773167 | NA | 1.998355 | 3.193768 | 2.688896 | 2.677566 | 2.676425 | 2.736637 | 6 | 3 | 2 | 15 |
| 349152 | NR_003561,NF | DPY19L2P2 | DPY19L2 pseudogene 2 | transcribed_ps | . | 21764 | . | . | . | 5.582676 | 5.067034 | 6.629875 | 5.496947 | 12.991118 | 11.712676 | 14.877247 | 12.568117 | 1.145578 | 697.704037 | 0.168660936 | 1.162545283 | 0.245014042 | 0.992733415 | 9.344593 | 9.541628 | 9.434666 | 9.352886 | 9.564754 | 9.405347 | 794 | 659 | 722 | 619 |
| 349667 | NM_178570 | RTN4RL2 | reticulon 4 receptor like 2 | protein_coding | NP_848665.1 | 23053 | 610462 | . | . | 4.981603 | 4.352195 | 6.167348 | 6.130431 | 11.592393 | 10.060294 | 13.839351 | 14.016501 | 1.298089 | 374.766637 | 0.189000439 | 1.991472086 | 0.046429011 | 0.654401867 | 8.349080 | 8.726060 | 8.502955 | 8.425119 | 8.620745 | 8.612124 | 402 | 327 | 371 | 394 |
| 352852 | gene-CYCSP45 | CYCSP45 | CYCS pseudogene 45 | pseudogene | . | 24419 | . | . | . | 0.221546 | 0.227873 | 0.130350 | 0.288776 | 0.515546 | 0.526738 | 0.292501 | 0.660253 | 1.067108 | 3.703508 | 1.564493575 | 0.059895269 | 0.952239048 | NA | 1.844134 | 1.932304 | 1.888489 | 1.889602 | 1.883355 | 1.896931 | 4 | 4 | 2 | 5 |
| 352909 | NM_00125671 | DNAAF3 | dynein axonemal assembly factor 3 | protein_coding | NP_001243643.1;NP_001243644.1;NP_001243645.1;NP_849159.2 | 30492 | 614566 | . | . | 1.903084 | 1.693147 | 1.466047 | 1.548203 | 4.428555 | 3.913786 | 3.289766 | 3.539781 | -1.174936 | 118.458691 | 0.298154213 | -0.78007539 | 0.435346504 | 0.998865404 | 6.999739 | 6.767397 | 6.936952 | 6.894239 | 6.855771 | 6.866891 | 156 | 130 | 93 | 104 |
| 352954 | NR_028038,NF | CASTOR3 | CASTOR family member 3 | transcribed_ps | . | 29954 | . | . | . | 1.127751 | 1.341327 | 1.687777 | 1.266409 | 2.624323 | 3.100538 | 3.787324 | 2.895492 | 1.205998 | 154.117010 | 0.276578024 | 0.977040186 | 0.328549258 | 0.998865404 | 7.125959 | 7.397086 | 7.201893 | 7.243859 | 7.352700 | 7.238249 | 152 | 159 | 167 | 136 |
| 353088 | NM_00100141 | ZNF429 | zinc finger protein 429 | protein_coding | NP_001001415.2;NP_001333841.1;NP_001333842.1;NP_001333843.1;NI | 20817 | . | . | . | 0.755592 | 0.624712 | 0.519858 | 0.299984 | 1.758296 | 1.444049 | 1.166545 | 0.685878 | -1.325955 | 46.810429 | 0.485952253 | -0.83759704 | 0.402257058 | 0.998865404 | 5.736193 | 5.333284 | 5.575306 | 5.564444 | 5.567101 | 5.472008 | 63 | 56 | 46 | 26 |
| 353091 | NM_00100178 | RAET1G | retinoic acid early transcript 1G | protein_coding | NP_001001788.2;XP_011534102.1;XP_016866320.1 | 16795 | 609244 | . | . | 0.225977 | 0.228154 | 0.140390 | 0.187561 | 0.525857 | 0.527389 | 0.315031 | 0.428837 | -1.245535 | 8.065020 | 1.067331285 | -0.29678291 | 0.766632253 | NA | 3.161369 | 2.844638 | 3.023057 | 3.017424 | 3.011119 | 3.013521 | 11 | 9 | 6 | 7 |
| 353116 | NM_00131924 | RILPL1 | Rab interacting lysosomal protein like 1 | protein_coding | NP_001306172.1;NP_001306173.1;NP_001306231.1;NP_847884.2;XP_00 | 26814 | 614092 | . | . | 3.690331 | 4.073336 | 4.421009 | 3.629029 | 8.587549 | 9.415702 | 9.920615 | 8.297343 | -1.033403 | 365.787698 | 0.19402165 | -0.244319 | 0.806983772 | 0.998865404 | 8.537949 | 8.491402 | 8.521710 | 8.524380 | 8.557337 | 8.448196 | 427 | 402 | 343 | 305 |
| 353139 | NM_178428 | LCE2A | late cornified envelope 2A | protein_coding | NP_848515.1 | 29469 | 612609 | . | . | 0.980496 | 0.891366 | 0.788106 | 1.323148 | 2.281655 | 2.060433 | 1.768488 | 3.025221 | 1.137647 | 19.603382 | 0.702174521 | 0.264966836 | 0.79103501 | NA | 4.198253 | 4.381963 | 4.288570 | 4.282082 | 4.276289 | 4.315516 | 22 | 19 | 14 | 24 |
| 353174 | NM_180990 | ZACN | zinc activated ion channel | protein_coding | NP_851321.2 | 29504 | 610935 | . | . | 0.004027 | 0.011136 | 0.006434 | 0.007810 | 0.009372 | 0.025742 | 0.014438 | 0.017856 | 1.234655 | 1.002957 | 2.868339486 | 0.106022226 | 0.915564717 | NA | -0.155866 | 0.148375 | -0.001261 | -0.000972 | 0.000053 | -0.000315 | 1 | 1 | 1 | 1 |
| 353194 | gene-KRT16P6 | KRT16P6 | keratin 16 pseudogene 6 | pseudogene | . | 50719 | . | . | . | 0.182618 | 0.094524 | 0.224745 | 0.090586 | 0.424960 | 0.218496 | 0.504321 | 0.207115 | 1.108372 | 7.986602 | 1.098629223 | 0.135115611 | 0.892520468 | NA | 2.917750 | 3.073227 | 2.998741 | 2.980217 | 3.006579 | 2.979602 | 11 | 6 | 10 | 5 |
| 353267 | gene-RAMACL | RAMACL | RNA guanine-7 methyltransferase activating subunit like (pseudog | pseudogene | . | 21234 | . | . | . | 3.418412 | 2.726429 | 3.218538 | 2.358076 | 7.954787 | 6.302263 | 7.222305 | 5.391461 | -1.117411 | 107.160818 | 0.327886114 | -0.48846184 | 0.625222745 | 0.998865404 | 6.820259 | 6.662709 | 6.796294 | 6.719131 | 6.769682 | 6.674295 | 145 | 108 | 102 | 80 |
| 353274 | NM_00136945 | ZNF445 | zinc finger protein 445 | protein_coding | NP_001356383.1;NP_852466.1 | 21018 | . | . | . | 1.079566 | 1.010979 | 1.081271 | 1.050663 | 2.512193 | 2.336924 | 2.426342 | 2.402217 | 1.003526 | 602.519165 | 0.158994777 | 0.031934827 | 0.974524025 | 0.998865404 | 9.232129 | 9.237592 | 9.257742 | 9.208479 | 9.249662 | 9.221826 | 718 | 625 | 536 | 554 |
| 353322 | NM_181726,XI | ANKRD37 | ankyrin repeat domain 37 | protein_coding | NP_859077.1;XP_016863665.1 | 29593 | . | . | . | 1.705578 | 1.413768 | 2.612526 | 2.119782 | 3.968952 | 3.267989 | 5.862433 | 4.846629 | 1.489985 | 50.116364 | 0.448671325 | 1.282225651 | 0.199763501 | 0.96621494 | 5.329726 | 5.907274 | 5.612930 | 5.581331 | 5.693183 | 5.647032 | 51 | 39 | 58 | 50 |
| 353355 | NM_00120700 | ZNF233 | zinc finger protein 233 | protein_coding | NP_001193934.1;NP_001317458.1;NP_861421.2;XP_016882247.1;XP_01 | 30946 | . | . | . | 0.256052 | 0.343030 | 0.385686 | 0.198415 | 0.595845 | 0.792931 | 0.865467 | 0.453653 | 1.425408 | 34.894579 | 0.526538109 | 0.971201439 | 0.331447977 | 0.998865404 | 4.847059 | 5.357829 | 5.078789 | 5.101857 | 5.144481 | 5.133272 | 30 | 34 | 37 | 37 |
| 353497 | NM_181808 | POLN | DNA polymerase nu | protein_coding | NP_861524.2 | 18870 | 610887 | . | . | 0.533020 | 0.566737 | 0.476633 | 0.474443 | 1.240358 | 1.310037 | 1.069551 | 1.084757 | -1.174239 | 52.348927 | 0.43191284 | -0.53651109 | 0.591605373 | 0.998865404 | 5.821414 | 5.589448 | 5.720177 | 5.733505 | 5.696876 | 5.695522 | 63 | 63 | 42 | 45 |
| 353500 | NM_181809,XI | BMP8A | bone morphogenetic protein 8a | protein_coding | NP_861525.2;XP_006710679.1;XP_011539683.1;XP_011539684.1;XP_01( | 21650 | . | . | . | 3.087990 | 2.876725 | 2.902198 | 2.650983 | 7.185880 | 6.649682 | 6.512447 | 6.061158 | -1.038084 | 272.473037 | 0.208055561 | -0.25917484 | 0.795500346 | 0.998865404 | 8.116642 | 8.062795 | 8.097129 | 8.102365 | 8.085871 | 8.074967 | 318 | 301 | 235 | 248 |
| 353512 | gene-PKD1P4 | PKD1P4 | polycystin 1, transient receptor potential channel interacting pseu | pseudogene | . | 30068 | . | . | . | 2.413892 | 2.671854 | 2.788303 | 2.954056 | 5.617227 | 6.176109 | 6.256872 | 6.754097 | 1.114218 | 541.947303 | 0.165256679 | 0.944178351 | 0.345078516 | 0.998865404 | 9.002137 | 9.157689 | 9.014559 | 9.074154 | 9.099113 | 9.130066 | 563 | 579 | 485 | 548 |
| 359744 | gene-MRPL35F | MRPL35P3 | mitochondrial ribosomal protein L35 pseudogene 3 | pseudogene | . | 23731 | . | . | . | 0.015135 | 0.052681 | 0.181315 | 0.096761 | 0.035221 | 0.121773 | 0.406865 | 0.221232 | 3.106129 | 1.844611 | 2.242171711 | 0.729256358 | 0.465844863 | NA | -0.155866 | 1.481107 | 0.852330 | 0.852687 | 0.864335 | 0.858436 | 1 | 1 | 3 | 2 |
| 359792 | gene-EIF4A1P2 | EIF4A1P2 | eukaryotic translation initiation factor 4A1 pseudogene 2 | pseudogene | . | 23946 | . | . | . | 0.316703 | 0.291980 | 0.148945 | 0.262026 | 0.736980 | 0.674925 | 0.334229 | 0.599091 | -1.438429 | 11.375671 | 0.916048409 | -0.57256144 | 0.566914672 | NA | 3.747897 | 3.219802 | 3.525891 | 3.521626 | 3.495304 | 3.516027 | 16 | 14 | 6 | 11 |
| 359800 | gene-VDAC1P6 | VDAC1P6 | voltage dependent anion channel 1 pseudogene 6 | pseudogene | . | 37481 | . | . | . | 1.406546 | 1.197880 | 1.389357 | 1.278210 | 3.273090 | 2.768952 | 3.117678 | 2.922475 | 1.011279 | 71.077514 | 0.376840234 | 0.042940184 | 0.965749216 | 0.998865404 | 6.142567 | 6.160023 | 6.169048 | 6.128994 | 6.162622 | 6.141133 | 88 | 70 | 65 | 64 |
| 359809 | NR_144431 | PEG13 | paternally expressed 13 | lncRNA | . | 19969 | . | . | . | 0.076384 | 0.056439 | 0.047120 | 0.038316 | 0.177748 | 0.130462 | 0.105736 | 0.087606 | -1.448257 | 10.188932 | 0.96146425 | -0.55573342 | 0.578393108 | NA | 3.590318 | 3.058876 | 3.371314 | 3.354466 | 3.350217 | 3.342501 | 16 | 11 | 8 | 7 |
| 359821 | NR_002208 | MRPL42P5 | mitochondrial ribosomal protein L42 pseudogene 5 | transcribed_ps | . | 29714 | . | . | . | 0.015112 | 0.021564 | 0.169093 | 0.003933 | 0.035167 | 0.049846 | 0.379439 | 0.008993 | 3.144341 | 1.865438 | 2.270779213 | 0.727837232 | 0.46671323 | NA | -0.155866 | 1.502474 | 0.861252 | 0.861611 | 0.878455 | 0.862426 | 1 | 1 | 4 | 1 |
| 359845 | NM_182705 | RFLNB | refilin B | protein_coding | NP_874364.1 | 28705 | 615928 | . | . | 5.582122 | 5.876837 | 4.582363 | 6.261734 | 12.989827 | 13.584571 | 10.282689 | 14.316710 | -1.071511 | 629.461159 | 0.178280266 | -0.55893324 | 0.576207283 | 0.998865404 | 9.347443 | 9.246747 | 9.303742 | 9.333013 | 9.167987 | 9.366218 | 734 | 718 | 449 | 655 |
| 359948 | NM_00107739 | IRF2BP2 | interferon regulatory factor 2 binding protein 2 | protein_coding | NP_001070865.1;NP_892017.2 | 21729 | 615332 | . | . | 10.553627 | 10.740515 | 10.398973 | 10.396452 | 24.558723 | 24.827184 | 23.334993 | 23.770254 | -1.039823 | 1521.305832 | 0.123740193 | -0.45529597 | 0.648896326 | 0.998865404 | 10.598986 | 10.542653 | 10.582165 | 10.589966 | 10.559368 | 10.552322 | 1777 | 1682 | 1303 | 1392 |
| 360023 | NM_194314,N | ZBTB41 | zinc finger and BTB domain containing 41 | protein_coding | NP_919290.2;XP_011507809.1;XP_016856694.1 | 24819 | . | . | . | 1.467247 | 1.437841 | 1.221019 | 1.017866 | 3.414344 | 3.323635 | 2.739933 | 2.327228 | -1.300747 | 336.065288 | 0.202626597 | -1.87211378 | 0.061190864 | 0.722512831 | 8.569289 | 8.191206 | 8.475378 | 8.443298 | 8.368516 | 8.253904 | 449 | 399 | 282 | 244 |
| 360132 | NR_003949,NF | FKBP9P1 | FKBP prolyl isomerase 9 pseudogene 1 | transcribed_ps | . | 23568 | . | . | . | 3.632673 | 4.158083 | 3.732134 | 3.965067 | 8.453378 | 9.611596 | 8.374800 | 9.065655 | -1.017835 | 297.419351 | 0.205035932 | -0.12438575 | 0.901009857 | 0.998865404 | 8.229381 | 8.203210 | 8.184437 | 8.255469 | 8.200110 | 8.221816 | 323 | 345 | 251 | 282 |
| 360177 | gene-CYCSP24 | CYCSP24 | CYCS pseudogene 24 | pseudogene | . | 24398 | . | . | . | 0.028532 | 0.020041 | 0.024288 | 0.014398 | 0.066394 | 0.046325 | 0.054501 | 0.032920 | 1.234655 | 1.002957 | 2.868339486 | 0.106022226 | 0.915564717 | NA | -0.155866 | 0.148375 | -0.001261 | -0.000972 | 0.000053 | -0.000315 | 1 | 1 | 1 | 1 |
| 360227 | gene-OTUD4P1 | OTUD4P1 | OTUD4 pseudogene 1 | pseudogene | . | 33912 | . | . | . | 0.046096 | 0.044486 | 0.034132 | 0.018823 | 0.107266 | 0.102831 | 0.076591 | 0.043037 | -1.679702 | 8.232996 | 1.077628522 | -0.69430742 | 0.487489437 | NA | 3.365657 | 2.622519 | 3.057262 | 3.056082 | 3.046382 | 3.029546 | 12 | 11 | 7 | 4 |
| 373156 | NM_00114367 | GSTK1 | glutathione S-transferase kappa 1 | protein_coding | NP_001137151.1;NP_001137152.1;NP_001137153.1;NP_057001.1 | 16906 | 602321 | . | . | 86.979794 | 90.270953 | 88.451729 | 99.749109 | 202.405558 | 208.665363 | 198.483107 | 228.064495 | 1.055403 | 3032.356292 | 0.116110887 | 0.670001548 | 0.502856803 | 0.998865404 | 11.526896 | 11.604507 | 11.532319 | 11.554327 | 11.544112 | 11.629016 | 3352 | 3227 | 2568 | 3070 |
| 373159 | gene-STK19B | STK19B | serine/threonine kinase 19B (pseudogene) | pseudogene | . | 21668 | . | . | . | 2.041846 | 1.617892 | 2.167367 | 1.749371 | 4.751459 | 3.739830 | 4.863509 | 3.999729 | 1.048457 | 35.260785 | 0.527480493 | 0.129422466 | 0.897023374 | 0.998865404 | 5.104211 | 5.174908 | 5.149777 | 5.118866 | 5.156690 | 5.125066 | 44 | 33 | 35 | 30 |
| 373856 | XM_01702916 | USP41 | ubiquitin specific peptidase 41 | protein_coding | XP_016884653.1 | 20070 | . | . | . | 0.775644 | 0.944467 | 0.845919 | 0.618573 | 1.804955 | 2.183177 | 1.898218 | 1.414294 | -1.166052 | 25.908364 | 0.612534518 | -0.36182713 | 0.717481219 | NA | 4.801350 | 4.580931 | 4.691915 | 4.714401 | 4.701699 | 4.671860 | 29 | 33 | 24 | 19 |
| 373861 | NR_024192,NF | H1-9P | H1.9 linker histone, pseudogene | transcribed_ps | . | 30616 | 608101 | . | . | 9.409607 | 10.996484 | 11.037806 | 8.029664 | 21.896539 | 25.418866 | 24.768516 | 18.358873 | -1.081209 | 307.636467 | 0.210732252 | -0.53454399 | 0.592965225 | 0.998865404 | 8.319867 | 8.208135 | 8.265824 | 8.301950 | 8.303208 | 8.176485 | 356 | 356 | 288 | 244 |
| 373863 | NM_194249 | DND1 | DND microRNA-mediated repression inhibitor 1 | protein_coding | NP_919225.1 | 23799 | 609385 | . | . | 0.482086 | 0.611752 | 0.668774 | 0.625039 | 1.121833 | 1.414093 | 1.500710 | 1.429078 | 1.171483 | 29.795389 | 0.567870212 | 0.402091464 | 0.687616715 | 0.998865404 | 4.778707 | 5.006356 | 4.869460 | 4.895889 | 4.906449 | 4.897166 | 28 | 33 | 29 | 29 |
| 374286 | NM_00128254 | CDRT1 | CMT1A duplicated region transcript 1 | protein_coding | NP_001269469.1;NP_006373.2 | 14379 | 604596 | . | . | 0.522280 | 0.579955 | 1.101249 | 0.663234 | 1.215366 | 1.340593 | 2.471170 | 1.516406 | 1.617876 | 164.863412 | 0.295600654 | 2.348104992 | 0.018869202 | 0.458615322 | 6.976092 | 7.671226 | 7.202848 | 7.267529 | 7.536751 | 7.314535 | 132 | 148 | 220 | 145 |
| 374291 | NM_00136360 | NDUFS7 | NADH:ubiquinone oxidoreductase core subunit S7 | protein_coding | NP_001350531.1;NP_077718.3;XP_016882257.2;XP_024307267.1 | 7714 | 601825 | . | . | 33.822603 | 32.919342 | 31.163608 | 35.475115 | 78.706591 | 76.094538 | 69.930233 | 81.109641 | 1.002788 | 879.191148 | 0.147177158 | 0.027293418 | 0.978225706 | 0.998865404 | 9.778241 | 9.781823 | 9.782737 | 9.774357 | 9.728334 | 9.830007 | 1017 | 942 | 711 | 884 |
| 374354 | NM_198514,XI | NHLRC2 | NHL repeat containing 2 | protein_coding | NP_940916.2;XP_011538071.1 | 24731 | 618277 | . | . | 1.647044 | 1.680692 | 1.524880 | 1.506520 | 3.832740 | 3.884995 | 3.421787 | 3.444479 | -1.101973 | 538.952216 | 0.163469619 | -0.85697415 | 0.391459169 | 0.998865404 | 9.142218 | 9.002424 | 9.115202 | 9.094004 | 9.056160 | 9.027683 | 661 | 600 | 456 | 470 |

| | | | | | | | | | | |
|---|---|---|---|---|---|---|---|---|---|---|
| 374378 | NM_00136346 | GALNT18 | polypeptide N-acetylgalactosaminyltransferase 18 | protein_coding | NP_001350393.1;NP_940918.2;XP_006718288.1;XP_011518370.1;XP_01 | 30488 | 615136 | . | . | [illegible] |
| 374383 | NM_00120243 | NCR3LG1 | natural killer cell cytotoxicity receptor 3 ligand 1 | protein_coding | NP_001189368.1;XP_011518374.1;XP_011518375.1;XP_011518376.1;XP | 42400 | 613714 | . | . | [illegible] |
| 374393 | NM_00114270 | FAM111B | family with sequence similarity 111 member B | protein_coding | NP_001136175.1;NP_001136176.1;NP_945185.1 | 24200 | 615584 | . | . | [illegible] |
| 374395 | NM_00136359 | TMEM179B | transmembrane protein 179B | protein_coding | NP_001350528.1;NP_001350529.1;NP_001350530.1;NP_955369.1 | 33744 | . | . | . | [illegible] |
| 374443 | NR_002814,N | LOC374443 | C-type lectin domain family 2 member D pseudogene | transcribed_ps | . | . | . | . | . | [illegible] |
| 374491 | NR_002815 | TPTE2P6 | transmembrane phosphoinositide 3-phosphatase and tensin hom | transcribed_ps | . | 42644 | . | . | . | [illegible] |
| 374500 | gene-THSD1P1 | THSD1P1 | thrombospondin type 1 domain containing 1 pseudogene 1 | pseudogene | . | 31375 | . | . | . | [illegible] |
| 374618 | NM_00128644 | TEX9 | testis expressed 9 | protein_coding | NP_001273378.1;NP_940926.1;XP_005254416.1;XP_005254418.1;XP_01 | 29585 | . | . | . | [illegible] |
| 374650 | NR_003246 | GOLGA6L5P | golgin A6 family like 5, pseudogene | transcribed_ps | . | 30472 | . | . | . | [illegible] |
| 374654 | NM_198525,X | KIF7 | kinesin family member 7 | protein_coding | NP_940927.2;XP_011519833.1 | 30497 | 611254 | . | . | [illegible] |
| 374655 | NM_198526,X | ZNF710 | zinc finger protein 710 | protein_coding | NP_940928.2;XP_005254963.1;XP_005254964.1;XP_011519842.1;XP_01 | 25352 | . | . | . | [illegible] |
| 374659 | NM_00128645 | HDDC3 | HD domain containing 3 | protein_coding | NP_001273380.1;NP_940929.1 | 30522 | . | . | . | [illegible] |
| 374666 | NR_003659 | WASH3P | WASP family homolog 3, pseudogene | transcribed_ps | . | 24362 | . | . | . | [illegible] |
| 374677 | gene-WASH4P | WASH4P | WASP family homolog 4, pseudogene | pseudogene | . | 14126 | . | . | . | [illegible] |
| 374786 | NM_00114505 | EFCAB5 | EF-hand calcium binding domain 5 | protein_coding | NP_001138525.2;NP_940931.3;XP_011523061.1;XP_011523065.1;XP_01 | 24801 | . | . | . | [illegible] |
| 374819 | NM_00130325 | LRRC37A3 | leucine rich repeat containing 37 member A3 | protein_coding | NP_001290184.1;NP_955372.2 | 32427 | 616557 | . | . | [illegible] |
| 374860 | NM_00136760 | ANKRD30B | ankyrin repeat domain 30B | protein_coding | NP_001354536.1;XP_011523966.1;XP_011523967.1;XP_011523968.1;XP | 24165 | 616565 | . | . | [illegible] |
| 374868 | NM_00130608 | ATP9B | ATPase phospholipid transporting 9B (putative) | protein_coding | NP_001293014.1;NP_940933.3;XP_011524265.1;XP_011524266.1;XP_01 | 13541 | 614446 | . | . | [illegible] |
| 374875 | NM_00126786 | HSD11B1L | hydroxysteroid 11-beta dehydrogenase 1 like | protein_coding | NP_001254797.1;NP_001254798.1;NP_001254799.1;NP_001254800.1;N | 30419 | . | . | . | [illegible] |
| 374879 | NM_198535 | ZNF699 | zinc finger protein 699 | protein_coding | NP_940937.1 | 24750 | 609571 | . | . | [illegible] |
| 374882 | NM_00114541 | TMEM205 | transmembrane protein 205 | protein_coding | NP_001138888.1;NP_001308041.1;NP_001308042.1;NP_001308043.1;N | 29631 | 613771 | . | . | [illegible] |
| 374887 | NM_00119032 | YJEFN3 | YjeF N-terminal domain containing 3 | protein_coding | NP_001177257.1;NP_940939.2;XP_005259960.1;XP_011526298.1;XP_01 | 24785 | 618607 | . | . | [illegible] |
| 374899 | NM_00103723 | ZNF829 | zinc finger protein 829 | protein_coding | NP_001032309.2;NP_001165450.1;XP_005258933.1;XP_011525235.1 | 34032 | . | . | . | [illegible] |
| 374900 | NM_00120483 | ZNF568 | zinc finger protein 568 | protein_coding | NP_001191764.1;NP_001191765.1;NP_001191766.1;NP_001191767.1;N | 25392 | 617566 | . | . | [illegible] |
| 374920 | NM_199341,X | ZSWIM9 | zinc finger SWIM-type containing 9 | protein_coding | NP_955373.3;XP_005259506.1;XP_006723267.1;XP_006723268.1;XP_01 | 34495 | . | . | . | [illegible] |
| 374928 | NM_00130433 | ZNF773 | zinc finger protein 773 | protein_coding | NP_001291263.1;NP_001291264.1;NP_001291265.1;NP_001291266.1;N | 30487 | . | . | . | [illegible] |
| 374969 | NM_199342,X | SVBP | small vasohibin binding protein | protein_coding | NP_955374.1;XP_016856715.1 | 29204 | 617853 | . | . | [illegible] |
| 374986 | NM_00127038 | MIGA1 | mitoguardin 1 | protein_coding | NP_001257313.1;NP_001350512.1;NP_001350513.1;NP_001350515.1;N | 24741 | 616773 | . | . | [illegible] |
| 375033 | NM_00108047 | PEAR1 | platelet endothelial aggregation receptor 1 | protein_coding | NP_001073940.1;NP_001340611.1;NP_001340612.1;XP_005245198.1;XP | 33631 | 610278 | . | . | [illegible] |
| 375035 | NM_199344 | SFT2D2 | SFT2 domain containing 2 | protein_coding | NP_955376.1 | 25140 | . | . | . | [illegible] |
| 375056 | NM_00130086 | MIA3 | MIA SH3 domain ER export factor 3 | protein_coding | NP_001287796.1;NP_001310991.1;NP_001310992.1;NP_001310993.1;N | 24008 | 613455 | . | . | [illegible] |
| 375057 | NM_00100366 | STUM | stum, mechanosensory transduction mediator homolog | protein_coding | NP_001003665.1;XP_011542485.1 | 30491 | . | . | . | [illegible] |
| 375061 | NM_198552 | FAM89A | family with sequence similarity 89 member A | protein_coding | NP_940954.1 | 25057 | . | . | . | [illegible] |
| 375133 | NR_003700 | PI4KAP2 | phosphatidylinositol 4-kinase alpha pseudogene 2 | transcribed_ps | . | 33577 | . | . | . | [illegible] |
| 375189 | NM_199346,X | PFN4 | profilin family member 4 | protein_coding | NP_955378.1;XP_005264349.1;XP_006712070.1;XP_011531125.1 | 31103 | . | . | . | [illegible] |
| 375190 | NM_00114571 | FAM228B | family with sequence similarity 228 member B | protein_coding | NP_001139182.1;NP_001278257.1 | 24736 | . | . | . | [illegible] |
| 375248 | NM_00135458 | ANKRD36 | ankyrin repeat domain 36 | protein_coding | NP_001341516.1;NP_940957.3;XP_006712577.1;XP_011509433.1;XP_01 | 24079 | . | . | . | [illegible] |
| 375260 | NR_024077 | WASH2P | WASP family homolog 2, pseudogene | transcribed_ps | . | 33145 | . | . | . | [illegible] |
| 375287 | NM_198557 | RBM43 | RNA binding motif protein 43 | protein_coding | NP_940959.1 | 24790 | . | . | . | [illegible] |
| 375295 | NR_040001 | LINC01116 | long intergenic non-protein coding RNA 1116 | lncRNA | . | 49259 | . | . | . | [illegible] |
| 375341 | NM_198562 | C3orf62 | chromosome 3 open reading frame 62 | protein_coding | NP_940964.1 | 24771 | . | . | . | [illegible] |
| 375346 | NM_198563 | STIMATE | STIM activating enhancer | protein_coding | NP_940965.1 | 30526 | 617189 | . | . | [illegible] |
| 375387 | NM_198565 | NRROS | negative regulator of reactive oxygen species | protein_coding | NP_940967.1 | 24613 | 615322 | . | . | [illegible] |
| 375444 | NM_198566,X | C5orf34 | chromosome 5 open reading frame 34 | protein_coding | NP_940968.1;XP_011512340.1;XP_016864934.1;XP_016864935.1 | 24738 | . | . | . | [illegible] |
| 375449 | NM_00116466 | MAST4 | microtubule associated serine/threonine kinase family member 4 | protein_coding | NP_001158136.1;NP_001277155.1;NP_001277156.1;NP_001277157.1;N | 19037 | 618002 | . | . | [illegible] |
| 375484 | NM_00130819 | SIMC1 | SUMO interacting motifs containing 1 | protein_coding | NP_001295124.1;NP_001295125.1;NP_001295129.1;NP_940969.3;XP_01 | 24779 | 618102 | . | . | [illegible] |
| 375513 | NR_132999,N | GUSBP4 | GUSB pseudogene 4 | transcribed_ps | . | 18220 | . | . | . | [illegible] |
| 375612 | NM_199000,X | LHFPL3 | LHFPL tetraspan subfamily member 3 | protein_coding | NP_945351.1;XP_005250384.1 | 6589 | 609719 | . | . | [illegible] |
| 375616 | NM_00136612 | KCP | kielin cysteine rich BMP regulator | protein_coding | NP_001353051.1;NP_955381.2;XP_016867673.1;XP_016867675.1;XP_01 | 17585 | 609344 | . | . | [illegible] |
| 375690 | NR_033266 | WASH5P | WASP family homolog 5, pseudogene | transcribed_ps | . | 33884 | . | . | . | [illegible] |
| 375704 | NM_198573 | ENHO | energy homeostasis associated | protein_coding | NP_940975.2 | 24838 | 618556 | . | . | [illegible] |
| 375743 | NM_00109966 | PTAR1 | protein prenyltransferase alpha subunit repeat containing 1 | protein_coding | NP_001093136.1;NP_001353864.1;NP_001353865.1;NP_001353866.1;N | 30449 | . | . | . | [illegible] |
| 375748 | NM_00101089 | ERCC6L2 | ERCC excision repair 6 like 2 | protein_coding | NP_001010895.2;NP_001362220.1;NP_001362221.1;NP_001362222.1;N | 26922 | 615667 | . | . | [illegible] |
| 375757 | NM_00104001 | SWI5 | SWI5 homologous recombination repair protein | protein_coding | NP_001035100.2;NP_001305018.2;NP_001305021.1;NP_001366196.1 | 31412 | 616528 | . | . | [illegible] |
| 375775 | NM_00109853 | PNPLA7 | patatin like phospholipase domain containing 7 | protein_coding | NP_001092007.2;NP_689499.4;XP_006717165.1;XP_006717167.1;XP_01 | 24768 | 612122 | . | . | [illegible] |
| 375790 | NM_00130527 | AGRN | agrin | protein_coding | NP_001292204.1;NP_001351656.1;NP_940978.2;XP_005244806.1;XP_01 | 329 | 103320 | . | . | [illegible] |
| 375791 | NM_199001 | CYSRT1 | cysteine rich tail 1 | protein_coding | NP_945352.4 | 30529 | . | . | . | [illegible] |
| 376267 | NM_00130815 | RAB15 | RAB15, member RAS oncogene family | protein_coding | NP_001295083.1;NP_001317111.1;NP_941959.1;XP_024305345.1 | 20150 | . | . | . | [illegible] |
| 376497 | NM_198580,X | SLC27A1 | solute carrier family 27 member 1 | protein_coding | NP_940982.1;XP_011526302.1;XP_011526303.1;XP_011526304.1;XP_01 | 10995 | 600691 | . | . | [illegible] |
| 376693 | NR_026667 | RPS10P7 | ribosomal protein S10 pseudogene 7 | transcribed_ps | . | 36423 | . | . | . | [illegible] |
| 376818 | gene-E2F6P2 | E2F6P2 | E2F transcription factor 6 pseudogene 2 | pseudogene | . | 51450 | . | . | . | [illegible] |
| 376940 | NM_198581,X | ZC3H6 | zinc finger CCCH-type containing 6 | protein_coding | NP_940983.2;XP_006712582.1 | 24762 | . | . | . | [illegible] |
| 377677 | NM_198584 | CA13 | carbonic anhydrase 13 | protein_coding | NP_940986.1 | 14914 | 611436 | . | . | [illegible] |
| 378108 | NM_00131781 | TRIM74 | tripartite motif containing 74 | protein_coding | NP_001304744.1;NP_942150.1;XP_011514489.1;XP_011514491.1;XP_01 | 17453 | 612550 | . | . | [illegible] |
| 378706 | NR_027260 | RN7SL2 | RNA component of signal recognition particle 7SL2 | scRNA | . | 23134 | 612179 | . | . | [illegible] |
| 378707 | NR_145670 | RN7SL3 | RNA component of signal recognition particle 7SL3 | scRNA | . | 23135 | 612180 | . | . | [illegible] |
| 378708 | NM_199294,N | CENPS | centromere protein S | protein_coding | NP_954988.1 | 23163 | 609130 | . | . | [illegible] |
| 378805 | NR_015431,N | LINC-PINT | long intergenic non-protein coding RNA, p53 induced transcript | lncRNA | . | 26885 | 618212 | . | . | [illegible] |
| 378884 | NM_198586 | NHLRC1 | NHL repeat containing E3 ubiquitin protein ligase 1 | protein_coding | NP_940988.2 | 21576 | 608072 | . | . | [illegible] |
| 378925 | NM_198085 | RNF148 | ring finger protein 148 | protein_coding | NP_932351.1 | 22411 | . | . | . | [illegible] |
| 378938 | NR_002819,N | MALAT1 | metastasis associated lung adenocarcinoma transcript 1 | lncRNA | . | 29665 | 607924 | . | . | [illegible] |
| 379025 | NR_029434,N | PSMA3-AS1 | PSMA3 antisense RNA 1 | lncRNA | . | 26445 | . | . | . | [illegible] |
| 379034 | NR_145429 | LINC01181 | long intergenic non-protein coding RNA 1181 | lncRNA | . | 25533 | . | . | . | [illegible] |
| 386593 | NR_027928 | CHKB-CPT1B | CHKB-CPT1B readthrough (NMD candidate) | lncRNA | . | 41998 | . | . | . | [illegible] |
| 386597 | NR_033997 | GRASLND | glycosaminoglycan regulatory associated long non-coding RNA | lncRNA | . | 30963 | . | . | . | [illegible] |
| 386610 | gene-E2F6P1 | E2F6P1 | E2F transcription factor 6 pseudogene 1 | pseudogene | . | 51449 | . | . | . | [illegible] |
| 386724 | NM_198722 | AMIGO3 | adhesion molecule with Ig like domain 3 | protein_coding | NP_942015.1 | 24075 | 615691 | . | . | [illegible] |
| 386758 | NR_037159,N | ZNF582-AS1 | ZNF582 antisense RNA 1 (head to head) | lncRNA | . | 25213 | . | . | . | [illegible] |
| 387032 | NM_00130450 | ZKSCAN4 | zinc finger with KRAB and SCAN domains 4 | protein_coding | NP_001291435.1;NP_061983.2;XP_005249152.1;XP_005249154.1;XP_00 | 13854 | 611643 | . | . | [illegible] |
| 387036 | NR_003504 | GUSBP2 | GUSB pseudogene 2 | transcribed_ps | . | 18792 | . | . | . | [illegible] |
| 387066 | NR_003038 | SNHG5 | small nucleolar RNA host gene 5 | lncRNA | . | 21026 | 613263 | . | . | [illegible] |
| 387101 | gene-RPL29P4 | RPL29P4 | ribosomal protein L29 pseudogene 4 | pseudogene | . | 21482 | . | . | . | [illegible] |
| 387103 | NM_00101250 | CENPW | centromere protein W | protein_coding | NP_001012525.1;NP_001273453.1;NP_001273454.1;XP_016866334.1 | 21488 | 611264 | . | . | [illegible] |
| 387104 | NM_00101227 | SOGA3 | SOGA family member 3 | protein_coding | NP_001012279.1 | 21494 | . | . | . | [illegible] |
| 387119 | NM_00104247 | CEP85L | centrosomal protein 85 like | protein_coding | NP_001035940.1;NP_001171506.1;NP_996804.2;XP_005267027.1;XP_00 | 21638 | 618865 | . | . | [illegible] |
| 387254 | NR_002594 | SLC7A5P2 | solute carrier family 7 member 5 pseudogene 2 | transcribed_ps | . | 24951 | . | . | . | [illegible] |
| 387263 | NM_00102986 | C6orf120 | chromosome 6 open reading frame 120 | protein_coding | NP_001025034.1;NP_001304271.1 | 21247 | 616987 | . | . | [illegible] |
| 387328 | gene-ZNF322P | ZNF322P1 | zinc finger protein 322 pseudogene 1 | pseudogene | . | 14003 | . | . | . | [illegible] |
| 387338 | NM_00125612 | NSUN4 | NOP2/Sun RNA methyltransferase 4 | protein_coding | NP_001243056.1;NP_001243057.1;NP_950245.2 | 31802 | 615394 | . | . | [illegible] |
| 387496 | NM_00133112 | RASL11A | RAS like family 11 member A | protein_coding | NP_001318055.1;NP_996563.1;XP_024305115.1 | 23802 | 612403 | . | . | [illegible] |
| 387509 | NM_207370,X | GPR153 | G protein-coupled receptor 153 | protein_coding | NP_997253.2;XP_011539736.1;XP_016856739.1;XP_016856740.1 | 23618 | 614269 | . | . | [illegible] |
| 387521 | NM_00116250 | TMEM189 | transmembrane protein 189 | protein_coding | NP_001155977.2;NP_954580.2 | 16735 | 610994 | . | . | [illegible] |
| 387522 | NM_199203 | TMEM189-UBE | TMEM189-UBE2V1 readthrough | protein_coding | NP_954673.2 | 33521 | . | . | . | [illegible] |
| 387553 | gene-DNM1P1 | DNM1P18 | dynamin 1 pseudogene 18 | pseudogene | . | 21135 | . | . | . | [illegible] |
| 387590 | NR_001591 | TPTEP1 | TPTE pseudogene 1 | transcribed_ps | . | 43648 | . | . | . | [illegible] |
| 387628 | NR_003674,N | FGF7P6 | fibroblast growth factor 7 pseudogene 6 | transcribed_ps | . | 27852 | . | . | . | [illegible] |
| 387640 | NM_207371,X | SKIDA1 | SKI/DACH domain containing 1 | protein_coding | NP_997254.3;XP_005252507.1;XP_011517782.1;XP_016871714.1;XP_01 | 32697 | . | . | . | [illegible] |
| 387680 | NM_00100575 | WASHC2A | WASH complex subunit 2A | protein_coding | NP_001005751.1;NP_001278327.1;NP_001317031.1;XP_005269863.2;XP | 23416 | . | . | . | [illegible] |
| 387707 | NM_00100173 | CC2D2B | coiled-coil and C2 domain containing 2B | protein_coding | NP_001001732.2;NP_001123918.2;NP_001153219.1;NP_001335937.1;XP | 31666 | . | . | . | [illegible] |
| 387723 | NM_00135504 | C10orf143 | chromosome 10 open reading frame 143 | protein_coding | NP_001341971.1;XP_024303775.1;XP_024303776.1;XP_024303777.1 | 48677 | . | . | . | [illegible] |
| 387753 | gene-RPL21P9 | RPL21P97 | ribosomal protein L21 pseudogene 97 | pseudogene | . | 36223 | . | . | . | [illegible] |
| 387763 | NM_00114503 | C11orf96 | chromosome 11 open reading frame 96 | protein_coding | NP_001138505.1 | 38675 | . | . | . | [illegible] |
| 387787 | NM_00114486 | LIPT2 | lipoyl(octanoyl) transferase 2 | protein_coding | NP_001138341.1;NP_001316870.1;NP_001316871.1 | 37216 | 617659 | . | . | [illegible] |
| 387841 | NR_003932 | RPL13AP20 | ribosomal protein L13a pseudogene 20 | transcribed_ps | . | 35709 | . | . | . | [illegible] |
| 387845 | gene-EEF1A1P | EEF1A1P16 | eukaryotic translation elongation factor 1 alpha 1 pseudogene 16 | pseudogene | . | 37889 | . | . | . | [illegible] |
| 387849 | NM_00102987 | REP15 | RAB15 effector protein | protein_coding | NP_001025045.3 | 33748 | 610848 | . | . | [illegible] |
| 387856 | NM_00101363 | CCDC184 | coiled-coil domain containing 184 | protein_coding | NP_001013657.3 | 33749 | . | . | . | [illegible] |
| 387867 | gene-RPSAP12 | RPSAP12 | ribosomal protein SA pseudogene 12 | pseudogene | . | 6504 | . | . | . | [illegible] |
| 387882 | NM_00114519 | C12orf75 | chromosome 12 open reading frame 75 | protein_coding | NP_001138671.1 | 35164 | . | . | . | [illegible] |
| 387885 | NM_00114487 | CFAP73 | cilia and flagella associated protein 73 | protein_coding | NP_001138344.1;XP_011536629.1;XP_024304746.1 | 37100 | . | . | . | [illegible] |
| 387893 | NM_00132450 | KMT5A | lysine methyltransferase 5A | protein_coding | NP_001311433.1;NP_001311434.1;NP_001311435.1;NP_001354315.1;N | 29489 | 607240 | . | . | [illegible] |
| 387914 | NM_00100753 | SHISA2 | shisa family member 2 | protein_coding | NP_001007539.1 | 20366 | 617324 | . | . | [illegible] |
| 387921 | NM_00101275 | NHLRC3 | NHL repeat containing 3 | protein_coding | NP_001012772.1;NP_001017370.1 | 33751 | . | . | . | [illegible] |
| 387933 | gene-HNRNPA | HNRNPA3P5 | heterogeneous nuclear ribonucleoprotein A3 pseudogene 5 | pseudogene | . | 39774 | . | . | . | [illegible] |
| 387934 | gene-FABP5P1 | FABP5P1 | fatty acid binding protein 5 pseudogene 1 | pseudogene | . | 31059 | . | . | . | [illegible] |
| 388021 | NM_00128638 | TMEM179 | transmembrane protein 179 | protein_coding | NP_001273318.1;NP_001273319.1;XP_011535047.1;XP_011535048.1;XP | 20137 | . | . | . | [illegible] |
| 388115 | NM_207380 | CCDC9B | coiled-coil domain containing 9B | protein_coding | NP_997263.2 | 33488 | . | . | . | [illegible] |
| 388152 | NR_027001 | GOLGA2P7 | GOLGA2 pseudogene 7 | transcribed_ps | . | 49516 | . | . | . | [illegible] |
| 388165 | NR_003661 | UBE2Q2P1 | ubiquitin conjugating enzyme E2 Q2 pseudogene 1 | transcribed_ps | . | 37439 | . | . | . | [illegible] |
| 388182 | NR_028139,N | SPATA41 | spermatogenesis associated 41 | lncRNA | . | 48613 | . | . | . | [illegible] |
| 388228 | NM_00102440 | SBK1 | SH3 domain binding kinase 1 | protein_coding | NP_001019572.1;XP_005255372.1 | 17699 | . | . | . | [illegible] |
| 388254 | gene-HERC2P5 | HERC2P5 | hect domain and RLD 2 pseudogene 5 | pseudogene | . | 4873 | . | . | . | [illegible] |
| 388272 | NM_00100143 | C16orf87 | chromosome 16 open reading frame 87 | protein_coding | NP_001001436.1;NP_001335589.1;NP_001335590.1;XP_024306039.1;XP | 33754 | . | . | . | [illegible] |
| 388275 | gene-HNRNPA | HNRNPA1P7 | heterogeneous nuclear ribonucleoprotein A1 pseudogene 7 | pseudogene | . | 31015 | . | . | . | [illegible] |
| 388284 | NM_00101298 | C16orf86 | chromosome 16 open reading frame 86 | protein_coding | NP_001013002.2;XP_005256009.1;XP_024306042.1 | 33755 | . | . | . | [illegible] |
| 388327 | NM_00110552 | C17orf100 | chromosome 17 open reading frame 100 | protein_coding | NP_001098990.1 | 34494 | . | . | . | [illegible] |
| 388339 | gene-RPS18P1 | RPS18P12 | ribosomal protein S18 pseudogene 12 | pseudogene | . | 35967 | . | . | . | [illegible] |
| 388341 | NM_00111356 | LRRC75A | leucine rich repeat containing 75A | protein_coding | NP_001107039.1;NP_997270.2;XP_011522147.1;XP_016880108.1;XP_01 | 32403 | . | . | . | [illegible] |
| 388344 | gene-RPL13P1 | RPL13P12 | ribosomal protein L13 pseudogene 12 | pseudogene | . | 35701 | . | . | . | [illegible] |
| 388363 | gene-RPS7P1 | RPS7P1 | ribosomal protein S7 pseudogene 1 | pseudogene | . | 17081 | . | . | . | [illegible] |
| 388389 | NM_00125839 | CCDC103 | coiled-coil domain containing 103 | protein_coding | NP_001245324.1;NP_001245325.1;NP_001245326.1;NP_001245327.1;N | 32700 | 614677 | . | . | [illegible] |
| 388403 | NM_00100540 | YPEL2 | yippee like 2 | protein_coding | NP_001005404.1;XP_016880110.1 | 18326 | 609723 | . | . | [illegible] |
| 388474 | gene-RPL7AP6 | RPL7AP66 | ribosomal protein L7a pseudogene 66 | pseudogene | . | 35682 | . | . | . | [illegible] |
| 388523 | NM_00126771 | ZNF728 | zinc finger protein 728 | protein_coding | NP_001254645.1 | 32463 | . | . | . | [illegible] |
| 388524 | NM_00135528 | RPSAP58 | ribosomal protein SA pseudogene 58 | protein_coding | NP_001342212.1;NP_001342216.1 | 36809 | . | . | . | [illegible] |
| 388531 | NM_207391 | RGS9BP | regulator of G protein signaling 9 binding protein | protein_coding | NP_997274.2 | 30304 | 607814 | . | . | [illegible] |
| 388536 | NM_00124280 | ZNF790 | zinc finger protein 790 | protein_coding | NP_001229729.1;NP_001229730.1;NP_001229731.1;NP_996777.2;XP_00 | 33114 | . | . | . | [illegible] |
| 388552 | NM_212550,X | BLOC1S3 | biogenesis of lysosomal organelles complex 1 subunit 3 | protein_coding | NP_997715.1 | 20914 | 609762 | . | . | [illegible] |
| 388556 | gene-RPS9P4 | RPS9P4 | ribosomal protein S9 pseudogene 4 | pseudogene | . | 36710 | . | . | . | [illegible] |
| 388558 | NM_00103988 | ZNF808 | zinc finger protein 808 | protein_coding | NP_001034975.2;NP_001308353.1;NP_001308354.1;NP_001350479.1;XP | 33230 | . | . | . | [illegible] |
| 388559 | NM_00131012 | ZNF888 | zinc finger protein 888 | protein_coding | NP_001297056.1;XP_005259508.1;XP_016882287.1;XP_016882288.1;XP | 38695 | . | . | . | [illegible] |
| 388561 | NM_00100840 | ZNF761 | zinc finger protein 761 | protein_coding | NP_001008401.3;NP_001276880.1;NP_001276881.1;NP_001276882.1 | 23179 | . | . | . | [illegible] |
| 388564 | NM_00119076 | TMEM238 | transmembrane protein 238 | protein_coding | NP_001177693.1 | 40042 | . | . | . | [illegible] |
| 388567 | NM_00102356 | ZNF749 | zinc finger protein 749 | protein_coding | NP_001018855.2;NP_001308881.1;NP_001308882.1;NP_001308883.1;XP | 32783 | . | . | . | [illegible] |
| 388569 | NM_207395,X | ZNF324B | zinc finger protein 324B | protein_coding | NP_997278.2;XP_005258972.1;XP_005258975.1 | 33107 | . | . | . | [illegible] |
| 388572 | XR_00295721 | LOC388572 | uncharacterized LOC388572 | lncRNA | . | . | . | . | . | [illegible] |
| 388574 | NR_029406 | RPL23AP87 | ribosomal protein L23a pseudogene 87 | transcribed_ps | . | 51578 | . | . | . | [illegible] |
| 388579 | gene-TUBB8P1 | TUBB8P11 | tubulin beta 8 class VIII pseudogene 11 | pseudogene | . | 32337 | . | . | . | [illegible] |
| 388581 | NM_00101498 | C1QTNF12 | C1q and TNF related 12 | protein_coding | NP_001014980.1;XP_011539737.1;XP_011539738.1;XP_011539739.1;XP | 32308 | 616593 | . | . | [illegible] |
| 388588 | NM_00116372 | SMIM1 | small integral membrane protein 1 (Vel blood group) | protein_coding | NP_001157196.1;NP_001275512.1;NP_001366619.1;NP_001366620.1 | 44204 | 615242 | . | . | [illegible] |
| 388591 | NM_207396,X | RNF207 | ring finger protein 207 | protein_coding | NP_997279.2;XP_011539741.1;XP_016856748.1 | 32947 | 616923 | . | . | [illegible] |
| 388605 | gene-HS6ST1P | HS6ST1P1 | heparan sulfate 6-O-sulfotransferase 1 pseudogene 1 | pseudogene | . | 31835 | . | . | . | [illegible] |
| 388610 | NM_00101364 | TRNP1 | TMF1 regulated nuclear protein 1 | protein_coding | NP_001013664.2;XP_005245924.1 | 34348 | 616824 | . | . | [illegible] |
| 388630 | NM_00119498 | TRABD2B | TraB domain containing 2B | protein_coding | NP_001181915.1;XP_006710703.1;XP_011539745.1;XP_011539746.1;XP | 44200 | 614913 | . | . | [illegible] |
| 388650 | NM_00100660 | DIPK1A | divergent protein kinase domain 1A | protein_coding | NP_001006606.2;NP_001239198.1;NP_001239199.1;NP_001239200.1;N | 32213 | 614542 | . | . | [illegible] |
| 388677 | NM_00136400 | NOTCH2NLA | notch 2 N-terminal like A | protein_coding | NP_001350935.1;NP_982283.2 | 31862 | 618023 | . | . | [illegible] |
| 388685 | NR_027468,N | LINC01138 | long intergenic non-protein coding RNA 1138 | lncRNA | . | 49454 | . | . | . | [illegible] |
| 388692 | NR_111933 | LINC02591 | long intergenic non-protein coding RNA 2591 | lncRNA | . | 53901 | . | . | . | [illegible] |
| 388695 | NM_00113654 | LYSMD1 | LysM domain containing 1 | protein_coding | NP_001130015.1;NP_997716.1 | 32070 | . | . | . | [illegible] |
| 388707 | gene-RPSAP18 | RPSAP18 | ribosomal protein SA pseudogene 18 | pseudogene | . | 36106 | . | . | . | [illegible] |
| 388722 | NM_00102459 | C1orf53 | chromosome 1 open reading frame 53 | protein_coding | NP_001019765.1 | 30003 | . | . | . | [illegible] |
| 388730 | NM_203376 | TMEM81 | transmembrane protein 81 | protein_coding | NP_976310.1 | 32349 | . | . | . | [illegible] |

| | | | | | | | | | | | | | | | | | | | | | | | | | | | | | | | | | |
|---|---|---|---|---|---|---|---|---|---|---|---|---|---|---|---|---|---|---|---|---|---|---|---|---|---|---|---|---|---|---|---|---|---|
| 388734 | gene-PRELID1F | PRELID1P5 | PRELID1 pseudogene 5 | pseudogene | . | 54770 | . | . | . | 0.177495 | 0.208219 | 0.048865 | 0.135674 | 0.413038 | 0.481307 | 0.109651 | 0.310203 | -1.992264 | 5.730381 | 1.297808251 | -0.76622166 | 0.44354442 | NA | 2.934350 | 1.932304 | 2.535821 | 2.542106 | 2.515101 | 2.528376 | 8 | 9 | 2 | 5 |
| 388753 | NM_00101298 | COA6 | cytochrome c oxidase assembly factor 6 | protein_coding | NP_001013003.1;NP_001193570.2;NP_001288662.1 | 18025 | 614772 | . | . | 21.364813 | 19.318946 | 22.599364 | 17.469993 | 49.716801 | 44.656606 | 50.712317 | 39.943063 | -1.069397 | 983.265180 | 0.15563026 | -0.62196841 | 0.533962642 | 0.998865404 | 9.988638 | 9.892639 | 10.011296 | 9.913414 | 9.987423 | 9.836367 | 1247 | 1024 | 913 | 798 |
| 388789 | NM_00134895 | SMIM26 | small integral membrane protein 26 | protein_coding | NP_001335886.1;NP_001335887.1 | 43430 | . | . | . | 29.825084 | 27.744105 | 28.764346 | 30.514784 | 69.404201 | 64.131744 | 64.546354 | 69.768431 | 1.020657 | 480.514971 | 0.170653344 | 0.172854464 | 0.862765825 | 0.998865404 | 8.893610 | 8.923114 | 8.922621 | 8.880460 | 8.899475 | 8.929460 | 566 | 496 | 412 | 466 |
| 388795 | NM_00114396 | EFCAB8 | EF-hand calcium binding domain 8 | protein_coding | NP_001137439.1;XP_011527423.1;XP_024307649.1;XP_024307650.1;XP_ | 34532 | . | . | . | 0.221690 | 0.158281 | 0.286167 | 0.102348 | 0.515882 | 0.365872 | 0.642150 | 0.234007 | 1.033154 | 50.894769 | 0.515273883 | 0.091322204 | 0.927236576 | 0.998865404 | 5.643098 | 5.695321 | 5.677527 | 5.621575 | 5.739385 | 5.564660 | 66 | 46 | 66 | 26 |
| 388796 | NR_015366,NF | SNHG17 | small nucleolar RNA host gene 17 | lncRNA | . | 48600 | . | . | . | 10.130185 | 10.710918 | 12.834863 | 10.939087 | 23.573355 | 24.758768 | 28.801060 | 25.010922 | 1.145108 | 386.642471 | 0.194432615 | 1.005405277 | 0.31470174 | 0.998865404 | 8.493448 | 8.689602 | 8.530380 | 8.569469 | 8.694350 | 8.560996 | 401 | 402 | 399 | 344 |
| 388815 | NR_027790,NF | MIR99AHG | mir-99a-let-7c cluster host gene | lncRNA | . | 1274 | 615964 | . | . | 0.214143 | 0.192082 | 0.164999 | 0.067858 | 0.498319 | 0.444005 | 0.370251 | 0.155150 | -1.732860 | 19.138235 | 0.736872005 | -1.07638105 | 0.281756874 | NA | 4.599021 | 3.811620 | 4.278118 | 4.284500 | 4.261087 | 4.214147 | 27 | 27 | 17 | 8 |
| 388907 | gene-RPL5P34 | RPL5P34 | ribosomal protein L5 pseudogene 34 | pseudogene | . | 36463 | . | . | . | 0.373218 | 0.336704 | 0.388313 | 0.166220 | 0.868494 | 0.778305 | 0.871363 | 0.380042 | -1.299976 | 10.315544 | 0.973381158 | -0.38883535 | 0.697397949 | NA | 3.540965 | 3.168566 | 3.375319 | 3.370519 | 3.376819 | 3.344867 | 14 | 12 | 11 | 5 |
| 388955 | gene-PRELID1F | PRELID1P6 | PRELID1 pseudogene 6 | pseudogene | . | 54771 | . | . | . | 1.813203 | 1.452129 | 1.451689 | 1.533024 | 4.219397 | 3.356661 | 3.257548 | 3.505078 | -1.108640 | 45.471906 | 0.465205252 | -0.31983849 | 0.749090768 | 0.998865404 | 5.579017 | 5.430994 | 5.536571 | 5.496956 | 5.497186 | 5.501425 | 61 | 46 | 37 | 41 |
| 388962 | NM_00103550 | BOLA3 | bolA family member 3 | protein_coding | NP_001030582.1;NP_997717.2 | 24415 | 613183 | . | . | 47.091411 | 44.183125 | 41.404695 | 54.804683 | 109.583647 | 102.131274 | 92.910942 | 125.304392 | 1.042074 | 792.749177 | 0.165513464 | 0.359231647 | 0.719421804 | 0.998865404 | 9.600993 | 9.659849 | 9.636213 | 9.590938 | 9.545597 | 9.731467 | 923 | 811 | 611 | 858 |
| 388963 | NM_00114505 | C2orf81 | chromosome 2 open reading frame 81 | protein_coding | NP_001138526.1;NP_001303693.1;NP_001303694.1;NP_001303695.1 | 34350 | . | . | . | 1.415881 | 1.171076 | 1.262023 | 1.111380 | 3.294812 | 2.706996 | 2.831943 | 2.541039 | -1.120454 | 84.645960 | 0.35005528 | -0.46873742 | 0.639257336 | 0.998865404 | 6.482357 | 6.319806 | 6.443968 | 6.393019 | 6.404584 | 6.370920 | 112 | 88 | 74 | 70 |
| 388969 | NM_00101364 | C2orf68 | chromosome 2 open reading frame 68 | protein_coding | NP_001013671.2;XP_005264362.1;XP_024308647.1 | 34353 | 618519 | . | . | 6.850351 | 7.167894 | 6.481237 | 5.789745 | 15.941049 | 16.568907 | 14.543707 | 13.237563 | -1.063482 | 624.765990 | 0.15918505 | -0.55781478 | 0.576970872 | 0.998865404 | 9.330668 | 9.242324 | 9.303909 | 9.310138 | 9.304415 | 9.225900 | 738 | 698 | 558 | 534 |
| 389053 | gene-HNRNPKF | HNRNPKP2 | heterogeneous nuclear ribonucleoprotein K pseudogene 2 | pseudogene | . | 42375 | . | . | . | 0.019134 | 0.003489 | 0.003925 | 0.040159 | 0.044525 | 0.008065 | 0.008807 | 0.091820 | 1.635035 | 1.753401 | 2.24714197 | 0.315654997 | 0.752264389 | NA | 0.413393 | 1.121007 | 0.804636 | 0.800773 | 0.802005 | 0.811322 | 2 | 1 | 1 | 3 |
| 389072 | NM_00108047 | PLEKHM3 | pleckstrin homology domain containing M3 | protein_coding | NP_001073944.1;XP_005246602.1;XP_006712584.1;XP_011509462.1;XP_ | 34006 | . | . | . | 1.126259 | 1.327398 | 1.155118 | 1.405238 | 2.620853 | 3.068337 | 2.592051 | 3.212909 | 1.018508 | 353.728792 | 0.201538337 | 0.131273392 | 0.895559039 | 0.998865404 | 8.453865 | 8.479025 | 8.410285 | 8.506309 | 8.416687 | 8.520075 | 370 | 410 | 282 | 365 |
| 389084 | NM_00134612 | SNORC | secondary ossification center associated regulator of chondrocyte | protein_coding | NP_001333049.1;NP_001333050.1;NP_001333051.1;NP_996778.1;XP_01 | 33763 | . | . | . | 0.428291 | 0.675400 | 0.942777 | 0.599482 | 0.996652 | 1.561218 | 2.115565 | 1.370643 | 1.279966 | 64.139889 | 0.419123884 | 0.849643438 | 0.395523354 | 0.998865404 | 5.813288 | 6.170907 | 5.940022 | 5.983622 | 6.074396 | 5.956591 | 58 | 67 | 78 | 51 |
| 389112 | gene-RPS27P4 | RPS27P4 | ribosomal protein S27 pseudogene 4 | pseudogene | . | 16694 | . | . | . | 1.999537 | 2.044205 | 1.243613 | 3.323013 | 4.653006 | 4.725272 | 2.790631 | 7.597674 | 1.120693 | 23.721936 | 0.678654097 | 0.242231469 | 0.808600812 | NA | 4.486115 | 4.645770 | 4.557285 | 4.555911 | 4.521543 | 4.612292 | 26 | 24 | 12 | 34 |
| 389119 | NM_00136628 | INKA1 | inka box actin regulator 1 | protein_coding | NP_001353210.1;NP_976248.2 | 32480 | . | . | . | 1.018537 | 0.702115 | 1.128337 | 1.230064 | 2.370177 | 1.622971 | 2.531956 | 2.812396 | 1.315730 | 33.038436 | 0.547620588 | 0.722879166 | 0.469754124 | 0.998865404 | 4.833848 | 5.231046 | 5.043264 | 4.998919 | 5.049834 | 5.064786 | 39 | 25 | 31 | 37 |
| 389125 | NM_205853 | MUSTN1 | musculoskeletal, embryonic nuclear protein 1 | protein_coding | NP_995325.4 | 22144 | 617195 | . | . | 1.339605 | 0.801809 | 0.796084 | 1.225603 | 3.117317 | 1.853418 | 1.786389 | 2.802195 | -1.070634 | 17.636758 | 0.747192886 | -0.13178088 | 0.895157611 | NA | 4.189060 | 4.090276 | 4.163319 | 4.125163 | 4.125724 | 4.152705 | 26 | 15 | 12 | 19 |
| 389136 | NM_00132049 | VGLL3 | vestigial like family member 3 | protein_coding | NP_001307422.1;NP_001307423.1;NP_057290.2;XP_006713201.1 | 24327 | 609980 | . | . | 0.277817 | 0.245914 | 0.401570 | 0.256387 | 0.646493 | 0.568440 | 0.901111 | 0.586199 | 1.225858 | 96.674305 | 0.348826972 | 0.842228436 | 0.3996601 | 0.998865404 | 6.439208 | 6.735706 | 6.573576 | 6.533814 | 6.681245 | 6.543345 | 107 | 87 | 113 | 78 |
| 389137 | gene-ARMC10I | ARMC10P1 | armadillo repeat containing 10 pseudogene 1 | pseudogene | . | 43646 | . | . | . | 3.293180 | 2.738277 | 2.949017 | 1.389654 | 7.663364 | 6.329649 | 6.617508 | 3.177278 | -1.404530 | 170.407451 | 0.324281393 | -1.51130447 | 0.130710896 | 0.882587711 | 7.635781 | 7.149048 | 7.509488 | 7.419785 | 7.450030 | 7.180881 | 251 | 194 | 168 | 85 |
| 389156 | gene-RPL21P3 | RPL21P39 | ribosomal protein L21 pseudogene 39 | pseudogene | . | 36127 | . | . | . | 0.331222 | 0.352035 | 0.213692 | 0.228692 | 0.770768 | 0.813745 | 0.479518 | 0.522878 | -1.417561 | 5.358218 | 1.304113678 | -0.38601767 | 0.699483584 | NA | 2.651489 | 2.148375 | 2.430947 | 2.432782 | 2.424234 | 2.422662 | 7 | 7 | 4 | 4 |
| 389168 | gene-SETP14 | SETP14 | SET pseudogene 14 | pseudogene | . | 42933 | . | . | . | 1.688108 | 1.449663 | 1.860543 | 1.413759 | 3.928297 | 3.350962 | 4.175004 | 3.232392 | 1.024714 | 90.614594 | 0.345240066 | 0.102019144 | 0.91874148 | 0.998865404 | 6.482827 | 6.520273 | 6.517463 | 6.472593 | 6.541107 | 6.461867 | 111 | 89 | 91 | 74 |
| 389170 | NM_00100431 | LEKR1 | leucine, glutamate and lysine rich 1 | protein_coding | NP_001004316.2;NP_001180212.1;XP_005247498.1;XP_005247499.1;XP | 33765 | 613536 | . | . | 0.965679 | 0.529165 | 0.868574 | 0.393140 | 2.247175 | 1.223187 | 1.949052 | 0.898869 | 1.164403 | 44.325713 | 0.489384492 | 0.448707366 | 0.65364277 | 0.998865404 | 5.354393 | 5.577160 | 5.490815 | 5.409532 | 5.491034 | 5.461867 | 59 | 33 | 45 | 41 |
| 389203 | NM_00114543 | SMIM20 | small integral membrane protein 20 | protein_coding | NP_001138904.1 | 37260 | 617465 | . | . | 0.682058 | 0.233499 | 0.165758 | 0.553292 | 1.587177 | 0.539743 | 0.371957 | 1.265036 | 1.364823 | 13.469166 | 0.93048328 | -0.35055579 | 0.725921629 | NA | 3.906576 | 3.577925 | 3.790647 | 3.723184 | 3.717518 | 3.770239 | 26 | 8 | 5 | 17 |
| 389223 | gene-EEF1A1P: | EEF1A1P35 | eukaryotic translation elongation factor 1 alpha 1 pseudogene 35 | pseudogene | . | 37913 | . | . | . | 0.022164 | 0.041404 | 0.020912 | 0.027935 | 0.051576 | 0.095708 | 0.046926 | 0.063870 | -1.369252 | 1.950107 | 2.123255725 | -0.21353427 | 0.830910261 | NA | 1.175403 | 0.715150 | 0.968658 | 0.973601 | 0.966115 | 0.970520 | 2 | 3 | 1 | 2 |
| 389247 | NR_033995 | LOC389247 | uncharacterized LOC389247 | lncRNA | . | | . | . | . | 0.496238 | 0.551405 | 0.238653 | 0.404214 | 1.154766 | 1.274598 | 0.535531 | 0.924187 | -1.529095 | 11.123580 | 0.924268951 | -0.66287825 | 0.507408533 | NA | 3.751025 | 3.134756 | 3.491080 | 3.494844 | 3.464336 | 3.480455 | 15 | 15 | 6 | 10 |
| 389289 | NM_00101427 | ANXA2R | annexin A2 receptor | protein_coding | NP_001014301.1;NP_001369281.1 | 33463 | 611296 | . | . | 1.411234 | 1.568743 | 1.629353 | 1.404199 | 3.283999 | 3.626222 | 3.656220 | 3.210536 | 1.084952 | 58.735814 | 0.423121089 | 0.278006895 | 0.781007066 | 0.998865404 | 5.816276 | 5.933673 | 5.826656 | 5.896353 | 5.902114 | 5.855670 | 54 | 71 | 59 | 51 |
| 389293 | gene-ISCA1P1 | ISCA1P1 | iron-sulfur cluster assembly 1 pseudogene 1 | pseudogene | . | 33263 | . | . | . | 0.031853 | 0.078608 | 0.126913 | 0.085829 | 0.074124 | 0.181705 | 0.284789 | 0.196238 | 1.641289 | 5.387238 | 1.319926079 | 0.541567758 | 0.588116308 | NA | 2.029593 | 2.742346 | 2.400760 | 2.414912 | 2.425786 | 2.413846 | 3 | 6 | 7 | 5 |
| 389333 | NM_00116154 | PROB1 | proline rich basic protein 1 | protein_coding | NP_001155018.1 | 41906 | . | . | . | 0.049530 | 0.071730 | 0.012577 | 0.015207 | 0.115259 | 0.165808 | 0.028222 | 0.034768 | -4.062607 | 5.610863 | 1.374008372 | -1.47190223 | 0.141047282 | NA | 3.170740 | 1.148375 | 2.513591 | 2.524196 | 2.488895 | 2.487825 | 9 | 11 | 2 | 2 |
| 389337 | NM_00100166 | ARHGEF37 | Rho guanine nucleotide exchange factor 37 | protein_coding | NP_001001669.2;XP_005268504.1;XP_005268505.1;XP_006714847.1;XP | 34430 | . | . | . | 0.164447 | 0.103510 | 0.166428 | 0.215452 | 0.382674 | 0.239268 | 0.373460 | 0.492606 | 1.437598 | 26.615203 | 0.609650409 | 0.858952351 | 0.390366805 | NA | 4.448287 | 4.972679 | 4.723385 | 4.687293 | 4.729635 | 4.759941 | 30 | 19 | 24 | 33 |
| 389342 | gene-RPL10P9 | RPL10P9 | ribosomal protein L10 pseudogene 9 | pseudogene | . | 35579 | . | . | . | 0.337776 | 0.396671 | 0.255919 | 0.469807 | 0.786018 | 0.916922 | 0.574276 | 1.074158 | 1.040584 | 9.146263 | 1.010537899 | 0.056794473 | 0.954708916 | NA | 3.166062 | 3.219802 | 3.191085 | 3.193781 | 3.183505 | 3.204547 | 10 | 10 | 6 | 11 |
| 389362 | NM_00112859 | PSMG4 | proteasome assembly chaperone 4 | protein_coding | NP_001122063.1;NP_001122064.1;NP_001129222.1;XP_011512893.1;XP | 21108 | 617550 | . | . | 8.421173 | 8.650289 | 7.976100 | 8.453946 | 19.596413 | 19.955529 | 17.898138 | 19.328942 | -1.032071 | 404.842978 | 0.180906787 | -0.25174599 | 0.801237405 | 0.998865404 | 8.683634 | 8.638450 | 8.667228 | 8.673497 | 8.675862 | 8.626801 | 471 | 446 | 361 | 358 |
| 389421 | NM_00100431 | LIN28B | lin-28 homolog B | protein_coding | NP_001004317.1;XP_006715540.2;XP_011534120.1 | 32207 | 611044 | . | . | 0.372117 | 0.276475 | 0.365819 | 0.231618 | 0.865932 | 0.639084 | 0.820887 | 0.529567 | -1.088726 | 53.560990 | 0.453081955 | -0.27068044 | 0.786638825 | 0.998865404 | 5.801271 | 5.682507 | 5.778340 | 5.715466 | 5.772656 | 5.687118 | 74 | 51 | 55 | 37 |
| 389432 | NM_00100300 | SAMD5 | sterile alpha motif domain containing 5 | protein_coding | NP_001025231.1;XP_016866339.1 | 21180 | . | . | . | 2.261236 | 2.011486 | 1.445347 | 1.397955 | 5.261990 | 4.649639 | 3.243317 | 3.196259 | -1.520173 | 351.638828 | 0.194042103 | -3.1139385 | 0.00184608 | 0.106540819 | 8.728422 | 8.124794 | 8.598683 | 8.520894 | 8.336976 | 8.314219 | 519 | 429 | 248 | 256 |
| 389435 | gene-RPL27AP | RPL27AP6 | ribosomal protein L27a pseudogene 6 | pseudogene | . | 35977 | . | . | . | 16.863190 | 17.319450 | 11.513106 | 18.562920 | 39.241337 | 40.034683 | 25.835077 | 42.441914 | -1.150405 | 253.840722 | 0.249326431 | -0.8107513 | 0.417508506 | 0.998865404 | 8.086225 | 7.882122 | 8.012940 | 8.022573 | 7.831078 | 8.051442 | 310 | 296 | 158 | 272 |
| 389473 | gene-LOC3894 | LOC389473 | chromosome 5 open reading frame 13 pseudogene | pseudogene | . | | . | . | . | 0.165374 | 0.426603 | 0.540151 | 0.326573 | 0.384832 | 0.986112 | 1.212083 | 0.746671 | 1.475966 | 22.496522 | 0.70390239 | 0.797922937 | 0.424915212 | NA | 4.186683 | 4.744590 | 4.417913 | 4.486914 | 4.515441 | 4.462787 | 12 | 28 | 29 | 19 |
| 389524 | NM_00100379 | GTF2IRD2B | GTF2I repeat domain containing 2B | protein_coding | NP_001003795.1;NP_001355230.1;NP_001355231.1 | 33125 | 608900 | . | . | 1.755387 | 2.263892 | 2.520528 | 1.967122 | 4.084858 | 5.233088 | 5.655990 | 4.497592 | 1.178589 | 138.667362 | 0.283097018 | 0.837383335 | 0.40237713 | 0.998865404 | 6.992595 | 7.228723 | 7.037829 | 7.119701 | 7.142199 | 7.139209 | 130 | 153 | 131 | 140 |
| 389538 | NR_002822,NF | CCZ1P-OR7E38 | CCZ1P-OR7E38P readthrough | transcribed_ps | . | | . | . | . | 17.275486 | 18.178840 | 17.374738 | 19.419369 | 40.200766 | 42.021200 | 38.988410 | 44.400080 | 1.029408 | 512.444900 | 0.168115825 | 0.248726784 | 0.803572129 | 0.998865404 | 8.980459 | 9.021752 | 8.979778 | 9.003192 | 8.978454 | 9.040558 | 571 | 555 | 430 | 511 |
| 389541 | NM_00100839 | LAMTOR4 | late endosomal/lysosomal adaptor, MAPK and MTOR activator 4 | protein_coding | NP_001008396.1;NP_001305165.1;NP_001305166.1;XP_024302528.1;XP | 33772 | 618834 | . | . | 64.022689 | 66.943837 | 61.178841 | 66.489304 | 148.983426 | 154.743679 | 137.283539 | 152.019887 | -1.037680 | 1211.085202 | 0.132685151 | -0.40216301 | 0.687564061 | 0.998865404 | 10.268668 | 10.215002 | 10.242076 | 10.269302 | 10.198145 | 10.256684 | 1395 | 1355 | 993 | 1158 |
| 389549 | NM_00102461 | FEZF1 | FEZ family zinc finger 1 | protein_coding | NP_001019784.2;NP_001153736.1;XP_005250394.1;XP_011514504.1 | 22788 | 613301 | . | . | 0.521510 | 0.451095 | 0.759437 | 0.489999 | 1.213576 | 1.042726 | 1.704155 | 1.120326 | 1.213288 | 61.099503 | 0.424724955 | 0.656711699 | 0.511366311 | 0.998865404 | 5.785505 | 6.066966 | 5.899781 | 5.903341 | 5.999066 | 5.884162 | 63 | 60 | 73 | 47 |
| 389599 | gene-STRADB | STRADBP1 | STE20-related kinase adaptor beta pseudogene 1 | pseudogene | . | 22478 | . | . | . | 0.017695 | 0.040028 | 0.082886 | 0.035054 | 0.041177 | 0.092526 | 0.185994 | 0.080147 | 1.981332 | 3.366748 | 1.651696735 | 0.597246919 | 0.550342538 | NA | 1.175403 | 2.161867 | 1.724130 | 1.729236 | 1.742621 | 1.731255 | 2 | 3 | 5 | 3 |
| 389634 | NR_024420 | LINC00937 | long intergenic non-protein coding RNA 937 | lncRNA | . | 48629 | . | . | . | 0.274168 | 0.044374 | 0.110630 | 0.087703 | 0.638000 | 0.102572 | 0.248250 | 0.200523 | -1.587301 | 2.859780 | 1.842571797 | -0.36176354 | 0.717528737 | NA | 1.808558 | 1.148375 | 1.540439 | 1.516128 | 1.522760 | 1.521949 | 7 | 1 | 2 | 2 |
| 389652 | NR_146077 | ASNSP1 | asparagine synthetase pseudogene 1 | transcribed_ps | . | 754 | . | . | . | 1.040128 | 0.701797 | 0.271629 | 0.668423 | 2.420422 | 1.622235 | 0.609527 | 1.528269 | -1.854082 | 31.162971 | 0.611789086 | -1.45590193 | 0.145419732 | 0.906189003 | 5.341075 | 4.445696 | 5.030128 | 4.965914 | 4.886703 | 4.959843 | 56 | 35 | 11 | 29 |
| 389668 | NM_00101172 | XKR9 | XK related 9 | protein_coding | NP_001011720.1;NP_001274187.1;NP_001274188.1;NP_001274189.1;XF | 20937 | . | . | . | 0.275792 | 0.358872 | 0.854597 | 0.175319 | 0.641779 | 0.829551 | 1.917690 | 0.400848 | 1.438556 | 33.920707 | 0.612116398 | 0.857061756 | 0.391410756 | 0.998865404 | 4.795304 | 5.324606 | 5.034695 | 5.027476 | 5.156680 | 4.996594 | 33 | 29 | 53 | 18 |
| 389672 | gene-RPSAP47 | RPSAP47 | ribosomal protein SA pseudogene 47 | pseudogene | . | 36162 | . | . | . | 18.302849 | 20.019638 | 15.871617 | 22.133169 | 42.591480 | 46.276295 | 35.615448 | 50.604862 | -1.022086 | 614.785775 | 0.183109164 | -0.17211636 | 0.863346054 | 0.998865404 | 9.280176 | 9.247519 | 9.240631 | 9.295182 | 9.141886 | 9.354076 | 687 | 698 | 444 | 661 |
| 389674 | gene-HNRNPA | HNRNPA1P4 | heterogeneous nuclear ribonucleoprotein A1 pseudogene 4 | pseudogene | . | 32234 | . | . | . | 2.597893 | 2.493911 | 2.682628 | 2.636207 | 6.045403 | 5.764788 | 6.019739 | 6.027375 | 1.030496 | 120.538897 | 0.293630898 | 0.147594769 | 0.882662583 | 0.998865404 | 6.891328 | 6.935051 | 6.915988 | 6.899346 | 6.923059 | 6.912940 | 140 | 125 | 108 | 113 |
| 389677 | NM_00137796 | RBM12B | RNA binding motif protein 12B | protein_coding | NP_001364889.1;NP_001364890.1;NP_001364891.1;NP_001364892.1;NI | 32310 | . | . | . | 3.545971 | 3.243645 | 3.388161 | 2.873753 | 8.251620 | 7.497833 | 7.602933 | 6.570492 | -1.098839 | 870.170267 | 0.153036465 | -0.88855001 | 0.374244968 | 0.998865404 | 9.831189 | 9.695951 | 9.831070 | 9.761275 | 9.787970 | 9.667429 | 1099 | 936 | 783 | 711 |
| 389741 | NR_015363,NF | GLIDR | glioblastoma down-regulated RNA | lncRNA | . | 48823 | . | . | . | 3.773525 | 4.090086 | 3.436535 | 3.766932 | 8.781150 | 9.454417 | 7.711485 | 8.612642 | -1.120385 | 182.264910 | 0.248084508 | -0.66104143 | 0.508585743 | 0.998865404 | 7.589950 | 7.425131 | 7.509807 | 7.557776 | 7.470187 | 7.499815 | 210 | 219 | 144 | 167 |
| 389765 | NR_029410 | LOC389765 | kinesin family member 27 pseudogene | transcribed_ps | . | | . | . | . | 0.393460 | 0.425148 | 0.356428 | 0.518993 | 0.915597 | 0.982748 | 0.799814 | 1.186617 | 1.037811 | 23.756108 | 0.638066793 | 0.083916188 | 0.933123069 | NA | 4.544574 | 4.595431 | 4.564411 | 4.570920 | 4.552441 | 4.589939 | 26 | 26 | 17 | 27 |
| 389787 | gene-TPT1P9 | TPT1P9 | tumor protein, translationally-controlled 1 pseudogene 9 | pseudogene | . | 49299 | . | . | . | 0.153778 | 0.540381 | 0.245945 | 0.245833 | 0.357848 | 1.249114 | 0.551893 | 0.562069 | -1.368208 | 7.885808 | 1.128421471 | -0.40081782 | 0.688554255 | NA | 3.189301 | 2.733338 | 2.959427 | 3.003314 | 2.971807 | 2.969492 | 5 | 15 | 6 | 6 |
| 389791 | NR_024425 | PTGES2-AS1 | PTGES2 antisense RNA 1 (head to head) | lncRNA | . | 48711 | . | . | . | 0.189859 | 0.159153 | 0.194220 | 0.144850 | 0.441811 | 0.367889 | 0.435825 | 0.331182 | -1.034196 | 11.886816 | 0.8875339 | -0.05465662 | 0.956412034 | NA | 3.593806 | 3.548418 | 3.575605 | 3.566956 | 3.578510 | 3.560429 | 15 | 12 | 12 | 9 |
| 389792 | NM_203434 | IER5L | immediate early response 5 like | protein_coding | NP_982258.2 | 23679 | . | . | . | 2.479743 | 2.894316 | 2.150220 | 2.253724 | 5.770464 | 6.690342 | 4.825032 | 5.152873 | -1.234161 | 206.995810 | 0.237179713 | -1.27974873 | 0.20063352 | 0.966265067 | 7.837588 | 7.533314 | 7.702607 | 7.774501 | 7.634996 | 7.650760 | 244 | 265 | 158 | 177 |
| 389813 | NM_00108048 | AJM1 | apical junction component 1 homolog | protein_coding | NP_001073951.2 | 37284 | . | . | . | 0.292334 | 0.332758 | 0.433307 | 0.328279 | 0.680273 | 0.769185 | 0.972328 | 0.750571 | 1.219789 | 34.913886 | 0.52931632 | 0.541517437 | 0.588154428 | 0.998865404 | 4.974755 | 5.262390 | 5.097211 | 5.113187 | 5.152520 | 5.112203 | 34 | 36 | 38 | 31 |
| 389840 | NM_00100167 | MAP3K15 | mitogen-activated protein kinase kinase kinase 15 | protein_coding | NP_001001671.3;XP_011543809.3;XP_011543810.3;XP_011543812.1;XP | 31689 | 300820 | . | . | 0.102278 | 0.062919 | 0.094374 | 0.205861 | 0.238006 | 0.145441 | 0.211771 | 0.470677 | 1.740191 | 18.909421 | 0.747469671 | 1.069268874 | 0.284948532 | NA | 3.787711 | 4.585454 | 4.214132 | 4.190181 | 4.208293 | 4.283008 | 19 | 12 | 13 | 31 |
| 389849 | gene-GAPDHP | GAPDHP65 | glyceraldehyde 3 phosphate dehydrogenase pseudogene 65 | pseudogene | . | 4143 | . | . | . | 0.089335 | 0.118148 | 0.114972 | 0.071914 | 0.207885 | 0.273105 | 0.257993 | 0.164424 | -1.270958 | 4.425644 | 1.431029408 | -0.24172537 | 0.808991909 | NA | 2.307819 | 1.963455 | 2.147718 | 2.153399 | 2.148878 | 2.142575 | 5 | 6 | 4 | 3 |
| 389856 | NM_00114507 | USP27X | ubiquitin specific peptidase 27 X-linked | protein_coding | NP_001138545.1 | 13486 | 300975 | . | . | 1.444270 | 1.333225 | 0.972682 | 1.549888 | 3.360877 | 3.081810 | 2.182670 | 3.543635 | -1.119740 | 127.463279 | 0.312781127 | -0.52165675 | 0.60190934 | 0.998865404 | 7.074169 | 6.908981 | 7.027764 | 6.994636 | 6.893005 | 7.043781 | 162 | 139 | 81 | 138 |
| 389866 | gene-SERBP1P | SERBP1P1 | SERPINE1 mRNA binding protein 1 pseudogene 1 | pseudogene | . | 32247 | . | . | . | 0.182232 | 0.130197 | 0.104896 | 0.289091 | 0.424060 | 0.300957 | 0.235384 | 0.660973 | 1.223752 | 14.854842 | 0.834028034 | 0.349282669 | 0.726877109 | NA | 3.741623 | 4.029739 | 3.891033 | 3.870852 | 3.863554 | 3.923700 | 18 | 12 | 8 | 22 |
| 389884 | gene-PA2G4P1 | PA2G4P1 | proliferation-associated 2G4 pseudogene 1 | pseudogene | . | 8551 | . | . | . | 0.118467 | 0.061480 | 0.045801 | 0.068133 | 0.275678 | 0.142113 | 0.102777 | 0.155777 | -1.487557 | 4.088173 | 1.503726114 | -0.38101697 | 0.703190655 | NA | 2.290730 | 1.715150 | 2.050610 | 2.039448 | 2.033341 | 2.042033 | 7 | 4 | 2 | 4 |
| 389895 | NM_00127156 | LOC389895 | chromosome 16 open reading frame 72-like | protein_coding | NP_001258489.1 | | . | . | . | 0.122545 | 0.063942 | 0.242358 | 0.135603 | 0.285167 | 0.147806 | 0.543845 | 0.310041 | 1.952820 | 5.918498 | 1.272998622 | 0.758491864 | 0.448156582 | NA | 1.998355 | 2.971139 | 2.542985 | 2.531557 | 2.565696 | 2.543696 | 6 | 3 | 9 | 5 |
| 389901 | gene-XRCC6P2 | XRCC6P2 | | pseudogene | . | 45184 | . | . | . | 0.489234 | 0.483050 | 0.237974 | 0.360000 | 1.138467 | 1.116592 | 0.534007 | 0.823097 | -1.652424 | 24.799041 | 0.631411919 | -1.14756082 | 0.251149912 | NA | 4.950620 | 4.223024 | 4.663814 | 4.660658 | 4.596722 | 4.626900 | 36 | 33 | 13 | 21 |
| 389903 | NM_00112982 | CSAG3 | CSAG family member 3 | protein_coding | NP_001123298.1;NP_001123300.1;NP_024308147.1 | 26237 | . | . | . | 8.389679 | 10.309316 | 11.708441 | 9.635697 | 19.523128 | 23.830444 | 26.273403 | 22.030878 | 1.156999 | 378.603461 | 0.207120434 | 1.015774257 | 0.309756807 | 0.998865404 | 8.455171 | 8.666206 | 8.472563 | 8.555230 | 8.685302 | 8.506864 | 375 | 406 | 406 | 324 |
| 389906 | NR_034031 | LOC389906 | zinc finger protein 839 pseudogene | transcribed_ps | . | | . | . | . | 1.186646 | 1.658047 | 0.687150 | 1.221832 | 2.761373 | 3.832650 | 1.541945 | 2.793573 | -1.498138 | 67.176381 | 0.42178426 | -1.38262734 | 0.16677916 | 0.945013445 | 6.333958 | 5.746391 | 6.066066 | 6.152652 | 5.965242 | 6.067739 | 78 | 101 | 34 | 64 |
| 390006 | gene-PPIAP19 | PPIAP19 | peptidylprolyl isomerase A pseudogene 19 | pseudogene | . | 31663 | . | . | . | 0.046409 | 0.040040 | 0.029651 | 0.040566 | 0.107996 | 0.092544 | 0.066535 | 0.092749 | -1.204659 | 1.220068 | 2.631544379 | -0.10207874 | 0.918694172 | NA | 0.413393 | 0.148375 | 0.294625 | 0.290739 | 0.291821 | 0.291432 | 1 | 1 | 1 | 1 |
| 390010 | NM_00114634 | NKX1-2 | NK1 homeobox 2 | protein_coding | NP_001139812.1 | 31652 | . | . | . | 0.160987 | 0.323764 | 0.090206 | 0.234782 | 0.374624 | 0.748394 | 0.202420 | 0.536801 | -1.518058 | 20.632421 | 0.744903296 | -0.808463 | 0.418824099 | NA | 4.639904 | 4.029739 | 4.345229 | 4.416475 | 4.315853 | 4.375054 | 19 | 36 | 8 | 22 |
| 390345 | gene-RPL10P1 | RPL10P13 | ribosomal protein L10 pseudogene 13 | pseudogene | . | 52346 | . | . | . | 0.823928 | 0.506967 | 0.130921 | 0.156812 | 1.917315 | 1.171877 | 0.293783 | 0.358533 | -3.820420 | 5.335450 | 1.403196849 | -1.37808984 | 0.168175553 | NA | 3.079678 | 1.148375 | 2.456458 | 2.438282 | 2.419756 | 2.418712 | 12 | 7 | 2 | 2 |
| 390354 | gene-RPL18AP | RPL18AP3 | ribosomal protein L18a pseudogene 3 | pseudogene | . | 31387 | . | . | . | 15.573434 | 14.237487 | 14.817169 | 18.678165 | 36.240017 | 32.910587 | 33.249294 | 42.705402 | 1.107187 | 307.885967 | 0.210306894 | 0.698498211 | 0.484865676 | 0.998865404 | 8.191265 | 8.337534 | 8.261274 | 8.209109 | 8.227547 | 8.350241 | 353 | 300 | 250 | 337 |
| 390363 | gene-CLIC1P1 | CLIC1P1 | chloride intracellular channel 1 pseudogene 1 | pseudogene | . | 32691 | . | . | . | 0.196206 | 0.131434 | 0.174449 | 0.271716 | 0.456579 | 0.303816 | 0.391457 | 0.621246 | 1.334073 | 6.249007 | 1.211341737 | 0.343286403 | 0.731383009 | NA | 2.421266 | 2.836248 | 2.638477 | 2.631748 | 2.637084 | 2.649445 | 7 | 5 | 5 | 8 |
| 390365 | NR_072991 | HERC2P10 | hect domain and RLD 2 pseudogene 10 | transcribed_ps | . | 39056 | . | . | . | 1.706674 | 2.516730 | 2.115032 | 2.547483 | 3.971500 | 5.817535 | 4.746071 | 5.824517 | 1.083363 | 44.597762 | 0.478929904 | 0.241197964 | 0.809401689 | 0.998865404 | 5.421369 | 5.534224 | 5.435748 | 5.499217 | 5.467413 | 5.495424 | 40 | 55 | 37 | 47 |
| 390595 | NM_00116369 | UBAP1L | ubiquitin associated protein 1 like | protein_coding | NP_001157164.1;XP_011519849.1;XP_016877661.1 | 40028 | . | . | . | 0.592281 | 1.015774 | 0.569406 | 0.521089 | 1.378262 | 2.348007 | 1.277730 | 1.191407 | -1.233810 | 214.477844 | 0.276160458 | -1.0976247 | 0.272368404 | 0.998865404 | 7.887678 | 7.585944 | 7.715370 | 7.845969 | 7.786276 | 7.572073 | 237 | 289 | 208 | 136 |
| 390627 | gene-CSPG4P1 | CSPG4P10 | chondroitin sulfate proteoglycan 4 pseudogene 10 | pseudogene | . | 48361 | . | . | . | 0.045168 | 0.006429 | 0.036930 | 0.009407 | 0.105108 | 0.014862 | 0.082869 | 0.021509 | -1.125035 | 4.171058 | 1.532172253 | -0.11093403 | 0.91166866 | NA | 2.137672 | 1.978782 | 2.072029 | 2.047999 | 2.065855 | 2.049677 | 8 | 2 | 5 | 2 |
| 390637 | NM_00101365 | GDPGP1 | GDP-D-glucose phosphorylase 1 | protein_coding | NP_001013679.2;NP_001309740.1 | 34360 | . | . | . | 0.488162 | 0.671493 | 0.577388 | 0.568255 | 1.135975 | 1.552186 | 1.295642 | 1.299245 | -1.015570 | 93.592868 | 0.340119912 | -0.04879584 | 0.961081994 | 0.998865404 | 6.557107 | 6.539401 | 6.505057 | 6.591633 | 6.547929 | 6.540050 | 92 | 117 | 82 | 86 |
| 390916 | NM_00110557 | NUDT19 | nudix hydrolase 19 | protein_coding | NP_001099040.1;XP_016882295.1 | 32036 | . | . | . | 5.921455 | 6.250182 | 6.327286 | 6.199592 | 13.779469 | 14.447574 | 14.198246 | 14.174628 | 1.019106 | 559.046086 | 0.16209211 | 0.168451398 | 0.866228176 | 0.998865404 | 9.113109 | 9.140409 | 9.100922 | 9.139248 | 9.146097 | 9.118941 | 619 | 615 | 501 | 518 |
| 390927 | NM_00101365 | ZNF793 | zinc finger protein 793 | protein_coding | NP_001013681.2;XP_005258984.1;XP_006723276.1;XP_024307279.1 | 33115 | . | . | . | 0.052880 | 0.037267 | 0.089063 | 0.103171 | 0.123055 | 0.086143 | 0.199856 | 0.235888 | 2.027076 | 16.881740 | 0.757744381 | 1.345308618 | 0.178525592 | NA | 3.478589 | 4.499300 | 4.042555 | 4.025794 | 4.075716 | 4.091992 | 15 | 10 | 18 | 23 |
| 390980 | NM_00102356 | ZNF805 | zinc finger protein 805 | protein_coding | NP_001018857.2;NP_001318550.1 | 23272 | . | . | . | 0.516607 | 0.458271 | 0.515950 | 0.446905 | 1.202165 | 1.059313 | 1.157776 | 1.021796 | -1.027908 | 154.098627 | 0.268949154 | -0.14765472 | 0.882615269 | 0.998865404 | 7.286780 | 7.248385 | 7.297159 | 7.246785 | 7.289647 | 7.232551 | 191 | 158 | 142 | 132 |
| 391040 | gene-BTF3L4P | BTF3L4P3 | basic transcription factor 3 like 4 pseudogene 3 | pseudogene | . | 39647 | . | . | . | 0.042205 | 0.077545 | 0.013735 | 0.037520 | 0.098212 | 0.179249 | 0.030821 | 0.085785 | -1.369252 | 1.950107 | 2.123255725 | -0.21353427 | 0.830910261 | NA | 1.175403 | 0.715150 | 0.968658 | 0.973601 | 0.966115 | 0.970520 | 2 | 3 | 1 | 2 |
| 391059 | NM_00101366 | FRRS1 | ferric chelate reductase 1 | protein_coding | NP_001013682.2;NP_001347970.1;XP_011539753.1;XP_011539754.1;XP | 27622 | 611578 | . | . | 2.539902 | 2.316058 | 1.971018 | 1.714115 | 5.910455 | 5.353672 | 4.422908 | 3.919120 | -1.037869 | 398.148152 | 0.188029416 | -0.2851908 | 0.775497968 | 0.998865404 | 8.663265 | 8.610577 | 8.661370 | 8.632465 | 8.675555 | 8.572098 | 478 | 427 | 370 | 334 |
| 391169 | gene-CDKN2A | CDKN2AIPNLP1 | CDKN2A interacting protein N-terminal like pseudogene 1 | pseudogene | . | 39854 | . | . | . | 0.079530 | 0.162025 | 0.155377 | 0.280650 | 0.185069 | 0.374527 | 0.348662 | 0.641672 | 1.705653 | 6.096028 | 1.239582667 | 0.621438178 | 0.534311359 | NA | 2.175403 | 2.940155 | 2.583839 | 2.593834 | 2.594797 | 2.611938 | 4 | 6 | 5 | 9 |
| 391256 | gene-KRT18P4 | KRT18P4 | keratin 18 pseudogene 4 | pseudogene | . | 16604 | . | . | . | 0.176059 | 0.164961 | 0.398875 | 0.296204 | 0.409696 | 0.381314 | 0.895064 | 0.677236 | 1.970464 | 11.319889 | 0.915306395 | 1.0690795 | 0.285033848 | NA | 2.928838 | 3.909275 | 3.466375 | 3.464761 | 3.507779 | 3.489118 | 9 | 8 | 15 | 12 |
| 391356 | NM_00101366 | PTRHD1 | peptidyl-tRNA hydrolase domain containing 1 | protein_coding | NP_001013685.1 | 33782 | 617342 | . | . | 14.831994 | 15.370568 | 16.007820 | 16.237427 | 34.514660 | 35.529758 | 35.921082 | 37.124950 | 1.052302 | 278.445940 | 0.206272741 | 0.356560205 | 0.721421073 | 0.998865404 | 8.084073 | 8.157500 | 8.099718 | 8.114720 | 8.133155 | 8.134505 | 308 | 297 | 248 | 268 |
| 391358 | gene-TRMT11 | TRMT112P6 | tRNA methyltransferase subunit 11-2 pseudogene 6 | pseudogene | . | 44025 | . | . | . | 0.562690 | 0.671869 | 0.893194 | 0.583690 | 1.309402 | 1.553056 | 2.004301 | 1.334538 | 1.148765 | 15.045115 | 0.852686879 | 0.234651156 | 0.814479501 | NA | 3.600756 | 3.803035 | 3.691252 | 3.698957 | 3.716926 | 3.688922 | 13 | 14 | 15 | 10 |
| 391485 | gene-KRT8P30 | KRT8P30 | keratin 8 pseudogene 30 | pseudogene | . | 33382 | . | . | . | 0.374919 | 0.296863 | 0.114459 | 0.112499 | 0.872452 | 0.686213 | 0.256844 | 0.257216 | -2.824453 | 6.352983 | 1.248525964 | -1.19979193 | 0.230220159 | NA | 3.229742 | 1.733338 | 2.700408 | 2.690719 | 2.669012 | 2.667612 | 12 | 9 | 3 | 3 |
| 391592 | gene-BZW1P1 | BZW1P1 | basic leucine zipper and W2 domains 1 pseudogene 1 | pseudogene | . | 31384 | . | . | . | 0.025573 | 0.060579 | 0.018627 | 0.038140 | 0.059508 | 0.140032 | 0.041798 | 0.087202 | -1.236989 | 3.683103 | 1.580948665 | -0.19408121 | 0.846112291 | NA | 2.029593 | 1.715150 | 1.878176 | 1.892094 | 1.877195 | 1.885962 | 3 | 6 | 2 | 4 |
| 391656 | gene-RPS15AP | RPS15AP17 | ribosomal protein S15a pseudogene 17 | pseudogene | . | 36888 | . | . | . | 0.205885 | 0.227981 | 0.070053 | 0.065645 | 0.479103 | 0.526988 | 0.157196 | 0.150088 | -3.240854 | 2.349347 | 2.037050328 | -0.83276002 | 0.404980085 | NA | 1.844134 | 0.148375 | 1.256048 | 1.257052 | 1.245413 | 1.244915 | 4 | 4 | 1 | 1 |
| 391712 | NM_00101241 | TRIM61 | tripartite motif containing 61 | protein_coding | NP_001012414.1 | 24339 | . | . | . | 0.536220 | 0.696340 | 0.525502 | 0.762951 | 1.247805 | 1.609621 | 1.179211 | 1.744398 | 1.044307 | 31.248505 | 0.563101896 | 0.111074195 | 0.911557512 | 0.998865404 | 4.935725 | 4.995095 | 4.945514 | 4.976085 | 4.944905 | 4.987523 | 31 | 37 | 23 | 35 |
| 392100 | gene-PSMC1P3 | PSMC1P3 | proteasome 26S subunit, ATPase 1 pseudogene 3 | pseudogene | . | 39778 | . | . | . | 0.027518 | 0.048069 | 0.020283 | 0.051382 | 0.064035 | 0.111114 | 0.045514 | 0.117479 | -1.032087 | 2.216774 | 2.002641667 | -0.02275192 | 0.981848159 | NA | 1.175403 | 1.121007 | 1.148069 | 1.155041 | 1.145613 | 1.154849 | 2 | 3 | 1 | 3 |
| 392232 | NR_033867 | LOC392232 | transient receptor potential cation channel subfamily A member 1 | transcribed_ps | . | | . | . | . | 0.118227 | 0.163677 | 0.058544 | 0.091644 | 0.275236 | 0.378346 | 0.131371 | 0.209533 | -1.836078 | 9.384754 | 1.009523385 | -0.86835783 | 0.38519847 | NA | 3.604218 | 2.724273 | 3.242741 | 3.258247 | 3.222311 | 3.229568 | 12 | 15 | 5 | 7 |
| 392256 | gene-RPS26P6 | RPS26P6 | ribosomal protein S26 pseudogene 6 | pseudogene | . | 31090 | . | . | . | 5.828068 | 6.101658 | 5.226029 | 7.102723 | 13.562155 | 14.104257 | 11.727058 | 16.239531 | 1.019327 | 85.105247 | 0.352412735 | 0.078365017 | 0.937537701 | 0.998865404 | 6.398224 | 6.424013 | 6.401299 | 6.413442 | 6.373652 | 6.450475 | 95 | 93 | 64 | 92 |
| 392281 | gene-GPS2P1 | GPS2P1 | G protein pathway suppressor 2 pseudogene 1 | pseudogene | . | 49622 | . | . | . | 0.165485 | 0.150234 | 0.158959 | 0.083524 | 0.385091 | 0.347373 | 0.356699 | 0.190968 | -1.304720 | 5.147359 | 1.33591402 | -0.28724888 | 0.773921261 | NA | 2.540965 | 2.161867 | 2.371045 | 2.368570 | 2.369294 | 2.357879 | 7 | 6 | 5 | 3 |
| 392352 | gene-PPIAP87 | PPIAP87 | peptidylprolyl isomerase A pseudogene 87 | pseudogene | . | 53711 | . | . | . | 0.017671 | 0.049024 | 0.013864 | 0.002354 | 0.041122 | 0.113322 | 0.031110 | 0.005381 | -1.225404 | 1.234643 | 2.621948942 | -0.11184694 | 0.910944771 | NA | 0.444631 | 0.148375 | 0.303453 | 0.308168 | 0.304847 | 0.304457 | 1 | 2 | 1 | 1 |
| 392617 | NM_00112863 | ELFN1 | extracellular leucine rich repeat and fibronectin type III domain co | protein_coding | NP_001122108.1;XP_006715788.1;XP_006715790.1;XP_011513699.1;XP | 33154 | 614964 | . | . | 0.146485 | 0.070394 | 0.046219 | 0.062936 | 0.340876 | 0.162718 | 0.103714 | 0.143897 | -1.378914 | 15.050449 | 0.795665529 | -0.5825715 | 0.560181793 | NA | 4.125927 | 3.660102 | 3.931339 | 3.924345 | 3.897386 | 3.916686 | 21 | 18 | 9 | 14 |
| 392636 | NM_00100432 | AGMO | alkylglycerol monooxygenase | protein_coding | NP_001004320.1;XP_006715793.1;XP_006715794.1;XP_011513704.1;XP | 33784 | 613738 | . | . | 0.671870 | 0.359507 | 0.565543 | 0.199016 | 1.560668 | 0.915157 | 1.269056 | 0.455548 | -1.044106 | 42.688790 | 0.513245392 | -0.12148297 | 0.903308502 | 0.998865404 | 5.444429 | 5.365566 | 5.438682 | 5.373041 | [illegible] | [illegible] | 64 | 34 | 44 | 31 |
| 399473 | NM_00104252 | SPRED3 | sprouty related EVH1 domain containing 3 | protein_coding | NP_001035987.1;XP_006723282.1;XP_011525280.1 | 31041 | 609293 | . | . | 0.217089 | 0.267998 | 0.225753 | 0.271370 | 0.505174 | 0.619489 | 0.506584 | 0.620455 | 1.018404 | 37.202537 | 0.512239677 | 0.051362576 | 0.959036606 | 0.998865404 | 5.205110 | 5.229445 | 5.199126 | 5.230338 | 5.205933 | 5.228972 | 38 | 44 | 30 | 38 |
| 399511 | gene-TCEA1P2 | TCEA1P2 | transcription elongation factor A1 pseudogene 2 | pseudogene | . | 29891 | . | . | . | 2.113962 | 1.581693 | 1.593379 | 1.232283 | 4.919277 | 3.656155 | 3.575496 | 2.817468 | -1.324058 | 133.628728 | 0.302181506 | -1.34014329 | 0.180198764 | 0.952541638 | 7.249356 | 6.846831 | 7.165045 | 7.046105 | 7.046968 | 6.962385 | 201 | 140 | 113 | 94 |
| 399512 | NM_00132087 | SLC25A35 | solute carrier family 25 member 35 | protein_coding | NP_001307799.1;NP_001307800.1;NP_001307801.1;NP_001307806.1;NI | 31921 | 610818 | . | . | 4.559730 | 5.354879 | 4.116135 | 5.546402 | 10.610679 | 12.378041 | 9.236489 | 12.681190 | -1.043770 | 428.031243 | 0.224109208 | -0.27577283 | 0.782722552 | 0.998865404 | 8.773000 | 8.709445 | 8.649927 | 8.836418 | 8.597667 | 8.831508 | 428 | 543 | 293 | 469 |
| 399664 | NM_00117411 | MEX3D | mex-3 RNA binding family member D | protein_coding | NP_001167589.1;NP_976049.3;XP_016882300.1 | 16734 | 611009 | . | . | 2.816934 | 2.697942 | 2.159870 | 2.535762 | 6.555121 | 6.236412 | 4.846685 | 5.797719 | -1.186877 | 228.186135 | 0.227240438 | -1.08770379 | 0.276725893 | 0.998865404 | 7.952641 | 7.704869 | 7.886008 | 7.861785 | 7.760745 | 7.822107 | 292 | 261 | 169 | 209 |
| 399665 | NM_00103525 | FAM102A | family with sequence similarity 102 member A | protein_coding | NP_001030331.1;NP_976050.1 | 31419 | 610891 | . | . | 6.924476 | 7.401414 | 9.657209 | 8.686859 | 16.113542 | 17.108699 | 21.670496 | 19.861471 | 1.225595 | 1108.718484 | 0.133656 | 2.195798476 | 0.028106371 | 0.539571956 | 9.960473 | 10.253979 | 10.019248 | 10.046338 | 10.199153 | 10.169053 | 1126 | 1095 | 1083 | 1122 |
| 399687 | NM_00134676 | MYO18A | myosin XVIIIA | protein_coding | NP_001333694.1;NP_001333695.1;NP_001333696.1;NP_001333697.1;NI | 31104 | 610067 | . | . | 9.765423 | 10.243142 | 10.816419 | 11.453132 | 22.724537 | 23.677479 | 24.271731 | 26.186225 | 1.102421 | 3278.308189 | 0.111157216 | 1.265550068 | 0.205674212 | 0.970176829 | 11.606738 | 11.747310 | 11.624773 | 11.649371 | 11.697730 | 11.736297 | 3537 | 3416 | 2920 | 3298 |
| 399694 | NM_203439,XI | SHC4 | SHC adaptor protein 4 | protein_coding | NP_976224.3;XP_005254432.1 | 16743 | 617372 | . | . | 3.552336 | 4.052891 | 3.462328 | 3.326714 | 8.266433 | 9.368440 | 7.769364 | 7.606136 | -1.134819 | 564.065551 | 0.165398562 | -1.10316708 | 0.269954614 | 0.998865404 | 9.228126 | 9.045539 | 9.136876 | 9.219723 | 9.112314 | 9.080362 | 648 | 687 | 471 | 483 |
| 399726 | NR_160800 | MIR1915HG | MIR1915 host gene | lncRNA | . | 31448 | . | . | . | 0.459794 | 0.624243 | 0.438959 | 0.558837 | 1.069960 | 1.442965 | 0.985012 | 1.277714 | 1.067098 | 62.188013 | 0.410294394 | -0.29617201 | 0.767098822 | 0.998865404 | 6.019256 | 5.895208 | [illegible] | [illegible] | 5.924979 | 5.970740 | 64 | 80 | 46 | 62 |
| 399744 | NR_024497 | LINC00999 | long intergenic non-protein coding RNA 999 | lncRNA | . | 38537 | . | . | . | 0.433161 | 0.487376 | 0.511698 | 0.480939 | 1.007983 | 1.126591 | 1.148236 | 1.099610 | 1.059685 | 78.630577 | 0.3579921 | 0.233624975 | 0.815276137 | 0.998865404 | 6.254627 | 6.338200 | 6.273083 | 6.302149 | 6.312388 | 6.292993 | 83 | 87 | 73 | 73 |
| 399761 | NR_003611 | BMS1P1 | BMS1 pseudogene 1 | transcribed_ps | . | 23649 | . | . | . | 0.352293 | 0.357342 | 0.543885 | 0.306710 | 0.819800 | 0.826012 | 1.220463 | 0.701257 | 1.175062 | 52.234192 | 0.459210712 | 0.506820096 | 0.612281055 | 0.998865404 | 5.584287 | 5.819946 | 5.680118 | 5.681939 | 5.766813 | 5.653755 | 55 | 52 | 63 | 38 |
| 399815 | NR_027282 | C10orf88B | C10orf88B (pseudogene) | transcribed_ps | . | 44080 | . | . | . | 1.324723 | 1.108078 | 1.731748 | 1.292796 | 3.082684 | 2.561373 | 3.885993 | 2.955824 | 1.222456 | 89.972020 | 0.349051401 | 0.83020171 | 0.406424749 | 0.998865404 | 6.338014 | 6.630188 | 6.480032 | 6.429166 | 6.555644 | 6.466471 | 102 | 79 | 99 | 79 |
| 399818 | NM_00130446 | EEF1AKMT2 | EEF1A lysine methyltransferase 2 | protein_coding | NP_001291396.1;NP_001291397.1;NP_997719.2 | 33787 | 617794 | . | . | 2.735160 | 2.511223 | 2.163325 | 2.326384 | 6.364831 | 5.804806 | 4.854437 | 5.319001 | -1.181987 | 270.167918 | 0.212101015 | -1.13774791 | 0.255225761 | 0.995760118 | 8.193263 | 7.952095 | 8.147952 | 8.089914 | 8.019062 | 8.047565 | 355 | 299 | 209 | 239 |
| 399821 | NR_033847 | EDRF1-DT | EDRF1 divergent transcript | lncRNA | . | 54128 | . | . | . | 0.056487 | 0.095439 | 0.102016 | 0.147061 | 0.131448 | 0.220611 | 0.228920 | 0.336237 | 1.464658 | 8.333629 | 1.06184233 | 0.518499087 | 0.604110097 | NA | 2.760365 | 3.306201 | 3.035438 | 3.050172 | 3.048893 | 3.065096 | 6 | 9 | 7 | 11 |
| 399844 | NR_028324 | LINC01002 | long intergenic non-protein coding RNA 1002 | lncRNA | . | 38538 | . | . | . | 0.200338 | 0.246076 | 0.331549 | 0.190986 | 0.466194 | 0.568816 | 0.743987 | 0.436667 | 1.146663 | 37.164832 | 0.532362419 | 0.366147468 | 0.714255039 | 0.998865404 | 5.113960 | 5.311047 | 5.182906 | 5.210350 | 5.256686 | 5.174181 | 36 | 41 | 44 | 27 |
| 399900 | XR_247232 | LOC399900 | uncharacterized LOC399900 | lncRNA | . | | . | . | . | 0.535278 | 0.441662 | 0.688604 | 0.583102 | 1.245613 | 1.020922 | 1.545208 | 1.333194 | 1.285712 | 28.115705 | 0.584776541 | 0.620009612 | 0.535251459 | 0.998865404 | 4.619518 | 4.984116 | 4.802048 | 4.782350 | 4.831958 | 4.809824 | 31 | 24 | 30 | 27 |
| 399909 | NM_032223,XI | PCNX3 | pecanex 3 | protein_coding | NP_115599.2;XP_005274044.1;XP_005274047.1;XP_006718612.1;XP_01 | 18760 | 617657 | . | . | 8.146736 | 8.414638 | 9.262317 | 7.995676 | 18.957791 | 19.450811 | 20.784368 | 18.281166 | 1.044292 | 1857.694713 | 0.123414804 | 0.506621021 | 0.612420756 | 0.998865404 | 10.827629 | 10.890286 | 10.825266 | 10.857409 | 10.911277 | 10.838883 | 2050 | 2001 | 1726 | 1698 |
| 399942 | gene-TUBAP2 | TUBAP2 | tubulin alpha pseudogene 2 | pseudogene | . | 14531 | . | . | . | 2.974227 | 2.785634 | 2.069864 | 2.802021 | 6.921149 | 6.439117 | 4.644715 | 6.406488 | -1.193583 | 134.088793 | 0.291673426 | -0.87529044 | 0.381415896 | 0.998865404 | 7.189697 | 6.932989 | 7.112126 | 7.084369 | 6.984369 | 7.080960 | 174 | 152 | 91 | 131 |
| 399959 | NR_024430,NF | MIR100HG | mir-100-let-7a-2-mir-125b-1 cluster host gene | lncRNA | . | 39522 | 615965 | . | . | 0.931881 | 0.681480 | 1.252278 | 0.766208 | 2.168525 | 1.575272 | 2.810076 | 1.751846 | 1.234458 | 88.779883 | 0.370627943 | 0.819898665 | 0.412273878 | 0.998865404 | 6.310360 | 6.617632 | 6.474609 | 6.383865 | 6.552511 | 6.422578 | 107 | 71 | 104 | 72 |
| 399965 | gene-KRT18P5 | KRT18P59 | keratin 18 pseudogene 59 | pseudogene | . | 48886 | . | . | . | 0.051004 | 0.137030 | 0.104997 | 0.062656 | 0.118689 | 0.316752 | 0.235611 | 0.143257 | -1.164597 | 4.223110 | 1.482550427 | -0.1482792 | 0.882122429 | NA | 2.184683 | 1.963455 | 2.067850 | 2.086146 | 2.077250 | 2.070955 | 3 | 7 | 4 | 3 |
| 399967 | NM_212555 | PATE2 | prostate and testis expressed 2 | protein_coding | NP_997720.1 | 32249 | . | . | . | 0.083386 | 0.109321 | 0.014946 | 0.068793 | 0.194044 | 0.252701 | 0.033539 | 0.157287 | -2.143057 | 4.278627 | 1.513563914 | -0.72654377 | 0.467505487 | NA | 2.548173 | 1.437399 | 2.114975 | 2.120795 | 2.096798 | 2.110542 | 6 | 7 | 1 | 4 |
| 399972 | NR_033839 | GSEC | G-quadruplex forming sequence containing lncRNA | lncRNA | . | 48645 | . | . | . | 0.995672 | 1.073987 | 1.095150 | 1.018035 | 2.316970 | 2.467585 | 2.457245 | 2.327615 | 1.014153 | 25.311996 | 0.610790001 | 0.017109436 | 0.986349223 | 0.998865404 | 4.651489 | 4.671937 | 4.656762 | 4.663699 | 4.665548 | 4.667646 | 28 | 28 | 23 | 23 |
| 399975 | NR_145484 | LOC399975 | uncharacterized LOC399975 | lncRNA | . | | . | . | . | 1.188119 | 1.145366 | 1.125840 | 0.915268 | 2.764802 | 2.647565 | 2.526353 | 2.092652 | -1.159845 | 123.529609 | 0.296396977 | -0.72177344 | 0.470433781 | 0.998865404 | 7.051052 | 6.838556 | 6.981633 | 6.965381 | 6.955932 | 6.887147 | 156 | 140 | 110 | 96 |
| 399979 | NM_00130108 | SNX19 | sorting nexin 19 | protein_coding | NP_001288018.1;NP_001334847.2;NP_001334848.2;NP_001334849.2;NI | 21532 | . | . | . | 2.764791 | 3.091178 | 3.331731 | 3.320860 | 6.433782 | 7.145395 | 7.476308 | 7.592752 | 1.129044 | 1680.612903 | 0.124757442 | 1.403541899 | 0.160455307 | 0.933525128 | 10.624541 | 10.799690 | 10.641595 | 10.687435 | 10.786230 | 10.732320 | 1764 | 1754 | 1603 | 1614 |
| 400011 | gene-SUPT16H | SUPT16HP1 | SPT16 homolog, facilitates chromatin remodeling subunit pseudog | pseudogene | . | 31388 | . | . | . | 0.118377 | 0.107045 | 0.158305 | 0.082290 | 0.275461 | 0.247440 | 0.355232 | 0.188146 | 1.038998 | 13.717273 | 0.839979033 | 0.065707334 | 0.947610833 | NA | 3.747897 | 3.807334 | 3.773910 | 3.769779 | 3.792662 | 3.755200 | 16 | 14 | 16 | 9 |
| 400027 | NR_028408 | LINC00938 | long intergenic non-protein coding RNA 938 | lncRNA | . | 48630 | . | . | . | 2.470384 | 2.297759 | 1.980590 | 2.387972 | 5.748685 | 5.311372 | 4.444387 | 5.459148 | -1.146222 | 157.063812 | 0.266494865 | -0.55840047 | 0.576407115 | 0.998865404 | 7.368178 | 7.218347 | 7.332789 | 7.298564 | 7.240429 | 7.307623 | 198 | 171 | 118 | 152 |
| 400043 | NR_026656 | LINC02381 | long intergenic non-protein coding RNA 2381 | lncRNA | . | 53304 | . | . | . | 4.325251 | 3.959613 | 5.446079 | 5.404739 | 10.065037 | 9.152822 | 12.220843 | 12.357294 | 1.287906 | 208.446665 | 0.233445851 | 1.563646939 | 0.117900505 | 0.857396579 | 7.509215 | 7.874761 | 7.658229 | 7.616388 | 7.758073 | 7.749637 | 220 | 187 | 206 | 218 |
| 400061 | gene-NSA2P2 | NSA2P2 | NSA2 pseudogene 2 | pseudogene | . | 54581 | . | . | . | 0.253405 | 0.147107 | 0.057519 | 0.132881 | 0.589685 | 0.340044 | 0.129071 | 0.303816 | -2.039890 | 3.316902 | 1.690653885 | -0.6083394 | 0.542962394 | NA | 2.156660 | 1.121007 | 1.753968 | 1.746851 | 1.735459 | 1.744512 | 6 | 4 | 1 | 3 |
| 400073 | NM_207435,N | C12orf76 | chromosome 12 open reading frame 76 | protein_coding | NP_997318.1;XP_016874786.1 | 33790 | . | . | . | 7.281689 | 7.551995 | 9.230745 | 7.017222 | 16.944792 | 17.456775 | 20.713524 | 16.044045 | 1.088116 | 426.808540 | 0.192886031 | 0.631629029 | 0.527629313 | 0.998865404 | 8.674772 | 8.797508 | 8.696734 | 8.718762 | 8.838285 | 8.673378 | 462 | 449 | 439 | 361 |
| 400154 | gene-DGKZP1 | DGKZP1 | diacylglycerol kinase zeta pseudogene 1 | pseudogene | . | 39263 | . | . | . | 0.065735 | 0.064899 | 0.041072 | 0.036664 | 0.152016 | 0.150099 | 0.092164 | 0.083268 | -1.718688 | 6.024125 | 1.241001524 | -0.62957839 | 0.528970467 | NA | 2.928838 | 2.439275 | 2.608017 | 2.606048 | 2.593439 | 2.591808 | 9 | 8 | 4 | 3 |
| 400156 | gene-RPS26P4 | RPS26P47 | ribosomal protein S26 pseudogene 47 | pseudogene | . | 36368 | . | . | . | 0.426949 | 0.562081 | 0.419012 | 0.347153 | 0.993527 | 1.299276 | 0.940250 | 0.793725 | -1.300288 | 6.375751 | 1.199853777 | -0.3157309 | 0.752206771 | NA | 2.849979 | 2.470304 | 2.674355 | 2.684794 | 2.673021 | 2.671097 | 7 | 9 | 5 | 5 |
| 400209 | gene-YTHDF2P | YTHDF2P1 | YTH domain family member 2 pseudogene 1 | pseudogene | . | 19865 | . | . | . | 0.852258 | 0.723930 | 0.782947 | 1.024107 | 1.983240 | 1.673444 | 1.756910 | 2.341498 | 1.125808 | 55.630417 | 0.426157218 | 0.405006895 | 0.685472443 | 0.998865404 | 5.709283 | 5.881202 | 5.798392 | 5.767668 | 5.781644 | 5.832083 | 65 | 52 | 45 | 62 |
| 400236 | NR_036500 | FOXN3-AS1 | FOXN3 antisense RNA 1 | lncRNA | . | 44290 | . | . | . | 0.354278 | 0.263754 | 0.068681 | 0.235452 | 0.824419 | 0.609678 | 0.154118 | 0.538334 | -1.966208 | 8.064730 | 1.128854182 | -0.8640764 | 0.38754592 | NA | 3.421266 | 2.437399 | 3.041756 | 3.028479 | 2.997949 | 3.025085 | 14 | 10 | 2 | 8 |
| 400242 | NR_015415 | DICER1-AS1 | DICER1 antisense RNA 1 | lncRNA | . | 43017 | . | . | . | 0.769187 | 0.487007 | 1.170956 | 0.800569 | 1.789929 | 1.125737 | 2.627591 | 1.830406 | 1.517967 | 20.778650 | 0.691689017 | 0.87053649 | 0.384007487 | NA | 4.042145 | 4.648638 | 4.360710 | 4.329944 | 4.400831 | 4.362067 | 23 | 15 | 26 | 19 |
| 400322 | NR_002824 | HERC2P2 | hect domain and RLD 2 pseudogene 2 | transcribed_ps | . | 4870 | . | . | . | 43.723763 | 48.256119 | 62.005768 | 50.098450 | 101.746994 | 111.546181 | 139.139145 | 114.544159 | 1.202773 | 9651.357827 | 0.126341759 | 2.108284525 | 0.035006386 | 0.583157098 | 13.097166 | 13.363584 | 13.112403 | 13.187974 | 13.400258 | 13.207220 | 9636 | 9881 | 10185 | 8782 |
| 400389 | gene-RPL12P3 | RPL12P35 | ribosomal protein L12 pseudogene 35 | pseudogene | . | 35969 | . | . | . | 0.008908 | 0.075625 | 0.102532 | 0.045789 | 0.020730 | 0.174810 | 0.230078 | 0.104691 | 1.236246 | 1.522137 | 2.383171416 | 0.128385813 | 0.897843658 | NA | 0.444631 | 0.751299 | 0.592746 | 0.597508 | 0.599417 | 0.593826 | 1 | 2 | 2 | 1 |
| 400410 | NR_037652,NF | ST20 | suppressor of tumorigenicity 20 | lncRNA | . | 33520 | . | . | . | 5.055412 | 4.988363 | 3.713114 | 5.489013 | 11.764152 | 11.530825 | 8.332121 | 12.549977 | -1.115058 | 97.692503 | 0.338955582 | -0.46353807 | 0.642978734 | 0.998865404 | 6.687585 | 6.528376 | 6.629286 | 6.622112 | 6.537824 | 6.645296 | 120 | 110 | 65 | 103 |
| 400451 | NM_207446,XI | FAM174B | family with sequence similarity 174 member B | protein_coding | NP_997329.2 | 34339 | . | . | . | 3.080381 | 3.329196 | 4.196103 | 4.236044 | 7.168172 | 7.695589 | 9.415931 | 9.685214 | 1.286486 | 295.497476 | 0.202472595 | 1.795400598 | 0.072589897 | 0.758556671 | 8.014059 | 8.377159 | 8.114825 | 8.148245 | 8.261626 | 8.270405 | 288 | 288 | 287 | 314 |
| 400464 | NR_135737 | LOC400464 | ubiquitin conjugating enzyme E2 Q2 pseudogene | transcribed_ps | . | | . | . | . | 1.703981 | 1.372271 | 1.272442 | 2.644106 | 3.965233 | 3.172065 | 2.855324 | 6.045435 | 1.265922 | 30.839252 | 0.588917286 | 0.577651647 | 0.56349933 | 0.998865404 | 4.767942 | 5.105724 | 4.937061 | 4.910763 | 4.907786 | 4.997009 | 35 | 26 | 20 | 43 |
| 400499 | NM_00137070 | LOC400499 | putative uncharacterized protein LOC400499 | protein_coding | NP_001357633.1;XP_016879418.1;XP_016879419.1;XP_016879420.1;XP | | . | . | . | 0.136928 | 0.154606 | 0.129457 | 0.168456 | 0.318637 | 0.357378 | 0.290498 | 0.385156 | -1.012178 | 49.974022 | 0.445170437 | -0.04127148 | 0.967079467 | 0.998865404 | 5.653161 | 5.632981 | 5.630961 | 5.656838 | 5.624217 | 5.658756 | 54 | 58 | 39 | 51 |
| 400506 | NM_00101299 | KNOP1 | lysine rich nucleolar protein 1 | protein_coding | NP_001013009.2;NP_001335456.1;NP_001335457.1;NP_001335458.1;NI | 34404 | . | . | . | 4.705213 | 4.650882 | 4.259810 | 4.798307 | 10.949221 | 10.750723 | 9.558889 | 10.970756 | -1.037962 | 900.303865 | 0.147886928 | -0.36347394 | 0.716250877 | 0.998865404 | 9.841171 | 9.786854 | 9.816522 | 9.837438 | 9.742676 | 9.854355 | 1041 | 1004 | 708 | 893 |
| 400508 | NR_026675 | CRYM-AS1 | CRYM antisense RNA 1 | lncRNA | . | 34405 | . | . | . | 0.254096 | 0.308546 | 0.353788 | 0.261527 | 0.591292 | 0.713219 | 0.793890 | 0.597951 | 1.103110 | 21.749254 | 0.663765299 | 0.21329406 | 0.831097606 | NA | 4.369732 | 4.515225 | 4.427041 | 4.440719 | 4.457312 | 4.427980 | 22 | 24 | 23 | 18 |
| 400555 | NR_146877 | ZC3H18-AS1 | ZC3H18 antisense RNA 1 (head to head) | lncRNA | . | 33799 | . | . | . | 0.783115 | 0.468955 | 1.127274 | 0.477147 | 1.822342 | 1.084010 | 2.529571 | 1.090941 | 1.258319 | 27.690682 | 0.644229106 | 0.514564267 | 0.60685754 | 0.998865404 | 4.612658 | 4.950290 | 4.786819 | 4.736475 | 4.836797 | 4.736072 | 35 | 20 | 38 | 17 |
| 400566 | NM_00101367 | C17orf97 | chromosome 17 open reading frame 97 | protein_coding | NP_001013694.4 | 33800 | . | . | . | 1.152655 | 1.312092 | 1.404669 | 1.585736 | 2.682275 | 3.032959 | 3.152037 | 3.625597 | 1.199165 | 78.507004 | 0.359608987 | 0.728653027 | 0.466213933 | 0.998865404 | 6.158490 | 6.419243 | 6.250645 | 6.282395 | 6.297184 | 6.329700 | 77 | 82 | 70 | 85 |
| 400569 | NM_00100168 | MED11 | mediator complex subunit 11 | protein_coding | NP_001001683.1;NP_001291929.1 | 32687 | 612383 | . | . | 12.618779 | 13.301821 | 14.650217 | 17.825750 | 29.364420 | 30.747754 | 32.874663 | 40.756458 | 1.247755 | 386.560410 | 0.190311364 | 1.677956998 | 0.093355494 | 0.809520218 | 8.426545 | 8.745016 | 8.511670 | 8.532540 | 8.594528 | 8.706410 | 389 | 378 | 339 | 439 |
| 400579 | NR_104343 | LINC02076 | long intergenic non-protein coding RNA 2076 | lncRNA | . | 52924 | . | . | . | 0.382888 | 0.394672 | 0.687351 | 0.425686 | 0.890995 | 0.912302 | 1.542396 | 0.973281 | 1.410128 | 28.116416 | 0.594279836 | 0.834331326 | 0.404094304 | 0.998865404 | 4.542770 | 5.041092 | 4.781602 | 4.780686 | 4.848325 | 4.791292 | 27 | 25 | 35 | 24 |
| 400604 | NR_038458 | TOB1-AS1 | TOB1 antisense RNA 1 | lncRNA | . | 44340 | . | . | . | 0.311717 | 0.240355 | 0.404925 | 0.202938 | 0.725378 | 0.555591 | 0.908640 | 0.463994 | 1.077844 | 13.471011 | 0.850632154 | 0.127139028 | 0.898830362 | NA | 3.694014 | 3.807334 | 3.752176 | 3.735760 | 3.766928 | 3.729478 | 17 | 12 | 16 | 9 |
| 400619 | NR_033876 | LINC00511 | long intergenic non-protein coding RNA 511 | lncRNA | . | 43564 | . | . | . | 5.057379 | 4.770142 | 4.735018 | 4.766860 | 11.768729 | 11.026396 | 10.625244 | 10.898861 | -1.050063 | 341.456603 | 0.191763592 | -0.36751713 | 0.713233313 | 0.998865404 | 8.450250 | 8.380012 | 8.448097 | 8.410206 | 8.402493 | 8.401239 | 416 | 365 | 290 | 312 |
| 400644 | NR_104164 | LINC01443 | long intergenic non-protein coding RNA 1443 | lncRNA | . | 50768 | . | . | . | 10.158397 | 10.869442 | 11.721042 | 13.180480 | 23.639008 | 25.125202 | 26.301678 | 30.135601 | 1.167600 | 669.093420 | 0.156396468 | 1.429358712 | 0.152901159 | 0.917207605 | 9.270262 | 9.493257 | 9.309431 | 9.349071 | 9.396582 | 9.471480 | 690 | 686 | 593 | 712 |
| 400652 | gene-RPS3AP4 | RPS3AP49 | RPS3A pseudogene 49 | pseudogene | . | 36810 | . | . | . | 0.020064 | 0.115697 | 0.012961 | 0.148949 | 0.046689 | 0.267439 | 0.029084 | 0.340553 | 1.194800 | 2.498016 | 1.945927483 | 0.131952036 | 0.89502223 | NA | 1.193905 | 1.437399 | 1.307800 | 1.321383 | 1.309640 | 1.323706 | 1 | 4 | 1 | 4 |
| 400657 | NR_024484 | LINC00909 | long intergenic non-protein coding RNA 909 | lncRNA | . | 44331 | . | . | . | 1.003305 | 0.676598 | 0.754535 | 0.863612 | 2.334732 | 1.563986 | 1.693154 | 1.974545 | -1.057815 | 89.719983 | 0.355362898 | -0.22818233 | 0.819504495 | 0.998865404 | 6.526943 | 6.446655 | 6.549609 | 6.433687 | 6.459801 | 6.494582 | 127 | 80 | 71 | 87 |
| 400658 | NR_164145 | ZNF516-DT | ZNF516 divergent transcript | lncRNA | . | 51248 | . | . | . | 0.268340 | 0.161378 | 0.071810 | 0.261572 | 0.624437 | 0.373031 | 0.161140 | 0.598053 | -1.244353 | 14.025822 | 0.878368097 | -0.35907005 | 0.719542687 | NA | 3.961318 | 3.640961 | 3.834846 | 3.803458 | 3.776997 | 3.834158 | 22 | 13 | 5 | 18 |
| 400673 | NM_00101792 | VMAC | vimentin type intermediate filament associated coiled-coil protein | protein_coding | NP_001017921.1 | 33803 | 617204 | . | . | 1.746392 | 1.796768 | 1.221160 | 1.318954 | 4.063926 | 4.153310 | 2.740248 | 3.015631 | -1.409549 | 106.546881 | 0.312278655 | -1.58587135 | 0.112768496 | 0.850211417 | 6.962063 | 6.466231 | 6.783489 | 6.790755 | 6.669566 | 6.689171 | 142 | 136 | 74 | 86 |
| 400684 | NR_046201 | LOC400684 | uncharacterized LOC400684 | lncRNA | . | | . | . | . | 1.210539 | 0.997723 | 1.174121 | 1.500662 | 2.816973 | 2.306280 | 2.634693 | 3.431085 | 1.216775 | 36.644905 | 0.517629861 | 0.546834419 | 0.58448631 | 0.998865404 | 5.047242 | 5.330007 | 5.190957 | 5.163340 | 5.187833 | 5.222676 | 42 | 32 | 31 | 42 |
| 400713 | NM_00114543 | ZNF880 | zinc finger protein 880 | protein_coding | NP_001138906.1;NP_016882301.1 | 37249 | . | . | . | 0.569200 | 0.776818 | 0.201165 | 0.797069 | 1.324550 | 1.795648 | 0.451408 | 1.822404 | -1.147374 | 47.136563 | 0.482092087 | -0.41140553 | 0.680775198 | 0.998865404 | 5.656500 | 5.453945 | 5.542090 | 5.595081 | 5.500819 | 5.586839 | 50 | 62 | 28 | 52 |
| 400720 | NM_00102455 | ZNF772 | zinc finger protein 772 | protein_coding | NP_001019767.1;NP_001137540.1;NP_001317542.1;XP_005259001.1;XI | 33106 | . | . | . | 0.601258 | 0.487732 | 0.537194 | 0.576091 | 1.399152 | 1.127415 | 1.205449 | 1.317163 | 1.060335 | 105.823749 | 0.329754536 | 0.256311807 | 0.797710081 | 0.998865404 | 6.683090 | 6.766737 | 6.746315 | 6.678796 | 6.677741 | 6.782768 | 131 | 99 | 79 | 119 |
| 400745 | NM_00101031 | SH2D5 | SH2 domain containing 5 | protein_coding | NP_001096630.1;NP_001096631.1;XP_011539761.1;XP_011539762.1;XP | 28819 | . | . | . | 1.857864 | 2.231826 | 1.757307 | 1.918696 | 4.323324 | 5.158964 | 3.943346 | 4.386870 | 1.003856 | 247.124079 | 0.217383088 | 0.025543717 | 0.979621279 | 0.998865404 | 7.946570 | 7.951610 | 7.940826 | 7.955204 | 7.930847 | 7.968585 | 280 | 270 | 207 | 241 |
| 400750 | gene-HSPA5P1 | HSPA5P1 | HSPA5 pseudogene 1 | pseudogene | . | 54532 | . | . | . | 0.235229 | 0.226168 | 0.088443 | 0.227259 | 0.547388 | 0.522797 | 0.198463 | 0.519601 | -1.451272 | 14.721302 | 0.830959048 | -0.64662372 | 0.517875505 | NA | 4.125927 | 3.582949 | 3.900885 | 3.893876 | 3.852097 | 3.895216 | 21 | 18 | 6 | 16 |
| 400752 | NR_024270 | LINC01144 | long intergenic non-protein coding RNA 1144 | lncRNA | . | 49460 | . | . | . | 0.662695 | 0.487018 | 0.393167 | 0.673844 | 1.542119 | 1.125763 | 0.882255 | 1.540663 | -1.088715 | 30.366589 | 0.581632325 | -0.21083211 | 0.833018279 | 0.998865404 | 4.985276 | 4.860868 | 4.949724 | 4.910244 | 4.890970 | 4.947803 | 42 | 29 | 19 | 34 |
| 400761 | NR_033981 | LRRC8C-DT | LRRC8C divergent transcript | lncRNA | . | 53731 | . | . | . | 1.066125 | 0.597938 | 0.432089 | 1.172077 | 2.480917 | 1.382161 | 0.969595 | 2.679815 | -1.034184 | 20.512344 | 0.735650753 | -0.06591767 | 0.947443376 | NA | 4.383691 | 4.332699 | 4.383235 | 4.330948 | 4.318329 | 4.389511 | 31 | 16 | 10 | 27 |
| 400793 | NM_00108537 | C1orf226 | chromosome 1 open reading frame 226 | protein_coding | NP_001078844.1;NP_001128712.1 | 34351 | . | . | . | 0.326365 | 0.388469 | 0.483988 | 0.597031 | 0.759465 | 0.897962 | 1.086054 | 1.365041 | 1.514408 | 58.161625 | 0.416918304 | 1.436143108 | 0.150961624 | 0.913627824 | 5.532907 | 6.129784 | 5.791099 | 5.820312 | 5.867736 | 5.914021 | 49 | 54 | 55 | 72 |
| 400798 | NR_033186 | C1orf220 | chromosome 1 putative open reading frame 220 | lncRNA | . | 33805 | . | . | . | 0.573822 | 0.481729 | 0.733791 | 0.385353 | 1.335308 | 1.113538 | 1.646605 | 0.881063 | 1.054171 | 44.137789 | 0.502024746 | 0.151603012 | 0.879500058 | 0.998865404 | 5.423228 | 5.503540 | 5.466598 | 5.435518 | 5.511796 | 5.405182 | 54 | 42 | 52 | 29 |
| 400818 | NM_00103767 | NBPF9 | NBPF member 9 | protein_coding | NP_001032764.2;NP_001264373.1 | 31991 | 613999 | . | . | 5.949828 | 6.684912 | 7.296871 | 6.964958 | 13.845493 | 15.452472 | 16.373968 | 15.924548 | 1.108883 | 1031.302500 | 0.140683545 | 1.059872916 | 0.289202419 | 0.998865404 | 9.933692 | 10.082953 | 9.944139 | 9.994928 | 10.085280 | 10.005151 | 1087 | 1092 | 994 | 962 |
| 400892 | NR_037566 | BCRP2 | BCR pseudogene 2 | transcribed_ps | . | 1015 | . | . | . | 0.040484 | 0.063610 | 0.059200 | 0.063790 | 0.094209 | 0.147037 | 0.132842 | 0.145848 | 1.102063 | 4.741867 | 1.38109312 | 0.101518922 | 0.919138538 | NA | 2.175403 | 2.312263 | 2.237617 | 2.246189 | 2.241736 | 2.245026 | 4 | 6 | 4 | 5 |
| 400916 | NM_00130133 | CHCHD10 | coiled-coil-helix-coiled-coil-helix domain containing 10 | protein_coding | NP_001288268.1;NP_998885.1 | 15559 | 615903 | . | . | 84.000255 | 83.362850 | 70.577137 | 95.669617 | 195.472042 | 192.696952 | 158.373051 | 218.737214 | -1.014493 | 1726.905948 | 0.147631938 | -0.1406099 | 0.888178125 | 0.998865404 | 10.764549 | 10.743320 | 10.755736 | 10.758981 | 10.630347 | 10.849841 | 1999 | 1881 | 1282 | 1832 |
| 400927 | NR_002821 | TPTEP2 | TPTE pseudogene 2 | transcribed_ps | . | 53828 | . | . | . | 2.837847 | 1.122739 | 1.248237 | 2.351310 | 6.603787 | 2.595263 | 2.801008 | 5.375991 | -1.108872 | 29.428735 | 0.64159367 | -0.23237833 | 0.816244179 | 0.998865404 | 4.951837 | 4.802613 | 4.932927 | 4.823017 | 4.833528 | 4.899717 | 51 | 19 | 17 | 34 |
| 400931 | NR_027033,NF | MIRLET7BHG | MIRLET7B host gene | lncRNA | . | 37189 | 618216 | . | . | 0.791764 | 0.882645 | 1.178157 | 0.953906 | 1.842467 | 2.040273 | 2.643751 | 2.180992 | 1.254791 | 140.272993 | 0.282994825 | 1.157077479 | 0.247240707 | 0.992733415 | 6.958722 | 7.286848 | 7.062964 | 7.099133 | 7.209310 | 7.121181 | 136 | 141 | 151 | 130 |
| 400932 | NR_033377 | LINC00898 | long intergenic non-protein coding RNA 898 | lncRNA | . | 48581 | . | . | . | 0.073956 | 0.105921 | 0.132736 | 0.081337 | 0.172098 | 0.244841 | 0.297857 | 0.185968 | 1.193600 | 13.347184 | 0.849448405 | 0.300570595 | 0.763741957 | NA | 3.604218 | 3.861274 | 3.716774 | 3.732650 | 3.751326 | 3.718505 | 12 | 15 | 16 | 10 |
| 400954 | NM_00103975 | EML6 | EMAP like 6 | protein_coding | NP_001034842.2;XP_016859587.1;XP_016859588.1;XP_016859589.1;XF | 35412 | . | . | . | 1.285598 | 1.498880 | 1.745651 | 1.160485 | 2.991639 | 3.464727 | 3.917190 | 2.653311 | 1.013496 | 306.236883 | 0.237168741 | 0.081547392 | 0.935006637 | 0.998865404 | 8.248337 | 8.268601 | 8.189509 | 8.302612 | 8.359599 | 8.138303 | 314 | 362 | 323 | 230 |
| 400960 | NR_033872 | PCBP1-AS1 | PCBP1 antisense RNA 1 | lncRNA | . | 42948 | . | . | . | 1.568638 | 1.953680 | 2.056221 | 2.088648 | 3.650019 | 4.614100 | [illegible] | 4.775446 | 1.159574 | 142.770110 | 0.278883646 | 0.765890311 | 0.443741542 | 0.998865404 | 7.084667 | 7.259963 | 7.126598 | 7.156216 | 7.179891 | 7.181903 | 136 | 158 | 133 | 144 |
| 400961 | NM_020459,XI | PAIP2B | poly(A) binding protein interacting protein 2B | protein_coding | NP_065192.1;XP_005264367.1;XP_005264368.1;XP_011531144.1;XP_01 | 29200 | 611018 | . | . | 0.463418 | 0.518926 | 0.443407 | 0.403767 | 1.078393 | 1.199521 | 0.994994 | 0.923167 | -1.164072 | 90.601228 | 0.337104347 | -0.65018355 | 0.515573664 | 0.998865404 | 6.606764 | 6.387857 | 6.505943 | 6.539679 | 6.494940 | 6.464274 | 106 | 111 | 77 | 74 |
| 400963 | gene-RPS2P17 | RPS2P17 | ribosomal protein S2 pseudogene 17 | pseudogene | . | 35764 | . | . | . | 0.171308 | 0.096133 | 0.008882 | 0.194660 | 0.398640 | 0.222216 | 0.019931 | 0.445068 | -1.174755 | 4.161902 | 1.537516593 | -0.15112651 | 0.87987592 | NA | 2.156660 | 1.916472 | 2.052971 | 2.045884 | 2.034663 | 2.058017 | 6 | 4 | 1 | 6 |
| 400966 | NM_00102445 | RGPD1 | RANBP2 like and GRIP domain containing 1 | protein_coding | NP_001019628.3;NP_001369273.1;XP_011531147.1;XP_011531151.1;XP | 32414 | 612704 | . | . | 0.273614 | 0.252661 | 0.288460 | 0.211921 | 0.636712 | 0.584037 | 0.647297 | 0.484531 | -1.076461 | 61.791750 | 0.40863747 | -0.26012287 | 0.794768995 | 0.998865404 | 6.000397 | 5.896415 | 5.966112 | 5.945662 | 5.972224 | 5.906581 | 77 | 66 | 60 | 47 |

| | | | | | | | | | |
|---|---|---|---|---|---|---|---|---|---|
| 400986 | NM_00131015 | ANKRD36C | ankyrin repeat domain 36C | protein_coding | NP_001297083.1 | 32946 | . | . | [illegible] |
| 400999 | NR_033871 | FLJ42351 | uncharacterized LOC400999 | lncRNA | . | . | . | . | [illegible] |
| 401014 | NR_033870 | TEX41 | testis expressed 41 | lncRNA | . | 48667 | . | . | [illegible] |
| 401019 | gene-RPS15P4 | RPS15P4 | ribosomal protein S15 pseudogene 4 | pseudogene | . | 35712 | . | . | [illegible] |
| 401021 | NR_148241 | LOC401021 | uncharacterized LOC401021 | lncRNA | . | . | . | . | [illegible] |
| 401022 | NR_033979,NF | HAGLR | HOXD antisense growth-associated long non-coding RNA | lncRNA | . | 43755 | 618209 | . | [illegible] |
| 401027 | NM_213608 | C2orf66 | chromosome 2 open reading frame 66 | protein_coding | NP_998773.2 | 33809 | . | . | [illegible] |
| 401074 | NR_040004,NF | LINC00960 | long intergenic non-protein coding RNA 960 | lncRNA | . | 48710 | . | . | [illegible] |
| 401093 | NR_027037,NF | MBNL1-AS1 | MBNL1 antisense RNA 1 | lncRNA | . | 44584 | . | . | [illegible] |
| 401097 | NM_00116821 | C3orf80 | chromosome 3 open reading frame 80 | protein_coding | NP_001161686.1 | 40048 | . | . | [illegible] |
| 401115 | NM_00114193 | C4orf48 | chromosome 4 open reading frame 48 | protein_coding | NP_001135408.3;NP_001161715.1;XP_011511771.1;XP_011511773.1;XP | 34437 | 614690 | . | [illegible] |
| 401149 | NR_037596 | LINC01061 | long intergenic non-protein coding RNA 1061 | lncRNA | . | 49084 | . | . | [illegible] |
| 401152 | NM_00100170 | C4orf3 | chromosome 4 open reading frame 3 | protein_coding | NP_001001701.2;NP_001163801.1 | 19225 | . | . | [illegible] |
| 401164 | NR_033869 | LINC01060 | long intergenic non-protein coding RNA 1060 | lncRNA | . | 49081 | . | . | [illegible] |
| 401206 | gene-RPS25P6 | RPS25P6 | ribosomal protein S25 pseudogene 6 | pseudogene | . | 35532 | . | . | [illegible] |
| 401232 | NR_026856 | LINC01011 | long intergenic non-protein coding RNA 1011 | lncRNA | . | 33812 | . | . | [illegible] |
| 401233 | NR_033884 | HTATSF1P2 | HIV-1 Tat specific factor 1 pseudogene 2 | transcribed_ps | . | 38586 | . | . | [illegible] |
| 401251 | NM_00103965 | SAPCD1 | suppressor APC domain containing 1 | protein_coding | NP_001034740.1 | 13938 | . | . | [illegible] |
| 401261 | NR_134654 | LOC401261 | uncharacterized LOC401261 | lncRNA | . | . | . | . | [illegible] |
| 401264 | NR_103446,NF | TRAM2-AS1 | TRAM2 antisense RNA 1 (head to head) | lncRNA | . | 48663 | . | . | [illegible] |
| 401265 | NM_00100376 | KLHL31 | kelch like family member 31 | protein_coding | NP_001003760.2;XP_005249164.1;XP_016866347.1 | 21353 | 610749 | . | [illegible] |
| 401303 | NR_023382 | ZNF815P | zinc finger protein 815, pseudogene | transcribed_ps | . | 22029 | . | . | [illegible] |
| 401316 | gene-EIF4HP1 | EIF4HP1 | eukaryotic translation initiation factor 4H pseudogene 1 | pseudogene | . | 49035 | . | . | [illegible] |
| 401320 | NR_038889 | GARS1-DT | GARS1 divergent transcript | lncRNA | . | 48951 | . | . | [illegible] |
| 401321 | NR_036501 | LINC00997 | long intergenic non-protein coding RNA 997 | lncRNA | . | 48952 | . | . | [illegible] |
| 401331 | NR_024116 | RASA4CP | RAS p21 protein activator 4C, pseudogene | transcribed_ps | . | 44185 | . | . | [illegible] |
| 401337 | NR_038371 | LINC01446 | long intergenic non-protein coding RNA 1446 | lncRNA | . | 50773 | . | . | [illegible] |
| 401375 | gene-GTF2IRD2 | GTF2IRD2P1 | GTF2I repeat domain containing 2 pseudogene 1 | pseudogene | . | 33127 | . | . | [illegible] |
| 401397 | NM_00135268 | SMIM30 | small integral membrane protein 30 | protein_coding | NP_001339616.1;NP_001339617.1 | 48953 | . | . | [illegible] |
| 401399 | NM_00111472 | PRRT4 | proline rich transmembrane protein 4 | protein_coding | NP_001108198.2;NP_001167635.1;XP_005250399.1;XP_005250400.1;XP | 37280 | . | . | [illegible] |
| 401427 | NM_00100532 | OR2A7 | olfactory receptor family 2 subfamily A member 7 | protein_coding | NP_001005328.1 | 8234 | . | . | [illegible] |
| 401428 | NR_002158 | OR2A20P | olfactory receptor family 2 subfamily A member 20 pseudogene | transcribed_ps | . | 15413 | . | . | [illegible] |
| 401431 | NR_027040 | ATP6V0E2-AS1 | ATP6V0E2 antisense RNA 1 | lncRNA | . | 44180 | . | . | [illegible] |
| 401466 | NM_00109967 | RBIS | ribosomal biogenesis factor | protein_coding | NP_001093140.1;NP_001093141.1;NP_001093142.1;NP_001093143.1;NI | 32235 | . | . | [illegible] |
| 401472 | NR_149020 | GASAL1 | growth arrest associated lncRNA 1 | lncRNA | . | 53461 | . | . | [illegible] |
| 401474 | NM_00110167 | SAMD12 | sterile alpha motif domain containing 12 | protein_coding | NP_001095146.1;NP_001336740.1;NP_001350203.1;NP_997389.2 | 31750 | 618073 | . | [illegible] |
| 401475 | gene-SRRM1P1 | SRRM1P1 | serine/arginine repetitive matrix 1 pseudogene 1 | pseudogene | . | 31861 | . | . | [illegible] |
| 401494 | NM_00101091 | HACD4 | 3-hydroxyacyl-CoA dehydratase 4 | protein_coding | NP_001010915.2;NP_001308812.1;NP_001308832.1;XP_011516178.1;XP | 20920 | 615941 | . | [illegible] |
| 401505 | NM_00100179 | TOMM5 | translocase of outer mitochondrial membrane 5 | protein_coding | NP_001001790.1;NP_001127956.1;NP_001127957.1 | 31369 | 616169 | . | [illegible] |
| 401509 | NR_003528 | ZNF658B | zinc finger protein 658B (pseudogene) | transcribed_ps | . | 32033 | . | . | [illegible] |
| 401525 | gene-CDK2AP2 | CDK2AP2P3 | cyclin dependent kinase 2 associated protein 2 pseudogene 3 | pseudogene | . | 38494 | . | . | [illegible] |
| 401541 | NM_00101226 | CENPP | centromere protein P | protein_coding | NP_001012267.1;NP_001273898.1;NP_001273900.1;XP_011516987.1;XP | 32933 | 611505 | . | [illegible] |
| 401548 | NM_00101299 | SNX30 | sorting nexin family member 30 | protein_coding | NP_001013012.1;XP_011516993.1;XP_016870205.1;XP_024303312.1 | 23685 | . | . | [illegible] |
| 401577 | NR_033380,NF | CD99P1 | CD99 molecule pseudogene 1 | transcribed_ps | . | 7083 | . | . | [illegible] |
| 401588 | NR_015378 | ZNF674-AS1 | ZNF674 antisense RNA 1 (head to head) | lncRNA | . | 44266 | . | . | [illegible] |
| 401612 | NM_00101275 | SLC25A53 | solute carrier family 25 member 53 | protein_coding | NP_001012773.2;XP_005262186.1;XP_011529254.1;XP_011529255.1;XP | 31894 | 300941 | . | [illegible] |
| 401647 | NM_00101091 | GOLGA7B | golgin A7 family member B | protein_coding | NP_001010917.1 | 31668 | 614189 | . | [illegible] |
| 401720 | NM_00101369 | FIGNL2 | fidgetin like 2 | protein_coding | NP_001013712.4;XP_005268927.1;XP_016874787.1;XP_016874788.1 | 13287 | . | . | [illegible] |
| 401805 | gene-LOC4018 | LOC401805 | ring finger protein 145 pseudogene | pseudogene | . | . | . | . | [illegible] |
| 401904 | gene-RPL23AP | RPL23AP2 | ribosomal protein L23a pseudogene 2 | pseudogene | . | 10319 | . | . | [illegible] |
| 401911 | gene-RPL29P3 | RPL29P33 | ribosomal protein L29 pseudogene 33 | pseudogene | . | 36923 | . | . | [illegible] |
| 401960 | gene-GAPDHP2 | GAPDHP27 | glyceraldehyde 3 phosphate dehydrogenase pseudogene 27 | pseudogene | . | 37779 | . | . | [illegible] |
| 401982 | gene-AKR1B1P | AKR1B1P1 | aldehyde reductase family 1 member B1 pseudogene 1 | pseudogene | . | 420 | . | . | [illegible] |
| 402055 | NM_00101369 | SRRD | SRR1 domain containing | protein_coding | NP_001013716.2;XP_011528480.1;XP_016884288.1 | 33910 | 602254 | . | [illegible] |
| 402057 | gene-RPS17P1 | RPS17P16 | ribosomal protein S17 pseudogene 16 | pseudogene | . | 36821 | . | . | [illegible] |
| 402096 | NR_157576 | ANAPC1P6 | ANAPC1 pseudogene 6 | transcribed_ps | . | 54712 | . | . | [illegible] |
| 402112 | gene-HNRNPA | HNRNPA1P39 | heterogeneous nuclear ribonucleoprotein A1 pseudogene 39 | pseudogene | . | 48768 | . | . | [illegible] |
| 402146 | gene-B3GAT3P | B3GAT3P1 | beta-1,3-glucuronyltransferase 3 pseudogene 1 | pseudogene | . | 35243 | . | . | [illegible] |
| 402175 | gene-NDUFB4F | NDUFB4P12 | NADH:ubiquinone oxidoreductase subunit B4 pseudogene 12 | pseudogene | . | 45260 | . | . | [illegible] |
| 402287 | gene-RPS3AP2 | RPS3AP21 | RPS3A pseudogene 21 | pseudogene | . | 36841 | . | . | [illegible] |
| 402483 | NR_024368 | LINC01000 | long intergenic non-protein coding RNA 1000 | lncRNA | . | 38541 | . | . | [illegible] |
| 402562 | gene-HNRNPA | HNRNPA1P8 | heterogeneous nuclear ribonucleoprotein A1 pseudogene 8 | pseudogene | . | 39126 | . | . | [illegible] |
| 402573 | NM_00100432 | C7orf61 | chromosome 7 open reading frame 61 | protein_coding | NP_001004323.1 | 22135 | . | . | [illegible] |
| 402665 | NM_00110137 | IGLON5 | IgLON family member 5 | protein_coding | NP_001094842.1 | 34550 | 618861 | . | [illegible] |
| 402682 | NM_00101507 | UFSP1 | UFM1 specific peptidase 1 (inactive) | protein_coding | NP_001015072.2 | 33821 | 611481 | . | [illegible] |
| 402778 | NM_00117082 | IFITM10 | interferon induced transmembrane protein 10 | protein_coding | NP_001164291.2 | 40022 | 618293 | . | [illegible] |
| 403313 | NM_203453 | PLPP6 | phospholipid phosphatase 6 | protein_coding | NP_982278.3 | 23682 | 611666 | . | [illegible] |
| 403323 | NR_122076,NF | LERFS | lncRNA negative regulator of fibroblast-like synoviocyte migration, | lncRNA | . | 51182 | . | . | [illegible] |
| 403341 | NM_00109927 | ZBTB34 | zinc finger and BTB domain containing 34 | protein_coding | NP_001092740.1;XP_005252046.1;XP_011517001.1 | 31446 | 611692 | . | [illegible] |
| 404093 | NM_00127187 | CUEDC1 | CUE domain containing 1 | protein_coding | NP_001258804.1;NP_001278954.1;XP_011523114.1;XP_016880129.1;XP | 31350 | . | . | [illegible] |
| 404201 | NR_015359,NF | WDFY3-AS2 | WDFY3 antisense RNA 2 | lncRNA | . | 21603 | . | . | [illegible] |
| 404217 | NM_206833 | CTXN1 | cortexin 1 | protein_coding | NP_996664.1 | 31108 | 600135 | . | [illegible] |
| 404281 | NM_206923 | YY2 | YY2 transcription factor | protein_coding | NP_996806.2 | 31684 | 300570 | . | [illegible] |
| 404550 | NM_206967,N | C16orf74 | chromosome 16 open reading frame 74 | protein_coding | NP_996850.1 | 23362 | . | . | [illegible] |
| 404636 | NM_00130311 | DENND10 | DENN domain containing 10 | protein_coding | NP_001290040.1;NP_001290041.1;NP_001290042.1;NP_996892.1;XP_00 | 31793 | . | . | [illegible] |
| 404663 | NR_033383 | LINC01194 | long intergenic non-protein coding RNA 1194 | lncRNA | . | 37171 | 617097 | . | [illegible] |
| 404672 | NM_207118,XI | GTF2H5 | general transcription factor IIH subunit 5 | protein_coding | NP_997001.1;XP_016866351.1 | 21157 | 608780 | . | [illegible] |
| 404734 | NM_020690 | ANKHD1-EIF4E | ANKHD1-EIF4EBP3 readthrough | protein_coding | NP_065741.3 | 33530 | . | . | [illegible] |
| 404785 | NM_00100535 | POTEG | POTE ankyrin domain family member G | protein_coding | NP_001005356.1 | 33896 | 608916 | . | [illegible] |
| 406884 | NR_029479,rn | MIRLET7B | microRNA let-7b | miRNA | . | 31479 | 611249 | . | [illegible] |
| 406886 | NR_029481,rn | MIRLET7D | microRNA let-7d | miRNA | . | 31481 | 612145 | . | [illegible] |
| 406891 | NR_029661,rn | MIRLET7I | microRNA let-7i | miRNA | . | 31486 | 612148 | . | [illegible] |
| 406929 | NR_029700,rn | MIR138-1 | microRNA 138-1 | miRNA | . | 31524 | 613394 | . | [illegible] |
| 406938 | NR_029701,rn | MIR146A | microRNA 146a | miRNA | . | 31533 | 610566 | . | [illegible] |
| 406962 | NR_029707,rn | MIR186 | microRNA 186 | miRNA | . | 31557 | . | . | [illegible] |
| 406975 | NR_029584,rn | MIR198 | microRNA 198 | miRNA | . | 31570 | . | . | [illegible] |
| 406993 | NR_029624,rn | MIR211 | microRNA 211 | miRNA | . | 31588 | 613753 | . | [illegible] |
| 407002 | NR_029633,rn | MIR219A1 | microRNA 219a-1 | miRNA | . | 31597 | 611500 | . | [illegible] |
| 407004 | NR_029494,rn | MIR22 | microRNA 22 | miRNA | . | 31599 | 612077 | . | [illegible] |
| 407006 | NR_029635,rn | MIR221 | microRNA 221 | miRNA | . | 31601 | 300568 | . | [illegible] |
| 407012 | NR_029496,rn | MIR24-1 | microRNA 24-1 | miRNA | . | 31607 | 609705 | . | [illegible] |
| 407014 | NR_029498,rn | MIR25 | microRNA 25 | miRNA | . | 31609 | 612150 | . | [illegible] |
| 407019 | NR_029665,rn | MIR27B | microRNA 27b | miRNA | . | 31614 | 610636 | . | [illegible] |
| 407021 | NR_029503,rn | MIR29A | microRNA 29a | miRNA | . | 31616 | 610782 | . | [illegible] |
| 407040 | NR_029610,rn | MIR34A | microRNA 34a | miRNA | . | 31635 | 611172 | . | [illegible] |
| 407043 | NR_029605,rn | MIR7-1 | microRNA 7-1 | miRNA | . | 31638 | 615239 | . | [illegible] |
| 407835 | NR_002144 | LOC407835 | mitogen-activated protein kinase kinase 2 pseudogene | transcribed_ps | . | . | . | . | [illegible] |
| 407975 | NR_027349,NF | MIR17HG | miR-17-92a-1 cluster host gene | lncRNA | . | 23564 | 609415 | . | [illegible] |
| 408050 | NM_00100406 | NOMO3 | NODAL modulator 3 | protein_coding | NP_001004067.1;XP_005255375.1 | 25242 | 609159 | . | [illegible] |
| 408186 | NR_153413 | OVOS | alpha-2-macroglobulin like 1 pseudogene | transcribed_ps | . | . | . | . | [illegible] |
| 414059 | NM_00100141 | TBC1D3B | TBC1 domain family member 3B | protein_coding | NP_001001417.6;XP_005258037.2 | 27011 | 610144 | . | [illegible] |
| 414060 | NM_00100141 | TBC1D3C | TBC1 domain family member 3C | protein_coding | NP_001001418.5;XP_006721967.1;XP_006721968.1;XP_006721970.1;XP | 24889 | 610806 | . | [illegible] |
| 414149 | NM_00103984 | ACBD7 | acyl-CoA binding domain containing 7 | protein_coding | NP_001034933.1 | 17715 | . | . | [illegible] |
| 414169 | gene-BANF1P2 | BANF1P2 | BANF1 pseudogene 2 | pseudogene | . | 23426 | . | . | [illegible] |
| 414189 | NM_00107766 | AGAP6 | ArfGAP with GTPase domain, ankyrin repeat and PH domain 6 | protein_coding | NP_001071133.2;NP_001352796.1 | 23466 | . | . | [illegible] |
| 414194 | NR_103829 | CCNYL2 | cyclin Y like 2 (pseudogene) | transcribed_ps | . | 23495 | . | . | [illegible] |
| 414196 | NR_126062 | CELF2-AS1 | CELF2 antisense RNA 1 | lncRNA | . | 23515 | . | . | [illegible] |
| 414200 | gene-NHP2P1 | NHP2P1 | NHP2 ribonucleoprotein pseudogene 1 | pseudogene | . | 23582 | . | . | [illegible] |
| 414212 | NR_111968 | GLUD1P2 | glutamate dehydrogenase 1 pseudogene 2 | transcribed_ps | . | 4337 | . | . | [illegible] |
| 414224 | NR_029396 | AGAP12P | ArfGAP with GTPase domain, ankyrin repeat and PH domain 12, ps | transcribed_ps | . | 23661 | . | . | [illegible] |
| 414235 | NM_00135538 | PRR26 | proline rich 26 | protein_coding | NP_001342317.1;NP_001342319.1;XP_011518067.1;XP_024303792.1;XP | 30724 | . | . | [illegible] |
| 414241 | NR_027632 | SHLD2P1 | shieldin complex subunit 2 pseudogene 1 | transcribed_ps | . | 31425 | . | . | [illegible] |
| 414245 | NR_038373,NF | DNAJC9-AS1 | DNAJC9 antisense RNA 1 | lncRNA | . | 31432 | . | . | [illegible] |
| 414325 | NM_00108155 | DEFB103A | defensin beta 103A | protein_coding | NP_001075020.1 | 15967 | . | . | [illegible] |
| 414328 | NM_00100155 | IDNK | IDNK gluconokinase | protein_coding | NP_001001551.2;NP_001243844.1;NP_001338464.1;XP_006717173.1;XP | 31367 | 611343 | . | [illegible] |
| 414761 | NR_135289,NF | HCG15 | HLA complex group 15 | lncRNA | . | 18361 | . | . | [illegible] |
| 414765 | NR_044997 | HCG25 | HLA complex group 25 | lncRNA | . | 20196 | . | . | [illegible] |
| 414777 | NR_024052,NF | HCG18 | HLA complex group 18 | lncRNA | . | 31337 | . | . | [illegible] |
| 414778 | NR_052012 | HCG17 | HLA complex group 17 | lncRNA | . | 31339 | . | . | [illegible] |
| 414918 | NM_00100179 | DENND6B | DENN domain containing 6B | protein_coding | NP_001001794.3;XP_005261971.1;XP_011528994.1;XP_011528995.1;XP | 32690 | . | . | [illegible] |
| 414919 | NM_00100179 | C8orf82 | chromosome 8 open reading frame 82 | protein_coding | NP_001001795.1 | 33826 | . | . | [illegible] |
| 415116 | NM_00100185 | PIM3 | Pim-3 proto-oncogene, serine/threonine kinase | protein_coding | NP_001001852.2 | 19310 | 610580 | . | [illegible] |
| 425054 | NM_00100188 | VCX3B | variable charge X-linked 3B | protein_coding | NP_001001888.3 | 31838 | 300981 | . | [illegible] |
| 431707 | NM_00100193 | LHX8 | LIM homeobox 8 | protein_coding | NP_001001933.1;NP_001243043.1;XP_016856805.1;XP_016856806.1 | 28838 | 604425 | . | [illegible] |
| 439921 | NM_00100852 | MXRA7 | matrix remodeling associated 7 | protein_coding | NP_001008528.1;NP_001008529.1;NP_001350698.1;NP_940932.2;XP_00 | 7541 | . | . | [illegible] |
| 439931 | NR_027051,NF | THAP7-AS1 | THAP7 antisense RNA 1 | lncRNA | . | 41013 | . | . | [illegible] |
| 439949 | NR_036502,NF | PRKCQ-AS1 | PRKCQ antisense RNA 1 | lncRNA | . | 44689 | . | . | [illegible] |
| 439953 | gene-PPIAP31 | PPIAP31 | peptidylprolyl isomerase A pseudogene 31 | pseudogene | . | 44962 | . | . | [illegible] |
| 439965 | NR_027634,NF | SHLD2P3 | shieldin complex subunit 2 pseudogene 3 | transcribed_ps | . | 34038 | . | . | [illegible] |
| 439990 | NR_038464 | LINC00857 | long intergenic non-protein coding RNA 857 | lncRNA | . | 45114 | . | . | [illegible] |
| 439992 | gene-RPS3AP5 | RPS3AP5 | RPS3A pseudogene 5 | pseudogene | . | 23744 | . | . | [illegible] |
| 439994 | NR_029408,NF | LINC00863 | long intergenic non-protein coding RNA 863 | lncRNA | . | 45162 | . | . | [illegible] |
| 440026 | NM_00116503 | TMEM41B | transmembrane protein 41B | protein_coding | NP_001158502.1;NP_055827.1 | 28948 | . | . | [illegible] |
| 440028 | NR_033972 | LINC02709 | long intergenic non-protein coding RNA 2709 | lncRNA | . | 54226 | . | . | [illegible] |
| 440043 | gene-PGAM1P | PGAM1P8 | phosphoglycerate mutase 1 pseudogene 8 | pseudogene | . | 42455 | . | . | [illegible] |
| 440055 | gene-RPS12P2 | RPS12P22 | ribosomal protein S12 pseudogene 22 | pseudogene | . | 35671 | . | . | [illegible] |
| 440078 | NR_026788 | FAM66C | family with sequence similarity 66 member C | lncRNA | . | 21644 | . | . | [illegible] |
| 440081 | NR_033399 | DDX12P | DEAD/H-box helicase 12, pseudogene | transcribed_ps | . | 2737 | 601151 | . | [illegible] |
| 440093 | NM_00101369 | H3-5 | H3.5 histone | protein_coding | NP_001013721.2 | 33164 | 616134 | . | [illegible] |
| 440101 | NR_026655 | FLJ12825 | uncharacterized LOC440101 | lncRNA | . | . | . | . | [illegible] |
| 440104 | NR_036476,NF | TMEM198B | transmembrane protein 198B (pseudogene) | transcribed_ps | . | 43629 | . | . | [illegible] |
| 440138 | NM_00100412 | ALG11 | ALG11 alpha-1,2-mannosyltransferase | protein_coding | NP_001004127.2 | 32456 | 613666 | . | [illegible] |
| 440145 | NM_00107177 | MZT1 | mitotic spindle organizing protein 1 | protein_coding | NP_001065243.1 | 33830 | 613448 | . | [illegible] |
| 440176 | gene-RPL12P6 | RPL12P6 | ribosomal protein L12 pseudogene 6 | pseudogene | . | 23538 | . | . | [illegible] |
| 440193 | NM_00108041 | CCDC88C | coiled-coil domain containing 88C | protein_coding | NP_001073883.2;XP_005267748.1;XP_011535098.1;XP_016876824.1;XP | 19967 | 611204 | . | [illegible] |
| 440248 | NR_036443 | HERC2P9 | hect domain and RLD 2 pseudogene 9 | transcribed_ps | . | 30495 | . | . | [illegible] |
| 440253 | NR_026589 | WHAMMP2 | WAS protein homolog associated with actin, golgi membranes anc | transcribed_ps | . | 32360 | . | . | [illegible] |
| 440270 | NM_00102356 | GOLGA8B | golgin A8 family member B | protein_coding | NP_001018861.3 | 31973 | 609619 | . | [illegible] |
| 440275 | NM_00101370 | EIF2AK4 | eukaryotic translation initiation factor 2 alpha kinase 4 | protein_coding | NP_001013725.2 | 19687 | 609280 | . | [illegible] |
| 440278 | NR_002318 | CATSPER2P1 | cation channel sperm associated 2 pseudogene 1 | transcribed_ps | . | 31054 | . | . | [illegible] |
| 440288 | NR_038448,NF | UBL7-AS1 | UBL7 antisense RNA 1 (head to head) | lncRNA | . | 48614 | . | . | [illegible] |
| 440292 | NR_157580 | COMMD4P1 | COMM domain containing 4 pseudogene 1 | transcribed_ps | . | 51909 | . | . | [illegible] |
| 440295 | NM_00129142 | GOLGA6L9 | golgin A6 family like 9 | protein_coding | NP_001278349.1;NP_937824.3;XP_011519903.1;XP_016877715.1 | 37229 | . | . | [illegible] |
| 440300 | NR_033738 | LOC440300 | chondroitin sulfate proteoglycan 4 pseudogene | transcribed_ps | . | . | . | . | [illegible] |
| 440307 | NR_104604 | TTLL13P | tubulin tyrosine ligase like 13, pseudogene | transcribed_ps | . | 32484 | . | . | [illegible] |
| 440313 | XR_00175171 | LOC440313 | protein enabled homolog | lncRNA | . | . | . | . | [illegible] |
| 440345 | NM_00131014 | NPIPB4 | nuclear pore complex interacting protein family member B4 | protein_coding | NP_001297077.1;XP_016878733.1;XP_016878736.1;XP_024306051.1;XP | 41985 | . | . | [illegible] |
| 440348 | NM_00130609 | NPIPB15 | nuclear pore complex interacting protein family member B15 | protein_coding | NP_001293023.1;XP_011521798.1;XP_011521800.1;XP_011521803.1;XP | 34409 | . | . | [illegible] |
| 440350 | gene-NPIPB7 | NPIPB7 | nuclear pore complex interacting protein family member B7 | pseudogene | . | 33832 | . | . | [illegible] |
| 440352 | NR_002939 | SNX29P2 | sorting nexin 29 pseudogene 2 | transcribed_ps | . | 31914 | . | . | [illegible] |
| 440353 | NM_00135540 | NPIPB12 | nuclear pore complex interacting protein family member B12 | protein_coding | NP_001342330.1 | 37491 | . | . | [illegible] |
| 440354 | NR_135305,NF | SMG1P2 | SMG1 pseudogene 2 | transcribed_ps | . | 49859 | . | . | [illegible] |

| | | | | | | | | | | | | | | | | | | | | | | | | | | | | | | | | | |
|---|---|---|---|---|---|---|---|---|---|---|---|---|---|---|---|---|---|---|---|---|---|---|---|---|---|---|---|---|---|---|---|---|---|
| 440356 | NR_015396,NF | CDIPTOSP | CDIP transferase opposite strand, pseudogene | lncRNA | . | 48609 | . | . | . | 4.403200 | 2.911102 | 3.629755 | 4.682632 | 10.246426 | 6.729142 | 8.145066 | 10.706283 | 1.103708 | 78.734090 | 0.378617779 | 0.375995353 | 0.706920351 | 0.998865404 | 6.225814 | 6.368493 | 6.330214 | 6.230602 | 6.274146 | 6.339394 | 103 | 65 | 63 | 87 |
| 440366 | gene-HERC2P8 | HERC2P8 | hect domain and RLD 2 pseudogene 8 | pseudogene | . | 38747 | . | . | . | 2.233801 | 2.529399 | 2.673123 | 2.037691 | 5.198148 | 5.846819 | 5.998409 | 4.658939 | -1.024663 | 291.643463 | 0.2204635 | -0.1594352 | 0.87332601 | 0.998865404 | 8.205095 | 8.170825 | 8.159350 | 8.222407 | 8.249886 | 8.101727 | 320 | 337 | 286 | 232 |
| 440375 | gene-RAB43P1 | RAB43P1 | RAB43 pseudogene 1 | pseudogene | . | 33153 | . | . | . | 4.154058 | 4.626941 | 4.979489 | 5.895262 | 9.666664 | 10.695381 | 11.173830 | 13.478816 | 1.221590 | 630.292200 | 0.164604006 | 1.754268183 | 0.079384565 | 0.776573458 | 9.148730 | 9.436680 | 9.192876 | 9.258238 | 9.304094 | 9.413519 | 621 | 643 | 555 | 701 |
| 440396 | gene-HNRNPA1 | HNRNPA1P16 | heterogeneous nuclear ribonucleoprotein A1 pseudogene 16 | pseudogene | . | 39134 | . | . | . | 1.978590 | 1.761684 | 2.677672 | 1.918878 | 4.604259 | 4.072212 | 6.008618 | 4.387287 | 1.213480 | 84.255518 | 0.359367799 | 0.776780817 | 0.437288096 | 0.998865404 | 6.249108 | 6.530583 | 6.378027 | 6.346393 | 6.462096 | 6.365496 | 93 | 77 | 94 | 72 |
| 440400 | NM_00100433 | RNASEK | ribonuclease K | protein_coding | NP_001004333.3 | 33911 | 617098 | . | . | 139.314517 | 128.416423 | 162.113320 | 173.792108 | 324.190608 | 296.840319 | 363.777576 | 397.355012 | 1.243854 | 2895.342213 | 0.11705161 | 2.689560052 | 0.007154627 | 0.26784811 | 11.333520 | 11.648372 | 11.439846 | 11.364967 | 11.557123 | 11.604615 | 3111 | 2654 | 2712 | 3095 |
| 440406 | gene-UBE2SP2 | UBE2SP2 | ubiquitin conjugating enzyme E2 S pseudogene 2 | pseudogene | . | 32196 | . | . | . | 1.694304 | 1.917292 | 2.055711 | 2.691050 | 3.942715 | 4.431907 | 4.612957 | 6.152765 | 1.286577 | 43.101306 | 0.478527782 | 0.759701536 | 0.44743301 | 0.998865404 | 5.237567 | 5.599155 | 5.393857 | 5.411067 | 5.421683 | 5.465355 | 41 | 43 | 37 | 51 |
| 440423 | NR_024187,NF | SUZ12P1 | SUZ12 pseudogene 1 | transcribed_ps | . | 32421 | . | . | . | 5.717693 | 7.635426 | 7.008801 | 6.335481 | 13.305306 | 17.649627 | 15.727548 | 14.485326 | 1.001426 | 368.508457 | 0.204563545 | 0.010046628 | 0.991984085 | 0.999587993 | 8.524071 | 8.527035 | 8.496935 | 8.546269 | 8.602337 | 8.436624 | 406 | 414 | 367 | 296 |
| 440434 | NR_036750 | NPEPPSP1 | NPEPPS pseudogene 1 | transcribed_ps | . | 44259 | . | . | . | 7.047050 | 6.576398 | 7.495124 | 6.814969 | 16.398775 | 15.201639 | 16.818842 | 15.581616 | 1.034009 | 359.843627 | 0.190887441 | 0.252759921 | 0.800453741 | 0.998865404 | 8.466518 | 8.515519 | 8.502503 | 8.458286 | 8.529621 | 8.469355 | 423 | 367 | 335 | 325 |
| 440435 | NM_00100433 | GPR179 | G protein-coupled receptor 179 | protein_coding | NP_001004334.3 | 31371 | 614515 | . | . | 0.035408 | 0.022629 | 0.066884 | 0.039725 | 0.082396 | 0.052307 | 0.150086 | 0.090826 | 1.843216 | 12.047855 | 0.902283485 | 0.977769295 | 0.328188441 | NA | 3.079678 | 3.967302 | 3.565015 | 3.547020 | 3.604046 | 3.571109 | 12 | 7 | 17 | 11 |
| 440456 | NR_024386 | PLEKHM1P1 | pleckstrin homology and RUN domain containing M1 pseudogene | transcribed_ps | . | 35411 | . | . | . | 1.931656 | 2.261088 | 2.260175 | 1.859313 | 4.495044 | 5.226605 | 5.071768 | 4.251097 | -1.030862 | 189.501709 | 0.251606777 | -0.17428556 | 0.861641044 | 0.998865404 | 7.587570 | 7.544239 | 7.536625 | 7.603507 | 7.600569 | 7.512123 | 205 | 223 | 179 | 157 |
| 440459 | gene-SLC16A6I | SLC16A6P1 | SLC16A6 pseudogene 1 | pseudogene | . | 48932 | . | . | . | 3.805519 | 4.086338 | 3.797933 | 4.020897 | 8.855599 | 9.445753 | 8.522452 | 9.193304 | -1.021159 | 214.042205 | 0.230288635 | -0.1311744 | 0.89563735 | 0.998865404 | 7.757023 | 7.726316 | 7.731116 | 7.761918 | 7.726515 | 7.747046 | 241 | 241 | 180 | 203 |
| 440465 | NR_026857 | BAIAP2-DT | BAIAP2 divergent transcript | lncRNA | . | 44342 | . | . | . | 2.480084 | 2.482214 | 2.503988 | 2.830285 | 5.771258 | 5.737749 | 5.618876 | 6.471110 | 1.062220 | 354.915321 | 0.190788667 | 0.456436873 | 0.648075855 | 0.998865404 | 8.427387 | 8.513976 | 8.455342 | 8.450900 | 8.454686 | 8.519310 | 398 | 370 | 300 | 362 |
| 440482 | NR_040113 | ANKRD20A5P | ankyrin repeat domain 20 family member A5, pseudogene | transcribed_ps | . | 33833 | . | . | . | 0.453240 | 0.466746 | 0.420712 | 0.330831 | 1.054709 | 1.078904 | 0.944066 | 0.756406 | -1.226754 | 48.875454 | 0.451234726 | -0.65342029 | 0.513485363 | 0.998865404 | 5.750243 | 5.456953 | 5.626878 | 5.632986 | 5.613402 | 5.572268 | 61 | 59 | 43 | 36 |
| 440498 | NM_00113618 | HSBP1L1 | heat shock factor binding protein 1 like 1 | protein_coding | NP_001129652.1 | 37243 | . | . | . | 1.504160 | 1.554054 | 2.435739 | 1.392989 | 3.500243 | 3.592265 | 5.465728 | 3.184903 | 1.227558 | 37.513962 | 0.528588644 | 0.55958657 | 0.575761468 | 0.998865404 | 5.072328 | 5.370958 | 5.200821 | 5.206286 | 5.276005 | 5.189196 | 38 | 37 | 46 | 28 |
| 440514 | gene-CSP3 | CSP3 | CS pseudogene 3 | pseudogene | . | 54780 | . | . | . | 0.386785 | 0.445573 | 0.279067 | 0.400235 | 0.900064 | 1.029962 | 0.626218 | 0.915089 | -1.238121 | 34.487253 | 0.536965025 | -0.57387761 | 0.566050624 | 0.998865404 | 5.255178 | 4.944065 | 5.114224 | 5.134640 | 5.073973 | 5.116587 | 41 | 44 | 22 | 34 |
| 440515 | NM_00109926 | ZNF506 | zinc finger protein 506 | protein_coding | NP_001092739.1;NP_001138876.1 | 23780 | . | . | . | 0.123119 | 0.221179 | 0.241883 | 0.192275 | 0.286502 | 0.511266 | 0.542777 | 0.439615 | 1.260240 | 20.440340 | 0.697571958 | 0.478371517 | 0.632385802 | NA | 4.177728 | 4.509889 | 4.310986 | 4.355772 | 4.363717 | 4.343304 | 15 | 25 | 22 | 19 |
| 440519 | NM_00135540 | ZNF724 | zinc finger protein 724 | protein_coding | NP_001342333.1;NP_001342334.1;NP_001342335.1;XP_024307288.1;XP | 32460 | . | . | . | 1.207299 | 1.380208 | 1.142550 | 0.862693 | 2.809433 | 3.190412 | 2.563848 | 1.972444 | -1.319586 | 101.833805 | 0.325450644 | -1.22932874 | 0.218948575 | 0.982008277 | 6.855724 | 6.456953 | 6.696399 | 6.723545 | 6.659118 | 6.589898 | 128 | 130 | 86 | 72 |
| 440525 | gene-PRR13P5 | PRR13P5 | proline rich 13 pseudogene 5 | pseudogene | . | 50618 | . | . | . | 4.841599 | 4.346233 | 3.977960 | 5.134938 | 11.266601 | 10.046513 | 8.926428 | 11.740425 | -1.022806 | 156.594732 | 0.272384593 | -0.11943768 | 0.90492861 | 0.998865404 | 7.307498 | 7.274093 | 7.317902 | 7.270565 | 7.236409 | 7.332310 | 193 | 161 | 118 | 163 |
| 440567 | NM_00108959 | UQCRHL | ubiquinol-cytochrome c reductase hinge protein like | protein_coding | NP_001083060.1 | 51714 | . | . | . | 2.146921 | 1.094762 | 1.886736 | 2.495755 | 4.995974 | 2.530592 | 4.233780 | 5.706248 | 1.339927 | 32.202190 | 0.577553623 | 0.73093521 | 0.464818728 | 0.998865404 | 4.781608 | 5.205535 | 5.017867 | 4.942765 | 5.000243 | 5.036801 | 42 | 20 | 28 | 39 |
| 440574 | NM_00103236 | MICOS10 | mitochondrial contact site and cristae organizing system subunit 1 | protein_coding | NP_001027535.1;NP_001191011.1;NP_001191012.1 | 32068 | 616574 | . | . | 15.573028 | 14.364407 | 14.375928 | 16.887755 | 36.239080 | 33.203969 | 32.259165 | 38.611842 | 1.032727 | 1785.320682 | 0.131648316 | 0.352899233 | 0.724163986 | 0.998865404 | 10.778633 | 10.824931 | 10.822730 | 10.752646 | 10.754271 | 10.869581 | 2113 | 1811 | 1462 | 1825 |
| 440584 | NR_033967 | SLC2A1-AS1 | SLC2A1 antisense RNA 1 | lncRNA | . | 44187 | . | . | . | 0.631335 | 0.499178 | 0.465224 | 0.609658 | 1.469143 | 1.153872 | 1.043949 | 1.393911 | -1.041420 | 19.692650 | 0.695357272 | -0.08420479 | 0.932893612 | NA | 4.328680 | 4.269892 | 4.313412 | 4.291788 | 4.290796 | 4.307665 | 26 | 19 | 15 | 20 |
| 440589 | gene-RPS2P8 | RPS2P8 | ribosomal protein S2 pseudogene 8 | pseudogene | . | 36207 | . | . | . | 0.029897 | 0.045869 | 0.113584 | 0.007797 | 0.069571 | 0.106029 | 0.254879 | 0.017827 | 1.252417 | 2.026741 | 2.090159777 | 0.155354231 | 0.876542064 | NA | 0.844134 | 1.175234 | 1.007230 | 1.007808 | 1.014998 | 1.004217 | 2 | 2 | 3 | 1 |
| 440590 | NM_00100433 | ZYG11A | zyg-11 family member A, cell cycle regulator | protein_coding | NP_001004339.2;NP_001294860.1;XP_011539782.1;XP_011539783.1;XP | 32058 | 618675 | . | . | 1.974753 | 2.419547 | 3.145426 | 2.249892 | 4.595333 | 5.592890 | 7.058243 | 5.144107 | 1.169491 | 281.970998 | 0.232137636 | 0.973044168 | 0.330531351 | 0.998865404 | 8.021651 | 8.248265 | 8.043537 | 8.133572 | 8.258809 | 8.076707 | 273 | 305 | 309 | 237 |
| 440595 | gene-EEF1A1P1 | EEF1A1P11 | eukaryotic translation elongation factor 1 alpha 1 pseudogene 11 | pseudogene | . | 3206 | . | . | . | 15.566810 | 15.841194 | 12.605130 | 17.458992 | 36.224609 | 36.617630 | 28.285549 | 39.917912 | -1.059981 | 837.179695 | 0.169668071 | -0.49530816 | 0.620382609 | 0.998865404 | 9.751210 | 9.666329 | 9.724079 | 9.730342 | 9.569508 | 9.789820 | 988 | 934 | 596 | 881 |
| 440600 | NR_036595 | RBM15-AS1 | RBM15 antisense RNA 1 | lncRNA | . | 53636 | . | . | . | 0.118016 | 0.345398 | 0.287085 | 0.177419 | 0.274627 | 0.798402 | 0.644212 | 0.405648 | -1.000928 | 19.558298 | 0.754992828 | -0.00177233 | 0.998585887 | NA | 4.290955 | 4.288462 | 4.236528 | 4.316555 | 4.296222 | 4.257992 | 12 | 31 | 21 | 14 |
| 440672 | NM_00135540 | NUDT4B | nudix hydrolase 4B | protein_coding | NP_001342336.1 | 18012 | . | . | . | 4.432635 | 3.328991 | 2.190103 | 2.237999 | 10.314924 | 7.695111 | 4.914527 | 5.116919 | -1.775729 | 347.754667 | 0.210880486 | -3.92834561 | 8.55322E-05 | 0.011137749 | 8.797241 | 7.969467 | 8.661105 | 8.473962 | 8.248913 | 8.251645 | 587 | 410 | 217 | 236 |
| 440686 | NM_00135540 | H3-2 | H3.2 histone (putative) | protein_coding | NP_001342338.1;NP_001359034.1 | 32060 | . | . | . | 3.751412 | 3.970343 | 4.873681 | 3.867371 | 8.729691 | 9.177626 | 10.936399 | 8.842284 | 1.138956 | 305.547145 | 0.206909291 | 0.907216815 | 0.364292153 | 0.998865404 | 8.157885 | 8.346459 | 8.214319 | 8.217351 | 8.339358 | 8.230453 | 328 | 309 | 311 | 275 |
| 440689 | NM_00102459 | H2BC18 | H2B clustered histone 18 | protein_coding | NP_001019770.1;NP_001154806.1 | 24700 | . | . | . | 0.718893 | 0.930328 | 0.748844 | 0.897388 | 1.672895 | 2.150496 | 1.680384 | 2.051771 | 1.127203 | 16.656991 | 0.74948324 | 0.23048838 | 0.817712291 | NA | 3.969395 | 4.141582 | 4.055656 | 4.048283 | 4.054029 | 4.067084 | 19 | 16 | 14 | 18 |
| 440730 | NM_00100434 | TRIM67 | tripartite motif containing 67 | protein_coding | NP_001004342.3;NP_001287818.1;XP_011542494.1;XP_016856812.1 | 31859 | 610584 | . | . | 0.026475 | 0.032264 | 0.027380 | 0.033779 | 0.061609 | 0.074580 | 0.061439 | 0.077232 | 1.034586 | 8.683313 | 1.027908499 | 0.047721396 | 0.961938282 | NA | 3.094525 | 3.141582 | 3.113535 | 3.120246 | 3.114833 | 3.121612 | 9 | 10 | 7 | 9 |
| 440732 | gene-RPS7P3 | RPS7P3 | ribosomal protein S7 pseudogene 3 | pseudogene | . | 36518 | . | . | . | 0.280718 | 0.173563 | 0.040687 | 0.381527 | 0.653243 | 0.401200 | 0.091301 | 0.872316 | -1.005721 | 4.867346 | 1.447767147 | -0.00568426 | 0.995464639 | NA | 2.290730 | 2.275501 | 2.291357 | 2.280232 | 2.269205 | 2.301951 | 7 | 4 | 1 | 8 |
| 440737 | gene-RPL35P1 | RPL35P1 | ribosomal protein L35 pseudogene 1 | pseudogene | . | 36695 | . | . | . | 0.116701 | 0.481491 | 0.127954 | 0.424392 | 0.271569 | 1.112987 | 0.287125 | 0.970323 | -1.043582 | 4.462699 | 1.495230047 | -0.04115989 | 0.967168431 | NA | 2.193905 | 2.121007 | 2.140947 | 2.167704 | 2.144405 | 2.162686 | 2 | 8 | 2 | 6 |
| 440804 | NM_00112863 | RIMBP3B | RIMS binding protein 3B | protein_coding | NP_001122107.1 | 33891 | 612700 | . | . | 0.005574 | 0.035516 | 0.011916 | 0.032516 | 0.012970 | 0.082096 | 0.026738 | 0.074344 | 1.066075 | 4.231012 | 1.521377408 | 0.060674142 | 0.951618725 | NA | 2.039857 | 2.121007 | 2.064348 | 2.086823 | 2.067728 | 2.086048 | 2 | 7 | 2 | 6 |
| 440823 | NR_003491,NF | MIAT | myocardial infarction associated transcript | lncRNA | . | 33425 | 611082 | . | . | 0.123172 | 0.125843 | 0.054472 | 0.110790 | 0.286627 | 0.290892 | 0.122234 | 0.253308 | -1.533978 | 33.293683 | 0.567934021 | -1.08688339 | 0.277088349 | 0.998865404 | 5.334945 | 4.712861 | 5.085572 | 5.088473 | 4.995030 | 5.066088 | 46 | 44 | 15 | 33 |
| 440836 | NM_00101444 | ODF3B | outer dense fiber of sperm tails 3B | protein_coding | NP_001014440.2;NP_001369736.1;NP_001369737.1 | 34388 | . | . | . | 0.326629 | 0.207011 | 0.235234 | 0.378203 | 0.760078 | 0.478515 | 0.527858 | 0.864716 | 1.099741 | 8.900001 | 1.026055427 | 0.133681312 | 0.893654583 | NA | 3.084644 | 3.219802 | 3.156417 | 3.146578 | 3.144727 | 3.165798 | 11 | 8 | 6 | 11 |
| 440854 | NM_00114512 | CAPN14 | calpain 14 | protein_coding | NP_001138594.1;NP_001308199.1;XP_011531165.1;XP_011531166.1;XP | 16664 | 610229 | . | . | 0.376811 | 0.296849 | 0.338719 | 0.232475 | 0.876855 | 0.686180 | 0.760076 | 0.531527 | -1.193996 | 40.259607 | 0.500471999 | -0.51111363 | 0.609271495 | 0.998865404 | 5.452135 | 5.199327 | 5.363736 | 5.325796 | 5.339549 | 5.295001 | 56 | 42 | 37 | 29 |
| 440900 | NR_034128,NF | LINC01191 | long intergenic non-protein coding RNA 1191 | lncRNA | . | 49595 | . | . | . | 0.474408 | 0.152552 | 0.299653 | 0.498674 | 1.103966 | 0.352630 | 0.672413 | 1.140160 | 1.312040 | 8.674144 | 1.081777235 | 0.362192273 | 0.717208353 | NA | 2.906576 | 3.300113 | 3.123684 | 3.089584 | 3.103894 | 3.129638 | 13 | 4 | 6 | 12 |
| 440905 | NR_026758 | FAR2P1 | fatty acyl-CoA reductase 2 pseudogene 1 | transcribed_ps | . | 49284 | . | . | . | 0.569066 | 0.505069 | 0.832786 | 0.496947 | 1.324241 | 1.167489 | 1.868748 | 1.136211 | 1.210034 | 85.137595 | 0.373806318 | 0.735802716 | 0.461850768 | 0.998865404 | 6.266033 | 6.544043 | 6.389450 | 6.358058 | 6.497486 | 6.349282 | 94 | 78 | 102 | 65 |
| 440918 | NR_147695 | FLJ46875 | uncharacterized LOC440918 | lncRNA | . | . | . | . | . | 0.202153 | 0.212972 | 0.174546 | 0.204438 | 0.470418 | 0.492295 | 0.391677 | 0.467424 | -1.096238 | 34.731309 | 0.525516332 | -0.25224824 | 0.800849194 | 0.998865404 | 5.183405 | 5.049834 | 5.123080 | 5.128880 | 5.105364 | 5.121351 | 41 | 40 | 27 | 33 |
| 440926 | NR_002315 | H3P6 | H3 histone pseudogene 6 | transcribed_ps | . | 42980 | . | . | . | 32.135204 | 27.408474 | 27.988508 | 28.841429 | 74.779945 | 63.355911 | 62.805401 | 65.942505 | -1.062864 | 960.722202 | 0.147094829 | -0.59795736 | 0.549868385 | 0.998865404 | 9.951146 | 9.863470 | 9.988859 | 9.865488 | 9.877587 | 9.891788 | 1235 | 979 | 802 | 882 |
| 440934 | NR_136638,NF | CT75 | cancer/testis associated transcript 75 | lncRNA | . | 37174 | . | . | . | 0.643376 | 0.687392 | 1.015665 | 0.772202 | 1.497161 | 1.588935 | 2.279121 | 1.765548 | 1.487410 | 25.115448 | 0.619724995 | 0.92428519 | 0.355337851 | NA | 4.334945 | 4.909275 | 4.616705 | 4.618807 | 4.675889 | 4.640946 | 23 | 22 | 30 | 24 |
| 440944 | NR_027007,NF | THUMPD3-AS1 | THUMPD3 antisense RNA 1 | lncRNA | . | 44478 | . | . | . | 4.548422 | 5.279880 | 4.581919 | 4.003577 | 10.584366 | 12.204680 | 10.281694 | 9.153703 | -1.142870 | 240.429014 | 0.228555496 | -0.84295209 | 0.399255239 | 0.998865404 | 8.002219 | 7.810340 | 7.914912 | 7.964222 | 7.923297 | 7.823164 | 281 | 290 | 216 | 188 |
| 440957 | NM_00112476 | SMIM4 | small integral membrane protein 4 | protein_coding | NP_001118239.1;XP_011532031.1 | 37257 | . | . | . | 8.295597 | 6.887921 | 7.196699 | 9.252443 | 19.304195 | 15.921739 | 16.149184 | 21.154612 | 1.068516 | 407.487503 | 0.192963837 | 0.495470445 | 0.620268079 | 0.998865404 | 8.622402 | 8.717263 | 8.668114 | 8.628337 | 8.613010 | 8.757712 | 469 | 411 | 322 | 442 |
| 440992 | gene-RPS20P14 | RPS20P14 | ribosomal protein S20 pseudogene 14 | pseudogene | . | 36692 | . | . | . | 0.332206 | 0.412658 | 0.427687 | 0.574996 | 0.773058 | 0.953878 | 0.959717 | 1.314659 | 1.225446 | 7.498226 | 1.106669262 | 0.265036069 | 0.790981676 | NA | 2.754145 | 3.044381 | 2.896057 | 2.902339 | 2.900115 | 2.912026 | 7 | 8 | 6 | 9 |
| 440993 | NR_122105 | MIR570HG | MIR570 host gene | lncRNA | . | 53743 | . | . | . | 0.849037 | 1.225850 | 1.583066 | 1.193472 | 1.975744 | 2.833606 | 3.552354 | 2.728732 | 1.342968 | 35.919207 | 0.531458513 | 0.800486109 | 0.423429208 | 0.998865404 | 4.938471 | 5.363679 | 5.111446 | 5.157005 | 5.198023 | 5.153270 | 29 | 39 | 41 | 33 |
| 441024 | NM_00100434 | MTHFD2L | methylenetetrahydrofolate dehydrogenase (NADP+ dependent) 2 | protein_coding | NP_001004346.2;NP_001138450.1;NP_001338239.1;NP_001338240.1;NI | 31865 | 614047 | . | . | 1.840810 | 1.304655 | 1.226628 | 1.489045 | 4.283640 | 3.015767 | 2.752519 | 3.404524 | -1.225921 | 110.146438 | 0.32125244 | -0.91474951 | 0.360323147 | 0.998865404 | 6.922265 | 6.629462 | 6.871450 | 6.753362 | 6.744714 | 6.751776 | 163 | 109 | 85 | 94 |
| 441027 | NM_00108050 | TMEM150C | transmembrane protein 150C | protein_coding | NP_001073975.1;NP_001340383.1;NP_001340384.1 | 37263 | 617292 | . | . | 0.932585 | 0.866295 | 0.497326 | 0.923589 | 2.170165 | 2.002479 | 1.115985 | 2.111676 | -1.278596 | 80.522004 | 0.388867197 | -0.91177803 | 0.361885578 | 0.998865404 | 6.499094 | 6.141415 | 6.371457 | 6.349491 | 6.223667 | 6.363455 | 108 | 94 | 43 | 86 |
| 441032 | gene-EEF1A1P9 | EEF1A1P9 | eukaryotic translation elongation factor 1 alpha 1 pseudogene 9 | pseudogene | . | 3204 | . | . | . | 0.712528 | 0.612523 | 0.362041 | 0.656128 | 1.658081 | 1.415873 | 0.812409 | 1.500158 | -1.300466 | 32.801027 | 0.561337115 | -0.67522394 | 0.499533498 | 0.998865404 | 5.214084 | 4.832035 | 5.066531 | 5.043648 | 4.990041 | 5.051139 | 46 | 37 | 18 | 34 |
| 441034 | gene-RPL7AP30 | RPL7AP30 | ribosomal protein L7a pseudogene 30 | pseudogene | . | 35757 | . | . | . | 0.651841 | 0.449156 | 0.124286 | 0.394275 | 1.516860 | 1.038244 | 0.278894 | 0.901463 | -2.199466 | 11.332076 | 0.965309023 | -1.17802023 | 0.238788543 | NA | 3.964015 | 2.819320 | 3.543099 | 3.519574 | 3.478402 | 3.509391 | 21 | 14 | 3 | 10 |
| 441050 | gene-PTGES3P3 | PTGES3P3 | prostaglandin E synthase 3 pseudogene 3 | pseudogene | . | 43823 | . | . | . | 0.052250 | 0.125707 | 0.030928 | 0.024731 | 0.121588 | 0.290577 | 0.069401 | 0.056544 | -2.452425 | 1.915126 | 2.205695884 | -0.58675761 | 0.55736651 | NA | 1.444631 | 0.148375 | 0.948324 | 0.957585 | 0.945780 | 0.945324 | 2 | 4 | 1 | 1 |
| 441054 | NM_00111435 | C4orf47 | chromosome 4 open reading frame 47 | protein_coding | NP_001107829.1;NP_001332936.1;XP_005263076.1;XP_011530291.1;XP | 34346 | . | . | . | 0.311577 | 0.801021 | 0.627796 | 0.317892 | 0.725052 | 1.851596 | 1.408754 | 0.726822 | 1.250861 | 27.973968 | 0.601760892 | 0.536626721 | 0.591525483 | 0.998865404 | 4.636526 | 4.957666 | 4.754995 | 4.811732 | 4.818113 | 4.800394 | 21 | 34 | 29 | 27 |
| 441073 | gene-RPL26P19 | RPL26P19 | ribosomal protein L26 pseudogene 19 | pseudogene | . | 36393 | . | . | . | 5.336926 | 5.746790 | 4.524796 | 6.741469 | 12.419245 | 13.283964 | 10.153512 | 15.413568 | -1.002240 | 91.376865 | 0.349364164 | -0.00924068 | 0.992627111 | 0.999862634 | 6.516559 | 6.510949 | 6.502496 | 6.521056 | 6.455663 | 6.564215 | 102 | 102 | 64 | 102 |
| 441087 | NR_149046 | LOC441087 | uncharacterized LOC441087 | lncRNA | . | . | . | . | . | 0.268566 | 0.107025 | 0.074909 | 0.312396 | 0.624965 | 0.247393 | 0.168093 | 0.714256 | 1.010690 | 7.559572 | 1.164024772 | 0.013179012 | 0.989484974 | NA | 2.912173 | 2.924410 | 2.932270 | 2.905500 | 2.901065 | 2.937321 | 12 | 5 | 3 | 11 |
| 441108 | NR_161242 | IRF1-AS1 | IRF1 antisense RNA 1 | lncRNA | . | 33838 | . | . | . | 0.279659 | 0.568050 | 0.179143 | 0.470565 | 0.650779 | 1.313074 | 0.401991 | 1.075892 | -1.269808 | 8.081670 | 1.100435163 | -0.31315848 | 0.754160264 | NA | 3.180050 | 2.827809 | 3.006769 | 3.004135 | 3.000971 | 3.022814 | 7 | 13 | 4 | 9 |
| 441114 | gene-RPLP0P9 | RPLP0P9 | ribosomal protein lateral stalk subunit P0 pseudogene | pseudogene | . | . | . | . | . | 53.290810 | 50.210827 | 44.402786 | 58.700932 | 124.009895 | 116.064575 | 99.638565 | 134.212692 | -1.017830 | 597.339815 | 0.176894868 | -0.14413102 | 0.885397011 | 0.998865404 | 9.235460 | 9.209237 | 9.249203 | 9.202982 | 9.122854 | 9.298589 | 718 | 628 | 446 | 629 |
| 441124 | NR_132119 | GTF2IP20 | general transcription factor IIi pseudogene 20 | transcribed_ps | . | 51732 | . | . | . | 3.786658 | 4.198052 | 4.624064 | 3.556339 | 8.811708 | 9.703986 | 10.376267 | 8.131146 | 1.012121 | 371.428060 | 0.202544534 | 0.085821722 | 0.931611398 | 0.998865404 | 8.527872 | 8.546008 | 8.502515 | 8.557478 | 8.612469 | 8.456464 | 405 | 417 | 369 | 303 |
| 441150 | NM_00100873 | C6orf226 | chromosome 6 open reading frame 226 | protein_coding | NP_001008739.1 | 34431 | . | . | . | 2.200107 | 2.338978 | 2.029465 | 2.716801 | 5.119740 | 5.406653 | 4.554061 | 6.211643 | 1.035479 | 40.609805 | 0.491833398 | 0.102266386 | 0.918545236 | 0.998865404 | 5.319341 | 5.367765 | 5.336898 | 5.343643 | 5.324327 | 5.366708 | 45 | 44 | 31 | 44 |
| 441151 | NM_00113756 | TMEM151B | transmembrane protein 151B | protein_coding | NP_001131032.1 | 21315 | . | . | . | 0.059577 | 0.038799 | 0.068003 | 0.053173 | 0.138637 | 0.089686 | 0.152596 | 0.121573 | 1.245648 | 9.018289 | 1.016390598 | 0.311786476 | 0.755202802 | NA | 3.003609 | 3.324312 | 3.169723 | 3.155579 | 3.177898 | 3.165088 | 11 | 7 | 10 | 8 |
| 441155 | NM_00127167 | LOC441155 | zinc finger CCCH-type domain-containing-like | protein_coding | NP_001258604.1;XP_016866378.1 | . | . | . | . | 1.696446 | 1.299504 | 1.271103 | 0.672483 | 3.947700 | 3.003862 | 2.852318 | 1.537554 | -1.552313 | 147.676296 | 0.325619921 | -1.94834473 | 0.051373728 | 0.672667836 | 7.487472 | 6.856716 | 7.329629 | 7.211825 | 7.201077 | 6.997248 | 234 | 168 | 132 | 75 |
| 441166 | gene-CNN2P9 | CNN2P9 | calponin 2 pseudogene 9 | pseudogene | . | 39534 | . | . | . | 0.052343 | 0.057047 | 0.046916 | 0.027374 | 0.121804 | 0.131867 | 0.105279 | 0.062587 | -1.606250 | 2.188044 | 2.021504645 | -0.33821176 | 0.735203614 | NA | 1.429096 | 0.751299 | 1.137918 | 1.138711 | 1.136419 | 1.130808 | 3 | 3 | 2 | 1 |
| 441172 | NR_033896 | FLJ46906 | uncharacterized LOC441172 | lncRNA | . | . | . | . | . | 8.686892 | 8.931846 | 10.586798 | 7.795949 | 20.214752 | 20.646362 | 23.756466 | 17.824512 | 1.029401 | 281.353497 | 0.223779889 | 0.186812288 | 0.851807812 | 0.998865404 | 8.114535 | 8.157623 | 8.119697 | 8.130906 | 8.218551 | 8.055382 | 316 | 302 | 287 | 226 |
| 441191 | NR_015449,NF | RNF216P1 | ring finger protein 216 pseudogene 1 | transcribed_ps | . | 33610 | . | . | . | 4.998129 | 5.568782 | 5.518570 | 6.038620 | 11.630855 | 12.872489 | 12.383510 | 13.806588 | 1.071066 | 395.336360 | 0.18683723 | 0.530125679 | 0.596024796 | 0.998865404 | 8.576996 | 8.675207 | 8.565673 | 8.642784 | 8.617420 | 8.672716 | 411 | 439 | 339 | 401 |
| 441194 | NR_002217 | PMS2CL | PMS2 C-terminal like pseudogene | transcribed_ps | . | 30061 | . | . | . | 3.188303 | 2.797027 | 3.735273 | 3.222538 | 7.419312 | 6.465453 | 8.381845 | 7.367950 | 1.144847 | 175.651745 | 0.256147489 | 0.761883488 | 0.446129546 | 0.998865404 | 7.354961 | 7.551499 | 7.452749 | 7.393973 | 7.514493 | 7.446434 | 202 | 164 | 176 | 162 |
| 441204 | NR_148499,NF | LOC441204 | uncharacterized LOC441204 | lncRNA | . | . | . | . | . | 0.297631 | 0.187032 | 0.013662 | 0.387399 | 0.692601 | 0.432332 | 0.030658 | 0.885742 | -1.313361 | 8.175923 | 1.1243015 | -0.34978476 | 0.72650024 | NA | 3.216230 | 2.819320 | 3.058831 | 3.024607 | 3.016060 | 3.047520 | 15 | 6 | 3 | 10 |
| 441208 | NR_003502 | ZNRF2P1 | zinc and ring finger 2 pseudogene 1 | transcribed_ps | . | 42792 | . | . | . | 0.208321 | 0.630840 | 0.553598 | 0.514292 | 0.484772 | 1.458216 | 1.242257 | 1.175867 | 1.260120 | 30.846932 | 0.622875995 | 0.535517453 | 0.592292092 | 0.998865404 | 4.772726 | 5.102572 | 4.857305 | 4.971200 | 4.949601 | 4.939803 | 16 | 44 | 31 | 31 |
| 441212 | NR_003500 | RP9P | RP9 pseudogene | transcribed_ps | . | 33969 | . | . | . | 6.625493 | 6.956649 | 8.148715 | 7.502709 | 15.417794 | 16.080606 | 18.285480 | 17.154053 | 1.135281 | 295.559295 | 0.203549246 | 0.899289273 | 0.368498599 | 0.998865404 | 8.112729 | 8.296058 | 8.159135 | 8.181486 | 8.262911 | 8.212723 | 312 | 305 | 286 | 281 |
| 441234 | NM_00115927 | ZNF716 | zinc finger protein 716 | protein_coding | NP_001152751.1 | 32458 | . | . | . | 0.252710 | 0.290487 | 0.237142 | 0.124019 | 0.588067 | 0.671473 | 0.532139 | 0.283554 | -1.517969 | 37.310436 | 0.538320902 | -1.11855677 | 0.263329275 | 0.996308385 | 5.489863 | 4.891560 | 5.236113 | 5.261008 | 5.225343 | 5.150642 | 48 | 52 | 34 | 19 |
| 441242 | NR_038378,NF | LOC441242 | uncharacterized LOC441242 | lncRNA | . | . | . | . | . | 0.729644 | 0.542827 | 0.494639 | 0.541072 | 1.697912 | 1.254769 | 1.109955 | 1.237097 | -1.220619 | 30.028310 | 0.571659459 | -0.50311904 | 0.614880583 | 0.998865404 | 5.044696 | 4.757542 | 4.943891 | 4.901249 | 4.891844 | 4.905544 | 44 | 30 | 22 | 27 |
| 441250 | NM_00114544 | TYW1B | tRNA-yW synthesizing protein 1 homolog B | protein_coding | NP_001138912.2 | 33908 | . | . | . | 0.727553 | 0.808027 | 0.444250 | 0.611976 | 1.693046 | 1.867791 | 0.996884 | 1.399210 | -1.455965 | 63.571594 | 0.404698245 | -1.33920966 | 0.180502428 | 0.952541638 | 6.237009 | 5.692544 | 6.019851 | 6.042293 | 5.919991 | 5.976406 | 83 | 85 | 38 | 56 |
| 441259 | gene-LOC4412 | LOC441259 | PMS1 homolog 2, mismatch repair system component pseudogen | pseudogene | . | . | . | . | . | 0.325282 | 0.406560 | 0.300441 | 0.301596 | 0.756943 | 0.939781 | 0.674182 | 0.689563 | -1.216522 | 8.598776 | 1.034543925 | -0.27332096 | 0.784606507 | NA | 3.238681 | 2.955730 | 3.107158 | 3.114028 | 3.104394 | 3.101762 | 10 | 11 | 7 | 7 |
| 441263 | NR_023383 | DTX2P1-UPK3E | DTX2P1-UPK3BP1-PMS2P11 readthrough, transcribed pseudogen | transcribed_ps | . | 42360 | . | . | . | 1.052607 | 1.010894 | 1.579459 | 1.340230 | 2.449459 | 2.336727 | 3.544261 | 3.064276 | 1.391782 | 69.145892 | 0.3817022 | 1.249490363 | 0.211485776 | 0.974996048 | 5.852816 | 6.330900 | 6.066419 | 6.057710 | 6.161478 | 6.117474 | 68 | 61 | 76 | 69 |
| 441294 | NM_00100874 | CTAGE15 | CTAGE family member 15 | protein_coding | NP_001008747.1 | 37295 | . | . | . | 0.140540 | 0.083035 | 0.169280 | 0.084221 | 0.327042 | 0.191939 | 0.379859 | 0.192561 | 1.082064 | 10.209627 | 0.972439107 | 0.117011413 | 0.906851009 | NA | 3.290730 | 3.410501 | 3.355929 | 3.334181 | 3.362260 | 3.334871 | 14 | 8 | 12 | 7 |
| 441295 | NR_002157 | OR2A9P | olfactory receptor family 2 subfamily A member 9 pseudogene | transcribed_ps | . | 8236 | . | . | . | 0.539582 | 0.645024 | 0.717359 | 0.876720 | 1.255630 | 1.491001 | 1.609732 | 2.004517 | 1.316972 | 21.855848 | 0.658358572 | 0.603356501 | 0.546271553 | NA | 4.238681 | 4.634191 | 4.424134 | 4.437540 | 4.448468 | 4.463584 | 20 | 22 | 20 | 25 |
| 441307 | NR_033963 | HRAT92 | heart tissue-associated transcript 92 | lncRNA | . | . | . | . | . | 0.109811 | 0.202063 | 0.214843 | 0.157478 | 0.255535 | 0.467077 | 0.482103 | 0.360054 | 1.198861 | 18.904049 | 0.728399504 | 0.359231267 | 0.719422088 | NA | 4.104339 | 4.365137 | 4.201022 | 4.245726 | 4.252666 | 4.223519 | 14 | 24 | 21 | 16 |
| 441308 | NM_00100550 | OR4F21 | olfactory receptor family 4 subfamily F member 21 | protein_coding | NP_001005504.1 | 19583 | . | . | . | 0.021044 | 0.008968 | 0.019083 | 0.010817 | 0.048970 | 0.020730 | 0.042821 | 0.024732 | 1.234655 | 1.002957 | 2.868339486 | 0.106022226 | 0.915564717 | NA | -0.155866 | 0.148375 | -0.001261 | -0.000972 | 0.000053 | -0.000315 | 1 | 1 | 1 | 1 |
| 441376 | NM_00102535 | AARD | alanine and arginine rich domain containing protein | protein_coding | NP_001020528.1 | 33842 | . | . | . | 2.003796 | 2.288763 | 1.829165 | 2.550802 | 4.662916 | 5.290579 | 4.104594 | 5.832105 | 1.008878 | 155.106866 | 0.279102284 | 0.045686176 | 0.963560383 | 0.998865404 | 7.271760 | 7.282458 | 7.247960 | 7.297349 | 7.212752 | 7.336230 | 167 | 177 | 114 | 169 |
| 441381 | NM_00102467 | LRRC24 | leucine rich repeat containing 24 | protein_coding | NP_001019849.2 | 28947 | . | . | . | 0.472349 | 0.724721 | 0.212210 | 0.617117 | 1.099175 | 1.675225 | 0.476192 | 1.410965 | -1.440036 | 28.620385 | 0.639881075 | -0.8221923 | 0.41096746 | 0.998865404 | 5.081051 | 4.547915 | 4.829031 | 4.885355 | 4.771347 | 4.858069 | 31 | 44 | 11 | 32 |
| 441389 | NR_038977 | LINC01239 | long intergenic non-protein coding RNA 1239 | lncRNA | . | 49796 | . | . | . | 2.735240 | 2.773353 | 4.387507 | 2.822920 | 6.365015 | 6.410731 | 9.845439 | 6.454272 | 1.292517 | 232.968438 | 0.254839903 | 1.452610189 | 0.146332016 | 0.907376824 | 7.666156 | 8.037939 | 7.784857 | 7.788778 | 8.018060 | 7.791816 | 233 | 220 | 279 | 192 |
| 441391 | gene-RBMXP2 | RBMXP2 | RBMX pseudogene 2 | pseudogene | . | 23284 | . | . | . | 0.140220 | 0.150238 | 0.078663 | 0.147256 | 0.326298 | 0.347283 | 0.176518 | 0.336684 | -1.282979 | 8.774233 | 1.033314626 | -0.34790758 | 0.727909595 | NA | 3.303566 | 2.940155 | 3.141983 | 3.144839 | 3.125291 | 3.142016 | 11 | 11 | 5 | 9 |
| 441394 | NR_003667 | SUGT1P1 | SUGT1 pseudogene 1 | transcribed_ps | . | 19384 | . | . | . | 0.789503 | 1.433318 | 1.540879 | 0.626863 | 1.837205 | 3.313178 | 3.457686 | 1.433247 | -1.031323 | 28.540376 | 0.644133506 | -0.06907929 | 0.944926508 | 0.998865404 | 4.855800 | 4.813758 | 4.788379 | 4.856425 | 4.866744 | 4.769560 | 24 | 40 | 35 | 15 |
| 441425 | XR_00174629 | ANKRD20A3P | ankyrin repeat domain 20 family member A3, pseudogene, transc | transcribed_ps | . | 31981 | . | . | . | 0.133470 | 0.117023 | 0.078679 | 0.131250 | 0.310590 | 0.270505 | 0.176553 | 0.300087 | -1.221735 | 17.070517 | 0.753376407 | -0.38351493 | 0.701338 | NA | 4.231982 | 3.940155 | 4.109907 | 4.099462 | 4.073825 | 4.106070 | 23 | 19 | 10 | 18 |
| 441459 | NM_00135343 | ANKRD18B | ankyrin repeat domain 18B | protein_coding | NP_001340361.1 | 23644 | . | . | . | 0.616906 | 0.640391 | 0.486844 | 0.625877 | 1.435565 | 1.480293 | 1.092463 | 1.430994 | -1.141407 | 70.227348 | 0.381618411 | -0.50000947 | 0.617068413 | 0.998865404 | 6.226996 | 6.034510 | 6.147052 | 6.154297 | 6.091677 | 6.145859 | 85 | 82 | 50 | 69 |
| 441481 | gene-GPX1P1 | GPX1P1 | glutathione peroxidase pseudogene 1 | pseudogene | . | 4560 | . | . | . | 1.249220 | 0.851486 | 0.845874 | 0.684893 | 2.906986 | 1.968247 | 1.898116 | 1.565927 | -1.374955 | 25.026448 | 0.629366049 | -0.72991542 | 0.465441884 | NA | 4.855648 | 4.399154 | 4.687869 | 4.639531 | 4.639268 | 4.623424 | 40 | 25 | 20 | 18 |
| 441502 | NR_002309 | RPS26P11 | ribosomal protein S26 pseudogene 11 | transcribed_ps | . | 31817 | . | . | . | 0.096951 | 0.189398 | 0.092193 | 0.138718 | 0.225608 | 0.437803 | 0.206879 | 0.317162 | 1.014447 | 2.735953 | 1.804031249 | 0.011470344 | 0.99084819 | NA | 1.444631 | 1.459419 | 1.445375 | 1.454766 | 1.448241 | 1.452296 | 2 | 4 | 2 | 3 |
| 441519 | NM_00101743 | CT45A3 | cancer/testis antigen family 45 member A3 | protein_coding | NP_001017435.1;NP_001357077.1;NP_001357078.1 | 33268 | 300794 | . | . | 1.631139 | 1.683467 | 1.078706 | 1.180995 | 3.795728 | 3.891408 | 2.420585 | 2.700205 | -1.459836 | 37.416318 | 0.509157221 | -1.07198052 | 0.283728782 | 0.998865404 | 5.473017 | 4.926775 | 5.251802 | 5.259153 | 5.199262 | 5.202785 | 50 | 49 | 26 | 29 |
| 441521 | NM_00100755 | CT45A5 | cancer/testis antigen family 45 member A5 | protein_coding | NP_001007552.2;NP_001165759.2;XP_011529643.1 | 33270 | 300796 | . | . | 1.444393 | 1.315637 | 0.993796 | 0.910661 | 3.361161 | 3.041155 | 2.230050 | 2.082119 | -1.459525 | 49.048679 | 0.448406883 | -1.21652638 | 0.223784469 | 0.982008277 | 5.862893 | 5.318300 | 5.661011 | 5.641636 | 5.589455 | 5.577628 | 70 | 60 | 36 | 36 |
| 441531 | NM_00102989 | PGAM4 | phosphoglycerate mutase family member 4 | protein_coding | NP_001025062.1 | 21731 | 300567 | . | . | 3.697567 | 3.973504 | 3.168636 | 4.204442 | 8.604390 | 9.184932 | 7.110328 | 9.612956 | -1.053222 | 198.496777 | 0.249302872 | -0.30007256 | 0.764121807 | 0.998865404 | 7.670682 | 7.594249 | 7.628301 | 7.658257 | 7.558309 | 7.678069 | 227 | 227 | 145 | 206 |
| 441549 | NM_00102995 | CDNF | cerebral dopamine neurotrophic factor | protein_coding | NP_001025125.2;XP_011517790.1 | 24913 | 611233 | . | . | 0.787769 | 0.357580 | 0.762665 | 0.494004 | 1.833171 | 0.826562 | 1.711399 | 1.129481 | 1.100067 | 26.560401 | 0.642571606 | 0.214127163 | 0.830447894 | NA | 4.657993 | 4.800881 | 4.757503 | 4.677122 | 4.749966 | 4.706397 | 40 | 17 | 29 | 21 |
| 441581 | NM_00108099 | FRG2B | FSHD region gene 2 family member B | protein_coding | NP_001074467.1 | 33518 | . | . | . | 0.225031 | 0.278746 | 0.333134 | 0.327902 | 0.523657 | 0.644335 | 0.747542 | 0.749710 | 1.264541 | 19.337427 | 0.697073808 | 0.485764425 | 0.6271342 | NA | 4.094525 | 4.432387 | 4.251296 | 4.264271 | 4.278022 | 4.276090 | 18 | 20 | 19 | 20 |
| 441666 | NR_024380 | LOC441666 | zinc finger protein 91 pseudogene | transcribed_ps | . | . | . | . | . | 0.238144 | 0.334359 | 0.439301 | 0.195036 | 0.554171 | 0.772885 | 0.985778 | 0.445927 | 1.095343 | 63.333383 | 0.459011027 | 0.286228692 | 0.774702969 | 0.998865404 | 5.916503 | 6.050189 | 5.920812 | 5.993577 | 6.061545 | 5.886530 | 58 | 76 | 80 | 38 |
| 441711 | gene-LOC4417 | LOC441711 | RING finger protein 145-like | pseudogene | . | . | . | . | . | 0.224068 | 0.169493 | 0.136670 | 0.125129 | 0.521416 | 0.391792 | 0.306684 | 0.286092 | -1.461443 | 10.904396 | 0.928791727 | -0.58936123 | 0.555618982 | NA | 3.694014 | 3.148375 | 3.470309 | 3.445547 | 3.457047 | 3.442466 | 17 | 12 | 8 | 8 |
| 441733 | NR_073405 | PRKXP1 | PRKX pseudogene 1 | transcribed_ps | . | 9442 | . | . | . | 0.219727 | 0.263775 | 0.394033 | 0.181004 | 0.511313 | 0.609727 | 0.884198 | 0.413844 | 1.175782 | 95.466226 | 0.389322032 | 0.600070187 | 0.548459461 | 0.998865404 | 6.454105 | 6.690093 | 6.508103 | 6.559141 | 6.690157 | 6.460632 | 92 | 103 | 123 | 61 |
| 441783 | gene-ACTG1P2 | ACTG1P24 | actin gamma 1 pseudogene 24 | pseudogene | . | 49661 | . | . | . | 0.180335 | 0.314343 | 0.182889 | 0.251918 | 0.419646 | 0.726617 | 0.410397 | 0.575980 | -1.092132 | 6.923660 | 1.159680259 | -0.10963963 | 0.912695179 | NA | 2.855800 | 2.724273 | 2.783247 | 2.802076 | 2.786189 | 2.793728 | 6 | 10 | 5 | 7 |
| 441876 | gene-RPS16P1 | RPS16P1 | ribosomal protein S16 pseudogene 1 | pseudogene | . | 36462 | . | . | . | 0.148491 | 0.214409 | 0.152928 | 0.094660 | 0.345545 | 0.495615 | 0.343166 | 0.216429 | -1.131926 | 2.973891 | 1.729253768 | -0.10338572 | 0.917656846 | NA | 1.658167 | 1.481107 | 1.569843 | 1.575066 | 1.573727 | 1.567785 | 3 | 4 | 3 | 2 |
| 441951 | NR_003604,NF | ZFAS1 | ZNFX1 antisense RNA 1 | lncRNA | . | 33101 | . | . | . | 89.961894 | 80.592244 | 88.244746 | 101.930343 | 209.345013 | 186.292579 | 198.018649 | 233.051624 | 1.071386 | 2546.470817 | 0.123464892 | 0.805723429 | 0.420402337 | 0.998865404 | 11.263749 | 11.363107 | 11.314015 | 11.255047 | 11.285011 | 11.393598 | 2935 | 2556 | 2138 | 2634 |
| 442075 | NR_103821 | EMC3-AS1 | EMC3 antisense RNA 1 | lncRNA | . | 49223 | . | . | . | 1.518852 | 1.335330 | 2.024544 | 1.777086 | 3.534431 | 3.086674 | 4.543018 | 4.063096 | 1.313018 | 243.789130 | 0.224694993 | 1.748535096 | 0.080371414 | 0.779933816 | 7.719034 | 8.113117 | 7.882908 | 7.820917 | 8.022607 | 7.948808 | 259 | 212 | 258 | 241 |
| 442112 | gene-SERBP1P | SERBP1P5 | SERPINE1 mRNA binding protein 1 pseudogene 5 | pseudogene | . | 44632 | . | . | . | 0.767565 | 0.610879 | 0.412699 | 0.319131 | 1.786156 | 1.412075 | 0.926085 | 0.729654 | -1.890937 | 32.116275 | 0.561680145 | -1.63634275 | 0.101767873 | 0.820390774 | 5.391727 | 4.475715 | 5.064920 | 5.025709 | 4.979035 | 4.953509 | 54 | 40 | 22 | 18 |
| 442155 | gene-LOC4421 | LOC442155 | transcription factor B2, mitochondrial pseudogene | pseudogene | . | . | . | . | . | 0.057629 | 0.100921 | 0.195724 | 0.304165 | 0.134106 | 0.233284 | 0.439199 | 0.695438 | 2.853317 | 7.793490 | 1.121657125 | 1.348576365 | 0.177473076 | NA | 2.019256 | 3.527725 | 2.917604 | 2.923398 | 2.944260 | 2.964923 | 4 | 5 | 8 | 13 |
| 442421 | NR_024496,NF | PTGER4P2-CDK | PTGER4P2-CDK2AP2P2 readthrough, transcribed pseudogene | transcribed_ps | . | 44749 | . | . | . | 0.121650 | 0.151968 | 0.214974 | 0.144941 | 0.283084 | 0.351282 | 0.482396 | 0.331390 | 1.427395 | 18.003960 | 0.730105227 | 0.703164993 | 0.481952932 | NA | 3.889947 | 4.404839 | 4.140698 | 4.149227 | 4.188894 | 4.155135 | 16 | 17 | 22 | 16 |
| 442429 | gene-EEF1DP2 | EEF1DP2 | eukaryotic translation elongation factor 1 delta pseudogene 2 | pseudogene | . | 33523 | . | . | . | 0.024724 | 0.021539 | 0.017184 | 0.011809 | 0.057534 | 0.049789 | 0.038561 | 0.026999 | 1.234655 | 1.002957 | 2.868339486 | 0.106022226 | 0.915564717 | NA | -0.155866 | 0.148375 | -0.001261 | -0.000972 | 0.000053 | -0.000315 | 1 | 1 | 1 | 1 |
| 442454 | NR_002308 | UQCRBP1 | ubiquinol-cytochrome c reductase binding protein pseudogene 1 | transcribed_ps | . | 12583 | . | . | . | 0.383370 | 0.427550 | 0.254308 | 0.266936 | 0.892118 | 0.988302 | 0.570659 | 0.610318 | -1.498545 | 8.977189 | 1.017774507 | -0.5733714 | 0.566393249 | NA | 3.429096 | 2.844638 | 3.179385 | 3.182444 | 3.163779 | 3.166090 | 12 | 12 | 6 | 7 |
| 442523 | NR_003551 | DPY19L2P4 | DPY19L2 pseudogene 4 | transcribed_ps | . | 22176 | . | . | . | 0.622277 | 0.528485 | 0.854401 | 0.359296 | 1.448064 | 1.221617 | 1.917251 | 0.821488 | 1.039193 | 37.439986 | 0.558239811 | 0.09935443 | 0.920856863 | 0.998865404 | 5.195961 | 5.256422 | 5.225580 | 5.199135 | 5.276406 | 5.155800 | 46 | 36 | 47 | 21 |
| 442578 | NR_040582 | STAG3L3 | stromal antigen 3-like 3 (pseudogene) | transcribed_ps | . | 33845 | . | . | . | 0.404111 | 0.480909 | 1.157572 | 0.442684 | 0.940382 | 1.111643 | 2.597558 | 1.012144 | 1.763792 | 42.304471 | 0.548101088 | 1.493666974 | 0.135262685 | 0.892805049 | 4.934350 | 5.755755 | 5.307797 | 5.301702 | 5.494405 | 5.318762 | 32 | 36 | 68 | 28 |
| 442582 | NR_040584 | STAG3L2 | stromal antigen 3-like 2 (pseudogene) | transcribed_ps | . | 33886 | . | . | . | 6.118652 | 6.567220 | 8.148101 | 6.012027 | 14.238355 | 15.180423 | 18.284103 | 13.745786 | 1.102240 | 392.135956 | 0.202947407 | 0.691992588 | 0.488941983 | 0.998865404 | 8.542782 | 8.684175 | 8.562672 | 8.599361 | 8.727211 | 8.542013 | 416 | 415 | 413 | 326 |
| 442673 | gene-TUBG1P | TUBG1P | tubulin gamma 1 pseudogene | pseudogene | . | 12418 | . | . | . | 0.035004 | 0.078808 | 0.049006 | 0.085263 | 0.081456 | 0.182169 | 0.109968 | 0.194943 | 1.069135 | 3.738910 | 1.550039158 | 0.062220578 | 0.950387176 | NA | 1.855800 | 1.947964 | 1.894546 | 1.904179 | 1.898695 | 1.902366 | 3 | 5 | 3 | 4 |
| 442775 | gene-RAC1P2 | RAC1P2 | Rac family small GTPase 1 pseudogene 2 | pseudogene | . | 31414 | . | . | . | 1.686614 | 1.594835 | 1.732545 | 1.865517 | 3.924819 | 3.686534 | 3.887781 | 4.265283 | 1.075543 | 50.658128 | 0.438185511 | 0.239774748 | 0.810504884 | 0.998865404 | 5.609306 | 5.714230 | 5.660635 | 5.648924 | 5.661510 | 5.675246 | 58 | 51 | 44 | 51 |
| 442777 | gene-TECRP1 | TECRP1 | trans-2,3-enoyl-CoA reductase pseudogene 1 | pseudogene | . | 44050 | . | . | . | 7.342976 | 7.999424 | 7.705109 | 8.123418 | 17.087408 | 18.491028 | 17.290041 | 18.573229 | 1.017297 | 134.901220 | 0.280074609 | 0.088334939 | 0.92961047 | 0.998865404 | 7.063617 | 7.087801 | 7.057539 | 7.086605 | 7.071369 | 7.085442 | 148 | 150 | 116 | 130 |
| 444882 | NM_00100292 | IGFL4 | IGF like family member 4 | protein_coding | NP_001002923.1 | 32931 | 610547 | . | . | 0.301527 | 0.266184 | 0.152445 | 0.276344 | 0.701666 | 0.615297 | 0.342081 | 0.631827 | -1.239846 | 24.566379 | 0.642517848 | -0.48272658 | 0.62928989 | NA | 4.766397 | 4.453945 | 4.649205 | 4.615120 | 4.586671 | 4.633840 | 36 | 25 | 14 | 26 |
| 445328 | NM_00100370 | ARHGEF35 | Rho guanine nucleotide exchange factor 35 | protein_coding | NP_001003702.2;NP_001355247.1 | 33846 | . | . | . | 0.686705 | 0.697050 | 0.700561 | 0.687416 | 1.597990 | 1.611263 | 1.572038 | 1.571696 | -1.016174 | 56.935189 | 0.414151539 | -0.05588971 | 0.955429672 | 0.998865404 | 5.842669 | 5.819737 | 5.832750 | 5.833246 | 5.833302 | 5.826124 | 66 | 62 | 50 | 52 |
| 445329 | NM_00101739 | SULT1A4 | sulfotransferase family 1A member 4 | protein_coding | NP_001017390.1 | 30004 | 615819 | . | . | 0.381999 | 0.237802 | 0.397011 | 0.578256 | 0.888927 | 0.549690 | 0.890881 | 1.322113 | 1.564396 | 17.119388 | 0.757565062 | 0.852211959 | 0.39409646 | NA | 3.738475 | 4.384843 | 4.084784 | 4.056638 | 4.088053 | 4.118491 | 19 | 11 | 15 | 23 |
| 445571 | NM_00129182 | CBWD3 | COBW domain containing 3 | protein_coding | NP_001278750.1;NP_001365042.1;NP_001365043.1;NP_001365044.1;NI | 18519 | 611080 | . | . | 8.940268 | 8.541902 | 10.608307 | 9.272045 | 20.804373 | 19.744994 | 23.804735 | 21.199436 | 1.139027 | 491.792308 | 0.173776928 | 1.080708509 | 0.279826797 | 0.998865404 | 8.844572 | 9.033085 | 8.908369 | 8.885928 | 9.026836 | 8.929901 | 539 | 487 | 490 | 454 |
| 445577 | NR_166069 | C9orf129 | chromosome 9 open reading frame 129 | transcribed_ps | . | 31116 | . | . | . | 0.150649 | 0.181769 | 0.123187 | 0.231558 | 0.350567 | 0.420166 | 0.276429 | 0.529429 | 1.060083 | 35.141624 | 0.548504268 | 0.153466469 | 0.878030431 | 0.998865404 | 5.094525 | 5.174582 | 5.115375 | 5.138995 | 5.093009 | 5.176095 | 36 | 40 | 22 | 44 |
| 445582 | NM_00108353 | POTEE | POTE ankyrin domain family member E | protein_coding | NP_001077007.1;XP_011509518.1;XP_016859648.1;XP_016859650.1 | 33895 | 608914 | . | . | 0.048755 | 0.041133 | 0.037104 | 0.066697 | 0.113455 | 0.095081 | 0.083260 | 0.152495 | 1.144771 | 7.196156 | 1.132244859 | 0.172276391 | 0.863220244 | NA | 2.747897 | 2.940155 | 2.844716 | 2.842605 | 2.839564 | 2.856514 | 8 | 7 | 5 | 9 |
| 445815 | NM_00100406 | PALM2AKAP2 | PALM2 and AKAP2 fusion | protein_coding | NP_001004065.2;NP_001032370.1;NP_001130034.1;NP_001185585.1;NI | 33529 | 604582 | . | . | 10.730962 | 11.513899 | 10.452722 | 11.139214 | 24.971389 | 26.614892 | 23.455604 | 25.468486 | -1.042007 | 2382.082088 | 0.117185738 | -0.50658756 | 0.61244424 | 0.998865404 | 11.247471 | 11.187929 | 11.211265 | 11.256734 | 11.179665 | 11.221987 | 2721 | 2697 | 1975 | 2245 |
| 446207 | gene-ACTBP10 | ACTBP10 | ACTB pseudogene 10 | pseudogene | . | 31819 | . | . | . | 0.007681 | 0.071880 | 0.013948 | 0.100927 | 0.017873 | 0.166154 | 0.031300 | 0.230759 | 1.385935 | 3.263036 | 1.764888514 | 0.266792733 | 0.78962875 | NA | 1.460000 | 1.916472 | 1.685462 | 1.703463 | 1.687568 | 1.711171 | 1 | 5 | 1 | 6 |
| 448831 | NM_00100521 | FRG2 | FSHD region gene 2 | protein_coding | NP_001005217.1;NP_001273749.1 | 19136 | 609032 | . | . | 0.712067 | 0.686920 | 0.798824 | 0.954863 | 1.657011 | 1.587846 | 1.792539 | 2.183180 | 1.220230 | 52.216766 | 0.433514388 | 0.662384693 | 0.507724711 | 0.998865404 | 5.556147 | 5.842545 | 5.685629 | 5.680416 | 5.704964 | 5.736860 | 55 | 50 | 46 | 58 |
| 450238 | gene-PHF5DP | PHF5DP | PHD finger protein 5D pseudogene | pseudogene | . | 23268 | . | . | . | 0.435870 | 0.381600 | 0.725697 | 0.393529 | 1.014288 | 0.882086 | 1.628443 | 0.899757 | 1.386680 | 14.432604 | 0.821760427 | 0.573932702 | 0.566013342 | NA | 3.593806 | 4.069653 | 3.833108 | 3.824606 | 3.870261 | 3.827784 | 15 | 12 | 19 | 11 |
| 474170 | NM_00100660 | LRRC37A2 | leucine rich repeat containing 37 member A2 | protein_coding | NP_001006608.2;XP_011523143.1;XP_011523144.1;XP_011523145.1;XP | 32404 | 616556 | . | . | 0.693080 | 0.664447 | 0.754905 | 0.544261 | 1.612827 | 1.535900 | 1.693984 | 1.244387 | -1.026176 | 126.274624 | 0.301075326 | -0.12381545 | 0.901461396 | 0.998865404 | 6.998130 | 6.962492 | 6.976663 | 6.987422 | 7.027228 | 6.915674 | 145 | 140 | 126 | 98 |
| 474338 | NR_002190 | SUMO1P3 | SUMO1 pseudogene 3 | transcribed_ps | . | 33150 | . | . | . | 0.191390 | 0.088621 | 0.039947 | 0.062962 | 0.445373 | 0.204852 | 0.089641 | 0.143955 | -2.094732 | 4.255727 | 1.500822087 | -0.71078735 | 0.477216021 | NA | 2.526439 | 1.459419 | 2.119918 | 2.100722 | 2.094682 | 2.098545 | 9 | 4 | 2 | 3 |
| 474343 | NM_00100668 | SPIN2B | spindlin family member 2B | protein_coding | NP_001006682.1;NP_001006683.1;NP_001006684.1;NP_001269390.1;NI | 33147 | 300517 | . | . | 2.675409 | 2.271635 | 2.516223 | 2.411619 | 6.225789 | 5.250985 | 5.646329 | 5.513879 | 1.043073 | 100.938427 | 0.323668731 | 0.187968728 | 0.850901166 | 0.998865404 | 6.625875 | 6.688117 | 6.679266 | 6.618285 | 6.676811 | 6.647369 | 125 | 96 | 94 | 92 |
| 474383 | NM_00100752 | F8A2 | coagulation factor VIII associated 2 | protein_coding | NP_001007524.1 | 31849 | . | . | . | 2.041292 | 2.400983 | 2.115654 | 2.780262 | 4.750171 | 5.549980 | 4.747466 | 6.356739 | 1.085915 | 81.503216 | 0.360431512 | 0.329914866 | 0.74146429 | 0.998865404 | 6.289230 | 6.405979 | 6.314047 | 6.355394 | 6.320668 | 6.391699 | 83 | 91 | 64 | 90 |
| 474384 | NM_00100752 | F8A3 | coagulation factor VIII associated 3 | protein_coding | NP_001007525.1 | 31850 | . | . | . | 2.015228 | 1.961176 | 1.913725 | 2.198534 | 4.689518 | 4.533347 | 4.294343 | 5.026687 | 1.025441 | 107.972961 | 0.310197651 | 0.11684291 | 0.90698454 | 0.998865404 | 6.736587 | 6.772245 | 6.755935 | 6.744355 | 6.737771 | 6.778198 | 125 | 113 | 89 | 109 |
| 492303 | NR_002830 | GEMIN8P4 | gem nuclear organelle associated protein 8 pseudogene 4 | transcribed_ps | . | 37979 | . | . | . | 3.529179 | 4.088948 | 3.288992 | 4.848752 | 8.212545 | 9.451786 | 7.380404 | 11.086095 | 1.057097 | 209.208732 | 0.257083663 | 0.311603909 | 0.755341563 | 0.998865404 | 7.669258 | 7.747285 | 7.658541 | 7.723162 | 7.626779 | 7.799697 | 218 | 235 | 152 | 239 |
| 492311 | NM_00100718 | IGIP | IgA inducing protein | protein_coding | NP_001007190.1 | 33847 | . | . | . | 0.935041 | 0.852455 | 0.971937 | 0.764127 | 2.175879 | 1.970489 | 2.180997 | 1.747087 | -1.046136 | 95.482916 | 0.333728915 | -0.19498033 | 0.84540835 | 0.998865404 | 6.608444 | 6.545204 | 6.597830 | 6.567027 | 6.603152 | 6.534176 | 118 | 100 | 91 | 77 |
| 493753 | NM_00100821 | COA5 | cytochrome c oxidase assembly factor 5 | protein_coding | NP_001008216.1 | 33848 | 613920 | . | . | 10.131248 | 9.496058 | 10.432539 | 9.191043 | 23.575830 | 21.950563 | 23.410315 | 21.014227 | -1.014005 | 541.785183 | 0.168729123 | -0.11891712 | 0.905341016 | 0.998865404 | 9.091197 | 9.071892 | 9.108847 | 9.061490 | 9.119334 | 9.031544 | 651 | 567 | 500 | 470 |
| 493754 | NR_111972,NF | GS1-124K5.11 | RAB guanine nucleotide exchange factor 1 pseudogene | transcribed_ps | . | . | . | . | . | 3.813980 | 4.431402 | 5.256577 | 5.033774 | 8.875289 | 10.243385 | 11.795606 | 11.509130 | 1.210904 | 279.585532 | 0.209653128 | 1.316863863 | 0.187884261 | 0.960264078 | 7.982705 | 8.258432 | 8.037374 | 8.104470 | 8.180734 | 8.162009 | 272 | 291 | 271 | 282 |
| 493812 | NR_026790 | HCG11 | HLA complex group 11 | lncRNA | . | 17707 | . | . | . | 0.090113 | 0.148867 | 0.180324 | 0.088440 | 0.209697 | 0.344112 | 0.404642 | 0.202208 | 1.109944 | 22.893821 | 0.680924079 | 0.221004342 | 0.825089052 | NA | 4.438825 | 4.590940 | 4.477844 | 4.522775 | 4.542844 | 4.477054 | 19 | 29 | 28 | 15 |
| 493856 | NM_00100838 | CISD2 | CDGSH iron sulfur domain 2 | protein_coding | NP_001008389.1 | 24212 | 611507 | . | . | 5.098024 | 4.123620 | 5.350265 | 4.037995 | 11.863310 | 9.531932 | 12.005838 | 9.232396 | 1.003558 | 852.827618 | 0.177334547 | 0.028894412 | 0.976948803 | 0.998865404 | 9.733028 | 9.739186 | 9.806663 | 9.648030 | 9.831491 | 9.624341 | 1088 | 817 | 851 | 685 |
| 493861 | NM_00100839 | EID3 | EP300 interacting inhibitor of differentiation 3 | protein_coding | NP_001008395.1 | 32961 | 612986 | . | . | 0.941770 | 0.978788 | 1.686148 | 0.533543 | 2.191536 | 2.262513 | 3.783666 | 1.219883 | 1.129073 | 47.820664 | 0.537948503 | 0.325568504 | 0.744750845 | 0.998865404 | 5.487053 | 5.666496 | 5.544740 | 5.549223 | 5.659930 | 5.467957 | 51 | 49 | 67 | 23 |
| 493869 | NM_00100839 | GPX8 | glutathione peroxidase 8 (putative) | protein_coding | NP_001008398.2;NP_001293126.1;NP_001293127.1;NP_001293130.1;XP | 33100 | 617172 | . | . | 26.716171 | 27.397963 | 22.174282 | 20.374222 | 62.169629 | 63.331619 | 49.758448 | 46.583238 | -1.285270 | 2795.219444 | 0.116312582 | -3.112917 | 0.001852481 | 0.106540819 | 11.618382 | 11.256472 | 11.535385 | 11.548349 | 11.374902 | 11.299206 | 3592 | 3419 | 2229 | 2184 |
| 493900 | NR_073431 | TMEM9B-AS1 | TMEM9B antisense RNA 1 | lncRNA | . | 19230 | . | . | . | 1.042413 | 0.730984 | 0.530778 | 0.996159 | 2.425738 | 1.689703 | 1.191051 | 2.277599 | -1.141509 | 27.574829 | 0.614930367 | -0.31050972 | 0.756173366 | 0.998865404 | 4.878555 | 4.685555 | 4.815487 | 4.771236 | 4.749002 | 4.807611 | 40 | 26 | 16 | 31 |
| 493911 | NM_00100848 | PHOSPHO2 | phosphatase, orphan 2 | protein_coding | NP_001008489.1;NP_001186214.1;NP_001186215.1;NP_001186216.1;NI | 28316 | . | . | . | 2.320149 | 2.314638 | 1.696355 | 2.080006 | 5.399084 | 5.350388 | 3.806572 | 4.755687 | -1.240562 | 80.970994 | 0.355740341 | -0.87421592 | 0.382000679 | 0.998865404 | 6.487053 | 6.174745 | 6.365035 | 6.371960 | 6.289787 | 6.333311 | 102 | 98 | 57 | 74 |
| 494115 | NM_00116253 | RBMXL1 | RBMX like 1 | protein_coding | NP_001156008.1;NP_062556.2 | 25073 | . | . | . | 4.565001 | 4.563670 | 4.289437 | 3.811642 | 10.622945 | 10.549128 | 9.625374 | 8.714867 | -1.136080 | 652.398688 | 0.155941543 | -1.18034378 | 0.23786351 | 0.992733415 | 9.438506 | 9.254874 | 9.395402 | 9.386515 | 9.336928 | 9.272372 | 804 | 744 | 557 | 545 |
| 494127 | NR_036691 | NFYCP2 | NFYC pseudogene 2 | transcribed_ps | . | 54924 | . | . | . | 0.245841 | 0.368292 | 0.049736 | 0.212799 | 0.572083 | 0.851323 | 0.111605 | 0.486541 | -2.182108 | 9.518980 | 1.06242796 | -1.05957558 | 0.28933773 | NA | 3.710171 | 2.572884 | 3.262987 | 3.286638 | 3.227575 | 3.259219 | 12 | 17 | 2 | 9 |
| 494143 | NM_00100870 | CHAC2 | ChaC glutathione specific gamma-glutamylcyclotransferase 2 | protein_coding | NP_001008708.1;NP_001333056.1 | 32363 | 617446 | . | . | 5.480290 | 4.899233 | 5.464295 | 5.052959 | 12.752860 | 11.324798 | 12.261720 | 11.552990 | 1.000651 | 214.918478 | 0.232632452 | 0.004034431 | 0.996780999 | 0.999862634 | 7.746723 | 7.748568 | 7.774513 | 7.718373 | 7.766720 | 7.727341 | 262 | 218 | 195 | 193 |
| 494470 | NM_00125675 | RNF165 | ring finger protein 165 | protein_coding | NP_001243687.1;NP_689683.2;XP_011524318.1;XP_016881277.1;XP_02 | 31696 | . | . | . | 0.191040 | 0.214241 | 0.192552 | 0.158864 | 0.444558 | 0.495229 | 0.432081 | 0.363225 | -1.166076 | 35.339335 | 0.521242364 | -0.4252569 | 0.670649407 | 0.998865404 | 5.249661 | 5.028259 | 5.163771 | 5.148446 | 5.133211 | 5.136380 | 46 | 39 | 28 | 31 |
| 494513 | NM_00104270 | PJVK | pejvakin | protein_coding | NP_001036167.1;NP_001340704.1;NP_001340705.1;NP_001340706.1;NI | 29502 | 610219 | . | . | 2.309688 | 2.181361 | 2.055153 | 1.659034 | 5.374741 | 5.042314 | 4.611701 | 3.793185 | -1.021095 | 137.700141 | 0.288560526 | -0.10436799 | 0.916877322 | 0.998865404 | 7.119654 | 7.090976 | 7.099228 | 7.114051 | 7.151032 | 7.044094 | 157 | 153 | 136 | 109 |
| 494514 | NM_00101271 | TYMSOS | TYMS opposite strand | protein_coding | NP_001012734.2 | 29553 | . | . | . | 1.869076 | 2.204660 | 2.131856 | 1.339033 | 4.349416 | 5.096169 | 4.783823 | 3.061539 | -1.194183 | 47.116513 | 0.47293817 | -0.54134798 | 0.588267755 | 0.998865404 | 5.679488 | 5.425686 | 5.554554 | 5.587226 | 5.576169 | 5.502088 | 54 | 60 | 46 | 31 |
| 494540 | gene-TBCAP1 | TBCAP1 | tubulin folding cofactor A pseudogene 1 | pseudogene | . | 42630 | . | . | . | 3.060226 | 3.072163 | 2.388324 | 2.261754 | 7.121272 | 7.101443 | 5.359329 | 5.171231 | -1.335074 | 49.786911 | 0.443213895 | -0.94067487 | 0.346871502 | 0.998865404 | 5.830973 | 5.414458 | 5.667473 | 5.664258 | 5.617245 | 5.609000 | 66 | 61 | 38 | 39 |
| 497048 | NR_122107 | KU-MEL-3 | uncharacterized LOC497048 | lncRNA | . | . | . | . | . | 2.076339 | 1.772167 | 3.051949 | 4.168748 | 4.831726 | 4.096444 | 6.848485 | 9.531347 | 1.846541 | 103.083234 | 0.330976417 | 2.673379223 | 0.007509131 | 0.2720434 | 6.178768 | 7.063112 | 6.582494 | 6.544295 | 6.694717 | 6.805249 | 90 | 72 | 99 | 144 |
| 497258 | NR_002832,NF | BDNF-AS | BDNF antisense RNA | lncRNA | . | 20608 | 611468 | . | . | 0.024426 | 0.027225 | 0.004435 | 0.031849 | 0.056841 | 0.062933 | 0.009952 | 0.072818 | -1.231830 | 2.433884 | 1.916745144 | -0.15693464 | 0.87529636 | NA | 1.429096 | 1.121007 | 1.290581 | 1.291393 | 1.284044 | 1.293236 | 3 | 3 | 1 | 3 |
| 497661 | NM_00103500 | C18orf32 | chromosome 18 open reading frame 32 | protein_coding | NP_001030177.1;NP_001186275.1 | 31690 | . | . | . | 1.866880 | 1.401656 | 1.870713 | 1.395318 | 4.344305 | 3.239989 | 4.197826 | 3.190230 | -1.010880 | 280.680914 | 0.237258495 | -0.06580079 | 0.947536426 | 0.998865404 | 8.139587 | 8.125955 | 8.204949 | 8.049836 | 8.198906 | 8.044860 | 372 | 260 | 279 | 223 |
| 503538 | NR_015380 | A1BG-AS1 | A1BG antisense RNA 1 | lncRNA | . | 37133 | . | . | . | 4.579208 | 5.387843 | 5.496779 | 4.179889 | 10.656004 | 12.454240 | 12.334613 | 9.556819 | -1.043438 | 326.188500 | 0.214880835 | -0.28548593 | 0.775271885 | 0.998865404 | 8.379549 | 8.318939 | 8.313384 | 8.400404 | 8.408585 | 8.254215 | 354 | 387 | 317 | 257 |
| 503569 | NR_033932 | RGMB-AS1 | RGMB antisense RNA 1 | lncRNA | . | 48666 | . | . | . | 0.285310 | 0.376281 | 0.304288 | 0.374732 | 0.663929 | 0.869789 | 0.682813 | 0.856781 | -1.002728 | 21.554332 | 0.66475188 | -0.00591198 | 0.995282949 | NA | 4.432996 | 4.426810 | 4.418602 | 4.440137 | 4.421134 | 4.437195 | 22 | 26 | 17 | 22 |
| 503637 | NR_122113 | DUXAP8 | double homeobox A pseudogene 8 | transcribed_ps | . | 32187 | . | . | . | 6.240156 | 6.224830 | 7.944502 | 7.652005 | 14.521099 | 14.388972 | 17.827234 | 17.495401 | 1.234059 | 460.700005 | 0.171880502 | 1.765242689 | 0.077522987 | 0.767975096 | 8.687891 | 8.991529 | 8.778974 | 8.773939 | 8.921179 | 8.891651 | 477 | 443 | 453 | 466 |
| 503638 | NR_122111,NF | DUXAP9 | double homeobox A pseudogene 9 | transcribed_ps | . | 32188 | . | . | . | 14.623268 | 16.072323 | 19.165818 | 15.784801 | 34.028949 | 37.151895 | 43.007541 | 36.090074 | 1.152718 | 928.268387 | 0.152876187 | 1.341213761 | 0.17985106 | 0.952541638 | 9.752044 | 9.957447 | 9.774535 | 9.829132 | 9.975418 | 9.829893 | 955 | 966 | 950 | 840 |
| 503639 | NR_110526 | DUXAP10 | double homeobox A pseudogene 10 | transcribed_ps | . | 32189 | . | . | . | 6.202594 | 6.803617 | 7.736740 | 6.656358 | 14.433691 | 15.726865 | 17.361021 | 15.218972 | 1.092044 | 511.922240 | 0.173768389 | 0.73103816 | 0.464755845 | 0.998865404 | 8.934676 | 9.062074 | 8.942282 | 8.995092 | 9.075498 | 8.972487 | 540 | 550 | 502 | 461 |
| 503693 | NR_024061 | LOH12CR2 | loss of heterozygosity on chromosome 12, region 2 | lncRNA | . | 26524 | . | . | . | 0.510548 | 0.634881 | 0.562372 | 0.688070 | 1.188065 | 1.467556 | 1.261947 | 1.573190 | 1.089381 | 29.108365 | 0.574938083 | 0.214821698 | 0.829906335 | 0.998865404 | 4.801350 | 4.922818 | 4.844691 | 4.867355 | 4.855332 | 4.876983 | 29 | 33 | 24 | 31 |
| 541465 | NM_00101743 | CT45A6 | cancer/testis antigen family 45 member A6 | protein_coding | NP_001017438.2;XP_005262482.2;XP_011529653.1 | 33271 | 300797 | . | . | 1.235665 | 1.230401 | 0.899469 | 0.910746 | 2.875446 | 2.844127 | 2.018383 | 2.082313 | -1.380427 | 33.606893 | 0.535290304 | -0.86890256 | 0.384900427 | 0.998865404 | 5.284382 | 4.819737 | 5.099080 | 5.091041 | 5.054709 | 5.050290 | 46 | 41 | 25 | 26 |
| 541466 | NM_00101741 | CT45A1 | cancer/testis antigen family 45 member A1 | protein_coding | NP_001017417.1;XP_005278198.1;XP_011529654.1 | 33267 | 300648 | . | . | 0.369656 | 0.571840 | 0.227127 | 0.531934 | 0.860207 | 1.321831 | 0.509668 | 1.216203 | -1.330014 | 13.419061 | 0.857696936 | -0.47970498 | 0.631437188 | NA | 3.939842 | 3.522505 | 3.743033 | 3.771261 | 3.725909 | 3.755240 | 14 | 20 | 7 | 14 |
| 541468 | NM_00101361 | LURAP1 | leucine rich adaptor protein 1 | protein_coding | NP_001013633.1 | 32327 | 616129 | . | . | 3.620578 | 3.356040 | 2.984138 | 3.478377 | 8.425234 | 7.757636 | 6.696319 | 7.952895 | -1.093339 | 194.136011 | 0.243088737 | -0.52960158 | 0.596388196 | 0.998865404 | 7.645962 | 7.534872 | 7.639326 | 7.601815 | 7.550940 | 7.610339 | 243 | 210 | 150 | 186 |
| 541471 | NR_015395,NF | MIR4435-2HG | MIR4435-2 host gene | lncRNA | . | 35163 | 617144 | . | . | 59.868834 | 61.892531 | 50.932880 | 64.535884 | 139.317228 | 143.067356 | 114.291912 | 147.553639 | -1.001891 | 2062.660367 | 0.132018383 | -0.02064882 | 0.983525794 | 0.998865404 | 11.011827 | 11.008753 | 11.006078 | 11.013909 | 10.928358 | 11.082627 | 2366 | 2239 | 1618 | 2119 |
| 541565 | NM_00101384 | C8orf58 | chromosome 8 open reading frame 58 | protein_coding | NP_001013864.1;NP_001185756.1;NP_775957.2 | 32233 | . | . | . | 9.295779 | 9.282155 | 10.673041 | 9.882033 | 21.631657 | 21.456119 | 23.949994 | 22.594096 | 1.090447 | 619.434444 | 0.157835508 | 0.791456458 | 0.428677675 | 0.998865404 | 9.210796 | 9.336099 | 9.248667 | 9.239670 | 9.331644 | 9.271835 | 687 | 635 | 587 | 579 |
| 541578 | NM_00101384 | EOLA2 | endothelium and lymphocyte associated ASCH domain 2 | protein_coding | NP_001013867.1;XP_005274755.1;XP_005274756.1;XP_005274757.1;XP | 17402 | . | . | . | 6.538221 | 6.820468 | 6.503794 | 7.738426 | 15.214706 | 15.765818 | 14.594326 | 17.692993 | -1.044564 | 395.009335 | 0.192424925 | -0.32688249 | 0.743756765 | 0.998865404 | 8.657279 | 8.593502 | 8.646788 | 8.628317 | 8.544013 | 8.675100 | 472 | 429 | 298 | 403 |
| 542767 | NM_00101444 | PCOTH | Pro-X-Gly collagen triple helix like repeat containing | protein_coding | NP_001014442.2;NP_001129288.1;NP_001335041.1;NP_001335042.1;NI | 39839 | 617122 | . | . | 4.124170 | 5.225421 | 5.416400 | 5.084317 | 9.597116 | 12.078793 | 12.154244 | 11.624690 | 1.115381 | 126.147002 | 0.296968238 | 0.530482152 | 0.59577768 | 0.998865404 | 6.898396 | 7.055266 | 6.910535 | 6.998329 | 7.007117 | 6.980384 | 120 | 145 | 120 | 120 |
| 548593 | NM_00101499 | SLX1A | SLX1 homolog A, structure-specific endonuclease subunit | protein_coding | NP_001014999.1;NP_001015000.1 | 20922 | 615822 | . | . | 14.774766 | 16.476397 | 13.394127 | 16.951011 | 34.381490 | 38.085928 | 30.056032 | 38.756468 | -1.012478 | 531.403217 | 0.17664714 | -0.10127562 | 0.91933167 | 0.998865404 | 9.063067 | 9.044198 | 9.038379 | 9.074087 | 8.974655 | 9.116819 | 599 | 593 | 408 | 550 |
| 548596 | NM_00101500 | CKMT1A | creatine kinase, mitochondrial 1A | protein_coding | NP_001015001.1;NP_001308855.1;NP_001308856.1;NP_001308857.1;NI | 31736 | 613415 | . | . | 1.201832 | 1.016191 | 1.568166 | 1.541545 | 2.796712 | 2.348972 | 3.518920 | 3.524562 | 1.383530 | 67.167574 | 0.386064866 | 1.213148396 | 0.22507309 | 0.982008277 | 5.816198 | 6.285233 | 6.038994 | 6.006783 | 6.094494 | 6.101249 | 70 | 56 | 67 | 74 |
| 548644 | NM_00109761 | POLR2J3 | RNA polymerase II subunit J3 | protein_coding | NP_001091084.2 | 33853 | . | . | . | 12.592464 | 13.528637 | 13.094825 | 16.115816 | 29.303183 | 31.272049 | 29.384409 | 36.846897 | 1.102639 | 724.036284 | 0.160975678 | 0.875662704 | 0.381213426 | 0.998865404 | 9.428126 | 9.568306 | 9.441294 | 9.485022 | 9.459325 | 9.597653 | 768 | 767 | 595 | 782 |
| 548645 | NM_00101588 | DNAJC25 | DnaJ heat shock protein family (Hsp40) member C25 | protein_coding | NP_001015882.2 | 34187 | . | . | . | 4.541818 | 4.401948 | 5.052091 | 4.668701 | 10.568995 | 10.175300 | 11.336747 | 10.674430 | 1.079633 | 347.518017 | 0.192273719 | 0.574913945 | 0.565349498 | 0.998865404 | 8.384274 | 8.495472 | 8.433829 | 8.401046 | 8.488208 | 8.433216 | 396 | 350 | 329 | 322 |
| 550112 | NR_015439 | UBA6-AS1 | UBA6 antisense RNA 1 (head to head) | lncRNA | . | 49083 | . | . | . | 2.040132 | 2.238757 | 2.207658 | 2.299923 | 4.747470 | 5.174986 | 4.953919 | 5.258501 | 1.041906 | 157.177135 | 0.262227168 | 0.225853449 | 0.821315404 | 0.998865404 | 7.266608 | 7.325290 | 7.270564 | 7.303640 | 7.296495 | 7.310616 | 170 | 173 | 137 | 153 |
| 550631 | NM_00101743 | CCDC157 | coiled-coil domain containing 157 | protein_coding | NP_001017437.2;NP_001305263.2;NP_001305264.1;XP_006724341.1;XP | 33854 | . | . | . | 2.234548 | 2.318401 | 2.056526 | 2.511600 | 5.199885 | 5.359087 | 4.614784 | 5.742476 | 1.001372 | 171.019434 | 0.256957458 | 0.007695251 | 0.993860139 | 0.999862634 | 7.417140 | 7.418892 | 7.443207 | 7.390759 | 7.390423 | 7.444613 | 209 | 173 | 139 | 171 |
| 550643 | NM_00134812 | NBDY | negative regulator of P-body association | protein_coding | NP_001335058.1;XP_024308162.1 | 50713 | 300992 | . | . | 3.651000 | 3.789876 | 3.595021 | 4.838784 | 8.496027 | 8.760470 | 8.067124 | 11.063306 | 1.107829 | 293.878549 | 0.21810708 | 0.677350793 | 0.498183411 | 0.998865404 | 8.124000 | 8.270438 | 8.160894 | 8.174954 | 8.141732 | 8.297252 | 317 | 305 | 230 | 331 |
| 550644 | gene-SMC3P1 | SMC3P1 | SMC3 pseudogene 1 | pseudogene | . | 54642 | . | . | . | 0.401114 | 0.207632 | 0.101547 | 0.260801 | 0.933409 | 0.479950 | 0.227869 | 0.596290 | -1.627488 | 8.589739 | 1.087945001 | -0.64584751 | 0.518378114 | NA | 3.413393 | 2.705970 | 3.135358 | 3.105818 | 3.088887 | 3.115607 | 16 | 8 | 3 | 9 |

| | | | | | | | | | | |
|---|---|---|---|---|---|---|---|---|---|---|
| 552889 | NM_00113626 | ATXN7L3B | ataxin 7 like 3B | protein_coding | NP_001129734.1 | 37931 | 615579 | . | . | [illegible] |
| 552891 | NM_004125 | DNAJC25-GNG | DNAJC25-GNG10 readthrough | protein_coding | NP_004116.2 | 37501 | . | . | . | [illegible] |
| 552900 | NM_00103182 | BOLA2 | bolA family member 2 | protein_coding | NP_001026997.1;NP_001307508.1 | 29488 | 613182 | . | . | [illegible] |
| 553103 | NR_110997 | MIR3936HG | MIR3936 host gene | lncRNA | . | 40538 | . | . | . | [illegible] |
| 553115 | NM_00135965 | PEF1 | penta-EF-hand domain containing 1 | protein_coding | NP_001346580.1;NP_036524.1;XP_011540047.1;XP_011540048.1;XP_01 | 30009 | 610033 | . | . | [illegible] |
| 553939 | gene-LOC5539 | LOC553939 | trimethyllysine hydroxylase, epsilon pseudogene | pseudogene | . | . | . | . | . | [illegible] |
| 554203 | NR_024582 | JPX | JPX transcript, XIST activator | lncRNA | . | 37191 | 300832 | . | . | [illegible] |
| 554206 | NR_038379 | LOC554206 | leucine carboxyl methyltransferase 1 pseudogene | transcribed_ps | . | . | . | . | . | [illegible] |
| 554224 | gene-PPIP5K1P | PPIP5K1P1 | diphosphoinositol pentakisphosphate kinase 1 pseudogene 1 | pseudogene | . | 32373 | . | . | . | [illegible] |
| 554225 | NR_146078 | STRCP1 | stereocilin pseudogene 1 | transcribed_ps | . | 33915 | . | . | . | [illegible] |
| 554236 | NR_002833 | DPY19L2P1 | DPY19L2 pseudogene 1 | transcribed_ps | . | 22305 | . | . | . | [illegible] |
| 554249 | NR_126049 | LOC554249 | uncharacterized LOC554249 | lncRNA | . | . | . | . | . | [illegible] |
| 554251 | NM_00102468 | FBXO48 | F-box protein 48 | protein_coding | NP_001019851.1;XP_005264464.1;XP_016859926.1 | 33857 | . | . | . | [illegible] |
| 554282 | NM_00128738 | FAM72C | family with sequence similarity 72 member C | protein_coding | NP_001274314.1;NP_001332996.1;NP_001332997.1;NP_001332999.1;NI | 30602 | 616853 | . | . | [illegible] |
| 554313 | NM_00103407 | H4C15 | H4 clustered histone 15 | protein_coding | NP_001029249.1;XP_024307476.1;XP_024307477.1;XP_024307478.1;XP | 29607 | . | . | . | [illegible] |
| 573971 | gene-DHFRP1 | DHFRP1 | dihydrofolate reductase pseudogene 1 | pseudogene | . | 2862 | . | . | . | [illegible] |
| 574029 | NR_002834 | DUSP5P1 | dual specificity phosphatase 5 pseudogene 1 | transcribed_ps | . | 32020 | . | . | . | [illegible] |
| 574036 | NR_024337 | SERTAD4-AS1 | SERTAD4 antisense RNA 1 | lncRNA | . | 32019 | . | . | . | [illegible] |
| 574040 | NR_002325 | SNORA6 | small nucleolar RNA, H/ACA box 6 | snoRNA | . | 32591 | . | . | . | [illegible] |
| 574042 | NR_002327 | SNORA10 | small nucleolar RNA, H/ACA box 10 | snoRNA | . | 32598 | . | . | . | [illegible] |
| 574407 | NR_073154,NF | OBSCN-AS1 | OBSCN antisense RNA 1 | lncRNA | . | 32047 | . | . | . | [illegible] |
| 574414 | NM_00119557 | PRR9 | proline rich 9 | protein_coding | NP_001182500.1 | 32057 | . | . | . | [illegible] |
| 574432 | NR_103539,NF | IBA57-DT | IBA57 divergent transcript | lncRNA | . | 32062 | . | . | . | [illegible] |
| 594837 | NR_002434 | SNORD101 | small nucleolar RNA, C/D box 101 | snoRNA | . | 32764 | . | . | . | [illegible] |
| 594838 | NR_002435 | SNORD100 | small nucleolar RNA, C/D box 100 | snoRNA | . | 32763 | . | . | . | [illegible] |
| 594839 | NR_002436 | SNORA33 | small nucleolar RNA, H/ACA box 33 | snoRNA | . | 32623 | . | . | . | [illegible] |
| 594842 | NR_002835 | HAS2-AS1 | HAS2 antisense RNA 1 | antisense_RNA | . | 34340 | 614353 | . | . | [illegible] |
| 595097 | NR_002440 | SNORD16 | small nucleolar RNA, C/D box 16 | snoRNA | . | 32712 | . | . | . | [illegible] |
| 595098 | NR_002441 | SNORD18A | small nucleolar RNA, C/D box 18A | snoRNA | . | 32714 | . | . | . | [illegible] |
| 595100 | NR_002443 | SNORD18C | small nucleolar RNA, C/D box 18C | snoRNA | . | 32716 | . | . | . | [illegible] |
| 595101 | NR_002453 | SMG1P5 | SMG1 pseudogene 5 | transcribed_ps | . | 49862 | . | . | . | [illegible] |
| 595135 | NR_002836 | PGM5P2 | phosphoglucomutase 5 pseudogene 2 | transcribed_ps | . | 18965 | . | . | . | [illegible] |
| 606495 | NM_00103167 | CYB5RL | cytochrome b5 reductase like | protein_coding | NP_001026842.2;NP_001340282.1;NP_001340283.1;NP_001340285.1 | 32220 | . | . | . | [illegible] |
| 606500 | NR_002450 | SNORD68 | small nucleolar RNA, C/D box 68 | snoRNA | . | 32729 | . | . | . | [illegible] |
| 606553 | NM_00103183 | C8orf49 | chromosome 8 open reading frame 49 | protein_coding | NP_001027009.2 | 32200 | . | . | . | [illegible] |
| 606724 | NR_002454 | LOC606724 | coronin 1A pseudogene | transcribed_ps | . | . | . | . | . | [illegible] |
| 613037 | NM_00132189 | NPIPB13 | nuclear pore complex interacting protein family, member B13 | protein_coding | NP_001308821.1 | 41989 | . | . | . | [illegible] |
| 613126 | NR_122109,NF | CYP51A1-AS1 | CYP51A1 antisense RNA 1 | lncRNA | . | 50694 | . | . | . | [illegible] |
| 613211 | NM_00130269 | DEFB134 | defensin beta 134 | protein_coding | NP_001289624.1;XP_016869213.1 | 32399 | . | . | . | [illegible] |
| 619189 | NM_00125803 | SERINC4 | serine incorporator 4 | protein_coding | NP_001244960.1;NP_001244961.1 | 32237 | 614550 | . | . | [illegible] |
| 619207 | NR_149718 | SCART1 | scavenger receptor family member expressed on T cells 1 | transcribed_ps | . | 32411 | . | . | . | [illegible] |
| 619208 | NM_00103356 | FAM229B | family with sequence similarity 229 member B | protein_coding | NP_001028736.1;XP_016866663.1;XP_016866664.1 | 33858 | . | . | . | [illegible] |
| 619279 | NM_00103372 | ZNF704 | zinc finger protein 704 | protein_coding | NP_001028895.1;NP_001354712.1;XP_016869214.1 | 32291 | . | . | . | [illegible] |
| 619349 | gene-SETP3 | SETP3 | SET pseudogene 3 | pseudogene | . | 31115 | . | . | . | [illegible] |
| 619351 | NR_026765 | LINC00589 | long intergenic non-protein coding RNA 589 | lncRNA | . | 32299 | . | . | . | [illegible] |
| 619383 | NR_002569 | SCARNA9 | small Cajal body-specific RNA 9 | guide_RNA | . | 32566 | . | . | . | [illegible] |
| 619443 | gene-RPS26P7 | RPS26P7 | ribosomal protein S26 pseudogene 7 | pseudogene | . | 20077 | . | . | . | [illegible] |
| 619498 | NR_002579 | SNORD74 | small nucleolar RNA, C/D box 74 | snoRNA | . | 32734 | . | . | . | [illegible] |
| 619499 | NR_002575 | SNORA27 | small nucleolar RNA, H/ACA box 27 | snoRNA | . | 32617 | . | . | . | [illegible] |
| 619505 | NR_002576 | SNORA21 | small nucleolar RNA, H/ACA box 21 | snoRNA | . | 32611 | . | . | . | [illegible] |
| 619519 | gene-TWF1P1 | TWF1P1 | twinfilin 1 pseudogene 1 | pseudogene | . | 21736 | . | . | . | [illegible] |
| 619562 | NR_002580 | SNORA3A | small nucleolar RNA, H/ACA box 3A | snoRNA | . | 32586 | . | . | . | [illegible] |
| 619565 | NR_002585 | SNORA52 | small nucleolar RNA, H/ACA box 52 | snoRNA | . | 32645 | . | . | . | [illegible] |
| 619567 | NR_002587 | SNORD2 | small nucleolar RNA, C/D box 2 | snoRNA | . | 32678 | . | . | . | [illegible] |
| 619568 | NR_002588 | SNORA4 | small nucleolar RNA, H/ACA box 4 | snoRNA | . | 32587 | . | . | . | [illegible] |
| 619569 | NR_002590 | SNORA41 | small nucleolar RNA, H/ACA box 41 | snoRNA | . | 32634 | . | . | . | [illegible] |
| 619570 | NR_002591 | SNORD95 | small nucleolar RNA, C/D box 95 | snoRNA | . | 32757 | . | . | . | [illegible] |
| 619571 | NR_002592 | SNORD96A | small nucleolar RNA, C/D box 96A | snoRNA | . | 32758 | . | . | . | [illegible] |
| 641293 | gene-RPL21P1 | RPL21P19 | ribosomal protein L21 pseudogene 19 | pseudogene | . | 35748 | . | . | . | [illegible] |
| 641298 | NR_027154 | SMG1P1 | SMG1 pseudogene 1 | transcribed_ps | . | 44186 | . | . | . | [illegible] |
| 641339 | NM_00103989 | ZNF674 | zinc finger protein 674 | protein_coding | NP_001034980.1;NP_001139763.1;NP_001177346.1;XP_011542243.1;XP | 17625 | 300573 | . | . | [illegible] |
| 641371 | NM_00103716 | ACOT1 | acyl-CoA thioesterase 1 | protein_coding | NP_001032238.1;XP_016877078.1;XP_016877079.1;XP_016877080.1;XP | 33128 | 614313 | . | . | [illegible] |
| 641467 | NR_038381 | TSC22D1-AS1 | TSC22D1 antisense RNA 1 | lncRNA | . | 43684 | . | . | . | [illegible] |
| 641518 | NR_029373,NF | LEF1-AS1 | LEF1 antisense RNA 1 | lncRNA | . | 40339 | . | . | . | [illegible] |
| 641638 | NR_002599,NF | SNHG6 | small nucleolar RNA host gene 6 | lncRNA | . | 32965 | 612215 | . | . | [illegible] |
| 641648 | NR_002598 | SNORD87 | small nucleolar RNA, C/D box 87 | snoRNA | . | 32746 | 612216 | . | . | [illegible] |
| 641649 | NM_00104259 | TMEM91 | transmembrane protein 91 | protein_coding | NP_001036060.1;NP_001092291.1;NP_001092292.1;NP_001092293.1;NI | 32393 | 618294 | . | . | [illegible] |
| 641808 | NR_146190 | FAM185BP | family with sequence similarity 185 member B, pseudogene | transcribed_ps | . | 21813 | . | . | . | [illegible] |
| 641977 | NR_024271 | SEPTIN7P2 | septin 7 pseudogene 2 | transcribed_ps | . | 32339 | 611563 | . | . | [illegible] |
| 642236 | NR_033907 | FRG1JP | FSHD region gene 1 family member J, pseudogene | transcribed_ps | . | 51768 | . | . | . | [illegible] |
| 642311 | gene-BMS1P15 | BMS1P15 | BMS1 pseudogene 15 | pseudogene | . | 49160 | . | . | . | [illegible] |
| 642361 | NR_029407 | LOC642361 | uncharacterized LOC642361 | lncRNA | . | . | . | . | . | [illegible] |
| 642366 | NR_046243 | LOC642366 | uncharacterized LOC642366 | lncRNA | . | . | . | . | . | [illegible] |
| 642395 | gene-PSPC1P1 | PSPC1P1 | paraspeckle component 1 pseudogene 1 | pseudogene | . | 42033 | . | . | . | [illegible] |
| 642402 | NR_111962 | GOLGA6L17P | golgin A6 family like 17, pseudogene | transcribed_ps | . | 49414 | . | . | . | [illegible] |
| 642460 | gene-ANKRD30 | ANKRD30BP1 | ankyrin repeat domain 30B pseudogene 1 | pseudogene | . | 19722 | . | . | . | [illegible] |
| 642475 | NM_00110087 | MROH6 | maestro heat like repeat family member 6 | protein_coding | NP_001094348.1;XP_006716678.1;XP_011515516.1;XP_011515517.1;XP | 27814 | . | . | . | [illegible] |
| 642489 | gene-FKBP1C | FKBP1C | FKBP prolyl isomerase 1C | pseudogene | . | 21376 | . | . | . | [illegible] |
| 642502 | gene-LOC6425 | LOC642502 | succinate dehydrogenase complex subunit C pseudogene | pseudogene | . | . | . | . | . | [illegible] |
| 642517 | NM_00119081 | AGAP9 | ArfGAP with GTPase domain, ankyrin repeat and PH domain 9 | protein_coding | NP_001177739.1 | 23463 | . | . | . | [illegible] |
| 642546 | gene-HK2P1 | HK2P1 | hexokinase 2 pseudogene 1 | pseudogene | . | 4924 | . | . | . | [illegible] |
| 642574 | NM_00138187 | IQANK1 | IQ motif and ankyrin repeat containing 1 | protein_coding | NP_001368803.1;NP_001368804.1;XP_016869616.1 | 49576 | . | . | . | [illegible] |
| 642585 | gene-VDAC1P1 | VDAC1P1 | voltage dependent anion channel 1 pseudogene 1 | pseudogene | . | 12671 | . | . | . | [illegible] |
| 642592 | gene-EIF5AP4 | EIF5AP4 | eukaryotic translation initiation factor 5A pseudogene 4 | pseudogene | . | 31442 | . | . | . | [illegible] |
| 642636 | NM_00113656 | RAD21L1 | RAD21 cohesin complex component like 1 | protein_coding | NP_001130038.2;XP_006723665.1;XP_006723666.1;XP_006723667.1;XP | 16271 | . | . | . | [illegible] |
| 642641 | gene-RPSAP54 | RPSAP54 | ribosomal protein SA pseudogene 54 | pseudogene | . | 36373 | . | . | . | [illegible] |
| 642658 | NM_00108051 | SCX | scleraxis bHLH transcription factor | protein_coding | NP_001073983.1;XP_006716679.1 | 32322 | 609067 | . | . | [illegible] |
| 642659 | gene-HNRNPA1 | HNRNPA1P48 | heterogeneous nuclear ribonucleoprotein A1 pseudogene 48 | pseudogene | . | 48778 | . | . | . | [illegible] |
| 642740 | gene-RPL9P11 | RPL9P11 | ribosomal protein L9 pseudogene 11 | pseudogene | . | 35621 | . | . | . | [illegible] |
| 642741 | gene-RPL3P7 | RPL3P7 | ribosomal protein L3 pseudogene 7 | pseudogene | . | 36797 | . | . | . | [illegible] |
| 642778 | NM_00127732 | NPIPA3 | nuclear pore complex interacting protein family member A3 | protein_coding | NP_001264252.1;XP_005255542.1;XP_011520878.2;XP_011520879.2;XP | 41978 | . | . | . | [illegible] |
| 642799 | NM_00127732 | NPIPA2 | nuclear pore complex interacting protein family member A2 | protein_coding | NP_001264253.1;XP_005255549.1;XP_006720980.2;XP_011520889.1;XP | 41979 | . | . | . | [illegible] |
| 642819 | NM_00135544 | ZNF487 | zinc finger protein 487 | protein_coding | NP_001342373.2;NP_001342374.2;NP_001342375.2;NP_001342376.1;NI | 23488 | . | . | . | [illegible] |
| 642826 | NR_024495 | BMS1P2 | BMS1 pseudogene 2 | transcribed_ps | . | 23650 | . | . | . | [illegible] |
| 642828 | gene-RPL6P10 | RPL6P10 | ribosomal protein L6 pseudogene 10 | pseudogene | . | 36575 | . | . | . | [illegible] |
| 642846 | NR_024374 | LOC642846 | DEAD/H (Asp-Glu-Ala-Asp/His) box polypeptide 11-like | lncRNA | . | . | . | . | . | [illegible] |
| 642852 | NR_026943 | LINC00205 | long intergenic non-protein coding RNA 205 | lncRNA | . | 16420 | . | . | . | [illegible] |
| 642909 | gene-CTBP2P4 | CTBP2P4 | CTBP2 pseudogene 4 | pseudogene | . | 45196 | . | . | . | [illegible] |
| 642924 | NR_033858 | LINC00535 | long intergenic non-protein coding RNA 535 | lncRNA | . | 43644 | . | . | . | [illegible] |
| 642946 | NR_027285,NF | FLVCR1-DT | FLVCR1 divergent transcript | lncRNA | . | 39077 | 610864 | . | . | [illegible] |
| 642954 | gene-RBBP4P1 | RBBP4P1 | RBBP4 pseudogene 1 | pseudogene | . | 42368 | . | . | . | [illegible] |
| 642956 | gene-FABP5P9 | FABP5P9 | fatty acid binding protein 5 pseudogene 9 | pseudogene | . | 31064 | . | . | . | [illegible] |
| 642969 | gene-LOC6429 | LOC642969 | phosphoglycerate mutase 1 (brain) pseudogene | pseudogene | . | . | . | . | . | [illegible] |
| 642975 | gene-LOC6429 | LOC642975 | transmembrane protein 230 pseudogene | pseudogene | . | . | . | . | . | [illegible] |
| 642987 | NM_00103976 | TMEM232 | transmembrane protein 232 | protein_coding | NP_001034852.3;XP_006714733.1;XP_011541854.1;XP_011541855.1;XP | 37270 | . | . | . | [illegible] |
| 643007 | gene-RPL36AP | RPL36AP8 | ribosomal protein L36a pseudogene 8 | pseudogene | . | 30973 | . | . | . | [illegible] |
| 643072 | NR_110587,NF | LOC643072 | uncharacterized LOC643072 | lncRNA | . | . | . | . | . | [illegible] |
| 643136 | NM_00135545 | ZC3H11B | zinc finger CCCH-type containing 11B | protein_coding | NP_001342386.1 | 25659 | . | . | . | [illegible] |
| 643155 | NM_00104824 | SMIM15 | small integral membrane protein 15 | protein_coding | NP_001041714.1 | 33861 | . | . | . | [illegible] |
| 643159 | gene-ATF4P3 | ATF4P3 | activating transcription factor 4 pseudogene 3 | pseudogene | . | 788 | . | . | . | [illegible] |
| 643180 | NR_033416 | CCT6P3 | chaperonin containing TCP1 subunit 6 pseudogene 3 | transcribed_ps | . | 35137 | . | . | . | [illegible] |
| 643224 | NR_027156 | TUBBP5 | tubulin beta pseudogene 5 | transcribed_ps | . | 23674 | . | . | . | [illegible] |
| 643232 | gene-NRBF2P3 | NRBF2P3 | nuclear receptor binding factor 2 pseudogene 3 | pseudogene | . | 44606 | . | . | . | [illegible] |
| 643246 | NM_00108548 | MAP1LC3B2 | microtubule associated protein 1 light chain 3 beta 2 | protein_coding | NP_001078950.1 | 34390 | . | . | . | [illegible] |
| 643253 | NR_003110 | CCT6P1 | chaperonin containing TCP1 subunit 6 pseudogene 1 | transcribed_ps | . | 33094 | . | . | . | [illegible] |
| 643284 | gene-RPS29P1 | RPS29P18 | ribosomal protein S29 pseudogene 18 | pseudogene | . | 37004 | . | . | . | [illegible] |
| 643300 | gene-HSPD1P1 | HSPD1P1 | heat shock protein family D (Hsp60) member 1 pseudogene 1 | pseudogene | . | 35133 | . | . | . | [illegible] |
| 643310 | gene-ST13P18 | ST13P18 | ST13, Hsp70 interacting protein pseudogene 18 | pseudogene | . | 38861 | . | . | . | [illegible] |
| 643314 | NM_015038 | KIAA0754 | KIAA0754 | protein_coding | NP_055853.1 | 29111 | . | . | . | [illegible] |
| 643339 | NR_040096 | LOC643339 | uncharacterized LOC643339 | lncRNA | . | . | . | . | . | [illegible] |
| 643343 | gene-RPL5P4 | RPL5P4 | ribosomal protein L5 pseudogene 4 | pseudogene | . | 36095 | . | . | . | [illegible] |
| 643382 | NM_00114668 | TMEM253 | transmembrane protein 253 | protein_coding | NP_001140155.1;XP_006720296.1;XP_011535380.1;XP_011535381.1;XP | 32545 | . | . | . | [illegible] |
| 643387 | NR_026923 | LOC643387 | TAR DNA binding protein pseudogene | transcribed_ps | . | . | . | . | . | [illegible] |
| 643412 | gene-BTF3P16 | BTF3P16 | basic transcription factor 3 pseudogene 16 | pseudogene | . | 38568 | . | . | . | [illegible] |
| 643427 | gene-RBMS2P1 | RBMS2P1 | RNA binding motif single stranded interacting protein 2 pseudoger | pseudogene | . | 30994 | . | . | . | [illegible] |
| 643446 | gene-RNPS1P1 | RNPS1P1 | RNA binding protein with serine rich domain 1 pseudogene 1 | pseudogene | . | 39525 | . | . | . | [illegible] |
| 643454 | gene-LOC6434 | LOC643454 | adaptor related protein complex 3 sigma 1 subunit pseudogene | pseudogene | . | . | . | . | . | [illegible] |
| 643531 | gene-RPL29P2 | RPL29P26 | ribosomal protein L29 pseudogene 26 | pseudogene | . | 35717 | . | . | . | [illegible] |
| 643535 | gene-KCTD9P2 | KCTD9P2 | potassium channel tetramerization domain containing 9 pseudoge | pseudogene | . | 29274 | . | . | . | [illegible] |
| 643562 | gene-LOC6435 | LOC643562 | Rho GTPase activating protein 21 pseudogene | pseudogene | . | . | . | . | . | [illegible] |
| 643576 | gene-LOC6435 | LOC643576 | phosphoglycerate mutase 1 (brain) pseudogene | pseudogene | . | . | . | . | . | [illegible] |
| 643586 | gene-PKMP1 | PKMP1 | pyruvate kinase M1/2 pseudogene 1 | pseudogene | . | 44243 | . | . | . | [illegible] |
| 643641 | NM_00109922 | ZNF862 | zinc finger protein 862 | protein_coding | NP_001092690.1;XP_006716148.1;XP_011514765.1;XP_011514766.1;XP | 34519 | . | . | . | [illegible] |
| 643668 | gene-PSMC1P9 | PSMC1P9 | proteasome 26S subunit, ATPase 1 pseudogene 9 | pseudogene | . | 39784 | . | . | . | [illegible] |
| 643689 | gene-HNRNPA3 | HNRNPA3P3 | heterogeneous nuclear ribonucleoprotein A3 pseudogene 3 | pseudogene | . | 39772 | . | . | . | [illegible] |
| 643699 | NM_00128249 | GOLGA8N | golgin A8 family member N | protein_coding | NP_001269423.1 | 44405 | . | . | . | [illegible] |
| 643707 | NM_00126753 | GOLGA6L4 | golgin A6 family like 4 | protein_coding | NP_001254465.2;XP_016877970.1;XP_016877971.1 | 27256 | . | . | . | [illegible] |
| 643719 | NR_027620 | SCGB1B2P | secretoglobin family 1B member 2, pseudogene | transcribed_ps | . | 20741 | . | . | . | [illegible] |
| 643723 | NR_038845 | LYPLAL1-DT | LYPLAL1 divergent transcript | lncRNA | . | 50560 | . | . | . | [illegible] |
| 643744 | gene-HMGN2P | HMGN2P4 | high mobility group nucleosomal binding domain 2 pseudogene 4 | pseudogene | . | 33567 | . | . | . | [illegible] |
| 643749 | NR_034108,NF | TRAF3IP2-AS1 | TRAF3IP2 antisense RNA 1 | antisense_RNA | . | 40005 | . | . | . | [illegible] |
| 643751 | gene-CDC42P6 | CDC42P6 | cell division cycle 42 pseudogene 6 | pseudogene | . | 44430 | . | . | . | [illegible] |
| 643752 | gene-RAP1BL | RAP1BL | RAP1B, member of RAS oncogene family pseudogene | pseudogene | . | . | . | . | . | [illegible] |
| 643783 | NR_125363 | USP46-DT | USP46 divergent transcript | lncRNA | . | 43991 | . | . | . | [illegible] |
| 643784 | gene-NAIPP3 | NAIPP3 | NAIP pseudogene 3 | pseudogene | . | 54493 | . | . | . | [illegible] |
| 643827 | NR_111893 | CNTNAP3P2 | CNTNAP3 pseudogene 2 | transcribed_ps | . | 49589 | . | . | . | [illegible] |
| 643836 | NM_00117263 | ZFP62 | ZFP62 zinc finger protein | protein_coding | NP_001166109.1;NP_001364868.1;NP_001364869.1;NP_001364870.1;NI | 23241 | 610281 | . | . | [illegible] |
| 643837 | NR_047519,NF | LINC01128 | long intergenic non-protein coding RNA 1128 | lncRNA | . | 49377 | . | . | . | [illegible] |
| 643853 | NM_00103977 | TMPPE | transmembrane protein with metallophosphoesterase domain | protein_coding | NP_001034859.2;NP_001129710.1 | 33865 | . | . | . | [illegible] |
| 643854 | NM_00114565 | CTAGE9 | CTAGE family member 9 | protein_coding | NP_001139131.1 | 37275 | . | . | . | [illegible] |
| 643866 | NM_00103977 | CBLN3 | cerebellin 3 precursor | protein_coding | NP_001034860.1 | 20146 | 612978 | . | . | [illegible] |
| 643872 | gene-HMGN2P | HMGN2P6 | high mobility group nucleosomal binding domain 2 pseudogene 6 | pseudogene | . | 33574 | . | . | . | [illegible] |
| 643873 | gene-EIF4BP7 | EIF4BP7 | eukaryotic translation initiation factor 4B pseudogene 7 | pseudogene | . | 37940 | . | . | . | [illegible] |
| 643911 | NM_00130896 | CRNDE | colorectal neoplasia differentially expressed | protein_coding | NP_001295892.1;XP_016879051.1;XP_016879052.1 | 37078 | 615624 | . | . | [illegible] |
| 643932 | gene-RPS3AP2 | RPS3AP20 | RPS3A pseudogene 20 | pseudogene | . | 36141 | . | . | . | [illegible] |
| 643988 | NM_00124265 | FNDC10 | fibronectin type III domain containing 10 | protein_coding | NP_001229588.1 | 42951 | . | . | . | [illegible] |
| 644019 | NM_00108545 | CBWD6 | COBW domain containing 6 | protein_coding | NP_001078926.1;XP_016870508.1;XP_016870509.1;XP_016870510.1;XP | 31978 | . | . | . | [illegible] |
| 644029 | gene-RPL7AP6 | RPL7AP62 | ribosomal protein L7a pseudogene 62 | pseudogene | . | 36890 | . | . | . | [illegible] |
| 644035 | gene-NONOP1 | NONOP1 | non-POU domain containing, octamer-binding pseudogene 1 | pseudogene | . | 42031 | . | . | . | [illegible] |
| 644037 | gene-HNRNPA1 | HNRNPA1P12 | heterogeneous nuclear ribonucleoprotein A1 pseudogene 12 | pseudogene | . | 39130 | . | . | . | [illegible] |
| 644060 | gene-LOC6440 | LOC644060 | insulin like growth factor 2 mRNA binding protein 2 pseudogene | pseudogene | . | . | . | . | . | [illegible] |
| 644063 | gene-HNRNPKI | HNRNPKP4 | heterogeneous nuclear ribonucleoprotein K pseudogene 4 | pseudogene | . | 42377 | . | . | . | [illegible] |

| | | | | | | | | | | | | | | | | | | | | | | | | | | | | | | | | | |
|---|---|---|---|---|---|---|---|---|---|---|---|---|---|---|---|---|---|---|---|---|---|---|---|---|---|---|---|---|---|---|---|---|---|
| 644068 | NR_077246 | RPS14P3 | ribosomal protein S14 pseudogene 3 | transcribed_ps | . | 35797 | . | . | . | 1.975987 | 2.423668 | 4.083527 | 1.550693 | 4.598203 | 5.602417 | 9.163317 | 3.545474 | 1.258857 | 66.100327 | 0.46599557 | 0.71269881 | 0.476032153 | 0.998865404 | 5.869382 | 6.204385 | 5.974295 | 6.014928 | 6.152629 | 5.930128 | 61 | 69 | 93 | 38 |
| 644079 | gene-BCRP1 | BCRP1 | BCR pseudogene 1 | pseudogene | . | 39073 | . | . | . | 0.056672 | 0.045585 | 0.051541 | 0.038589 | 0.131879 | 0.105371 | 0.115656 | 0.088229 | -1.072000 | 5.182339 | 1.322372303 | -0.07585255 | 0.939536404 | NA | 2.421266 | 2.324312 | 2.378555 | 2.371759 | 2.376811 | 2.370200 | 7 | 5 | 5 | 4 |
| 644096 | NM_00104263 | SDHAF1 | succinate dehydrogenase complex assembly factor 1 | protein_coding | NP_001036096.2 | 33867 | 612848 | . | . | 7.792237 | 9.765778 | 7.359498 | 11.302045 | 18.132856 | 22.574032 | 16.514502 | 25.840782 | 1.046854 | 317.479103 | 0.235973029 | 0.279945788 | 0.779519097 | 0.998865404 | 8.278113 | 8.342211 | 8.230258 | 8.348900 | 8.196139 | 8.423940 | 319 | 371 | 224 | 367 |
| 644101 | gene-CBX3P9 | CBX3P9 | chromobox 3 pseudogene 9 | pseudogene | . | 42881 | . | . | . | 0.095152 | 0.204474 | 0.342180 | 0.040490 | 0.221422 | 0.472652 | 0.767842 | 0.092576 | 1.252754 | 4.608064 | 1.505874269 | 0.215890216 | 0.829073324 | NA | 2.029593 | 2.359867 | 2.174962 | 2.189061 | 2.204746 | 2.168390 | 3 | 6 | 8 | 1 |
| 644110 | gene-LOC6441 | LOC644110 | hyperpolarization activated cyclic nucleotide gated potassium and | pseudogene | . | . | . | . | . | 0.373905 | 0.398458 | 0.571357 | 0.329961 | 0.870092 | 0.921054 | 1.282107 | 0.754417 | 1.157952 | 42.156717 | 0.500663631 | 0.422591043 | 0.672593663 | 0.998865404 | 5.286539 | 5.500888 | 5.373495 | 5.380172 | 5.444700 | 5.354179 | 44 | 43 | 50 | 31 |
| 644128 | NR_003572 | RPL23AP53 | ribosomal protein L23a pseudogene 53 | transcribed_ps | . | 35921 | . | . | . | 0.921097 | 0.710238 | 0.893493 | 0.673270 | 2.143430 | 1.641747 | 2.004973 | 1.539351 | -1.057758 | 144.706264 | 0.292425238 | -0.27702847 | 0.781758249 | 0.998865404 | 7.215753 | 7.137144 | 7.235244 | 7.130073 | 7.214409 | 7.108257 | 194 | 139 | 140 | 113 |
| 644131 | gene-CCT8P1 | CCT8P1 | chaperonin containing TCP1 subunit 8 pseudogene 1 | pseudogene | . | 35144 | . | . | . | 0.150911 | 0.164794 | 0.288462 | 0.039828 | 0.351177 | 0.380929 | 0.647299 | 0.091061 | 1.030285 | 10.049167 | 1.042308583 | 0.04129576 | 0.967060113 | NA | 3.303566 | 3.354002 | 3.313740 | 3.316694 | 3.346534 | 3.285471 | 11 | 11 | 15 | 3 |
| 644153 | gene-DNM1P3 | DNM1P31 | dynamin 1 pseudogene 31 | pseudogene | . | 35178 | . | . | . | 0.244677 | 0.277437 | 0.640376 | 0.445992 | 0.569372 | 0.641308 | 1.436985 | 1.019709 | 1.964419 | 7.988544 | 1.085280156 | 0.897558439 | 0.369421005 | NA | 2.429096 | 3.404839 | 2.966562 | 2.968401 | 2.999826 | 2.982111 | 6 | 6 | 11 | 8 |
| 644157 | gene-DND1P1 | DND1P1 | DND microRNA-mediated repression inhibitor 1 pseudogene 1 | pseudogene | . | 31444 | . | . | . | 0.230881 | 0.121191 | 0.280796 | 0.054516 | 0.537269 | 0.280139 | 0.630099 | 0.124645 | -1.104417 | 8.911273 | 1.092699911 | -0.13112932 | 0.895673007 | NA | 3.220748 | 3.087437 | 3.163645 | 3.137531 | 3.169556 | 3.123177 | 14 | 7 | 12 | 3 |
| 644165 | NR_024494 | BCRP3 | BCR pseudogene 3 | transcribed_ps | . | 1016 | . | . | . | 0.113290 | 0.115218 | 0.166004 | 0.147130 | 0.263631 | 0.266331 | 0.372507 | 0.336395 | 1.440494 | 6.571903 | 1.176576929 | 0.447538777 | 0.654486096 | NA | 2.429096 | 2.955730 | 2.701555 | 2.703288 | 2.714444 | 2.711990 | 6 | 6 | 7 | 7 |
| 644191 | gene-RPS26P8 | RPS26P8 | ribosomal protein S26 pseudogene 8 | pseudogene | . | 31329 | . | . | . | 4.412173 | 4.822114 | 3.803913 | 4.878545 | 10.267308 | 11.146533 | 8.535871 | 11.154214 | -1.062985 | 63.123921 | 0.400250073 | -0.22016476 | 0.825742838 | 0.998865404 | 6.024366 | 5.934463 | 5.978147 | 5.996257 | 5.949000 | 5.997905 | 72 | 73 | 47 | 64 |
| 644192 | NR_102743,NF | NR2F2-AS1 | NR2F2 antisense RNA 1 | lncRNA | . | 44222 | . | . | . | 7.185516 | 7.447694 | 7.552325 | 8.339481 | 16.720990 | 17.215679 | 16.947201 | 19.067232 | 1.072902 | 269.968852 | 0.211254548 | 0.480551254 | 0.630835467 | 0.998865404 | 8.025318 | 8.126216 | 8.053810 | 8.060760 | 8.063162 | 8.122894 | 298 | 283 | 229 | 277 |
| 644215 | NR_157582 | LOC644215 | uncharacterized LOC644215 | lncRNA | . | . | . | . | . | 0.755434 | 0.696173 | 0.563289 | 0.556731 | 1.757925 | 1.609236 | 1.264004 | 1.272900 | -1.327313 | 16.095060 | 0.763801595 | -0.53483553 | 0.592763594 | NA | 4.198253 | 3.790061 | 4.026192 | 4.019489 | 4.003294 | 4.003377 | 22 | 19 | 12 | 13 |
| 644236 | gene-RPL7AP4 | RPL7AP43 | ribosomal protein L7a pseudogene 43 | pseudogene | . | 35775 | . | . | . | 0.786581 | 1.157777 | 1.706804 | 1.785740 | 1.830405 | 2.676253 | 3.830018 | 4.082882 | 1.769112 | 36.804842 | 0.522222527 | 1.576005062 | 0.115024637 | 0.855274326 | 4.733907 | 5.554573 | 5.120792 | 5.162778 | 5.218748 | 5.226909 | 25 | 34 | 40 | 45 |
| 644237 | gene-GAPDHP3 | GAPDHP33 | glyceraldehyde 3 phosphate dehydrogenase pseudogene 33 | pseudogene | . | 37785 | . | . | . | 0.868030 | 0.784417 | 1.416332 | 0.667607 | 2.019943 | 1.813215 | 3.178207 | 1.526404 | 1.250380 | 37.820209 | 0.545173461 | 0.591310533 | 0.554312374 | 0.998865404 | 5.068573 | 5.395155 | 5.217948 | 5.201394 | 5.299017 | 5.182782 | 41 | 34 | 50 | 25 |
| 644242 | NR_036540 | LINC00622 | long intergenic non-protein coding RNA 622 | lncRNA | . | 44251 | . | . | . | 8.290194 | 7.114074 | 12.890814 | 6.813270 | 19.291620 | 16.444502 | 28.926613 | 15.577730 | 1.261093 | 435.525688 | 0.250749059 | 1.334701151 | 0.181974177 | 0.953687323 | 8.588715 | 8.924965 | 8.719921 | 8.625364 | 8.997576 | 8.594460 | 479 | 382 | 555 | 313 |
| 644246 | NR_034172 | KANSL1-AS1 | KANSL1 antisense RNA 1 | lncRNA | . | 43740 | . | . | . | 0.613310 | 0.649979 | 0.908447 | 0.680369 | 1.427198 | 1.502455 | 2.038528 | 1.555582 | 1.237670 | 12.056311 | 0.877494364 | 0.3505738 | 0.725908115 | NA | 3.429096 | 3.737849 | 3.578369 | 3.581654 | 3.598117 | 3.584466 | 12 | 12 | 13 | 11 |
| 644248 | NR_045481 | SNCA-AS1 | SNCA antisense RNA 1 | lncRNA | . | 50600 | . | . | . | 0.179218 | 0.205928 | 0.094497 | 0.397972 | 0.417047 | 0.476013 | 0.212047 | 0.909916 | 1.316239 | 6.221929 | 1.244159818 | 0.318625416 | 0.750010578 | NA | 2.429096 | 2.819320 | 2.627874 | 2.629577 | 2.620443 | 2.652325 | 6 | 6 | 3 | 10 |
| 644253 | gene-MAPK8IP | MAPK8IP1P1 | mitogen-activated protein kinase 8 interacting protein 1 pseudoge | pseudogene | . | 6885 | . | . | . | 0.006657 | 0.015703 | 0.002382 | 0.009694 | 0.015491 | 0.036298 | 0.005344 | 0.022165 | 1.234655 | 1.002957 | 2.868339486 | 0.106022226 | 0.915564717 | NA | -0.155866 | 0.148375 | -0.001261 | -0.000972 | 0.000053 | -0.000315 | 1 | 1 | 1 | 1 |
| 644261 | gene-LOC6442 | LOC644261 | solute carrier family 25 member 53 pseudogene | pseudogene | . | . | . | . | . | 0.136584 | 0.099617 | 0.216598 | 0.153863 | 0.317836 | 0.230270 | 0.486040 | 0.351790 | 1.616620 | 5.841863 | 1.253289013 | 0.552929536 | 0.580311662 | NA | 2.156660 | 2.852980 | 2.532348 | 2.525326 | 2.545007 | 2.537844 | 6 | 4 | 7 | 6 |
| 644295 | gene-KIAA114 | KIAA1143P1 | KIAA1143 pseudogene 1 | pseudogene | . | 54591 | . | . | . | 0.297691 | 0.193745 | 0.252778 | 0.317559 | 0.692739 | 0.447849 | 0.567227 | 0.726060 | 1.099746 | 8.422475 | 1.047549307 | 0.130944555 | 0.895819168 | NA | 3.003609 | 3.141582 | 3.078697 | 3.064538 | 3.071834 | 3.078637 | 11 | 7 | 7 | 9 |
| 644315 | gene-RPS7P11 | RPS7P11 | ribosomal protein S7 pseudogene 11 | pseudogene | . | 35841 | . | . | . | 2.554974 | 2.793160 | 1.932114 | 2.748195 | 5.945529 | 6.456515 | 4.335607 | 6.283422 | -1.160108 | 54.707407 | 0.431736405 | -0.49627425 | 0.619700934 | 0.998865404 | 5.877915 | 5.661290 | 5.780751 | 5.797757 | 5.730715 | 5.788334 | 65 | 66 | 37 | 55 |
| 644330 | gene-TPM3P5 | TPM3P5 | tropomyosin 3 pseudogene 5 | pseudogene | . | 39168 | . | . | . | 0.185359 | 0.209392 | 0.111085 | 0.318909 | 0.431338 | 0.484018 | 0.249272 | 0.729148 | 1.095368 | 14.077741 | 0.850706199 | 0.154477948 | 0.877232897 | NA | 3.751025 | 3.876918 | 3.807574 | 3.811544 | 3.786666 | 3.842986 | 15 | 15 | 7 | 20 |
| 644353 | NM_00114397 | ZCCHC18 | zinc finger CCHC-type containing 18 | protein_coding | NP_001137450.1;XP_011529314.1 | 32459 | . | . | . | 0.089019 | 0.501297 | 1.059051 | 0.110225 | 0.207151 | 1.158772 | 2.376479 | 0.252016 | 1.831181 | 40.861042 | 0.556441389 | 1.568493454 | 0.116766016 | 0.857396579 | 4.849979 | 5.724718 | 5.247170 | 5.285964 | 5.440929 | 5.272819 | 28 | 36 | 66 | 28 |
| 644390 | gene-HNRNPLF | HNRNPLP2 | heterogeneous nuclear ribonucleoprotein L pseudogene 2 | pseudogene | . | 48747 | . | . | . | 0.501361 | 0.359718 | 0.188869 | 0.285865 | 1.166687 | 0.831505 | 0.423817 | 0.653596 | -1.777444 | 19.852144 | 0.709489304 | -1.16957984 | 0.242170093 | NA | 4.667953 | 3.836248 | 4.357895 | 4.322904 | 4.282097 | 4.305019 | 34 | 23 | 10 | 16 |
| 644397 | gene-LRRC37A | LRRC37A17P | leucine rich repeat containing 37 member A17, pseudogene | pseudogene | . | 48365 | . | . | . | 0.081407 | 0.139253 | 0.102934 | 0.111496 | 0.189436 | 0.321889 | 0.230980 | 0.254923 | -1.033582 | 16.877019 | 0.755891755 | -0.06304155 | 0.949733418 | NA | 4.101892 | 4.051647 | 4.057559 | 4.094142 | 4.071622 | 4.075660 | 15 | 23 | 14 | 16 |
| 644435 | gene-AGGF1P2 | AGGF1P2 | angiogenic factor with G-patch and FHA domains 1 pseudogene 2 | pseudogene | . | 38052 | . | . | . | 0.206456 | 0.279960 | 0.239101 | 0.017827 | 0.480432 | 0.647140 | 0.536535 | 0.040760 | -1.944399 | 7.517955 | 1.21305235 | -0.79083479 | 0.429040405 | NA | 3.307819 | 2.359867 | 2.917091 | 2.928059 | 2.918929 | 2.882735 | 10 | 12 | 8 | 1 |
| 644456 | gene-LOC6444 | LOC644456 | IK cytokine, down-regulator of HLA II pseudogene | pseudogene | . | . | . | . | . | 0.151042 | 0.105566 | 0.109679 | 0.166506 | 0.351482 | 0.244022 | 0.246115 | 0.380696 | 1.028853 | 8.134981 | 1.068085913 | 0.038421356 | 0.969351734 | NA | 3.003609 | 3.044381 | 3.031606 | 3.017439 | 3.019666 | 3.031481 | 11 | 7 | 6 | 9 |
| 644462 | gene-LOC6444 | LOC644462 | STE20 related adaptor beta pseudogene | pseudogene | . | . | . | . | . | 0.401790 | 0.493008 | 0.386621 | 0.346139 | 0.934982 | 1.139611 | 0.867566 | 0.791406 | -1.243621 | 27.998947 | 0.586847485 | -0.5359943 | 0.591962492 | 0.998865404 | 4.956050 | 4.641433 | 4.807583 | 4.834210 | 4.802350 | 4.790115 | 32 | 37 | 23 | 22 |
| 644464 | gene-RPSAP61 | RPSAP61 | ribosomal protein SA pseudogene 61 | pseudogene | . | 36760 | . | . | . | 0.095005 | 0.078731 | 0.085706 | 0.050879 | 0.221080 | 0.181989 | 0.192321 | 0.116330 | -1.123618 | 2.959315 | 1.731081282 | -0.09713705 | 0.922617562 | NA | 1.644780 | 1.481107 | 1.568690 | 1.565383 | 1.568406 | 1.562470 | 4 | 3 | 3 | 2 |
| 644498 | gene-HMGN2P | HMGN2P7 | high mobility group nucleosomal binding domain 2 pseudogene 7 | pseudogene | . | 33575 | . | . | . | 0.191037 | 0.085854 | 0.153764 | 0.124128 | 0.444551 | 0.198454 | 0.345041 | 0.283804 | -1.042482 | 5.651541 | 1.280806605 | -0.04686352 | 0.962622001 | NA | 2.526439 | 2.470304 | 2.510053 | 2.490820 | 2.500326 | 2.498470 | 9 | 4 | 5 | 5 |
| 644511 | NR_026715 | RPL13AP6 | ribosomal protein L13a pseudogene 6 | transcribed_ps | . | 23737 | . | . | . | 0.042215 | 0.008097 | 0.020749 | 0.031005 | 0.098237 | 0.018718 | 0.046560 | 0.070889 | -1.204659 | 1.220068 | 2.631544379 | -0.10207874 | 0.918694172 | NA | 0.413393 | 0.148375 | 0.294625 | 0.290739 | 0.291821 | 0.291432 | 2 | 1 | 1 | 1 |
| 644524 | NM_033176 | NKX2-4 | NK2 homeobox 4 | protein_coding | NP_149416.1 | 7837 | 607808 | . | . | 0.557325 | 0.707010 | 0.474238 | 0.381476 | 1.296919 | 1.634286 | 1.064177 | 0.872200 | -1.469601 | 29.492486 | 0.577043616 | -0.96253478 | 0.335781039 | 0.998865404 | 5.133136 | 4.578418 | 4.890849 | 4.921522 | 4.871541 | 4.850487 | 36 | 42 | 23 | 20 |
| 644538 | NM_00116343 | SMIM10 | small integral membrane protein 10 | protein_coding | NP_001156910.1 | 41913 | . | . | . | 3.889495 | 3.455553 | 3.120348 | 3.104479 | 9.051014 | 7.987666 | 7.001970 | 7.098022 | -1.195571 | 157.365789 | 0.264160762 | -0.97553975 | 0.329292596 | 0.998865404 | 7.420801 | 7.163718 | 7.360117 | 7.304973 | 7.265954 | 7.257673 | 210 | 173 | 126 | 133 |
| 644596 | NM_00134825 | SMIM10L2B | small integral membrane protein 10 like 2B | protein_coding | NP_001335184.1 | 34500 | . | . | . | 0.102119 | 0.077689 | 0.106853 | 0.220600 | 0.237635 | 0.179581 | 0.239774 | 0.504377 | 1.743610 | 11.629150 | 0.914705634 | 0.876869807 | 0.380557357 | NA | 3.084644 | 3.885075 | 3.517795 | 3.508122 | 3.521865 | 3.559847 | 11 | 8 | 9 | 18 |
| 644604 | gene-EEF1A1P1 | EEF1A1P12 | eukaryotic translation elongation factor 1 alpha 1 pseudogene 12 | pseudogene | . | 3195 | . | . | . | 0.178670 | 0.216567 | 0.106180 | 0.270693 | 0.415771 | 0.500605 | 0.238264 | 0.618907 | -1.081616 | 10.788050 | 0.957550303 | -0.11820601 | 0.905904429 | NA | 3.489863 | 3.370388 | 3.430888 | 3.438245 | 3.410852 | 3.450473 | 12 | 13 | 5 | 14 |
| 644616 | gene-ILF2P2 | ILF2P2 | interleukin enhancer binding factor 2 pseudogene 2 | pseudogene | . | 44696 | . | . | . | 0.024749 | 0.100733 | 0.298373 | 0.037031 | 0.057591 | 0.232848 | 0.669540 | 0.084668 | 2.341036 | 6.095088 | 1.369094513 | 0.896320207 | 0.37008177 | NA | 1.867373 | 3.094489 | 2.546540 | 2.565049 | 2.606019 | 2.549442 | 2 | 6 | 13 | 2 |
| 644619 | NR_027392 | INTS4P2 | integrator complex subunit 4 pseudogene 2 | transcribed_ps | . | 22351 | . | . | . | 0.319771 | 0.399128 | 0.369310 | 0.486976 | 0.744120 | 0.922602 | 0.828722 | 1.113414 | 1.172175 | 21.961343 | 0.659553001 | 0.347490746 | 0.728222669 | NA | 4.339107 | 4.565787 | 4.437059 | 4.454573 | 4.448228 | 4.472649 | 21 | 24 | 18 | 25 |
| 644634 | NR_145442 | LOC644634 | uncharacterized LOC644634 | lncRNA | . | . | . | . | . | 0.230437 | 0.331471 | 0.235859 | 0.477332 | 0.536237 | 0.766211 | 0.529260 | 1.091364 | 1.217160 | 13.966417 | 0.840088173 | 0.337487005 | 0.735749806 | NA | 3.658167 | 3.936235 | 3.782300 | 3.802304 | 3.783107 | 3.825201 | 12 | 16 | 9 | 19 |
| 644637 | gene-NF1P6 | NF1P6 | neurofibromin 1 pseudogene 6 | pseudogene | . | 7771 | . | . | . | 0.218347 | 0.392920 | 0.848696 | 0.345619 | 0.508102 | 0.908251 | 1.904449 | 0.790217 | 1.880532 | 24.113914 | 0.692694661 | 1.315356742 | 0.188390037 | NA | 4.064940 | 4.976875 | 4.505984 | 4.547462 | 4.640645 | 4.534329 | 14 | 23 | 39 | 17 |
| 644656 | NR_036539 | LOC644656 | uncharacterized LOC644656 | lncRNA | . | . | . | . | . | 3.016806 | 3.502464 | 3.713106 | 3.239289 | 7.020231 | 8.096102 | 8.332102 | 7.406249 | 1.050324 | 120.228666 | 0.298838518 | 0.237031444 | 0.812632396 | 0.998865404 | 6.873655 | 6.944744 | 6.875725 | 6.921660 | 6.940529 | 6.890353 | 126 | 135 | 115 | 107 |
| 644660 | NR_033886 | RAD21-AS1 | RAD21 antisense RNA 1 | lncRNA | . | 32158 | . | . | . | 0.065861 | 0.117287 | 0.141255 | 0.042722 | 0.153262 | 0.271114 | 0.316971 | 0.097679 | 1.052153 | 6.038991 | 1.264416888 | 0.058006184 | 0.953743703 | NA | 2.555346 | 2.632244 | 2.577125 | 2.591563 | 2.598958 | 2.572359 | 5 | 8 | 8 | 3 |
| 644714 | NR_033947 | LIMD1-AS1 | LIMD1 antisense RNA 1 | lncRNA | . | 44107 | . | . | . | 0.020326 | 0.012931 | 0.013881 | 0.064195 | 0.047300 | 0.029891 | 0.031148 | 0.146775 | 1.815038 | 2.518421 | 1.923208224 | 0.447169498 | 0.654752683 | NA | 0.844134 | 1.696731 | 1.319806 | 1.320421 | 1.317443 | 1.336353 | 2 | 2 | 1 | 5 |
| 644739 | gene-WASF4P | WASF4P | WASP family member 4, pseudogene | pseudogene | . | 20801 | . | . | . | 0.052539 | 0.073657 | 0.040400 | 0.136021 | 0.122262 | 0.170261 | 0.090656 | 0.310996 | 1.392013 | 10.740013 | 0.982489546 | 0.485676728 | 0.627196386 | NA | 3.170740 | 3.640961 | 3.401849 | 3.412922 | 3.393795 | 3.451701 | 9 | 11 | 5 | 18 |
| 644745 | gene-HSPD1P4 | HSPD1P4 | heat shock protein family D (Hsp60) member 1 pseudogene 4 | pseudogene | . | 35146 | . | . | . | 0.210351 | 0.251676 | 0.125350 | 0.134082 | 0.489496 | 0.581760 | 0.281282 | 0.306561 | -1.816114 | 13.241663 | 0.850273606 | -1.01244476 | 0.311325448 | NA | 4.094525 | 3.232645 | 3.744303 | 3.756828 | 3.716462 | 3.717840 | 18 | 20 | 8 | 9 |
| 644794 | NR_136637 | LINC02604 | long intergenic non-protein coding RNA 2604 | lncRNA | . | 53972 | . | . | . | 1.882572 | 2.225969 | 3.091722 | 2.279581 | 4.380822 | 5.145427 | 6.937734 | 5.211991 | 1.293088 | 161.357341 | 0.276779365 | 1.339767522 | 0.180320938 | 0.952541638 | 7.136396 | 7.507974 | 7.238953 | 7.300449 | 7.438913 | 7.304984 | 149 | 164 | 183 | 144 |
| 644815 | NM_00103999 | FAM83G | family with sequence similarity 83 member G | protein_coding | NP_001035088.2;XP_016880442.1 | 32554 | 615886 | . | . | 2.504436 | 2.674557 | 2.140298 | 2.621960 | 5.827927 | 6.182359 | 4.802767 | 5.994801 | -1.096441 | 406.561058 | 0.187119062 | -0.70985967 | 0.477791164 | 0.998865404 | 8.732679 | 8.598876 | 8.676310 | 8.710851 | 8.585522 | 8.689097 | 476 | 472 | 307 | 396 |
| 644820 | gene-EEF1B2P3 | EEF1B2P3 | eukaryotic translation elongation factor 1 beta 2 pseudogene 3 | pseudogene | . | 3210 | . | . | . | 0.707393 | 1.353754 | 1.071474 | 1.626034 | 1.646132 | 3.129263 | 2.404356 | 3.717734 | 1.306408 | 28.663584 | 0.607986036 | 0.634234744 | 0.525927636 | 0.998865404 | 4.638216 | 5.019022 | 4.781990 | 4.846434 | 4.822032 | 4.869286 | 20 | 35 | 23 | 36 |
| 644850 | gene-PDCL3P5 | PDCL3P5 | PDCL3 pseudogene 5 | pseudogene | . | 44506 | . | . | . | 0.013701 | 0.126970 | 0.041177 | 0.174672 | 0.031884 | 0.293497 | 0.092401 | 0.399366 | 1.595113 | 3.550530 | 1.668534548 | 0.403742695 | 0.68640194 | NA | 1.460000 | 2.121007 | 1.802903 | 1.820961 | 1.810270 | 1.828731 | 1 | 5 | 2 | 6 |
| 644873 | NR_015360,NF | LINC01184 | long intergenic non-protein coding RNA 1184 | lncRNA | . | 49565 | . | . | . | 8.608944 | 8.778300 | 8.195992 | 9.655589 | 20.033366 | 20.291435 | 18.391573 | 22.076362 | 1.068668 | 586.351539 | 0.159468626 | 0.60083475 | 0.547950057 | 0.998865404 | 9.147017 | 9.242643 | 9.179985 | 9.165601 | 9.195207 | 9.237558 | 660 | 605 | 510 | 586 |
| 644877 | gene-CACYBPP | CACYBPP2 | calcyclin binding protein pseudogene 2 | pseudogene | . | 45123 | . | . | . | 0.425335 | 0.238606 | 0.064506 | 0.246758 | 0.989772 | 0.551550 | 0.144750 | 0.564183 | -2.041589 | 8.267265 | 1.125100394 | -0.91520076 | 0.360086244 | NA | 3.474811 | 2.437399 | 3.084327 | 3.059041 | 3.032790 | 3.059884 | 16 | 9 | 2 | 8 |
| 644890 | NM_00108083 | MEIG1 | meiosis/spermiogenesis associated 1 | protein_coding | NP_001074305.1;XP_024303904.1 | 23429 | 614174 | . | . | 0.538259 | 0.341944 | 0.617906 | 0.519242 | 1.252550 | 0.790419 | 1.386562 | 1.187185 | 1.132005 | 10.932993 | 0.922973906 | 0.193808728 | 0.846325653 | NA | 3.357474 | 3.538109 | 3.454220 | 3.436801 | 3.451020 | 3.452089 | 14 | 9 | 10 | 11 |
| 644907 | gene-RPL32P1 | RPL32P18 | ribosomal protein L32 pseudogene 18 | pseudogene | . | 36301 | . | . | . | 0.270773 | 0.269954 | 0.138948 | 0.027125 | 0.630101 | 0.624011 | 0.311795 | 0.062019 | -2.679779 | 3.085637 | 1.76917515 | -0.80382895 | 0.421495771 | NA | 2.166062 | 0.751299 | 1.645501 | 1.646763 | 1.636091 | 1.630421 | 5 | 5 | 2 | 1 |
| 644914 | gene-H3P16 | H3P16 | H3 histone pseudogene 16 | pseudogene | . | 42982 | . | . | . | 5.240489 | 4.150990 | 4.443222 | 3.291289 | 12.194832 | 9.595201 | 9.970460 | 7.525141 | -1.235014 | 135.475992 | 0.300033145 | -1.01497882 | 0.310115922 | 0.998865404 | 7.225000 | 6.923012 | 7.163106 | 7.067035 | 7.090415 | 6.984689 | 193 | 142 | 122 | 96 |
| 644928 | gene-RPS26P1 | RPS26P15 | ribosomal protein S26 pseudogene 15 | pseudogene | . | 35680 | . | . | . | 0.456089 | 0.299122 | 0.459413 | 0.789221 | 1.061339 | 0.691433 | 1.030910 | 1.804460 | 1.609494 | 7.553611 | 1.117543845 | 0.614389353 | 0.53895807 | NA | 2.533720 | 3.219802 | 2.905626 | 2.894893 | 2.905605 | 2.926894 | 8 | 5 | 6 | 11 |
| 644961 | NR_033926 | ACTG1P20 | actin gamma 1 pseudogene 20 | transcribed_ps | . | 51500 | . | . | . | 0.401765 | 0.277640 | 0.369882 | 0.302628 | 0.934924 | 0.641777 | 0.830005 | 0.691922 | -1.008879 | 28.421793 | 0.589067404 | -0.02165067 | 0.982726616 | 0.998865404 | 4.833848 | 4.823987 | 4.850178 | 4.805536 | 4.837826 | 4.816169 | 39 | 25 | 27 | 24 |
| 644962 | NR_077215 | TNRC18P1 | trinucleotide repeat containing 18 pseudogene 1 | transcribed_ps | . | 43881 | . | . | . | 0.188886 | 0.107244 | 0.154999 | 0.198515 | 0.439545 | 0.247900 | 0.347812 | 0.453880 | 1.176514 | 9.677523 | 0.987298542 | 0.237535587 | 0.812241315 | NA | 3.151937 | 3.387716 | 3.279822 | 3.257754 | 3.270332 | 3.281379 | 13 | 7 | 8 | 11 |
| 644972 | gene-RPS3AP2 | RPS3AP26 | RPS3A pseudogene 26 | pseudogene | . | 36513 | . | . | . | 16.369823 | 11.990900 | 7.915517 | 16.991735 | 38.093250 | 27.717505 | 17.762192 | 38.849586 | -1.156904 | 356.287002 | 0.568360412 | -0.36995655 | 0.711414863 | 0.998865404 | 8.578346 | 8.367769 | 8.591711 | 8.400919 | 8.197788 | 8.598040 | 510 | 347 | 184 | 421 |
| 644988 | gene-ABHD17A | ABHD17AP6 | ABHD17A pseudogene 6 | pseudogene | . | 34044 | . | . | . | 0.820232 | 0.688580 | 0.915538 | 0.650941 | 1.908714 | 1.591683 | 2.054440 | 1.488298 | 1.036499 | 35.534125 | 0.529738729 | 0.097629935 | 0.922226153 | 0.998865404 | 5.123517 | 5.178231 | 5.158938 | 5.131828 | 5.174495 | 5.125839 | 44 | 34 | 37 | 28 |
| 645018 | gene-RPS2P20 | RPS2P20 | ribosomal protein S2 pseudogene 20 | pseudogene | . | 35992 | . | . | . | 0.040350 | 0.102191 | 0.016829 | 0.092759 | 0.093897 | 0.236220 | 0.037764 | 0.212082 | -1.242626 | 2.448460 | 1.925493808 | -0.16275906 | 0.87070814 | NA | 1.444631 | 1.121007 | 1.292286 | 1.301626 | 1.289916 | 1.299116 | 2 | 4 | 1 | 3 |
| 645086 | gene-LOC6450 | LOC645086 | chromosome 11 open reading frame 58 pseudogene | pseudogene | . | . | . | . | . | 0.037688 | 0.072630 | 0.048673 | 0.023396 | 0.087701 | 0.167887 | 0.109221 | 0.053492 | -1.620424 | 2.202620 | 2.029505069 | -0.34312378 | 0.731505345 | NA | 1.444631 | 0.751299 | 1.140280 | 1.149605 | 1.142955 | 1.137337 | 2 | 4 | 2 | 1 |
| 645139 | gene-LOC6451 | LOC645139 | poly(A) binding protein interacting protein 1 pseudogene | pseudogene | . | . | . | . | . | 3.341465 | 2.735917 | 2.449944 | 1.308840 | 7.775727 | 6.324193 | 5.497602 | 2.992506 | -1.645559 | 105.710186 | 0.358390706 | -2.00501103 | 0.044961867 | 0.649185583 | 7.037417 | 6.322959 | 6.825200 | 6.746889 | 6.707779 | 6.549496 | 167 | 127 | 91 | 52 |
| 645158 | NR_033754 | CBX3P2 | chromobox 3 pseudogene 2 | transcribed_ps | . | 42874 | . | . | . | 0.383135 | 0.392635 | 0.358753 | 0.108538 | 0.891570 | 0.907593 | 0.805032 | 0.248158 | -1.640254 | 14.102491 | 0.862497437 | -0.82773455 | 0.407820846 | NA | 4.128334 | 3.421760 | 3.833148 | 3.833977 | 3.826896 | 3.780305 | 20 | 19 | 14 | 5 |
| 645161 | gene-RPL12P8 | RPL12P8 | ribosomal protein L12 pseudogene 8 | pseudogene | . | 23735 | . | . | . | 0.155587 | 0.207413 | 0.200421 | 0.074695 | 0.362057 | 0.479445 | 0.449738 | 0.170780 | -1.070014 | 3.478495 | 1.609933276 | -0.06064235 | 0.951644048 | NA | 1.844134 | 1.751299 | 1.796180 | 1.797278 | 1.801166 | 1.790082 | 4 | 4 | 4 | 2 |
| 645166 | NR_027354,NF | LSP1P5 | LSP1 pseudogene 5 | transcribed_ps | . | 50697 | . | . | . | 3.597766 | 3.354155 | 3.036545 | 3.255358 | 8.372151 | 7.753282 | 6.813917 | 7.442989 | -1.079739 | 69.017998 | 0.378965642 | -0.29206614 | 0.770236057 | 0.998865404 | 6.163131 | 6.052552 | 6.126448 | 6.111433 | 6.098163 | 6.103592 | 85 | 75 | 57 | 63 |
| 645173 | gene-RPS2P48 | RPS2P48 | ribosomal protein S2 pseudogene 48 | pseudogene | . | 35985 | . | . | . | 0.036257 | 0.001634 | 0.007599 | 0.050331 | 0.084371 | 0.003778 | 0.017052 | 0.115076 | 1.233060 | 1.486734 | 2.400765875 | 0.125894475 | 0.899815456 | NA | 0.413393 | 0.715150 | 0.572744 | 0.568868 | 0.570025 | 0.574493 | 2 | 1 | 1 | 2 |
| 645174 | gene-RPL21P4 | RPL21P46 | ribosomal protein L21 pseudogene 46 | pseudogene | . | 35687 | . | . | . | 0.068886 | 0.036234 | 0.136382 | 0.021762 | 0.160301 | 0.083757 | 0.306037 | 0.049757 | 1.256691 | 1.507561 | 2.39030677 | 0.137902842 | 0.890317204 | NA | 0.413393 | 0.751299 | 0.586366 | 0.582479 | 0.588807 | 0.583226 | 2 | 1 | 2 | 1 |
| 645206 | NR_038840 | LINC00693 | long intergenic non-protein coding RNA 693 | lncRNA | . | 44526 | . | . | . | 0.488107 | 0.559630 | 0.208110 | 0.218873 | 1.135846 | 1.293609 | 0.466992 | 0.500426 | -2.396748 | 19.735544 | 0.710770384 | -1.77424161 | 0.076023194 | NA | 4.799841 | 3.538109 | 4.333961 | 4.348502 | 4.272786 | 4.273077 | 30 | 32 | 10 | 11 |
| 645212 | NR_034170,NF | EIF3J-DT | EIF3J divergent transcript | lncRNA | . | 48616 | . | . | . | 1.842070 | 2.587664 | 2.679197 | 1.804690 | 4.286574 | 5.981501 | 6.012042 | 4.126207 | 1.013990 | 120.636979 | 0.32056002 | 0.062524658 | 0.950145026 | 0.998865404 | 6.903989 | 6.924991 | 6.859450 | 6.949725 | 6.973494 | 6.842996 | 120 | 146 | 126 | 92 |
| 645233 | gene-TDGP1 | TDGP1 | thymine-DNA glycosylase pseudogene 1 | pseudogene | . | 49891 | . | . | . | 0.105897 | 0.019347 | 0.060056 | 0.044865 | 0.246428 | 0.044722 | 0.134763 | 0.102578 | -1.231886 | 5.572833 | 1.355867751 | -0.2219015 | 0.824390561 | NA | 2.617627 | 2.324312 | 2.502218 | 2.462265 | 2.480434 | 2.473808 | 12 | 2 | 5 | 4 |
| 645292 | gene-HMGB1P | HMGB1P6 | high mobility group box 1 pseudogene 6 | pseudogene | . | 4998 | . | . | . | 16.570780 | 15.262781 | 13.261582 | 13.394137 | 38.560883 | 35.280598 | 29.758604 | 30.624104 | -1.211218 | 930.510476 | 0.141641776 | -1.95181456 | 0.050960225 | 0.672475152 | 9.993406 | 9.717147 | 9.960373 | 9.892386 | 9.788567 | 9.788140 | 1227 | 1050 | 732 | 789 |
| 645296 | gene-RPL17P3 | RPL17P39 | ribosomal protein L17 pseudogene 39 | pseudogene | . | 36427 | . | . | . | 2.499722 | 1.951155 | 1.281208 | 2.401257 | 5.816957 | 4.510181 | 2.874995 | 5.490189 | -1.206641 | 33.859370 | 0.558034974 | -0.48562584 | 0.627232471 | 0.998865404 | 5.211816 | 4.938196 | 5.113689 | 5.076487 | 5.035210 | 5.103889 | 48 | 35 | 19 | 37 |
| 645312 | gene-GLTPP1 | GLTPP1 | glycolipid transfer protein pseudogene 1 | pseudogene | . | 33991 | . | . | . | 0.300106 | 0.293250 | 0.290896 | 0.179484 | 0.698360 | 0.677861 | 0.652762 | 0.410369 | -1.231822 | 19.902701 | 0.698041241 | -0.43091175 | 0.666532542 | NA | 4.455972 | 4.158506 | 4.327240 | 4.321665 | 4.323517 | 4.290109 | 26 | 23 | 19 | 13 |
| 645313 | gene-SCDP1 | SCDP1 | stearoyl-CoA desaturase pseudogene 1 | pseudogene | . | 10572 | . | . | . | 0.030152 | 0.055405 | 0.062149 | 0.034897 | 0.070165 | 0.128071 | 0.139460 | 0.079788 | 1.156849 | 7.806968 | 1.099835263 | 0.191120407 | 0.84843126 | NA | 2.855800 | 3.066070 | 2.944583 | 2.963533 | 2.967816 | 2.950539 | 6 | 10 | 9 | 6 |
| 645317 | gene-CHCHD2F | CHCHD2P6 | coiled-coil-helix-coiled-coil-helix domain containing 2 pseudogene | pseudogene | . | 39590 | . | . | . | 0.210969 | 0.435596 | 0.564864 | 0.940481 | 0.490934 | 1.006899 | 1.267539 | 2.150297 | 2.242793 | 13.904094 | 0.852478583 | 1.366950839 | 0.171640715 | NA | 3.104339 | 4.263648 | 3.739122 | 3.762830 | 3.779754 | 3.815885 | 7 | 12 | 13 | 22 |
| 645323 | NR_015436,NF | LINC00461 | long intergenic non-protein coding RNA 461 | lncRNA | . | 42810 | 616611 | . | . | 1.328188 | 1.455141 | 1.319142 | 0.980513 | 3.090749 | 3.363623 | 2.960116 | 2.241826 | -1.121115 | 158.059125 | 0.274569403 | -0.60070223 | 0.548038331 | 0.998865404 | 7.383518 | 7.220521 | 7.343718 | 7.305380 | 7.334365 | 7.220634 | 201 | 172 | 149 | 119 |
| 645332 | NR_024249 | FAM86C2P | family with sequence similarity 86 member C2, pseudogene | transcribed_ps | . | 42392 | . | . | . | 1.157021 | 1.261884 | 1.155020 | 1.309811 | 2.692435 | 2.916902 | 2.591831 | 2.994727 | 1.006562 | 71.582088 | 0.373758909 | 0.025247339 | 0.979857678 | 0.998865404 | 6.157312 | 6.165730 | 6.150852 | 6.170003 | 6.148892 | 6.175036 | 79 | 80 | 59 | 71 |
| 645345 | gene-CHCHD2F | CHCHD2P9 | coiled-coil-helix-coiled-coil-helix domain containing 2 pseudogene | pseudogene | . | 23676 | . | . | . | 6.980975 | 5.945803 | 5.201108 | 7.058293 | 16.245016 | 13.743990 | 11.671136 | 16.137947 | -1.068227 | 151.451393 | 0.281062604 | -0.33877992 | 0.73477553 | 0.998865404 | 7.289980 | 7.193840 | 7.287123 | 7.220418 | 7.172827 | 7.281193 | 195 | 155 | 109 | 157 |
| 645369 | NM_00108020 | TMEM200C | transmembrane protein 200C | protein_coding | NP_001073678.1;XP_011524031.1 | 37208 | . | . | . | 0.431999 | 0.428836 | 0.439849 | 0.422804 | 1.005279 | 0.991273 | 0.987008 | 0.966691 | 1.181095 | 61.423309 | 0.41096722 | 0.584293172 | 0.559023085 | 0.998865404 | 5.816276 | 6.055266 | 5.880696 | 5.950555 | 5.960591 | 5.943889 | 54 | 71 | 60 | 60 |
| 645387 | gene-RPL6P27 | RPL6P27 | ribosomal protein L6 pseudogene 27 | pseudogene | . | 36133 | . | . | . | 20.014994 | 20.225336 | 15.352449 | 22.774551 | 46.575718 | 46.751774 | 34.450451 | 52.071304 | -1.070588 | 561.684700 | 0.193966063 | -0.50732535 | 0.611926551 | 0.998865404 | 9.182537 | 9.082979 | 9.150815 | 9.152866 | 8.977504 | 9.222031 | 668 | 628 | 382 | 605 |
| 645426 | NM_00120705 | TMEM191C | transmembrane protein 191C | protein_coding | NP_001193981.1 | 33601 | . | . | . | 0.594481 | 0.755594 | 1.038366 | 1.057959 | 1.383382 | 1.746590 | 2.330064 | 2.418897 | 1.504379 | 35.477469 | 0.523402434 | 1.125650671 | 0.260313411 | 0.995760118 | 4.825128 | 5.413048 | 5.095088 | 5.121874 | 5.164603 | 5.165560 | 29 | 34 | 37 | 40 |
| 645431 | NR_024334 | FUT8-AS1 | FUT8 antisense RNA 1 | lncRNA | . | 44294 | . | . | . | 0.030617 | 0.134535 | 0.308378 | 0.327131 | 0.071247 | 0.310984 | 0.691991 | 0.747946 | 3.465481 | 13.188920 | 0.917088423 | 1.955161294 | 0.050564035 | NA | 2.569585 | 4.353326 | 3.634828 | 3.667002 | 3.711922 | 3.720104 | 3 | 10 | 17 | 20 |
| 645432 | NM_00108052 | ARRDC5 | arrestin domain containing 5 | protein_coding | NP_001073992.2;NP_001354118.1;XP_005259673.1;XP_011526483.1;XP | 31407 | . | . | . | 0.031824 | 0.004798 | 0.000575 | 0.016044 | 0.074056 | 0.011090 | 0.001289 | 0.036682 | 1.234655 | 1.002957 | 2.868339486 | 0.106022226 | 0.915564717 | NA | -0.155866 | 0.148375 | -0.001261 | -0.000972 | 0.000053 | -0.000315 | 1 | 1 | 1 | 1 |
| 645441 | gene-RPL17P6 | RPL17P6 | ribosomal protein L17 pseudogene 6 | pseudogene | . | 36520 | . | . | . | 13.171231 | 11.990872 | 11.768597 | 14.271255 | 30.649998 | 27.717438 | 26.408390 | 32.629528 | 1.022241 | 243.884034 | 0.225234781 | 0.140897561 | 0.887950869 | 0.998865404 | 7.914358 | 7.945576 | 7.948724 | 7.897888 | 7.889107 | 7.977817 | 292 | 247 | 195 | 252 |
| 645453 | gene-RLIMP3 | RLIMP3 | ring finger protein, LIM domain interacting pseudogene 3 | pseudogene | . | 39684 | . | . | . | 0.025098 | 0.022055 | 0.016186 | 0.043141 | 0.058405 | 0.050980 | 0.036322 | 0.098636 | -1.021598 | 2.202198 | 2.006095747 | -0.01536684 | 0.987739514 | NA | 1.156660 | 1.121007 | 1.145060 | 1.141477 | 1.138432 | 1.147659 | 3 | 2 | 1 | 3 |
| 645455 | NR_003135 | CEP170P1 | centrosomal protein 170 pseudogene 1 | transcribed_ps | . | 28364 | . | . | . | 5.057396 | 6.097347 | 6.023260 | 6.191428 | 11.768770 | 14.094289 | 13.516020 | 14.155965 | 1.079786 | 197.707841 | 0.242081319 | 0.457473246 | 0.647330928 | 0.998865404 | 7.571193 | 7.681164 | 7.568266 | 7.645740 | 7.638301 | 7.645955 | 200 | 223 | 177 | 194 |
| 645468 | gene-LOC6454 | LOC645468 | insulin like growth factor 2 mRNA binding protein 3 pseudogene | pseudogene | . | . | . | . | . | 0.006631 | 0.010351 | 0.009125 | 0.009103 | 0.015430 | 0.023926 | 0.020477 | 0.020812 | -1.225404 | 1.234643 | 2.621948492 | -0.11184694 | 0.910944771 | NA | 0.444631 | 0.148375 | 0.303453 | 0.308168 | 0.304847 | 0.304457 | 1 | 2 | 1 | 1 |
| 645513 | NR_037630 | LOC645513 | septin 7 pseudogene | transcribed_ps | . | . | . | . | . | 3.267046 | 2.761282 | 3.430832 | 3.744709 | 7.602551 | 6.382827 | 7.698687 | 8.561833 | 1.174835 | 108.418444 | 0.312264857 | 0.744426602 | 0.45661843 | 0.998865404 | 6.639350 | 6.872198 | 6.757705 | 6.702966 | 6.769544 | 6.795196 | 125 | 98 | 98 | 114 |
| 645529 | gene-ACRP1 | ACRP1 | ACR pseudogene 1 | pseudogene | . | 54745 | . | . | . | 0.138590 | 0.066139 | 0.062712 | 0.161092 | 0.322504 | 0.152884 | 0.140723 | 0.368318 | 1.059782 | 3.190579 | 1.688200453 | 0.049619648 | 0.960425489 | NA | 1.631267 | 1.715150 | 1.681404 | 1.669604 | 1.671959 | 1.680780 | 5 | 2 | 2 | 4 |
| 645548 | gene-HSPD1P6 | HSPD1P6 | heat shock protein family D (Hsp60) member 1 pseudogene 6 | pseudogene | . | 5267 | . | . | . | 0.327296 | 0.308235 | 0.158130 | 0.270468 | 0.761631 | 0.712498 | 0.354840 | 0.618394 | -1.484667 | 19.097390 | 0.716882308 | -0.79530433 | 0.426436493 | NA | 4.513800 | 3.940155 | 4.279418 | 4.274028 | 4.229002 | 4.260977 | 27 | 24 | 10 | 18 |
| 645553 | NR_135234 | IGFL2-AS1 | IGFL2 antisense RNA 1 | lncRNA | . | 52559 | . | . | . | 1.436231 | 1.341275 | 1.982698 | 2.317881 | 3.342167 | 3.100417 | 4.449118 | 5.299560 | 1.523875 | 90.968243 | 0.336696065 | 1.805024612 | 0.071070831 | 0.751131579 | 6.171846 | 6.779267 | 6.441347 | 6.424330 | 6.529644 | 6.574272 | 86 | 75 | 89 | 110 |
| 645579 | gene-EIF4BP6 | EIF4BP6 | eukaryotic translation initiation factor 4B pseudogene 6 | pseudogene | . | 37939 | . | . | . | 1.523887 | 1.330832 | 1.345603 | 1.442234 | 3.546147 | 3.076277 | 3.019494 | 3.297496 | -1.041302 | 169.242034 | 0.256848447 | -0.22732483 | 0.820171165 | 0.998865404 | 7.431727 | 7.373575 | 7.441303 | 7.380411 | 7.382758 | 7.406202 | 213 | 173 | 140 | 160 |
| 645591 | NR_034141,NF | LHFPL3-AS1 | LHFPL3 antisense RNA 1 | lncRNA | . | 40341 | . | . | . | 11.304892 | 12.069958 | 10.261708 | 13.487393 | 26.306949 | 27.900250 | 23.026973 | 30.837322 | -1.030924 | 409.278473 | 0.19889044 | -0.22091716 | 0.82515694 | 0.998865404 | 8.699407 | 8.654115 | 8.658513 | 8.708439 | 8.575235 | 8.748172 | 459 | 467 | 299 | 433 |
| 645619 | gene-AK4P3 | AK4P3 | adenylate kinase 4 pseudogene 3 | pseudogene | . | 21596 | . | . | . | 1.109289 | 0.894659 | 1.161335 | 1.071255 | 2.581362 | 2.068044 | 2.606002 | 2.449298 | 1.097722 | 77.816417 | 0.364166047 | 0.369371216 | 0.71185105 | 0.998865404 | 6.212383 | 6.348417 | 6.295414 | 6.238476 | 6.302764 | 6.279783 | 95 | 71 | 74 | 73 |
| 645644 | NR_024492 | FLJ42627 | uncharacterized LOC645644 | lncRNA | . | . | . | . | . | 0.129663 | 0.171863 | 0.249570 | 0.154107 | 0.301730 | 0.397270 | 0.560028 | 0.352348 | 1.320711 | 39.733099 | 0.516113444 | 0.777570705 | 0.436822137 | 0.998865404 | 5.096985 | 5.499560 | 5.257959 | 5.296545 | 5.360411 | 5.278009 | 34 | 42 | 49 | 32 |
| 645668 | gene-TUFMP1 | TUFMP1 | Tu translation elongation factor, mitochondrial pseudogene 1 | pseudogene | . | 12421 | . | . | . | 2.484474 | 3.012266 | 2.811679 | 2.296447 | 5.781473 | 6.962988 | 6.309326 | 5.250553 | -1.086356 | 135.595639 | 0.289267561 | -0.41310059 | 0.679532915 | 0.998865404 | 7.141438 | 7.022443 | 7.061391 | 7.129922 | 7.102471 | 7.027940 | 148 | 166 | 125 | 109 |
| 645676 | NR_147963,NF | ASH1L-AS1 | ASH1L antisense RNA 1 | lncRNA | . | 44146 | . | . | . | 1.365089 | 0.781316 | 0.956683 | 1.089577 | 3.176619 | 1.806046 | 2.146769 | 2.491189 | -1.044017 | 40.040387 | 0.51525237 | -0.12061092 | 0.903999223 | 0.998865404 | 5.353365 | 5.292767 | 5.368000 | 5.279602 | 5.309011 | 5.327101 | 60 | 32 | 32 | 39 |
| 645687 | NR_026796,NF | LINC00520 | long intergenic non-protein coding RNA 520 | lncRNA | . | 19843 | . | . | . | 13.257645 | 11.433361 | 15.298853 | 18.995354 | 30.851087 | 26.428724 | 34.330185 | 43.430621 | 1.368673 | 926.871760 | 0.158563384 | 2.855498586 | 0.004296931 | 0.193526306 | 9.612231 | 10.064863 | 9.777964 | 9.673934 | 9.873939 | 10.028867 | 969 | 781 | 836 | 1107 |
| 645691 | gene-HNRNPA1 | HNRNPA1P35 | heterogeneous nuclear ribonucleoprotein A1 pseudogene 35 | pseudogene | . | 48764 | . | . | . | 7.049213 | 6.179858 | 6.150823 | 6.578290 | 16.403811 | 14.285018 | 13.802267 | 15.040477 | -1.051925 | 247.817616 | 0.220491137 | -0.33122437 | 0.74047501 | 0.998865404 | 7.989089 | 7.916262 | 8.000505 | 7.929219 | 7.926345 | 7.954183 | 313 | 255 | 204 | 233 |
| 645693 | gene-EEF1A1P2 | EEF1A1P22 | eukaryotic translation elongation factor 1 alpha 1 pseudogene 22 | pseudogene | . | 37895 | . | . | . | 0.020238 | 0.053231 | 0.046918 | 0.027356 | 0.047095 | 0.123046 | 0.105283 | 0.062547 | 1.235624 | 2.525094 | 1.867936317 | 0.163410023 | 0.870195612 | NA | 1.175403 | 1.481107 | 1.324448 | 1.329469 | 1.332382 | 1.326463 | 2 | 3 | 3 | 2 |
| 645700 | NR_034163 | ZNF890P | zinc finger protein 890, pseudogene | transcribed_ps | . | 38691 | . | . | . | 0.057495 | 0.162440 | 0.096197 | 0.185603 | 0.133793 | 0.375487 | 0.215863 | 0.424359 | 1.305754 | 6.791087 | 1.189073998 | 0.323683256 | 0.746177848 | NA | 2.562483 | 2.940155 | 2.740052 | 2.762951 | 2.751223 | 2.768257 | 4 | 9 | 5 | 9 |
| 645784 | NR_015424 | ANKRD36BP2 | ankyrin repeat domain 36B pseudogene 2 | transcribed_ps | . | 33607 | . | . | . | 1.337692 | 1.417278 | 1.147633 | 1.108126 | 3.112865 | 3.276103 | 2.575254 | 2.533598 | -1.230162 | 67.925229 | 0.382512483 | -0.78127579 | 0.434640308 | 0.998865404 | 6.227558 | 5.928749 | 6.104267 | 6.117558 | 6.067124 | 6.058592 | 84 | 83 | 54 | 56 |
| 645811 | NM_00114398 | CCDC154 | coiled-coil domain containing 154 | protein_coding | NP_001137452.1 | 34454 | 618740 | . | . | 0.186293 | 0.196285 | 0.262396 | 0.195455 | 0.433512 | 0.453721 | 0.588809 | 0.446885 | 1.197104 | 14.798516 | 0.795811104 | 0.326143851 | 0.744315521 | NA | 3.751025 | 4.011955 | 3.873978 | 3.878005 | 3.897007 | 3.877988 | 15 | 15 | 16 | 13 |
| 645899 | gene-RPS28P4 | RPS28P4 | ribosomal protein S28 pseudogene 4 | pseudogene | . | 36208 | . | . | . | 1.377567 | 1.821828 | 2.389498 | 2.520623 | 3.205654 | 4.211237 | 5.361965 | 5.763104 | 1.481427 | 24.556688 | 0.624347605 | 0.908127928 | 0.363810637 | NA | 4.307819 | 4.873224 | 4.576707 | 4.598236 | 4.624761 | 4.629591 | 20 | 24 | 25 | 28 |
| 645931 | gene-FEN1P1 | FEN1P1 | flap structure-specific endonuclease 1 pseudogene 1 | pseudogene | . | 3651 | . | . | . | 0.109917 | 0.065020 | 0.162466 | 0.036528 | 0.255782 | 0.150297 | 0.364569 | 0.083517 | 1.081741 | 6.499868 | 1.22943355 | 0.092201625 | 0.926537847 | NA | 2.638039 | 2.760196 | 2.698670 | 2.683786 | 2.709187 | 2.677615 | 9 | 5 | 9 | 3 |
| 645949 | NR_038837 | LINC01822 | long intergenic non-protein coding RNA 1822 | lncRNA | . | 52627 | . | . | . | 0.074937 | 0.111454 | 0.139622 | 0.146162 | 0.174382 | 0.257630 | 0.313307 | 0.334183 | 1.582033 | 8.143596 | 1.06672418 | 0.620385271 | 0.535004167 | NA | 2.658167 | 3.318300 | 3.000346 | 3.010810 | 3.023722 | 3.020625 | 6 | 8 | 9 | 9 |
| 645958 | NR_077247 | RPS18P9 | ribosomal protein S18 pseudogene 9 | transcribed_ps | . | 36483 | . | . | . | 0.387413 | 0.372925 | 0.233364 | 0.542701 | 0.901525 | 0.862033 | 0.523662 | 1.240821 | 1.040784 | 12.330591 | 0.887806619 | 0.064959235 | 0.948206457 | NA | 3.597285 | 3.650563 | 3.622924 | 3.622439 | 3.605401 | 3.644000 | 14 | 13 | 7 | 16 |
| 645965 | gene-LOC6459 | LOC645965 | adipose differentiation-related protein pseudogene | pseudogene | . | . | . | . | . | 0.130500 | 0.113008 | 0.107855 | 0.119443 | 0.303678 | 0.261223 | 0.242023 | 0.273093 | 1.004159 | 7.167427 | 1.127206955 | 0.005311571 | 0.995762 | NA | 2.838265 | 2.844638 | 2.845094 | 2.838919 | 2.840897 | 2.843391 | 9 | 7 | 6 | 7 |
| 645968 | gene-RPS3AP2 | RPS3AP25 | RPS3A pseudogene 25 | pseudogene | . | 36801 | . | . | . | 0.220982 | 0.215729 | 0.086963 | 0.096272 | 0.514234 | 0.498668 | 0.195143 | 0.220114 | -1.890214 | 4.804058 | 1.390306576 | -0.66068137 | 0.508816673 | NA | 2.651489 | 1.733338 | 2.280421 | 2.282212 | 2.268487 | 2.267229 | 7 | 7 | 3 | 3 |
| 645969 | gene-SELENOTI | SELENOTP1 | selenoprotein T pseudogene 1 | pseudogene | . | 53722 | . | . | . | 0.070886 | 0.023196 | 0.050008 | 0.005409 | 0.164956 | 0.053618 | 0.112217 | 0.012367 | -1.569340 | 4.353188 | 1.532138176 | -0.42434701 | 0.671312759 | NA | 2.405477 | 1.769039 | 2.141056 | 2.117422 | 2.130914 | 2.109910 | 9 | 3 | 5 | 1 |
| 645979 | gene-RPS26P3 | RPS26P31 | ribosomal protein S26 pseudogene 31 | pseudogene | . | 36614 | . | . | . | 0.480871 | 0.562995 | 0.163154 | 0.294527 | 1.119008 | 1.301389 | 0.366113 | 0.673402 | -2.317952 | 5.463714 | 1.331409912 | -0.91095214 | 0.362320592 | NA | 2.934350 | 1.715150 | 2.469781 | 2.476042 | 2.448989 | 2.457518 | 8 | 9 | 2 | 4 |
| 646024 | NR_024045 | RAET1K | retinoic acid early transcript 1K pseudogene | transcribed_ps | . | 16797 | . | . | . | 0.371855 | 0.641283 | 0.340223 | 0.799938 | 0.865323 | 1.482353 | 0.763451 | 1.828963 | 1.121102 | 32.052231 | 0.603080934 | 0.273459334 | 0.784500148 | 0.998865404 | 4.920675 | 5.079653 | 4.952497 | 5.016363 | 4.947489 | 5.049429 | 26 | 41 | 18 | 44 |
| 646050 | gene-AIDAP1 | AIDAP1 | AIDA pseudogene 1 | pseudogene | . | 54663 | . | . | . | 0.025871 | 0.077593 | 0.009876 | 0.009075 | 0.060203 | 0.179359 | 0.022161 | 0.020749 | -2.452425 | 1.915126 | 2.205695884 | -0.58675761 | 0.55736651 | NA | 1.444631 | 0.148375 | 0.948324 | 0.957585 | 0.945780 | 0.945324 | 2 | 4 | 1 | 1 |
| 646104 | gene-RPL32P1 | RPL32P16 | ribosomal protein L32 pseudogene 16 | pseudogene | . | 36086 | . | . | . | 0.347850 | 0.542616 | 0.000220 | 0.616098 | 0.809461 | 1.254282 | 0.000494 | 1.408634 | -1.288317 | 6.790797 | 1.267223733 | -0.28841581 | 0.773028466 | NA | 2.939842 | 2.562749 | 2.763178 | 2.777842 | 2.741726 | 2.783561 | 7 | 10 | 1 | 10 |
| 646192 | gene-NOP56P1 | NOP56P1 | NOP56 ribonucleoprotein pseudogene 1 | pseudogene | . | 13962 | . | . | . | 2.559300 | 1.643124 | 3.874172 | 3.974974 | 5.955597 | 3.798155 | 8.693530 | 9.088305 | 1.809193 | 45.066852 | 0.480161018 | 1.781374082 | 0.074851359 | 0.765313565 | 5.002297 | 5.860037 | 5.451741 | 5.394611 | 5.520260 | 5.525269 | 45 | 27 | 50 | 55 |
| 646195 | gene-RPS28P7 | RPS28P7 | ribosomal protein S28 pseudogene 7 | pseudogene | . | 35787 | . | . | . | 32.726109 | 34.548264 | 34.869316 | 37.845486 | 76.154999 | 79.859856 | 78.245735 | 86.529205 | 1.066124 | 409.184259 | 0.180225726 | 0.512551795 | 0.608264881 | 0.998865404 | 8.629926 | 8.721825 | 8.643085 | 8.669730 | 8.672887 | 8.716031 | 446 | 437 | 354 | 410 |
| 646202 | gene-TLK2P1 | TLK2P1 | tousled like kinase 2 pseudogene 1 | pseudogene | . | 18048 | . | . | . | 0.116120 | 0.159473 | 0.083532 | 0.099549 | 0.270216 | 0.368629 | 0.187444 | 0.227608 | -1.493147 | 12.393626 | 0.873515929 | -0.66210143 | 0.507906218 | NA | 3.892781 | 3.312263 | 3.639409 | 3.655532 | 3.622962 | 3.629068 | 15 | 18 | 8 | 10 |
| 646214 | NR_027053 | LOC646214 | p21 (RAC1) activated kinase 2 pseudogene | transcribed_ps | . | . | . | . | . | 1.023061 | 1.070787 | 1.553102 | 0.872399 | 2.380704 | 2.475172 | 3.485116 | 1.994637 | 1.144315 | 290.186078 | 0.258382393 | 0.752701497 | 0.45162931 | 0.998865404 | 8.079449 | 8.275561 | 8.121850 | 8.140359 | 8.346717 | 8.035398 | 306 | 297 | 346 | 208 |
| 646270 | gene-ELL2P1 | ELL2P1 | elongation factor for RNA polymerase II 2 pseudogene 1 | pseudogene | . | 39343 | . | . | . | 0.015614 | 0.033610 | 0.031008 | 0.015355 | 0.036334 | 0.077691 | 0.069581 | 0.035106 | -1.131926 | 2.973891 | 1.729253768 | -0.10338572 | 0.917656846 | NA | 1.658167 | 1.481107 | 1.569843 | 1.575066 | 1.573727 | 1.567785 | 3 | 4 | 3 | 2 |
| 646278 | NR_037599 | PDCD6IPP2 | PDCD6IP pseudogene 2 | transcribed_ps | . | 49873 | . | . | . | 0.400949 | 0.467516 | 0.384988 | 0.376449 | 0.933024 | 1.080684 | 0.863903 | 0.860706 | -1.172146 | 27.065951 | 0.59360932 | -0.38603298 | 0.699472241 | 0.998865404 | 4.868663 | 4.639023 | 4.759188 | 4.778144 | 4.752797 | 4.749542 | 31 | 34 | 22 | 23 |
| 646294 | gene-RPS2P29 | RPS2P29 | ribosomal protein S2 pseudogene 29 | pseudogene | . | 36011 | . | . | . | 0.120482 | 0.177000 | 0.204569 | 0.110374 | 0.280365 | 0.409144 | 0.459047 | 0.252357 | 1.017348 | 5.000631 | 1.358013481 | 0.018271916 | 0.985421932 | NA | 2.307819 | 2.336262 | 2.313958 | 2.319705 | 2.325426 | 2.308943 | 5 | 6 | 6 | 3 |
| 646309 | gene-NAMPTP | NAMPTP1 | nicotinamide phosphoribosyltransferase pseudogene 1 | pseudogene | . | 17633 | . | . | . | 1.878214 | 1.990815 | 1.867893 | 1.477735 | 4.370681 | 4.601858 | 4.191499 | 3.378666 | -1.170195 | 141.094206 | 0.282419608 | -0.80287792 | 0.422045301 | 0.998865404 | 7.248861 | 7.023367 | 7.158827 | 7.177475 | 7.151663 | 7.065085 | 171 | 168 | 127 | 107 |
| 646347 | gene-LOC6463 | LOC646347 | spermine synthase pseudogene | pseudogene | . | . | . | . | . | 0.297626 | 0.220900 | 0.153919 | 0.375181 | 0.692589 | 0.510620 | 0.345390 | 0.857806 | 1.034994 | 11.796836 | 0.9181667 | 0.054045134 | 0.956899205 | NA | 3.537347 | 3.582949 | 3.565604 | 3.552775 | 3.538993 | 3.582604 | 15 | 11 | 6 | 16 |
| 646405 | NR_026730 | TPTE2P1 | transmembrane phosphoinositide 3-phosphatase and tensin hom | transcribed_ps | . | 35196 | . | . | . | 0.051704 | 0.072430 | 0.132577 | 0.057641 | 0.120317 | 0.167425 | 0.297499 | 0.131789 | 1.503113 | 16.362596 | 0.794860214 | 0.739694519 | 0.459485374 | NA | 3.706954 | 4.297658 | 3.990560 | 4.006783 | 4.059498 | 3.993601 | 13 | 16 | 24 | 11 |
| 646450 | NR_161301 | ARIH2OS | ARIH2 opposite strand lncRNA | lncRNA | . | 34425 | . | . | . | 0.688718 | 0.530671 | 0.722229 | 0.415620 | 1.602676 | 1.226670 | 1.620662 | 0.950265 | -1.063069 | 29.936449 | 0.585940505 | -0.15058784 | 0.880300853 | 0.998865404 | 4.945169 | 4.861274 | 4.920335 | 4.887821 | 4.926951 | 4.865645 | 40 | 29 | 32 | 20 |
| 646476 | gene-RPL7AP3 | RPL7AP39 | ribosomal protein L7a pseudogene 39 | pseudogene | . | 35881 | . | . | . | 2.175989 | 2.436643 | 2.943316 | 3.338252 | 5.063616 | 5.632410 | 6.604715 | 7.632517 | 1.345702 | 73.183634 | 0.370890879 | 1.154944714 | 0.248113067 | 0.992733415 | 5.964086 | 6.391300 | 6.136180 | 6.160715 | 6.206239 | 6.236090 | 68 | 71 | 69 | 83 |
| 646483 | gene-RPL6P19 | RPL6P19 | ribosomal protein L6 pseudogene 19 | pseudogene | . | 36754 | . | . | . | 1.822706 | 1.793174 | 1.494425 | 2.057486 | 4.241511 | 4.145001 | 3.353446 | 4.704196 | -1.023279 | 51.809597 | 0.440245418 | -0.07541184 | 0.939887037 | 0.998865404 | 5.712495 | 5.677589 | 5.699504 | 5.695444 | 5.665877 | 5.719617 | 61 | 56 | 38 | 55 |
| 646517 | gene-LOC6465 | LOC646517 | WT1 associated protein pseudogene | pseudogene | . | . | . | . | . | 0.407159 | 0.540732 | 0.348605 | 0.645702 | 0.947476 | 1.249927 | 0.782258 | 1.476322 | 1.052905 | 13.383527 | 0.845542395 | 0.087960894 | 0.929907757 | NA | 3.706954 | 3.776970 | 3.731372 | 3.747340 | 3.728023 | 3.756657 | 13 | 16 | 9 | 16 |
| 646531 | gene-YBX1P2 | YBX1P2 | Y-box binding protein 1 pseudogene 2 | pseudogene | . | 22635 | . | . | . | 0.250407 | 0.201393 | 0.144373 | 0.254646 | 0.582707 | 0.465530 | 0.323968 | 0.582219 | -1.112204 | 11.017662 | 0.925727585 | -0.16573072 | 0.868368868 | NA | 3.537347 | 3.381963 | 3.474021 | 3.461142 | 3.452144 | 3.472594 | 15 | 11 | 7 | 12 |
| 646555 | gene-ARID3BP | ARID3BP1 | ARID3B pseudogene 1 | pseudogene | . | 54553 | . | . | . | 0.040969 | 0.065083 | 0.055320 | 0.049156 | 0.095337 | 0.150442 | 0.124135 | 0.112389 | -1.061742 | 7.449091 | 1.113902058 | -0.07759548 | 0.938149836 | NA | 2.939842 | 2.852980 | 2.889028 | 2.903806 | 2.898069 | 2.890790 | 7 | 10 | 7 | 6 |
| 646609 | gene-CLIC4P1 | CLIC4P1 | chloride intracellular channel 4 pseudogene 1 | pseudogene | . | 49797 | . | . | . | 0.164655 | 0.139042 | 0.161802 | 0.034267 | 0.383158 | 0.321402 | 0.363080 | 0.078348 | -1.430486 | 3.428938 | 1.65997574 | -0.31115212 | 0.755684977 | NA | 2.008843 | 1.502474 | 1.783939 | 1.780895 | 1.784763 | 1.768843 | 5 | 4 | 4 | 1 |
| 646612 | gene-EEF1A1P2 | EEF1A1P25 | eukaryotic translation elongation factor 1 alpha 1 pseudogene 25 | pseudogene | . | 37898 | . | . | . | 0.759661 | 0.728251 | 0.491887 | 0.840607 | 1.767762 | 1.683384 | 1.103781 | 1.921949 | -1.135988 | 32.823796 | 0.555776255 | -0.33097379 | 0.740664286 | 0.998865404 | 5.127131 | 4.940155 | 5.049047 | 5.043605 | 4.999161 | 5.059329 | 41 | 37 | 20 | 36 |
| 646658 | NM_00110557 | SYNDIG1L | synapse differentiation inducing 1 like | protein_coding | NP_001099049.1;XP_016877089.1 | 32388 | 609999 | . | . | 0.039050 | 0.028397 | 0.030322 | 0.010374 | 0.090871 | 0.065641 | 0.068041 | 0.023719 | -1.477981 | 4.866116 | 1.396863018 | -0.40349556 | 0.686583701 | NA | 2.533720 | 1.978782 | 2.296576 | 2.285700 | 2.290659 | 2.274450 | 8 | 5 | 5 | 2 |
| 646660 | gene-RPS20P2 | RPS20P26 | ribosomal protein S20 pseudogene 26 | pseudogene | . | 36980 | . | . | . | 0.178856 | 0.389199 | 0.185230 | 0.295855 | 0.416206 | 0.899651 | 0.415650 | 0.676437 | -1.129525 | 4.686060 | 1.400898652 | -0.12543149 | 0.900181963 | NA | 2.316288 | 2.134756 | 2.223529 | 2.237674 | 2.223842 | 2.232207 | 4 | 7 | 3 | 5 |
| 646719 | NR_046262 | NIPBL-DT | NIPBL divergent transcript | lncRNA | . | 51293 | . | . | . | 1.253671 | 0.875432 | 1.376797 | 0.713111 | 2.917343 | 2.023601 | 3.089492 | 1.630444 | -1.034798 | 175.908835 | 0.321438946 | -0.15352619 | 0.877983338 | 0.998865404 | 7.481626 | 7.435371 | 7.527226 | 7.369166 | 7.562419 | 7.292694 | 243 | 158 | 199 | 110 |
| 646762 | NR_024278 | LOC646762 | uncharacterized LOC646762 | lncRNA | . | . | . | . | . | 2.828917 | 3.207129 | 3.353299 | 3.497362 | 6.583005 | 7.413422 | 7.524705 | 7.996303 | 1.119597 | 580.666727 | 0.162899078 | 1.000490427 | 0.317073228 | 0.998865404 | 9.098028 | 9.260532 | 9.102924 | 9.179378 | 9.205563 | 9.226648 | 594 | 626 | 525 | 584 |
| 646766 | gene-RPL35P2 | RPL35P2 | ribosomal protein L35 pseudogene 2 | pseudogene | . | 35789 | . | . | . | 0.110022 | 0.076387 | 0.033921 | 0.247164 | 0.256026 | 0.176571 | 0.076117 | 0.565111 | 1.516129 | 2.251754 | 2.002462329 | 0.29982714 | 0.764309013 | NA | 0.844134 | 1.437399 | 1.163414 | 1.164008 | 1.160961 | 1.175061 | 2 | 2 | 1 | 4 |
| 646780 | gene-PHKA1P1 | PHKA1P1 | phosphorylase kinase regulatory subunit alpha 1 pseudogene | pseudogene | . | 33919 | . | . | . | 0.062480 | 0.107567 | 0.095854 | 0.044575 | 0.145393 | 0.248645 | 0.215094 | 0.101915 | -1.131926 | 2.973891 | 1.729253768 | -0.10338572 | 0.917656846 | NA | 1.658167 | 1.481107 | 1.569843 | 1.575066 | 1.573727 | 1.567785 | 3 | 4 | 3 | 2 |
| 646802 | NR_103761 | CYP4F62P | cytochrome P450 family 4 subfamily F member 62, pseudogene | transcribed_ps | . | 39956 | . | . | . | 0.066117 | 0.056825 | 0.066443 | 0.054924 | 0.153856 | 0.131353 | 0.149096 | 0.125577 | 1.015669 | 4.965229 | 1.346923932 | 0.01665281 | 0.986713594 | NA | 2.299300 | 2.324312 | 2.311711 | 2.308997 | 2.313995 | 2.307397 | 6 | 5 | 5 | 4 |
| 646808 | gene-LAGE3P1 | LAGE3P1 | L antigen family member 3 pseudogene 1 | pseudogene | . | 49909 | . | . | . | 0.190926 | 0.251662 | 0.223999 | 0.373280 | 0.444292 | 0.581729 | 0.502647 | 0.853459 | 1.222121 | 5.492312 | 1.289276996 | 0.224457361 | 0.822401441 | NA | 2.307819 | 2.592943 | 2.448470 | 2.454248 | 2.449997 | 2.462756 | 5 | 6 | 4 | 7 |
| 646819 | gene-RPS15AP | RPS15AP19 | ribosomal protein S15a pseudogene 19 | pseudogene | . | 36603 | . | . | . | 0.799468 | 1.271025 | 0.993645 | 0.927382 | 1.860394 | 2.938032 | 2.229711 | 2.120348 | -1.064079 | 14.877356 | 0.799326484 | -0.11210149 | 0.910742938 | NA | 3.939842 | 3.848815 | 3.881314 | 3.909704 | 3.893926 | 3.889292 | 14 | 20 | 13 | 13 |
| 646839 | gene-TPM3P8 | TPM3P8 | tropomyosin 3 pseudogene 8 | pseudogene | . | 39171 | . | . | . | 0.573018 | 0.557881 | 0.483896 | 0.531022 | 1.333437 | 1.289567 | 1.085849 | 1.214119 | -1.127005 | 34.667177 | 0.525652067 | -0.32815187 | 0.742796838 | 0.998865404 | 5.199399 | 5.026416 | 5.128154 | 5.123227 | 5.103402 | 5.115236 | 43 | 39 | 27 | 32 |
| 646851 | NM_00101364 | FAM227A | family with sequence similarity 227 member A | protein_coding | NP_001013669.1;NP_001277959.1;XP_006724354.2;XP_006724355.2;XP | 44197 | . | . | . | 1.266523 | 1.429324 | 1.361202 | 1.354383 | 2.947249 | 3.303945 | 3.054500 | 3.096637 | 1.095942 | 222.791219 | 0.227903101 | 0.579948465 | 0.561949371 | 0.998865404 | 7.731670 | 7.864377 | 7.802821 | 7.748337 | 7.822593 | 7.816368 | 259 | 216 | 204 | 217 |
| 646864 | NM_00134972 | ZNF723 | zinc finger protein 723 | protein_coding | NP_001336655.1;XP_024307420.1;XP_024307421.1 | 32286 | . | . | . | 0.056196 | 0.093843 | 0.105492 | 0.064695 | 0.130770 | 0.216921 | 0.236721 | 0.147918 | 1.390349 | 8.158593 | 1.103285617 | 0.430937765 | 0.666513623 | NA | 2.772726 | 3.245375 | 2.991795 | 3.023342 | 3.028417 | 3.010900 | 4 | 11 | 10 | 7 |
| 646890 | gene-AIDAP2 | AIDAP2 | AIDA pseudogene 2 | pseudogene | . | 54664 | . | . | . | 0.069664 | 0.121521 | 0.204859 | 0.028795 | 0.162112 | 0.280902 | 0.459697 | 0.065837 | 1.254241 | 4.088884 | 1.569974973 | 0.208165181 | 0.835099995 | NA | 1.855800 | 2.188478 | 2.006873 | 2.016579 | 2.031466 | 2.000171 | 3 | 5 | 7 | 1 |
| 646903 | NR_036538 | LOC646903 | uncharacterized LOC646903 | lncRNA | . | . | . | . | . | 0.842695 | 0.949175 | 0.803965 | 0.612730 | 1.960986 | 2.194061 | 1.804075 | 1.400933 | -1.284543 | 49.535532 | 0.450731424 | -0.80148589 | 0.422850409 | 0.998865404 | 5.799086 | 5.439328 | 5.643251 | 5.662959 | 5.629925 | 5.584719 | 61 | 63 | 43 | 35 |
| 646962 | NM_00103979 | HRCT1 | histidine rich carboxyl terminus 1 | protein_coding | NP_001034881.1 | 33872 | . | . | . | 2.852785 | 2.415402 | 1.854030 | 2.426953 | 6.638547 | 5.583311 | 4.160390 | 5.548939 | -1.249397 | 70.011761 | 0.386308751 | -0.83154181 | 0.405667616 | 0.998865404 | 6.281682 | 5.959409 | 6.178329 | 6.134521 | 6.075765 | 6.131184 | 97 | 77 | 47 | 66 |
| 647000 | gene-TUBBP2 | TUBBP2 | tubulin beta pseudogene 2 | pseudogene | . | 12415 | . | . | . | 0.114064 | 0.102552 | 0.119962 | 0.107985 | 0.265431 | 0.237053 | 0.269191 | 0.246895 | 1.124984 | 5.231896 | 1.310758984 | 0.129622743 | 0.896864909 | NA | 2.299300 | 2.470304 | 2.384811 | 2.382116 | 2.387186 | 2.385366 | 6 | 5 | 5 | 5 |
| 647009 | gene-RWDD4P | RWDD4P2 | RWD domain containing 4 pseudogene 2 | pseudogene | . | 23754 | . | . | . | 0.960421 | 0.883046 | 0.720418 | 0.485494 | 2.234939 | 2.041200 | 1.616598 | 1.110025 | -1.562527 | 61.354464 | 0.417073644 | -1.54380664 | 0.122635181 | 0.866753766 | 6.224186 | 5.583440 | 5.995198 | 5.971252 | 5.922673 | 5.852651 | 90 | 77 | 50 | 36 |
| 647030 | gene-EEF1B2P6 | EEF1B2P6 | eukaryotic translation elongation factor 1 beta 2 pseudogene 6 | pseudogene | . | 51453 | . | . | . | 1.205444 | 1.308802 | 1.397573 | 2.226097 | 2.805117 | 3.025355 | 3.136114 | 5.089706 | 1.422905 | 36.929741 | 0.526068928 | 0.96724937 | 0.333419387 | 0.998865404 | 4.931597 | 5.437671 | 5.166201 | 5.174758 | 5.182925 | 5.258354 | 34 | 34 | 29 | 50 |
| 647042 | NM_00116446 | GOLGA6L10 | golgin A6 family like 10 | protein_coding | NP_001157937.2 | 37228 | . | . | . | 0.396226 | 0.340009 | 0.503520 | 0.297035 | 0.922034 | 0.785947 | 1.129885 | 0.679135 | 1.074339 | 53.879866 | 0.449565869 | 0.230109573 | 0.818006627 | 0.998865404 | 5.697260 | 5.804111 | 5.751474 | 5.723835 | 5.800926 | 5.700003 | 64 | 52 | 61 | 39 |
| 647070 | NR_148933 | LOC647070 | uncharacterized LOC647070 | lncRNA | . | . | . | . | . | 0.193480 | 0.202455 | 0.205327 | 0.227113 | 0.450236 | 0.467983 | 0.460749 | 0.519268 | 1.095636 | 29.604645 | 0.56594095 | 0.232830579 | 0.815892966 | 0.998865404 | 4.820670 | 4.951852 | 4.878564 | 4.882564 | 4.887328 | 4.895692 | 32 | 31 | 26 | 30 |
| 647076 | gene-BRD7P2 | BRD7P2 | bromodomain containing 7 pseudogene 2 | pseudogene | . | 33541 | . | . | . | 0.015989 | 0.198768 | 0.123600 | 0.018763 | 0.037206 | 0.459462 | 0.277354 | 0.042899 | -1.382037 | 6.742796 | 1.329880445 | -0.3510062 | 0.7255837 | NA | 2.966989 | 2.502474 | 2.713884 | 2.770304 | 2.748375 | 2.716925 | 2 | 15 | 8 | 2 |
| 647081 | gene-SDHDP2 | SDHDP2 | succinate dehydrogenase complex subunit D pseudogene 2 | pseudogene | . | 10685 | . | . | . | 0.044684 | 0.082302 | 0.028428 | 0.029013 | 0.103982 | 0.190244 | 0.063792 | 0.066334 | -1.422882 | 2.686397 | 1.818605976 | -0.27978332 | 0.779643752 | NA | 1.658167 | 1.148375 | 1.430406 | 1.435592 | 1.429080 | 1.428283 | 3 | 4 | 2 | 2 |
| 647087 | NM_00113092 | STMP1 | short transmembrane mitochondrial protein 1 | protein_coding | NP_001124401.1 | 41909 | 618755 | . | . | 11.118413 | 9.807963 | 8.653400 | 11.050623 | 25.875003 | 22.671547 | 19.417978 | 25.269536 | -1.077702 | 779.113017 | 0.165899944 | -0.65074711 | 0.515209747 | 0.998865404 | 9.658866 | 9.550476 | 9.676279 | 9.580736 | 9.493278 | 9.654375 | 993 | 814 | 576 | 785 |
| 647099 | gene-RPL23AP | RPL23AP42 | ribosomal protein L23a pseudogene 42 | pseudogene | . | 35942 | . | . | . | 40.390514 | 40.132828 | 40.386726 | 44.318172 | 93.990387 | 92.768829 | 90.626640 | 101.328239 | 1.037293 | 696.205042 | 0.152235724 | 0.346980241 | 0.728606163 | 0.998865404 | 9.416852 | 9.469406 | 9.435515 | 9.425222 | 9.426696 | 9.483368 | 793 | 732 | 591 | 692 |
| 647121 | NR_003955 | EMBP1 | embigin pseudogene 1 | transcribed_ps | . | 38661 | . | . | . | 0.582631 | 0.474933 | 0.389033 | 0.290812 | 1.355806 | 1.097829 | 0.872980 | 0.664906 | -1.569215 | 57.831821 | 0.426353835 | -1.52465577 | 0.127344979 | 0.874064017 | 6.141376 | 5.494237 | 5.923168 | 5.869995 | 5.830074 | 5.780586 | 90 | 68 | 45 | 36 |
| 647135 | NM_00127187 | SRGAP2B | SLIT-ROBO Rho GTPase activating protein 2B | protein_coding | NP_001258799.2;NP_001317613.1;NP_001353209.1;NP_001355253.1;XF | 35237 | 614703 | . | . | 4.776810 | 5.455883 | 5.396380 | 5.137423 | 11.115833 | 12.611521 | 12.109316 | 11.746106 | 1.016948 | 1110.184446 | 0.138349731 | 0.175255034 | 0.860879243 | 0.998865404 | 10.104454 | 10.128612 | 10.066029 | 10.151549 | 10.141434 | 10.102503 | 1195 | 1256 | 999 | 1022 |
| 647167 | gene-EEF1A1P1 | EEF1A1P14 | eukaryotic translation elongation factor 1 alpha 1 pseudogene 14 | pseudogene | . | 3197 | . | . | . | 0.052446 | 0.065358 | 0.030962 | 0.099688 | 0.122043 | 0.151077 | 0.069479 | 0.227926 | 1.067108 | 3.703508 | 1.564493575 | 0.059895269 | 0.952239048 | NA | 1.844134 | 1.932304 | 1.888489 | 1.889602 | 1.883355 | 1.896931 | 4 | 4 | 2 | 5 |
| 647276 | gene-RPL4P4 | RPL4P4 | ribosomal protein L4 pseudogene 4 | pseudogene | . | 36179 | . | . | . | 3.136240 | 2.292581 | 2.268594 | 3.282216 | 7.298160 | 5.299404 | 5.090658 | 7.504396 | 1.009022 | 123.176901 | 0.31919836 | 0.040595427 | 0.96761843 | 0.998865404 | 6.938363 | 6.950786 | 6.994256 | 6.880674 | 6.881356 | 6.999536 | 164 | 111 | 89 | 136 |
| 647309 | NM_00114668 | GMNC | geminin coiled-coil domain containing | protein_coding | NP_001140158.1 | 40049 | 614448 | . | . | 0.095173 | 0.069125 | 0.024030 | 0.080180 | 0.221473 | 0.159785 | 0.053923 | 0.183321 | -1.568352 | 8.401780 | 1.083807868 | -0.5990445 | 0.549143213 | NA | 3.361571 | 2.705970 | 3.092650 | 3.083320 | 3.057876 | 3.084643 | 13 | 10 | 3 | 9 |
| 647310 | NM_00119508 | TEX22 | testis expressed 22 | protein_coding | NP_001182011.1;XP_006720297.1 | 40026 | . | . | . | 0.117327 | 0.078129 | 0.076069 | 0.140731 | 0.273024 | 0.180600 | 0.170696 | 0.321765 | 1.111882 | 9.867556 | 0.989020227 | 0.154702167 | 0.877056121 | NA | 3.225252 | 3.376188 | 3.308341 | 3.290585 | 3.288983 | 3.319173 | 13 | 8 | 6 | 13 |

| | | | | | | | | | | | | | | | | | | | | | | | | | | | | | | | | | |
|---|---|---|---|---|---|---|---|---|---|---|---|---|---|---|---|---|---|---|---|---|---|---|---|---|---|---|---|---|---|---|---|---|---|
| 647319 | gene-VEZF1P1 | VEZF1P1 | vascular endothelial zinc finger 1 pseudogene 1 | pseudogene | . | 32320 | . | . | . | 0.107417 | 0.029196 | 0.031334 | 0.008480 | 0.249964 | 0.067488 | 0.070314 | 0.019389 | -2.984929 | 5.600043 | 1.41585904 | -1.11430334 | 0.265149077 | NA | 3.064677 | 1.502474 | 2.529463 | 2.486348 | 2.490785 | 2.474727 | 15 | 4 | 4 | 1 |
| 647323 | NR_033944 | LINC02026 | long intergenic non-protein coding RNA 2026 | lncRNA | . | 52861 | . | . | . | 0.069289 | 0.059479 | 0.141032 | 0.129183 | 0.161238 | 0.137488 | 0.316472 | 0.295362 | 1.925772 | 5.891420 | 1.250543577 | 0.756020834 | 0.449636705 | NA | 2.008843 | 2.955730 | 2.536601 | 2.533718 | 2.553436 | 2.551042 | 5 | 4 | 7 | 7 |
| 647859 | NR_026578 | OCLNP1 | OCLN pseudogene 1 | transcribed_ps | . | 53916 | . | . | . | 0.361481 | 0.467826 | 1.171728 | 0.316429 | 0.841182 | 1.081400 | 2.629324 | 0.723478 | 1.737697 | 11.116124 | 0.976601741 | 0.816276025 | 0.414342283 | NA | 3.019256 | 3.820154 | 3.428940 | 3.439945 | 3.493499 | 3.427442 | 8 | 10 | 19 | 6 |
| 647946 | NR_024391 | MIR924HG | MIR924 host gene | lncRNA | . | 44332 | . | . | . | 0.355052 | 0.190524 | 0.194179 | 0.191309 | 0.826220 | 0.440406 | 0.435731 | 0.437406 | -1.404817 | 15.972625 | 0.788220601 | -0.62213823 | 0.53385098 | NA | 4.220748 | 3.733338 | 4.037041 | 3.987326 | 3.991520 | 3.987044 | 28 | 14 | 12 | 12 |
| 647979 | NR_027451 | NORAD | non-coding RNA activated by DNA damage | lncRNA | . | 44311 | 617037 | . | . | 39.229156 | 37.216541 | 40.478657 | 34.947578 | 91.287857 | 86.027702 | 90.832924 | 79.903488 | -1.027632 | 6321.963849 | 0.117041437 | -0.33598586 | 0.736881531 | 0.998865404 | 12.645614 | 12.606434 | 12.662207 | 12.610455 | 12.677628 | 12.545578 | 7604 | 6703 | 5848 | 5388 |
| 648000 | gene-RPL7P23 | RPL7P23 | ribosomal protein L7 pseudogene 23 | pseudogene | . | 35658 | . | . | . | 1.159088 | 1.235474 | 0.799028 | 1.169157 | 2.697244 | 2.855853 | 1.792996 | 2.673138 | -1.204753 | 29.120710 | 0.5789464 | -0.46418278 | 0.642516799 | 0.998865404 | 4.993221 | 4.721998 | 4.875262 | 4.879919 | 4.837500 | 4.873557 | 36 | 35 | 19 | 29 |
| 648987 | NR_034085,NF | LOC648987 | uncharacterized LOC648987 | lncRNA | . | . | . | . | . | 3.446278 | 2.984599 | 2.845184 | 4.612341 | 8.019634 | 6.899033 | 6.384511 | 10.545571 | 1.167931 | 233.454269 | 0.227805847 | 0.983094215 | 0.325561071 | 0.998865404 | 7.751004 | 7.974353 | 7.840098 | 7.810175 | 7.854387 | 7.944491 | 256 | 225 | 199 | 257 |
| 649395 | NR_029404 | YWHAEP1 | tyrosine 3-monooxygenase/tryptophan 5-monooxygenase activati | transcribed_ps | . | 49442 | . | . | . | 0.075051 | 0.015588 | 0.047276 | 0.016143 | 0.174647 | 0.036031 | 0.106085 | 0.036910 | -1.177196 | 2.446386 | 1.967286461 | -0.11963392 | 0.904773146 | NA | 1.397517 | 1.175234 | 1.300148 | 1.283886 | 1.295548 | 1.284795 | 5 | 1 | 3 | 1 |
| 649446 | NR_024101 | DLGAP1-AS1 | DLGAP1 antisense RNA 1 | lncRNA | . | 31676 | . | . | . | 4.557506 | 4.996526 | 4.182544 | 3.847999 | 10.605504 | 11.549694 | 9.385508 | 8.797992 | -1.204115 | 136.833300 | 0.28001692 | -0.9569873 | 0.338573664 | 0.998865404 | 7.223934 | 6.956214 | 7.113239 | 7.145770 | 7.077274 | 7.045162 | 165 | 168 | 113 | 111 |
| 650368 | NR_024248 | TSSC2 | tumor suppressing subtransferable candidate 2 (pseudogene) | transcribed_ps | . | 12384 | 608999 | . | . | 0.417157 | 0.320415 | 0.276188 | 0.285433 | 0.970742 | 0.740653 | 0.619758 | 0.652609 | -1.259013 | 15.617112 | 0.778548669 | -0.42681131 | 0.669516771 | NA | 4.121102 | 3.790061 | 3.988046 | 3.965286 | 3.961178 | 3.961304 | 23 | 16 | 12 | 13 |
| 650655 | NR_003574 | ABCA17P | ATP binding cassette subfamily A member 17, pseudogene | transcribed_ps | . | 32972 | . | . | . | 0.182149 | 0.120505 | 0.132816 | 0.092095 | 0.423869 | 0.278552 | 0.298036 | 0.210563 | -1.317765 | 18.049344 | 0.736078696 | -0.54082977 | 0.588624921 | NA | 4.357474 | 3.963455 | 4.203132 | 4.169889 | 4.177306 | 4.153563 | 28 | 18 | 16 | 12 |
| 650669 | NR_044995 | GAS6-AS1 | GAS6 antisense RNA 1 | lncRNA | . | 39826 | . | . | . | 0.239979 | 0.120082 | 0.129901 | 0.097387 | 0.558440 | 0.277575 | 0.291493 | 0.222663 | -1.371066 | 4.613603 | 1.415513638 | -0.32164865 | 0.747718885 | NA | 2.413393 | 1.963455 | 2.222348 | 2.207129 | 2.211307 | 2.204999 | 8 | 4 | 4 | 3 |
| 650788 | gene-RPS28P9 | RPS28P9 | ribosomal protein S28 pseudogene 9 | pseudogene | . | 36193 | . | . | . | 0.704656 | 1.438232 | 0.885105 | 1.701452 | 1.639764 | 3.324537 | 1.986151 | 3.890168 | 1.213509 | 14.514748 | 0.836527365 | 0.333742182 | 0.738574116 | NA | 3.716583 | 3.989431 | 3.825183 | 3.865521 | 3.839088 | 3.876236 | 10 | 19 | 10 | 19 |
| 650794 | NR_038939,NF | MIPEPP3 | mitochondrial intermediate peptidase pseudogene 3 | transcribed_ps | . | 39458 | . | . | . | 0.842098 | 0.912647 | 1.060751 | 1.389829 | 1.959599 | 2.109623 | 2.380294 | 3.177680 | 1.293548 | 53.457950 | 0.426700609 | 0.870245016 | 0.38416652 | 0.998865404 | 5.542770 | 5.914070 | 5.709896 | 5.708081 | 5.757002 | 5.760913 | 54 | 50 | 52 | 57 |
| 650901 | gene-RPS2P12 | RPS2P12 | ribosomal protein S2 pseudogene 12 | pseudogene | . | 36546 | . | . | . | 0.039957 | 0.132940 | 0.082586 | 0.026786 | 0.092981 | 0.307296 | 0.185321 | 0.061242 | -1.414555 | 2.721799 | 1.855026378 | -0.26972566 | 0.787371318 | NA | 1.671431 | 1.175234 | 1.437784 | 1.451514 | 1.445759 | 1.434966 | 2 | 5 | 3 | 1 |
| 651302 | NR_103448 | ZNF192P1 | zinc finger protein 192 pseudogene 1 | transcribed_ps | . | 18777 | . | . | . | 0.442085 | 0.284493 | 0.853173 | 0.446980 | 1.028751 | 0.657619 | 1.914495 | 1.021967 | 1.778182 | 26.647881 | 0.637227213 | 1.303151117 | 0.192523175 | NA | 4.258362 | 5.094145 | 4.698839 | 4.661337 | 4.783058 | 4.699482 | 27 | 16 | 39 | 22 |
| 651430 | NR_046264 | FLJ38576 | uncharacterized LOC651430 | lncRNA | . | . | . | . | . | 0.033631 | 0.013021 | 0.101114 | 0.048422 | 0.078260 | 0.030099 | 0.226897 | 0.110712 | 2.300109 | 4.410936 | 1.469160464 | 0.81795173 | 0.41338475 | NA | 1.413393 | 2.622519 | 2.114691 | 2.107038 | 2.135254 | 2.118528 | 4 | 2 | 7 | 4 |
| 651714 | gene-MUC20P | MUC20P1 | mucin 20, cell surface associated pseudogene 1 | pseudogene | . | 51921 | . | . | . | 0.021747 | 0.063063 | 0.167264 | 0.117395 | 0.050607 | 0.145774 | 0.375335 | 0.268409 | 2.463951 | 4.706755 | 1.427043167 | 0.911656896 | 0.361949361 | NA | 1.444631 | 2.742346 | 2.196176 | 2.205796 | 2.225279 | 2.213361 | 2 | 4 | 7 | 5 |
| 651746 | NM_00116444 | ANKRD33B | ankyrin repeat domain 33B | protein_coding | NP_001157912.1;XP_005248400.1;XP_005248401.1 | 35240 | . | . | . | 0.439342 | 0.565636 | 0.477996 | 0.603090 | 1.022366 | 1.307492 | 1.072609 | 1.378892 | 1.090578 | 154.328821 | 0.283495959 | 0.441249092 | 0.659032676 | 0.998865404 | 7.207104 | 7.330007 | 7.188002 | 7.304090 | 7.235324 | 7.326529 | 145 | 183 | 124 | 168 |
| 652276 | NR_015441 | LOC652276 | potassium channel tetramerization domain containing 5 pseudoge | transcribed_ps | . | . | . | . | . | 2.912453 | 3.495984 | 4.242537 | 3.850111 | 6.777397 | 8.081123 | 9.520131 | 8.802822 | 1.247732 | 306.681661 | 0.207298062 | 1.540331271 | 0.123479625 | 0.869586359 | 8.092232 | 8.411353 | 8.146417 | 8.236961 | 8.342203 | 8.281956 | 287 | 320 | 312 | 302 |
| 652959 | gene-BTF3P5 | BTF3P5 | basic transcription factor 3 pseudogene 5 | pseudogene | . | 38565 | . | . | . | 0.384389 | 0.091264 | 0.154428 | 0.021030 | 0.894488 | 0.210962 | 0.346533 | 0.048082 | -2.353652 | 4.717025 | 1.508551739 | -0.81860035 | 0.413014466 | NA | 2.722634 | 1.502474 | 2.272817 | 2.237343 | 2.245851 | 2.229932 | 12 | 3 | 4 | 1 |
| 652965 | NR_002918 | SNORA48 | small nucleolar RNA, H/ACA box 48 | snoRNA | . | 32641 | . | . | . | 9.783055 | 4.948168 | 6.368901 | 6.201982 | 22.765574 | 11.437911 | 14.291630 | 14.180095 | -1.163206 | 29.316602 | 0.599573556 | -0.36376977 | 0.716029941 | 0.998865404 | 4.977288 | 4.761978 | 4.922621 | 4.838520 | 4.865378 | 4.861489 | 48 | 23 | 24 | 25 |
| 652966 | NR_002604 | SNORD10 | small nucleolar RNA, C/D box 10 | snoRNA | . | 32706 | . | . | . | 4.848212 | 6.960842 | 7.509260 | 3.640093 | 11.281987 | 16.090298 | 16.850561 | 8.322639 | -1.062183 | 27.149969 | 0.628004033 | -0.13858601 | 0.889777298 | 0.998865404 | 4.804363 | 4.720166 | 4.739898 | 4.777515 | 4.788779 | 4.714135 | 27 | 35 | 31 | 16 |
| 652968 | NM_00103766 | CASTOR1 | cytosolic arginine sensor for mTORC1 subunit 1 | protein_coding | NP_001032755.1 | 34423 | 617034 | . | . | 5.876196 | 6.129847 | 5.989962 | 6.720768 | 13.674150 | 14.169415 | 13.441299 | 15.366239 | 1.046295 | 289.544902 | 0.205489776 | 0.317723911 | 0.750694372 | 0.998865404 | 8.144962 | 8.209601 | 8.155160 | 8.174558 | 8.160745 | 8.216087 | 320 | 311 | 244 | 292 |
| 653061 | NM_00135546 | GOLGA8S | golgin A8 family member S | protein_coding | NP_001342394.1;XP_024305798.1;XP_024305799.1 | 44409 | . | . | . | 0.028747 | 0.010241 | 0.016411 | 0.018671 | 0.066896 | 0.023672 | 0.036825 | 0.042690 | 1.089774 | 3.695183 | 1.576587396 | 0.078669071 | 0.937295847 | NA | 1.820514 | 1.947964 | 1.893116 | 1.877225 | 1.884826 | 1.888494 | 6 | 2 | 3 | 4 |
| 653073 | NM_00128247 | GOLGA8J | golgin A8 family member J | protein_coding | NP_001269401.1;XP_006720714.1;XP_011520252.1 | 38650 | . | . | . | 0.156145 | 0.103240 | 0.083245 | 0.065778 | 0.363356 | 0.238645 | 0.186800 | 0.150394 | -1.574500 | 15.729280 | 0.780354939 | -0.83922565 | 0.401342694 | NA | 4.264940 | 3.612728 | 4.002958 | 3.988642 | 3.972378 | 3.958715 | 24 | 19 | 12 | 10 |
| 653075 | NM_00135546 | GOLGA8T | golgin A8 family member T | protein_coding | NP_001342398.1;XP_024305800.1;XP_024305801.1;XP_024305802.1;XP | 44410 | . | . | . | 0.324833 | 0.284249 | 0.357782 | 0.239233 | 0.755899 | 0.657054 | 0.802854 | 0.546978 | -1.026606 | 46.831799 | 0.468542784 | -0.08085154 | 0.935560015 | 0.998865404 | 5.566740 | 5.531882 | 5.563624 | 5.534196 | 5.576679 | 5.511021 | 59 | 47 | 48 | 35 |
| 653079 | gene-RPL21P9 | RPL21P98 | ribosomal protein L21 pseudogene 98 | pseudogene | . | 36397 | . | . | . | 0.099446 | 0.030209 | 0.064926 | 0.124724 | 0.231414 | 0.069830 | 0.145693 | 0.285166 | 1.232222 | 1.970512 | 2.132558215 | 0.141267792 | 0.887658392 | NA | 0.820514 | 1.121007 | 0.982345 | 0.974334 | 0.975623 | 0.984888 | 3 | 1 | 1 | 3 |
| 653080 | NR_146391 | GUSBP16 | GUSB pseudogene 16 | transcribed_ps | . | 54498 | . | . | . | 1.882080 | 2.182564 | 2.942680 | 2.067407 | 4.379677 | 5.045095 | 6.603288 | 4.726880 | 1.216951 | 168.922911 | 0.275307754 | 1.028923414 | 0.303515665 | 0.998865404 | 7.251071 | 7.535386 | 7.320752 | 7.377348 | 7.504979 | 7.351062 | 163 | 176 | 190 | 143 |
| 653082 | NM_0013513C | ZDHHC11B | zinc finger DHHC-type containing 11B | protein_coding | NP_001338232.1;XP_016865599.1;XP_016865600.1;XP_016865601.1;XP | 32962 | . | . | . | 9.711261 | 9.878812 | 4.510974 | 5.318205 | 22.598506 | 22.835318 | 10.122494 | 12.159448 | -1.975649 | 1054.356840 | 0.139477506 | -7.04290251 | 1.88276E-12 | 5.19755E-09 | 10.451422 | 9.468643 | 10.241552 | 10.228432 | 9.700386 | 9.771721 | 1625 | 1499 | 582 | 701 |
| 653113 | NR_024254 | FAM86FP | family with sequence similarity 86 member F, pseudogene | transcribed_ps | . | 42357 | . | . | . | 0.553019 | 0.456902 | 0.709131 | 0.986381 | 1.286899 | 1.056148 | 1.591268 | 2.255243 | 1.647156 | 45.554487 | 0.472012144 | 1.525336433 | 0.127175205 | 0.874064017 | 5.105434 | 5.824839 | 5.462456 | 5.439333 | 5.505016 | 5.565515 | 43 | 34 | 42 | 61 |
| 653115 | gene-HNRNPA | HNRNPA3P6 | heterogeneous nuclear ribonucleoprotein A3 pseudogene 6 | pseudogene | . | 48495 | . | . | . | 6.278072 | 5.723287 | 5.799128 | 4.732688 | 14.609332 | 13.229634 | 13.013074 | 10.820729 | -1.156849 | 228.750429 | 0.232882563 | -0.90260197 | 0.366737181 | 0.998865404 | 7.938228 | 7.729490 | 7.896166 | 7.843869 | 7.847678 | 7.751257 | 297 | 251 | 204 | 178 |
| 653121 | NM_00104044 | ZBTB8A | zinc finger and BTB domain containing 8A | protein_coding | NP_001035531.2;NP_001278425.1 | 24172 | 618742 | . | . | 1.023846 | 1.146377 | 0.984252 | 0.952688 | 2.382530 | 2.649902 | 2.208632 | 2.178207 | -1.152348 | 225.228829 | 0.227271504 | -0.90014293 | 0.368044194 | 0.998865404 | 7.914046 | 7.709183 | 7.819404 | 7.875873 | 7.785688 | 7.775263 | 262 | 275 | 184 | 194 |
| 653122 | gene-LSM12P1 | LSM12P1 | LSM12 pseudogene 1 | pseudogene | . | 32453 | . | . | . | 4.395130 | 4.769110 | 4.465880 | 4.691441 | 10.227647 | 11.024012 | 10.021307 | 10.726421 | -1.013540 | 286.185126 | 0.205410536 | -0.09445896 | 0.924744581 | 0.998865404 | 8.170709 | 8.150832 | 8.145039 | 8.183113 | 8.146229 | 8.167849 | 320 | 322 | 242 | 272 |
| 653125 | NM_00128249 | GOLGA8K | golgin A8 family member K | protein_coding | NP_001269422.1 | 38652 | . | . | . | 0.173474 | 0.117664 | 0.178120 | 0.168130 | 0.403681 | 0.271985 | 0.399697 | 0.384408 | 1.172087 | 26.150732 | 0.610123092 | 0.375463953 | 0.70731545 | NA | 4.588571 | 4.819737 | 4.715794 | 4.676715 | 4.715449 | 4.711567 | 33 | 21 | 25 | 26 |
| 653156 | gene-RPL21P5 | RPL21P53 | ribosomal protein L21 pseudogene 53 | pseudogene | . | 36574 | . | . | . | 5.283704 | 4.272891 | 4.559727 | 5.328738 | 12.295396 | 9.876979 | 10.231895 | 12.183527 | 1.021713 | 86.254540 | 0.349409213 | 0.088693078 | 0.929323833 | 0.998865404 | 6.414924 | 6.445966 | 6.456728 | 6.395619 | 6.411404 | 6.453306 | 109 | 82 | 70 | 88 |
| 653160 | NR_037869 | LOC653160 | uncharacterized LOC653160 | lncRNA | . | . | . | . | . | 0.202492 | 0.240983 | 0.158544 | 0.128747 | 0.471208 | 0.557044 | 0.355768 | 0.294364 | -1.429731 | 11.467303 | 0.905234974 | -0.56973426 | 0.568857949 | NA | 3.754145 | 3.239024 | 3.526558 | 3.538408 | 3.518657 | 3.510603 | 14 | 16 | 9 | 8 |
| 653162 | NR_026890 | RPSAP9 | ribosomal protein SA pseudogene 9 | pseudogene | . | 31463 | . | . | . | 4.012633 | 3.696468 | 2.995601 | 4.089825 | 9.337562 | 8.544551 | 6.722043 | 9.350898 | -1.104476 | 166.266823 | 0.269061131 | -0.53282478 | 0.59415488 | 0.998865404 | 7.447856 | 7.303235 | 7.415696 | 7.377237 | 7.295992 | 7.412142 | 210 | 180 | 117 | 170 |
| 653175 | gene-HAUS6P1 | HAUS6P1 | HAUS augmin like complex subunit 6 pseudogene 1 | pseudogene | . | 43764 | . | . | . | 0.061920 | 0.115062 | 0.054573 | 0.011501 | 0.144090 | 0.265970 | 0.122459 | 0.026297 | -2.618477 | 4.068190 | 1.57057194 | -0.8842178 | 0.37657865 | NA | 2.555346 | 1.175234 | 2.034729 | 2.048895 | 2.030722 | 2.019876 | 5 | 8 | 3 | 1 |
| 653188 | NR_027386 | GUSBP3 | GUSB pseudogene 3 | transcribed_ps | . | 37301 | . | . | . | 0.938295 | 1.076378 | 1.283120 | 1.098233 | 2.183451 | 2.488095 | 2.879283 | 2.510979 | 1.167547 | 53.570962 | 0.429797305 | 0.519966657 | 0.603086815 | 0.998865404 | 5.627196 | 5.850899 | 5.710468 | 5.736414 | 5.768871 | 5.738309 | 53 | 57 | 54 | 50 |
| 653189 | gene-BTF3L4P | BTF3L4P2 | basic transcription factor 3 like 4 pseudogene 2 | pseudogene | . | 39646 | . | . | . | 0.508821 | 0.520717 | 0.278238 | 0.474989 | 1.184049 | 1.203662 | 0.624359 | 1.086006 | -1.371605 | 14.328735 | 0.818007433 | -0.55728715 | 0.577331253 | NA | 4.052320 | 3.592943 | 3.855814 | 3.856467 | 3.824414 | 3.848694 | 19 | 18 | 8 | 14 |
| 653199 | gene-TCAF1P1 | TCAF1P1 | TRPM8 channel associated factor 1 pseudogene 1 | pseudogene | . | 33604 | . | . | . | 6.184433 | 6.593812 | 5.164566 | 6.782176 | 14.391429 | 15.241892 | 11.589135 | 15.506639 | -1.084133 | 679.110801 | 0.17029179 | -0.68436893 | 0.493742232 | 0.998865404 | 9.465062 | 9.347552 | 9.413150 | 9.452086 | 9.286834 | 9.460700 | 791 | 784 | 493 | 690 |
| 653214 | gene-PPIAP22 | PPIAP22 | peptidylprolyl isomerase A pseudogene 22 | pseudogene | . | 17236 | . | . | . | 35.758606 | 34.456913 | 33.794483 | 40.609882 | 83.211746 | 79.646592 | 75.833832 | 92.849678 | 1.045508 | 833.776132 | 0.152562965 | 0.420832345 | 0.673877511 | 0.998865404 | 9.671280 | 9.735048 | 9.702317 | 9.670407 | 9.654511 | 9.777807 | 960 | 860 | 677 | 868 |
| 653226 | gene-SRP9P1 | SRP9P1 | signal recognition particle 9 pseudogene 1 | pseudogene | . | 23402 | . | . | . | 5.889180 | 4.969784 | 4.600102 | 4.085029 | 13.704363 | 11.487877 | 10.322495 | 9.339933 | -1.269837 | 224.627641 | 0.233945244 | -1.47317912 | 0.140702745 | 0.90099803 | 7.972880 | 7.629522 | 7.908691 | 7.817403 | 7.778231 | 7.722549 | 315 | 247 | 183 | 174 |
| 653232 | gene-RPL15P3 | RPL15P3 | ribosomal protein L15 pseudogene 3 | pseudogene | . | 21538 | . | . | . | 36.808304 | 35.137451 | 28.806295 | 41.325760 | 85.654442 | 81.221794 | 64.640495 | 94.486435 | -1.041245 | 781.311452 | 0.177667789 | -0.32819265 | 0.742766008 | 0.998865404 | 9.639003 | 9.579900 | 9.639609 | 9.600968 | 9.463754 | 9.705803 | 943 | 837 | 550 | 842 |
| 653234 | NR_160521 | AGAP10P | ArfGAP with GTPase domain, ankyrin repeat and PH domain 10, p | transcribed_ps | . | 23659 | . | . | . | 0.507744 | 0.590541 | 0.645879 | 0.554391 | 1.181542 | 1.365061 | 1.449335 | 1.267549 | 1.087947 | 27.690362 | 0.587486507 | 0.206997265 | 0.836011995 | 0.998865404 | 4.729157 | 4.850899 | 4.774069 | 4.792738 | 4.801595 | 4.784282 | 28 | 31 | 27 | 25 |
| 653236 | gene-RPL23AP | RPL23AP49 | ribosomal protein L23a pseudogene 49 | pseudogene | . | 35689 | . | . | . | 2.950548 | 1.463776 | 2.408635 | 1.638375 | 6.866047 | 3.383584 | 5.404908 | 3.745950 | -1.117410 | 35.359770 | 0.564073272 | -0.28393244 | 0.776462158 | 0.998865404 | 5.219620 | 5.064275 | 5.194250 | 5.096141 | 5.156896 | 5.104130 | 57 | 27 | 35 | 25 |
| 653238 | NR_033417 | GTF2H2B | general transcription factor IIH subunit 2B (pseudogene) | transcribed_ps | . | 31393 | . | . | . | 4.048594 | 3.738738 | 4.113784 | 3.289272 | 9.421244 | 8.642260 | 9.231213 | 7.520528 | -1.066766 | 518.435582 | 0.179982886 | -0.51807062 | 0.604408995 | 0.998865404 | 9.063356 | 8.971214 | 9.064673 | 9.007423 | 9.065614 | 8.919743 | 643 | 552 | 487 | 416 |
| 653259 | NR_165820 | AGAP14P | ArfGAP with GTPase domain, ankyrin repeat and PH domain 14, p | transcribed_ps | . | 23660 | . | . | . | 0.249795 | 0.317458 | 0.491719 | 0.241612 | 0.581283 | 0.733819 | 1.103404 | 0.552418 | 1.294062 | 12.400034 | 0.88384443 | 0.420783224 | 0.673913388 | NA | 3.432996 | 3.807334 | 3.608245 | 3.619766 | 3.646563 | 3.609095 | 11 | 13 | 16 | 9 |
| 653268 | NR_126580 | AGAP7P | ArfGAP with GTPase domain, ankyrin repeat and PH domain 7, ps | transcribed_ps | . | 23465 | . | . | . | 1.127686 | 1.257330 | 1.531969 | 0.908369 | 2.624172 | 2.906374 | 3.437694 | 2.076878 | 1.008554 | 91.173649 | 0.36246212 | 0.033902489 | 0.972954908 | 0.998865404 | 6.503273 | 6.517781 | 6.486955 | 6.517424 | 6.574274 | 6.430003 | 99 | 103 | 100 | 64 |
| 653269 | NM_0012774C | POTEI | POTE ankyrin domain family member I | protein_coding | NP_001264335.1;NP_001358855.1;XP_011509972.1;XP_016860221.1;XP | 37093 | . | . | . | 0.028187 | 0.041831 | 0.010288 | 0.019609 | 0.065593 | 0.096693 | 0.023085 | 0.044833 | -2.325463 | 5.478290 | 1.334935418 | -0.91204258 | 0.361746303 | NA | 2.939842 | 1.715150 | 2.467986 | 2.482516 | 2.451242 | 2.459774 | 7 | 10 | 2 | 4 |
| 653303 | gene-LOC6533 | LOC653303 | proprotein convertase subtilisin/kexin type 7 pseudogene | pseudogene | . | . | . | . | . | 2.712652 | 3.222735 | 3.405418 | 3.313497 | 6.312452 | 7.449497 | 7.641659 | 7.575917 | 1.114388 | 109.290098 | 0.310234614 | 0.503658309 | 0.61450151 | 0.998865404 | 6.692067 | 6.847772 | 6.724230 | 6.777934 | 6.792237 | 6.780409 | 109 | 121 | 102 | 106 |
| 653308 | NM_00107951 | ASAH2B | N-acylsphingosine amidohydrolase 2B | protein_coding | NP_001072984.1;NP_001308886.1;NP_001308887.1;NP_001308888.1;NI | 23456 | . | . | . | 0.515270 | 0.440059 | 0.500779 | 0.330070 | 1.199057 | 1.017212 | 1.123732 | 0.754667 | -1.165517 | 73.612914 | 0.388351469 | -0.56899421 | 0.569360072 | 0.998865404 | 6.306701 | 6.088857 | 6.237754 | 6.197459 | 6.228403 | 6.129107 | 98 | 79 | 72 | 50 |
| 653314 | gene-RPL19P9 | RPL19P9 | ribosomal protein L19 pseudogene 9 | pseudogene | . | 35918 | . | . | . | 0.162505 | 0.067433 | 0.055689 | 0.369235 | 0.378157 | 0.155875 | 0.124965 | 0.844212 | 1.750018 | 4.257246 | 1.51820269 | 0.531793193 | 0.594869248 | NA | 1.631267 | 2.437399 | 2.081420 | 2.069663 | 2.072308 | 2.100162 | 5 | 2 | 2 | 8 |
| 653319 | NM_00104071 | KIAA0895L | KIAA0895 like | protein_coding | NP_001035805.1;NP_001356609.1;NP_001356610.1;NP_001356611.1;NI | 34408 | . | . | . | 4.549220 | 5.017535 | 5.253745 | 4.549714 | 10.586219 | 11.598260 | 11.789251 | 10.402381 | 1.009931 | 521.245556 | 0.172657015 | 0.082575668 | 0.934188949 | 0.998865404 | 9.018501 | 9.033085 | 8.990452 | 9.051291 | 9.074789 | 8.978376 | 569 | 586 | 490 | 454 |
| 653333 | NM_00113761 | FAM86B2 | family with sequence similarity 86 member B2 | protein_coding | NP_001131082.1 | 32222 | 616123 | . | . | 0.396211 | 0.370302 | 0.459255 | 0.265627 | 0.922000 | 0.855969 | 1.030554 | 0.607324 | -1.071025 | 17.321668 | 0.745991205 | -0.13269916 | 0.894431311 | NA | 4.161369 | 4.066070 | 4.120982 | 4.110281 | 4.126980 | 4.093791 | 22 | 18 | 18 | 12 |
| 653354 | gene-ARMCX7I | ARMCX7P | armadillo repeat containing X-linked 7, pseudogene | pseudogene | . | 49917 | . | . | . | 4.188572 | 4.396856 | 4.943725 | 4.754632 | 9.746981 | 10.163529 | 11.093575 | 10.870901 | 1.119954 | 112.701544 | 0.302584109 | 0.540145034 | 0.589097015 | 0.998865404 | 6.732284 | 6.895812 | 6.788747 | 6.801711 | 6.840268 | 6.824659 | 120 | 117 | 106 | 109 |
| 653364 | gene-HSPB1P2 | HSPB1P2 | heat shock protein family B (small) member 1 pseudogene 2 | pseudogene | . | 5252 | . | . | . | 0.128546 | 0.076908 | 0.092905 | 0.082428 | 0.299133 | 0.177776 | 0.208476 | 0.188463 | -1.204527 | 2.440135 | 1.90164038 | -0.14117614 | 0.887730791 | NA | 1.413393 | 1.148375 | 1.296738 | 1.289023 | 1.291174 | 1.290401 | 4 | 2 | 2 | 2 |
| 653381 | NR_146393,NF | SORD2P | sorbitol dehydrogenase 2, pseudogene | transcribed_ps | . | 49919 | . | . | . | 6.040959 | 5.810010 | 6.555464 | 6.325539 | 14.057559 | 13.430099 | 14.710271 | 14.462592 | 1.071449 | 526.219420 | 0.164970997 | 0.603520165 | 0.546162705 | 0.998865404 | 8.988692 | 9.088595 | 9.031419 | 9.000841 | 9.074913 | 9.046188 | 599 | 535 | 484 | 499 |
| 653390 | NR_003369 | RRN3P2 | RRN3 pseudogene 2 | transcribed_ps | . | 37619 | . | . | . | 0.679784 | 0.863868 | 1.071844 | 0.862938 | 1.581885 | 1.996870 | 2.405188 | 1.973006 | 1.235067 | 65.369221 | 0.397957207 | 0.765382796 | 0.444043634 | 0.998865404 | 5.870101 | 6.174908 | 5.975084 | 6.021856 | 6.073378 | 6.018866 | 60 | 70 | 70 | 60 |
| 653404 | NM_00108547 | FOXD4L6 | forkhead box D4 like 6 | protein_coding | NP_001078945.1 | 31986 | . | . | . | 0.009500 | 0.060475 | 0.013019 | 0.034468 | 0.022108 | 0.139791 | 0.029215 | 0.078807 | -1.253864 | 2.463036 | 1.965225217 | -0.16607795 | 0.868095609 | NA | 1.460000 | 1.121007 | 1.292751 | 1.310635 | 1.294575 | 1.303782 | 1 | 5 | 1 | 3 |
| 653406 | gene-NAIPP4 | NAIPP4 | NAIP pseudogene 4 | pseudogene | . | 54494 | . | . | . | 0.415257 | 0.520832 | 0.814800 | 0.523793 | 0.966321 | 1.203927 | 1.828388 | 1.197591 | 1.416203 | 83.744140 | 0.37270566 | 1.346982309 | 0.17798593 | 0.951511445 | 6.114567 | 6.617632 | 6.295891 | 6.349729 | 6.480125 | 6.349848 | 71 | 83 | 104 | 72 |
| 653440 | gene-WASH6P, | WASH6P | WASP family homolog 6, pseudogene | pseudogene | . | 31685 | . | . | . | 14.134564 | 12.725538 | 15.862621 | 13.598045 | 32.891708 | 29.415648 | 35.595265 | 31.090315 | 1.081179 | 688.448675 | 0.162112678 | 0.694611963 | 0.487298513 | 0.998865404 | 9.369374 | 9.482808 | 9.435969 | 9.358506 | 9.506384 | 9.393749 | 805 | 673 | 673 | 616 |
| 653441 | gene-PHC1P1 | PHC1P1 | polyhomeotic homolog 1 pseudogene 1 | pseudogene | . | 34502 | . | . | . | 1.519484 | 1.589071 | 1.664268 | 1.422223 | 3.535902 | 3.673208 | 3.734570 | 3.251744 | -1.021044 | 260.233426 | 0.217472044 | -0.13815879 | 0.890114927 | 0.998865404 | 8.038251 | 8.008925 | 8.018125 | 8.037535 | 8.058894 | 7.974128 | 297 | 289 | 243 | 221 |
| 653458 | gene-BMS1P9 | BMS1P9 | BMS1 pseudogene 9 | pseudogene | . | 49155 | . | . | . | 0.094358 | 0.134409 | 0.103911 | 0.112581 | 0.219574 | 0.310692 | 0.233174 | 0.257404 | -1.218521 | 3.689354 | 1.556720669 | -0.18316111 | 0.854671607 | NA | 2.019256 | 1.733338 | 1.884830 | 1.890282 | 1.884801 | 1.883646 | 4 | 5 | 3 | 3 |
| 653464 | NM_00127187 | SRGAP2C | SLIT-ROBO Rho GTPase activating protein 2C | protein_coding | NP_001258801.1;NP_001316913.1 | 30584 | 614704 | . | . | 6.920344 | 7.240163 | 7.218314 | 7.510581 | 16.103927 | 16.735962 | 16.197688 | 17.172050 | 1.025875 | 1595.747231 | 0.123462487 | 0.298512558 | 0.765311993 | 0.998865404 | 10.621544 | 10.658255 | 10.615590 | 10.644000 | 10.637847 | 10.661448 | 1781 | 1731 | 1384 | 1538 |
| 653479 | NR_033934 | MRPL45P2 | mitochondrial ribosomal protein L45 pseudogene 2 | transcribed_ps | . | 29716 | . | . | . | 2.120591 | 2.137732 | 2.013491 | 2.200438 | 4.934702 | 4.941464 | 4.518215 | 5.031040 | -1.027407 | 92.431096 | 0.331385611 | -0.11771251 | 0.906295452 | 0.998865404 | 6.549923 | 6.510419 | 6.534977 | 6.534329 | 6.515167 | 6.538337 | 108 | 101 | 76 | 89 |
| 653483 | NR_027906 | AFDN-DT | AFDN divergent transcript | lncRNA | . | 21236 | . | . | . | 0.051393 | 0.046662 | 0.167168 | 0.100077 | 0.119593 | 0.107862 | 0.375119 | 0.228815 | 2.492754 | 7.041394 | 1.170709358 | 1.125591611 | 0.260338421 | NA | 2.008843 | 3.330299 | 2.778034 | 2.775227 | 2.815226 | 2.792780 | 5 | 4 | 11 | 7 |
| 653489 | NM_00114401 | RGPD3 | RANBP2 like and GRIP domain containing 3 | protein_coding | NP_001137485.1;XP_016860227.1;XP_016860228.1;XP_016860229.1 | 32416 | 612706 | . | . | 0.828524 | 0.824554 | 0.960236 | 0.637399 | 1.928010 | 1.905994 | 2.154743 | 1.457337 | -1.033508 | 181.317231 | 0.271084584 | -0.17540563 | 0.860760919 | 0.998865404 | 7.524961 | 7.479424 | 7.510710 | 7.498493 | 7.573346 | 7.400270 | 215 | 196 | 187 | 133 |
| 653503 | gene-LOC6535 | LOC653503 | G protein subunit gamma 10 pseudogene | pseudogene | . | . | . | . | . | 3.571479 | 1.338936 | 1.569872 | 0.949372 | 8.310979 | 3.095011 | 3.522748 | 2.170625 | -1.973486 | 62.972800 | 0.499561574 | -1.96321418 | 0.049621293 | 0.664973499 | 6.383638 | 5.407673 | 6.140288 | 5.880977 | 5.910897 | 5.831477 | 140 | 49 | 46 | 30 |
| 653510 | gene-CBWD4P | CBWD4P | COBW domain containing 4, pseudogene | pseudogene | . | 18520 | . | . | . | 1.736501 | 2.061927 | 2.537486 | 1.571123 | 4.040910 | 4.766237 | 5.694046 | 3.592186 | 1.068751 | 77.232882 | 0.38525781 | 0.248990614 | 0.803368042 | 0.998865404 | 6.221430 | 6.319201 | 6.234153 | 6.275398 | 6.331704 | 6.208545 | 79 | 87 | 86 | 57 |
| 653513 | NR_037182 | LOC653513 | phosphodiesterase 4D interacting protein-like | transcribed_ps | . | . | . | . | . | 2.999914 | 2.815842 | 2.364380 | 3.770359 | 6.980924 | 6.508946 | 5.305600 | 8.620478 | 1.041099 | 81.288437 | 0.372074737 | 0.156170755 | 0.875898428 | 0.998865404 | 6.316705 | 6.372708 | 6.345821 | 6.327165 | 6.286600 | 6.404464 | 95 | 83 | 56 | 95 |
| 653519 | NM_00109761 | GPR89A | G protein-coupled receptor 89A | protein_coding | NP_001091081.1;NP_001091082.2;XP_006711555.1;XP_006711556.1;XP | 31984 | 612821 | . | . | 9.343022 | 9.289913 | 10.534402 | 7.516412 | 21.741592 | 21.474048 | 23.638890 | 17.185376 | -1.087369 | 604.988059 | 0.186968804 | -0.64632064 | 0.518071723 | 0.998865404 | 9.299380 | 9.179662 | 9.272949 | 9.255081 | 9.317236 | 9.086768 | 735 | 671 | 593 | 448 |
| 653550 | NM_00120525 | TP53TG3C | TP53 target 3C | protein_coding | NP_001192188.1;XP_006721137.1;XP_011544222.1;XP_016879080.1 | 42962 | . | . | . | 0.889613 | 0.686419 | 0.627815 | 0.560134 | 2.070166 | 1.586690 | 1.408797 | 1.280679 | -1.250265 | 47.482973 | 0.460592696 | -0.6996096 | 0.484172793 | 0.998865404 | 5.720410 | 5.400577 | 5.609929 | 5.568830 | 5.568645 | 5.531615 | 67 | 51 | 41 | 35 |
| 653553 | gene-HSPB1P1 | HSPB1P1 | heat shock protein family B (small) member 1 pseudogene 1 | pseudogene | . | 5251 | . | . | . | 0.588101 | 0.723900 | 0.265920 | 0.377184 | 1.368535 | 1.673328 | 0.596716 | 0.862387 | -1.939051 | 6.466671 | 1.211746109 | -0.78840832 | 0.430457907 | NA | 3.094525 | 2.134756 | 2.709726 | 2.716244 | 2.690278 | 2.698416 | 9 | 10 | 3 | 5 |
| 653566 | gene-SPCS2P4 | SPCS2P4 | signal peptidase complex subunit 2 pseudogene 4 | pseudogene | . | 45237 | . | . | . | 6.339538 | 5.357408 | 5.690427 | 5.237918 | 14.752363 | 12.383889 | 12.769153 | 11.975878 | -1.086714 | 248.609912 | 0.22335683 | -0.53713098 | 0.591177145 | 0.998865404 | 8.015959 | 7.897073 | 8.021250 | 7.931523 | 7.958505 | 7.913177 | 324 | 255 | 217 | 213 |
| 653567 | NM_00109884 | TMEM236 | transmembrane protein 236 | protein_coding | NP_001092314.1;XP_011517928.1;XP_011517929.1;XP_011517930.1;XP | 23473 | . | . | . | 0.274054 | 0.172128 | 0.325823 | 0.231893 | 0.637734 | 0.397882 | 0.731138 | 0.530196 | -1.167702 | 33.871450 | 0.563193285 | -0.39715079 | 0.691256271 | 0.998865404 | 5.190212 | 4.965003 | 5.123184 | 5.060593 | 5.039046 | 5.103139 | 51 | 31 | 20 | 37 |
| 653583 | NM_198850,XI | PHLDB3 | pleckstrin homology like domain family B member 3 | protein_coding | NP_942147.3;XP_005259226.1;XP_005259227.1;XP_005259228.1;XP_01 | 30499 | . | . | . | 1.959621 | 2.233509 | 1.570084 | 1.280214 | 4.560116 | 5.162854 | 3.523223 | 2.927058 | -1.104537 | 145.061427 | 0.271559842 | -0.52821267 | 0.597351739 | 0.998865404 | 7.250214 | 7.107288 | 7.215002 | 7.184338 | 7.168386 | 7.155412 | 182 | 158 | 122 | 127 |
| 653631 | gene-LOC6536 | LOC653631 | axin interactor, dorsalization associated pseudogene | pseudogene | . | . | . | . | . | 3.219115 | 2.953444 | 2.943379 | 2.942704 | 7.491012 | 6.827018 | 6.604858 | 6.728143 | -1.062088 | 204.260029 | 0.235439393 | -0.36911057 | 0.712045309 | 0.998865404 | 7.716858 | 7.630373 | 7.710638 | 7.665020 | 7.662563 | 7.658822 | 254 | 216 | 173 | 185 |
| 653635 | NR_024540 | WASH7P | WASP family homolog 7, pseudogene | transcribed_ps | . | 38034 | . | . | . | 13.503942 | 14.098948 | 18.141588 | 14.176060 | 31.424229 | 32.590347 | 40.709194 | 32.411877 | 1.154908 | 825.694066 | 0.16255495 | 1.278200033 | 0.201178905 | 0.967101044 | 9.581513 | 9.789896 | 9.620575 | 9.644822 | 9.823280 | 9.638715 | 867 | 841 | 868 | 724 |
| 653639 | gene-LYPLA2P1 | LYPLA2P1 | LYPLA2 pseudogene 1 | pseudogene | . | 21069 | . | . | . | 0.713212 | 0.743696 | 0.707748 | 0.813644 | 1.659674 | 1.719086 | 1.588166 | 1.860301 | 1.041224 | 38.434685 | 0.500065772 | 0.11654662 | 0.90721934 | 0.998865404 | 5.235335 | 5.292767 | 5.259442 | 5.262094 | 5.257652 | 5.275889 | 43 | 41 | 32 | 39 |
| 653645 | NR_002924 | TBC1D3P1-DH | TBC1D3P1-DHX40P1 readthrough, transcribed pseudogene | transcribed_ps | . | 42362 | . | . | . | 0.354466 | 0.746111 | 0.280061 | 0.644776 | 0.824858 | 1.724669 | 0.628448 | 1.474204 | -1.200648 | 30.692795 | 0.628248967 | -0.41991869 | 0.674544851 | 0.998865404 | 5.068704 | 4.798300 | 4.893542 | 4.989981 | 4.877902 | 4.961609 | 25 | 49 | 15 | 36 |
| 653650 | gene-PDPK2P | PDPK2P | 3-phosphoinositide dependent protein kinase 2, pseudogene | pseudogene | . | 49897 | . | . | . | 1.496052 | 2.061294 | 2.317843 | 2.090747 | 3.481374 | 4.764773 | 5.201172 | 4.780244 | 1.231650 | 71.866563 | 0.382161997 | 0.786557545 | 0.43154092 | 0.998865404 | 6.009565 | 6.309384 | 6.098889 | 6.168344 | 6.199571 | 6.169847 | 63 | 80 | 73 | 70 |
| 653653 | NR_027408 | LOC653653 | adaptor related protein complex 1 sigma 2 subunit pseudogene | pseudogene | . | . | . | . | . | 0.302694 | 0.182079 | 0.227918 | 0.143617 | 0.704383 | 0.420883 | 0.511442 | 0.328363 | -1.291986 | 8.309208 | 1.068687947 | -0.34583538 | 0.729466437 | NA | 3.225252 | 2.861274 | 3.071029 | 3.053175 | 3.061123 | 3.044070 | 13 | 8 | 8 | 5 |
| 653658 | gene-RPS23P8 | RPS23P8 | ribosomal protein S23 pseudogene 8 | pseudogene | . | 35982 | . | . | . | 15.696208 | 13.126665 | 12.442182 | 10.918590 | 36.525723 | 30.342875 | 27.919895 | 24.964060 | -1.252219 | 166.091471 | 0.263764135 | -1.23021624 | 0.21861614 | 0.982008277 | 7.528334 | 7.205288 | 7.459122 | 7.377669 | 7.352587 | 7.301114 | 232 | 181 | 137 | 129 |
| 653659 | NM_0010798C | TMEM183B | transmembrane protein 183B | protein_coding | NP_001073277.1 | 33205 | 611365 | . | . | 3.569779 | 3.146606 | 3.493243 | 3.900077 | 8.307023 | 7.273520 | 7.838737 | 8.917063 | 1.087346 | 187.690604 | 0.246516197 | 0.490071269 | 0.624083468 | 0.998865404 | 7.490516 | 7.611379 | 7.560786 | 7.502680 | 7.547567 | 7.589545 | 221 | 181 | 162 | 192 |
| 653665 | gene-RPL21P9 | RPL21P93 | ribosomal protein L21 pseudogene 93 | pseudogene | . | 35646 | . | . | . | 0.458408 | 0.453417 | 1.213885 | 0.695845 | 1.066735 | 1.048093 | 2.723923 | 1.590966 | 2.031092 | 12.918660 | 0.869007643 | 1.176348415 | 0.239455676 | NA | 3.089593 | 4.114763 | 3.649845 | 3.648554 | 3.706742 | 3.668872 | 10 | 9 | 19 | 12 |
| 653689 | NM_00108084 | GSTT2B | glutathione S-transferase theta 2B | protein_coding | NP_001074312.1;NP_001350733.1 | 33437 | . | . | . | 9.419545 | 8.305324 | 14.642812 | 10.467278 | 21.919668 | 19.198130 | 32.858043 | 23.932186 | 1.395447 | 368.893955 | 0.213714074 | 2.249393365 | 0.02448748 | 0.51603159 | 8.265897 | 8.748118 | 8.445886 | 8.377440 | 8.704984 | 8.494213 | 377 | 311 | 438 | 334 |
| 653691 | NR_110549 | TCAF2P1 | TRPM8 channel associated factor 2 pseudogene 1 | transcribed_ps | . | 33603 | . | . | . | 0.127437 | 0.119270 | 0.189119 | 0.186706 | 0.296550 | 0.275698 | 0.424378 | 0.426881 | 1.469880 | 11.621669 | 0.892388709 | 0.622708518 | 0.533476084 | NA | 3.234218 | 3.790061 | 3.520792 | 3.519631 | 3.539582 | 3.540072 | 11 | 10 | 12 | 13 |
| 653698 | gene-LOC6536 | LOC653698 | solute carrier family 25 member 33 pseudogene | pseudogene | . | . | . | . | . | 0.828288 | 0.965642 | 0.945681 | 1.149777 | 1.927460 | 2.232125 | 2.122080 | 2.628829 | 1.150806 | 38.149133 | 0.505141247 | 0.4011633 | 0.688299899 | 0.998865404 | 5.149693 | 5.350483 | 5.229645 | 5.249946 | 5.245431 | 5.275767 | 38 | 41 | 32 | 42 |
| 653702 | gene-RPL7P9 | RPL7P9 | ribosomal protein L7 pseudogene 9 | pseudogene | . | 37028 | . | . | . | 22.011787 | 20.293991 | 17.237160 | 22.084774 | 51.222332 | 46.910469 | 38.679688 | 50.494213 | -1.091130 | 530.091790 | 0.179316714 | -0.70168185 | 0.482877594 | 0.998865404 | 9.111953 | 8.985472 | 9.105726 | 9.045885 | 8.943468 | 9.091139 | 666 | 570 | 389 | 531 |
| 653712 | NR_034179 | LOC653712 | intraflagellar transport 122 homolog (Chlamydomonas) pseudoge | transcribed_ps | . | . | . | . | . | 0.383315 | 0.331940 | 0.678968 | 0.530255 | 0.891989 | 0.767294 | 1.523584 | 1.212364 | 1.627153 | 17.056389 | 0.747626086 | 0.939440611 | 0.347504571 | NA | 3.697260 | 4.401999 | 4.064785 | 4.056703 | 4.108106 | 4.083355 | 16 | 13 | 21 | 17 |
| 653720 | NM_00128246 | GOLGA8M | golgin A8 family member M | protein_coding | NP_001269397.1;XP_005268341.1;XP_005268342.1;XP_006720717.1;XP | 44404 | . | . | . | 0.035255 | 0.022027 | 0.036986 | 0.032639 | 0.082041 | 0.050917 | 0.082997 | 0.074624 | 1.243304 | 7.510728 | 1.106317816 | 0.283985773 | 0.776421283 | NA | 2.741623 | 3.058876 | 2.905417 | 2.894943 | 2.911307 | 2.903785 | 9 | 6 | 8 | 7 |
| 653737 | gene-RPL21P1 | RPL21P119 | ribosomal protein L21 pseudogene 119 | pseudogene | . | 35614 | . | . | . | 6.101172 | 5.280231 | 5.796406 | 6.543021 | 14.197678 | 12.205490 | 13.006967 | 14.959840 | 1.065111 | 107.597881 | 0.313057533 | 0.290693375 | 0.771285839 | 0.998865404 | 6.703284 | 6.794293 | 6.761718 | 6.714019 | 6.739749 | 6.776522 | 129 | 104 | 91 | 110 |
| 653758 | gene-TPM3P6 | TPM3P6 | tropomyosin 3 pseudogene 6 | pseudogene | . | 39169 | . | . | . | 1.207419 | 1.311698 | 1.171070 | 1.410454 | 2.809714 | 3.032049 | 2.627847 | 3.224836 | 1.025131 | 43.604522 | 0.472556912 | 0.075774232 | 0.939598717 | 0.998865404 | 5.429096 | 5.463510 | 5.438401 | 5.449338 | 5.433519 | 5.462282 | 48 | 48 | 35 | 45 |
| 653784 | NM_00108536 | MZT2A | mitotic spindle organizing protein 2A | protein_coding | NP_001078834.1;XP_005263799.1;XP_016860233.1;XP_024308816.1;XP | 33187 | 613449 | . | . | 43.078182 | 43.767155 | 35.851742 | 61.430094 | 100.244695 | 101.169740 | 80.450268 | 140.452615 | -1.016996 | 1431.545865 | 0.147577018 | -0.16475403 | 0.869137586 | 0.998865404 | 10.495727 | 10.470882 | 10.483520 | 10.492436 | 10.371860 | 10.568497 | 1653 | 1567 | 1076 | 1501 |
| 653789 | gene-RPL23AP | RPL23AP25 | ribosomal protein L23a pseudogene 25 | pseudogene | . | 35672 | . | . | . | 0.817745 | 0.511170 | 0.688953 | 0.926967 | 1.902927 | 1.181592 | 1.545990 | 2.119400 | 1.187216 | 12.665568 | 0.865892429 | 0.285926969 | 0.774934057 | NA | 3.533720 | 3.781347 | 3.665915 | 3.644972 | 3.655372 | 3.674652 | 16 | 10 | 10 | 15 |
| 653805 | gene-RPS27P8 | RPS27P8 | ribosomal protein S27 pseudogene 8 | pseudogene | . | 36022 | . | . | . | 2.872738 | 2.288724 | 1.266788 | 2.002691 | 6.684979 | 5.290488 | 2.842636 | 4.578915 | -1.599422 | 22.854764 | 0.660025931 | -1.02650092 | 0.304631974 | NA | 4.814705 | 4.134756 | 4.554655 | 4.528322 | 4.482174 | 4.512939 | 36 | 27 | 12 | 20 |
| 653820 | NM_00110091 | FAM72B | family with sequence similarity 72 member B | protein_coding | NP_001094380.1;NP_001307078.1 | 24805 | 614711 | . | . | 7.412538 | 7.260636 | 5.915273 | 5.402884 | 17.249281 | 16.783286 | 13.273698 | 12.353051 | -1.334009 | 475.884182 | 0.17176248 | -2.42060072 | 0.015494886 | 0.417236976 | 9.087214 | 8.671937 | 8.990066 | 8.962821 | 8.819018 | 8.774575 | 639 | 575 | 368 | 368 |
| 653857 | NM_00116445 | ACTR3C | actin related protein 3C | protein_coding | NP_001157930.1;NP_001157931.1;NP_001337956.1;NP_001337957.1;NI | 37282 | . | . | . | 3.810027 | 4.336133 | 1.814152 | 2.980289 | 8.866084 | 10.023163 | 4.070907 | 6.814076 | -1.582886 | 200.920464 | 0.238845243 | -2.77400294 | 0.005537118 | 0.227466779 | 7.943839 | 7.281837 | 7.763406 | 7.717725 | 7.543826 | 7.527376 | 296 | 254 | 138 | 143 |
| 654029 | gene-RPS10P1 | RPS10P15 | ribosomal protein S10 pseudogene 15 | pseudogene | . | 35860 | . | . | . | 0.169618 | 0.121154 | 0.169645 | 0.208729 | 0.394708 | 0.280053 | 0.380678 | 0.477233 | 1.233975 | 3.492648 | 1.593779309 | 0.190310891 | 0.84906552 | NA | 1.644780 | 1.947964 | 1.801217 | 1.797945 | 1.801116 | 1.804812 | 4 | 3 | 3 | 4 |
| 654254 | NM_0011376C | ZNF732 | zinc finger protein 732 | protein_coding | NP_001131080.1 | 37138 | . | . | . | 0.303470 | 0.383975 | 0.199170 | 0.267084 | 0.706187 | 0.887575 | 0.446931 | 0.610655 | -1.453722 | 21.631231 | 0.671254407 | -0.80409431 | 0.421342511 | NA | 4.681128 | 4.138173 | 4.442495 | 4.468239 | 4.411019 | 4.432759 | 26 | 31 | 13 | 19 |
| 654319 | NR_002919 | SNORA5A | small nucleolar RNA, H/ACA box 5A | snoRNA | . | 32588 | . | . | . | 3.495145 | 3.897757 | 3.618123 | 3.866754 | 8.133346 | 9.009839 | 8.118963 | 8.840874 | 1.021213 | 15.886142 | 0.76567872 | 0.039550988 | 0.968451102 | NA | 3.974755 | 4.004486 | 3.983843 | 3.992383 | 3.989770 | 3.989365 | 17 | 18 | 14 | 15 |
| 654320 | NR_002920 | SNORA8 | small nucleolar RNA, H/ACA box 8 | snoRNA | . | 32596 | . | . | . | 6.416821 | 5.564470 | 6.683529 | 2.073667 | 14.932207 | 12.862521 | 14.997644 | 4.741193 | -1.365510 | 23.295007 | 0.699094544 | -0.64288803 | 0.520296755 | NA | 4.746331 | 4.303756 | 4.559872 | 4.544289 | 4.563263 | 4.478943 | 33 | 27 | 26 | 9 |
| 654321 | NR_002921 | SNORA75 | small nucleolar RNA, H/ACA box 75 | snoRNA | . | 32661 | . | . | . | 1.815887 | 3.177718 | 2.483106 | 1.031932 | 4.225644 | 7.345438 | 5.572019 | 2.359390 | -1.344808 | 9.854668 | 0.998740593 | -0.42793937 | 0.66869526 | NA | 3.497331 | 3.073227 | 3.294004 | 3.317635 | 3.306453 | 3.279409 | 10 | 15 | 10 | 5 |
| 654322 | NR_002922 | SNORA13 | small nucleolar RNA, H/ACA box 13 | snoRNA | . | 32601 | . | . | . | 0.402214 | 0.753166 | 1.946561 | 0.861726 | 0.935968 | 1.740976 | 4.368026 | 1.970234 | 2.476473 | 4.727581 | 1.436908312 | 0.910487441 | 0.362565502 | NA | 1.444631 | 2.751299 | 2.199185 | 2.208814 | 2.233355 | 2.211534 | 2 | 4 | 8 | 4 |
| 654342 | NR_027438 | LSP1P4 | LSP1 pseudogene 4 | transcribed_ps | . | 53915 | . | . | . | 4.596441 | 4.443746 | 4.218962 | 5.227477 | 10.696109 | 10.271918 | 9.467229 | 11.952004 | 1.038981 | 87.669499 | 0.343137089 | 0.160780392 | 0.872266369 | 0.998865404 | 6.426654 | 6.480843 | 6.453356 | 6.442377 | 6.430726 | 6.485623 | 101 | 91 | 70 | 92 |
| 654350 | gene-RPL9P25 | RPL9P25 | ribosomal protein L9 pseudogene 25 | pseudogene | . | 36112 | . | . | . | 0.098255 | 0.332365 | 0.267838 | 0.384320 | 0.228644 | 0.768276 | 0.601021 | 0.878701 | 1.547419 | 6.363117 | 1.229354575 | 0.512353087 | 0.608403918 | NA | 2.324708 | 2.947964 | 2.639391 | 2.662207 | 2.659666 | 2.667006 | 3 | 8 | 6 | 8 |
| 654364 | NM_00101813 | NME1-NME2 | NME1-NME2 readthrough | protein_coding | NP_001018146.1 | 33531 | . | . | . | 31.880235 | 15.925945 | 26.044379 | 28.092237 | 74.186615 | 36.813532 | 58.442831 | 64.229563 | 1.105941 | 900.166851 | 0.50168416 | 0.28957349 | 0.77214255 | 0.998865404 | 9.739517 | 9.884918 | 9.981467 | 9.479788 | 9.817304 | 9.849846 | 1322 | 606 | 804 | 906 |
| 654365 | gene-RPS27P3 | RPS27P3 | ribosomal protein S27 pseudogene 3 | pseudogene | . | 17445 | . | . | . | 4.356562 | 4.376356 | 3.525849 | 4.022041 | 10.137898 | 10.116143 | 7.911904 | 9.195918 | -1.173926 | 44.409386 | 0.468098702 | -0.49421519 | 0.621154214 | 0.998865404 | 5.584287 | 5.351955 | 5.487587 | 5.489040 | 5.450676 | 5.472087 | 55 | 52 | 33 | 41 |
| 654388 | gene-RPL32P2 | RPL32P2 | ribosomal protein L32 pseudogene 2 | pseudogene | . | 31033 | . | . | . | 0.091156 | 0.077335 | 0.182013 | 0.090336 | 0.212123 | 0.178763 | 0.408432 | 0.206543 | 1.552428 | 2.293408 | 1.956121338 | 0.324380053 | 0.74565032 | NA | 0.844134 | 1.481107 | 1.182896 | 1.183495 | 1.190762 | 1.184854 | 2 | 2 | 3 | 2 |
| 654427 | gene-PCNPP1 | PCNPP1 | PEST containing nuclear protein pseudogene 1 | pseudogene | . | 32440 | . | . | . | 0.164227 | 0.059362 | 0.265037 | 0.138064 | 0.382164 | 0.137219 | 0.594735 | 0.315666 | 1.658208 | 11.197876 | 0.961886464 | 0.758535679 | 0.448130362 | NA | 3.069695 | 3.807334 | 3.472502 | 3.437815 | 3.498465 | 3.461031 | 14 | 5 | 16 | 9 |
| 654433 | NR_015377,NF | PAX8-AS1 | PAX8 antisense RNA 1 | lncRNA | . | 49271 | . | . | . | 5.368441 | 5.964665 | 6.015711 | 5.842911 | 12.492583 | 13.787592 | 13.499082 | 13.359122 | 1.036735 | 383.181134 | 0.185395129 | 0.280735031 | 0.778913638 | 0.998865404 | 8.555835 | 8.607469 | 8.539845 | 8.600394 | 8.587830 | 8.595392 | 411 | 427 | 337 | 368 |
| 654434 | NR_027058 | SNHG20 | small nucleolar RNA host gene 20 | lncRNA | . | 33099 | . | . | . | 3.931830 | 3.961498 | 5.615907 | 3.967393 | 9.149529 | 9.157179 | 12.601933 | 9.070972 | 1.197993 | 295.766159 | 0.225524368 | 1.1556178 | 0.247837524 | 0.992733415 | 8.071390 | 8.333360 | 8.151220 | 8.150633 | 8.340620 | 8.145365 | 310 | 290 | 330 | 249 |
| 654466 | NR_003670 | FGF7P3 | fibroblast growth factor 7 pseudogene 3 | transcribed_ps | . | 26671 | . | . | . | 0.097603 | 0.070764 | 0.048672 | 0.067379 | 0.227126 | 0.163573 | 0.109219 | 0.154054 | -1.382128 | 11.150236 | 0.920067484 | -0.50745305 | 0.611836972 | NA | 3.694014 | 3.226238 | 3.501495 | 3.484997 | 3.471874 | 3.483036 | 17 | 12 | 7 | 10 |
| 654483 | NM_00103918 | BOLA2B | bolA family member 2B | protein_coding | NP_001034271.2;NP_001307536.2 | 32479 | . | . | . | 72.595014 | 63.111539 | 60.479417 | 71.725304 | 168.931588 | 145.885166 | 135.714065 | 163.991385 | -1.028461 | 766.849679 | 0.163195463 | -0.2480934 | 0.804062144 | 0.998865404 | 9.603021 | 9.562292 | 9.644509 | 9.534618 | 9.504497 | 9.634082 | 965 | 774 | 596 | 775 |
| 654841 | NR_102371 | MFF-DT | MFF divergent transcript | lncRNA | . | 41067 | . | . | . | 0.137447 | 0.175761 | 0.131721 | 0.164577 | 0.319845 | 0.406279 | 0.295578 | 0.376285 | -1.086361 | 12.085173 | 0.878174745 | -0.13608172 | 0.891756677 | NA | 3.654832 | 3.532926 | 3.592759 | 3.604518 | 3.589004 | 3.599489 | 13 | 15 | 9 | 12 |
| 664709 | NR_002944 | HNRNPA1P10 | heterogeneous nuclear ribonucleoprotein A1 pseudogene 10 | transcribed_ps | . | 39128 | . | . | . | 7.625631 | 7.424681 | 8.644411 | 9.053889 | 17.745155 | 17.162481 | 19.397808 | 20.700644 | 1.159922 | 345.493564 | 0.190387979 | 1.124168841 | 0.260941405 | 0.995760118 | 8.321550 | 8.535552 | 8.397089 | 8.377500 | 8.460790 | 8.481437 | 375 | 339 | 317 | 354 |
| 677768 | NR_003002 | SCARNA13 | small Cajal body-specific RNA 13 | guide_RNA | . | 32570 | . | . | . | 0.014776 | 0.041463 | 0.031043 | 0.022831 | 0.034385 | 0.095843 | 0.069660 | 0.052201 | 1.234655 | 1.002957 | 2.868339486 | 0.106022226 | 0.915564717 | NA | -0.155866 | 0.148375 | -0.001261 | -0.000972 | 0.000053 | -0.000315 | 1 | 1 | 1 | 1 |
| 677769 | NR_003003 | SCARNA17 | small Cajal body-specific RNA 17 | guide_RNA | . | 32574 | 615645 | . | . | 1.775671 | 0.976863 | 1.639568 | 0.611299 | 4.132061 | 2.258063 | 3.679143 | 1.397663 | -1.233960 | 16.918412 | 0.801710618 | -0.3783111 | 0.705199503 | NA | 4.220748 | 3.925185 | 4.108394 | 4.058629 | 4.096282 | 4.039957 | 28 | 14 | 19 | 8 |
| 677770 | NR_003004 | SCARNA22 | small Cajal body-specific RNA 22 | guide_RNA | . | 32580 | . | . | . | 0.810059 | 0.556963 | 0.934503 | 0.807071 | 1.885040 | 1.287444 | 2.096998 | 1.845272 | 1.244163 | 3.513475 | 1.59175418 | 0.198005305 | 0.843040913 | NA | 1.644780 | 1.963455 | 1.806945 | 1.803669 | 1.811949 | 1.805707 | 4 | 3 | 4 | 3 |
| 677771 | NR_003006 | SCARNA6 | small Cajal body-specific RNA 6 | guide_RNA | . | 32562 | 615641 | . | . | 0.246254 | 0.050800 | 0.155560 | 0.379263 | 0.573042 | 0.117426 | 0.349071 | 0.867139 | 1.553018 | 2.258005 | 1.98954141 | 0.31920654 | 0.749569898 | NA | 0.820514 | 1.459419 | 1.170516 | 1.162497 | 1.169025 | 1.173137 | 3 | 1 | 2 | 7 |
| 677775 | NR_003008 | SCARNA5 | small Cajal body-specific RNA 5 | guide_RNA | . | 32561 | 615640 | . | . | 0.185554 | 0.162729 | 0.528202 | 0.113558 | 0.431791 | 0.376154 | 1.185269 | 0.259637 | 1.570810 | 2.314234 | 1.987698758 | 0.327770168 | 0.743085445 | NA | 0.844134 | 1.502474 | 1.190054 | 1.190655 | 1.203070 | 1.187104 | 2 | 2 | 4 | 1 |
| 677777 | NR_003010 | SCARNA12 | small Cajal body-specific RNA 12 | guide_RNA | . | 32569 | 615642 | . | . | 10.936441 | 10.689479 | 15.564215 | 11.253801 | 25.449547 | 24.709208 | 34.925648 | 25.730478 | 1.217018 | 102.394125 | 0.32910352 | 0.860976649 | 0.389250904 | 0.998865404 | 6.528358 | 6.813549 | 6.643959 | 6.634804 | 6.752486 | 6.645079 | 108 | 98 | 114 | 88 |
| 677778 | NR_003011 | SCARNA15 | small Cajal body-specific RNA 15 | guide_RNA | . | 32572 | 612675 | . | . | 1.356058 | 1.047809 | 1.254255 | 0.183730 | 3.155603 | 2.422058 | 2.814511 | 0.420076 | -1.584271 | 4.382339 | 1.500556848 | -0.44238166 | 0.658213049 | NA | 2.421266 | 1.769039 | 2.142651 | 2.135809 | 2.140642 | 2.119613 | 7 | 5 | 5 | 1 |
| 677781 | NR_003013 | SCARNA16 | small Cajal body-specific RNA 16 | guide_RNA | . | 32573 | . | . | . | 1.093346 | 1.575047 | 0.486472 | 0.926974 | 2.544262 | 3.640791 | 1.091628 | 2.119415 | -1.639398 | 6.516227 | 1.205611083 | -0.59193932 | 0.554159116 | NA | 3.019256 | 2.300113 | 2.714122 | 2.724697 | 2.698706 | 2.711599 | 8 | 10 | 3 | 6 |
| 677792 | NR_003026 | SNORA1 | small nucleolar RNA, H/ACA box 1 | snoRNA | . | 32557 | . | . | . | 2.256928 | 2.544384 | 2.963279 | 0.221723 | 5.251966 | 5.881459 | 6.649513 | 0.506943 | -1.518101 | 8.597546 | 1.163323291 | -0.51771301 | 0.60465852 | NA | 3.369732 | 2.777828 | 3.103723 | 3.110765 | 3.116632 | 3.065535 | 11 | 12 | 11 | 1 |
| 677793 | NR_002950 | SNORA2A | small nucleolar RNA, H/ACA box 2A | snoRNA | . | 32584 | . | . | . | 0.557992 | 0.535290 | 0.621158 | 0.221136 | 1.298471 | 1.237347 | 1.393860 | 0.505601 | -1.198555 | 2.475538 | 1.907561801 | -0.13697918 | 0.89104725 | NA | 1.429096 | 1.175234 | 1.307104 | 1.307921 | 1.310815 | 1.300040 | 3 | 3 | 3 | 1 |
| 677795 | NR_002990 | SNORA5B | small nucleolar RNA, H/ACA box 5B | snoRNA | . | 32589 | . | . | . | 0.723680 | 0.618669 | 2.219301 | 1.835292 | 1.684034 | 1.430082 | 4.980047 | 4.196177 | 2.659867 | 5.730117 | 1.289011743 | 1.094911939 | 0.273555218 | NA | 1.644780 | 3.058876 | 2.484709 | 2.481580 | 2.510736 | 2.503327 | 4 | 3 | 8 | 7 |
| 677796 | NR_002991 | SNORA5C | small nucleolar RNA, H/ACA box 5C | snoRNA | . | 32590 | 611335 | . | . | 3.544117 | 2.682406 | 4.894045 | 4.295542 | 8.247306 | 6.200501 | 10.982095 | 9.821248 | 1.441994 | 16.915623 | 0.74857498 | 0.705427809 | 0.480544043 | NA | 3.790753 | 4.321310 | 4.065575 | 4.049661 | 4.091633 | 4.076464 | 18 | 13 | 19 | 17 |
| 677797 | NR_002992 | SNORA7B | small nucleolar RNA, H/ACA box 7B | snoRNA | . | 32593 | . | . | . | 0.292566 | 0.443805 | 1.061672 | 0.464106 | 0.680813 | 1.025875 | 2.382362 | 1.061124 | 1.488926 | 2.812587 | 1.788565834 | 0.321079512 | 0.748150136 | NA | 1.175403 | 1.751299 | 1.472828 | 1.477882 | 1.485999 | 1.474918 | 2 | 3 | 4 | 2 |
| 677798 | NR_002952 | SNORA9 | small nucleolar RNA, H/ACA box 9 | snoRNA | . | 32597 | . | . | . | 2.091102 | 3.874054 | 5.941553 | 5.392238 | 4.866080 | 8.955050 | 13.332672 | 12.328712 | 1.821541 | 18.483428 | 0.731298342 | 1.18304534 | 0.236791179 | NA | 3.713381 | 4.575901 | 4.149567 | 4.182250 | 4.218075 | 4.206628 | 11 | 18 | 22 | 21 |
| 677799 | NR_002953 | SNORA11 | small nucleolar RNA, H/ACA box 11 | snoRNA | . | 32599 | 300662 | . | . | 2.768894 | 1.188481 | 2.490615 | 0.458396 | 6.443331 | 2.747228 | 5.588868 | 1.048069 | -1.430909 | 7.550440 | 1.189937587 | -0.43441894 | 0.663984243 | NA | 3.147197 | 2.641905 | 2.936430 | 2.905998 | 2.927237 | 2.890912 | 14 | 6 | 9 | 2 |
| 677802 | NR_002956 | SNORA14B | small nucleolar RNA, H/ACA box 14B | snoRNA | . | 32603 | . | . | . | 0.034948 | 0.030752 | 0.152497 | 0.080530 | 0.081325 | 0.071084 | 0.342199 | 0.184123 | 1.234655 | 1.002957 | 2.868339486 | 0.106022226 | 0.915564717 | NA | -0.155866 | 0.148375 | -0.001261 | -0.000972 | 0.000053 | -0.000315 | 1 | 1 | 1 | 1 |
| 677804 | NR_002958 | SNORA17A | small nucleolar RNA, H/ACA box 17A | snoRNA | . | 32607 | . | . | . | 0.014545 | 0.015941 | 0.019239 | 0.019209 | 0.033848 | 0.036848 | 0.043173 | 0.043919 | 1.234655 | 1.002957 | 2.868339486 | 0.106022226 | 0.915564717 | NA | -0.155866 | 0.148375 | -0.001261 | -0.000972 | 0.000053 | -0.000315 | 1 | 1 | 1 | 1 |
| 677805 | NR_002959 | SNORA18 | small nucleolar RNA, H/ACA box 18 | snoRNA | . | 32608 | . | . | . | 7.330073 | 6.234131 | 8.792095 | 6.809854 | 17.057381 | 14.410473 | 19.729206 | 15.569921 | 1.125922 | 30.436322 | 0.565861809 | 0.302383707 | 0.762359573 | 0.998865404 | 4.838265 | 5.011955 | 4.927282 | 4.905213 | 4.948351 | 4.913012 | 36 | 28 | 32 | 26 |
| 677806 | NR_002960 | SNORA20 | small nucleolar RNA, H/ACA box 20 | snoRNA | . | 32610 | . | . | . | 0.715410 | 0.903524 | 1.339903 | 0.595300 | 1.664788 | 2.088537 | 3.006704 | 1.361083 | 1.102192 | 4.264341 | 1.457581144 | 0.096306883 | 0.923276845 | NA | 2.019256 | 2.161867 | 2.082773 | 2.088287 | 2.093105 | 2.081722 | 4 | 5 | 5 | 3 |
| 677807 | NR_002961 | SNORA22 | small nucleolar RNA, H/ACA box 22 | snoRNA | . | 32612 | . | . | . | 0.415507 | 0.951421 | 0.472761 | 0.888585 | 0.966901 | 2.199252 | 1.060862 | 2.031645 | -1.097167 | 3.451417 | 1.619995521 | -0.08258222 | 0.934183741 | NA | 1.855800 | 1.715150 | 1.784522 | 1.794114 | 1.783459 | 1.792257 | 3 | 5 | 2 | 4 |
| 677809 | NR_002963 | SNORA24 | small nucleolar RNA, H/ACA box 24 | snoRNA | . | 32614 | . | . | . | 5.331527 | 5.256227 | 10.471205 | 6.393974 | 12.406685 | 12.150002 | 23.497078 | 14.619062 | 1.567076 | 28.796767 | 0.591606282 | 1.095450801 | 0.273319189 | 0.998865404 | 4.486115 | 5.136800 | 4.805237 | 4.804155 | 4.889160 | 4.823203 | 26 | 24 | 38 | 25 |
| 677810 | NR_003016 | SNORA26 | small nucleolar RNA, H/ACA box 26 | snoRNA | . | 32616 | 614626 | . | . | 2.642060 | 5.995525 | 5.435741 | 2.295117 | 6.148183 | 13.858926 | 12.197646 | 5.247512 | -1.109623 | 15.972366 | 0.820483197 | -0.18290361 | 0.854873654 | NA | 4.069957 | 3.921224 | 3.962815 | 4.019245 | 4.007518 | 3.960468 | 12 | 25 | 18 | 9 |
| 677811 | NR_002964 | SNORA28 | small nucleolar RNA, H/ACA box 28 | snoRNA | . | 32618 | . | . | . | 0.392067 | 0.442966 | 0.305154 | 0.280505 | 0.912357 | 1.023936 | 0.684756 | 0.641342 | 1.234663 | 2.005914 | 2.070995667 | 0.146845664 | 0.883253841 | NA | 0.844134 | 1.148375 | 0.998092 | 0.998667 | 1.000707 | 0.999974 | 2 | 2 | 2 | 2 |
| 677812 | NR_002965 | SNORA29 | small nucleolar RNA, H/ACA box 29 | snoRNA | . | 32619 | . | . | . | 0.356760 | 0.887565 | 0.261561 | 0.639716 | 0.830194 | 2.051646 | 0.586934 | 1.462634 | -1.453250 | 2.680146 | 1.862965018 | -0.2894755 | 0.772217523 | NA | 1.671431 | 1.121007 | 1.422771 | 1.436450 | 1.420486 | 1.429652 | 2 | 5 | 1 | 3 |
| 677814 | NR_002967 | SNORA31 | small nucleolar RNA, H/ACA box 31 | snoRNA | . | 32621 | . | . | . | 0.834874 | 1.131103 | 0.649228 | 0.594926 | 1.942787 | 2.614596 | 1.456849 | 1.360227 | -1.218521 | 3.689354 | 1.556720669 | -0.18316111 | 0.854671607 | NA | 2.019256 | 1.733338 | 1.884830 | 1.890282 | 1.884801 | 1.883646 | 4 | 5 | 3 | 3 |
| 677821 | NR_002972 | SNORA71E | small nucleolar RNA, H/ACA box 71E | snoRNA | . | 32632 | . | . | . | 0.274938 | 0.412555 | 0.568125 | 0.534899 | 0.639791 | 0.953640 | 1.274856 | 1.222982 | 1.852159 | 2.560074 | 1.861050267 | 0.477799129 | 0.632793181 | NA | 0.844134 | 1.733338 | 1.338676 | 1.339295 | 1.346639 | 1.345588 | 2 | 2 | 3 | 3 |
| 677822 | NR_002973 | SNORA40 | small nucleolar RNA, H/ACA box 40 | snoRNA | . | 32633 | . | . | . | 11.439449 | 8.336213 | 10.974008 | 7.000647 | 26.620068 | 19.269529 | 24.625353 | 16.006147 | -1.114531 | 37.705651 | 0.526435526 | -0.29716235 | 0.766342562 | 0.998865404 | 5.310890 | 5.158506 | 5.267009 | 5.216735 | 5.256468 | 5.195049 | 53 | 36 | 38 | 26 |
| 677824 | NR_002975 | SNORA17B | small nucleolar RNA, H/ACA box 17B | snoRNA | . | 32636 | . | . | . | 0.004662 | 0.005781 | 0.005471 | 0.011261 | 0.010849 | 0.013363 | 0.012276 | 0.025748 | 1.234655 | 1.002957 | 2.868339486 | 0.106022226 | 0.915564717 | NA | -0.155866 | 0.148375 | -0.001261 | -0.000972 | 0.000053 | -0.000315 | 1 | 1 | 1 | 1 |
| 677825 | NR_002976 | SNORA44 | small nucleolar RNA, H/ACA box 44 | snoRNA | . | 32637 | . | . | . | 3.167808 | 5.096292 | 3.242274 | 5.629297 | 7.371618 | 11.780307 | 7.275568 | 12.870719 | 1.062110 | 18.119143 | 0.741213475 | 0.11728532 | 0.906633958 | NA | 4.137922 | 4.219802 | 4.157267 | 4.189977 | 4.158149 | 4.198404 | 16 | 23 | 12 | 22 |
| 677826 | NR_002977 | SNORA38 | small nucleolar RNA, H/ACA box 38 | snoRNA | . | 32638 | 611333 | . | . | 5.027345 | 2.271569 | 4.438941 | 5.605533 | 11.698839 | 5.250833 | 9.960855 | 12.816385 | 1.346931 | 18.225767 | 0.74441358 | 0.577200439 | 0.563804061 | NA | 3.955908 | 4.387716 | 4.192470 | 4.145266 | 4.181640 | 4.202777 | 24 | 11 | 16 | 22 |
| 677828 | NR_003014 | SNORA47 | small nucleolar RNA, H/ACA box 47 | snoRNA | . | 32640 | . | . | . | 0.547839 | 0.183066 | 0.371492 | 0.402819 | 1.274845 | 0.423164 | 0.833617 | 0.920999 | 1.250132 | 1.991338 | 2.100718025 | 0.153318985 | 0.87814673 | NA | 0.820514 | 1.148375 | 0.993554 | 0.985525 | 0.991973 | 0.991243 | 3 | 1 | 2 | 2 |
| 677831 | NR_002981 | SNORA51 | small nucleolar RNA, H/ACA box 51 | snoRNA | . | 32644 | . | . | . | 4.162196 | 3.685311 | 3.345596 | 4.674016 | 9.685601 | 8.518762 | 7.507421 | 10.686583 | -1.002601 | 16.530800 | 0.756141545 | -0.00495546 | 0.996046134 | NA | 4.049783 | 4.044381 | 4.051575 | 4.044433 | 4.036439 | 4.059103 | 20 | 17 | 12 | 18 |
| 677832 | NR_003015 | SNORA53 | small nucleolar RNA, H/ACA box 53 | snoRNA | . | 32646 | . | . | . | 0.477083 | 0.870695 | 0.443816 | 0.334907 | 1.110192 | 2.012650 | 0.995910 | 0.765725 | -1.766046 | 4.601523 | 1.423994021 | -0.5762125 | 0.564471557 | NA | 2.555346 | 1.733338 | 2.207672 | 2.221907 | 2.203838 | 2.202598 | 5 | 8 | 3 | 3 |
| 677834 | NR_002983 | SNORA55 | small nucleolar RNA, H/ACA box 55 | snoRNA | . | 32649 | . | . | . | 1.241138 | 2.335044 | 2.165033 | 2.467119 | 2.888180 | 5.397559 | 4.858273 | 5.640775 | 1.294054 | 9.322432 | 1.101708085 | 0.371263635 | 0.710441181 | NA | 3.024434 | 3.393446 | 3.198224 | 3.217391 | 3.217825 | 3.219315 | 7 | 11 | 9 | 10 |
| 677835 | NR_002984 | SNORA56 | small nucleolar RNA, H/ACA box 56 | snoRNA | . | 32650 | . | . | . | 0.010169 | 0.005521 | 0.001922 | 0.000656 | 0.023665 | 0.012763 | 0.004313 | 0.001499 | 1.234655 | 1.002957 | 2.868339486 | 0.106022226 | 0.915564717 | NA | -0.155866 | 0.148375 | -0.001261 | -0.000972 | 0.000053 | -0.000315 | 1 | 1 | 1 | 1 |
| 677837 | NR_002986 | SNORA60 | small nucleolar RNA, H/ACA box 60 | snoRNA | . | 32654 | . | . | . | 1.097744 | 0.591833 | 0.344644 | 0.517237 | 2.554496 | 1.368049 | 0.773372 | 1.182600 | -1.452968 | 3.372709 | 1.642924592 | -0.32807522 | 0.742854792 | NA | 1.998355 | 1.459419 | 1.771805 | 1.760323 | 1.758360 | 1.762326 | 6 | 3 | 2 | 3 |
| 677838 | NR_002987 | SNORA61 | small nucleolar RNA, H/ACA box 61 | snoRNA | . | 32655 | . | . | . | 0.036971 | 0.028061 | 0.032314 | 0.046401 | 0.086034 | 0.064864 | 0.072511 | 0.106091 | 1.234655 | 1.002957 | 2.868339486 | 0.106022226 | 0.915564717 | NA | -0.155866 | 0.148375 | -0.001261 | -0.000972 | 0.000053 | -0.000315 | 1 | 1 | 1 | 1 |
| 677839 | NR_003017 | SNORA71C | small nucleolar RNA, H/ACA box 71C | snoRNA | . | 32656 | . | . | . | 1.147101 | 1.943477 | 0.525396 | 1.031914 | 2.669351 | 4.492435 | 1.178972 | 2.359348 | -1.885125 | 5.527846 | 1.326652642 | -0.68944954 | 0.490540414 | NA | 2.855800 | 1.932304 | 2.473691 | 2.492327 | 2.461012 | 2.474323 | 6 | 10 | 2 | 5 |
| 677840 | NR_003018 | SNORA71D | small nucleolar RNA, H/ACA box 71D | snoRNA | . | 32657 | . | . | . | 0.011866 | 0.240540 | 0.413948 | 0.243681 | 0.027614 | 0.556019 | 0.928887 | 0.557147 | 1.236246 | 1.522137 | 2.383171416 | 0.128385813 | 0.897843658 | NA | 0.444631 | 0.751299 | 0.592746 | 0.597508 | 0.599417 | 0.593826 | 1 | 2 | 2 | 1 |
| 677842 | NR_002995 | SNORA50C | small nucleolar RNA, H/ACA box 50C | snoRNA | . | 32662 | . | . | . | 4.245820 | 2.869097 | 3.262623 | 2.856080 | 9.880197 | 6.632046 | 7.321231 | 6.530088 | -1.188512 | 13.954498 | 0.825968595 | -0.30165331 | 0.762916365 | NA | 3.920530 | 3.674293 | 3.824032 | 3.796393 | 3.804258 | 3.795335 | 21 | 13 | 12 | 11 |
| 677843 | NR_003019 | SNORA77 | small nucleolar RNA, H/ACA box 77 | snoRNA | . | 32663 | . | . | . | 0.720538 | 0.547562 | 0.703075 | 0.252553 | 1.676720 | 1.265715 | 1.577679 | 0.577432 | -1.393479 | 2.692648 | 1.841362901 | -0.25996586 | 0.794890106 | NA | 1.644780 | 1.175234 | 1.435544 | 1.432221 | 1.435169 | 1.424396 | 4 | 3 | 3 | 1 |
| 677844 | NR_003020 | SNORA78 | small nucleolar RNA, H/ACA box 78 | snoRNA | . | 32664 | . | . | . | 0.793004 | 1.207409 | 0.588202 | 0.910543 | 1.845353 | 2.790981 | 1.319908 | 2.081849 | -1.167760 | 4.187707 | 1.469228883 | -0.15228666 | 0.878960843 | NA | 2.175403 | 1.947964 | 2.064565 | 2.074374 | 2.064708 | 2.068325 | 4 | 6 | 3 | 4 |
| 677847 | NR_002989 | SNORA81 | small nucleolar RNA, H/ACA box 81 | snoRNA | . | 32667 | 611334 | . | . | 6.580270 | 4.966776 | 5.130190 | 2.496621 | 15.312559 | 11.480927 | 11.511997 | 5.708227 | -1.534949 | 26.940344 | 0.63046173 | -0.98053675 | 0.326821237 | 0.998865404 | 5.025590 | 4.413324 | 4.793445 | 4.754099 | 4.757632 | 4.696491 | 43 | 30 | 25 | 13 |
| 677848 | NR_004395 | SNORD1A | small nucleolar RNA, C/D box 1A | snoRNA | . | 32556 | . | . | . | 2.142876 | 0.928906 | 3.142261 | 1.511682 | 4.986561 | 2.147208 | 7.051141 | 3.456281 | 1.533856 | 5.076844 | 1.357295535 | 0.454700618 | 0.649324636 | NA | 1.998355 | 2.622519 | 2.332395 | 2.320927 | 2.344801 | 2.328081 | 6 | 3 | 7 | 4 |

| | | | | | | | | | | | | | | | | | | | | | | | | | | | | | | | | | |
|---|---|---|---|---|---|---|---|---|---|---|---|---|---|---|---|---|---|---|---|---|---|---|---|---|---|---|---|---|---|---|---|---|---|
| 677849 | NR_004396 | SNORD1B | small nucleolar RNA, C/D box 1B | snoRNA | . | 32676 | . | . | . | 1.424953 | 0.963249 | 3.419143 | 1.260849 | 3.315924 | 2.226593 | 7.672458 | 2.882782 | 1.881961 | 5.147227 | 1.356351197 | 0.67256683 | 0.50122291 | NA | 1.832372 | 2.751299 | 2.341373 | 2.334034 | 2.362997 | 2.341214 | 5 | 3 | 8 | 4 |
| 677850 | NR_004397 | SNORD1C | small nucleolar RNA, C/D box 1C | snoRNA | . | 32677 | . | . | . | 3.309287 | 4.263144 | 5.932194 | 4.057659 | 7.700847 | 9.854447 | 13.311671 | 9.277356 | 1.292796 | 11.355424 | 0.908380823 | 0.407862562 | 0.68337458 | NA | 3.307819 | 3.678992 | 3.484817 | 3.496111 | 3.512417 | 3.494150 | 10 | 12 | 13 | 10 |
| 678655 | NR_015382 | CD27-AS1 | CD27 antisense RNA 1 | lncRNA | . | 43896 | . | . | . | 4.155381 | 3.766866 | 4.225392 | 3.398765 | 9.669743 | 8.707279 | 9.481658 | 7.770872 | -1.054469 | 287.023444 | 0.216224954 | -0.35387414 | 0.72343321 | 0.998865404 | 8.202085 | 8.126988 | 8.205250 | 8.148057 | 8.206820 | 8.088121 | 357 | 301 | 271 | 232 |
| 684959 | NR_003028 | SNORA25 | small nucleolar RNA, H/ACA box 25 | snoRNA | . | 32615 | . | . | . | 4.081742 | 6.914243 | 6.654177 | 3.037067 | 9.498383 | 15.982582 | 14.931780 | 6.943892 | -1.133759 | 22.143509 | 0.701464804 | -0.25819438 | 0.796256899 | NA | 4.555346 | 4.376754 | 4.439625 | 4.492167 | 4.486955 | 4.421181 | 20 | 32 | 25 | 12 |
| 692057 | NR_003030 | SNORD12 | small nucleolar RNA, C/D box 12 | snoRNA | . | 32708 | . | . | . | 4.160680 | 7.402668 | 7.018539 | 2.340994 | 9.682073 | 17.111599 | 15.749399 | 5.352405 | -1.191436 | 15.409881 | 0.835817649 | -0.30234099 | 0.76239213 | NA | 4.064940 | 3.815893 | 3.923192 | 3.963670 | 3.959769 | 3.903399 | 14 | 23 | 18 | 7 |
| 692058 | NR_003031 | SNORD11 | small nucleolar RNA, C/D box 11 | snoRNA | . | 32707 | . | . | . | 1.055983 | 1.454795 | 1.769893 | 0.367653 | 2.457315 | 3.362824 | 3.971587 | 0.840594 | -1.196175 | 3.731008 | 1.605927474 | -0.16092174 | 0.872155043 | NA | 2.019256 | 1.769039 | 1.893674 | 1.899141 | 1.903813 | 1.882757 | 4 | 5 | 5 | 1 |
| 692063 | NR_003032 | SNORA32 | small nucleolar RNA, H/ACA box 32 | snoRNA | . | 32622 | . | . | . | 6.175907 | 5.311898 | 3.958426 | 0.876289 | 14.371590 | 12.278689 | 8.882593 | 2.003531 | **-2.342493** | 15.980273 | 0.835773177 | -1.469352 | 0.141737339 | NA | 4.482357 | 3.264261 | 4.036635 | 4.019409 | 3.996024 | 3.949539 | 28 | 22 | 13 | 4 |
| 692072 | NR_003033 | SNORD5 | small nucleolar RNA, C/D box 5 | snoRNA | . | 32702 | . | . | . | 4.258470 | 1.991912 | 6.044555 | 1.428977 | 9.909633 | 4.604393 | 13.563807 | 3.267185 | 1.192188 | 8.280347 | 1.117107511 | 0.227025197 | 0.820404149 | NA | 2.912173 | 3.175234 | 3.047995 | 3.021122 | 3.061704 | 3.020075 | 12 | 5 | 12 | 4 |
| 692073 | NR_003035 | SNORA16A | small nucleolar RNA, H/ACA box 16A | snoRNA | . | 32605 | . | . | . | 9.363158 | 7.251844 | 9.162815 | 6.970401 | 21.788454 | 16.762962 | 20.561090 | 15.936993 | -1.040970 | 34.607512 | 0.536768341 | -0.10792074 | 0.914058566 | 0.998865404 | 5.140184 | 5.085321 | 5.133060 | 5.095167 | 5.128539 | 5.088607 | 46 | 33 | 34 | 27 |
| 692075 | NR_003036 | SNORD6 | small nucleolar RNA, C/D box 6 | snoRNA | . | 32703 | . | . | . | 2.414093 | 0.939194 | 3.450388 | 0.343142 | 5.617694 | 2.170989 | 7.742570 | 0.784554 | 1.020234 | 4.493954 | 1.511550839 | 0.019119767 | 0.984745563 | NA | 2.147197 | 2.188478 | 2.166098 | 2.150501 | 2.174165 | 2.142963 | 7 | 3 | 7 | 1 |
| 692085 | NR_003042 | SNORD45C | small nucleolar RNA, C/D box 45C | snoRNA | . | 32720 | . | . | . | 2.480872 | 1.250349 | 1.765198 | 0.941442 | 5.773091 | 2.890237 | 3.961053 | 2.152496 | -1.371066 | 4.613603 | 1.415513638 | -0.32164865 | 0.747718885 | NA | 2.413393 | 1.963455 | 2.222348 | 2.207129 | 2.211307 | 2.204999 | 8 | 4 | 4 | 3 |
| 692086 | NR_003045 | SNORD17 | small nucleolar RNA, C/D box 17 | snoRNA | . | 32713 | . | . | . | 5.685484 | 8.015715 | 4.550270 | 6.256196 | 13.230355 | 18.528685 | 10.210673 | 14.304049 | -1.276220 | 45.789497 | 0.477140994 | -0.73746864 | 0.46083741 | 0.998865404 | 5.683585 | 5.328512 | 5.503306 | 5.569779 | 5.473899 | 5.516223 | 49 | 65 | 30 | 43 |
| 692087 | NR_003043 | SNORD49B | small nucleolar RNA, C/D box 49B | snoRNA | . | 32721 | . | . | . | 10.966494 | 11.872782 | 15.449190 | 9.991860 | 25.519480 | 27.444468 | 34.667534 | 22.845201 | 1.086595 | 18.746634 | 0.717758 | 0.166928452 | 0.867426346 | NA | 4.166062 | 4.288462 | 4.217506 | 4.222686 | 4.245617 | 4.207438 | 20 | 20 | 21 | 14 |
| 692088 | NR_003044 | SNORD50B | small nucleolar RNA, C/D box 50B | snoRNA | . | 32722 | 613264 | . | . | 5.039611 | 8.911998 | 11.099006 | 2.962624 | 11.727382 | 20.600483 | 24.905848 | 6.773687 | 1.031802 | 16.111059 | 0.837931891 | 0.053902228 | 0.957013062 | NA | 3.985415 | 4.034132 | 3.973343 | 4.013908 | 4.032838 | 3.957633 | 13 | 22 | 22 | 7 |
| 692089 | NR_003047 | SNORD19 | small nucleolar RNA, C/D box 19 | snoRNA | . | 32717 | . | . | . | 1.174321 | 1.785731 | 0.452913 | 2.550993 | 2.732693 | 4.127799 | 1.016323 | 5.832542 | -1.066221 | 3.914367 | 1.570963747 | -0.05888515 | 0.953043589 | NA | 2.019256 | 1.916472 | 1.969412 | 1.974872 | 1.959240 | 1.982658 | 4 | 5 | 1 | 6 |
| 692090 | NR_003046 | SNORD59B | small nucleolar RNA, C/D box 59B | snoRNA | . | 32723 | . | . | . | 0.895213 | 1.758623 | 4.189153 | 1.093481 | 2.083198 | 4.065136 | 9.400337 | 2.500114 | 1.869173 | 5.197205 | 1.374937507 | 0.65632057 | 0.511617885 | NA | 1.855800 | 2.760196 | 2.341172 | 2.350983 | 2.376219 | 2.344531 | 3 | 5 | 9 | 3 |
| 692091 | NR_003048 | SNORD23 | small nucleolar RNA, C/D box 23 | snoRNA | . | 32718 | . | . | . | 11.249715 | 9.202596 | 7.167233 | 11.329940 | 26.178547 | 21.272211 | 16.083063 | 25.904560 | -1.121286 | 33.803852 | 0.545385078 | -0.30282195 | 0.762025558 | 0.998865404 | 5.160193 | 4.993209 | 5.101622 | 5.074748 | 5.047164 | 5.098082 | 45 | 35 | 22 | 36 |
| 692093 | NR_003050 | SNORD62B | small nucleolar RNA, C/D box 62B | snoRNA | . | 23031 | . | . | . | 1.126633 | 1.663483 | 1.068748 | 1.169078 | 2.621722 | 3.845215 | 2.398241 | 2.672959 | -1.218521 | 3.689354 | 1.556720669 | -0.18316111 | 0.854671607 | NA | 2.019256 | 1.733338 | 1.884830 | 1.890282 | 1.884801 | 1.883646 | 4 | 5 | 3 | 3 |
| 692099 | NR_024241 | FAM86DP | family with sequence similarity 86 member D, pseudogene | transcribed_ps | . | 32659 | . | . | . | 9.635351 | 10.302587 | 9.125526 | 10.622355 | 22.421858 | 23.814892 | 20.477417 | 24.286751 | -1.022950 | 775.012758 | 0.152800519 | -0.21424044 | 0.830359561 | 0.998865404 | 9.614664 | 9.581296 | 9.584334 | 9.626211 | 9.539104 | 9.637391 | 876 | 871 | 619 | 769 |
| 692106 | NR_003054 | SNORD65 | small nucleolar RNA, C/D box 65 | snoRNA | . | 32726 | . | . | . | 26.014427 | 19.939236 | 21.444378 | 15.803354 | 60.536640 | 46.090439 | 48.120560 | 36.132496 | -1.242284 | 47.993828 | 0.463481454 | -0.67531316 | 0.499476824 | 0.998865404 | 5.731451 | 5.421486 | 5.628806 | 5.577836 | 5.589360 | 5.540130 | 69 | 50 | 43 | 34 |
| 692108 | NR_003056 | SNORD67 | small nucleolar RNA, C/D box 67 | snoRNA | . | 32728 | . | . | . | 0.410626 | 0.256706 | 0.610308 | 0.574996 | 0.955543 | 0.593388 | 1.369513 | 1.314659 | 1.659343 | 1.774228 | 2.21400588 | 0.329995761 | 0.741403165 | NA | 0.413393 | 1.148375 | 0.817239 | 0.813367 | 0.819776 | 0.819065 | 2 | 1 | 2 | 2 |
| 692109 | NR_003057 | SNORD69 | small nucleolar RNA, C/D box 69 | snoRNA | . | 32730 | . | . | . | 1.988496 | 2.083002 | 2.929477 | 2.295044 | 4.627313 | 4.814952 | 6.573662 | 5.247345 | 1.340584 | 6.305236 | 1.200698396 | 0.352179518 | 0.724703637 | NA | 2.429096 | 2.852980 | 2.643888 | 2.645602 | 2.656687 | 2.649491 | 6 | 6 | 7 | 6 |
| 692158 | NR_004390 | SNORA57 | small nucleolar RNA, H/ACA box 57 | snoRNA | . | 32651 | . | . | . | 0.572728 | 0.150038 | 1.092622 | 0.391403 | 1.332762 | 0.346819 | 2.451812 | 0.894897 | 1.776638 | 3.070930 | 1.763842668 | 0.470081381 | 0.638296876 | NA | 1.137672 | 1.978782 | 1.603704 | 1.591525 | 1.613664 | 1.597457 | 4 | 1 | 5 | 2 |
| 692195 | NR_003941 | SNORD75 | small nucleolar RNA, C/D box 75 | snoRNA | . | 32735 | . | . | . | 1.737660 | 1.880043 | 2.068943 | 1.229594 | 4.043606 | 4.345803 | 4.642649 | 2.811320 | 1.084682 | 3.745161 | 1.542159506 | 0.07604373 | 0.939384307 | NA | 1.844134 | 1.963455 | 1.900254 | 1.901373 | 1.905335 | 1.899077 | 4 | 4 | 4 | 3 |
| 692196 | NR_003942 | SNORD76 | small nucleolar RNA, C/D box 76 | snoRNA | . | 32736 | . | . | . | 31.693678 | 31.986835 | 35.235104 | 24.867237 | 73.752487 | 73.939003 | 79.066551 | 56.855976 | -1.070694 | 78.954611 | 0.36826183 | -0.26759859 | 0.789008316 | 0.998865404 | 6.350383 | 6.253908 | 6.309712 | 6.309340 | 6.334531 | 6.247153 | 94 | 88 | 78 | 59 |
| 692197 | NR_003943 | SNORD77 | small nucleolar RNA, C/D box 77 | snoRNA | . | 32737 | . | . | . | 5.489810 | 4.805020 | 8.908750 | 3.063772 | 12.775013 | 11.107018 | 19.990978 | 7.004951 | 1.156052 | 12.573839 | 0.894872241 | 0.23378336 | 0.815153168 | NA | 3.540965 | 3.755755 | 3.640215 | 3.635552 | 3.671312 | 3.619689 | 14 | 12 | 17 | 7 |
| 692198 | NR_003944 | SNORD78 | small nucleolar RNA, C/D box 78 | snoRNA | . | 32738 | . | . | . | 10.556127 | 9.915418 | 11.678474 | 8.821401 | 24.564539 | 22.919930 | 26.206158 | 20.169085 | 1.002877 | 21.095559 | 0.671295847 | 0.006174352 | 0.995073611 | NA | 4.395729 | 4.401999 | 4.400098 | 4.394373 | 4.409768 | 4.385039 | 25 | 22 | 21 | 17 |
| 692199 | NR_003065 | SNORD84 | small nucleolar RNA, C/D box 84 | snoRNA | . | 32743 | . | . | . | 5.435272 | 4.594067 | 2.483483 | 3.242256 | 12.648102 | 10.619392 | 5.572864 | 7.413033 | -1.679626 | 10.343984 | 0.956093767 | -0.78249689 | 0.433922611 | NA | 3.697260 | 2.947964 | 3.395144 | 3.386672 | 3.364243 | 3.371112 | 16 | 13 | 6 | 8 |
| 692201 | NR_004399 | SNORD86 | small nucleolar RNA, C/D box 86 | snoRNA | . | 32745 | . | . | . | 0.007237 | 0.008097 | 0.008998 | 0.008516 | 0.016840 | 0.018717 | 0.020191 | 0.019470 | 1.234655 | 1.002957 | 2.868339486 | 0.106022226 | 0.915564717 | NA | -0.155866 | 0.148375 | -0.001261 | -0.000972 | 0.000053 | -0.000315 | 1 | 1 | 1 | 1 |
| 692202 | NR_003067 | SNORD88A | small nucleolar RNA, C/D box 88A | snoRNA | . | 32747 | . | . | . | 0.433313 | 0.956980 | 2.204658 | 0.272391 | 1.008337 | 2.212103 | 4.947189 | 0.622790 | 1.459205 | 3.352594 | 1.712794051 | 0.318300287 | 0.750257166 | NA | 1.444631 | 1.993948 | 1.716624 | 1.726128 | 1.740176 | 1.713936 | 2 | 4 | 6 | 1 |
| 692203 | NR_003068 | SNORD88B | small nucleolar RNA, C/D box 88B | snoRNA | . | 32748 | . | . | . | 12.048350 | 11.411083 | 10.679757 | 12.586607 | 28.037006 | 26.377232 | 23.965065 | 28.777779 | -1.010850 | 36.077145 | 0.516121093 | -0.03016497 | 0.975935487 | 0.998865404 | 5.181089 | 5.164892 | 5.178593 | 5.170030 | 5.163060 | 5.182433 | 43 | 38 | 29 | 36 |
| 692204 | NR_003069 | SNORD88C | small nucleolar RNA, C/D box 88C | snoRNA | . | 32749 | . | . | . | 4.473815 | 3.485710 | 4.968031 | 0.693360 | 10.410752 | 8.057376 | 11.148118 | 1.585285 | -1.381735 | 10.812246 | 1.03614121 | -0.45020976 | 0.652559197 | NA | 3.644780 | 3.188478 | 3.442979 | 3.430321 | 3.451250 | 3.390357 | 16 | 12 | 14 | 2 |
| 692205 | NR_003070 | SNORD89 | small nucleolar RNA, C/D box 89 | snoRNA | . | 32750 | . | . | . | 0.003453 | 0.012151 | 0.007639 | 0.018244 | 0.008036 | 0.028087 | 0.017142 | 0.041712 | 1.234655 | 1.002957 | 2.868339486 | 0.106022226 | 0.915564717 | NA | -0.155866 | 0.148375 | -0.001261 | -0.000972 | 0.000053 | -0.000315 | 1 | 1 | 1 | 1 |
| 692210 | NR_003075 | SNORD93 | small nucleolar RNA, C/D box 93 | snoRNA | . | 32755 | . | . | . | 5.122986 | 2.409599 | 6.061841 | 1.112894 | 11.921400 | 5.569896 | 13.602595 | 2.544501 | -1.034426 | 9.198767 | 1.084131736 | -0.04504135 | 0.96407435 | NA | 3.220748 | 3.181871 | 3.205769 | 3.179657 | 3.216624 | 3.165339 | 14 | 7 | 13 | 3 |
| 692211 | NR_003076 | SNORD98 | small nucleolar RNA, C/D box 98 | snoRNA | . | 32761 | . | . | . | 0.032589 | 0.438443 | 0.267807 | 0.522396 | 0.075835 | 1.013480 | 0.600950 | 1.194396 | 1.834012 | 1.269624 | 2.598200026 | 0.336772769 | 0.736288203 | NA | -0.155866 | 0.715150 | 0.332517 | 0.332827 | 0.333928 | 0.338438 | 1 | 1 | 1 | 2 |
| 692212 | NR_003077 | SNORD99 | small nucleolar RNA, C/D box 99 | snoRNA | . | 32762 | . | . | . | 10.238320 | 12.099103 | 18.629644 | 12.314525 | 23.824993 | 27.967619 | 41.804386 | 28.155697 | 1.377300 | 33.679533 | 0.546716695 | 0.844756778 | 0.398246651 | 0.998865404 | 4.823643 | 5.286924 | 5.030373 | 5.049594 | 5.112815 | 5.051357 | 30 | 33 | 41 | 29 |
| 692213 | NR_003078 | SNORD110 | small nucleolar RNA, C/D box 110 | snoRNA | . | 32775 | . | . | . | 9.276011 | 7.735592 | 5.624597 | 4.552356 | 21.585659 | 17.881168 | 12.621431 | 10.408422 | -1.683745 | 16.395187 | 0.767410513 | -0.97949373 | 0.327336087 | NA | 4.361571 | 3.612728 | 4.067593 | 4.049832 | 4.029717 | 4.016012 | 26 | 20 | 12 | 10 |
| 692223 | NR_004403 | SNORD97 | small nucleolar RNA, C/D box 97 | snoRNA | . | 32760 | . | . | . | 1.193592 | 1.448096 | 2.367307 | 1.652883 | 2.777538 | 3.347339 | 5.312168 | 3.779121 | 1.507146 | 7.883180 | 1.085058917 | 0.545425769 | 0.585460721 | NA | 2.651489 | 3.245375 | 2.958563 | 2.960591 | 2.982685 | 2.965194 | 7 | 7 | 10 | 7 |
| 692225 | NR_004378 | SNORD94 | small nucleolar RNA, C/D box 94 | snoRNA | . | 32756 | . | . | . | 0.962231 | 1.423755 | 1.005003 | 1.780834 | 2.239151 | 3.291073 | 2.255198 | 4.071666 | 1.217569 | 5.990665 | 1.244693342 | 0.22817186 | 0.819512637 | NA | 2.436885 | 2.715150 | 2.570991 | 2.581111 | 2.572686 | 2.590127 | 5 | 7 | 4 | 8 |
| 692227 | NR_004380 | SNORD104 | small nucleolar RNA, C/D box 104 | snoRNA | . | 32768 | . | . | . | 26.317360 | 25.619251 | 23.975941 | 20.326900 | 61.241577 | 59.220051 | 53.801319 | 46.475044 | -1.199470 | 60.418547 | 0.4062672 | -0.64587462 | 0.518360556 | 0.998865404 | 6.041573 | 5.780468 | 5.940643 | 5.933334 | 5.915339 | 5.880927 | 77 | 70 | 52 | 47 |
| 692233 | NR_003140 | SNORD117 | small nucleolar RNA, C/D box 117 | snoRNA | . | 32742 | . | . | . | 0.402214 | 0.394252 | 1.267192 | 0.257357 | 0.935968 | 0.911332 | 2.843541 | 0.588416 | 1.252417 | 2.026741 | 2.090159777 | 0.155354231 | 0.876542064 | NA | 0.844134 | 1.175234 | 1.007230 | 1.007808 | 1.014998 | 1.004217 | 2 | 2 | 3 | 1 |
| 692234 | NR_004054 | SNORD103A | small nucleolar RNA, C/D box 103A | snoRNA | . | 32766 | . | . | . | 1.163252 | 0.424544 | 0.549602 | 0.831992 | 2.706934 | 0.981353 | 1.233291 | 1.902252 | 1.031783 | 2.706802 | 1.80851132 | 0.024959558 | 0.980087222 | NA | 1.413393 | 1.459419 | 1.442085 | 1.434379 | 1.436604 | 1.440654 | 4 | 2 | 2 | 3 |
| 692235 | NR_033295 | SNORD103B | small nucleolar RNA, C/D box 103B | snoRNA | . | 32767 | . | . | . | 1.217325 | 1.069823 | 1.219862 | 2.101763 | 2.832765 | 2.472944 | 2.737336 | 4.805430 | 1.375493 | 4.257668 | 1.459684053 | 0.315101379 | 0.752684682 | NA | 1.844134 | 2.300113 | 2.081326 | 2.082483 | 2.081487 | 2.094711 | 4 | 4 | 3 | 6 |
| 692247 | NR_147607,NF | LMNTD2-AS1 | LMNTD2 antisense RNA 1 | lncRNA | . | 41204 | . | . | . | 0.897601 | 0.593172 | 1.381977 | 0.497054 | 2.088754 | 1.371144 | 3.101117 | 1.136456 | 1.258281 | 15.111013 | 0.83269672 | 0.398049254 | 0.69059388 | NA | 3.738475 | 4.076793 | 3.910850 | 3.882516 | 3.941827 | 3.880675 | 19 | 11 | 21 | 9 |
| 692312 | NM_00104066 | PPAN-P2RY11 | PPAN-P2RY11 readthrough | protein_coding | NP_001035754.1;NP_001185619.1 | 33526 | . | . | . | 3.372220 | 5.057461 | 4.275669 | 2.507324 | 7.847292 | 11.690548 | 9.594478 | 5.732697 | -1.256548 | 363.076868 | 0.258672806 | -1.27367623 | 0.202778183 | 0.968254058 | 8.659104 | 8.330486 | 8.426430 | 8.664698 | 8.553741 | 8.267954 | 375 | 521 | 353 | 223 |
| 693120 | NR_030361,rn | MIR33B | microRNA 33b | miRNA | . | 32791 | 613486 | . | . | 0.172327 | 0.312132 | 0.262470 | 0.243083 | 0.401014 | 0.721509 | 0.588976 | 0.555782 | 1.221536 | 2.504267 | 1.872041618 | 0.154214919 | 0.877440279 | NA | 1.175403 | 1.459419 | 1.316408 | 1.321418 | 1.319196 | 1.323280 | 2 | 3 | 2 | 3 |
| 693130 | NR_030382,rn | MIR548D1 | microRNA 548d-1 | miRNA | . | 32801 | . | . | . | 0.240594 | 0.130222 | 0.348175 | 0.141502 | 0.559871 | 0.301014 | 0.781295 | 0.323527 | -1.056394 | 1.724672 | 2.259675635 | -0.03502612 | 0.972058914 | NA | 0.820514 | 0.751299 | 0.790628 | 0.782592 | 0.788957 | 0.783375 | 3 | 1 | 2 | 1 |
| 693140 | NR_030282,rn | MIR555 | microRNA 555 | miRNA | . | 32811 | . | . | . | 0.000274 | 0.000326 | 0.000079 | 0.008679 | 0.000639 | 0.000753 | 0.000177 | 0.019842 | 1.234663 | 2.005914 | 2.070995667 | 0.146845664 | 0.883253841 | NA | 0.844134 | 1.148375 | 0.998092 | 0.998667 | 1.000707 | 0.999974 | 2 | 2 | 2 | 2 |
| 693149 | NR_030290,rn | MIR564 | microRNA 564 | miRNA | . | 32820 | . | . | . | 0.126869 | 0.307796 | 0.001168 | 0.146182 | 0.295230 | 0.711485 | 0.002620 | 0.334228 | 1.234663 | 2.005914 | 2.070995667 | 0.146845664 | 0.883253841 | NA | 0.844134 | 1.148375 | 0.998092 | 0.998667 | 1.000707 | 0.999974 | 2 | 2 | 2 | 2 |
| 693155 | NR_030296,rn | MIR570 | microRNA 570 | miRNA | . | 32826 | 614538 | . | . | 2.640361 | 3.585355 | 4.948524 | 1.865092 | 6.144227 | 8.287708 | 11.104345 | 4.264311 | 1.099695 | 12.294670 | 0.888876116 | 0.154243439 | 0.877417793 | NA | 3.548173 | 3.688346 | 3.603994 | 3.615647 | 3.633440 | 3.596113 | 12 | 14 | 15 | 8 |
| 693158 | NR_030299,rn | MIR573 | microRNA 573 | miRNA | . | 32829 | . | . | . | 0.366713 | 0.310110 | 0.697880 | 0.904911 | 0.853355 | 0.716834 | 1.566024 | 2.068970 | 1.729504 | 3.043852 | 1.713699129 | 0.461199977 | 0.644655138 | NA | 1.156660 | 1.947964 | 1.593764 | 1.590220 | 1.597682 | 1.601437 | 3 | 2 | 3 | 4 |
| 693174 | NR_030318,rn | MIR589 | microRNA 589 | miRNA | . | 32845 | . | . | . | 0.156095 | 0.561903 | 0.548647 | 0.157777 | 0.363238 | 1.298865 | 1.231149 | 0.360739 | 1.236277 | 3.044274 | 1.727664661 | 0.177118846 | 0.859415039 | NA | 1.444631 | 1.751299 | 1.589015 | 1.598466 | 1.602247 | 1.591161 | 2 | 4 | 4 | 2 |
| 693175 | NR_030321,rn | MIR590 | microRNA 590 | miRNA | . | 32846 | 615070 | . | . | 1.194158 | 1.575824 | 1.602334 | 0.862312 | 2.778855 | 3.642589 | 3.595592 | 1.971575 | 1.045330 | 5.982762 | 1.231979154 | 0.051915284 | 0.958596195 | NA | 2.548173 | 2.612728 | 2.574413 | 2.580407 | 2.582075 | 2.575150 | 6 | 7 | 6 | 5 |
| 693196 | NR_030342,rn | MIR611 | microRNA 611 | miRNA | . | 32867 | . | . | . | 0.787533 | 0.148723 | 0.223399 | 0.542800 | 1.832621 | 0.343781 | 0.501300 | 1.241046 | -1.007340 | 2.223025 | 1.975467281 | -0.0053412 | 0.995738358 | NA | 1.156660 | 1.148375 | 1.155078 | 1.151489 | 1.153584 | 1.152831 | 3 | 2 | 2 | 2 |
| 693197 | NR_030343,rn | MIR612 | microRNA 612 | miRNA | . | 32868 | . | . | . | 0.000801 | 0.002824 | 0.000558 | 0.004878 | 0.001865 | 0.006528 | 0.001252 | 0.011152 | 1.234663 | 2.005914 | 2.070995667 | 0.146845664 | 0.883253841 | NA | 0.844134 | 1.148375 | 0.998092 | 0.998667 | 1.000707 | 0.999974 | 2 | 2 | 2 | 2 |
| 693201 | NR_030346,rn | MIR616 | microRNA 616 | miRNA | . | 32872 | 614489 | . | . | 0.530471 | 1.101420 | 3.214582 | 1.400806 | 1.234428 | 2.545982 | 7.213429 | 3.202778 | **2.108210** | 7.020989 | 1.175003786 | 0.915757387 | 0.359794158 | NA | 2.175403 | 3.251698 | 2.771301 | 2.781403 | 2.812643 | 2.785420 | 4 | 6 | 11 | 6 |
| 693206 | NR_030352,rn | MIR621 | microRNA 621 | miRNA | . | 32877 | . | . | . | 0.006033 | 0.004344 | 0.007130 | 0.006910 | 0.014040 | 0.010043 | 0.016000 | 0.015799 | 1.234663 | 2.005914 | 2.070995667 | 0.146845664 | 0.883253841 | NA | 0.844134 | 1.148375 | 0.998092 | 0.998667 | 1.000707 | 0.999974 | 2 | 2 | 2 | 2 |
| 693216 | NR_030360,rn | MIR631 | microRNA 631 | miRNA | . | 32887 | . | . | . | 0.210743 | 0.304855 | 0.060726 | 0.178243 | 0.490407 | 0.704687 | 0.136268 | 0.407532 | -1.089958 | 1.718421 | 2.237283933 | -0.0555461 | 0.955703407 | NA | 0.844134 | 0.715150 | 0.784731 | 0.785283 | 0.782101 | 0.786536 | 2 | 2 | 1 | 2 |
| 693217 | NR_030362,rn | MIR632 | microRNA 632 | miRNA | . | 32888 | . | . | . | 0.000295 | 0.000221 | 0.000247 | 0.000124 | 0.000686 | 0.000512 | 0.000554 | 0.000282 | 1.234663 | 2.005914 | 2.070995667 | 0.146845664 | 0.883253841 | NA | 0.844134 | 1.148375 | 0.998092 | 0.998667 | 1.000707 | 0.999974 | 2 | 2 | 2 | 2 |
| 693220 | NR_030365,rn | MIR635 | microRNA 635 | miRNA | . | 32891 | . | . | . | 3.743002 | 5.477076 | 3.237328 | 3.041525 | 8.710119 | 12.660506 | 7.264470 | 6.954086 | -1.381352 | 13.200299 | 0.846248401 | -0.55076137 | 0.581797267 | NA | 3.937099 | 3.470304 | 3.724794 | 3.745041 | 3.718396 | 3.714625 | 15 | 19 | 10 | 10 |
| 693221 | NR_030366,rn | MIR636 | microRNA 636 | miRNA | . | 32892 | . | . | . | 0.549619 | 0.551300 | 0.476695 | 0.741455 | 1.278985 | 1.274353 | 1.069690 | 1.695249 | 1.234665 | 3.008871 | 1.708874417 | 0.17796512 | 0.858750369 | NA | 1.429096 | 1.733338 | 1.582415 | 1.583274 | 1.586316 | 1.585224 | 3 | 3 | 3 | 3 |
| 693222 | NR_030367,rn | MIR637 | microRNA 637 | miRNA | . | 32893 | . | . | . | 0.790727 | 1.037114 | 1.252555 | 1.063045 | 1.840055 | 2.397334 | 2.810696 | 2.430526 | 1.235205 | 4.531008 | 1.409796152 | 0.21616645 | 0.828858004 | NA | 2.019256 | 2.324312 | 2.168104 | 2.173640 | 2.178531 | 2.171959 | 4 | 5 | 5 | 4 |
| 693224 | NR_030369,rn | MIR639 | microRNA 639 | miRNA | . | 32895 | . | . | . | 0.000249 | 0.000243 | 0.000757 | 0.000748 | 0.000581 | 0.000562 | 0.001698 | 0.001711 | -1.089958 | 1.718421 | 2.237283933 | -0.0555461 | 0.955703407 | NA | 0.844134 | 0.715150 | 0.784731 | 0.785283 | 0.782101 | 0.786536 | 2 | 2 | 1 | 2 |
| 693226 | NR_030371,rn | MIR641 | microRNA 641 | miRNA | . | 32897 | . | . | . | 0.000867 | 0.007686 | 0.001266 | 0.000760 | 0.002019 | 0.017766 | 0.002840 | 0.001737 | 1.234663 | 2.005914 | 2.070995667 | 0.146845664 | 0.883253841 | NA | 0.844134 | 1.148375 | 0.998092 | 0.998667 | 1.000707 | 0.999974 | 2 | 2 | 2 | 2 |
| 693232 | NR_030377,rn | MIR647 | microRNA 647 | miRNA | . | 32903 | . | . | . | 3.384269 | 5.012807 | 4.338925 | 3.200790 | 7.875334 | 11.587327 | 9.736420 | 7.318227 | -1.092281 | 13.396872 | 0.843919959 | -0.15089644 | 0.880057412 | NA | 3.805868 | 3.678992 | 3.732196 | 3.756360 | 3.748330 | 3.729959 | 13 | 18 | 13 | 10 |
| 723790 | NM_00104087 | H2AC19 | H2A clustered histone 19 | protein_coding | NP_001035807.1 | 29668 | . | . | . | 1.056458 | 1.207435 | 0.923340 | 1.445502 | 2.458421 | 2.791040 | 2.071949 | 3.304968 | 1.054238 | 19.814663 | 0.69691912 | 0.109340105 | 0.912932738 | NA | 4.271488 | 4.344579 | 4.300453 | 4.309873 | 4.292075 | 4.326906 | 21 | 22 | 14 | 23 |
| 724028 | NR_030395,rn | MIR658 | microRNA 658 | miRNA | . | 32914 | . | . | . | 0.000247 | 0.000195 | 0.000246 | 0.000522 | 0.000574 | 0.000451 | 0.000552 | 0.001194 | 1.234663 | 2.005914 | 2.070995667 | 0.146845664 | 0.883253841 | NA | 0.844134 | 1.148375 | 0.998092 | 0.998667 | 1.000707 | 0.999974 | 2 | 2 | 2 | 2 |
| 724087 | NR_046101 | LINC01545 | long intergenic non-protein coding RNA 1545 | lncRNA | . | 17986 | . | . | . | 0.256722 | 0.303323 | 0.601890 | 0.504196 | 0.597403 | 0.701146 | 1.350623 | 1.152784 | 1.907612 | 7.196446 | 1.137275903 | 0.819297771 | 0.412616544 | NA | 2.307819 | 3.239024 | 2.817505 | 2.823433 | 2.844762 | 2.837013 | 5 | 6 | 9 | 8 |
| 724102 | NR_003141,NF | SNHG4 | small nucleolar RNA host gene 4 | lncRNA | . | 32964 | . | . | . | 21.310111 | 23.149821 | 30.126026 | 20.619827 | 49.589505 | 53.511849 | 67.601930 | 47.144785 | 1.125200 | 786.738825 | 0.18391544 | 0.925325048 | 0.354796854 | 0.998865404 | 9.531765 | 9.702659 | 9.543921 | 9.596779 | 9.781791 | 9.507704 | 820 | 829 | 863 | 632 |
| 727676 | NR_033294 | SNORD118 | small nucleolar RNA, C/D box 118 | snoRNA | . | 32952 | 616663 | . | . | 0.004738 | 0.009832 | 0.005457 | 0.015951 | 0.011026 | 0.022727 | 0.012246 | 0.036469 | 1.234655 | 1.002957 | 2.868339486 | 0.106022226 | 0.915564717 | NA | -0.155866 | 0.148375 | -0.001261 | -0.000972 | 0.000053 | -0.000315 | 1 | 1 | 1 | 1 |
| 727708 | NR_001290 | SNORD116-19 | small nucleolar RNA, C/D box 116-19 | snoRNA | . | 33085 | . | . | . | 0.000238 | 0.015729 | 0.019514 | 0.018367 | 0.000554 | 0.036359 | 0.043788 | 0.041993 | 1.234655 | 1.002957 | 2.868339486 | 0.106022226 | 0.915564717 | NA | -0.155866 | 0.148375 | -0.001261 | -0.000972 | 0.000053 | -0.000315 | 1 | 1 | 1 | 1 |
| 727751 | NR_102747,NF | LOC727751 | golgin A2 pseudogene | transcribed_ps | . | . | . | . | . | 4.043204 | 4.197052 | 3.971915 | 3.788749 | 9.408702 | 9.701674 | 8.912862 | 8.662525 | -1.077891 | 383.880388 | 0.183669772 | -0.58915999 | 0.555753954 | 0.998865404 | 8.637494 | 8.529512 | 8.600220 | 8.612068 | 8.579399 | 8.544881 | 454 | 434 | 331 | 336 |
| 727758 | NR_033770,NF | ROCK1P1 | Rho associated coiled-coil containing protein kinase 1 pseudogene | transcribed_ps | . | 37832 | . | . | . | 0.518785 | 0.353103 | 0.586388 | 0.285628 | 1.207233 | 0.816212 | 1.315837 | 0.653054 | -1.005937 | 36.498375 | 0.558531305 | -0.01528911 | 0.98780153 | 0.999197585 | 5.191364 | 5.188155 | 5.208934 | 5.153580 | 5.229265 | 5.129583 | 50 | 32 | 43 | 22 |
| 727768 | gene-TEKT4P1 | TEKT4P1 | tektin 4 pseudogene 1 | pseudogene | . | 37711 | . | . | . | 0.183054 | 0.121082 | 0.132561 | 0.161886 | 0.425975 | 0.279887 | 0.297464 | 0.370132 | -1.064981 | 10.806803 | 0.930291256 | -0.09763291 | 0.92222379 | NA | 3.478589 | 3.387716 | 3.446176 | 3.429063 | 3.429166 | 3.440078 | 15 | 10 | 8 | 11 |
| 727795 | gene-HMGN2P | HMGN2P5 | high mobility group nucleosomal binding domain 2 pseudogene 5 | pseudogene | . | 33568 | . | . | . | 4.249521 | 4.006072 | 3.599149 | 3.906638 | 9.888810 | 9.260215 | 8.076386 | 8.932064 | -1.115665 | 147.335370 | 0.270050082 | -0.58472147 | 0.558735013 | 0.998865404 | 7.279817 | 7.121777 | 7.238140 | 7.210602 | 7.170043 | 7.194574 | 185 | 162 | 117 | 135 |
| 727800 | NM_031297 | RNF208 | ring finger protein 208 | protein_coding | NP_112587.2 | 25420 | . | . | . | 1.526588 | 1.099474 | 1.122014 | 1.610890 | 3.552434 | 2.541482 | 2.517768 | 3.683108 | 1.036006 | 45.381372 | 0.480646752 | 0.106173324 | 0.915444836 | 0.998865404 | 5.478589 | 5.529027 | 5.527694 | 5.469552 | 5.474417 | 5.534888 | 60 | 40 | 33 | 51 |
| 727831 | gene-RPL13AP | RPL13AP8 | ribosomal protein L13a pseudogene 8 | pseudogene | . | 36966 | . | . | . | 0.073120 | 0.052466 | 0.045490 | 0.046116 | 0.170152 | 0.121277 | 0.102077 | 0.105439 | -1.204527 | 2.440135 | 1.90164038 | -0.14117614 | 0.887730791 | NA | 1.413393 | 1.148375 | 1.296738 | 1.289023 | 1.291174 | 1.290401 | 4 | 2 | 2 | 2 |
| 727851 | NM_00116446 | RGPD8 | RANBP2 like and GRIP domain containing 8 | protein_coding | NP_001157935.1;XP_005263804.1;XP_011510035.1;XP_016860314.1;XP_ | 9849 | 602752 | . | . | 3.164093 | 3.309260 | 3.885804 | 3.040074 | 7.362977 | 7.649501 | 8.719632 | 6.950769 | 1.055508 | 763.206783 | 0.164880615 | 0.472693664 | 0.636431747 | 0.998865404 | 9.536098 | 9.614692 | 9.541015 | 9.566998 | 9.677149 | 9.499190 | 839 | 816 | 768 | 642 |
| 727856 | NR_110561,NF | DDX11L16 | DEAD/H-box helicase 11 like 16 (pseudogene) | transcribed_ps | . | 37115 | . | . | . | 0.241153 | 0.213495 | 0.239461 | 0.300964 | 0.561171 | 0.493504 | 0.537343 | 0.688117 | 1.233969 | 13.970594 | 0.81863592 | 0.370502329 | 0.711008242 | NA | 3.644780 | 3.947964 | 3.800498 | 3.788051 | 3.800108 | 3.814065 | 16 | 12 | 12 | 16 |
| 727866 | NM_00109968 | FAM156B | family with sequence similarity 156 member B | protein_coding | NP_001093154.1;NP_001308107.1;NP_001308108.1;NP_001308109.1;NI | 31962 | . | . | . | 7.258447 | 7.497205 | 9.234858 | 7.233465 | 16.890705 | 17.330125 | 20.722753 | 16.538457 | 1.063649 | 862.210259 | 0.159817285 | 0.557028349 | 0.57750806 | 0.998865404 | 9.706363 | 9.796035 | 9.721675 | 9.732071 | 9.856691 | 9.677917 | 955 | 908 | 869 | 730 |
| 727884 | gene-LOC7278 | LOC727884 | insulin like growth factor 2 mRNA binding protein 2 pseudogene | pseudogene | . | . | . | . | . | 0.015362 | 0.008477 | 0.006016 | 0.013235 | 0.035748 | 0.019595 | 0.013500 | 0.030261 | 1.233060 | 1.486734 | 2.400765875 | 0.125894475 | 0.899815456 | NA | 0.413393 | 0.715150 | 0.572744 | 0.568868 | 0.570025 | 0.574493 | 2 | 1 | 1 | 2 |
| 727897 | NM_002458 | MUC5B | mucin 5B, oligomeric mucus/gel-forming | protein_coding | NP_002449.2 | 7516 | 600770 | . | . | 0.335841 | 0.412236 | 0.747542 | 0.461942 | 0.781517 | 0.952901 | 1.677462 | 1.056177 | 1.596405 | 273.330590 | 0.255625666 | 2.639900931 | **0.008293027** | 0.289062237 | 7.717499 | 8.393120 | 7.893575 | 7.983921 | 8.310445 | 8.033835 | 219 | 249 | 363 | 239 |
| 727909 | NM_00135547 | GOLGA8Q | golgin A8 family member Q | protein_coding | NP_001342405.1 | 44408 | . | . | . | 0.256874 | 0.266334 | 0.216761 | 0.199178 | 0.597756 | 0.615642 | 0.486406 | 0.455396 | -1.266323 | 14.461598 | 0.804743735 | -0.42329673 | 0.67207878 | NA | 4.014059 | 3.674293 | 3.862271 | 3.866811 | 3.854248 | 3.845283 | 18 | 18 | 12 | 11 |
| 727910 | NM_00116440 | TLCD2 | TLC domain containing 2 | protein_coding | NP_001157879.1 | 33522 | . | . | . | 0.568683 | 0.640484 | 0.408815 | 0.560726 | 1.323350 | 1.480506 | 0.917369 | 1.282034 | -1.257572 | 100.717902 | 0.330283962 | -1.00107942 | 0.316788413 | 0.998865404 | 6.811042 | 6.478150 | 6.669068 | 6.706626 | 6.576555 | 6.657935 | 122 | 128 | 66 | 96 |
| 727914 | gene-BMS1P16 | BMS1P16 | BMS1 pseudogene 16 | pseudogene | . | 49161 | . | . | . | 0.217759 | 0.428067 | 0.500465 | 0.515926 | 0.506734 | 0.989496 | 1.123029 | 1.179603 | 1.566652 | 14.506713 | 0.820107132 | 0.789755865 | 0.429670359 | NA | 3.501050 | 4.144983 | 3.815058 | 3.847260 | 3.857299 | 3.861300 | 9 | 16 | 15 | 17 |
| 727919 | gene-NDUFAF2 | NDUFAF2P1 | NDUFAF2 pseudogene 1 | pseudogene | . | 54722 | . | . | . | 0.310502 | 0.283996 | 0.224810 | 0.414540 | 0.722551 | 0.656469 | 0.504466 | 0.947797 | -1.020375 | 6.641996 | 1.180246235 | -0.02465506 | 0.980330105 | NA | 2.747897 | 2.715150 | 2.734976 | 2.732828 | 2.724607 | 2.741931 | 8 | 7 | 4 | 8 |
| 727936 | NM_00108039 | GXYLT2 | glucoside xylosyltransferase 2 | protein_coding | NP_001073862.1;XP_011532369.1;XP_011532370.1;XP_016862612.1;XP_ | 33383 | 613322 | . | . | 6.962029 | 7.562341 | 5.919971 | 6.145521 | 16.200928 | 17.480692 | 13.284243 | 14.051004 | -1.234242 | 664.272972 | 0.153290259 | -1.98071881 | **0.047622816** | 0.659753773 | 9.519575 | 9.215720 | 9.427357 | 9.457366 | 9.291315 | 9.308550 | 827 | 809 | 510 | 565 |
| 727947 | gene-LOC7279 | LOC727947 | ubiquinol-cytochrome c reductase binding protein pseudogene | pseudogene | . | . | . | . | . | 0.284746 | 0.492138 | 0.079261 | 0.072788 | 0.662616 | 1.137599 | 0.177859 | 0.166420 | **-5.301224** | 3.493202 | 1.793052019 | -1.3420277 | 0.179587015 | NA | 2.555346 | 0.148375 | 1.826351 | 1.840398 | 1.812047 | 1.811440 | 5 | 8 | 1 | 1 |
| 727956 | NR_003265 | SDHAP2 | succinate dehydrogenase complex flavoprotein subunit A pseudog | transcribed_ps | . | 27408 | . | . | . | 6.161878 | 5.654534 | 6.507177 | 5.836409 | 14.338943 | 13.070708 | 14.601915 | 13.344255 | 1.030160 | 435.429671 | 0.18025204 | 0.237825248 | 0.812016635 | 0.998865404 | 8.744294 | 8.787967 | 8.784401 | 8.727034 | 8.809518 | 8.737974 | 517 | 441 | 407 | 390 |
| 727957 | NM_00109928 | MROH1 | maestro heat like repeat family member 1 | protein_coding | NP_001092750.2;NP_001092751.2;NP_001275743.1;NP_115826.3;XP_01 | 26958 | . | . | . | 7.236559 | 6.996737 | 8.036056 | 7.645392 | 16.839770 | 16.173273 | 18.032676 | 17.480277 | 1.096253 | 1087.891499 | 0.137820327 | 0.961984151 | 0.336057565 | 0.998865404 | 10.019331 | 10.152247 | 10.052475 | 10.050128 | 10.161395 | 10.076057 | 1197 | 1118 | 1046 | 1006 |
| 727991 | gene-SLC35E1F | SLC35E1P1 | solute carrier family 35 member E1 pseudogene 1 | pseudogene | . | 42046 | . | . | . | 0.464544 | 0.593221 | 0.572840 | 0.649081 | 1.081012 | 1.371258 | 1.285437 | 1.484047 | 1.143982 | 63.155172 | 0.401037388 | 0.483905849 | 0.628452689 | 0.998865404 | 5.881486 | 6.073770 | 5.936651 | 5.986623 | 5.977548 | 6.005585 | 60 | 71 | 55 | 67 |
| 728005 | NR_165819 | AGAP13P | ArfGAP with GTPase domain, ankyrin repeat and PH domain 13, p: | transcribed_ps | . | 23499 | . | . | . | 0.524365 | 0.457636 | 0.700431 | 0.465190 | 1.220220 | 1.057847 | 1.571746 | 1.063601 | 1.178559 | 18.535353 | 0.71960894 | 0.329378521 | 0.741869601 | NA | 4.087121 | 4.327309 | 4.205128 | 4.194246 | 4.229332 | 4.195700 | 21 | 17 | 21 | 15 |
| 728024 | NR_003671 | LOC728024 | chromosome X open reading frame 56 pseudogene | transcribed_ps | . | . | . | . | . | 0.357936 | 0.370688 | 0.610471 | 0.388420 | 0.832932 | 0.856863 | 1.369880 | 0.888076 | 1.338063 | 20.464922 | 0.688824908 | 0.609946571 | 0.541897201 | NA | 4.128334 | 4.550984 | 4.331526 | 4.332767 | 4.378530 | 4.335389 | 20 | 19 | 25 | 17 |
| 728034 | NR_146105,NF | BMS1P14 | BMS1 pseudogene 14 | transcribed_ps | . | 49159 | . | . | . | 1.103620 | 1.093939 | 0.991112 | 0.753732 | 2.568171 | 2.528691 | 2.224028 | 1.723318 | -1.267400 | 145.663116 | 0.28316699 | -1.20731613 | 0.227310421 | 0.982008277 | 7.346539 | 7.006448 | 7.227253 | 7.228667 | 7.190132 | 7.082414 | 187 | 176 | 129 | 102 |
| 728039 | NR_027292 | SSR4P1 | signal sequence receptor subunit 4 pseudogene 1 | transcribed_ps | . | 23131 | . | . | . | 0.507891 | 0.502003 | 0.367483 | 0.587310 | 1.181883 | 1.160401 | 0.824621 | 1.342815 | -1.065109 | 35.166760 | 0.53487508 | -0.17013585 | 0.864903303 | 0.998865404 | 5.182248 | 5.088511 | 5.141721 | 5.140326 | 5.102641 | 5.161103 | 42 | 39 | 23 | 39 |
| 728047 | NM_00127730 | GOLGA8O | golgin A8 family member O | protein_coding | NP_001264237.1;XP_011520290.1;XP_011520291.1;XP_024305810.1 | 44406 | . | . | . | 0.165864 | 0.171441 | 0.200232 | 0.125030 | 0.385972 | 0.396294 | 0.449313 | 0.285866 | 1.006934 | 26.117017 | 0.613897577 | 0.016239152 | 0.987043601 | NA | 4.700498 | 4.713310 | 4.703328 | 4.702890 | 4.727497 | 4.679576 | 30 | 28 | 28 | 19 |
| 728066 | NR_034169 | FAM133DP | family with sequence similarity 133 member D, pseudogene | transcribed_ps | . | 44192 | . | . | . | 1.647325 | 1.173308 | 1.539167 | 1.210806 | 3.833393 | 2.712154 | 3.453847 | 2.768365 | -1.040371 | 88.407247 | 0.357497398 | -0.1597156 | 0.873105109 | 0.998865404 | 6.493037 | 6.438635 | 6.515650 | 6.418885 | 6.491265 | 6.422212 | 121 | 81 | 85 | 71 |
| 728116 | NM_00114572 | ZBTB8B | zinc finger and BTB domain containing 8B | protein_coding | NP_001139192.1 | 37057 | . | . | . | 0.252766 | 0.217885 | 0.181337 | 0.259142 | 0.588196 | 0.503651 | 0.406915 | 0.592497 | -1.088575 | 91.341335 | 0.348190744 | -0.35165048 | 0.725100405 | 0.998865404 | 6.573775 | 6.449961 | 6.547397 | 6.501555 | 6.452587 | 6.545939 | 118 | 95 | 63 | 96 |
| 728118 | NM_00109933 | NUTM2A | NUT family member 2A | protein_coding | NP_001092808.1;XP_011538424.1;XP_016872096.1 | 23438 | . | . | . | 0.036534 | 0.107668 | 0.031705 | 0.026653 | 0.085016 | 0.248879 | 0.071144 | 0.060939 | **-2.379977** | 10.341072 | 1.015585325 | -1.23175044 | 0.218042323 | NA | 3.864489 | 2.612728 | 3.357538 | 3.417850 | 3.354388 | 3.347084 | 9 | 23 | 6 | 5 |
| 728121 | gene-CSPG4P1 | CSPG4P12 | chondroitin sulfate proteoglycan 4 pseudogene 12 | pseudogene | . | 48362 | . | . | . | 0.055776 | 0.022213 | 0.030260 | 0.044116 | 0.129792 | 0.051346 | 0.067903 | 0.100865 | -1.127606 | 5.847825 | 1.270095999 | -0.13641764 | 0.891491131 | NA | 2.631267 | 2.459419 | 2.564915 | 2.541695 | 2.546198 | 2.554126 | 10 | 4 | 4 | 6 |
| 728130 | NM_00100961 | NUTM2D | NUT family member 2D | protein_coding | NP_001009610.1;NP_001369233.1 | 23447 | . | . | . | 0.078176 | 0.196336 | 0.046607 | 0.068651 | 0.181917 | 0.453840 | 0.104584 | 0.156962 | -1.959489 | 13.035372 | 0.907670981 | -1.06919517 | 0.284981733 | NA | 4.111656 | 3.134756 | 3.686654 | 3.754885 | 3.676316 | 3.692330 | 11 | 27 | 6 | 10 |
| 728138 | gene-LOC7281 | LOC728138 | KIAA2013 pseudogene | pseudogene | . | . | . | . | . | 2.237550 | 1.952570 | 1.717498 | 2.038500 | 5.206872 | 4.513453 | 3.854015 | 4.660788 | -1.131379 | 144.064528 | 0.278304797 | -0.6398813 | 0.522249771 | 0.998865404 | 7.257046 | 7.078581 | 7.222733 | 7.164047 | 7.117585 | 7.175518 | 189 | 153 | 108 | 137 |
| 728139 | gene-RPL7AP7( | RPL7AP70 | ribosomal protein L7a pseudogene 70 | pseudogene | . | 36752 | . | . | . | 0.186293 | 0.045232 | 0.112352 | 0.134327 | 0.433510 | 0.104555 | 0.252115 | 0.307123 | 1.089774 | 3.695183 | 1.576587396 | 0.078669071 | 0.937295847 | NA | 1.820514 | 1.947964 | 1.893116 | 1.877225 | 1.884826 | 1.888494 | 6 | 2 | 3 | 4 |
| 728162 | gene-ST13P11 | ST13P11 | ST13, Hsp70 interacting protein pseudogene 11 | pseudogene | . | 38746 | . | . | . | 0.042820 | 0.025185 | 0.044812 | 0.109726 | 0.099645 | 0.058217 | 0.100556 | 0.250875 | 1.714765 | 3.023025 | 1.735050114 | 0.44840823 | 0.653858604 | NA | 1.156660 | 1.932304 | 1.585902 | 1.582362 | 1.584678 | 1.598382 | 3 | 2 | 2 | 5 |
| 728178 | NR_168439,NF | LOC728178 | uncharacterized LOC728178 | lncRNA | . | . | . | . | . | 0.179846 | 0.003915 | 0.018524 | 0.008814 | 0.418509 | 0.009050 | 0.041567 | 0.020153 | **-4.352887** | 2.956951 | 1.980247933 | -1.07156907 | 0.283913634 | NA | 2.264711 | 0.148375 | 1.609711 | 1.573635 | 1.575172 | 1.574620 | 10 | 1 | 1 | 1 |
| 728190 | NR_024397 | NUTM2A-AS1 | NUTM2A antisense RNA 1 | lncRNA | . | 45161 | . | . | . | 6.983221 | 7.213732 | 6.377502 | 7.659425 | 16.250240 | 16.674864 | 14.310930 | 17.512363 | -1.025638 | 181.070777 | 0.250509351 | -0.14578694 | 0.884089584 | 0.998865404 | 7.519060 | 7.481516 | 7.501075 | 7.510230 | 7.457860 | 7.530281 | 209 | 200 | 142 | 182 |
| 728202 | gene-RPL36AP | RPL36AP7 | ribosomal protein L36a pseudogene 7 | pseudogene | . | 31328 | . | . | . | 0.309718 | 0.312725 | 0.120658 | 0.261442 | 0.720727 | 0.722877 | 0.270753 | 0.597756 | -1.362299 | 3.885638 | 1.529316846 | -0.29166198 | 0.770545078 | NA | 2.166062 | 1.715150 | 1.967471 | 1.968798 | 1.958296 | 1.967024 | 5 | 5 | 2 | 4 |
| 728229 | NM_00124231 | TMEM191B | transmembrane protein 191B | protein_coding | NP_001229242.1;XP_011544462.1;XP_024308033.1 | 33600 | . | . | . | 0.744654 | 0.777278 | 0.737079 | 0.809336 | 1.732839 | 1.796712 | 1.653983 | 1.850450 | 1.102091 | 61.776037 | 0.398583255 | 0.351853046 | 0.724948474 | 0.998865404 | 5.877199 | 6.017349 | 5.930902 | 5.941665 | 5.959987 | 5.954876 | 66 | 65 | 57 | 60 |
| 728233 | NR_003563 | PI4KAP1 | phosphatidylinositol 4-kinase alpha pseudogene 1 | transcribed_ps | . | 33576 | . | . | . | 11.874170 | 12.187314 | 13.946537 | 10.789179 | 27.631680 | 28.171520 | 31.295626 | 24.668175 | 1.013742 | 897.406352 | 0.160791744 | 0.122463184 | 0.902532201 | 0.998865404 | 9.799401 | 9.819762 | 9.795743 | 9.808501 | 9.902973 | 9.712409 | 1017 | 970 | 890 | 735 |
| 728239 | NM_00109880 | MAGED4 | MAGE family member D4 | protein_coding | NP_001092270.1;NP_001258990.1;NP_001258991.1;NP_001258992.1 | 23793 | 300702 | . | . | 6.007392 | 6.041910 | 6.612000 | 5.972978 | 13.979448 | 13.966145 | 14.837137 | 13.656506 | 1.029954 | 487.860640 | 0.170786385 | 0.249315352 | 0.803116857 | 0.998865404 | 8.908649 | 8.951681 | 8.920945 | 8.919589 | 8.971138 | 8.905693 | 554 | 518 | 453 | 440 |
| 728262 | NR_146164 | FAM157A | family with sequence similarity 157 member A | lncRNA | . | 34079 | . | . | . | 0.258320 | 0.296943 | 0.338719 | 0.272983 | 0.601122 | 0.686396 | 0.760076 | 0.624142 | 1.098034 | 63.164792 | 0.400638463 | 0.336769048 | 0.736291007 | 0.998865404 | 5.911621 | 6.047288 | 5.953109 | 5.980370 | 6.012458 | 5.961900 | 65 | 69 | 64 | 55 |
| 728294 | NM_00128724 | D2HGDH | D-2-hydroxyglutarate dehydrogenase | protein_coding | NP_001274178.1;NP_001339753.1;NP_689996.4;XP_011510036.1;XP_01 | 28358 | 609186 | . | . | 13.163848 | 12.335734 | 13.747313 | 12.549929 | 30.632817 | 28.514598 | 30.848574 | 28.693921 | 1.014408 | 905.649572 | 0.149236016 | 0.138287198 | 0.890013443 | 0.998865404 | 9.812126 | 9.833413 | 9.847317 | 9.784675 | 9.879624 | 9.771440 | 1080 | 928 | 853 | 791 |
| 728300 | gene-KRT18P5 | KRT18P5 | keratin 18 pseudogene 5 | pseudogene | . | 19259 | . | . | . | 0.650940 | 1.445283 | 1.983675 | 0.584567 | 1.514765 | 3.340836 | 4.451309 | 1.336544 | 1.207554 | 51.617618 | 0.571176267 | 0.476364227 | 0.633814915 | 0.998865404 | 5.546469 | 5.820154 | 5.565416 | 5.699914 | 5.775939 | 5.553425 | 34 | 69 | 76 | 24 |
| 728323 | NR_130699,NF | LINC01881 | long intergenic non-protein coding RNA 1881 | lncRNA | . | 52700 | . | . | . | 1.850995 | 1.852316 | 1.504590 | 1.564804 | 4.307341 | 4.281712 | 3.376257 | 3.577739 | -1.193348 | 77.027765 | 0.360300176 | -0.70778333 | 0.47907982 | 0.998865404 | 6.389273 | 6.134069 | 6.292529 | 6.289531 | 6.242147 | 6.249972 | 97 | 90 | 60 | 67 |
| 728324 | gene-CHCHD2F | CHCHD2P2 | coiled-coil-helix-coiled-coil-helix domain containing 2 pseudogene | pseudogene | . | 39586 | . | . | . | 0.476484 | 0.509789 | 0.287023 | 0.349514 | 1.108797 | 1.178399 | 0.644071 | 0.799122 | -1.566642 | 9.909763 | 0.972698183 | -0.66585413 | 0.505504341 | NA | 3.597285 | 2.947964 | 3.326506 | 3.325859 | 3.303294 | 3.310212 | 14 | 13 | 6 | 8 |
| 728325 | gene-HMGN1P | HMGN1P21 | high mobility group nucleosome binding domain 1 pseudogene 21 | pseudogene | . | 39365 | . | . | . | 0.025110 | 0.043014 | 0.033005 | 0.004621 | 0.058431 | 0.099429 | 0.074063 | 0.010566 | -1.070164 | 1.739247 | 2.228774873 | -0.04389516 | 0.964987971 | NA | 0.844134 | 0.751299 | 0.796508 | 0.797062 | 0.799027 | 0.793436 | 2 | 2 | 2 | 1 |
| 728340 | NM_00109872 | GTF2H2C | GTF2H2 family member C | protein_coding | NP_001092198.1;NP_001362927.1;NP_001362928.1;NP_001362929.1;NI | 31394 | . | . | . | 5.688470 | 7.102917 | 6.273270 | 6.867984 | 13.237304 | 16.418709 | 14.077038 | 15.702834 | -1.049413 | 759.766256 | 0.146959188 | -0.47348283 | 0.635868751 | 0.998865404 | 9.603687 | 9.534303 | 9.593241 | 9.578813 | 9.565430 | 9.539286 | 905 | 831 | 657 | 682 |
| 728350 | gene-EIF2S2P4 | EIF2S2P4 | eukaryotic translation initiation factor 2 subunit 2 beta pseudogen | pseudogene | . | 37626 | . | . | . | 1.569058 | 1.414970 | 1.006350 | 1.285922 | 3.651263 | 3.270767 | 2.258221 | 2.940107 | -1.325666 | 57.683244 | 0.419149868 | -0.97033741 | 0.331878338 | 0.998865404 | 6.039657 | 5.631770 | 5.893616 | 5.867116 | 5.801631 | 5.843407 | 80 | 67 | 38 | 52 |
| 728377 | NR_033942 | ARHGEF34P | Rho guanine nucleotide exchange factor 34, pseudogene | transcribed_ps | . | 38086 | . | . | . | 0.782262 | 1.008020 | 0.908780 | 1.113775 | 1.820356 | 2.330083 | 2.039275 | 2.546515 | 1.119501 | 125.350402 | 0.300894411 | 0.541237608 | 0.588343819 | 0.998865404 | 6.887145 | 7.048019 | 6.903491 | 6.986156 | 6.950080 | 7.019259 | 120 | 143 | 104 | 136 |
| 728378 | NM_00109977 | POTEF | POTE ankyrin domain family member F | protein_coding | NP_001093241.1;XP_016860322.1;XP_016860323.1;XP_016860324.1 | 33905 | . | . | . | 0.090602 | 0.118938 | 0.117231 | 0.151934 | 0.210834 | 0.274930 | 0.263062 | 0.347379 | 1.262595 | 16.785255 | 0.751984639 | 0.44733854 | 0.654630644 | NA | 3.892781 | 4.226238 | 4.046950 | 4.063412 | 4.060997 | 4.082984 | 15 | 18 | 14 | 20 |
| 728380 | gene-RPL7P26 | RPL7P26 | ribosomal protein L7 pseudogene 26 | pseudogene | . | 36630 | . | . | . | 0.026282 | 0.010258 | 0.119620 | 0.157736 | 0.061160 | 0.023712 | 0.268425 | 0.360646 | **3.704896** | 2.111278 | 2.128970011 | 0.887486872 | 0.374816831 | NA | -0.155866 | 1.733338 | 1.045261 | 1.045639 | 1.057359 | 1.056349 | 1 | 1 | 3 | 3 |
| 728388 | gene-DNM1P3( | DNM1P30 | dynamin 1 pseudogene 30 | pseudogene | . | 35177 | . | . | . | 0.139463 | 0.123611 | 0.475285 | 0.325266 | 0.324535 | 0.285732 | 1.066527 | 0.743683 | **2.597582** | 8.865178 | 1.044675171 | 1.318275211 | 0.187411536 | NA | 2.299300 | 3.678992 | 3.105661 | 3.103211 | 3.149060 | 3.130893 | 6 | 5 | 13 | 10 |
| 728392 | NM_00116237 | LOC728392 | uncharacterized LOC728392 | protein_coding | NP_001155843.1 | . | . | . | . | 1.571718 | 1.481121 | 2.733935 | 2.167974 | 3.657453 | 3.423678 | 6.134871 | 4.956815 | 1.578928 | 93.026432 | 0.338380186 | 1.947352419 | 0.051492498 | 0.672667836 | 6.171846 | 6.832345 | 6.463082 | 6.446016 | 6.624885 | 6.546221 | 86 | 75 | 111 | 94 |
| 728402 | NR_027338 | TPI1P3 | triosephosphate isomerase 1 pseudogene 3 | transcribed_ps | . | 38070 | . | . | . | 0.092476 | 0.228343 | 0.105673 | 0.161396 | 0.215195 | 0.527825 | 0.237127 | 0.369012 | -1.232535 | 6.152390 | 1.238135294 | -0.24361488 | 0.807529107 | NA | 2.766559 | 2.459419 | 2.613407 | 2.636282 | 2.615158 | 2.623067 | 5 | 10 | 4 | 6 |
| 728407 | NR_029388 | PARGP1 | poly(ADP-ribose) glycohydrolase pseudogene 1 | transcribed_ps | . | 45008 | . | . | . | 1.317336 | 1.152681 | 0.990875 | 0.935729 | 3.065496 | 2.664475 | 2.223495 | 2.139434 | -1.299062 | 50.282769 | 0.442740352 | -0.85257768 | 0.393893544 | 0.998865404 | 5.828014 | 5.451470 | 5.692524 | 5.662941 | 5.633877 | 5.625400 | 70 | 57 | 39 | 40 |
| 728411 | NR_027026,NF | GUSBP1 | GUSB pseudogene 1 | transcribed_ps | . | 13670 | . | . | . | 2.852391 | 3.595338 | 3.960091 | 3.935034 | 6.637630 | 8.310784 | 8.886331 | 8.996990 | 1.166566 | 150.997953 | 0.272347516 | 0.816119925 | 0.414431549 | 0.998865404 | 7.123580 | 7.344724 | 7.163641 | 7.242080 | 7.253939 | 7.273391 | 144 | 166 | 138 | 156 |
| 728424 | gene-LOC7284 | LOC728424 | hyperpolarization activated cyclic nucleotide gated potassium and | pseudogene | . | . | . | . | . | 0.584521 | 0.427736 | 0.746950 | 0.380189 | 1.360203 | 0.988732 | 1.676134 | 0.869257 | 1.098786 | 53.441498 | 0.473964587 | 0.286752577 | 0.774301777 | 0.998865404 | 5.667953 | 5.808407 | 5.749513 | 5.687731 | 5.801239 | 5.668931 | 68 | 46 | 65 | 35 |
| 728431 | NR_038842 | LINC01137 | long intergenic non-protein coding RNA 1137 | lncRNA | . | 49453 | . | . | . | 1.094210 | 0.807779 | 0.507321 | 0.862305 | 2.546272 | 1.867218 | 1.138414 | 1.971557 | -1.392760 | 36.760936 | 0.536919351 | -0.89016423 | 0.373377711 | 0.998865404 | 5.420284 | 4.940155 | 5.251121 | 5.198710 | 5.147854 | 5.207460 | 57 | 39 | 20 | 36 |
| 728433 | gene-FGF7P7 | FGF7P7 | fibroblast growth factor 7 pseudogene 7 | pseudogene | . | 33589 | . | . | . | 0.021971 | 0.015006 | 0.005565 | 0.010275 | 0.051127 | 0.034687 | 0.012487 | 0.023492 | **-2.014451** | 1.668864 | 2.31231603 | -0.43695888 | 0.662141169 | NA | 1.156660 | 0.148375 | 0.757225 | 0.753619 | 0.750445 | 0.750013 | 3 | 2 | 1 | 1 |
| 728448 | NR_003929,NF | PPIEL | peptidylprolyl isomerase E like pseudogene | transcribed_ps | . | 33195 | . | . | . | 2.057916 | 2.740461 | 2.406919 | 2.698115 | 4.788855 | 6.334697 | 5.401056 | 6.168918 | 1.044501 | 207.853860 | 0.241950973 | 0.259614952 | 0.795160802 | 0.998865404 | 7.668247 | 7.729945 | 7.631961 | 7.739294 | 7.695391 | 7.718437 | 207 | 245 | 180 | 204 |
| 728468 | gene-BCRP6 | BCRP6 | BCR pseudogene 6 | pseudogene | . | 39074 | . | . | . | 0.038705 | 0.064698 | 0.040446 | 0.038377 | 0.090067 | 0.149551 | 0.090759 | 0.087745 | -1.220295 | 4.923997 | 1.35824335 | -0.2114715 | 0.83251937 | NA | 2.436885 | 2.148375 | 2.298926 | 2.308949 | 2.300263 | 2.298731 | 5 | 7 | 4 | 4 |
| 728481 | gene-RPL23AP | RPL23AP21 | ribosomal protein L23a pseudogene 21 | pseudogene | . | 35827 | . | . | . | 1.232458 | 1.281204 | 1.694255 | 1.967924 | 2.867981 | 2.961561 | 3.801858 | 4.499424 | 1.441733 | 25.726778 | 0.607308134 | 0.869088113 | 0.384798938 | NA | 4.397726 | 4.924798 | 4.655880 | 4.658207 | 4.688927 | 4.702165 | 24 | 23 | 25 | 30 |
| 728485 | NR_164115,NF | LOC728485 | uncharacterized LOC728485 | lncRNA | . | . | . | . | . | 0.176678 | 0.203137 | 0.218919 | 0.280635 | 0.411137 | 0.469559 | 0.491248 | 0.641638 | 1.182508 | 13.228474 | 0.839303108 | 0.288155816 | 0.773227468 | NA | 3.600756 | 3.840449 | 3.713502 | 3.721212 | 3.719891 | 3.734242 | 13 | 14 | 11 | 15 |
| 728489 | NM_00108084 | DNLZ | DNL-type zinc finger | protein_coding | NP_001074318.1 | 33879 | . | . | . | 1.429227 | 1.296364 | 1.329233 | 1.480673 | 3.325870 | 2.996604 | 2.982761 | 3.385382 | 1.019105 | 134.634680 | 0.28205034 | 0.096801562 | 0.922883984 | 0.998865404 | 7.059209 | 7.086475 | 7.089188 | 7.048599 | 7.059998 | 7.091793 | 162 | 136 | 113 | 133 |
| 728492 | NM_00117808 | SERF1B | small EDRK-rich factor 1B | protein_coding | NP_001171558.1;NP_075267.1 | 10756 | . | . | . | 11.630249 | 11.298949 | 12.898491 | 13.159416 | 27.064068 | 26.118027 | 28.943840 | 30.087442 | 1.180128 | 417.277562 | 0.177481066 | 1.346305348 | 0.178204062 | 0.951511445 | 8.580360 | 8.819471 | 8.655233 | 8.646893 | 8.758490 | 8.742707 | 444 | 410 | 398 | 418 |
| 728498 | NM_00128249 | GOLGA8H | golgin A8 family member H | protein_coding | NP_001269419.1 | 37443 | . | . | . | 0.233450 | 0.223178 | 0.222311 | 0.177561 | 0.543249 | 0.515886 | 0.498859 | 0.405972 | -1.162332 | 35.437218 | 0.52400804 | -0.41415743 | 0.678758819 | 0.998865404 | 5.250766 | 5.035606 | 5.162560 | 5.154396 | 5.153055 | 5.121791 | 45 | 40 | 32 | 27 |
| 728501 | gene-RPL21P7! | RPL21P75 | ribosomal protein L21 pseudogene 75 | pseudogene | . | 36460 | . | . | . | 5.272918 | 4.002479 | 5.250520 | 5.004138 | 12.270296 | 9.251909 | 11.782016 | 11.441365 | 1.093194 | 86.925204 | 0.349421343 | 0.367891423 | 0.712954196 | 0.998865404 | 6.375209 | 6.505266 | 6.464132 | 6.387251 | 6.458244 | 6.444131 | 109 | 77 | 81 | 83 |
| 728517 | gene-RPL23AP | RPL23AP24 | ribosomal protein L23a pseudogene 24 | pseudogene | . | 36176 | . | . | . | 0.209140 | 0.122305 | 0.144096 | 0.345653 | 0.486678 | 0.282714 | 0.323348 | 0.790294 | 1.434070 | 3.240136 | 1.679368828 | 0.309709143 | 0.75678215 | NA | 1.413393 | 1.932304 | 1.693199 | 1.685516 | 1.687889 | 1.701542 | 4 | 2 | 2 | 5 |
| 728519 | gene-NAIPP2 | NAIPP2 | NAIP pseudogene 2 | pseudogene | . | 16042 | . | . | . | 0.509589 | 0.487659 | 0.821292 | 0.373638 | 1.185835 | 1.127246 | 1.842954 | 0.854278 | 1.177713 | 92.297022 | 0.394321959 | 0.598466457 | 0.549528734 | 0.998865404 | 6.403647 | 6.642872 | 6.492936 | 6.478757 | 6.642283 | 6.410890 | 100 | 89 | 120 | 58 |
| 728532 | gene-DYNC1I2I | DYNC1I2P1 | dynein cytoplasmic 1 intermediate chain 2 pseudogene 1 | pseudogene | . | 45012 | . | . | . | 0.291147 | 0.240022 | 0.292454 | 0.353630 | 0.677511 | 0.554823 | 0.656259 | 0.808532 | 1.176532 | 20.920910 | 0.672366373 | 0.348827732 | 0.727218642 | NA | 4.264940 | 4.499300 | 4.383873 | 4.369853 | 4.383300 | 4.399114 | 24 | 19 | 18 | 23 |
| 728535 | gene-NAIPP1 | NAIPP1 | NAIP pseudogene 1 | pseudogene | . | 54492 | . | . | . | 0.060415 | 0.112873 | 0.102475 | 0.071615 | 0.140588 | 0.260910 | 0.229950 | 0.163740 | -1.086933 | 3.472244 | 1.606184845 | -0.07487456 | 0.940314518 | NA | 1.855800 | 1.733338 | 1.790869 | 1.800475 | 1.794924 | 1.793788 | 3 | 5 | 3 | 3 |
| 728537 | NM_00135136 | C2orf92 | chromosome 2 open reading frame 92 | protein_coding | NP_001338297.1;XP_024308873.1;XP_024308874.1;XP_024308876.1;XP_ | 49272 | . | . | . | 1.450255 | 1.922927 | 3.859157 | 1.982502 | 3.374801 | 4.444928 | 8.659836 | 4.532754 | 1.615131 | 104.212391 | 0.339439428 | 2.037627947 | **0.041587155** | 0.63293663 | 6.316233 | 7.008316 | 6.568266 | 6.637435 | 6.817072 | 6.683252 | 80 | 97 | 133 | 98 |
| 728553 | gene-RPS8P8 | RPS8P8 | ribosomal protein S8 pseudogene 8 | pseudogene | . | 36213 | . | . | . | 0.128172 | 0.031242 | 0.051084 | 0.283108 | 0.298261 | 0.072218 | 0.114631 | 0.647293 | 1.973582 | 3.275116 | 1.719558537 | 0.570388466 | 0.568414251 | NA | 1.137672 | 2.121007 | 1.702196 | 1.690080 | 1.696883 | 1.715359 | 4 | 1 | 2 | 6 |
| 728554 | NR_003615 | LOC728554 | THO complex 3 pseudogene | transcribed_ps | . | . | . | . | . | 16.777510 | 16.506567 | 14.066642 | 16.856615 | 39.041954 | 38.155670 | 31.565140 | 38.540646 | -1.091592 | 913.926546 | 0.148679876 | -0.85037279 | 0.39511786 | 0.998865404 | 9.898002 | 9.771076 | 9.875136 | 9.857177 | 9.740527 | 9.861467 | 1112 | 1017 | 695 | 889 |
| 728558 | NR_038444,NF | ENTPD1-AS1 | ENTPD1 antisense RNA 1 | lncRNA | . | 45203 | . | . | . | 0.389493 | 0.357608 | 0.609690 | 0.369946 | 0.906365 | 0.826627 | 1.368127 | 0.845836 | 1.346455 | 12.616723 | 0.863484462 | 0.497016238 | 0.6191776 | NA | 3.425187 | 3.857133 | 3.642618 | 3.637734 | 3.668184 | 3.644840 | 13 | 11 | 15 | 11 |
| 728562 | gene-RPSAP45 | RPSAP45 | ribosomal protein SA pseudogene 45 | pseudogene | . | 35857 | . | . | . | 0.088585 | 0.059343 | 0.017805 | 0.028943 | 0.206141 | 0.137175 | 0.039955 | 0.066174 | **-2.824587** | 2.117661 | 2.1147751 | -0.70836839 | 0.478716515 | NA | 1.644780 | 0.148375 | 1.107057 | 1.103720 | 1.096335 | 1.095860 | 4 | 3 | 1 | 1 |
| 728564 | gene-GMPSP1 | GMPSP1 | guanine monophosphate synthase pseudogene 1 | pseudogene | . | 39428 | . | . | . | 0.323650 | 0.060710 | 0.300271 | 0.271083 | 0.753148 | 0.140333 | 0.673800 | 0.619798 | 1.253769 | 4.971058 | 1.388964774 | 0.234903065 | 0.81428397 | NA | 2.137672 | 2.470304 | 2.318474 | 2.294460 | 2.312557 | 2.310763 | 8 | 2 | 5 | 5 |
| 728568 | NM_00113557 | C12orf73 | chromosome 12 open reading frame 73 | protein_coding | NP_001129042.1;XP_011537020.1;XP_016875405.1;XP_016875406.1 | 34450 | 618812 | . | . | 4.836553 | 3.916085 | 4.455954 | 3.377811 | 11.254857 | 9.052205 | 9.999031 | 7.722962 | -1.091053 | 257.123144 | 0.238645489 | -0.5268109 | 0.598324915 | 0.998865404 | 8.066849 | 7.943131 | 8.073944 | 7.971357 | 8.057072 | 7.896664 | 339 | 261 | 249 | 193 |
| 728576 | gene-RPL15P7 | RPL15P7 | ribosomal protein L15 pseudogene 7 | pseudogene | . | 36808 | . | . | . | 1.084254 | 0.412187 | 0.212645 | 0.376313 | 2.523103 | 0.952789 | 0.477168 | 0.860394 | **-2.464396** | 16.152102 | 0.837478739 | -1.55375167 | 0.120243663 | NA | 4.522785 | 3.219802 | 4.086980 | 4.000428 | 3.975260 | 3.995564 | 38 | 14 | 6 | 11 |
| 728577 | NM_00120138 | CNTNAP3B | contactin associated protein family member 3B | protein_coding | NP_001188309.2 | 32035 | . | . | . | 1.341560 | 1.114638 | 1.190260 | 1.163560 | 3.121866 | 2.576535 | 2.670909 | 2.660342 | -1.059106 | 287.252117 | 0.211876714 | -0.39101716 | 0.695784554 | 0.998865404 | 8.206603 | 8.124579 | 8.231395 | 8.126704 | 8.159329 | 8.140907 | 373 | 288 | 247 | 257 |
| 728589 | gene-NAP1L4P | NAP1L4P1 | nucleosome assembly protein 1 like 4 pseudogene 1 | pseudogene | . | 39740 | . | . | . | 0.994671 | 1.270988 | 1.166776 | 0.762182 | 2.314641 | 2.937946 | 2.618211 | 1.742639 | -1.182728 | 77.283389 | 0.384357899 | -0.62992863 | 0.528741277 | 0.998865404 | 6.387318 | 6.146846 | 6.255594 | 6.320957 | 6.295956 | 6.193276 | 85 | 101 | 75 | 52 |
| 728591 | NM_00114498 | CCDC169 | coiled-coil domain containing 169 | protein_coding | NP_001138453.1;NP_001138454.1;NP_001138455.1;NP_001138456.1;NI | 34361 | . | . | . | 0.081434 | 0.032754 | 0.029450 | 0.023150 | 0.189500 | 0.075711 | 0.066086 | 0.052929 | **-2.079474** | 8.517704 | 1.092836941 | -0.96649232 | 0.333797887 | NA | 3.522785 | 2.470304 | 3.134946 | 3.089348 | 3.086677 | 3.084558 | 19 | 7 | 5 | 5 |
| 728599 | gene-CIAPIN1P | CIAPIN1P | cytokine induced apoptosis inhibitor 1 pseudogene | pseudogene | . | 31087 | . | . | . | 0.021872 | 0.021494 | 0.031077 | 0.102994 | 0.050898 | 0.049685 | 0.069737 | 0.235483 | **2.431989** | 3.072581 | 1.752427149 | 0.731634934 | 0.46439142 | NA | 0.844134 | 2.121007 | 1.597058 | 1.597717 | 1.600049 | 1.618600 | 2 | 2 | 2 | 6 |
| 728609 | NR_003263 | SDHAP3 | succinate dehydrogenase complex flavoprotein subunit A pseudog | transcribed_ps | . | 18781 | . | . | . | 0.431292 | 0.400763 | 0.243911 | 0.405194 | 1.003635 | 0.926381 | 0.547330 | 0.926428 | -1.295513 | 13.865362 | 0.830469347 | -0.44977405 | 0.652873374 | NA | 3.969395 | 3.592943 | 3.810125 | 3.802588 | 3.778560 | 3.802897 | 19 | 16 | 8 | 14 |
| 728621 | NM_00108085 | CCDC30 | coiled-coil domain containing 30 | protein_coding | NP_001074319.1;NP_001342153.1;NP_001342155.1;NP_001342156.1;XI | 26103 | . | . | . | 0.360046 | 0.307829 | 0.390332 | 0.209647 | 0.837843 | 0.711560 | 0.875893 | 0.479331 | -1.068159 | 51.280766 | 0.449358709 | -0.21169295 | 0.832346585 | 0.998865404 | 5.725983 | 5.633218 | 5.680927 | 5.686991 | 5.707706 | 5.634019 | 60 | 58 | 52 | 37 |
| 728632 | gene-HMGN2P | HMGN2P3 | high mobility group nucleosomal binding domain 2 pseudogene 3 | pseudogene | . | 33566 | . | . | . | 10.058338 | 10.808909 | 9.014646 | 9.198656 | 23.406164 | 24.985279 | 20.228601 | 21.031633 | -1.160219 | 370.553010 | 0.186237027 | -1.15120826 | 0.249646569 | 0.993339639 | 8.636871 | 8.422224 | 8.556145 | 8.593653 | 8.486897 | 8.491389 | 444 | 443 | 297 | 323 |
| 728635 | NM_00127786 | DHRS4L1 | dehydrogenase/reductase 4 like 1 | protein_coding | NP_001264793.1;XP_006720309.1 | 19732 | 615195 | . | . | 0.641727 | 0.549738 | 0.391531 | 0.518322 | 1.493325 | 1.270743 | 0.878583 | 1.185082 | -1.343910 | 22.617116 | 0.651053326 | -0.65499427 | 0.512471445 | NA | 4.697260 | 4.269892 | 4.525574 | 4.509790 | 4.481692 | 4.498249 | 32 | 26 | 15 | 20 |
| 728641 | gene-FABP5P7 | FABP5P7 | fatty acid binding protein 5 pseudogene 7 | pseudogene | . | 31070 | . | . | . | 9.199907 | 7.645446 | 7.577513 | 5.903001 | 21.408562 | 17.672791 | 17.003719 | 13.496511 | -1.263513 | 153.424120 | 0.279118438 | -1.20894995 | 0.226682073 | 0.982008277 | 7.419328 | 7.083993 | 7.343684 | 7.261381 | 7.256653 | 7.164557 | 216 | 167 | 133 | 111 |
| 728642 | NM_00131389 | CDK11A | cyclin dependent kinase 11A | protein_coding | NP_001300825.1;NP_001300911.1;NP_076916.2;NP_277071.2 | 1730 | 116951 | . | . | 20.036267 | 22.153799 | 26.420833 | 23.242427 | 46.625218 | 51.209499 | 59.287585 | 53.141045 | 1.160999 | 2112.423078 | 0.128448926 | 1.676672066 | 0.093606626 | 0.810572308 | 10.932927 | 11.148401 | 10.943278 | 11.016877 | 11.155498 | 11.041280 | 2146 | 2208 | 2113 | 1978 |
| 728658 | NR_026712 | RPL13AP5 | ribosomal protein L13a pseudogene 5 | transcribed_ps | . | 23736 | . | . | . | 4.301573 | 3.499255 | 3.236129 | 5.238713 | 10.009937 | 8.088685 | 7.261780 | 11.977694 | 1.073033 | 83.775211 | 0.373419702 | 0.272333135 | 0.785365881 | 0.998865404 | 6.337494 | 6.437671 | 6.402477 | 6.345303 | 6.328476 | 6.454513 | 103 | 78 | 58 | 100 |
| 728661 | NM_00111078 | SLC35E2B | solute carrier family 35 member E2B | protein_coding | NP_001104251.1;NP_001277193.1;XP_011540374.1;XP_024305278.1;XP | 33941 | . | . | . | 7.892534 | 8.005320 | 10.421710 | 9.149017 | 18.366248 | 18.504655 | 23.386015 | 20.918139 | 1.225124 | 1637.639804 | 0.128073651 | 2.287178474 | **0.022185408** | 0.492741101 | 10.523342 | 10.816587 | 10.606823 | 10.576099 | 10.795056 | 10.703005 | 1729 | 1556 | 1659 | 1593 |
| 728666 | gene-PRELID1F | PRELID1P1 | PRELID1 pseudogene 1 | pseudogene | . | 43886 | . | . | . | 0.316045 | 0.193022 | 0.266352 | 0.303493 | 0.735451 | 0.446180 | 0.597686 | 0.693901 | 1.099746 | 8.422475 | 1.047549307 | 0.130944555 | 0.895819168 | NA | 3.003609 | 3.141582 | 3.078697 | 3.064538 | 3.071834 | 3.078637 | 11 | 7 | 7 | 9 |

| | | | | | | | | | | | | | | | | | | | | | | | | | | | | | | | | | |
|---|---|---|---|---|---|---|---|---|---|---|---|---|---|---|---|---|---|---|---|---|---|---|---|---|---|---|---|---|---|---|---|---|---|
| 728673 | NR_126047 | LOC728673 | uncharacterized LOC728673 | lncRNA | . | . | . | . | . | 0.856684 | 1.029807 | 1.045440 | 1.131264 | 1.993538 | 2.380445 | 2.345938 | 2.586501 | 1.131880 | 71.850986 | 0.374798467 | 0.476844102 | 0.633473138 | 0.998865404 | 6.075449 | 6.252966 | 6.128065 | 6.168804 | 6.169843 | 6.187520 | 71 | 79 | 64 | 74 |
| 728688 | gene-LOC7286 | LOC728688 | ubiquitin like with PHD and ring finger domains 1 pseudogene | pseudogene | . | . | . | . | . | 3.706073 | 3.840300 | 3.225822 | 3.701686 | 8.624184 | 8.877027 | 7.238652 | 8.463465 | -1.103663 | 315.385753 | 0.200144898 | -0.71098276 | 0.477094921 | 0.998865404 | 8.370722 | 8.227681 | 8.319910 | 8.335035 | 8.239181 | 8.306577 | 376 | 362 | 244 | 299 |
| 728689 | NM_00109966 | EIF3CL | eukaryotic translation initiation factor 3 subunit C like | protein_coding | NP_001093131.1;NP_001304785.1;NP_001304786.1;XP_016879109.1;XP | 26347 | . | . | . | 65.432371 | 67.994816 | 56.603078 | 69.078756 | 152.263814 | 157.173063 | 127.015673 | 157.940382 | -1.079266 | 5888.312186 | 0.120854663 | -0.91060729 | 0.362502328 | 0.998865404 | 12.577690 | 12.467483 | 12.543327 | 12.564325 | 12.407031 | 12.566031 | 6952 | 6678 | 4457 | 5814 |
| 728698 | gene-EIF4A1P4 | EIF4A1P4 | eukaryotic translation initiation factor 4A1 pseudogene 4 | pseudogene | . | 37923 | . | . | . | 0.040154 | 0.086437 | 0.026937 | 0.059481 | 0.093441 | 0.199802 | 0.060446 | 0.135997 | 1.014447 | 2.735953 | 1.804031249 | 0.011470344 | 0.99084819 | NA | 1.444631 | 1.459419 | 1.445375 | 1.454766 | 1.448241 | 1.452296 | 2 | 4 | 2 | 3 |
| 728715 | NR_144634 | LOC728715 | ovostatin homolog 2 | lncRNA | . | . | . | . | . | 182.545807 | 185.377838 | 244.735367 | 219.429459 | 424.791626 | 428.509216 | 549.179077 | 501.699371 | 1.244424 | 5777.099154 | 0.111324476 | 2.833860166 | **0.004598945** | 0.200249966 | 12.329735 | 12.645302 | 12.393622 | 12.399406 | 12.630322 | 12.528115 | 5907 | 5574 | 5903 | 5648 |
| 728723 | NR_024398 | ZBED3-AS1 | ZBED3 antisense RNA 1 | lncRNA | . | 44188 | . | . | . | 0.315711 | 0.368353 | 0.425826 | 0.556480 | 0.734672 | 0.851465 | 0.955541 | 1.272324 | 1.397514 | 15.606419 | 0.776217029 | 0.622071912 | 0.533894583 | NA | 3.703730 | 4.184531 | 3.941255 | 3.949294 | 3.958916 | 3.976582 | 14 | 15 | 14 | 19 |
| 728730 | NR_037875 | MAP4K3-DT | MAP4K3 divergent transcript | lncRNA | . | 54056 | . | . | . | 0.942680 | 0.725639 | 0.568554 | 0.846064 | 2.193654 | 1.677346 | 1.275818 | 1.934424 | -1.197431 | 60.998748 | 0.421913883 | -0.61610229 | 0.537827014 | 0.998865404 | 6.055419 | 5.794187 | 5.981405 | 5.917293 | 5.873284 | 5.947760 | 87 | 62 | 39 | 62 |
| 728734 | NM_00131013 | NPIPB8 | nuclear pore complex interacting protein family member B8 | protein_coding | NP_001297065.1;XP_016879111.1;XP_016879112.1;XP_016879113.1;XP | 37490 | . | . | . | 0.724051 | 1.130871 | 1.386062 | 0.921733 | 1.684896 | 2.614058 | 3.110283 | 2.107434 | 1.194794 | 58.355616 | 0.431551451 | 0.59497441 | 0.551860577 | 0.998865404 | 5.732325 | 5.989801 | 5.808071 | 5.867985 | 5.916876 | 5.832209 | 52 | 66 | 66 | 48 |
| 728739 | NR_037629 | LOC728739 | programmed cell death 2 pseudogene | transcribed_ps | . | . | . | . | . | 0.355863 | 0.274393 | 0.220771 | 0.317765 | 0.828109 | 0.634273 | 0.495403 | 0.726533 | -1.215358 | 6.094087 | 1.227907856 | -0.22915497 | 0.818748471 | NA | 2.741623 | 2.459419 | 2.620667 | 2.610140 | 2.606076 | 2.613978 | 9 | 6 | 4 | 6 |
| 728741 | NM_00128252 | NPIPB6 | nuclear pore complex interacting protein family member B6 | protein_coding | NP_001269453.1;XP_005255798.1;XP_005255799.1;XP_005255800.1;XP | 37454 | . | . | . | 0.423585 | 0.569002 | 0.657095 | 0.308709 | 0.985701 | 1.315274 | 1.474503 | 0.705829 | -1.033043 | 30.144265 | 0.591465166 | -0.07929518 | 0.936797836 | 0.998865404 | 4.935725 | 4.891560 | 4.893951 | 4.924399 | 4.941857 | 4.867362 | 31 | 37 | 34 | 19 |
| 728743 | NR_027237 | LOC728743 | zinc finger protein pseudogene | transcribed_ps | . | . | . | . | . | 0.861810 | 0.812648 | 0.871930 | 1.094002 | 2.005468 | 1.878473 | 1.956586 | 2.501305 | 1.157178 | 96.093362 | 0.330707595 | 0.636849325 | 0.524222994 | 0.998865404 | 6.477694 | 6.687420 | 6.570471 | 6.551778 | 6.570900 | 6.637080 | 106 | 93 | 80 | 107 |
| 728752 | NR_036504 | LOC728752 | uncharacterized LOC728752 | lncRNA | . | . | . | . | . | 0.522645 | 0.437818 | 0.679829 | 0.745192 | 1.216217 | 1.012035 | 1.525517 | 1.703793 | 1.466940 | 26.468897 | 0.601221231 | 0.919478239 | 0.357845483 | NA | 4.423228 | 4.976501 | 4.704710 | 4.687799 | 4.731169 | 4.744041 | 27 | 21 | 26 | 31 |
| 728758 | NR_003571 | PIN4P1 | peptidylprolyl cis/trans isomerase, NIMA-interacting 4 pseudogen | transcribed_ps | . | 44193 | . | . | . | 0.092976 | 0.122916 | 0.268061 | 0.117620 | 0.216360 | 0.284126 | 0.601521 | 0.268924 | 1.866049 | 11.628631 | 0.926221336 | 0.971675777 | 0.331211871 | NA | 3.019256 | 3.921224 | 3.495145 | 3.506183 | 3.555103 | 3.508010 | 8 | 10 | 18 | 9 |
| 728769 | NR_105014 | SCAMP1-AS1 | SCAMP1 antisense RNA 1 | lncRNA | . | 50544 | . | . | . | 3.004156 | 2.802628 | 2.835923 | 3.263838 | 6.990794 | 6.478401 | 6.363729 | 7.462378 | 1.034153 | 143.894632 | 0.274390116 | 0.176571834 | 0.85984472 | 0.998865404 | 7.144649 | 7.192689 | 7.175999 | 7.147597 | 7.149914 | 7.198773 | 169 | 147 | 119 | 146 |
| 728770 | gene-DNM1P5 | DNM1P5 | dynamin 1 pseudogene 5 | pseudogene | . | 21122 | . | . | . | 0.022326 | 0.165950 | 0.116299 | 0.101345 | 0.051953 | 0.383600 | 0.260971 | 0.231713 | 1.097765 | 4.791846 | 1.423817317 | 0.094513289 | 0.924701427 | NA | 2.193905 | 2.324312 | 2.236276 | 2.263149 | 2.255151 | 2.248555 | 2 | 8 | 5 | 4 |
| 728776 | gene-HMGN1P | HMGN1P37 | high mobility group nucleosome binding domain 1 pseudogene 37 | pseudogene | . | 39421 | . | . | . | 0.400935 | 0.195033 | 0.322184 | 0.132896 | 0.932992 | 0.450827 | 0.722972 | 0.303850 | -1.288925 | 10.257242 | 0.998185593 | -0.36683399 | 0.71374285 | NA | 3.526439 | 3.168566 | 3.382107 | 3.344742 | 3.367886 | 3.335965 | 18 | 8 | 11 | 5 |
| 728806 | NR_033799 | NSFP1 | N-ethylmaleimide-sensitive factor pseudogene 1 | transcribed_ps | . | 31405 | . | . | . | 4.912755 | 6.161301 | 5.180026 | 6.528283 | 11.432181 | 14.242123 | 11.623827 | 14.926144 | 1.048495 | 104.524299 | 0.323961835 | 0.210890393 | 0.832972801 | 0.998865404 | 6.674187 | 6.740441 | 6.663381 | 6.730848 | 6.678294 | 6.745557 | 105 | 122 | 83 | 111 |
| 728825 | gene-LOC7288 | LOC728825 | small ubiquitin-like modifier 2 | pseudogene | . | . | . | . | . | 3.615541 | 3.480561 | 3.276812 | 3.127667 | 8.413514 | 8.045474 | 7.353072 | 7.151040 | -1.129105 | 107.302916 | 0.309885382 | -0.56530535 | 0.571866092 | 0.998865404 | 6.830234 | 6.655574 | 6.773170 | 6.757670 | 6.734907 | 6.718691 | 134 | 120 | 90 | 92 |
| 728833 | NM_00134594 | FAM72D | family with sequence similarity 72 member D | protein_coding | NP_001332871.1;NP_997301.2;XP_011508264.1;XP_011508265.1;XP_02 | 33593 | 614712 | . | . | 3.945438 | 4.338722 | 3.030271 | 3.148384 | 9.181195 | 10.029151 | 6.799839 | 7.198404 | -1.366592 | 268.996981 | 0.211975066 | -2.12563832 | **0.033533386** | 0.575719787 | 8.279439 | 7.828333 | 8.118588 | 8.170429 | 7.976670 | 7.993055 | 341 | 351 | 194 | 217 |
| 728841 | NM_00103750 | NBPF8 | NBPF member 8 | protein_coding | NP_001032590.2 | 31990 | 613998 | . | . | 13.161186 | 13.672669 | 16.816955 | 14.068795 | 30.626623 | 31.604985 | 37.736757 | 32.166628 | 1.133674 | 3255.695943 | 0.125833666 | 1.438455404 | 0.150304882 | 0.913261272 | 11.575311 | 11.756504 | 11.600549 | 11.626394 | 11.792652 | 11.634773 | 3458 | 3345 | 3294 | 2936 |
| 728851 | gene-HMGN1P | HMGN1P36 | high mobility group nucleosome binding domain 1 pseudogene 36 | pseudogene | . | 39380 | . | . | . | 9.567365 | 7.016682 | 7.845386 | 3.999010 | 22.263649 | 16.219374 | 17.604820 | 9.143262 | -1.419030 | 266.711497 | 0.288868907 | -1.74787001 | 0.080486539 | 0.779933816 | 8.288487 | 7.786303 | 8.211947 | 8.032843 | 8.088861 | 7.781750 | 417 | 284 | 255 | 139 |
| 728855 | NR_024510,NF | LINC00623 | long intergenic non-protein coding RNA 623 | lncRNA | . | 44252 | . | . | . | 3.849012 | 3.765558 | 3.024826 | 3.866027 | 8.956808 | 8.704257 | 6.787623 | 8.839210 | -1.108071 | 396.247165 | 0.192294552 | -0.76991258 | 0.441351754 | 0.998865404 | 8.702800 | 8.553871 | 8.670799 | 8.647258 | 8.531632 | 8.660716 | 489 | 441 | 290 | 392 |
| 728888 | NM_00131013 | NPIPB11 | nuclear pore complex interacting protein family member B11 | protein_coding | NP_001297066.2;XP_024306188.1;XP_024306189.1;XP_024306190.1 | 37453 | . | . | . | 2.640385 | 2.965309 | 4.771719 | 3.051288 | 6.144286 | 6.854445 | 10.707597 | 6.976408 | 1.388057 | 410.914995 | 0.223879163 | 2.11304356 | **0.03459704** | 0.580952481 | 8.426313 | 8.900319 | 8.531869 | 8.592586 | 8.895425 | 8.602188 | 375 | 391 | 511 | 345 |
| 728911 | NM_152582,XI | CT45A2 | cancer/testis antigen family 45 member A2 | protein_coding | NP_689795.4;XP_006724846.1;XP_011529691.1 | 28400 | 300793 | . | . | 0.498535 | 0.628677 | 0.390391 | 0.326528 | 1.160113 | 1.453217 | 0.876027 | 0.746568 | -1.620142 | 18.922777 | 0.715347302 | -0.97312228 | 0.330492533 | NA | 4.548173 | 3.852980 | 4.253709 | 4.275247 | 4.233697 | 4.219407 | 24 | 28 | 14 | 12 |
| 728913 | gene-RCN1P2 | RCN1P2 | reticulocalbin 1 pseudogene 2 | pseudogene | . | 39204 | . | . | . | 6.310275 | 6.107935 | 4.917152 | 4.365703 | 14.684269 | 14.118767 | 11.033947 | 9.981661 | -1.357571 | 361.490499 | 0.19130616 | -2.30534989 | **0.021146985** | 0.4816697 | 8.701195 | 8.260981 | 8.589940 | 8.564506 | 8.436507 | 8.367477 | 489 | 440 | 284 | 269 |
| 728917 | gene-ABHD17A | ABHD17AP1 | ABHD17A pseudogene 1 | pseudogene | . | 28394 | . | . | . | 0.280052 | 0.306822 | 0.273603 | 0.297837 | 0.651693 | 0.709234 | 0.613957 | 0.680969 | -1.082519 | 13.809845 | 0.820062723 | -0.13949262 | 0.889060889 | NA | 3.844134 | 3.728813 | 3.789476 | 3.793615 | 3.784201 | 3.789339 | 16 | 16 | 11 | 13 |
| 728920 | NR_146087 | PDE4DIPP1 | phosphodiesterase 4D interacting protein pseudogene 1 | transcribed_ps | . | 50867 | . | . | . | 0.020946 | 0.141723 | 0.094159 | 0.056682 | 0.048741 | 0.327600 | 0.211291 | 0.129597 | -1.066154 | 5.774398 | 1.339202654 | -0.06900771 | 0.94498348 | NA | 2.576652 | 2.481107 | 2.499814 | 2.539486 | 2.524033 | 2.512314 | 2 | 11 | 6 | 4 |
| 728927 | NM_00117090 | ZNF736 | zinc finger protein 736 | protein_coding | NP_001164376.1;NP_001281184.1 | 32467 | . | . | . | 0.734149 | 0.893404 | 0.908630 | 0.768751 | 1.708394 | 2.065144 | 2.038939 | 1.757659 | 1.026486 | 190.941408 | 0.255637474 | 0.147526849 | 0.882716189 | 0.998865404 | 7.557733 | 7.595986 | 7.525844 | 7.607640 | 7.631774 | 7.523565 | 197 | 222 | 189 | 159 |
| 728931 | gene-AMD1P3 | AMD1P3 | adenosylmethionine decarboxylase 1 pseudogene 3 | pseudogene | . | 51608 | . | . | . | 0.420398 | 0.064146 | 0.230112 | 0.371173 | 0.978284 | 0.148277 | 0.516364 | 0.848642 | 1.211335 | 16.504561 | 0.869048362 | 0.318277295 | 0.750274606 | NA | 3.898138 | 4.177906 | 4.072180 | 3.980089 | 4.022595 | 4.058817 | 29 | 5 | 12 | 21 |
| 728933 | gene-RPL23AP | RPL23AP60 | ribosomal protein L23a pseudogene 60 | pseudogene | . | 35966 | . | . | . | 0.112714 | 0.111374 | 0.132199 | 0.093909 | 0.262291 | 0.257445 | 0.296650 | 0.214712 | -1.007340 | 2.223025 | 1.975467281 | -0.0053412 | 0.995738358 | NA | 1.156660 | 1.148375 | 1.155078 | 1.151489 | 1.153584 | 1.152831 | 3 | 2 | 2 | 2 |
| 728937 | gene-RPS26P2 | RPS26P25 | ribosomal protein S26 pseudogene 25 | pseudogene | . | 35730 | . | . | . | 12.272170 | 14.085109 | 9.822182 | 15.988586 | 28.557844 | 32.558361 | 22.040691 | 36.556004 | -1.035294 | 180.464601 | 0.280743369 | -0.17824501 | 0.858530562 | 0.998865404 | 7.521579 | 7.469087 | 7.469445 | 7.525957 | 7.380059 | 7.576695 | 198 | 211 | 118 | 205 |
| 728963 | NR_026768 | RPS15AP10 | ribosomal protein S15a pseudogene 10 | transcribed_ps | . | 37001 | . | . | . | 0.102125 | 0.062030 | 0.187499 | 0.037918 | 0.237649 | 0.143384 | 0.420743 | 0.086696 | 1.269574 | 3.035949 | 1.770619904 | 0.194477046 | 0.845802367 | NA | 1.413393 | 1.769039 | 1.591721 | 1.583993 | 1.601652 | 1.580582 | 4 | 2 | 5 | 1 |
| 728975 | NR_145677 | LOC728975 | uncharacterized LOC728975 | lncRNA | . | . | . | . | . | 0.334712 | 0.381703 | 0.310104 | 0.208307 | 0.778889 | 0.882323 | 0.695865 | 0.476268 | -1.403711 | 18.087242 | 0.732413268 | -0.66799117 | 0.504139226 | NA | 4.399719 | 3.913269 | 4.187117 | 4.196731 | 4.180114 | 4.151771 | 23 | 24 | 16 | 11 |
| 728979 | gene-RPL10AP | RPL10AP9 | ribosomal protein L10a pseudogene 9 | pseudogene | . | 35947 | . | . | . | 0.109147 | 0.413589 | 0.273860 | 0.167683 | 0.253989 | 0.956030 | 0.614535 | 0.383387 | -1.061773 | 5.759822 | 1.30614911 | -0.06620678 | 0.9472132 | NA | 2.569585 | 2.481107 | 2.503288 | 2.534527 | 2.523314 | 2.511600 | 3 | 10 | 6 | 4 |
| 728989 | NR_024442 | LOC728989 | phosphodiesterase 4D interacting protein pseudogene | transcribed_ps | . | . | . | . | . | 0.572538 | 0.617612 | 0.949990 | 0.732128 | 1.332319 | 1.427638 | 2.131751 | 1.673923 | 1.372890 | 24.505480 | 0.625450656 | 0.731018859 | 0.464767634 | NA | 4.367696 | 4.826107 | 4.586384 | 4.592440 | 4.636467 | 4.606144 | 23 | 23 | 28 | 23 |
| 728992 | gene-RPL7AP1 | RPL7AP11 | ribosomal protein L7a pseudogene 11 | pseudogene | . | 35242 | . | . | . | 0.248245 | 0.268601 | 0.112528 | 0.182331 | 0.577676 | 0.620883 | 0.252509 | 0.416878 | -1.630830 | 5.786188 | 1.267580534 | -0.55665605 | 0.57776245 | NA | 2.844134 | 2.134756 | 2.547117 | 2.549177 | 2.531452 | 2.539668 | 8 | 8 | 3 | 5 |
| 729009 | gene-FTH1P20 | FTH1P20 | ferritin heavy chain 1 pseudogene 20 | pseudogene | . | 37639 | . | . | . | 1.654946 | 1.655817 | 1.797690 | 1.808436 | 3.851126 | 3.827495 | 4.033963 | 4.134774 | 1.063252 | 46.717105 | 0.454525372 | 0.194672541 | 0.84564931 | 0.998865404 | 5.500953 | 5.589448 | 5.538819 | 5.539860 | 5.550581 | 5.549586 | 52 | 49 | 42 | 45 |
| 729013 | NR_034137 | ZBED5-AS1 | ZBED5 antisense RNA 1 | lncRNA | . | 48646 | . | . | . | 1.338171 | 2.107763 | 1.949037 | 1.335882 | 3.113978 | 4.872190 | 4.373582 | 3.054335 | -1.072381 | 58.904766 | 0.438831683 | -0.22973976 | 0.818293994 | 0.998865404 | 5.929600 | 5.829281 | 5.832048 | 5.927366 | 5.904150 | 5.826798 | 55 | 80 | 59 | 43 |
| 729046 | gene-RPL17P5 | RPL17P50 | ribosomal protein L17 pseudogene 50 | pseudogene | . | 45098 | . | . | . | 0.590971 | 0.395027 | 0.436487 | 0.215968 | 1.375213 | 0.913122 | 0.979465 | 0.493784 | -1.395749 | 8.042542 | 1.095967627 | -0.43891737 | 0.660721411 | NA | 3.225252 | 2.751299 | 3.025396 | 3.007529 | 3.015406 | 2.993606 | 13 | 8 | 8 | 4 |
| 729082 | NR_026757,NF | OIP5-AS1 | OIP5 antisense RNA 1 | lncRNA | . | 43563 | . | . | . | 5.640364 | 4.833456 | 5.138590 | 4.250101 | 13.125358 | 11.172753 | 11.530848 | 9.717351 | -1.136021 | 1364.692830 | 0.149428364 | -1.23128972 | 0.218214525 | 0.982008277 | 10.503115 | 10.319786 | 10.522747 | 10.391046 | 10.433716 | 10.284132 | 1812 | 1434 | 1222 | 1079 |
| 729086 | gene-LOC7290 | LOC729086 | VOPP1, WBP1/VOPP1 family member pseudogene | pseudogene | . | . | . | . | . | 5.556520 | 6.095527 | 4.697198 | 5.122542 | 12.930251 | 14.090083 | 10.540377 | 11.712085 | -1.201489 | 489.752927 | 0.171439119 | -1.54470551 | 0.122417511 | 0.866527641 | 9.062524 | 8.797106 | 8.962999 | 9.017373 | 8.852028 | 8.897650 | 590 | 601 | 372 | 433 |
| 729092 | NM_00114400 | AGAP5 | ArfGAP with GTPase domain, ankyrin repeat and PH domain 5 | protein_coding | NP_001137472.1 | 23467 | . | . | . | 0.504134 | 0.651653 | 0.468092 | 0.406191 | 1.173142 | 1.506326 | 1.050386 | 0.928708 | -1.333276 | 94.532123 | 0.339000195 | -1.22411509 | 0.220908812 | 0.982008277 | 6.755313 | 6.340427 | 6.560135 | 6.641085 | 6.537833 | 6.496681 | 109 | 131 | 76 | 70 |
| 729148 | gene-NUS1P1 | NUS1P1 | NUS1 pseudogene 1 | pseudogene | . | 38472 | . | . | . | 2.169363 | 1.841260 | 1.815390 | 1.332096 | 5.048197 | 4.256155 | 4.073681 | 3.045679 | -1.293002 | 216.180517 | 0.249991859 | -1.48294466 | 0.138089114 | 0.899796454 | 7.928647 | 7.560060 | 7.852624 | 7.764105 | 7.755374 | 7.619167 | 305 | 240 | 190 | 149 |
| 729162 | NR_146578,NF | FAM239B | family with sequence similarity 239 member B | transcribed_ps | . | 53415 | . | . | . | 0.227129 | 0.287128 | 0.230176 | 0.204151 | 0.528539 | 0.663709 | 0.516508 | 0.466766 | -1.340599 | 39.669206 | 0.501216051 | -0.84370409 | 0.398834779 | 0.998865404 | 5.505589 | 5.083550 | 5.314542 | 5.349895 | 5.303536 | 5.271982 | 47 | 54 | 33 | 28 |
| 729163 | gene-FABP5P2 | FABP5P2 | fatty acid binding protein 5 pseudogene 2 | pseudogene | . | 31060 | . | . | . | 0.205763 | 0.064635 | 0.077425 | 0.043218 | 0.478819 | 0.149407 | 0.173739 | 0.098813 | -1.852709 | 2.390579 | 1.968149219 | -0.45201679 | 0.651256889 | NA | 1.631267 | 0.751299 | 1.277471 | 1.265665 | 1.267797 | 1.262184 | 5 | 2 | 2 | 1 |
| 729188 | gene-RPS26P5 | RPS26P58 | ribosomal protein S26 pseudogene 58 | pseudogene | . | 36571 | . | . | . | 1.681055 | 1.483689 | 1.108957 | 1.996380 | 3.911884 | 3.429613 | 2.488467 | 4.564487 | -1.039055 | 22.366123 | 0.664998777 | -0.08311557 | 0.933759647 | NA | 4.511958 | 4.453945 | 4.492198 | 4.479309 | 4.458176 | 4.505674 | 28 | 23 | 14 | 26 |
| 729209 | gene-RPSAP26 | RPSAP26 | ribosomal protein SA pseudogene 26 | pseudogene | . | 36195 | . | . | . | 0.251570 | 0.311286 | 0.522273 | 0.366125 | 0.585414 | 0.719552 | 1.171965 | 0.837101 | 1.533930 | 11.965391 | 0.88829509 | 0.694850506 | 0.487148993 | NA | 3.238681 | 3.857133 | 3.553073 | 3.560207 | 3.590545 | 3.567218 | 10 | 11 | 15 | 11 |
| 729212 | gene-RPL13AP | RPL13AP26 | ribosomal protein L13a pseudogene 26 | pseudogene | . | 35492 | . | . | . | 1.152780 | 1.077925 | 0.688965 | 0.935478 | 2.682566 | 2.491672 | 1.546018 | 2.138860 | -1.344125 | 18.725228 | 0.713680781 | -0.59784122 | 0.549945885 | NA | 4.425187 | 3.996978 | 4.248369 | 4.238754 | 4.211184 | 4.229104 | 26 | 22 | 12 | 17 |
| 729217 | gene-LOC7292 | LOC729217 | EBP like pseudogene | pseudogene | . | . | . | . | . | 0.143767 | 0.103092 | 0.074619 | 0.077045 | 0.334551 | 0.238301 | 0.167443 | 0.176154 | -1.609374 | 2.888932 | 1.766430981 | -0.38863673 | 0.697544891 | NA | 1.832372 | 1.148375 | 1.547205 | 1.539788 | 1.537694 | 1.536879 | 5 | 3 | 2 | 2 |
| 729218 | NR_103825,NF | LOC729218 | uncharacterized LOC729218 | transcribed_ps | . | . | . | . | . | 2.102983 | 2.228299 | 2.562679 | 2.413285 | 4.893729 | 5.150812 | 5.750576 | 5.517691 | 1.133736 | 415.037349 | 0.17939172 | 1.009438729 | 0.312764272 | 0.998865404 | 8.603644 | 8.784864 | 8.641980 | 8.670974 | 8.753000 | 8.710436 | 437 | 430 | 397 | 399 |
| 729220 | NR_164135 | FLJ45513 | uncharacterized LOC729220 | lncRNA | . | . | . | . | . | 0.315824 | 0.256760 | 0.675609 | 0.314701 | 0.734936 | 0.593513 | 1.516047 | 0.719527 | 1.624070 | 11.760783 | 0.911713382 | 0.767361967 | 0.442866335 | NA | 3.161369 | 3.865403 | 3.528908 | 3.523494 | 3.571871 | 3.529602 | 11 | 9 | 17 | 9 |
| 729234 | NR_003698 | FAHD2CP | fumarylacetoacetate hydrolase domain containing 2C, pseudogen | transcribed_ps | . | 44135 | . | . | . | 4.241801 | 3.311408 | 3.180477 | 4.832992 | 9.870845 | 7.654467 | 7.136899 | 11.050062 | 1.042149 | 103.908333 | 0.339423781 | 0.175478193 | 0.860703905 | 0.998865404 | 6.669607 | 6.728135 | 6.727558 | 6.648511 | 6.638420 | 6.761237 | 132 | 96 | 74 | 119 |
| 729236 | gene-RPS13P2 | RPS13P2 | ribosomal protein S13 pseudogene 2 | pseudogene | . | 36364 | . | . | . | 0.862202 | 0.487356 | 0.433144 | 0.035948 | 2.006379 | 1.126546 | 0.971963 | 0.082191 | **-2.560997** | 8.055176 | 1.189466925 | -1.14059944 | 0.254036651 | NA | 3.530084 | 2.188478 | 3.046692 | 3.017464 | 3.016151 | 2.985002 | 17 | 9 | 7 | 1 |
| 729262 | NM_00127849 | NUTM2B | NUT family member 2B | protein_coding | NP_001265424.1 | 23445 | . | . | . | 0.023354 | 0.027346 | 0.061633 | 0.039387 | 0.054345 | 0.063211 | 0.138303 | 0.090054 | 1.768494 | 4.369705 | 1.449320338 | 0.567522094 | 0.57035952 | NA | 1.658167 | 2.481107 | 2.102996 | 2.108355 | 2.122659 | 2.111009 | 3 | 4 | 6 | 4 |
| 729264 | NM_00124372 | TP53TG3D | TP53 target 3D | protein_coding | NP_001230651.1;XP_016879123.1;XP_016879126.1 | 44657 | . | . | . | 1.898156 | 1.633710 | 2.006597 | 0.905732 | 4.417087 | 3.776392 | 4.502746 | 2.070850 | -1.150155 | 92.680532 | 0.391008545 | -0.51617279 | 0.60573373 | 0.998865404 | 6.629658 | 6.431964 | 6.591436 | 6.493066 | 6.592436 | 6.403984 | 132 | 90 | 101 | 53 |
| 729277 | gene-RPL23AP | RPL23AP47 | ribosomal protein L23a pseudogene 47 | pseudogene | . | 36576 | . | . | . | 0.125821 | 0.228245 | 0.094991 | 0.097502 | 0.292790 | 0.527599 | 0.213158 | 0.222926 | -1.422882 | 2.686397 | 1.818605976 | -0.27978332 | 0.779643752 | NA | 1.658167 | 1.148375 | 1.430406 | 1.435592 | 1.429080 | 1.428283 | 3 | 4 | 2 | 2 |
| 729288 | NR_160540 | ZNF286B | zinc finger protein 286B (pseudogene) | transcribed_ps | . | 33241 | . | . | . | 0.761524 | 0.889082 | 1.131875 | 1.025512 | 1.772098 | 2.055152 | 2.539895 | 2.344711 | 1.291339 | 155.475573 | 0.267455112 | 1.37917552 | 0.167840644 | 0.945638095 | 7.084399 | 7.453185 | 7.193694 | 7.247818 | 7.345629 | 7.298601 | 145 | 157 | 161 | 155 |
| 729291 | NR_148936 | LOC729291 | uncharacterized LOC729291 | lncRNA | . | . | . | . | . | 0.452107 | 0.472954 | 0.502630 | 0.308089 | 1.052071 | 1.093253 | 1.127888 | 0.704410 | -1.151701 | 69.626772 | 0.39979764 | -0.50967378 | 0.610280022 | 0.998865404 | 6.218609 | 6.017531 | 6.129471 | 6.139713 | 6.153317 | 6.045992 | 84 | 82 | 70 | 46 |
| 729299 | gene-PMS2P8 | PMS2P8 | PMS1 homolog 2, mismatch repair system component pseudogen | pseudogene | . | 33516 | . | . | . | 0.412141 | 0.346061 | 0.490646 | 0.386811 | 0.959070 | 0.799935 | 1.100995 | 0.884397 | 1.117578 | 9.481661 | 0.986836499 | 0.162515194 | 0.870900164 | NA | 3.161369 | 3.324312 | 3.242058 | 3.236505 | 3.250380 | 3.237541 | 11 | 9 | 10 | 8 |
| 729317 | gene-VDAC2P5 | VDAC2P5 | VDAC2 pseudogene 5 | pseudogene | . | 54753 | . | . | . | 4.657181 | 4.984584 | 4.380518 | 5.370051 | 10.837452 | 11.522090 | 9.829756 | 12.277984 | -1.002913 | 182.301111 | 0.251488294 | -0.01668626 | 0.986686907 | 0.999197585 | 7.512904 | 7.507450 | 7.496911 | 7.523193 | 7.467135 | 7.548806 | 204 | 203 | 143 | 187 |
| 729338 | NR_024041 | CETN4P | centrin 4, pseudogene | transcribed_ps | . | 35450 | . | . | . | 0.947665 | 0.649627 | 1.098554 | 1.095710 | 2.205255 | 1.501643 | 2.465123 | 2.505211 | 1.344075 | 18.815751 | 0.713080016 | 0.598268535 | 0.549660768 | NA | 4.003609 | 4.432387 | 4.228697 | 4.201644 | 4.239818 | 4.237930 | 22 | 14 | 19 | 20 |
| 729355 | NM_00109968 | TP53TG3B | TP53 target 3B | protein_coding | NP_001093157.1;XP_006721147.1;XP_011544235.1;XP_016879128.1 | 37202 | . | . | . | 0.959205 | 0.742914 | 0.821282 | 0.543501 | 2.232110 | 1.717279 | 1.842933 | 1.242650 | -1.237986 | 43.624413 | 0.490105869 | -0.62842429 | 0.529726029 | 0.998865404 | 5.591191 | 5.286924 | 5.487960 | 5.438115 | 5.458422 | 5.396932 | 63 | 45 | 41 | 29 |
| 729362 | gene-RPL36AP | RPL36AP37 | ribosomal protein L36a pseudogene 37 | pseudogene | . | 36969 | . | . | . | 7.126060 | 9.247688 | 5.421264 | 11.464762 | 16.582634 | 21.376448 | 12.165159 | 26.212816 | 1.014660 | 103.702405 | 0.374854963 | 0.056010903 | 0.955333121 | 0.998865404 | 6.687628 | 6.704931 | 6.643197 | 6.722027 | 6.573920 | 6.790720 | 104 | 125 | 59 | 132 |
| 729375 | NR_024252 | FAM86HP | family with sequence similarity 86 member H, pseudogene | transcribed_ps | . | 42359 | . | . | . | 0.660099 | 0.791799 | 1.306740 | 0.934155 | 1.536078 | 1.830280 | 2.932286 | 2.135834 | 1.524353 | 61.307440 | 0.414471603 | 1.467404294 | 0.142266107 | 0.903146012 | 5.601622 | 6.210572 | 5.858617 | 5.891719 | 6.005291 | 5.924695 | 51 | 57 | 75 | 58 |
| 729389 | gene-RPS6P25 | RPS6P25 | ribosomal protein S6 pseudogene 25 | pseudogene | . | 35948 | . | . | . | 0.277936 | 0.219779 | 0.404236 | 0.103195 | 0.646768 | 0.508028 | 0.907093 | 0.235942 | -1.061206 | 6.963240 | 1.183747283 | -0.07240176 | 0.942282183 | NA | 2.838265 | 2.760196 | 2.798748 | 2.792546 | 2.809409 | 2.777836 | 9 | 7 | 9 | 3 |
| 729402 | gene-RPL21P1 | RPL21P16 | ribosomal protein L21 pseudogene 16 | pseudogene | . | 31396 | . | . | . | 11.378744 | 10.016980 | 8.625706 | 10.344584 | 26.478806 | 23.154699 | 19.355833 | 23.651661 | -1.145447 | 183.464958 | 0.252344497 | -0.77636392 | 0.437534142 | 0.998865404 | 7.614206 | 7.417838 | 7.578578 | 7.516349 | 7.454037 | 7.523452 | 241 | 197 | 136 | 174 |
| 729416 | gene-RPLP1P6 | RPLP1P6 | ribosomal protein lateral stalk subunit P1 pseudogene 6 | pseudogene | . | 35809 | . | . | . | 0.627072 | 0.614406 | 0.492544 | 0.407421 | 1.459222 | 1.420228 | 1.105255 | 0.931520 | -1.360039 | 12.441108 | 0.867335401 | -0.51150715 | 0.60899598 | NA | 3.841203 | 3.399154 | 3.651811 | 3.647831 | 3.637338 | 3.628980 | 17 | 15 | 10 | 9 |
| 729438 | NM_00114506 | CASTOR2 | cytosolic arginine sensor for mTORC1 subunit 2 | protein_coding | NP_001138536.1;XP_016868063.1 | 37073 | 617033 | . | . | 0.569526 | 0.724837 | 0.961036 | 0.844254 | 1.325310 | 1.675494 | 2.156538 | 1.930287 | 1.286540 | 176.608572 | 0.250425969 | 1.451514264 | 0.146636713 | 0.90884489 | 7.271488 | 7.634553 | 7.380410 | 7.425344 | 7.512654 | 7.507002 | 168 | 176 | 174 | 185 |
| 729440 | NM_00126772 | CCDC61 | coiled-coil domain containing 61 | protein_coding | NP_001254652.1;XP_005259248.1;XP_005259249.1;XP_011525559.1 | 33629 | . | . | . | 5.354951 | 5.649943 | 4.889935 | 6.330025 | 12.461192 | 13.060095 | 10.972872 | 14.472852 | -1.017487 | 349.316595 | 0.20083805 | -0.12453296 | 0.900893304 | 0.998865404 | 8.461459 | 8.435205 | 8.423458 | 8.479905 | 8.380736 | 8.499881 | 386 | 399 | 269 | 359 |
| 729453 | gene-PMS2P6 | PMS2P6 | PMS1 homolog 2, mismatch repair system component pseudogen | pseudogene | . | 33514 | . | . | . | 0.366010 | 0.404527 | 0.430772 | 0.319938 | 0.851720 | 0.935083 | 0.966639 | 0.731500 | -1.109843 | 9.390873 | 0.992687278 | -0.15146281 | 0.879610644 | NA | 3.303566 | 3.155137 | 3.232259 | 3.235164 | 3.235628 | 3.223017 | 11 | 11 | 9 | 7 |
| 729480 | gene-RPL23AP | RPL23AP5 | ribosomal protein L23a pseudogene 5 | pseudogene | . | 14194 | . | . | . | 0.205619 | 0.679939 | 0.452979 | 0.494748 | 0.478485 | 1.571709 | 1.016472 | 1.131183 | 1.015102 | 8.243262 | 1.088667203 | 0.019862957 | 0.984152696 | NA | 3.034734 | 3.051647 | 3.019057 | 3.054748 | 3.036581 | 3.038707 | 5 | 13 | 7 | 8 |
| 729494 | gene-ARPC3P3 | ARPC3P3 | actin related protein 2/3 complex subunit 3 pseudogene 3 | pseudogene | . | 39432 | . | . | . | 0.657483 | 0.378048 | 0.480486 | 0.074682 | 1.529989 | 0.873874 | 1.078197 | 0.170750 | -1.857944 | 11.322811 | 1.015101461 | -0.88041153 | 0.37863641 | NA | 3.875693 | 2.993948 | 3.531907 | 3.499973 | 3.511341 | 3.460253 | 21 | 12 | 12 | 2 |
| 729495 | gene-ABHD17A | ABHD17AP4 | ABHD17A pseudogene 4 | pseudogene | . | 34042 | . | . | . | 0.172747 | 0.293398 | 0.262348 | 0.120256 | 0.401990 | 0.678202 | 0.588700 | 0.274950 | -1.187618 | 10.408828 | 0.967752238 | -0.25633733 | 0.797690372 | NA | 3.497331 | 3.251698 | 3.368760 | 3.392448 | 3.386285 | 3.359041 | 10 | 15 | 11 | 6 |
| 729515 | NM_018452 | TMEM242 | transmembrane protein 242 | protein_coding | NP_060922.2 | 17206 | . | . | . | 1.552397 | 1.281192 | 1.172604 | 1.395594 | 3.612492 | 2.961534 | 2.631290 | 3.190861 | -1.115230 | 182.641081 | 0.255904807 | -0.61484348 | 0.538658093 | 0.998865404 | 7.589468 | 7.431971 | 7.579711 | 7.489073 | 7.454597 | 7.521012 | 244 | 187 | 138 | 175 |
| 729522 | NR_024035,NF | AACSP1 | acetoacetyl-CoA synthetase pseudogene 1 | transcribed_ps | . | 18226 | . | . | . | 0.255251 | 0.240366 | 0.505011 | 0.346908 | 0.593979 | 0.555617 | 1.133229 | 0.793163 | 1.656076 | 30.368038 | 0.571396157 | 1.273667355 | 0.202781331 | 0.968254058 | 4.513800 | 5.243790 | 4.878840 | 4.874105 | 4.963323 | 4.910415 | 27 | 24 | 39 | 29 |
| 729524 | gene-FEM1AP4 | FEM1AP4 | fem-1 homolog A pseudogene 4 | pseudogene | . | 39832 | . | . | . | 0.019350 | 0.031240 | 0.039072 | 0.012164 | 0.045028 | 0.072212 | 0.087676 | 0.027811 | -1.003532 | 2.258427 | 1.991984473 | -0.00255353 | 0.997962579 | NA | 1.175403 | 1.175234 | 1.166206 | 1.171202 | 1.174041 | 1.163259 | 2 | 3 | 3 | 1 |
| 729530 | gene-CNOT6LP | CNOT6LP1 | CNOT6L pseudogene 1 | pseudogene | . | 32355 | . | . | . | 0.088218 | 0.038245 | 0.012569 | 0.033960 | 0.205288 | 0.088406 | 0.028203 | 0.077646 | **-2.420444** | 5.622522 | 1.341758376 | -0.95044813 | 0.341884598 | NA | 2.993083 | 1.715150 | 2.531212 | 2.500533 | 2.490577 | 2.499071 | 13 | 5 | 2 | 4 |
| 729533 | NM_00112316 | FAM72A | family with sequence similarity 72 member A | protein_coding | NP_001116640.1;NP_001304830.1;XP_006711589.1;XP_011508267.1;XP | 24044 | 614710 | . | . | 4.515633 | 4.417022 | 2.927322 | 2.801506 | 10.508065 | 10.210147 | 6.568829 | 6.405309 | -1.394925 | 325.613134 | 0.196979385 | -2.43775427 | **0.014778817** | 0.406358577 | 8.567306 | 8.086917 | 8.449876 | 8.410455 | 8.229274 | 8.264351 | 451 | 396 | 228 | 264 |
| 729540 | NM_00103786 | RGPD6 | RANBP2 like and GRIP domain containing 6 | protein_coding | NP_001032955.1;NP_001116835.1;XP_011510068.1;XP_011510071.1;XP | 32419 | 612709 | . | . | 3.508490 | 3.681636 | 4.013963 | 3.223908 | 8.164403 | 8.510266 | 9.007217 | 7.371083 | 1.008486 | 773.699191 | 0.167830233 | 0.07263566 | 0.942096049 | 0.998865404 | 9.589120 | 9.602109 | 9.589919 | 9.589754 | 9.689668 | 9.492726 | 887 | 831 | 771 | 626 |
| 729557 | gene-LOC7295 | LOC729557 | density regulated re-initiation and release factor pseudogene | pseudogene | . | . | . | . | . | 0.067091 | 0.073345 | 0.011075 | 0.044162 | 0.156124 | 0.169539 | 0.024853 | 0.100972 | -1.620137 | 1.451754 | 2.440017378 | -0.28529139 | 0.775420906 | NA | 0.844134 | 0.148375 | 0.548030 | 0.548560 | 0.545314 | 0.544903 | 2 | 2 | 1 | 1 |
| 729564 | gene-HMGN2P | HMGN2P41 | high mobility group nucleosomal binding domain 2 pseudogene 4 | pseudogene | . | 39412 | . | . | . | 20.097914 | 18.961847 | 17.803413 | 18.134499 | 46.768673 | 43.831161 | 39.950344 | 41.462376 | -1.102936 | 698.087044 | 0.151508548 | -0.93294121 | 0.350850321 | 0.998865404 | 9.516139 | 9.374933 | 9.502931 | 9.456527 | 9.410185 | 9.415505 | 871 | 764 | 575 | 625 |
| 729595 | gene-HMGB3P | HMGB3P22 | high mobility group box 3 pseudogene 22 | pseudogene | . | 39314 | . | . | . | 0.139827 | 0.038106 | 0.285612 | 0.108183 | 0.325382 | 0.088085 | 0.640905 | 0.247347 | 1.776638 | 3.070930 | 1.763842668 | 0.470081381 | 0.638296876 | NA | 1.137672 | 1.978782 | 1.603704 | 1.591525 | 1.613664 | 1.597457 | 4 | 1 | 5 | 2 |
| 729603 | NR_003288 | CHP1P2 | CHP1 pseudogene 2 | transcribed_ps | . | 54896 | . | . | . | 0.848256 | 0.864570 | 1.181241 | 0.696039 | 1.973927 | 1.998492 | 2.650670 | 1.591410 | 1.072992 | 87.924212 | 0.368260487 | 0.275997055 | 0.782550328 | 0.998865404 | 6.405136 | 6.509359 | 6.438309 | 6.441767 | 6.531282 | 6.382769 | 97 | 92 | 100 | 63 |
| 729614 | NR_024279 | FLJ37453 | uncharacterized LOC729614 | lncRNA | . | . | . | . | . | 0.995250 | 1.179478 | 1.054865 | 1.013878 | 2.315988 | 2.726415 | 2.367086 | 2.318110 | -1.068049 | 90.505577 | 0.337419598 | -0.28148378 | 0.77833937 | 0.998865404 | 6.546825 | 6.451470 | 6.484739 | 6.532520 | 6.497001 | 6.483240 | 99 | 109 | 78 | 80 |
| 729621 | gene-RPL36AP | RPL36AP46 | ribosomal protein L36a pseudogene 46 | pseudogene | . | 35812 | . | . | . | 0.113963 | 0.064064 | 0.105911 | 0.165183 | 0.265197 | 0.148087 | 0.237662 | 0.377671 | 1.659343 | 1.774228 | 2.21400588 | 0.329995761 | 0.741403165 | NA | 0.413393 | 1.148375 | 0.817239 | 0.813367 | 0.819776 | 0.819065 | 2 | 1 | 2 | 2 |
| 729646 | gene-RPL31P4 | RPL31P4 | ribosomal protein L31 pseudogene 4 | pseudogene | . | 23546 | . | . | . | 0.537526 | 0.414715 | 0.155753 | 0.574492 | 1.250844 | 0.958633 | 0.349504 | 1.313507 | -1.313680 | 6.284119 | 1.249846439 | -0.31492957 | 0.752815126 | NA | 2.838265 | 2.437399 | 2.665683 | 2.659471 | 2.641056 | 2.668489 | 9 | 7 | 2 | 8 |
| 729652 | NR_135198 | LOC729652 | uncharacterized LOC729652 | lncRNA | . | . | . | . | . | 0.106841 | 0.086872 | 0.095813 | 0.176617 | 0.248623 | 0.200809 | 0.215001 | 0.403813 | 1.387519 | 12.260076 | 0.884014967 | 0.534502046 | 0.592994234 | NA | 3.361571 | 3.832035 | 3.605164 | 3.596046 | 3.601437 | 3.634771 | 13 | 10 | 9 | 17 |
| 729673 | gene-PPIAP18 | PPIAP18 | peptidylprolyl isomerase A pseudogene 18 | pseudogene | . | 32454 | . | . | . | 0.350175 | 0.918089 | 0.677590 | 0.871730 | 0.814873 | 2.122203 | 1.520491 | 1.993106 | 1.216628 | 16.591467 | 0.789454505 | 0.358334021 | 0.720093361 | NA | 3.906869 | 4.184531 | 4.006987 | 4.063560 | 4.040900 | 4.058466 | 10 | 23 | 14 | 19 |
| 729674 | gene-PSMC2P1 | PSMC2P1 | PSMC2 pseudogene 1 | pseudogene | . | 54571 | . | . | . | 0.070446 | 0.014688 | 0.171463 | 0.117781 | 0.163930 | 0.033952 | 0.384759 | 0.269293 | **3.259855** | 4.712584 | 1.466468407 | 1.162526198 | 0.24502179 | NA | 1.137672 | 2.852980 | 2.204072 | 2.191983 | 2.224760 | 2.217665 | 4 | 1 | 7 | 6 |
| 729678 | NR_027183,NF | LINC00847 | long intergenic non-protein coding RNA 847 | lncRNA | . | 45050 | . | . | . | 12.432434 | 12.169038 | 11.620761 | 12.443344 | 28.930784 | 28.129278 | 26.076652 | 28.450229 | -1.038001 | 714.273681 | 0.150496642 | -0.35753162 | 0.720693857 | 0.998865404 | 9.507044 | 9.453118 | 9.506502 | 9.479556 | 9.447642 | 9.486730 | 854 | 770 | 592 | 676 |
| 729681 | NR_147704 | LOC729681 | uncharacterized LOC729681 | lncRNA | . | . | . | . | . | 0.367968 | 0.191954 | 0.418843 | 0.153138 | 0.856276 | 0.443709 | 0.939871 | 0.350132 | 1.022604 | 16.645363 | 0.80750165 | 0.03993525 | 0.968144748 | NA | 4.037030 | 4.076793 | 4.070555 | 4.023616 | 4.078923 | 4.017840 | 25 | 12 | 21 | 9 |
| 729683 | NR_046273 | LOC729683 | uncharacterized LOC729683 | lncRNA | . | . | . | . | . | 0.285136 | 0.277993 | 0.180753 | 0.307338 | 0.663523 | 0.642593 | 0.405605 | 0.702691 | -1.160638 | 19.574784 | 0.704105747 | -0.30523624 | 0.76018623 | NA | 4.395729 | 4.177906 | 4.302909 | 4.297102 | 4.269713 | 4.305319 | 25 | 22 | 12 | 21 |
| 729708 | gene-TPI1P1 | TPI1P1 | triosephosphate isomerase 1 pseudogene 1 | pseudogene | . | 35449 | . | . | . | 1.451373 | 1.921861 | 1.401306 | 2.290363 | 3.377405 | 4.442467 | 3.144490 | 5.236643 | 1.082873 | 70.263554 | 0.403253439 | 0.284842478 | 0.775764827 | 0.998865404 | 6.077942 | 6.189318 | 6.085881 | 6.151760 | 6.081911 | 6.191943 | 67 | 83 | 49 | 84 |
| 729727 | gene-RPL12P1 | RPL12P1 | ribosomal protein L12 pseudogene 1 | pseudogene | . | 13976 | . | . | . | 0.019756 | 0.053731 | 0.073758 | 0.058177 | 0.045974 | 0.124201 | 0.165510 | 0.133015 | 1.633456 | 1.788804 | 2.208363229 | 0.320566613 | 0.748538842 | NA | 0.444631 | 1.148375 | 0.822034 | 0.826821 | 0.828805 | 0.828093 | 1 | 2 | 2 | 2 |
| 729732 | NR_047662 | LOC729732 | uncharacterized LOC729732 | lncRNA | . | . | . | . | . | 4.954792 | 4.659501 | 6.733371 | 5.251307 | 11.530004 | 10.770646 | 15.109489 | 12.006489 | 1.228894 | 669.986254 | 0.171284202 | 1.73606152 | 0.082552957 | 0.78412713 | 9.231157 | 9.529429 | 9.331754 | 9.286152 | 9.535603 | 9.355949 | 716 | 626 | 725 | 604 |
| 729737 | NR_039983 | LOC729737 | uncharacterized LOC729737 | lncRNA | . | . | . | . | . | 0.521254 | 0.599500 | 0.765780 | 0.660673 | 1.212980 | 1.385771 | 1.718387 | 1.510550 | 1.258155 | 109.070940 | 0.31206056 | 1.061682765 | 0.288379726 | 0.998865404 | 6.593851 | 6.925389 | 6.704606 | 6.744262 | 6.824576 | 6.771515 | 104 | 111 | 114 | 105 |
| 729747 | NM_00108040 | ZNF878 | zinc finger protein 878 | protein_coding | NP_001073873.2;XP_016882684.1;XP_016882685.1 | 37246 | . | . | . | 0.132647 | 0.139607 | 0.183802 | 0.175978 | 0.308675 | 0.322708 | 0.412446 | 0.402352 | 1.237586 | 11.537132 | 0.894790733 | 0.343687541 | 0.731081282 | NA | 3.365657 | 3.674293 | 3.517953 | 3.516997 | 3.532688 | 3.523933 | 12 | 11 | 12 | 11 |
| 729759 | NM_00100522 | OR4F29 | olfactory receptor family 4 subfamily F member 29 | protein_coding | NP_001005221.2 | 31275 | . | . | . | 0.062143 | 0.014234 | 0.033460 | 0.029788 | 0.144610 | 0.032903 | 0.075083 | 0.068107 | -1.598769 | 1.437178 | 2.472395132 | -0.2738078 | 0.784232327 | NA | 0.820514 | 0.148375 | 0.540170 | 0.532159 | 0.533298 | 0.532889 | 3 | 1 | 1 | 1 |
| 729764 | gene-RPL23AP | RPL23AP45 | ribosomal protein L23a pseudogene 45 | pseudogene | . | 36896 | . | . | . | 0.112714 | 0.105754 | 0.171892 | 0.093909 | 0.262291 | 0.244455 | 0.385720 | 0.214712 | 1.247923 | 2.510518 | 1.870625777 | 0.170814096 | 0.864369945 | NA | 1.156660 | 1.481107 | 1.322319 | 1.318739 | 1.326048 | 1.320134 | 3 | 2 | 3 | 2 |
| 729769 | gene-UQCRHP2 | UQCRHP2 | ubiquinol-cytochrome c reductase hinge protein pseudogene 2 | pseudogene | . | 38045 | . | . | . | 1.251890 | 1.188452 | 2.315599 | 2.089593 | 2.913199 | 2.747161 | 5.196137 | 4.777607 | 1.790816 | 29.387138 | 0.573162067 | 1.466630072 | 0.142476715 | 0.903146012 | 4.395729 | 5.237432 | 4.827904 | 4.822679 | 4.902495 | 4.883705 | 25 | 22 | 35 | 33 |
| 729798 | gene-RPL23AP | RPL23AP63 | ribosomal protein L23a pseudogene 63 | pseudogene | . | 35502 | . | . | . | 0.026431 | 0.024989 | 0.055833 | 0.017676 | 0.061506 | 0.057763 | 0.125288 | 0.040413 | 1.870326 | 1.290451 | 2.585246895 | 0.349401681 | 0.726787772 | NA | -0.155866 | 0.751299 | 0.348480 | 0.348793 | 0.355090 | 0.349503 | 1 | 1 | 2 | 1 |
| 729799 | NR_026952 | SEC14L1P1 | SEC14 like 1 pseudogene 1 | transcribed_ps | . | 44189 | . | . | . | 0.495518 | 0.727349 | 0.579155 | 0.594727 | 1.153091 | 1.681299 | 1.299607 | 1.359773 | -1.051098 | 48.819352 | 0.457803692 | -0.15704675 | 0.875208005 | 0.998865404 | 5.645708 | 5.572116 | 5.576019 | 5.645964 | 5.601073 | 5.604104 | 47 | 64 | 41 | 45 |
| 729806 | gene-ZNF731P | ZNF731P | zinc finger protein 731, pseudogene | pseudogene | . | 32471 | . | . | . | 10.432743 | 8.479399 | 12.056468 | 6.928330 | 24.277420 | 19.600512 | 27.054365 | 15.840803 | -1.009099 | 200.330170 | 0.286462627 | -0.04561785 | 0.963614845 | 0.998865404 | 7.651445 | 7.641008 | 7.685967 | 7.588426 | 7.749463 | 7.504205 | 256 | 194 | 221 | 136 |
| 729830 | NM_00110997 | FAM160A1 | family with sequence similarity 160 member A1 | protein_coding | NP_001103447.1;NP_001335623.1;XP_005263256.1;XP_011530522.1;XP | 34237 | . | . | . | 0.327550 | 0.300886 | 0.414747 | 0.362649 | 0.762222 | 0.695510 | 0.930681 | 0.829153 | 1.214319 | 126.905663 | 0.290563782 | 0.964153282 | 0.334969083 | 0.998865404 | 6.840102 | 7.121430 | 6.965444 | 6.929707 | 7.039928 | 6.992065 | 139 | 117 | 129 | 122 |
| 729839 | gene-LOC7298 | LOC729839 | DTW domain containing 2 pseudogene | pseudogene | . | . | . | . | . | 0.661363 | 0.326502 | 0.359484 | 0.143590 | 1.539019 | 0.754723 | 0.806671 | 0.328302 | -1.985749 | 13.369667 | 0.894552549 | -1.10634477 | 0.268577321 | NA | 4.149569 | 3.168566 | 3.788139 | 3.737628 | 3.740508 | 3.708351 | 27 | 13 | 11 | 5 |
| 729852 | NM_00130234 | UMAD1 | UBAP1-MVB12-associated (UMA) domain containing 1 | protein_coding | NP_001289277.1;NP_001289278.1;NP_001289279.1 | 48955 | . | . | . | 4.739305 | 3.563650 | 4.025860 | 4.201617 | 11.028558 | 8.237535 | 9.033914 | 9.606498 | -1.025390 | 290.030104 | 0.220881926 | -0.16376635 | 0.869915084 | 0.998865404 | 8.197651 | 8.162250 | 8.261203 | 8.101402 | 8.162580 | 8.179756 | 388 | 270 | 245 | 273 |
| 729857 | NM_00107817 | RGPD2 | RANBP2 like and GRIP domain containing 2 | protein_coding | NP_001071638.2;XP_016860334.1;XP_016860335.1;XP_016860336.1;XP | 32415 | 612705 | . | . | 0.356369 | 0.373487 | 0.405243 | 0.255580 | 0.829283 | 0.863330 | 0.909351 | 0.584354 | -1.092994 | 83.451482 | 0.368657915 | -0.34797868 | 0.727856196 | 0.998865404 | 6.444530 | 6.318448 | 6.378522 | 6.402727 | 6.422884 | 6.307159 | 96 | 98 | 85 | 58 |
| 729859 | gene-WBP1P2 | WBP1P2 | WBP1 pseudogene 2 | pseudogene | . | 54568 | . | . | . | 0.106994 | 0.109686 | 0.075904 | 0.036909 | 0.248978 | 0.253543 | 0.170326 | 0.084389 | -1.611705 | 3.639798 | 1.582737814 | -0.43506102 | 0.663518135 | NA | 2.166062 | 1.481107 | 1.874045 | 1.875353 | 1.869876 | 1.863894 | 5 | 5 | 3 | 2 |
| 729870 | NR_136614 | LOC729870 | uncharacterized LOC729870 | lncRNA | . | . | . | . | . | 0.268317 | 0.263702 | 0.350873 | 0.416106 | 0.624385 | 0.609558 | 0.787348 | 0.951377 | 1.466398 | 11.600842 | 0.894340109 | 0.617524017 | 0.536889145 | NA | 3.234218 | 3.785711 | 3.518881 | 3.517719 | 3.532781 | 3.542747 | 11 | 10 | 11 | 14 |
| 729873 | NM_00112339 | TBC1D3 | TBC1 domain family member 3 | protein_coding | NP_001116863.3 | 19031 | 607741 | . | . | 0.751820 | 0.535077 | 0.765983 | 0.575151 | 1.749516 | 1.236855 | 1.718843 | 1.315015 | 1.027599 | 46.605520 | 0.472020593 | 0.083210301 | 0.933684322 | 0.998865404 | 5.521050 | 5.563496 | 5.566691 | 5.505715 | 5.565796 | 5.516377 | 62 | 41 | 47 | 38 |
| 729877 | NM_00112339 | TBC1D3H | TBC1 domain family member 3H | protein_coding | NP_001116864.3;XP_006722118.1;XP_006722119.1;XP_011523474.1;XP | 30708 | 610811 | . | . | 0.342731 | 0.322425 | 0.448986 | 0.382966 | 0.797549 | 0.745300 | 1.007513 | 0.875607 | 1.210226 | 44.591409 | 0.467133479 | 0.589287728 | 0.555668276 | 0.998865404 | 5.333903 | 5.610270 | 5.459203 | 5.455603 | 5.504067 | 5.474808 | 47 | 43 | 46 | 42 |
| 729903 | gene-RPS16P1 | RPS16P10 | ribosomal protein S16 pseudogene 10 | pseudogene | . | 35739 | . | . | . | 0.053846 | 0.038221 | 0.081949 | 0.102850 | 0.125302 | 0.088349 | 0.183892 | 0.235155 | 1.659343 | 1.774228 | 2.21400588 | 0.329995761 | 0.741403165 | NA | 0.413393 | 1.148375 | 0.817239 | 0.813367 | 0.819776 | 0.819065 | 2 | 1 | 2 | 2 |
| 729920 | NM_00110141 | CRPPA | CDP-L-ribitol pyrophosphorylase A | protein_coding | NP_001094887.1;NP_001094896.1;NP_001355126.1 | 37276 | 614631 | . | . | 0.138783 | 0.215432 | 0.271601 | 0.165177 | 0.322954 | 0.497980 | 0.609464 | 0.377657 | 1.236660 | 35.855918 | 0.547298536 | 0.559929762 | 0.575527488 | 0.998865404 | 5.002435 | 5.309532 | 5.107919 | 5.164551 | 5.202838 | 5.128717 | 29 | 42 | 43 | 28 |
| 729927 | gene-RPS20P2 | RPS20P27 | ribosomal protein S20 pseudogene 27 | pseudogene | . | 36154 | . | . | . | 4.673809 | 4.023438 | 4.650688 | 4.789975 | 10.876145 | 9.300356 | 10.436008 | 10.951708 | 1.072131 | 73.210849 | 0.370928782 | 0.270891968 | 0.786474123 | 0.998865404 | 6.142567 | 6.243635 | 6.203836 | 6.163900 | 6.197802 | 6.204648 | 88 | 70 | 65 | 72 |
| 729948 | gene-CDC26P1 | CDC26P1 | cell division cycle 26 pseudogene 1 | pseudogene | . | 22770 | . | . | . | 2.864408 | 3.037377 | 3.525641 | 2.826139 | 6.665595 | 7.021032 | 7.911436 | 6.461631 | 1.066793 | 67.259215 | 0.387767361 | 0.240557356 | 0.809898205 | 0.998865404 | 6.023720 | 6.118058 | 6.055407 | 6.067307 | 6.101407 | 6.050649 | 73 | 72 | 67 | 58 |
| 729967 | NM_00114545 | MORN2 | MORN repeat containing 2 | protein_coding | NP_001138922.1 | 30166 | . | . | . | 17.886732 | 17.168314 | 16.910715 | 17.025145 | 41.623161 | 39.685326 | 37.947151 | 38.925968 | -1.046480 | 398.617227 | 0.180862096 | -0.3624044 | 0.71704985 | 0.998865404 | 8.671177 | 8.605803 | 8.667203 | 8.637290 | 8.625880 | 8.625025 | 481 | 429 | 339 | 365 |
| 729970 | NR_033998,NF | LOC729970 | hCG2028352-like | lncRNA | . | . | . | . | . | 0.567975 | 1.171144 | 1.227919 | 0.738190 | 1.321702 | 2.707151 | 2.755414 | 1.687784 | 1.107556 | 28.163193 | 0.621547398 | 0.237118235 | 0.812565066 | 0.998865404 | 4.740215 | 4.887505 | 4.761943 | 4.833634 | 4.838210 | 4.781236 | 21 | 38 | 32 | 21 |
| 729975 | NR_034159 | LINC01530 | long intergenic non-protein coding RNA 1530 | lncRNA | . | 44320 | . | . | . | 0.179254 | 0.505424 | 0.349103 | 0.432855 | 0.417131 | 1.168310 | 0.783376 | 0.989672 | 1.157279 | 20.078286 | 0.730177557 | 0.288610732 | 0.772879282 | NA | 4.220984 | 4.426810 | 4.274156 | 4.346684 | 4.315420 | 4.331708 | 12 | 29 | 17 | 22 |
| 729978 | NM_00135551 | NPIPB2 | nuclear pore complex interacting protein family member B2 | protein_coding | NP_001342443.1;XP_024306192.1;XP_024306193.1;XP_024306194.1;XP | 37451 | . | . | . | 1.011261 | 1.242777 | 1.958872 | 1.125842 | 2.353246 | 2.872736 | 4.395652 | 2.574103 | 1.252909 | 64.768320 | 0.416713644 | 0.780587832 | 0.435044951 | 0.998865404 | 5.844134 | 6.171741 | 5.972269 | 5.986268 | 6.090702 | 5.965521 | 64 | 64 | 79 | 50 |
| 729991 | NM_00114578 | BORCS8 | BLOC-1 related complex subunit 8 | protein_coding | NP_001139255.1;NP_001139256.1 | 37247 | 616601 | . | . | 16.620324 | 18.812461 | 14.586042 | 16.571081 | 38.676177 | 43.485843 | 32.730655 | 37.887809 | -1.150037 | 521.373036 | 0.171652503 | -1.17493476 | 0.240020817 | 0.992733415 | 9.123865 | 8.921380 | 9.028465 | 9.107346 | 8.944072 | 9.012992 | 604 | 638 | 398 | 480 |
| 729998 | gene-LOC7299 | LOC729998 | eukaryotic translation elongation factor 1 gamma pseudogene | pseudogene | . | . | . | . | . | 18.165852 | 18.660606 | 15.115158 | 19.134998 | 42.272682 | 43.134827 | 33.917976 | 43.749901 | -1.090100 | 782.270286 | 0.159881844 | -0.7784505 | 0.436303475 | 0.998865404 | 9.672775 | 9.547555 | 9.633260 | 9.646630 | 9.499242 | 9.653355 | 931 | 889 | 578 | 780 |
| 730010 | gene-BRI3P1 | BRI3P1 | brain protein I3 pseudogene 1 | pseudogene | . | 33533 | . | . | . | 1.460630 | 1.361247 | 0.678297 | 1.410278 | 3.398946 | 3.146585 | 1.522078 | 3.224432 | -1.355332 | 28.369554 | 0.606505945 | -0.72323508 | 0.469535471 | 0.998865404 | 5.030746 | 4.587955 | 4.851404 | 4.841496 | 4.779205 | 4.843027 | 39 | 34 | 14 | 30 |
| 730029 | gene-RPSAP19 | RPSAP19 | ribosomal protein SA pseudogene 19 | pseudogene | . | 36508 | . | . | . | 11.231901 | 10.030210 | 7.717630 | 11.738555 | 26.137094 | 23.185280 | 17.318138 | 26.838810 | -1.108818 | 327.925678 | 0.226904156 | -0.65676269 | 0.511333518 | 0.998865404 | 8.430400 | 8.280140 | 8.415306 | 8.346397 | 8.210080 | 8.426286 | 421 | 350 | 216 | 350 |
| 730041 | gene-RHOQP2 | RHOQP2 | ras homolog family member Q pseudogene 2 | pseudogene | . | 37836 | . | . | . | 0.470075 | 0.365808 | 0.233036 | 0.245783 | 1.093885 | 0.845581 | 0.522926 | 0.561954 | -1.776775 | 30.707405 | 0.568739667 | -1.45806773 | 0.144821874 | 0.906189003 | 5.296092 | 4.467590 | 4.997719 | 4.957448 | 4.907860 | 4.909417 | 51 | 37 | 19 | 21 |
| 730051 | NM_00114498 | ZNF814 | zinc finger protein 814 | protein_coding | NP_001138461.1;XP_016882686.1 | 33258 | . | . | . | 0.368720 | 0.302349 | 0.410071 | 0.295276 | 0.858025 | 0.698893 | 0.920188 | 0.675113 | 1.049455 | 67.358068 | 0.398270535 | 0.174858245 | 0.86119102 | 0.998865404 | 6.037097 | 6.109551 | 6.089486 | 6.038006 | 6.112892 | 6.036138 | 84 | 63 | 70 | 54 |
| 730052 | gene-UBE2V1P | UBE2V1P2 | ubiquitin conjugating enzyme E2 V1 pseudogene 2 | pseudogene | . | 32648 | . | . | . | 0.378603 | 0.287286 | 0.236057 | 0.248413 | 0.881025 | 0.664075 | 0.529706 | 0.567966 | -1.370627 | 6.563710 | 1.184217007 | -0.38408144 | 0.700918087 | NA | 2.923304 | 2.470304 | 2.730318 | 2.720083 | 2.716895 | 2.714956 | 10 | 7 | 5 | 5 |
| 730069 | gene-NRBF2P6 | NRBF2P6 | NRBF2 pseudogene 6 | pseudogene | . | 54797 | . | . | . | 1.076480 | 0.703585 | 1.083324 | 0.870395 | 2.505012 | 1.626368 | 2.430947 | 1.990055 | 1.092748 | 49.438091 | 0.462897267 | 0.27643564 | 0.782213486 | 0.998865404 | 5.560516 | 5.691610 | 5.652560 | 5.575196 | 5.651463 | 5.609764 | 66 | 40 | 50 | 43 |
| 730087 | NM_00124403 | ZNF726 | zinc finger protein 726 | protein_coding | NP_001230967.1;NP_001335616.1;NP_001335617.1;NP_001335618.1 | 32462 | . | . | . | 4.058995 | 4.500936 | 5.796999 | 4.065911 | 9.445449 | 10.404117 | 13.008297 | 9.296223 | 1.121830 | 282.310548 | 0.232330937 | 0.713871558 | 0.475306606 | 0.998865404 | 8.055327 | 8.222132 | 8.066834 | 8.138994 | 8.255112 | 8.063999 | 285 | 307 | 306 | 230 |
| 730092 | NR_003370 | RRN3P1 | RRN3 pseudogene 1 | transcribed_ps | . | 30548 | . | . | . | 0.311480 | 0.401838 | 0.474136 | 0.320181 | 0.724827 | 0.928867 | 1.063947 | 0.732056 | 1.087275 | 19.259563 | 0.707815355 | 0.170548321 | 0.864578936 | NA | 4.205110 | 4.327309 | 4.250283 | 4.267404 | 4.282406 | 4.248740 | 19 | 22 | 21 | 15 |
| 730094 | NM_00116457 | MOSMO | modulator of smoothened | protein_coding | NP_001158051.1 | 27087 | . | . | . | 4.654910 | 4.415454 | 4.344889 | 3.541070 | 10.832167 | 10.206520 | 9.749804 | 8.096236 | -1.166052 | 591.309771 | 0.171019356 | -1.29594991 | 0.194992744 | 0.96621494 | 9.313887 | 9.093224 | 9.275353 | 9.233869 | 9.220050 | 9.082256 | 755 | 666 | 526 | 457 |
| 730098 | NM_00132003 | LOC730098 | uncharacterized LOC730098 | protein_coding | NP_001306966.1;NP_001306967.1;XP_016870595.1 | . | . | . | . | 5.667552 | 6.909685 | 4.324394 | 5.298495 | 13.188626 | 15.972045 | 9.703814 | 12.114382 | -1.296556 | 247.117899 | 0.227888038 | -1.64416257 | 0.100142626 | 0.815618258 | 8.124920 | 7.748738 | 7.963426 | 8.050570 | 7.837873 | 7.919900 | 295 | 326 | 170 | 220 |
| 730101 | NR_024403,NF | LOC730101 | uncharacterized LOC730101 | lncRNA | . | . | . | . | . | 3.605697 | 3.091879 | 3.375447 | 3.186484 | 8.390605 | 7.147018 | 7.574407 | 7.285516 | -1.030361 | 376.811961 | 0.191146389 | -0.22574303 | 0.821401289 | 0.998865404 | 8.578699 | 8.536393 | 8.611161 | 8.517735 | 8.569599 | 8.526448 | 475 | 380 | 335 | 335 |
| 730102 | NR_037167,NF | CRYZL2P | crystallin zeta like 2, pseudogene | transcribed_ps | . | 52164 | . | . | . | 0.251082 | 0.364405 | 0.297625 | 0.206418 | 0.584279 | 0.842340 | 0.667863 | 0.471949 | -1.233424 | 26.119645 | 0.616818768 | -0.49069284 | 0.623643709 | NA | 4.849979 | 4.548418 | 4.698201 | 4.735757 | 4.711747 | 4.677472 | 28 | 36 | 24 | 18 |
| 730112 | NM_00109995 | FAM166B | family with sequence similarity 166 member B | protein_coding | NP_001093421.1;NP_001157782.1;NP_001274167.1;NP_001274168.1;XI | 34242 | . | . | . | 0.994229 | 0.710875 | 0.984354 | 0.584303 | 2.313611 | 1.643222 | 2.208861 | 1.335939 | -1.105764 | 29.346886 | 0.589718957 | -0.24595452 | 0.805717441 | 0.998865404 | 4.943804 | 4.803035 | 4.898752 | 4.858800 | 4.892885 | 4.840418 | 41 | 28 | 30 | 20 |
| 730138 | gene-PFN1P1 | PFN1P1 | profilin 1 pseudogene 1 | pseudogene | . | 42989 | . | . | . | 0.253842 | 0.267985 | 0.163098 | 0.229253 | 0.590702 | 0.619460 | 0.365988 | 0.524159 | -1.303139 | 6.340349 | 1.202737085 | -0.31760106 | 0.750787571 | NA | 2.844134 | 2.459419 | 2.672855 | 2.674958 | 2.662404 | 2.670281 | 8 | 8 | 4 | 6 |
| 730153 | gene-NPIPB10I | NPIPB10P | nuclear pore complex interacting protein family, member B10, pse | pseudogene | . | 37455 | . | . | . | 1.106668 | 0.691779 | 1.306839 | 1.005274 | 2.575263 | 1.599078 | 2.932509 | 2.298439 | 1.268819 | 28.814519 | 0.592910904 | 0.579321911 | 0.562371984 | 0.998865404 | 4.664641 | 5.011955 | 4.851336 | 4.801537 | 4.871987 | 4.836714 | 36 | 21 | 32 | 26 |
| 730183 | NM_00125693 | LOC730183 | uncharacterized LOC730183 | protein_coding | NP_001243861.1 | . | . | . | . | 1.297475 | 1.287607 | 1.606258 | 0.907095 | 3.019276 | 2.976362 | 3.604397 | 2.073965 | -1.038699 | 21.615161 | 0.673111936 | -0.08138036 | 0.935139466 | NA | 4.459800 | 4.407673 | 4.427594 | 4.437595 | 4.450395 | 4.407420 | 24 | 25 | 23 | 15 |
| 730202 | NR_122121 | LOC730202 | uncharacterized LOC730202 | lncRNA | . | . | . | . | . | 0.828440 | 1.132196 | 0.761492 | 1.541867 | 1.927813 | 2.617122 | 1.708765 | 3.525295 | 1.165543 | 22.874742 | 0.6736492 | 0.328067115 | 0.742860917 | NA | 4.403699 | 4.619599 | 4.489529 | 4.514961 | 4.482038 | 4.550862 | 21 | 26 | 14 | 31 |
| 730234 | NR_146388 | LOC730234 | uncharacterized LOC730234 | lncRNA | . | . | . | . | . | 0.966630 | 0.710238 | 0.416366 | 0.553476 | 2.249388 | 1.641747 | 0.934312 | 1.265457 | -1.617313 | 11.549476 | 0.908332213 | -0.76359576 | 0.445108175 | NA | 3.835322 | 3.141582 | 3.559551 | 3.539506 | 3.522350 | 3.528813 | 19 | 13 | 7 | 9 |
| 730257 | gene-LOC7302 | LOC730257 | . | pseudogene | . | . | . | . | . | 0.585367 | 0.424959 | 0.368608 | 0.676204 | 1.362172 | 0.982312 | 0.827145 | 1.546061 | 1.063292 | 17.014156 | 0.760590345 | 0.116405894 | 0.907330864 | NA | 4.044696 | 4.131331 | 4.096982 | 4.073995 | 4.069474 | 4.110250 | 22 | 15 | 11 | 21 |
| 730268 | NR_160651 | ANAPC1P4 | ANAPC1 pseudogene 4 | transcribed_ps | . | 54710 | . | . | . | 3.869529 | 3.750434 | 4.099930 | 4.282488 | 9.004554 | 8.669296 | 9.200124 | 9.791401 | 1.085811 | 279.409601 | 0.206384384 | 0.575491232 | 0.564959119 | 0.998865404 | 8.065626 | 8.184405 | 8.114228 | 8.092542 | 8.138064 | 8.154776 | 315 | 283 | 249 | 277 |
| 730338 | NR_134575 | LOC730338 | uncharacterized LOC730338 | lncRNA | . | . | . | . | . | 0.220784 | 0.274143 | 0.250330 | 0.081600 | 0.513773 | 0.633693 | 0.561733 | 0.186569 | -1.432839 | 13.002363 | 0.893607809 | -0.58065394 | 0.561473712 | NA | 3.934350 | 3.421760 | 3.703111 | 3.715367 | 3.712062 | 3.665459 | 16 | 18 | 14 | 5 |
| 730747 | gene-RPL14P3 | RPL14P3 | ribosomal protein L14 pseudogene 3 | pseudogene | . | 36638 | . | . | . | 0.227643 | 0.107409 | 0.054309 | 0.110854 | 0.529734 | 0.248280 | 0.121868 | 0.253453 | -1.829845 | 3.085216 | 1.747177154 | -0.49893145 | 0.61782768 | NA | 1.998355 | 1.121007 | 1.649647 | 1.638179 | 1.631048 | 1.640128 | 6 | 3 | 1 | 3 |
| 730755 | NM_00116525 | KRTAP2-3 | keratin associated protein 2-3 | protein_coding | NP_001158724.1 | 18906 | . | . | . | 0.220528 | 0.320782 | 0.056288 | 0.116216 | 0.513178 | 0.741501 | 0.126308 | 0.265714 | **-2.780921** | 5.211623 | 1.378544392 | -1.07037748 | 0.284449433 | NA | 2.939842 | 1.459419 | 2.398514 | 2.413011 | 2.381696 | 2.385465 | 7 | 10 | 2 | 3 |
| 731157 | NR_105024 | LOC731157 | uncharacterized LOC731157 | lncRNA | . | . | . | . | . | 0.534370 | 0.638843 | 0.516513 | 0.482567 | 1.243501 | 1.476715 | 1.159040 | 1.103331 | -1.216522 | 8.598776 | 1.034543925 | -0.27332096 | 0.784606507 | NA | 3.238681 | 2.955730 | 3.107158 | 3.114028 | 3.104394 | 3.101762 | 10 | 11 | 7 | 7 |
| 731223 | NR_038421 | LINC01220 | long intergenic non-protein coding RNA 1220 | lncRNA | . | 49664 | . | . | . | 0.484592 | 0.459819 | 0.336292 | 0.280816 | 1.127664 | 1.062892 | 0.754629 | 0.642053 | -1.523198 | 6.311619 | 1.210134287 | -0.5016828 | 0.615890663 | NA | 2.928838 | 2.324312 | 2.671386 | 2.669434 | 2.661916 | 2.655222 | 9 | 8 | 5 | 4 |
| 731275 | NR_029401 | LINC01347 | long intergenic non-protein coding RNA 1347 | lncRNA | . | 50566 | . | . | . | 0.304234 | 0.221121 | 0.338589 | 0.211227 | 0.707964 | 0.511131 | 0.759785 | 0.482946 | 1.040274 | 32.804422 | 0.561370676 | 0.101473307 | 0.919174746 | 0.998865404 | 5.004919 | 5.066070 | 5.049446 | 5.006534 | 5.063541 | 5.002169 | 43 | 29 | 36 | 24 |
| 731605 | gene-BCLAF1P2 | BCLAF1P2 | . | pseudogene | . | 51807 | . | . | . | 2.270018 | 2.923640 | 2.668452 | 2.841166 | 5.282424 | 6.758126 | 5.987929 | 6.495988 | 1.048649 | 244.713386 | 0.228607522 | 0.299779535 | 0.764345327 | 0.998865404 | 7.900843 | 7.968267 | 7.858974 | 7.978738 | 7.930766 | 7.957115 | 242 | 289 | 212 | 241 |
| 731688 | gene-RPL21P1 | RPL21P120 | ribosomal protein L21 pseudogene 120 | pseudogene | . | 35743 | . | . | . | 0.108132 | 0.058751 | 0.093799 | 0.120130 | 0.251627 | 0.135806 | 0.210483 | 0.274662 | 1.233700 | 2.489691 | 1.874770521 | 0.161615342 | 0.871608775 | NA | 1.156660 | 1.459419 | 1.314228 | 1.310654 | 1.312823 | 1.316904 | 3 | 2 | 2 | 3 |
| 732229 | gene-LOC7322 | LOC732229 | . | pseudogene | . | . | . | . | . | 2.600127 | 2.736889 | 2.595951 | 2.611419 | 6.050602 | 6.326443 | 5.825239 | 5.970700 | -1.044365 | 58.829488 | 0.40791365 | -0.15352683 | 0.877982834 | 0.998865404 | 5.909524 | 5.846728 | 5.879724 | 5.887319 | 5.874296 | 5.874574 | 68 | 66 | 50 | 54 |
| 735301 | NR_003142 | SNHG9 | small nucleolar RNA host gene 9 | lncRNA | . | 33102 | . | . | . | 9.615834 | 7.976817 | 8.755428 | 10.862741 | 22.376444 | 18.438770 | 19.646929 | 24.836367 | 1.106183 | 68.232274 | 0.388362477 | 0.374881406 | 0.707748668 | 0.998865404 | 6.017890 | 6.163216 | 6.100646 | 6.055284 | 6.077314 | 6.125793 | 82 | 63 | 56 | 74 |
| 751071 | NM_00104322 | CSKMT | citrate synthase lysine methyltransferase | protein_coding | NP_001036694.1;XP_005274289.1 | 33113 | 617897 | . | . | 1.637813 | 1.535121 | 1.462593 | 1.338200 | 3.811257 | 3.548499 | 3.282016 | 3.059635 | -1.096289 | 68.561721 | 0.381032072 | -0.3480758 | 0.727783257 | 0.998865404 | 6.163717 | 6.031937 | 6.115104 | 6.106274 | 6.101246 | 6.076901 | 84 | 76 | 60 | 58 |
| 751580 | NR_130733,NF | LINC00106 | long intergenic non-protein coding RNA 106 | lncRNA | . | 31843 | . | . | . | 3.040306 | 2.660044 | 3.755320 | 3.602093 | 7.074918 | 6.148812 | 8.426830 | 8.235757 | 1.284205 | 39.892425 | 0.491203152 | 0.734676212 | 0.462536711 | 0.998865404 | 5.125927 | 5.487556 | 5.298881 | 5.286561 | 5.335199 | 5.327135 | 42 | 36 | 40 | 41 |
| 751603 | gene-LOC7516 | LOC751603 | tropomyosin 3 pseudogene | pseudogene | . | . | . | . | . | 0.023590 | 0.017092 | 0.025167 | 0.016735 | 0.054895 | 0.039509 | 0.056474 | 0.038262 | 1.234663 | 2.005914 | 2.070995667 | 0.146845664 | 0.883253841 | NA | 0.844134 | 1.148375 | 0.998092 | 0.998667 | 1.000707 | 0.999974 | 2 | 2 | 2 | 2 |
| 767557 | NR_003189 | SYS1-DBNDD2 | SYS1-DBNDD2 readthrough (NMD candidate) | lncRNA | . | 33535 | . | . | . | 1.095711 | 1.760713 | 2.048710 | 2.181164 | 2.549765 | 4.069966 | 4.597247 | 4.986973 | 1.464100 | 74.487558 | 0.386549049 | 1.422883855 | 0.154769827 | 0.923600487 | 5.919286 | 6.466911 | 6.105038 | 6.204241 | 6.243121 | 6.257810 | 54 | 80 | 75 | 85 |
| 767558 | NM_00112861 | LUZP6 | leucine zipper protein 6 | protein_coding | NP_001122091.2 | 33955 | 611050 | . | . | 7.682522 | 4.546929 | 6.960569 | 4.775867 | 17.877544 | 10.510429 | 15.619314 | 10.919451 | -1.057025 | 707.325047 | 0.464836588 | -0.17212496 | 0.863339293 | 0.998865404 | 9.505532 | 9.425826 | 9.640127 | 9.270485 | 9.555262 | 9.294255 | 1055 | 580 | 712 | 522 |
| 767811 | NR_003238 | H2BW4P | H2B.W histone 4, pseudogene | transcribed_ps | . | 25757 | . | . | . | 0.224422 | 0.142077 | 0.288011 | 0.259372 | 0.522240 | 0.328418 | 0.646287 | 0.593023 | 1.528262 | 7.328598 | 1.122671973 | 0.545031634 | 0.585731761 | NA | 2.533720 | 3.148375 | 2.862746 | 2.851986 | 2.872660 | 2.869886 | 8 | 5 | 8 | 8 |
| 768206 | NM_00107762 | PRCD | photoreceptor disc component | protein_coding | NP_001071088.1;XP_016880502.1;XP_016880503.1;XP_016880504.1 | 32528 | 610598 | . | . | 0.141056 | 0.279268 | 0.291240 | 0.249083 | 0.328244 | 0.645540 | 0.653535 | 0.569497 | 1.009184 | 45.910727 | 0.491858802 | 0.026814796 | 0.978607452 | 0.998865404 | 5.514817 | 5.526677 | 5.458273 | 5.563203 | 5.530120 | 5.497037 | 37 | 64 | 44 | 39 |
| 768211 | NM_00108539 | RELL1 | RELT like 1 | protein_coding | NP_001078868.1;NP_001078869.1;XP_016864079.1 | 27379 | 611212 | . | . | 12.480717 | 11.792197 | 12.889958 | 11.662988 | 29.043140 | 27.258190 | 28.924693 | 26.666034 | -1.001306 | 1350.878770 | 0.130250391 | -0.0144557 | 0.988466426 | 0.999197585 | 10.400468 | 10.398896 | 10.416739 | 10.382629 | 10.430168 | 10.366722 | 1591 | 1426 | 1223 | 1213 |
| 768213 | NR_030407,rn | MIR671 | microRNA 671 | miRNA | . | 33134 | 615245 | . | . | 0.010247 | 0.009216 | 0.015703 | 0.008178 | 0.023846 | 0.021301 | 0.035236 | 0.018698 | 1.234665 | 3.008871 | 1.708874417 | 0.17796512 | 0.858750369 | NA | 1.429096 | 1.733338 | 1.582415 | 1.583274 | 1.586316 | 1.585224 | 3 | 3 | 3 | 3 |
| 768220 | NR_030527,rn | MIR765 | microRNA 765 | miRNA | . | 33141 | . | . | . | 1.153121 | 1.945525 | 0.913512 | 0.565590 | 2.683361 | 4.497168 | 2.049895 | 1.293156 | -1.738748 | 5.337813 | 1.326875536 | -0.60144981 | 0.54754043 | NA | 2.760365 | 1.963455 | 2.421394 | 2.435827 | 2.418766 | 2.412395 | 6 | 9 | 4 | 3 |
| 780779 | gene-USP10P1 | USP10P1 | ubiquitin specific peptidase 10 pseudogene 1 | pseudogene | . | 31752 | . | . | . | 0.013933 | 0.002070 | 0.012500 | 0.003385 | 0.032422 | 0.004785 | 0.028051 | 0.007740 | 1.256691 | 1.507561 | 2.39030677 | 0.137902842 | 0.890317204 | NA | 0.413393 | 0.751299 | 0.586366 | 0.582479 | 0.588807 | 0.583226 | 2 | 1 | 2 | 1 |

| | | | | | | | | | | | | | | | | | | | | | | | | | | | | | | | | | |
|---|---|---|---|---|---|---|---|---|---|---|---|---|---|---|---|---|---|---|---|---|---|---|---|---|---|---|---|---|---|---|---|---|---|
| 780813 | gene-PAICSP4 | PAICSP4 | phosphoribosylaminoimidazole carboxylase, phosphoribosylaminc | pseudogene | . | 38097 | . | . | . | 0.091447 | 0.136820 | 0.054729 | 0.179590 | 0.212800 | 0.316266 | 0.122810 | 0.410612 | -1.015935 | 7.134098 | 1.169849924 | -0.01949715 | 0.98444451 | NA | 2.849979 | 2.819320 | 2.829972 | 2.840464 | 2.818782 | 2.850486 | 7 | 9 | 3 | 10 |
| 780851 | NR_006880 | SNORD3A | small nucleolar RNA, C/D box 3A | snoRNA | . | 33189 | 180710 | . | . | 3.669605 | 4.689815 | 4.442181 | 4.887111 | 8.539323 | 10.840717 | 9.968125 | 11.173800 | 1.113961 | 30.434218 | 0.562835833 | 0.276631637 | 0.78206297 | 0.998865404 | 4.848520 | 5.002612 | 4.902491 | 4.932808 | 4.926998 | 4.935021 | 29 | 35 | 27 | 31 |
| 780852 | NR_003924 | SNORD3B-2 | small nucleolar RNA, C/D box 3B-2 | snoRNA | . | 33190 | . | . | . | 0.156714 | 0.249919 | 0.763104 | 0.179582 | 0.364681 | 0.577699 | 1.712384 | 0.410593 | 2.188957 | 2.868395 | 1.799190675 | 0.62819551 | 0.529875871 | NA | 0.844134 | 1.978782 | 1.491365 | 1.492010 | 1.509713 | 1.493473 | 2 | 2 | 5 | 2 |
| 780853 | NR_006881 | SNORD3C | small nucleolar RNA, C/D box 3C | snoRNA | . | 33191 | . | . | . | 0.107663 | 0.270112 | 0.332567 | 0.229921 | 0.250536 | 0.624376 | 0.746270 | 0.525688 | 1.633456 | 1.788804 | 2.208363229 | 0.320566613 | 0.748538842 | NA | 0.444631 | 1.148375 | 0.822034 | 0.826821 | 0.828805 | 0.828093 | 1 | 2 | 2 | 2 |
| 780854 | NR_006882 | SNORD3D | small nucleolar RNA, C/D box 3D | snoRNA | . | 33192 | . | . | . | 0.155877 | 0.253980 | 0.789659 | 0.180110 | 0.362732 | 0.587085 | 1.771973 | 0.411801 | 2.188957 | 2.868395 | 1.799190675 | 0.62819551 | 0.529875871 | NA | 0.844134 | 1.978782 | 1.491365 | 1.492010 | 1.509713 | 1.493473 | 2 | 2 | 5 | 2 |
| 790955 | NM_00108537 | UQCC3 | ubiquinol-cytochrome c reductase complex assembly factor 3 | protein_coding | NP_001078841.1 | 34399 | 616097 | . | . | 3.496040 | 3.685691 | 3.588389 | 3.251012 | 8.135429 | 8.519640 | 8.052241 | 7.433054 | -1.066656 | 210.824746 | 0.232999323 | -0.39955184 | 0.689486636 | 0.998865404 | 7.765492 | 7.672821 | 7.723889 | 7.744917 | 7.728603 | 7.680173 | 245 | 240 | 187 | 181 |
| 791126 | gene-LOC7911 | LOC791126 | WDR45-like pseudogene | pseudogene | . | . | . | . | . | 0.073363 | 0.038503 | 0.049214 | 0.300221 | 0.170718 | 0.089001 | 0.110436 | 0.686420 | 2.966072 | 6.144741 | 1.330722677 | 1.178723267 | 0.238508388 | NA | 1.631267 | 3.200321 | 2.585635 | 2.573983 | 2.582257 | 2.632939 | 5 | 2 | 3 | 14 |
| 100009676 | NR_024407 | ZBTB11-AS1 | ZBTB11 antisense RNA 1 | lncRNA | . | 48573 | . | . | . | 0.435337 | 0.642082 | 0.261829 | 0.421279 | 1.013047 | 1.484201 | 0.587537 | 0.963205 | -1.608253 | 38.719373 | 0.526264722 | -1.30256514 | 0.192723265 | 0.96621494 | 5.579110 | 4.889137 | 5.274583 | 5.341161 | 5.212236 | 5.263689 | 45 | 61 | 20 | 34 |
| 100033413 | NR_003316 | SNORD116-1 | small nucleolar RNA, C/D box 116-1 | snoRNA | . | 33067 | 605436 | . | . | 1.024194 | 0.614769 | 1.094779 | 1.415020 | 2.383342 | 1.421066 | 2.456652 | 3.235276 | 1.233975 | 3.492648 | 1.593779309 | 0.190310891 | 0.84906552 | NA | 1.644780 | 1.947964 | 1.801217 | 1.797945 | 1.801116 | 1.804812 | 4 | 3 | 3 | 4 |
| 100033414 | NR_003317 | SNORD116-2 | small nucleolar RNA, C/D box 116-2 | snoRNA | . | 33068 | . | . | . | 0.416431 | 1.432442 | 0.747470 | 0.580158 | 0.969051 | 3.311155 | 1.677300 | 1.326463 | -1.433603 | 2.700973 | 1.834732622 | -0.28322691 | 0.777002898 | NA | 1.671431 | 1.148375 | 1.430998 | 1.444703 | 1.433848 | 1.433049 | 2 | 5 | 2 | 2 |
| 100033416 | NR_003319 | SNORD116-4 | small nucleolar RNA, C/D box 116-4 | snoRNA | . | 33070 | . | . | . | 0.004717 | 0.001673 | 0.001729 | 0.001515 | 0.010976 | 0.003866 | 0.003879 | 0.003463 | 1.234655 | 1.002957 | 2.868339486 | 0.106022226 | 0.915564717 | NA | -0.155866 | 0.148375 | -0.001261 | -0.000972 | 0.000053 | -0.000315 | 1 | 1 | 1 | 1 |
| 100033418 | NR_003321 | SNORD116-6 | small nucleolar RNA, C/D box 116-6 | snoRNA | . | 33072 | . | . | . | 1.186414 | 1.052129 | 1.700143 | 2.058855 | 2.760834 | 2.432042 | 3.815072 | 4.707326 | 1.509134 | 5.049766 | 1.337874017 | 0.443779403 | 0.657202081 | NA | 2.008843 | 2.602870 | 2.324842 | 2.321904 | 2.331291 | 2.334281 | 5 | 4 | 5 | 6 |
| 100033424 | NR_003327 | SNORD116-12 | small nucleolar RNA, C/D box 116-12 | snoRNA | . | 33078 | . | . | . | 0.566186 | 0.059376 | 0.490843 | 0.339492 | 1.317539 | 0.137251 | 1.101439 | 0.776208 | 1.256691 | 1.507561 | 2.39030677 | 0.137902842 | 0.890317204 | NA | 0.413393 | 0.751299 | 0.586366 | 0.582479 | 0.588807 | 0.583226 | 2 | 1 | 2 | 1 |
| 100033425 | NR_003328 | SNORD116-13 | small nucleolar RNA, C/D box 116-13 | snoRNA | . | 33079 | . | . | . | 2.128280 | 3.021944 | 1.484217 | 3.304871 | 4.952596 | 6.985358 | 3.330540 | 7.556194 | -1.137113 | 7.603721 | 1.115770547 | -0.16614081 | 0.86804614 | NA | 3.019256 | 2.827809 | 2.926120 | 2.936799 | 2.916087 | 2.937990 | 8 | 10 | 4 | 9 |
| 100033426 | NR_003329 | SNORD116-14 | small nucleolar RNA, C/D box 116-14 | snoRNA | . | 33080 | . | . | . | 3.450834 | 2.350053 | 1.838714 | 1.800885 | 8.030231 | 5.432253 | 4.126022 | 4.117509 | -1.611439 | 7.229617 | 1.137542664 | -0.6051198 | 0.545099397 | NA | 3.156660 | 2.470304 | 2.876816 | 2.862905 | 2.855642 | 2.853644 | 12 | 8 | 5 | 5 |
| 100033430 | NR_003333 | SNORD116-18 | small nucleolar RNA, C/D box 116-18 | snoRNA | . | 33084 | . | . | . | 0.746205 | 0.115628 | 0.198675 | 0.449006 | 1.736449 | 0.267278 | 0.445820 | 1.026598 | -1.076094 | 1.703845 | 2.268349467 | -0.04664372 | 0.962797187 | NA | 0.820514 | 0.715150 | 0.778739 | 0.770724 | 0.771939 | 0.776369 | 3 | 1 | 1 | 2 |
| 100033431 | NR_003334 | SNORD116-20 | small nucleolar RNA, C/D box 116-20 | snoRNA | . | 33086 | . | . | . | 3.379697 | 1.859417 | 2.687951 | 3.873129 | 7.864694 | 4.298126 | 6.031684 | 8.855450 | 1.237076 | 8.941233 | 1.031286202 | 0.297622491 | 0.765991307 | NA | 2.998355 | 3.306201 | 3.164246 | 3.141812 | 3.153534 | 3.169627 | 12 | 6 | 7 | 11 |
| 100033432 | NR_003335 | SNORD116-21 | small nucleolar RNA, C/D box 116-21 | snoRNA | . | 33087 | . | . | . | 8.089207 | 4.412600 | 6.408232 | 9.111527 | 18.823917 | 10.199923 | 14.379887 | 20.832424 | 1.238302 | 20.876761 | 0.696587148 | 0.442677525 | 0.657999001 | NA | 4.220748 | 4.530328 | 4.393626 | 4.344122 | 4.373278 | 4.402494 | 28 | 14 | 17 | 25 |
| 100033433 | NR_003336 | SNORD116-22 | small nucleolar RNA, C/D box 116-22 | snoRNA | . | 33088 | . | . | . | 1.312391 | 2.596934 | 2.317871 | 4.665277 | 3.053988 | 6.002929 | 5.201235 | 10.666602 | 1.645524 | 8.362358 | 1.081642483 | 0.664310723 | 0.50649146 | NA | 2.664814 | 3.376188 | 3.032595 | 3.051468 | 3.045177 | 3.075682 | 5 | 9 | 6 | 13 |
| 100033435 | NR_003338 | SNORD116-24 | small nucleolar RNA, C/D box 116-24 | snoRNA | . | 33090 | . | . | . | 0.464563 | 0.571888 | 0.845341 | 0.558519 | 1.081058 | 1.321944 | 1.896922 | 1.276988 | 1.552428 | 2.293408 | 1.956121338 | 0.324380053 | 0.74565032 | NA | 0.844134 | 1.481107 | 1.182896 | 1.183495 | 1.190762 | 1.184854 | 2 | 2 | 3 | 2 |
| 100033439 | NR_003341 | SNORD116-27 | small nucleolar RNA, C/D box 116-27 | snoRNA | . | 33093 | . | . | . | 0.728784 | 0.093752 | 0.514217 | 0.051106 | 1.695909 | 0.216712 | 1.153887 | 0.116848 | -1.056394 | 1.724672 | 2.259675635 | -0.03502612 | 0.972058914 | NA | 0.820514 | 0.751299 | 0.790628 | 0.782592 | 0.788957 | 0.783375 | 3 | 1 | 2 | 1 |
| 100033819 | NR_030533,rn | MIR675 | microRNA 675 | miRNA | . | 33351 | 615509 | . | . | 0.011213 | 0.017225 | 0.006434 | 0.013736 | 0.026091 | 0.039817 | 0.014437 | 0.031408 | 1.234665 | 3.008871 | 1.708874417 | 0.17796512 | 0.858750369 | NA | 1.429096 | 1.733338 | 1.582415 | 1.583274 | 1.586316 | 1.585224 | 3 | 3 | 3 | 3 |
| 100033820 | NR_003361 | SNORD116-28 | small nucleolar RNA, C/D box 116-28 | snoRNA | . | 33347 | . | . | . | 2.544420 | 0.789668 | 4.000469 | 3.103041 | 5.920970 | 1.825354 | 8.976936 | 7.094734 | 2.088995 | 8.211483 | 1.100955051 | 0.965351804 | 0.334368633 | NA | 2.405477 | 3.475715 | 3.017374 | 2.993815 | 3.038359 | 3.025369 | 9 | 3 | 11 | 9 |
| 100036574 | gene-HMGN1P | HMGN1P3 | high mobility group nucleosome binding domain 1 pseudogene 3 | pseudogene | . | 23343 | . | . | . | 0.951398 | 0.730482 | 0.764205 | 0.617771 | 2.213942 | 1.688541 | 1.714853 | 1.412459 | -1.233137 | 29.123235 | 0.579579144 | -0.52164286 | 0.601919019 | 0.998865404 | 5.006229 | 4.706421 | 4.895780 | 4.860059 | 4.864064 | 4.842701 | 42 | 30 | 25 | 22 |
| 100036575 | gene-HMGN1P | HMGN1P4 | high mobility group nucleosome binding domain 1 pseudogene 4 | pseudogene | . | 39336 | . | . | . | 1.468400 | 0.446357 | 0.674052 | 0.170091 | 3.417027 | 1.031774 | 1.512554 | 0.388893 | -2.269954 | 26.726572 | 0.757772512 | -1.56070961 | 0.118592283 | NA | 5.210562 | 4.037796 | 4.837682 | 4.678723 | 4.716803 | 4.631360 | 65 | 19 | 23 | 6 |
| 100037417 | NM_00108439 | DDTL | D-dopachrome tautomerase like | protein_coding | NP_001077862.1;XP_011528118.1 | 33446 | . | . | . | 14.307762 | 13.425805 | 14.223527 | 15.961364 | 33.294750 | 31.034350 | 31.917184 | 36.493765 | 1.022847 | 996.932120 | 0.144625158 | 0.225346506 | 0.821709724 | 0.998865404 | 9.945162 | 9.977361 | 9.963774 | 9.940717 | 9.909831 | 10.024463 | 1153 | 1047 | 808 | 1019 |
| 100048912 | NR_003529,NF | CDKN2B-AS1 | CDKN2B antisense RNA 1 | lncRNA | . | 34341 | 613149 | . | . | 0.277610 | 0.298372 | 0.391543 | 0.312865 | 0.646010 | 0.689699 | 0.878613 | 0.715330 | 1.236512 | 35.593525 | 0.518596251 | 0.590586815 | 0.554797302 | 0.998865404 | 4.991900 | 5.298886 | 5.134580 | 5.132228 | 5.170658 | 5.155489 | 37 | 34 | 36 | 35 |
| 100049076 | NR_033968 | GUSBP9 | GUSB pseudogene 9 | transcribed_ps | . | 42323 | . | . | . | 1.382095 | 2.113076 | 1.451172 | 1.671718 | 3.216193 | 4.884471 | 3.256389 | 3.822186 | -1.133770 | 31.824468 | 0.560980861 | -0.32287807 | 0.746787584 | 0.998865404 | 5.081051 | 4.897226 | 4.970416 | 5.026707 | 4.974398 | 4.991580 | 31 | 44 | 24 | 30 |
| 100093630 | NR_003584,NF | SNHG8 | small nucleolar RNA host gene 8 | lncRNA | . | 33098 | . | . | . | 46.194173 | 48.883689 | 45.545662 | 44.107288 | 107.495735 | 112.996842 | 102.203147 | 100.846075 | -1.079154 | 783.236930 | 0.146230849 | -0.75155223 | 0.45232038 | 0.998865404 | 9.667243 | 9.557272 | 9.620085 | 9.658515 | 9.592192 | 9.579750 | 911 | 901 | 661 | 700 |
| 100093631 | NR_003580 | GTF2IP4 | general transcription factor IIi pseudogene 4 | transcribed_ps | . | 51716 | . | . | . | 11.001663 | 11.102663 | 13.098724 | 13.192740 | 25.601320 | 25.664301 | 29.393516 | 30.163635 | 1.172943 | 1370.194583 | 0.126332619 | 1.82164524 | 0.068508832 | 0.741130004 | 10.300511 | 10.530651 | 10.355268 | 10.355582 | 10.478233 | 10.476377 | 1452 | 1361 | 1288 | 1385 |
| 100101267 | NM_00109941 | POM121C | POM121 transmembrane nucleoporin C | protein_coding | NP_001092885.2 | 34005 | 615754 | . | . | 7.867836 | 7.997212 | 8.221855 | 8.459625 | 18.308778 | 18.485914 | 18.449602 | 19.341928 | 1.039613 | 1465.394237 | 0.124973743 | 0.448468396 | 0.65381519 | 0.998865404 | 10.488824 | 10.544780 | 10.500085 | 10.503520 | 10.519746 | 10.543551 | 1651 | 1554 | 1279 | 1422 |
| 100101440 | NR_130940 | PMS2P7 | PMS1 homolog 2, mismatch repair system component pseudogen | transcribed_ps | . | 33515 | . | . | . | 2.328927 | 1.355773 | 2.407982 | 2.349886 | 5.419510 | 3.133931 | 5.403442 | 5.372735 | 1.271676 | 44.473960 | 0.488596996 | 0.709646122 | 0.477923611 | 0.998865404 | 5.289655 | 5.639023 | 5.486949 | 5.404661 | 5.489699 | 5.484482 | 57 | 31 | 44 | 46 |
| 100101467 | NM_00111273 | ZSCAN30 | zinc finger and SCAN domain containing 30 | protein_coding | NP_001106205.1;NP_001159484.1;NP_001275640.1;XP_005258240.1;XF | 33517 | . | . | . | 1.155185 | 0.564115 | 1.089593 | 1.242035 | 2.688162 | 1.303978 | 2.445016 | 2.839763 | 1.351009 | 133.105489 | 0.333547656 | 1.301274115 | 0.193164644 | 0.96621494 | 6.822041 | 7.258001 | 7.086244 | 6.871537 | 7.078526 | 7.109385 | 172 | 83 | 128 | 149 |
| 100101478 | NR_130172 | H2BW3P | H2B.W histone 3, pseudogene | transcribed_ps | . | 44390 | . | . | . | 0.314191 | 0.142077 | 0.628012 | 0.311948 | 0.731136 | 0.328418 | 1.409241 | 0.713231 | 1.925668 | 11.051149 | 0.965237632 | 0.979405633 | 0.327379595 | NA | 2.912173 | 3.865403 | 3.441492 | 3.414749 | 3.480245 | 3.437978 | 12 | 5 | 17 | 9 |
| 100101490 | gene-MCTS2P | MCTS2P | MCTS family member 2, pseudogene | pseudogene | . | 49760 | . | . | . | 7.210799 | 7.124601 | 6.253594 | 7.623973 | 16.779825 | 16.468834 | 14.032885 | 17.431307 | -1.046285 | 166.487298 | 0.260221646 | -0.25084671 | 0.801932625 | 0.998865404 | 7.412015 | 7.345761 | 7.392611 | 7.384372 | 7.331200 | 7.407052 | 198 | 182 | 128 | 167 |
| 100101917 | gene-ELF2P1 | ELF2P1 | ELF2 pseudogene 1 | pseudogene | . | 33539 | . | . | . | 0.191492 | 0.073966 | 0.212787 | 0.186484 | 0.445609 | 0.170975 | 0.477489 | 0.426373 | 1.476055 | 12.067970 | 0.90892316 | 0.618035528 | 0.536551919 | NA | 3.282109 | 3.848815 | 3.591949 | 3.553827 | 3.595797 | 3.591394 | 16 | 6 | 13 | 13 |
| 100113373 | gene-HMGN2P | HMGN2P17 | high mobility group nucleosomal binding domain 2 pseudogene 1 | pseudogene | . | 39381 | . | . | . | 0.367357 | 0.481091 | 0.365201 | 0.299049 | 0.854854 | 1.112064 | 0.819501 | 0.683741 | -1.270038 | 14.490749 | 0.807955614 | -0.42684516 | 0.669492112 | NA | 4.019256 | 3.674293 | 3.856068 | 3.876520 | 3.855857 | 3.846887 | 16 | 20 | 12 | 11 |
| 100113378 | NR_003684 | SNORD119 | small nucleolar RNA, C/D box 119 | snoRNA | . | 33558 | . | . | . | 2.553041 | 1.875759 | 3.799166 | 3.352787 | 5.941031 | 4.335901 | 8.525218 | 7.665750 | 1.548119 | 9.117401 | 1.006562427 | 0.626406071 | 0.531048629 | NA | 2.838265 | 3.470304 | 3.172870 | 3.166812 | 3.189189 | 3.185760 | 9 | 7 | 10 | 10 |
| 100113380 | NR_003686 | SNORD125 | small nucleolar RNA, C/D box 125 | snoRNA | . | 33564 | . | . | . | 2.311383 | 1.508563 | 2.338242 | 1.579885 | 5.378684 | 3.487111 | 5.246947 | 3.612220 | 1.071153 | 6.458214 | 1.198243817 | 0.082758037 | 0.934043936 | NA | 2.638039 | 2.742346 | 2.694829 | 2.679965 | 2.695389 | 2.683418 | 9 | 5 | 7 | 5 |
| 100113381 | NR_003687 | SNORD19B | small nucleolar RNA, C/D box 19B | snoRNA | . | 33570 | . | . | . | 5.464487 | 4.908579 | 7.185882 | 4.243159 | 12.716086 | 11.346400 | 16.124910 | 9.701479 | 1.103176 | 16.494458 | 0.765312783 | 0.185104533 | 0.853147046 | NA | 3.969395 | 4.114763 | 4.038648 | 4.031252 | 4.060668 | 4.022780 | 19 | 16 | 19 | 12 |
| 100113386 | NR_027287 | UCKL1-AS1 | UCKL1 antisense RNA 1 | lncRNA | . | 31967 | . | . | . | 0.023973 | 0.049592 | 0.206214 | 0.104530 | 0.055787 | 0.114635 | 0.462738 | 0.238996 | 3.607467 | 11.460945 | 0.972233662 | 1.90384907 | 0.056929844 | NA | 2.316288 | 4.165221 | 3.437313 | 3.452231 | 3.529118 | 3.476580 | 4 | 7 | 21 | 11 |
| 100113393 | NR_003695 | SNORD12B | small nucleolar RNA, C/D box 12B | snoRNA | . | 33573 | . | . | . | 4.684872 | 4.982455 | 8.464910 | 3.331480 | 10.901889 | 11.517168 | 18.995012 | 7.617033 | 1.180461 | 17.644854 | 0.769842396 | 0.310908024 | 0.755870544 | NA | 4.014059 | 4.257993 | 4.119893 | 4.124648 | 4.169325 | 4.098884 | 18 | 18 | 24 | 10 |
| 100113407 | NM_00110082 | TMEM170B | transmembrane protein 170B | protein_coding | NP_001094299.1 | 34244 | . | . | . | 0.404807 | 0.228866 | 0.446909 | 0.180313 | 0.942002 | 0.529035 | 1.002852 | 0.412264 | -1.022626 | 88.010466 | 0.439528642 | -0.07343861 | 0.941457096 | 0.998865404 | 6.473391 | 6.445682 | 6.519597 | 6.356436 | 6.545324 | 6.310031 | 131 | 69 | 108 | 47 |
| 100124516 | NR_003701 | SNORD58C | small nucleolar RNA, C/D box 58C | snoRNA | . | 33613 | . | . | . | 5.331250 | 8.277411 | 10.228758 | 3.012028 | 12.406036 | 19.133604 | 22.953037 | 6.886644 | -1.068803 | 19.252503 | 0.771994636 | -0.12434783 | 0.901039881 | NA | 4.312060 | 4.220434 | 4.238705 | 4.275592 | 4.293059 | 4.208789 | 18 | 26 | 25 | 8 |
| 100124533 | NR_003703 | SCARNA27 | small Cajal body-specific RNA 27 | guide_RNA | . | 33614 | . | . | . | 1.037575 | 0.394016 | 0.496965 | 1.162874 | 2.414479 | 0.910785 | 1.115175 | 2.658772 | 1.232559 | 3.457246 | 1.634513507 | 0.18455436 | 0.853578586 | NA | 1.631267 | 1.932304 | 1.793133 | 1.781341 | 1.783769 | 1.797375 | 5 | 2 | 2 | 5 |
| 100124534 | NR_003704 | SNORA84 | small nucleolar RNA, H/ACA box 84 | snoRNA | . | 33615 | . | . | . | 0.583707 | 0.297184 | 0.681783 | 0.756938 | 1.358310 | 0.686955 | 1.529902 | 1.730649 | 1.488840 | 2.777185 | 1.782378592 | 0.322147257 | 0.747341139 | NA | 1.156660 | 1.733338 | 1.464780 | 1.461218 | 1.468604 | 1.467534 | 3 | 2 | 3 | 3 |
| 100124536 | NR_003706 | SNORA38B | small nucleolar RNA, H/ACA box 38B | snoRNA | . | 33617 | . | . | . | 0.128195 | 0.209190 | 1.154036 | 0.493917 | 0.298315 | 0.483553 | 2.589624 | 1.129283 | 4.380364 | 2.419598 | 2.050691654 | 1.039186408 | 0.298718051 | NA | -0.155866 | 1.978782 | 1.229833 | 1.230235 | 1.252391 | 1.236103 | 1 | 1 | 5 | 2 |
| 100125288 | NM_00110316 | ZGLP1 | zinc finger GATA like protein 1 | protein_coding | NP_001096637.1 | 37245 | 611639 | . | . | 0.326001 | 0.326990 | 0.660612 | 0.410385 | 0.758617 | 0.755851 | 1.482394 | 0.938296 | 1.622068 | 27.057520 | 0.608435842 | 1.146931192 | 0.25141006 | 0.993760899 | 4.365657 | 5.066070 | 4.714335 | 4.712743 | 4.796834 | 4.735214 | 24 | 22 | 36 | 24 |
| 100125387 | gene-HNRNPRI | HNRNPRP1 | heterogeneous nuclear ribonucleoprotein R pseudogene 1 | pseudogene | . | 44907 | . | . | . | 0.015231 | 0.026264 | 0.039751 | 0.007268 | 0.035442 | 0.060712 | 0.089200 | 0.016618 | -1.003532 | 2.258427 | 1.991984473 | -0.00255353 | 0.997962579 | NA | 1.175403 | 1.175234 | 1.166206 | 1.171202 | 1.174041 | 1.163259 | 2 | 3 | 3 | 1 |
| 100125556 | NR_024250,NF | FAM86JP | family with sequence similarity 86 member J, pseudogene | transcribed_ps | . | 44097 | . | . | . | 1.522398 | 1.493276 | 2.105427 | 2.822394 | 3.542682 | 3.451774 | 4.724517 | 6.453066 | 1.611353 | 124.553451 | 0.299003192 | 2.301891037 | 0.021341318 | 0.484497055 | 6.576422 | 7.263725 | 6.862213 | 6.851578 | 6.971374 | 7.070972 | 112 | 101 | 117 | 162 |
| 100126311 | NR_030599,rn | MIR147B | microRNA 147b | miRNA | . | 33655 | . | . | . | 0.146466 | 0.322526 | 0.221791 | 0.221902 | 0.340833 | 0.745531 | 0.497692 | 0.507352 | -1.215015 | 2.454711 | 1.887245685 | -0.1488804 | 0.881648006 | NA | 1.429096 | 1.148375 | 1.299640 | 1.300454 | 1.298230 | 1.297456 | 3 | 3 | 2 | 2 |
| 100126321 | NR_030627,rn | MIR922 | microRNA 922 | miRNA | . | 33672 | . | . | . | 0.011568 | 0.004485 | 0.009255 | 0.009420 | 0.026919 | 0.010368 | 0.020768 | 0.021537 | 1.234663 | 2.005914 | 2.070995667 | 0.146852664 | 0.883253841 | NA | 0.844134 | 1.148375 | 0.998092 | 0.998667 | 1.000707 | 0.999974 | 2 | 2 | 2 | 2 |
| 100126332 | NR_030641,rn | MIR943 | microRNA 943 | miRNA | . | 33689 | . | . | . | 0.452736 | 0.000304 | 0.189678 | 0.206722 | 1.053534 | 0.000703 | 0.425632 | 0.472645 | 1.250132 | 1.991338 | 2.100718025 | 0.153318985 | 0.87814673 | NA | 0.820514 | 1.148375 | 0.993554 | 0.985525 | 0.991973 | 0.991243 | 3 | 1 | 2 | 2 |
| 100126338 | NR_030633,rn | MIR937 | microRNA 937 | miRNA | . | 33680 | . | . | . | 0.006419 | 0.370958 | 0.533759 | 0.264618 | 0.014938 | 0.857487 | 1.197740 | 0.605020 | 1.830661 | 2.574650 | 1.882789928 | 0.463336015 | 0.643123535 | NA | 0.867373 | 1.733338 | 1.340114 | 1.349386 | 1.352319 | 1.351266 | 1 | 3 | 3 | 3 |
| 100126351 | NR_030635,rn | MIR939 | microRNA 939 | miRNA | . | 33682 | . | . | . | 3.897191 | 3.774746 | 3.277091 | 5.981183 | 9.068922 | 8.725497 | 7.353696 | 13.675265 | 1.182054 | 13.193072 | 0.843969501 | 0.285905883 | 0.774950208 | NA | 3.597285 | 3.836248 | 3.714301 | 3.713874 | 3.711820 | 3.735591 | 14 | 13 | 10 | 16 |
| 100126784 | NR_015384 | LOC10012678 | uncharacterized LOC100126784 | lncRNA | . | . | . | . | . | 0.762184 | 0.853207 | 0.456294 | 0.625224 | 1.773633 | 1.972227 | 1.023910 | 1.429500 | -1.501721 | 79.247141 | 0.366666526 | -1.59986498 | 0.109628541 | 0.843391194 | 6.572938 | 5.983930 | 6.342629 | 6.373187 | 6.221932 | 6.283850 | 104 | 108 | 47 | 68 |
| 100126793 | NR_004431,NF | GHRLOS | ghrelin opposite strand/antisense RNA | lncRNA | . | 33885 | 618445 | . | . | 0.116801 | 0.263001 | 0.207231 | 0.104157 | 0.271801 | 0.607939 | 0.465019 | 0.238144 | -1.238705 | 10.655090 | 0.969219832 | -0.31864076 | 0.749998938 | NA | 3.558919 | 3.251698 | 3.396015 | 3.431990 | 3.417653 | 3.390395 | 9 | 17 | 11 | 6 |
| 100127888 | NR_024470 | SLCO4A1-AS1 | SLCO4A1 antisense RNA 1 | lncRNA | . | 40537 | . | . | . | 2.242579 | 2.840378 | 3.137948 | 3.524611 | 5.218575 | 6.565661 | 7.041464 | 8.058602 | 1.291734 | 130.379234 | 0.292174595 | 1.264001441 | 0.20622951 | 0.970482251 | 6.831011 | 7.198758 | 6.932198 | 7.010010 | 7.042921 | 7.083902 | 116 | 137 | 121 | 145 |
| 100127922 | gene-SUMO2P | SUMO2P6 | SUMO2 pseudogene 6 | pseudogene | . | 39016 | . | . | . | 0.251936 | 0.060086 | 0.083864 | 0.037194 | 0.586266 | 0.138893 | 0.188188 | 0.085039 | -2.113398 | 2.607689 | 1.911185074 | -0.56486668 | 0.572164446 | NA | 1.820514 | 0.751299 | 1.407921 | 1.392061 | 1.394249 | 1.388624 | 6 | 2 | 2 | 1 |
| 100127924 | gene-RPS15AP | RPS15AP14 | ribosomal protein S15a pseudogene 14 | pseudogene | . | 36629 | . | . | . | 0.147765 | 0.176503 | 0.330421 | 0.166107 | 0.343855 | 0.407996 | 0.741456 | 0.379784 | 1.658520 | 3.583858 | 1.589233434 | 0.459275663 | 0.646036222 | NA | 1.429096 | 2.161867 | 1.822455 | 1.823360 | 1.836777 | 1.825414 | 3 | 3 | 5 | 3 |
| 100127952 | gene-ATF4P4 | ATF4P4 | activating transcription factor 4 pseudogene 4 | pseudogene | . | 787 | . | . | . | 0.377015 | 0.164927 | 0.205720 | 0.282923 | 0.877329 | 0.381236 | 0.461631 | 0.646870 | -1.127537 | 11.695650 | 0.924032549 | -0.18741211 | 0.851337528 | NA | 3.631267 | 3.459419 | 3.574990 | 3.530036 | 3.538815 | 3.554238 | 20 | 8 | 8 | 12 |
| 100127983 | NM_00119097 | C8orf88 | chromosome 8 open reading frame 88 | protein_coding | NP_001177901.1;NP_001350204.1 | 44672 | . | . | . | 6.419250 | 5.939270 | 5.347449 | 4.183964 | 14.937860 | 13.728886 | 11.999520 | 9.566138 | -1.315445 | 178.191809 | 0.258994728 | -1.52725363 | 0.126697955 | 0.874064017 | 7.660784 | 7.267003 | 7.550556 | 7.510287 | 7.462680 | 7.363771 | 243 | 209 | 151 | 126 |
| 100127993 | gene-RPL15P5 | RPL15P5 | ribosomal protein L15 pseudogene 5 | pseudogene | . | 35746 | . | . | . | 1.031782 | 0.915198 | 0.529123 | 1.204343 | 2.400999 | 2.115520 | 1.187337 | 2.753587 | -1.138348 | 29.793676 | 0.602787157 | -0.31012911 | 0.75646278 | 0.998865404 | 4.989254 | 4.798300 | 4.913236 | 4.895836 | 4.846292 | 4.930045 | 39 | 32 | 15 | 36 |
| 100128010 | gene-SUMO2P | SUMO2P17 | SUMO2 pseudogene 17 | pseudogene | . | 49358 | . | . | . | 3.094556 | 2.210284 | 1.973670 | 2.798334 | 7.201160 | 5.109170 | 4.428859 | 6.398058 | -1.127990 | 46.256619 | 0.474675185 | -0.36604884 | 0.714328634 | 0.998865404 | 5.616092 | 5.441824 | 5.573345 | 5.510418 | 5.496594 | 5.545781 | 66 | 44 | 32 | 47 |
| 100128012 | gene-ISCA1P4 | ISCA1P4 | iron-sulfur cluster assembly 1 pseudogene 4 | pseudogene | . | 38025 | . | . | . | 0.407297 | 0.278391 | 0.070983 | 0.155734 | 0.947797 | 0.643513 | 0.159285 | 0.356067 | -2.754035 | 5.167896 | 1.379837477 | -1.0592166 | 0.289501145 | NA | 2.923304 | 1.459419 | 2.401834 | 2.391566 | 2.372949 | 2.376716 | 10 | 7 | 2 | 3 |
| 100128025 | NR_040250 | WWTR1-AS1 | WWTR1 antisense RNA 1 | lncRNA | . | 41035 | . | . | . | 0.419992 | 0.120440 | 0.441791 | 0.288240 | 0.977339 | 0.278403 | 0.991367 | 0.659026 | 1.374625 | 17.304043 | 0.808869194 | 0.567506593 | 0.570370049 | NA | 3.861296 | 4.327309 | 4.122817 | 4.050623 | 4.127601 | 4.093898 | 26 | 7 | 21 | 15 |
| 100128060 | gene-RPS7P10 | RPS7P10 | ribosomal protein S7 pseudogene 10 | pseudogene | . | 39903 | . | . | . | 0.224900 | 0.501649 | 0.288191 | 0.301675 | 0.523351 | 1.159585 | 0.646691 | 0.689744 | -1.225692 | 7.407860 | 1.131913755 | -0.25938096 | 0.79534132 | NA | 3.029593 | 2.733338 | 2.877903 | 2.905179 | 2.886023 | 2.883750 | 6 | 12 | 6 | 6 |
| 100128062 | gene-SLC2A3P | SLC2A3P1 | solute carrier family 2 member 3 pseudogene 1 | pseudogene | . | 11008 | . | . | . | 0.031904 | 0.033421 | 0.032899 | 0.029569 | 0.074241 | 0.077255 | 0.073824 | 0.067607 | -1.070164 | 1.739247 | 2.228774873 | -0.04389516 | 0.964987971 | NA | 0.844134 | 0.751299 | 0.796508 | 0.797062 | 0.799027 | 0.793436 | 2 | 2 | 2 | 1 |
| 100128071 | NM_00116767 | FAM229A | family with sequence similarity 229 member A | protein_coding | NP_001161148.1 | 44652 | . | . | . | 1.632119 | 1.259918 | 2.244256 | 1.331825 | 3.798007 | 2.912356 | 5.036047 | 3.045059 | 1.213912 | 35.631453 | 0.543317365 | 0.514734351 | 0.606738665 | 0.998865404 | 5.006229 | 5.289999 | 5.149208 | 5.113821 | 5.197130 | 5.118818 | 42 | 30 | 43 | 27 |
| 100128076 | NR_015444 | LOC10012807 | protein tyrosine phosphatase pseudogene | transcribed_ps | . | . | . | . | . | 0.821167 | 1.175757 | 1.180454 | 1.242101 | 1.910888 | 2.717814 | 2.648905 | 2.839915 | 1.210399 | 69.715284 | 0.389195138 | 0.707826734 | 0.479052862 | 0.998865404 | 5.980095 | 6.253753 | 6.053307 | 6.132452 | 6.133744 | 6.144337 | 60 | 80 | 65 | 73 |
| 100128079 | NR_147898 | LOC10012807 | uncharacterized LOC100128079 | lncRNA | . | . | . | . | . | 0.123945 | 0.128628 | 0.184711 | 0.035342 | 0.288426 | 0.297328 | 0.414486 | 0.080805 | 1.001795 | 4.502278 | 1.443995283 | 0.001792081 | 0.998570127 | NA | 2.166062 | 2.175015 | 2.163703 | 2.165080 | 2.175005 | 2.153702 | 5 | 5 | 6 | 2 |
| 100128108 | NR_168283 | LOC10012810 | ubiquitin-conjugating enzyme E2Q2 pseudogene | transcribed_ps | . | . | . | . | . | 2.600481 | 1.789669 | 1.784809 | 2.627985 | 6.051427 | 4.136899 | 4.005059 | 6.008574 | -1.016377 | 37.126451 | 0.52797968 | -0.04438855 | 0.964594685 | 0.998865404 | 5.226376 | 5.202274 | 5.241747 | 5.186940 | 5.185609 | 5.238426 | 51 | 33 | 26 | 41 |
| 100128140 | gene-RPS4XP1 | RPS4XP17 | ribosomal protein S4X pseudogene 17 | pseudogene | . | 36789 | . | . | . | 0.014469 | 0.082863 | 0.081280 | 0.213423 | 0.033670 | 0.191542 | 0.182390 | 0.487965 | 2.404518 | 3.087157 | 1.773701975 | 0.713619117 | 0.475462733 | NA | 0.867373 | 2.121007 | 1.597567 | 1.606871 | 1.604792 | 1.623355 | 1 | 3 | 2 | 6 |
| 100128191 | NR_027157 | TMPO-AS1 | TMPO antisense RNA 1 | lncRNA | . | 44158 | . | . | . | 1.694402 | 1.691520 | 1.824343 | 1.760491 | 3.942944 | 3.910025 | 4.093774 | 4.025153 | 1.041455 | 172.054099 | 0.251335069 | 0.233155287 | 0.815640825 | 0.998865404 | 7.396977 | 7.455859 | 7.419480 | 7.415552 | 7.443559 | 7.425438 | 195 | 181 | 156 | 161 |
| 100128288 | NR_024447 | LOC10012828 | uncharacterized LOC100128288 | lncRNA | . | . | . | . | . | 0.028905 | 0.007889 | 0.015230 | 0.017368 | 0.067264 | 0.018235 | 0.034176 | 0.039711 | -1.076094 | 1.703845 | 2.268349467 | -0.04664372 | 0.962797187 | NA | 0.820514 | 0.715150 | 0.778739 | 0.770724 | 0.771939 | 0.776369 | 3 | 1 | 1 | 2 |
| 100128292 | NR_024585 | DLG5-AS1 | DLG5 antisense RNA 1 | lncRNA | . | 45109 | . | . | . | 0.056415 | 0.026705 | 0.044588 | 0.154266 | 0.131281 | 0.061730 | 0.100055 | 0.352711 | 2.205143 | 7.098142 | 1.209847634 | 0.942988215 | 0.345686931 | NA | 2.147197 | 3.287859 | 2.804971 | 2.789559 | 2.798772 | 2.844151 | 7 | 3 | 4 | 14 |
| 100128340 | NR_149047 | LOC10012834 | uncharacterized LOC100128340 | lncRNA | . | . | . | . | . | 1.268508 | 1.552714 | 1.864213 | 1.392624 | 2.951871 | 3.589167 | 4.183239 | 3.184069 | 1.137283 | 29.808313 | 0.573200321 | 0.32378046 | 0.746104251 | 0.998865404 | 4.801350 | 4.987909 | 4.872779 | 4.895498 | 4.919299 | 4.879675 | 29 | 33 | 32 | 25 |
| 100128361 | NR_036505 | LOC10012836 | uncharacterized LOC100128361 | lncRNA | . | . | . | . | . | 0.020562 | 0.013843 | 0.028167 | 0.022236 | 0.047848 | 0.031999 | 0.063206 | 0.050841 | 1.234663 | 2.005914 | 2.070995667 | 0.146852664 | 0.883253841 | NA | 0.844134 | 1.148375 | 0.998092 | 0.998667 | 1.000707 | 0.999974 | 2 | 2 | 2 | 2 |
| 100128385 | NR_024376 | FAM225B | family with sequence similarity 225 member B | lncRNA | . | 21865 | . | . | . | 0.220211 | 0.220444 | 0.197851 | 0.217493 | 0.512440 | 0.509566 | 0.443971 | 0.497272 | -1.069520 | 39.464852 | 0.493376245 | -0.19653168 | 0.844194038 | 0.998865404 | 5.350383 | 5.252966 | 5.308933 | 5.308707 | 5.293535 | 5.303568 | 47 | 44 | 32 | 37 |
| 100128494 | NR_104178 | LOC10012849 | uncharacterized LOC100128494 | lncRNA | . | . | . | . | . | 0.053723 | 0.011967 | 0.025123 | 0.046362 | 0.125015 | 0.027662 | 0.056376 | 0.106001 | 1.017298 | 4.879848 | 1.419531655 | 0.01742964 | 0.986093863 | NA | 2.273436 | 2.300113 | 2.302135 | 2.274146 | 2.282045 | 2.295142 | 9 | 2 | 3 | 6 |
| 100128525 | gene-UQCRFS1 | UQCRFS1P1 | ubiquinol-cytochrome c reductase, Rieske iron-sulfur polypeptide | pseudogene | . | 12588 | . | . | . | 5.756876 | 4.990384 | 3.886329 | 5.973320 | 13.396485 | 11.535495 | 8.720810 | 13.657288 | -1.104966 | 163.074266 | 0.284259914 | -0.50658564 | 0.612445583 | 0.998865404 | 7.420310 | 7.274793 | 7.398085 | 7.334165 | 7.242916 | 7.402638 | 212 | 171 | 107 | 175 |
| 100128535 | gene-RPS18P1 | RPS18P14 | ribosomal protein S18 pseudogene 14 | pseudogene | . | 36386 | . | . | . | 0.269893 | 0.569528 | 0.137234 | 0.128599 | 0.628051 | 1.316490 | 0.307950 | 0.294026 | -3.471614 | 4.959532 | 1.446610154 | -1.24125128 | 0.214512935 | NA | 2.945312 | 1.148375 | 2.322986 | 2.345708 | 2.310154 | 2.309144 | 6 | 11 | 2 | 2 |
| 100128553 | NM_198495 | CTAGE4 | CTAGE family member 4 | protein_coding | NP_940897.2 | 24772 | 608910 | . | . | 0.467237 | 0.368502 | 0.557804 | 0.313952 | 1.087281 | 0.851808 | 1.251696 | 0.717815 | 1.024729 | 35.315362 | 0.547249372 | 0.064398277 | 0.948653102 | 0.998865404 | 5.122310 | 5.161867 | 5.151616 | 5.117207 | 5.176357 | 5.097883 | 45 | 33 | 40 | 24 |
| 100128594 | NR_146556,NF | TMSB15B-AS1 | TMSB15B antisense RNA 1 | lncRNA | . | 53858 | . | . | . | 0.311566 | 0.323348 | 0.505352 | 0.441381 | 0.725027 | 0.747434 | 1.133995 | 1.009165 | 1.407469 | 15.654324 | 0.772552456 | 0.638278273 | 0.523292553 | NA | 3.700498 | 4.194412 | 3.948406 | 3.948346 | 3.976454 | 3.966294 | 15 | 14 | 17 | 16 |
| 100128640 | NR_028389 | ACVR2B-AS1 | ACVR2B antisense RNA 1 | lncRNA | . | 44161 | . | . | . | 0.721759 | 0.604436 | 0.705891 | 0.610349 | 1.679563 | 1.397181 | 1.584000 | 1.395491 | -1.012345 | 52.464165 | 0.43535217 | -0.04066024 | 0.96756676 | 0.998865404 | 5.721207 | 5.705270 | 5.730607 | 5.696484 | 5.726184 | 5.696258 | 66 | 52 | 49 | 45 |
| 100128665 | gene-FYTTD1P | FYTTD1P1 | forty-two-three domain containing 1 pseudogene 1 | pseudogene | . | 38035 | . | . | . | 0.103501 | 0.161416 | 0.284255 | 0.189203 | 0.240850 | 0.373120 | 0.637860 | 0.432591 | 1.778453 | 15.683765 | 0.793282494 | 1.04707082 | 0.295066917 | NA | 3.497331 | 4.327309 | 3.920330 | 3.944449 | 3.987306 | 3.953722 | 10 | 15 | 21 | 15 |
| 100128675 | NR_024561,NF | HPN-AS1 | HPN antisense RNA 1 | lncRNA | . | 47041 | . | . | . | 2.134514 | 2.237835 | 2.810631 | 3.321450 | 4.967103 | 5.172856 | 6.306973 | 7.594102 | 1.500317 | 158.345899 | 0.263473459 | 2.221353502 | 0.026327028 | 0.531274802 | 6.985014 | 7.569966 | 7.186973 | 7.231101 | 7.383142 | 7.358563 | 137 | 145 | 170 | 173 |
| 100128682 | NR_040046 | LINC01531 | long intergenic non-protein coding RNA 1531 | lncRNA | . | 51270 | . | . | . | 1.334494 | 1.313834 | 0.870463 | 0.828743 | 3.105422 | 3.036986 | 1.953293 | 1.894822 | -1.577645 | 112.770502 | 0.305090091 | -2.15599412 | 0.031084124 | 0.560017015 | 7.108789 | 6.451470 | 6.889872 | 6.879335 | 6.744769 | 6.730286 | 161 | 147 | 78 | 80 |
| 100128731 | NM_00113469 | OST4 | oligosaccharyltransferase complex subunit 4, non-catalytic | protein_coding | NP_001128165.1 | 32483 | . | . | . | 207.332565 | 189.856522 | 177.223511 | 210.851089 | 482.471405 | 438.861847 | 397.684418 | 482.085968 | -1.038419 | 2703.577710 | 0.127605299 | -0.4262277 | 0.669941938 | 0.998865404 | 11.427661 | 11.373132 | 11.453508 | 11.374569 | 11.315840 | 11.448115 | 3325 | 2829 | 2118 | 2690 |
| 100128750 | NR_046205 | RBPMS-AS1 | RBPMS antisense RNA 1 | lncRNA | . | 48721 | . | . | . | 0.310503 | 0.206306 | 1.067475 | 0.277568 | 0.722553 | 0.476886 | 2.395383 | 0.634625 | 2.455689 | 13.905679 | 0.908631911 | 1.42646097 | 0.153735325 | NA | 3.003609 | 4.306796 | 3.735900 | 3.721859 | 3.827211 | 3.732281 | 11 | 7 | 27 | 8 |
| 100128771 | gene-EIF4BP3 | EIF4BP3 | eukaryotic translation initiation factor 4B pseudogene 3 | pseudogene | . | 37936 | . | . | . | 0.714278 | 0.757959 | 0.599429 | 0.885947 | 1.662155 | 1.752057 | 1.345101 | 2.025614 | 1.000836 | 46.193036 | 0.467527239 | 0.002579573 | 0.997941801 | 0.999862634 | 5.530179 | 5.529027 | 5.524833 | 5.532923 | 5.498397 | 5.558787 | 52 | 51 | 33 | 51 |
| 100128775 | gene-NEDD8P1 | NEDD8P1 | NEDD8 pseudogene 1 | pseudogene | . | 54555 | . | . | . | 0.219899 | 0.206909 | 0.089860 | 0.318427 | 0.511714 | 0.478279 | 0.201642 | 0.728045 | -1.027040 | 4.418971 | 1.448485533 | -0.02657402 | 0.978799495 | NA | 2.166062 | 2.121007 | 2.145780 | 2.147148 | 2.136769 | 2.155024 | 5 | 5 | 2 | 6 |
| 100128782 | NR_023389,NF | LINC00476 | long intergenic non-protein coding RNA 476 | lncRNA | . | 27858 | . | . | . | 3.333414 | 3.498728 | 3.912808 | 3.549721 | 7.756990 | 8.087465 | 8.780229 | 8.116015 | 1.094500 | 161.299928 | 0.261465231 | 0.49823873 | 0.618315788 | 0.998865404 | 7.266882 | 7.397373 | 7.293718 | 7.331493 | 7.372384 | 7.325694 | 169 | 174 | 155 | 149 |
| 100128822 | NR_027387 | LINC01003 | long intergenic non-protein coding RNA 1003 | lncRNA | . | 48957 | . | . | . | 1.727986 | 1.227911 | 1.668174 | 1.867737 | 4.021095 | 2.838372 | 3.743335 | 4.270360 | 1.180120 | 72.429855 | 0.383815323 | 0.622523067 | 0.533597981 | 0.998865404 | 6.053521 | 6.293533 | 6.193711 | 6.111112 | 6.181552 | 6.205875 | 90 | 59 | 65 | 77 |
| 100128834 | gene-CLK3P2 | CLK3P2 | CDC like kinase 3 pseudogene 2 | pseudogene | . | 49786 | . | . | . | 0.101493 | 0.012039 | 0.016987 | 0.256548 | 0.236178 | 0.027829 | 0.038117 | 0.586566 | 2.241427 | 4.271400 | 1.617311691 | 0.719970925 | 0.471542898 | NA | 1.397517 | 2.562749 | 2.078960 | 2.062783 | 2.064694 | 2.107131 | 5 | 1 | 1 | 10 |
| 100128881 | NR_036480 | VPS9D1-AS1 | VPS9D1 antisense RNA 1 | lncRNA | . | 48915 | . | . | . | 0.162255 | 0.272814 | 0.302876 | 0.352345 | 0.377575 | 0.630621 | 0.679645 | 0.805595 | 1.443549 | 15.439287 | 0.789221523 | 0.671067033 | 0.502177828 | NA | 3.661494 | 4.187832 | 3.911746 | 3.940007 | 3.946148 | 3.954565 | 11 | 17 | 15 | 18 |
| 100128890 | NR_027423 | FAM66B | family with sequence similarity 66 member B | lncRNA | . | 28890 | . | . | . | 0.108913 | 0.057722 | 0.082536 | 0.073161 | 0.253444 | 0.133426 | 0.185209 | 0.167273 | -1.083920 | 11.534768 | 0.911783634 | -0.12750689 | 0.89853922 | NA | 3.583318 | 3.470304 | 3.547396 | 3.514362 | 3.528701 | 3.525072 | 18 | 9 | 10 | 10 |
| 100128927 | NM_00113760 | ZBTB42 | zinc finger and BTB domain containing 42 | protein_coding | NP_001131073.1;NP_001357271.1 | 32550 | 613915 | . | . | 0.891635 | 0.790020 | 0.743926 | 0.930119 | 2.074870 | 1.826167 | 1.669349 | 2.126608 | -1.009741 | 95.262533 | 0.333922498 | -0.04188068 | 0.966593818 | 0.998865404 | 6.581215 | 6.566421 | 6.593187 | 6.566238 | 6.538519 | 6.604417 | 117 | 97 | 73 | 99 |
| 100128936 | gene-RPL10AP | RPL10AP6 | ribosomal protein L10a pseudogene 6 | pseudogene | . | 36504 | . | . | . | 0.288753 | 0.196113 | 0.269888 | 0.475009 | 0.671940 | 0.453323 | 0.605621 | 1.086052 | 1.516950 | 7.286944 | 1.132054839 | 0.531045886 | 0.595386982 | NA | 2.533720 | 3.134756 | 2.856448 | 2.845699 | 2.856341 | 2.872981 | 8 | 5 | 6 | 10 |
| 100128961 | gene-GAPDHP | GAPDHP62 | glyceraldehyde 3 phosphate dehydrogenase pseudogene 62 | pseudogene | . | 4156 | . | . | . | 1.152527 | 0.966642 | 1.173999 | 1.547302 | 2.681978 | 2.234435 | 2.634420 | 3.537722 | 1.273712 | 45.216049 | 0.470956558 | 0.741128275 | 0.458615667 | 0.998865404 | 5.314065 | 5.662477 | 5.486274 | 5.457793 | 5.487670 | 5.538351 | 50 | 39 | 38 | 54 |
| 100128966 | NR_146628 | LOC10012896 | uncharacterized LOC100128966 | lncRNA | . | . | . | . | . | 0.481963 | 0.760249 | 0.406060 | 0.471432 | 1.121548 | 1.757349 | 0.911186 | 1.077873 | -1.359689 | 9.792187 | 0.982185974 | -0.45131662 | 0.65176137 | NA | 3.497331 | 3.051647 | 3.289154 | 3.312750 | 3.286784 | 3.288753 | 10 | 15 | 7 | 8 |
| 100129028 | gene-RPLP1P4 | RPLP1P4 | ribosomal protein lateral stalk subunit P1 pseudogene 4 | pseudogene | . | 36939 | . | . | . | 0.473506 | 1.062792 | 0.867280 | 1.422737 | 1.101868 | 2.456692 | 1.946150 | 3.252919 | 1.517633 | 15.960118 | 0.804625049 | 0.747955107 | 0.454487236 | NA | 3.668126 | 4.263648 | 3.945766 | 3.990289 | 3.978752 | 4.014554 | 9 | 19 | 13 | 22 |
| 100129034 | NR_027406 | LOC10012903 | uncharacterized LOC100129034 | lncRNA | . | . | . | . | . | 2.344332 | 1.818859 | 1.765346 | 1.855902 | 5.455357 | 4.204374 | 3.961386 | 4.243300 | -1.167564 | 346.923338 | 0.205254637 | -1.08890141 | 0.276197364 | 0.998865404 | 8.545641 | 8.322701 | 8.552906 | 8.399346 | 8.380989 | 8.401687 | 486 | 351 | 273 | 306 |
| 100129075 | NR_027123 | KTN1-AS1 | KTN1 antisense RNA 1 | lncRNA | . | 19842 | . | . | . | 0.879625 | 0.968115 | 0.777174 | 0.882593 | 2.046922 | 2.237841 | 1.743956 | 2.017945 | -1.132891 | 41.822523 | 0.481607836 | -0.37376621 | 0.708578261 | 0.998865404 | 5.473964 | 5.292767 | 5.390431 | 5.404969 | 5.368782 | 5.386720 | 49 | 50 | 32 | 39 |
| 100129141 | gene-RPL21P3 | RPL21P37 | ribosomal protein L21 pseudogene 37 | pseudogene | . | 35704 | . | . | . | 0.194022 | 0.138893 | 0.117530 | 0.104861 | 0.451497 | 0.321056 | 0.263734 | 0.239753 | -1.412257 | 2.671821 | 1.820869166 | -0.27349696 | 0.784471225 | NA | 1.644780 | 1.148375 | 1.428682 | 1.425364 | 1.423205 | 1.422411 | 4 | 3 | 2 | 2 |
| 100129195 | NR_103455,NF | ZSCAN16-AS1 | ZSCAN16 antisense RNA 1 | lncRNA | . | 48982 | . | . | . | 1.359440 | 1.292736 | 0.978465 | 1.280155 | 3.163473 | 2.988218 | 2.195646 | 2.926922 | -1.171364 | 77.175301 | 0.366577452 | -0.62248643 | 0.533622061 | 0.998865404 | 6.380267 | 6.150749 | 6.300084 | 6.282145 | 6.220874 | 6.279815 | 99 | 87 | 54 | 75 |
| 100129196 | NR_034182 | MATN1-AS1 | MATN1 antisense RNA 1 | lncRNA | . | 40364 | . | . | . | 0.840353 | 0.798722 | 0.303993 | 0.551011 | 1.955536 | 1.846281 | 0.682152 | 1.269172 | -1.941450 | 40.339783 | 0.516681643 | -1.85246414 | 0.063959202 | 0.731304863 | 5.736193 | 4.774776 | 5.388503 | 5.377304 | 5.252258 | 5.313226 | 63 | 56 | 17 | 33 |
| 100129203 | NR_110295 | LOC10012920 | uncharacterized LOC100129203 | lncRNA | . | . | . | . | . | 0.320755 | 0.366419 | 0.328271 | 0.311567 | 0.746411 | 0.846995 | 0.736630 | 0.712360 | -1.081149 | 20.760583 | 0.673840832 | -0.16704982 | 0.867330852 | NA | 4.431047 | 4.318004 | 4.372756 | 4.386496 | 4.375947 | 4.369689 | 23 | 25 | 18 | 18 |
| 100129215 | XR_109334 | LOC10012921 | uncharacterized LOC100129215 | lncRNA | . | . | . | . | . | 0.071377 | 0.131391 | 0.252440 | 0.237992 | 0.166098 | 0.303716 | 0.566468 | 0.544141 | 2.388306 | 14.541561 | 0.827176656 | 1.518403176 | 0.128912792 | NA | 3.104339 | 4.356363 | 3.798098 | 3.821995 | 3.863257 | 3.861976 | 7 | 12 | 18 | 19 |
| 100129224 | gene-SOWAHC | SOWAHCP4 | . | pseudogene | . | 54508 | . | . | . | 0.082121 | 0.146618 | 0.357039 | 0.126730 | 0.191098 | 0.338915 | 0.801185 | 0.289753 | 1.991514 | 10.141897 | 1.005848585 | 0.988086649 | 0.323110214 | NA | 2.760365 | 3.755755 | 3.292060 | 3.306981 | 3.355132 | 3.303353 | 6 | 9 | 17 | 7 |
| 100129250 | NM_00134911 | SMIM27 | small integral membrane protein 27 | protein_coding | NP_001336047.1;NP_001336048.1;XP_024303136.1 | 31420 | . | . | . | 0.839428 | 0.960812 | 1.451982 | 0.699247 | 1.953385 | 2.220960 | 3.258205 | 1.598746 | -1.195550 | 26.805535 | 0.597071388 | -0.4315641 | 0.666058258 | NA | 4.867222 | 4.610270 | 4.749961 | 4.761439 | 4.744400 | 4.727871 | 32 | 33 | 23 | 21 |
| 100129269 | NR_034126 | LINC01160 | long intergenic non-protein coding RNA 1160 | lncRNA | . | 49525 | . | . | . | 0.231159 | 0.078583 | 0.455712 | 0.100361 | 0.537917 | 0.181648 | 1.022606 | 0.229464 | 1.767350 | 11.064072 | 1.021494331 | 0.804299614 | 0.421223961 | NA | 2.993083 | 3.824402 | 3.439892 | 3.409114 | 3.488842 | 3.413319 | 13 | 5 | 20 | 5 |
| 100129272 | gene-TOMM20 | TOMM20P4 | TOMM20 pseudogene 4 | pseudogene | . | 50522 | . | . | . | 0.297135 | 0.299844 | 0.461973 | 0.212017 | 0.691445 | 0.693103 | 1.036655 | 0.484753 | 1.101565 | 8.015754 | 1.086429071 | 0.128452094 | 0.897791207 | NA | 2.928838 | 3.073227 | 2.995512 | 2.993670 | 3.011459 | 2.984465 | 9 | 8 | 10 | 5 |
| 100129361 | NM_00127159 | SMIM10L1 | small integral membrane protein 10 like 1 | protein_coding | NP_001258521.1 | 49847 | . | . | . | 3.658882 | 3.656328 | 4.003904 | 3.436758 | 8.514369 | 8.451767 | 8.984645 | 7.857738 | 1.003404 | 555.113645 | 0.168095083 | 0.029169033 | 0.976729779 | 0.998865404 | 9.113852 | 9.119421 | 9.117569 | 9.111349 | 9.168144 | 9.062784 | 641 | 595 | 523 | 479 |
| 100129387 | NR_024490 | GABPB1-AS1 | GABPB1 antisense RNA 1 | lncRNA | . | 44157 | . | . | . | 1.140609 | 1.355556 | 1.662946 | 1.631130 | 2.654244 | 3.133428 | 3.731604 | 3.729386 | 1.300367 | 187.124706 | 0.247066255 | 1.53367366 | 0.125109957 | 0.873049916 | 7.346308 | 7.724668 | 7.447094 | 7.514932 | 7.603468 | 7.588663 | 172 | 190 | 187 | 195 |
| 100129396 | NM_00134816 | FAM106C | family with sequence similarity 106 member C | protein_coding | NP_001335091.1 | 38396 | . | . | . | 0.258445 | 0.332802 | 0.346229 | 0.250349 | 0.601413 | 0.769286 | 0.776928 | 0.572394 | 1.002763 | 16.678661 | 0.755504192 | 0.005268304 | 0.995796521 | NA | 4.057381 | 4.062477 | 4.048531 | 4.065236 | 4.068973 | 4.045102 | 17 | 20 | 17 | 13 |
| 100129405 | NR_024117 | MSTO2P | misato family member 2, pseudogene | transcribed_ps | . | 26329 | . | . | . | 8.096533 | 8.865922 | 9.745714 | 8.526494 | 18.840963 | 20.493975 | 21.869101 | 19.494816 | 1.062034 | 612.405467 | 0.163740312 | 0.530292068 | 0.595909446 | 0.998865404 | 9.214121 | 9.301251 | 9.208955 | 9.263255 | 9.323630 | 9.227483 | 656 | 667 | 588 | 549 |
| 100129455 | XR_241351 | LOC10012945 | uncharacterized LOC100129455 | lncRNA | . | . | . | . | . | 0.483414 | 0.499649 | 0.679946 | 0.820793 | 1.124925 | 1.154962 | 1.525778 | 1.876646 | 1.501241 | 45.969147 | 0.461683568 | 1.269604571 | 0.20422552 | 0.969317911 | 5.200543 | 5.785711 | 5.474725 | 5.477577 | 5.529009 | 5.562173 | 42 | 40 | 44 | 56 |
| 100129460 | NR_036680 | DPY19L1P1 | DPY19L1 pseudogene 1 | transcribed_ps | . | 22395 | . | . | . | 0.277560 | 0.290047 | 0.280798 | 0.339476 | 0.645894 | 0.670457 | 0.630103 | 0.776171 | 1.073410 | 18.123320 | 0.718677097 | 0.142206541 | 0.886916865 | NA | 4.128334 | 4.229445 | 4.175463 | 4.176558 | 4.175192 | 4.187793 | 20 | 19 | 15 | 19 |
| 100129461 | NR_126015,NF | LYRM4-AS1 | LYRM4 antisense RNA 1 | lncRNA | . | 52055 | . | . | . | 1.049056 | 0.900649 | 1.411722 | 1.536824 | 2.441195 | 2.081890 | 3.167864 | 3.513765 | 1.526585 | 50.265973 | 0.444156962 | 1.374081457 | 0.169416395 | 0.946181982 | 5.314065 | 5.924798 | 5.614043 | 5.585755 | 5.666161 | 5.689573 | 50 | 39 | 50 | 60 |
| 100129482 | NR_026777 | ZNF37BP | zinc finger protein 37B, pseudogene | transcribed_ps | . | 13103 | . | . | . | 2.925401 | 3.104506 | 4.973627 | 3.061502 | 6.807528 | 7.176203 | 11.160674 | 6.999760 | 1.315202 | 954.400909 | 0.19586705 | 2.018123772 | 0.043578372 | 0.642641319 | 9.686938 | 10.082883 | 9.761473 | 9.797464 | 10.147799 | 9.777946 | 925 | 912 | 1172 | 770 |
| 100129503 | XR_243769 | LOC10012950 | uncharacterized LOC100129503 | lncRNA | . | . | . | . | . | 0.135479 | 0.204568 | 0.304639 | 0.362987 | 0.315265 | 0.472868 | 0.683602 | 0.829926 | 1.868097 | 13.559141 | 0.839594523 | 1.073814815 | 0.282905689 | NA | 3.243129 | 4.141582 | 3.719117 | 3.734627 | 3.756307 | 3.769732 | 9 | 12 | 14 | 18 |
| 100129518 | NR_037166 | SOD2-OT1 | SOD2 overlapping transcript 1 | lncRNA | . | 53445 | . | . | . | 0.125693 | 0.411305 | 0.374437 | 0.399493 | 0.292492 | 0.950750 | 0.840226 | 0.913393 | 1.445204 | 17.805573 | 0.784859119 | 0.676902692 | 0.498467696 | NA | 3.867373 | 4.393446 | 4.088585 | 4.157669 | 4.150267 | 4.153186 | 8 | 24 | 18 | 20 |
| 100129534 | NR_024489 | LOC10012953 | small nuclear ribonucleoprotein polypeptide N pseudogene | transcribed_ps | . | . | . | . | . | 0.189036 | 0.407052 | 0.397786 | 0.103798 | 0.439895 | 0.940918 | 0.892620 | 0.237321 | -1.180163 | 19.710047 | 0.792716763 | -0.30147789 | 0.76305011 | NA | 4.413600 | 4.178556 | 4.259493 | 4.324087 | 4.322131 | 4.232628 | 16 | 31 | 25 | 7 |
| 100129543 | NM_00127740 | ZNF730 | zinc finger protein 730 | protein_coding | NP_001264332.1;XP_016881603.1;XP_016881604.1 | 32470 | . | . | . | 2.217859 | 2.188164 | 2.158173 | 2.206386 | 5.161047 | 5.058041 | 4.842877 | 5.044642 | -1.097028 | 131.841146 | 0.283531385 | -0.47120288 | 0.637495865 | 0.998865404 | 7.107570 | 6.974685 | 7.074354 | 7.044698 | 7.037722 | 7.014642 | 165 | 143 | 113 | 114 |
| 100129550 | NR_024618 | LINC02035 | long intergenic non-protein coding RNA 2035 | lncRNA | . | 52875 | . | . | . | 0.171224 | 0.095138 | 0.062815 | 0.033138 | 0.398446 | 0.219917 | 0.140955 | 0.075765 | -2.625324 | 17.197556 | 0.799199783 | -1.74236203 | 0.081445115 | NA | 4.636349 | 3.251698 | 4.169456 | 4.107088 | 4.086285 | 4.058596 | 37 | 19 | 11 | 6 |
| 100129577 | gene-LOC1001 | LOC10012957 | mitochondrial carrier 1 pseudogene | pseudogene | . | . | . | . | . | 0.053534 | 0.057228 | 0.189910 | 0.046087 | 0.124575 | 0.132285 | 0.426152 | 0.105373 | 2.295365 | 4.446339 | 1.473797331 | 0.813357123 | 0.416013335 | NA | 1.429096 | 2.632244 | 2.117236 | 2.118208 | 2.147076 | 2.120413 | 3 | 3 | 8 | 3 |
| 100129617 | NR_045112 | LOC10012961 | uncharacterized LOC100129617 | lncRNA | . | . | . | . | . | 0.140227 | 0.069353 | 0.054842 | 0.123660 | 0.326313 | 0.160312 | 0.123064 | 0.282733 | -1.217920 | 6.058684 | 1.254325506 | -0.22675066 | 0.820617634 | NA | 2.735321 | 2.448451 | 2.617090 | 2.598237 | 2.593413 | 2.611100 | 10 | 5 | 3 | 7 |
| 100129662 | NR_038405 | FGF13-AS1 | FGF13 antisense RNA 1 | lncRNA | . | 44264 | . | . | . | 0.197932 | 0.115818 | 0.407309 | 0.275498 | 0.460596 | 0.267718 | 0.913990 | 0.629893 | 1.930923 | 7.188121 | 1.149307135 | 0.825967554 | 0.408822498 | NA | 2.290730 | 3.245375 | 2.823218 | 2.812157 | 2.847155 | 2.829691 | 7 | 4 | 10 | 7 |
| 100129716 | NR_027435 | ARRDC3-AS1 | ARRDC3 antisense RNA 1 | lncRNA | . | 44145 | . | . | . | 0.387579 | 0.326396 | 0.259344 | 0.331073 | 0.901913 | 0.754480 | 0.581961 | 0.756960 | -1.235138 | 25.878079 | 0.610321388 | -0.49919891 | 0.61763926 | NA | 4.838265 | 4.532926 | 4.719392 | 4.697091 | 4.676038 | 4.695677 | 36 | 28 | 18 | 24 |
| 100129722 | NR_038389 | STPG3-AS1 | STPG3 antisense RNA 1 | lncRNA | . | 51176 | . | . | . | 0.715461 | 0.556872 | 0.317704 | 0.609843 | 1.664907 | 1.287236 | 0.712919 | 1.394334 | -1.392762 | 29.496338 | 0.596613665 | -0.80110274 | 0.423072171 | 0.998865404 | 5.102988 | 4.622041 | 4.923738 | 4.885287 | 4.835751 | 4.894543 | 45 | 32 | 15 | 30 |
| 100129726 | NR_027251 | LINC01126 | long intergenic non-protein coding RNA 1126 | lncRNA | . | 49275 | . | . | . | 0.673783 | 0.569656 | 1.054256 | 0.801848 | 1.567920 | 1.316785 | 2.365719 | 1.833330 | 1.463204 | 40.227691 | 0.497265607 | 1.104301171 | 0.269462515 | 0.998865404 | 5.028170 | 5.579675 | 5.300339 | 5.276433 | 5.369352 | 5.324112 | 41 | 32 | 47 | 39 |
| 100129781 | NR_132983,NF | MIR193BHG | MIR193B host gene | lncRNA | . | 51945 | . | . | . | 0.325795 | 0.438789 | 0.654433 | 0.115784 | 0.758139 | 1.014281 | 1.468529 | 0.264727 | -1.018341 | 10.295429 | 1.035126327 | -0.02533136 | 0.979790659 | NA | 3.373794 | 3.354002 | 3.343547 | 3.358957 | 3.380446 | 3.319381 | 10 | 13 | 15 | 3 |
| 100129794 | NR_033240 | SLC25A21-AS1 | SLC25A21 antisense RNA 1 | lncRNA | . | 44298 | . | . | . | 0.431161 | 0.557418 | 0.244876 | 0.405737 | 1.003329 | 1.288497 | 0.549496 | 0.927670 | -1.537350 | 23.006350 | 0.660383177 | -0.9395235 | 0.347462034 | NA | 4.802857 | 4.177906 | 4.532557 | 4.562324 | 4.489291 | 4.524462 | 28 | 34 | 12 | 21 |
| 100129800 | NR_146880 | ZNF807P | zinc finger protein 807, pseudogene | transcribed_ps | . | 33229 | . | . | . | 0.114774 | 0.042621 | 0.072607 | 0.032192 | 0.267084 | 0.098521 | 0.162927 | 0.073602 | -1.489653 | 5.552428 | 1.321202371 | -0.43519137 | 0.663423528 | NA | 2.728992 | 2.161867 | 2.496812 | 2.469588 | 2.479041 | 2.467621 | 11 | 4 | 5 | 3 |
| 100129842 | NM_00115929 | ZNF737 | zinc finger protein 737 | protein_coding | NP_001152765.1;XP_005259754.1;XP_006722660.1;XP_011525900.1;XP | 32468 | 603984 | . | . | 0.610599 | 0.696489 | 0.463831 | 0.679754 | 1.420888 | 1.609965 | 1.040822 | 1.554177 | 1.039577 | 117.802522 | 0.297343882 | 0.188324187 | 0.850622526 | 0.998865404 | 6.851724 | 6.908177 | 6.888196 | 6.857598 | 6.887896 | 6.884399 | 139 | 119 | 105 | 112 |
| 100129858 | NR_033939 | SCOC-AS1 | SCOC antisense RNA 1 | lncRNA | . | 50601 | . | . | . | 0.271442 | 0.237102 | 0.210416 | 0.180162 | 0.631657 | 0.548073 | 0.472166 | 0.411920 | -1.298721 | 14.664133 | 0.800047532 | -0.47133598 | 0.637400825 | NA | 4.049783 | 3.674293 | 3.889897 | 3.882637 | 3.874206 | 3.865228 | 20 | 17 | 12 | 11 |
| 100129905 | gene-RPS27P1 | RPS27P19 | ribosomal protein S27 pseudogene 19 | pseudogene | . | 36094 | . | . | . | 1.785350 | 1.173164 | 0.775285 | 0.645740 | 4.154583 | 2.711822 | 1.739716 | 1.476408 | -1.981564 | 12.403764 | 0.890335023 | -1.10816635 | 0.267789981 | NA | 4.042145 | 3.058876 | 3.673622 | 3.642509 | 3.626391 | 3.618511 | 23 | 14 | 8 | 7 |
| 100129917 | NR_036511,NF | LOC10012991 | uncharacterized LOC100129917 | lncRNA | . | . | . | . | . | 1.452724 | 1.903687 | 1.638156 | 1.246415 | 3.380548 | 4.400456 | 3.675973 | 2.849779 | -1.033536 | 73.362181 | 0.372017491 | -0.12791928 | 0.898212856 | 0.998865404 | 6.220302 | 6.173244 | 6.185538 | 6.213417 | 6.211692 | 6.171716 | 81 | 85 | 68 | 62 |
| 100129961 | NR_036549 | CCNT2-AS1 | CCNT2 antisense RNA 1 | lncRNA | . | 40130 | . | . | . | 0.912813 | 0.795199 | 0.906470 | 0.610330 | 2.124153 | 1.838138 | 2.034092 | 1.395446 | -1.138799 | 35.737213 | 0.530615053 | -0.35338894 | 0.723796879 | 0.998865404 | 5.248555 | 5.064275 | 5.178723 | 5.156240 | 5.175349 | 5.122544 | 47 | 38 | 35 | 25 |
| 100130009 | gene-HMGA1P | HMGA1P8 | high mobility group AT-hook 1 pseudogene 8 | pseudogene | . | 39183 | . | . | . | 0.448890 | 0.506981 | 0.559106 | 0.615760 | 1.044585 | 1.171909 | 1.254617 | 1.407862 | 1.211584 | 31.780055 | 0.548335867 | 0.504973115 | 0.613577719 | 0.998865404 | 4.845597 | 5.121346 | 4.965637 | 4.981065 | 4.992021 | 5.003546 | 31 | 33 | 29 | 34 |
| 100130027 | NR_147501 | LOC10013002 | uncharacterized LOC100130027 | lncRNA | . | . | . | . | . | 0.781807 | 0.624229 | 0.580687 | 0.464344 | 1.819298 | 1.442934 | 1.303045 | 1.061669 | -1.370945 | 84.256025 | 0.358244928 | -1.27055809 | 0.203885905 | 0.969101675 | 6.605424 | 6.152605 | 6.471622 | 6.400553 | 6.378680 | 6.323797 | 125 | 93 | 69 | 59 |
| 100130086 | NM_00116441 | HSFX2 | heat shock transcription factor family, X-linked 2 | protein_coding | NP_001157887.1 | 32701 | . | . | . | 0.053000 | 0.080326 | 0.121128 | 0.076866 | 0.123334 | 0.185678 | 0.271809 | 0.175745 | 1.481702 | 5.604348 | 1.28298601 | 0.442136823 | 0.6583902 | NA | 2.175403 | 2.742346 | 2.464234 | 2.474203 | 2.485124 | 2.473179 | 4 | 6 | 7 | 5 |
| 100130105 | gene-TRIM51C | TRIM51CP | tripartite motif-containing 51C, pseudogene | pseudogene | . | 43968 | . | . | . | 0.702974 | 0.444450 | 0.299843 | 0.746223 | 1.635850 | 1.027366 | 0.672840 | 1.706151 | -1.110179 | 27.048976 | 0.646415211 | -0.23327459 | 0.81554819 | 0.998865404 | 4.832372 | 4.678533 | 4.787139 | 4.735012 | 4.706319 | 4.791647 | 40 | 24 | 13 | 34 |
| 100130108 | gene-ZNF849P | ZNF849P | zinc finger protein 849, pseudogene | pseudogene | . | 34389 | . | . | . | 0.053895 | 0.196896 | 0.021283 | 0.010739 | 0.125415 | 0.455133 | 0.047759 | 0.024554 | -4.930232 | 3.290668 | 1.852860038 | -1.22421768 | 0.214156259 | NA | 2.452336 | 0.148375 | 1.727172 | 1.753664 | 1.720982 | 1.720395 | 3 | 9 | 1 | 1 |
| 100130109 | gene-SNRPEP4 | SNRPEP4 | SNRPE pseudogene 4 | pseudogene | . | 43446 | . | . | . | 0.573389 | 0.217481 | 0.425120 | 0.362288 | 1.334300 | 0.502716 | 0.953957 | 0.828328 | -1.016772 | 6.156145 | 1.237251678 | -0.01939494 | 0.984526043 | NA | 2.631267 | 2.612728 | 2.633578 | 2.610331 | 2.624994 | 2.618066 | 10 | 4 | 6 | 5 |
| 100130183 | gene-RPS24P1 | RPS24P19 | ribosomal protein S24 pseudogene 19 | pseudogene | . | 35683 | . | . | . | 1.672660 | 1.572329 | 1.144853 | 1.243947 | 3.892348 | 3.634509 | 2.569017 | 2.844137 | -1.349700 | 23.388809 | 0.637372248 | -0.67878571 | 0.49727365 | NA | 4.747897 | 4.315285 | 4.570318 | 4.562377 | 4.536076 | 4.538736 | 32 | 28 | 17 | 19 |
| 100130211 | gene-EEF1A1P | EEF1A1P29 | eukaryotic translation elongation factor 1 alpha 1 pseudogene 29 | pseudogene | . | 37904 | . | . | . | 0.037232 | 0.026936 | 0.015661 | 0.012154 | 0.086641 | 0.062264 | 0.035143 | 0.027789 | -1.620137 | 1.451754 | 2.440017378 | -0.28529139 | 0.775420906 | NA | 0.844134 | 0.148375 | 0.548030 | 0.548560 | 0.545314 | 0.544903 | 2 | 2 | 1 | 1 |
| 100130239 | gene-UBA52P6 | UBA52P6 | ubiquitin A-52 residue ribosomal protein fusion product 1 pseudog | pseudogene | . | 36763 | . | . | . | 0.422828 | 0.743644 | 0.841216 | 0.354626 | 0.983938 | 1.718966 | 1.887663 | 0.810810 | 1.061967 | 9.798728 | 1.001129982 | 0.086640804 | 0.930957031 | NA | 3.247564 | 3.336262 | 3.270774 | 3.294278 | 3.301147 | 3.269000 | 8 | 13 | 12 | 6 |
| 100130275 | NR_033910 | TFAP2A-AS1 | TFAP2A antisense RNA 1 | lncRNA | . | 40579 | . | . | . | 1.146192 | 0.911234 | 1.148562 | 0.925003 | 2.667237 | 2.106358 | 2.577341 | 2.114909 | -1.016861 | 42.560785 | 0.483917369 | -0.04984805 | 0.960243475 | 0.998865404 | 5.422247 | 5.400577 | 5.430398 | 5.392254 | 5.426952 | 5.390263 | 55 | 41 | 41 | 35 |
| 100130283 | NR_147908 | LOC10013028 | uncharacterized LOC100130283 | lncRNA | . | . | . | . | . | 0.541600 | 0.839398 | 0.612092 | 0.507076 | 1.260325 | 1.940306 | 1.373516 | 1.159268 | -1.245419 | 18.713859 | 0.720516367 | -0.43945032 | 0.660335272 | NA | 4.375822 | 4.058876 | 4.215940 | 4.252753 | 4.224450 | 4.209763 | 19 | 27 | 16 | 14 |
| 100130289 | gene-SNRPGP1 | SNRPGP10 | small nuclear ribonucleoprotein polypeptide G pseudogene 10 | pseudogene | . | 39329 | . | . | . | 0.221448 | 0.304384 | 0.191135 | 0.225658 | 0.515318 | 0.703596 | 0.428902 | 0.515941 | -1.079982 | 3.457668 | 1.600358864 | -0.06936412 | 0.94469979 | NA | 1.844134 | 1.733338 | 1.790916 | 1.792011 | 1.790805 | 1.789671 | 4 | 4 | 3 | 3 |
| 100130311 | NM_00114553 | C17orf107 | chromosome 17 open reading frame 107 | protein_coding | NP_001139008.1 | 37238 | . | . | . | 0.231317 | 0.365776 | 0.212942 | 0.246701 | 0.538283 | 0.845508 | 0.477835 | 0.564053 | -1.302367 | 26.725145 | 0.611652568 | -0.62312425 | 0.533202877 | NA | 4.919286 | 4.535520 | 4.725840 | 4.782147 | 4.720319 | 4.730935 | 27 | 40 | 19 | 23 |
| 100130357 | NR_134859,NF | LOC10013035 | uncharacterized LOC100130357 | lncRNA | . | . | . | . | . | 1.041019 | 1.186162 | 1.254011 | 0.862505 | 2.422494 | 2.741866 | 2.813963 | 1.972016 | -1.027247 | 61.543548 | 0.434086801 | -0.08934403 | 0.928808505 | 0.998865404 | 5.962808 | 5.924002 | 5.881194 | 5.992449 | 5.968549 | 5.895214 | 54 | 84 | 62 | 47 |
| 100130370 | NR_164137,NF | LOC10013037 | uncharacterized LOC100130370 | lncRNA | . | . | . | . | . | 10.581110 | 11.692836 | 9.746915 | 10.037484 | 24.622679 | 27.028516 | 21.871794 | 22.949515 | -1.143479 | 395.546969 | 0.183203578 | -1.05582238 | 0.29104937 | 0.998865404 | 8.721349 | 8.527559 | 8.636637 | 8.694521 | 8.583460 | 8.589885 | 463 | 477 | 319 | 348 |
| 100130383 | gene-HIGD1AP | HIGD1AP12 | HIG1 hypoxia inducible domain family member 1A pseudogene 12 | pseudogene | . | 43007 | . | . | . | 0.037985 | 0.007823 | 0.008494 | 0.016509 | 0.088393 | 0.018084 | 0.019060 | 0.037745 | 1.234655 | 1.002957 | 2.868339486 | 0.106022226 | 0.915564717 | NA | -0.155866 | 0.148375 | -0.001261 | -0.000972 | 0.000053 | -0.000315 | 1 | 1 | 1 | 1 |
| 100130388 | gene-EEF1E1P1 | EEF1E1P1 | eukaryotic translation elongation factor 1 epsilon 1 pseudogene 1 | pseudogene | . | 51459 | . | . | . | 0.338247 | 0.426980 | 0.670382 | 0.682342 | 0.787115 | 0.986983 | 1.504319 | 1.560093 | 1.752496 | 16.090486 | 0.767266312 | 1.054928534 | 0.291458011 | NA | 3.548173 | 4.356363 | 3.968047 | 3.979962 | 4.012937 | 4.011523 | 12 | 14 | 18 | 19 |
| 100130394 | gene-LOC1001 | LOC10013039 | translation machinery associated 7 homolog (S. cerevisiae) pseudc | pseudogene | . | . | . | . | . | 4.770169 | 4.622986 | 3.736087 | 3.567666 | 11.100381 | 10.686239 | 8.383671 | 8.157090 | -1.290729 | 50.109385 | 0.441598209 | -0.83375709 | 0.404417878 | 0.998865404 | 5.819181 | 5.451470 | 5.675613 | 5.669016 | 5.629652 | 5.621184 | 66 | 60 | 39 | 40 |
| 100130418 | NR_015352,NF | CECR7 | cat eye syndrome chromosome region, candidate 7 | lncRNA | . | 1845 | . | . | . | 6.502438 | 6.483162 | 5.976835 | 6.280774 | 15.131442 | 14.986119 | 13.411843 | 14.360243 | -1.067425 | 768.729719 | 0.146674874 | -0.64179152 | 0.521008559 | 0.998865404 | 9.632709 | 9.538416 | 9.612721 | 9.605034 | 9.546471 | 9.579657 | 919 | 852 | 631 | 714 |
| 100130447 | gene-RPL12P1 | RPL12P16 | ribosomal protein L12 pseudogene 16 | pseudogene | . | 36926 | . | . | . | 0.207033 | 1.719942 | 1.833356 | 2.066375 | 0.481775 | 3.975723 | 4.113998 | 4.724520 | 2.010939 | 28.991075 | 0.737520719 | 1.366564467 | 0.171761861 | 0.947194954 | 4.271716 | 5.273004 | 4.710906 | 4.844200 | 4.852687 | 4.867956 | 5 | 37 | 32 | 38 |
| 100130460 | NR_103765 | CAND1.11 | uncharacterized LOC100130460 | lncRNA | . | . | . | . | . | 0.224719 | 0.096862 | 0.214895 | 0.177127 | 0.522931 | 0.223901 | 0.482219 | 0.404981 | 1.150933 | 6.205701 | 1.224445671 | 0.165629398 | 0.86844861 | NA | 2.526439 | 2.733338 | 2.638648 | 2.619424 | 2.634118 | 2.631953 | 9 | 4 | 6 | 6 |
| 100130476 | NR_049793 | WAKMAR2 | wound and keratinocyte migration associated lncRNA 2 | lncRNA | . | 53754 | 618508 | . | . | 0.134461 | 0.162886 | 0.004678 | 0.264230 | 0.312896 | 0.376518 | 0.010498 | 0.604130 | -1.093388 | 9.007149 | 1.153355951 | -0.11167857 | 0.911078271 | NA | 3.238681 | 3.100135 | 3.167817 | 3.174698 | 3.134976 | 3.199537 | 10 | 11 | 1 | 15 |

| | | | | | | | | | | | | | | | | | | | | | | | | | | | | | | | | | |
|---|---|---|---|---|---|---|---|---|---|---|---|---|---|---|---|---|---|---|---|---|---|---|---|---|---|---|---|---|---|---|---|---|---|
| 100130522 | NR_028339,NF | PARD6G-AS1 | PARD6G antisense RNA 1 | lncRNA | . | 44109 | . | . | . | 0.201055 | 0.126701 | 0.229222 | 0.241970 | 0.467864 | 0.292876 | 0.514369 | 0.553236 | 1.395038 | 21.340423 | 0.674657521 | 0.711922651 | 0.476512675 | NA | 4.154300 | 4.636609 | 4.409721 | 4.375583 | 4.419450 | 4.425626 | 25 | 15 | 21 | 24 |
| 100130557 | NR_024567 | NFYC-AS1 | NFYC antisense RNA 1 | lncRNA | . | 49451 | . | . | . | 1.073638 | 1.388405 | 1.741170 | 1.035220 | 2.498400 | 3.209361 | 3.907135 | 2.366908 | 1.117458 | 130.166992 | 0.324653663 | 0.493514028 | 0.62164943 | 0.998865404 | 6.941175 | 7.102743 | 6.947759 | 7.035438 | 7.121959 | 6.934307 | 124 | 149 | 150 | 96 |
| 100130561 | gene-HMGB1P | HMGB1P10 | high mobility group box 1 pseudogene 10 | pseudogene | . | 4994 | . | . | . | 3.786659 | 3.367495 | 2.051680 | 2.462911 | 8.811710 | 7.784115 | 4.603911 | 5.631153 | -1.605309 | 108.525011 | 0.315972245 | -2.16111019 | 0.030686828 | 0.556596966 | 7.063584 | 6.379942 | 6.853777 | 6.804895 | 6.658073 | 6.702932 | 164 | 135 | 66 | 85 |
| 100130562 | gene-RPS2P5 | RPS2P5 | ribosomal protein S2 pseudogene 5 | pseudogene | . | 31386 | . | . | . | 45.053612 | 47.096233 | 41.198597 | 53.936619 | 104.841599 | 108.865044 | 92.448456 | 123.319664 | 1.018196 | 1374.021156 | 0.149040895 | 0.174553743 | 0.861430298 | 0.998865404 | 10.411411 | 10.436854 | 10.401373 | 10.428972 | 10.325279 | 10.523060 | 1540 | 1496 | 1050 | 1467 |
| 100130581 | NR_027412,NF | LINC00910 | long intergenic non-protein coding RNA 910 | lncRNA | . | 44361 | . | . | . | 0.574583 | 0.267369 | 0.344934 | 0.574200 | 1.337079 | 0.618037 | 0.774021 | 1.312841 | 1.096364 | 40.882427 | 0.542547341 | 0.244636144 | 0.806738175 | 0.998865404 | 5.285348 | 5.418403 | 5.392206 | 5.280851 | 5.312183 | 5.391499 | 61 | 27 | 28 | 50 |
| 100130624 | gene-RPL15P2 | RPL15P22 | ribosomal protein L15 pseudogene 22 | pseudogene | . | 36933 | . | . | . | 0.213180 | 0.215854 | 0.245303 | 0.065880 | 0.496079 | 0.498957 | 0.550454 | 0.150626 | -1.372606 | 4.663582 | 1.417171851 | -0.32241518 | 0.747138182 | NA | 2.429096 | 1.978782 | 2.225615 | 2.227202 | 2.227796 | 2.211581 | 6 | 6 | 5 | 2 |
| 100130673 | NR_038454 | LOC10013067 | phosphoribosyl pyrophosphate synthetase 2 pseudogene | transcribed_ps | . | . | . | . | . | 0.067806 | 0.028309 | 0.253900 | 0.172860 | 0.157787 | 0.065438 | 0.569745 | 0.395223 | 3.956381 | 6.627578 | 1.248867833 | 1.588784136 | 0.112109136 | NA | 1.413393 | 3.404839 | 2.679126 | 2.671622 | 2.720472 | 2.702786 | 4 | 2 | 11 | 8 |
| 100130691 | NR_026966 | LOC10013069 | uncharacterized LOC100130691 | lncRNA | . | . | . | . | . | 0.071824 | 0.081536 | 0.120561 | 0.073097 | 0.167137 | 0.188475 | 0.270535 | 0.167128 | 1.198522 | 14.335565 | 0.816919114 | 0.319807023 | 0.749114623 | NA | 3.703730 | 3.967302 | 3.824144 | 3.832103 | 3.855758 | 3.822775 | 14 | 15 | 17 | 11 |
| 100130705 | NM_00119515 | ATP6V1FNB | ATP6V1F neighbor | protein_coding | NP_001182079.1 | 52392 | . | . | . | 0.249828 | 0.135946 | 0.238591 | 0.244963 | 0.581360 | 0.314246 | 0.535390 | 0.560079 | 1.244027 | 26.927832 | 0.616307489 | 0.511136884 | 0.60925521 | 0.998865404 | 4.583318 | 4.901253 | 4.763090 | 4.700963 | 4.756546 | 4.756652 | 36 | 18 | 26 | 28 |
| 100130717 | NR_024482,NF | HDHD5-AS1 | HDHD5 antisense RNA 1 | lncRNA | . | 1842 | . | . | . | 0.088070 | 0.220552 | 0.567809 | 0.305320 | 0.204944 | 0.509816 | 1.274149 | 0.698078 | 1.999574 | 5.422218 | 1.313203924 | 0.761262233 | 0.446500453 | NA | 1.855800 | 2.852980 | 2.407310 | 2.417124 | 2.432373 | 2.425230 | 3 | 5 | 7 | 6 |
| 100130742 | NM_00112989 | LRRC69 | leucine rich repeat containing 69 | protein_coding | NP_001123362.1;NP_001341399.1 | 34303 | . | . | . | 0.484874 | 0.229123 | 0.535023 | 0.377494 | 1.128324 | 0.529629 | 1.200575 | 0.863096 | 1.281575 | 18.757870 | 0.730345753 | 0.490066443 | 0.624086883 | NA | 4.037030 | 4.399154 | 4.235316 | 4.188531 | 4.239659 | 4.224191 | 25 | 12 | 20 | 18 |
| 100130744 | NR_046285 | LOC10013074 | uncharacterized LOC100130744 | lncRNA | . | . | . | . | . | 0.385373 | 0.181843 | 0.490768 | 0.015131 | 0.896779 | 0.420338 | 1.101268 | 0.034596 | -1.114303 | 22.335675 | 0.901859368 | -0.17313258 | 0.862547213 | NA | 4.553369 | 4.405394 | 4.494114 | 4.418884 | 4.522824 | 4.363638 | 37 | 16 | 35 | 2 |
| 100130746 | gene-LOC1001 | LOC10013074 | AKT interacting protein pseudogene | pseudogene | . | . | . | . | . | 0.055978 | 0.102181 | 0.127375 | 0.155289 | 0.130262 | 0.236196 | 0.285825 | 0.355049 | 1.683119 | 4.847231 | 1.376290116 | 0.54576923 | 0.585224577 | NA | 1.855800 | 2.602870 | 2.254057 | 2.263809 | 2.268790 | 2.271806 | 3 | 5 | 5 | 6 |
| 100130758 | gene-METTL15 | METTL15P1 | methyltransferase like 15 pseudogene 1 | pseudogene | . | 31926 | . | . | . | 0.468016 | 0.458170 | 0.412324 | 0.385054 | 1.089092 | 1.059080 | 0.925242 | 0.880379 | -1.174170 | 49.387539 | 0.444065502 | -0.52163719 | 0.601922968 | 0.998865404 | 5.736982 | 5.505927 | 5.644226 | 5.640240 | 5.619331 | 5.606531 | 62 | 57 | 41 | 41 |
| 100130776 | NR_027032 | AGAP2-AS1 | AGAP2 antisense RNA 1 | lncRNA | . | 48633 | . | . | . | 3.813217 | 3.975685 | 3.700446 | 4.259517 | 8.873512 | 9.189973 | 8.303694 | 9.738878 | 1.010397 | 188.428186 | 0.244542266 | 0.061021126 | 0.951342384 | 0.998865404 | 7.550820 | 7.564897 | 7.547185 | 7.563278 | 7.531293 | 7.587385 | 212 | 206 | 154 | 189 |
| 100130855 | NR_034080 | USP3-AS1 | USP3 antisense RNA 1 | lncRNA | . | 44140 | . | . | . | 0.241210 | 0.110823 | 0.409726 | 0.147148 | 0.561306 | 0.256171 | 0.919414 | 0.336436 | 1.538331 | 12.978777 | 0.910828375 | 0.682198385 | 0.495113516 | NA | 3.349244 | 3.978782 | 3.682780 | 3.648950 | 3.720276 | 3.659084 | 16 | 7 | 20 | 8 |
| 100130890 | NM_00119513 | TSTD3 | thiosulfate sulfurtransferase like domain containing 3 | protein_coding | NP_001182060.1;XP_016865630.1;XP_024302060.1 | 40910 | . | . | . | 3.888789 | 4.151578 | 4.403441 | 3.909777 | 9.049374 | 9.596560 | 9.881194 | 8.939244 | -1.026376 | 234.448534 | 0.237640921 | -0.1580494 | 0.87441787 | 0.998865404 | 7.890974 | 7.855058 | 7.892090 | 7.860594 | 7.934129 | 7.788552 | 282 | 248 | 232 | 184 |
| 100130892 | gene-RPL7P32 | RPL7P32 | ribosomal protein L7 pseudogene 32 | pseudogene | . | 35692 | . | . | . | 3.050371 | 2.429140 | 2.511898 | 3.376567 | 7.098339 | 5.615066 | 5.636627 | 7.720119 | 1.063593 | 73.715971 | 0.380412095 | 0.233815066 | 0.815128553 | 0.998865404 | 6.159017 | 6.247439 | 6.222752 | 6.164537 | 6.174648 | 6.242854 | 92 | 68 | 57 | 81 |
| 100130932 | gene-SNRPGP1 | SNRPGP15 | small nuclear ribonucleoprotein polypeptide G pseudogene 15 | pseudogene | . | 49371 | . | . | . | 0.658434 | 0.423821 | 0.298989 | 0.346031 | 1.532204 | 0.979680 | 0.670924 | 0.791158 | -1.759672 | 8.379302 | 1.066920656 | -0.76416812 | 0.444767062 | NA | 3.417335 | 2.602870 | 3.097240 | 3.075810 | 3.064628 | 3.067263 | 15 | 9 | 5 | 6 |
| 100130935 | NR_073432 | CSAG4 | CSAG family member 4 (pseudogene) | transcribed_ps | . | 20923 | . | . | . | 1.263263 | 1.143191 | 1.825935 | 0.977102 | 2.939665 | 2.642538 | 4.097345 | 2.234027 | 1.151077 | 31.340878 | 0.57600876 | 0.352398181 | 0.724539666 | 0.998865404 | 4.862893 | 5.069653 | 4.957511 | 4.946831 | 5.008361 | 4.929706 | 35 | 30 | 38 | 22 |
| 100130950 | NR_034082,NF | LOC10013095 | uncharacterized LOC100130950 | lncRNA | . | . | . | . | . | 0.382254 | 0.622011 | 0.645207 | 0.534462 | 0.889521 | 1.437807 | 1.447825 | 1.221984 | 1.287172 | 35.525928 | 0.52625609 | 0.692066842 | 0.488895353 | 0.998865404 | 4.957404 | 5.321310 | 5.107440 | 5.141881 | 5.173841 | 5.145978 | 31 | 38 | 38 | 34 |
| 100130954 | NR_034016,NF | LINC01502 | long intergenic non-protein coding RNA 1502 | lncRNA | . | 51183 | . | . | . | 0.131603 | 0.074052 | 0.105635 | 0.143081 | 0.306245 | 0.171174 | 0.237043 | 0.327139 | 1.176514 | 9.677523 | 0.987298542 | 0.237535587 | 0.812241315 | NA | 3.151937 | 3.387716 | 3.279822 | 3.257754 | 3.270332 | 3.281379 | 13 | 7 | 8 | 11 |
| 100130958 | NM_00112997 | SYCE1L | synaptonemal complex central element protein 1 like | protein_coding | NP_001123451.1;NP_001335853.1 | 37236 | . | . | . | 0.376089 | 0.576403 | 0.559172 | 0.198477 | 0.875174 | 1.332381 | 1.254766 | 0.453794 | -1.259480 | 14.588632 | 0.84058171 | -0.39594937 | 0.692142377 | NA | 4.021847 | 3.692999 | 3.855507 | 3.883952 | 3.878424 | 3.831593 | 15 | 21 | 16 | 7 |
| 100130967 | NR_160976,NF | C6orf99 | chromosome 6 open reading frame 99 | lncRNA | . | 21179 | . | . | . | 0.994389 | 0.811111 | 0.858623 | 0.753155 | 2.313985 | 1.874920 | 1.926725 | 1.721998 | -1.139318 | 24.348329 | 0.628413842 | -0.29943658 | 0.764606952 | NA | 4.695638 | 4.509889 | 4.625013 | 4.601730 | 4.610529 | 4.589924 | 33 | 25 | 22 | 19 |
| 100130987 | NR_024469 | LOC10013098 | uncharacterized LOC100130987 | lncRNA | . | . | . | . | . | 1.494698 | 1.225138 | 1.419935 | 1.270030 | 3.478223 | 2.831962 | 3.186293 | 2.903774 | -1.030228 | 36.565521 | 0.515210995 | -0.08339045 | 0.933541078 | 0.998865404 | 5.212950 | 5.171577 | 5.209388 | 5.179492 | 5.197585 | 5.182910 | 47 | 36 | 33 | 32 |
| 100131017 | NM_00127855 | ZNF316 | zinc finger protein 316 | protein_coding | NP_001265488.1;XP_006715693.1;XP_024302386.1;XP_024302387.1 | 13843 | . | . | . | 5.425192 | 5.463608 | 6.354499 | 5.377243 | 12.624644 | 12.629376 | 14.259311 | 12.294425 | 1.123052 | 1087.229790 | 0.138622188 | 1.207777745 | 0.227132764 | 0.982008277 | 10.000126 | 10.167882 | 10.037973 | 10.043332 | 10.172095 | 10.079585 | 1175 | 1109 | 1062 | 1012 |
| 100131067 | NR_034121,NF | CKMT2-AS1 | CKMT2 antisense RNA 1 | lncRNA | . | 48997 | . | . | . | 6.805506 | 7.018731 | 8.268763 | 8.243435 | 15.836692 | 16.224114 | 18.554867 | 18.847636 | 1.204209 | 594.155067 | 0.158834913 | 1.687828601 | 0.091444136 | 0.805877675 | 9.074255 | 9.342670 | 9.158614 | 9.137521 | 9.290383 | 9.251311 | 631 | 572 | 581 | 591 |
| 100131089 | NR_040059,NF | SRP14-AS1 | SRP14 antisense RNA1 (head to head) | lncRNA | . | 48619 | . | . | . | 2.697541 | 4.199648 | 2.649841 | 2.855929 | 6.277289 | 9.707673 | 5.946166 | 6.529741 | -1.108651 | 114.758910 | 0.314236645 | -0.47354404 | 0.635825091 | 0.998865404 | 6.915807 | 6.765188 | 6.803009 | 6.908977 | 6.804379 | 6.841395 | 118 | 150 | 89 | 108 |
| 100131187 | NM_00111320 | TSTD1 | thiosulfate sulfurtransferase like domain containing 1 | protein_coding | NP_001106676.1;NP_001106677.1;NP_001106678.1 | 35410 | 616041 | . | . | 12.681734 | 9.548693 | 7.414193 | 13.736722 | 29.510917 | 22.072228 | 16.637235 | 31.407378 | -1.077365 | 176.847118 | 0.3069498 | -0.35024116 | 0.726157717 | 0.998865404 | 7.519928 | 7.410723 | 7.525731 | 7.418251 | 7.308837 | 7.559142 | 237 | 174 | 103 | 208 |
| 100131193 | NR_024580 | CCDC183-AS1 | CCDC183 antisense RNA 1 | lncRNA | . | 44105 | . | . | . | 1.398161 | 2.111277 | 2.669342 | 1.880540 | 3.253578 | 4.880312 | 5.989926 | 4.299631 | 1.281250 | 150.277205 | 0.304903269 | 1.172674394 | 0.240926404 | 0.992733415 | 7.041640 | 7.399224 | 7.092728 | 7.237178 | 7.335200 | 7.187680 | 122 | 170 | 173 | 130 |
| 100131205 | NR_026911 | RPL21P28 | ribosomal protein L21 pseudogene 28 | transcribed_ps | . | 36258 | . | . | . | 1.059381 | 1.336537 | 1.392097 | 1.538891 | 2.465224 | 3.089465 | 3.123824 | 3.518491 | 1.229054 | 24.058467 | 0.629231941 | 0.472874645 | 0.636302616 | NA | 4.432996 | 4.728813 | 4.562046 | 4.583744 | 4.588545 | 4.598530 | 22 | 26 | 22 | 26 |
| 100131211 | NM_00114264 | NEMP2 | nuclear envelope integral membrane protein 2 | protein_coding | NP_001136117.1;XP_005246269.1;XP_005246270.1;XP_011508760.1;XP | 33700 | 616497 | . | . | 4.784857 | 4.396586 | 4.620226 | 5.243012 | 11.134560 | 10.162899 | 10.367652 | 11.987523 | 1.010597 | 727.641509 | 0.153394703 | 0.099145071 | 0.921023087 | 0.998865404 | 9.499495 | 9.514633 | 9.535417 | 9.469488 | 9.479376 | 9.540240 | 873 | 744 | 608 | 716 |
| 100131213 | NR_024421,NF | ZNF503-AS2 | ZNF503 antisense RNA 2 | lncRNA | . | 23525 | . | . | . | 1.895818 | 1.644176 | 2.355065 | 2.125060 | 4.411648 | 3.800587 | 5.284698 | 4.858698 | 1.238186 | 142.272071 | 0.276192972 | 1.115989581 | 0.264426595 | 0.996598404 | 6.989585 | 7.298886 | 7.126786 | 7.084805 | 7.206555 | 7.166148 | 155 | 129 | 144 | 140 |
| 100131234 | NR_040073 | MIR181A1HG | MIR181A1 host gene | lncRNA | . | 48659 | . | . | . | 0.424308 | 0.381786 | 0.373460 | 0.066470 | 0.987383 | 0.882516 | 0.838033 | 0.151975 | -1.591286 | 6.570383 | 1.230709142 | -0.54455856 | 0.586057161 | NA | 3.008843 | 2.348113 | 2.728637 | 2.722665 | 2.725205 | 2.698858 | 10 | 8 | 7 | 2 |
| 100131261 | gene-PNRC2P1 | PNRC2P1 | proline rich nuclear receptor coactivator 2 pseudogene 1 | pseudogene | . | 51933 | . | . | . | 2.102680 | 1.938065 | 1.993159 | 1.544574 | 4.893024 | 4.479925 | 4.472591 | 3.531484 | -1.154416 | 141.277174 | 0.284133058 | -0.72910427 | 0.46593788 | 0.998865404 | 7.241405 | 7.036064 | 7.185243 | 7.149452 | 7.158840 | 7.066357 | 182 | 156 | 129 | 107 |
| 100131294 | gene-LOC1001 | LOC10013129 | RAB13, member RAS oncogene family pseudogene | pseudogene | . | . | . | . | . | 0.900996 | 1.047784 | 0.830275 | 0.585287 | 2.096655 | 2.421999 | 1.863113 | 1.338189 | -1.376997 | 29.105748 | 0.582067837 | -0.79290677 | 0.427832144 | 0.998865404 | 5.074826 | 4.615182 | 4.873036 | 4.892646 | 4.862661 | 4.828167 | 36 | 39 | 25 | 19 |
| 100131378 | NM_00116669 | C11orf91 | chromosome 11 open reading frame 91 | protein_coding | NP_001160164.1;XP_016872542.1 | 34444 | . | . | . | 0.831811 | 0.613043 | 0.210528 | 0.679910 | 1.935658 | 1.417076 | 0.472420 | 1.554534 | -1.318669 | 21.536682 | 0.700016038 | -0.57010408 | 0.568607108 | NA | 4.616092 | 4.213338 | 4.461556 | 4.426285 | 4.389798 | 4.447833 | 33 | 22 | 10 | 24 |
| 100131390 | NM_00114525 | SP9 | Sp9 transcription factor | protein_coding | NP_001138722.1 | 30690 | . | . | . | 0.043707 | 0.029100 | 0.084019 | 0.054279 | 0.101709 | 0.067265 | 0.188536 | 0.124103 | 1.562015 | 4.572240 | 1.41581461 | 0.454443717 | 0.649509493 | NA | 1.832372 | 2.481107 | 2.180260 | 2.172895 | 2.191611 | 2.179966 | 5 | 3 | 6 | 4 |
| 100131434 | NR_027455 | LINC00893 | long intergenic non-protein coding RNA 893 | lncRNA | . | 44265 | . | . | . | 0.521657 | 0.506921 | 0.458315 | 0.507927 | 1.213916 | 1.171770 | 1.028445 | 1.161314 | -1.072711 | 37.227251 | 0.507518401 | -0.19952206 | 0.841854385 | 0.998865404 | 5.268218 | 5.166566 | 5.226908 | 5.222526 | 5.209115 | 5.219886 | 45 | 41 | 30 | 35 |
| 100131454 | NR_024120 | DBIL5P | diazepam binding inhibitor-like 5, pseudogene | transcribed_ps | . | 38519 | . | . | . | 0.323628 | 0.318294 | 0.324008 | 0.345199 | 0.753096 | 0.735752 | 0.727063 | 0.789256 | 1.019511 | 31.223953 | 0.55143522 | 0.050555029 | 0.959680098 | 0.998865404 | 4.950620 | 4.978409 | 4.964558 | 4.961765 | 4.963712 | 4.967459 | 36 | 33 | 27 | 30 |
| 100131564 | NR_034089 | CCDC18-AS1 | CCDC18 antisense RNA 1 | lncRNA | . | 52262 | . | . | . | 2.262941 | 1.907561 | 1.882405 | 1.572346 | 5.265957 | 4.409412 | 4.224063 | 3.594983 | -1.226249 | 240.342793 | 0.231629968 | -1.27035393 | 0.203958586 | 0.969101675 | 8.047877 | 7.755204 | 8.000465 | 7.907008 | 7.898766 | 7.810073 | 332 | 260 | 206 | 183 |
| 100131572 | gene-RPS27P9 | RPS27P9 | ribosomal protein S27 pseudogene 9 | pseudogene | . | 36964 | . | . | . | 0.405303 | 0.174526 | 0.953157 | 0.656714 | 0.943156 | 0.403425 | 2.138857 | 1.501498 | 2.493067 | 6.234721 | 1.23919918 | 1.063526808 | 0.287543115 | NA | 1.832372 | 3.155137 | 2.608878 | 2.601614 | 2.635878 | 2.623430 | 5 | 3 | 9 | 7 |
| 100131609 | gene-HNRNPA1 | HNRNPA1P2 | heterogeneous nuclear ribonucleoprotein A1 pseudogene 2 | pseudogene | . | 13958 | . | . | . | 0.040823 | 0.015697 | 0.022405 | 0.063571 | 0.094998 | 0.036285 | 0.050275 | 0.145347 | 1.635035 | 1.753401 | 2.24714197 | 0.315654997 | 0.752264389 | NA | 0.413393 | 1.121007 | 0.804636 | 0.800773 | 0.802005 | 0.811322 | 2 | 1 | 1 | 3 |
| 100131655 | NR_040024 | ZNF236-DT | ZNF236 divergent transcript | lncRNA | . | 53779 | . | . | . | 0.231833 | 0.169449 | 0.067582 | 0.295712 | 0.539485 | 0.391689 | 0.151651 | 0.676110 | -1.039817 | 8.304187 | 1.102830163 | -0.05107733 | 0.959263898 | NA | 3.084644 | 3.022362 | 3.061141 | 3.051281 | 3.034194 | 3.074988 | 11 | 8 | 3 | 12 |
| 100131691 | NR_027334 | MZF1-AS1 | MZF1 antisense RNA 1 | lncRNA | . | 51271 | . | . | . | 0.995459 | 1.149899 | 1.490745 | 1.044757 | 2.316474 | 2.658043 | 3.345189 | 2.388713 | 1.163521 | 95.798484 | 0.342956988 | 0.637099455 | 0.524060064 | 0.998865404 | 6.467850 | 6.687648 | 6.531813 | 6.571008 | 6.650983 | 6.538960 | 95 | 102 | 106 | 79 |
| 100131693 | gene-EIF4EP2 | EIF4EP2 | eukaryotic translation initiation factor 4E pseudogene 2 | pseudogene | . | 32428 | . | . | . | 1.778602 | 1.680891 | 1.350970 | 1.307184 | 4.138881 | 3.885454 | 3.031538 | 2.988721 | -1.306246 | 77.894687 | 0.359640322 | -1.07170003 | 0.283854785 | 0.998865404 | 6.463090 | 6.078224 | 6.327826 | 6.308195 | 6.254897 | 6.244774 | 105 | 92 | 60 | 62 |
| 100131713 | gene-RPL29P1 | RPL29P11 | ribosomal protein L29 pseudogene 11 | pseudogene | . | 36905 | . | . | . | 1.485635 | 1.153906 | 0.983275 | 1.196175 | 3.457133 | 2.667306 | 2.206441 | 2.734912 | -1.245112 | 24.145082 | 0.63281504 | -0.49979176 | 0.617221717 | NA | 4.743194 | 4.426810 | 4.623183 | 4.592660 | 4.578246 | 4.594026 | 35 | 25 | 17 | 22 |
| 100131801 | NM_00117115 | PET100 | PET100 cytochrome c oxidase chaperone | protein_coding | NP_001164626.1 | 40038 | 614770 | . | . | 26.101237 | 24.196759 | 20.981811 | 26.822740 | 60.738649 | 55.931891 | 47.082572 | 61.327009 | -1.068871 | 514.216072 | 0.180179786 | -0.53328618 | 0.593835495 | 0.998865404 | 9.053800 | 8.957040 | 9.052276 | 8.998854 | 8.907977 | 9.053343 | 637 | 550 | 382 | 520 |
| 100131827 | NM_00112822 | ZNF717 | zinc finger protein 717 | protein_coding | NP_001121695.1;NP_001277137.1;NP_001277138.1;NP_001277139.1;NI | 29448 | 618405 | . | . | 3.068031 | 2.663760 | 3.375748 | 3.137830 | 7.139433 | 6.157402 | 7.575079 | 7.174275 | 1.256817 | 280.932556 | 0.212242119 | 1.553765546 | 0.120240352 | 0.860382361 | 7.959193 | 8.290040 | 8.091008 | 8.040974 | 8.226806 | 8.143644 | 301 | 255 | 293 | 271 |
| 100131828 | gene-CIR1P2 | CIR1P2 | CIR1 pseudogene 2 | pseudogene | . | 44012 | . | . | . | 0.796244 | 0.748468 | 0.541653 | 0.456314 | 1.852892 | 1.730117 | 1.215454 | 1.043307 | -1.558100 | 30.534311 | 0.565270905 | -1.13182555 | 0.257707784 | 0.995760118 | 5.216348 | 4.578418 | 4.966145 | 4.957501 | 4.918849 | 4.897749 | 44 | 39 | 23 | 20 |
| 100131863 | gene-RPS18P5 | RPS18P5 | ribosomal protein S18 pseudogene 5 | pseudogene | . | 36287 | . | . | . | 1.654541 | 2.229706 | 1.891333 | 2.799984 | 3.850186 | 5.154065 | 4.244095 | 6.401831 | 1.206143 | 36.713607 | 0.527462381 | 0.512644198 | 0.60820023 | 0.998865404 | 5.058644 | 5.325518 | 5.159238 | 5.197482 | 5.175065 | 5.235061 | 33 | 41 | 28 | 45 |
| 100131897 | NM_00112989 | INSYN2B | inhibitory synaptic factor family member 2B | protein_coding | NP_001123363.1;NP_001333233.1;XP_016864397.1 | 37271 | . | . | . | 0.591135 | 0.606589 | 0.458196 | 0.737621 | 1.375596 | 1.402158 | 1.028177 | 1.686483 | -1.019007 | 106.988955 | 0.334287331 | -0.0812588 | 0.935236141 | 0.998865404 | 6.756051 | 6.726433 | 6.737509 | 6.745139 | 6.661615 | 6.804013 | 123 | 118 | 71 | 122 |
| 100131905 | gene-RPS27P2 | RPS27P21 | ribosomal protein S27 pseudogene 21 | pseudogene | . | 35826 | . | . | . | 0.244999 | 0.173505 | 0.167031 | 0.129431 | 0.401065 | 0.570124 | 0.374814 | 0.295928 | -1.204527 | 2.440135 | 1.90164038 | -0.14117614 | 0.887730791 | NA | 1.413393 | 1.148375 | 1.296738 | 1.289023 | 1.291174 | 1.290401 | 4 | 2 | 2 | 2 |
| 100131997 | NR_103833 | FAM27E3 | family with sequence similarity 27 member E3 | lncRNA | . | 28655 | . | . | . | 2.475496 | 3.064038 | 3.717009 | 2.348291 | 5.760580 | 7.082661 | 8.340860 | 5.369088 | 1.082665 | 136.835776 | 0.310340009 | 0.369231151 | 0.711955439 | 0.998865404 | 7.037164 | 7.153110 | 7.034908 | 7.109092 | 7.183914 | 7.013030 | 136 | 156 | 152 | 103 |
| 100131998 | NR_027460 | RRN3P3 | RRN3 pseudogene 3 | transcribed_ps | . | 37620 | . | . | . | 2.439176 | 2.209937 | 3.100756 | 2.772637 | 5.676064 | 5.108369 | 6.958007 | 6.339304 | 1.244448 | 204.494567 | 0.238004272 | 1.325632889 | 0.184961327 | 0.958025663 | 7.508984 | 7.825525 | 7.642108 | 7.595992 | 7.746566 | 7.690381 | 221 | 186 | 209 | 200 |
| 100132057 | NR_168361,NF | LOC10013205 | phosphodiesterase 4D interacting protein pseudogene | transcribed_ps | . | . | . | . | . | 0.094623 | 0.282566 | 0.118342 | 0.036300 | 0.220191 | 0.653164 | 0.265555 | 0.082995 | -2.324830 | 7.485048 | 1.196190786 | -1.01750068 | 0.308915308 | NA | 3.385915 | 2.175234 | 2.897925 | 2.937562 | 2.901958 | 2.880568 | 7 | 16 | 6 | 2 |
| 100132062 | NR_028325 | LOC10013206 | uncharacterized LOC100132062 | lncRNA | . | . | . | . | . | 0.896993 | 1.036921 | 1.441379 | 1.183299 | 2.087340 | 2.396890 | 3.234413 | 2.705473 | 1.340507 | 155.656209 | 0.271361791 | 1.557990204 | 0.11923557 | 0.858706505 | 7.055169 | 7.478348 | 7.187625 | 7.236834 | 7.371160 | 7.284268 | 143 | 153 | 171 | 150 |
| 100132074 | NM_00129128 | FOXO6 | forkhead box O6 | protein_coding | NP_001278210.2 | 24814 | 611457 | . | . | 0.090409 | 0.038577 | 0.087803 | 0.140301 | 0.210386 | 0.089173 | 0.197027 | 0.320781 | 1.679989 | 8.310306 | 1.084550513 | 0.690103599 | 0.49012904 | NA | 2.631267 | 3.381963 | 3.046547 | 3.023392 | 3.043649 | 3.064499 | 10 | 4 | 7 | 12 |
| 100132101 | NR_036470 | HERC2P7 | hect domain and RLD 2 pseudogene 7 | transcribed_ps | . | 4875 | . | . | . | 0.742196 | 0.783798 | 1.207394 | 0.648194 | 1.727121 | 1.811784 | 2.709358 | 1.482018 | 1.192904 | 46.785186 | 0.486446886 | 0.523135988 | 0.600879623 | 0.998865404 | 5.413496 | 5.670989 | 5.516898 | 5.524431 | 5.607067 | 5.492844 | 48 | 47 | 58 | 33 |
| 100132111 | NR_024237,NF | C2CD4D-AS1 | C2CD4D and THEM5 antisense RNA 1 | lncRNA | . | 54045 | . | . | . | 0.394120 | 0.297324 | 0.136860 | 0.399277 | 0.917132 | 0.687278 | 0.307109 | 0.912900 | -1.317905 | 22.729671 | 0.69902917 | -0.5697135 | 0.568872033 | NA | 4.694014 | 4.290932 | 4.536609 | 4.505599 | 4.456479 | 4.532364 | 34 | 24 | 9 | 27 |
| 100132153 | gene-TUBB2BP | TUBB2BP1 | tubulin beta 2B class IIb pseudogene 1 | pseudogene | . | 42332 | . | . | . | 5.262115 | 5.609645 | 4.733618 | 5.279335 | 12.245158 | 12.966947 | 10.622102 | 12.070574 | -1.098843 | 134.520840 | 0.281238928 | -0.4835235 | 0.628724077 | 0.998865404 | 7.138456 | 7.001675 | 7.079363 | 7.099179 | 7.037848 | 7.071593 | 158 | 156 | 106 | 126 |
| 100132247 | NM_00113586 | NPIPB5 | nuclear pore complex interacting protein family member B5 | protein_coding | NP_001129337.1 | 37233 | . | . | . | 27.975361 | 32.915657 | 49.756596 | 31.377472 | 65.099815 | 76.086021 | 111.652351 | 71.740860 | 1.315997 | 4220.444498 | 0.351078793 | 1.128398348 | 0.259151725 | 0.995760118 | 11.831527 | 12.227717 | 11.845352 | 11.970458 | 12.319503 | 11.919330 | 3874 | 4235 | 5136 | 3456 |
| 100132249 | NR_135128 | LOC10013224 | uncharacterized LOC100132249 | lncRNA | . | . | . | . | . | 0.178191 | 0.151800 | 0.371277 | 0.222679 | 0.414659 | 0.350892 | 0.833134 | 0.509128 | 1.792362 | 30.002127 | 0.586418663 | 1.435599578 | 0.151116315 | 0.913648048 | 4.423228 | 5.268639 | 4.857490 | 4.840671 | 4.955186 | 4.880590 | 27 | 21 | 42 | 27 |
| 100132273 | NR_034118 | NDUFA6-DT | NDUFA6 divergent transcript | lncRNA | . | 45273 | . | . | . | 1.129160 | 1.275319 | 1.125314 | 1.408856 | 2.627602 | 2.947956 | 2.525173 | 3.221182 | 1.044258 | 64.240434 | 0.396914411 | 0.157410179 | 0.874921592 | 0.998865404 | 5.974755 | 6.035427 | 5.985343 | 6.012860 | 5.984989 | 6.032234 | 68 | 72 | 51 | 68 |
| 100132287 | NR_028322 | LOC10013228 | uncharacterized LOC100132287 | lncRNA | . | . | . | . | . | 1.460607 | 1.744541 | 2.268184 | 1.855435 | 3.398893 | 4.032584 | 5.089739 | 4.242232 | 1.271082 | 249.648788 | 0.229017653 | 1.511052232 | 0.130775144 | 0.882587711 | 7.780218 | 8.126558 | 7.853332 | 7.933050 | 8.067822 | 7.956363 | 232 | 257 | 269 | 234 |
| 100132288 | NR_038327,NF | TEKT4P2 | tektin 4 pseudogene 2 | transcribed_ps | . | 40046 | . | . | . | 15.503823 | 14.338664 | 15.392508 | 13.138483 | 36.078031 | 33.144460 | 34.540341 | 30.039579 | -1.055947 | 680.581662 | 0.161047897 | -0.48766276 | 0.625788736 | 0.998865404 | 9.448993 | 9.371251 | 9.455256 | 9.399723 | 9.445628 | 9.333047 | 837 | 724 | 624 | 569 |
| 100132341 | NR_024034 | CLUHP3 | clustered mitochondria homolog pseudogene 3 | transcribed_ps | . | 28447 | . | . | . | 1.767109 | 1.567633 | 1.969087 | 1.410164 | 4.112136 | 3.623653 | 4.418574 | 3.224173 | 1.000198 | 94.043954 | 0.343428835 | 0.000832157 | 0.999336035 | 0.999862634 | 6.553910 | 6.556615 | 6.570443 | 6.532879 | 6.600406 | 6.500962 | 115 | 95 | 96 | 73 |
| 100132352 | NR_034006,NF | FRG1HP | FSHD region gene 1 family member H, pseudogene | transcribed_ps | . | 51767 | . | . | . | 6.904224 | 6.174091 | 8.323074 | 7.976577 | 16.066412 | 14.271690 | 18.676736 | 18.237493 | 1.231536 | 336.094289 | 0.19520853 | 1.539169449 | 0.123762934 | 0.871137062 | 8.234343 | 8.535421 | 8.358971 | 8.296390 | 8.456013 | 8.433599 | 367 | 306 | 329 | 341 |
| 100132356 | NR_034127 | LOC10013235 | uncharacterized LOC100132356 | lncRNA | . | . | . | . | . | 0.317273 | 0.301738 | 0.576048 | 0.221457 | 0.738306 | 0.697482 | 1.292636 | 0.506334 | 1.251236 | 20.211179 | 0.72419127 | 0.446502753 | 0.655234124 | NA | 4.163717 | 4.491830 | 4.314420 | 4.311784 | 4.371066 | 4.291615 | 21 | 19 | 28 | 12 |
| 100132380 | gene-AGGF1P1 | AGGF1P1 | angiogenic factor with G-patch and FHA domains 1 pseudogene 1 | pseudogene | . | 38051 | . | . | . | 0.055629 | 0.070744 | 0.049758 | 0.017017 | 0.129452 | 0.163528 | 0.111654 | 0.038907 | -1.719630 | 5.344064 | 1.339561378 | -0.58384643 | 0.559323635 | NA | 2.754145 | 1.978782 | 2.426213 | 2.432314 | 2.424534 | 2.408294 | 7 | 8 | 5 | 2 |
| 100132403 | NR_146178 | FAM157B | family with sequence similarity 157 member B | lncRNA | . | 34080 | . | . | . | 0.205057 | 0.349777 | 0.326514 | 0.285548 | 0.477176 | 0.808526 | 0.732687 | 0.652872 | 1.092119 | 34.027245 | 0.549300285 | 0.23144027 | 0.816972783 | 0.998865404 | 5.024434 | 5.150069 | 5.041889 | 5.109423 | 5.097436 | 5.078723 | 28 | 44 | 33 | 31 |
| 100132406 | NM_00103970 | NBPF10 | NBPF member 10 | protein_coding | NP_001034792.4;NP_001289300.1 | 31992 | 614000 | . | . | 1.407287 | 1.555950 | 1.785188 | 1.429460 | 3.274815 | 3.596646 | 4.005911 | 3.268289 | 1.069666 | 627.369138 | 0.171501986 | 0.566528525 | 0.571034548 | 0.998865404 | 9.243502 | 9.341186 | 9.236355 | 9.297611 | 9.387637 | 9.231945 | 666 | 684 | 629 | 538 |
| 100132570 | gene-LOC1001 | LOC10013257 | chromosome 17 open reading frame 58 pseudogene | pseudogene | . | . | . | . | . | 0.361076 | 0.214888 | 0.181668 | 0.196050 | 0.840238 | 0.496723 | 0.407657 | 0.448244 | -1.563371 | 12.039504 | 0.895037431 | -0.72026053 | 0.4713646 | NA | 3.875693 | 3.232645 | 3.623734 | 3.591874 | 3.583926 | 3.585414 | 21 | 12 | 8 | 9 |
| 100132678 | gene-RPL7AP4 | RPL7AP45 | ribosomal protein L7a pseudogene 45 | pseudogene | . | 35935 | . | . | . | 0.016409 | 0.004599 | 0.006754 | 0.004318 | 0.038183 | 0.010631 | 0.015156 | 0.009874 | 1.234655 | 1.002957 | 2.868339486 | 0.106022226 | 0.915564717 | NA | -0.155866 | 0.148375 | -0.001261 | -0.000972 | 0.000053 | -0.000315 | 1 | 1 | 1 | 1 |
| 100132707 | NR_024476,NF | PAXIP1-AS2 | PAXIP1 antisense RNA 2 | lncRNA | . | 48958 | . | . | . | 1.071164 | 1.115953 | 1.353152 | 1.394524 | 2.492644 | 2.579574 | 3.036432 | 3.188414 | 1.197473 | 63.606362 | 0.394678871 | 0.65874598 | 0.510058902 | 0.998865404 | 5.854998 | 6.115460 | 5.960858 | 5.971556 | 6.020294 | 5.991946 | 65 | 64 | 64 | 61 |
| 100132742 | gene-RPL17P7 | RPL17P7 | ribosomal protein L17 pseudogene 7 | pseudogene | . | 36005 | . | . | . | 18.746492 | 16.934793 | 15.196490 | 19.192114 | 43.623856 | 39.145535 | 34.100487 | 43.880489 | -1.052597 | 283.991577 | 0.216481875 | -0.34161519 | 0.732640504 | 0.998865404 | 8.186539 | 8.111904 | 8.191500 | 8.131903 | 8.076823 | 8.190205 | 354 | 297 | 214 | 288 |
| 100132774 | NR_033827 | KDM4A-AS1 | KDM4A antisense RNA 1 | lncRNA | . | 40528 | . | . | . | 1.449637 | 1.137823 | 1.025478 | 0.960551 | 3.373366 | 2.630130 | 2.301143 | 2.196186 | -1.320014 | 56.536085 | 0.423441671 | -0.94594597 | 0.344176143 | 0.998865404 | 6.006883 | 5.607807 | 5.877403 | 5.821508 | 5.800605 | 5.786867 | 83 | 61 | 44 | 44 |
| 100132815 | NR_103741,NF | IPO5P1 | importin 5 pseudogene 1 | transcribed_ps | . | 49687 | . | . | . | 1.349309 | 1.244141 | 1.244262 | 1.160576 | 3.139898 | 2.875886 | 2.792088 | 2.653519 | -1.085947 | 213.755168 | 0.231434769 | -0.5139854 | 0.607262201 | 0.998865404 | 7.797744 | 7.679464 | 7.778980 | 7.738314 | 7.735013 | 7.705930 | 267 | 230 | 184 | 186 |
| 100132832 | NR_028058 | PMS2P9 | PMS1 homolog 2, mismatch repair system component pseudogen | transcribed_ps | . | 9135 | . | . | . | 0.205683 | 0.447604 | 0.555791 | 0.489017 | 0.478634 | 1.034656 | 1.247180 | 1.118079 | 1.566540 | 31.398465 | 0.588388621 | 1.100601676 | 0.271070056 | 0.998865404 | 4.614556 | 5.259255 | 4.886971 | 4.963252 | 4.993517 | 4.970184 | 18 | 36 | 36 | 33 |
| 100132863 | gene-HMGB1P | HMGB1P37 | high mobility group box 1 pseudogene 37 | pseudogene | . | 39184 | . | . | . | 0.026334 | 0.016752 | 0.019509 | 0.006121 | 0.061280 | 0.038723 | 0.043778 | 0.013995 | 1.234655 | 1.002957 | 2.868339486 | 0.106022226 | 0.915564717 | NA | -0.155866 | 0.148375 | -0.001261 | -0.000972 | 0.000053 | -0.000315 | 1 | 1 | 1 | 1 |
| 100132891 | NR_033651,NF | MSC-AS1 | MSC antisense RNA 1 | lncRNA | . | 48724 | . | . | . | 1.233573 | 1.414855 | 1.475166 | 1.543994 | 2.870576 | 3.270500 | 3.310229 | 3.530158 | 1.132940 | 203.990431 | 0.236690108 | 0.760790873 | 0.446781987 | 0.998865404 | 7.579943 | 7.759207 | 7.615672 | 7.665759 | 7.681723 | 7.713548 | 208 | 218 | 182 | 210 |
| 100132948 | NR_027421 | FAM27C | family with sequence similarity 27 member C | lncRNA | . | 23668 | . | . | . | 4.211616 | 4.087174 | 5.894633 | 5.797875 | 9.800603 | 9.447687 | 13.227385 | 13.256153 | 1.392302 | 109.874950 | 0.3063636 | 1.558513995 | 0.119111453 | 0.858706505 | 6.521098 | 6.998950 | 6.721595 | 6.709757 | 6.827719 | 6.817156 | 108 | 97 | 113 | 118 |
| 100133005 | NR_146066 | RASA4DP | RAS p21 protein activator 4CD, pseudogene | transcribed_ps | . | 44226 | . | . | . | 0.507406 | 0.444275 | 0.452577 | 0.607538 | 1.180753 | 1.026961 | 1.015570 | 1.389064 | 1.101405 | 75.633039 | 0.373411354 | 0.37316658 | 0.709024468 | 0.998865404 | 6.170095 | 6.308477 | 6.244056 | 6.207794 | 6.212828 | 6.286685 | 89 | 72 | 59 | 85 |
| 100133036 | NR_026759 | FAM95B1 | family with sequence similarity 95 member B1 | lncRNA | . | 32318 | . | . | . | 0.203311 | 0.273672 | 0.324257 | 0.233655 | 0.473114 | 0.632606 | 0.727624 | 0.534225 | 1.143937 | 13.038864 | 0.85005618 | 0.228228625 | 0.819468509 | NA | 3.604218 | 3.798723 | 3.687277 | 3.703123 | 3.712141 | 3.693593 | 12 | 15 | 14 | 11 |
| 100133091 | NR_029411 | LOC10013309 | uncharacterized LOC100133091 | lncRNA | . | . | . | . | . | 0.525135 | 0.858429 | 0.961076 | 0.962670 | 1.222012 | 1.984298 | 2.156626 | 2.201030 | 1.368591 | 69.485829 | 0.398675275 | 1.135487649 | 0.256171054 | 0.995760118 | 5.875842 | 6.326414 | 6.017584 | 6.117032 | 6.143303 | 6.142278 | 52 | 78 | 70 | 75 |
| 100133121 | NR_027422 | FAM27B | family with sequence similarity 27 member B | lncRNA | . | 23667 | . | . | . | 8.799605 | 11.216546 | 11.592412 | 13.907855 | 20.477043 | 25.927549 | 26.013037 | 31.798656 | 1.259574 | 249.481173 | 0.230694169 | 1.44319358 | 0.148965948 | 0.909660767 | 7.787620 | 8.118970 | 7.839418 | 7.950051 | 7.965126 | 8.055269 | 225 | 266 | 221 | 283 |
| 100133161 | NR_028326,NF | LINC01001 | long intergenic non-protein coding RNA 1001 | lncRNA | . | 38540 | . | . | . | 0.463287 | 0.520421 | 0.760902 | 0.656617 | 1.078088 | 1.202977 | 1.707442 | 1.501276 | 1.328610 | 90.462998 | 0.338274222 | 1.211792643 | 0.225591765 | 0.982008277 | 6.279574 | 6.689864 | 6.436140 | 6.462572 | 6.554304 | 6.505342 | 85 | 88 | 97 | 89 |
| 100133163 | gene-BCRP7 | BCRP7 | BCR pseudogene 7 | pseudogene | . | 39075 | . | . | . | 0.013360 | 0.015527 | 0.013734 | 0.036803 | 0.031089 | 0.035890 | 0.030818 | 0.084147 | 1.833800 | 2.539248 | 1.876543826 | 0.466195476 | 0.641075587 | NA | 0.844134 | 1.715150 | 1.330022 | 1.330639 | 1.332827 | 1.341772 | 2 | 2 | 2 | 4 |
| 100133165 | gene-LOC1001 | LOC10013316 | Dexi homolog (mouse) pseudogene | pseudogene | . | . | . | . | . | 0.075014 | 0.026954 | 0.027004 | 0.098747 | 0.174561 | 0.062306 | 0.060596 | 0.225773 | 1.232559 | 3.457246 | 1.634513507 | 0.18455436 | 0.853578586 | NA | 1.631267 | 1.932304 | 1.793133 | 1.781341 | 1.783769 | 1.797375 | 5 | 2 | 2 | 5 |
| 100133172 | NR_026789 | FAM66A | family with sequence similarity 66 member A | lncRNA | . | 30444 | . | . | . | 0.213702 | 0.239175 | 0.214671 | 0.201458 | 0.497292 | 0.552864 | 0.481716 | 0.460610 | -1.133991 | 5.912379 | 1.237552025 | -0.14658695 | 0.883458053 | NA | 2.651489 | 2.470304 | 2.566431 | 2.568309 | 2.564936 | 2.563055 | 7 | 7 | 5 | 5 |
| 100133204 | NM_00135064 | C9orf147 | chromosome 9 open reading frame 147 | protein_coding | NP_001337578.1 | 31438 | . | . | . | 0.157352 | 0.062822 | 0.068451 | 0.148457 | 0.366165 | 0.145216 | 0.153602 | 0.339430 | -1.030099 | 12.208156 | 0.918986206 | -0.04655476 | 0.962868092 | NA | 3.631267 | 3.587955 | 3.632810 | 3.587888 | 3.591848 | 3.625875 | 20 | 8 | 7 | 15 |
| 100133205 | NR_026775 | LINC00240 | long intergenic non-protein coding RNA 240 | lncRNA | . | 18772 | . | . | . | 0.400596 | 0.167475 | 0.101572 | 0.324177 | 0.932204 | 0.387125 | 0.227925 | 0.741191 | -1.313361 | 8.175923 | 1.1243015 | -0.34978476 | 0.72650024 | NA | 3.216230 | 2.819320 | 3.058831 | 3.024607 | 3.016060 | 3.047520 | 15 | 6 | 3 | 10 |
| 100133220 | NM_00131015 | GOLGA6L3 | golgin A6 family like 3 | protein_coding | NP_001297082.1 | 37441 | . | . | . | 0.230556 | 0.311308 | 0.108209 | 0.114689 | 0.536514 | 0.719602 | 0.242819 | 0.262222 | -2.302764 | 10.983657 | 0.945964016 | -1.27210611 | 0.203335422 | NA | 3.937099 | 2.733338 | 3.473963 | 3.493990 | 3.447234 | 3.444650 | 15 | 19 | 6 | 6 |
| 100133265 | gene-BMS1P7 | BMS1P7 | BMS1 pseudogene 7 | pseudogene | . | 23655 | . | . | . | 0.265967 | 0.330880 | 0.572340 | 0.199176 | 0.618915 | 0.764843 | 1.284315 | 0.455391 | 1.344223 | 13.732138 | 0.866204047 | 0.492692836 | 0.62222964 | NA | 3.548173 | 3.978782 | 3.748245 | 3.760011 | 3.801873 | 3.740666 | 12 | 14 | 20 | 8 |
| 100133273 | gene-RPL21P8 | RPL21P80 | ribosomal protein L21 pseudogene 80 | pseudogene | . | 35697 | . | . | . | 0.115148 | 0.060504 | 0.017251 | 0.195854 | 0.267955 | 0.139858 | 0.038711 | 0.447798 | 1.219351 | 2.468865 | 1.915493108 | 0.149368098 | 0.881263183 | NA | 1.156660 | 1.437399 | 1.304555 | 1.300986 | 1.298008 | 1.312051 | 3 | 2 | 1 | 4 |
| 100133315 | NM_00137584 | LOC10013331 | XRCC1 N-terminal domain containing 1-like | protein_coding | NP_001362776.1;NP_001362777.1 | . | . | . | . | 0.368175 | 0.209603 | 0.189579 | 0.417313 | 0.856760 | 0.484506 | 0.425410 | 0.954137 | 1.052036 | 24.889602 | 0.617806292 | 0.118458738 | 0.905704187 | NA | 4.600756 | 4.674293 | 4.625854 | 4.640252 | 4.646571 | 4.629966 | 26 | 28 | 24 | 22 |
| 100133612 | NR_024455 | LINC01134 | long intergenic non-protein coding RNA 1134 | lncRNA | . | 49449 | . | . | . | 0.831097 | 0.797748 | 1.771268 | 0.534578 | 1.933997 | 1.844030 | 3.974674 | 1.222249 | 1.402893 | 60.379953 | 0.50715094 | 0.963037143 | 0.335528884 | 0.998865404 | 5.648977 | 6.141231 | 5.847620 | 5.839929 | 6.036126 | 5.779926 | 59 | 53 | 94 | 31 |
| 100133669 | NR_026913 | LY6E-DT | LY6E divergent transcript | lncRNA | . | 53728 | . | . | . | 1.236353 | 1.885566 | 1.252435 | 1.317823 | 2.877045 | 4.358571 | 2.810428 | 3.013047 | -1.230503 | 31.473264 | 0.562922236 | -0.53159683 | 0.595005264 | 0.998865404 | 5.118809 | 4.817608 | 4.960088 | 5.016314 | 4.960322 | 4.964833 | 32 | 45 | 24 | 27 |
| 100133941 | NM_00129173 | CD24 | CD24 molecule | protein_coding | NP_001278666.1;NP_001278667.1;NP_001278668.1;NP_001346013.1;NI | 1645 | 600074 | . | . | 0.264069 | 0.123299 | 0.212051 | 0.183190 | 0.614500 | 0.285011 | 0.475836 | 0.418841 | -1.060334 | 14.276973 | 0.819056552 | -0.10319087 | 0.917811484 | NA | 3.875693 | 3.794399 | 3.853555 | 3.821746 | 3.838685 | 3.829513 | 21 | 12 | 13 | 12 |
| 100133957 | NR_028119 | UXT-AS1 | UXT antisense RNA 1 | lncRNA | . | 49239 | . | . | . | 0.160597 | 0.175016 | 0.031708 | 0.078752 | 0.373716 | 0.404558 | 0.071151 | 0.180057 | -2.178033 | 2.616014 | 1.88223668 | -0.59664442 | 0.550744806 | NA | 1.844134 | 0.715150 | 1.406131 | 1.407157 | 1.395562 | 1.399878 | 4 | 4 | 1 | 2 |
| 100133985 | NR_024444 | LINC01816 | long intergenic non-protein coding RNA 1816 | lncRNA | . | 52621 | . | . | . | 0.519023 | 0.944377 | 0.982771 | 1.562570 | 1.207788 | 2.182969 | 2.205309 | 3.572631 | 1.669735 | 21.094421 | 0.698399078 | 1.059020659 | 0.289590371 | NA | 3.985415 | 4.719719 | 4.339364 | 4.380170 | 4.381551 | 4.427623 | 13 | 22 | 18 | 30 |
| 100133991 | NR_024434,NF | MAP3K14-AS1 | MAP3K14 antisense RNA 1 | lncRNA | . | 44359 | . | . | . | 0.218814 | 0.375108 | 0.167732 | 0.235750 | 0.509189 | 0.867077 | 0.376384 | 0.539014 | -1.343693 | 9.055897 | 1.02200911 | -0.41702517 | 0.676659999 | NA | 3.377846 | 2.947964 | 3.175862 | 3.199310 | 3.172305 | 3.179323 | 9 | 14 | 6 | 8 |
| 100134229 | NR_024451 | KDM7A-DT | KDM7A divergent transcript | lncRNA | . | 48959 | . | . | . | 0.800789 | 0.648116 | 0.890383 | 0.808692 | 1.863468 | 1.498149 | 1.997993 | 1.848977 | 1.158736 | 58.853384 | 0.413776729 | 0.513686883 | 0.607470924 | 0.998865404 | 5.767942 | 5.982216 | 5.882587 | 5.836110 | 5.902262 | 5.878573 | 70 | 52 | 58 | 56 |
| 100134368 | NR_024453 | LOC10013436 | uncharacterized LOC100134368 | lncRNA | . | . | . | . | . | 0.406015 | 0.318856 | 0.611455 | 0.467784 | 0.944813 | 0.737049 | 1.372088 | 1.069534 | 1.455310 | 22.490819 | 0.655429774 | 0.825910691 | 0.408854756 | NA | 4.193664 | 4.737849 | 4.474078 | 4.451993 | 4.510828 | 4.485257 | 24 | 17 | 26 | 22 |
| 100134713 | NR_024454 | NDUFB2-AS1 | NDUFB2 antisense RNA 1 | lncRNA | . | 40396 | . | . | . | 0.318200 | 0.810587 | 0.306148 | 0.698709 | 0.740465 | 1.873707 | 0.686988 | 1.597516 | -1.140287 | 28.091357 | 0.664612511 | -0.28497348 | 0.775664463 | 0.998865404 | 4.906869 | 4.710567 | 4.755148 | 4.859772 | 4.752968 | 4.833549 | 20 | 46 | 14 | 34 |
| 100134868 | NR_004846 | LOC10013486 | uncharacterized LOC100134868 | lncRNA | . | . | . | . | . | 1.217476 | 1.243234 | 1.383579 | 1.390383 | 2.833117 | 2.873791 | 3.104711 | 3.178945 | 1.111679 | 124.056246 | 0.289471206 | 0.527653559 | 0.597739812 | 0.998865404 | 6.876484 | 7.029179 | 6.930947 | 6.937695 | 6.972418 | 6.970043 | 134 | 128 | 114 | 122 |
| 100134869 | NR_004847 | UBE2Q2P2 | ubiquitin conjugating enzyme E2 Q2 pseudogene 2 | transcribed_ps | . | 37440 | . | . | . | 0.851620 | 0.983303 | 0.830475 | 0.576863 | 1.981755 | 2.272949 | 1.863561 | 1.318929 | -1.310237 | 39.232911 | 0.506900085 | -0.7690437 | 0.44186737 | 0.998865404 | 5.474910 | 5.087090 | 5.303346 | 5.324718 | 5.297407 | 5.248787 | 48 | 51 | 35 | 26 |
| 100134934 | NM_00111332 | TEN1 | TEN1 subunit of CST complex | protein_coding | NP_001106795.2 | 37242 | 613130 | . | . | 3.045228 | 4.450932 | 4.239151 | 5.148005 | 7.086370 | 10.288529 | 9.512532 | 11.770302 | 1.234320 | 128.371771 | 0.307972195 | 0.986182559 | 0.324043536 | 0.998865404 | 6.845636 | 7.147021 | 6.892618 | 7.018049 | 6.997218 | 7.066041 | 108 | 147 | 112 | 145 |
| 100134938 | NM_00111440 | UPK3BL1 | uroplakin 3B like 1 | protein_coding | NP_001107875.1 | 37278 | . | . | . | 5.107440 | 4.099146 | 10.169310 | 3.933527 | 11.885224 | 9.475361 | 22.819635 | 8.993541 | 1.511324 | 244.039289 | 0.673757789 | 0.884313157 | 0.376527188 | 0.998865404 | 7.602204 | 8.198545 | 7.829386 | 7.728013 | 8.201474 | 7.708593 | 249 | 186 | 369 | 153 |
| 100137047 | NM_00111463 | JMJD7 | jumonji domain containing 7 | protein_coding | NP_001108104.1 | 34397 | . | . | . | 4.314303 | 3.117854 | 2.263599 | 5.591103 | 10.039560 | 7.207060 | 5.079450 | 12.783393 | 1.041348 | 168.314634 | 0.352949578 | 0.165609835 | 0.868464007 | 0.998865404 | 7.366369 | 7.423107 | 7.433163 | 7.296339 | 7.194237 | 7.544162 | 221 | 149 | 87 | 228 |
| 100137049 | NM_00111463 | PLA2G4B | phospholipase A2 group IVB | protein_coding | NP_001108105.1 | 9036 | 606088 | . | . | 3.847926 | 0.073507 | 0.099608 | 3.700920 | 8.954282 | 0.169914 | 0.223518 | 8.461714 | -1.051236 | 164.991333 | 2.441221547 | -0.02952871 | 0.976442923 | 0.998865404 | 7.401869 | 7.329721 | 7.525915 | 6.572085 | 6.611344 | 7.478441 | 382 | 7 | 8 | 293 |
| 100142659 | NM_00127850 | CTAGE8 | CTAGE family member 8 | protein_coding | NP_001265436.1 | 37294 | . | . | . | 0.418976 | 0.337394 | 0.560196 | 0.275778 | 0.974975 | 0.779902 | 1.257064 | 0.630533 | 1.087359 | 39.228368 | 0.534515617 | 0.226051071 | 0.821161699 | 0.998865404 | 5.229742 | 5.355183 | 5.293909 | 5.260682 | 5.342317 | 5.234877 | 48 | 36 | 48 | 25 |
| 100144602 | NR_034138 | EPHA5-AS1 | EPHA5 antisense RNA 1 | lncRNA | . | 50602 | . | . | . | 0.235616 | 0.220792 | 0.327743 | 0.382600 | 0.548288 | 0.510370 | 0.735446 | 0.874770 | 1.481642 | 11.102489 | 0.913637013 | 0.62081192 | 0.534723379 | NA | 3.161369 | 3.728813 | 3.457572 | 3.452127 | 3.471318 | 3.476735 | 11 | 9 | 11 | 13 |
| 100144603 | NR_021492,NF | CHKB-DT | CHKB divergent transcript | lncRNA | . | 40146 | . | . | . | 2.290999 | 1.706764 | 1.637474 | 1.872146 | 5.331250 | 3.945261 | 3.674444 | 4.280440 | -1.126139 | 38.738278 | 0.507450637 | -0.33773712 | 0.735561297 | 0.998865404 | 5.358499 | 5.187832 | 5.311943 | 5.258813 | 5.260493 | 5.275230 | 55 | 37 | 30 | 36 |
| 100147744 | NR_023317 | RNU7-1 | RNA, U7 small nuclear 1 | snRNA | . | 34033 | 617876 | . | . | 0.007657 | 0.014041 | 0.003253 | 0.000411 | 0.017818 | 0.032456 | 0.007299 | 0.000941 | 1.234655 | 1.002957 | 2.868339486 | 0.106022226 | 0.915564717 | NA | -0.155866 | 0.148375 | -0.001261 | -0.000972 | 0.000053 | -0.000315 | 1 | 1 | 1 | 1 |
| 100147773 | NR_125374 | TMCO1-AS1 | TMCO1 antisense RNA 1 | lncRNA | . | 54046 | . | . | . | 0.109401 | 0.156315 | 0.198094 | 0.235791 | 0.254581 | 0.361328 | 0.444518 | 0.539107 | 1.576751 | 12.184154 | 0.879444476 | 0.747011102 | 0.455056859 | NA | 3.243129 | 3.897226 | 3.575773 | 3.591183 | 3.602834 | 3.612453 | 9 | 12 | 12 | 15 |
| 100151640 | gene-EXOSC3P | EXOSC3P1 | exosome component 3 pseudogene 1 | pseudogene | . | 33989 | . | . | . | 0.259230 | 0.273578 | 0.244284 | 0.312319 | 0.603240 | 0.632388 | 0.548167 | 0.714079 | -1.015512 | 6.662823 | 1.168730264 | -0.01900087 | 0.984840415 | NA | 2.747897 | 2.724273 | 2.738749 | 2.736602 | 2.733426 | 2.740989 | 8 | 7 | 5 | 7 |
| 100151642 | gene-ATG12P1 | ATG12P1 | autophagy related 12 pseudogene 1 | pseudogene | . | 21466 | . | . | . | 0.103476 | 0.053163 | 0.069174 | 0.086579 | 0.240792 | 0.122890 | 0.155224 | 0.197952 | -1.007340 | 2.223025 | 1.975467281 | -0.0053412 | 0.995738358 | NA | 1.156660 | 1.148375 | 1.155078 | 1.151489 | 1.153584 | 1.152831 | 3 | 2 | 2 | 2 |
| 100151683 | NR_023343 | RNU4ATAC | RNA, U4atac small nuclear (U12-dependent splicing) | snRNA | . | 34016 | 601428 | . | . | 1.056033 | 1.231518 | 1.346433 | 1.127091 | 2.457431 | 2.846709 | 3.021356 | 2.576959 | 1.119830 | 5.246471 | 1.310503281 | 0.124593545 | 0.900845339 | NA | 2.307819 | 2.470304 | 2.383652 | 2.389415 | 2.390160 | 2.388338 | 5 | 6 | 5 | 5 |
| 100151684 | NR_023344 | RNU6ATAC | RNA, U6atac small nuclear (U12-dependent splicing) | snRNA | . | 34017 | 601429 | . | . | 0.882113 | 1.205577 | 0.588825 | 0.277259 | 2.052712 | 2.786746 | 1.321306 | 0.633920 | -2.694437 | 3.100213 | 1.771949195 | -0.80701188 | 0.419659637 | NA | 2.175403 | 0.751299 | 1.645950 | 1.655621 | 1.640648 | 1.634973 | 4 | 6 | 2 | 1 |
| 100158257 | gene-NF1P7 | NF1P7 | neurofibromin 1 pseudogene 7 | pseudogene | . | 34230 | . | . | . | 0.078528 | 0.088490 | 0.105265 | 0.114304 | 0.182738 | 0.204548 | 0.236212 | 0.261342 | 1.234668 | 5.014785 | 1.337965904 | 0.227302363 | 0.820188636 | NA | 2.166062 | 2.470304 | 2.318123 | 2.319540 | 2.324556 | 2.322755 | 5 | 5 | 5 | 5 |
| 100158258 | gene-NEK2P3 | NEK2P3 | NEK2 pseudogene 3 | pseudogene | . | 37817 | . | . | . | 0.055720 | 0.085790 | 0.108268 | 0.092366 | 0.129662 | 0.198308 | 0.242949 | 0.211183 | 1.335067 | 6.319812 | 1.204375098 | 0.346165065 | 0.729218667 | NA | 2.436885 | 2.852980 | 2.641987 | 2.652155 | 2.658930 | 2.651730 | 5 | 7 | 7 | 6 |
| 100169751 | NR_023363 | RNA5S1 | RNA, 5S ribosomal 1 | rRNA | . | 34362 | . | . | . | 0.022782 | 0.024520 | 0.091092 | 0.083942 | 0.053014 | 0.056679 | 0.204408 | 0.191924 | 1.234655 | 1.002957 | 2.868339486 | 0.106022226 | 0.915564717 | NA | -0.155866 | 0.148375 | -0.001261 | -0.000972 | 0.000053 | -0.000315 | 1 | 1 | 1 | 1 |
| 100169753 | NR_023364 | RNA5S2 | RNA, 5S ribosomal 2 | rRNA | . | 34363 | . | . | . | 0.022782 | 0.047098 | 0.152224 | 0.081957 | 0.053014 | 0.108870 | 0.341586 | 0.187385 | 1.234655 | 1.002957 | 2.868339486 | 0.106022226 | 0.915564717 | NA | -0.155866 | 0.148375 | -0.001261 | -0.000972 | 0.000053 | -0.000315 | 1 | 1 | 1 | 1 |
| 100169754 | NR_023365 | RNA5S3 | RNA, 5S ribosomal 3 | rRNA | . | 34364 | . | . | . | 0.022782 | 0.047098 | 0.093816 | 0.080823 | 0.053014 | 0.108870 | 0.210520 | 0.184792 | 1.234655 | 1.002957 | 2.868339486 | 0.106022226 | 0.915564717 | NA | -0.155866 | 0.148375 | -0.001261 | -0.000972 | 0.000053 | -0.000315 | 1 | 1 | 1 | 1 |
| 100169755 | NR_023366 | RNA5S4 | RNA, 5S ribosomal 4 | rRNA | . | 34365 | . | . | . | 0.022782 | 0.024520 | 0.063856 | 0.027792 | 0.053014 | 0.056679 | 0.143290 | 0.063543 | 1.234655 | 1.002957 | 2.868339486 | 0.106022226 | 0.915564717 | NA | -0.155866 | 0.148375 | -0.001261 | -0.000972 | 0.000053 | -0.000315 | 1 | 1 | 1 | 1 |
| 100169756 | NR_023367 | RNA5S5 | RNA, 5S ribosomal 5 | rRNA | . | 34366 | . | . | . | 0.022782 | 0.022578 | 0.030566 | 0.080823 | 0.053014 | 0.052190 | 0.068589 | 0.184792 | 1.234655 | 1.002957 | 2.868339486 | 0.106022226 | 0.915564717 | NA | -0.155866 | 0.148375 | -0.001261 | -0.000972 | 0.000053 | -0.000315 | 1 | 1 | 1 | 1 |
| 100169757 | NR_023368 | RNA5S6 | RNA, 5S ribosomal 6 | rRNA | . | 34367 | . | . | . | 0.468945 | 0.635583 | 1.143646 | 2.069914 | 1.091255 | 1.469180 | 2.566309 | 4.732614 | 2.444539 | 4.629699 | 1.430455622 | 0.901504836 | 0.367319964 | NA | 1.429096 | 2.715150 | 2.183400 | 2.184383 | 2.192983 | 2.210672 | 3 | 3 | 4 | 8 |
| 100169758 | NR_023369 | RNA5S7 | RNA, 5S ribosomal 7 | rRNA | . | 34368 | . | . | . | 0.043759 | 0.087156 | 0.150711 | 0.300320 | 0.101830 | 0.201465 | 0.338190 | 0.686647 | 1.834012 | 1.269624 | 2.598200026 | 0.336772769 | 0.736288203 | NA | -0.155866 | 0.715150 | 0.332517 | 0.332827 | 0.333928 | 0.338438 | 1 | 1 | 1 | 2 |
| 100169759 | NR_023370 | RNA5S8 | RNA, 5S ribosomal 8 | rRNA | . | 34369 | . | . | . | 0.043759 | 0.085214 | 0.119540 | 0.298052 | 0.101830 | 0.196975 | 0.268244 | 0.681460 | 1.834012 | 1.269624 | 2.598200026 | 0.336772769 | 0.736288203 | NA | -0.155866 | 0.715150 | 0.332517 | 0.332827 | 0.333928 | 0.338438 | 1 | 1 | 1 | 2 |
| 100169761 | NR_023372 | RNA5S10 | RNA, 5S ribosomal 10 | rRNA | . | 34371 | . | . | . | 0.043759 | 0.087156 | 0.208514 | 0.270544 | 0.101830 | 0.201465 | 0.467898 | 0.618566 | 1.234655 | 1.002957 | 2.868339486 | 0.106022226 | 0.915564717 | NA | -0.155866 | 0.148375 | -0.001261 | -0.000972 | 0.000053 | -0.000315 | 1 | 1 | 1 | 1 |
| 100169762 | NR_023373 | RNA5S11 | RNA, 5S ribosomal 11 | rRNA | . | 34372 | . | . | . | 0.066541 | 0.109734 | 0.205790 | 0.239916 | 0.154843 | 0.253655 | 0.461787 | 0.548540 | 1.234655 | 1.002957 | 2.868339486 | 0.106022226 | 0.915564717 | NA | -0.155866 | 0.148375 | -0.001261 | -0.000972 | 0.000053 | -0.000315 | 1 | 1 | 1 | 1 |
| 100169763 | NR_023374 | RNA5S12 | RNA, 5S ribosomal 12 | rRNA | . | 34373 | . | . | . | 0.066541 | 0.085214 | 0.117421 | 0.300320 | 0.154843 | 0.196975 | 0.263490 | 0.686647 | 1.834012 | 1.269624 | 2.598200026 | 0.336772769 | 0.736288203 | NA | -0.155866 | 0.715150 | 0.332517 | 0.332827 | 0.333928 | 0.338438 | 1 | 1 | 1 | 2 |
| 100169764 | NR_023375 | RNA5S13 | RNA, 5S ribosomal 13 | rRNA | . | 34374 | . | . | . | 0.066541 | 0.062636 | 0.086855 | 0.184616 | 0.154843 | 0.144785 | 0.194901 | 0.422103 | 1.234655 | 1.002957 | 2.868339486 | 0.106022226 | 0.915564717 | NA | -0.155866 | 0.148375 | -0.001261 | -0.000972 | 0.000053 | -0.000315 | 1 | 1 | 1 | 1 |
| 100169765 | NR_023376 | RNA5S14 | RNA, 5S ribosomal 14 | rRNA | . | 34375 | . | . | . | 0.066541 | 0.109734 | 0.205790 | 0.213259 | 0.154843 | 0.253655 | 0.461787 | 0.487590 | 1.234655 | 1.002957 | 2.868339486 | 0.106022226 | 0.915564717 | NA | -0.155866 | 0.148375 | -0.001261 | -0.000972 | 0.000053 | -0.000315 | 1 | 1 | 1 | 1 |
| 100169767 | NR_023378 | RNA5S16 | RNA, 5S ribosomal 16 | rRNA | . | 34377 | . | . | . | 0.043759 | 0.067977 | 0.116816 | 0.240767 | 0.101830 | 0.157131 | 0.262131 | 0.550485 | 1.234655 | 1.002957 | 2.868339486 | 0.106022226 | 0.915564717 | NA | -0.155866 | 0.148375 | -0.001261 | -0.000972 | 0.000053 | -0.000315 | 1 | 1 | 1 | 1 |
| 100169768 | NR_023379 | RNA5S17 | RNA, 5S ribosomal 17 | rRNA | . | 34378 | . | . | . | 0.066542 | 0.109734 | 0.144658 | 0.215527 | 0.154845 | 0.253655 | 0.324607 | 0.492777 | 1.234655 | 1.002957 | 2.868339486 | 0.106022226 | 0.915564717 | NA | -0.155866 | 0.148375 | -0.001261 | -0.000972 | 0.000053 | -0.000315 | 1 | 1 | 1 | 1 |
| 100170217 | NR_024230 | SNAR-B2 | small NF90 (ILF3) associated RNA B2 | snRNA | . | 34318 | . | . | . | 0.977243 | 0.646265 | 0.247175 | 0.450851 | 2.274086 | 1.493870 | 0.554653 | 1.030818 | -2.163369 | 2.601438 | 1.891323799 | -0.58862454 | 0.556113169 | NA | 1.832372 | 0.715150 | 1.403689 | 1.396279 | 1.389020 | 1.393333 | 5 | 3 | 1 | 2 |
| 100170224 | NR_024231 | SNAR-B1 | small NF90 (ILF3) associated RNA B1 | snRNA | . | 34317 | . | . | . | 0.979518 | 0.462667 | 0.247175 | 0.449898 | 2.279379 | 1.069475 | 0.554653 | 1.028638 | -1.884343 | 2.369752 | 1.973624408 | -0.46313857 | 0.643265045 | NA | 1.631267 | 0.715150 | 1.268913 | 1.257131 | 1.254154 | 1.258492 | 5 | 2 | 1 | 2 |
| 100170229 | NM_00114564 | SRRM5 | serine/arginine repetitive matrix 5 | protein_coding | NP_001139113.1 | 37248 | . | . | . | 0.243491 | 0.336841 | 0.309591 | 0.349857 | 0.566614 | 0.778624 | 0.694714 | 0.799906 | 1.129518 | 23.967679 | 0.632769914 | 0.277679613 | 0.781258312 | NA | 4.493602 | 4.667215 | 4.558364 | 4.587869 | 4.580220 | 4.590498 | 22 | 28 | 21 | 25 |
| 100170841 | NM_00113067 | EPOP | elongin BC and polycomb repressive complex 2 associated protein | protein_coding | NP_001124149.1 | 34493 | 617795 | . | . | 1.634483 | 1.979857 | 2.078646 | 2.062439 | 3.803509 | 4.576528 | 4.664422 | 4.715522 | 1.127009 | 196.971709 | 0.242872834 | 0.710245434 | 0.477551951 | 0.998865404 | 7.533361 | 7.705213 | 7.550845 | 7.630724 | 7.648779 | 7.641379 | 194 | 218 | 183 | 194 |
| 100174949 | NR_148401,NF | TRIM59-IFT80 | TRIM59-IFT80 protein | lncRNA | . | . | . | . | . | 0.485879 | 0.490829 | 1.597781 | 0.622081 | 1.130660 | 1.134574 | 3.585373 | 1.422313 | 1.959322 | 140.789774 | 0.358506414 | 2.706659091 | 0.006796401 | 0.261867098 | 6.570728 | 7.543203 | 6.949190 | 6.952971 | 7.365347 | 7.005343 | 109 | 103 | 227 | 105 |
| 100188893 | NM_00113449 | TOMM6 | translocase of outer mitochondrial membrane 6 | protein_coding | NP_001127965.1;NP_001369223.1 | 34528 | 616168 | . | . | 98.019392 | 96.167492 | 95.451618 | 113.845483 | 228.095161 | 222.295475 | 214.190666 | 260.294167 | 1.063351 | 1806.087595 | 0.13076427 | 0.677688686 | 0.497969101 | 0.998865404 | 10.773807 | 10.862145 | 10.803872 | 10.782151 | 10.774375 | 10.904261 | 2044 | 1863 | 1486 | 1888 |
| 100188947 | NR_024467 | HECTD2-AS1 | HECTD2 antisense RNA 1 | lncRNA | . | 48679 | . | . | . | 1.697347 | 1.450693 | 1.771146 | 1.277308 | 3.949797 | 3.353341 | 3.974399 | 2.920414 | -1.054503 | 49.714480 | 0.452400489 | -0.16923761 | 0.865609749 | 0.998865404 | 5.672083 | 5.598158 | 5.652831 | 5.624706 | 5.657485 | 5.599886 | 63 | 51 | 49 | 38 |
| 100189119 | rna-TRK-CTT3- | TRK-CTT3-1 | tRNA-Lys | tRNA | . | 34681 | . | . | . | 0.257976 | 0.092553 | 0.075243 | 0.286735 | 0.600320 | 0.213941 | 0.168844 | 0.655586 | 1.234655 | 1.002957 | 2.868339486 | 0.106022226 | 0.915564717 | NA | -0.155866 | 0.148375 | -0.001261 | -0.000972 | 0.000053 | -0.000315 | 1 | 1 | 1 | 1 |
| 100189122 | rna-TRK-TTT3-1 | TRK-TTT3-1 | tRNA-Lys | tRNA | . | 34684 | . | . | . | 0.665503 | 0.579465 | 0.321039 | 0.244430 | 1.548652 | 1.339460 | 0.720401 | 0.558861 | -1.620137 | 1.451754 | 2.440017378 | -0.28529139 | 0.775420906 | NA | 0.844134 | 0.148375 | 0.548030 | 0.548560 | 0.545314 | 0.544903 | 2 | 2 | 1 | 1 |
| 100189123 | rna-TRW-CCA1 | TRW-CCA1-1 | tRNA-Trp | tRNA | . | 34685 | . | . | . | 0.424559 | 0.005228 | 0.006085 | 0.018227 | 0.987967 | 0.012085 | 0.013653 | 0.041673 | -1.204659 | 1.220068 | 2.631544379 | -0.10207874 | 0.918694172 | NA | 0.413393 | 0.148375 | 0.294625 | 0.290739 | 0.291821 | 0.291432 | 2 | 1 | 1 | 1 |
| 100189169 | rna-TRL-AAG3- | TRL-AAG3-1 | tRNA-Leu | tRNA | . | 34731 | . | . | . | 0.113166 | 0.103890 | 0.977981 | 0.343142 | 0.263343 | 0.240146 | 2.194561 | 0.784554 | 2.507047 | 1.577944 | 2.402382155 | 0.551947588 | 0.580984263 | NA | -0.155866 | 1.175234 | 0.628257 | 0.628593 | 0.640169 | 0.629356 | 1 | 1 | 3 | 1 |
| 100189310 | rna-TRI-AAT5-3 | TRI-AAT5-3 | tRNA-Ile | tRNA | . | 34877 | . | . | . | 0.726586 | 0.254060 | 0.158350 | 0.199394 | 1.690796 | 0.587271 | 0.355333 | 0.455890 | -1.204659 | 1.220068 | 2.631544379 | -0.10207874 | 0.918694172 | NA | 0.413393 | 0.148375 | 0.294625 | 0.290739 | 0.291821 | 0.291432 | 2 | 1 | 1 | 1 |
| 100190938 | NR_024461,NF | RAMP2-AS1 | RAMP2 antisense RNA 1 | lncRNA | . | 44358 | . | . | . | 0.455211 | 0.633137 | 0.832127 | 0.533230 | 1.059294 | 1.463525 | 1.867270 | 1.219167 | 1.178421 | 60.424433 | 0.439397249 | 0.53904514 | 0.589855708 | 0.998865404 | 5.793110 | 6.031199 | 5.851044 | 5.921153 | 5.979641 | 5.860528 | 53 | 70 | 72 | 45 |
| 100190939 | NR_024458 | TPT1-AS1 | TPT1 antisense RNA 1 | lncRNA | . | 43686 | . | . | . | 0.683787 | 0.726229 | 0.879907 | 0.629496 | 1.591199 | 1.678711 | 1.974486 | 1.439267 | 1.060208 | 35.158206 | 0.529867673 | 0.159185071 | 0.873523066 | 0.998865404 | 5.092061 | 5.178231 | 5.123325 | 5.132386 | 5.159991 | 5.111329 | 38 | 38 | 37 | 28 |
| 100190952 | gene-NDUFB1F | NDUFB1P1 | NADH:ubiquinone oxidoreductase subunit B1 pseudogene 1 | pseudogene | . | 30893 | . | . | . | 0.343722 | 0.519582 | 0.461164 | 0.432145 | 0.799856 | 1.201037 | 1.034838 | 0.988048 | -1.017658 | 4.475201 | 1.420029653 | -0.01778304 | 0.985811938 | NA | 2.175403 | 2.148375 | 2.155464 | 2.165310 | 2.160790 | 2.159299 | 4 | 6 | 4 | 4 |
| 100190986 | NR_024456 | LOC10019098 | uncharacterized LOC100190986 | lncRNA | . | . | . | . | . | 1.206720 | 1.308236 | 2.006615 | 1.357039 | 2.808087 | 3.024046 | 4.502785 | 3.102708 | 1.320744 | 113.092546 | 0.322485474 | 1.244553883 | 0.213295781 | 0.976167662 | 6.605901 | 7.008783 | 6.750546 | 6.773628 | 6.920978 | 6.781763 | 108 | 109 | 134 | 97 |
| 100191040 | NM_00113600 | C2CD4D | C2 calcium dependent domain containing 4D | protein_coding | NP_001129475.1;XP_011507357.1;XP_016855478.1 | 37210 | . | . | . | 0.492715 | 0.412590 | 0.530862 | 0.697818 | 1.146567 | 0.953720 | 1.191240 | 1.595478 | 1.373448 | 37.136875 | 0.514067006 | 0.890550861 | 0.373170171 | 0.998865404 | 4.968052 | 5.425412 | 5.194312 | 5.173500 | 5.210632 | 5.248667 | 39 | 31 | 33 | 45 |
| 100192204 | NR_036506 | PPIAP30 | peptidylprolyl isomerase A pseudogene 30 | transcribed_ps | . | 44878 | . | . | . | 0.099537 | 0.094791 | 0.080724 | 0.221453 | 0.231626 | 0.219113 | 0.181143 | 0.506326 | 1.222837 | 2.988044 | 1.723505885 | 0.168396441 | 0.866271408 | NA | 1.429096 | 1.715150 | 1.575076 | 1.575932 | 1.573846 | 1.582711 | 3 | 3 | 2 | 4 |
| 100192386 | NR_024389 | FLJ16779 | uncharacterized LOC100192386 | lncRNA | . | . | . | . | . | 0.199607 | 0.161013 | 0.195625 | 0.178130 | 0.464493 | 0.372189 | 0.438977 | 0.407274 | 1.024648 | 44.342916 | 0.471047101 | 0.074573614 | 0.940553969 | 0.998865404 | 5.452135 | 5.488895 | 5.486219 | 5.448513 | 5.479855 | 5.463330 | 56 | 42 | 41 | 40 |
| 100216336 | gene-RPS24P8 | RPS24P8 | ribosomal protein S24 pseudogene 8 | pseudogene | . | 37016 | . | . | . | 1.705059 | 1.440176 | 1.266808 | 1.246976 | 3.967743 | 3.329032 | 2.842680 | 2.851063 | -1.244730 | 22.981244 | 0.643444572 | -0.49084692 | 0.623534719 | NA | 4.671258 | 4.356363 | 4.545905 | 4.526205 | 4.516196 | 4.514211 | 32 | 25 | 18 | 19 |
| 100216545 | NR_024586 | KMT2E-AS1 | KMT2E antisense RNA 1 | lncRNA | . | 40845 | . | . | . | 0.955946 | 0.662076 | 1.069395 | 1.484168 | 2.224526 | 1.530418 | 2.399691 | 3.393373 | 1.557673 | 117.425573 | 0.326185326 | 1.960210919 | 0.049971143 | 0.667071261 | 6.520592 | 7.160278 | 6.831817 | 6.725761 | 6.867527 | 6.986983 | 125 | 81 | 105 | 155 |
| 100216546 | NR_039981 | LINC01004 | long intergenic non-protein coding RNA 1004 | lncRNA | . | 48961 | . | . | . | 0.677358 | 0.672843 | 1.102131 | 0.592402 | 1.576239 | 1.555306 | 2.473150 | 1.354458 | 1.236113 | 223.669526 | 0.283462089 | 1.078842 | 0.280658168 | 0.998865404 | 7.643142 | 7.950924 | 7.739635 | 7.732771 | 7.979228 | 7.672267 | 232 | 214 | 281 | 161 |
| 100240708 | gene-GAPDHP4 | GAPDHP43 | glyceraldehyde 3 phosphate dehydrogenase pseudogene 43 | pseudogene | . | 37800 | . | . | . | 0.299697 | 0.366572 | 0.776198 | 0.313730 | 0.697408 | 0.847347 | 1.741767 | 0.717307 | 1.611513 | 18.213458 | 0.762468099 | 0.90287865 | 0.366590303 | NA | 3.799841 | 4.491830 | 4.141331 | 4.149762 | 4.221449 | 4.141456 | 15 | 16 | 28 | 12 |
| 100268168 | NR_026682,NF | LOC10026816 | uncharacterized LOC100268168 | lncRNA | . | . | . | . | . | 0.562152 | 0.743783 | 0.682735 | 0.520950 | 1.308150 | 1.719287 | 1.532039 | 1.191090 | -1.082876 | 25.550777 | 0.622801865 | -0.18443826 | 0.853669658 | NA | 4.730742 | 4.617632 | 4.663569 | 4.689682 | 4.689756 | 4.646421 | 27 | 32 | 26 | 18 |
| 100270746 | NR_026776 | LOC10027074 | uncharacterized LOC100270746 | lncRNA | . | . | . | . | . | 0.692914 | 0.423621 | 0.473978 | 0.273128 | 1.612439 | 0.979220 | 1.063592 | 0.624476 | -1.445866 | 14.325009 | 0.833703118 | -0.6380375 | 0.52344927 | NA | 4.079678 | 3.553545 | 3.871587 | 3.836723 | 3.845343 | 3.817658 | 24 | 14 | 13 | 8 |

| | | | | | | | | | | | | | | | | | | | | | | | | | | | | | | | | | |
|---|---|---|---|---|---|---|---|---|---|---|---|---|---|---|---|---|---|---|---|---|---|---|---|---|---|---|---|---|---|---|---|---|---|
| 100270804 | NR_026885 | LINC00653 | long intergenic non-protein coding RNA 653 | lncRNA | . | 44308 | . | . | . | 0.568189 | 0.515705 | 0.571125 | 0.639671 | 1.322199 | 1.192074 | 1.281587 | 1.462531 | 1.107445 | 36.792476 | 0.51071903 | 0.288289797 | 0.773124917 | 0.998865404 | 5.125927 | 5.273004 | 5.198514 | 5.186025 | 5.199815 | 5.214140 | 42 | 36 | 32 | 38 |
| 100270853 | gene-RPL7P19 | RPL7P19 | ribosomal protein L7 pseudogene 19 | pseudogene | . | 36268 | . | . | . | 0.671259 | 0.851269 | 0.315024 | 0.556105 | 1.562047 | 1.967748 | 0.706904 | 1.271469 | -1.780757 | 12.982899 | 0.868614339 | -0.9584125 | 0.337854795 | NA | 4.057381 | 3.219802 | 3.713314 | 3.729701 | 3.679407 | 3.699935 | 17 | 20 | 6 | 11 |
| 100270854 | gene-RPL8P3 | RPL8P3 | ribosomal protein L8 pseudogene 3 | pseudogene | . | 36407 | . | . | . | 0.053524 | 0.263414 | 0.060848 | 0.133559 | 0.124552 | 0.608893 | 0.136542 | 0.305366 | -1.320345 | 3.199326 | 1.703056615 | -0.235409 | 0.813891303 | NA | 1.867373 | 1.459419 | 1.671597 | 1.689660 | 1.674619 | 1.678618 | 2 | 6 | 2 | 3 |
| 100270862 | gene-RPS3P4 | RPS3P4 | ribosomal protein S3 pseudogene 4 | pseudogene | . | 36616 | . | . | . | 0.140493 | 0.129666 | 0.126353 | 0.578830 | 0.326933 | 0.299729 | 0.283532 | 1.323425 | **2.421587** | 6.124336 | 1.295976927 | 0.984549043 | 0.32484563 | NA | 1.844134 | 3.114084 | 2.581490 | 2.582786 | 2.582272 | 2.628303 | 4 | 4 | 3 | 13 |
| 100270896 | gene-RPL7AP1 | RPL7AP16 | ribosomal protein L7a pseudogene 16 | pseudogene | . | 35898 | . | . | . | 0.279184 | 0.515841 | 0.471888 | 0.481500 | 0.649672 | 1.192389 | 1.058902 | 1.100892 | 1.224492 | 12.577565 | 0.874622545 | 0.334067946 | 0.738328287 | NA | 3.501050 | 3.790061 | 3.623511 | 3.655559 | 3.650656 | 3.651059 | 9 | 16 | 12 | 13 |
| 100270918 | gene-RPSAP41 | RPSAP41 | ribosomal protein SA pseudogene 41 | pseudogene | . | 36663 | . | . | . | 0.535445 | 0.746725 | 0.949937 | 0.257139 | 1.246001 | 1.726088 | 2.131632 | 0.587917 | -1.065583 | 15.744857 | 0.842351746 | -0.10879364 | 0.913366169 | NA | 4.019256 | 3.933075 | 3.957398 | 3.977982 | 4.000241 | 3.925042 | 16 | 20 | 21 | 6 |
| 100271027 | gene-RPL39P1 | RPL39P19 | ribosomal protein L39 pseudogene 19 | pseudogene | . | 36346 | . | . | . | 0.091966 | 0.076737 | 0.171893 | 0.112130 | 0.214008 | 0.177382 | 0.385722 | 0.256371 | 1.234663 | 2.005914 | 2.070995667 | 0.146845664 | 0.883253841 | NA | 0.844134 | 1.148375 | 0.998092 | 0.998667 | 1.000707 | 0.999974 | 2 | 2 | 2 | 2 |
| 100271033 | gene-RPL4P6 | RPL4P6 | ribosomal protein L4 pseudogene 6 | pseudogene | . | 35611 | . | . | . | 0.461214 | 0.334203 | 0.361594 | 0.587409 | 1.073264 | 0.772525 | 0.811406 | 1.343042 | 1.169860 | 19.342544 | 0.712607698 | 0.317616982 | 0.750775491 | NA | 4.156660 | 4.381963 | 4.277309 | 4.250970 | 4.257329 | 4.296584 | 24 | 16 | 14 | 24 |
| 100271039 | gene-RPL7AP2 | RPL7AP22 | ribosomal protein L7a pseudogene 22 | pseudogene | . | 36592 | . | . | . | 0.102780 | 0.391901 | 0.471710 | 0.796595 | 0.239173 | 0.905896 | 1.058503 | 1.821320 | **2.495780** | 12.698602 | 0.924451863 | 1.427322143 | 0.153487061 | NA | 2.867373 | 4.177906 | 3.596196 | 3.632502 | 3.644346 | 3.681295 | 4 | 12 | 12 | 21 |
| 100271063 | gene-RPS14P1 | RPS14P1 | ribosomal protein S14 pseudogene 1 | pseudogene | . | 35572 | . | . | . | 0.452877 | 0.367619 | 0.044279 | 0.152142 | 1.053863 | 0.849767 | 0.099362 | 0.347854 | **-3.266098** | 4.663292 | 1.484545012 | -1.15022989 | 0.250049197 | NA | 2.838265 | 1.121007 | 2.254468 | 2.248182 | 2.224382 | 2.233268 | 9 | 7 | 1 | 3 |
| 100271082 | gene-RPS20P1 | RPS20P11 | ribosomal protein S20 pseudogene 11 | pseudogene | . | 36084 | . | . | . | 1.125773 | 1.595462 | 0.183452 | 1.090662 | 2.619720 | 3.687983 | 0.411661 | 2.493670 | **-2.031903** | 16.442350 | 0.855385311 | -1.19575522 | 0.231792097 | NA | 4.465522 | 3.431841 | 4.049947 | 4.082649 | 3.986935 | 4.049138 | 21 | 28 | 3 | 17 |
| 100271090 | gene-RPS21P1 | RPS21P1 | ribosomal protein S21 pseudogene 1 | pseudogene | . | 36401 | . | . | . | 0.186727 | 0.212250 | 0.108587 | 0.116045 | 0.434522 | 0.490625 | 0.243666 | 0.265324 | **-2.036119** | 1.683440 | 2.306917194 | -0.44467227 | 0.656556613 | NA | 1.175403 | 0.148375 | 0.762508 | 0.767419 | 0.759880 | 0.759446 | 2 | 3 | 1 | 1 |
| 100271234 | gene-RPS20P2 | RPS20P21 | ribosomal protein S20 pseudogene 21 | pseudogene | . | 35993 | . | . | . | 0.000175 | 0.001436 | 0.002610 | 0.003665 | 0.000406 | 0.003320 | 0.005857 | 0.008380 | 1.234655 | 1.002957 | 2.868339486 | 0.106022226 | 0.915564717 | NA | -0.155866 | 0.148375 | -0.001261 | -0.000972 | 0.000053 | -0.000315 | 1 | 1 | 1 | 1 |
| 100271238 | gene-RPS26P3 | RPS26P37 | ribosomal protein S26 pseudogene 37 | pseudogene | . | 36875 | . | . | . | 0.125872 | 0.158280 | 0.039294 | 0.187240 | 0.292910 | 0.365871 | 0.088174 | 0.428102 | -1.032087 | 2.216774 | 2.002641667 | -0.02275192 | 0.981848159 | NA | 1.175403 | 1.121007 | 1.148069 | 1.153041 | 1.145613 | 1.154849 | 2 | 3 | 1 | 3 |
| 100271273 | gene-RPL12P2 | RPL12P28 | ribosomal protein L12 pseudogene 28 | pseudogene | . | 36818 | . | . | . | 0.174469 | 0.013346 | 0.387816 | 0.285228 | 0.405996 | 0.030850 | 0.870248 | 0.652140 | **2.261266** | 3.604263 | 1.630894347 | 0.72177021 | 0.470435768 | NA | 1.137672 | 2.324312 | 1.830974 | 1.818829 | 1.841122 | 1.834610 | 4 | 1 | 5 | 4 |
| 100271332 | gene-RPL36AP | RPL36AP21 | ribosomal protein L36a pseudogene 21 | pseudogene | . | 37041 | . | . | . | 0.124646 | 0.050116 | 0.053582 | 0.131793 | 0.290055 | 0.115846 | 0.120236 | 0.301329 | 1.233060 | 1.486734 | 2.400765875 | 0.125894475 | 0.899815456 | NA | 0.413393 | 0.715150 | 0.572744 | 0.568868 | 0.570025 | 0.574493 | 2 | 1 | 1 | 2 |
| 100271350 | gene-RPS10P1 | RPS10P19 | ribosomal protein S10 pseudogene 19 | pseudogene | . | 35907 | . | . | . | 5.518628 | 4.562750 | 3.999476 | 4.869433 | 12.842074 | 10.547002 | 8.974709 | 11.133380 | -1.155027 | 115.890678 | 0.308174963 | -0.67470283 | 0.499864583 | 0.998865404 | 6.956995 | 6.748738 | 6.914551 | 6.844985 | 6.803658 | 6.860119 | 157 | 121 | 85 | 110 |
| 100271374 | gene-RPS27AP | RPS27AP5 | ribosomal protein S27a pseudogene 5 | pseudogene | . | 36867 | . | . | . | 0.179112 | 0.308680 | 0.236876 | 0.281950 | 0.416800 | 0.713527 | 0.531542 | 0.644644 | 1.102063 | 4.741867 | 1.38109312 | 0.101518922 | 0.919138538 | NA | 2.175403 | 2.312263 | 2.236317 | 2.246189 | 2.241736 | 2.245026 | 4 | 6 | 4 | 5 |
| 100271391 | gene-RPL21P1 | RPL21P133 | ribosomal protein L21 pseudogene 133 | pseudogene | . | 36003 | . | . | . | 0.250024 | 0.289036 | 0.346568 | 0.193630 | 0.581816 | 0.668119 | 0.777688 | 0.442713 | 1.034696 | 5.484409 | 1.288032471 | 0.038202669 | 0.969526093 | NA | 2.429096 | 2.481107 | 2.450875 | 2.452526 | 2.458371 | 2.446685 | 6 | 6 | 6 | 4 |
| 100271392 | gene-RPL31P6 | RPL31P63 | ribosomal protein L31 pseudogene 63 | pseudogene | . | 35666 | . | . | . | 0.103420 | 0.044788 | 0.274228 | 0.182342 | 0.240663 | 0.103529 | 0.615360 | 0.416904 | **2.914566** | 2.615882 | 1.896469644 | 0.813765289 | 0.415779423 | NA | 0.413393 | 1.963455 | 1.359184 | 1.355355 | 1.372337 | 1.366135 | 2 | 1 | 4 | 3 |
| 100271479 | gene-RPL31P4 | RPL31P46 | ribosomal protein L31 pseudogene 46 | pseudogene | . | 35559 | . | . | . | 0.177899 | 0.194747 | 0.331504 | 0.184475 | 0.413979 | 0.450166 | 0.743886 | 0.421780 | 1.246534 | 3.029698 | 1.717790234 | 0.185076135 | 0.85316932 | NA | 1.429096 | 1.751299 | 1.588467 | 1.589328 | 1.597487 | 1.586411 | 3 | 3 | 4 | 2 |
| 100271489 | gene-RPL32P2 | RPL32P24 | ribosomal protein L32 pseudogene 24 | pseudogene | . | 36722 | . | . | . | 0.242450 | 0.201988 | 0.710012 | 0.145747 | 0.564191 | 0.466904 | 1.593247 | 0.333232 | 1.817694 | 5.687233 | 1.317302595 | 0.654450607 | 0.512821539 | NA | 2.008843 | 2.877719 | 2.478432 | 2.475525 | 2.510212 | 2.473542 | 5 | 4 | 10 | 3 |
| 100271524 | gene-RPL39P3 | RPL39P35 | ribosomal protein L39 pseudogene 35 | pseudogene | . | 35815 | . | . | . | 0.056678 | 0.034397 | 0.015955 | 0.067273 | 0.131892 | 0.079510 | 0.035803 | 0.153811 | 1.234655 | 1.002957 | 2.868339486 | 0.106022226 | 0.915564717 | NA | -0.155866 | 0.148375 | -0.001261 | -0.000972 | 0.000053 | -0.000315 | 1 | 1 | 1 | 1 |
| 100271526 | gene-RPL39P3 | RPL39P36 | ribosomal protein L39 pseudogene 36 | pseudogene | . | 36076 | . | . | . | 0.136804 | 0.286467 | 0.177186 | 0.343994 | 0.318349 | 0.662182 | 0.397601 | 0.786501 | 1.044475 | 3.219731 | 1.666442807 | 0.037672219 | 0.969949027 | NA | 1.658167 | 1.715150 | 1.683853 | 1.689092 | 1.682707 | 1.691538 | 3 | 4 | 2 | 4 |
| 100271528 | gene-RPL39P3 | RPL39P37 | ribosomal protein L39 pseudogene 37 | pseudogene | . | 36282 | . | . | . | 0.000235 | 0.001492 | 0.069600 | 0.001413 | 0.000547 | 0.003448 | 0.156180 | 0.003231 | 1.234655 | 1.002957 | 2.868339486 | 0.106022226 | 0.915564717 | NA | -0.155866 | 0.148375 | -0.001261 | -0.000972 | 0.000053 | -0.000315 | 1 | 1 | 1 | 1 |
| 100271548 | gene-RPS10P2 | RPS10P26 | ribosomal protein S10 pseudogene 26 | pseudogene | . | 36321 | . | . | . | 0.521684 | 0.596373 | 0.544737 | 0.917414 | 1.213981 | 1.378542 | 1.222375 | 2.097557 | 1.279247 | 11.236714 | 0.918262669 | 0.386920805 | 0.698814839 | NA | 3.303566 | 3.655341 | 3.477753 | 3.480789 | 3.476768 | 3.506070 | 11 | 11 | 8 | 15 |
| 100271566 | gene-RPS20P3 | RPS20P35 | ribosomal protein S20 pseudogene 35 | pseudogene | . | 36929 | . | . | . | 1.096975 | 0.816693 | 0.837113 | 1.196074 | 2.552707 | 1.887823 | 1.878458 | 2.734682 | 1.027364 | 16.284539 | 0.764690991 | 0.050932848 | 0.959379031 | NA | 4.006229 | 4.044381 | 4.034368 | 4.015155 | 4.015303 | 4.037994 | 21 | 15 | 12 | 18 |
| 100271594 | gene-RPS3AP3 | RPS3AP38 | RPS3A pseudogene 38 | pseudogene | . | 36145 | . | . | . | 0.109509 | 0.075071 | 0.312387 | 0.116499 | 0.254833 | 0.173531 | 0.700987 | 0.266361 | 1.791252 | 4.375956 | 1.464111527 | 0.574387903 | 0.565705337 | NA | 1.644780 | 2.491830 | 2.106828 | 2.103600 | 2.127354 | 2.105791 | 4 | 3 | 7 | 3 |
| 100271603 | gene-RPL13AP | RPL13AP24 | ribosomal protein L13a pseudogene 24 | pseudogene | . | 36209 | . | . | . | 0.085332 | 0.069401 | 0.229608 | 0.139236 | 0.198570 | 0.160423 | 0.515233 | 0.318346 | **2.070415** | 4.138019 | 1.492770801 | 0.703336237 | 0.481846233 | NA | 1.429096 | 2.481107 | 2.022586 | 2.023534 | 2.042192 | 2.030551 | 3 | 3 | 6 | 4 |
| 100271626 | gene-RPL23AP | RPL23AP79 | ribosomal protein L23a pseudogene 79 | pseudogene | . | 36226 | . | . | . | 4.081128 | 4.045833 | 4.103174 | 4.582798 | 9.496951 | 9.352124 | 9.207402 | 10.478024 | 1.046285 | 69.693618 | 0.377415304 | 0.172956366 | 0.862685726 | 0.998865404 | 6.090211 | 6.154972 | 6.118597 | 6.115317 | 6.116273 | 6.139023 | 79 | 73 | 59 | 70 |
| 100271645 | gene-RPL36AP | RPL36AP42 | ribosomal protein L36a pseudogene 42 | pseudogene | . | 36277 | . | . | . | 48.827297 | 48.780380 | 50.402992 | 50.984325 | 113.623123 | 112.758034 | 113.102859 | 116.569595 | 1.025218 | 586.452933 | 0.158193767 | 0.227131224 | 0.820321705 | 0.998865404 | 9.177782 | 9.213737 | 9.190534 | 9.184372 | 9.203467 | 9.204339 | 670 | 622 | 516 | 557 |
| 100271646 | gene-RPL36AP | RPL36AP43 | ribosomal protein L36a pseudogene 43 | pseudogene | . | 36791 | . | . | . | 0.191862 | 0.503167 | 0.347150 | 0.343142 | 0.446471 | 1.163094 | 0.778995 | 0.784554 | -1.022918 | 4.489776 | 1.434937436 | -0.02278153 | 0.981824542 | NA | 2.184683 | 2.148375 | 2.153818 | 2.172145 | 2.163314 | 2.161822 | 3 | 7 | 4 | 4 |
| 100271700 | gene-NONOP2 | NONOP2 | non-POU domain containing, octamer-binding pseudogene 2 | pseudogene | . | 42032 | . | . | . | 0.095096 | 0.149191 | 0.203276 | 0.184912 | 0.221292 | 0.344863 | 0.456145 | 0.422779 | 1.490804 | 10.668690 | 0.937440107 | 0.614535557 | 0.538861484 | NA | 3.099441 | 3.674293 | 3.387067 | 3.402248 | 3.417753 | 3.409056 | 8 | 11 | 12 | 11 |
| 100271835 | NR_027145 | LIMS3-LOC440 | LIMS3-LOC440895 readthrough | lncRNA | . | . | . | . | . | 0.184572 | 0.180790 | 0.254328 | 0.323908 | 0.429507 | 0.417905 | 0.570704 | 0.740578 | 1.582311 | 24.289178 | 0.627992552 | 1.054205311 | 0.291788929 | NA | 4.234218 | 4.895208 | 4.568826 | 4.566666 | 4.599994 | 4.626727 | 22 | 20 | 23 | 31 |
| 100271836 | NR_027155 | SMG1P3 | SMG1 pseudogene 3 | transcribed_ps | . | 49860 | . | . | . | 2.801260 | 3.225335 | 3.730617 | 2.777551 | 6.518648 | 7.455505 | 8.371395 | 6.350541 | 1.064774 | 493.643974 | 0.192996488 | 0.469163882 | 0.6389525 | 0.998865404 | 8.901001 | 8.992212 | 8.880095 | 8.963688 | 9.054602 | 8.861355 | 514 | 550 | 510 | 405 |
| 100271874 | NR_027347 | ZNRF2P2 | zinc and ring finger 2 pseudogene 2 | transcribed_ps | . | 42793 | . | . | . | 0.378067 | 0.720093 | 0.433345 | 0.649413 | 0.879777 | 1.664527 | 0.972413 | 1.484805 | -1.019726 | 18.351251 | 0.739872923 | -0.03808885 | 0.969616842 | NA | 4.214202 | 4.181222 | 4.170601 | 4.219116 | 4.180294 | 4.206879 | 15 | 26 | 13 | 20 |
| 100271927 | NM_00127733 | RASA4B | RAS p21 protein activator 4B | protein_coding | NP_001264264.1;NP_001354696.1;XP_011513992.1;XP_016867115.1;XP | 35202 | . | . | . | 0.818312 | 0.796648 | 0.792026 | 1.151138 | 1.904247 | 1.841488 | 1.777285 | 2.631942 | -1.007016 | 134.619815 | 0.284275383 | -0.03547983 | 0.971697126 | 0.998865404 | 7.077877 | 7.067599 | 7.098479 | 7.048462 | 7.047423 | 7.094460 | 166 | 136 | 109 | 134 |
| 100272146 | gene-NFE2L3P | NFE2L3P2 | nuclear factor, erythroid 2 like 3 pseudogene 2 | pseudogene | . | 44191 | . | . | . | 0.107883 | 0.049965 | 0.037721 | 0.011626 | 0.251049 | 0.115496 | 0.084646 | 0.026581 | **-3.365610** | 4.892904 | 1.479888282 | -1.18310828 | 0.236766238 | NA | 2.912173 | 1.175234 | 2.330618 | 2.303860 | 2.298766 | 2.287899 | 12 | 5 | 3 | 1 |
| 100272147 | NM_00101802 | CMC4 | C-X9-C motif containing 4 | protein_coding | NP_001018024.1 | 35428 | . | . | . | 6.589534 | 6.655333 | 6.996648 | 6.234842 | 15.334117 | 15.384102 | 15.700275 | 14.255225 | -1.014556 | 182.095628 | 0.247584802 | -0.08420565 | 0.932892927 | 0.998865404 | 7.518601 | 7.498434 | 7.510836 | 7.511418 | 7.530889 | 7.478342 | 211 | 198 | 167 | 159 |
| 100272217 | NR_027440 | LOC100272217 | uncharacterized LOC100272217 | lncRNA | . | . | . | . | . | 0.543953 | 0.899758 | 1.267391 | 0.436247 | 1.265801 | 2.079831 | 2.843990 | 0.997427 | 1.165250 | 53.368234 | 0.52336582 | 0.421579075 | 0.673332282 | 0.998865404 | 5.622171 | 5.845048 | 5.651001 | 5.741690 | 5.819483 | 5.624665 | 43 | 66 | 74 | 28 |
| 100272228 | NR_027456 | LINC00894 | long intergenic non-protein coding RNA 894 | lncRNA | . | 48579 | . | . | . | 0.357932 | 0.504580 | 0.299816 | 0.338514 | 0.832923 | 1.166358 | 0.672777 | 0.773972 | -1.363516 | 37.869552 | 0.51544929 | -0.86784698 | 0.385478108 | 0.998865404 | 5.450313 | 5.000737 | 5.237438 | 5.293700 | 5.212553 | 5.228605 | 42 | 55 | 26 | 32 |
| 100286844 | NR_027499,NF | BCDIN3D-AS1 | BCDIN3D antisense RNA 1 | lncRNA | . | 44113 | . | . | . | 0.210673 | 0.155611 | 0.273753 | 0.146926 | 0.490244 | 0.359701 | 0.614294 | 0.335927 | -1.015176 | 18.660023 | 0.749196056 | -0.02900453 | 0.976860981 | NA | 4.229742 | 4.213972 | 4.224547 | 4.206297 | 4.246522 | 4.180770 | 24 | 18 | 23 | 10 |
| 100286925 | NR_147620 | PPM1F-AS1 | PPM1F antisense RNA 1 | lncRNA | . | 40888 | . | . | . | 0.058765 | 0.031673 | 0.067379 | 0.141279 | 0.136748 | 0.073213 | 0.151198 | 0.323018 | **2.356217** | 8.942753 | 1.063311957 | 1.162850051 | 0.244890346 | NA | 2.413393 | 3.650563 | 3.131807 | 3.116799 | 3.137245 | 3.176496 | 8 | 4 | 7 | 16 |
| 100286979 | NR_037931 | ANAPC1P1 | anaphase promoting complex subunit 1 pseudogene 1 | transcribed_ps | . | 44150 | . | . | . | 0.262837 | 0.207808 | 0.324383 | 0.358966 | 0.611631 | 0.480358 | 0.727907 | 0.820733 | 1.441592 | 16.369365 | 0.757811365 | 0.696298964 | 0.486241622 | NA | 3.744764 | 4.273004 | 4.018113 | 4.006075 | 4.033739 | 4.041771 | 17 | 13 | 16 | 19 |
| 100287015 | NR_040040 | LOC100287015 | uncharacterized LOC100287015 | lncRNA | . | . | . | . | . | 1.044573 | 0.994844 | 1.212455 | 1.311152 | 2.430764 | 2.299628 | 2.720713 | 2.997793 | 1.219639 | 85.986085 | 0.342792201 | 0.835649352 | 0.403352196 | 0.998865404 | 6.275780 | 6.562100 | 6.402042 | 6.386095 | 6.437313 | 6.460048 | 92 | 81 | 79 | 92 |
| 100287029 | NR_045117 | DDX11L10 | DEAD/H-box helicase 11 like 10 (pseudogene) | transcribed_ps | . | 14125 | . | . | . | 0.044611 | 0.075352 | 0.213268 | 0.076819 | 0.103812 | 0.174180 | 0.478568 | 0.175637 | **2.176703** | 5.751365 | 1.310280585 | 0.856415632 | 0.391767921 | NA | 1.855800 | 2.978782 | 2.481933 | 2.491792 | 2.522167 | 2.490298 | 3 | 5 | 10 | 4 |
| 100287042 | NR_036520 | LOC100287042 | uncharacterized LOC100287042 | lncRNA | . | . | . | . | . | 0.441861 | 0.337295 | 0.297168 | 0.268378 | 1.028229 | 0.779672 | 0.666836 | 0.613613 | -1.382411 | 27.706594 | 0.592819937 | -0.78807586 | 0.430652333 | 0.998865404 | 5.006229 | 4.540693 | 4.831176 | 4.795396 | 4.780916 | 4.773490 | 42 | 30 | 21 | 21 |
| 100287046 | gene-GUSBP13 | GUSBP13 | GUSB pseudogene 13 | pseudogene | . | 54495 | . | . | . | 1.495418 | 1.354199 | 1.905348 | 1.464341 | 3.479899 | 3.130291 | 4.275545 | 3.348041 | 1.167877 | 33.010972 | 0.542504051 | 0.412694491 | 0.679830462 | 0.998865404 | 4.927457 | 5.153450 | 5.036730 | 5.022800 | 5.067886 | 5.032131 | 37 | 31 | 35 | 29 |
| 100287102 | NR_046018 | DDX11L1 | DEAD/H-box helicase 11 like 1 (pseudogene) | transcribed_ps | . | 37102 | . | . | . | 0.139645 | 0.164779 | 0.359649 | 0.311661 | 0.324959 | 0.380894 | 0.807042 | 0.712575 | **2.081083** | 13.158249 | 0.851752087 | 1.241364153 | 0.214471255 | NA | 3.094525 | 4.151760 | 3.673394 | 3.680442 | 3.724917 | 3.710362 | 9 | 10 | 17 | 15 |
| 100287171 | NM_00137809 | WASHC1 | WASH complex subunit 1 | protein_coding | NP_001365019.1;NP_878908.4;XP_011515961.1;XP_011515962.1;XP_01 | 24361 | 613632 | . | . | 4.261528 | 5.556880 | 5.003094 | 4.398864 | 9.916751 | 12.844976 | 11.226798 | 10.057483 | 1.075275 | 889.691156 | 0.150831369 | 0.694189476 | 0.487563392 | 0.998865404 | 9.743687 | 9.848723 | 9.742289 | 9.794792 | 9.880015 | 9.758715 | 951 | 959 | 869 | 792 |
| 100287177 | NR_130728,NF | EML2-AS1 | EML2 antisense RNA 1 | lncRNA | . | 48331 | . | . | . | 0.633657 | 0.506971 | 0.953180 | 0.762006 | 1.474547 | 1.171887 | 2.138908 | 1.742236 | 1.467540 | 24.279623 | 0.629870308 | 0.878592704 | 0.379622153 | NA | 4.297162 | 4.852980 | 4.580368 | 4.562758 | 4.622807 | 4.596855 | 25 | 19 | 28 | 24 |
| 100287178 | NM_00125685 | USP17L11 | ubiquitin specific peptidase 17 like family member 11 | protein_coding | NP_001243783.1 | 44439 | . | . | . | 0.003461 | 0.016038 | 0.015723 | 0.034135 | 0.008053 | 0.037072 | 0.035282 | 0.078045 | 1.834012 | 1.269624 | 2.598200026 | 0.336772769 | 0.736288203 | NA | -0.155866 | 0.715150 | 0.332517 | 0.332827 | 0.333928 | 0.338438 | 1 | 1 | 1 | 2 |
| 100287205 | NM_00125685 | USP17L12 | ubiquitin specific peptidase 17 like family member 12 | protein_coding | NP_001243782.1 | 44440 | . | . | . | 0.003461 | 0.012106 | 0.014781 | 0.024772 | 0.008053 | 0.027984 | 0.033167 | 0.056638 | 1.834012 | 1.269624 | 2.598200026 | 0.336772769 | 0.736288203 | NA | -0.155866 | 0.715150 | 0.332517 | 0.332827 | 0.333928 | 0.338438 | 1 | 1 | 1 | 2 |
| 100287226 | NM_00124268 | ZNF729 | zinc finger protein 729 | protein_coding | NP_001229609.1 | 32464 | . | . | . | 0.285676 | 0.335023 | 0.267684 | 0.190585 | 0.664779 | 0.774421 | 0.600676 | 0.435750 | -1.358038 | 32.795102 | 0.549364142 | -0.8036994 | 0.4215706 | 0.998865404 | 5.238681 | 4.798723 | 5.043868 | 5.067739 | 5.033624 | 4.998648 | 40 | 44 | 28 | 22 |
| 100287227 | NR_027954 | TIPARP-AS1 | TIPARP antisense RNA 1 | lncRNA | . | 41028 | . | . | . | 0.287784 | 0.201199 | 0.328221 | 0.169341 | 0.669684 | 0.465080 | 0.736519 | 0.387179 | 1.000996 | 20.111355 | 0.710106023 | 0.002022998 | 0.998385883 | NA | 4.326585 | 4.333284 | 4.339659 | 4.310212 | 4.350725 | 4.298734 | 27 | 18 | 23 | 13 |
| 100287238 | NM_00125685 | USP17L13 | ubiquitin specific peptidase 17 like family member 13 | protein_coding | NP_001243784.1 | 44441 | . | . | . | 0.008429 | 0.004036 | 0.001793 | 0.022416 | 0.019616 | 0.010763 | 0.004023 | 0.051253 | 1.834012 | 1.269624 | 2.598200026 | 0.336772769 | 0.736288203 | NA | -0.155866 | 0.715150 | 0.332517 | 0.332827 | 0.333928 | 0.338438 | 1 | 1 | 1 | 2 |
| 100287270 | gene-USP17L1 | USP17L14P | ubiquitin specific peptidase 17 like family member 14, pseudogene | pseudogene | . | 44442 | . | . | . | 0.006699 | 0.013388 | 0.000896 | 0.014734 | 0.015589 | 0.030946 | 0.002012 | 0.033687 | 1.234655 | 1.002957 | 2.868339486 | 0.106022226 | 0.915564717 | NA | -0.155866 | 0.148375 | -0.001261 | -0.000972 | 0.000053 | -0.000315 | 1 | 1 | 1 | 1 |
| 100287275 | gene-PCOTHP1 | PCOTHP1 | Pro-X-Gly collagen triple helix like repeat containing pseudogene 1 | pseudogene | . | 39840 | . | . | . | 0.239421 | 0.469597 | 0.324924 | 0.202986 | 0.557142 | 1.085495 | 0.729121 | 0.464102 | -1.380277 | 6.642840 | 1.197146727 | -0.38838802 | 0.697728906 | NA | 2.945312 | 2.481107 | 2.725140 | 2.748155 | 2.733005 | 2.721249 | 6 | 11 | 6 | 4 |
| 100287290 | XR_00174050 | LOC100287290 | cytokine receptor CRL2, transcript variant X9 | misc_RNA | . | . | . | . | . | 0.059396 | 0.043972 | 0.036387 | 0.028115 | 0.138218 | 0.101643 | 0.081652 | 0.064281 | -1.322441 | 18.093071 | 0.728097097 | -0.55377759 | 0.579731069 | NA | 4.363616 | 3.963455 | 4.195647 | 4.185741 | 4.181074 | 4.157319 | 25 | 21 | 16 | 12 |
| 100287314 | NR_040245 | LINC00941 | long intergenic non-protein coding RNA 941 | lncRNA | . | 48635 | . | . | . | 2.525799 | 2.361137 | 2.086190 | 2.544415 | 5.877638 | 5.457874 | 4.681350 | 5.817502 | -1.067940 | 140.857005 | 0.279919172 | -0.33877917 | 0.734776095 | 0.998865404 | 7.185110 | 7.089480 | 7.165375 | 7.135391 | 7.090418 | 7.159988 | 174 | 151 | 107 | 140 |
| 100287327 | NM_00125685 | USP17L17 | ubiquitin specific peptidase 17 like family member 17 | protein_coding | NP_001243786.1 | 44445 | . | . | . | 0.001730 | 0.005794 | 0.000896 | 0.013053 | 0.004027 | 0.013394 | 0.002012 | 0.029845 | 1.234655 | 1.002957 | 2.868339486 | 0.106022226 | 0.915564717 | NA | -0.155866 | 0.148375 | -0.001261 | -0.000972 | 0.000053 | -0.000315 | 1 | 1 | 1 | 1 |
| 100287369 | gene-HSPE1P4 | HSPE1P4 | heat shock protein family E (Hsp10) member 1 pseudogene 4 | pseudogene | . | 49323 | . | . | . | 0.088733 | 0.329431 | 0.306428 | 0.253287 | 0.206485 | 0.761495 | 0.687615 | 0.579110 | 1.376459 | 4.328473 | 1.47435995 | 0.312652338 | 0.75454481 | NA | 1.867373 | 2.324312 | 2.089088 | 2.107348 | 2.107843 | 2.101278 | 2 | 6 | 5 | 4 |
| 100287425 | gene-LOC1002 | LOC100287425 | short transmembrane mitochondrial protein 1 pseudogene | pseudogene | . | . | . | . | . | 0.095834 | 0.081431 | 0.052634 | 0.166991 | 0.223010 | 0.188232 | 0.118110 | 0.381806 | 1.216986 | 1.985087 | 2.102786642 | 0.134731979 | 0.892823788 | NA | 0.844134 | 1.121007 | 0.986974 | 0.987547 | 0.984432 | 0.993706 | 2 | 2 | 1 | 3 |
| 100287482 | NM_00119524 | SMKR1 | small lysine rich protein 1 | protein_coding | NP_001182172.1;XP_024302388.1 | 43561 | . | . | . | 0.532905 | 0.184015 | 0.345022 | 0.333935 | 1.240092 | 0.425358 | 0.774220 | 0.763503 | -1.040151 | 11.288506 | 0.939333477 | -0.06046104 | 0.951788444 | NA | 3.522785 | 3.470304 | 3.519442 | 3.473993 | 3.496776 | 3.493169 | 19 | 7 | 10 | 10 |
| 100287497 | NR_136590,NF | LOC100287497 | septin 7 pseudogene | transcribed_ps | . | . | . | . | . | 2.145401 | 1.801349 | 2.529999 | 2.604234 | 4.992435 | 4.163898 | 5.677245 | 5.954272 | 1.246742 | 134.150110 | 0.283783803 | 1.121146367 | 0.262225565 | 0.995760118 | 6.899379 | 7.218426 | 7.049593 | 6.992008 | 7.105909 | 7.097512 | 149 | 118 | 131 | 138 |
| 100287551 | gene-HSPA8P8 | HSPA8P8 | heat shock protein family A (Hsp70) member 8 pseudogene 8 | pseudogene | . | 44923 | . | . | . | 2.490042 | 1.751891 | 1.825743 | 0.219508 | 5.794430 | 4.049575 | 4.096914 | 0.501879 | **-2.088453** | 111.752881 | 1.264074633 | -0.84048427 | 0.400636916 | 0.998865404 | 7.239429 | 6.177889 | 6.937627 | 6.789592 | 6.800614 | 6.420156 | 205 | 134 | 112 | 15 |
| 100287569 | NR_027345,NF | LINC00173 | long intergenic non-protein coding RNA 173 | lncRNA | . | 33791 | . | . | . | 1.535274 | 1.214348 | 1.039531 | 0.885417 | 3.572645 | 2.807017 | 2.332678 | 2.024402 | -1.402545 | 38.767175 | 0.508551203 | -0.9596805 | 0.337216041 | 0.998865404 | 5.499094 | 5.013816 | 5.309297 | 5.296064 | 5.274296 | 5.230087 | 54 | 47 | 33 | 25 |
| 100287596 | NR_051986 | DDX11L5 | DEAD/H-box helicase 11 like 5 (pseudogene) | transcribed_ps | . | 37106 | . | . | . | 0.091607 | 0.188244 | 0.153790 | 0.204353 | 0.213173 | 0.435135 | 0.345101 | 0.467230 | 1.219196 | 8.530334 | 1.053979542 | 0.271285666 | 0.78617133 | NA | 2.945312 | 3.226238 | 3.070870 | 3.094077 | 3.084377 | 3.095873 | 6 | 11 | 7 | 10 |
| 100287653 | gene-LOC1002 | LOC100287653 | PDGFA associated protein 1 pseudogene | pseudogene | . | . | . | . | . | 0.178566 | 0.310554 | 0.371785 | 0.396074 | 0.415531 | 0.717859 | 0.834274 | 0.905576 | 1.602850 | 7.638992 | 1.105503304 | 0.615683227 | 0.538103609 | NA | 2.555346 | 3.232645 | 2.905905 | 2.920474 | 2.928287 | 2.930208 | 5 | 8 | 8 | 9 |
| 100287765 | NR_038988,NF | LINC00630 | long intergenic non-protein coding RNA 630 | lncRNA | . | 44263 | . | . | . | 0.191864 | 0.125856 | 0.368201 | 0.146103 | 0.446477 | 0.290920 | 0.826231 | 0.334050 | 1.627623 | 27.601703 | 0.632696468 | 1.110748714 | 0.266676528 | 0.998274218 | 4.389723 | 5.097659 | 4.749942 | 4.721061 | 4.839151 | 4.738143 | 28 | 19 | 41 | 20 |
| 100287808 | NR_149004 | LOC100287808 | uncharacterized LOC100287808 | lncRNA | . | . | . | . | . | 0.670564 | 0.229094 | 0.683589 | 0.445824 | 1.560430 | 0.529561 | 1.533954 | 1.019323 | 1.231548 | 43.407323 | 0.563323488 | 0.533392414 | 0.593761968 | 0.998865404 | 5.278863 | 5.584693 | 5.478324 | 5.323352 | 5.476547 | 5.398217 | 67 | 21 | 51 | 35 |
| 100287837 | NR_147696 | LOC100287837 | uncharacterized LOC100287837 | lncRNA | . | . | . | . | . | 0.148035 | 0.084197 | 0.036666 | 0.201342 | 0.344484 | 0.194625 | 0.082277 | 0.460345 | 1.031537 | 5.870303 | 1.298011959 | 0.034510512 | 0.97247006 | NA | 2.533720 | 2.572884 | 2.559869 | 2.549066 | 2.539091 | 2.571329 | 8 | 5 | 2 | 9 |
| 100287887 | gene-RPL13AP | RPL13AP25 | ribosomal protein L13a pseudogene 25 | pseudogene | . | 36981 | . | . | . | 2.939655 | 3.515826 | 2.117769 | 4.965225 | 6.840699 | 8.126989 | 4.752212 | 11.352397 | 1.078388 | 69.976157 | 0.438796396 | 0.24812592 | 0.804036892 | 0.998865404 | 6.075449 | 6.180232 | 6.087810 | 6.128684 | 6.026042 | 6.218306 | 71 | 79 | 38 | 95 |
| 100287896 | NM_00131924 | LOC100287896 | uncharacterized LOC100287896 | protein_coding | NP_001306169.1;XP_016872544.1 | . | . | . | . | 1.733478 | 1.886487 | 1.492301 | 1.071724 | 4.033874 | 4.360701 | 3.348680 | 2.450369 | -1.426341 | 116.004026 | 0.30931173 | -1.6563205 | 0.097656931 | 0.813553488 | 7.090828 | 6.580303 | 6.909923 | 6.914032 | 6.834652 | 6.752111 | 156 | 148 | 95 | 77 |
| 100287898 | NM_00124267 | TTC34 | tetratricopeptide repeat domain 34 | protein_coding | NP_001229601.2;XP_016855479.1 | 34297 | . | . | . | 0.137384 | 0.238033 | 0.173753 | 0.156377 | 0.319698 | 0.550225 | 0.389896 | 0.357538 | -1.152527 | 31.649433 | 0.569318372 | -0.35972846 | 0.719050206 | 0.998865404 | 5.083534 | 4.877319 | 4.953777 | 5.024695 | 4.978552 | 4.965299 | 29 | 46 | 27 | 26 |
| 100287932 | NM_006327,N | TIMM23 | translocase of inner mitochondrial membrane 23 | protein_coding | NP_006318.1 | 17312 | 605034 | . | . | 48.877711 | 50.401713 | 44.139571 | 47.605229 | 113.740450 | 116.505810 | 99.047915 | 108.843707 | -1.096048 | 1774.382997 | 0.122090237 | -1.08371752 | 0.278490064 | 0.998865404 | 10.857832 | 10.725330 | 10.819519 | 10.836571 | 10.730812 | 10.780425 | 2114 | 2024 | 1425 | 1638 |
| 100288069 | NR_168328 | LOC100288069 | uncharacterized LOC100288069 | lncRNA | . | . | . | . | . | 2.058054 | 2.219319 | 2.767729 | 2.346576 | 4.789177 | 5.130055 | 6.210703 | 5.365167 | 1.178626 | 520.758994 | 0.173616156 | 1.36568806 | 0.172036894 | 0.947431715 | 8.900816 | 9.138361 | 8.945190 | 8.987271 | 9.127672 | 9.012841 | 532 | 533 | 533 | 482 |
| 100288073 | gene-LOC1002 | LOC100288073 | heat shock protein family A (Hsp70) member 5 pseudogene | pseudogene | . | . | . | . | . | 1.452411 | 1.839925 | 0.982644 | 1.376428 | 3.379819 | 4.253069 | 2.205024 | 3.147040 | -1.408527 | 106.093877 | 0.330584953 | -1.49488813 | 0.134943642 | 0.891755273 | 6.956388 | 6.459419 | 6.737477 | 6.817188 | 6.625279 | 6.713682 | 127 | 149 | 64 | 96 |
| 100288142 | NM_00127826 | NBPF20 | NBPF member 20 | protein_coding | NP_001265196.1 | 32000 | 614007 | . | . | 0.822815 | 0.788443 | 1.014473 | 0.807803 | 1.914725 | 1.822520 | 2.276447 | 1.846945 | 1.116231 | 494.078432 | 0.18178873 | 0.872638899 | 0.382859931 | 0.998865404 | 8.866573 | 9.026206 | 8.925187 | 8.893951 | 9.052410 | 8.900763 | 551 | 491 | 507 | 431 |
| 100288152 | NR_125375 | SLC9A3-AS1 | SLC9A3 antisense RNA 1 | lncRNA | . | 40550 | . | . | . | 4.070812 | 4.397021 | 3.917336 | 4.810452 | 9.472947 | 10.163911 | 8.790389 | 10.998528 | 1.015649 | 239.183664 | 0.226798071 | 0.098772738 | 0.921318715 | 0.998865404 | 7.891346 | 7.912526 | 7.879994 | 7.913976 | 7.855928 | 7.951053 | 264 | 265 | 189 | 248 |
| 100288165 | gene-HMGN2P | HMGN2P15 | high mobility group nucleosomal binding domain 2 pseudogene 1 | pseudogene | . | 39020 | . | . | . | 0.517748 | 0.733593 | 0.829832 | 0.662358 | 1.204821 | 1.695732 | 1.862119 | 1.514403 | 1.163561 | 25.839790 | 0.614441459 | 0.355683062 | 0.722077931 | NA | 4.578230 | 4.796563 | 4.660270 | 4.693831 | 4.706156 | 4.680270 | 23 | 30 | 27 | 23 |
| 100288175 | NR_148960 | LOC100288175 | uncharacterized LOC100288175 | lncRNA | . | . | . | . | . | 1.859811 | 1.721099 | 2.212770 | 1.664771 | 4.327856 | 3.978397 | 4.965392 | 3.806301 | 1.069603 | 57.030866 | 0.422429404 | 0.229803164 | 0.818244726 | 0.998865404 | 5.783216 | 5.882421 | 5.831504 | 5.814967 | 5.867432 | 5.805679 | 66 | 57 | 59 | 47 |
| 100288181 | NR_038925 | LNCOC1 | lncRNA associated with ovarian cancer 1 | lncRNA | . | 53947 | . | . | . | 0.227295 | 0.200638 | 0.164102 | 0.258453 | 0.528926 | 0.463783 | 0.368241 | 0.590922 | -1.014331 | 38.181460 | 0.514866045 | -0.03987109 | 0.968195899 | 0.998865404 | 5.266033 | 5.243479 | 5.264886 | 5.246273 | 5.222721 | 5.283411 | 47 | 39 | 26 | 43 |
| 100288198 | NR_040671 | HMGN3-AS1 | HMGN3 antisense RNA 1 | lncRNA | . | 48984 | . | . | . | 0.312861 | 0.477734 | 0.286146 | 0.235306 | 0.728041 | 1.104303 | 0.642104 | 0.537999 | -1.498637 | 31.234229 | 0.571825404 | -1.02068023 | 0.307405963 | 0.998865404 | 5.227616 | 4.643839 | 4.956453 | 5.019947 | 4.949329 | 4.927944 | 34 | 49 | 24 | 21 |
| 100288203 | NR_146103,NF | LOC100288203 | WASP homolog associated with actin, golgi membranes and micro | transcribed_ps | . | . | . | . | . | 1.624914 | 1.651567 | 1.948094 | 1.554733 | 3.781242 | 3.817672 | 4.371467 | 3.554714 | 1.081619 | 118.183492 | 0.303894922 | 0.372472474 | 0.709541103 | 0.998865404 | 6.826572 | 6.940932 | 6.861058 | 6.875051 | 6.935884 | 6.851750 | 128 | 125 | 120 | 101 |
| 100288332 | NM_00127732 | NPIPA5 | nuclear pore complex interacting protein family member A5 | protein_coding | NP_001264254.1;NP_001338129.1;XP_011520631.1;XP_011520638.1;XP | 41980 | . | . | . | 22.754222 | 23.970076 | 34.311124 | 21.428988 | 52.950009 | 55.407904 | 76.993166 | 48.994834 | 1.161660 | 960.072787 | 0.185550421 | 1.16511892 | 0.24397086 | 0.992733415 | 9.794472 | 10.011383 | 9.833566 | 9.851314 | 10.100540 | 9.778058 | 1010 | 970 | 1091 | 759 |
| 100288380 | NR_027470 | ULK4P2 | ULK4 pseudogene 2 | transcribed_ps | . | 15776 | . | . | . | 0.336308 | 0.471263 | 0.657266 | 0.494161 | 0.782603 | 1.089345 | 1.474885 | 1.129842 | 1.432397 | 13.698388 | 0.831465443 | 0.623515391 | 0.532945891 | NA | 3.493602 | 4.011955 | 3.746279 | 3.762056 | 3.784983 | 3.766013 | 11 | 14 | 16 | 13 |
| 100288413 | NM_00124269 | ERVMER34-1 | endogenous retrovirus group MER34 member 1, envelope | protein_coding | NP_001229619.1;NP_078810.1 | 42970 | . | . | . | 1.617685 | 1.051824 | 1.192547 | 1.095383 | 3.764420 | 2.431337 | 2.676040 | 2.504463 | -1.261387 | 128.349182 | 0.303521969 | -1.10374427 | 0.269704086 | 0.998865404 | 7.161044 | 6.827591 | 7.105952 | 6.972662 | 6.969923 | 6.948256 | 193 | 128 | 102 | 103 |
| 100288428 | NR_033378 | LMCD1-AS1 | LMCD1 antisense RNA 1 | lncRNA | . | 44477 | . | . | . | 0.175301 | 0.187446 | 0.134228 | 0.169893 | 0.407933 | 0.433290 | 0.301204 | 0.388440 | -1.198946 | 11.516437 | 0.898847659 | -0.29122465 | 0.770879509 | NA | 3.651489 | 3.387716 | 3.532036 | 3.535648 | 3.519219 | 3.530055 | 14 | 14 | 8 | 11 |
| 100288436 | gene-SLC4A1AI | SLC4A1APP1 | solute carrier family 4 member 1 adaptor protein pseudogene 1 | pseudogene | . | 49811 | . | . | . | 0.109788 | 0.074912 | 0.024091 | 0.111801 | 0.255481 | 0.173162 | 0.054059 | 0.255620 | -1.431344 | 2.650995 | 1.84938096 | -0.27975329 | 0.779666793 | NA | 1.644780 | 1.121007 | 1.420364 | 1.417051 | 1.409777 | 1.418929 | 4 | 3 | 1 | 3 |
| 100288486 | NR_034090,NF | DDX11L9 | DEAD/H-box helicase 11 like 9 (pseudogene) | transcribed_ps | . | 37109 | . | . | . | 0.127608 | 0.107102 | 0.153712 | 0.207547 | 0.296948 | 0.247572 | 0.344926 | 0.474531 | 1.496705 | 7.806124 | 1.087556986 | 0.534951285 | 0.592683544 | NA | 2.644780 | 3.226238 | 2.953131 | 2.946747 | 2.958208 | 2.969788 | 8 | 6 | 7 | 10 |
| 100288520 | NM_00125689 | USP17L15 | ubiquitin specific peptidase 17 like family member 15 | protein_coding | NP_001243823.2 | 44443 | . | . | . | 0.004976 | 0.015982 | 0.006676 | 0.024652 | 0.011579 | 0.036943 | 0.014981 | 0.056363 | 1.834012 | 1.269624 | 2.598200026 | 0.336772769 | 0.736288203 | NA | -0.155866 | 0.715150 | 0.332517 | 0.332827 | 0.333928 | 0.338438 | 1 | 1 | 1 | 2 |
| 100288615 | NR_036650 | WHAMMP1 | WAS protein homolog associated with actin, golgi membranes and | transcribed_ps | . | 32361 | . | . | . | 1.595495 | 1.869301 | 1.986188 | 1.608631 | 3.712782 | 4.320973 | 4.456948 | 3.677943 | 1.027860 | 86.597618 | 0.348834092 | 0.113646458 | 0.909518058 | 0.998865404 | 6.415961 | 6.456268 | 6.408569 | 6.450295 | 6.467845 | 6.405716 | 91 | 99 | 85 | 73 |
| 100288637 | NR_038253,NF | LOC100288637 | OTU deubiquitinase 7A pseudogene | transcribed_ps | . | . | . | . | . | 1.350504 | 1.298819 | 1.595070 | 1.354943 | 3.142678 | 3.002280 | 3.579291 | 3.097916 | 1.099645 | 247.913797 | 0.221190255 | 0.619544814 | 0.535557509 | 0.998865404 | 7.883034 | 8.021055 | 7.940136 | 7.914344 | 8.014201 | 7.933581 | 279 | 248 | 244 | 224 |
| 100288690 | gene-FAM21FP | FAM21FP | . | pseudogene | . | 45011 | . | . | . | 0.137825 | 0.363930 | 0.190276 | 0.294615 | 0.320724 | 0.841239 | 0.426973 | 0.673602 | -1.052067 | 16.374488 | 0.803507671 | -0.0911343 | 0.92738588 | NA | 4.072459 | 3.993209 | 3.996346 | 4.060655 | 4.011650 | 4.039199 | 11 | 26 | 11 | 18 |
| 100288695 | NM_00120528 | LIMS4 | LIM zinc finger domain containing 4 | protein_coding | NP_001192217.1;NP_001358269.1;XP_016858592.1;XP_016858593.1;XP | 39941 | . | . | . | 0.605914 | 0.631459 | 0.598299 | 0.754816 | 1.409987 | 1.459646 | 1.342565 | 1.725798 | 1.063178 | 43.862442 | 0.472779122 | 0.186944298 | 0.851704307 | 0.998865404 | 5.410533 | 5.497971 | 5.455673 | 5.442013 | 5.440621 | 5.477198 | 51 | 44 | 35 | 47 |
| 100288713 | gene-PMS2P10 | PMS2P10 | PMS1 homolog 2, mismatch repair system component pseudogen | pseudogene | . | 9124 | . | . | . | 0.354370 | 0.490952 | 0.383082 | 0.271395 | 0.824633 | 1.134858 | 0.859625 | 0.620513 | -1.330136 | 9.082975 | 1.017198023 | -0.40461506 | 0.685760489 | NA | 3.373794 | 2.963455 | 3.183162 | 3.198433 | 3.185516 | 3.173147 | 10 | 13 | 8 | 6 |
| 100288730 | NR_029383 | PAN3-AS1 | PAN3 antisense RNA 1 | lncRNA | . | 39932 | . | . | . | 0.493463 | 0.599512 | 0.757834 | 0.563157 | 1.148309 | 1.385800 | 1.700557 | 1.287591 | 1.178643 | 41.647514 | 0.489819233 | 0.484111605 | 0.628306666 | 0.998865404 | 5.256278 | 5.494237 | 5.346397 | 5.374345 | 5.410970 | 5.361733 | 40 | 45 | 45 | 36 |
| 100288778 | NR_130745 | WASH8P | WAS protein family homolog 8, pseudogene | transcribed_ps | . | 53913 | . | . | . | 11.369511 | 11.622448 | 14.104360 | 11.304282 | 26.457317 | 26.865808 | 31.649775 | 25.845896 | 1.090258 | 707.378350 | 0.164691963 | 0.75698664 | 0.449057866 | 0.998865404 | 9.402300 | 9.527655 | 9.428713 | 9.437879 | 9.570212 | 9.409460 | 774 | 735 | 716 | 612 |
| 100288842 | NR_027442 | B3GNT10 | UDP-GlcNAc:betaGal beta-1,3-N-acetylglucosaminyltransferase 10 | transcribed_ps | . | 53652 | . | . | . | 1.203914 | 1.071502 | 1.049573 | 0.902270 | 2.801557 | 2.476826 | 2.355211 | 2.062933 | -1.181623 | 118.596576 | 0.301111107 | -0.79960363 | 0.423940483 | 0.998865404 | 7.004626 | 6.765297 | 6.939795 | 6.894667 | 6.885597 | 6.836035 | 157 | 130 | 102 | 94 |
| 100288846 | NR_038935 | LOC100288846 | uncharacterized LOC100288846 | lncRNA | . | . | . | . | . | 0.331845 | 0.342707 | 0.541759 | 0.173369 | 0.772218 | 0.792184 | 1.215691 | 0.396387 | 1.025386 | 15.014396 | 0.838625591 | 0.043127319 | 0.965600042 | NA | 3.887108 | 3.929136 | 3.897465 | 3.897761 | 3.931534 | 3.865724 | 17 | 16 | 20 | 7 |
| 100289019 | NR_033374 | SLC25A25-AS1 | SLC25A25 antisense RNA 1 | lncRNA | . | 27844 | . | . | . | 0.233789 | 0.185643 | 0.348171 | 0.182289 | 0.544035 | 0.429123 | 0.781286 | 0.416782 | 1.249417 | 29.194066 | 0.598748716 | 0.536544981 | 0.591581957 | 0.998865404 | 4.695638 | 5.021237 | 4.854384 | 4.831350 | 4.907742 | 4.829002 | 33 | 25 | 37 | 21 |
| 100289027 | gene-CDK2AP2 | CDK2AP2P1 | cyclin dependent kinase 2 associated protein 2 pseudogene 1 | pseudogene | . | 38492 | . | . | . | 0.191465 | 0.402384 | 0.478843 | 0.110762 | 0.445546 | 0.930129 | 1.074510 | 0.253245 | 1.012256 | 7.294461 | 1.17614978 | 0.014942201 | 0.988078292 | NA | 2.855800 | 2.877719 | 2.846436 | 2.865353 | 2.874443 | 2.837849 | 6 | 10 | 10 | 3 |
| 100289034 | gene-RPL23AP | RPL23AP88 | ribosomal protein L23a pseudogene 88 | pseudogene | . | 51579 | . | . | . | 0.110647 | 0.121129 | 0.132199 | 0.093909 | 0.257479 | 0.279994 | 0.296650 | 0.214712 | -1.215015 | 2.454711 | 1.887245685 | -0.1488804 | 0.881648006 | NA | 1.429096 | 1.148375 | 1.299640 | 1.300454 | 1.298230 | 1.297456 | 3 | 3 | 2 | 2 |
| 100289092 | NR_046287,NF | ATP2A1-AS1 | ATP2A1 antisense RNA 1 | lncRNA | . | 51370 | . | . | . | 3.670509 | 6.192038 | 3.391371 | 4.284094 | 8.541424 | 14.313175 | 7.610139 | 9.795073 | -1.286001 | 75.056807 | 0.396896447 | -0.91432406 | 0.360546594 | 0.998865404 | 6.401265 | 6.035427 | 6.182705 | 6.322552 | 6.168718 | 6.215223 | 73 | 114 | 51 | 68 |
| 100289097 | NR_132315 | FRG1CP | FSHD region gene 1 family member C, pseudogene | transcribed_ps | . | 51762 | . | . | . | 11.097891 | 10.655224 | 12.685822 | 12.887277 | 25.825249 | 24.630026 | 28.466618 | 29.465225 | 1.160404 | 312.127952 | 0.197758353 | 1.085298061 | 0.277789664 | 0.998865404 | 8.174599 | 8.389401 | 8.255209 | 8.228957 | 8.321946 | 8.324530 | 341 | 304 | 291 | 315 |
| 100289098 | NR_134570,NF | GS1-124K5.4 | uncharacterized LOC100289098 | lncRNA | . | . | . | . | . | 1.636532 | 2.486072 | 1.660477 | 1.503548 | 3.808276 | 5.746667 | 3.726062 | 3.437683 | -1.211452 | 34.958010 | 0.532569125 | -0.51962786 | 0.603322974 | 0.998865404 | 5.259576 | 4.982216 | 5.115480 | 5.164567 | 5.120890 | 5.107055 | 37 | 48 | 29 | 28 |
| 100289124 | NR_103714 | FAM27E2 | family with sequence similarity 27 member E2 | lncRNA | . | 32013 | . | . | . | 0.240786 | 0.421510 | 0.263658 | 0.351952 | 0.560318 | 0.974339 | 0.591641 | 0.804696 | -1.109420 | 18.035028 | 0.739476435 | -0.20258414 | 0.839460085 | NA | 4.247564 | 4.093800 | 4.152714 | 4.197264 | 4.158391 | 4.176093 | 16 | 26 | 13 | 18 |
| 100289170 | gene-COX20P2 | COX20P2 | COX20 pseudogene 2 | pseudogene | . | 43772 | . | . | . | 0.345207 | 0.097919 | 0.246836 | 0.081973 | 0.803311 | 0.226344 | 0.553892 | 0.187421 | -1.186152 | 3.666454 | 1.607991748 | -0.15316549 | 0.878267774 | NA | 1.987791 | 1.751299 | 1.887026 | 1.867054 | 1.879715 | 1.868644 | 7 | 2 | 4 | 2 |
| 100289178 | NR_040077 | GNG12-AS1 | GNG12, DIRAS3 and WLS antisense RNA 1 | lncRNA | . | 43938 | 615406 | . | . | 0.050639 | 0.099996 | 0.095845 | 0.104553 | 0.117838 | 0.231145 | 0.215073 | 0.239047 | 1.224804 | 6.530671 | 1.187254179 | 0.246409692 | 0.805365104 | NA | 2.555346 | 2.844638 | 2.691015 | 2.705476 | 2.702994 | 2.705559 | 5 | 8 | 6 | 7 |
| 100289230 | NR_036530 | LOC100289230 | uncharacterized LOC100289230 | lncRNA | . | . | . | . | . | 1.340698 | 0.878999 | 1.115503 | 0.922044 | 3.119860 | 2.031845 | 2.503156 | 2.108144 | -1.110817 | 62.669065 | 0.418227159 | -0.36253281 | 0.716953911 | 0.998865404 | 6.042145 | 5.893385 | 6.027226 | 5.927718 | 5.976659 | 5.934003 | 92 | 56 | 57 | 50 |
| 100289274 | NR_132117 | DNAJC3-DT | DNAJC3 divergent transcript | lncRNA | . | 39808 | . | . | . | 0.766703 | 0.656205 | 0.539203 | 0.464682 | 1.784149 | 1.516847 | 1.209957 | 1.062440 | -1.436022 | 55.407781 | 0.426751517 | -1.22337626 | 0.221187616 | 0.982008277 | 6.028815 | 5.508570 | 5.845882 | 5.809573 | 5.770980 | 5.741704 | 81 | 65 | 43 | 39 |
| 100289279 | XM_01154567 | CNTNAP3C | contactin associated protein family member 3C, transcript variant | protein_coding | XP_011543974.1;XP_011543975.1;XP_011543976.1;XP_016870869.1 | 53878 | . | . | . | 1.438258 | 1.677584 | 1.644389 | 1.590528 | 3.346885 | 3.877811 | 3.689961 | 3.636554 | -1.052981 | 106.996279 | 0.316573296 | -0.23526694 | 0.814001556 | 0.998865404 | 6.778324 | 6.703541 | 6.712685 | 6.782533 | 6.746539 | 6.716836 | 113 | 131 | 95 | 93 |
| 100289341 | NR_027447 | MAN1B1-DT | MAN1B1 divergent transcript | lncRNA | . | 48715 | . | . | . | 0.825499 | 0.502776 | 0.355426 | 0.930077 | 1.920971 | 1.162189 | 0.797566 | 2.126511 | -1.052057 | 38.564148 | 0.574660912 | -0.12740255 | 0.898621795 | 0.998865404 | 5.306645 | 5.230733 | 5.304540 | 5.225500 | 5.191734 | 5.320599 | 57 | 32 | 18 | 51 |
| 100289361 | NR_034032 | LOC100289361 | uncharacterized LOC100289361 | lncRNA | . | . | . | . | . | 1.898315 | 1.959635 | 1.532333 | 1.920606 | 4.417457 | 4.529785 | 3.438511 | 4.391238 | -1.133067 | 35.874742 | 0.519966679 | -0.3466245 | 0.72887344 | 0.998865404 | 5.252974 | 5.071091 | 5.172469 | 5.178651 | 5.143585 | 5.172177 | 43 | 42 | 26 | 35 |
| 100289373 | NR_036531,NF | UBAC2-AS1 | UBAC2 antisense RNA 1 | lncRNA | . | 42502 | . | . | . | 0.667015 | 0.835061 | 0.630435 | 0.688169 | 1.552171 | 1.930283 | 1.414677 | 1.573417 | -1.119051 | 40.420748 | 0.490472384 | -0.33085684 | 0.740752626 | 0.998865404 | 5.416453 | 5.252966 | 5.332403 | 5.360649 | 5.324156 | 5.334116 | 45 | 50 | 32 | 37 |
| 100289388 | NR_102280,NF | KCTD21-AS1 | KCTD21 antisense RNA 1 | lncRNA | . | 48674 | . | . | . | 1.458951 | 1.343702 | 1.497837 | 1.871097 | 3.395039 | 3.106028 | 3.361102 | 4.278042 | 1.174681 | 52.559777 | 0.434674353 | 0.534352504 | 0.593097673 | 0.998865404 | 5.595547 | 5.826958 | 5.704500 | 5.689361 | 5.705775 | 5.749532 | 58 | 50 | 44 | 59 |
| 100289495 | NR_040022 | LOC100289495 | uncharacterized LOC100289495 | lncRNA | . | . | . | . | . | 0.074801 | 0.157251 | 0.186559 | 0.241640 | 0.174066 | 0.363494 | 0.418634 | 0.552481 | 1.828661 | 20.470168 | 0.709613912 | 1.22712871 | 0.219774219 | NA | 3.858702 | 4.724273 | 4.287614 | 4.332617 | 4.347308 | 4.375692 | 11 | 21 | 20 | 28 |
| 100289509 | NR_045118 | KCNIP2-AS1 | KCNIP2 antisense RNA 1 | lncRNA | . | 48680 | . | . | . | 0.195347 | 0.345884 | 0.208274 | 0.137241 | 0.454580 | 0.799526 | 0.467361 | 0.313786 | -1.541677 | 7.091636 | 1.162054209 | -0.53741117 | 0.590983628 | NA | 3.104339 | 2.481107 | 2.823332 | 2.846453 | 2.827249 | 2.815467 | 7 | 12 | 6 | 4 |
| 100289561 | NM_00124271 | LOC100289561 | uncharacterized LOC100289561 | protein_coding | NP_001229642.1;NP_001334808.1;NP_001334809.1 | . | . | . | . | 4.502533 | 5.209752 | 4.184402 | 4.731326 | 10.477578 | 12.042576 | 9.389676 | 10.817617 | -1.087092 | 164.256376 | 0.261667887 | -0.46040936 | 0.645222418 | 0.998865404 | 7.419379 | 7.297666 | 7.347866 | 7.404229 | 7.316863 | 7.367118 | 184 | 197 | 128 | 157 |
| 100289574 | NR_109773 | HERC2P4 | hect domain and RLD 2 pseudogene 4 | transcribed_ps | . | 4872 | . | . | . | 3.009630 | 3.317539 | 3.858450 | 2.676637 | 7.003532 | 7.668641 | 8.658251 | 6.119812 | 1.020647 | 271.826330 | 0.234305819 | 0.125836085 | 0.899861677 | 0.998865404 | 8.071094 | 8.101826 | 8.052893 | 8.098862 | 8.177329 | 7.987338 | 296 | 303 | 283 | 210 |
| 100289635 | NM_00116471 | ZNF605 | zinc finger protein 605 | protein_coding | NP_001158187.1;NP_899061.1 | 28068 | . | . | . | 1.772011 | 1.757187 | 2.131888 | 1.466680 | 4.123543 | 4.061816 | 4.783894 | 3.353390 | 1.006456 | 518.660352 | 0.197237008 | 0.047072672 | 0.962455308 | 0.998865404 | 9.013400 | 9.023874 | 9.017004 | 9.006293 | 9.129577 | 8.888078 | 600 | 553 | 538 | 396 |
| 100289640 | gene-PPATP1 | PPATP1 | phosphoribosyl pyrophosphate amidotransferase pseudogene 1 | pseudogene | . | 9239 | . | . | . | 0.049419 | 0.106771 | 0.070307 | 0.031661 | 0.115000 | 0.246806 | 0.157768 | 0.072388 | -1.482605 | 4.173553 | 1.498374381 | -0.37916728 | 0.704563648 | NA | 2.316288 | 1.751299 | 2.059301 | 2.073402 | 2.064515 | 2.053396 | 4 | 7 | 4 | 2 |
| 100289678 | NM_00119522 | ZNF783 | zinc finger family member 783 | protein_coding | NP_001182149.1;XP_011514003.1 | 27222 | . | . | . | 4.888658 | 5.606846 | 7.470063 | 5.525573 | 11.376109 | 12.960478 | 16.762604 | 12.633565 | 1.207982 | 797.381058 | 0.17249014 | 1.580371734 | 0.114021761 | 0.852965542 | 9.496118 | 9.769227 | 9.526851 | 9.599163 | 9.800257 | 9.582265 | 788 | 820 | 873 | 695 |
| 100293516 | NM_00120481 | ZNF587B | zinc finger protein 587B | protein_coding | NP_001191747.1;NP_001363152.1 | 37142 | . | . | . | 2.894120 | 3.183506 | 4.093802 | 2.692984 | 6.734736 | 7.358815 | 9.186374 | 6.157189 | 1.111063 | 569.381130 | 0.207515209 | 0.732191155 | 0.464051902 | 0.998865404 | 9.074731 | 9.227717 | 9.096722 | 9.121744 | 9.321973 | 9.011839 | 609 | 593 | 642 | 432 |
| 100294145 | NR_037177,NF | LOC100294145 | uncharacterized LOC100294145 | lncRNA | . | . | . | . | . | 2.936311 | 2.673189 | 2.359269 | 2.628809 | 6.832914 | 6.179197 | 5.294131 | 6.010460 | -1.140603 | 304.471933 | 0.203751359 | -0.93151112 | 0.351589234 | 0.998865404 | 8.342023 | 8.152059 | 8.313413 | 8.255036 | 8.187739 | 8.239030 | 393 | 332 | 235 | 280 |
| 100294362 | NR_029376 | RNF213-AS1 | RNF213 antisense RNA 1 | lncRNA | . | 54402 | . | . | . | 0.042959 | 0.049066 | 0.055391 | 0.100456 | 0.099967 | 0.113419 | 0.124296 | 0.229681 | 1.597494 | 7.582763 | 1.111546909 | 0.607991074 | 0.543193394 | NA | 2.548173 | 3.219802 | 2.901884 | 2.908003 | 2.910082 | 2.931381 | 6 | 7 | 6 | 11 |
| 100302090 | gene-DDX11L8 | DDX11L8 | DEAD/H-box helicase 11 like 8 (pseudogene) | pseudogene | . | 37101 | . | . | . | 0.037380 | 0.105470 | 0.097770 | 0.055946 | 0.086984 | 0.243798 | 0.219393 | 0.127914 | 1.226593 | 3.542627 | 1.608010717 | 0.183243119 | 0.854607263 | NA | 1.671431 | 1.963455 | 1.806711 | 1.820556 | 1.820104 | 1.813852 | 2 | 5 | 4 | 3 |
| 100302128 | NR_031632,rn | MIR1302-3 | microRNA 1302-3 | miRNA | . | 35295 | . | . | . | 0.321884 | 0.034059 | 0.344957 | 0.213842 | 0.749036 | 0.078728 | 0.774073 | 0.488925 | -1.056394 | 1.724672 | 2.259675635 | -0.03502612 | 0.972058914 | NA | 0.820514 | 0.751299 | 0.790628 | 0.782592 | 0.788957 | 0.783375 | 3 | 1 | 2 | 1 |
| 100302132 | NR_031593,rn | MIR1182 | microRNA 1182 | miRNA | . | 35263 | . | . | . | 0.028880 | 0.110094 | 0.071809 | 0.158357 | 0.067204 | 0.254487 | 0.161137 | 0.362066 | 1.234663 | 2.005914 | 2.070995667 | 0.146845664 | 0.883253841 | NA | 0.844134 | 1.148375 | 0.998092 | 0.998667 | 1.000707 | 0.999974 | 2 | 2 | 2 | 2 |
| 100302133 | NR_031619,rn | MIR1287 | microRNA 1287 | miRNA | . | 35280 | . | . | . | 0.000849 | 0.001293 | 0.001875 | 0.002473 | 0.001975 | 0.002988 | 0.004207 | 0.005654 | 1.234665 | 3.008871 | 1.708874417 | 0.17796512 | 0.858750369 | NA | 1.429096 | 1.733338 | 1.582415 | 1.583274 | 1.586316 | 1.585224 | 3 | 3 | 3 | 3 |
| 100302137 | NR_031735,rn | MIR1914 | microRNA 1914 | miRNA | . | 35398 | . | . | . | 0.140214 | 0.057481 | 0.054772 | 0.133597 | 0.326282 | 0.132870 | 0.122905 | 0.305454 | 1.234665 | 3.008871 | 1.708874417 | 0.17796512 | 0.858750369 | NA | 1.429096 | 1.733338 | 1.582415 | 1.583274 | 1.586316 | 1.585224 | 3 | 3 | 3 | 3 |
| 100302138 | NR_031699,rn | MIR1292 | microRNA 1292 | miRNA | . | 35364 | 614155 | . | . | 0.965246 | 1.123236 | 1.881609 | 0.216460 | 2.246168 | 2.596409 | 4.222276 | 0.494910 | 1.001795 | 4.502278 | 1.443995283 | 0.001792081 | 0.998570127 | NA | 2.166062 | 2.175234 | 2.163703 | 2.165080 | 2.175005 | 2.153702 | 5 | 5 | 6 | 2 |
| 100302143 | NR_031650,rn | MIR1248 | microRNA 1248 | miRNA | . | 35314 | . | . | . | 0.235319 | 0.115944 | 0.147675 | 0.051905 | 0.547596 | 0.268009 | 0.331379 | 0.118674 | 1.234663 | 2.005914 | 2.070995667 | 0.146845664 | 0.883253841 | NA | 0.844134 | 1.148375 | 0.998092 | 0.998667 | 1.000707 | 0.999974 | 2 | 2 | 2 | 2 |
| 100302149 | NR_031651,rn | MIR1249 | microRNA 1249 | miRNA | . | 35315 | . | . | . | 0.950050 | 0.384396 | 2.882250 | 3.490972 | 2.210805 | 0.888549 | 6.467685 | 7.981691 | **2.323724** | 5.926400 | 1.261494038 | 0.96428433 | 0.334903395 | NA | 1.832372 | 3.051647 | 2.541498 | 2.534222 | 2.558357 | 2.560723 | 5 | 3 | 7 | 8 |
| 100302156 | NR_031598,rn | MIR1229 | hsa-miR-1229-3p | . | . | 33924 | . | . | . | 2.320581 | 1.533434 | 4.102516 | 1.871061 | 5.400086 | 3.544602 | 9.205928 | 4.277958 | 1.352243 | 6.311487 | 1.210515872 | 0.359643626 | 0.719113655 | NA | 2.421266 | 2.861274 | 2.647847 | 2.641107 | 2.661528 | 2.644542 | 7 | 5 | 8 | 5 |
| 100302174 | NR_031707,rn | MIR1307 | microRNA 1307 | miRNA | . | 35372 | . | . | . | 0.020003 | 0.064438 | 0.050793 | 0.013402 | 0.046549 | 0.148951 | 0.113977 | 0.030643 | 1.234665 | 3.008871 | 1.708874417 | 0.17796512 | 0.858750369 | NA | 1.429096 | 1.733338 | 1.582415 | 1.583274 | 1.586316 | 1.585224 | 3 | 3 | 3 | 3 |
| 100302183 | NR_031726,rn | MIR1825 | microRNA 1825 | miRNA | . | 35389 | . | . | . | 0.005511 | 0.004311 | 0.006427 | 0.005592 | 0.012826 | 0.009965 | 0.014422 | 0.012785 | 1.234663 | 2.005914 | 2.070995667 | 0.146845664 | 0.883253841 | NA | 0.844134 | 1.148375 | 0.998092 | 0.998667 | 1.000707 | 0.999974 | 2 | 2 | 2 | 2 |
| 100302185 | NR_031609,rn | MIR1204 | microRNA 1204 | miRNA | . | 37059 | . | . | . | 0.629031 | 1.171284 | 1.144293 | 1.203863 | 1.463782 | 2.707476 | 2.567760 | 2.752490 | 1.406178 | 3.794718 | 1.535154763 | 0.320344955 | 0.748706848 | NA | 1.658167 | 2.148375 | 1.909779 | 1.915081 | 1.919062 | 1.917633 | 3 | 4 | 4 | 4 |
| 100302197 | NR_031706,rn | MIR1306 | microRNA 1306 | miRNA | . | 35371 | . | . | . | 0.001187 | 0.003601 | 0.004824 | 0.002509 | 0.002762 | 0.008322 | 0.010825 | 0.005736 | 1.234665 | 3.008871 | 1.708874417 | 0.17796512 | 0.858750369 | NA | 1.429096 | 1.733338 | 1.582415 | 1.583274 | 1.586316 | 1.585224 | 3 | 3 | 3 | 3 |
| 100302210 | NR_031730,rn | MIR1909 | microRNA 1909 | miRNA | . | 35393 | 615201 | . | . | 0.001609 | 0.003746 | 0.001567 | 0.000806 | 0.003744 | 0.008660 | 0.003516 | 0.001842 | 1.234663 | 2.005914 | 2.070995667 | 0.146845664 | 0.883253841 | NA | 0.844134 | 1.148375 | 0.998092 | 0.998667 | 1.000707 | 0.999974 | 2 | 2 | 2 | 2 |
| 100302213 | NR_031592,rn | MIR1181 | microRNA 1181 | miRNA | . | 35262 | . | . | . | 0.004973 | 0.004833 | 0.005661 | 0.000959 | 0.011572 | 0.011171 | 0.012704 | 0.002193 | 1.234663 | 2.005914 | 2.070995667 | 0.146845664 | 0.883253841 | NA | 0.844134 | 1.148375 | 0.998092 | 0.998667 | 1.000707 | 0.999974 | 2 | 2 | 2 | 2 |
| 100302218 | NR_031616,rn | MIR1285-1 | microRNA 1285-1 | miRNA | . | 35277 | . | . | . | 0.168965 | 0.236344 | 0.221100 | 0.543737 | 0.393187 | 0.546319 | 0.496144 | 1.243190 | 1.833800 | 2.539248 | 1.876543826 | 0.466195476 | 0.641075587 | NA | 0.844134 | 1.715150 | 1.330022 | 1.330639 | 1.332827 | 1.341772 | 2 | 2 | 2 | 4 |
| 100302224 | NR_031747,rn | MIR2110 | microRNA 2110 | miRNA | . | 37071 | . | . | . | 0.000373 | 0.000789 | 0.001255 | 0.000085 | 0.000869 | 0.001823 | 0.002815 | 0.000196 | 1.234663 | 2.005914 | 2.070995667 | 0.146845664 | 0.883253841 | NA | 0.844134 | 1.148375 | 0.998092 | 0.998667 | 1.000707 | 0.999974 | 2 | 2 | 2 | 2 |
| 100302237 | NR_031694,rn | MIR1281 | microRNA 1281 | miRNA | . | 35359 | . | . | . | 0.763947 | 3.096780 | 2.440772 | 1.411988 | 1.777735 | 7.158345 | 5.477022 | 3.228342 | 1.103469 | 4.777270 | 1.39527179 | 0.101805687 | 0.918910911 | NA | 2.184683 | 2.324312 | 2.238701 | 2.257075 | 2.253377 | 2.246785 | 3 | 7 | 5 | 4 |
| 100302240 | NR_031639,rn | MIR1304 | microRNA 1304 | miRNA | . | 35302 | . | . | . | 5.139423 | 4.692519 | 6.083266 | 6.302659 | 11.959649 | 10.846969 | 13.650673 | 14.410282 | 1.234399 | 18.017825 | 0.71977523 | 0.422088188 | 0.672960649 | NA | 4.011454 | 4.315285 | 4.160484 | 4.157295 | 4.173536 | 4.176671 | 19 | 17 | 17 | 19 |
| 100302258 | NR_031715,rn | MIR1469 | hsa-miR-1469 | . | . | 35378 | . | . | . | 0.885920 | 1.361818 | 0.439407 | 0.835709 | 2.061571 | 3.147906 | 0.986018 | 1.910750 | -1.215015 | 2.454711 | 1.887245685 | -0.1488804 | 0.881648006 | NA | 1.429096 | 1.148375 | 1.299640 | 1.300454 | 1.298230 | 1.297456 | 3 | 3 | 2 | 2 |
| 100302274 | NR_031589,rn | MIR1178 | microRNA 1178 | miRNA | . | 35259 | . | . | . | 0.000038 | 0.273595 | 0.068506 | 0.000022 | 0.000088 | 0.632428 | 0.153726 | 0.000051 | -1.641549 | 1.466329 | 2.458262141 | -0.29087928 | 0.771143647 | NA | 0.867373 | 0.148375 | 0.553736 | 0.562835 | 0.555209 | 0.554796 | 1 | 3 | 1 | 1 |
| 100302285 | NR_036052,rn | MIR1244-1 | microRNA 1244-1 | miRNA | . | 35310 | . | . | . | 0.001083 | 0.000654 | 0.001055 | 0.000415 | 0.002521 | 0.001512 | 0.002368 | 0.000949 | 1.256691 | 1.507561 | 2.39030677 | 0.137902842 | 0.890317204 | NA | 0.413393 | 0.751299 | 0.586366 | 0.582479 | 0.588807 | 0.583226 | 2 | 1 | 2 | 1 |
| 100302401 | NR_027982 | RASAL2-AS1 | RASAL2 antisense RNA 1 | lncRNA | . | 44170 | . | . | . | 0.589747 | 0.760973 | 0.830075 | 0.758655 | 1.372366 | 1.759023 | 1.862664 | 1.734574 | 1.165809 | 54.077640 | 0.429637625 | 0.515158988 | 0.606441925 | 0.998865404 | 5.642344 | 5.863137 | 5.715237 | 5.758204 | 5.776504 | 5.757527 | 51 | 60 | 53 | 52 |
| 100302650 | NR_028308 | BABAM2-AS1 | BABAM2 antisense RNA 1 | lncRNA | . | 44171 | . | . | . | 0.036396 | 0.157129 | 0.065209 | 0.122032 | 0.084695 | 0.363211 | 0.146328 | 0.279012 | -1.104674 | 5.198989 | 1.36579238 | -0.10515577 | 0.916252205 | NA | 2.452336 | 2.300113 | 2.363406 | 2.390293 | 2.368047 | 2.381140 | 3 | 9 | 3 | 6 |
| 100302666 | gene-CSPG4P1 | CSPG4P13 | chondroitin sulfate proteoglycan 4 pseudogene 13 | pseudogene | . | 49195 | . | . | . | 0.192053 | 0.157123 | 0.163974 | 0.176337 | 0.446916 | 0.363196 | 0.367953 | 0.403174 | -1.045182 | 27.112590 | 0.593576102 | -0.10740714 | 0.91446599 | 0.998865404 | 4.792272 | 4.728813 | 4.775663 | 4.753132 | 4.754694 | 4.764387 | 35 | 27 | 22 | 26 |
| 100302692 | NR_028379 | FTX | FTX transcript, XIST regulator | lncRNA | . | 37190 | 300936 | . | . | 0.183568 | 0.284715 | 0.353489 | 0.301154 | 0.427171 | 0.658132 | 0.793220 | 0.688553 | 1.388628 | 21.014905 | 0.680147689 | 0.696407385 | 0.48617374 | NA | 4.137922 | 4.610270 | 4.351568 | 4.384516 | 4.405147 | 4.388955 | 16 | 23 | 23 | 21 |
| 100302736 | NM_00116446 | TMED7-TICAM | TMED7-TICAM2 readthrough | protein_coding | NP_001157940.1;NP_001157941.1 | 33945 | . | . | . | 5.280069 | 5.648952 | 4.595744 | 5.027069 | 12.286936 | 13.057805 | 10.312716 | 11.493797 | -1.153214 | 600.285962 | 0.160449825 | -1.28177638 | 0.199921103 | 0.96621494 | 9.328924 | 9.122727 | 9.252943 | 9.296196 | 9.152561 | 9.207186 | 717 | 716 | 464 | 545 |
| 100302739 | NR_028370 | PCNA-AS1 | PCNA antisense RNA 1 | lncRNA | . | 37184 | . | . | . | 0.086603 | 0.132357 | 0.105029 | 0.148337 | 0.201529 | 0.305949 | 0.235682 | 0.339156 | 1.234663 | 2.005914 | 2.070995667 | 0.146845664 | 0.883253841 | NA | 0.844134 | 1.148375 | 0.998092 | 0.998667 | 1.000707 | 0.999974 | 2 | 2 | 2 | 2 |
| 100303728 | NR_028443,NF | SLC25A5-AS1 | SLC25A5 antisense RNA 1 | lncRNA | . | 43438 | . | . | . | 0.492508 | 0.499984 | 0.731581 | 0.988614 | 1.146085 | 1.155736 | 1.641647 | 2.260350 | 1.644049 | 45.562811 | 0.472238913 | 1.518835401 | 0.128803933 | 0.878399546 | 5.107875 | 5.823779 | 5.455429 | 5.446892 | 5.500847 | 5.569262 | 41 | 36 | 41 | 62 |
| 100303755 | NM_00116481 | PET117 | PET117 cytochrome c oxidase chaperone | protein_coding | NP_001158283.1 | 40045 | 614771 | . | . | 3.573103 | 2.526234 | 2.845856 | 2.649961 | 8.314757 | 5.839504 | 6.386019 | 6.058822 | -1.126719 | 104.823780 | 0.328925758 | -0.52330272 | 0.600763612 | 0.998865404 | 6.794507 | 6.624109 | 6.784897 | 6.669107 | 6.702386 | 6.679887 | 150 | 99 | 89 | 89 |
| 100306951 | NR_028514 | PITPNA-AS1 | PITPNA antisense RNA 1 | lncRNA | . | 44116 | . | . | . | 5.916412 | 5.794834 | 4.176080 | 4.607035 | 13.767734 | 13.395018 | 9.371002 | 10.533438 | -1.349499 | 88.935382 | 0.338740746 | -1.27656308 | 0.201756542 | 0.967296129 | 6.674967 | 6.242048 | 6.520189 | 6.512552 | 6.422425 | 6.442731 | 119 | 109 | 63 | 74 |
| 100310756 | NR_147504 | LOC100310756 | uncharacterized LOC100310756 | lncRNA | . | . | . | . | . | 2.077130 | 2.117757 | 2.296908 | 2.060272 | 4.833567 | 4.895290 | 5.154195 | 4.710566 | 1.028378 | 97.339445 | 0.324518239 | 0.124401428 | 0.900997443 | 0.998865404 | 6.584287 | 6.625327 | 6.598333 | 6.601063 | 6.625729 | 6.590145 | 110 | 104 | 91 | 87 |
| 100310840 | gene-VDAC1P8 | VDAC1P8 | voltage dependent anion channel 1 pseudogene 8 | pseudogene | . | 37483 | . | . | . | 0.090869 | 0.056833 | 0.032266 | 0.076154 | 0.211457 | 0.131372 | 0.072404 | 0.174116 | -1.355150 | 3.871062 | 1.535090604 | -0.28561979 | 0.775169348 | NA | 2.156660 | 1.715150 | 1.967195 | 1.960089 | 1.953915 | 1.962640 | 6 | 4 | 2 | 4 |
| 100313843 | NR_031756,rn | MIR711 | microRNA 711 | miRNA | . | 37316 | . | . | . | 0.000325 | 0.000336 | 0.000323 | 0.000607 | 0.000758 | 0.000776 | 0.000724 | 0.001388 | 1.234663 | 2.005914 | 2.070995667 | 0.146845664 | 0.883253841 | NA | 0.844134 | 1.148375 | 0.998092 | 0.998667 | 1.000707 | 0.999974 | 2 | 2 | 2 | 2 |
| 100313887 | NR_031754,rn | MIR2277 | microRNA 2277 | miRNA | . | 37314 | . | . | . | 0.002787 | 0.000657 | 0.001843 | 0.005305 | 0.006485 | 0.001518 | 0.004136 | 0.012130 | 1.234665 | 3.008871 | 1.708874417 | 0.17796512 | 0.858750369 | NA | 1.429096 | 1.733338 | 1.582415 | 1.583274 | 1.586316 | 1.585224 | 3 | 3 | 3 | 3 |

| | | | | | | | | | | | | | | | | | | | | | | | | | | | | | | | | | |
|---|---|---|---|---|---|---|---|---|---|---|---|---|---|---|---|---|---|---|---|---|---|---|---|---|---|---|---|---|---|---|---|---|---|
| 100313892 | NR_031580,rn | MIR761 | microRNA 761 | miRNA | . | 37305 | . | . | . | 0.001709 | 0.001446 | 0.001751 | 0.003252 | 0.003975 | 0.003343 | 0.003931 | 0.007435 | 1.234663 | 2.005914 | 2.070995667 | 0.146845664 | 0.883253841 | NA | 0.844134 | 1.148375 | 0.998092 | 0.998667 | 1.000707 | 0.999974 | 2 | 2 | 2 | 2 |
| 100313968 | gene-LOC1003 | LOC10031396 | riboflavin kinase pseudogene | pseudogene | . | . | . | . | . | 0.088962 | 0.133125 | 0.047114 | 0.024037 | 0.207017 | 0.307724 | 0.105723 | 0.054958 | -3.231877 | 3.549010 | 1.685819399 | -1.00388699 | 0.31543309 | NA | 2.436885 | 0.751299 | 1.844977 | 1.854827 | 1.835736 | 1.830028 | 5 | 7 | 2 | 1 |
| 100316868 | NR_037843 | HOTTIP | HOXA distal transcript antisense RNA | lncRNA | . | 37461 | 614060 | . | . | 0.085419 | 0.062845 | 0.055497 | 0.056197 | 0.198773 | 0.145268 | 0.124533 | 0.128489 | -1.345335 | 9.719309 | 0.980900585 | -0.43629861 | 0.66262009 | NA | 3.478589 | 3.051647 | 3.301486 | 3.284323 | 3.279207 | 3.281179 | 15 | 10 | 7 | 8 |
| 100316904 | NM_00116868 | SAP25 | Sin3A associated protein 25 | protein_coding | NP_001162153.2;NP_001335606.1;NP_001335609.1 | 41908 | . | . | . | 1.445260 | 1.519354 | 1.365851 | 0.981907 | 3.363179 | 3.512054 | 3.064931 | 2.245014 | -1.276226 | 54.436629 | 0.432141704 | -0.81427813 | 0.415485633 | 0.998865404 | 5.930908 | 5.580931 | 5.786383 | 5.793943 | 5.768785 | 5.715067 | 69 | 67 | 48 | 38 |
| 100329108 | gene-GCSHP5 | GCSHP5 | glycine cleavage system protein H pseudogene 5 | pseudogene | . | 44195 | . | . | . | 4.280085 | 3.306639 | 2.903990 | 0.950822 | 9.959934 | 7.643444 | 6.516470 | 2.173941 | -1.979382 | 102.260461 | 0.423097799 | -2.32818473 | 0.019902297 | 0.467054728 | 7.084334 | 6.104662 | 6.798070 | 6.695301 | 6.651896 | 6.417961 | 177 | 127 | 90 | 32 |
| 100329109 | NR_033248 | GCSHP3 | glycine cleavage system protein H pseudogene 3 | transcribed_ps | . | 43929 | . | . | . | 0.005109 | 0.002823 | 0.005020 | 0.000808 | 0.011889 | 0.006526 | 0.011265 | 0.001847 | 1.234655 | 1.002957 | 2.868339486 | 0.106022226 | 0.915564717 | NA | -0.155866 | 0.148375 | -0.001261 | -0.000972 | 0.000053 | -0.000315 | 1 | 1 | 1 | 1 |
| 100335030 | NR_033267 | LOC10033503( | FGFR1 oncogene partner 2 pseudogene | transcribed_ps | . | . | . | . | . | 0.283634 | 0.107023 | 0.179618 | 0.146503 | 0.660028 | 0.247389 | 0.403058 | 0.334962 | -1.197320 | 18.278113 | 0.764088764 | -0.34002446 | 0.733838105 | NA | 4.313954 | 4.058876 | 4.234413 | 4.158111 | 4.190204 | 4.175570 | 33 | 12 | 16 | 14 |
| 100379224 | NR_033341 | LOC10037922 | uncharacterized LOC100379224 | lncRNA | . | . | . | . | . | 0.045145 | 0.172964 | 0.111457 | 0.055378 | 0.105055 | 0.399813 | 0.250106 | 0.126616 | -1.246345 | 10.698817 | 1.007600725 | -0.31530737 | 0.752528289 | NA | 3.569585 | 3.251698 | 3.383747 | 3.444202 | 3.417630 | 3.390356 | 6 | 20 | 11 | 6 |
| 100379251 | NR_037806 | NPPA-AS1 | NPPA antisense RNA 1 | antisense_RNA | . | 37635 | . | . | . | 3.180541 | 1.967112 | 4.814234 | 3.169115 | 7.401250 | 4.547068 | 10.803002 | 7.245803 | 1.523799 | 170.321839 | 0.308309652 | 1.97098159 | 0.048725982 | 0.661973154 | 7.074728 | 7.685372 | 7.380066 | 7.204750 | 7.564510 | 7.368616 | 192 | 111 | 217 | 152 |
| 100379571 | NR_037803 | BACE1-AS | BACE1 antisense RNA | antisense_RNA | . | 37125 | 614263 | . | . | 0.601488 | 0.113576 | 1.050771 | 0.288081 | 1.399686 | 0.262537 | 2.357899 | 0.658663 | 1.723003 | 13.818357 | 0.96372346 | 0.81447117 | 0.415375081 | NA | 3.336810 | 4.132003 | 3.770192 | 3.711637 | 3.814745 | 3.730131 | 19 | 4 | 24 | 7 |
| 100379667 | gene-NEK2P2 | NEK2P2 | NEK2 pseudogene 2 | pseudogene | . | 37816 | . | . | . | 0.083009 | 0.114587 | 0.136761 | 0.046802 | 0.193166 | 0.264873 | 0.306888 | 0.107006 | -1.043217 | 5.736922 | 1.279932587 | -0.04768946 | 0.961963737 | NA | 2.548173 | 2.491830 | 2.513117 | 2.519095 | 2.525709 | 2.504160 | 6 | 7 | 7 | 3 |
| 100381270 | NM_00117410 | ZBED6 | zinc finger BED-type containing 6 | protein_coding | NP_001167579.1 | 33273 | 613512 | . | . | 1.680692 | 1.955566 | 1.962573 | 2.118653 | 3.911039 | 4.520378 | 4.403956 | 4.844049 | 1.108133 | 214.028880 | 0.233682054 | 0.633898028 | 0.526147372 | 0.998865404 | 7.666178 | 7.813392 | 7.681318 | 7.747890 | 7.746703 | 7.778869 | 217 | 235 | 189 | 218 |
| 100418737 | gene-KRT8P33 | KRT8P33 | keratin 8 pseudogene 33 | pseudogene | . | 39867 | . | . | . | 0.250278 | 0.337307 | 0.383373 | 0.415817 | 0.582407 | 0.779701 | 0.860277 | 0.950717 | 1.366064 | 19.169872 | 0.703438528 | 0.639750014 | 0.522335135 | NA | 4.019256 | 4.467590 | 4.229389 | 4.250184 | 4.263900 | 4.266414 | 16 | 20 | 19 | 21 |
| 100418744 | gene-KRT8P46 | KRT8P46 | keratin 8 pseudogene 46 | pseudogene | . | 39880 | . | . | . | 0.185011 | 0.115562 | 0.160391 | 0.122267 | 0.430528 | 0.267128 | 0.359914 | 0.279548 | -1.017055 | 8.393745 | 1.053119078 | -0.02316724 | 0.981516872 | NA | 3.079678 | 3.058876 | 3.078629 | 3.060466 | 3.072720 | 3.065137 | 12 | 7 | 8 | 7 |
| 100418748 | gene-KRT8P36 | KRT8P36 | keratin 8 pseudogene 36 | pseudogene | . | 39870 | . | . | . | 0.163091 | 0.110284 | 0.231209 | 0.152248 | 0.379519 | 0.254927 | 0.518825 | 0.348096 | 1.316662 | 9.305782 | 1.004313492 | 0.395180543 | 0.692709651 | NA | 3.003609 | 3.404839 | 3.211684 | 3.197552 | 3.224862 | 3.207111 | 11 | 7 | 11 | 8 |
| 100418781 | gene-KRT8P39 | KRT8P39 | keratin 8 pseudogene 39 | pseudogene | . | 39873 | . | . | . | 0.109013 | 0.224698 | 0.217998 | 0.035719 | 0.253677 | 0.519400 | 0.489181 | 0.081667 | -1.231517 | 8.227457 | 1.155228603 | -0.2600664 | 0.794812557 | NA | 3.180050 | 2.885872 | 3.020521 | 3.047988 | 3.049577 | 3.003235 | 7 | 13 | 11 | 2 |
| 100418843 | gene-PHBP13 | PHBP13 | prohibitin pseudogene 13 | pseudogene | . | 39292 | . | . | . | 0.532215 | 0.392335 | 0.361607 | 0.152435 | 1.238487 | 0.906899 | 0.811436 | 0.348523 | -1.744795 | 12.298690 | 0.908987391 | -0.88346359 | 0.376985846 | NA | 3.964015 | 3.168566 | 3.650731 | 3.627173 | 3.625647 | 3.593665 | 21 | 14 | 11 | 5 |
| 100418902 | gene-TUBBP10 | TUBBP10 | tubulin beta class I pseudogene 10 | pseudogene | . | 42344 | . | . | . | 0.200648 | 0.302811 | 0.423453 | 0.302404 | 0.466916 | 0.699961 | 0.950217 | 0.691411 | 1.473743 | 17.296822 | 0.749394791 | 0.746582509 | 0.45531561 | NA | 3.805868 | 4.365137 | 4.072595 | 4.097065 | 4.127859 | 4.098773 | 13 | 18 | 21 | 16 |
| 100418910 | gene-ACTG1P1 | ACTG1P14 | actin gamma 1 pseudogene 14 | pseudogene | . | 49751 | . | . | . | 0.076523 | 0.102628 | 0.211248 | 0.098541 | 0.178072 | 0.237229 | 0.474035 | 0.225301 | 1.517177 | 5.105995 | 1.344812288 | 0.447192103 | 0.654736363 | NA | 2.019256 | 2.622519 | 2.330836 | 2.336427 | 2.351572 | 2.334837 | 4 | 5 | 7 | 4 |
| 100418943 | gene-PRDX2P3 | PRDX2P3 | peroxiredoxin 2 pseudogene 3 | pseudogene | . | 44968 | . | . | . | 0.111669 | 0.142586 | 0.195888 | 0.070634 | 0.259859 | 0.329594 | 0.439567 | 0.161497 | 1.093564 | 3.765988 | 1.559157961 | 0.082761252 | 0.93404138 | NA | 1.844134 | 1.978782 | 1.904604 | 1.905725 | 1.914771 | 1.898564 | 4 | 4 | 5 | 2 |
| 100419029 | gene-NF1P4 | NF1P4 | neurofibromin 1 pseudogene 4 | pseudogene | . | 7769 | . | . | . | 0.170347 | 0.096850 | 0.174932 | 0.128625 | 0.396404 | 0.223874 | 0.392543 | 0.294086 | 1.099750 | 7.944949 | 1.082845079 | 0.12668085 | 0.899193003 | NA | 2.917750 | 3.058876 | 2.995201 | 2.976696 | 2.993147 | 2.985596 | 11 | 6 | 8 | 7 |
| 100419041 | gene-SNX18P1 | SNX18P12 | sorting nexin 18 pseudogene 12 | pseudogene | . | 39620 | . | . | . | 0.701959 | 0.766599 | 1.009577 | 0.488225 | 1.633488 | 1.772029 | 2.265462 | 1.116269 | 1.037197 | 13.731849 | 0.839542116 | 0.062760427 | 0.949957279 | NA | 3.751025 | 3.807334 | 3.771127 | 3.775082 | 3.793828 | 3.756359 | 15 | 15 | 16 | 9 |
| 100419075 | gene-AK3P3 | AK3P3 | adenylate kinase 3 pseudogene 3 | pseudogene | . | 39062 | . | . | . | 0.835162 | 0.847472 | 1.144719 | 1.064288 | 1.943456 | 1.958969 | 2.568717 | 2.433367 | 1.310953 | 25.396401 | 0.609969033 | 0.640385847 | 0.521921782 | NA | 4.457887 | 4.848815 | 4.644452 | 4.646946 | 4.678156 | 4.669801 | 25 | 24 | 26 | 26 |
| 100419185 | gene-PDLIM1P | PDLIM1P4 | PDZ and LIM domain 1 pseudogene 4 | pseudogene | . | 48947 | . | . | . | 0.723828 | 0.596839 | 0.579727 | 0.692854 | 1.684377 | 1.379621 | 1.300890 | 1.584128 | -1.036019 | 24.153275 | 0.628185824 | -0.08126568 | 0.93523067 | NA | 4.619518 | 4.568322 | 4.606976 | 4.587060 | 4.585903 | 4.601074 | 31 | 24 | 19 | 24 |
| 100419583 | NR_125376 | LOC10041958 | ring finger protein 4 pseudogene | transcribed_ps | . | . | . | . | . | 5.210801 | 5.750566 | 5.281332 | 5.214462 | 12.125748 | 13.292689 | 11.851154 | 11.922248 | -1.057925 | 344.088591 | 0.192245177 | -0.42257378 | 0.67260626 | 0.998865404 | 8.466780 | 8.385343 | 8.415870 | 8.467915 | 8.416583 | 8.404205 | 389 | 399 | 294 | 310 |
| 100419620 | gene-LOC1004 | LOC10041962( | RAN binding protein 1 pseudogene | pseudogene | . | . | . | . | . | 0.203956 | 0.257526 | 0.406921 | 0.083815 | 0.474613 | 0.595282 | 0.913119 | 0.191632 | 1.005250 | 6.781532 | 1.231886712 | 0.006132376 | 0.995107102 | NA | 2.754145 | 2.769039 | 2.747539 | 2.753778 | 2.771268 | 2.729859 | 7 | 8 | 10 | 2 |
| 100419673 | gene-LOC1004 | LOC10041967 | PHD finger protein 10 pseudogene | pseudogene | . | . | . | . | . | 0.365359 | 0.161759 | 0.379925 | 0.081762 | 0.850205 | 0.373914 | 0.852542 | 0.186939 | -1.181286 | 8.609204 | 1.114277974 | -0.21570756 | 0.82921571 | NA | 3.216230 | 2.986385 | 3.120937 | 3.086580 | 3.117922 | 3.076480 | 15 | 6 | 11 | 3 |
| 100419709 | gene-ZNF887P | ZNF887P | zinc finger protein 887, pseudogene | pseudogene | . | 38700 | . | . | . | 0.748910 | 0.849166 | 0.958956 | 0.664320 | 1.742744 | 1.962885 | 2.151870 | 1.518889 | -1.002600 | 18.403333 | 0.720388654 | -0.00520063 | 0.99585052 | NA | 4.202828 | 4.200961 | 4.194809 | 4.203992 | 4.213499 | 4.184681 | 20 | 21 | 19 | 14 |
| 100419798 | gene-LOC1004 | LOC10041979 | gon-4 like pseudogene | pseudogene | . | . | . | . | . | 0.580000 | 0.823995 | 0.464131 | 0.698923 | 1.349684 | 1.904700 | 1.041496 | 1.598003 | -1.216996 | 23.785814 | 0.64628764 | -0.4383879 | 0.661105118 | NA | 4.708563 | 4.421211 | 4.563866 | 4.601155 | 4.546264 | 4.580211 | 25 | 33 | 15 | 24 |
| 100419814 | gene-ANKRD4 | ANKRD49P1 | ANKRD49 pseudogene 1 | pseudogene | . | 54619 | . | . | . | 0.105176 | 0.172915 | 0.207081 | 0.145786 | 0.244748 | 0.399700 | 0.464685 | 0.333323 | 1.229505 | 5.548541 | 1.288054024 | 0.231416997 | 0.816990862 | NA | 2.316288 | 2.612728 | 2.454436 | 2.468700 | 2.470251 | 2.463353 | 4 | 7 | 6 | 5 |
| 100419892 | gene-LOC1004 | LOC10041989 | zinc finger protein 592 pseudogene | pseudogene | . | . | . | . | . | 0.254793 | 0.231659 | 0.324931 | 0.387881 | 0.592914 | 0.535490 | 0.729136 | 0.886845 | 1.314372 | 7.771144 | 1.084462749 | 0.363658162 | 0.716113289 | NA | 2.747897 | 3.141582 | 2.949294 | 2.947220 | 2.954360 | 2.961243 | 8 | 7 | 7 | 9 |
| 100419903 | gene-RWDD4P | RWDD4P1 | RWD domain containing 4 pseudogene 1 | pseudogene | . | 23753 | . | . | . | 0.028371 | 0.058018 | 0.107854 | 0.048749 | 0.066021 | 0.134112 | 0.242022 | 0.111458 | 2.054720 | 2.076297 | 2.080611993 | 0.499344301 | 0.617536851 | NA | 0.444631 | 1.481107 | 1.028646 | 1.033470 | 1.040700 | 1.034794 | 1 | 2 | 3 | 2 |
| 100419906 | gene-ANKRD11 | ANKRD11P2 | ANKRD11 pseudogene 2 | pseudogene | . | 54738 | . | . | . | 3.372533 | 3.450306 | 4.125769 | 3.472227 | 7.848022 | 7.975536 | 9.258107 | 7.938835 | 1.098423 | 154.835492 | 0.26774661 | 0.505827282 | 0.612977904 | 0.998865404 | 7.204795 | 7.341168 | 7.250904 | 7.255076 | 7.326441 | 7.252631 | 169 | 160 | 154 | 138 |
| 100419920 | gene-LOC1004 | LOC10041992( | ankyrin repeat domain 33B pseudogene | pseudogene | . | . | . | . | . | 1.960271 | 1.845708 | 2.325466 | 2.721003 | 4.561631 | 4.266436 | 5.218278 | 6.221250 | 1.302749 | 118.060560 | 0.299335996 | 1.27468588 | 0.202420447 | 0.968125233 | 6.680265 | 7.061402 | 6.841210 | 6.820925 | 6.894356 | 6.945547 | 122 | 107 | 108 | 134 |
| 100419947 | gene-HMGB1P | HMGB1P21 | high mobility group box 1 pseudogene 21 | pseudogene | . | 39106 | . | . | . | 0.050647 | 0.151084 | 0.136323 | 0.051491 | 0.117858 | 0.349238 | 0.305905 | 0.117728 | 1.056751 | 3.275960 | 1.678573033 | 0.047442362 | 0.962160667 | NA | 1.671431 | 1.751299 | 1.696027 | 1.709843 | 1.709321 | 1.698229 | 2 | 5 | 4 | 2 |
| 100420003 | gene-E2F3P2 | E2F3P2 | E2F transcription factor 3 pseudogene 2 | pseudogene | . | 3117 | . | . | . | 0.182471 | 0.253556 | 0.310828 | 0.243999 | 0.424617 | 0.586107 | 0.697489 | 0.557874 | 1.234186 | 34.221323 | 0.538755538 | 0.563446208 | 0.57313108 | 0.998865404 | 4.937099 | 5.240615 | 5.055378 | 5.093445 | 5.120894 | 5.084876 | 30 | 38 | 37 | 31 |
| 100420054 | gene-LOC1004 | LOC10042005 | NSFL1 cofactor pseudogene | pseudogene | . | . | . | . | . | 0.026723 | 0.120760 | 0.103080 | 0.049903 | 0.062186 | 0.279141 | 0.231308 | 0.114096 | -1.083834 | 3.507646 | 1.637532976 | -0.07092593 | 0.943456705 | NA | 1.867373 | 1.751299 | 1.795173 | 1.813325 | 1.808532 | 1.797431 | 2 | 6 | 4 | 2 |
| 100420063 | gene-HMGB1P | HMGB1P24 | high mobility group box 1 pseudogene 24 | pseudogene | . | 39115 | . | . | . | 0.145597 | 0.125302 | 0.181963 | 0.255957 | 0.338811 | 0.289642 | 0.408321 | 0.585215 | 1.642558 | 5.316433 | 1.30947547 | 0.546740961 | 0.584556709 | NA | 2.008843 | 2.724273 | 2.396405 | 2.393486 | 2.402945 | 2.410684 | 5 | 4 | 5 | 7 |
| 100420067 | gene-TMEM23 | TMEM231P1 | TMEM231 pseudogene 1 | pseudogene | . | 53870 | . | . | . | 0.169372 | 0.298908 | 0.279471 | 0.202871 | 0.394135 | 0.690939 | 0.627126 | 0.463840 | -1.012895 | 6.733628 | 1.173659783 | -0.01574899 | 0.987434645 | NA | 2.760365 | 2.742346 | 2.740583 | 2.755184 | 2.753500 | 2.741501 | 6 | 9 | 7 | 5 |
| 100420109 | gene-WBP1P1 | WBP1P1 | WBP1 pseudogene 1 | pseudogene | . | 54567 | . | . | . | 0.333022 | 0.131903 | 0.010360 | 0.058248 | 0.774955 | 0.304899 | 0.023247 | 0.133177 | -4.247082 | 5.734268 | 1.402656069 | -1.48751498 | 0.136878851 | NA | 3.216230 | 1.121007 | 2.570748 | 2.536407 | 2.517099 | 2.525901 | 15 | 6 | 1 | 3 |
| 100420110 | gene-SCYL2P1 | SCYL2P1 | SCYL2 pseudogene 1 | pseudogene | . | 54685 | . | . | . | 0.116396 | 0.063376 | 0.170978 | 0.015925 | 0.270858 | 0.146496 | 0.383670 | 0.036411 | 1.032071 | 9.507087 | 1.111577547 | 0.040970626 | 0.967319313 | NA | 3.220748 | 3.276716 | 3.246285 | 3.220164 | 3.266885 | 3.201039 | 14 | 7 | 15 | 2 |
| 100420159 | gene-RRM2P3 | RRM2P3 | ribonucleotide reductase M2 polypeptide pseudogene 3 | pseudogene | . | 10455 | . | . | . | 0.165994 | 0.187960 | 0.255517 | 0.086570 | 0.386276 | 0.434478 | 0.573372 | 0.197931 | -1.039362 | 17.217016 | 0.788251242 | -0.07066101 | 0.943667557 | NA | 4.130737 | 4.080349 | 4.093900 | 4.102852 | 4.130067 | 4.059624 | 19 | 20 | 22 | 8 |
| 100420174 | gene-LOC1004 | LOC10042017 | centrosomal protein 170kDa pseudogene | pseudogene | . | . | . | . | . | 0.384864 | 0.916166 | 0.821023 | 0.437996 | 0.895594 | 2.117758 | 1.842351 | 1.001425 | -1.041934 | 8.003673 | 1.1031202 | -0.05372427 | 0.957154844 | NA | 3.029593 | 2.971139 | 2.980879 | 3.008258 | 3.004148 | 2.982100 | 6 | 12 | 9 | 5 |
| 100420178 | gene-E2F6P3 | E2F6P3 | E2F transcription factor 6 pseudogene 3 | pseudogene | . | 51451 | . | . | . | 0.161661 | 0.174695 | 0.239480 | 0.230044 | 0.376193 | 0.403815 | 0.537385 | 0.525968 | 1.370625 | 14.378449 | 0.805060557 | 0.564968718 | 0.572095041 | NA | 3.600756 | 4.055266 | 3.824191 | 3.831987 | 3.850275 | 3.845204 | 13 | 14 | 15 | 15 |
| 100420253 | gene-CTBP2P8 | CTBP2P8 | CTBP2 pseudogene 8 | pseudogene | . | 45200 | . | . | . | 0.149435 | 0.110616 | 0.183859 | 0.065736 | 0.347741 | 0.255694 | 0.412575 | 0.150299 | -1.042858 | 7.447018 | 1.133530941 | -0.05341123 | 0.957404254 | NA | 2.923304 | 2.869520 | 2.898902 | 2.888689 | 2.905668 | 2.878890 | 10 | 7 | 9 | 4 |
| 100420258 | gene-PLA2G12 | PLA2G12AP1 | phospholipase A2 group XIIA pseudogene 1 | pseudogene | . | 39248 | . | . | . | 0.063663 | 0.574623 | 0.449063 | 0.212582 | 0.148146 | 1.328267 | 1.007683 | 0.486043 | 1.091320 | 9.135316 | 1.145420381 | 0.110067771 | 0.912355628 | NA | 3.128586 | 3.251698 | 3.137056 | 3.202009 | 3.187192 | 3.159940 | 2 | 17 | 11 | 6 |
| 100420259 | gene-GDI2P2 | GDI2P2 | GDP dissociation inhibitor 2 pseudogene 2 | pseudogene | . | 37976 | . | . | . | 0.467183 | 0.770753 | 0.625317 | 1.032781 | 1.087154 | 1.781629 | 1.403193 | 2.361330 | 1.354667 | 14.324293 | 0.824341005 | 0.531259228 | 0.595239155 | NA | 3.607672 | 4.040734 | 3.810674 | 3.834733 | 3.825395 | 3.857706 | 11 | 16 | 11 | 19 |
| 100420300 | gene-FTH1P23 | FTH1P23 | ferritin heavy chain 1 pseudogene 23 | pseudogene | . | 37641 | . | . | . | 1.471737 | 2.292252 | 1.605370 | 2.189010 | 3.424793 | 5.298642 | 3.602404 | 5.004910 | -1.002945 | 51.318598 | 0.457517785 | -0.0092744 | 0.992600208 | 0.999862634 | 5.685221 | 5.677589 | 5.636976 | 5.716978 | 5.649990 | 5.703819 | 47 | 67 | 38 | 55 |
| 100420307 | gene-NTAN1P2 | NTAN1P2 | N-terminal asparagine amidase pseudogene 2 | pseudogene | . | 45262 | . | . | . | 0.132257 | 0.441023 | 0.289399 | 0.180188 | 0.307768 | 1.019445 | 0.649402 | 0.411977 | -1.250852 | 9.428771 | 1.042956325 | -0.30961163 | 0.756856314 | NA | 3.389932 | 3.066070 | 3.211018 | 3.259166 | 3.234694 | 3.217337 | 6 | 17 | 9 | 6 |
| 100420326 | gene-LOC1004 | LOC10042032( | ferritin heavy chain like 17 pseudogene | pseudogene | . | . | . | . | . | 0.409396 | 0.003266 | 0.059572 | 0.490872 | 0.952682 | 0.007549 | 0.133679 | 1.122322 | 1.330022 | 6.612001 | 1.366717533 | 0.301049887 | 0.763376453 | NA | 2.504372 | 2.916472 | 2.734585 | 2.690278 | 2.697916 | 2.744038 | 12 | 1 | 2 | 12 |
| 100420341 | gene-SEC63P1 | SEC63P1 | SEC63 homolog, protein translocation regulator pseudogene 1 | pseudogene | . | 41953 | . | . | . | 0.519069 | 0.274599 | 0.521540 | 0.111428 | 1.207895 | 0.634747 | 1.170320 | 0.254767 | -1.272987 | 16.693399 | 0.85205878 | -0.40867777 | 0.682776146 | NA | 4.220748 | 3.881801 | 4.085576 | 4.035771 | 4.082680 | 4.002932 | 28 | 14 | 21 | 5 |
| 100420401 | gene-BTBD10P | BTBD10P2 | BTB domain containing 10 pseudogene 2 | pseudogene | . | 51541 | . | . | . | 0.152801 | 0.231002 | 0.405298 | 0.013271 | 0.355575 | 0.533971 | 0.909478 | 0.030342 | -1.023336 | 4.018501 | 1.562816923 | -0.02129443 | 0.983010785 | NA | 2.019256 | 1.993948 | 1.994726 | 2.000224 | 2.010028 | 1.983860 | 4 | 5 | 6 | 1 |
| 100420429 | gene-LOC1004 | LOC10042042 | ubiquitin specific peptidase 15 pseudogene | pseudogene | . | . | . | . | . | 0.318694 | 0.117591 | 0.229557 | 0.149282 | 0.741615 | 0.271818 | 0.515119 | 0.341316 | -1.092797 | 4.669411 | 1.419095876 | -0.09021596 | 0.928115605 | NA | 2.282109 | 2.161867 | 2.233991 | 2.214380 | 2.227989 | 2.216609 | 8 | 3 | 5 | 3 |
| 100420464 | gene-LOC1004 | LOC10042046 | salvador family WW domain containing protein 1 pseudogene | pseudogene | . | . | . | . | . | 0.193788 | 0.155237 | 0.070458 | 0.026968 | 0.450953 | 0.358838 | 0.158106 | 0.061659 | -2.665017 | 3.071062 | 1.775354898 | -0.79654205 | 0.425717043 | NA | 2.156660 | 0.751299 | 1.644074 | 1.636941 | 1.630580 | 1.624916 | 6 | 4 | 2 | 1 |
| 100420528 | gene-LOC1004 | LOC10042052 | CTD nuclear envelope phosphatase 1 pseudogene | pseudogene | . | . | . | . | . | 0.537536 | 0.988539 | 0.763128 | 0.888347 | 1.250869 | 2.285053 | 1.712437 | 2.031101 | 1.062883 | 28.430800 | 0.598637934 | 0.146971501 | 0.883154515 | 0.998865404 | 4.786347 | 4.871172 | 4.788057 | 4.852048 | 4.820995 | 4.834277 | 23 | 38 | 24 | 29 |
| 100420650 | gene-PGAM1P | PGAM1P7 | phosphoglycerate mutase 1 pseudogene 7 | pseudogene | . | 42454 | . | . | . | 0.194536 | 0.217193 | 0.499414 | 0.083230 | 0.452693 | 0.502050 | 1.120670 | 0.190296 | 1.437899 | 7.678993 | 1.162708735 | 0.450639046 | 0.65224972 | NA | 2.651489 | 3.181871 | 2.913117 | 2.915130 | 2.951918 | 2.900460 | 7 | 7 | 13 | 3 |
| 100420889 | gene-ANAPC1F | ANAPC1P3 | ANAPC1 pseudogene 3 | pseudogene | . | 54709 | . | . | . | 1.171546 | 0.473903 | 0.832425 | 0.614909 | 2.726235 | 1.095448 | 1.867937 | 1.405915 | -1.081119 | 8.097927 | 1.099248492 | -0.10236597 | 0.918466193 | NA | 3.069695 | 2.963455 | 3.035647 | 3.000916 | 3.021682 | 3.009394 | 14 | 5 | 8 | 6 |
| 100420981 | gene-LOC1004 | LOC10042098 | nucleosome assembly protein 1 like 1 pseudogene | pseudogene | . | . | . | . | . | 0.264433 | 0.142347 | 0.473847 | 0.372101 | 0.615347 | 0.329041 | 1.063299 | 0.850765 | 1.981567 | 11.290738 | 0.927069685 | 1.06425806 | 0.287211811 | NA | 2.917750 | 3.909275 | 3.471047 | 3.452706 | 3.504312 | 3.485659 | 11 | 6 | 15 | 12 |
| 100421096 | gene-NCAPGP1 | NCAPGP1 | non-SMC condensin I complex subunit G pseudogene 1 | pseudogene | . | 51904 | . | . | . | 0.132765 | 0.093671 | 0.224596 | 0.034612 | 0.308950 | 0.216524 | 0.503987 | 0.079136 | 1.183389 | 7.327368 | 1.209512789 | 0.200845228 | 0.840819597 | NA | 2.741623 | 2.993948 | 2.859931 | 2.849416 | 2.885490 | 2.834179 | 9 | 6 | 12 | 2 |
| 100421130 | gene-LOC1004 | LOC10042113( | LSM14A, SCD6 homolog A (S. cerevisiae) pseudogene | pseudogene | . | . | . | . | . | 0.148425 | 0.256096 | 0.590251 | 0.312533 | 0.345390 | 0.591978 | 1.324506 | 0.714571 | 2.149402 | 13.537374 | 0.871060499 | 1.267346117 | 0.205031557 | NA | 3.104339 | 4.207482 | 3.696998 | 3.720738 | 3.775784 | 3.728427 | 7 | 12 | 21 | 12 |
| 100421131 | gene-RCC2P6 | RCC2P6 | regulator of chromosome condensation 2 pseudogene 6 | pseudogene | . | 42382 | . | . | . | 0.132386 | 0.276821 | 0.111289 | 0.147515 | 0.308068 | 0.639885 | 0.249729 | 0.337274 | -1.564150 | 7.800849 | 1.112928194 | -0.5798925 | 0.561987115 | NA | 3.251986 | 2.602870 | 2.958129 | 2.989601 | 2.957271 | 2.959972 | 7 | 14 | 5 | 6 |
| 100421239 | gene-FANCD2P | FANCD2P1 | FANCD2 pseudogene 1 | pseudogene | . | 44487 | . | . | . | 1.016400 | 0.223213 | 0.917431 | 0.709448 | 2.365203 | 0.515967 | 2.058688 | 1.622068 | 1.166232 | 6.695730 | 1.210604231 | 0.183259632 | 0.854594307 | NA | 2.624463 | 2.852980 | 2.752222 | 2.720595 | 2.744772 | 2.737559 | 11 | 3 | 7 | 6 |
| 100421296 | gene-SRSF11P1 | SRSF11P1 | serine and arginine rich splicing factor 11 pseudogene 1 | pseudogene | . | 49884 | . | . | . | 4.411591 | 4.314551 | 7.133069 | 5.304048 | 10.265953 | 9.973278 | 16.006401 | 12.127076 | 1.404353 | 208.585479 | 0.247920007 | 1.976064286 | 0.048147497 | 0.660542298 | 7.438021 | 7.929339 | 7.616973 | 7.603515 | 7.838379 | 7.688858 | 203 | 184 | 244 | 194 |
| 100421306 | gene-LOC1004 | LOC10042130( | RecQ like helicase pseudogene | pseudogene | . | . | . | . | . | 0.972032 | 1.301709 | 1.253038 | 1.181339 | 2.261958 | 3.008959 | 2.811781 | 2.700992 | 1.072809 | 18.208701 | 0.721732706 | 0.140485555 | 0.888276364 | NA | 4.135531 | 4.235838 | 4.168080 | 4.192917 | 4.188966 | 4.183130 | 17 | 22 | 17 | 17 |
| 100421353 | gene-HNRNPA | HNRNPA1P15 | heterogeneous nuclear ribonucleoprotein A1 pseudogene 15 | pseudogene | . | 39133 | . | . | . | 0.028401 | 0.043012 | 0.023539 | 0.018887 | 0.066089 | 0.099424 | 0.052821 | 0.043183 | -1.620137 | 1.451754 | 2.440017378 | -0.28529139 | 0.775420906 | NA | 0.844134 | 0.148375 | 0.548030 | 0.548560 | 0.545314 | 0.544903 | 2 | 2 | 1 | 1 |
| 100421358 | gene-LOC1004 | LOC10042135 | CREB regulated transcription coactivator 2 pseudogene | pseudogene | . | . | . | . | . | 0.180428 | 0.216586 | 0.328794 | 0.122715 | 0.419862 | 0.500647 | 0.737804 | 0.280574 | 1.122647 | 14.342238 | 0.843549186 | 0.197860237 | 0.843154415 | NA | 3.754145 | 3.925185 | 3.821726 | 3.833815 | 3.862780 | 3.806340 | 14 | 16 | 19 | 8 |
| 100421372 | NM_00137878 | LOC10042137 | zinc finger and SCAN domain containing 29 | protein_coding | NP_001365714.1;NP_001365719.1 | . | . | . | . | 4.343071 | 3.685813 | 4.099571 | 3.986375 | 10.106506 | 8.519924 | 9.199318 | 9.114372 | -1.016380 | 571.966562 | 0.167494657 | -0.13994307 | 0.888704969 | 0.998865404 | 9.171109 | 9.148375 | 9.213094 | 9.112838 | 9.177839 | 9.128976 | 714 | 575 | 512 | 512 |
| 100421374 | gene-MPRIPP1 | MPRIPP1 | myosin phosphatase Rho interacting protein pseudogene 1 | pseudogene | . | 44594 | . | . | . | 0.176224 | 0.221796 | 0.113365 | 0.145264 | 0.410080 | 0.512691 | 0.254388 | 0.332129 | -1.518443 | 20.129920 | 0.693811703 | -0.86852555 | 0.385106689 | NA | 4.602488 | 3.996978 | 4.342185 | 4.363893 | 4.309218 | 4.326992 | 25 | 29 | 12 | 17 |
| 100421378 | gene-HNRNPA | HNRNPA1P77 | heterogeneous nuclear ribonucleoprotein A1 pseudogene 77 | pseudogene | . | 48807 | . | . | . | 0.105729 | 0.261907 | 0.379458 | 0.269978 | 0.246036 | 0.605410 | 0.851492 | 0.617272 | 1.699509 | 7.961888 | 1.103393268 | 0.69342296 | 0.488044158 | NA | 2.562483 | 3.324312 | 2.955163 | 2.978222 | 2.991769 | 2.979000 | 4 | 9 | 10 | 8 |
| 100421384 | gene-HNRNPA | HNRNPA1P38 | heterogeneous nuclear ribonucleoprotein A1pseudogene 38 | pseudogene | . | 48767 | . | . | . | 0.952105 | 0.534676 | 0.634932 | 1.018386 | 2.215589 | 1.235927 | 1.424768 | 2.328418 | 1.100448 | 29.853107 | 0.598839829 | 0.229720691 | 0.818308816 | 0.998865404 | 4.829417 | 4.966926 | 4.922827 | 4.855857 | 4.873041 | 4.929034 | 42 | 22 | 21 | 36 |
| 100421395 | gene-HNRNPA | HNRNPA3P12 | heterogeneous nuclear ribonucleoprotein A3 pseudogene 12 | pseudogene | . | 48758 | . | . | . | 0.117502 | 0.178558 | 0.142665 | 0.059092 | 0.273432 | 0.412745 | 0.320137 | 0.135107 | -1.496353 | 4.909843 | 1.395423789 | -0.41668419 | 0.676909419 | NA | 2.555346 | 1.978782 | 2.293763 | 2.308058 | 2.300142 | 2.283908 | 5 | 8 | 5 | 2 |
| 100421398 | gene-HNRNPA | HNRNPA1P54 | heterogeneous nuclear ribonucleoprotein A1 pseudogene 54 | pseudogene | . | 48784 | . | . | . | 0.055309 | 0.058977 | 0.040968 | 0.070807 | 0.128705 | 0.136328 | 0.091931 | 0.161891 | 1.222837 | 2.988044 | 1.723505885 | 0.168396441 | 0.866271408 | NA | 1.429096 | 1.715150 | 1.575076 | 1.575932 | 1.573846 | 1.582711 | 3 | 3 | 2 | 4 |
| 100421409 | gene-LOC1004 | LOC10042140 | glucosamine-phosphate N-acetyltransferase 1 pseudogene | pseudogene | . | . | . | . | . | 0.124729 | 0.211799 | 0.301370 | 0.261131 | 0.290250 | 0.489582 | 0.676266 | 0.597046 | 1.608485 | 7.659819 | 1.105490188 | 0.620270116 | 0.535079966 | NA | 2.555346 | 3.239024 | 2.908426 | 2.923004 | 2.935800 | 2.928019 | 5 | 8 | 9 | 8 |
| 100421436 | NR_168330 | UBE2FP1 | UBE2F pseudogene 1 | transcribed_ps | . | 44535 | . | . | . | 1.355240 | 1.562434 | 1.949330 | 1.254850 | 3.153698 | 3.611637 | 4.374241 | 2.869066 | 1.088473 | 126.277857 | 0.315158568 | 0.388075285 | 0.697960321 | 0.998865404 | 6.917163 | 7.041092 | 6.934144 | 6.980347 | 7.063905 | 6.904719 | 130 | 139 | 140 | 96 |
| 100421438 | gene-LOC1004 | LOC10042143 | DnaJ heat shock protein family (Hsp40) member C8 pseudogene | pseudogene | . | . | . | . | . | 0.094957 | 0.079865 | 0.041196 | 0.047897 | 0.220970 | 0.184612 | 0.092442 | 0.109510 | -1.412257 | 2.671821 | 1.820869166 | -0.27349696 | 0.784471225 | NA | 1.644780 | 1.148375 | 1.428682 | 1.425364 | 1.423205 | 1.422411 | 4 | 3 | 2 | 2 |
| 100421490 | gene-LOC1004 | LOC10042149( | DnaJ heat shock protein family (Hsp40) member C9 pseudogene | pseudogene | . | . | . | . | . | 0.316571 | 0.365075 | 0.273366 | 0.388467 | 0.736673 | 0.843888 | 0.613425 | 0.888183 | -1.042703 | 10.114240 | 0.959293734 | -0.06288887 | 0.949854997 | NA | 3.369732 | 3.306201 | 3.336828 | 3.343977 | 3.330535 | 3.346482 | 11 | 12 | 7 | 11 |
| 100421515 | gene-BMPR1AI | BMPR1AP1 | bone morphogenetic protein receptor type 1A pseudogene 1 | pseudogene | . | 21742 | . | . | . | 0.193104 | 0.191543 | 0.243573 | 0.047300 | 0.449360 | 0.442759 | 0.546570 | 0.108147 | -1.261522 | 10.876378 | 1.001703099 | -0.33459585 | 0.737929974 | NA | 3.597285 | 3.270502 | 3.443097 | 3.442507 | 3.459292 | 3.403160 | 14 | 13 | 14 | 3 |
| 100421533 | gene-EIF4A1P1 | EIF4A1P10 | eukaryotic translation initiation factor 4A1 pseudogene 10 | pseudogene | . | 37930 | . | . | . | 1.084406 | 1.499578 | 0.950887 | 1.770975 | 2.523457 | 3.466342 | 2.133762 | 4.049125 | 1.042951 | 57.442223 | 0.450951136 | 0.134539743 | 0.892975786 | 0.998865404 | 5.815530 | 5.871997 | 5.801278 | 5.864274 | 5.780248 | 5.900207 | 55 | 70 | 36 | 71 |
| 100421542 | gene-DNAJA1P | DNAJA1P6 | DnaJ heat shock protein family (Hsp40) member A1 pseudogene 6 | pseudogene | . | 39342 | . | . | . | 0.032891 | 0.080993 | 0.346126 | 0.200706 | 0.076539 | 0.187219 | 0.776696 | 0.458891 | 4.582753 | 8.859217 | 1.119311557 | 1.962111887 | 0.049749459 | NA | 1.671431 | 3.861274 | 3.068589 | 3.083046 | 3.144156 | 3.111060 | 2 | 5 | 16 | 10 |
| 100421561 | gene-LOC1004 | LOC10042156 | family with sequence similarity 133 member B pseudogene | pseudogene | . | . | . | . | . | 0.736733 | 0.589399 | 0.862095 | 0.408702 | 1.714409 | 1.362421 | 1.934515 | 0.934448 | -1.046759 | 40.686504 | 0.527879988 | -0.12489495 | 0.90060672 | 0.998865404 | 5.376728 | 5.315581 | 5.360152 | 5.324780 | 5.387854 | 5.279931 | 53 | 40 | 47 | 24 |
| 100421569 | gene-LOC1004 | LOC10042156 | glyoxylate reductase 1 homolog (Arabidopsis) pseudogene | pseudogene | . | . | . | . | . | 0.018140 | 0.072538 | 0.032388 | 0.074383 | 0.042211 | 0.167675 | 0.072677 | 0.170067 | 1.206200 | 3.500973 | 1.630637355 | 0.165867226 | 0.86826144 | NA | 1.671431 | 1.932304 | 1.794260 | 1.808066 | 1.797393 | 1.811022 | 2 | 5 | 2 | 5 |
| 100421577 | NR_038275 | FAM21EP | family with sequence similarity 21 member E, pseudogene | transcribed_ps | . | 45010 | . | . | . | 0.226774 | 0.168716 | 0.130276 | 0.242816 | 0.527712 | 0.389994 | 0.292335 | 0.555170 | -1.051849 | 13.486527 | 0.847901821 | -0.08601022 | 0.931458297 | NA | 3.790753 | 3.715150 | 3.764778 | 3.748692 | 3.736914 | 3.770402 | 18 | 13 | 8 | 16 |
| 100421579 | gene-EIF4A2P2 | EIF4A2P2 | eukaryotic translation initiation factor 4A2 pseudogene 2 | pseudogene | . | 45099 | . | . | . | 0.065238 | 0.183769 | 0.113514 | 0.288034 | 0.151812 | 0.424791 | 0.254722 | 0.658557 | 1.571106 | 5.823110 | 1.289679786 | 0.505381849 | 0.613290665 | NA | 2.184683 | 2.827809 | 2.515225 | 2.533684 | 2.525185 | 2.547414 | 3 | 7 | 4 | 9 |
| 100421582 | gene-PAIP1P1 | PAIP1P1 | poly(A) binding protein interacting protein 1 pseudogene 1 | pseudogene | . | 18240 | . | . | . | 0.301185 | 0.172737 | 0.205786 | 0.070803 | 0.700870 | 0.399290 | 0.461777 | 0.161883 | -1.805102 | 8.658892 | 1.084696481 | -0.78554728 | 0.432132762 | NA | 3.474811 | 2.632244 | 3.142862 | 3.117505 | 3.121285 | 3.094703 | 16 | 9 | 8 | 3 |
| 100421614 | gene-LOC1004 | LOC10042161 | actin related protein 2 pseudogene | pseudogene | . | . | . | . | . | 0.026727 | 0.018288 | 0.037640 | 0.056968 | 0.062194 | 0.042274 | 0.084463 | 0.130250 | 2.061309 | 2.040895 | 2.090979391 | 0.499077532 | 0.617724765 | NA | 0.413393 | 1.459419 | 1.015206 | 1.011351 | 1.017838 | 1.021985 | 2 | 1 | 2 | 3 |
| 100421693 | gene-NSA2P7 | NSA2P7 | NSA2 pseudogene 7 | pseudogene | . | 54588 | . | . | . | 0.445799 | 0.534484 | 0.194678 | 0.303123 | 1.037391 | 1.235484 | 0.436851 | 0.693056 | -1.819679 | 10.085642 | 0.974609807 | -0.88618448 | 0.375518128 | NA | 3.703730 | 2.836248 | 3.352152 | 3.359777 | 3.324048 | 3.335923 | 14 | 15 | 5 | 8 |
| 100421724 | gene-BNIP3P5 | BNIP3P5 | BCL2 interacting protein 3 pseudogene 5 | pseudogene | . | 39658 | . | . | . | 0.408040 | 1.058930 | 1.067096 | 0.473496 | 0.949526 | 2.447763 | 2.394532 | 1.082592 | 1.028425 | 16.596910 | 0.8127788 | 0.049751756 | 0.960320212 | NA | 4.032166 | 4.073227 | 4.007973 | 4.068500 | 4.066279 | 4.014419 | 11 | 25 | 20 | 10 |
| 100421788 | gene-KMT5AP | KMT5AP2 | KMT5A pseudogene 2 | pseudogene | . | 54727 | . | . | . | 0.105884 | 0.121986 | 0.123924 | 0.165370 | 0.246397 | 0.281975 | 0.278082 | 0.378099 | 1.228632 | 5.513138 | 1.280223022 | 0.232032061 | 0.816513112 | NA | 2.307819 | 2.602870 | 2.452722 | 2.458507 | 2.459322 | 2.462261 | 5 | 6 | 5 | 6 |
| 100421796 | gene-EEF1A1P | EEF1A1P19 | eukaryotic translation elongation factor 1 alpha 1 pseudogene 19 | pseudogene | . | 37892 | . | . | . | 4.502714 | 4.677640 | 3.036349 | 4.266798 | 10.478000 | 10.812575 | 6.813478 | 9.755528 | -1.270943 | 211.873620 | 0.247330262 | -1.39853428 | 0.161952687 | 0.935295382 | 7.890460 | 7.542761 | 7.770701 | 7.785739 | 7.596474 | 7.733409 | 269 | 260 | 136 | 203 |
| 100421854 | gene-EEF1A1P | EEF1A1P4 | eukaryotic translation elongation factor 1 alpha 1 pseudogene 4 | pseudogene | . | 3199 | . | . | . | 0.592941 | 0.596322 | 0.292150 | 0.605481 | 1.379799 | 1.378426 | 0.655576 | 1.384360 | -1.330723 | 27.451556 | 0.612587266 | -0.67290105 | 0.501010243 | 0.998865404 | 4.972078 | 4.555601 | 4.797127 | 4.797913 | 4.735389 | 4.795098 | 36 | 34 | 14 | 29 |
| 100421976 | gene-LOC1004 | LOC10042197 | interferon stimulated exonuclease gene 20 like 2 pseudogene | pseudogene | . | . | . | . | . | 0.222157 | 0.022096 | 0.339625 | 0.108082 | 0.516969 | 0.051075 | 0.762108 | 0.247116 | 1.732199 | 8.426942 | 1.16544507 | 0.680087452 | 0.496449089 | NA | 2.617627 | 3.421760 | 3.061386 | 3.021296 | 3.084897 | 3.038212 | 12 | 2 | 14 | 5 |
| 100422094 | gene-KCTD5P1 | KCTD5P1 | KCTD5 pseudogene 1 | pseudogene | . | 54736 | . | . | . | 0.031635 | 0.053744 | 0.131495 | 0.060830 | 0.073616 | 0.124232 | 0.295071 | 0.139080 | 1.870564 | 2.580901 | 1.868819978 | 0.483445811 | 0.628779227 | NA | 0.844134 | 1.751299 | 1.345797 | 1.346419 | 1.358911 | 1.347831 | 2 | 2 | 4 | 2 |
| 100422338 | gene-G3BP1P1 | G3BP1P1 | G3BP1 pseudogene 1 | pseudogene | . | 54590 | . | . | . | 0.111165 | 0.249676 | 0.357274 | 0.045129 | 0.258685 | 0.577138 | 0.801712 | 0.103183 | 1.135447 | 10.682457 | 1.0523909 | 0.174136725 | 0.861758012 | NA | 3.320504 | 3.507767 | 3.374124 | 3.410170 | 3.433005 | 3.362225 | 7 | 15 | 17 | 3 |
| 100422343 | gene-UPF3AP3 | UPF3AP3 | UPF3A pseudogene 3 | pseudogene | . | 38395 | . | . | . | 0.144303 | 0.068725 | 0.292267 | 0.150125 | 0.335799 | 0.158862 | 0.655840 | 0.343243 | 2.009721 | 6.689766 | 1.206072103 | 0.834937802 | 0.403752729 | NA | 2.147197 | 3.161867 | 2.719778 | 2.704280 | 2.743565 | 2.721349 | 7 | 3 | 10 | 6 |
| 100422368 | gene-LOC1004 | LOC10042236 | UPF3B, regulator of nonsense mediated mRNA decay pseudogene | pseudogene | . | . | . | . | . | 0.251582 | 0.150932 | 0.216497 | 0.276929 | 0.585442 | 0.348885 | 0.485813 | 0.633164 | 1.238434 | 6.970721 | 1.151826538 | 0.26785054 | 0.788814366 | NA | 2.638039 | 2.947964 | 2.802222 | 2.787396 | 2.797946 | 2.805195 | 9 | 5 | 6 | 8 |
| 100422414 | gene-LOC1004 | LOC10042241 | proteasome (prosome, macropain) 26S subunit, non-ATPase, 7 ps | pseudogene | . | . | . | . | . | 0.197892 | 0.278747 | 0.208485 | 0.083907 | 0.460502 | 0.644337 | 0.467834 | 0.191844 | -1.607997 | 5.126954 | 1.365632868 | -0.50179283 | 0.615813256 | NA | 2.658167 | 1.978782 | 2.361483 | 2.371668 | 2.363821 | 2.347584 | 6 | 8 | 5 | 2 |
| 100422495 | gene-LOC1004 | LOC10042249 | protein tyrosine phosphatase non-receptor type 4 pseudogene | pseudogene | . | . | . | . | . | 0.042912 | 0.095985 | 0.045432 | 0.048198 | 0.099859 | 0.221874 | 0.101948 | 0.110200 | -1.428313 | 5.387370 | 1.315024922 | -0.39110465 | 0.695719881 | NA | 2.664814 | 2.148375 | 2.427402 | 2.445975 | 2.428895 | 2.427321 | 5 | 9 | 4 | 4 |
| 100422558 | NR_135312 | SMG1P6 | SMG1 pseudogene 6 | transcribed_ps | . | 49863 | . | . | . | 1.357880 | 1.346502 | 2.042216 | 1.402688 | 3.159842 | 3.112499 | 4.582674 | 3.207080 | 1.250305 | 150.386323 | 0.287115171 | 1.12247833 | 0.261659116 | 0.995760118 | 7.061414 | 7.385484 | 7.180790 | 7.175421 | 7.340098 | 7.184305 | 155 | 143 | 173 | 127 |
| 100422581 | gene-DFFBP1 | DFFBP1 | DNA fragmentation factor subunit beta pseudogene 1 | pseudogene | . | 49815 | . | . | . | 0.820164 | 0.757412 | 0.897995 | 1.167386 | 1.908556 | 1.750792 | 2.015074 | 2.669089 | 1.307017 | 33.553439 | 0.53785835 | 0.718178331 | 0.472647337 | 0.998865404 | 4.862893 | 5.248230 | 5.049694 | 5.039165 | 5.061933 | 5.097860 | 35 | 30 | 29 | 40 |
| 100422627 | gene-LOC1004 | LOC10042262 | TATA-box binding protein associated factor 4b pseudogene | pseudogene | . | . | . | . | . | 0.198652 | 0.071683 | 0.445499 | 0.121862 | 0.462272 | 0.165699 | 0.999686 | 0.278622 | 2.029925 | 10.089845 | 1.04436463 | 0.978035964 | 0.328056537 | NA | 2.728992 | 3.760196 | 3.303419 | 3.276229 | 3.350733 | 3.289444 | 11 | 4 | 18 | 6 |
| 100422666 | gene-YWHABP | YWHABP2 | tyrosine 3-monooxygenase/tryptophan 5-monooxygenase activati | pseudogene | . | 50295 | . | . | . | 0.072782 | 0.180511 | 0.204497 | 0.431995 | 0.169366 | 0.417260 | 0.458886 | 0.987705 | 2.298065 | 7.464765 | 1.162904415 | 1.03226001 | 0.30195033 | NA | 2.184683 | 3.376188 | 2.856350 | 2.874964 | 2.877005 | 2.907701 | 3 | 7 | 6 | 13 |
| 100422691 | gene-ITGB1P1 | ITGB1P1 | integrin subunit beta 1 pseudogene 1 | pseudogene | . | 3781 | . | . | . | 3.082609 | 2.625932 | 2.367994 | 1.004002 | 7.173357 | 6.069958 | 5.313710 | 2.295532 | -1.716382 | 76.167174 | 0.427808893 | -1.82177334 | 0.068489385 | 0.741130004 | 6.586868 | 5.812690 | 6.329401 | 6.276973 | 6.247611 | 6.083880 | 120 | 95 | 69 | 31 |
| 100422712 | gene-THAP12P | THAP12P7 | THAP domain containing 12 pseudogene 7 | pseudogene | . | 39574 | . | . | . | 0.077065 | 0.086711 | 0.067988 | 0.047332 | 0.179333 | 0.200437 | 0.152563 | 0.108218 | -1.469222 | 7.546262 | 1.108359946 | -0.50078741 | 0.616520745 | NA | 3.166062 | 2.612728 | 2.926385 | 2.928963 | 2.918300 | 2.911263 | 10 | 10 | 6 | 5 |
| 100422717 | gene-LOC1004 | LOC10042271 | pterin-4 alpha-carbinolamine dehydratase 2 pseudogene | pseudogene | . | . | . | . | . | 0.518520 | 0.108254 | 0.113982 | 0.154690 | 1.206617 | 0.250233 | 0.255773 | 0.353681 | -2.243898 | 4.458261 | 1.50024945 | -0.77720862 | 0.4370357 | NA | 2.624463 | 1.459419 | 2.193446 | 2.161973 | 2.160314 | 2.164152 | 11 | 3 | 2 | 3 |
| 100422832 | NR_036159,rn | MIR3191 | microRNA 3191 | miRNA | . | 38222 | . | . | . | 1.123028 | 2.345654 | 0.852233 | 2.258882 | 2.613331 | 5.422085 | 1.912387 | 5.164665 | 1.030432 | 5.955684 | 1.247516659 | 0.034668175 | 0.972344339 | NA | 2.555346 | 2.592943 | 2.565179 | 2.579570 | 2.566864 | 2.579559 | 5 | 8 | 4 | 7 |
| 100422860 | NR_036251,rn | MIR4292 | microRNA 4292 | miRNA | . | 38348 | . | . | . | 2.292111 | 1.407063 | 1.796750 | 1.322102 | 5.333838 | 3.252489 | 4.031857 | 3.022829 | -1.072000 | 5.182339 | 1.322372303 | -0.07585255 | 0.939536404 | NA | 2.421266 | 2.324312 | 2.378555 | 2.371759 | 2.376811 | 2.370200 | 7 | 5 | 5 | 4 |
| 100422872 | NR_036263,rn | MIR1244-3 | microRNA 1244-3 | miRNA | . | 38390 | . | . | . | 1.693369 | 1.865351 | 0.696500 | 0.127952 | 3.940540 | 4.311840 | 1.562927 | 0.292547 | -1.934468 | 4.088594 | 1.509517358 | -0.63062329 | 0.528286867 | NA | 2.429096 | 1.481107 | 2.046486 | 2.048025 | 2.038381 | 2.032368 | 6 | 6 | 3 | 2 |
| 100422885 | NR_036262,rn | MIR1244-2 | microRNA 1244-2 | miRNA | . | 38321 | . | . | . | 78.889275 | 41.716774 | 45.003222 | 13.666140 | 183.578581 | 96.430193 | 100.985927 | 31.246006 | -2.057110 | 120.746261 | 0.885795363 | -1.17478519 | 0.240080667 | 0.992733415 | 7.343871 | 6.304517 | 7.111868 | 6.845262 | 6.870325 | 6.598394 | 245 | 121 | 105 | 35 |
| 100422886 | NR_036143,rn | MIR3179-2 | microRNA 3179-2 | miRNA | . | 38325 | . | . | . | 0.249739 | 0.448730 | 0.335908 | 0.498909 | 0.581153 | 1.037259 | 0.753767 | 1.140697 | 1.221536 | 2.504267 | 1.872041618 | 0.154214919 | 0.877440279 | NA | 1.175403 | 1.459419 | 1.316408 | 1.321418 | 1.319196 | 1.323280 | 2 | 3 | 2 | 3 |
| 100422894 | NR_036213,rn | MIR4260 | microRNA 4260 | miRNA | . | 38179 | . | . | . | 0.000019 | 0.000017 | 0.000032 | 0.000037 | 0.000045 | 0.00004 | 0.000071 | 0.000085 | 1.234655 | 1.002957 | 2.868339486 | 0.106022226 | 0.915564717 | NA | -0.155866 | 0.148375 | -0.001261 | -0.000972 | 0.000053 | -0.000315 | 1 | 1 | 1 | 1 |
| 100422910 | NR_036055,rn | MIR2861 | microRNA 2861 | miRNA | . | 38221 | 613405 | . | . | 0.413945 | 0.009792 | 0.302596 | 0.595192 | 0.963265 | 0.022634 | 0.679017 | 1.360836 | 1.232222 | 1.970512 | 2.132558215 | 0.141267792 | 0.887658392 | NA | 0.820514 | 1.121007 | 0.982345 | 0.974334 | 0.975623 | 0.984888 | 3 | 1 | 1 | 3 |
| 100422936 | NR_036108,rn | MIR3153 | microRNA 3153 | miRNA | . | 38289 | . | . | . | 2.154298 | 0.693536 | 1.543450 | 1.201703 | 5.013139 | 1.603139 | 3.463457 | 2.747551 | 1.025364 | 4.936078 | 1.374632832 | 0.026287866 | 0.979027733 | NA | 2.282109 | 2.324312 | 2.311881 | 2.292277 | 2.305965 | 2.299376 | 8 | 3 | 5 | 4 |
| 100422945 | NR_036220,rn | MIR4326 | microRNA 4326 | miRNA | . | 38314 | . | . | . | 1.446202 | 0.469453 | 0.376777 | 0.614123 | 3.365371 | 1.085161 | 0.845475 | 1.404119 | -1.401566 | 2.657246 | 1.841602615 | -0.2644653 | 0.791421401 | NA | 1.631267 | 1.148375 | 1.425810 | 1.414002 | 1.416209 | 1.415417 | 5 | 2 | 2 | 2 |
| 100422960 | NR_036140,rn | MIR3179-1 | microRNA 3179-1 | miRNA | . | 38353 | . | . | . | 0.083130 | 0.445757 | 0.297475 | 0.409445 | 0.193447 | 1.030388 | 0.667527 | 0.936148 | -1.017609 | 2.237600 | 1.972119472 | -0.01276997 | 0.989811317 | NA | 1.175403 | 1.148375 | 1.158018 | 1.163002 | 1.160711 | 1.159957 | 2 | 3 | 2 | 2 |
| 100422961 | NR_036271,rn | MIR4315-2 | microRNA 4315-2 | miRNA | . | 38356 | . | . | . | 0.358332 | 0.116948 | 0.285988 | 0.138075 | 0.833854 | 0.270331 | 0.641749 | 0.315693 | 1.234663 | 2.005914 | 2.070995667 | 0.146845664 | 0.883253841 | NA | 0.844134 | 1.148375 | 0.998092 | 0.998667 | 1.000707 | 0.999974 | 2 | 2 | 2 | 2 |
| 100422971 | NR_036197,rn | MIR4312 | microRNA 4312 | miRNA | . | 38376 | . | . | . | 0.132802 | 0.077304 | 0.106001 | 0.054180 | 0.309036 | 0.178693 | 0.237863 | 0.123877 | -1.204659 | 1.220068 | 2.631544379 | -0.10207874 | 0.918694172 | NA | 0.413393 | 0.148375 | 0.294625 | 0.290739 | 0.291821 | 0.291432 | 2 | 1 | 1 | 1 |
| 100422986 | NR_036072,rn | MIR3125 | microRNA 3125 | miRNA | . | 38219 | . | . | . | 0.118970 | 1.733305 | 1.715284 | 0.808137 | 0.276848 | 4.006612 | 3.849046 | 1.847708 | 1.396524 | 3.844696 | 1.602632163 | 0.300655489 | 0.763677214 | NA | 1.684574 | 2.161867 | 1.912109 | 1.934584 | 1.934947 | 1.923562 | 1 | 6 | 5 | 3 |
| 100423004 | NR_036199,rn | MIR4315-1 | microRNA 4315-1 | miRNA | . | 38342 | . | . | . | 0.000588 | 0.000747 | 0.001741 | 0.000438 | 0.001369 | 0.001726 | 0.003906 | 0.001002 | 2.469696 | 1.557117 | 2.386421563 | 0.546564555 | 0.584677926 | NA | -0.155866 | 1.148375 | 0.616459 | 0.616793 | 0.623168 | 0.622477 | 1 | 1 | 2 | 2 |
| 100423006 | NR_036145,rn | MIR3179-3 | microRNA 3179-3 | miRNA | . | 38214 | . | . | . | 0.284939 | 0.641171 | 0.533130 | 0.729439 | 0.663064 | 1.482097 | 1.196329 | 1.667776 | 1.474528 | 2.791761 | 1.780166124 | 0.314719634 | 0.752974536 | NA | 1.175403 | 1.733338 | 1.466303 | 1.471348 | 1.474336 | 1.473263 | 2 | 3 | 3 | 3 |
| 100423014 | NR_036163,rn | MIR3196 | microRNA 3196 | miRNA | . | 38198 | . | . | . | 1.167019 | 1.382741 | 1.933885 | 2.555735 | 2.715700 | 3.196267 | 4.339582 | 5.843385 | 1.693200 | 4.832655 | 1.369616787 | 0.554719208 | 0.579086736 | NA | 1.844134 | 2.602870 | 2.255277 | 2.256480 | 2.265820 | 2.268836 | 4 | 4 | 5 | 6 |
| 100423020 | NR_036212,rn | MIR4258 | microRNA 4258 | miRNA | . | 38281 | . | . | . | 1.544342 | 2.467375 | 2.750066 | 2.040161 | 3.593748 | 5.703447 | 6.171067 | 4.664586 | 1.230871 | 7.554455 | 1.106883667 | 0.270741783 | 0.786589639 | NA | 2.760365 | 3.058876 | 2.899253 | 2.913932 | 2.917443 | 2.909911 | 6 | 9 | 8 | 7 |
| 100423031 | NR_036207,rn | MIR4321 | microRNA 4321 | miRNA | . | 38244 | . | . | . | 0.089757 | 0.204734 | 0.167854 | 0.058766 | 0.208869 | 0.473252 | 0.376659 | 0.134362 | 1.234663 | 2.005914 | 2.070995667 | 0.146845664 | 0.883253841 | NA | 0.844134 | 1.148375 | 0.998092 | 0.998667 | 1.000707 | 0.999974 | 2 | 2 | 2 | 2 |
| 100423037 | NR_036137,rn | MIR3176 | microRNA 3176 | miRNA | . | 38345 | . | . | . | 2.215746 | 0.893964 | 1.152885 | 1.960623 | 5.156132 | 2.066436 | 2.587040 | 4.482733 | -1.048405 | 5.630714 | 1.286383674 | -0.0530141 | 0.957720673 | NA | 2.526439 | 2.459419 | 2.506208 | 2.486990 | 2.491440 | 2.499396 | 9 | 4 | 4 | 6 |
| 100462836 | gene-EIF3EP3 | EIF3EP3 | eukaryotic translation initiation factor 3 subunit E pseudogene 3 | pseudogene | . | 37920 | . | . | . | 1.478213 | 0.671968 | 0.950773 | 0.848052 | 3.439862 | 1.553285 | 2.133506 | 1.938971 | -1.226820 | 9.262187 | 1.02129699 | -0.28877352 | 0.772754694 | NA | 3.349244 | 3.058876 | 3.234938 | 3.201060 | 3.213504 | 3.205862 | 16 | 7 | 8 | 7 |
| 100462880 | gene-ELOCP19 | ELOCP19 | elongin C pseudogene 19 | pseudogene | . | 38155 | . | . | . | 0.628704 | 1.236543 | 0.919339 | 1.389525 | 1.463021 | 2.858324 | 2.062970 | 3.176984 | 1.260388 | 14.303888 | 0.832539427 | 0.401023961 | 0.688402483 | NA | 3.664814 | 3.993209 | 3.804983 | 3.841258 | 3.823718 | 3.851542 | 10 | 18 | 11 | 18 |
| 100462977 | NM_00119045 | MTRNR2L1 | MT-RNR2 like 1 | protein_coding | NP_001177381.1 | 37155 | 616985 | . | . | 0.221680 | 0.088032 | 0.212686 | 0.255095 | 0.515858 | 0.203491 | 0.477260 | 0.583244 | 1.450465 | 9.768311 | 1.000670905 | 0.536155753 | 0.59185091 | NA | 2.993083 | 3.532926 | 3.287673 | 3.256951 | 3.283135 | 3.293884 | 13 | 5 | 9 | 12 |
| 100462981 | NM_00119047 | MTRNR2L2 | MT-RNR2 like 2 | protein_coding | NP_001177399.1 | 37156 | . | . | . | 0.490927 | 0.282817 | 0.656887 | 0.356470 | 1.142407 | 0.653745 | 1.474035 | 0.815027 | 1.297435 | 14.333070 | 0.83181263 | 0.451618738 | 0.651543669 | NA | 3.638039 | 4.019385 | 3.837536 | 3.808829 | 3.858126 | 3.820409 | 18 | 10 | 18 | 11 |
| 100463486 | NM_00119070 | MTRNR2L8 | MT-RNR2 like 8 | protein_coding | NP_001177631.1 | 37165 | . | . | . | 0.690187 | 0.382501 | 0.719994 | 0.676550 | 1.606092 | 0.884168 | 1.615645 | 1.546851 | 1.294701 | 25.511481 | 0.62968132 | 0.591757324 | 0.554013104 | NA | 4.472918 | 4.848815 | 4.679354 | 4.623927 | 4.683344 | 4.674974 | 33 | 17 | 26 | 26 |
| 100463487 | NM_00119070 | MTRNR2L9 | MT-RNR2 like 9 | protein_coding | NP_001177635.1 | 37166 | . | . | . | 0.154116 | 0.044093 | 0.253528 | 0.127098 | 0.358635 | 0.101923 | 0.568909 | 0.290593 | 1.892882 | 2.566326 | 1.896119385 | 0.485509686 | 0.627314843 | NA | 0.820514 | 1.751299 | 1.343198 | 1.335165 | 1.352083 | 1.341014 | 3 | 1 | 4 | 2 |
| 100499177 | NR_034075,NF | THAP9-AS1 | THAP9 antisense RNA 1 | lncRNA | . | 44172 | . | . | . | 9.530920 | 8.696585 | 8.922709 | 7.856931 | 22.178845 | 20.102548 | 20.022303 | 17.963940 | -1.086881 | 620.980728 | 0.164219498 | -0.73190954 | 0.464223786 | 0.998865404 | 9.336864 | 9.217476 | 9.342044 | 9.266017 | 9.293019 | 9.203398 | 786 | 659 | 553 | 520 |
| 100499466 | NR_027418 | LINC00674 | long intergenic non-protein coding RNA 674 | lncRNA | . | 44355 | . | . | . | 2.252893 | 2.398239 | 2.510048 | 2.223965 | 5.242575 | 5.543636 | 5.632476 | 5.084831 | 1.004430 | 818.560649 | 0.149655613 | 0.042608431 | 0.966013674 | 0.998865404 | 9.673599 | 9.680284 | 9.655182 | 9.693593 | 9.722757 | 9.630736 | 915 | 905 | 760 | 719 |
| 100499467 | NR_036488,NF | LINC00673 | long intergenic non-protein coding RNA 673 | lncRNA | . | 44354 | 617079 | . | . | 21.572172 | 23.264707 | 22.842165 | 25.614186 | 50.199332 | 53.777412 | 51.257156 | 58.563794 | 1.067699 | 1705.560436 | 0.127672587 | 0.740215413 | 0.459169297 | 0.998865404 | 10.688182 | 10.782342 | 10.681393 | 10.736319 | 10.717614 | 10.801517 | 1833 | 1843 | 1450 | 1739 |
| 100499484 | NR_036526 | SUGT1P4-STRA | SUGT1P4-STRA6LP readthrough | transcribed_ps | . | 53834 | . | . | . | 1.271333 | 1.672146 | 1.883540 | 1.353190 | 2.958444 | 3.865241 | 4.226609 | 3.093908 | 1.083186 | 155.265313 | 0.28856353 | 0.399497444 | 0.689526707 | 0.998865404 | 7.219485 | 7.335372 | 7.202364 | 7.306137 | 7.352063 | 7.217863 | 149 | 182 | 164 | 126 |
| 100500802 | NR_037456,rn | MIR3685 | microRNA 3685 | miRNA | . | 38886 | . | . | . | 1.267208 | 0.639379 | 0.997072 | 0.404808 | 2.948843 | 1.477955 | 2.237400 | 0.925544 | -1.123618 | 2.959315 | 1.731081282 | -0.09713705 | 0.922617562 | NA | 1.644780 | 1.481107 | 1.568690 | 1.565383 | 1.568406 | 1.562470 | 4 | 3 | 3 | 2 |
| 100500804 | NR_037427,rn | MIR3654 | hsa-miR-3654 | . | . | 38896 | . | . | . | 0.003470 | 0.004490 | 0.007509 | 0.005026 | 0.008075 | 0.010378 | 0.016850 | 0.011492 | 1.234663 | 2.005914 | 2.070995667 | 0.146845664 | 0.883253841 | NA | 0.844134 | 1.148375 | 0.998092 | 0.998667 | 1.000707 | 0.999974 | 2 | 2 | 2 | 2 |
| 100500808 | NR_037481,rn | MIR3917 | microRNA 3917 | miRNA | . | 38911 | . | . | . | 0.333394 | 0.352797 | 0.317532 | 0.134097 | 0.775822 | 0.815505 | 0.712532 | 0.306599 | -1.215015 | 2.454711 | 1.887245685 | -0.1488804 | 0.881648006 | NA | 1.429096 | 1.148375 | 1.299640 | 1.300454 | 1.298230 | 1.297456 | 3 | 3 | 2 | 2 |
| 100500810 | NR_037415,rn | MIR3620 | microRNA 3620 | miRNA | . | 38917 | . | . | . | 0.072551 | 1.678357 | 0.634632 | 0.366582 | 0.168830 | 3.879598 | 1.424096 | 0.838147 | 1.064030 | 3.788889 | 1.618225908 | 0.055331888 | 0.955874058 | NA | 1.878854 | 1.963455 | 1.896759 | 1.923488 | 1.914439 | 1.908167 | 1 | 7 | 4 | 3 |
| 100500820 | NR_037428,rn | MIR3655 | hsa-miR-3655 | . | . | 38960 | . | . | . | 0.003792 | 0.004053 | 0.011037 | 0.004610 | 0.008823 | 0.009368 | 0.024769 | 0.010542 | 1.234663 | 2.005914 | 2.070995667 | 0.146845664 | 0.883253841 | NA | 0.844134 | 1.148375 | 0.998092 | 0.998667 | 1.000707 | 0.999974 | 2 | 2 | 2 | 2 |
| 100500827 | NR_037408,rn | MIR3614 | microRNA 3614 | miRNA | . | 38995 | . | . | . | 0.000882 | 0.000423 | 0.000742 | 0.002379 | 0.002052 | 0.000977 | 0.001664 | 0.005439 | 1.234665 | 3.008871 | 1.708874417 | 0.17796512 | 0.858750369 | NA | 1.429096 | 1.733338 | 1.582415 | 1.583274 | 1.586316 | 1.585224 | 3 | 3 | 3 | 3 |
| 100500832 | NR_037431,rn | MIR3658 | microRNA 3658 | miRNA | . | 38963 | . | . | . | 0.008840 | 0.003754 | 0.006200 | 0.002861 | 0.020571 | 0.008677 | 0.013911 | 0.006539 | 1.234663 | 2.005914 | 2.070995667 | 0.146845664 | 0.883253841 | NA | 0.844134 | 1.148375 | 0.998092 | 0.998667 | 1.000707 | 0.999974 | 2 | 2 | 2 | 2 |
| 100500842 | NR_037425,rn | MIR3652 | hsa-miR-3652 | . | . | 38894 | . | . | . | 0.000055 | 0.000054 | 0.000040 | 0.000056 | 0.000128 | 0.000125 | 0.00009 | 0.000128 | -1.620137 | 1.451754 | 2.440017378 | -0.28529139 | 0.775420906 | NA | 0.844134 | 0.148375 | 0.548030 | 0.548560 | 0.545314 | 0.544903 | 2 | 2 | 1 | 1 |
| 100500847 | NR_037409,rn | MIR3615 | microRNA 3615 | miRNA | . | 38905 | . | . | . | 0.000690 | 0.001325 | 0.013803 | 0.090327 | 0.001606 | 0.003063 | 0.030975 | 0.206521 | 1.234663 | 2.005914 | 2.070995667 | 0.146845664 | 0.883253841 | NA | 0.844134 | 1.148375 | 0.998092 | 0.998667 | 1.000707 | 0.999974 | 2 | 2 | 2 | 2 |
| 100500849 | NR_037480,rn | MIR3916 | microRNA 3916 | miRNA | . | 38912 | . | . | . | 5.996747 | 5.518861 | 6.793643 | 5.795548 | 13.954678 | 12.757095 | 15.244740 | 13.250831 | 1.087853 | 19.174182 | 0.700193816 | 0.173500284 | 0.862258205 | NA | 4.198253 | 4.321310 | 4.258460 | 4.251938 | 4.269688 | 4.254381 | 22 | 19 | 19 | 17 |
| 100500853 | NR_037400,rn | MIR3605 | microRNA 3605 | miRNA | . | 38922 | . | . | . | 0.867596 | 2.282748 | 1.110103 | 0.532889 | 2.018933 | 5.276676 | 2.491038 | 1.218387 | -1.357858 | 5.906549 | 1.264379839 | -0.34905074 | 0.727051219 | NA | 2.766559 | 2.324312 | 2.555203 | 2.578067 | 2.561918 | 2.555247 | 5 | 10 | 5 | 4 |
| 100500860 | NR_037412,rn | MIR3618 | microRNA 3618 | miRNA | . | 38937 | . | . | . | 0.000991 | 0.000764 | 0.002366 | 0.002235 | 0.002304 | 0.001768 | 0.005308 | 0.005111 | 1.234663 | 2.005914 | 2.070995667 | 0.146845664 | 0.883253841 | NA | 0.844134 | 1.148375 | 0.998092 | 0.998667 | 1.000707 | 0.999974 | 2 | 2 | 2 | 2 |
| 100500862 | NR_037421,rn | MIR3648-1 | microRNA 3648-1 | miRNA | . | 38941 | . | . | . | 0.000975 | 0.000818 | 0.000698 | 0.000854 | 0.002269 | 0.001891 | 0.001565 | 0.001951 | 1.234663 | 2.005914 | 2.070995667 | 0.146845664 | 0.883253841 | NA | 0.844134 | 1.148375 | 0.998092 | 0.998667 | 1.000707 | 0.999974 | 2 | 2 | 2 | 2 |
| 100500865 | NR_037500,rn | MIR3936 | microRNA 3936 | miRNA | . | 38947 | . | . | . | 0.343019 | 0.254485 | 1.811293 | 2.148622 | 0.798218 | 0.588253 | 4.064489 | 4.912567 | 3.712057 | 5.260493 | 1.376023207 | 1.375135801 | 0.169089347 | NA | 1.156660 | 3.051647 | 2.354103 | 2.350706 | 2.379184 | 2.381627 | 3 | 2 | 7 | 8 |
| 100500888 | NR_037505,rn | MIR3940 | microRNA 3940 | miRNA | . | 38893 | . | . | . | 0.778956 | 0.749905 | 0.682892 | 0.158821 | 1.812664 | 1.733438 | 1.532390 | 0.363124 | -1.152593 | 4.193958 | 1.463795295 | -0.13996666 | 0.888686332 | NA | 2.166062 | 1.963455 | 2.070107 | 2.071460 | 2.071194 | 2.064908 | 5 | 5 | 4 | 3 |
| 100500914 | NR_037404,rn | MIR3610 | microRNA 3610 | miRNA | . | 38942 | . | . | . | 0.000328 | 0.087060 | 0.024749 | 0.026575 | 0.000763 | 0.201243 | 0.055535 | 0.060759 | 1.234663 | 2.005914 | 2.070995667 | 0.146845664 | 0.883253841 | NA | 0.844134 | 1.148375 | 0.998092 | 0.998667 | 1.000707 | 0.999974 | 2 | 2 | 2 | 2 |
| 100500918 | NR_037424,rn | MIR3651 | microRNA 3651 | miRNA | . | 38897 | . | . | . | 0.054471 | 0.507503 | 0.142106 | 0.179566 | 0.126756 | 1.173115 | 0.318882 | 0.410556 | -1.017609 | 2.237600 | 1.972119472 | -0.01276997 | 0.989811317 | NA | 1.175403 | 1.148375 | 1.158018 | 1.163002 | 1.160711 | 1.159957 | 2 | 3 | 2 | 2 |
| 100500934 | gene-RBX1P2 | RBX1P2 | RBX1 pseudogene 2 | pseudogene | . | 54741 | . | . | . | 0.563186 | 0.423364 | 0.651969 | 0.627369 | 1.310558 | 0.978625 | 1.462999 | 1.434404 | 1.275600 | 17.273210 | 0.739102312 | 0.475138971 | 0.634687923 | NA | 3.923304 | 4.276108 | 4.105925 | 4.086457 | 4.114886 | 4.113615 | 20 | 14 | 17 | 18 |
| 100505483 | NR_038926 | PRKAG2-AS1 | PRKAG2 antisense RNA 1 | lncRNA | . | 40468 | . | . | . | 1.197943 | 0.970600 | 0.883346 | 0.873954 | 2.787662 | 2.243585 | 1.982202 | 1.998193 | -1.262879 | 24.418422 | 0.626522537 | -0.53743617 | 0.590966368 | NA | 4.767942 | 4.432387 | 4.638065 | 4.611476 | 4.602529 | 4.600191 | 35 | 26 | 19 | 20 |
| 100505498 | XR_923410 | LOC10050549 | uncharacterized LOC100505498 | lncRNA | . | . | . | . | . | 0.273868 | 0.194876 | 0.189095 | 0.304275 | 0.637301 | 0.450464 | 0.424323 | 0.695689 | 1.046822 | 63.861436 | 0.422153542 | 0.156380975 | 0.875732733 | 0.998865404 | 5.964015 | 6.028999 | 6.024715 | 5.949336 | 5.946694 | 6.046715 | 84 | 56 | 44 | 75 |
| 100505501 | NR_149030,NF | LOC10050550 | uncharacterized LOC100505501 | lncRNA | . | . | . | . | . | 0.296917 | 0.247719 | 0.332936 | 0.337099 | 0.690937 | 0.572614 | 0.747098 | 0.770737 | 1.198772 | 31.461336 | 0.55177152 | 0.474032787 | 0.635476527 | 0.998865404 | 4.838265 | 5.100822 | 4.971128 | 4.949131 | 4.983660 | 4.982456 | 36 | 28 | 30 | 32 |
| 100505506 | gene-BANF1P3 | BANF1P3 | BANF1 pseudogene 3 | pseudogene | . | 43883 | . | . | . | 5.639420 | 4.872602 | 4.739542 | 6.173668 | 13.123161 | 11.263236 | 10.635395 | 14.115359 | 1.026192 | 119.121509 | 0.305816405 | 0.12197106 | 0.902921938 | 0.998865404 | 6.877877 | 6.914471 | 6.916897 | 6.863333 | 6.856907 | 6.939881 | 146 | 117 | 92 | 127 |
| 100505532 | NR_103854 | LINC01301 | long intergenic non-protein coding RNA 1301 | lncRNA | . | 50464 | . | . | . | 0.215682 | 0.185948 | 0.387095 | 0.183535 | 0.501901 | 0.429826 | 0.868629 | 0.419630 | 1.400762 | 26.880121 | 0.629067956 | 0.77290795 | 0.439576859 | NA | 4.482357 | 4.973054 | 4.721648 | 4.704928 | 4.793299 | 4.705315 | 28 | 22 | 37 | 19 |
| 100505534 | XR_109178,XR | GPR176-DT | GPR176-DT, transcript variant X2 | lncRNA | . | 54389 | . | . | . | 0.028344 | 0.024054 | 0.038223 | 0.040740 | 0.065958 | 0.055602 | 0.085772 | 0.093146 | 1.384537 | 4.278495 | 1.446462025 | 0.324518665 | 0.745545394 | NA | 1.844134 | 2.312263 | 2.086858 | 2.088018 | 2.092125 | 2.095468 | 4 | 4 | 4 | 5 |
| 100505538 | NR_038991 | RBM26-AS1 | RBM26 antisense RNA 1 | lncRNA | . | 39805 | . | . | . | 0.394834 | 0.467826 | 0.739918 | 0.508472 | 0.918795 | 1.081401 | 1.660354 | 1.162561 | 1.417909 | 41.324775 | 0.496183239 | 1.015279787 | 0.309972477 | 0.998865404 | 5.094525 | 5.599398 | 5.315791 | 5.339811 | 5.415355 | 5.349522 | 36 | 40 | 50 | 37 |
| 100505549 | NR_164148 | LOC10050554 | uncharacterized LOC100505549 | lncRNA | . | . | . | . | . | 0.743382 | 0.520819 | 0.470062 | 0.558307 | 1.729881 | 1.203895 | 1.054805 | 1.276502 | -1.248015 | 35.336550 | 0.532484642 | -0.60027138 | 0.548325386 | 0.998865404 | 5.293950 | 4.974591 | 5.187123 | 5.133023 | 5.119336 | 5.140033 | 53 | 35 | 25 | 32 |
| 100505573 | NM_00130126 | INAFM2 | InaF motif containing 2 | protein_coding | NP_001288197.1;XP_011519451.1 | 35165 | . | . | . | 6.212699 | 6.067427 | 4.969067 | 5.200829 | 14.457206 | 14.025130 | 11.150443 | 11.891077 | -1.230684 | 521.408512 | 0.164995152 | -1.81496323 | 0.069529561 | 0.744724233 | 9.168226 | 8.868807 | 9.100488 | 9.077020 | 8.952687 | 8.959827 | 674 | 610 | 404 | 441 |
| 100505591 | NM_00119554 | LRRC3C | leucine rich repeat containing 3C | protein_coding | NP_001182474.1;XP_011522460.1;XP_016879492.1 | 40034 | . | . | . | 0.101742 | 0.134358 | 0.096205 | 0.330742 | 0.236757 | 0.310575 | 0.215882 | 0.756202 | 1.560771 | 8.060289 | 1.098529015 | 0.584653421 | 0.558780779 | NA | 2.658167 | 3.293999 | 2.987306 | 2.997742 | 2.990620 | 3.026159 | 6 | 8 | 5 | 13 |
| 100505592 | NR_110428,NF | GAPLINC | gastric adenocarcinoma associated, positive CD44 regulator, long | lncRNA | . | 51308 | . | . | . | 1.975770 | 2.802063 | 1.802226 | 4.150516 | 4.597698 | 6.477094 | 4.044143 | 9.489661 | 1.244079 | 41.408799 | 0.53448242 | 0.589501991 | 0.555524577 | 0.998865404 | 5.208527 | 5.518578 | 5.314796 | 5.368017 | 5.311127 | 5.438338 | 35 | 47 | 25 | 59 |

| | | | | | | | | |
|---|---|---|---|---|---|---|---|---|
| 100505616 | NR_110096,NF | SNHG21 | small nucleolar RNA host gene 21 | lncRNA | . | 50284 | . | [illegible] |
| 100505636 | NR_102739,NF | MALINC1 | mitosis associated long intergenic non-coding RNA 1 | lncRNA | . | 49009 | . | [illegible] |
| 100505641 | NR_046251,NF | FGD5-AS1 | FGD5 antisense RNA 1 | lncRNA | . | 40410 | . | [illegible] |
| 100505648 | NR_040058 | RAD51-AS1 | RAD51 antisense RNA 1 | lncRNA | . | 48621 | . | [illegible] |
| 100505666 | NR_040772,NF | DCST1-AS1 | DCST1 antisense RNA 1 | lncRNA | . | 41147 | . | [illegible] |
| 100505678 | NR_040093 | STARD4-AS1 | STARD4 antisense RNA 1 | lncRNA | . | 44117 | . | [illegible] |
| 100505681 | NR_038452 | NAPA-AS1 | NAPA antisense RNA 1 | lncRNA | . | 44118 | . | [illegible] |
| 100505683 | NR_109884 | LINC01431 | long intergenic non-protein coding RNA 1431 | lncRNA | . | 50744 | . | [illegible] |
| 100505687 | NR_038301,NF | LINC00888 | long intergenic non-protein coding RNA 888 | lncRNA | . | 48575 | . | [illegible] |
| 100505692 | NR_103838 | FTO-IT1 | FTO intronic transcript 1 | lncRNA | . | 43758 | . | [illegible] |
| 100505696 | NR_046084 | SH3BP5-AS1 | SH3BP5 antisense RNA 1 | lncRNA | . | 44501 | . | [illegible] |
| 100505715 | NR_038334 | LOC100505715 | uncharacterized LOC100505715 | lncRNA | . | . | . | [illegible] |
| 100505728 | XR_922172 | LOC100505728 | uncharacterized LOC100505728 | misc_RNA | . | . | . | [illegible] |
| 100505738 | NR_039989 | MIR4458HG | MIR4458 host gene | lncRNA | . | 49008 | . | [illegible] |
| 100505761 | NR_038937,NF | RPARP-AS1 | RPARP antisense RNA 1 | lncRNA | . | 45238 | . | [illegible] |
| 100505771 | NR_132417,NF | MHENCR | melanoma highly expressed competing endogenous lncRNA for m | lncRNA | . | 53110 | . | [illegible] |
| 100505783 | NR_038337,NF | OSER1-DT | OSER1 divergent transcript | lncRNA | . | 48585 | . | [illegible] |
| 100505785 | NR_145471 | KIRREL1-IT1 | KIRREL1 intronic transcript 1 | lncRNA | . | 41412 | . | [illegible] |
| 100505806 | NR_045196 | SNHG18 | small nucleolar RNA host gene 18 | lncRNA | . | 49007 | . | [illegible] |
| 100505812 | NR_040599 | CARD8-AS1 | CARD8 antisense RNA 1 | lncRNA | . | 51408 | . | [illegible] |
| 100505826 | NR_038341 | STK4-AS1 | STK4 antisense RNA 1 (head to head) | lncRNA | . | 43811 | . | [illegible] |
| 100505839 | NR_038940 | SH3PXD2A-AS1 | SH3PXD2A antisense RNA 1 | lncRNA | . | 45242 | . | [illegible] |
| 100505854 | NR_038361,NF | APTR | Alu-mediated CDKN1A/p21 transcriptional regulator | lncRNA | . | 44173 | 616048 | [illegible] |
| 100505865 | NR_046242 | LINC00920 | long intergenic non-protein coding RNA 920 | lncRNA | . | 48611 | . | [illegible] |
| 100505876 | NM_00132237 | CEBPZOS | CEBPZ opposite strand | protein_coding | NP_001309302.1;NP_001309303.1;NP_001309304.1;NP_001309305.1;XF | 49288 | . | [illegible] |
| 100505881 | NR_038343,NF | MAGI2-AS3 | MAGI2 antisense RNA 3 | lncRNA | . | 40862 | . | [illegible] |
| 100505894 | NR_039993,NF | TMEM161B-AS | TMEM161B antisense RNA 1 | lncRNA | . | 43839 | . | [illegible] |
| 100505915 | NR_125434 | LOC100505915 | uncharacterized LOC100505915 | lncRNA | . | . | . | [illegible] |
| 100505938 | NR_125740 | LOC100505938 | uncharacterized LOC100505938 | lncRNA | . | . | . | [illegible] |
| 100505942 | NR_104656 | LOC100505942 | uncharacterized LOC100505942 | lncRNA | . | . | . | [illegible] |
| 100505975 | NR_038351 | ENO1-AS1 | ENO1 antisense RNA 1 | lncRNA | . | 40214 | . | [illegible] |
| 100505989 | NM_00135288 | SMIM31 | small integral membrane protein 31 | protein_coding | NP_001339814.1 | 49638 | . | [illegible] |
| 100505994 | NR_103548,NF | LUCAT1 | lung cancer associated transcript 1 | lncRNA | . | 48498 | 618190 | [illegible] |
| 100506007 | NR_110434,NF | RTCA-AS1 | RTCA antisense RNA 1 | lncRNA | . | 50573 | . | [illegible] |
| 100506022 | NR_104622,NF | LINC02606 | long intergenic non-protein coding RNA 2606 | lncRNA | . | 54047 | . | [illegible] |
| 100506023 | NR_037845 | LOC100506023 | uncharacterized LOC100506023 | lncRNA | . | . | . | [illegible] |
| 100506046 | NR_037605 | GAS5-AS1 | GAS5 antisense RNA 1 | lncRNA | . | 44119 | . | [illegible] |
| 100506054 | NR_038429,NF | RNASEH1-AS1 | RNASEH1 antisense RNA 1 | lncRNA | . | 49289 | . | [illegible] |
| 100506055 | gene-MXRA7P1 | MXRA7P1 | MXRA7 pseudogene 1 | pseudogene | . | 54643 | . | [illegible] |
| 100506060 | NR_033959 | SMG1P7 | SMG1 pseudogene 7 | transcribed_ps | . | 49864 | . | [illegible] |
| 100506076 | NR_103732,NF | LOC100506076 | uncharacterized LOC100506076 | lncRNA | . | . | . | [illegible] |
| 100506083 | NR_039997 | LOC100506083 | uncharacterized LOC100506083 | lncRNA | . | . | . | [illegible] |
| 100506084 | NM_00103908 | ARL17B | ADP ribosylation factor like GTPase 17B | protein_coding | NP_001034172.3;NP_001096624.1;NP_001317169.1;NP_001339698.1;NI | 32387 | . | [illegible] |
| 100506100 | NR_046240 | LOC100506100 | uncharacterized LOC100506100 | lncRNA | . | . | . | [illegible] |
| 100506115 | NM_00127865 | PELATON | plaque enriched lncRNA in atherosclerotic and inflammatory bowe | protein_coding | NP_001265584.1;XP_006723743.1 | 50328 | . | [illegible] |
| 100506123 | NR_130704 | LOC100506123 | uncharacterized LOC100506123 | lncRNA | . | . | . | [illegible] |
| 100506124 | NR_045375 | LOC100506124 | uncharacterized LOC100506124 | lncRNA | . | . | . | [illegible] |
| 100506127 | NM_00128245 | GVQW3 | GVQW motif containing 3 | protein_coding | NP_001269385.1;NP_001292154.1;NP_001334813.1;NP_001334814.1;XF | 51239 | . | [illegible] |
| 100506144 | NM_00119515 | TMEM35B | transmembrane protein 35B | protein_coding | NP_001182085.1 | 40021 | . | [illegible] |
| 100506161 | NR_110613,NF | TAF1A-AS1 | TAF1A antisense RNA 1 | lncRNA | . | 40573 | . | [illegible] |
| 100506164 | NM_016153 | HSFX1 | heat shock transcription factor family, X-linked 1 | protein_coding | NP_057237.1 | 29603 | . | [illegible] |
| 100506190 | NR_038955 | LINC00963 | long intergenic non-protein coding RNA 963 | lncRNA | . | 48716 | . | [illegible] |
| 100506205 | gene-TMEM18 | TMEM185AP1 | transmembrane protein 185A pseudogene 1 | pseudogene | . | 31717 | . | [illegible] |
| 100506207 | NR_038979,NF | LOC100506207 | uncharacterized LOC100506207 | lncRNA | . | . | . | [illegible] |
| 100506211 | NR_038262 | MIR210HG | MIR210 host gene | lncRNA | . | 39524 | . | [illegible] |
| 100506215 | NR_038879,NF | PAXBP1-AS1 | PAXBP1 antisense RNA 1 | lncRNA | . | 39603 | . | [illegible] |
| 100506233 | NR_038903 | RAB30-DT | RAB30 divergent transcript | lncRNA | . | 48672 | . | [illegible] |
| 100506235 | XR_245000 | LOC100506235 | uncharacterized LOC100506235 | lncRNA | . | . | . | [illegible] |
| 100506258 | NR_103855 | LOC100506258 | uncharacterized LOC100506258 | lncRNA | . | . | . | [illegible] |
| 100506282 | NR_149008,NF | LOC100506282 | uncharacterized LOC100506282 | lncRNA | . | . | . | [illegible] |
| 100506286 | NR_135594 | TBC1D8-AS1 | TBC1D8 antisense RNA 1 | lncRNA | . | 52782 | . | [illegible] |
| 100506299 | NR_110153,NF | ID2-AS1 | ID2 antisense RNA 1 | lncRNA | . | 51103 | . | [illegible] |
| 100506302 | NR_126344 | LOC100506302 | uncharacterized LOC100506302 | lncRNA | . | . | . | [illegible] |
| 100506305 | NR_038904 | LINC00958 | long intergenic non-protein coding RNA 958 | lncRNA | . | 48671 | 618335 | [illegible] |
| 100506311 | NR_038366,NF | HOTAIRM1 | HOXA transcript antisense RNA, myeloid-specific 1 | lncRNA | . | 37117 | . | [illegible] |
| 100506325 | NR_103856,NF | SP2-AS1 | SP2 antisense RNA 1 | lncRNA | . | 51341 | . | [illegible] |
| 100506330 | gene-WBP1LP2 | WBP1LP2 | WBP1L pseudogene 2 | pseudogene | . | 43949 | . | [illegible] |
| 100506343 | NR_038951,NF | INKA2-AS1 | INKA2 antisense RNA 1 | lncRNA | . | 49446 | . | [illegible] |
| 100506358 | NR_149012 | LOC100506358 | uncharacterized LOC100506358 | lncRNA | . | . | . | [illegible] |
| 100506365 | NR_110438,NF | OTUD6B-AS1 | OTUD6B antisense RNA 1 (head to head) | lncRNA | . | 50466 | . | [illegible] |
| 100506368 | NR_038905 | FZD4-DT | FZD4 divergent transcript | lncRNA | . | 54219 | . | [illegible] |
| 100506380 | NR_026865,NF | LINC01006 | long intergenic non-protein coding RNA 1006 | lncRNA | . | 48971 | 610242 | [illegible] |
| 100506392 | NR_103743,NF | SLC16A1-AS1 | SLC16A1 antisense RNA 1 | lncRNA | . | 49445 | . | [illegible] |
| 100506394 | NR_044993 | GAS6-DT | GAS6 divergent transcript | lncRNA | . | 43694 | . | [illegible] |
| 100506411 | XR_001750801 | LOC100506411 | uncharacterized LOC100506411, transcript variant X6 | lncRNA | . | . | . | [illegible] |
| 100506428 | NR_038892,NF | CBR3-AS1 | CBR3 antisense RNA 1 | lncRNA | . | 43664 | . | [illegible] |
| 100506433 | NR_039996 | LINC00648 | long intergenic non-protein coding RNA 648 | lncRNA | . | 44302 | . | [illegible] |
| 100506436 | NR_040012 | SPAG5-AS1 | SPAG5 antisense RNA 1 | lncRNA | . | 41140 | . | [illegible] |
| 100506444 | NR_125918 | LOC100506444 | uncharacterized LOC100506444 | lncRNA | . | . | . | [illegible] |
| 100506451 | NR_038227,NF | RASSF8-AS1 | RASSF8 antisense RNA 1 | lncRNA | . | 48637 | . | [illegible] |
| 100506457 | NR_110196,NF | MIR3681HG | MIR3681 host gene | lncRNA | . | 52001 | . | [illegible] |
| 100506465 | NR_110025,NF | LINC01234 | long intergenic non-protein coding RNA 1234 | lncRNA | . | 49757 | . | [illegible] |
| 100506469 | NR_038396 | TMEM147-AS1 | TMEM147 antisense RNA 1 | lncRNA | . | 51273 | . | [illegible] |
| 100506472 | NR_040535 | LOC100506472 | uncharacterized LOC100506472 | lncRNA | . | . | . | [illegible] |
| 100506473 | XR_427149 | LOC100506473 | uncharacterized LOC100506473 | lncRNA | . | . | . | [illegible] |
| 100506474 | NR_038434 | LOC100506474 | uncharacterized LOC100506474 | lncRNA | . | . | . | [illegible] |
| 100506476 | NR_109995 | LOC100506476 | uncharacterized LOC100506476 | lncRNA | . | . | . | [illegible] |
| 100506495 | NR_103553,NF | LIFR-AS1 | LIFR antisense RNA 1 | lncRNA | . | 43600 | . | [illegible] |
| 100506516 | NR_108081 | LINC01176 | long intergenic non-protein coding RNA 1176 | lncRNA | . | 49548 | . | [illegible] |
| 100506540 | NM_00135557 | SPTY2D1OS | SPTY2D1 opposite strand | protein_coding | NP_001342499.1 | 44122 | . | [illegible] |
| 100506571 | NR_158166 | LINC02809 | long intergenic non-protein coding RNA 2809 | lncRNA | . | 54341 | . | [illegible] |
| 100506581 | NM_00119512 | C16orf95 | chromosome 16 open reading frame 95 | protein_coding | NP_001182053.1;NP_001182054.1;NP_001243846.1 | 40033 | . | [illegible] |
| 100506599 | NR_038969 | PPP1R26-AS1 | PPP1R26 antisense RNA 1 | lncRNA | . | 48717 | . | [illegible] |
| 100506603 | NR_104183,NF | VASH1-AS1 | VASH1 antisense RNA 1 | lncRNA | . | 53215 | . | [illegible] |
| 100506606 | NR_103860 | LOC100506606 | uncharacterized LOC100506606 | lncRNA | . | . | . | [illegible] |
| 100506637 | NR_109996,NF | PRKAR2A-AS1 | PRKAR2A antisense RNA 1 | lncRNA | . | 40471 | . | [illegible] |
| 100506639 | NR_102752 | LOC100506639 | uncharacterized LOC100506639 | lncRNA | . | . | . | [illegible] |
| 100506649 | NR_038924 | PXN-AS1 | PXN antisense RNA 1 | lncRNA | . | 44123 | . | [illegible] |
| 100506658 | NM_00120525 | OCLN | occludin | protein_coding | NP_001192183.1;NP_001192184.1;NP_002529.1;XP_016864402.1;XP_01 | 8104 | 602876 | [illegible] |
| 100506660 | NR_038927 | DDX11-AS1 | DDX11 antisense RNA 1 | lncRNA | . | 44176 | . | [illegible] |
| 100506668 | NR_038854 | NRAV | negative regulator of antiviral response | lncRNA | . | 48588 | 616207 | [illegible] |
| 100506672 | gene-LOC1005 | LOC100506672 | cytochrome c oxidase subunit 7C pseudogene | pseudogene | . | . | . | [illegible] |
| 100506674 | NR_109862,NF | MRPS30-DT | MRPS30 divergent transcript | lncRNA | . | 53420 | . | [illegible] |
| 100506685 | gene-STMN1P1 | STMN1P1 | stathmin 1 pseudogene 1 | pseudogene | . | 44062 | . | [illegible] |
| 100506686 | NR_040051,NF | IQCH-AS1 | IQCH antisense RNA 1 | lncRNA | . | 44104 | . | [illegible] |
| 100506691 | NR_135044 | LOC100506691 | uncharacterized LOC100506691 | lncRNA | . | . | . | [illegible] |
| 100506696 | NR_046325,NF | PCAT6 | prostate cancer associated transcript 6 | lncRNA | . | 43714 | . | [illegible] |
| 100506697 | NR_038995 | LINC00327 | long intergenic non-protein coding RNA 327 | lncRNA | . | 42009 | . | [illegible] |
| 100506710 | NR_036592 | EBLN3P | endogenous Bornavirus like nucleoprotein 3, pseudogene | transcribed_ps | . | 50682 | . | [illegible] |
| 100506713 | NR_040089 | ALOX12-AS1 | ALOX12 antisense RNA 1 | lncRNA | . | 51342 | . | [illegible] |
| 100506714 | NR_038956,NF | NUP50-DT | NUP50 divergent transcript | lncRNA | . | 50502 | . | [illegible] |
| 100506730 | NR_037847 | LOC100506730 | uncharacterized LOC100506730 | lncRNA | . | . | . | [illegible] |
| 100506746 | NR_038841 | AFF1-AS1 | AFF1 antisense RNA 1 | lncRNA | . | 53447 | . | [illegible] |
| 100506749 | NR_131900 | AGAP1-IT1 | AGAP1 intronic transcript 1 | lncRNA | . | 41427 | . | [illegible] |
| 100506753 | NR_126338,NF | MAILR | macrophage interferon regulatory lncRNA | lncRNA | . | 51558 | . | [illegible] |
| 100506757 | NR_038998 | LINC00629 | long intergenic non-protein coding RNA 629 | lncRNA | . | 44262 | . | [illegible] |
| 100506759 | NR_002163 | OR7E37P | olfactory receptor family 7 subfamily E member 37 pseudogene | transcribed_ps | . | 8410 | . | [illegible] |
| 100506776 | NR_040085 | TRG-AS1 | T cell receptor gamma locus antisense RNA 1 | lncRNA | . | 48974 | . | [illegible] |
| 100506779 | NR_038410,NF | TSPOAP1-AS1 | TSPOAP1, SUPT4H1 and RNF43 antisense RNA 1 | lncRNA | . | 44148 | . | [illegible] |
| 100506783 | NR_038435 | HOXD-AS2 | HOXD cluster antisense RNA 2 | lncRNA | . | 43756 | . | [illegible] |
| 100506804 | NR_040081 | TMEM30A-DT | TMEM30A divergent transcript | lncRNA | . | 48985 | . | [illegible] |
| 100506826 | NR_038266,NF | MYLK-AS1 | MYLK antisense RNA 1 | lncRNA | . | 42440 | . | [illegible] |
| 100506827 | NR_131958,NF | LINC02475 | long intergenic non-protein coding RNA 2475 | lncRNA | . | 53418 | . | [illegible] |
| 100506834 | NR_049794 | GAS1RR | GAS1 adjacent regulatory RNA | lncRNA | . | 52261 | . | [illegible] |
| 100506844 | NR_038269 | GIHCG | GIHCG inhibitor of miR-200b/200a/429 expression | lncRNA | . | 52649 | . | [illegible] |
| 100506866 | NR_038271,NF | TTN-AS1 | TTN antisense RNA 1 | lncRNA | . | 44124 | . | [illegible] |
| 100506881 | NR_125364 | MKLN1-AS | MKLN1 antisense RNA | lncRNA | . | 40374 | . | [illegible] |
| 100506885 | NR_134649,NF | LNC-LBCS | lncRNA bladder and prostate cancer suppressor, hnRNPK interacti | lncRNA | . | 54418 | . | [illegible] |
| 100506901 | XR_001755918 | LL0XNC01-250I | uncharacterized LL0XNC01-250H12.3, transcript variant X1 | lncRNA | . | . | . | [illegible] |
| 100506922 | XR_001739698 | MAP3K2-DT | MAP3K2 divergent transcript, transcript variant X4 | lncRNA | . | 54088 | . | [illegible] |
| 100506930 | NR_038278,NF | LINC00665 | long intergenic non-protein coding RNA 665 | lncRNA | . | 44323 | . | [illegible] |
| 100506963 | NR_038280 | ELOA-AS1 | ELOA antisense RNA 1 | lncRNA | . | 50582 | . | [illegible] |
| 100506965 | NR_146168 | PWAR6 | Prader Willi/Angelman region RNA 6 | lncRNA | . | 49129 | . | [illegible] |
| 100506990 | NR_040091,NF | LOC100506990 | uncharacterized LOC100506990 | lncRNA | . | . | . | [illegible] |
| 100506994 | NR_038281,NF | PTPRG-AS1 | PTPRG antisense RNA 1 | lncRNA | . | 44638 | . | [illegible] |
| 100507002 | NR_110801 | LOC100507002 | uncharacterized LOC100507002 | lncRNA | . | . | . | [illegible] |
| 100507032 | NR_037893 | TMCC1-AS1 | TMCC1 antisense RNA 1 (head to head) | lncRNA | . | 49060 | . | [illegible] |
| 100507046 | gene-HSPE1P3 | HSPE1P3 | heat shock protein family E (Hsp10) member 1 pseudogene 3 | pseudogene | . | 49322 | . | [illegible] |
| 100507050 | NM_00119552 | TPBGL | trophoblast glycoprotein like | protein_coding | NP_001182457.1 | 44159 | . | [illegible] |
| 100507053 | NR_037884 | LOC100507053 | uncharacterized LOC100507053 | lncRNA | . | . | . | [illegible] |
| 100507062 | NR_038286 | PSMD6-AS2 | PSMD6 antisense RNA 2 | lncRNA | . | 44125 | . | [illegible] |
| 100507065 | NR_120431,NF | LOC100507065 | uncharacterized LOC100507065 | lncRNA | . | . | . | [illegible] |
| 100507118 | NR_051984 | PRC1-AS1 | PRC1 antisense RNA 1 | lncRNA | . | 48587 | . | [illegible] |
| 100507124 | gene-HMGN1P | HMGN1P25 | high mobility group nucleosome binding domain 1 pseudogene 25 | pseudogene | . | 39369 | . | [illegible] |
| 100507135 | NR_045180,NF | SPART-AS1 | SPART antisense RNA 1 | lncRNA | . | 39933 | . | [illegible] |
| 100507171 | NR_045634,NF | BOLA3-AS1 | BOLA3 divergent transcript | lncRNA | . | 42922 | . | [illegible] |
| 100507178 | NR_037868 | SLFNL1-AS1 | SLFNL1 antisense RNA 1 | lncRNA | . | 44126 | . | [illegible] |
| 100507194 | NR_038295,NF | LINC02525 | long intergenic non-protein coding RNA 2525 | lncRNA | . | 53545 | . | [illegible] |
| 100507217 | NR_037600,NF | CHASERR | long intergenic non-protein coding RNA 1578 | lncRNA | . | 48626 | . | [illegible] |
| 100507246 | NR_038108,NF | SNHG16 | small nucleolar RNA host gene 16 | lncRNA | . | 44352 | . | [illegible] |
| 100507254 | NR_038981,NF | LINC01013 | long intergenic non-protein coding RNA 1013 | lncRNA | . | 48987 | . | [illegible] |
| 100507266 | NR_037888 | STX18-AS1 | STX18 antisense RNA 1 (head to head) | lncRNA | . | 48877 | . | [illegible] |
| 100507290 | NM_00119560 | ZNF865 | zinc finger protein 865 | protein_coding | NP_001182534.1 | 38705 | . | [illegible] |
| 100507291 | NR_121608,NF | LOC100507291 | uncharacterized LOC100507291 | lncRNA | . | . | . | [illegible] |
| 100507297 | NR_047573,NF | TMEM44-AS1 | TMEM44 antisense RNA 1 | lncRNA | . | 44272 | . | [illegible] |
| 100507299 | NR_039990 | SMC5-AS1 | SMC5 antisense RNA 1 (head to head) | lncRNA | . | 48718 | . | [illegible] |
| 100507303 | NR_132114 | SNHG19 | small nucleolar RNA host gene 19 | lncRNA | . | 49574 | . | [illegible] |
| 100507316 | NR_120682,NF | MINCR | MYC-induced long non-coding RNA | lncRNA | . | 51653 | . | [illegible] |
| 100507321 | NR_040023 | ERVK13-1 | endogenous retrovirus group K13 member 1 | lncRNA | . | 27548 | . | [illegible] |
| 100507331 | NR_038357 | ZSWIM8-AS1 | ZSWIM8 antisense RNA 1 | lncRNA | . | 45103 | . | [illegible] |
| 100507347 | NR_108060,NF | VIM-AS1 | VIM antisense RNA 1 | lncRNA | . | 44879 | . | [illegible] |
| 100507373 | NR_045214 | LOC100507373 | uncharacterized LOC100507373 | lncRNA | . | . | . | [illegible] |

| | | | | | | | | | | | |
|---|---|---|---|---|---|---|---|---|---|---|---|
| 100507381 | NR_120503 | KCNQ5-IT1 | KCNQ5 intronic transcript 1 | lncRNA | . | 41354 | . | . | . | . | [illegible] |
| 100507391 | NR_037891 | LINC01968 | long intergenic non-protein coding RNA 1968 | lncRNA | . | 52794 | . | . | . | . | [illegible] |
| 100507398 | NR_103812 | INTS6-AS1 | INTS6 antisense RNA 1 | lncRNA | . | 42691 | . | . | . | . | [illegible] |
| 100507419 | NR_123723 | MMP25-AS1 | MMP25 antisense RNA 1 | lncRNA | . | 51372 | . | . | . | . | [illegible] |
| 100507436 | NM_00117751 | MICA | MHC class I polypeptide-related sequence A | protein_coding | NP_001170990.1;NP_001276081.1;NP_001276082.1;NP_001276083.1 | 7090 | 600169 | . | . | MICA | [illegible] |
| 100507437 | NR_125384,NF | LOC100507437 | uncharacterized LOC100507437 | lncRNA | . | . | . | . | . | . | [illegible] |
| 100507458 | NR_110900 | ZNF213-AS1 | ZNF213 antisense RNA 1 (head to head) | lncRNA | . | 50505 | . | . | . | . | [illegible] |
| 100507459 | NR_109990 | NRSN2-AS1 | NRSN2 antisense RNA 1 | lncRNA | . | 51222 | . | . | . | . | [illegible] |
| 100507462 | NR_161203,NF | CCDC28A-AS1 | CCDC28A antisense RNA 1 | lncRNA | . | 51715 | . | . | . | . | [illegible] |
| 100507463 | NR_037173,NF | PSMB8-AS1 | PSMB8 antisense RNA 1 (head to head) | lncRNA | . | 39758 | . | . | . | . | [illegible] |
| 100507468 | NR_108105,NF | CT66 | cancer/testis associated transcript 66 | lncRNA | . | 54198 | . | . | . | . | [illegible] |
| 100507472 | NR_045387 | LOC100507472 | uncharacterized LOC100507472 | lncRNA | . | . | . | . | . | . | [illegible] |
| 100507475 | NR_046452 | IDH1-AS1 | IDH1 antisense RNA 1 | lncRNA | . | 40292 | . | . | . | . | [illegible] |
| 100507477 | NR_121622,NF | LOC100507477 | uncharacterized LOC100507477 | lncRNA | . | . | . | . | . | . | [illegible] |
| 100507487 | NR_125882 | LINC02615 | long intergenic non-protein coding RNA 2615 | lncRNA | . | 53402 | . | . | . | . | [illegible] |
| 100507495 | NR_040047,NF | SDCBP2-AS1 | SDCBP2 antisense RNA 1 | lncRNA | . | 44314 | . | . | . | . | [illegible] |
| 100507501 | NR_039999 | LINC01569 | long intergenic non-protein coding RNA 1569 | lncRNA | . | 51380 | . | . | . | . | [illegible] |
| 100507506 | NR_103764 | ECI2-DT | ECI2 divergent transcript | lncRNA | . | 54425 | . | . | . | . | [illegible] |
| 100507507 | NR_161353 | LOC100507507 | uncharacterized LOC100507507 | lncRNA | . | . | . | . | . | . | [illegible] |
| 100507524 | NR_037901 | ARHGEF26-AS1 | ARHGEF26 antisense RNA 1 | lncRNA | . | 41048 | . | . | . | . | [illegible] |
| 100507526 | gene-SMG1P4 | SMG1P4 | SMG1 pseudogene 4 | pseudogene | . | 49861 | . | . | . | . | [illegible] |
| 100507540 | NR_110952,NF | LINC01504 | long intergenic non-protein coding RNA 1504 | lncRNA | . | 51185 | . | . | . | . | [illegible] |
| 100507547 | NR_037169,NF | LOC100507547 | uncharacterized LOC100507547 | lncRNA | . | . | . | . | . | . | [illegible] |
| 100507551 | NR_147835 | LOC100507551 | uncharacterized LOC100507551 | lncRNA | . | . | . | . | . | . | [illegible] |
| 100507554 | XR_108428,XR | RPL37A-DT | RPL37A divergent transcript, transcript variant X3 | lncRNA | . | 40510 | . | . | . | . | [illegible] |
| 100507561 | gene-LOC1005 | LOC100507561 | . | pseudogene | . | . | . | . | . | . | [illegible] |
| 100507564 | NR_038953 | LOC100507564 | uncharacterized LOC100507564 | lncRNA | . | . | . | . | . | . | [illegible] |
| 100507567 | NR_038237 | RAB11B-AS1 | RAB11B antisense RNA 1 | lncRNA | . | 44178 | . | . | . | . | [illegible] |
| 100507599 | NR_144529 | LOC100507599 | uncharacterized LOC100507599 | lncRNA | . | . | . | . | . | . | [illegible] |
| 100507602 | NR_102759,NF | TRIM52-AS1 | TRIM52 antisense RNA 1 (head to head) | lncRNA | . | 49006 | . | . | . | . | [illegible] |
| 100507607 | NM_00128725 | NPIPB9 | nuclear pore complex interacting protein family member B9 | protein_coding | NP_001274179.1;NP_001274180.1;XP_003118767.2;XP_011543992.1;XP | 41987 | . | . | . | . | [illegible] |
| 100507617 | gene-PPP1R14 | PPP1R14BP3 | protein phosphatase 1 regulatory inhibitor subunit 14B pseudoger | pseudogene | . | 16330 | . | . | . | . | [illegible] |
| 100507634 | NR_038320 | LOC100507634 | uncharacterized LOC100507634 | lncRNA | . | . | . | . | . | . | [illegible] |
| 100507642 | NR_108064,NF | LOC100507642 | uncharacterized LOC100507642 | lncRNA | . | . | . | . | . | . | [illegible] |
| 100507661 | NR_109989 | LINC02082 | long intergenic non-protein coding RNA 2082 | lncRNA | . | 52931 | . | . | . | . | [illegible] |
| 100507670 | NR_135595 | ZNF687-AS1 | ZNF687 antisense RNA 1 | lncRNA | . | 54048 | . | . | . | . | [illegible] |
| 100507747 | NM_00136545 | C13orf46 | chromosome 13 open reading frame 46 | protein_coding | NP_001352384.1;XP_006720058.3;XP_011533160.2;XP_011533161.2;XP | 53786 | . | . | . | . | [illegible] |
| 100509894 | NR_046226,NF | CPB2-AS1 | CPB2 antisense RNA 1 | lncRNA | . | 39898 | . | . | . | . | [illegible] |
| 100510707 | NM_00135557 | TBC1D3J | TBC1 domain family member 3J | protein_coding | NP_001342500.1;XP_024306292.1 | 51247 | . | . | . | . | [illegible] |
| 100526694 | NM_00119881 | MSANTD3-TME | MSANTD3-TMEFF1 readthrough | protein_coding | NP_001185741.1 | 38838 | . | . | . | . | [illegible] |
| 100526737 | NM_00119884 | RBM14-RBM4 | RBM14-RBM4 readthrough | protein_coding | NP_001185774.1;NP_001185775.1 | 38840 | . | . | . | . | [illegible] |
| 100526739 | NM_00124376 | CENPS-CORT | CENPS-CORT readthrough | protein_coding | NP_001230697.1;NP_001257446.1;NP_940946.1;NP_950171.2 | 38843 | . | . | . | . | [illegible] |
| 100526740 | NM_00119887 | ATP5MF-PTCD1 | ATP5MF-PTCD1 readthrough | protein_coding | NP_001185808.1 | 38844 | . | . | . | . | [illegible] |
| 100526767 | NM_00119895 | RNF103-CHMP | RNF103-CHMP3 readthrough | protein_coding | NP_001185883.1 | 38847 | . | . | . | . | [illegible] |
| 100526820 | NR_037593 | CAHM | colon adenocarcinoma hypermethylated | lncRNA | . | 42860 | 615930 | . | . | . | [illegible] |
| 100526830 | NR_037608 | SLX1A-SULT1A3 | SLX1A-SULT1A3 readthrough (NMD candidate) | lncRNA | . | 44437 | . | . | . | . | [illegible] |
| 100526831 | NR_037609 | SLX1B-SULT1A4 | SLX1B-SULT1A4 readthrough (NMD candidate) | lncRNA | . | 48353 | . | . | . | . | [illegible] |
| 100526832 | NM_00119929 | PHOSPHO2-KLI | PHOSPHO2-KLHL23 readthrough | protein_coding | NP_001186219.1 | . | . | . | . | . | [illegible] |
| 100526833 | NR_037611,NF | SEPT5-GP1BB | SEPT5-GP1BB readthrough | lncRNA | . | . | . | . | . | . | [illegible] |
| 100526835 | NM_00111280 | FPGT-TNNI3K | FPGT-TNNI3K readthrough | protein_coding | NP_001106279.2;NP_001186256.2 | 42952 | . | . | . | . | [illegible] |
| 100526836 | NR_037616 | BLOC1S5-TXND | BLOC1S5-TXNDC5 readthrough (NMD candidate) | lncRNA | . | 42001 | . | . | . | . | [illegible] |
| 100526837 | NR_037618 | EEF1E1-BLOC1S | EEF1E1-BLOC1S5 readthrough (NMD candidate) | lncRNA | . | 49187 | . | . | . | . | [illegible] |
| 100526842 | NM_00119935 | RPL17-C18orf3 | RPL17-C18orf32 readthrough | protein_coding | NP_001186284.1;NP_001186285.1 | 44661 | . | . | . | . | [illegible] |
| 100527943 | NM_00119953 | TGIF2-RAB5IF | TGIF2-RAB5IF readthrough | protein_coding | NP_001186464.1 | 44664 | . | . | . | . | [illegible] |
| 100527963 | NM_00119966 | PMF1-BGLAP | PMF1-BGLAP readthrough | protein_coding | NP_001186590.1;NP_001186591.1;NP_001186592.1;NP_001186593.1 | 42953 | . | . | . | . | [illegible] |
| 100527964 | NR_037642 | LHX4-AS1 | LHX4 antisense RNA 1 | lncRNA | . | 40982 | . | . | . | . | [illegible] |
| 100527978 | NM_00119969 | TLCD4-RWDD3 | TLCD4-RWDD3 readthrough | protein_coding | NP_001186620.1 | 49388 | . | . | . | . | [illegible] |
| 100528020 | NM_00125840 | FAM187A | family with sequence similarity 187 member A | protein_coding | NP_001245329.1 | 35153 | . | . | . | . | [illegible] |
| 100528021 | NM_00119976 | ST20-MTHFS | ST20-MTHFS readthrough | protein_coding | NP_001186689.1 | 44655 | . | . | . | . | [illegible] |
| 100528022 | NR_037658 | BLOC1S1-RDH5 | BLOC1S1-RDH5 readthrough | lncRNA | . | . | . | . | . | . | [illegible] |
| 100528031 | NR_037661 | FKBP1A-SDCBP | FKBP1A-SDCBP2 readthrough (NMD candidate) | lncRNA | . | 41997 | . | . | . | . | [illegible] |
| 100528062 | NM_00119981 | ARMCX5-GPRA | ARMCX5-GPRASP2 readthrough | protein_coding | NP_001186747.1;NP_001337197.1;NP_001337198.1;NP_001337199.1 | 42000 | . | . | . | . | [illegible] |
| 100528064 | NM_00119982 | NEDD8-MDP1 | NEDD8-MDP1 readthrough | protein_coding | NP_001186752.1 | 39551 | . | . | . | . | [illegible] |
| 100529063 | NM_00119986 | BCL2L2-PABPN | BCL2L2-PABPN1 readthrough | protein_coding | NP_001186793.2 | 42959 | . | . | . | . | [illegible] |
| 100529067 | NR_037673 | SERF2-C15ORF | SERF2-C15orf63 readthrough | lncRNA | . | . | . | . | . | . | [illegible] |
| 100529097 | NM_00119997 | RPL36A-HNRNI | RPL36A-HNRNPH2 readthrough | protein_coding | NP_001186902.2;NP_001186903.2 | 48349 | . | . | . | . | [illegible] |
| 100529145 | NR_037709 | TEN1-CDK3 | TEN1-CDK3 readthrough (NMD candidate) | lncRNA | . | 44420 | . | . | . | . | [illegible] |
| 100529209 | NR_037717 | RNASEK-C17or | RNASEK-C17orf49 readthrough | lncRNA | . | 44419 | . | . | . | . | [illegible] |
| 100529211 | NR_037719 | TMEM256-PLS | TMEM256-PLSCR3 readthrough (NMD candidate) | lncRNA | . | 49186 | . | . | . | . | [illegible] |
| 100529215 | NM_00117265 | ZNF559-ZNF17 | ZNF559-ZNF177 readthrough | protein_coding | NP_001166121.1;NP_001189354.1 | 42964 | . | . | . | . | [illegible] |
| 100529239 | NM_00120247 | RPS10-NUDT3 | RPS10-NUDT3 readthrough | protein_coding | NP_001189399.1 | 49181 | . | . | . | . | [illegible] |
| 100529241 | NM_00120248 | HSPE1-MOB4 | HSPE1-MOB4 readthrough | protein_coding | NP_001189414.1 | 49184 | . | . | . | . | [illegible] |
| 100529251 | NM_00120250 | CKLF-CMTM1 | CKLF-CMTM1 readthrough | protein_coding | NP_001189438.1;NP_001191027.1;NP_001191028.1 | 39977 | . | . | . | . | [illegible] |
| 100529261 | NM_00120255 | CHURC1-FNTB | CHURC1-FNTB readthrough | protein_coding | NP_001189487.1;NP_001189488.1 | 42960 | . | . | . | . | [illegible] |
| 100529262 | NR_037775 | MIA-RAB4B | MIA-RAB4B readthrough (NMD candidate) | lncRNA | . | 48352 | . | . | . | . | [illegible] |
| 100529264 | NR_037791 | RAB4B-EGLN2 | RAB4B-EGLN2 readthrough (NMD candidate) | lncRNA | . | 44465 | . | . | . | . | [illegible] |
| 100529855 | NR_037802 | ZNF625-ZNF20 | ZNF625-ZNF20 readthrough (NMD candidate) | lncRNA | . | 48368 | . | . | . | . | [illegible] |
| 100532724 | NR_037804 | NPHP3-ACAD1 | NPHP3-ACAD11 readthrough (NMD candidate) | lncRNA | . | 48351 | . | . | . | . | [illegible] |
| 100532731 | NM_00120406 | COMMD3-BMI | COMMD3-BMI1 readthrough | protein_coding | NP_001190991.1 | 48326 | . | . | . | . | [illegible] |
| 100532732 | NR_037846 | MSH5-SAPCD1 | MSH5-SAPCD1 readthrough (NMD candidate) | lncRNA | . | 41994 | . | . | . | . | [illegible] |
| 100532735 | NR_037849 | INO80B-WBP1 | INO80B-WBP1 readthrough (NMD candidate) | lncRNA | . | 49199 | . | . | . | . | [illegible] |
| 100532736 | NM_00120408 | MICOS10-NBL1 | MICOS10-NBL1 readthrough | protein_coding | NP_001191017.1;NP_001191018.1 | 48338 | . | . | . | . | [illegible] |
| 100532737 | NR_037853 | ATP6V1G2-DD | ATP6V1G2-DDX39B readthrough (NMD candidate) | lncRNA | . | 41999 | . | . | . | . | [illegible] |
| 100532742 | NR_037859 | PIR-FIGF | PIR-FIGF readthrough | lncRNA | . | . | . | . | . | . | [illegible] |
| 100532746 | NR_037861 | PPT2-EGFL8 | PPT2-EGFL8 readthrough (NMD candidate) | lncRNA | . | 48343 | . | . | . | . | [illegible] |
| 100533106 | NM_00120418 | ZHX1-C8orf76 | ZHX1-C8orf76 readthrough | protein_coding | NP_001191109.1 | 42975 | . | . | . | . | [illegible] |
| 100533107 | NR_037882 | RTEL1-TNFRSF | RTEL1-TNFRSF6B readthrough (NMD candidate) | lncRNA | . | 44095 | . | . | . | . | [illegible] |
| 100533111 | NR_037894 | ZNF670-ZNF69 | ZNF670-ZNF695 readthrough (NMD candidate) | lncRNA | . | 49200 | . | . | . | . | [illegible] |
| 100533179 | NR_037904 | UBE2F-SCLY | UBE2F-SCLY readthrough (NMD candidate) | lncRNA | . | 48339 | . | . | . | . | [illegible] |
| 100533182 | NR_039978 | SEC24B-AS1 | SEC24B antisense RNA 1 | lncRNA | . | 44003 | . | . | . | . | [illegible] |
| 100533195 | NR_037915 | FAM24B-CUZD | FAM24B-CUZD1 readthrough | lncRNA | . | . | . | . | . | . | [illegible] |
| 100533464 | NR_037918 | PRH1-PRR4 | PRH1-PRR4 readthrough | lncRNA | . | . | . | . | . | . | [illegible] |
| 100533496 | NM_00120447 | TVP23C-CDRT4 | TVP23C-CDRT4 readthrough | protein_coding | NP_001191407.1 | 42961 | . | . | . | . | [illegible] |
| 100533619 | gene-TMPOP2 | TMPOP2 | thymopoietin pseudogene 2 | pseudogene | . | 50860 | . | . | . | . | [illegible] |
| 100533673 | gene-OSBPL9P | OSBPL9P6 | oxysterol binding protein like 9 pseudogene 6 | pseudogene | . | 48736 | . | . | . | . | [illegible] |
| 100533789 | NR_126038 | LRRC37A8P | leucine rich repeat containing 37 member A8, pseudogene | transcribed_ps | . | 43793 | . | . | . | . | [illegible] |
| 100533800 | gene-BMPR1A | BMPR1AP2 | bone morphogenetic protein receptor type 1A pseudogene 2 | pseudogene | . | 21743 | . | . | . | . | [illegible] |
| 100533814 | gene-TCAF2C | TCAF2C | TRPM8 channel associated factor 2C | pseudogene | . | 53641 | . | . | . | . | [illegible] |
| 100533955 | NR_037926 | SENP3-EIF4A1 | SENP3-EIF4A1 readthrough (NMD candidate) | lncRNA | . | 49182 | . | . | . | . | [illegible] |
| 100533975 | NR_037929,NF | SLMO2-ATP5E | SLMO2-ATP5E readthrough | lncRNA | . | . | . | . | . | . | [illegible] |
| 100533997 | NM_00120481 | MAGEA10-MA | MAGEA10-MAGEA5 readthrough | protein_coding | NP_001191740.1 | . | . | . | . | . | [illegible] |
| 100534012 | NM_00120484 | TNFAIP8L2-SCN | TNFAIP8L2-SCNM1 readthrough | protein_coding | NP_001191777.1 | . | . | . | . | . | [illegible] |
| 100534593 | NR_037945 | STX16-NPEPL1 | STX16-NPEPL1 readthrough (NMD candidate) | lncRNA | . | 41993 | . | . | . | . | [illegible] |
| 100534595 | NR_037946 | HNRNPUL2-BS | HNRNPUL2-BSCL2 readthrough (NMD candidate) | lncRNA | . | 49189 | . | . | . | . | [illegible] |
| 100534599 | NM_00120489 | ISY1-RAB43 | ISY1-RAB43 readthrough | protein_coding | NP_001191819.1 | 42969 | . | . | . | . | [illegible] |
| 100534611 | NR_037950 | TM4SF19-TCTE | TM4SF19-TCTEX1D2 readthrough (NMD candidate) | lncRNA | . | 49190 | . | . | . | . | [illegible] |
| 100616112 | NR_039956,rn | MIR4793 | microRNA 4793 | miRNA | . | 41538 | . | . | . | . | [illegible] |
| 100616113 | NR_039828,rn | MIR4680 | microRNA 4680 | miRNA | . | 41541 | . | . | . | . | [illegible] |
| 100616115 | NR_039679,rn | MIR4469 | microRNA 4469 | miRNA | . | 41556 | . | . | . | . | [illegible] |
| 100616117 | NR_039797,rn | MIR4653 | microRNA 4653 | miRNA | . | 41562 | . | . | . | . | [illegible] |
| 100616122 | NR_039749,rn | MIR4523 | microRNA 4523 | miRNA | . | 41583 | . | . | . | . | [illegible] |
| 100616124 | NR_039790,rn | MIR4647 | microRNA 4647 | miRNA | . | 41594 | . | . | . | . | [illegible] |
| 100616139 | NR_039895,rn | MIR4741 | microRNA 4741 | miRNA | . | 41661 | . | . | . | . | [illegible] |
| 100616142 | NR_039663,rn | MIR4458 | microRNA 4458 | miRNA | . | 41674 | . | . | . | . | [illegible] |
| 100616143 | NR_039920,rn | MIR4763 | microRNA 4763 | miRNA | . | 41677 | . | . | . | . | [illegible] |
| 100616146 | NR_039759,rn | MIR4534 | microRNA 4534 | miRNA | . | 41692 | . | . | . | . | [illegible] |
| 100616167 | NR_039873,rn | MIR4722 | hsa-miR-4722-3p | . | . | 41782 | . | . | . | . | [illegible] |
| 100616193 | NR_039658,rn | MIR4453 | microRNA 4453 | miRNA | . | 41892 | . | . | . | . | [illegible] |
| 100616194 | NR_039689,rn | MIR4477B | microRNA 4477b | miRNA | . | 41898 | . | . | . | . | [illegible] |
| 100616210 | NR_039890,rn | MIR4737 | microRNA 4737 | miRNA | . | 41669 | . | . | . | . | [illegible] |
| 100616211 | NR_039858,rn | MIR4709 | microRNA 4709 | miRNA | . | 41690 | . | . | . | . | [illegible] |
| 100616248 | NR_039877,rn | MIR4724 | microRNA 4724 | miRNA | . | 41590 | . | . | . | . | [illegible] |
| 100616256 | NR_039872,rn | MIR4721 | microRNA 4721 | miRNA | . | 41609 | . | . | . | . | [illegible] |
| 100616262 | NR_039850,rn | MIR4701 | microRNA 4701 | miRNA | . | 41627 | . | . | . | . | [illegible] |
| 100616269 | NR_039782,rn | MIR4639 | hsa-miR-4639-5p | . | . | 41645 | . | . | . | . | [illegible] |
| 100616270 | NR_039795,rn | MIR4651 | microRNA 4651 | miRNA | . | 41647 | . | . | . | . | [illegible] |
| 100616284 | NR_039709,rn | MIR4489 | microRNA 4489 | miRNA | . | 41685 | . | . | . | . | [illegible] |
| 100616292 | NR_039839,rn | MIR4690 | hsa-miR-4690-5p | . | . | 41707 | . | . | . | . | [illegible] |
| 100616314 | NR_039905,rn | MIR4750 | microRNA 4750 | miRNA | . | 41765 | . | . | . | . | [illegible] |
| 100616318 | NR_039810,rn | MIR4664 | microRNA 4664 | miRNA | . | 41777 | . | . | . | . | [illegible] |
| 100616329 | NR_039849,rn | MIR4700 | microRNA 4700 | miRNA | . | 41806 | . | . | . | . | [illegible] |
| 100616343 | NR_039824,rn | MIR4677 | microRNA 4677 | miRNA | . | 41844 | . | . | . | . | [illegible] |
| 100616345 | NR_039624,rn | MIR4426 | microRNA 4426 | miRNA | . | 41848 | . | . | . | . | [illegible] |
| 100616361 | NR_039934,rn | MIR4775 | microRNA 4775 | miRNA | . | 41885 | . | . | . | . | [illegible] |
| 100616364 | NR_039946,rn | MIR4785 | microRNA 4785 | miRNA | . | 41889 | . | . | . | . | [illegible] |
| 100616367 | NR_039677,rn | MIR4467 | microRNA 4467 | miRNA | . | 41896 | . | . | . | . | [illegible] |
| 100616378 | NR_039945,rn | MIR4784 | microRNA 4784 | miRNA | . | 41580 | . | . | . | . | [illegible] |
| 100616387 | NR_039891,rn | MIR3064 | microRNA 3064 | miRNA | . | 41652 | . | . | . | . | [illegible] |
| 100616393 | NR_039801,rn | MIR4657 | microRNA 4657 | miRNA | . | 41706 | . | . | . | . | [illegible] |
| 100616394 | NR_039646,rn | MIR4444-1 | microRNA 4444-1 | miRNA | . | 41710 | . | . | . | . | [illegible] |
| 100616396 | NR_039862,rn | MIR4712 | microRNA 4712 | miRNA | . | 41732 | . | . | . | . | [illegible] |
| 100616400 | NR_039851,rn | MIR3198-2 | microRNA 3198-2 | miRNA | . | 41766 | . | . | . | . | [illegible] |
| 100616403 | NR_039840,rn | MIR4691 | microRNA 4691 | miRNA | . | 41796 | . | . | . | . | [illegible] |
| 100616408 | NR_039969,rn | MIR5047 | microRNA 5047 | miRNA | . | 41834 | . | . | . | . | [illegible] |
| 100616414 | NR_039918,rn | MIR4761 | microRNA 4761 | miRNA | . | 41591 | . | . | . | . | [illegible] |
| 100616417 | NR_039949,rn | MIR4786 | microRNA 4786 | miRNA | . | 41751 | . | . | . | . | [illegible] |
| 100616424 | NR_039856,rn | MIR4707 | microRNA 4707 | miRNA | . | 41531 | . | . | . | . | [illegible] |
| 100616439 | NR_039802,rn | MIR4658 | microRNA 4658 | miRNA | . | 41673 | . | . | . | . | [illegible] |
| 100616453 | NR_039835,rn | MIR4687 | microRNA 4687 | miRNA | . | 41712 | . | . | . | . | [illegible] |
| 100616477 | NR_039644,rn | MIR4442 | microRNA 4442 | miRNA | . | 41718 | . | . | . | . | [illegible] |
| 100616480 | NR_039697,rn | MIR4479 | microRNA 4479 | miRNA | . | 41767 | . | . | . | . | [illegible] |
| 100616483 | NR_039906,rn | MIR4751 | microRNA 4751 | miRNA | . | 41819 | . | . | . | . | [illegible] |
| 100616487 | NR_039742,rn | MIR4517 | microRNA 4517 | miRNA | . | 41878 | . | . | . | . | [illegible] |
| 100616493 | NR_039643,rn | MIR4441 | microRNA 4441 | miRNA | . | 41728 | . | . | . | . | [illegible] |
| 100616500 | NR_039831,rn | MIR4683 | microRNA 4683 | miRNA | . | 41883 | . | . | . | . | [illegible] |
| 100616668 | NR_038258,NF | TPTE2P5 | transmembrane phosphoinositide 3-phosphatase and tensin hom | transcribed_ps | . | 42356 | . | . | . | . | [illegible] |
| 100630918 | NR_038942 | PARTICL | promoter of MAT2A antisense radiation-induced circulating long n | lncRNA | . | 50886 | 616350 | . | . | . | [illegible] |
| 100630923 | NR_038967 | LOC100630923 | LOC100289561-PRKRIP1 readthrough | lncRNA | . | . | . | . | . | . | [illegible] |
| 100642175 | NR_131221 | SPRY4-IT1 | SPRY4 intronic transcript 1 | lncRNA | . | 42394 | 617617 | . | . | . | [illegible] |
| 100652736 | NR_103767 | LINC01311 | long intergenic non-protein coding RNA 1311 | lncRNA | . | 50503 | . | . | . | . | [illegible] |
| 100652739 | NR_045126,NF | RAET1E-AS1 | RAET1E antisense RNA 1 | lncRNA | . | 48994 | . | . | . | . | [illegible] |
| 100652748 | NM_00129011 | TIMM23B | translocase of inner mitochondrial membrane 23 homolog B | protein_coding | NP_001277046.1;NP_001277047.1;NP_001352800.1 | 23581 | . | . | . | . | [illegible] |

| | | | | | | | | | | |
|---|---|---|---|---|---|---|---|---|---|---|
| 100652758 | NR_132345,NF | LOC100652758 | uncharacterized LOC100652758 | lncRNA | . | . | . | . | . | [illegible] |
| 100652768 | NR_045215 | LOC100652768 | uncharacterized LOC100652768 | lncRNA | . | . | . | . | . | [illegible] |
| 100652772 | NR_073113 | NNT-AS1 | NNT antisense RNA 1 | lncRNA | . | 49005 | . | . | . | [illegible] |
| 100652781 | NR_045011 | SNX29P1 | sorting nexin 29 pseudogene 1 | transcribed_ps | . | 31913 | . | . | . | [illegible] |
| 100652853 | NR_046368 | LINC01023 | long intergenic non-protein coding RNA 1023 | lncRNA | . | 49004 | . | . | . | [illegible] |
| 100652883 | NR_045985 | ALOX15P1 | arachidonate 15-lipoxygenase pseudogene 1 | transcribed_ps | . | 13742 | . | . | . | [illegible] |
| 100653233 | NR_120505 | LOC100653233 | uncharacterized LOC100653233 | lncRNA | . | . | . | . | . | [illegible] |
| 100653395 | gene-LOC1006 | LOC100653395 | methionine adenosyltransferase 2A pseudogene | pseudogene | . | . | . | . | . | [illegible] |
| 100653515 | NM_00124354 | CEP295NL | CEP295 N-terminal like | protein_coding | NP_001230469.1;NP_001230470.1 | 44659 | . | . | . | [illegible] |
| 100689074 | NR_138047 | POU6F2-AS2 | POU6F2 antisense RNA 2 | lncRNA | . | 21887 | . | . | . | [illegible] |
| 100750246 | NR_047116 | HIF1A-AS1 | HIF1A antisense RNA 1 | lncRNA | . | 43014 | 614528 | . | . | [illegible] |
| 100750325 | NR_045407,NF | RCAN3AS | RCAN3 antisense RNA | lncRNA | . | 39009 | . | . | . | [illegible] |
| 100775104 | NR_046220 | KLHL7-DT | KLHL7 divergent transcript | lncRNA | . | 43431 | . | . | . | [illegible] |
| 100847004 | NR_049820,rna | MIR5188 | microRNA 5188 | miRNA | . | 43502 | . | . | . | [illegible] |
| 100847010 | NR_049844,rna | MIR5581 | hsa-miR-5581-3p | . | . | 43527 | . | . | . | [illegible] |
| 100847012 | NR_049800,rna | MIR5004 | microRNA 5004 | miRNA | . | 43532 | . | . | . | [illegible] |
| 100847023 | NR_049814,rna | MIR5091 | microRNA 5091 | miRNA | . | 43478 | . | . | . | [illegible] |
| 100847026 | NR_049803,rna | MIR5006 | microRNA 5006 | miRNA | . | 43495 | . | . | . | [illegible] |
| 100847028 | NR_049852,rna | MIR5587 | hsa-miR-5587-5p | . | . | 43508 | . | . | . | [illegible] |
| 100847037 | NR_049797,rna | MIR5001 | microRNA 5001 | miRNA | . | 43462 | . | . | . | [illegible] |
| 100847046 | NR_049808,rna | MIR5010 | microRNA 5010 | miRNA | . | 43484 | . | . | . | [illegible] |
| 100847070 | NR_049828,rna | MIR5196 | microRNA 5196 | miRNA | . | 43542 | . | . | . | [illegible] |
| 100847073 | NR_049813,rna | MIR5090 | microRNA 5090 | miRNA | . | 43547 | . | . | . | [illegible] |
| 100847079 | NR_049825,rna | MIR5193 | microRNA 5193 | miRNA | . | 43534 | . | . | . | [illegible] |
| 100847090 | NR_049819,rna | MIR5187 | microRNA 5187 | miRNA | . | 43523 | . | . | . | [illegible] |
| 100859930 | NR_045680 | HEIH | hepatocellular carcinoma up-regulated EZH2-associated long non- | lncRNA | . | 45049 | 618615 | . | . | [illegible] |
| 100861412 | NM_00125614 | FSBP | fibrinogen silencer binding protein | protein_coding | NP_001243070.1 | 43653 | 616306 | . | . | [illegible] |
| 100861430 | gene-PDCD6IPI | PDCD6IPP1 | PDCD6IP pseudogene 1 | pseudogene | . | 49872 | . | . | . | [illegible] |
| 100861541 | NR_047482 | DOCK9-DT | DOCK9 divergent transcript | lncRNA | . | 43696 | . | . | . | [illegible] |
| 100861548 | NR_046507 | PINK1-AS | PINK1 antisense RNA | antisense_RNA | . | 38872 | . | . | . | [illegible] |
| 100861555 | NR_047495 | LINC00565 | long intergenic non-protein coding RNA 565 | lncRNA | . | 43709 | . | . | . | [illegible] |
| 100862671 | NM_00125682 | TMEM265 | transmembrane protein 265 | protein_coding | NP_001243758.1 | 51241 | . | . | . | [illegible] |
| 100862682 | gene-LOC1008 | LOC100862682 | nucleus accumbens associated 1, BEN and BTB (POZ) domain cont | pseudogene | . | . | . | . | . | [illegible] |
| 100862692 | NR_046400 | DIP2A-IT1 | DIP2A intronic transcript 1 | lncRNA | . | 41430 | . | . | . | [illegible] |
| 100862728 | NR_046420 | UPK1A-AS1 | UPK1A antisense RNA 1 | lncRNA | . | 40603 | . | . | . | [illegible] |
| 100873757 | gene-RNU6-36 | RNU6-36P | RNA, U6 small nuclear 36, pseudogene | pseudogene | . | 34280 | . | . | . | [illegible] |
| 100873896 | gene-HMGB1P | HMGB1P41 | high mobility group box 1 pseudogene 41 | pseudogene | . | 39188 | . | . | . | [illegible] |
| 100873949 | NR_046696 | IPO9-AS1 | IPO9 antisense RNA 1 | lncRNA | . | 40892 | . | . | . | [illegible] |
| 100873979 | NR_109804 | RBMS3-AS3 | RBMS3 antisense RNA 3 | lncRNA | . | 39989 | . | . | . | [illegible] |
| 100874032 | NR_046734 | PRRT3-AS1 | PRRT3 antisense RNA 1 | lncRNA | . | 41151 | . | . | . | [illegible] |
| 100874034 | NR_046745 | UBXN7-AS1 | UBXN7 antisense RNA 1 | lncRNA | . | 41227 | . | . | . | [illegible] |
| 100874054 | NR_051960 | FALEC | focally amplified long non-coding RNA in epithelial cancer | lncRNA | . | 43713 | 616092 | . | . | [illegible] |
| 100874058 | NR_049718 | COX10-AS1 | COX10 antisense RNA 1 | lncRNA | . | 38873 | . | . | . | [illegible] |
| 100874069 | NR_046542 | STK24-AS1 | STK24 antisense RNA 1 | lncRNA | . | 39935 | . | . | . | [illegible] |
| 100874094 | NR_046658 | ZNF197-AS1 | ZNF197 antisense RNA 1 | lncRNA | . | 40620 | . | . | . | [illegible] |
| 100874194 | NR_046533 | CLDN10-AS1 | CLDN10 antisense RNA 1 | lncRNA | . | 39907 | . | . | . | [illegible] |
| 100874207 | NR_046683 | ST3GAL6-AS1 | ST3GAL6 antisense RNA 1 | lncRNA | . | 40828 | . | . | . | [illegible] |
| 100874211 | NR_046706 | MID1IP1-AS1 | MID1IP1 antisense RNA 1 | lncRNA | . | 40932 | . | . | . | [illegible] |
| 100874214 | NR_046724,NF | TM4SF19-AS1 | TM4SF19 antisense RNA 1 | lncRNA | . | 41085 | . | . | . | [illegible] |
| 100874246 | NR_046748 | ARHGAP31-AS1 | ARHGAP31 antisense RNA 1 | lncRNA | . | 41235 | . | . | . | [illegible] |
| 100874252 | gene-USP32P4 | USP32P4 | ubiquitin specific peptidase 32 pseudogene 4 | pseudogene | . | 43577 | . | . | . | [illegible] |
| 100874260 | NR_144562 | UBE2CP5 | ubiquitin conjugating enzyme E2 C pseudogene 5 | transcribed_ps | . | 43554 | . | . | . | [illegible] |
| 100874278 | NR_046817,NF | ERI3-IT1 | ERI3 intronic transcript 1 | lncRNA | . | 41431 | . | . | . | [illegible] |
| 100874291 | NR_046762 | ALMS1-IT1 | ALMS1 intronic transcript 1 | lncRNA | . | 41305 | . | . | . | [illegible] |
| 100874296 | NR_046784 | KCNH1-IT1 | KCNH1 intronic transcript 1 | lncRNA | . | 41351 | . | . | . | [illegible] |
| 100874353 | NR_046793 | RPS6KA2-IT1 | RPS6KA2 intronic transcript 1 | lncRNA | . | 41378 | . | . | . | [illegible] |
| 100874362 | NR_102279 | HOXB-AS1 | HOXB cluster antisense RNA 1 | lncRNA | . | 43744 | 618066 | . | . | [illegible] |
| 100874363 | NR_047504 | HOXC-AS1 | HOXC cluster antisense RNA 1 | lncRNA | . | 43749 | . | . | . | [illegible] |
| 100874365 | NR_047506 | HOXC-AS3 | HOXC cluster antisense RNA 3 | lncRNA | . | 43751 | . | . | . | [illegible] |
| 100874366 | NR_047507 | HOXC13-AS | HOXC13 antisense RNA | lncRNA | . | 43753 | . | . | . | [illegible] |
| 100874381 | gene-NPIPP1 | NPIPP1 | nuclear pore complex interacting protein pseudogene 1 | pseudogene | . | 35407 | . | . | . | [illegible] |
| 100874391 | gene-OSTCP7 | OSTCP7 | oligosaccharyltransferase complex subunit pseudogene 7 | pseudogene | . | 42870 | . | . | . | [illegible] |
| 100874417 | gene-TIMM8BI | TIMM8BP2 | translocase of inner mitochondrial membrane 8B pseudogene 2 | pseudogene | . | 39212 | . | . | . | [illegible] |
| 100874467 | gene-HMGN2P | HMGN2P21 | high mobility group nucleosomal binding domain 2 pseudogene 2: | pseudogene | . | 39386 | . | . | . | [illegible] |
| 100885776 | NR_047679 | UGDH-AS1 | UGDH antisense RNA 1 | lncRNA | . | 40601 | . | . | . | [illegible] |
| 100885778 | NR_047687 | NALCN-AS1 | NALCN antisense RNA 1 | lncRNA | . | 42743 | . | . | . | [illegible] |
| 100885799 | gene-ENO1P4 | ENO1P4 | enolase 1 pseudogene 4 | pseudogene | . | 37945 | . | . | . | [illegible] |
| 100885865 | gene-HMGN1P | HMGN1P29 | high mobility group nucleosome binding domain 1 pseudogene 29 | pseudogene | . | 39373 | . | . | . | [illegible] |
| 100887068 | gene-KRT8P52 | KRT8P52 | keratin 8 pseudogene 52 | pseudogene | . | 54654 | . | . | . | [illegible] |
| 100887750 | NR_002816,NF | MRPS31P5 | mitochondrial ribosomal protein S31 pseudogene 5 | transcribed_ps | . | 39253 | . | . | . | [illegible] |
| 100996246 | NR_146452 | LINC02361 | long intergenic non-protein coding RNA 2361 | lncRNA | . | 53283 | . | . | . | [illegible] |
| 100996255 | NR_102756 | LINC02256 | long intergenic non-protein coding RNA 2256 | lncRNA | . | 53157 | . | . | . | [illegible] |
| 100996259 | NM_00134816 | FAM106B | family with sequence similarity 106 member B | protein_coding | NP_001335090.1 | 32450 | . | . | . | [illegible] |
| 100996294 | XR_002958103 | LOC100996294 | uncharacterized LOC100996294 | lncRNA | . | . | . | . | . | [illegible] |
| 100996301 | NR_121634,NF | FOXD3-AS1 | FOXD3 antisense RNA 1 | lncRNA | . | 40241 | . | . | . | [illegible] |
| 100996333 | gene-LOC1009 | LOC100996333 | G protein-coupled receptor 125 pseudogene | pseudogene | . | . | . | . | . | [illegible] |
| 100996379 | XR_001751424 | LOC100996379 | E3 ubiquitin-protein ligase HERC2-like, transcript variant X2 | misc_RNA | . | . | . | . | . | [illegible] |
| 100996419 | NR_134283 | LOC100996419 | uncharacterized LOC100996419 | lncRNA | . | . | . | . | . | [illegible] |
| 100996425 | NR_131947,NF | ARLNC1 | androgen receptor regulated long noncoding RNA 1 | lncRNA | . | 53032 | 618053 | . | . | [illegible] |
| 100996437 | NR_110037 | LOC100996437 | uncharacterized LOC100996437 | lncRNA | . | . | . | . | . | [illegible] |
| 100996438 | gene-MED15P( | MED15P6 | . | pseudogene | . | 48655 | . | . | . | [illegible] |
| 100996442 | XR_001737578 | LOC100996442 | uncharacterized LOC100996442, transcript variant X14 | misc_RNA | . | . | . | . | . | [illegible] |
| 100996465 | NR_130768 | LCA10 | lung carcinoma-associated 10 | lncRNA | . | . | . | . | . | [illegible] |
| 100996492 | NM_00135288 | CTXND1 | cortexin domain containing 1 | protein_coding | NP_001339817.1 | 50507 | . | . | . | [illegible] |
| 100996511 | NR_110616 | LINC01355 | long intergenic non-protein coding RNA 1355 | lncRNA | . | 50584 | . | . | . | [illegible] |
| 100996517 | NR_158170 | SEC22B3P | SEC22 homolog B3, pseudogene | transcribed_ps | . | 53891 | . | . | . | [illegible] |
| 100996539 | NR_108075 | ITPR1-DT | ITPR1 divergent transcript | lncRNA | . | 44470 | . | . | . | [illegible] |
| 100996583 | NR_121638 | LOC100996583 | uncharacterized LOC100996583 | lncRNA | . | . | . | . | . | [illegible] |
| 100996645 | NR_120607 | LINCR-0003 | uncharacterized LincR-0003 | lncRNA | . | . | . | . | . | [illegible] |
| 100996660 | XR_001752953 | LOC100996660 | uncharacterized LOC100996660, transcript variant X2 | lncRNA | . | . | . | . | . | [illegible] |
| 100996664 | NR_135816 | LOC100996664 | uncharacterized LOC100996664 | lncRNA | . | . | . | . | . | [illegible] |
| 100996702 | NR_103746 | LINC01356 | long intergenic non-protein coding RNA 1356 | lncRNA | . | 50587 | . | . | . | [illegible] |
| 100996712 | NR_120535 | SRGAP2D | SLIT-ROBO Rho GTPase activating protein 2D (pseudogene) | transcribed_ps | . | 43932 | 614705 | . | . | [illegible] |
| 100996716 | NR_158171 | SEC22B2P | SEC22 homolog B2, pseudogene | transcribed_ps | . | 53890 | . | . | . | [illegible] |
| 100996717 | NM_00136401 | NOTCH2NLC | notch 2 N-terminal like C | protein_coding | NP_001350941.1;NP_001350942.1 | 53924 | 618025 | . | . | [illegible] |
| 100996720 | NR_160510 | LOC100996720 | uncharacterized LOC100996720 | lncRNA | . | . | . | . | . | [illegible] |
| 100996723 | gene-LOC1009 | LOC100996723 | . | pseudogene | . | . | . | . | . | [illegible] |
| 100996724 | NR_144516,NF | LOC100996724 | phosphodiesterase 4D interacting protein-like | transcribed_ps | . | . | . | . | . | [illegible] |
| 100996732 | XR_159162,XR | LINC02802 | long intergenic non-protein coding RNA 2802, transcript variant X5 | lncRNA | . | 54326 | . | . | . | [illegible] |
| 100996738 | NR_145479 | HERC2P11 | hect domain and RLD 2 pseudogene 11 | transcribed_ps | . | 49882 | . | . | . | [illegible] |
| 100996740 | NR_148945 | LOC100996740 | uncharacterized LOC100996740 | lncRNA | . | . | . | . | . | [illegible] |
| 100996756 | XR_159169 | LOC100996756 | uncharacterized LOC100996756 | lncRNA | . | . | . | . | . | [illegible] |
| 100996763 | NM_00136400 | NOTCH2NLB | notch 2 N-terminal like B | protein_coding | NP_001350936.1;NP_001350937.1;XP_006711763.1;XP_011508558.1;XP | 53923 | 618024 | . | . | [illegible] |
| 100996842 | NM_00132280 | LOC100996842 | uncharacterized LOC100996842 | protein_coding | NP_001309729.1;XP_006721686.1 | . | . | . | . | [illegible] |
| 100996928 | NM_00124458 | FMC1-LUC7L2 | FMC1-LUC7L2 readthrough | protein_coding | NP_001231513.1 | 44671 | . | . | . | [illegible] |
| 100996939 | NM_032906 | PYURF | PIGY upstream reading frame | protein_coding | NP_116295.1 | 44317 | . | . | . | [illegible] |
| 101059906 | XM_00395993 | LOC101059906 | collagen alpha-2(IV) chain-like | protein_coding | XP_003959982.2 | . | . | . | . | [illegible] |
| 101059918 | NM_00128248 | GOLGA8R | golgin A8 family member R | protein_coding | NP_001269413.1 | 44407 | . | . | . | [illegible] |
| 101059938 | NM_00128250 | NPIPA7 | nuclear pore complex interacting protein family member A7 | protein_coding | NP_001269436.1;XP_005255792.1;XP_011520641.1;XP_016878322.1;XP | 41982 | . | . | . | [illegible] |
| 101059953 | NM_00128251 | NPIPA8 | nuclear pore complex interacting protein family member A8 | protein_coding | NP_001269440.1 | 41983 | . | . | . | [illegible] |
| 101059954 | XR_171071 | LOC101059954 | uncharacterized LOC101059954 | lncRNA | . | . | . | . | . | [illegible] |
| 101060048 | gene-LOC1010 | LOC101060048 | . | pseudogene | . | . | . | . | . | [illegible] |
| 101060104 | gene-LOC1010 | LOC101060104 | . | pseudogene | . | . | . | . | . | [illegible] |
| 101060146 | NR_110646 | CHMP1B2P | charged multivesicular body protein 1B2, pseudogene | transcribed_ps | . | 49380 | . | . | . | [illegible] |
| 101060179 | XM_00395993 | LOC101060179 | uncharacterized LOC101060179 | protein_coding | XP_003959981.1 | . | . | . | . | [illegible] |
| 101060195 | NR_134638 | FCF1P2 | FCF1 pseudogene 2 | transcribed_ps | . | 44614 | . | . | . | [illegible] |
| 101060200 | NM_00127729 | ZNF891 | zinc finger protein 891 | protein_coding | NP_001264220.1;XP_016874155.1;XP_016874156.1;XP_016874157.1 | 38709 | . | . | . | [illegible] |
| 101060211 | NM_00129154 | CT45A7 | cancer/testis antigen family 45 member A7 | protein_coding | NP_001278472.1;XP_003960131.1;XP_011529531.1 | 51260 | . | . | . | [illegible] |
| 101060212 | gene-LOC1010 | LOC101060212 | . | pseudogene | . | . | . | . | . | [illegible] |
| 101060226 | NM_00135136 | NBPF19 | NBPF member 19 | protein_coding | NP_001338294.1 | 31999 | 614006 | . | . | [illegible] |
| 101060264 | NR_126415,NF | FOXP4-AS1 | FOXP4 antisense RNA 1 | lncRNA | . | 50332 | . | . | . | [illegible] |
| 101060321 | NM_00129146 | TBC1D3G | TBC1 domain family member 3G | protein_coding | NP_001278391.1;XP_005276971.1 | 29860 | 610810 | . | . | [illegible] |
| 101060341 | XM_00527179 | LOC101060341 | putative uncharacterized protein FLJ44672, transcript variant X1 | protein_coding | XP_005271850.1;XP_016868414.1;XP_016868416.1;XP_024302801.1 | . | . | . | . | [illegible] |
| 101060351 | NM_00129146 | TBC1D3K | TBC1 domain family member 3K | protein_coding | NP_001278393.1;XP_006722298.1;XP_006722299.1;XP_006722300.1;XP | 51245 | . | . | . | [illegible] |
| 101060376 | NM_00129145 | TBC1D3L | TBC1 domain family member 3L | protein_coding | NP_001278388.1;NP_001356428.1;NP_001356429.1;NP_001356430.1;NI | 51246 | . | . | . | [illegible] |
| 101060389 | NM_00129146 | TBC1D3D | TBC1 domain family member 3D | protein_coding | NP_001278394.1;XP_006722311.1;XP_006722312.1;XP_006722313.1;XP | 28944 | 610807 | . | . | [illegible] |
| 101060684 | NM_00135137 | NBPF26 | NBPF member 26 | protein_coding | NP_001338301.1 | 49571 | . | . | . | [illegible] |
| 101060691 | NR_120611,NF | NUTM2B-AS1 | NUTM2B antisense RNA 1 | lncRNA | . | 51204 | 618639 | . | . | [illegible] |
| 101101772 | NR_073453 | LINC00601 | long intergenic non-protein coding RNA 601 | lncRNA | . | 43916 | . | . | . | [illegible] |
| 101232810 | gene-LOC1012 | LOC101232810 | stathmin 3 pseudogene | pseudogene | . | . | . | . | . | [illegible] |
| 101234262 | gene-COX5BP6 | COX5BP6 | cytochrome c oxidase subunit 5B pseudogene 6 | pseudogene | . | 2275 | . | . | . | [illegible] |
| 101241892 | NR_103844 | NPTN-IT1 | NPTN intronic transcript 1 | lncRNA | . | 45091 | 615176 | . | . | [illegible] |
| 101241901 | gene-LOC1012 | LOC101241901 | chromosome 4 open reading frame 46 pseudogene | pseudogene | . | . | . | . | . | [illegible] |
| 101243545 | NR_102265 | LINC02067 | long intergenic non-protein coding RNA 2067 | lncRNA | . | 52913 | . | . | . | [illegible] |
| 101243555 | NR_103745 | PTENP1-AS | PTENP1 antisense RNA | lncRNA | . | 48773 | . | . | . | [illegible] |
| 101290501 | gene-TBCAP3 | TBCAP3 | tubulin folding cofactor A pseudogene 3 | pseudogene | . | 42632 | . | . | . | [illegible] |
| 101340251 | NR_102369 | SNORD124 | small nucleolar RNA, C/D box 124 | snoRNA | . | 33563 | . | . | . | [illegible] |
| 101340252 | NR_102370 | SNORD121B | small nucleolar RNA, C/D box 121B | snoRNA | . | 33569 | . | . | . | [illegible] |
| 101409254 | NR_102423 | LINC00681 | long intergenic non-protein coding RNA 681 | lncRNA | . | 44423 | . | . | . | [illegible] |
| 101409258 | NR_102429 | OSGEPL1-AS1 | OSGEPL1 antisense RNA 1 | lncRNA | . | 41009 | . | . | . | [illegible] |
| 101410533 | NR_102684 | YTHDF3-AS1 | YTHDF3 antisense RNA 1 (head to head) | lncRNA | . | 48728 | . | . | . | [illegible] |
| 101410535 | NR_102697 | LAMTOR5-AS1 | LAMTOR5 antisense RNA 1 | lncRNA | . | 40823 | . | . | . | [illegible] |
| 101410538 | NM_00135500 | MMP24OS | MMP24 opposite strand | protein_coding | NP_001341932.1;NP_001341933.1 | 44421 | . | . | . | [illegible] |
| 101559451 | NR_103482 | LOC101559451 | uncharacterized LOC101559451 | lncRNA | . | . | . | . | . | [illegible] |
| 101593348 | NR_103497 | LOC101593348 | uncharacterized LOC101593348 | lncRNA | . | . | . | . | . | [illegible] |
| 101669762 | NR_103783 | BLACAT1 | bladder cancer associated transcript 1 | lncRNA | . | 48597 | 615480 | . | . | [illegible] |
| 101730217 | NR_103546 | SPECC1L-ADOR | SPECC1L-ADORA2A readthrough (NMD candidate) | lncRNA | . | 49185 | . | . | . | [illegible] |
| 101735302 | NR_103720 | STAG3L5P | stromal antigen 3-like 5 pseudogene | transcribed_ps | . | 48896 | . | . | . | [illegible] |
| 101752334 | NR_103728 | PVRIG2P | poliovirus receptor related immunoglobulin domain containing 2, | transcribed_ps | . | 48897 | . | . | . | [illegible] |
| 101752399 | NR_036569,NF | STAG3L5P-PVR | STAG3L5P-PVRIG2P-PILRB readthrough | lncRNA | . | 48898 | . | . | . | [illegible] |
| 101752400 | NR_103792 | CAPN10-DT | CAPN10 divergent transcript | lncRNA | . | 48839 | . | . | . | [illegible] |
| 101867536 | NR_109833 | PRNCR1 | prostate cancer associated non-coding RNA 1 | lncRNA | . | 48942 | 615452 | . | . | [illegible] |
| 101926888 | NR_109885,NF | RALY-AS1 | RALY antisense RNA 1 | lncRNA | . | 50743 | . | . | . | [illegible] |
| 101926889 | NR_109994 | LOC101926889 | uncharacterized LOC101926889 | lncRNA | . | . | . | . | . | [illegible] |
| 101926895 | NR_120372 | CRTC3-AS1 | CRTC3 antisense RNA 1 | lncRNA | . | 51433 | . | . | . | [illegible] |
| 101926907 | XR_246327 | LOC101926907 | uncharacterized LOC101926907 | lncRNA | . | . | . | . | . | [illegible] |
| 101926911 | NR_110104 | VPS33B-DT | VPS33B divergent transcript | lncRNA | . | 51413 | . | . | . | [illegible] |
| 101926913 | NR_110185 | LOC101926913 | uncharacterized LOC101926913 | lncRNA | . | . | . | . | . | [illegible] |
| 101926917 | NR_136286 | ANKRD18CP | ankyrin repeat domain 18C, pseudogene | transcribed_ps | . | 43601 | . | . | . | [illegible] |
| 101926918 | NR_135112 | GTF2IP12 | general transcription factor IIi pseudogene 12 | transcribed_ps | . | 51723 | . | . | . | [illegible] |
| 101926935 | NR_110001 | LOC101926935 | uncharacterized LOC101926935 | lncRNA | . | . | . | . | . | [illegible] |
| 101926941 | NR_120664 | SPRY4-AS1 | SPRY4 antisense RNA 1 | lncRNA | . | 53465 | . | . | . | [illegible] |

| | | | | | | | | | | | | | | | | | | | | | | | | | | | | | | | | | |
|---|---|---|---|---|---|---|---|---|---|---|---|---|---|---|---|---|---|---|---|---|---|---|---|---|---|---|---|---|---|---|---|---|---|
| 101926948 | NR_135130 | LOC10192694 | uncharacterized LOC101926948 | lncRNA | . | . | . | . | . | 0.219263 | 0.062087 | 0.179435 | 0.133865 | 0.510234 | 0.143517 | 0.402646 | 0.306067 | 1.132228 | 4.704391 | 1.426797835 | 0.125571176 | 0.900071381 | NA | 2.137672 | 2.324312 | 2.241337 | 2.217316 | 2.235334 | 2.228762 | 8 | 2 | 5 | 4 |
| 101926963 | NR_110054,N | PRKAR1B-AS1 | PRKAR1B antisense RNA 1 | lncRNA | . | 40470 | . | . | . | 2.215718 | 2.421536 | 1.754399 | 2.287235 | 5.156067 | 5.597489 | 3.936819 | 5.229492 | -1.179776 | 72.072091 | 0.37810998 | -0.6308026 | 0.528169603 | 0.998865404 | 6.286539 | 6.046201 | 6.186871 | 6.197711 | 6.122905 | 6.180515 | 88 | 86 | 50 | 70 |
| 101926967 | XR_00175290 | LOC10192696 | uncharacterized LOC101926967, transcript variant X1 | lncRNA | . | . | . | . | . | 0.409396 | 0.499266 | 0.682840 | 0.343849 | 0.952682 | 1.154075 | 1.532274 | 0.786169 | 1.137831 | 22.618407 | 0.673369786 | 0.276647516 | 0.782050776 | NA | 4.401710 | 4.590940 | 4.474718 | 4.492411 | 4.528106 | 4.462331 | 22 | 25 | 28 | 15 |
| 101926977 | XR_00174600 | LOC10192697 | uncharacterized LOC101926977, transcript variant X1 | lncRNA | . | . | . | . | . | 0.245790 | 0.309145 | 0.266108 | 0.350109 | 0.571962 | 0.714604 | 0.597138 | 0.800483 | 1.139279 | 13.460161 | 0.833332939 | 0.225746074 | 0.821398921 | NA | 3.654832 | 3.840449 | 3.737999 | 3.749864 | 3.744460 | 3.758784 | 13 | 15 | 11 | 15 |
| 101927009 | NR_135195 | LINC02177 | long intergenic non-protein coding RNA 2177 | lncRNA | . | 53039 | . | . | . | 0.049301 | 0.097475 | 0.100566 | 0.057514 | 0.114725 | 0.225318 | 0.225667 | 0.131499 | 1.058110 | 13.053861 | 0.875168951 | 0.093113385 | 0.925813484 | NA | 3.664814 | 3.746830 | 3.677889 | 3.714117 | 3.715595 | 3.682919 | 10 | 18 | 15 | 9 |
| 101927021 | NR_110065,N | ZFAND2A-DT | ZFAND2A divergent transcript | lncRNA | . | 41187 | . | . | . | 2.369872 | 2.875148 | 3.010382 | 2.546917 | 5.514789 | 6.646033 | 6.755212 | 5.823224 | 1.072281 | 192.987126 | 0.250117341 | 0.402544037 | 0.687283686 | 0.998865404 | 7.541238 | 7.641734 | 7.523814 | 7.620731 | 7.633043 | 7.574893 | 191 | 223 | 185 | 175 |
| 101927027 | NR_110204,N | CHROMR | cholesterol induced regulator of metabolism RNA | lncRNA | . | 54059 | . | . | . | 6.200853 | 5.943883 | 6.801730 | 7.087935 | 14.429641 | 13.739550 | 15.262885 | 16.205721 | 1.136892 | 346.070400 | 0.19076469 | 0.970279904 | 0.331906991 | 0.998865404 | 8.339196 | 8.524689 | 8.410683 | 8.380877 | 8.480427 | 8.456451 | 383 | 340 | 327 | 338 |
| 101927045 | NR_110032 | LOC10192704 | uncharacterized LOC101927045 | lncRNA | . | . | . | . | . | 1.184851 | 1.301775 | 1.481871 | 1.249972 | 2.757198 | 3.009112 | 3.325275 | 2.857914 | 1.088307 | 86.274168 | 0.345330008 | 0.353532689 | 0.723689127 | 0.998865404 | 6.368205 | 6.490901 | 6.404222 | 6.428036 | 6.463465 | 6.415106 | 91 | 93 | 85 | 77 |
| 101927056 | NR_125395,N | LINC02614 | long intergenic non-protein coding RNA 2614 | lncRNA | . | 54072 | . | . | . | 0.085719 | 0.128795 | 0.401837 | 0.445740 | 0.199473 | 0.297716 | 0.901711 | 1.019132 | 3.507213 | 7.098853 | 1.185750427 | 1.526733618 | 0.126827264 | NA | 1.658167 | 3.464871 | 2.776931 | 2.782503 | 2.812622 | 2.819053 | 3 | 4 | 9 | 11 |
| 101927060 | XR_00175310 | LOC10192706 | uncharacterized LOC101927060, transcript variant X25 | lncRNA | . | . | . | . | . | 1.629292 | 1.923472 | 1.899283 | 1.640344 | 3.791427 | 4.446192 | 4.261937 | 3.750453 | 1.043152 | 249.360252 | 0.223773976 | 0.272372094 | 0.785335928 | 0.998865404 | 7.931270 | 7.992261 | 7.907443 | 7.989531 | 8.002584 | 7.937093 | 258 | 285 | 236 | 223 |
| 101927086 | XR_00174670 | KLF9-DT | KLF9 divergent transcript, transcript variant X7 | lncRNA | . | 54815 | . | . | . | 1.805373 | 1.961416 | 0.701129 | 1.422885 | 4.201176 | 4.533901 | 1.573312 | 3.253258 | -1.746311 | 49.691077 | 0.472923766 | -1.70071981 | 0.088995614 | 0.802216049 | 5.983353 | 5.174582 | 5.677070 | 5.688027 | 5.540868 | 5.622291 | 71 | 70 | 22 | 44 |
| 101927096 | NR_109873 | CLMAT3 | colorectal liver metastasis associated transcript 3 | lncRNA | . | 52287 | . | . | . | 1.669260 | 1.802075 | 1.163030 | 1.494223 | 3.884436 | 4.165576 | 2.609806 | 3.416363 | -1.310485 | 94.901042 | 0.334303538 | -1.16690485 | 0.243248798 | 0.992733415 | 6.751025 | 6.359150 | 6.596803 | 6.617662 | 6.497379 | 6.557841 | 120 | 120 | 63 | 86 |
| 101927111 | NR_109992,N | SUCLG2-AS1 | SUCLG2 antisense RNA 1 (head to head) | lncRNA | . | 49643 | . | . | . | 0.748044 | 0.509152 | 0.487795 | 0.641325 | 1.740729 | 1.176927 | 1.094596 | 1.466315 | -1.057206 | 20.393538 | 0.687494997 | -0.11673826 | 0.907067469 | NA | 4.389723 | 4.309235 | 4.368331 | 4.339284 | 4.338455 | 4.359662 | 28 | 19 | 15 | 21 |
| 101927117 | NR_109889 | PLCG1-AS1 | PLCG1 antisense RNA 1 | lncRNA | . | 40450 | . | . | . | 0.099882 | 0.099212 | 0.206550 | 0.173571 | 0.232430 | 0.229333 | 0.463493 | 0.396849 | 1.908847 | 8.472321 | 1.05101073 | 0.887432989 | 0.374845829 | NA | 2.540965 | 3.475715 | 3.055127 | 3.052876 | 3.084396 | 3.071383 | 7 | 6 | 11 | 9 |
| 101927125 | NR_120508,N | ELFN1-AS1 | ELFN1 antisense RNA 1 | lncRNA | . | 39071 | . | . | . | 2.552970 | 2.384387 | 2.666569 | 3.527182 | 5.940865 | 5.511617 | 5.983703 | 8.064485 | 1.187788 | 68.550860 | 0.38441607 | 0.645855017 | 0.518373254 | 0.998865404 | 5.970066 | 6.217540 | 6.083522 | 6.066264 | 6.089409 | 6.140563 | 75 | 65 | 58 | 77 |
| 101927151 | NR_110687,N | LOC10192715 | uncharacterized LOC101927151 | lncRNA | . | . | . | . | . | 2.806822 | 3.630951 | 3.075307 | 4.277058 | 6.531591 | 8.393108 | 6.900901 | 9.778987 | 1.134869 | 191.624474 | 0.266142717 | 0.685817132 | 0.492828431 | 0.998865404 | 7.489186 | 7.669462 | 7.487318 | 7.604173 | 7.531724 | 7.669797 | 179 | 220 | 150 | 220 |
| 101927152 | NR_110000 | SAMMSON | survival associated mitochondrial melanoma specific oncogenic nc | lncRNA | . | 49644 | 616895 | . | . | 0.766415 | 0.947064 | 0.891886 | 0.863019 | 1.783479 | 2.189182 | 2.001367 | 1.973191 | 1.009666 | 55.122093 | 0.424144416 | 0.032719128 | 0.97389857 | 0.998865404 | 5.777940 | 5.791147 | 5.762151 | 5.801943 | 5.787594 | 5.780719 | 57 | 65 | 49 | 51 |
| 101927174 | NR_144486,N | CHL1-AS2 | CHL1 antisense RNA 2 | lncRNA | . | 40147 | . | . | . | 0.931436 | 0.666525 | 1.253780 | 0.705237 | 2.167489 | 1.540702 | 2.813446 | 1.612440 | 1.220446 | 20.322082 | 0.705167925 | 0.407575325 | 0.683585483 | NA | 4.191364 | 4.483795 | 4.342204 | 4.312086 | 4.370014 | 4.317894 | 25 | 16 | 25 | 15 |
| 101927204 | NR_110184 | FAM111A-DT | FAM111A divergent transcript | lncRNA | . | 53752 | . | . | . | 0.908555 | 0.834337 | 0.745896 | 0.953530 | 2.114245 | 1.928607 | 1.673768 | 2.180132 | -1.044772 | 60.869249 | 0.406733037 | -0.15535523 | 0.876541281 | 0.998865404 | 5.959362 | 5.895208 | 5.943365 | 5.922404 | 5.900327 | 5.946190 | 75 | 64 | 46 | 62 |
| 101927209 | XR_00174168 | LOC10192720 | uncharacterized LOC101927209, transcript variant X3 | lncRNA | . | . | . | . | . | 5.629669 | 4.546081 | 5.199630 | 6.913030 | 13.100471 | 10.508468 | 11.667820 | 15.805822 | 1.089353 | 55.502152 | 0.429056276 | 0.287774514 | 0.773519347 | 0.998865404 | 5.731451 | 5.854854 | 5.808725 | 5.758207 | 5.779177 | 5.822261 | 69 | 50 | 45 | 60 |
| 101927233 | NR_109879 | SEMA6A-AS1 | SEMA6A antisense RNA 1 | lncRNA | . | 51110 | . | . | . | 0.128329 | 0.097893 | 0.053232 | 0.096699 | 0.298628 | 0.226284 | 0.119451 | 0.221091 | -1.461556 | 3.387285 | 1.629878023 | -0.33591778 | 0.73693287 | NA | 2.008843 | 1.459419 | 1.773175 | 1.770137 | 1.763831 | 1.767798 | 5 | 4 | 2 | 3 |
| 101927245 | NR_160672,N | LOC10192724 | uncharacterized LOC101927245 | lncRNA | . | . | . | . | . | 0.179575 | 0.236231 | 0.245594 | 0.355086 | 0.417877 | 0.546060 | 0.551105 | 0.811862 | -1.308546 | 8.246728 | 1.068154521 | -0.36321038 | 0.716447733 | NA | 3.225252 | 2.836248 | 3.063912 | 3.046084 | 3.039079 | 3.051164 | 13 | 8 | 5 | 8 |
| 101927249 | NR_046681 | PDZRN3-AS1 | PDZRN3 antisense RNA 1 | lncRNA | . | 40814 | . | . | . | 0.311610 | 0.310144 | 0.264297 | 0.195589 | 0.725130 | 0.716913 | 0.593076 | 0.447191 | -1.359163 | 24.917619 | 0.622381336 | -0.71133037 | 0.476879537 | NA | 4.842669 | 4.401999 | 4.657169 | 4.657241 | 4.638279 | 4.613371 | 33 | 31 | 21 | 17 |
| 101927263 | XR_932655 | LOC10192726 | uncharacterized LOC101927263 | lncRNA | . | . | . | . | . | 0.048318 | 0.034436 | 0.116513 | 0.104393 | 0.112437 | 0.079599 | 0.261453 | 0.238682 | 2.483531 | 6.213894 | 1.237367269 | 1.060633115 | 0.288856667 | NA | 1.832372 | 3.148375 | 2.606046 | 2.598786 | 2.628030 | 2.625352 | 5 | 3 | 8 | 8 |
| 101927278 | NR_134451,N | LOC10192727 | uncharacterized LOC101927278 | lncRNA | . | . | . | . | . | 0.561765 | 0.644818 | 0.819924 | 0.979708 | 1.307251 | 1.490526 | 1.839885 | 2.239986 | 1.429764 | 23.447946 | 0.636735934 | 0.810032102 | 0.417921726 | NA | 4.271488 | 4.785711 | 4.520495 | 4.530146 | 4.550674 | 4.569316 | 21 | 22 | 22 | 28 |
| 101927279 | NR_110514 | EP300-AS1 | EP300 antisense RNA 1 | lncRNA | . | 50504 | . | . | . | 1.382084 | 0.838635 | 0.513137 | 0.420088 | 3.216166 | 1.938542 | 1.151464 | 0.960481 | -2.362576 | 34.516703 | 0.564873494 | -2.19582055 | 0.02810479 | 0.539571956 | 5.598931 | 4.362218 | 5.204651 | 5.109126 | 5.057346 | 5.036346 | 70 | 39 | 20 | 17 |
| 101927288 | NR_047502,N | MANEA-DT | MANEA divergent transcript | lncRNA | . | 43732 | . | . | . | 0.822681 | 1.158058 | 1.114435 | 0.772346 | 1.914413 | 2.676903 | 2.500761 | 1.765877 | -1.154127 | 18.531729 | 0.722398267 | -0.28627082 | 0.774670707 | NA | 4.312060 | 4.104320 | 4.198089 | 4.234868 | 4.210425 | 4.200280 | 18 | 26 | 16 | 15 |
| 101927293 | XR_00174418 | LOC10192729 | uncharacterized LOC101927293, transcript variant X6 | lncRNA | . | . | . | . | . | 1.091771 | 1.120728 | 0.976865 | 1.403537 | 2.540595 | 2.590615 | 2.192058 | 3.209022 | 1.085299 | 36.023279 | 0.518784634 | 0.227633423 | 0.819931234 | 0.998865404 | 5.111529 | 5.227843 | 5.156921 | 5.169736 | 5.159025 | 5.190635 | 38 | 39 | 29 | 39 |
| 101927300 | XR_00174756 | LOC10192730 | uncharacterized LOC101927300, transcript variant X2 | lncRNA | . | . | . | . | . | 0.754493 | 0.812888 | 0.930275 | 0.643569 | 1.755735 | 1.879027 | 2.087511 | 1.471444 | -1.023482 | 125.609139 | 0.304422373 | -0.10999936 | 0.912409881 | 0.998865404 | 6.988627 | 6.956793 | 6.962879 | 6.983699 | 7.023850 | 6.902978 | 142 | 141 | 127 | 96 |
| 101927322 | NR_164150 | LOC10192732 | uncharacterized LOC101927322 | lncRNA | . | . | . | . | . | 0.353329 | 0.233203 | 0.223102 | 0.253713 | 0.822210 | 0.539059 | 0.500634 | 0.580084 | -1.233313 | 10.490580 | 0.945657645 | -0.3199248 | 0.74902534 | NA | 3.533720 | 3.232645 | 3.410565 | 3.389517 | 3.389543 | 3.391176 | 16 | 10 | 8 | 9 |
| 101927330 | XR_923203,X | LOC10192733 | uncharacterized LOC101927330, transcript variant X1 | lncRNA | . | . | . | . | . | 1.058757 | 1.105462 | 1.493003 | 1.339171 | 2.463770 | 2.555324 | 3.350255 | 3.061856 | 1.295655 | 53.051228 | 0.429556318 | 0.869925779 | 0.384340967 | 0.998865404 | 5.530179 | 5.904267 | 5.692917 | 5.701321 | 5.758546 | 5.735518 | 52 | 51 | 55 | 53 |
| 101927345 | XM_01154400 | LOC10192734 | putative ankyrin repeat domain-containing protein 20A12 pseudo | protein_coding | XP_011542311.1 | . | . | . | . | 1.197971 | 1.249742 | 1.385978 | 0.858101 | 2.787727 | 2.888837 | 3.110093 | 1.961947 | -1.072354 | 67.741087 | 0.412860436 | -0.24410403 | 0.807150251 | 0.998865404 | 6.130137 | 6.032117 | 6.072905 | 6.094948 | 6.128818 | 6.002561 | 77 | 79 | 73 | 44 |
| 101927352 | NR_110383,N | PLS3-AS1 | PLS3 antisense RNA 1 | lncRNA | . | 50343 | . | . | . | 0.452553 | 0.313502 | 0.231256 | 0.288054 | 1.053110 | 0.724673 | 0.518932 | 0.658601 | -1.461297 | 21.016272 | 0.683830074 | -0.80027051 | 0.423554087 | NA | 4.641413 | 4.093800 | 4.433996 | 4.395060 | 4.372874 | 4.390221 | 34 | 22 | 13 | 18 |
| 101927365 | NR_125843 | LOC10192736 | uncharacterized LOC101927365 | lncRNA | . | . | . | . | . | 0.992745 | 1.292450 | 1.246407 | 1.086834 | 2.310159 | 2.987556 | 2.796902 | 2.484918 | 1.007518 | 37.054580 | 0.515132777 | 0.020975483 | 0.983265213 | 0.998865404 | 5.206250 | 5.216890 | 5.189182 | 5.227603 | 5.221275 | 5.197916 | 37 | 45 | 35 | 32 |
| 101927400 | XR_00173964 | LOC10192740 | uncharacterized LOC101927400, transcript variant X5 | lncRNA | . | . | . | . | . | 0.167689 | 0.209846 | 0.151955 | 0.110925 | 0.390219 | 0.485068 | 0.340983 | 0.253617 | -1.334527 | 11.018506 | 0.921328653 | -0.45187786 | 0.651356976 | NA | 3.654832 | 3.239024 | 3.466182 | 3.477859 | 3.462094 | 3.454073 | 13 | 15 | 9 | 8 |
| 101927402 | NR_136167 | LOC10192740 | uncharacterized LOC101927402 | lncRNA | . | . | . | . | . | 0.183989 | 0.186529 | 0.210924 | 0.231495 | 0.428150 | 0.431170 | 0.473307 | 0.529285 | 1.187392 | 13.234725 | 0.836155263 | 0.296351473 | 0.76696168 | NA | 3.597285 | 3.844638 | 3.718163 | 3.717738 | 3.725445 | 3.730370 | 14 | 13 | 12 | 14 |
| 101927405 | XR_245566 | LOC10192740 | uncharacterized LOC101927405 | lncRNA | . | . | . | . | . | 0.046579 | 0.058305 | 0.037648 | 0.043901 | 0.108391 | 0.134774 | 0.084481 | 0.100375 | -1.256899 | 6.859528 | 1.155225536 | -0.28554533 | 0.775226383 | NA | 2.934350 | 2.602870 | 2.782313 | 2.788708 | 2.777068 | 2.779861 | 8 | 9 | 5 | 6 |
| 101927413 | XR_242419,X | LOC10192741 | uncharacterized LOC101927413, transcript variant X2 | lncRNA | . | . | . | . | . | 0.057364 | 0.223511 | 0.027747 | 0.389321 | 0.133488 | 0.516655 | 0.062264 | 0.890137 | 1.216734 | 3.970175 | 1.523074266 | 0.185817133 | 0.852588168 | NA | 1.844134 | 2.121007 | 1.984946 | 1.986080 | 1.979899 | 1.998241 | 4 | 4 | 2 | 6 |
| 101927415 | NR_110049 | TMED2-DT | TMED2 divergent transcript | lncRNA | . | 53346 | . | . | . | 0.308285 | 0.288176 | 0.502176 | 0.216020 | 0.717391 | 0.666131 | 1.126868 | 0.493904 | 1.199104 | 24.173451 | 0.662931518 | 0.395149656 | 0.692732444 | NA | 4.455972 | 4.722444 | 4.576389 | 4.571058 | 4.632765 | 4.549147 | 26 | 23 | 32 | 15 |
| 101927420 | NR_136252,N | LOC10192742 | uncharacterized LOC101927420 | lncRNA | . | . | . | . | . | 1.576274 | 2.191066 | 1.651429 | 2.806410 | 3.668055 | 5.064749 | 3.705758 | 6.416524 | 1.151456 | 51.290123 | 0.462765066 | 0.439659265 | 0.66018391 | 0.998865404 | 5.577349 | 5.776970 | 5.633398 | 5.686608 | 5.637958 | 5.733925 | 47 | 59 | 36 | 64 |
| 101927423 | XM_03475180 | PANO1 | proapoptotic nucleolar protein 1 | protein_coding | XP_034607700.1 | 51237 | . | . | . | 0.412174 | 0.266763 | 0.534111 | 0.208782 | 0.959147 | 0.616634 | 1.198529 | 0.477356 | 1.075660 | 28.700022 | 0.635499781 | 0.165572838 | 0.868493124 | 0.998865404 | 4.786188 | 4.897621 | 4.852360 | 4.799750 | 4.883867 | 4.783069 | 39 | 23 | 37 | 16 |
| 101927441 | XR_00295825 | LOC10192744 | uncharacterized LOC101927441, transcript variant X1 | lncRNA | . | . | . | . | . | 0.131045 | 0.072558 | 0.112774 | 0.131679 | 0.304947 | 0.167721 | 0.253061 | 0.301068 | 1.233852 | 11.964680 | 0.884101771 | 0.342912356 | 0.731664396 | NA | 3.421266 | 3.724273 | 3.578290 | 3.565181 | 3.575416 | 3.590216 | 14 | 10 | 10 | 14 |
| 101927481 | NR_126336 | LINC01909 | long intergenic non-protein coding RNA 1909 | lncRNA | . | 52728 | . | . | . | 0.377750 | 0.133002 | 0.222631 | 0.289882 | 0.879039 | 0.307439 | 0.499578 | 0.662780 | -1.070902 | 6.836206 | 1.189666424 | -0.0830705 | 0.933795485 | NA | 2.820514 | 2.724273 | 2.791781 | 2.760609 | 2.770407 | 2.777940 | 12 | 4 | 5 | 7 |
| 101927482 | NR_110225,N | HECW2-AS1 | HECW2 antisense RNA 1 | lncRNA | . | 54062 | . | . | . | 17.580028 | 17.262533 | 14.302102 | 14.660461 | 40.909447 | 39.903118 | 32.093504 | 33.519404 | -1.218885 | 706.463113 | 0.149983821 | -1.90394915 | 0.056916807 | 0.708295137 | 9.600191 | 9.314643 | 9.538314 | 9.518570 | 9.388425 | 9.396957 | 906 | 826 | 550 | 601 |
| 101927506 | NR_168395 | LOC10192750 | uncharacterized LOC101927506 | lncRNA | . | . | . | . | . | 0.303375 | 0.312833 | 0.368837 | 0.281228 | 0.705966 | 0.723127 | 0.827660 | 0.642994 | 1.020051 | 23.116049 | 0.64170756 | 0.044633576 | 0.964399379 | NA | 4.515640 | 4.545847 | 4.526669 | 4.529210 | 4.542336 | 4.517193 | 26 | 25 | 23 | 19 |
| 101927509 | NR_168372 | LOC10192750 | uncharacterized LOC101927509 | lncRNA | . | . | . | . | . | 0.048086 | 0.065472 | 0.112851 | 0.079169 | 0.111897 | 0.151342 | 0.253234 | 0.181011 | 1.716531 | 11.011991 | 0.926940728 | 0.840933909 | 0.40038496 | NA | 3.019256 | 3.798723 | 3.426799 | 3.437782 | 3.467343 | 3.448889 | 8 | 10 | 14 | 11 |
| 101927510 | XR_00175294 | LOC10192751 | uncharacterized LOC101927510, transcript variant X2 | lncRNA | . | . | . | . | . | 0.296159 | 0.210284 | 0.303002 | 0.270674 | 0.689174 | 0.486081 | 0.679927 | 0.618862 | 1.093259 | 31.298091 | 0.559008728 | 0.230112165 | 0.818004613 | 0.998865404 | 4.901103 | 5.031937 | 4.979735 | 4.939553 | 4.977917 | 4.964114 | 40 | 27 | 30 | 29 |
| 101927512 | XR_00175471 | LINC01275 | long intergenic non-protein coding RNA 1275, transcript variant X | lncRNA | . | 50331 | . | . | . | 0.549735 | 0.770104 | 0.292398 | 0.937230 | 1.279256 | 1.780130 | 0.656132 | 2.142865 | -1.086232 | 21.584424 | 0.721220719 | -0.16545808 | 0.868583441 | NA | 4.493602 | 4.367480 | 4.417543 | 4.446847 | 4.381802 | 4.466259 | 22 | 28 | 9 | 29 |
| 101927523 | NR_120624 | LINC02620 | long intergenic non-protein coding RNA 2620 | lncRNA | . | 54097 | . | . | . | 0.864819 | 0.833180 | 0.617531 | 1.207428 | 2.012470 | 1.925932 | 1.385721 | 2.760642 | 1.062716 | 28.153831 | 0.606876622 | 0.144602558 | 0.885024677 | 0.998865404 | 4.772565 | 4.856726 | 4.810876 | 4.807087 | 4.777960 | 4.852684 | 32 | 29 | 17 | 36 |
| 101927547 | NR_046837 | ANKRD44-IT1 | ANKRD44 intronic transcript 1 | lncRNA | . | 41477 | . | . | . | 0.069190 | 0.010718 | 0.009591 | 0.022148 | 0.161008 | 0.024776 | 0.021523 | 0.050640 | -1.598769 | 1.437178 | 2.472395132 | -0.2738078 | 0.784232327 | NA | 0.820514 | 0.148375 | 0.540170 | 0.532159 | 0.533298 | 0.532889 | 3 | 1 | 1 | 1 |
| 101927550 | NR_110102 | LOC10192755 | uncharacterized LOC101927550 | lncRNA | . | . | . | . | . | 0.213702 | 0.116710 | 0.139251 | 0.169599 | 0.497292 | 0.269779 | 0.312474 | 0.387768 | -1.101908 | 17.376342 | 0.753290413 | -0.18585595 | 0.852557727 | NA | 4.186752 | 4.048019 | 4.146663 | 4.100709 | 4.110048 | 4.123314 | 27 | 14 | 13 | 17 |
| 101927556 | XR_00175222 | LOC10192755 | uncharacterized LOC101927556, transcript variant X4 | lncRNA | . | . | . | . | . | 0.042676 | 0.032667 | 0.152532 | 0.096864 | 0.099309 | 0.075512 | 0.342277 | 0.221470 | 2.682315 | 11.543540 | 0.943561108 | 1.508623641 | 0.131394989 | NA | 2.644780 | 4.073227 | 3.474449 | 3.468294 | 3.543996 | 3.492026 | 8 | 6 | 20 | 10 |
| 101927559 | NR_125739 | CEBPB-AS1 | CEBPB antisense RNA 1 | lncRNA | . | 51226 | . | . | . | 0.363132 | 0.360815 | 0.458819 | 0.490103 | 0.845023 | 0.834039 | 1.029576 | 1.120563 | 1.281210 | 39.360743 | 0.493980851 | 0.723727327 | 0.469233153 | 0.998865404 | 5.109094 | 5.466231 | 5.274942 | 5.273385 | 5.305472 | 5.318155 | 40 | 37 | 37 | 43 |
| 101927572 | NM_00129000 | LOC10192757 | uncharacterized LOC101927572 | protein_coding | NP_001276985.1;NP_001334836.1 | . | . | . | . | 0.511619 | 0.952878 | 1.104198 | 1.216104 | 1.190559 | 2.202620 | 2.477788 | 2.780477 | 1.494558 | 20.301094 | 0.70205468 | 0.825745829 | 0.408948289 | NA | 4.027016 | 4.602870 | 4.290664 | 4.335559 | 4.342152 | 4.353111 | 13 | 23 | 20 | 24 |
| 101927587 | NR_119375 | LINC01725 | long intergenic non-protein coding RNA 1725 | lncRNA | . | 52513 | . | . | . | 0.071466 | 0.118950 | 0.096924 | 0.106975 | 0.166303 | 0.274958 | 0.217495 | 0.244585 | 1.059717 | 10.220025 | 0.955383348 | 0.087586512 | 0.930205323 | NA | 3.312060 | 3.393446 | 3.339275 | 3.358704 | 3.350985 | 3.352386 | 9 | 13 | 9 | 10 |
| 101927597 | NR_110175,N | NCK1-DT | NCK1 divergent transcript | lncRNA | . | 49645 | . | . | . | 2.505910 | 1.539222 | 1.989892 | 2.656288 | 5.831355 | 3.557981 | 4.465260 | 6.073288 | 1.111752 | 69.531598 | 0.407510941 | 0.375044139 | 0.707627641 | 0.998865404 | 6.040868 | 6.194251 | 6.155679 | 6.043899 | 6.092871 | 6.159157 | 94 | 54 | 55 | 78 |
| 101927599 | NR_110703,N | ZNF529-AS1 | ZNF529 antisense RNA 1 | lncRNA | . | 51275 | . | . | . | 2.057585 | 2.174299 | 1.923475 | 2.244550 | 4.788084 | 5.025992 | 4.316223 | 5.131898 | -1.027073 | 84.521127 | 0.346250039 | -0.11130409 | 0.911375215 | 0.998865404 | 6.420827 | 6.381384 | 6.400501 | 6.409874 | 6.381419 | 6.414132 | 97 | 94 | 68 | 83 |
| 101927603 | XR_00174652 | LOC10192760 | uncharacterized LOC101927603 | lncRNA | . | . | . | . | . | 0.015116 | 0.013958 | 0.117674 | 0.046215 | 0.035176 | 0.032265 | 0.264057 | 0.105665 | 4.690549 | 10.232553 | 1.051127339 | 2.121300365 | 0.033896534 | NA | 1.844134 | 4.076793 | 3.270667 | 3.272290 | 3.362505 | 3.300267 | 4 | 4 | 21 | 9 |
| 101927610 | XR_00174529 | LOC10192761 | uncharacterized LOC101927610, transcript variant X1 | lncRNA | . | . | . | . | . | 0.108080 | 0.143186 | 0.246128 | 0.131919 | 0.251507 | 0.330980 | 0.552304 | 0.301616 | 1.459269 | 22.163228 | 0.679674452 | 0.802216413 | 0.422427784 | NA | 4.170740 | 4.717884 | 4.425269 | 4.446477 | 4.503037 | 4.437232 | 18 | 22 | 30 | 17 |
| 101927612 | NR_108047,N | RNF139-AS1 | RNF139 antisense RNA 1 (head to head) | lncRNA | . | 48940 | . | . | . | 0.491326 | 0.560521 | 0.437314 | 0.839163 | 1.143336 | 1.295671 | 0.981320 | 1.918646 | 1.080072 | 42.554372 | 0.480404084 | 0.231320986 | 0.817065445 | 0.998865404 | 5.355528 | 5.464871 | 5.384801 | 5.420066 | 5.404735 | 5.425383 | 42 | 49 | 36 | 44 |
| 101927613 | XR_00174143 | LOC10192761 | uncharacterized LOC101927613 | lncRNA | . | . | . | . | . | 0.049015 | 0.114665 | 0.066903 | 0.067506 | 0.114060 | 0.265053 | 0.150129 | 0.154344 | -1.117001 | 6.439883 | 1.206684569 | -0.13228822 | 0.894756327 | NA | 2.766559 | 2.602870 | 2.675786 | 2.698708 | 2.682683 | 2.685524 | 5 | 10 | 5 | 6 |
| 101927621 | NR_110707,N | LINC01534 | long intergenic non-protein coding RNA 1534 | lncRNA | . | 51281 | . | . | . | 0.843870 | 0.839405 | 0.974983 | 1.361529 | 1.963721 | 1.940322 | 2.187834 | 3.112972 | 1.348384 | 27.353988 | 0.59216113 | 0.728233773 | 0.466470495 | 0.998865404 | 4.542770 | 4.972679 | 4.750628 | 4.749673 | 4.768246 | 4.798474 | 27 | 25 | 24 | 33 |
| 101927631 | NR_110007,N | ADNP-AS1 | ADNP antisense RNA 1 | lncRNA | . | 51227 | . | . | . | 0.309290 | 0.491042 | 0.672200 | 0.914465 | 0.719730 | 1.135066 | 1.508397 | 2.090815 | 1.853941 | 19.921836 | 0.703797204 | 1.265415046 | 0.205722585 | NA | 3.805868 | 4.692544 | 4.260631 | 4.285236 | 4.307244 | 4.340234 | 13 | 18 | 19 | 28 |
| 101927632 | XR_242320,X | PPP1R35-AS1 | PPP1R35 antisense RNA 1, transcript variant X2 | lncRNA | . | 41194 | . | . | . | 1.736150 | 1.126579 | 1.808934 | 1.593507 | 4.040092 | 2.604137 | 4.059198 | 3.643364 | 1.177875 | 36.893693 | 0.522910899 | 0.451677341 | 0.651501444 | 0.998865404 | 5.080921 | 5.319806 | 5.216289 | 5.159777 | 5.221749 | 5.202230 | 47 | 29 | 37 | 35 |
| 101927655 | NM_00128993 | ZASP | ZO-2 associated speckle protein | protein_coding | NP_001276862.1 | . | 617677 | . | . | 0.064466 | 0.033548 | 2.241061 | 0.047031 | 0.150015 | 0.077547 | 5.028877 | 0.107531 | 19.532895 | 22.922561 | 1.997440679 | 2.146663934 | 0.031820046 | NA | 1.156660 | 5.446776 | 4.214444 | 4.212058 | 4.560738 | 4.217059 | 3 | 2 | 74 | 2 |
| 101927673 | XR_247234 | LOC10192767 | uncharacterized LOC101927673 | lncRNA | . | . | . | . | . | 0.576640 | 0.731914 | 0.728311 | 0.544646 | 1.341866 | 1.691852 | 1.634308 | 1.245268 | -1.049797 | 42.187785 | 0.488438233 | -0.14353902 | 0.885864495 | 0.998865404 | 5.432996 | 5.363679 | 5.380556 | 5.419436 | 5.414555 | 5.369649 | 44 | 52 | 41 | 33 |
| 101927699 | NR_155739,N | LOC10192769 | uncharacterized LOC101927699 | lncRNA | . | . | . | . | . | 0.620444 | 0.189105 | 0.236117 | 0.192299 | 1.443798 | 0.437123 | 0.529839 | 0.439668 | -1.313952 | 10.901900 | 0.967305895 | -0.40722655 | 0.683841604 | NA | 3.627869 | 3.239024 | 3.480811 | 3.427727 | 3.445476 | 3.437479 | 21 | 7 | 9 | 8 |
| 101927740 | NR_109890 | LINC02202 | long intergenic non-protein coding RNA 2202 | lncRNA | . | 53068 | . | . | . | 0.119830 | 0.057869 | 0.024941 | 0.178942 | 0.278850 | 0.133767 | 0.055967 | 0.409130 | 1.089813 | 8.787543 | 1.13718468 | 0.109112632 | 0.913113156 | NA | 3.074695 | 3.193768 | 3.143400 | 3.117049 | 3.103556 | 3.163095 | 13 | 6 | 2 | 15 |
| 101927750 | NR_110273,N | ATP6V1B1-AS1 | ATP6V1B1 antisense RNA 1 | lncRNA | . | 51118 | . | . | . | 9.756619 | 8.306878 | 8.905615 | 8.751446 | 22.704051 | 19.201721 | 19.983942 | 20.009142 | -1.057065 | 256.798043 | 0.217900452 | -0.36743192 | 0.713296868 | 0.998865404 | 8.043614 | 7.964277 | 8.056810 | 7.977702 | 7.998801 | 7.981322 | 327 | 263 | 221 | 230 |
| 101927751 | NR_135827,N | LOC10192775 | uncharacterized LOC101927751 | lncRNA | . | . | . | . | . | 0.769000 | 0.870321 | 1.379774 | 0.873365 | 1.789497 | 2.011788 | 3.096172 | 1.996844 | 1.378789 | 113.213293 | 0.312574867 | 1.482529137 | 0.138199557 | 0.899796454 | 6.572054 | 7.036522 | 6.746863 | 6.766895 | 6.910937 | 6.811824 | 106 | 106 | 130 | 106 |
| 101927752 | NR_134302,N | LOC10192775 | uncharacterized LOC101927752 | lncRNA | . | . | . | . | . | 7.053228 | 7.723159 | 7.228002 | 7.869448 | 16.413151 | 17.852429 | 16.219428 | 17.992557 | 1.011038 | 297.423782 | 0.203527872 | 0.077811626 | 0.937977899 | 0.998865404 | 8.208766 | 8.223946 | 8.189486 | 8.236046 | 8.200268 | 8.237031 | 326 | 333 | 251 | 290 |
| 101927755 | NR_110815,N | RNFT1-DT | RNFT1 divergent transcript | lncRNA | . | 51346 | . | . | . | 1.250344 | 1.128717 | 1.151974 | 1.445235 | 2.909603 | 2.609080 | 2.584996 | 3.304359 | 1.066171 | 17.105788 | 0.73982909 | 0.124946725 | 0.900565727 | NA | 4.049783 | 4.141582 | 4.096992 | 4.089883 | 4.091639 | 4.104644 | 20 | 17 | 14 | 18 |
| 101927759 | NR_152848 | LOC10192775 | uncharacterized LOC101927759 | lncRNA | . | . | . | . | . | 0.829028 | 0.758871 | 1.013619 | 0.993683 | 1.929181 | 1.754165 | 2.274532 | 2.271939 | 1.236076 | 20.528343 | 0.675856885 | 0.452413695 | 0.650970983 | NA | 4.198253 | 4.504604 | 4.348974 | 4.342537 | 4.365255 | 4.362984 | 22 | 19 | 20 | 21 |
| 101927765 | NR_125961,N | COA6-AS1 | COA6 antisense RNA 1 | lncRNA | . | 40825 | . | . | . | 3.369348 | 3.699858 | 5.306918 | 4.206225 | 7.840610 | 8.552388 | 11.908569 | 9.617034 | 1.329281 | 77.416654 | 0.366004881 | 1.121967724 | 0.261876163 | 0.995760118 | 6.054220 | 6.465685 | 6.217019 | 6.238989 | 6.333152 | 6.267462 | 73 | 75 | 86 | 73 |
| 101927789 | NR_160547,N | LOC10192778 | FAU, ubiquitin like and ribosomal protein S30 fusion pseudogene | transcribed_ps | . | . | . | . | . | 1.028213 | 1.023297 | 1.496639 | 1.433600 | 2.392693 | 2.365397 | 3.358413 | 3.277757 | 1.134718 | 23.980181 | 0.627865716 | 0.290402868 | 0.77150805 | NA | 4.489863 | 4.671937 | 4.566947 | 4.581026 | 4.590403 | 4.582795 | 24 | 26 | 23 | 23 |
| 101927809 | NR_134605 | LOC10192780 | uncharacterized LOC101927809 | lncRNA | . | . | . | . | . | 0.885685 | 0.850599 | 1.076989 | 1.003376 | 2.061026 | 1.966197 | 2.416732 | 2.294100 | 1.236404 | 18.038651 | 0.719621588 | 0.425432407 | 0.670521486 | NA | 4.011454 | 4.318300 | 4.161572 | 4.158383 | 4.179347 | 4.173295 | 19 | 17 | 18 | 18 |
| 101927811 | NR_110119,N | LOC10192781 | uncharacterized LOC101927811 | lncRNA | . | . | . | . | . | 0.604732 | 0.554814 | 0.676071 | 0.424367 | 1.407235 | 1.282476 | 1.517085 | 0.970264 | 1.000915 | 16.151157 | 0.767445629 | 0.001718965 | 0.998628465 | NA | 4.011454 | 4.015675 | 4.012792 | 4.009481 | 4.025291 | 3.996869 | 19 | 17 | 17 | 12 |
| 101927824 | NR_160681,N | LOC10192782 | uncharacterized LOC101927824 | lncRNA | . | . | . | . | . | 0.462151 | 0.208390 | 0.577648 | 0.296276 | 1.075445 | 0.481703 | 1.296224 | 0.677399 | 1.256996 | 15.559388 | 0.815272961 | 0.404747914 | 0.685662819 | NA | 3.781608 | 4.118228 | 3.961039 | 3.920808 | 3.979730 | 3.932563 | 21 | 10 | 20 | 11 |
| 101927825 | XR_00174810 | LOC10192782 | uncharacterized LOC101927825, transcript variant X7 | lncRNA | . | . | . | . | . | 0.446912 | 0.576220 | 0.568316 | 0.654474 | 1.039984 | 1.331957 | 1.275285 | 1.496376 | 1.804149 | 43.713921 | 0.489759388 | 1.738238936 | 0.082168721 | 0.783811235 | 4.961318 | 5.814408 | 5.409755 | 5.352112 | 5.460976 | 5.494213 | 44 | 26 | 45 | 57 |
| 101927837 | XR_00175224 | LOC10192783 | uncharacterized LOC101927837, transcript variant X1 | lncRNA | . | . | . | . | . | 0.963103 | 1.235218 | 0.911820 | 0.632191 | 2.241181 | 2.855262 | 2.046098 | 1.445428 | -1.513629 | 48.169001 | 0.4579953 | -1.30571586 | 0.191649201 | 0.964394432 | 5.857901 | 5.260824 | 5.607440 | 5.643750 | 5.568761 | 5.536423 | 61 | 68 | 37 | 32 |
| 101927858 | XR_242051 | RNF32-AS1 | RNF32 antisense RNA 1 | lncRNA | . | 40798 | . | . | . | 0.138177 | 0.244547 | 0.202669 | 0.107291 | 0.321543 | 0.565282 | 0.454782 | 0.245309 | -1.213258 | 23.518628 | 0.67082766 | -0.41573478 | 0.677604088 | NA | 4.687671 | 4.410501 | 4.530991 | 4.587300 | 4.566442 | 4.514321 | 22 | 35 | 24 | 14 |
| 101927861 | XR_940252 | LINC01293 | long intergenic non-protein coding RNA 1293 | lncRNA | . | 50362 | . | . | . | 13.747653 | 15.135458 | 13.706513 | 11.536550 | 31.991354 | 34.986286 | 30.757019 | 26.376949 | -1.159542 | 1276.928247 | 0.14265046 | -1.49704896 | 0.134380519 | 0.891755273 | 10.421140 | 10.207912 | 10.336319 | 10.403845 | 10.323803 | 10.188746 | 1510 | 1544 | 1122 | 1008 |
| 101927862 | NR_109894 | CTB-178M22.2 | uncharacterized LOC101927862 | lncRNA | . | . | . | . | . | 0.670209 | 0.850079 | 1.118348 | 0.473643 | 1.559603 | 1.964997 | 2.509540 | 1.082928 | 1.066571 | 13.030961 | 0.868806296 | 0.107020202 | 0.914772953 | NA | 3.654832 | 3.751299 | 3.688595 | 3.700445 | 3.719036 | 3.676894 | 13 | 15 | 16 | 8 |
| 101927864 | XR_934153 | LOC10192786 | uncharacterized LOC101927864 | lncRNA | . | . | . | . | . | 0.200556 | 0.165215 | 0.171119 | 0.199942 | 0.466701 | 0.381902 | 0.383986 | 0.457143 | 1.008808 | 26.930460 | 0.595993083 | 0.021227934 | 0.983063831 | 0.998865404 | 4.744764 | 4.757542 | 4.762995 | 4.739463 | 4.744890 | 4.758885 | 34 | 26 | 22 | 27 |
| 101927868 | XR_246196 | GRK5-IT1 | GRK5 intronic transcript 1 | lncRNA | . | 49478 | . | . | . | 0.264708 | 0.306799 | 0.346830 | 0.220359 | 0.615987 | 0.709179 | 0.778277 | 0.503824 | -1.014983 | 17.400376 | 0.740527095 | -0.0289727 | 0.976886362 | NA | 4.130737 | 4.111291 | 4.114640 | 4.123601 | 4.132218 | 4.103590 | 19 | 20 | 18 | 13 |
| 101927886 | NR_131972 | RFX5-AS1 | RFX5 antisense RNA 1 | lncRNA | . | 40503 | . | . | . | 0.603824 | 0.528212 | 1.112274 | 0.689066 | 1.405123 | 1.220987 | 2.495911 | 1.575468 | 1.592310 | 10.449506 | 0.95343881 | 0.703895367 | 0.481497937 | NA | 3.008843 | 3.683676 | 3.363316 | 3.357580 | 3.395526 | 3.367758 | 10 | 8 | 14 | 9 |
| 101927888 | XR_241961,X | LOC10192788 | uncharacterized LOC101927888, transcript variant X2 | lncRNA | . | . | . | . | . | 0.194606 | 0.323175 | 0.199871 | 0.233182 | 0.452856 | 0.747032 | 0.448505 | 0.533143 | -1.181930 | 28.331819 | 0.596085699 | -0.40454637 | 0.68581099 | 0.998865404 | 4.941211 | 4.697185 | 4.802208 | 4.862176 | 4.806365 | 4.820541 | 27 | 41 | 21 | 26 |
| 101927895 | NR_135114,N | EMC1-AS1 | EMC1 antisense RNA 1 | lncRNA | . | 54050 | . | . | . | 0.165850 | 0.069930 | 0.008070 | 0.057371 | 0.385940 | 0.161646 | 0.018110 | 0.131171 | -1.619811 | 2.152642 | 2.042559916 | -0.3406635 | 0.733356919 | NA | 1.413393 | 0.715150 | 1.124741 | 1.117036 | 1.113993 | 1.118361 | 4 | 2 | 1 | 2 |
| 101927897 | XR_00175351 | LOC10192789 | uncharacterized LOC101927897 | lncRNA | . | . | . | . | . | 0.122076 | 0.063038 | 0.116243 | 0.093209 | 0.284076 | 0.145714 | 0.260846 | 0.213112 | 1.112039 | 25.811609 | 0.632727091 | 0.24213877 | 0.808672638 | NA | 4.609216 | 4.766401 | 4.711505 | 4.645843 | 4.701076 | 4.679726 | 37 | 18 | 26 | 23 |
| 101927919 | NR_110854 | LOC10192791 | uncharacterized LOC101927919 | lncRNA | . | . | . | . | . | 0.465822 | 0.360463 | 0.555169 | 0.567946 | 1.083987 | 0.833226 | 1.245783 | 1.298540 | 1.313517 | 25.331847 | 0.61339477 | 0.641406449 | 0.521258644 | NA | 4.452135 | 4.846728 | 4.652781 | 4.632066 | 4.670675 | 4.671152 | 28 | 21 | 25 | 27 |
| 101927943 | XR_00295820 | SETBP1-DT | SETBP1 divergent transcript | lncRNA | . | 54819 | . | . | . | 0.163517 | 0.246073 | 0.267299 | 0.154608 | 0.380511 | 0.568810 | 0.599811 | 0.353494 | 1.031230 | 13.740173 | 0.836229804 | 0.053055603 | 0.957687605 | NA | 3.757258 | 3.803035 | 3.764319 | 3.784441 | 3.790182 | 3.762152 | 13 | 17 | 15 | 10 |
| 101927957 | NR_126330,N | LINC01572 | long intergenic non-protein coding RNA 1572 | lncRNA | . | 51385 | . | . | . | 0.727017 | 0.621687 | 0.849080 | 0.673165 | 1.691800 | 1.437059 | 1.905308 | 1.539113 | -1.046041 | 46.045775 | 0.462955975 | -0.14027186 | 0.888445198 | 0.998865404 | 5.556147 | 5.492904 | 5.530736 | 5.525214 | 5.540548 | 5.499405 | 55 | 50 | 44 | 37 |
| 101927963 | XR_00174612 | LOC10192796 | uncharacterized LOC101927963, transcript variant X9 | lncRNA | . | . | . | . | . | 1.049591 | 1.297359 | 1.111164 | 1.541419 | 2.442441 | 2.998904 | 2.493420 | 3.524271 | 1.148880 | 37.644529 | 0.509946618 | 0.392644315 | 0.694582204 | 0.998865404 | 5.131937 | 5.330007 | 5.208714 | 5.232565 | 5.223581 | 5.258329 | 37 | 41 | 31 | 42 |
| 101927974 | NR_144536,N | LOC10192797 | uncharacterized LOC101927974 | lncRNA | . | . | . | . | . | 2.017442 | 1.716267 | 2.374661 | 2.369438 | 4.694672 | 3.967230 | 5.328668 | 5.417439 | 1.319756 | 54.458411 | 0.426065642 | 0.939458267 | 0.347495509 | 0.998865404 | 5.552567 | 5.953793 | 5.748728 | 5.716174 | 5.787367 | 5.787023 | 59 | 46 | 54 | 58 |
| 101927979 | XR_00175276 | LOC10192797 | uncharacterized LOC101927979 | lncRNA | . | . | . | . | . | 1.052423 | 1.292512 | 1.265841 | 1.439582 | 2.449033 | 2.987700 | 2.840512 | 3.291433 | 1.138259 | 136.970419 | 0.28485478 | 0.655872944 | 0.511905878 | 0.998865404 | 7.002072 | 7.187420 | 7.033988 | 7.105021 | 7.094063 | 7.140673 | 133 | 152 | 119 | 145 |
| 101927989 | NR_136504 | LOC10192798 | uncharacterized LOC101927989 | lncRNA | . | . | . | . | . | 0.365540 | 0.127121 | 0.300679 | 0.352945 | 0.850628 | 0.293846 | 0.674715 | 0.806966 | 1.300865 | 11.205489 | 0.940654082 | 0.403412019 | 0.686645147 | NA | 3.282109 | 3.664848 | 3.492855 | 3.454723 | 3.481826 | 3.492114 | 16 | 6 | 10 | 13 |
| 101927999 | XR_00175231 | LOC10192799 | putative uncharacterized protein FLJ44672 | transcribed_ps | . | . | . | . | . | 0.423814 | 0.492978 | 0.581048 | 0.439326 | 0.986232 | 1.139542 | 1.303856 | 1.004467 | 1.097888 | 33.483091 | 0.540883159 | 0.249094316 | 0.803287826 | 0.998865404 | 4.995859 | 5.131667 | 5.044822 | 5.064473 | 5.086915 | 5.046907 | 34 | 37 | 35 | 28 |
| 101928000 | NR_135657,N | ZNF232-AS1 | ZNF232 antisense RNA 1 | lncRNA | . | 40623 | . | . | . | 21.493692 | 20.844699 | 20.598436 | 21.546874 | 50.016706 | 48.183457 | 46.222293 | 49.264369 | -1.018460 | 312.556573 | 0.197693295 | -0.13348703 | 0.893808218 | 0.998865404 | 8.301168 | 8.274751 | 8.303443 | 8.282719 | 8.274528 | 8.291704 | 370 | 334 | 265 | 295 |
| 101928005 | XR_948815 | LOC10192800 | uncharacterized LOC101928005 | lncRNA | . | . | . | . | . | 0.286593 | 0.520936 | 0.335432 | 0.424443 | 0.666913 | 1.204167 | 0.752699 | 0.970439 | -1.044216 | 22.559785 | 0.6687604 | -0.09333787 | 0.925635146 | NA | 4.528454 | 4.462148 | 4.465845 | 4.522215 | 4.480170 | 4.500513 | 19 | 32 | 17 | 23 |
| 101928020 | NR_110291 | KANSL1L-AS1 | KANSL1L antisense RNA 1 | lncRNA | . | 41139 | . | . | . | 0.946208 | 1.056210 | 1.813818 | 1.094873 | 2.201866 | 2.441478 | 4.070155 | 2.503297 | 1.437907 | 59.165781 | 0.432508934 | 1.211467149 | 0.225716417 | 0.982008277 | 5.599889 | 6.125824 | 5.821203 | 5.840552 | 5.964541 | 5.846074 | 53 | 55 | 76 | 49 |
| 101928041 | gene-LOC1019 | LOC10192804 | . | pseudogene | . | . | . | . | . | 0.349472 | 0.314834 | 0.457243 | 0.075427 | 0.813237 | 0.727753 | 1.026039 | 0.172455 | -1.243414 | 31.148228 | 0.685597756 | -0.45844159 | 0.646635218 | 0.998865404 | 5.106655 | 4.799146 | 4.966483 | 4.949727 | 5.005185 | 4.849847 | 42 | 35 | 41 | 8 |
| 101928042 | NR_135833 | LOC10192804 | uncharacterized LOC101928042 | lncRNA | . | . | . | . | . | 0.107592 | 0.315604 | 0.068622 | 0.252654 | 0.250371 | 0.729533 | 0.153985 | 0.577663 | -1.063293 | 3.949770 | 1.534504604 | -0.05769897 | 0.953988412 | NA | 2.029593 | 1.932304 | 1.975490 | 1.989440 | 1.974599 | 1.988147 | 3 | 6 | 2 | 5 |
| 101928055 | XR_00175574 | LOC10192805 | uncharacterized LOC101928055, transcript variant X1 | lncRNA | . | . | . | . | . | 0.173731 | 0.343976 | 0.261480 | 0.116678 | 0.404281 | 0.795114 | 0.586752 | 0.266772 | -1.315238 | 10.353021 | 0.970441096 | -0.40736516 | 0.68373981 | NA | 3.555346 | 3.161867 | 3.362975 | 3.390758 | 3.375581 | 3.353255 | 10 | 16 | 10 | 6 |
| 101928062 | NR_120459,N | PRANCR | progenitor renewal associated non-coding RNA | lncRNA | . | 51126 | . | . | . | 0.904074 | 0.767393 | 1.164921 | 1.205041 | 2.103817 | 1.773863 | 2.614048 | 2.755183 | 1.480184 | 27.788499 | 0.584853121 | 0.967382003 | 0.333353103 | 0.998865404 | 4.486115 | 5.051647 | 4.765047 | 4.763917 | 4.804588 | 4.812571 | 26 | 24 | 28 | 32 |
| 101928068 | NR_111907 | LOC10192806 | uncharacterized LOC101928068 | lncRNA | . | . | . | . | . | 0.119958 | 0.125757 | 0.164852 | 0.116919 | 0.279147 | 0.290693 | 0.369924 | 0.267320 | 1.124644 | 20.500035 | 0.681839604 | 0.248545394 | 0.803712452 | NA | 4.269309 | 4.440712 | 4.347128 | 4.348758 | 4.372662 | 4.343324 | 22 | 21 | 22 | 17 |
| 101928069 | NR_120586 | LOC10192806 | uncharacterized LOC101928069 | lncRNA | . | . | . | . | . | 0.739564 | 0.744219 | 0.565364 | 0.708493 | 1.720997 | 1.720295 | 1.268660 | 1.619884 | -1.180446 | 18.628056 | 0.712501257 | -0.33590397 | 0.736943284 | NA | 4.334945 | 4.093800 | 4.228274 | 4.229976 | 4.207148 | 4.224766 | 23 | 22 | 13 | 18 |
| 101928077 | XR_927131 | LOC10192807 | uncharacterized LOC101928077 | lncRNA | . | . | . | . | . | 0.155936 | 0.228982 | 0.209442 | 0.122506 | 0.362870 | 0.529301 | 0.469982 | 0.280095 | -1.160204 | 9.657962 | 0.992704283 | -0.21595352 | 0.829023978 | NA | 3.373794 | 3.161867 | 3.265500 | 3.280833 | 3.277898 | 3.255604 | 10 | 13 | 10 | 6 |
| 101928079 | NR_104131,N | SLC44A3-AS1 | SLC44A3 antisense RNA 1 | lncRNA | . | 49057 | . | . | . | 0.695355 | 1.020809 | 0.652802 | 0.922344 | 1.618118 | 2.359647 | 1.464869 | 2.108829 | 1.033504 | 41.231015 | 0.498417776 | 0.095389284 | 0.924005628 | 0.998865404 | 5.343257 | 5.387716 | 5.329350 | 5.389307 | 5.345853 | 5.383841 | 38 | 52 | 32 | 44 |
| 101928085 | NR_121674,N | ERICH6-AS1 | ERICH6 antisense RNA 1 | lncRNA | . | 41205 | . | . | . | 0.590228 | 0.892355 | 0.802095 | 0.455697 | 1.373485 | 2.062719 | 1.799879 | 1.041898 | -1.214679 | 14.813645 | 0.808910345 | -0.34685525 | 0.728700066 | NA | 4.021847 | 3.742346 | 3.879653 | 3.908101 | 3.893095 | 3.869830 | 15 | 21 | 14 | 10 |
| 101928095 | NR_168401 | LOC10192809 | uncharacterized LOC101928095 | lncRNA | . | . | . | . | . | 0.203895 | 0.102123 | 0.309882 | 0.261959 | 0.474472 | 0.236063 | 0.695367 | 0.598938 | 1.698227 | 6.613134 | 1.185992633 | 0.644211187 | 0.519438497 | NA | 2.290730 | 3.058876 | 2.710664 | 2.699577 | 2.724456 | 2.716995 | 7 | 4 | 8 | 7 |
| 101928096 | XR_00174383 | LOC10192809 | uncharacterized LOC101928096 | lncRNA | . | . | . | . | . | 0.055621 | 0.030170 | 0.066850 | 0.049872 | 0.129431 | 0.069739 | 0.150009 | 0.114027 | 1.353628 | 31.401249 | 0.581331024 | 0.751433584 | 0.452391758 | 0.998865404 | 4.735321 | 5.176571 | 4.975891 | 4.907744 | 5.000306 | 4.960178 | 40 | 20 | 36 | 29 |
| 101928099 | NR_121566,N | PCAT7 | prostate cancer associated transcript 7 | lncRNA | . | 48824 | . | . | . | 1.570385 | 1.471362 | 1.491262 | 1.635462 | 3.654349 | 3.401118 | 3.346350 | 3.739291 | 1.022814 | 92.879338 | 0.330934192 | 0.098340433 | 0.921661973 | 0.998865404 | 6.521098 | 6.553294 | 6.539995 | 6.527568 | 6.527342 | 6.553180 | 108 | 97 | 78 | 92 |
| 101928100 | NR_120430 | KLRK1-AS1 | KLRK1 antisense RNA 1 | lncRNA | . | 54868 | . | . | . | 0.337258 | 0.290331 | 0.232521 | 0.371697 | 0.784813 | 0.671113 | 0.521770 | 0.849842 | -1.046367 | 27.527636 | 0.600931722 | -0.10881288 | 0.91335091 | 0.998865404 | 4.816198 | 4.748628 | 4.795341 | 4.776737 | 4.756035 | 4.805358 | 35 | 28 | 18 | 31 |
| 101928103 | NR_110292 | SNHG31 | small nucleolar RNA host gene 31 | lncRNA | . | 54196 | . | . | . | 0.140486 | 0.287068 | 0.102172 | 0.168201 | 0.326917 | 0.663571 | 0.229271 | 0.384572 | -1.551673 | 12.648212 | 0.888870117 | -0.71306756 | 0.475803955 | NA | 3.945312 | 3.306201 | 3.652331 | 3.696450 | 3.642888 | 3.658536 | 12 | 22 | 7 | 11 |
| 101928104 | NR_110825 | LINC01482 | long intergenic non-protein coding RNA 1482 | lncRNA | . | 51128 | . | . | . | 0.062803 | 0.232681 | 0.302649 | 0.187222 | 0.146145 | 0.537854 | 0.679137 | 0.428062 | 1.678269 | 16.492934 | 0.833297216 | 0.896407991 | 0.370034901 | NA | 3.624820 | 4.368050 | 3.967781 | 4.033206 | 4.056413 | 4.017928 | 6 | 21 | 22 | 15 |
| 101928111 | NR_135575,N | HDAC4-AS1 | HDAC4 antisense RNA 1 | lncRNA | . | 54064 | . | . | . | 1.282660 | 1.038639 | 2.281873 | 2.115042 | 2.984802 | 2.400860 | 5.120457 | 4.835793 | 1.813346 | 25.063264 | 0.645285263 | 1.330658764 | 0.183301317 | NA | 4.151937 | 5.015675 | 4.616697 | 4.574861 | 4.682167 | 4.629548 | 26 | 14 | 34 | 24 |
| 101928113 | NR_110569 | ST3GAL5-AS1 | ST3GAL5 antisense RNA 1 (head to head) | lncRNA | . | 51129 | . | . | . | 0.440211 | 0.316657 | 0.620151 | 0.619255 | 1.024389 | 0.731967 | 1.391602 | 1.415853 | 1.611802 | 17.483937 | 0.738790804 | 0.93216491 | 0.351251307 | NA | 3.741623 | 4.432387 | 4.108492 | 4.088434 | 4.134718 | 4.132941 | 18 | 12 | 19 | 20 |
| 101928118 | NR_125948,N | LOC10192811 | uncharacterized LOC101928118 | lncRNA | . | . | . | . | . | 0.092816 | 0.131971 | 0.037315 | 0.146665 | 0.215988 | 0.305056 | 0.083733 | 0.335331 | -1.239759 | 8.530044 | 1.087147048 | -0.28520505 | 0.775487049 | NA | 3.243129 | 2.924410 | 3.092098 | 3.107171 | 3.073292 | 3.109419 | 9 | 12 | 3 | 11 |
| 101928120 | NM_00132172 | LOC10192812 | uncharacterized LOC101928120 | protein_coding | NP_001308655.1 | . | . | . | . | 0.174243 | 0.197517 | 0.334035 | 0.354943 | 0.405471 | 0.456569 | 0.749565 | 0.811536 | 1.772660 | 9.952804 | 0.966685303 | 0.854379615 | 0.392894687 | NA | 2.844134 | 3.669578 | 3.287827 | 3.290200 | 3.313422 | 3.314329 | 8 | 8 | 11 | 12 |
| 101928123 | NR_135248 | LOC10192812 | uncharacterized LOC101928123 | lncRNA | . | . | . | . | . | 0.556606 | 1.000765 | 0.624064 | 1.224635 | 1.295246 | 2.313312 | 1.400382 | 2.799983 | 1.173809 | 45.640656 | 0.517255614 | 0.446970704 | 0.654896212 | 0.998865404 | 5.394834 | 5.620820 | 5.439034 | 5.530658 | 5.454097 | 5.569696 | 35 | 58 | 29 | 61 |
| 101928126 | NR_134561 | LOC10192812 | uncharacterized LOC101928126 | lncRNA | . | . | . | . | . | 0.832993 | 0.482600 | 0.154999 | 0.312580 | 1.938408 | 1.115552 | 0.347812 | 0.714678 | -2.736733 | 16.371708 | 0.816752346 | -1.77832956 | 0.075349745 | NA | 4.585071 | 3.127898 | 4.098554 | 4.043461 | 3.993007 | 4.018220 | 35 | 19 | 5 | 11 |
| 101928143 | NR_135251 | LOC10192814 | uncharacterized LOC101928143 | lncRNA | . | . | . | . | . | 0.282871 | 0.285287 | 0.075015 | 0.347829 | 0.658252 | 0.659454 | 0.168332 | 0.795269 | -1.279773 | 5.583232 | 1.309829524 | -0.27170576 | 0.785848274 | NA | 2.651489 | 2.287859 | 2.489483 | 2.491332 | 2.472709 | 2.495532 | 7 | 7 | 2 | 7 |
| 101928146 | XR_00174396 | LOC10192814 | uncharacterized LOC101928146, transcript variant X1 | lncRNA | . | . | . | . | . | 0.195207 | 0.390111 | 0.254333 | 0.276966 | 0.454255 | 0.901759 | 0.570716 | 0.633251 | -1.057424 | 9.680440 | 0.989464028 | -0.08141216 | 0.935114178 | NA | 3.316288 | 3.232645 | 3.260245 | 3.287851 | 3.270894 | 3.272611 | 8 | 14 | 8 | 9 |
| 101928153 | NR_110531,N | ITGA9-AS1 | ITGA9 antisense RNA 1 | lncRNA | . | 49668 | . | . | . | 0.709682 | 0.686400 | 1.000433 | 1.114807 | 1.651459 | 1.586644 | 2.244942 | 2.548874 | 1.542167 | 15.935144 | 0.766973848 | 0.814837173 | 0.41516552 | NA | 3.648138 | 4.273004 | 3.971498 | 3.967293 | 3.994876 | 4.002955 | 15 | 13 | 16 | 19 |
| 101928156 | XR_923872 | LOC10192815 | uncharacterized LOC101928156 | lncRNA | . | . | . | . | . | 0.612145 | 0.537388 | 0.522435 | 0.381269 | 1.424486 | 1.242197 | 1.172330 | 0.871727 | -1.309115 | 10.273891 | 0.953492544 | -0.4075453 | 0.683607529 | NA | 3.540965 | 3.155137 | 3.372923 | 3.368126 | 3.364629 | 3.351966 | 14 | 12 | 9 | 7 |
| 101928161 | NR_110294 | NCKAP5-AS2 | NCKAP5 antisense RNA 2 | lncRNA | . | 54065 | . | . | . | 0.553058 | 0.870206 | 1.466808 | 1.158671 | 1.286988 | 2.011519 | 3.291474 | 2.649164 | 1.826712 | 17.220055 | 0.754033252 | 1.152799733 | 0.248992594 | NA | 3.607672 | 4.475715 | 4.052763 | 4.077060 | 4.120746 | 4.096014 | 11 | 16 | 22 | 18 |
| 101928195 | NR_135597 | LOC10192819 | methylenetetrahydrofolate dehydrogenase (NADP+ dependent) 1 | transcribed_ps | . | . | . | . | . | 0.011312 | 0.041151 | 0.071573 | 0.145501 | 0.026324 | 0.095124 | 0.160607 | 0.332671 | 2.426831 | 2.322137 | 1.993060038 | 0.641763705 | 0.521026619 | NA | 0.444631 | 1.715150 | 1.191101 | 1.195936 | 1.198067 | 1.207063 | 1 | 2 | 2 | 4 |
| 101928216 | XR_00173794 | LOC10192821 | uncharacterized LOC101928216, transcript variant X3 | lncRNA | . | . | . | . | . | 2.096933 | 2.617444 | 2.317772 | 3.499724 | 4.879648 | 6.050337 | 5.201015 | 8.001704 | 1.185408 | 129.083834 | 0.315741326 | 0.777166944 | 0.437060282 | 0.998865404 | 6.885723 | 7.128412 | 6.941226 | 7.003098 | 6.945219 | 7.114706 | 124 | 139 | 94 | 161 |
| 101928222 | NR_110235 | LOC10192822 | uncharacterized LOC101928222 | lncRNA | . | . | . | . | . | 0.504566 | 0.156585 | 0.371279 | 0.366715 | 1.174146 | 0.361954 | 0.833140 | 0.838450 | 1.077764 | 13.314825 | 0.881982461 | 0.12249808 | 0.902504567 | NA | 3.677674 | 3.790061 | 3.757019 | 3.700156 | 3.733156 | 3.733489 | 22 | 7 | 12 | 13 |
| 101928230 | XR_243478,X | LOC10192823 | uncharacterized LOC101928230, transcript variant X4 | lncRNA | . | . | . | . | . | 0.469420 | 0.661835 | 0.582530 | 0.374958 | 1.092360 | 1.529862 | 1.307179 | 0.857296 | -1.172336 | 29.683098 | 0.58812452 | -0.39002953 | 0.696514711 | 0.998865404 | 5.001122 | 4.773010 | 4.872922 | 4.921857 | 4.902425 | 4.854357 | 30 | 41 | 29 | 20 |
| 101928236 | XR_00174097 | LOC10192823 | uncharacterized LOC101928236, transcript variant X7 | lncRNA | . | . | . | . | . | 0.014273 | 0.025857 | 0.041891 | 0.070403 | 0.033213 | 0.059770 | 0.094003 | 0.160968 | 2.572604 | 14.493235 | 0.838452168 | 1.62588807 | 0.103973468 | NA | 3.024434 | 4.381963 | 3.792181 | 3.811720 | 3.837844 | 3.877927 | 7 | 11 | 14 | 24 |
| 101928238 | NR_134920,N | ZNF426-DT | ZNF426 divergent transcript | lncRNA | . | 53777 | . | . | . | 0.483488 | 0.244693 | 0.574531 | 0.561304 | 1.125097 | 0.565618 | 1.289232 | 1.283355 | 1.543357 | 14.123440 | 0.832557685 | 0.751985632 | 0.4520597 | NA | 3.471023 | 4.100822 | 3.812825 | 3.779513 | 3.823004 | 3.822504 | 17 | 8 | 15 | 16 |
| 101928243 | NR_110199,N | DARS-AS1 | DARS antisense RNA 1 | lncRNA | . | 40170 | . | . | . | 0.426822 | 0.578742 | 1.184418 | 1.312831 | 0.993232 | 1.337788 | 2.657800 | 3.001630 | 2.272050 | 24.276833 | 0.636251861 | 1.860890175 | 0.062759688 | NA | 3.892781 | 5.074662 | 4.527112 | 4.544159 | 4.608761 | 4.621363 | 15 | 18 | 28 | 33 |
| 101928274 | XR_00174934 | LOC10192827 | uncharacterized LOC101928274, transcript variant X7 | lncRNA | . | . | . | . | . | 1.761445 | 1.029782 | 2.407945 | 2.061608 | 4.098958 | 2.380387 | 5.403357 | 4.713624 | 1.606056 | 74.272576 | 0.395014415 | 1.73037323 | 0.083563614 | 0.785005898 | 5.830895 | 6.517652 | 6.186409 | 6.082130 | 6.278230 | 6.226296 | 82 | 46 | 87 | 78 |
| 101928295 | NR_110729 | LOC10192829 | uncharacterized LOC101928295 | lncRNA | . | . | . | . | . | 0.450675 | 0.299698 | 0.320734 | 0.402033 | 1.048738 | 0.692766 | 0.719718 | 0.919200 | -1.049119 | 37.840263 | 0.519809057 | -0.13308482 | 0.894126298 | 0.998865404 | 5.275723 | 5.207163 | 5.276955 | 5.212078 | 5.224192 | 5.251545 | 54 | 33 | 29 | 38 |
| 101928304 | NR_125855,N | LOC10192830 | uncharacterized LOC101928304 | lncRNA | . | . | . | . | . | 15.194299 | 12.988782 | 6.205883 | 11.619194 | 35.357760 | 30.024155 | 13.925820 | 26.565904 | -1.594354 | 203.946503 | 0.288066166 | -2.33617214 | 0.019482272 | 0.462516353 | 7.970552 | 7.295222 | 7.800761 | 7.713482 | 7.426010 | 7.655758 | 313 | 248 | 96 | 191 |
| 101928344 | XR_00295843 | LOC10192834 | uncharacterized LOC101928344, transcript variant X1 | lncRNA | . | . | . | . | . | 0.441045 | 0.491550 | 0.579252 | 0.506834 | 1.026330 | 1.136239 | 1.299822 | 1.158817 | 1.202785 | 37.095257 | 0.510043219 | 0.522266558 | 0.60148475 | 0.998865404 | 5.073577 | 5.340427 | 5.188149 | 5.200888 | 5.233118 | 5.209311 | 37 | 38 | 38 | 35 |
| 101928346 | XR_00295743 | KDM2B-DT | KDM2B divergent transcript, transcript variant X1 | lncRNA | . | 53287 | . | . | . | 1.157555 | 0.746514 | 1.619292 | 1.623038 | 2.693677 | 1.725601 | 3.633644 | 3.710883 | 1.544641 | 16.416848 | 0.770778812 | 0.813815011 | 0.415750934 | NA | 3.687500 | 4.318300 | 4.026980 | 3.994539 | 4.044289 | 4.038365 | 19 | 10 | 18 | 18 |
| 101928361 | XR_244865 | LOC10192836 | uncharacterized LOC101928361 | lncRNA | . | . | . | . | . | 0.087723 | 0.065430 | 0.047184 | 0.083882 | 0.204135 | 0.151245 | 0.105879 | 0.191786 | -1.153040 | 14.152434 | 0.830589281 | -0.24734516 | 0.804641112 | NA | 3.923304 | 3.715150 | 3.840441 | 3.820826 | 3.805088 | 3.838472 | 20 | 14 | 8 | 16 |
| 101928369 | XR_244325,X | LOC10192836 | uncharacterized LOC101928369, transcript variant X2 | lncRNA | . | . | . | . | . | 0.130485 | 0.127077 | 0.147725 | 0.332936 | 0.303644 | 0.293745 | 0.331492 | 0.761219 | 1.759228 | 9.854922 | 0.994537982 | 0.81941778 | 0.412548094 | NA | 2.838265 | 3.650563 | 3.279015 | 3.273016 | 3.280677 | 3.319739 | 9 | 7 | 7 | 16 |
| 101928378 | NR_110730 | PTOV1-AS2 | PTOV1 antisense RNA 2 | lncRNA | . | 51284 | . | . | . | 6.328393 | 7.414813 | 8.852185 | 6.381801 | 14.726431 | 17.139671 | 19.864046 | 14.591231 | 1.093175 | 248.906007 | 0.239519232 | 0.536593271 | 0.591548593 | 0.998865404 | 7.893294 | 8.022719 | 7.892733 | 7.967860 | 8.056557 | 7.887413 | 253 | 276 | 264 | 203 |
| 101928403 | NR_135540 | EIF2AK3-DT | EIF2AK3 divergent transcript | lncRNA | . | 54066 | . | . | . | 0.597780 | 1.144105 | 0.942719 | 1.415914 | 1.391059 | 2.644650 | 2.115434 | 3.237319 | 1.336750 | 42.785342 | 0.517762794 | 0.808729732 | 0.418670622 | 0.998865404 | 5.197227 | 5.611258 | 5.333603 | 5.426711 | 5.391958 | 5.464166 | 29 | 52 | 34 | 55 |
| 101928421 | XR_00174516 | LOC10192842 | uncharacterized LOC101928421, transcript variant X4 | lncRNA | . | . | . | . | . | 0.036394 | 0.062608 | 0.012157 | 0.022941 | 0.084691 | 0.144721 | 0.027280 | 0.052452 | -2.391934 | 11.173690 | 0.952817027 | -1.3204819 | 0.186674175 | NA | 3.980095 | 2.715150 | 3.498348 | 3.522391 | 3.461746 | 3.478483 | 15 | 20 | 4 | 8 |
| 101928433 | NR_136240 | ATXN1-AS1 | ATXN1 antisense RNA 1 | lncRNA | . | 40515 | . | . | . | 0.286745 | 0.303773 | 0.426362 | 0.228035 | 0.667267 | 0.702184 | 0.956745 | 0.521375 | 1.093926 | 46.553632 | 0.48490372 | 0.267095583 | 0.789395569 | 0.998865404 | 5.473017 | 5.605582 | 5.520237 | 5.528062 | 5.591598 | 5.485197 | 50 | 49 | 55 | 32 |
| 101928451 | XR_927951,X | LOC10192845 | uncharacterized LOC101928451, transcript variant X1 | lncRNA | . | . | . | . | . | 2.164949 | 2.737801 | 2.670419 | 2.009470 | 5.037927 | 6.328551 | 5.992343 | 4.594417 | -1.041358 | 260.729066 | 0.238728507 | -0.24490923 | 0.806526719 | 0.998865404 | 8.055028 | 7.997208 | 7.970059 | 8.089856 | 8.086376 | 7.929905 | 271 | 320 | 257 | 202 |
| 101928462 | XR_00175101 | LOC10192846 | uncharacterized LOC101928462, transcript variant X8 | lncRNA | . | . | . | . | . | 0.086785 | 0.066121 | 0.069983 | 0.070971 | 0.201950 | 0.152841 | 0.157040 | 0.162267 | -1.064029 | 10.842205 | 0.922842911 | -0.09702317 | 0.922707995 | NA | 3.482357 | 3.393446 | 3.446186 | 3.437230 | 3.438062 | 3.439400 | 14 | 11 | 9 | 10 |
| 101928464 | NR_134927,N | ZNF433-AS1 | ZNF433 and ZNF878 antisense RNA 1 | lncRNA | . | 53776 | . | . | . | 3.947792 | 3.014265 | 4.252936 | 3.420765 | 9.186674 | 6.967609 | 9.543467 | 7.821172 | 1.104783 | 79.426038 | 0.373135062 | 0.385283394 | 0.700027463 | 0.998865404 | 6.236393 | 6.382966 | 6.331188 | 6.253461 | 6.356394 | 6.280253 | 100 | 69 | 83 | 67 |
| 101928487 | XR_241416,X | LOC10192848 | uncharacterized LOC101928487, transcript variant X1 | lncRNA | . | . | . | . | . | 1.212036 | 0.867664 | 1.467332 | 1.028612 | 2.820457 | 2.005644 | 3.292649 | 2.351800 | 1.167758 | 30.483805 | 0.575838474 | 0.388548358 | 0.697610275 | 0.998865404 | 4.811713 | 5.039266 | 4.934214 | 4.893233 | 4.956823 | 4.908451 | 38 | 25 | 34 | 25 |
| 101928489 | NR_110889 | LOC10192848 | uncharacterized LOC101928489 | lncRNA | . | . | . | . | . | 1.604236 | 1.422231 | 1.146253 | 2.173656 | 3.733124 | 3.287550 | 2.572158 | 4.969806 | 1.085025 | 97.869180 | 0.364213388 | 0.323240935 | 0.746512782 | 0.998865404 | 6.553910 | 6.669347 | 6.611391 | 6.573882 | 6.521964 | 6.704770 | 115 | 95 | 62 | 124 |
| 101928492 | NR_126023 | OR2A1-AS1 | OR2A1 antisense RNA 1 | lncRNA | . | 49168 | . | . | . | 0.499668 | 0.577451 | 0.357576 | 0.467253 | 1.162749 | 1.334805 | 0.802391 | 1.068319 | -1.308696 | 14.651631 | 0.801126561 | -0.48447964 | 0.628045509 | NA | 4.054853 | 3.664848 | 3.881463 | 3.890067 | 3.863805 | 3.873719 | 18 | 19 | 10 | 13 |
| 101928504 | NR_135252 | LOC10192850 | uncharacterized LOC101928504 | lncRNA | . | . | . | . | . | 0.193581 | 0.431624 | 0.237798 | 0.410229 | 0.450471 | 0.997718 | 0.533613 | 0.937940 | 1.051732 | 20.898824 | 0.714026133 | 0.101911791 | 0.918826691 | NA | 4.351521 | 4.418403 | 4.343549 | 4.407775 | 4.358760 | 4.402201 | 15 | 30 | 14 | 25 |
| 101928513 | NR_110573 | CATIP-AS1 | CATIP antisense RNA 1 | lncRNA | . | 41080 | . | . | . | 0.874066 | 0.863548 | 1.062152 | 2.149817 | 2.033986 | 1.996130 | 2.383438 | 4.915302 | 1.798328 | 21.281537 | 0.694680635 | 1.218770556 | 0.22293129 | NA | 3.928838 | 4.772579 | 4.369341 | 4.366126 | 4.382521 | 4.455234 | 18 | 16 | 16 | 34 |
| 101928527 | NM_00130842 | PIGBOS1 | PIGB opposite strand 1 | protein_coding | NP_001295350.1;NP_001295351.1;NP_001295352.1;XP_024305583.1 | 50696 | 618809 | . | . | 5.494388 | 5.308384 | 4.976019 | 5.272415 | 12.785668 | 12.270567 | 11.166039 | 12.054751 | -1.075523 | 146.957247 | 0.269353969 | -0.38996613 | 0.696561595 | 0.998865404 | 7.250766 | 7.145832 | 7.224291 | 7.203099 | 7.182431 | 7.189133 | 180 | 160 | 122 | 134 |
| 101928530 | NR_110919 | LOC10192853 | uncharacterized LOC101928530 | lncRNA | . | . | . | . | . | 0.199546 | 0.195297 | 0.305491 | 0.868915 | 0.464352 | 0.451437 | 0.685513 | 1.986671 | 2.889525 | 6.965990 | 1.207336908 | 1.267941487 | 0.204818847 | NA | 1.844134 | 3.370388 | 2.759324 | 2.760684 | 2.770581 | 2.811034 | 4 | 4 | 5 | 14 |
| 101928545 | NR_120511,N | SEPTIN7-DT | SEPTIN7 divergent transcript | lncRNA | . | 51153 | . | . | . | 0.248550 | 0.711141 | 0.464211 | 0.385290 | 0.578386 | 1.643833 | 1.041676 | 0.880920 | -1.143656 | 11.933038 | 0.924282821 | -0.20951716 | 0.834044545 | NA | 3.671431 | 3.475715 | 3.547693 | 3.600022 | 3.573783 | 3.560560 | 8 | 20 | 11 | 9 |
| 101928578 | NR_110410 | RAP2C-AS1 | RAP2C antisense RNA 1 | lncRNA | . | 40957 | . | . | . | 0.058383 | 0.048330 | 0.110112 | 0.064778 | 0.135859 | 0.111717 | 0.247088 | 0.148107 | 1.603633 | 8.156098 | 1.073925027 | 0.634442994 | 0.525791758 | NA | 2.644780 | 3.330299 | 3.008614 | 3.002240 | 3.033653 | 3.011206 | 8 | 6 | 11 | 7 |
| 101928589 | XM_00526223 | LOC10192858 | uncharacterized LOC101928589 | protein_coding | XP_005262296.1 | . | . | . | . | 0.118749 | 0.008846 | 0.299547 | 0.027231 | 0.276334 | 0.020447 | 0.672176 | 0.062261 | 2.325920 | 7.318911 | 1.339804913 | 0.908939574 | 0.363382023 | NA | 2.128083 | 3.359867 | 2.832857 | 2.800254 | 2.878003 | 2.806907 | 9 | 1 | 16 | 2 |
| 101928605 | NR_126022 | ARHGEF35-AS1 | ARHGEF35 antisense RNA 1 | lncRNA | . | 41292 | . | . | . | 0.653366 | 0.550187 | 0.683487 | 0.757163 | 1.520409 | 1.271781 | 1.533726 | 1.731163 | 1.178209 | 20.941737 | 0.670702381 | 0.352758224 | 0.724269705 | NA | 4.264940 | 4.501955 | 4.385026 | 4.371005 | 4.389139 | 4.395894 | 24 | 19 | 19 | 22 |
| 101928607 | NR_109964 | TRPM2-AS | TRPM2 antisense RNA | lncRNA | . | 50758 | . | . | . | 3.697587 | 3.714776 | 4.449092 | 4.111384 | 8.604439 | 8.586871 | 9.983633 | 9.400189 | 1.138555 | 109.312185 | 0.307400764 | 0.608987998 | 0.542532395 | 0.998865404 | 6.675379 | 6.863137 | 6.748984 | 6.747559 | 6.805455 | 6.774568 | 118 | 110 | 106 | 104 |
| 101928649 | NR_109909 | CTC-338M12.4 | uncharacterized LOC101928649 | lncRNA | . | . | . | . | . | 7.661592 | 6.861142 | 7.937840 | 6.056980 | 17.828838 | 15.859837 | 17.812283 | 13.848564 | -1.051480 | 225.811702 | 0.239728852 | -0.30209837 | 0.762577072 | 0.998865404 | 7.853889 | 7.783198 | 7.856735 | 7.799180 | 7.866911 | 7.738049 | 282 | 235 | 218 | 178 |
| 101928650 | NR_125958 | GORAB-AS1 | GORAB antisense RNA 1 | lncRNA | . | 54051 | . | . | . | 1.731432 | 3.040891 | 3.282955 | 3.925571 | 4.029114 | 7.029156 | 7.366857 | 8.975352 | 1.502235 | 59.331166 | 0.435237024 | 1.348944533 | 0.177354783 | 0.950321455 | 5.569492 | 6.153284 | 5.780499 | 5.878834 | 5.896801 | 5.935398 | 40 | 65 | 57 | 72 |
| 101928659 | NR_135180 | LOC10192865 | uncharacterized LOC101928659 | lncRNA | . | . | . | . | . | 0.775233 | 0.298885 | 0.455676 | 1.639554 | 1.803998 | 0.690885 | 1.022524 | 3.748644 | 1.729463 | 10.267894 | 1.031234634 | 0.766386163 | 0.443446563 | NA | 2.912173 | 3.701358 | 3.346072 | 3.319401 | 3.325746 | 3.388242 | 12 | 5 | 5 | 19 |

| | | | | | | | | | | | | | | | | | | | | | | | | | | | | | | | | | |
|---|---|---|---|---|---|---|---|---|---|---|---|---|---|---|---|---|---|---|---|---|---|---|---|---|---|---|---|---|---|---|---|---|---|
| 101928689 | NR_109928 | LINC01424 | long intergenic non-protein coding RNA 1424 | lncRNA | . | 40558 | . | . | . | 0.715370 | 0.613515 | 0.655757 | 0.305741 | 1.664696 | 1.418168 | 1.471500 | 0.699040 | -1.369279 | 25.188885 | 0.643889803 | -0.70418326 | 0.481318654 | NA | 4.861447 | 4.413324 | 4.679143 | 4.660602 | 4.667643 | 4.606323 | 36 | 29 | 25 | 13 |
| 101928705 | XR_242929,XR | LOC10192870 | uncharacterized LOC101928705, transcript variant X1 | lncRNA | . | . | . | . | . | 0.075900 | 0.082108 | 0.070175 | 0.042658 | 0.176623 | 0.189796 | 0.157470 | 0.097533 | -1.410266 | 21.495353 | 0.674978287 | -0.73479058 | 0.462467043 | NA | 4.651489 | 4.158506 | 4.439744 | 4.446444 | 4.429020 | 4.395553 | 28 | 28 | 19 | 13 |
| 101928707 | XR_00175211 | CD2BP2-DT | CD2BP2 divergent transcript, transcript variant X4 | lncRNA | . | 53029 | . | . | . | 1.284850 | 1.929849 | 1.505700 | 1.748106 | 2.989896 | 4.460931 | 3.378748 | 3.996838 | -1.032071 | 61.185346 | 0.412347961 | -0.11044724 | 0.912054692 | 0.998865404 | 5.958757 | 5.911078 | 5.899560 | 5.972052 | 5.916342 | 5.942804 | 60 | 78 | 49 | 60 |
| 101928716 | XR_242166 | SUGCT-AS1 | SUGCT antisense RNA 1 | lncRNA | . | 40745 | . | . | . | 2.283910 | 1.890623 | 1.502921 | 1.683681 | 5.314753 | 4.370259 | 3.372511 | 3.849536 | -1.330635 | 173.771898 | 0.259185622 | -1.59003857 | 0.111826111 | 0.848562841 | 7.632416 | 7.220363 | 7.541455 | 7.450206 | 7.357994 | 7.396172 | 251 | 193 | 123 | 147 |
| 101928725 | NR_135474 | LOC10192872 | uncharacterized LOC101928725 | lncRNA | . | . | . | . | . | 0.165010 | 0.102057 | 0.091808 | 0.224173 | 0.383985 | 0.235910 | 0.206015 | 0.512545 | 1.228038 | 6.929068 | 1.176043161 | 0.251993288 | 0.801046252 | NA | 2.638039 | 2.932304 | 2.794510 | 2.779700 | 2.780157 | 2.806855 | 9 | 5 | 4 | 10 |
| 101928728 | NR_135804 | LOC10192872 | uncharacterized LOC101928728 | lncRNA | . | . | . | . | . | 0.279932 | 0.418160 | 0.194134 | 0.189794 | 0.651414 | 0.966596 | 0.435631 | 0.433941 | -1.791985 | 19.324091 | 0.715115238 | -1.17681566 | 0.239269091 | NA | 4.633141 | 3.790061 | 4.277696 | 4.318575 | 4.251964 | 4.251774 | 23 | 32 | 12 | 13 |
| 101928736 | NR_110940,NF | MIR762HG | MIR762 host gene | lncRNA | . | 51386 | . | . | . | 0.832991 | 0.515544 | 1.102133 | 0.751633 | 1.938404 | 1.191705 | 2.473155 | 1.718519 | 1.263648 | 37.278225 | 0.531137368 | 0.635606924 | 0.525032655 | 0.998865404 | 5.039590 | 5.380804 | 5.229935 | 5.158160 | 5.244791 | 5.212433 | 48 | 26 | 40 | 35 |
| 101928755 | gene-LOC1019 | LOC10192875 | . | pseudogene | . | . | . | . | . | 0.019019 | 0.030882 | 0.036264 | 0.009921 | 0.044259 | 0.071386 | 0.081375 | 0.022682 | -1.070164 | 1.739247 | 2.228774873 | -0.04389516 | 0.964987971 | NA | 0.844134 | 0.751299 | 0.796508 | 0.797062 | 0.799027 | 0.793436 | 2 | 2 | 2 | 1 |
| 101928762 | XR_00175211 | LOC10192876 | uncharacterized LOC101928762, transcript variant X1 | lncRNA | . | . | . | . | . | 1.137186 | 0.719329 | 1.548557 | 1.082891 | 2.646279 | 1.662762 | 3.474916 | 2.475902 | 1.399147 | 62.019437 | 0.429007179 | 1.129462156 | 0.258702925 | 0.995760118 | 5.689946 | 6.178231 | 5.946103 | 5.857976 | 6.017359 | 5.932987 | 73 | 43 | 74 | 56 |
| 101928764 | NM_00136545 | LOC10192876 | coiled-coil domain-containing protein 144B | protein_coding | NP_001352383.1;XP_016876377.1;XP_016876378.1;XP_016876379.1;XP_ | . | . | . | . | 0.226935 | 0.298417 | 0.339641 | 0.155017 | 0.528088 | 0.689805 | 0.762145 | 0.354429 | -1.285107 | 34.443851 | 0.550184062 | -0.65775922 | 0.510692862 | 0.998865404 | 5.274751 | 4.915262 | 5.101286 | 5.139636 | 5.116942 | 5.055574 | 39 | 47 | 33 | 21 |
| 101928768 | XR_241720 | LOC10192876 | uncharacterized LOC101928768 | lncRNA | . | . | . | . | . | 2.064448 | 1.784413 | 0.847657 | 1.479565 | 4.804056 | 4.124752 | 1.902117 | 3.382850 | -1.672898 | 53.551318 | 0.452127103 | -1.64190411 | 0.100609878 | 0.815978072 | 6.067946 | 5.322518 | 5.807990 | 5.771983 | 5.653648 | 5.728465 | 83 | 67 | 26 | 47 |
| 101928784 | NR_120339,NF | RORA-AS1 | RORA antisense RNA 1 | lncRNA | . | 51410 | . | . | . | 0.255552 | 0.321904 | 0.220519 | 0.463244 | 0.594678 | 0.744096 | 0.494838 | 1.059152 | 1.064027 | 14.351081 | 0.821151977 | 0.109036228 | 0.913173756 | NA | 3.799841 | 3.885075 | 3.835756 | 3.843862 | 3.824799 | 3.862321 | 15 | 16 | 9 | 18 |
| 101928788 | gene-ARL4AP5 | ARL4AP5 | . | pseudogene | . | 52374 | . | . | . | 0.078673 | 0.203527 | 0.271223 | 0.261615 | 0.183075 | 0.470462 | 0.608616 | 0.598151 | 1.732242 | 7.421881 | 1.131595913 | 0.700462418 | 0.483638568 | NA | 2.444631 | 3.232645 | 2.858556 | 2.877282 | 2.885043 | 2.886988 | 4 | 8 | 8 | 9 |
| 101928820 | NR_134633,NF | LINC02535 | long intergenic non-protein coding RNA 2535 | lncRNA | . | 53569 | . | . | . | 0.099645 | 0.085383 | 0.095669 | 0.066023 | 0.231877 | 0.197367 | 0.214678 | 0.150955 | -1.115889 | 7.637050 | 1.095650786 | -0.14438354 | 0.885197614 | NA | 3.008843 | 2.852980 | 2.939371 | 2.933467 | 2.936284 | 2.928997 | 10 | 8 | 7 | 6 |
| 101928824 | XR_00175561 | LOC10192882 | uncharacterized LOC101928824, transcript variant X2 | lncRNA | . | . | . | . | . | 1.855489 | 1.836352 | 2.569286 | 2.392024 | 4.317799 | 4.244809 | 5.765404 | 5.469079 | 1.235858 | 28.587402 | 0.576271274 | 0.530153964 | 0.596005186 | 0.998865404 | 4.676202 | 4.982216 | 4.817959 | 4.821369 | 4.850941 | 4.837489 | 29 | 28 | 29 | 28 |
| 101928837 | NR_120561,NF | LOC10192883 | uncharacterized LOC101928837 | lncRNA | . | . | . | . | . | 1.728747 | 1.024651 | 0.826020 | 1.245370 | 4.022866 | 2.368527 | 1.853565 | 2.847391 | -1.347573 | 13.540393 | 0.853931146 | -0.50397917 | 0.614276015 | NA | 3.958615 | 3.527725 | 3.792319 | 3.752926 | 3.745348 | 3.765141 | 23 | 12 | 8 | 13 |
| 101928847 | NR_120563 | LOC10192884 | uncharacterized LOC101928847 | lncRNA | . | . | . | . | . | 0.495814 | 0.737844 | 0.551121 | 0.505246 | 1.153779 | 1.705559 | 1.236699 | 1.155184 | -1.178303 | 35.022142 | 0.53547963 | -0.44205293 | 0.658450903 | 0.998865404 | 5.244239 | 5.006356 | 5.109734 | 5.169591 | 5.122339 | 5.112721 | 35 | 49 | 29 | 29 |
| 101928865 | NR_120564 | LINC02728 | long intergenic non-protein coding RNA 2728 | lncRNA | . | 54245 | . | . | . | 0.565642 | 0.434374 | 0.891600 | 0.441989 | 1.316272 | 1.004074 | 2.000724 | 1.010556 | 1.293788 | 30.273657 | 0.596824145 | 0.622630857 | 0.533527129 | 0.998865404 | 4.719612 | 5.095903 | 4.903404 | 4.876688 | 4.966431 | 4.874474 | 34 | 25 | 40 | 21 |
| 101928868 | XR_00174442 | LOC10192886 | uncharacterized LOC101928868 | lncRNA | . | . | . | . | . | 0.763217 | 0.708919 | 0.564285 | 0.571707 | 1.776038 | 1.638698 | 1.266239 | 1.307139 | -1.299348 | 24.160080 | 0.62720063 | -0.60234002 | 0.546947826 | NA | 4.771026 | 4.393446 | 4.617125 | 4.605578 | 4.584128 | 4.586463 | 33 | 28 | 18 | 20 |
| 101928891 | NR_110537 | LINC01637 | long intergenic non-protein coding RNA 1637 | lncRNA | . | 52424 | . | . | . | 0.196032 | 0.504747 | 0.164649 | 0.446837 | 0.456175 | 1.166745 | 0.369469 | 1.021639 | -1.082689 | 7.415762 | 1.155764421 | -0.09917095 | 0.921002538 | NA | 2.950762 | 2.827809 | 2.873669 | 2.905059 | 2.875841 | 2.897795 | 5 | 12 | 4 | 9 |
| 101928895 | XR_946815,XR | LINC01786 | long intergenic non-protein coding RNA 1786, transcript variant X | lncRNA | . | 52575 | . | . | . | 2.109444 | 1.184119 | 0.675701 | 1.509545 | 4.908763 | 2.737144 | 1.516254 | 3.451395 | -1.535165 | 56.140249 | 0.48106971 | -1.28545564 | 0.19863311 | 0.96621494 | 6.088293 | 5.467324 | 5.905494 | 5.784090 | 5.702394 | 5.815968 | 98 | 55 | 25 | 56 |
| 101928933 | XR_00173831 | LOC10192893 | uncharacterized LOC101928933, transcript variant X3 | lncRNA | . | . | . | . | . | 0.016237 | 0.043366 | 0.036768 | 0.106064 | 0.037783 | 0.100241 | 0.082507 | 0.242503 | 2.413261 | 5.409294 | 1.359992485 | 0.934552064 | 0.350019193 | NA | 1.671431 | 2.932304 | 2.397545 | 2.411553 | 2.411592 | 2.438658 | 2 | 5 | 4 | 10 |
| 101928953 | XR_00174613 | LOC10192895 | uncharacterized LOC101928953, transcript variant X1 | lncRNA | . | . | . | . | . | 1.996964 | 1.881842 | 2.588925 | 1.309042 | 4.647018 | 4.349960 | 5.809474 | 2.992975 | 1.008368 | 76.972628 | 0.410084868 | 0.029314971 | 0.976613387 | 0.998865404 | 6.258420 | 6.274085 | 6.265536 | 6.242759 | 6.341452 | 6.163079 | 92 | 79 | 91 | 47 |
| 101928963 | XR_244663 | LOC10192896 | uncharacterized LOC101928963 | lncRNA | . | . | . | . | . | 0.387098 | 0.627486 | 0.409658 | 0.726508 | 0.900792 | 1.450462 | 0.919262 | 1.661075 | 1.120004 | 11.946349 | 0.900621939 | 0.181545473 | 0.855939444 | NA | 3.497331 | 3.655341 | 3.559759 | 3.583546 | 3.563028 | 3.592233 | 10 | 15 | 8 | 15 |
| 101929004 | XR_942763,XR | LOC10192900 | uncharacterized LOC101929004, transcript variant X4 | lncRNA | . | . | . | . | . | 0.208744 | 0.123826 | 0.191029 | 0.170631 | 0.485755 | 0.286229 | 0.428664 | 0.390128 | 1.046069 | 11.388041 | 0.911495196 | 0.071287714 | 0.943168773 | NA | 3.474811 | 3.543272 | 3.519429 | 3.494160 | 3.513370 | 3.504862 | 16 | 9 | 11 | 10 |
| 101929007 | XR_00175404 | LOC10192900 | WAS/WASL-interacting protein family member 1, transcript varian | lncRNA | . | . | . | . | . | 0.058950 | 0.025797 | 0.031156 | 0.026569 | 0.137179 | 0.059631 | 0.069913 | 0.060746 | -1.401185 | 10.628982 | 0.965934602 | -0.50380941 | 0.614395315 | NA | 3.631267 | 3.148375 | 3.443995 | 3.399014 | 3.407522 | 3.404469 | 20 | 8 | 8 | 8 |
| 101929008 | XR_00174519 | GTF2IP13 | general transcription factor IIi pseudogene 13, transcript variant X | transcribed_ps | . | 51725 | . | . | . | 0.882335 | 1.205410 | 0.861621 | 1.252979 | 2.053230 | 2.786359 | 1.933452 | 2.864788 | -1.133894 | 45.411373 | 0.477739704 | -0.37946572 | 0.704342055 | 0.998865404 | 5.593897 | 5.410224 | 5.474433 | 5.551416 | 5.483775 | 5.500551 | 44 | 63 | 35 | 42 |
| 101929018 | XR_00174105 | LOC10192901 | uncharacterized LOC101929018 | lncRNA | . | . | . | . | . | 0.006105 | 0.030570 | 0.111772 | 0.000005 | 0.014207 | 0.070664 | 0.250812 | 0.000013 | 2.501663 | 2.384617 | 2.02142424 | 0.654433344 | 0.512832658 | NA | 0.444631 | 1.769039 | 1.214508 | 1.219375 | 1.237000 | 1.215813 | 1 | 2 | 5 | 1 |
| 101929054 | NR_135307 | LOC10192905 | uncharacterized LOC101929054 | lncRNA | . | . | . | . | . | 0.354737 | 0.234796 | 0.223108 | 0.601800 | 0.825488 | 0.542741 | 0.500648 | 1.375944 | 1.377609 | 12.687624 | 0.901724962 | 0.51253583 | 0.60827605 | NA | 3.417335 | 3.876918 | 3.659024 | 3.637786 | 3.637614 | 3.694189 | 15 | 9 | 7 | 20 |
| 101929066 | NR_125429,NF | LOC10192906 | uncharacterized LOC101929066 | lncRNA | . | . | . | . | . | 0.409960 | 0.488586 | 0.475020 | 1.017034 | 0.953992 | 1.129389 | 1.065932 | 2.325327 | 1.965173 | 24.591115 | 0.625565379 | 1.55804027 | 0.119223702 | NA | 4.052320 | 5.026416 | 4.566391 | 4.567693 | 4.627437 | 4.640212 | 19 | 18 | 27 | 32 |
| 101929072 | NR_148222,NF | MICA-AS1 | MICA antisense RNA 1 | lncRNA | . | 53631 | . | . | . | 2.096143 | 1.722287 | 2.992513 | 2.021917 | 4.877811 | 3.981144 | 6.715113 | 4.622873 | 1.285754 | 94.576627 | 0.350602408 | 1.034263031 | 0.301013215 | 0.998865404 | 6.369170 | 6.734578 | 6.542072 | 6.486057 | 6.648808 | 6.523948 | 105 | 80 | 111 | 80 |
| 101929074 | NR_126366,NF | PIK3CD-AS2 | PIK3CD antisense RNA 2 | lncRNA | . | 51334 | . | . | . | 20.399912 | 17.761550 | 17.891838 | 23.997794 | 47.471431 | 41.056616 | 40.148765 | 54.868103 | 1.044283 | 352.944550 | 0.209241817 | 0.298760124 | 0.765123078 | 0.998865404 | 8.432093 | 8.493842 | 8.483605 | 8.411071 | 8.386319 | 8.552186 | 423 | 349 | 268 | 387 |
| 101929086 | NR_108090 | LINC01447 | long intergenic non-protein coding RNA 1447 | lncRNA | . | 50783 | . | . | . | 2.665648 | 2.871176 | 2.572709 | 3.178884 | 6.203073 | 6.636852 | 5.773085 | 7.268140 | 1.027186 | 243.280029 | 0.225576027 | 0.171547878 | 0.863792987 | 0.998865404 | 7.907617 | 7.945088 | 7.901421 | 7.935552 | 7.881765 | 7.979352 | 267 | 268 | 193 | 254 |
| 101929099 | NR_110786 | LOC10192909 | uncharacterized LOC101929099 | lncRNA | . | . | . | . | . | 0.127845 | 0.145088 | 0.165295 | 0.156558 | 0.297502 | 0.335378 | 0.370916 | 0.357951 | 1.234670 | 9.026614 | 1.007130212 | 0.301971883 | 0.762673497 | NA | 3.014059 | 3.318300 | 3.163682 | 3.166181 | 3.175021 | 3.171849 | 9 | 9 | 9 | 9 |
| 101929125 | NR_109859 | LINC01730 | long intergenic non-protein coding RNA 1730 | lncRNA | . | 52518 | . | . | . | 1.093656 | 1.154222 | 1.284240 | 1.878535 | 2.544983 | 2.668036 | 2.881797 | 4.295046 | 1.377877 | 30.370762 | 0.568874918 | 0.812915648 | 0.416266422 | 0.998865404 | 4.676202 | 5.136466 | 4.895035 | 4.898544 | 4.910518 | 4.960624 | 29 | 28 | 25 | 39 |
| 101929141 | NR_164151 | LOC10192914 | coiled-coil domain-containing protein 144A-like | transcribed_ps | . | . | . | . | . | 0.242015 | 0.369650 | 0.343121 | 0.224282 | 0.563180 | 0.854462 | 0.769954 | 0.512795 | -1.066981 | 12.681409 | 0.870116826 | -0.10749631 | 0.914395255 | NA | 3.710171 | 3.617632 | 3.651102 | 3.675136 | 3.671012 | 3.648021 | 12 | 17 | 13 | 9 |
| 101929145 | NR_132103,NF | PPP3CB-AS1 | PPP3CB antisense RNA 1 (head to head) | lncRNA | . | 50750 | . | . | . | 0.488442 | 0.716760 | 0.969380 | 0.668899 | 1.136625 | 1.656823 | 2.175260 | 1.529358 | 1.352026 | 67.874490 | 0.408106873 | 1.06619778 | 0.286334246 | 0.998865404 | 5.850708 | 6.286154 | 5.997631 | 6.070634 | 6.152476 | 6.054394 | 55 | 73 | 81 | 59 |
| 101929147 | NR_125974,NF | WARS2-AS1 | WARS2 antisense RNA 1 | lncRNA | . | 40612 | . | . | . | 4.837426 | 4.818313 | 5.963418 | 5.723274 | 11.256889 | 11.137746 | 13.381738 | 13.085589 | 1.169179 | 317.053794 | 0.196797651 | 1.145823527 | 0.251868178 | 0.994172589 | 8.191280 | 8.417063 | 8.262526 | 8.261439 | 8.363914 | 8.331130 | 337 | 315 | 306 | 311 |
| 101929185 | XR_00175271 | LOC10192918 | uncharacterized LOC101929185 | lncRNA | . | . | . | . | . | 0.025916 | 0.034911 | 0.042015 | 0.058560 | 0.060307 | 0.080697 | 0.094281 | 0.133891 | 1.529550 | 9.104899 | 1.013429435 | 0.604982497 | 0.545190627 | NA | 2.849979 | 3.459419 | 3.163093 | 3.173769 | 3.177673 | 3.193465 | 7 | 9 | 8 | 12 |
| 101929188 | NR_134628 | LOC10192918 | uncharacterized LOC101929188 | lncRNA | . | . | . | . | . | 0.285106 | 0.430124 | 0.103372 | 0.384600 | 0.663454 | 0.994251 | 0.231965 | 0.879343 | -1.466760 | 20.819958 | 0.740655472 | -0.74614059 | 0.455582493 | NA | 4.633141 | 4.072527 | 4.375800 | 4.416604 | 4.326337 | 4.399076 | 23 | 32 | 7 | 24 |
| 101929200 | XR_00174258 | LOC10192920 | uncharacterized LOC101929200, transcript variant X3 | lncRNA | . | . | . | . | . | 0.325084 | 0.623327 | 0.238001 | 0.289968 | 0.756485 | 1.440848 | 0.534067 | 0.662979 | -1.622849 | 31.162164 | 0.60637758 | -1.15196984 | 0.249333468 | 0.992945717 | 5.270513 | 4.568322 | 4.920269 | 5.041637 | 4.915918 | 4.930558 | 27 | 58 | 19 | 24 |
| 101929201 | XR_00295621 | LOC10192920 | uncharacterized LOC101929201 | lncRNA | . | . | . | . | . | 0.108356 | 0.060915 | 0.150651 | 0.106116 | 0.252150 | 0.140807 | 0.338056 | 0.242621 | 1.465361 | 12.622552 | 0.874467538 | 0.630390786 | 0.528438938 | NA | 3.353365 | 3.909275 | 3.649701 | 3.624133 | 3.667314 | 3.648606 | 15 | 8 | 15 | 12 |
| 101929212 | NR_104064,NF | SMIM2-AS1 | SMIM2 antisense RNA 1 | lncRNA | . | 42674 | . | . | . | 0.300315 | 0.412880 | 0.798228 | 0.752848 | 0.698846 | 0.954391 | 1.791201 | 1.721297 | 2.066600 | 21.386382 | 0.67370706 | 1.554472897 | 0.120071657 | NA | 3.802857 | 4.848815 | 4.356323 | 4.373048 | 4.432046 | 4.423946 | 14 | 17 | 26 | 26 |
| 101929215 | NR_109930 | THBS4-AS1 | THBS4 antisense RNA 1 | lncRNA | . | 40583 | . | . | . | 0.116442 | 0.053538 | 0.077929 | 0.027603 | 0.270964 | 0.123756 | 0.174870 | 0.063112 | -1.499490 | 6.288719 | 1.247437951 | -0.4685379 | 0.639399976 | NA | 2.912173 | 2.336262 | 2.676104 | 2.649255 | 2.659587 | 2.643092 | 12 | 5 | 6 | 3 |
| 101929230 | XR_00173932 | LOC10192923 | uncharacterized LOC101929230, transcript variant X8 | lncRNA | . | . | . | . | . | 0.193443 | 0.237862 | 0.093074 | 0.227785 | 0.450150 | 0.549830 | 0.208856 | 0.520806 | -1.240294 | 21.462833 | 0.687959875 | -0.45159912 | 0.651557803 | NA | 4.572938 | 4.257378 | 4.430690 | 4.440960 | 4.389538 | 4.442816 | 26 | 27 | 11 | 24 |
| 101929234 | NR_110304 | LOC10192923 | uncharacterized LOC101929234 | lncRNA | . | . | . | . | . | 0.316098 | 0.398242 | 0.485488 | 0.437181 | 0.735574 | 0.920553 | 1.089422 | 0.999563 | 1.303471 | 18.138186 | 0.721489176 | 0.529957259 | 0.596141565 | NA | 3.977427 | 4.359293 | 4.154852 | 4.171538 | 4.189077 | 4.178317 | 16 | 19 | 19 | 18 |
| 101929243 | NR_134625,NF | LOC10192924 | uncharacterized LOC101929243 | lncRNA | . | . | . | . | . | 2.447705 | 1.962391 | 2.973440 | 1.288098 | 5.695911 | 4.536154 | 6.672314 | 2.945082 | 1.163468 | 29.011646 | 0.588265511 | 0.371314713 | 0.710403141 | 0.998865404 | 4.743194 | 4.965380 | 4.857998 | 4.827704 | 4.886696 | 4.834079 | 35 | 25 | 33 | 23 |
| 101929248 | XR_00295668 | LOC10192924 | uncharacterized LOC101929248, transcript variant X1 | lncRNA | . | . | . | . | . | 0.047505 | 0.042154 | 0.031194 | 0.044326 | 0.110546 | 0.097441 | 0.069999 | 0.101346 | -1.146685 | 10.050108 | 0.960355551 | -0.20562147 | 0.837086606 | NA | 3.425187 | 3.226238 | 3.338226 | 3.333189 | 3.323939 | 3.335233 | 13 | 11 | 7 | 10 |
| 101929280 | XR_243327 | LOC10192928 | uncharacterized LOC101929280 | lncRNA | . | . | . | . | . | 0.023493 | 0.040244 | 0.066669 | 0.040361 | 0.054668 | 0.093027 | 0.149603 | 0.092280 | 1.645385 | 3.598434 | 1.59960535 | 0.449126741 | 0.653340231 | NA | 1.444631 | 2.161867 | 1.822271 | 1.831784 | 1.840816 | 1.829444 | 2 | 4 | 5 | 3 |
| 101929289 | XR_246494 | LOC10192928 | uncharacterized LOC101929289 | lncRNA | . | . | . | . | . | 0.673230 | 0.414757 | 0.353979 | 0.823542 | 1.566633 | 0.958728 | 0.794318 | 1.882931 | 1.087098 | 7.882468 | 1.098307151 | 0.109697745 | 0.912649089 | NA | 2.917750 | 3.037079 | 2.986453 | 2.967976 | 2.969413 | 2.990942 | 11 | 6 | 5 | 10 |
| 101929295 | NR_110303 | LOC10192929 | uncharacterized LOC101929295 | lncRNA | . | . | . | . | . | 0.089418 | 0.274386 | 0.292035 | 0.380735 | 0.208080 | 0.634256 | 0.655319 | 0.870505 | 1.847647 | 16.156692 | 0.809173266 | 1.094560796 | 0.273709099 | NA | 3.508460 | 4.387716 | 3.948011 | 3.996915 | 4.003916 | 4.025443 | 7 | 18 | 16 | 22 |
| 101929302 | NR_121586 | RFX3-AS1 | RFX3 antisense RNA 1 | lncRNA | . | 51197 | . | . | . | 0.253242 | 0.303557 | 0.131087 | 0.088493 | 0.589306 | 0.701686 | 0.294155 | 0.202329 | -2.656179 | 7.118424 | 1.178118278 | -1.19627424 | 0.231589559 | NA | 3.369732 | 1.963455 | 2.853968 | 2.860867 | 2.831799 | 2.825293 | 11 | 12 | 4 | 3 |
| 101929307 | XR_925771,XR | LOC10192930 | uncharacterized LOC101929307, transcript variant X1 | lncRNA | . | . | . | . | . | 1.519960 | 1.011003 | 1.310605 | 1.611616 | 3.537011 | 2.336980 | 2.940960 | 3.684771 | -1.001841 | 89.496489 | 0.35474933 | -0.00747809 | 0.9940334 | 0.999862634 | 6.484657 | 6.482860 | 6.535759 | 6.424383 | 6.463899 | 6.497999 | 123 | 78 | 73 | 89 |
| 101929351 | NR_121597,NF | CDC37L1-DT | CDC37L1 divergent transcript | lncRNA | . | 49735 | . | . | . | 0.966220 | 0.796970 | 0.898829 | 1.047544 | 2.248434 | 1.842231 | 2.016946 | 2.395084 | 1.110454 | 9.440008 | 0.986297278 | 0.153250128 | 0.878201028 | NA | 3.161369 | 3.312263 | 3.237996 | 3.232446 | 3.236436 | 3.242852 | 11 | 9 | 8 | 10 |
| 101929356 | XR_00173973 | LOC10192935 | uncharacterized LOC101929356, transcript variant X3 | lncRNA | . | . | . | . | . | 0.260557 | 0.170166 | 0.254929 | 0.247171 | 0.606327 | 0.393347 | 0.572051 | 0.565128 | 1.160219 | 52.170157 | 0.43747449 | 0.490079048 | 0.624077964 | 0.998865404 | 5.592935 | 5.809269 | 5.704328 | 5.668659 | 5.732002 | 5.697754 | 61 | 47 | 53 | 48 |
| 101929378 | NR_110249,NF | LINC01876 | long intergenic non-protein coding RNA 1876 | lncRNA | . | 52695 | . | . | . | 0.117443 | 0.064445 | 0.105377 | 0.018749 | 0.273296 | 0.148968 | 0.236463 | 0.042867 | -1.576598 | 5.104054 | 1.41755721 | -0.46334276 | 0.6431187 | NA | 2.638039 | 1.993948 | 2.366339 | 2.351425 | 2.361461 | 2.335364 | 9 | 5 | 6 | 1 |
| 101929400 | XM_02444736 | LOC10192940 | uncharacterized LOC101929400, transcript variant X1 | protein_coding | XP_024303128.1;XP_024303129.1;XP_024303130.1 | . | . | . | . | 0.271838 | 0.192909 | 0.247028 | 0.266413 | 0.632579 | 0.445918 | 0.554323 | 0.609120 | 1.117465 | 14.657328 | 0.798823703 | 0.20058134 | 0.84102595 | NA | 3.790753 | 3.951852 | 3.876272 | 3.860234 | 3.873110 | 3.877622 | 18 | 13 | 13 | 15 |
| 101929402 | NR_126027 | EXTL3-AS1 | EXTL3 antisense RNA 1 | lncRNA | . | 27985 | . | . | . | 0.298824 | 0.273043 | 0.425807 | 0.362939 | 0.695377 | 0.631151 | 0.955498 | 0.829817 | 1.309230 | 8.794927 | 1.020374198 | 0.380956715 | 0.703235367 | NA | 2.928838 | 3.318300 | 3.126217 | 3.124432 | 3.137485 | 3.134333 | 9 | 8 | 9 | 9 |
| 101929450 | NR_125814,NF | LINC02099 | long intergenic non-protein coding RNA 2099 | lncRNA | . | 52953 | . | . | . | 1.538792 | 1.299738 | 1.726504 | 1.976135 | 3.580832 | 3.004403 | 3.874225 | 4.518195 | 1.237176 | 134.254732 | 0.282849351 | 1.085564922 | 0.277671526 | 0.998865404 | 6.907423 | 7.213981 | 7.033548 | 7.015024 | 7.074339 | 7.128954 | 142 | 126 | 120 | 149 |
| 101929467 | NR_125775 | LURAP1L-AS1 | LURAP1L antisense RNA 1 | lncRNA | . | 49761 | . | . | . | 0.454499 | 0.617076 | 0.847190 | 0.476546 | 1.057638 | 1.426398 | 1.901070 | 1.089566 | 1.206068 | 17.899019 | 0.744433253 | 0.363109499 | 0.716523089 | NA | 4.019256 | 4.291534 | 4.134256 | 4.154985 | 4.182385 | 4.134995 | 16 | 20 | 22 | 13 |
| 101929475 | XR_00174741 | ITGB1-DT | ITGB1 divergent transcript, transcript variant X2 | lncRNA | . | 53718 | . | . | . | 0.223624 | 0.273264 | 0.103957 | 0.159504 | 0.520382 | 0.631662 | 0.233275 | 0.364687 | -1.876226 | 21.350120 | 0.685366306 | -1.32459659 | 0.185304987 | NA | 4.801350 | 3.889137 | 4.436797 | 4.458943 | 4.379755 | 4.407000 | 29 | 33 | 10 | 17 |
| 101929479 | NR_158179,NF | LOC10192947 | golgin A2 pseudogene | transcribed_ps | . | . | . | . | . | 3.627636 | 4.361145 | 3.888461 | 4.588591 | 8.441657 | 10.080982 | 8.725594 | 10.491267 | 1.047087 | 242.409676 | 0.22651656 | 0.293053427 | 0.769481314 | 0.998865404 | 7.888369 | 7.953503 | 7.864070 | 7.950325 | 7.899157 | 7.961171 | 249 | 278 | 202 | 247 |
| 101929506 | XR_00174751 | LOC10192950 | uncharacterized LOC101929506 | lncRNA | . | . | . | . | . | 0.005799 | 0.013945 | 0.016514 | 0.030669 | 0.013494 | 0.032234 | 0.037057 | 0.070120 | 2.190250 | 3.591761 | 1.609506468 | 0.702759375 | 0.482205719 | NA | 1.175403 | 2.300113 | 1.820133 | 1.825246 | 1.828451 | 1.841807 | 2 | 3 | 3 | 6 |
| 101929511 | NR_135639 | LINC01970 | long intergenic non-protein coding RNA 1970 | lncRNA | . | 52796 | . | . | . | 0.120783 | 0.058589 | 0.151329 | 0.230342 | 0.281067 | 0.135432 | 0.339579 | 0.526649 | 2.161918 | 8.430246 | 1.073970412 | 1.035700363 | 0.300341953 | NA | 2.413393 | 3.527725 | 3.052231 | 3.037181 | 3.062506 | 3.083039 | 8 | 4 | 8 | 13 |
| 101929523 | NR_134621 | LOC10192952 | uncharacterized LOC101929523 | lncRNA | . | . | . | . | . | 0.662565 | 0.690702 | 0.536020 | 0.637600 | 1.541816 | 1.596588 | 1.202813 | 1.457797 | -1.170578 | 20.367304 | 0.680578527 | -0.33386546 | 0.738481085 | NA | 4.457887 | 4.229445 | 4.356703 | 4.358889 | 4.338075 | 4.350442 | 25 | 24 | 15 | 19 |
| 101929530 | gene-FANCD2P | FANCD2P2 | FANCD2 pseudogene 2 | pseudogene | . | 44488 | . | . | . | 0.181992 | 0.241555 | 0.180176 | 0.217448 | 0.423503 | 0.558366 | 0.404311 | 0.497170 | -1.048617 | 7.912042 | 1.078291573 | -0.06351561 | 0.949355933 | NA | 3.019256 | 2.947964 | 2.979983 | 2.990700 | 2.980097 | 2.987242 | 8 | 10 | 6 | 8 |
| 101929536 | NR_135805 | LOC10192953 | uncharacterized LOC101929536 | lncRNA | . | . | . | . | . | 0.281374 | 0.480425 | 0.355775 | 0.225797 | 0.654769 | 1.110524 | 0.798348 | 0.516258 | -1.336684 | 13.039418 | 0.866862953 | -0.48295851 | 0.629125193 | NA | 3.898433 | 3.481107 | 3.696029 | 3.728247 | 3.707203 | 3.684381 | 13 | 20 | 12 | 8 |
| 101929540 | XR_00174743 | LOC10192954 | uncharacterized LOC101929540, transcript variant X4 | lncRNA | . | . | . | . | . | 0.995693 | 1.099510 | 1.007455 | 1.257468 | 2.317017 | 2.541568 | 2.260698 | 2.875052 | 1.111716 | 34.556950 | 0.528857377 | 0.288901926 | 0.77265643 | 0.998865404 | 5.033316 | 5.184531 | 5.099056 | 5.104185 | 5.099931 | 5.132052 | 37 | 36 | 28 | 38 |
| 101929541 | NR_125982,NF | LINC02608 | long intergenic non-protein coding RNA 2608 | lncRNA | . | 54052 | . | . | . | 0.614742 | 0.811036 | 0.483614 | 0.842837 | 1.430530 | 1.874746 | 1.085216 | 1.927047 | -1.106073 | 11.559742 | 0.908930332 | -0.16001939 | 0.872865796 | NA | 3.604218 | 3.453945 | 3.527155 | 3.542857 | 3.517339 | 3.542344 | 12 | 15 | 7 | 13 |
| 101929566 | XR_243337,XR | LOC10192956 | uncharacterized LOC101929566, transcript variant X2 | lncRNA | . | . | . | . | . | 0.371897 | 0.395782 | 0.288430 | 0.459805 | 0.865419 | 0.914867 | 0.647230 | 1.051288 | -1.060739 | 20.437265 | 0.681056674 | -0.12490868 | 0.900595843 | NA | 4.395729 | 4.309235 | 4.360204 | 4.354447 | 4.341583 | 4.362791 | 25 | 22 | 15 | 21 |
| 101929580 | NR_135738 | GTF2IP23 | general transcription factor IIi pseudogene 23 | transcribed_ps | . | 51735 | . | . | . | 0.874228 | 1.060835 | 1.434490 | 0.573071 | 2.034364 | 2.452169 | 3.218955 | 1.310259 | 1.033098 | 32.036780 | 0.593204105 | 0.079191497 | 0.936880308 | 0.998865404 | 4.976092 | 5.026777 | 4.976849 | 5.000072 | 5.043597 | 4.938884 | 33 | 37 | 40 | 18 |
| 101929609 | XR_246317 | LOC10192960 | uncharacterized LOC101929609 | lncRNA | . | . | . | . | . | 0.255708 | 0.273363 | 0.491692 | 0.303662 | 0.595042 | 0.631891 | 1.103342 | 0.694288 | 1.512913 | 18.059768 | 0.731211518 | 0.816903373 | 0.41398365 | NA | 3.844134 | 4.443477 | 4.142944 | 4.147370 | 4.195794 | 4.157395 | 16 | 16 | 23 | 16 |
| 101929626 | NR_125987 | LOC10192962 | uncharacterized LOC101929626 | lncRNA | . | . | . | . | . | 0.704755 | 0.587513 | 0.453906 | 0.634670 | 1.639993 | 1.358063 | 1.018551 | 1.451098 | -1.217073 | 7.328730 | 1.119650443 | -0.25312863 | 0.800168815 | NA | 3.008843 | 2.724273 | 2.884552 | 2.878636 | 2.871377 | 2.878859 | 10 | 8 | 5 | 7 |
| 101929638 | XR_00175547 | LOC10192963 | uncharacterized LOC101929638, transcript variant X12 | lncRNA | . | . | . | . | . | 0.831799 | 1.116738 | 1.102705 | 1.377984 | 1.935630 | 2.581390 | 2.474439 | 3.150597 | 1.206594 | 27.263622 | 0.592041524 | 0.457638298 | 0.647212324 | 0.998865404 | 4.628047 | 4.897226 | 4.744748 | 4.763159 | 4.766091 | 4.783712 | 26 | 29 | 24 | 30 |
| 101929655 | NR_120365 | TMC3-AS1 | TMC3 antisense RNA 1 | lncRNA | . | 51424 | . | . | . | 6.119011 | 6.821319 | 6.951899 | 8.221081 | 14.239190 | 15.767784 | 15.599859 | 18.796526 | 1.156644 | 236.192591 | 0.226380477 | 0.927396772 | 0.353720566 | 0.998865404 | 7.775639 | 7.984451 | 7.820844 | 7.867523 | 7.875231 | 7.953727 | 240 | 248 | 203 | 256 |
| 101929690 | NR_120367 | LINC01583 | long intergenic non-protein coding RNA 1583 | lncRNA | . | 51425 | . | . | . | 0.032097 | 0.077443 | 0.121348 | 0.037281 | 0.074691 | 0.179012 | 0.272302 | 0.085238 | 1.489097 | 5.660577 | 1.328353907 | 0.432442992 | 0.665419478 | NA | 2.184683 | 2.760196 | 2.466458 | 2.484970 | 2.501590 | 2.469926 | 3 | 7 | 9 | 3 |
| 101929693 | NR_110861 | RARA-AS1 | RARA antisense RNA 1 | lncRNA | . | 49577 | . | . | . | 0.456208 | 0.360617 | 0.079793 | 0.446463 | 1.061614 | 0.833582 | 0.179053 | 1.020784 | -1.529790 | 19.160389 | 0.808492983 | -0.75861328 | 0.448083927 | NA | 4.537347 | 3.916472 | 4.290893 | 4.266345 | 4.199496 | 4.287651 | 30 | 22 | 4 | 24 |
| 101929703 | XR_00295836 | LOC10192970 | uncharacterized LOC101929703 | transcribed_ps | . | . | . | . | . | 0.020028 | 0.020026 | 0.186034 | 0.053043 | 0.046607 | 0.046291 | 0.417455 | 0.121277 | 5.018843 | 5.414183 | 1.43717729 | 1.619392923 | 0.105362748 | NA | 0.844134 | 3.175234 | 2.370912 | 2.371751 | 2.425464 | 2.383478 | 2 | 2 | 12 | 4 |
| 101929704 | XR_241754 | LOC10192970 | uncharacterized LOC101929704 | lncRNA | . | . | . | . | . | 0.636776 | 0.342945 | 1.019458 | 0.779349 | 1.481803 | 0.792733 | 2.287634 | 1.781888 | 1.787464 | 31.620169 | 0.580206245 | 1.44416588 | 0.14869232 | 0.909660767 | 4.500856 | 5.343390 | 4.954695 | 4.895790 | 5.018150 | 4.977574 | 34 | 17 | 40 | 33 |
| 101929709 | NR_125822 | LOC10192970 | uncharacterized LOC101929709 | lncRNA | . | . | . | . | . | 1.431310 | 1.380161 | 1.996290 | 1.463338 | 3.330718 | 3.190304 | 4.479617 | 3.345750 | 1.209854 | 78.759029 | 0.367048805 | 0.748765336 | 0.453998656 | 0.998865404 | 6.154362 | 6.431131 | 6.273017 | 6.261386 | 6.359070 | 6.272339 | 84 | 75 | 87 | 68 |
| 101929723 | NR_110262,NF | LINC01914 | long intergenic non-protein coding RNA 1914 | lncRNA | . | 52733 | . | . | . | 8.782528 | 10.647066 | 5.636780 | 5.476254 | 20.437305 | 24.611172 | 12.648770 | 12.520803 | -1.767547 | 190.082549 | 0.250110876 | -3.28553568 | 0.001017886 | 0.068536035 | 7.923723 | 7.101697 | 7.625978 | 7.718633 | 7.435388 | 7.418489 | 254 | 286 | 122 | 126 |
| 101929745 | NR_134272 | LINC02119 | long intergenic non-protein coding RNA 2119 | lncRNA | . | 52975 | . | . | . | 1.902242 | 1.634271 | 1.566923 | 1.245742 | 4.426596 | 3.777691 | 3.516130 | 2.848241 | -1.269773 | 127.969412 | 0.297505549 | -1.15819961 | 0.246782586 | 0.992733415 | 7.160781 | 6.818245 | 7.069934 | 7.007246 | 6.993059 | 6.914724 | 178 | 142 | 110 | 93 |
| 101929748 | NR_136301 | LOC10192974 | uncharacterized LOC101929748 | lncRNA | . | . | . | . | . | 0.308272 | 0.594903 | 0.338819 | 0.461839 | 0.717361 | 1.375146 | 0.760300 | 1.055941 | -1.138505 | 36.527177 | 0.548867927 | -0.34095707 | 0.733135898 | 0.998865404 | 5.283416 | 5.092039 | 5.145973 | 5.241884 | 5.155681 | 5.197034 | 31 | 55 | 25 | 37 |
| 101929759 | XR_929025 | LOC10192975 | uncharacterized LOC101929759 | lncRNA | . | . | . | . | . | 0.430399 | 0.948973 | 0.835173 | 0.618082 | 1.001557 | 2.193593 | 1.874104 | 1.413170 | 1.052431 | 29.314951 | 0.610488437 | 0.120764383 | 0.903877662 | 0.998865404 | 4.836948 | 4.909275 | 4.816350 | 4.902850 | 4.884020 | 4.848918 | 21 | 42 | 30 | 24 |
| 101929762 | NR_125931 | LOC10192976 | uncharacterized LOC101929762 | lncRNA | . | . | . | . | . | 0.015281 | 0.134103 | 0.082888 | 0.103140 | 0.035560 | 0.309985 | 0.185999 | 0.235817 | 1.211526 | 5.556865 | 1.337483281 | 0.206975182 | 0.836029241 | NA | 2.333078 | 2.602870 | 2.445557 | 2.476759 | 2.464686 | 2.467626 | 2 | 9 | 5 | 6 |
| 101929767 | NR_110868 | LOC10192976 | uncharacterized LOC101929767 | lncRNA | . | . | . | . | . | 1.528570 | 1.388202 | 2.080742 | 1.912859 | 3.557044 | 3.208892 | 4.669124 | 4.373525 | 1.358093 | 38.012702 | 0.503717659 | 0.876646775 | 0.380678524 | 0.998865404 | 5.010149 | 5.452842 | 5.224305 | 5.211158 | 5.270823 | 5.254812 | 39 | 33 | 40 | 39 |
| 101929780 | NR_104217 | NBPF25P | NBPF member 25, pseudogene | transcribed_ps | . | 45046 | . | . | . | 3.041305 | 3.246816 | 4.181063 | 3.349455 | 7.077242 | 7.505160 | 9.382183 | 7.658132 | 1.180652 | 412.065685 | 0.191267256 | 1.252611126 | 0.210347264 | 0.972954982 | 8.561588 | 8.801879 | 8.614268 | 8.646945 | 8.799060 | 8.656832 | 423 | 419 | 433 | 370 |
| 101929796 | NR_157049 | NOTCH2NLR | notch 2 N-terminal like R (pseudogene) | transcribed_ps | . | 53925 | 618026 | . | . | 5.641540 | 6.295278 | 5.601416 | 5.038277 | 13.128095 | 14.551816 | 12.569414 | 11.519421 | -1.136732 | 199.654923 | 0.24024017 | -0.7696145 | 0.4415286 | 0.998865404 | 7.730722 | 7.546106 | 7.644614 | 7.693013 | 7.636291 | 7.586075 | 232 | 241 | 172 | 165 |
| 101929817 | gene-LOC1019 | LOC10192981 | . | pseudogene | . | . | . | . | . | 0.356797 | 0.409696 | 0.814765 | 0.513188 | 0.830280 | 0.947031 | 1.828309 | 1.173344 | 1.693308 | 33.286411 | 0.554835459 | 1.369495624 | 0.170844402 | 0.946389416 | 4.626345 | 5.387998 | 4.995716 | 5.006896 | 5.106612 | 5.032203 | 27 | 28 | 45 | 30 |
| 101929823 | XR_00175271 | LOC10192982 | uncharacterized LOC101929823, transcript variant X1 | lncRNA | . | . | . | . | . | 0.552423 | 0.629503 | 0.641020 | 0.565201 | 1.285509 | 1.455123 | 1.438429 | 1.292265 | -1.056627 | 33.679507 | 0.535617041 | -0.1483641 | 0.882055434 | 0.998865404 | 5.112745 | 5.033773 | 5.066833 | 5.086809 | 5.080733 | 5.058156 | 37 | 40 | 31 | 28 |
| 101929828 | XR_942193,XR | LOC10192982 | uncharacterized LOC101929828, transcript variant X1 | lncRNA | . | . | . | . | . | 0.071612 | 0.022686 | 0.104281 | 0.057419 | 0.166645 | 0.052440 | 0.234003 | 0.131282 | 1.761293 | 3.050103 | 1.748313667 | 0.467098531 | 0.640429385 | NA | 1.137672 | 1.963455 | 1.598347 | 1.586185 | 1.603199 | 1.596985 | 4 | 1 | 4 | 3 |
| 101929898 | XR_925675 | LOC10192989 | uncharacterized LOC101929898 | lncRNA | . | . | . | . | . | 0.635388 | 0.637232 | 0.237453 | 0.575030 | 1.478574 | 1.472991 | 0.532838 | 1.314737 | -1.515463 | 18.571959 | 0.741476786 | -0.80887077 | 0.418589481 | NA | 4.486115 | 3.881002 | 4.237355 | 4.235671 | 4.180784 | 4.226984 | 26 | 24 | 8 | 19 |
| 101929950 | NR_164156 | LOC10192995 | puromycin-sensitive aminopeptidase-like protein | transcribed_ps | . | . | . | . | . | 3.066968 | 2.987879 | 3.418667 | 3.178841 | 7.136961 | 6.906615 | 7.671390 | 7.268042 | 1.078142 | 76.375905 | 0.361732821 | 0.300074631 | 0.764120229 | 0.998865404 | 6.199399 | 6.308626 | 6.247817 | 6.239702 | 6.274143 | 6.251478 | 86 | 78 | 72 | 71 |
| 101929951 | XR_930805 | MACORIS | macrophage enriched lincRNA repressor of IFN-gamma signaling | lncRNA | . | 53963 | . | . | . | 0.640779 | 0.513045 | 0.471142 | 1.136523 | 1.491120 | 1.185927 | 1.057229 | 2.598524 | 1.334045 | 17.176979 | 0.776916013 | 0.535202181 | 0.592510058 | NA | 3.881411 | 4.293999 | 4.091414 | 4.075827 | 4.070581 | 4.138114 | 19 | 14 | 10 | 26 |
| 101929959 | XM_01151809 | LOC10192995 | ribosome biogenesis protein BMS1 homolog, transcript variant X6 | protein_coding | XP_011516396.1;XP_016870865.1;XP_016870866.1;XP_016870867.1;XP_ | . | . | . | . | 0.160476 | 0.059242 | 0.140797 | 0.145186 | 0.373435 | 0.136940 | 0.315947 | 0.331952 | 1.406731 | 8.519514 | 1.072197629 | 0.459193617 | 0.646095134 | NA | 2.820514 | 3.318300 | 3.091853 | 3.060674 | 3.090910 | 3.087784 | 12 | 4 | 9 | 9 |
| 101929964 | XR_427768 | LOC10192996 | uncharacterized LOC101929964 | lncRNA | . | . | . | . | . | 0.182799 | 0.059418 | 0.152672 | 0.046390 | 0.425381 | 0.137348 | 0.342591 | 0.106065 | -1.013826 | 3.953947 | 1.559807483 | -0.01269991 | 0.989867212 | NA | 1.987791 | 1.978782 | 1.989930 | 1.969938 | 1.987760 | 1.971580 | 7 | 2 | 5 | 2 |
| 101929977 | XR_427677,XR | LOC10192997 | uncharacterized LOC101929977, transcript variant X1 | lncRNA | . | . | . | . | . | 0.279637 | 0.096303 | 0.185504 | 0.473283 | 0.650727 | 0.222610 | 0.416266 | 1.082106 | 1.451539 | 27.282340 | 0.622336298 | 0.863814999 | 0.387689527 | 0.998865404 | 4.476701 | 5.013451 | 4.757499 | 4.717623 | 4.741808 | 4.815420 | 31 | 19 | 20 | 39 |
| 101930026 | XR_00174651 | LOC10193002 | uncharacterized LOC101930026, transcript variant X2 | lncRNA | . | . | . | . | . | 0.383653 | 0.572177 | 0.518882 | 0.503675 | 0.892777 | 1.322611 | 1.164357 | 1.151592 | 1.113129 | 49.968772 | 0.449182774 | 0.344227679 | 0.730675068 | 0.998865404 | 5.564169 | 5.717660 | 5.600003 | 5.656502 | 5.653448 | 5.643409 | 46 | 59 | 47 | 48 |
| 101930071 | NR_126041 | LOC10193007 | uncharacterized LOC101930071 | lncRNA | . | . | . | . | . | 3.404095 | 2.932922 | 1.597244 | 4.197131 | 7.921469 | 6.779580 | 3.584168 | 9.596241 | -1.102756 | 28.089409 | 0.631538174 | -0.22344352 | 0.823190338 | 0.998865404 | 4.882837 | 4.737408 | 4.826816 | 4.804936 | 4.757006 | 4.850453 | 37 | 29 | 13 | 36 |
| 101930085 | NR_136255 | LOC10193008 | uncharacterized LOC101930085 | lncRNA | . | . | . | . | . | 1.176663 | 1.292410 | 1.480469 | 1.192668 | 2.738143 | 2.987464 | 3.322129 | 2.726893 | 1.067224 | 168.963247 | 0.262134677 | 0.358070556 | 0.720290513 | 0.998865404 | 7.352443 | 7.447134 | 7.365682 | 7.401717 | 7.457158 | 7.362423 | 180 | 184 | 169 | 145 |
| 101930090 | gene-LOC1019 | LOC10193009 | contactin associated protein family member 3B pseudogene | pseudogene | . | . | . | . | . | 0.406045 | 0.498330 | 0.355793 | 0.346698 | 0.944883 | 1.151912 | 0.798390 | 0.792684 | -1.283562 | 19.596455 | 0.695399758 | -0.51790752 | 0.604522792 | NA | 4.461710 | 4.100822 | 4.296776 | 4.314363 | 4.285489 | 4.284518 | 23 | 26 | 15 | 16 |
| 101930100 | XR_00175494 | LOC10193010 | uncharacterized LOC101930100, transcript variant X1 | lncRNA | . | . | . | . | . | 4.205128 | 6.719722 | 5.897362 | 4.414055 | 9.785508 | 15.532937 | 13.233510 | 10.092211 | -1.029429 | 184.110487 | 0.267602532 | -0.1563674 | 0.875743433 | 0.998865404 | 7.544822 | 7.503741 | 7.475121 | 7.574334 | 7.578979 | 7.443266 | 191 | 224 | 183 | 143 |
| 101930105 | NR_146580,NF | FAM239A | family with sequence similarity 239 member A | transcribed_ps | . | 53414 | . | . | . | 0.361891 | 0.372642 | 0.281703 | 0.295365 | 0.842135 | 0.861379 | 0.632133 | 0.675317 | -1.291441 | 51.316951 | 0.43658304 | -0.84515707 | 0.398023149 | 0.998865404 | 5.854271 | 5.484875 | 5.705474 | 5.709086 | 5.655621 | 5.663155 | 66 | 63 | 38 | 43 |
| 101930112 | XR_933895 | LOC10193011 | uncharacterized LOC101930112 | lncRNA | . | . | . | . | . | 0.552928 | 0.584186 | 0.636577 | 0.419211 | 1.286686 | 1.350372 | 1.428459 | 0.958477 | -1.098970 | 32.155718 | 0.554941317 | -0.24534481 | 0.806189458 | 0.998865404 | 5.072328 | 4.938580 | 5.009076 | 5.014242 | 5.023389 | 4.975093 | 38 | 37 | 32 | 23 |
| 101930129 | XR_00174808 | LOC10193012 | uncharacterized LOC101930129, transcript variant X1 | lncRNA | . | . | . | . | . | 0.117237 | 0.077496 | 0.168806 | 0.070895 | 0.272814 | 0.179136 | 0.378796 | 0.162093 | 1.187007 | 8.793275 | 1.05683662 | 0.234026807 | 0.814964168 | NA | 3.003609 | 3.257993 | 3.130440 | 3.116266 | 3.148339 | 3.111475 | 11 | 7 | 12 | 5 |
| 101930275 | XR_428319,XR | LOC10193027 | uncharacterized LOC101930275, transcript variant X2 | lncRNA | . | . | . | . | . | 0.040127 | 0.037904 | 0.071010 | 0.056780 | 0.093376 | 0.087617 | 0.159346 | 0.129820 | 1.430357 | 7.111909 | 1.141777763 | 0.452255867 | 0.651084665 | NA | 2.548173 | 3.066070 | 2.810096 | 2.816189 | 2.833142 | 2.815895 | 6 | 7 | 9 | 6 |
| 101930307 | XM_01701541 | LOC10193030 | uncharacterized LOC101930307, transcript variant X4 | protein_coding | XP_016870904.1;XP_016870906.1;XP_016870907.1;XP_016870908.1;XP_ | . | . | . | . | 1.581245 | 1.642742 | 1.855405 | 1.338946 | 3.679621 | 3.797273 | 4.163475 | 3.061340 | 1.001324 | 132.466462 | 0.300036814 | 0.006362403 | 0.994923571 | 0.999862634 | 7.047593 | 7.051371 | 7.041811 | 7.048255 | 7.110418 | 6.976244 | 151 | 144 | 137 | 101 |
| 101930370 | NR_134241,NF | LOC10193037 | uncharacterized LOC101930370 | lncRNA | . | . | . | . | . | 0.883791 | 0.733091 | 0.926289 | 0.967025 | 2.056618 | 1.694570 | 2.078566 | 2.210986 | 1.152019 | 56.772488 | 0.417744674 | 0.488729871 | 0.625032953 | 0.998865404 | 5.721207 | 5.925786 | 5.826577 | 5.792704 | 5.831064 | 5.846172 | 66 | 52 | 51 | 59 |
| 101930420 | XM_01700760 | LOC10193042 | DNA primase large subunit-like | protein_coding | XP_016863098.1 | . | . | . | . | 0.333391 | 0.185811 | 0.137522 | 0.389077 | 0.775814 | 0.429511 | 0.308596 | 0.889579 | -1.027246 | 12.700258 | 0.903901798 | -0.0429054 | 0.965776945 | NA | 3.687500 | 3.645770 | 3.683823 | 3.651265 | 3.641930 | 3.689894 | 19 | 10 | 6 | 17 |
| 101930452 | NR_120479 | LINC02367 | long intergenic non-protein coding RNA 2367 | lncRNA | . | 53290 | . | . | . | 1.159481 | 1.100378 | 2.215243 | 1.064600 | 2.698161 | 2.543572 | 4.970942 | 2.434081 | 1.429995 | 189.541116 | 0.314036076 | 1.643154023 | 0.10035107 | 0.815618258 | 7.284112 | 7.802333 | 7.468356 | 7.444147 | 7.771132 | 7.427733 | 185 | 163 | 263 | 135 |
| 101954266 | NR_104075 | RNVU1-14 | RNA, variant U1 small nuclear 14 | snRNA | . | 48319 | . | . | . | 1.191576 | 1.167862 | 1.042733 | 1.267953 | 2.772847 | 2.699565 | 2.339863 | 2.899024 | -1.015512 | 6.662823 | 1.168730264 | -0.01900087 | 0.984840415 | NA | 2.747897 | 2.724273 | 2.738749 | 2.736602 | 2.733426 | 2.740989 | 8 | 7 | 5 | 7 |
| 101954267 | NR_104076 | RNVU1-15 | RNA, variant U1 small nuclear 15 | snRNA | . | 48320 | . | . | . | 0.276794 | 0.109272 | 0.338254 | 0.420089 | 0.644111 | 0.252586 | 0.759032 | 0.960485 | 1.659343 | 1.774228 | 2.21400588 | 0.329995761 | 0.741403165 | NA | 0.413393 | 1.148375 | 0.817239 | 0.813367 | 0.819776 | 0.819065 | 2 | 1 | 2 | 2 |
| 101954271 | NR_104080 | RNU6-9 | RNA, U6 small nuclear 9 | snRNA | . | 34269 | . | . | . | 0.350729 | 1.203212 | 0.723243 | 0.510082 | 0.816161 | 2.781279 | 1.622936 | 1.166241 | -1.140274 | 2.988466 | 1.7459779 | -0.10846704 | 0.913625222 | NA | 1.671431 | 1.481107 | 1.569974 | 1.583738 | 1.578049 | 1.572101 | 2 | 5 | 3 | 2 |
| 101954278 | NR_104088 | RNU6-8 | RNA, U6 small nuclear 8 | snRNA | . | 34285 | . | . | . | 1.987039 | 3.324667 | 1.673498 | 2.187132 | 4.623922 | 7.685116 | 3.755282 | 5.000618 | -1.363402 | 7.821254 | 1.095486662 | -0.40823084 | 0.68310421 | NA | 3.175403 | 2.724273 | 2.966931 | 2.986022 | 2.962012 | 2.969445 | 8 | 12 | 5 | 7 |
| 102060282 | NR_109831 | RASSF1-AS1 | RASSF1 antisense RNA 1 | lncRNA | . | 49091 | . | . | . | 0.863414 | 0.708188 | 1.361333 | 0.969209 | 2.009199 | 1.637008 | 3.054792 | 2.215982 | 1.498264 | 24.587943 | 0.630323018 | 0.925384929 | 0.354765715 | NA | 4.297162 | 4.883439 | 4.595492 | 4.577897 | 4.646948 | 4.607675 | 25 | 19 | 30 | 23 |
| 102157401 | NR_109828 | PCBP2-OT1 | PCBP2 overlapping transcript 1 | lncRNA | . | 49150 | . | . | . | 0.371768 | 0.094595 | 0.292348 | 0.240655 | 0.865119 | 0.218660 | 0.656019 | 0.550229 | 1.253769 | 4.971058 | 1.388964774 | 0.234903065 | 0.81428397 | NA | 2.137672 | 2.470304 | 2.318474 | 2.294460 | 2.312557 | 2.310763 | 8 | 2 | 5 | 5 |
| 102157402 | NM_00101589 | AK6 | adenylate kinase 6 | protein_coding | NP_001015891.1;NP_057367.1 | 49151 | . | . | . | 12.432416 | 12.312812 | 12.363448 | 8.211438 | 28.930743 | 28.461618 | 27.743219 | 18.774480 | -1.221218 | 590.014639 | 0.197236878 | -1.46180087 | 0.143795786 | 0.906161673 | 9.341007 | 9.053952 | 9.268347 | 9.254744 | 9.253300 | 8.988822 | 754 | 693 | 558 | 395 |
| 102238594 | NR_108100 | SPACA6P-AS | SPACA6P antisense RNA | lncRNA | . | 49383 | . | . | . | 0.133933 | 0.200980 | 0.244110 | 0.207457 | 0.311668 | 0.464573 | 0.547776 | 0.474327 | 1.337088 | 24.760067 | 0.627613518 | 0.667758891 | 0.504287511 | NA | 4.405684 | 4.823987 | 4.588596 | 4.621954 | 4.645738 | 4.624253 | 20 | 27 | 27 | 24 |
| 102288414 | NM_00128608 | C11orf98 | chromosome 11 open reading frame 98 | protein_coding | NP_001273015.1 | 51238 | . | . | . | 33.824939 | 32.209731 | 33.639684 | 38.891906 | 78.712019 | 74.454240 | 75.486479 | 88.921720 | 1.097550 | 717.737648 | 0.156390541 | 0.858663897 | 0.390525974 | 0.998865404 | 9.418800 | 9.552719 | 9.471024 | 9.434899 | 9.463477 | 9.568461 | 809 | 719 | 604 | 757 |
| 102388891 | gene-MCRIP2P | MCRIP2P1 | MAPK regulated corepressor interacting protein 2 pseudogene 1 | pseudogene | . | 49629 | . | . | . | 0.224312 | 0.357467 | 0.266581 | 0.036487 | 0.521982 | 0.826302 | 0.598199 | 0.083423 | -1.940262 | 5.828264 | 1.341276612 | -0.71294152 | 0.475881945 | NA | 2.939842 | 1.993948 | 2.546523 | 2.561135 | 2.550016 | 2.523877 | 7 | 10 | 6 | 1 |
| 102464826 | NR_106721,rn | MIR6073 | microRNA 6073 | miRNA | . | 50128 | . | . | . | 0.001387 | 0.008118 | 0.005455 | 0.009239 | 0.003227 | 0.018766 | 0.012241 | 0.021125 | 1.234663 | 2.005914 | 2.070995667 | 0.146845664 | 0.883253841 | NA | 0.844134 | 1.148375 | 0.998092 | 0.998667 | 1.000707 | 0.999974 | 2 | 2 | 2 | 2 |
| 102464833 | NR_106732,rn | MIR6084 | microRNA 6084 | miRNA | . | 50235 | . | . | . | 0.002823 | 0.001933 | 0.004470 | 0.003077 | 0.006568 | 0.004469 | 0.010031 | 0.007035 | 1.234663 | 2.005914 | 2.070995667 | 0.146845664 | 0.883253841 | NA | 0.844134 | 1.148375 | 0.998092 | 0.998667 | 1.000707 | 0.999974 | 2 | 2 | 2 | 2 |
| 102465133 | NR_106740,rn | MIR6125 | microRNA 6125 | miRNA | . | 49931 | . | . | . | 0.000353 | 0.000620 | 0.001383 | 0.000286 | 0.000821 | 0.001433 | 0.003105 | 0.000653 | 1.234663 | 2.005914 | 2.070995667 | 0.146845664 | 0.883253841 | NA | 0.844134 | 1.148375 | 0.998092 | 0.998667 | 1.000707 | 0.999974 | 2 | 2 | 2 | 2 |
| 102465136 | NR_106744,rn | MIR378J | microRNA 378j | miRNA | . | 49979 | . | . | . | 0.438871 | 0.424721 | 0.371046 | 0.471260 | 1.021270 | 0.981763 | 0.832618 | 1.077480 | 1.234665 | 3.008871 | 1.708874417 | 0.17796512 | 0.858750369 | NA | 1.429096 | 1.733338 | 1.582415 | 1.583274 | 1.586316 | 1.585224 | 3 | 3 | 3 | 3 |
| 102465248 | NR_106756,rn | MIR6501 | microRNA 6501 | miRNA | . | 50033 | . | . | . | 0.004552 | 0.002510 | 0.002350 | 0.004939 | 0.010594 | 0.005801 | 0.005276 | 0.011293 | 1.234665 | 3.008871 | 1.708874417 | 0.17796512 | 0.858750369 | NA | 1.429096 | 1.733338 | 1.582415 | 1.583274 | 1.586316 | 1.585224 | 3 | 3 | 3 | 3 |
| 102465252 | NR_106761,rn | MIR6506 | microRNA 6506 | miRNA | . | 50158 | . | . | . | 0.000843 | 0.003330 | 0.001452 | 0.002015 | 0.001959 | 0.007699 | 0.003259 | 0.004608 | 1.234665 | 3.008871 | 1.708874417 | 0.17796512 | 0.858750369 | NA | 1.429096 | 1.733338 | 1.582415 | 1.583274 | 1.586316 | 1.585224 | 3 | 3 | 3 | 3 |
| 102465256 | NR_106768,rn | MIR6513 | microRNA 6513 | miRNA | . | 50247 | . | . | . | 0.440604 | 0.000002 | 0.000003 | 0.387668 | 1.025302 | 0.000005 | 0.000006 | 0.886358 | 1.232222 | 1.970512 | 2.132558215 | 0.141267792 | 0.887658392 | NA | 0.820514 | 1.121007 | 0.982345 | 0.974334 | 0.975623 | 0.984888 | 3 | 1 | 1 | 3 |
| 102465428 | NR_106774,rn | MIR6717 | microRNA 6717 | miRNA | . | 50000 | . | . | . | 0.000095 | 0.000031 | 0.720758 | 0.001244 | 0.000220 | 0.000073 | 1.617362 | 0.002843 | 2.507047 | 1.577944 | 2.402382155 | 0.551947588 | 0.580984263 | NA | -0.155866 | 1.175234 | 0.628257 | 0.628593 | 0.640169 | 0.629356 | 1 | 1 | 3 | 1 |
| 102465429 | NR_106775,rn | MIR6511B1 | microRNA 6511b-1 | miRNA | . | 50228 | . | . | . | 0.000006 | 0.000012 | 0.000015 | 0.000008 | 0.000013 | 0.000027 | 0.000035 | 0.000018 | 1.234655 | 1.002957 | 2.868339486 | 0.106022226 | 0.915564717 | NA | -0.155866 | 0.148375 | -0.001261 | -0.000972 | 0.000053 | -0.000315 | 1 | 1 | 1 | 1 |
| 102465433 | NR_106782,rn | MIR6724-1 | microRNA 6724-1 | miRNA | . | 50837 | . | . | . | 0.023837 | 0.036331 | 0.152694 | 0.147835 | 0.055469 | 0.083980 | 0.342642 | 0.338006 | 1.234663 | 2.005914 | 2.070995667 | 0.146845664 | 0.883253841 | NA | 0.844134 | 1.148375 | 0.998092 | 0.998667 | 1.000707 | 0.999974 | 2 | 2 | 2 | 2 |
| 102465435 | NR_106785,rn | MIR6727 | hsa-miR-6727-3p | . | . | 50171 | . | . | . | 0.628382 | 1.785815 | 1.587793 | 1.259152 | 1.462272 | 4.127991 | 3.562963 | 2.878902 | 1.406178 | 3.794718 | 1.535154763 | 0.320344955 | 0.748706848 | NA | 1.658167 | 2.148375 | 1.909779 | 1.915081 | 1.919062 | 1.917633 | 3 | 4 | 4 | 4 |
| 102465436 | NR_106786,rn | MIR6728 | hsa-miR-6728-3p | . | . | 49961 | . | . | . | 0.928712 | 0.742858 | 1.758864 | 0.356051 | 2.161153 | 1.717149 | 3.946840 | 0.814068 | 1.416461 | 3.815544 | 1.540627593 | 0.326029967 | 0.744401682 | NA | 1.658167 | 2.161867 | 1.914543 | 1.919852 | 1.928928 | 1.917557 | 3 | 4 | 5 | 3 |
| 102465439 | NR_106791,rn | MIR6733 | hsa-miR-6733-3p | . | . | 50239 | . | . | . | 1.376624 | 1.405448 | 0.603446 | 0.405802 | 3.203462 | 3.248754 | 1.354115 | 0.927816 | -1.079982 | 3.457668 | 1.600358864 | -0.06936412 | 0.94469979 | NA | 1.844134 | 1.733338 | 1.790916 | 1.792011 | 1.790805 | 1.789671 | 4 | 4 | 3 | 3 |
| 102465442 | NR_106796,rn | MIR6738 | hsa-miR-6738-3p | . | . | 49939 | . | . | . | 0.253198 | 0.091799 | 1.268734 | 0.098296 | 0.589204 | 0.212197 | 2.847002 | 0.224742 | 1.870564 | 2.580901 | 1.868819978 | 0.483445811 | 0.628779227 | NA | 0.844134 | 1.751299 | 1.345797 | 1.346419 | 1.358911 | 1.347831 | 2 | 2 | 4 | 2 |
| 102465444 | NR_106800,rn | MIR6742 | hsa-miR-6742-3p | . | . | 50090 | . | . | . | 0.290915 | 0.515621 | 0.390577 | 1.318964 | 0.676972 | 1.191882 | 0.876444 | 3.015655 | 1.474528 | 2.791761 | 1.780166124 | 0.314719634 | 0.752974536 | NA | 1.175403 | 1.733338 | 1.466303 | 1.471348 | 1.474336 | 1.473263 | 2 | 3 | 3 | 3 |
| 102465446 | NR_106804,rn | MIR6746 | hsa-miR-6746-3p | . | . | 49925 | . | . | . | 2.538247 | 1.051242 | 0.591194 | 0.967577 | 5.906607 | 2.429994 | 1.326621 | 2.212252 | -1.155700 | 4.158556 | 1.470708199 | -0.14194993 | 0.887119558 | NA | 2.156660 | 1.947964 | 2.065350 | 2.058249 | 2.057237 | 2.060853 | 6 | 4 | 3 | 4 |
| 102465448 | NR_106806,rn | MIR6748 | microRNA 6748 | miRNA | . | 50141 | . | . | . | 0.946271 | 1.529454 | 2.478582 | 2.604054 | 2.202012 | 3.535400 | 5.561866 | 5.953861 | 1.676756 | 6.621459 | 1.177900279 | 0.633052533 | 0.526699338 | NA | 2.307819 | 3.051647 | 2.704608 | 2.710487 | 2.721666 | 2.723956 | 5 | 6 | 7 | 8 |
| 102465451 | NR_106811,rn | MIR6753 | hsa-miR-6753-5p | . | . | 50255 | . | . | . | 1.243069 | 2.762151 | 3.295811 | 3.348404 | 2.892673 | 6.384835 | 7.395703 | 7.655726 | 1.516075 | 14.219219 | 0.82795878 | 0.725085607 | 0.468399513 | NA | 3.501050 | 4.097315 | 3.788525 | 3.820701 | 3.825865 | 3.834695 | 9 | 16 | 14 | 17 |
| 102465452 | NR_106813,rn | MIR6755 | microRNA 6755 | miRNA | . | 50224 | . | . | . | 1.067497 | 1.020479 | 1.380337 | 0.272572 | 2.484108 | 2.358882 | 3.097435 | 0.623203 | -1.005372 | 4.481452 | 1.421774944 | -0.00543691 | 0.995661997 | NA | 2.166062 | 2.161867 | 2.160490 | 2.161865 | 2.166738 | 2.155350 | 5 | 5 | 5 | 3 |
| 102465453 | NR_106814,rn | MIR6756 | hsa-miR-6756-3p | . | . | 49974 | . | . | . | 1.080695 | 0.963428 | 2.314342 | 1.527542 | 2.514822 | 2.227009 | 5.193319 | 3.492545 | 1.543408 | 4.565989 | 1.404110616 | 0.445918993 | 0.655655762 | NA | 1.844134 | 2.470304 | 2.175928 | 2.177111 | 2.186379 | 2.184618 | 4 | 4 | 5 | 5 |
| 102465454 | NR_106816,rn | MIR6758 | microRNA 6758 | miRNA | . | 50038 | . | . | . | 2.596021 | 4.534798 | 4.447515 | 1.791930 | 6.041047 | 10.482389 | 9.980096 | 4.097036 | -1.010484 | 8.992055 | 1.029735372 | -0.01461178 | 0.988341897 | NA | 3.175403 | 3.161867 | 3.154048 | 3.173290 | 3.174365 | 3.152089 | 8 | 12 | 10 | 6 |
| 102465463 | NR_106830,rn | MIR6772 | microRNA 6772 | miRNA | . | 50064 | . | . | . | 3.596152 | 4.956075 | 3.883916 | 3.660588 | 8.368394 | 11.456190 | 8.715395 | 8.369499 | -1.038462 | 10.170469 | 0.955111171 | -0.05700705 | 0.954539582 | NA | 3.373794 | 3.318300 | 3.338483 | 3.353851 | 3.346129 | 3.342859 | 10 | 13 | 9 | 9 |
| 102465464 | NR_106833,rn | MIR6775 | hsa-miR-6775-3p | . | . | 50100 | . | . | . | 1.467886 | 0.533949 | 1.031701 | 0.387540 | 3.415832 | 1.234248 | 2.315108 | 0.886064 | 1.091603 | 3.730586 | 1.550327645 | 0.081562453 | 0.93499466 | NA | 1.832372 | 1.963455 | 1.899807 | 1.892406 | 1.900732 | 1.894479 | 5 | 3 | 4 | 3 |
| 102465466 | NR_106835,rn | MIR6777 | hsa-miR-6777-3p | . | . | 50173 | . | . | . | 2.280013 | 2.152299 | 2.403182 | 3.388064 | 5.305685 | 4.975135 | 5.392671 | 7.746407 | 1.340107 | 6.269833 | 1.204593662 | 0.350614734 | 0.725877401 | NA | 2.421266 | 2.844638 | 2.642191 | 2.635459 | 2.645842 | 2.648431 | 7 | 5 | 6 | 7 |
| 102465474 | NR_106849,rn | MIR6791 | hsa-miR-6791-3p | . | . | 50097 | . | . | . | 0.989665 | 0.452443 | 1.056338 | 0.414972 | 2.302989 | 1.045841 | 2.370390 | 0.948786 | 1.244163 | 3.513475 | 1.59175418 | 0.198005305 | 0.843040913 | NA | 1.644780 | 1.963455 | 1.806945 | 1.803669 | 1.811949 | 1.805707 | 4 | 3 | 4 | 3 |
| 102465478 | NR_106855,rn | MIR6797 | microRNA 6797 | miRNA | . | 50169 | . | . | . | 0.936402 | 1.157373 | 1.956207 | 0.882895 | 2.179044 | 2.675320 | 4.389672 | 2.018634 | 1.241819 | 5.035612 | 1.342131562 | 0.23280467 | 0.815913086 | NA | 2.166062 | 2.481107 | 2.321700 | 2.323119 | 2.333191 | 2.321524 | 5 | 5 | 6 | 4 |
| 102465480 | NR_106858,rn | MIR6800 | microRNA 6800 | miRNA | . | 50042 | . | . | . | 0.228762 | 0.233795 | 0.000001 | 0.002114 | 0.532338 | 0.540428 | 0.000002 | 0.004834 | -2.014451 | 1.668864 | 2.31231603 | -0.43695888 | 0.662141169 | NA | 1.156660 | 0.148375 | 0.757225 | 0.753619 | 0.750445 | 0.750013 | 3 | 2 | 1 | 1 |
| 102465481 | NR_106860,rn | MIR6802 | hsa-miR-6802-3p | . | . | 50154 | . | . | . | 1.690181 | 0.824295 | 0.719478 | 0.792919 | 3.933120 | 1.905395 | 1.614491 | 1.812916 | -1.073044 | 3.443092 | 1.608707002 | -0.06322457 | 0.949587677 | NA | 1.832372 | 1.733338 | 1.790084 | 1.782678 | 1.785831 | 1.784699 | 5 | 3 | 3 | 3 |
| 102465483 | NR_106863,rn | MIR6805 | microRNA 6805 | miRNA | . | 50215 | . | . | . | 0.000109 | 0.000042 | 0.000256 | 0.000063 | 0.000253 | 0.000098 | 0.000574 | 0.000144 | 1.234663 | 2.005914 | 2.070995667 | 0.146845664 | 0.883253841 | NA | 0.844134 | 1.148375 | 0.998092 | 0.998667 | 1.000707 | 0.999974 | 2 | 2 | 2 | 2 |
| 102465487 | NR_106870,rn | MIR6812 | microRNA 6812 | miRNA | . | 50116 | . | . | . | 0.223083 | 0.198599 | 0.303945 | 0.436552 | 0.519122 | 0.459069 | 0.682044 | 0.998122 | 1.234665 | 3.008871 | 1.708874417 | 0.17796512 | 0.858750369 | NA | 1.429096 | 1.733338 | 1.582415 | 1.583274 | 1.586316 | 1.585224 | 3 | 3 | 3 | 3 |
| 102465488 | NR_106872,rn | MIR6814 | hsa-miR-6814-3p | . | . | 50145 | . | . | . | 1.521446 | 1.041598 | 1.710982 | 0.836061 | 3.540468 | 2.407701 | 3.839393 | 1.911554 | 1.120373 | 4.733543 | 1.380832867 | 0.118754081 | 0.905470189 | NA | 2.156660 | 2.324312 | 2.242200 | 2.235117 | 2.244405 | 2.237825 | 6 | 4 | 5 | 4 |
| 102465490 | NR_106874,rn | MIR6816 | microRNA 6816 | miRNA | . | 50256 | . | . | . | 0.875222 | 0.660495 | 2.024856 | 0.862892 | 2.036676 | 1.526763 | 4.543719 | 1.972900 | 1.552526 | 4.586815 | 1.408219862 | 0.450652422 | 0.652240078 | NA | 1.844134 | 2.481107 | 2.179860 | 2.181044 | 2.195385 | 2.183733 | 4 | 4 | 6 | 4 |
| 102465491 | NR_106877,rn | MIR6819 | hsa-miR-6819-3p | . | . | 50155 | . | . | . | 0.628052 | 0.430701 | 0.503048 | 0.123987 | 1.461503 | 0.995585 | 1.128826 | 0.283482 | -1.123618 | 2.959315 | 1.731081282 | -0.09713705 | 0.922617562 | NA | 1.644780 | 1.481107 | 1.568690 | 1.565383 | 1.568406 | 1.562470 | 4 | 3 | 3 | 2 |
| 102465492 | NR_106878,rn | MIR6820 | microRNA 6820 | miRNA | . | 50095 | . | . | . | 0.657402 | 1.229177 | 2.870523 | 2.276515 | 1.529800 | 2.841299 | 6.441371 | 5.204981 | 1.782960 | 5.639329 | 1.277144333 | 0.653234273 | 0.513605258 | NA | 2.019256 | 2.852980 | 2.470525 | 2.476158 | 2.491447 | 2.484292 | 4 | 5 | 7 | 6 |
| 102465493 | NR_106879,rn | MIR6821 | hsa-miR-6821-5p | . | . | 49980 | . | . | . | 4.194480 | 4.772867 | 4.929760 | 3.947128 | 9.760728 | 11.032696 | 11.062238 | 9.024639 | -1.032880 | 10.176720 | 0.954451278 | -0.0488993 | 0.960999547 | NA | 3.369732 | 3.324312 | 3.343027 | 3.350189 | 3.351567 | 3.338682 | 11 | 12 | 10 | 8 |
| 102465495 | NR_106882,rn | MIR6824 | hsa-miR-6824-3p | . | . | 49946 | . | . | . | 0.004255 | 0.006683 | 0.006599 | 0.017641 | 0.009901 | 0.015449 | 0.014808 | 0.040334 | 1.234665 | 3.008871 | 1.708874417 | 0.17796512 | 0.858750369 | NA | 1.429096 | 1.733338 | 1.582415 | 1.583274 | 1.586316 | 1.585224 | 3 | 3 | 3 | 3 |
| 102465500 | NR_106891,rn | MIR6833 | microRNA 6833 | miRNA | . | 50245 | . | . | . | 0.977171 | 0.129065 | 0.746067 | 0.389521 | 2.273918 | 0.298338 | 1.674151 | 0.890593 | 1.091603 | 3.730586 | 1.550327645 | 0.081562453 | 0.93499466 | NA | 1.832372 | 1.963455 | 1.899807 | 1.892406 | 1.900732 | 1.894479 | 5 | 3 | 4 | 3 |
| 102465501 | NR_106892,rn | MIR6834 | hsa-miR-6834-5p | . | . | 50108 | . | . | . | 1.254346 | 1.193379 | 3.504610 | 1.262791 | 2.918913 | 2.758547 | 7.864242 | 2.887222 | 1.743483 | 6.164760 | 1.233303377 | 0.650263393 | 0.515522099 | NA | 2.166062 | 2.971139 | 2.598433 | 2.599939 | 2.625365 | 2.603355 | 5 | 5 | 9 | 5 |
| 102465502 | NR_106893,rn | MIR6835 | microRNA 6835 | miRNA | . | 49963 | . | . | . | 3.730109 | 3.827416 | 4.596923 | 1.990699 | 8.680118 | 8.847242 | 10.315362 | 4.551497 | -1.054302 | 9.180014 | 1.011388087 | -0.07542901 | 0.939873375 | NA | 3.234218 | 3.161867 | 3.198220 | 3.196891 | 3.206441 | 3.184174 | 11 | 10 | 10 | 6 |
| 102465503 | NR_106895,rn | MIR6836 | hsa-miR-6836-3p | . | . | 50067 | . | . | . | 0.423393 | 1.131688 | 0.136011 | 0.646075 | 0.985254 | 2.615948 | 0.305206 | 1.477171 | -1.144474 | 2.953064 | 1.732307522 | -0.11238452 | 0.91051853 | NA | 1.658167 | 1.459419 | 1.563031 | 1.568244 | 1.561794 | 1.565819 | 3 | 4 | 2 | 3 |
| 102465504 | NR_106897,rn | MIR6838 | hsa-miR-6838-3p | . | . | 50050 | . | . | . | 0.035114 | 0.053192 | 0.085923 | 0.024009 | 0.081712 | 0.122955 | 0.192808 | 0.054894 | 1.234665 | 3.008871 | 1.708874417 | 0.17796512 | 0.858750369 | NA | 1.429096 | 1.733338 | 1.582415 | 1.583274 | 1.586316 | 1.585224 | 3 | 3 | 3 | 3 |
| 102465510 | NR_106906,rn | MIR6847 | microRNA 6847 | miRNA | . | 50022 | . | . | . | 0.470437 | 1.233527 | 1.236237 | 0.766156 | 1.094727 | 2.851351 | 2.774080 | 1.751726 | 1.235205 | 4.531008 | 1.409796152 | 0.21616645 | 0.828858004 | NA | 2.019256 | 2.324312 | 2.168104 | 2.173640 | 2.178531 | 2.171959 | 4 | 5 | 5 | 4 |

| | | | | | | | | | | | | | | | | | | | | | | | | | | | | | | | | | | |
|---|---|---|---|---|---|---|---|---|---|---|---|---|---|---|---|---|---|---|---|---|---|---|---|---|---|---|---|---|---|---|---|---|---|---|
| 102465511 | NR_106907,rn | MIR6848 | microRNA 6848 | miRNA | . | | 50176 | . | . | . | 0.644869 | 1.195086 | 2.203942 | 2.380771 | 1.500637 | 2.762496 | 4.945583 | 5.443351 | 1.774780 | 5.618502 | 1.277655733 | 0.647780304 | 0.517127056 | NA | 2.019256 | 2.844638 | 2.467041 | 2.472668 | 2.482909 | 2.485573 | 4 | 5 | 6 | 7 |
| 102465512 | NR_106910,rn | MIR6851 | hsa-miR-6851-3p | . | . | | 50083 | . | . | . | 0.496439 | 0.235747 | 0.254807 | 0.137367 | 1.155234 | 0.544938 | 0.571778 | 0.314072 | -1.215015 | 2.454711 | 1.887245685 | -0.1488804 | 0.881648006 | NA | 1.429096 | 1.148375 | 1.299640 | 1.300454 | 1.298230 | 1.297456 | 3 | 3 | 2 | 2 |
| 102465513 | NR_106911,rn | MIR6852 | microRNA 6852 | miRNA | . | | 49993 | . | . | . | 0.000029 | 0.000031 | 0.000036 | 0.000037 | 0.000067 | 0.000071 | 0.000081 | 0.000086 | 1.234655 | 1.002957 | 2.868339486 | 0.106022226 | 0.915564717 | NA | -0.155866 | 0.148375 | -0.001261 | -0.000972 | 0.000053 | -0.000315 | 1 | 1 | 1 | 1 |
| 102465517 | NR_106917,rn | MIR6858 | hsa-miR-6858-5p | . | . | | 50250 | . | . | . | 0.148799 | 1.041888 | 0.720220 | 0.444978 | 0.346263 | 2.408371 | 1.616153 | 1.017390 | 1.224497 | 3.023447 | 1.718828416 | 0.169993295 | 0.86501541 | NA | 1.444631 | 1.733338 | 1.582994 | 1.592430 | 1.591096 | 1.590002 | 2 | 4 | 3 | 3 |
| 102465519 | NR_106921,rn | MIR6861 | microRNA 6861 | miRNA | . | | 50129 | . | . | . | 0.000007 | 0.000015 | 0.000022 | 0.000024 | 0.000017 | 0.000035 | 0.000048 | 0.000054 | 1.234655 | 1.002957 | 2.868339486 | 0.106022226 | 0.915564717 | NA | -0.155866 | 0.148375 | -0.001261 | -0.000972 | 0.000053 | -0.000315 | 1 | 1 | 1 | 1 |
| 102465524 | NR_106929,rn | MIR6869 | microRNA 6869 | miRNA | . | | 50056 | . | . | . | 2.619583 | 4.731799 | 1.625975 | 2.787717 | 6.095878 | 10.937766 | 3.648640 | 6.373783 | -1.363402 | 7.821254 | 1.095486662 | -0.40823084 | 0.68310421 | NA | 3.175403 | 2.724273 | 2.966931 | 2.986022 | 2.962012 | 2.969445 | 8 | 12 | 5 | 7 |
| 102465526 | NR_106932,rn | MIR6872 | microRNA 6872 | miRNA | . | | 50037 | . | . | . | 0.001085 | 0.000613 | 0.000811 | 0.001373 | 0.002527 | 0.001417 | 0.001818 | 0.003141 | 1.234665 | 3.008871 | 1.708874417 | 0.17796512 | 0.858750369 | NA | 1.429096 | 1.733338 | 1.582415 | 1.583274 | 1.586316 | 1.585224 | 3 | 3 | 3 | 3 |
| 102465536 | NR_106950,rn | MIR6890 | hsa-miR-6890-3p | . | . | | 49990 | . | . | . | 0.954840 | 1.122586 | 1.533441 | 1.338823 | 2.221953 | 2.594907 | 3.440996 | 3.061060 | -1.090094 | 3.436841 | 1.614238885 | -0.07709675 | 0.93854658 | NA | 1.844134 | 1.715150 | 1.784541 | 1.785633 | 1.779321 | 1.788114 | 4 | 4 | 2 | 4 |
| 102465537 | NR_106951,rn | MIR6891 | microRNA 6891 | miRNA | . | | 50243 | . | . | . | 0.240559 | 0.705200 | 0.956143 | 0.176752 | 0.559789 | 1.630103 | 2.145559 | 0.404121 | 1.086663 | 3.780564 | 1.565154252 | 0.076608921 | 0.938934662 | NA | 1.855800 | 1.978782 | 1.904222 | 1.913881 | 1.918573 | 1.902356 | 3 | 5 | 5 | 2 |
| 102465665 | NR_106957,rn | MIR7107 | microRNA 7107 | miRNA | . | | 50006 | . | . | . | 0.000026 | 0.000003 | 0.351494 | 0.000017 | 0.000061 | 0.000006 | 0.788744 | 0.000039 | 1.234663 | 2.005914 | 2.070995667 | 0.146845664 | 0.883253841 | NA | 0.844134 | 1.148375 | 0.998092 | 0.998667 | 1.000707 | 0.999974 | 2 | 2 | 2 | 2 |
| 102465666 | NR_106959,rn | MIR7109 | microRNA 7109 | miRNA | . | | 50122 | . | . | . | 0.283933 | 0.865871 | 0.632974 | 0.568123 | 0.660723 | 2.001498 | 1.420376 | 1.298945 | 1.054438 | 3.240557 | 1.652230047 | 0.046285504 | 0.963082693 | NA | 1.658167 | 1.733338 | 1.690660 | 1.695907 | 1.694638 | 1.693524 | 3 | 4 | 3 | 3 |
| 102465667 | NR_106960,rn | MIR7110 | microRNA 7110 | miRNA | . | | 50024 | . | . | . | 3.830403 | 4.152895 | 5.002722 | 2.419574 | 8.913506 | 9.599600 | 11.225965 | 5.532067 | -1.022873 | 12.441398 | 0.878373894 | -0.03714413 | 0.970370088 | NA | 3.651489 | 3.622519 | 3.632193 | 3.635874 | 3.648887 | 3.616399 | 14 | 14 | 14 | 8 |
| 102465668 | NR_106961,rn | MIR7111 | microRNA 7111 | miRNA | . | | 50073 | . | . | . | 3.165245 | 4.492007 | 4.864723 | 2.234107 | 7.365658 | 10.383475 | 10.916300 | 5.108021 | -1.017669 | 10.716726 | 0.943270865 | -0.02678867 | 0.978628291 | NA | 3.432996 | 3.410501 | 3.412379 | 3.423766 | 3.430860 | 3.403435 | 11 | 13 | 12 | 7 |
| 102465669 | NR_106963,rn | MIR7113 | microRNA 7113 | miRNA | . | | 49947 | . | . | . | 5.161455 | 3.924570 | 7.054578 | 5.220378 | 12.010920 | 9.071820 | 15.830270 | 11.935776 | 1.351200 | 12.097543 | 0.880155769 | 0.493369048 | 0.621751848 | NA | 3.361571 | 3.798723 | 3.584967 | 3.575821 | 3.605565 | 3.587072 | 13 | 10 | 14 | 11 |
| 102465694 | NR_106981,rn | MIR7161 | hsa-miR-7161-5p | . | . | | 49985 | . | . | . | 0.002471 | 0.004291 | 0.008823 | 0.001252 | 0.005751 | 0.009917 | 0.019796 | 0.002862 | 1.234665 | 3.008871 | 1.708874417 | 0.17796512 | 0.858750369 | NA | 1.429096 | 1.733338 | 1.582415 | 1.583274 | 1.586316 | 1.585224 | 3 | 3 | 3 | 3 |
| 102465801 | NR_106990,rn | MIR7703 | hsa-miR-7703 | . | . | | 50271 | . | . | . | 0.234513 | 0.829456 | 0.341104 | 1.281589 | 0.545719 | 1.917324 | 0.765429 | 2.930201 | 1.221536 | 2.504267 | 1.872041618 | 0.154214919 | 0.877440279 | NA | 1.175403 | 1.459419 | 1.316408 | 1.321418 | 1.319196 | 1.323280 | 2 | 3 | 2 | 3 |
| 102465834 | NR_106998,rn | MIR7844 | microRNA 7844 | miRNA | . | | 50044 | . | . | . | 0.004578 | 0.000803 | 0.007221 | 0.002579 | 0.010651 | 0.001857 | 0.016203 | 0.005895 | 1.234663 | 2.005914 | 2.070995667 | 0.146845664 | 0.883253841 | NA | 0.844134 | 1.148375 | 0.998092 | 0.998667 | 1.000707 | 0.999974 | 2 | 2 | 2 | 2 |
| 102465909 | NR_107062,rn | MIR6859-2 | hsa-miR-6859-3p | . | . | | 49954 | . | . | . | 11.691889 | 10.956785 | 16.260882 | 11.235190 | 27.207505 | 25.327102 | 36.488946 | 25.687930 | 1.242446 | 22.617564 | 0.65365705 | 0.479125335 | 0.631849467 | NA | 4.332860 | 4.648638 | 4.484563 | 4.478702 | 4.521328 | 4.482510 | 24 | 21 | 26 | 19 |
| 102465910 | NR_107063,rn | MIR6859-3 | microRNA 6859-3 | miRNA | . | | 50248 | . | . | . | 6.958016 | 6.883723 | 8.734323 | 6.062619 | 16.191588 | 15.912034 | 19.599568 | 13.861454 | 1.072390 | 17.203670 | 0.741130488 | 0.136049121 | 0.891782449 | NA | 4.052320 | 4.155137 | 4.098354 | 4.099194 | 4.115882 | 4.091822 | 19 | 18 | 18 | 14 |
| 102465976 | NR_106865,rn | MIR6807 | hsa-miR-6807-5p | . | . | | 50191 | . | . | . | 3.631439 | 2.627161 | 5.311690 | 5.483775 | 8.450507 | 6.072800 | 11.919278 | 12.538001 | 1.692201 | 11.424963 | 0.905433551 | 0.83816276 | 0.401939299 | NA | 3.084644 | 3.844638 | 3.494024 | 3.484327 | 3.512763 | 3.517875 | 11 | 8 | 12 | 14 |
| 102465978 | NR_106909,rn | MIR6850 | microRNA 6850 | miRNA | . | | 50093 | . | . | . | 2.709950 | 3.588006 | 2.993843 | 3.035907 | 6.306164 | 8.293839 | 6.718096 | 6.941240 | 1.023981 | 8.199535 | 1.057914703 | 0.032317857 | 0.974218569 | NA | 3.019256 | 3.051647 | 3.028367 | 3.039112 | 3.033572 | 3.035697 | 8 | 10 | 7 | 8 |
| 102466190 | NR_106779,rn | MIR6721 | microRNA 6721 | miRNA | . | | 50049 | . | . | . | 0.334286 | 0.055776 | 0.629255 | 0.114481 | 0.777898 | 0.128929 | 1.412030 | 0.261747 | 1.683953 | 1.795055 | 2.231855449 | 0.336873073 | 0.736212585 | NA | 0.413393 | 1.175234 | 0.827582 | 0.823701 | 0.835285 | 0.824501 | 2 | 1 | 3 | 1 |
| 102466192 | NR_106808,rn | MIR6750 | hsa-miR-6750-3p | . | . | | 49965 | . | . | . | 0.344893 | 0.671259 | 0.462735 | 0.300813 | 0.802580 | 1.551645 | 1.038364 | 0.687774 | 1.054438 | 3.240557 | 1.652230047 | 0.046285504 | 0.963082693 | NA | 1.658167 | 1.733338 | 1.690660 | 1.695907 | 1.694638 | 1.693524 | 3 | 4 | 3 | 3 |
| 102466193 | NR_106815,rn | MIR6757 | microRNA 6757 | miRNA | . | | 50160 | . | . | . | 0.201763 | 1.362365 | 1.178863 | 0.661650 | 0.469509 | 3.149168 | 2.645335 | 1.512785 | 1.396326 | 3.809293 | 1.552872665 | 0.310158077 | 0.756440749 | NA | 1.671431 | 2.148375 | 1.908976 | 1.922846 | 1.922466 | 1.921035 | 2 | 5 | 4 | 4 |
| 102466205 | NR_106947,rn | MIR6887 | microRNA 6887 | miRNA | . | | 50065 | . | . | . | 29.039248 | 29.950720 | 45.378131 | 33.337768 | 67.575524 | 69.232431 | 101.827215 | 76.222852 | 1.311053 | 72.077321 | 0.380830188 | 1.025984793 | 0.304898752 | 0.998865404 | 5.962063 | 6.354305 | 6.125757 | 6.131029 | 6.233451 | 6.153446 | 71 | 68 | 82 | 65 |
| 102466270 | NR_106799,rn | MIR6741 | hsa-miR-6741-3p | . | . | | 50004 | . | . | . | 0.736029 | 1.047336 | 1.285416 | 1.790892 | 1.712769 | 2.420963 | 2.884435 | 4.094661 | 1.384537 | 4.278495 | 1.446462025 | 0.324518665 | 0.745545394 | NA | 1.844134 | 2.312263 | 2.086858 | 2.088018 | 2.092125 | 2.095468 | 4 | 4 | 4 | 5 |
| 102466616 | NR_106748,rn | MIR6132 | microRNA 6132 | miRNA | . | | 50272 | . | . | . | 0.000358 | 0.018990 | 0.025942 | 0.000010 | 0.000833 | 0.043896 | 0.058214 | 0.000023 | -1.070164 | 1.739247 | 2.228774873 | -0.04389516 | 0.964987971 | NA | 0.844134 | 0.751299 | 0.796508 | 0.797062 | 0.799027 | 0.793436 | 2 | 2 | 2 | 1 |
| 102466657 | NR_106760,rn | MIR6505 | microRNA 6505 | miRNA | . | | 50104 | . | . | . | 0.318053 | 0.317004 | 0.662498 | 0.858596 | 0.740125 | 0.732768 | 1.486627 | 1.963077 | 1.646324 | 3.563032 | 1.584094361 | 0.454043688 | 0.649797382 | NA | 1.429096 | 2.148375 | 1.817378 | 1.818280 | 1.826590 | 1.825182 | 3 | 3 | 4 | 4 |
| 102466658 | NR_106765,rn | MIR6510 | microRNA 6510 | miRNA | . | | 50101 | . | . | . | 0.000002 | 0.000008 | 0.006789 | 0.394090 | 0.000006 | 0.000019 | 0.015234 | 0.901041 | 2.432439 | 1.536291 | 2.42076221 | 0.52975204 | 0.596283861 | NA | -0.155866 | 1.121007 | 0.602056 | 0.602388 | 0.603562 | 0.612940 | 1 | 1 | 1 | 3 |
| 102466719 | NR_106773,rn | MIR6716 | microRNA 6716 | miRNA | . | | 50048 | . | . | . | 0.442234 | 0.981128 | 0.655490 | 0.338589 | 1.029096 | 2.267921 | 1.470902 | 0.774143 | -1.040207 | 3.976848 | 1.501826248 | -0.0378681 | 0.969792848 | NA | 2.019256 | 1.963455 | 1.987477 | 1.992960 | 1.992634 | 1.986359 | 4 | 5 | 4 | 3 |
| 102466723 | NR_106792,rn | MIR6734 | hsa-miR-6734-3p | . | . | | 50219 | . | . | . | 1.873345 | 1.730481 | 3.448850 | 3.551044 | 4.359350 | 4.000085 | 7.739117 | 8.119043 | 1.684070 | 6.606883 | 1.177794842 | 0.638440942 | 0.523186687 | NA | 2.299300 | 3.051647 | 2.706388 | 2.703786 | 2.719297 | 2.721588 | 6 | 5 | 7 | 8 |
| 102466724 | NR_106797,rn | MIR6739 | microRNA 6739 | miRNA | . | | 50165 | . | . | . | 0.105185 | 1.481571 | 2.175833 | 0.721506 | 0.244771 | 3.424717 | 4.882506 | 1.649636 | 1.691201 | 4.903460 | 1.405704797 | 0.539265496 | 0.589703674 | NA | 1.867373 | 2.622519 | 2.259555 | 2.277899 | 2.288657 | 2.271910 | 2 | 6 | 7 | 4 |
| 102466729 | NR_106817,rn | MIR6759 | hsa-miR-6759-3p | . | . | | 50063 | . | . | . | 0.004394 | 0.005355 | 0.004714 | 0.018836 | 0.010225 | 0.012378 | 0.010578 | 0.043067 | 1.234665 | 3.008871 | 1.708874417 | 0.17796512 | 0.858750369 | NA | 1.429096 | 1.733338 | 1.582415 | 1.583274 | 1.586316 | 1.585224 | 3 | 3 | 3 | 3 |
| 102466733 | NR_106836,rn | MIR6778 | hsa-miR-6778-3p | . | . | | 50183 | . | . | . | 0.676749 | 1.150275 | 1.781541 | 0.607100 | 1.574823 | 2.658912 | 3.997728 | 1.388062 | 1.243590 | 4.032655 | 1.495177637 | 0.210349977 | 0.833394532 | NA | 1.844134 | 2.161867 | 2.001326 | 2.002467 | 2.011588 | 2.000215 | 4 | 4 | 5 | 3 |
| 102466734 | NR_106841,rn | MIR6783 | hsa-miR-6783-3p | . | . | | 50159 | . | . | . | 2.591221 | 2.847332 | 1.966460 | 1.029087 | 6.029877 | 6.581736 | 4.412679 | 2.352885 | -1.164080 | 5.414026 | 1.29429889 | -0.16935084 | 0.865520687 | NA | 2.540965 | 2.324312 | 2.441780 | 2.439324 | 2.440117 | 2.433489 | 7 | 6 | 5 | 4 |
| 102466736 | NR_106847,rn | MIR6789 | hsa-miR-6789-3p | . | . | | 50094 | . | . | . | 1.096170 | 2.920320 | 3.580732 | 1.479109 | 2.550834 | 6.750449 | 8.035061 | 3.381808 | 1.320081 | 7.862775 | 1.097182031 | 0.36514115 | 0.715006045 | NA | 2.760365 | 3.161867 | 2.951397 | 2.966117 | 2.979620 | 2.957372 | 6 | 9 | 10 | 6 |
| 102466740 | NR_106866,rn | MIR6808 | hsa-miR-6808-3p | . | . | | 50046 | . | . | . | 1.150976 | 0.286172 | 2.618370 | 1.017510 | 2.678369 | 0.661499 | 5.875546 | 2.326417 | 1.646324 | 3.563032 | 1.584094361 | 0.454043688 | 0.649797382 | NA | 1.429096 | 2.148375 | 1.817378 | 1.818280 | 1.826590 | 1.825182 | 3 | 3 | 4 | 4 |
| 102466747 | NR_106899,rn | MIR6840 | microRNA 6840 | miRNA | . | | 50138 | . | . | . | 4.798758 | 4.170259 | 6.481500 | 4.795686 | 11.166909 | 9.639741 | 14.544299 | 10.964766 | 1.194955 | 13.261803 | 0.838503553 | 0.306446645 | 0.759264598 | NA | 3.593806 | 3.852980 | 3.723623 | 3.715058 | 3.736618 | 3.722686 | 15 | 12 | 14 | 12 |
| 102466748 | NR_106904,rn | MIR6845 | hsa-miR-6845-3p | . | . | | 49956 | . | . | . | 4.177436 | 6.356561 | 5.946394 | 3.336579 | 9.721067 | 14.693476 | 13.343536 | 7.628690 | 1.000793 | 9.475832 | 0.995330916 | 0.001148537 | 0.999083601 | NA | 3.243129 | 3.245375 | 3.232882 | 3.248082 | 3.249281 | 3.231721 | 9 | 12 | 10 | 7 |
| 102466751 | NR_106918,rn | MIR6859-1 | hsa-miR-6859-3p | . | . | | 50039 | . | . | . | 17.417391 | 13.504039 | 28.864513 | 18.068752 | 40.530987 | 31.215189 | 64.771125 | 41.312056 | 1.494979 | 36.278343 | 0.531784885 | 1.090902863 | 0.275315627 | 0.998865404 | 4.860000 | 5.443477 | 5.149877 | 5.124458 | 5.228522 | 5.158472 | 37 | 28 | 46 | 32 |
| 102466754 | NR_106933,rn | MIR6873 | hsa-miR-6873-3p | . | . | | 50231 | . | . | . | 1.622219 | 2.779873 | 0.209645 | 0.828189 | 3.774972 | 6.425800 | 0.470437 | 1.893555 | -1.962877 | 4.082343 | 1.513755166 | -0.64275228 | 0.520384855 | NA | 2.436885 | 1.459419 | 2.040875 | 2.050791 | 2.031792 | 2.035682 | 5 | 7 | 2 | 3 |
| 102466755 | NR_106935,rn | MIR6875 | microRNA 6875 | miRNA | . | | 50015 | . | . | . | 1.637035 | 1.542242 | 1.324105 | 1.914487 | 3.809448 | 3.564960 | 2.971254 | 4.377246 | 1.044033 | 5.947359 | 1.232384315 | 0.050445116 | 0.959767685 | NA | 2.540965 | 2.602870 | 2.572624 | 2.570204 | 2.571137 | 2.574026 | 7 | 6 | 5 | 6 |
| 102466759 | NR_106954,rn | MIR6894 | hsa-miR-6894-3p | . | . | | 49938 | . | . | . | 0.844777 | 1.642916 | 1.363428 | 0.846080 | 1.965829 | 3.797674 | 3.059492 | 1.934460 | 1.094380 | 4.243514 | 1.451691029 | 0.089629449 | 0.928581681 | NA | 2.019256 | 2.148375 | 2.078515 | 2.084022 | 2.083765 | 2.082295 | 4 | 5 | 4 | 4 |
| 102466854 | NR_106992,rn | MIR7705 | microRNA 7705 | miRNA | . | | 50143 | . | . | . | 0.000017 | 0.000005 | 0.000009 | 0.000004 | 0.000040 | 0.000013 | 0.000019 | 0.000009 | 1.234663 | 2.005914 | 2.070995667 | 0.146845664 | 0.883253841 | NA | 0.844134 | 1.148375 | 0.998092 | 0.998667 | 1.000707 | 0.999974 | 2 | 2 | 2 | 2 |
| 102466864 | NR_106997,rn | MIR6516 | microRNA 6516 | miRNA | . | | 50233 | . | . | . | 0.143645 | 0.182111 | 0.238806 | 0.054352 | 0.334269 | 0.420957 | 0.535876 | 0.124270 | 1.234665 | 3.008871 | 1.708874417 | 0.17796512 | 0.858750369 | NA | 1.429096 | 1.733338 | 1.582415 | 1.583274 | 1.586316 | 1.585224 | 3 | 3 | 3 | 3 |
| 102466877 | NR_107039,rn | MIR8072 | microRNA 8072 | miRNA | . | | 50174 | . | . | . | 0.000163 | 0.000169 | 0.000393 | 0.000048 | 0.000378 | 0.000389 | 0.000882 | 0.000109 | 1.234663 | 2.005914 | 2.070995667 | 0.146845664 | 0.883253841 | NA | 0.844134 | 1.148375 | 0.998092 | 0.998667 | 1.000707 | 0.999974 | 2 | 2 | 2 | 2 |
| 102466911 | NR_106843,rn | MIR6785 | microRNA 6785 | miRNA | . | | 50206 | . | . | . | 0.000449 | 0.000543 | 0.000707 | 0.000003 | 0.001045 | 0.001255 | 0.001587 | 0.000008 | -1.198555 | 2.475538 | 1.907561801 | -0.13697918 | 0.89104725 | NA | 1.429096 | 1.175234 | 1.307104 | 1.307921 | 1.310815 | 1.300040 | 3 | 3 | 3 | 1 |
| 102466912 | NR_106931,rn | MIR6871 | microRNA 6871 | miRNA | . | | 49964 | . | . | . | 0.586191 | 1.828610 | 1.465292 | 0.769707 | 1.364091 | 4.226913 | 3.288074 | 1.759844 | 1.109089 | 4.762694 | 1.38020612 | 0.108226518 | 0.91381601 | NA | 2.175403 | 2.324312 | 2.240500 | 2.250382 | 2.250998 | 2.244409 | 4 | 6 | 5 | 4 |
| 102466982 | NR_106787,rn | MIR6729 | microRNA 6729 | miRNA | . | | 50103 | . | . | . | 1.555357 | 2.853451 | 2.505221 | 1.609424 | 3.619380 | 6.595881 | 5.621646 | 3.679757 | -1.012895 | 6.733628 | 1.173659783 | -0.01574899 | 0.987434645 | NA | 2.760365 | 2.742346 | 2.740583 | 2.755184 | 2.753500 | 2.741501 | 6 | 9 | 7 | 5 |
| 102466983 | NR_106824,rn | MIR6766 | microRNA 6766 | miRNA | . | | 49941 | . | . | . | 0.680922 | 0.725109 | 0.269859 | 1.014944 | 1.584536 | 1.676122 | 0.605555 | 2.320549 | -1.090094 | 3.436841 | 1.614238885 | -0.07709675 | 0.93854658 | NA | 1.844134 | 1.715150 | 1.784541 | 1.785633 | 1.779321 | 1.788114 | 4 | 4 | 2 | 4 |
| 102466985 | NR_106896,rn | MIR6837 | microRNA 6837 | miRNA | . | | 50125 | . | . | . | 0.142912 | 0.786816 | 0.833961 | 0.502193 | 0.332562 | 1.818760 | 1.871383 | 1.148204 | 1.434830 | 3.310940 | 1.652984007 | 0.315114584 | 0.752674656 | NA | 1.444631 | 1.963455 | 1.707928 | 1.717404 | 1.721258 | 1.715020 | 2 | 4 | 4 | 3 |
| 102467073 | NR_104613 | HRAT5 | heart tissue-associated transcript 5 | lncRNA | . | . | | . | . | . | 0.349037 | 0.577274 | 1.122321 | 0.927969 | 0.812222 | 1.334395 | 2.518456 | 2.121692 | 2.154170 | 19.935050 | 0.708198818 | 1.563306761 | 0.117980459 | NA | 3.661494 | 4.766401 | 4.246180 | 4.274906 | 4.333704 | 4.312532 | 11 | 17 | 26 | 23 |
| 102467147 | NR_104664 | LINC01948 | long intergenic non-protein coding RNA 1948 | lncRNA | . | | 52772 | . | . | . | 0.366959 | 0.321968 | 0.249519 | 0.148556 | 0.853928 | 0.744244 | 0.559913 | 0.339655 | -1.707528 | 6.044952 | 1.24758069 | -0.61872457 | 0.536097823 | NA | 2.928838 | 2.161867 | 2.611025 | 2.609056 | 2.601472 | 2.590008 | 9 | 8 | 5 | 3 |
| 102546294 | NR_105057 | LOC10254629 | uncharacterized LOC102546294 | lncRNA | . | . | | . | . | . | 0.051207 | 0.094142 | 0.144556 | 0.187094 | 0.119161 | 0.217613 | 0.324379 | 0.427769 | 2.332593 | 8.277689 | 1.081965836 | 1.12936505 | 0.25874387 | NA | 2.316288 | 3.532926 | 3.005827 | 3.020328 | 3.037605 | 3.048567 | 4 | 7 | 9 | 12 |
| 102606465 | NR_109849 | LOC10260646 | uncharacterized LOC102606465 | lncRNA | . | . | | . | . | . | 1.044383 | 0.810600 | 1.153233 | 1.086877 | 2.430321 | 1.873738 | 2.587822 | 2.485014 | 1.183389 | 42.919760 | 0.479084604 | 0.507058523 | 0.612113756 | 0.998865404 | 5.296092 | 5.540693 | 5.425141 | 5.385690 | 5.439421 | 5.426793 | 51 | 37 | 42 | 42 |
| 102723099 | NR_130107 | GHET1 | gastric carcinoma proliferation enhancing transcript 1 | lncRNA | . | | 49425 | . | . | . | 0.461530 | 0.327408 | 0.332895 | 0.244509 | 1.073999 | 0.756819 | 0.747007 | 0.559041 | -1.355435 | 20.952851 | 0.683120155 | -0.64228284 | 0.520689556 | NA | 4.590318 | 4.155137 | 4.420612 | 4.388904 | 4.390305 | 4.366083 | 32 | 22 | 18 | 14 |
| 102723166 | NR_110370 | STAM-AS1 | STAM antisense RNA 1 (head to head) | lncRNA | . | | 49568 | . | . | . | 0.294061 | 0.574247 | 0.657187 | 0.423100 | 0.684292 | 1.327398 | 1.474708 | 0.967369 | 1.216806 | 40.656089 | 0.529922494 | 0.534227938 | 0.593183842 | 0.998865404 | 5.197227 | 5.479761 | 5.268084 | 5.361182 | 5.380891 | 5.310808 | 29 | 52 | 47 | 33 |
| 102723305 | XR_00295872 | LOC10272330 | uncharacterized LOC102723305, transcript variant X2 | lncRNA | . | . | | . | . | . | 0.908535 | 0.566312 | 1.273767 | 0.854693 | 2.114200 | 1.309055 | 2.858297 | 1.954155 | 1.493459 | 25.568000 | 0.631776343 | 0.915920943 | 0.359708362 | NA | 4.355421 | 4.938580 | 4.661405 | 4.620760 | 4.706930 | 4.658767 | 29 | 17 | 32 | 23 |
| 102723324 | XR_00174655 | LOC10272332 | uncharacterized LOC102723324, transcript variant X3 | lncRNA | . | . | | . | . | . | 0.848335 | 0.883193 | 1.052482 | 0.932369 | 1.974110 | 2.041540 | 2.361737 | 2.131753 | 1.142957 | 38.476628 | 0.50023224 | 0.385363486 | 0.699968131 | 0.998865404 | 5.166062 | 5.359293 | 5.247515 | 5.257047 | 5.282102 | 5.262328 | 40 | 40 | 38 | 36 |
| 102723334 | XR_428139,XR | LOC10272333 | uncharacterized LOC102723334, transcript variant X3 | lncRNA | . | . | | . | . | . | 0.515236 | 0.304653 | 0.373445 | 0.264414 | 1.198974 | 0.704219 | 0.838000 | 0.604549 | -1.450999 | 10.896071 | 0.943679503 | -0.56909895 | 0.569288988 | NA | 3.687500 | 3.155137 | 3.475296 | 3.442630 | 3.447676 | 3.434984 | 19 | 10 | 9 | 7 |
| 102723341 | XR_00174432 | LOC10272334 | uncharacterized LOC102723341, transcript variant X3 | lncRNA | . | . | | . | . | . | 0.121607 | 0.151243 | 0.080428 | 0.203577 | 0.282985 | 0.349606 | 0.180478 | 0.465455 | 1.088230 | 17.831228 | 0.75752289 | 0.161029884 | 0.872069864 | NA | 4.096985 | 4.213338 | 4.138792 | 4.159567 | 4.125975 | 4.184690 | 17 | 21 | 10 | 24 |
| 102723342 | XR_429419 | LOC10272334 | uncharacterized LOC102723342 | lncRNA | . | . | | . | . | . | 0.054918 | 0.152713 | 0.352668 | 0.135277 | 0.127796 | 0.353002 | 0.791376 | 0.309294 | 2.209044 | 16.764068 | 0.831397156 | 1.375302201 | 0.169037774 | NA | 3.385915 | 4.528236 | 3.981281 | 4.021909 | 4.097658 | 4.013358 | 7 | 16 | 29 | 12 |
| 102723356 | NR_125796,NF | ZFPM2-AS1 | ZFPM2 antisense RNA 1 | lncRNA | . | | 50698 | . | . | . | 3.172867 | 3.211135 | 4.502044 | 3.146742 | 7.383393 | 7.422682 | 10.102456 | 7.194653 | 1.158044 | 108.454075 | 0.324427479 | 0.652503286 | 0.514076554 | 0.998865404 | 6.650233 | 6.863754 | 6.728119 | 6.731903 | 6.836651 | 6.713996 | 115 | 109 | 120 | 89 |
| 102723360 | NM_00132297 | LOC10272336 | uncharacterized LOC102723360 | protein_coding | NP_001309904.1;XP_006724005.1;XP_006724007.1;XP_011544411.1;XP | . | | . | . | . | 2.880720 | 3.479505 | 3.941688 | 3.551578 | 6.703555 | 8.043031 | 8.845035 | 8.120261 | 1.012737 | 324.022370 | 0.20461005 | 0.089240697 | 0.928890623 | 0.998865404 | 8.330276 | 8.349559 | 8.336584 | 8.332435 | 8.400533 | 8.279127 | 372 | 346 | 315 | 272 |
| 102723390 | NR_146067 | FRG1EP | FSHD region gene 1 family member E, pseudogene | transcribed_ps | . | | 51764 | . | . | . | 1.949226 | 2.106831 | 2.352676 | 2.131035 | 4.535929 | 4.870035 | 5.279337 | 4.872359 | 1.088451 | 256.389138 | 0.216117208 | 0.565785173 | 0.571539827 | 0.998865404 | 7.939617 | 8.062164 | 7.961309 | 7.995744 | 8.048546 | 7.993592 | 273 | 274 | 245 | 237 |
| 102723409 | XR_00174385 | LOC10272340 | uncharacterized LOC102723409, transcript variant X6 | lncRNA | . | . | | . | . | . | 0.028784 | 0.019085 | 0.063116 | 0.017467 | 0.066981 | 0.044116 | 0.141629 | 0.039936 | 1.396178 | 9.634929 | 1.014365153 | 0.474663808 | 0.635026619 | NA | 3.003609 | 3.491830 | 3.254006 | 3.239874 | 3.281902 | 3.239996 | 11 | 7 | 14 | 6 |
| 102723446 | XR_00174520 | LOC10272344 | uncharacterized LOC102723446, transcript variant X1 | lncRNA | . | . | | . | . | . | 0.022333 | 0.090099 | 0.027341 | 0.043561 | 0.051969 | 0.208269 | 0.061352 | 0.099597 | -1.400213 | 8.595020 | 1.100894155 | -0.44113809 | 0.659113029 | NA | 3.328899 | 2.836248 | 3.079995 | 3.132106 | 3.087584 | 3.099661 | 5 | 17 | 5 | 8 |
| 102723458 | XR_00174175 | LOC10272345 | uncharacterized LOC102723458, transcript variant X12 | lncRNA | . | . | | . | . | . | 1.931677 | 2.269195 | 2.663367 | 1.905914 | 4.495093 | 5.245346 | 5.976517 | 4.357647 | 1.088641 | 144.401274 | 0.277999883 | 0.440747174 | 0.659396041 | 0.998865404 | 7.110920 | 7.234321 | 7.142000 | 7.166648 | 7.227257 | 7.144643 | 154 | 154 | 145 | 126 |
| 102723471 | NR_135668,NF | LHX1-DT | LHX1 divergent transcript | lncRNA | . | | 53778 | . | . | . | 2.640312 | 3.077405 | 3.082717 | 2.866650 | 6.144115 | 7.113559 | 6.917528 | 6.554254 | 1.024052 | 133.709786 | 0.284804239 | 0.120395164 | 0.904170122 | 0.998865404 | 7.045717 | 7.080001 | 7.029682 | 7.083591 | 7.084927 | 7.046003 | 141 | 153 | 124 | 120 |
| 102723508 | NR_110456,NF | KANTR | KDM5C adjacent transcript | lncRNA | . | | 49510 | 301019 | . | . | 0.154501 | 0.352341 | 0.198507 | 0.242063 | 0.359531 | 0.814450 | 0.445443 | 0.553450 | -1.198953 | 39.410280 | 0.531470958 | -0.49254924 | 0.622331123 | 0.998865404 | 5.427245 | 5.161538 | 5.254659 | 5.360437 | 5.263106 | 5.299304 | 34 | 61 | 27 | 38 |
| 102723510 | NR_120306 | LCMT1-AS1 | LCMT1 antisense RNA 1 | lncRNA | . | | 51177 | . | . | . | 0.765445 | 0.317852 | 0.848679 | 1.091708 | 1.782222 | 0.734729 | 1.904411 | 2.496061 | 1.780410 | 14.187150 | 0.839494615 | 0.991322147 | 0.321528302 | NA | 3.349244 | 4.184531 | 3.811769 | 3.778095 | 3.821093 | 3.838925 | 16 | 7 | 14 | 19 |
| 102723534 | NR_160687,NF | LOC10272353 | HECT and RLD domain containing E3 ubiquitin protein ligase 2 pse | transcribed_ps | . | . | | . | . | . | 2.026571 | 1.904584 | 2.785371 | 1.934103 | 4.715915 | 4.402532 | 6.250291 | 4.422096 | 1.185618 | 159.654893 | 0.28088591 | 0.874517813 | 0.381836324 | 0.998865404 | 7.189697 | 7.437317 | 7.289194 | 7.262227 | 7.419781 | 7.263045 | 174 | 152 | 179 | 132 |
| 102723539 | XR_00174817 | LOC10272353 | uncharacterized LOC102723539 | lncRNA | . | . | | . | . | . | 0.150012 | 0.192877 | 0.396614 | 0.233504 | 0.349084 | 0.445843 | 0.889989 | 0.533878 | 1.822244 | 31.148345 | 0.579953234 | 1.492734683 | 0.135506651 | 0.893217909 | 4.463617 | 5.330299 | 4.885601 | 4.911799 | 5.011111 | 4.932270 | 22 | 27 | 44 | 28 |
| 102723547 | XM_00672485 | CSAG2 | CSAG family member 2, transcript variant X1 | protein_coding | XP_006724920.1;XP_016885512.1 | | 16847 | . | . | . | 6.094207 | 6.537029 | 7.712500 | 7.046639 | 14.181468 | 15.110636 | 17.306626 | 16.111302 | 1.216148 | 207.761786 | 0.235862741 | 1.196963924 | 0.231320617 | 0.986686796 | 7.550372 | 7.833348 | 7.643857 | 7.651764 | 7.772289 | 7.701998 | 214 | 204 | 213 | 198 |
| 102723553 | NM_00131369 | SMIM11B | small integral membrane protein 11B | protein_coding | NP_001300621.1;XP_016883693.1;XP_016883694.1 | | 51846 | . | . | . | 4.122700 | 3.898899 | 4.842604 | 5.242811 | 9.593689 | 9.012478 | 10.866661 | 11.987064 | 1.188032 | 144.662330 | 0.270614175 | 0.91855502 | 0.358328368 | 0.998865404 | 7.046957 | 7.295447 | 7.141396 | 7.138174 | 7.200534 | 7.210201 | 153 | 142 | 135 | 149 |
| 102723564 | XR_00175143 | LOC10272356 | uncharacterized LOC102723564, transcript variant X2 | lncRNA | . | . | | . | . | . | 5.866776 | 5.559759 | 7.271663 | 6.659301 | 13.652232 | 12.851631 | 16.317401 | 15.225701 | 1.196669 | 476.279416 | 0.173504103 | 1.492901858 | 0.135462879 | 0.893217909 | 8.759993 | 9.019667 | 8.853217 | 8.814969 | 8.981033 | 8.911150 | 515 | 453 | 476 | 460 |
| 102723566 | NR_136302 | LOC10272356 | uncharacterized LOC102723566 | lncRNA | . | . | | . | . | . | 0.206365 | 0.180623 | 0.276363 | 0.134291 | 0.480221 | 0.417518 | 0.620151 | 0.307040 | 1.034452 | 13.218920 | 0.862019625 | 0.056688596 | 0.954793259 | NA | 3.697260 | 3.751299 | 3.721983 | 3.713655 | 3.740585 | 3.698463 | 16 | 13 | 16 | 8 |
| 102723570 | XR_430342 | CDC27P9 | cell division cycle 27 pseudogene 9 | transcribed_ps | . | | 51759 | . | . | . | 0.292221 | 0.379946 | 0.564291 | 0.427067 | 0.680009 | 0.878263 | 1.266253 | 0.976438 | 1.433923 | 30.693140 | 0.568048799 | 0.915357568 | 0.360003946 | 0.998865404 | 4.656500 | 5.176571 | 4.889976 | 4.920063 | 4.969120 | 4.929100 | 25 | 31 | 36 | 29 |
| 102723574 | gene-HAUS6P3 | HAUS6P3 | HAUS augmin like complex subunit 6 pseudogene 3 | pseudogene | . | | 50772 | . | . | . | 0.205204 | 0.161124 | 0.256880 | 0.180428 | 0.477517 | 0.372445 | 0.576432 | 0.412527 | 1.179595 | 27.770752 | 0.596408897 | 0.399543771 | 0.689492578 | 0.998865404 | 4.669607 | 4.911273 | 4.793354 | 4.766302 | 4.820246 | 4.776458 | 33 | 24 | 31 | 23 |
| 102723584 | XR_430343 | CDC27P10 | cell division cycle 27 pseudogene 10 | transcribed_ps | . | | 51760 | . | . | . | 0.293949 | 0.393194 | 0.538917 | 0.410972 | 0.684031 | 0.908886 | 1.209315 | 0.939640 | 1.367243 | 29.851486 | 0.574221688 | 0.785880308 | 0.431937612 | 0.998865404 | 4.656500 | 5.107809 | 4.855037 | 4.885081 | 4.925309 | 4.889759 | 25 | 31 | 34 | 28 |
| 102723591 | XR_430558,XR | LOC10272359 | uncharacterized LOC102723591, transcript variant X1 | lncRNA | . | . | | . | . | . | 2.032002 | 1.792650 | 1.085504 | 1.408089 | 4.728552 | 4.143790 | 2.435840 | 3.219427 | -1.525059 | 41.874580 | 0.483896797 | -1.25825455 | 0.208299711 | 0.970482251 | 5.660576 | 5.051647 | 5.432631 | 5.414156 | 5.353558 | 5.360385 | 61 | 52 | 28 | 32 |
| 102723603 | XR_430344 | CDC27P11 | cell division cycle 27 pseudogene 11 | transcribed_ps | . | | 51761 | . | . | . | 0.355292 | 0.437296 | 0.565852 | 0.374745 | 0.826780 | 1.010828 | 1.269755 | 0.856810 | 1.169315 | 52.223764 | 0.449270429 | 0.50228935 | 0.615464 | 0.998865404 | 5.588663 | 5.815684 | 5.665127 | 5.701238 | 5.753174 | 5.671681 | 50 | 57 | 59 | 42 |
| 102723619 | NR_146497 | ATXN2-AS | ATXN2 antisense RNA | lncRNA | . | | 51838 | . | . | . | 0.174247 | 0.453381 | 0.621909 | 0.224957 | 0.405481 | 1.048009 | 1.395545 | 0.514337 | 1.341597 | 6.376041 | 1.235039831 | 0.343269156 | 0.731395983 | NA | 2.444631 | 2.869520 | 2.644056 | 2.662711 | 2.675182 | 2.648363 | 4 | 8 | 9 | 4 |
| 102723626 | gene-TUBB8P8 | TUBB8P8 | tubulin beta 8 class VIII pseudogene 8 | pseudogene | . | | 42346 | . | . | . | 0.105435 | 0.110408 | 0.151682 | 0.157208 | 0.245352 | 0.255214 | 0.340369 | 0.359438 | 1.481659 | 5.568946 | 1.275118504 | 0.444832318 | 0.656440938 | NA | 2.166062 | 2.733338 | 2.462655 | 2.464114 | 2.474331 | 2.472221 | 5 | 5 | 6 | 6 |
| 102723631 | NM_00129152 | CT45A10 | cancer/testis antigen family 45 member A10 | protein_coding | NP_001278456.1;NP_001278457.1;NP_001278458.1;NP_001278459.1 | | 51263 | . | . | . | 4.407587 | 4.328064 | 2.859046 | 2.931114 | 10.256637 | 10.004513 | 6.415616 | 6.701642 | -1.517281 | 137.170171 | 0.279108997 | -2.15503123 | 0.031159391 | 0.560017015 | 7.369451 | 6.767948 | 7.175472 | 7.163428 | 7.015758 | 7.019750 | 193 | 176 | 94 | 103 |
| 102723648 | XR_00174410 | LOC10272364 | uncharacterized LOC102723648, transcript variant X4 | lncRNA | . | . | | . | . | . | 1.943073 | 2.458205 | 1.976732 | 1.093614 | 4.521610 | 5.682249 | 4.435730 | 2.500421 | -1.454007 | 56.248078 | 0.453483104 | -1.19085847 | 0.233709149 | 0.988893316 | 6.057381 | 5.520401 | 5.818015 | 5.871195 | 5.819949 | 5.717302 | 68 | 80 | 52 | 30 |
| 102723660 | gene-SOWAHC | SOWAHCP3 | . | pseudogene | . | | 54507 | . | . | . | 0.051175 | 0.192083 | 0.228561 | 0.185670 | 0.119085 | 0.444008 | 0.512885 | 0.424511 | 1.725030 | 10.551114 | 0.992238963 | 0.792773938 | 0.427909543 | NA | 2.956191 | 3.737849 | 3.346766 | 3.386917 | 3.398853 | 3.385277 | 4 | 13 | 13 | 11 |
| 102723665 | NR_135921 | LOC10272366 | uncharacterized LOC102723665 | lncRNA | . | . | | . | . | . | 0.630807 | 0.530910 | 0.425913 | 0.454508 | 1.467915 | 1.227222 | 0.955736 | 1.039178 | -1.318043 | 29.489147 | 0.571427353 | -0.69719678 | 0.485679652 | 0.998865404 | 5.067319 | 4.669578 | 4.913857 | 4.889642 | 4.868674 | 4.869562 | 42 | 33 | 22 | 24 |
| 102723670 | XR_00175077 | LOC10272367 | uncharacterized LOC102723670, transcript variant X1 | lncRNA | . | . | | . | . | . | 0.086781 | 0.063252 | 0.088545 | 0.225336 | 0.201944 | 0.146210 | 0.198691 | 0.515204 | 1.613967 | 9.342415 | 1.002143523 | 0.68913434 | 0.490738732 | NA | 2.838265 | 3.527725 | 3.207264 | 3.201227 | 3.213749 | 3.234138 | 9 | 7 | 8 | 13 |
| 102723672 | NR_134325 | LOC10272367 | uncharacterized LOC102723672 | lncRNA | . | . | | . | . | . | 0.375340 | 0.805881 | 0.961154 | 0.429991 | 0.873432 | 1.862830 | 2.156801 | 0.983123 | 1.183706 | 11.889601 | 0.924634042 | 0.263143131 | 0.792440261 | NA | 3.444631 | 3.688346 | 3.536376 | 3.572494 | 3.581908 | 3.544561 | 8 | 16 | 15 | 8 |
| 102723678 | XR_00174649 | LOC10272367 | uncharacterized LOC102723678, transcript variant X3 | lncRNA | . | . | | . | . | . | 0.018528 | 0.035288 | 0.060570 | 0.049122 | 0.043116 | 0.081571 | 0.135918 | 0.112311 | 2.458823 | 3.900081 | 1.546624225 | 0.839226336 | 0.401342312 | NA | 1.175403 | 2.470304 | 1.932259 | 1.937407 | 1.950937 | 1.949236 | 2 | 3 | 5 | 5 |
| 102723680 | NM_00129154 | CT45A9 | cancer/testis antigen family 45 member A9 | protein_coding | NP_001278469.1;NP_001308200.1;XP_006724859.1 | | 51262 | . | . | . | 0.558433 | 0.596976 | 0.371110 | 0.331232 | 1.299497 | 1.379937 | 0.832762 | 0.757323 | -1.680102 | 20.072461 | 0.693142548 | -1.07993494 | 0.280171155 | NA | 4.653161 | 3.905270 | 4.345792 | 4.359995 | 4.314885 | 4.305074 | 27 | 29 | 14 | 13 |
| 102723694 | XR_00295672 | LOC10272369 | uncharacterized LOC102723694, transcript variant X1 | lncRNA | . | . | | . | . | . | 1.256275 | 1.143862 | 1.835078 | 0.885859 | 2.923404 | 2.644088 | 4.117861 | 2.025412 | 1.119363 | 401.323504 | 0.520467036 | 0.312561909 | 0.754613522 | 0.998865404 | 8.564766 | 8.727870 | 8.625117 | 8.565771 | 8.857466 | 8.421805 | 458 | 388 | 499 | 257 |
| 102723696 | XR_00174087 | LOC10272369 | uncharacterized LOC102723696, transcript variant X5 | lncRNA | . | . | | . | . | . | 0.385945 | 0.445195 | 0.286122 | 0.261888 | 0.898111 | 1.029091 | 0.642049 | 0.598775 | -1.416783 | 18.746476 | 0.710421308 | -0.70749386 | 0.479259627 | NA | 4.457887 | 3.955730 | 4.245642 | 4.247726 | 4.221793 | 4.216617 | 25 | 24 | 14 | 14 |
| 102723722 | XM_01702917 | LOC10272372 | uncharacterized LOC102723722, transcript variant X1 | protein_coding | XP_016884663.1 | . | | . | . | . | 1.142327 | 0.825163 | 0.935759 | 0.819232 | 2.658243 | 1.907402 | 2.099817 | 1.873077 | -1.134047 | 26.037598 | 0.611516407 | -0.29677147 | 0.766640987 | NA | 4.789233 | 4.610270 | 4.728850 | 4.691217 | 4.704944 | 4.688468 | 37 | 25 | 23 | 21 |
| 102723728 | XM_00672099 | LOC10272372 | nodal modulator 3-like | protein_coding | XP_006721059.1 | . | | . | . | . | 84.818596 | 83.606277 | 80.349220 | 76.587654 | 197.376358 | 193.259659 | 180.301315 | 175.108566 | -1.088198 | 3957.365023 | 0.109410399 | -1.11452973 | 0.265051999 | 0.996598404 | 12.009959 | 11.888118 | 11.993884 | 11.974540 | 11.937876 | 11.890039 | 4804 | 4399 | 3392 | 3450 |
| 102723737 | NM_00129153 | CT45A8 | cancer/testis antigen family 45 member A8 | protein_coding | NP_001278464.1;XP_006724862.1;XP_011529543.1 | | 51261 | . | . | . | 0.803967 | 0.863471 | 0.548879 | 0.487837 | 1.870862 | 1.995953 | 1.231669 | 1.115383 | -1.591897 | 22.280910 | 0.656494653 | -1.02170995 | 0.306918199 | NA | 4.774103 | 4.104320 | 4.501548 | 4.504937 | 4.465972 | 4.455574 | 31 | 30 | 16 | 15 |
| 102723765 | XR_00174824 | LOC10272376 | uncharacterized LOC102723765, transcript variant X8 | lncRNA | . | . | | . | . | . | 2.164290 | 1.939356 | 1.155072 | 1.682537 | 5.036393 | 4.482908 | 2.591949 | 3.846918 | -1.500240 | 75.643294 | 0.379019291 | -1.5439671 | 0.122596302 | 0.866753766 | 6.505078 | 5.917857 | 6.307584 | 6.270996 | 6.150279 | 6.225088 | 111 | 92 | 43 | 67 |
| 102723775 | NR_120595,NF | LINC01315 | long intergenic non-protein coding RNA 1315 | lncRNA | . | | 50513 | . | . | . | 0.807883 | 1.016909 | 0.949026 | 0.959445 | 1.879979 | 2.350632 | 2.129585 | 2.193658 | 1.097449 | 30.053020 | 0.563690806 | 0.237992147 | 0.811887185 | 0.998865404 | 4.841203 | 4.974591 | 4.905910 | 4.898801 | 4.902994 | 4.924218 | 34 | 30 | 25 | 32 |
| 102723777 | gene-LOC1027 | LOC10272377 | . | pseudogene | . | . | | . | . | . | 0.329041 | 0.384799 | 0.490745 | 0.431527 | 0.765692 | 0.889479 | 1.101218 | 0.986634 | 1.266443 | 19.358253 | 0.697585396 | 0.488517257 | 0.625183505 | NA | 4.094525 | 4.435167 | 4.252212 | 4.265191 | 4.283619 | 4.272565 | 18 | 20 | 20 | 19 |
| 102723780 | NR_132320 | FRG1FP | FSHD region gene 1 family member F, pseudogene | transcribed_ps | . | | 51765 | . | . | . | 0.582127 | 0.455673 | 0.570693 | 0.740832 | 1.354633 | 1.053309 | 1.280618 | 1.693824 | 1.233893 | 18.949977 | 0.708879336 | 0.427741602 | 0.668839258 | NA | 4.084644 | 4.387716 | 4.241127 | 4.222356 | 4.238158 | 4.259214 | 22 | 16 | 16 | 22 |
| 102723811 | XR_00295844 | LOC10272381 | uncharacterized LOC102723811 | lncRNA | . | . | | . | . | . | 0.048617 | 0.182367 | 0.288039 | 0.143430 | 0.113135 | 0.421550 | 0.646352 | 0.327935 | 1.726848 | 6.193911 | 1.253562803 | 0.628720706 | 0.52953192 | NA | 2.184683 | 2.971139 | 2.594235 | 2.612790 | 2.629550 | 2.607523 | 3 | 7 | 9 | 5 |
| 102723814 | XR_430332 | LOC10272381 | uncharacterized LOC102723814 | lncRNA | . | . | | . | . | . | 0.155546 | 0.125218 | 0.166830 | 0.101353 | 0.361963 | 0.289447 | 0.374361 | 0.231732 | -1.068534 | 10.385506 | 0.954737985 | -0.100167 | 0.920211746 | NA | 3.421266 | 3.330299 | 3.383509 | 3.370294 | 3.385029 | 3.362529 | 14 | 10 | 11 | 7 |
| 102723834 | XR_00173849 | LOC10272383 | uncharacterized LOC102723834, transcript variant X1 | lncRNA | . | . | | . | . | . | 0.256943 | 0.566190 | 0.058528 | 0.205598 | 0.597918 | 1.308775 | 0.131336 | 0.470076 | -2.925893 | 20.337710 | 0.785186608 | -1.97262266 | 0.04853857 | NA | 4.924835 | 3.364566 | 4.337407 | 4.423650 | 4.272407 | 4.320689 | 23 | 44 | 4 | 15 |
| 102723859 | NM_00129146 | TBC1D3E | TBC1 domain family member 3E | protein_coding | NP_001278395.1;XP_006722317.1 | | 27071 | 610808 | . | . | 2.461880 | 2.730614 | 3.169017 | 3.226574 | 5.728896 | 6.311937 | 7.111181 | 7.377179 | 1.215910 | 221.348625 | 0.227703553 | 1.238611848 | 0.215489278 | 0.981973755 | 7.642532 | 7.924104 | 7.716251 | 7.760423 | 7.827939 | 7.834001 | 219 | 226 | 210 | 229 |
| 102723870 | XR_426677 | LOC10272387 | uncharacterized LOC102723870 | lncRNA | . | . | | . | . | . | 0.599104 | 0.378879 | 0.571470 | 0.632344 | 1.394140 | 0.875797 | 1.282362 | 1.445779 | 1.241007 | 9.481239 | 0.991747032 | 0.314102984 | 0.753442824 | NA | 3.079678 | 3.393446 | 3.245519 | 3.227409 | 3.244898 | 3.246368 | 12 | 7 | 9 | 10 |
| 102723872 | XR_00295652 | LOC10272387 | uncharacterized LOC102723872, transcript variant X1 | lncRNA | . | . | | . | . | . | 0.188653 | 0.218219 | 0.248067 | 0.150809 | 0.439004 | 0.504423 | 0.556655 | 0.344807 | -1.082548 | 16.867042 | 0.759378133 | -0.15068954 | 0.880220624 | NA | 4.130737 | 4.019385 | 4.071961 | 4.080886 | 4.089378 | 4.051599 | 19 | 20 | 18 | 11 |
| 102723897 | XR_00173755 | WASH9P | WAS protein family homolog 9, pseudogene | transcribed_ps | . | | 53981 | . | . | . | 14.424350 | 15.311575 | 17.885582 | 15.360954 | 33.566055 | 35.393391 | 40.134731 | 35.120995 | 1.103015 | 863.989916 | 0.15042652 | 0.940341203 | 0.347042571 | 0.998865404 | 9.682217 | 9.824040 | 9.698923 | 9.734993 | 9.844169 | 9.727223 | 922 | 909 | 852 | 781 |
| 102723922 | XR_00174391 | LOC10272392 | uncharacterized LOC102723922, transcript variant X3 | lncRNA | . | . | | . | . | . | 0.083517 | 0.053700 | 0.059276 | 0.135457 | 0.194347 | 0.124130 | 0.133013 | 0.309706 | 1.913714 | 7.054836 | 1.219171304 | 0.76804229 | 0.442462061 | NA | 2.273436 | 3.213338 | 2.808848 | 2.780885 | 2.799535 | 2.830575 | 9 | 2 | 5 | 12 |
| 102723927 | NR_110592 | LINC01238 | long intergenic non-protein coding RNA 1238 | lncRNA | . | | 49795 | . | . | . | 0.310412 | 0.312844 | 0.508524 | 0.500640 | 0.722341 | 0.723153 | 1.141114 | 1.144654 | 1.594953 | 27.914690 | 0.584242842 | 1.152798343 | 0.248993165 | 0.992733415 | 4.427143 | 5.100822 | 4.764714 | 4.763388 | 4.816981 | 4.816050 | 25 | 23 | 30 | 32 |
| 102723961 | XR_430026,XR | LOC10272396 | uncharacterized LOC102723961, transcript variant X10 | lncRNA | . | . | | . | . | . | 0.033634 | 0.049975 | 0.037416 | 0.033257 | 0.078269 | 0.115520 | 0.083961 | 0.076039 | -1.083473 | 6.929911 | 1.148960728 | -0.10066775 | 0.919814216 | NA | 2.849979 | 2.733338 | 2.788849 | 2.799342 | 2.792718 | 2.790487 | 7 | 9 | 6 | 6 |
| 102723981 | gene-RPL23AP | RPL23AP84 | ribosomal protein L23a pseudogene 84 | pseudogene | . | | 51575 | . | . | . | 0.112714 | 0.111374 | 0.132199 | 0.093909 | 0.262291 | 0.257445 | 0.296650 | 0.214712 | -1.007340 | 2.223025 | 1.975467281 | -0.0053412 | 0.995738358 | NA | 1.156660 | 1.148375 | 1.155078 | 1.151489 | 1.153584 | 1.152831 | 3 | 2 | 2 | 2 |
| 102723983 | XR_429136 | LOC10272398 | uncharacterized LOC102723983 | lncRNA | . | . | | . | . | . | 2.132471 | 1.914912 | 2.352337 | 1.729607 | 4.962348 | 4.426406 | 5.278576 | 3.954541 | -1.006696 | 49.098129 | 0.452873085 | -0.02126121 | 0.983037285 | 0.998865404 | 5.621228 | 5.613955 | 5.627114 | 5.605205 | 5.641424 | 5.587844 | 60 | 50 | 49 | 39 |
| 102723989 | XR_428560,XR | LOC10272398 | uncharacterized LOC102723989, transcript variant X2 | lncRNA | . | . | | . | . | . | 1.431606 | 1.640121 | 1.411328 | 1.652013 | 3.331405 | 3.791214 | 3.166980 | 3.777131 | -1.019177 | 44.032914 | 0.47090801 | -0.05819585 | 0.953592627 | 0.998865404 | 5.474910 | 5.445966 | 5.451219 | 5.472819 | 5.446444 | 5.471225 | 48 | 51 | 35 | 44 |
| 102723996 | NM_00136377 | LOC10272399 | ICOS ligand | protein_coding | NP_001350699.1;XP_006723962.1;XP_011544380.1;XP_011544381.1 | . | | . | . | . | 0.166674 | 0.181000 | 0.153118 | 0.147728 | 0.387857 | 0.418390 | 0.343592 | 0.337763 | -1.118309 | 17.000846 | 0.741227739 | -0.21763709 | 0.827711882 | NA | 4.166062 | 4.004486 | 4.090790 | 4.095853 | 4.085419 | 4.084917 | 20 | 20 | 14 | 15 |
| 102724009 | NR_120665 | KIF1C-AS1 | KIF1C antisense RNA 1 | lncRNA | . | | 40324 | . | . | . | 0.425389 | 0.219089 | 0.357346 | 0.796817 | 0.989898 | 0.506433 | 0.801874 | 1.821826 | 1.754688 | 8.570722 | 1.070075685 | 0.758090851 | 0.448396596 | NA | 2.638039 | 3.448451 | 3.084379 | 3.069650 | 3.080615 | 3.115677 | 9 | 5 | 6 | 14 |
| 102724023 | NM_00136375 | GATD3B | glutamine amidotransferase like class 1 domain containing 3B | protein_coding | NP_001350687.1;NP_001350689.1;NP_001350690.1 | | 53816 | . | . | . | 12.685601 | 12.837985 | 12.124569 | 14.063709 | 29.519912 | 29.675574 | 27.207183 | 32.155003 | 1.015751 | 636.531481 | 0.159456941 | 0.141401434 | 0.887552821 | 0.998865404 | 9.303026 | 9.325066 | 9.308932 | 9.307759 | 9.272347 | 9.363106 | 728 | 681 | 520 | 642 |
| 102724055 | XR_00175176 | LOC10272405 | uncharacterized LOC102724055, transcript variant X2 | lncRNA | . | . | | . | . | . | 0.806261 | 0.713053 | 0.552116 | 0.615785 | 1.876202 | 1.648253 | 1.238931 | 1.407919 | -1.132274 | 65.807954 | 0.392998193 | -0.4560411 | 0.64836042 | 0.998865404 | 6.127733 | 5.946990 | 6.056977 | 6.053890 | 6.000708 | 6.052958 | 81 | 75 | 47 | 65 |
| 102724064 | NR_135761 | LOC10272406 | uncharacterized LOC102724064 | lncRNA | . | . | | . | . | . | 0.484311 | 1.349568 | 1.411820 | 1.540299 | 1.127011 | 3.119586 | 3.168084 | 3.521711 | 1.539127 | 21.901938 | 0.70300692 | 0.884930174 | 0.376194292 | NA | 4.111656 | 4.728813 | 4.381149 | 4.450106 | 4.450466 | 4.460716 | 11 | 27 | 22 | 26 |
| 102724078 | XR_00175146 | LOC10272407 | uncharacterized LOC102724078, transcript variant X2 | lncRNA | . | . | | . | . | . | 0.170031 | 0.267820 | 0.272606 | 0.147242 | 0.395669 | 0.619079 | 0.611720 | 0.336652 | -1.052914 | 45.359514 | 0.504885152 | -0.14733479 | 0.882867773 | 0.998865404 | 5.539252 | 5.466498 | 5.458911 | 5.536325 | 5.537751 | 5.438804 | 42 | 61 | 50 | 29 |
| 102724081 | XR_00173976 | LOC10272408 | uncharacterized LOC102724081, transcript variant X2 | lncRNA | . | . | | . | . | . | 0.040987 | 0.021710 | 0.021466 | 0.054668 | 0.095378 | 0.050184 | 0.048169 | 0.124991 | -1.103401 | 6.346178 | 1.215547826 | -0.11678479 | 0.907030595 | NA | 2.735321 | 2.592943 | 2.680537 | 2.661680 | 2.661989 | 2.674606 | 10 | 5 | 4 | 7 |
| 102724093 | XR_429585 | LOC10272409 | golgin subfamily A member 6-like protein 4 | transcribed_ps | . | . | | . | . | . | 0.184570 | 0.161940 | 0.244648 | 0.170499 | 0.429501 | 0.374333 | 0.548984 | 0.389825 | 1.177433 | 27.799903 | 0.592879652 | 0.397458368 | 0.691029481 | 0.998865404 | 4.672908 | 4.911273 | 4.787579 | 4.775737 | 4.821821 | 4.778026 | 31 | 26 | 31 | 23 |
| 102724101 | NM_00133006 | TP53TG3E | TP53 target 3 family member E | protein_coding | NP_001316990.1;XP_006721180.1;XP_011544334.1;XP_016879440.1 | | 51816 | . | . | . | 1.024863 | 0.801999 | 0.815732 | 0.577071 | 2.384899 | 1.853858 | 1.830479 | 1.319406 | -1.220716 | 52.454453 | 0.443169141 | -0.64925053 | 0.516176457 | 0.998865404 | 5.848444 | 5.563496 | 5.755810 | 5.706866 | 5.718021 | 5.668491 | 74 | 55 | 47 | 38 |
| 102724105 | NR_125345 | ZNF528-AS1 | ZNF528 antisense RNA 1 | lncRNA | . | | 51305 | . | . | . | 0.839438 | 1.006372 | 0.782666 | 1.647707 | 1.953407 | 2.326274 | 1.756280 | 3.767286 | 1.289391 | 44.177238 | 0.509435999 | 0.71979608 | 0.471650559 | 0.998865404 | 5.272576 | 5.635164 | 5.419525 | 5.444132 | 5.408508 | 5.536759 | 41 | 45 | 28 | 63 |
| 102724117 | XR_429586 | LOC10272411 | golgin subfamily A member 6-like protein 4 | transcribed_ps | . | . | | . | . | . | 0.239524 | 0.237843 | 0.335392 | 0.195615 | 0.557381 | 0.549785 | 0.752610 | 0.447251 | 1.095463 | 35.261918 | 0.542300433 | 0.242560283 | 0.808346052 | 0.998865404 | 5.071077 | 5.205854 | 5.127481 | 5.125499 | 5.177563 | 5.099121 | 39 | 36 | 41 | 25 |
| 102724122 | XR_427722 | LOC10272412 | uncharacterized LOC102724122 | lncRNA | . | . | | . | . | . | 3.786835 | 3.272857 | 2.614487 | 2.467817 | 8.812120 | 7.565356 | 5.866833 | 5.642369 | -1.411002 | 359.239394 | 0.194570527 | -2.55290279 | 0.010682931 | 0.333896806 | 8.715558 | 8.219636 | 8.622960 | 8.528001 | 8.398253 | 8.362101 | 521 | 419 | 268 | 270 |
| 102724127 | NM_00133006 | TP53TG3F | TP53 target 3 family member F | protein_coding | NP_001316995.1;XP_006721181.1;XP_011544335.1;XP_016879445.1 | | 51817 | . | . | . | 0.937386 | 0.797758 | 1.077536 | 0.600117 | 2.181335 | 1.844052 | 2.417960 | 1.372098 | -1.195903 | 56.226692 | 0.435096631 | -0.5932028 | 0.553045436 | 0.998865404 | 5.934966 | 5.680165 | 5.856032 | 5.802175 | 5.827604 | 5.757981 | 79 | 58 | 53 | 39 |
| 102724135 | NR_158192 | LOC10272413 | uncharacterized LOC102724135 | transcribed_ps | . | . | | . | . | . | 4.449051 | 4.605465 | 4.461332 | 4.664124 | 10.353124 | 10.645739 | 10.011100 | 10.663964 | -1.005584 | 364.936323 | 0.186480523 | -0.04307983 | 0.965637898 | 0.998865404 | 8.515640 | 8.507350 | 8.505604 | 8.520600 | 8.500052 | 8.519606 | 416 | 400 | 311 | 347 |
| 102724146 | XR_00295740 | LOC10272414 | uncharacterized LOC102724146, transcript variant X1 | lncRNA | . | . | | . | . | . | 0.085398 | 0.073251 | 0.226652 | 0.145219 | 0.198724 | 0.169322 | 0.508600 | 0.332027 | 2.350945 | 6.753900 | 1.187338459 | 1.038660011 | 0.298962885 | NA | 2.008843 | 3.245375 | 2.721830 | 2.719004 | 2.753964 | 2.736502 | 5 | 4 | 10 | 7 |
| 102724149 | XR_429273 | LOC10272414 | uncharacterized LOC102724149 | lncRNA | . | . | | . | . | . | 0.241542 | 0.381500 | 0.281510 | 0.508510 | 0.562077 | 0.881854 | 0.631701 | 1.162647 | 1.287059 | 18.042377 | 0.745475692 | 0.488383146 | 0.625278477 | NA | 3.982758 | 4.341619 | 4.139171 | 4.171794 | 4.152722 | 4.196987 | 14 | 21 | 13 | 24 |
| 102724159 | NM_00136823 | LOC10272415 | periodic tryptophan protein 2 homolog | protein_coding | NP_001355167.1 | . | | . | . | . | 5.304486 | 5.663676 | 4.752754 | 5.120751 | 12.343756 | 13.091842 | 10.665043 | 11.707989 | -1.122686 | 519.173516 | 0.166996963 | -0.99974344 | 0.317434683 | 0.998865404 | 9.101369 | 8.933921 | 9.036682 | 9.074961 | 8.959477 | 9.003867 | 614 | 610 | 410 | 475 |
| 102724194 | XR_427194,XR | LOC10272419 | uncharacterized LOC102724194, transcript variant X2 | lncRNA | . | . | | . | . | . | 0.073820 | 0.069003 | 0.151881 | 0.129197 | 0.171781 | 0.159504 | 0.340816 | 0.295393 | 2.011194 | 5.407643 | 1.306422518 | 0.771612725 | 0.440343835 | NA | 1.844134 | 2.852980 | 2.408846 | 2.410093 | 2.429707 | 2.422568 | 4 | 4 | 7 | 6 |
| 102724200 | XM_01154608 | LOC10272420 | trafficking protein particle complex subunit 10-like | protein_coding | XP_011544384.3 | . | | . | . | . | 1.767437 | 1.803469 | 2.466884 | 2.134434 | 4.112899 | 4.168799 | 5.535615 | 4.880128 | 1.270831 | 205.230177 | 0.237301404 | 1.457101551 | 0.14508835 | 0.906189003 | 7.497490 | 7.843956 | 7.615891 | 7.620821 | 7.763474 | 7.690829 | 207 | 196 | 215 | 199 |
| 102724219 | NM_00132204 | LOC10272421 | uncharacterized LOC102724219 | protein_coding | NP_001308973.1;XP_011544385.1;XP_011544386.1;XP_011544387.2;XP | . | | . | . | . | 3.928923 | 3.752784 | 4.014891 | 3.423325 | 9.142765 | 8.674729 | 9.009298 | 7.827028 | 1.024623 | 327.211165 | 0.206038443 | 0.170323209 | 0.86475596 | 0.998865404 | 8.335843 | 8.372085 | 8.352307 | 8.337487 | 8.423394 | 8.289355 | 377 | 344 | 323 | 273 |
| 102724224 | NR_110599 | LINC01117 | long intergenic non-protein coding RNA 1117 | lncRNA | . | | 49260 | . | . | . | 0.836267 | 0.913512 | 1.299509 | 0.630315 | 1.946027 | 2.111623 | 2.916060 | 1.441140 | 1.094055 | 19.307467 | 0.726074266 | 0.178610686 | 0.858243403 | NA | 4.202828 | 4.336262 | 4.254047 | 4.263292 | 4.296134 | 4.234934 | 20 | 21 | 24 | 12 |
| 102724233 | XR_427040,XR | LINC02576 | long intergenic non-protein coding RNA 2576, transcript variant X | lncRNA | . | | 53748 | . | . | . | 0.113894 | 0.026248 | 0.035505 | 0.072412 | 0.265036 | 0.060674 | 0.079671 | 0.165562 | 1.040789 | 2.692226 | 1.853674999 | 0.031115311 | 0.975177579 | NA | 1.397517 | 1.459419 | 1.438737 | 1.422501 | 1.429119 | 1.433167 | 5 | 1 | 2 | 3 |
| 102724236 | gene-LOC1027 | LOC10272423 | . | pseudogene | . | . | | . | . | . | 0.026485 | 0.021000 | 0.084055 | 0.005644 | 0.061632 | 0.048542 | 0.188618 | 0.012904 | 1.683953 | 1.795055 | 2.231855449 | 0.336873073 | 0.736212585 | NA | 0.413393 | 1.175234 | 0.827582 | 0.823701 | 0.835285 | 0.824501 | 2 | 1 | 3 | 1 |
| 102724238 | NR_126048 | LOC10272423 | uncharacterized LOC102724238 | lncRNA | . | . | | . | . | . | 1.102616 | 1.040254 | 0.944997 | 0.969108 | 2.565832 | 2.404594 | 2.120546 | 2.215750 | -1.125464 | 72.320746 | 0.371003967 | -0.459616 | 0.645791872 | 0.998865404 | 6.258969 | 6.088684 | 6.200286 | 6.183534 | 6.162566 | 6.163734 | 91 | 80 | 59 | 64 |
| 102724301 | NR_120587 | LOC10272430 | uncharacterized LOC102724301 | lncRNA | . | . | | . | . | . | 1.279592 | 0.698725 | 1.446535 | 1.072074 | 2.977662 | 1.615135 | 3.245982 | 2.451170 | 1.248655 | 19.010383 | 0.723495246 | 0.442814828 | 0.657899677 | NA | 4.077189 | 4.401999 | 4.253632 | 4.211033 | 4.262697 | 4.238075 | 25 | 13 | 21 | 17 |
| 102724316 | NR_003930,NF | SVIL-AS1 | SVIL antisense RNA 1 | lncRNA | . | | 51219 | . | . | . | 8.690468 | 11.654998 | 10.018231 | 11.242170 | 20.223073 | 26.941053 | 22.480622 | 25.703884 | 1.007716 | 677.054989 | 0.175298009 | 0.06326263 | 0.949557373 | 0.998865404 | 9.398014 | 9.408226 | 9.296637 | 9.489556 | 9.370176 | 9.434231 | 668 | 830 | 563 | 667 |
| 102724322 | XR_00174665 | LOC10272432 | uncharacterized LOC102724322, transcript variant X2 | lncRNA | . | . | | . | . | . | 0.090651 | 0.105532 | 0.056696 | 0.118244 | 0.210948 | 0.243943 | 0.127223 | 0.270353 | -1.330063 | 10.205160 | 0.970428644 | -0.4240338 | 0.671541166 | NA | 3.544574 | 3.127898 | 3.361579 | 3.364911 | 3.337410 | 3.363264 | 13 | 13 | 5 | 11 |
| 102724329 | XR_426856,XR | LOC10272432 | uncharacterized LOC102724329, transcript variant X1 | lncRNA | . | . | | . | . | . | 0.140313 | 0.213288 | 0.147086 | 0.105941 | 0.326514 | 0.493025 | 0.330056 | 0.242221 | -1.389348 | 13.971992 | 0.832264159 | -0.57002092 | 0.568663509 | NA | 4.021847 | 3.548418 | 3.801132 | 3.829500 | 3.804686 | 3.786482 | 15 | 21 | 12 | 9 |
| 102724330 | XR_00174589 | LOC10272433 | uncharacterized LOC102724330, transcript variant X1 | lncRNA | . | . | | . | . | . | 0.660910 | 0.988583 | 1.296656 | 0.470952 | 1.537964 | 2.285153 | 2.909659 | 1.076776 | -1.171972 | 79.823607 | 0.368523056 | -0.62123077 | 0.534447796 | 0.998865404 | 6.427631 | 6.200961 | 6.337082 | 6.337540 | 6.340454 | 6.249204 | 99 | 93 | 76 | 56 |
| 102724334 | XM_00672391 | LOC10272433 | histone H2B type F-S-like | protein_coding | XP_006723981.1 | . | | . | . | . | 0.250081 | 0.126347 | 0.191811 | 0.123605 | 0.581949 | 0.292057 | 0.430418 | 0.282608 | 1.043482 | 2.727629 | 1.804920749 | 0.034021556 | 0.972859962 | NA | 1.413393 | 1.481107 | 1.449528 | 1.441809 | 1.449161 | 1.443240 | 4 | 2 | 3 | 2 |
| 102724351 | XR_00174757 | LOC10272435 | uncharacterized LOC102724351, transcript variant X2 | lncRNA | . | . | | . | . | . | 0.580164 | 0.582462 | 0.663882 | 0.222519 | 1.350066 | 1.346386 | 1.489732 | 0.508763 | -1.422915 | 27.054033 | 0.615621963 | -0.82656226 | 0.408485217 | 0.998865404 | 4.987929 | 4.483795 | 4.789749 | 4.764798 | 4.764524 | 4.712353 | 40 | 31 | 25 | 15 |
| 102724360 | NR_134912,NF | HOMER3-AS1 | HOMER3 antisense RNA 1 | lncRNA | . | | 53775 | . | . | . | 0.261847 | 0.044315 | 0.356661 | 0.334412 | 0.609328 | 0.102437 | 0.800337 | 0.764592 | 2.091485 | 8.176081 | 1.130675516 | 0.941497121 | 0.346450164 | NA | 2.397517 | 3.470304 | 3.016209 | 2.984198 | 3.028171 | 3.024826 | 10 | 2 | 10 | 10 |
| 102724364 | XR_922650 | SEC22B4P | SEC22 homolog B4, pseudogene | transcribed_ps | . | | 53892 | . | . | . | 1.200966 | 1.062481 | 1.204887 | 0.941589 | 2.794697 | 2.455973 | 2.703731 | 2.152832 | -1.071527 | 249.227167 | 0.230273197 | -0.43282684 | 0.665140575 | 0.998865404 | 8.009496 | 7.911474 | 8.008984 | 7.942146 | 8.001698 | 7.878186 | 316 | 260 | 236 | 197 |
| 102724378 | NR_160696,NF | LOC10272437 | uncharacterized LOC102724378 | lncRNA | . | . | | . | . | . | 1.995160 | 1.837695 | 2.050910 | 2.106572 | 4.642821 | 4.247913 | 4.602184 | 4.816426 | 1.075842 | 57.614696 | 0.412592349 | 0.255617697 | 0.798246052 | 0.998865404 | 5.794547 | 5.900248 | 5.849420 | 5.829729 | 5.850896 | 5.858508 | 67 | 57 | 51 | 57 |
| 102724397 | XR_430055,XR | LOC10272439 | uncharacterized LOC102724397, transcript variant X1 | lncRNA | . | . | | . | . | . | 0.978302 | 0.710260 | 0.908405 | 0.839411 | 2.276550 | 1.641797 | 2.038435 | 1.919215 | 1.030971 | 26.746678 | 0.601378559 | 0.07317046 | 0.941670473 | NA | 4.718015 | 4.764191 | 4.755420 | 4.721069 | 4.747979 | 4.735427 | 35 | 24 | 25 | 24 |
| 102724428 | NM_00132064 | SIK1B | salt inducible kinase 1B (putative) | protein_coding | NP_001307572.1;XP_016883697.1 | | 52389 | . | . | . | 0.579842 | 0.621193 | 0.507236 | 0.453090 | 1.349315 | 1.435915 | 1.138222 | 1.035936 | -1.254986 | 80.654253 | 0.354909847 | -0.92325053 | 0.355876664 | 0.998865404 | 6.487990 | 6.160863 | 6.354198 | 6.372713 | 6.318444 | 6.289309 | 100 | 100 | 66 | 63 |
| 102724430 | gene-KRT8P50 | KRT8P50 | keratin 8 pseudogene 50 | pseudogene | . | | 48346 | . | . | . | 0.062861 | 0.020366 | 0.086921 | 0.055443 | 0.146279 | 0.047077 | 0.195047 | 0.126764 | 1.671915 | 3.569283 | 1.602038051 | 0.462848637 | 0.643472866 | NA | 1.413393 | 2.161867 | 1.821784 | 1.814090 | 1.831907 | 1.820551 | 4 | 2 | 5 | 3 |
| 102724450 | NR_125997 | RNF207-AS1 | RNF207 antisense RNA 1 | lncRNA | . | | 41286 | . | . | . | 13.218796 | 10.694854 | 12.496509 | 11.708195 | 30.760681 | 24.721632 | 28.041803 | 26.769398 | -1.005439 | 142.187645 | 0.280639122 | -0.02788627 | 0.977752859 | 0.998865404 | 7.154922 | 7.148375 | 7.192551 | 7.108721 | 7.163364 | 7.135478 | 182 | 137 | 128 | 128 |
| 102724474 | XM_01153753 | LOC10272447 | WAS/WASL-interacting protein family member 2-like | protein_coding | XP_011535834.1 | . | | . | . | . | 0.208057 | 0.232294 | 0.296891 | 0.341295 | 0.484158 | 0.536957 | 0.666215 | 0.780330 | 1.405971 | 11.348751 | 0.903497258 | 0.544071039 | 0.586392589 | NA | 3.238681 | 3.728813 | 3.484633 | 3.491716 | 3.502491 | 3.507884 | 10 | 11 | 11 | 13 |
| 102724479 | XR_00174053 | LOC10272447 | uncharacterized LOC102724479, transcript variant X1 | lncRNA | . | . | | . | . | . | 1.963848 | 1.053223 | 0.817700 | 2.049879 | 4.569956 | 2.434571 | 1.834894 | 4.686806 | -1.086787 | 63.120714 | 0.408119297 | -0.29419975 | 0.768605274 | 0.998865404 | 6.039657 | 5.917857 | 6.002415 | 5.976170 | 5.932344 | 6.008240 | 80 | 67 | 43 | 67 |
| 102724484 | NR_120522 | CHN2-AS1 | CHN2 antisense RNA 1 | lncRNA | . | | 40149 | . | . | . | 0.174945 | 0.174489 | 0.136261 | 0.193778 | 0.407104 | 0.403339 | 0.305767 | 0.443050 | -1.064509 | 7.378286 | 1.115912755 | -0.08082017 | 0.935584965 | NA | 2.928838 | 2.836248 | 2.887681 | 2.885802 | 2.878545 | 2.890750 | 9 | 8 | 5 | 8 |
| 102724497 | XR_00173954 | LOC10272449 | uncharacterized LOC102724497, transcript variant X1 | lncRNA | . | . | | . | . | . | 0.070940 | 0.254477 | 0.188537 | 0.169363 | 0.165081 | 0.588235 | 0.423071 | 0.387227 | 1.231406 | 11.011701 | 0.914719976 | 0.328304929 | 0.742681118 | NA | 3.303566 | 3.602870 | 3.450133 | 3.453154 | 3.458942 | 3.464641 | 11 | 11 | 10 | 12 |
| 102724515 | NR_125792 | LINC01291 | long intergenic non-protein coding RNA 1291 | lncRNA | . | | 50358 | . | . | . | 23.632633 | 26.157272 | 17.479305 | 17.784653 | 54.994110 | 60.463703 | 39.223053 | 40.662491 | -1.429753 | 328.076893 | 0.196585273 | -2.62362648 | 0.008699914 | 0.293930247 | 8.592980 | 8.076889 | 8.418129 | 8.473143 | 8.248290 | 8.250802 | 424 | 436 | 234 | 254 |
| 102724517 | XR_00175282 | LOC10272451 | uncharacterized LOC102724517 | lncRNA | . | . | | . | . | . | 1.332461 | 1.108742 | 1.377127 | 0.963880 | 3.100691 | 2.562907 | 3.090234 | 2.203796 | -1.062072 | 78.606319 | 0.372927596 | -0.23297276 | 0.815782556 | 0.998865404 | 6.338014 | 6.253908 | 6.326987 | 6.275623 | 6.328737 | 6.241491 | 102 | 79 | 78 | 59 |
| 102724528 | XR_00173829 | LOC10272452 | uncharacterized LOC102724528, transcript variant X3 | lncRNA | . | . | | . | . | . | 0.093888 | 0.140309 | 0.268879 | 0.157289 | 0.218482 | 0.324331 | 0.603356 | 0.359623 | 1.751529 | 7.448959 | 1.124141958 | 0.719317228 | 0.471945483 | NA | 2.436885 | 3.245375 | 2.865900 | 2.876172 | 2.898192 | 2.880708 | 5 | 7 | 10 | 7 |
| 102724532 | NR_135480 | LOC10272453 | uncharacterized LOC102724532 | lncRNA | . | . | | . | . | . | 4.833886 | 4.527539 | 2.894321 | 4.370173 | 11.248652 | 10.465608 | 6.494772 | 9.991882 | -1.299153 | 68.899263 | 0.394192757 | -0.95783386 | 0.338146543 | 0.998865404 | 6.283842 | 5.904070 | 6.147050 | 6.127397 | 6.035349 | 6.114838 | 93 | 81 | 42 | 67 |
| 102724542 | XR_00173957 | LOC10272454 | uncharacterized LOC102724542, transcript variant X6 | lncRNA | . | . | | . | . | . | 4.314555 | 4.674045 | 5.821436 | 7.417227 | 10.040147 | 10.804267 | 13.063134 | 16.958608 | 1.349409 | 126.452781 | 0.289478813 | 1.493468272 | 0.135314654 | 0.892805049 | 6.750634 | 7.182131 | 6.912317 | 6.928436 | 7.003093 | 7.047483 | 121 | 119 | 119 | 144 |
| 102724545 | XR_00175446 | LOC10272454 | uncharacterized LOC102724545, transcript variant X3 | lncRNA | . | . | | . | . | . | 4.674273 | 4.675190 | 5.227613 | 5.646450 | 10.877226 | 10.806910 | 11.730614 | 12.909936 | 1.158108 | 275.647232 | 0.207327607 | 1.021426702 | 0.30705232 | 0.998865404 | 7.996978 | 8.208625 | 8.070145 | 8.060636 | 8.130530 | 8.152559 | 297 | 273 | 252 | 283 |

| | | | | | | | | | | | | | | | | | | | | | | | | | | | | | | | | | | |
|---|---|---|---|---|---|---|---|---|---|---|---|---|---|---|---|---|---|---|---|---|---|---|---|---|---|---|---|---|---|---|---|---|---|---|
| 102724553 | XR_00175152 | LOC10272455 | uncharacterized LOC102724553, transcript variant X1 | lncRNA | . | . | | . | . | . | 0.287294 | 0.104626 | 0.301835 | 0.195978 | 0.668544 | 0.241847 | 0.677308 | 0.448080 | 1.253467 | 6.997799 | 1.166845076 | 0.279321164 | 0.779998366 | NA | 2.631267 | 2.963455 | 2.808661 | 2.785419 | 2.810323 | 2.798090 | 10 | 4 | 8 | 6 |
| 102724560 | NM_00132107 | CBSL | cystathionine beta-synthase like | protein_coding | NP_001308002.1;NP_001340935.1;NP_001340936.1;NP_001340937.1;N | | 51829 | . | . | . | 2.689470 | 3.556172 | 1.721854 | 3.065871 | 6.258509 | 8.220250 | 3.863791 | 7.009750 | -1.307674 | 222.625079 | 0.28256963 | -1.36958615 | 0.170816126 | 0.946389416 | 7.980244 | 7.590451 | 7.784530 | 7.910849 | 7.591635 | 7.838395 | 255 | 306 | 120 | 232 |
| 102724571 | NR_132738 | LNCTAM34A | long non coding transcriptional activator of miR34a | lncRNA | . | | 52548 | . | . | . | 3.104455 | 2.376106 | 2.162801 | 2.706676 | 7.224193 | 5.492474 | 4.853263 | 6.188492 | -1.139474 | 79.610445 | 0.367440311 | -0.51264914 | 0.608196775 | 0.998865404 | 6.406077 | 6.217540 | 6.369340 | 6.292793 | 6.272251 | 6.322490 | 111 | 79 | 58 | 77 |
| 102724579 | NR_136323 | LOC10272457 | uncharacterized LOC102724579 | lncRNA | . | . | | . | . | . | 0.266090 | 0.203200 | 0.245959 | 0.036120 | 0.619203 | 0.469706 | 0.551924 | 0.082585 | -1.393479 | 2.692648 | 1.841362901 | -0.25996586 | 0.794890106 | NA | 1.644780 | 1.175234 | 1.435544 | 1.432221 | 1.435169 | 1.424396 | 4 | 3 | 3 | 1 |
| 102724580 | NR_136300 | LOC10272458 | methylenetetrahydrofolate dehydrogenase (NADP+ dependent) 1 | transcribed_ps | . | . | | . | . | . | 0.054862 | 0.129708 | 0.100188 | 0.098549 | 0.127666 | 0.299825 | 0.224820 | 0.225320 | 1.144258 | 7.779890 | 1.106929037 | 0.175632178 | 0.860582921 | NA | 2.861598 | 3.051647 | 2.936999 | 2.964290 | 2.954390 | 2.956562 | 5 | 11 | 7 | 8 |
| 102724594 | NM_00132064 | U2AF1L5 | U2 small nuclear RNA auxiliary factor 1 like 5 | protein_coding | NP_001307575.1;NP_001307577.1;NP_001307579.1;NP_001307580.1;X | | 51830 | . | . | . | 22.247720 | 21.410186 | 19.622620 | 21.728776 | 51.771358 | 49.490606 | 44.032591 | 49.680264 | -1.068449 | 639.971552 | 0.157058358 | -0.60817044 | 0.54307444 | 0.998865404 | 9.368942 | 9.273198 | 9.360430 | 9.326812 | 9.269218 | 9.328209 | 780 | 696 | 514 | 606 |
| 102724596 | NR_110882,N | LOC10272459 | uncharacterized LOC102724596 | lncRNA | . | . | | . | . | . | 0.755242 | 0.439581 | 0.595823 | 0.308705 | 1.757479 | 1.016112 | 1.337008 | 0.705818 | -1.443643 | 10.952300 | 0.956475347 | -0.55381847 | 0.579703088 | NA | 3.690761 | 3.168566 | 3.476023 | 3.451410 | 3.462043 | 3.430097 | 18 | 11 | 11 | 5 |
| 102724642 | XM_02445328 | LOC10272464 | anaphase-promoting complex subunit 1-like | protein_coding | XP_024309048.1 | . | | . | . | . | 1.799028 | 1.723485 | 2.053500 | 1.876004 | 4.186413 | 3.983914 | 4.607995 | 4.289259 | 1.099058 | 147.666510 | 0.269835661 | 0.505002401 | 0.61355715 | 0.998865404 | 7.136066 | 7.273080 | 7.195167 | 7.176395 | 7.240869 | 7.203331 | 166 | 148 | 141 | 138 |
| 102724660 | XR_428245,XR | LOC10272466 | uncharacterized LOC102724660, transcript variant X4 | lncRNA | . | . | | . | . | . | 0.973518 | 0.685326 | 0.969761 | 0.793164 | 2.265417 | 1.584162 | 2.176115 | 1.813476 | -1.001450 | 49.529855 | 0.459624243 | -0.0045465 | 0.996372429 | 0.999862634 | 5.629658 | 5.630795 | 5.656346 | 5.597164 | 5.655284 | 5.597675 | 66 | 45 | 50 | 39 |
| 102724668 | NR_132360 | DPY19L1P2 | DPY19L1 pseudogene 2 | transcribed_ps | . | | 22851 | . | . | . | 0.740378 | 0.861997 | 1.113472 | 0.754266 | 1.722889 | 1.992545 | 2.498598 | 1.724539 | 1.164445 | 32.107972 | 0.556047716 | 0.395006595 | 0.692838021 | 0.998865404 | 4.889947 | 5.111291 | 4.980412 | 4.995992 | 5.033802 | 4.980988 | 32 | 34 | 36 | 26 |
| 102724699 | NR_125409 | LINC02043 | long intergenic non-protein coding RNA 2043 | lncRNA | . | | 52883 | . | . | . | 0.313468 | 0.372968 | 0.214696 | 0.442787 | 0.729453 | 0.862133 | 0.481771 | 1.012379 | -1.054595 | 19.682222 | 0.716378299 | -0.10705068 | 0.91474877 | NA | 4.339107 | 4.257378 | 4.291917 | 4.309268 | 4.269098 | 4.322669 | 21 | 24 | 11 | 24 |
| 102724701 | XR_00175510 | LOC10272470 | uncharacterized LOC102724701, transcript variant X6 | lncRNA | . | . | | . | . | . | 5.772083 | 5.721312 | 6.109233 | 4.871456 | 13.431871 | 13.225067 | 13.708937 | 11.138009 | -1.005041 | 539.112391 | 0.181832159 | -0.03989368 | 0.968177887 | 0.998865404 | 9.077590 | 9.071288 | 9.071086 | 9.075553 | 9.156059 | 8.976556 | 620 | 585 | 533 | 434 |
| 102724715 | XR_00295813 | LOC10272471 | uncharacterized LOC102724715, transcript variant X1 | lncRNA | . | . | | . | . | . | 0.140410 | 0.343844 | 0.233491 | 0.140286 | 0.326739 | 0.794812 | 0.523946 | 0.320747 | -1.174530 | 10.936332 | 0.961304571 | -0.24142614 | 0.809224858 | NA | 3.562483 | 3.330299 | 3.426717 | 3.470874 | 3.452507 | 3.429955 | 8 | 18 | 11 | 7 |
| 102724748 | XR_00174244 | LOC10272474 | uncharacterized LOC102724748, transcript variant X1 | lncRNA | . | . | | . | . | . | 0.567137 | 0.361867 | 0.729530 | 0.186294 | 1.319752 | 0.836472 | 1.637044 | 0.425939 | -1.024049 | 57.734344 | 0.54433121 | -0.06298533 | 0.949778189 | 0.998865404 | 5.865705 | 5.836866 | 5.874206 | 5.778710 | 5.935243 | 5.694523 | 82 | 49 | 79 | 22 |
| 102724774 | NR_131251 | LINC01594 | long intergenic non-protein coding RNA 1594 | lncRNA | . | | 51584 | . | . | . | 0.164831 | 0.211062 | 0.421922 | 0.240088 | 0.383568 | 0.487880 | 0.946780 | 0.548932 | 1.650085 | 5.372662 | 1.308034072 | 0.55238618 | 0.580683797 | NA | 2.019256 | 2.742346 | 2.402555 | 2.408167 | 2.423384 | 2.411450 | 4 | 5 | 7 | 5 |
| 102724805 | XR_940344,XR | LOC10272480 | uncharacterized LOC102724805, transcript variant X2 | lncRNA | . | . | | . | . | . | 0.359177 | 0.444160 | 0.432261 | 0.535760 | 0.835820 | 1.026696 | 0.969980 | 1.224951 | 1.189647 | 31.486310 | 0.553989063 | 0.452235476 | 0.651099354 | 0.998865404 | 4.847059 | 5.095559 | 4.949912 | 4.972801 | 4.970988 | 4.995580 | 30 | 34 | 27 | 35 |
| 102724814 | NR_110555 | LOC10272481 | uncharacterized LOC102724814 | lncRNA | . | . | | . | . | . | 1.905594 | 1.953743 | 2.675472 | 1.481508 | 4.434395 | 4.516164 | 6.003681 | 3.387292 | 1.061424 | 203.058379 | 0.285318614 | 0.301420723 | 0.763093694 | 0.998865404 | 7.621037 | 7.709121 | 7.637263 | 7.644095 | 7.793275 | 7.524698 | 225 | 214 | 235 | 139 |
| 102724826 | NR_126465,N | ZNF337-AS1 | ZNF337 antisense RNA 1 | lncRNA | . | | 40759 | . | . | . | 1.647961 | 1.671317 | 1.967643 | 1.819013 | 3.834874 | 3.863323 | 4.415333 | 4.158955 | 1.166253 | 51.072946 | 0.440456419 | 0.503750704 | 0.614436573 | 0.998865404 | 5.558826 | 5.781561 | 5.647330 | 5.662666 | 5.704671 | 5.662679 | 52 | 53 | 53 | 46 |
| 102724827 | NR_110543 | MIATNB | MIAT neighbor | lncRNA | . | | 50731 | . | . | . | 0.624076 | 0.753121 | 0.814896 | 1.071026 | 1.452251 | 1.740873 | 1.828603 | 2.448775 | 1.347185 | 43.221245 | 0.478964409 | 0.89766233 | 0.369365598 | 0.998865404 | 5.203970 | 5.631770 | 5.388776 | 5.413059 | 5.427940 | 5.471211 | 39 | 43 | 38 | 52 |
| 102724832 | XR_00174171 | LOC10272483 | uncharacterized LOC102724832, transcript variant X2 | lncRNA | . | . | | . | . | . | 0.089302 | 0.033187 | 0.000025 | 0.000009 | 0.207810 | 0.076714 | 0.000057 | 0.00002 | -3.218571 | 2.334771 | 2.046315186 | -0.82412528 | 0.409868376 | NA | 1.832372 | 0.148375 | 1.252857 | 1.245446 | 1.238133 | 1.237637 | 5 | 3 | 1 | 1 |
| 102724843 | NM_00132204 | LOC10272484 | uncharacterized LOC102724843 | protein_coding | NP_001308976.1;XP_011544404.1;XP_011544405.1;XP_016883710.1;XP | . | | . | . | . | 6.247564 | 6.517755 | 7.406915 | 5.952951 | 14.538340 | 15.066084 | 16.620903 | 13.610714 | 1.117203 | 829.656901 | 0.177360951 | 0.901500933 | 0.367322038 | 0.998865404 | 9.613805 | 9.774466 | 9.642456 | 9.658452 | 9.850799 | 9.591748 | 892 | 855 | 898 | 674 |
| 102724851 | XR_00174405 | LOC10272485 | uncharacterized LOC102724851, transcript variant X8 | lncRNA | . | . | | . | . | . | 0.033400 | 0.028463 | 0.023635 | 0.023491 | 0.077724 | 0.065794 | 0.053037 | 0.053712 | -1.215204 | 11.003508 | 0.918596862 | -0.3061176 | 0.759515108 | NA | 3.593806 | 3.312263 | 3.473181 | 3.464491 | 3.456254 | 3.462500 | 15 | 12 | 8 | 10 |
| 102724862 | NM_00129146 | TBC1D3I | TBC1 domain family member 3I | protein_coding | NP_001278392.1;XP_006722286.1;XP_006722287.1;XP_006722289.1;XP | | 32709 | . | . | . | 0.467106 | 0.491643 | 0.674152 | 0.606932 | 1.086975 | 1.136454 | 1.512778 | 1.387677 | 1.346932 | 64.259503 | 0.393309622 | 1.092465098 | 0.274628723 | 0.998865404 | 5.774871 | 6.204880 | 5.956892 | 5.970440 | 6.044371 | 6.015537 | 61 | 61 | 68 | 65 |
| 102724880 | XR_00174579 | LOC10272488 | uncharacterized LOC102724880, transcript variant X3 | lncRNA | . | . | | . | . | . | 1.264028 | 0.503394 | 1.199710 | 0.442567 | 2.941444 | 1.163617 | 2.692115 | 1.011876 | 1.108581 | 45.615355 | 0.493116006 | 0.301580569 | 0.762971822 | 0.998865404 | 5.432894 | 5.585944 | 5.528352 | 5.459040 | 5.553923 | 5.468039 | 60 | 37 | 52 | 34 |
| 102724883 | XR_00174766 | LOC10272488 | uncharacterized LOC102724883 | lncRNA | . | . | | . | . | . | 0.249655 | 0.163710 | 0.144869 | 0.217016 | 0.580956 | 0.378424 | 0.325082 | 0.496182 | -1.124976 | 21.024043 | 0.686388989 | -0.24751908 | 0.804506524 | NA | 4.476701 | 4.306201 | 4.420950 | 4.380875 | 4.375440 | 4.405706 | 31 | 19 | 14 | 22 |
| 102724917 | XR_00175069 | LINC02327 | long intergenic non-protein coding RNA 2327, transcript variant X3 | lncRNA | . | | 53247 | . | . | . | 1.119942 | 0.897891 | 1.476813 | 1.797672 | 2.606150 | 2.075516 | 3.313927 | 4.110163 | 1.420071 | 40.593181 | 0.489730427 | 1.033145318 | 0.301535898 | 0.998865404 | 5.067319 | 5.574640 | 5.318044 | 5.294454 | 5.367083 | 5.354404 | 42 | 33 | 43 | 43 |
| 102724933 | NR_120492 | LINC02412 | long intergenic non-protein coding RNA 2412 | lncRNA | . | | 53341 | . | . | . | 0.210344 | 0.154402 | 0.050519 | 0.035423 | 0.489479 | 0.356907 | 0.113364 | 0.080990 | -3.426594 | 4.901229 | 1.440895084 | -1.23310513 | 0.217536543 | NA | 2.923304 | 1.148375 | 2.327952 | 2.317677 | 2.299010 | 2.298006 | 10 | 7 | 2 | 2 |
| 102724945 | XR_00175094 | LOC10272494 | uncharacterized LOC102724945, transcript variant X2 | lncRNA | . | . | | . | . | . | 1.579835 | 1.885244 | 1.073479 | 1.231552 | 3.676339 | 4.357825 | 2.408854 | 2.815798 | -1.524428 | 50.232084 | 0.446183547 | -1.36326896 | 0.17279776 | 0.947431715 | 5.923377 | 5.313775 | 5.670711 | 5.707361 | 5.605652 | 5.618641 | 64 | 71 | 33 | 39 |
| 102724951 | NM_00132203 | LOC10272495 | uncharacterized LOC102724951 | protein_coding | NP_001308965.1;XP_011544407.1;XP_011544408.1;XP_011544409.2;XP | . | | . | . | . | 6.109933 | 5.766501 | 5.798669 | 5.058121 | 14.218065 | 13.329525 | 13.012045 | 11.564794 | 1.019324 | 502.314624 | 0.185183549 | 0.149112417 | 0.881464928 | 0.998865404 | 8.958028 | 8.986725 | 8.976003 | 8.950753 | 9.059568 | 8.884553 | 584 | 526 | 502 | 410 |
| 102724965 | XM_02445327 | LOC10272496 | uncharacterized LOC102724965, transcript variant X1 | protein_coding | XP_024309045.1;XP_024309046.1 | . | | . | . | . | 0.463252 | 0.349854 | 0.623877 | 0.415915 | 1.078006 | 0.808703 | 1.399962 | 0.950940 | 1.245798 | 40.660392 | 0.503208446 | 0.630096846 | 0.528631223 | 0.998865404 | 5.176444 | 5.496901 | 5.338003 | 5.300796 | 5.386122 | 5.320488 | 47 | 34 | 47 | 34 |
| 102725021 | NR_157593,N | LOC10272502 | uncharacterized LOC102725021 | lncRNA | . | . | | . | . | . | 0.369735 | 0.459037 | 0.449172 | 0.410470 | 0.860388 | 1.061083 | 1.007929 | 0.938491 | 1.005386 | 19.906717 | 0.69646954 | 0.011126837 | 0.991122252 | NA | 4.312060 | 4.318300 | 4.293384 | 4.330267 | 4.315554 | 4.309360 | 18 | 26 | 18 | 18 |
| 102725044 | XR_00175066 | LOC10272504 | uncharacterized LOC102725044, transcript variant X5 | lncRNA | . | . | | . | . | . | 1.007054 | 0.869357 | 1.062429 | 1.049602 | 2.343455 | 2.009559 | 2.384059 | 2.399791 | 1.218037 | 36.288162 | 0.512989107 | 0.554705255 | 0.579096281 | 0.998865404 | 5.032032 | 5.316794 | 5.164005 | 5.161891 | 5.192360 | 5.189654 | 38 | 35 | 35 | 37 |
| 102725068 | NR_149132 | MICB-DT | MICB divergent transcript | lncRNA | . | | 53632 | . | . | . | 0.293587 | 0.143321 | 0.071691 | 0.203427 | 0.683188 | 0.331293 | 0.160871 | 0.465112 | -1.625807 | 11.443269 | 0.963369355 | -0.72781607 | 0.466726188 | NA | 3.826455 | 3.121007 | 3.556777 | 3.512767 | 3.493192 | 3.528417 | 22 | 10 | 4 | 12 |
| 102725121 | NR_148357 | LOC10272512 | DEAD/H-box helicase 11 pseudogene | transcribed_ps | . | . | | . | . | . | 0.288902 | 0.308478 | 0.710353 | 0.512910 | 0.672286 | 0.713060 | 1.594011 | 1.172708 | 2.005241 | 22.941847 | 0.654030239 | 1.534754295 | 0.124844194 | NA | 3.931597 | 4.936619 | 4.461911 | 4.466821 | 4.548333 | 4.508993 | 17 | 17 | 31 | 24 |
| 102725180 | XM_01702755 | LOC10272518 | collagen alpha-1(VII) chain-like, transcript variant X1 | protein_coding | XP_016883040.1 | . | | . | . | . | 0.171453 | 0.276038 | 0.314735 | 0.320525 | 0.398977 | 0.638073 | 0.706257 | 0.732841 | 1.406727 | 22.278278 | 0.664566368 | 0.740847591 | 0.458785856 | NA | 4.211935 | 4.701810 | 4.430064 | 4.471007 | 4.483957 | 4.480837 | 16 | 25 | 23 | 24 |
| 102725191 | NR_136261,N | LOC10272519 | uncharacterized LOC102725191 | lncRNA | . | . | | . | . | . | 0.217618 | 0.597439 | 0.493644 | 0.729255 | 0.506406 | 1.381007 | 1.107722 | 1.667354 | 1.526331 | 11.462861 | 0.930579178 | 0.655578431 | 0.512095407 | NA | 3.184683 | 3.790061 | 3.476180 | 3.512209 | 3.515310 | 3.515819 | 6 | 14 | 12 | 13 |
| 102725509 | NR_110731 | LINC01232 | long intergenic non-protein coding RNA 1232 | lncRNA | . | | 49755 | . | . | . | 0.510713 | 0.718144 | 0.857320 | 0.708003 | 1.188449 | 1.660021 | 1.923800 | 1.618764 | 1.280334 | 40.071511 | 0.500432822 | 0.712424716 | 0.476201815 | 0.998865404 | 5.135531 | 5.491568 | 5.273388 | 5.319133 | 5.350844 | 5.318090 | 34 | 44 | 43 | 38 |
| 102800310 | NR_110457 | HAGLROS | HAGLR opposite strand lncRNA | lncRNA | . | | 50646 | . | . | . | 0.589660 | 0.344826 | 0.421139 | 0.794377 | 1.372164 | 0.797080 | 0.945025 | 1.816249 | 1.285566 | 11.676764 | 0.917635706 | 0.394932019 | 0.692893059 | NA | 3.353365 | 3.715150 | 3.547141 | 3.521574 | 3.530367 | 3.564141 | 15 | 8 | 8 | 16 |
| 102800317 | NM_00128991 | TPTEP2-CSNK1 | TPTEP2-CSNK1E readthrough | protein_coding | NP_001276841.1 | | 53829 | . | . | . | 0.224376 | 0.648075 | 0.988377 | 0.205896 | 0.522133 | 1.498054 | 2.217890 | 0.470756 | 1.353880 | 55.217475 | 1.181323466 | 0.370008408 | 0.711376226 | 0.998865404 | 5.551858 | 5.989226 | 5.604565 | 5.782814 | 5.894103 | 5.602173 | 28 | 75 | 91 | 21 |
| 102929163 | gene-LOC1029 | LOC10292916 | ADP ribosylation factor like GTPase 17A pseudogene | pseudogene | | . | | . | . | . | 0.067761 | 0.071141 | 0.065282 | 0.072035 | 0.157683 | 0.164445 | 0.146492 | 0.164698 | -1.012468 | 4.460625 | 1.415462877 | -0.01262893 | 0.989923838 | NA | 2.166062 | 2.148375 | 2.156438 | 2.157811 | 2.157616 | 2.156126 | 5 | 5 | 4 | 4 |
| 103021295 | NR_110982 | LOC10302129 | uncharacterized LOC103021295 | lncRNA | . | . | | . | . | . | 4.238369 | 3.716908 | 3.902155 | 1.963085 | 9.862861 | 8.591800 | 8.756324 | 4.488361 | -1.380790 | 50.588629 | 0.474317589 | -0.98139714 | 0.326396937 | 0.998865404 | 5.872900 | 5.411913 | 5.700736 | 5.671573 | 5.676729 | 5.573614 | 72 | 59 | 49 | 27 |
| 103091865 | NR_111942 | BRWD1-AS2 | BRWD1 antisense RNA 2 | lncRNA | . | | 16423 | . | . | . | 0.148503 | 0.040177 | 0.119763 | 0.735256 | 0.345571 | 0.092870 | 0.268744 | 1.681077 | 3.966226 | 8.782679 | 1.184485565 | 1.678168959 | 0.09331412 | NA | 1.820514 | 3.810781 | 3.077731 | 3.062129 | 3.076166 | 3.157692 | 6 | 2 | 4 | 22 |
| 103091866 | NR_111954,N | LINC01145 | long intergenic non-protein coding RNA 1145 | lncRNA | . | | 49462 | . | . | . | 0.466601 | 0.608844 | 0.589944 | 0.739395 | 1.085799 | 1.407367 | 1.323818 | 1.690538 | 1.281208 | 81.575698 | 0.360294173 | 0.992256372 | 0.32107248 | 0.998865404 | 6.161430 | 6.516868 | 6.277068 | 6.339965 | 6.350569 | 6.398468 | 72 | 87 | 73 | 93 |
| 103344926 | NR_120606 | LINCR-0002 | uncharacterized LincR-0002 | lncRNA | . | . | | . | . | . | 4.891859 | 4.933865 | 5.734621 | 5.807100 | 11.383557 | 11.404851 | 12.868323 | 13.277246 | 1.161469 | 197.200894 | 0.236974004 | 0.911284721 | 0.362145375 | 0.998865404 | 7.511522 | 7.727451 | 7.586364 | 7.587101 | 7.652849 | 7.654659 | 210 | 197 | 184 | 199 |
| 103344928 | NR_145481 | KIAA1614-AS1 | KIAA1614 antisense RNA 1 | lncRNA | . | | 50644 | . | . | . | 0.696644 | 0.369067 | 0.572072 | 0.767569 | 1.621119 | 0.853115 | 1.283712 | 1.754954 | 1.238386 | 27.847482 | 0.610309391 | 0.505417279 | 0.613265785 | 0.998865404 | 4.636349 | 4.946016 | 4.811578 | 4.750140 | 4.788104 | 4.822799 | 37 | 19 | 23 | 33 |
| 103344931 | NR_120684 | LOC10334493 | uncharacterized LOC103344931 | lncRNA | . | . | | . | . | . | 0.534460 | 0.455576 | 0.674032 | 0.457658 | 1.243711 | 1.053084 | 1.512509 | 1.046381 | 1.119250 | 70.669299 | 0.392996493 | 0.413570851 | 0.679188424 | 0.998865404 | 6.057947 | 6.223339 | 6.141940 | 6.103564 | 6.198105 | 6.099487 | 83 | 66 | 78 | 56 |
| 103352539 | NR_121647 | LINC01410 | long intergenic non-protein coding RNA 1410 | lncRNA | . | | 50702 | . | . | . | 0.678888 | 0.702219 | 1.058057 | 0.959316 | 1.579801 | 1.623210 | 2.374249 | 2.193362 | 1.438879 | 76.596503 | 0.362542111 | 1.447956182 | 0.147629306 | 0.90884489 | 5.972747 | 6.498101 | 6.196968 | 6.205663 | 6.308472 | 6.277592 | 71 | 69 | 83 | 80 |
| 103504727 | NR_128715,rn | MIR6724-2 | microRNA 6724-2 | miRNA | . | | 50832 | . | . | . | 0.024865 | 0.037047 | 0.204869 | 0.160135 | 0.057861 | 0.085636 | 0.459721 | 0.366130 | 1.234663 | 2.005914 | 2.070995667 | 0.146845664 | 0.883253841 | NA | 0.844134 | 1.148375 | 0.998092 | 0.998667 | 1.000707 | 0.999974 | 2 | 2 | 2 | 2 |
| 103504729 | NR_128709,rn | MIR3179-4 | microRNA 3179-4 | miRNA | . | | 50838 | . | . | . | 0.388724 | 0.654891 | 0.483449 | 0.764803 | 0.904577 | 1.513812 | 1.084846 | 1.748629 | 1.234665 | 3.008871 | 1.708874417 | 0.17796512 | 0.858750369 | NA | 1.429096 | 1.733338 | 1.582415 | 1.583274 | 1.586316 | 1.585224 | 3 | 3 | 3 | 3 |
| 103504731 | NR_128711,rn | MIR3648-2 | microRNA 3648-2 | miRNA | . | | 50843 | . | . | . | 1.787103 | 1.606548 | 1.654861 | 2.130635 | 4.158663 | 3.713607 | 3.713458 | 4.871445 | 1.078174 | 11.158428 | 0.910210117 | 0.119301873 | 0.9050362 | NA | 3.425187 | 3.532926 | 3.480015 | 3.475046 | 3.475954 | 3.486541 | 13 | 11 | 9 | 12 |
| 103504733 | NR_128717,rn | MIR6724-4 | microRNA 6724-4 | miRNA | . | | 50830 | . | . | . | 0.072102 | 0.109991 | 0.363364 | 0.327308 | 0.167784 | 0.254251 | 0.815376 | 0.748352 | 1.234663 | 2.005914 | 2.070995667 | 0.146845664 | 0.883253841 | NA | 0.844134 | 1.148375 | 0.998092 | 0.998667 | 1.000707 | 0.999974 | 2 | 2 | 2 | 2 |
| 103504738 | NR_128720,rn | MIR6859-4 | hsa-miR-6859-3p | . | . | | 50840 | . | . | . | 11.432351 | 7.340686 | 15.280315 | 6.491267 | 26.603551 | 16.968324 | 34.288584 | 14.841512 | 1.155790 | 18.266612 | 0.752374557 | 0.277627106 | 0.781298622 | NA | 4.079678 | 4.294599 | 4.193421 | 4.158691 | 4.215298 | 4.158740 | 24 | 14 | 23 | 12 |
| 103504739 | NR_128716,rn | MIR6724-3 | microRNA 6724-3 | miRNA | . | | 50842 | . | . | . | 0.048298 | 0.073378 | 0.255338 | 0.245607 | 0.112393 | 0.169616 | 0.572971 | 0.561550 | 1.234663 | 2.005914 | 2.070995667 | 0.146845664 | 0.883253841 | NA | 0.844134 | 1.148375 | 0.998092 | 0.998667 | 1.000707 | 0.999974 | 2 | 2 | 2 | 2 |
| 103625684 | NR_125730 | RNU6-2 | RNA, U6 small nuclear 2 | snRNA | . | | 34270 | . | . | . | 1.547033 | 1.197722 | 0.706702 | 2.809458 | 3.600010 | 2.768587 | 1.585819 | 6.423492 | 1.221732 | 5.940686 | 1.26581468 | 0.228254738 | 0.81944821 | NA | 2.421266 | 2.705970 | 2.569190 | 2.562451 | 2.557571 | 2.584772 | 7 | 5 | 3 | 9 |
| 103689912 | NR_125749,N | TBX2-AS1 | TBX2 antisense RNA 1 | lncRNA | . | | 50355 | . | . | . | 2.370863 | 2.152339 | 2.656874 | 2.296001 | 5.517095 | 4.975229 | 5.961949 | 5.249533 | 1.077102 | 57.249628 | 0.415638403 | 0.257806432 | 0.7965563 | 0.998865404 | 5.783979 | 5.892374 | 5.834211 | 5.824330 | 5.861743 | 5.827017 | 65 | 58 | 56 | 51 |
| 103752588 | NR_125801 | PACERR | PTGS2 antisense NFKB1 complex-mediated expression regulator R | lncRNA | . | | 50552 | 617650 | . | . | 0.424712 | 0.231153 | 0.519493 | 0.280399 | 0.988323 | 0.534320 | 1.165726 | 0.641099 | 1.191141 | 9.760830 | 0.995437448 | 0.253500662 | 0.799881349 | NA | 3.151937 | 3.410501 | 3.286996 | 3.264892 | 3.297170 | 3.269789 | 13 | 7 | 12 | 7 |
| 103908604 | NR_147619 | LOC10390860 | uncharacterized LOC103908604 | lncRNA | . | . | | . | . | . | 0.423067 | 0.243834 | 0.496544 | 0.495070 | 0.984495 | 0.563633 | 1.114230 | 1.131920 | 1.421125 | 7.545708 | 1.109312455 | 0.457069945 | 0.647620772 | NA | 2.638039 | 3.148375 | 2.909653 | 2.894843 | 2.915555 | 2.912763 | 9 | 5 | 8 | 8 |
| 103908605 | NR_126050 | LOC10390860 | uncharacterized LOC103908605 | lncRNA | . | . | | . | . | . | 2.734493 | 4.038260 | 3.128821 | 3.267303 | 6.363278 | 9.334620 | 7.020984 | 7.470301 | -1.063406 | 54.241703 | 0.438077567 | -0.2024583 | 0.839558453 | 0.998865404 | 5.805868 | 5.715375 | 5.724258 | 5.803791 | 5.749955 | 5.755519 | 52 | 72 | 45 | 50 |
| 104310353 | NR_126356 | LINC01399 | long intergenic non-protein coding RNA 1399 | lncRNA | . | | 50680 | . | . | . | 1.072563 | 0.625139 | 0.944847 | 0.506241 | 2.495898 | 1.445038 | 2.120209 | 1.157460 | -1.181584 | 16.442538 | 0.786154484 | -0.30620268 | 0.759450331 | NA | 4.151937 | 3.917252 | 4.064692 | 4.022377 | 4.050547 | 4.012936 | 26 | 14 | 17 | 10 |
| 104326193 | NR_126376 | DLX2-DT | DLX2 divergent transcript | lncRNA | . | | 50638 | . | . | . | 0.402214 | 0.378792 | 0.232431 | 0.477510 | 0.935968 | 0.875594 | 0.521568 | 1.091769 | -1.105008 | 11.509764 | 0.918053148 | -0.15691593 | 0.875311106 | NA | 3.597285 | 3.448451 | 3.530076 | 3.529536 | 3.507339 | 3.541873 | 14 | 13 | 6 | 14 |
| 104355135 | NR_126378 | LINC00431 | long intergenic non-protein coding RNA 431 | lncRNA | . | | 42766 | . | . | . | 0.258255 | 0.139083 | 0.385363 | 0.080753 | 0.600970 | 0.321497 | 0.864744 | 0.184633 | 1.150331 | 5.279800 | 1.367054442 | 0.147798436 | 0.882501843 | NA | 2.290730 | 2.502474 | 2.393600 | 2.382425 | 2.406934 | 2.375527 | 7 | 4 | 8 | 2 |
| 104355287 | NR_126403 | LINC01317 | long intergenic non-protein coding RNA 1317 | lncRNA | . | | 50523 | . | . | . | 6.530190 | 6.040594 | 6.924068 | 7.339781 | 15.196019 | 13.963104 | 15.537408 | 16.781538 | 1.118903 | 161.757562 | 0.259146782 | 0.625453759 | 0.531673288 | 0.998865404 | 7.254323 | 7.416501 | 7.328825 | 7.296478 | 7.348230 | 7.368261 | 183 | 158 | 145 | 164 |
| 104355295 | NR_126411 | SGMS1-AS1 | SGMS1 antisense RNA 1 | lncRNA | . | | 49683 | . | . | . | 0.342193 | 0.411487 | 0.318021 | 0.317125 | 0.796297 | 0.951170 | 0.713630 | 0.725068 | -1.191220 | 28.517705 | 0.578953189 | -0.43602857 | 0.662816004 | 0.998865404 | 4.954694 | 4.701810 | 4.835895 | 4.855152 | 4.827157 | 4.823561 | 33 | 36 | 23 | 24 |
| 104355426 | NR_126418 | LENG8-AS1 | LENG8 antisense RNA 1 | lncRNA | . | | 40705 | . | . | . | 1.678762 | 2.146174 | 2.202706 | 2.417223 | 3.906549 | 4.960978 | 4.942808 | 5.526692 | 1.198473 | 59.325341 | 0.411737136 | 0.634379562 | 0.525833144 | 0.998865404 | 5.754924 | 6.014563 | 5.842772 | 5.889317 | 5.895522 | 5.913292 | 55 | 65 | 54 | 63 |
| 104472715 | NR_146177 | SNHG14 | small nucleolar RNA host gene 14 | lncRNA | . | | 37462 | 616259 | . | . | 11.465152 | 12.009668 | 11.793537 | 12.184583 | 26.679878 | 27.760885 | 26.464355 | 27.858603 | 1.007368 | 7118.385457 | 0.103464122 | 0.10236418 | 0.918467615 | 0.998865404 | 12.792049 | 12.802600 | 12.778004 | 12.810078 | 12.790826 | 12.809806 | 8012 | 7798 | 6143 | 6773 |
| 104472848 | NR_126453 | IRAIN | IGF1R antisense imprinted non-protein coding RNA | lncRNA | . | | 50365 | . | . | . | 0.097851 | 0.156965 | 0.142385 | 0.133755 | 0.227703 | 0.362832 | 0.319508 | 0.305815 | 1.053235 | 22.698477 | 0.654866234 | 0.114263068 | 0.909029258 | NA | 4.467424 | 4.540693 | 4.477388 | 4.518391 | 4.506567 | 4.499491 | 20 | 29 | 21 | 21 |
| 104968399 | NR_130143,N | LOC10496839 | uncharacterized LOC104968399 | lncRNA | . | . | | . | . | . | 0.376185 | 0.122384 | 0.225412 | 0.290331 | 0.875399 | 0.282895 | 0.505817 | 0.663807 | 1.323198 | 7.741992 | 1.099842842 | 0.367352032 | 0.713356447 | NA | 2.735321 | 3.141582 | 2.952377 | 2.933550 | 2.949305 | 2.956187 | 10 | 5 | 7 | 9 |
| 105221694 | NR_130765,N | BISPR | BST2 interferon stimulated positive regulator | lncRNA | . | | 51290 | . | . | . | 0.297276 | 0.093393 | 0.169614 | 0.230045 | 0.691774 | 0.215883 | 0.380608 | 0.525970 | 1.021172 | 12.501901 | 0.909062555 | 0.033249961 | 0.973475257 | NA | 3.627869 | 3.660102 | 3.668045 | 3.615004 | 3.633161 | 3.652765 | 21 | 7 | 9 | 14 |
| 105274304 | NR_130916 | LOC10527430 | uncharacterized LOC105274304 | lncRNA | . | . | | . | . | . | 0.222355 | 0.151953 | 0.087016 | 0.170881 | 0.517428 | 0.351245 | 0.195262 | 0.390699 | -1.333541 | 8.962192 | 1.0293692 | -0.40341465 | 0.686643208 | NA | 3.357474 | 2.940155 | 3.184830 | 3.167344 | 3.156290 | 3.172978 | 14 | 9 | 5 | 9 |
| 105369147 | NR_148213,N | LOC10536914 | uncharacterized LOC105369147 | lncRNA | . | . | | . | . | . | 1.683754 | 1.908856 | 1.456650 | 2.614786 | 3.918165 | 4.412407 | 3.268679 | 5.978396 | 1.150784 | 170.463384 | 0.27752831 | 0.730078301 | 0.465342324 | 0.998865404 | 7.309355 | 7.510290 | 7.384676 | 7.369955 | 7.343893 | 7.520285 | 186 | 168 | 126 | 206 |
| 105369149 | XR_00174847 | LOC10536914 | uncharacterized LOC105369149, transcript variant X1 | lncRNA | . | . | | . | . | . | 0.221020 | 0.211035 | 0.095112 | 0.358734 | 0.514323 | 0.487816 | 0.213429 | 0.820204 | 1.067108 | 3.703508 | 1.564493575 | 0.059895269 | 0.952239048 | NA | 1.844134 | 1.932304 | 1.888489 | 1.889602 | 1.883355 | 1.896931 | 4 | 4 | 2 | 5 |
| 105369151 | XR_931426 | LOC10536915 | uncharacterized LOC105369151 | lncRNA | . | . | | . | . | . | 3.896237 | 3.815103 | 4.742297 | 3.131807 | 9.066705 | 8.818781 | 10.641577 | 7.160504 | 1.006907 | 955.791509 | 0.183300063 | 0.054173092 | 0.956797259 | 0.998865404 | 9.895152 | 9.905932 | 9.903072 | 9.881009 | 10.039172 | 9.732489 | 1113 | 1012 | 1009 | 711 |
| 105369154 | NR_123722 | PKD1P6-NPIPP | PKD1P6-NPIPP1 readthrough | transcribed_ps | . | . | | . | . | . | 13.050104 | 14.066918 | 18.460455 | 13.080777 | 30.368128 | 32.516312 | 41.424728 | 29.907644 | 1.148272 | 1325.955011 | 0.165355173 | 1.206275472 | 0.227711292 | 0.982008277 | 10.269384 | 10.469326 | 10.287450 | 10.337500 | 10.545825 | 10.272866 | 1374 | 1376 | 1449 | 1096 |
| 105369161 | XR_938320 | LOC10536916 | uncharacterized LOC105369161 | lncRNA | . | . | | . | . | . | 0.039651 | 0.021883 | 0.041790 | 0.035193 | 0.092269 | 0.050582 | 0.093777 | 0.080464 | 1.197969 | 12.713472 | 0.867633242 | 0.300347081 | 0.763912425 | NA | 3.530084 | 3.794399 | 3.673188 | 3.644070 | 3.673319 | 3.664257 | 17 | 9 | 13 | 12 |
| 105369164 | XR_00173934 | LOC10536916 | uncharacterized LOC105369164, transcript variant X2 | lncRNA | . | . | | . | . | . | 0.366922 | 0.255084 | 0.162820 | 0.500967 | 0.853843 | 0.589639 | 0.365363 | 1.145401 | -1.058759 | 11.248927 | 0.934710087 | -0.08812782 | 0.929775086 | NA | 3.533720 | 3.448451 | 3.505622 | 3.484629 | 3.474923 | 3.509482 | 16 | 10 | 6 | 14 |
| 105369165 | XR_00173946 | LOC10536916 | uncharacterized LOC105369165, transcript variant X2 | lncRNA | . | . | | . | . | . | 0.432621 | 0.264913 | 0.265557 | 0.231368 | 1.006726 | 0.612359 | 0.595903 | 0.528996 | -1.759314 | 8.429280 | 1.057930156 | -0.77038476 | 0.441071688 | NA | 3.425187 | 2.612728 | 3.096537 | 3.091400 | 3.076795 | 3.069694 | 13 | 11 | 6 | 5 |
| 105369180 | XR_949578 | LOC10536918 | uncharacterized LOC105369180 | lncRNA | . | . | | . | . | . | 0.347253 | 0.405233 | 0.437058 | 0.363032 | 0.808071 | 0.936713 | 0.980744 | 0.830030 | 1.054050 | 84.942907 | 0.350186978 | 0.216866005 | 0.828312771 | 0.998865404 | 6.369732 | 6.446101 | 6.377334 | 6.418932 | 6.438247 | 6.386556 | 88 | 96 | 83 | 74 |
| 105369187 | NR_135033 | LOC10536918 | uncharacterized LOC105369187 | lncRNA | . | . | | . | . | . | 0.544732 | 0.710283 | 0.271229 | 0.321371 | 1.267615 | 1.641852 | 0.608631 | 0.734776 | -2.065523 | 11.018638 | 0.937087007 | -1.11676657 | 0.264094148 | NA | 3.892781 | 2.844638 | 3.478682 | 3.494677 | 3.451950 | 3.454065 | 15 | 18 | 6 | 7 |
| 105369199 | NR_160532,N | LOC10536919 | NBPF member 6 pseudogene | transcribed_ps | . | . | | . | . | . | 0.292285 | 0.390631 | 0.277246 | 0.183201 | 0.680159 | 0.902962 | 0.622131 | 0.418867 | -1.392555 | 12.021885 | 0.898546259 | -0.53167462 | 0.594951374 | NA | 3.805868 | 3.330299 | 3.584822 | 3.608862 | 3.590828 | 3.568242 | 13 | 18 | 11 | 7 |
| 105369201 | NR_160702,N | LOC10536920 | uncharacterized LOC105369201 | lncRNA | . | . | | . | . | . | 5.692885 | 5.574301 | 7.070750 | 9.134000 | 13.247576 | 12.885245 | 15.866557 | 20.883806 | 1.123360 | 262.747168 | 0.215706738 | 0.778000046 | 0.436568986 | 0.998865404 | 7.951657 | 8.118580 | 7.970789 | 8.038077 | 8.043058 | 8.084882 | 266 | 285 | 232 | 271 |
| 105369214 | XM_01151419 | LOC10536921 | uncharacterized LOC105369214 | protein_coding | XP_011512492.1 | . | | . | . | . | 0.049147 | 0.082912 | 0.084091 | 0.098239 | 0.114367 | 0.191654 | 0.188698 | 0.224612 | 1.347887 | 5.296028 | 1.311069465 | 0.328510142 | 0.742525977 | NA | 2.175403 | 2.602870 | 2.388885 | 2.398813 | 2.399569 | 2.402533 | 4 | 6 | 5 | 6 |
| 105369225 | XR_00175290 | LOC10536922 | uncharacterized LOC105369225, transcript variant X1 | lncRNA | . | . | | . | . | . | 0.305377 | 0.162077 | 0.156241 | 0.116336 | 0.710623 | 0.374649 | 0.350600 | 0.265989 | -1.605757 | 6.514154 | 1.199396266 | -0.56966425 | 0.56890544 | NA | 3.003609 | 2.324312 | 2.725265 | 2.711029 | 2.707834 | 2.701129 | 11 | 7 | 5 | 4 |
| 105369228 | XR_00174241 | LOC10536922 | uncharacterized LOC105369228 | lncRNA | . | . | | . | . | . | 0.341121 | 0.500096 | 0.512902 | 0.303233 | 0.793803 | 1.155994 | 1.150937 | 0.693306 | -1.039953 | 41.599355 | 0.512937488 | -0.11018608 | 0.912261801 | 0.998865404 | 5.405684 | 5.350772 | 5.344054 | 5.403982 | 5.408552 | 5.326171 | 40 | 54 | 45 | 28 |
| 105369250 | XR_00174157 | LOC10536925 | uncharacterized LOC105369250, transcript variant X2 | lncRNA | . | . | | . | . | . | 0.086902 | 0.087366 | 0.204516 | 0.243612 | 0.202224 | 0.201950 | 0.458928 | 0.556989 | 2.279063 | 10.247779 | 0.977103778 | 1.216289034 | 0.223874837 | NA | 2.644780 | 3.832035 | 3.322757 | 3.316537 | 3.338578 | 3.372283 | 8 | 6 | 9 | 17 |
| 105369299 | XR_937652 | LOC10536929 | uncharacterized LOC105369299 | lncRNA | . | . | | . | . | . | 0.154074 | 0.167861 | 0.270842 | 0.228498 | 0.358536 | 0.388018 | 0.607761 | 0.522434 | 1.446499 | 3.296365 | 1.642758879 | 0.324189341 | 0.745794654 | NA | 1.429096 | 1.963455 | 1.707764 | 1.708646 | 1.716881 | 1.710648 | 3 | 3 | 4 | 3 |
| 105369312 | XR_950123 | LOC10536931 | uncharacterized LOC105369312 | lncRNA | . | . | | . | . | . | 0.029081 | 0.052094 | 0.044136 | 0.028886 | 0.067672 | 0.120417 | 0.099040 | 0.066044 | -1.079491 | 8.704561 | 1.042201933 | -0.10588303 | 0.915675155 | NA | 3.175403 | 3.066070 | 3.110482 | 3.129692 | 3.125771 | 3.108459 | 8 | 12 | 9 | 6 |
| 105369313 | XR_00174821 | LOC10536931 | uncharacterized LOC105369313 | lncRNA | . | . | | . | . | . | 0.540282 | 0.653422 | 0.982492 | 0.364080 | 1.257258 | 1.510414 | 2.204684 | 0.832426 | 1.116862 | 93.844169 | 0.421129815 | 0.378626019 | 0.704965598 | 0.998865404 | 6.468800 | 6.631032 | 6.485848 | 6.536987 | 6.674450 | 6.394213 | 93 | 104 | 126 | 50 |
| 105369315 | XR_00174822 | LOC10536931 | uncharacterized LOC105369315, transcript variant X1 | lncRNA | . | . | | . | . | . | 0.031116 | 0.054704 | 0.046956 | 0.034008 | 0.072409 | 0.126450 | 0.105370 | 0.077754 | -1.022965 | 15.200252 | 0.800498996 | -0.04091987 | 0.967359773 | NA | 3.942580 | 3.909275 | 3.904024 | 3.940458 | 3.930302 | 3.911445 | 13 | 21 | 15 | 12 |
| 105369339 | XR_950186,XR | LOC10536933 | uncharacterized LOC105369339, transcript variant X2 | lncRNA | . | . | | . | . | . | 0.397081 | 0.503740 | 0.842682 | 0.109456 | 0.924025 | 1.164418 | 1.890955 | 0.250258 | 1.023666 | 7.286136 | 1.200167306 | 0.02811741 | 0.977568508 | NA | 2.844134 | 2.885872 | 2.851910 | 2.854089 | 2.876621 | 2.830260 | 8 | 8 | 11 | 2 |
| 105369340 | NR_135086,N | PPP1R14B-AS1 | PPP1R14B antisense RNA 1 | lncRNA | . | | 54233 | . | . | . | 2.102902 | 3.204073 | 1.758752 | 3.157138 | 4.893538 | 7.406359 | 3.946589 | 7.218422 | -1.086601 | 38.945642 | 0.529469854 | -0.22630646 | 0.820963078 | 0.998865404 | 5.344293 | 5.219802 | 5.252143 | 5.319057 | 5.236278 | 5.309808 | 37 | 53 | 24 | 44 |
| 105369347 | XR_00174826 | RELA-DT | RELA divergent transcript, transcript variant X3 | lncRNA | . | | 54185 | . | . | . | 1.618626 | 1.458558 | 1.794430 | 1.122063 | 3.766610 | 3.371520 | 4.026648 | 2.565463 | 1.233457 | 85.583022 | 0.35220471 | 0.859464836 | 0.390084112 | 0.998865404 | 6.258969 | 6.563496 | 6.390814 | 6.374588 | 6.480883 | 6.398174 | 91 | 80 | 94 | 76 |
| 105369352 | XR_00174827 | LOC10536935 | uncharacterized LOC105369352, transcript variant X6 | lncRNA | . | . | | . | . | . | 0.365598 | 0.710291 | 0.472380 | 0.314479 | 0.850761 | 1.641871 | 1.060005 | 0.719018 | -1.267055 | 20.236418 | 0.710422337 | -0.48067033 | 0.630750823 | NA | 4.499192 | 4.158506 | 4.318970 | 4.371384 | 4.342005 | 4.308563 | 19 | 31 | 19 | 13 |
| 105369362 | XR_950242,XR | LOC10536936 | uncharacterized LOC105369362, transcript variant X1 | lncRNA | . | . | | . | . | . | 0.314371 | 0.366054 | 0.491406 | 0.303452 | 0.731554 | 0.846151 | 1.102700 | 0.693807 | 1.067188 | 16.649088 | 0.749838477 | 0.125111894 | 0.900434967 | NA | 4.008843 | 4.104320 | 4.058975 | 4.047732 | 4.063017 | 4.053007 | 20 | 16 | 16 | 15 |
| 105369370 | XR_950269 | LOC10536937 | uncharacterized LOC105369370 | lncRNA | . | . | | . | . | . | 5.132875 | 5.804722 | 4.068109 | 4.567183 | 11.944413 | 13.417875 | 9.128718 | 10.442322 | -1.283297 | 814.180372 | 0.15156674 | -2.37423587 | 0.017585313 | 0.445376286 | 9.838221 | 9.477727 | 9.707060 | 9.790424 | 9.537682 | 9.608726 | 994 | 1044 | 587 | 704 |
| 105369383 | XR_950297 | LOC10536938 | uncharacterized LOC105369383 | lncRNA | . | . | | . | . | . | 0.037180 | 0.050837 | 0.033995 | 0.023026 | 0.086520 | 0.117512 | 0.076284 | 0.052647 | -1.422882 | 2.686397 | 1.818605976 | -0.27978332 | 0.779643752 | NA | 1.658167 | 1.148375 | 1.430406 | 1.435592 | 1.429080 | 1.428283 | 3 | 4 | 2 | 2 |
| 105369401 | XR_950342 | LOC10536940 | uncharacterized LOC105369401 | lncRNA | . | . | | . | . | . | 6.215874 | 6.365664 | 4.957591 | 5.635342 | 14.464594 | 14.714518 | 11.124691 | 12.884541 | -1.203254 | 359.573443 | 0.190447087 | -1.40165593 | 0.161018018 | 0.933525128 | 8.617744 | 8.350151 | 8.536446 | 8.545030 | 8.402340 | 8.466128 | 449 | 427 | 267 | 324 |
| 105369403 | XR_00174831 | LOC10536940 | uncharacterized LOC105369403 | lncRNA | . | . | | . | . | . | 0.075510 | 0.072286 | 0.222150 | 0.066882 | 0.175716 | 0.167093 | 0.498497 | 0.152918 | 1.852337 | 13.450764 | 0.891041956 | 0.998097105 | 0.318232274 | NA | 3.234218 | 4.128572 | 3.703908 | 3.702860 | 3.775621 | 3.700250 | 11 | 10 | 23 | 8 |
| 105369435 | XR_947901,XR | LOC10536943 | uncharacterized LOC105369435, transcript variant X1 | lncRNA | . | . | | . | . | . | 0.333823 | 0.322565 | 0.424895 | 0.294896 | 0.776819 | 0.745623 | 0.953451 | 0.674244 | 1.087418 | 97.805922 | 0.338195481 | 0.357504558 | 0.720714115 | 0.998865404 | 6.549025 | 6.672053 | 6.599805 | 6.587912 | 6.673569 | 6.561516 | 110 | 99 | 105 | 78 |
| 105369477 | XR_00174836 | LOC10536947 | uncharacterized LOC105369477, transcript variant X4 | lncRNA | . | . | | . | . | . | 0.044305 | 0.087312 | 0.063532 | 0.030843 | 0.103101 | 0.201826 | 0.142563 | 0.070520 | -1.390232 | 16.714618 | 0.781621197 | -0.60812855 | 0.543102218 | NA | 4.280172 | 3.807334 | 4.051939 | 4.092491 | 4.067621 | 4.030081 | 17 | 26 | 16 | 9 |
| 105369486 | NR_135100 | LINC02732 | long intergenic non-protein coding RNA 2732 | lncRNA | . | | 54249 | . | . | . | 14.737249 | 14.395392 | 12.178647 | 16.089539 | 34.294186 | 33.275597 | 27.328531 | 36.786819 | -1.044725 | 599.701512 | 0.175709766 | -0.35924517 | 0.71941169 | 0.998865404 | 9.259724 | 9.195769 | 9.251397 | 9.230060 | 9.120536 | 9.295160 | 717 | 651 | 442 | 623 |
| 105369490 | NR_149131 | LINC02550 | long intergenic non-protein coding RNA 2550 | lncRNA | . | | 53585 | . | . | . | 1.035275 | 1.098612 | 1.196932 | 1.691568 | 2.409128 | 2.539492 | 2.685881 | 3.867568 | 1.330495 | 18.811574 | 0.709916278 | 0.58029885 | 0.561713103 | NA | 4.014059 | 4.424013 | 4.213494 | 4.218323 | 4.225922 | 4.251402 | 18 | 18 | 16 | 23 |
| 105369547 | XR_948132,XR | CCDC15-DT | CCDC15 divergent transcript, transcript variant X1 | lncRNA | . | | 54193 | . | . | . | 0.209964 | 0.312865 | 0.642908 | 0.288802 | 0.488594 | 0.723203 | 1.442667 | 0.660312 | 1.366022 | 14.933031 | 0.798488769 | 0.563540948 | 0.573066585 | NA | 3.658167 | 4.107809 | 3.870206 | 3.890319 | 3.909992 | 3.890771 | 12 | 16 | 17 | 14 |
| 105369557 | XR_948146 | HP11014 | uncharacterized HP11014 | lncRNA | . | . | | . | . | . | 0.019247 | 0.148057 | 0.104559 | 0.206047 | 0.044789 | 0.342240 | 0.234626 | 0.471103 | 1.841041 | 9.746092 | 1.086806381 | 0.81019226 | 0.417829685 | NA | 2.784982 | 3.655341 | 3.225463 | 3.274146 | 3.261204 | 3.290951 | 2 | 13 | 8 | 15 |
| 105369617 | XR_00174897 | LOC10536961 | uncharacterized LOC105369617, transcript variant X3 | lncRNA | . | . | | . | . | . | 0.131910 | 0.197987 | 0.106490 | 0.118617 | 0.306961 | 0.457656 | 0.238960 | 0.271204 | -1.412354 | 10.009298 | 0.969560087 | -0.5137399 | 0.607433855 | NA | 3.551764 | 3.051647 | 3.324937 | 3.344521 | 3.318627 | 3.320575 | 11 | 15 | 7 | 8 |
| 105369647 | XR_00174899 | LOC10536964 | uncharacterized LOC105369647, transcript variant X2 | lncRNA | . | . | | . | . | . | 7.921907 | 6.010146 | 9.331768 | 10.921976 | 18.434605 | 13.892722 | 20.940218 | 24.971801 | 1.300914 | 173.911940 | 0.26229373 | 1.446947973 | 0.147911496 | 0.909078163 | 7.240878 | 7.618861 | 7.362861 | 7.391333 | 7.425719 | 7.546257 | 168 | 169 | 148 | 209 |
| 105369691 | NR_135030,N | LOC10536969 | uncharacterized LOC105369691 | lncRNA | . | . | | . | . | . | 0.504297 | 0.244665 | 0.419977 | 0.171109 | 1.173519 | 0.565555 | 0.942417 | 0.391220 | -1.282959 | 8.315459 | 1.09130868 | -0.32939847 | 0.741854522 | NA | 3.220748 | 2.869520 | 3.073697 | 3.047599 | 3.064752 | 3.037979 | 14 | 7 | 9 | 4 |
| 105369698 | XR_00174904 | LOC10536969 | uncharacterized LOC105369698 | lncRNA | . | . | | . | . | . | 0.097258 | 0.036442 | 0.175389 | 0.049849 | 0.226324 | 0.084238 | 0.393568 | 0.113973 | 1.632344 | 8.189427 | 1.140557987 | 0.619823997 | 0.535373668 | NA | 2.631267 | 3.348113 | 3.014076 | 2.990824 | 3.045618 | 2.994112 | 10 | 4 | 14 | 4 |
| 105369760 | XR_944935,XR | LOC10536976 | uncharacterized LOC105369760, transcript variant X1 | lncRNA | . | . | | . | . | . | 0.156526 | 0.237726 | 0.346255 | 0.934046 | 0.364243 | 0.549514 | 0.776988 | 2.135586 | 2.093694 | 9.743597 | 0.994474415 | 1.071974186 | 0.283731627 | NA | 2.658167 | 3.719719 | 3.245695 | 3.256274 | 3.269585 | 3.294227 | 6 | 8 | 9 | 15 |
| 105369779 | XR_944987 | LOC10536977 | uncharacterized LOC105369779 | lncRNA | . | . | | . | . | . | 3.801126 | 3.386925 | 4.419890 | 4.616523 | 8.845377 | 7.829030 | 9.918106 | 10.555132 | 1.239116 | 183.856358 | 0.246966596 | 1.252442544 | 0.210408652 | 0.972954982 | 7.359295 | 7.668988 | 7.495582 | 7.443938 | 7.554998 | 7.570723 | 201 | 166 | 174 | 194 |
| 105369803 | XR_00295741 | LOC10536980 | uncharacterized LOC105369803, transcript variant X1 | lncRNA | . | . | | . | . | . | 0.426846 | 0.398361 | 0.507848 | 0.596520 | 0.993288 | 0.920830 | 1.139596 | 1.363871 | 1.329713 | 20.850817 | 0.671877244 | 0.611889714 | 0.540610728 | NA | 4.161369 | 4.573379 | 4.368283 | 4.357802 | 4.389300 | 4.386764 | 22 | 18 | 21 | 22 |
| 105369807 | NR_146529 | LINC02454 | long intergenic non-protein coding RNA 2454 | lncRNA | . | | 53387 | . | . | . | 2.143691 | 1.310126 | 1.428180 | 3.425808 | 4.988457 | 3.028415 | 3.204795 | 7.832704 | 1.390534 | 39.843352 | 0.5591751 | 0.850607818 | 0.394987245 | 0.998865404 | 5.059773 | 5.533970 | 5.307222 | 5.239278 | 5.252521 | 5.392250 | 48 | 27 | 24 | 61 |
| 105369812 | XR_00174918 | LOC10536981 | uncharacterized LOC105369812, transcript variant X1 | lncRNA | . | . | | . | . | . | 0.047014 | 0.106242 | 0.072960 | 0.043445 | 0.109404 | 0.245582 | 0.163720 | 0.099331 | -1.375387 | 8.628349 | 1.070557361 | -0.4295313 | 0.667536624 | NA | 3.320504 | 2.861274 | 3.094744 | 3.130476 | 3.109119 | 3.092022 | 7 | 15 | 8 | 5 |
| 105369816 | XR_00174918 | LOC10536981 | uncharacterized LOC105369816, transcript variant X4 | lncRNA | . | . | | . | . | . | 0.154930 | 0.165104 | 0.194664 | 0.238849 | 0.360527 | 0.381644 | 0.436822 | 0.546101 | 1.269930 | 17.316937 | 0.734001216 | 0.469684491 | 0.63858045 | NA | 3.931597 | 4.276108 | 4.097428 | 4.101984 | 4.118073 | 4.116797 | 17 | 17 | 17 | 18 |
| 105369844 | XR_00174921 | LOC10536984 | uncharacterized LOC105369844, transcript variant X1 | lncRNA | . | . | | . | . | . | 0.078222 | 0.070975 | 0.130787 | 0.011084 | 0.182025 | 0.164062 | 0.293484 | 0.025343 | 1.015487 | 9.017059 | 1.119709025 | 0.019801843 | 0.984201447 | NA | 3.156660 | 3.188478 | 3.165790 | 3.151903 | 3.189424 | 3.128403 | 12 | 8 | 14 | 2 |
| 105369850 | XR_00174922 | LOC10536985 | uncharacterized LOC105369850, transcript variant X2 | lncRNA | . | . | | . | . | . | 0.129860 | 0.355796 | 0.353830 | 0.178393 | 0.302190 | 0.822439 | 0.793982 | 0.407873 | 1.373667 | 14.470080 | 0.821136311 | 0.557803342 | 0.57697868 | NA | 3.607672 | 4.066070 | 3.820421 | 3.844534 | 3.868846 | 3.835705 | 11 | 16 | 18 | 12 |
| 105369865 | NR_146533,N | PPP1R12A-AS1 | PPP1R12A antisense RNA 1 | lncRNA | . | | 51903 | . | . | . | 0.287137 | 0.215587 | 0.486433 | 0.332203 | 0.668179 | 0.498339 | 1.091543 | 0.759543 | 1.547027 | 16.508480 | 0.76573025 | 0.822088562 | 0.411026495 | NA | 3.694014 | 4.327309 | 4.024435 | 4.008202 | 4.063662 | 4.030084 | 17 | 12 | 21 | 15 |
| 105369893 | NR_135018 | LINC02392 | long intergenic non-protein coding RNA 2392 | lncRNA | . | | 53319 | . | . | . | 0.209899 | 0.209540 | 0.444921 | 0.070659 | 0.488444 | 0.484361 | 0.998389 | 0.161552 | 1.170212 | 6.837340 | 1.235058146 | 0.18361035 | 0.854319143 | NA | 2.651489 | 2.885872 | 2.754709 | 2.756659 | 2.783410 | 2.737010 | 7 | 7 | 11 | 2 |
| 105369957 | XR_00174929 | LOC10536995 | uncharacterized LOC105369957, transcript variant X6 | lncRNA | . | . | | . | . | . | 0.105278 | 0.073049 | 0.167074 | 0.091250 | 0.244987 | 0.168857 | 0.374908 | 0.208633 | 1.435362 | 10.372871 | 0.945545141 | 0.551443828 | 0.581329463 | NA | 3.089593 | 3.612728 | 3.357706 | 3.356245 | 3.380157 | 3.366821 | 10 | 9 | 12 | 10 |
| 105370088 | XR_945573 | LOC10537008 | uncharacterized LOC105370088 | lncRNA | . | . | | . | . | . | 0.362309 | 0.328482 | 0.823038 | 0.678611 | 0.843108 | 0.759301 | 1.846874 | 1.551563 | 2.092145 | 11.067376 | 0.925641981 | 1.150534079 | 0.249923966 | NA | 2.838265 | 3.905270 | 3.434704 | 3.428772 | 3.471207 | 3.462028 | 9 | 7 | 14 | 13 |
| 105370135 | XR_941799 | LOC10537013 | uncharacterized LOC105370135 | lncRNA | . | . | | . | . | . | 0.145618 | 0.178506 | 0.181153 | 0.087906 | 0.338859 | 0.412624 | 0.406503 | 0.200986 | -1.157574 | 9.643386 | 0.989882155 | -0.21326171 | 0.831122838 | NA | 3.369732 | 3.161867 | 3.268266 | 3.275388 | 3.276633 | 3.254344 | 11 | 12 | 10 | 6 |
| 105370148 | XR_00174980 | LOC10537014 | uncharacterized LOC105370148, transcript variant X1 | lncRNA | . | . | | . | . | . | 0.043129 | 0.037954 | 0.067011 | 0.048126 | 0.100363 | 0.087733 | 0.150370 | 0.110034 | 1.244163 | 3.513475 | 1.59175418 | 0.198005305 | 0.843040913 | NA | 1.644780 | 1.963455 | 1.806945 | 1.803669 | 1.811949 | 1.805707 | 4 | 3 | 4 | 3 |
| 105370203 | NR_135318 | LOC10537020 | uncharacterized LOC105370203 | lncRNA | . | . | | . | . | . | 1.022970 | 1.084882 | 1.253973 | 0.903172 | 2.380493 | 2.507752 | 2.813879 | 2.064995 | 1.037238 | 22.400585 | 0.653188956 | 0.080753746 | 0.935637793 | NA | 4.457887 | 4.512525 | 4.479918 | 4.482230 | 4.499157 | 4.469603 | 25 | 24 | 23 | 18 |
| 105370322 | XR_931655 | LOC10537032 | uncharacterized LOC105370322 | lncRNA | . | . | | . | . | . | 0.021311 | 0.030772 | 0.045578 | 0.073350 | 0.049592 | 0.071132 | 0.102276 | 0.167707 | 2.045617 | 4.096365 | 1.495516147 | 0.690421152 | 0.489929379 | NA | 1.429096 | 2.459419 | 2.012929 | 2.013872 | 2.022329 | 2.030512 | 3 | 3 | 4 | 6 |
| 105370324 | XR_931663,XR | LOC10537032 | uncharacterized LOC105370324, transcript variant X1 | lncRNA | . | . | | . | . | . | 0.448631 | 0.424084 | 0.490285 | 0.449138 | 1.043982 | 0.980289 | 1.100185 | 1.026900 | 1.056716 | 59.900622 | 0.406015056 | 0.196022243 | 0.844592756 | 0.998865404 | 5.863615 | 5.944256 | 5.903195 | 5.890890 | 5.922372 | 5.895015 | 69 | 61 | 57 | 54 |
| 105370327 | XR_00295749 | LOC10537032 | uncharacterized LOC105370327, transcript variant X1 | lncRNA | . | . | | . | . | . | 0.009444 | 0.013567 | 0.011394 | 0.021545 | 0.021977 | 0.031360 | 0.025567 | 0.049261 | 1.395892 | 3.773891 | 1.543982731 | 0.311653565 | 0.755303821 | NA | 1.658167 | 2.134756 | 1.904007 | 1.909301 | 1.908172 | 1.916665 | 3 | 4 | 3 | 5 |
| 105370333 | NR_132422 | PCCA-DT | PCCA divergent transcript | lncRNA | . | | 53266 | . | . | . | 8.606799 | 9.178730 | 8.135621 | 10.669755 | 20.028376 | 21.217045 | 18.256098 | 24.395128 | 1.044199 | 287.432847 | 0.217371015 | 0.287051271 | 0.774073063 | 0.998865404 | 8.136231 | 8.197285 | 8.138671 | 8.168070 | 8.104950 | 8.242799 | 315 | 312 | 222 | 311 |
| 105370413 | XR_00175066 | LOC10537041 | uncharacterized LOC105370413, transcript variant X1 | lncRNA | . | . | | . | . | . | 11.871988 | 12.292250 | 15.937344 | 15.876327 | 27.626605 | 28.414089 | 35.762940 | 36.299335 | 1.321901 | 1347.036551 | 0.131650107 | 3.058215735 | 0.002226592 | 0.119586193 | 10.180322 | 10.582839 | 10.240305 | 10.313598 | 10.509820 | 10.470783 | 1270 | 1314 | 1367 | 1402 |
| 105370447 | XR_943739 | LOC10537044 | uncharacterized LOC105370447 | lncRNA | . | . | | . | . | . | 2.386436 | 2.104888 | 2.074354 | 2.005262 | 5.553335 | 4.865544 | 4.654790 | 4.584793 | -1.110788 | 104.593197 | 0.315292851 | -0.48076874 | 0.63068087 | 0.998865404 | 6.782070 | 6.631282 | 6.746951 | 6.703638 | 6.698921 | 6.686332 | 135 | 111 | 88 | 91 |
| 105370452 | XR_943752 | LOC10537045 | uncharacterized LOC105370452 | lncRNA | . | . | | . | . | . | 5.674561 | 5.092332 | 7.246034 | 6.713205 | 13.204934 | 11.771154 | 16.259890 | 15.348948 | 1.277913 | 411.417311 | 0.183334327 | 1.92974891 | 0.053637957 | 0.686359128 | 8.496314 | 8.850874 | 8.635023 | 8.568682 | 8.775933 | 8.722046 | 440 | 367 | 419 | 414 |
| 105370492 | XR_943849 | LOC10537049 | uncharacterized LOC105370492 | lncRNA | . | . | | . | . | . | 0.261414 | 0.393679 | 1.002560 | 0.615638 | 0.608321 | 0.910006 | 2.249714 | 1.407583 | 2.347235 | 14.343626 | 0.837644058 | 1.469552703 | 0.141682938 | NA | 3.099441 | 4.330299 | 3.780064 | 3.795525 | 3.859646 | 3.816811 | 8 | 11 | 22 | 14 |
| 105370504 | XR_00175096 | LOC10537050 | uncharacterized LOC105370504, transcript variant X9 | lncRNA | . | . | | . | . | . | 0.061604 | 0.039223 | 0.047448 | 0.093679 | 0.143355 | 0.090666 | 0.106473 | 0.214185 | 1.336626 | 17.233208 | 0.755659045 | 0.553947336 | 0.579614888 | NA | 3.884262 | 4.300113 | 4.093477 | 4.085905 | 4.086262 | 4.135423 | 18 | 15 | 12 | 24 |
| 105370532 | XR_00175079 | LOC10537053 | uncharacterized LOC105370532, transcript variant X1 | lncRNA | . | . | | . | . | . | 1.005901 | 1.276552 | 0.811208 | 0.660039 | 2.340774 | 2.950806 | 1.820327 | 1.509101 | -1.567653 | 10.696032 | 0.940798319 | -0.6894213 | 0.490558182 | NA | 3.706954 | 3.058876 | 3.426348 | 3.442101 | 3.417243 | 3.409483 | 13 | 16 | 8 | 7 |
| 105370543 | XR_00175098 | LOC10537054 | uncharacterized LOC105370543, transcript variant X4 | lncRNA | . | . | | . | . | . | 0.207028 | 0.191647 | 0.148308 | 0.225385 | 0.481763 | 0.443001 | 0.332800 | 0.515314 | -1.140329 | 5.891552 | 1.242811549 | -0.15243699 | 0.878842286 | NA | 2.651489 | 2.459419 | 2.562767 | 2.564642 | 2.556224 | 2.564156 | 7 | 7 | 4 | 6 |
| 105370560 | XR_944020,XR | LOC10537056 | uncharacterized LOC105370560, transcript variant X1 | lncRNA | . | . | | . | . | . | 0.295826 | 0.447394 | 0.231314 | 0.235394 | 0.688398 | 1.034172 | 0.519063 | 0.538201 | -1.543307 | 19.696121 | 0.705327729 | -0.88756627 | 0.374774103 | NA | 4.579989 | 3.951852 | 4.299322 | 4.340084 | 4.282354 | 4.286401 | 22 | 31 | 13 | 15 |
| 105370567 | XR_944031 | LOC10537056 | uncharacterized LOC105370567 | lncRNA | . | . | | . | . | . | 0.122290 | 0.180663 | 0.331110 | 0.156237 | 0.284573 | 0.417610 | 0.743001 | 0.357217 | 1.572687 | 37.159029 | 0.559990897 | 1.166503149 | 0.243411076 | 0.992733415 | 4.851436 | 5.506187 | 5.133435 | 5.179425 | 5.282226 | 5.157828 | 27 | 37 | 54 | 27 |
| 105370594 | XR_944073 | LOC10537059 | uncharacterized LOC105370594 | lncRNA | . | . | | . | . | . | 0.077756 | 0.047136 | 0.071852 | 0.061952 | 0.180942 | 0.108957 | 0.161235 | 0.141645 | 1.011829 | 7.173678 | 1.134087183 | 0.014959477 | 0.98806451 | NA | 2.832372 | 2.852980 | 2.849158 | 2.834634 | 2.845910 | 2.838658 | 10 | 6 | 7 | 6 |
| 105370624 | XR_944148,XR | LOC10537062 | uncharacterized LOC105370624, transcript variant X2 | lncRNA | . | . | | . | . | . | 0.125303 | 0.128448 | 0.069630 | 0.040560 | 0.291585 | 0.296913 | 0.156248 | 0.092735 | -2.214103 | 7.174232 | 1.164151668 | -0.98502838 | 0.324610129 | NA | 3.303566 | 2.161867 | 2.863154 | 2.865890 | 2.845983 | 2.834446 | 11 | 11 | 5 | 3 |
| 105370687 | NR_158216 | LOC10537068 | uncharacterized LOC105370687 | lncRNA | . | . | | . | . | . | 0.229817 | 0.246659 | 0.179727 | 0.262434 | 0.534793 | 0.570163 | 0.403302 | 0.600023 | -1.063045 | 13.052728 | 0.847875909 | -0.10402769 | 0.917147373 | NA | 3.751025 | 3.660102 | 3.707742 | 3.711645 | 3.696385 | 3.715940 | 15 | 15 | 9 | 14 |
| 105370697 | NR_135205 | LOC10537069 | uncharacterized LOC105370697 | lncRNA | . | . | | . | . | . | 0.054722 | 0.124209 | 0.056518 | 0.113416 | 0.127339 | 0.287114 | 0.126824 | 0.259312 | 1.026278 | 5.970260 | 1.261048588 | 0.029675309 | 0.976326004 | NA | 2.562483 | 2.592943 | 2.562676 | 2.585479 | 2.568504 | 2.581203 | 4 | 9 | 4 | 7 |
| 105370714 | XR_948836 | LOC10537071 | uncharacterized LOC105370714, transcript variant X2 | lncRNA | . | . | | . | . | . | 0.871979 | 0.860777 | 0.957848 | 1.125834 | 2.029131 | 1.989725 | 2.149383 | 2.574085 | 1.180241 | 81.568897 | 0.352556791 | 0.678136672 | 0.497685041 | 0.998865404 | 6.225873 | 6.464191 | 6.326523 | 6.321960 | 6.347070 | 6.388988 | 87 | 80 | 71 | 89 |
| 105370751 | XR_00175176 | LOC10537075 | uncharacterized LOC105370751, transcript variant X1 | lncRNA | . | . | | . | . | . | 0.327285 | 0.452423 | 0.646342 | 0.504140 | 0.761606 | 1.045796 | 1.450373 | 1.152656 | 1.394404 | 12.407937 | 0.870694539 | 0.550880815 | 0.58171538 | NA | 3.373794 | 3.852980 | 3.606421 | 3.621979 | 3.639181 | 3.625293 | 10 | 13 | 14 | 12 |
| 105370754 | XR_00175147 | LOC10537075 | uncharacterized LOC105370754, transcript variant X2 | lncRNA | . | . | | . | . | . | 1.189359 | 0.668826 | 1.978158 | 0.382692 | 2.767689 | 1.546022 | 4.438929 | 0.874979 | 1.219819 | 13.378439 | 0.924760265 | 0.309990728 | 0.756568007 | NA | 3.586822 | 3.881801 | 3.729981 | 3.705080 | 3.768283 | 3.688256 | 17 | 10 | 21 | 5 |
| 105370849 | XM_01152229 | LOC10537084 | golgin subfamily A member 2-like, transcript variant X1 | protein_coding | XP_011520597.1;XP_016878276.1;XP_016878277.1 | . | | . | . | . | 5.015152 | 5.678194 | 6.942977 | 4.529286 | 11.670463 | 13.125396 | 15.579840 | 10.355675 | 1.097840 | 1068.008695 | 0.180159812 | 0.747487376 | 0.45476942 | 0.998865404 | 9.991508 | 10.126739 | 9.981533 | 10.054632 | 10.229290 | 9.924235 | 1114 | 1153 | 1169 | 836 |
| 105370850 | XR_932340 | LOC10537085 | uncharacterized LOC105370850 | lncRNA | . | . | | . | . | . | 1.648106 | 1.746472 | 2.298319 | 1.371692 | 3.835211 | 4.037048 | 5.157360 | 3.136210 | 1.066437 | 172.589536 | 0.289800253 | 0.320215391 | 0.748805057 | 0.998865404 | 7.383041 | 7.477806 | 7.396896 | 7.419701 | 7.538952 | 7.320057 | 187 | 185 | 195 | 124 |
| 105370852 | XR_932347 | LOC10537085 | uncharacterized LOC105370852 | lncRNA | . | . | | . | . | . | 1.382856 | 1.559220 | 1.851153 | 1.265761 | 3.217963 | 3.604207 | 4.153934 | 2.894013 | 1.045874 | 313.908776 | 0.227424401 | 0.284528179 | 0.776005642 | 0.998865404 | 8.260904 | 8.326748 | 8.245565 | 8.305242 | 8.400890 | 8.189072 | 334 | 349 | 333 | 243 |
| 105370876 | XR_932420,XR | LOC10537087 | uncharacterized LOC105370876, transcript variant X1 | lncRNA | . | . | | . | . | . | 0.244704 | 0.425714 | 0.372134 | 1.023381 | 0.569438 | 0.984056 | 0.835058 | 2.339839 | 1.827190 | 8.901943 | 1.040303285 | 0.83593553 | 0.403191173 | NA | 2.658167 | 3.522505 | 3.122909 | 3.133419 | 3.136554 | 3.166615 | 6 | 8 | 7 | 14 |
| 105370877 | XR_932423 | LOC10537087 | uncharacterized LOC105370877 | lncRNA | . | . | | . | . | . | 0.044041 | 0.056473 | 0.025761 | 0.070628 | 0.102486 | 0.130540 | 0.057808 | 0.161483 | -1.019977 | 13.790670 | 0.853781395 | -0.03342435 | 0.973336193 | NA | 3.802857 | 3.768175 | 3.777213 | 3.793320 | 3.760173 | 3.807586 | 14 | 17 | 7 | 18 |
| 105370888 | NR_135678 | TMEM202-AS1 | TMEM202 antisense RNA 1 | lncRNA | . | | 53265 | . | . | . | 0.186188 | 0.303913 | 0.295577 | 0.090349 | 0.433268 | 0.702509 | 0.663267 | 0.206572 | -1.245377 | 5.701941 | 1.310129278 | -0.24164195 | 0.809057613 | NA | 2.658167 | 2.348113 | 2.505500 | 2.515754 | 2.518068 | 2.491700 | 6 | 8 | 7 | 2 |
| 105370910 | XR_932500 | LOC10537091 | uncharacterized LOC105370910 | lncRNA | . | . | | . | . | . | 0.799243 | 0.885766 | 0.654142 | 0.574683 | 1.859871 | 2.047487 | 1.467877 | 1.313944 | -1.394572 | 47.810570 | 0.454291947 | -1.05619892 | 0.290877344 | 0.998865404 | 5.799086 | 5.319806 | 5.599658 | 5.619283 | 5.560893 | 5.540832 | 61 | 63 | 37 | 35 |
| 105370941 | NR_135665 | LOC10537094 | uncharacterized LOC105370941 | lncRNA | . | . | | . | . | . | 0.531169 | 0.423954 | 0.685111 | 0.356428 | 1.236051 | 0.979989 | 1.537369 | 0.814930 | 1.073896 | 42.671460 | 0.512456567 | 0.200708031 | 0.840926879 | 0.998865404 | 5.360548 | 5.467856 | 5.419470 | 5.380629 | 5.463718 | 5.356897 | 53 | 39 | 51 | 28 |
| 105370953 | XR_00175174 | LOC10537095 | uncharacterized LOC105370953 | lncRNA | . | . | | . | . | . | 0.249495 | 0.134378 | 0.554007 | 0.259964 | 0.580586 | 0.310621 | 1.243176 | 0.594378 | 2.096011 | 12.411982 | 0.913999694 | 1.168103605 | 0.242764976 | NA | 2.998355 | 4.073227 | 3.598825 | 3.576511 | 3.652216 | 3.600317 | 12 | 6 | 20 | 10 |
| 105371008 | NR_146563,N | LINC02251 | long intergenic non-protein coding RNA 2251 | lncRNA | . | | 53149 | . | . | . | 0.148319 | 0.271774 | 0.250539 | 0.489390 | 0.345144 | 0.628218 | 0.562202 | 1.118931 | 1.680261 | 20.554415 | 0.712368933 | 1.050979689 | 0.293267918 | NA | 3.942580 | 4.685555 | 4.303624 | 4.340382 | 4.336202 | 4.396013 | 13 | 21 | 16 | 31 |
| 105371017 | XR_00175170 | LOC10537101 | uncharacterized LOC105371017, transcript variant X3 | lncRNA | . | . | | . | . | . | 0.068742 | 0.024641 | 0.060355 | 0.043368 | 0.159965 | 0.056960 | 0.135434 | 0.099156 | 1.099042 | 7.432020 | 1.145244617 | 0.11896668 | 0.905301754 | NA | 2.820514 | 2.963455 | 2.903857 | 2.872644 | 2.897616 | 2.885364 | 12 | 4 | 8 | 6 |
| 105371019 | XR_00175170 | LOC10537101 | uncharacterized LOC105371019, transcript variant X2 | lncRNA | . | . | | . | . | . | 0.126309 | 0.066525 | 0.139765 | 0.103467 | 0.293925 | 0.153776 | 0.313629 | 0.236564 | 1.193379 | 11.226738 | 0.921335631 | 0.276829067 | 0.781911361 | NA | 3.353365 | 3.612728 | 3.492363 | 3.466732 | 3.495054 | 3.481677 | 15 | 8 | 12 | 10 |
| 105371026 | XR_00175172 | LOC10537102 | uncharacterized LOC105371026, transcript variant X8 | lncRNA | . | . | | . | . | . | 0.105222 | 0.373947 | 0.214299 | 0.282085 | 0.244855 | 0.864396 | 0.480880 | 0.644954 | 1.121834 | 29.301605 | 0.600912556 | 0.276013251 | 0.782537888 | 0.998865404 | 4.789392 | 4.951852 | 4.817300 | 4.896618 | 4.867254 | 4.875680 | 21 | 40 | 26 | 30 |
| 105371036 | XR_932985,XR | LOC10537103 | uncharacterized LOC105371036, transcript variant X2 | lncRNA | . | . | | . | . | . | 0.618468 | 0.521155 | 0.474116 | 0.631662 | 1.439201 | 1.204672 | 1.063902 | 1.444220 | -1.010908 | 31.581118 | 0.553646296 | -0.02826942 | 0.977447271 | 0.998865404 | 4.989254 | 4.972679 | 4.990876 | 4.973566 | 4.965908 | 4.995650 | 39 | 32 | 24 | 33 |
| 105371038 | NR_135177 | LOC10537103 | uncharacterized LOC105371038 | lncRNA | . | . | | . | . | . | 0.689886 | 0.635722 | 1.175866 | 0.888762 | 1.605392 | 1.469501 | 2.638609 | 2.032048 | 1.536983 | 14.223397 | 0.813307705 | 0.762443162 | 0.445795554 | NA | 3.486115 | 4.107809 | 3.805422 | 3.804855 | 3.841028 | 3.821848 | 13 | 12 | 17 | 14 |
| 105371044 | XR_932996 | LOC10537104 | uncharacterized LOC105371044 | lncRNA | . | . | | . | . | . | 0.026726 | 0.012107 | 0.084393 | 0.037109 | 0.062193 | 0.027985 | 0.189375 | 0.084845 | 2.751459 | 4.193826 | 1.512979257 | 0.965113488 | 0.334487972 | NA | 1.156660 | 2.622519 | 2.034317 | 2.030839 | 2.059048 | 2.042312 | 3 | 2 | 7 | 4 |
| 105371082 | XR_933070,XR | LOC10537108 | uncharacterized LOC105371082, transcript variant X2 | lncRNA | . | . | | . | . | . | 0.201095 | 0.212083 | 0.133229 | 0.168683 | 0.467957 | 0.490241 | 0.298963 | 0.385674 | -1.273089 | 9.994300 | 0.968213045 | -0.35976927 | 0.719019686 | NA | 3.486115 | 3.134756 | 3.332065 | 3.331217 | 3.312805 | 3.329065 | 13 | 12 | 6 | 10 |
| 105371083 | NR_135174 | LOC10537108 | uncharacterized LOC105371083 | lncRNA | . | . | | . | . | . | 0.270259 | 0.116270 | 0.589480 | 0.152135 | 0.628904 | 0.268763 | 1.322777 | 0.347838 | 1.844242 | 6.948532 | 1.219957338 | 0.723818795 | 0.469176989 | NA | 2.282109 | 3.175234 | 2.773720 | 2.754125 | 2.803329 | 2.761599 | 8 | 3 | 12 | 4 |
| 105371094 | XR_933105 | LOC10537109 | uncharacterized LOC105371094 | lncRNA | . | . | | . | . | . | 0.456931 | 0.322278 | 0.423031 | 0.454198 | 1.063297 | 0.744961 | 0.949269 | 1.038469 | 1.093368 | 23.801517 | 0.637258017 | 0.202083286 | 0.839851614 | NA | 4.506418 | 4.636609 | 4.584569 | 4.548655 | 4.572630 | 4.578579 | 31 | 20 | 21 | 24 |
| 105371115 | XR_00175210 | LOC10537111 | uncharacterized LOC105371115, transcript variant X3 | lncRNA | . | . | | . | . | . | 0.298830 | 0.186514 | 0.264154 | 0.374236 | 0.695389 | 0.431135 | 0.592754 | 0.855645 | 1.240360 | 26.971559 | 0.601327745 | 0.516787035 | 0.605304832 | 0.998865404 | 4.588571 | 4.901253 | 4.755723 | 4.716695 | 4.760114 | 4.760214 | 33 | 21 | 26 | 28 |

| | | | | | | | | | | | | | | | | | | | | | | | | | | | | | | | | | |
|---|---|---|---|---|---|---|---|---|---|---|---|---|---|---|---|---|---|---|---|---|---|---|---|---|---|---|---|---|---|---|---|---|---|
| 105371159 | XR_001752365 | LOC10537115 | uncharacterized LOC105371159, transcript variant X2 | lncRNA | . | . | . | . | . | 0.078409 | 0.120886 | 0.202553 | 0.159541 | 0.182460 | 0.279434 | 0.454523 | 0.364772 | 1.788356 | 32.076320 | 0.56304101 | 1.489472705 | 0.136362936 | 0.89431118 | 4.524804 | 5.362218 | 4.924362 | 4.966103 | 5.036743 | 5.000323 | 21 | 30 | 40 | 34 |
| 105371184 | NR_136337 | LINC02867 | long intergenic non-protein coding RNA 2867 | transcribed_ps | . | 54519 | . | . | . | 0.185639 | 0.120211 | 0.098195 | 0.230522 | 0.431989 | 0.277872 | 0.220346 | 0.527061 | 1.073915 | 3.688932 | 1.572624472 | 0.065419025 | 0.947840377 | NA | 1.832372 | 1.932304 | 1.887985 | 1.880601 | 1.878713 | 1.892282 | 5 | 3 | 2 | 5 |
| 105371196 | XR_001737752 | LOC10537119 | uncharacterized LOC105371196 | lncRNA | . | . | . | . | . | 0.176109 | 0.282669 | 0.299896 | 0.154135 | 0.409812 | 0.653401 | 0.672957 | 0.352411 | 1.017348 | 5.000631 | 1.358013481 | 0.018271916 | 0.985421932 | NA | 2.307819 | 2.336262 | 2.313958 | 2.319705 | 2.325426 | 2.308943 | 5 | 6 | 6 | 3 |
| 105371204 | XR_942166,XR | LOC10537120 | uncharacterized LOC105371204, transcript variant X2 | lncRNA | . | . | . | . | . | 0.900546 | 1.005165 | 1.205697 | 0.894236 | 2.095607 | 2.323483 | 2.705549 | 2.044564 | 1.084585 | 67.905007 | 0.391774597 | 0.299005548 | 0.764935812 | 0.998865404 | 6.025012 | 6.143450 | 6.059014 | 6.083670 | 6.126178 | 6.053175 | 71 | 74 | 71 | 56 |
| 105371214 | XR_001737751 | LOC10537121 | uncharacterized LOC105371214 | lncRNA | . | . | . | . | . | 0.238996 | 0.352945 | 0.542209 | 0.252323 | 0.556153 | 0.815848 | 1.216702 | 0.576906 | 1.331132 | 7.364422 | 1.136334463 | 0.363144714 | 0.716496784 | NA | 2.658167 | 3.073227 | 2.858209 | 2.868620 | 2.886283 | 2.859275 | 6 | 8 | 10 | 5 |
| 105371222 | XR_922062 | LOC10537122 | uncharacterized LOC105371222 | lncRNA | . | . | . | . | . | 0.209640 | 0.243476 | 0.262262 | 0.222921 | 0.487841 | 0.562806 | 0.588508 | 0.509683 | 1.077466 | 18.200376 | 0.718555054 | 0.149803252 | 0.880919844 | NA | 4.130737 | 4.239024 | 4.175634 | 4.184648 | 4.193449 | 4.178396 | 19 | 20 | 18 | 16 |
| 105371224 | XR_001737711 | LOC10537122 | uncharacterized LOC105371224, transcript variant X2 | lncRNA | . | . | . | . | . | 0.045850 | 0.036489 | 0.065630 | 0.038414 | 0.106692 | 0.084347 | 0.147272 | 0.087829 | 1.216713 | 19.832054 | 0.705784283 | 0.400956975 | 0.688451801 | NA | 4.159017 | 4.446236 | 4.302375 | 4.283915 | 4.333095 | 4.285558 | 23 | 17 | 24 | 15 |
| 105371226 | XR_922067 | LINC01731 | long intergenic non-protein coding RNA 1731 | lncRNA | . | 52519 | . | . | . | 0.145195 | 0.048575 | 0.175018 | 0.059020 | 0.337876 | 0.112284 | 0.392734 | 0.134942 | 1.161978 | 5.749002 | 1.315170554 | 0.164680053 | 0.869195818 | NA | 2.405477 | 2.632244 | 2.524249 | 2.500583 | 2.529553 | 2.502999 | 9 | 3 | 8 | 3 |
| 105371228 | XR_001737719 | LINC02804 | long intergenic non-protein coding RNA 2804, transcript variant X | lncRNA | . | 54338 | . | . | . | 0.160335 | 0.228293 | 0.220779 | 0.130793 | 0.373106 | 0.527710 | 0.495422 | 0.299042 | -1.022964 | 39.962264 | 0.498305928 | -0.06573242 | 0.947590859 | 0.998865404 | 5.335987 | 5.304979 | 5.315017 | 5.325355 | 5.340853 | 5.291924 | 45 | 45 | 40 | 31 |
| 105371240 | XR_933516 | LOC10537124 | uncharacterized LOC105371240 | lncRNA | . | . | . | . | . | 0.222200 | 0.237559 | 0.245233 | 0.220522 | 0.517068 | 0.549128 | 0.550296 | 0.504198 | 1.003416 | 31.013515 | 0.554726249 | 0.008867732 | 0.992924667 | 0.999862634 | 4.951979 | 4.957666 | 4.951440 | 4.956038 | 4.963050 | 4.945195 | 35 | 34 | 29 | 27 |
| 105371267 | NR_136518,NF | LOC10537126 | p53-regulated lncRNA 1 | lncRNA | . | . | . | . | . | 0.266411 | 0.439745 | 0.440728 | 0.152405 | 0.619948 | 1.016491 | 0.988980 | 0.348456 | -1.190212 | 30.805187 | 0.629780171 | -0.39889776 | 0.689968543 | 0.998865404 | 5.063683 | 4.815893 | 4.909212 | 4.976921 | 4.973581 | 4.867570 | 29 | 45 | 36 | 14 |
| 105371271 | XR_933590 | LOC10537127 | uncharacterized LOC105371271 | lncRNA | . | . | . | . | . | 0.028767 | 0.009098 | 0.053279 | 0.026117 | 0.066943 | 0.021031 | 0.119557 | 0.059712 | 1.792857 | 4.340553 | 1.469052581 | 0.573335663 | 0.566417443 | NA | 1.631267 | 2.481107 | 2.103399 | 2.091597 | 2.114680 | 2.103044 | 5 | 2 | 6 | 4 |
| 105371286 | XR_933619,XR | LOC10537128 | uncharacterized LOC105371286, transcript variant X1 | lncRNA | . | . | . | . | . | 0.258204 | 0.377952 | 0.054401 | 0.306111 | 0.600851 | 0.873652 | 0.122075 | 0.699886 | -1.707115 | 11.770734 | 0.959387761 | -0.80422142 | 0.421269112 | NA | 3.895610 | 3.114084 | 3.563865 | 3.587889 | 3.526306 | 3.571128 | 14 | 19 | 3 | 13 |
| 105371288 | XR_001737726 | LOC10537128 | uncharacterized LOC105371288, transcript variant X3 | lncRNA | . | . | . | . | . | 0.185974 | 0.184248 | 0.153786 | 0.174264 | 0.432767 | 0.425897 | 0.345092 | 0.398433 | -1.105427 | 6.396156 | 1.191194435 | -0.12139337 | 0.903379466 | NA | 2.747897 | 2.602870 | 2.682244 | 2.680078 | 2.676835 | 2.679676 | 8 | 7 | 5 | 6 |
| 105371312 | XR_002958598 | LOC10537131 | uncharacterized LOC105371312 | lncRNA | . | . | . | . | . | 3.639834 | 4.290743 | 4.046873 | 4.394042 | 8.470043 | 9.918244 | 9.081065 | 10.046454 | 1.050042 | 138.166620 | 0.280348427 | 0.251285618 | 0.801593294 | 0.998865404 | 7.075138 | 7.144558 | 7.069783 | 7.128667 | 7.105428 | 7.129894 | 143 | 157 | 119 | 137 |
| 105371354 | XR_001752380 | LOC10537135 | uncharacterized LOC105371354 | lncRNA | . | . | . | . | . | 0.181450 | 0.222353 | 0.163651 | 0.154085 | 0.422241 | 0.513979 | 0.367228 | 0.352297 | -1.253772 | 13.739883 | 0.825407512 | -0.39529 | 0.692628878 | NA | 3.934350 | 3.607807 | 3.783594 | 3.795897 | 3.778304 | 3.774245 | 16 | 18 | 11 | 11 |
| 105371366 | XR_001752382 | LOC10537136 | uncharacterized LOC105371366, transcript variant X6 | lncRNA | . | . | . | . | . | 0.064956 | 0.100301 | 0.059878 | 0.081523 | 0.151155 | 0.231851 | 0.134364 | 0.186395 | -1.166259 | 29.284798 | 0.581194287 | -0.38177888 | 0.702625384 | 0.998865404 | 4.980095 | 4.755319 | 4.858447 | 4.903636 | 4.851758 | 4.874375 | 30 | 40 | 21 | 28 |
| 105371374 | NR_146503 | ATP2C2-AS1 | ATP2C2 antisense RNA 1 | lncRNA | . | 53167 | . | . | . | 0.075561 | 0.126987 | 0.218147 | 0.074855 | 0.175834 | 0.293537 | 0.489516 | 0.171147 | 1.415242 | 12.007467 | 0.927607153 | 0.540151968 | 0.589092234 | NA | 3.312060 | 3.815893 | 3.548163 | 3.567781 | 3.604043 | 3.547568 | 9 | 13 | 18 | 7 |
| 105371416 | XR_001752310 | LINC02166 | long intergenic non-protein coding RNA 2166, transcript variant X | lncRNA | . | 53027 | . | . | . | 0.551655 | 0.804137 | 0.623395 | 1.212762 | 1.283724 | 1.858799 | 1.398878 | 2.772836 | 1.461869 | 17.157285 | 0.746618015 | 0.73372795 | 0.46311456 | NA | 3.802857 | 4.347534 | 4.069225 | 4.085591 | 4.092131 | 4.118175 | 14 | 17 | 15 | 22 |
| 105371423 | XR_001752318 | LINC02193 | long intergenic non-protein coding RNA 2193 | lncRNA | . | 53055 | . | . | . | 0.092512 | 0.076819 | 0.172244 | 0.115651 | 0.215278 | 0.177571 | 0.386510 | 0.264422 | 1.675589 | 15.555079 | 0.786145005 | 0.94724006 | 0.343516467 | NA | 3.537347 | 4.285383 | 3.933287 | 3.920650 | 3.975484 | 3.946618 | 15 | 11 | 20 | 15 |
| 105371450 | XR_001738241 | THBS3-AS1 | THBS3 antisense RNA 1, transcript variant X1 | lncRNA | . | 40582 | . | . | . | 0.097276 | 0.166895 | 0.408043 | 0.342601 | 0.226365 | 0.385786 | 0.915638 | 0.783317 | 2.739185 | 15.193183 | 0.816036017 | 1.781473864 | 0.074835071 | NA | 3.024434 | 4.475715 | 3.852080 | 3.871833 | 3.936524 | 3.911724 | 7 | 11 | 22 | 18 |
| 105371452 | XR_922171 | LOC10537145 | uncharacterized LOC105371452 | lncRNA | . | . | . | . | . | 0.510318 | 0.295193 | 0.351791 | 0.285961 | 1.187532 | 0.682351 | 0.789409 | 0.653816 | -1.258856 | 4.396493 | 1.440664735 | -0.23052804 | 0.817681476 | NA | 2.290730 | 1.963455 | 2.148233 | 2.137057 | 2.141181 | 2.134886 | 7 | 4 | 4 | 3 |
| 105371453 | XR_922175 | LOC10537145 | uncharacterized LOC105371453 | lncRNA | . | . | . | . | . | 0.126311 | 0.125672 | 0.086353 | 0.089911 | 0.293931 | 0.290497 | 0.193773 | 0.205571 | -1.434706 | 32.331018 | 0.545651571 | -0.95437225 | 0.339895249 | 0.998865404 | 5.251870 | 4.731077 | 5.041763 | 5.040596 | 4.994966 | 4.995388 | 44 | 41 | 23 | 25 |
| 105371480 | XR_001753073 | LOC10537148 | uncharacterized LOC105371480 | lncRNA | . | . | . | . | . | 1.096322 | 1.282776 | 1.480677 | 1.414604 | 2.551187 | 2.965194 | 3.322596 | 3.234325 | 1.201917 | 117.741461 | 0.299723783 | 0.88527199 | 0.376009953 | 0.998865404 | 6.740919 | 7.005889 | 6.818598 | 6.867022 | 6.915150 | 6.895132 | 114 | 124 | 115 | 117 |
| 105371490 | XR_934156 | LOC10537149 | uncharacterized LOC105371490 | lncRNA | . | . | . | . | . | 0.429998 | 0.346152 | 0.381864 | 0.169859 | 1.000624 | 0.800145 | 0.856893 | 0.388362 | -1.412949 | 11.465229 | 0.920505504 | -0.54177772 | 0.587971642 | NA | 3.744764 | 3.251698 | 3.536648 | 3.524317 | 3.526728 | 3.499479 | 17 | 13 | 11 | 6 |
| 105371492 | XR_934158,XR | LOC10537149 | uncharacterized LOC105371492, transcript variant X3 | lncRNA | . | . | . | . | . | 0.433840 | 0.811670 | 0.928107 | 0.950780 | 1.009565 | 1.876212 | 2.082644 | 2.173846 | 1.535201 | 29.209709 | 0.592860862 | 1.043124752 | 0.296890519 | 0.998865404 | 4.528454 | 5.143283 | 4.799607 | 4.856567 | 4.870853 | 4.886830 | 19 | 32 | 29 | 35 |
| 105371525 | XR_001752781 | LOC10537152 | uncharacterized LOC105371525, transcript variant X2 | lncRNA | . | . | . | . | . | 0.793453 | 1.130078 | 0.802204 | 1.500399 | 1.846399 | 2.612226 | 1.800122 | 3.430484 | 1.147015 | 40.501362 | 0.519132936 | 0.381183056 | 0.703067421 | 0.998865404 | 5.239794 | 5.433505 | 5.306547 | 5.338044 | 5.292847 | 5.392626 | 39 | 45 | 26 | 53 |
| 105371531 | XR_934222 | LOC10537153 | uncharacterized LOC105371531 | lncRNA | . | . | . | . | . | 0.041470 | 0.047373 | 0.158964 | 0.074660 | 0.096503 | 0.109505 | 0.356710 | 0.170702 | 2.701151 | 9.980172 | 1.004820745 | 1.426696665 | 0.153667347 | NA | 2.429096 | 3.865403 | 3.264919 | 3.266902 | 3.328041 | 3.285696 | 6 | 6 | 17 | 9 |
| 105371554 | XR_001753079 | LOC10537155 | uncharacterized LOC105371554, transcript variant X3 | lncRNA | . | . | . | . | . | 0.420209 | 0.538189 | 0.564750 | 0.200766 | 0.977844 | 1.244047 | 1.267282 | 0.459028 | -1.330230 | 40.634590 | 0.55735114 | -0.73862838 | 0.460132688 | 0.998865404 | 5.533815 | 5.126853 | 5.339001 | 5.374663 | 5.375547 | 5.245116 | 48 | 55 | 45 | 17 |
| 105371564 | NR_131243 | SMCR2 | Smith-Magenis syndrome chromosome region, candidate 2 | lncRNA | . | 17914 | . | . | . | 0.140337 | 0.170316 | 0.333722 | 0.247014 | 0.326569 | 0.393694 | 0.748862 | 0.564768 | 1.939228 | 4.636372 | 1.408501212 | 0.678368243 | 0.497538239 | NA | 1.658167 | 2.612728 | 2.186518 | 2.191898 | 2.206274 | 2.199435 | 3 | 4 | 6 | 5 |
| 105371586 | XR_001737762 | FAM78B-AS1 | FAM78B antisense RNA 1 | lncRNA | . | 40639 | . | . | . | 0.105832 | 0.171381 | 0.090114 | 0.140271 | 0.246275 | 0.396155 | 0.202213 | 0.320713 | -1.210143 | 25.533386 | 0.632080024 | -0.43535297 | 0.663306239 | NA | 4.807371 | 4.527725 | 4.657126 | 4.709595 | 4.644745 | 4.682421 | 25 | 37 | 16 | 26 |
| 105371592 | NR_136412 | LOC10537159 | uncharacterized LOC105371592 | lncRNA | . | . | . | . | . | 3.591844 | 3.363945 | 3.637049 | 4.439163 | 8.358370 | 7.775910 | 8.161432 | 10.149619 | 1.141603 | 52.258129 | 0.434310545 | 0.439917297 | 0.659997008 | 0.998865404 | 5.609306 | 5.799591 | 5.698117 | 5.686488 | 5.699333 | 5.735658 | 58 | 51 | 44 | 57 |
| 105371622 | XR_922296 | LOC10537162 | uncharacterized LOC105371622 | lncRNA | . | . | . | . | . | 0.201879 | 0.305476 | 0.344091 | 0.179759 | 0.469781 | 0.706120 | 0.772129 | 0.410998 | 1.012495 | 54.518991 | 0.471431052 | 0.038001223 | 0.969686707 | 0.998865404 | 5.758813 | 5.778494 | 5.714918 | 5.795094 | 5.818904 | 5.693616 | 50 | 70 | 63 | 35 |
| 105371630 | XR_922307 | LOC10537163 | uncharacterized LOC105371630 | lncRNA | . | . | . | . | . | 0.284121 | 0.338701 | 0.585163 | 0.248435 | 0.661161 | 0.782924 | 1.313089 | 0.568017 | 1.299686 | 11.901681 | 0.90627867 | 0.417270672 | 0.676480437 | NA | 3.369732 | 3.751299 | 3.550425 | 3.557723 | 3.588602 | 3.546452 | 11 | 12 | 16 | 8 |
| 105371637 | XR_922334 | LOC10537163 | uncharacterized LOC105371637 | lncRNA | . | . | . | . | . | 0.084898 | 0.056850 | 0.019279 | 0.040828 | 0.197561 | 0.131412 | 0.043262 | 0.093348 | -2.163723 | 10.405753 | 0.983001018 | -1.13277202 | 0.257310002 | NA | 3.832372 | 2.715150 | 3.420831 | 3.392782 | 3.364625 | 3.381429 | 20 | 12 | 4 | 8 |
| 105371675 | XR_922396 | LOC10537167 | uncharacterized LOC105371675 | lncRNA | . | . | . | . | . | 1.841284 | 1.716280 | 1.877508 | 1.799448 | 4.284743 | 3.967257 | 4.213074 | 4.114223 | 1.015880 | 146.711010 | 0.270210168 | 0.084120785 | 0.9329604 | 0.998865404 | 7.185110 | 7.208462 | 7.208158 | 7.178367 | 7.209410 | 7.189413 | 174 | 151 | 132 | 135 |
| 105371692 | XR_001738393 | LOC10537169 | uncharacterized LOC105371692, transcript variant X2 | lncRNA | . | . | . | . | . | 0.023854 | 0.026839 | 0.028721 | 0.030444 | 0.055510 | 0.062039 | 0.064449 | 0.069606 | 1.171140 | 9.742077 | 0.970077039 | 0.234944127 | 0.8142521 | NA | 3.166062 | 3.393446 | 3.276053 | 3.278794 | 3.283567 | 3.285012 | 10 | 10 | 9 | 10 |
| 105371703 | NR_135673 | LOC10537170 | uncharacterized LOC105371703 | lncRNA | . | . | . | . | . | 6.628016 | 9.452944 | 7.857514 | 6.905406 | 15.423666 | 21.850903 | 17.632034 | 15.788390 | -1.105377 | 184.590031 | 0.266206749 | -0.54295455 | 0.587161093 | 0.998865404 | 7.598850 | 7.453871 | 7.464763 | 7.620103 | 7.530371 | 7.472637 | 185 | 245 | 163 | 153 |
| 105371711 | XR_934639 | LOC10537171 | uncharacterized LOC105371711 | lncRNA | . | . | . | . | . | 0.348204 | 0.275688 | 0.741088 | 0.567138 | 0.810284 | 0.637266 | 1.662980 | 1.296694 | 2.072262 | 8.240635 | 1.070593916 | 0.981890748 | 0.326153674 | NA | 2.421266 | 3.475715 | 3.014342 | 3.007725 | 3.043552 | 3.030552 | 7 | 5 | 11 | 9 |
| 105371716 | XR_001752822 | LOC10537171 | uncharacterized LOC105371716 | lncRNA | . | . | . | . | . | 0.037681 | 0.018857 | 0.014610 | 0.025687 | 0.087685 | 0.043588 | 0.032784 | 0.058729 | -1.487557 | 4.088173 | 1.503726114 | -0.38101697 | 0.703190655 | NA | 2.290730 | 1.715150 | 2.050610 | 2.039448 | 2.033341 | 2.042033 | 7 | 4 | 2 | 4 |
| 105371722 | XR_934657 | LOC10537172 | uncharacterized LOC105371722 | lncRNA | . | . | . | . | . | 0.947504 | 0.540779 | 0.615856 | 1.007981 | 2.204882 | 1.250036 | 1.381963 | 2.304629 | 1.007635 | 4.909000 | 1.369565717 | 0.008011823 | 0.993607558 | NA | 2.290730 | 2.300113 | 2.303276 | 2.292132 | 2.291320 | 2.304426 | 7 | 4 | 3 | 6 |
| 105371734 | XR_934677 | LOC10537173 | uncharacterized LOC105371734 | lncRNA | . | . | . | . | . | 0.397284 | 0.385881 | 0.180771 | 0.494264 | 0.924496 | 0.891982 | 0.405644 | 1.130077 | -1.212038 | 13.873265 | 0.855367979 | -0.32434525 | 0.745676665 | NA | 3.928838 | 3.645770 | 3.805100 | 3.801436 | 3.767519 | 3.815302 | 18 | 16 | 6 | 17 |
| 105371743 | NR_135479 | SNHG30 | small nucleolar RNA host gene 30 | lncRNA | . | 52836 | . | . | . | 9.866202 | 9.790821 | 9.553204 | 9.549758 | 22.959059 | 22.631920 | 21.437113 | 21.834389 | -1.042529 | 249.838691 | 0.215304655 | -0.27908281 | 0.780181273 | 0.998865404 | 7.994524 | 7.934560 | 7.979519 | 7.971331 | 7.957604 | 7.951736 | 296 | 273 | 214 | 228 |
| 105371761 | XR_934734 | LOC10537176 | uncharacterized LOC105371761 | lncRNA | . | . | . | . | . | 0.607534 | 0.276809 | 0.442327 | 0.406657 | 1.413755 | 0.639857 | 0.992568 | 0.929772 | -1.081734 | 11.520193 | 0.922256975 | -0.12290061 | 0.902185805 | NA | 3.579805 | 3.470304 | 3.548835 | 3.507679 | 3.526252 | 3.522626 | 19 | 8 | 10 | 10 |
| 105371763 | NR_158157 | LOC10537176 | uncharacterized LOC105371763 | lncRNA | . | . | . | . | . | 0.833705 | 0.774195 | 0.734249 | 0.742280 | 1.940067 | 1.789586 | 1.647634 | 1.697135 | -1.100329 | 66.247064 | 0.386545421 | -0.35684186 | 0.721210193 | 0.998865404 | 6.116929 | 5.979362 | 6.071194 | 6.052378 | 6.040596 | 6.039338 | 83 | 72 | 55 | 59 |
| 105371781 | XR_001752888 | LOC10537178 | uncharacterized LOC105371781, transcript variant X1 | lncRNA | . | . | . | . | . | 0.030489 | 0.078305 | 0.058539 | 0.029212 | 0.070949 | 0.181005 | 0.131359 | 0.066790 | -1.015711 | 4.510603 | 1.440876329 | -0.01560899 | 0.987546337 | NA | 2.184683 | 2.161867 | 2.157842 | 2.176189 | 2.172422 | 2.161022 | 3 | 7 | 5 | 3 |
| 105371814 | NR_135674 | LOC10537181 | uncharacterized LOC105371814 | lncRNA | . | . | . | . | . | 0.423490 | 0.313502 | 0.611596 | 0.270576 | 0.985478 | 0.724674 | 1.372404 | 0.618641 | 1.191775 | 9.796233 | 0.998788318 | 0.253418677 | 0.799944697 | NA | 3.156660 | 3.416142 | 3.286122 | 3.272294 | 3.305174 | 3.268162 | 12 | 8 | 13 | 6 |
| 105371818 | XR_001752932 | LOC10537181 | uncharacterized LOC105371818, transcript variant X1 | lncRNA | . | . | . | . | . | 1.182136 | 0.977852 | 1.418580 | 1.045499 | 2.750879 | 2.260350 | 3.183253 | 2.390409 | 1.122120 | 116.751330 | 0.314538262 | 0.528480707 | 0.597165734 | 0.998865404 | 6.780541 | 6.949127 | 6.873773 | 6.809355 | 6.932406 | 6.826606 | 139 | 107 | 124 | 98 |
| 105371824 | NR_135675 | LOC10537182 | uncharacterized LOC105371824 | lncRNA | . | . | . | . | . | 0.440435 | 0.482249 | 0.145295 | 0.482208 | 1.024909 | 1.114739 | 0.326038 | 1.102512 | -1.368940 | 7.750449 | 1.120072859 | -0.40449092 | 0.685851754 | NA | 3.166062 | 2.705970 | 2.964968 | 2.967557 | 2.941890 | 2.968774 | 10 | 10 | 3 | 9 |
| 105371825 | XR_001752940 | LOC10537182 | uncharacterized LOC105371825 | lncRNA | . | . | . | . | . | 0.048872 | 0.046037 | 0.111854 | 0.088496 | 0.113726 | 0.106417 | 0.250997 | 0.202337 | 2.140736 | 7.736031 | 1.101601024 | 0.996828028 | 0.318847991 | NA | 2.299300 | 3.399154 | 2.921224 | 2.918697 | 2.949476 | 2.941453 | 6 | 5 | 10 | 9 |
| 105371841 | XR_934884 | LOC10537184 | uncharacterized LOC105371841 | lncRNA | . | . | . | . | . | 0.051895 | 0.120738 | 0.101510 | 0.075131 | 0.120763 | 0.279090 | 0.227785 | 0.171778 | 1.103469 | 4.777270 | 1.39527179 | 0.101805687 | 0.918910911 | NA | 2.184683 | 2.324312 | 2.238701 | 2.257075 | 2.253377 | 2.246785 | 3 | 7 | 5 | 4 |
| 105371903 | XR_934990 | LOC10537190 | uncharacterized LOC105371903 | lncRNA | . | . | . | . | . | 0.055363 | 0.041018 | 0.025693 | 0.053635 | 0.128833 | 0.094815 | 0.057653 | 0.122629 | -1.164984 | 4.137729 | 1.490722843 | -0.14778737 | 0.88251058 | NA | 2.156660 | 1.932304 | 2.059624 | 2.052531 | 2.046422 | 2.059916 | 6 | 4 | 2 | 5 |
| 105371927 | XR_935047 | LINC01971 | long intergenic non-protein coding RNA 1971 | lncRNA | . | 52797 | . | . | . | 0.417487 | 0.344264 | 0.578224 | 0.225823 | 0.971510 | 0.795782 | 1.297518 | 0.516318 | 1.072337 | 6.493617 | 1.200672597 | 0.083918644 | 0.933121116 | NA | 2.644780 | 2.751299 | 2.695991 | 2.689514 | 2.705619 | 2.683845 | 8 | 6 | 8 | 4 |
| 105371932 | XM_01702549 | LOC10537193 | uncharacterized LOC105371932 | protein_coding | XP_016880981.1 | . | . | . | . | 0.717819 | 0.964552 | 0.633832 | 0.827462 | 1.670394 | 2.229606 | 1.422301 | 1.891893 | -1.165359 | 29.270222 | 0.578810802 | -0.38142679 | 0.702886589 | 0.998865404 | 4.978762 | 4.755319 | 4.862043 | 4.899848 | 4.851678 | 4.874294 | 31 | 39 | 21 | 28 |
| 105371941 | XR_933959 | LOC10537194 | uncharacterized LOC105371941 | lncRNA | . | . | . | . | . | 0.503776 | 0.335991 | 0.892089 | 0.939702 | 1.172307 | 0.776657 | 2.001822 | 2.148518 | 2.079114 | 19.230955 | 0.712174491 | 1.482738943 | 0.138143784 | NA | 3.641413 | 4.699499 | 4.226160 | 4.205827 | 4.270970 | 4.277110 | 17 | 11 | 22 | 25 |
| 105371960 | XR_935094 | LOC10537196 | uncharacterized LOC105371960 | lncRNA | . | . | . | . | . | 0.691998 | 0.414035 | 0.406551 | 0.403935 | 1.610308 | 0.957061 | 0.912290 | 0.923550 | -1.474348 | 4.108999 | 1.49076593 | -0.37569768 | 0.707141664 | NA | 2.290730 | 1.733338 | 2.055975 | 2.044800 | 2.043780 | 2.042586 | 7 | 4 | 3 | 3 |
| 105371967 | XR_002958220 | LOC10537196 | uncharacterized LOC105371967, transcript variant X1 | lncRNA | . | . | . | . | . | 0.682327 | 0.919786 | 0.850582 | 0.867608 | 1.587803 | 2.126126 | 1.908681 | 1.983682 | 1.133700 | 53.957730 | 0.438842104 | 0.41253696 | 0.679945897 | 0.998865404 | 5.658912 | 5.842751 | 5.751153 | 5.719726 | 5.796165 | 5.722754 | 63 | 50 | 59 | 44 |
| 105371998 | NR_136514 | LOC10537199 | uncharacterized LOC105371998 | lncRNA | . | . | . | . | . | 0.239656 | 0.197460 | 0.596868 | 0.470977 | 0.557690 | 0.456437 | 1.339354 | 1.076832 | 2.288945 | 13.970752 | 0.833276924 | 1.433716653 | 0.151653138 | NA | 3.084644 | 4.282298 | 3.761387 | 3.751848 | 3.814530 | 3.794988 | 11 | 8 | 19 | 16 |
| 105372007 | XR_935178 | LOC10537200 | uncharacterized LOC105372007 | lncRNA | . | . | . | . | . | 0.316315 | 0.129848 | 0.276283 | 0.319793 | 0.736077 | 0.300149 | 0.619971 | 0.731169 | 1.244548 | 15.419852 | 0.808152366 | 0.390548023 | 0.696131352 | NA | 3.778546 | 4.097315 | 3.957748 | 3.909516 | 3.944159 | 3.952845 | 22 | 9 | 14 | 17 |
| 105372061 | XR_001753404 | LOC10537206 | uncharacterized LOC105372061, transcript variant X6 | lncRNA | . | . | . | . | . | 0.054874 | 0.039981 | 0.082494 | 0.099726 | 0.127694 | 0.092419 | 0.185116 | 0.228012 | 1.713457 | 7.861931 | 1.088882719 | 0.713493181 | 0.475540631 | NA | 2.533720 | 3.312263 | 2.959689 | 2.948966 | 2.969779 | 2.976377 | 8 | 5 | 8 | 10 |
| 105372098 | XR_002958192 | LOC10537209 | uncharacterized LOC105372098, transcript variant X2 | lncRNA | . | . | . | . | . | 0.132252 | 0.183733 | 0.165972 | 0.113611 | 0.307754 | 0.424706 | 0.372437 | 0.259759 | -1.076493 | 18.998435 | 0.703490572 | -0.15115832 | 0.879850826 | NA | 4.299300 | 4.194412 | 4.256758 | 4.246661 | 4.250938 | 4.240549 | 24 | 20 | 17 | 16 |
| 105372105 | XR_935449 | LOC10537210 | uncharacterized LOC105372105 | lncRNA | . | . | . | . | . | 0.046911 | 0.047806 | 0.061412 | 0.039997 | 0.109163 | 0.110505 | 0.137808 | 0.091449 | 1.071109 | 10.232527 | 0.953281953 | 0.103961572 | 0.917199845 | NA | 3.303566 | 3.404839 | 3.348356 | 3.351323 | 3.361817 | 3.344016 | 11 | 11 | 11 | 8 |
| 105372117 | XR_935468 | LOC10537211 | uncharacterized LOC105372117 | lncRNA | . | . | . | . | . | 0.162579 | 0.138683 | 0.303634 | 0.213762 | 0.378327 | 0.320573 | 0.681345 | 0.488741 | 1.680244 | 44.400980 | 0.480540687 | 1.557975323 | 0.119239098 | 0.858706505 | 5.048513 | 5.799803 | 5.419335 | 5.399355 | 5.529850 | 5.459986 | 41 | 33 | 57 | 43 |
| 105372118 | XR_001753450 | LOC10537211 | uncharacterized LOC105372118 | lncRNA | . | . | . | . | . | 0.109317 | 0.066523 | 0.161586 | 0.095862 | 0.254385 | 0.153770 | 0.362595 | 0.219177 | 1.441305 | 7.082758 | 1.147930876 | 0.459414123 | 0.645936808 | NA | 2.533720 | 3.066070 | 2.813551 | 2.802767 | 2.828356 | 2.811121 | 8 | 5 | 9 | 6 |
| 105372130 | XR_001753451 | LOC10537213 | uncharacterized LOC105372130, transcript variant X7 | lncRNA | . | . | . | . | . | 0.030433 | 0.064603 | 0.028485 | 0.051029 | 0.070818 | 0.149333 | 0.063919 | 0.116673 | -1.195011 | 6.636167 | 1.189857468 | -0.21601241 | 0.82897807 | NA | 2.855800 | 2.592943 | 2.725324 | 2.744113 | 2.723127 | 2.735728 | 6 | 10 | 4 | 7 |
| 105372141 | XR_001753464 | LOC10537214 | uncharacterized LOC105372141, transcript variant X2 | lncRNA | . | . | . | . | . | 0.154398 | 0.227652 | 0.264366 | 0.336584 | 0.359290 | 0.526228 | 0.593230 | 0.769560 | 1.491956 | 12.899617 | 0.855156956 | 0.674969268 | 0.49969529 | NA | 3.373794 | 3.947964 | 3.660429 | 3.676012 | 3.683640 | 3.697737 | 10 | 13 | 12 | 16 |
| 105372165 | XR_935576 | LOC10537216 | uncharacterized LOC105372165 | lncRNA | . | . | . | . | . | 0.202291 | 0.107406 | 0.107015 | 0.144570 | 0.470740 | 0.248274 | 0.240139 | 0.330541 | -1.251780 | 17.971866 | 0.747511616 | -0.43341233 | 0.664715246 | NA | 4.320284 | 3.996978 | 4.204927 | 4.152056 | 4.152608 | 4.170602 | 30 | 15 | 12 | 17 |
| 105372212 | XR_935657 | LOC10537221 | uncharacterized LOC105372212 | lncRNA | . | . | . | . | . | 0.067497 | 0.359406 | 0.179250 | 0.275215 | 0.157069 | 0.830784 | 0.402232 | 0.629246 | 1.071003 | 12.770093 | 0.946086514 | 0.104601714 | 0.916691855 | NA | 3.628225 | 3.719719 | 3.623142 | 3.695895 | 3.651962 | 3.676174 | 5 | 22 | 9 | 15 |
| 105372233 | XR_936223,XR | LOC10537223 | uncharacterized LOC105372233, transcript variant X1 | lncRNA | . | . | . | . | . | 0.419519 | 0.524518 | 0.187487 | 0.209663 | 0.976239 | 1.212448 | 0.420715 | 0.479369 | -1.738748 | 5.337813 | 1.326875536 | -0.60144981 | 0.54754043 | NA | 2.760365 | 1.963455 | 2.421394 | 2.435827 | 2.418766 | 2.412395 | 6 | 9 | 4 | 3 |
| 105372234 | XR_936226 | LOC10537223 | uncharacterized LOC105372234 | lncRNA | . | . | . | . | . | 0.388293 | 0.623854 | 0.329378 | 0.739348 | 0.903574 | 1.442067 | 0.739114 | 1.690432 | 1.052721 | 6.937814 | 1.168177378 | 0.06345173 | 0.949406796 | NA | 2.760365 | 2.827809 | 2.784807 | 2.799401 | 2.782705 | 2.804722 | 6 | 9 | 4 | 9 |
| 105372253 | XR_936280 | LOC10537225 | uncharacterized LOC105372253 | lncRNA | . | . | . | . | . | 0.319819 | 0.323509 | 0.180745 | 0.160133 | 0.744232 | 0.747805 | 0.405587 | 0.366126 | -1.921238 | 6.472922 | 1.204835937 | -0.78187907 | 0.434285649 | NA | 3.089593 | 2.148375 | 2.714790 | 2.713068 | 2.696358 | 2.694686 | 10 | 9 | 4 | 4 |
| 105372295 | XR_936359,XR | LOC10537229 | uncharacterized LOC105372295, transcript variant X2 | lncRNA | . | . | . | . | . | 0.650465 | 0.618548 | 0.451274 | 0.591444 | 1.513660 | 1.429802 | 1.012645 | 1.352266 | -1.217390 | 18.346814 | 0.717120023 | -0.39573808 | 0.692298262 | NA | 4.332860 | 4.048019 | 4.211922 | 4.205813 | 4.186923 | 4.200092 | 24 | 21 | 13 | 17 |
| 105372296 | XR_936361 | LOC10537229 | uncharacterized LOC105372296 | lncRNA | . | . | . | . | . | 0.732361 | 0.700937 | 0.649582 | 1.430971 | 1.704233 | 1.620246 | 1.457644 | 3.271745 | 1.425223 | 32.538788 | 0.580938242 | 0.879935003 | 0.378894521 | 0.998865404 | 4.747897 | 5.255806 | 4.991374 | 4.983923 | 4.978075 | 5.086466 | 32 | 28 | 21 | 49 |
| 105372308 | XR_001754062 | LOC10537230 | uncharacterized LOC105372308 | lncRNA | . | . | . | . | . | 0.628541 | 0.703294 | 1.043686 | 0.761173 | 1.462641 | 1.625695 | 2.342001 | 1.740331 | 1.321852 | 15.640170 | 0.776906257 | 0.518158569 | 0.604347639 | NA | 3.751025 | 4.155137 | 3.947235 | 3.951320 | 3.980006 | 3.956012 | 15 | 15 | 18 | 14 |
| 105372310 | XR_001754063 | LOC10537231 | uncharacterized LOC105372310, transcript variant X6 | lncRNA | . | . | . | . | . | 0.996643 | 0.983452 | 1.110430 | 0.709278 | 2.319229 | 2.273294 | 2.491775 | 1.621681 | -1.091447 | 117.171539 | 0.322266284 | -0.39173161 | 0.695256534 | 0.998865404 | 6.932938 | 6.809374 | 6.900945 | 6.864139 | 6.921853 | 6.779627 | 148 | 125 | 119 | 82 |
| 105372352 | XR_002958393 | LOC10537235 | uncharacterized LOC105372352, transcript variant X6 | lncRNA | . | . | . | . | . | 0.312556 | 0.499042 | 0.508659 | 0.499735 | 0.727331 | 1.153557 | 1.141415 | 1.142584 | -1.146463 | 23.053279 | 0.644073459 | -0.30615961 | 0.759483124 | NA | 4.622936 | 4.424013 | 4.538522 | 4.533799 | 4.510489 | 4.535447 | 29 | 26 | 16 | 23 |
| 105372353 | XR_935896 | LOC10537235 | uncharacterized LOC105372353 | lncRNA | . | . | . | . | . | 0.546474 | 0.844139 | 0.861232 | 0.922661 | 1.271668 | 1.951264 | 1.932580 | 2.109554 | 1.263978 | 33.697544 | 0.544823328 | 0.620331131 | 0.535039803 | 0.998865404 | 4.897022 | 5.232645 | 5.024054 | 5.077032 | 5.079008 | 5.085441 | 27 | 39 | 32 | 36 |
| 105372401 | XR_935968 | LOC10537240 | uncharacterized LOC105372401 | lncRNA | . | . | . | . | . | 0.210702 | 0.115099 | 0.137481 | 0.115766 | 0.490312 | 0.266056 | 0.308504 | 0.264685 | -1.325374 | 14.822940 | 0.813154617 | -0.49978112 | 0.617229204 | NA | 4.077189 | 3.674293 | 3.922909 | 3.880165 | 3.888227 | 3.879244 | 25 | 13 | 12 | 11 |
| 105372411 | XR_001753942 | LOC10537241 | uncharacterized LOC105372411, transcript variant X2 | lncRNA | . | . | . | . | . | 0.078159 | 0.036681 | 0.094643 | 0.099796 | 0.181880 | 0.084791 | 0.212375 | 0.228172 | 1.617170 | 5.806461 | 1.269262177 | 0.546357857 | 0.584819973 | NA | 2.147197 | 2.844638 | 2.529670 | 2.514159 | 2.533164 | 2.535804 | 7 | 3 | 6 | 7 |
| 105372419 | NR_134887 | GEMIN7-AS1 | GEMIN7 antisense RNA 1 | lncRNA | . | 53773 | . | . | . | 0.698457 | 0.601987 | 1.067093 | 0.666860 | 1.625337 | 1.391519 | 2.394526 | 1.524696 | 1.328778 | 15.631845 | 0.783682925 | 0.523298681 | 0.600766418 | NA | 3.744764 | 4.158506 | 3.953529 | 3.941431 | 3.983129 | 3.949844 | 17 | 13 | 19 | 13 |
| 105372421 | XR_001753960 | LOC10537242 | uncharacterized LOC105372421, transcript variant X4 | lncRNA | . | . | . | . | . | 0.042842 | 0.144979 | 0.254244 | 0.234729 | 0.099694 | 0.335125 | 0.570516 | 0.536681 | 2.106598 | 4.938441 | 1.394311221 | 0.770929082 | 0.440748968 | NA | 1.671431 | 2.742346 | 2.267119 | 2.281126 | 2.296267 | 2.284340 | 2 | 5 | 7 | 5 |
| 105372462 | XR_936075,XR | LOC10537246 | uncharacterized LOC105372462, transcript variant X1 | lncRNA | . | . | . | . | . | 0.135270 | 0.153347 | 0.079206 | 0.175602 | 0.314779 | 0.354469 | 0.177735 | 0.401494 | -1.078198 | 10.808876 | 0.942144449 | -0.11529285 | 0.908213005 | NA | 3.489863 | 3.376188 | 3.433962 | 3.441323 | 3.418926 | 3.448983 | 12 | 13 | 6 | 13 |
| 105372470 | XR_936090 | LOC10537247 | uncharacterized LOC105372470 | lncRNA | . | . | . | . | . | 0.058676 | 0.042102 | 0.200785 | 0.210133 | 0.136541 | 0.097320 | 0.450556 | 0.480444 | 3.386552 | 3.935062 | 1.568607645 | 1.121897516 | 0.261906017 | NA | 0.844134 | 2.602870 | 1.940157 | 1.940884 | 1.958872 | 1.961999 | 2 | 2 | 5 | 6 |
| 105372480 | NR_144444 | LOC10537248 | uncharacterized LOC105372480 | lncRNA | . | . | . | . | . | 2.409007 | 2.638047 | 2.670460 | 3.229877 | 5.605858 | 6.097963 | 5.992435 | 7.384730 | 1.151028 | 60.319262 | 0.406582024 | 0.499095197 | 0.617712321 | 0.998865404 | 5.810293 | 6.011772 | 5.887819 | 5.904712 | 5.905916 | 5.946338 | 62 | 63 | 51 | 66 |
| 105372481 | NR_144445 | ERVK3-1 | endogenous retrovirus group K3 member 1 | lncRNA | . | 30466 | . | . | . | 27.269337 | 28.485353 | 23.204695 | 29.112700 | 63.456863 | 65.845169 | 52.070663 | 66.562729 | -1.080417 | 1185.285796 | 0.146496255 | -0.76171125 | 0.446232359 | 0.998865404 | 10.266024 | 10.153834 | 10.224802 | 10.251480 | 10.097047 | 10.256659 | 1393 | 1352 | 884 | 1183 |
| 105372482 | XR_936129 | LOC10537248 | uncharacterized LOC105372482 | lncRNA | . | . | . | . | . | 0.701327 | 0.681186 | 0.642929 | 0.616906 | 1.632017 | 1.574593 | 1.442713 | 1.410481 | -1.115684 | 55.657372 | 0.419305447 | -0.37664331 | 0.706438696 | 0.998865404 | 5.875051 | 5.717660 | 5.815197 | 5.806076 | 5.793860 | 5.783511 | 69 | 62 | 47 | 48 |
| 105372486 | XR_936132 | LOC10537248 | uncharacterized LOC105372486 | lncRNA | . | . | . | . | . | 1.718883 | 1.458608 | 1.476371 | 1.865586 | 3.999911 | 3.371638 | 3.312933 | 4.265441 | 1.033228 | 16.803718 | 0.749637761 | 0.062907812 | 0.949839914 | NA | 4.047242 | 4.093800 | 4.077219 | 4.062151 | 4.063133 | 4.080956 | 21 | 16 | 13 | 18 |
| 105372493 | NR_136535 | LOC10537249 | uncharacterized LOC105372493 | lncRNA | . | . | . | . | . | 1.203868 | 1.909418 | 0.928151 | 1.024331 | 2.801451 | 4.413706 | 2.082744 | 2.342011 | -1.492281 | 8.290877 | 1.065054906 | -0.54224346 | 0.5876508 | NA | 3.312060 | 2.733338 | 3.053327 | 3.072561 | 3.049521 | 3.047170 | 9 | 13 | 6 | 6 |
| 105372553 | XR_937308 | LOC10537255 | uncharacterized LOC105372553 | lncRNA | . | . | . | . | . | 0.590403 | 1.209454 | 1.645224 | 1.203817 | 1.373893 | 2.795707 | 3.691835 | 2.752386 | 1.547699 | 33.466859 | 0.573694697 | 1.098362441 | 0.272046256 | 0.998865404 | 4.716583 | 5.344869 | 4.977124 | 5.053290 | 5.097837 | 5.048740 | 20 | 38 | 41 | 32 |
| 105372558 | XR_001754526 | LOC10537255 | uncharacterized LOC105372558, transcript variant X9 | lncRNA | . | . | . | . | . | 0.265350 | 0.205277 | 0.262683 | 0.184729 | 0.617480 | 0.474506 | 0.589454 | 0.422359 | 1.152455 | 17.178274 | 0.78639495 | 0.260314257 | 0.794621376 | NA | 3.993083 | 4.204225 | 4.117879 | 4.058811 | 4.117977 | 4.079988 | 26 | 10 | 20 | 13 |
| 105372579 | XR_001754564 | LOC10537257 | uncharacterized LOC105372579 | lncRNA | . | . | . | . | . | 0.378459 | 0.335093 | 0.170355 | 0.250969 | 0.880690 | 0.774583 | 0.382272 | 0.573811 | -1.714112 | 61.945618 | 0.416303174 | -1.86753635 | 0.061826725 | 0.723723143 | 6.290786 | 5.510949 | 6.024563 | 5.989400 | 5.860081 | 5.923287 | 96 | 79 | 32 | 51 |
| 105372582 | NR_135006 | LOC10537258 | uncharacterized LOC105372582 | lncRNA | . | . | . | . | . | 0.090273 | 0.107551 | 0.091318 | 0.059640 | 0.210069 | 0.248609 | 0.204916 | 0.136359 | -1.314448 | 10.303042 | 0.954071931 | -0.4134456 | 0.679280174 | NA | 3.548173 | 3.155137 | 3.367835 | 3.379318 | 3.367496 | 3.354827 | 12 | 14 | 9 | 7 |
| 105372609 | XR_936703,XR | LOC10537260 | uncharacterized LOC105372609, transcript variant X2 | lncRNA | . | . | . | . | . | 0.199791 | 0.486113 | 0.452099 | 0.607398 | 0.464921 | 1.123672 | 1.014498 | 1.388741 | 1.471324 | 10.635361 | 0.963451458 | 0.578249159 | 0.563095914 | NA | 3.109221 | 3.660102 | 3.374130 | 3.405903 | 3.398253 | 3.418160 | 6 | 13 | 9 | 14 |
| 105372692 | XR_001754688 | LOC10537269 | uncharacterized LOC105372692, transcript variant X1 | lncRNA | . | . | . | . | . | 0.153748 | 0.245037 | 0.320837 | 0.159944 | 0.357777 | 0.566414 | 0.719948 | 0.365693 | 1.239835 | 14.602232 | 0.806328952 | 0.38464249 | 0.700502309 | NA | 3.703730 | 4.015675 | 3.849470 | 3.857447 | 3.881165 | 3.852789 | 14 | 15 | 17 | 12 |
| 105372704 | XM_01152912 | LOC10537270 | uncharacterized LOC105372704 | protein_coding | XP_011527425.1 | . | . | . | . | 0.232334 | 0.316218 | 0.222467 | 0.302146 | 0.540650 | 0.730953 | 0.499210 | 0.690821 | -1.035003 | 14.589440 | 0.802993292 | -0.06181185 | 0.950712669 | NA | 3.892781 | 3.840449 | 3.859920 | 3.876222 | 3.858816 | 3.873008 | 15 | 18 | 11 | 15 |
| 105372705 | XR_936960 | LOC10537270 | uncharacterized LOC105372705 | lncRNA | . | . | . | . | . | 0.588833 | 0.379387 | 0.735294 | 0.450607 | 1.370239 | 0.876969 | 1.649978 | 1.030259 | 1.188863 | 29.774592 | 0.592393583 | 0.421312049 | 0.673527233 | 0.998865404 | 4.763303 | 5.017531 | 4.901086 | 4.852141 | 4.927848 | 4.866503 | 38 | 23 | 35 | 23 |
| 105372709 | XR_936967 | LOC10537270 | uncharacterized LOC105372709 | lncRNA | . | . | . | . | . | 0.516740 | 0.856605 | 0.775838 | 0.565524 | 1.202476 | 1.980080 | 1.740957 | 1.293005 | -1.034065 | 70.227282 | 0.410876109 | -0.11761947 | 0.906369173 | 0.998865404 | 6.157962 | 6.109551 | 6.067111 | 6.188835 | 6.159493 | 6.082710 | 62 | 96 | 70 | 54 |
| 105372717 | XR_936988,XR | LOC10537271 | uncharacterized LOC105372717, transcript variant X3 | lncRNA | . | . | . | . | . | 0.035121 | 0.079638 | 0.066921 | 0.060555 | 0.081728 | 0.184088 | 0.150169 | 0.138452 | -1.099721 | 5.184413 | 1.346074248 | -0.10187919 | 0.918852567 | NA | 2.444631 | 2.300113 | 2.365952 | 2.384411 | 2.366434 | 2.379523 | 4 | 8 | 3 | 6 |
| 105372730 | XR_001754941 | LINC01666 | long intergenic non-protein coding RNA 1666 | lncRNA | . | 52454 | . | . | . | 1.093830 | 0.252715 | 1.244601 | 0.612679 | 2.545386 | 0.584163 | 2.792849 | 1.400817 | 1.316975 | 12.788032 | 0.935755694 | 0.424499687 | 0.671201433 | NA | 3.459600 | 3.865403 | 3.683510 | 3.625520 | 3.691107 | 3.648905 | 20 | 5 | 17 | 9 |
| 105372731 | gene-CDRT15P | CDRT15P9 | . | pseudogene | . | 54917 | . | . | . | 0.150402 | 0.173399 | 0.299027 | 0.167994 | 0.349991 | 0.400820 | 0.671008 | 0.384098 | 1.430357 | 7.111909 | 1.141777763 | 0.452255867 | 0.651084665 | NA | 2.548173 | 3.066070 | 2.810096 | 2.816189 | 2.833142 | 2.815895 | 6 | 7 | 9 | 6 |
| 105372791 | XR_001755012 | LOC10537279 | uncharacterized LOC105372791, transcript variant X3 | lncRNA | . | . | . | . | . | 0.262367 | 0.468243 | 0.142755 | 0.357535 | 0.610540 | 1.082365 | 0.320337 | 0.817462 | -1.326051 | 32.394276 | 0.600608377 | -0.67787399 | 0.497851594 | 0.998865404 | 5.209663 | 4.796139 | 5.002235 | 5.062223 | 4.949068 | 5.041660 | 34 | 48 | 14 | 37 |
| 105372795 | NR_131189 | LOC10537279 | uncharacterized LOC105372795 | lncRNA | . | . | . | . | . | 0.144571 | 0.046624 | 0.318763 | 0.148016 | 0.336423 | 0.107773 | 0.715294 | 0.338420 | 2.205588 | 5.686811 | 1.31870708 | 0.865365505 | 0.386838208 | NA | 1.820514 | 2.971139 | 2.483551 | 2.467676 | 2.506202 | 2.484203 | 6 | 2 | 9 | 5 |
| 105372817 | XR_937764,XR | LOC10537281 | uncharacterized LOC105372817, transcript variant X1 | lncRNA | . | . | . | . | . | 0.397587 | 2.145288 | 1.113903 | 2.096867 | 0.925201 | 4.958929 | 2.499566 | 4.794238 | -1.031420 | 17.840396 | 0.7709893 | -0.05789 | 0.953836252 | NA | 4.182369 | 4.131331 | 4.122075 | 4.182379 | 4.129848 | 4.170560 | 13 | 27 | 11 | 21 |
| 105372824 | XM_01152981 | LOC10537282 | uncharacterized LOC105372824 | protein_coding | XP_011528113.1 | . | . | . | . | 0.983595 | 0.820539 | 1.499423 | 0.935360 | 2.288866 | 1.896713 | 3.364662 | 2.138589 | 1.316409 | 36.803906 | 0.529369651 | 0.749206819 | 0.45373256 | 0.998865404 | 4.987929 | 5.387998 | 5.182792 | 5.158310 | 5.246560 | 5.172353 | 40 | 31 | 45 | 30 |
| 105372836 | XM_01152981 | LOC10537283 | uncharacterized LOC105372836, transcript variant X1 | protein_coding | XP_011528114.1 | . | . | . | . | 0.087012 | 0.128741 | 0.091620 | 0.050405 | 0.202480 | 0.297590 | 0.205593 | 0.115246 | -1.344985 | 3.941867 | 1.529916775 | -0.27948559 | 0.779872196 | NA | 2.175403 | 1.751299 | 1.977228 | 1.987021 | 1.982366 | 1.971259 | 4 | 6 | 4 | 2 |
| 105372840 | NR_146912 | LINC01694 | long intergenic non-protein coding RNA 1694 | lncRNA | . | 52481 | . | . | . | 0.174008 | 0.115125 | 0.108695 | 0.165777 | 0.404924 | 0.266117 | 0.243908 | 0.379028 | -1.070911 | 30.141154 | 0.583546914 | -0.16937529 | 0.865501455 | 0.998865404 | 4.962667 | 4.862935 | 4.943078 | 4.892323 | 4.885350 | 4.933458 | 43 | 27 | 20 | 33 |
| 105372860 | XR_938003 | LOC10537286 | uncharacterized LOC105372860 | lncRNA | . | . | . | . | . | 0.121731 | 0.090256 | 0.221366 | 0.216935 | 0.283274 | 0.208630 | 0.496740 | 0.495997 | 2.008078 | 19.483468 | 0.70414115 | 1.428428079 | 0.153168682 | NA | 3.694014 | 4.701810 | 4.244840 | 4.228814 | 4.294176 | 4.291280 | 17 | 12 | 23 | 24 |
| 105372876 | XR_922480 | DYRK3-AS1 | DYRK3 antisense RNA 1 | lncRNA | . | 40199 | . | . | . | 0.415698 | 0.423808 | 0.309554 | 0.311991 | 0.967346 | 0.979652 | 0.694630 | 0.713331 | -1.350772 | 13.654925 | 0.827377824 | -0.52428785 | 0.600078349 | NA | 3.972078 | 3.538109 | 3.784832 | 3.785243 | 3.766859 | 3.767676 | 18 | 17 | 10 | 11 |
| 105372879 | XR_922483 | LOC10537287 | uncharacterized LOC105372879 | lncRNA | . | . | . | . | . | 0.575134 | 0.601269 | 0.224578 | 0.505124 | 1.338360 | 1.389861 | 0.503948 | 1.154907 | -1.525791 | 10.387290 | 0.961410403 | -0.63402393 | 0.526065203 | NA | 3.651489 | 3.037079 | 3.391943 | 3.395472 | 3.363909 | 3.385087 | 14 | 14 | 5 | 10 |
| 105372888 | XR_001738429 | LOC10537288 | uncharacterized LOC105372888 | lncRNA | . | . | . | . | . | 0.040331 | 0.079725 | 0.035627 | 0.088108 | 0.093851 | 0.184287 | 0.079946 | 0.201448 | 1.044475 | 3.219731 | 1.666442807 | 0.037672219 | 0.969949027 | NA | 1.658167 | 1.715150 | 1.683853 | 1.689092 | 1.682707 | 1.691538 | 3 | 4 | 2 | 4 |
| 105372898 | XR_922535 | LOC10537289 | uncharacterized LOC105372898 | lncRNA | . | . | . | . | . | 1.836912 | 1.908738 | 1.221800 | 1.189157 | 4.274570 | 4.412133 | 2.741684 | 2.718866 | -1.575115 | 218.160904 | 0.229490446 | -2.85614302 | 0.004288219 | 0.193526306 | 8.060013 | 7.404700 | 7.855055 | 7.869142 | 7.658818 | 7.643134 | 303 | 292 | 150 | 156 |
| 105372899 | XR_922537 | LOC10537289 | uncharacterized LOC105372899 | lncRNA | . | . | . | . | . | 0.856290 | 1.313304 | 0.578968 | 1.205636 | 1.992622 | 3.035761 | 1.299187 | 2.756543 | -1.163674 | 7.119944 | 1.149717835 | -0.19020912 | 0.849145262 | NA | 2.939842 | 2.715150 | 2.829202 | 2.843926 | 2.823077 | 2.840342 | 7 | 10 | 4 | 8 |
| 105372902 | XR_922552 | LOC10537290 | uncharacterized LOC105372902 | lncRNA | . | . | . | . | . | 0.244732 | 0.334298 | 0.431082 | 0.206507 | 0.569502 | 0.772744 | 0.967335 | 0.472153 | 1.105002 | 12.813850 | 0.877400262 | 0.164177513 | 0.869591403 | NA | 3.604218 | 3.751299 | 3.660162 | 3.676000 | 3.694545 | 3.652399 | 12 | 15 | 16 | 8 |
| 105372906 | XR_922576,XR | LOC10537290 | uncharacterized LOC105372906, transcript variant X1 | lncRNA | . | . | . | . | . | 0.752945 | 0.728115 | 0.447559 | 0.394174 | 1.752135 | 1.683070 | 1.004309 | 0.901232 | -1.722235 | 11.332787 | 0.914899496 | -0.85723286 | 0.391316204 | NA | 3.841203 | 3.058876 | 3.524900 | 3.520852 | 3.500305 | 3.492500 | 17 | 15 | 8 | 7 |
| 105372926 | XR_001738472 | LOC10537292 | uncharacterized LOC105372926, transcript variant X2 | lncRNA | . | . | . | . | . | 0.172671 | 0.206946 | 0.163463 | 0.146306 | 0.401813 | 0.478366 | 0.366807 | 0.334511 | -1.254178 | 33.125479 | 0.540430711 | -0.60459625 | 0.545447299 | 0.998865404 | 5.203970 | 4.877319 | 5.054212 | 5.078014 | 5.043156 | 5.029815 | 39 | 43 | 27 | 26 |
| 105372935 | XR_938017 | FAM247A | family with sequence similarity 247 member A | lncRNA | . | 54926 | . | . | . | 0.560592 | 0.548834 | 0.321953 | 0.421442 | 1.304522 | 1.268655 | 0.722454 | 0.963576 | -1.500016 | 17.918976 | 0.72963273 | -0.80174271 | 0.422701803 | NA | 4.427143 | 3.840449 | 4.185492 | 4.183563 | 4.147017 | 4.160834 | 25 | 23 | 11 | 15 |
| 105372942 | XR_951230 | FAM247D | family with sequence similarity 247 member D | lncRNA | . | 54929 | . | . | . | 0.410263 | 0.347942 | 0.548976 | 0.271087 | 0.954700 | 0.804282 | 1.231886 | 0.619807 | 1.058782 | 15.224834 | 0.801460364 | 0.102819232 | 0.918106441 | NA | 3.884262 | 3.971139 | 3.924733 | 3.917017 | 3.945571 | 3.903207 | 18 | 15 | 18 | 10 |
| 105372952 | XR_938055,XR | LOC10537295 | uncharacterized LOC105372952, transcript variant X1 | lncRNA | . | . | . | . | . | 0.194372 | 0.362937 | 0.409044 | 0.346521 | 0.452311 | 0.838945 | 0.917884 | 0.792280 | 1.315976 | 14.674689 | 0.816958738 | 0.484887522 | 0.627756134 | NA | 3.664814 | 4.058876 | 3.837601 | 3.873950 | 3.880592 | 3.866149 | 10 | 18 | 16 | 14 |
| 105372956 | XR_002958463 | LOC10537295 | uncharacterized LOC105372956, transcript variant X2 | lncRNA | . | . | . | . | . | 0.318585 | 0.122122 | 0.564799 | 0.311046 | 0.741361 | 0.282291 | 1.267392 | 0.711169 | 1.426407 | 3.800969 | 1.541606122 | 0.332371436 | 0.739608805 | NA | 1.644780 | 2.161867 | 1.914567 | 1.911306 | 1.924759 | 1.913395 | 4 | 3 | 5 | 3 |
| 105372976 | XR_938114 | LOC10537297 | uncharacterized LOC105372976 | lncRNA | . | . | . | . | . | 0.060448 | 0.126544 | 0.053150 | 0.036333 | 0.140665 | 0.292513 | 0.119266 | 0.083070 | -2.098296 | 16.400600 | 0.802435328 | -1.33246662 | 0.182706903 | NA | 4.473116 | 3.404839 | 4.026424 | 4.090455 | 4.018021 | 3.999764 | 17 | 32 | 11 | 8 |
| 105372988 | XR_938153 | LOC10537298 | uncharacterized LOC105372988 | lncRNA | . | . | . | . | . | 0.026775 | 0.066534 | 0.056947 | 0.021253 | 0.062306 | 0.153795 | 0.127788 | 0.048591 | -1.076899 | 3.493070 | 1.615701837 | -0.06615256 | 0.947256369 | NA | 1.855800 | 1.751299 | 1.796109 | 1.805729 | 1.805270 | 1.794177 | 3 | 5 | 4 | 2 |
| 105372989 | XR_001755486 | LOC10537298 | uncharacterized LOC105372989, transcript variant X1 | lncRNA | . | . | . | . | . | 0.211975 | 0.130656 | 0.543608 | 0.259338 | 0.493276 | 0.302018 | 1.219842 | 0.592946 | 1.984989 | 11.346967 | 0.929626238 | 1.064009571 | 0.287324364 | NA | 2.923304 | 3.917252 | 3.470591 | 3.460606 | 3.517539 | 3.479937 | 10 | 7 | 17 | 10 |
| 105373017 | XR_938209,XR | LOC10537301 | uncharacterized LOC105373017, transcript variant X5 | lncRNA | . | . | . | . | . | 0.758361 | 0.580619 | 0.841306 | 0.614349 | 1.764737 | 1.342125 | 1.887868 | 1.404635 | -1.336344 | 11.681917 | 0.906297702 | -0.46153865 | 0.644412196 | NA | 3.738475 | 3.324312 | 3.571370 | 3.542941 | 3.548880 | 3.535913 | 19 | 11 | 10 | 8 |
| 105373027 | XR_001755536 | LOC10537302 | uncharacterized LOC105373027, transcript variant X2 | lncRNA | . | . | . | . | . | 0.448445 | 0.276265 | 0.662857 | 0.694772 | 1.043550 | 0.638601 | 1.487432 | 1.588513 | 1.729578 | 18.263111 | 0.727193778 | 1.086945391 | 0.277060944 | NA | 3.741623 | 4.532926 | 4.166323 | 4.146324 | 4.188061 | 4.208561 | 18 | 12 | 18 | 24 |
| 105373038 | XR_001755554 | LOC10537303 | uncharacterized LOC105373038 | lncRNA | . | . | . | . | . | 0.105297 | 0.194329 | 0.107014 | 0.106512 | 0.245030 | 0.449200 | 0.240136 | 0.243527 | -1.291443 | 7.624970 | 1.110621733 | -0.33223235 | 0.739713821 | NA | 3.104339 | 2.733338 | 2.924499 | 2.947676 | 2.928595 | 2.926303 | 7 | 12 | 6 | 6 |
| 105373041 | XR_949167 | LOC10537304 | uncharacterized LOC105373041 | lncRNA | . | . | . | . | . | 0.221230 | 0.249758 | 0.595252 | 0.272096 | 0.514812 | 0.577326 | 1.335728 | 0.622114 | 1.870773 | 5.161802 | 1.349094562 | 0.669808174 | 0.502980082 | NA | 1.844134 | 2.751299 | 2.340554 | 2.341783 | 2.366365 | 2.344572 | 4 | 4 | 8 | 4 |
| 105373043 | XR_938274 | LOC10537304 | uncharacterized LOC105373043 | lncRNA | . | . | . | . | . | 0.225875 | 0.133114 | 0.236912 | 0.283429 | 0.525619 | 0.307699 | 0.531623 | 0.648028 | 1.429987 | 7.041104 | 1.143026134 | 0.451434996 | 0.651676066 | NA | 2.533720 | 3.051647 | 2.808549 | 2.797777 | 2.813374 | 2.815617 | 8 | 5 | 7 | 8 |
| 105373063 | XR_001755587 | LOC10537306 | uncharacterized LOC105373063 | lncRNA | . | . | . | . | . | 0.166318 | 0.116650 | 0.126947 | 0.148513 | 0.387029 | 0.269641 | 0.284865 | 0.339558 | -1.050677 | 22.637389 | 0.651791317 | -0.10941971 | 0.9128696 | NA | 4.535535 | 4.464871 | 4.519943 | 4.487992 | 4.493763 | 4.505019 | 31 | 21 | 18 | 22 |
| 105373064 | NR_146602 | LOC10537306 | uncharacterized LOC105373064 | lncRNA | . | . | . | . | . | 1.306611 | 1.367996 | 1.676559 | 1.775393 | 3.040538 | 3.162183 | 3.762149 | 4.059225 | 1.269810 | 50.891624 | 0.437029772 | 0.788533446 | 0.430384743 | 0.998865404 | 5.487053 | 5.831186 | 5.639012 | 5.643628 | 5.679646 | 5.692271 | 51 | 49 | 48 | 55 |
| 105373077 | XR_001755593 | LOC10537307 | uncharacterized LOC105373077 | lncRNA | . | . | . | . | . | 0.645144 | 0.433525 | 0.492298 | 0.555293 | 1.501276 | 1.002112 | 1.104702 | 1.269611 | -1.027779 | 24.628728 | 0.627930465 | -0.06295348 | 0.949803552 | NA | 4.641413 | 4.602870 | 4.643333 | 4.604550 | 4.616065 | 4.626554 | 34 | 22 | 20 | 24 |
| 105373095 | XR_938352 | LOC10537309 | uncharacterized LOC105373095 | lncRNA | . | . | . | . | . | 0.298275 | 0.169183 | 0.369164 | 0.279526 | 0.694097 | 0.391073 | 0.828393 | 0.639102 | 1.353896 | 49.760301 | 0.474106923 | 0.92197886 | 0.356539602 | 0.998865404 | 5.399615 | 5.840655 | 5.640642 | 5.549733 | 5.680445 | 5.622494 | 62 | 33 | 57 | 46 |
| 105373102 | XM_01154618 | LOC10537310 | uncharacterized LOC105373102, transcript variant X1 | protein_coding | XP_011544488.1 | . | . | . | . | 1.428152 | 1.159936 | 2.307454 | 1.275806 | 3.323368 | 2.681244 | 5.177860 | 2.916978 | 1.328513 | 65.625809 | 0.424867438 | 0.964565107 | 0.334762683 | 0.998865404 | 5.814705 | 6.228157 | 6.010055 | 5.964362 | 6.121061 | 5.977854 | 72 | 54 | 84 | 50 |
| 105373105 | XR_001755765 | LOC10537310 | uncharacterized LOC105373105, transcript variant X4 | lncRNA | . | . | . | . | . | 0.088431 | 0.078760 | 0.109705 | 0.045662 | 0.205780 | 0.182060 | 0.246176 | 0.104400 | -1.412210 | 5.343643 | 1.307168744 | -0.3809413 | 0.703246805 | NA | 2.644780 | 2.148375 | 2.431886 | 2.425361 | 2.421100 | 2.419531 | 8 | 6 | 4 | 4 |
| 105373106 | XR_001738492 | CNIH3-AS1 | CNIH3 antisense RNA 1, transcript variant X1 | lncRNA | . | 41163 | . | . | . | 0.160590 | 0.253720 | 0.169855 | 0.086788 | 0.373699 | 0.586486 | 0.381151 | 0.198430 | -2.131577 | 6.886316 | 1.182352995 | -0.92351551 | 0.355738623 | NA | 3.229742 | 2.134756 | 2.812491 | 2.802825 | 2.781226 | 2.789308 | 12 | 9 | 3 | 5 |
| 105373113 | NR_146920 | LINC01703 | long intergenic non-protein coding RNA 1703 | lncRNA | . | 52491 | . | . | . | 0.744216 | 0.454290 | 0.834503 | 0.826732 | 1.731822 | 1.050112 | 1.872600 | 1.890225 | 1.371817 | 20.070799 | 0.695448164 | 0.655819363 | 0.511940356 | NA | 4.079678 | 4.538109 | 4.323326 | 4.288746 | 4.331842 | 4.334016 | 24 | 14 | 20 | 22 |
| 105373117 | XR_001738505 | LOC10537311 | uncharacterized LOC105373117 | lncRNA | . | . | . | . | . | 0.064267 | 0.046114 | 0.153754 | 0.044174 | 0.149552 | 0.106594 | 0.345019 | 0.100999 | 1.765277 | 9.277343 | 1.058749314 | 0.774398925 | 0.438694919 | NA | 2.741623 | 3.568818 | 3.182600 | 3.172201 | 3.228230 | 3.171782 | 9 | 6 | 16 | 5 |
| 105373148 | XR_001755813 | LOC10537314 | uncharacterized LOC105373148, transcript variant X6 | lncRNA | . | . | . | . | . | 0.624445 | 0.809858 | 1.164876 | 0.857851 | 1.453109 | 1.872022 | 2.613947 | 1.961375 | 1.493635 | 88.545860 | 0.367316471 | 1.575829241 | 0.115065161 | 0.855274326 | 6.149100 | 6.729601 | 6.375939 | 6.405008 | 6.577591 | 6.422256 | 77 | 81 | 115 | 75 |
| 105373150 | XR_950537,XR | LOC10537315 | uncharacterized LOC105373150, transcript variant X1 | lncRNA | . | . | . | . | . | 0.395839 | 0.325761 | 0.413674 | 0.238307 | 0.921133 | 0.753011 | 0.928273 | 0.544860 | -1.161431 | 22.903248 | 0.657345943 | -0.32844709 | 0.742573641 | NA | 4.619518 | 4.407673 | 4.533912 | 4.513904 | 4.530917 | 4.487934 | 31 | 24 | 23 | 15 |
| 105373159 | NR_149312,NF | LOC10537315 | uncharacterized LOC105373159 | lncRNA | . | . | . | . | . | 1.238651 | 1.565145 | 1.085595 | 1.245647 | 2.882393 | 3.617902 | 2.436044 | 2.848023 | -1.219793 | 163.691797 | 0.267659547 | -1.07089772 | 0.284215419 | 0.998865404 | 7.491734 | 7.203578 | 7.342460 | 7.440426 | 7.287668 | 7.335225 | 184 | 216 | 120 | 147 |
| 105373174 | XR_949021 | LOC10537317 | uncharacterized LOC105373174 | lncRNA | . | . | . | . | . | 0.469792 | 0.695704 | 0.454506 | 0.347872 | 1.093225 | 1.608151 | 1.019899 | 0.795368 | -1.412862 | 20.796539 | 0.686812211 | -0.72599217 | 0.46784357 | NA | 4.605946 | 4.107809 | 4.375360 | 4.412413 | 4.373869 | 4.354335 | 23 | 31 | 17 | 14 |
| 105373195 | XR_001755844 | LOC10537319 | uncharacterized LOC105373195, transcript variant X2 | lncRNA | . | . | . | . | . | 0.108045 | 0.073896 | 0.159337 | 0.094045 | 0.251424 | 0.170814 | 0.357547 | 0.215024 | 1.334512 | 15.133492 | 0.801338131 | 0.519521989 | 0.603396783 | NA | 3.694014 | 4.114763 | 3.908168 | 3.891848 | 3.937625 | 3.899762 | 17 | 12 | 19 | 12 |
| 105373212 | XR_949297 | LINC02768 | long intergenic non-protein coding RNA 2768 | lncRNA | . | 54288 | . | . | . | 0.140024 | 0.194990 | 0.248074 | 0.163479 | 0.325843 | 0.450729 | 0.556671 | 0.373775 | 1.192684 | 13.845537 | 0.832724766 | 0.30527787 | 0.76015453 | NA | 3.658167 | 3.913269 | 3.767871 | 3.787906 | 3.802564 | 3.774366 | 12 | 16 | 16 | 11 |
| 105373215 | XR_001738537 | LOC10537321 | uncharacterized LOC105373215, transcript variant X6 | lncRNA | . | . | . | . | . | 0.214962 | 0.240706 | 0.062967 | 0.032999 | 0.500224 | 0.556402 | 0.141296 | 0.075447 | -2.513785 | 7.840139 | 1.124339063 | -1.18279362 | 0.236890948 | NA | 3.486115 | 2.161867 | 2.997092 | 2.996098 | 2.972225 | 2.960645 | 13 | 12 | 5 | 3 |
| 105373218 | XR_001738550 | LOC10537321 | uncharacterized LOC105373218, transcript variant X3 | lncRNA | . | . | . | . | . | 0.167587 | 0.299096 | 0.304572 | 0.053387 | 0.389981 | 0.691374 | 0.683451 | 0.122062 | 1.318998 | 22.377035 | 0.747887139 | 0.534094392 | 0.59327623 | NA | 4.267126 | 4.672399 | 4.448085 | 4.441907 | 4.533634 | 4.403791 | 23 | 20 | 36 | 9 |
| 105373249 | gene-LOC1053 | LOC10537324 | . | pseudogene | . | . | . | . | . | 0.060548 | 0.040555 | 0.086886 | 0.113426 | 0.140898 | 0.093744 | 0.194970 | 0.259335 | 1.729504 | 3.043852 | 1.713699129 | 0.461199977 | 0.644655138 | NA | 1.156660 | 1.947964 | 1.593764 | 1.590220 | 1.597682 | 1.601437 | 3 | 2 | 3 | 4 |
| 105373255 | XR_949335 | LOC10537325 | uncharacterized LOC105373255 | lncRNA | . | . | . | . | . | 0.402457 | 0.457609 | 0.822082 | 0.679940 | 0.936535 | 1.057783 | 1.844727 | 1.554602 | 1.698520 | 24.261160 | 0.62994924 | 1.21323829 | 0.225038729 | NA | 4.168403 | 4.932690 | 4.551524 | 4.564943 | 4.621463 | 4.599681 | 19 | 21 | 29 | 26 |
| 105373264 | XR_949348 | LOC10537326 | uncharacterized LOC105373264 | lncRNA | . | . | . | . | . | 0.410674 | 0.557862 | 0.598241 | 0.281370 | 0.955656 | 1.289524 | 1.342436 | 0.643318 | -1.140191 | 14.391927 | 0.833040389 | -0.22721035 | 0.820260182 | NA | 3.937099 | 3.751299 | 3.837001 | 3.857370 | 3.859875 | 3.817727 | 15 | 19 | 16 | 8 |
| 105373296 | XR_938468 | LOC10537329 | uncharacterized LOC105373296 | lncRNA | . | . | . | . | . | 1.408879 | 1.287226 | 1.424052 | 1.177963 | 3.278520 | 2.975480 | 3.195530 | 2.693273 | -1.046238 | 36.348833 | 0.516442594 | -0.12626977 | 0.899518382 | 0.998865404 | 5.215216 | 5.151760 | 5.194576 | 5.179051 | 5.193972 | 5.166622 | 45 | 38 | 34 | 30 |
| 105373311 | NR_158569 | LOC10537331 | uncharacterized LOC105373311 | lncRNA | . | . | . | . | . | 0.194962 | 0.123443 | 0.140499 | 0.048789 | 0.453686 | 0.285345 | 0.315275 | 0.111551 | -1.647760 | 10.398140 | 0.989580059 | -0.72809297 | 0.466556679 | NA | 3.690761 | 2.978782 | 3.405029 | 3.380401 | 3.386018 | 3.354256 | 18 | 11 | 10 | 4 |
| 105373354 | XR_001739245 | TMEM18-DT | TMEM18 divergent transcript, transcript variant X1 | lncRNA | . | 54481 | . | . | . | 0.133179 | 0.390211 | 0.150585 | 0.358580 | 0.309913 | 0.901989 | 0.337909 | 0.819851 | -1.300063 | 10.824296 | 0.950105546 | -0.39846214 | 0.690289559 | NA | 3.614556 | 3.232645 | 3.421544 | 3.461558 | 3.428489 | 3.430097 | 9 | 18 | 8 | 9 |
| 105373366 | XR_938518,XR | IDS2 | uncharacterized IDS2, transcript variant X1 | lncRNA | . | . | . | . | . | 0.194912 | 0.889658 | 0.844408 | 0.211366 | 0.453567 | 2.056484 | 1.894827 | 0.483264 | 1.040762 | 23.461845 | 0.645547751 | 0.089289323 | 0.928851979 | NA | 4.522976 | 4.580931 | 4.529005 | 4.562377 | 4.564492 | 4.534747 | 22 | 29 | 24 | 19 |
| 105373368 | NR_136578 | MAGEA4-AS1 | MAGEA4 antisense RNA 1 | lncRNA | . | 53860 | . | . | . | 9.317341 | 10.347928 | 13.706641 | 11.309812 | 21.681835 | 23.919699 | 30.757305 | 25.858540 | 1.257834 | 146.536576 | 0.276343253 | 1.197573016 | 0.231083286 | 0.986164394 | 7.019870 | 7.351364 | 7.124529 | 7.161046 | 7.269907 | 7.190493 | 142 | 147 | 156 | 138 |
| 105373370 | XR_001755975 | LOC10537337 | uncharacterized LOC105373370, transcript variant X2 | lncRNA | . | . | . | . | . | 3.524148 | 4.302124 | 3.230910 | 3.595304 | 8.200836 | 9.944552 | 7.250069 | 8.220235 | -1.159796 | 57.295403 | 0.416056977 | -0.51404349 | 0.607221585 | 0.998865404 | 5.943875 | 5.728813 | 5.841212 | 5.871600 | 5.817601 | 5.834652 | 66 | 71 | 44 | 52 |
| 105373383 | NR_131982 | LOC10537338 | uncharacterized LOC105373383 | lncRNA | . | . | . | . | . | 1.523782 | 0.890814 | 0.885717 | 2.423676 | 3.545904 | 2.059158 | 1.987524 | 5.541447 | 1.353782 | 25.094006 | 0.678932788 | 0.643651285 | 0.519801586 | NA | 4.415365 | 4.850491 | 4.644623 | 4.596380 | 4.599949 | 4.702217 | 31 | 17 | 14 | 39 |
| 105373428 | XR_922797 | LOC10537342 | uncharacterized LOC105373428 | lncRNA | . | . | . | . | . | 0.477029 | 0.279598 | 0.744042 | 0.227544 | 1.110065 | 0.646304 | 1.669608 | 0.520251 | 1.262550 | 15.636444 | 0.83957931 | 0.400606646 | 0.688709752 | NA | 3.784663 | 4.128572 | 3.958603 | 3.926421 | 3.995111 | 3.919968 | 20 | 11 | 23 | 8 |
| 105373456 | XR_001739303 | LOC10537345 | uncharacterized LOC105373456, transcript variant X2 | lncRNA | . | . | . | . | . | 0.191530 | 0.145821 | 0.333897 | 0.277194 | 0.445698 | 0.337072 | 0.749256 | 0.633772 | 1.486473 | 22.813715 | 0.652958284 | 0.875848939 | 0.381112161 | NA | 4.195961 | 4.770810 | 4.487645 | 4.473459 | 4.537320 | 4.498292 | 23 | 18 | 28 | 21 |
| 105373463 | XR_001739316 | LAPTM4A-DT | LAPTM4A divergent transcript, transcript variant X1 | lncRNA | . | 54382 | . | . | . | 0.106461 | 0.098277 | 0.039052 | 0.147885 | 0.247739 | 0.227171 | 0.087633 | 0.338121 | -1.126908 | 4.636082 | 1.417744254 | -0.12158066 | 0.903231132 | NA | 2.299300 | 2.121007 | 2.220259 | 2.217532 | 2.207226 | 2.225437 | 6 | 5 | 2 | 6 |
| 105373490 | XR_923077 | LOC10537349 | uncharacterized LOC105373490 | lncRNA | . | . | . | . | . | 0.054736 | 0.020881 | 0.113987 | 0.078814 | 0.127372 | 0.048268 | 0.255783 | 0.180198 | 2.327297 | 10.343588 | 0.987857878 | 1.233633958 | 0.217339333 | NA | 2.631267 | 3.857133 | 3.339795 | 3.316699 | 3.376869 | 3.353602 | 10 | 4 | 15 | 11 |
| 105373507 | NR_146956 | LINC01849 | long intergenic non-protein coding RNA 1849 | lncRNA | . | 52665 | . | . | . | 0.281792 | 0.330381 | 0.262987 | 0.836911 | 0.655742 | 0.763690 | 0.590136 | 1.913498 | 1.784230 | 10.613691 | 0.983084861 | 0.849674181 | 0.395506257 | NA | 2.934350 | 3.763758 | 3.375373 | 3.382052 | 3.376344 | 3.433860 | 8 | 9 | 6 | 19 |
| 105373511 | XR_001739613 | LOC10537351 | uncharacterized LOC105373511, transcript variant X2 | lncRNA | . | . | . | . | . | 0.225542 | 0.184988 | 0.358724 | 0.179654 | 0.524845 | 0.427608 | 0.804966 | 0.410758 | 1.053421 | 11.500499 | 0.933773525 | 0.080407097 | 0.935913482 | NA | 3.482357 | 3.563745 | 3.518908 | 3.509970 | 3.540125 | 3.493405 | 14 | 11 | 15 | 6 |
| 105373629 | XR_923354 | LOC10537362 | uncharacterized LOC105373629 | lncRNA | . | . | . | . | . | 0.081675 | 0.369943 | 0.092231 | 0.171999 | 0.190061 | 0.855140 | 0.206964 | 0.393256 | -2.207935 | 2.645165 | 1.906394295 | -0.59940248 | 0.548904522 | NA | 1.867373 | 0.715150 | 1.407548 | 1.425490 | 1.405258 | 1.409579 | 2 | 6 | 1 | 2 |
| 105373637 | XR_923368 | LOC10537363 | uncharacterized LOC105373637 | lncRNA | . | . | . | . | . | 0.036916 | 0.049884 | 0.082926 | 0.040908 | 0.085906 | 0.115309 | 0.186083 | 0.093531 | 1.441283 | 13.760868 | 0.849278007 | 0.620943463 | 0.534636823 | NA | 3.493602 | 4.023086 | 3.748055 | 3.763848 | 3.801030 | 3.753890 | 11 | 14 | 19 | 10 |

| | | | | | | | | | | | | | | | | | | | | | | | | | | | | | | | | | |
|---|---|---|---|---|---|---|---|---|---|---|---|---|---|---|---|---|---|---|---|---|---|---|---|---|---|---|---|---|---|---|---|---|---|
| 105373645 | XR_923378 | LOC105373645 | uncharacterized LOC105373645 | lncRNA | . | . | . | . | . | 1.659402 | 1.385400 | 2.084804 | 1.438623 | 3.861497 | 3.202415 | 4.678239 | 3.289241 | 1.149687 | 31.747599 | 0.558576906 | 0.360274916 | 0.718641556 | 0.998865404 | 4.882837 | 5.087090 | 4.984892 | 4.963178 | 5.015737 | 4.967272 | 37 | 29 | 35 | 26 |
| 105373656 | NR_136320 | LOC105373656 | uncharacterized LOC105373656 | lncRNA | . | . | . | . | . | 0.841868 | 1.026007 | 1.336628 | 0.678281 | 1.959062 | 2.371661 | 2.999353 | 1.550810 | 1.081728 | 10.772534 | 0.943474572 | 0.120128001 | 0.904381751 | NA | 3.369732 | 3.486479 | 3.417238 | 3.424450 | 3.440602 | 3.408295 | 11 | 12 | 13 | 7 |
| 105373690 | XR_001739737 | LOC105373690 | uncharacterized LOC105373690, transcript variant X2 | lncRNA | . | . | . | . | . | 0.207054 | 0.556024 | 1.043492 | 0.183167 | 0.481823 | 1.285274 | 2.341565 | 0.418790 | 1.623484 | 11.937795 | 1.018120102 | 0.686651283 | 0.492302503 | NA | 3.184683 | 3.885872 | 3.511550 | 3.547726 | 3.598675 | 3.508854 | 6 | 14 | 22 | 4 |
| 105373716 | XR_002959431 | LOC105373716 | uncharacterized LOC105373716, transcript variant X2 | lncRNA | . | . | . | . | . | 0.213372 | 0.284889 | 0.104293 | 0.116191 | 0.496525 | 0.658535 | 0.234030 | 0.265657 | -2.191985 | 20.318855 | 0.700170788 | -1.61708831 | 0.105859221 | NA | 4.802857 | 3.669578 | 4.365290 | 4.395017 | 4.316242 | 4.316224 | 28 | 34 | 11 | 12 |
| 105373730 | XR_001739760 | LOC105373730 | uncharacterized LOC105373730, transcript variant X4 | lncRNA | . | . | . | . | . | 1.037233 | 1.115393 | 1.467548 | 1.556653 | 2.413685 | 2.578280 | 3.293136 | 3.559102 | 1.221683 | 117.694720 | 0.303409394 | 0.952081268 | 0.341055768 | 0.998865404 | 6.726777 | 7.016512 | 6.827499 | 6.848715 | 6.943930 | 6.866147 | 118 | 118 | 125 | 108 |
| 105373748 | XR_923592 | LOC105373748 | uncharacterized LOC105373748 | lncRNA | . | . | . | . | . | 0.864249 | 1.095446 | 0.811941 | 0.677455 | 2.011142 | 2.532172 | 1.821971 | 1.548920 | -1.319055 | 37.979515 | 0.510541305 | -0.78251112 | 0.433914252 | 0.998865404 | 5.432996 | 5.033773 | 5.248340 | 5.287002 | 5.238601 | 5.215819 | 44 | 52 | 31 | 28 |
| 105373754 | XR_923600 | LOC105373754 | uncharacterized LOC105373754 | lncRNA | . | . | . | . | . | 0.083539 | 0.041015 | 0.117456 | 0.029063 | 0.194398 | 0.094808 | 0.263567 | 0.066449 | 1.056112 | 5.496911 | 1.346552897 | 0.058492297 | 0.953356498 | NA | 2.413393 | 2.502474 | 2.456939 | 2.441701 | 2.466238 | 2.434850 | 8 | 4 | 8 | 2 |
| 105373759 | NR_157853 | LOC105373759 | uncharacterized LOC105373759 | lncRNA | . | . | . | . | . | 0.201276 | 0.502881 | 1.411089 | 1.069242 | 0.468379 | 1.162433 | 3.166444 | 2.444694 | 3.354514 | 25.652822 | 0.665962503 | 2.621925104 | 0.008743464 | NA | 3.562483 | 5.303458 | 4.550157 | 4.596304 | 4.710750 | 4.670101 | 8 | 18 | 39 | 32 |
| 105373774 | XR_923645 | LOC105373774 | uncharacterized LOC105373774 | lncRNA | . | . | . | . | . | 0.124811 | 0.066279 | 0.139520 | 0.040923 | 0.290441 | 0.153208 | 0.313079 | 0.093565 | -1.060071 | 22.057736 | 0.745053934 | -0.11295957 | 0.910062606 | NA | 4.500856 | 4.424561 | 4.482360 | 4.422772 | 4.495249 | 4.397084 | 34 | 17 | 29 | 9 |
| 105373785 | XR_001739147 | LOC105373785 | uncharacterized LOC105373785, transcript variant X2 | lncRNA | . | . | . | . | . | 0.411282 | 0.702429 | 0.565220 | 0.293221 | 0.957070 | 1.623697 | 1.268337 | 0.670415 | -1.184919 | 34.588343 | 0.549160603 | -0.44575084 | 0.655777236 | 0.998865404 | 5.229860 | 4.984116 | 5.080286 | 5.158341 | 5.108129 | 5.085613 | 32 | 51 | 30 | 27 |
| 105373794 | XR_001739152 | LOC105373794 | uncharacterized LOC105373794 | lncRNA | . | . | . | . | . | 0.073143 | 0.043821 | 0.049441 | 0.007064 | 0.170207 | 0.101295 | 0.110945 | 0.016152 | -1.974526 | 10.081917 | 1.052952384 | -0.93214703 | 0.351260544 | NA | 3.738475 | 2.769039 | 3.365039 | 3.336548 | 3.342079 | 3.300799 | 19 | 11 | 10 | 2 |
| 105373884 | XR_923918,XR | LOC105373884 | uncharacterized LOC105373884, transcript variant X1 | lncRNA | . | . | . | . | . | 0.072615 | 0.202349 | 0.353694 | 0.160435 | 0.168977 | 0.467738 | 0.793680 | 0.366814 | 1.542997 | 9.238185 | 1.042235272 | 0.600378312 | 0.548254139 | NA | 2.861598 | 3.486479 | 3.166403 | 3.193888 | 3.213943 | 3.181613 | 5 | 11 | 13 | 7 |
| 105373886 | XR_923922 | LINC02832 | long intergenic non-protein coding RNA 2832 | lncRNA | . | 54366 | . | . | . | 0.702453 | 0.482902 | 1.074186 | 0.812803 | 1.634638 | 1.116250 | 2.410441 | 1.858378 | 1.543081 | 14.194245 | 0.819131113 | 0.763996812 | 0.444869143 | NA | 3.478589 | 4.107809 | 3.810716 | 3.793767 | 3.838353 | 3.819180 | 15 | 10 | 17 | 14 |
| 105373887 | XR_923923 | LOC105373887 | uncharacterized LOC105373887 | lncRNA | . | . | . | . | . | 0.510175 | 0.520972 | 1.033814 | 0.619299 | 1.187197 | 1.204250 | 2.319849 | 1.415954 | 1.541073 | 11.971642 | 0.893625107 | 0.698206568 | 0.485048019 | NA | 3.234218 | 3.861274 | 3.556686 | 3.555544 | 3.594916 | 3.562122 | 11 | 10 | 16 | 10 |
| 105373945 | XR_001739947 | LOC105373945 | uncharacterized LOC105373945, transcript variant X1 | lncRNA | . | . | . | . | . | 0.043425 | 0.024450 | 0.099439 | 0.036078 | 0.101052 | 0.056518 | 0.223138 | 0.082487 | 1.614572 | 8.197751 | 1.098175054 | 0.62936401 | 0.529110774 | NA | 2.644780 | 3.342199 | 3.010694 | 3.004313 | 3.045550 | 3.003720 | 8 | 6 | 13 | 5 |
| 105373957 | XR_001739962 | LOC105373957 | uncharacterized LOC105373957 | lncRNA | . | . | . | . | . | 1.127247 | 1.452400 | 0.766434 | 1.616076 | 2.623151 | 3.357287 | 1.719857 | 3.694965 | -1.092321 | 53.800615 | 0.471649118 | -0.27011051 | 0.787075237 | 0.998865404 | 5.814036 | 5.682048 | 5.728114 | 5.777962 | 5.669359 | 5.798755 | 57 | 68 | 29 | 65 |
| 105373958 | XR_001739964 | LOC105373958 | uncharacterized LOC105373958, transcript variant X2 | lncRNA | . | . | . | . | . | 0.316477 | 0.350598 | 0.359666 | 0.535246 | 0.736456 | 0.810423 | 0.807080 | 1.223776 | 1.263243 | 17.219054 | 0.746150005 | 0.451828567 | 0.651392488 | NA | 3.928838 | 4.263648 | 4.094320 | 4.090865 | 4.091934 | 4.127505 | 18 | 16 | 13 | 22 |
| 105373976 | XR_924075 | LOC105373976 | uncharacterized LOC105373976 | lncRNA | . | . | . | . | . | 0.341770 | 0.141330 | 0.410330 | 0.346574 | 0.795313 | 0.326690 | 0.920770 | 0.792400 | 1.485042 | 17.628591 | 0.765626482 | 0.745145982 | 0.456183474 | NA | 3.823488 | 4.399154 | 4.138812 | 4.086697 | 4.150456 | 4.135018 | 23 | 9 | 20 | 18 |
| 105373984 | NR_136172 | LINC01918 | long intergenic non-protein coding RNA 1918 | lncRNA | . | 52737 | . | . | . | 20.538050 | 20.606634 | 18.687790 | 22.361448 | 47.792885 | 47.633156 | 41.934856 | 51.126793 | -1.015255 | 284.674033 | 0.209954652 | -0.10403388 | 0.917142461 | 0.998865404 | 8.164450 | 8.141795 | 8.157814 | 8.155756 | 8.104314 | 8.191260 | 331 | 309 | 225 | 287 |
| 105374002 | XR_924264 | LOC105374002 | uncharacterized LOC105374002 | lncRNA | . | . | . | . | . | 0.269437 | 0.147296 | 0.252631 | 0.197197 | 0.626991 | 0.340481 | 0.566896 | 0.450868 | 1.130204 | 5.217320 | 1.319545931 | 0.13382156 | 0.893543678 | NA | 2.290730 | 2.470304 | 2.385394 | 2.374246 | 2.383656 | 2.381838 | 7 | 4 | 5 | 5 |
| 105374010 | XR_001740453 | LOC105374010 | uncharacterized LOC105374010, transcript variant X1 | lncRNA | . | . | . | . | . | 0.007027 | 0.007643 | 0.015096 | 0.015750 | 0.016351 | 0.017666 | 0.033874 | 0.036010 | 2.058005 | 4.117192 | 1.489335242 | 0.699135083 | 0.484467615 | NA | 1.429096 | 2.470304 | 2.018221 | 2.019166 | 2.032732 | 2.031011 | 3 | 3 | 5 | 5 |
| 105374013 | XM_01151334 | LOC105374013 | endogenous retrovirus group K member 5 Gag polyprotein | protein_coding | XP_011511643.1 | . | . | . | . | 0.830076 | 0.914987 | 0.811256 | 0.949942 | 1.931621 | 2.115034 | 1.820435 | 2.171929 | -1.005846 | 109.632240 | 0.309739526 | -0.02714994 | 0.978340141 | 0.998865404 | 6.781345 | 6.771695 | 6.762273 | 6.791755 | 6.751555 | 6.798305 | 121 | 124 | 88 | 110 |
| 105374027 | XR_924311 | LOC105374027 | uncharacterized LOC105374027 | lncRNA | . | . | . | . | . | 0.072201 | 0.040343 | 0.091924 | 0.127471 | 0.168015 | 0.093255 | 0.206275 | 0.291448 | 1.910413 | 7.125641 | 1.149698837 | 0.812286327 | 0.416627354 | NA | 2.290730 | 3.226238 | 2.815104 | 2.804060 | 2.824061 | 2.835735 | 7 | 4 | 7 | 10 |
| 105374042 | NR_147411 | LOC105374042 | uncharacterized LOC105374042 | lncRNA | . | . | . | . | . | 0.955097 | 0.645840 | 1.082663 | 1.543874 | 2.222551 | 1.492889 | 2.429465 | 3.529885 | 1.618047 | 35.593358 | 0.537059339 | 1.292695063 | 0.196116546 | 0.96621494 | 4.764851 | 5.459419 | 5.125914 | 5.084957 | 5.144623 | 5.199006 | 37 | 24 | 32 | 48 |
| 105374080 | XR_924427 | LOC105374080 | uncharacterized LOC105374080 | lncRNA | . | . | . | . | . | 0.104119 | 0.053343 | 0.138204 | 0.035257 | 0.242289 | 0.123304 | 0.310126 | 0.080611 | 1.156727 | 5.763578 | 1.295089261 | 0.162188149 | 0.871157692 | NA | 2.413393 | 2.632244 | 2.524116 | 2.508892 | 2.533506 | 2.506941 | 8 | 4 | 8 | 3 |
| 105374085 | XR_924442 | LOC105374085 | uncharacterized LOC105374085 | lncRNA | . | . | . | . | . | 0.400459 | 0.462407 | 0.405087 | 0.671397 | 0.931885 | 1.068874 | 0.909004 | 1.535069 | 1.222747 | 20.951713 | 0.684574042 | 0.423804977 | 0.671708048 | NA | 4.238681 | 4.525118 | 4.368297 | 4.381637 | 4.368848 | 4.416161 | 20 | 22 | 15 | 27 |
| 105374101 | XR_924469 | LOC105374101 | uncharacterized LOC105374101 | lncRNA | . | . | . | . | . | 0.794357 | 1.125547 | 0.837946 | 0.849272 | 1.848502 | 2.601753 | 1.880327 | 1.941760 | -1.145735 | 18.994680 | 0.710658491 | -0.27618546 | 0.782405623 | NA | 4.343257 | 4.144983 | 4.236587 | 4.269503 | 4.240439 | 4.244015 | 19 | 26 | 15 | 17 |
| 105374104 | NR_146653,NF | LINC02021 | long intergenic non-protein coding RNA 2021 | lncRNA | . | 52856 | . | . | . | 0.740239 | 0.635675 | 0.847124 | 0.835574 | 1.722567 | 1.469391 | 1.900921 | 1.910440 | 1.134386 | 44.926518 | 0.467766219 | 0.388894671 | 0.697354064 | 0.998865404 | 5.394730 | 5.578418 | 5.483528 | 5.466282 | 5.514781 | 5.477525 | 51 | 43 | 46 | 40 |
| 105374128 | XR_001740930 | LOC105374128 | uncharacterized LOC105374128, transcript variant X1 | lncRNA | . | . | . | . | . | 0.558995 | 0.396091 | 0.502148 | 0.405781 | 1.300805 | 0.915583 | 1.126806 | 0.927769 | -1.091745 | 53.928131 | 0.432928586 | -0.29251064 | 0.769896225 | 0.998865404 | 5.813957 | 5.689282 | 5.787468 | 5.734775 | 5.757167 | 5.731223 | 73 | 53 | 48 | 45 |
| 105374145 | XR_001740949 | LOC105374145 | uncharacterized LOC105374145, transcript variant X2 | lncRNA | . | . | . | . | . | 0.016840 | 0.029416 | 0.044478 | 0.011130 | 0.039187 | 0.067998 | 0.099806 | 0.025448 | 1.065794 | 6.558171 | 1.229338796 | 0.074778761 | 0.940390741 | NA | 2.664814 | 2.760196 | 2.691717 | 2.710472 | 2.718693 | 2.687076 | 5 | 9 | 9 | 3 |
| 105374152 | XR_924575 | LOC105374152 | uncharacterized LOC105374152 | lncRNA | . | . | . | . | . | 0.111443 | 0.048318 | 0.167925 | 0.092396 | 0.259331 | 0.111690 | 0.376820 | 0.211252 | 1.590572 | 19.556509 | 0.744271524 | 0.899598648 | 0.368333877 | NA | 3.912173 | 4.588444 | 4.277371 | 4.225930 | 4.317875 | 4.261146 | 24 | 10 | 27 | 16 |
| 105374162 | XR_924588 | LOC105374162 | uncharacterized LOC105374162 | lncRNA | . | . | . | . | . | 0.021178 | 0.008094 | 0.004764 | 0.016650 | 0.049281 | 0.018709 | 0.010690 | 0.038067 | -1.272675 | 4.340264 | 1.480088783 | -0.23502885 | 0.814186339 | NA | 2.282109 | 1.932304 | 2.137398 | 2.117841 | 2.116139 | 2.129589 | 8 | 3 | 2 | 5 |
| 105374164 | XR_001740973 | LOC105374164 | uncharacterized LOC105374164, transcript variant X3 | lncRNA | . | . | . | . | . | 0.045570 | 0.148542 | 0.161584 | 0.048903 | 0.106043 | 0.343363 | 0.362590 | 0.111812 | 1.132254 | 5.836456 | 1.314085624 | 0.136366454 | 0.891531591 | NA | 2.452336 | 2.632244 | 2.515627 | 2.542688 | 2.545752 | 2.519133 | 3 | 9 | 8 | 3 |
| 105374167 | XR_924597 | LOC105374167 | uncharacterized LOC105374167, transcript variant X1 | lncRNA | . | . | . | . | . | 0.064093 | 0.099521 | 0.168615 | 0.086546 | 0.149147 | 0.230047 | 0.378367 | 0.197876 | 1.604069 | 9.994880 | 0.984892339 | 0.692193204 | 0.488816006 | NA | 2.939842 | 3.622519 | 3.286123 | 3.301144 | 3.330474 | 3.297992 | 7 | 10 | 14 | 8 |
| 105374179 | XR_924615 | LOC105374179 | uncharacterized LOC105374179 | lncRNA | . | . | . | . | . | 0.045436 | 0.090770 | 0.154698 | 0.074596 | 0.105731 | 0.209819 | 0.347139 | 0.170555 | 1.574418 | 12.822043 | 0.887507599 | 0.737817058 | 0.460625626 | NA | 3.316288 | 3.971139 | 3.634996 | 3.662931 | 3.695209 | 3.652803 | 8 | 14 | 18 | 10 |
| 105374187 | XR_001740993 | LOC105374187 | uncharacterized LOC105374187, transcript variant X1 | lncRNA | . | . | . | . | . | 0.052346 | 0.048361 | 0.052189 | 0.039705 | 0.121811 | 0.111788 | 0.117111 | 0.090781 | -1.135568 | 13.051076 | 0.846336176 | -0.21671554 | 0.828430033 | NA | 3.793789 | 3.612728 | 3.715159 | 3.707078 | 3.710581 | 3.697087 | 17 | 14 | 12 | 10 |
| 105374249 | XR_002959682 | LOC105374249 | uncharacterized LOC105374249, transcript variant X3 | lncRNA | . | . | . | . | . | 0.196275 | 0.356810 | 0.257828 | 0.199232 | 0.456739 | 0.824783 | 0.578558 | 0.455520 | -1.175570 | 17.549599 | 0.745117837 | -0.31318535 | 0.754139846 | NA | 4.245349 | 4.011955 | 4.119375 | 4.156063 | 4.135318 | 4.116146 | 17 | 25 | 16 | 13 |
| 105374254 | XR_924792,XR | LOC105374254 | uncharacterized LOC105374254, transcript variant X1 | lncRNA | . | . | . | . | . | 0.318283 | 0.130852 | 0.195005 | 0.228075 | 0.740657 | 0.302471 | 0.437587 | 0.521466 | -1.050133 | 7.355386 | 1.13521804 | -0.06216635 | 0.95043036 | NA | 2.912173 | 2.844638 | 2.894432 | 2.867609 | 2.878233 | 2.880706 | 12 | 5 | 6 | 7 |
| 105374292 | XR_001741084 | LOC105374292 | uncharacterized LOC105374292, transcript variant X1 | lncRNA | . | . | . | . | . | 0.388747 | 0.052664 | 0.163518 | 0.380109 | 0.904630 | 0.121734 | 0.366930 | 0.869074 | 1.184045 | 10.061344 | 1.067710735 | 0.228268167 | 0.81943777 | NA | 3.202589 | 3.448451 | 3.349742 | 3.290823 | 3.310770 | 3.345500 | 18 | 3 | 6 | 14 |
| 105374296 | XR_001741088 | LOC105374296 | uncharacterized LOC105374296 | lncRNA | . | . | . | . | . | 1.285462 | 1.103769 | 1.178015 | 1.135126 | 2.991324 | 2.551411 | 2.643431 | 2.595330 | -1.048163 | 89.591882 | 0.338850257 | -0.20027356 | 0.841266637 | 0.998865404 | 6.518348 | 6.451470 | 6.515298 | 6.468249 | 6.485583 | 6.471859 | 114 | 91 | 78 | 80 |
| 105374298 | XR_002959696 | LOC105374298 | uncharacterized LOC105374298, transcript variant X1 | lncRNA | . | . | . | . | . | 0.606068 | 0.616825 | 0.910056 | 0.424421 | 1.410346 | 1.425819 | 2.042139 | 0.970387 | 1.079299 | 113.393557 | 0.364173759 | 0.302315071 | 0.762411891 | 0.998865404 | 6.767596 | 6.880582 | 6.795770 | 6.798281 | 6.938032 | 6.691300 | 125 | 118 | 140 | 70 |
| 105374299 | XM_01151336 | LOC105374299 | uncharacterized LOC105374299 | protein_coding | XP_011511664.1 | . | . | . | . | 0.113999 | 0.063014 | 0.080717 | 0.075096 | 0.265281 | 0.145660 | 0.181126 | 0.171699 | -1.138679 | 7.103295 | 1.14295911 | -0.16392585 | 0.869789521 | NA | 2.917750 | 2.733338 | 2.842608 | 2.824084 | 2.830335 | 2.828092 | 11 | 6 | 6 | 6 |
| 105374306 | XR_924896 | LOC105374306 | uncharacterized LOC105374306 | lncRNA | . | . | . | . | . | 0.552145 | 0.728711 | 0.857128 | 0.562425 | 1.284863 | 1.684449 | 1.923371 | 1.285918 | 1.088840 | 11.769240 | 0.897093245 | 0.136877507 | 0.891127618 | NA | 3.493602 | 3.617632 | 3.541378 | 3.557007 | 3.565043 | 3.542091 | 11 | 14 | 13 | 9 |
| 105374312 | NR_153417 | LOC105374312 | uncharacterized LOC105374312 | lncRNA | . | . | . | . | . | 0.179271 | 0.175624 | 0.353422 | 0.302665 | 0.417171 | 0.405962 | 0.793069 | 0.692007 | 1.646324 | 3.563032 | 1.584094361 | 0.454043688 | 0.649797382 | NA | 1.429096 | 2.148375 | 1.817378 | 1.818280 | 1.826590 | 1.825182 | 3 | 3 | 4 | 4 |
| 105374333 | XR_939851 | LOC105374333 | uncharacterized LOC105374333 | lncRNA | . | . | . | . | . | 0.476481 | 0.487315 | 0.218476 | 0.608844 | 1.108792 | 1.126450 | 0.490254 | 1.392050 | -1.165635 | 11.222271 | 0.939985454 | -0.23523381 | 0.814027262 | NA | 3.597285 | 3.370388 | 3.495542 | 3.494983 | 3.467734 | 3.507268 | 14 | 13 | 5 | 14 |
| 105374352 | XR_925056 | LOC105374352 | uncharacterized LOC105374352 | lncRNA | . | . | . | . | . | 0.094920 | 0.110899 | 0.047250 | 0.125911 | 0.220884 | 0.256348 | 0.106027 | 0.287880 | -1.120833 | 6.369078 | 1.218328943 | -0.13507949 | 0.892549022 | NA | 2.754145 | 2.582949 | 2.672673 | 2.678854 | 2.661231 | 2.683634 | 7 | 8 | 3 | 8 |
| 105374363 | XR_001739353 | LOC105374363 | uncharacterized LOC105374363 | lncRNA | . | . | . | . | . | 2.214112 | 2.295154 | 3.873286 | 1.718212 | 5.152330 | 5.305351 | 8.691541 | 3.928488 | 1.222759 | 115.771949 | 0.370372004 | 0.783374209 | 0.433407397 | 0.998865404 | 6.701307 | 6.994143 | 6.793993 | 6.804051 | 6.995351 | 6.718636 | 118 | 114 | 154 | 73 |
| 105374366 | NR_134664,NF | LINC02482 | long intergenic non-protein coding RNA 2482 | lncRNA | . | 53458 | . | . | . | 4.832459 | 4.392077 | 4.116969 | 6.574464 | 11.245329 | 10.152482 | 9.238359 | 15.031730 | 1.147503 | 73.475376 | 0.388901201 | 0.510408432 | 0.609765351 | 0.998865404 | 6.097535 | 6.294149 | 6.189258 | 6.164217 | 6.152182 | 6.266372 | 83 | 70 | 53 | 90 |
| 105374369 | XR_939869,XR | LOC105374369 | uncharacterized LOC105374369, transcript variant X2 | lncRNA | . | . | . | . | . | 1.201018 | 0.548607 | 0.880681 | 0.406163 | 2.794818 | 1.268130 | 1.976222 | 0.928643 | -1.264128 | 7.558342 | 1.124563822 | -0.30068791 | 0.763652486 | NA | 3.074695 | 2.742346 | 2.937996 | 2.911536 | 2.922883 | 2.910858 | 13 | 6 | 7 | 5 |
| 105374407 | XR_925212 | LOC105374407 | uncharacterized LOC105374407 | lncRNA | . | . | . | . | . | 0.047125 | 0.096456 | 0.078954 | 0.111078 | 0.109663 | 0.222961 | 0.177170 | 0.253967 | 1.321397 | 18.379849 | 0.745096909 | 0.539612799 | 0.589464088 | NA | 3.988068 | 4.384843 | 4.154400 | 4.203034 | 4.185594 | 4.215897 | 12 | 23 | 15 | 23 |
| 105374419 | XR_002959786 | LOC105374419 | uncharacterized LOC105374419, transcript variant X4 | lncRNA | . | . | . | . | . | 0.261064 | 0.195183 | 0.138688 | 0.178266 | 0.607508 | 0.451174 | 0.311212 | 0.407583 | -1.346981 | 5.827420 | 1.254842717 | -0.34245694 | 0.732007044 | NA | 2.741623 | 2.312263 | 2.558676 | 2.548136 | 2.544005 | 2.547171 | 9 | 6 | 4 | 5 |
| 105374445 | XR_001741684 | LOC105374445 | uncharacterized LOC105374445, transcript variant X2 | lncRNA | . | . | . | . | . | 0.100559 | 0.099062 | 0.144460 | 0.141842 | 0.234005 | 0.228985 | 0.324165 | 0.324306 | 1.687524 | 11.474941 | 0.902076858 | 0.836855687 | 0.402673691 | NA | 3.094525 | 3.848815 | 3.490596 | 3.497524 | 3.522329 | 3.517970 | 9 | 10 | 13 | 13 |
| 105374571 | XR_940033 | LOC105374571 | uncharacterized LOC105374571 | lncRNA | . | . | . | . | . | 0.662680 | 0.593228 | 0.420329 | 0.582808 | 1.542084 | 1.371274 | 0.943206 | 1.332522 | -1.300311 | 9.495947 | 0.99033519 | -0.38255364 | 0.702050753 | NA | 3.425187 | 3.044381 | 3.261034 | 3.255964 | 3.241610 | 3.253259 | 13 | 11 | 6 | 9 |
| 105374584 | XR_940058 | LOC105374584 | uncharacterized LOC105374584 | lncRNA | . | . | . | . | . | 0.055076 | 0.045914 | 0.049932 | 0.049998 | 0.128164 | 0.106133 | 0.112046 | 0.114314 | -1.054136 | 13.065230 | 0.843637962 | -0.09015902 | 0.928160845 | NA | 3.744764 | 3.669578 | 3.716641 | 3.704407 | 3.707211 | 3.707823 | 17 | 13 | 11 | 12 |
| 105374612 | XR_925680,XR | LOC105374612 | uncharacterized LOC105374612, transcript variant X1 | lncRNA | . | . | . | . | . | 0.020745 | 0.025739 | 0.026646 | 0.026885 | 0.048275 | 0.059496 | 0.059793 | 0.061469 | 1.225237 | 3.507224 | 1.592557788 | 0.184019254 | 0.85399835 | NA | 1.658167 | 1.947964 | 1.801559 | 1.806830 | 1.805631 | 1.809330 | 3 | 4 | 3 | 4 |
| 105374727 | NR_134263 | LOC105374727 | uncharacterized LOC105374727 | lncRNA | . | . | . | . | . | 0.572042 | 0.272324 | 0.147780 | 0.419131 | 1.331166 | 0.629489 | 0.331614 | 0.958292 | -1.321088 | 5.055727 | 1.381512501 | -0.29078714 | 0.771214118 | NA | 2.526439 | 2.121007 | 2.358611 | 2.339413 | 2.333566 | 2.351688 | 9 | 4 | 2 | 6 |
| 105374748 | XR_001742653 | LOC105374748 | uncharacterized LOC105374748, transcript variant X2 | lncRNA | . | . | . | . | . | 0.208034 | 0.194054 | 0.229787 | 0.133579 | 0.484102 | 0.448564 | 0.515635 | 0.305414 | 1.018609 | 12.722640 | 0.865416941 | 0.030736365 | 0.97547979 | NA | 3.654832 | 3.683676 | 3.658661 | 3.670483 | 3.679421 | 3.651594 | 13 | 15 | 14 | 9 |
| 105374749 | XR_001742656 | C5orf34-AS1 | C5orf34 antisense RNA 1, transcript variant X1 | lncRNA | . | 41061 | . | . | . | 0.739222 | 0.523468 | 0.936079 | 0.299241 | 1.720200 | 1.210022 | 2.100533 | 0.684180 | 1.003969 | 15.700708 | 0.81434827 | 0.007017024 | 0.994401271 | NA | 3.966708 | 3.978782 | 3.972593 | 3.957147 | 3.995172 | 3.934077 | 20 | 15 | 20 | 8 |
| 105374775 | XR_001739502 | LOC105374775 | uncharacterized LOC105374775, transcript variant X1 | lncRNA | . | . | . | . | . | 0.340602 | 0.230852 | 0.237664 | 0.414462 | 0.792593 | 0.533625 | 0.533312 | 0.947617 | 1.130783 | 23.248587 | 0.660023054 | 0.268660452 | 0.788190988 | NA | 4.448287 | 4.624479 | 4.548025 | 4.511745 | 4.516281 | 4.567073 | 30 | 19 | 16 | 29 |
| 105374786 | XR_001739524 | LOC105374786 | uncharacterized LOC105374786, transcript variant X4 | lncRNA | . | . | . | . | . | 0.042735 | 0.062397 | 0.052020 | 0.022570 | 0.099446 | 0.144233 | 0.116731 | 0.051604 | -1.203695 | 6.185718 | 1.2372718 | -0.21617063 | 0.828849835 | NA | 2.754145 | 2.491830 | 2.626774 | 2.632955 | 2.635396 | 2.613843 | 7 | 8 | 7 | 3 |
| 105374800 | XR_940239 | LOC105374800 | uncharacterized LOC105374800 | lncRNA | . | . | . | . | . | 0.088550 | 0.168781 | 0.111652 | 0.003433 | 0.206059 | 0.390144 | 0.250544 | 0.007849 | -1.899591 | 6.579130 | 1.286327779 | -0.71963696 | 0.471748547 | NA | 3.104339 | 2.188478 | 2.714991 | 2.738068 | 2.723720 | 2.692523 | 7 | 2 | 7 | 1 |
| 105374809 | XR_002959406 | LOC105374809 | uncharacterized LOC105374809, transcript variant X1 | lncRNA | . | . | . | . | . | 2.239365 | 2.407484 | 2.707316 | 3.032781 | 5.211096 | 5.565008 | 6.075139 | 6.934094 | 1.227616 | 261.927007 | 0.217172264 | 1.362328096 | 0.173094365 | 0.947431715 | 7.878093 | 8.172913 | 7.952707 | 7.996031 | 8.037853 | 8.118250 | 259 | 265 | 232 | 291 |
| 105374811 | XR_004643303 | LOC105374811 | uncharacterized LOC105374811, transcript variant X3 | lncRNA | . | . | . | . | . | 0.742886 | 0.833294 | 0.739606 | 0.353993 | 1.728726 | 1.926195 | 1.659655 | 0.809363 | -1.896132 | 31.615300 | 0.589443164 | -1.5659859 | 0.117351918 | 0.857396579 | 5.369732 | 4.451739 | 5.006298 | 5.030581 | 4.972620 | 4.906516 | 44 | 48 | 26 | 13 |
| 105374888 | XR_926394 | NQO2-AS1 | NQO2 antisense RNA 1 | lncRNA | . | 40407 | . | . | . | 0.179254 | 0.397532 | 0.620347 | 0.385897 | 0.417131 | 0.918913 | 1.392041 | 0.882308 | 1.744531 | 7.463535 | 1.135530354 | 0.707016962 | 0.479555937 | NA | 2.444631 | 3.245375 | 2.863256 | 2.882006 | 2.899725 | 2.882234 | 4 | 8 | 10 | 7 |
| 105374897 | XR_926412 | LOC105374897 | uncharacterized LOC105374897 | lncRNA | . | . | . | . | . | 0.009699 | 0.015628 | 0.010501 | 0.008058 | 0.022570 | 0.036124 | 0.023565 | 0.018424 | -1.017609 | 2.237600 | 1.972119472 | -0.01276997 | 0.989811317 | NA | 1.175403 | 1.148375 | 1.158018 | 1.163002 | 1.160711 | 1.159957 | 2 | 3 | 2 | 2 |
| 105374898 | XR_001743791 | LOC105374898 | uncharacterized LOC105374898, transcript variant X2 | lncRNA | . | . | . | . | . | 0.005324 | 0.183586 | 0.147224 | 0.002357 | 0.012389 | 0.424368 | 0.330367 | 0.005388 | -1.210489 | 5.030205 | 1.492472079 | -0.18465361 | 0.853500732 | NA | 2.460000 | 2.188478 | 2.300208 | 2.335624 | 2.329218 | 2.297956 | 2 | 10 | 7 | 1 |
| 105374937 | XR_926497 | LOC105374937 | uncharacterized LOC105374937 | lncRNA | . | . | . | . | . | 1.056176 | 0.840045 | 1.391069 | 0.807628 | 2.457765 | 1.941802 | 3.121518 | 1.846545 | 1.145898 | 52.791818 | 0.457711985 | 0.429261662 | 0.667732814 | 0.998865404 | 5.618662 | 5.818882 | 5.720882 | 5.678595 | 5.777399 | 5.672271 | 63 | 47 | 62 | 39 |
| 105374945 | XR_001744005 | LOC105374945 | uncharacterized LOC105374945, transcript variant X1 | lncRNA | . | . | . | . | . | 0.125878 | 0.188979 | 0.149946 | 0.294334 | 0.292922 | 0.436834 | 0.336475 | 0.672959 | 1.330766 | 5.758978 | 1.265991016 | 0.325639372 | 0.74469722 | NA | 2.307819 | 2.715150 | 2.513936 | 2.519736 | 2.515553 | 2.533030 | 5 | 6 | 4 | 8 |
| 105374952 | NR_134616,NF | LOC105374952 | uncharacterized LOC105374952 | lncRNA | . | . | . | . | . | 0.693879 | 0.670064 | 0.679958 | 1.088114 | 1.614684 | 1.548884 | 1.525806 | 2.487844 | 1.289154 | 30.229996 | 0.573980841 | 0.638390916 | 0.523219243 | 0.998865404 | 4.724393 | 5.088511 | 4.897850 | 4.893971 | 4.896842 | 4.955999 | 31 | 28 | 23 | 39 |
| 105374956 | NR_149116 | MIR548A1HG | MIR548A1 host gene | lncRNA | . | 53539 | . | . | . | 0.137794 | 0.100947 | 0.270339 | 0.062693 | 0.320652 | 0.233344 | 0.606633 | 0.143340 | 1.428952 | 8.148195 | 1.115705232 | 0.461552998 | 0.644401907 | NA | 2.741623 | 3.264261 | 3.007802 | 2.997339 | 3.038526 | 2.991925 | 9 | 6 | 13 | 4 |
| 105374985 | XR_926612 | LOC105374985 | uncharacterized LOC105374985 | lncRNA | . | . | . | . | . | 0.204131 | 0.295284 | 0.318984 | 0.252363 | 0.475021 | 0.682561 | 0.715791 | 0.576998 | 1.126748 | 16.777774 | 0.75310093 | 0.228608602 | 0.819173138 | NA | 3.980095 | 4.151760 | 4.046698 | 4.071281 | 4.075038 | 4.060268 | 15 | 20 | 17 | 15 |
| 105374997 | XR_926658 | LOC105374997 | uncharacterized LOC105374997 | lncRNA | . | . | . | . | . | 0.232463 | 0.324459 | 0.306181 | 0.123945 | 0.540950 | 0.750002 | 0.687061 | 0.283385 | -1.300124 | 43.898048 | 0.528882018 | -0.71594247 | 0.474026873 | 0.998865404 | 5.631445 | 5.256422 | 5.438022 | 5.497961 | 5.483493 | 5.362496 | 48 | 62 | 47 | 21 |
| 105375001 | XR_926664 | LOC105375001 | uncharacterized LOC105375001 | lncRNA | . | . | . | . | . | 0.021398 | 0.027012 | 0.115654 | 0.091339 | 0.049794 | 0.062440 | 0.259524 | 0.208835 | 3.932936 | 12.204716 | 0.921682175 | 2.143479258 | 0.032074635 | NA | 2.307819 | 4.282298 | 3.533326 | 3.539676 | 3.611403 | 3.591787 | 5 | 6 | 19 | 16 |
| 105375013 | NR_138037 | HCG20 | HLA complex group 20 | lncRNA | . | 31334 | . | . | . | 14.821259 | 12.531214 | 16.974224 | 18.752077 | 34.489681 | 28.966463 | 38.089668 | 42.874401 | 1.286935 | 590.463613 | 0.168653301 | 2.157916119 | 0.030934354 | 0.558665013 | 9.012161 | 9.376327 | 9.167108 | 9.055164 | 9.248024 | 9.309970 | 647 | 508 | 552 | 651 |
| 105375022 | XR_926705 | LOC105375022 | uncharacterized LOC105375022 | lncRNA | . | . | . | . | . | 0.450468 | 0.393220 | 0.549722 | 0.241532 | 1.048258 | 0.908946 | 1.233559 | 0.552235 | -1.051266 | 10.925512 | 0.946274282 | -0.07622339 | 0.939241373 | NA | 3.482357 | 3.416142 | 3.451296 | 3.442329 | 3.462699 | 3.425689 | 14 | 11 | 13 | 6 |
| 105375044 | XR_926767 | LOC105375044 | uncharacterized LOC105375044 | lncRNA | . | . | . | . | . | 0.150340 | 0.049181 | 0.378325 | 0.040106 | 0.349848 | 0.113684 | 0.848951 | 0.091699 | 1.991292 | 5.966402 | 1.367964732 | 0.726410989 | 0.467586857 | NA | 1.987791 | 2.993948 | 2.546946 | 2.526935 | 2.580406 | 2.528925 | 7 | 2 | 12 | 2 |
| 105375101 | XR_001744179 | LOC105375101 | uncharacterized LOC105375101 | lncRNA | . | . | . | . | . | 0.200743 | 0.210283 | 0.238261 | 0.174005 | 0.467138 | 0.486078 | 0.534651 | 0.397842 | -1.018550 | 10.681324 | 0.934139782 | -0.02838599 | 0.977354295 | NA | 3.429096 | 3.404839 | 3.413998 | 3.417185 | 3.423582 | 3.405761 | 12 | 12 | 11 | 8 |
| 105375119 | XR_001745042 | LOC105375119 | uncharacterized LOC105375119 | lncRNA | . | . | . | . | . | 0.147150 | 0.118027 | 0.403658 | 0.042186 | 0.342424 | 0.272825 | 0.905797 | 0.096453 | 1.632226 | 8.260232 | 1.180525006 | 0.59875128 | 0.549338756 | NA | 2.644780 | 3.359867 | 3.010545 | 3.004155 | 3.059981 | 2.989016 | 8 | 6 | 16 | 2 |
| 105375210 | XR_927138 | TAX1BP1-AS1 | TAX1BP1 antisense RNA 1 | lncRNA | . | 40576 | . | . | . | 0.029289 | 0.051606 | 0.101880 | 0.051281 | 0.068157 | 0.119290 | 0.228615 | 0.117247 | 1.488926 | 2.812587 | 1.788565834 | 0.321079512 | 0.748150136 | NA | 1.175403 | 1.751299 | 1.472828 | 1.477882 | 1.485999 | 1.474918 | 2 | 3 | 4 | 2 |
| 105375213 | XR_927142 | LOC105375213 | uncharacterized LOC105375213 | lncRNA | . | . | . | . | . | 0.114828 | 0.077370 | 0.184830 | 0.037010 | 0.267210 | 0.178844 | 0.414754 | 0.084620 | 1.154299 | 16.358581 | 0.851484936 | 0.243124452 | 0.80790899 | NA | 3.920530 | 4.135426 | 4.022392 | 3.994790 | 4.064391 | 3.970648 | 21 | 13 | 25 | 6 |
| 105375215 | XR_927145 | FKBP14-AS1 | FKBP14 antisense RNA 1 | lncRNA | . | 40990 | . | . | . | 0.197658 | 0.246166 | 0.206575 | 0.218724 | 0.459957 | 0.569025 | 0.463548 | 0.500088 | -1.063112 | 7.413689 | 1.109555193 | -0.07957582 | 0.936574631 | NA | 2.934350 | 2.844638 | 2.888809 | 2.895252 | 2.888758 | 2.891228 | 8 | 9 | 6 | 7 |
| 105375259 | XR_927225 | LOC105375259 | uncharacterized LOC105375259 | lncRNA | . | . | . | . | . | 0.471661 | 0.322807 | 0.424232 | 0.464093 | 1.097575 | 0.746182 | 0.951963 | 1.061093 | 1.053064 | 9.158765 | 1.004040776 | 0.07429332 | 0.940776992 | NA | 3.156660 | 3.232645 | 3.200610 | 3.186777 | 3.194950 | 3.196714 | 12 | 8 | 8 | 9 |
| 105375270 | XR_927251 | LOC105375270 | uncharacterized LOC105375270 | lncRNA | . | . | . | . | . | 0.142342 | 0.191309 | 0.263007 | 0.197202 | 0.331235 | 0.442219 | 0.590179 | 0.450879 | 1.342428 | 12.141270 | 0.881490032 | 0.481962097 | 0.629832872 | NA | 3.373794 | 3.798723 | 3.576612 | 3.592152 | 3.609289 | 3.590784 | 10 | 13 | 14 | 11 |
| 105375310 | XR_001739612 | LOC105375310 | uncharacterized LOC105375310 | lncRNA | . | . | . | . | . | 0.081257 | 0.027580 | 0.084911 | 0.057895 | 0.189088 | 0.063752 | 0.190538 | 0.132369 | 1.291579 | 18.755797 | 0.767069992 | 0.48122872 | 0.630353951 | NA | 4.029324 | 4.404839 | 4.242247 | 4.171276 | 4.244562 | 4.210702 | 28 | 9 | 22 | 16 |
| 105375334 | XR_001744935 | LOC105375334 | uncharacterized LOC105375334, transcript variant X1 | lncRNA | . | . | . | . | . | 0.589882 | 0.481483 | 0.703505 | 0.427719 | 1.372680 | 1.112969 | 1.578644 | 0.977927 | -1.039765 | 55.315524 | 0.456797433 | -0.12315694 | 0.90198282 | 0.998865404 | 5.815452 | 5.763302 | 5.806108 | 5.766557 | 5.837399 | 5.716198 | 71 | 55 | 62 | 35 |
| 105375384 | XR_927722 | LOC105375384 | uncharacterized LOC105375384 | lncRNA | . | . | . | . | . | 0.195987 | 0.327644 | 0.265324 | 0.293328 | 0.456069 | 0.757365 | 0.595379 | 0.670659 | 1.097625 | 13.741825 | 0.835266009 | 0.1608884 | 0.872181299 | NA | 3.713381 | 3.844638 | 3.756875 | 3.789079 | 3.776279 | 3.781158 | 11 | 18 | 12 | 14 |
| 105375434 | XR_001744972 | LOC105375434 | uncharacterized LOC105375434, transcript variant X3 | lncRNA | . | . | . | . | . | 0.172143 | 0.158680 | 0.276857 | 0.242773 | 0.400584 | 0.366796 | 0.621259 | 0.555071 | 1.521604 | 16.973504 | 0.74307602 | 0.814981011 | 0.41508318 | NA | 3.751025 | 4.356363 | 4.057531 | 4.061703 | 4.090691 | 4.089201 | 15 | 15 | 18 | 19 |
| 105375436 | XR_927837,XR | LOC105375436 | uncharacterized LOC105375436, transcript variant X1 | lncRNA | . | . | . | . | . | 0.353739 | 0.482621 | 0.353886 | 0.252561 | 0.823165 | 1.115601 | 0.794110 | 0.577450 | -1.390936 | 59.252350 | 0.427850813 | -1.11266931 | 0.265850479 | 0.997969584 | 6.106718 | 5.632007 | 5.884269 | 5.952951 | 5.882872 | 5.817186 | 68 | 85 | 51 | 38 |
| 105375508 | XR_001744984 | LOC105375508 | uncharacterized LOC105375508, transcript variant X4 | lncRNA | . | . | . | . | . | 0.748666 | 0.522488 | 0.551492 | 1.025051 | 1.742176 | 1.207753 | 1.237532 | 2.343656 | 1.198425 | 17.659526 | 0.735917266 | 0.354848569 | 0.72270304 | NA | 4.006229 | 4.266773 | 4.141934 | 4.122798 | 4.132886 | 4.159186 | 21 | 15 | 14 | 21 |
| 105375513 | XR_002956515 | LOC105375513 | uncharacterized LOC105375513, transcript variant X2 | lncRNA | . | . | . | . | . | 0.096665 | 0.083336 | 0.046787 | 0.098818 | 0.224944 | 0.192634 | 0.104988 | 0.225936 | -1.260755 | 13.612849 | 0.847441103 | -0.39446807 | 0.693235498 | NA | 3.926074 | 3.587955 | 3.784286 | 3.772626 | 3.747708 | 3.781542 | 19 | 15 | 7 | 15 |
| 105375519 | XR_001745364 | LOC105375519 | uncharacterized LOC105375519, transcript variant X3 | lncRNA | . | . | . | . | . | 3.494970 | 2.152133 | 3.616326 | 3.618398 | 8.132941 | 4.974752 | 8.114930 | 8.273033 | 1.337681 | 175.624519 | 0.271884287 | 1.543795673 | 0.12263784 | 0.866753766 | 7.230273 | 7.651753 | 7.454562 | 7.304303 | 7.518153 | 7.492289 | 206 | 131 | 180 | 183 |
| 105375521 | XR_928021 | LOC105375521 | uncharacterized LOC105375521 | lncRNA | . | . | . | . | . | 0.139004 | 0.159933 | 0.101121 | 0.189799 | 0.323468 | 0.369692 | 0.226912 | 0.433954 | -1.052422 | 11.811833 | 0.899798632 | -0.08192165 | 0.934709016 | NA | 3.600756 | 3.522505 | 3.560899 | 3.568499 | 3.547181 | 3.576737 | 13 | 14 | 7 | 14 |
| 105375523 | XR_001745378 | LOC105375523 | uncharacterized LOC105375523, transcript variant X5 | lncRNA | . | . | . | . | . | 0.460025 | 0.702939 | 0.459586 | 0.401979 | 1.070498 | 1.624876 | 1.031297 | 0.919077 | -1.181140 | 40.713684 | 0.501108978 | -0.47929731 | 0.631727136 | 0.998865404 | 5.461710 | 5.223339 | 5.342548 | 5.374414 | 5.360862 | 5.303593 | 46 | 52 | 39 | 28 |
| 105375532 | XR_928043 | LOC105375532 | uncharacterized LOC105375532 | lncRNA | . | . | . | . | . | 0.027581 | 0.087926 | 0.111901 | 0.045925 | 0.064182 | 0.203245 | 0.251103 | 0.105002 | 1.362217 | 15.593821 | 0.851319804 | 0.523841986 | 0.600388448 | NA | 3.722967 | 4.168566 | 3.902215 | 3.958990 | 3.981867 | 3.920593 | 8 | 21 | 22 | 10 |
| 105375536 | XR_001744989 | LOC105375536 | uncharacterized LOC105375536, transcript variant X2 | lncRNA | . | . | . | . | . | 0.174718 | 0.068001 | 0.177367 | 0.097315 | 0.406576 | 0.157187 | 0.398007 | 0.222498 | 1.107497 | 7.951200 | 1.101226442 | 0.133763023 | 0.893589968 | NA | 2.912173 | 3.066070 | 2.998244 | 2.971398 | 2.997124 | 2.979869 | 12 | 5 | 9 | 6 |
| 105375545 | NR_133932 | TMEM139-AS1 | TMEM139 antisense RNA 1 | lncRNA | . | 40988 | . | . | . | 0.440712 | 0.260575 | 0.215331 | 0.405849 | 1.025555 | 0.602330 | 0.483197 | 0.927925 | -1.083150 | 6.844531 | 1.170865692 | -0.09841676 | 0.92160137 | NA | 2.832372 | 2.715150 | 2.787008 | 2.772496 | 2.768627 | 2.785911 | 10 | 6 | 4 | 8 |
| 105375563 | XR_928157,XR | REPIN1-AS1 | REPIN1 antisense RNA 1, transcript variant X3 | lncRNA | . | 41201 | . | . | . | 0.205990 | 0.075944 | 0.131218 | 0.279048 | 0.479348 | 0.175547 | 0.294448 | 0.638011 | 1.375031 | 9.459991 | 1.024663584 | 0.448405267 | 0.653860741 | NA | 2.993083 | 3.453945 | 3.244344 | 3.213632 | 3.229797 | 3.255094 | 13 | 5 | 7 | 13 |
| 105375591 | XR_928234 | LOC105375591 | uncharacterized LOC105375591 | lncRNA | . | . | . | . | . | 0.732047 | 0.949936 | 0.589127 | 0.848546 | 1.703504 | 2.195819 | 1.321984 | 1.940099 | -1.172304 | 41.243359 | 0.497969082 | -0.46056517 | 0.645110603 | 0.998865404 | 5.477744 | 5.245065 | 5.357406 | 5.399731 | 5.327444 | 5.379354 | 45 | 54 | 27 | 42 |
| 105375666 | XR_928448 | VPS13B-DT | VPS13B divergent transcript | lncRNA | . | 54375 | . | . | . | 0.843873 | 0.524792 | 0.705648 | 0.558302 | 1.963727 | 1.213081 | 1.583453 | 1.276491 | -1.087340 | 8.127079 | 1.073213251 | -0.11256249 | 0.910377426 | NA | 3.079678 | 2.963455 | 3.033811 | 3.015634 | 3.027817 | 3.015519 | 12 | 7 | 8 | 6 |
| 105375674 | XR_928458,XR | LOC105375674 | uncharacterized LOC105375674, transcript variant X2 | lncRNA | . | . | . | . | . | 1.728360 | 1.942315 | 1.548384 | 1.552565 | 4.021963 | 4.489749 | 3.474528 | 3.549754 | -1.203144 | 190.763607 | 0.243002559 | -1.09796976 | 0.272217695 | 0.998865404 | 7.703124 | 7.435794 | 7.589541 | 7.639929 | 7.529636 | 7.537682 | 227 | 237 | 149 | 164 |
| 105375675 | XR_928460,XR | LOC105375675 | uncharacterized LOC105375675, transcript variant X1 | lncRNA | . | . | . | . | . | 0.246278 | 0.106113 | 0.207674 | 0.223130 | 0.573099 | 0.245285 | 0.466013 | 0.510160 | 1.190806 | 10.672155 | 0.960169856 | 0.262389808 | 0.793020929 | NA | 3.282109 | 3.538109 | 3.426397 | 3.388237 | 3.415205 | 3.416299 | 16 | 6 | 10 | 11 |
| 105375700 | XR_001746028 | LOC105375700 | uncharacterized LOC105375700, transcript variant X2 | lncRNA | . | . | . | . | . | 0.343875 | 0.149996 | 0.240466 | 0.183352 | 0.800211 | 0.346722 | 0.539598 | 0.419213 | 1.053602 | 5.953610 | 1.238589718 | 0.060818682 | 0.951503612 | NA | 2.533720 | 2.612728 | 2.576957 | 2.566126 | 2.576413 | 2.569495 | 8 | 5 | 6 | 5 |
| 105375721 | XR_001746041 | LOC105375721 | uncharacterized LOC105375721, transcript variant X4 | lncRNA | . | . | . | . | . | 0.276067 | 0.155812 | 0.124944 | 0.166706 | 0.642418 | 0.360165 | 0.280372 | 0.381152 | 1.152243 | 23.166835 | 0.640390715 | 0.319250258 | 0.749536749 | NA | 4.429096 | 4.631770 | 4.521305 | 4.527446 | 4.526054 | 4.549994 | 24 | 24 | 19 | 26 |
| 105375737 | XR_001746070 | LOC105375737 | uncharacterized LOC105375737, transcript variant X1 | lncRNA | . | . | . | . | . | 0.180237 | 0.505914 | 0.472989 | 0.597493 | 0.419420 | 1.169443 | 1.061373 | 1.366098 | 1.673901 | 4.826404 | 1.385828044 | 0.536296092 | 0.591753931 | NA | 1.855800 | 2.592943 | 2.249208 | 2.258950 | 2.258842 | 2.271713 | 3 | 5 | 4 | 7 |
| 105375740 | XR_928608 | LOC105375740 | uncharacterized LOC105375740 | lncRNA | . | . | . | . | . | 0.034145 | 0.203536 | 0.125161 | 0.059800 | 0.079457 | 0.470483 | 0.280857 | 0.136726 | -1.284369 | 8.959570 | 1.10971196 | -0.32536373 | 0.7449058 | NA | 3.333078 | 2.971139 | 3.126490 | 3.186984 | 3.158272 | 3.136178 | 4 | 18 | 9 | 5 |
| 105375846 | XR_001745916 | LOC105375846 | uncharacterized LOC105375846, transcript variant X1 | lncRNA | . | . | . | . | . | 0.073464 | 0.075170 | 0.102818 | 0.030522 | 0.170954 | 0.173758 | 0.230722 | 0.069785 | -1.129624 | 7.194926 | 1.165729084 | -0.15084322 | 0.88009939 | NA | 2.928838 | 2.760196 | 2.845692 | 2.844064 | 2.856696 | 2.825122 | 9 | 8 | 9 | 3 |
| 105375884 | XR_929015 | LOC105375884 | uncharacterized LOC105375884 | lncRNA | . | . | . | . | . | 0.895687 | 0.778226 | 0.647959 | 0.767676 | 2.084301 | 1.798904 | 1.454002 | 1.755201 | -1.215210 | 22.007017 | 0.657329844 | -0.42779964 | 0.668796998 | NA | 4.593806 | 4.312263 | 4.480496 | 4.464142 | 4.448462 | 4.460366 | 30 | 24 | 16 | 20 |
| 105375892 | XR_929032 | LOC105375892 | uncharacterized LOC105375892 | lncRNA | . | . | . | . | . | 0.364790 | 0.205263 | 0.572354 | 0.222900 | 0.848881 | 0.474475 | 1.284345 | 0.509634 | 1.386459 | 15.938092 | 0.820759819 | 0.574351762 | 0.565729788 | NA | 3.735321 | 4.213972 | 3.984601 | 3.948211 | 4.021209 | 3.955380 | 20 | 10 | 23 | 10 |
| 105375921 | XR_001745976 | LOC105375921 | uncharacterized LOC105375921 | lncRNA | . | . | . | . | . | 0.904394 | 0.387225 | 0.920455 | 0.160653 | 2.104561 | 0.895087 | 2.065474 | 0.367314 | -1.084900 | 4.690237 | 1.440974544 | -0.08158513 | 0.934976627 | NA | 2.282109 | 2.175234 | 2.237092 | 2.217461 | 2.236120 | 2.214848 | 8 | 3 | 6 | 2 |
| 105375942 | XR_929381 | LOC105375942 | uncharacterized LOC105375942 | lncRNA | . | . | . | . | . | 0.023541 | 0.006582 | 0.039933 | 0.021256 | 0.054781 | 0.015215 | 0.089608 | 0.048598 | 1.976352 | 4.628047 | 1.435114189 | 0.684851565 | 0.493437593 | NA | 1.631267 | 2.622519 | 2.190873 | 2.179079 | 2.207306 | 2.190587 | 5 | 2 | 7 | 4 |
| 105375957 | XR_929435,XR | LOC105375957 | uncharacterized LOC105375957, transcript variant X1 | lncRNA | . | . | . | . | . | 0.876292 | 0.920274 | 0.839342 | 0.454603 | 2.039167 | 2.127255 | 1.883458 | 1.039396 | -1.231697 | 8.092098 | 1.078775072 | -0.2786934 | 0.780480125 | NA | 3.156660 | 2.861274 | 3.029094 | 3.015199 | 3.023096 | 3.006050 | 12 | 8 | 8 | 5 |
| 105375980 | XR_929488,XR | LOC105375980 | uncharacterized LOC105375980, transcript variant X1 | lncRNA | . | . | . | . | . | 0.193674 | 0.173252 | 0.135326 | 0.297579 | 0.450687 | 0.400480 | 0.303668 | 0.680380 | 1.157530 | 17.843308 | 0.747714311 | 0.282259897 | 0.777744233 | NA | 4.049783 | 4.257378 | 4.151770 | 4.144711 | 4.132179 | 4.186042 | 20 | 17 | 11 | 24 |
| 105375983 | XR_001746628 | LOC105375983 | uncharacterized LOC105375983, transcript variant X1 | lncRNA | . | . | . | . | . | 2.540472 | 1.963118 | 1.827073 | 2.346615 | 5.911784 | 4.537834 | 4.099899 | 5.365256 | -1.100004 | 121.022486 | 0.306653001 | -0.44841769 | 0.653851779 | 0.998865404 | 6.986272 | 6.848713 | 6.980487 | 6.886456 | 6.871389 | 6.931347 | 165 | 119 | 91 | 118 |
| 105376033 | XR_001746665 | LOC105376033 | uncharacterized LOC105376033 | lncRNA | . | . | . | . | . | 0.078756 | 0.124043 | 0.315174 | 0.131026 | 0.183269 | 0.286731 | 0.707241 | 0.299575 | 2.174621 | 8.598933 | 1.073248424 | 1.044272628 | 0.296359273 | NA | 2.436885 | 3.558654 | 3.058043 | 3.068416 | 3.110384 | 3.073142 | 5 | 7 | 14 | 7 |
| 105376034 | XR_929594 | LOC105376034 | uncharacterized LOC105376034 | lncRNA | . | . | . | . | . | 0.033644 | 0.084493 | 0.106797 | 0.054866 | 0.078291 | 0.195309 | 0.239649 | 0.125445 | 1.355548 | 5.352257 | 1.333713351 | 0.329063175 | 0.742107937 | NA | 2.184683 | 2.622519 | 2.393565 | 2.412024 | 2.418553 | 2.401804 | 3 | 7 | 7 | 4 |
| 105376037 | XR_929599 | LOC105376037 | uncharacterized LOC105376037 | lncRNA | . | . | . | . | . | 0.125167 | 0.129994 | 0.182924 | 0.136578 | 0.291270 | 0.300486 | 0.410476 | 0.312270 | 1.240730 | 11.557958 | 0.897804292 | 0.34661154 | 0.728883175 | NA | 3.365657 | 3.678992 | 3.519283 | 3.518327 | 3.538880 | 3.520609 | 12 | 11 | 13 | 10 |
| 105376057 | gene-LOC1053 | LOC105376057 | protein FRG1B-like | pseudogene | . | . | . | . | . | 1.285963 | 0.834339 | 0.771316 | 0.956217 | 2.992489 | 1.928612 | 1.730812 | 2.186277 | -1.231038 | 12.909888 | 0.859691217 | -0.34881694 | 0.727226743 | NA | 3.832372 | 3.532926 | 3.714346 | 3.686334 | 3.683502 | 3.693885 | 20 | 12 | 9 | 12 |
| 105376060 | gene-LOC1053 | LOC105376060 | . | pseudogene | . | . | . | . | . | 0.154389 | 0.189578 | 0.279002 | 0.143514 | 0.359270 | 0.438218 | 0.626072 | 0.328128 | 1.193139 | 24.694705 | 0.647086146 | 0.393706233 | 0.693797939 | NA | 4.491734 | 4.749066 | 4.594960 | 4.616817 | 4.658419 | 4.588199 | 23 | 27 | 31 | 17 |
| 105376063 | XR_929667,XR | LOC105376063 | uncharacterized LOC105376063, transcript variant X1 | lncRNA | . | . | . | . | . | 0.297538 | 0.319155 | 0.412127 | 0.379913 | 0.692384 | 0.737742 | 0.924801 | 0.868626 | 1.299973 | 48.513969 | 0.44739125 | 0.845974981 | 0.397566701 | 0.998865404 | 5.398723 | 5.777184 | 5.564268 | 5.575292 | 5.622134 | 5.611797 | 47 | 47 | 49 | 50 |
| 105376074 | XR_929905 | LOC105376074 | uncharacterized LOC105376074 | lncRNA | . | . | . | . | . | 0.041294 | 0.045258 | 0.017518 | 0.033272 | 0.096093 | 0.104616 | 0.039309 | 0.076074 | -1.427143 | 5.337392 | 1.311725711 | -0.39118671 | 0.69565923 | NA | 2.651489 | 2.134756 | 2.426873 | 2.428705 | 2.415100 | 2.423372 | 7 | 7 | 3 | 5 |
| 105376076 | XR_929908,XR | LOC105376076 | uncharacterized LOC105376076, transcript variant X1 | lncRNA | . | . | . | . | . | 0.106321 | 0.182831 | 0.192384 | 0.048399 | 0.247413 | 0.422621 | 0.431703 | 0.110657 | -1.336579 | 5.933627 | 1.291231352 | -0.32414444 | 0.745828683 | NA | 2.760365 | 2.348113 | 2.562553 | 2.577080 | 2.575191 | 2.548820 | 6 | 9 | 7 | 2 |
| 105376114 | NR_135839,NF | LOC105376114 | uncharacterized LOC105376114 | lncRNA | . | . | . | . | . | 1.084997 | 1.607976 | 1.156647 | 1.735901 | 2.524833 | 3.716910 | 2.595483 | 3.968931 | -1.008409 | 55.683414 | 0.458648831 | -0.02633962 | 0.978986451 | 0.998865404 | 5.807371 | 5.790934 | 5.746099 | 5.838862 | 5.741412 | 5.840120 | 50 | 74 | 36 | 65 |
| 105376154 | XR_001746842 | LOC105376154 | uncharacterized LOC105376154, transcript variant X1 | lncRNA | . | . | . | . | . | 0.067629 | 0.270035 | 0.158383 | 0.206872 | 0.157375 | 0.624199 | 0.355407 | 0.472989 | -1.153006 | 2.967640 | 1.749036955 | -0.11743631 | 0.906514309 | NA | 1.671431 | 1.459419 | 1.563193 | 1.576934 | 1.566137 | 1.570164 | 2 | 5 | 2 | 3 |
| 105376156 | XR_001746848 | LOC105376156 | uncharacterized LOC105376156, transcript variant X2 | lncRNA | . | . | . | . | . | 0.725235 | 0.555335 | 0.542133 | 0.566749 | 1.687651 | 1.283679 | 1.216532 | 1.295805 | -1.183036 | 42.862275 | 0.479798863 | -0.50540735 | 0.613272756 | 0.998865404 | 5.537347 | 5.295830 | 5.458584 | 5.414881 | 5.409030 | 5.410168 | 60 | 44 | 34 | 37 |
| 105376177 | XR_930174,XR | LOC105376177 | uncharacterized LOC105376177, transcript variant X3 | lncRNA | . | . | . | . | . | 0.469585 | 0.696826 | 0.272903 | 0.446841 | 1.092744 | 1.610745 | 0.612386 | 1.021649 | -1.497041 | 16.475547 | 0.766728568 | -0.7592178 | 0.44772228 | NA | 4.305694 | 3.719719 | 4.055121 | 4.068287 | 4.021646 | 4.045323 | 21 | 23 | 9 | 15 |
| 105376190 | XR_930191 | LOC105376190 | uncharacterized LOC105376190 | lncRNA | . | . | . | . | . | 0.913769 | 0.675748 | 0.433251 | 0.686285 | 2.126377 | 1.562021 | 0.972204 | 1.569109 | -1.378087 | 6.542883 | 1.189228609 | -0.38904758 | 0.697240952 | NA | 2.923304 | 2.459419 | 2.727015 | 2.716786 | 2.708565 | 2.716413 | 10 | 7 | 4 | 6 |
| 105376204 | XM_01701540 | LOC105376204 | uncharacterized LOC105376204 | protein_coding | XP_016870898.1 | . | . | . | . | 0.055600 | 0.021856 | 0.062252 | 0.015905 | 0.129382 | 0.050521 | 0.139693 | 0.036366 | -1.019421 | 9.667968 | 1.056440159 | -0.02626756 | 0.979043933 | NA | 3.282109 | 3.264261 | 3.283927 | 3.245667 | 3.287019 | 3.240542 | 16 | 6 | 13 | 4 |
| 105376266 | XR_930332 | LOC105376266 | uncharacterized LOC105376266 | lncRNA | . | . | . | . | . | 0.133855 | 0.129585 | 0.184602 | 0.135656 | 0.311487 | 0.299541 | 0.414243 | 0.310160 | 1.217240 | 26.341898 | 0.604687199 | 0.469026166 | 0.639050933 | NA | 4.569400 | 4.855058 | 4.704742 | 4.699990 | 4.740880 | 4.706014 | 28 | 25 | 29 | 23 |
| 105376281 | XR_930372 | LOC105376281 | uncharacterized LOC105376281 | lncRNA | . | . | . | . | . | 0.046640 | 0.036771 | 0.062835 | 0.056949 | 0.108533 | 0.084998 | 0.141001 | 0.130208 | 1.488840 | 2.777185 | 1.782378592 | 0.322147257 | 0.747341139 | NA | 1.156660 | 1.733338 | 1.464780 | 1.461218 | 1.468604 | 1.467534 | 3 | 2 | 3 | 3 |
| 105376306 | NR_135119 | LOC105376306 | uncharacterized LOC105376306 | lncRNA | . | . | . | . | . | 0.542901 | 0.228418 | 0.586028 | 0.416363 | 1.263352 | 0.527999 | 1.315030 | 0.951964 | 1.251082 | 8.000756 | 1.089526312 | 0.296621116 | 0.766755788 | NA | 2.826455 | 3.155137 | 3.001416 | 2.978566 | 3.004328 | 2.991807 | 11 | 5 | 9 | 7 |
| 105376333 | XR_930463 | LOC105376333 | uncharacterized LOC105376333 | transcribed_ps | . | . | . | . | . | 0.369985 | 0.420592 | 0.671688 | 0.322977 | 0.860971 | 0.972217 | 1.507250 | 0.738448 | 1.245373 | 24.735936 | 0.647019578 | 0.489286772 | 0.624638685 | NA | 4.459800 | 4.779589 | 4.599892 | 4.610086 | 4.663984 | 4.589331 | 24 | 25 | 32 | 17 |
| 105376358 | XR_930568 | LOC105376358 | uncharacterized LOC105376358 | lncRNA | . | . | . | . | . | 0.349480 | 0.146878 | 0.141250 | 0.182027 | 0.813255 | 0.339516 | 0.316961 | 0.416183 | -1.551021 | 9.836885 | 1.002240325 | -0.631803 | 0.527515615 | NA | 3.579805 | 2.947964 | 3.334475 | 3.293340 | 3.291747 | 3.298665 | 19 | 8 | 6 | 8 |
| 105376367 | XR_001747333 | LOC105376367 | uncharacterized LOC105376367, transcript variant X3 | lncRNA | . | . | . | . | . | 0.471393 | 0.678128 | 0.127931 | 0.322849 | 1.096951 | 1.567523 | 0.287073 | 0.738156 | -2.506277 | 19.212186 | 0.749946495 | -1.76752035 | 0.077141124 | NA | 4.781768 | 3.448451 | 4.280788 | 4.321757 | 4.214345 | 4.248045 | 26 | 35 | 6 | 14 |
| 105376373 | XR_001747338 | LOC105376373 | uncharacterized LOC105376373, transcript variant X1 | lncRNA | . | . | . | . | . | 0.185361 | 0.276916 | 0.261053 | 0.153407 | 0.431342 | 0.640104 | 0.585795 | 0.350746 | -1.119082 | 15.388633 | 0.799238616 | -0.20308845 | 0.839065898 | NA | 4.021847 | 3.861274 | 3.929745 | 3.958250 | 3.952911 | 3.920040 | 15 | 21 | 16 | 10 |
| 105376486 | XR_930807 | LOC105376486 | uncharacterized LOC105376486 | lncRNA | . | . | . | . | . | 0.121265 | 0.275111 | 0.393026 | 0.293118 | 0.282190 | 0.635931 | 0.881940 | 0.670180 | 1.697528 | 20.791834 | 0.710913246 | 1.073879486 | 0.282876699 | NA | 3.948040 | 4.708721 | 4.307670 | 4.360770 | 4.395227 | 4.365184 | 11 | 23 | 26 | 21 |
| 105376494 | XR_930820 | LOC105376494 | uncharacterized LOC105376494 | lncRNA | . | . | . | . | . | 0.081773 | 0.035577 | 0.084422 | 0.089791 | 0.190289 | 0.082237 | 0.189441 | 0.205297 | 1.368121 | 5.252301 | 1.327545888 | 0.340625565 | 0.733385481 | NA | 2.147197 | 2.602870 | 2.391574 | 2.376044 | 2.389841 | 2.392805 | 7 | 3 | 5 | 6 |
| 105376512 | XR_930968 | LOC105376512 | uncharacterized LOC105376512 | lncRNA | . | . | . | . | . | 0.328340 | 0.140849 | 0.135294 | 0.109392 | 0.764061 | 0.325578 | 0.303595 | 0.250111 | -1.923290 | 17.842082 | 0.778891559 | -1.21143455 | 0.225728905 | NA | 4.551576 | 3.612728 | 4.222225 | 4.139945 | 4.140464 | 4.126591 | 38 | 15 | 12 | 10 |
| 105376517 | XR_002957212 | LOC105376517 | uncharacterized LOC105376517 | lncRNA | . | . | . | . | . | 0.577986 | 0.773058 | 0.383795 | 0.564896 | 1.344997 | 1.786958 | 0.861225 | 1.291566 | -1.397081 | 20.446144 | 0.691580624 | -0.69755481 | 0.485455649 | NA | 4.576468 | 4.090276 | 4.358657 | 4.384166 | 4.329540 | 4.356195 | 24 | 29 | 12 | 19 |
| 105376567 | XR_001748140 | LOC105376567 | uncharacterized LOC105376567, transcript variant X8 | lncRNA | . | . | . | . | . | 0.674318 | 0.896042 | 0.483614 | 0.979585 | 1.569162 | 2.071241 | 1.085216 | 2.239704 | 1.046058 | 64.624255 | 0.398285926 | 0.163104969 | 0.870435788 | 0.998865404 | 5.982023 | 6.045291 | 6.008274 | 6.006983 | 5.984317 | 6.050120 | 73 | 68 | 49 | 71 |
| 105376577 | XR_931096 | LOC105376577 | uncharacterized LOC105376577 | lncRNA | . | . | . | . | . | 0.637128 | 0.396852 | 0.524787 | 0.269924 | 1.482624 | 0.917341 | 1.177607 | 0.617149 | -1.285011 | 22.977519 | 0.678359658 | -0.53331673 | 0.593814347 | NA | 4.689131 | 4.333284 | 4.558644 | 4.504938 | 4.533863 | 4.481874 | 37 | 21 | 23 | 13 |
| 105376579 | XR_001748146 | LOC105376579 | uncharacterized LOC105376579, transcript variant X1 | lncRNA | . | . | . | . | . | 0.095938 | 0.148069 | 0.390389 | 0.177669 | 0.223252 | 0.342268 | 0.876023 | 0.406219 | 2.386356 | 9.173920 | 1.054156208 | 1.190344988 | 0.233910825 | NA | 2.436885 | 3.692999 | 3.143950 | 3.154370 | 3.206181 | 3.159189 | 5 | 7 | 16 | 7 |
| 105376580 | XR_001748148 | CSRP3-AS1 | CSRP3 and E2F8 antisense RNA 1, transcript variant X1 | lncRNA | . | 54183 | . | . | . | 0.068083 | 0.105798 | 0.218259 | 0.071457 | 0.158432 | 0.244557 | 0.489766 | 0.163377 | 1.582956 | 15.163355 | 0.841723724 | 0.787219631 | 0.431153307 | NA | 3.551764 | 4.217207 | 3.873453 | 3.893507 | 3.950156 | 3.874905 | 11 | 15 | 24 | 9 |
| 105376634 | XR_931202 | LOC105376634 | uncharacterized LOC105376634 | lncRNA | . | . | . | . | . | 3.426947 | 4.560393 | 3.815465 | 3.212985 | 7.974646 | 10.541553 | 8.561794 | 7.346108 | -1.137779 | 48.662780 | 0.456644885 | -0.40780018 | 0.683420385 | 0.998865404 | 5.694911 | 5.508570 | 5.586385 | 5.642684 | 5.606080 | 5.577031 | 51 | 64 | 43 | 39 |
| 105376678 | XM_01152090 | LOC105376678 | ovostatin-like | protein_coding | XP_011519205.1 | . | . | . | . | 0.350918 | 0.497795 | 0.564545 | 0.551507 | 0.816602 | 1.150675 | 1.266822 | 1.260956 | 1.315180 | 15.108488 | 0.78795366 | 0.501628671 | 0.615928748 | NA | 3.706954 | 4.100822 | 3.893087 | 3.909200 | 3.919377 | 3.918790 | 13 | 16 | 15 | 16 |
| 105376689 | XR_001737883 | LOC105376689 | uncharacterized LOC105376689, transcript variant X4 | lncRNA | . | . | . | . | . | 0.862612 | 0.802221 | 1.190023 | 0.597359 | 2.007333 | 1.854369 | 2.670375 | 1.365793 | -1.114024 | 79.012654 | 0.382293571 | -0.40748727 | 0.683650144 | 0.998865404 | 6.378246 | 6.225750 | 6.334252 | 6.292518 | 6.343865 | 6.223541 | 103 | 83 | 81 | 53 |
| 105376736 | NR_135061,NF | EXOSC10-AS1 | EXOSC10 antisense RNA 1 | lncRNA | . | 40456 | . | . | . | 1.839602 | 2.369870 | 3.344520 | 2.246262 | 4.280830 | 5.478062 | 7.505005 | 5.135810 | 1.396079 | 38.280924 | 0.517708361 | 0.929829644 | 0.352459295 | 0.998865404 | 4.997177 | 5.479761 | 5.206354 | 5.233698 | 5.303980 | 5.233977 | 33 | 38 | 47 | 33 |
| 105376748 | XR_001739154 | LOC105376748 | uncharacterized LOC105376748, transcript variant X1 | lncRNA | . | . | . | . | . | 0.293884 | 0.269839 | 0.312867 | 0.340593 | 0.683879 | 0.623744 | 0.702066 | 0.778725 | 1.157704 | 23.179337 | 0.637449741 | 0.33142534 | 0.740323227 | NA | 4.425187 | 4.636609 | 4.529645 | 4.520286 | 4.536132 | 4.542139 | 26 | 22 | 21 | 24 |
| 105376752 | gene-NPIPA9 | NPIPA9 | nuclear pore complex interacting protein family, member A9 | pseudogene | . | 41984 | . | . | . | 2.395941 | 2.393724 | 3.409789 | 1.989602 | 5.575453 | 5.533202 | 7.651469 | 4.548989 | 1.110601 | 92.335149 | 0.362306207 | 0.417716206 | 0.676154622 | 0.998865404 | 6.449781 | 6.603730 | 6.505078 | 6.503278 | 6.609996 | 6.452987 | 101 | 94 | 107 | 67 |
| 105376781 | XR_001752320 | LOC105376781 | uncharacterized LOC105376781, transcript variant X1 | lncRNA | . | . | . | . | . | 0.641795 | 0.799723 | 1.037574 | 0.748194 | 1.493484 | 1.848593 | 2.328283 | 1.710656 | 1.020844 | 18.236201 | 0.735378782 | 0.04047175 | 0.967717028 | NA | 4.173073 | 4.204225 | 4.167747 | 4.196589 | 4.202715 | 4.164653 | 17 | 23 | 20 | 13 |
| 105376805 | NR_135056,NF | LOC105376805 | uncharacterized LOC105376805 | lncRNA | . | . | . | . | . | 15.715262 | 15.155574 | 14.035641 | 19.485281 | 36.570062 | 35.032790 | 31.495573 | 44.550781 | 1.057862 | 240.945770 | 0.234856765 | 0.345535768 | 0.729691625 | 0.998865404 | 7.872021 | 7.952000 | 7.909004 | 7.882728 | 7.846681 | 7.994978 | 277 | 246 | 183 | 267 |
| 105376843 | NR_132278 | SNHG25 | small nucleolar RNA host gene 25 | lncRNA | . | 51534 | . | . | . | 7.312581 | 5.224105 | 5.670914 | 2.674038 | 17.016676 | 12.075751 | 12.725367 | 6.113870 | -1.478164 | 18.721924 | 0.744022755 | -0.75778159 | 0.448581747 | NA | 4.478589 | 3.921224 | 4.258648 | 4.226129 | 4.235024 | 4.188024 | 30 | 20 | 18 | 15 |
| 105376845 | XR_001737923 | LOC105376845 | uncharacterized LOC105376845, transcript variant X3 | lncRNA | . | . | . | . | . | 0.198374 | 0.436784 | 0.484078 | 0.197038 | 0.461625 | 1.009645 | 1.086257 | 0.450504 | 1.156580 | 14.652632 | 0.842504408 | 0.249096066 | 0.803286473 | NA | 3.763465 | 3.974966 | 3.837095 | 3.873489 | 3.890242 | 3.838450 | 11 | 19 | 19 | 9 |
| 105376859 | XR_947064,XR | LOC105376859 | uncharacterized LOC105376859, transcript variant X1 | lncRNA | . | . | . | . | . | 0.122760 | 0.140996 | 0.051893 | 0.128758 | 0.285667 | 0.325918 | 0.116447 | 0.294389 | -1.310430 | 6.319522 | 1.218371574 | -0.3201328 | 0.748867659 | NA | 2.844134 | 2.448451 | 2.668864 | 2.670965 | 2.653357 | 2.671021 | 8 | 8 | 3 | 7 |
| 105376861 | XR_947068 | LOC105376861 | uncharacterized LOC105376861 | lncRNA | . | . | . | . | . | 0.164817 | 0.217673 | 0.255112 | 0.299418 | 0.383536 | 0.503160 | 0.572464 | 0.684583 | 1.337373 | 5.779805 | 1.254252371 | 0.334383714 | 0.738090027 | NA | 2.307819 | 2.724273 | 2.518296 | 2.524102 | 2.524987 | 2.532666 | 5 | 6 | 5 | 7 |
| 105376878 | XR_001737933 | RUNX3-AS1 | RUNX3 antisense RNA 1, transcript variant X3 | lncRNA | . | 40513 | . | . | . | 0.158770 | 0.330304 | 0.069589 | 0.098292 | 0.369464 | 0.763512 | 0.156157 | 0.224732 | -1.731893 | 39.878226 | 0.52461118 | -1.51035578 | 0.130952665 | 0.882587711 | 5.661494 | 4.864999 | 5.305918 | 5.399766 | 5.251858 | 5.290399 | 44 | 68 | 21 | 32 |
| 105376912 | XR_936215 | LOC105376912 | uncharacterized LOC105376912, transcript variant X2 | lncRNA | . | . | . | . | . | 0.090045 | 0.102625 | 0.213529 | 0.056390 | 0.209538 | 0.237222 | 0.479153 | 0.128930 | 1.252417 | 2.026741 | 2.090159777 | 0.155354231 | 0.876542064 | NA | 0.844134 | 1.175234 | 1.007230 | 1.007808 | 1.014998 | 1.004217 | 2 | 2 | 3 | 1 |
| 105376959 | XR_940608 | LOC105376959 | uncharacterized LOC105376959 | lncRNA | . | . | . | . | . | 0.142806 | 0.203701 | 0.167557 | 0.192194 | 0.332316 | 0.470864 | 0.375994 | 0.439430 | -1.018500 | 11.152599 | 0.913328993 | -0.02895549 | 0.97690009 | NA | 3.493602 | 3.464871 | 3.471405 | 3.486967 | 3.475339 | 3.481299 | 11 | 14 | 9 | 11 |
| 105376987 | XR_001740620 | LOC105376987 | uncharacterized LOC105376987, transcript variant X3 | lncRNA | . | . | . | . | . | 0.407603 | 0.258708 | 0.501699 | 0.665729 | 0.948508 | 0.598015 | 1.125798 | 1.522110 | 1.600026 | 13.900079 | 0.828615466 | 0.818347701 | 0.413158676 | NA | 3.417335 | 4.097315 | 3.782886 | 3.761685 | 3.796079 | 3.804932 | 15 | 9 | 14 | 17 |
| 105376993 | XR_940651,XR | LOC105376993 | uncharacterized LOC105376993, transcript variant X2 | lncRNA | . | . | . | . | . | 0.245301 | 0.133089 | 0.238180 | 0.127486 | 0.570826 | 0.307640 | 0.534471 | 0.291482 | -1.101006 | 15.764129 | 0.777872955 | -0.17846393 | 0.858358643 | NA | 4.044696 | 3.909275 | 3.993778 | 3.970694 | 3.985118 | 3.966248 | 22 | 15 | 15 | 12 |
| 105377016 | XR_001740630 | LOC105377016 | uncharacterized LOC105377016, transcript variant X1 | lncRNA | . | . | . | . | . | 0.008095 | 0.010257 | 0.020604 | 0.021907 | 0.018837 | 0.023710 | 0.046236 | 0.050087 | 2.280548 | 5.169705 | 1.343937686 | 0.88499657 | 0.376158481 | NA | 1.658167 | 2.844638 | 2.339168 | 2.344590 | 2.359107 | 2.361823 | 3 | 4 | 6 | 7 |
| 105377023 | XR_940698,XR | LOC105377023 | uncharacterized LOC105377023, transcript variant X2 | lncRNA | . | . | . | . | . | 0.400625 | 0.427463 | 0.479839 | 0.338971 | 0.932271 | 0.988099 | 1.076744 | 0.775017 | 1.043628 | 46.886215 | 0.466898919 | 0.131949871 | 0.895023942 | 0.998865404 | 5.519313 | 5.582186 | 5.526608 | 5.558869 | 5.580687 | 5.519068 | 48 | 54 | 49 | 37 |
| 105377116 | XR_940895,XR | LOC105377116 | uncharacterized LOC105377116, transcript variant X1 | lncRNA | . | . | . | . | . | 0.141704 | 0.120647 | 0.077606 | 0.084162 | 0.329751 | 0.278882 | 0.174145 | 0.192427 | -1.618390 | 18.808245 | 0.714046407 | -0.97270945 | 0.330697725 | NA | 4.539157 | 3.844638 | 4.266513 | 4.249718 | 4.218238 | 4.222628 | 29 | 23 | 12 | 14 |

| | | | | | | | | | | | | | | | | | | | | | | | | | | | | | | | | | |
|---|---|---|---|---|---|---|---|---|---|---|---|---|---|---|---|---|---|---|---|---|---|---|---|---|---|---|---|---|---|---|---|---|---|
| 105377134 | XR_937583 | LOC105377134 | uncharacterized LOC105377134 | lncRNA | . | . | . | . | . | 0.220991 | 0.188301 | 0.162822 | 0.244010 | 0.514255 | 0.435265 | 0.365368 | 0.557900 | -1.016013 | 19.953066 | 0.69379205 | -0.03303479 | 0.973646843 | NA | 4.330772 | 4.306201 | 4.327022 | 4.313215 | 4.303488 | 4.333900 | 25 | 20 | 14 | 22 |
| 105377182 | XR_937989 | FAM247C | family with sequence similarity 247 member C | lncRNA | . | 54928 | . | . | . | 0.374962 | 0.201601 | 0.483402 | 0.315982 | 0.872553 | 0.466010 | 1.084739 | 0.722455 | 1.299383 | 13.309286 | 0.857449496 | 0.440640104 | 0.659473565 | NA | 3.530084 | 3.913269 | 3.732511 | 3.703402 | 3.747222 | 3.719066 | 17 | 9 | 16 | 11 |
| 105377198 | XR_941033 | LOC105377198 | uncharacterized LOC105377198 | lncRNA | . | . | . | . | . | 0.627240 | 0.689250 | 0.660140 | 0.607976 | 1.459614 | 1.593232 | 1.481335 | 1.390065 | -1.038437 | 18.074186 | 0.719710283 | -0.07560461 | 0.939733672 | NA | 4.202828 | 4.148375 | 4.172709 | 4.181866 | 4.177177 | 4.171584 | 20 | 21 | 16 | 16 |
| 105377203 | XR_938280 | LOC105377203 | uncharacterized LOC105377203 | lncRNA | . | . | . | . | . | 0.580225 | 0.672112 | 0.332269 | 0.242335 | 1.350208 | 1.553616 | 0.745602 | 0.554070 | -2.085851 | 10.309425 | 0.971257626 | -1.09202372 | 0.274822675 | NA | 3.799841 | 2.742346 | 3.385677 | 3.393483 | 3.363657 | 3.351441 | 15 | 16 | 7 | 5 |
| 105377205 | XR_001755602 | LOC105377205 | uncharacterized LOC105377205, transcript variant X1 | lncRNA | . | . | . | . | . | 0.192388 | 0.120244 | 0.118063 | 0.197604 | 0.447693 | 0.277947 | 0.264930 | 0.451796 | 1.008159 | 14.762402 | 0.808587308 | 0.014498607 | 0.98843219 | NA | 3.878555 | 3.889137 | 3.896056 | 3.872309 | 3.870652 | 3.898717 | 20 | 13 | 10 | 17 |
| 105377236 | XR_938659,XR | LOC105377236 | uncharacterized LOC105377236, transcript variant X1 | lncRNA | . | . | . | . | . | 0.083671 | 0.095286 | 0.131566 | 0.217411 | 0.194705 | 0.220259 | 0.295230 | 0.497084 | 1.931438 | 12.499955 | 0.878978415 | 1.080430952 | 0.279950319 | NA | 3.094525 | 4.040734 | 3.606392 | 3.613384 | 3.628671 | 3.661272 | 9 | 10 | 11 | 19 |
| 105377265 | XR_001741714 | LOC105377265 | uncharacterized LOC105377265, transcript variant X1 | lncRNA | . | . | . | . | . | 3.997238 | 3.832806 | 1.798903 | 2.121414 | 9.301737 | 8.859701 | 4.036685 | 4.850362 | -1.684223 | 55.993949 | 0.442009165 | -1.70150968 | 0.088847321 | 0.802216049 | 6.136066 | 5.380522 | 5.862851 | 5.849978 | 5.713632 | 5.791506 | 83 | 74 | 27 | 49 |
| 105377294 | XR_001741738 | LOC105377294 | uncharacterized LOC105377294, transcript variant X1 | lncRNA | . | . | . | . | . | 0.101140 | 0.070043 | 0.043766 | 0.073472 | 0.235358 | 0.161907 | 0.098211 | 0.167984 | -1.421455 | 7.959235 | 1.095427149 | -0.46316949 | 0.643242881 | NA | 3.225252 | 2.715150 | 3.015533 | 2.997703 | 2.985611 | 3.002733 | 13 | 8 | 4 | 8 |
| 105377319 | XR_938955 | LOC105377319 | uncharacterized LOC105377319 | lncRNA | . | . | . | . | . | 0.007521 | 0.010688 | 0.022029 | 0.016502 | 0.017503 | 0.024705 | 0.049433 | 0.037730 | 2.170210 | 2.847568 | 1.783980699 | 0.6265958 | 0.530924222 | NA | 0.844134 | 1.963455 | 1.485636 | 1.486278 | 1.498845 | 1.492632 | 2 | 2 | 4 | 3 |
| 105377348 | NR_131185,NR | UBE2D3-AS1 | UBE2D3 antisense RNA 1 | lncRNA | . | 54095 | . | . | . | 0.891441 | 1.065334 | 1.415837 | 0.847326 | 2.074420 | 2.462568 | 3.177099 | 1.937311 | 1.146213 | 31.378354 | 0.571631321 | 0.344408891 | 0.730538804 | 0.998865404 | 4.868663 | 5.067862 | 4.944921 | 4.964136 | 5.006108 | 4.936073 | 31 | 34 | 37 | 23 |
| 105377359 | XR_001741788 | LOC105377359 | uncharacterized LOC105377359, transcript variant X2 | lncRNA | . | . | . | . | . | 0.253904 | 0.297520 | 0.362673 | 0.181628 | 0.590845 | 0.687732 | 0.813829 | 0.415271 | -1.019612 | 29.399650 | 0.599294305 | -0.04675438 | 0.962708986 | 0.998865404 | 4.889947 | 4.865403 | 4.864473 | 4.879906 | 4.908533 | 4.829625 | 32 | 34 | 34 | 18 |
| 105377372 | NR_161159 | NEUROG2-AS1 | NEUROG2 and ZGRF1 antisense RNA 1 | lncRNA | . | 40656 | . | . | . | 0.459772 | 0.447583 | 0.266817 | 0.331991 | 1.069908 | 1.034608 | 0.598729 | 0.759057 | -1.548230 | 7.742546 | 1.094712193 | -0.57605988 | 0.564574711 | NA | 3.234218 | 2.602870 | 2.968719 | 2.967296 | 2.951701 | 2.954402 | 11 | 10 | 5 | 6 |
| 105377486 | XR_001741443 | FAM160A1-DT | FAM160A1 divergent transcript, transcript variant X2 | lncRNA | . | 53123 | . | . | . | 0.108023 | 0.140240 | 0.190756 | 0.105548 | 0.251373 | 0.324171 | 0.428054 | 0.241322 | 1.115475 | 9.531639 | 0.993579444 | 0.158676866 | 0.873923465 | NA | 3.170740 | 3.330299 | 3.238605 | 3.249614 | 3.259940 | 3.237452 | 9 | 11 | 11 | 7 |
| 105377527 | XR_939429 | LOC105377527 | uncharacterized LOC105377527 | lncRNA | . | . | . | . | . | 0.209063 | 0.309949 | 0.277257 | 0.399530 | 0.486497 | 0.716461 | 0.622156 | 0.913478 | 1.289686 | 18.567808 | 0.724369587 | 0.50667425 | 0.612383401 | NA | 4.021847 | 4.384843 | 4.183095 | 4.211803 | 4.202441 | 4.232705 | 15 | 21 | 15 | 23 |
| 105377540 | XR_939462,XR | SAP30-DT | SAP30 divergent transcript, transcript variant X1 | lncRNA | . | 54424 | . | . | . | 1.521778 | 1.539026 | 1.930995 | 2.324059 | 3.541240 | 3.557529 | 4.333097 | 5.313685 | 1.374179 | 54.799187 | 0.425243781 | 1.078370245 | 0.28086856 | 0.998865404 | 5.529269 | 5.986776 | 5.735231 | 5.736825 | 5.777937 | 5.819230 | 53 | 50 | 50 | 65 |
| 105377548 | XR_002959809 | LOC105377548 | uncharacterized LOC105377548, transcript variant X3 | lncRNA | . | . | . | . | . | 0.123573 | 0.124792 | 0.183664 | 0.052040 | 0.287559 | 0.288461 | 0.412136 | 0.118983 | -1.081932 | 18.170417 | 0.773611005 | -0.14685578 | 0.883245854 | NA | 4.236451 | 4.128572 | 4.175548 | 4.180862 | 4.208848 | 4.133798 | 21 | 21 | 23 | 8 |
| 105377567 | XR_001741467 | LOC105377567 | uncharacterized LOC105377567, transcript variant X3 | lncRNA | . | . | . | . | . | 0.148947 | 0.099673 | 0.181268 | 0.149557 | 0.346604 | 0.230398 | 0.406761 | 0.341944 | 1.141397 | 12.434303 | 0.864443285 | 0.220720769 | 0.825309859 | NA | 3.537347 | 3.728813 | 3.636966 | 3.624155 | 3.635177 | 3.640452 | 15 | 11 | 11 | 13 |
| 105377576 | XR_001741930 | LOC105377576 | uncharacterized LOC105377576, transcript variant X1 | lncRNA | . | . | . | . | . | 0.114841 | 0.060309 | 0.171629 | 0.062031 | 0.267239 | 0.139408 | 0.385129 | 0.141826 | 1.205237 | 22.563860 | 0.706027 | 0.381454265 | 0.702866201 | NA | 4.351306 | 4.627390 | 4.501106 | 4.444640 | 4.529547 | 4.450325 | 31 | 15 | 30 | 14 |
| 105377621 | NR_136202 | LOC105377621 | uncharacterized LOC105377621 | lncRNA | . | . | . | . | . | 0.626031 | 1.146265 | 1.276388 | 1.212646 | 1.456800 | 2.649642 | 2.864177 | 2.772570 | 1.380279 | 25.820905 | 0.625050712 | 0.743875346 | 0.456951892 | NA | 4.440763 | 4.903263 | 4.634871 | 4.687816 | 4.700156 | 4.691544 | 18 | 30 | 27 | 27 |
| 105377623 | NR_136203 | LOC105377623 | uncharacterized LOC105377623 | lncRNA | . | . | . | . | . | 0.214618 | 0.166376 | 0.206008 | 0.496711 | 0.499425 | 0.384585 | 0.462276 | 1.135670 | 1.734475 | 7.315674 | 1.14589712 | 0.69334228 | 0.488094776 | NA | 2.421266 | 3.213338 | 2.854394 | 2.847735 | 2.853348 | 2.884359 | 7 | 5 | 5 | 12 |
| 105377626 | XR_939842 | LOC105377626 | uncharacterized LOC105377626 | lncRNA | . | . | . | . | . | 0.171325 | 0.212319 | 0.146111 | 0.317322 | 0.398680 | 0.490785 | 0.327870 | 0.725518 | 1.136043 | 7.189905 | 1.147821382 | 0.160318925 | 0.872629851 | NA | 2.754145 | 2.932304 | 2.838795 | 2.845046 | 2.832677 | 2.859351 | 7 | 8 | 4 | 10 |
| 105377651 | NR_134682 | LOC105377651 | uncharacterized LOC105377651 | lncRNA | . | . | . | . | . | 7.671864 | 8.709192 | 3.184807 | 3.718370 | 17.852741 | 20.131689 | 7.146615 | 8.501611 | -2.387698 | 73.899757 | 0.38045045 | -3.30035256 | 0.000965634 | 0.066311699 | 6.703326 | 5.445966 | 6.272817 | 6.309635 | 6.075095 | 6.097796 | 113 | 119 | 35 | 44 |
| 105377663 | XR_001741694 | LOC105377663 | uncharacterized LOC105377663, transcript variant X3 | lncRNA | . | . | . | . | . | 0.561178 | 0.218587 | 0.518697 | 0.180569 | 1.305884 | 0.505273 | 1.163939 | 0.412849 | 1.112666 | 12.171814 | 0.91719965 | 0.167924289 | 0.866642842 | NA | 3.522785 | 3.683676 | 3.618585 | 3.573109 | 3.615558 | 3.587783 | 19 | 7 | 14 | 9 |
| 105377750 | XR_001743016 | LOC105377750 | uncharacterized LOC105377750, transcript variant X3 | lncRNA | . | . | . | . | . | 0.335894 | 0.943127 | 0.126357 | 0.336912 | 0.781640 | 2.180080 | 0.283542 | 0.770308 | -2.284047 | 9.047704 | 1.03897785 | -1.14688937 | 0.251427348 | NA | 3.654832 | 2.459419 | 3.197833 | 3.209324 | 3.168367 | 3.175939 | 13 | 15 | 4 | 6 |
| 105377763 | NR_134259,NR | LOC105377763 | uncharacterized LOC105377763 | lncRNA | . | . | . | . | . | 0.550370 | 0.877876 | 0.425382 | 0.923886 | 1.280734 | 2.029248 | 0.954545 | 2.112355 | -1.112295 | 19.676393 | 0.721396719 | -0.21283714 | 0.831453998 | NA | 4.375822 | 4.216574 | 4.283221 | 4.320065 | 4.268094 | 4.317309 | 19 | 27 | 11 | 23 |
| 105377769 | XR_941333,XR | LOC105377769 | uncharacterized LOC105377769, transcript variant X1 | lncRNA | . | . | . | . | . | 1.834065 | 1.628867 | 1.748567 | 2.076207 | 4.267944 | 3.765197 | 3.923734 | 4.746999 | 1.105613 | 18.375411 | 0.715308645 | 0.202495451 | 0.839529415 | NA | 4.125927 | 4.269892 | 4.197784 | 4.190988 | 4.193738 | 4.210759 | 21 | 18 | 15 | 20 |
| 105377771 | XR_001743047 | LOC105377771 | uncharacterized LOC105377771, transcript variant X2 | lncRNA | . | . | . | . | . | 1.578555 | 1.747033 | 2.099751 | 1.838573 | 3.673363 | 4.038347 | 4.711781 | 4.203679 | 1.181987 | 56.333571 | 0.418659986 | 0.576158923 | 0.564507767 | 0.998865404 | 5.690031 | 5.931705 | 5.786087 | 5.802245 | 5.843441 | 5.812439 | 57 | 58 | 57 | 53 |
| 105377805 | XM_01153485 | LOC105377805 | basic salivary proline-rich protein 4-like, transcript variant X1 | protein_coding | XP_011533156.1;XP_016876396.1;XP_016876397.1 | . | . | . | . | 0.056160 | 0.034407 | 0.051127 | 0.052312 | 0.130686 | 0.079534 | 0.114727 | 0.119605 | 1.180127 | 9.698350 | 0.98404517 | 0.242816284 | 0.80814772 | NA | 3.151937 | 3.393446 | 3.282168 | 3.260091 | 3.277625 | 3.279073 | 13 | 7 | 9 | 10 |
| 105377806 | XR_001750048 | LOC105377806 | uncharacterized LOC105377806 | lncRNA | . | . | . | . | . | 0.116180 | 0.128353 | 0.187249 | 0.166901 | 0.270356 | 0.296694 | 0.420182 | 0.381599 | 1.417366 | 36.892169 | 0.511250895 | 0.984276917 | 0.324979376 | 0.998865404 | 4.931597 | 5.435167 | 5.165959 | 5.174494 | 5.230212 | 5.210201 | 34 | 34 | 40 | 38 |
| 105377826 | XR_001752723 | LOC105377826 | uncharacterized LOC105377826, transcript variant X10 | lncRNA | . | . | . | . | . | 2.199081 | 2.180375 | 3.755385 | 3.088820 | 5.117354 | 5.040032 | 8.426976 | 7.062220 | 1.488589 | 59.842473 | 0.409921321 | 1.400136883 | 0.161472332 | 0.933525128 | 5.587789 | 6.161703 | 5.832767 | 5.862197 | 5.948981 | 5.913134 | 51 | 56 | 67 | 62 |
| 105377887 | XR_942770 | LOC105377887 | uncharacterized LOC105377887 | lncRNA | . | . | . | . | . | 0.299304 | 0.180511 | 0.461223 | 0.230812 | 0.696493 | 0.417260 | 1.034971 | 0.527724 | 1.348644 | 6.816091 | 1.174683559 | 0.367340891 | 0.713364757 | NA | 2.533720 | 2.971139 | 2.759758 | 2.748957 | 2.774473 | 2.752475 | 8 | 5 | 9 | 5 |
| 105377929 | XR_942843 | LOC105377929 | uncharacterized LOC105377929 | lncRNA | . | . | . | . | . | 1.117360 | 0.838819 | 1.672173 | 0.431407 | 2.600143 | 1.938969 | 3.752307 | 0.986361 | 1.056185 | 22.529723 | 0.734717187 | 0.107336797 | 0.914521793 | NA | 4.450212 | 4.536027 | 4.489475 | 4.460786 | 4.534151 | 4.423140 | 29 | 20 | 32 | 9 |
| 105377940 | XR_942864 | LOC105377940 | uncharacterized LOC105377940 | lncRNA | . | . | . | . | . | 0.119181 | 0.181835 | 0.167570 | 0.262702 | 0.277338 | 0.420320 | 0.376022 | 0.600637 | 1.330890 | 11.553359 | 0.909505697 | 0.453423515 | 0.650243807 | NA | 3.312060 | 3.719719 | 3.505964 | 3.525494 | 3.518098 | 3.542451 | 9 | 13 | 9 | 15 |
| 105378029 | XR_943077 | LOC105378029 | uncharacterized LOC105378029 | lncRNA | . | . | . | . | . | 0.257597 | 0.218124 | 0.531576 | 0.166390 | 0.599438 | 0.504204 | 1.192842 | 0.380430 | 1.447692 | 54.810163 | 0.522026155 | 1.022468179 | 0.306559366 | 0.998865404 | 5.482357 | 6.020493 | 5.718981 | 5.688659 | 5.888813 | 5.651837 | 56 | 44 | 86 | 29 |
| 105378035 | XR_001744402 | LOC105378035 | uncharacterized LOC105378035, transcript variant X4 | lncRNA | . | . | . | . | . | 0.108630 | 0.037178 | 0.403682 | 0.088516 | 0.252787 | 0.085938 | 0.905850 | 0.202382 | 2.170914 | 14.752873 | 0.883464895 | 1.265814069 | 0.205579657 | NA | 3.211697 | 4.339233 | 3.847899 | 3.805492 | 3.909186 | 3.838616 | 16 | 5 | 25 | 11 |
| 105378077 | XR_943166 | LOC105378077 | uncharacterized LOC105378077 | lncRNA | . | . | . | . | . | 0.045598 | 0.037196 | 0.032084 | 0.033988 | 0.106108 | 0.085980 | 0.071995 | 0.077709 | -1.231861 | 12.959866 | 0.848617855 | -0.35450501 | 0.722960448 | NA | 3.838265 | 3.538109 | 3.711808 | 3.699836 | 3.693598 | 3.694478 | 18 | 14 | 10 | 11 |
| 105378083 | XR_001744428 | LOC105378083 | uncharacterized LOC105378083, transcript variant X2 | lncRNA | . | . | . | . | . | 0.266334 | 0.146525 | 0.142579 | 0.398401 | 0.619769 | 0.338699 | 0.319944 | 0.910895 | 1.225855 | 14.840266 | 0.839545884 | 0.349936901 | 0.726386053 | NA | 3.738475 | 4.029739 | 3.893311 | 3.865063 | 3.861946 | 3.922086 | 19 | 11 | 8 | 22 |
| 105378114 | XR_002956378 | LOC105378114 | uncharacterized LOC105378114 | lncRNA | . | . | . | . | . | 0.039045 | 0.061659 | 0.021038 | 0.065974 | 0.090860 | 0.142527 | 0.047209 | 0.150842 | -1.071449 | 5.668190 | 1.288127328 | -0.07729331 | 0.938390213 | NA | 2.555346 | 2.448451 | 2.497307 | 2.511659 | 2.493822 | 2.511602 | 5 | 8 | 3 | 7 |
| 105378120 | XR_943243,XR | LOC105378120 | uncharacterized LOC105378120, transcript variant X3 | lncRNA | . | . | . | . | . | 0.099157 | 0.246566 | 0.456473 | 0.097681 | 0.230742 | 0.569949 | 1.024312 | 0.223336 | 1.581702 | 24.015645 | 0.747952926 | 0.884384877 | 0.376488484 | NA | 4.216466 | 4.879762 | 4.495705 | 4.553316 | 4.639264 | 4.496390 | 14 | 27 | 41 | 11 |
| 105378130 | XR_943270,XR | LOC105378130 | uncharacterized LOC105378130, transcript variant X1 | lncRNA | . | . | . | . | . | 0.036925 | 0.045320 | 0.096570 | 0.041168 | 0.085926 | 0.104758 | 0.216701 | 0.094126 | 1.640271 | 11.291581 | 0.936891554 | 0.762023954 | 0.446045708 | NA | 3.094525 | 3.811620 | 3.462002 | 3.468924 | 3.512966 | 3.465981 | 9 | 10 | 17 | 8 |
| 105378171 | XR_943570 | LOC105378171 | uncharacterized LOC105378171 | lncRNA | . | . | . | . | . | 0.063355 | 0.097227 | 0.298154 | 0.132845 | 0.147429 | 0.224745 | 0.669048 | 0.303735 | 2.484275 | 6.305526 | 1.256979386 | 1.044428213 | 0.296287316 | NA | 1.855800 | 3.168566 | 2.609446 | 2.619351 | 2.654840 | 2.622796 | 3 | 5 | 11 | 5 |
| 105378174 | XR_943577 | LOC105378174 | uncharacterized LOC105378174 | lncRNA | . | . | . | . | . | 0.063421 | 0.085654 | 0.240828 | 0.132961 | 0.147583 | 0.197993 | 0.540412 | 0.304000 | 2.483428 | 5.498852 | 1.324621337 | 0.99072301 | 0.321820852 | NA | 1.658167 | 2.971139 | 2.419035 | 2.424493 | 2.454219 | 2.432191 | 3 | 4 | 9 | 5 |
| 105378179 | XR_001750763 | LOC105378179 | uncharacterized LOC105378179, transcript variant X2 | lncRNA | . | . | . | . | . | 0.260007 | 0.283828 | 0.156834 | 0.226184 | 0.605047 | 0.656081 | 0.351930 | 0.517142 | -1.464082 | 32.466895 | 0.561986694 | -0.97866455 | 0.327745755 | 0.998865404 | 5.271488 | 4.717436 | 5.039332 | 5.056108 | 4.972066 | 5.025364 | 42 | 44 | 17 | 31 |
| 105378217 | NR_147190,NR | STK32A-AS1 | STK32A antisense RNA 1 | lncRNA | . | 53468 | . | . | . | 1.713541 | 1.776221 | 1.758417 | 1.786225 | 3.987479 | 4.105813 | 3.945834 | 4.083991 | 1.000195 | 104.551961 | 0.313025668 | 0.000898817 | 0.999282848 | 0.999862634 | 6.708119 | 6.708034 | 6.698866 | 6.716880 | 6.704861 | 6.711055 | 117 | 116 | 90 | 99 |
| 105378260 | XM_01702852 | LOC105378260 | histone-lysine N-methyltransferase 2C, transcript variant X3 | protein_coding | XP_016884009.1;XP_016884010.1 | . | . | . | . | 0.452133 | 0.921187 | 0.402761 | 0.527024 | 1.052133 | 2.129365 | 0.903784 | 1.204978 | -1.451145 | 32.493424 | 0.578874495 | -0.92799356 | 0.353410908 | 0.998865404 | 5.267240 | 4.726544 | 4.990248 | 5.089789 | 4.984388 | 5.002537 | 30 | 55 | 21 | 27 |
| 105378367 | NR_132791 | LOC105378367 | uncharacterized LOC105378367 | lncRNA | . | . | . | . | . | 0.414725 | 0.226530 | 0.332823 | 0.532950 | 0.965082 | 0.523635 | 0.746845 | 1.218527 | 1.346188 | 11.480481 | 0.914569117 | 0.468941505 | 0.639111448 | NA | 3.290730 | 3.719719 | 3.518929 | 3.497285 | 3.510977 | 3.535314 | 14 | 8 | 9 | 15 |
| 105378383 | XR_946104 | LOC105378383 | uncharacterized LOC105378383 | lncRNA | . | . | . | . | . | 0.041145 | 0.104424 | 0.055203 | 0.146857 | 0.095745 | 0.241380 | 0.123873 | 0.335772 | 1.345980 | 9.036590 | 1.069745406 | 0.400709149 | 0.688634274 | NA | 2.950762 | 3.370388 | 3.143424 | 3.174987 | 3.151177 | 3.191174 | 5 | 12 | 5 | 14 |
| 105378421 | XR_001747542 | LOC105378421 | uncharacterized LOC105378421, transcript variant X4 | lncRNA | . | . | . | . | . | 0.019593 | 0.053029 | 0.081453 | 0.076621 | 0.045593 | 0.122579 | 0.182778 | 0.175184 | 1.958750 | 10.087741 | 1.003262551 | 0.966779356 | 0.333654347 | NA | 2.772726 | 3.737849 | 3.282190 | 3.313913 | 3.334188 | 3.320639 | 4 | 11 | 13 | 11 |
| 105378507 | XR_946354 | LOC105378507 | uncharacterized LOC105378507 | lncRNA | . | . | . | . | . | 0.131452 | 0.142659 | 0.063599 | 0.084582 | 0.305894 | 0.329764 | 0.142715 | 0.193387 | -1.858404 | 5.519521 | 1.300202808 | -0.68763465 | 0.491682881 | NA | 2.844134 | 1.947964 | 2.481628 | 2.483667 | 2.465885 | 2.469351 | 8 | 8 | 3 | 4 |
| 105378512 | XR_946358 | LOC105378512 | uncharacterized LOC105378512 | lncRNA | . | . | . | . | . | 0.750825 | 1.213243 | 1.187308 | 0.873159 | 1.747201 | 2.804466 | 2.664284 | 1.996374 | 1.038585 | 114.185275 | 0.339362799 | 0.160946289 | 0.872135704 | 0.998865404 | 6.807746 | 6.862206 | 6.738324 | 6.894430 | 6.882494 | 6.778806 | 99 | 149 | 117 | 92 |
| 105378516 | XR_001747608 | LOC105378516 | uncharacterized LOC105378516, transcript variant X2 | lncRNA | . | . | . | . | . | 0.040089 | 0.019908 | 0.026300 | 0.030356 | 0.093288 | 0.046018 | 0.059017 | 0.069405 | 1.126185 | 4.718967 | 1.397067092 | 0.122716929 | 0.902331257 | NA | 2.147197 | 2.324312 | 2.242091 | 2.226536 | 2.240182 | 2.233606 | 7 | 3 | 5 | 4 |
| 105378525 | XR_946382,XR | LOC105378525 | uncharacterized LOC105378525, transcript variant X3 | lncRNA | . | . | . | . | . | 0.056891 | 0.056757 | 0.082299 | 0.055204 | 0.132388 | 0.131197 | 0.184676 | 0.126217 | 1.164185 | 7.777816 | 1.090175924 | 0.201178586 | 0.840558935 | NA | 2.844134 | 3.066070 | 2.949579 | 2.951794 | 2.964592 | 2.947326 | 8 | 8 | 9 | 6 |
| 105378526 | XR_946386 | LOC105378526 | uncharacterized LOC105378526 | lncRNA | . | . | . | . | . | 0.045847 | 0.049739 | 0.133643 | 0.089338 | 0.106688 | 0.114974 | 0.299892 | 0.204260 | 2.357599 | 15.079494 | 0.806889165 | 1.533442462 | 0.125166873 | NA | 3.166062 | 4.404839 | 3.859192 | 3.862306 | 3.930782 | 3.897081 | 10 | 10 | 22 | 16 |
| 105378550 | XR_946449 | LOC105378550 | uncharacterized LOC105378550 | lncRNA | . | . | . | . | . | 1.795144 | 2.244431 | 2.547672 | 1.856581 | 4.177374 | 5.188103 | 5.716901 | 4.244852 | 1.076854 | 119.163422 | 0.314267906 | 0.33991065 | 0.733923816 | 0.998865404 | 6.841974 | 6.949614 | 6.841410 | 6.913720 | 6.957682 | 6.845987 | 118 | 137 | 125 | 97 |
| 105378560 | XR_946467 | LOC105378560 | uncharacterized LOC105378560 | lncRNA | . | . | . | . | . | 0.015530 | 0.007896 | 0.020930 | 0.009456 | 0.036140 | 0.018251 | 0.046966 | 0.021621 | 1.253092 | 5.525640 | 1.296761534 | 0.251004315 | 0.801810773 | NA | 2.290730 | 2.622519 | 2.461300 | 2.450150 | 2.469719 | 2.452993 | 7 | 4 | 7 | 4 |
| 105378561 | XR_001747657 | LOC105378561 | uncharacterized LOC105378561, transcript variant X2 | lncRNA | . | . | . | . | . | 0.169561 | 0.098218 | 0.186984 | 0.236243 | 0.394576 | 0.227036 | 0.419587 | 0.540142 | 1.110003 | 22.103923 | 0.661621509 | 0.227568098 | 0.819982023 | NA | 4.387715 | 4.540693 | 4.476928 | 4.440184 | 4.471895 | 4.464866 | 29 | 18 | 21 | 21 |
| 105378577 | NR_160731 | LOC105378577 | uncharacterized LOC105378577 | lncRNA | . | . | . | . | . | 0.383244 | 0.214176 | 0.185325 | 0.202358 | 0.891825 | 0.495078 | 0.415865 | 0.462668 | -1.549273 | 9.150573 | 1.020164788 | -0.61910688 | 0.535845952 | NA | 3.474811 | 2.844638 | 3.222758 | 3.197442 | 3.191440 | 3.193732 | 16 | 9 | 6 | 7 |
| 105378580 | XR_002958522 | LINC01409 | long intergenic non-protein coding RNA 1409, transcript variant X4 | lncRNA | . | 50701 | . | . | . | 0.125858 | 0.125376 | 0.069692 | 0.086294 | 0.292876 | 0.289812 | 0.156388 | 0.197301 | -1.616502 | 8.695947 | 1.036629584 | -0.66839262 | 0.503883007 | NA | 3.425187 | 2.733338 | 3.139786 | 3.134665 | 3.120123 | 3.117737 | 13 | 11 | 6 | 6 |
| 105378593 | XR_946826 | LOC105378593 | uncharacterized LOC105378593 | lncRNA | . | . | . | . | . | 0.184579 | 0.035339 | 0.056828 | 0.187380 | 0.429523 | 0.081688 | 0.127520 | 0.428423 | 1.097214 | 4.158134 | 1.525336187 | 0.08774815 | 0.930076849 | NA | 1.987791 | 2.121007 | 2.066342 | 2.046420 | 2.049033 | 2.067309 | 7 | 2 | 2 | 6 |
| 105378616 | NR_146730,NR | LINC01715 | long intergenic non-protein coding RNA 1715 | lncRNA | . | 52502 | . | . | . | 0.342345 | 0.130052 | 0.233157 | 0.644632 | 0.796651 | 0.300620 | 0.523199 | 1.473875 | 1.378855 | 8.919984 | 1.086250483 | 0.426670597 | 0.66961927 | NA | 2.906576 | 3.370388 | 3.161013 | 3.125949 | 3.136288 | 3.176220 | 13 | 4 | 5 | 14 |
| 105378622 | NR_146732 | LINC01778 | long intergenic non-protein coding RNA 1778 | lncRNA | . | 52568 | . | . | . | 0.469066 | 0.406651 | 0.491836 | 0.238181 | 1.091537 | 0.939993 | 1.103666 | 0.544573 | -1.165295 | 13.309840 | 0.853324291 | -0.25863053 | 0.795920328 | NA | 3.838265 | 3.622519 | 3.744228 | 3.732261 | 3.745464 | 3.712970 | 18 | 14 | 14 | 8 |
| 105378642 | XR_001737967 | LOC105378642 | uncharacterized LOC105378642, transcript variant X3 | lncRNA | . | . | . | . | . | 0.065152 | 0.032616 | 0.049060 | 0.096771 | 0.151611 | 0.075392 | 0.110088 | 0.221255 | 1.481598 | 5.498141 | 1.311943316 | 0.432300558 | 0.665522982 | NA | 2.147197 | 2.715150 | 2.455766 | 2.440262 | 2.449053 | 2.466556 | 7 | 3 | 4 | 8 |
| 105378662 | XR_001737994 | LOC105378662 | uncharacterized LOC105378662, transcript variant X2 | lncRNA | . | . | . | . | . | 0.598622 | 1.025317 | 1.093806 | 0.719642 | 1.393018 | 2.370068 | 2.454471 | 1.645378 | 1.086614 | 11.783815 | 0.902070552 | 0.132849603 | 0.894312325 | NA | 3.497331 | 3.617632 | 3.538146 | 3.561960 | 3.565848 | 3.542892 | 10 | 15 | 13 | 9 |
| 105378663 | NR_135048 | LOC105378663 | uncharacterized LOC105378663 | lncRNA | . | . | . | . | . | 0.656795 | 1.303269 | 1.851398 | 1.064444 | 1.528389 | 3.012564 | 4.154484 | 2.433726 | 1.459350 | 18.370584 | 0.749008334 | 0.728063901 | 0.466574471 | NA | 3.901250 | 4.446236 | 4.147161 | 4.187868 | 4.220577 | 4.173005 | 12 | 21 | 24 | 15 |
| 105378669 | XR_001738013 | LOC105378669 | uncharacterized LOC105378669, transcript variant X1 | lncRNA | . | . | . | . | . | 0.127485 | 0.133007 | 0.165801 | 0.141109 | 0.296663 | 0.307451 | 0.372052 | 0.322628 | 1.168043 | 8.759947 | 1.022801912 | 0.219097462 | 0.826574127 | NA | 3.014059 | 3.239024 | 3.122285 | 3.124765 | 3.133537 | 3.125682 | 9 | 9 | 9 | 8 |
| 105378672 | XR_947234 | LOC105378672 | uncharacterized LOC105378672 | lncRNA | . | . | . | . | . | 1.472924 | 1.449665 | 1.958029 | 1.373988 | 3.427556 | 3.350965 | 4.393760 | 3.141459 | 1.141379 | 24.013510 | 0.632648884 | 0.301553916 | 0.762992143 | NA | 4.486115 | 4.678992 | 4.574930 | 4.573572 | 4.604491 | 4.570055 | 26 | 24 | 26 | 20 |
| 105378687 | XR_001738025 | LOC105378687 | uncharacterized LOC105378687, transcript variant X1 | lncRNA | . | . | . | . | . | 1.379517 | 0.885632 | 1.809886 | 1.208798 | 3.210193 | 2.047181 | 4.061332 | 2.763773 | 1.459397 | 42.364817 | 0.506654563 | 1.076419151 | 0.281739841 | 0.998865404 | 5.104211 | 5.653422 | 5.371926 | 5.341258 | 5.467474 | 5.365042 | 44 | 33 | 56 | 34 |
| 105378688 | XR_947283 | LOC105378688 | uncharacterized LOC105378688 | lncRNA | . | . | . | . | . | 0.395929 | 0.571086 | 0.767000 | 0.382931 | 0.921344 | 1.320091 | 1.721126 | 0.875527 | 1.182009 | 20.676625 | 0.705449974 | 0.341966813 | 0.732375868 | NA | 4.243129 | 4.486479 | 4.336607 | 4.365689 | 4.395946 | 4.334688 | 18 | 24 | 26 | 14 |
| 105378698 | XR_947298 | LOC105378698 | uncharacterized LOC105378698 | lncRNA | . | . | . | . | . | 0.366548 | 0.824301 | 0.750727 | 0.327298 | 0.852973 | 1.905410 | 1.684609 | 0.748328 | -1.112866 | 14.218543 | 0.859503718 | -0.17949868 | 0.857546152 | NA | 3.904062 | 3.751299 | 3.799987 | 3.848392 | 3.838735 | 3.796566 | 11 | 22 | 16 | 8 |
| 105378701 | XR_001738039 | LOC105378701 | uncharacterized LOC105378701, transcript variant X1 | lncRNA | . | . | . | . | . | 0.059233 | 0.246983 | 0.151315 | 0.326410 | 0.137838 | 0.570912 | 0.339546 | 0.746299 | 1.557649 | 13.912551 | 0.88423502 | 0.723077136 | 0.469632496 | NA | 3.448488 | 4.079653 | 3.746265 | 3.790653 | 3.767257 | 3.822800 | 7 | 17 | 9 | 22 |
| 105378730 | XR_947362 | LOC105378730 | uncharacterized LOC105378730 | lncRNA | . | . | . | . | . | 1.274924 | 0.937819 | 1.416689 | 0.896079 | 2.966800 | 2.167810 | 3.179008 | 2.048779 | 1.030838 | 49.431032 | 0.470106101 | 0.093207375 | 0.925738815 | 0.998865404 | 5.603263 | 5.651032 | 5.648025 | 5.588419 | 5.666080 | 5.580405 | 65 | 44 | 54 | 36 |
| 105378732 | NR_131923 | LINC01771 | long intergenic non-protein coding RNA 1771 | lncRNA | . | 52561 | . | . | . | 1.074876 | 1.627837 | 1.198658 | 0.993116 | 2.501281 | 3.762818 | 2.689753 | 2.270642 | -1.240955 | 22.403213 | 0.660785584 | -0.4713333 | 0.637402736 | NA | 4.633141 | 4.321310 | 4.474646 | 4.515646 | 4.482970 | 4.467462 | 23 | 32 | 19 | 17 |
| 105378756 | XR_001738082 | LOC105378756 | uncharacterized LOC105378756, transcript variant X2 | lncRNA | . | . | . | . | . | 0.368229 | 0.155101 | 0.249960 | 0.228296 | 0.856884 | 0.358522 | 0.560905 | 0.521973 | -1.169362 | 31.300037 | 0.589168459 | -0.3831183 | 0.701632048 | 0.998865404 | 5.074695 | 4.852980 | 5.020741 | 4.927000 | 4.968293 | 4.941958 | 52 | 24 | 28 | 24 |
| 105378819 | XR_947544 | LOC105378819 | uncharacterized LOC105378819 | lncRNA | . | . | . | . | . | 0.211398 | 0.143053 | 0.183459 | 0.465202 | 0.491933 | 0.330673 | 0.411677 | 1.063629 | 1.788622 | 8.066118 | 1.10786544 | 0.757175733 | 0.448944587 | NA | 2.533720 | 3.370388 | 2.994364 | 2.983671 | 2.989473 | 3.029625 | 8 | 5 | 5 | 14 |
| 105378828 | NR_135837,NR | CLCA4-AS1 | CLCA4 antisense RNA 1 | lncRNA | . | 54055 | . | . | . | 0.352673 | 0.566868 | 0.286036 | 0.380141 | 0.820684 | 1.310340 | 0.641856 | 0.869149 | -1.288626 | 13.222777 | 0.858040342 | -0.42635924 | 0.669846097 | NA | 3.898433 | 3.527725 | 3.717817 | 3.750012 | 3.709514 | 3.729380 | 13 | 20 | 8 | 13 |
| 105378852 | XR_001737796 | LINC02787 | long intergenic non-protein coding RNA 2787, transcript variant X3 | lncRNA | . | 54308 | . | . | . | 0.739687 | 0.982884 | 1.318304 | 1.519514 | 1.721282 | 2.271980 | 2.958235 | 3.474190 | 1.639592 | 30.245151 | 0.565260489 | 1.26196152 | 0.206962633 | 0.970482251 | 4.519313 | 5.231046 | 4.865482 | 4.883822 | 4.926493 | 4.941717 | 24 | 27 | 31 | 37 |
| 105378853 | NR_135038,NR | LINC02609 | long intergenic non-protein coding RNA 2609 | lncRNA | . | 27140 | . | . | . | 1.441513 | 1.756739 | 2.224824 | 1.672686 | 3.354462 | 4.060783 | 4.992441 | 3.824402 | 1.183743 | 150.106544 | 0.274455445 | 0.886686647 | 0.375247629 | 0.998865404 | 7.102746 | 7.346644 | 7.168631 | 7.212910 | 7.296900 | 7.214780 | 149 | 157 | 156 | 137 |
| 105378854 | NR_146610 | LINC01763 | long intergenic non-protein coding RNA 1763 | lncRNA | . | 52554 | . | . | . | 0.207043 | 0.255098 | 0.551886 | 0.354360 | 0.481798 | 0.589670 | 1.238416 | 0.810202 | 1.918744 | 13.144095 | 0.857015151 | 1.097019655 | 0.2726328 | NA | 3.170740 | 4.111291 | 3.672167 | 3.683452 | 3.728434 | 3.699931 | 9 | 11 | 18 | 13 |
| 105378890 | XR_001738175 | LOC105378890 | uncharacterized LOC105378890, transcript variant X2 | lncRNA | . | . | . | . | . | 0.398446 | 0.370897 | 0.319001 | 0.542202 | 0.927200 | 0.857345 | 0.715828 | 1.239683 | 1.055422 | 29.436379 | 0.578007493 | 0.134634498 | 0.892900864 | 0.998865404 | 4.841203 | 4.916862 | 4.878807 | 4.871664 | 4.857516 | 4.905314 | 34 | 30 | 21 | 34 |
| 105378912 | XR_947716,XR | LINC01357 | long intergenic non-protein coding RNA 1357, transcript variant X1 | lncRNA | . | 50588 | . | . | . | 8.940224 | 10.512327 | 10.166109 | 13.557296 | 20.804267 | 24.299717 | 22.812454 | 30.997143 | 1.218891 | 78.733669 | 0.366762949 | 0.778620977 | 0.436203018 | 0.998865404 | 6.150285 | 6.433641 | 6.246268 | 6.287371 | 6.280538 | 6.354832 | 75 | 83 | 65 | 92 |
| 105378913 | XR_001738195 | LOC105378913 | uncharacterized LOC105378913, transcript variant X2 | lncRNA | . | . | . | . | . | 0.100900 | 0.128167 | 0.162816 | 0.112571 | 0.234799 | 0.296264 | 0.365355 | 0.257381 | 1.126252 | 5.267298 | 1.314790513 | 0.130461513 | 0.896201302 | NA | 2.307819 | 2.481107 | 2.387070 | 2.392839 | 2.398631 | 2.386952 | 5 | 6 | 6 | 4 |
| 105378936 | XR_947756 | LOC105378936 | uncharacterized LOC105378936 | lncRNA | . | . | . | . | . | 3.533089 | 3.935774 | 3.815912 | 3.717365 | 8.221643 | 9.097717 | 8.562795 | 8.499313 | -1.005335 | 467.537954 | 0.173439682 | -0.04425681 | 0.964699696 | 0.998865404 | 8.872848 | 8.865021 | 8.838470 | 8.900881 | 8.878215 | 8.855279 | 513 | 531 | 413 | 429 |
| 105378941 | XR_001737691 | LOC105378941 | uncharacterized LOC105378941 | lncRNA | . | . | . | . | . | 0.086976 | 0.080699 | 0.049572 | 0.128502 | 0.202397 | 0.186539 | 0.111237 | 0.293805 | -1.089958 | 1.718421 | 2.237283933 | -0.0555461 | 0.955703407 | NA | 0.844134 | 0.715150 | 0.784731 | 0.785283 | 0.782101 | 0.786536 | 2 | 2 | 1 | 2 |
| 105378948 | NR_168432,NR | LOC105378948 | uncharacterized LOC105378948 | lncRNA | . | . | . | . | . | 0.301866 | 0.365555 | 0.387208 | 0.322666 | 0.702454 | 0.844998 | 0.868883 | 0.737738 | -1.060821 | 14.848204 | 0.792915018 | -0.10742826 | 0.914449239 | NA | 3.934350 | 3.848815 | 3.887973 | 3.900361 | 3.892696 | 3.888064 | 16 | 18 | 13 | 13 |
| 105378949 | NR_168263,NR | LOC105378949 | collagen alpha-2(VIII) chain | lncRNA | . | . | . | . | . | 0.501123 | 0.506310 | 1.363533 | 0.877326 | 1.166135 | 1.170358 | 3.059728 | 2.005903 | 2.078654 | 25.557729 | 0.622248152 | 1.696509317 | 0.089789467 | NA | 4.052320 | 5.109551 | 4.612406 | 4.613763 | 4.709218 | 4.665151 | 19 | 18 | 35 | 27 |
| 105378954 | XR_002958577 | LOC105378954 | uncharacterized LOC105378954 | transcribed_ps | . | . | . | . | . | 0.143971 | 0.264347 | 0.193648 | 0.333627 | 0.335027 | 0.611050 | 0.434541 | 0.762800 | 1.261176 | 13.784709 | 0.847412176 | 0.395049759 | 0.692806166 | NA | 3.611118 | 3.940155 | 3.754339 | 3.786490 | 3.768027 | 3.800787 | 10 | 17 | 10 | 18 |
| 105378988 | XR_001742413 | LOC105378988 | uncharacterized LOC105378988 | lncRNA | . | . | . | . | . | 0.126863 | 0.114225 | 0.059337 | 0.143625 | 0.295215 | 0.264036 | 0.133150 | 0.328382 | -1.207284 | 12.659449 | 0.884038141 | -0.30741266 | 0.75852929 | NA | 3.793789 | 3.517267 | 3.675322 | 3.667232 | 3.641221 | 3.680131 | 17 | 14 | 6 | 15 |
| 105379016 | XR_948416,XR | LOC105379016 | uncharacterized LOC105379016, transcript variant X1 | lncRNA | . | . | . | . | . | 0.143025 | 0.074548 | 0.101886 | 0.068820 | 0.332823 | 0.172322 | 0.228630 | 0.157348 | -1.197683 | 6.121164 | 1.237852115 | -0.21024047 | 0.833479994 | NA | 2.735321 | 2.481107 | 2.627774 | 2.608882 | 2.619191 | 2.607485 | 10 | 5 | 6 | 4 |
| 105379020 | XR_948429 | LOC105379020 | uncharacterized LOC105379020 | lncRNA | . | . | . | . | . | 0.016296 | 0.019415 | 0.005395 | 0.029396 | 0.037922 | 0.044879 | 0.012107 | 0.067210 | 1.212988 | 1.501310 | 2.393488391 | 0.116384722 | 0.907347643 | NA | 0.444631 | 0.715150 | 0.579258 | 0.584004 | 0.580748 | 0.585221 | 1 | 2 | 1 | 2 |
| 105379021 | XR_948435,XR | LOC105379021 | uncharacterized LOC105379021, transcript variant X1 | lncRNA | . | . | . | . | . | 0.016191 | 0.034196 | 0.012635 | 0.019893 | 0.037677 | 0.079046 | 0.028353 | 0.045484 | -1.032087 | 2.216774 | 2.002641667 | -0.02275192 | 0.981848159 | NA | 1.175403 | 1.121007 | 1.148069 | 1.153041 | 1.145613 | 1.154849 | 2 | 3 | 1 | 3 |
| 105379028 | XR_001742725 | LOC105379028 | uncharacterized LOC105379028, transcript variant X7 | lncRNA | . | . | . | . | . | 0.431895 | 0.640942 | 0.396780 | 0.261315 | 1.005037 | 1.481567 | 0.890361 | 0.597465 | -1.396020 | 66.682007 | 0.398154064 | -1.20887705 | 0.226710086 | 0.982008277 | 6.279033 | 5.799803 | 6.085090 | 6.104825 | 6.053243 | 5.983463 | 86 | 87 | 57 | 43 |
| 105379052 | XR_001742772 | LOC105379052 | uncharacterized LOC105379052, transcript variant X2 | lncRNA | . | . | . | . | . | 0.154650 | 0.137856 | 0.094060 | 0.136848 | 0.359877 | 0.318661 | 0.211069 | 0.312886 | -1.215275 | 25.646392 | 0.613762308 | -0.45829242 | 0.646742368 | NA | 4.814705 | 4.532926 | 4.707199 | 4.680992 | 4.663770 | 4.683432 | 36 | 27 | 18 | 24 |
| 105379110 | XR_948649,XR | LOC105379110 | uncharacterized LOC105379110, transcript variant X1 | lncRNA | . | . | . | . | . | 0.293767 | 0.199528 | 0.170211 | 0.356937 | 0.683608 | 0.461218 | 0.381950 | 0.816094 | 1.109169 | 9.882132 | 0.983318583 | 0.15201452 | 0.879175483 | NA | 3.229742 | 3.376188 | 3.306428 | 3.296911 | 3.291061 | 3.321256 | 12 | 9 | 6 | 13 |
| 105379130 | XR_001742844 | LOC105379130 | uncharacterized LOC105379130, transcript variant X3 | lncRNA | . | . | . | . | . | 0.068557 | 0.152930 | 0.458166 | 0.128610 | 0.159535 | 0.353505 | 1.028112 | 0.294053 | 2.380897 | 8.423476 | 1.125606129 | 1.111850205 | 0.266202561 | NA | 2.316288 | 3.568818 | 3.015822 | 3.030379 | 3.082062 | 3.025463 | 4 | 7 | 16 | 5 |
| 105379152 | NR_146771 | LOC105379152 | uncharacterized LOC105379152 | lncRNA | . | . | . | . | . | 0.762073 | 0.979324 | 0.894164 | 1.400044 | 1.773374 | 2.263751 | 2.006477 | 3.201035 | 1.294166 | 61.867049 | 0.419982829 | 0.885804974 | 0.375722629 | 0.998865404 | 5.754924 | 6.123764 | 5.889602 | 5.936354 | 5.919016 | 6.017456 | 55 | 65 | 48 | 79 |
| 105379154 | XR_948723 | LOC105379154 | uncharacterized LOC105379154 | lncRNA | . | . | . | . | . | 0.467451 | 0.529898 | 0.322182 | 0.530354 | 1.087777 | 1.224883 | 0.722966 | 1.212592 | -1.116682 | 11.046814 | 0.92213671 | -0.17266223 | 0.862916934 | NA | 3.544574 | 3.381963 | 3.469267 | 3.472661 | 3.455306 | 3.475761 | 13 | 13 | 7 | 12 |
| 105379173 | XR_002956227 | LOC105379173 | uncharacterized LOC105379173 | lncRNA | . | . | . | . | . | 0.056715 | 0.083658 | 0.093993 | 0.048647 | 0.131979 | 0.193379 | 0.210917 | 0.111225 | 1.013731 | 14.995643 | 0.809978553 | 0.024291051 | 0.980620452 | NA | 3.895610 | 3.917252 | 3.887691 | 3.912074 | 3.919466 | 3.881835 | 14 | 19 | 17 | 10 |
| 105379176 | NR_134248 | LOC105379176 | uncharacterized LOC105379176 | lncRNA | . | . | . | . | . | 0.060099 | 0.098243 | 0.122466 | 0.159641 | 0.139853 | 0.227093 | 0.274809 | 0.365000 | 1.747510 | 4.328051 | 1.451691788 | 0.554732405 | 0.579077708 | NA | 1.658167 | 2.459419 | 2.093865 | 2.099211 | 2.103335 | 2.111485 | 3 | 4 | 4 | 6 |
| 105379183 | NR_134247 | LOC105379183 | uncharacterized LOC105379183 | lncRNA | . | . | . | . | . | 0.251645 | 0.551876 | 0.380781 | 0.204577 | 0.585588 | 1.275685 | 0.854462 | 0.467741 | -1.380277 | 6.642840 | 1.197146727 | -0.38838802 | 0.697728906 | NA | 2.945312 | 2.481107 | 2.725140 | 2.748155 | 2.733005 | 2.721249 | 6 | 11 | 6 | 4 |
| 105379191 | XR_948811 | CTB-1I21.1 | uncharacterized CTB-1I21.1 | lncRNA | . | . | . | . | . | 0.258303 | 0.425926 | 0.370317 | 0.353790 | 0.601081 | 0.984548 | 0.830980 | 0.808899 | 1.052811 | 16.217363 | 0.768693098 | 0.09658732 | 0.923054126 | NA | 3.982758 | 4.055266 | 3.997378 | 4.029899 | 4.019990 | 4.014774 | 14 | 21 | 15 | 15 |
| 105379194 | NR_134244,NR | LOC105379194 | uncharacterized LOC105379194 | lncRNA | . | . | . | . | . | 1.366964 | 0.972727 | 0.780698 | 1.254125 | 3.180980 | 2.248503 | 1.751865 | 2.867407 | -1.159474 | 20.751125 | 0.69015169 | -0.30930955 | 0.757086069 | NA | 4.478589 | 4.263648 | 4.400250 | 4.367875 | 4.353582 | 4.388640 | 30 | 20 | 13 | 22 |
| 105379220 | gene-LOC1053 | LOC105379220 | . | pseudogene | . | . | . | . | . | 1.561385 | 1.393074 | 1.992921 | 1.834574 | 3.633406 | 3.220152 | 4.472058 | 4.194536 | 1.271417 | 74.741336 | 0.366789816 | 0.944511651 | 0.344908251 | 0.998865404 | 6.039657 | 6.387139 | 6.200132 | 6.174453 | 6.258743 | 6.235613 | 80 | 67 | 76 | 75 |
| 105379231 | XR_002956685 | LOC105379231 | uncharacterized LOC105379231, transcript variant X1 | lncRNA | . | . | . | . | . | 1.119170 | 0.929942 | 0.492453 | 0.828011 | 2.604354 | 2.149604 | 1.105053 | 1.893148 | -1.358298 | 14.276683 | 0.828194591 | -0.53344901 | 0.593722801 | NA | 4.039590 | 3.597915 | 3.869162 | 3.830187 | 3.823543 | 3.838359 | 24 | 13 | 9 | 13 |
| 105379241 | XR_948955,XR | LOC105379241 | uncharacterized LOC105379241, transcript variant X2 | lncRNA | . | . | . | . | . | 0.662492 | 0.446613 | 0.481441 | 0.339152 | 1.541646 | 1.032365 | 1.080340 | 0.775432 | -1.320117 | 10.981030 | 0.932319651 | -0.429752 | 0.667376051 | NA | 3.641413 | 3.245375 | 3.477076 | 3.456354 | 3.462123 | 3.444506 | 17 | 11 | 10 | 7 |
| 105379242 | XR_002956695 | LOC105379242 | uncharacterized LOC105379242, transcript variant X3 | lncRNA | . | . | . | . | . | 0.463526 | 0.380683 | 0.409792 | 0.291876 | 1.078645 | 0.879967 | 0.919560 | 0.667340 | -1.216774 | 28.956103 | 0.579558767 | -0.48840771 | 0.625261079 | 0.998865404 | 4.989254 | 4.708721 | 4.877272 | 4.859802 | 4.860613 | 4.830388 | 39 | 32 | 26 | 21 |
| 105379250 | XR_948993 | LOC105379250 | uncharacterized LOC105379250 | lncRNA | . | . | . | . | . | 0.021829 | 0.020092 | 0.060968 | 0.019812 | 0.050797 | 0.046444 | 0.136811 | 0.045297 | 1.670938 | 3.604685 | 1.61049105 | 0.459895863 | 0.64559097 | NA | 1.429096 | 2.175234 | 1.826469 | 1.827376 | 1.845887 | 1.824546 | 3 | 3 | 6 | 2 |
| 105379251 | XR_001746684 | LOC105379251 | uncharacterized LOC105379251, transcript variant X1 | lncRNA | . | . | . | . | . | 0.423360 | 0.485799 | 0.386627 | 0.408055 | 0.985175 | 1.122946 | 0.867579 | 0.932970 | -1.127762 | 104.130638 | 0.314332721 | -0.55184476 | 0.581054719 | 0.998865404 | 6.786648 | 6.612607 | 6.710935 | 6.732257 | 6.676979 | 6.691172 | 123 | 123 | 83 | 94 |
| 105379252 | NR_136310 | LOC105379252 | family with sequence similarity 95 member B1-like | lncRNA | . | . | . | . | . | 0.322442 | 0.228837 | 0.290085 | 0.188730 | 0.750337 | 0.528967 | 0.650941 | 0.431508 | -1.128752 | 13.042751 | 0.858521254 | -0.2035222 | 0.838726893 | NA | 3.787711 | 3.617632 | 3.720761 | 3.696570 | 3.713222 | 3.690240 | 19 | 12 | 13 | 9 |
| 105379273 | XR_002958840 | LOC105379273 | uncharacterized LOC105379273 | lncRNA | . | . | . | . | . | 7.266521 | 7.181098 | 8.260918 | 7.530513 | 16.909492 | 16.599430 | 18.537262 | 17.217625 | 1.070024 | 31.083476 | 0.552874039 | 0.176609752 | 0.859814933 | 0.998865404 | 4.908124 | 5.006356 | 4.953778 | 4.950773 | 4.965405 | 4.956038 | 35 | 32 | 29 | 29 |
| 105379278 | XR_001749751 | LOC105379278 | uncharacterized LOC105379278, transcript variant X2 | lncRNA | . | . | . | . | . | 0.162950 | 0.194732 | 0.221733 | 0.287251 | 0.379190 | 0.450131 | 0.497563 | 0.656765 | 1.413598 | 7.049429 | 1.142049881 | 0.437258845 | 0.661923638 | NA | 2.548173 | 3.044381 | 2.801859 | 2.807938 | 2.809882 | 2.821855 | 6 | 7 | 6 | 9 |
| 105379283 | XR_949295 | LOC105379283 | uncharacterized LOC105379283 | lncRNA | . | . | . | . | . | 0.618269 | 0.855662 | 0.507001 | 0.771924 | 1.438737 | 1.977902 | 1.137695 | 1.764912 | -1.172436 | 22.102374 | 0.668810493 | -0.34316002 | 0.731478082 | NA | 4.578230 | 4.344579 | 4.457525 | 4.490808 | 4.442226 | 4.476763 | 23 | 30 | 14 | 23 |
| 105379326 | XR_001745836 | LOC105379326 | uncharacterized LOC105379326, transcript variant X1 | lncRNA | . | . | . | . | . | 0.178469 | 0.039193 | 0.127705 | 0.140067 | 0.415304 | 0.090597 | 0.286566 | 0.320247 | 1.357524 | 6.226106 | 1.269056112 | 0.347484811 | 0.728227127 | NA | 2.397517 | 2.844638 | 2.643542 | 2.611502 | 2.634953 | 2.637544 | 10 | 2 | 6 | 7 |
| 105379362 | XR_949653,XR | LOC105379362 | uncharacterized LOC105379362, transcript variant X1 | lncRNA | . | . | . | . | . | 2.970414 | 2.830260 | 3.078290 | 2.424489 | 6.912275 | 6.542275 | 6.907594 | 5.543307 | -1.065718 | 531.533592 | 0.179367311 | -0.51194254 | 0.608691227 | 0.998865404 | 9.098686 | 9.007920 | 9.090442 | 9.053532 | 9.106688 | 8.949993 | 649 | 575 | 503 | 423 |
| 105379383 | XR_949689 | LOC105379383 | uncharacterized LOC105379383 | lncRNA | . | . | . | . | . | 0.052129 | 0.067236 | 0.077448 | 0.037014 | 0.121306 | 0.155419 | 0.173791 | 0.084628 | -1.072619 | 8.710812 | 1.047938977 | -0.09651069 | 0.923114983 | NA | 3.170740 | 3.073227 | 3.114704 | 3.125646 | 3.130857 | 3.103845 | 9 | 11 | 10 | 5 |
| 105379395 | XR_949714 | LOC105379395 | uncharacterized LOC105379395 | lncRNA | . | . | . | . | . | 0.245367 | 0.083792 | 0.319958 | 0.287448 | 0.570979 | 0.193688 | 0.717975 | 0.657215 | 1.783013 | 24.634160 | 0.675318605 | 1.235441967 | 0.216666062 | NA | 4.142442 | 4.982216 | 4.610131 | 4.536262 | 4.638524 | 4.625269 | 30 | 10 | 29 | 28 |
| 105379427 | gene-ZNF73P | ZNF73P | zinc finger protein 73, pseudogene | pseudogene | . | 13143 | 194547 | . | . | 0.051287 | 0.074889 | 0.059551 | 0.057706 | 0.119346 | 0.173110 | 0.133631 | 0.131937 | -1.138198 | 5.926954 | 1.240250569 | -0.15057607 | 0.880310142 | NA | 2.658167 | 2.470304 | 2.564687 | 2.574947 | 2.567296 | 2.565412 | 6 | 8 | 5 | 5 |
| 105379428 | XR_001755412 | LOC105379428 | uncharacterized LOC105379428, transcript variant X1 | lncRNA | . | . | . | . | . | 1.985638 | 2.362899 | 1.480745 | 2.708244 | 4.620662 | 5.461948 | 3.322749 | 6.192078 | 1.074306 | 27.019144 | 0.610777637 | 0.169301146 | 0.865559775 | 0.998865404 | 4.705343 | 4.804765 | 4.737785 | 4.760127 | 4.727681 | 4.785494 | 27 | 31 | 18 | 33 |
| 105379444 | XR_950677 | LOC105379444 | uncharacterized LOC105379444 | lncRNA | . | . | . | . | . | 1.611498 | 1.760224 | 2.434352 | 2.456763 | 3.750022 | 4.068835 | 5.462615 | 5.617096 | 1.435674 | 39.908943 | 0.491607154 | 1.061270409 | 0.28856703 | 0.998865404 | 5.034600 | 5.555851 | 5.274600 | 5.287336 | 5.336467 | 5.336166 | 36 | 37 | 41 | 44 |
| 105379477 | NR_134504 | LOC105379477 | uncharacterized LOC105379477 | lncRNA | . | . | . | . | . | 0.364935 | 0.380256 | 0.423256 | 0.275802 | 0.849219 | 0.878979 | 0.949775 | 0.630590 | -1.060970 | 21.811444 | 0.664903106 | -0.12841451 | 0.897820952 | NA | 4.487990 | 4.404839 | 4.445057 | 4.451285 | 4.459537 | 4.425667 | 25 | 25 | 22 | 16 |
| 105379482 | gene-LOC1053 | LOC105379482 | . | pseudogene | . | . | . | . | . | 0.260008 | 0.090756 | 0.196620 | 0.161466 | 0.605050 | 0.209786 | 0.441209 | 0.369174 | 1.007521 | 27.070544 | 0.656794596 | 0.016459104 | 0.986868128 | 0.999197585 | 4.750861 | 4.766401 | 4.803583 | 4.694167 | 4.761726 | 4.740326 | 46 | 15 | 26 | 23 |
| 105379487 | XR_951086 | LINC01670 | long intergenic non-protein coding RNA 1670 | lncRNA | . | 52458 | . | . | . | 0.571738 | 0.539326 | 0.440159 | 0.379642 | 1.330459 | 1.246676 | 0.987705 | 0.868006 | -1.334838 | 14.895819 | 0.794085921 | -0.52471018 | 0.599784677 | NA | 4.089593 | 3.674293 | 3.911679 | 3.908509 | 3.896064 | 3.887069 | 20 | 18 | 12 | 11 |
| 105379490 | XR_951092 | LOC105379490 | uncharacterized LOC105379490 | lncRNA | . | . | . | . | . | 0.101887 | 0.103841 | 0.253733 | 0.161374 | 0.237095 | 0.240033 | 0.569369 | 0.368962 | 1.861419 | 5.140976 | 1.338462045 | 0.669725756 | 0.50303263 | NA | 1.844134 | 2.742346 | 2.337800 | 2.339027 | 2.358569 | 2.346645 | 4 | 4 | 7 | 5 |
| 105379493 | XR_001755117 | CH507-145C22 | uncharacterized CH507-145C22.1, transcript variant X2 | lncRNA | . | . | . | . | . | 1.178039 | 0.954298 | 1.389258 | 0.868069 | 2.741344 | 2.205902 | 3.117454 | 1.984736 | 1.051580 | 122.498004 | 0.328957738 | 0.220569192 | 0.825427891 | 0.998865404 | 6.898322 | 6.973917 | 6.955507 | 6.881506 | 7.020940 | 6.844569 | 152 | 115 | 137 | 88 |
| 105379499 | XR_001755120 | LOC105379499 | uncharacterized LOC105379499, transcript variant X1 | lncRNA | . | . | . | . | . | 1.830275 | 1.726804 | 1.929392 | 1.153473 | 4.259125 | 3.991586 | 4.329499 | 2.637279 | -1.107319 | 140.833881 | 0.322050815 | -0.45667092 | 0.647907594 | 0.998865404 | 7.208183 | 7.063913 | 7.154244 | 7.141387 | 7.213974 | 6.992811 | 173 | 157 | 152 | 87 |
| 105379506 | XR_951149 | LOC105379506 | uncharacterized LOC105379506 | lncRNA | . | . | . | . | . | 38.094383 | 35.644138 | 55.384583 | 34.689674 | 88.647194 | 82.393021 | 124.281395 | 79.313828 | 1.204913 | 1030.077412 | 0.191489912 | 1.404404291 | 0.160198494 | 0.933525128 | 9.867374 | 10.137110 | 9.953572 | 9.899462 | 10.227714 | 9.870424 | 1116 | 970 | 1209 | 808 |
| 105379507 | XR_951150 | LOC105379507 | uncharacterized LOC105379507 | lncRNA | . | . | . | . | . | 42.974098 | 39.257946 | 60.556015 | 39.764679 | 100.002495 | 90.746506 | 135.885941 | 90.917213 | 1.203318 | 1147.782915 | 0.182075388 | 1.466525897 | 0.142505071 | 0.903146012 | 10.024538 | 10.292322 | 10.119498 | 10.046681 | 10.373293 | 10.045797 | 1259 | 1068 | 1322 | 926 |
| 105379508 | XR_951151 | LOC105379508 | uncharacterized LOC105379508 | lncRNA | . | . | . | . | . | 45.034225 | 42.071907 | 64.800209 | 41.570148 | 104.796501 | 97.251091 | 145.409775 | 95.045212 | 1.205218 | 1095.862649 | 0.185955034 | 1.448168149 | 0.14757003 | 0.90884489 | 9.956509 | 10.226573 | 10.043699 | 9.987966 | 10.312074 | 9.970205 | 1188 | 1031 | 1274 | 873 |
| 105379516 | XR_951179 | LOC105379516 | uncharacterized LOC105379516 | lncRNA | . | . | . | . | . | 0.097081 | 0.160100 | 0.130969 | 0.097708 | 0.225911 | 0.370078 | 0.293890 | 0.223397 | -1.127348 | 10.654668 | 0.943034674 | -0.18337926 | 0.85450045 | NA | 3.497331 | 3.324312 | 3.402744 | 3.426444 | 3.415455 | 3.402537 | 10 | 15 | 10 | 8 |
| 105379623 | XR_001742418 | LOC105379623 | uncharacterized LOC105379623, transcript variant X1 | lncRNA | . | . | . | . | . | 0.013990 | 0.034259 | 0.011178 | 0.025185 | 0.032556 | 0.079192 | 0.025083 | 0.057583 | 1.609291 | 1.767977 | 2.241358656 | 0.306254063 | 0.759411212 | NA | 0.444631 | 1.121007 | 0.809537 | 0.814310 | 0.811122 | 0.820450 | 1 | 2 | 1 | 3 |
| 105379695 | NR_149095 | IER3-AS1 | IER3 antisense RNA 1 | lncRNA | . | 53629 | . | . | . | 0.410927 | 0.251305 | 0.136960 | 0.280484 | 0.956243 | 0.580902 | 0.307334 | 0.641292 | -1.416373 | 5.308240 | 1.323580529 | -0.37942662 | 0.704371085 | NA | 2.638039 | 2.134756 | 2.428160 | 2.413292 | 2.408274 | 2.416542 | 9 | 5 | 3 | 5 |
| 105379719 | XR_001745081 | LOC105379719 | uncharacterized LOC105379719 | lncRNA | . | . | . | . | . | 0.441274 | 0.481146 | 0.306595 | 0.424200 | 1.026863 | 1.112191 | 0.687991 | 0.969882 | -1.263857 | 5.624885 | 1.270626224 | -0.26587959 | 0.790331944 | NA | 2.651489 | 2.312263 | 2.498537 | 2.500393 | 2.491909 | 2.495098 | 7 | 7 | 4 | 5 |
| 105379752 | XM_01701293 | LOC105379752 | putative uncharacterized protein FLJ46204 | protein_coding | XP_016868426.1 | . | . | . | . | 0.033475 | 0.042715 | 0.016494 | 0.008442 | 0.077897 | 0.098738 | 0.037012 | 0.019302 | -2.955379 | 6.634647 | 1.240001581 | -1.26075911 | 0.20739565 | NA | 3.307819 | 1.751299 | 2.750258 | 2.761118 | 2.731925 | 2.720664 | 10 | 12 | 4 | 2 |
| 105379807 | NR_144466 | LOC105379807 | uncharacterized LOC105379807 | lncRNA | . | . | . | . | . | 0.652601 | 1.334611 | 1.099981 | 1.872122 | 1.518629 | 3.085014 | 2.468325 | 4.280384 | 1.468852 | 21.214493 | 0.709591928 | 0.781702045 | 0.434389699 | NA | 4.106782 | 4.655341 | 4.348955 | 4.401666 | 4.381695 | 4.437145 | 13 | 25 | 16 | 30 |
| 105416157 | NR_131157 | NKILA | NF-kappaB interacting lncRNA | lncRNA | . | 51599 | . | . | . | 0.126695 | 0.184418 | 0.028791 | 0.170647 | 0.294825 | 0.426290 | 0.064605 | 0.390164 | -1.501459 | 10.873141 | 0.992055847 | -0.5910607 | 0.554479748 | NA | 3.710171 | 3.114084 | 3.444358 | 3.468157 | 3.414371 | 3.459351 | 12 | 17 | 3 | 13 |
| 105447645 | NR_131188 | LOC105447645 | uncharacterized LOC105447645 | lncRNA | . | . | . | . | . | 0.133442 | 0.048294 | 0.146579 | 0.066470 | 0.310524 | 0.111635 | 0.328918 | 0.151976 | 1.133496 | 4.739794 | 1.408037939 | 0.128390785 | 0.897839723 | NA | 2.147197 | 2.336262 | 2.245528 | 2.229959 | 2.248662 | 2.232205 | 7 | 3 | 6 | 3 |
| 105447646 | NR_148381 | PNKY | long intergenic non-protein coding RNA PNKY | lncRNA | . | . | 616328 | . | . | 0.174606 | 0.054044 | 0.286610 | 0.135937 | 0.406315 | 0.124924 | 0.643144 | 0.310804 | 1.942650 | 8.470247 | 1.104311388 | 0.867532637 | 0.385650237 | NA | 2.519121 | 3.486479 | 3.062725 | 3.035109 | 3.089508 | 3.057219 | 10 | 3 | 13 | 7 |
| 105747689 | NR_131917 | NABP1-OT1 | uncharacterized LOC105747689 | lncRNA | . | 55015 | . | . | . | 0.349828 | 0.443155 | 0.513359 | 0.158101 | 0.814064 | 1.024372 | 1.151962 | 0.361480 | -1.199718 | 22.308989 | 0.716616367 | -0.36657674 | 0.713934762 | NA | 4.602488 | 4.345159 | 4.468414 | 4.490398 | 4.505889 | 4.416920 | 25 | 29 | 27 | 9 |
| 106144532 | NR_132369,NR | ARF4-AS1 | ARF4 antisense RNA 1 | lncRNA | . | 51593 | . | . | . | 1.474074 | 1.042411 | 1.227971 | 1.325918 | 3.430231 | 2.409580 | 2.755532 | 3.031553 | 1.044816 | 20.519728 | 0.683151979 | 0.092584444 | 0.926233702 | NA | 4.326585 | 4.390584 | 4.371149 | 4.341766 | 4.354470 | 4.366229 | 27 | 18 | 17 | 21 |
| 106144535 | NR_132374 | LINC01037 | long intergenic non-protein coding RNA 1037 | lncRNA | . | 49025 | . | . | . | 0.383004 | 0.872956 | 0.854143 | 0.637727 | 0.891267 | 2.017877 | 1.916672 | 1.458087 | 1.162496 | 9.322854 | 1.020368784 | 0.212889063 | 0.831413493 | NA | 3.109221 | 3.324312 | 3.192736 | 3.224419 | 3.221377 | 3.208534 | 6 | 13 | 10 | 8 |
| 106146148 | NR_132385 | PSLNR | prostate enriched lncRNA | lncRNA | . | 54405 | . | . | . | 0.524002 | 1.115657 | 1.082518 | 0.603094 | 1.219375 | 2.578891 | 2.429139 | 1.378904 | 1.026978 | 30.345408 | 0.614652644 | 0.062483066 | 0.950178147 | 0.998865404 | 4.904062 | 4.942494 | 4.862833 | 4.953290 | 4.944821 | 4.878788 | 22 | 44 | 34 | 21 |
| 106478913 | gene-RNU6-5P | RNU6-5P | RNA, U6 small nuclear 5, pseudogene | pseudogene | . | 34249 | . | . | . | 0.746054 | 0.425239 | 0.234047 | 0.414630 | 1.736098 | 0.982960 | 0.525195 | 0.948003 | -1.355045 | 1.935531 | 2.127533516 | -0.20603216 | 0.836765791 | NA | 1.156660 | 0.715150 | 0.964669 | 0.961073 | 0.957962 | 0.962362 | 3 | 2 | 1 | 2 |
| 106478914 | gene-RNU6-4P | RNU6-4P | RNA, U6 small nuclear 4, pseudogene | pseudogene | . | 34253 | . | . | . | 0.317561 | 0.573226 | 0.357548 | 0.162939 | 0.738978 | 1.325037 | 0.802328 | 0.372540 | -1.345114 | 1.970934 | 2.116122047 | -0.20212829 | 0.839816435 | NA | 1.175403 | 0.751299 | 0.979035 | 0.983991 | 0.981632 | 0.976028 | 2 | 3 | 2 | 1 |
| 106478919 | gene-RNU6-3P | RNU6-3P | RNA, U6 small nuclear 3, pseudogene | pseudogene | . | 34275 | . | . | . | 0.579335 | 0.620616 | 0.310912 | 0.812806 | 1.348135 | 1.434581 | 0.697677 | 1.858384 | -1.231830 | 2.433884 | 1.916745144 | -0.15693464 | 0.87529636 | NA | 1.429096 | 1.121007 | 1.290581 | 1.291393 | 1.284044 | 1.293236 | 3 | 3 | 1 | 3 |
| 106479023 | NR_160940,NR | H3P4 | H3 histone pseudogene 4 | transcribed_ps | . | 43797 | . | . | . | 1.597631 | 1.701292 | 2.082805 | 1.486872 | 3.717751 | 3.932613 | 4.673754 | 3.399558 | 1.055567 | 71.716211 | 0.374864373 | 0.208124656 | 0.835131636 | 0.998865404 | 6.124120 | 6.203250 | 6.176276 | 6.135958 | 6.176676 | 6.162180 | 87 | 69 | 66 | 67 |
| 106479130 | gene-RN7SKP8 | RN7SKP80 | RN7SK pseudogene 80 | pseudogene | . | 45804 | . | . | . | 0.179255 | 0.099764 | 0.210420 | 0.180534 | 0.417133 | 0.230610 | 0.472175 | 0.412769 | 1.659343 | 1.774228 | 2.21400588 | 0.329995761 | 0.741403165 | NA | 0.413393 | 1.148375 | 0.817239 | 0.813367 | 0.819776 | 0.819065 | 2 | 1 | 2 | 2 |
| 106479413 | gene-RN7SL52 | RN7SL521P | RNA, 7SL, cytoplasmic 521, pseudogene | pseudogene | . | 46537 | . | . | . | 0.522653 | 0.891262 | 0.870623 | 0.634930 | 1.216236 | 2.060192 | 1.953652 | 1.451692 | 1.066916 | 7.000295 | 1.147954231 | 0.081402485 | 0.935121873 | NA | 2.760365 | 2.852980 | 2.794792 | 2.809417 | 2.807801 | 2.800553 | 6 | 9 | 7 | 6 |
| 106479524 | NR_144555 | RN7SL832P | RNA, 7SL, cytoplasmic 832, pseudogene | transcribed_ps | . | 46848 | . | . | . | 0.128946 | 0.301639 | 0.286702 | 0.328674 | 0.300062 | 0.697253 | 0.643350 | 0.751474 | 1.410675 | 17.488928 | 0.765760239 | 0.648225282 | 0.516839252 | NA | 3.861598 | 4.353426 | 4.075849 | 4.128666 | 4.124495 | 4.132169 | 10 | 22 | 17 | 20 |
| 106479836 | gene-RNU6-61 | RNU6-611P | RNA, U6 small nuclear 611, pseudogene | pseudogene | . | 47574 | . | . | . | 0.697190 | 0.747480 | 0.210284 | 0.886734 | 1.622390 | 1.727833 | 0.471871 | 2.027413 | -1.231830 | 2.433884 | 1.916745144 | -0.15693464 | 0.87529636 | NA | 1.429096 | 1.121007 | 1.290581 | 1.291393 | 1.284044 | 1.293236 | 3 | 3 | 1 | 3 |
| 106480341 | gene-SEPTIN14 | SEPTIN14P8 | septin 14 pseudogene 8 | pseudogene | . | 51692 | . | . | . | 0.034008 | 0.029937 | 0.058318 | 0.060408 | 0.079138 | 0.069200 | 0.130864 | 0.138115 | 1.234655 | 1.002957 | 2.868339486 | 0.106022226 | 0.915564717 | NA | -0.155866 | 0.148375 | -0.001261 | -0.000972 | 0.000053 | -0.000315 | 1 | 1 | 1 | 1 |
| 106480343 | gene-SEPTIN14 | SEPTIN14P12 | septin 14 pseudogene 12 | pseudogene | . | 51699 | . | . | . | 0.111327 | 0.171616 | 0.184538 | 0.409062 | 0.259063 | 0.396698 | 0.414097 | 0.935271 | 1.758244 | 5.576849 | 1.298692836 | 0.626888515 | 0.530732316 | NA | 2.019256 | 2.827809 | 2.458064 | 2.463680 | 2.463772 | 2.486036 | 4 | 5 | 4 | 9 |
| 106480348 | gene-NF1P10 | NF1P10 | neurofibromin 1 pseudogene 10 | pseudogene | . | 51737 | . | . | . | 0.512534 | 0.464299 | 0.445668 | 0.595873 | 1.192687 | 1.073248 | 1.000067 | 1.362394 | 1.019606 | 13.093959 | 0.844082369 | 0.033186443 | 0.973525909 | NA | 3.697260 | 3.724273 | 3.714976 | 3.706658 | 3.704590 | 3.719251 | 16 | 13 | 10 | 14 |
| 106480443 | gene-PKMP4 | PKMP4 | pyruvate kinase M1/2 pseudogene 4 | pseudogene | . | 44246 | . | . | . | 0.148747 | 0.104528 | 0.070186 | 0.178720 | 0.346141 | 0.241622 | 0.157494 | 0.408622 | 1.052081 | 3.205155 | 1.667970995 | 0.043913111 | 0.964973664 | NA | 1.644780 | 1.715150 | 1.683106 | 1.679819 | 1.677798 | 1.686624 | 4 | 3 | 2 | 4 |
| 106480492 | gene-RN7SL18 | RN7SL181P | RNA, 7SL, cytoplasmic 181, pseudogene | pseudogene | . | 46197 | . | . | . | 0.549997 | 0.022254 | 0.628340 | 0.217064 | 1.279866 | 0.051442 | 1.409976 | 0.496290 | 1.273595 | 3.505151 | 1.675357258 | 0.208258073 | 0.835027467 | NA | 1.617627 | 1.978782 | 1.808391 | 1.788008 | 1.810173 | 1.793992 | 6 | 1 | 5 | 2 |
| 106480521 | gene-RN7SL60 | RN7SL608P | RNA, 7SL, cytoplasmic 608, pseudogene | pseudogene | . | 46624 | . | . | . | 1.114223 | 0.924438 | 0.860993 | 0.863391 | 2.592843 | 2.136881 | 1.932044 | 1.974040 | -1.210460 | 8.569624 | 1.037087504 | -0.26570168 | 0.790468973 | NA | 3.229742 | 2.955730 | 3.111371 | 3.101770 | 3.100577 | 3.097949 | 12 | 9 | 7 | 7 |
| 106480738 | NR_133663 | OSTN-AS1 | OSTN antisense RNA 1 | lncRNA | . | 41250 | . | . | . | 0.594618 | 0.355645 | 0.344854 | 0.356468 | 1.383701 | 0.822090 | 0.773842 | 0.815023 | -1.392805 | 7.264598 | 1.133608593 | -0.42165628 | 0.673275918 | NA | 3.079678 | 2.602870 | 2.882090 | 2.863907 | 2.860906 | 2.863654 | 12 | 7 | 5 | 6 |
| 106480792 | gene-CHCHD4P | CHCHD4P3 | coiled-coil-helix-coiled-coil-helix domain containing 4 pseudogene | pseudogene | . | 44491 | . | . | . | 0.425311 | 0.540409 | 0.322413 | 0.514213 | 0.989715 | 1.249181 | 0.723486 | 1.175688 | -1.155567 | 10.058432 | 0.969288548 | -0.21520972 | 0.829603811 | NA | 3.432996 | 3.219802 | 3.330570 | 3.341881 | 3.319290 | 3.340204 | 11 | 13 | 6 | 11 |
| 106480951 | gene-RN7SL11 | RN7SL113P | RNA, 7SL, cytoplasmic 113, pseudogene | pseudogene | . | 46129 | . | . | . | 0.167045 | 0.297687 | 0.160435 | 0.150340 | 0.388720 | 0.688116 | 0.360012 | 0.343734 | -1.225524 | 2.469287 | 1.896160988 | -0.15473285 | 0.877031932 | NA | 1.444631 | 1.148375 | 1.301291 | 1.310651 | 1.304063 | 1.303286 | 2 | 4 | 2 | 2 |
| 106481054 | gene-RN7SL48 | RN7SL481P | RNA, 7SL, cytoplasmic 481, pseudogene | pseudogene | . | 46497 | . | . | . | 0.217607 | 0.160772 | 0.716287 | 0.052167 | 0.506380 | 0.371632 | 1.607327 | 0.119273 | 1.774673 | 3.106332 | 1.770289815 | 0.467467804 | 0.640165222 | NA | 1.156660 | 1.993948 | 1.609751 | 1.606193 | 1.629022 | 1.602777 | 3 | 2 | 6 | 1 |
| 106481111 | gene-RN7SL68 | RN7SL689P | RNA, 7SL, cytoplasmic 689, pseudogene | pseudogene | . | 46705 | . | . | . | 0.281258 | 0.818536 | 0.336393 | 0.069792 | 0.654498 | 1.892082 | 0.754857 | 0.159571 | -2.629382 | 4.082765 | 1.582025684 | -0.88160609 | 0.377989859 | NA | 2.562483 | 1.175234 | 2.032994 | 2.055527 | 2.033103 | 2.022250 | 4 | 9 | 3 | 1 |
| 106481526 | gene-RNU6-10 | RNU6-1099P | RNA, U6 small nuclear 1099, pseudogene | pseudogene | . | 48062 | . | . | . | 1.658380 | 1.499558 | 0.979980 | 1.218973 | 3.859118 | 3.466295 | 2.199046 | 2.787036 | -1.507010 | 4.839039 | 1.373456333 | -0.43080322 | 0.666611457 | NA | 2.540965 | 1.947964 | 2.288759 | 2.286269 | 2.276818 | 2.280358 | 7 | 6 | 3 | 4 |
| 106481624 | gene-RNVU1-3 | RNVU1-32 | RNA, variant U1 small nuclear 32 | pseudogene | . | 48448 | . | . | . | 0.161429 | 0.832908 | 0.471128 | 0.537729 | 0.375651 | 1.925303 | 1.057197 | 1.229454 | 1.214180 | 3.038023 | 1.759389422 | 0.159136332 | 0.873561464 | NA | 1.460000 | 1.733338 | 1.582567 | 1.600591 | 1.594894 | 1.593798 | 1 | 5 | 3 | 3 |
| 106481673 | gene-HSPE1P1 | HSPE1P11 | heat shock protein family E (Hsp10) member 1 pseudogene 11 | pseudogene | . | 49330 | . | . | . | 0.730990 | 0.878426 | 1.117005 | 0.788031 | 1.701044 | 2.030521 | 2.506528 | 1.801739 | 1.155170 | 14.546425 | 0.806265882 | 0.258109461 | 0.79632243 | NA | 3.754145 | 3.963455 | 3.846088 | 3.858184 | 3.873001 | 3.849396 | 14 | 16 | 16 | 12 |

| | | | | | | | | | | | | | | | | | | | | | | | | | | | | | | | | | |
|---|---|---|---|---|---|---|---|---|---|---|---|---|---|---|---|---|---|---|---|---|---|---|---|---|---|---|---|---|---|---|---|---|---|
| 106481687 | gene-PDIA3P2 | PDIA3P2 | protein disulfide isomerase family A member 3 pseudogene 2 | pseudogene | . | 49403 | . | . | . | 11.803831 | 13.043636 | 9.659758 | 10.960519 | 27.468000 | 30.150949 | 21.676214 | 25.059923 | -1.226503 | 96.549659 | 0.327399198 | -0.899669 | 0.368296423 | 0.998865404 | 6.733470 | 6.437807 | 6.609133 | 6.638049 | 6.547560 | 6.578507 | 117 | 120 | 71 | 86 |
| 106481733 | gene-SEPTIN14 | SEPTIN14P13 | septin 14 pseudogene 13 | pseudogene | . | 51700 | . | . | . | 0.104139 | 0.054960 | 0.200504 | 0.111970 | 0.242336 | 0.127043 | 0.449924 | 0.256006 | **2.086101** | 2.061721 | 2.085192682 | 0.50873437 | 0.610938427 | NA | 0.413393 | 1.481107 | 1.025095 | 1.021232 | 1.032895 | 1.026997 | 2 | 1 | 3 | 2 |
| 106481953 | gene-RNU1-39 | RNU1-39P | RNA, U1 small nuclear 39, pseudogene | pseudogene | . | 48381 | . | . | . | 0.002430 | 0.002113 | 0.001502 | 0.033734 | 0.005654 | 0.004884 | 0.003371 | 0.077130 | 1.234655 | 1.002957 | 2.868339486 | 0.106022226 | 0.915564717 | NA | -0.155866 | 0.148375 | -0.001261 | -0.000972 | 0.000053 | -0.000315 | 1 | 1 | 1 | 1 |
| 106631777 | NR_145819 | RNA45SN1 | RNA, 45S pre-ribosomal N1 | rRNA | . | 53514 | . | . | . | 35.229092 | 29.814592 | 40.180408 | 36.490406 | 81.979553 | 68.917763 | 90.163673 | 83.430977 | 1.162251 | 14731.986079 | 0.11932408 | 1.817916351 | 0.069076919 | 0.743155658 | 13.734094 | 13.951084 | 13.849024 | 13.700587 | 13.951632 | 13.859223 | 17063 | 13417 | 14505 | 14058 |
| 106631781 | NR_145820 | RNA18SN1 | RNA, 18S ribosomal N1 | rRNA | . | 53515 | . | . | . | 66.899040 | 46.890732 | 57.341007 | 50.649105 | 155.676804 | 108.390030 | 128.671570 | 115.803162 | -1.069071 | 3230.904260 | 0.157945328 | -0.6100703 | 0.541815239 | 0.998865404 | 11.704953 | 11.608892 | 11.819302 | 11.518726 | 11.678317 | 11.568929 | 4536 | 2954 | 2898 | 2732 |
| 106632260 | NR_145821 | RNA5-8SN1 | RNA, 5.8S ribosomal N1 | rRNA | . | 53517 | . | . | . | 0.145811 | 0.109607 | 0.206910 | 0.127657 | 0.339309 | 0.253361 | 0.464300 | 0.291873 | 1.234655 | 1.002957 | 2.868339486 | 0.106022226 | 0.915564717 | NA | -0.155866 | 0.148375 | -0.001261 | -0.000972 | 0.000053 | -0.000315 | 1 | 1 | 1 | 1 |
| 106632264 | NR_145822 | RNA28SN1 | RNA, 28S ribosomal N1 | rRNA | . | 53516 | . | . | . | 112.897865 | 68.433266 | 93.103844 | 115.971878 | 262.717957 | 158.186554 | 208.922318 | 265.155884 | 1.135205 | 15399.286538 | 0.390596557 | 0.468393067 | 0.639503525 | 0.998865404 | 13.816199 | 13.999156 | 14.037588 | 13.606912 | 13.858821 | 14.040627 | 20749 | 11685 | 12753 | 16953 |
| 106632270 | gene-DUX4L35 | DUX4L35 | double homeobox 4 like 35 (pseudogene) | pseudogene | . | 51773 | . | . | . | 0.433979 | 0.576276 | 0.595419 | 0.623868 | 1.009886 | 1.332087 | 1.336103 | 1.426401 | 1.186497 | 14.266834 | 0.809703314 | 0.304690609 | 0.760601803 | NA | 3.706954 | 3.951852 | 3.816793 | 3.832841 | 3.833206 | 3.837760 | 13 | 16 | 13 | 15 |
| 106633800 | NR_132753 | SNORD133 | small nucleolar RNA, C/D box 133 | snoRNA | . | 50414 | . | . | . | 0.475102 | 0.455595 | 0.620479 | 0.147742 | 1.105582 | 1.053128 | 1.392338 | 0.337795 | -1.274956 | 3.211828 | 1.706060013 | -0.20541333 | 0.837249206 | NA | 1.844134 | 1.502474 | 1.683379 | 1.684455 | 1.688269 | 1.672344 | 4 | 4 | 4 | 1 |
| 106633803 | NR_132755 | SNORD134 | small nucleolar RNA, C/D box 134 | snoRNA | . | 50415 | . | . | . | 0.224678 | 0.483644 | 1.175433 | 0.739368 | 0.522834 | 1.117964 | 2.637637 | 1.690476 | **2.473804** | 3.920908 | 1.549808182 | 0.843156827 | 0.39914074 | NA | 1.175403 | 2.481107 | 1.936871 | 1.942026 | 1.960660 | 1.949017 | 2 | 3 | 6 | 4 |
| 106633804 | NR_132756 | SNORD135 | small nucleolar RNA, C/D box 135 | snoRNA | . | 50416 | . | . | . | 1.193618 | 1.272946 | 0.976493 | 0.992825 | 2.777597 | 2.942471 | 2.191221 | 2.269977 | -1.303068 | 3.170174 | 1.676119146 | -0.22785485 | 0.819759083 | NA | 1.844134 | 1.459419 | 1.671887 | 1.672959 | 1.666582 | 1.670577 | 4 | 4 | 2 | 3 |
| 106633805 | NR_132757 | SNORD136 | small nucleolar RNA, C/D box 136 | snoRNA | . | 50417 | . | . | . | 0.033420 | 0.167861 | 0.328819 | 0.119049 | 0.077770 | 0.388018 | 0.737860 | 0.272193 | 1.870326 | 1.290451 | 2.585246895 | 0.349401681 | 0.726787772 | NA | -0.155866 | 0.751299 | 0.348480 | 0.348793 | 0.355090 | 0.349503 | 1 | 1 | 2 | 1 |
| 106633806 | NR_132758 | SNORD142 | small nucleolar RNA, C/D box 142 | snoRNA | . | 50389 | . | . | . | 0.411496 | 0.399208 | 0.242245 | 0.265715 | 0.957567 | 0.922786 | 0.543591 | 0.607527 | -1.215015 | 2.454711 | 1.887245685 | -0.1488804 | 0.881648006 | NA | 1.429096 | 1.148375 | 1.299640 | 1.300454 | 1.298230 | 1.297456 | 3 | 3 | 2 | 2 |
| 106633814 | NR_132768 | SNORA86 | small nucleolar RNA, H/ACA box 86 | snoRNA | . | 50390 | . | . | . | 0.003065 | 0.003282 | 0.004346 | 0.003674 | 0.007133 | 0.007586 | 0.009753 | 0.008400 | 1.234655 | 1.002957 | 2.868339486 | 0.106022226 | 0.915564717 | NA | -0.155866 | 0.148375 | -0.001261 | -0.000972 | 0.000053 | -0.000315 | 1 | 1 | 1 | 1 |
| 106635525 | NR_132774 | SNORA92 | small nucleolar RNA, H/ACA box 92 | snoRNA | . | 50396 | . | . | . | 1.606006 | 0.986877 | 2.156253 | 1.152788 | 3.737243 | 2.281210 | 4.838569 | 2.635714 | 1.251769 | 12.054238 | 0.898009468 | 0.360763116 | 0.718276538 | NA | 3.417335 | 3.746830 | 3.587564 | 3.566229 | 3.604996 | 3.572374 | 15 | 9 | 15 | 9 |
| 106635532 | NR_132782 | SNORA100 | small nucleolar RNA, H/ACA box 100 | snoRNA | . | 50404 | . | . | . | 0.002079 | 0.000990 | 0.001532 | 0.000479 | 0.004838 | 0.002288 | 0.003439 | 0.001096 | 1.234655 | 1.002957 | 2.868339486 | 0.106022226 | 0.915564717 | NA | -0.155866 | 0.148375 | -0.001261 | -0.000972 | 0.000053 | -0.000315 | 1 | 1 | 1 | 1 |
| 106635533 | NR_132783 | SNORA103 | small nucleolar RNA, H/ACA box 103 | snoRNA | . | 51395 | . | . | . | 7.232668 | 8.028900 | 11.529739 | 3.326472 | 16.830717 | 18.559162 | 25.872400 | 7.605583 | -1.042475 | 20.716337 | 0.737188986 | -0.08140782 | 0.935117634 | NA | 4.399719 | 4.345159 | 4.358906 | 4.368701 | 4.403439 | 4.314765 | 23 | 24 | 27 | 9 |
| 106635534 | NR_132784 | SNORA104 | small nucleolar RNA, H/ACA box 104 | snoRNA | . | 51396 | . | . | . | 25.775455 | 22.011675 | 16.278360 | 8.492486 | 59.980541 | 50.880970 | 36.528164 | 19.417059 | -1.932800 | 79.477667 | 0.396851557 | -2.39558651 | **0.016593798** | 0.431581156 | 6.709283 | 5.763302 | 6.406254 | 6.354072 | 6.272188 | 6.151063 | 130 | 104 | 62 | 35 |
| 106635539 | NR_132787 | SNORA105A | small nucleolar RNA, H/ACA box 105A | snoRNA | . | 51397 | . | . | . | 0.016608 | 0.005178 | 0.015651 | 0.005937 | 0.038648 | 0.011969 | 0.035121 | 0.013575 | 1.234655 | 1.002957 | 2.868339486 | 0.106022226 | 0.915564717 | NA | -0.155866 | 0.148375 | -0.001261 | -0.000972 | 0.000053 | -0.000315 | 1 | 1 | 1 | 1 |
| 106635540 | NR_132788 | SNORA105B | small nucleolar RNA, H/ACA box 105B | snoRNA | . | 51398 | . | . | . | 0.005236 | 0.004069 | 0.005968 | 0.003111 | 0.012184 | 0.009405 | 0.013393 | 0.007114 | 1.234655 | 1.002957 | 2.868339486 | 0.106022226 | 0.915564717 | NA | -0.155866 | 0.148375 | -0.001261 | -0.000972 | 0.000053 | -0.000315 | 1 | 1 | 1 | 1 |
| 106635541 | NR_132789 | SNORA105C | small nucleolar RNA, H/ACA box 105C | snoRNA | . | 51399 | . | . | . | 0.093124 | 0.007001 | 0.024742 | 0.000811 | 0.216703 | 0.016183 | 0.055520 | 0.001855 | 1.234655 | 1.002957 | 2.868339486 | 0.106022226 | 0.915564717 | NA | -0.155866 | 0.148375 | -0.001261 | -0.000972 | 0.000053 | -0.000315 | 1 | 1 | 1 | 1 |
| 106635549 | NR_132974 | SNORD131 | small nucleolar RNA, C/D box 131 | snoRNA | . | 50412 | . | . | . | 0.734670 | 0.301805 | 0.933018 | 0.394849 | 1.709608 | 0.697635 | 2.093666 | 0.902775 | 1.257030 | 3.015122 | 1.731267741 | 0.190623073 | 0.848820912 | NA | 1.413393 | 1.751299 | 1.586906 | 1.579188 | 1.591739 | 1.580672 | 4 | 2 | 4 | 2 |
| 106635683 | NR_132980 | SNORD141A | small nucleolar RNA, C/D box 141A | snoRNA | . | 51406 | . | . | . | 0.000696 | 0.000521 | 0.001013 | 0.000577 | 0.001620 | 0.001205 | 0.002274 | 0.001319 | 1.234655 | 1.002957 | 2.868339486 | 0.106022226 | 0.915564717 | NA | -0.155866 | 0.148375 | -0.001261 | -0.000972 | 0.000053 | -0.000315 | 1 | 1 | 1 | 1 |
| 106635684 | NR_132981 | SNORD141B | small nucleolar RNA, C/D box 141B | snoRNA | . | 51407 | . | . | . | 0.022397 | 0.005174 | 0.032621 | 0.057947 | 0.052118 | 0.011960 | 0.073200 | 0.132490 | 1.234655 | 1.002957 | 2.868339486 | 0.106022226 | 0.915564717 | NA | -0.155866 | 0.148375 | -0.001261 | -0.000972 | 0.000053 | -0.000315 | 1 | 1 | 1 | 1 |
| 106660606 | NR_132988 | LOC10666060 | uncharacterized LOC106660606 | lncRNA | . | . | . | . | . | 0.437326 | 0.169686 | 0.294919 | 0.483063 | 1.017676 | 0.392236 | 0.661789 | 1.104466 | 1.239059 | 5.448584 | 1.316786994 | 0.234848032 | 0.814326686 | NA | 2.282109 | 2.592943 | 2.450686 | 2.431118 | 2.439885 | 2.452633 | 8 | 3 | 4 | 7 |
| 106660610 | NR_132993 | MIR5689HG | MIR5689 host gene | lncRNA | . | 52007 | . | . | . | 0.358800 | 0.627499 | 0.174376 | 0.259455 | 0.834942 | 1.450493 | 0.391294 | 0.593213 | **-2.088395** | 7.680909 | 1.12448512 | -0.94478331 | 0.344769516 | NA | 3.377846 | 2.312263 | 2.949454 | 2.972733 | 2.935455 | 2.938434 | 9 | 14 | 4 | 5 |
| 106660612 | NR_125727,NF | LINC00680 | long intergenic non-protein coding RNA 680 | lncRNA | . | 44417 | . | . | . | 2.318530 | 2.678944 | 3.616281 | 3.258008 | 5.395314 | 6.192499 | 8.114829 | 7.449048 | 1.289070 | 200.445264 | 0.239994392 | 1.52641269 | 0.126907119 | 0.874064017 | 7.452185 | 7.818725 | 7.560391 | 7.601457 | 7.725350 | 7.664545 | 192 | 198 | 209 | 198 |
| 106660614 | gene-EGLN1P1 | EGLN1P1 | egl-9 family hypoxia inducible factor 1 pseudogene 1 | pseudogene | . | 44481 | . | . | . | 0.166174 | 0.339087 | 0.595610 | 0.228939 | 0.386694 | 0.783815 | 1.336532 | 0.523441 | 1.592273 | 27.032419 | 0.674283639 | 0.995260843 | 0.319609417 | 0.998865404 | 4.381886 | 5.053812 | 4.668829 | 4.730188 | 4.808842 | 4.689751 | 16 | 30 | 42 | 17 |
| 106699570 | NR_133569 | LOC10669957 | uncharacterized LOC106699570 | lncRNA | . | . | . | . | . | 0.173632 | 0.206792 | 0.310007 | 0.187776 | 0.404048 | 0.478010 | 0.695646 | 0.429329 | 1.272175 | 20.444517 | 0.693389152 | 0.500868469 | 0.616463695 | NA | 4.168403 | 4.517781 | 4.326490 | 4.339682 | 4.377353 | 4.329713 | 19 | 21 | 25 | 16 |
| 106707243 | NM_00131689 | PRH1-TAS2R14 | PRH1-TAS2R14 readthrough | protein_coding | NP_001303822.1 | . | . | . | . | 0.072218 | 0.145068 | 0.302120 | 0.471793 | 0.168053 | 0.335331 | 0.677949 | 1.078699 | **3.303911** | 13.595642 | 0.881286733 | 1.956429038 | 0.050414634 | NA | 2.664814 | 4.381963 | 3.687735 | 3.707182 | 3.741911 | 3.782485 | 5 | 9 | 14 | 24 |
| 106736475 | NR_133914 | PCF11-AS1 | PCF11 antisense RNA 1 | lncRNA | . | 52263 | . | . | . | 0.080035 | 0.091769 | 0.274359 | 0.038116 | 0.186244 | 0.212128 | 0.615653 | 0.087147 | 1.741682 | 8.036870 | 1.159002118 | 0.690663939 | 0.489776756 | NA | 2.548173 | 3.354002 | 2.964817 | 2.970993 | 3.017649 | 2.956346 | 6 | 7 | 15 | 3 |
| 106865373 | NM_00131774 | GET1-SH3BGR | GET1-SH3BGR readthrough | protein_coding | NP_001304673.1;NP_001337229.1 | 54635 | . | . | . | 1.924326 | 0.361228 | 0.696331 | 0.102432 | 4.477985 | 0.834994 | 1.562545 | 0.234200 | **-2.383783** | 37.281578 | 0.704727059 | -1.77835225 | 0.075346019 | 0.766002767 | 5.712327 | 4.468122 | 5.347060 | 5.128105 | 5.180938 | 5.058701 | 93 | 26 | 32 | 7 |
| 106903081 | gene-DUSP8P5 | DUSP8P5 | dual specificity phosphatase 8 pseudogene 5 | pseudogene | . | 45003 | . | . | . | 0.452624 | 0.564798 | 0.527600 | 0.454160 | 1.053276 | 1.305557 | 1.183919 | 1.038383 | -1.044507 | 59.000249 | 0.413901633 | -0.15178003 | 0.879360436 | 0.998865404 | 5.913716 | 5.850899 | 5.862154 | 5.908774 | 5.891860 | 5.861103 | 62 | 72 | 54 | 50 |
| 107075116 | NR_132748 | MIR3142HG | MIR3142 host gene | lncRNA | . | 51944 | . | . | . | 26.325691 | 24.447308 | 44.005661 | 26.496206 | 61.260967 | 56.511044 | 98.747429 | 60.580425 | 1.370862 | 2189.429928 | 0.368553348 | 1.234783866 | 0.216910953 | 0.982008277 | 10.850867 | 11.306056 | 10.977833 | 10.914262 | 11.390303 | 10.965361 | 2214 | 1911 | 2759 | 1773 |
| 107080638 | NM_00131880 | TBC1D7-LOC1C | TBC1D7-LOC100130357 readthrough | protein_coding | NP_001305738.1 | . | . | . | . | 0.226490 | 0.067831 | 0.130358 | 0.070213 | 0.527052 | 0.156794 | 0.292521 | 0.160533 | -1.159559 | 7.866663 | 1.121139998 | -0.19049864 | 0.848918405 | NA | 3.074695 | 2.869520 | 2.989899 | 2.963418 | 2.984768 | 2.958002 | 13 | 6 | 9 | 4 |
| 107105263 | gene-SEPTIN14 | SEPTIN14P2 | septin 14 pseudogene 2 | pseudogene | . | 51686 | . | . | . | 0.225160 | 0.303992 | 0.722405 | 0.388813 | 0.523957 | 0.702692 | 1.621055 | 0.888976 | **2.069473** | 33.210357 | 0.57656473 | 1.819853973 | 0.068781246 | 0.741130004 | 4.434941 | 5.485138 | 4.956877 | 4.987138 | 5.116215 | 5.012216 | 21 | 27 | 51 | 29 |
| 107105265 | gene-SEPTIN14 | SEPTIN14P3 | septin 14 pseudogene 3 | pseudogene | . | 51687 | . | . | . | 0.054221 | 0.076968 | 0.079976 | 0.064961 | 0.126175 | 0.177914 | 0.179464 | 0.148526 | 1.045330 | 5.982762 | 1.231979154 | 0.051915284 | 0.958596195 | NA | 2.548173 | 2.612728 | 2.574413 | 2.580407 | 2.582075 | 2.575150 | 6 | 7 | 6 | 5 |
| 107105266 | gene-SEPTIN14 | SEPTIN14P4 | septin 14 pseudogene 4 | pseudogene | . | 51688 | . | . | . | 0.036130 | 0.138859 | 0.063014 | 0.049511 | 0.084076 | 0.320979 | 0.141401 | 0.113200 | -1.547350 | 6.384498 | 1.253716361 | -0.50234606 | 0.615424115 | NA | 2.956191 | 2.324312 | 2.658487 | 2.698091 | 2.669482 | 2.662773 | 4 | 13 | 5 | 4 |
| 107105271 | gene-SEPTIN14 | SEPTIN14P19 | septin 14 pseudogene 19 | pseudogene | . | 51707 | . | . | . | 0.051708 | 0.030334 | 0.058798 | 0.036018 | 0.120326 | 0.070119 | 0.131942 | 0.082352 | 1.251430 | 4.018079 | 1.503132973 | 0.215268804 | 0.82955775 | NA | 1.832372 | 2.161867 | 2.001207 | 1.993815 | 2.007304 | 1.995939 | 5 | 3 | 5 | 3 |
| 107105276 | gene-SEPTIN14 | SEPTIN14P5 | septin 14 pseudogene 5 | pseudogene | . | 51689 | . | . | . | 0.045896 | 0.048096 | 0.097693 | 0.040339 | 0.106801 | 0.111176 | 0.219221 | 0.092231 | 1.369025 | 5.323106 | 1.313648056 | 0.344954579 | 0.730128518 | NA | 2.166062 | 2.622519 | 2.397111 | 2.398552 | 2.413736 | 2.397003 | 5 | 5 | 7 | 4 |
| 107105278 | gene-SEPTIN14 | SEPTIN14P6 | septin 14 pseudogene 6 | pseudogene | . | 51690 | . | . | . | 0.036333 | 0.032737 | 0.044338 | 0.025630 | 0.084549 | 0.075673 | 0.099493 | 0.058599 | 1.072156 | 3.246808 | 1.662992442 | 0.060442512 | 0.951803201 | NA | 1.644780 | 1.751299 | 1.695592 | 1.692298 | 1.700502 | 1.689427 | 4 | 3 | 4 | 2 |
| 107105280 | gene-SEPTIN14 | SEPTIN14P7 | septin 14 pseudogene 7 | pseudogene | . | 51691 | . | . | . | 0.047149 | 0.032795 | 0.002266 | 0.003218 | 0.109717 | 0.075807 | 0.005084 | 0.007357 | **-3.218571** | 2.334771 | 2.046315186 | -0.82412528 | 0.409868376 | NA | 1.832372 | 0.148375 | 1.252857 | 1.245446 | 1.238133 | 1.237637 | 5 | 3 | 1 | 1 |
| 107105352 | gene-SEPTIN14 | SEPTIN14P10 | septin 14 pseudogene 10 | pseudogene | . | 51697 | . | . | . | 0.008251 | 0.011646 | 0.039873 | 0.004555 | 0.019201 | 0.026920 | 0.089473 | 0.010413 | 1.657496 | 1.809630 | 2.226257051 | 0.32745803 | 0.743321482 | NA | 0.444631 | 1.175234 | 0.832289 | 0.837090 | 0.844244 | 0.833445 | 1 | 2 | 3 | 1 |
| 107105353 | gene-SEPTIN14 | SEPTIN14P11 | septin 14 pseudogene 11 | pseudogene | . | 51698 | . | . | . | 0.031387 | 0.057515 | 0.052901 | 0.029245 | 0.073038 | 0.132949 | 0.118709 | 0.066864 | -1.046143 | 3.991423 | 1.512902382 | -0.04301703 | 0.965687959 | NA | 2.029593 | 1.963455 | 1.986518 | 2.000503 | 1.995847 | 1.989568 | 3 | 6 | 4 | 3 |
| 107105354 | gene-SEPTIN14 | SEPTIN14P14 | septin 14 pseudogene 14 | pseudogene | . | 51701 | . | . | . | 0.071088 | 0.156697 | 0.174918 | 0.120554 | 0.165424 | 0.362212 | 0.392511 | 0.275632 | 1.289535 | 10.900798 | 0.949656643 | 0.386298936 | 0.699275288 | NA | 3.251986 | 3.617632 | 3.412226 | 3.444096 | 3.451924 | 3.428993 | 7 | 14 | 13 | 9 |
| 107126282 | gene-SEPTIN14 | SEPTIN14P15 | septin 14 pseudogene 15 | pseudogene | . | 51702 | . | . | . | 0.037267 | 0.074816 | 0.053148 | 0.081402 | 0.086721 | 0.172940 | 0.119262 | 0.186115 | 1.216502 | 5.506887 | 1.297811257 | 0.217858176 | 0.827539615 | NA | 2.316288 | 2.592943 | 2.446633 | 2.460872 | 2.452311 | 2.465075 | 4 | 7 | 4 | 7 |
| 107126283 | gene-SEPTIN14 | SEPTIN14P16 | septin 14 pseudogene 16 | pseudogene | . | 51704 | . | . | . | 0.050641 | 0.039601 | 0.055523 | 0.065067 | 0.117843 | 0.091540 | 0.124592 | 0.148767 | 1.234129 | 4.495606 | 1.411117721 | 0.215072734 | 0.829710612 | NA | 2.008843 | 2.312263 | 2.164373 | 2.161400 | 2.165561 | 2.168877 | 5 | 4 | 4 | 5 |
| 107126284 | gene-SEPTIN14 | SEPTIN14P18 | septin 14 pseudogene 18 | pseudogene | . | 51705 | . | . | . | 0.052378 | 0.034100 | 0.099745 | 0.053389 | 0.121885 | 0.078823 | 0.223825 | 0.122066 | 1.721953 | 4.859733 | 1.383986919 | 0.566512248 | 0.57104561 | NA | 1.832372 | 2.622519 | 2.263289 | 2.255936 | 2.279785 | 2.263062 | 5 | 3 | 7 | 4 |
| 107126285 | gene-SEPTIN14 | SEPTIN14P20 | septin 14 pseudogene 20 | pseudogene | . | 51706 | . | . | . | 0.041871 | 0.032707 | 0.079733 | 0.020592 | 0.097435 | 0.075605 | 0.178919 | 0.047082 | 1.437025 | 3.821796 | 1.563200556 | 0.33462445 | 0.737908399 | NA | 1.644780 | 2.175234 | 1.918354 | 1.915090 | 1.933625 | 1.912305 | 4 | 3 | 6 | 2 |
| 107126290 | gene-SEPTIN14 | SEPTIN14P23 | septin 14 pseudogene 23 | pseudogene | . | 51709 | . | . | . | 0.005200 | 0.007259 | 0.004920 | 0.003213 | 0.012101 | 0.016779 | 0.011040 | 0.007346 | 1.234655 | 1.002957 | 2.868339486 | 0.106022226 | 0.915564717 | NA | -0.155866 | 0.148375 | -0.001261 | -0.000972 | 0.000053 | -0.000315 | 1 | 1 | 1 | 1 |
| 107133486 | gene-CDK2AP2 | CDK2AP2P2 | cyclin dependent kinase 2 associated protein 2 pseudogene 2 | pseudogene | . | 38493 | . | . | . | 0.040426 | 0.110873 | 0.093101 | 0.116755 | 0.094072 | 0.256288 | 0.208916 | 0.266945 | 1.224497 | 3.023447 | 1.718828416 | 0.169993295 | 0.86501541 | NA | 1.444631 | 1.733338 | 1.582994 | 1.592430 | 1.591096 | 1.590002 | 2 | 4 | 3 | 3 |
| 107133515 | NR_015341 | SMURF2P1-LRF | SMURF2P1-LRRC37BP1 readthrough transcribed pseudogene | transcribed_ps | . | . | . | . | . | 2.999011 | 2.852111 | 3.388174 | 3.414679 | 6.978822 | 6.592783 | 7.602965 | 7.807258 | 1.146353 | 453.300002 | 0.173161925 | 1.137959063 | 0.255137577 | 0.995760118 | 8.722313 | 8.919592 | 8.797246 | 8.760957 | 8.864763 | 8.862318 | 501 | 442 | 422 | 453 |
| 107133516 | gene-SMURF2F | SMURF2P1 | SMAD specific E3 ubiquitin protein ligase 2 pseudogene 1 | pseudogene | . | 44402 | . | . | . | 1.732986 | 2.123843 | 1.570927 | 1.399149 | 4.032729 | 4.909359 | 3.525115 | 3.198987 | -1.328563 | 30.348294 | 0.565115783 | -0.7252785 | 0.468281193 | 0.998865404 | 5.113960 | 4.704117 | 4.928860 | 4.955920 | 4.914313 | 4.901731 | 36 | 41 | 24 | 23 |
| 107133522 | gene-DHX40P1 | DHX40P1 | DEAH-box helicase 40 pseudogene 1 | pseudogene | . | 20687 | . | . | . | 1.606775 | 1.519037 | 1.962017 | 1.108027 | 3.739032 | 3.511321 | 4.402709 | 2.533373 | -1.032144 | 67.541886 | 0.412234835 | -0.11072453 | 0.911834795 | 0.998865404 | 6.098763 | 6.056346 | 6.081651 | 6.068892 | 6.128919 | 6.002899 | 81 | 72 | 74 | 45 |
| 107178919 | gene-RNY3P15 | RNY3P15 | RNY3 pseudogene 15 | pseudogene | . | 50890 | . | . | . | 0.000334 | 0.000660 | 0.000374 | 0.000012 | 0.000777 | 0.001526 | 0.000838 | 0.000027 | 1.234655 | 1.002957 | 2.868339486 | 0.106022226 | 0.915564717 | NA | -0.155866 | 0.148375 | -0.001261 | -0.000972 | 0.000053 | -0.000315 | 1 | 1 | 1 | 1 |
| 107282092 | NM_00132062 | BOLA2-SMG1P | BOLA2-SMG1P6 readthrough | protein_coding | NP_001307551.1;NP_001307552.1;NP_001307553.1;NP_001307554.1;N | 53563 | . | . | . | 14.453970 | 11.435394 | 12.866964 | 9.961110 | 33.634983 | 26.433427 | 28.873092 | 22.774892 | -1.144612 | 732.525091 | 0.179436361 | -1.0859466 | 0.277502619 | 0.998865404 | 9.610349 | 9.416621 | 9.637174 | 9.472088 | 9.548134 | 9.373335 | 1004 | 746 | 671 | 558 |
| 107983949 | XR_932444 | LOC10798394 | uncharacterized LOC107983949 | lncRNA | . | . | . | . | . | 0.360898 | 0.106341 | 0.492861 | 0.109300 | 0.839825 | 0.245812 | 1.105966 | 0.249901 | 1.346206 | 9.345362 | 1.100345027 | 0.38978578 | 0.69669496 | NA | 2.987791 | 3.427357 | 3.218594 | 3.179424 | 3.239170 | 3.182895 | 14 | 4 | 15 | 4 |
| 107983950 | XR_933884 | LOC10798395 | uncharacterized LOC107983950 | lncRNA | . | . | . | . | . | 0.261428 | 0.291417 | 0.516056 | 0.324243 | 0.608354 | 0.673623 | 1.158015 | 0.741343 | 1.528188 | 14.763403 | 0.805318221 | 0.759726953 | 0.447417814 | NA | 3.544574 | 4.158506 | 3.854959 | 3.858617 | 3.900185 | 3.866908 | 13 | 13 | 19 | 13 |
| 107983971 | XR_242293,XR | LOC10798397 | uncharacterized LOC107983971, transcript variant X1 | lncRNA | . | . | . | . | . | 1.343094 | 1.168598 | 1.518803 | 1.603982 | 3.125435 | 2.701266 | 3.408151 | 3.667315 | 1.362675 | 25.900583 | 0.605281976 | 0.737575329 | 0.460772552 | NA | 4.454055 | 4.901253 | 4.677683 | 4.664738 | 4.703995 | 4.704181 | 27 | 22 | 26 | 28 |
| 107983979 | XR_00174142 | LOC10798397 | uncharacterized LOC107983979, transcript variant X10 | lncRNA | . | . | . | . | . | 0.246285 | 0.465918 | 0.505099 | 0.441175 | 0.573116 | 1.076989 | 1.133426 | 1.008695 | 1.264090 | 16.372283 | 0.7701624 | 0.438997143 | 0.660663607 | NA | 3.855800 | 4.191126 | 3.998288 | 4.034788 | 4.033697 | 4.032753 | 12 | 20 | 16 | 17 |
| 107983993 | NM_00136350 | UPK3BL2 | uroplakin 3B like 2 | protein_coding | NP_001350435.1 | 53444 | . | . | . | 4.766788 | 4.478804 | 6.404907 | 3.459928 | 11.092512 | 10.352955 | 14.372427 | 7.910714 | 1.051562 | 277.186351 | 0.267387288 | 0.271270935 | 0.786182659 | 0.998865404 | 8.076959 | 8.151503 | 8.107449 | 8.070269 | 8.262542 | 7.943066 | 322 | 281 | 322 | 186 |
| 107983998 | XM_01152701 | LOC10798399 | uncharacterized LOC107983998, transcript variant X2 | protein_coding | XP_011525320.1;XP_011525323.1 | . | . | . | . | 1.873733 | 1.913320 | 2.131013 | 1.620842 | 4.360254 | 4.422725 | 4.781933 | 3.705864 | 1.005099 | 85.906447 | 0.34829953 | 0.021066895 | 0.983192293 | 0.998865404 | 6.420336 | 6.429040 | 6.420338 | 6.423879 | 6.455790 | 6.390194 | 98 | 93 | 84 | 71 |
| 107984002 | XR_001747662 | LINC02646 | long intergenic non-protein coding RNA 2646, transcript variant X | lncRNA | . | 54130 | . | . | . | 0.132892 | 0.181855 | 0.041403 | 0.178290 | 0.309244 | 0.420367 | 0.092906 | 0.407638 | -1.305302 | 10.718088 | 0.955887938 | -0.40212224 | 0.687594067 | NA | 3.604218 | 3.213338 | 3.424426 | 3.440048 | 3.404395 | 3.434816 | 12 | 15 | 5 | 12 |
| 107984011 | XR_00174670 | LOC10798401 | uncharacterized LOC107984011 | lncRNA | . | . | . | . | . | 13.565982 | 15.272016 | 9.944662 | 10.009816 | 31.568600 | 35.301949 | 22.315533 | 22.886255 | -1.464520 | 960.935294 | 0.142259901 | -3.86917049 | **0.000109206** | **0.012952189** | 10.157500 | 9.606867 | 9.986242 | 10.070272 | 9.753451 | 9.750978 | 1243 | 1300 | 679 | 730 |
| 107984018 | XR_00173847 | LOC10798401 | uncharacterized LOC107984018, transcript variant X2 | lncRNA | . | . | . | . | . | 0.021051 | 0.030209 | 0.075586 | 0.006840 | 0.048988 | 0.069829 | 0.169613 | 0.015639 | 1.670225 | 3.640087 | 1.667184083 | 0.443887728 | 0.657123758 | NA | 1.444631 | 2.188478 | 1.829218 | 1.838756 | 1.857956 | 1.826597 | 2 | 4 | 7 | 1 |
| 107984034 | XR_00174529 | LOC10798403 | uncharacterized LOC107984034 | lncRNA | . | . | . | . | . | 0.779163 | 0.770853 | 0.352463 | 0.730484 | 1.813145 | 1.781862 | 0.790918 | 1.670164 | -1.361044 | 9.706807 | 0.989700859 | -0.4493419 | 0.653185037 | NA | 3.486115 | 3.037079 | 3.292360 | 3.291493 | 3.268028 | 3.289298 | 13 | 12 | 5 | 10 |
| 107984035 | NR_148348 | LOC10798403 | uncharacterized LOC107984035 | lncRNA | . | . | . | . | . | 0.275026 | 0.301611 | 0.392964 | 0.359152 | 0.639996 | 0.697188 | 0.881800 | 0.821158 | 1.280411 | 27.634001 | 0.585708877 | 0.608846497 | 0.54262619 | 0.998865404 | 4.599021 | 4.955730 | 4.764783 | 4.771692 | 4.800523 | 4.791593 | 27 | 27 | 28 | 28 |
| 107984036 | NR_159963,NF | LOC10798403 | uncharacterized LOC107984036 | lncRNA | . | . | . | . | . | 0.453386 | 0.175258 | 0.213182 | 0.096392 | 1.055048 | 0.405118 | 0.478374 | 0.220388 | -1.999090 | 17.602493 | 0.811809585 | -1.23100734 | 0.21832012 | NA | 4.549782 | 3.558654 | 4.203804 | 4.113614 | 4.127834 | 4.090296 | 39 | 14 | 14 | 7 |
| 107984037 | XR_00175541 | LOC10798403 | uncharacterized LOC107984037 | lncRNA | . | . | . | . | . | 0.207385 | 0.263109 | 0.238380 | 0.667034 | 0.482593 | 0.608188 | 0.534919 | 1.525094 | 1.890835 | 5.843515 | 1.278524369 | 0.718816027 | 0.472254281 | NA | 2.019256 | 2.932304 | 2.522311 | 2.527947 | 2.528108 | 2.555057 | 4 | 5 | 4 | 10 |
| 107984041 | XR_00295638 | LOC10798404 | uncharacterized LOC107984041, transcript variant X3 | lncRNA | . | . | . | . | . | 0.000050 | 0.000029 | 0.000710 | 0.014079 | 0.000116 | 0.000068 | 0.001592 | 0.032189 | 1.234655 | 1.002957 | 2.868339486 | 0.106022226 | 0.915564717 | NA | -0.155866 | 0.148375 | -0.001261 | -0.000972 | 0.000053 | -0.000315 | 1 | 1 | 1 | 1 |
| 107984063 | XR_00295939 | LOC10798406 | uncharacterized LOC107984063, transcript variant X4 | lncRNA | . | . | . | . | . | 1.339564 | 1.334198 | 1.135217 | 1.413269 | 3.117218 | 3.084058 | 2.547393 | 3.231271 | -1.041187 | 86.684167 | 0.347008652 | -0.16780421 | 0.866737313 | 0.998865404 | 6.467374 | 6.407395 | 6.426334 | 6.459428 | 6.403145 | 6.459689 | 96 | 101 | 66 | 88 |
| 107984088 | XR_00175446 | LOC10798408 | uncharacterized LOC107984088, transcript variant X2 | lncRNA | . | . | . | . | . | 2.759923 | 2.719768 | 2.313683 | 3.861586 | 6.422453 | 6.286867 | 5.191840 | 8.829061 | 1.042866 | 66.328100 | 0.394196323 | 0.153611651 | 0.87791595 | 0.998865404 | 6.021779 | 6.080715 | 6.049060 | 6.041820 | 6.020139 | 6.088826 | 76 | 69 | 50 | 73 |
| 107984120 | XR_00174420 | LOC10798412 | uncharacterized LOC107984120 | lncRNA | . | . | . | . | . | 0.081095 | 0.096821 | 0.061319 | 0.056549 | 0.188712 | 0.223805 | 0.137598 | 0.129292 | -1.527348 | 23.811372 | 0.636528324 | -0.95993959 | 0.33708562 | NA | 4.847059 | 4.235838 | 4.586100 | 4.608527 | 4.559339 | 4.553087 | 30 | 34 | 17 | 17 |
| 107984126 | XM_01701538 | LOC10798412 | zinc finger protein 658B-like | protein_coding | XP_016870876.1 | . | . | . | . | 0.193391 | 0.132105 | 0.140517 | 0.163643 | 0.450030 | 0.305367 | 0.315316 | 0.374150 | -1.039615 | 7.868315 | 1.083527528 | -0.05172815 | 0.958745308 | NA | 3.003609 | 2.947964 | 2.985822 | 2.971641 | 2.973800 | 2.980941 | 11 | 7 | 6 | 8 |
| 107984129 | XR_00174901 | LOC10798412 | uncharacterized LOC107984129 | lncRNA | . | . | . | . | . | 0.225504 | 0.256515 | 0.453429 | 0.145132 | 0.524756 | 0.592947 | 1.017480 | 0.331828 | 1.253019 | 8.106963 | 1.10629996 | 0.294141542 | 0.768649748 | NA | 2.844134 | 3.175234 | 3.000405 | 3.002645 | 3.030301 | 2.988620 | 8 | 8 | 12 | 4 |
| 107984133 | gene-LOC1079 | LOC10798413 | . | pseudogene | . | . | . | . | . | 0.033491 | 0.058142 | 0.092622 | 0.027000 | 0.077935 | 0.134397 | 0.207842 | 0.061733 | 1.426913 | 3.836371 | 1.562203198 | 0.328317003 | 0.74267199 | NA | 1.658167 | 2.175234 | 1.918311 | 1.923628 | 1.937784 | 1.916451 | 3 | 4 | 6 | 2 |
| 107984138 | XM_01702396 | LOC10798413 | serine/threonine-protein kinase SMG1-like, transcript variant X1 | protein_coding | XP_016879454.1;XP_016879455.1 | . | . | . | . | 0.679024 | 0.792217 | 1.141687 | 0.715913 | 1.580116 | 1.831244 | 2.561913 | 1.636850 | 1.240586 | 73.031987 | 0.390827733 | 0.795801871 | 0.426147204 | 0.998865404 | 6.025658 | 6.338346 | 6.137047 | 6.168313 | 6.262132 | 6.146133 | 70 | 75 | 86 | 59 |
| 107984140 | XR_00175396 | LOC10798414 | uncharacterized LOC107984140, transcript variant X2 | lncRNA | . | . | . | . | . | 0.157598 | 0.049509 | 0.066994 | 0.131799 | 0.366738 | 0.114443 | 0.150332 | 0.301341 | -1.076037 | 6.815379 | 1.200894634 | -0.08804115 | 0.929843972 | NA | 2.820514 | 2.715150 | 2.788091 | 2.756938 | 2.761691 | 2.778967 | 12 | 4 | 4 | 8 |
| 107984224 | XR_00174742 | LOC10798422 | uncharacterized LOC107984224 | lncRNA | . | . | . | . | . | 0.283088 | 0.279651 | 0.238481 | 0.163539 | 0.658757 | 0.646425 | 0.535145 | 0.373912 | -1.441084 | 12.910732 | 0.857841149 | -0.61451332 | 0.538876177 | NA | 3.928838 | 3.404839 | 3.706662 | 3.702931 | 3.693318 | 3.675384 | 18 | 16 | 11 | 8 |
| 107984227 | gene-LOC1079 | LOC10798422 | . | pseudogene | . | . | . | . | . | 0.137389 | 0.264267 | 0.452769 | 0.200330 | 0.319709 | 0.610864 | 1.016000 | 0.458030 | 1.614675 | 13.628716 | 0.873895101 | 0.790992074 | 0.428948617 | NA | 3.381886 | 4.073227 | 3.716666 | 3.748818 | 3.786555 | 3.734650 | 8 | 15 | 20 | 10 |
| 107984243 | XR_00174750 | LOC10798424 | uncharacterized LOC107984243 | lncRNA | . | . | . | . | . | 1.049419 | 1.008075 | 1.026233 | 0.723745 | 2.442042 | 2.330211 | 2.302836 | 1.654756 | -1.168263 | 36.194335 | 0.522738287 | -0.42921206 | 0.667768905 | 0.998865404 | 5.284382 | 5.062477 | 5.192681 | 5.184773 | 5.188572 | 5.144340 | 46 | 41 | 34 | 26 |
| 107984247 | XR_00174751 | LOC10798424 | uncharacterized LOC107984247, transcript variant X1 | lncRNA | . | . | . | . | . | 0.061306 | 0.055338 | 0.123975 | 0.133016 | 0.142661 | 0.127917 | 0.278197 | 0.304126 | **2.058695** | 10.275279 | 0.956851796 | 1.088705805 | 0.276283641 | NA | 2.747897 | 3.790061 | 3.328521 | 3.326619 | 3.359136 | 3.359755 | 8 | 7 | 12 | 13 |
| 107984251 | XR_00174754 | LOC10798425 | uncharacterized LOC107984251 | lncRNA | . | . | . | . | . | 0.195352 | 0.308972 | 0.091503 | 0.324735 | 0.454593 | 0.714202 | 0.205331 | 0.742469 | -1.177075 | 15.412051 | 0.837842151 | -0.2807293 | 0.778918034 | NA | 4.062425 | 3.819320 | 3.933884 | 3.966312 | 3.908500 | 3.969505 | 15 | 22 | 6 | 20 |
| 107984260 | XR_00174756 | LOC10798426 | uncharacterized LOC107984260 | lncRNA | . | . | . | . | . | 0.581708 | 0.983004 | 0.730784 | 0.571904 | 1.353659 | 2.272257 | 1.639858 | 1.307591 | -1.173425 | 5.443177 | 1.300174921 | -0.17745717 | 0.859149305 | NA | 2.555346 | 2.324312 | 2.438719 | 2.453070 | 2.445284 | 2.438648 | 5 | 8 | 5 | 4 |
| 107984261 | XR_00174756 | LOC10798426 | uncharacterized LOC107984261 | lncRNA | . | . | . | . | . | 0.102407 | 0.042209 | 0.162596 | 0.122557 | 0.238305 | 0.097569 | 0.364860 | 0.280211 | 1.936600 | 14.326265 | 0.848850813 | 1.123313922 | 0.261304193 | NA | 3.282109 | 4.242203 | 3.819592 | 3.781407 | 3.852918 | 3.828734 | 16 | 6 | 19 | 15 |
| 107984262 | XR_00174756 | LOC10798426 | uncharacterized LOC107984262 | lncRNA | . | . | . | . | . | 0.978739 | 0.879438 | 1.133757 | 0.999804 | 2.277567 | 2.032861 | 2.544117 | 2.285933 | 1.138336 | 70.833483 | 0.377112256 | 0.495678417 | 0.620121317 | 0.998865404 | 6.049148 | 6.237432 | 6.141716 | 6.112798 | 6.174821 | 6.141948 | 81 | 67 | 70 | 66 |
| 107984280 | XR_00174764 | LINC02666 | long intergenic non-protein coding RNA 2666, transcript variant X | lncRNA | . | 54152 | . | . | . | 0.009558 | 0.006558 | 0.025549 | 0.010410 | 0.022242 | 0.015160 | 0.057331 | 0.023802 | **2.223171** | 9.361879 | 1.041135601 | 1.107078328 | 0.268260064 | NA | 2.533720 | 3.692999 | 3.188105 | 3.177443 | 3.237919 | 3.190972 | 8 | 5 | 16 | 7 |
| 107984285 | XR_00174767 | LOC10798428 | uncharacterized LOC107984285 | lncRNA | . | . | . | . | . | 0.339950 | 0.318181 | 0.306951 | 0.327712 | 0.791077 | 0.735490 | 0.688788 | 0.749274 | -1.056125 | 31.819477 | 0.547335601 | -0.14393379 | 0.885552753 | 0.998865404 | 5.030746 | 4.951852 | 5.001356 | 4.991657 | 4.985466 | 4.993638 | 39 | 34 | 26 | 30 |
| 107984289 | gene-LOC1079 | LOC10798428 | Ras homolog, mTORC1 binding pseudogene | pseudogene | . | . | . | . | . | 1.827904 | 1.550420 | 0.966213 | 1.002201 | 4.253609 | 3.583865 | 2.168153 | 2.291414 | -1.735179 | 56.739438 | 0.423027461 | -1.87951024 | 0.060174857 | 0.719735192 | 6.169511 | 5.375024 | 5.900374 | 5.856726 | 5.769171 | 5.768934 | 90 | 71 | 36 | 39 |
| 107984293 | XR_00174770 | LOC10798429 | uncharacterized LOC107984293 | lncRNA | . | . | . | . | . | 0.330697 | 0.234757 | 0.435344 | 0.548528 | 0.769545 | 0.542651 | 0.976900 | 1.254145 | 1.709952 | 51.962591 | 0.448961134 | 1.723882337 | 0.084729087 | 0.788085288 | 5.260558 | 6.035427 | 5.656564 | 5.602786 | 5.706322 | 5.755037 | 52 | 34 | 51 | 68 |
| 107984316 | XR_00174814 | LOC10798431 | uncharacterized LOC107984316 | lncRNA | . | . | . | . | . | 1.636021 | 1.672161 | 2.103988 | 1.688551 | 3.807088 | 3.865276 | 4.721288 | 3.860672 | 1.131265 | 154.891589 | 0.270112607 | 0.658749606 | 0.510056573 | 0.998865404 | 7.182827 | 7.361853 | 7.243227 | 7.249481 | 7.339340 | 7.249162 | 166 | 158 | 159 | 137 |
| 107984333 | XR_00174824 | LOC10798433 | uncharacterized LOC107984333 | lncRNA | . | . | . | . | . | 0.230881 | 0.271693 | 0.202045 | 0.129360 | 0.537269 | 0.628031 | 0.453383 | 0.295766 | -1.532987 | 14.124970 | 0.82749212 | -0.74483503 | 0.456371454 | NA | 4.094525 | 3.481107 | 3.829948 | 3.842553 | 3.821830 | 3.798967 | 18 | 20 | 12 | 8 |
| 107984345 | NM_00136920 | SMIM38 | small integral membrane protein 38 | protein_coding | NP_001356130.1 | 54074 | . | . | . | 0.089057 | 0.085398 | 0.075889 | 0.074897 | 0.207239 | 0.197401 | 0.170291 | 0.171243 | -1.213493 | 8.584200 | 1.034325131 | -0.26990148 | 0.787236049 | NA | 3.234218 | 2.955730 | 3.109436 | 3.108069 | 3.102648 | 3.100018 | 11 | 10 | 7 | 7 |
| 107984346 | XR_00174828 | LOC10798434 | uncharacterized LOC107984346 | lncRNA | . | . | . | . | . | 0.548382 | 0.913823 | 1.099037 | 0.646448 | 1.276108 | 2.112344 | 2.466207 | 1.478026 | 1.174174 | 19.646591 | 0.715230578 | 0.323875684 | 0.746032154 | NA | 4.175403 | 4.407673 | 4.261234 | 4.298082 | 4.314425 | 4.271467 | 16 | 24 | 23 | 15 |
| 107984370 | XR_00174805 | LOC10798437 | uncharacterized LOC107984370 | lncRNA | . | . | . | . | . | 1.287372 | 1.314682 | 0.594300 | 1.302509 | 2.995767 | 3.038945 | 1.333591 | 2.978033 | -1.371849 | 23.993888 | 0.660035879 | -0.69105628 | 0.489530171 | NA | 4.796818 | 4.335679 | 4.603404 | 4.603230 | 4.540756 | 4.602436 | 32 | 30 | 11 | 26 |
| 107984377 | XR_00174833 | LOC10798437 | uncharacterized LOC107984377 | lncRNA | . | . | . | . | . | 0.489518 | 0.296425 | 0.363766 | 0.417098 | 1.139128 | 0.685199 | 0.816280 | 0.953645 | -1.046239 | 15.238276 | 0.792511241 | -0.08228549 | 0.93441969 | NA | 3.961318 | 3.897226 | 3.947889 | 3.916502 | 3.924674 | 3.933945 | 22 | 13 | 12 | 15 |
| 107984384 | XR_00174837 | LOC10798438 | uncharacterized LOC107984384 | lncRNA | . | . | . | . | . | 0.825428 | 0.687496 | 1.070363 | 0.819482 | 1.920805 | 1.589177 | 2.401864 | 1.873649 | 1.241838 | 33.101760 | 0.543171119 | 0.575282242 | 0.565100429 | 0.998865404 | 4.882837 | 5.197690 | 5.038808 | 5.017180 | 5.074300 | 5.038439 | 37 | 29 | 36 | 30 |
| 107984387 | XR_00174838 | LOC10798438 | uncharacterized LOC107984387 | lncRNA | . | . | . | . | . | 0.069831 | 0.057934 | 0.077521 | 0.071669 | 0.162500 | 0.133917 | 0.173956 | 0.163862 | 1.161178 | 15.464001 | 0.776895265 | 0.277500689 | 0.781395677 | NA | 3.838265 | 4.055266 | 3.948569 | 3.936736 | 3.955249 | 3.950094 | 18 | 14 | 15 | 15 |
| 107984390 | XR_00174839 | LOC10798439 | uncharacterized LOC107984390, transcript variant X2 | lncRNA | . | . | . | . | . | 0.024691 | 0.018419 | 0.042109 | 0.051448 | 0.057458 | 0.042577 | 0.094491 | 0.117629 | **2.113075** | 4.867636 | 1.378538024 | 0.782962859 | 0.433648923 | NA | 1.644780 | 2.724273 | 2.261588 | 2.258404 | 2.272152 | 2.279954 | 4 | 3 | 5 | 7 |
| 107984396 | XR_00174841 | CCDC84-DT | CCDC84 divergent transcript | lncRNA | . | 54215 | . | . | . | 0.298883 | 0.562185 | 0.640602 | 0.458169 | 0.695512 | 1.299517 | 1.437492 | 1.047548 | 1.263492 | 19.458210 | 0.718513513 | 0.469603655 | 0.638638213 | NA | 4.104339 | 4.440712 | 4.239181 | 4.283924 | 4.295619 | 4.266305 | 14 | 24 | 22 | 17 |
| 107984397 | XR_00174841 | LOC10798439 | uncharacterized LOC107984397 | lncRNA | . | . | . | . | . | 0.323446 | 0.196379 | 0.179552 | 0.156879 | 0.752672 | 0.453938 | 0.402910 | 0.358684 | -1.406830 | 5.329067 | 1.315982612 | -0.37420577 | 0.708251232 | NA | 2.638039 | 2.148375 | 2.432264 | 2.417385 | 2.417427 | 2.415859 | 9 | 5 | 4 | 4 |
| 107984421 | XR_00174849 | LOC10798442 | uncharacterized LOC107984421, transcript variant X2 | lncRNA | . | . | . | . | . | 0.642620 | 0.441235 | 0.891443 | 0.662921 | 1.495404 | 1.019935 | 2.000372 | 1.515689 | 1.422033 | 45.926715 | 0.478582544 | 1.061373698 | 0.288520105 | 0.998865404 | 5.243013 | 5.754424 | 5.506538 | 5.448868 | 5.566859 | 5.509232 | 52 | 33 | 54 | 43 |
| 107984433 | XR_00174929 | LOC10798443 | uncharacterized LOC107984433 | lncRNA | . | . | . | . | . | 0.086206 | 0.076945 | 0.058237 | 0.038401 | 0.200606 | 0.177861 | 0.130683 | 0.087799 | -1.610905 | 10.168527 | 0.964840029 | -0.71293854 | 0.475883789 | NA | 3.648138 | 2.963455 | 3.365078 | 3.360505 | 3.347909 | 3.335479 | 15 | 13 | 8 | 6 |
| 107984449 | XM_02444928 | LOC10798444 | uncharacterized LOC107984449, transcript variant X1 | protein_coding | XP_024305057.1;XP_024305058.1 | . | . | . | . | 0.533148 | 0.540176 | 0.727525 | 0.955609 | 1.240656 | 1.248641 | 1.632546 | 2.184886 | 1.548903 | 46.249967 | 0.463272481 | 1.362582588 | 0.1730141 | 0.947431715 | 5.182248 | 5.812271 | 5.481000 | 5.480179 | 5.531190 | 5.579724 | 42 | 39 | 43 | 59 |
| 107984455 | XR_00174941 | LOC10798445 | uncharacterized LOC107984455 | lncRNA | . | . | . | . | . | 0.774545 | 0.447944 | 1.439464 | 0.339842 | 1.802399 | 1.035442 | 3.230116 | 0.777009 | 1.446613 | 15.278857 | 0.86823697 | 0.613517822 | 0.539534002 | NA | 3.638039 | 4.178556 | 3.911863 | 3.883159 | 3.965295 | 3.876133 | 18 | 10 | 25 | 7 |
| 107984459 | XR_00174943 | LOC10798445 | uncharacterized LOC107984459 | lncRNA | . | . | . | . | . | 0.016457 | 0.010592 | 0.007534 | 0.007060 | 0.038296 | 0.024483 | 0.016906 | 0.016141 | **-2.014346** | 3.337728 | 1.664788034 | -0.60687087 | 0.543936609 | NA | 2.156660 | 1.148375 | 1.760619 | 1.753493 | 1.747193 | 1.746336 | 6 | 4 | 2 | 2 |
| 107984461 | XR_00174944 | LOC10798446 | uncharacterized LOC107984461, transcript variant X2 | lncRNA | . | . | . | . | . | 1.649840 | 1.546550 | 1.705550 | 1.780874 | 3.839247 | 3.574921 | 3.827206 | 4.071757 | 1.080679 | 55.910851 | 0.417913504 | 0.267849508 | 0.788815161 | 0.998865404 | 5.747897 | 5.860037 | 5.802689 | 5.789093 | 5.809447 | 5.813672 | 64 | 56 | 50 | 55 |
| 107984485 | XR_00174900 | LOC10798448 | uncharacterized LOC107984485 | lncRNA | . | . | . | . | . | 0.528406 | 0.487334 | 0.810467 | 0.514779 | 1.229622 | 1.126493 | 1.818665 | 1.176981 | 1.285889 | 76.821714 | 0.382287926 | 0.94893308 | 0.342654649 | 0.998865404 | 6.069199 | 6.434609 | 6.230300 | 6.208142 | 6.342519 | 6.220446 | 81 | 69 | 92 | 63 |
| 107984489 | XR_00174899 | LOC10798448 | uncharacterized LOC107984489 | lncRNA | . | . | . | . | . | 0.541728 | 0.524814 | 0.745420 | 0.411318 | 1.260623 | 1.213132 | 1.672700 | 0.940430 | 1.069593 | 49.196302 | 0.471558773 | 0.205832582 | 0.836921689 | 0.998865404 | 5.569400 | 5.669809 | 5.610953 | 5.602242 | 5.670256 | 5.564580 | 56 | 50 | 57 | 34 |
| 107984495 | XR_00174896 | LOC10798449 | uncharacterized LOC107984495, transcript variant X1 | lncRNA | . | . | . | . | . | 0.902046 | 0.916978 | 1.138127 | 1.342615 | 2.099099 | 2.119635 | 2.553925 | 3.069732 | 1.350672 | 26.870211 | 0.596358341 | 0.727210016 | 0.467097311 | NA | 4.515640 | 4.947964 | 4.723115 | 4.725880 | 4.744356 | 4.770449 | 26 | 25 | 24 | 32 |
| 107984504 | XR_00174908 | CPNE8-AS1 | CPNE8 antisense RNA 1 | lncRNA | . | 53937 | . | . | . | 1.955864 | 1.990112 | 1.530109 | 1.364582 | 4.551376 | 4.600233 | 3.433519 | 3.119955 | -1.378749 | 69.255524 | 0.381163788 | -1.21564494 | 0.224120209 | 0.982008277 | 6.326640 | 5.864169 | 6.150320 | 6.152339 | 6.089744 | 6.062453 | 92 | 87 | 54 | 51 |
| 107984529 | XR_00174920 | LOC10798452 | uncharacterized LOC107984529 | lncRNA | . | . | . | . | . | 0.550952 | 0.407551 | 0.845324 | 0.580930 | 1.282088 | 0.942072 | 1.896884 | 1.328227 | 1.452618 | 13.156307 | 0.849335482 | 0.634208513 | 0.525944752 | NA | 3.421266 | 3.963455 | 3.702877 | 3.689819 | 3.729411 | 3.705871 | 14 | 10 | 16 | 12 |
| 107984551 | XR_00174996 | LOC10798455 | uncharacterized LOC107984551 | lncRNA | . | . | . | . | . | 7.615652 | 9.158854 | 4.351830 | 8.756968 | 17.721933 | 21.171101 | 9.765382 | 20.021769 | -1.286854 | 123.308429 | 0.344416108 | -1.05642199 | 0.290775462 | 0.998865404 | 7.118172 | 6.750752 | 6.947598 | 7.014009 | 6.786514 | 6.990981 | 146 | 163 | 63 | 134 |
| 107984578 | XR_00174986 | LOC10798457 | uncharacterized LOC107984578, transcript variant X1 | lncRNA | . | . | . | . | . | 0.172848 | 0.210409 | 0.420219 | 0.149232 | 0.402225 | 0.486369 | 0.942960 | 0.341201 | 1.137831 | 22.618407 | 0.673369786 | 0.276647516 | 0.782050776 | NA | 4.401710 | 4.590940 | 4.474718 | 4.492411 | 4.528106 | 4.462331 | 22 | 25 | 28 | 15 |
| 107984583 | XR_00175003 | LOC10798458 | uncharacterized LOC107984583 | lncRNA | . | . | . | . | . | 0.051296 | 0.088940 | 0.079243 | 0.030517 | 0.119368 | 0.205588 | 0.177820 | 0.069773 | -1.310628 | 10.373847 | 0.982890304 | -0.39705139 | 0.691329568 | NA | 3.555346 | 3.168566 | 3.363991 | 3.391787 | 3.381495 | 3.349514 | 10 | 16 | 11 | 5 |
| 107984638 | XM_02444979 | LOC10798463 | arf-GAP with GTPase, ANK repeat and PH domain-containing prote | protein_coding | XP_024305566.1 | . | . | . | . | 0.116116 | 0.228454 | 0.058753 | 0.152758 | 0.270207 | 0.528082 | 0.131841 | 0.349262 | -1.403292 | 6.580360 | 1.21274852 | -0.40306425 | 0.686900956 | NA | 2.945312 | 2.448451 | 2.715331 | 2.738295 | 2.708063 | 2.725706 | 6 | 11 | 3 | 7 |
| 107984643 | XR_00175062 | LOC10798464 | uncharacterized LOC107984643 | lncRNA | . | . | . | . | . | 0.186718 | 0.170063 | 0.227598 | 0.491004 | 0.434501 | 0.393108 | 0.510723 | 1.122623 | **2.070179** | 8.922348 | 1.043378291 | 1.006112285 | 0.31436156 | NA | 2.540965 | 3.587955 | 3.127860 | 3.125646 | 3.137395 | 3.172060 | 7 | 6 | 7 | 15 |
| 107984655 | XR_00175079 | LOC10798465 | uncharacterized LOC107984655 | lncRNA | . | . | . | . | . | 0.593768 | 0.448533 | 0.658887 | 0.437965 | 1.381722 | 1.036805 | 1.478523 | 1.001356 | 1.023304 | 17.629567 | 0.742103926 | 0.044784854 | 0.964278798 | NA | 4.121102 | 4.158506 | 4.147225 | 4.124543 | 4.154180 | 4.120850 | 23 | 16 | 19 | 13 |
| 107984658 | XR_00175090 | LOC10798465 | uncharacterized LOC107984658 | lncRNA | . | . | . | . | . | 2.661076 | 2.519690 | 2.536316 | 2.816084 | 6.192434 | 5.824375 | 5.691420 | 6.438641 | 1.017933 | 173.048960 | 0.252420895 | 0.101585659 | 0.919085563 | 0.998865404 | 7.422273 | 7.447688 | 7.444882 | 7.417050 | 7.419674 | 7.456995 | 204 | 179 | 145 | 171 |
| 107984659 | XR_00175090 | LOC10798465 | uncharacterized LOC107984659 | lncRNA | . | . | . | . | . | 0.498000 | 0.391061 | 0.495341 | 0.414763 | 1.158866 | 0.903956 | 1.111531 | 0.948306 | 1.019425 | 42.862433 | 0.481475243 | 0.057646967 | 0.954029838 | 0.998865404 | 5.406572 | 5.436556 | 5.439175 | 5.397375 | 5.435798 | 5.407245 | 55 | 40 | 41 | 37 |
| 107984660 | XR_00175063 | LOC10798466 | uncharacterized LOC107984660, transcript variant X8 | lncRNA | . | . | . | . | . | 0.109034 | 0.244425 | 0.077597 | 0.162091 | 0.253727 | 0.564999 | 0.174125 | 0.370601 | -1.377627 | 11.744368 | 0.940619356 | -0.49136293 | 0.62316978 | NA | 3.769646 | 3.300113 | 3.534209 | 3.586350 | 3.531521 | 3.556859 | 9 | 21 | 6 | 12 |
| 107984667 | XR_00175066 | LOC10798466 | uncharacterized LOC107984667 | lncRNA | . | . | . | . | . | 9.760788 | 8.612059 | 13.377614 | 11.790252 | 22.713755 | 19.907158 | 30.018978 | 26.957010 | 1.346908 | 199.059318 | 0.242273148 | 1.773417833 | 0.076159496 | 0.767975096 | 7.405606 | 7.836454 | 7.585700 | 7.531543 | 7.724124 | 7.660718 | 208 | 171 | 212 | 200 |
| 107984740 | XR_00175171 | LOC10798474 | uncharacterized LOC107984740 | lncRNA | . | . | . | . | . | 0.041234 | 0.032241 | 0.037017 | 0.026759 | 0.095953 | 0.074526 | 0.083066 | 0.061182 | -1.072000 | 5.182339 | 1.322372303 | -0.07585255 | 0.939536404 | NA | 2.421266 | 2.324312 | 2.378555 | 2.371759 | 2.376811 | 2.370200 | 7 | 5 | 5 | 4 |
| 107984749 | XR_00175163 | LOC10798474 | uncharacterized LOC107984749, transcript variant X1 | lncRNA | . | . | . | . | . | 0.051614 | 0.031039 | 0.017560 | 0.013387 | 0.120107 | 0.071748 | 0.039404 | 0.030609 | **-2.211347** | 3.554839 | 1.628026365 | -0.70325993 | 0.481893779 | NA | 2.290730 | 1.148375 | 1.855928 | 1.844755 | 1.838523 | 1.837645 | 7 | 4 | 2 | 2 |
| 107984751 | XM_01702280 | LOC10798475 | uncharacterized LOC107984751, transcript variant X1 | protein_coding | XP_016878292.1;XP_016878293.1 | . | . | . | . | 0.007964 | 0.012701 | 0.010000 | 0.015980 | 0.018532 | 0.029360 | 0.022439 | 0.036537 | 1.221395 | 5.008534 | 1.348182172 | 0.214013852 | 0.830536255 | NA | 2.175403 | 2.459419 | 2.312483 | 2.322378 | 2.317994 | 2.326047 | 4 | 6 | 4 | 6 |
| 107984754 | XR_00175159 | LOC10798475 | uncharacterized LOC107984754 | lncRNA | . | . | . | . | . | 1.103125 | 1.238208 | 1.719307 | 1.102169 | 2.567017 | 2.862174 | 3.858075 | 2.519978 | 1.191180 | 243.178045 | 0.253113738 | 0.997145035 | 0.318694116 | 0.998865404 | 7.793449 | 8.047150 | 7.845454 | 7.896177 | 8.064873 | 7.837756 | 242 | 252 | 281 | 193 |
| 107984760 | XR_00175148 | LOC10798476 | uncharacterized LOC107984760 | lncRNA | . | . | . | . | . | 1.143007 | 1.078667 | 0.388343 | 0.858949 | 2.659824 | 2.493387 | 0.871429 | 1.963885 | -1.766922 | 9.103090 | 1.028825644 | -0.79822921 | 0.424737491 | NA | 3.540965 | 2.715150 | 3.209902 | 3.205044 | 3.176468 | 3.193445 | 14 | 12 | 4 | 8 |
| 107984761 | gene-LOC1079 | LOC10798476 | . | pseudogene | . | . | . | . | . | 0.024428 | 0.128214 | 0.040048 | 0.025865 | 0.056844 | 0.296372 | 0.089867 | 0.059137 | -1.225404 | 1.234643 | 2.621948492 | -0.11184694 | 0.910944771 | NA | 0.444631 | 0.148375 | 0.303453 | 0.308168 | 0.304847 | 0.304457 | 1 | 2 | 1 | 1 |
| 107984773 | NR_147199,NF | LINC01852 | long intergenic non-protein coding RNA 1852 | lncRNA | . | 52668 | . | . | . | 0.386661 | 0.221903 | 0.089384 | 0.546630 | 0.899775 | 0.512938 | 0.200575 | 1.249804 | -1.220988 | 12.046564 | 0.971299242 | -0.29656082 | 0.766801831 | NA | 3.728992 | 3.437399 | 3.622434 | 3.569950 | 3.558924 | 3.612340 | 22 | 8 | 4 | 16 |
| 107984813 | NR_158165 | LOC10798481 | uncharacterized LOC107984813 | lncRNA | . | . | . | . | . | 0.096543 | 0.133318 | 0.107534 | 0.077102 | 0.224659 | 0.308170 | 0.241303 | 0.176285 | -1.454615 | 3.422687 | 1.626206149 | -0.33245338 | 0.739546934 | NA | 2.019256 | 1.481107 | 1.779562 | 1.784988 | 1.779442 | 1.773473 | 4 | 5 | 3 | 2 |
| 107984814 | XM_01702392 | LOC10798481 | uncharacterized LOC107984814, transcript variant X1 | protein_coding | XP_016879409.1 | . | . | . | . | 0.336663 | 0.249603 | 0.471142 | 0.170927 | 0.783428 | 0.576968 | 1.057230 | 0.390805 | 1.099507 | 15.020225 | 0.833005495 | 0.164292751 | 0.869500688 | NA | 3.835322 | 3.978782 | 3.905701 | 3.885765 | 3.931971 | 3.870859 | 19 | 13 | 20 | 8 |
| 107984817 | XM_01702397 | LOC10798481 | uncharacterized LOC107984817 | protein_coding | XP_016879461.1 | . | . | . | . | 0.091358 | 0.082127 | 0.115190 | 0.061215 | 0.212593 | 0.189840 | 0.258483 | 0.139961 | 1.026618 | 7.728260 | 1.10027347 | 0.034446008 | 0.972521496 | NA | 2.928838 | 2.971139 | 2.946563 | 2.944703 | 2.957478 | 2.935459 | 9 | 8 | 9 | 5 |
| 107984836 | XR_00175236 | LOC10798483 | uncharacterized LOC107984836 | lncRNA | . | . | . | . | . | 0.412323 | 0.695493 | 0.689998 | 0.810877 | 0.959493 | 1.607663 | 1.548335 | 1.853974 | 1.350935 | 40.860661 | 0.506156937 | 0.857359376 | 0.391246303 | 0.998865404 | 5.121228 | 5.552014 | 5.281554 | 5.352954 | 5.352913 | 5.377103 | 30 | 47 | 38 | 47 |
| 107984862 | XM_01702396 | LOC10798486 | uncharacterized LOC107984862 | protein_coding | XP_016879458.1 | . | . | . | . | 0.015524 | 0.058461 | 0.066895 | 0.037106 | 0.036125 | 0.135134 | 0.150111 | 0.084839 | 1.230352 | 5.583943 | 1.314109443 | 0.227584438 | 0.819969319 | NA | 2.324708 | 2.622519 | 2.454920 | 2.477683 | 2.479964 | 2.463209 | 3 | 8 | 7 | 4 |
| 107984865 | XR_00175208 | LOC10798486 | uncharacterized LOC107984865 | lncRNA | . | . | . | . | . | 0.076814 | 0.051706 | 0.056494 | 0.015080 | 0.178749 | 0.119521 | 0.126771 | 0.034478 | -1.764153 | 7.004604 | 1.202401884 | -0.68111672 | 0.49579764 | NA | 3.156660 | 2.348113 | 2.828664 | 2.814724 | 2.817375 | 2.791030 | 12 | 8 | 7 | 2 |
| 107984875 | XR_00175211 | LOC10798487 | uncharacterized LOC107984875 | lncRNA | . | . | . | . | . | 3.291985 | 3.776872 | 4.044706 | 3.658144 | 7.660584 | 8.730408 | 9.076203 | 8.363911 | 1.078253 | 129.318990 | 0.288179661 | 0.377178253 | 0.70604114 | 0.998865404 | 6.959398 | 7.068135 | 6.975526 | 7.021721 | 7.045006 | 7.006225 | 134 | 143 | 123 | 119 |
| 107984890 | XR_00175229 | LOC10798489 | uncharacterized LOC107984890 | lncRNA | . | . | . | . | . | 0.120362 | 0.074135 | 0.176627 | 0.384225 | 0.280088 | 0.171367 | 0.396345 | 0.878484 | **2.596597** | 7.189351 | 1.178464729 | 1.168148747 | 0.24274677 | NA | 1.998355 | 3.376188 | 2.815996 | 2.804681 | 2.824071 | 2.854790 | 6 | 3 | 6 | 13 |
| 107984896 | gene-LOC1079 | LOC10798489 | . | pseudogene | . | . | . | . | . | 0.011154 | 0.007705 | 0.002010 | 0.019674 | 0.025955 | 0.017810 | 0.004510 | 0.044982 | 1.234655 | 1.002957 | 2.868339486 | 0.106022226 | 0.915564717 | NA | -0.155866 | 0.148375 | -0.001261 | -0.000972 | 0.000053 | -0.000315 | 1 | 1 | 1 | 1 |
| 107984910 | XR_00173787 | LOC10798491 | uncharacterized LOC107984910 | lncRNA | . | . | . | . | . | 0.121984 | 0.124695 | 0.103519 | 0.264118 | 0.283861 | 0.288237 | 0.232293 | 0.603875 | 1.452185 | 11.517535 | 0.926841285 | 0.580709384 | 0.561436336 | NA | 3.234218 | 3.768175 | 3.508176 | 3.507009 | 3.502304 | 3.550138 | 11 | 10 | 7 | 18 |
| 107984913 | XR_00173788 | LOC10798491 | uncharacterized LOC107984913 | lncRNA | . | . | . | . | . | 0.223360 | 0.185703 | 0.534321 | 0.295788 | 0.519769 | 0.429261 | 1.199002 | 0.676285 | 1.979290 | 19.356337 | 0.723807413 | 1.36083612 | 0.173565484 | NA | 3.697260 | 4.686013 | 4.226093 | 4.218150 | 4.306687 | 4.245342 | 16 | 13 | 29 | 17 |
| 107984921 | NR_147210,NF | LINC01772 | long intergenic non-protein coding RNA 1772 | lncRNA | . | 52562 | . | . | . | 0.014203 | 0.032208 | 0.050001 | 0.025874 | 0.033052 | 0.074451 | 0.112202 | 0.059158 | 1.446856 | 3.331767 | 1.669192382 | 0.319269252 | 0.749522347 | NA | 1.444631 | 1.978782 | 1.712848 | 1.722338 | 1.731295 | 1.715068 | 2 | 4 | 5 | 2 |
| 107984948 | XR_00173802 | LOC10798494 | uncharacterized LOC107984948, transcript variant X1 | lncRNA | . | . | . | . | . | 0.168976 | 0.215969 | 0.352274 | 0.349277 | 0.393214 | 0.499221 | 0.790492 | 0.798580 | 1.798531 | 8.178576 | 1.065322664 | 0.794894419 | 0.426674921 | NA | 2.548173 | 3.393446 | 3.004191 | 3.010363 | 3.027591 | 3.029195 | 6 | 7 | 9 | 10 |
| 107984950 | XR_00173803 | LOC10798495 | uncharacterized LOC107984950 | lncRNA | . | . | . | . | . | 0.466784 | 0.667194 | 0.582536 | 0.840753 | 1.086225 | 1.542248 | 1.307195 | 1.922281 | 1.231340 | 65.817662 | 0.411027929 | 0.730436334 | 0.465123516 | 0.998865404 | 5.884337 | 6.181222 | 5.969116 | 6.045192 | 6.011751 | 6.096896 | 56 | 75 | 52 | 80 |
| 107984952 | XR_00173803 | LOC10798495 | uncharacterized LOC107984952 | lncRNA | . | . | . | . | . | 0.558116 | 0.501984 | 0.788003 | 0.570192 | 1.298758 | 1.160359 | 1.768255 | 1.303677 | 1.256700 | 41.952917 | 0.487395726 | 0.676330686 | 0.498830714 | 0.998865404 | 5.215216 | 5.547133 | 5.372215 | 5.356992 | 5.427544 | 5.374155 | 45 | 38 | 47 | 37 |
| 107984958 | XR_00173805 | DHCR24-DT | DHCR24 divergent transcript | lncRNA | . | 53969 | . | . | . | 0.631707 | 0.876207 | 0.772961 | 1.121631 | 1.470009 | 2.025392 | 1.734502 | 2.564475 | 1.240774 | 68.075668 | 0.4028711 | 0.772554822 | 0.439785891 | 0.998865404 | 5.926838 | 6.234884 | 6.019853 | 6.089314 | 6.060908 | 6.148168 | 59 | 76 | 54 | 83 |
| 107984969 | XR_00175309 | LOC10798496 | uncharacterized LOC107984969 | lncRNA | . | . | . | . | . | 0.243574 | 0.084623 | 0.309444 | 0.357212 | 0.566807 | 0.195610 | 0.694383 | 0.816722 | 1.948698 | 15.014229 | 0.831697744 | 1.157284094 | 0.24715631 | NA | 3.345111 | 4.312263 | 3.890202 | 3.848171 | 3.905477 | 3.918175 | 17 | 6 | 16 | 20 |
| 107984984 | XR_00175296 | LOC10798498 | uncharacterized LOC107984984, transcript variant X1 | lncRNA | . | . | . | . | . | 0.163856 | 0.104236 | 0.082906 | 0.109871 | 0.381300 | 0.240947 | 0.186038 | 0.251207 | -1.294582 | 6.311197 | 1.211990228 | -0.30733451 | 0.758588772 | NA | 2.832372 | 2.459419 | 2.675235 | 2.660699 | 2.656697 | 2.664571 | 10 | 6 | 4 | 6 |
| 107984993 | XR_00175294 | LOC10798499 | uncharacterized LOC107984993 | lncRNA | . | . | . | . | . | 0.024566 | 0.015917 | 0.018078 | 0.025392 | 0.057166 | 0.036792 | 0.040567 | 0.058056 | 1.068406 | 6.914914 | 1.154321747 | 0.082697809 | 0.934091827 | NA | 2.741623 | 2.836248 | 2.794289 | 2.783799 | 2.784985 | 2.797248 | 9 | 6 | 5 | 8 |
| 107985000 | XR_00175297 | LOC10798500 | uncharacterized LOC107985000 | lncRNA | . | . | . | . | . | 0.257502 | 0.104410 | 0.223894 | 0.123454 | 0.599219 | 0.241348 | 0.502413 | 0.282264 | -1.013454 | 16.287064 | 0.811299124 | -0.0237649 | 0.981040139 | NA | 4.031897 | 4.019385 | 4.051032 | 3.988209 | 4.037763 | 4.000042 | 27 | 10 | 18 | 11 |
| 107985005 | XR_00175299 | LOC10798500 | uncharacterized LOC107985005 | lncRNA | . | . | . | . | . | 0.331515 | 0.304155 | 0.280310 | 0.097553 | 0.771449 | 0.703068 | 0.629007 | 0.223043 | -1.681093 | 7.517533 | 1.137241336 | -0.65896267 | 0.509919739 | NA | 3.234218 | 2.491830 | 2.924448 | 2.923008 | 2.917322 | 2.895743 | 11 | 10 | 7 | 3 |
| 107985010 | XR_00175303 | LOC10798501 | uncharacterized LOC107985010 | lncRNA | . | . | . | . | . | 0.149321 | 0.249055 | 0.403334 | 0.394117 | 0.347476 | 0.575701 | 0.905069 | 0.901101 | **2.148599** | 5.688885 | 1.285576988 | 0.85828879 | 0.390733018 | NA | 1.855800 | 2.955730 | 2.474550 | 2.484384 | 2.499703 | 2.497326 | 3 | 5 | 7 | 7 |
| 107985012 | XR_00175282 | LOC10798501 | uncharacterized LOC107985012 | lncRNA | . | . | . | . | . | 0.237721 | 0.255057 | 0.202298 | 0.144874 | 0.553186 | 0.589575 | 0.453951 | 0.331237 | -1.411711 | 26.853572 | 0.604765237 | -0.82254183 | 0.410768592 | NA | 4.973417 | 4.478414 | 4.763349 | 4.771483 | 4.746902 | 4.712735 | 35 | 35 | 23 | 17 |
| 107985031 | XR_00175286 | LOC10798503 | uncharacterized LOC107985031 | lncRNA | . | . | . | . | . | 1.600845 | 1.761513 | 1.959600 | 1.138009 | 3.725231 | 4.071815 | 4.397285 | 2.601923 | -1.093977 | 29.238057 | 0.589284158 | -0.21989878 | 0.825949984 | 0.998865404 | 4.931597 | 4.805186 | 4.867886 | 4.876008 | 4.891486 | 4.830177 | 34 | 34 | 31 | 19 |
| 107985034 | XR_00175301 | LOC10798503 | uncharacterized LOC107985034 | lncRNA | . | . | . | . | . | 0.476369 | 0.801995 | 0.985741 | 0.567742 | 1.108529 | 1.853848 | 2.211974 | 1.298075 | 1.209164 | 22.459178 | 0.675533872 | 0.405620473 | 0.685021484 | NA | 4.345328 | 4.620077 | 4.448418 | 4.489422 | 4.512726 | 4.460437 | 18 | 27 | 27 | 17 |
| 107985040 | XR_00175290 | LOC10798504 | uncharacterized LOC107985040 | lncRNA | . | . | . | . | . | 0.924910 | 0.872137 | 0.905936 | 0.496017 | 2.152303 | 2.015985 | 2.032894 | 1.134083 | -1.296569 | 50.036410 | 0.468750162 | -0.79935645 | 0.424083758 | 0.998865404 | 5.818436 | 5.447614 | 5.672169 | 5.658899 | 5.663967 | 5.569311 | 67 | 59 | 49 | 29 |
| 107985041 | XR_00173812 | LOC10798504 | uncharacterized LOC107985041 | lncRNA | . | . | . | . | . | 0.449029 | 0.586918 | 0.595955 | 0.764548 | 1.044908 | 1.356687 | 1.337306 | 1.748049 | 1.276255 | 37.365228 | 0.515380422 | 0.682829513 | 0.494714579 | 0.998865404 | 5.038444 | 5.387716 | 5.181499 | 5.216098 | 5.215094 | 5.253496 | 33 | 40 | 32 | 44 |
| 107985046 | XR_00173812 | LOC10798504 | uncharacterized LOC107985046 | lncRNA | . | . | . | . | . | 0.056199 | 0.075024 | 0.236165 | 0.101880 | 0.130778 | 0.173421 | 0.529947 | 0.232937 | **2.554675** | 16.019031 | 0.817091863 | 1.656043384 | 0.097713033 | NA | 3.170740 | 4.525629 | 3.927160 | 3.938644 | 4.030722 | 3.955560 | 9 | 11 | 28 | 13 |
| 107985082 | XM_01702551 | LOC10798508 | leucine-rich repeat-containing protein 37A3-like | protein_coding | XP_016880999.1 | . | . | . | . | 0.527360 | 0.430344 | 0.443153 | 0.591679 | 1.227188 | 0.994760 | 0.994423 | 1.352803 | 1.071849 | 17.574989 | 0.734284658 | 0.13632499 | 0.891564369 | NA | 4.084644 | 4.184531 | 4.141600 | 4.122750 | 4.128709 | 4.146129 | 22 | 16 | 14 | 19 |
| 107985099 | gene-DNAJB6P | DNAJB6P6 | . | pseudogene | . | 54759 | . | . | . | 0.439972 | 0.398685 | 0.728962 | 0.572805 | 1.023832 | 0.921578 | 1.635770 | 1.309650 | 1.535839 | 23.845402 | 0.634227651 | 0.976032329 | 0.329048444 | NA | 4.231982 | 4.852980 | 4.547246 | 4.537192 | 4.597185 | 4.571250 | 23 | 19 | 28 | 24 |
| 107985115 | NR_168396,NF | LOC10798511 | LOC107985115 | lncRNA | . | . | . | . | . | 2.220885 | 1.757000 | 2.646428 | 2.066947 | 5.168091 | 4.061386 | 5.938510 | 4.725829 | 1.223703 | 163.564656 | 0.272536868 | 1.068675191 | 0.285216054 | 0.998865404 | 7.199655 | 7.492904 | 7.344130 | 7.262862 | 7.437892 | 7.332823 | 187 | 142 | 176 | 148 |
| 107985140 | XR_00175346 | WDR7-OT1 | WDR7 overlapping transcript 1 | lncRNA | . | 45131 | . | . | . | 0.023277 | 0.024216 | 0.036966 | 0.031002 | 0.054166 | 0.055977 | 0.082951 | 0.070883 | 1.340584 | 6.305236 | 1.200698396 | 0.352179518 | 0.724703637 | NA | 2.429096 | 2.852980 | 2.643888 | 2.645602 | 2.656687 | 2.649491 | 6 | 6 | 7 | 6 |
| 107985147 | XR_00175344 | LOC10798514 | uncharacterized LOC107985147 | lncRNA | . | . | . | . | . | 0.002237 | 0.011499 | 0.004039 | 0.005310 | 0.005206 | 0.026581 | 0.009064 | 0.012140 | -1.641802 | 2.932659 | 1.783481861 | -0.40105847 | 0.688377074 | NA | 1.867373 | 1.148375 | 1.549439 | 1.567461 | 1.552366 | 1.551546 | 2 | 6 | 2 | 2 |

| | | | | | | | | | | | | | | | | | | | | | | | | | | | | | | | | | |
|---|---|---|---|---|---|---|---|---|---|---|---|---|---|---|---|---|---|---|---|---|---|---|---|---|---|---|---|---|---|---|---|---|---|
| 107985162 | XR_001753518 | LOC107985162 | uncharacterized LOC107985162 | lncRNA | . | . | . | . | . | 0.238807 | 0.098265 | 0.281942 | 0.113841 | 0.555713 | 0.227144 | 0.632671 | 0.260283 | 1.159106 | 17.240755 | 0.809679776 | 0.263082358 | 0.792487102 | NA | 3.993083 | 4.213972 | 4.118175 | 4.059065 | 4.132168 | 4.066417 | 26 | 10 | 23 | 10 |
| 107985163 | XR_001753519 | LOC107985163 | uncharacterized LOC107985163 | lncRNA | . | . | . | . | . | 0.175621 | 0.220608 | 0.298833 | 0.103312 | 0.408677 | 0.509946 | 0.670573 | 0.236210 | 1.007628 | 11.761337 | 0.925026996 | 0.011851319 | 0.990544237 | NA | 3.548173 | 3.563745 | 3.541972 | 3.553587 | 3.571246 | 3.524509 | 12 | 14 | 15 | 6 |
| 107985184 | NR_160737 | LOC107985184 | uncharacterized LOC107985184 | lncRNA | . | . | . | . | . | 0.081713 | 0.133139 | 0.220041 | 0.147947 | 0.190150 | 0.307758 | 0.493765 | 0.338262 | 1.643847 | 12.548703 | 0.880084644 | 0.814780834 | 0.415197774 | NA | 3.247564 | 3.963455 | 3.607594 | 3.631325 | 3.658195 | 3.634659 | 8 | 13 | 16 | 12 |
| 107985194 | XR_001738201 | LINC02798 | long intergenic non-protein coding RNA 2798 | lncRNA | . | 54323 | . | . | . | 0.022908 | 0.043738 | 0.050074 | 0.052005 | 0.053307 | 0.101103 | 0.112364 | 0.118903 | 1.534105 | 4.580564 | 1.410961501 | 0.437572199 | 0.661696428 | NA | 1.855800 | 2.470304 | 2.174880 | 2.184610 | 2.189519 | 2.187757 | 3 | 5 | 5 | 5 |
| 107985200 | XR_001738207 | LOC107985200 | uncharacterized LOC107985200, transcript variant X1 | lncRNA | . | . | . | . | . | 3.366068 | 2.635383 | 2.386354 | 2.586573 | 7.832979 | 6.091806 | 5.354908 | 5.913891 | -1.241015 | 52.006597 | 0.440270566 | -0.70756607 | 0.479214769 | 0.998865404 | 5.847713 | 5.536815 | 5.749931 | 5.694310 | 5.675908 | 5.687063 | 75 | 54 | 39 | 45 |
| 107985201 | XR_001738209 | LOC107985201 | uncharacterized LOC107985201 | lncRNA | . | . | . | . | . | 0.205644 | 0.175129 | 0.203573 | 0.207747 | 0.478542 | 0.404820 | 0.456812 | 0.474990 | 1.092961 | 12.182212 | 0.870488189 | 0.14732172 | 0.88287809 | NA | 3.540965 | 3.669578 | 3.605093 | 3.600418 | 3.607183 | 3.607877 | 14 | 12 | 11 | 12 |
| 107985209 | XR_001738246 | LOC107985209 | uncharacterized LOC107985209, transcript variant X2 | lncRNA | . | . | . | . | . | 0.184780 | 0.159000 | 0.274265 | 0.180638 | 0.429992 | 0.367536 | 0.615444 | 0.413007 | 1.336631 | 14.124284 | 0.820166837 | 0.510386058 | 0.609781023 | NA | 3.593806 | 4.015675 | 3.806375 | 3.797858 | 3.833976 | 3.805622 | 15 | 12 | 17 | 12 |
| 107985216 | XR_001738261 | LOC107985216 | uncharacterized LOC107985216, transcript variant X3 | lncRNA | . | . | . | . | . | 0.279216 | 0.429610 | 0.405878 | 0.179955 | 0.649746 | 0.993064 | 0.910778 | 0.411445 | -1.228265 | 72.523093 | 0.443890644 | -0.66823165 | 0.503985736 | 0.998865404 | 6.319922 | 6.025855 | 6.133514 | 6.244858 | 6.224491 | 6.048392 | 73 | 104 | 79 | 37 |
| 107985228 | XR_001738304 | LINC02803 | long intergenic non-protein coding RNA 2803 | lncRNA | . | 54328 | . | . | . | 0.024967 | 0.027979 | 0.052614 | 0.008698 | 0.058100 | 0.064674 | 0.118065 | 0.019887 | 1.103415 | 7.573630 | 1.188002771 | 0.119507402 | 0.904873376 | NA | 2.844134 | 2.993948 | 2.903528 | 2.905728 | 2.933241 | 2.881929 | 8 | 8 | 12 | 2 |
| 107985246 | NR_160746,NF | LOC107985246 | uncharacterized LOC107985246 | lncRNA | . | . | . | . | . | 1.465222 | 1.780293 | 1.459108 | 1.709187 | 3.409632 | 4.115225 | 3.274195 | 3.907853 | -1.063944 | 105.209549 | 0.320793339 | -0.27875436 | 0.780433344 | 0.998865404 | 6.761488 | 6.671348 | 6.682586 | 6.766135 | 6.713819 | 6.697508 | 109 | 132 | 91 | 93 |
| 107985263 | XR_001738465 | LOC107985263 | uncharacterized LOC107985263 | lncRNA | . | . | . | . | . | 0.154557 | 0.092704 | 0.131313 | 0.058205 | 0.359660 | 0.214290 | 0.294663 | 0.133078 | -1.261839 | 7.593745 | 1.125024112 | -0.29824028 | 0.76551978 | NA | 3.079678 | 2.751299 | 2.939000 | 2.920796 | 2.932836 | 2.911050 | 12 | 7 | 8 | 4 |
| 107985289 | XR_001753939 | LOC107985289 | uncharacterized LOC107985289 | lncRNA | . | . | . | . | . | 0.050916 | 0.021704 | 0.119446 | 0.079986 | 0.118484 | 0.050169 | 0.268034 | 0.182879 | 2.860223 | 6.003035 | 1.283700493 | 1.181060274 | 0.23757878 | NA | 1.631267 | 3.155137 | 2.552345 | 2.540634 | 2.579291 | 2.566853 | 5 | 2 | 9 | 7 |
| 107985295 | XR_001753826 | LOC107985295 | uncharacterized LOC107985295, transcript variant X1 | lncRNA | . | . | . | . | . | 0.078282 | 0.057947 | 0.055367 | 0.210988 | 0.182165 | 0.133947 | 0.124242 | 0.482400 | 1.828037 | 7.561514 | 1.153468883 | 0.754502425 | 0.45054759 | NA | 2.421266 | 3.287859 | 2.898361 | 2.891722 | 2.892332 | 2.937603 | 7 | 5 | 4 | 14 |
| 107985297 | XR_001753847 | LOC107985297 | uncharacterized LOC107985297 | lncRNA | . | . | . | . | . | 0.352907 | 0.260115 | 0.360328 | 0.413176 | 0.821228 | 0.601267 | 0.808565 | 0.944678 | 1.237841 | 17.469493 | 0.737017468 | 0.417665176 | 0.676191937 | NA | 3.964015 | 4.273004 | 4.126216 | 4.102919 | 4.126641 | 4.134559 | 21 | 14 | 16 | 19 |
| 107985303 | XR_001753834 | LOC107985303 | uncharacterized LOC107985303 | lncRNA | . | . | . | . | . | 1.136683 | 1.474701 | 1.463537 | 1.063219 | 2.645107 | 3.408836 | 3.284133 | 2.430925 | -1.041455 | 45.449169 | 0.474098588 | -0.12360392 | 0.901628887 | 0.998865404 | 5.534723 | 5.477065 | 5.485022 | 5.527605 | 5.525925 | 5.472535 | 47 | 56 | 45 | 35 |
| 107985307 | XR_001738485 | LINC01705 | long intergenic non-protein coding RNA 1705 | lncRNA | . | 52493 | . | . | . | 0.706971 | 0.531481 | 0.841068 | 0.300927 | 1.645150 | 1.228541 | 1.887331 | 0.688035 | -1.069929 | 16.871220 | 0.794966984 | -0.12266595 | 0.902371631 | NA | 4.121102 | 4.030460 | 4.083375 | 4.060630 | 4.099412 | 4.033665 | 23 | 16 | 21 | 8 |
| 107985336 | XR_001753890 | LOC107985336 | uncharacterized LOC107985336 | lncRNA | . | . | . | . | . | 0.983181 | 0.876407 | 1.300242 | 0.882175 | 2.287902 | 2.025854 | 2.917704 | 2.016988 | 1.155480 | 169.392955 | 0.278087593 | 0.749738132 | 0.453412437 | 0.998865404 | 7.295049 | 7.505726 | 7.390302 | 7.340611 | 7.506314 | 7.338586 | 192 | 159 | 189 | 137 |
| 107985338 | XR_001738495 | LINC02813 | long intergenic non-protein coding RNA 2813, transcript variant X | lncRNA | . | 54345 | . | . | . | 0.023593 | 0.150399 | 0.131510 | 0.049617 | 0.054902 | 0.347653 | 0.295105 | 0.113444 | 1.076313 | 7.604855 | 1.194408473 | 0.088828396 | 0.92921829 | NA | 2.873125 | 2.978782 | 2.887015 | 2.931095 | 2.927598 | 2.895777 | 3 | 13 | 10 | 4 |
| 107985354 | XR_001738506 | LOC107985354 | uncharacterized LOC107985354 | lncRNA | . | . | . | . | . | 0.243272 | 0.236376 | 0.233626 | 0.094707 | 0.566105 | 0.546394 | 0.524250 | 0.216537 | -1.388434 | 10.028051 | 0.982375579 | -0.48195289 | 0.629839412 | NA | 3.540965 | 3.073227 | 3.337645 | 3.332829 | 3.334173 | 3.307147 | 14 | 12 | 10 | 5 |
| 107985355 | NR_153418 | LOC107985355 | uncharacterized LOC107985355 | lncRNA | . | . | . | . | . | 0.619505 | 0.733604 | 0.857710 | 0.998436 | 1.441612 | 1.695757 | 1.924677 | 2.282806 | 1.314955 | 15.073085 | 0.786807831 | 0.502045325 | 0.615635639 | NA | 3.703730 | 4.097315 | 3.894834 | 3.902838 | 3.912336 | 3.921067 | 14 | 15 | 14 | 17 |
| 107985360 | XR_001738520 | LOC107985360 | uncharacterized LOC107985360 | lncRNA | . | . | . | . | . | 0.186490 | 0.143846 | 0.120825 | 0.084215 | 0.433971 | 0.332506 | 0.271128 | 0.192548 | -1.557467 | 11.374019 | 0.91510487 | -0.69850057 | 0.484864204 | NA | 3.790753 | 3.155137 | 3.532537 | 3.516329 | 3.508959 | 3.496230 | 18 | 13 | 9 | 7 |
| 107985367 | XR_001738546 | LOC107985367 | uncharacterized LOC107985367 | lncRNA | . | . | . | . | . | 0.300334 | 0.219853 | 0.369703 | 0.123223 | 0.698889 | 0.508201 | 0.829604 | 0.281735 | -1.051226 | 17.895003 | 0.782589896 | -0.09209573 | 0.926621984 | NA | 4.193664 | 4.128572 | 4.165943 | 4.143558 | 4.187695 | 4.112665 | 24 | 17 | 23 | 8 |
| 107985369 | XR_001738560 | LOC107985369 | uncharacterized LOC107985369 | lncRNA | . | . | . | . | . | 0.021553 | 0.044557 | 0.052251 | 0.109730 | 0.050154 | 0.102996 | 0.117249 | 0.250886 | 2.258938 | 5.128052 | 1.365082818 | 0.861225711 | 0.389113742 | NA | 1.658167 | 2.827809 | 2.329385 | 2.334794 | 2.339131 | 2.361493 | 3 | 4 | 4 | 9 |
| 107985373 | XR_001738572 | LOC107985373 | uncharacterized LOC107985373 | lncRNA | . | . | . | . | . | 0.375728 | 0.193097 | 0.424546 | 0.202924 | 0.874334 | 0.446352 | 0.952670 | 0.463963 | 1.108372 | 7.986602 | 1.098629223 | 0.135115611 | 0.892520468 | NA | 2.917750 | 3.073227 | 2.998741 | 2.980217 | 3.006579 | 2.979602 | 11 | 6 | 10 | 5 |
| 107985388 | XM_01702818 | LOC107985388 | pre-mRNA-splicing factor cwc22-like | protein_coding | XP_016883675.1 | . | . | . | . | 0.182040 | 0.227613 | 0.218909 | 0.205203 | 0.423615 | 0.526138 | 0.491226 | 0.469173 | 1.056319 | 12.947654 | 0.846921431 | 0.093333631 | 0.925638515 | NA | 3.654832 | 3.733338 | 3.684942 | 3.696775 | 3.696109 | 3.691871 | 13 | 15 | 12 | 12 |
| 107985393 | XR_001754582 | LOC107985393 | uncharacterized LOC107985393, transcript variant X2 | lncRNA | . | . | . | . | . | 0.050445 | 0.245220 | 0.276060 | 0.141187 | 0.117388 | 0.566836 | 0.619472 | 0.322808 | 1.385461 | 4.349300 | 1.484829981 | 0.316781226 | 0.751409611 | NA | 1.867373 | 2.336262 | 2.092822 | 2.111102 | 2.116665 | 2.100173 | 2 | 6 | 6 | 3 |
| 107985395 | XR_001754478 | PANK2-AS1 | PANK2 antisense RNA 1 | lncRNA | . | 40732 | . | . | . | 1.044876 | 0.797153 | 1.235020 | 1.079781 | 2.431469 | 1.842655 | 2.771349 | 2.468790 | 1.241174 | 25.036450 | 0.619142132 | 0.503446474 | 0.614650404 | NA | 4.480474 | 4.794399 | 4.641429 | 4.616974 | 4.660046 | 4.643072 | 29 | 21 | 26 | 24 |
| 107985400 | XR_001754565 | LOC107985400 | uncharacterized LOC107985400, transcript variant X1 | lncRNA | . | . | . | . | . | 0.073778 | 0.107962 | 0.108212 | 0.056556 | 0.171684 | 0.249558 | 0.242825 | 0.129308 | -1.046096 | 11.453017 | 0.918148166 | -0.07081133 | 0.943547917 | NA | 3.548173 | 3.486479 | 3.508794 | 3.520380 | 3.528334 | 3.496016 | 12 | 14 | 13 | 7 |
| 107985423 | XR_001754452 | LOC107985423 | uncharacterized LOC107985423 | lncRNA | . | . | . | . | . | 0.250899 | 0.426659 | 0.431284 | 0.430411 | 0.583853 | 0.986241 | 0.967789 | 0.984083 | 1.265779 | 63.454065 | 0.418941808 | 0.811630625 | 0.417003613 | 0.998865404 | 5.808872 | 6.146680 | 5.895951 | 6.002456 | 6.004198 | 6.001619 | 48 | 76 | 62 | 66 |
| 107985433 | XM_01702819 | LOC107985433 | putative uncharacterized protein LOC401522 | protein_coding | XP_016883681.1 | . | . | . | . | 0.092416 | 0.094188 | 0.338543 | 0.078693 | 0.215055 | 0.217720 | 0.759681 | 0.179923 | 2.133538 | 7.048067 | 1.201613752 | 0.909816364 | 0.362919363 | NA | 2.166062 | 3.264261 | 2.775334 | 2.776906 | 2.822363 | 2.775631 | 5 | 5 | 13 | 4 |
| 107985440 | XR_001754508 | LOC107985440 | uncharacterized LOC107985440, transcript variant X1 | lncRNA | . | . | . | . | . | 0.705632 | 0.635183 | 0.220415 | 0.294233 | 1.642035 | 1.468255 | 0.494606 | 0.672728 | -2.009522 | 19.095870 | 0.715925847 | -1.40636423 | 0.159615996 | NA | 4.672908 | 3.664848 | 4.294875 | 4.282567 | 4.229787 | 4.239248 | 31 | 26 | 10 | 13 |
| 107985469 | XR_001737616 | LOC107985469 | uncharacterized LOC107985469 | lncRNA | . | . | . | . | . | 0.021730 | 0.053769 | 0.027254 | 0.058353 | 0.050568 | 0.124289 | 0.061157 | 0.133418 | 1.036824 | 3.234306 | 1.683534089 | 0.030989169 | 0.975278178 | NA | 1.671431 | 1.715150 | 1.683658 | 1.697432 | 1.686697 | 1.695533 | 2 | 5 | 2 | 4 |
| 107985522 | XR_001737690 | LOC107985522 | uncharacterized LOC107985522 | lncRNA | . | . | . | . | . | 0.136462 | 0.040626 | 0.048266 | 0.056204 | 0.317554 | 0.093909 | 0.108308 | 0.128504 | -1.600469 | 6.449600 | 1.243341424 | -0.54570272 | 0.585270304 | NA | 2.987791 | 2.312263 | 2.722553 | 2.683554 | 2.688200 | 2.691296 | 14 | 4 | 4 | 5 |
| 107985523 | XR_001737693 | LOC107985523 | uncharacterized LOC107985523 | lncRNA | . | . | . | . | . | 0.638515 | 1.227389 | 0.722978 | 0.608248 | 1.485850 | 2.837163 | 1.622341 | 1.390688 | -1.360810 | 7.856656 | 1.100259856 | -0.40396423 | 0.686239022 | NA | 3.180050 | 2.733338 | 2.966626 | 2.993987 | 2.970797 | 2.968484 | 7 | 13 | 6 | 6 |
| 107985524 | NR_168400 | LOC107985524 | uncharacterized LOC107985524 | lncRNA | . | . | . | . | . | 0.236491 | 0.124669 | 0.323430 | 0.130993 | 0.550324 | 0.288179 | 0.725768 | 0.299499 | 1.364294 | 6.317738 | 1.235068193 | 0.36285784 | 0.716711082 | NA | 2.413393 | 2.869520 | 2.650763 | 2.635560 | 2.665318 | 2.638538 | 8 | 4 | 9 | 4 |
| 107985532 | XM_01702916 | LOC107985532 | uncharacterized LOC107985532 | protein_coding | XP_016884654.1 | . | . | . | . | 1.770641 | 1.604450 | 1.857377 | 1.804109 | 4.120354 | 3.708759 | 4.167899 | 4.124881 | 1.066807 | 121.210501 | 0.294138911 | 0.317194234 | 0.751096225 | 0.998865404 | 6.873617 | 6.967594 | 6.926439 | 6.891185 | 6.938003 | 6.924411 | 142 | 120 | 111 | 115 |
| 107985544 | NR_146603 | LOC107985544 | uncharacterized LOC107985544 | lncRNA | . | . | . | . | . | 0.031364 | 0.034075 | 0.040981 | 0.017387 | 0.072984 | 0.078766 | 0.091960 | 0.039753 | -1.206689 | 4.930248 | 1.36136082 | -0.1991048 | 0.842180764 | NA | 2.429096 | 2.161867 | 2.303870 | 2.305477 | 2.306142 | 2.294735 | 6 | 6 | 5 | 3 |
| 107985551 | XR_001755557 | LOC107985551 | uncharacterized LOC107985551 | lncRNA | . | . | . | . | . | 1.187198 | 1.340334 | 0.689207 | 0.416207 | 2.762657 | 3.098243 | 1.546560 | 0.951608 | -2.154356 | 7.889695 | 1.103931612 | -1.00301173 | 0.315855199 | NA | 3.429096 | 2.324312 | 3.001213 | 3.004189 | 2.980301 | 2.973486 | 12 | 12 | 5 | 4 |
| 107985558 | XR_001755455 | LOC107985558 | uncharacterized LOC107985558 | lncRNA | . | . | . | . | . | 0.677211 | 0.581409 | 0.880992 | 0.821570 | 1.575898 | 1.343952 | 1.976920 | 1.878423 | 1.329103 | 113.580694 | 0.305212008 | 1.344812351 | 0.178685841 | 0.951511445 | 6.607151 | 7.018740 | 6.794866 | 6.747953 | 6.878519 | 6.852394 | 121 | 97 | 117 | 117 |
| 107985564 | XR_001755582 | LOC107985564 | uncharacterized LOC107985564 | lncRNA | . | . | . | . | . | 0.129308 | 0.383852 | 0.319655 | 0.336100 | 0.300906 | 0.887290 | 0.717296 | 0.768454 | 1.220007 | 6.545247 | 1.200796003 | 0.238916412 | 0.811170399 | NA | 2.562483 | 2.844638 | 2.688363 | 2.711254 | 2.704497 | 2.707062 | 4 | 9 | 6 | 7 |
| 107985565 | XR_001755630 | LOC107985565 | uncharacterized LOC107985565 | lncRNA | . | . | . | . | . | 0.869551 | 1.009778 | 0.395239 | 0.833766 | 2.023481 | 2.334147 | 0.886905 | 1.906307 | -1.497198 | 15.718430 | 0.798788499 | -0.72893473 | 0.466041592 | NA | 4.238681 | 3.650563 | 3.986591 | 3.999564 | 3.946903 | 3.984974 | 20 | 22 | 7 | 16 |
| 107985567 | XR_001755555 | LOC107985567 | protein LLP homolog pseudogene | transcribed_ps | . | . | . | . | . | 0.992045 | 0.938268 | 0.798624 | 0.648605 | 2.308530 | 2.168848 | 1.792089 | 1.482959 | -1.353065 | 86.876898 | 0.346717186 | -1.25817555 | 0.208328276 | 0.970482251 | 6.641879 | 6.207322 | 6.492180 | 6.469968 | 6.423205 | 6.371874 | 119 | 104 | 71 | 62 |
| 107985595 | XR_001737760 | LOC107985595 | uncharacterized LOC107985595 | lncRNA | . | . | . | . | . | 0.023589 | 0.005085 | 0.051994 | 0.000348 | 0.054894 | 0.011753 | 0.116673 | 0.000796 | 1.912449 | 2.587152 | 1.934465599 | 0.483555537 | 0.628701337 | NA | 0.820514 | 1.769039 | 1.348859 | 1.340813 | 1.362872 | 1.341752 | 3 | 1 | 5 | 1 |
| 107985638 | XR_001755998 | LOC107985638 | uncharacterized LOC107985638 | lncRNA | . | . | . | . | . | 1.282129 | 0.770025 | 1.303531 | 1.637744 | 2.983567 | 1.779947 | 2.925085 | 3.744506 | 1.398360 | 33.072995 | 0.559006803 | 0.865347996 | 0.386847815 | 0.998865404 | 4.784663 | 5.269892 | 5.045586 | 4.985660 | 5.044180 | 5.075829 | 40 | 22 | 30 | 40 |
| 107985665 | XR_001755896 | LOC107985665 | uncharacterized LOC107985665, transcript variant X1 | lncRNA | . | . | . | . | . | 0.084445 | 0.100322 | 0.062433 | 0.103301 | 0.196506 | 0.231899 | 0.140098 | 0.236185 | -1.061020 | 5.674441 | 1.265769002 | -0.06751 | 0.9461757 | NA | 2.548173 | 2.459419 | 2.503664 | 2.509625 | 2.501138 | 2.509100 | 6 | 7 | 4 | 6 |
| 107985675 | XR_001755782 | LOC107985675 | uncharacterized LOC107985675, transcript variant X8 | lncRNA | . | . | . | . | . | 0.178731 | 0.057358 | 0.099965 | 0.114476 | 0.415915 | 0.132586 | 0.224321 | 0.261736 | -1.098818 | 20.348159 | 0.70897312 | -0.19175874 | 0.847931188 | NA | 4.411418 | 4.279207 | 4.381086 | 4.317309 | 4.346887 | 4.336202 | 33 | 15 | 18 | 17 |
| 107985678 | XM_01703000 | LOC107985678 | uncharacterized LOC107985678 | protein_coding | XP_016885496.1 | . | . | . | . | 0.135974 | 0.218204 | 0.151863 | 0.185303 | 0.316418 | 0.504389 | 0.340777 | 0.423674 | -1.039322 | 12.401818 | 0.874960068 | -0.06359386 | 0.949293624 | NA | 3.661494 | 3.602870 | 3.617820 | 3.645829 | 3.627056 | 3.632614 | 11 | 17 | 10 | 12 |
| 107985721 | XR_001737606 | LOC107985721 | uncharacterized LOC107985721 | lncRNA | . | . | . | . | . | 0.013059 | 0.043966 | 0.059756 | 0.011622 | 0.030389 | 0.101629 | 0.134090 | 0.026571 | 1.066915 | 3.296787 | 1.717757441 | 0.054399537 | 0.956616848 | NA | 1.671431 | 1.769039 | 1.700452 | 1.714288 | 1.718854 | 1.697761 | 2 | 5 | 5 | 1 |
| 107985730 | XR_001737835 | MIR1302-2HG | MIR1302-2 host gene, transcript variant X2 | lncRNA | . | 52482 | . | . | . | 0.092156 | 0.162364 | 0.175770 | 0.057185 | 0.214450 | 0.375312 | 0.394424 | 0.130748 | -1.003532 | 2.258427 | 1.991984473 | -0.00255353 | 0.997962579 | NA | 1.175403 | 1.175234 | 1.166206 | 1.171202 | 1.174041 | 1.163259 | 2 | 3 | 3 | 1 |
| 107985733 | XR_001737880 | LOC107985733 | uncharacterized LOC107985733 | lncRNA | . | . | . | . | . | 20.476145 | 21.786345 | 22.558575 | 22.947392 | 47.648827 | 50.360107 | 50.620789 | 52.466484 | 1.062796 | 358.186489 | 0.188009276 | 0.467345231 | 0.6402529 | 0.998865404 | 8.440082 | 8.527722 | 8.450454 | 8.481726 | 8.499309 | 8.502775 | 389 | 385 | 320 | 347 |
| 107985736 | XR_001737913 | LOC107985736 | uncharacterized LOC107985736 | lncRNA | . | . | . | . | . | 0.108824 | 0.132133 | 0.174856 | 0.153305 | 0.253238 | 0.305431 | 0.392371 | 0.350513 | 1.333586 | 13.108825 | 0.842561121 | 0.492915108 | 0.622072572 | NA | 3.489863 | 3.905270 | 3.692850 | 3.700394 | 3.717752 | 3.708430 | 12 | 13 | 14 | 13 |
| 107985767 | XR_001739092 | LOC107985767 | uncharacterized LOC107985767 | lncRNA | . | . | . | . | . | 0.301648 | 0.298691 | 0.283623 | 0.422329 | 0.701947 | 0.690439 | 0.636442 | 0.965605 | 1.129885 | 11.923448 | 0.884808603 | 0.1991117 | 0.842175369 | NA | 3.486115 | 3.660102 | 3.571189 | 3.570470 | 3.567370 | 3.587074 | 13 | 12 | 9 | 14 |
| 107985770 | NR_164153,NF | LOC107985770 | MANTIS | lncRNA | . | . | . | . | . | 0.101295 | 0.040495 | 0.026928 | 0.152350 | 0.235718 | 0.093606 | 0.060425 | 0.348330 | 1.335132 | 11.347195 | 1.011857496 | 0.412095524 | 0.68026941 | NA | 3.282109 | 3.696731 | 3.505593 | 3.467547 | 3.464823 | 3.536489 | 16 | 6 | 4 | 20 |
| 107985780 | XR_001739138 | LOC107985780 | uncharacterized LOC107985780 | lncRNA | . | . | . | . | . | 0.186526 | 0.203323 | 0.109707 | 0.083899 | 0.434053 | 0.469991 | 0.246180 | 0.191826 | -2.026032 | 58.019628 | 0.424645386 | -2.39884075 | 0.016447066 | 0.430775811 | 6.279033 | 5.262390 | 5.911130 | 5.930608 | 5.800364 | 5.759072 | 86 | 87 | 38 | 31 |
| 107985788 | XR_001739177 | LOC107985788 | uncharacterized LOC107985788 | lncRNA | . | . | . | . | . | 0.094699 | 0.262875 | 0.163122 | 0.050466 | 0.220367 | 0.607648 | 0.366042 | 0.115384 | -1.815599 | 3.170596 | 1.735886308 | -0.49568087 | 0.620119588 | NA | 2.029593 | 1.175234 | 1.665242 | 1.679110 | 1.669183 | 1.658378 | 3 | 6 | 3 | 1 |
| 107985853 | XR_001739290 | LOC107985853 | uncharacterized LOC107985853 | lncRNA | . | . | . | . | . | 0.099301 | 0.078836 | 0.183752 | 0.028430 | 0.231078 | 0.182233 | 0.412334 | 0.065003 | 1.130853 | 9.066615 | 1.0860626 | 0.163352267 | 0.870241083 | NA | 3.084644 | 3.270502 | 3.168900 | 3.159028 | 3.196593 | 3.140361 | 11 | 8 | 14 | 3 |
| 107985864 | XR_001739395 | LOC107985864 | uncharacterized LOC107985864 | lncRNA | . | . | . | . | . | 0.487354 | 0.740447 | 0.793185 | 0.258266 | 1.134092 | 1.711576 | 1.779883 | 0.590495 | -1.173123 | 15.899909 | 0.820853939 | -0.28062806 | 0.778995693 | NA | 4.099441 | 3.873625 | 3.974718 | 4.003337 | 4.008246 | 3.947161 | 16 | 22 | 19 | 7 |
| 107985873 | XR_001739426 | LOC107985873 | uncharacterized LOC107985873 | lncRNA | . | . | . | . | . | 0.326249 | 0.554640 | 0.381018 | 0.180063 | 0.759195 | 1.282076 | 0.854993 | 0.411692 | -1.562147 | 10.766837 | 0.966490522 | -0.66584212 | 0.505512018 | NA | 3.713381 | 3.073227 | 3.422475 | 3.454409 | 3.431181 | 3.404107 | 11 | 18 | 10 | 5 |
| 107985875 | XR_002959382 | LOC107985875 | uncharacterized LOC107985875 | lncRNA | . | . | . | . | . | 0.292329 | 0.141751 | 0.254092 | 0.312652 | 0.680261 | 0.327663 | 0.570175 | 0.714841 | 1.290570 | 13.211403 | 0.861703823 | 0.427070815 | 0.669327749 | NA | 3.526439 | 3.897226 | 3.729288 | 3.692056 | 3.720821 | 3.730311 | 18 | 8 | 12 | 15 |
| 107985877 | XR_001739444 | LOC107985877 | uncharacterized LOC107985877 | lncRNA | . | . | . | . | . | 0.978243 | 1.082514 | 1.089681 | 0.785775 | 2.276411 | 2.502280 | 2.445212 | 1.796582 | -1.066389 | 14.370678 | 0.808826144 | -0.11465265 | 0.908720445 | NA | 3.889947 | 3.798723 | 3.842067 | 3.850345 | 3.851451 | 3.832838 | 16 | 17 | 14 | 11 |
| 107985897 | XR_001739535 | LOC107985897 | uncharacterized LOC107985897 | lncRNA | . | . | . | . | . | 0.049759 | 0.089586 | 0.160691 | 0.108076 | 0.115792 | 0.207082 | 0.360586 | 0.247104 | 2.004134 | 10.865263 | 0.948396069 | 1.05755283 | 0.290259357 | NA | 2.855800 | 3.857133 | 3.394057 | 3.413276 | 3.447676 | 3.424373 | 6 | 10 | 15 | 11 |
| 107985911 | NR_160761,NF | LOC107985911 | lymphocyte specific protein 1 pseudogene | transcribed_ps | . | . | . | . | . | 5.914820 | 5.464691 | 4.527802 | 6.271004 | 13.764027 | 12.631878 | 10.160255 | 14.337904 | -1.099325 | 189.990994 | 0.261330667 | -0.52277895 | 0.601128087 | 0.998865404 | 7.637050 | 7.499235 | 7.619638 | 7.556316 | 7.473663 | 7.614857 | 245 | 200 | 131 | 198 |
| 107985915 | XR_001739592 | LOC107985915 | uncharacterized LOC107985915, transcript variant X3 | lncRNA | . | . | . | . | . | 0.369309 | 0.699370 | 0.652499 | 0.321744 | 0.859396 | 1.616625 | 1.464190 | 0.735630 | 1.066420 | 29.850810 | 0.604125494 | 0.153570205 | 0.877948631 | 0.998865404 | 4.851436 | 4.946398 | 4.862282 | 4.907574 | 4.933631 | 4.850414 | 27 | 37 | 36 | 19 |
| 107985929 | XR_001739626 | LOC107985929 | uncharacterized LOC107985929 | lncRNA | . | . | . | . | . | 0.485617 | 0.185568 | 0.334032 | 0.281628 | 1.130051 | 0.428949 | 0.749558 | 0.643909 | -1.119053 | 8.813391 | 1.04840913 | -0.15478552 | 0.876990405 | NA | 3.216230 | 3.058876 | 3.159853 | 3.125556 | 3.142184 | 3.134574 | 15 | 6 | 8 | 7 |
| 107985930 | XR_001739628 | LOC107985930 | uncharacterized LOC107985930 | lncRNA | . | . | . | . | . | 0.084606 | 0.154010 | 0.108385 | 0.042933 | 0.196882 | 0.356000 | 0.243213 | 0.098161 | -1.533357 | 7.112463 | 1.177015386 | -0.52394722 | 0.600315249 | NA | 3.104339 | 2.491830 | 2.825357 | 2.848493 | 2.834270 | 2.812683 | 7 | 12 | 7 | 3 |
| 107985939 | NR_158150 | LOC107985939 | uncharacterized LOC107985939 | lncRNA | . | . | . | . | . | 0.296997 | 0.290056 | 0.385168 | 0.257024 | 0.691125 | 0.670478 | 0.864306 | 0.587656 | 1.072417 | 20.450479 | 0.68497969 | 0.147254303 | 0.8829313 | NA | 4.301434 | 4.404839 | 4.348288 | 4.346097 | 4.369984 | 4.336161 | 23 | 21 | 22 | 16 |
| 107985942 | XR_001739683 | LOC107985942 | uncharacterized LOC107985942 | lncRNA | . | . | . | . | . | 0.490890 | 0.665145 | 0.620799 | 0.419078 | 1.142322 | 1.537512 | 1.393056 | 0.958173 | -1.107055 | 27.017106 | 0.60815379 | -0.24126648 | 0.80934859 | 0.998865404 | 4.826611 | 4.681336 | 4.741540 | 4.775397 | 4.768870 | 4.725411 | 28 | 35 | 27 | 19 |
| 107985944 | XR_001739707 | LOC107985944 | uncharacterized LOC107985944 | lncRNA | . | . | . | . | . | 0.977630 | 1.115445 | 1.327521 | 0.857780 | 2.274986 | 2.578400 | 2.978918 | 1.961213 | 1.032362 | 76.285443 | 0.381888897 | 0.120318631 | 0.904230746 | 0.998865404 | 6.229240 | 6.277036 | 6.227498 | 6.259457 | 6.304599 | 6.195776 | 81 | 86 | 82 | 57 |
| 107985963 | gene-LOC1079 | LOC107985963 | . | pseudogene | . | . | . | . | . | 0.176576 | 0.044480 | 0.366185 | 0.200329 | 0.410901 | 0.102817 | 0.821708 | 0.458028 | 2.648900 | 8.071007 | 1.143506864 | 1.229020462 | 0.219064134 | NA | 2.137672 | 3.553545 | 2.980055 | 2.956106 | 3.014954 | 2.987385 | 8 | 2 | 13 | 8 |
| 107985986 | XR_001739885 | LOC107985986 | uncharacterized LOC107985986 | lncRNA | . | . | . | . | . | 0.695121 | 1.065378 | 1.766714 | 1.087809 | 1.617575 | 2.462669 | 3.964455 | 2.487147 | 1.614734 | 21.238557 | 0.687899703 | 1.004937494 | 0.314926949 | NA | 4.021847 | 4.713310 | 4.354813 | 4.383734 | 4.435214 | 4.387333 | 15 | 21 | 28 | 19 |
| 107986016 | XR_001740435 | LOC107986016 | uncharacterized LOC107986016 | lncRNA | . | . | . | . | . | 0.005525 | 0.011983 | 0.021620 | 0.010577 | 0.012856 | 0.027698 | 0.048514 | 0.024182 | 1.488926 | 2.812587 | 1.788565834 | 0.321079512 | 0.748150136 | NA | 1.175403 | 1.751299 | 1.472828 | 1.477882 | 1.485999 | 1.474918 | 2 | 3 | 4 | 2 |
| 107986024 | XR_001740495 | LOC107986024 | uncharacterized LOC107986024 | lncRNA | . | . | . | . | . | 0.036717 | 0.238550 | 0.090315 | 0.165206 | 0.085442 | 0.551419 | 0.202664 | 0.377724 | -1.110638 | 10.427129 | 1.047309691 | -0.14455051 | 0.885065775 | NA | 3.460000 | 3.300113 | 3.339372 | 3.408006 | 3.356690 | 3.382240 | 4 | 20 | 6 | 12 |
| 107986031 | XR_001740547 | LOC107986031 | uncharacterized LOC107986031 | lncRNA | . | . | . | . | . | 0.436221 | 0.392600 | 0.500714 | 0.356986 | 1.015104 | 0.907513 | 1.123588 | 0.816206 | -1.006099 | 15.631977 | 0.77774981 | -0.01127935 | 0.991000575 | NA | 3.969395 | 3.963455 | 3.968685 | 3.961241 | 3.976291 | 3.952649 | 19 | 16 | 16 | 12 |
| 107986043 | XR_001740556 | LOC107986043 | uncharacterized LOC107986043, transcript variant X2 | lncRNA | . | . | . | . | . | 1.666326 | 2.140571 | 1.334907 | 1.475648 | 3.877610 | 4.948026 | 2.995493 | 3.373894 | -1.357611 | 25.376392 | 0.617568458 | -0.71420427 | 0.475100874 | NA | 4.870101 | 4.426810 | 4.670918 | 4.697219 | 4.644606 | 4.660273 | 30 | 35 | 17 | 22 |
| 107986063 | XR_001740603 | XPC-AS1 | XPC antisense RNA 1 | lncRNA | . | 55014 | . | . | . | 0.116349 | 0.161039 | 0.252098 | 0.055231 | 0.270749 | 0.372250 | 0.565701 | 0.126279 | 1.062047 | 6.045242 | 1.28857603 | 0.06739863 | 0.94626436 | NA | 2.548173 | 2.641905 | 2.580621 | 2.586631 | 2.603296 | 2.566859 | 6 | 7 | 9 | 2 |
| 107986064 | XR_001740605 | LOC107986064 | uncharacterized LOC107986064, transcript variant X2 | lncRNA | . | . | . | . | . | 0.174307 | 0.357532 | 0.150825 | 0.391049 | 0.405620 | 0.826450 | 0.338446 | 0.894086 | 1.020488 | 28.206305 | 0.613857193 | 0.047664258 | 0.96198382 | 0.998865404 | 4.805868 | 4.829924 | 4.789715 | 4.834831 | 4.779012 | 4.849630 | 26 | 36 | 17 | 35 |
| 107986068 | XR_001740619 | LOC107986068 | uncharacterized LOC107986068 | lncRNA | . | . | . | . | . | 0.167069 | 0.166352 | 0.305498 | 0.313619 | 0.388777 | 0.384530 | 0.685529 | 0.717054 | 1.816192 | 10.724075 | 0.932893152 | 0.922845994 | 0.356087468 | NA | 2.928838 | 3.790061 | 3.395478 | 3.393825 | 3.422123 | 3.422698 | 9 | 8 | 12 | 13 |
| 107986077 | XR_001740668 | LOC107986077 | uncharacterized LOC107986077 | lncRNA | . | . | . | . | . | 0.078796 | 0.062070 | 0.077374 | 0.113065 | 0.183361 | 0.143477 | 0.173624 | 0.258509 | 1.474352 | 2.756358 | 1.797441179 | 0.31159909 | 0.755345226 | NA | 1.156660 | 1.715150 | 1.456815 | 1.453257 | 1.455501 | 1.464401 | 3 | 2 | 2 | 4 |
| 107986084 | XM_01700761 | LOC107986084 | Sjoegren syndrome nuclear autoantigen 1 homolog | protein_coding | XP_016863107.1 | . | . | . | . | 0.870416 | 1.082891 | 1.300204 | 0.897018 | 2.025495 | 2.503150 | 2.917620 | 2.050924 | 1.122862 | 41.079200 | 0.497849059 | 0.335805618 | 0.737017454 | 0.998865404 | 5.273664 | 5.442095 | 5.328785 | 5.360320 | 5.393229 | 5.331686 | 40 | 46 | 45 | 33 |
| 107986100 | NR_151705,NF | LINC02018 | long intergenic non-protein coding RNA 2018 | lncRNA | . | 52853 | . | . | . | 0.308284 | 0.309717 | 0.369642 | 0.351539 | 0.717391 | 0.715925 | 0.829466 | 0.803752 | -1.029872 | 16.797889 | 0.746720486 | -0.05686899 | 0.954649559 | NA | 4.092061 | 4.048019 | 4.069653 | 4.074527 | 4.063282 | 4.076621 | 19 | 19 | 13 | 17 |
| 107986119 | XR_001740863 | LOC107986119 | uncharacterized LOC107986119, transcript variant X2 | lncRNA | . | . | . | . | . | 0.727697 | 0.758373 | 1.327397 | 0.896372 | 1.693381 | 1.753012 | 2.978642 | 2.049449 | 1.540214 | 57.083101 | 0.438615379 | 1.420677003 | 0.155410681 | 0.924159366 | 5.488927 | 6.113896 | 5.761100 | 5.779939 | 5.914236 | 5.799291 | 49 | 51 | 75 | 49 |
| 107986127 | XR_001740893 | LOC107986127 | uncharacterized LOC107986127 | lncRNA | . | . | . | . | . | 0.266641 | 0.225271 | 0.275886 | 0.194342 | 0.620485 | 0.520724 | 0.619080 | 0.444339 | -1.058746 | 45.253855 | 0.474018384 | -0.173742 | 0.86206823 | 0.998865404 | 5.539157 | 5.459686 | 5.517870 | 5.488067 | 5.521639 | 5.464185 | 58 | 46 | 45 | 34 |
| 107986148 | XR_001740978 | LOC107986148 | uncharacterized LOC107986148 | lncRNA | . | . | . | . | . | 0.079110 | 0.034721 | 0.013091 | 0.057688 | 0.184093 | 0.080259 | 0.029375 | 0.131896 | -1.321088 | 5.055727 | 1.381512501 | -0.29078714 | 0.771214118 | NA | 2.526439 | 2.121007 | 2.358611 | 2.339413 | 2.333566 | 2.351688 | 9 | 4 | 2 | 6 |
| 107986153 | XR_001741020 | LOC107986153 | uncharacterized LOC107986153 | lncRNA | . | . | . | . | . | 0.380507 | 0.482654 | 0.362576 | 0.386880 | 0.885455 | 1.115676 | 0.813611 | 0.884556 | -1.165616 | 40.112849 | 0.494668569 | -0.44695169 | 0.654909944 | 0.998865404 | 5.432996 | 5.210413 | 5.318946 | 5.357710 | 5.309742 | 5.320005 | 44 | 52 | 31 | 36 |
| 107986160 | XR_001741040 | LOC107986160 | uncharacterized LOC107986160 | lncRNA | . | . | . | . | . | 0.052519 | 0.057987 | 0.046991 | 0.056795 | 0.122214 | 0.134040 | 0.105447 | 0.129855 | -1.033538 | 12.358091 | 0.865254846 | -0.05500303 | 0.956136053 | NA | 3.651489 | 3.602870 | 3.627208 | 3.630882 | 3.624475 | 3.630032 | 14 | 14 | 10 | 12 |
| 107986163 | NR_160768 | LOC107986163 | uncharacterized LOC107986163 | lncRNA | . | . | . | . | . | 0.287453 | 0.500934 | 0.318239 | 0.220886 | 0.668915 | 1.157932 | 0.714118 | 0.505029 | -1.435232 | 20.341913 | 0.707417653 | -0.73688348 | 0.461193208 | NA | 4.583502 | 4.062477 | 4.331321 | 4.387478 | 4.341150 | 4.317100 | 20 | 33 | 17 | 13 |
| 107986175 | XM_01700762 | LOC107986175 | uncharacterized LOC107986175 | protein_coding | XP_016863111.1 | . | . | . | . | 0.023410 | 0.079277 | 0.047752 | 0.040201 | 0.054477 | 0.183253 | 0.107154 | 0.091914 | -1.225524 | 2.469287 | 1.896160988 | -0.15473285 | 0.877031932 | NA | 1.444631 | 1.148375 | 1.301291 | 1.310651 | 1.304063 | 1.303286 | 2 | 4 | 2 | 2 |
| 107986176 | gene-LOC1079 | LOC107986176 | . | pseudogene | . | . | . | . | . | 0.038265 | 0.063983 | 0.311702 | 0.045654 | 0.089044 | 0.147901 | 0.699451 | 0.104383 | 2.924706 | 5.344222 | 1.439655909 | 1.0754596 | 0.28216901 | NA | 1.444631 | 2.993948 | 2.357289 | 2.366990 | 2.411707 | 2.360011 | 2 | 4 | 12 | 2 |
| 107986182 | XR_001741379 | LOC107986182 | uncharacterized LOC107986182, transcript variant X4 | lncRNA | . | . | . | . | . | 0.184790 | 0.192629 | 0.147306 | 0.285089 | 0.430014 | 0.445269 | 0.330549 | 0.651824 | 1.133286 | 39.173435 | 0.523316149 | 0.344938258 | 0.730140788 | 0.998865404 | 5.200543 | 5.377634 | 5.275694 | 5.278209 | 5.246570 | 5.343248 | 42 | 40 | 25 | 51 |
| 107986197 | XR_001741447 | LOC107986197 | uncharacterized LOC107986197 | lncRNA | . | . | . | . | . | 0.068815 | 0.042644 | 0.050854 | 0.053977 | 0.160136 | 0.098574 | 0.114115 | 0.123413 | -1.091740 | 8.106252 | 1.070230862 | -0.11831924 | 0.905814715 | NA | 3.079678 | 2.955730 | 3.031623 | 3.013456 | 3.020664 | 3.018077 | 12 | 7 | 7 | 7 |
| 107986209 | gene-H3P14 | H3P14 | . | pseudogene | . | 43805 | . | . | . | 0.195491 | 0.145885 | 0.191230 | 0.054041 | 0.454915 | 0.337219 | 0.429114 | 0.123558 | -1.198555 | 2.475538 | 1.907561801 | -0.13697918 | 0.89104725 | NA | 1.429096 | 1.175234 | 1.307104 | 1.307921 | 1.310815 | 1.300040 | 3 | 3 | 3 | 1 |
| 107986211 | XM_02445431 | LOC107986211 | translation initiation factor IF-2 | protein_coding | XP_024310085.1 | . | . | . | . | 0.064566 | 0.068756 | 0.107641 | 0.110831 | 0.150247 | 0.158932 | 0.241544 | 0.253401 | 1.603931 | 17.527664 | 0.732404994 | 0.930649315 | 0.352034996 | NA | 3.751025 | 4.432387 | 4.099887 | 4.104095 | 4.137918 | 4.136138 | 15 | 15 | 19 | 20 |
| 107986235 | XR_001741519 | LOC107986235 | uncharacterized LOC107986235 | lncRNA | . | . | . | . | . | 0.190903 | 0.337900 | 0.429216 | 0.351112 | 0.444238 | 0.781072 | 0.963149 | 0.802775 | 1.394845 | 8.108615 | 1.071142664 | 0.448217218 | 0.653996438 | NA | 2.760365 | 3.339024 | 2.996097 | 3.010828 | 3.019447 | 3.011635 | 6 | 9 | 9 | 8 |
| 107986246 | XR_001741543 | LOC107986246 | uncharacterized LOC107986246 | lncRNA | . | . | . | . | . | 0.091792 | 0.076310 | 0.172954 | 0.093610 | 0.213603 | 0.176393 | 0.388103 | 0.214027 | 1.383700 | 4.803926 | 1.375150259 | 0.340712422 | 0.733320086 | NA | 2.008843 | 2.481107 | 2.252885 | 2.249927 | 2.264306 | 2.252650 | 5 | 4 | 6 | 4 |
| 107986250 | XR_001741551 | LOC107986250 | uncharacterized LOC107986250 | lncRNA | . | . | . | . | . | 0.330972 | 0.409772 | 0.427833 | 0.411771 | 0.770185 | 0.947206 | 0.960046 | 0.941465 | 1.174450 | 10.261257 | 0.947035457 | 0.244960012 | 0.806487394 | NA | 3.238681 | 3.470304 | 3.347750 | 3.354753 | 3.360355 | 3.356829 | 10 | 11 | 10 | 10 |
| 107986298 | XR_001741783 | LOC107986298 | uncharacterized LOC107986298, transcript variant X2 | lncRNA | . | . | . | . | . | 0.187567 | 0.309754 | 0.253106 | 0.221560 | 0.436476 | 0.716011 | 0.567963 | 0.506571 | 1.043303 | 24.452300 | 0.635831056 | 0.096187023 | 0.923372037 | NA | 4.581746 | 4.641433 | 4.580449 | 4.629250 | 4.615446 | 4.603435 | 21 | 32 | 23 | 22 |
| 107986303 | XR_001741792 | LOC107986303 | uncharacterized LOC107986303 | lncRNA | . | . | . | . | . | 0.258131 | 0.246917 | 0.090310 | 0.385885 | 0.600683 | 0.570759 | 0.202653 | 0.882281 | -1.048621 | 12.218555 | 0.939455349 | -0.07290759 | 0.941879652 | NA | 3.648138 | 3.572884 | 3.613816 | 3.609381 | 3.577374 | 3.639878 | 15 | 13 | 4 | 18 |
| 107986334 | XR_002959825 | LOC107986334 | uncharacterized LOC107986334 | lncRNA | . | . | . | . | . | 0.023657 | 0.054976 | 0.042130 | 0.014739 | 0.055052 | 0.127079 | 0.094537 | 0.033698 | -1.372709 | 6.663666 | 1.212343035 | -0.37697705 | 0.706190659 | NA | 2.945312 | 2.491830 | 2.727297 | 2.750328 | 2.740169 | 2.718592 | 6 | 11 | 7 | 3 |
| 107986361 | XR_001742438 | LOC107986361 | uncharacterized LOC107986361 | lncRNA | . | . | . | . | . | 0.081500 | 0.026932 | 0.034694 | 0.016013 | 0.189653 | 0.062254 | 0.077852 | 0.036611 | -2.242786 | 5.432489 | 1.374368554 | -0.84787454 | 0.396507846 | NA | 2.906576 | 1.751299 | 2.478079 | 2.443020 | 2.447394 | 2.436239 | 13 | 4 | 4 | 2 |
| 107986363 | XR_001742454 | LOC107986363 | uncharacterized LOC107986363 | lncRNA | . | . | . | . | . | 0.056502 | 0.079008 | 0.076725 | 0.039891 | 0.131482 | 0.182631 | 0.172169 | 0.091206 | -1.160115 | 19.315924 | 0.719788139 | -0.29768162 | 0.765946174 | NA | 4.373794 | 4.161867 | 4.260017 | 4.289111 | 4.283610 | 4.240962 | 20 | 26 | 20 | 12 |
| 107986381 | XR_001742513 | LOC107986381 | uncharacterized LOC107986381 | lncRNA | . | . | . | . | . | 52.299862 | 53.053448 | 45.310783 | 61.685638 | 121.703934 | 122.635429 | 101.676079 | 141.036880 | 1.001382 | 1055.028216 | 0.160595732 | 0.012405957 | 0.990101733 | 0.999289967 | 10.042411 | 10.043720 | 10.038047 | 10.042077 | 9.925723 | 10.144483 | 1211 | 1141 | 782 | 1136 |
| 107986389 | XR_001742526 | LINC02163 | long intergenic non-protein coding RNA 2163 | lncRNA | . | 53024 | . | . | . | 2.417079 | 1.866656 | 3.633456 | 0.911065 | 5.624642 | 4.314859 | 8.153369 | 2.083041 | 1.077691 | 22.312613 | 0.736624663 | 0.146538959 | 0.883495937 | NA | 4.421266 | 4.536027 | 4.472261 | 4.447278 | 4.520619 | 4.409584 | 28 | 20 | 32 | 9 |
| 107986411 | XR_001742630 | LOC107986411 | uncharacterized LOC107986411 | lncRNA | . | . | . | . | . | 0.779428 | 0.607756 | 0.788119 | 0.519663 | 1.813760 | 1.404855 | 1.768516 | 1.188147 | -1.036890 | 13.611487 | 0.835499299 | -0.06255255 | 0.950122814 | NA | 3.790753 | 3.742346 | 3.774641 | 3.758532 | 3.775969 | 3.752775 | 18 | 13 | 14 | 10 |
| 107986451 | XR_001742893 | LOC107986451 | uncharacterized LOC107986451 | lncRNA | . | . | . | . | . | 0.155514 | 0.181334 | 0.243140 | 0.128517 | 0.361888 | 0.419162 | 0.545598 | 0.293838 | 1.140655 | 5.771902 | 1.262142153 | 0.150428314 | 0.880426704 | NA | 2.429096 | 2.622519 | 2.520125 | 2.521798 | 2.532740 | 2.515998 | 6 | 6 | 7 | 4 |
| 107986453 | NM_00136519 | LOC107986453 | uncharacterized LOC107986453 | protein_coding | NP_001352125.1;XP_016865619.1;XP_016865621.1;XP_016865622.1;XP_ | . | . | . | . | 0.473595 | 0.552071 | 0.288002 | 0.635147 | 1.102075 | 1.276138 | 0.646269 | 1.452188 | -1.173789 | 57.020478 | 0.470038509 | -0.4918168 | 0.622848857 | 0.998865404 | 5.946605 | 5.710567 | 5.815468 | 5.871936 | 5.736717 | 5.880926 | 62 | 75 | 28 | 68 |
| 107986465 | XR_001742939 | LOC107986465 | uncharacterized LOC107986465 | lncRNA | . | . | . | . | . | 0.537560 | 0.648037 | 0.522131 | 0.576553 | 1.250923 | 1.497966 | 1.171648 | 1.318220 | -1.086342 | 70.342170 | 0.379354573 | -0.31495095 | 0.752798898 | 0.998865404 | 6.195447 | 6.074662 | 6.124044 | 6.168031 | 6.116419 | 6.136442 | 77 | 86 | 56 | 66 |
| 107986467 | XR_001742942 | LOC107986467 | uncharacterized LOC107986467 | lncRNA | . | . | . | . | . | 0.768064 | 1.161489 | 1.407927 | 1.774747 | 1.787315 | 2.684835 | 3.159347 | 4.057749 | 1.605235 | 26.402951 | 0.612234951 | 1.115232868 | 0.264750645 | NA | 4.343257 | 5.022724 | 4.666401 | 4.699773 | 4.718768 | 4.748772 | 19 | 26 | 25 | 34 |
| 107986468 | XR_001742943 | LOC107986468 | uncharacterized LOC107986468 | lncRNA | . | . | . | . | . | 0.437108 | 0.795823 | 0.078982 | 0.071233 | 1.017167 | 1.839580 | 0.177233 | 0.162865 | -6.945687 | 4.405372 | 1.676923759 | -1.66740876 | 0.095433153 | NA | 2.945312 | 0.148375 | 2.158599 | 2.181198 | 2.140557 | 2.139864 | 6 | 11 | 1 | 1 |
| 107986476 | XR_001742987 | LOC107986476 | uncharacterized LOC107986476, transcript variant X1 | lncRNA | . | . | . | . | . | 0.047013 | 0.041653 | 0.065464 | 0.017902 | 0.109402 | 0.096283 | 0.146899 | 0.040931 | -1.169589 | 9.151284 | 1.040945982 | -0.2171122 | 0.82812091 | NA | 3.299300 | 3.080349 | 3.196760 | 3.191417 | 3.205851 | 3.169152 | 12 | 10 | 11 | 4 |
| 107986477 | XR_001742993 | LOC107986477 | uncharacterized LOC107986477 | lncRNA | . | . | . | . | . | 0.476098 | 0.394205 | 0.652710 | 0.639164 | 1.107901 | 0.911223 | 1.464662 | 1.461372 | 1.452551 | 25.220100 | 0.614817283 | 0.876015076 | 0.381021838 | NA | 4.361571 | 4.901253 | 4.636633 | 4.619367 | 4.666526 | 4.666767 | 26 | 20 | 26 | 28 |
| 107986488 | XR_001743015 | LOC107986488 | uncharacterized LOC107986488 | lncRNA | . | . | . | . | . | 0.112033 | 0.146829 | 0.123108 | 0.045416 | 0.260707 | 0.339401 | 0.276252 | 0.103837 | -1.559686 | 12.946556 | 0.892678215 | -0.71835071 | 0.472541071 | NA | 3.977427 | 3.342199 | 3.698571 | 3.714875 | 3.702701 | 3.660938 | 16 | 19 | 13 | 5 |
| 107986496 | XR_001743050 | TRIM7-AS1 | TRIM7 antisense RNA 1 | lncRNA | . | 40764 | . | . | . | 0.055983 | 0.026807 | 0.052635 | 0.060228 | 0.130275 | 0.061965 | 0.118112 | 0.137703 | 1.345721 | 6.255258 | 1.215938715 | 0.352303757 | 0.724610471 | NA | 2.413393 | 2.844638 | 2.643126 | 2.627953 | 2.642677 | 2.645267 | 8 | 4 | 6 | 7 |
| 107986513 | XR_001743787 | LOC107986513 | uncharacterized LOC107986513 | lncRNA | . | . | . | . | . | 0.005214 | 0.015181 | 0.012454 | 0.002690 | 0.012133 | 0.035091 | 0.027947 | 0.006151 | -1.254719 | 5.731092 | 1.33258195 | -0.24566154 | 0.805944246 | NA | 2.671431 | 2.348113 | 2.500639 | 2.527692 | 2.521491 | 2.495102 | 4 | 10 | 7 | 2 |
| 107986515 | XR_001743790 | LOC107986515 | uncharacterized LOC107986515 | lncRNA | . | . | . | . | . | 0.007513 | 0.021205 | 0.011983 | 0.010502 | 0.017482 | 0.049016 | 0.026890 | 0.024012 | -1.225524 | 2.469287 | 1.896160988 | -0.15473285 | 0.877031932 | NA | 1.444631 | 1.148375 | 1.301291 | 1.310651 | 1.304063 | 1.303286 | 2 | 4 | 2 | 2 |
| 107986556 | XR_001743927 | LOC107986556 | uncharacterized LOC107986556, transcript variant X1 | lncRNA | . | . | . | . | . | 0.083808 | 0.029659 | 0.180726 | 0.195213 | 0.195025 | 0.068557 | 0.405544 | 0.446332 | 2.658488 | 6.486812 | 1.237905158 | 1.139510355 | 0.254490352 | NA | 1.820514 | 3.239024 | 2.668547 | 2.652733 | 2.691464 | 2.683766 | 6 | 2 | 9 | 8 |
| 107986558 | XR_001743932 | LOC107986558 | uncharacterized LOC107986558 | lncRNA | . | . | . | . | . | 0.090902 | 0.326531 | 0.243215 | 0.328396 | 0.211532 | 0.754791 | 0.545768 | 0.750838 | 1.372849 | 28.105958 | 0.67007814 | 0.682268506 | 0.495069184 | 0.998865404 | 4.569585 | 5.020874 | 4.715330 | 4.830428 | 4.791458 | 4.830234 | 12 | 40 | 24 | 35 |
| 107986593 | XR_001744107 | LOC107986593 | uncharacterized LOC107986593 | lncRNA | . | . | . | . | . | 0.221933 | 0.167919 | 0.171055 | 0.282350 | 0.516448 | 0.388152 | 0.383843 | 0.645560 | 1.112624 | 9.902958 | 0.972408685 | 0.158334227 | 0.874193439 | NA | 3.229742 | 3.381963 | 3.309438 | 3.299918 | 3.299058 | 3.319669 | 12 | 9 | 7 | 12 |
| 107986594 | XR_001744111 | LOC107986594 | uncharacterized LOC107986594 | lncRNA | . | . | . | . | . | 0.314097 | 0.341608 | 0.200674 | 0.183216 | 0.730917 | 0.789643 | 0.450307 | 0.418903 | -1.693772 | 16.418088 | 0.762563137 | -0.99695331 | 0.318787174 | NA | 4.367696 | 3.607807 | 4.058434 | 4.063969 | 4.027054 | 4.022778 | 23 | 23 | 11 | 11 |
| 107986596 | XM_01701161 | LOC107986596 | arf-GAP with GTPase, ANK repeat and PH domain-containing prot | protein_coding | XP_016867102.1 | . | . | . | . | 0.204013 | 0.105997 | 0.166543 | 0.185725 | 0.474746 | 0.245016 | 0.373718 | 0.424639 | 1.123132 | 14.613601 | 0.817879359 | 0.204831018 | 0.837704152 | NA | 3.781608 | 3.951852 | 3.882747 | 3.842537 | 3.867958 | 3.872473 | 21 | 10 | 13 | 15 |
| 107986597 | XR_001744125 | LOC107986597 | uncharacterized LOC107986597 | lncRNA | . | . | . | . | . | 0.697682 | 0.538120 | 0.965859 | 0.448286 | 1.623534 | 1.243889 | 2.167360 | 1.024952 | 1.072337 | 6.493617 | 1.200672597 | 0.083918644 | 0.933121116 | NA | 2.644780 | 2.751299 | 2.695991 | 2.689514 | 2.705619 | 2.683845 | 8 | 6 | 8 | 4 |
| 107986613 | XR_001744210 | LOC107986613 | uncharacterized LOC107986613 | lncRNA | . | . | . | . | . | 0.053350 | 0.174543 | 0.116373 | 0.072848 | 0.124147 | 0.403464 | 0.261138 | 0.166559 | -1.182962 | 5.472329 | 1.334986123 | -0.18157781 | 0.855914061 | NA | 2.569585 | 2.324312 | 2.433470 | 2.464651 | 2.448346 | 2.441704 | 3 | 10 | 5 | 4 |
| 107986630 | XR_001744287 | LOC107986630 | uncharacterized LOC107986630 | lncRNA | . | . | . | . | . | 0.081731 | 0.413179 | 0.166810 | 0.117391 | 0.190191 | 0.955081 | 0.374317 | 0.268400 | -1.824503 | 7.079556 | 1.221068208 | -0.71044628 | 0.477427431 | NA | 3.193905 | 2.324312 | 2.802876 | 2.854880 | 2.814082 | 2.807320 | 4 | 16 | 5 | 4 |
| 107986644 | XR_001744345 | LOC107986644 | uncharacterized LOC107986644 | lncRNA | . | . | . | . | . | 0.119579 | 0.107137 | 0.066993 | 0.161312 | 0.278265 | 0.247652 | 0.150330 | 0.368820 | -1.020375 | 6.641996 | 1.180246235 | -0.02465506 | 0.980330105 | NA | 2.747897 | 2.715150 | 2.734976 | 2.732828 | 2.724607 | 2.741931 | 8 | 7 | 4 | 8 |
| 107986649 | XR_001744379 | LOC107986649 | uncharacterized LOC107986649, transcript variant X1 | lncRNA | . | . | . | . | . | 0.268876 | 0.274418 | 0.205611 | 0.170856 | 0.625685 | 0.634330 | 0.461385 | 0.390642 | -1.412555 | 88.334613 | 0.343426207 | -1.45098871 | 0.146783004 | 0.90884489 | 6.692024 | 6.195232 | 6.526831 | 6.494565 | 6.435062 | 6.394730 | 125 | 106 | 69 | 63 |
| 107986655 | XR_001744396 | LOC107986655 | uncharacterized LOC107986655, transcript variant X1 | lncRNA | . | . | . | . | . | 0.071717 | 0.058001 | 0.046747 | 0.088195 | 0.166888 | 0.134073 | 0.104899 | 0.201647 | -1.041354 | 10.064261 | 0.970963127 | -0.06020827 | 0.951989761 | NA | 3.361571 | 3.300113 | 3.338816 | 3.329568 | 3.319548 | 3.345087 | 13 | 10 | 6 | 12 |
| 107986727 | gene-LOC1079 | LOC107986727 | . | pseudogene | . | . | . | . | . | 0.008124 | 0.026190 | 0.030497 | 0.000126 | 0.018904 | 0.060540 | 0.068433 | 0.000287 | 1.236246 | 1.522137 | 2.383171416 | 0.128385813 | 0.897843658 | NA | 0.444631 | 0.751299 | 0.592746 | 0.597508 | 0.599417 | 0.593826 | 1 | 2 | 2 | 1 |
| 107986742 | XR_001745024 | LOC107986742 | uncharacterized LOC107986742 | lncRNA | . | . | . | . | . | 0.284842 | 0.197474 | 0.265883 | 0.445416 | 0.662838 | 0.456471 | 0.596635 | 1.018391 | 1.353992 | 10.492099 | 0.950825896 | 0.45983099 | 0.645637537 | NA | 3.156660 | 3.592943 | 3.385696 | 3.371944 | 3.380454 | 3.405273 | 12 | 8 | 8 | 14 |
| 107986785 | XR_001745175 | LOC107986785 | uncharacterized LOC107986785 | lncRNA | . | . | . | . | . | 0.094410 | 0.147497 | 0.131353 | 0.065569 | 0.219695 | 0.340945 | 0.294753 | 0.149917 | -1.211317 | 17.324585 | 0.766147977 | -0.36099658 | 0.718102006 | NA | 4.245349 | 3.971139 | 4.099409 | 4.136096 | 4.124724 | 4.082311 | 17 | 25 | 18 | 10 |
| 107986798 | XR_001745214 | LOC107986798 | uncharacterized LOC107986798 | lncRNA | . | . | . | . | . | 0.063008 | 0.022946 | 0.073775 | 0.040206 | 0.146623 | 0.053041 | 0.165549 | 0.091925 | 1.257223 | 4.522683 | 1.433731616 | 0.230336237 | 0.817830504 | NA | 1.998355 | 2.336262 | 2.172403 | 2.160913 | 2.179583 | 2.163123 | 6 | 3 | 6 | 3 |
| 107986800 | XM_01701292 | LOC107986800 | putative uncharacterized protein FLJ44672, transcript variant X2 | protein_coding | XP_016868417.1;XP_016868418.1 | . | . | . | . | 0.239382 | 0.039638 | 0.096006 | 0.051077 | 0.557053 | 0.091624 | 0.215434 | 0.116782 | -1.197014 | 7.291254 | 1.165804426 | -0.22254165 | 0.823892255 | NA | 2.987791 | 2.733338 | 2.890588 | 2.851529 | 2.866444 | 2.864186 | 14 | 4 | 6 | 6 |
| 107986820 | XR_001745270 | LOC107986820 | uncharacterized LOC107986820, transcript variant X1 | lncRNA | . | . | . | . | . | 0.134012 | 0.174953 | 0.249004 | 0.178758 | 0.311852 | 0.404410 | 0.558760 | 0.408708 | 1.081261 | 33.251827 | 0.548786763 | 0.205388523 | 0.837268586 | 0.998865404 | 4.997177 | 5.111291 | 5.031052 | 5.058066 | 5.081034 | 5.028200 | 33 | 38 | 36 | 26 |
| 107986827 | NR_147989 | CZ1P-ASNS | CZ1P-ASNS readthrough | lncRNA | . | . | . | . | . | 0.191532 | 0.405995 | 0.016866 | 0.137465 | 0.445703 | 0.938476 | 0.037846 | 0.314297 | -3.630595 | 21.355375 | 0.819837523 | -2.26899326 | 0.023268736 | NA | 5.068704 | 3.193768 | 4.405667 | 4.502763 | 4.323095 | 4.381492 | 25 | 49 | 2 | 15 |
| 107986847 | XR_001745359 | LOC107986847 | uncharacterized LOC107986847 | lncRNA | . | . | . | . | . | 0.051106 | 0.118180 | 0.240616 | 0.267914 | 0.118925 | 0.273179 | 0.539935 | 0.612553 | 2.807888 | 14.626098 | 0.834499609 | 1.784884726 | 0.074280017 | NA | 2.945312 | 4.429601 | 3.797170 | 3.821021 | 3.866630 | 3.874356 | 6 | 11 | 18 | 21 |
| 107986850 | XR_001745375 | LOC107986850 | uncharacterized LOC107986850 | lncRNA | . | . | . | . | . | 0.294256 | 0.148097 | 0.188127 | 0.134351 | 0.684745 | 0.342332 | 0.422152 | 0.307178 | -1.408073 | 14.057921 | 0.843766931 | -0.5851406 | 0.558453181 | NA | 4.037030 | 3.548418 | 3.849309 | 3.802360 | 3.814394 | 3.796213 | 25 | 12 | 12 | 9 |
| 107986852 | XR_001745381 | LOC107986852 | uncharacterized LOC107986852, transcript variant X1 | lncRNA | . | . | . | . | . | 3.861370 | 3.371444 | 4.614297 | 4.105697 | 8.985566 | 7.793246 | 10.354347 | 9.387190 | 1.191416 | 50.978819 | 0.439769456 | 0.574568508 | 0.565583155 | 0.998865404 | 5.539157 | 5.793316 | 5.665558 | 5.636086 | 5.694882 | 5.672519 | 58 | 46 | 51 | 49 |
| 107986853 | XR_001745384 | LOC107986853 | uncharacterized LOC107986853 | lncRNA | . | . | . | . | . | 0.110577 | 0.096612 | 0.379973 | 0.168300 | 0.257318 | 0.223323 | 0.852649 | 0.384798 | 2.580743 | 12.833991 | 0.896378499 | 1.525902971 | 0.127034031 | NA | 2.838265 | 4.210731 | 3.625781 | 3.619956 | 3.700940 | 3.644225 | 9 | 7 | 22 | 11 |
| 107986858 | XR_001745420 | LOC107986858 | uncharacterized LOC107986858 | lncRNA | . | . | . | . | . | 0.050563 | 0.156012 | 0.054724 | 0.109060 | 0.117663 | 0.360630 | 0.122799 | 0.249352 | -1.241112 | 7.380782 | 1.161054771 | -0.26840497 | 0.788387616 | NA | 3.034734 | 2.715150 | 2.868229 | 2.903768 | 2.870388 | 2.887631 | 5 | 13 | 4 | 8 |
| 107986859 | XR_001745421 | LOC107986859 | uncharacterized LOC107986859 | lncRNA | . | . | . | . | . | 0.223478 | 0.246818 | 0.388301 | 0.310350 | 0.520042 | 0.570532 | 0.871337 | 0.709578 | 1.462531 | 12.161675 | 0.875378796 | 0.626548533 | 0.530955214 | NA | 3.303566 | 3.852980 | 3.582422 | 3.585527 | 3.611077 | 3.597207 | 11 | 11 | 14 | 12 |
| 107986862 | XR_001745427 | LOC107986862 | uncharacterized LOC107986862 | lncRNA | . | . | . | . | . | 0.311358 | 0.347604 | 0.505575 | 0.208642 | 0.724544 | 0.803503 | 1.134495 | 0.477036 | 1.081467 | 28.094913 | 0.626852974 | 0.18024832 | 0.856957625 | 0.998865404 | 4.752586 | 4.869520 | 4.788655 | 4.803636 | 4.852156 | 4.755603 | 29 | 31 | 36 | 16 |
| 107986871 | XR_002956669 | LOC107986871 | ubiquitin-conjugating enzyme E2 variant 1 pseudogene | transcribed_ps | . | . | . | . | . | 0.133025 | 0.160207 | 0.261353 | 0.090512 | 0.309556 | 0.370326 | 0.586469 | 0.206945 | 1.206356 | 28.312314 | 0.643087557 | 0.42086886 | 0.673850846 | 0.998865404 | 4.679488 | 4.954173 | 4.784858 | 4.803565 | 4.873534 | 4.754569 | 27 | 30 | 40 | 15 |
| 107986874 | XR_001745685 | LOC107986874 | uncharacterized LOC107986874 | lncRNA | . | . | . | . | . | 0.051577 | 0.036683 | 0.067497 | 0.104675 | 0.120021 | 0.084794 | 0.151460 | 0.239327 | 1.846010 | 12.687914 | 0.874833967 | 1.01094681 | 0.312041888 | NA | 3.156660 | 4.040734 | 3.641167 | 3.627548 | 3.651380 | 3.683932 | 12 | 8 | 11 | 19 |
| 107986876 | NR_160769 | LOC107986876 | uncharacterized LOC107986876 | lncRNA | . | . | . | . | . | 0.168608 | 0.154257 | 0.076103 | 0.206014 | 0.392357 | 0.356571 | 0.170774 | 0.471028 | -1.127141 | 9.272164 | 1.029994981 | -0.16764002 | 0.866866485 | NA | 3.299300 | 3.121007 | 3.221274 | 3.215960 | 3.195720 | 3.231341 | 12 | 10 | 4 | 12 |
| 107986898 | XR_001745731 | LOC107986898 | uncharacterized LOC107986898 | lncRNA | . | . | . | . | . | 0.134346 | 0.340283 | 0.267815 | 0.168907 | 0.312630 | 0.786580 | 0.600969 | 0.386186 | -1.085934 | 5.226067 | 1.332906201 | -0.08923083 | 0.928898464 | NA | 2.444631 | 2.324312 | 2.374294 | 2.392788 | 2.384933 | 2.378311 | 4 | 8 | 5 | 4 |
| 107986910 | XM_01701415 | LOC107986910 | uncharacterized LOC107986910, transcript variant X4 | protein_coding | XP_016869640.1;XP_016869641.1;XP_016869642.1 | . | . | . | . | 2.901904 | 3.313864 | 3.804977 | 2.846819 | 6.752849 | 7.660145 | 8.538257 | 6.508912 | 1.069699 | 103.226169 | 0.323169105 | 0.30078711 | 0.763576838 | 0.998865404 | 6.639816 | 6.737849 | 6.656960 | 6.693899 | 6.733509 | 6.658203 | 108 | 114 | 104 | 88 |
| 107986925 | XR_001745828 | LOC107986925 | uncharacterized LOC107986925, transcript variant X1 | lncRNA | . | . | . | . | . | 9.418409 | 8.940829 | 9.468426 | 9.366309 | 21.917024 | 20.667127 | 21.246874 | 21.414953 | 1.010276 | 1191.093292 | 0.131143139 | 0.112468418 | 0.91045201 | 0.998865404 | 10.210651 | 10.225453 | 10.230296 | 10.197719 | 10.216728 | 10.226922 | 1394 | 1251 | 1034 | 1130 |
| 107986935 | XR_001745860 | LOC107986935 | uncharacterized LOC107986935 | lncRNA | . | . | . | . | . | 0.112000 | 0.085979 | 0.142139 | 0.175003 | 0.260628 | 0.198744 | 0.318956 | 0.400123 | 1.410871 | 7.539457 | 1.103028596 | 0.450202627 | 0.652564342 | NA | 2.644780 | 3.141582 | 2.905372 | 2.898970 | 2.910361 | 2.917271 | 8 | 6 | 7 | 9 |
| 107986939 | XR_001745885 | LOC107986939 | uncharacterized LOC107986939 | lncRNA | . | . | . | . | . | 0.026439 | 0.022367 | 0.058109 | 0.052254 | 0.061525 | 0.051701 | 0.130395 | 0.119472 | 2.240255 | 3.618839 | 1.595365712 | 0.729402146 | 0.465755705 | NA | 1.156660 | 2.324312 | 1.832024 | 1.828510 | 1.846379 | 1.839861 | 3 | 2 | 5 | 4 |
| 107986942 | XR_001745895 | LOC107986942 | uncharacterized LOC107986942 | lncRNA | . | . | . | . | . | 0.027116 | 0.074040 | 0.023461 | 0.021985 | 0.063100 | 0.171147 | 0.052647 | 0.050266 | -1.773381 | 4.616099 | 1.43640747 | -0.57539547 | 0.565023865 | NA | 2.562483 | 1.733338 | 2.205653 | 2.228270 | 2.205944 | 2.204703 | 4 | 9 | 3 | 3 |
| 107986985 | XR_001746138 | LOC107986985 | uncharacterized LOC107986985 | lncRNA | . | . | . | . | . | 1.891434 | 1.477202 | 1.230603 | 1.535743 | 4.401444 | 3.414619 | 2.761437 | 3.511295 | -1.230890 | 61.495871 | 0.410610581 | -0.72989254 | 0.46545587 | 0.998865404 | 6.084644 | 5.784621 | 5.995499 | 5.935015 | 5.900819 | 5.940443 | 88 | 64 | 43 | 57 |
| 107986991 | XR_001746442 | LOC107986991 | uncharacterized LOC107986991 | lncRNA | . | . | . | . | . | 1.146227 | 1.065897 | 1.462486 | 1.168036 | 2.667317 | 2.463869 | 3.281777 | 2.670576 | 1.176463 | 38.025758 | 0.506390248 | 0.462995352 | 0.643367699 | 0.998865404 | 5.125927 | 5.362218 | 5.238147 | 5.225717 | 5.274041 | 5.237598 | 42 | 36 | 40 | 34 |
| 107986997 | XR_001746463 | LOC107986997 | uncharacterized LOC107986997 | lncRNA | . | . | . | . | . | 0.147216 | 0.148638 | 0.421427 | 0.056674 | 0.342578 | 0.343583 | 0.945670 | 0.129579 | 1.587125 | 34.298470 | 0.704041769 | 0.94655776 | 0.343864177 | 0.998865404 | 4.725983 | 5.396862 | 5.013252 | 5.017195 | 5.191893 | 4.949004 | 30 | 29 | 64 | 10 |
| 107986998 | XR_001746464 | LOC107986998 | uncharacterized LOC107986998 | lncRNA | . | . | . | . | . | 0.043737 | 0.009601 | 0.105629 | 0.112336 | 0.101778 | 0.022193 | 0.237028 | 0.256844 | 3.120142 | 3.653820 | 1.63463852 | 1.004265982 | 0.315250426 | NA | 0.820514 | 2.470304 | 1.840605 | 1.832638 | 1.854992 | 1.853314 | 3 | 1 | 5 | 5 |
| 107987000 | XR_001746475 | LOC107987000 | uncharacterized LOC107987000, transcript variant X4 | lncRNA | . | . | . | . | . | 0.136619 | 0.135922 | 0.107998 | 0.139387 | 0.317918 | 0.314189 | 0.242344 | 0.318692 | -1.176016 | 16.223324 | 0.762265664 | -0.30685787 | 0.758951557 | NA | 4.133136 | 3.897226 | 4.019369 | 4.036143 | 4.011891 | 4.021068 | 18 | 21 | 12 | 15 |
| 107987001 | XR_001746484 | FAM27E4 | family with sequence similarity 27 member E4, transcript variant X | lncRNA | . | 44415 | . | . | . | 1.893557 | 2.082783 | 2.248924 | 1.738081 | 4.406386 | 4.814446 | 5.046521 | 3.973915 | 1.030405 | 107.161073 | 0.313618848 | 0.137782299 | 0.890412475 | 0.998865404 | 6.721647 | 6.765297 | 6.719592 | 6.754161 | 6.771002 | 6.721541 | 115 | 120 | 102 | 94 |
| 107987007 | XR_001746507 | LOC107987007 | uncharacterized LOC107987007, transcript variant X3 | lncRNA | . | . | . | . | . | 0.240627 | 0.184395 | 0.092033 | 0.228504 | 0.559948 | 0.426238 | 0.206520 | 0.522446 | -1.211978 | 15.143311 | 0.805040722 | -0.34453286 | 0.730445585 | NA | 4.054853 | 3.772579 | 3.925686 | 3.934320 | 3.898392 | 3.936166 | 18 | 19 | 8 | 17 |
| 107987041 | XR_001746588 | LOC107987041 | uncharacterized LOC107987041 | lncRNA | . | . | . | . | . | 0.142124 | 0.202014 | 0.297310 | 0.092964 | 0.330728 | 0.466965 | 0.667155 | 0.212552 | -1.070164 | 1.739247 | 2.228774873 | -0.04389516 | 0.964987971 | NA | 0.844134 | 0.751299 | 0.796508 | 0.797062 | 0.799027 | 0.793436 | 2 | 2 | 2 | 1 |
| 107987042 | XM_01701539 | LOC107987042 | phosphatidylinositol 4,5-bisphosphate 5-phosphatase A-like | protein_coding | XP_016870882.1 | . | . | . | . | 0.012589 | 0.029647 | 0.131922 | 0.052178 | 0.029294 | 0.068530 | 0.296030 | 0.119299 | 3.728483 | 5.337549 | 1.388024093 | 1.367835601 | 0.171363541 | NA | 1.175403 | 3.073227 | 2.361919 | 2.367194 | 2.406393 | 2.379308 | 2 | 3 | 10 | 5 |
| 107987057 | XR_001746639 | LOC107987057 | uncharacterized LOC107987057 | lncRNA | . | . | . | . | . | 0.027551 | 0.019925 | 0.017292 | 0.033965 | 0.064112 | 0.046057 | 0.038803 | 0.077657 | -1.016871 | 6.627420 | 1.185292646 | -0.02036291 | 0.983753873 | NA | 2.741623 | 2.715150 | 2.736303 | 2.725804 | 2.721874 | 2.739194 | 9 | 6 | 4 | 8 |
| 107987063 | XR_001746653 | LOC107987063 | uncharacterized LOC107987063 | lncRNA | . | . | . | . | . | 1.499186 | 1.210876 | 0.824562 | 0.717966 | 3.488668 | 2.798994 | 1.850293 | 1.641544 | -1.770839 | 43.014420 | 0.484907248 | -1.70018758 | 0.089095648 | 0.802216049 | 5.780158 | 4.957666 | 5.493449 | 5.449397 | 5.389711 | 5.371141 | 70 | 53 | 29 | 27 |
| 107987072 | XR_001746688 | LOC107987072 | uncharacterized LOC107987072 | lncRNA | . | . | . | . | . | 3.952108 | 3.986424 | 4.753719 | 4.519300 | 9.196718 | 9.214798 | 10.667208 | 10.332842 | 1.155240 | 148.820078 | 0.267742809 | 0.777584815 | 0.436813816 | 0.998865404 | 7.109398 | 7.317924 | 7.186028 | 7.186292 | 7.253976 | 7.229914 | 159 | 149 | 143 | 145 |
| 107987075 | XR_001746692 | LOC107987075 | uncharacterized LOC107987075, transcript variant X1 | misc_RNA | . | . | . | . | . | 0.626403 | 0.906250 | 0.474022 | 0.378662 | 1.457664 | 2.094835 | 1.063691 | 0.865766 | -1.353607 | 7.827505 | 1.087071699 | -0.40182189 | 0.687815111 | NA | 3.170740 | 2.733338 | 2.972336 | 2.983185 | 2.968374 | 2.966064 | 9 | 11 | 6 | 6 |
| 107987095 | XR_001746837 | LOC107987095 | uncharacterized LOC107987095 | lncRNA | . | . | . | . | . | 0.974367 | 0.606118 | 1.524894 | 1.256265 | 2.267393 | 1.401070 | 3.421818 | 2.872300 | 1.726268 | 69.245578 | 0.403968761 | 1.949796159 | 0.05120042 | 0.672475152 | 5.664641 | 6.455584 | 6.069169 | 5.980891 | 6.180869 | 6.125584 | 72 | 42 | 84 | 74 |

| | | | | | | | | | | | | | | | | | | | | | | | | | | | | | | | | | |
|---|---|---|---|---|---|---|---|---|---|---|---|---|---|---|---|---|---|---|---|---|---|---|---|---|---|---|---|---|---|---|---|---|---|
| 107987097 | XR_001746839 | LOC107987097 | uncharacterized LOC107987097, transcript variant X3 | lncRNA | . | . | . | . | . | 0.059926 | 0.109125 | 0.100695 | 0.063838 | 0.139450 | 0.252247 | 0.225955 | 0.145958 | -1.015426 | 17.957032 | 0.743773009 | -0.0296924 | 0.976312375 | NA | 4.177728 | 4.155137 | 4.139195 | 4.183802 | 4.172560 | 4.148454 | 15 | 25 | 18 | 14 |
| 107987102 | XR_001746847 | LOC107987102 | uncharacterized LOC107987102 | lncRNA | . | . | . | . | . | 0.042164 | 0.044405 | 0.040400 | 0.121298 | 0.098117 | 0.102644 | 0.090657 | 0.277335 | 1.674670 | 4.791002 | 1.398226315 | 0.532014868 | 0.594715698 | NA | 1.844134 | 2.582949 | 2.244755 | 2.245952 | 2.245095 | 2.267798 | 4 | 4 | 3 | 8 |
| 107987111 | XR_001746874 | LOC107987111 | uncharacterized LOC107987111, transcript variant X1 | lncRNA | . | . | . | . | . | 0.450530 | 0.713220 | 0.492513 | 0.261286 | 1.048400 | 1.648640 | 1.105185 | 0.597399 | -1.620183 | 10.963120 | 0.943930251 | -0.7375091 | 0.460812811 | NA | 3.760365 | 3.066070 | 3.454251 | 3.482175 | 3.454124 | 3.436697 | 12 | 18 | 9 | 6 |
| 107987116 | XR_001746897 | LOC107987116 | uncharacterized LOC107987116, transcript variant X2 | lncRNA | . | . | . | . | . | 0.858720 | 0.787019 | 0.751087 | 0.979656 | 1.998277 | 1.819228 | 1.685417 | 2.239868 | 1.031789 | 14.111492 | 0.81272158 | 0.055551326 | 0.955699241 | NA | 3.796818 | 3.840449 | 3.817792 | 3.817834 | 3.812604 | 3.826846 | 16 | 15 | 11 | 15 |
| 107987118 | XR_001746900 | LOC107987118 | uncharacterized LOC107987118 | lncRNA | . | . | . | . | . | 0.481680 | 0.285376 | 0.433934 | 0.312190 | 1.120889 | 0.659659 | 0.973735 | 0.713786 | -1.042219 | 41.441644 | 0.510407487 | -0.1168834 | 0.906952452 | 0.998865404 | 5.400611 | 5.344869 | 5.413483 | 5.329422 | 5.389894 | 5.340911 | 61 | 34 | 41 | 32 |
| 107987126 | XR_001746926 | LOC107987126 | uncharacterized LOC107987126 | lncRNA | . | . | . | . | . | 0.371988 | 0.815198 | 0.445271 | 0.642871 | 0.865632 | 1.884367 | 0.999175 | 1.469849 | -1.108182 | 28.433006 | 0.622877265 | -0.2379189 | 0.811943996 | 0.998865404 | 4.904062 | 4.750862 | 4.783242 | 4.873051 | 4.796988 | 4.837408 | 22 | 44 | 19 | 30 |
| 107987135 | XR_001746960 | LOC107987135 | uncharacterized LOC107987135 | transcribed_ps | . | . | . | . | . | 0.038062 | 0.059181 | 0.073698 | 0.054251 | 0.088571 | 0.136800 | 0.165377 | 0.124038 | 1.315256 | 7.841948 | 1.090170043 | 0.362643613 | 0.716871126 | NA | 2.760365 | 3.155137 | 2.949621 | 2.964331 | 2.972882 | 2.960353 | 6 | 9 | 9 | 7 |
| 107987141 | XR_001746979 | LOC107987141 | uncharacterized LOC107987141 | lncRNA | . | . | . | . | . | 1.144505 | 1.385366 | 1.173847 | 1.562635 | 2.663310 | 3.202336 | 2.634080 | 3.572778 | 1.069955 | 91.148254 | 0.344735777 | 0.282969213 | 0.777200437 | 0.998865404 | 6.461710 | 6.557003 | 6.470369 | 6.523887 | 6.477781 | 6.554443 | 92 | 104 | 71 | 100 |
| 107987142 | NR_165240,NF | LOC107987142 | uncharacterized LOC107987142 | lncRNA | . | . | . | . | . | 0.525946 | 0.683241 | 0.859064 | 0.780522 | 1.223896 | 1.579342 | 1.927715 | 1.784572 | -1.116369 | 22.961129 | 0.646579232 | -0.24562164 | 0.805975137 | NA | 4.597285 | 4.440712 | 4.526893 | 4.525874 | 4.530445 | 4.501012 | 28 | 26 | 22 | 17 |
| 107987147 | XR_001747447 | LOC107987147 | uncharacterized LOC107987147 | lncRNA | . | . | . | . | . | 0.199497 | 0.360815 | 0.386267 | 0.227206 | 0.464237 | 0.834040 | 0.866772 | 0.519479 | 1.075856 | 43.678016 | 0.51683596 | 0.204097022 | 0.838277681 | 0.998865404 | 5.394834 | 5.500888 | 5.383171 | 5.475297 | 5.486748 | 5.396078 | 35 | 58 | 50 | 31 |
| 107987150 | XR_001747529 | LOC107987150 | uncharacterized LOC107987150 | lncRNA | . | . | . | . | . | 0.228860 | 0.157409 | 0.247129 | 0.154457 | 0.532568 | 0.363858 | 0.554550 | 0.353148 | 1.026083 | 20.357195 | 0.695772274 | 0.05339089 | 0.957420464 | NA | 4.326585 | 4.368050 | 4.357453 | 4.328029 | 4.363997 | 4.325684 | 27 | 18 | 22 | 15 |
| 107987158 | XM_01701863 | LOC107987158 | uncharacterized LOC107987158, transcript variant X1 | protein_coding | XP_016874124.1;XP_016874125.1;XP_016874126.1;XP_016874127.1 | . | . | . | . | 0.061623 | 0.055213 | 0.112990 | 0.035327 | 0.143400 | 0.127627 | 0.253546 | 0.080771 | 1.213373 | 14.389721 | 0.86071199 | 0.324176475 | 0.745804431 | NA | 3.697260 | 3.982589 | 3.830233 | 3.821966 | 3.872763 | 3.802210 | 16 | 13 | 21 | 7 |
| 107987170 | XR_001748946 | LOC107987170 | uncharacterized LOC107987170, transcript variant X2 | lncRNA | . | . | . | . | . | 0.087069 | 0.047659 | 0.276153 | 0.075967 | 0.202612 | 0.110167 | 0.619680 | 0.173689 | **2.112546** | 4.165096 | 1.545851521 | 0.697985932 | 0.485185992 | NA | 1.413393 | 2.502474 | 2.028595 | 2.020915 | 2.054123 | 2.022601 | 4 | 2 | 8 | 2 |
| 107987172 | XR_001748990 | LOC107987172 | uncharacterized LOC107987172, transcript variant X1 | lncRNA | . | . | . | . | . | 3.144769 | 4.347093 | 3.369942 | 3.204731 | 7.318005 | 10.048501 | 7.562050 | 7.327236 | -1.156507 | 89.036467 | 0.348886973 | -0.60126799 | 0.547661508 | 0.998865404 | 6.577789 | 6.367180 | 6.446849 | 6.540253 | 6.461869 | 6.445375 | 93 | 119 | 74 | 75 |
| 107987175 | XM_01702029 | LOC107987175 | uncharacterized LOC107987175 | protein_coding | XP_016875783.1 | . | . | . | . | 0.026380 | 0.022340 | 0.000716 | 0.012520 | 0.061386 | 0.051639 | 0.001607 | 0.028624 | -1.204659 | 1.220068 | 2.631544379 | -0.10207874 | 0.918694172 | NA | 0.413393 | 0.148375 | 0.294625 | 0.290739 | 0.291821 | 0.291432 | 2 | 1 | 1 | 1 |
| 107987181 | XR_001749348 | LOC107987181 | uncharacterized LOC107987181 | lncRNA | . | . | . | . | . | 0.411540 | 0.350662 | 0.555987 | 0.821027 | 0.957670 | 0.810572 | 1.247620 | 1.877180 | 1.634156 | 3.542205 | 1.593114532 | 0.444754905 | 0.656496886 | NA | 1.429096 | 2.134756 | 1.811221 | 1.812122 | 1.815310 | 1.823837 | 3 | 3 | 3 | 5 |
| 107987222 | XR_001751474 | LOC107987222 | uncharacterized LOC107987222 | lncRNA | . | . | . | . | . | 3.066953 | 2.672370 | 4.041836 | 2.169831 | 7.136924 | 6.177302 | 9.069763 | 4.961061 | 1.066570 | 47.665840 | 0.483287674 | 0.192387051 | 0.847439035 | 0.998865404 | 5.525622 | 5.622519 | 5.573862 | 5.547477 | 5.625613 | 5.515634 | 57 | 46 | 56 | 32 |
| 107987234 | XR_001752091 | LOC107987234 | uncharacterized LOC107987234 | lncRNA | . | . | . | . | . | 0.024150 | 0.023090 | 0.015085 | 0.025242 | 0.056197 | 0.053375 | 0.033849 | 0.057712 | -1.222666 | 3.653952 | 1.57137831 | -0.18457097 | 0.853565556 | NA | 2.008843 | 1.715150 | 1.878723 | 1.875702 | 1.869460 | 1.878218 | 5 | 4 | 2 | 4 |
| 107987245 | XR_001752780 | LOC107987245 | uncharacterized LOC107987245 | lncRNA | . | . | . | . | . | 0.436733 | 0.498548 | 0.655463 | 0.257520 | 1.016296 | 1.152415 | 1.470840 | 0.588790 | -1.030963 | 49.632210 | 0.505091423 | -0.08709949 | 0.930592432 | 0.998865404 | 5.653161 | 5.612968 | 5.610132 | 5.636136 | 5.692101 | 5.536793 | 54 | 58 | 61 | 26 |
| 107987250 | XR_001753042 | LOC107987250 | uncharacterized LOC107987250 | lncRNA | . | . | . | . | . | 0.043941 | 0.038470 | 0.084432 | 0.046704 | 0.102252 | 0.088925 | 0.189463 | 0.106783 | 1.482287 | 8.892388 | 1.032190944 | 0.550115647 | 0.582240055 | NA | 2.838265 | 3.410501 | 3.135058 | 3.128977 | 3.161128 | 3.133742 | 9 | 7 | 12 | 7 |
| 107987256 | XR_001753380 | LOC107987256 | uncharacterized LOC107987256 | lncRNA | . | . | . | . | . | 0.067522 | 0.102318 | 0.057693 | 0.058754 | 0.157126 | 0.236511 | 0.129462 | 0.134335 | -1.357142 | 3.921040 | 1.519987402 | -0.28985247 | 0.771929104 | NA | 2.175403 | 1.733338 | 1.972586 | 1.982366 | 1.972638 | 1.971460 | 4 | 6 | 3 | 3 |
| 107987258 | XR_001753445 | LOC107987258 | uncharacterized LOC107987258 | lncRNA | . | . | . | . | . | 0.060691 | 0.041536 | 0.103446 | 0.041710 | 0.141229 | 0.096013 | 0.232130 | 0.095364 | 1.388102 | 20.272816 | 0.727392266 | 0.650423846 | 0.515418478 | NA | 4.082163 | 4.561202 | 4.323591 | 4.296904 | 4.377084 | 4.297695 | 23 | 15 | 29 | 13 |
| 107987261 | XR_001753508 | LOC107987261 | uncharacterized LOC107987261 | lncRNA | . | . | . | . | . | 11.031267 | 12.415820 | 13.587190 | 12.299229 | 25.670212 | 28.699726 | 30.489262 | 28.120728 | 1.089209 | 1967.674852 | 0.127956058 | 0.963461395 | 0.335316029 | 0.998865404 | 10.879295 | 11.002623 | 10.862279 | 10.948726 | 11.017038 | 10.929435 | 2050 | 2144 | 1882 | 1818 |
| 107987272 | XR_001754025 | LOC107987272 | uncharacterized LOC107987272 | lncRNA | . | . | . | . | . | 0.762148 | 0.651440 | 1.136686 | 0.684164 | 1.773550 | 1.505833 | 2.550690 | 1.564260 | 1.259157 | 50.072382 | 0.464879199 | 0.715149153 | 0.474516884 | 0.998865404 | 5.468275 | 5.804111 | 5.626507 | 5.599615 | 5.705152 | 5.603783 | 55 | 44 | 61 | 39 |
| 107987280 | XR_001754572 | LOC107987280 | uncharacterized LOC107987280 | lncRNA | . | . | . | . | . | 4.168135 | 3.986931 | 5.796059 | 3.662569 | 9.699421 | 9.215969 | 13.006188 | 8.374029 | 1.143825 | 148.972893 | 0.296901756 | 0.65296508 | 0.513778791 | 0.998865404 | 7.117535 | 7.313619 | 7.193011 | 7.173827 | 7.322925 | 7.140142 | 164 | 146 | 170 | 115 |
| 107987290 | XM_02445214 | LOC107987290 | uncharacterized LOC107987290 | protein_coding | XP_024307912.1 | . | . | . | . | 0.799422 | 0.724579 | 0.786623 | 1.049181 | 1.860287 | 1.674898 | 1.765158 | 2.398828 | 1.204990 | 22.688887 | 0.647415538 | 0.415531501 | 0.677752859 | NA | 4.363616 | 4.631770 | 4.496420 | 4.486793 | 4.497247 | 4.521238 | 25 | 21 | 19 | 26 |
| 107987293 | XR_001754938 | LOC107987293 | uncharacterized LOC107987293 | lncRNA | . | . | . | . | . | 4.156023 | 4.004974 | 5.425955 | 3.935336 | 9.671236 | 9.257677 | 12.175685 | 8.997678 | 1.130807 | 352.891675 | 0.210973175 | 0.840638819 | 0.400550305 | 0.998865404 | 8.370963 | 8.549670 | 8.434471 | 8.408816 | 8.582465 | 8.392728 | 390 | 349 | 379 | 294 |
| 107987294 | XR_001754939 | LOC107987294 | uncharacterized LOC107987294, transcript variant X1 | lncRNA | . | . | . | . | . | 5.593771 | 5.654664 | 7.183008 | 5.467374 | 13.016934 | 13.071009 | 16.118463 | 12.500501 | 1.107126 | 419.477634 | 0.200682274 | 0.73160296 | 0.464410941 | 0.998865404 | 8.636543 | 8.784562 | 8.682224 | 8.670050 | 8.832714 | 8.632970 | 463 | 425 | 446 | 346 |
| 107987295 | NR_146322 | LINC02246 | long intergenic non-protein coding RNA 2246 | lncRNA | . | 53135 | . | . | . | 0.081438 | 0.053366 | 0.097810 | 0.039269 | 0.189509 | 0.123357 | 0.219483 | 0.089784 | 1.011224 | 18.413761 | 0.75852663 | 0.02122862 | 0.983063284 | NA | 4.191364 | 4.213972 | 4.209430 | 4.179195 | 4.227543 | 4.161796 | 25 | 16 | 23 | 10 |
| 107987305 | gene-CTBP2P10 | CTBP2P10 | . | pseudogene | . | 54512 | . | . | . | 0.024254 | 0.034361 | 0.022543 | 0.012728 | 0.056441 | 0.079428 | 0.050585 | 0.029101 | -1.614875 | 5.842417 | 1.268770627 | -0.54495464 | 0.585784716 | NA | 2.849979 | 2.161867 | 2.552293 | 2.562681 | 2.550782 | 2.539323 | 7 | 9 | 5 | 3 |
| 107987307 | gene-CTBP2P11 | CTBP2P11 | . | pseudogene | . | 54513 | . | . | . | 0.016255 | 0.021848 | 0.023633 | 0.003904 | 0.037826 | 0.050504 | 0.053031 | 0.008926 | -1.462062 | 4.179804 | 1.529828488 | -0.35821271 | 0.720184137 | NA | 2.307819 | 1.769039 | 2.063891 | 2.069561 | 2.070026 | 2.048978 | 5 | 6 | 5 | 1 |
| 107987308 | XR_001755123 | LOC107987308 | uncharacterized LOC107987308, transcript variant X1 | lncRNA | . | . | . | . | . | 0.046976 | 0.043969 | 0.040681 | 0.045797 | 0.109314 | 0.101635 | 0.091287 | 0.104708 | 1.091603 | 3.730586 | 1.550327645 | 0.081562453 | 0.93499466 | NA | 1.832372 | 1.963455 | 1.899807 | 1.892406 | 1.900732 | 1.894479 | 5 | 3 | 4 | 3 |
| 107987309 | XR_001755125 | LOC107987309 | uncharacterized LOC107987309, transcript variant X1 | lncRNA | . | . | . | . | . | 0.031801 | 0.031442 | 0.029491 | 0.028744 | 0.074003 | 0.072679 | 0.066177 | 0.065722 | -1.136075 | 2.938488 | 1.734160894 | -0.10613692 | 0.915473716 | NA | 1.644780 | 1.459419 | 1.561840 | 1.558537 | 1.556447 | 1.560469 | 4 | 3 | 2 | 3 |
| 107987323 | XR_001755413 | LOC107987323 | uncharacterized LOC107987323, transcript variant X2 | lncRNA | . | . | . | . | . | 4.256707 | 6.226103 | 5.193558 | 4.973758 | 9.905531 | 14.391918 | 11.654194 | 11.371905 | -1.012841 | 77.921888 | 0.372794473 | -0.04937743 | 0.960618511 | 0.998865404 | 6.294062 | 6.273780 | 6.233468 | 6.329143 | 6.270615 | 6.287636 | 74 | 100 | 65 | 75 |
| 107987324 | XR_001755418 | LOC107987324 | uncharacterized LOC107987324 | lncRNA | . | . | . | . | . | 0.108858 | 0.107429 | 0.194172 | 0.066321 | 0.253318 | 0.248327 | 0.435717 | 0.151636 | 1.250253 | 7.083180 | 1.164141431 | 0.27678793 | 0.78194295 | NA | 2.651489 | 2.978782 | 2.808312 | 2.810281 | 2.832164 | 2.800379 | 7 | 7 | 10 | 4 |
| 107987326 | XR_001755583 | LOC107987326 | uncharacterized LOC107987326 | lncRNA | . | . | . | . | . | 0.342876 | 0.507277 | 0.865377 | 0.416401 | 0.797886 | 1.172593 | 1.941880 | 0.952051 | 1.479965 | 40.848383 | 0.5332715 | 1.060553679 | 0.288892783 | 0.998865404 | 5.041002 | 5.608049 | 5.273185 | 5.322985 | 5.419477 | 5.296064 | 31 | 42 | 57 | 30 |
| 107987330 | XR_001755780 | FAM239C | family with sequence similarity 239 member C, transcript variant X | lncRNA | . | 53416 | . | . | . | 0.212372 | 0.241781 | 0.199777 | 0.100117 | 0.494198 | 0.558887 | 0.448293 | 0.228905 | -1.426258 | 14.917068 | 0.81082668 | -0.63174409 | 0.52755411 | NA | 4.130737 | 3.622519 | 3.907372 | 3.916158 | 3.905230 | 3.872680 | 19 | 20 | 14 | 8 |
| 108783654 | NR_144402 | LOC108783654 | uncharacterized LOC108783654 | lncRNA | . | . | . | . | . | 6.933606 | 6.988525 | 6.145115 | 7.989561 | 16.134785 | 16.154287 | 13.789461 | 18.267183 | 1.001384 | 517.551292 | 0.180175138 | 0.011070672 | 0.991167062 | 0.999444868 | 9.015018 | 9.016098 | 9.013850 | 9.013262 | 8.931841 | 9.090544 | 596 | 558 | 394 | 546 |
| 108903150 | NR_144447 | ZNF8-ERVK3-1 | ZNF8-ERVK3-1 readthrough | lncRNA | . | . | . | . | . | 0.523400 | 0.709722 | 0.947834 | 0.606811 | 1.217972 | 1.640555 | 2.126912 | 1.387401 | 1.250189 | 102.423440 | 0.349604689 | 0.921457554 | 0.356811592 | 0.998865404 | 6.507902 | 6.830865 | 6.584519 | 6.673052 | 6.769301 | 6.623288 | 89 | 113 | 121 | 83 |
| 109136578 | NR_144569 | MASCRNA | MALAT1-associated small cytoplasmic RNA | scRNA | . | . | . | . | . | 0.000035 | 0.000038 | 0.000100 | 0.000038 | 0.000081 | 0.000087 | 0.000224 | 0.000087 | 1.234655 | 1.002957 | 2.868339486 | 0.106022226 | 0.915564717 | NA | -0.155866 | 0.148375 | -0.001261 | -0.000972 | 0.000053 | -0.000315 | 1 | 1 | 1 | 1 |
| 109136579 | NR_145459 | TALAM1 | TALAM1 transcript, MALAT1 antisense RNA | antisense_RNA | . | 54476 | . | . | . | 0.021653 | 0.012348 | 0.023762 | 0.004201 | 0.050387 | 0.028542 | 0.053321 | 0.009605 | -1.238550 | 4.438146 | 1.501285215 | -0.20559201 | 0.83710962 | NA | 2.290730 | 1.993948 | 2.154819 | 2.143620 | 2.157849 | 2.131733 | 7 | 4 | 6 | 1 |
| 109286551 | gene-BPTFP1 | BPTFP1 | bromodomain PHD finger transcription factor pseudogene 1 | pseudogene | . | 52869 | . | . | . | 0.056725 | 0.032262 | 0.063295 | 0.062829 | 0.132000 | 0.074574 | 0.142032 | 0.143651 | -1.204659 | 1.220068 | 2.631544379 | -0.10207874 | 0.918694172 | NA | 0.413393 | 0.148375 | 0.294625 | 0.290739 | 0.291821 | 0.291432 | 2 | 1 | 1 | 1 |
| 109286552 | gene-KPNA2P3 | KPNA2P3 | karyopherin subunit alpha 2 pseudogene 3 | pseudogene | . | 52870 | . | . | . | 0.590611 | 0.462253 | 0.125866 | 0.507388 | 1.374375 | 1.068517 | 0.282439 | 1.160083 | -1.593340 | 12.700812 | 0.909224349 | -0.73915114 | 0.459815228 | NA | 3.966708 | 3.287859 | 3.694439 | 3.678897 | 3.638897 | 3.683126 | 20 | 15 | 4 | 14 |
| 109286553 | NR_026899 | ARHGAP27P1-I | ARHGAP27P1-BPTFP1-KPNA2P3 readthrough, transcribed pseudo | transcribed_ps | . | 52873 | . | . | . | 1.152400 | 1.120318 | 2.033182 | 1.768857 | 2.681683 | 2.589665 | 4.562402 | 4.044282 | 1.648378 | 135.931058 | 0.282570417 | 2.55174241 | **0.010718573** | 0.333896806 | 6.681085 | 7.402995 | 6.977008 | 6.968172 | 7.184615 | 7.124231 | 120 | 109 | 158 | 147 |
| 109616966 | NR_145985 | SNORA31B | small nucleolar RNA, H/ACA box 31B | snoRNA | . | 52201 | . | . | . | 0.000027 | 0.000176 | 0.000004 | 0.000007 | 0.000063 | 0.000406 | 0.000009 | 0.000016 | 1.234655 | 1.002957 | 2.868339486 | 0.106022226 | 0.915564717 | NA | -0.155866 | 0.148375 | -0.001261 | -0.000972 | 0.000053 | -0.000315 | 1 | 1 | 1 | 1 |
| 109616975 | NR_145749 | SNORA74D | small nucleolar RNA, H/ACA box 74D | snoRNA | . | 52219 | . | . | . | 0.832060 | 0.458995 | 0.844259 | 0.111998 | 1.936238 | 1.060987 | 1.894493 | 0.256070 | -1.266475 | 3.197252 | 1.714541063 | -0.19878122 | 0.842433886 | NA | 1.832372 | 1.502474 | 1.682158 | 1.674730 | 1.682901 | 1.666988 | 5 | 3 | 4 | 1 |
| 109616976 | NR_145735 | SNORA79B | small nucleolar RNA, H/ACA box 79B | snoRNA | . | 52222 | . | . | . | 0.896246 | 2.276614 | 0.116289 | 0.878723 | 2.085601 | 5.262496 | 0.260948 | 2.009096 | **-2.827041** | 5.219948 | 1.42906209 | -1.04914471 | 0.294111523 | NA | 2.950762 | 1.437399 | 2.388907 | 2.419917 | 2.375151 | 2.388790 | 5 | 12 | 1 | 4 |
| 109616991 | NR_145744 | SNORD19C | small nucleolar RNA, C/D box 19C | snoRNA | . | 52224 | . | . | . | 1.238373 | 0.149071 | 2.688998 | 1.346962 | 2.881744 | 0.344583 | 6.034034 | 3.079669 | **2.019024** | 3.337596 | 1.689537975 | 0.599961532 | 0.548531873 | NA | 1.137672 | 2.161867 | 1.722113 | 1.709950 | 1.732167 | 1.720818 | 4 | 1 | 5 | 3 |
| 109616995 | NR_145774 | SNORD70B | small nucleolar RNA, C/D box 70B | snoRNA | . | 52233 | . | . | . | 2.103387 | 2.471473 | 0.986257 | 4.593534 | 4.894668 | 5.712920 | 2.213133 | 10.502568 | 1.115052 | 6.166122 | 1.273655981 | 0.123354604 | 0.901826303 | NA | 2.548173 | 2.696731 | 2.617816 | 2.623809 | 2.605286 | 2.642214 | 6 | 7 | 2 | 10 |
| 109617000 | NR_145776 | SNORA11F | small nucleolar RNA, H/ACA box 11F | snoRNA | . | 52189 | . | . | . | 1.339067 | 3.217553 | 2.034042 | 2.002558 | 3.116064 | 7.437519 | 4.564331 | 4.578611 | -1.072368 | 9.196663 | 1.020893452 | -0.0987372 | 0.921346933 | NA | 3.251986 | 3.148375 | 3.183299 | 3.214965 | 3.197875 | 3.194938 | 7 | 14 | 8 | 8 |
| 109617002 | NR_145761 | SNORA21B | small nucleolar RNA, H/ACA box 21B | snoRNA | . | 52195 | . | . | . | 3.058847 | 2.491492 | 4.212480 | 2.374038 | 7.118064 | 5.759195 | 9.452683 | 5.427955 | 1.200377 | 13.303457 | 0.84814934 | 0.310662438 | 0.756057255 | NA | 3.593806 | 3.861274 | 3.725395 | 3.716825 | 3.747983 | 3.715179 | 15 | 12 | 16 | 10 |
| 109617006 | NR_145766 | SNORA63B | small nucleolar RNA, H/ACA box 63B | snoRNA | . | 52209 | . | . | . | 6.901876 | 6.977309 | 4.801414 | 4.584383 | 16.060949 | 16.128361 | 10.774233 | 10.481647 | -1.475852 | 23.318848 | 0.639815879 | -0.87767116 | 0.380122197 | NA | 4.796818 | 4.235838 | 4.566338 | 4.566125 | 4.532091 | 4.525875 | 32 | 30 | 17 | 17 |
| 109617007 | NR_145763 | SNORA63E | small nucleolar RNA, H/ACA box 63E | snoRNA | . | 52212 | . | . | . | 1.128661 | 1.172818 | 3.091920 | 3.800367 | 2.626441 | 2.711021 | 6.938179 | 8.689088 | **2.768970** | 10.142705 | 0.978433687 | 1.501736592 | 0.13316515 | NA | 2.429096 | 3.897226 | 3.294472 | 3.296466 | 3.333333 | 3.343199 | 6 | 6 | 12 | 15 |
| 109617010 | NR_145765 | SNORA77B | small nucleolar RNA, H/ACA box 77B | snoRNA | . | 52221 | . | . | . | 0.000305 | 0.000576 | 0.004325 | 0.000945 | 0.000711 | 0.001332 | 0.009705 | 0.002160 | 1.234655 | 1.002957 | 2.868339486 | 0.106022226 | 0.915564717 | NA | -0.155866 | 0.148375 | -0.001261 | -0.000972 | 0.000053 | -0.000315 | 1 | 1 | 1 | 1 |
| 109617013 | NR_145752 | SNORD63B | small nucleolar RNA, C/D box 63B | snoRNA | . | 52230 | . | . | . | 3.749995 | 4.084425 | 5.547949 | 1.936640 | 8.726394 | 9.441332 | 12.449436 | 4.427896 | -1.053114 | 9.215416 | 1.024054896 | -0.0729073 | 0.941879884 | NA | 3.238681 | 3.168566 | 3.196961 | 3.203888 | 3.214127 | 3.182173 | 10 | 11 | 11 | 5 |
| 109617023 | NR_145714 | SNORA58B | small nucleolar RNA, H/ACA box 58B | snoRNA | . | 52208 | . | . | . | 2.008852 | 2.246375 | 1.497049 | 2.563476 | 4.674680 | 5.192595 | 3.359334 | 5.861083 | -1.010259 | 9.377949 | 0.999626907 | -0.0147314 | 0.988246471 | NA | 3.238681 | 3.219802 | 3.226696 | 3.233622 | 3.219196 | 3.240206 | 10 | 11 | 6 | 11 |
| 109623451 | NR_145781 | SNORD157 | small nucleolar RNA, C/D box 157 | snoRNA | . | 51871 | . | . | . | 0.585999 | 1.183667 | 2.089407 | 0.908318 | 1.363643 | 2.736098 | 4.688569 | 2.076762 | 1.488926 | 2.812587 | 1.788565834 | 0.321079512 | 0.748150136 | NA | 1.175403 | 1.751299 | 1.472828 | 1.477882 | 1.485999 | 1.474918 | 2 | 3 | 4 | 2 |
| 109623452 | NR_145780 | SNORD159 | small nucleolar RNA, C/D box 159 | snoRNA | . | 51873 | . | . | . | 4.429278 | 3.008908 | 4.247742 | 3.451033 | 10.307113 | 6.955225 | 9.531811 | 7.890376 | -1.003663 | 8.912925 | 1.021457839 | -0.00516459 | 0.995879269 | NA | 3.156660 | 3.155137 | 3.161907 | 3.148052 | 3.161109 | 3.148533 | 12 | 8 | 9 | 7 |
| 109623456 | NR_145806 | SNORD167 | small nucleolar RNA, C/D box 167 | snoRNA | . | 51881 | . | . | . | 0.172704 | 0.010121 | 0.003889 | 0.002030 | 0.401889 | 0.023394 | 0.008727 | 0.004641 | 1.234655 | 1.002957 | 2.868339486 | 0.106022226 | 0.915564717 | NA | -0.155866 | 0.148375 | -0.001261 | -0.000972 | 0.000053 | -0.000315 | 1 | 1 | 1 | 1 |
| 109623460 | NR_145802 | SNORD138 | small nucleolar RNA, C/D box 138 | snoRNA | . | 50420 | . | . | . | 1.438303 | 0.363699 | 1.389177 | 0.343142 | 3.346989 | 0.840706 | 3.117273 | 0.784554 | 1.017451 | 2.229276 | 2.035134528 | 0.012264417 | 0.990214656 | NA | 1.137672 | 1.175234 | 1.159059 | 1.146887 | 1.158517 | 1.147760 | 4 | 1 | 3 | 1 |
| 109623464 | NR_145807 | SNORA119 | small nucleolar RNA, H/ACA box 119 | snoRNA | . | 51857 | . | . | . | 0.197117 | 0.330476 | 0.905290 | 0.710114 | 0.458699 | 0.763911 | 2.031444 | 1.623591 | **2.170210** | 2.847568 | 1.783980699 | 0.6265958 | 0.530924222 | NA | 0.844134 | 1.963455 | 1.485636 | 1.486278 | 1.498845 | 1.492632 | 2 | 2 | 4 | 3 |
| 109623469 | NR_145805 | SNORD175 | small nucleolar RNA, C/D box 175 | snoRNA | . | 51889 | . | . | . | 1.546608 | 2.651970 | 3.076968 | 0.924578 | 3.599021 | 6.130147 | 6.904628 | 2.113938 | -1.203501 | 4.965651 | 1.387001101 | -0.19267279 | 0.847215238 | NA | 2.436885 | 2.175234 | 2.305509 | 2.315553 | 2.316952 | 2.295643 | 5 | 7 | 6 | 2 |
| 109623476 | NR_145793 | SNORD160 | small nucleolar RNA, C/D box 160 | snoRNA | . | 51874 | . | . | . | 0.695413 | 0.696163 | 0.180584 | 0.319640 | 1.618254 | 1.609212 | 0.405226 | 0.730818 | -1.620137 | 1.451754 | 2.440017378 | -0.28529139 | 0.775420906 | NA | 0.844134 | 0.148375 | 0.548030 | 0.548560 | 0.545314 | 0.544903 | 2 | 2 | 1 | 1 |
| 109703458 | NM_00134871 | HTD2 | hydroxyacyl-thioester dehydratase type 2 | protein_coding | NP_001335641.1;NP_001335642.1;NP_001335643.1;NP_001335644.1 | 53111 | . | . | . | 1.903139 | 1.736777 | 2.153172 | 1.923609 | 4.428680 | 4.014635 | 4.831654 | 4.398103 | 1.087810 | 196.385804 | 0.241511381 | 0.502778066 | 0.615120317 | 0.998865404 | 7.555029 | 7.677468 | 7.613513 | 7.579083 | 7.664010 | 7.603392 | 225 | 195 | 190 | 179 |
| 109729141 | NR_146292,NF | TDRKH-AS1 | TDRKH antisense RNA 1 | lncRNA | . | 40578 | . | . | . | 0.528253 | 0.682153 | 0.973095 | 0.849475 | 1.229267 | 1.576827 | 2.183598 | 1.942224 | 1.542527 | 25.137926 | 0.616391813 | 1.014445221 | 0.310370356 | NA | 4.305694 | 4.930721 | 4.610290 | 4.624035 | 4.668204 | 4.655160 | 21 | 23 | 28 | 27 |
| 109729146 | NR_146294 | POC1B-AS1 | POC1B antisense RNA 1 | lncRNA | . | 52949 | . | . | . | 0.135973 | 0.218285 | 0.242581 | 0.269435 | 0.316414 | 0.504576 | 0.544344 | 0.616031 | 1.337551 | 11.595013 | 0.90032373 | 0.466047634 | 0.641181405 | NA | 3.312060 | 3.728813 | 3.510410 | 3.529957 | 3.532392 | 3.537761 | 9 | 13 | 11 | 13 |
| 109729173 | NR_145448 | TFAP2A-AS2 | TFAP2A antisense RNA 2 | lncRNA | . | 52289 | 617136 | . | . | 0.347342 | 0.324593 | 0.428619 | 0.394802 | 0.808279 | 0.750311 | 0.961809 | 0.902668 | 1.206807 | 40.523352 | 0.48748077 | 0.556320021 | 0.577992103 | 0.998865404 | 5.198253 | 5.470304 | 5.327504 | 5.315688 | 5.356959 | 5.344839 | 44 | 38 | 40 | 40 |
| 109729180 | NR_146320 | SNHG26 | small nucleolar RNA host gene 26 | lncRNA | . | 53131 | . | . | . | 2.100101 | 2.080552 | 2.014256 | 1.884585 | 4.887022 | 4.809289 | 4.519931 | 4.308879 | -1.088348 | 48.595309 | 0.447046742 | -0.27321521 | 0.784687792 | 0.998865404 | 5.662238 | 5.540693 | 5.613582 | 5.608980 | 5.602457 | 5.589512 | 59 | 54 | 42 | 42 |
| 109729182 | NR_147193 | DM1-AS | DM1 locus antisense RNA | lncRNA | . | 53125 | . | . | . | 0.285929 | 0.103304 | 0.368100 | 0.152652 | 0.665369 | 0.238791 | 0.826006 | 0.349020 | 1.376014 | 5.798558 | 1.291282512 | 0.35661882 | 0.721377186 | NA | 2.282109 | 2.751299 | 2.530344 | 2.510729 | 2.539752 | 2.517994 | 8 | 3 | 8 | 4 |
| 109731405 | gene-PDXDC2P | PDXDC2P | pyridoxal dependent decarboxylase domain containing 2, pseudog | pseudogene | . | 27559 | . | . | . | 0.106079 | 0.074530 | 0.103942 | 0.096206 | 0.246849 | 0.172280 | 0.233243 | 0.219964 | 1.158198 | 7.223234 | 1.125532463 | 0.188250765 | 0.850680079 | NA | 2.741623 | 2.955730 | 2.852685 | 2.842200 | 2.853499 | 2.850988 | 9 | 6 | 7 | 7 |
| 109864279 | NR_146144 | RNA45SN2 | RNA, 45S pre-ribosomal N2 | rRNA | . | 53518 | . | . | . | 30.194065 | 25.915844 | 36.333195 | 34.222591 | 70.262833 | 59.905632 | 81.530632 | 78.245888 | 1.239742 | 13128.137118 | 0.115092971 | 2.693817442 | **0.007063884** | 0.266401081 | 13.517005 | 13.827112 | 13.646010 | 13.510494 | 13.794560 | 13.733530 | 14585 | 11631 | 13081 | 13148 |
| 109864280 | NR_146146 | RNA18SN2 | RNA, 18S ribosomal N2 | rRNA | . | 53519 | . | . | . | 77.342728 | 55.368015 | 63.586285 | 55.997059 | 179.979721 | 127.985657 | 142.685791 | 128.030624 | -1.125869 | 3675.986920 | 0.153658593 | -1.11311397 | 0.265659483 | 0.997577536 | 11.926777 | 11.756004 | 12.020090 | 11.733679 | 11.843332 | 11.731096 | 5244 | 3488 | 3214 | 3020 |
| 109864281 | NR_146147 | RNA5-8SN2 | RNA, 5.8S ribosomal N2 | rRNA | . | 53521 | . | . | . | 0.849756 | 0.760644 | 1.070985 | 0.686339 | 1.977416 | 1.758262 | 2.403260 | 1.569234 | 1.114700 | 4.748118 | 1.375504002 | 0.113889582 | 0.909325324 | NA | 2.166062 | 2.324312 | 2.241668 | 2.243064 | 2.248009 | 2.241424 | 5 | 5 | 5 | 4 |
| 109864282 | NR_146148 | RNA28SN2 | RNA, 28S ribosomal N2 | rRNA | . | 53520 | . | . | . | 93.887993 | 54.826267 | 73.124069 | 88.589973 | 218.481201 | 126.733368 | 164.088287 | 202.550430 | 1.070608 | 12219.280686 | 0.407696349 | 0.241430994 | 0.809221093 | 0.998865404 | 13.526814 | 13.625250 | 13.742664 | 13.279939 | 13.514399 | 13.670864 | 17214 | 9340 | 9993 | 12919 |
| 109910379 | NR_146151 | RNA45SN3 | RNA, 45S pre-ribosomal N3 | rRNA | . | 53522 | . | . | . | 41.361370 | 32.662766 | 44.463722 | 35.277718 | 96.249603 | 75.501442 | 99.775314 | 80.658318 | 1.062374 | 15943.348997 | 0.142775648 | 0.611394561 | 0.540938402 | 0.998865404 | 13.916320 | 14.003691 | 14.029672 | 13.820788 | 14.081408 | 13.873158 | 19970 | 14652 | 16001 | 13548 |
| 109910380 | NR_146152 | RNA18SN3 | RNA, 18S ribosomal N3 | rRNA | . | 53523 | . | . | . | 62.846573 | 47.084946 | 55.012840 | 55.850784 | 146.246552 | 108.838959 | 123.447212 | 127.696190 | -1.006385 | 3215.457870 | 0.142492026 | -0.06444443 | 0.948616353 | 0.998865404 | 11.655296 | 11.646307 | 11.765538 | 11.521434 | 11.643646 | 11.647413 | 4262 | 2967 | 2781 | 3012 |
| 109910381 | NR_146153 | RNA5-8SN3 | RNA, 5.8S ribosomal N3 | rRNA | . | 53525 | . | . | . | 0.246059 | 0.224828 | 0.337806 | 0.272973 | 0.572590 | 0.519699 | 0.758027 | 0.624121 | 1.234663 | 2.005914 | 2.070995667 | 0.146845664 | 0.883253841 | NA | 0.844134 | 1.148375 | 0.998092 | 0.998667 | 1.000707 | 0.999974 | 2 | 2 | 2 | 2 |
| 109910382 | NR_146154 | RNA28SN3 | RNA, 28S ribosomal N3 | rRNA | . | 53524 | . | . | . | 81.344658 | 48.752407 | 66.595268 | 97.542328 | 189.292374 | 112.693375 | 149.437851 | 223.018921 | 1.241694 | 11574.443657 | 0.401495282 | 0.777867387 | 0.436647196 | 0.998865404 | 13.334065 | 13.646376 | 13.581299 | 13.146253 | 13.398244 | 13.721284 | 14917 | 8307 | 9103 | 14228 |
| 109951030 | NR_146231 | PKD1P3-NPIPA | PKD1P3-NPIPA1 readthrough | transcribed_ps | . | . | . | . | . | 11.821568 | 12.322559 | 15.694532 | 12.576652 | 27.509277 | 28.484146 | 35.218079 | 28.755018 | 1.155369 | 3006.085365 | 0.132816283 | 1.568737513 | 0.116709114 | 0.857396579 | 11.445615 | 11.654193 | 11.476247 | 11.502792 | 11.699280 | 11.507524 | 3159 | 3059 | 3126 | 2673 |
| 110006322 | NR_146331 | PKD1P5-LOC10 | PKD1P5-LOC105376752 readthrough | transcribed_ps | . | . | . | . | . | 4.645393 | 4.890157 | 5.940300 | 4.752206 | 10.810021 | 11.303817 | 13.329860 | 10.865355 | 1.106101 | 1850.755511 | 0.139895611 | 1.039941142 | 0.298367247 | 0.998865404 | 10.779153 | 10.924962 | 10.793561 | 10.827283 | 10.978931 | 10.793343 | 1980 | 1937 | 1887 | 1611 |
| 110006323 | NR_146336 | PKD1P4-NPIPA | PKD1P4-NPIPA8 readthrough | transcribed_ps | . | . | . | . | . | 4.377130 | 5.136453 | 6.242777 | 3.792123 | 10.185761 | 11.873141 | 14.008611 | 8.670238 | 1.041678 | 1082.046210 | 0.197914318 | 0.297652179 | 0.765968646 | 0.998865404 | 10.049507 | 10.108973 | 9.997187 | 10.109656 | 10.255662 | 9.881142 | 1128 | 1230 | 1199 | 778 |
| 110006325 | NR_146339 | PPIP5K1P1-CAT | PPIP5K1P1-CATSPER2 readthrough | transcribed_ps | . | . | . | . | . | 1.362164 | 1.743422 | 2.304746 | 2.104360 | 3.169811 | 4.029998 | 5.171783 | 4.811369 | 1.400164 | 309.410277 | 0.211494926 | 2.296014668 | **0.021675043** | 0.48805973 | 8.010526 | 8.495639 | 8.104734 | 8.224593 | 8.377976 | 8.319136 | 262 | 311 | 330 | 321 |
| 110006327 | NR_146419 | ANKRD20A4-A | ANKRD20A4-ANKRD20A20P readthrough | lncRNA | . | . | . | . | . | 0.318138 | 0.236604 | 0.323206 | 0.289210 | 0.740320 | 0.546921 | 0.725265 | 0.661245 | 1.097235 | 40.260573 | 0.496898236 | 0.269417497 | 0.787608423 | 0.998865404 | 5.261655 | 5.397729 | 5.344465 | 5.297359 | 5.345541 | 5.325524 | 51 | 35 | 39 | 37 |
| 110091775 | NM_00135033 | CFAP298-TCP1 | CFAP298-TCP10L readthrough | protein_coding | NP_001337267.1 | 54636 | . | . | . | 0.129961 | 0.463644 | 0.284505 | 0.251416 | 0.302425 | 1.071733 | 0.638421 | 0.574834 | -1.117982 | 65.261607 | 0.509533785 | -0.31577307 | 0.752174762 | 0.998865404 | 6.107456 | 5.944256 | 5.866132 | 6.141323 | 6.001689 | 5.973854 | 35 | 116 | 57 | 54 |
| 110091777 | NR_146655,NF | LOC110091777 | uncharacterized LOC110091777 | lncRNA | . | . | . | . | . | 0.025354 | 0.010817 | 0.008927 | 0.035016 | 0.059001 | 0.025004 | 0.020032 | 0.080059 | 1.383419 | 4.207690 | 1.51920085 | 0.308213542 | 0.757919849 | NA | 1.820514 | 2.287859 | 2.074329 | 2.058470 | 2.061100 | 2.084167 | 6 | 2 | 2 | 7 |
| 110440226 | gene-CCZ1P | CCZ1P | CCZ1 homolog, vacuolar protein trafficking and biogenesis associa | pseudogene | . | . | . | . | . | 0.371789 | 0.815203 | 1.034291 | 0.189752 | 0.865169 | 1.884377 | 2.320920 | 0.433846 | 1.070049 | 9.875784 | 1.070774185 | 0.091220357 | 0.927317499 | NA | 3.251986 | 3.354002 | 3.268378 | 3.300195 | 3.317425 | 3.256296 | 7 | 14 | 15 | 3 |
| 110599564 | NM_032331 | EEF1AKMT4 | EEF1A lysine methyltransferase 4 | protein_coding | NP_115707.2 | 53611 | . | . | . | 6.322003 | 5.981445 | 5.388736 | 4.865521 | 14.711561 | 13.826378 | 12.092168 | 11.124434 | -1.214974 | 171.919862 | 0.254452679 | -1.10403848 | 0.269576445 | 0.998865404 | 7.558802 | 7.278820 | 7.479055 | 7.450395 | 7.406581 | 7.360467 | 224 | 197 | 143 | 137 |
| 110599588 | NM_00135349 | ASDURF | ASNSD1 upstream reading frame | protein_coding | NP_001340422.1;NP_001340423.1 | 53619 | . | . | . | 2.602876 | 1.602178 | 1.390017 | 4.385047 | 6.056999 | 3.703505 | 3.119157 | 10.025890 | 1.347749 | 51.143065 | 0.533483414 | 0.807057528 | 0.41963334 | 0.998865404 | 5.446359 | 5.874871 | 5.664183 | 5.584250 | 5.574159 | 5.778941 | 62 | 36 | 26 | 82 |
| 110806272 | NR_149073 | BTG3-AS1 | BTG3 antisense RNA 1 | lncRNA | . | 53145 | . | . | . | 0.536322 | 0.177720 | 0.618195 | 0.726800 | 1.248043 | 0.410809 | 1.387211 | 1.661741 | 1.821661 | 39.283840 | 0.569174513 | 1.520191828 | 0.128462771 | 0.876939477 | 4.796660 | 5.666032 | 5.282514 | 5.166104 | 5.302074 | 5.329479 | 48 | 15 | 41 | 51 |
| 110806273 | NR_149072 | SLC5A4-AS1 | SLC5A4 antisense RNA 1 | lncRNA | . | 53163 | . | . | . | 0.122069 | 0.133515 | 0.085076 | 0.153472 | 0.284061 | 0.308625 | 0.190909 | 0.350895 | -1.057125 | 19.190120 | 0.71245002 | -0.11249376 | 0.910431914 | NA | 4.303566 | 4.219802 | 4.262738 | 4.268286 | 4.240771 | 4.280845 | 22 | 22 | 12 | 22 |
| 110841581 | NR_149049 | CAMTA1-DT | CAMTA1 divergent transcript | lncRNA | . | 53020 | . | . | . | 0.423735 | 0.774836 | 0.986443 | 0.396603 | 0.986049 | 1.791069 | 2.213550 | 0.906786 | 1.136554 | 11.608358 | 0.932558979 | 0.198021206 | 0.843028472 | NA | 3.440763 | 3.627390 | 3.508231 | 3.536116 | 3.549599 | 3.507557 | 9 | 15 | 15 | 7 |
| 111089941 | NM_00135500 | MMP24-AS1-E | MMP24-AS1-EDEM2 readthrough | protein_coding | NP_001341937.1 | . | . | . | . | 0.079247 | 0.069313 | 0.063652 | 0.234613 | 0.184410 | 0.160220 | 0.142834 | 0.536415 | 1.966137 | 8.614027 | 1.082605901 | 0.900940864 | 0.367619763 | NA | 2.540965 | 3.512009 | 3.078678 | 3.076443 | 3.078055 | 3.127326 | 7 | 6 | 5 | 16 |
| 111216284 | NR_151723 | BNC2-AS1 | BNC2 antisense RNA 1 | lncRNA | . | 53727 | . | . | . | 0.376874 | 0.253949 | 1.168711 | 0.966491 | 0.877002 | 0.587013 | 2.622554 | 2.209767 | **3.232086** | 9.454319 | 1.027465613 | 1.647223734 | 0.099512067 | NA | 2.156660 | 3.852980 | 3.190727 | 3.183942 | 3.239217 | 3.225492 | 6 | 4 | 14 | 12 |
| 111240474 | NM_00135650 | CRYZL2P-SEC16 | CRYZL2P-SEC16B readthrough | protein_coding | NP_001343434.1;NP_001343435.1 | 53757 | . | . | . | 0.148957 | 0.266523 | 0.172637 | 0.055168 | 0.346629 | 0.616081 | 0.387393 | 0.126135 | -1.823050 | 27.004482 | 0.675129823 | -1.28324052 | 0.199407824 | 0.96621494 | 5.122436 | 4.261131 | 4.737545 | 4.815330 | 4.752071 | 4.672681 | 29 | 48 | 25 | 9 |
| 111822955 | NR_153405 | ATRIP-TREX1 | ATRIP-TREX1 readthrough | lncRNA | . | . | . | . | . | 0.827417 | 0.156826 | 1.095601 | 0.308227 | 1.925433 | 0.362511 | 2.458496 | 0.704724 | 1.405752 | 80.752326 | 1.293018973 | 0.379996433 | 0.703948063 | 0.998865404 | 6.068376 | 6.560690 | 6.371287 | 6.037838 | 6.462900 | 6.125470 | 130 | 23 | 128 | 39 |
| 112267857 | XM_01153628 | LOC112267857 | uncharacterized LOC112267857, transcript variant X2 | protein_coding | XP_011534584.1;XP_016866834.1 | . | . | . | . | 0.364246 | 0.345701 | 0.365388 | 0.203869 | 0.847615 | 0.799104 | 0.819920 | 0.466123 | -1.054302 | 9.180014 | 1.011388087 | -0.07542901 | 0.939873375 | NA | 3.234218 | 3.161867 | 3.198220 | 3.196891 | 3.206441 | 3.184174 | 11 | 10 | 10 | 6 |
| 112267861 | XR_002956938 | LOC112267861 | uncharacterized LOC112267861, transcript variant X5 | lncRNA | . | . | . | . | . | 0.167402 | 0.276189 | 0.281997 | 0.068018 | 0.389552 | 0.638422 | 0.632792 | 0.155515 | -1.210829 | 7.434937 | 1.136919727 | -0.24275653 | 0.80819401 | NA | 3.024434 | 2.751299 | 2.885220 | 2.904216 | 2.899227 | 2.877414 | 7 | 11 | 8 | 4 |
| 112267866 | XR_002957444 | LOC112267866 | uncharacterized LOC112267866, transcript variant X1 | lncRNA | . | . | . | . | . | 0.117392 | 0.095279 | 0.009422 | 0.048999 | 0.273175 | 0.220243 | 0.021143 | 0.112030 | **-2.049260** | 4.964939 | 1.42229392 | -0.72777037 | 0.466754163 | NA | 2.741623 | 1.696731 | 2.339350 | 2.328804 | 2.309286 | 2.327703 | 9 | 6 | 1 | 5 |
| 112267872 | XR_001754053 | LOC112267872 | uncharacterized LOC112267872, transcript variant X5 | lncRNA | . | . | . | . | . | 0.142501 | 1.191534 | 2.367480 | 1.957800 | 0.331605 | 2.754285 | 5.312555 | 4.476276 | 1.494815 | 23.680765 | 0.653158362 | 0.887942735 | 0.374571555 | NA | 4.247564 | 4.826107 | 4.508166 | 4.553281 | 4.585371 | 4.554989 | 16 | 26 | 28 | 23 |
| 112267877 | XR_001739359 | LOC112267877 | uncharacterized LOC112267877, transcript variant X8 | lncRNA | . | . | . | . | . | 0.318493 | 0.363091 | 0.499811 | 0.454806 | 0.741149 | 0.839300 | 1.121561 | 1.039864 | 1.299373 | 52.721563 | 0.442558162 | 0.853707934 | 0.393266837 | 0.998865404 | 5.518396 | 5.897423 | 5.671744 | 5.697384 | 5.771096 | 5.697558 | 49 | 53 | 61 | 46 |
| 112267904 | XM_02444878 | LOC112267904 | uncharacterized LOC112267904 | protein_coding | XP_024304555.1 | . | . | . | . | 0.041705 | 0.011775 | 0.051760 | 0.011583 | 0.097048 | 0.027218 | 0.116148 | 0.026483 | 1.182730 | 6.758210 | 1.250460191 | 0.193625114 | 0.846469432 | NA | 2.624463 | 2.877719 | 2.757780 | 2.726091 | 2.765216 | 2.728675 | 11 | 3 | 10 | 3 |
| 112267908 | gene-LOC1122 | LOC112267908 | . | pseudogene | . | . | . | . | . | 5.912230 | 6.402047 | 5.305197 | 6.309942 | 13.758002 | 14.798617 | 11.904707 | 14.426931 | -1.073210 | 160.774964 | 0.263216402 | -0.38725501 | 0.69856743 | 0.998865404 | 7.379536 | 7.276420 | 7.328341 | 7.357654 | 7.282961 | 7.345142 | 185 | 186 | 124 | 157 |
| 112267911 | gene-LOC1122 | LOC112267911 | . | pseudogene | . | . | . | . | . | 1.067110 | 1.059681 | 0.863588 | 0.878258 | 2.483208 | 2.449500 | 1.937866 | 2.008033 | -1.227792 | 38.222438 | 0.501671865 | -0.59015924 | 0.555083894 | 0.998865404 | 5.396728 | 5.100822 | 5.275685 | 5.272154 | 5.244401 | 5.242709 | 49 | 45 | 30 | 32 |
| 112267918 | gene-LOC1122 | LOC112267918 | . | pseudogene | . | . | . | . | . | 0.776730 | 0.831837 | 1.467524 | 1.077750 | 1.807484 | 1.922828 | 3.293082 | 2.464147 | 1.557098 | 44.781889 | 0.474484215 | 1.346430621 | 0.178163681 | 0.951511445 | 5.129536 | 5.769770 | 5.426511 | 5.436238 | 5.536128 | 5.474272 | 39 | 39 | 55 | 43 |
| 112267932 | XR_002956215 | LOC112267932 | uncharacterized LOC112267932 | lncRNA | . | . | . | . | . | 0.772772 | 0.854357 | 1.006830 | 0.710699 | 1.798272 | 1.974884 | 2.259296 | 1.624930 | 1.049476 | 27.163280 | 0.596541507 | 0.116789161 | 0.907027133 | 0.998865404 | 4.727571 | 4.798723 | 4.752091 | 4.763165 | 4.780240 | 4.745457 | 29 | 30 | 28 | 22 |
| 112267936 | XR_002956231 | LOC112267936 | uncharacterized LOC112267936 | lncRNA | . | . | . | . | . | 0.381651 | 0.564955 | 0.414727 | 0.367667 | 0.888118 | 1.305919 | 0.930636 | 0.840627 | -1.187932 | 23.672837 | 0.641837211 | -0.38709492 | 0.698685935 | NA | 4.684403 | 4.435167 | 4.552376 | 4.593470 | 4.561911 | 4.550569 | 24 | 33 | 20 | 19 |
| 112267938 | XR_002956241 | LOC112267938 | uncharacterized LOC112267938 | lncRNA | . | . | . | . | . | 0.070432 | 0.025329 | 0.041967 | 0.034724 | 0.163899 | 0.058549 | 0.094173 | 0.079391 | -1.159628 | 6.590366 | 1.211306726 | -0.17638995 | 0.859987603 | NA | 2.820514 | 2.612728 | 2.739739 | 2.708542 | 2.723293 | 2.716336 | 12 | 4 | 6 | 5 |
| 112267939 | XM_02444628 | LOC112267939 | uncharacterized LOC112267939 | protein_coding | XP_024302052.1 | . | . | . | . | 0.021056 | 0.022627 | 0.029424 | 0.002577 | 0.048998 | 0.052302 | 0.066026 | 0.005891 | -1.198555 | 2.475538 | 1.907561801 | -0.13697918 | 0.89104725 | NA | 1.429096 | 1.175234 | 1.307104 | 1.307921 | 1.310815 | 1.300040 | 3 | 3 | 3 | 1 |
| 112267941 | XR_002956242 | LOC112267941 | uncharacterized LOC112267941 | lncRNA | . | . | . | . | . | 0.377818 | 0.329616 | 0.453153 | 0.261576 | 0.879199 | 0.761921 | 1.016862 | 0.598062 | -1.002285 | 109.582129 | 0.342905973 | -0.00960185 | 0.992338953 | 0.999810065 | 6.775983 | 6.775755 | 6.791280 | 6.745166 | 6.850231 | 6.677812 | 135 | 110 | 121 | 75 |
| 112267942 | XR_002956243 | LOC112267942 | uncharacterized LOC112267942, transcript variant X2 | lncRNA | . | . | . | . | . | 0.342755 | 0.385565 | 0.447364 | 0.722946 | 0.797605 | 0.891251 | 1.003871 | 1.652930 | 1.480037 | 16.139753 | 0.767188617 | 0.737280879 | 0.460951558 | NA | 3.703730 | 4.266773 | 3.985793 | 3.993867 | 4.003611 | 4.030130 | 14 | 15 | 14 | 21 |
| 112267943 | XR_002956245 | LOC112267943 | uncharacterized LOC112267943 | lncRNA | . | . | . | . | . | 0.353751 | 0.347479 | 0.490972 | 0.206962 | 0.823193 | 0.803214 | 1.101726 | 0.473194 | -1.030235 | 41.572307 | 0.53017285 | -0.08105675 | 0.935396823 | 0.998865404 | 5.396728 | 5.358116 | 5.372054 | 5.368663 | 5.425841 | 5.301268 | 49 | 45 | 50 | 23 |
| 112267965 | XR_002956368 | LOC112267965 | uncharacterized LOC112267965 | lncRNA | . | . | . | . | . | 0.118779 | 0.039081 | 0.062469 | 0.042450 | 0.276404 | 0.090338 | 0.140178 | 0.097058 | -1.590783 | 8.643895 | 1.083902335 | -0.61789397 | 0.536645232 | NA | 3.405477 | 2.742346 | 3.147376 | 3.101473 | 3.113061 | 3.100977 | 18 | 6 | 7 | 5 |
| 112267967 | XR_002956372 | LOC112267967 | uncharacterized LOC112267967 | lncRNA | . | . | . | . | . | 0.025105 | 0.017308 | 0.027592 | 0.016985 | 0.058421 | 0.040007 | 0.061917 | 0.038835 | 1.091603 | 3.730586 | 1.550327645 | 0.081562453 | 0.93499466 | NA | 1.832372 | 1.963455 | 1.899807 | 1.892406 | 1.900732 | 1.894479 | 5 | 3 | 4 | 3 |
| 112267983 | NR_158201 | LOC112267983 | uncharacterized LOC112267983 | lncRNA | . | . | . | . | . | 0.277530 | 0.179905 | 0.313735 | 0.176721 | 0.645824 | 0.415858 | 0.704012 | 0.404052 | 1.056494 | 14.676503 | 0.820825284 | 0.096591473 | 0.923050829 | NA | 3.832372 | 3.917252 | 3.882503 | 3.854520 | 3.890686 | 3.853096 | 20 | 12 | 17 | 10 |
| 112267986 | XR_002956514 | LOC112267986 | uncharacterized LOC112267986 | lncRNA | . | . | . | . | . | 0.562287 | 0.424282 | 0.386386 | 0.519016 | 1.308464 | 0.980748 | 0.867038 | 1.186668 | -1.121493 | 15.197045 | 0.788745466 | -0.20972544 | 0.833881972 | NA | 4.006229 | 3.840449 | 3.941942 | 3.922670 | 3.917721 | 3.931832 | 21 | 15 | 11 | 15 |
| 112267987 | XR_002956520 | LOC112267987 | uncharacterized LOC112267987, transcript variant X1 | lncRNA | . | . | . | . | . | 0.359799 | 0.613114 | 0.782070 | 0.590838 | 0.837267 | 1.417240 | 1.754943 | 1.350880 | 1.408794 | 15.235100 | 0.798649881 | 0.619120107 | 0.535837238 | NA | 3.661494 | 4.155137 | 3.891637 | 3.919905 | 3.940272 | 3.916285 | 11 | 17 | 18 | 14 |
| 112267993 | XR_002956543 | LOC112267993 | uncharacterized LOC112267993 | lncRNA | . | . | . | . | . | 0.071960 | 0.125911 | 0.083212 | 0.052733 | 0.167454 | 0.291049 | 0.186725 | 0.120569 | -1.351039 | 7.862907 | 1.09572583 | -0.39614752 | 0.691996201 | NA | 3.175403 | 2.742346 | 2.971940 | 2.991055 | 2.977028 | 2.964965 | 8 | 12 | 7 | 5 |
| 112268014 | XR_002956659 | LOC112268014 | uncharacterized LOC112268014 | lncRNA | . | . | . | . | . | 0.261668 | 0.513723 | 1.096583 | 0.670991 | 0.608912 | 1.187493 | 2.460700 | 1.534140 | **2.276434** | 33.367350 | 0.58406636 | 2.031919698 | **0.042161786** | 0.63521717 | 4.349460 | 5.534478 | 4.940120 | 4.997911 | 5.115188 | 5.032188 | 16 | 29 | 50 | 33 |
| 112268021 | XR_002956687 | LOC112268021 | uncharacterized LOC112268021, transcript variant X1 | lncRNA | . | . | . | . | . | 0.024484 | 0.027623 | 0.042967 | 0.011455 | 0.056975 | 0.063852 | 0.096416 | 0.026191 | 1.517154 | 5.141398 | 1.373556358 | 0.43781782 | 0.661518352 | NA | 2.029593 | 2.632244 | 2.331582 | 2.345724 | 2.361555 | 2.334924 | 3 | 6 | 8 | 3 |
| 112268024 | XR_002956699 | LOC112268024 | uncharacterized LOC112268024 | lncRNA | . | . | . | . | . | 0.106367 | 0.066644 | 0.093838 | 0.074710 | 0.247521 | 0.154050 | 0.210571 | 0.170816 | -1.058238 | 12.573128 | 0.868670539 | -0.09401061 | 0.925100731 | NA | 3.690761 | 3.612728 | 3.666114 | 3.641576 | 3.657478 | 3.644018 | 18 | 11 | 12 | 10 |
| 112268035 | XR_002956842 | LOC112268035 | uncharacterized LOC112268035 | lncRNA | . | . | . | . | . | 0.014492 | 0.011912 | 0.028839 | 0.016839 | 0.033722 | 0.027536 | 0.064714 | 0.038500 | 1.665368 | 7.153141 | 1.147088389 | 0.641486008 | 0.521206969 | NA | 2.421266 | 3.161867 | 2.818721 | 2.812030 | 2.842631 | 2.820407 | 7 | 5 | 10 | 6 |
| 112268037 | XR_002956860 | LOC112268037 | uncharacterized LOC112268037 | lncRNA | . | . | . | . | . | 0.191935 | 0.221883 | 0.223121 | 0.284778 | 0.446640 | 0.512892 | 0.500677 | 0.651112 | 1.228951 | 18.509927 | 0.713071026 | 0.417107913 | 0.676599477 | NA | 4.054853 | 4.350483 | 4.193259 | 4.202132 | 4.205601 | 4.222303 | 18 | 19 | 16 | 21 |
| 112268047 | XR_002956917 | LOC112268047 | uncharacterized LOC112268047, transcript variant X1 | lncRNA | . | . | . | . | . | 0.162560 | 0.194113 | 0.119065 | 0.092553 | 0.378283 | 0.448701 | 0.267178 | 0.211610 | -1.643998 | 24.196036 | 0.632087947 | -1.13466623 | 0.25651519 | NA | 4.910922 | 4.194412 | 4.619001 | 4.630451 | 4.581390 | 4.570649 | 33 | 34 | 17 | 16 |
| 112268063 | XR_002957089 | LOC112268063 | uncharacterized LOC112268063, transcript variant X4 | lncRNA | . | . | . | . | . | 0.549905 | 0.561072 | 0.703572 | 0.555388 | 1.279653 | 1.296944 | 1.578797 | 1.269831 | 1.316430 | 41.981224 | 0.482965573 | 0.821239378 | 0.411509931 | 0.998865404 | 5.178768 | 5.577160 | 5.373198 | 5.350522 | 5.420326 | 5.391302 | 45 | 36 | 45 | 41 |
| 112268064 | XR_002957094 | LOC112268064 | uncharacterized LOC112268064 | lncRNA | . | . | . | . | . | 0.214340 | 0.051228 | 0.075963 | 0.188949 | 0.498779 | 0.118416 | 0.170458 | 0.432009 | -1.042978 | 9.472203 | 1.07526193 | -0.056459 | 0.954976159 | NA | 3.273436 | 3.213338 | 3.270343 | 3.215813 | 3.226259 | 3.256819 | 18 | 4 | 5 | 12 |
| 112268070 | XR_002957218 | LOC112268070 | uncharacterized LOC112268070, transcript variant X7 | lncRNA | . | . | . | . | . | 1.114158 | 1.115218 | 1.517849 | 1.655239 | 2.592690 | 2.577880 | 3.406011 | 3.784505 | 1.368010 | 24.407597 | 0.62184548 | 0.726995048 | 0.467228989 | NA | 4.365657 | 4.817608 | 4.587047 | 4.585308 | 4.615198 | 4.620300 | 24 | 22 | 24 | 27 |
| 112268076 | XM_02444878 | LOC112268076 | uncharacterized LOC112268076, transcript variant X4 | protein_coding | XP_024304552.1 | . | . | . | . | 0.063699 | 0.037421 | 0.011846 | 0.040311 | 0.148231 | 0.086500 | 0.026583 | 0.092166 | -1.501375 | 4.067346 | 1.545785722 | -0.37927902 | 0.704480678 | NA | 2.290730 | 1.696731 | 2.044312 | 2.033161 | 2.021954 | 2.040512 | 7 | 4 | 1 | 5 |
| 112268086 | XR_002957277 | LOC112268086 | uncharacterized LOC112268086 | lncRNA | . | . | . | . | . | 0.169943 | 0.058173 | 0.097270 | 0.092456 | 0.395464 | 0.134470 | 0.218270 | 0.211389 | -1.199355 | 10.951456 | 0.958412131 | -0.27363874 | 0.784362261 | NA | 3.576283 | 3.318300 | 3.482968 | 3.433694 | 3.451468 | 3.448138 | 20 | 7 | 9 | 9 |
| 112268089 | XR_002957397 | LOC112268089 | uncharacterized LOC112268089 | lncRNA | . | . | . | . | . | 0.172677 | 0.216976 | 0.253806 | 0.196364 | 0.401827 | 0.501550 | 0.569534 | 0.448964 | 1.143696 | 26.042325 | 0.608140039 | 0.318518765 | 0.750091463 | NA | 4.602488 | 4.796563 | 4.679402 | 4.701521 | 4.717760 | 4.691867 | 25 | 29 | 27 | 23 |
| 112268093 | XR_002957409 | LOC112268093 | uncharacterized LOC112268093 | lncRNA | . | . | . | . | . | 2.297998 | 2.035126 | 2.247700 | 2.290507 | 5.347536 | 4.704285 | 5.043774 | 5.236971 | 1.034698 | 231.710370 | 0.224955566 | 0.218750336 | 0.826844536 | 0.998865404 | 7.831138 | 7.880792 | 7.877728 | 7.815744 | 7.861432 | 7.866018 | 279 | 230 | 204 | 222 |
| 112268097 | XR_002957414 | MYG1-AS1 | MYG1 antisense RNA 1 | lncRNA | . | 54810 | . | . | . | 0.655229 | 0.658908 | 0.707369 | 1.097196 | 1.524745 | 1.523096 | 1.587315 | 2.508607 | 1.349961 | 26.309378 | 0.609971331 | 0.709734994 | 0.477868488 | NA | 4.486115 | 4.916862 | 4.694594 | 4.693377 | 4.701930 | 4.750164 | 26 | 24 | 21 | 34 |
| 112268098 | XR_002957416 | LOC112268098 | uncharacterized LOC112268098 | lncRNA | . | . | . | . | . | 0.325362 | 0.392344 | 0.495249 | 0.341035 | 0.757131 | 0.906921 | 1.111324 | 0.779736 | 1.114700 | 4.748118 | 1.375504002 | 0.113889582 | 0.909325324 | NA | 2.166062 | 2.324312 | 2.241668 | 2.243064 | 2.248009 | 2.241424 | 5 | 5 | 5 | 4 |
| 112268105 | NR_157805,NF | SBNO1-AS1 | SBNO1 antisense RNA 1 | lncRNA | . | 54355 | . | . | . | 3.510773 | 1.410341 | 3.285018 | 3.259087 | 8.169712 | 3.260067 | 7.371485 | 7.451515 | 1.278854 | 41.392210 | 0.535053225 | 0.663208159 | 0.507197247 | 0.998865404 | 5.180967 | 5.539401 | 5.398485 | 5.277945 | 5.381285 | 5.376907 | 59 | 23 | 41 | 43 |
| 112268113 | gene-LOC1122 | LOC112268113 | . | pseudogene | . | . | . | . | . | 0.023890 | 0.026246 | 0.033301 | 0.009214 | 0.055593 | 0.060668 | 0.074726 | 0.021066 | 1.001795 | 4.502278 | 1.443995283 | 0.001792081 | 0.998570127 | NA | 2.166062 | 2.175234 | 2.163703 | 2.165080 | 2.175005 | 2.153702 | 5 | 5 | 6 | 2 |
| 112268124 | NR_160776 | LOC112268124 | uncharacterized LOC112268124 | lncRNA | . | . | . | . | . | 0.296490 | 0.329239 | 0.351723 | 0.261644 | 0.689944 | 0.761051 | 0.789257 | 0.598219 | -1.030295 | 22.576464 | 0.650097431 | -0.06623221 | 0.947192951 | NA | 4.517478 | 4.475715 | 4.491702 | 4.501911 | 4.506397 | 4.481435 | 25 | 26 | 22 | 18 |
| 112268131 | XM_02444979 | LOC112268131 | uncharacterized LOC112268131, transcript variant X7 | protein_coding | XP_024305567.1;XP_024305568.1 | . | . | . | . | 0.137568 | 0.225243 | 0.168353 | 0.243481 | 0.320128 | 0.520659 | 0.377781 | 0.556688 | -1.019982 | 16.113651 | 0.788258413 | -0.03621086 | 0.971114223 | NA | 4.027016 | 3.993209 | 3.985141 | 4.029584 | 3.992405 | 4.019977 | 13 | 23 | 11 | 18 |
| 112268133 | XR_002957606 | LOC112268133 | uncharacterized LOC112268133 | lncRNA | . | . | . | . | . | 0.063109 | 0.016797 | 0.084390 | 0.056221 | 0.146858 | 0.038828 | 0.189369 | 0.128544 | 1.762385 | 9.150308 | 1.063969934 | 0.768375911 | 0.442263887 | NA | 2.722634 | 3.548418 | 3.184087 | 3.148507 | 3.198090 | 3.180144 | 12 | 3 | 12 | 9 |
| 112268148 | XR_002957712 | LOC112268148 | uncharacterized LOC112268148, transcript variant X8 | lncRNA | . | . | . | . | . | 0.329315 | 0.281236 | 0.405102 | 0.530667 | 0.766330 | 0.650089 | 0.909038 | 1.213305 | 1.446233 | 12.603799 | 0.858776138 | 0.619835443 | 0.535366132 | NA | 3.365657 | 3.897226 | 3.638171 | 3.637288 | 3.653238 | 3.662805 | 12 | 11 | 12 | 15 |
| 112268151 | XR_002957720 | LOC112268151 | uncharacterized LOC112268151 | lncRNA | . | . | . | . | . | 0.088663 | 0.028213 | 0.110635 | 0.101810 | 0.206322 | 0.065216 | 0.248263 | 0.232777 | 1.752123 | 18.252712 | 0.772989171 | 1.046721003 | 0.295228274 | NA | 3.725816 | 4.540693 | 4.181254 | 4.120512 | 4.197765 | 4.191000 | 23 | 7 | 21 | 21 |
| 112268155 | XR_002957735 | LOC112268155 | vesicle-associated membrane protein-associated protein A pseudo | transcribed_ps | . | . | . | . | . | 0.025958 | 0.010203 | 0.025072 | 0.032083 | 0.060406 | 0.023584 | 0.056262 | 0.073355 | 1.375174 | 4.747697 | 1.389328296 | 0.330817656 | 0.740782224 | NA | 1.998355 | 2.459419 | 2.244817 | 2.233361 | 2.241959 | 2.250038 | 6 | 3 | 4 | 6 |
| 112268170 | XR_002957880 | LOC112268170 | uncharacterized LOC112268170 | transcribed_ps | . | . | . | . | . | 0.580253 | 0.381081 | 0.698321 | 0.787518 | 1.350273 | 0.880885 | 1.567012 | 1.800566 | 1.535771 | 28.248954 | 0.59237309 | 1.044887946 | 0.296074762 | 0.998865404 | 4.476701 | 5.097315 | 4.803920 | 4.764131 | 4.825002 | 4.841344 | 31 | 19 | 28 | 34 |
| 112268177 | XR_002957914 | LOC112268177 | uncharacterized LOC112268177 | lncRNA | . | . | . | . | . | 0.045612 | 0.029155 | 0.185965 | 0.058868 | 0.106142 | 0.067394 | 0.417299 | 0.134594 | **3.153234** | 7.426348 | 1.199030378 | 1.381809953 | 0.167030062 | NA | 1.832372 | 3.497162 | 2.838485 | 2.831283 | 2.895500 | 2.843715 | 5 | 3 | 15 | 5 |
| 112268179 | XR_002957933 | LOC112268179 | uncharacterized LOC112268179 | lncRNA | . | . | . | . | . | 0.129904 | 0.156994 | 0.135127 | 0.141727 | 0.302293 | 0.362898 | 0.303220 | 0.324043 | 1.000941 | 7.182003 | 1.125027953 | 0.001205684 | 0.999038003 | NA | 2.844134 | 2.844638 | 2.843352 | 2.845521 | 2.843223 | 2.845716 | 8 | 8 | 6 | 7 |
| 112268184 | XM_02445105 | LOC112268184 | uncharacterized LOC112268184 | protein_coding | XP_024306822.1 | . | . | . | . | 0.231163 | 0.206652 | 0.105959 | 0.227710 | 0.537925 | 0.477686 | 0.237769 | 0.520632 | -1.334410 | 19.146525 | 0.729848558 | -0.57025732 | 0.568503186 | NA | 4.454055 | 4.033413 | 4.281412 | 4.268089 | 4.226124 | 4.276235 | 27 | 22 | 9 | 21 |
| 112268190 | XR_002958108 | LOC112268190 | uncharacterized LOC112268190 | lncRNA | . | . | . | . | . | 0.103602 | 0.040087 | 0.116527 | 0.137040 | 0.241085 | 0.092662 | 0.261483 | 0.313327 | 1.617170 | 5.806461 | 1.269262177 | 0.546357857 | 0.584819973 | NA | 2.147197 | 2.844638 | 2.529670 | 2.514159 | 2.533164 | 2.535804 | 7 | 3 | 6 | 7 |
| 112268192 | gene-LOC1122 | LOC112268192 | . | pseudogene | . | . | . | . | . | 0.066264 | 0.170887 | 0.070200 | 0.083311 | 0.154200 | 0.395014 | 0.157527 | 0.190480 | -1.433783 | 5.401945 | 1.329715414 | -0.39093046 | 0.695848638 | NA | 2.671431 | 2.148375 | 2.424803 | 2.451754 | 2.430431 | 2.428855 | 4 | 10 | 4 | 4 |
| 112268193 | gene-LOC1122 | LOC112268193 | . | pseudogene | . | . | . | . | . | 0.230710 | 0.220260 | 0.568208 | 0.118895 | 0.536871 | 0.509140 | 1.275042 | 0.271839 | 1.498972 | 11.788283 | 0.965024305 | 0.605139025 | 0.545086626 | NA | 3.234218 | 3.824402 | 3.525286 | 3.524125 | 3.582462 | 3.506975 | 11 | 10 | 20 | 5 |
| 112268202 | XR_002958144 | LOC112268202 | uncharacterized LOC112268202 | lncRNA | . | . | . | . | . | 0.041406 | 0.256913 | 0.257691 | 0.240663 | 0.096354 | 0.593866 | 0.578251 | 0.550247 | 1.590730 | 17.175068 | 0.839034183 | 0.798166047 | 0.424774138 | NA | 3.732491 | 4.396303 | 4.020822 | 4.102461 | 4.099605 | 4.093364 | 5 | 24 | 19 | 19 |
| 112268203 | XR_002958145 | LOC112268203 | uncharacterized LOC112268203 | lncRNA | . | . | . | . | . | 0.516123 | 0.212106 | 0.470313 | 0.399186 | 1.201040 | 0.490291 | 1.055369 | 0.912690 | 1.252317 | 5.490238 | 1.307232634 | 0.248310521 | 0.803894159 | NA | 2.282109 | 2.612728 | 2.458597 | 2.438999 | 2.457891 | 2.451008 | 8 | 3 | 6 | 5 |
| 112268218 | XR_002958243 | LOC112268218 | uncharacterized LOC112268218 | lncRNA | . | . | . | . | . | 0.021355 | 0.076824 | 0.020392 | 0.028962 | 0.049695 | 0.177582 | 0.045759 | 0.066217 | -1.641802 | 2.932659 | 1.783481861 | -0.40105847 | 0.688377074 | NA | 1.867373 | 1.148375 | 1.549439 | 1.567461 | 1.552366 | 1.551546 | 2 | 6 | 2 | 2 |
| 112268219 | NR_168005 | LOC112268219 | uncharacterized LOC112268219 | lncRNA | . | . | . | . | . | 1.229184 | 1.004384 | 1.243278 | 0.964506 | 2.860363 | 2.321679 | 2.789880 | 2.205229 | -1.024402 | 92.794023 | 0.342386832 | -0.10158728 | 0.919084275 | 0.998865404 | 6.552119 | 6.519618 | 6.567040 | 6.506605 | 6.565862 | 6.493458 | 119 | 91 | 90 | 75 |
| 112268225 | XR_002958304 | LOC112268225 | uncharacterized LOC112268225 | lncRNA | . | . | . | . | . | 0.012601 | 0.054857 | 0.044528 | 0.128159 | 0.029324 | 0.126804 | 0.099920 | 0.293020 | **2.504829** | 8.777694 | 1.136208041 | 1.165906269 | 0.243652344 | NA | 2.333078 | 3.645770 | 3.073468 | 3.104953 | 3.098730 | 3.147646 | 2 | 9 | 6 | 17 |
| 112268230 | XR_002958326 | LOC112268230 | uncharacterized LOC112268230, transcript variant X2 | lncRNA | . | . | . | . | . | 0.481948 | 0.440171 | 1.043511 | 0.306765 | 1.121514 | 1.017476 | 2.341608 | 0.701381 | 1.370542 | 11.697073 | 0.942643405 | 0.482415956 | 0.629510489 | NA | 3.299300 | 3.760196 | 3.525065 | 3.519876 | 3.568697 | 3.507519 | 12 | 10 | 18 | 6 |
| 112268235 | XR_002958352 | LOC112268235 | uncharacterized LOC112268235 | lncRNA | . | . | . | . | . | 0.022982 | 0.030446 | 0.036415 | 0.029947 | 0.053479 | 0.070378 | 0.081713 | 0.068470 | 1.235159 | 10.064973 | 0.960867344 | 0.317106031 | 0.751163149 | NA | 3.170740 | 3.475715 | 3.314122 | 3.325172 | 3.335632 | 3.322530 | 9 | 11 | 11 | 9 |
| 112268254 | XR_002958451 | LOC112268254 | uncharacterized LOC112268254 | lncRNA | . | . | . | . | . | 0.682140 | 0.681403 | 1.110944 | 0.629330 | 1.587368 | 1.575093 | 2.492927 | 1.438888 | 1.245164 | 53.398320 | 0.454975729 | 0.695279668 | 0.486880054 | 0.998865404 | 5.570285 | 5.889534 | 5.705879 | 5.704226 | 5.804263 | 5.687735 | 55 | 51 | 66 | 40 |
| 112268256 | XR_002958489 | LOC112268256 | uncharacterized LOC112268256 | lncRNA | . | . | . | . | . | 1.300144 | 1.171288 | 1.759240 | 0.883020 | 3.025488 | 2.707485 | 3.947684 | 2.018920 | 1.057918 | 66.404571 | 0.427379666 | 0.190059899 | 0.849262193 | 0.998865404 | 6.010149 | 6.095024 | 6.048047 | 6.024715 | 6.122139 | 5.970633 | 78 | 66 | 79 | 43 |
| 112268259 | XR_002958497 | LOC112268259 | uncharacterized LOC112268259 | lncRNA | . | . | . | . | . | 4.256510 | 4.303800 | 5.893358 | 5.714845 | 9.905073 | 9.948426 | 13.224524 | 13.066315 | 1.338364 | 244.299371 | 0.217564041 | 1.932626425 | 0.053282235 | 0.683508073 | 7.706912 | 8.127548 | 7.849061 | 7.851854 | 8.004252 | 7.984169 | 240 | 226 | 248 | 257 |
| 112268260 | XM_02445196 | LOC112268260 | uncharacterized LOC112268260, transcript variant X6 | protein_coding | XP_024307730.1;XP_024307731.1 | . | . | . | . | 0.191524 | 0.189801 | 0.195206 | 0.188783 | 0.445685 | 0.438732 | 0.438036 | 0.431631 | 1.095697 | 12.203039 | 0.870752421 | 0.151419138 | 0.879645094 | NA | 3.540965 | 3.674293 | 3.606639 | 3.601963 | 3.613601 | 3.604799 | 14 | 12 | 12 | 11 |
| 112268269 | NR_165198,NF | LOC112268269 | uncharacterized LOC112268269 | lncRNA | . | . | . | . | . | 3.795974 | 3.638552 | 5.307806 | 3.713125 | 8.833388 | 8.410674 | 11.910561 | 8.489621 | 1.165978 | 470.208174 | 0.19919299 | 1.112189858 | 0.266056528 | 0.998274218 | 8.761544 | 8.984186 | 8.820100 | 8.822130 | 9.024972 | 8.796461 | 499 | 469 | 519 | 390 |
| 112268271 | XM_02445202 | LOC112268271 | translation initiation factor IF-2 | protein_coding | XP_024307795.1 | . | . | . | . | 1.158734 | 1.152951 | 1.648398 | 1.264983 | 2.696422 | 2.665097 | 3.698958 | 2.892232 | 1.241990 | 302.606635 | 0.214488367 | 1.457669627 | 0.144931628 | 0.906189003 | 8.075884 | 8.389685 | 8.177531 | 8.170242 | 8.363646 | 8.212434 | 313 | 289 | 331 | 272 |
| 112268272 | XR_002958600 | LOC112268272 | uncharacterized LOC112268272, transcript variant X3 | lncRNA | . | . | . | . | . | 0.072485 | 0.031057 | 0.039464 | 0.048152 | 0.168675 | 0.071791 | 0.088555 | 0.110093 | -1.265622 | 8.736757 | 1.046650593 | -0.32469966 | 0.745408389 | NA | 3.286426 | 2.947964 | 3.151377 | 3.121416 | 3.123795 | 3.130834 | 15 | 7 | 6 | 8 |
| 112268275 | XR_002958629 | LINC02799 | long intergenic non-protein coding RNA 2799 | lncRNA | . | 54324 | . | . | . | 0.183330 | 0.282219 | 0.313900 | 0.182591 | 0.426617 | 0.652361 | 0.704383 | 0.417472 | 1.126252 | 5.267298 | 1.314790513 | 0.130461513 | 0.896201302 | NA | 2.307819 | 2.481107 | 2.387070 | 2.392839 | 2.398631 | 2.386952 | 5 | 6 | 6 | 4 |

| | | | | | | | | | | | | | | | | | | | | | | | | | | | | | | | | | |
|---|---|---|---|---|---|---|---|---|---|---|---|---|---|---|---|---|---|---|---|---|---|---|---|---|---|---|---|---|---|---|---|---|---|
| 112268288 | XR_002958732 | LOC112268288 | uncharacterized LOC112268288, transcript variant X1 | lncRNA | . | . | . | . | . | 0.037121 | 0.093433 | 0.036027 | 0.029729 | 0.086382 | 0.215975 | 0.080845 | 0.067971 | -1.814482 | 7.050405 | 1.18511069 | -0.72529715 | 0.468269751 | NA | 3.184683 | 2.324312 | 2.810776 | 2.846273 | 2.813752 | 2.806994 | 6 | 14 | 5 | 4 |
| 112268291 | XR_002958737 | LOC112268291 | uncharacterized LOC112268291 | lncRNA | . | . | . | . | . | 0.045914 | 0.055383 | 0.180547 | 0.107176 | 0.106843 | 0.128019 | 0.405142 | 0.245045 | **2.754437** | 7.610130 | 1.134564701 | 1.288386125 | 0.197611586 | NA | 2.019256 | 3.481107 | 2.880015 | 2.885808 | 2.926519 | 2.903856 | 4 | 5 | 12 | 8 |
| 112268292 | XR_002958738 | LOC112268292 | uncharacterized LOC112268292, transcript variant X1 | lncRNA | . | . | . | . | . | 0.352627 | 0.247740 | 0.567907 | 0.221995 | 0.820578 | 0.572662 | 1.274367 | 0.507565 | 1.284491 | 23.470068 | 0.688422188 | 0.524672902 | 0.599810598 | NA | 4.357474 | 4.724718 | 4.540516 | 4.507499 | 4.593611 | 4.501045 | 28 | 18 | 33 | 14 |
| 112268294 | XR_002958749 | LOC112268294 | uncharacterized LOC112268294, transcript variant X1 | lncRNA | . | . | . | . | . | 0.555452 | 0.392995 | 0.218879 | 0.311543 | 1.292560 | 0.908425 | 0.491158 | 0.712306 | -1.945305 | 4.053192 | 1.515469114 | -0.6334646 | 0.526430288 | NA | 2.421266 | 1.459419 | 2.041460 | 2.034627 | 2.024211 | 2.028099 | 7 | 5 | 2 | 3 |
| 112268301 | XR_002958812 | LOC112268301 | uncharacterized LOC112268301 | lncRNA | . | . | . | . | . | 0.252091 | 0.245816 | 0.325190 | 0.141919 | 0.586627 | 0.568214 | 0.729716 | 0.324481 | -1.084212 | 44.426259 | 0.516029806 | -0.22604691 | 0.821164935 | 0.998865404 | 5.528358 | 5.416142 | 5.474314 | 5.468468 | 5.519733 | 5.391490 | 54 | 49 | 52 | 24 |
| 112268302 | XR_002958814 | LOC112268302 | uncharacterized LOC112268302 | lncRNA | . | . | . | . | . | 0.052984 | 0.073163 | 0.143278 | 0.048799 | 0.123297 | 0.169119 | 0.321511 | 0.111573 | 1.426913 | 3.836371 | 1.562203198 | 0.328317003 | 0.74267199 | NA | 1.658167 | 2.175234 | 1.918311 | 1.923628 | 1.937784 | 1.916451 | 3 | 4 | 6 | 2 |
| 112268418 | XR_002959404 | LOC112268418 | uncharacterized LOC112268418 | lncRNA | . | . | . | . | . | 0.052429 | 0.029376 | 0.149803 | 0.057778 | 0.122004 | 0.067903 | 0.336153 | 0.132102 | **2.512512** | 6.276374 | 1.257012866 | 1.057372033 | 0.29034183 | NA | 1.832372 | 3.168566 | 2.612769 | 2.605497 | 2.649745 | 2.617724 | 5 | 3 | 11 | 5 |
| 112268420 | XR_002959408 | LOC112268420 | uncharacterized LOC112268420 | lncRNA | . | . | . | . | . | 0.073921 | 0.089324 | 0.339047 | 0.115745 | 0.172018 | 0.206477 | 0.760813 | 0.264637 | **2.676009** | 42.199001 | 0.573355402 | 2.476793357 | **0.013256863** | 0.381141357 | 4.519313 | 5.941708 | 5.244559 | 5.263967 | 5.509491 | 5.290379 | 24 | 27 | 80 | 29 |
| 112268421 | XR_002959410 | LOC112268421 | uncharacterized LOC112268421 | lncRNA | . | . | . | . | . | 0.003884 | 0.032746 | 0.037528 | 0.012850 | 0.009038 | 0.075693 | 0.084212 | 0.029381 | 1.345315 | 5.906839 | 1.339762179 | 0.319418031 | 0.74940954 | NA | 2.333078 | 2.760196 | 2.521824 | 2.553151 | 2.561219 | 2.529552 | 2 | 9 | 9 | 3 |
| 112268422 | XR_002959415 | LOC112268422 | uncharacterized LOC112268422 | lncRNA | . | . | . | . | . | 0.104842 | 0.014213 | 0.023922 | 0.021331 | 0.243971 | 0.032854 | 0.053681 | 0.048772 | **-2.476979** | 7.688131 | 1.234459561 | -1.06004424 | 0.289124476 | NA | 3.451934 | 2.148375 | 2.997474 | 2.922826 | 2.932163 | 2.930372 | 22 | 3 | 4 | 4 |
| 112268423 | XR_002959416 | LOC112268423 | uncharacterized LOC112268423 | lncRNA | . | . | . | . | . | 0.101267 | 0.308802 | 0.387063 | 0.386619 | 0.235653 | 0.713810 | 0.868558 | 0.883959 | 1.829620 | 19.292175 | 0.751855847 | 1.159190755 | 0.246378439 | NA | 3.772726 | 4.639023 | 4.192165 | 4.253427 | 4.273173 | 4.270523 | 8 | 22 | 22 | 23 |
| 112268435 | gene-PPIAP61 | PPIAP61 | peptidylprolyl isomerase A pseudogene 61 | pseudogene | . | 53685 | . | . | . | 0.036176 | 0.130687 | 0.190781 | 0.069349 | 0.084183 | 0.302089 | 0.428107 | 0.158558 | 1.236277 | 3.044274 | 1.727664661 | 0.177118846 | 0.859415039 | NA | 1.444631 | 1.751299 | 1.589015 | 1.598466 | 1.602247 | 1.591161 | 2 | 4 | 4 | 2 |
| 112268450 | XR_002959658 | LOC112268450 | uncharacterized LOC112268450 | lncRNA | . | . | . | . | . | 0.013400 | 0.061521 | 0.104600 | 0.058159 | 0.031182 | 0.142210 | 0.234720 | 0.132974 | **2.044542** | 16.674931 | 0.843505956 | 1.223201469 | 0.221253611 | NA | 3.456173 | 4.483795 | 3.970953 | 4.032608 | 4.078071 | 4.025486 | 5 | 19 | 25 | 15 |
| 112268458 | XR_002959697 | LOC112268458 | uncharacterized LOC112268458, transcript variant X1 | lncRNA | . | . | . | . | . | 0.602679 | 0.711170 | 0.882111 | 0.562470 | 1.402458 | 1.643900 | 1.979433 | 1.286021 | 1.074184 | 93.721089 | 0.354812846 | 0.29097196 | 0.771072765 | 0.998865404 | 6.496816 | 6.601875 | 6.512424 | 6.551668 | 6.619147 | 6.482565 | 97 | 104 | 104 | 70 |
| 112268469 | XR_002959800 | LOC112268469 | uncharacterized LOC112268469 | lncRNA | . | . | . | . | . | 0.526856 | 0.533244 | 0.411749 | 0.763583 | 1.226016 | 1.232618 | 0.923952 | 1.745842 | 1.077470 | 23.251082 | 0.654648986 | 0.164436111 | 0.869387838 | NA | 4.486115 | 4.590451 | 4.534298 | 4.532898 | 4.512736 | 4.568252 | 26 | 24 | 15 | 29 |
| 112441434 | NM_00136534 | SHLD3 | shieldin complex subunit 3 | protein_coding | NP_001352270.1 | 53826 | 618030 | . | . | 2.512853 | 1.774232 | 1.962603 | 1.443431 | 5.847513 | 4.101216 | 4.404024 | 3.300235 | -1.277686 | 93.067210 | 0.355172295 | -0.99538607 | 0.319548531 | 0.998865404 | 6.704855 | 6.354305 | 6.625759 | 6.513647 | 6.539427 | 6.459477 | 141 | 93 | 82 | 65 |
| 112543491 | NR_156734 | LOC112543491 | linc-UFC1 | lncRNA | . | . | . | . | . | 0.110228 | 0.038705 | 0.061490 | 0.014743 | 0.256505 | 0.089468 | 0.137981 | 0.033708 | -1.805704 | 5.502872 | 1.355749281 | -0.62884907 | 0.529447873 | NA | 2.820514 | 1.978782 | 2.489160 | 2.457988 | 2.467414 | 2.451201 | 12 | 4 | 5 | 2 |
| 112637020 | NR_157279 | SREBF2-AS1 | SREBF2 antisense RNA 1 | lncRNA | . | 53953 | . | . | . | 0.066590 | 0.075182 | 0.055983 | 0.029060 | 0.154958 | 0.173787 | 0.125624 | 0.066442 | -1.656032 | 12.623660 | 0.882562024 | -0.82456629 | 0.409617868 | NA | 3.974755 | 3.251698 | 3.671081 | 3.679386 | 3.661543 | 3.634241 | 17 | 18 | 11 | 6 |
| 112694756 | NM_00136530 | LOC112694756 | uncharaterized LOC112694756 | protein_coding | NP_001352233.1;NP_001352234.1;NP_001352236.1 | . | . | . | . | 45.678004 | 36.190074 | 8.873032 | 35.662314 | 106.294595 | 83.654978 | 19.910828 | 81.537642 | -1.892453 | 2514.434646 | 0.954448048 | -0.96417773 | 0.334956826 | 0.998865404 | 11.684000 | 10.763691 | 11.552202 | 11.340045 | 10.297482 | 11.317464 | 4248 | 3120 | 574 | 2641 |
| 112806053 | NR_157393 | TBILA | TGF-beta induced lncRNA | lncRNA | . | 53943 | . | . | . | 0.031176 | 0.016704 | 0.024496 | 0.054028 | 0.072548 | 0.038613 | 0.054969 | 0.123529 | 1.474352 | 2.756358 | 1.797441179 | 0.31159909 | 0.755345226 | NA | 1.156660 | 1.715150 | 1.456815 | 1.453257 | 1.455501 | 1.464401 | 3 | 2 | 2 | 4 |
| 113218477 | NR_158654,NF | TIMM23B-AGA | TIMM23B-AGAP6 readthrough (NMD candidate) | lncRNA | . | 45009 | . | . | . | 2.054249 | 4.243038 | 3.419832 | 1.482805 | 4.780321 | 9.807972 | 7.674004 | 3.390255 | -1.337663 | 284.788659 | 0.731277542 | -0.57394651 | 0.566003997 | 0.998865404 | 8.348312 | 7.928792 | 7.978424 | 8.369203 | 8.209563 | 7.830797 | 249 | 470 | 293 | 141 |
| 113218482 | NR_162090,rn | MIR9898 | microRNA 9898 | miRNA | . | 53990 | . | . | . | 0.000054 | 0.000067 | 0.000174 | 0.000078 | 0.000126 | 0.000155 | 0.000389 | 0.000177 | 1.234655 | 1.002957 | 2.868339486 | 0.106022226 | 0.915564717 | NA | -0.155866 | 0.148375 | -0.001261 | -0.000972 | 0.000053 | -0.000315 | 1 | 1 | 1 | 1 |
| 113218486 | NR_162101,rn | MIR10392 | hsa-miR-10392-3p | . | . | 54001 | . | . | . | 0.001166 | 0.001867 | 0.001967 | 0.001293 | 0.002712 | 0.004315 | 0.004413 | 0.002955 | 1.234665 | 3.008871 | 1.708874417 | 0.17796512 | 0.858750369 | NA | 1.429096 | 1.733338 | 1.582415 | 1.583274 | 1.586316 | 1.585224 | 3 | 3 | 3 | 3 |
| 113218487 | NR_162103,rn | MIR10394 | hsa-miR-10394-5p | . | . | 54003 | . | . | . | 0.113156 | 0.054876 | 0.014933 | 0.064261 | 0.263316 | 0.126848 | 0.033509 | 0.146926 | 1.234665 | 3.008871 | 1.708874417 | 0.17796512 | 0.858750369 | NA | 1.429096 | 1.733338 | 1.582415 | 1.583274 | 1.586316 | 1.585224 | 3 | 3 | 3 | 3 |
| 113218488 | NR_162111,rn | MIR10396B | hsa-miR-10396b-5p | . | . | 54006 | . | . | . | 0.000081 | 0.000083 | 0.000021 | 0.000004 | 0.000191 | 0.000193 | 0.000048 | 0.00001 | 1.034628 | 2.742204 | 1.790997379 | 0.027421635 | 0.978123443 | NA | 1.429096 | 1.481107 | 1.451689 | 1.452527 | 1.455495 | 1.449568 | 3 | 3 | 3 | 2 |
| 113218500 | NR_162102,rn | MIR10393 | hsa-miR-10393-5p | . | . | 54002 | . | . | . | 0.107053 | 0.016249 | 0.021233 | 0.007547 | 0.249117 | 0.037559 | 0.047647 | 0.017256 | 1.234665 | 3.008871 | 1.708874417 | 0.17796512 | 0.858750369 | NA | 1.429096 | 1.733338 | 1.582415 | 1.583274 | 1.586316 | 1.585224 | 3 | 3 | 3 | 3 |
| 113218502 | NR_162106,rn | MIR10397 | hsa-miR-10397-5p | . | . | 54007 | . | . | . | 0.211477 | 0.001054 | 0.276113 | 0.000931 | 0.492114 | 0.002435 | 0.619587 | 0.002129 | 1.234665 | 3.008871 | 1.708874417 | 0.17796512 | 0.858750369 | NA | 1.429096 | 1.733338 | 1.582415 | 1.583274 | 1.586316 | 1.585224 | 3 | 3 | 3 | 3 |
| 113218503 | NR_162108,rn | MIR10399 | hsa-miR-10399-3p | . | . | 54009 | . | . | . | 0.006949 | 0.003266 | 0.003733 | 0.002479 | 0.016172 | 0.007551 | 0.008377 | 0.005667 | 1.234665 | 3.008871 | 1.708874417 | 0.17796512 | 0.858750369 | NA | 1.429096 | 1.733338 | 1.582415 | 1.583274 | 1.586316 | 1.585224 | 3 | 3 | 3 | 3 |
| 113218510 | NR_162113,rn | MIR10523 | hsa-miR-10523-5p | . | . | 54013 | . | . | . | 1.100502 | 0.673010 | 0.419458 | 3.100434 | 2.560913 | 1.555694 | 0.941253 | 7.088775 | 1.395256 | 3.738489 | 1.570130403 | 0.306044729 | 0.75957059 | NA | 1.644780 | 2.121007 | 1.897160 | 1.893908 | 1.892025 | 1.910409 | 4 | 3 | 2 | 6 |
| 113219453 | NR_162133,rn | MIR12119 | microRNA 12119 | miRNA | . | 54027 | . | . | . | 0.012732 | 0.011182 | 0.017215 | 0.011778 | 0.029629 | 0.025847 | 0.038629 | 0.026930 | 1.234663 | 2.005914 | 2.070995667 | 0.146845664 | 0.883253841 | NA | 0.844134 | 1.148375 | 0.998092 | 0.998667 | 1.000707 | 0.999974 | 2 | 2 | 2 | 2 |
| 113219455 | NR_162136,rn | MIR12122 | microRNA 12122 | miRNA | . | 54030 | . | . | . | 1.928911 | 2.817620 | 4.011132 | 2.064969 | 4.488657 | 6.513053 | 9.000863 | 4.721306 | 1.309733 | 9.370336 | 1.006887163 | 0.386610044 | 0.699044921 | NA | 3.019256 | 3.410501 | 3.206132 | 3.216991 | 3.236414 | 3.209011 | 8 | 10 | 12 | 7 |
| 113219467 | NR_162149,rn | MIR12136 | microRNA 12136 | miRNA | . | 54043 | . | . | . | 1.642253 | 2.434159 | 3.770444 | 3.326475 | 3.821591 | 5.626668 | 8.460768 | 7.605590 | 1.614486 | 7.645243 | 1.09877429 | 0.628950417 | 0.529381518 | NA | 2.548173 | 3.239024 | 2.910909 | 2.917042 | 2.934134 | 2.926355 | 6 | 7 | 9 | 8 |
| 113219470 | NR_162141,rn | MIR12127 | microRNA 12127 | miRNA | . | 54035 | . | . | . | 0.005350 | 0.006661 | 0.005569 | 0.002109 | 0.012449 | 0.015396 | 0.012498 | 0.004822 | 1.234663 | 2.005914 | 2.070995667 | 0.146845664 | 0.883253841 | NA | 0.844134 | 1.148375 | 0.998092 | 0.998667 | 1.000707 | 0.999974 | 2 | 2 | 2 | 2 |
| 113455421 | NM_00100284 | DERPC | DERPC proline and glycine rich nuclear protein | protein_coding | NP_001002847.1;NP_001035234.2;NP_001353531.1;NP_001353532.1;NI | 54084 | . | . | . | 11.304711 | 12.056260 | 10.138299 | 15.282741 | 26.306525 | 27.868584 | 22.750053 | 34.942173 | 1.108078 | 926.283418 | 0.183885385 | 0.805170089 | 0.420721534 | 0.998865404 | 9.779815 | 9.927050 | 9.802338 | 9.819763 | 9.719946 | 10.027837 | 1000 | 960 | 670 | 1103 |
| 113523640 | NR_159947,NF | DCXR-DT | DCXR divergent transcript | lncRNA | . | 54083 | . | . | . | 0.713699 | 0.660530 | 0.357767 | 0.585273 | 1.660806 | 1.526846 | 0.802818 | 1.338157 | -1.580855 | 12.777868 | 0.864398321 | -0.76435238 | 0.444657279 | NA | 3.969395 | 3.306201 | 3.699527 | 3.691927 | 3.662729 | 3.678338 | 19 | 16 | 7 | 11 |
| 113939912 | NR_160414,NF | BMS1P2-AGAP | BMS1P2-AGAP9 readthrough | transcribed_ps | . | . | . | . | . | 0.271136 | 0.354105 | 0.820954 | 0.226084 | 0.630945 | 0.818529 | 1.842197 | 0.516913 | 1.641789 | 50.011386 | 0.559271222 | 1.278930383 | 0.200921572 | 0.96631375 | 5.240907 | 5.959011 | 5.530859 | 5.570531 | 5.760419 | 5.508219 | 38 | 46 | 85 | 25 |
| 113939925 | NR_160425,NF | BMS1P4-AGAP | BMS1P4-AGAP5 readthough | transcribed_ps | . | . | . | . | . | 0.969230 | 0.581101 | 1.335126 | 0.982052 | 2.255438 | 1.343242 | 2.995986 | 2.245345 | 1.474069 | 136.620870 | 0.317444438 | 1.763472495 | 0.077820832 | 0.76835547 | 6.785425 | 7.348120 | 7.078482 | 6.918287 | 7.195473 | 7.090683 | 158 | 90 | 160 | 133 |
| 114004351 | NR_160518,NF | PARGP1-AGAP | PARGP1-AGAP4 readthrough | transcribed_ps | . | . | . | . | . | 0.910712 | 1.128988 | 1.577685 | 0.764281 | 2.119264 | 2.609705 | 3.540277 | 1.747438 | 1.133920 | 150.803549 | 0.333933157 | 0.542980309 | 0.58714336 | 0.998865404 | 7.142034 | 7.325209 | 7.153415 | 7.232175 | 7.373722 | 7.092231 | 146 | 168 | 188 | 98 |
| 114118903 | NM_00136831 | ARHGAP11A-S | ARHGAP11A-SCG5 readthrough | protein_coding | NP_001355248.1 | . | . | . | . | 0.042298 | 0.023775 | 0.027859 | 0.043171 | 0.098430 | 0.054957 | 0.062516 | 0.098706 | 1.082770 | 3.709759 | 1.552098074 | 0.07391741 | 0.941076103 | NA | 1.832372 | 1.947964 | 1.894413 | 1.887021 | 1.890247 | 1.893916 | 5 | 3 | 3 | 4 |
| 114841035 | NM_00137041 | LOC114841035 | Uncharacterized protein LOC114841035 | protein_coding | NP_001357343.1 | . | . | . | . | 3.457672 | 4.764260 | 4.871304 | 3.778432 | 8.046145 | 11.012801 | 10.931063 | 8.638937 | 1.038797 | 217.727799 | 0.254860332 | 0.215464029 | 0.829405554 | 0.998865404 | 7.738599 | 7.793640 | 7.675793 | 7.821785 | 7.828476 | 7.706504 | 208 | 266 | 218 | 181 |
| 114841040 | NR_163486 | LOC114841040 | uncharacterized LOC114841040 | lncRNA | . | . | . | . | . | 0.206111 | 0.054726 | 0.338479 | 0.146221 | 0.479628 | 0.126501 | 0.759537 | 0.334317 | 1.719259 | 8.420691 | 1.118158781 | 0.699173683 | 0.484443495 | NA | 2.624463 | 3.416142 | 3.060105 | 3.028459 | 3.082797 | 3.045771 | 11 | 3 | 13 | 6 |
| 115253422 | NM_00137136 | DUS4L-BCAP29 | DUS4L-BCAP29 readthrough | protein_coding | NP_001358293.1;NP_001358294.1;NP_001358295.1;NP_001358296.1 | 54422 | . | . | . | 1.161037 | 2.541231 | 2.998483 | 2.136474 | 2.701781 | 5.874170 | 6.728509 | 4.884792 | 1.385564 | 466.082549 | 0.598807454 | 0.785684041 | 0.432052615 | 0.998865404 | 8.610184 | 9.080532 | 8.476011 | 8.928225 | 9.038903 | 8.810878 | 281 | 580 | 552 | 420 |
| 115308161 | NR_163974 | LOC115308161 | uncharacterized LOC115308161 | lncRNA | . | . | . | . | . | 0.282811 | 0.214223 | 0.493677 | 0.636110 | 0.658112 | 0.495186 | 1.107797 | 1.454390 | **2.209869** | 17.197963 | 0.753720521 | 1.517752439 | 0.129076819 | NA | 3.421266 | 4.565787 | 4.061090 | 4.048310 | 4.098313 | 4.123333 | 14 | 10 | 18 | 25 |
| 115482706 | gene-H3P47 | H3P47 | H3 histone pseudogene 47 | pseudogene | . | 54475 | . | . | . | 0.318355 | 0.311454 | 0.447045 | 0.632553 | 0.740825 | 0.719939 | 1.003156 | 1.446257 | 1.591513 | 9.861595 | 0.971692881 | 0.689929239 | 0.490238687 | NA | 2.928838 | 3.597915 | 3.282849 | 3.281139 | 3.294461 | 3.309843 | 9 | 8 | 9 | 13 |
| 115694671 | NR_164166 | LINC01297 | long intergenic non-protein coding RNA 1297 | lncRNA | . | 50443 | . | . | . | 0.122689 | 0.109637 | 0.111269 | 0.118234 | 0.285502 | 0.253430 | 0.249685 | 0.270328 | -1.002996 | 13.851076 | 0.818552833 | -0.00527326 | 0.995792563 | NA | 3.793789 | 3.790061 | 3.796360 | 3.788330 | 3.792027 | 3.792310 | 17 | 14 | 12 | 13 |
| 115801414 | NR_164308,NF | LINC01297-DU | LINC01297-DUXAP10-NBEAP6 readthrough | lncRNA | . | . | . | . | . | 0.530553 | 0.336162 | 0.592253 | 0.805499 | 1.234618 | 0.777052 | 1.329000 | 1.841679 | 1.579707 | 122.005197 | 0.30955729 | 2.130970458 | **0.033091576** | 0.575268292 | 6.563187 | 7.223505 | 6.877589 | 6.784682 | 6.952118 | 7.024791 | 126 | 86 | 118 | 153 |
| 115804232 | NR_036442,NF | CEROX1 | cytoplasmic endogenous regulator of oxidative phosphorylation 1 | lncRNA | . | 53928 | . | . | . | 0.976733 | 1.095601 | 1.137645 | 0.805354 | 2.272898 | 2.532530 | 2.552844 | 1.841347 | 1.140792 | 98.341716 | 0.323474804 | 0.587481022 | 0.556880693 | 0.998865404 | 6.521098 | 6.712052 | 6.602575 | 6.590351 | 6.655676 | 6.616322 | 108 | 97 | 97 | 92 |
| 115830329 | NR_164368 | ZNNT1 | ZNF706 neighboring transcript 1 | lncRNA | . | 54438 | . | . | . | 0.612446 | 0.802849 | 0.836735 | 0.202489 | 1.425186 | 1.855822 | 1.877608 | 0.462967 | -1.369497 | 66.288825 | 0.516366147 | -0.87853561 | 0.379653123 | 0.998865404 | 6.257928 | 5.808617 | 6.030466 | 6.096517 | 6.104648 | 5.864409 | 77 | 93 | 78 | 21 |
| 116412618 | NR_165159,NF | PDCD6-AHRR | PDCD6-AHRR readthrough (NMD candidate) | lncRNA | . | 54724 | . | . | . | 0.014978 | 0.159080 | 0.014425 | 0.050542 | 0.034854 | 0.367721 | 0.032370 | 0.115559 | **-2.520036** | 11.544882 | 1.097917418 | -1.21452132 | 0.224548715 | NA | 4.050049 | 2.705970 | 3.471627 | 3.588817 | 3.474237 | 3.501071 | 4 | 32 | 3 | 9 |
| 116804918 | NM_00137876 | TBCEL-TECTA | TBCEL-TECTA readthrough | protein_coding | NP_001365690.1 | 54857 | . | . | . | 0.025289 | 0.047690 | 0.067519 | 0.048109 | 0.058848 | 0.110238 | 0.151511 | 0.109995 | 1.567679 | 12.780389 | 0.875226748 | 0.741099296 | 0.458633237 | NA | 3.316288 | 3.963455 | 3.633728 | 3.661644 | 3.684379 | 3.660833 | 8 | 14 | 16 | 12 |
